# Genomic Signature of Sexual Reproduction in the Bdelloid Rotifer *Macrotrachella quadricornifera*

**DOI:** 10.1101/2020.08.06.239590

**Authors:** Veronika N. Laine, Timothy Sackton, Matthew Meselson

## Abstract

Bdelloid rotifers, common freshwater invertebrates of ancient origin and worldwide distribution have long been thought to be entirely asexual, being the principal exception to the view that in eukaryotes the loss of sex leads to early extinction. That bdelloids are facultatively sexual is shown by a study of allele sharing within a group of closely related bdelloids of the species *Macrotrachella quadricornifera*, making it likely that sexual reproduction is essential for long-term success in all eukaryotes.

## INTRODUCTION

Nearly all eukaryotes reproduce sexually, either constitutively or facultatively. That sexual reproduction may be essential for evolution was first suggested by Weismann (Weismann 1887) who, noting that, in comparison with asexuals, “in sexual reproduction twice as many individuals are required to produce any number of descendants” proposed that the compensating advantage of sexual reproduction is as a “source of individual variability, furnishing material for the operation of natural selection”. More specific hypotheses for the evolutionary benefit of sexual reproduction put forward since then include (*i*) bringing favorable mutations in different individuals together in the same lineage (Fisher 1930; Muller 1932) and reconstituting lines less loaded with detrimental mutations when, without sexual reproduction, recurring mutation and the continuing stochastic loss of less-loaded lines might drive the population to extinction (Muller 1964), and, more generally, weakening linkage, allowing selection to operate more nearly independently on linked loci (Hill and Robertson 1966); (*ii*) producing genotypes resistant to co-evolving biological antagonists (Hamilton 1980; Lively 2010); and (*iii*) purging synergistically-acting deleterious mutations from effectively infinite populations (Kondrashov 1988). An apparent challenge to all hypotheses for the evolutionary advantage of sexual reproduction is posed by a few groups of eukaryotes that have been designated “ancient asexuals” (Judson and Normark 1996; Schon *et al.* 2009). Of these, the most extensively studied are the rotifers of Class Bdelloidia.

## BDELLOID ROTIFERS

First described nearly 350 years ago (van Leewenhoeck 1677, 1702), bdelloid rotifers are minute freshwater invertebrates commonly found in lakes, ponds and streams and ephemerally aquatic habitats such as temporary pools and the water films on lichens and mosses (Figure 1). For review see: (Mark Welch *et al.* 2009). Characterized by their ciliated head and bilateral ovaries, bdelloids are classified in 4 families, 19 genera and some 500 morphospecies. The bdelloid radiation began tens of millions of years ago, as shown by the synonymous site difference between families and by the presence of bdelloid remains in ancient amber. Although typically only several tenths of a millimeter in size and containing only *ca*. 1,000 nuclei, mostly in syncytial tissues, bdelloids have ganglia, muscles, digestive, excretory, reproductive and secretory systems; photosensitive and tactile sensory organs; and structures for crawling, feeding and swimming. Bdelloids are degenerate tetraploids, descended from an ancient tetraploid ancestor (Mark Welch *et al.* 2008; Hur *et al.* 2009; Flot *et al.* 2013; Nowell *et al.* 2018).

**Figure 1.**
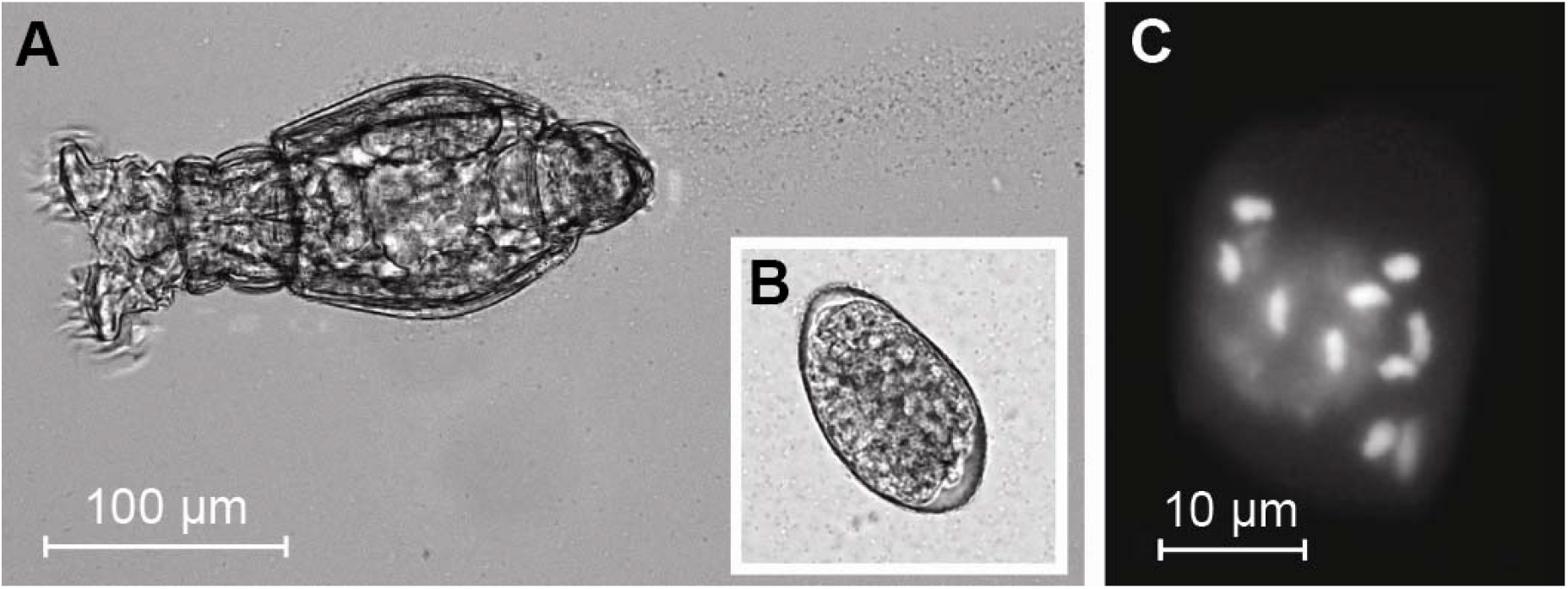
The bdelloid *M. quadricornifera*. (A) A feeding adult. Water and food particles are swept into the mouth by the motion of a pair of ciliated rings, the corona. The animal has attached itself to the glass slide by an adhesive secreted by its pedal gland, somewhat obscuring the tail. (B) An unhatched egg shown at the same scale. (C) A metaphase nucleus, showing the 10 chromosomes characteristic of the species.

The only observed means of bdelloid reproduction is from eggs produced in well-differentiated ovaries, with no reduction in chromosome number (Hsu 1956a; b). A few days after deposition a young bdelloid emerges and a few days later commences egg-laying, producing up to 32 eggs over a period of up to a few weeks during which there is little death, after which the death rate increases more or less exponentially (Meadow and Barrows Jr. 1971; Ricci 1983). Depending on species and conditions, the time from egg deposition to death is about a month (Ricci 1983; Ricci and Fascio 1995). Bdelloids are eutelic, with no cell division after eclosion except in the germ line.

Bdelloids are extremophiles, being able to survive prolonged desiccation, some species more than others, as well as starvation and extremes of temperature, and to resume reproduction upon restoration of favorable conditions, highly unusual abilities probably apomorphic to the Class (Ricci 1998; Ricci and Caprioli 2005; Ricci and Perletti 2006; Nowell *et al.* 2018). Bdelloids have a highly effective system of anti-oxidant protection, as manifested by their extreme resistance to ionizing radiation and to IR-induced protein carbonylation (Gladyshev and Meselson 2008; Krisko *et al.* 2012), apparently an adaptation to avoid the oxidative damage caused by desiccation, as known in other systems (França *et al.* 2007; Fredrickson *et al.* 2008).

Although bdelloids have been systematically studied ever since adequate microscopes became available (Ehrenberg 1838; Hudson and Gosse 1886) there is no confirmed observation of males. It has been estimated that contemporary rotifer workers have examined some 500,000 bdelloids from a variety of habitats and from laboratory culture without ever having seen males or hermaphrodites (Birky 2010). The only report to the contrary, in a treatise otherwise devoted to observations of males and sexual periods of the facultatively sexual rotifers of Class Monogononta, is a hesitant account of having twice seen a single male among many bdelloids of species *Rotaria rotatoria* “present in almost incredible numbers” beneath the ice of a frozen lake in Denmark in November 1923 (Wesenberg-Lund 1930). Sampling conducted there in January 2019 found few bdelloids and no males (Martin Sorensen, personal communication) nor was there any ice, the winter of that year having been among the warmest on record.

Despite the failure to document the existence of males, it may be that bdelloids reproduce sexually only rarely and under conditions not adequately investigated—a possibility made less implausible by estimates that outcrossing of *Saccharomyces cerevisiae* in the field may occur as seldom as once in 25-50,000 generations (Ruderfer *et al.* 2006; Magwene *et al.* 2011), owing to the repression of meiosis which, however, can be relieved in the laboratory by growth in specific media.

In the following, we review observations once interpreted as evidence for bdelloid asexuality but now known to have other explanations. We also summarize recent findings strongly suggestive of bdelloid sex. We then present a study of allele sharing in the bdelloid *Macrotrachella quadricornifera* with results in exact accord with expectations for facultative sexual reproduction and not explicable by horizontal gene transfer (HGT) or parasexuality. We also discuss the relation between bdelloid life history and population structure and implications for how bdelloid males and mating might be discovered.

## MATERIALS AND METHODS

### Sample collection and 10x sequencing

*M. quadrifornifera* isolates MA, MM and CR are from a group of 29 individuals morphologically identified as belonging to Family Philodinidae collected from ground moss in northeast United States in the autumn of 2011 (Signorovitch *et al.* 2015). Six of the 29, including MA, MM and CR, each collected at a site distant from the others, belong to the same mitochondrial clade (Lasek-Nesselquist 2012). Cultures of MA, MM and CR were twice established from single eggs, fed *E. coli* and maintained in 0.24u Millipore-filtered spring water at 20 °C in 100X20 mm plastic Petri dishes with continuous gentle rotation. Washed rotifers were shipped under dry ice in February 2016 to HudsonAlpha (Huntsville AL) for DNA extraction, library preparation, 10x Illumina sequencing at 56x coverage and provision to us of fastq files.

### Nanopore sequencing

Washed flash-frozen rotifers of isolate MA were digested for 17 h at 58 °C in 100 mM EDTA, 50 mM tris pH 9.0, 1% sodium sarcosyl, 1 mg/ml freshly dissolved proteinase K. DNA was isolated with a Qiagen MagAttract HMW DNA Kit and its size distribution analyzed with an Agilent Technologies 4200 TapeStation. DNA purity was verified by determining 260/280 and 260/230 ratios with a Nanodrop ND-1000 and its concentration was determined with a Qubit 3.0 fluorometer. DNA was prepared for sequencing with an Oxford Nanopore Ligation Sequencing Kit 1D without shearing and 1.1 μg (48μl) of the ligated DNA was loaded into the DNA repair end-end preparation step. Flow cells were prepared following the protocol from the same kit. DNA libraries were quantified with the Qubit and 420 ng of DNA was loaded for each sequencing run. Base calling for Nanopore reads was done with Albacore 2.3.4 and the results summarized with Nanoplot 1.20.0 (de Coster *et al.* 2018). Reads longer than 10kb were selected with fastp 0.19.5 (Chen *et al.* 2018) and aligned to the NCBI-nt database with Blastn (Altschul *et al.* 1990) using an e-value cutoff of 1e-25, removing reads with a best hit to a non-Animalia sequence, leaving 25,640 reads of length 10-92 kb. Statistics for the Nanopore reads are given in Table S1.

### Assembly of 10x reads

Scaffolds assembled from the 10x reads of each of the three isolates were obtained with Supernova 2.1.1 (Weisenfeld *et al.* 2017) using default parameters. Genome size, needed as an input parameter for Supernova, was estimated with Jellyfish 2.2.5 (Marçais and Kingsford 2011), based on the distribution of k-mers. Scaffolds were aligned to the NCBI nt database with Blastn with settings as above and non-Animalia scaffolds were removed. The mean assembly size was 328 Mb, range 352-371. Assembly statistics are presented in Table S2. Phased sequences (megabubbles) were then obtained from the 10x scaffolds with Supernova mkoutput with style=megabubbles.

### Alignment of 10x megabubbles to Nanopore reads

In order to select and align groups of homologous megabubbles from 2 or all 3 of the isolates, we aligned the Nanopore reads longer than 10 kb that did not have a significant Blast hit to non-Animalia sequences to the megabubbles from the 10x assemblies with Minimap 2 2.15-r905 (Li 2018). From the resulting alignments longer than 8kb, those in which megabubbles of at least two of the three isolates aligned to the same Nanopore read were chosen for analysis with Samtools faidx (Li *et al.* 2009). These alignments of 10x megabubbles, comprising groups of either 6 or 4 homologous sequences, were then realigned among themselves with Clustal Omega 1.2.3 (Sievers *et al.* 2011), and trimmed with Gblocks 0.91b (Talavera and Castresana 2007) to remove indels and alignment disruptions caused by repeats of unequal length, processes that neither add nor rearrange sequences. When two or more alignments overlapped, only the longest was retained. Lastly, 1 kb was removed from the ends of each alignment using EMBOSS seqret (Madeira *et al.* 2019) and alignments shorter than 1 kb were discarded. Pairwise SNP differences between homologs in each alignment were obtained with snp-dists 0.6.3 (*https://github.com/tseemann/snp-dists*). The workflow is depicted in Figure S1.

In five alignments MM or CR differed by more than 10% from both homologs of MA and were rejected. This may occur when, owing to a deletion in MM or CR occurring since their divergence from MA, there is no MM or CR sequence homologous to the Nanopore read, leaving only the homeologous sequence to align with it. This left 1,117 non-overlapping genomic regions for analysis, none of which contain homeologous sequences (Figure S2). That trimming neither adds nor rearranges sequences is confirmed by manual inspection of 5 randomly chosen regions of MA-MM sharing (regions 10, 104,142, 179, 262, total length 62,172 bp) in untrimmed MA-MM-CR alignments revealing only a single departure from perfect alignment, a one base deletion in a homopolymer run of As.

The sequence accuracy of the 10x assemblies was determined by comparison with published sequences of the four regions of MA, MA, and CR of total length 19.37 kb sequenced by Signorovitch *et al*. 2015, revealing a difference of 31 substitutions overall, or a 99.84% match (Table S3). Comparison of mitochondrial sequences in the 10x assemblies identified by blast searches against published mitochondrial sequences of MA, MM and CR (Lasek-Nesselquist 2012) revealed a perfect or near-perfect match for each isolate.

### Test for contamination

It might be asked if, despite precautions taken against it, contamination of MM DNA with MA DNA or the reverse in the DNA sent to HudsonAlpha for sequencing or occurring there, has mimicked allele sharing. Although it is most unlikely that contamination could be so massive and of the particular frequency required to mimic the MA-MM sharing we observe in half of the 622 MA-MM alignments, a test was conducted to rule out the possibility. All 10x Illumina reads sequenced from MA, MM, and CR were aligned to each of the 1,177 alignments. Using each haplotype in turn as a reference, bwa mem (Li and Durbin 2009) and GATK were used to produce variant calls (McKenna *et al.* 2010), implemented in a Snakemake (Molder *et al.* 2021) pipeline developed by the Harvard Informatics group (*https://github.com/harvardinformatics/shortRead_mapping_variantCalling*). The variant calls were then processed with vcftools (Danecek *et al.* 2011) to generate counts of reads at each variable nucleotide position supporting each allele. The fraction of Illumina reads supporting the alternate allele (the nucleotide that differs from the reference haplotype) shows a characteristic tri-modal pattern for all three isolates, with peaks at 0 (reference homozygote), 0.5 (heterozygote), and 1 (alternate homozygote), as shown for MA and MM in Figure S3. Such a pattern is expected for true variable positions in a diploid. Contamination would instead produce a pattern in which heterozygous positions are supported by a fraction of reads that departs from ½, and is determined by the proportion of the total DNA that was from a contaminating source, for which no evidence is seen in any of the three assemblies.

## PREVIOUS STUDIES

### Heterozygosity

In sexuals, heterozygosity caused by mutation is limited by haploid drift. The finding of much greater synonymous difference between gene copies in bdelloids than in monogononts was therefore initially interpreted as evidence for asexuality (Mark Welch and Meselson 2000, 2001). Continued investigation, however, showed that bdelloids are degenerate tetraploids, and that the highly diverged gene copies are homeologs, not homologs, with many genes present in only one or the other pair of homologs (Mark Welch *et al.* 2008; Hur *et al.* 2009). Bdelloid silent-site heterozygosity, the difference between homologs, lies within the range known for sexuals, providing no evidence for asexuality. Moreover, in asexual *Daphnia pulex* and *Saccharomyces cerevisiae* the frequency with which a nucleotide site is covered by a tract of homozygosity, as may result from germline crossing-over at the four-strand stage of mitosis or from certain processes of DNA damage repair, is much greater than the frequency of nucleotide substitution (Omilian et al. 2006; Xu et al. 2011; St. Charles and Petes 2013; Flynn et al. 2017). In sexuals, heterozygosity lost by such processes may be regained by outcrossing. But if bdelloids are ancient asexuals and if loss of heterozygosity were more frequent than substitution, the absence of outcrossing would be manifested as unusually *low* heterozygosity, the opposite of what had been thought (Magwene *et al.* 2011; Hartfield *et al.* 2018). A wide range of heterozygosity values could therefore be consistent with either sexual or asexual reproduction.

In a finite population, computer simulations demonstrate that small populations are driven to extinction by a Muller’s ratchet-like process of element accumulation, but that large populations can be cured of vertically transmitted TEs, even with excision rates well below transposition rates.

### Paucity of retrotransposons

Sexual reproduction allows vertically transmitted deleterious transposable elements to proliferate in populations, whereas under some theoretical models, the loss of sex may eventually free a population of such elements or drive it to extinction (Hickey 1982; Dolgin and Charlesworth 2006). As a test for asexuality, bdelloids, monogonont rotifers and sexually-reproducing animals of 23 other animal phyla were examined for genomic sequences coding for reverse transcriptases of LINE-like retrotransposons. These were found to be abundant in all the sexually-reproducing taxa but were not detected in bdelloids (Arkhipova and Meselson 2000, 2005a; b). Nevertheless, although bdelloids are nearly devoid of LINE-like retrotransposons, later work showed that they are not entirely absent (Gladyshev *et al.* 2007; Gladyshev and Arkhipova 2010) and that bdelloids have particularly effective retrotransposon silencing systems (Rodriguez and Arkhipova 2016). The paucity of LINE-like retrotransposons is therefore non-evidentiary as regards bdelloid sexuality.

### Genome structure

A draft genome sequence of the bdelloid, *Adineta vaga*, with numerous breaks in the colinearity of homologous regions and individual scaffolds containing genes in direct or palindromic repeats but no copy elsewhere in the genome was initially taken as evidence that bdelloids lack homologous chromosome pairs and had therefore evolved ameiotically (Flot *et al.* 2013). But subsequent genomic sequencing of three other bdelloid species, including *Adineta ricciae*, a close relative of *A. vaga*, found that the unusual genomic features that had been interpreted as evidence for ameiotic evolution are largely absent, suggesting that their apparent presence in *A. vaga* resulted from mis-assembly (Nowell *et al.* 2018) as later shown to be the case by the demonstration in *A. vaga* of homologous chromosome pairs (Simion *et al.* 2020).

### Allele sharing

A finding of two individuals closely related with respect to a given genomic region but more distantly related with respect to its homolog, a form of phylogenetic noncongruence known as allele sharing, would mean that recently in their ancestry the region had undergone some form of genetic exchange between individuals. A striking example of allele sharing among isolates MA, MM and CR was found by (Signorovitch *et al.* 2015, 2016) in each of the four genomic regions examined, 2.4 to 9.7 kb in length. At each region, MA identically shared a homolog with MM while the other homolog of MA was identical to a homolog of CR in two regions and nearly so in the other two. That these observations were evidence for sexual reproduction was disputed and attributed instead to HGT on the basis of observations in *A. vaga* (Debortoli *et al.* 2016; Flot *et al.* 2018). Subsequent analysis, however, showed that what had been interpreted as HGT could be explained as the result of cross-contamination among isolates (Wilson *et al.* 2018), leaving no agreed evidence for homologous HGT in bdelloids.

### Meiosis-associated genes

A survey of the genomes of four bdelloid species, including *A. vaga*, belonging to two bdelloid families for 11 genes associated with meiosis found all but one, *red1*, to be present in each species (Nowell *et al.* 2018). But neither was *red1* found in *Drosophila melanogaster*, a sexual species known to lack it. Although five of these genes had not been found in the draft assembly of the *A. vaga* genome (Flot *et al.* 2013) their subsequent detection and conservation in this and the three other bdelloid species tested suggests that bdelloids engage in meiosis, although it cannot be excluded that they are retained instead for other functions.

### Population Genetics

A recent population genetics survey of 11 individuals of the bdelloid rotifer A. vaga collected at diverse sites in the wild found a number of features consistent with sexual reproduction, including genotype proportions in Hardy-Weinberg equilibrium, indicating some form of genetic exchange between individuals. Recombination either between or within individuals is indicated by a decay of linkage disequilibrium with increasing physical distance along scaffolds and by signatures of reciprocal recombination (Vakhrusheva *et al.* 2020). Phylogenetic analysis of certain genomic segments in a group of three closely-related individuals revealed incongruence possibly resulting from sexual reproduction between their ancestors and members of a distinctly different population. Although the authors caution that neither sexual reproduction nor HGT by themselves provide a simple explanation for their data, their observations nevertheless provide strong evidence for recombination and some form of genetic exchange.

## RESULTS

### Alignments

As described in METHODS, we obtained alignments of phased sequences from 1,177 non-overlapping genomic regions of *M. quadricornifera* isolates MA, MM and CR. Of these, 331 alignments are with all three isolates, 291 with MA and MM, 110 with MA and CR and 445 with MM and CR, having a mean length of 12,490 bp (range 2,051 - 32,937 bp) and altogether covering 14.7 Mb, approximately 4% of the *ca.* 360 Mb genome (Table 2). Matrices giving pair-wise differences between homologous sequences, phylograms and plots of the spatial distribution of differences between homologs (“tic” plots) for four representative regions are given in Table 1 and, for all 1,177 regions, in Supplemental Material Tables 4-6.

**Table 1.**
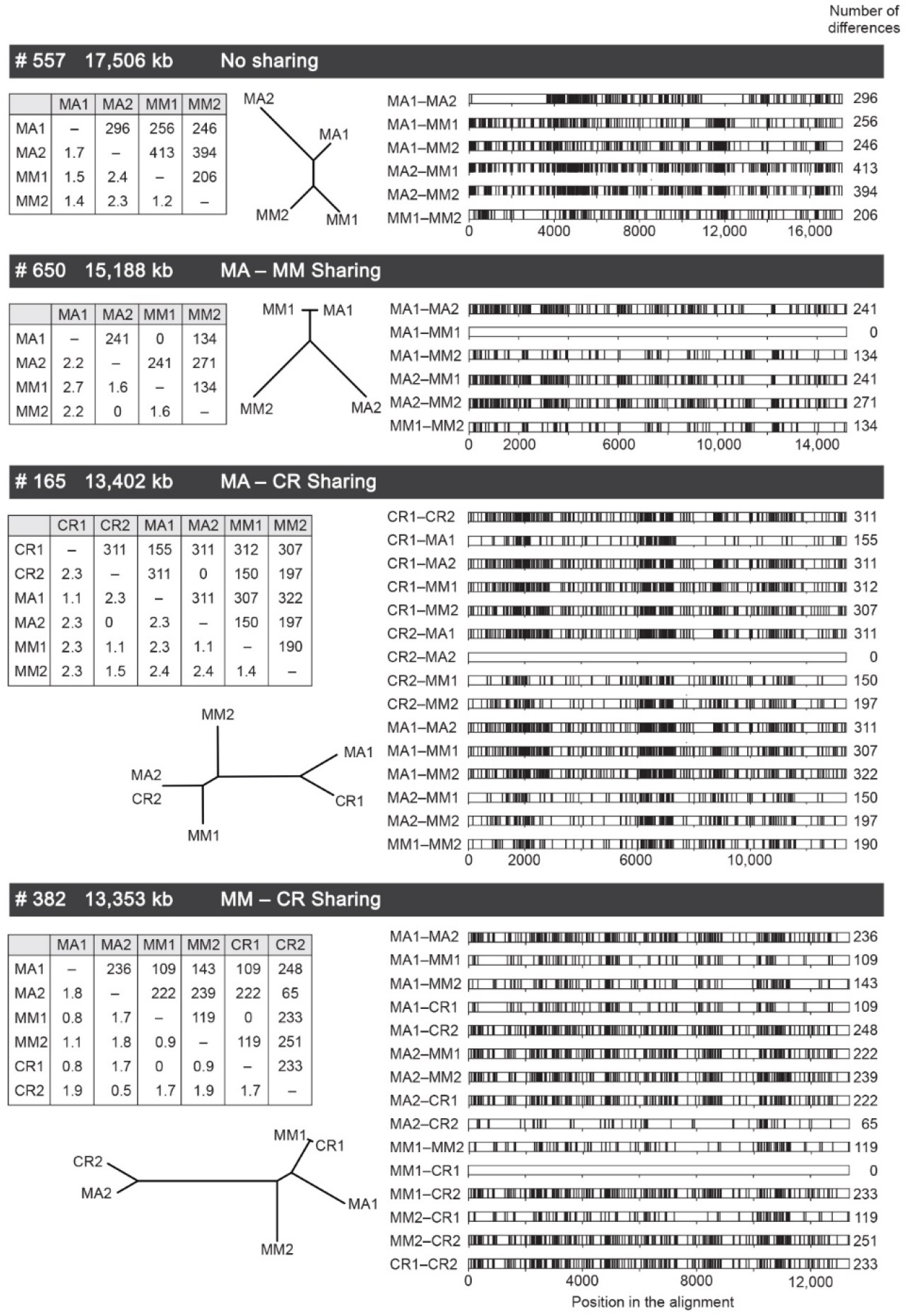
Representative difference matrices, phylograms and tic plots for four alignments. In the difference matrices, the number of alignments is given above the diagonal and the percent of the corresponding genotype is given below. Vertical lines in the tic plots represent sites of nucleotide difference. Allele numbers (1 or 2) are arbitrary.

**Table 2.** Summary statistics for the 1,177 non-overlapping alignments examined. Identical MA-MM sharing occurs in 315 of the 622 alignments in which MA and MM are included. The more frequent homozygosity and lower heterozygosity in alignments of of isolate MM may reflect more frequent occurrence of homozygosing events along its clonal lineage.

| Summary Statistics |  |  |  |  |  |  |  |  |  |  |
| --- | --- | --- | --- | --- | --- | --- | --- | --- | --- | --- |
| Alignment | Size Range (kb)<br># of Alignments | Identical Sharing<br>Number / Proportion |  |  | Homozygous<br>Number / Proportion |  |  | Average Heterozygosity |  |  |
|  |  | MA-MM | MA-CR | MM-CR | MA | MM | CR | MA | MM | CR |
| MA-MM | 2,149-32,937 | 151 |  |  | 12 | 8 |  | 0.019 | 0.010 |  |
|  | 291 | 0.519 |  |  | 0.041 | 0.028 |  |  |  |  |
| MA-CR | 2,051-29,791 |  | 1 |  | 1 |  | 0 | 0.018 |  | 0.018 |
|  | 110 |  | 0.009 |  | 0.009 |  | 0 |  |  |  |
| MM-CR | 4,511-31,052 |  |  | 7 |  | 18 | 12 |  | 0.010 | 0.020 |
|  | 445 |  |  | 0.016 |  | 0.04 | 0.027 |  |  |  |
| MA-MM-CR | 2,705-29,510 | 164 | 11 | 11 | 1 | 4 | 3 | 0.018 | 0.009 | 0.018 |
|  | 331 | 0.495 | 0.033 | 0.033 | 0.003 | 0.012 | 0.009 |  |  |  |
| All alignments | 2,051-32,937 | 315 | 12 | 18 | 14 | 30 | 15 | 0.018 | 0.010 | 0.019 |
|  | 1,177 | 0.506 | 0.027 | 0.023 | 0.019 | 0.028 | 0.017 |  |  |  |

### Allele sharing

Half of the MA-MM and MA-MM-CR alignments, 315 of 622, comprise a discrete class in which a homolog of MA is identical to a homolog of MM (Table 2; Fig. 2B). The frequencies of identical MA-MM sharing in the MA-MM and MA-MM-CR alignments considered separately are 0.519 and 0.495, respectively or 0.506 overall (S.E. = 0.02). MA and MM also share identical homologs with CR, but in a much smaller proportion of the alignments (Fig. 2A, C). CR shares identical homologs with MA in 12 of 441 alignments, and with MM in 18 of 776 alignments, or 2.7% and 2.3% respectively. In alignments without identical sharing the differences between the homolog of MA most similar to a homolog of MM form a broad distribution with a mean of 0.96 SNPs per 100 bp (S.E. = 0.61), Fig. 2B. Most or all of the regions identically shared between MA and MM must be considerably longer than the alignments in which we find them, as shown in a plot of the frequency of identical MA-MM sharing against alignment length in consecutive intervals each comprising 76-79 alignments (Fig. 3). The frequency of identical sharing is not significantly different from 50 percent in even the longest alignments (18-33 kb).

**Figure 2.**
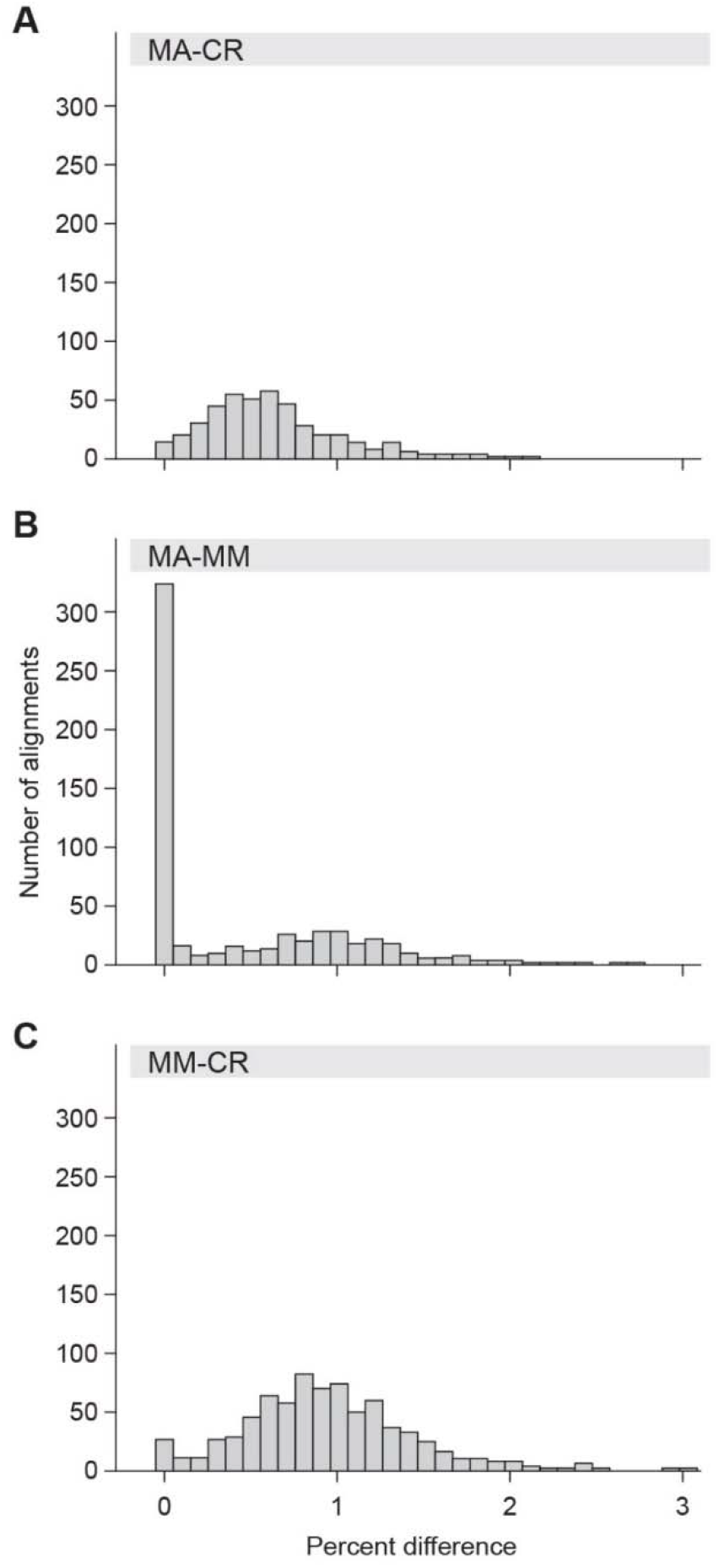
Histograms showing the distribution of divergence between the most similar homologs in each of the three pairs of isolates. (A) MA-CR, (B) MA-MM, (C) MM-CR. Alignments with identical or very nearly identical MA-MM sharing form a discrete class constituting half of the regions (panel B) showing that isolates MA and MM identically share ¼ of their genomes inherited from their recent ancewstors. Bin size = 0.05 percent difference for the first bar in panel B, otherwise 0.1 percent.

**Figure 3.**
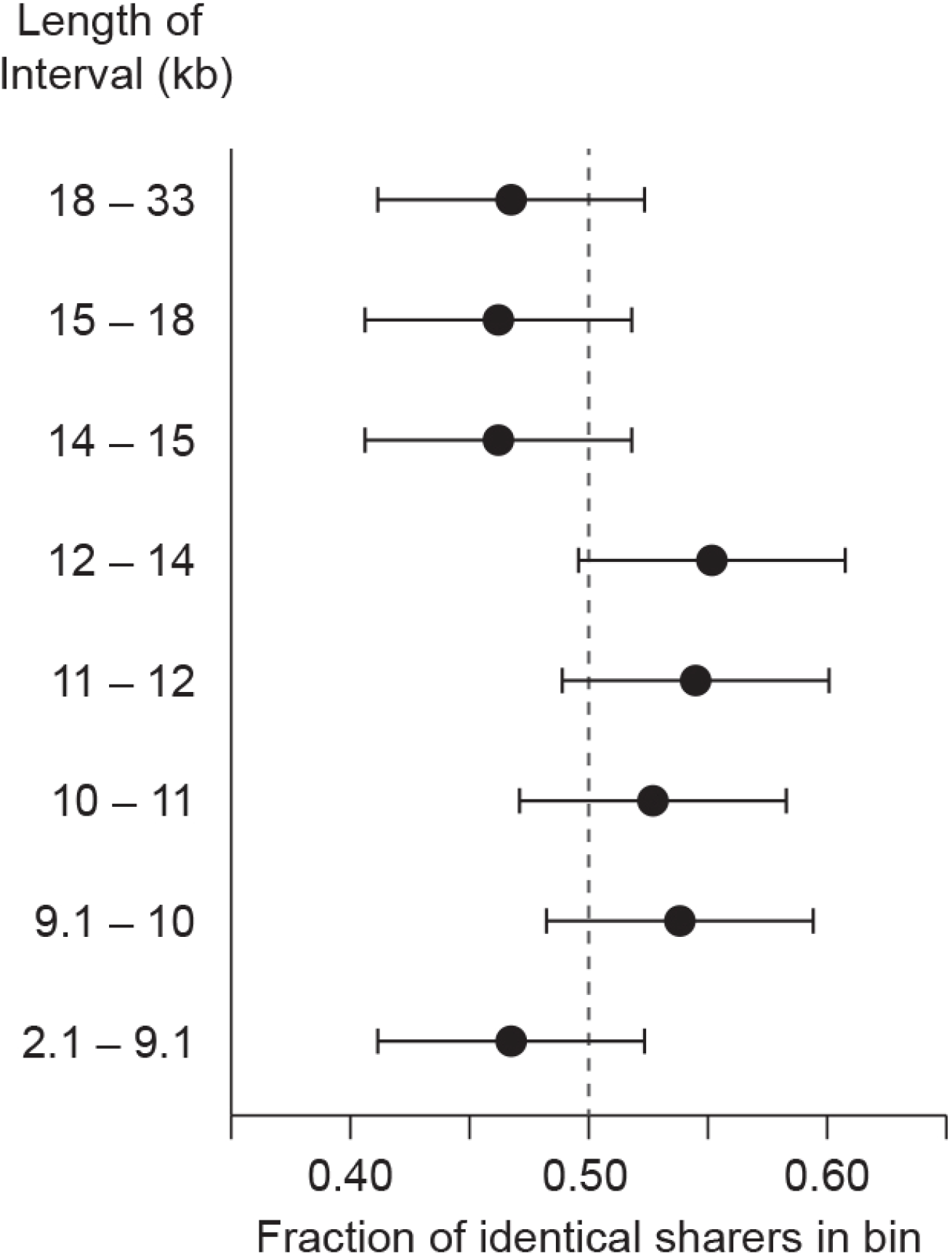
The mean frequency of identical MM-MA sharing in each of 8 bins of alignments of increasing length, each bin containing 76-79 alignments. Error bars represent standard errors. It may be seen that the identically shared segments must generally be longer than the longest regions examined.

### Genealogy

MA and MM identically share a homolog in a discrete class amounting to half of the 622 MA-MM and MA-MM-CR alignments while their other homologs, within and between MA and MM, are substantially diverged. MA and MM therefore identically share ¼ of their genomes as segments at least the length of the alignments, a degree of relationship expected for grandchild and grandparent, half siblings, aunt or uncle and nephew or niece from a large panmictic sexually-reproducing population. The near equality of MA-CR and MM-CR sharing frequencies indicates that CR is equidistant from MA and MM and therefore that MA and MM are not grandparent and grandchild or aunt/uncle-nephew/niece but rather half siblings or double first cousins.

Inspection of tic plots for the triple alignments reveals a few with interior regions of MA-MM identity covering much but not all of the alignment (Supplemental Material Table 4). In these regions there is substantial divergence from CR, showing that such identity is not the result of extreme conservation but instead reflects more remote relationships between MA and MM in addition to their relation as half siblings or double first cousins. Similarly, alignments in which CR is identical to MA or MM over much but not all the alignment are likely to reflect remote relationships between CR and MA and between CR and MM. In general, more distant relations will be manifested as shorter regions of identity by descent, owing to the crossing-over that occurs at each meiosis. For individuals related as half-sibs or double first cousins the regions of identity by descent from their common grandparents, assuming one cross-over per meiosis in each arm of the 10 chromosomes of *M. quadricornifera*, will average several Mbp in length, far longer than our longest alignments, consistent with the observation that over the region examined the frequency of identically shared regions does not fall off with length (Fig. 3).

### Homozygosity

In each of the three isolates there are a few regions that are entirely homozygous (Table 2). No more frequent in the shorter half of the alignments than in the longer half, they must generally be longer than the alignments in which they occur. These regions may be identical by descent or may have arisen by conversion or by germ-line crossing-over at the four-strand stage of mitosis. That these processes may have occurred more frequently in MM than in MA and CR is suggested by its higher frequency of homozygous regions and lower mean heterozygosity (Table 2 and Figure S2). The infrequency of homozygous regions suggests, for each isolate, that its parents cannot have been closely related, as expected for individuals from a large panmictic population. The few alignments in which conversion or mitotic crossing-over may have erased evidence of sharing are not included in the totals given above or in Table 2.

### Recombination

The finding that the pattern of sharing was the same in every one of the four regions examined by (Signorovitch *et al.* 2015), with MA sharing one of its homologs with MM and its other homolog with CR, suggested that the regions may not have recombined and therefore that entire haplotypes may have passed through meiosis intact, as in certain plants (Holsinger and Ellstrand 1984), an interpretation made plausible at the time by the earlier but subsequently disproven report that *A. vaga* has genomic features incompatible with meiotic recombination (Flot *et al.* 2016). In contrast, this pattern of sharing occurs in only 4 of our 331 much longer MM-MA-CR alignments. It may therefore be that the pattern encountered in the earlier study resulted from identity by descent from remote ancestors, in which case such regions would generally be shorter than the present alignments and therefore not counted as allele sharers. That entire haplotypes are not conserved and that recombination definitely occurs is seen in the observation that MA and MM share only a quarter of their genomes and share considerably less with CR and in the existence of numerous alignments in which sharing extends over only part of the region.

## DISCUSSION

We find that MA and MM identically share homologs in 315 or just half (0.506, S.E. = 0.02) of the 622 MA-MM and MA-MM-CR alignments, therefore sharing ¼ of their genome in segments at least as long as our alignments and possibly much longer. This degree of sharing is consistent with sexual reproduction, in which the proportion of the genome shared by two individuals inherited from their closest common ancestors, their coefficient of relationship, is a power of ½.

In all of biology there are only three known modes of homologous genetic exchange between individuals: sexual reproduction, horizontal gene transfer including its various forms in prokaryotes, and parasexuality. For the observed allele sharing to have occurred by homologous HGT would require massive horizontal transfer of long DNA segments between MA and MM, displacing a homolog coincidentally in exactly half (0.5, S.E. = 0.02) of the 622 alignments containing MA and MM, a degree of consanguinity consistent with sexual reproduction, yet never displacing both, there being no alignment in which MA and MM are of the form a/b and a/b.

Nor can the observed allele sharing be explained as the result of parasexuality. In the parasexual cycle, known only in certain fungi and protozoans, nuclei from two individuals fuse, forming nuclei of doubled ploidy that during subsequent generations undergo occasional mis-division, only rarely yielding viable diploids. In bdelloids, this would require nuclei from different individuals, sequestered in ovaries within the body of the animal, somehow to undergo fusion, followed by a series of random chromosome losses to give viable segregants, all having ten chromosomes and with MA-MM identical sharing in exactly half of the 1,177 genomic regions we examined.

The finding in *A. vaga* of meiosis-related genes (Nowell *et al.* 2018), of homologous chromosome pairs (Simion *et al.* 2020), and of Hardy-Weinberg equilibrium and decay of linkage disequilibrium with increasing physical distance and phylogenetic noncongruence (Vakhrusheva *et al.* 2020), although susceptible of other explanations, are in entire agreement with our conclusion from the pattern of allele sharing that bdelloids are facultatively sexual.

### Generations since the MA-MM sharing event

The number of clonal generations since the MA-MM sharing event may be estimated from the frequency of substitution differences between shared homologs, a few of which differ by one or more substitutions; from the number of generations that would cause mutational reduction of the frequency of identical sharers to fall significantly below the observed value of 0.5; from the frequency of homozygosity; and from the mitochondrial difference between MA and MM.

In addition to the 315 alignments in which MA and MM share identical homologs there are 6 in which they share homologs that differ by a single substitution. For a mean alignment length of 15 kb this is a frequency of 1.3X10^−6^ per bp. Substitution rates measured in accumulation experiments with *Caenorhabditis elegans*, asexual *D. pulex*, and *D. melanogaster range* from 2.3 to 5.5X10^−9^ per generation (Flynn *et al.* 2017). Taking a substitution rate of 4X10^−9^ and assuming a Poisson distribution of the few nucleotide substitution differences between shared homologs, this suggests that the shared homologs may be separated by 100-200 generations.

If there were as many as 2,000 generations separating the shared homologs of MA and MM and again assuming a substitution rate of 4X10^−9^ per generation, the expected number of substitutions in regions 18-33 kb in length, the longest interval in Figure 3, would be 0.14 – 0.26, reducing the proportion of identical sharers to 0.43 – 0.39, substantially less than the observed value of 0.5, suggesting that the number of generations between the shared homologs is no more than about 1,000.

As tracts of homozygosity arising in a genetically diverse population are generally erased by outcrossing, the frequency of substitutions in such tracts will increase with the number of clonal generations since the last outcross. Assuming that, as in daphnia and yeast, the likelihood of a site being covered by a tract of homozygosity to be about 4X10^−5^ per generation (Omilian et al. 2006; Xu et al. 2011; St. Charles and Petes 2013; Flynn et al. 2017) and considering that the total length of MA regions is 13.4 Mbp of which perfectly homozygous regions comprise some 151 Kbp, or about 1.1%, it appears that there have been some 340-500 generations from when the sharing event occurred to MA and 550-800 generations to MM.

A fourth estimate of the number of generations since the MA-MM sharing event may be obtained by assuming that their mitochondria descend from a common mother or maternal grandmother. Taking the difference of 5 substitutions or 2.5 × 10^−5^ between their 14 kb mitochondria (Lasek-Nesselquist 2012) and a mitochondrial mutation rate of 1.5 × 10^−7^ (Xu *et al.* 2012; Flynn *et al.* 2017), suggests that the shared homologs separated some 170 generations ago. These various estimates agree in suggesting that the shared homologs of MA and MM are separated by no more than about a thousand clonal generations.

### Abundance of close relatives in the sampled population

Isolates MA, MM and CR were collected at widely separated sites as part of a collection of only 29 individuals. What aspects of bdelloid life history could make finding relatives as close as MA and MM in so small and widely dispersed a sample of what must be an enormous population? It must be that the sampled population is largely made up of relatively few, very large, widely dispersed clones descended from recent crossing. Such an unusual population structure would result if sexual periods occur only rarely, during a population bloom, with multiple rounds of mating among two or more founding types, producing large numbers of closely related individuals. It may be that males are produced and mating occurs only when particular mating types are present together, causing one or both to produce haploid eggs and haploid males. At some stage, from fertilization to zygote development, selfing may be prevented, allowing mating only between different types, thereby avoiding the production of homozygous progeny. Such mating, followed by wide dispersion and extensive clonal reproduction would give rise to very large, widely dispersed clones of the products of recent crossing. Meanwhile, lines that fail to outcross would suffer loss of heterozygosity caused by conversion and mitotic crossing-over, and rapid clonal erosion, as seen in asexual *Daphnia pulex* (Tucker *et al.* 2013), driving them to extinction unless revived by timely outcrossing.

On this picture, field observations intended to detect males and mating should be made during population blooms, as may require specific external stimuli, and in sizeable bodies of water should it be that different but compatible types must be present in order to initiate mixis. Further, by analogy with monogononts, the appearance of bdelloid males may be confined to only a short interval during a population bloom, therefor requiring frequent sampling for their detection (Wesenberg-Lund 1930).

### Bdelloid life history - Eluding the Red Queen

It may be asked if there is a relation between the infrequency of bdelloid outcrossing and bdelloid life history. A distinctive feature of the latter is the ability of bdelloids to withstand desiccation and resume reproduction upon rehydration, an ability not present in most of the fungi and other organisms that infect, parasitize, prey on or compete with bdelloids (Wilson and Sherman 2010). In habitats that undergo desiccation and rehydration the population of desiccation-intolerant antagonists will be greatly reduced at each episode of desiccation while bdelloids will resume reproduction upon rehydration. Bdelloids gain additional freedom from having to co-evolve with biological antagonists by their ability to survive prolonged starvation, extremes of temperature and exposure to toxic conditions lethal to other taxa (Ricci and Perletti 2006; Aguilera *et al.* 2007). Further, owing to their small mass when desiccated, about 10ug, once airborne, even if associated with a small amount of adhering material, bdelloids may be transported by wind or vectors over considerable distances (Fontaneto *et al.* 2008), transferring to an environment where antagonists may be less abundant or less antagonistic. The combination of anhydrobiosis and resistance to conditions inimical to other taxa and dispersibility by wind, birds or other vectors therefore affords substantial protection from what would otherwise be co-evolving biotic antagonists, reducing the need for frequent recombination by largely eluding the “Red Queen” (Ladle *et al.* 1993; Wilson and Sherman 2010; Wilson 2011).

Although bdelloids may be substantially freed of the need for sexual reproduction to keep up with biological antagonists, other benefits of sex apparently maintain it. In addition to the benefit of recombination in facilitating adaptive selection and limiting the load of deleterious mutation, a further and more immediate benefit may be the restoration of lost heterozygosity. An example of this may be seen in the suggestion of (Tucker *et al.* 2013) that the early extinction of asexual *Daphnia pulex* may result from the uncovering of preexisting recessive deleterious alleles caused by conversion and deletion. Whether it is the masking of deleterious recessive alleles and/or some other benefit of heterozygosity, its restoration by occasional outcrossing may constitute an important benefit maintaining sex in bdelloid rotifers and more generally.

## Acknowledgements

We thank Nicole El-Ali and Claire Hartmann for Nanopore sequencing, Jae Hur for the script used in generating tic plots, Janet Montgomery for overall editing, and Irina Arkhipova, Timothy Barraclough, Brian and Deborah Charlesworth, Antoine Hout, Paul Simion and Karine van Doninck for critical reading of the manuscript. This work was supported by Oxford Nanopore, the Harvard Faculty of Arts and Sciences and by an anonymous donor.

## Supplemental Material

is available for this paper.

## Correspondence

and requests for materials should be addressed to Matthew Meselson.

## Author Contributions

VL and TS: Genome assembly and data analysis. MM: Rotifer culturing DNA preparation and data analysis. MM wrote the manuscript, which was edited and approved by all authors.

