## Supplemental Materials for "Genomic Signature of Sexual Reproduction in the Bdelloid Rotifer *Macrotrachella quadricornifera*"

**SUPLEMENTAL MATERIAL**

**Sexual Reproduction in Bdelloid Rotifers**

This Supplemental Material includes:

Figure S1: Workflow

Figure S2: Histograms showing heterozygosity for MA, MM and CR

Figure S3: Reads supporting the alternate allele in MA and MM.

Table S1: Statistics for Nanopore reads of isolate MA.

Table S2: 10x assembly statistics.

Table S3: Comparison of Genbank and 10x sequences.

Table S4: tic plots.

Table S5: Phylograms.

Table S6: Difference matrices.

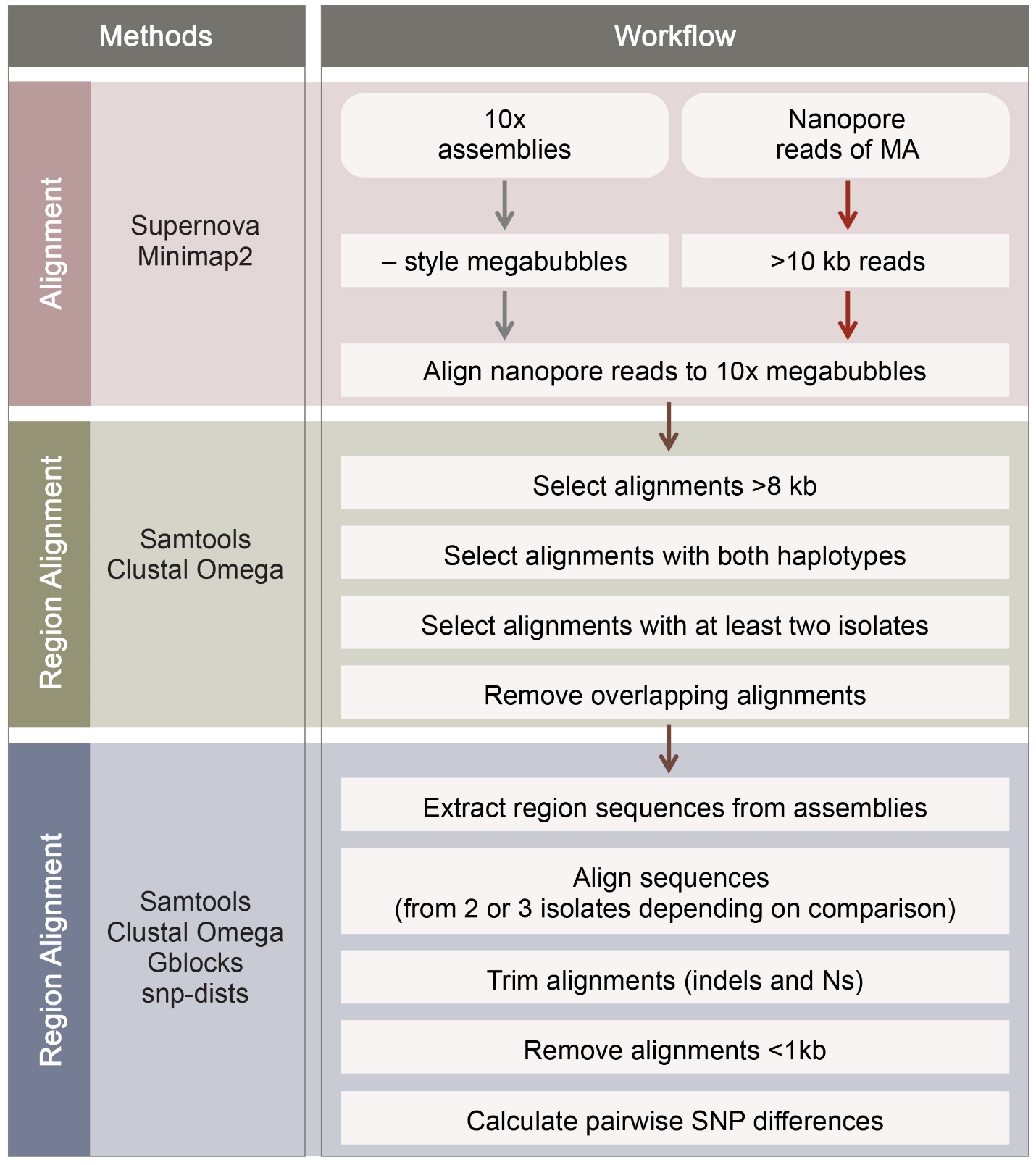

**Figure S1**. Workflow

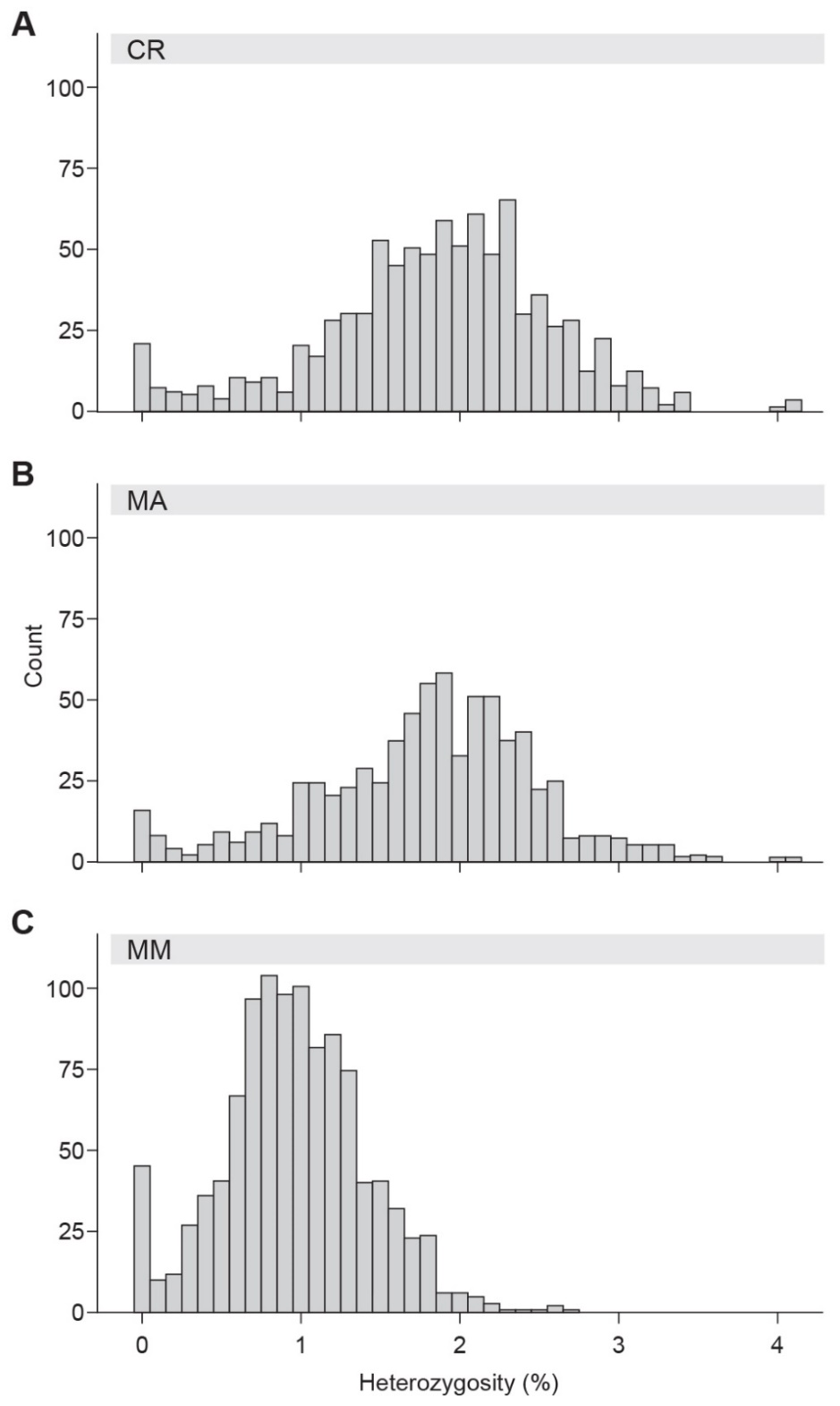

**Figure S2.** Histograms showing heterozygosity for MA (732 regions), MM (1067 regions) and CR (886 regions). Bin size = 0.1 percent difference.

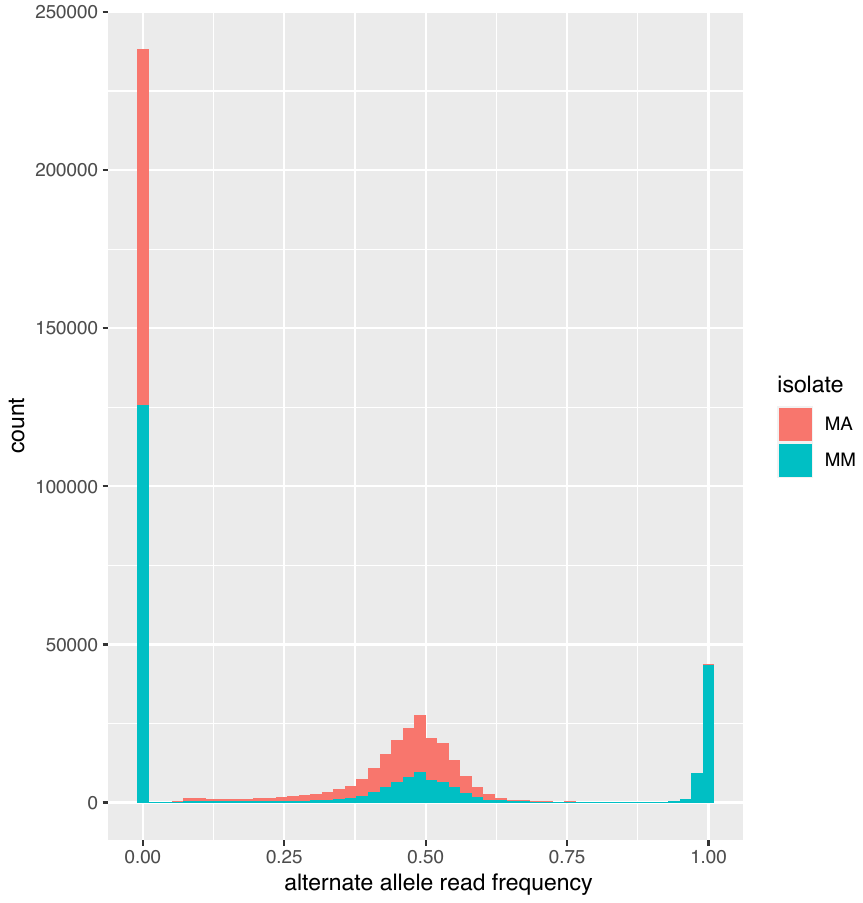

**Figure S3.** Reads supporting the alternate alleles in MA and MM. Using phased sequences from our 1,177 regions as a reference, we aligned 10x Illumina reads from MA or MM, and counted the number of Illumina reads at each variable nucleotide site supporting either the reference nucleotide or the non-reference nucleotide. We summarize these as the proportion of reads supporting the (alternate) non-reference nucleotide, as show in the X axis in this histogram. As expected, we find a tri-modal distribution with peaks representing invariant positions (alternate allele read frequency = 0), heterozygous positions (alternate allele read frequency distributed around 0.50), and positions where the isolate is homozygous for a different nucleotide than the reference (alternate allele read frequency = 1). Variable positions representing contamination would be expected to be supported by a proportion of reads with a mode at the fraction of the total DNA represented by contamination, which is extremely unlikely to be coincidentally 0.50. Results using a MA reference haplotype are plotted, but similar results are obtained with MM reference haplotypes.

**Table S1.** Statistics for Nanopore reads of isolate MA after removing non-animalia sequences.

| Total reads | **821,623** |
| --- | --- |
| Total bases | **1924.941 Mb** |
| Mean length | **2342 bp** |
| Read N50 | **4090 bp** |
| Maximum read length | **91,488 bp** |
| Reads longer than 10 kb | **25,640** |
| Estimated coverage | **5.46x** |
| % GC | **35.31%** |

**Table S2.** 10x assembly statistics.

|  | CR | MA | MM |
| --- | --- | --- | --- |
| Reads | **217.68 M** | **217.67 M** | **217.67 M** |
| Raw coverage | **54.67x** | **56.68x** | **51.51x** |
| Assembly size | **370.55 Mb** | **352.28 Mb** | **262.19 Mb** |
| Number of scaffolds (>10 kb) | **5,650** | **5,780** | **5,630** |
| Contig N50 | **41.31 Kb** | **39.12 Kb** | **35.20 Kb** |
| Scaffold N50 | **275.58 Kb** | **224.37 Kb** | **90.61 Kb** |
| % GC | **29.76%** | **30.52%** | **29.97%** |

**Table S3.** Comparison of Genbank and 10x sequences.

**hisA (4,205 bp)**

| Genbank Region | Differences from 10x | Differences per base pair |
| --- | --- | --- |
| CR1_KR133439 | 1 | 0.00024 |
| CR2_KR133440 | 2 | 0.00048 |
| MM1_KR133443 | 1 | 0.00024 |
| MM2_KR133444 | 0 | 0 |
| MA1_KR133441 | N/A | N/A |
| MA2_KR133442 | 1 | 0.00024 |

**hisB (9,819 bp)**

| Genbank Region | Differences from 10x | Differences per base pair |
| --- | --- | --- |
| CR1_KR133445 | 3 | 0.0003 |
| CR2_KR133446 | 1 | 0.0001 |
| MM1_KR133453 | 1 | 0.0001 |
| MM2_KR133454 | 10 | 0.001 |
| MA1_KR133451 | 0 | 0 |
| MA2_KR133452 | 3 | 0.0003 |

**hspA (2,635 bp)**

| Genbank Region | Differences from 10x | Differences per base pair |
| --- | --- | --- |
| CR1_KR133457 | 1 | 0.00038 |
| CR2_KR133458 | 2 | 0.00076 |
| MM1_KR133461 | 1 | 0.00038 |
| MM2_KR133462 | 1 | 0.00038 |
| MA1_KR133459 | 1 | 0.00038 |
| MA2_KR133460 | 2 | 0.00076 |

**hspB (2,711 bp)**

| Genbank Region | Differences from 10x | Differences per base pair |
| --- | --- | --- |
| CR1_KR133463 | 0 | 0 |
| CR2_KR133464 | N/A | N/A |
| MM1_KR133467 | 0 | 0 |
| MM2_KR133468 | 0 | 0 |
| MA1_KR133465 | 0 | 0 |
| MA2_KR133466 | N/A | N/A |

The three regions marked N/A were not assembled in the Supernova assembly and therefore could not be compared to 10x sequences. There are 31 SNPs in the 19.37 kb sequenced, a 99.84% match.

**Table S4.** TIC Plots.

In order of alignment (region) number

<https://github.com/tsackton/rotifer-outcrossing/tree/main/tables>

**Table S5.** Phylograms.

In order of alignment (region) number.

<https://github.com/tsackton/rotifer-outcrossing/tree/main/tables>

**Table S6.** Difference matrices.

In order of alignment (region) number.

<https://github.com/tsackton/rotifer-outcrossing/tree/main/tables>
