## Supplementary material for "Genomic Signature of Sexual Reproduction in the Bdelloid Rotifer *Macrotrachella quadricornifera*": Table S4

### Region: 15



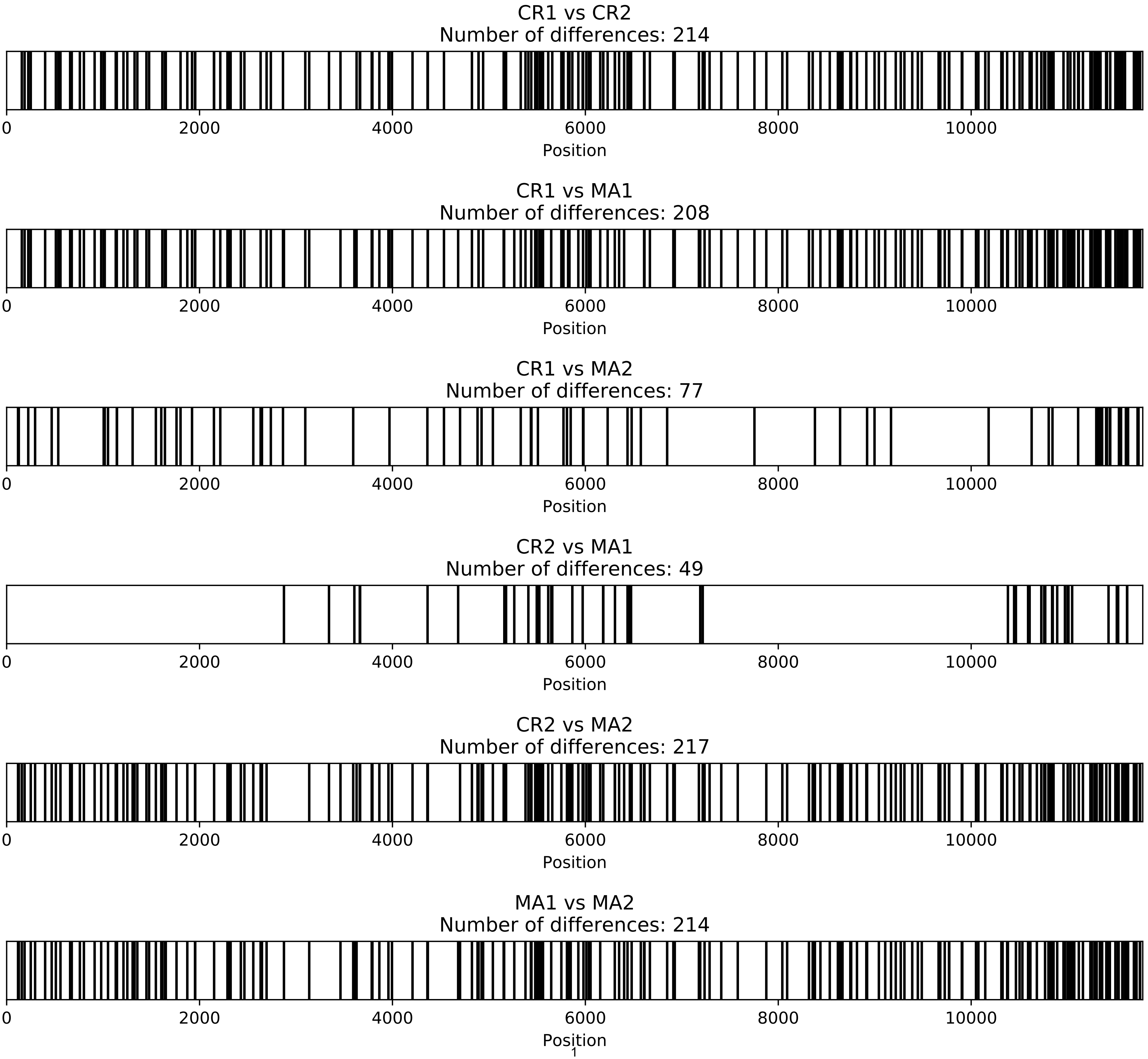
CR1 vs CR2
Number of differences: 214
CR1 vs MA1
Number of differences: 208
CR1 vs MA2
Number of differences: 77
CR2 vs MA1
Number of differences: 49
CR2 vs MA2
Number of differences: 217
MA1 vs MA2
Number of differences: 214
0
2000
4000
6000
8000
10000
Position

### Region: 18



CR1 vs CR2
Number of differences: 263

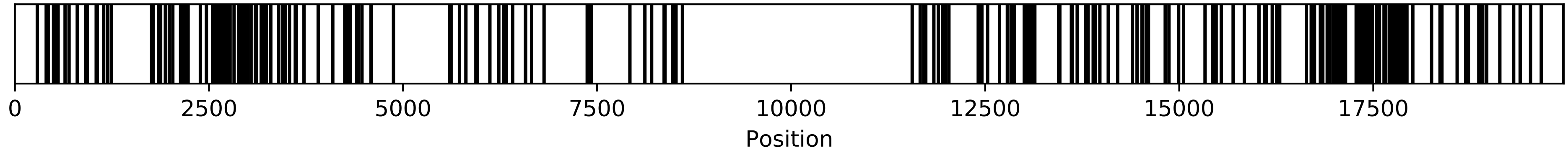


CR1 vs MA1
Number of differences: 275

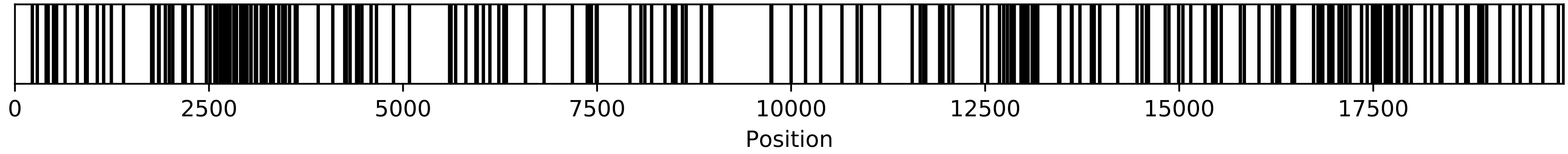


CR1 vs MA2
Number of differences: 96

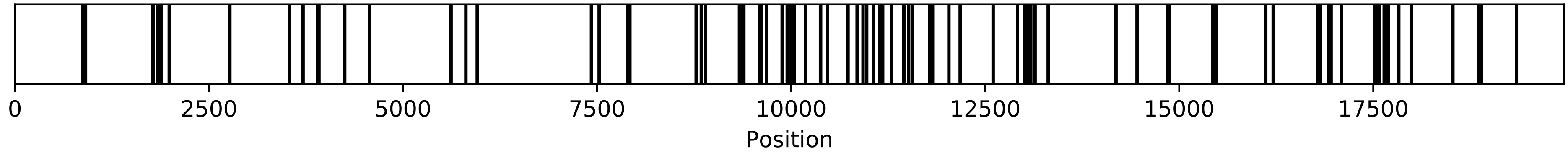


CR2 vs MA1
Number of differences: 168

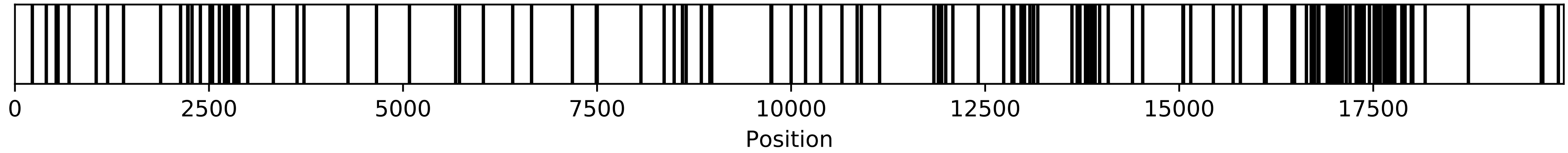


CR2 vs MA2
Number of differences: 302

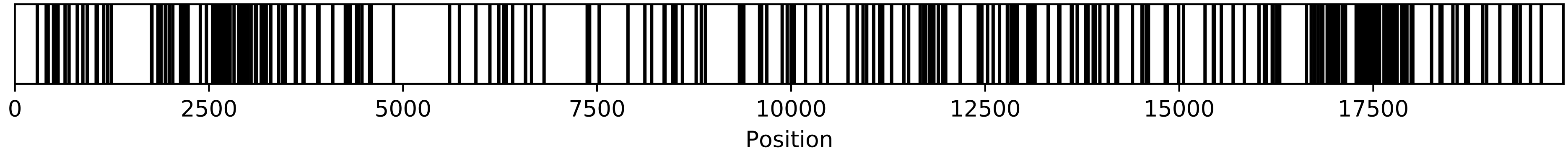


MA1 vs MA2
Number of differences: 295

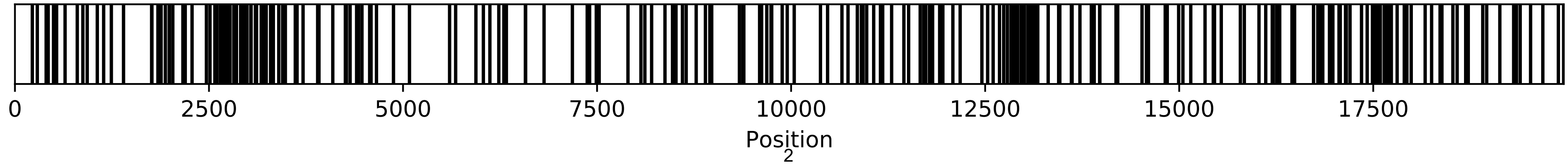

### Region: 20



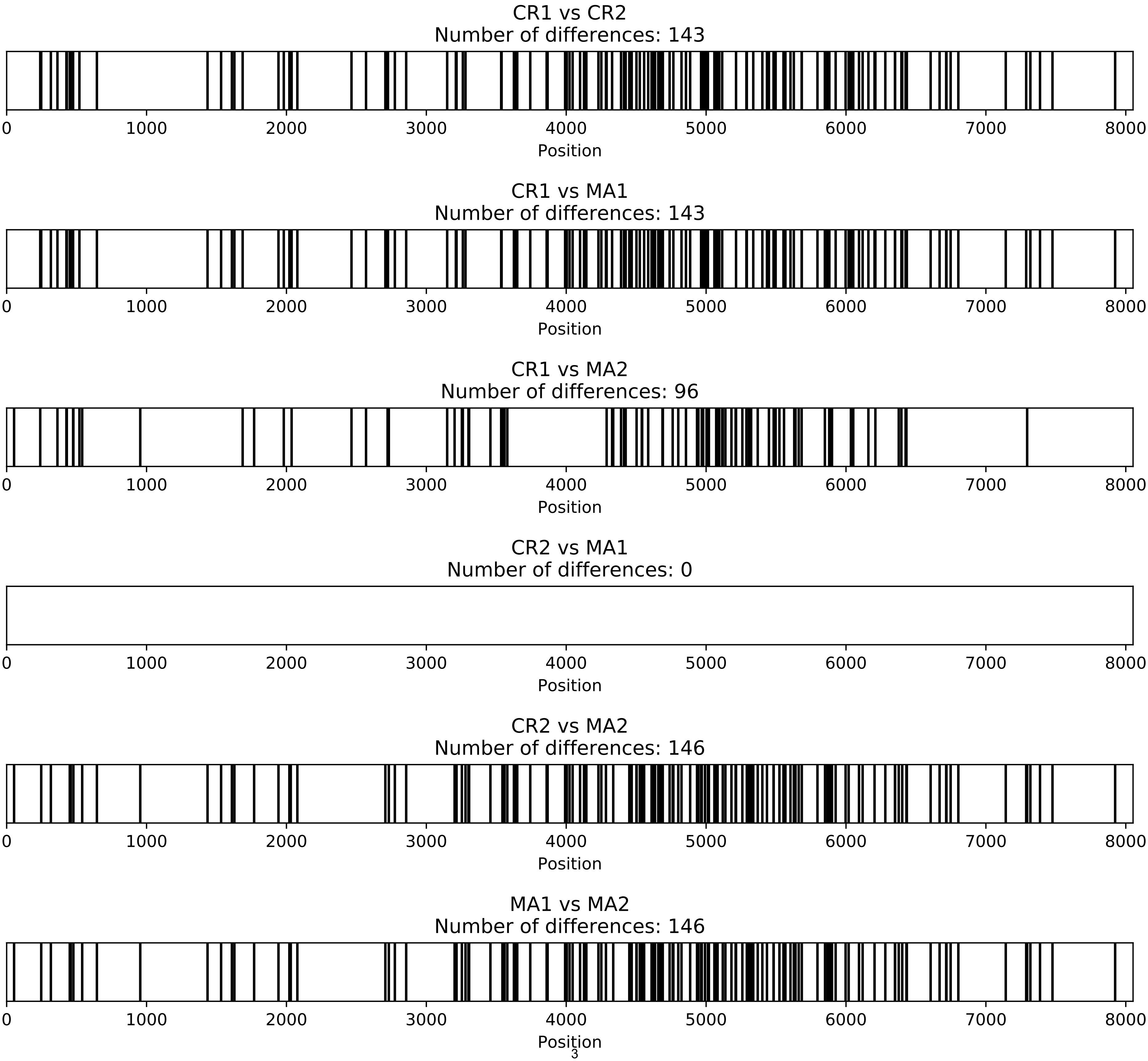
CR1 vs CR2
Number of differences: 143
Position
CR1 vs MA1
Number of differences: 143
Position
CR1 vs MA2
Number of differences: 96
Position
CR2 vs MA1
Number of differences: 0
Position
CR2 vs MA2
Number of differences: 146
Position
MA1 vs MA2
Number of differences: 146
Position
0
1000
2000
3000
4000
5000
6000
7000
8000

### Region: 22



CR1 vs CR2
Number of differences: 146

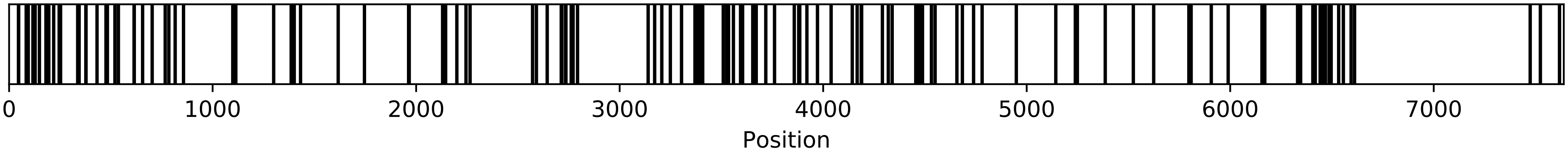


CR1 vs MA1
Number of differences: 149

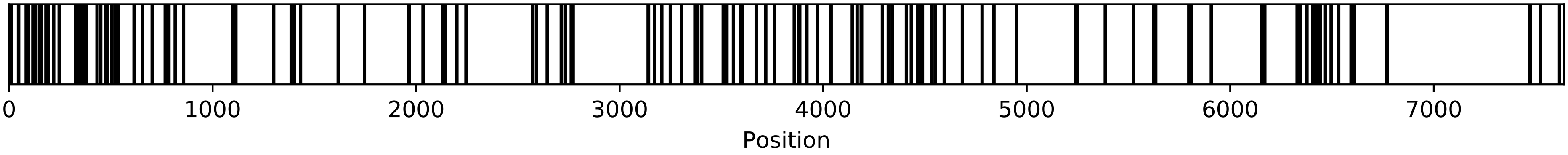


CR1 vs MA2
Number of differences: 84

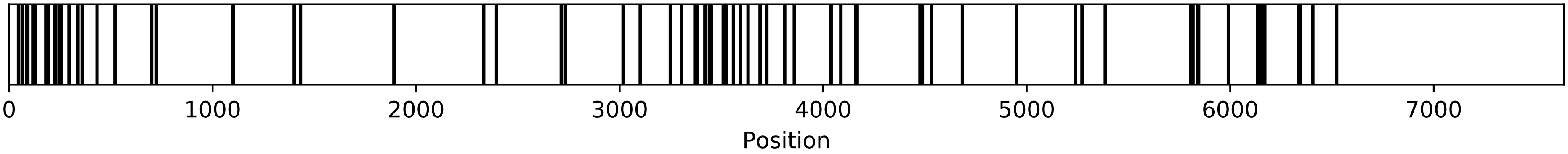


CR2 vs MA1
Number of differences: 41

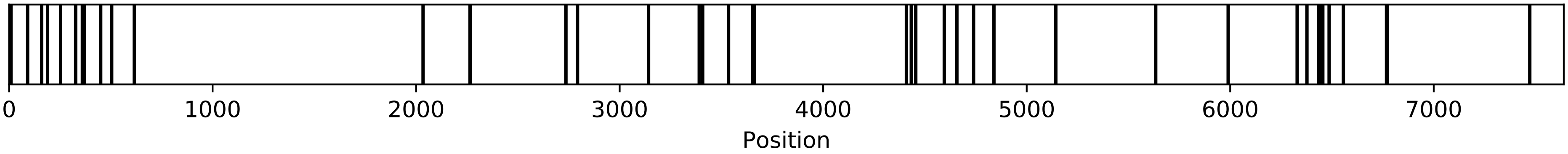


CR2 vs MA2
Number of differences: 149

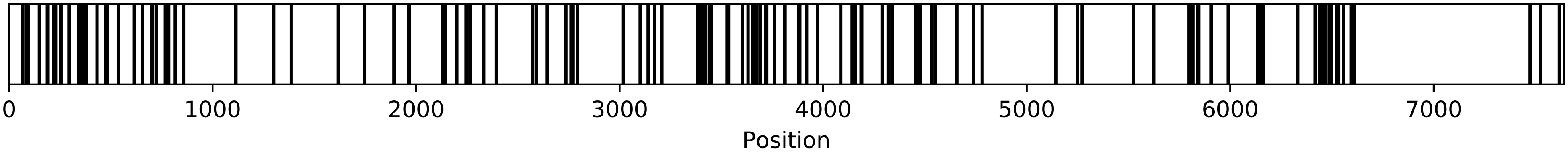


MA1 vs MA2
Number of differences: 150

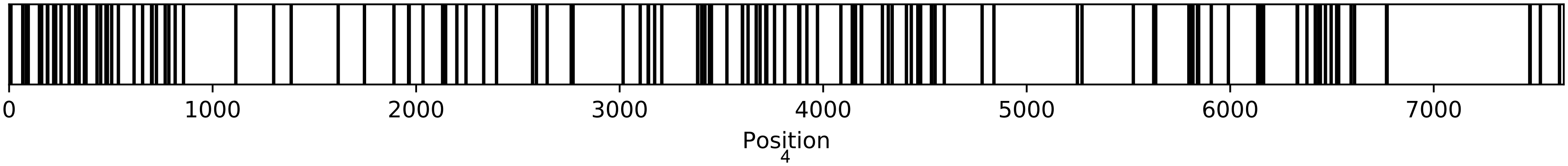

### Region: 29



## CR1 vs CR2
Number of differences: 52

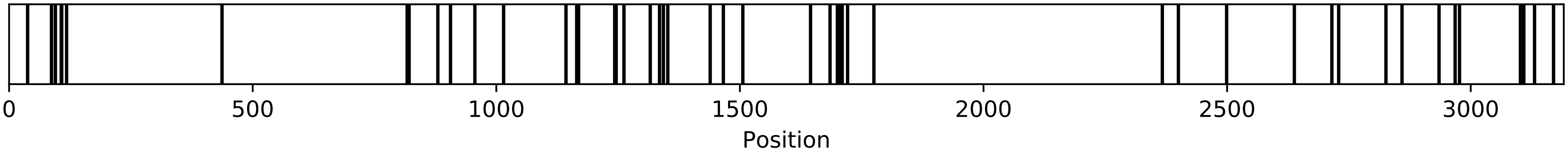


## CR1 vs MA1
Number of differences: 27

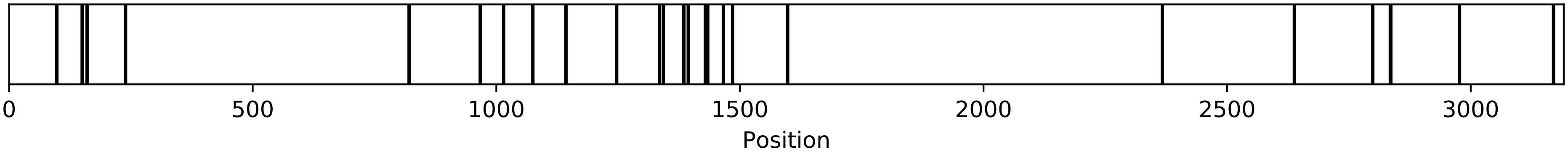


## CR1 vs MA2
Number of differences: 63

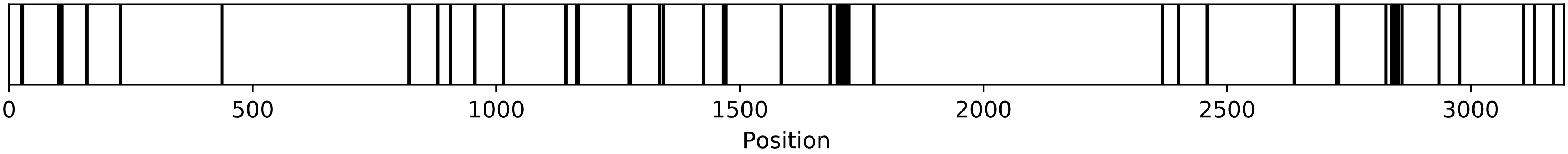


## CR2 vs MA1
Number of differences: 59

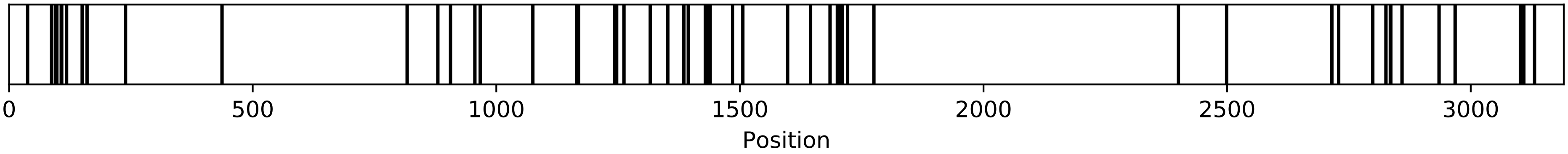


## CR2 vs MA2
Number of differences: 45

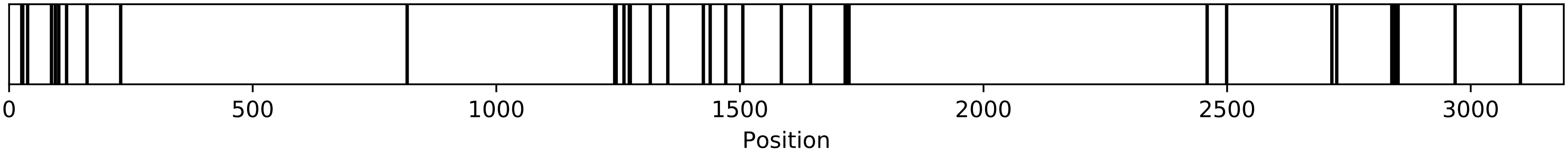


## MA1 vs MA2
Number of differences: 68

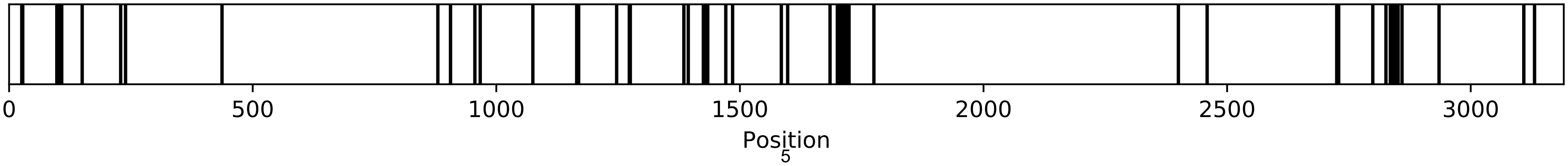

### Region: 31

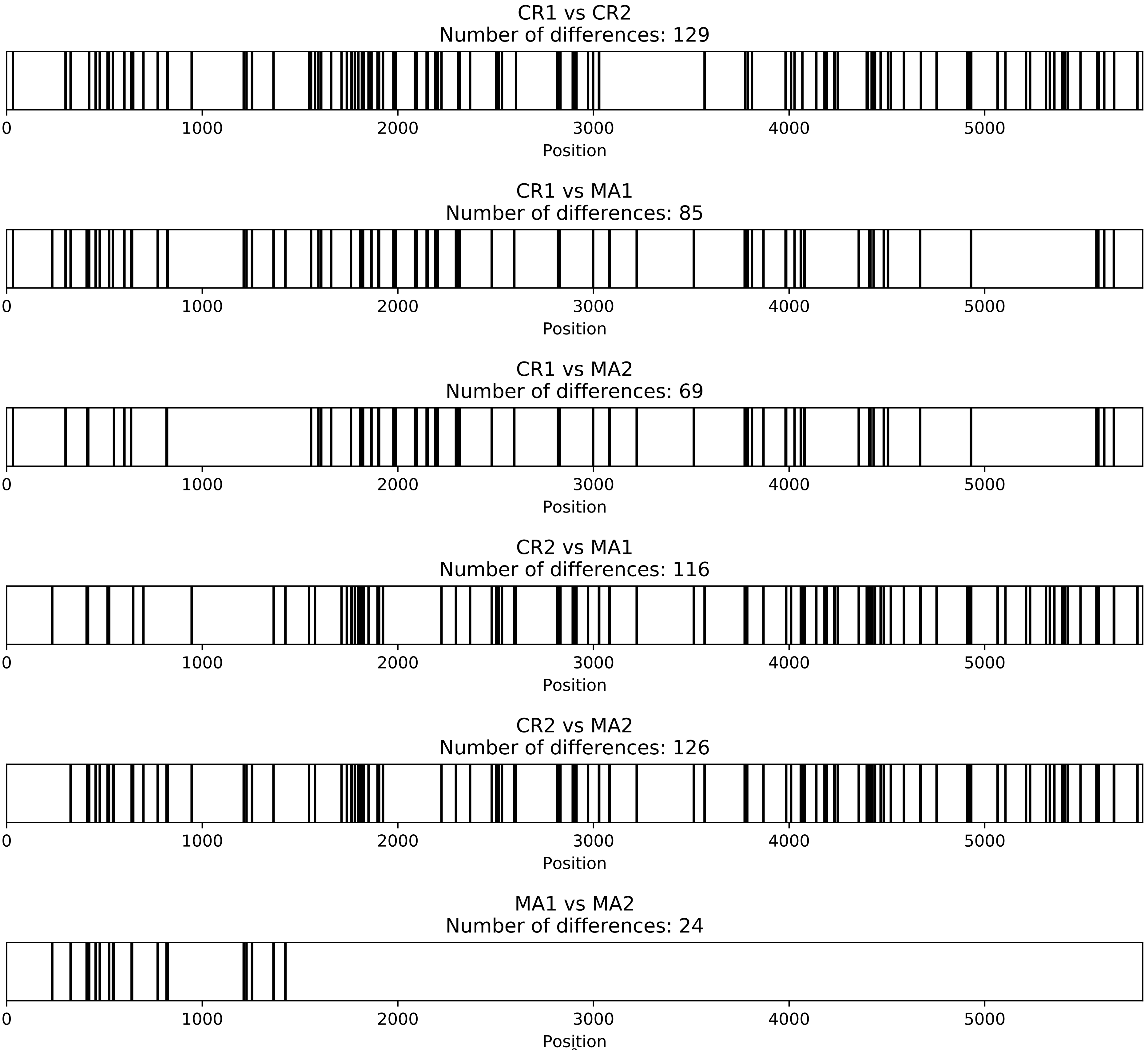
CR1 vs CR2
Number of differences: 129
0 1000 2000 3000 4000 5000
Position
CR1 vs MA1
Number of differences: 85
0 1000 2000 3000 4000 5000
Position
CR1 vs MA2
Number of differences: 69
0 1000 2000 3000 4000 5000
Position
CR2 vs MA1
Number of differences: 116
0 1000 2000 3000 4000 5000
Position
CR2 vs MA2
Number of differences: 126
0 1000 2000 3000 4000 5000
Position
MA1 vs MA2
Number of differences: 24
0 1000 2000 3000 4000 5000
Position

### Region: 42



CR1 vs CR2
Number of differences: 238

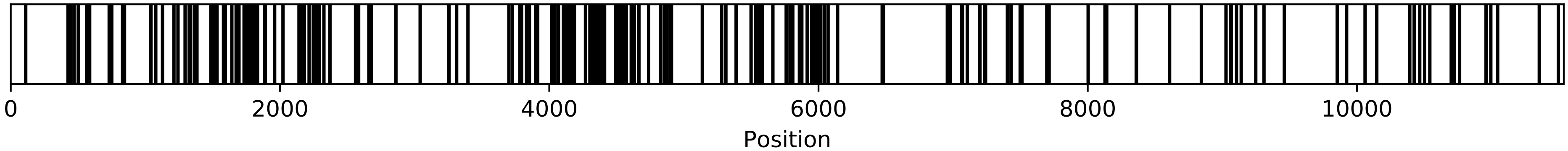


CR1 vs MA1
Number of differences: 69

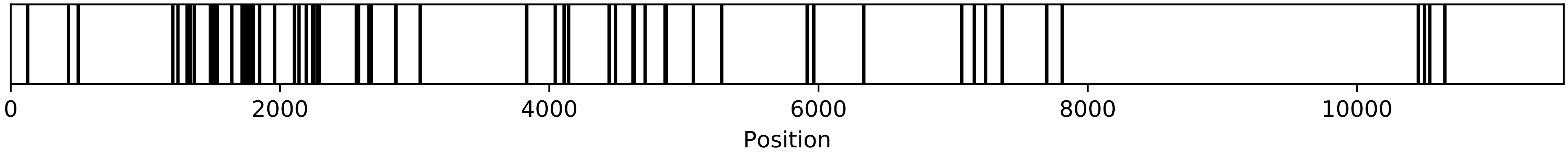


CR1 vs MA2
Number of differences: 237

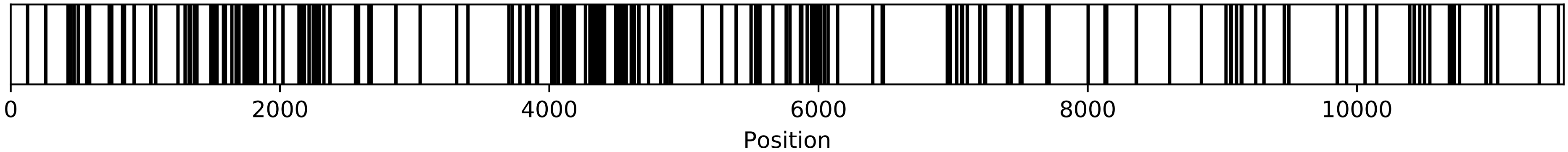


CR2 vs MA1
Number of differences: 223

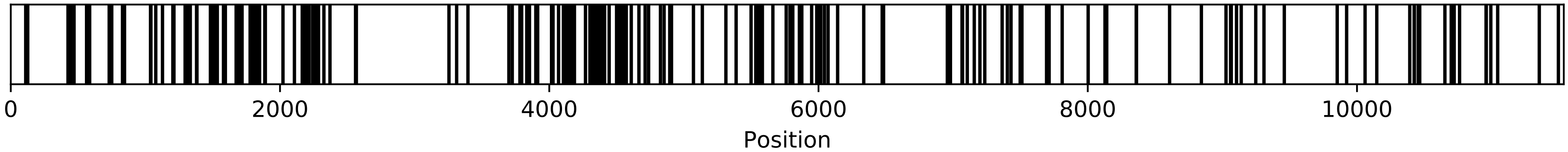


CR2 vs MA2
Number of differences: 25

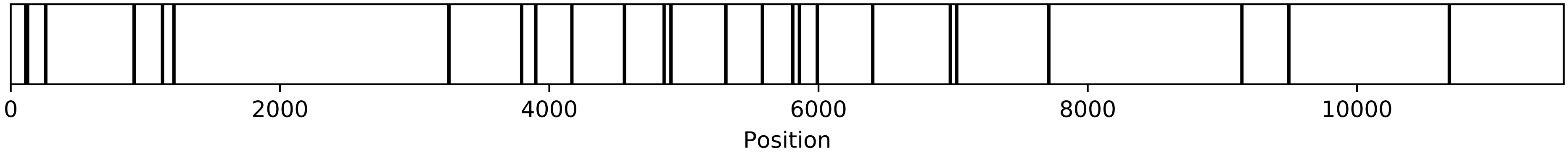


MA1 vs MA2
Number of differences: 222

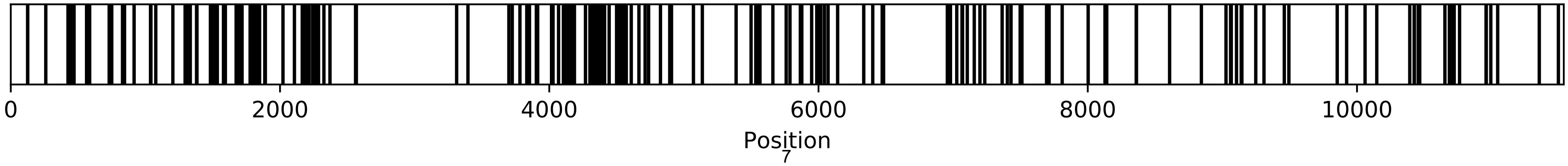

### Region: 44



CR1 vs CR2
Number of differences: 162

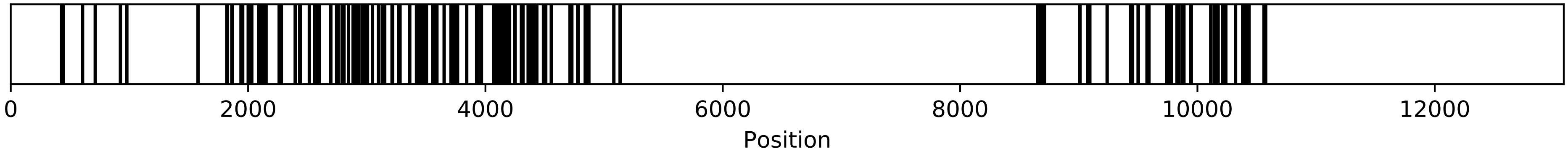


CR1 vs MA1
Number of differences: 221

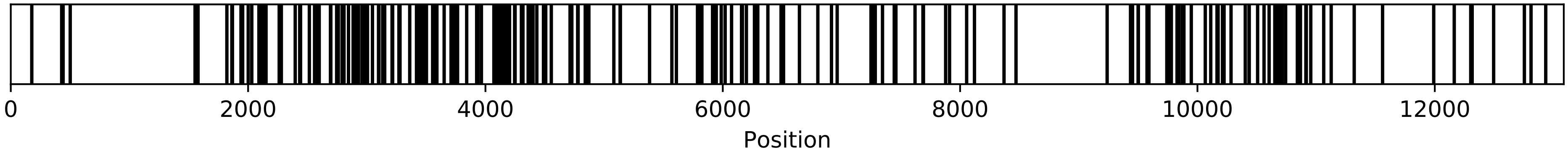


CR1 vs MA2
Number of differences: 102

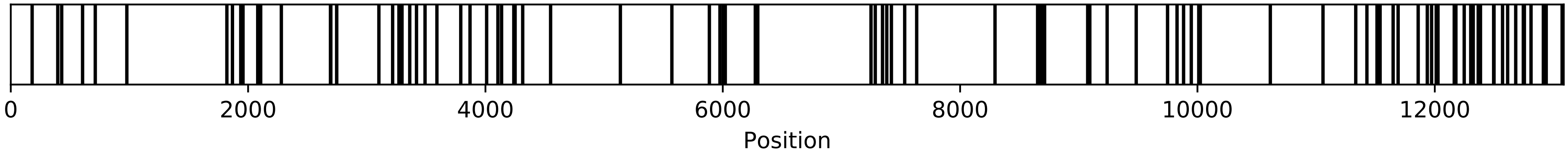


CR2 vs MA1
Number of differences: 105

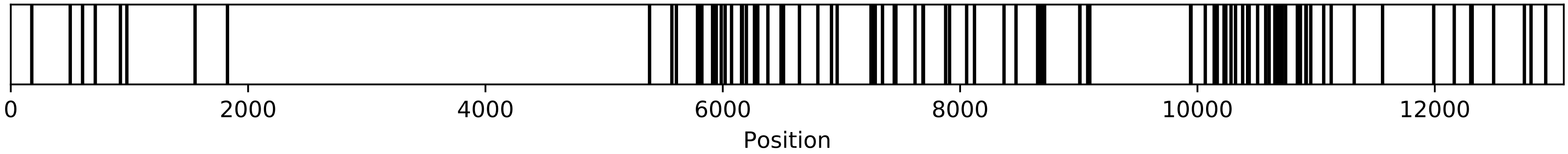


CR2 vs MA2
Number of differences: 192

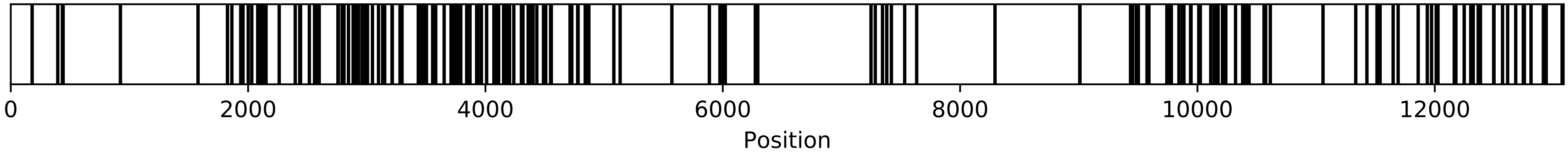


MA1 vs MA2
Number of differences: 245

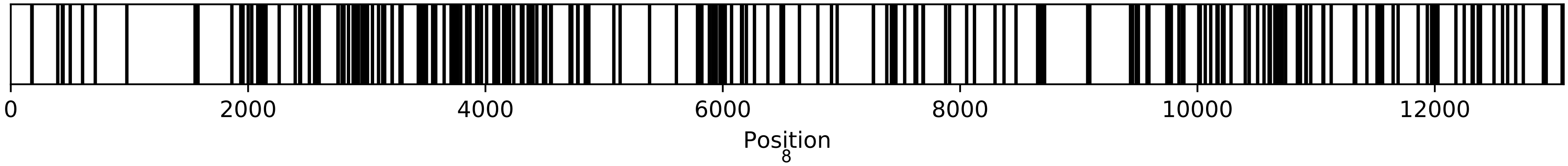

### Region: 48



CR1 vs CR2
Number of differences: 549

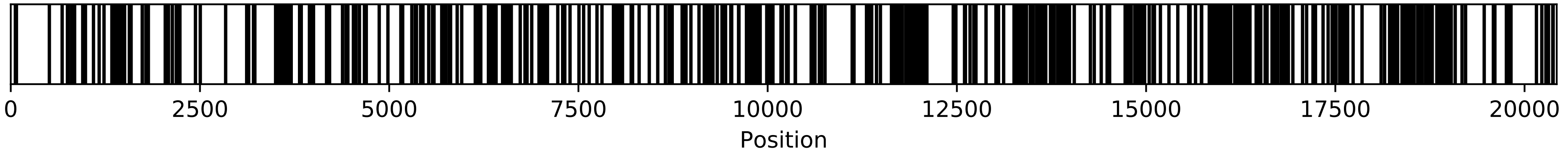


CR1 vs MA1
Number of differences: 192

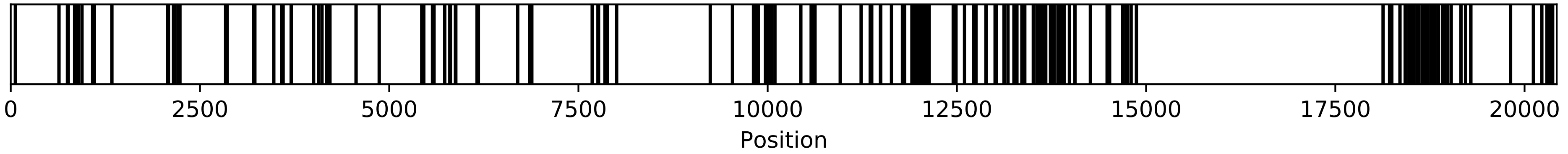


CR1 vs MA2
Number of differences: 537

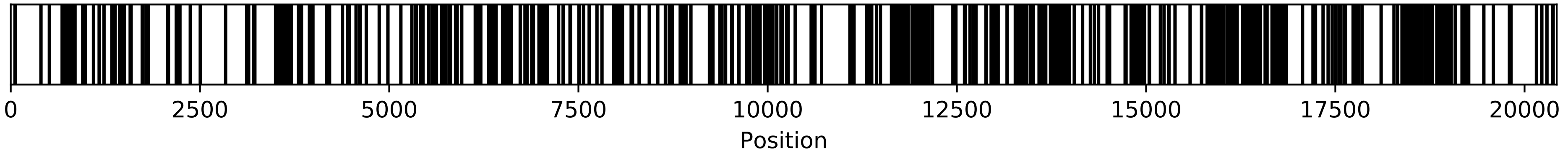


CR2 vs MA1
Number of differences: 553

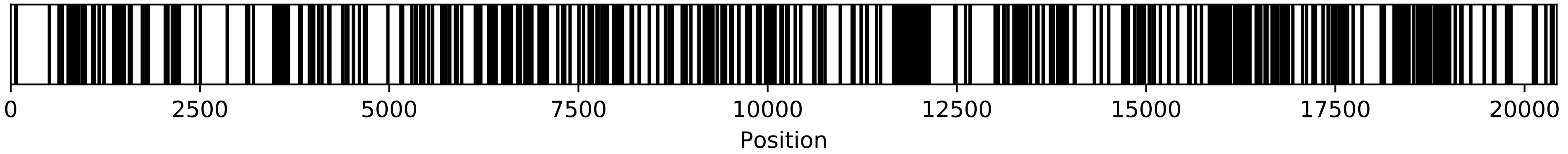


CR2 vs MA2
Number of differences: 205

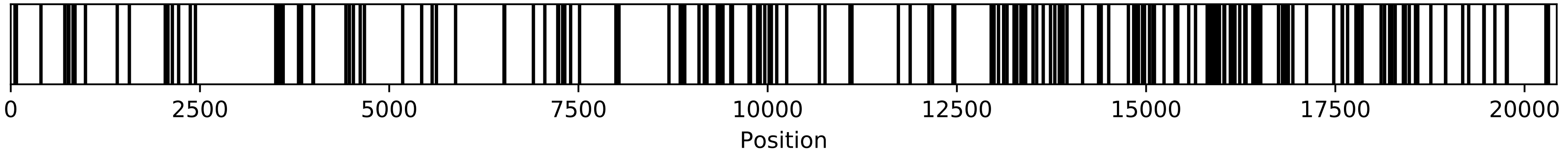


MA1 vs MA2
Number of differences: 548

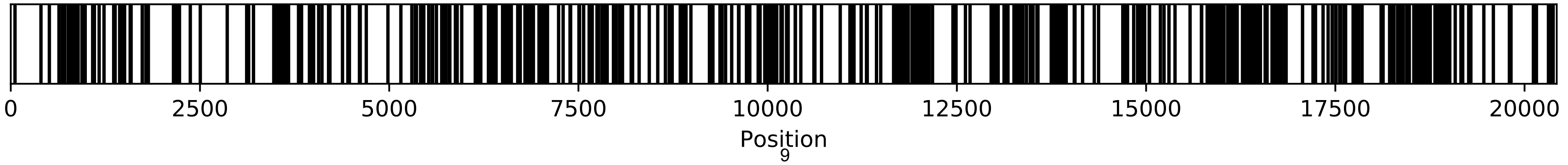

### Region: 49



CR1 vs CR2
Number of differences: 281

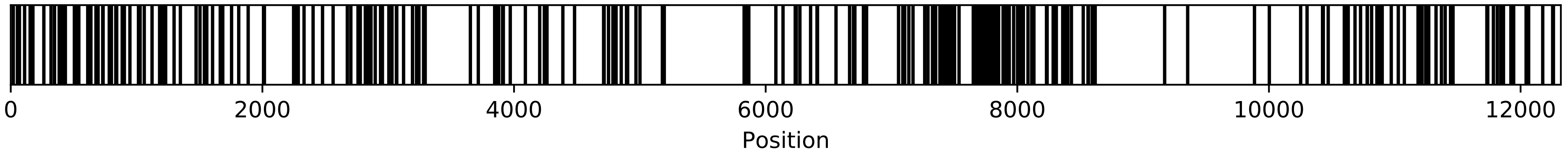


CR1 vs MA1
Number of differences: 274

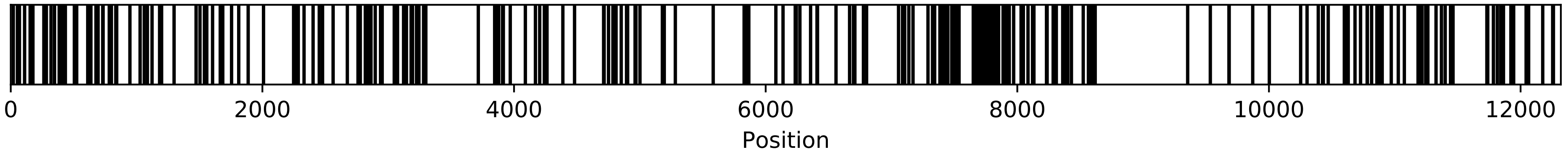


CR1 vs MA2
Number of differences: 117

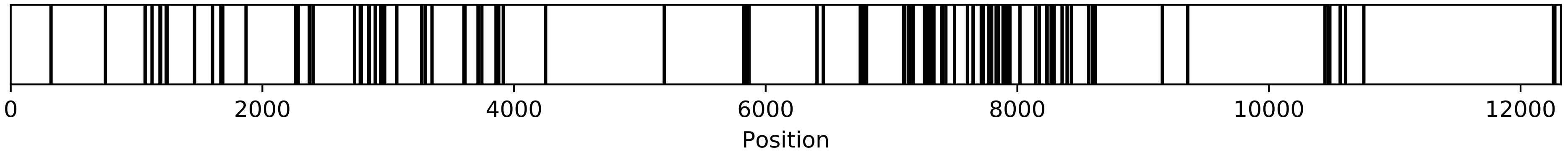


CR2 vs MA1
Number of differences: 65

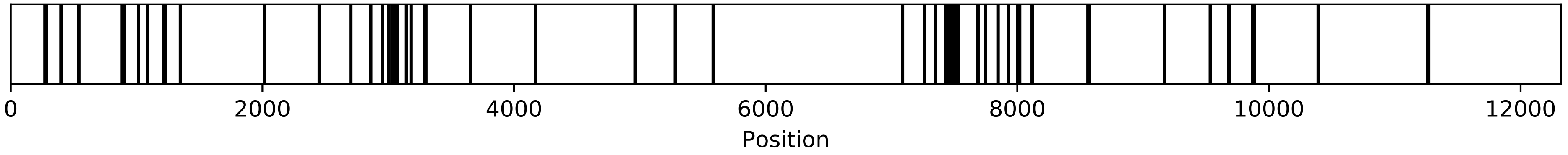


CR2 vs MA2
Number of differences: 292

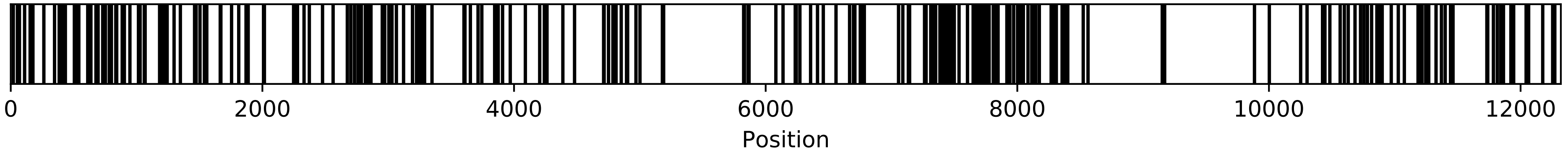


MA1 vs MA2
Number of differences: 285

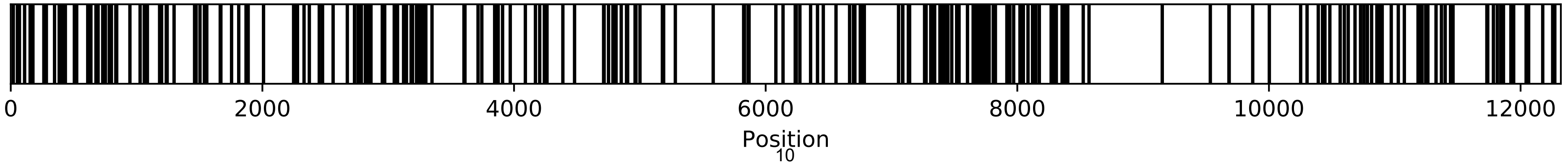

### Region: 55



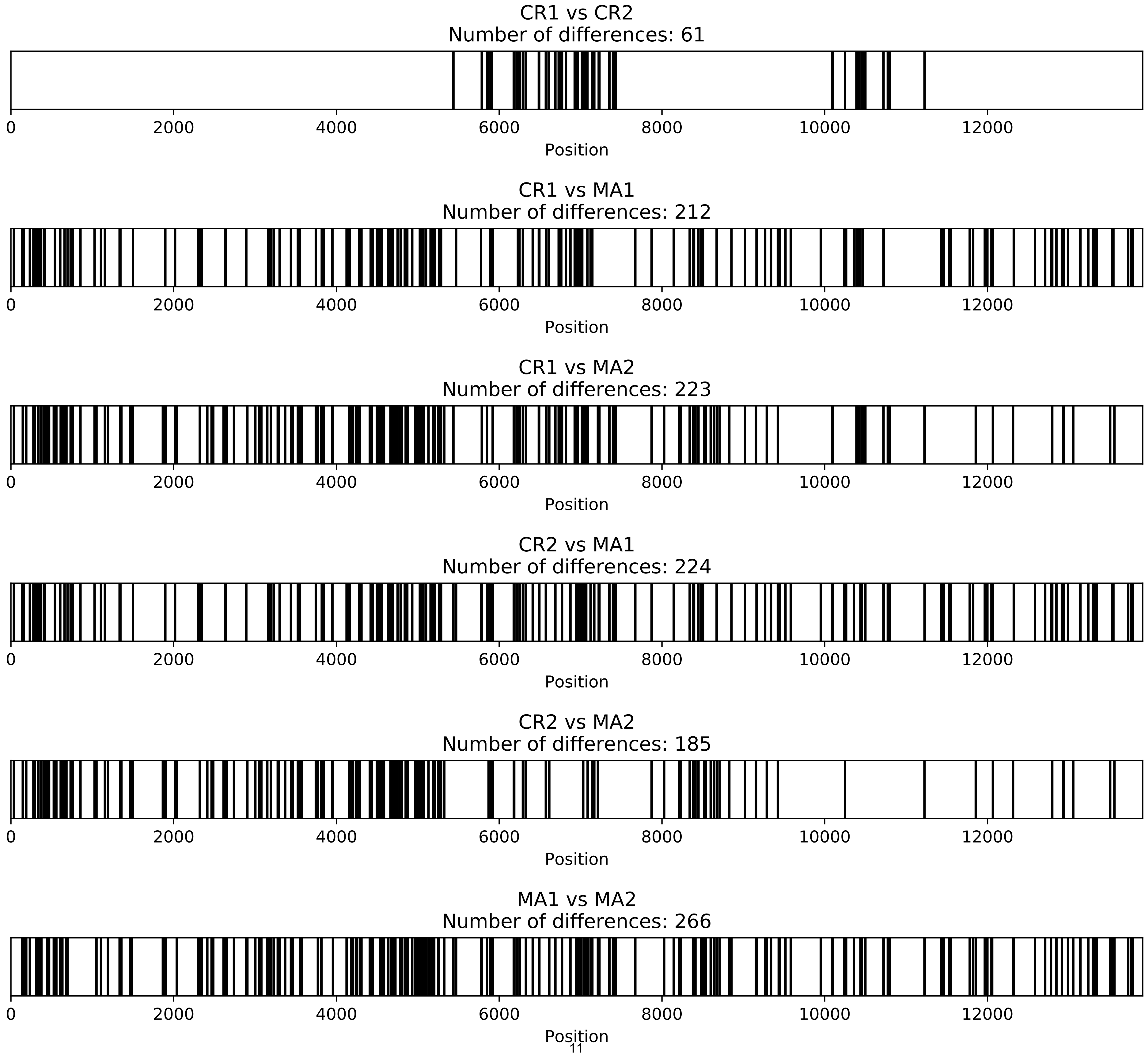
CR1 vs CR2
Number of differences: 61
CR1 vs MA1
Number of differences: 212
CR1 vs MA2
Number of differences: 223
CR2 vs MA1
Number of differences: 224
CR2 vs MA2
Number of differences: 185
MA1 vs MA2
Number of differences: 266
0
2000
4000
6000
8000
10000
12000
Position

### Region: 59



CR1 vs CR2
Number of differences: 144

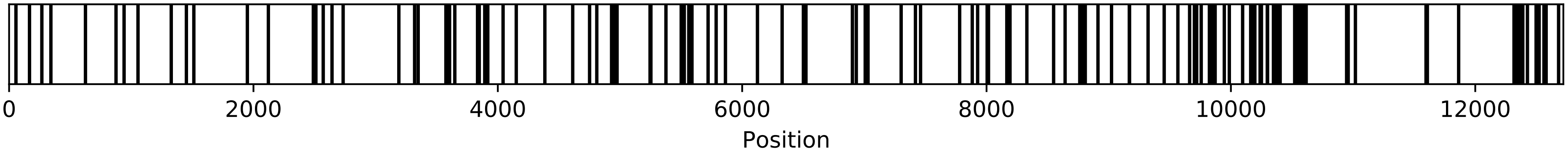


CR1 vs MA1
Number of differences: 48

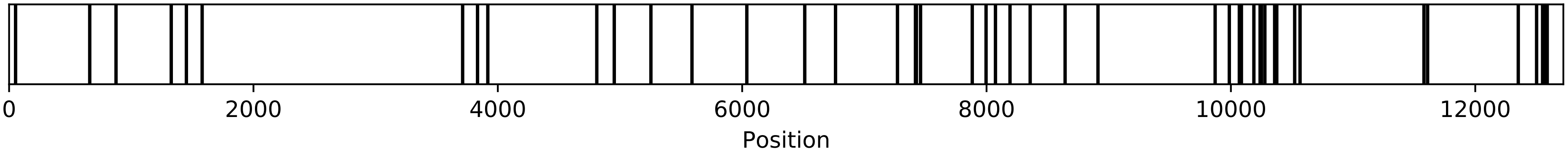


CR1 vs MA2
Number of differences: 152

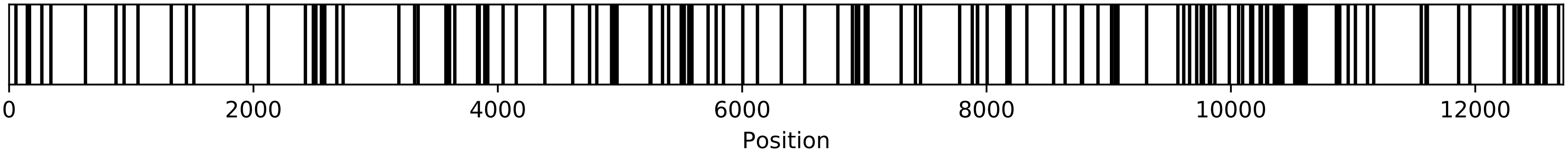


CR2 vs MA1
Number of differences: 145

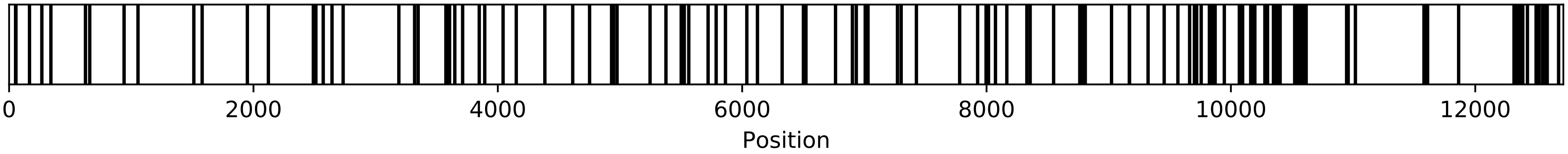


CR2 vs MA2
Number of differences: 62

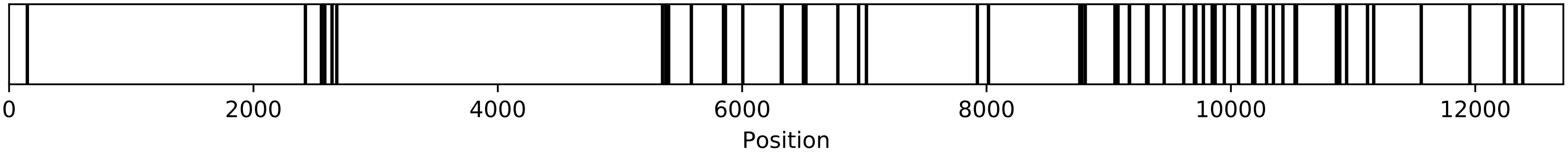


MA1 vs MA2
Number of differences: 154

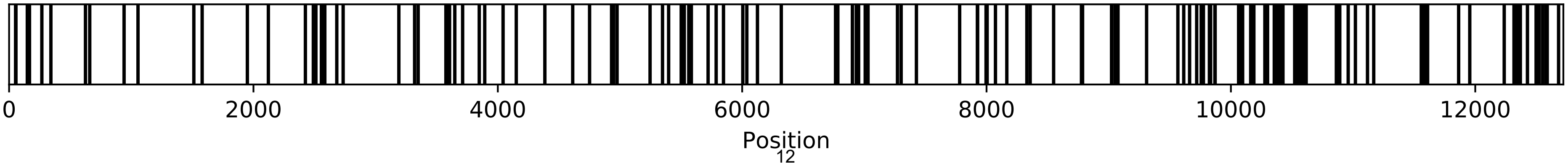

### Region: 73



CR1 vs CR2
Number of differences: 141

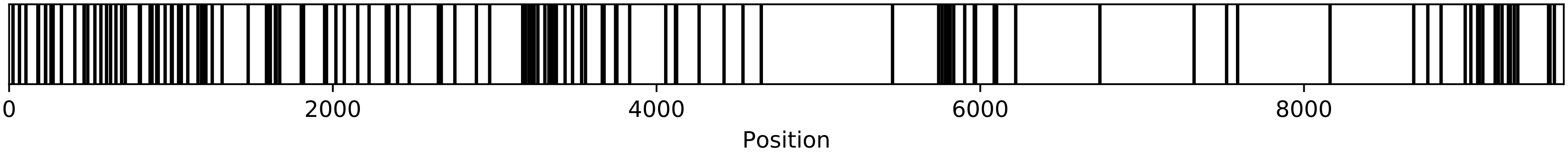


CR1 vs MA1
Number of differences: 147

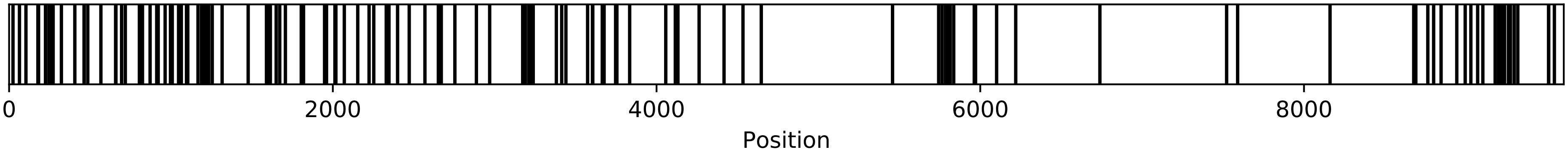


CR1 vs MA2
Number of differences: 40

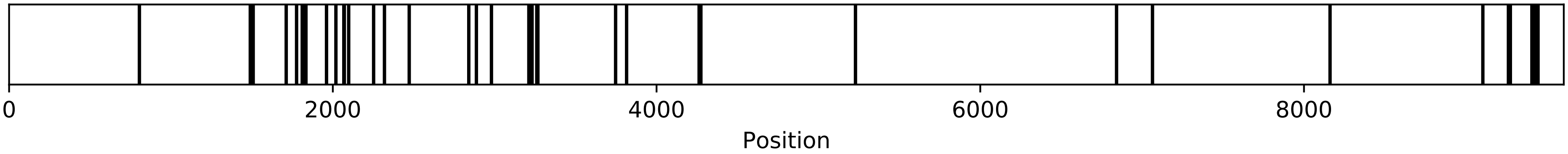


CR2 vs MA1
Number of differences: 55

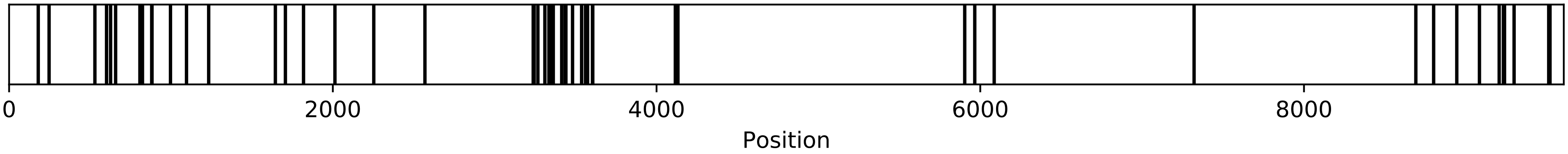


CR2 vs MA2
Number of differences: 147

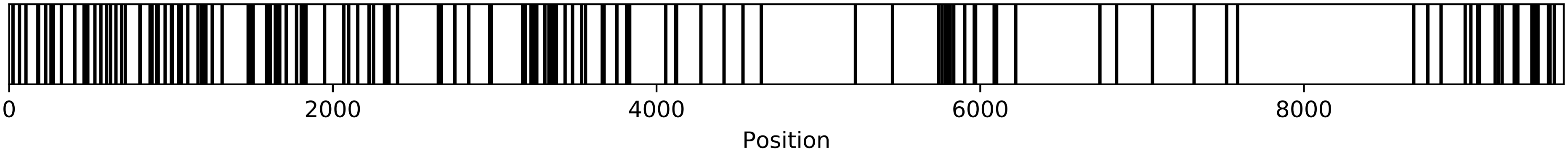


MA1 vs MA2
Number of differences: 154

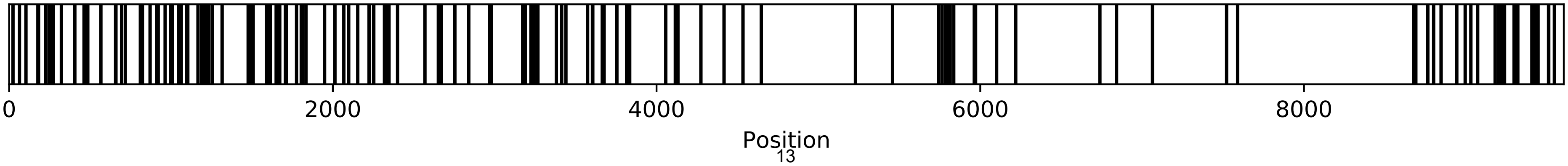

### Region: 84



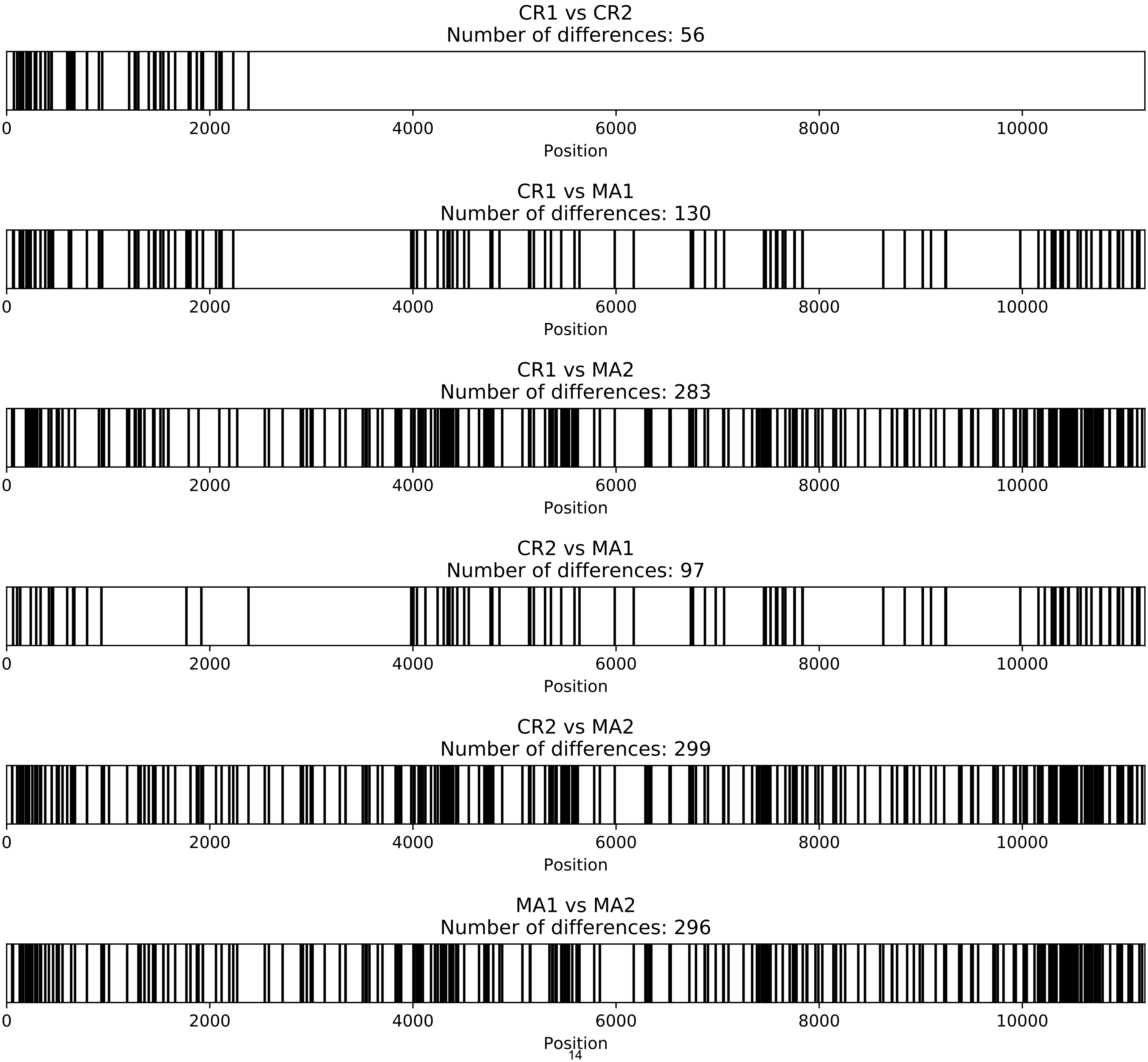
CR1 vs CR2
Number of differences: 56
CR1 vs MA1
Number of differences: 130
CR1 vs MA2
Number of differences: 283
CR2 vs MA1
Number of differences: 97
CR2 vs MA2
Number of differences: 299
MA1 vs MA2
Number of differences: 296
0
2000
4000
6000
8000
10000
Position

### Region: 109



CR1 vs CR2
Number of differences: 227

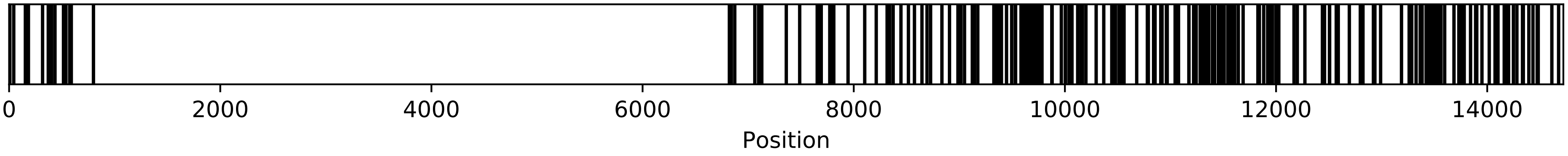


CR1 vs MA1
Number of differences: 288

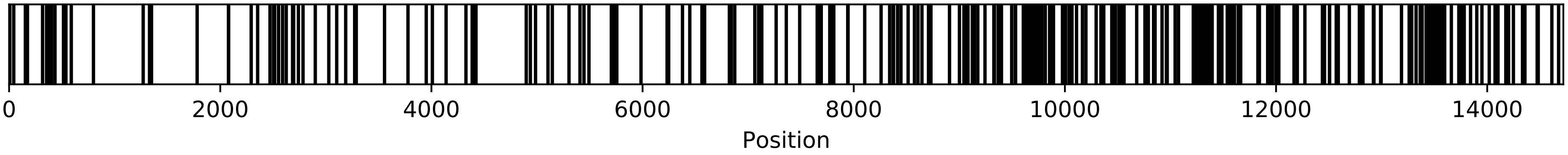


CR1 vs MA2
Number of differences: 297

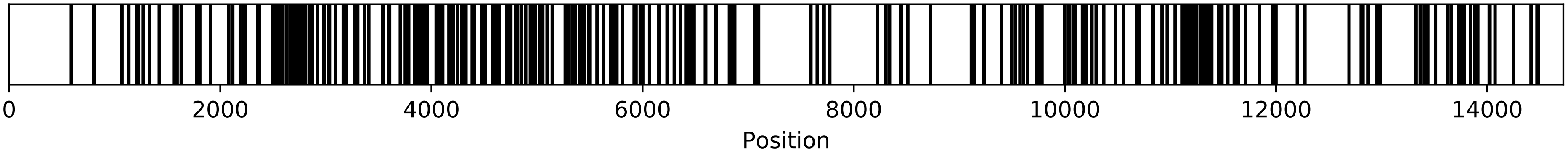


CR2 vs MA1
Number of differences: 155

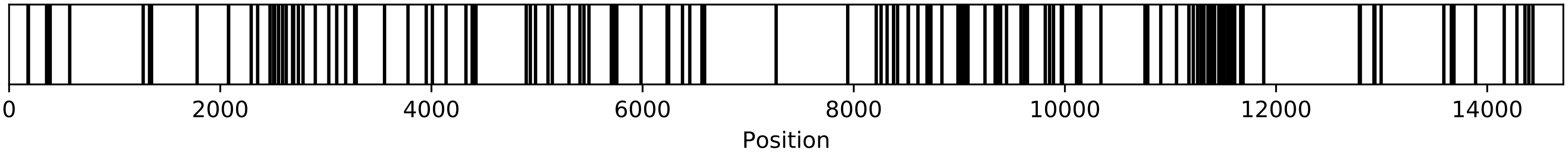


CR2 vs MA2
Number of differences: 390

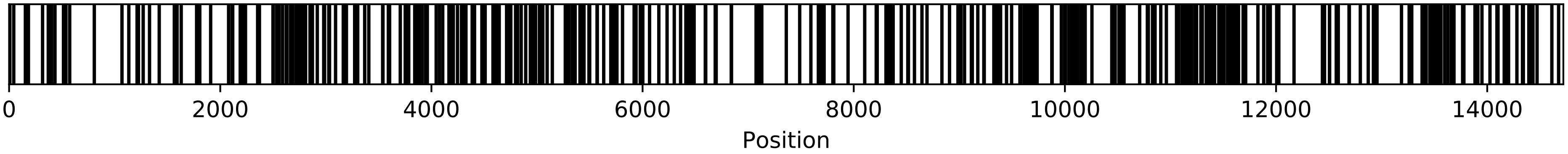


MA1 vs MA2
Number of differences: 388

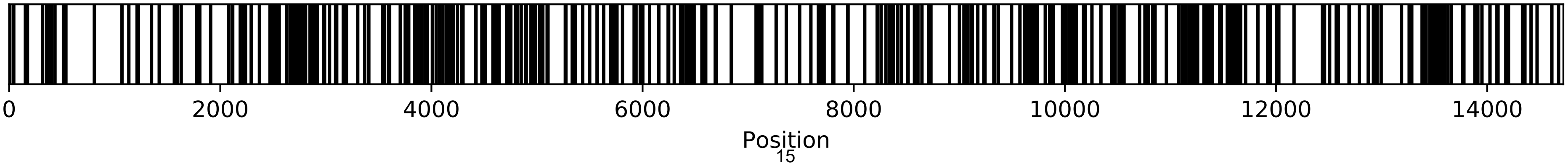

### Region: 111



CR1 vs CR2
Number of differences: 212

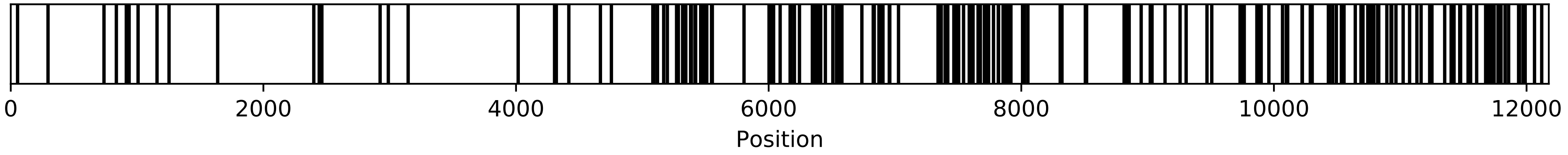


CR1 vs MA1
Number of differences: 165

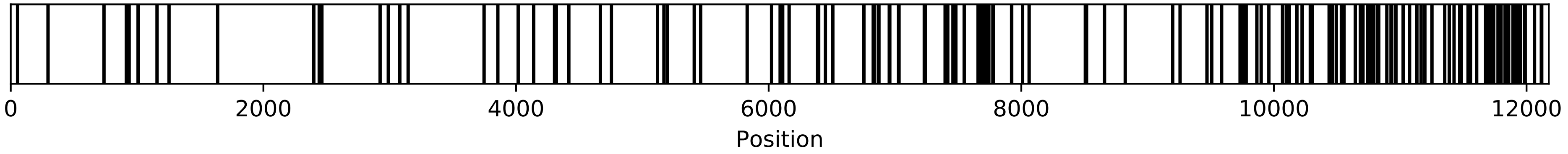


CR1 vs MA2
Number of differences: 133

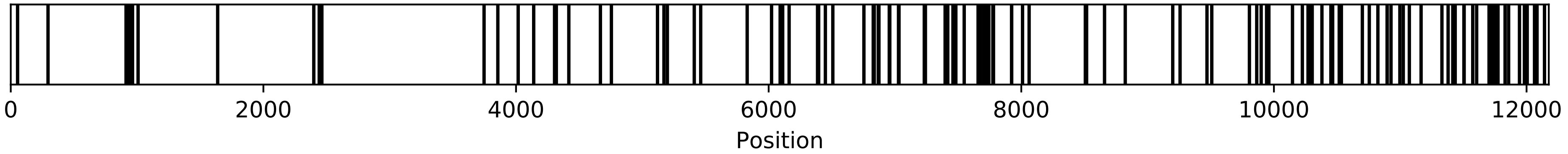


CR2 vs MA1
Number of differences: 134

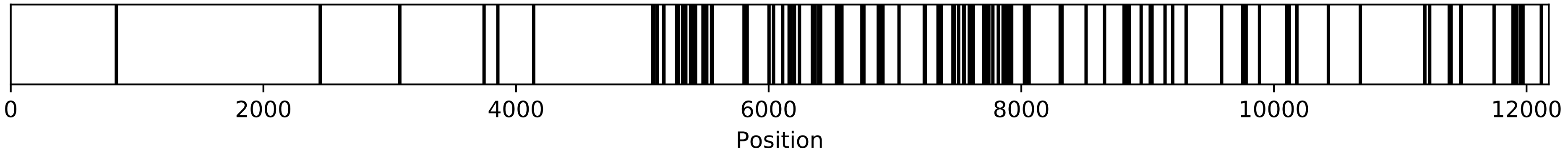


CR2 vs MA2
Number of differences: 203

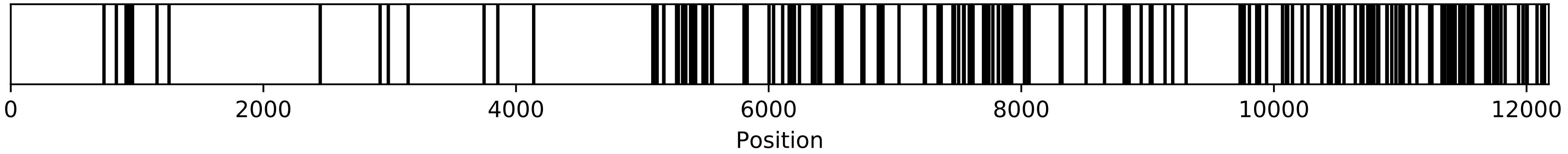


MA1 vs MA2
Number of differences: 101

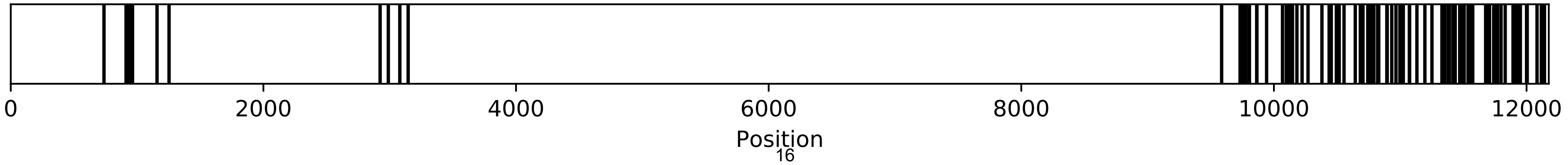

### Region: 122



CR1 vs CR2
Number of differences: 133

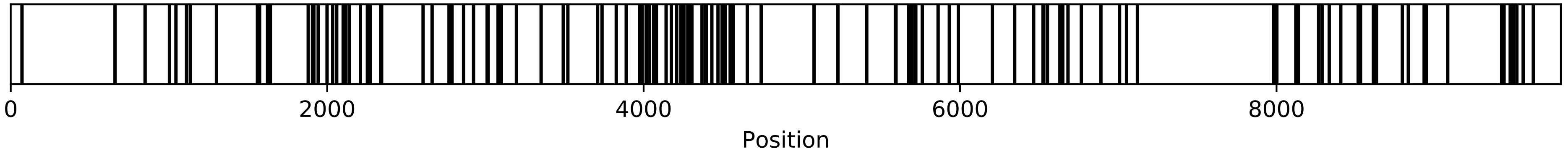


CR1 vs MA1
Number of differences: 91

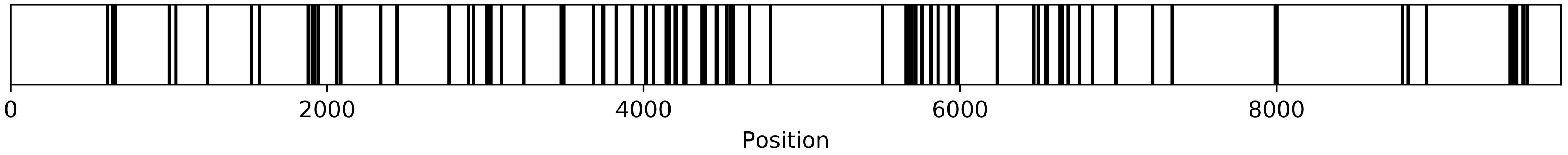


CR1 vs MA2
Number of differences: 134

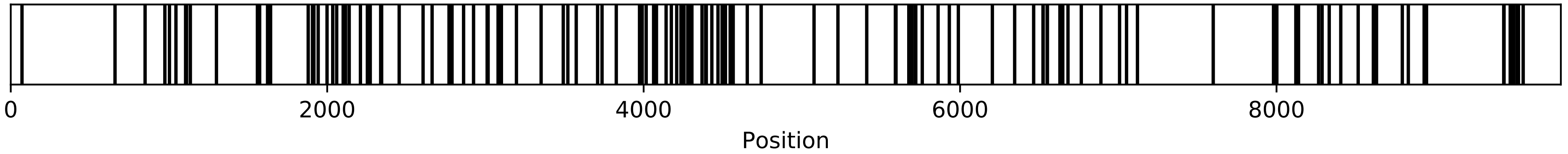


CR2 vs MA1
Number of differences: 133

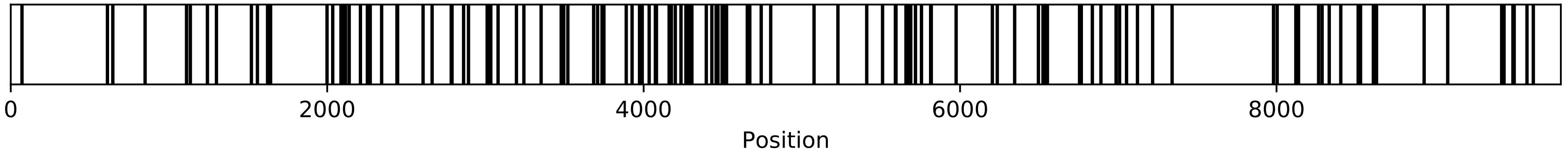


CR2 vs MA2
Number of differences: 13

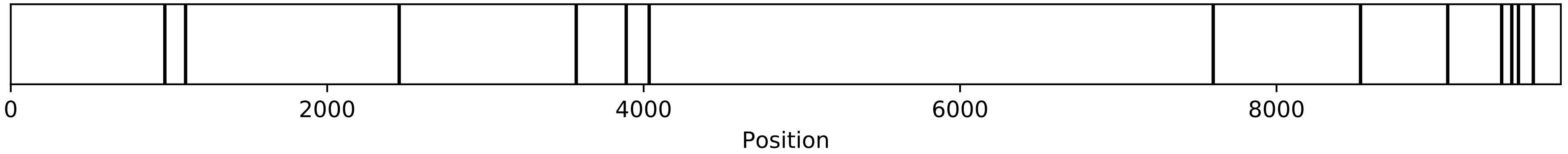


MA1 vs MA2
Number of differences: 134

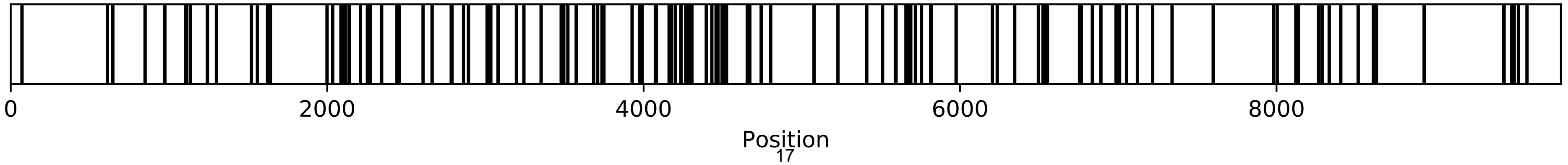

### Region: 127



CR1 vs CR2
Number of differences: 65

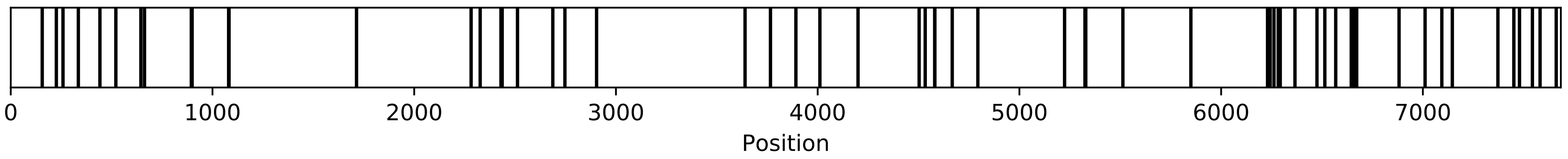


CR1 vs MA1
Number of differences: 75

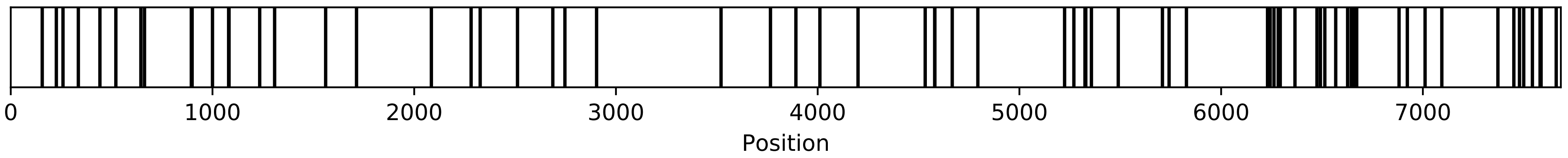


CR1 vs MA2
Number of differences: 24

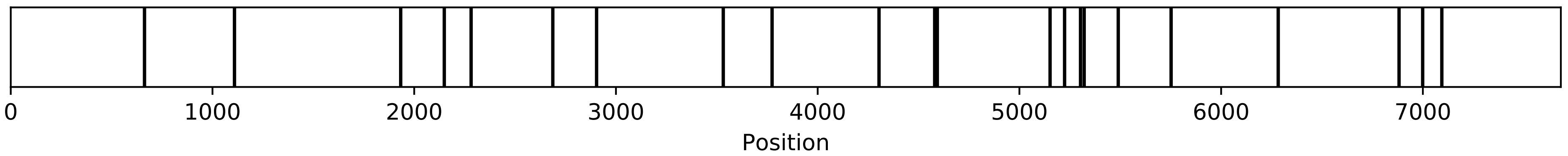


CR2 vs MA1
Number of differences: 26

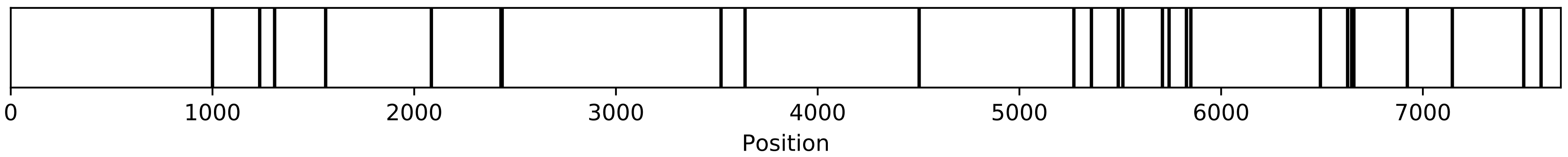


CR2 vs MA2
Number of differences: 69

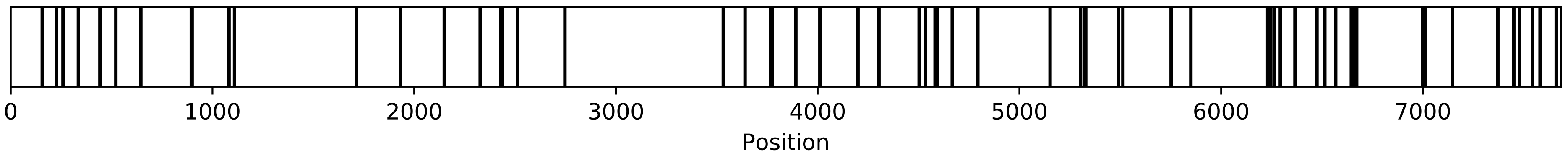


MA1 vs MA2
Number of differences: 77

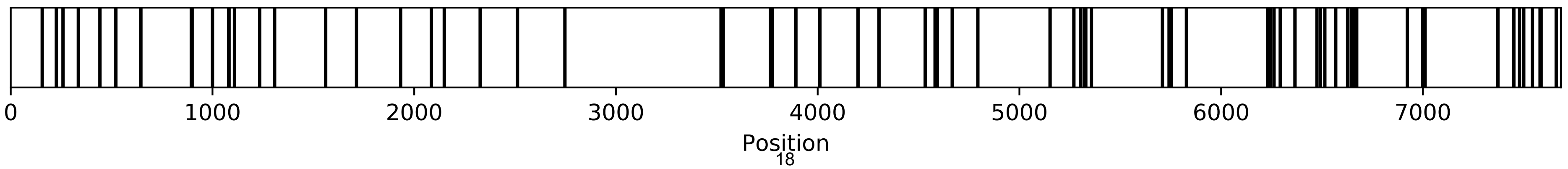

### Region: 129



CR1 vs CR2
Number of differences: 307

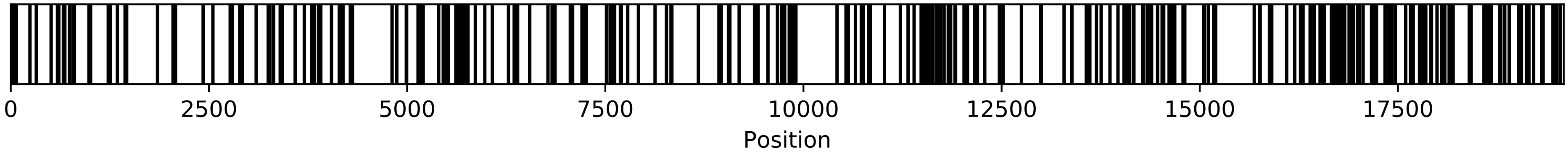


CR1 vs MA1
Number of differences: 311

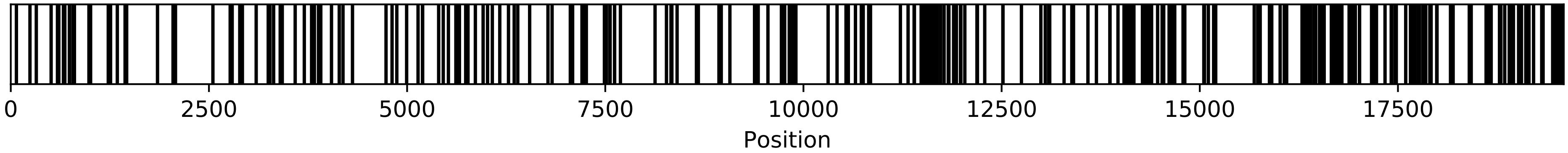


CR1 vs MA2
Number of differences: 64

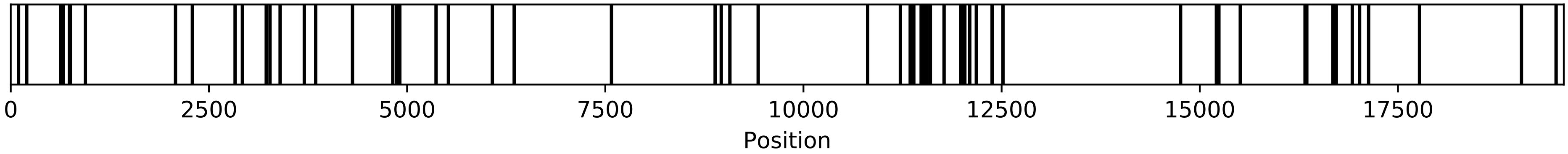


CR2 vs MA1
Number of differences: 119

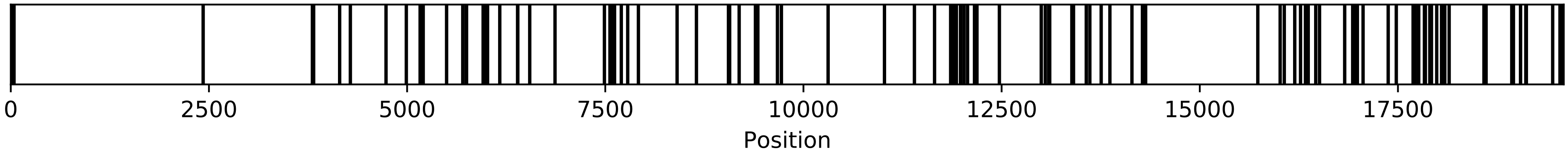


CR2 vs MA2
Number of differences: 318

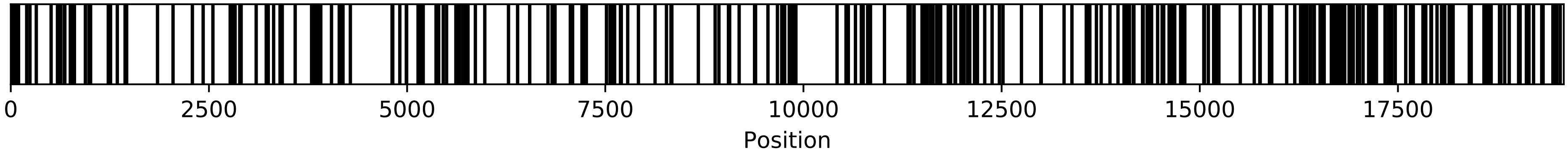


MA1 vs MA2
Number of differences: 320

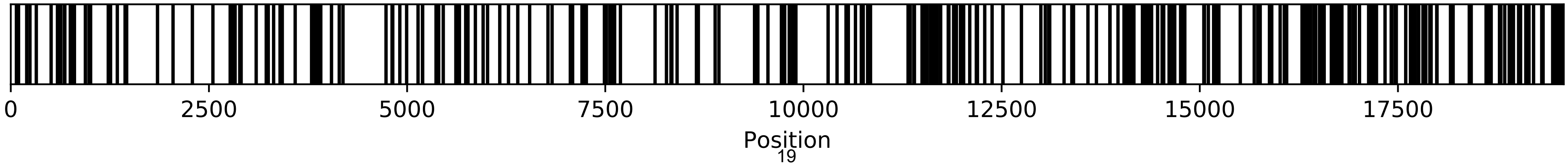

### Region: 132



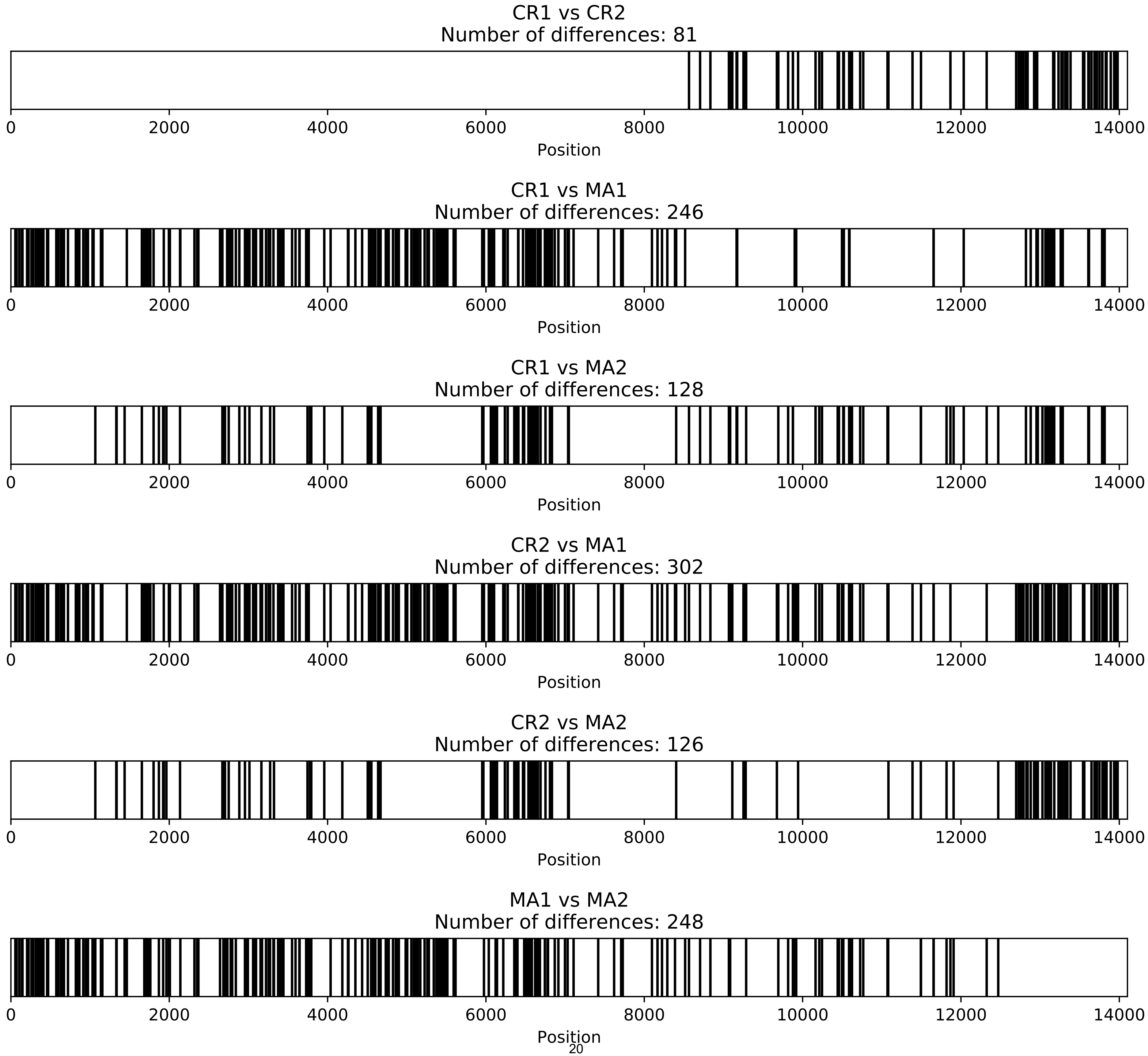
CR1 vs CR2
Number of differences: 81
CR1 vs MA1
Number of differences: 246
CR1 vs MA2
Number of differences: 128
CR2 vs MA1
Number of differences: 302
CR2 vs MA2
Number of differences: 126
MA1 vs MA2
Number of differences: 248
0
2000
4000
6000
8000
10000
12000
14000
Position

### Region: 142



CR1 vs CR2
Number of differences: 104

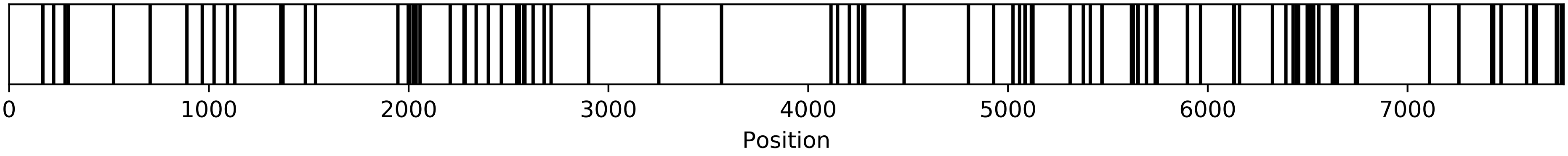


CR1 vs MA1
Number of differences: 36

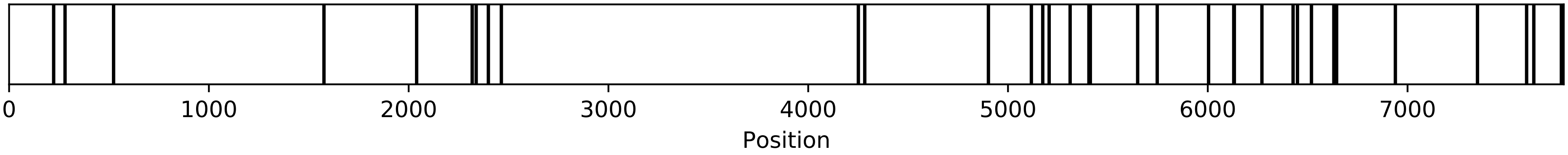


CR1 vs MA2
Number of differences: 60

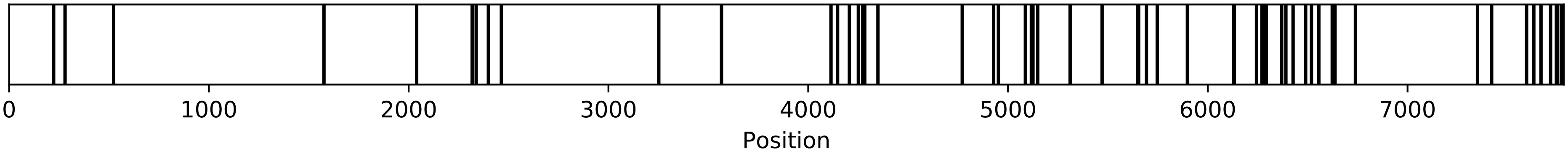


CR2 vs MA1
Number of differences: 96

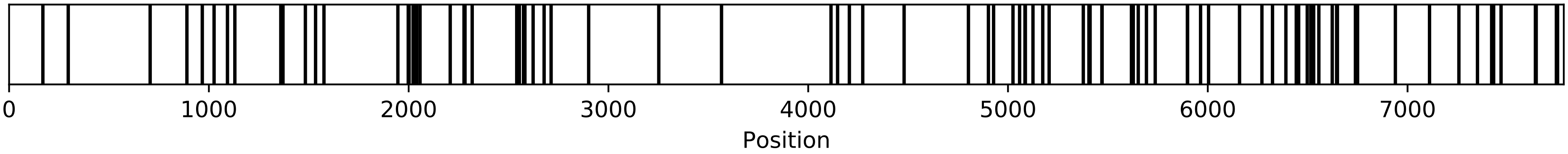


CR2 vs MA2
Number of differences: 84

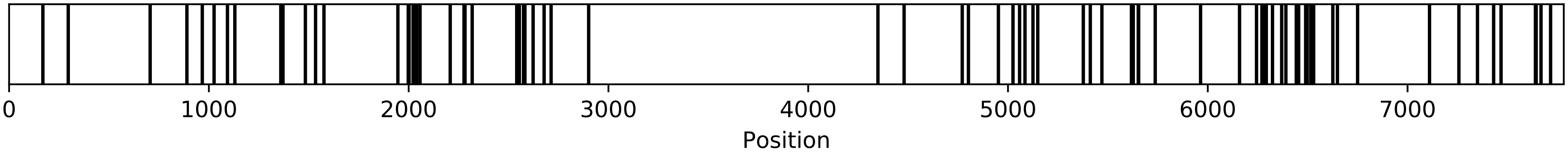


MA1 vs MA2
Number of differences: 44

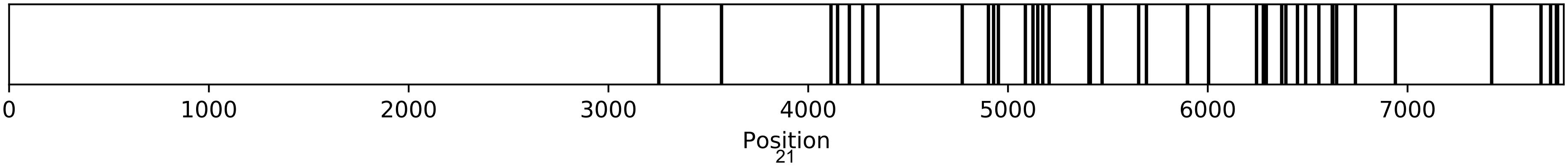

### Region: 144



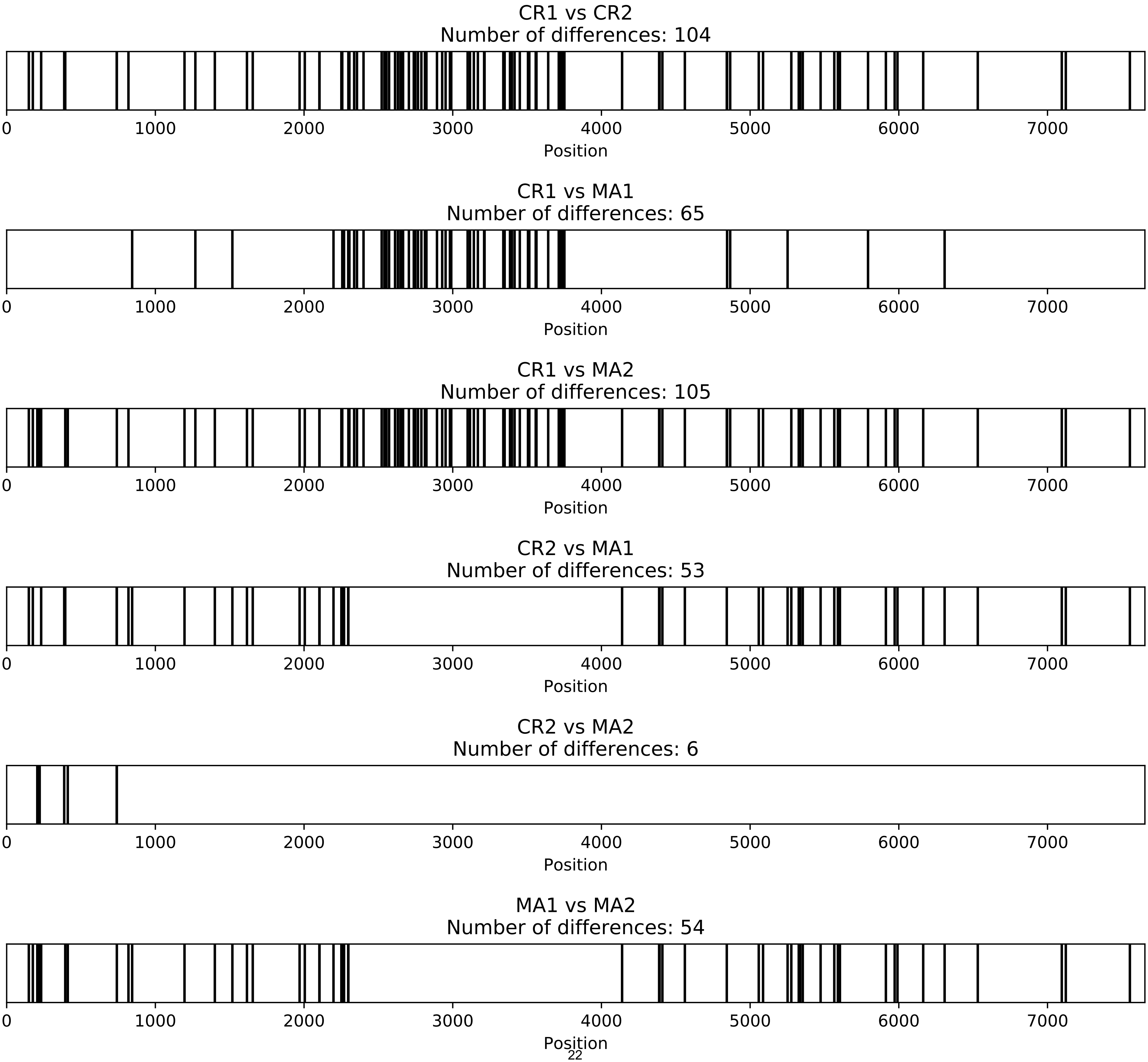
CR1 vs CR2
Number of differences: 104
0 1000 2000 3000 4000 5000 6000 7000
Position
CR1 vs MA1
Number of differences: 65
0 1000 2000 3000 4000 5000 6000 7000
Position
CR1 vs MA2
Number of differences: 105
0 1000 2000 3000 4000 5000 6000 7000
Position
CR2 vs MA1
Number of differences: 53
0 1000 2000 3000 4000 5000 6000 7000
Position
CR2 vs MA2
Number of differences: 6
0 1000 2000 3000 4000 5000 6000 7000
Position
MA1 vs MA2
Number of differences: 54
0 1000 2000 3000 4000 5000 6000 7000
Position

### Region: 153



CR1 vs CR2
Number of differences: 570

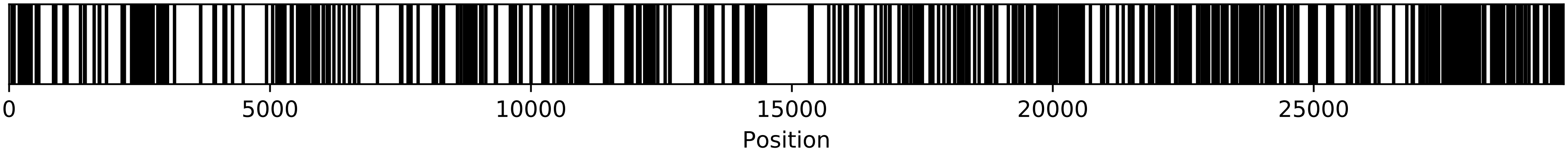


CR1 vs MA1
Number of differences: 568

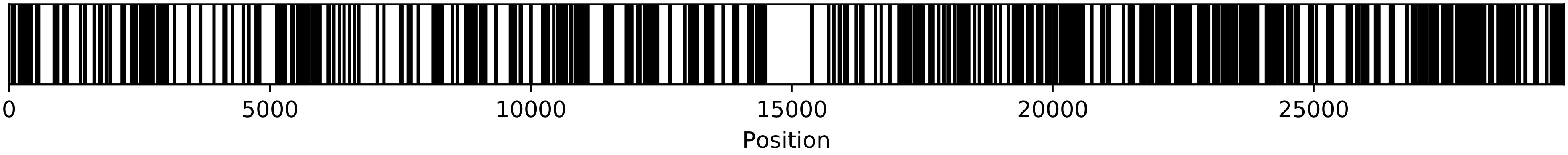


CR1 vs MA2
Number of differences: 90

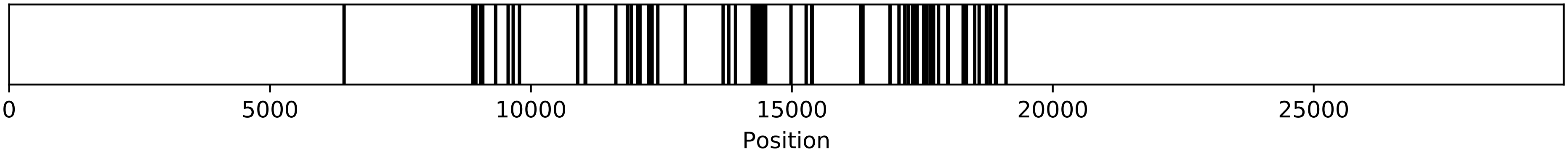


CR2 vs MA1
Number of differences: 161

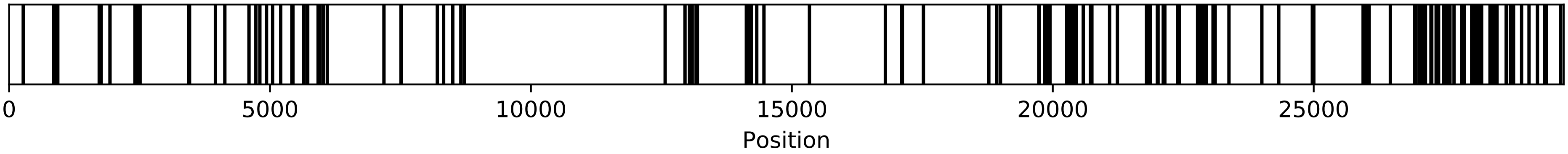


CR2 vs MA2
Number of differences: 567

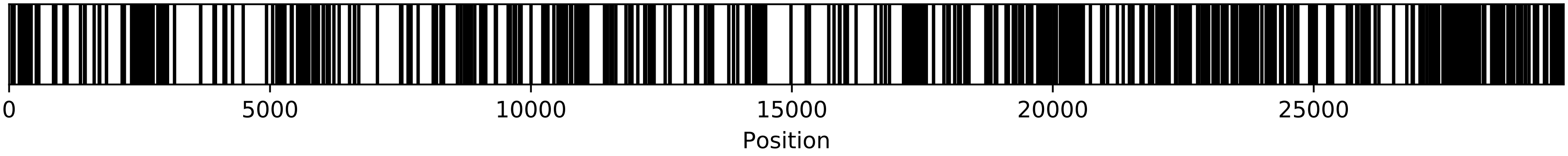


MA1 vs MA2
Number of differences: 565

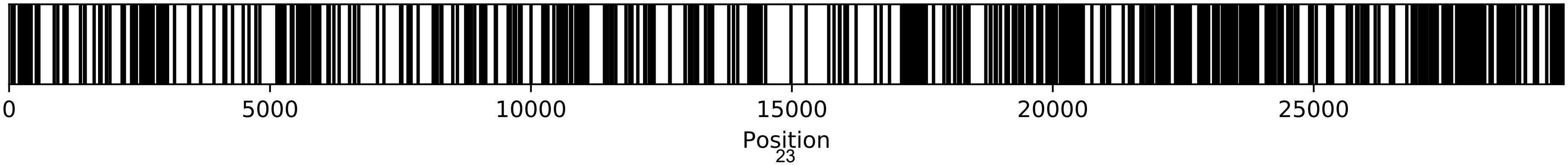

### Region: 161



CR1 vs CR2
Number of differences: 216

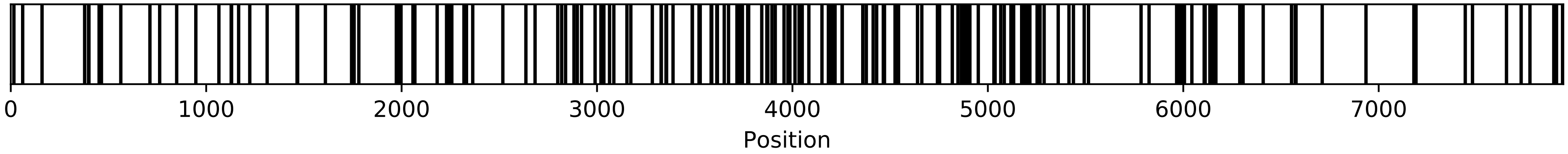


CR1 vs MA1
Number of differences: 109

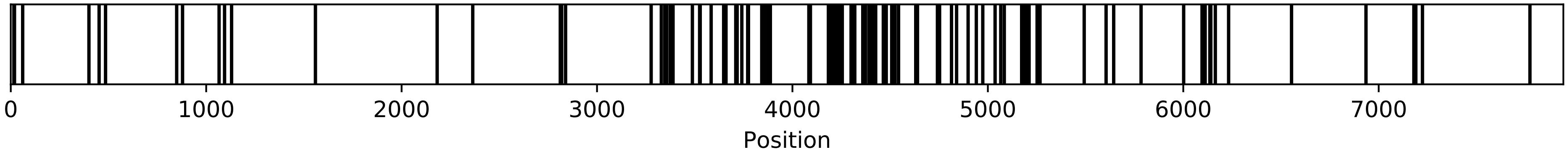


CR1 vs MA2
Number of differences: 195

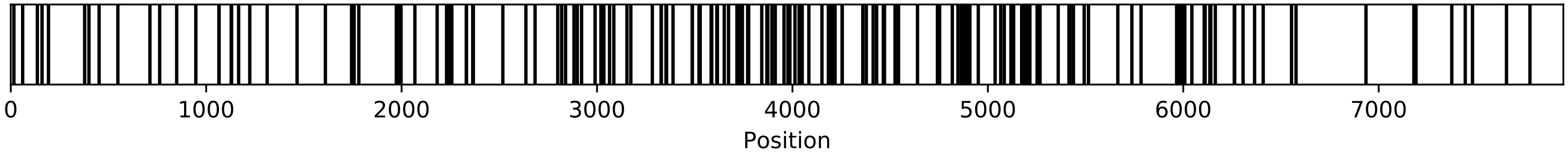


CR2 vs MA1
Number of differences: 204

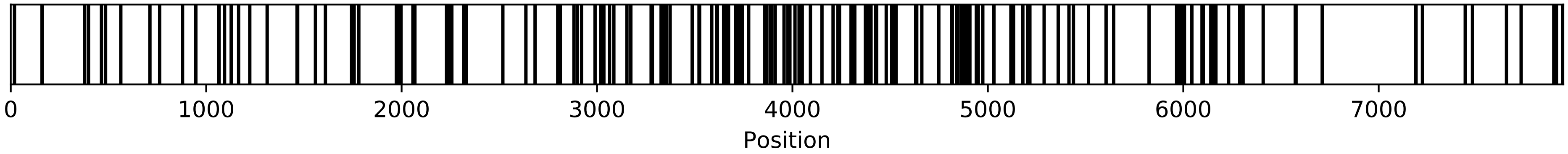


CR2 vs MA2
Number of differences: 55

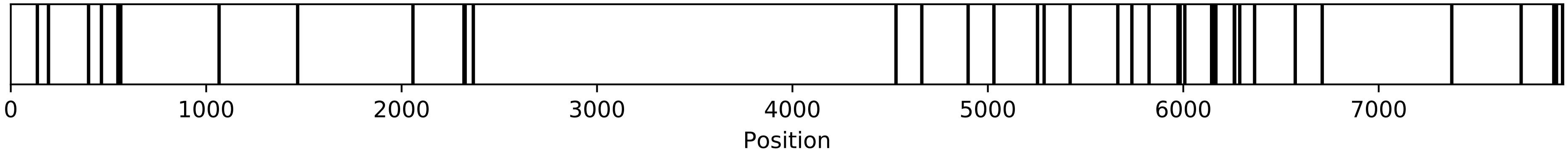


MA1 vs MA2
Number of differences: 183

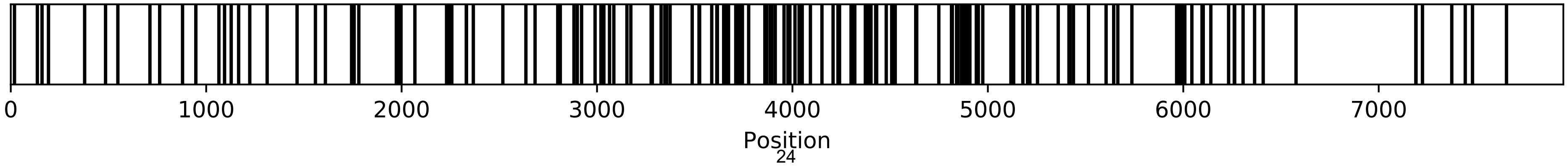

### Region: 165



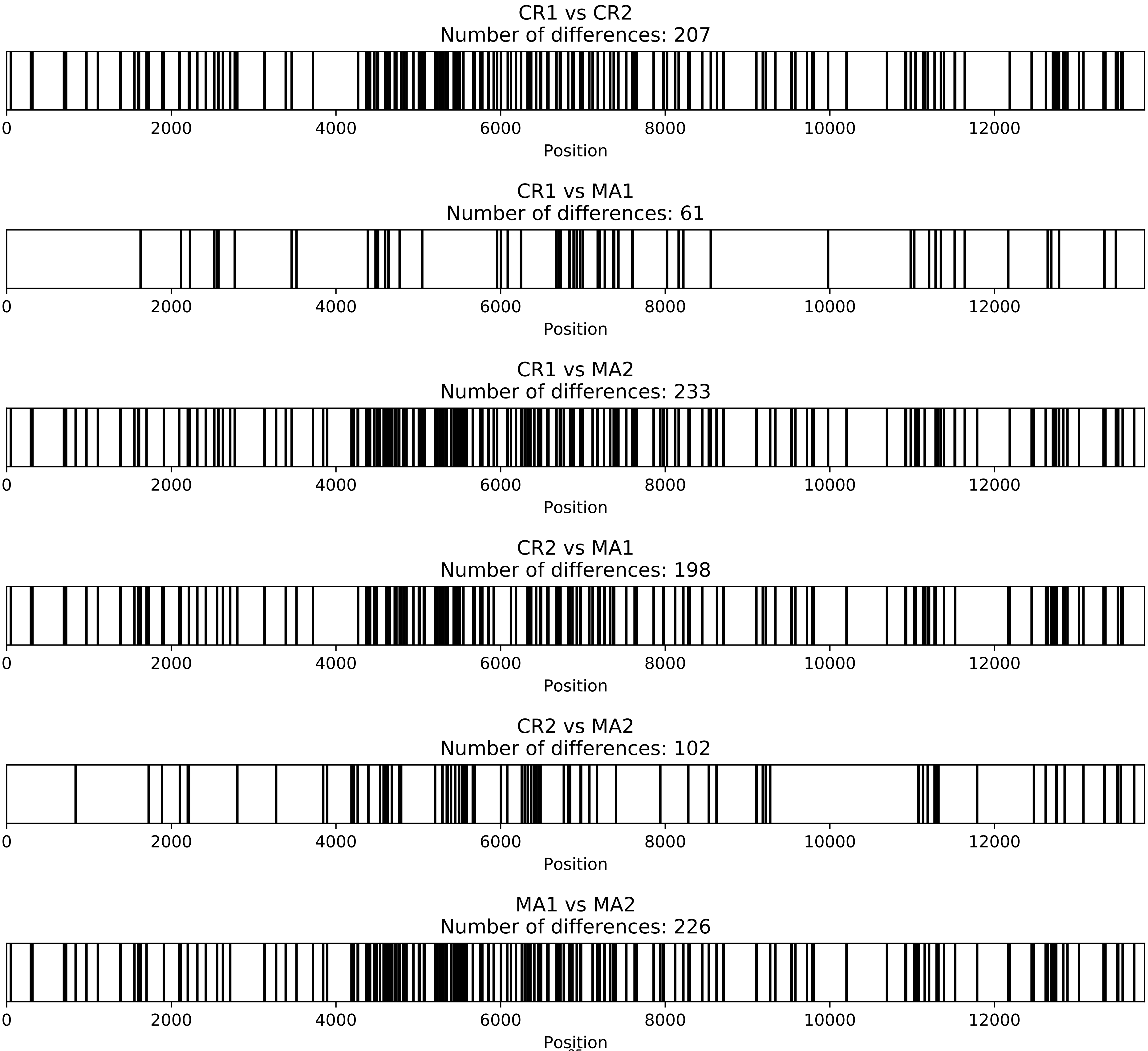
CR1 vs CR2
Number of differences: 207
CR1 vs MA1
Number of differences: 61
CR1 vs MA2
Number of differences: 233
CR2 vs MA1
Number of differences: 198
CR2 vs MA2
Number of differences: 102
MA1 vs MA2
Number of differences: 226
0
2000
4000
6000
8000
10000
12000
Position

### Region: 181



CR1 vs CR2
Number of differences: 162

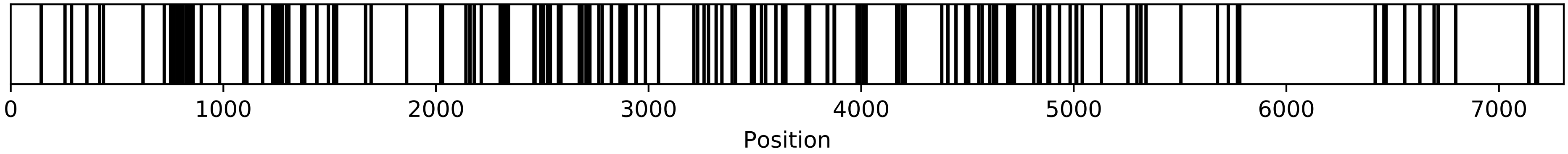


CR1 vs MA1
Number of differences: 133

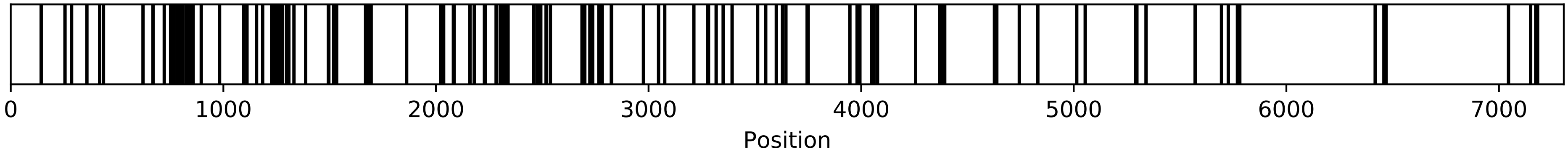


CR1 vs MA2
Number of differences: 112

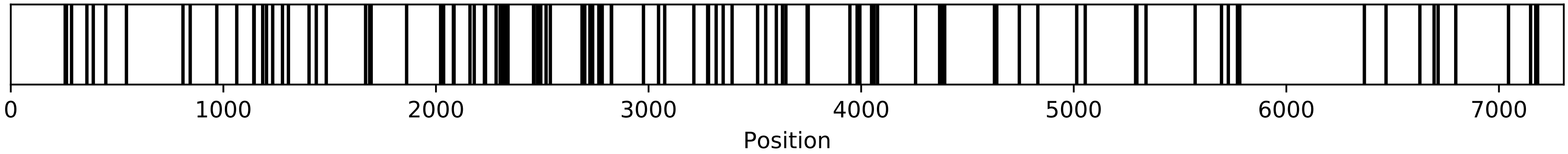


CR2 vs MA1
Number of differences: 135

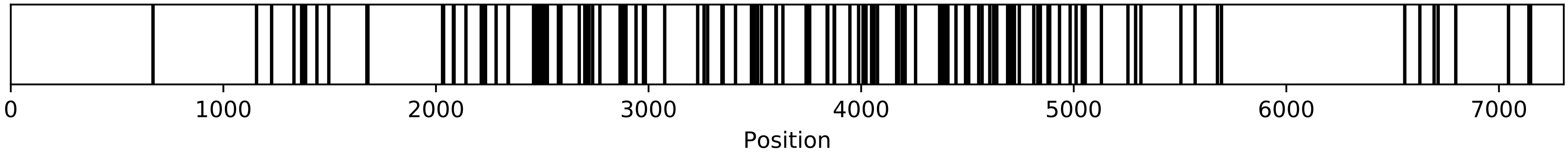


CR2 vs MA2
Number of differences: 172

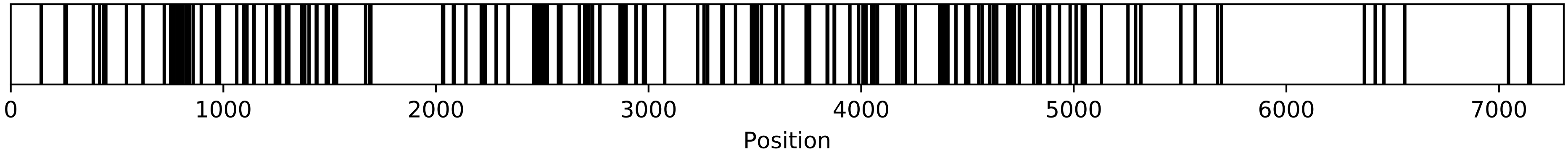


MA1 vs MA2
Number of differences: 65

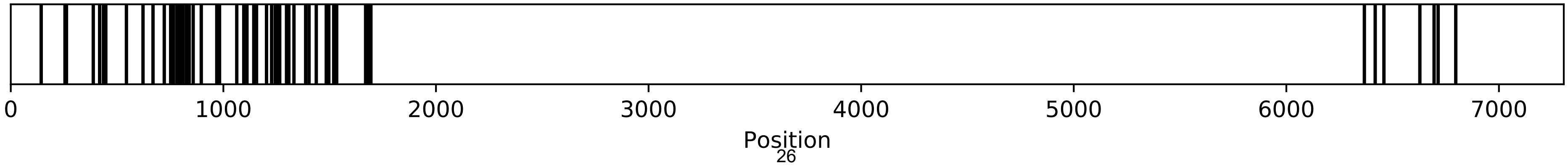

### Region: 184



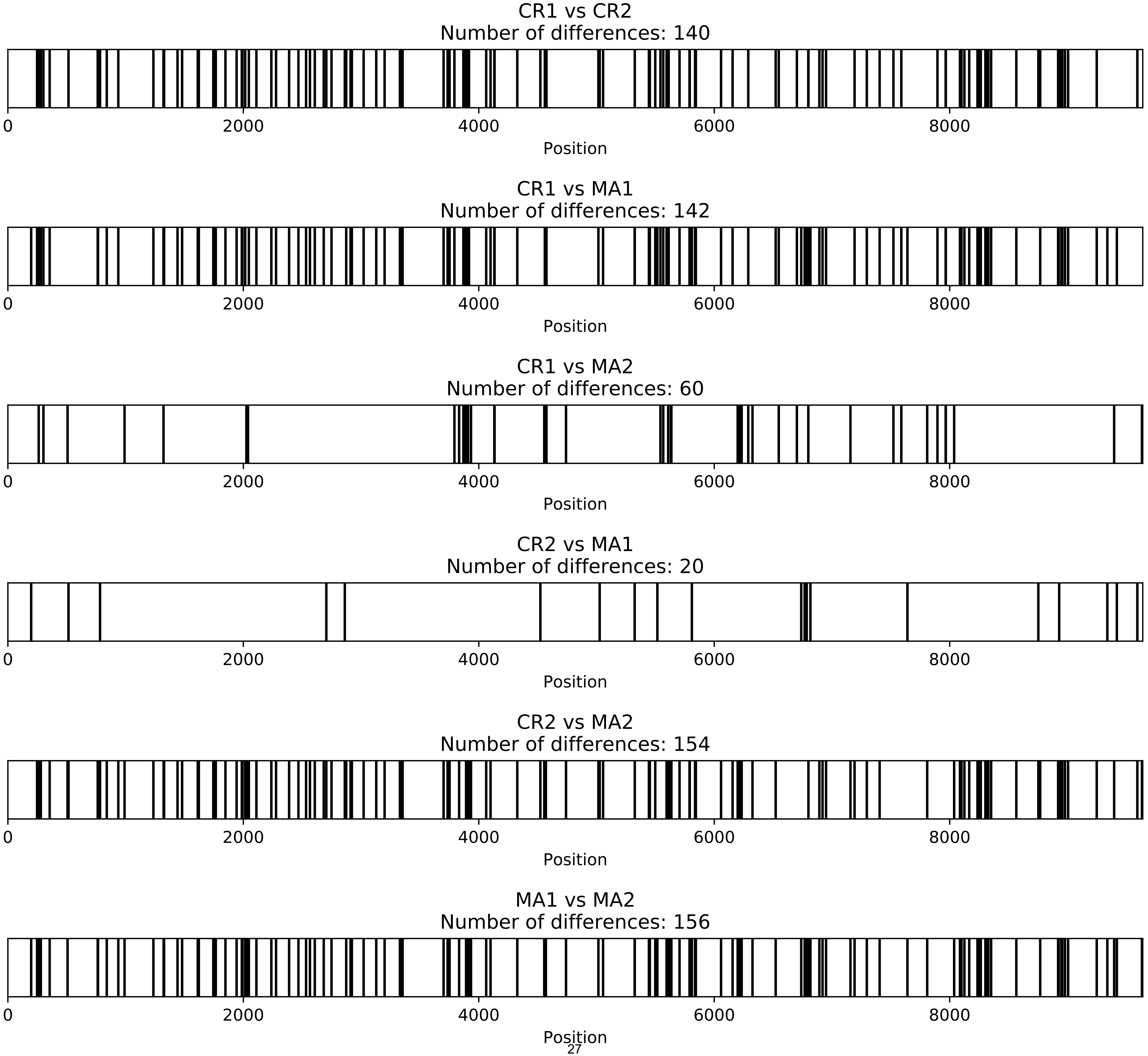
CR1 vs CR2
Number of differences: 140
0
2000
4000
6000
8000
Position
CR1 vs MA1
Number of differences: 142
0
2000
4000
6000
8000
Position
CR1 vs MA2
Number of differences: 60
0
2000
4000
6000
8000
Position
CR2 vs MA1
Number of differences: 20
0
2000
4000
6000
8000
Position
CR2 vs MA2
Number of differences: 154
0
2000
4000
6000
8000
Position
MA1 vs MA2
Number of differences: 156
0
2000
4000
6000
8000
Position

### Region: 185



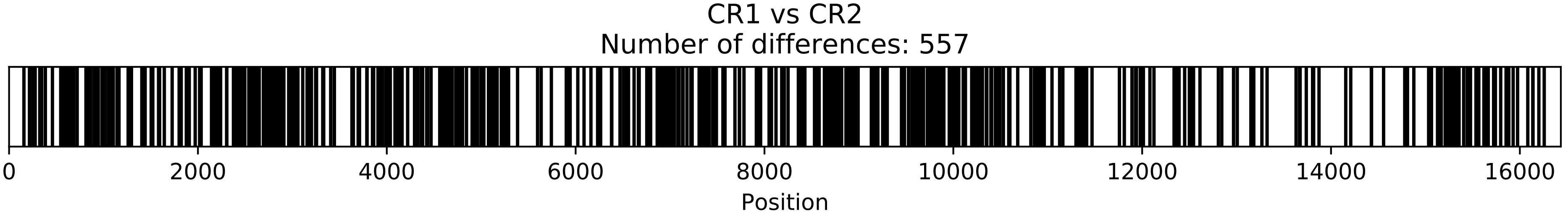
CR1 vs CR2
Number of differences: 557
0
2000
4000
6000
8000
10000
12000
14000
16000
Position

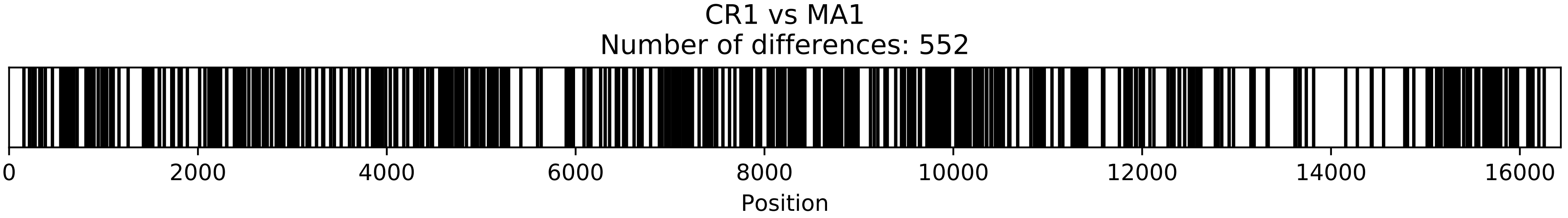
CR1 vs MA1
Number of differences: 552
0
2000
4000
6000
8000
10000
12000
14000
16000
Position

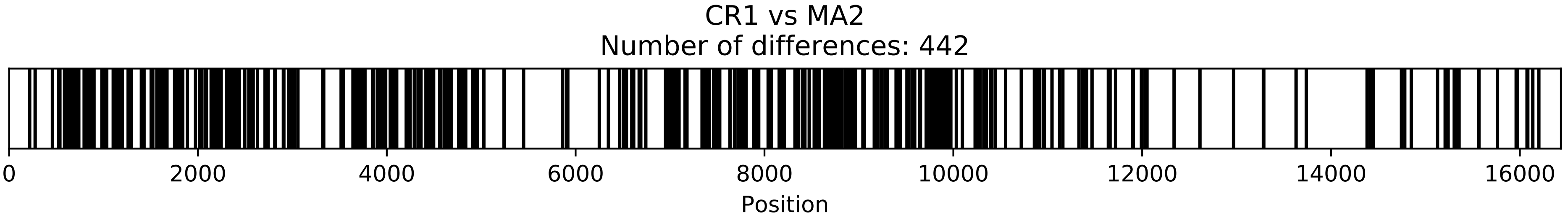
CR1 vs MA2
Number of differences: 442
0
2000
4000
6000
8000
10000
12000
14000
16000
Position

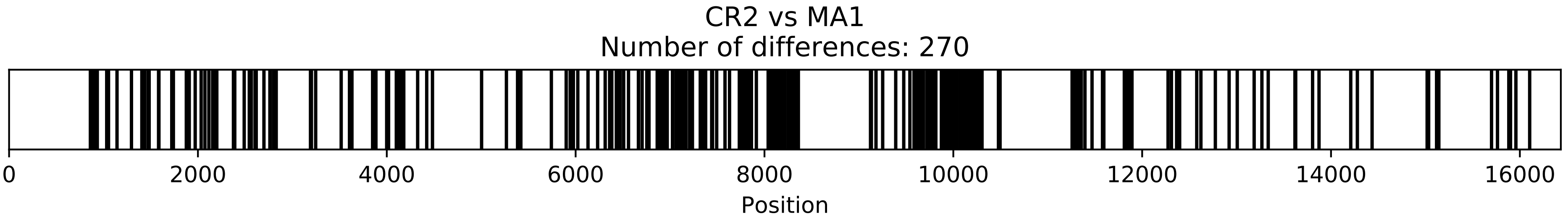
CR2 vs MA1
Number of differences: 270
0
2000
4000
6000
8000
10000
12000
14000
16000
Position

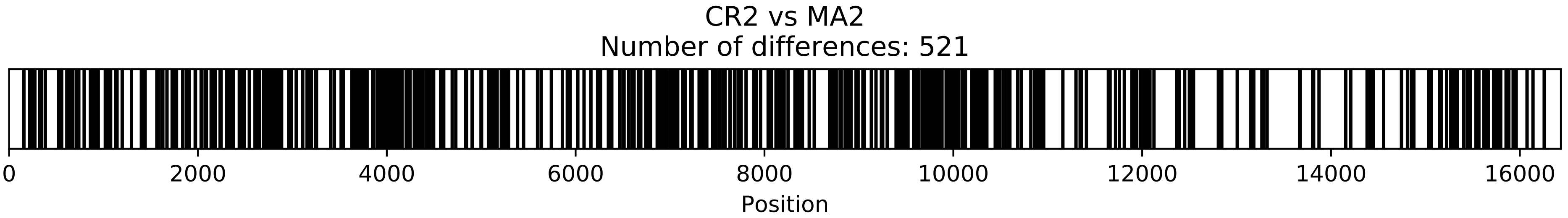
CR2 vs MA2
Number of differences: 521
0
2000
4000
6000
8000
10000
12000
14000
16000
Position

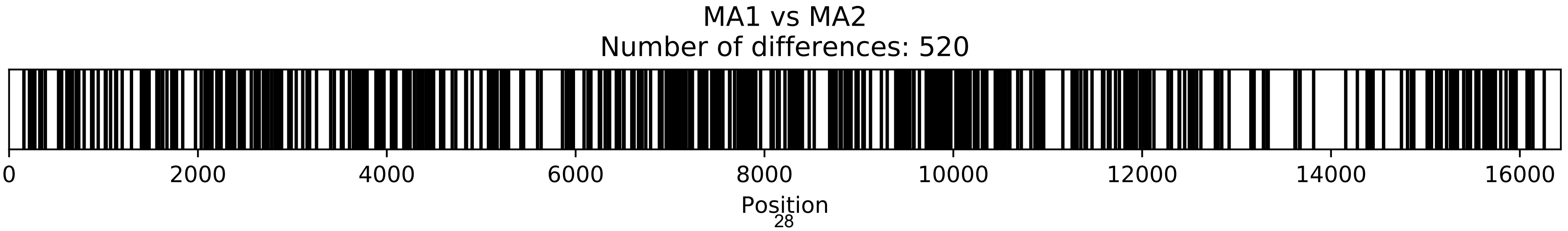
MA1 vs MA2
Number of differences: 520
0
2000
4000
6000
8000
10000
12000
14000
16000
Position

### Region: 195



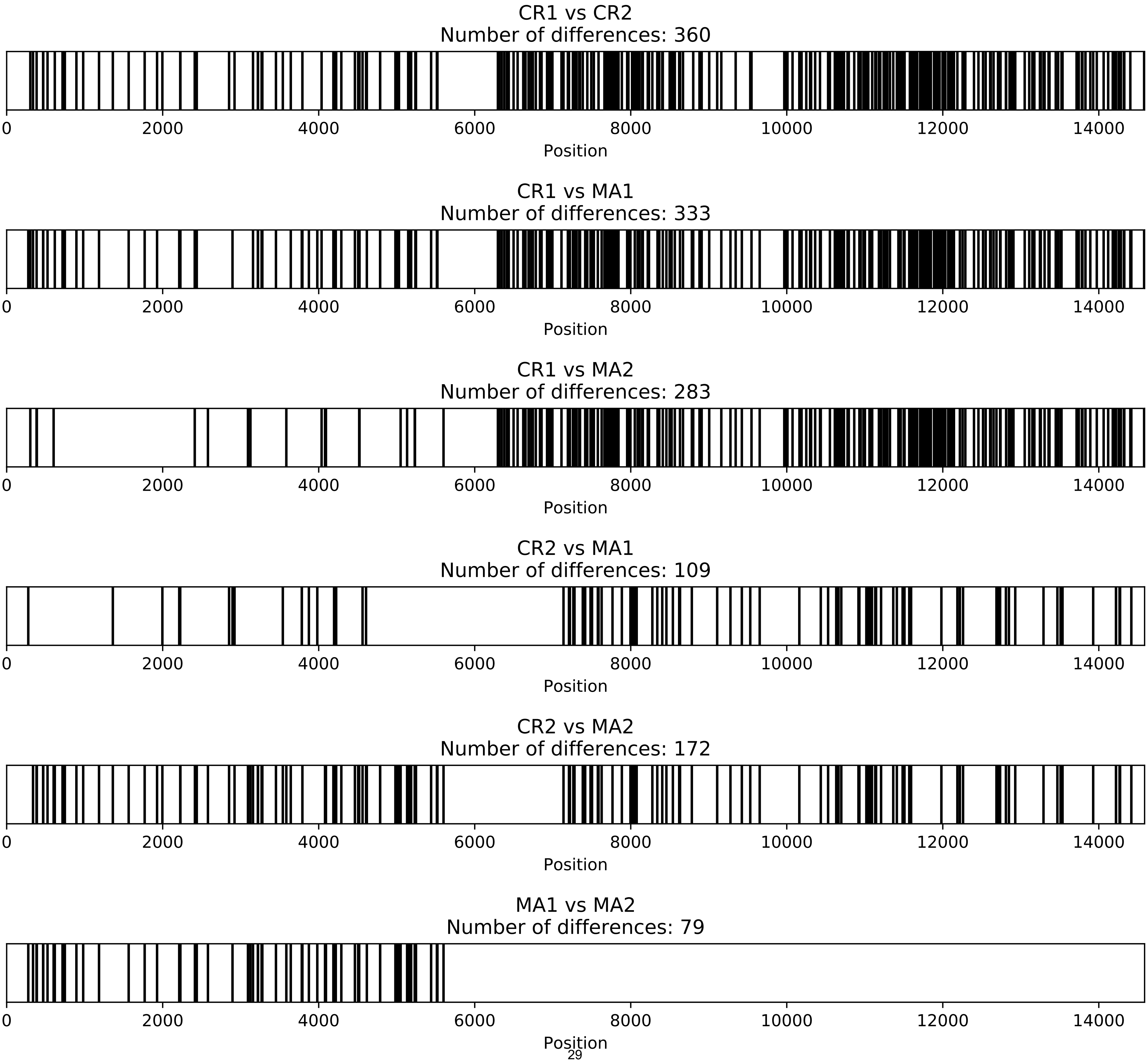
CR1 vs CR2
Number of differences: 360
CR1 vs MA1
Number of differences: 333
CR1 vs MA2
Number of differences: 283
CR2 vs MA1
Number of differences: 109
CR2 vs MA2
Number of differences: 172
MA1 vs MA2
Number of differences: 79
0
2000
4000
6000
8000
10000
12000
14000
Position

### Region: 205



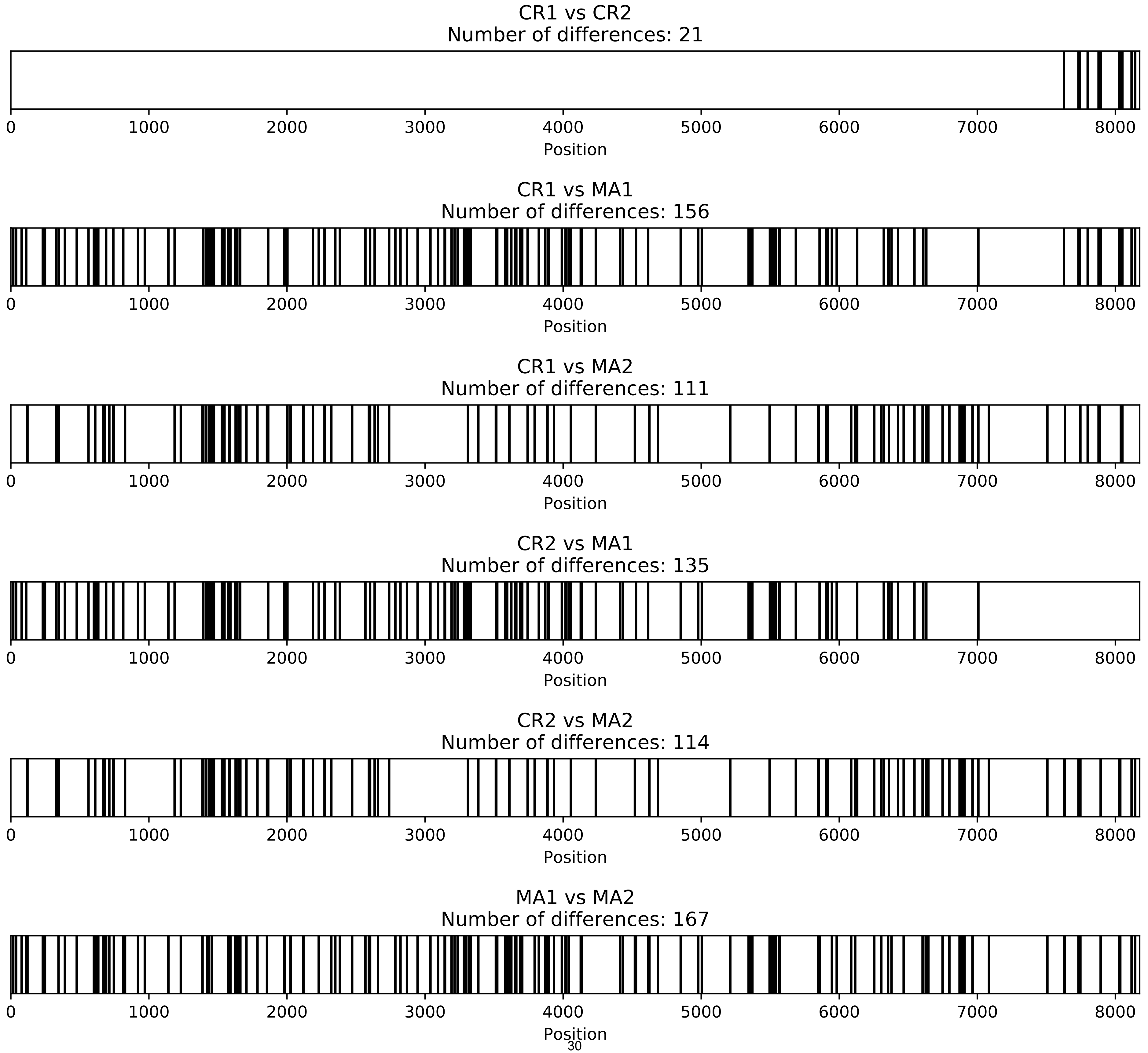
CR1 vs CR2
Number of differences: 21
CR1 vs MA1
Number of differences: 156
CR1 vs MA2
Number of differences: 111
CR2 vs MA1
Number of differences: 135
CR2 vs MA2
Number of differences: 114
MA1 vs MA2
Number of differences: 167
0
1000
2000
3000
4000
5000
6000
7000
8000
Position

### Region: 207



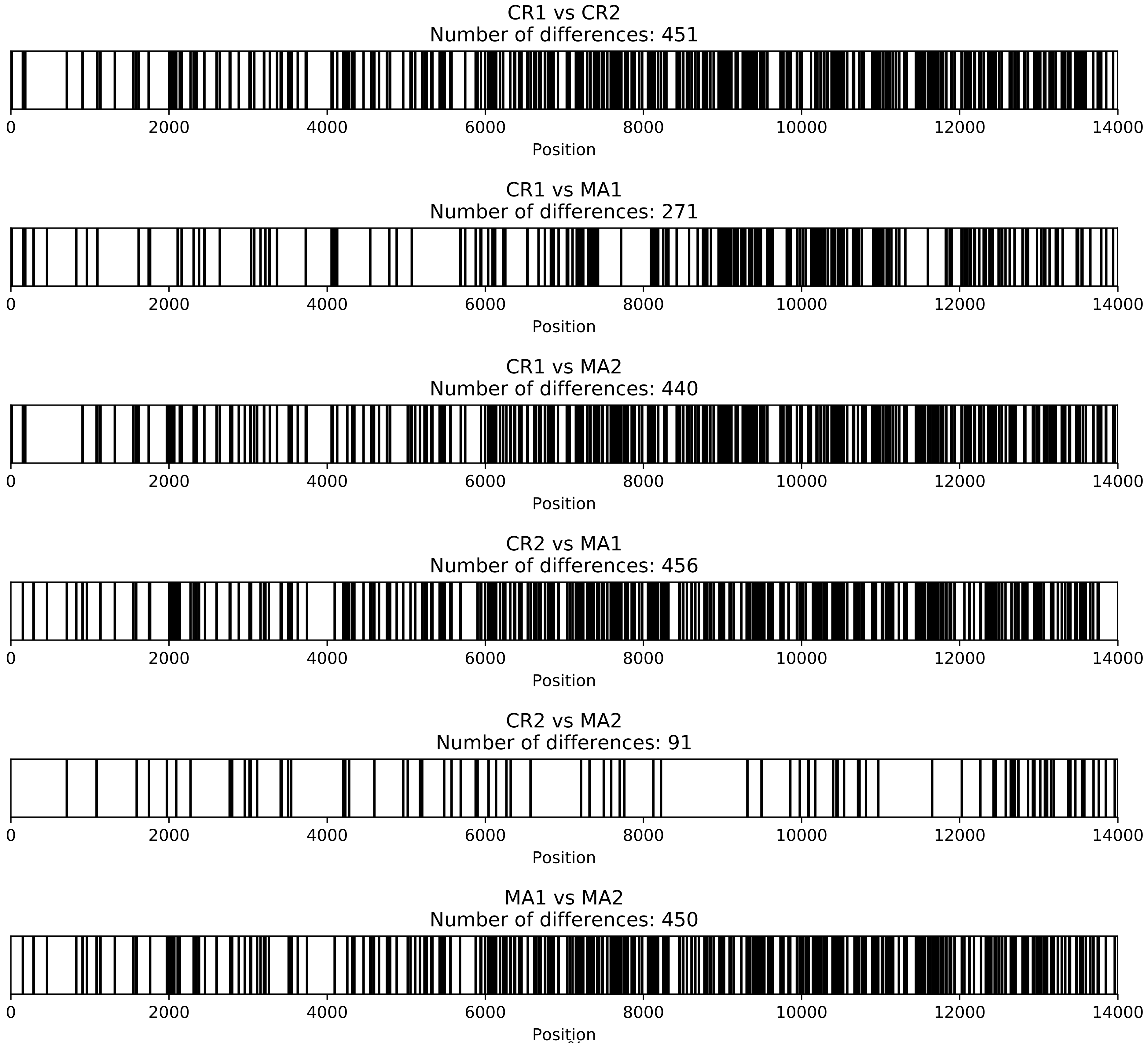
CR1 vs CR2
Number of differences: 451
CR1 vs MA1
Number of differences: 271
CR1 vs MA2
Number of differences: 440
CR2 vs MA1
Number of differences: 456
CR2 vs MA2
Number of differences: 91
MA1 vs MA2
Number of differences: 450
0
2000
4000
6000
8000
10000
12000
14000
Position

### Region: 218

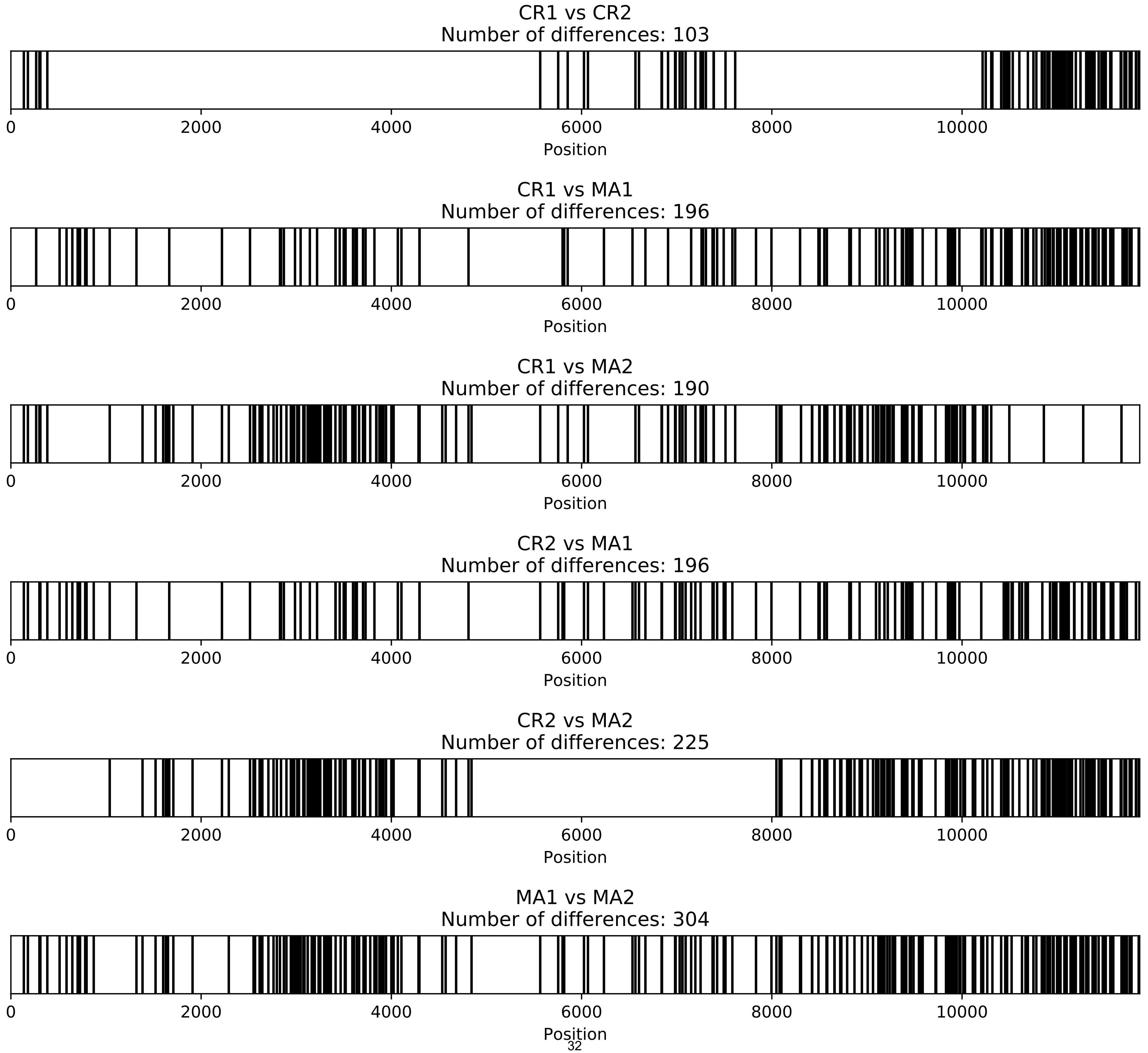
CR1 vs CR2
Number of differences: 103
0
2000
4000
6000
8000
10000
Position
CR1 vs MA1
Number of differences: 196
Position
CR1 vs MA2
Number of differences: 190
Position
CR2 vs MA1
Number of differences: 196
Position
CR2 vs MA2
Number of differences: 225
Position
MA1 vs MA2
Number of differences: 304
Position

### Region: 221



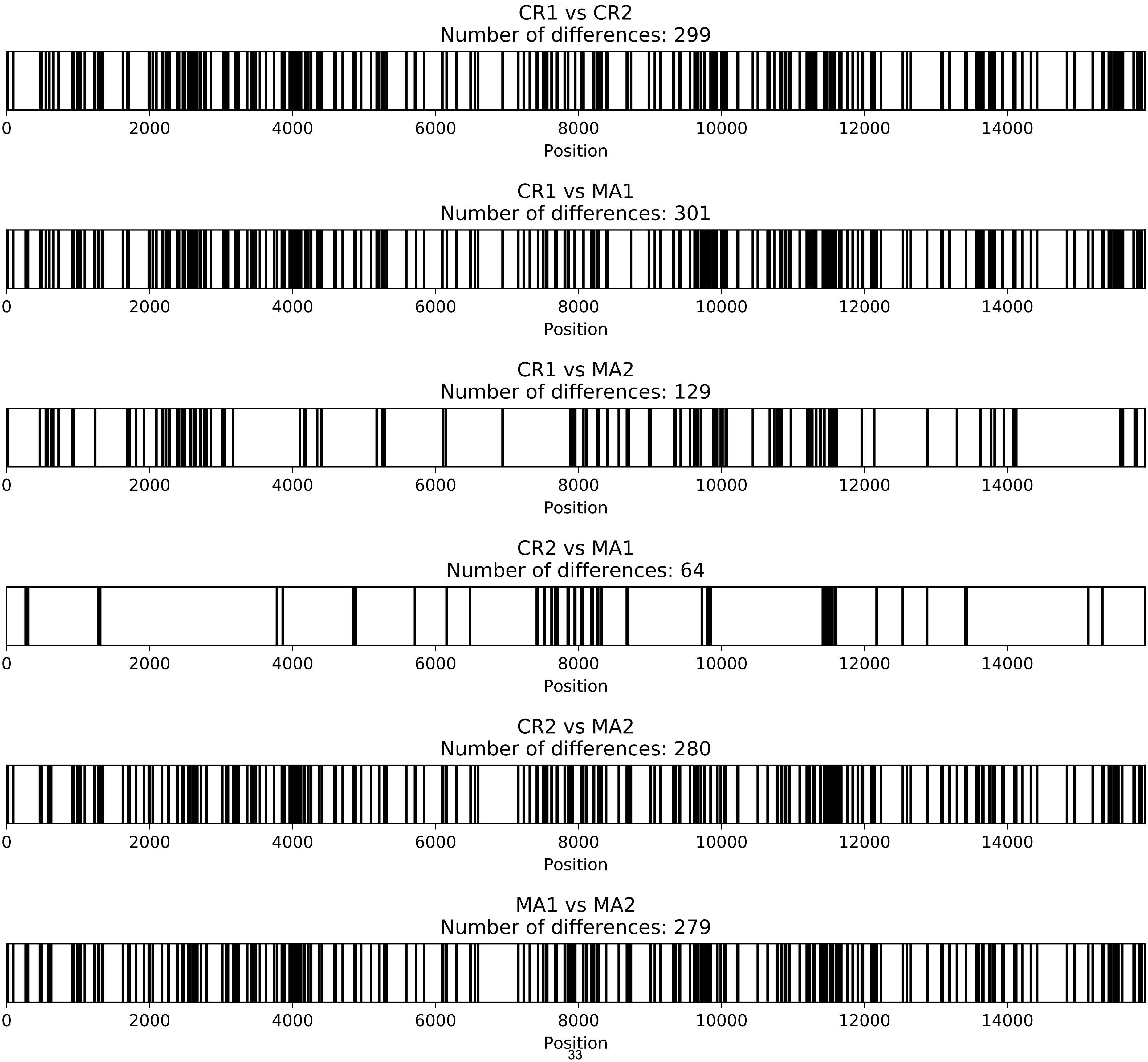
CR1 vs CR2
Number of differences: 299
CR1 vs MA1
Number of differences: 301
CR1 vs MA2
Number of differences: 129
CR2 vs MA1
Number of differences: 64
CR2 vs MA2
Number of differences: 280
MA1 vs MA2
Number of differences: 279
0
2000
4000
6000
8000
10000
12000
14000
Position

### Region: 225

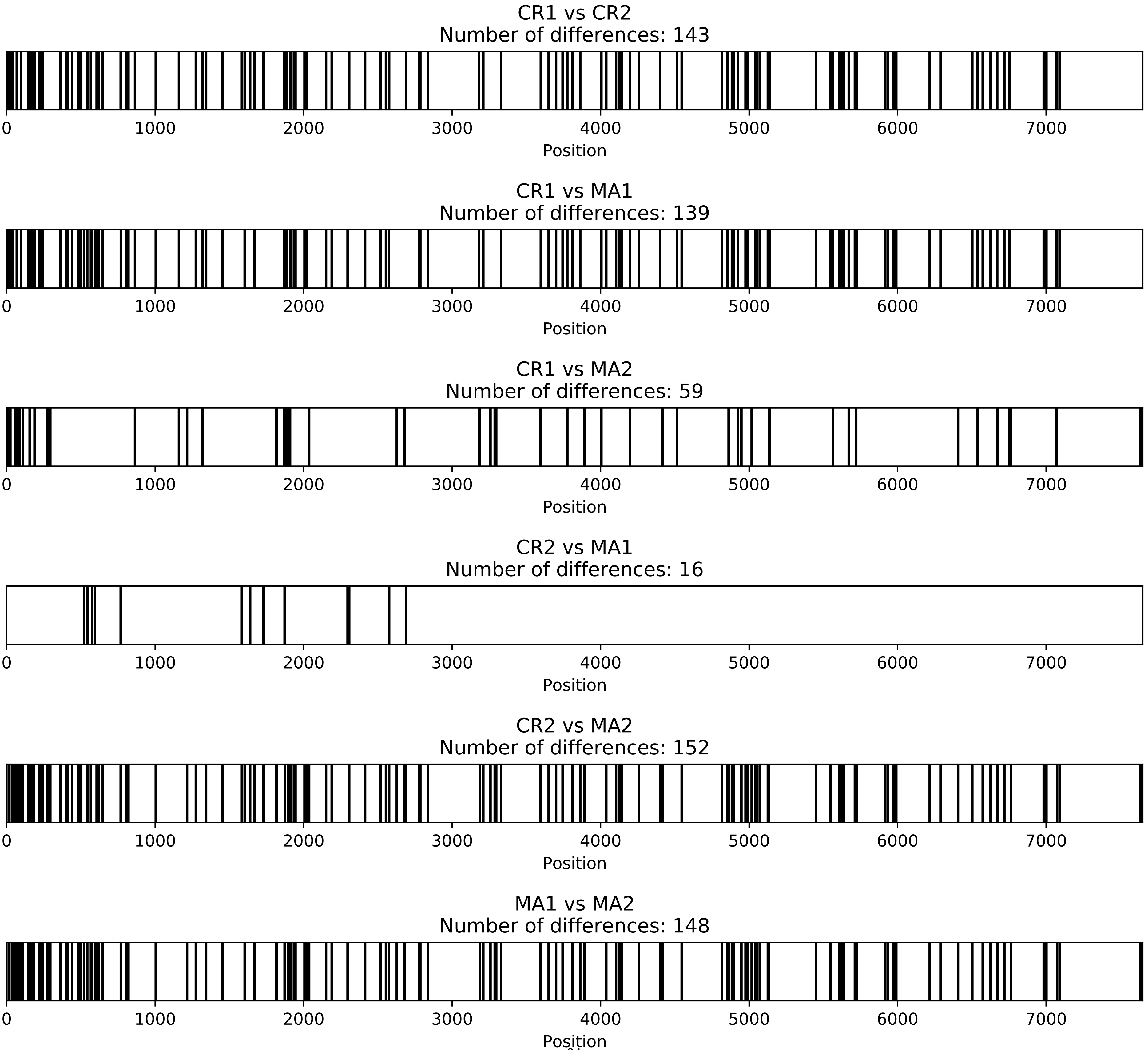
CR1 vs CR2
Number of differences: 143
0 1000 2000 3000 4000 5000 6000 7000
Position
CR1 vs MA1
Number of differences: 139
0 1000 2000 3000 4000 5000 6000 7000
Position
CR1 vs MA2
Number of differences: 59
0 1000 2000 3000 4000 5000 6000 7000
Position
CR2 vs MA1
Number of differences: 16
0 1000 2000 3000 4000 5000 6000 7000
Position
CR2 vs MA2
Number of differences: 152
0 1000 2000 3000 4000 5000 6000 7000
Position
MA1 vs MA2
Number of differences: 148
0 1000 2000 3000 4000 5000 6000 7000
Position

### Region: 234



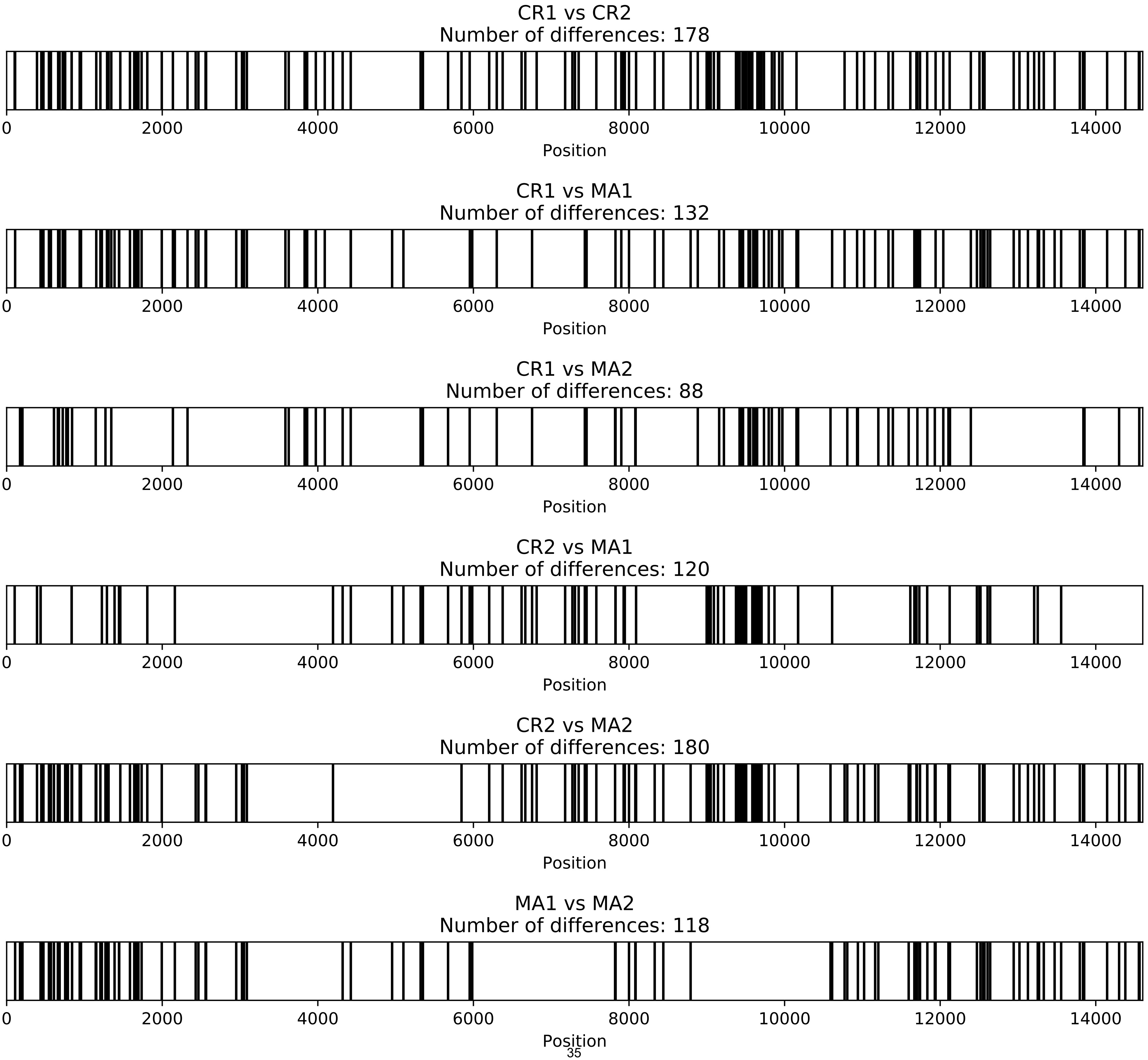
CR1 vs CR2
Number of differences: 178
0 2000 4000 6000 8000 10000 12000 14000
Position
CR1 vs MA1
Number of differences: 132
0 2000 4000 6000 8000 10000 12000 14000
Position
CR1 vs MA2
Number of differences: 88
0 2000 4000 6000 8000 10000 12000 14000
Position
CR2 vs MA1
Number of differences: 120
0 2000 4000 6000 8000 10000 12000 14000
Position
CR2 vs MA2
Number of differences: 180
0 2000 4000 6000 8000 10000 12000 14000
Position
MA1 vs MA2
Number of differences: 118
0 2000 4000 6000 8000 10000 12000 14000
Position

### Region: 240



CR1 vs CR2
Number of differences: 429

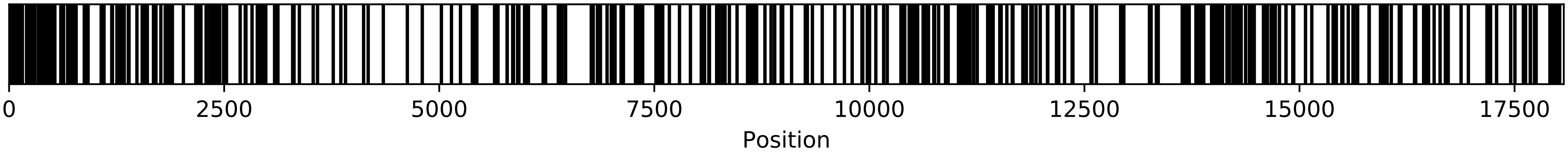


CR1 vs MA1
Number of differences: 423

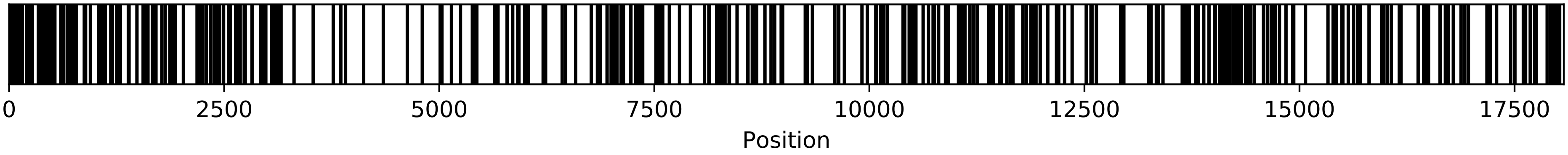


CR1 vs MA2
Number of differences: 204

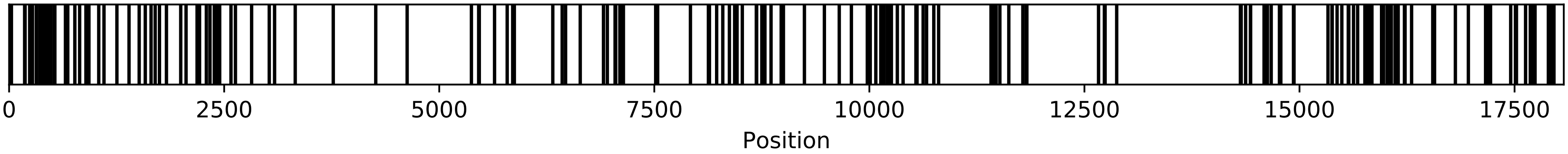


CR2 vs MA1
Number of differences: 137

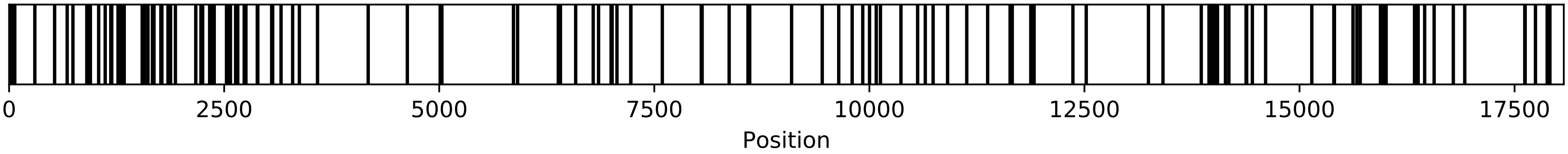


CR2 vs MA2
Number of differences: 461

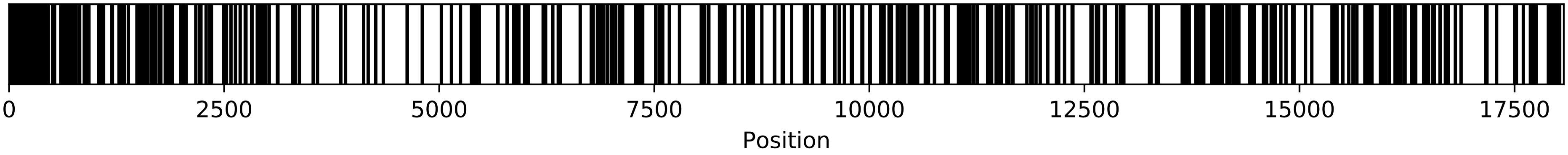


MA1 vs MA2
Number of differences: 455

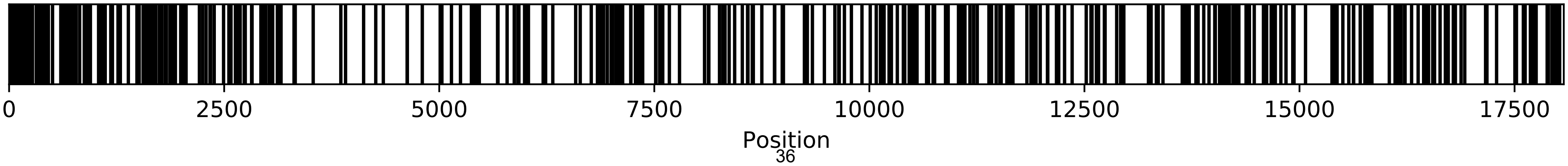

### Region: 241



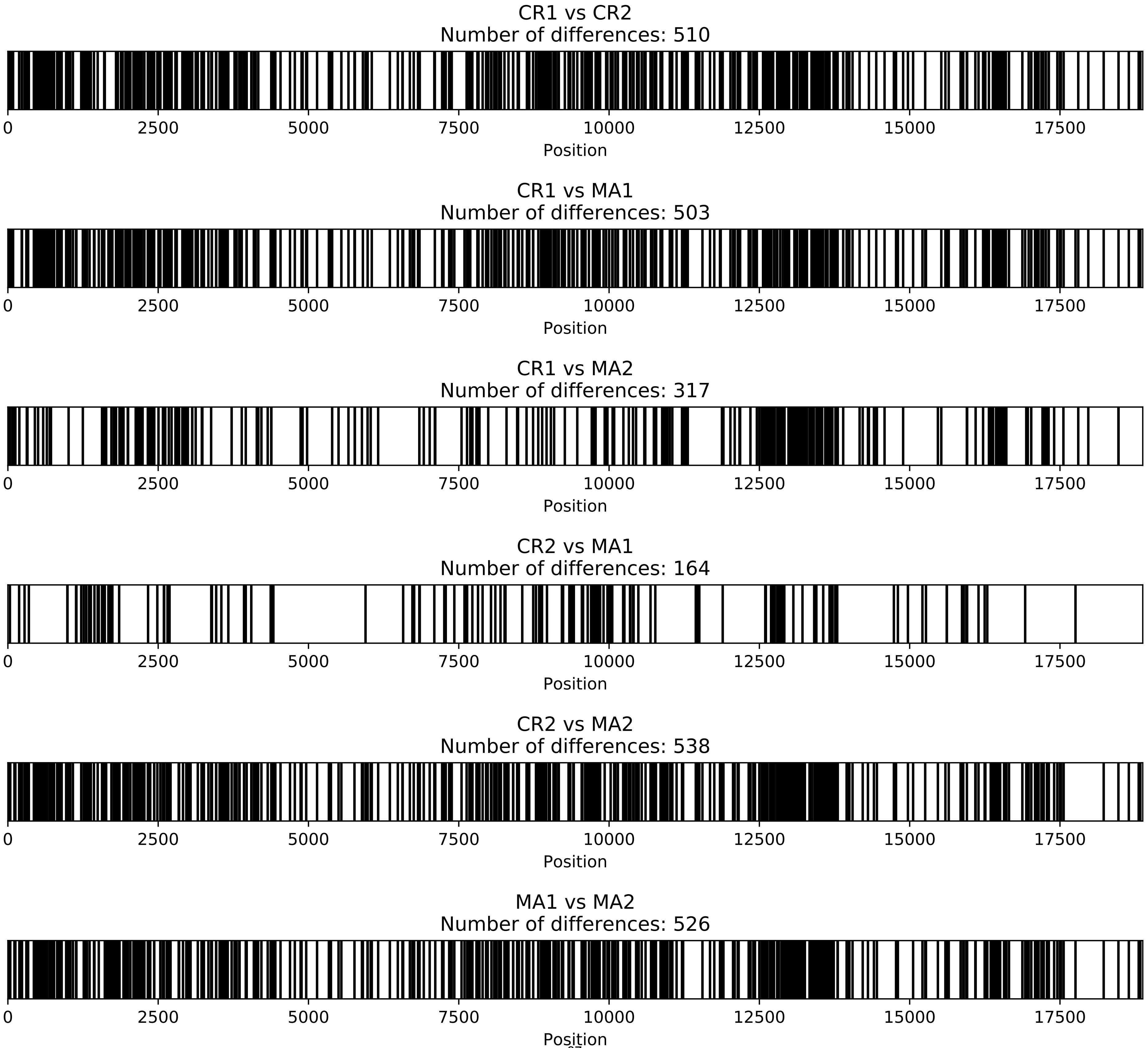
CR1 vs CR2
Number of differences: 510
CR1 vs MA1
Number of differences: 503
CR1 vs MA2
Number of differences: 317
CR2 vs MA1
Number of differences: 164
CR2 vs MA2
Number of differences: 538
MA1 vs MA2
Number of differences: 526
0
2500
5000
7500
10000
12500
15000
17500
Position

### Region: 249



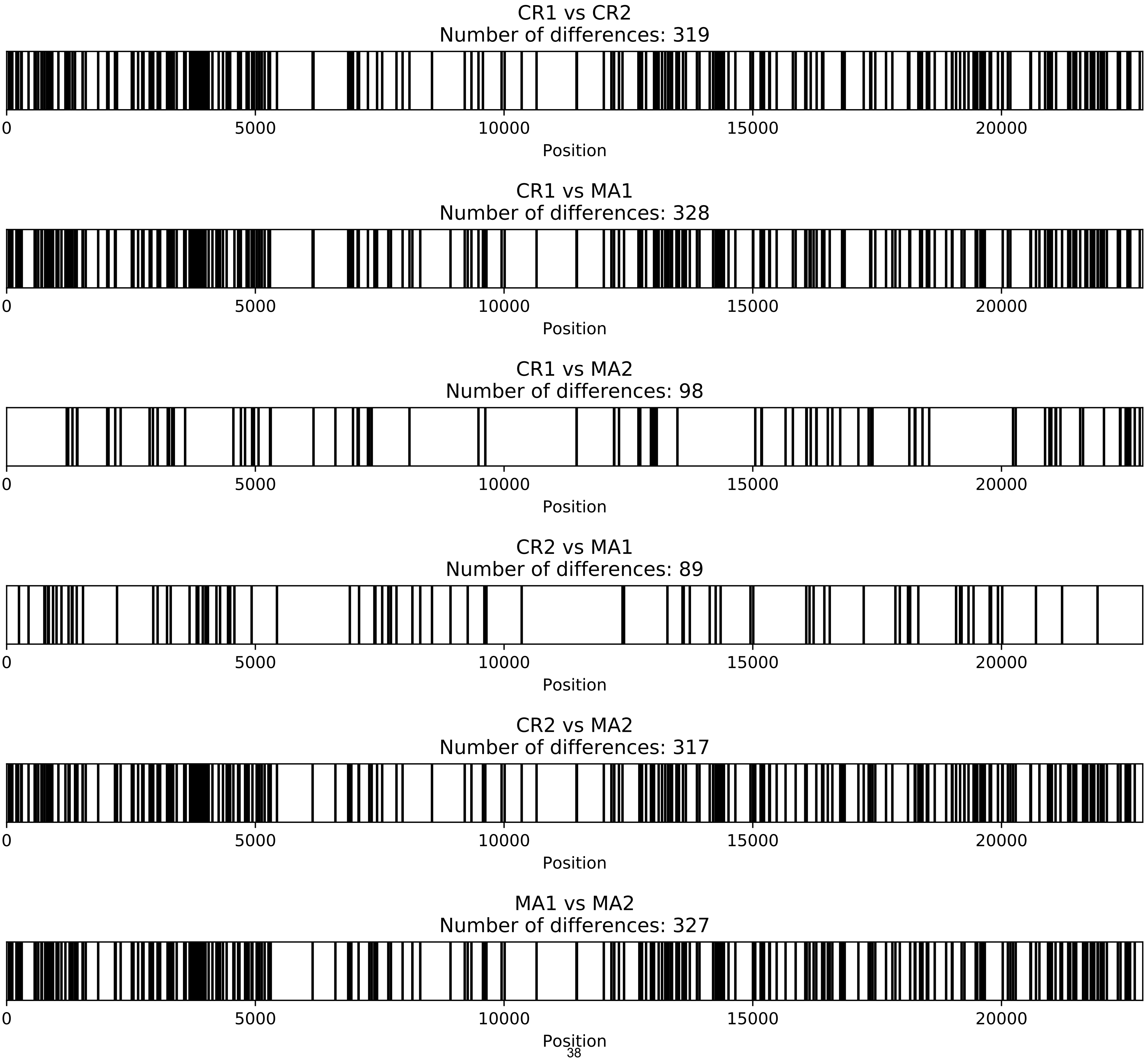
CR1 vs CR2
Number of differences: 319
CR1 vs MA1
Number of differences: 328
CR1 vs MA2
Number of differences: 98
CR2 vs MA1
Number of differences: 89
CR2 vs MA2
Number of differences: 317
MA1 vs MA2
Number of differences: 327
0
5000
10000
15000
20000
Position

### Region: 251



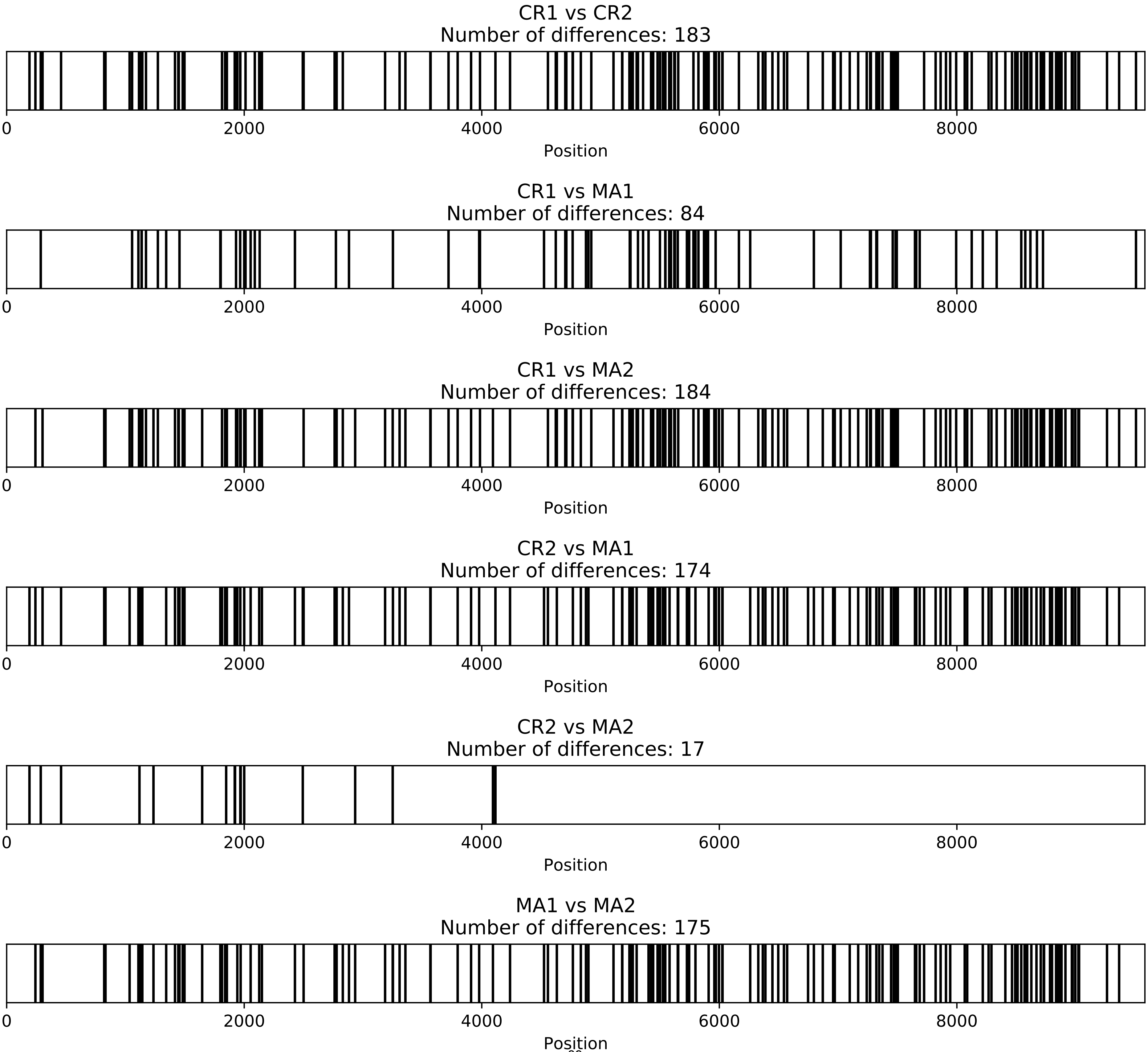
CR1 vs CR2
Number of differences: 183
CR1 vs MA1
Number of differences: 84
CR1 vs MA2
Number of differences: 184
CR2 vs MA1
Number of differences: 174
CR2 vs MA2
Number of differences: 17
MA1 vs MA2
Number of differences: 175
0
2000
4000
6000
8000
Position

### Region: 263



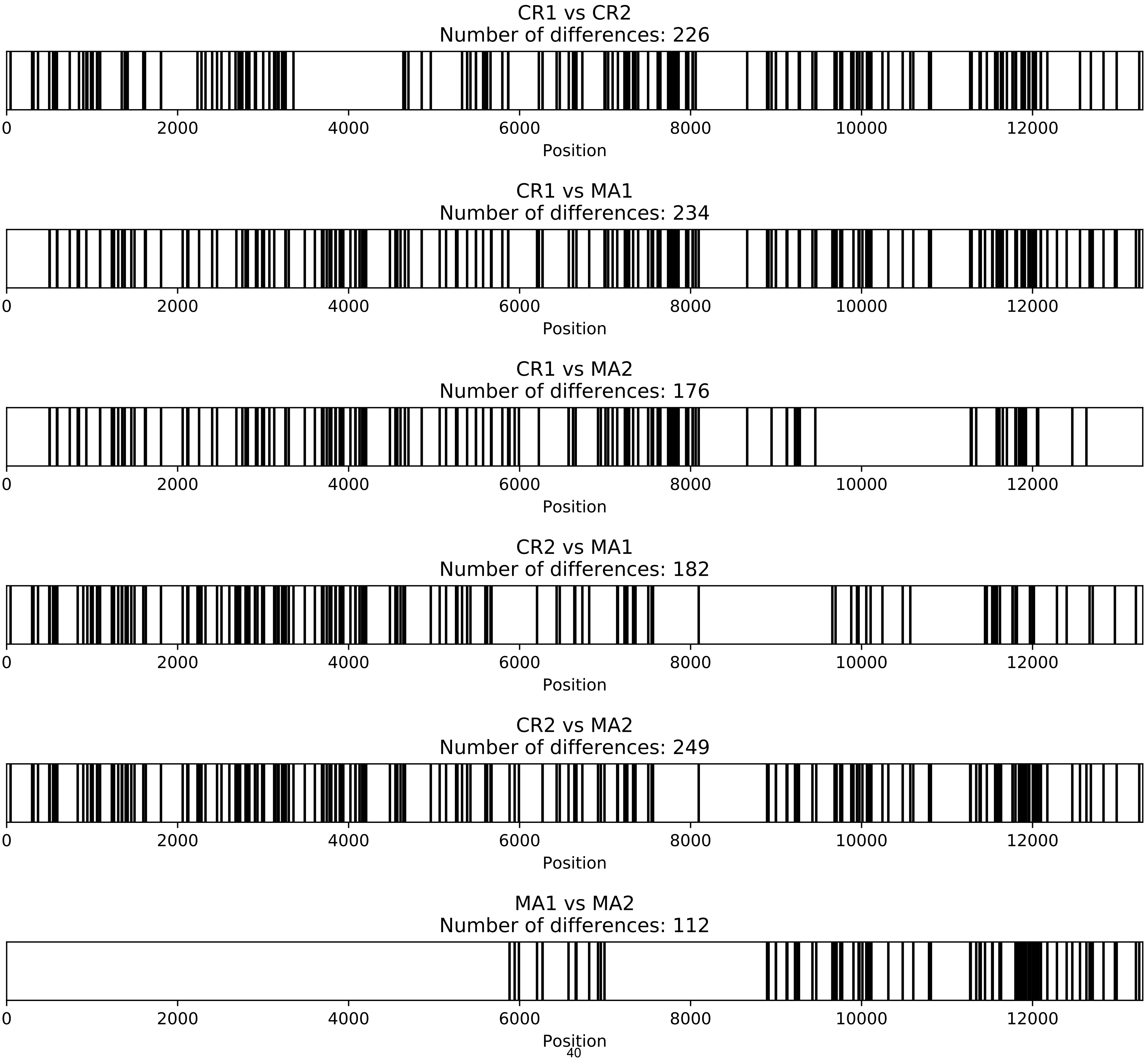
CR1 vs CR2
Number of differences: 226
CR1 vs MA1
Number of differences: 234
CR1 vs MA2
Number of differences: 176
CR2 vs MA1
Number of differences: 182
CR2 vs MA2
Number of differences: 249
MA1 vs MA2
Number of differences: 112
0
2000
4000
6000
8000
10000
12000
Position

### Region: 271



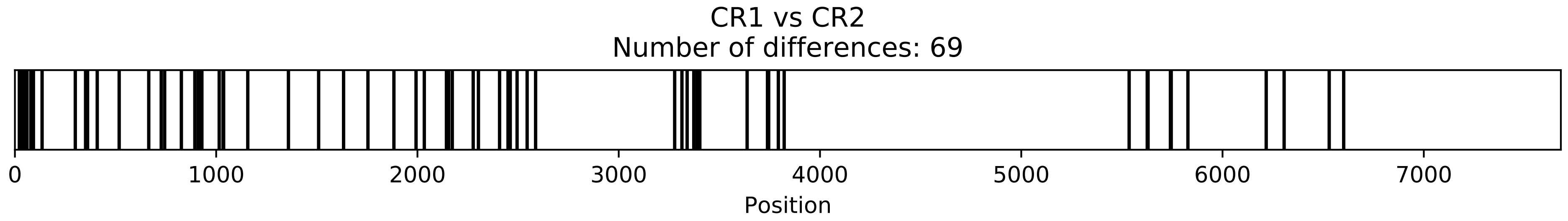
CR1 vs CR2
Number of differences: 69
0
1000
2000
3000
4000
5000
6000
7000
Position

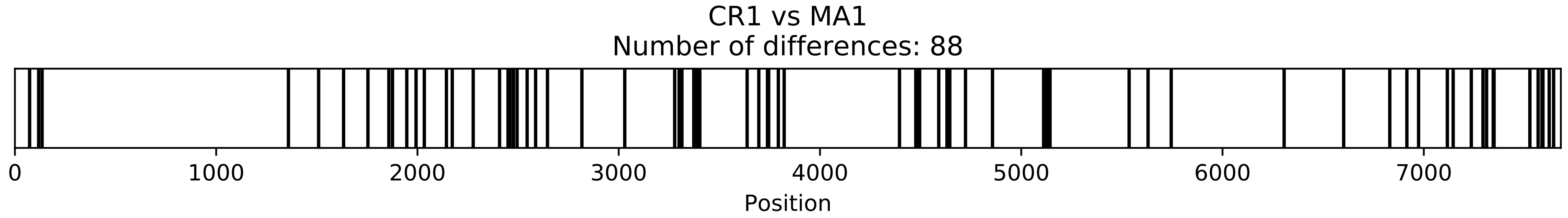
CR1 vs MA1
Number of differences: 88
0
1000
2000
3000
4000
5000
6000
7000
Position

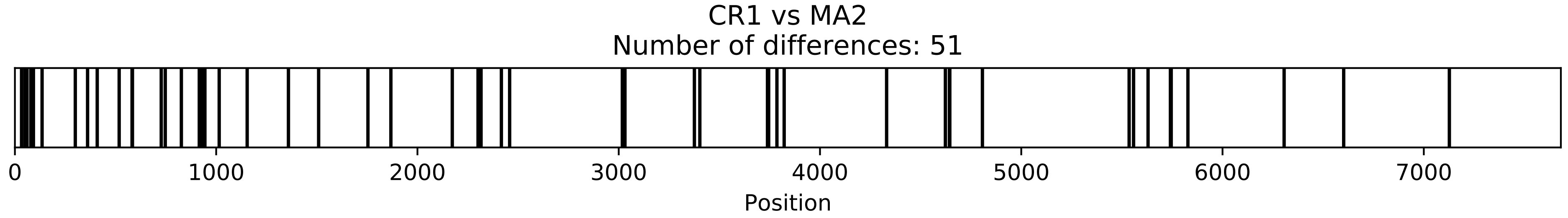
CR1 vs MA2
Number of differences: 51
0
1000
2000
3000
4000
5000
6000
7000
Position

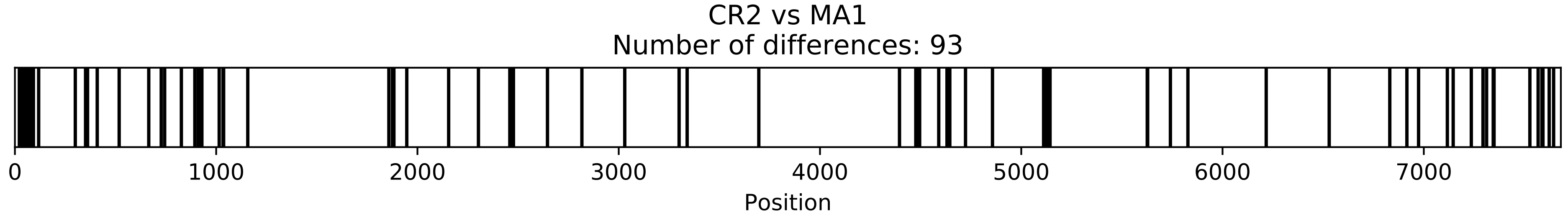
CR2 vs MA1
Number of differences: 93
0
1000
2000
3000
4000
5000
6000
7000
Position

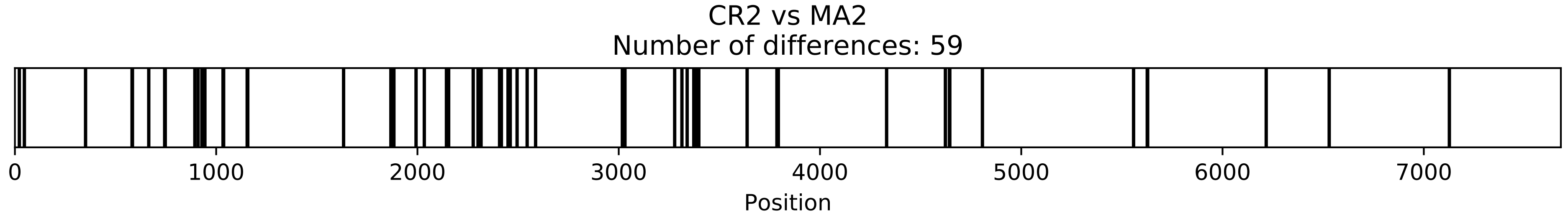
CR2 vs MA2
Number of differences: 59
0
1000
2000
3000
4000
5000
6000
7000
Position

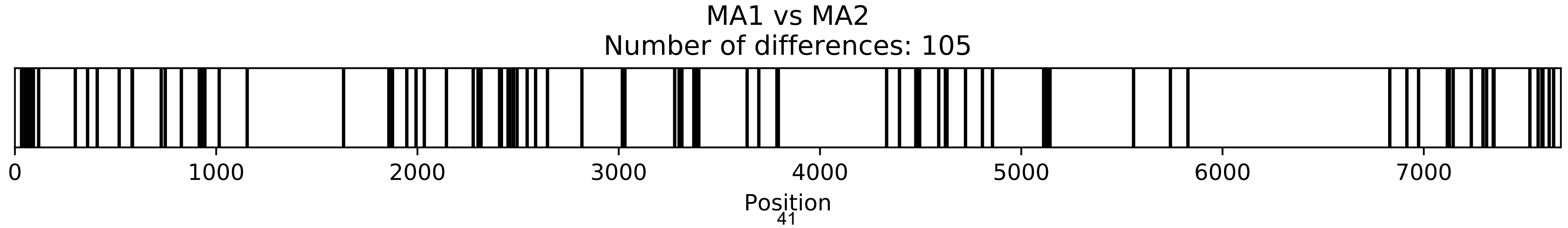
MA1 vs MA2
Number of differences: 105
0
1000
2000
3000
4000
5000
6000
7000
Position

### Region: 274



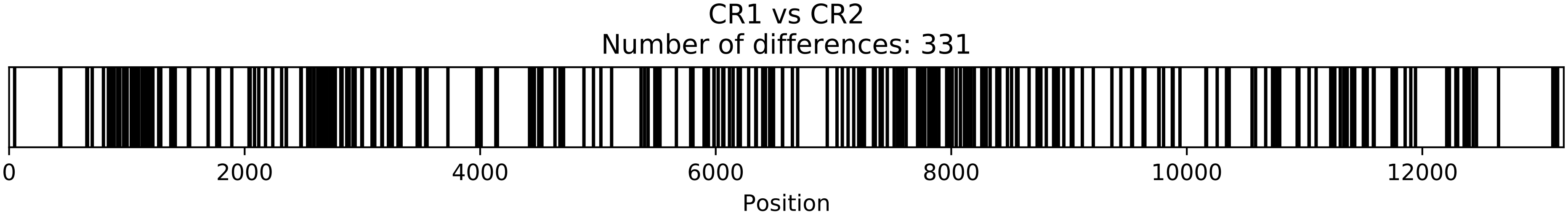
CR1 vs CR2
Number of differences: 331
0
2000
4000
6000
8000
10000
12000
Position

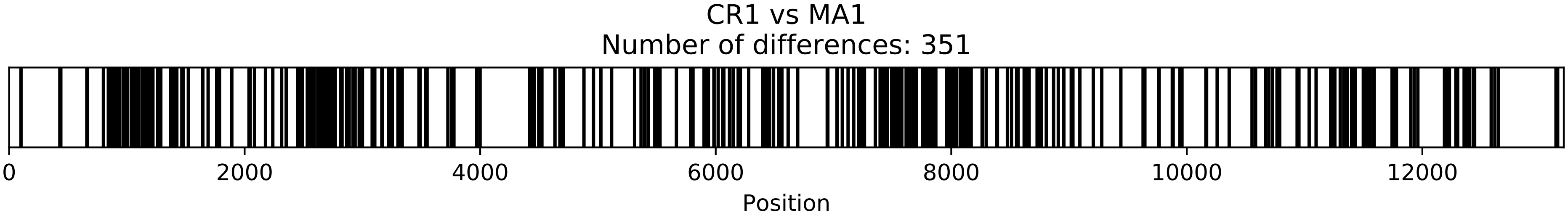
CR1 vs MA1
Number of differences: 351
0
2000
4000
6000
8000
10000
12000
Position

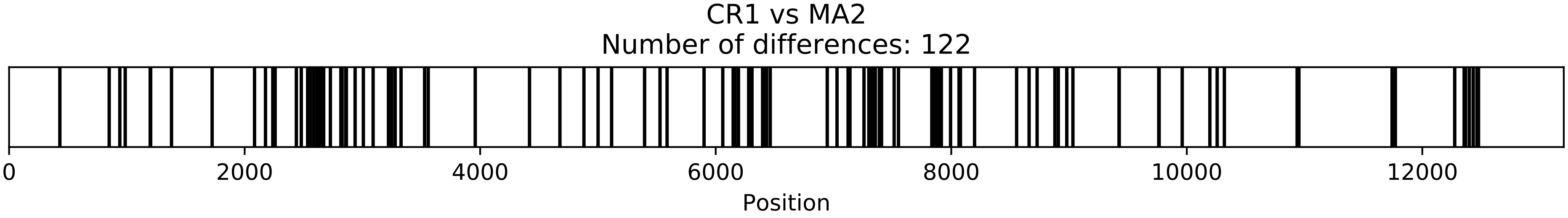
CR1 vs MA2
Number of differences: 122
0
2000
4000
6000
8000
10000
12000
Position

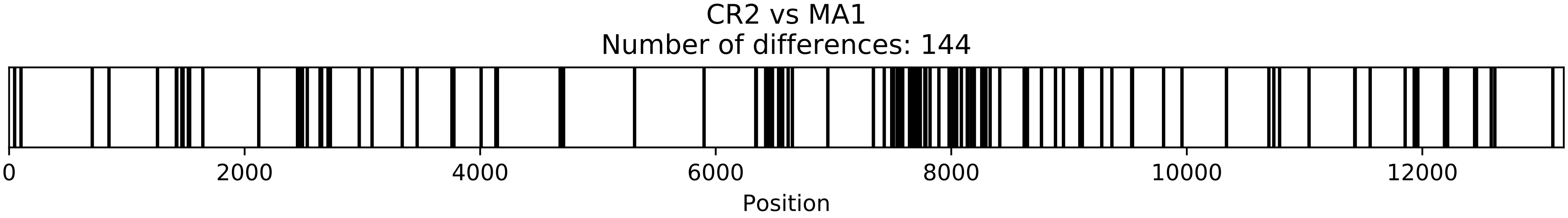
CR2 vs MA1
Number of differences: 144
0
2000
4000
6000
8000
10000
12000
Position

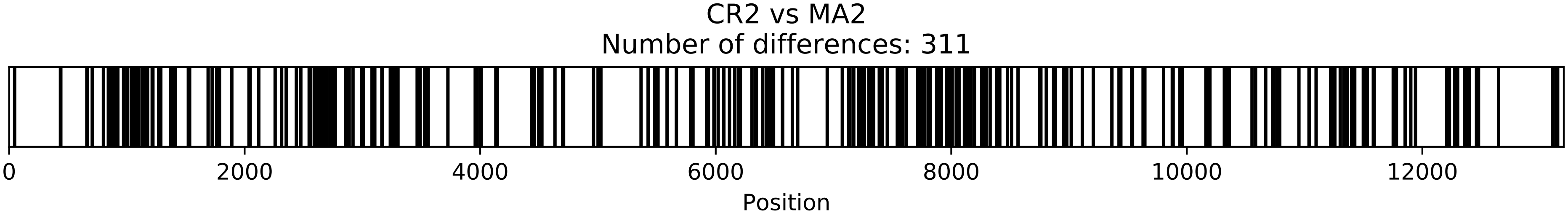
CR2 vs MA2
Number of differences: 311
0
2000
4000
6000
8000
10000
12000
Position

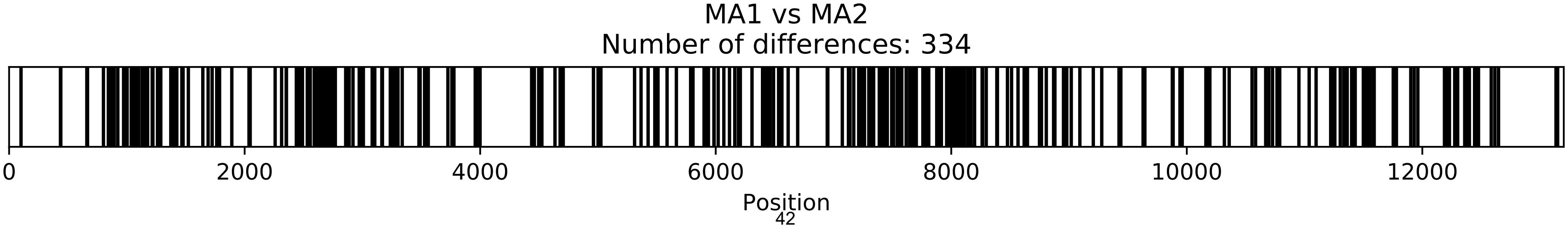
MA1 vs MA2
Number of differences: 334
0
2000
4000
6000
8000
10000
12000
Position

### Region: 278



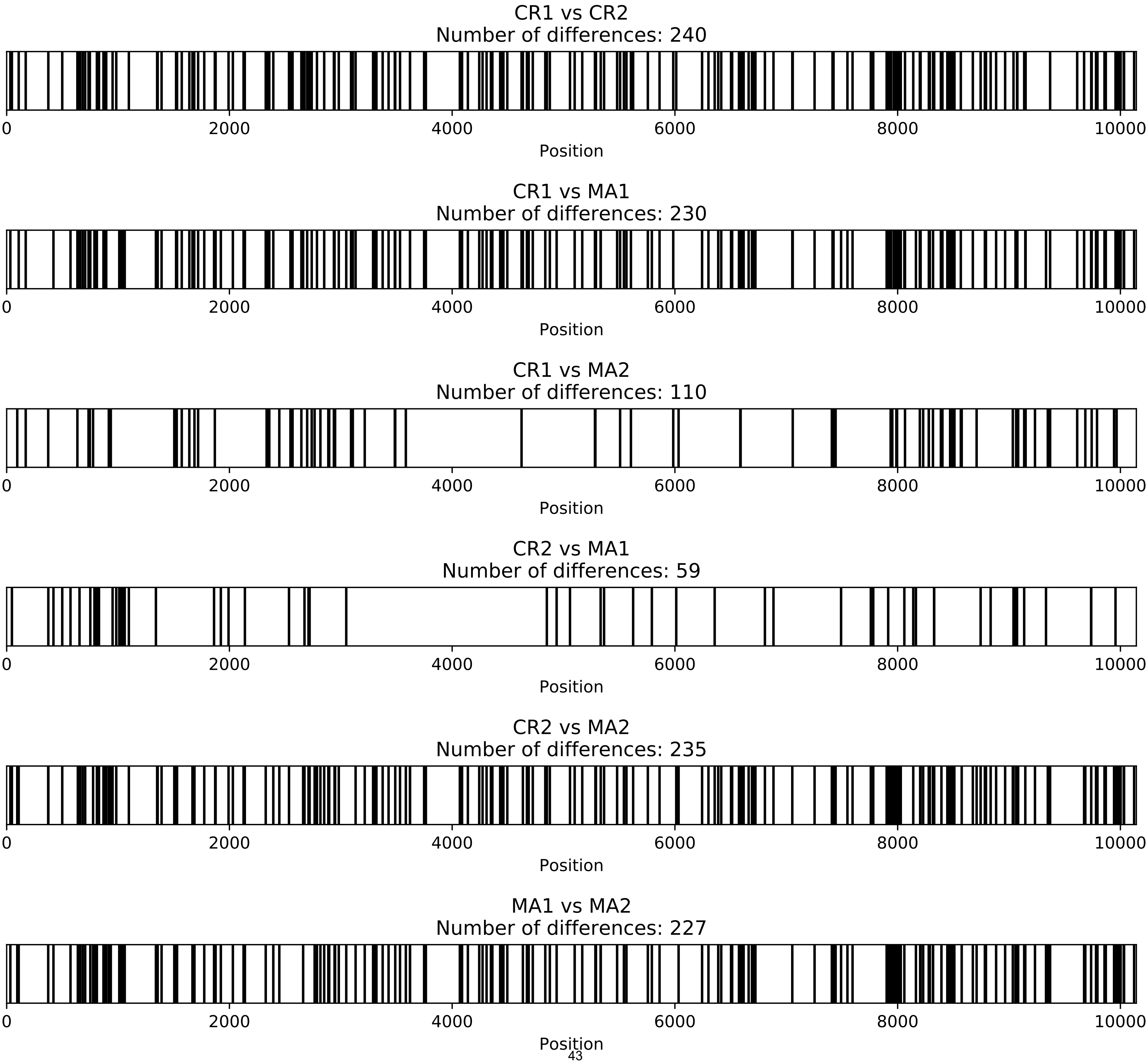
CR1 vs CR2
Number of differences: 240
0
2000
4000
6000
8000
10000
Position
CR1 vs MA1
Number of differences: 230
Position
CR1 vs MA2
Number of differences: 110
Position
CR2 vs MA1
Number of differences: 59
Position
CR2 vs MA2
Number of differences: 235
Position
MA1 vs MA2
Number of differences: 227
Position

### Region: 284



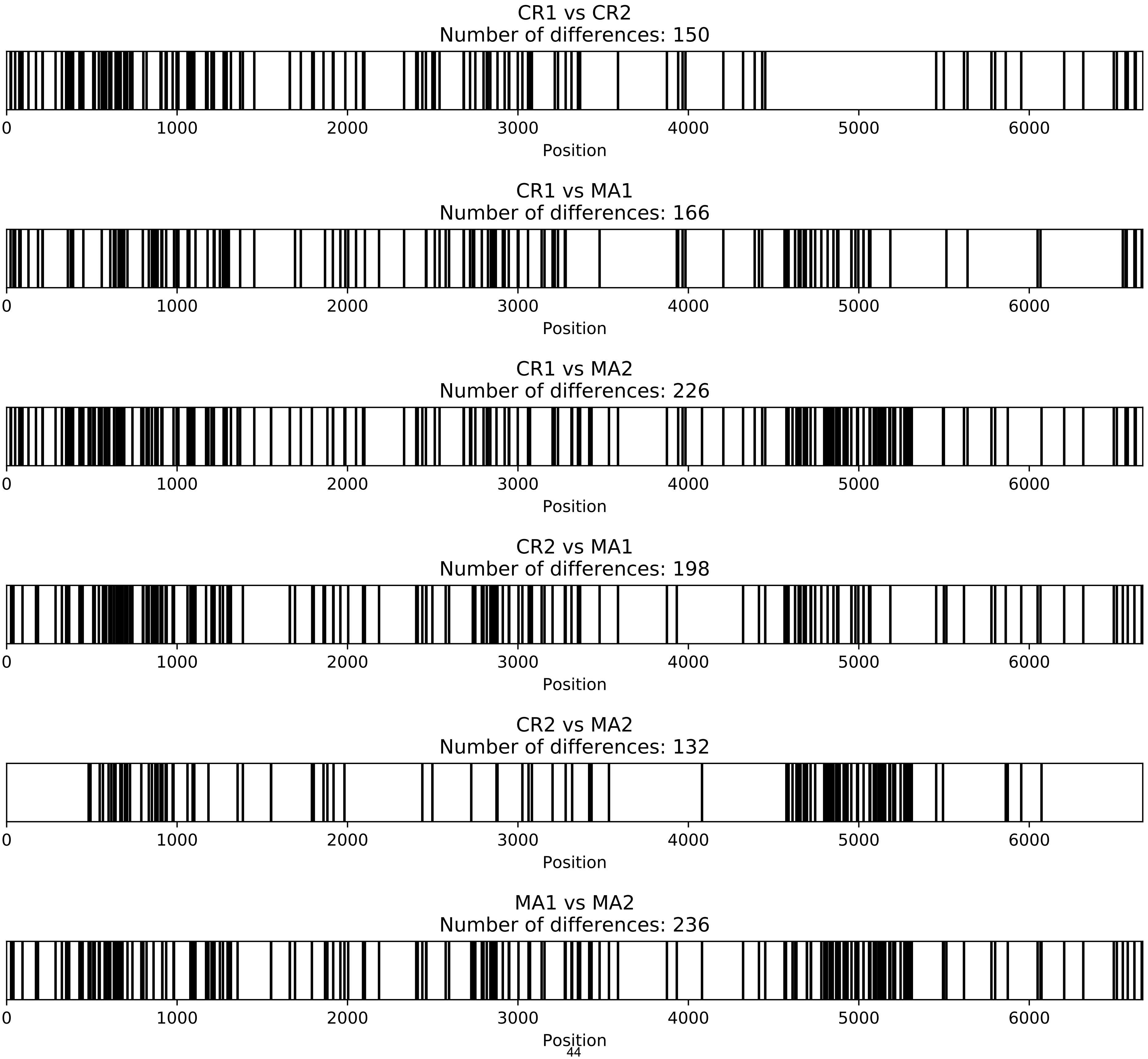
CR1 vs CR2
Number of differences: 150
CR1 vs MA1
Number of differences: 166
CR1 vs MA2
Number of differences: 226
CR2 vs MA1
Number of differences: 198
CR2 vs MA2
Number of differences: 132
MA1 vs MA2
Number of differences: 236
0
1000
2000
3000
4000
5000
6000
Position

### Region: 287



CR1 vs CR2
Number of differences: 352

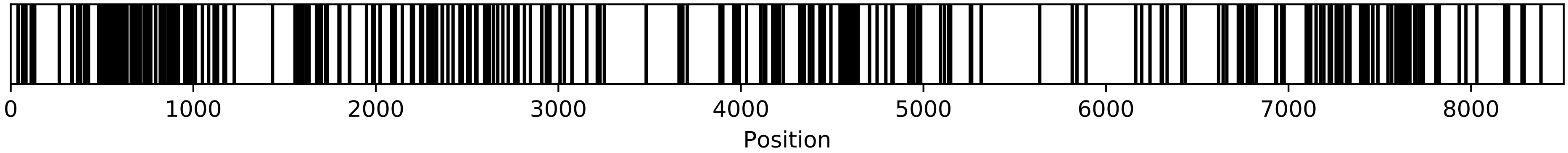


CR1 vs MA1
Number of differences: 153

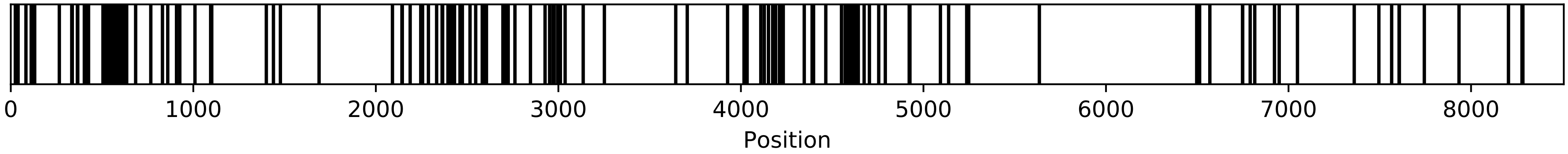


CR1 vs MA2
Number of differences: 328

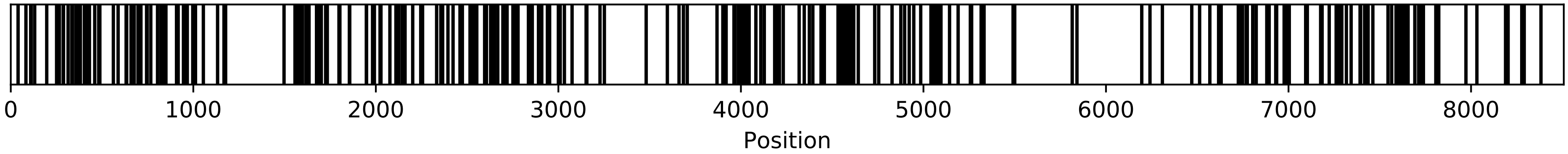


CR2 vs MA1
Number of differences: 334

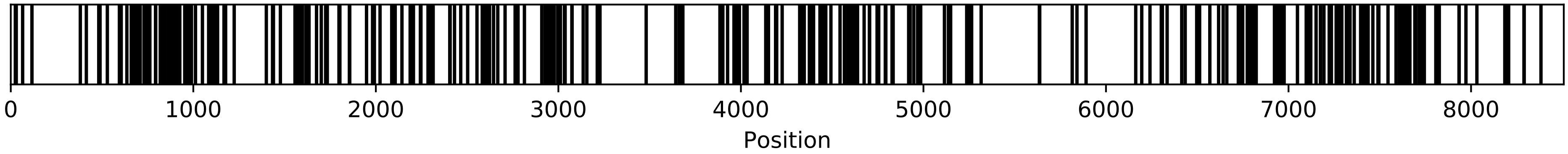


CR2 vs MA2
Number of differences: 320

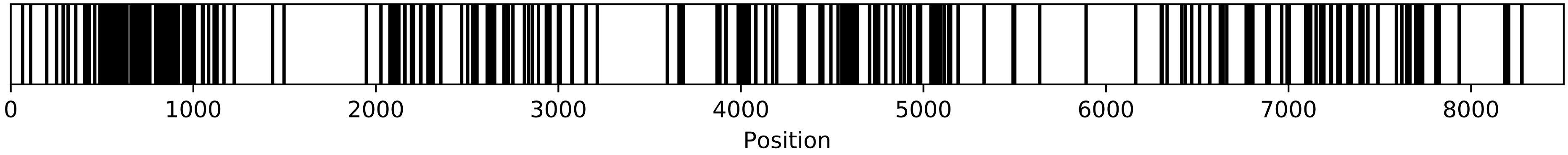


MA1 vs MA2
Number of differences: 349

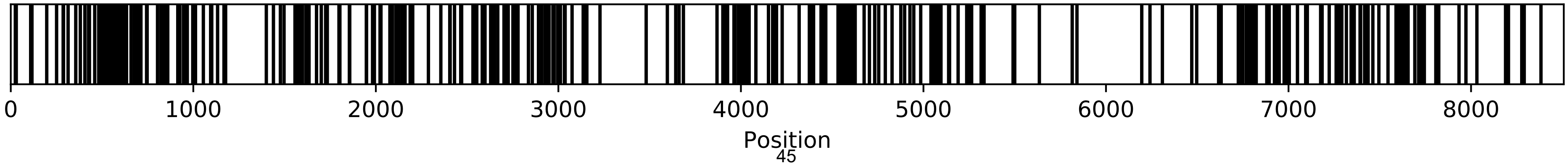

### Region: 290



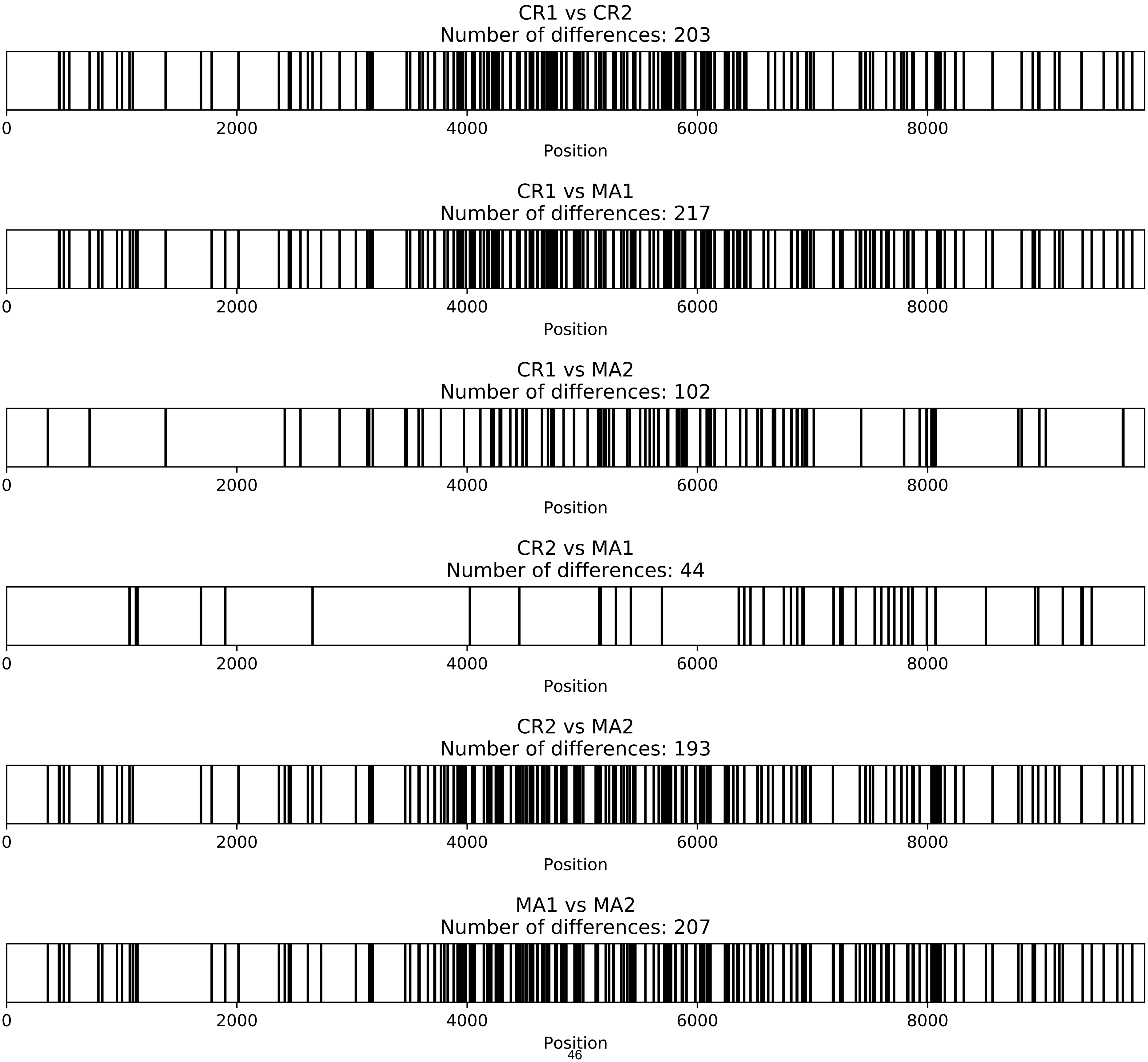
CR1 vs CR2
Number of differences: 203
CR1 vs MA1
Number of differences: 217
CR1 vs MA2
Number of differences: 102
CR2 vs MA1
Number of differences: 44
CR2 vs MA2
Number of differences: 193
MA1 vs MA2
Number of differences: 207
0
2000
4000
6000
8000
Position

### Region: 296



CR1 vs CR2
Number of differences: 176

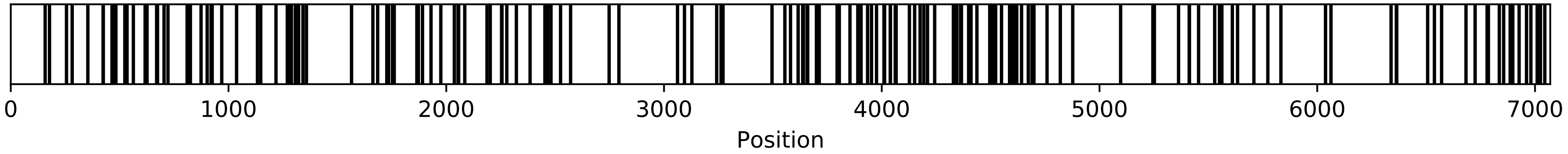


CR1 vs MA1
Number of differences: 65

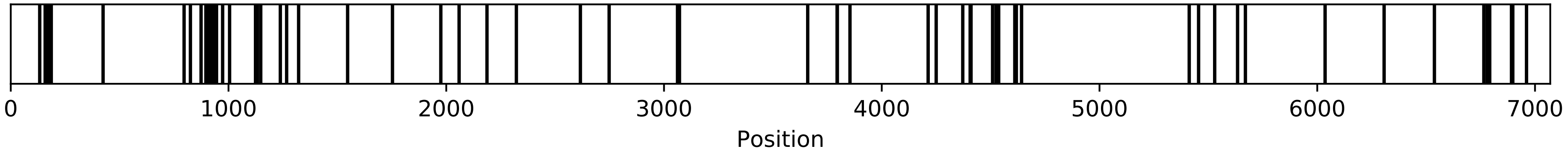


CR1 vs MA2
Number of differences: 177

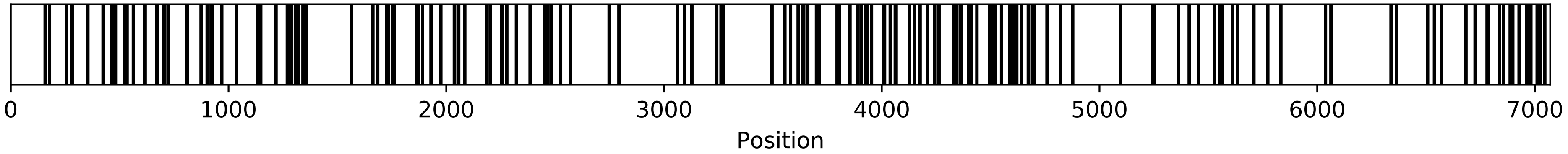


CR2 vs MA1
Number of differences: 178

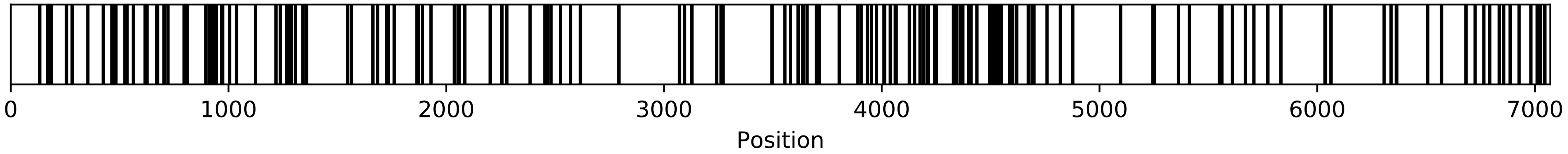


CR2 vs MA2
Number of differences: 11

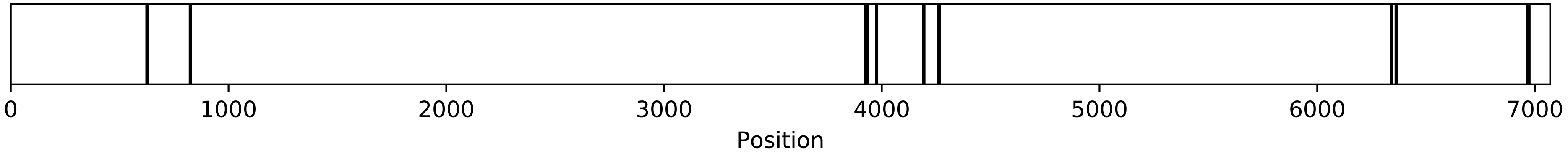


MA1 vs MA2
Number of differences: 181

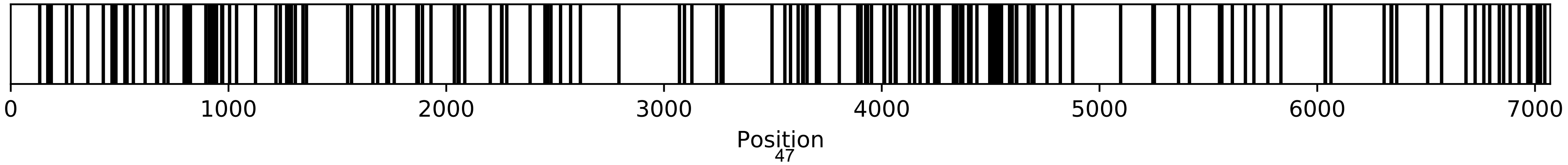

### Region: 298



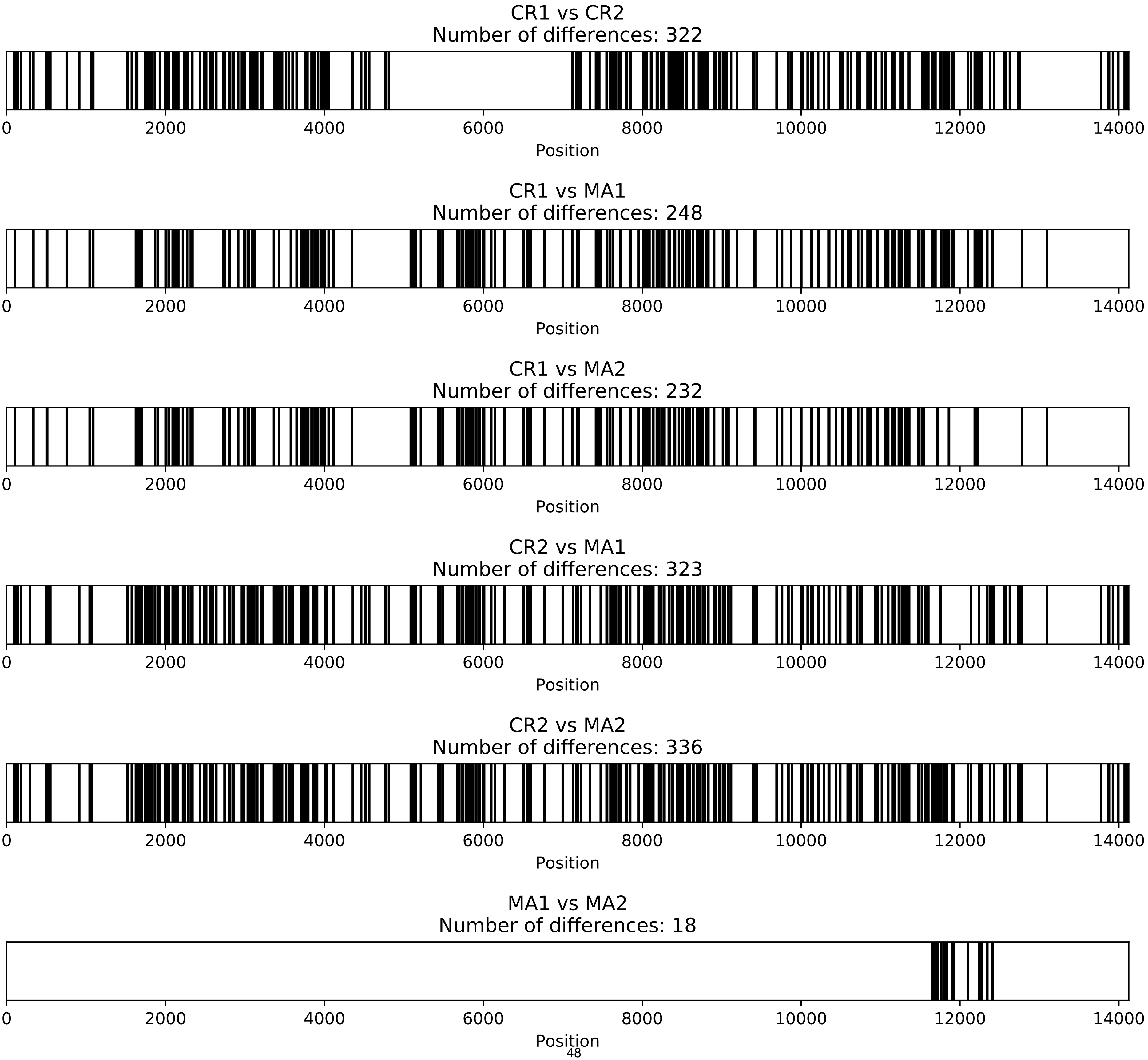
CR1 vs CR2
Number of differences: 322
0 2000 4000 6000 8000 10000 12000 14000
Position
CR1 vs MA1
Number of differences: 248
0 2000 4000 6000 8000 10000 12000 14000
Position
CR1 vs MA2
Number of differences: 232
0 2000 4000 6000 8000 10000 12000 14000
Position
CR2 vs MA1
Number of differences: 323
0 2000 4000 6000 8000 10000 12000 14000
Position
CR2 vs MA2
Number of differences: 336
0 2000 4000 6000 8000 10000 12000 14000
Position
MA1 vs MA2
Number of differences: 18
0 2000 4000 6000 8000 10000 12000 14000
Position

### Region: 299



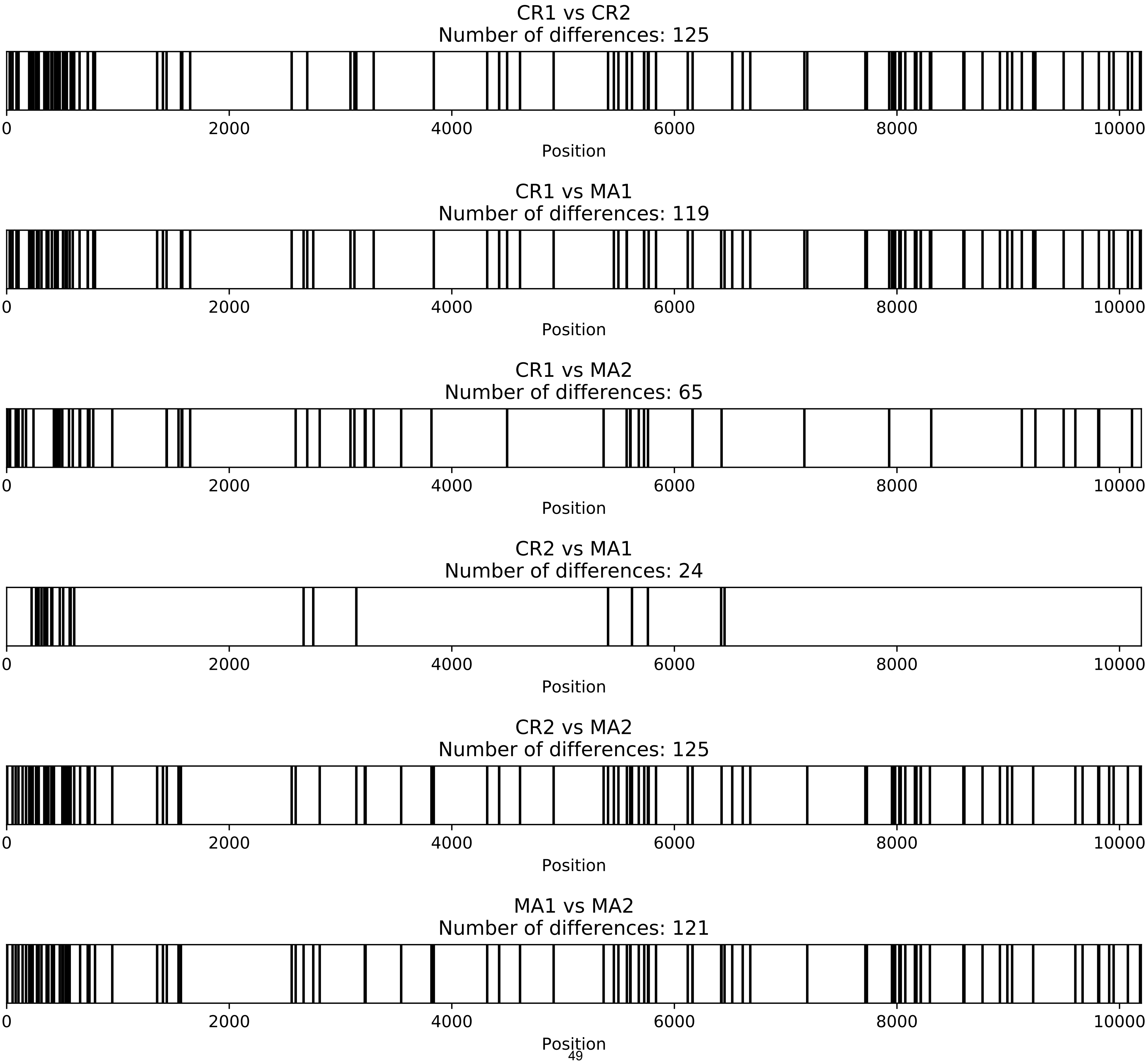
CR1 vs CR2
Number of differences: 125
CR1 vs MA1
Number of differences: 119
CR1 vs MA2
Number of differences: 65
CR2 vs MA1
Number of differences: 24
CR2 vs MA2
Number of differences: 125
MA1 vs MA2
Number of differences: 121
0
2000
4000
6000
8000
10000
Position

### Region: 303



CR1 vs CR2
Number of differences: 185

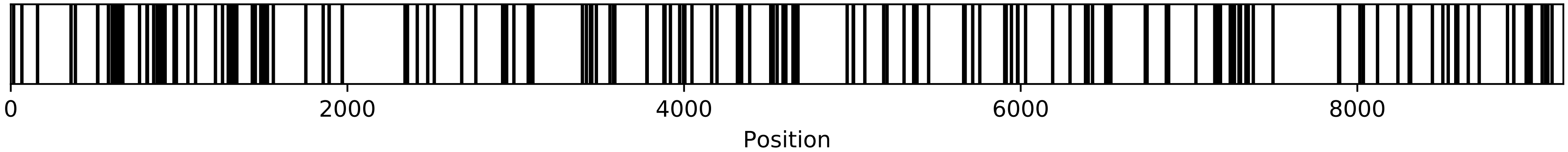


CR1 vs MA1
Number of differences: 178

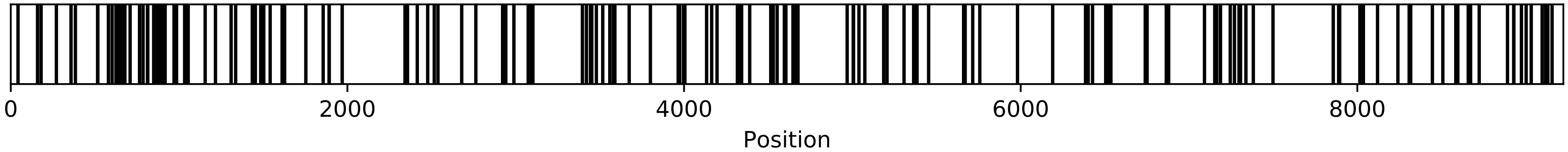


CR1 vs MA2
Number of differences: 105

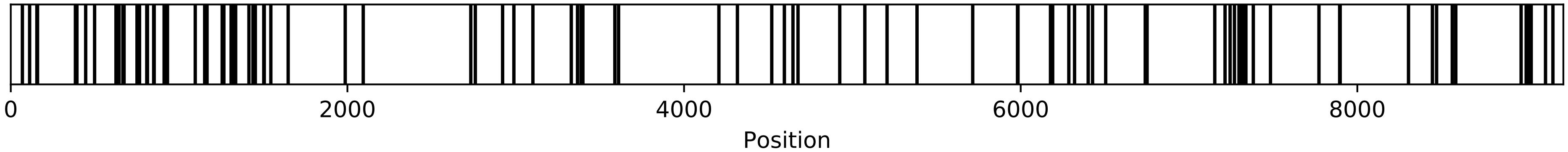


CR2 vs MA1
Number of differences: 72

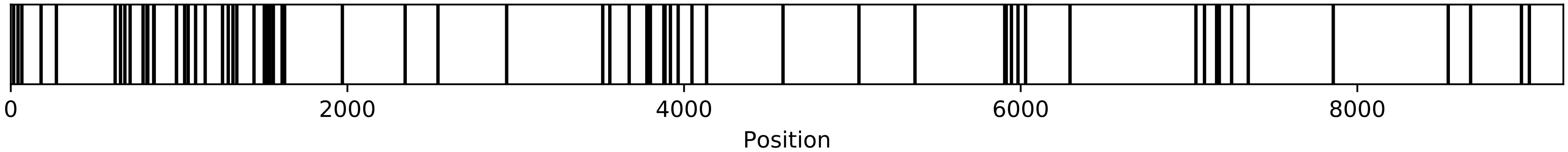


CR2 vs MA2
Number of differences: 187

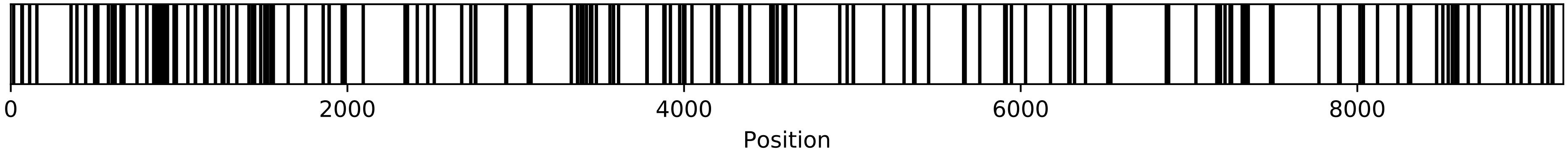


MA1 vs MA2
Number of differences: 185

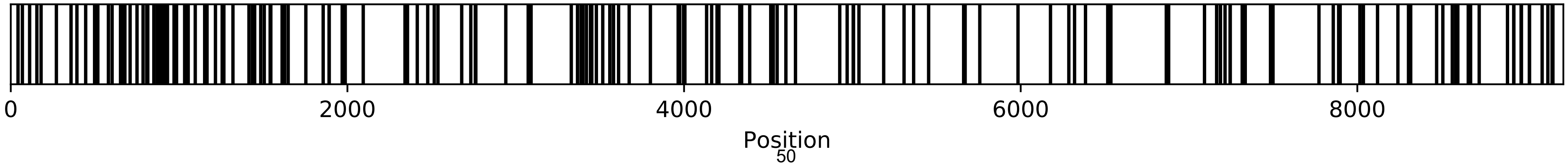

### Region: 309



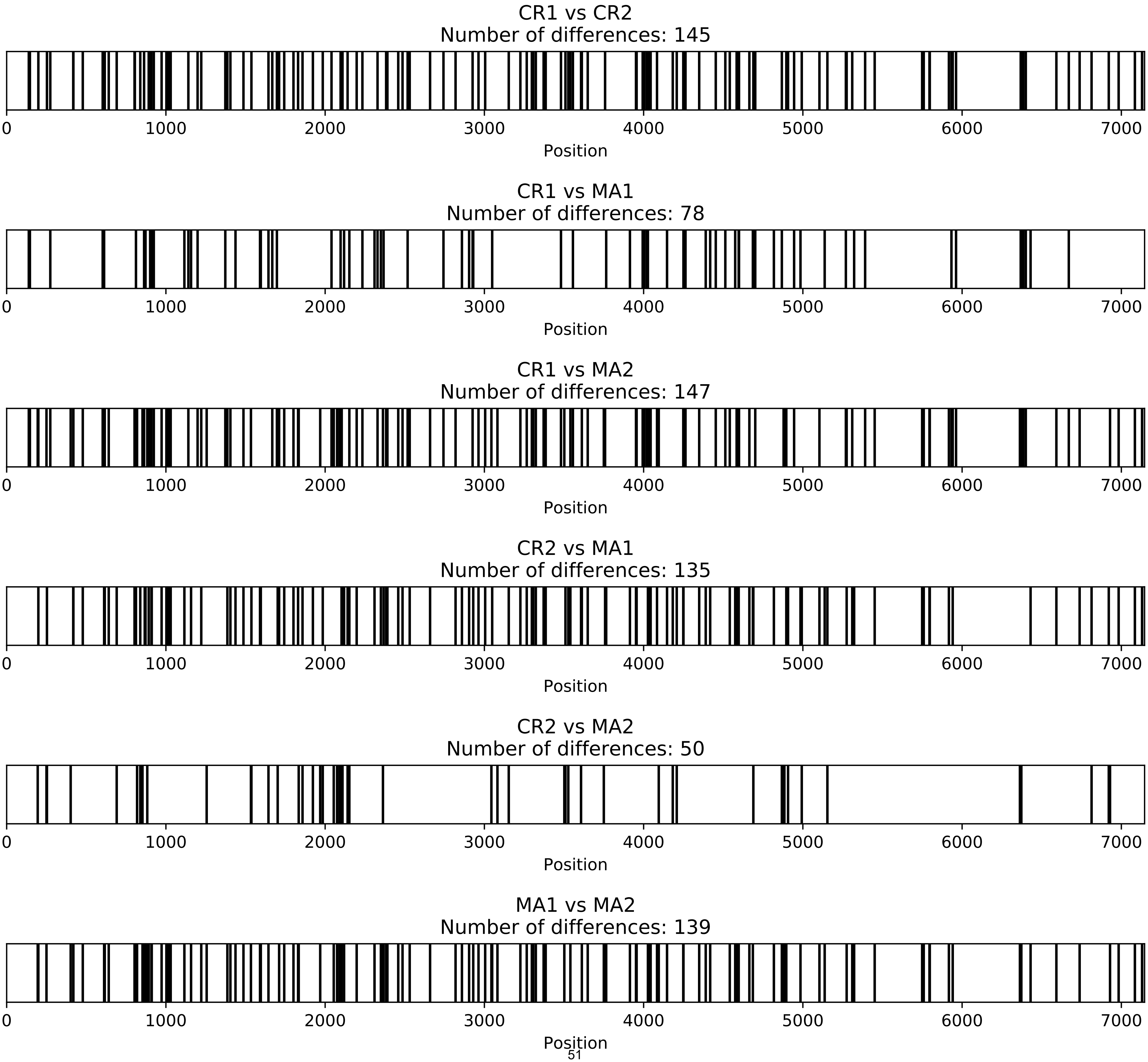
CR1 vs CR2
Number of differences: 145
CR1 vs MA1
Number of differences: 78
CR1 vs MA2
Number of differences: 147
CR2 vs MA1
Number of differences: 135
CR2 vs MA2
Number of differences: 50
MA1 vs MA2
Number of differences: 139
Position
0
1000
2000
3000
4000
5000
6000
7000

### Region: 312



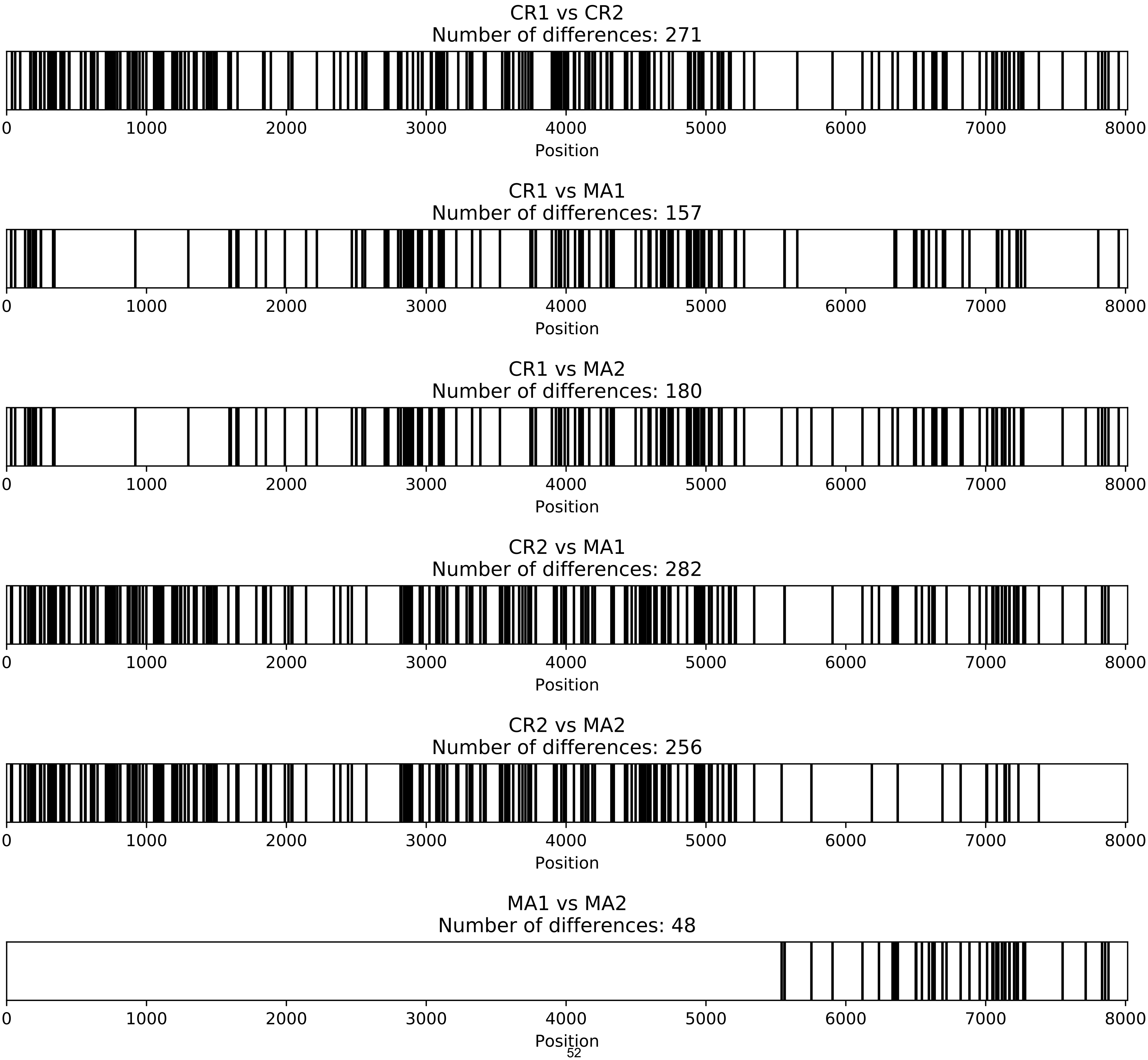
CR1 vs CR2
Number of differences: 271
CR1 vs MA1
Number of differences: 157
CR1 vs MA2
Number of differences: 180
CR2 vs MA1
Number of differences: 282
CR2 vs MA2
Number of differences: 256
MA1 vs MA2
Number of differences: 48
0 1000 2000 3000 4000 5000 6000 7000 8000
Position

### Region: 313



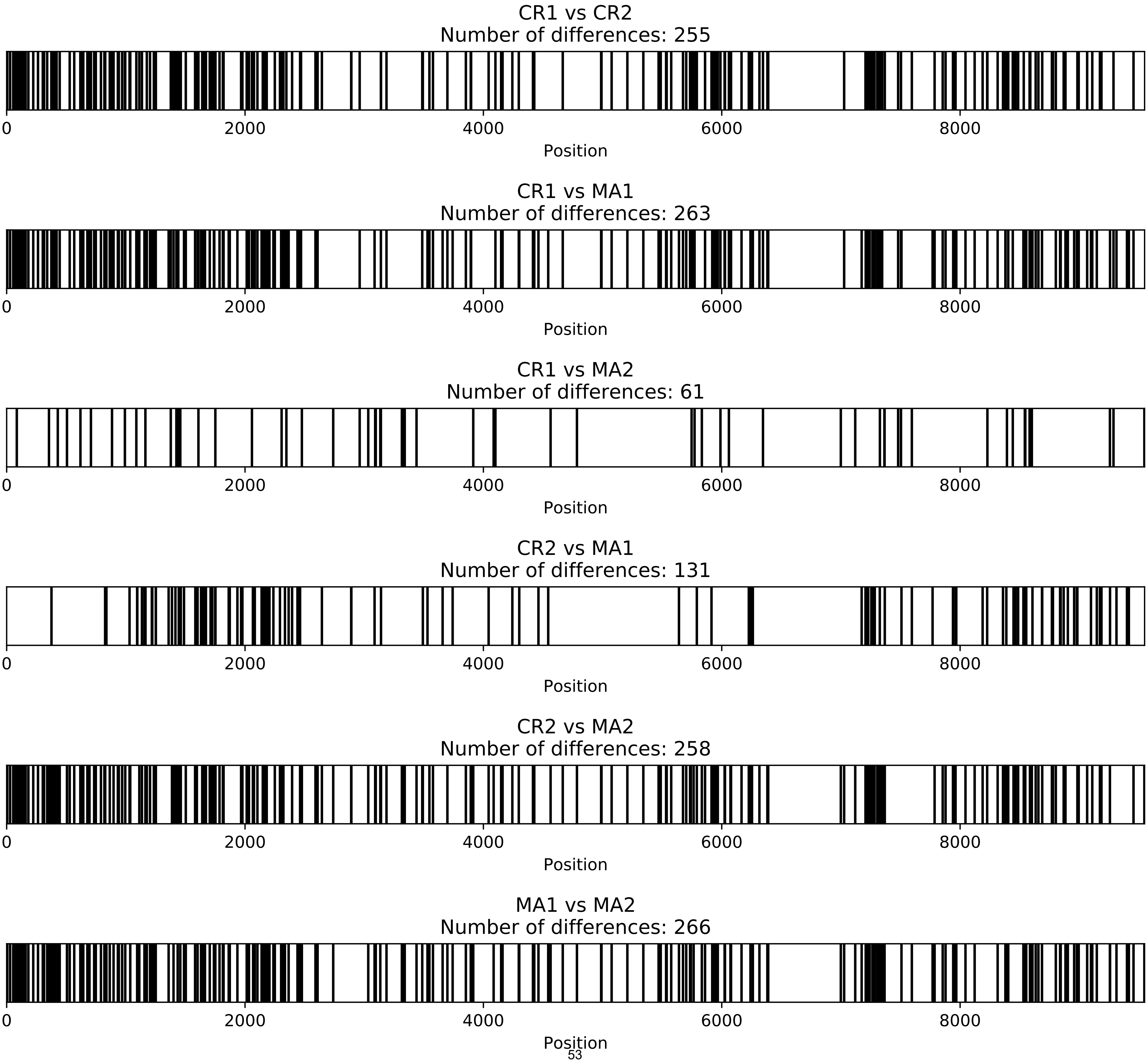
CR1 vs CR2
Number of differences: 255
CR1 vs MA1
Number of differences: 263
CR1 vs MA2
Number of differences: 61
CR2 vs MA1
Number of differences: 131
CR2 vs MA2
Number of differences: 258
MA1 vs MA2
Number of differences: 266
0
2000
4000
6000
8000
Position

### Region: 329



CR1 vs CR2
Number of differences: 184

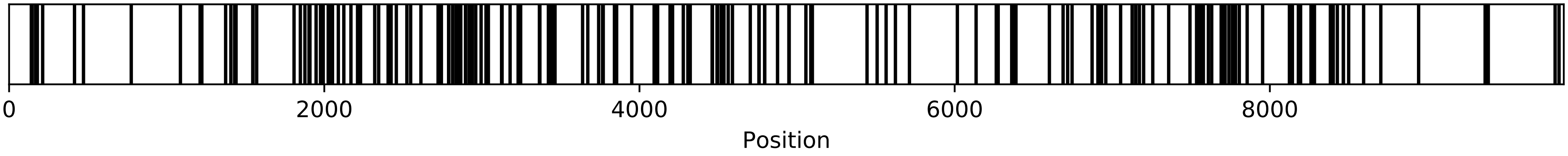


CR1 vs MA1
Number of differences: 28

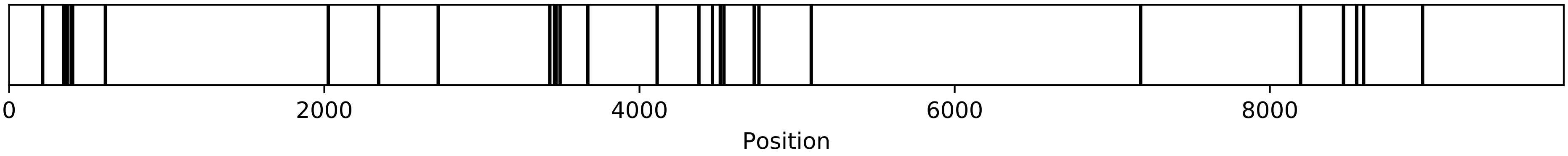


CR1 vs MA2
Number of differences: 197

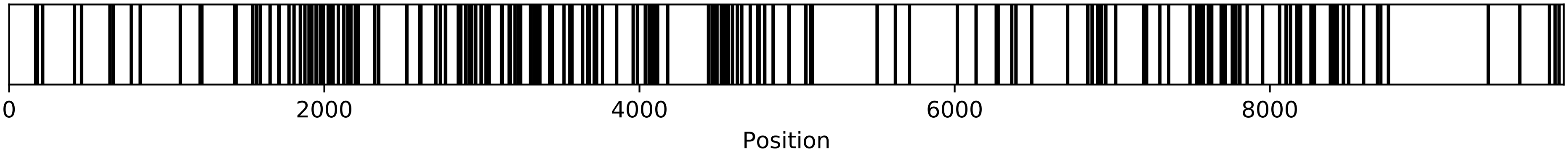


CR2 vs MA1
Number of differences: 186

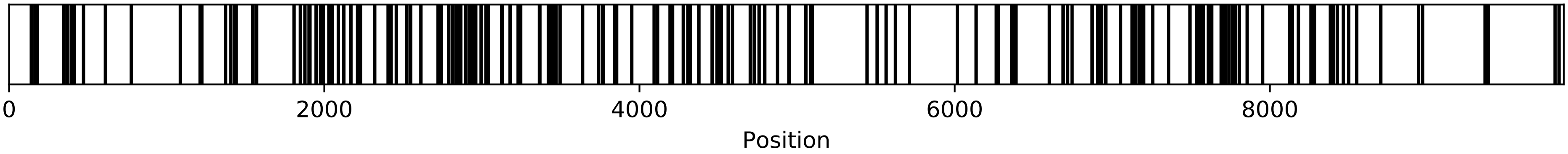


CR2 vs MA2
Number of differences: 129

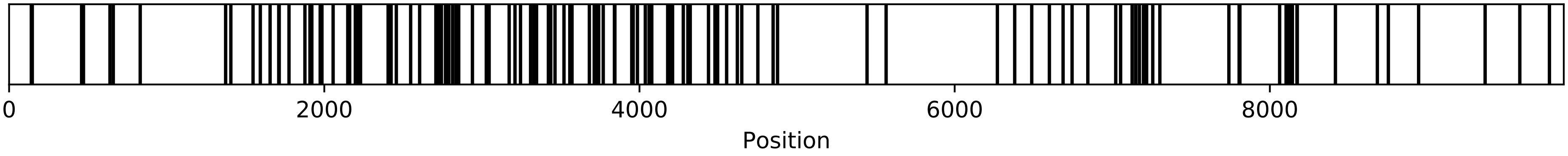


MA1 vs MA2
Number of differences: 199

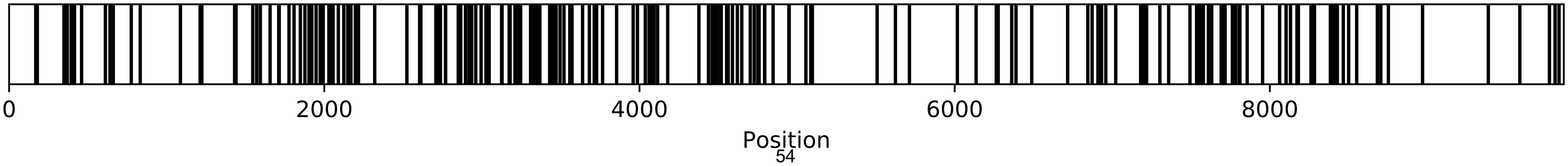

### Region: 332

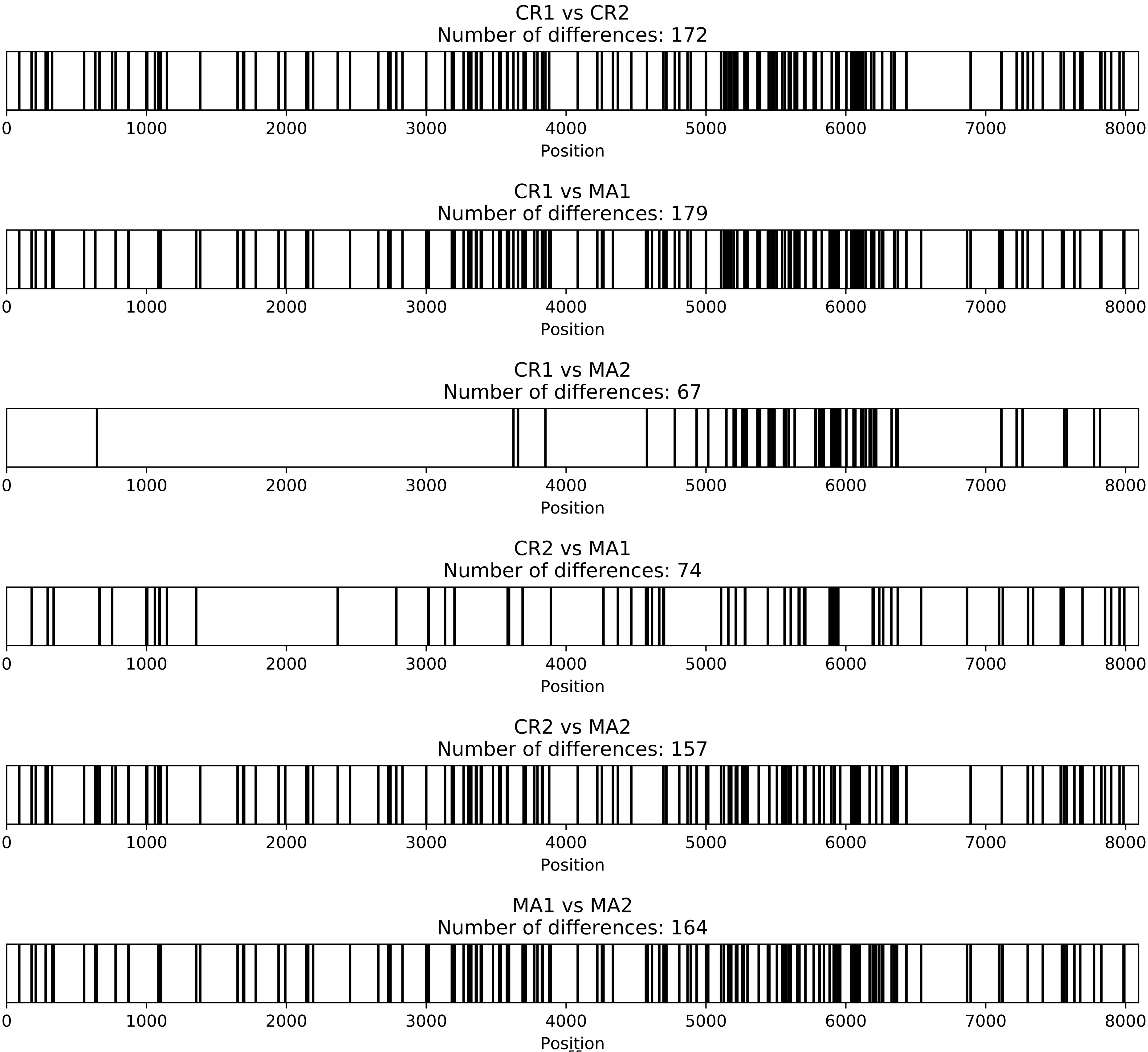
CR1 vs CR2
Number of differences: 172
CR1 vs MA1
Number of differences: 179
CR1 vs MA2
Number of differences: 67
CR2 vs MA1
Number of differences: 74
CR2 vs MA2
Number of differences: 157
MA1 vs MA2
Number of differences: 164
0
1000
2000
3000
4000
5000
6000
7000
8000
Position

### Region: 334



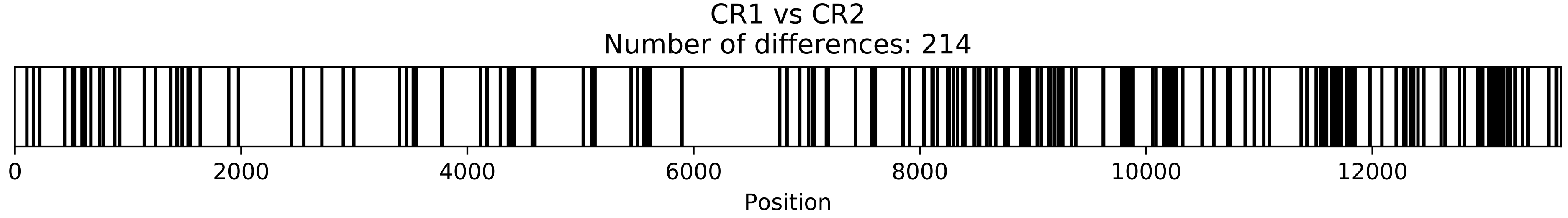
CR1 vs CR2
Number of differences: 214
0
2000
4000
6000
8000
10000
12000
Position

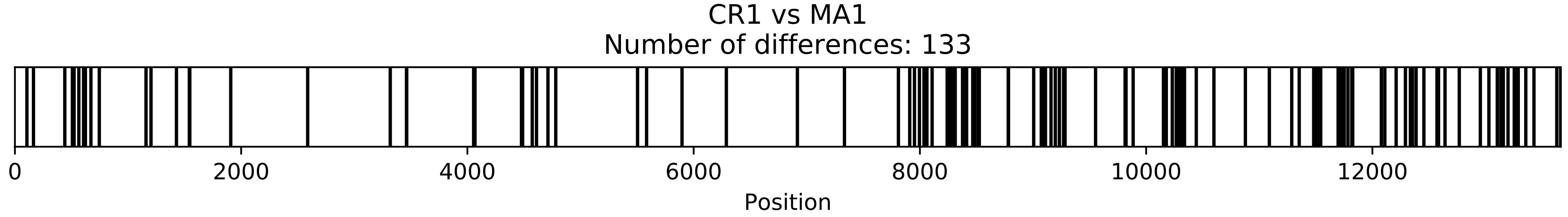
CR1 vs MA1
Number of differences: 133
0
2000
4000
6000
8000
10000
12000
Position

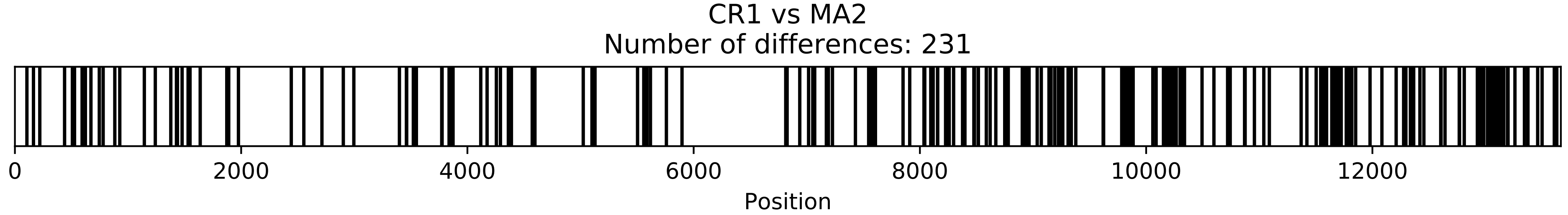
CR1 vs MA2
Number of differences: 231
0
2000
4000
6000
8000
10000
12000
Position

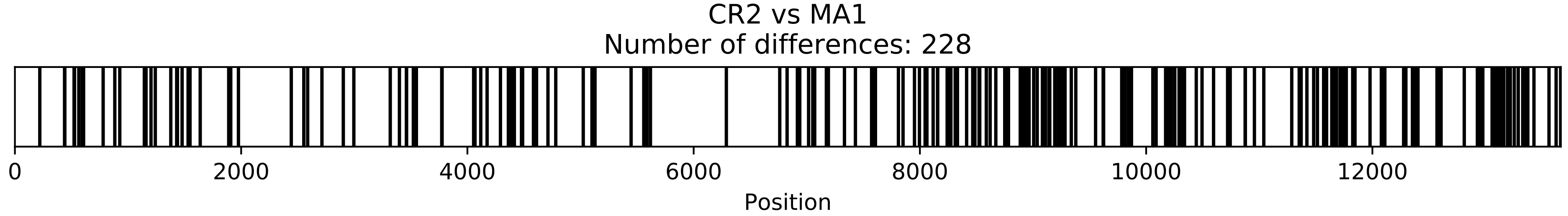
CR2 vs MA1
Number of differences: 228
0
2000
4000
6000
8000
10000
12000
Position

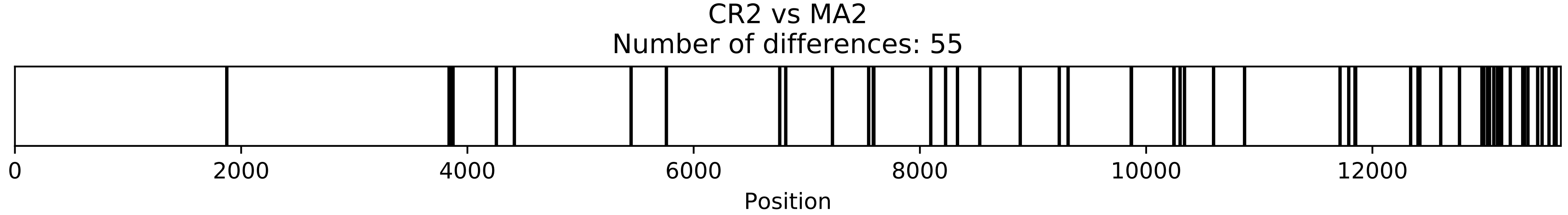
CR2 vs MA2
Number of differences: 55
0
2000
4000
6000
8000
10000
12000
Position

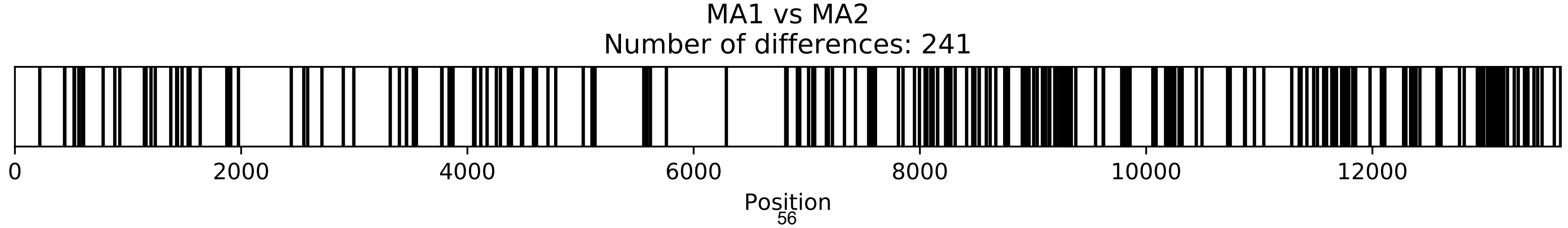
MA1 vs MA2
Number of differences: 241
0
2000
4000
6000
8000
10000
12000
Position

### Region: 340



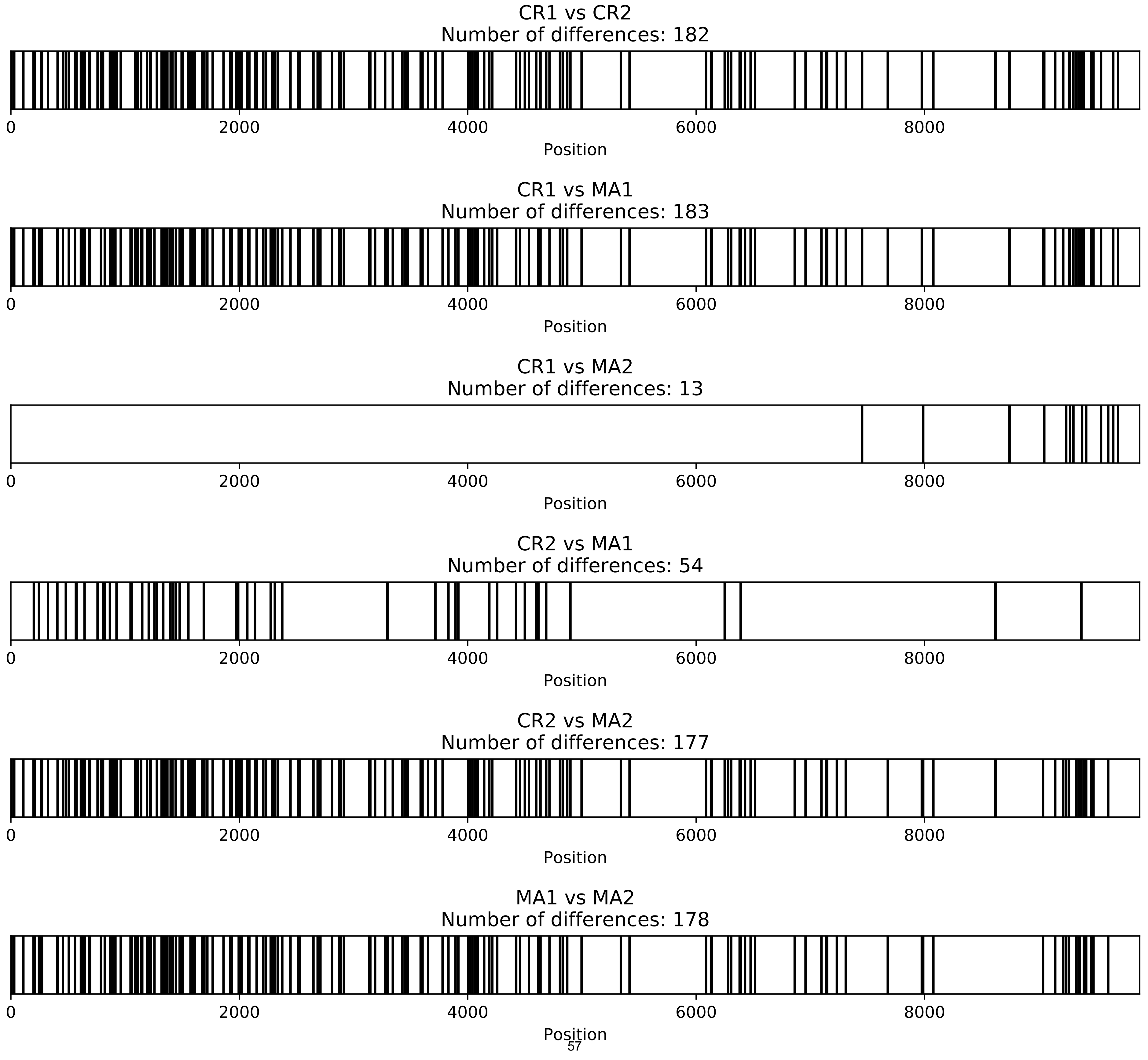
CR1 vs CR2
Number of differences: 182
0
2000
4000
6000
8000
Position
CR1 vs MA1
Number of differences: 183
Position
CR1 vs MA2
Number of differences: 13
Position
CR2 vs MA1
Number of differences: 54
Position
CR2 vs MA2
Number of differences: 177
Position
MA1 vs MA2
Number of differences: 178
Position

### Region: 341



CR1 vs CR2
Number of differences: 135

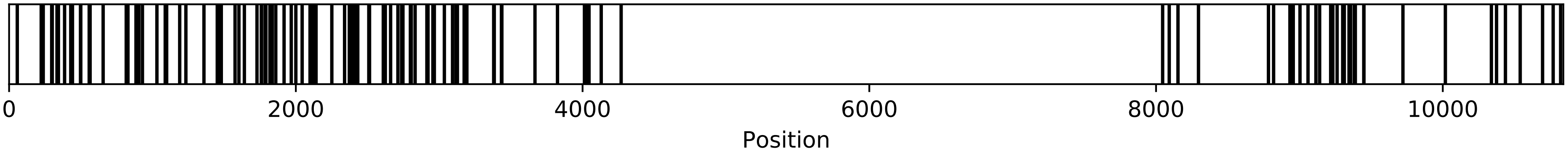


CR1 vs MA1
Number of differences: 159

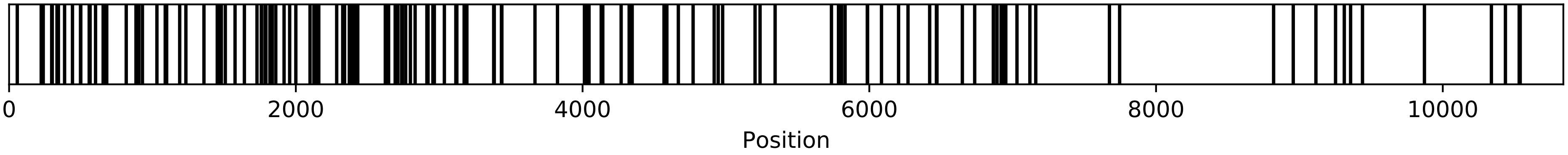


CR1 vs MA2
Number of differences: 79

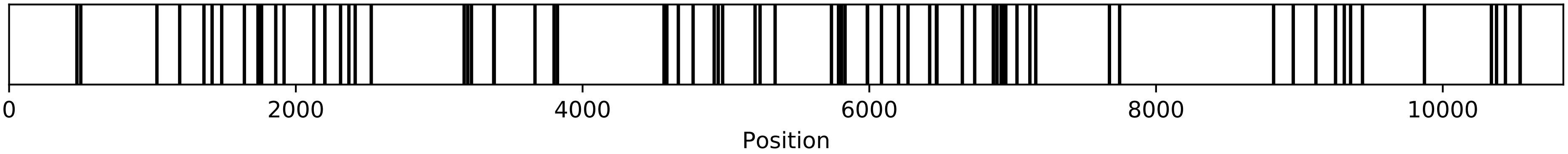


CR2 vs MA1
Number of differences: 113

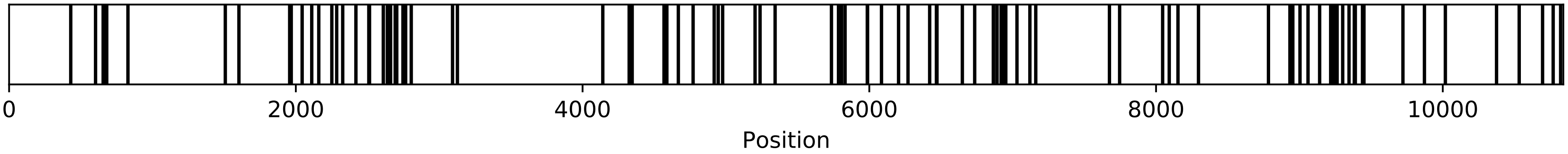


CR2 vs MA2
Number of differences: 166

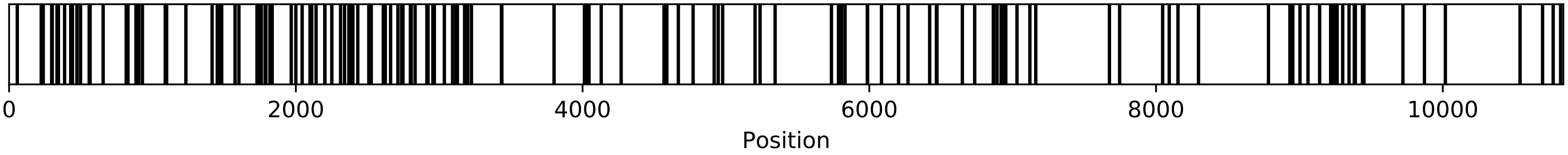


MA1 vs MA2
Number of differences: 110

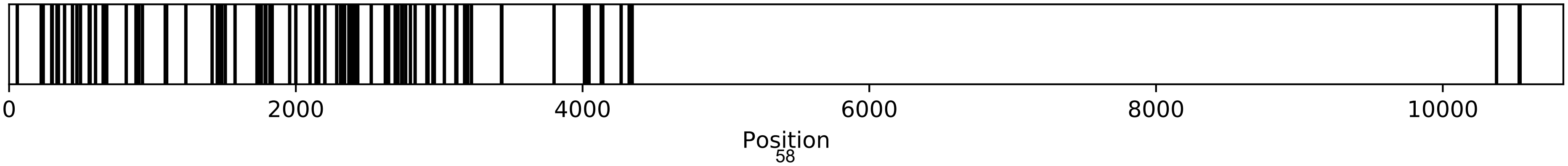

### Region: 344



CR1 vs CR2
Number of differences: 193

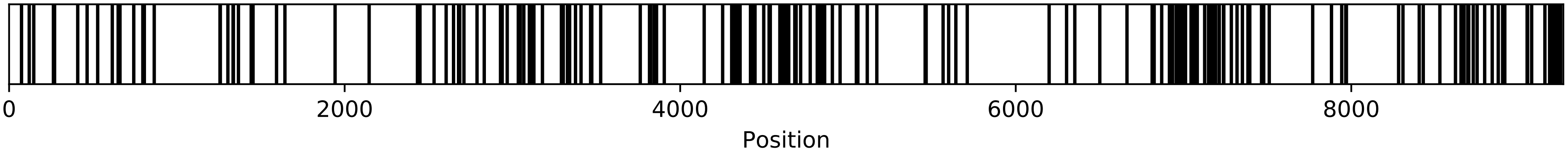


CR1 vs MA1
Number of differences: 110

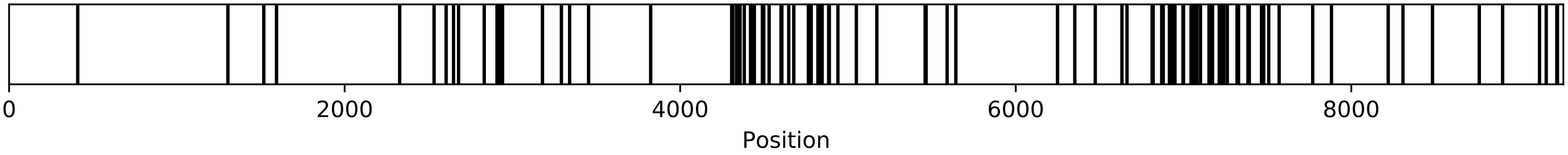


CR1 vs MA2
Number of differences: 194

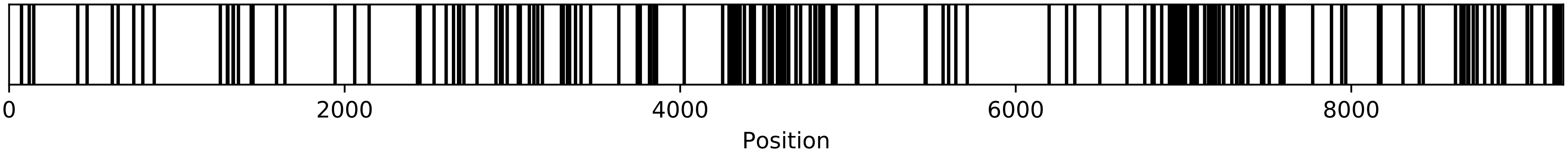


CR2 vs MA1
Number of differences: 193

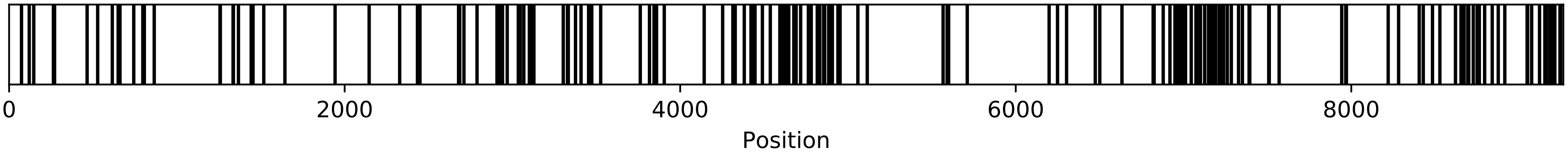


CR2 vs MA2
Number of differences: 71

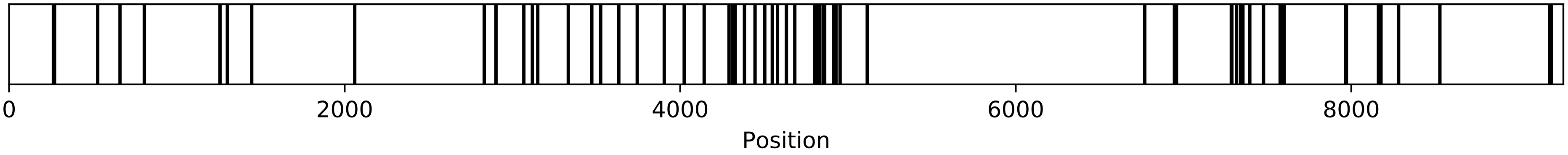


MA1 vs MA2
Number of differences: 195

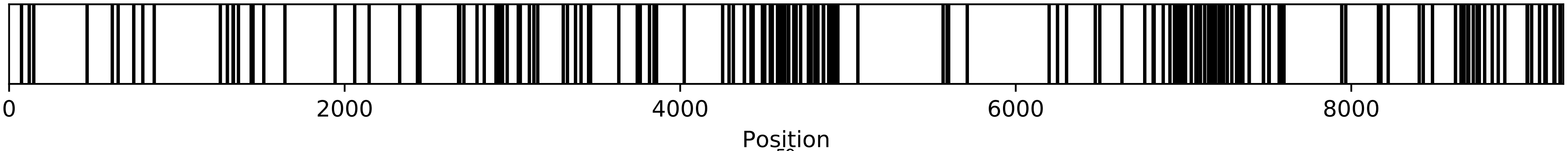

### Region: 346



CR1 vs CR2
Number of differences: 247

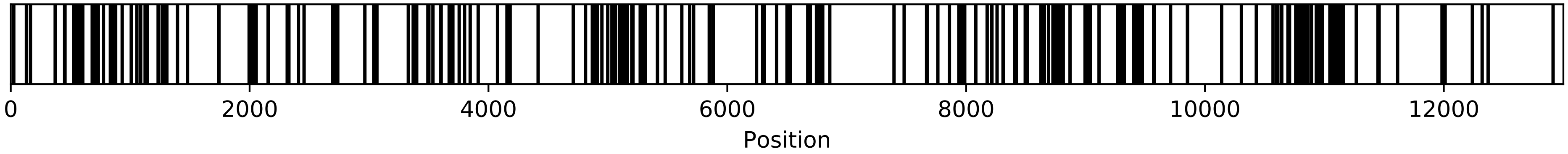


CR1 vs MA1
Number of differences: 100

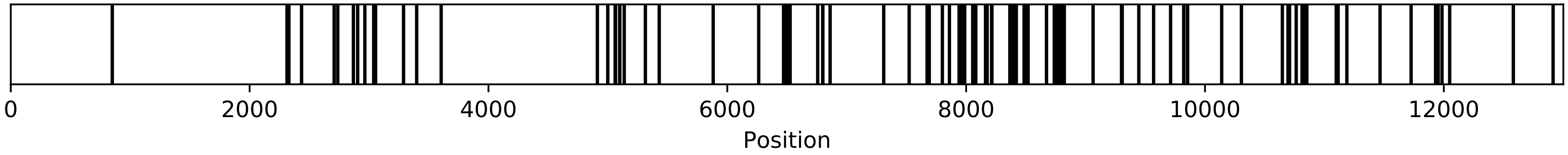


CR1 vs MA2
Number of differences: 243

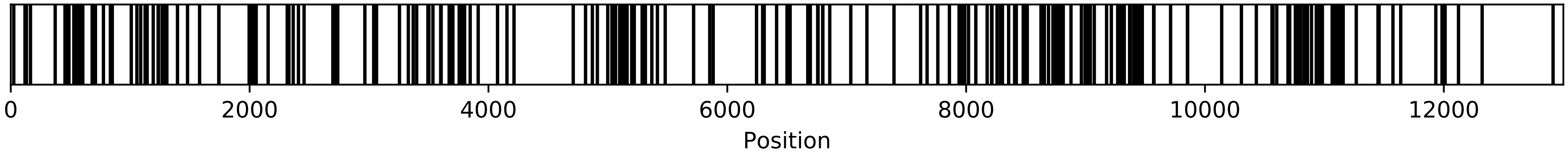


CR2 vs MA1
Number of differences: 243

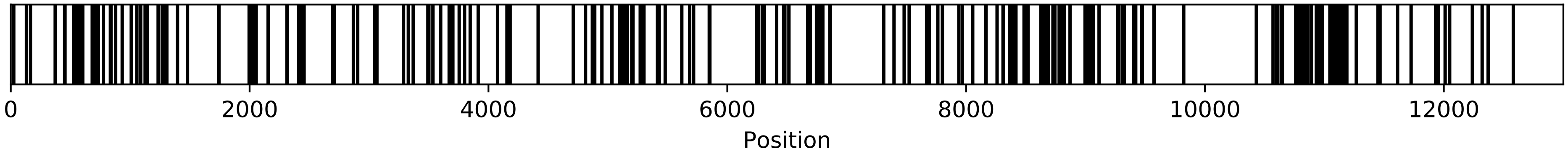


CR2 vs MA2
Number of differences: 90

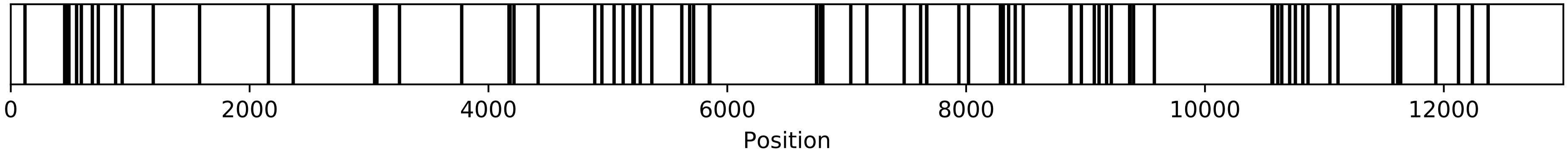


MA1 vs MA2
Number of differences: 237

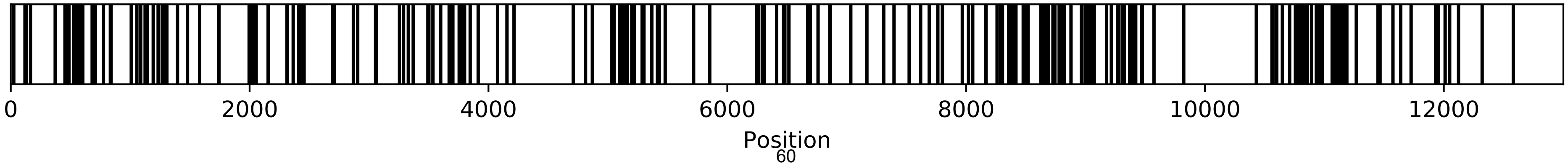

### Region: 349



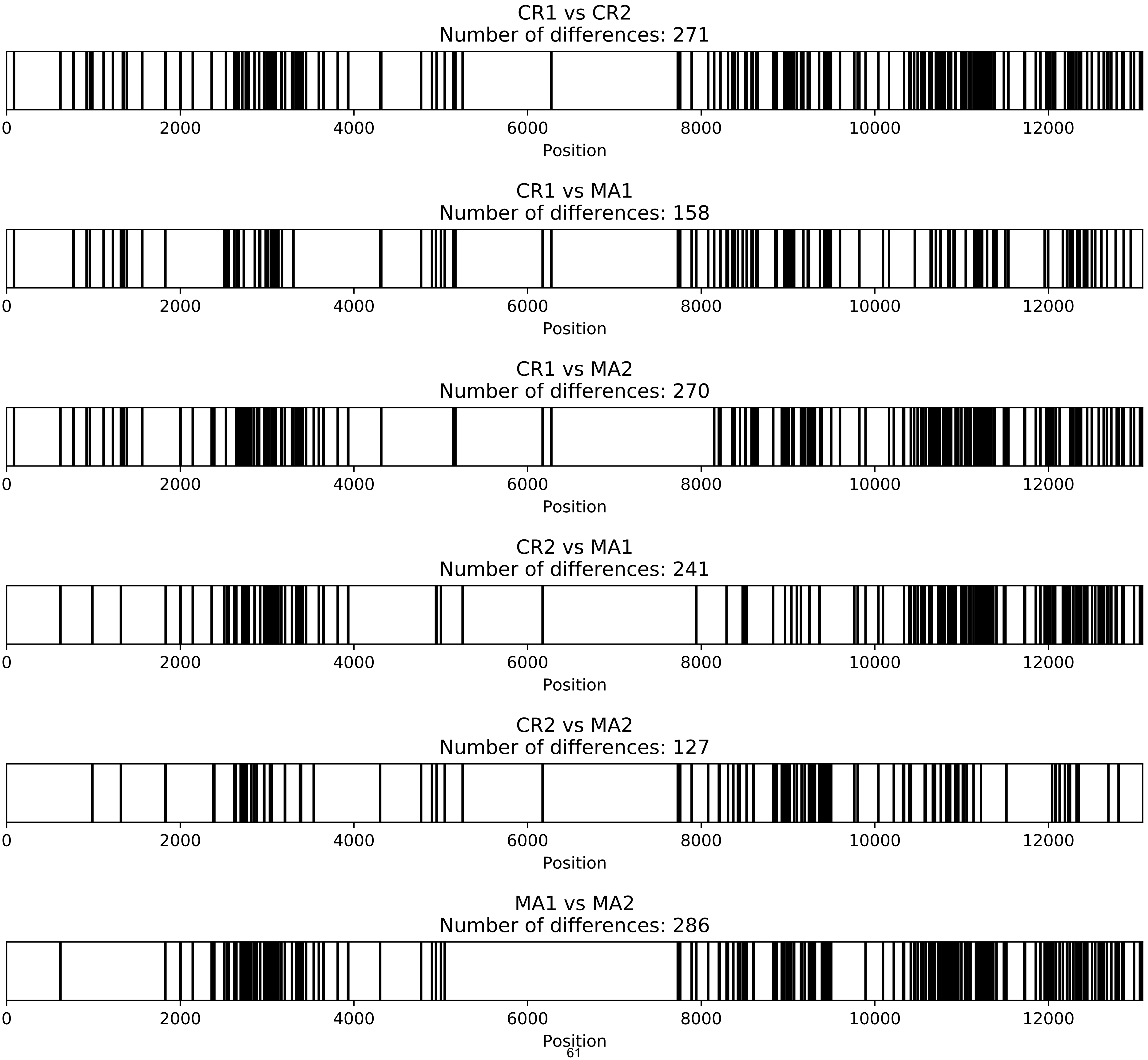
CR1 vs CR2
Number of differences: 271
CR1 vs MA1
Number of differences: 158
CR1 vs MA2
Number of differences: 270
CR2 vs MA1
Number of differences: 241
CR2 vs MA2
Number of differences: 127
MA1 vs MA2
Number of differences: 286
0
2000
4000
6000
8000
10000
12000
Position

### Region: 353



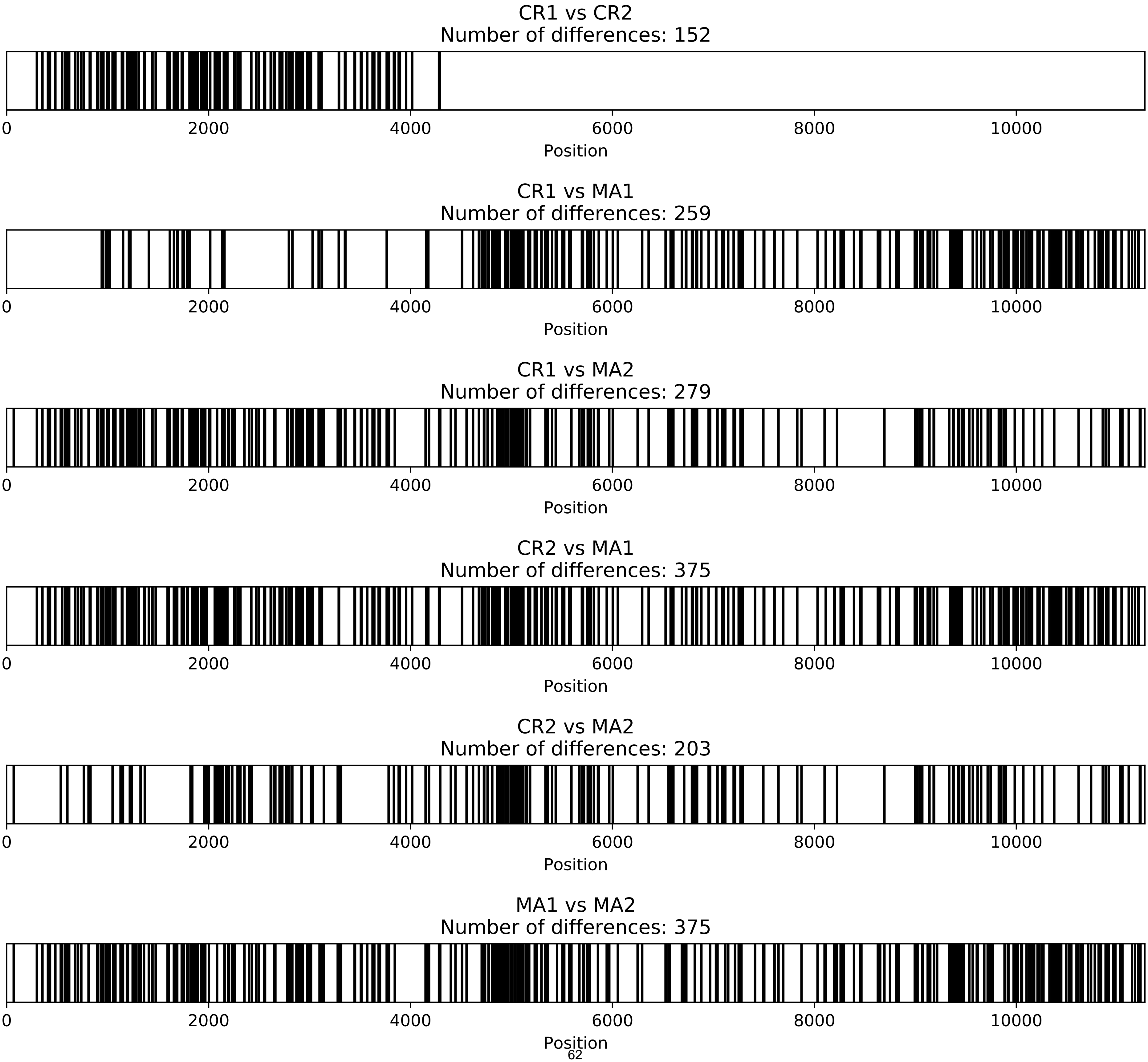
CR1 vs CR2
Number of differences: 152
CR1 vs MA1
Number of differences: 259
CR1 vs MA2
Number of differences: 279
CR2 vs MA1
Number of differences: 375
CR2 vs MA2
Number of differences: 203
MA1 vs MA2
Number of differences: 375
0
2000
4000
6000
8000
10000
Position

### Region: 360



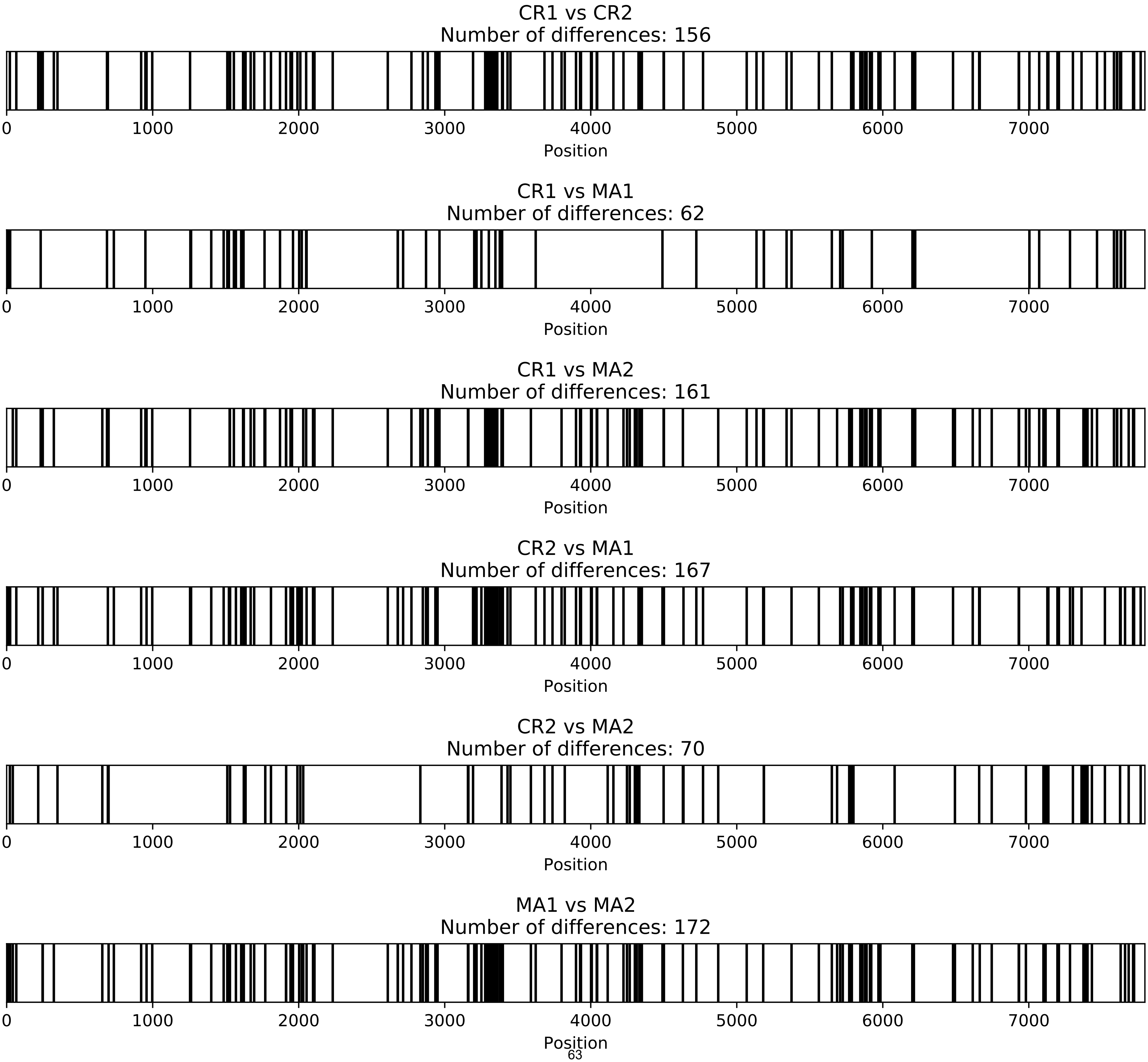
CR1 vs CR2
Number of differences: 156
CR1 vs MA1
Number of differences: 62
CR1 vs MA2
Number of differences: 161
CR2 vs MA1
Number of differences: 167
CR2 vs MA2
Number of differences: 70
MA1 vs MA2
Number of differences: 172
0
1000
2000
3000
4000
5000
6000
7000
Position

### Region: 361



CR1 vs CR2
Number of differences: 130

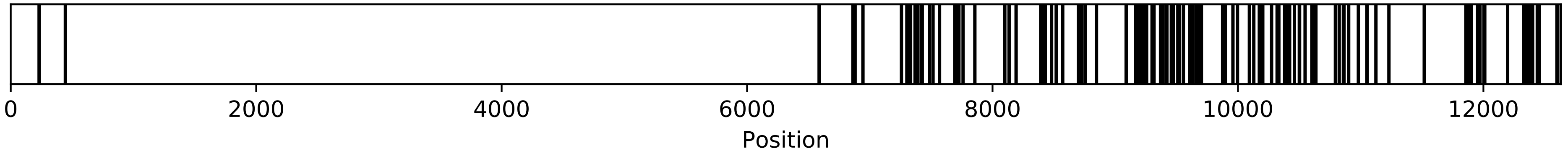


CR1 vs MA1
Number of differences: 194

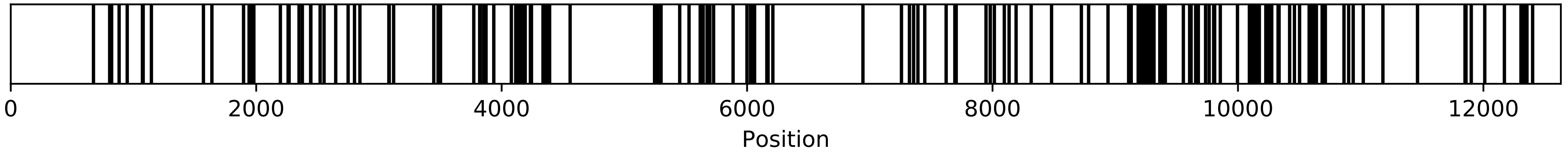


CR1 vs MA2
Number of differences: 156

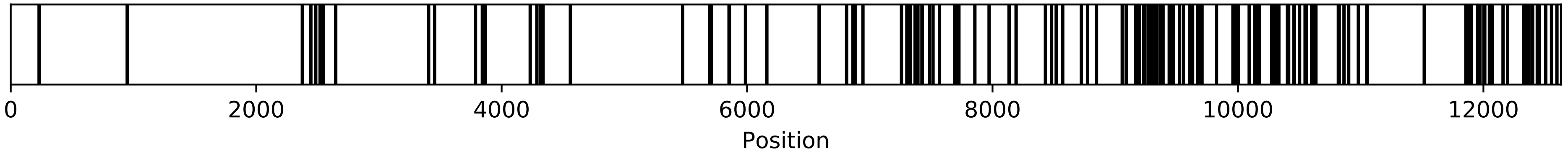


CR2 vs MA1
Number of differences: 240

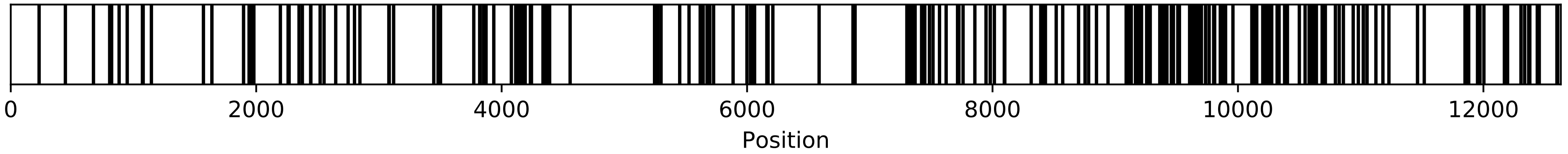


CR2 vs MA2
Number of differences: 102

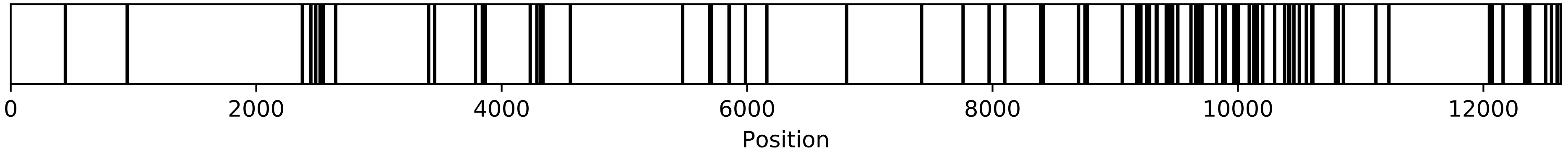


MA1 vs MA2
Number of differences: 241

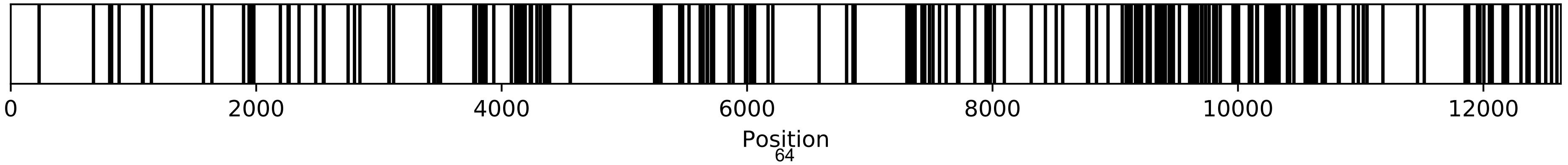

### Region: 365



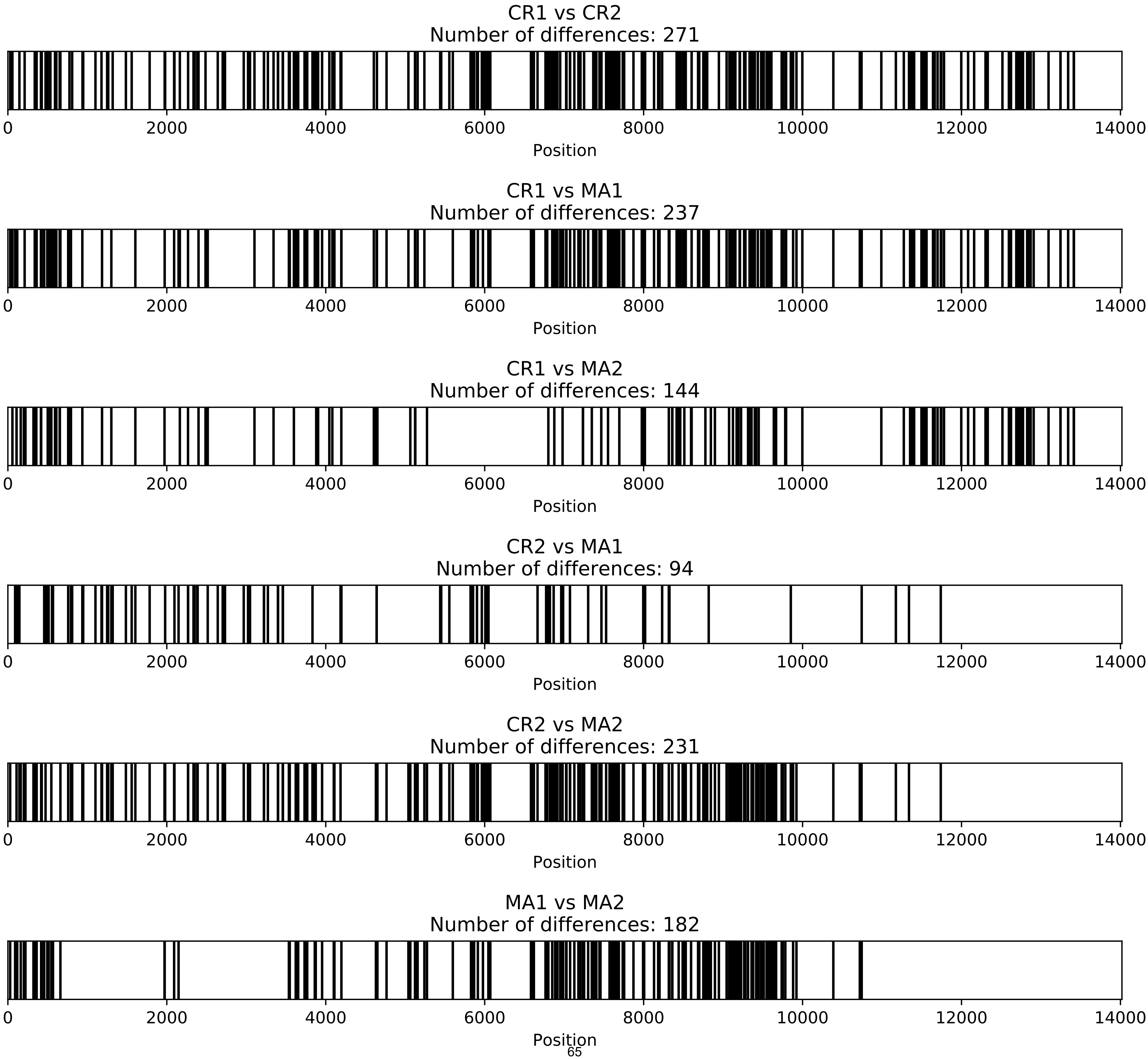
CR1 vs CR2
Number of differences: 271
CR1 vs MA1
Number of differences: 237
CR1 vs MA2
Number of differences: 144
CR2 vs MA1
Number of differences: 94
CR2 vs MA2
Number of differences: 231
MA1 vs MA2
Number of differences: 182
0
2000
4000
6000
8000
10000
12000
14000
Position

### Region: 366



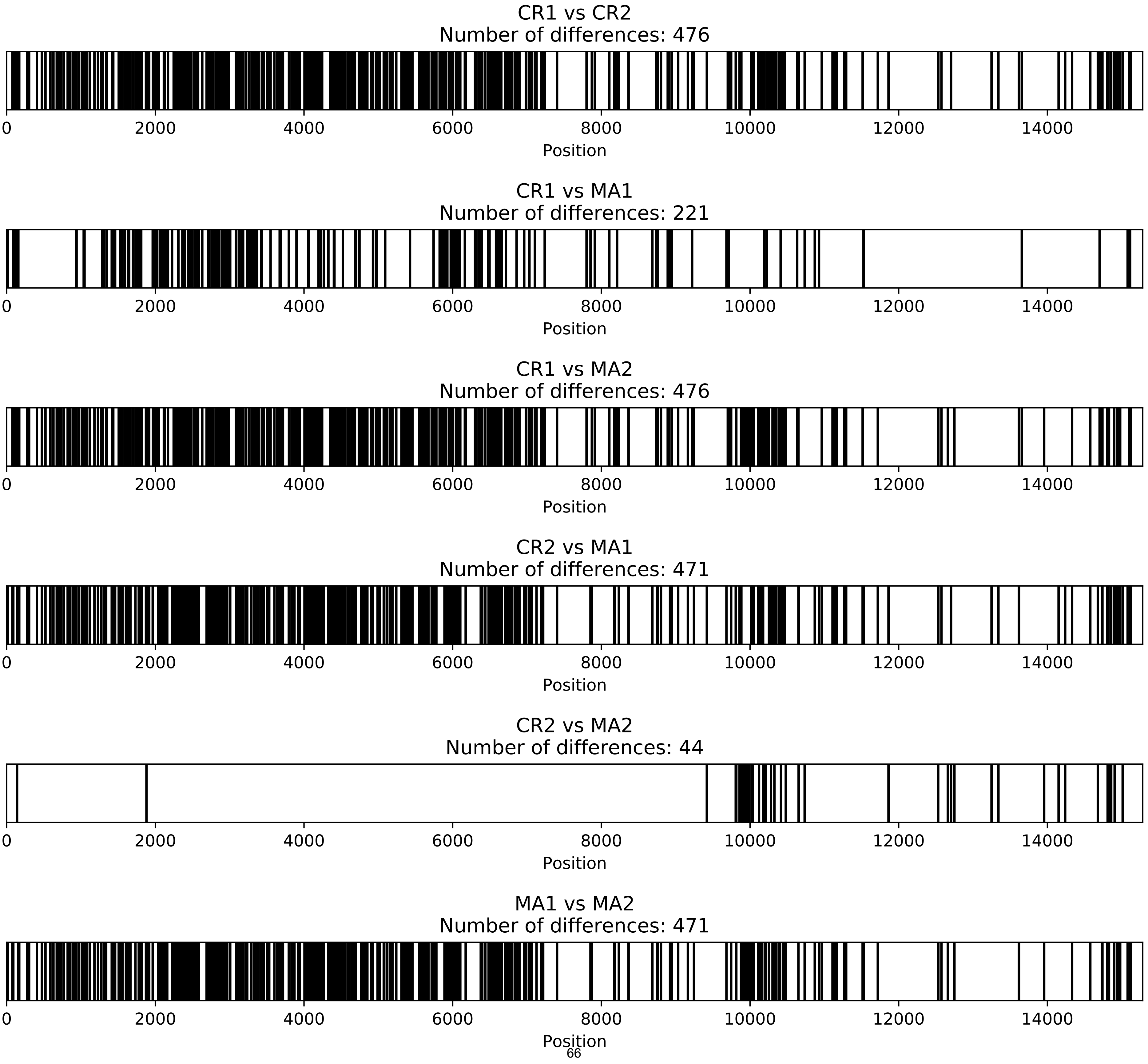
CR1 vs CR2
Number of differences: 476
CR1 vs MA1
Number of differences: 221
CR1 vs MA2
Number of differences: 476
CR2 vs MA1
Number of differences: 471
CR2 vs MA2
Number of differences: 44
MA1 vs MA2
Number of differences: 471
Position
0
2000
4000
6000
8000
10000
12000
14000

### Region: 372



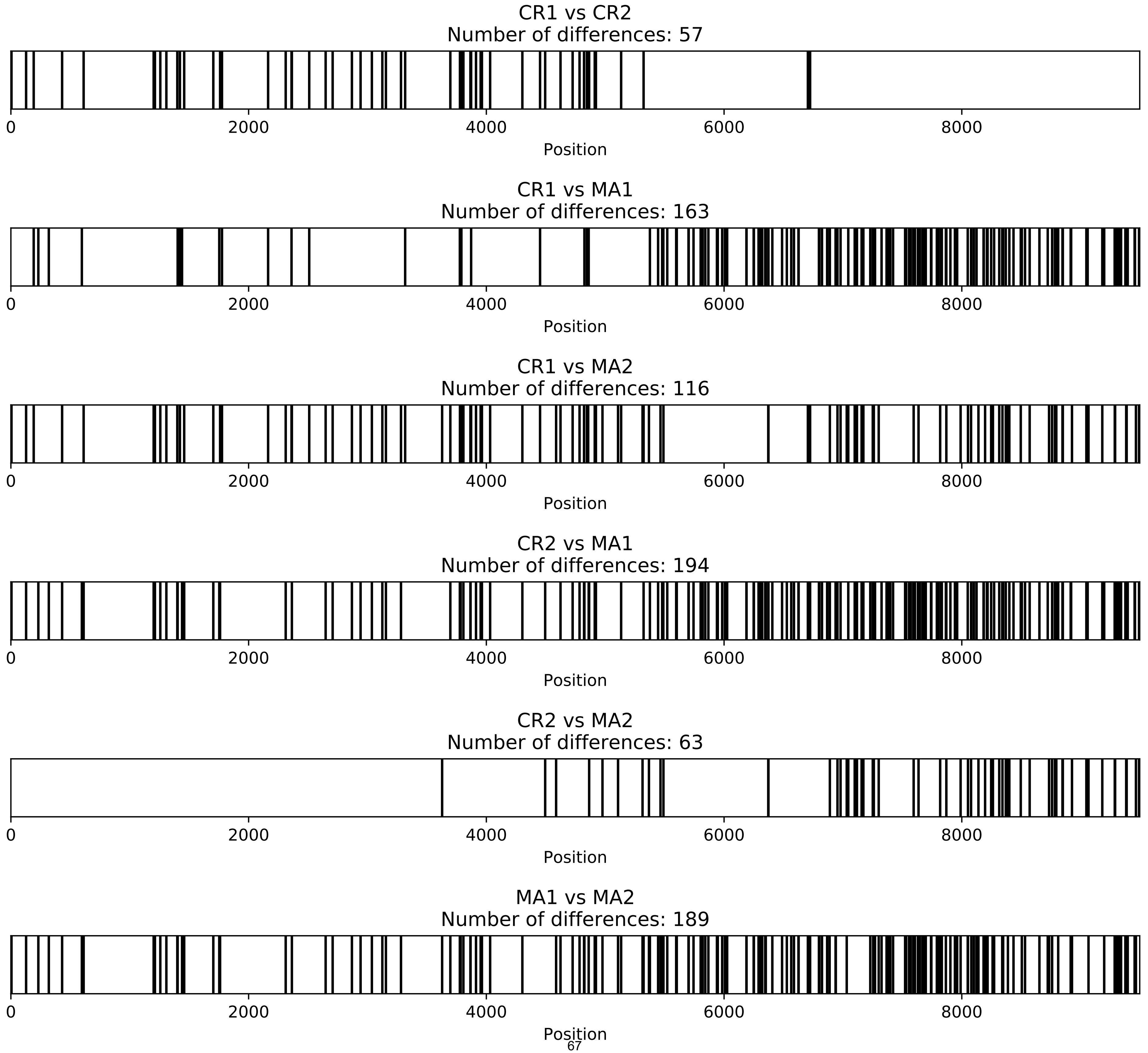
CR1 vs CR2
Number of differences: 57
0
2000
4000
6000
8000
Position
CR1 vs MA1
Number of differences: 163
0
2000
4000
6000
8000
Position
CR1 vs MA2
Number of differences: 116
0
2000
4000
6000
8000
Position
CR2 vs MA1
Number of differences: 194
0
2000
4000
6000
8000
Position
CR2 vs MA2
Number of differences: 63
0
2000
4000
6000
8000
Position
MA1 vs MA2
Number of differences: 189
0
2000
4000
6000
8000
Position

### Region: 374



CR1 vs CR2
Number of differences: 142

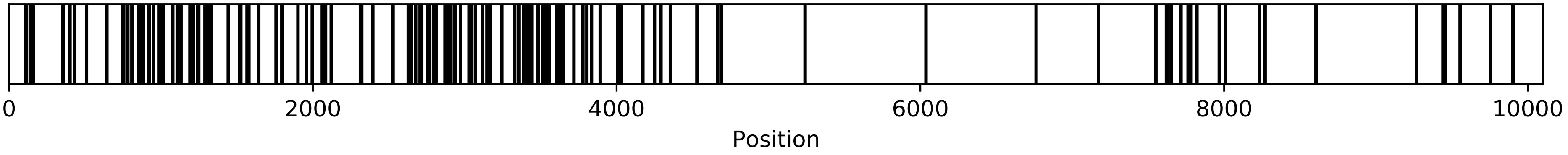


CR1 vs MA1
Number of differences: 107

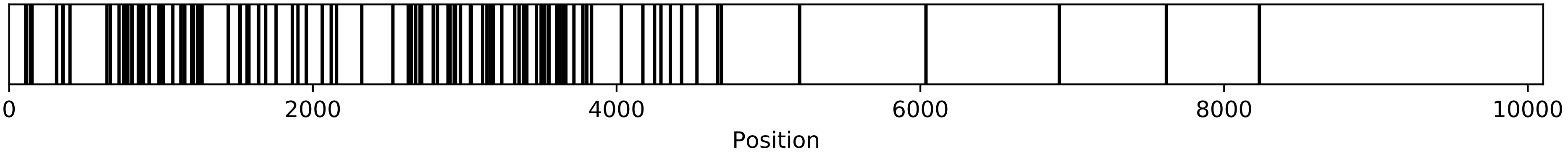


CR1 vs MA2
Number of differences: 62

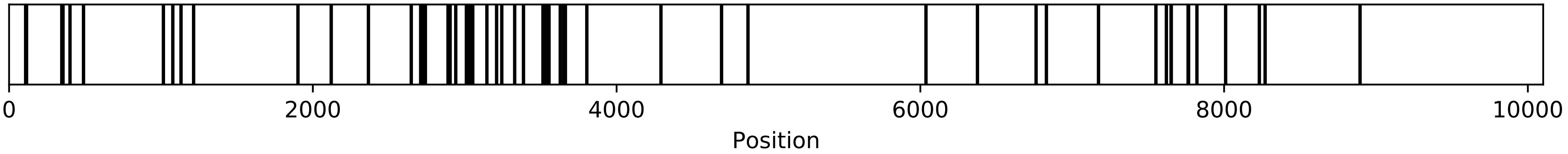


CR2 vs MA1
Number of differences: 84

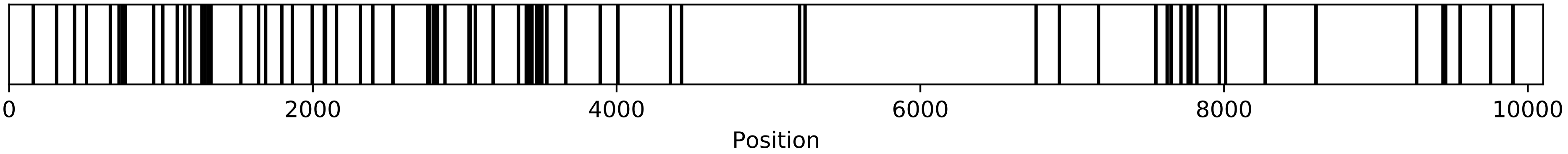


CR2 vs MA2
Number of differences: 114

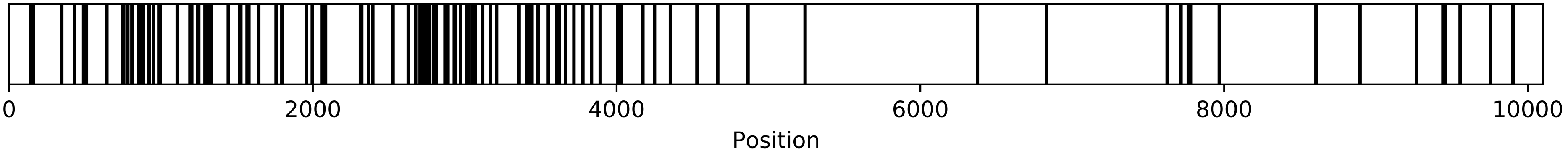


MA1 vs MA2
Number of differences: 104

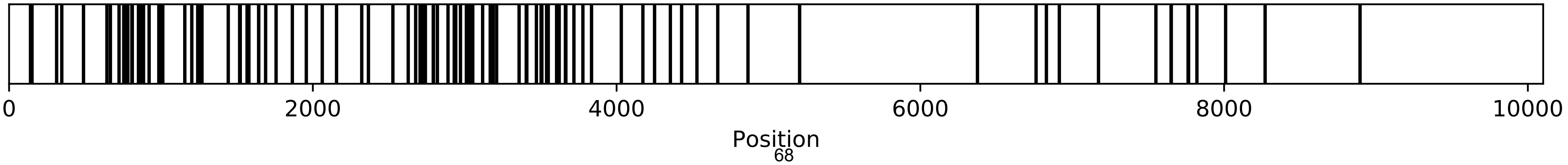

### Region: 377



CR1 vs CR2
Number of differences: 1

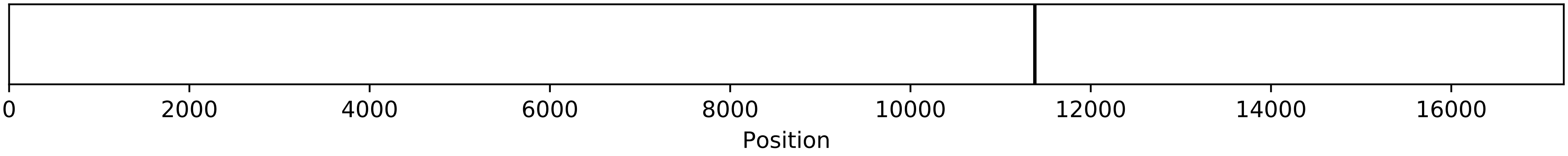


CR1 vs MA1
Number of differences: 353

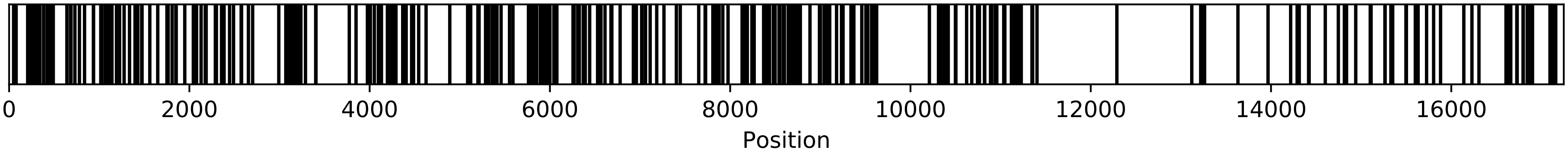


CR1 vs MA2
Number of differences: 331

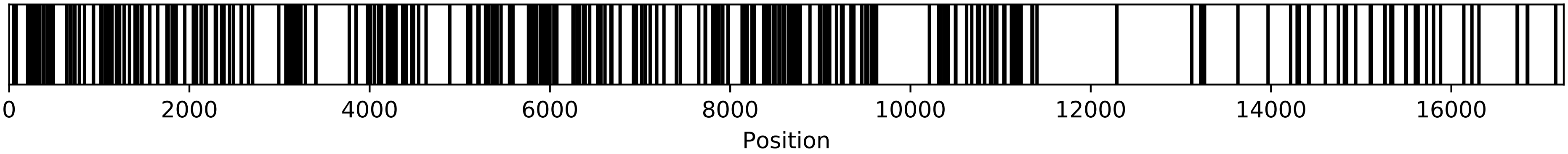


CR2 vs MA1
Number of differences: 354

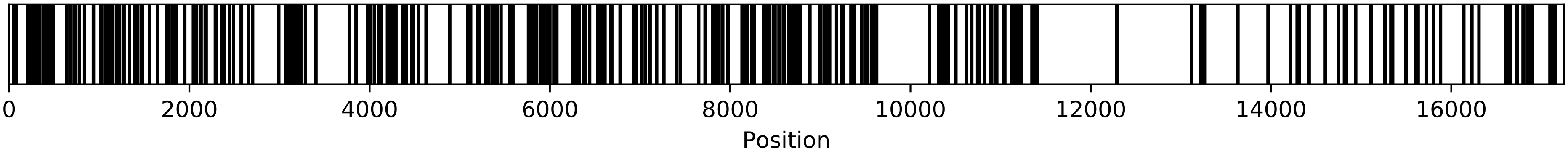


CR2 vs MA2
Number of differences: 332

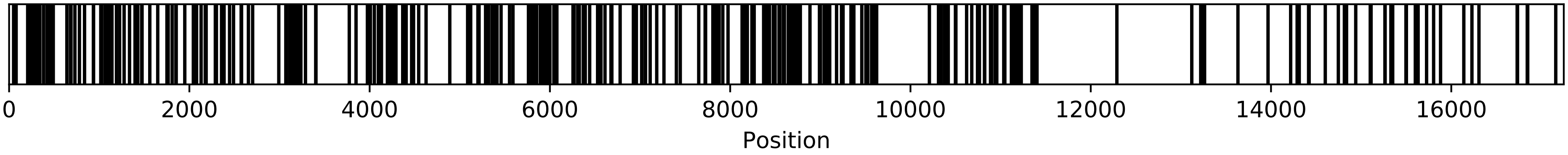


MA1 vs MA2
Number of differences: 26

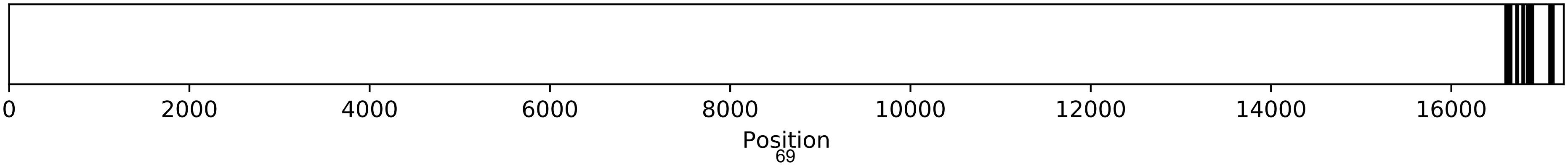

### Region: 389



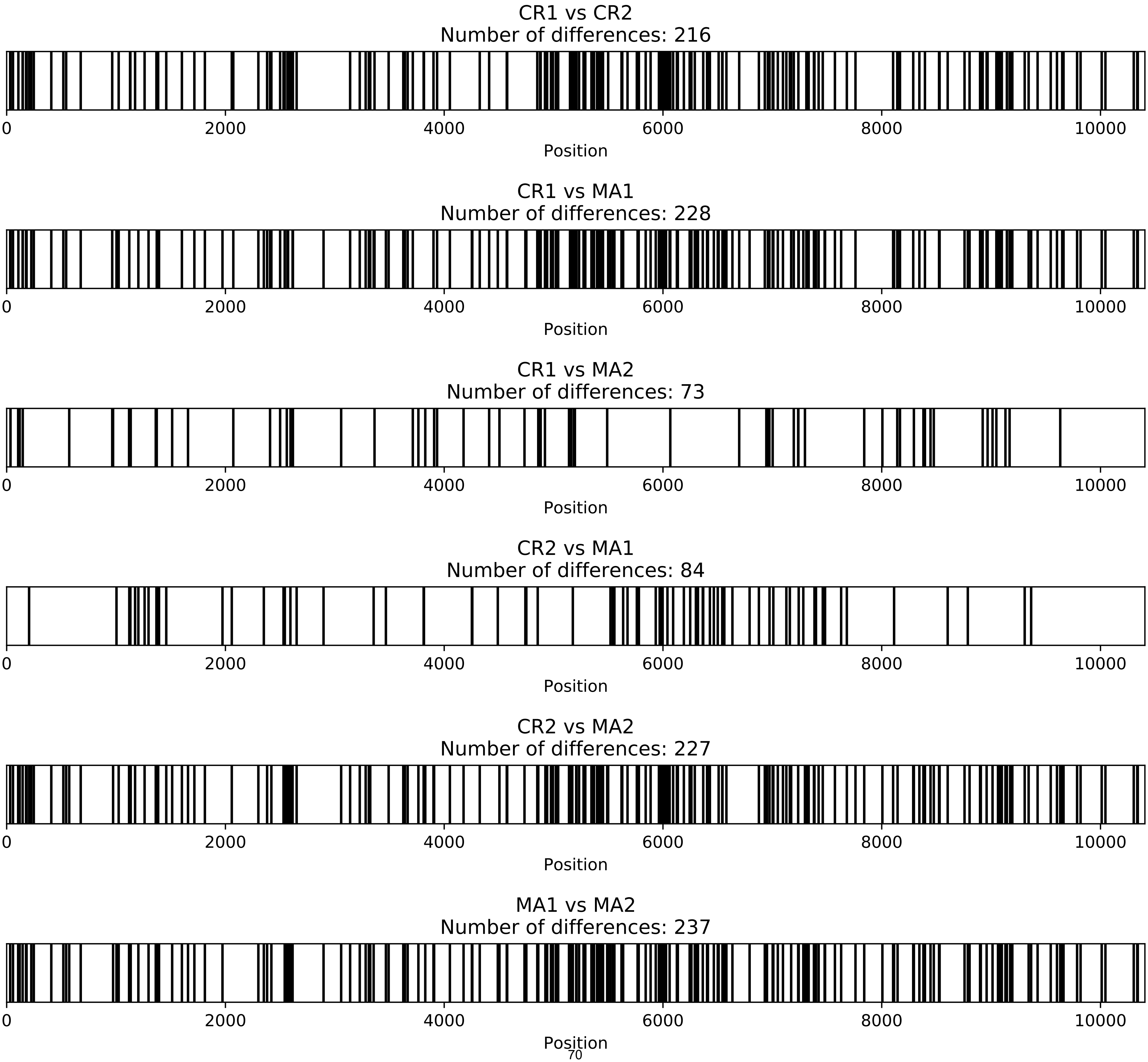
CR1 vs CR2
Number of differences: 216
0
2000
4000
6000
8000
10000
Position
CR1 vs MA1
Number of differences: 228
0
2000
4000
6000
8000
10000
Position
CR1 vs MA2
Number of differences: 73
0
2000
4000
6000
8000
10000
Position
CR2 vs MA1
Number of differences: 84
0
2000
4000
6000
8000
10000
Position
CR2 vs MA2
Number of differences: 227
0
2000
4000
6000
8000
10000
Position
MA1 vs MA2
Number of differences: 237
0
2000
4000
6000
8000
10000
Position

### Region: 410



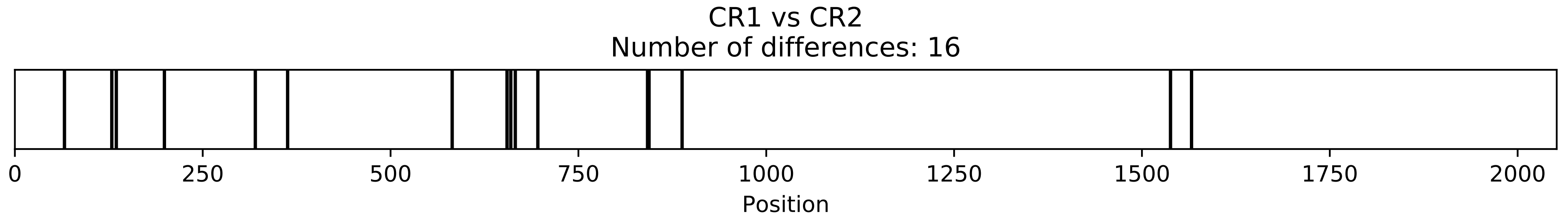
CR1 vs CR2
Number of differences: 16
0
250
500
750
1000
1250
1500
1750
2000
Position

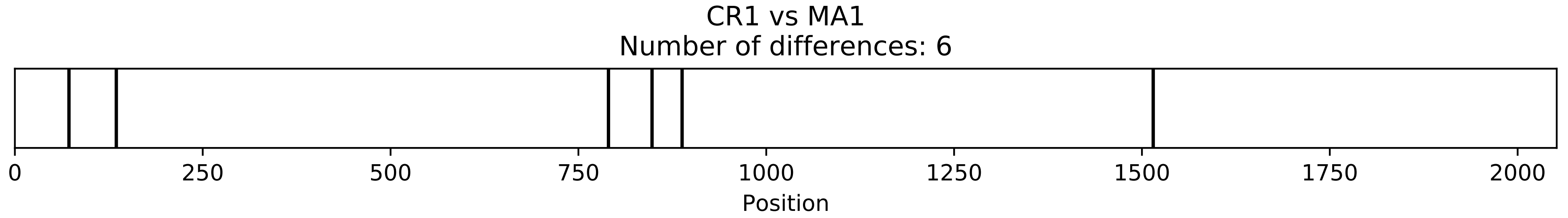
CR1 vs MA1
Number of differences: 6
0
250
500
750
1000
1250
1500
1750
2000
Position

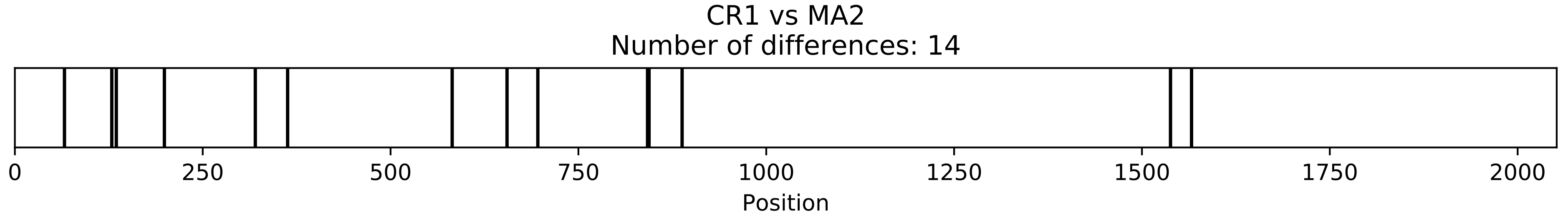
CR1 vs MA2
Number of differences: 14
0
250
500
750
1000
1250
1500
1750
2000
Position

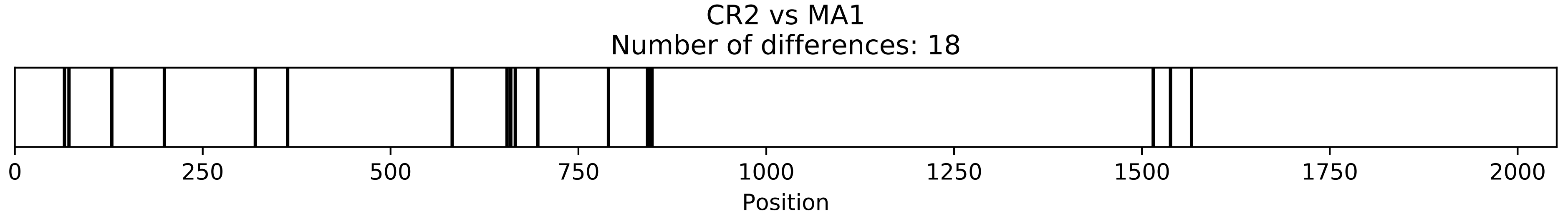
CR2 vs MA1
Number of differences: 18
0
250
500
750
1000
1250
1500
1750
2000
Position

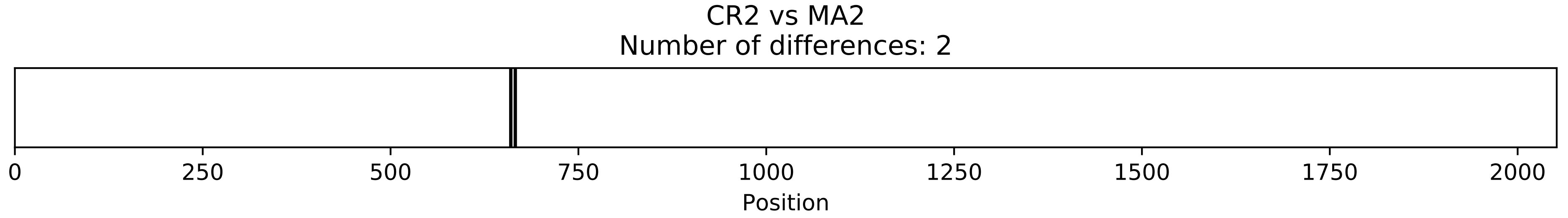
CR2 vs MA2
Number of differences: 2
0
250
500
750
1000
1250
1500
1750
2000
Position

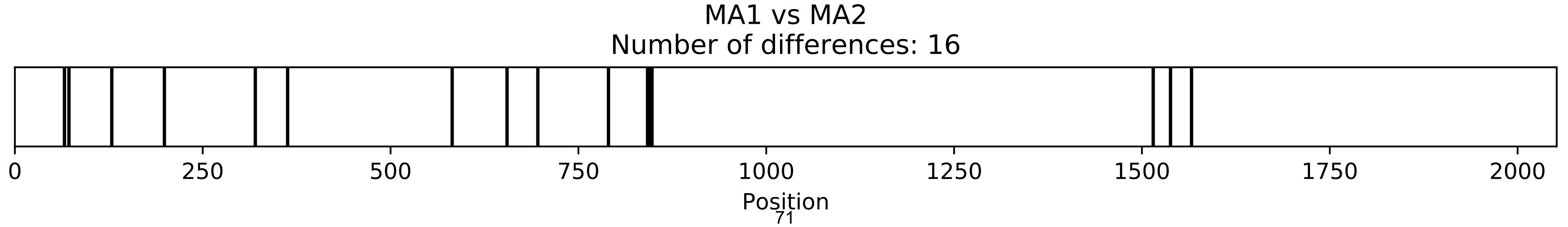
MA1 vs MA2
Number of differences: 16
0
250
500
750
1000
1250
1500
1750
2000
Position

### Region: 417



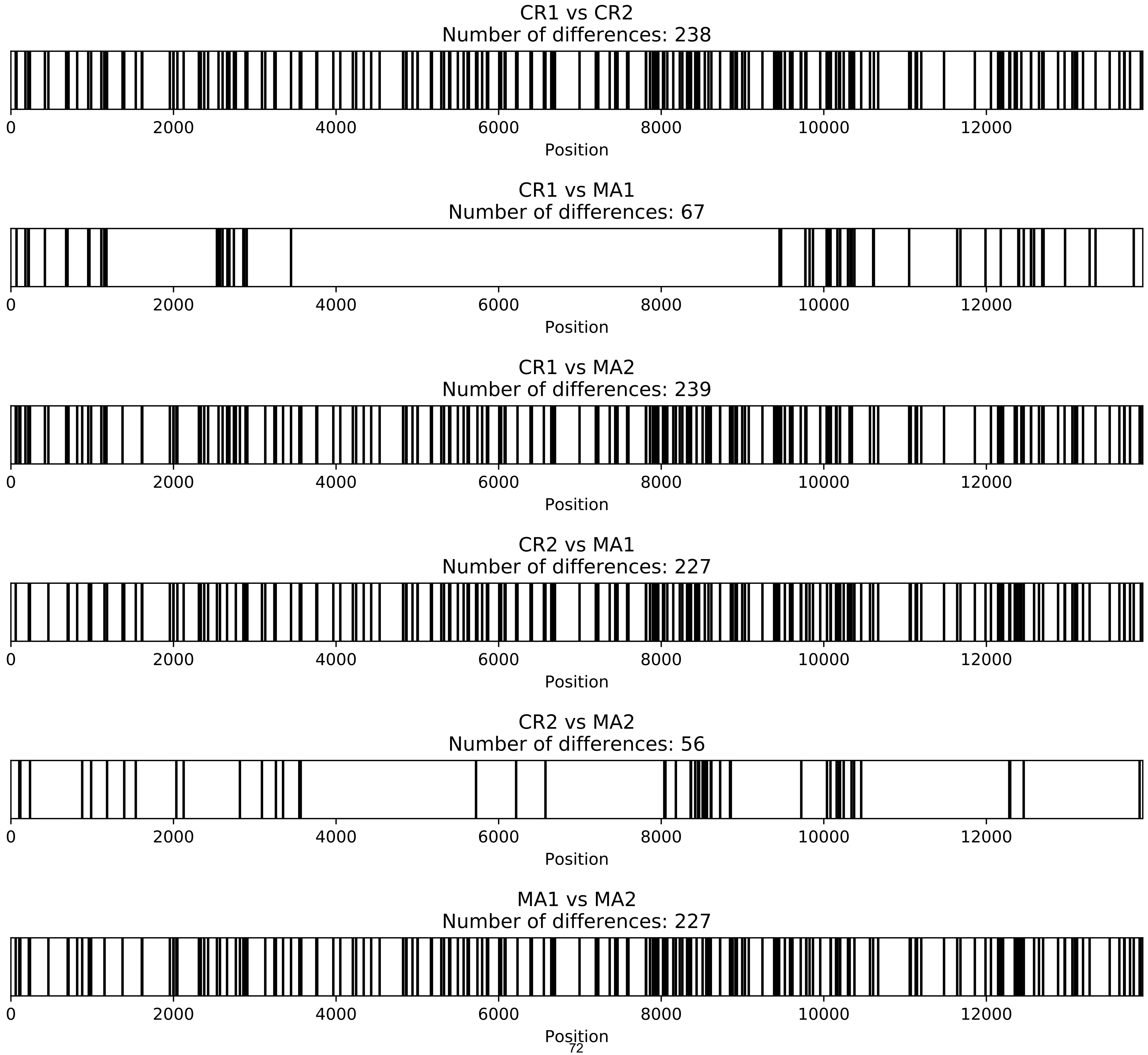
CR1 vs CR2
Number of differences: 238
CR1 vs MA1
Number of differences: 67
CR1 vs MA2
Number of differences: 239
CR2 vs MA1
Number of differences: 227
CR2 vs MA2
Number of differences: 56
MA1 vs MA2
Number of differences: 227
0
2000
4000
6000
8000
10000
12000
Position

### Region: 421



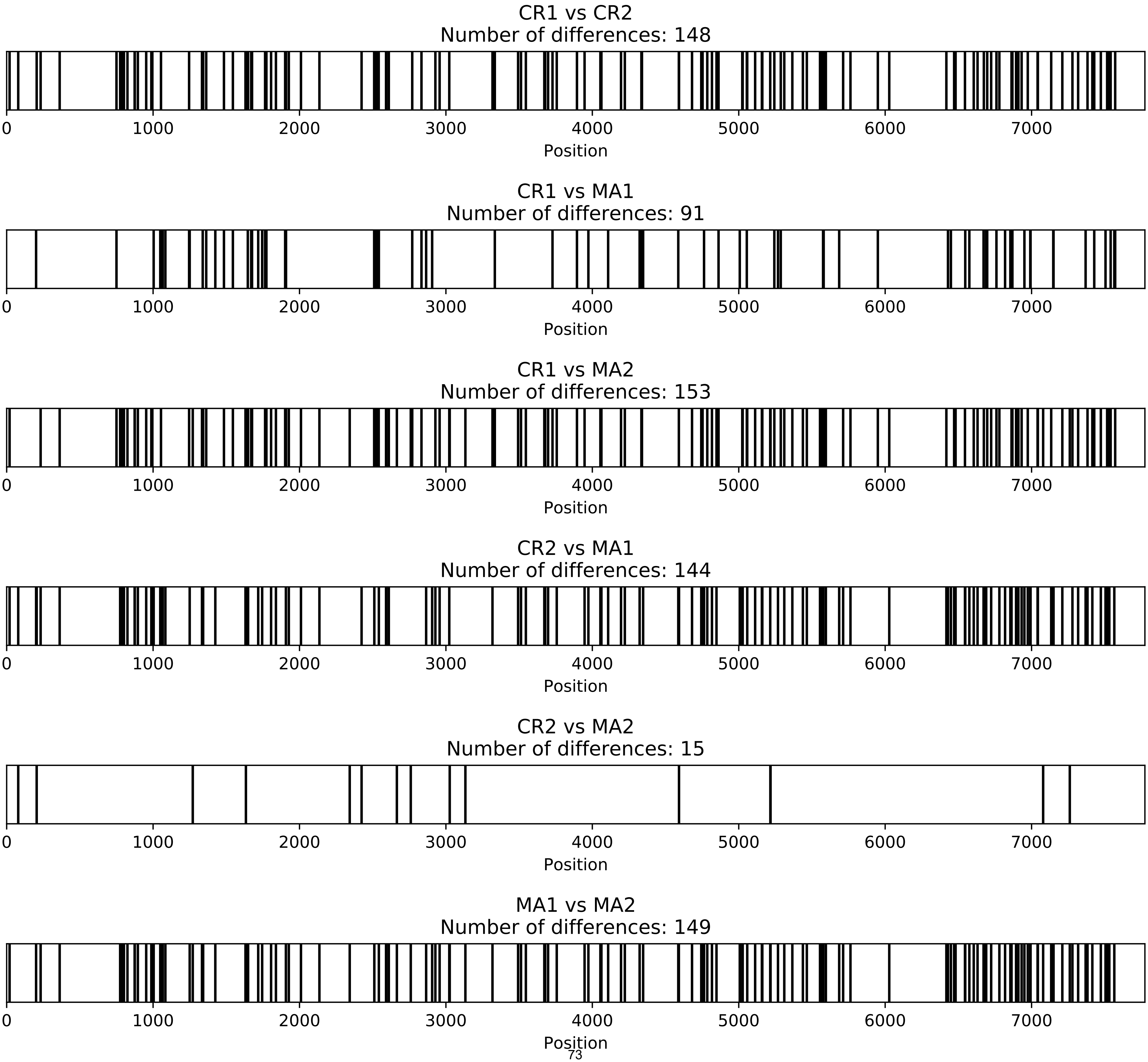
CR1 vs CR2
Number of differences: 148
CR1 vs MA1
Number of differences: 91
CR1 vs MA2
Number of differences: 153
CR2 vs MA1
Number of differences: 144
CR2 vs MA2
Number of differences: 15
MA1 vs MA2
Number of differences: 149
0
1000
2000
3000
4000
5000
6000
7000
Position

### Region: 422



CR1 vs CR2
Number of differences: 84

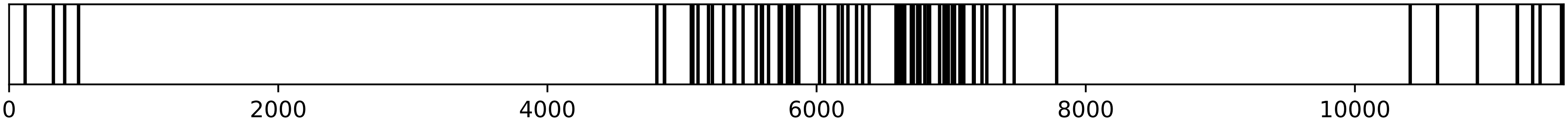


CR1 vs MA1
Number of differences: 145

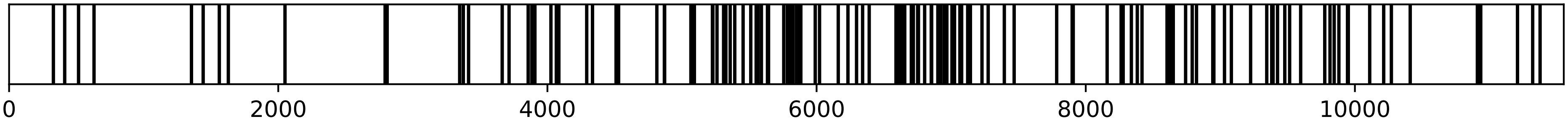


CR1 vs MA2
Number of differences: 73

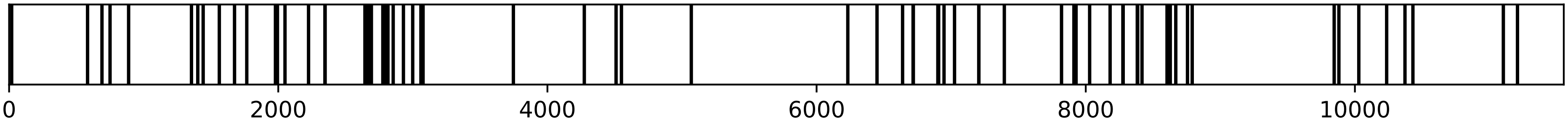


CR2 vs MA1
Number of differences: 125

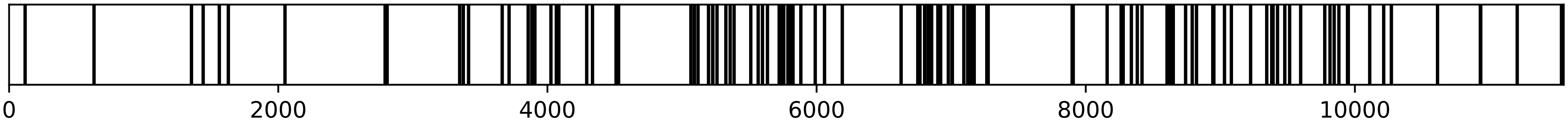


CR2 vs MA2
Number of differences: 141

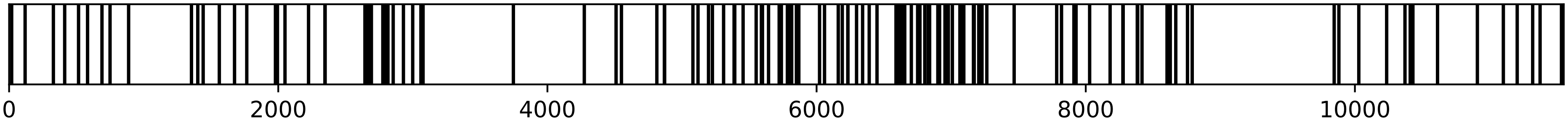


MA1 vs MA2
Number of differences: 172

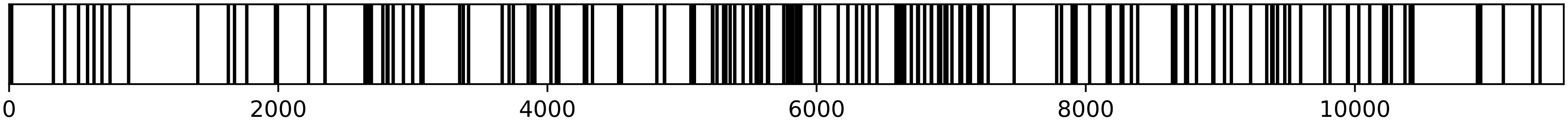

### Region: 442



CR1 vs CR2
Number of differences: 226

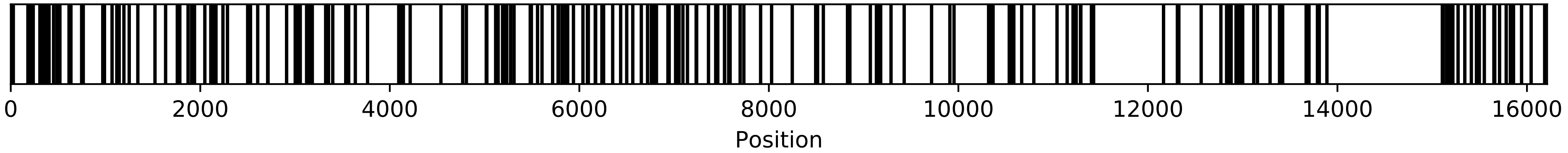


CR1 vs MA1
Number of differences: 114

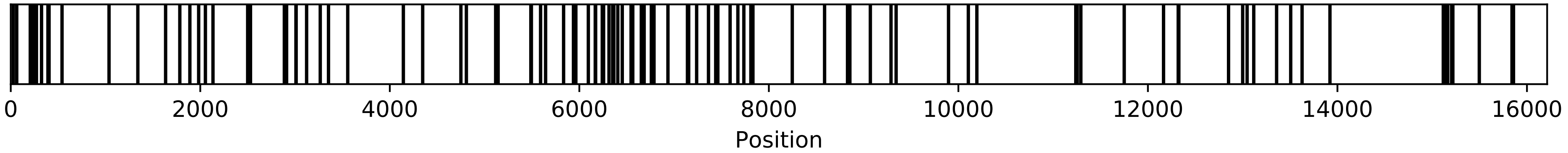


CR1 vs MA2
Number of differences: 241

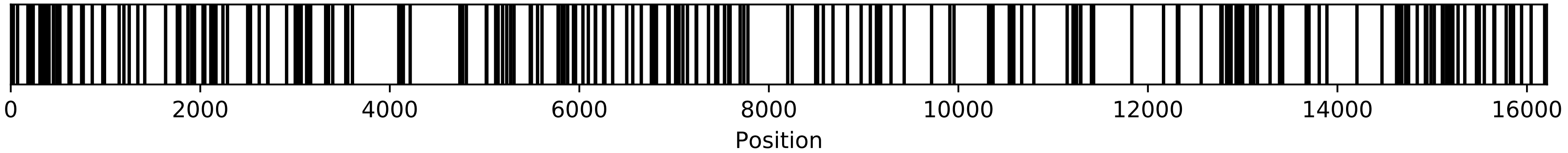


CR2 vs MA1
Number of differences: 218

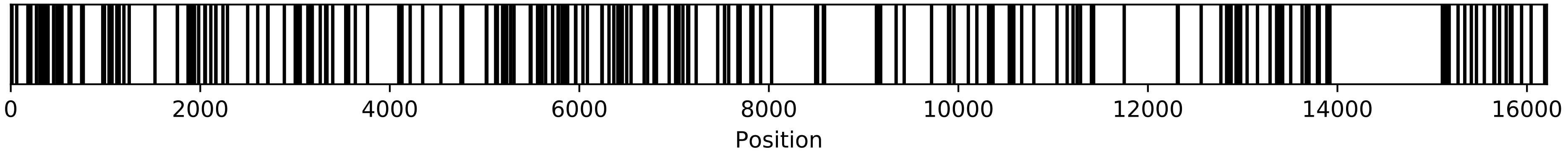


CR2 vs MA2
Number of differences: 70

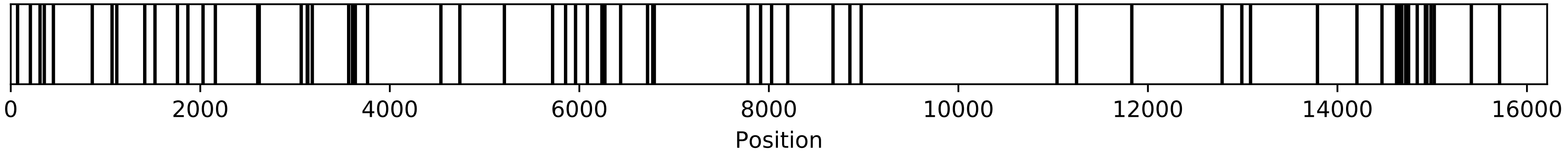


MA1 vs MA2
Number of differences: 234

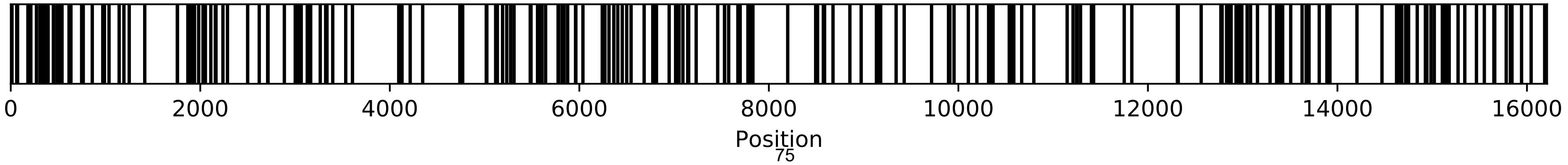

### Region: 450



CR1 vs CR2
Number of differences: 556

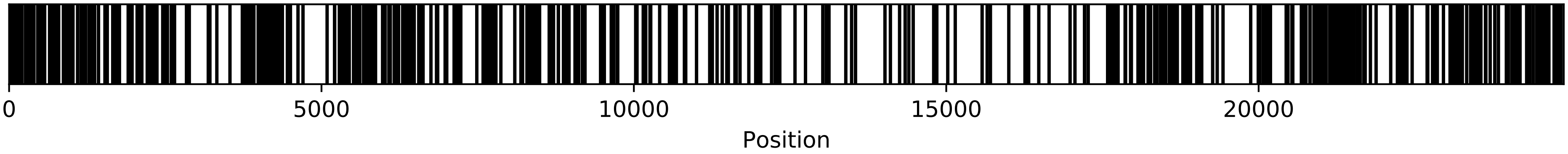


CR1 vs MA1
Number of differences: 295

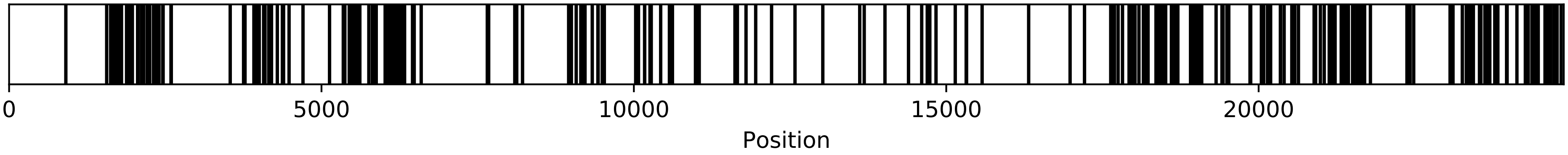


CR1 vs MA2
Number of differences: 556

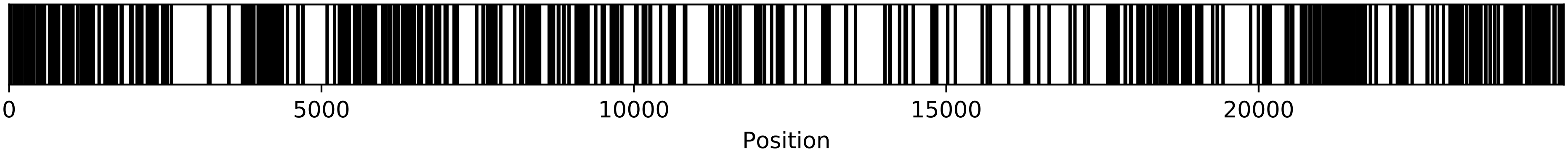


CR2 vs MA1
Number of differences: 587

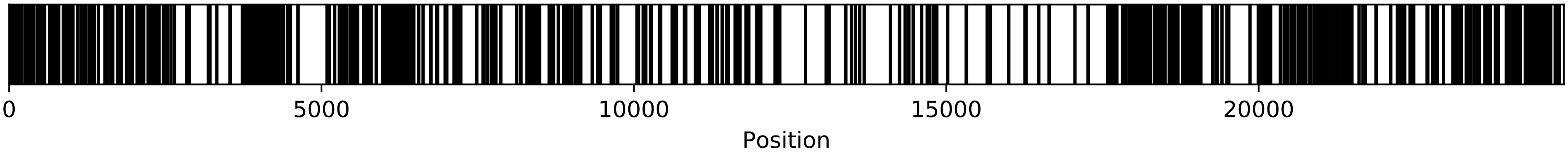


CR2 vs MA2
Number of differences: 141

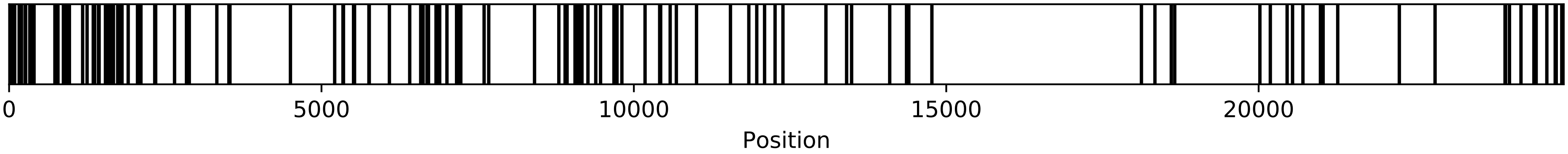


MA1 vs MA2
Number of differences: 587

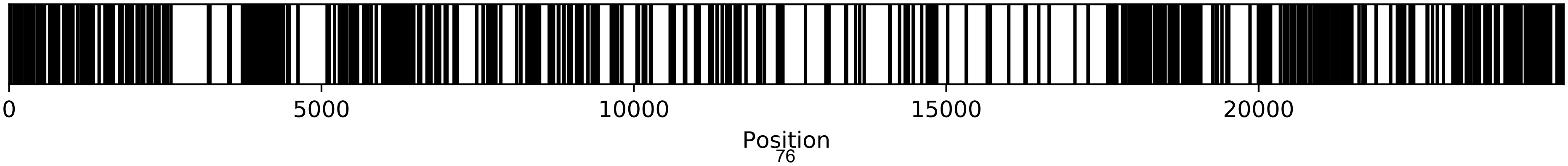

### Region: 471

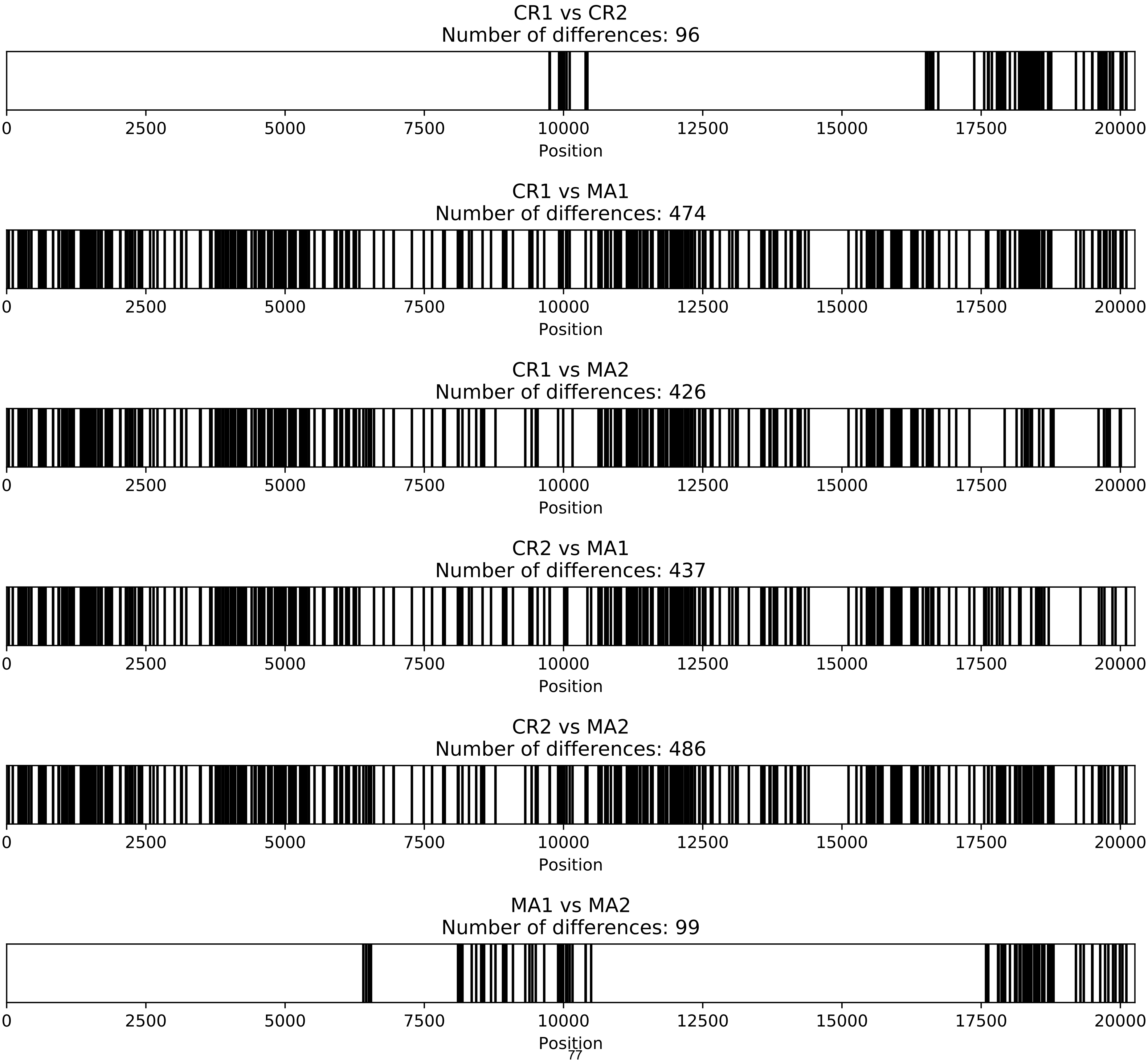
CR1 vs CR2
Number of differences: 96
CR1 vs MA1
Number of differences: 474
CR1 vs MA2
Number of differences: 426
CR2 vs MA1
Number of differences: 437
CR2 vs MA2
Number of differences: 486
MA1 vs MA2
Number of differences: 99
0
2500
5000
7500
10000
12500
15000
17500
20000
Position

### Region: 492



CR1 vs CR2
Number of differences: 301

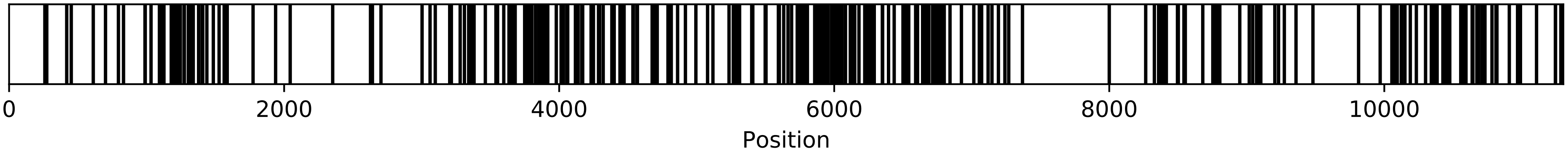


CR1 vs MA1
Number of differences: 138

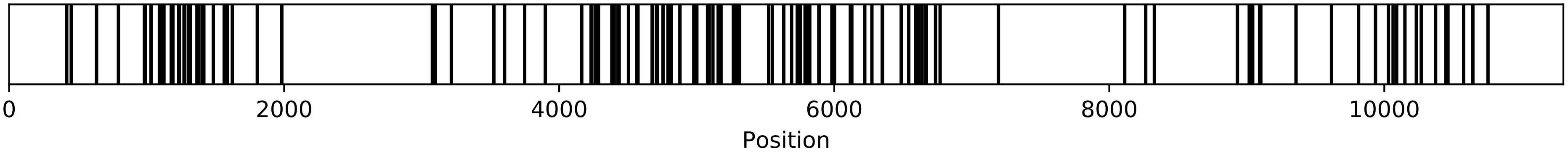


CR1 vs MA2
Number of differences: 300

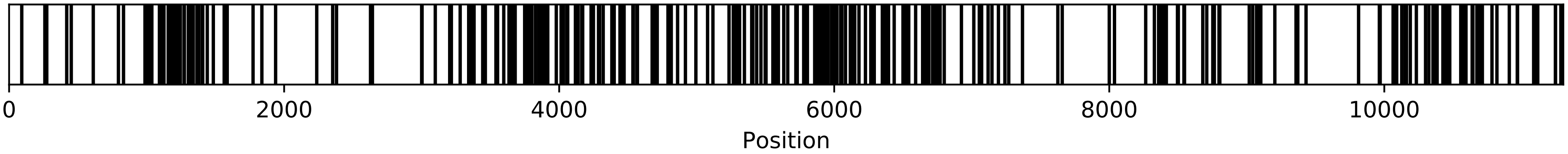


CR2 vs MA1
Number of differences: 279

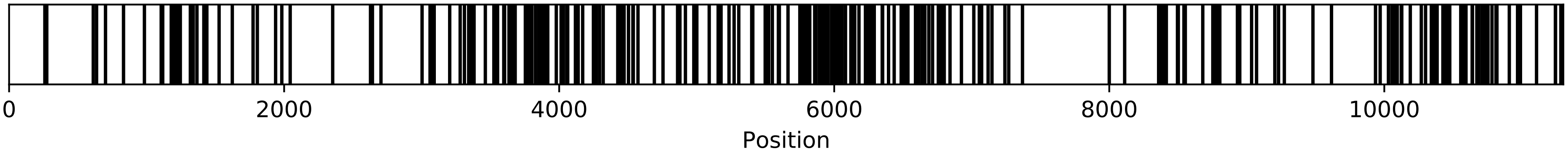


CR2 vs MA2
Number of differences: 86

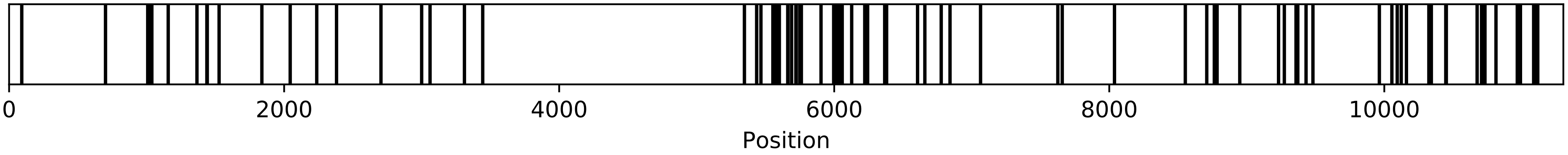


MA1 vs MA2
Number of differences: 283

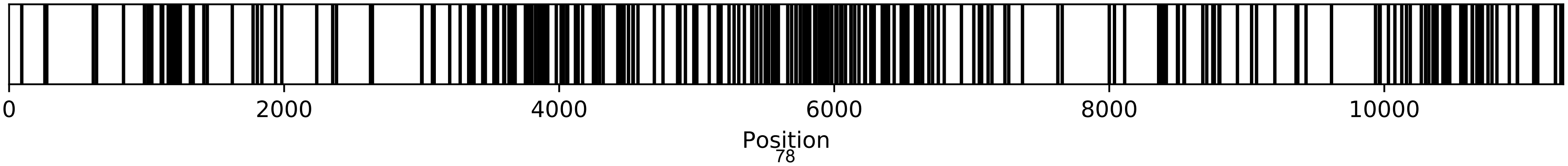

### Region: 494



## CR1 vs CR2
Number of differences: 136

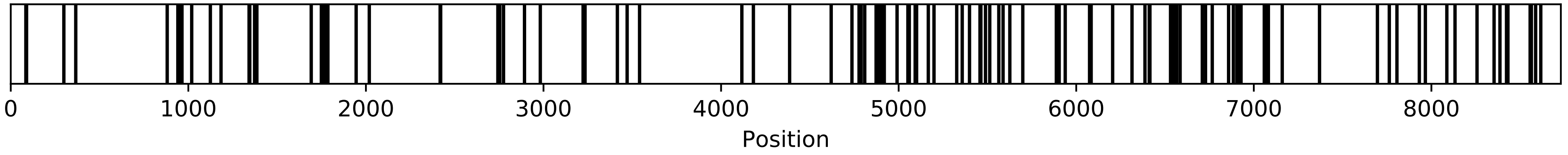


## CR1 vs MA1
Number of differences: 45

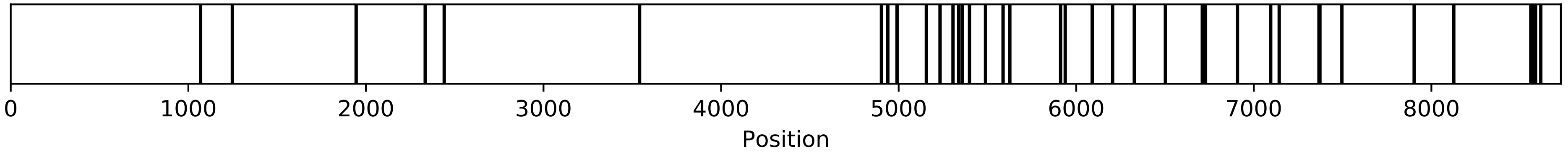


## CR1 vs MA2
Number of differences: 83

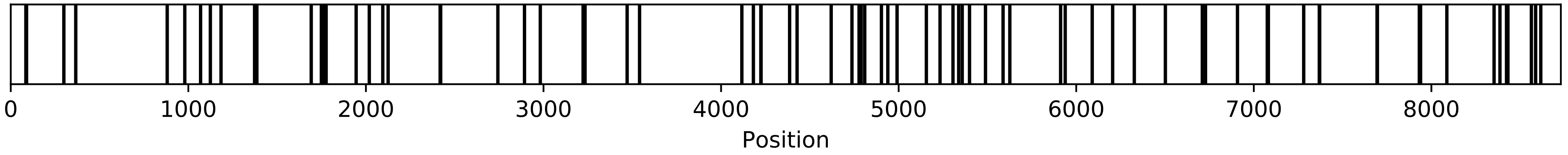


## CR2 vs MA1
Number of differences: 145

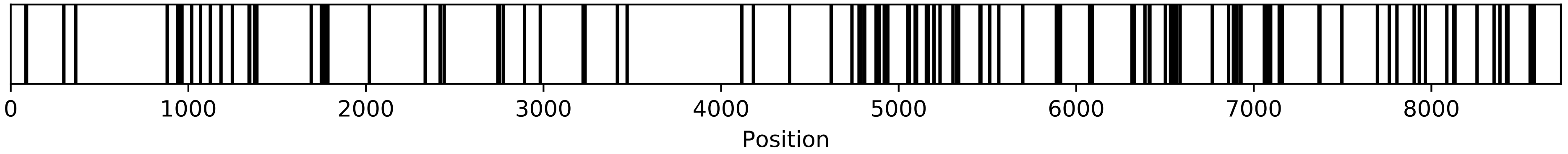


## CR2 vs MA2
Number of differences: 97

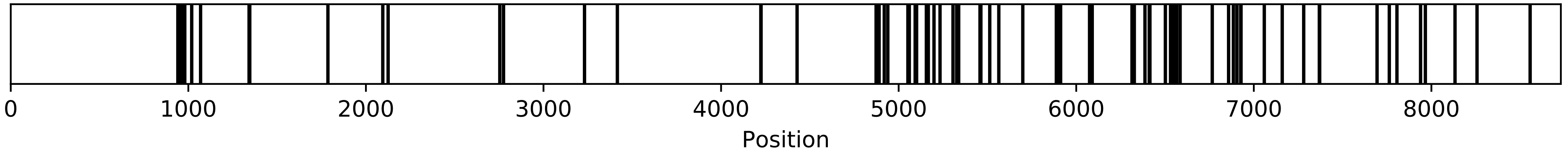


## MA1 vs MA2
Number of differences: 70

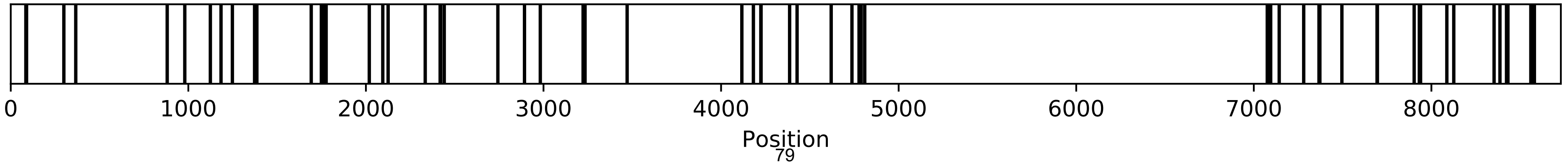

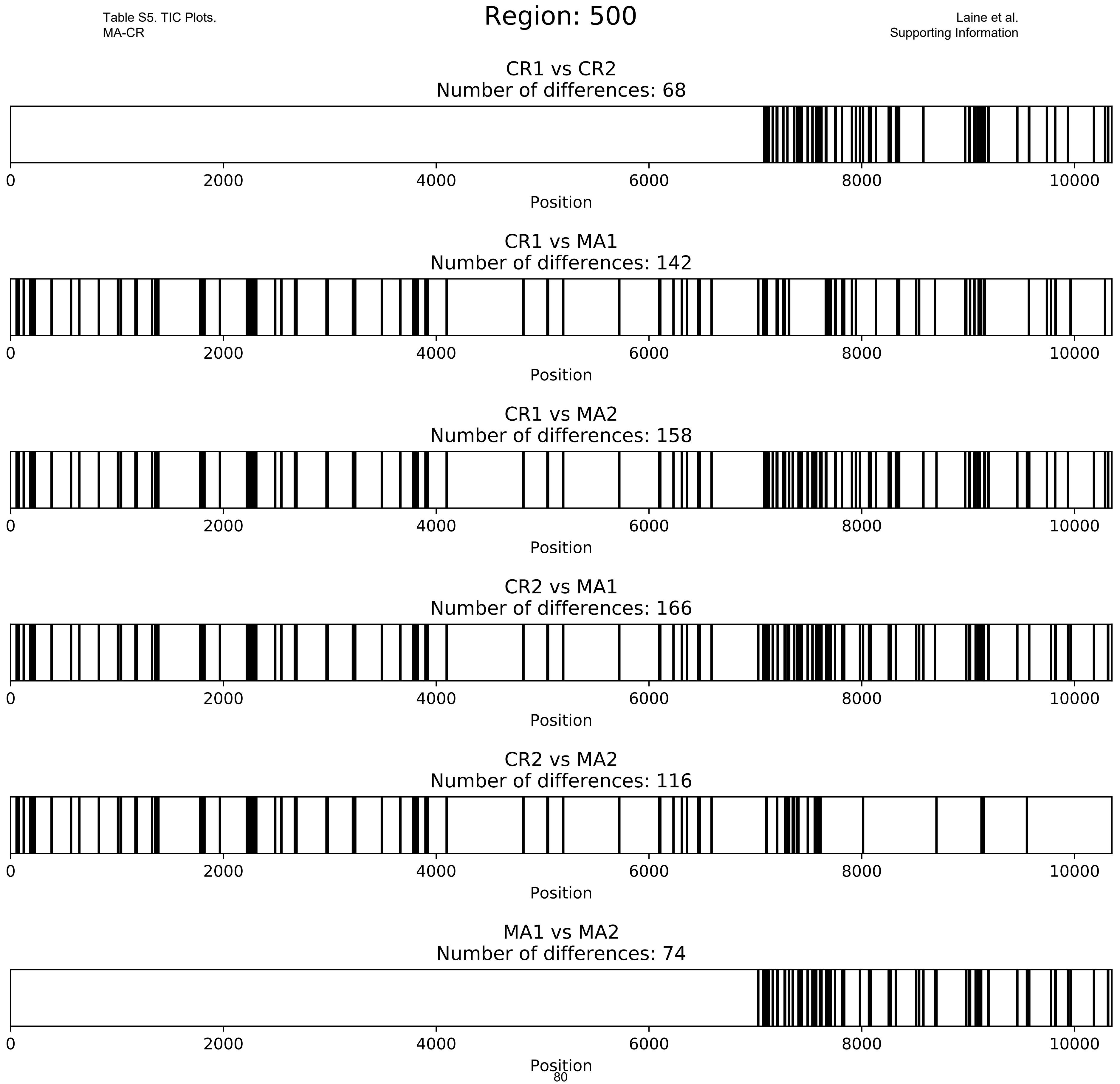
Region: 500
CR1 vs CR2
Number of differences: 68
CR1 vs MA1
Number of differences: 142
CR1 vs MA2
Number of differences: 158
CR2 vs MA1
Number of differences: 166
CR2 vs MA2
Number of differences: 116
MA1 vs MA2
Number of differences: 74
0
2000
4000
6000
8000
10000
Position

### Region: 502



## CR1 vs CR2
Number of differences: 166

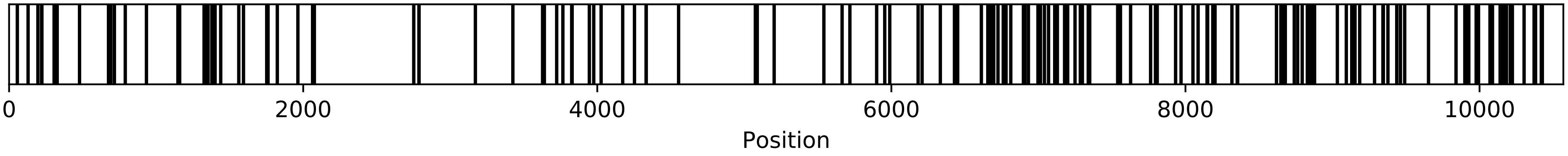


## CR1 vs MA1
Number of differences: 77

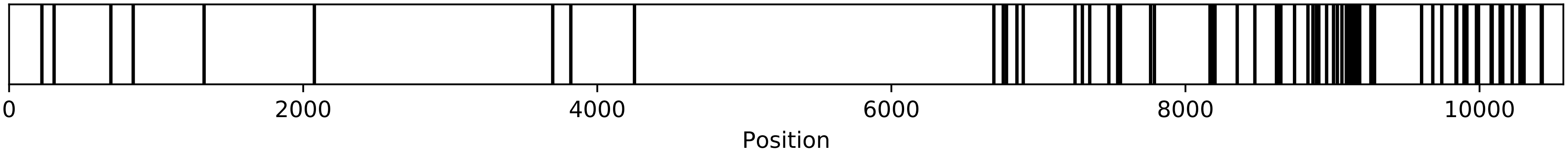


## CR1 vs MA2
Number of differences: 166

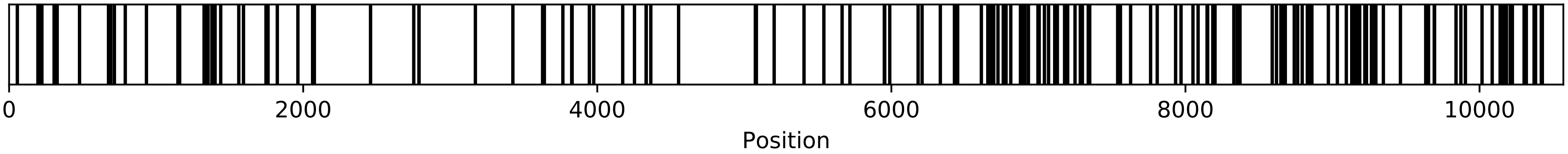


## CR2 vs MA1
Number of differences: 180

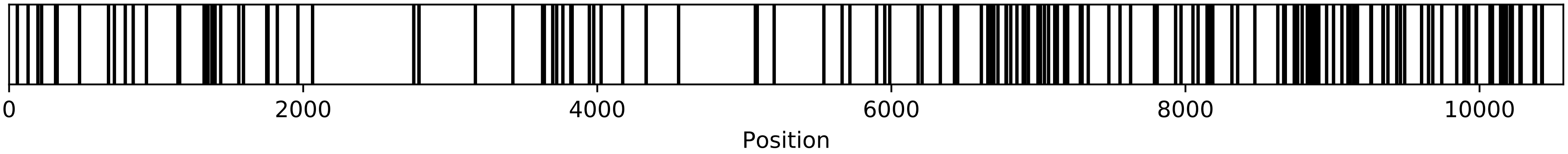


## CR2 vs MA2
Number of differences: 48

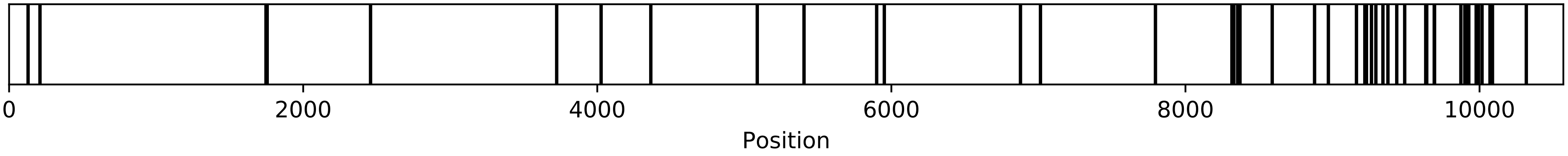


## MA1 vs MA2
Number of differences: 182

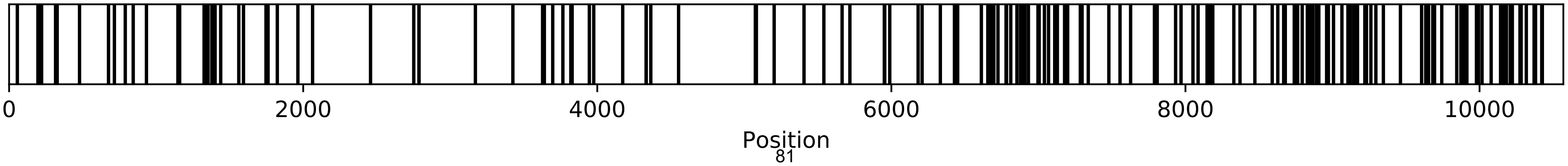

### Region: 506



CR1 vs CR2
Number of differences: 167

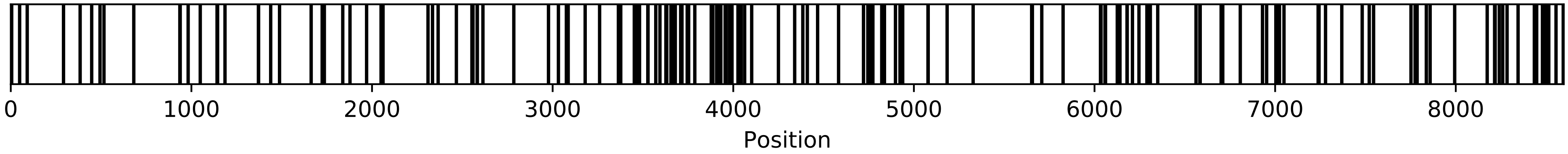


CR1 vs MA1
Number of differences: 148

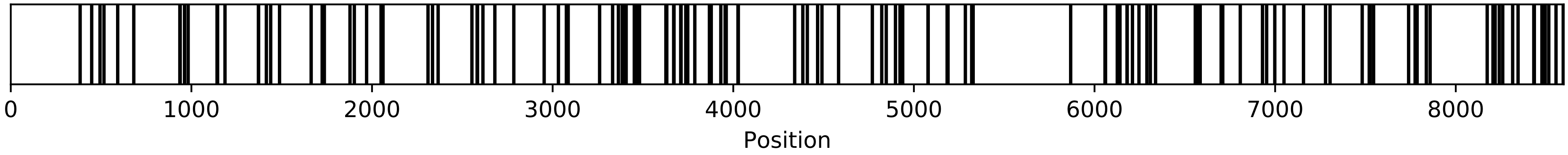


CR1 vs MA2
Number of differences: 48

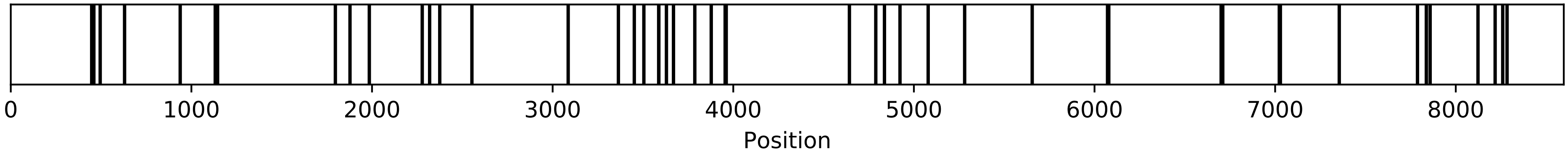


CR2 vs MA1
Number of differences: 103

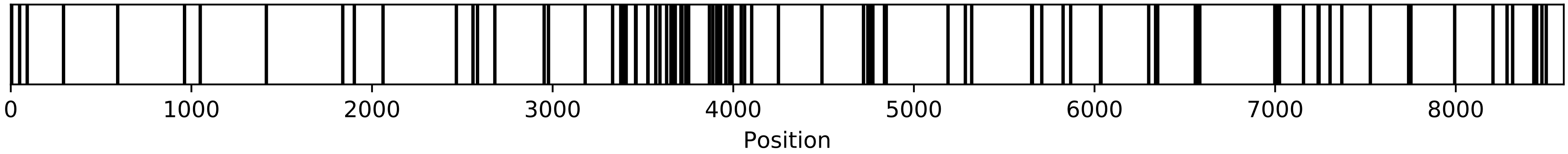


CR2 vs MA2
Number of differences: 168

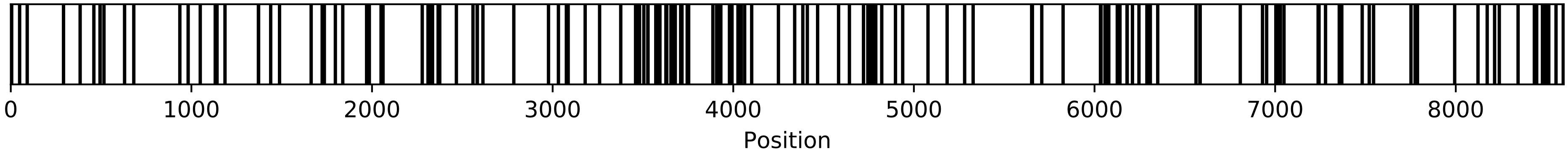


MA1 vs MA2
Number of differences: 155

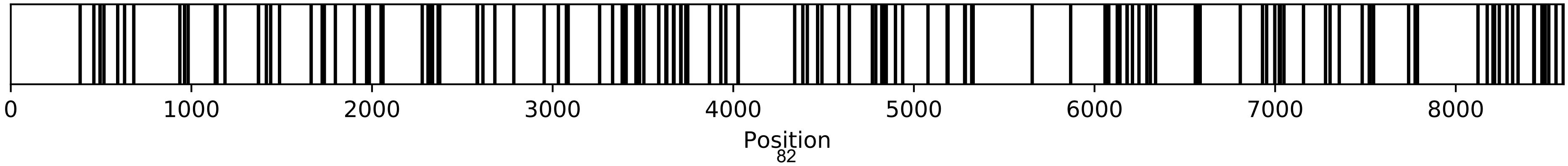

### Region: 511



## CR1 vs CR2
Number of differences: 317

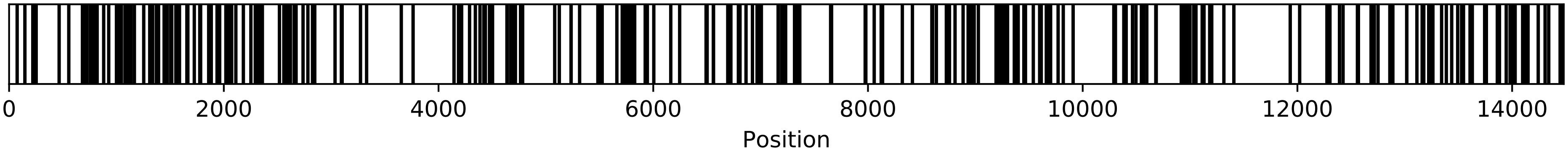


## CR1 vs MA1
Number of differences: 114

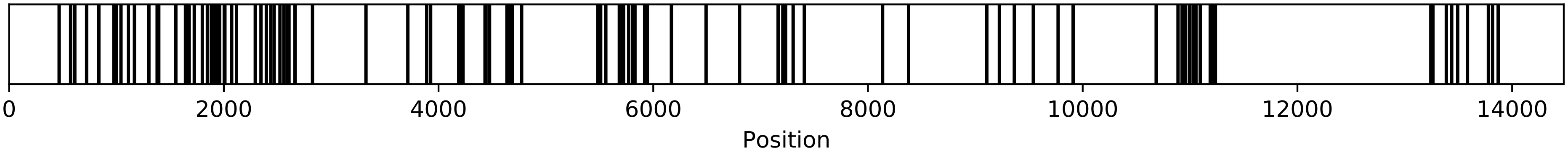


## CR1 vs MA2
Number of differences: 306

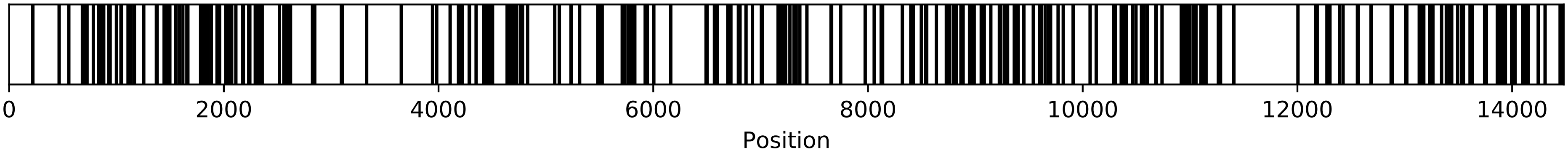


## CR2 vs MA1
Number of differences: 314

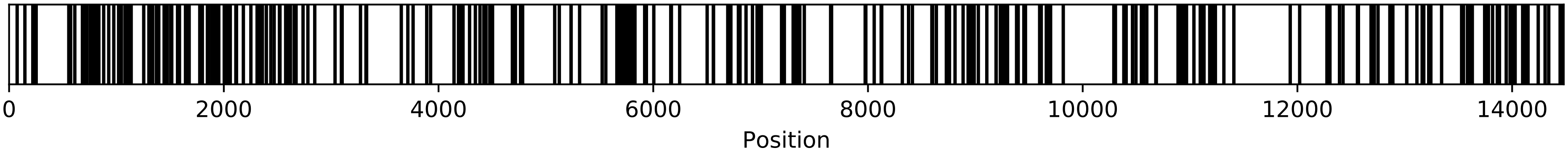


## CR2 vs MA2
Number of differences: 180

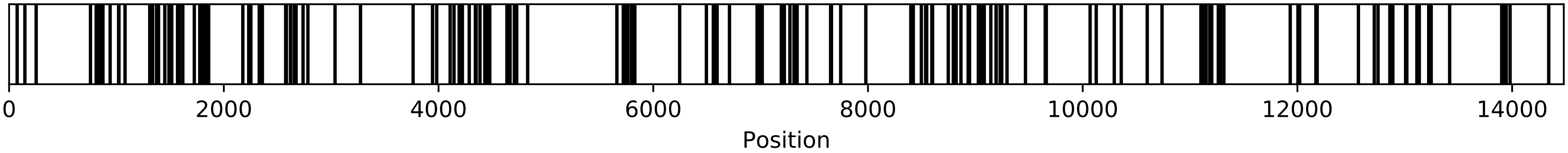


## MA1 vs MA2
Number of differences: 306

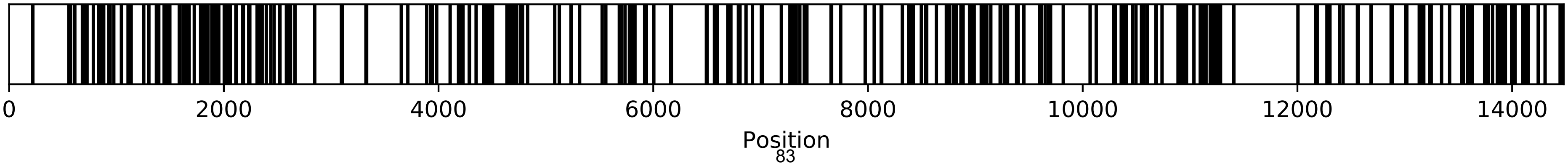

### Region: 515



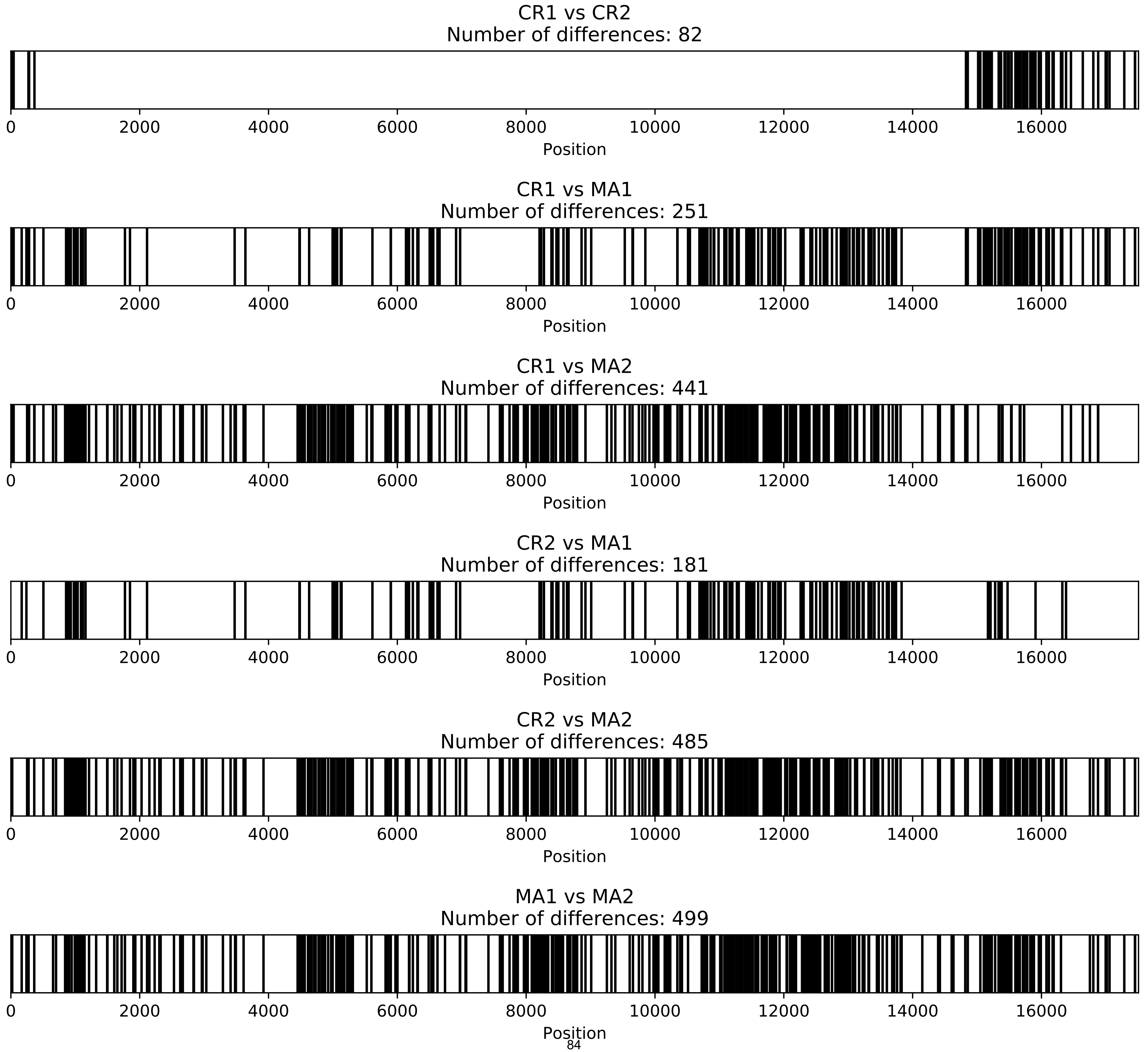
CR1 vs CR2
Number of differences: 82
CR1 vs MA1
Number of differences: 251
CR1 vs MA2
Number of differences: 441
CR2 vs MA1
Number of differences: 181
CR2 vs MA2
Number of differences: 485
MA1 vs MA2
Number of differences: 499
0
2000
4000
6000
8000
10000
12000
14000
16000
Position

### Region: 521



## CR1 vs CR2
Number of differences: 133

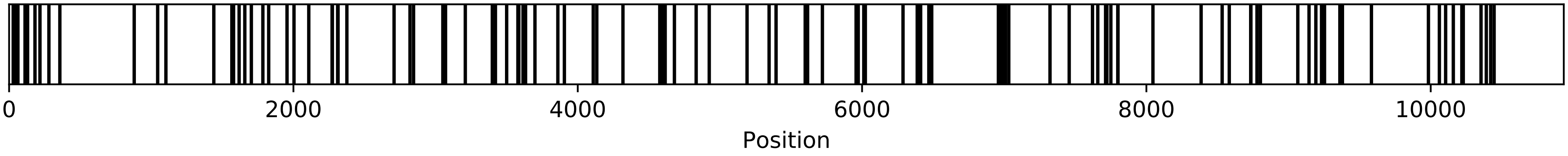


## CR1 vs MA1
Number of differences: 118

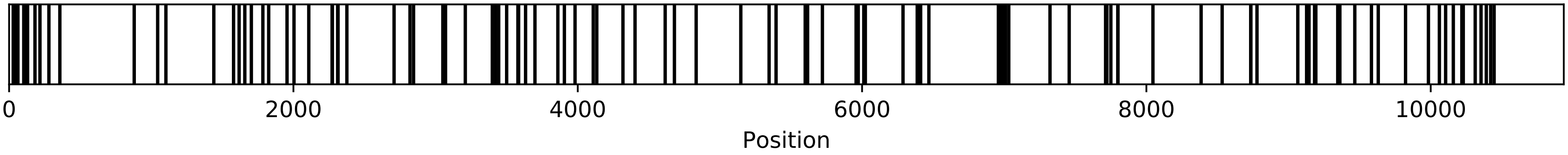


## CR1 vs MA2
Number of differences: 41

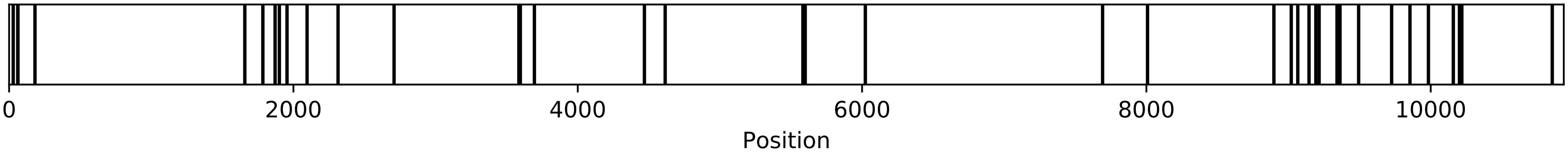


## CR2 vs MA1
Number of differences: 50

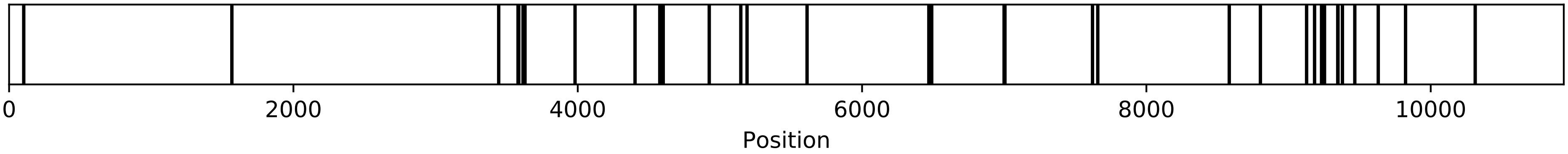


## CR2 vs MA2
Number of differences: 140

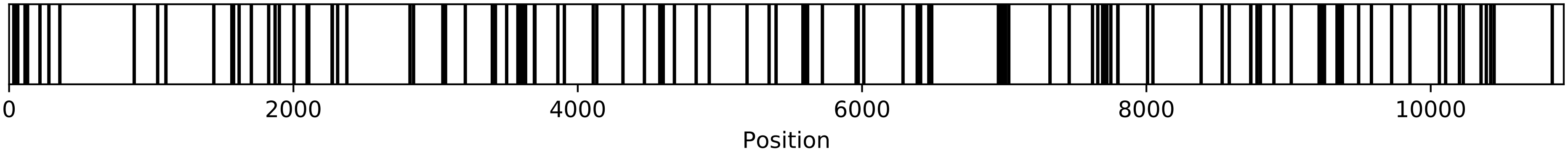


## MA1 vs MA2
Number of differences: 125

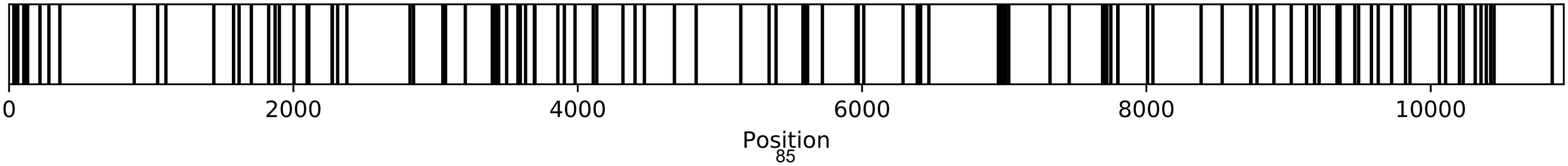

### Region: 527



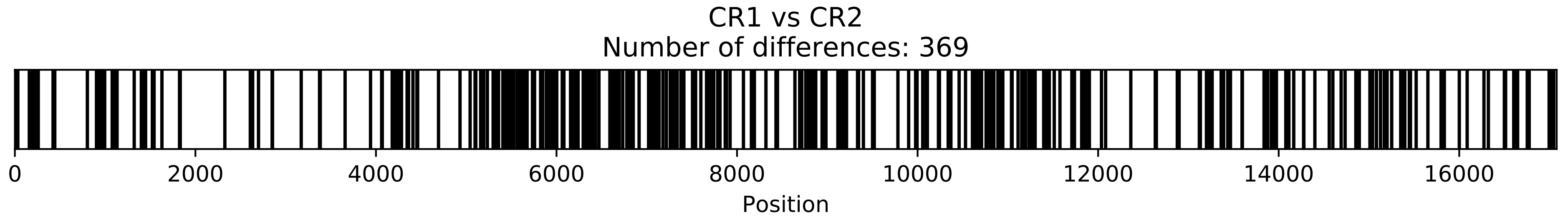
CR1 vs CR2
Number of differences: 369
0
2000
4000
6000
8000
10000
12000
14000
16000
Position

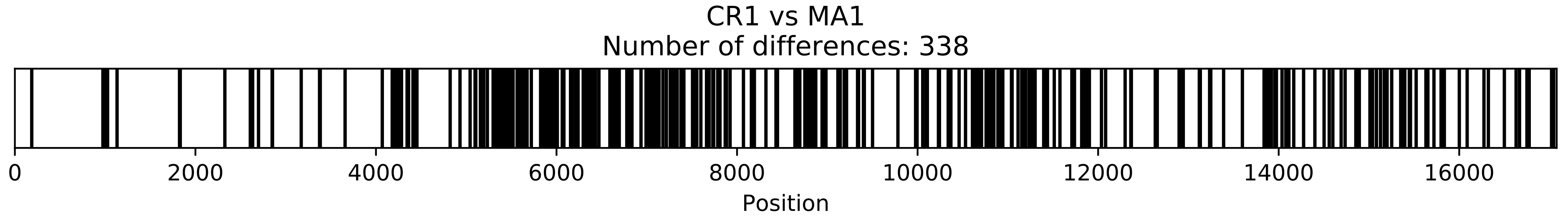
CR1 vs MA1
Number of differences: 338
0
2000
4000
6000
8000
10000
12000
14000
16000
Position

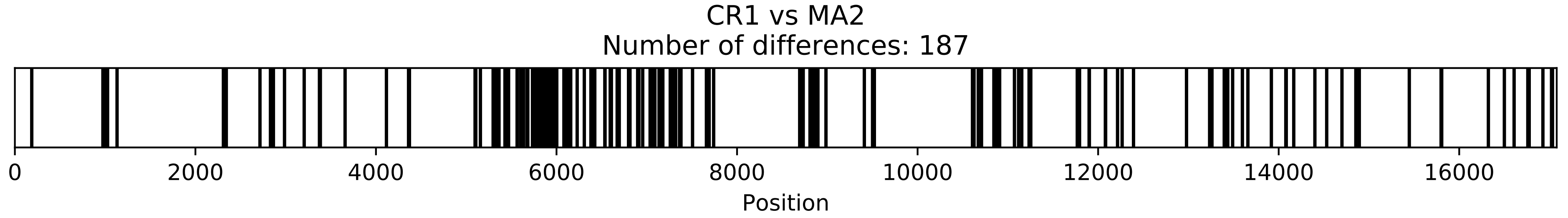
CR1 vs MA2
Number of differences: 187
0
2000
4000
6000
8000
10000
12000
14000
16000
Position

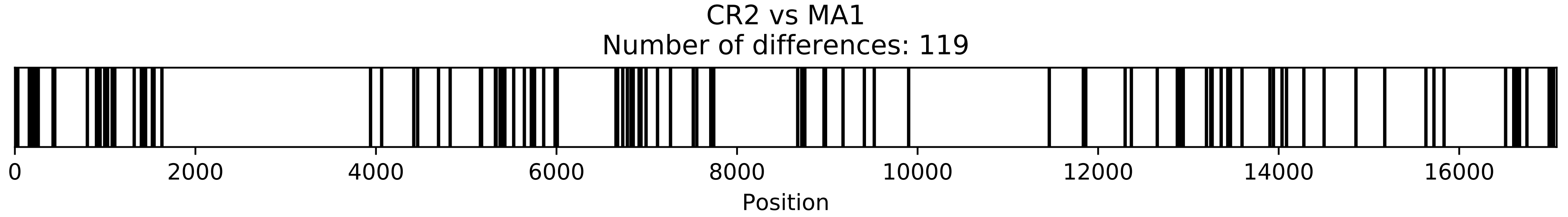
CR2 vs MA1
Number of differences: 119
0
2000
4000
6000
8000
10000
12000
14000
16000
Position

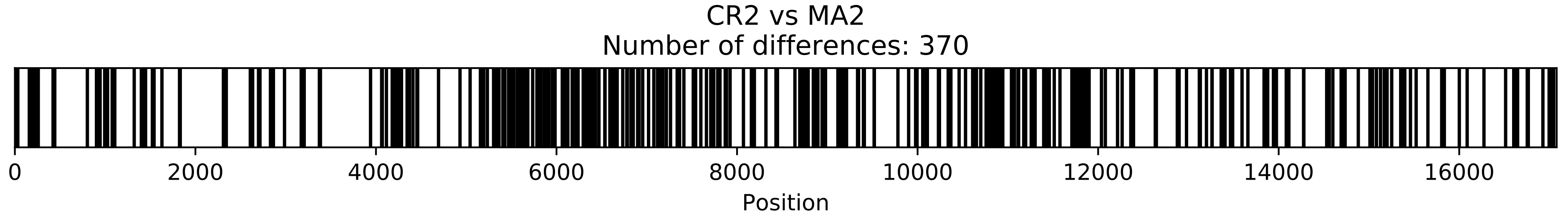
CR2 vs MA2
Number of differences: 370
0
2000
4000
6000
8000
10000
12000
14000
16000
Position

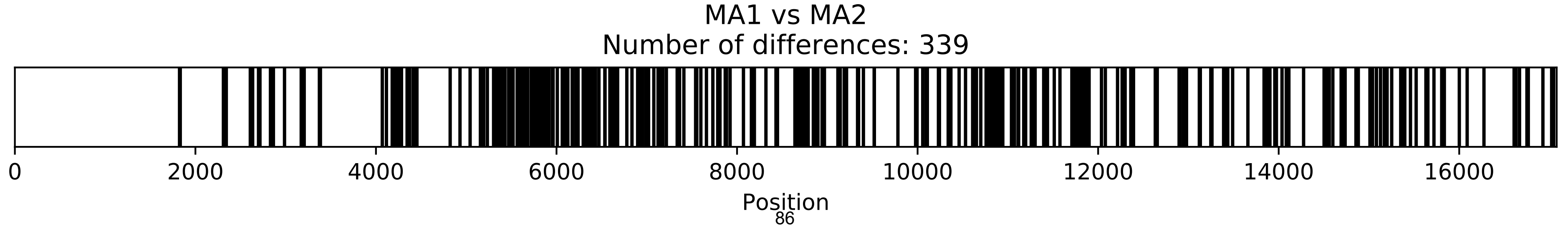
MA1 vs MA2
Number of differences: 339
0
2000
4000
6000
8000
10000
12000
14000
16000
Position

### Region: 539



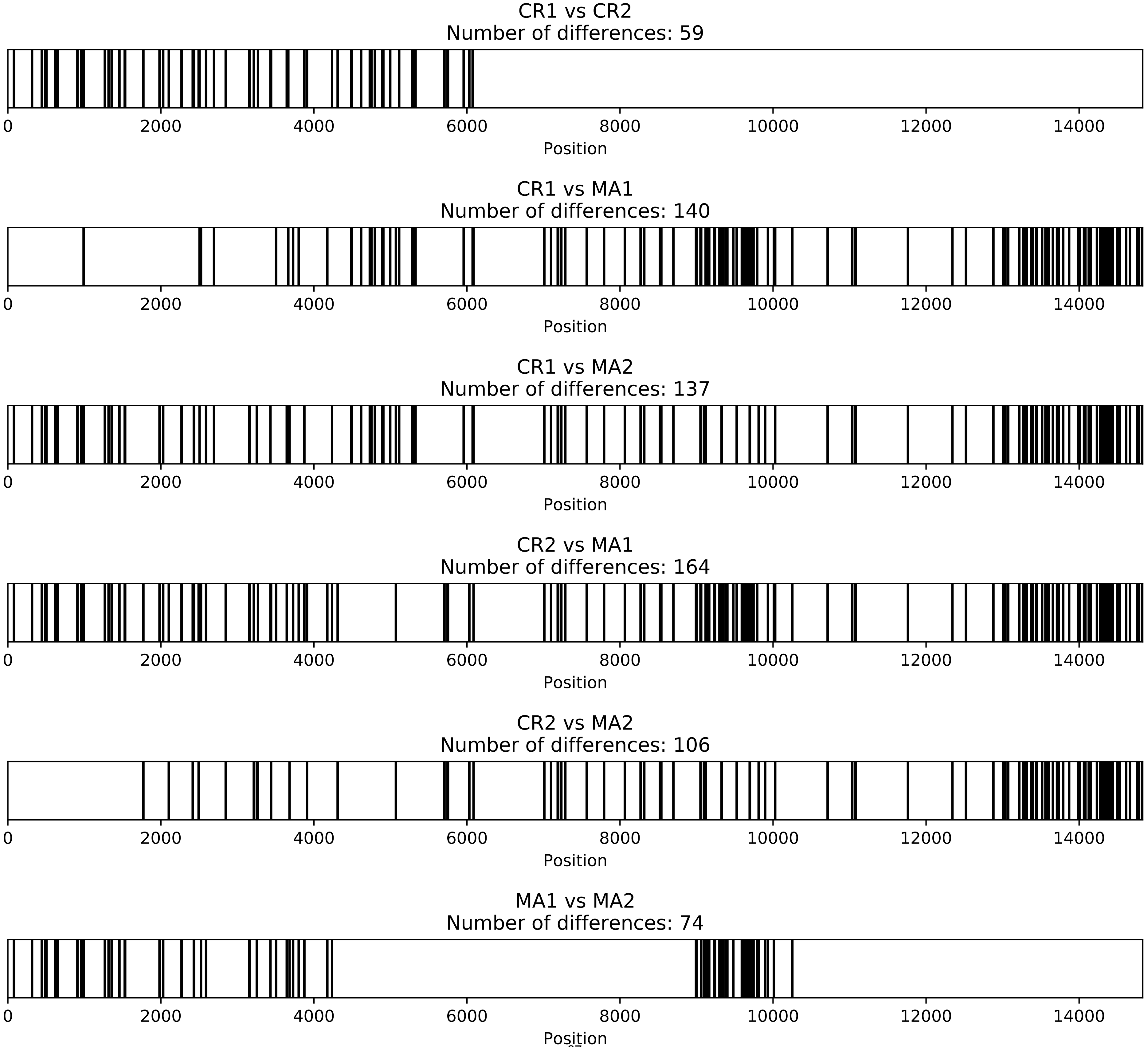
CR1 vs CR2
Number of differences: 59
CR1 vs MA1
Number of differences: 140
CR1 vs MA2
Number of differences: 137
CR2 vs MA1
Number of differences: 164
CR2 vs MA2
Number of differences: 106
MA1 vs MA2
Number of differences: 74
0
2000
4000
6000
8000
10000
12000
14000
Position

### Region: 562



CR1 vs CR2
Number of differences: 213

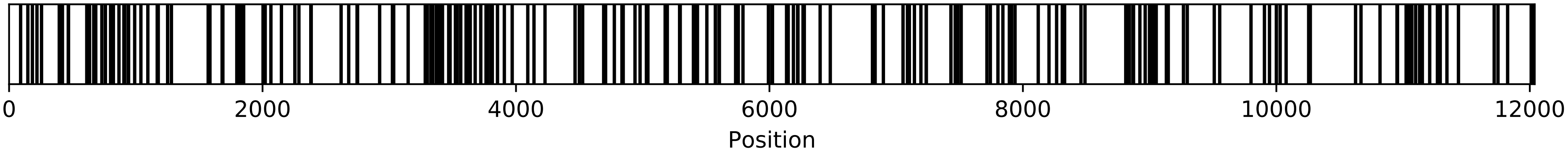


CR1 vs MA1
Number of differences: 64

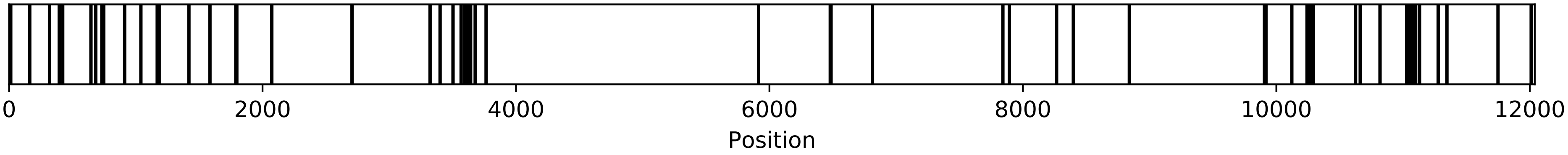


CR1 vs MA2
Number of differences: 216

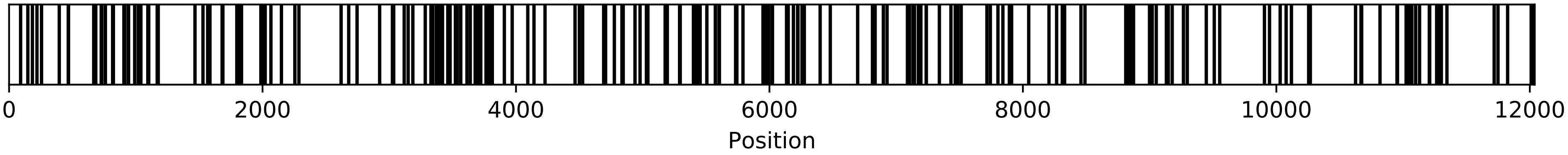


CR2 vs MA1
Number of differences: 211

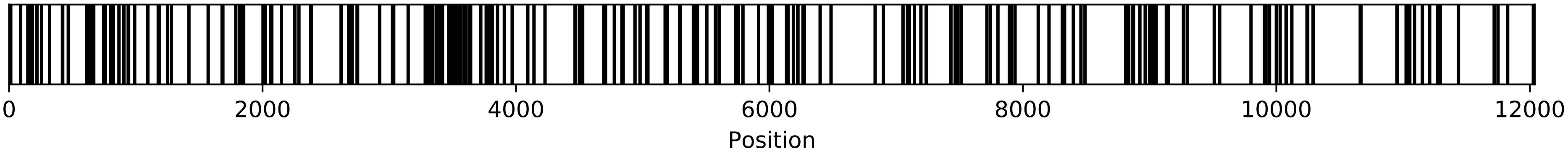


CR2 vs MA2
Number of differences: 68

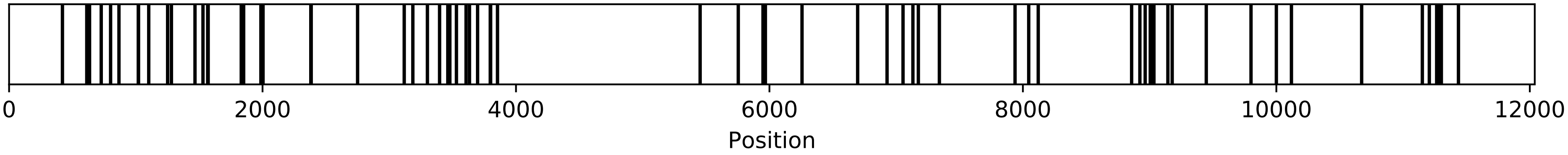


MA1 vs MA2
Number of differences: 214

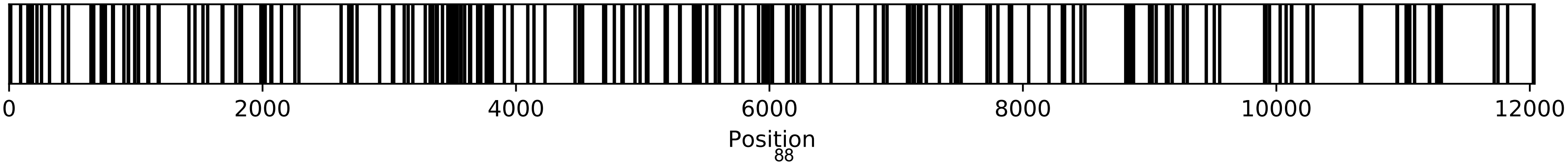

### Region: 563



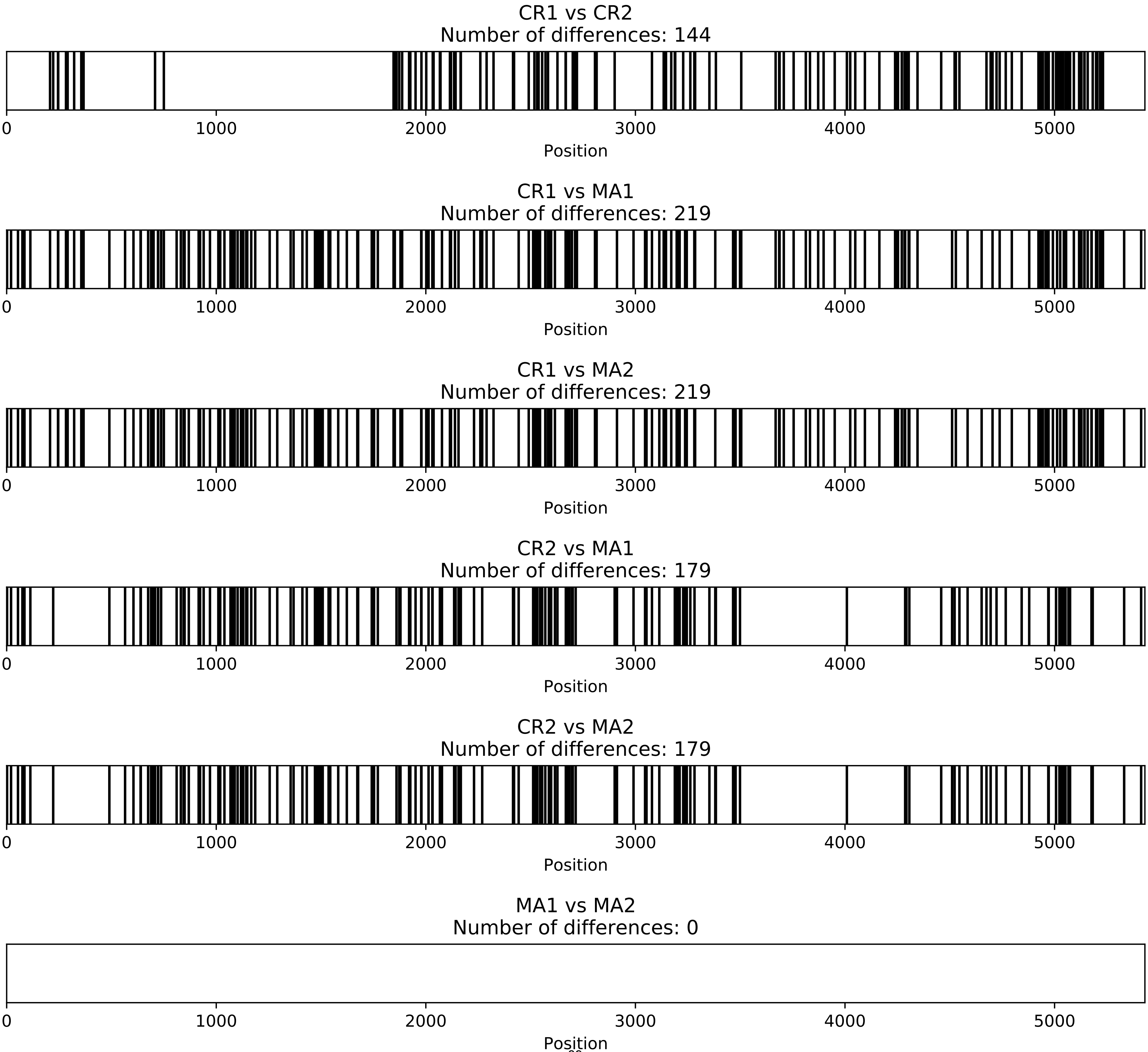
CR1 vs CR2
Number of differences: 144
CR1 vs MA1
Number of differences: 219
CR1 vs MA2
Number of differences: 219
CR2 vs MA1
Number of differences: 179
CR2 vs MA2
Number of differences: 179
MA1 vs MA2
Number of differences: 0
0
1000
2000
3000
4000
5000
Position

### Region: 566



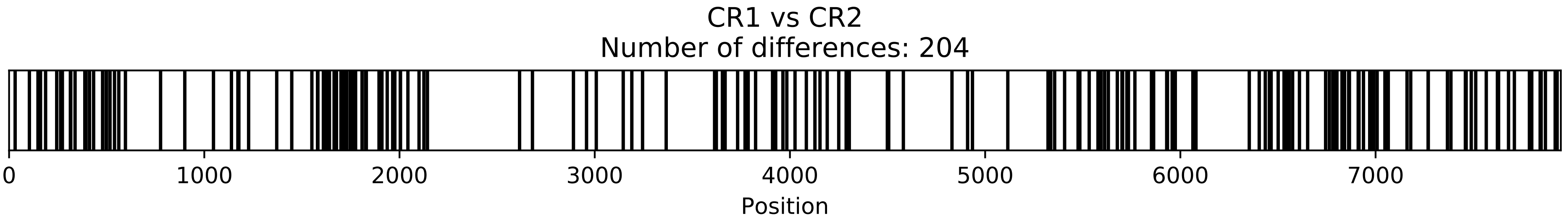
CR1 vs CR2
Number of differences: 204
0
1000
2000
3000
4000
5000
6000
7000
Position

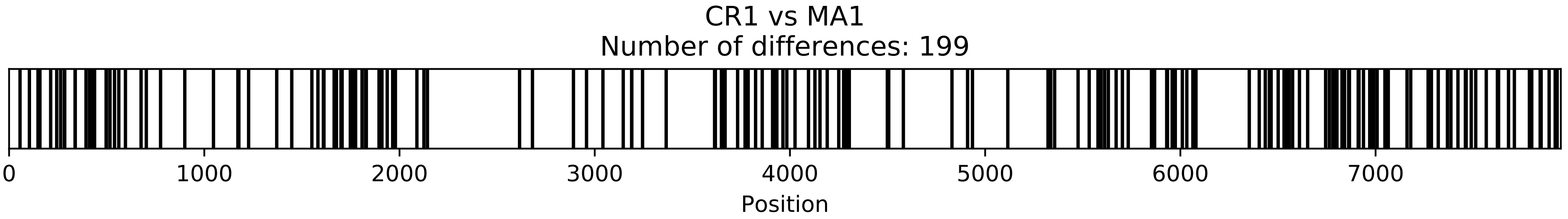
CR1 vs MA1
Number of differences: 199
0
1000
2000
3000
4000
5000
6000
7000
Position

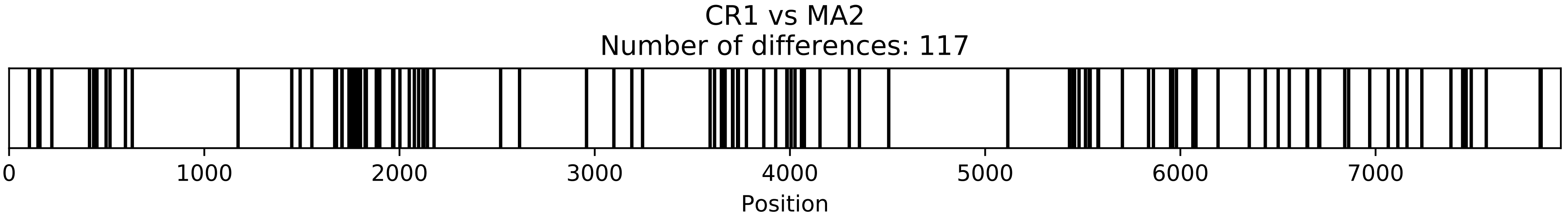
CR1 vs MA2
Number of differences: 117
0
1000
2000
3000
4000
5000
6000
7000
Position

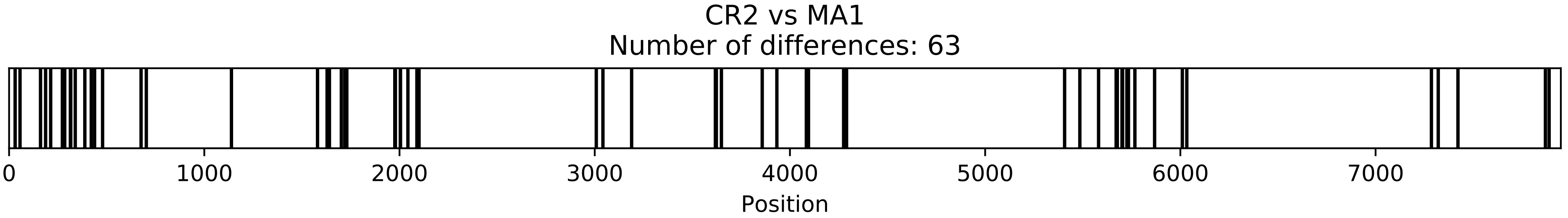
CR2 vs MA1
Number of differences: 63
0
1000
2000
3000
4000
5000
6000
7000
Position

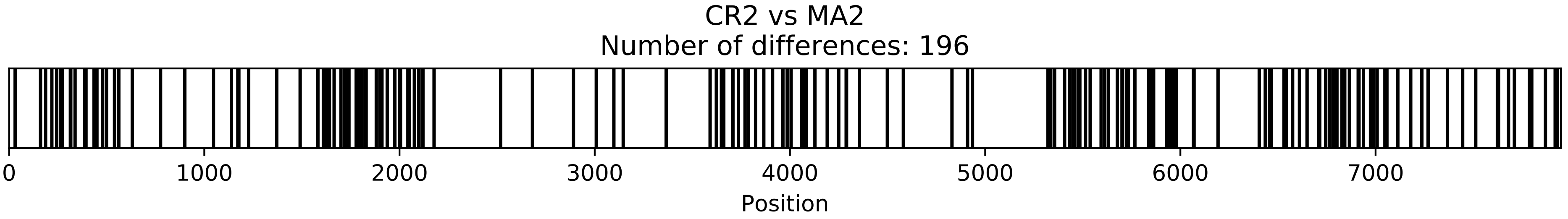
CR2 vs MA2
Number of differences: 196
0
1000
2000
3000
4000
5000
6000
7000
Position

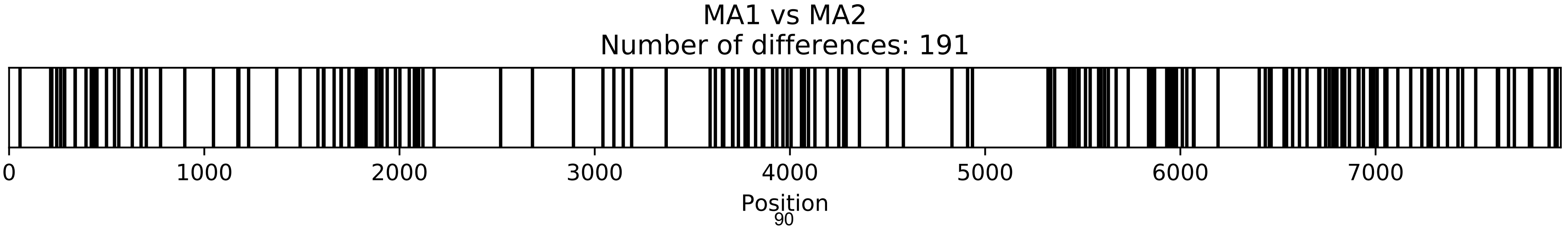
MA1 vs MA2
Number of differences: 191
0
1000
2000
3000
4000
5000
6000
7000
Position

### Region: 573



CR1 vs CR2
Number of differences: 216

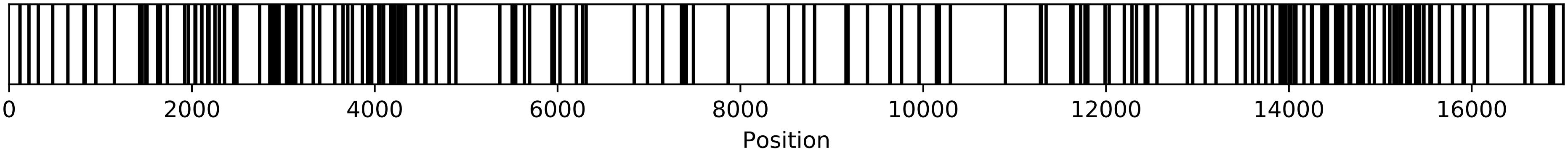


CR1 vs MA1
Number of differences: 119

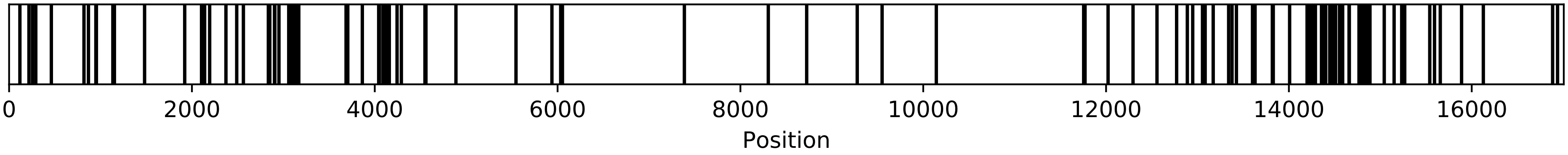


CR1 vs MA2
Number of differences: 218

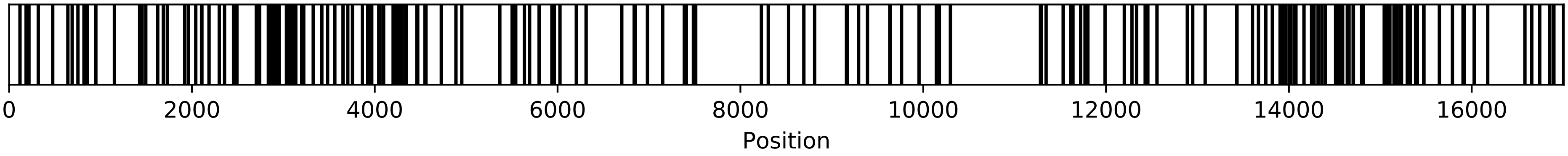


CR2 vs MA1
Number of differences: 233

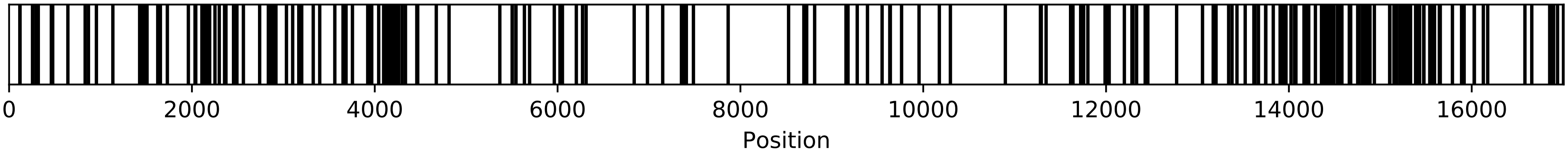


CR2 vs MA2
Number of differences: 81

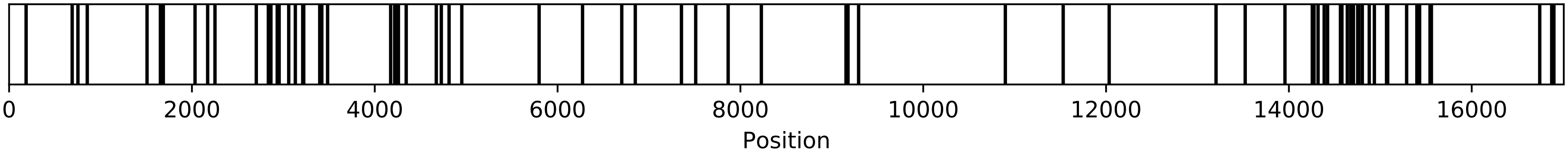


MA1 vs MA2
Number of differences: 238

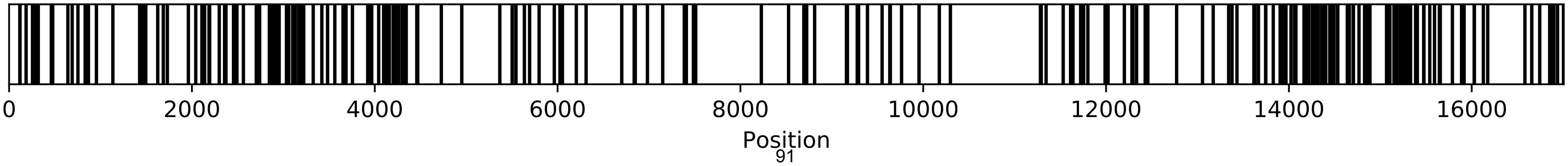

### Region: 590



CR1 vs CR2
Number of differences: 145

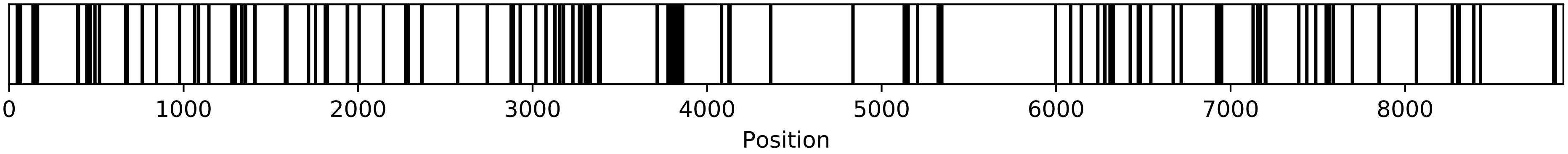


CR1 vs MA1
Number of differences: 94

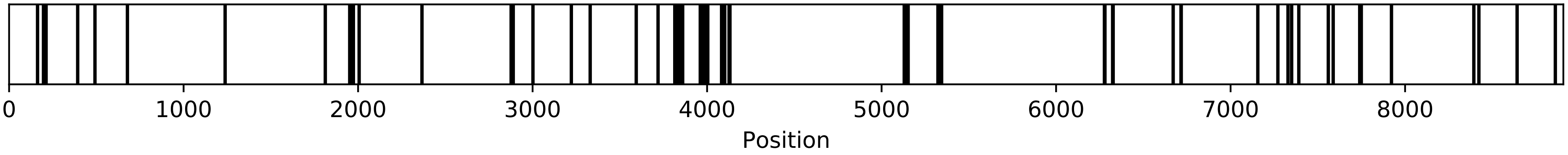


CR1 vs MA2
Number of differences: 158

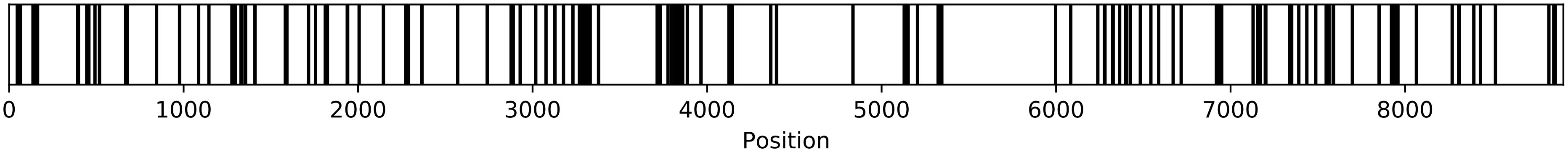


CR2 vs MA1
Number of differences: 150

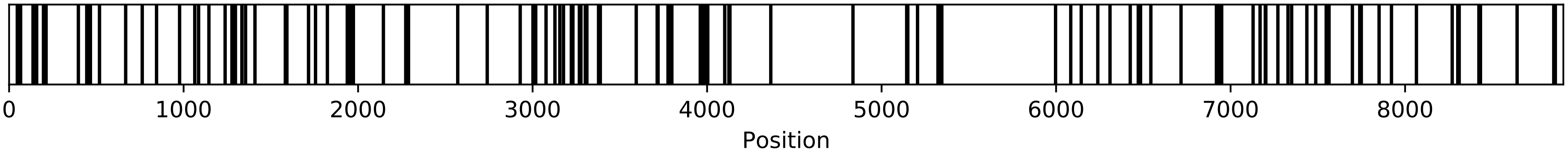


CR2 vs MA2
Number of differences: 48

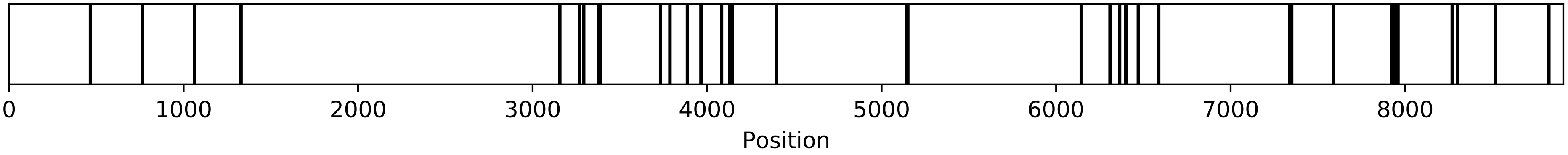


MA1 vs MA2
Number of differences: 155

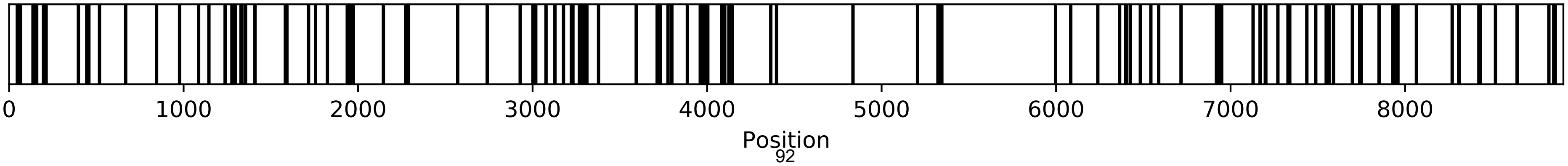

### Region: 594



CR1 vs CR2
Number of differences: 303

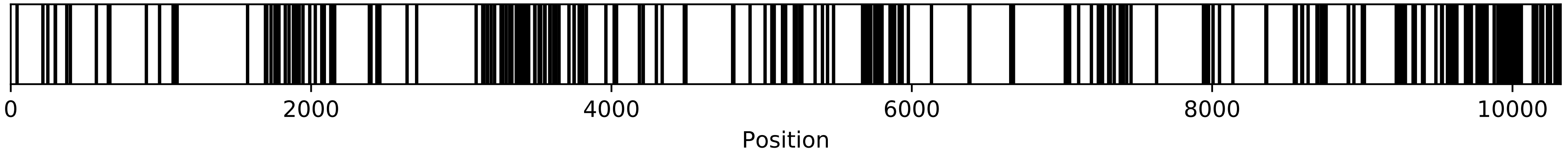


CR1 vs MA1
Number of differences: 312

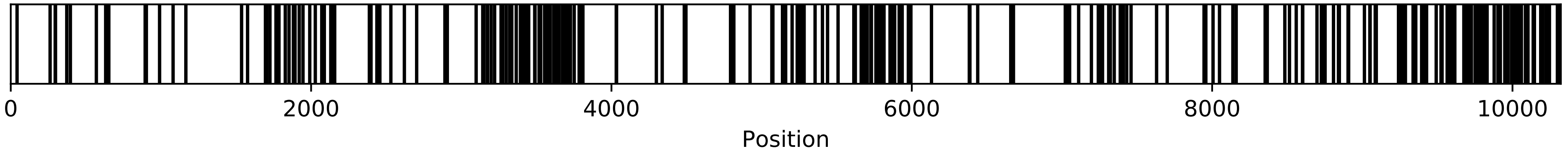


CR1 vs MA2
Number of differences: 99

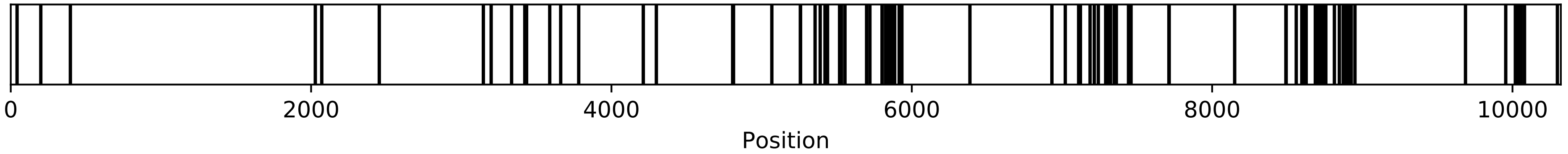


CR2 vs MA1
Number of differences: 174

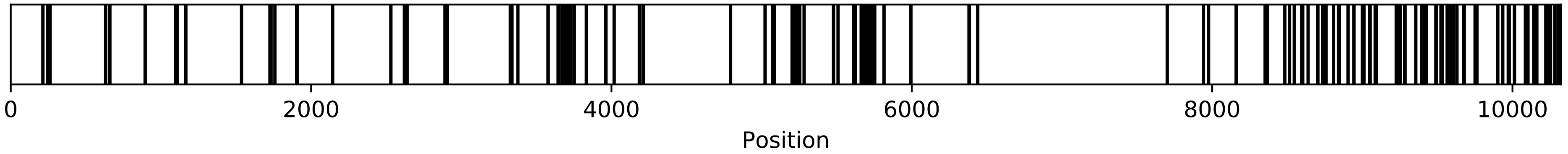


CR2 vs MA2
Number of differences: 318

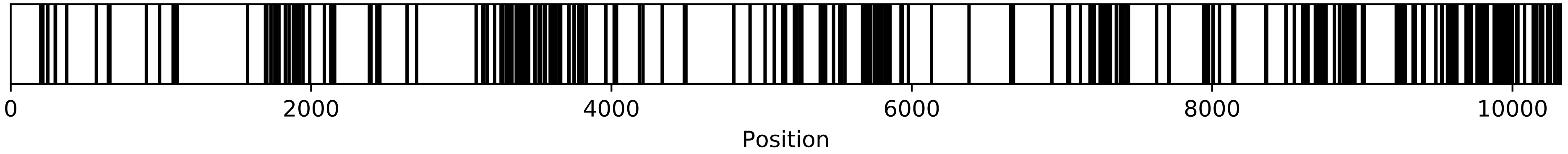


MA1 vs MA2
Number of differences: 329

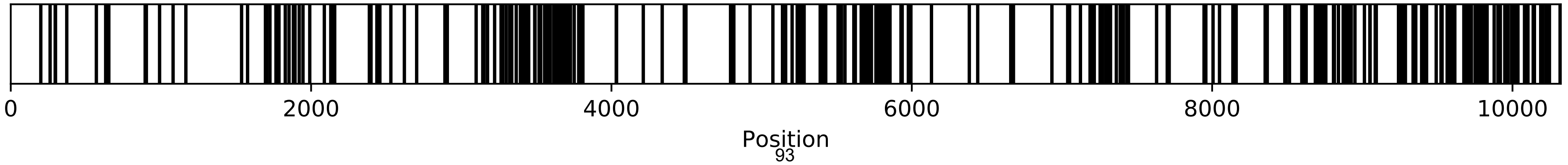

### Region: 604



CR1 vs CR2
Number of differences: 392

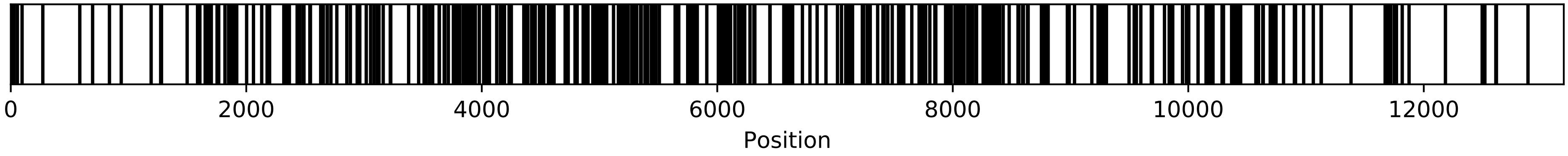


CR1 vs MA1
Number of differences: 408

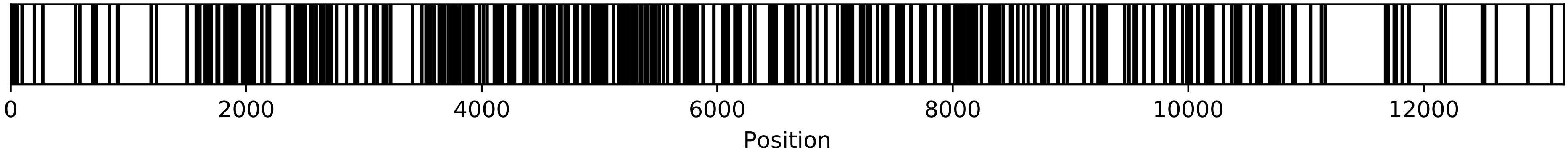


CR1 vs MA2
Number of differences: 173

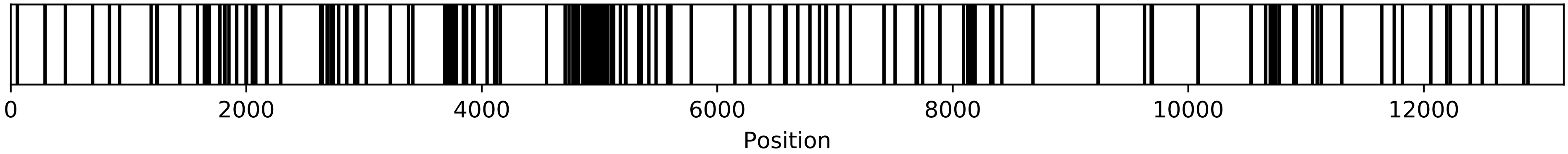


CR2 vs MA1
Number of differences: 254

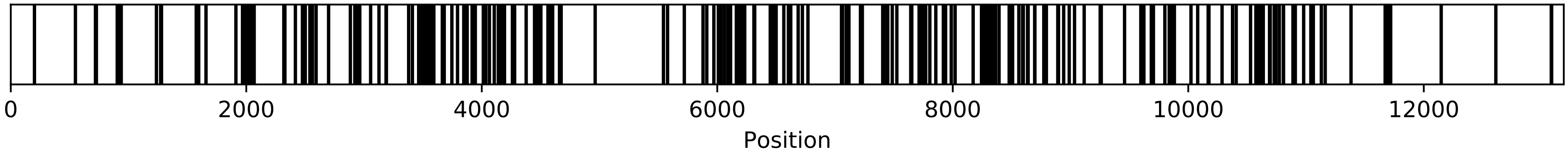


CR2 vs MA2
Number of differences: 392

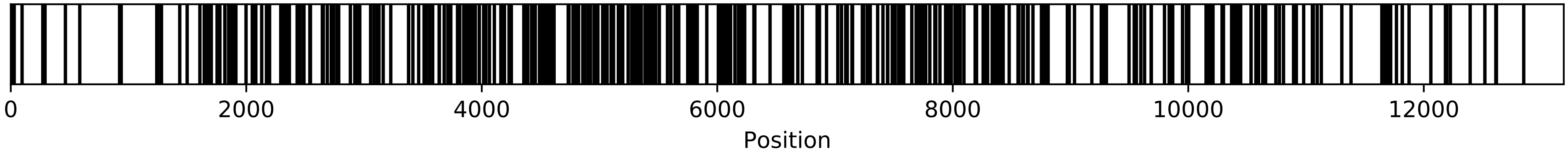


MA1 vs MA2
Number of differences: 409

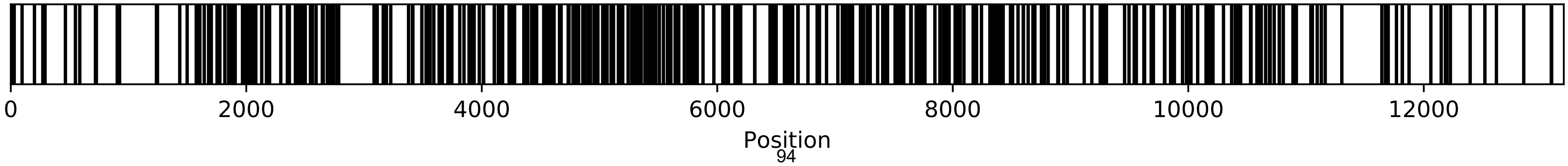

### Region: 617



## CR1 vs CR2
Number of differences: 204

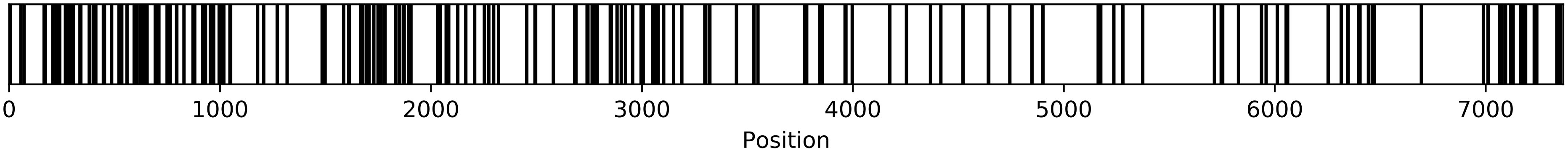


## CR1 vs MA1
Number of differences: 104

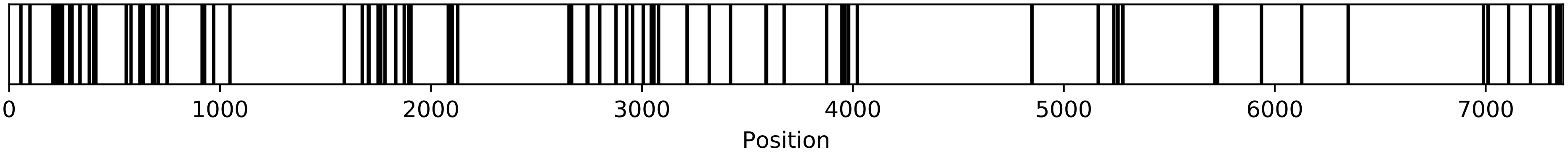


## CR1 vs MA2
Number of differences: 209

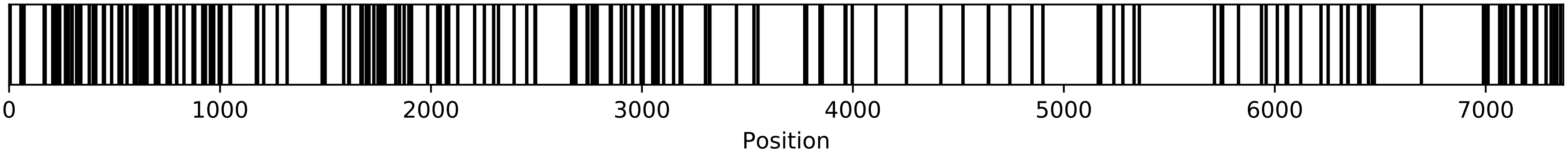


## CR2 vs MA1
Number of differences: 217

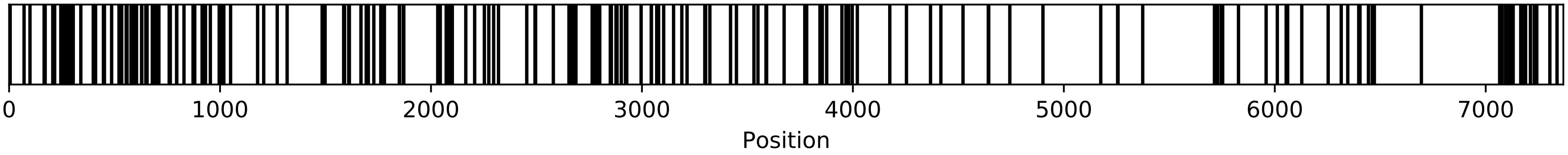


## CR2 vs MA2
Number of differences: 36

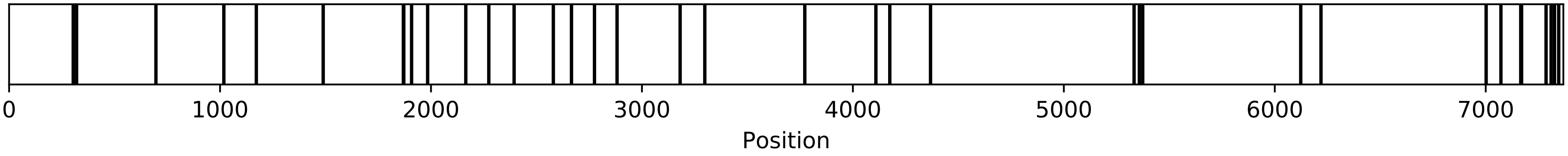


## MA1 vs MA2
Number of differences: 224

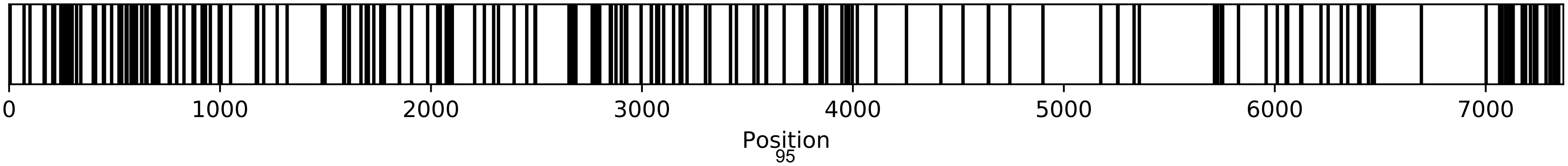

### Region: 620



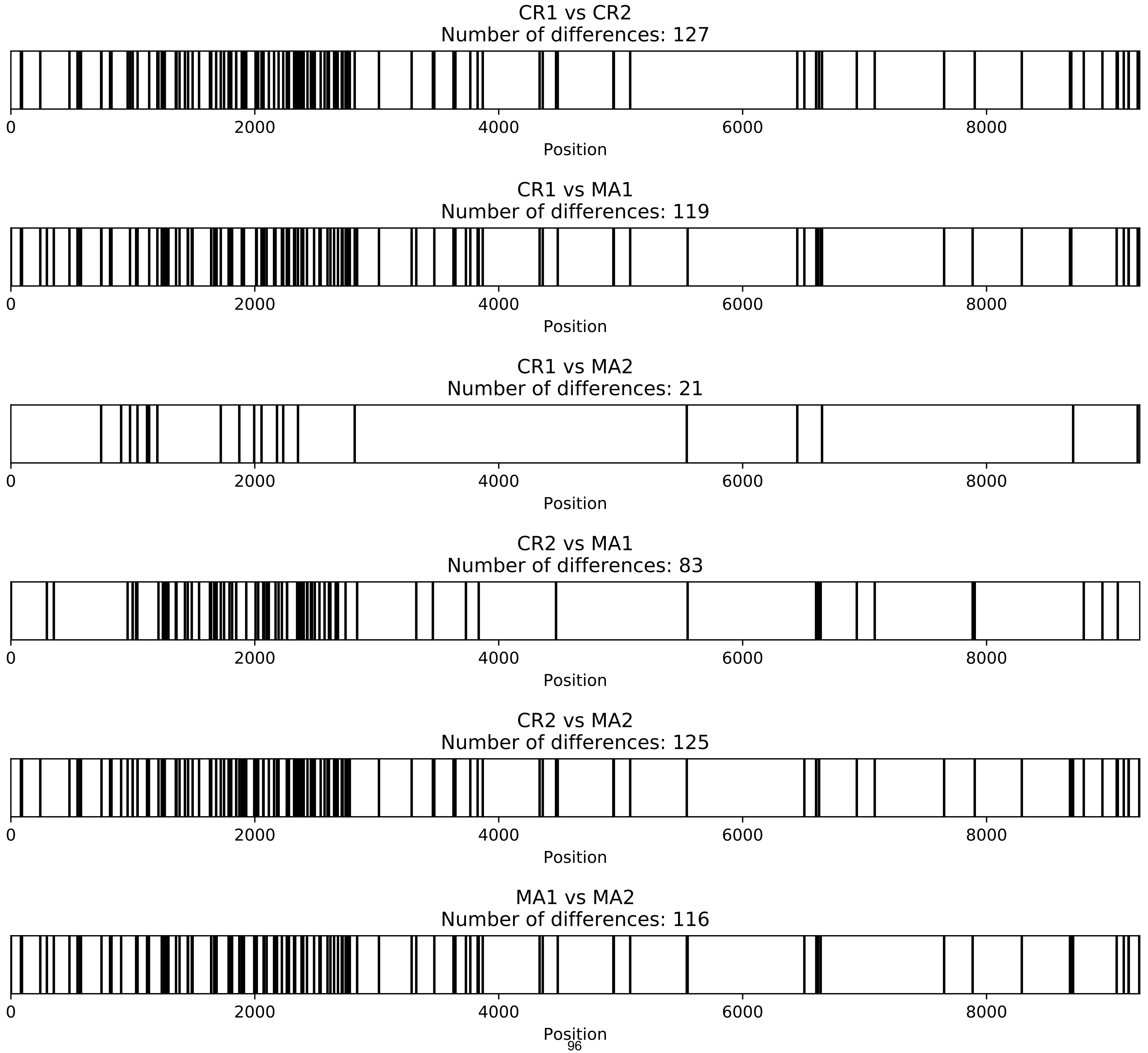
CR1 vs CR2
Number of differences: 127
Position
CR1 vs MA1
Number of differences: 119
Position
CR1 vs MA2
Number of differences: 21
Position
CR2 vs MA1
Number of differences: 83
Position
CR2 vs MA2
Number of differences: 125
Position
MA1 vs MA2
Number of differences: 116
0
2000
4000
6000
8000
Position

### Region: 622



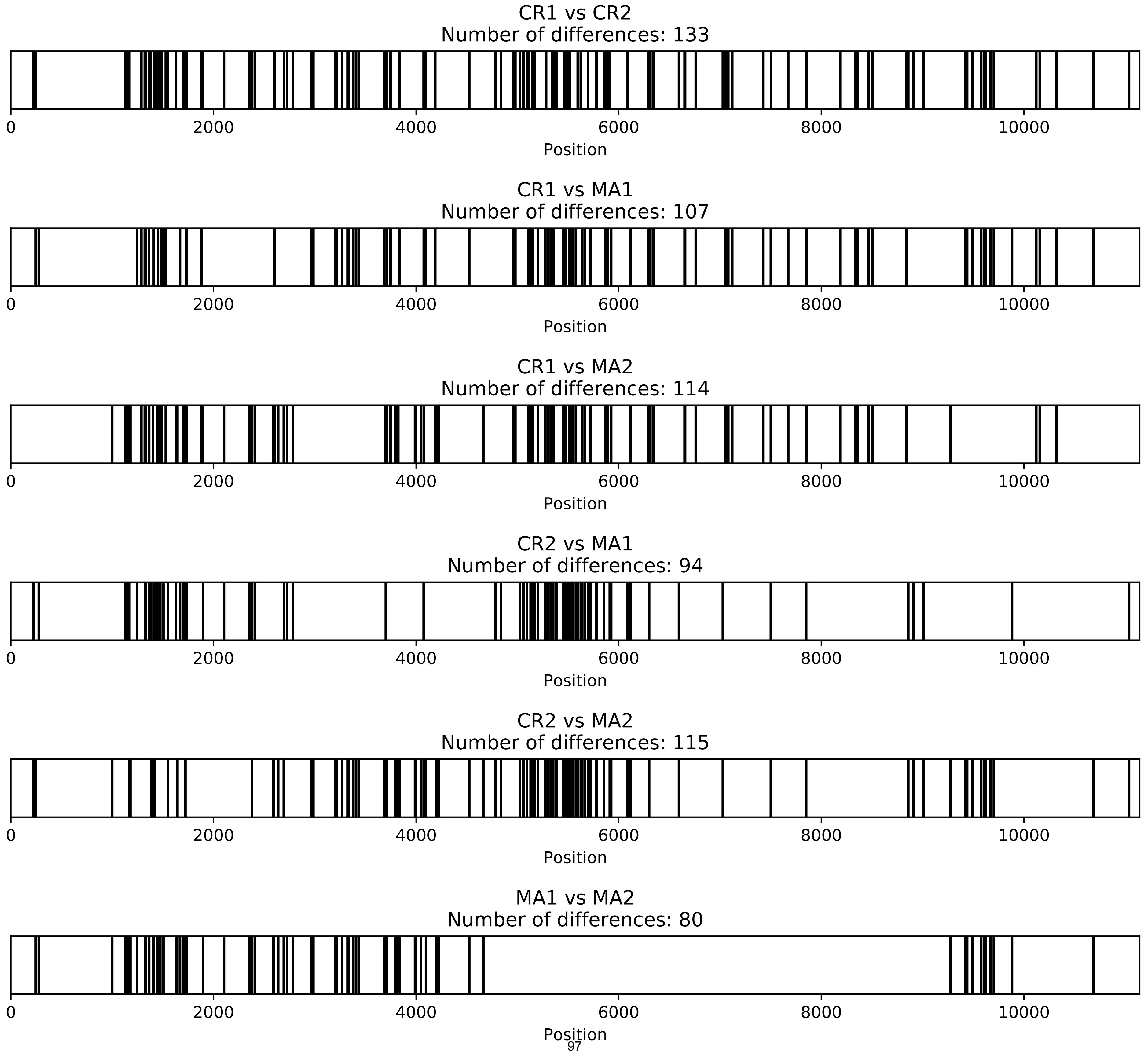
CR1 vs CR2
Number of differences: 133
0
2000
4000
6000
8000
10000
Position
CR1 vs MA1
Number of differences: 107
0
2000
4000
6000
8000
10000
Position
CR1 vs MA2
Number of differences: 114
0
2000
4000
6000
8000
10000
Position
CR2 vs MA1
Number of differences: 94
0
2000
4000
6000
8000
10000
Position
CR2 vs MA2
Number of differences: 115
0
2000
4000
6000
8000
10000
Position
MA1 vs MA2
Number of differences: 80
0
2000
4000
6000
8000
10000
Position

### Region: 636



CR1 vs CR2
Number of differences: 360

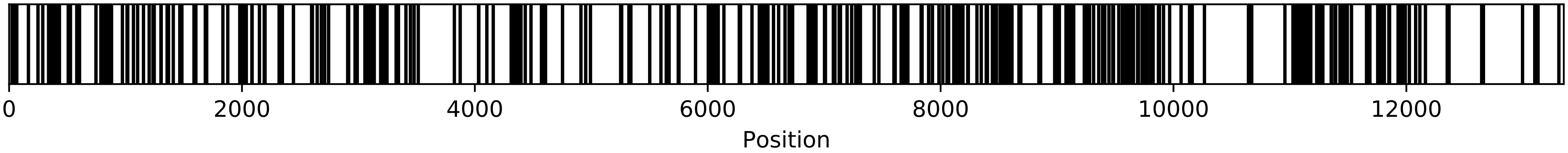


CR1 vs MA1
Number of differences: 224

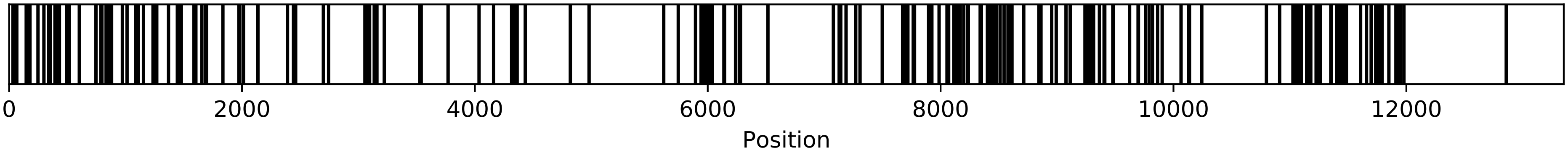


CR1 vs MA2
Number of differences: 294

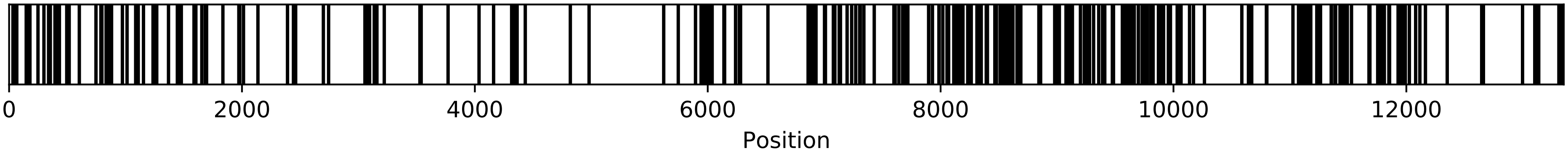


CR2 vs MA1
Number of differences: 326

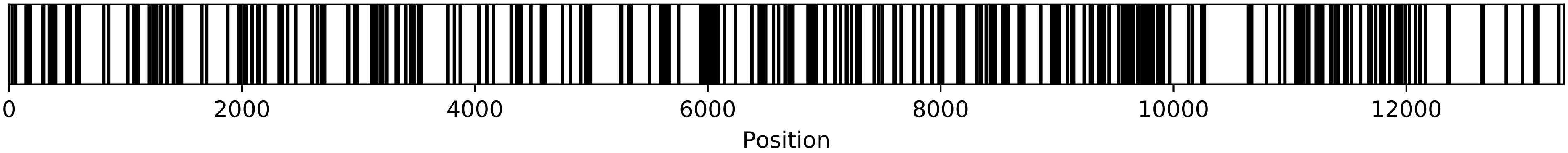


CR2 vs MA2
Number of differences: 207

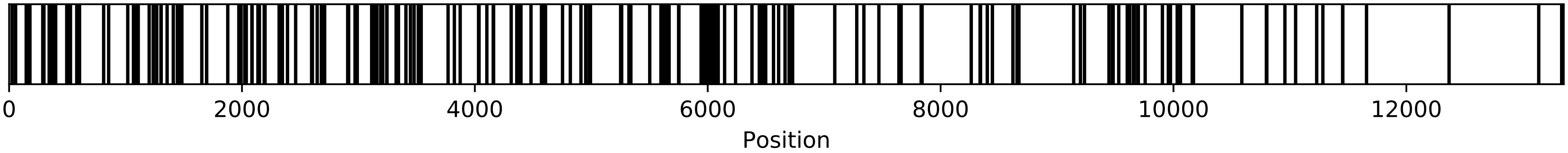


MA1 vs MA2
Number of differences: 180

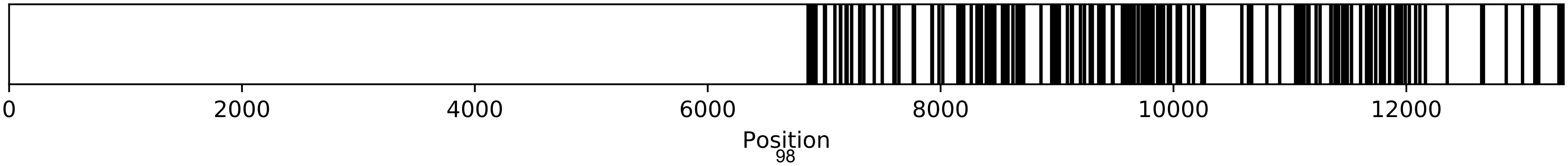

### Region: 637



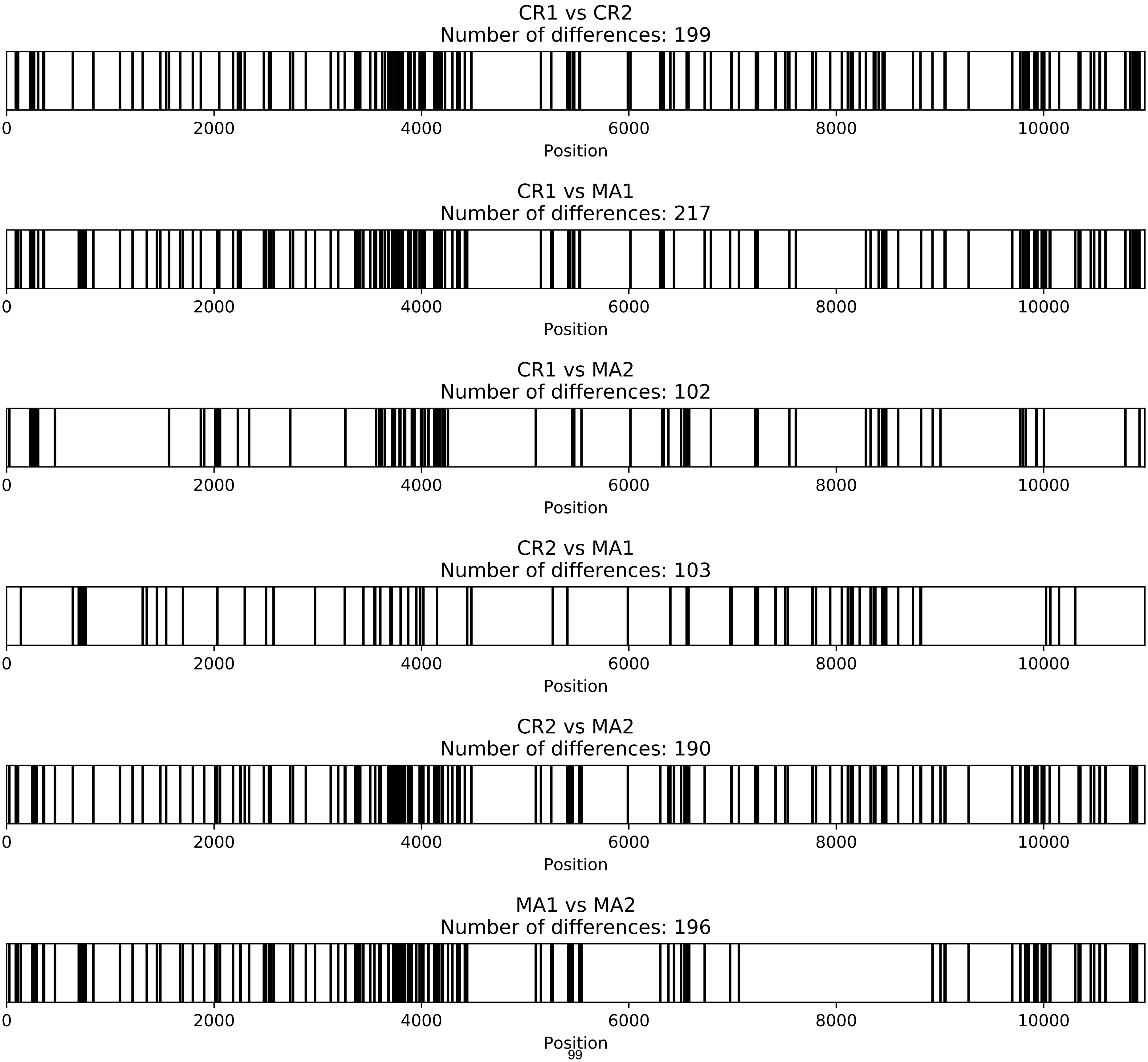
CR1 vs CR2
Number of differences: 199
CR1 vs MA1
Number of differences: 217
CR1 vs MA2
Number of differences: 102
CR2 vs MA1
Number of differences: 103
CR2 vs MA2
Number of differences: 190
MA1 vs MA2
Number of differences: 196
0
2000
4000
6000
8000
10000
Position

### Region: 643



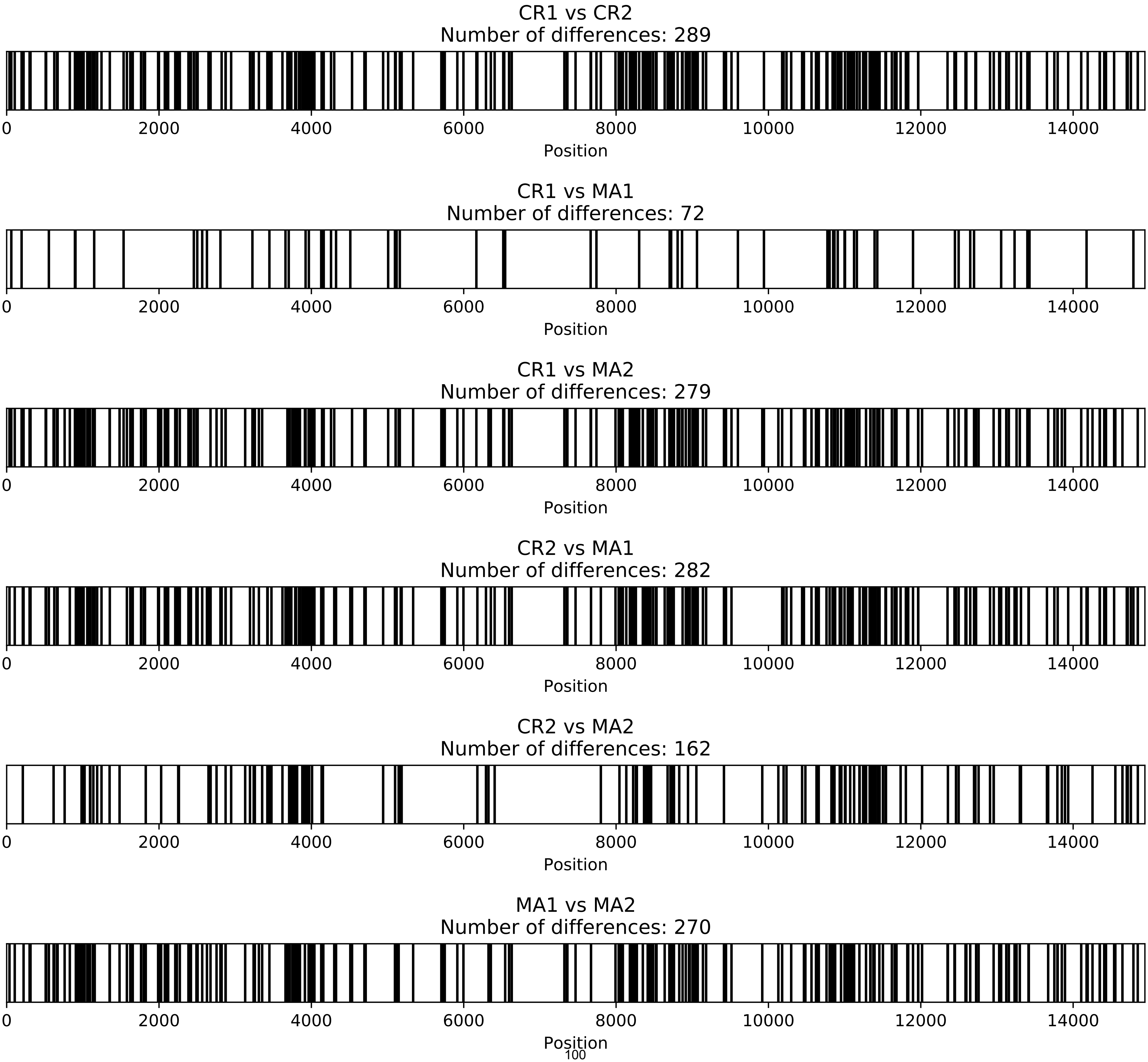
CR1 vs CR2
Number of differences: 289
CR1 vs MA1
Number of differences: 72
CR1 vs MA2
Number of differences: 279
CR2 vs MA1
Number of differences: 282
CR2 vs MA2
Number of differences: 162
MA1 vs MA2
Number of differences: 270
0 2000 4000 6000 8000 10000 12000 14000
Position

### Region: 647



CR1 vs CR2
Number of differences: 308

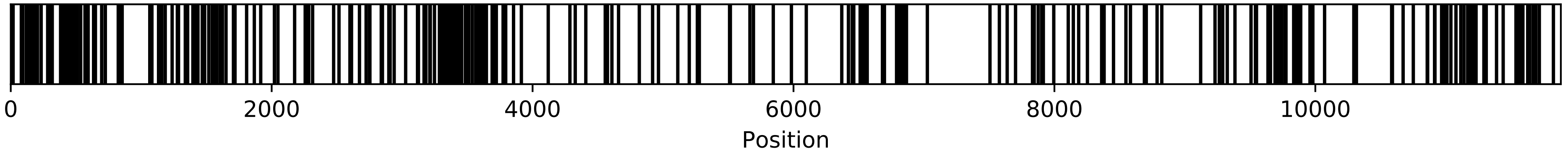


CR1 vs MA1
Number of differences: 155

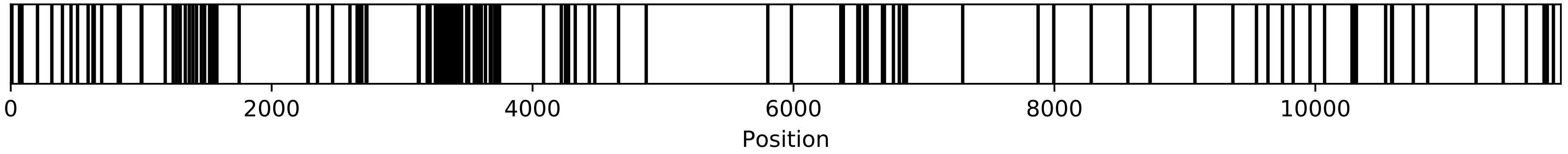


CR1 vs MA2
Number of differences: 291

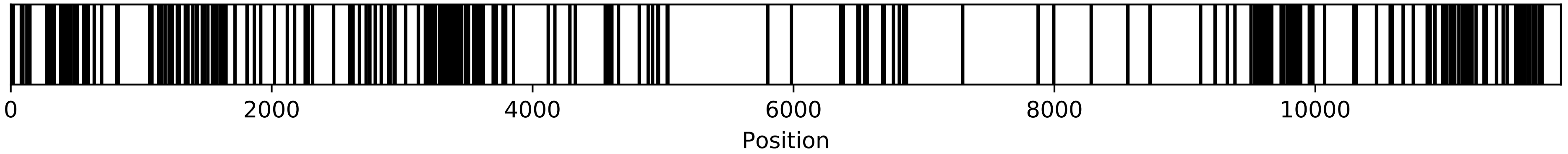


CR2 vs MA1
Number of differences: 304

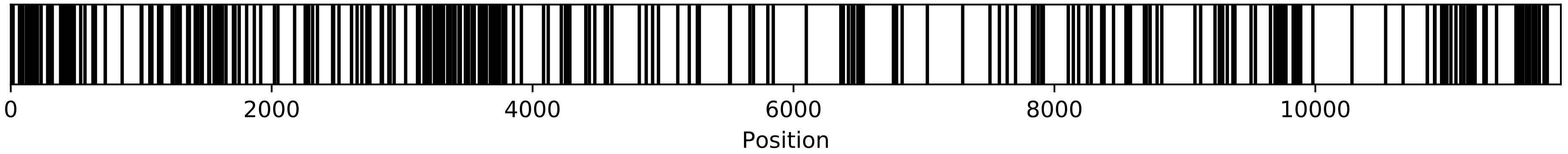


CR2 vs MA2
Number of differences: 205

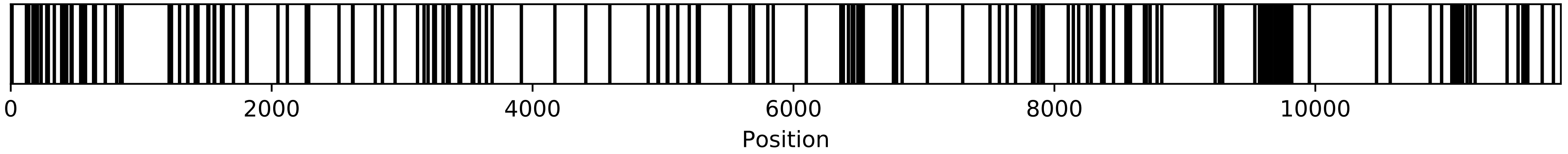


MA1 vs MA2
Number of differences: 268

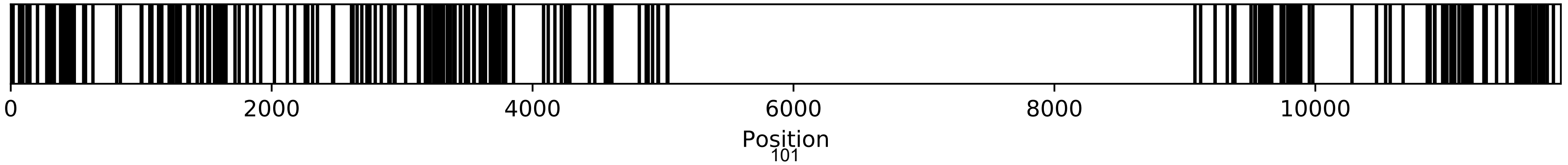

### Region: 653



CR1 vs CR2
Number of differences: 179

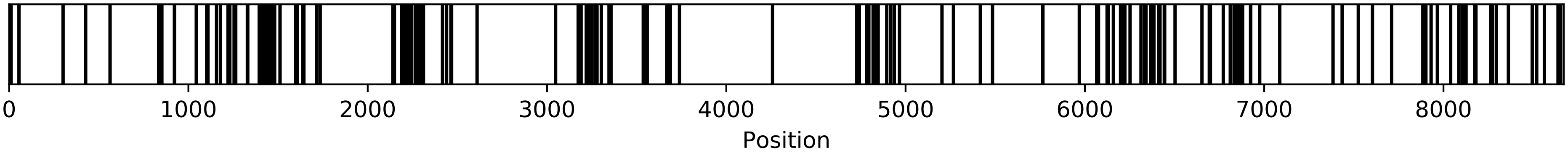


CR1 vs MA1
Number of differences: 113

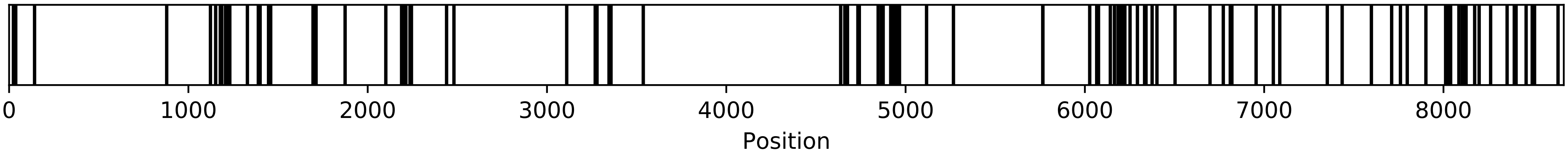


CR1 vs MA2
Number of differences: 120

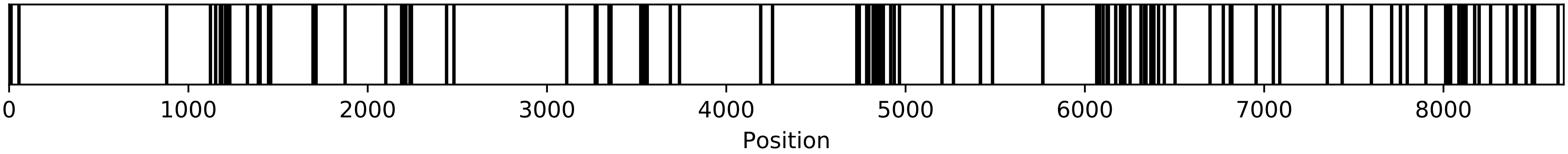


CR2 vs MA1
Number of differences: 190

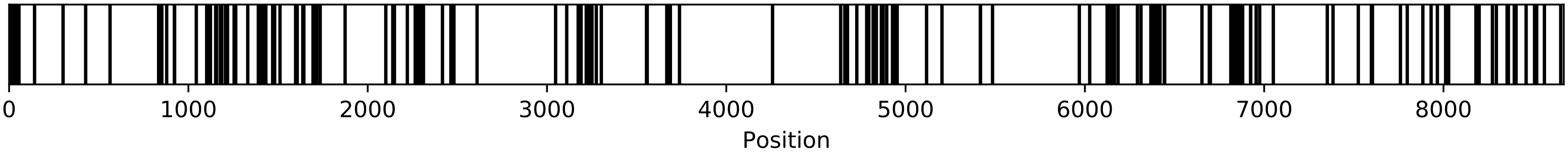


CR2 vs MA2
Number of differences: 144

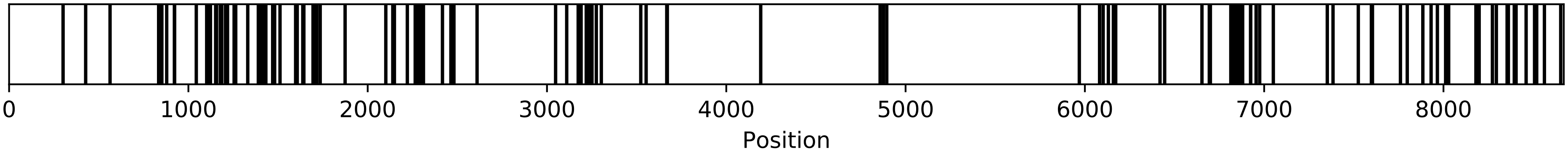


MA1 vs MA2
Number of differences: 63

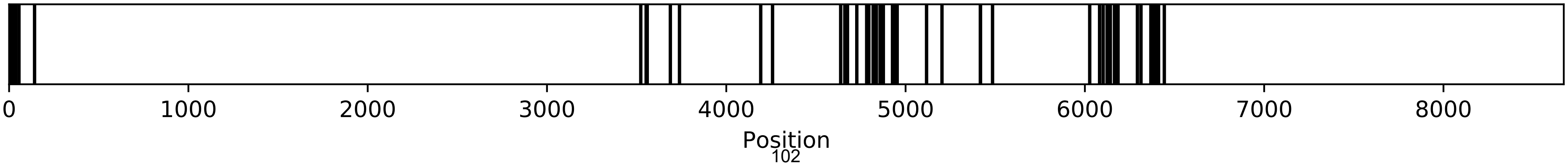

### Region: 656



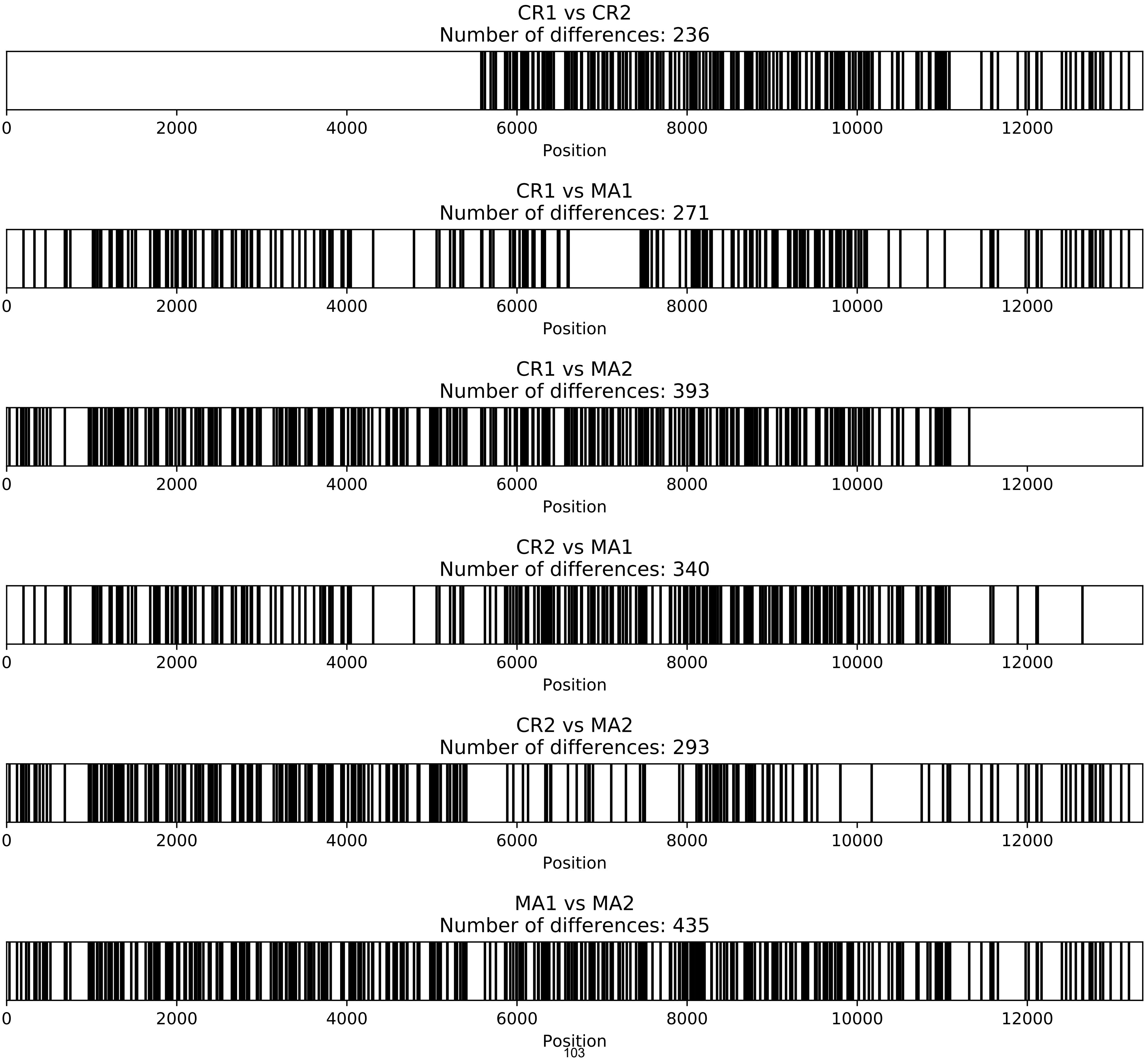
CR1 vs CR2
Number of differences: 236
CR1 vs MA1
Number of differences: 271
CR1 vs MA2
Number of differences: 393
CR2 vs MA1
Number of differences: 340
CR2 vs MA2
Number of differences: 293
MA1 vs MA2
Number of differences: 435
0 2000 4000 6000 8000 10000 12000
Position

### Region: 660



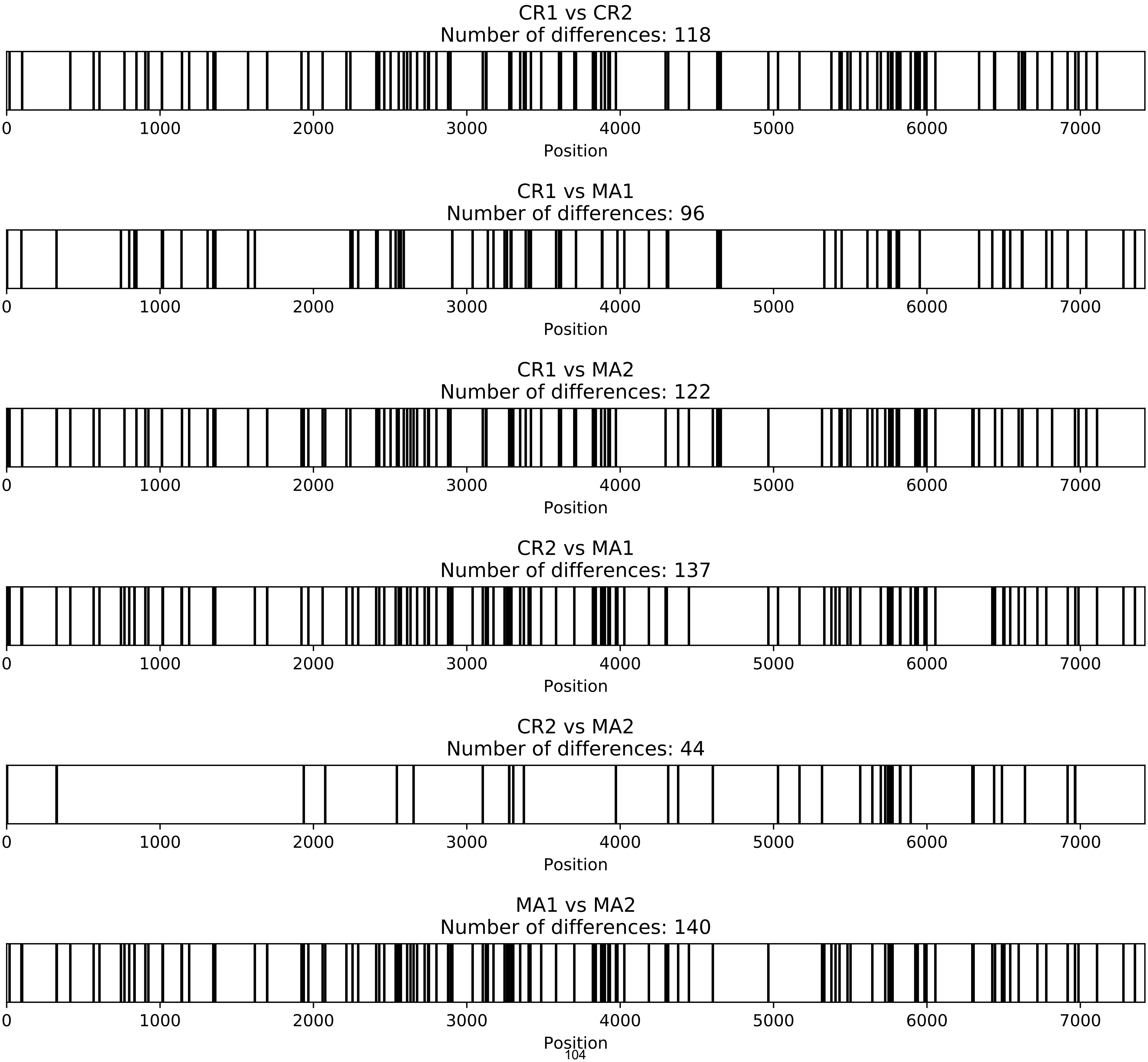
CR1 vs CR2
Number of differences: 118
CR1 vs MA1
Number of differences: 96
CR1 vs MA2
Number of differences: 122
CR2 vs MA1
Number of differences: 137
CR2 vs MA2
Number of differences: 44
MA1 vs MA2
Number of differences: 140
0
1000
2000
3000
4000
5000
6000
7000
Position

### Region: 664



## CR1 vs CR2
Number of differences: 211

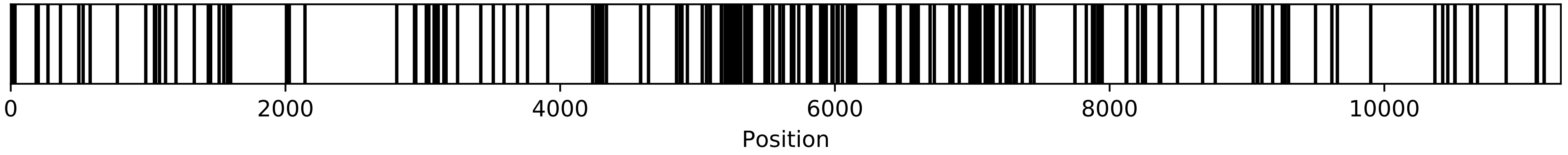


## CR1 vs MA1
Number of differences: 217

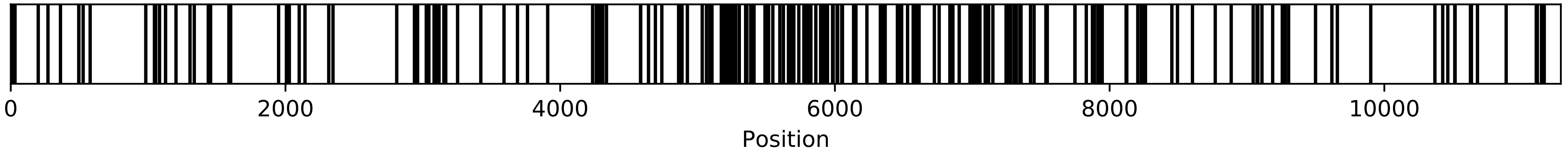


## CR1 vs MA2
Number of differences: 124

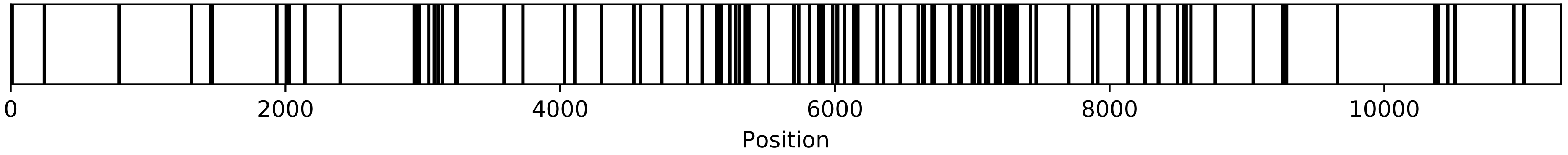


## CR2 vs MA1
Number of differences: 76

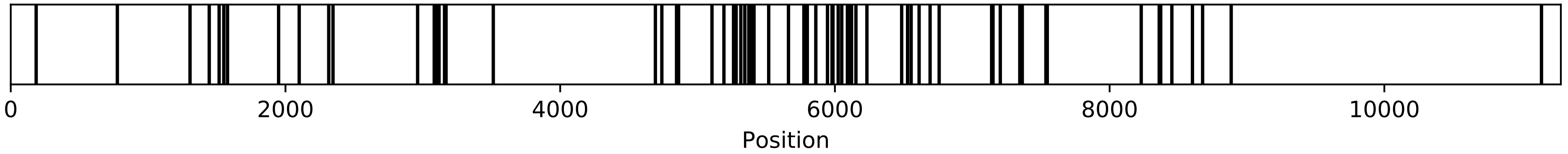


## CR2 vs MA2
Number of differences: 202

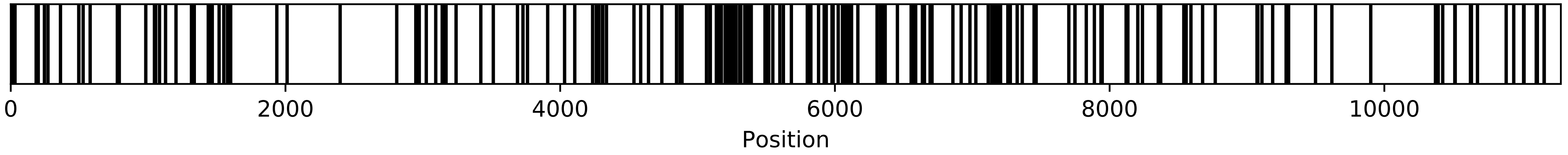


## MA1 vs MA2
Number of differences: 210

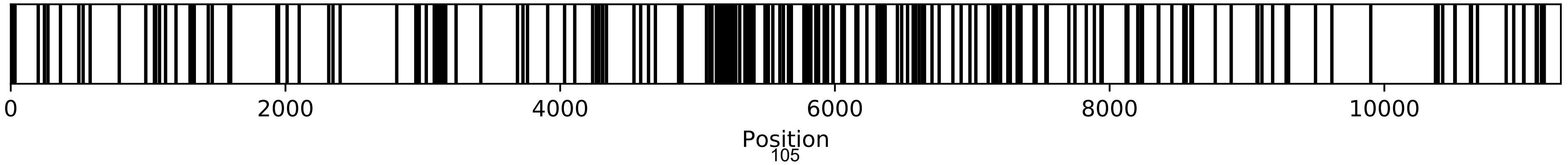

### Region: 665



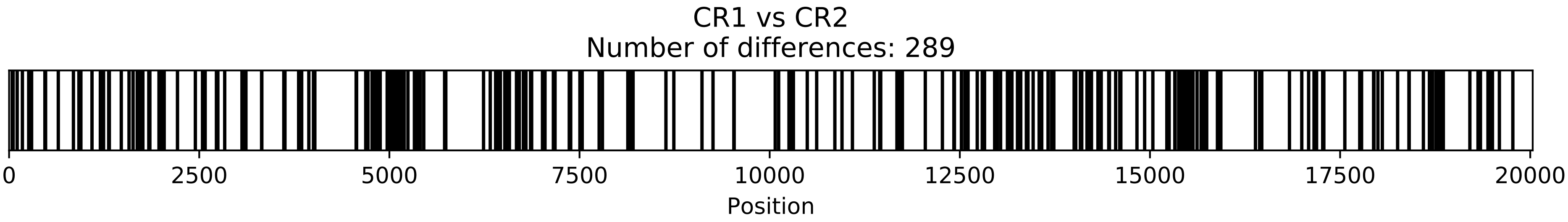
CR1 vs CR2
Number of differences: 289
0
2500
5000
7500
10000
12500
15000
17500
20000
Position

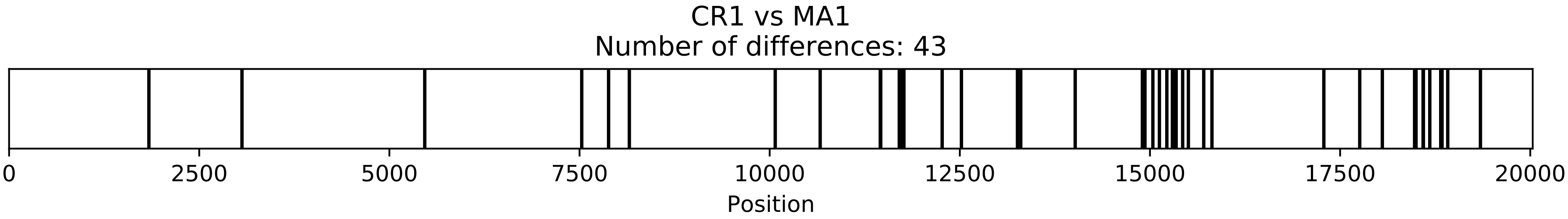
CR1 vs MA1
Number of differences: 43
0
2500
5000
7500
10000
12500
15000
17500
20000
Position

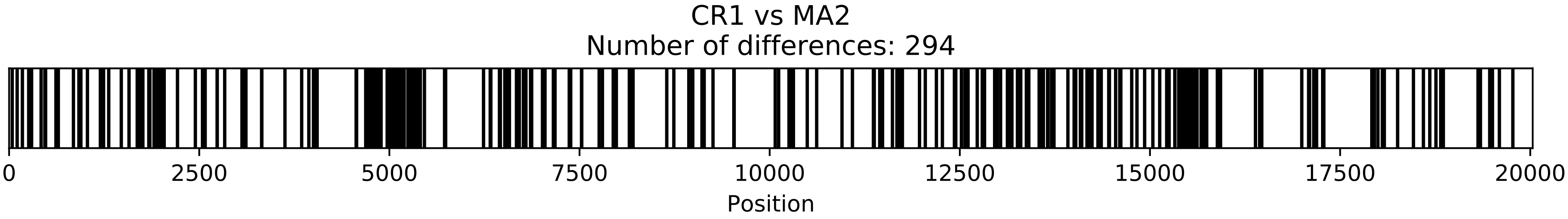
CR1 vs MA2
Number of differences: 294
0
2500
5000
7500
10000
12500
15000
17500
20000
Position

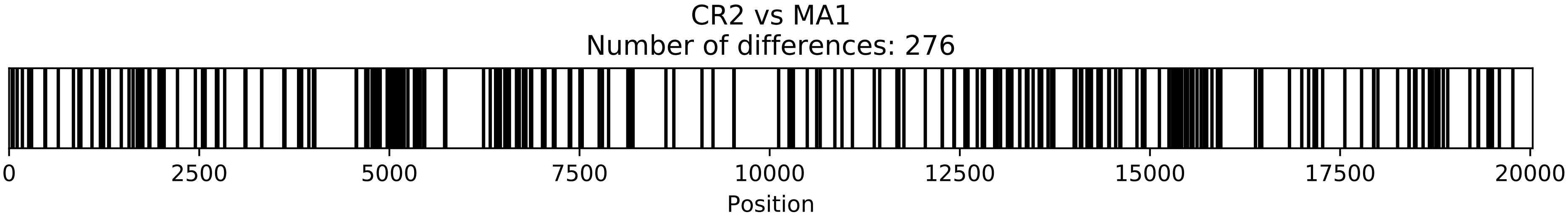
CR2 vs MA1
Number of differences: 276
0
2500
5000
7500
10000
12500
15000
17500
20000
Position

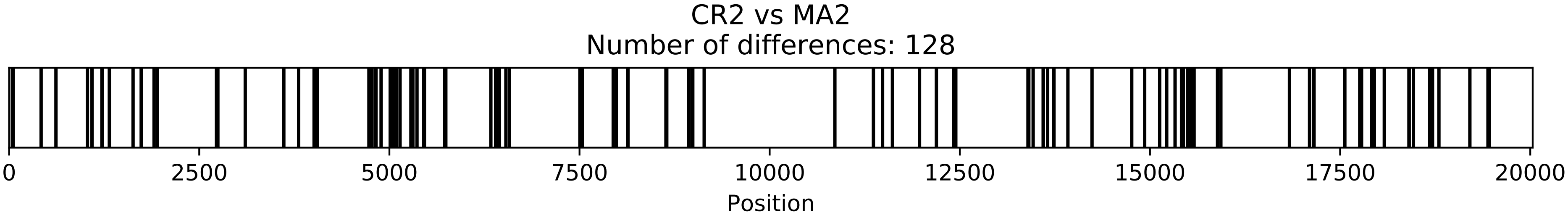
CR2 vs MA2
Number of differences: 128
0
2500
5000
7500
10000
12500
15000
17500
20000
Position

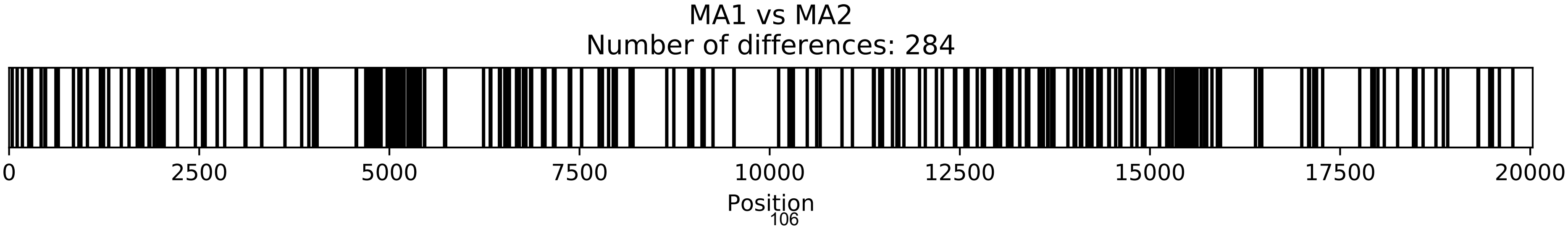
MA1 vs MA2
Number of differences: 284
0
2500
5000
7500
10000
12500
15000
17500
20000
Position

### Region: 666



CR1 vs CR2
Number of differences: 184

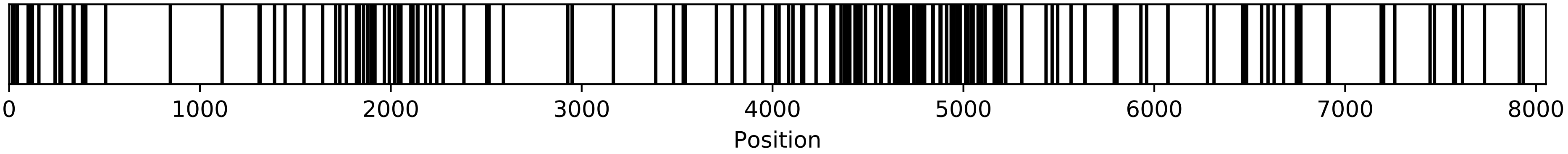


CR1 vs MA1
Number of differences: 174

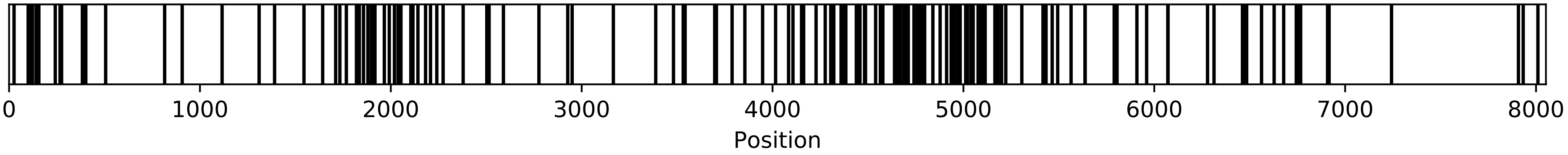


CR1 vs MA2
Number of differences: 114

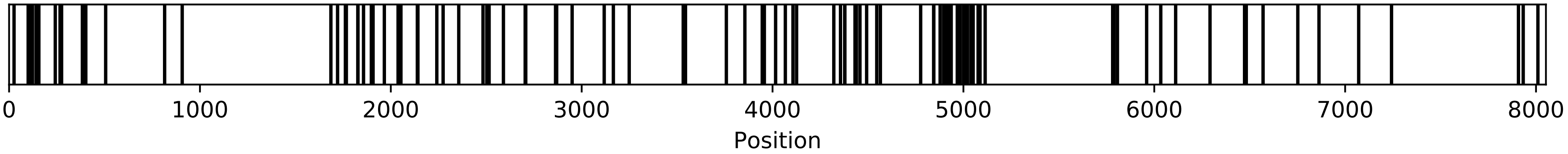


CR2 vs MA1
Number of differences: 63

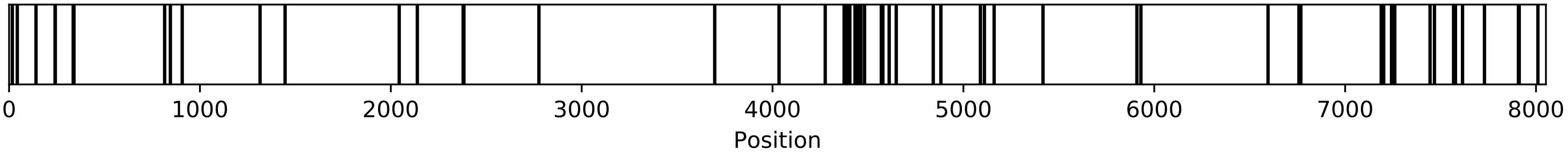


CR2 vs MA2
Number of differences: 171

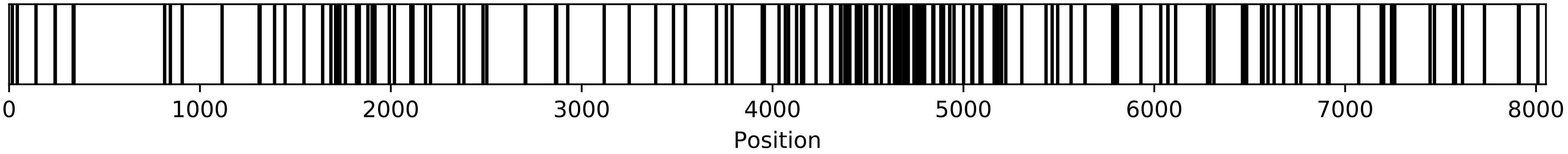


MA1 vs MA2
Number of differences: 150

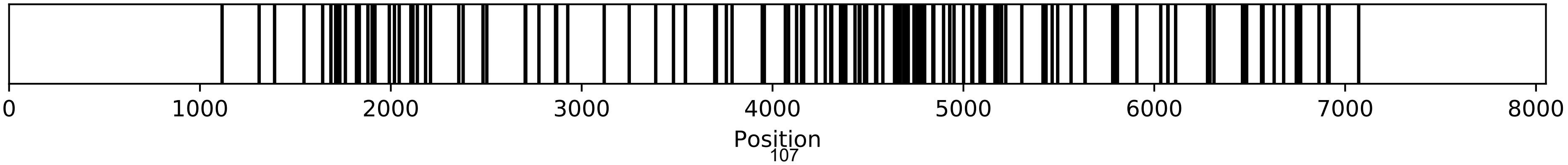

### Region: 675



CR1 vs CR2
Number of differences: 191

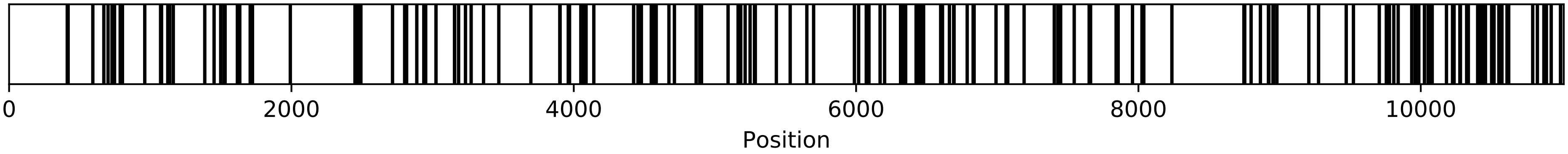


CR1 vs MA1
Number of differences: 180

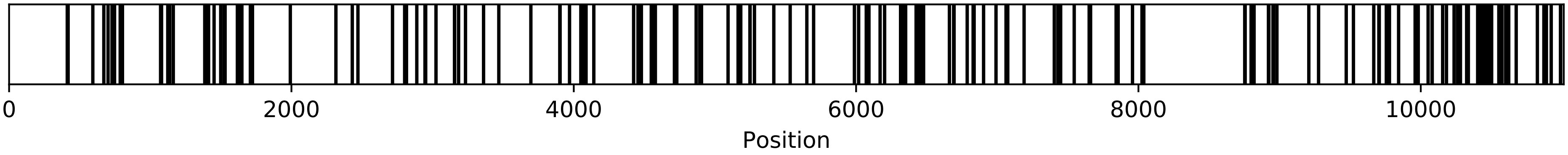


CR1 vs MA2
Number of differences: 48

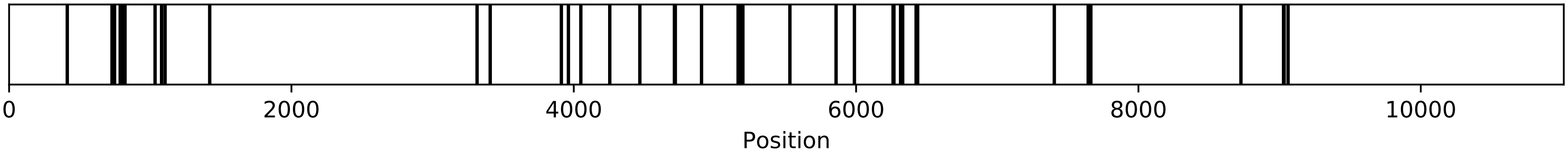


CR2 vs MA1
Number of differences: 54

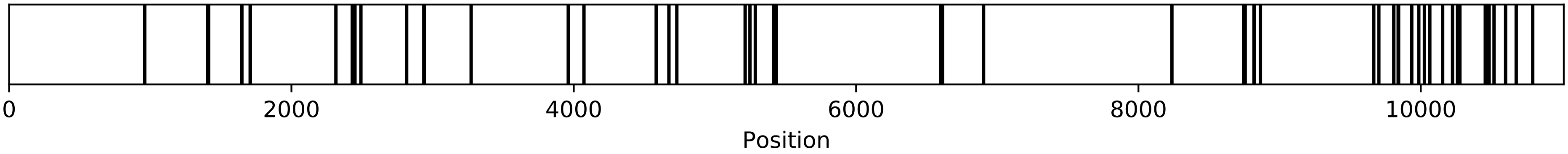


CR2 vs MA2
Number of differences: 202

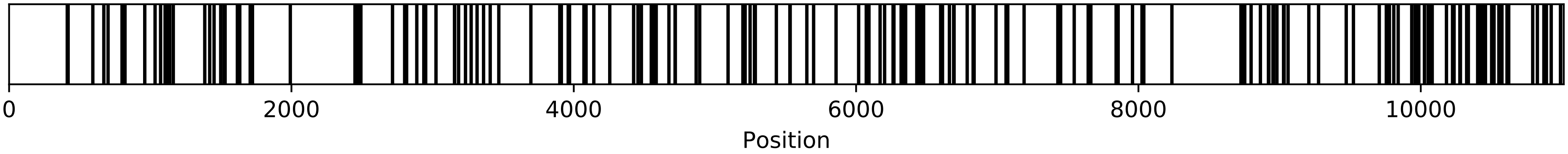


MA1 vs MA2
Number of differences: 191

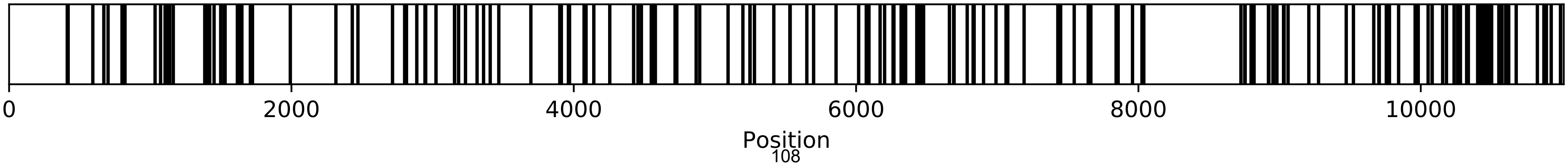

### Region: 680



## CR1 vs CR2
Number of differences: 275

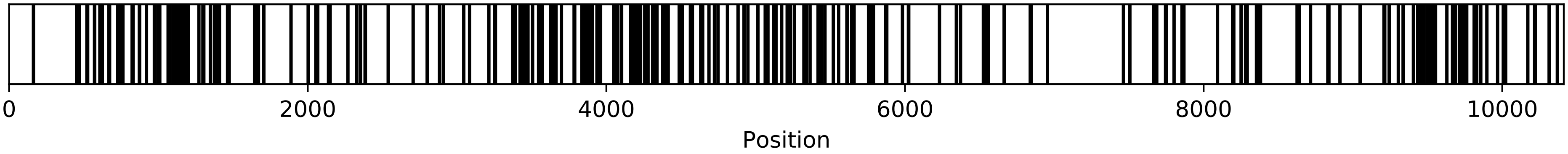


## CR1 vs MA1
Number of differences: 291

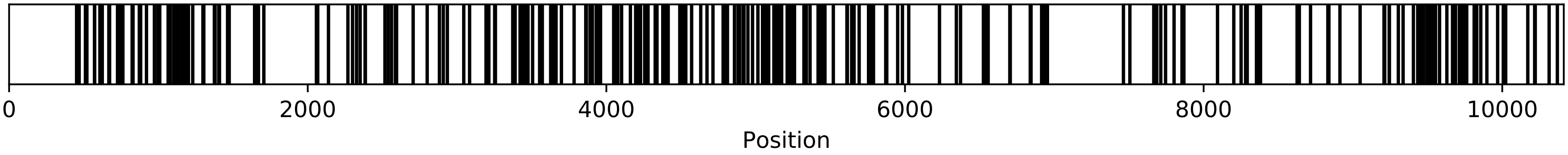


## CR1 vs MA2
Number of differences: 131

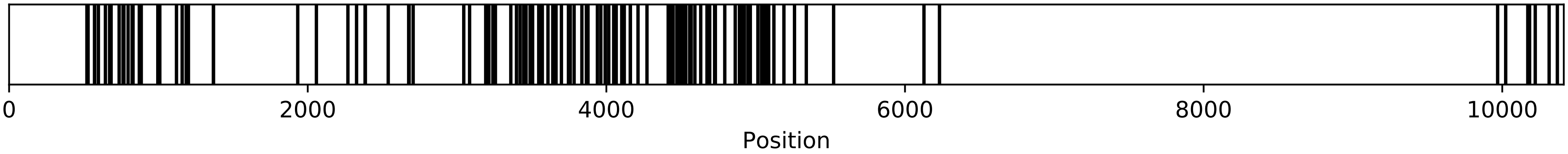


## CR2 vs MA1
Number of differences: 101

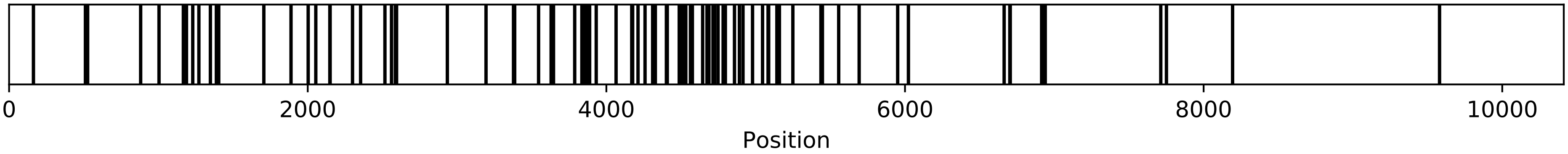


## CR2 vs MA2
Number of differences: 281

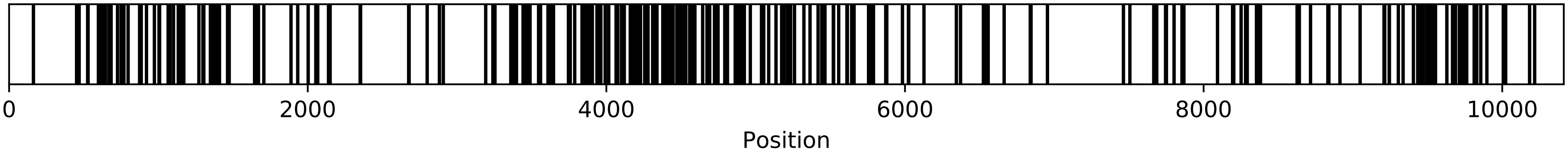


## MA1 vs MA2
Number of differences: 293

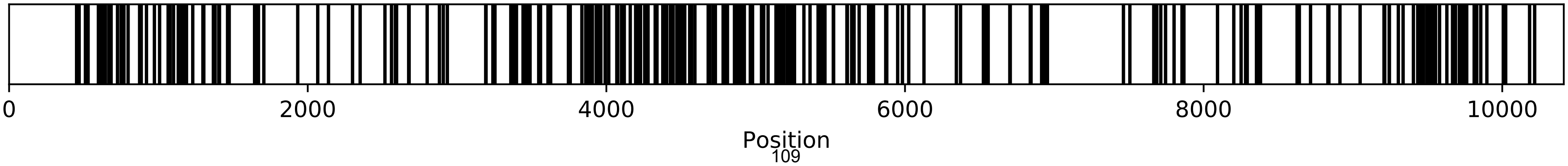

### Region: 683



CR1 vs CR2
Number of differences: 116

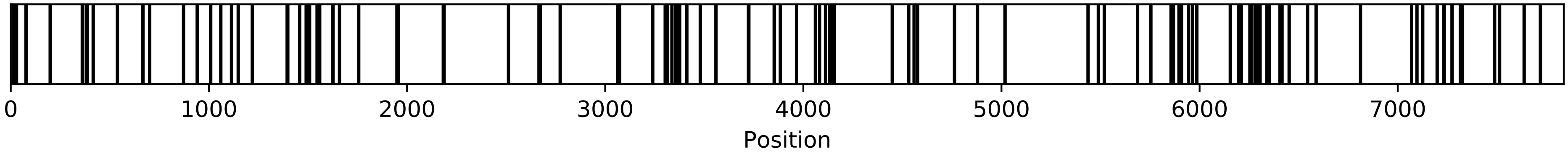


CR1 vs MA1
Number of differences: 91

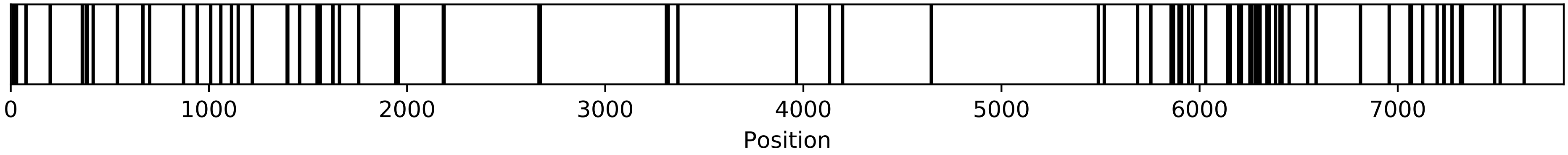


CR1 vs MA2
Number of differences: 29

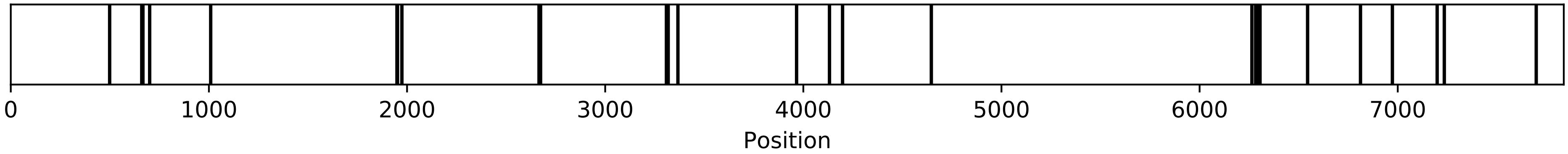


CR2 vs MA1
Number of differences: 53

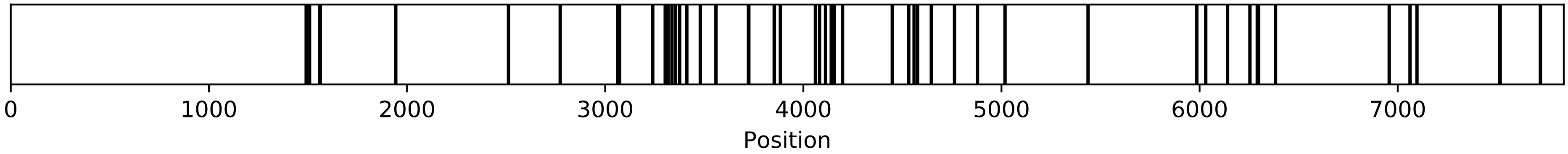


CR2 vs MA2
Number of differences: 111

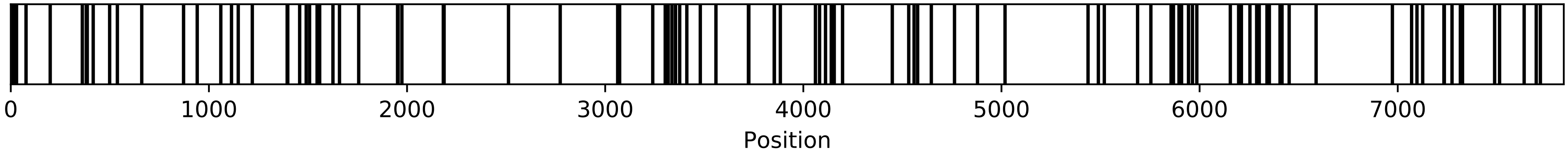


MA1 vs MA2
Number of differences: 80

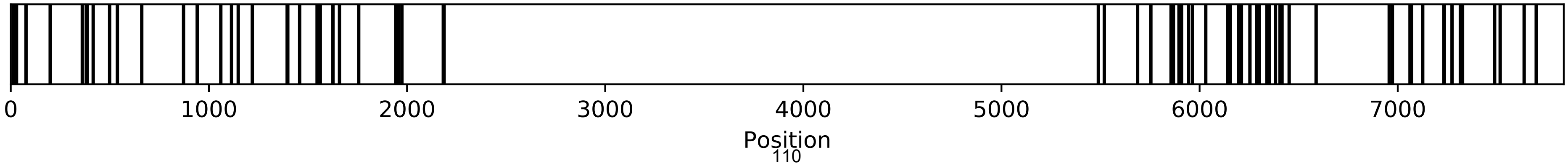

### Region: 5



## MA1 vs MA2
Number of differences: 526

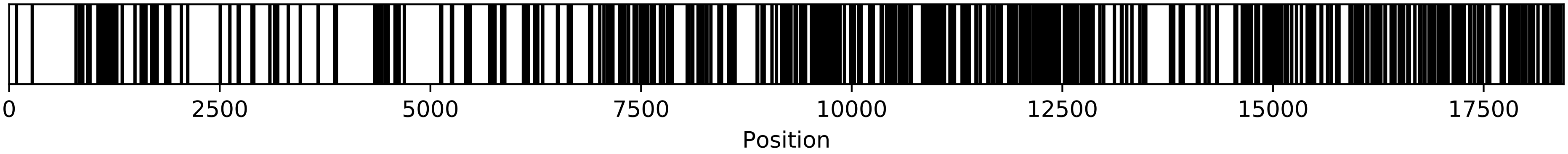


## MA1 vs MM1
Number of differences: 525

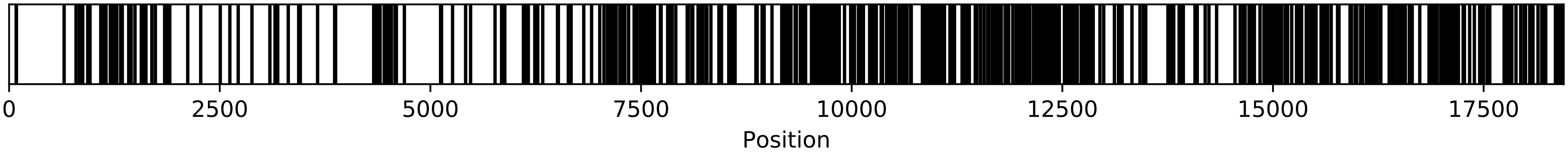


## MA1 vs MM2
Number of differences: 538

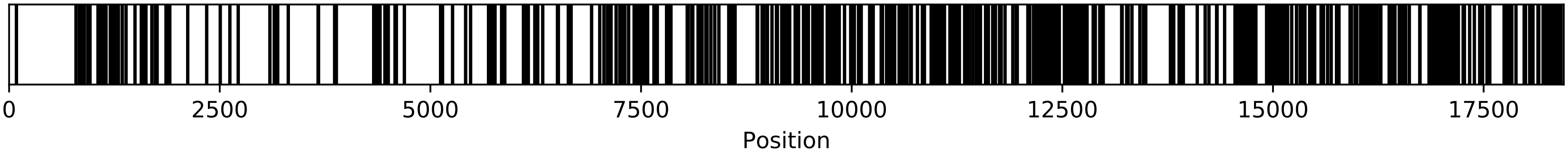


## MA2 vs MM1
Number of differences: 201

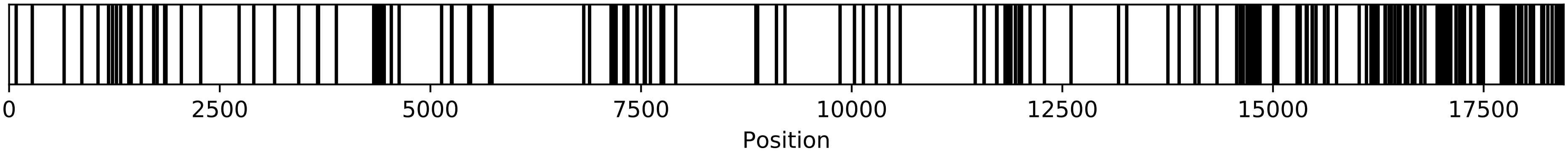


## MA2 vs MM2
Number of differences: 297

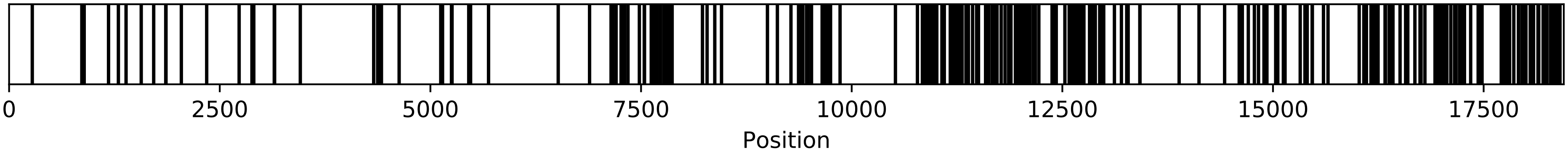


## MM1 vs MM2
Number of differences: 286

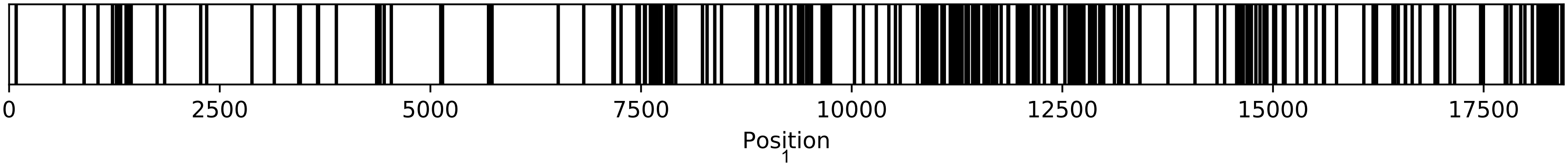

### Region: 12

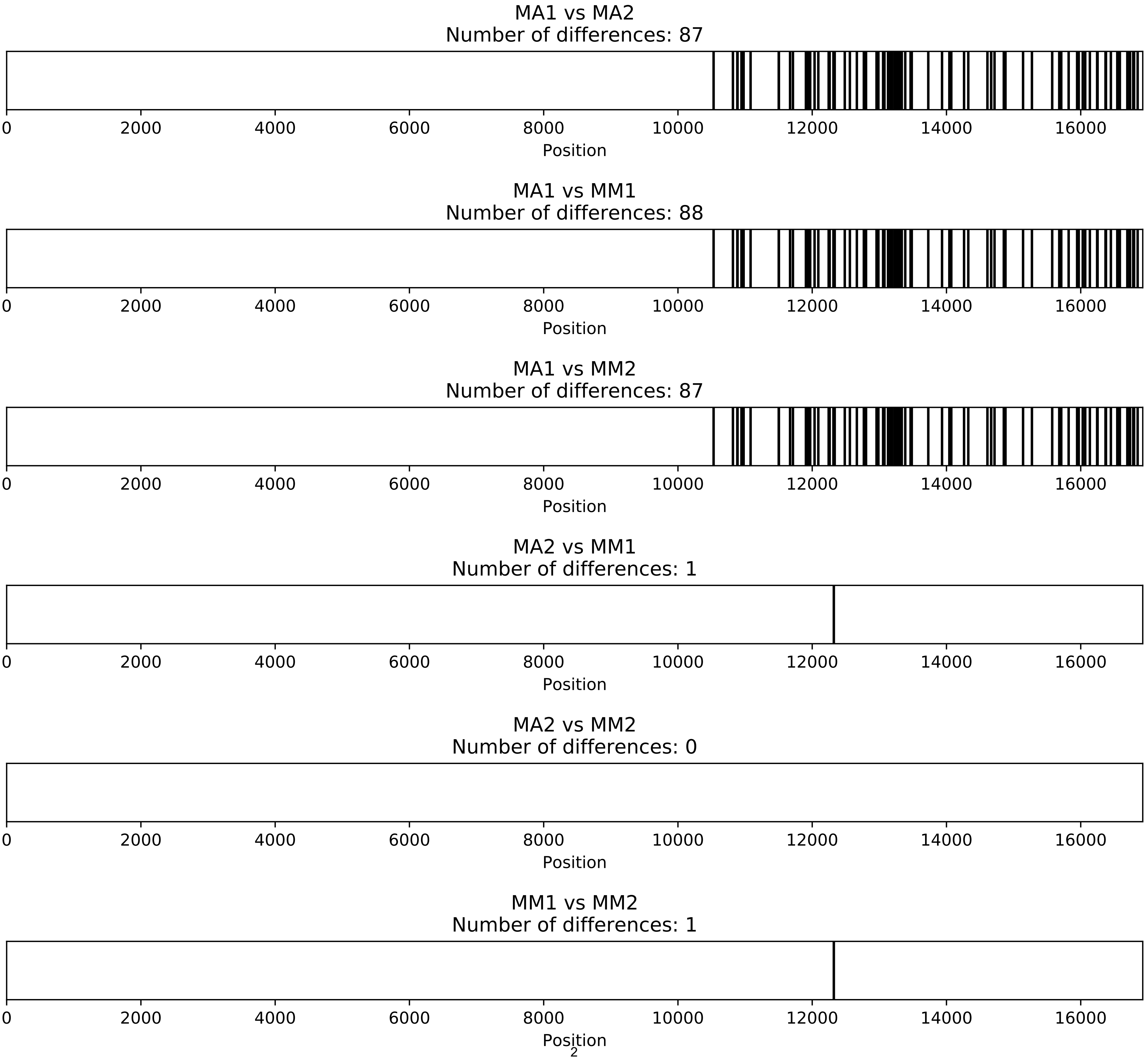
MA1 vs MA2
Number of differences: 87
MA1 vs MM1
Number of differences: 88
MA1 vs MM2
Number of differences: 87
MA2 vs MM1
Number of differences: 1
MA2 vs MM2
Number of differences: 0
MM1 vs MM2
Number of differences: 1
0
2000
4000
6000
8000
10000
12000
14000
16000
Position

### Region: 15



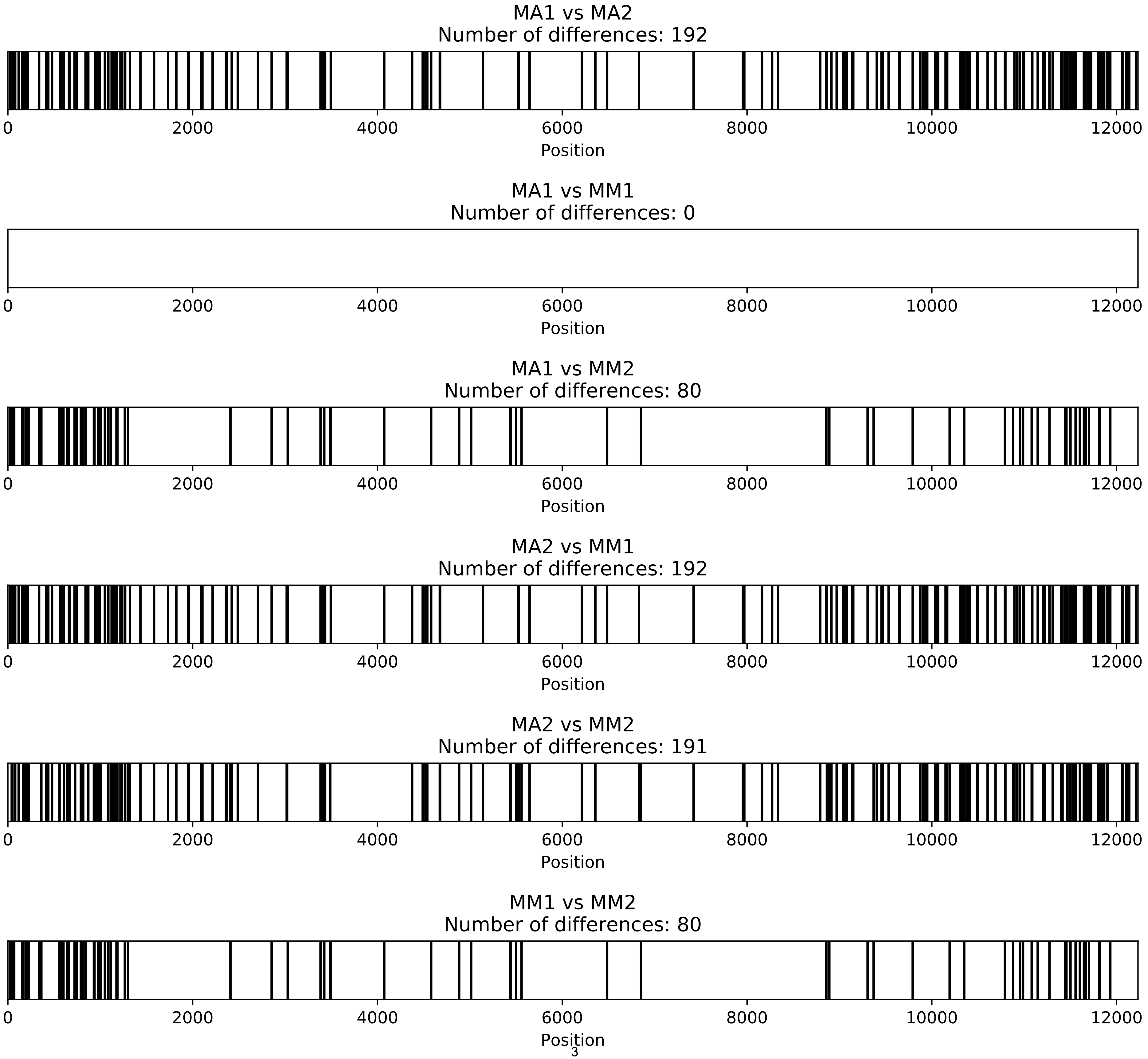
MA1 vs MA2
Number of differences: 192
0
2000
4000
6000
8000
10000
12000
Position
MA1 vs MM1
Number of differences: 0
0
2000
4000
6000
8000
10000
12000
Position
MA1 vs MM2
Number of differences: 80
0
2000
4000
6000
8000
10000
12000
Position
MA2 vs MM1
Number of differences: 192
0
2000
4000
6000
8000
10000
12000
Position
MA2 vs MM2
Number of differences: 191
0
2000
4000
6000
8000
10000
12000
Position
MM1 vs MM2
Number of differences: 80
0
2000
4000
6000
8000
10000
12000
Position

### Region: 20



MA1 vs MA2
Number of differences: 59

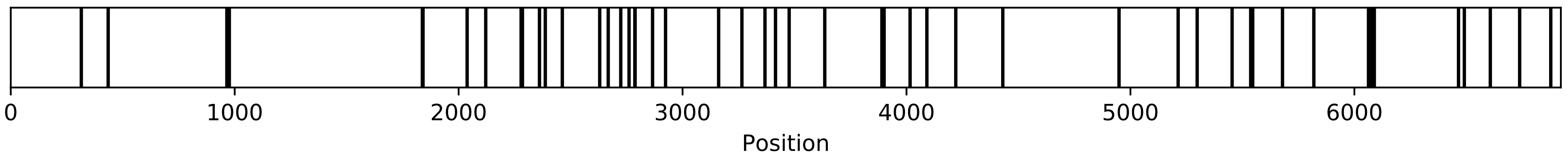


MA1 vs MM1
Number of differences: 62

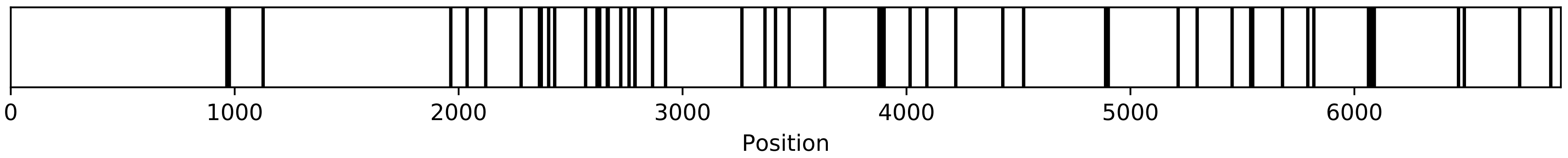


MA1 vs MM2
Number of differences: 59

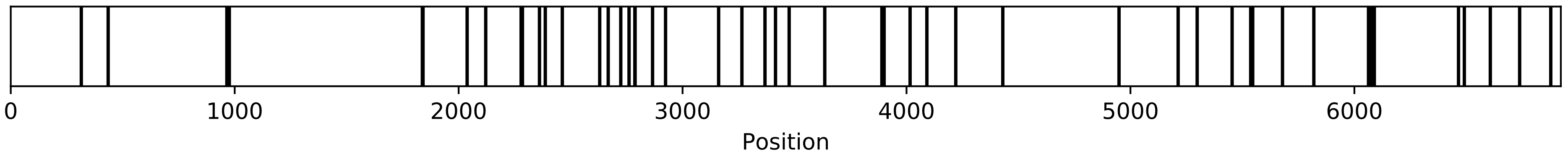


MA2 vs MM1
Number of differences: 24

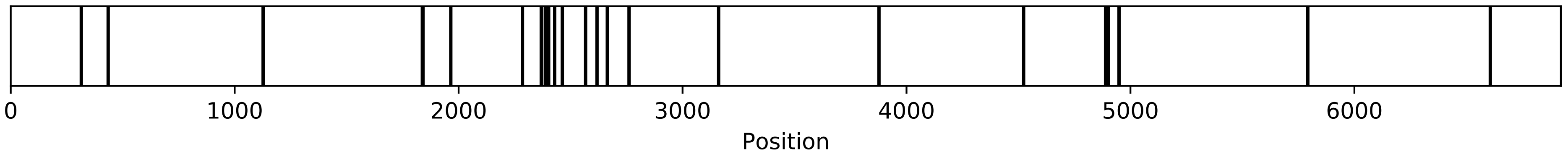


MA2 vs MM2
Number of differences: 0

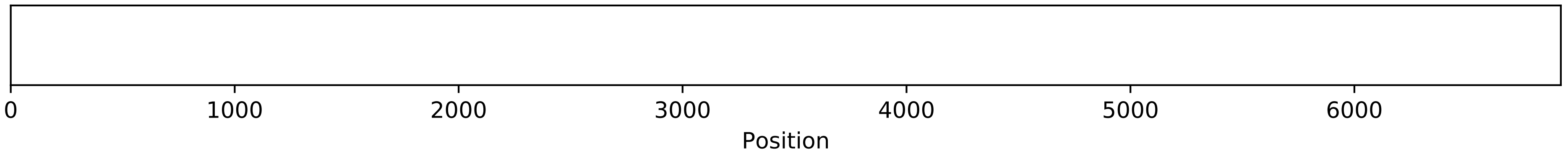


MM1 vs MM2
Number of differences: 24

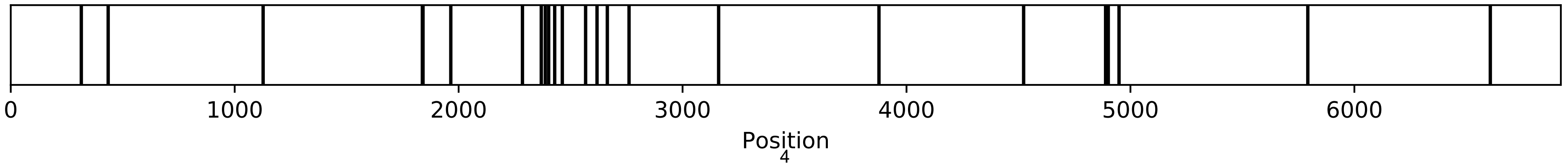

### Region: 22



MA1 vs MA2
Number of differences: 193

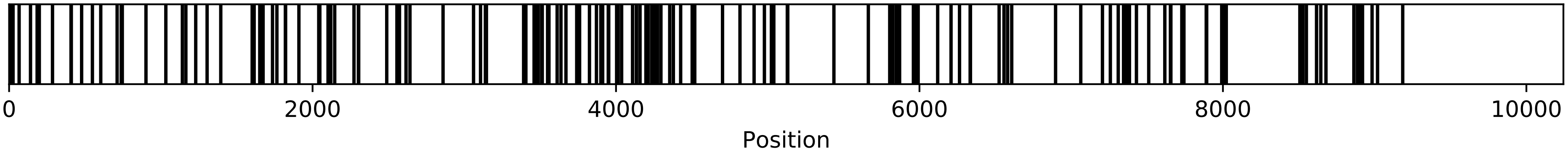


MA1 vs MM1
Number of differences: 193

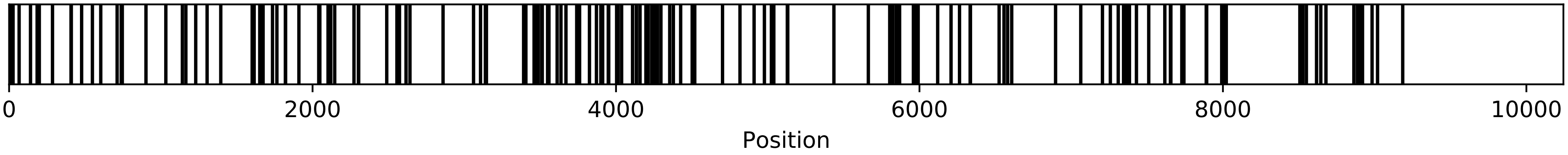


MA1 vs MM2
Number of differences: 235

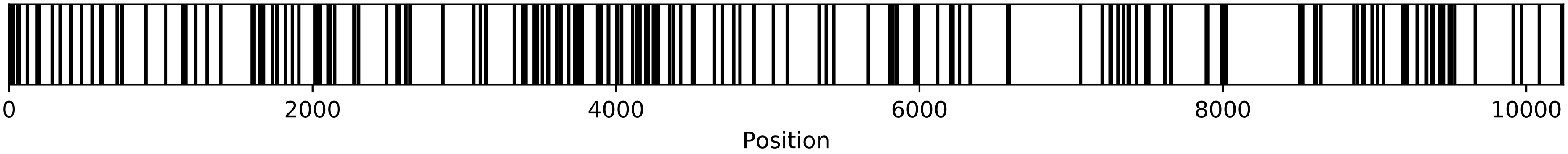


MA2 vs MM1
Number of differences: 0

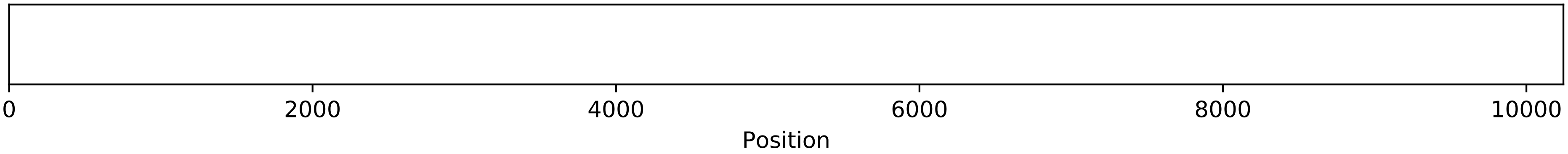


MA2 vs MM2
Number of differences: 117

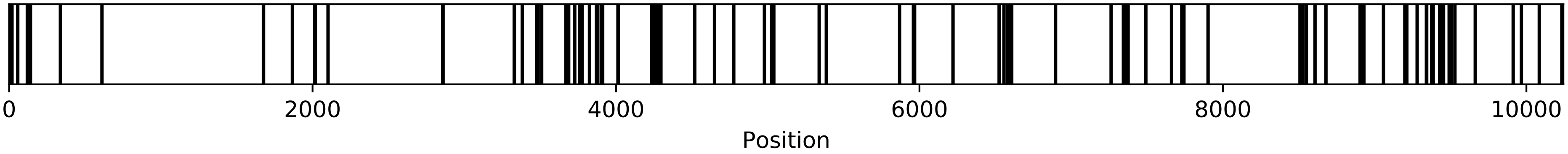


MM1 vs MM2
Number of differences: 117

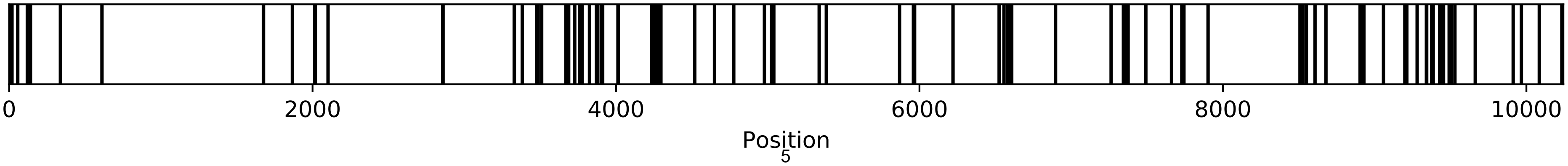

### Region: 24



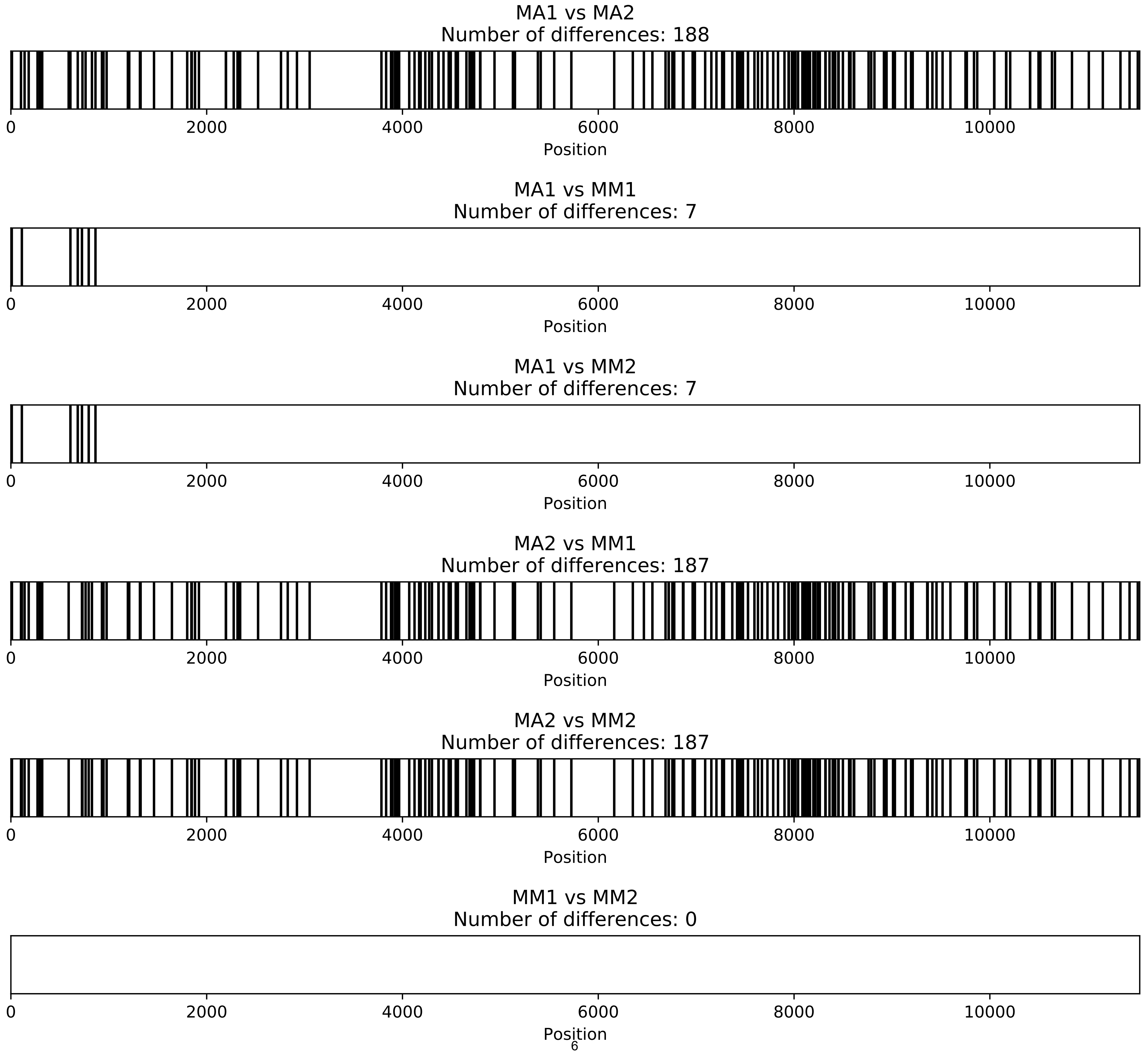
MA1 vs MA2
Number of differences: 188
MA1 vs MM1
Number of differences: 7
MA1 vs MM2
Number of differences: 7
MA2 vs MM1
Number of differences: 187
MA2 vs MM2
Number of differences: 187
MM1 vs MM2
Number of differences: 0
0
2000
4000
6000
8000
10000
Position

### Region: 26



MA1 vs MA2
Number of differences: 96

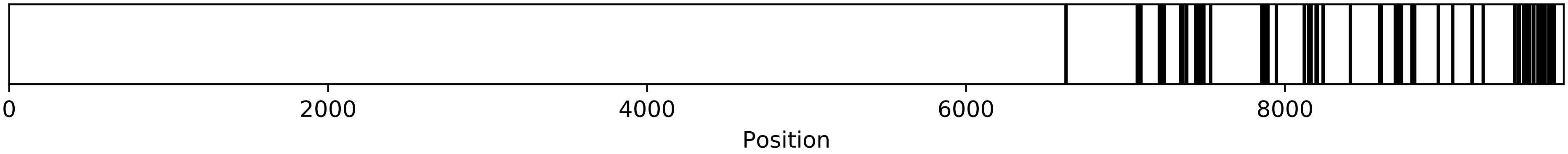


MA1 vs MM1
Number of differences: 232

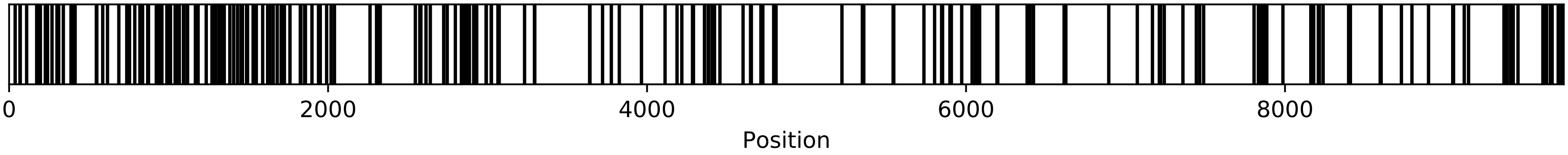


MA1 vs MM2
Number of differences: 153

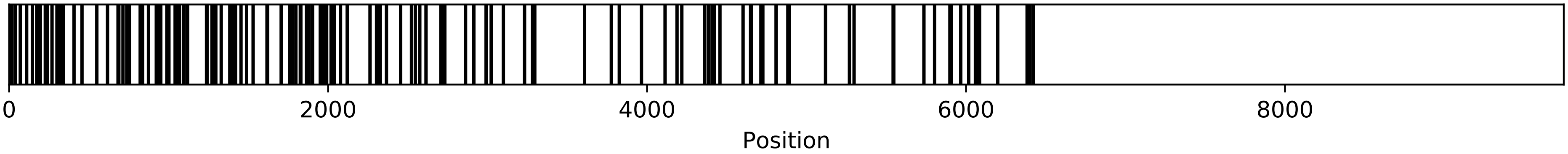


MA2 vs MM1
Number of differences: 286

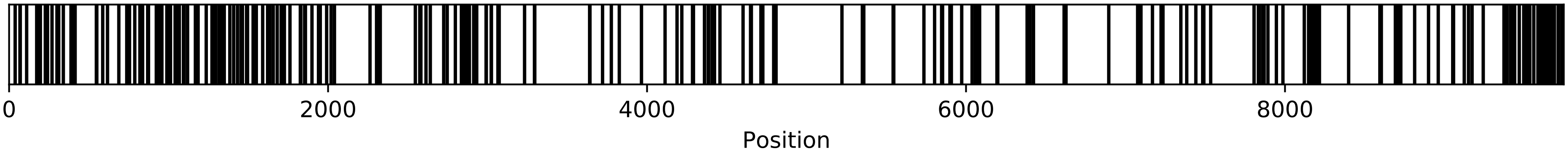


MA2 vs MM2
Number of differences: 249

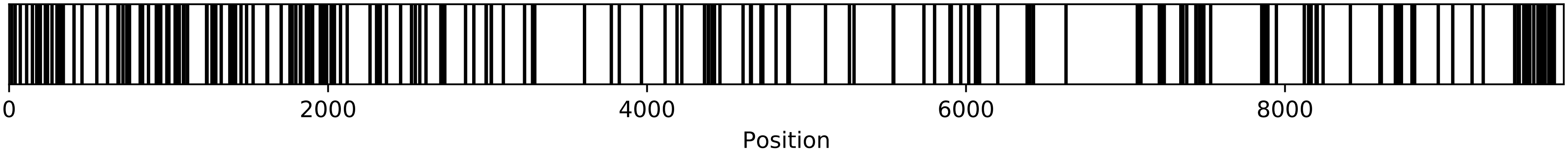


MM1 vs MM2
Number of differences: 177

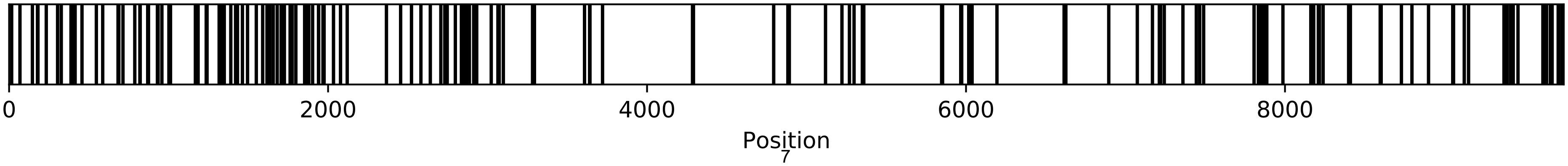

### Region: 27



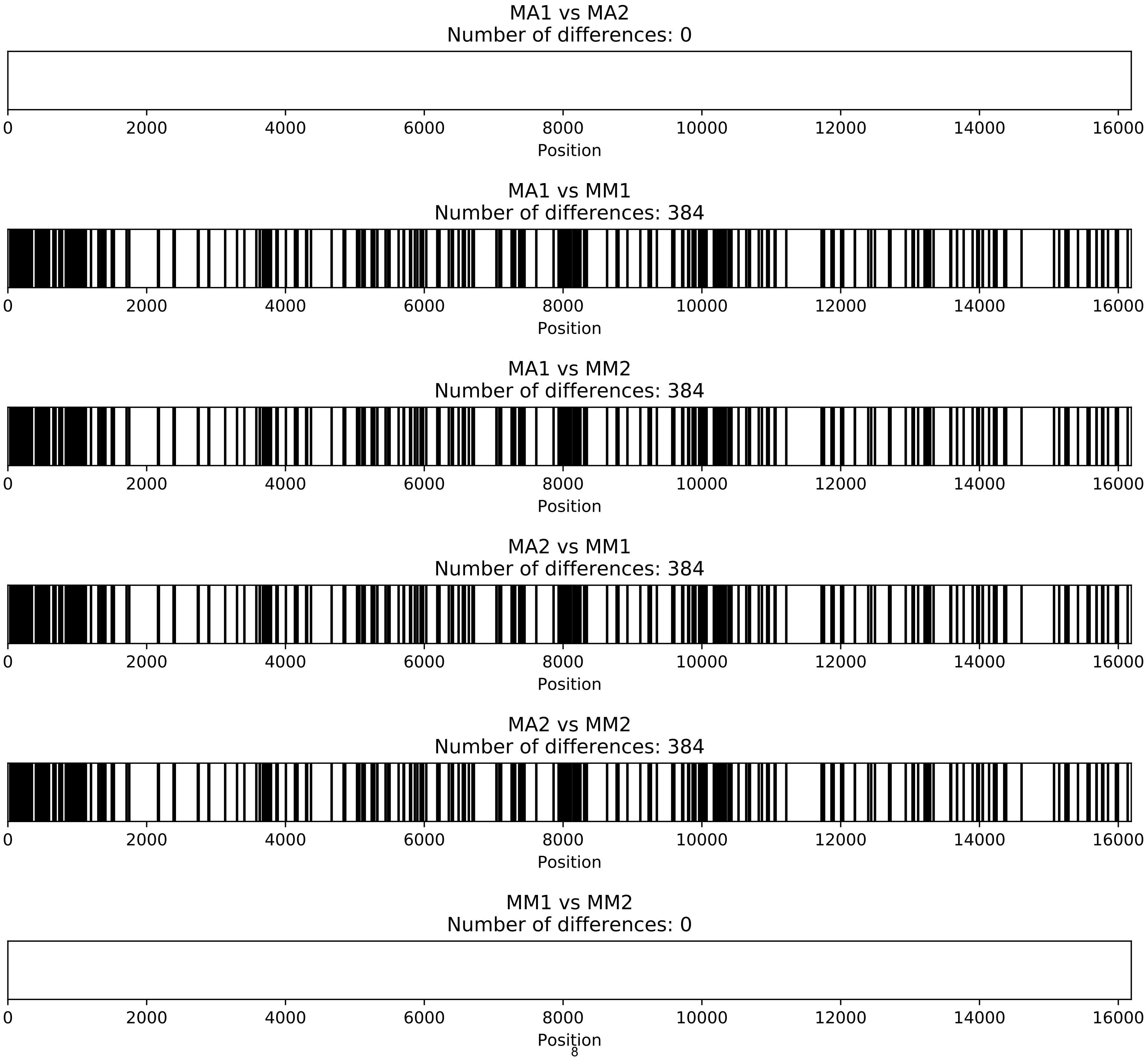
MA1 vs MA2
Number of differences: 0
MA1 vs MM1
Number of differences: 384
MA1 vs MM2
Number of differences: 384
MA2 vs MM1
Number of differences: 384
MA2 vs MM2
Number of differences: 384
MM1 vs MM2
Number of differences: 0
0 2000 4000 6000 8000 10000 12000 14000 16000
Position

### Region: 31



MA1 vs MA2
Number of differences: 295

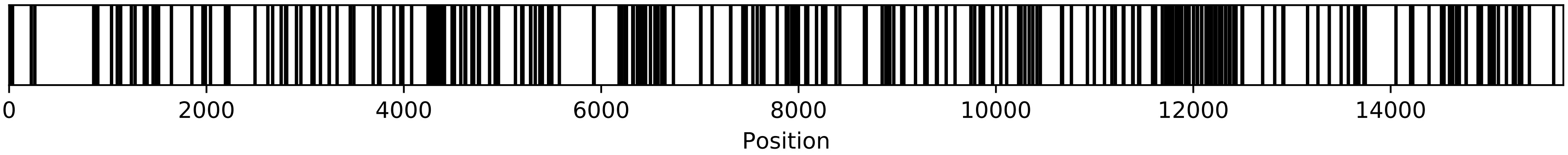


MA1 vs MM1
Number of differences: 295

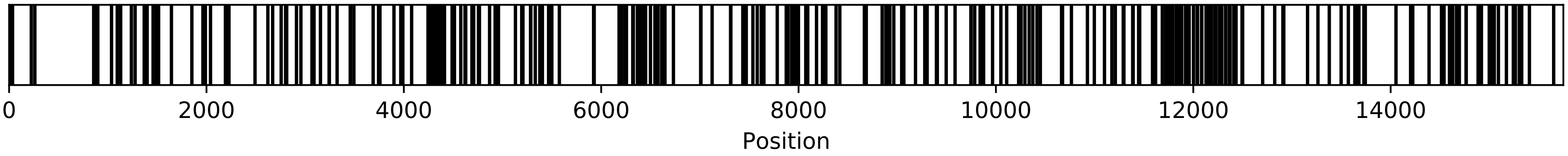


MA1 vs MM2
Number of differences: 311

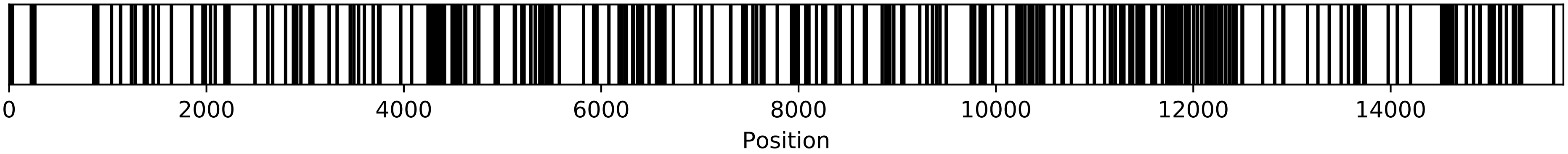


MA2 vs MM1
Number of differences: 0

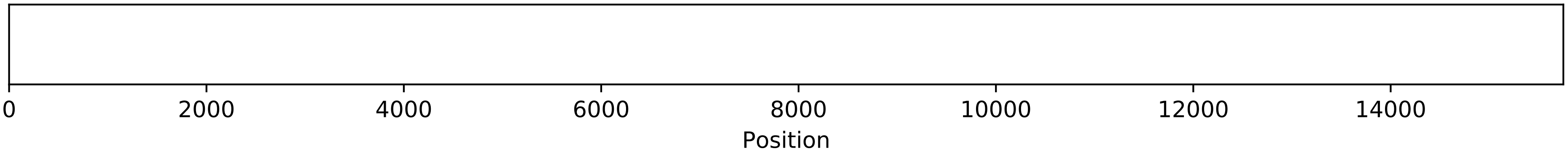


MA2 vs MM2
Number of differences: 118

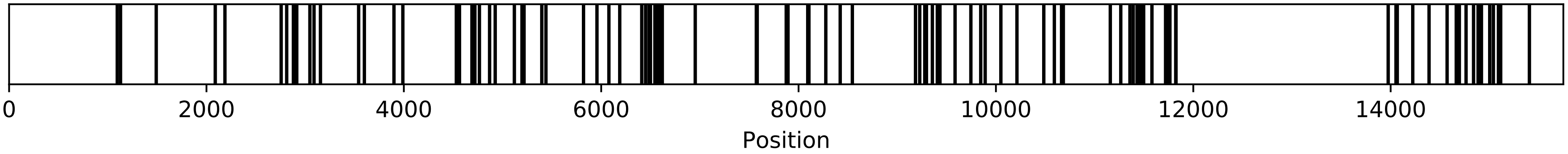


MM1 vs MM2
Number of differences: 118

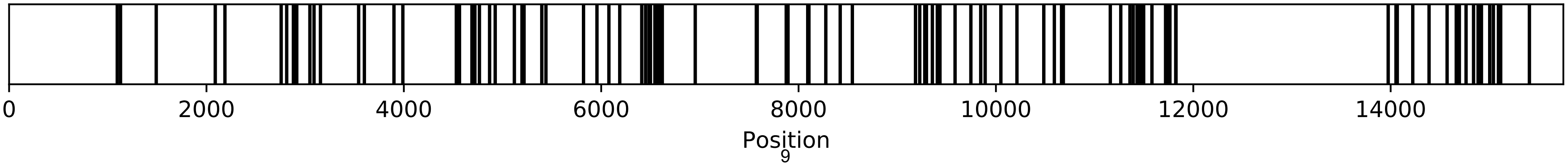

### Region: 32



MA1 vs MA2
Number of differences: 115

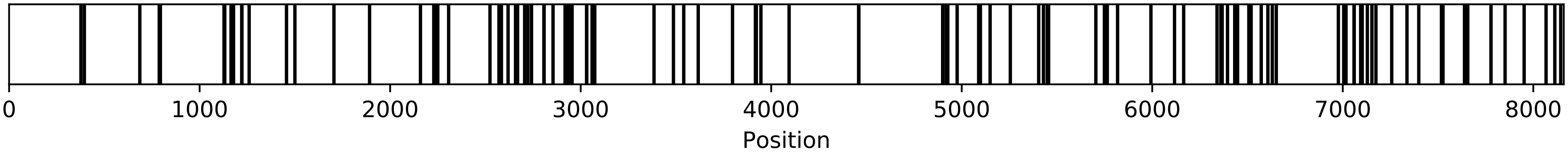


MA1 vs MM1
Number of differences: 57

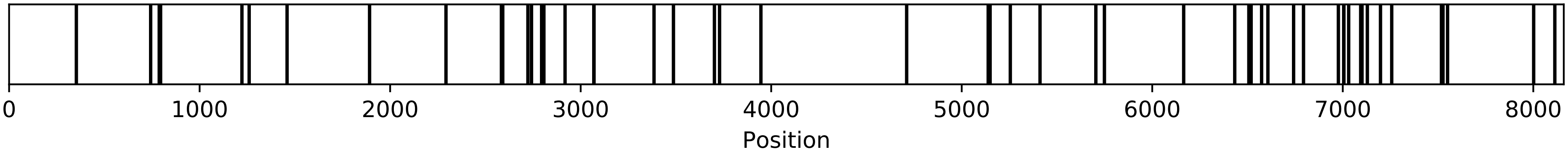


MA1 vs MM2
Number of differences: 0

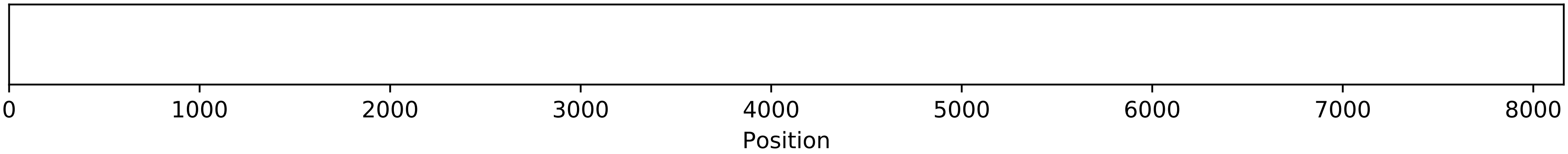


MA2 vs MM1
Number of differences: 103

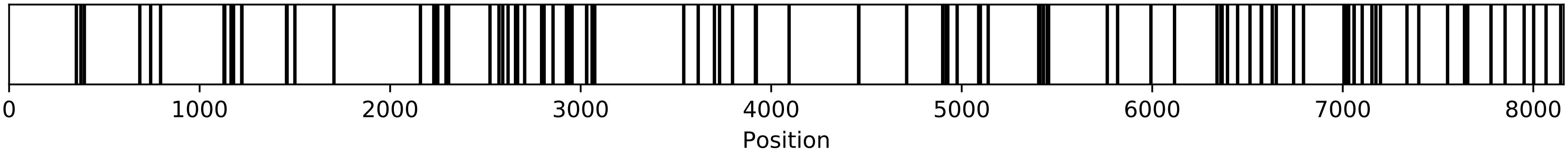


MA2 vs MM2
Number of differences: 115

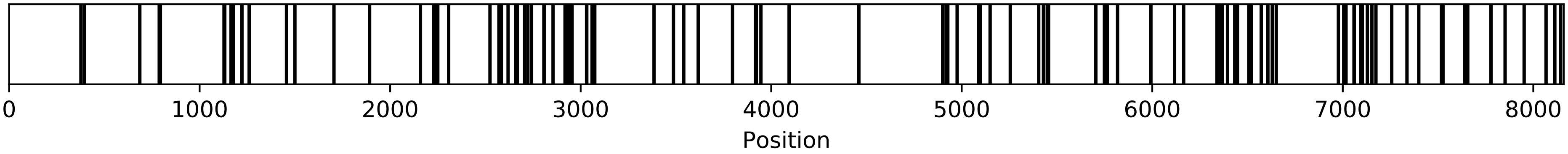


MM1 vs MM2
Number of differences: 57

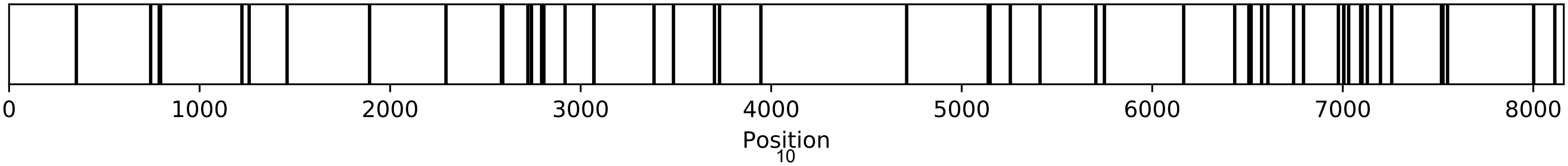

### Region: 33



MA1 vs MA2
Number of differences: 222

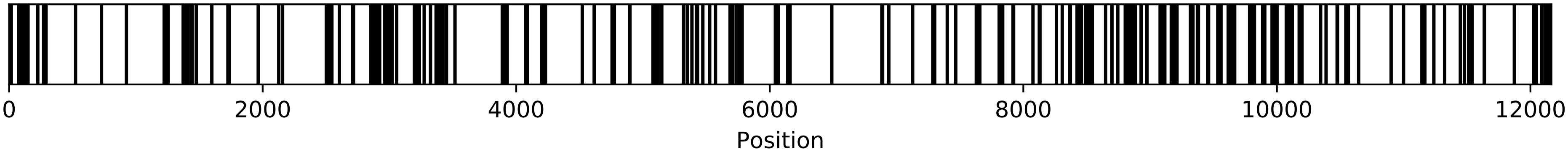


MA1 vs MM1
Number of differences: 233

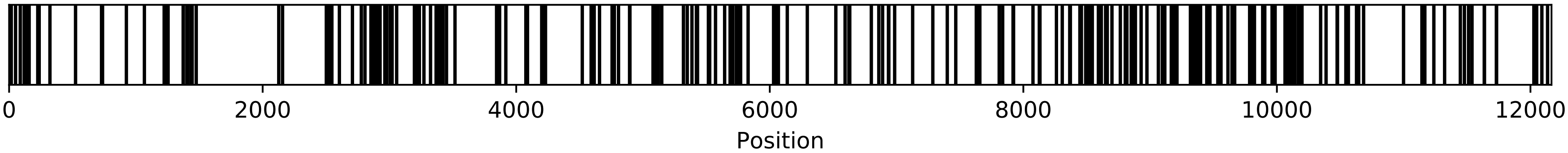


MA1 vs MM2
Number of differences: 245

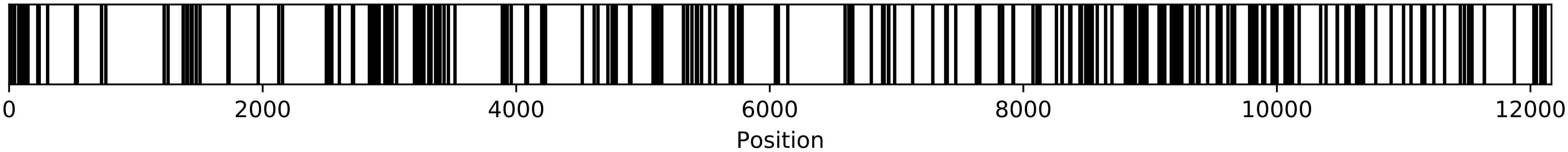


MA2 vs MM1
Number of differences: 112

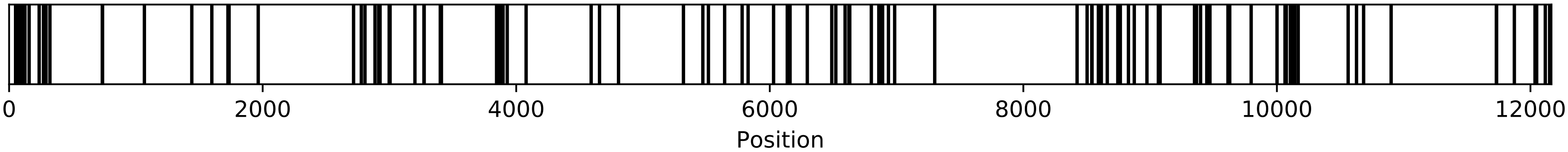


MA2 vs MM2
Number of differences: 102

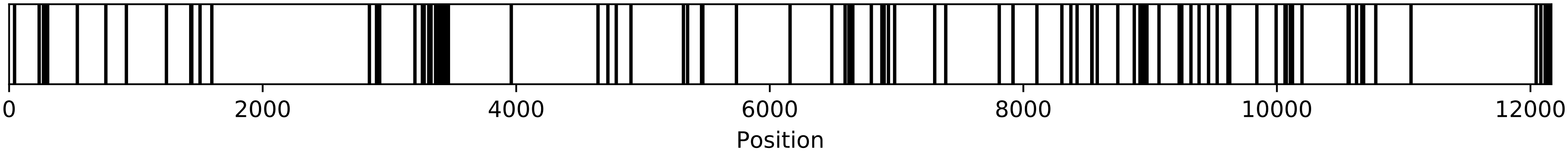


MM1 vs MM2
Number of differences: 128

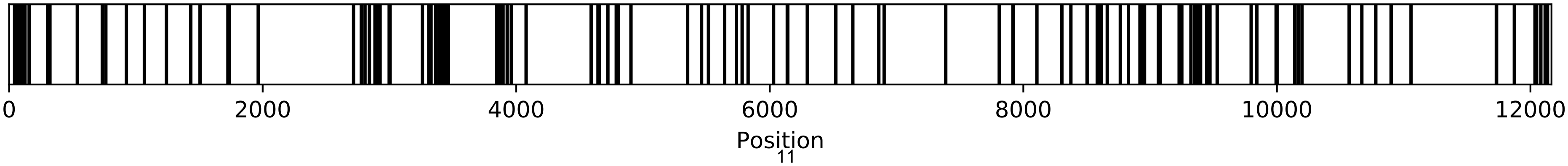

### Region: 49



MA1 vs MA2
Number of differences: 198

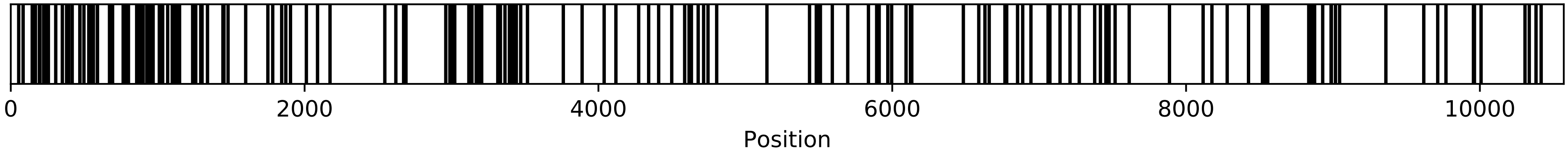


MA1 vs MM1
Number of differences: 79

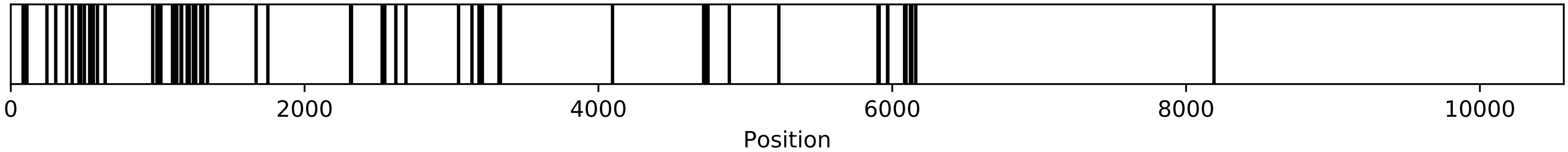


MA1 vs MM2
Number of differences: 7

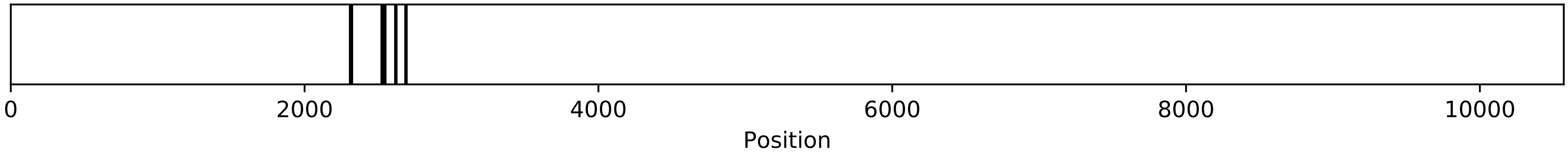


MA2 vs MM1
Number of differences: 208

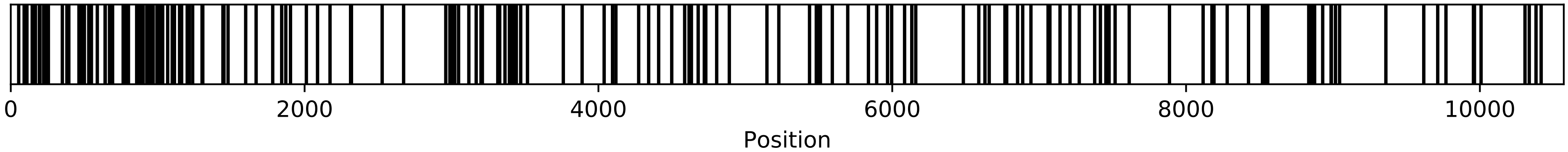


MA2 vs MM2
Number of differences: 199

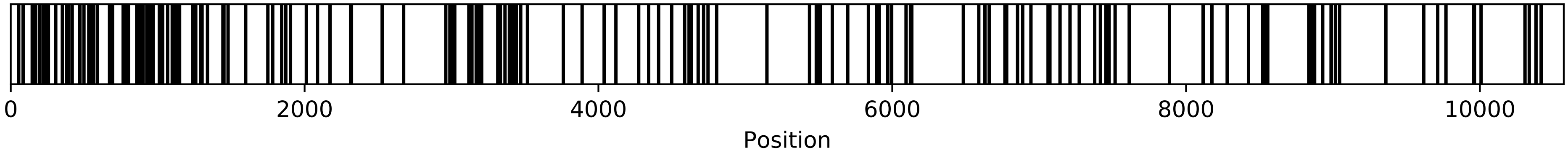


MM1 vs MM2
Number of differences: 72

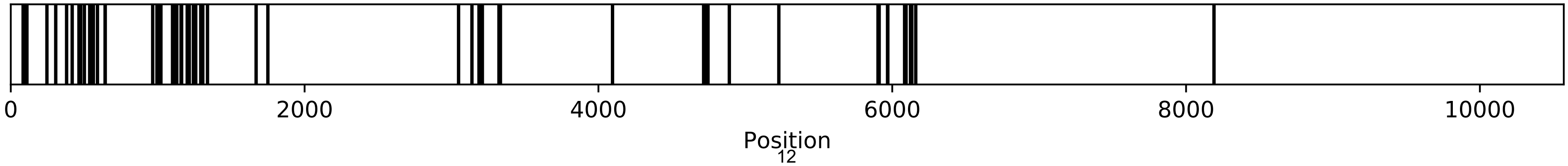

### Region: 50



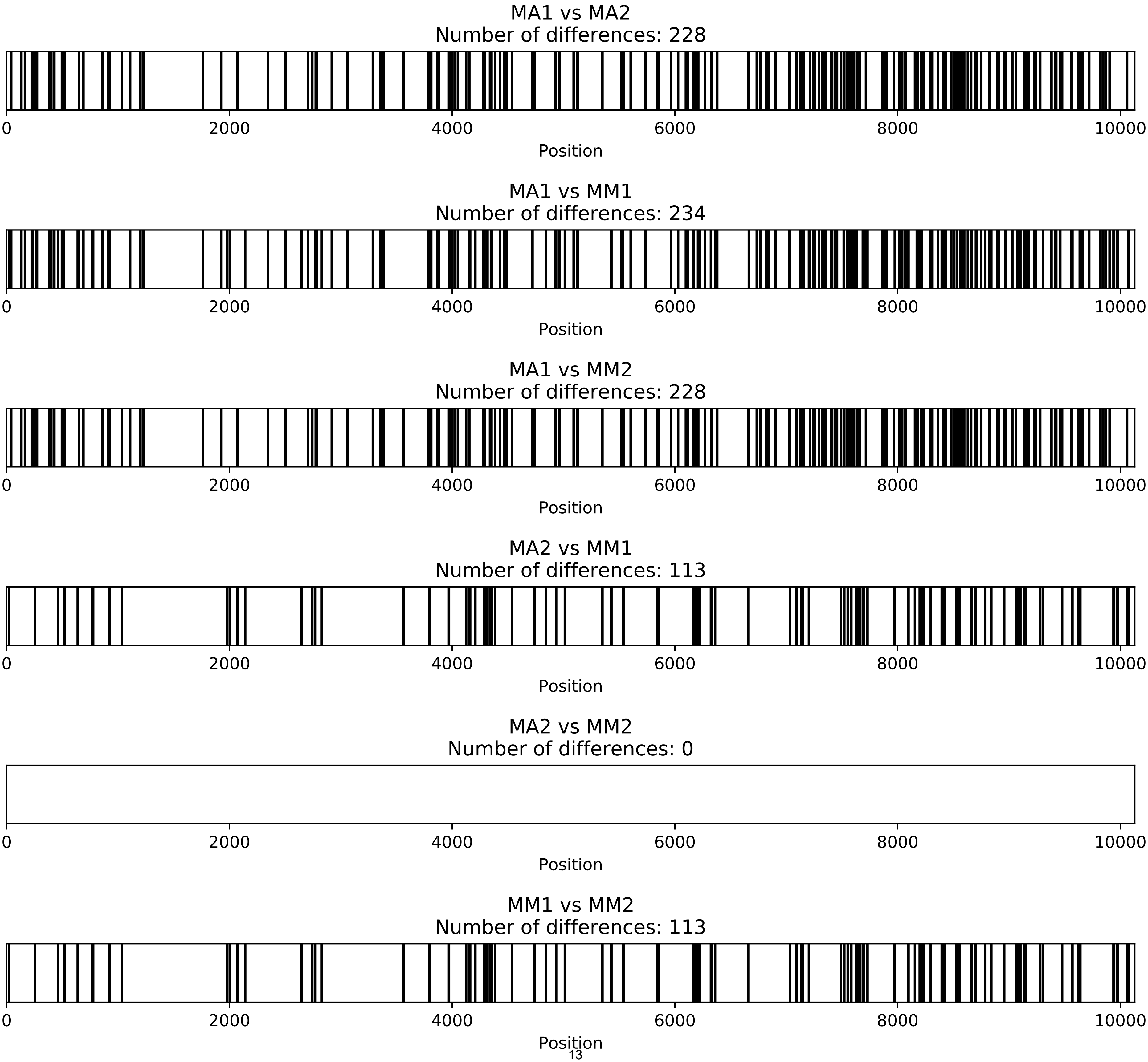
MA1 vs MA2
Number of differences: 228
MA1 vs MM1
Number of differences: 234
MA1 vs MM2
Number of differences: 228
MA2 vs MM1
Number of differences: 113
MA2 vs MM2
Number of differences: 0
MM1 vs MM2
Number of differences: 113
0
2000
4000
6000
8000
10000
Position

### Region: 59



MA1 vs MA2
Number of differences: 90

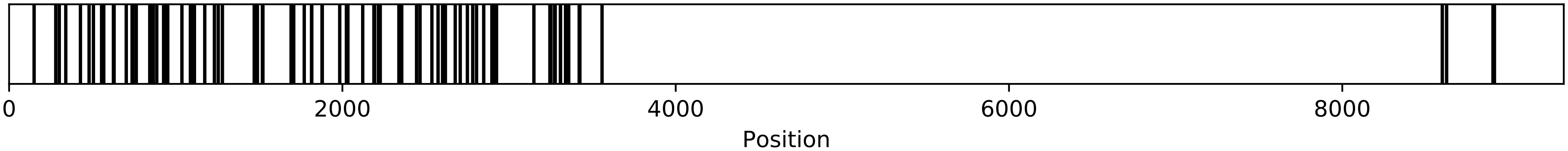


MA1 vs MM1
Number of differences: 211

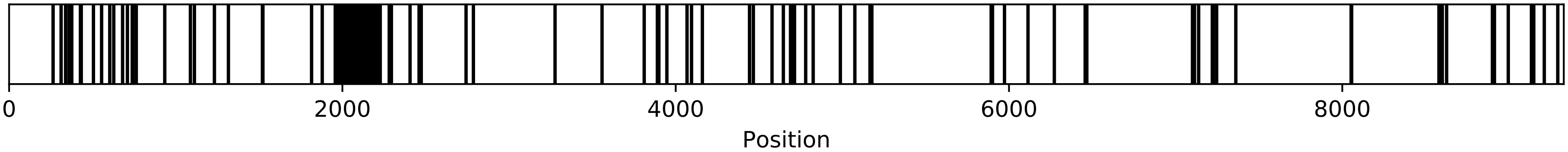


MA1 vs MM2
Number of differences: 9

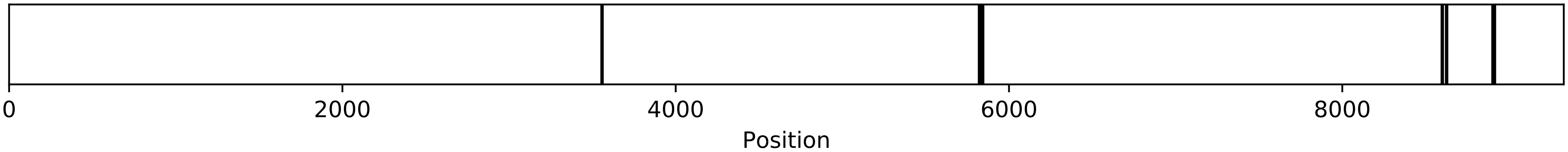


MA2 vs MM1
Number of differences: 250

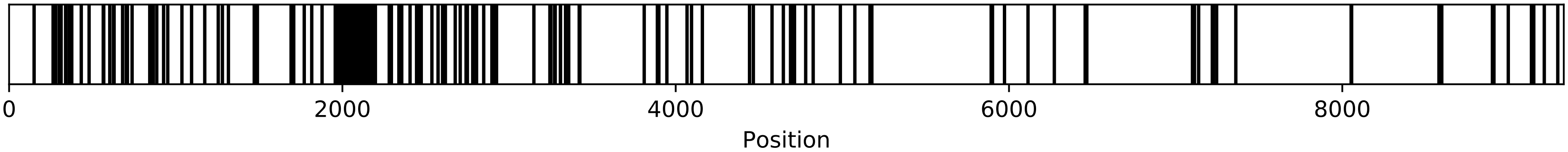


MA2 vs MM2
Number of differences: 85

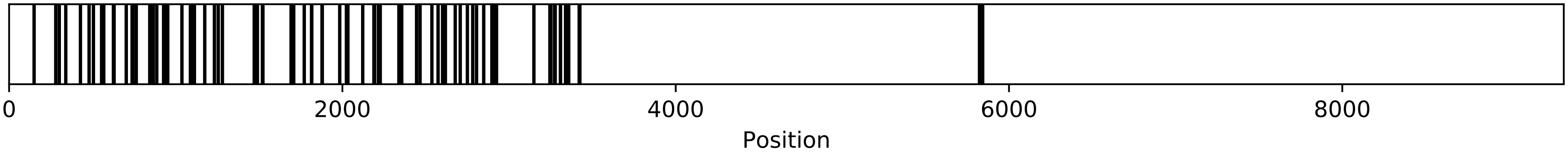


MM1 vs MM2
Number of differences: 210

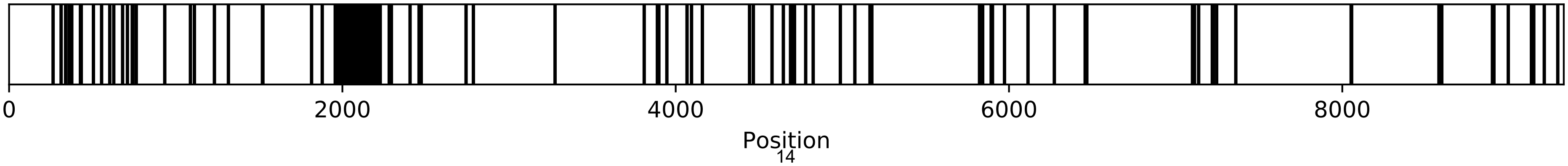

### Region: 61



MA1 vs MA2
Number of differences: 371

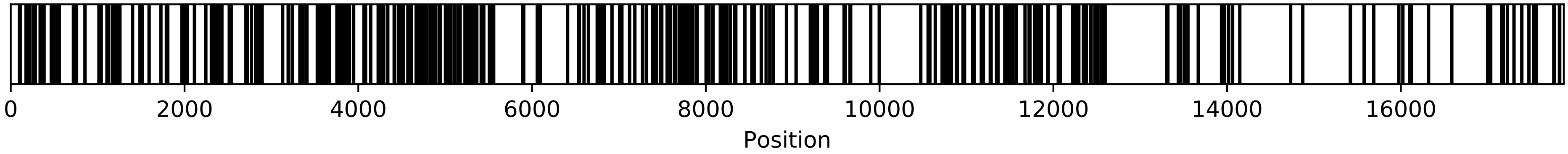


MA1 vs MM1
Number of differences: 371

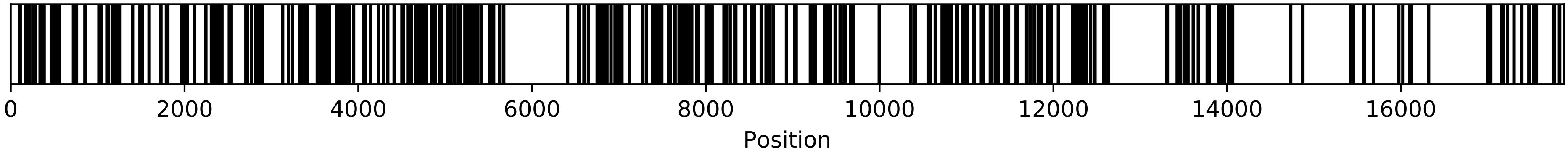


MA1 vs MM2
Number of differences: 373

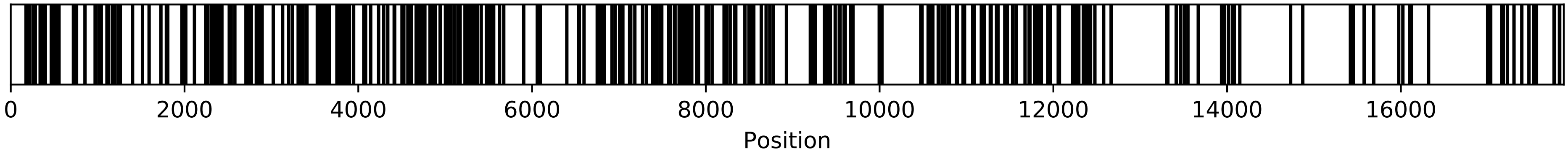


MA2 vs MM1
Number of differences: 120

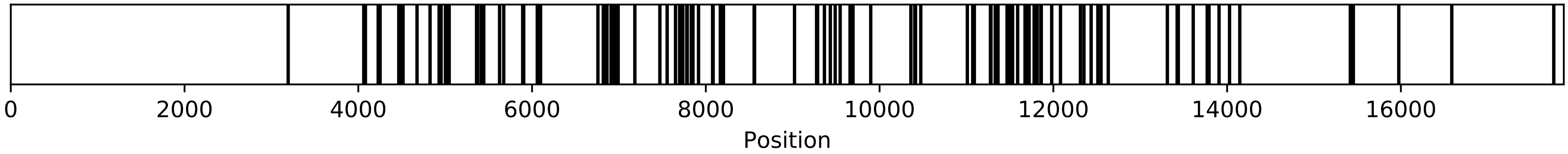


MA2 vs MM2
Number of differences: 153

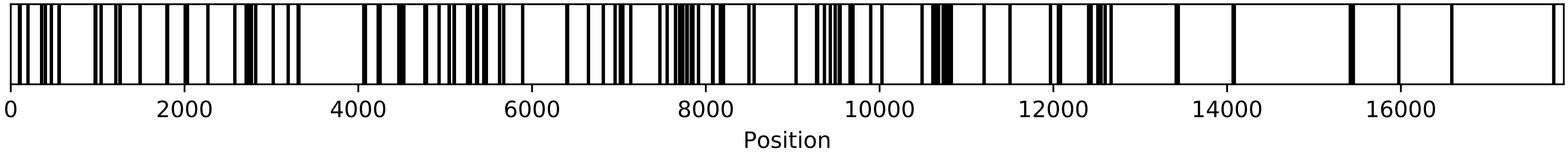


MM1 vs MM2
Number of differences: 150

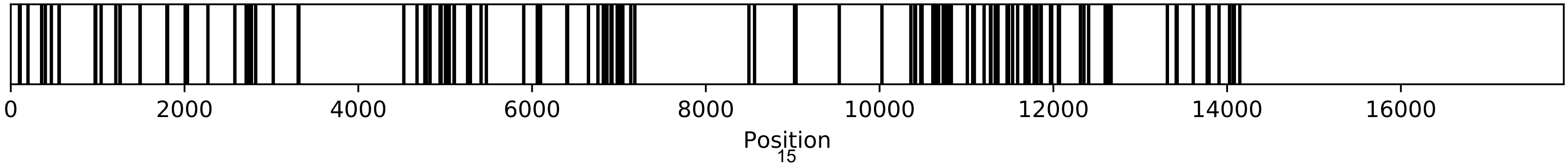

### Region: 65



MA1 vs MA2
Number of differences: 440

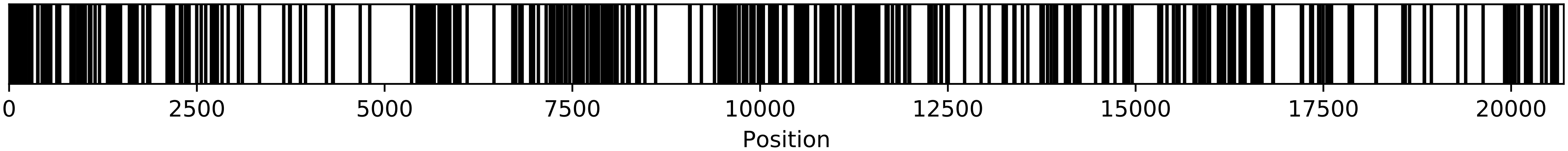


MA1 vs MM1
Number of differences: 236

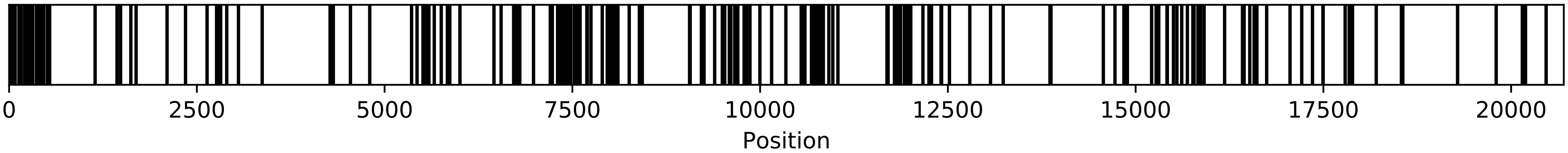


MA1 vs MM2
Number of differences: 229

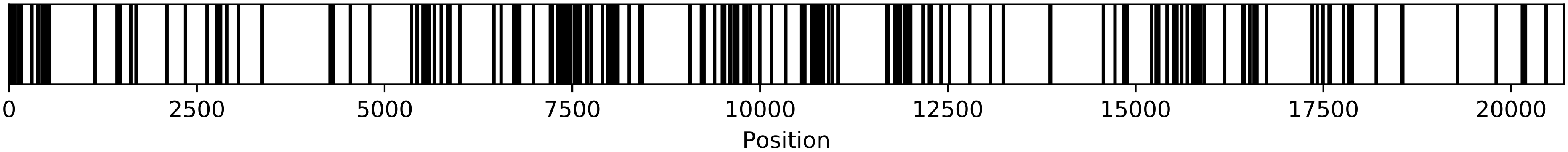


MA2 vs MM1
Number of differences: 442

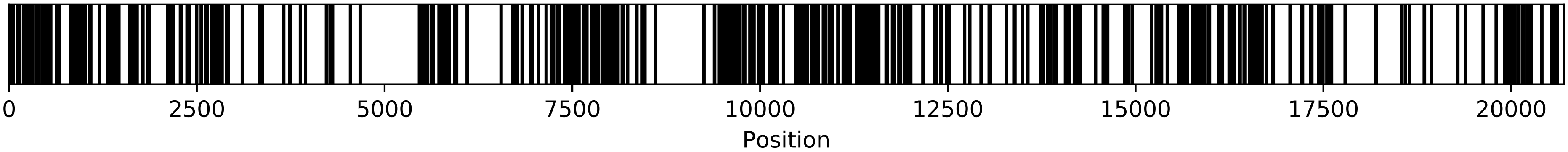


MA2 vs MM2
Number of differences: 437

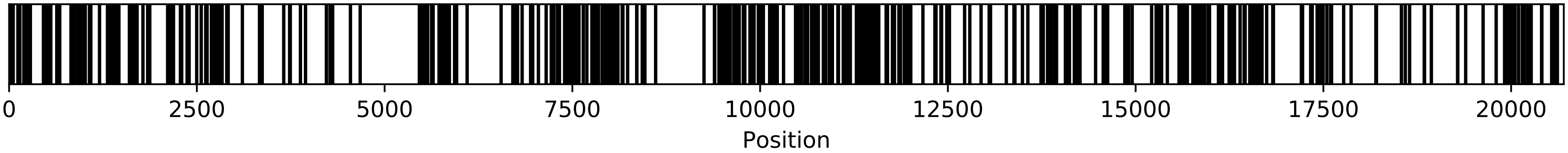


MM1 vs MM2
Number of differences: 27

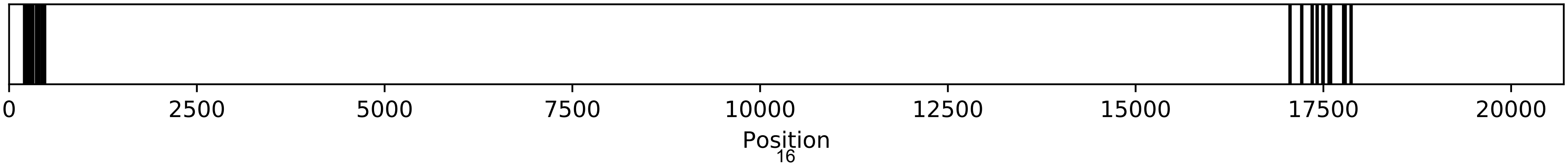

### Region: 67



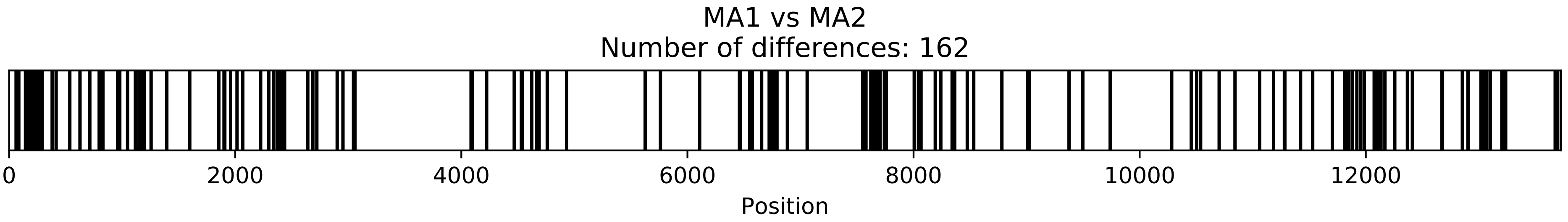
MA1 vs MA2
Number of differences: 162
0
2000
4000
6000
8000
10000
12000
Position

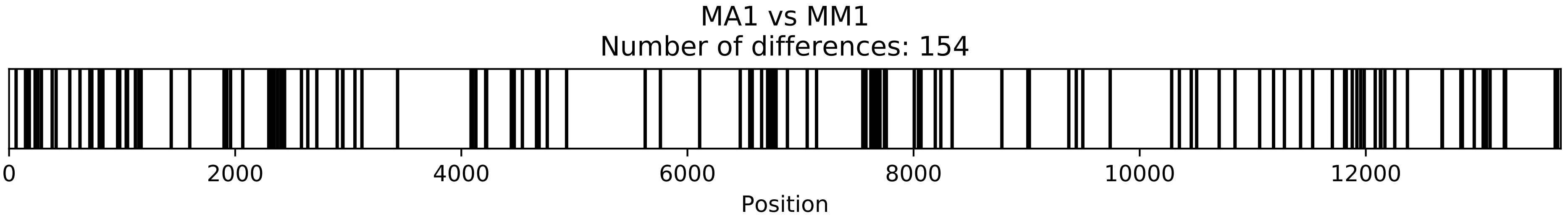
MA1 vs MM1
Number of differences: 154
0
2000
4000
6000
8000
10000
12000
Position

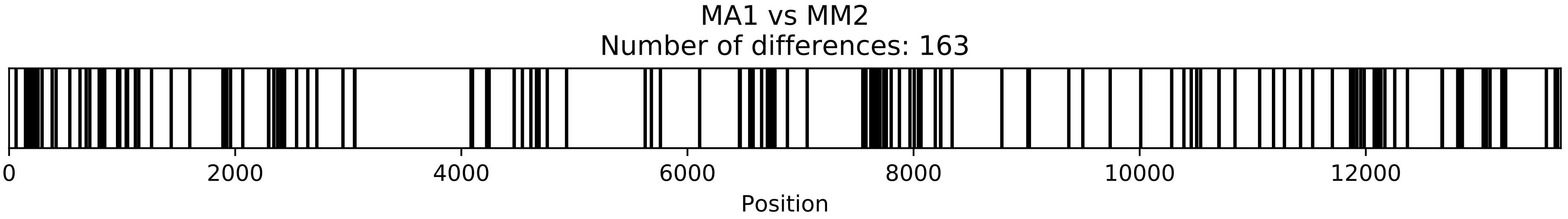
MA1 vs MM2
Number of differences: 163
0
2000
4000
6000
8000
10000
12000
Position

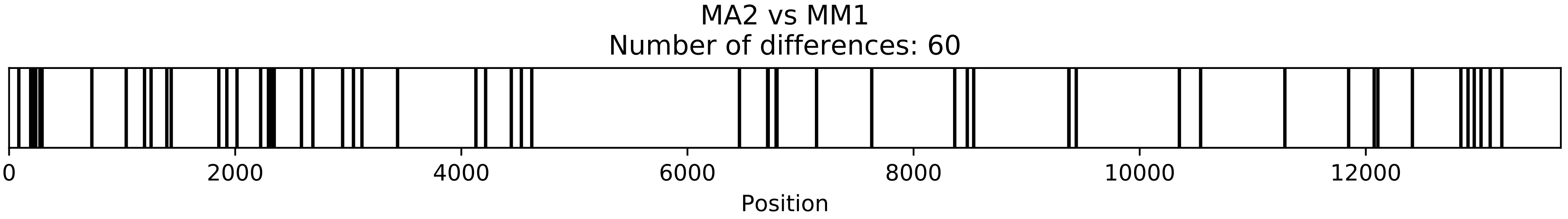
MA2 vs MM1
Number of differences: 60
0
2000
4000
6000
8000
10000
12000
Position

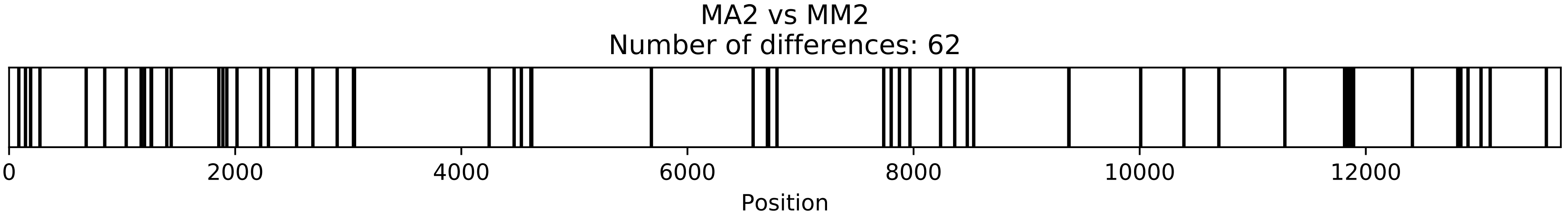
MA2 vs MM2
Number of differences: 62
0
2000
4000
6000
8000
10000
12000
Position

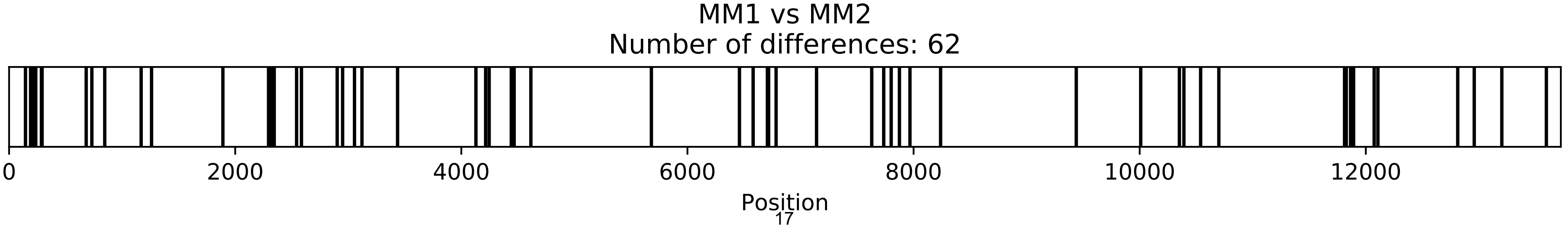
MM1 vs MM2
Number of differences: 62
0
2000
4000
6000
8000
10000
12000
Position

### Region: 71



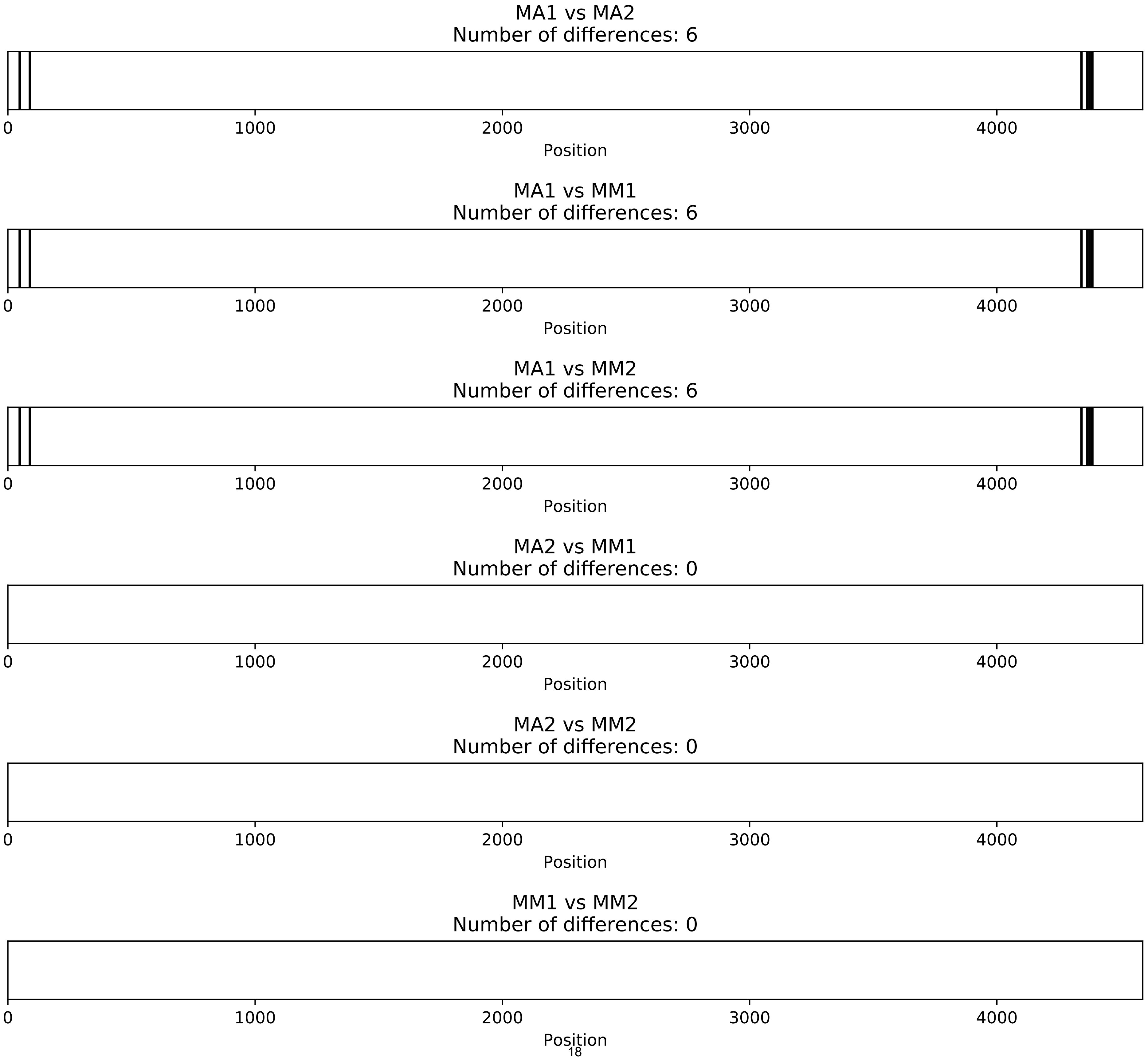
MA1 vs MA2
Number of differences: 6
0
1000
2000
3000
4000
Position
MA1 vs MM1
Number of differences: 6
Position
MA1 vs MM2
Number of differences: 6
Position
MA2 vs MM1
Number of differences: 0
Position
MA2 vs MM2
Number of differences: 0
Position
MM1 vs MM2
Number of differences: 0
Position

### Region: 72



MA1 vs MA2
Number of differences: 209

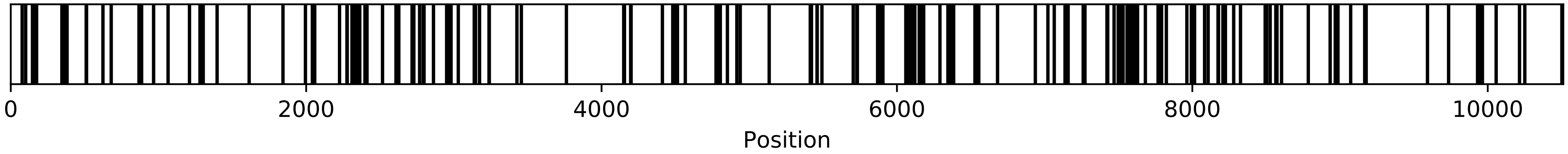


MA1 vs MM1
Number of differences: 209

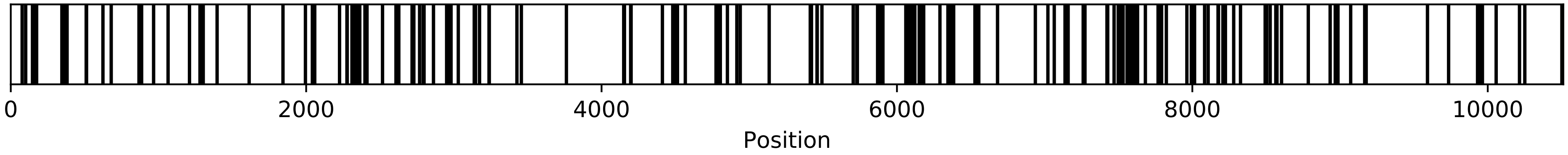


MA1 vs MM2
Number of differences: 198

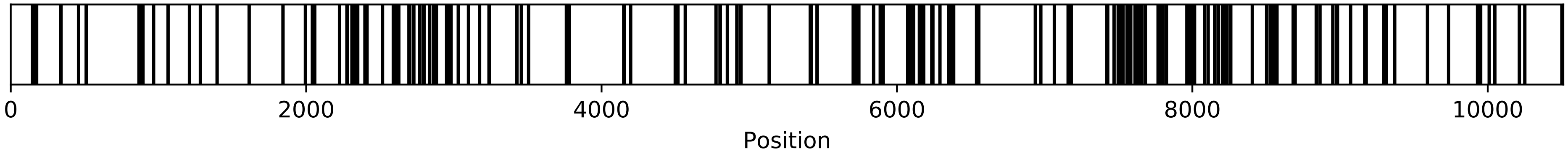


MA2 vs MM1
Number of differences: 0

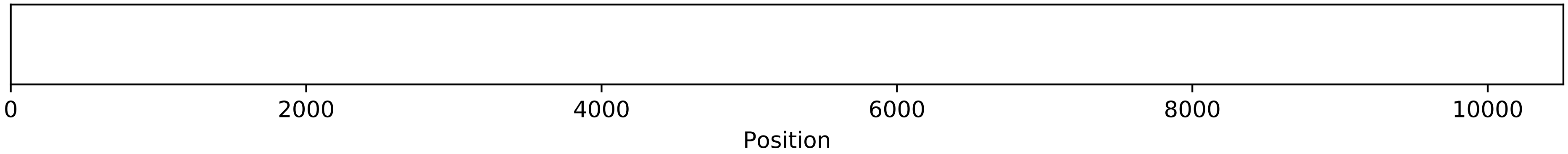


MA2 vs MM2
Number of differences: 129

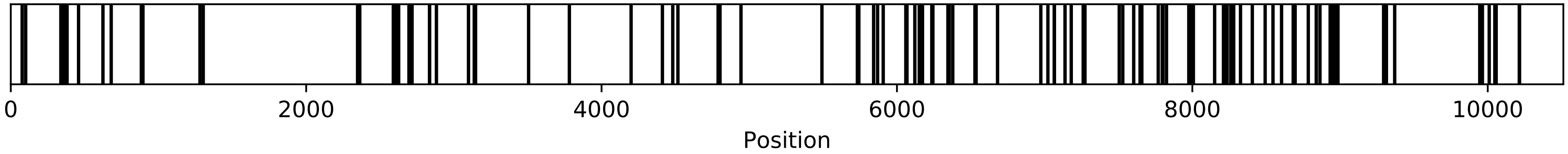


MM1 vs MM2
Number of differences: 129

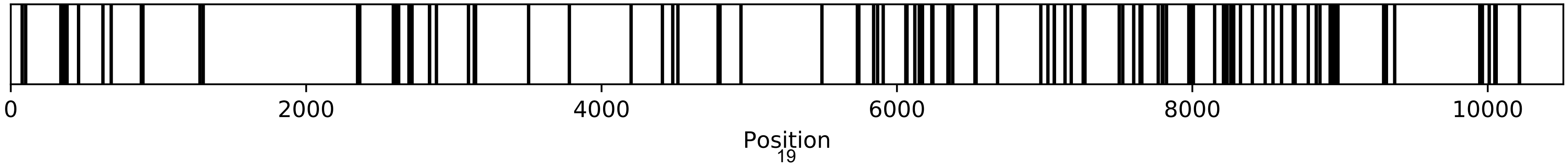

### Region: 83



MA1 vs MA2
Number of differences: 279

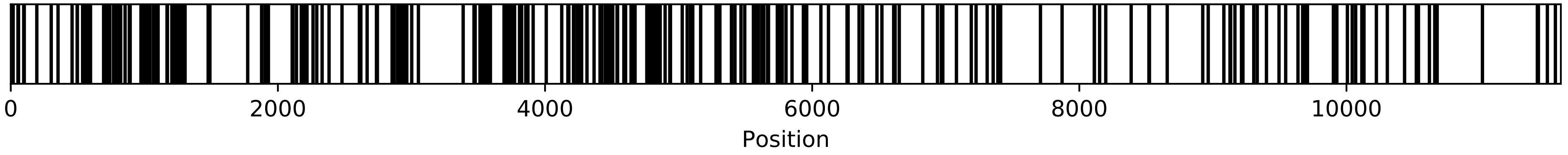


MA1 vs MM1
Number of differences: 271

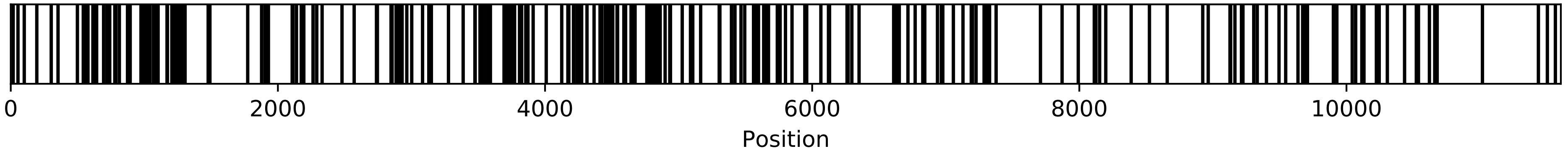


MA1 vs MM2
Number of differences: 279

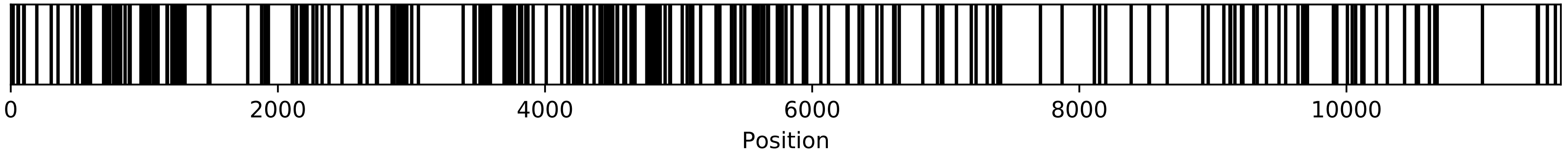


MA2 vs MM1
Number of differences: 93

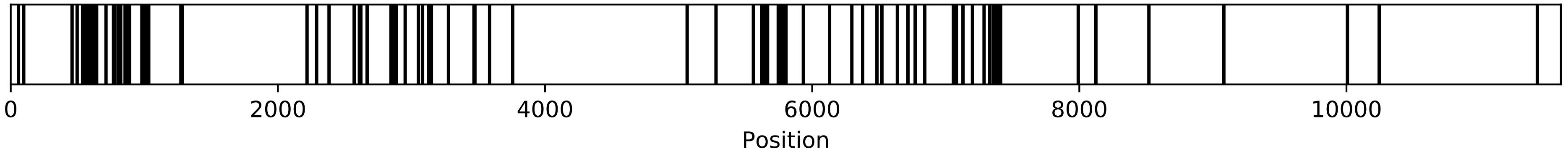


MA2 vs MM2
Number of differences: 0

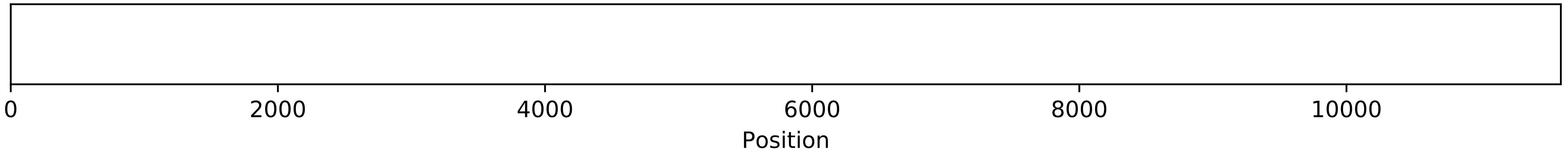


MM1 vs MM2
Number of differences: 93

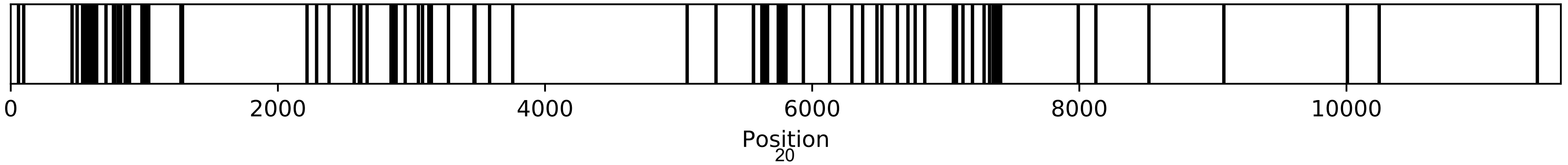

### Region: 87



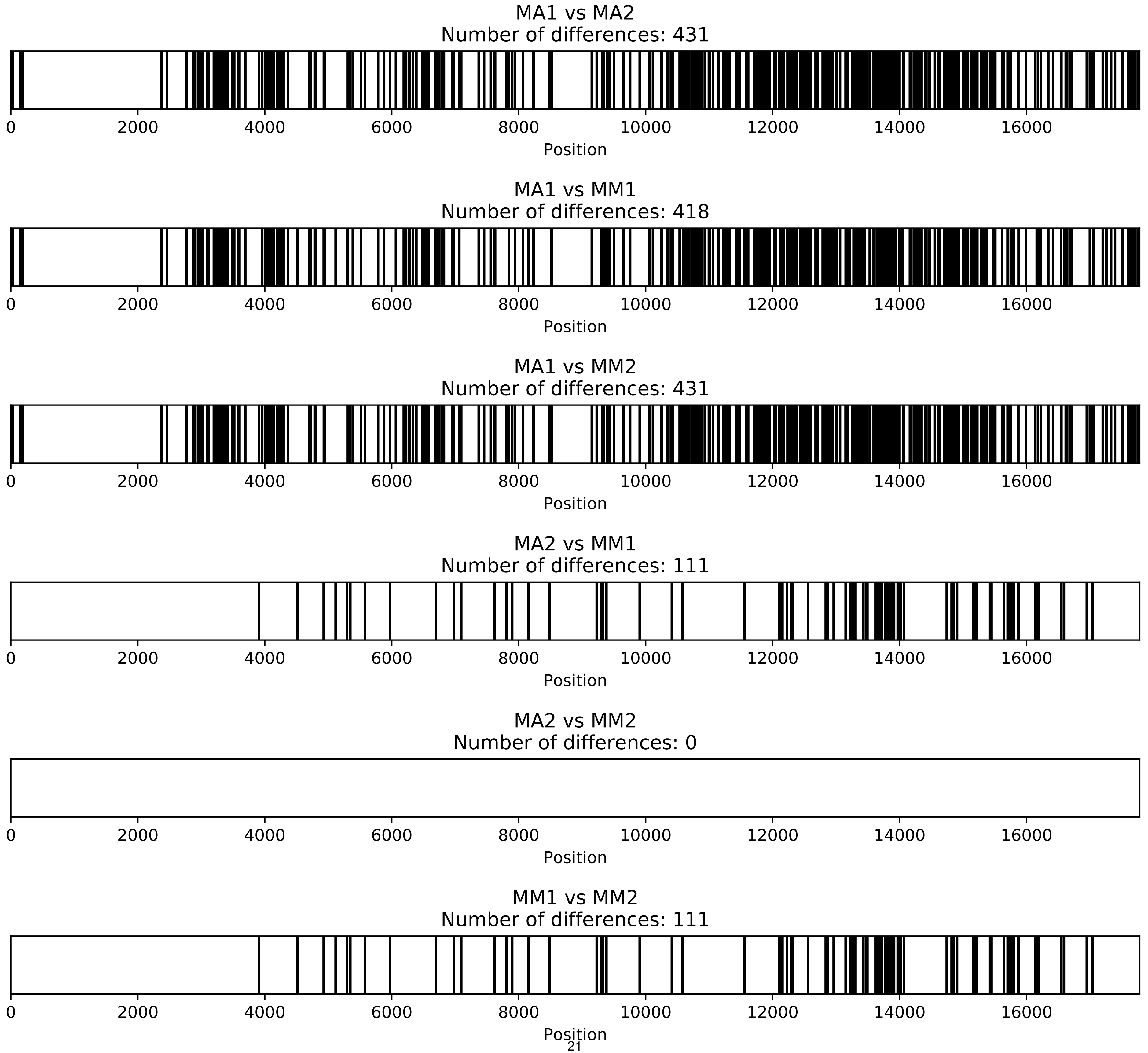
MA1 vs MA2
Number of differences: 431
0
2000
4000
6000
8000
10000
12000
14000
16000
Position
MA1 vs MM1
Number of differences: 418
Position
MA1 vs MM2
Number of differences: 431
Position
MA2 vs MM1
Number of differences: 111
Position
MA2 vs MM2
Number of differences: 0
Position
MM1 vs MM2
Number of differences: 111
Position

### Region: 92



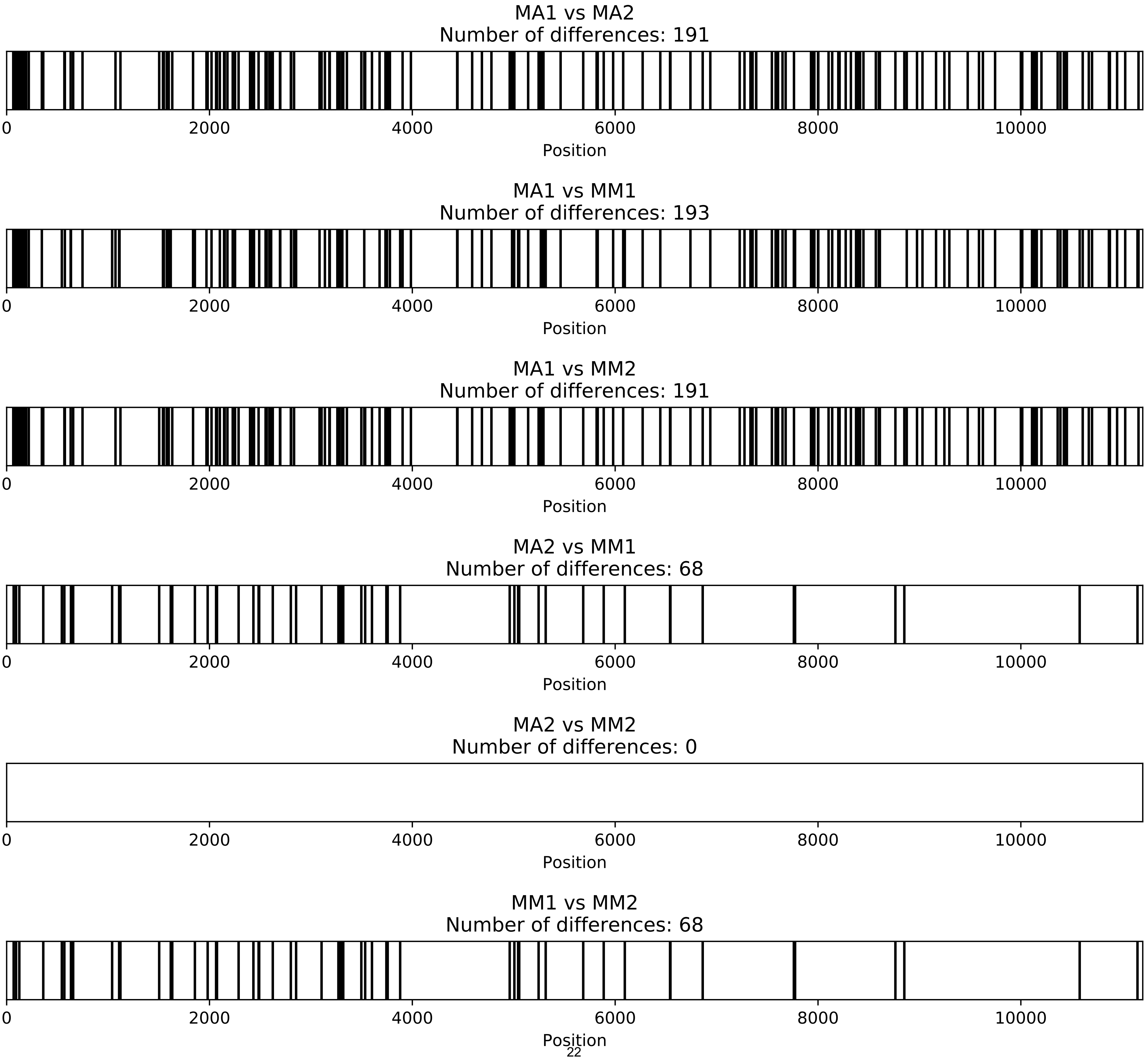
MA1 vs MA2
Number of differences: 191
0 2000 4000 6000 8000 10000
Position
MA1 vs MM1
Number of differences: 193
0 2000 4000 6000 8000 10000
Position
MA1 vs MM2
Number of differences: 191
0 2000 4000 6000 8000 10000
Position
MA2 vs MM1
Number of differences: 68
0 2000 4000 6000 8000 10000
Position
MA2 vs MM2
Number of differences: 0
0 2000 4000 6000 8000 10000
Position
MM1 vs MM2
Number of differences: 68
0 2000 4000 6000 8000 10000
Position

### Region: 95



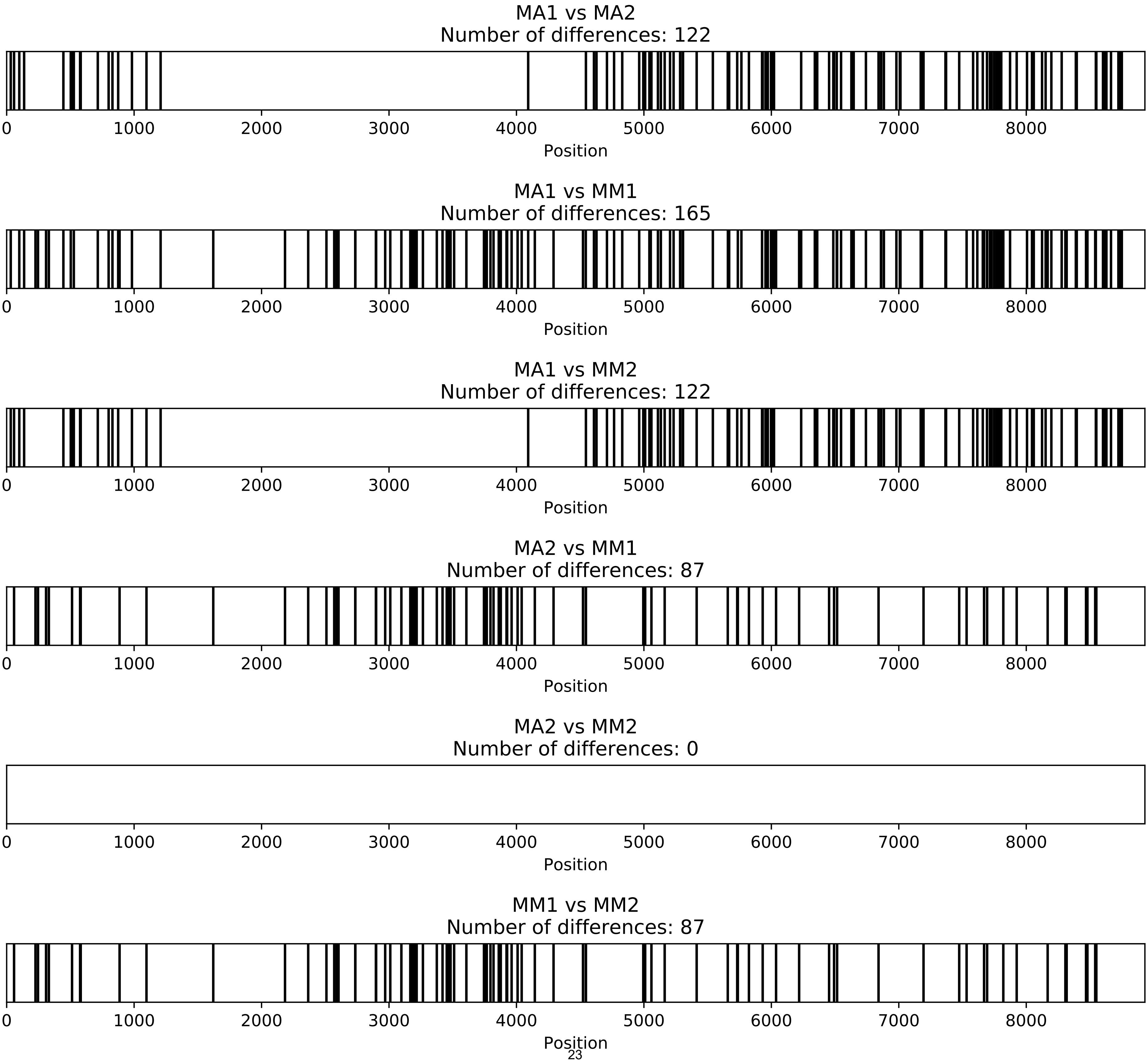
MA1 vs MA2
Number of differences: 122
0
1000
2000
3000
4000
5000
6000
7000
8000
Position
MA1 vs MM1
Number of differences: 165
Position
MA1 vs MM2
Number of differences: 122
Position
MA2 vs MM1
Number of differences: 87
Position
MA2 vs MM2
Number of differences: 0
Position
MM1 vs MM2
Number of differences: 87
Position

### Region: 101



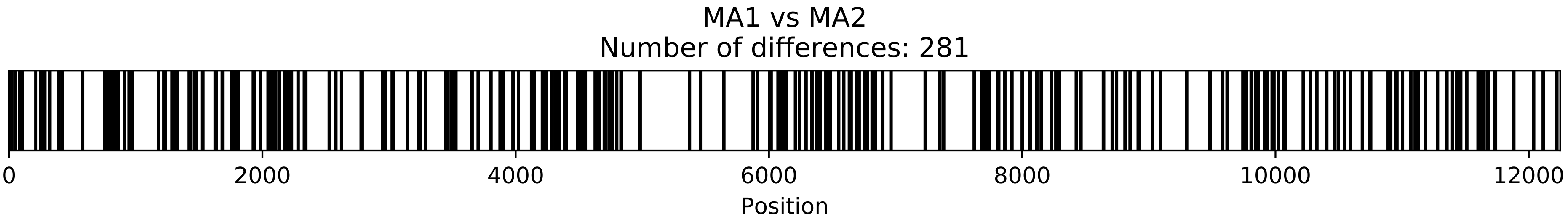
MA1 vs MA2
Number of differences: 281
0
2000
4000
6000
8000
10000
12000
Position

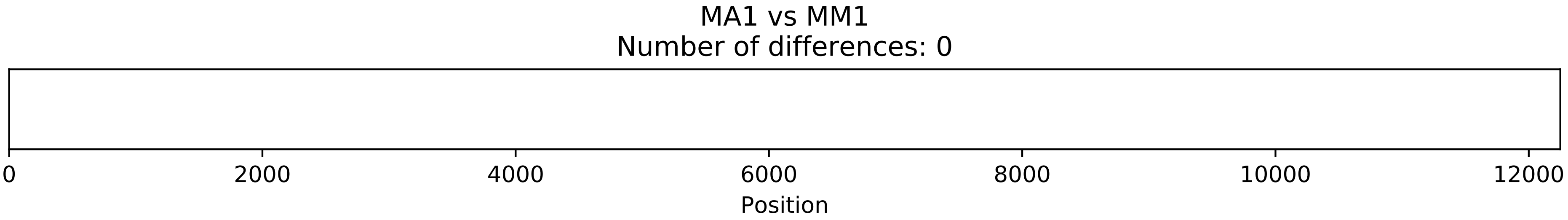
MA1 vs MM1
Number of differences: 0
0
2000
4000
6000
8000
10000
12000
Position

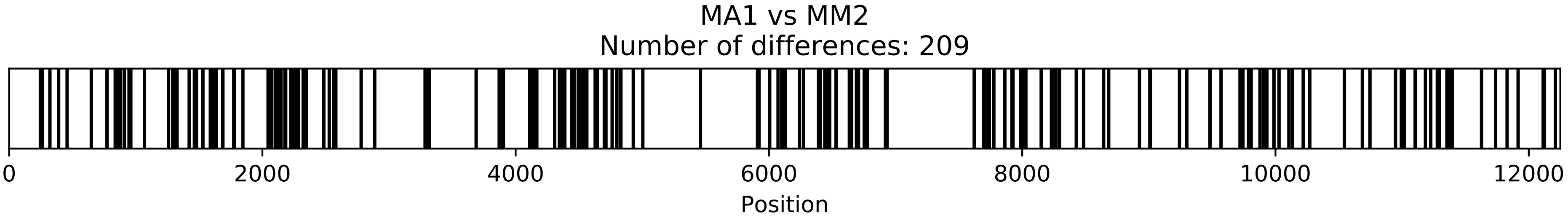
MA1 vs MM2
Number of differences: 209
0
2000
4000
6000
8000
10000
12000
Position

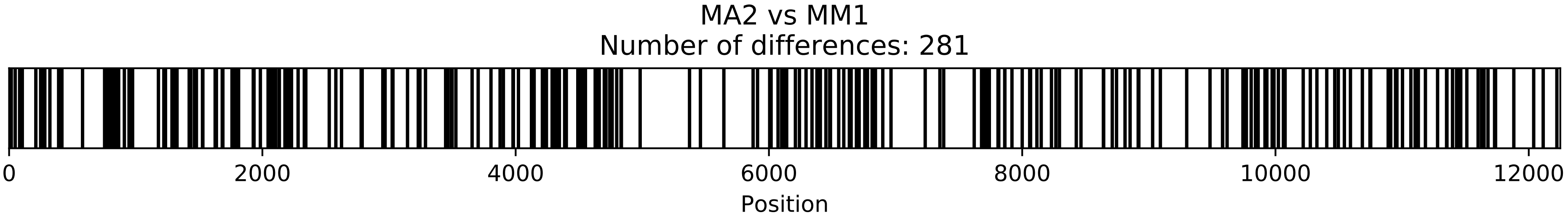
MA2 vs MM1
Number of differences: 281
0
2000
4000
6000
8000
10000
12000
Position

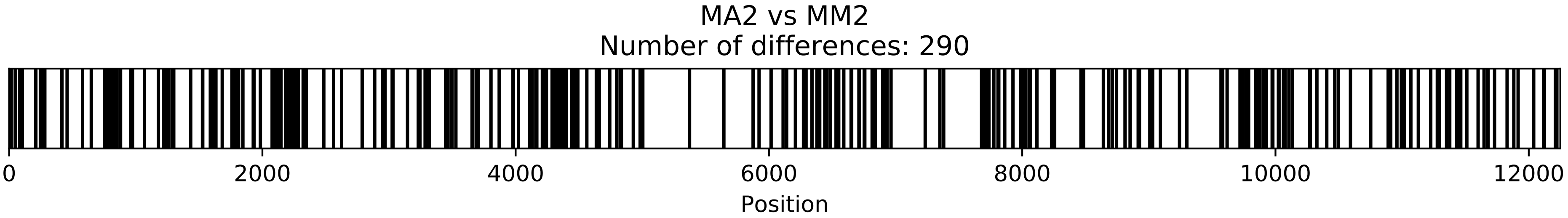
MA2 vs MM2
Number of differences: 290
0
2000
4000
6000
8000
10000
12000
Position

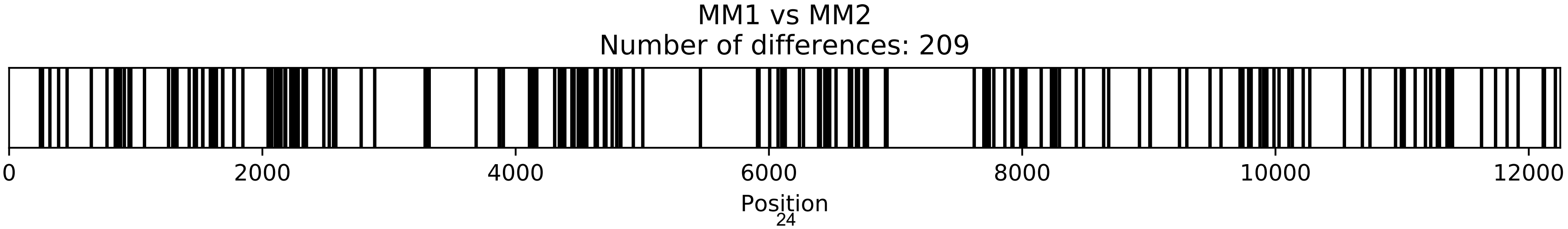
MM1 vs MM2
Number of differences: 209
0
2000
4000
6000
8000
10000
12000
Position

### Region: 102



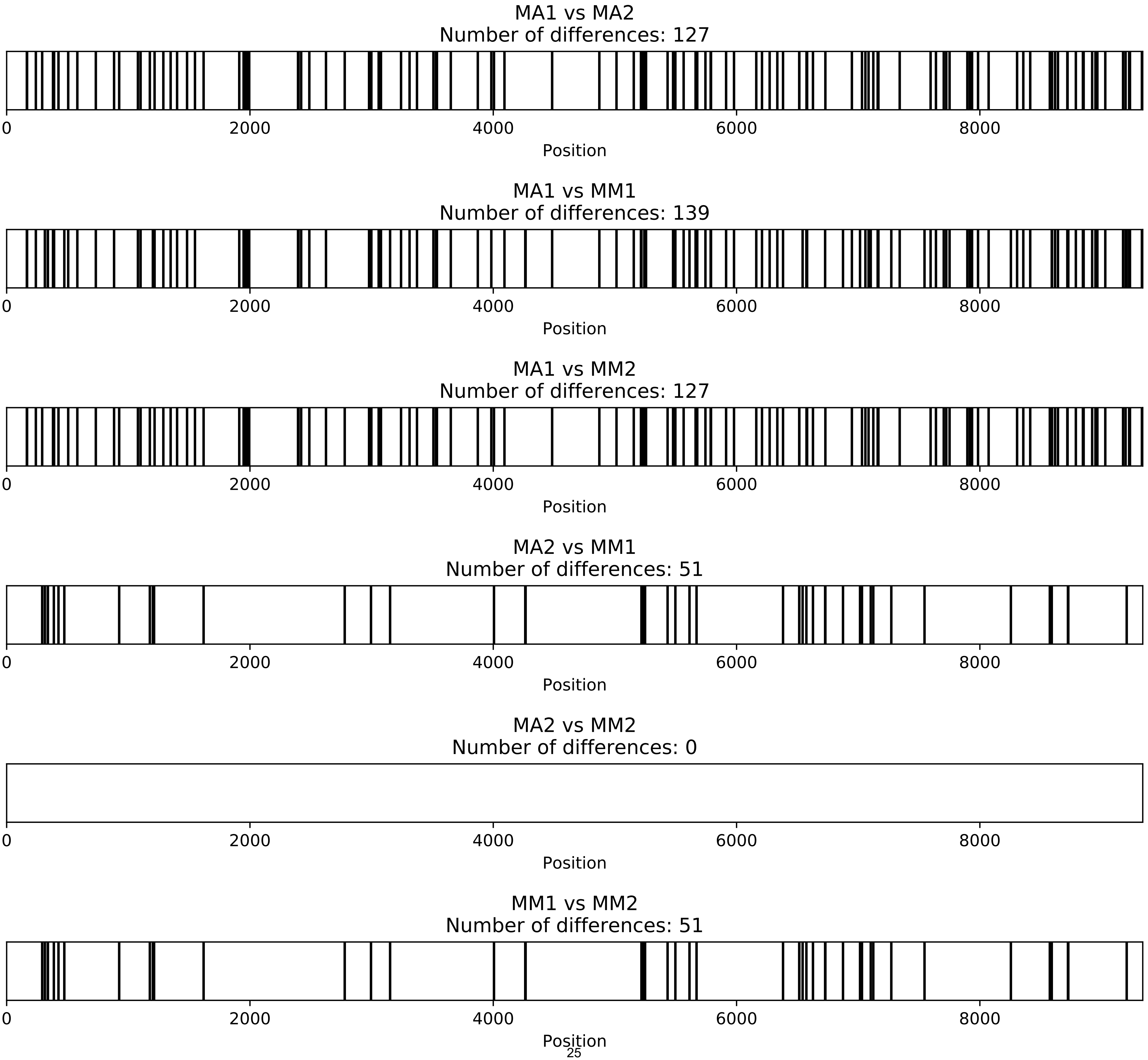
MA1 vs MA2
Number of differences: 127
MA1 vs MM1
Number of differences: 139
MA1 vs MM2
Number of differences: 127
MA2 vs MM1
Number of differences: 51
MA2 vs MM2
Number of differences: 0
MM1 vs MM2
Number of differences: 51
0
2000
4000
6000
8000
Position

### Region: 107



MA1 vs MA2
Number of differences: 236

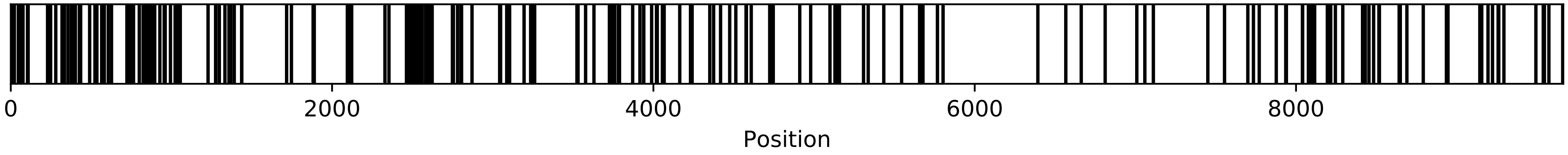


MA1 vs MM1
Number of differences: 118

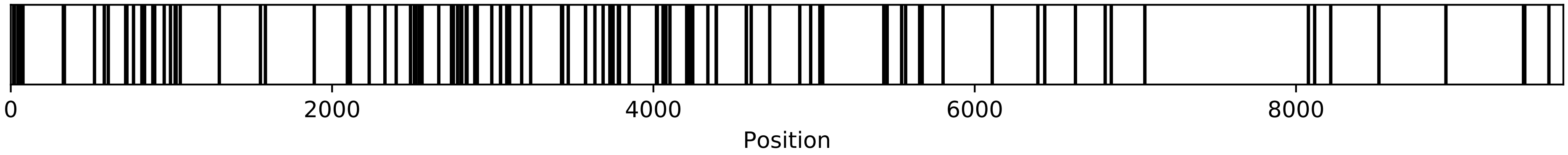


MA1 vs MM2
Number of differences: 122

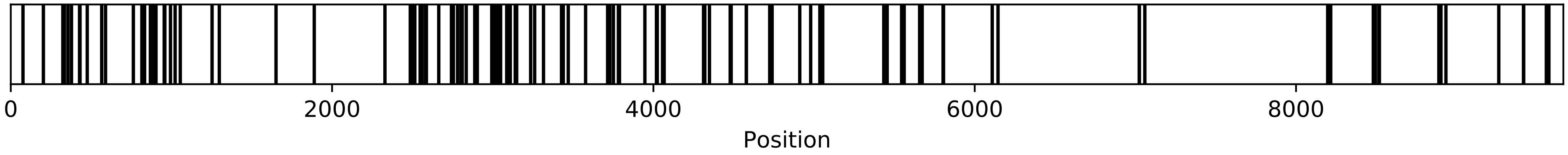


MA2 vs MM1
Number of differences: 236

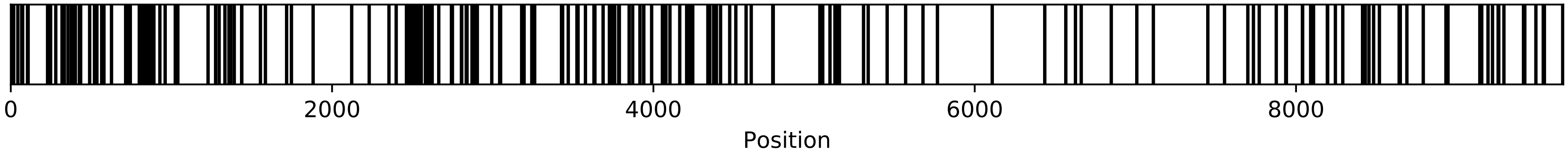


MA2 vs MM2
Number of differences: 253

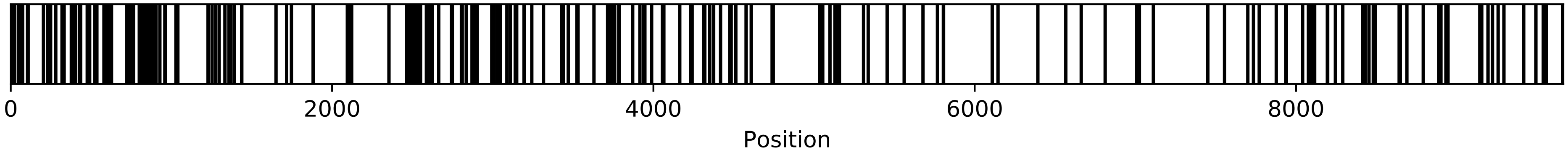


MM1 vs MM2
Number of differences: 116

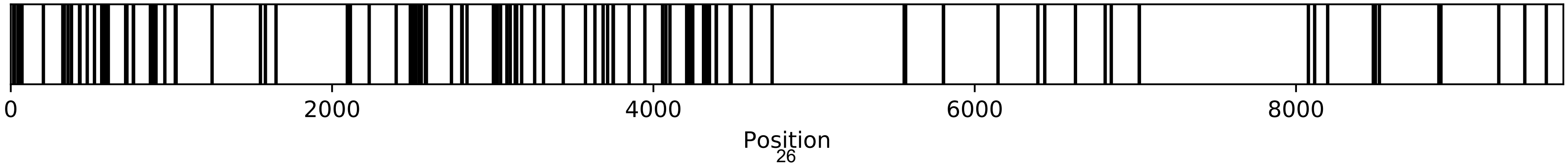

### Region: 108



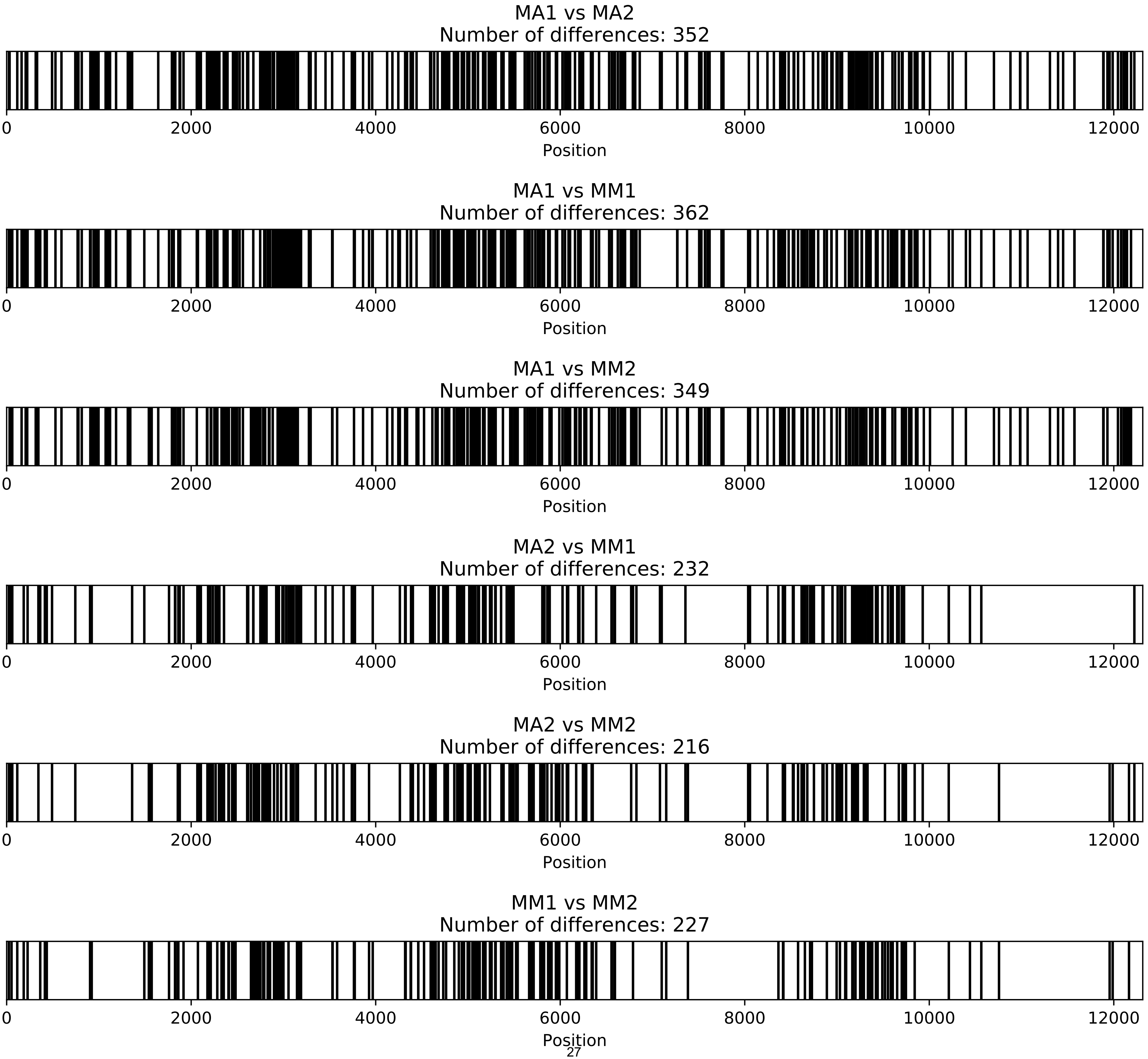
MA1 vs MA2
Number of differences: 352
MA1 vs MM1
Number of differences: 362
MA1 vs MM2
Number of differences: 349
MA2 vs MM1
Number of differences: 232
MA2 vs MM2
Number of differences: 216
MM1 vs MM2
Number of differences: 227
0
2000
4000
6000
8000
10000
12000
Position

### Region: 110



## MA1 vs MA2
Number of differences: 111

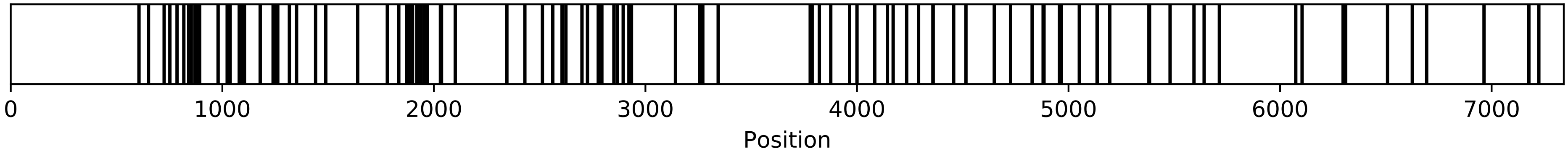


## MA1 vs MM1
Number of differences: 0

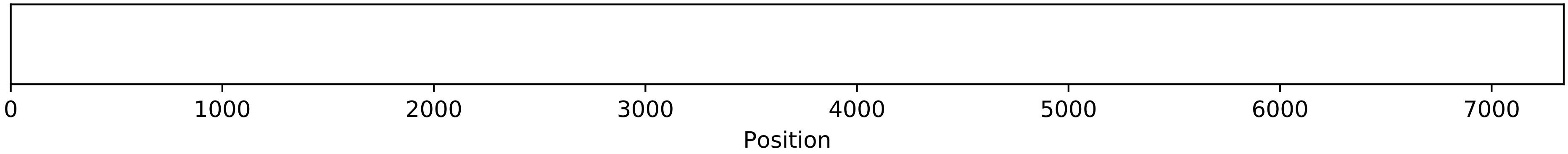


## MA1 vs MM2
Number of differences: 52

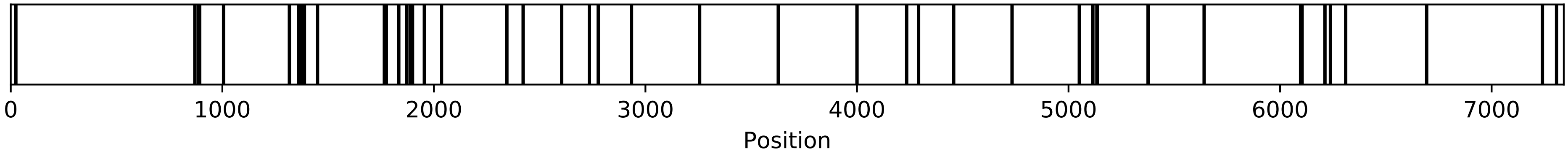


## MA2 vs MM1
Number of differences: 111

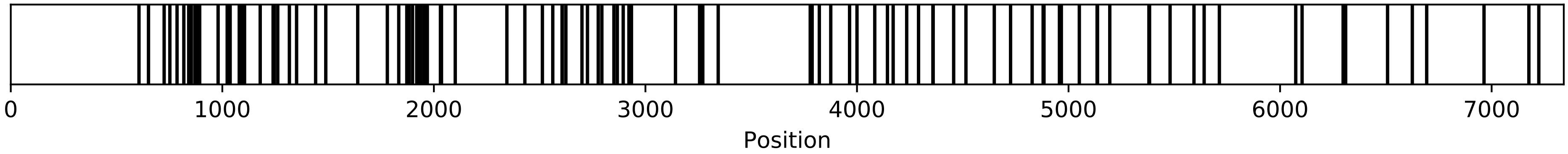


## MA2 vs MM2
Number of differences: 120

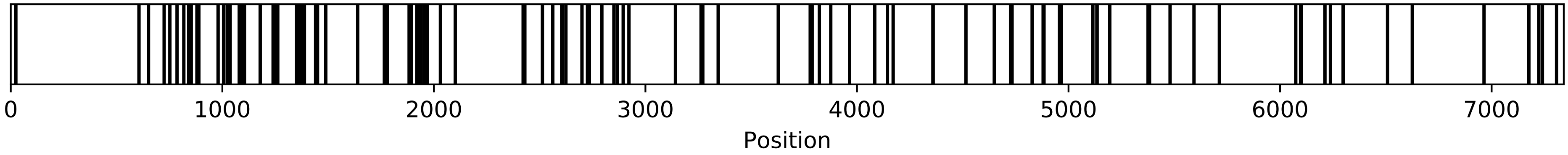


## MM1 vs MM2
Number of differences: 52

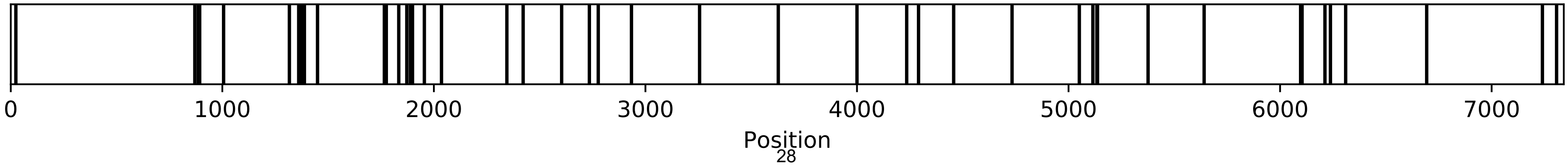

### Region: 112

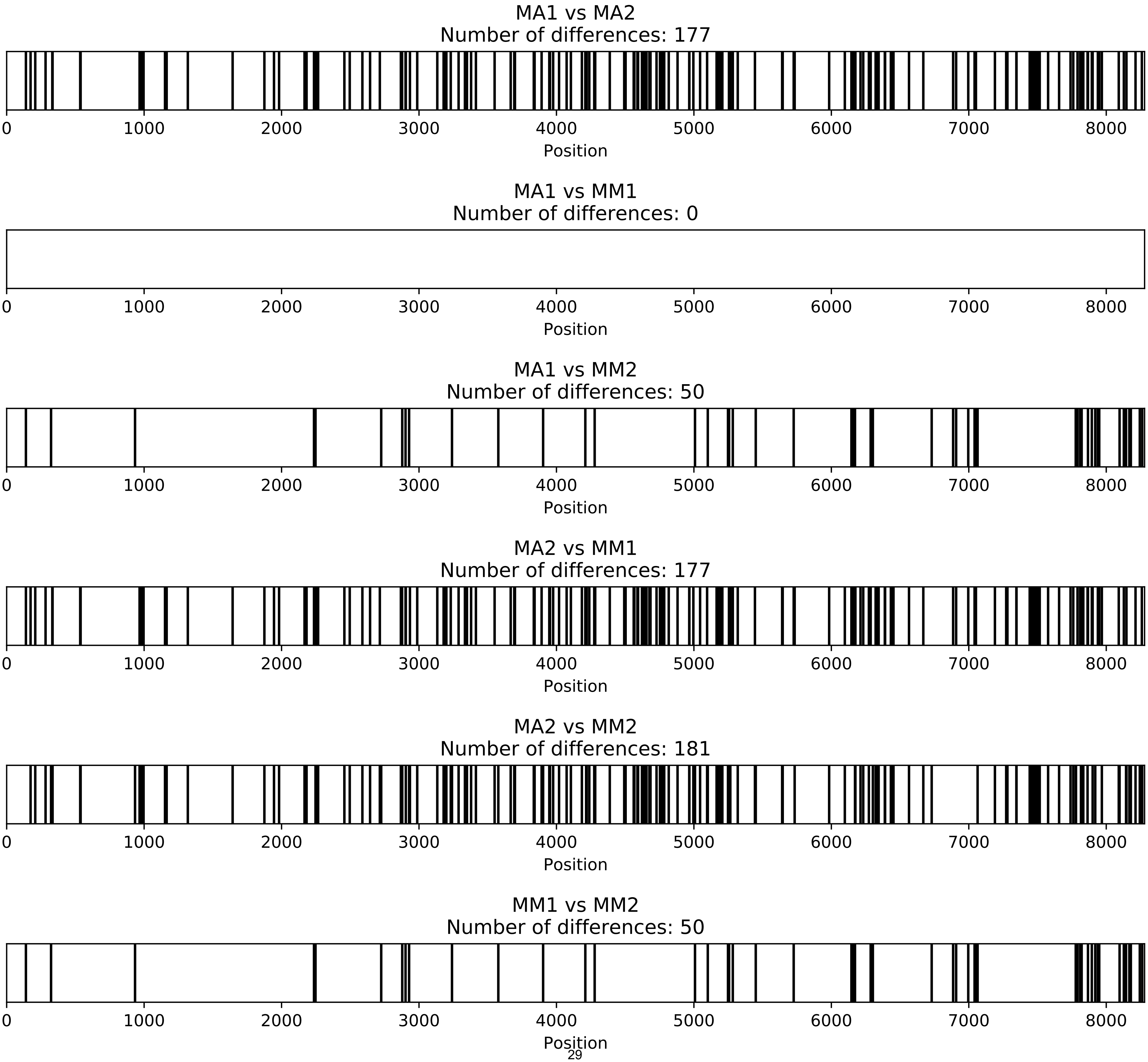
MA1 vs MA2
Number of differences: 177
MA1 vs MM1
Number of differences: 0
MA1 vs MM2
Number of differences: 50
MA2 vs MM1
Number of differences: 177
MA2 vs MM2
Number of differences: 181
MM1 vs MM2
Number of differences: 50
0
1000
2000
3000
4000
5000
6000
7000
8000
Position

### Region: 121



MA1 vs MA2
Number of differences: 23

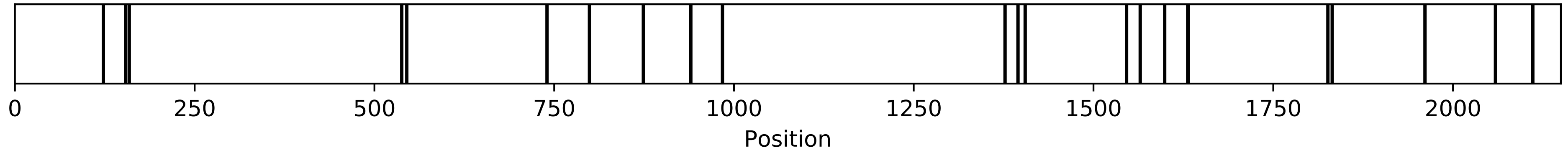


MA1 vs MM1
Number of differences: 13

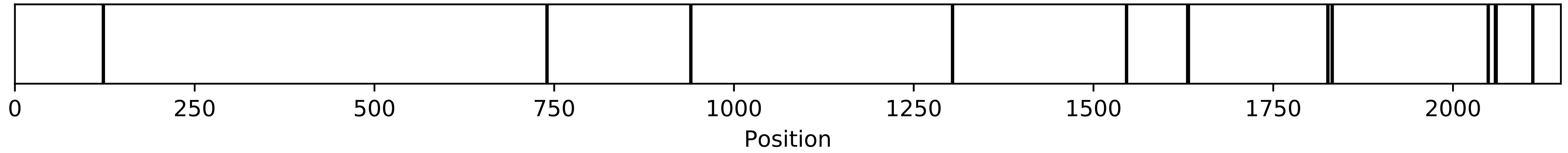


MA1 vs MM2
Number of differences: 12

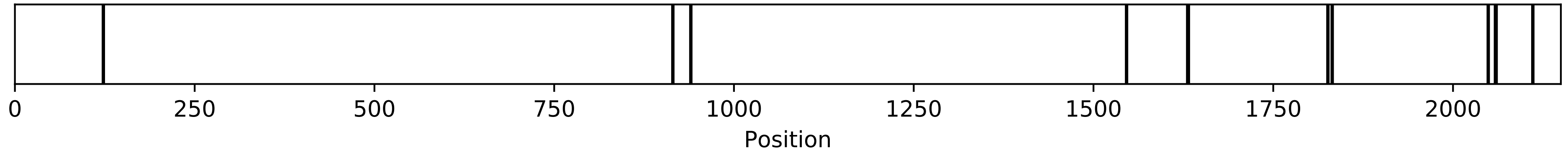


MA2 vs MM1
Number of differences: 16

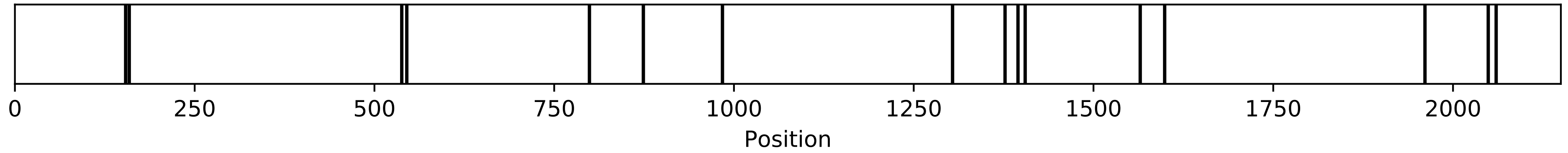


MA2 vs MM2
Number of differences: 17

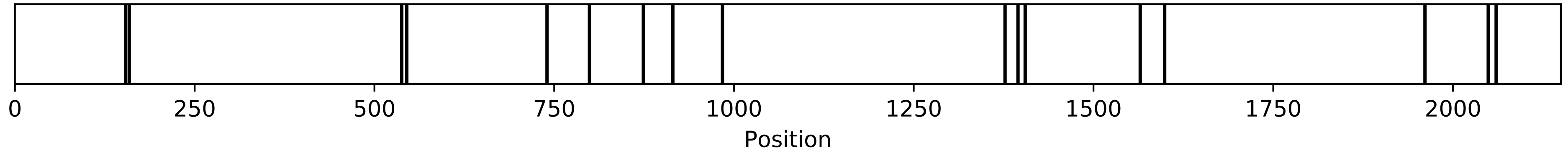


MM1 vs MM2
Number of differences: 3

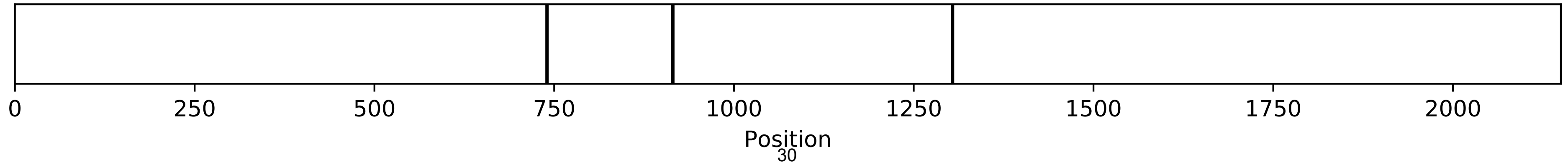

### Region: 125



MA1 vs MA2
Number of differences: 219

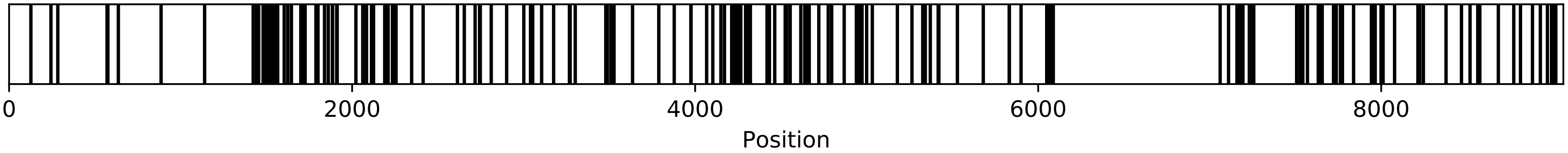


MA1 vs MM1
Number of differences: 219

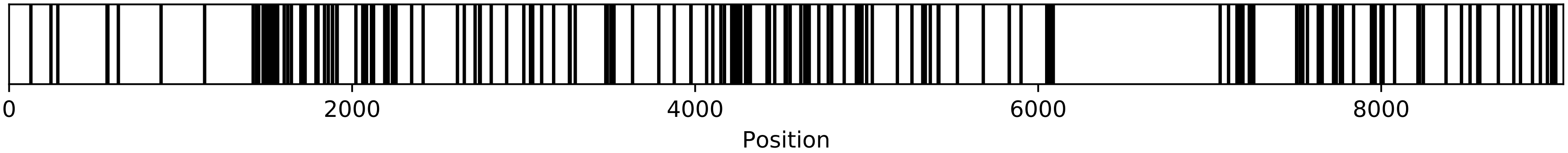


MA1 vs MM2
Number of differences: 217

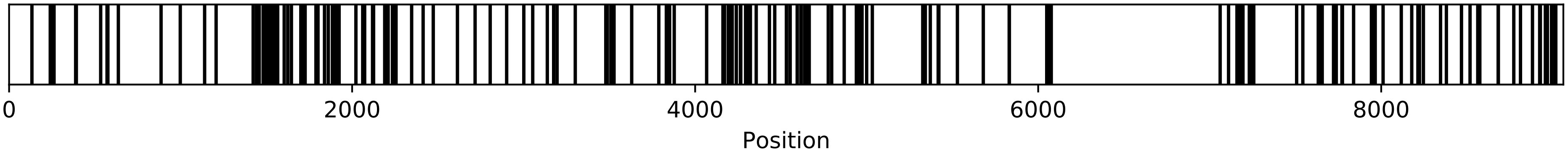


MA2 vs MM1
Number of differences: 0

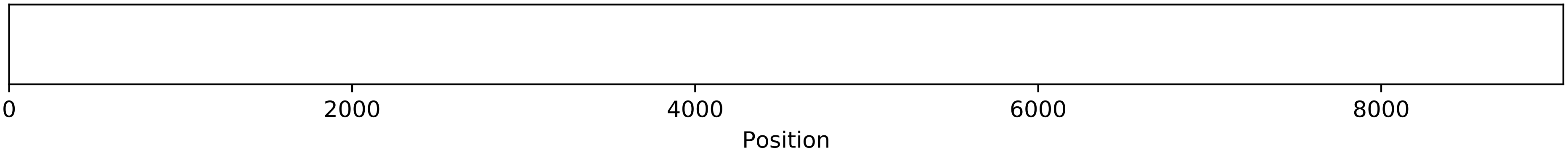


MA2 vs MM2
Number of differences: 93

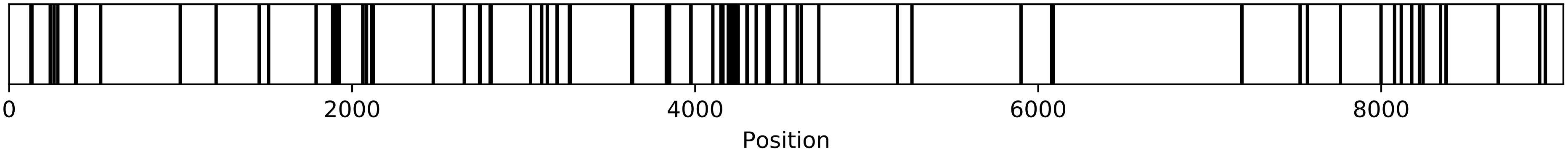


MM1 vs MM2
Number of differences: 93

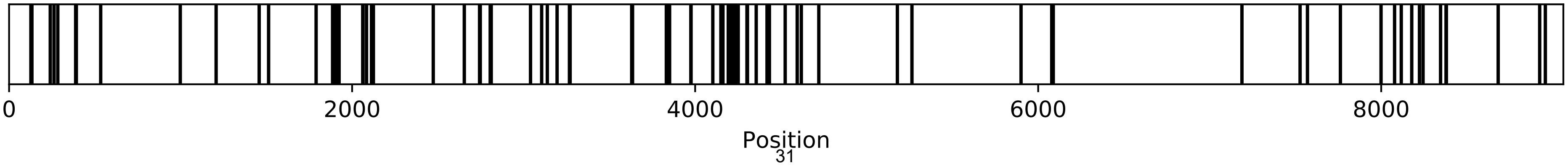

### Region: 127



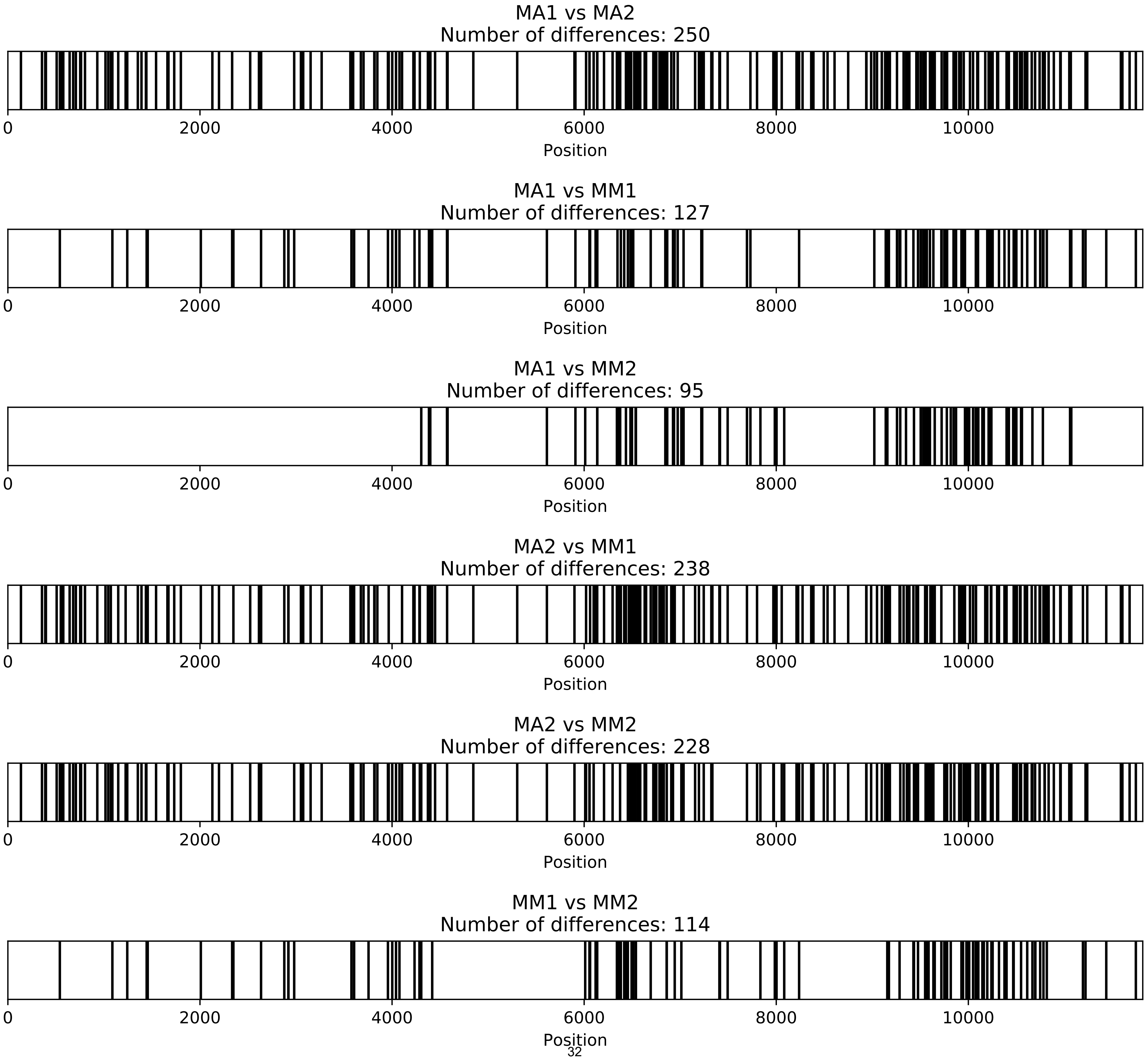
MA1 vs MA2
Number of differences: 250
MA1 vs MM1
Number of differences: 127
MA1 vs MM2
Number of differences: 95
MA2 vs MM1
Number of differences: 238
MA2 vs MM2
Number of differences: 228
MM1 vs MM2
Number of differences: 114
0
2000
4000
6000
8000
10000
Position

### Region: 131



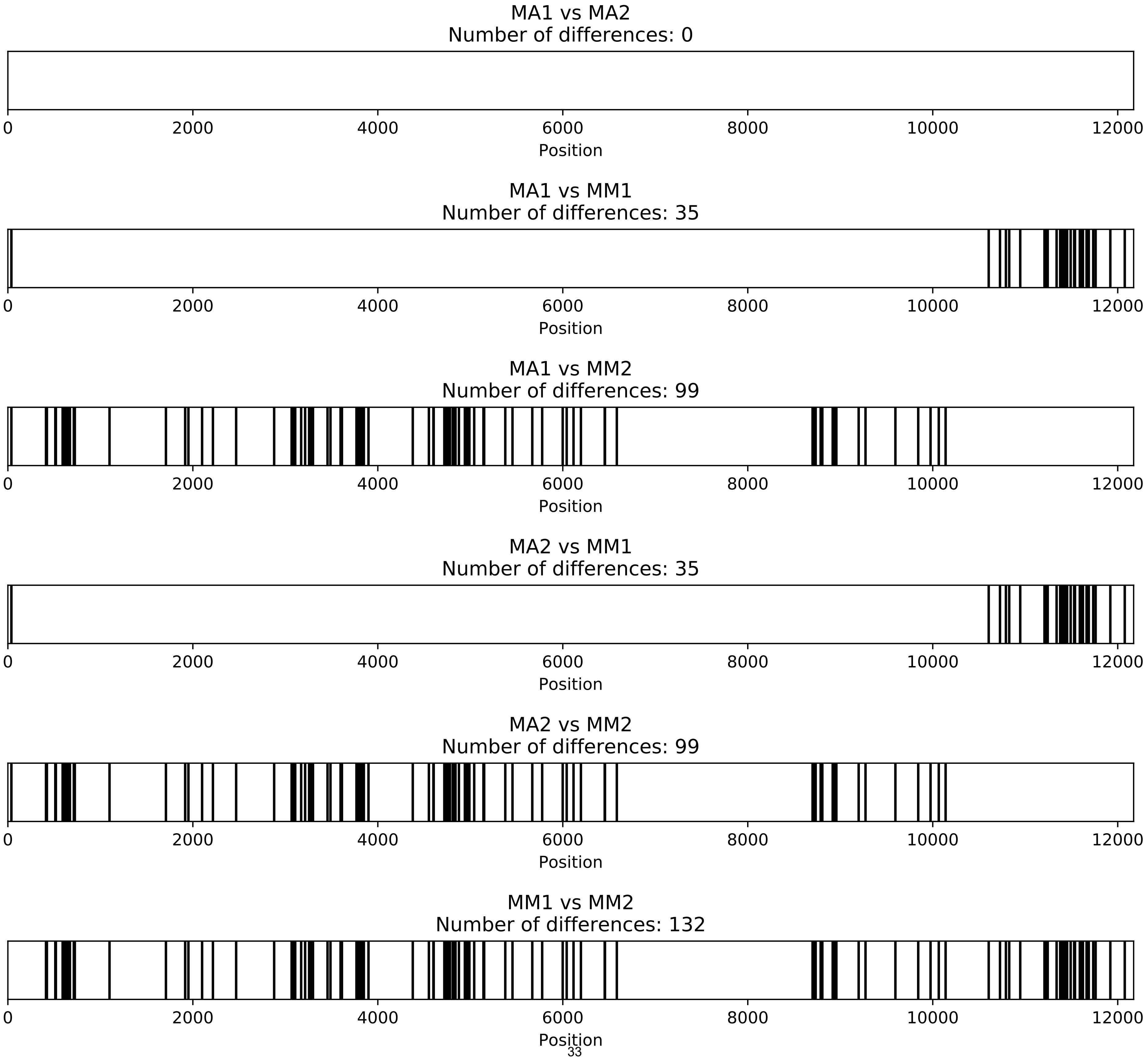
MA1 vs MA2
Number of differences: 0
MA1 vs MM1
Number of differences: 35
MA1 vs MM2
Number of differences: 99
MA2 vs MM1
Number of differences: 35
MA2 vs MM2
Number of differences: 99
MM1 vs MM2
Number of differences: 132
0
2000
4000
6000
8000
10000
12000
Position

### Region: 133



MA1 vs MA2
Number of differences: 66

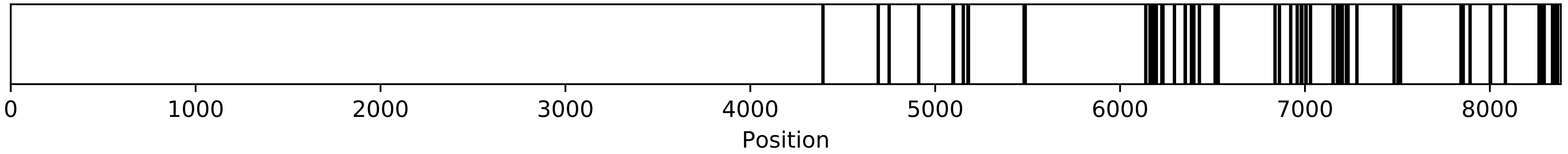


MA1 vs MM1
Number of differences: 60

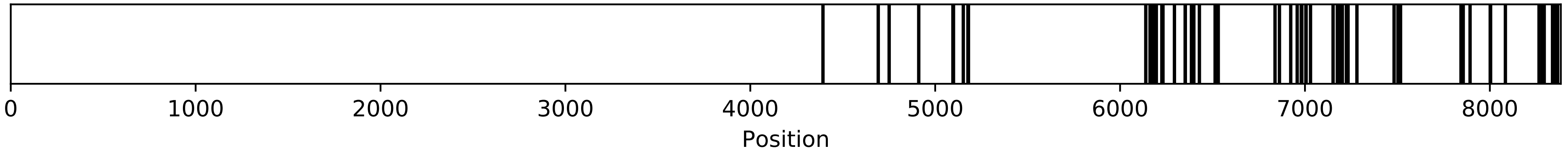


MA1 vs MM2
Number of differences: 103

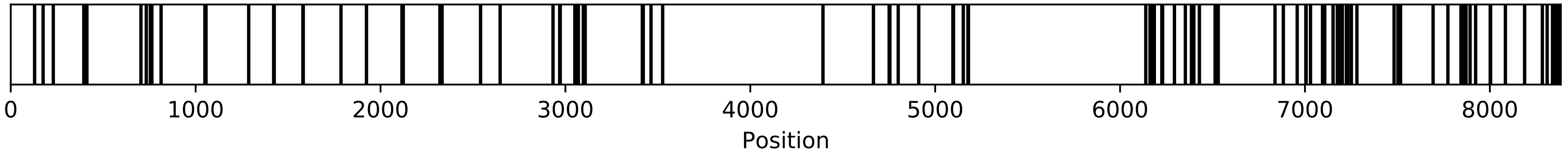


MA2 vs MM1
Number of differences: 6

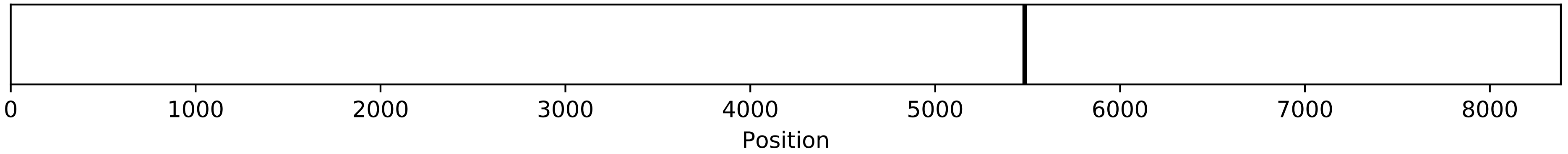


MA2 vs MM2
Number of differences: 76

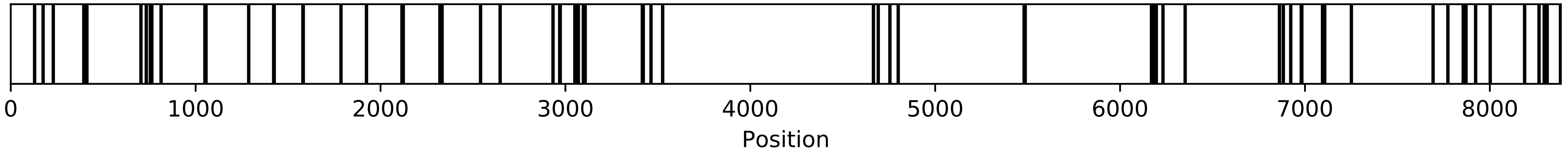


MM1 vs MM2
Number of differences: 70

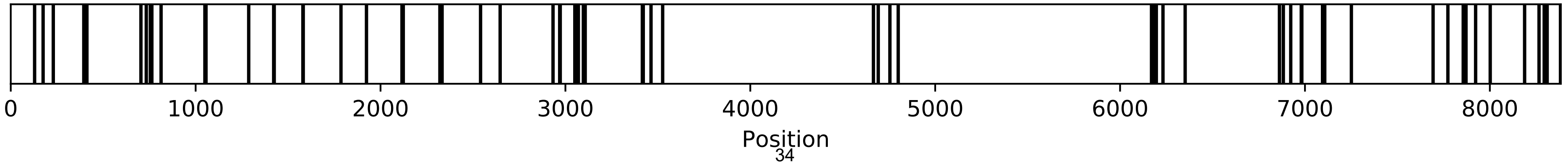

### Region: 141



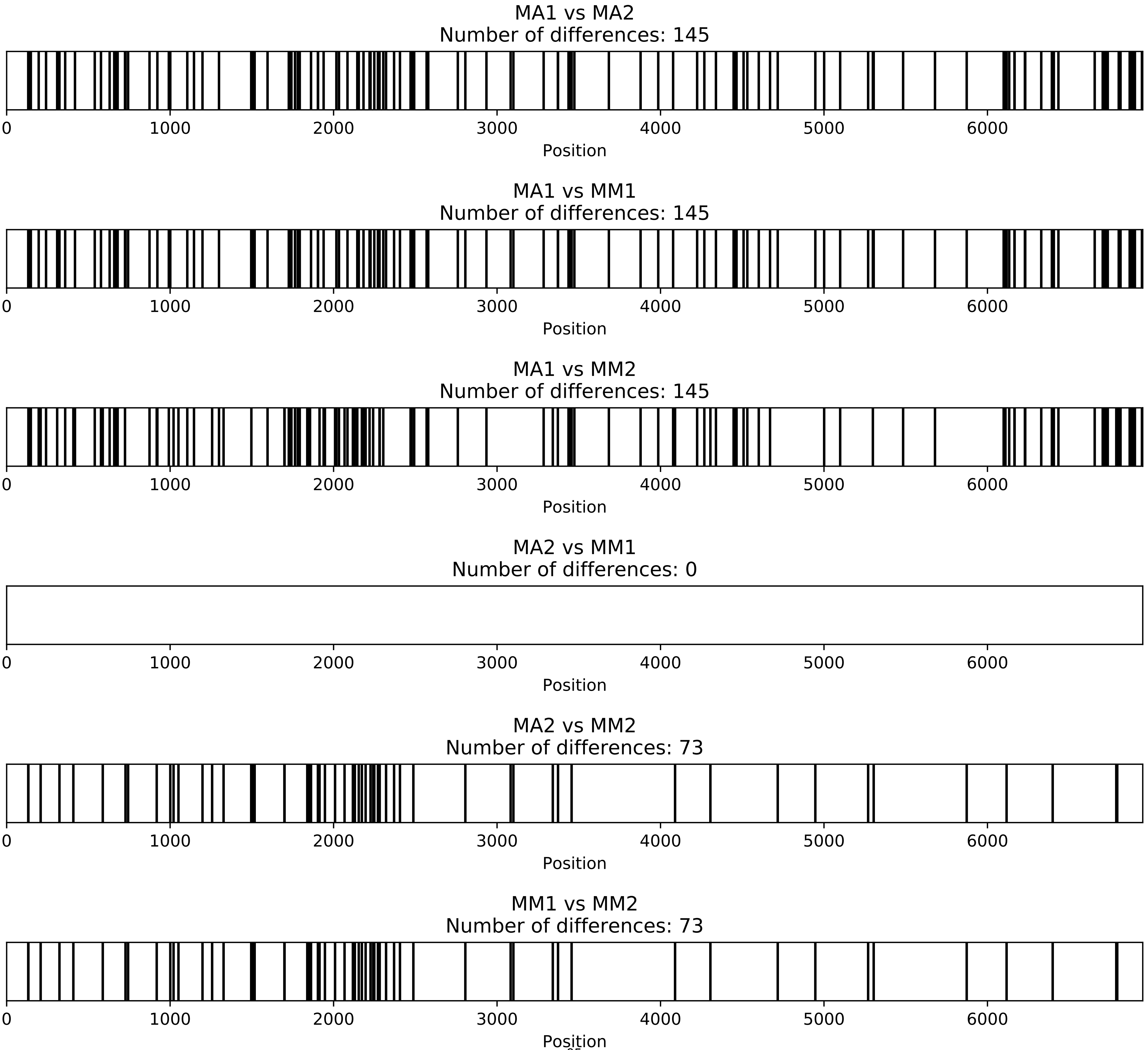
MA1 vs MA2
Number of differences: 145
MA1 vs MM1
Number of differences: 145
MA1 vs MM2
Number of differences: 145
MA2 vs MM1
Number of differences: 0
MA2 vs MM2
Number of differences: 73
MM1 vs MM2
Number of differences: 73
0
1000
2000
3000
4000
5000
6000
Position

### Region: 146



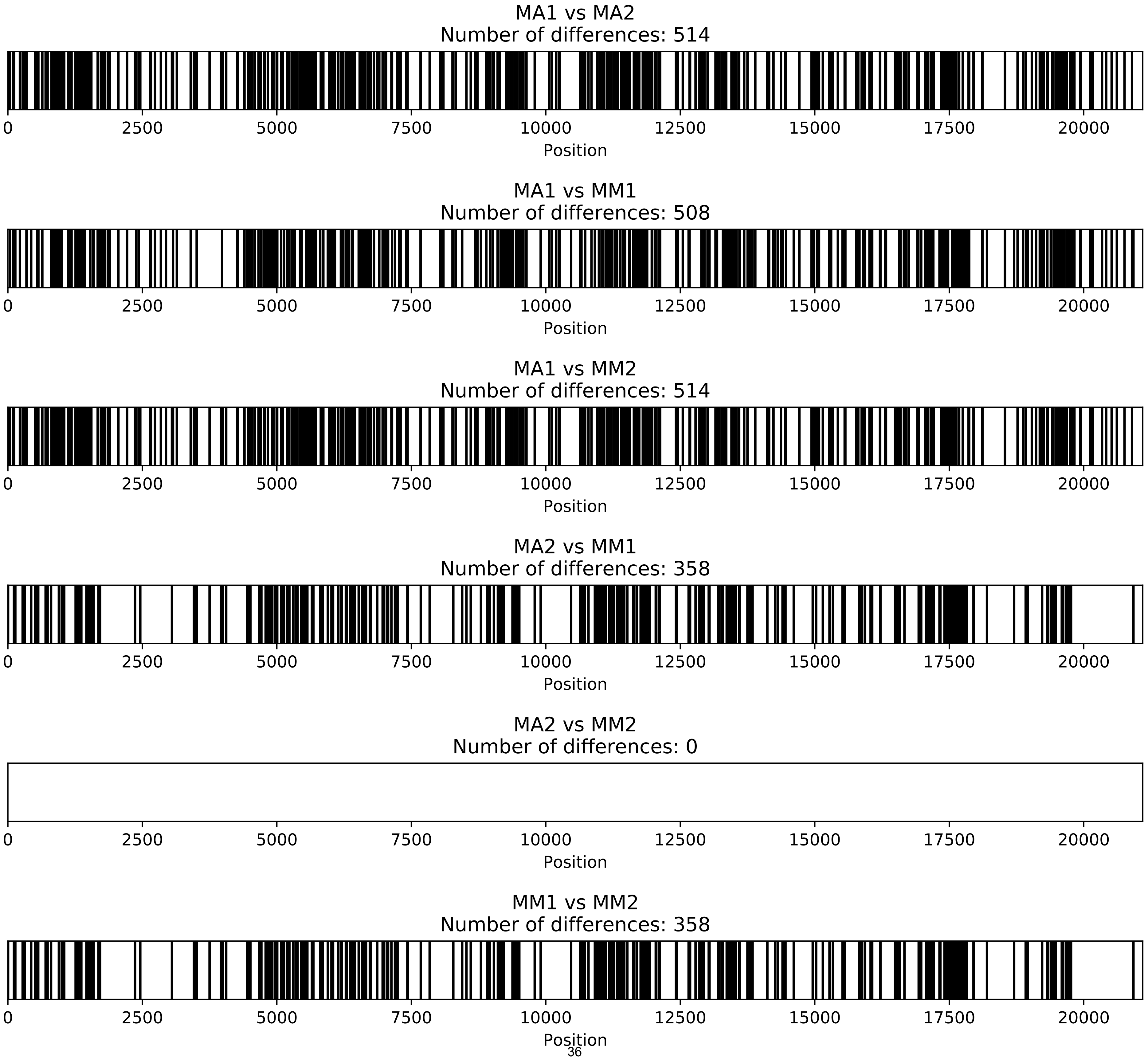
MA1 vs MA2
Number of differences: 514
0 2500 5000 7500 10000 12500 15000 17500 20000
Position
MA1 vs MM1
Number of differences: 508
0 2500 5000 7500 10000 12500 15000 17500 20000
Position
MA1 vs MM2
Number of differences: 514
0 2500 5000 7500 10000 12500 15000 17500 20000
Position
MA2 vs MM1
Number of differences: 358
0 2500 5000 7500 10000 12500 15000 17500 20000
Position
MA2 vs MM2
Number of differences: 0
0 2500 5000 7500 10000 12500 15000 17500 20000
Position
MM1 vs MM2
Number of differences: 358
0 2500 5000 7500 10000 12500 15000 17500 20000
Position

### Region: 147



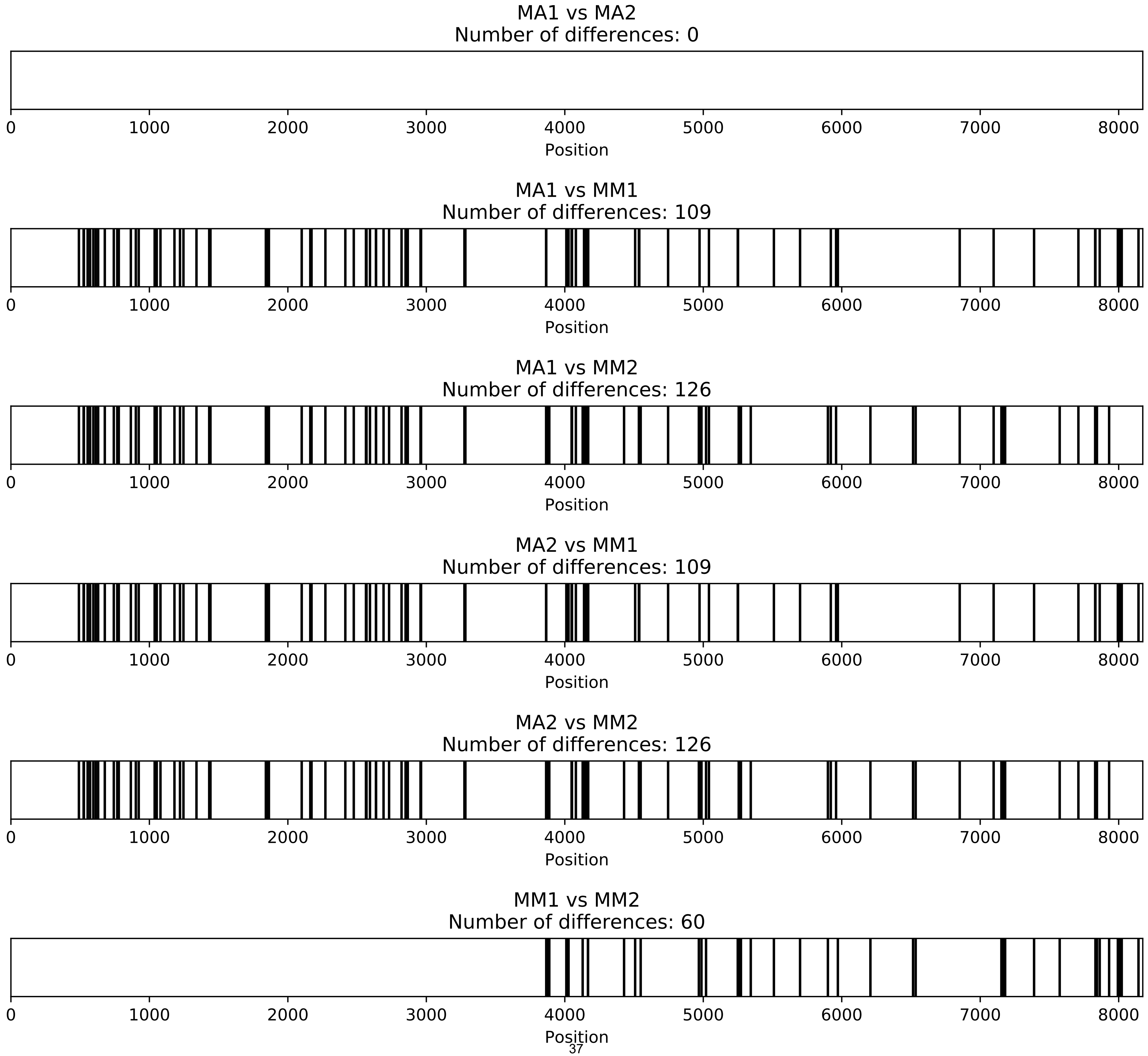
MA1 vs MA2
Number of differences: 0
MA1 vs MM1
Number of differences: 109
MA1 vs MM2
Number of differences: 126
MA2 vs MM1
Number of differences: 109
MA2 vs MM2
Number of differences: 126
MM1 vs MM2
Number of differences: 60
0
1000
2000
3000
4000
5000
6000
7000
8000
Position

### Region: 149



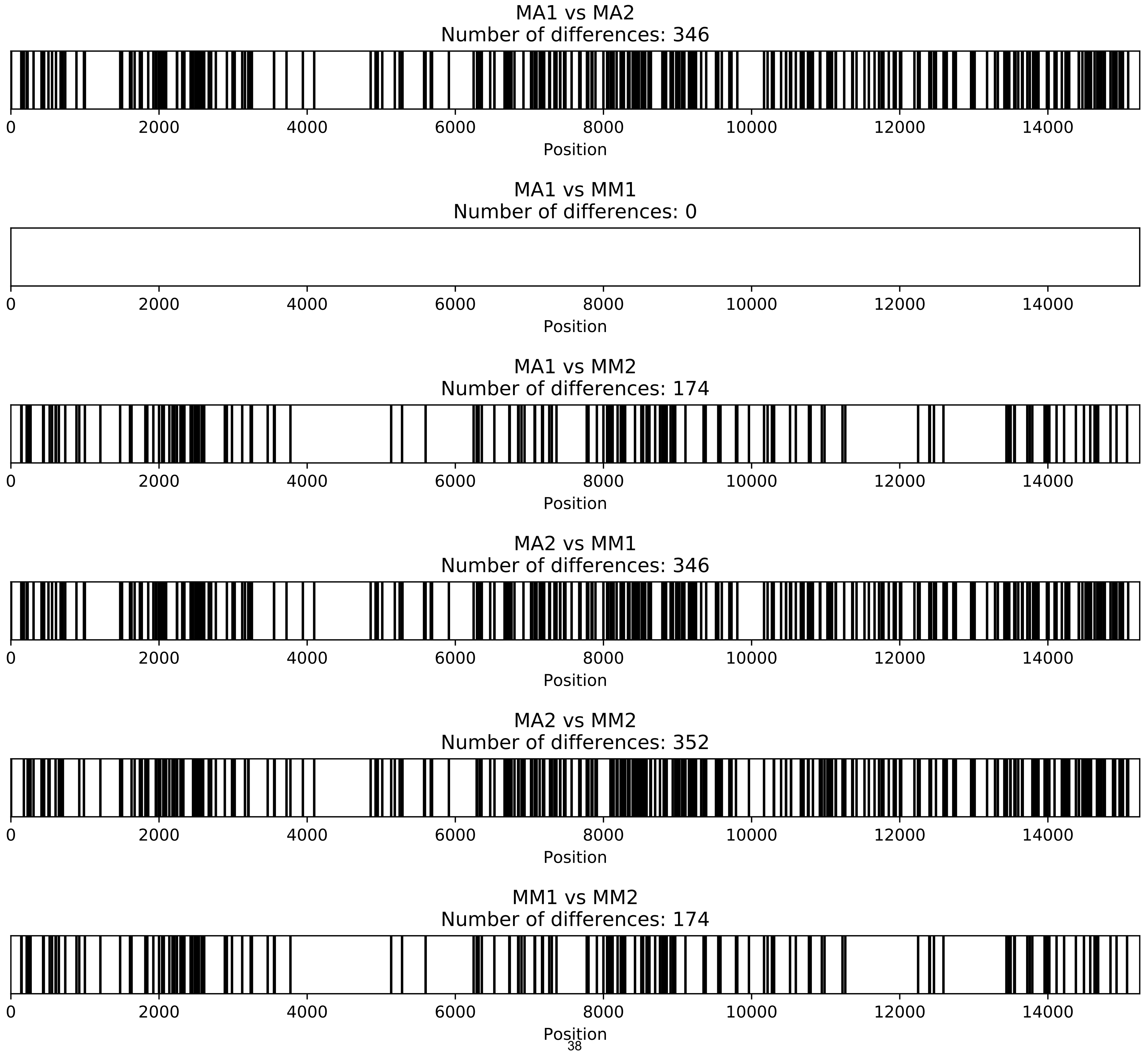
MA1 vs MA2
Number of differences: 346
0 2000 4000 6000 8000 10000 12000 14000
Position
MA1 vs MM1
Number of differences: 0
0 2000 4000 6000 8000 10000 12000 14000
Position
MA1 vs MM2
Number of differences: 174
0 2000 4000 6000 8000 10000 12000 14000
Position
MA2 vs MM1
Number of differences: 346
0 2000 4000 6000 8000 10000 12000 14000
Position
MA2 vs MM2
Number of differences: 352
0 2000 4000 6000 8000 10000 12000 14000
Position
MM1 vs MM2
Number of differences: 174
0 2000 4000 6000 8000 10000 12000 14000
Position

### Region: 150



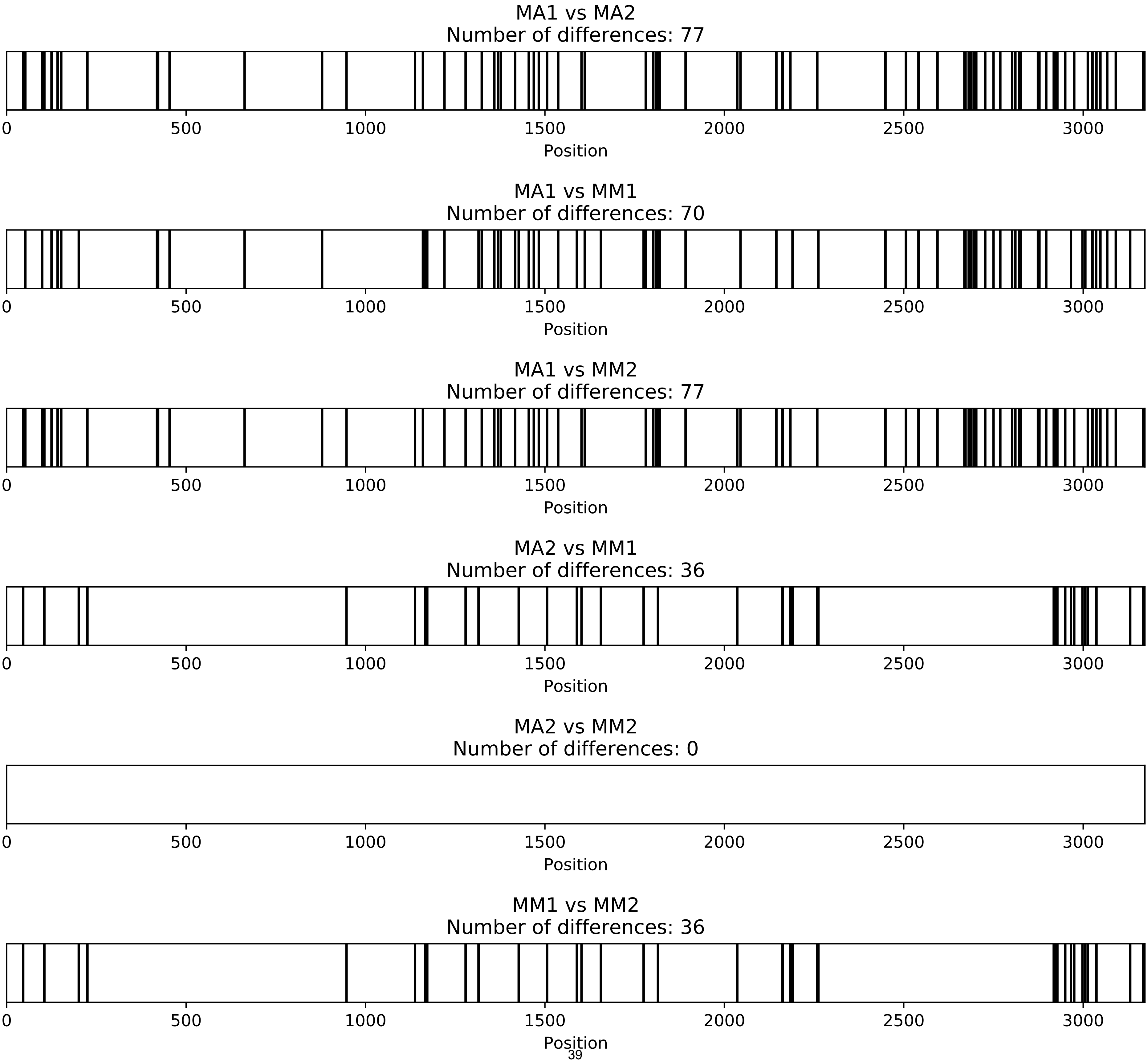
MA1 vs MA2
Number of differences: 77
0
500
1000
1500
2000
2500
3000
Position
MA1 vs MM1
Number of differences: 70
0
500
1000
1500
2000
2500
3000
Position
MA1 vs MM2
Number of differences: 77
0
500
1000
1500
2000
2500
3000
Position
MA2 vs MM1
Number of differences: 36
0
500
1000
1500
2000
2500
3000
Position
MA2 vs MM2
Number of differences: 0
0
500
1000
1500
2000
2500
3000
Position
MM1 vs MM2
Number of differences: 36
0
500
1000
1500
2000
2500
3000
Position

### Region: 153



MA1 vs MA2
Number of differences: 149

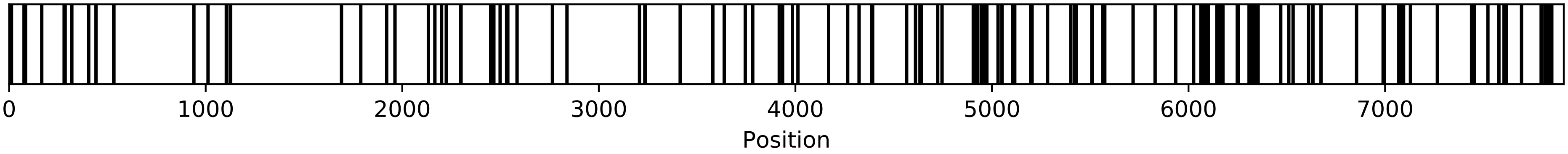


MA1 vs MM1
Number of differences: 151

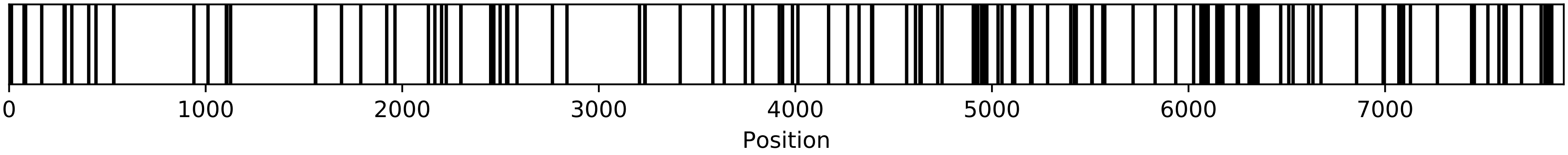


MA1 vs MM2
Number of differences: 146

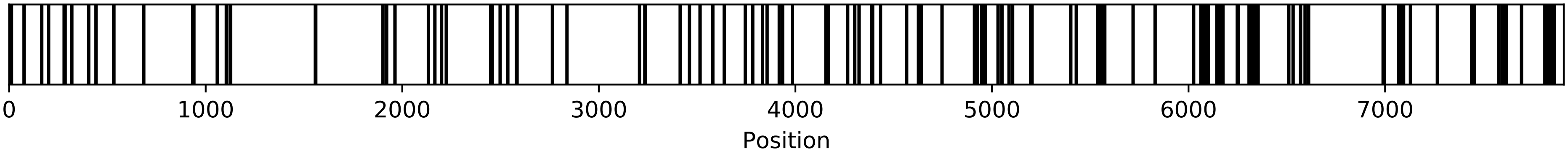


MA2 vs MM1
Number of differences: 2

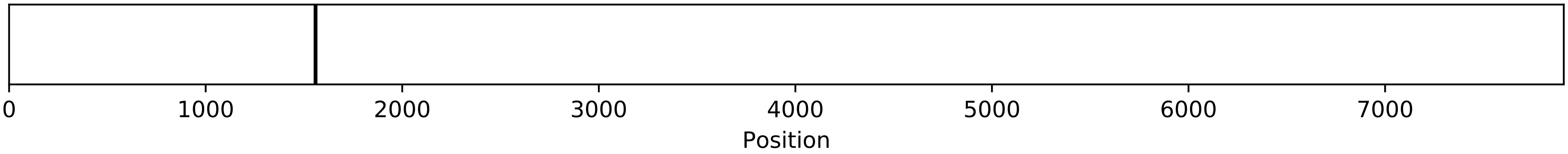


MA2 vs MM2
Number of differences: 63

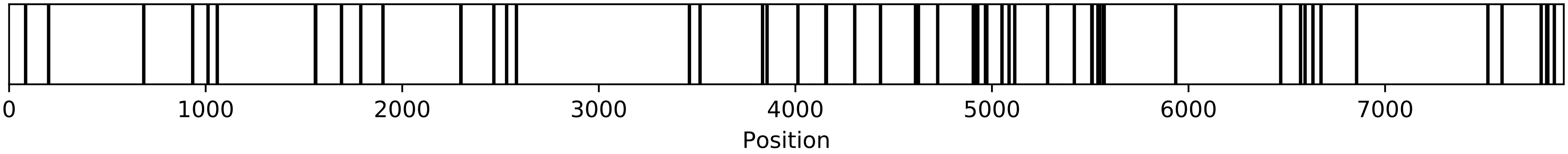


MM1 vs MM2
Number of differences: 61

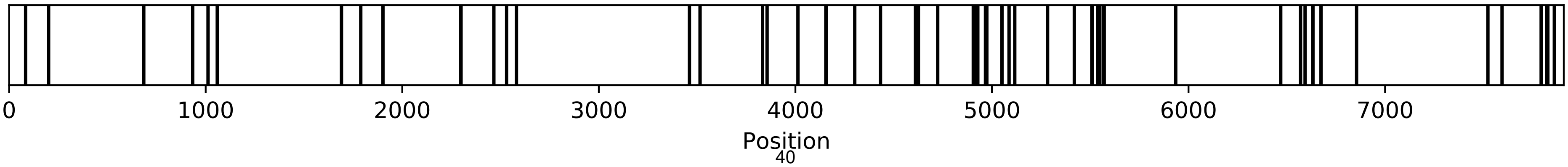

### Region: 158



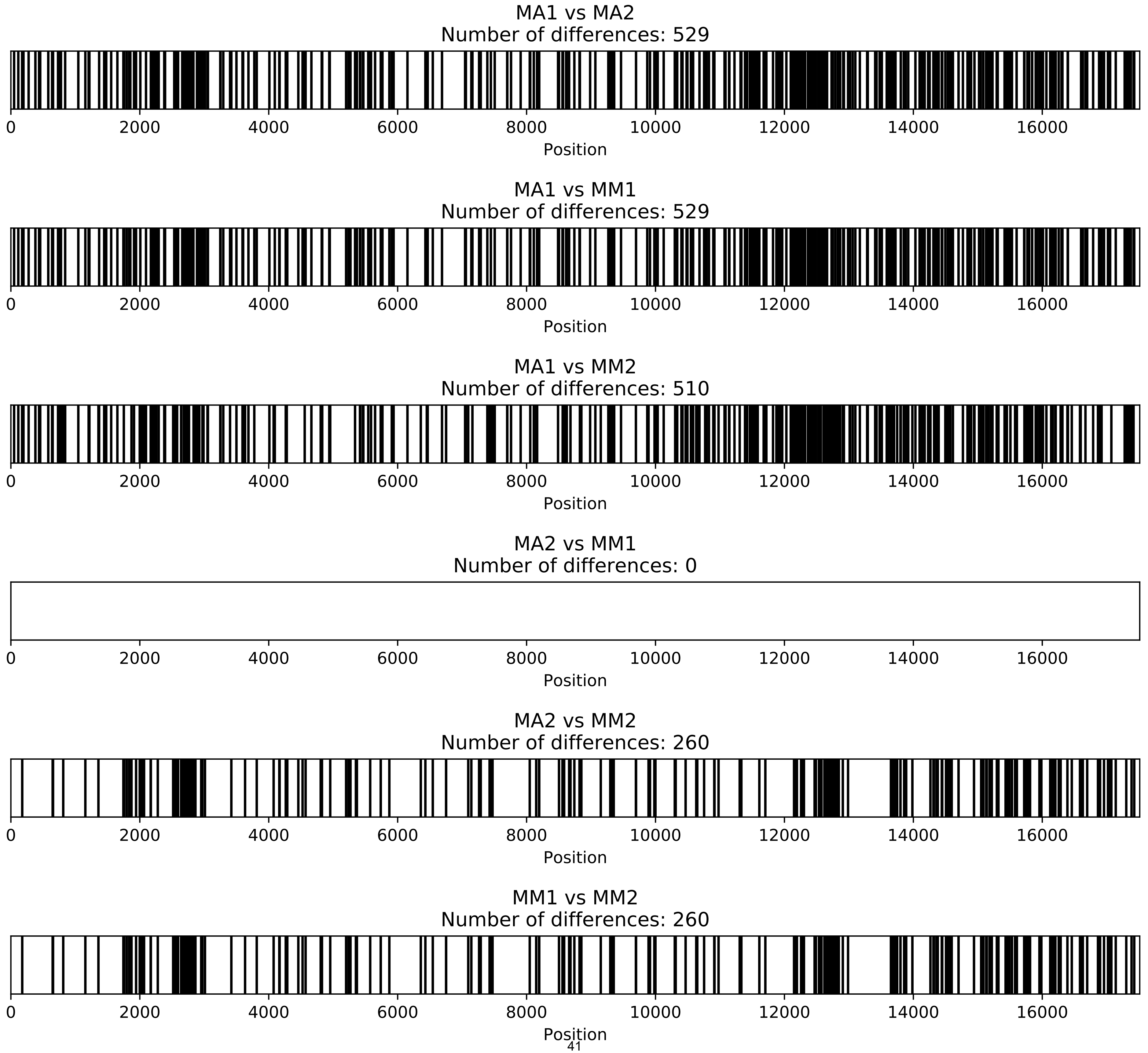
MA1 vs MA2
Number of differences: 529
MA1 vs MM1
Number of differences: 529
MA1 vs MM2
Number of differences: 510
MA2 vs MM1
Number of differences: 0
MA2 vs MM2
Number of differences: 260
MM1 vs MM2
Number of differences: 260
0 2000 4000 6000 8000 10000 12000 14000 16000
Position

### Region: 165



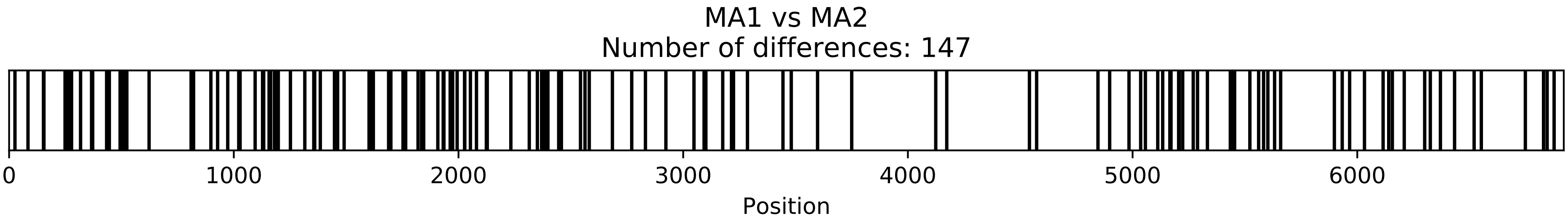
MA1 vs MA2
Number of differences: 147
0
1000
2000
3000
4000
5000
6000
Position

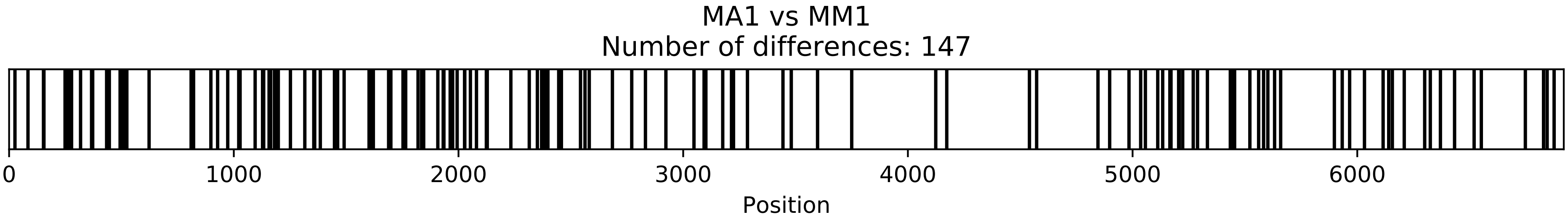
MA1 vs MM1
Number of differences: 147
0
1000
2000
3000
4000
5000
6000
Position

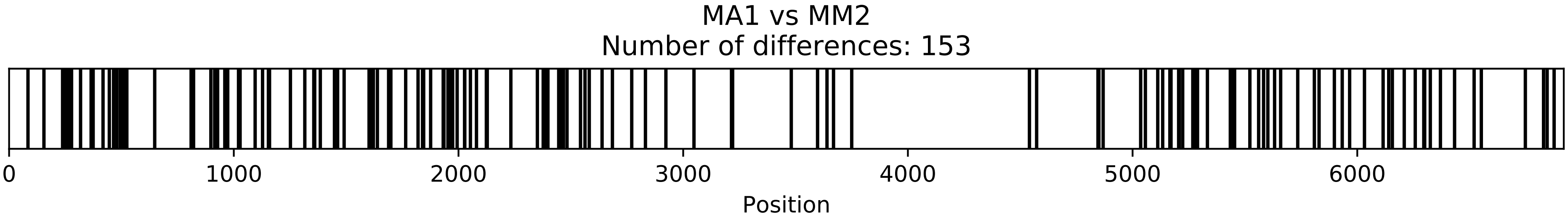
MA1 vs MM2
Number of differences: 153
0
1000
2000
3000
4000
5000
6000
Position

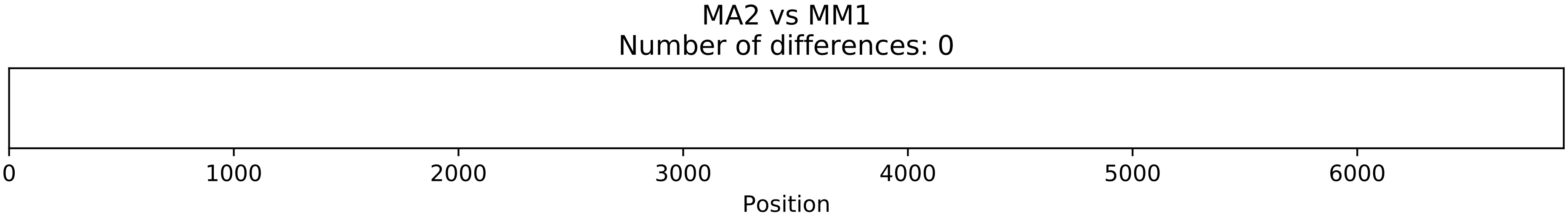
MA2 vs MM1
Number of differences: 0
0
1000
2000
3000
4000
5000
6000
Position

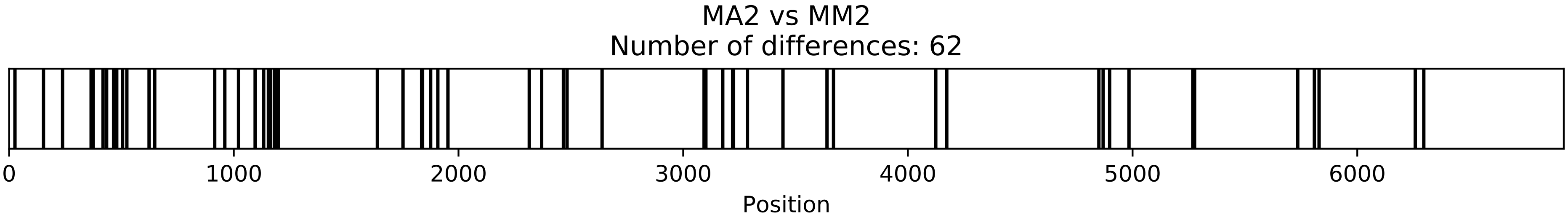
MA2 vs MM2
Number of differences: 62
0
1000
2000
3000
4000
5000
6000
Position

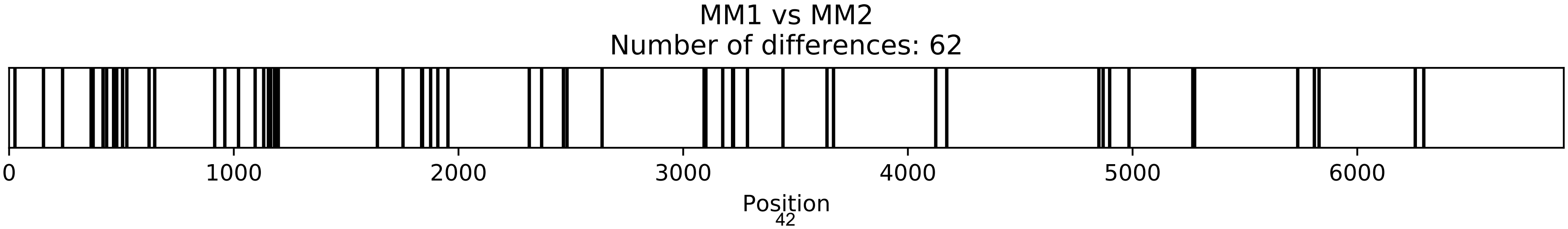
MM1 vs MM2
Number of differences: 62
0
1000
2000
3000
4000
5000
6000
Position

### Region: 167



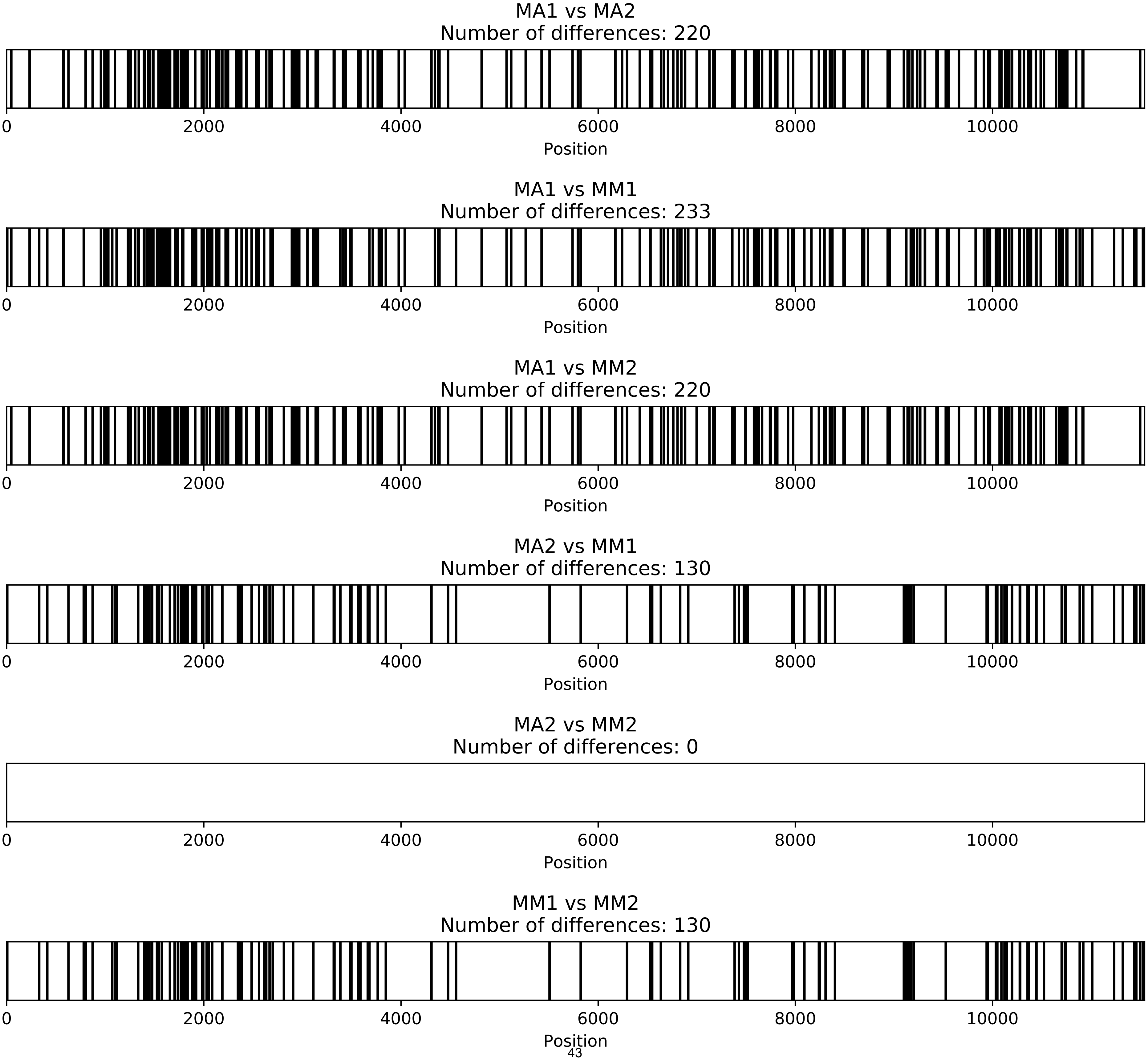
MA1 vs MA2
Number of differences: 220
MA1 vs MM1
Number of differences: 233
MA1 vs MM2
Number of differences: 220
MA2 vs MM1
Number of differences: 130
MA2 vs MM2
Number of differences: 0
MM1 vs MM2
Number of differences: 130
0
2000
4000
6000
8000
10000
Position

### Region: 173



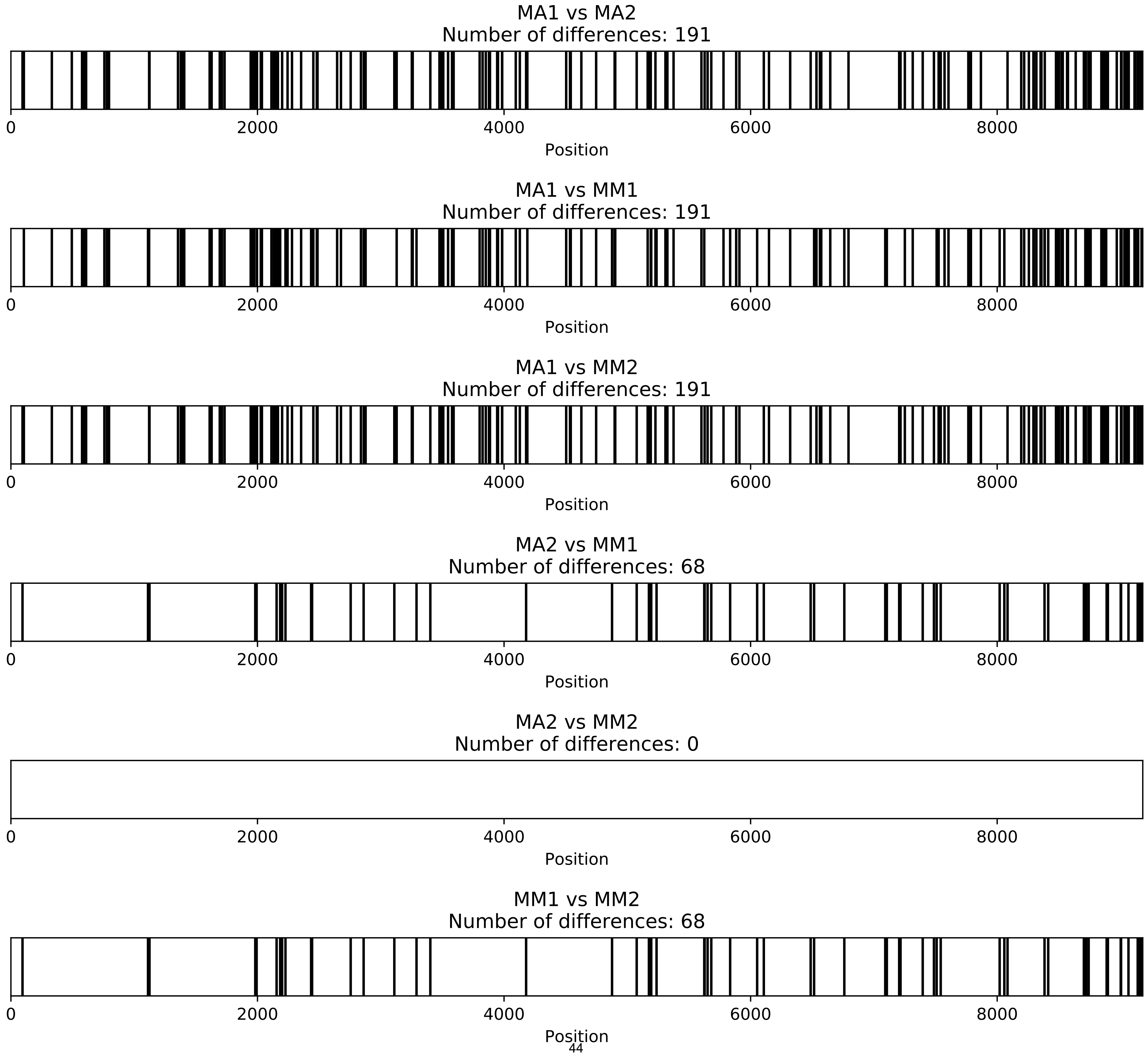
MA1 vs MA2
Number of differences: 191
MA1 vs MM1
Number of differences: 191
MA1 vs MM2
Number of differences: 191
MA2 vs MM1
Number of differences: 68
MA2 vs MM2
Number of differences: 0
MM1 vs MM2
Number of differences: 68
0
2000
4000
6000
8000
Position

### Region: 176



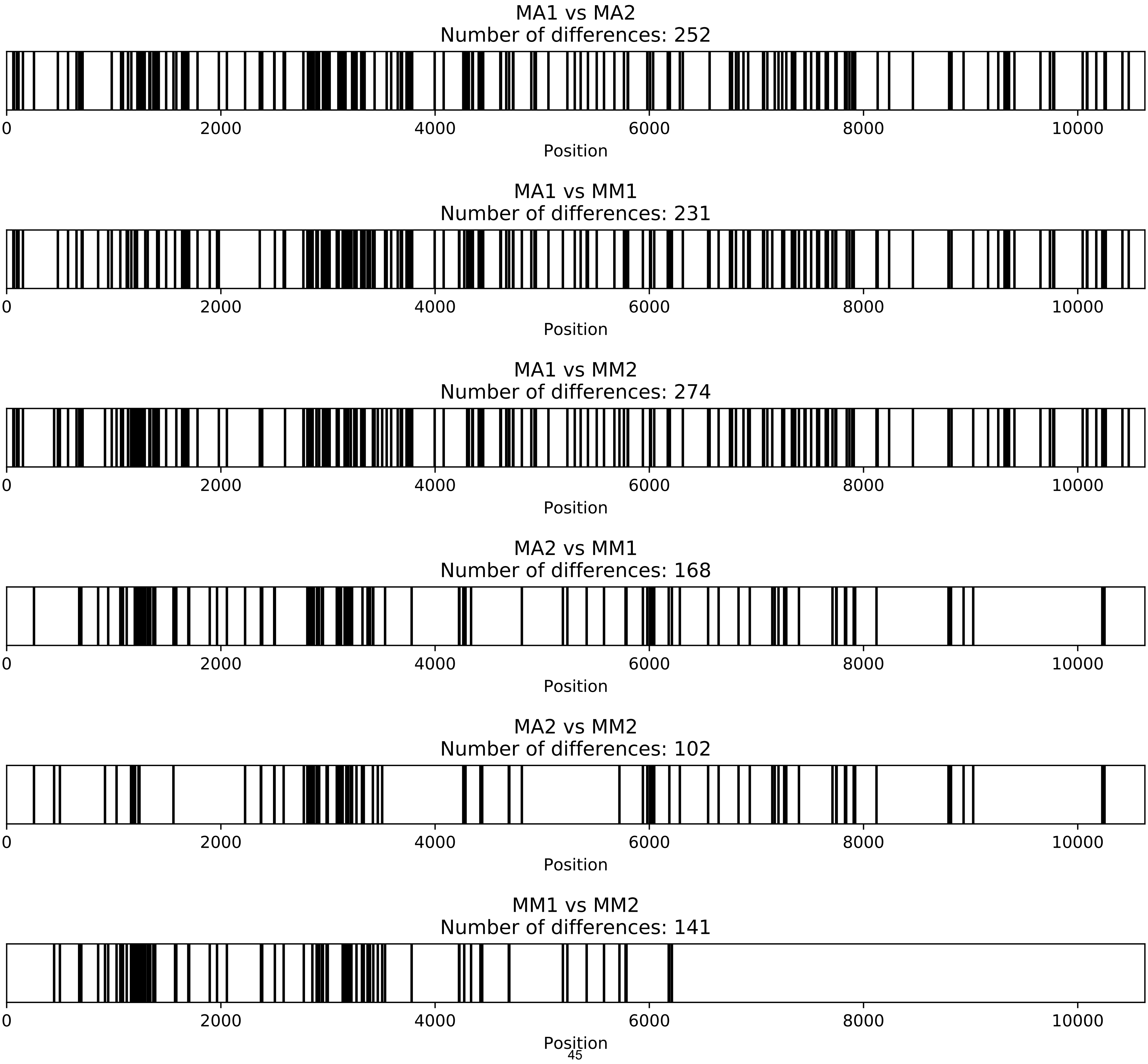
MA1 vs MA2
Number of differences: 252
0 2000 4000 6000 8000 10000
Position
MA1 vs MM1
Number of differences: 231
0 2000 4000 6000 8000 10000
Position
MA1 vs MM2
Number of differences: 274
0 2000 4000 6000 8000 10000
Position
MA2 vs MM1
Number of differences: 168
0 2000 4000 6000 8000 10000
Position
MA2 vs MM2
Number of differences: 102
0 2000 4000 6000 8000 10000
Position
MM1 vs MM2
Number of differences: 141
0 2000 4000 6000 8000 10000
Position

### Region: 178



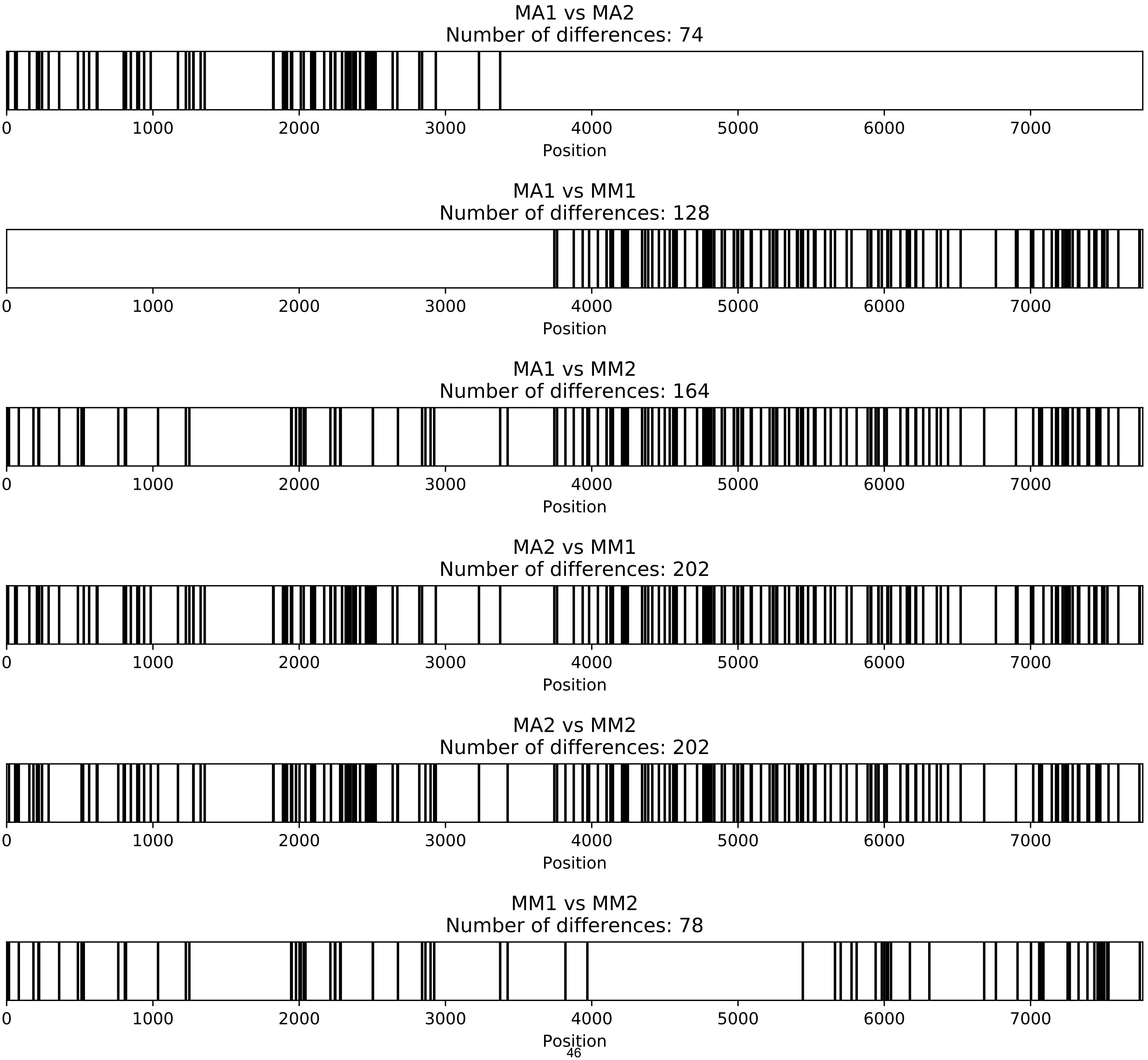
MA1 vs MA2
Number of differences: 74
MA1 vs MM1
Number of differences: 128
MA1 vs MM2
Number of differences: 164
MA2 vs MM1
Number of differences: 202
MA2 vs MM2
Number of differences: 202
MM1 vs MM2
Number of differences: 78
0
1000
2000
3000
4000
5000
6000
7000
Position

### Region: 179



MA1 vs MA2
Number of differences: 261

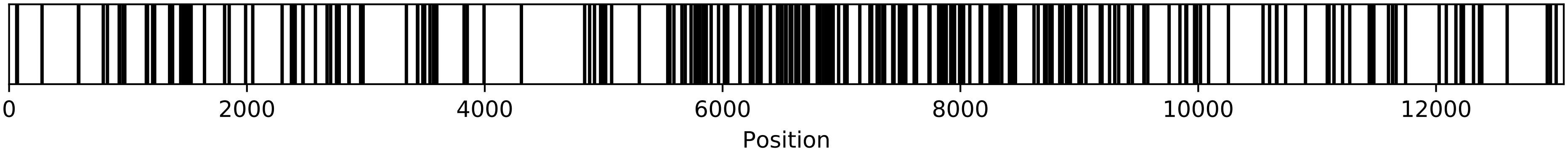


MA1 vs MM1
Number of differences: 155

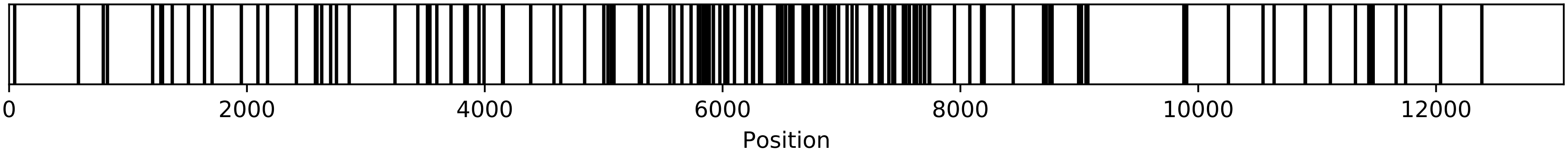


MA1 vs MM2
Number of differences: 0

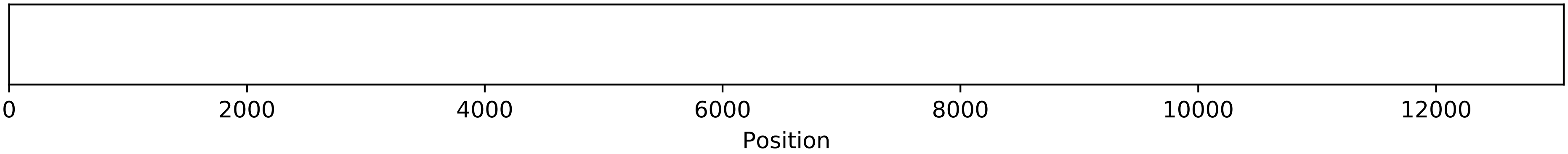


MA2 vs MM1
Number of differences: 275

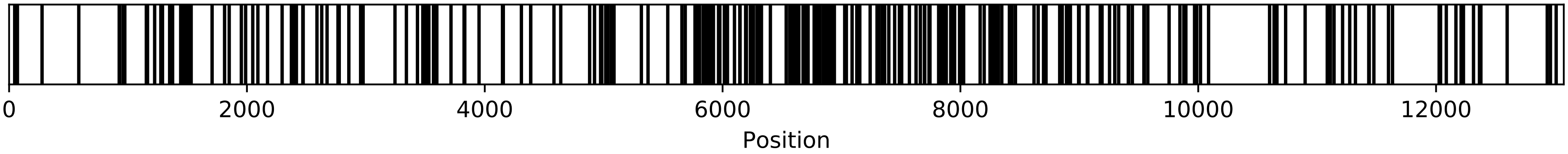


MA2 vs MM2
Number of differences: 261

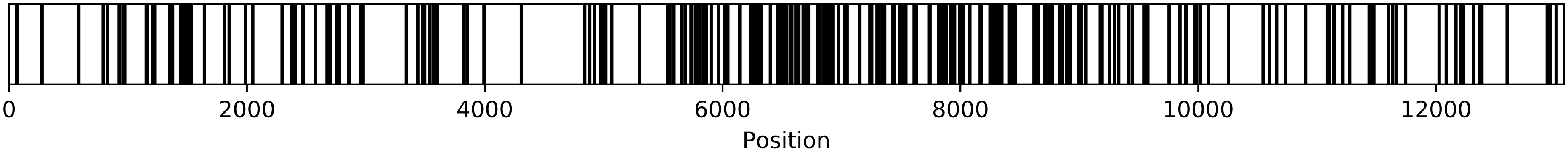


MM1 vs MM2
Number of differences: 155

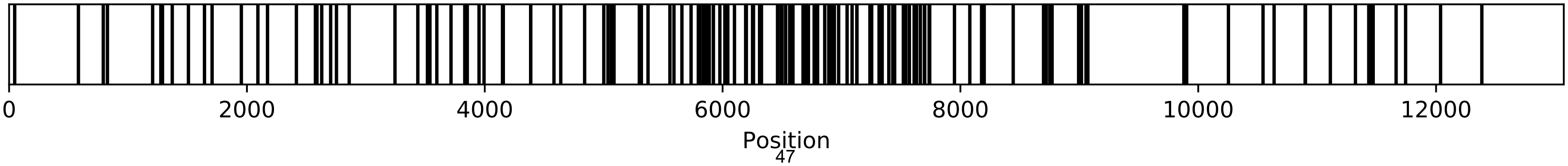

### Region: 181



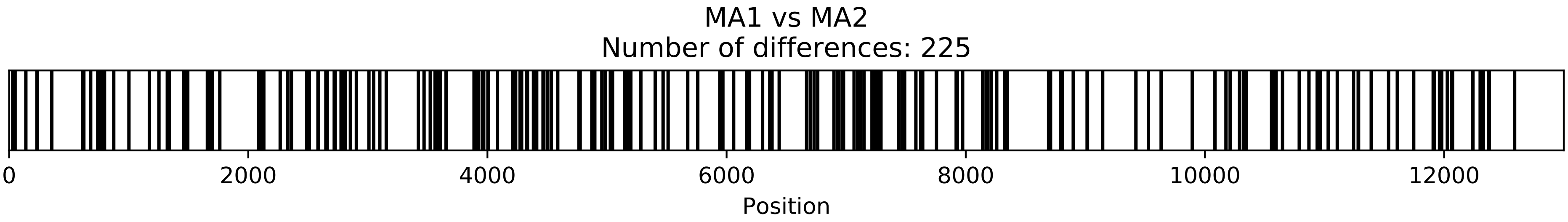
MA1 vs MA2
Number of differences: 225
0
2000
4000
6000
8000
10000
12000
Position

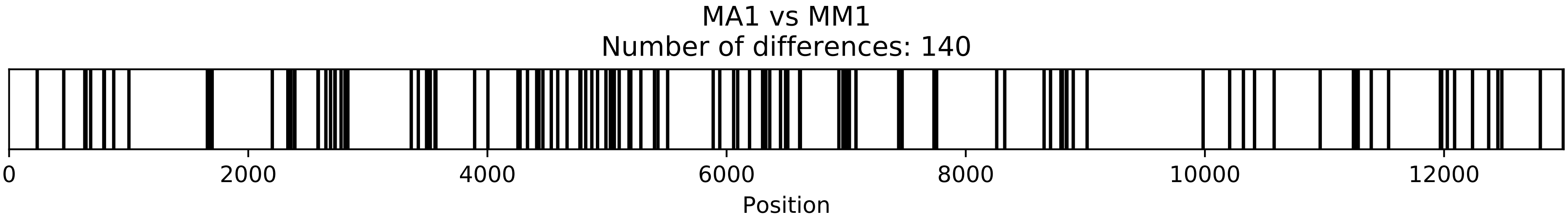
MA1 vs MM1
Number of differences: 140
0
2000
4000
6000
8000
10000
12000
Position

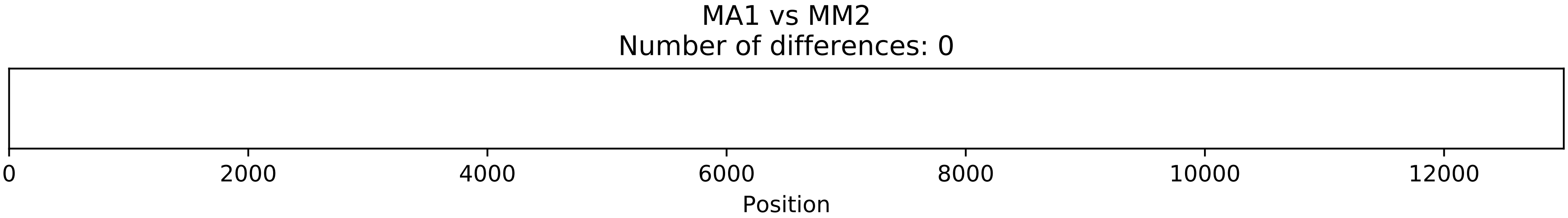
MA1 vs MM2
Number of differences: 0
0
2000
4000
6000
8000
10000
12000
Position

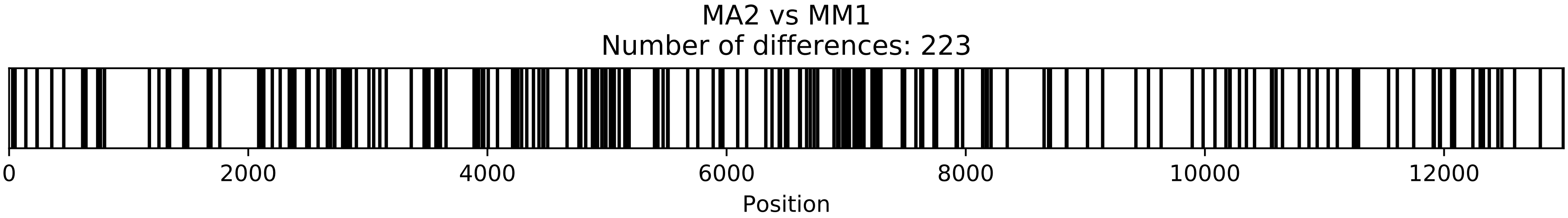
MA2 vs MM1
Number of differences: 223
0
2000
4000
6000
8000
10000
12000
Position

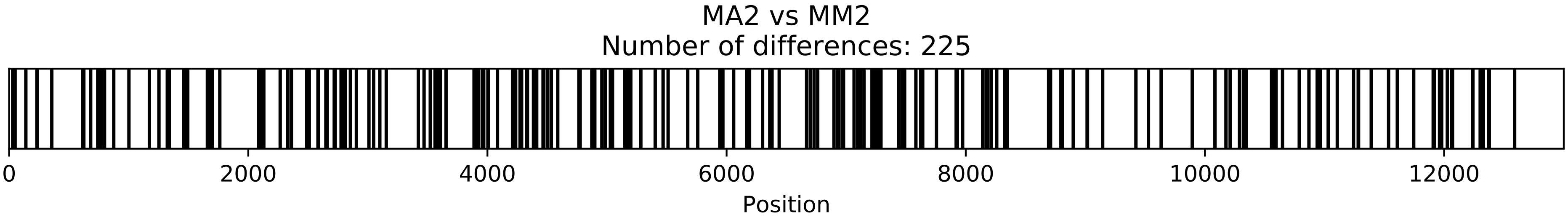
MA2 vs MM2
Number of differences: 225
0
2000
4000
6000
8000
10000
12000
Position

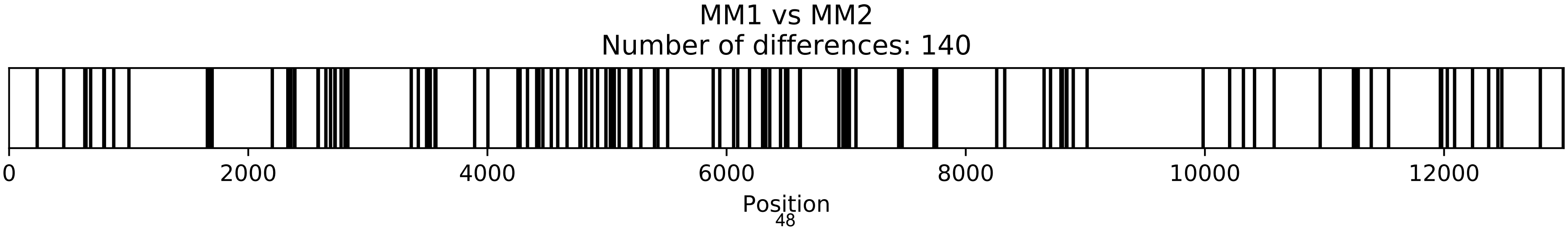
MM1 vs MM2
Number of differences: 140
0
2000
4000
6000
8000
10000
12000
Position

### Region: 183

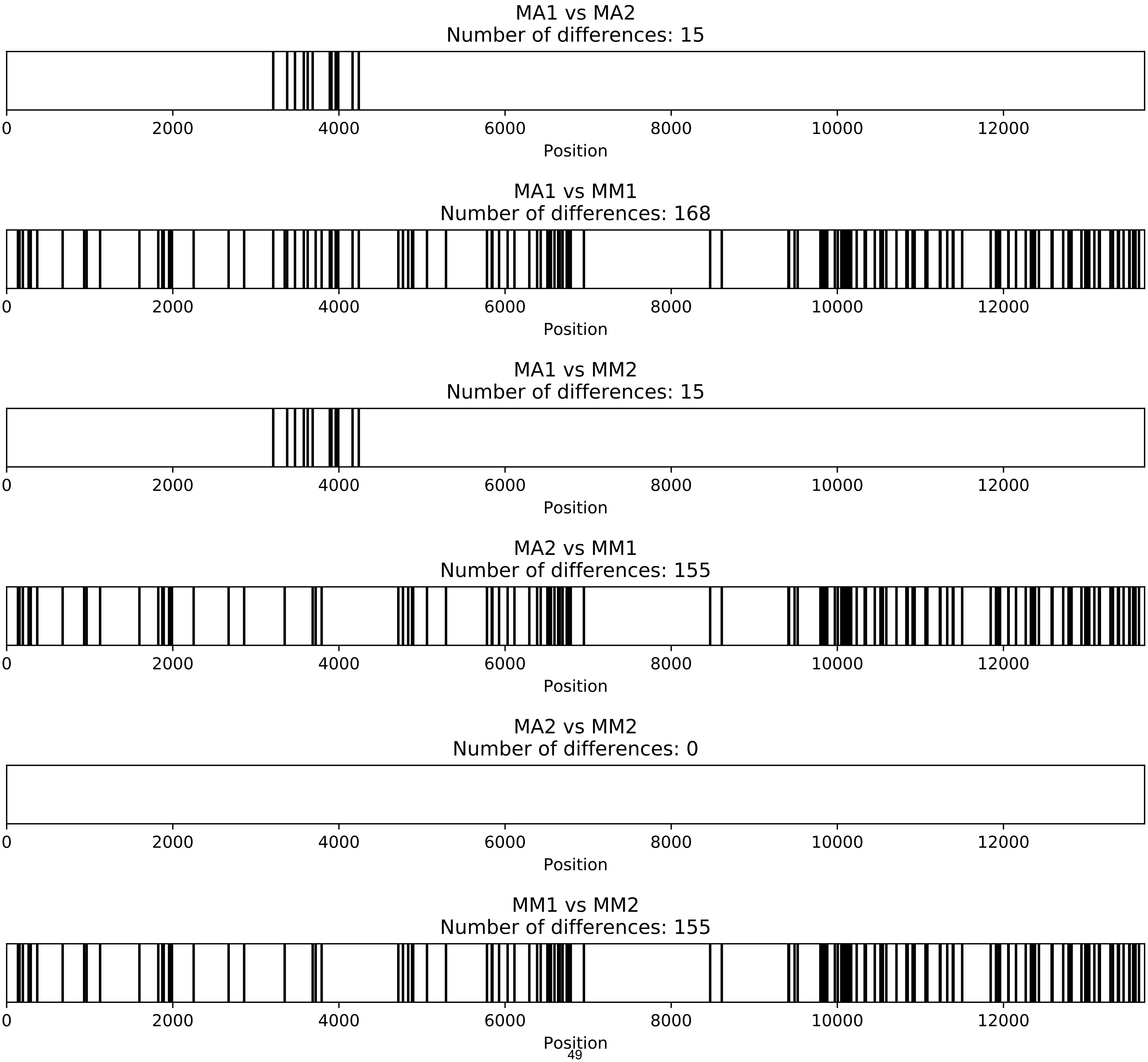
MA1 vs MA2
Number of differences: 15
MA1 vs MM1
Number of differences: 168
MA1 vs MM2
Number of differences: 15
MA2 vs MM1
Number of differences: 155
MA2 vs MM2
Number of differences: 0
MM1 vs MM2
Number of differences: 155
0
2000
4000
6000
8000
10000
12000
Position

### Region: 184



MA1 vs MA2
Number of differences: 296

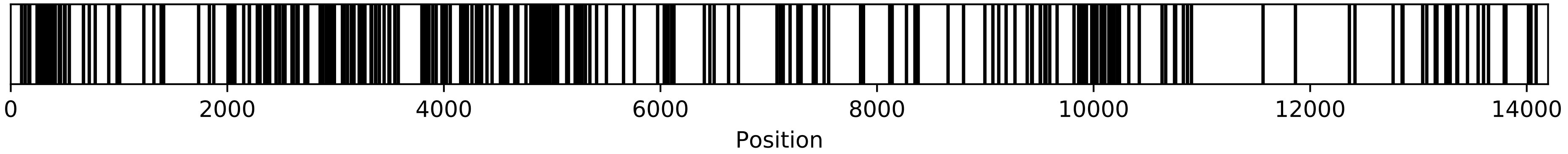


MA1 vs MM1
Number of differences: 296

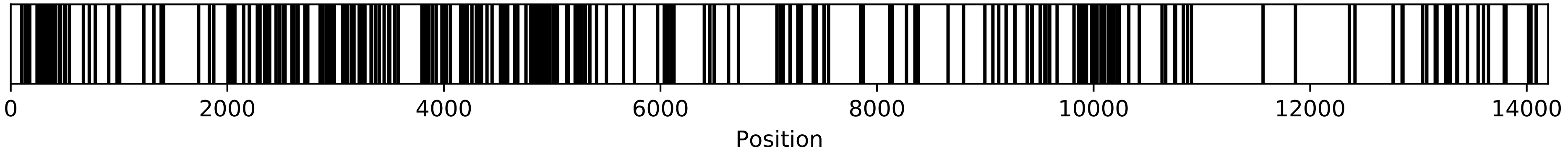


MA1 vs MM2
Number of differences: 293

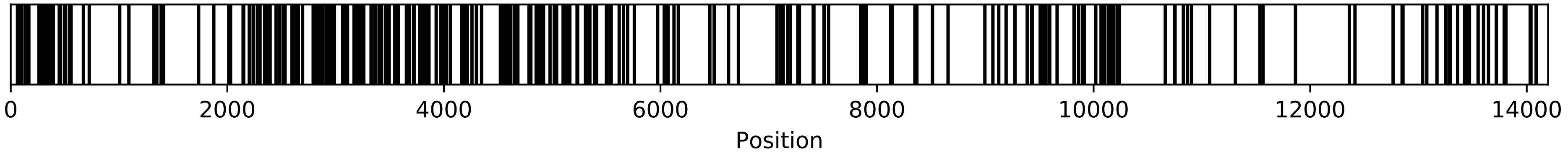


MA2 vs MM1
Number of differences: 0

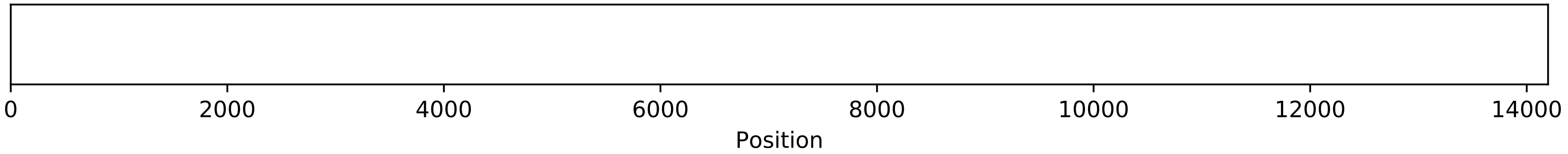


MA2 vs MM2
Number of differences: 168

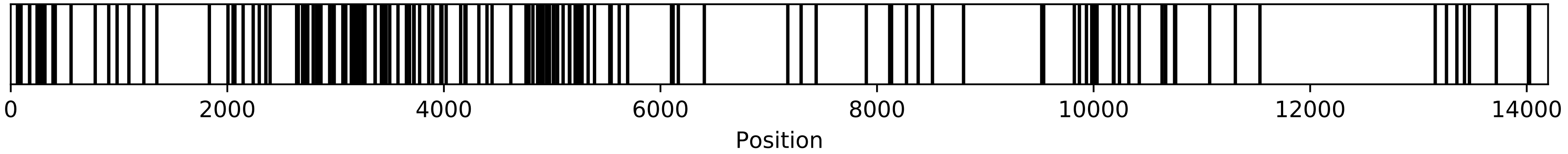


MM1 vs MM2
Number of differences: 168

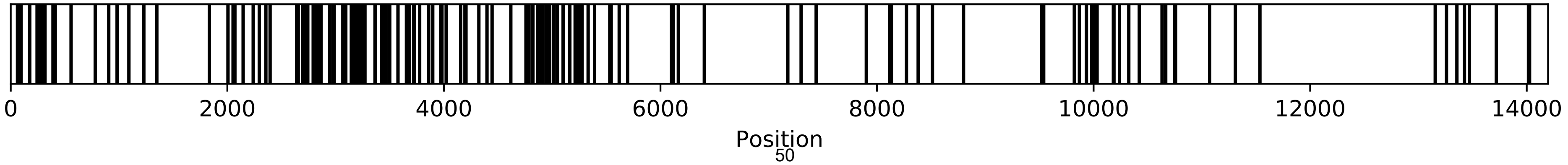

### Region: 186



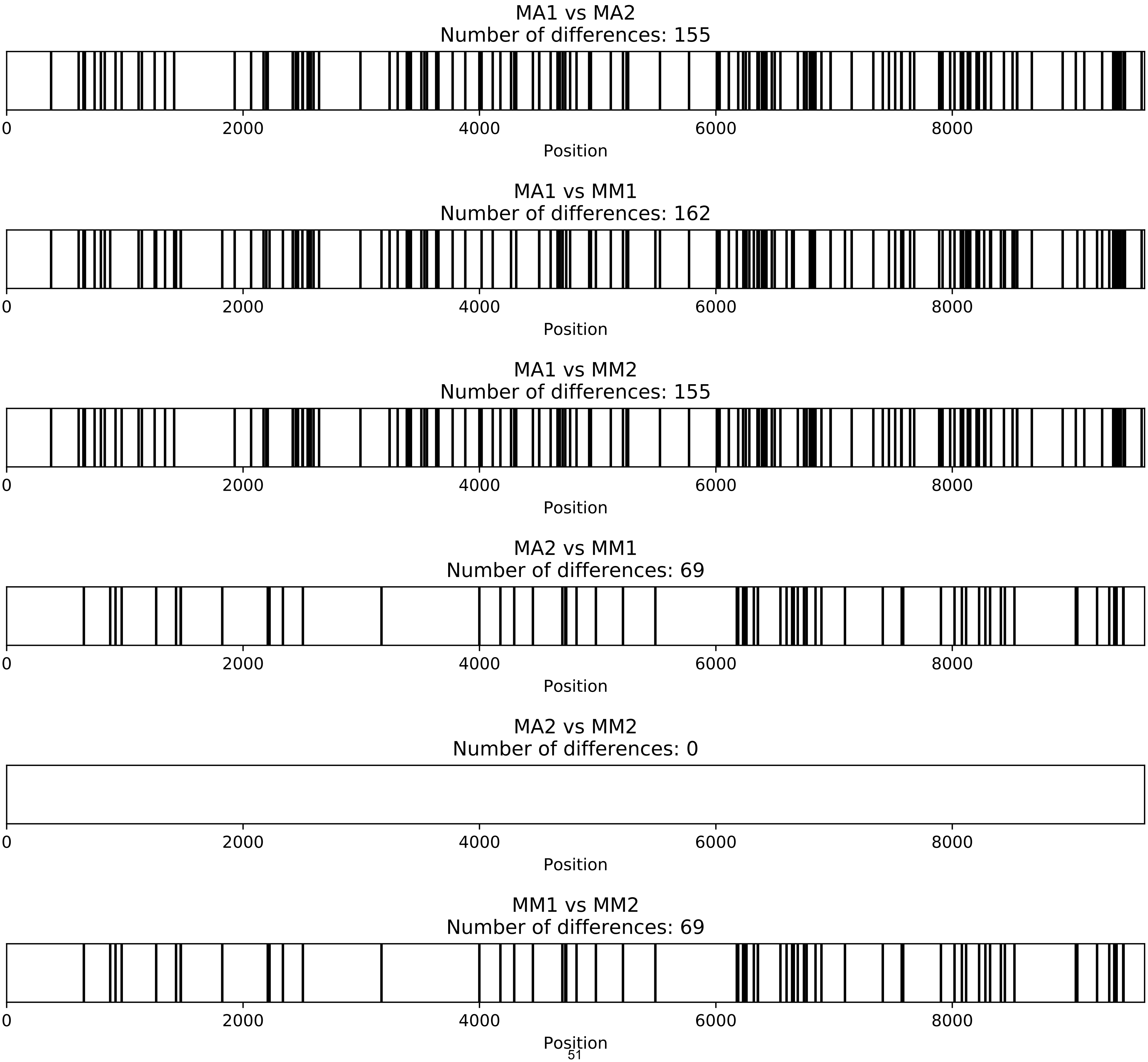
MA1 vs MA2
Number of differences: 155
MA1 vs MM1
Number of differences: 162
MA1 vs MM2
Number of differences: 155
MA2 vs MM1
Number of differences: 69
MA2 vs MM2
Number of differences: 0
MM1 vs MM2
Number of differences: 69
0
2000
4000
6000
8000
Position

### Region: 189



MA1 vs MA2
Number of differences: 229

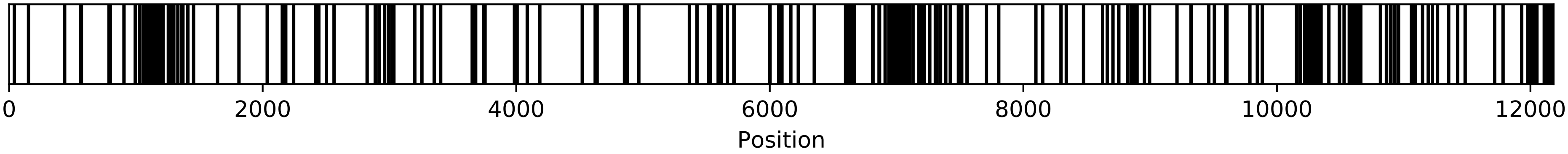


MA1 vs MM1
Number of differences: 220

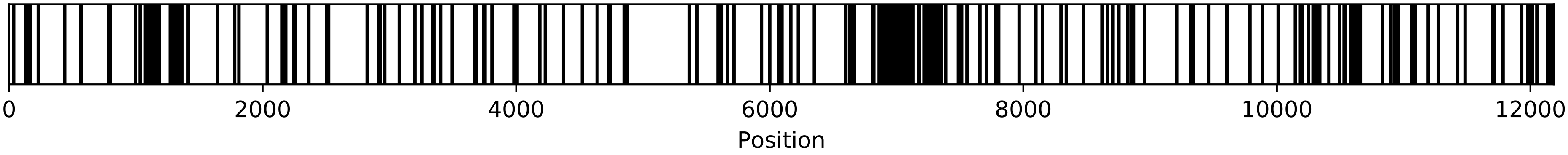


MA1 vs MM2
Number of differences: 229

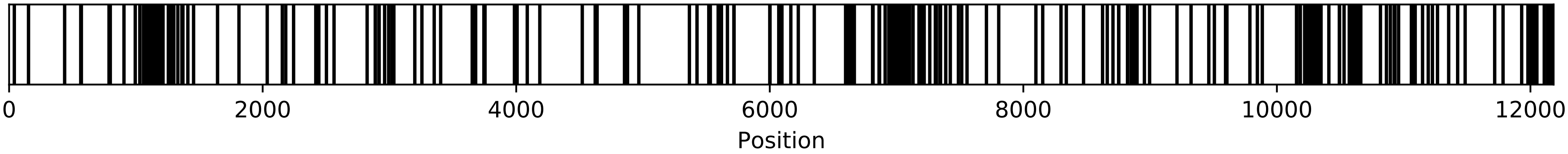


MA2 vs MM1
Number of differences: 142

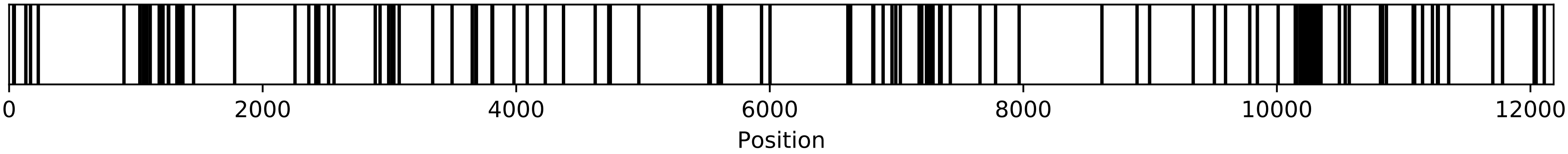


MA2 vs MM2
Number of differences: 0

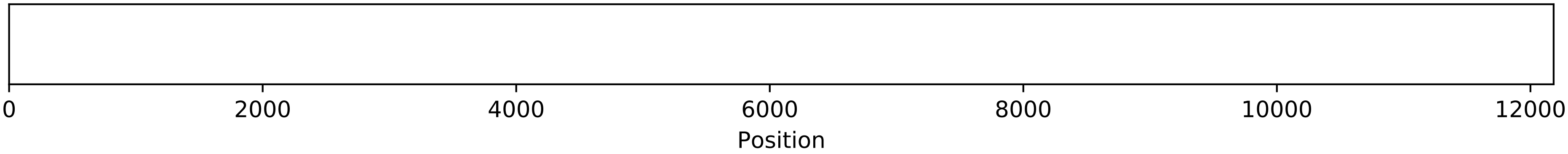


MM1 vs MM2
Number of differences: 142

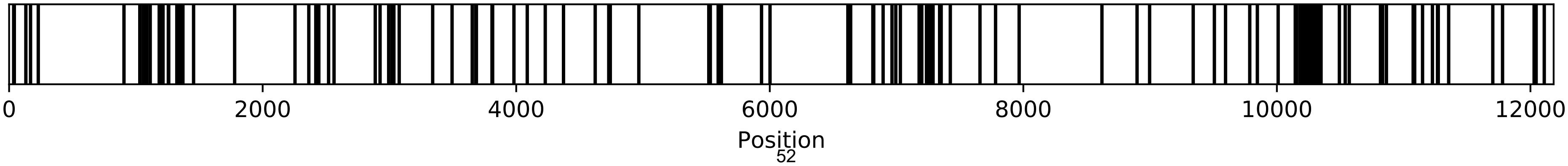

### Region: 191



MA1 vs MA2
Number of differences: 243

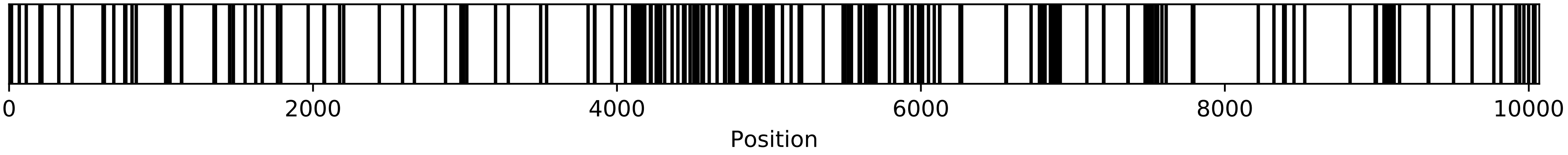


MA1 vs MM1
Number of differences: 108

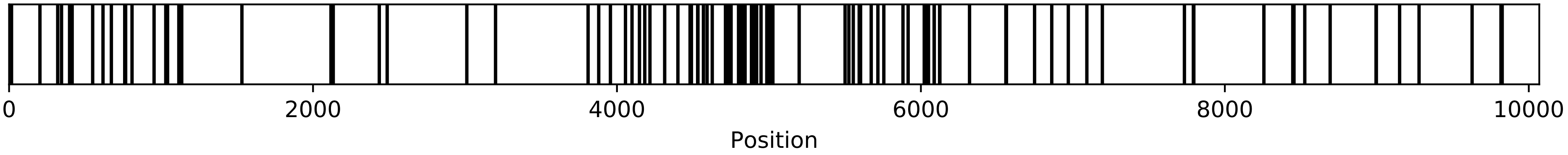


MA1 vs MM2
Number of differences: 115

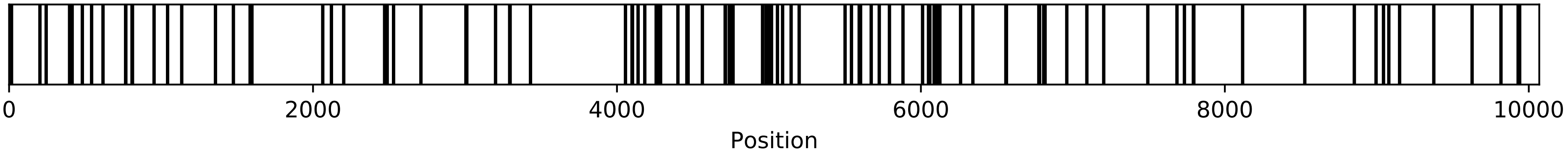


MA2 vs MM1
Number of differences: 235

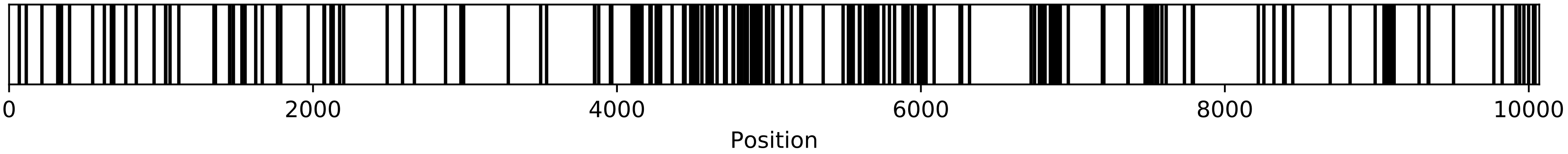


MA2 vs MM2
Number of differences: 246

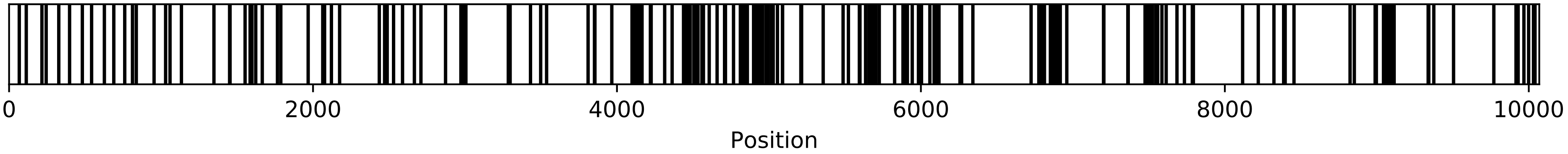


MM1 vs MM2
Number of differences: 130

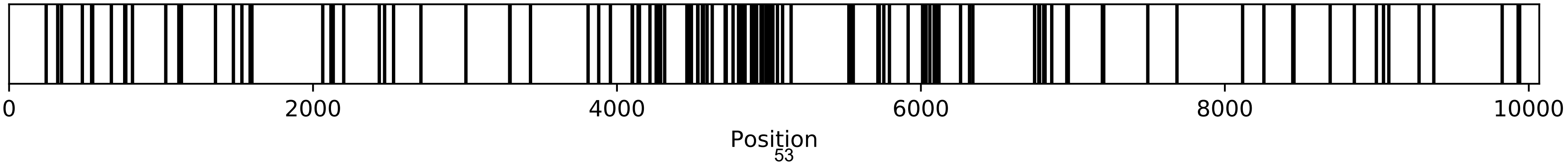

### Region: 196



MA1 vs MA2
Number of differences: 167

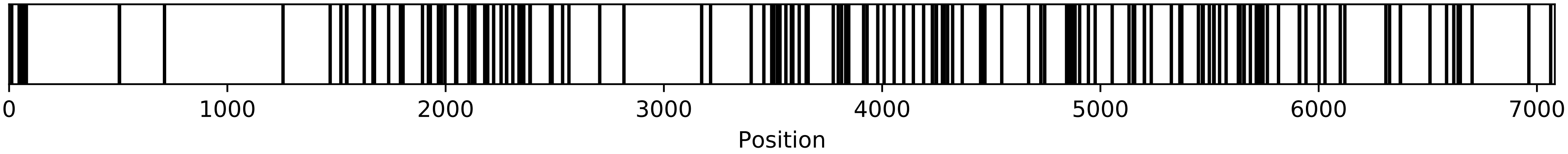


MA1 vs MM1
Number of differences: 195

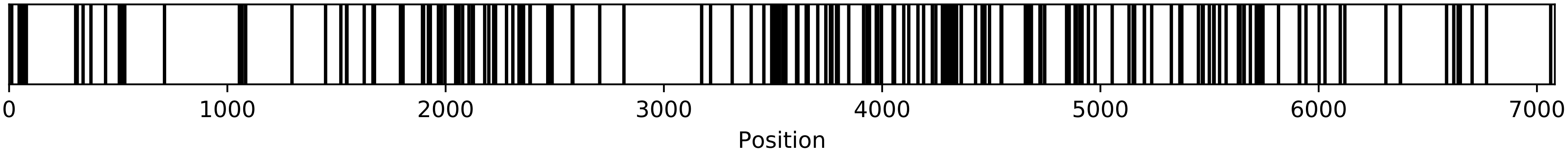


MA1 vs MM2
Number of differences: 179

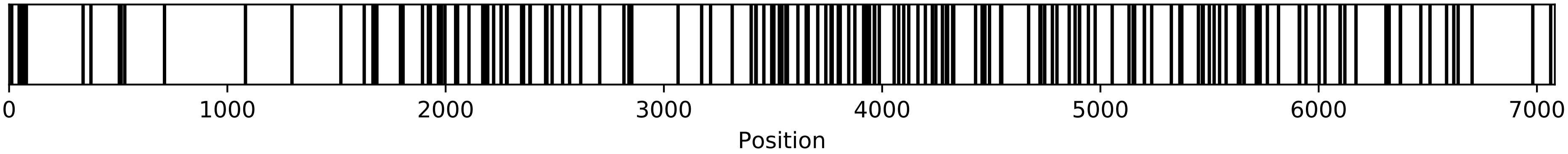


MA2 vs MM1
Number of differences: 92

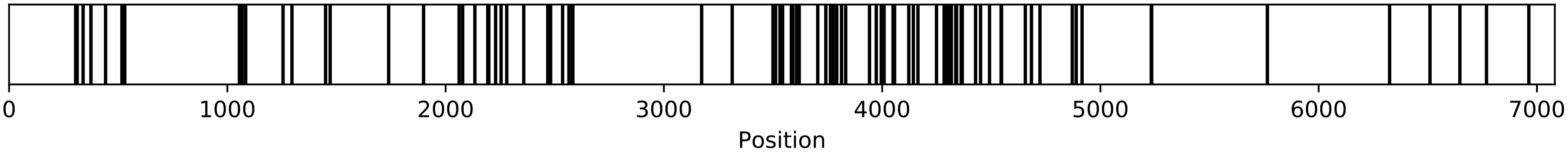


MA2 vs MM2
Number of differences: 97

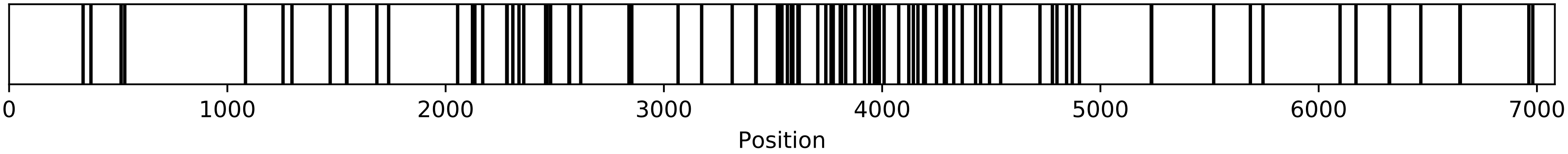


MM1 vs MM2
Number of differences: 106

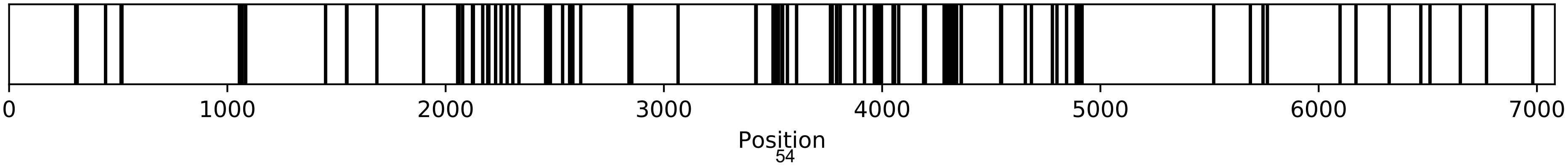

### Region: 198



MA1 vs MA2
Number of differences: 403

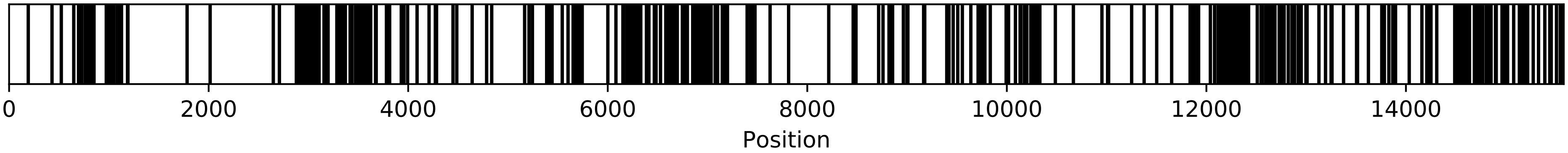


MA1 vs MM1
Number of differences: 411

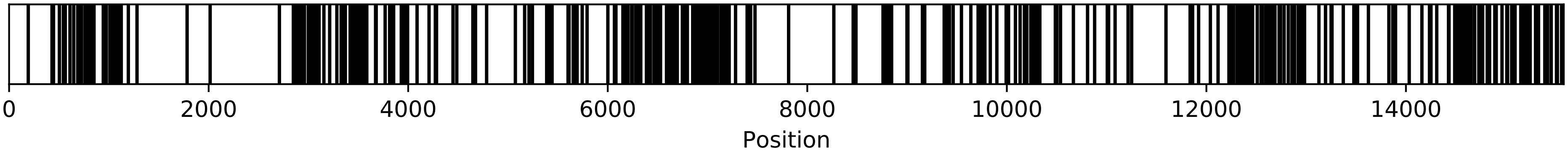


MA1 vs MM2
Number of differences: 403

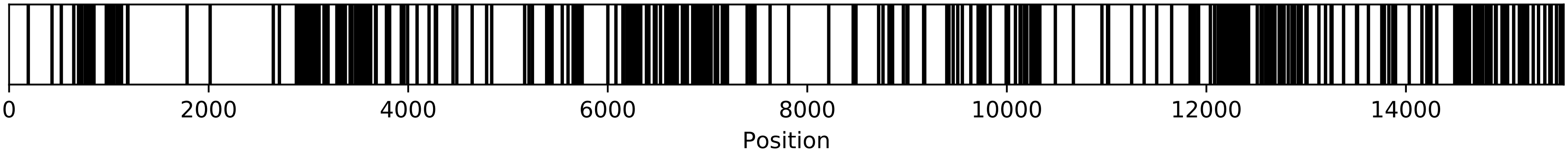


MA2 vs MM1
Number of differences: 270

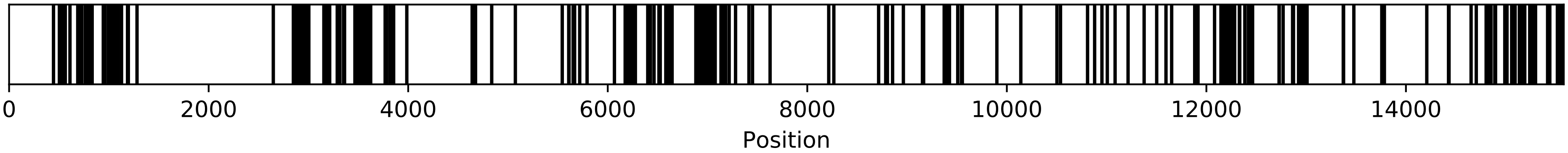


MA2 vs MM2
Number of differences: 0

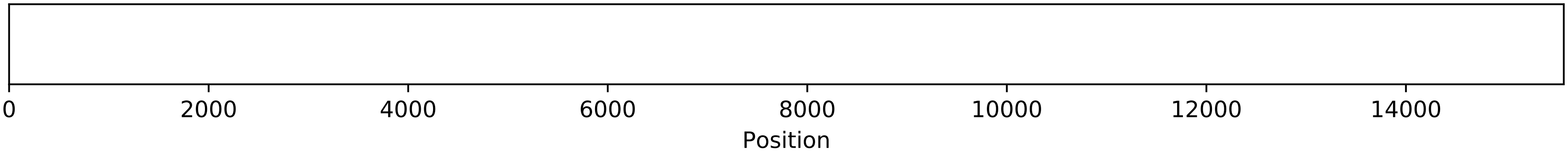


MM1 vs MM2
Number of differences: 270

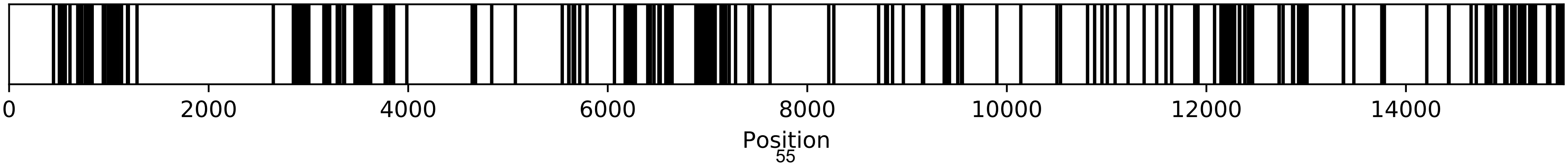

### Region: 199



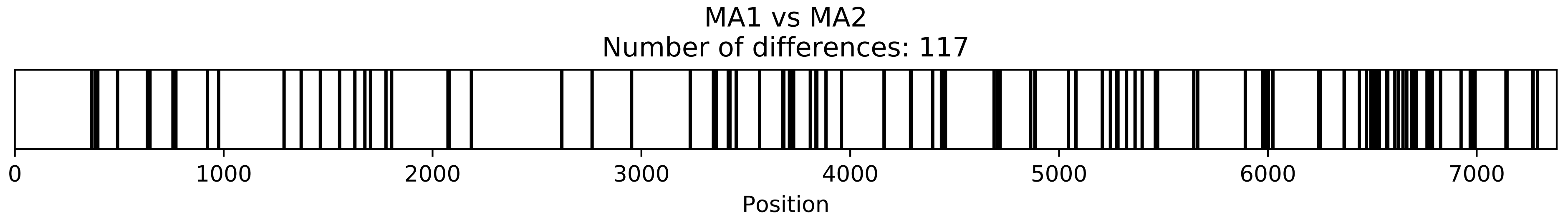
MA1 vs MA2
Number of differences: 117
0
1000
2000
3000
4000
5000
6000
7000
Position

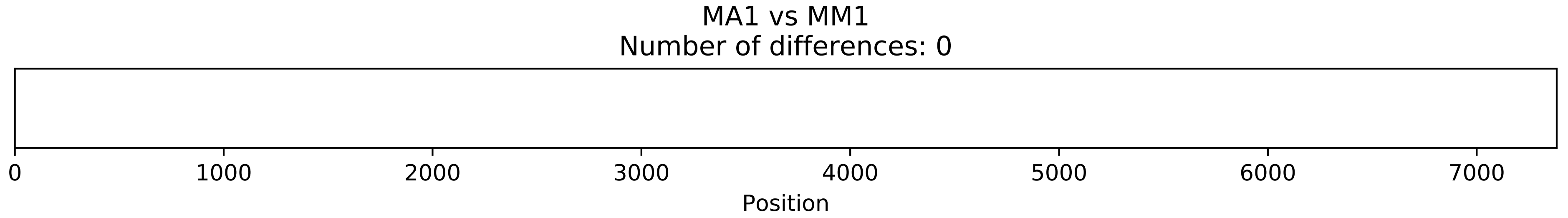
MA1 vs MM1
Number of differences: 0
0
1000
2000
3000
4000
5000
6000
7000
Position

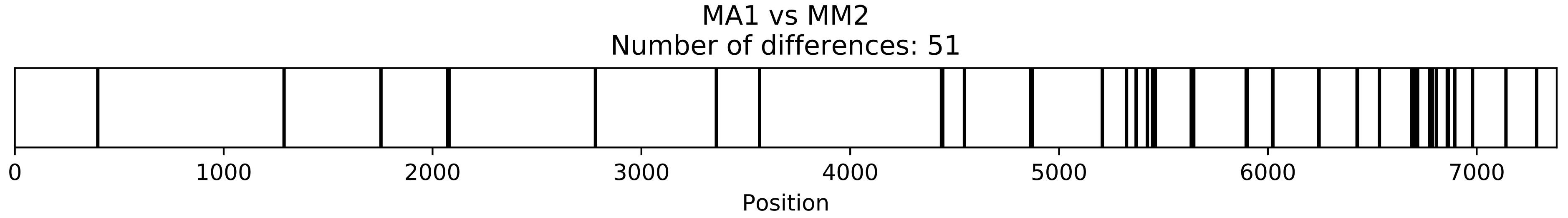
MA1 vs MM2
Number of differences: 51
0
1000
2000
3000
4000
5000
6000
7000
Position

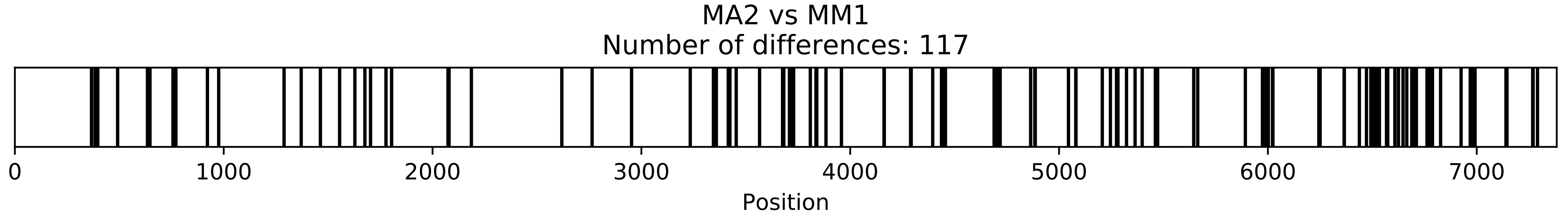
MA2 vs MM1
Number of differences: 117
0
1000
2000
3000
4000
5000
6000
7000
Position

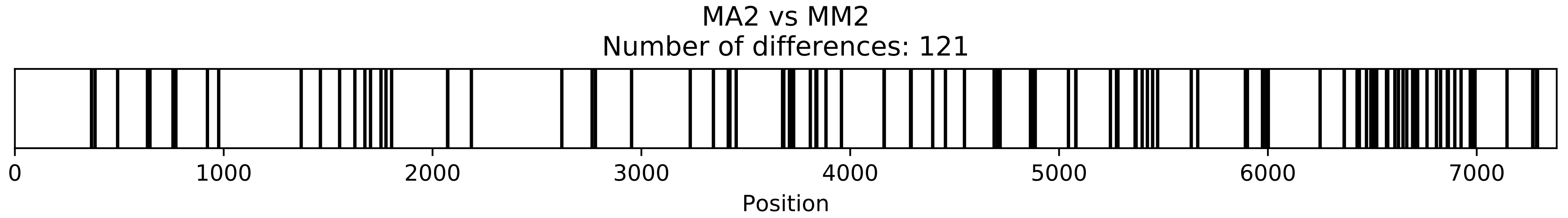
MA2 vs MM2
Number of differences: 121
0
1000
2000
3000
4000
5000
6000
7000
Position

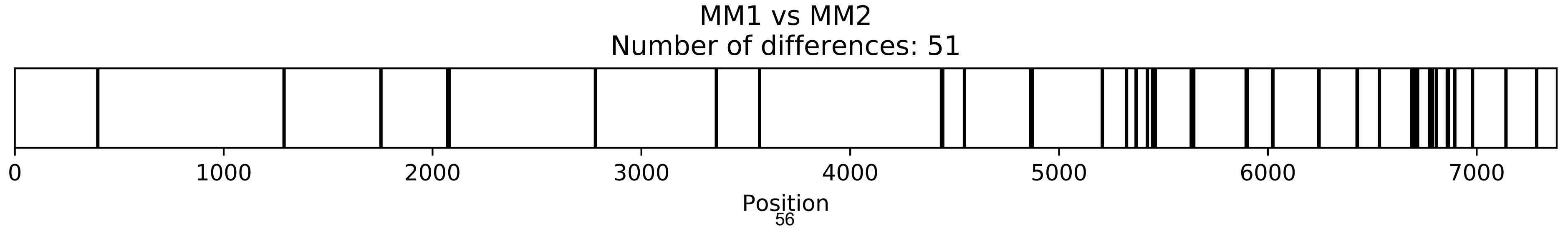
MM1 vs MM2
Number of differences: 51
0
1000
2000
3000
4000
5000
6000
7000
Position

### Region: 203



MA1 vs MA2
Number of differences: 445

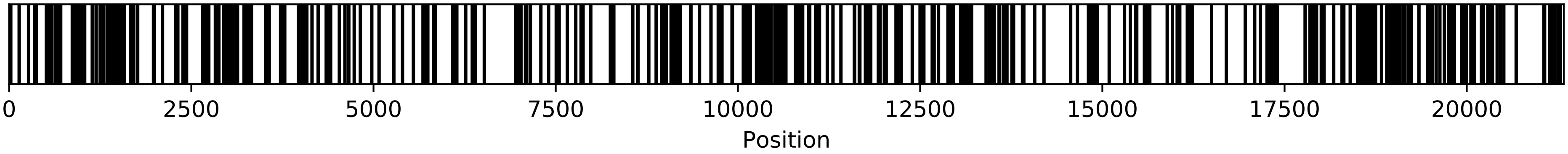


MA1 vs MM1
Number of differences: 244

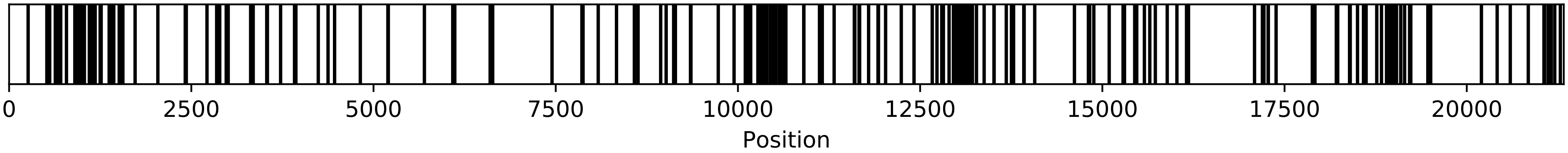


MA1 vs MM2
Number of differences: 241

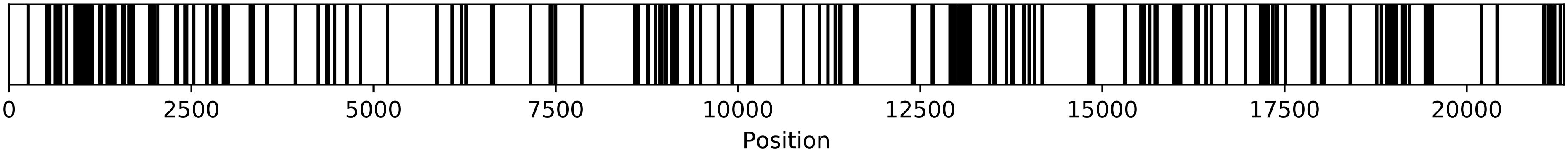


MA2 vs MM1
Number of differences: 471

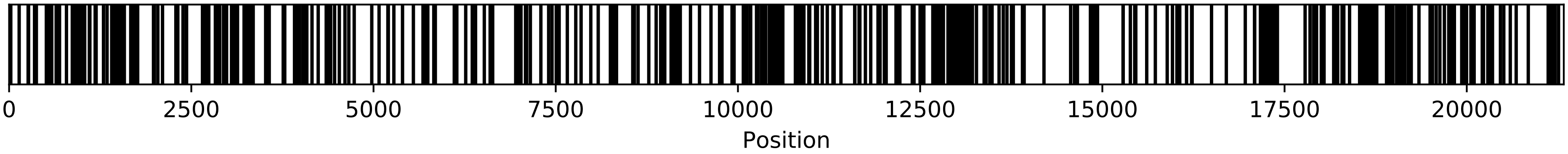


MA2 vs MM2
Number of differences: 461

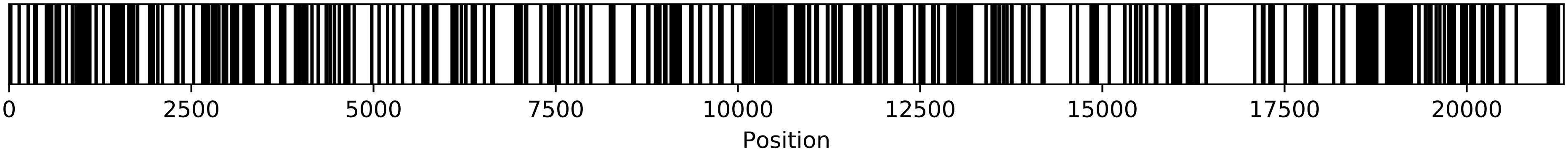


MM1 vs MM2
Number of differences: 235

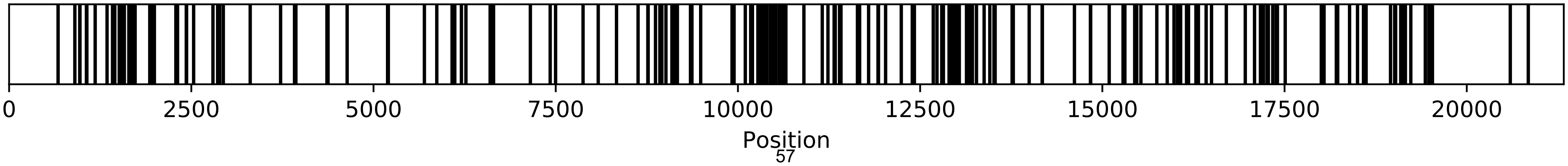

### Region: 206



MA1 vs MA2
Number of differences: 265

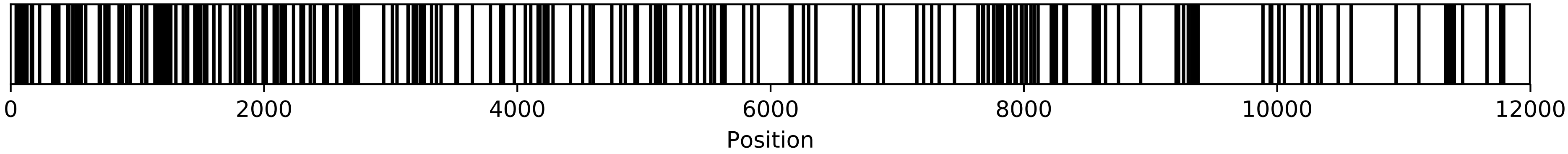


MA1 vs MM1
Number of differences: 265

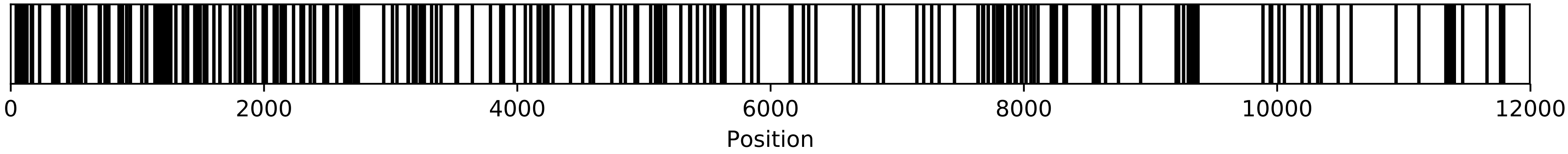


MA1 vs MM2
Number of differences: 265

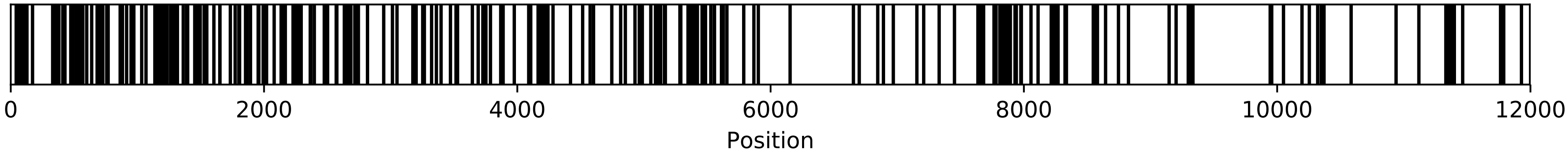


MA2 vs MM1
Number of differences: 0

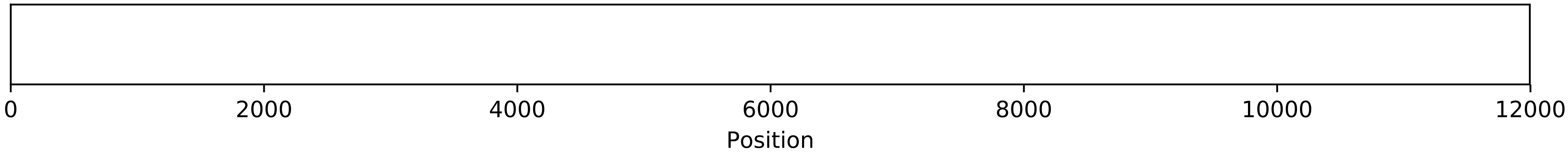


MA2 vs MM2
Number of differences: 144

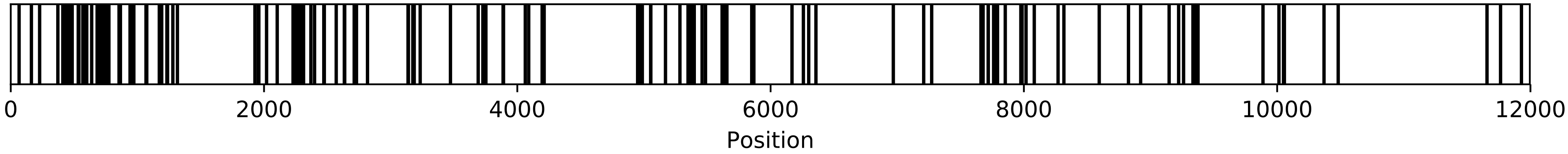


MM1 vs MM2
Number of differences: 144

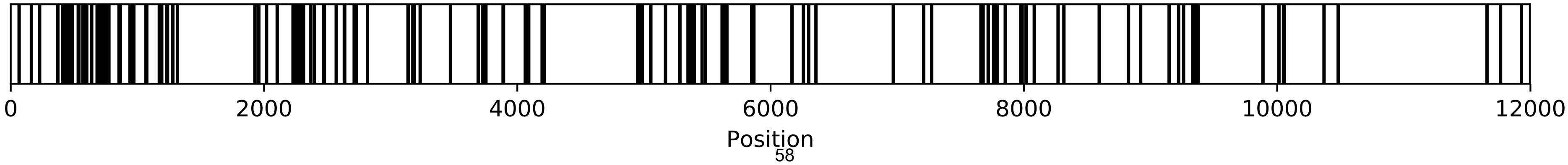

### Region: 212



MA1 vs MA2
Number of differences: 196

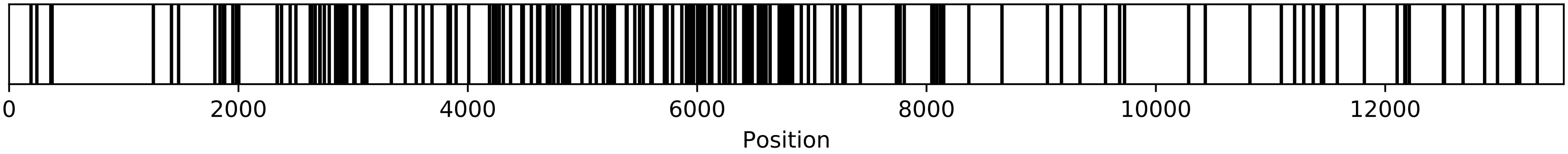


MA1 vs MM1
Number of differences: 200

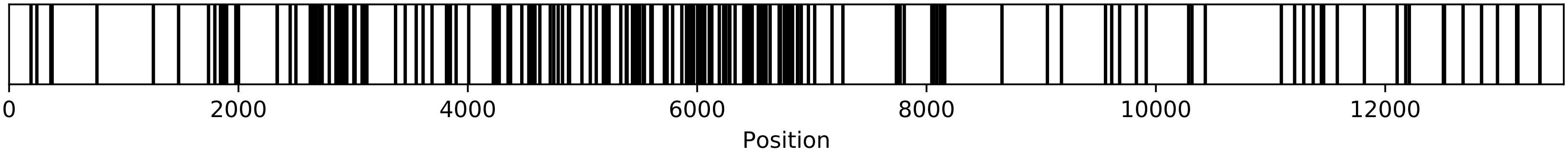


MA1 vs MM2
Number of differences: 196

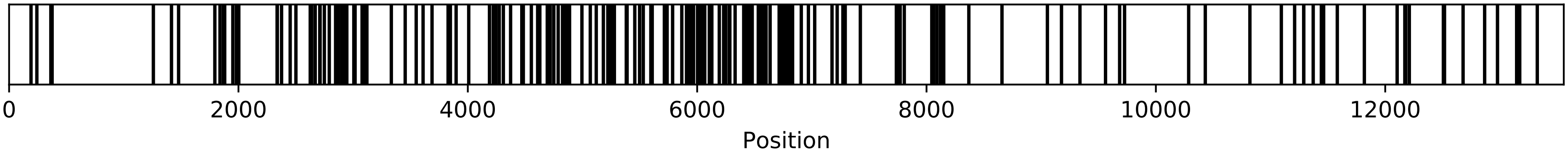


MA2 vs MM1
Number of differences: 78

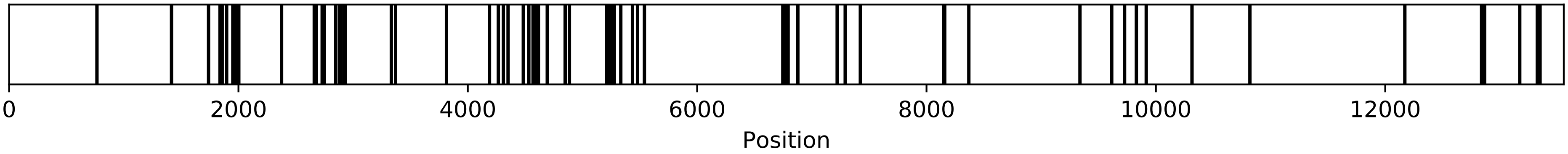


MA2 vs MM2
Number of differences: 0

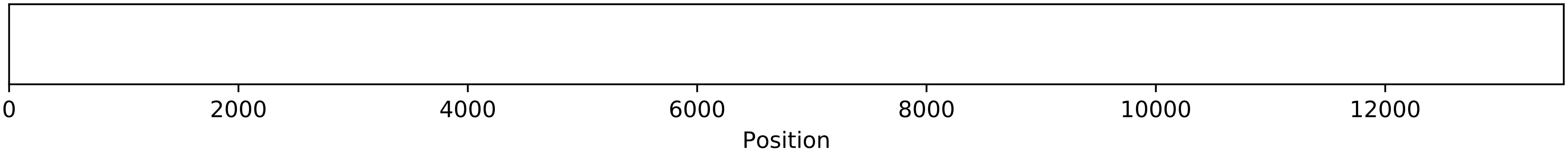


MM1 vs MM2
Number of differences: 78

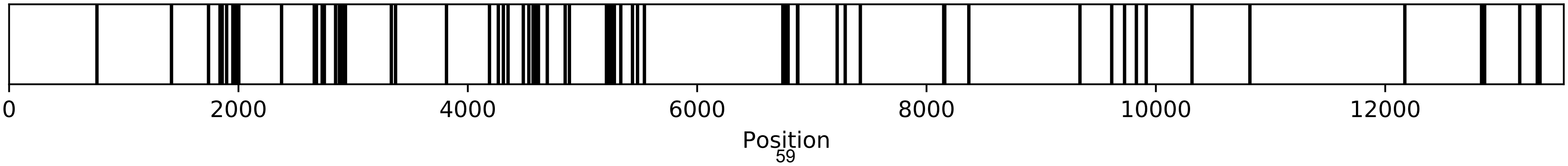

### Region: 215



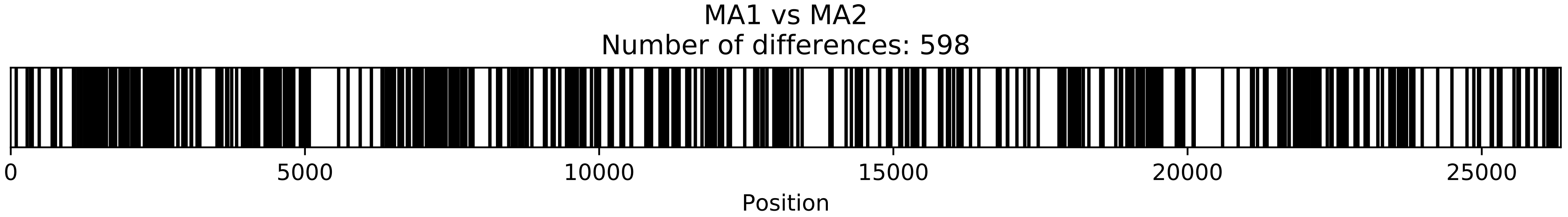
MA1 vs MA2
Number of differences: 598
0
5000
10000
15000
20000
25000
Position

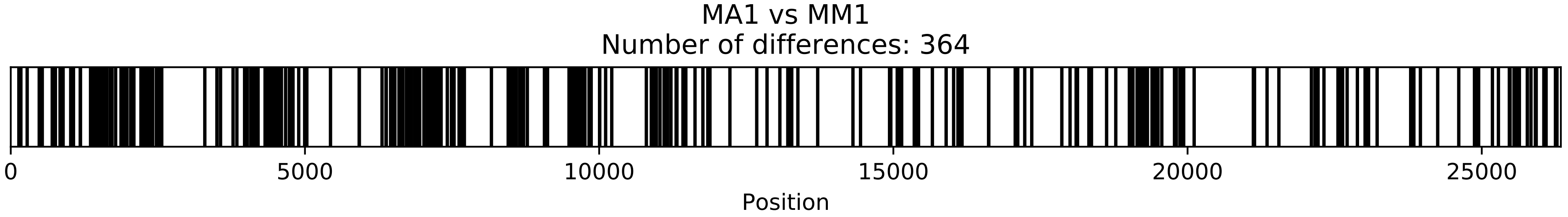
MA1 vs MM1
Number of differences: 364
0
5000
10000
15000
20000
25000
Position

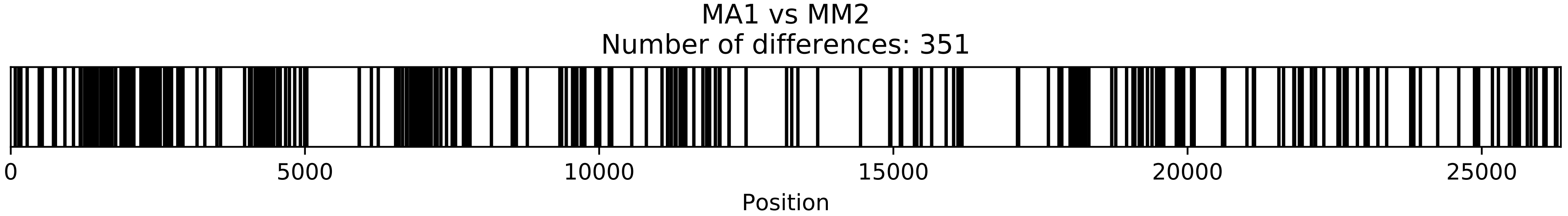
MA1 vs MM2
Number of differences: 351
0
5000
10000
15000
20000
25000
Position

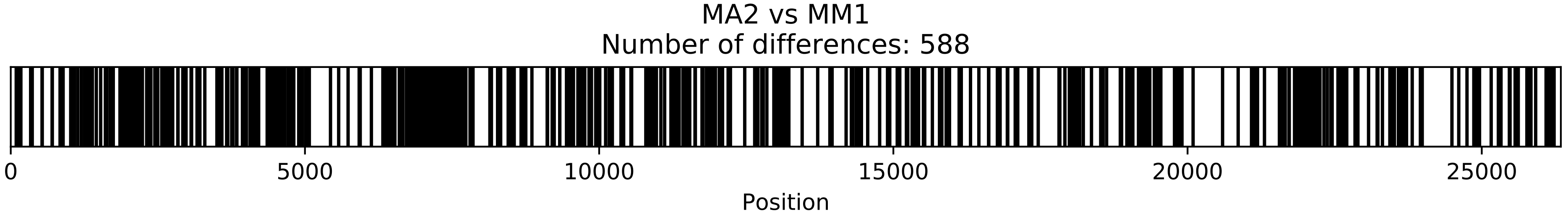
MA2 vs MM1
Number of differences: 588
0
5000
10000
15000
20000
25000
Position

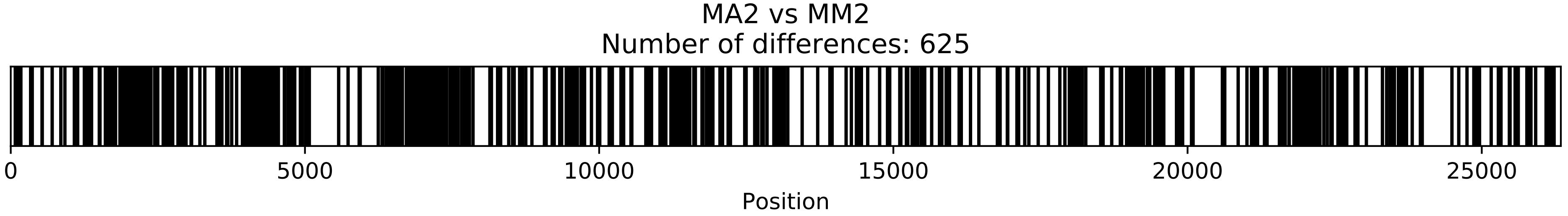
MA2 vs MM2
Number of differences: 625
0
5000
10000
15000
20000
25000
Position

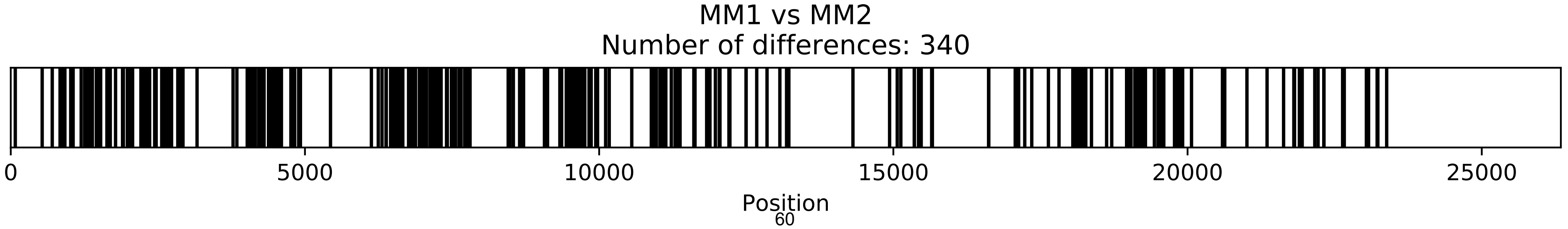
MM1 vs MM2
Number of differences: 340
0
5000
10000
15000
20000
25000
Position

### Region: 218



MA1 vs MA2
Number of differences: 291

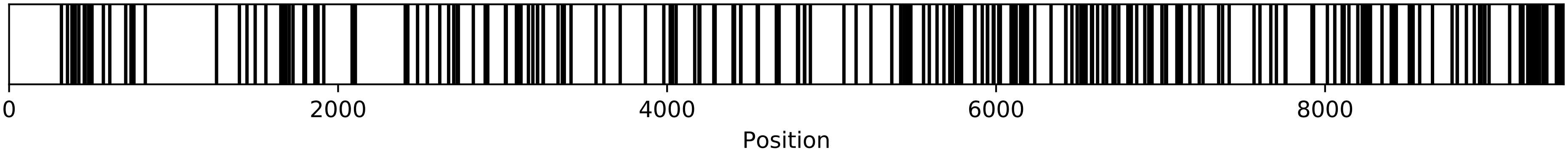


MA1 vs MM1
Number of differences: 198

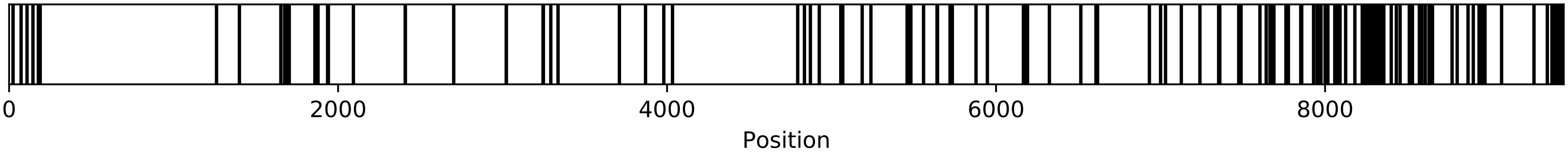


MA1 vs MM2
Number of differences: 194

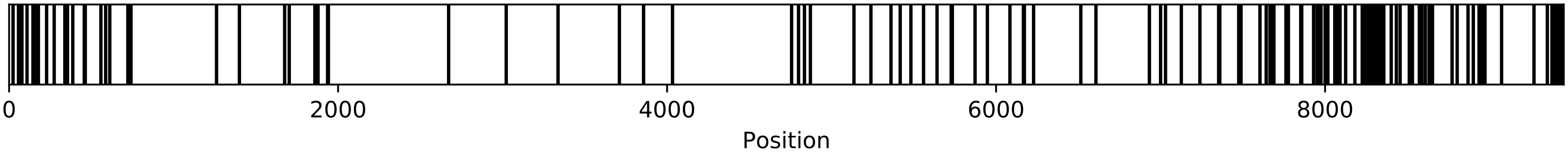


MA2 vs MM1
Number of differences: 323

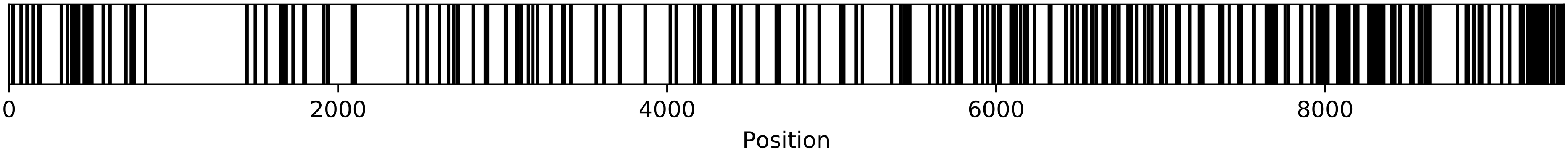


MA2 vs MM2
Number of differences: 327

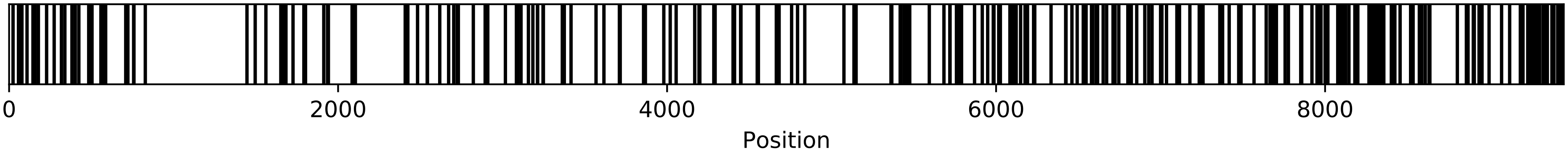


MM1 vs MM2
Number of differences: 56

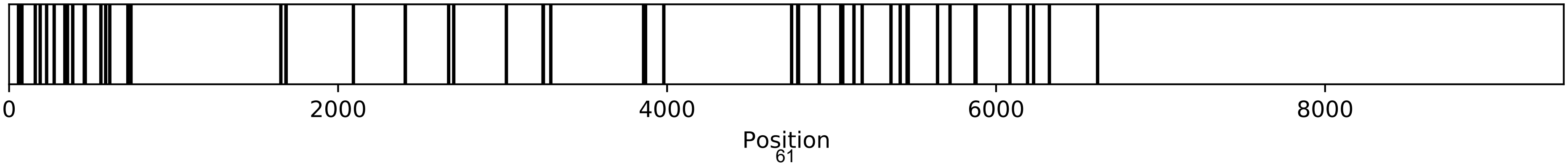

### Region: 219



MA1 vs MA2
Number of differences: 0

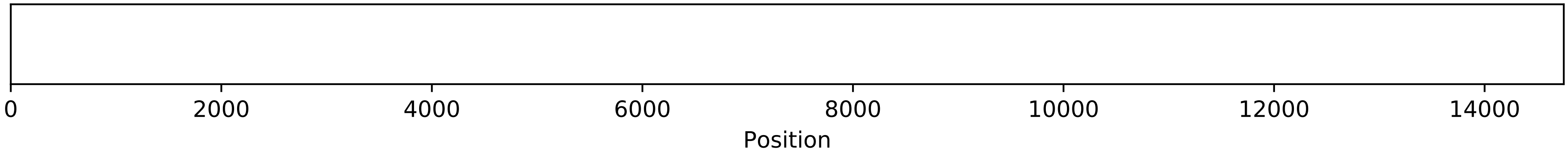


MA1 vs MM1
Number of differences: 734

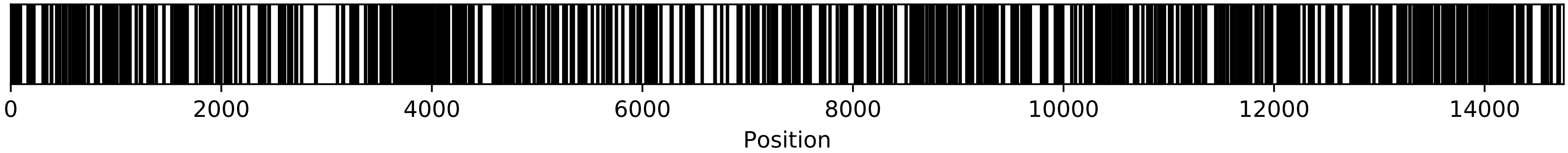


MA1 vs MM2
Number of differences: 734

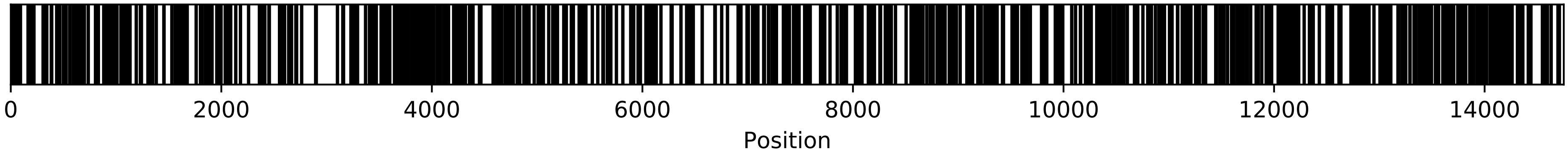


MA2 vs MM1
Number of differences: 734

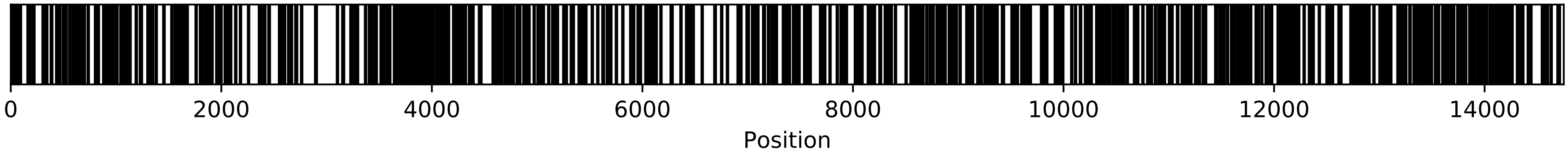


MA2 vs MM2
Number of differences: 734

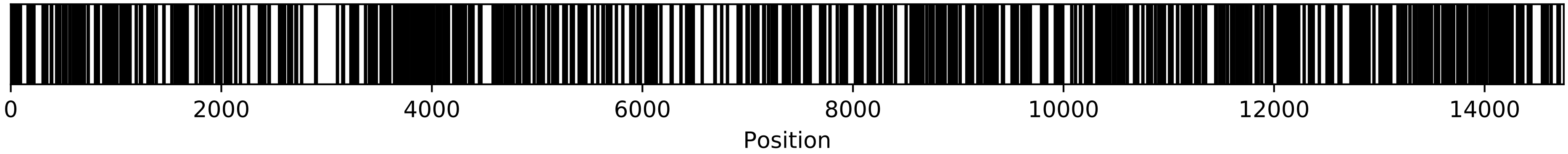


MM1 vs MM2
Number of differences: 0

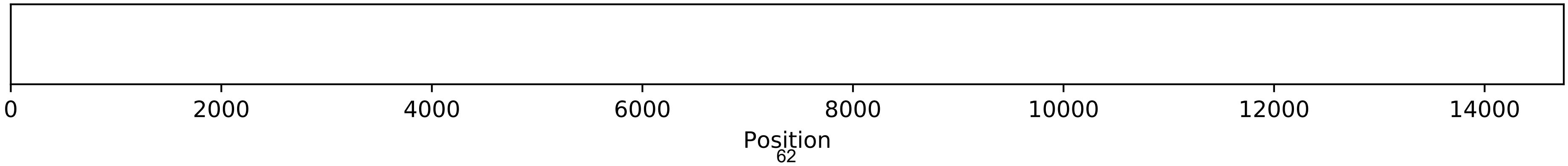

### Region: 223



## MA1 vs MA2
Number of differences: 277

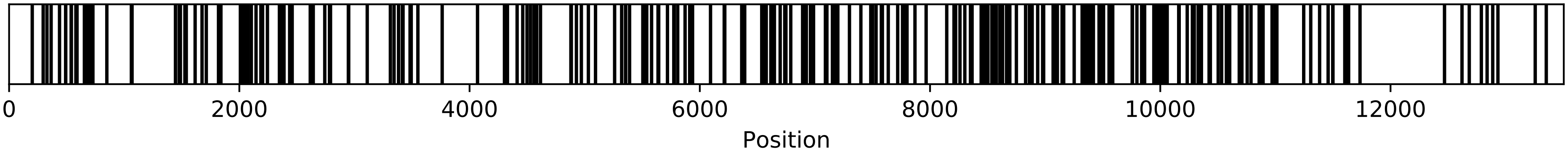


## MA1 vs MM1
Number of differences: 109

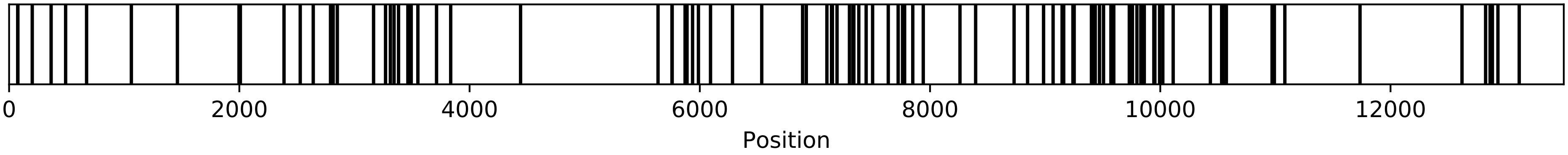


## MA1 vs MM2
Number of differences: 128

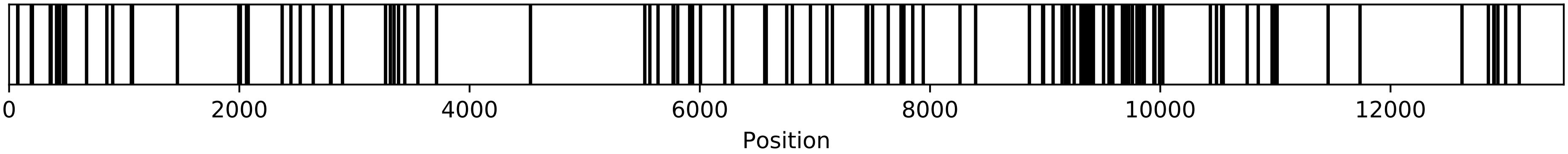


## MA2 vs MM1
Number of differences: 283

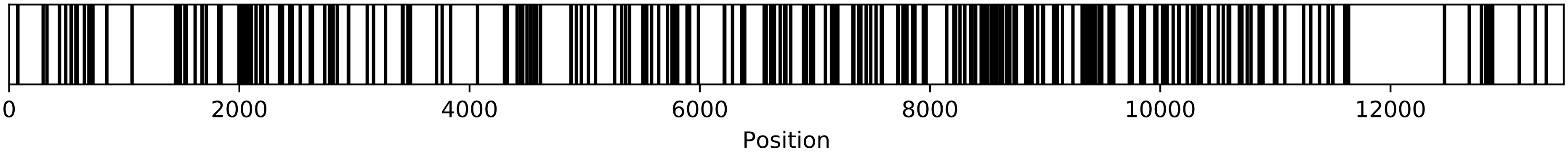


## MA2 vs MM2
Number of differences: 283

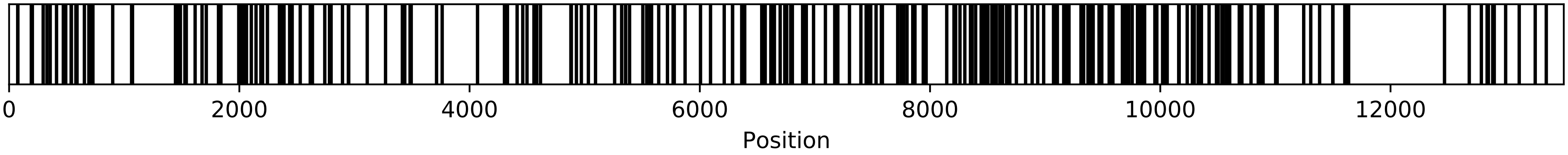


## MM1 vs MM2
Number of differences: 121

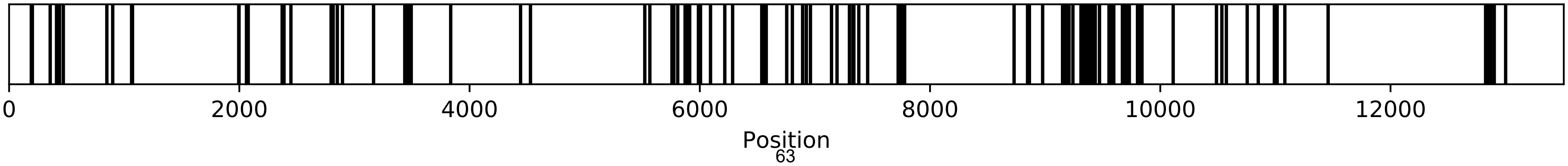

### Region: 225



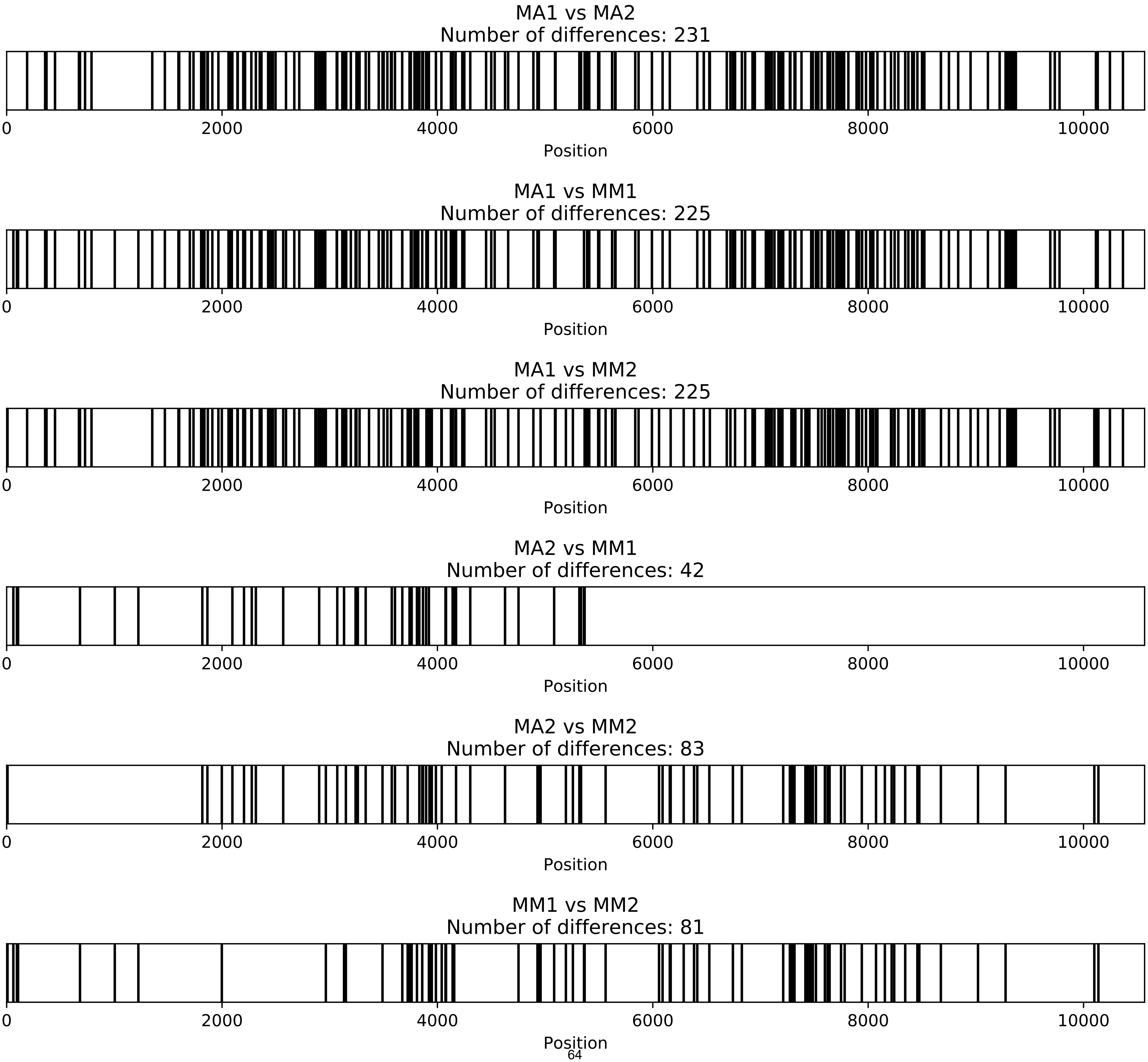
MA1 vs MA2
Number of differences: 231
MA1 vs MM1
Number of differences: 225
MA1 vs MM2
Number of differences: 225
MA2 vs MM1
Number of differences: 42
MA2 vs MM2
Number of differences: 83
MM1 vs MM2
Number of differences: 81
0
2000
4000
6000
8000
10000
Position

### Region: 226



MA1 vs MA2
Number of differences: 0

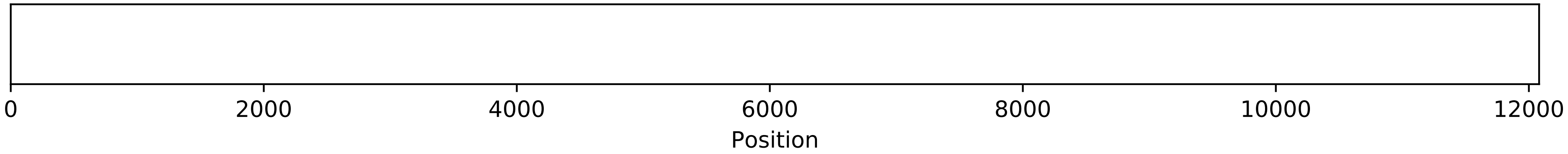


MA1 vs MM1
Number of differences: 117

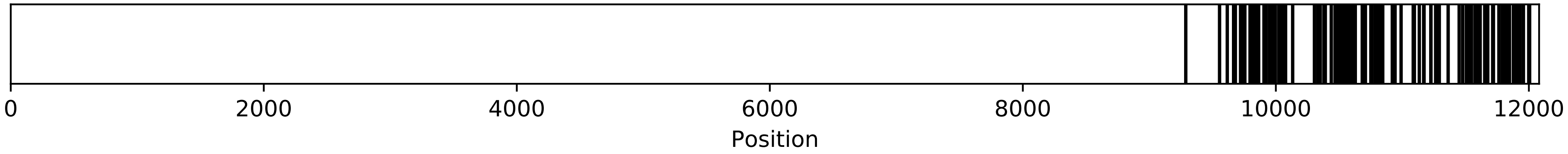


MA1 vs MM2
Number of differences: 259

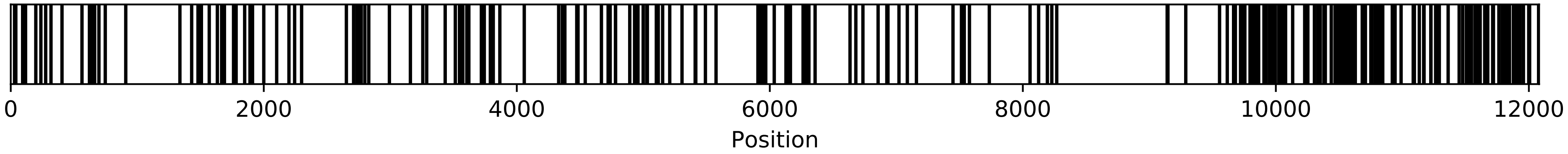


MA2 vs MM1
Number of differences: 117

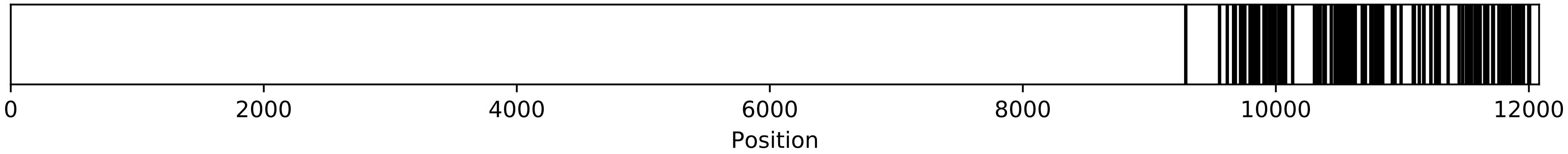


MA2 vs MM2
Number of differences: 259

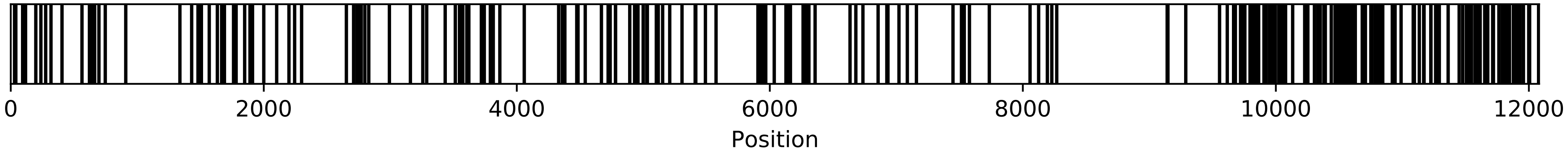


MM1 vs MM2
Number of differences: 142

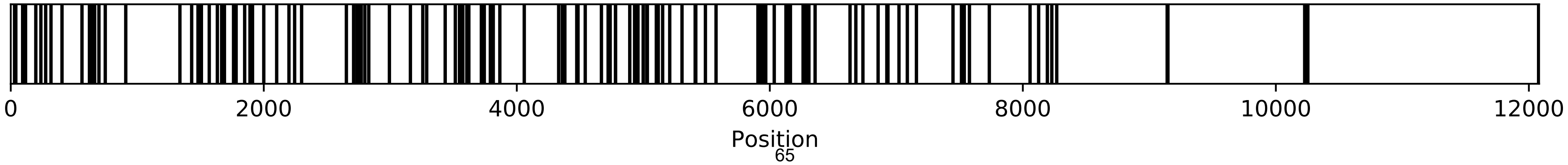

### Region: 227



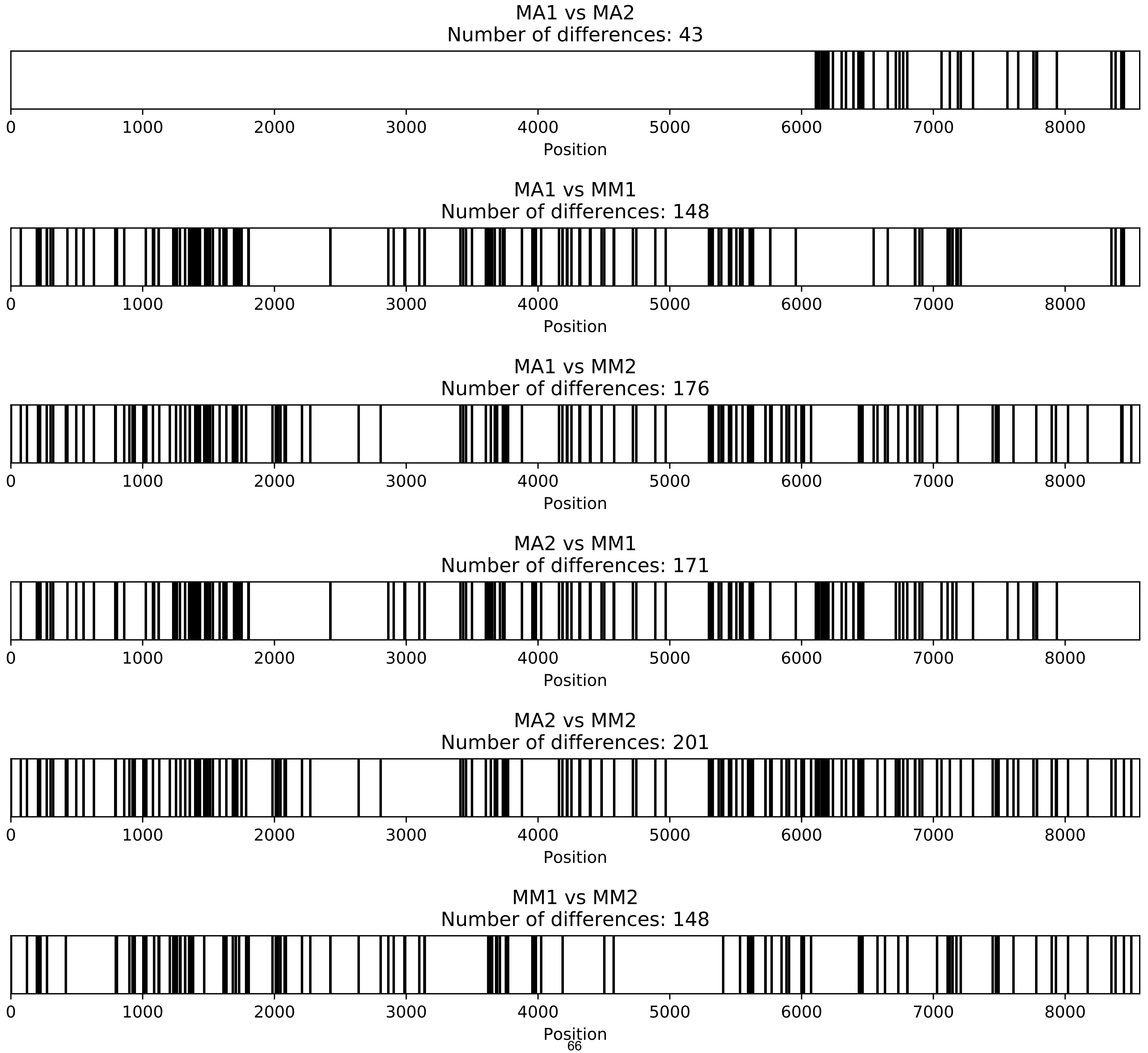
MA1 vs MA2
Number of differences: 43
MA1 vs MM1
Number of differences: 148
MA1 vs MM2
Number of differences: 176
MA2 vs MM1
Number of differences: 171
MA2 vs MM2
Number of differences: 201
MM1 vs MM2
Number of differences: 148
0
1000
2000
3000
4000
5000
6000
7000
8000
Position

### Region: 228



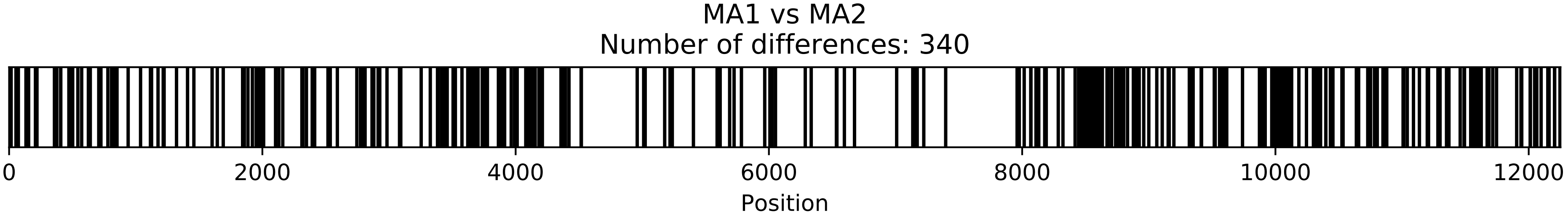
MA1 vs MA2
Number of differences: 340
0
2000
4000
6000
8000
10000
12000
Position

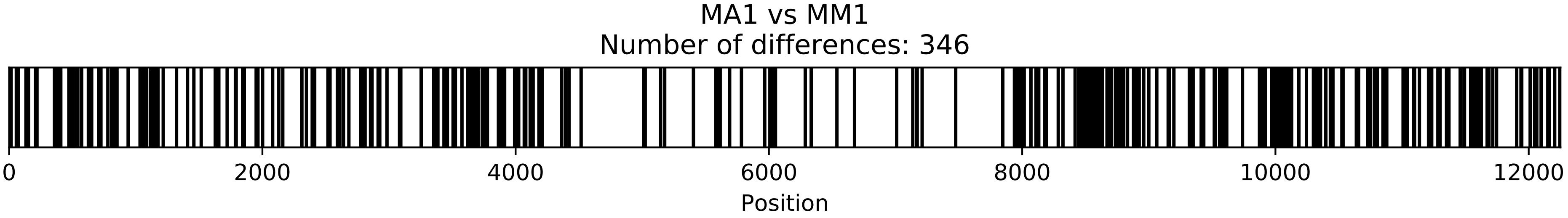
MA1 vs MM1
Number of differences: 346
0
2000
4000
6000
8000
10000
12000
Position

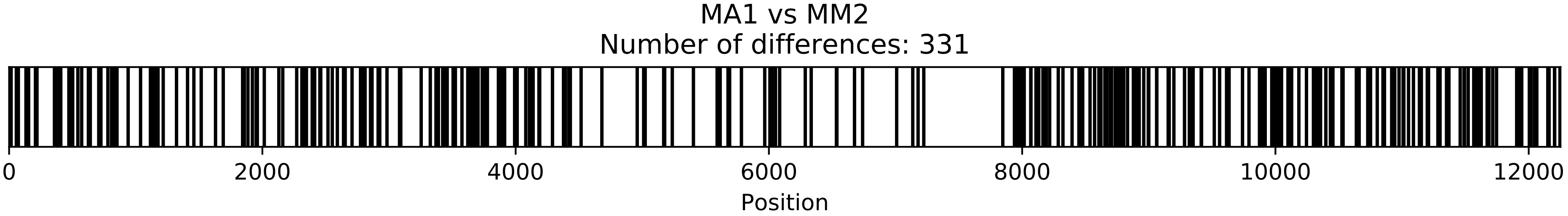
MA1 vs MM2
Number of differences: 331
0
2000
4000
6000
8000
10000
12000
Position

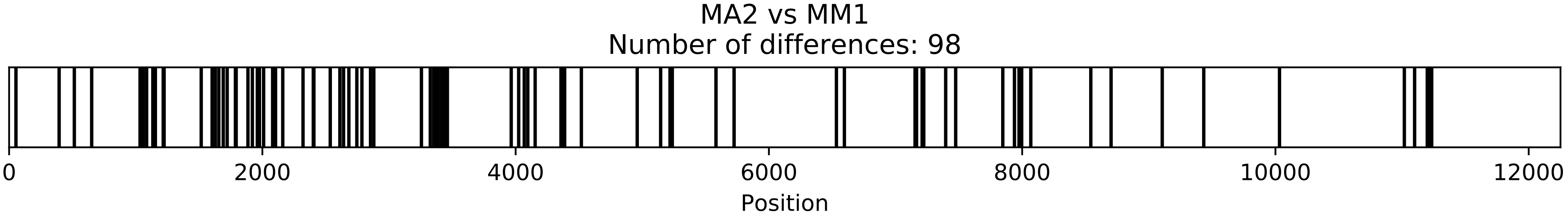
MA2 vs MM1
Number of differences: 98
0
2000
4000
6000
8000
10000
12000
Position

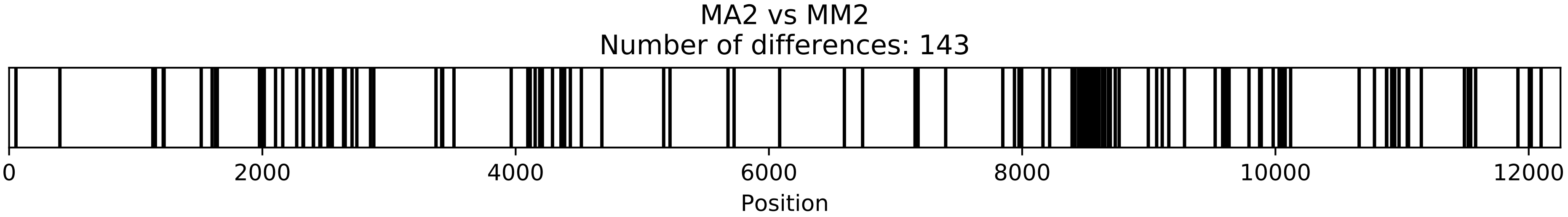
MA2 vs MM2
Number of differences: 143
0
2000
4000
6000
8000
10000
12000
Position

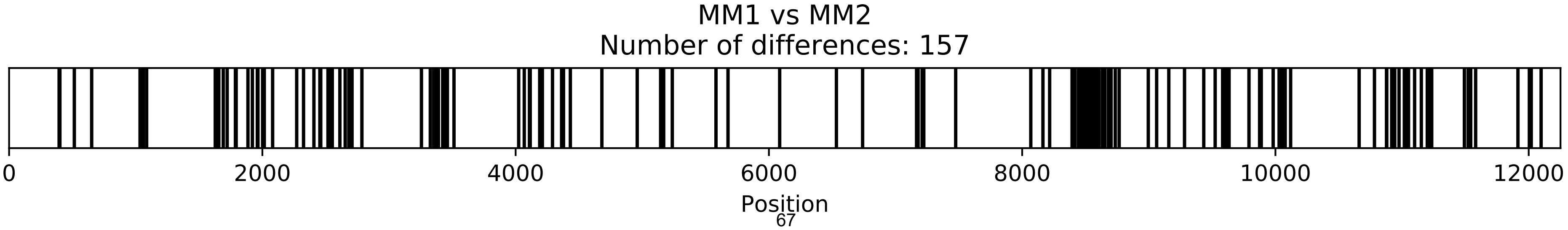
MM1 vs MM2
Number of differences: 157
0
2000
4000
6000
8000
10000
12000
Position

### Region: 237



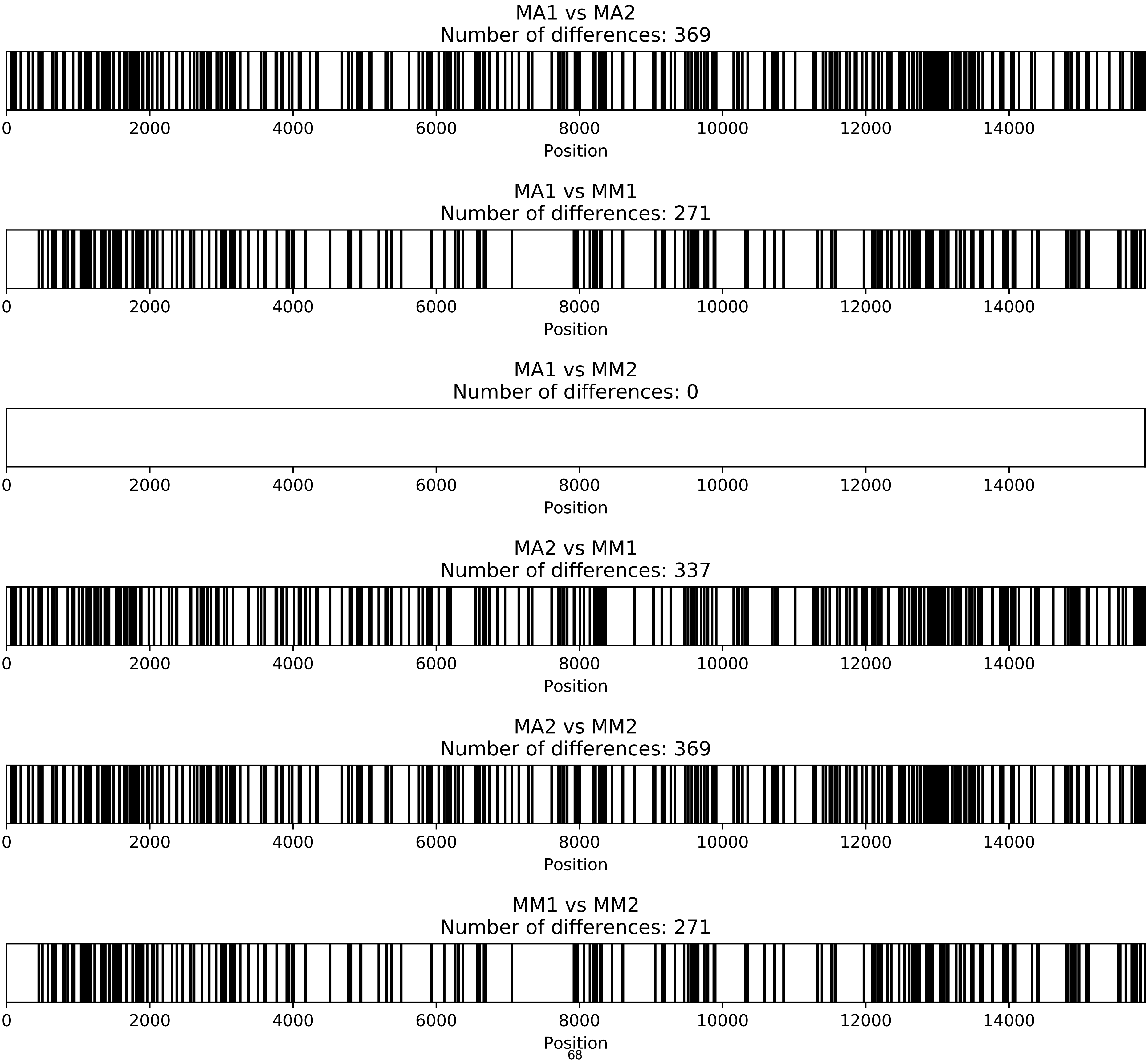
MA1 vs MA2
Number of differences: 369
0
2000
4000
6000
8000
10000
12000
14000
Position
MA1 vs MM1
Number of differences: 271
Position
MA1 vs MM2
Number of differences: 0
Position
MA2 vs MM1
Number of differences: 337
Position
MA2 vs MM2
Number of differences: 369
Position
MM1 vs MM2
Number of differences: 271
Position

### Region: 238



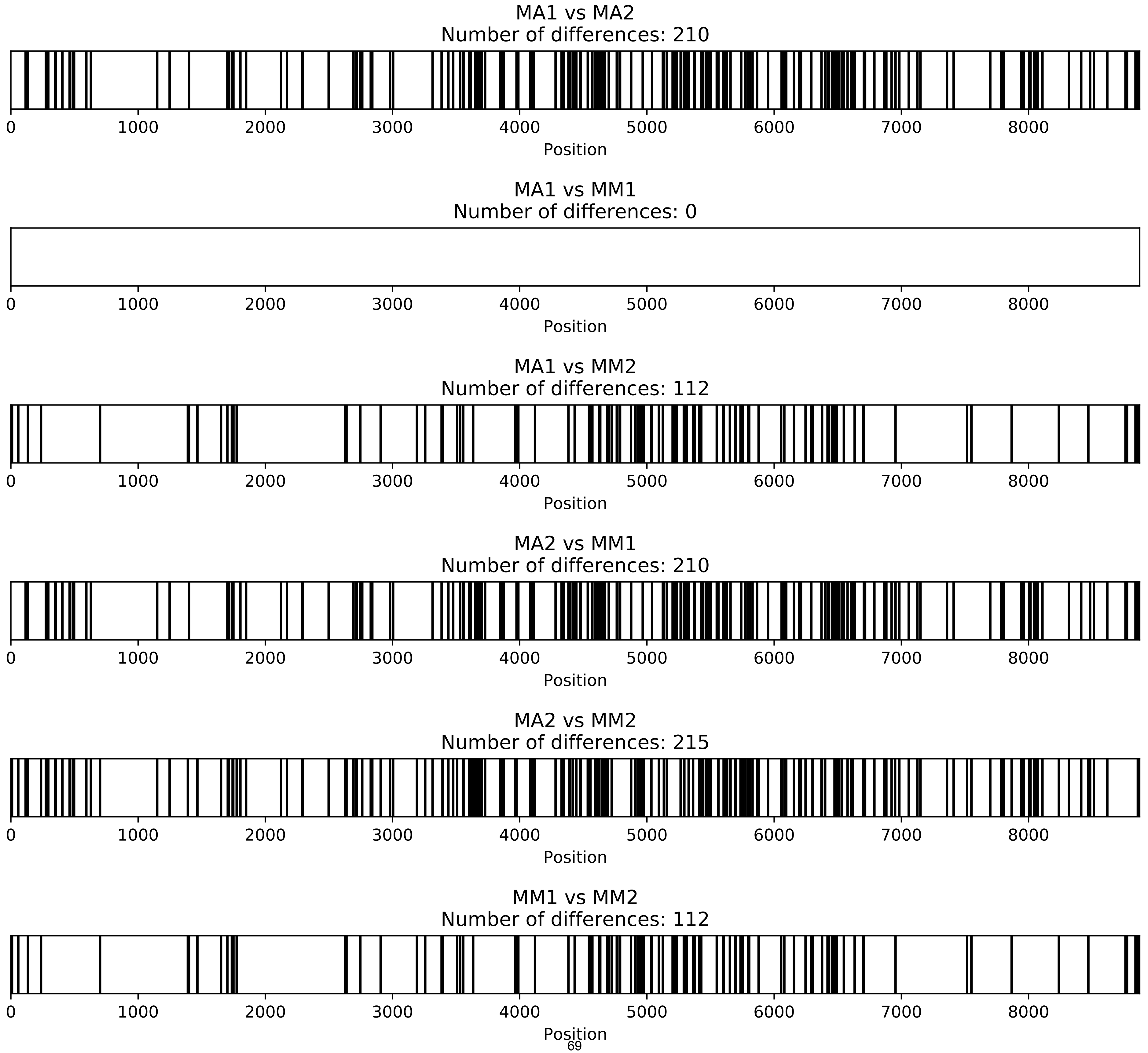
MA1 vs MA2
Number of differences: 210
0
1000
2000
3000
4000
5000
6000
7000
8000
Position
MA1 vs MM1
Number of differences: 0
0
1000
2000
3000
4000
5000
6000
7000
8000
Position
MA1 vs MM2
Number of differences: 112
0
1000
2000
3000
4000
5000
6000
7000
8000
Position
MA2 vs MM1
Number of differences: 210
0
1000
2000
3000
4000
5000
6000
7000
8000
Position
MA2 vs MM2
Number of differences: 215
0
1000
2000
3000
4000
5000
6000
7000
8000
Position
MM1 vs MM2
Number of differences: 112
0
1000
2000
3000
4000
5000
6000
7000
8000
Position

### Region: 240



MA1 vs MA2
Number of differences: 271

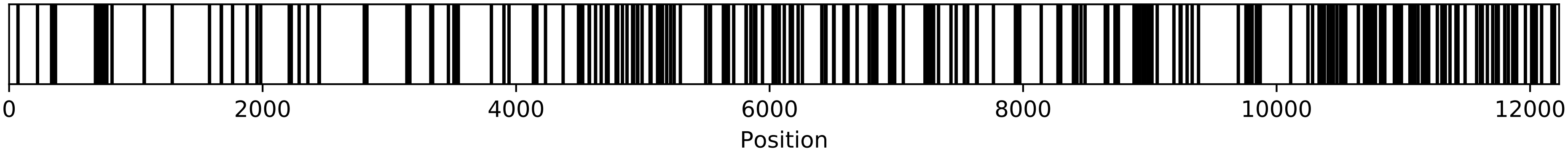


MA1 vs MM1
Number of differences: 130

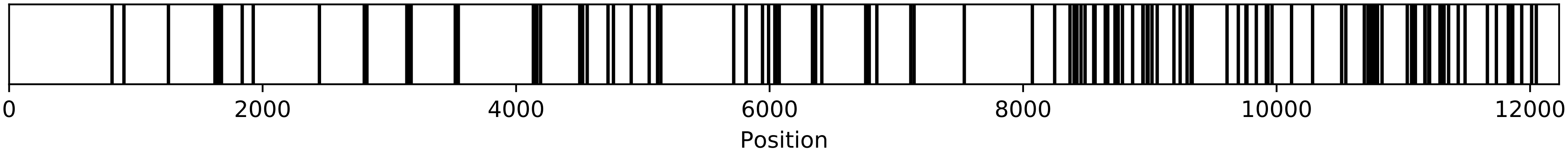


MA1 vs MM2
Number of differences: 0

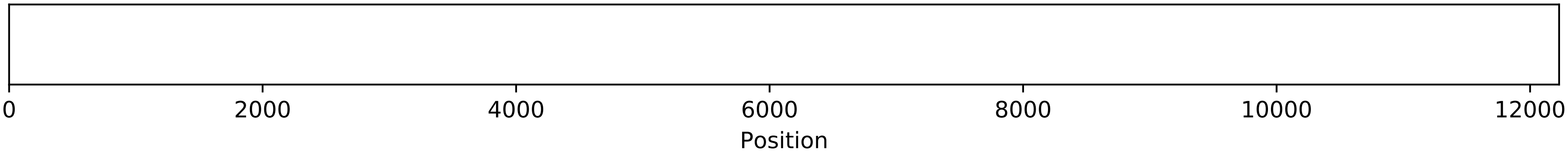


MA2 vs MM1
Number of differences: 275

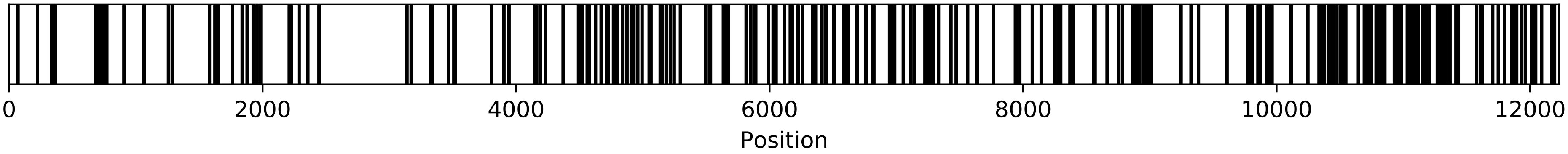


MA2 vs MM2
Number of differences: 271

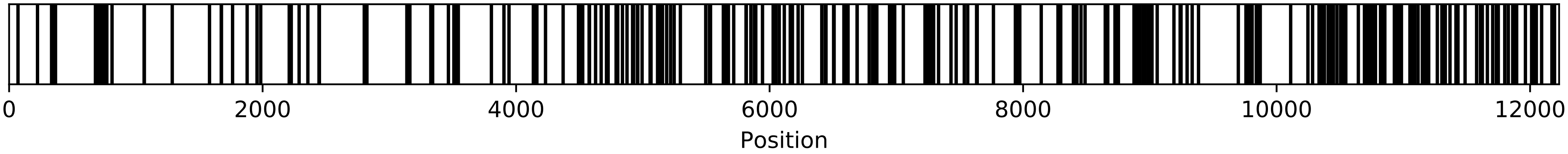


MM1 vs MM2
Number of differences: 130

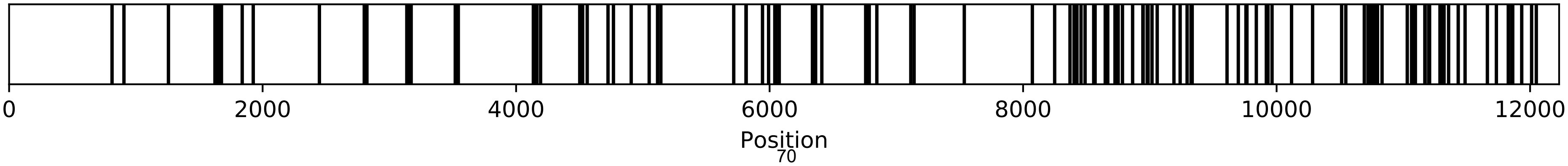

### Region: 248



MA1 vs MA2
Number of differences: 189

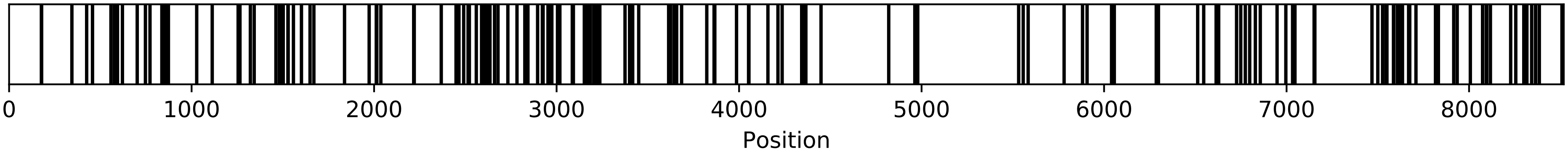


MA1 vs MM1
Number of differences: 200

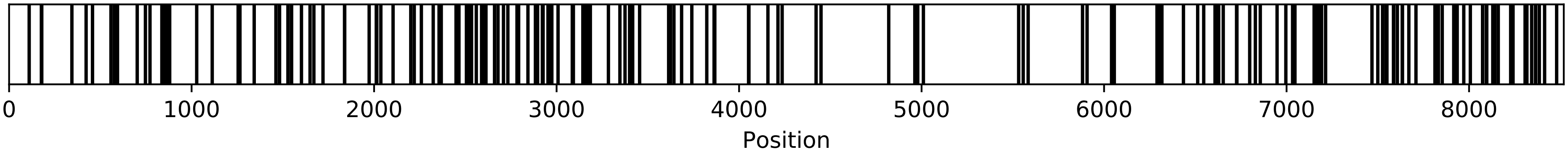


MA1 vs MM2
Number of differences: 212

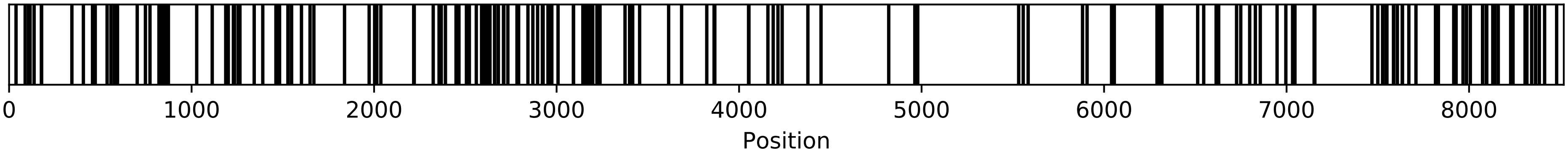


MA2 vs MM1
Number of differences: 99

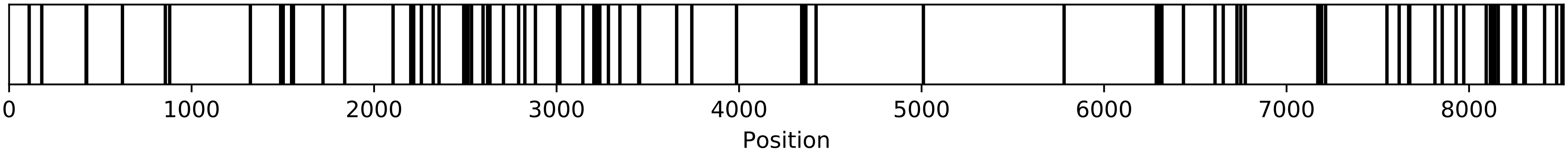


MA2 vs MM2
Number of differences: 110

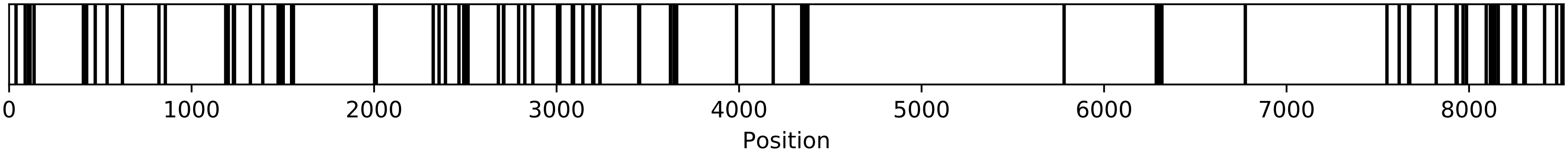


MM1 vs MM2
Number of differences: 76

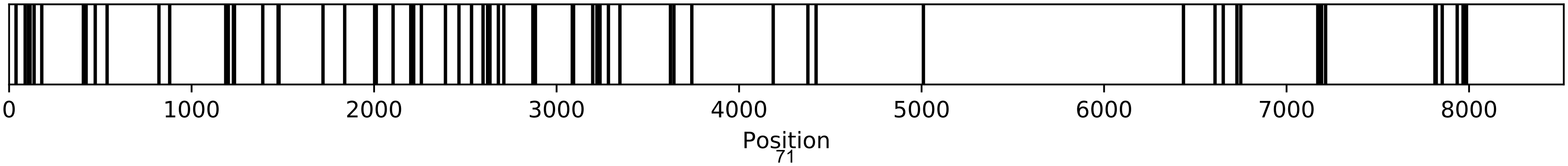

### Region: 251



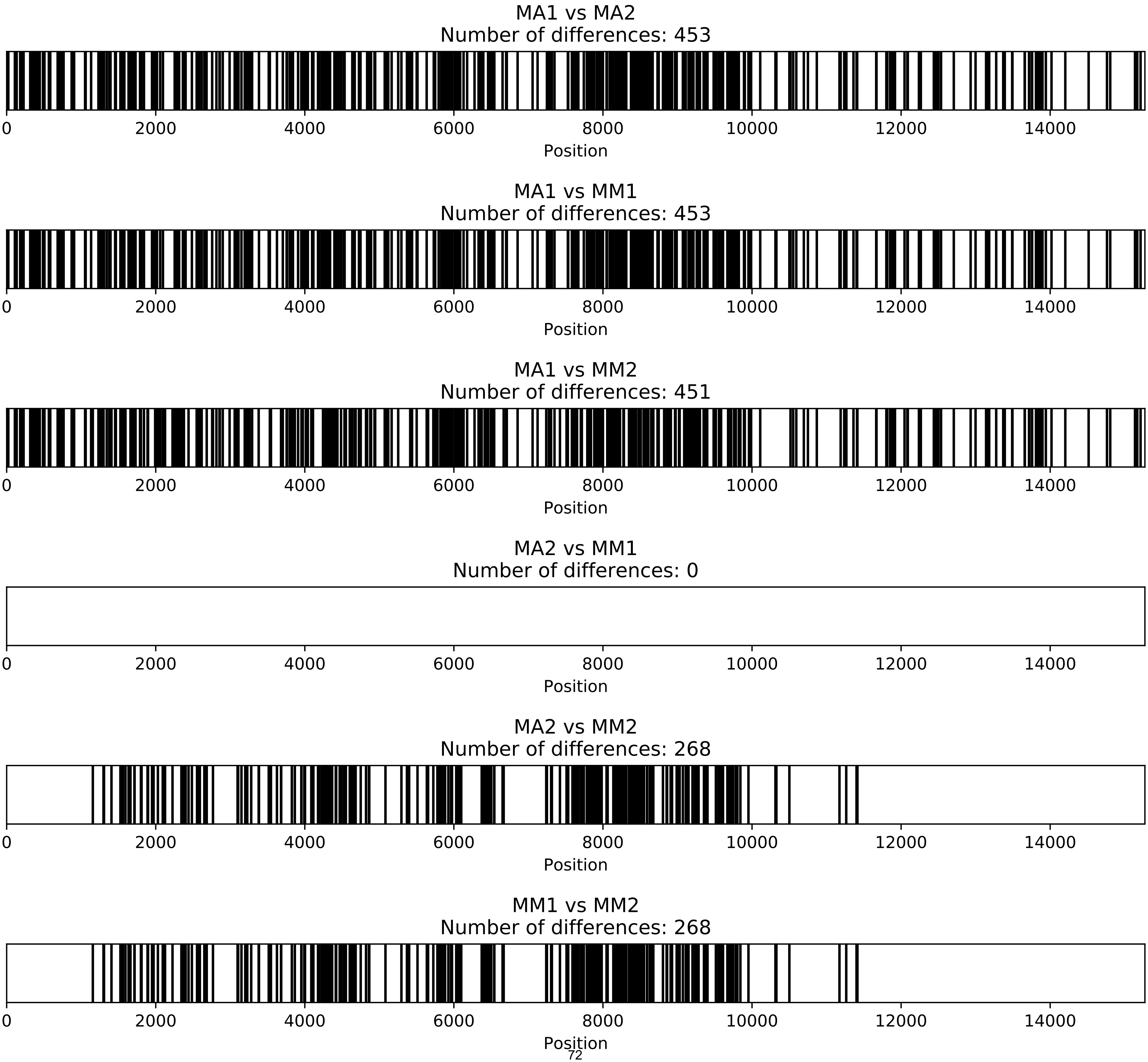
MA1 vs MA2
Number of differences: 453
MA1 vs MM1
Number of differences: 453
MA1 vs MM2
Number of differences: 451
MA2 vs MM1
Number of differences: 0
MA2 vs MM2
Number of differences: 268
MM1 vs MM2
Number of differences: 268
0
2000
4000
6000
8000
10000
12000
14000
Position

### Region: 256



MA1 vs MA2
Number of differences: 188

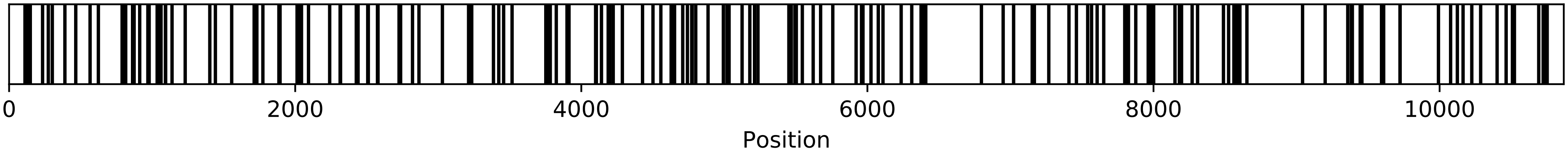


MA1 vs MM1
Number of differences: 183

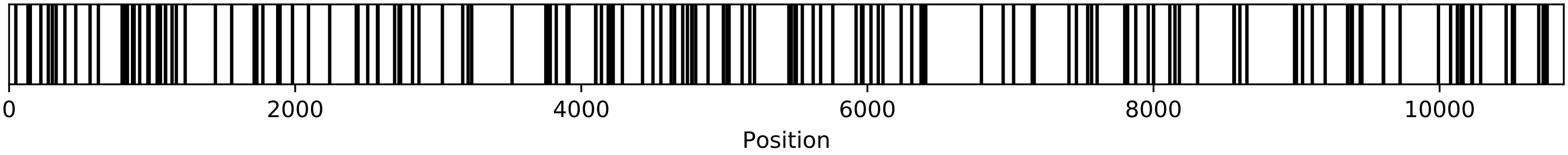


MA1 vs MM2
Number of differences: 188

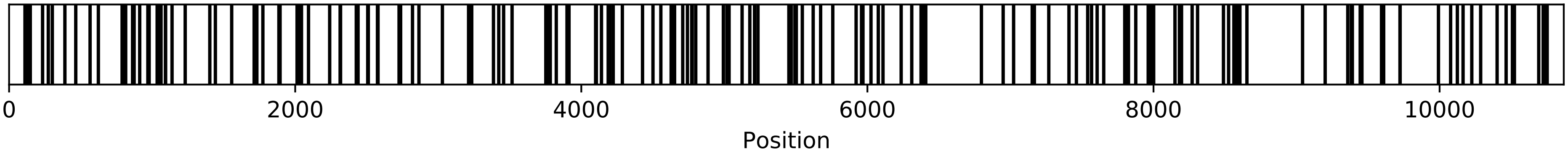


MA2 vs MM1
Number of differences: 64

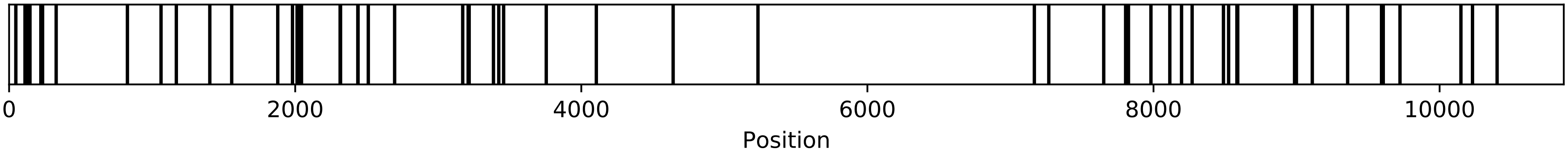


MA2 vs MM2
Number of differences: 0

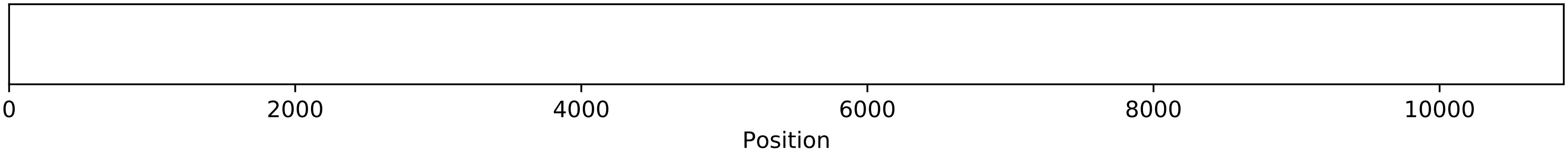


MM1 vs MM2
Number of differences: 64

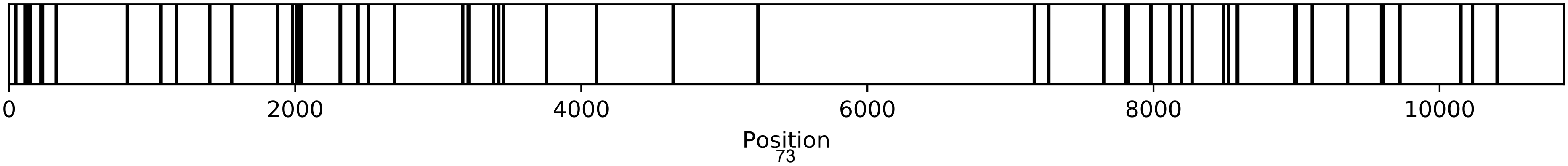

### Region: 261



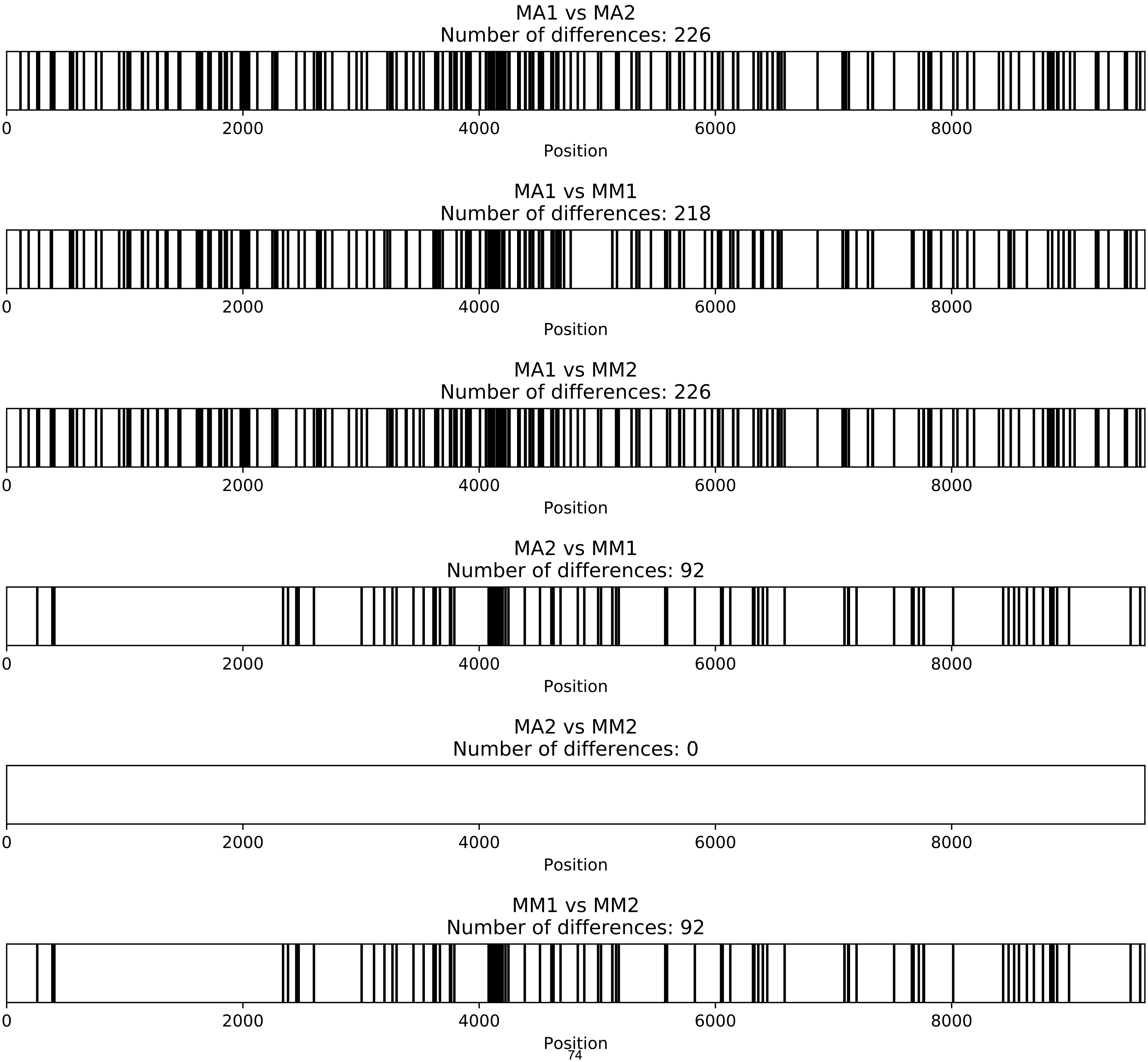
MA1 vs MA2
Number of differences: 226
MA1 vs MM1
Number of differences: 218
MA1 vs MM2
Number of differences: 226
MA2 vs MM1
Number of differences: 92
MA2 vs MM2
Number of differences: 0
MM1 vs MM2
Number of differences: 92
0
2000
4000
6000
8000
Position

### Region: 264



MA1 vs MA2
Number of differences: 220

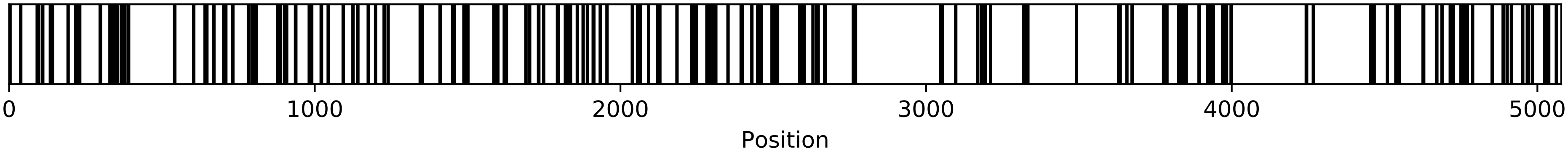


MA1 vs MM1
Number of differences: 237

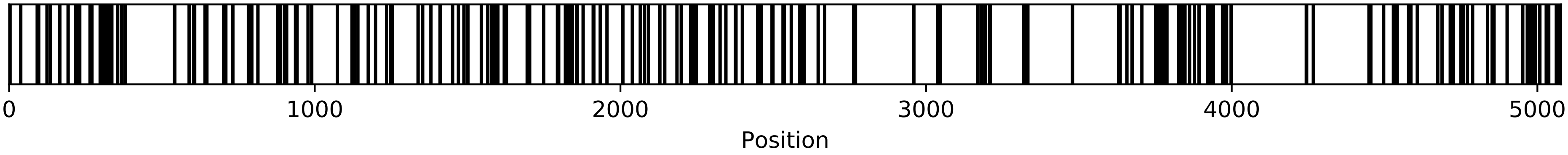


MA1 vs MM2
Number of differences: 233

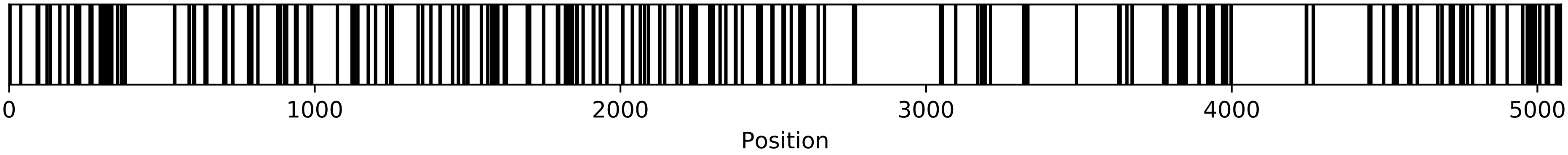


MA2 vs MM1
Number of differences: 167

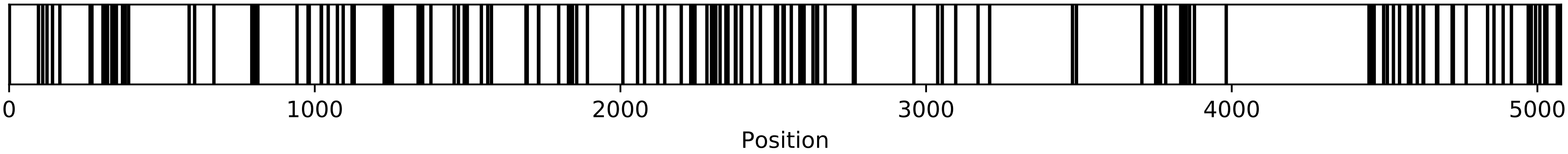


MA2 vs MM2
Number of differences: 140

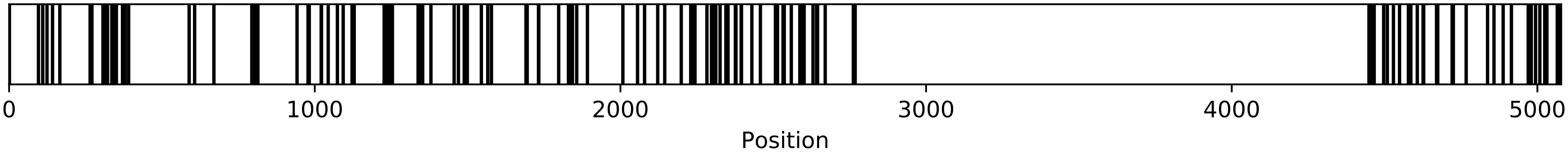


MM1 vs MM2
Number of differences: 27

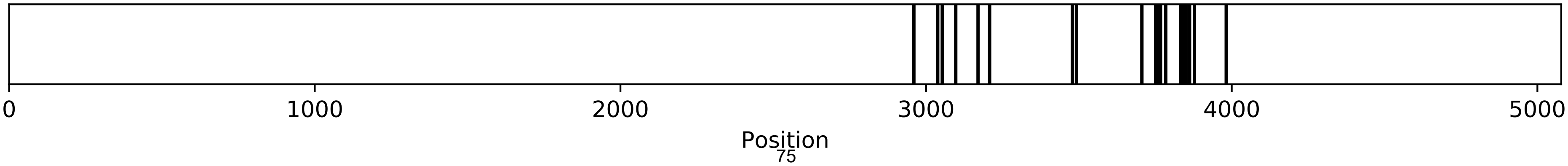

### Region: 267



MA1 vs MA2
Number of differences: 114

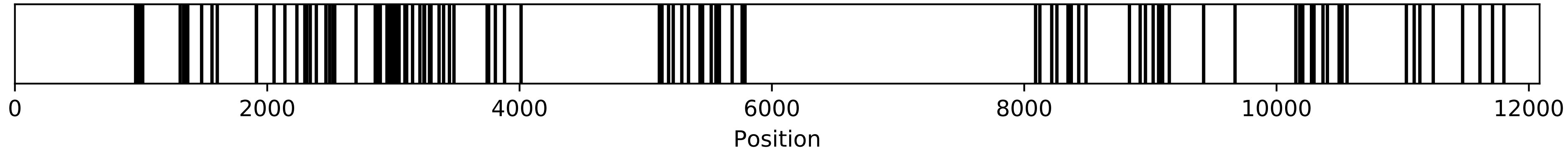


MA1 vs MM1
Number of differences: 109

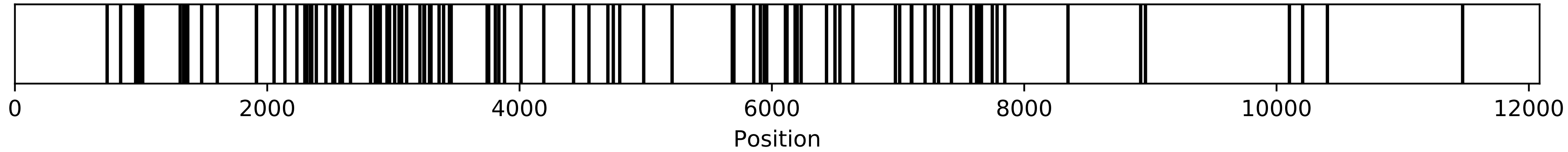


MA1 vs MM2
Number of differences: 95

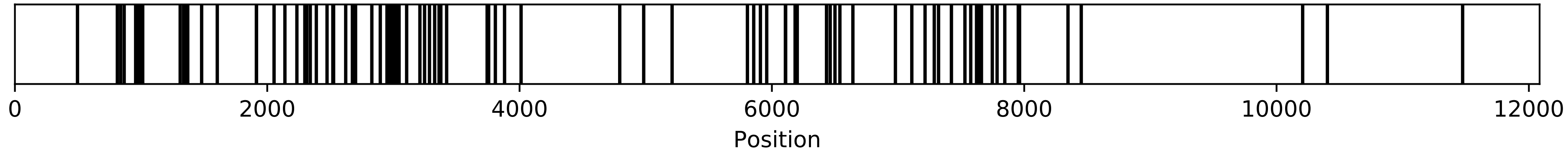


MA2 vs MM1
Number of differences: 121

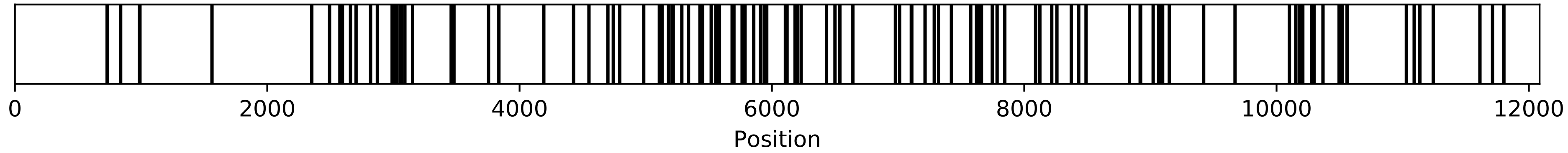


MA2 vs MM2
Number of differences: 123

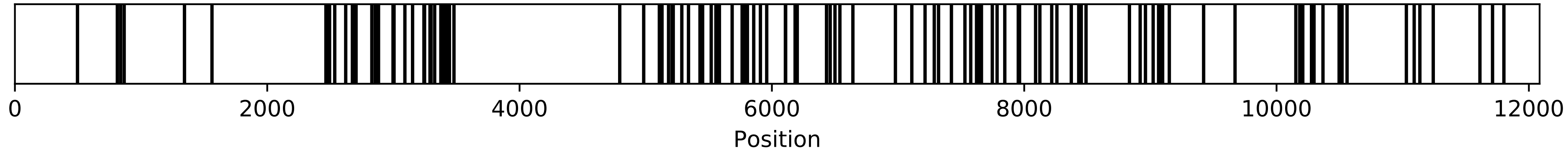


MM1 vs MM2
Number of differences: 67

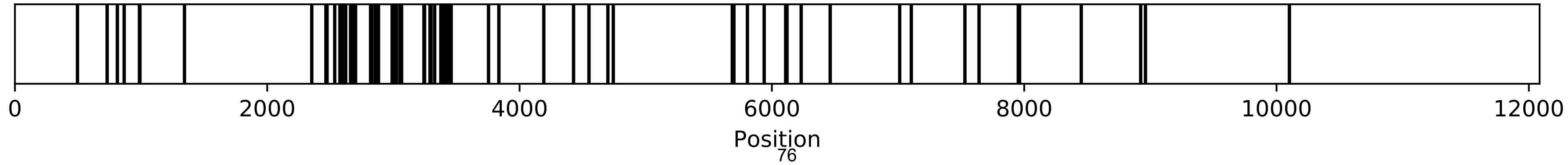

### Region: 272



MA1 vs MA2
Number of differences: 164

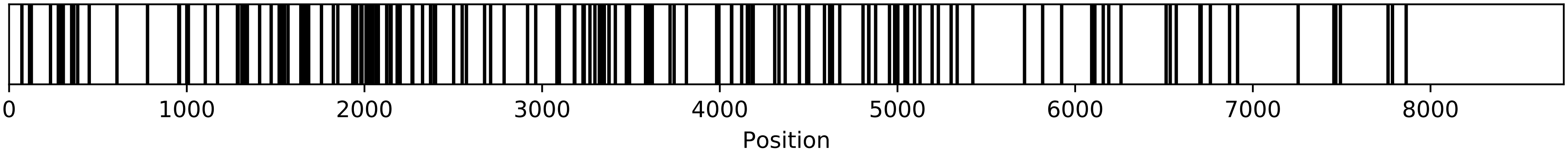


MA1 vs MM1
Number of differences: 169

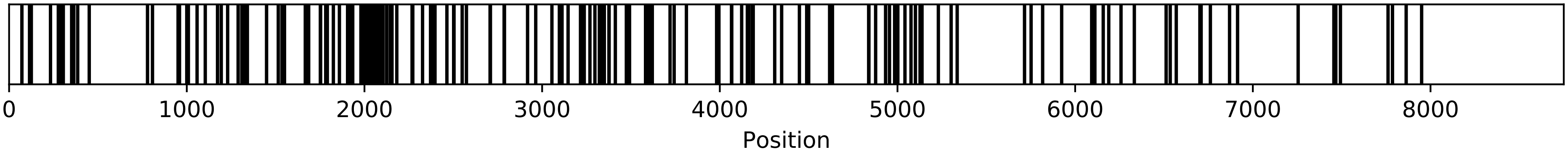


MA1 vs MM2
Number of differences: 164

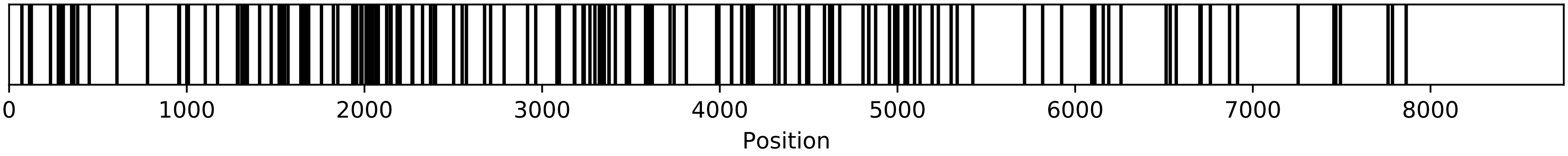


MA2 vs MM1
Number of differences: 75

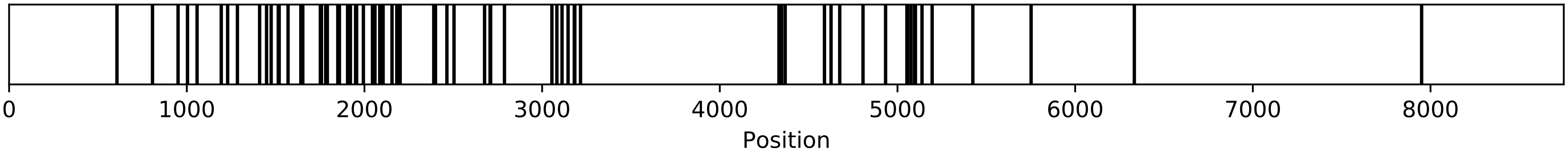


MA2 vs MM2
Number of differences: 0

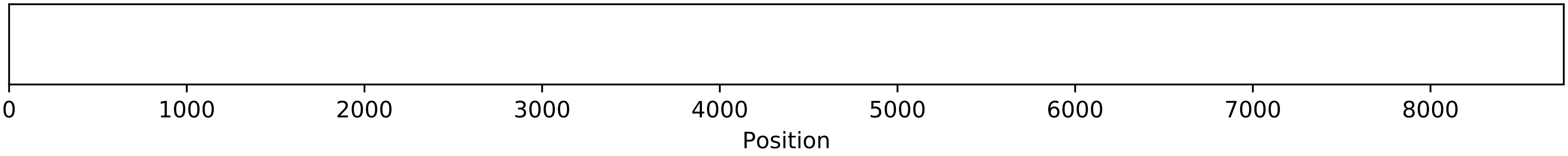


MM1 vs MM2
Number of differences: 75

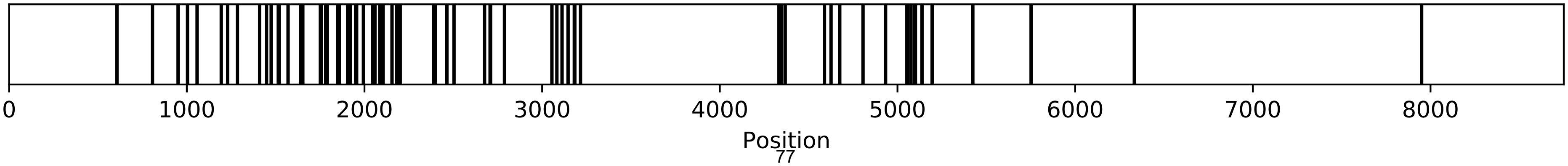

### Region: 277



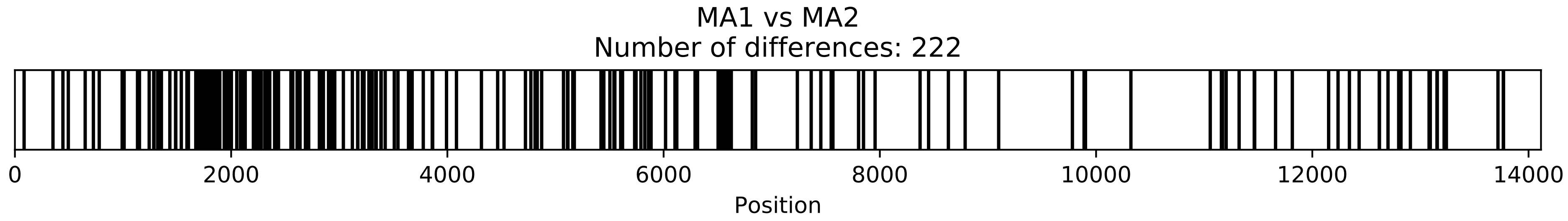
MA1 vs MA2
Number of differences: 222
0
2000
4000
6000
8000
10000
12000
14000
Position

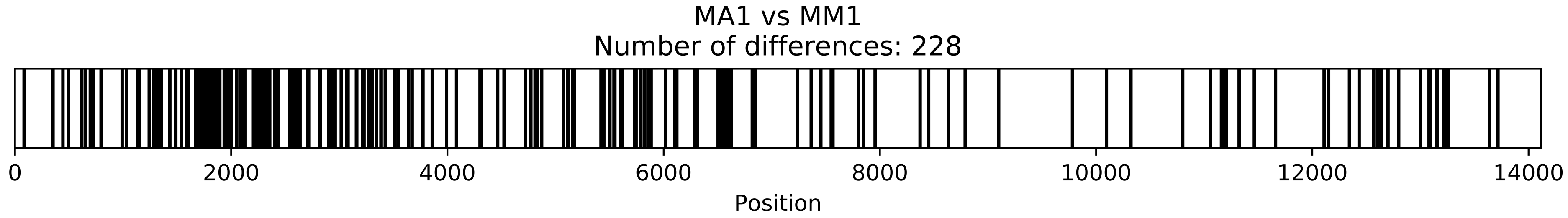
MA1 vs MM1
Number of differences: 228
0
2000
4000
6000
8000
10000
12000
14000
Position

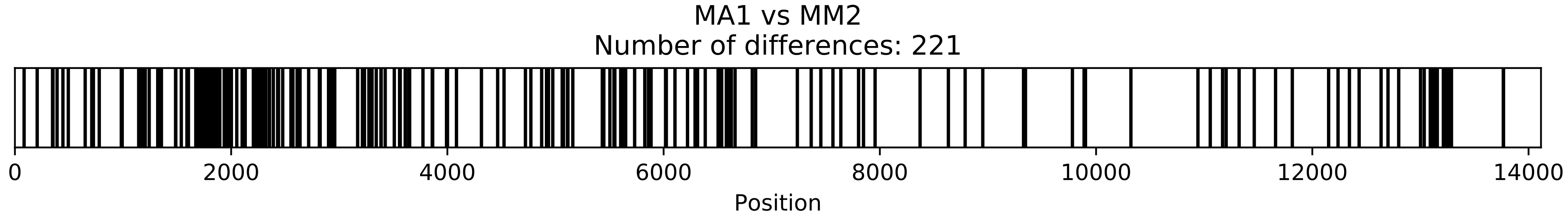
MA1 vs MM2
Number of differences: 221
0
2000
4000
6000
8000
10000
12000
14000
Position

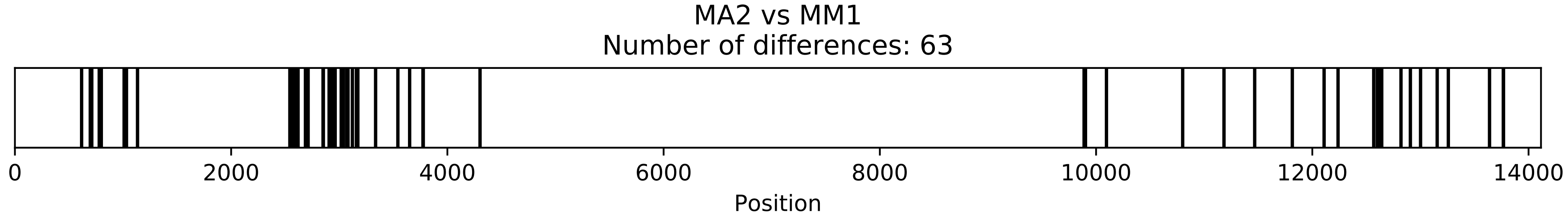
MA2 vs MM1
Number of differences: 63
0
2000
4000
6000
8000
10000
12000
14000
Position

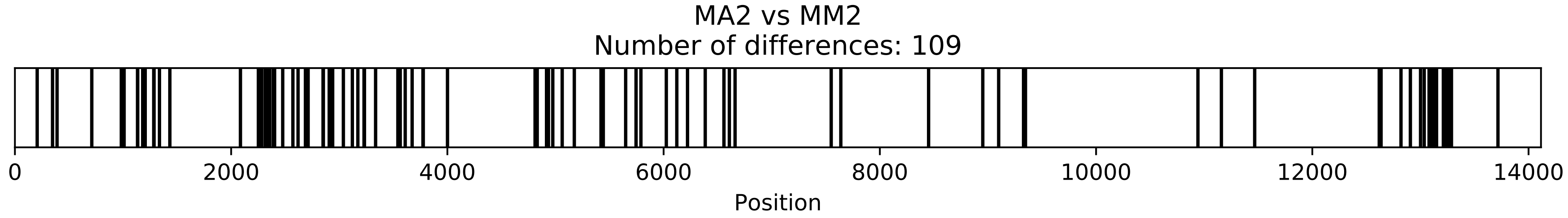
MA2 vs MM2
Number of differences: 109
0
2000
4000
6000
8000
10000
12000
14000
Position

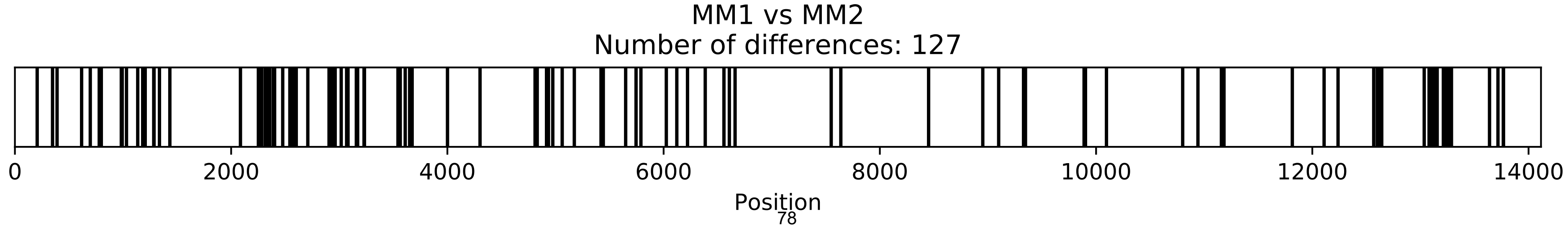
MM1 vs MM2
Number of differences: 127
0
2000
4000
6000
8000
10000
12000
14000
Position

### Region: 280

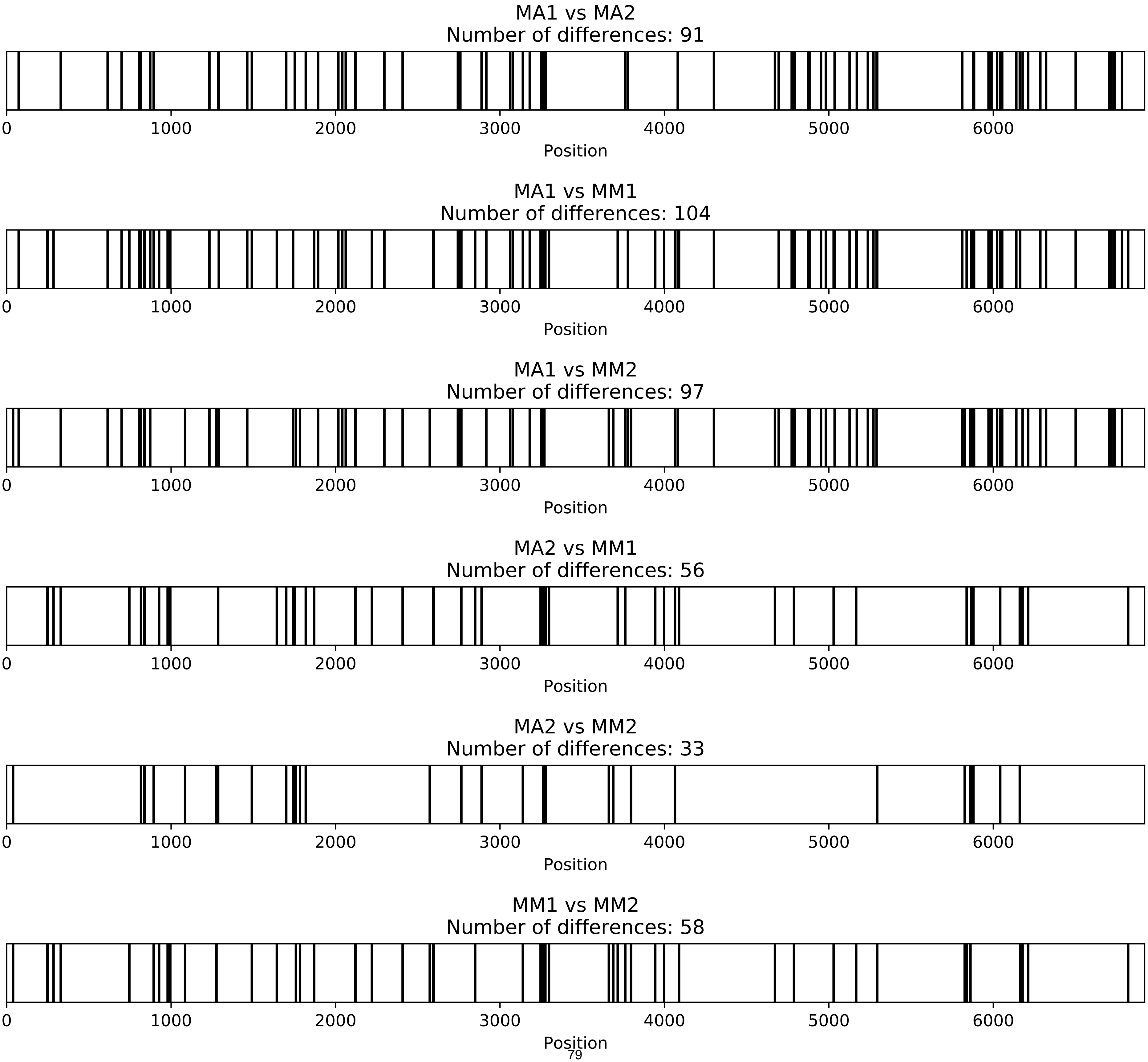
MA1 vs MA2
Number of differences: 91
0 1000 2000 3000 4000 5000 6000
Position
MA1 vs MM1
Number of differences: 104
0 1000 2000 3000 4000 5000 6000
Position
MA1 vs MM2
Number of differences: 97
0 1000 2000 3000 4000 5000 6000
Position
MA2 vs MM1
Number of differences: 56
0 1000 2000 3000 4000 5000 6000
Position
MA2 vs MM2
Number of differences: 33
0 1000 2000 3000 4000 5000 6000
Position
MM1 vs MM2
Number of differences: 58
0 1000 2000 3000 4000 5000 6000
Position

### Region: 282



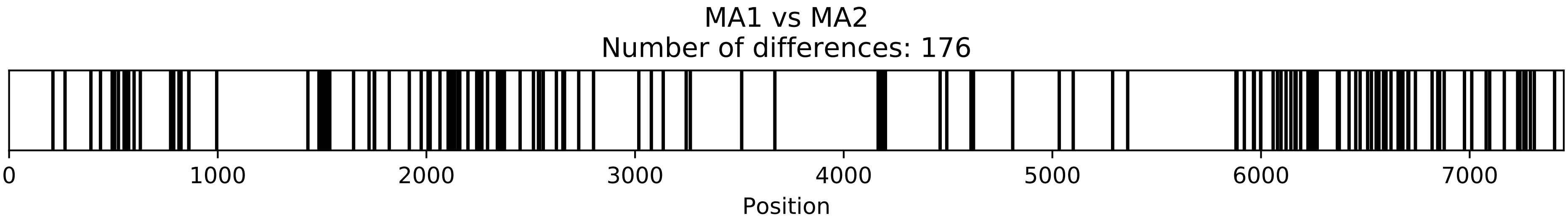
MA1 vs MA2
Number of differences: 176
0
1000
2000
3000
4000
5000
6000
7000
Position

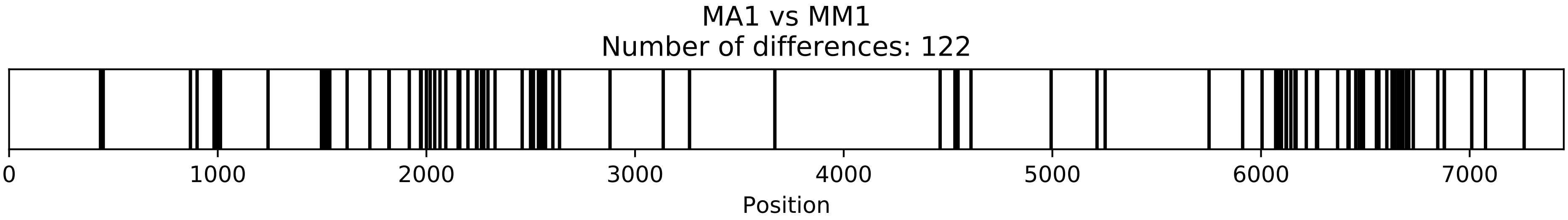
MA1 vs MM1
Number of differences: 122
0
1000
2000
3000
4000
5000
6000
7000
Position

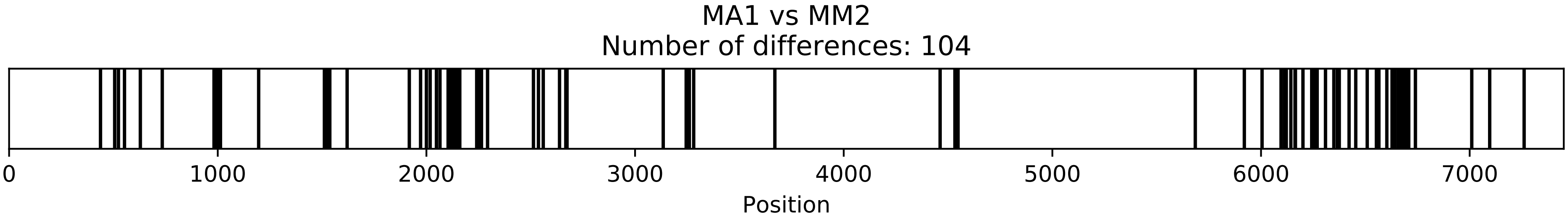
MA1 vs MM2
Number of differences: 104
0
1000
2000
3000
4000
5000
6000
7000
Position

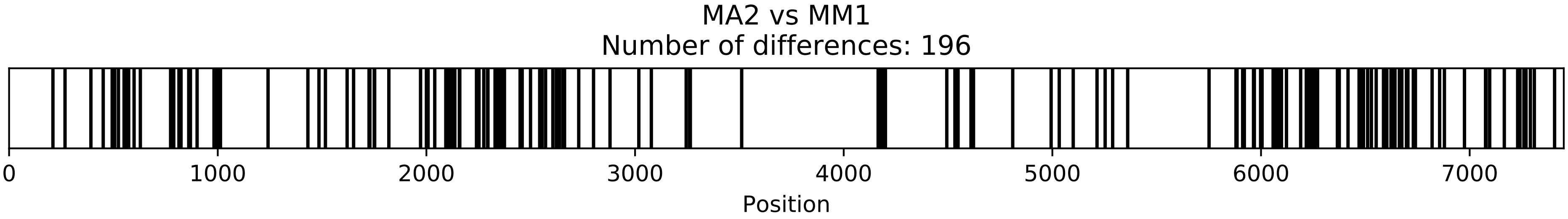
MA2 vs MM1
Number of differences: 196
0
1000
2000
3000
4000
5000
6000
7000
Position

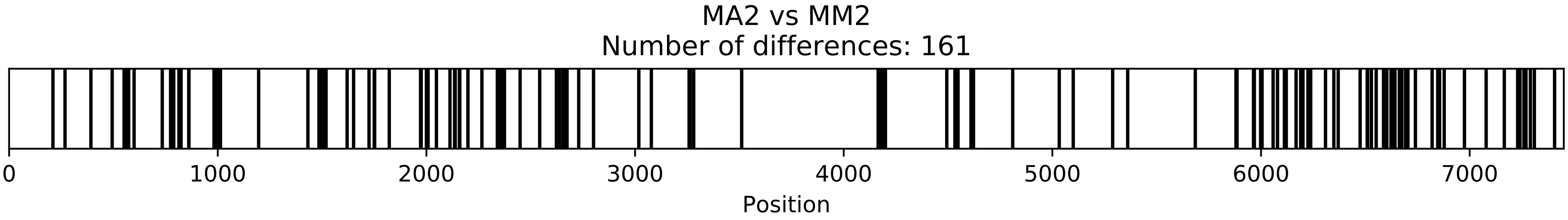
MA2 vs MM2
Number of differences: 161
0
1000
2000
3000
4000
5000
6000
7000
Position

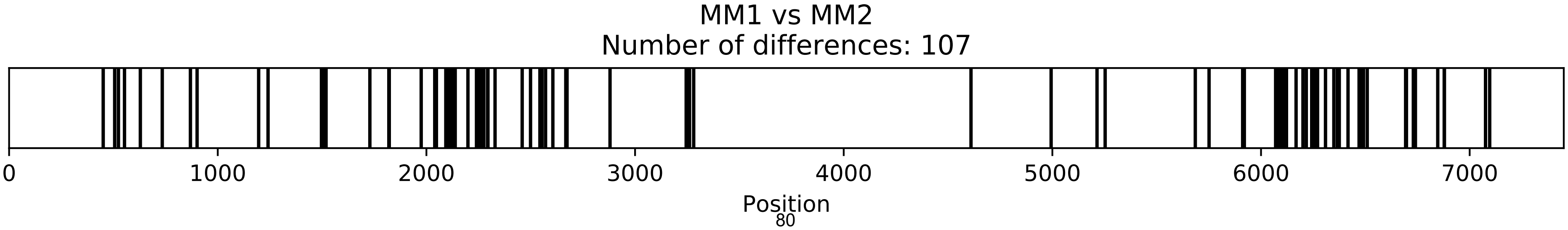
MM1 vs MM2
Number of differences: 107
0
1000
2000
3000
4000
5000
6000
7000
Position

### Region: 291



MA1 vs MA2
Number of differences: 367

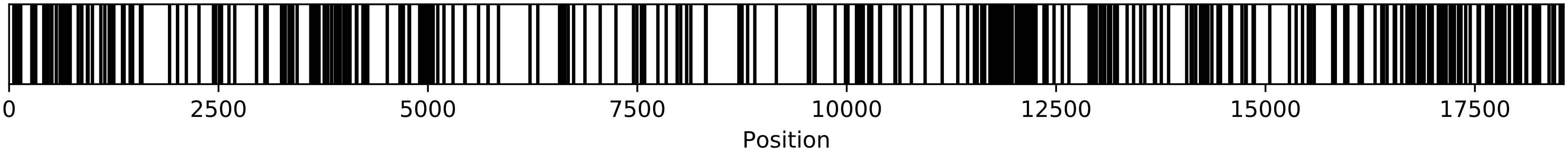


MA1 vs MM1
Number of differences: 386

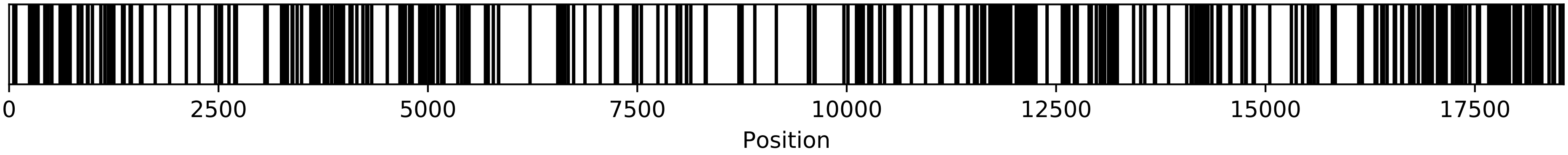


MA1 vs MM2
Number of differences: 371

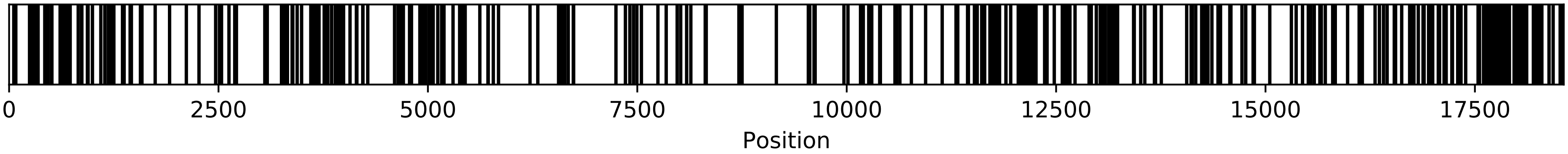


MA2 vs MM1
Number of differences: 171

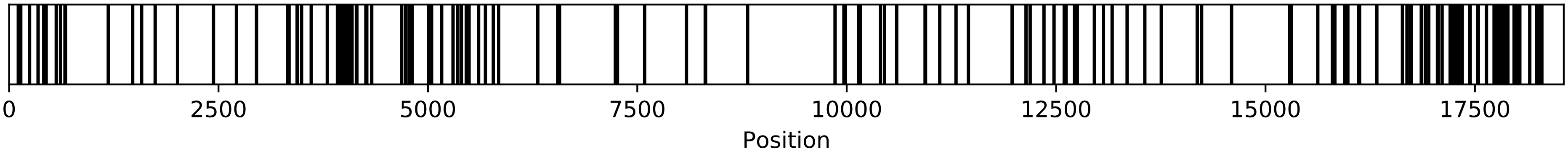


MA2 vs MM2
Number of differences: 193

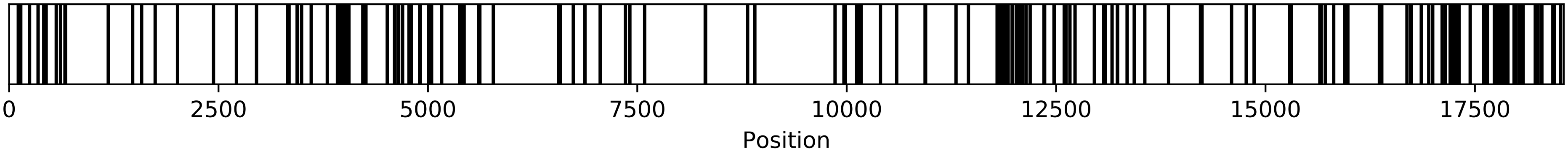


MM1 vs MM2
Number of differences: 149

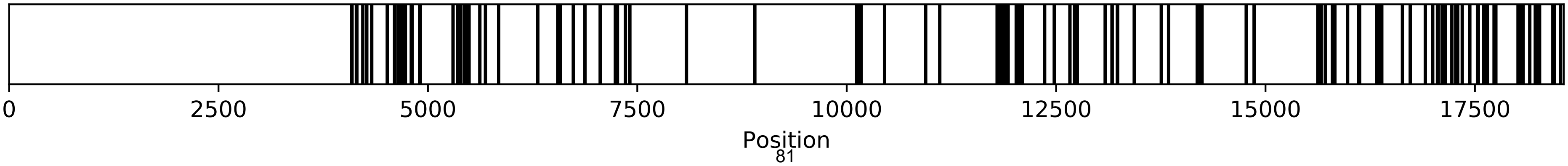

### Region: 293



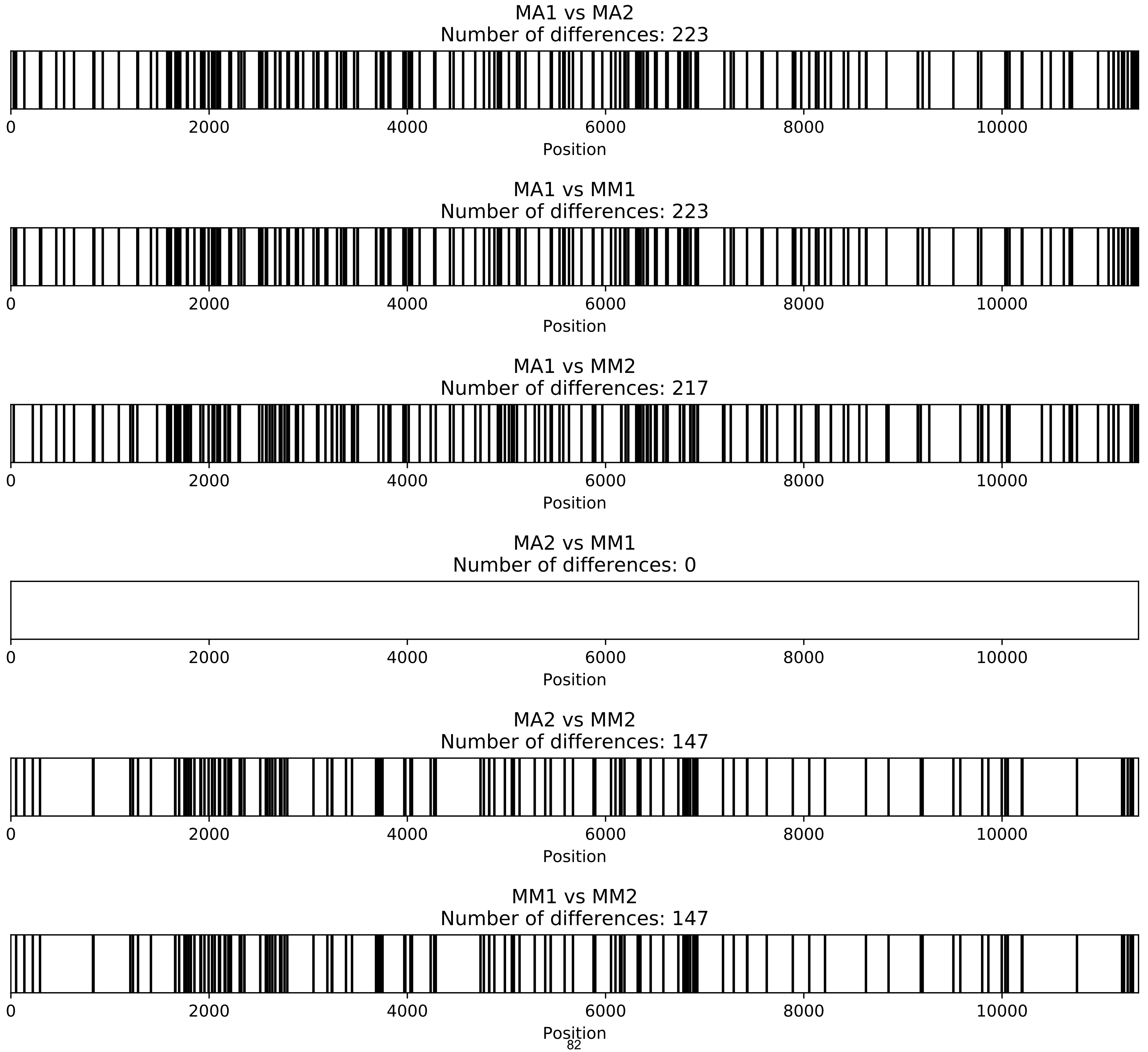
MA1 vs MA2
Number of differences: 223
MA1 vs MM1
Number of differences: 223
MA1 vs MM2
Number of differences: 217
MA2 vs MM1
Number of differences: 0
MA2 vs MM2
Number of differences: 147
MM1 vs MM2
Number of differences: 147
0
2000
4000
6000
8000
10000
Position

### Region: 295



MA1 vs MA2
Number of differences: 303

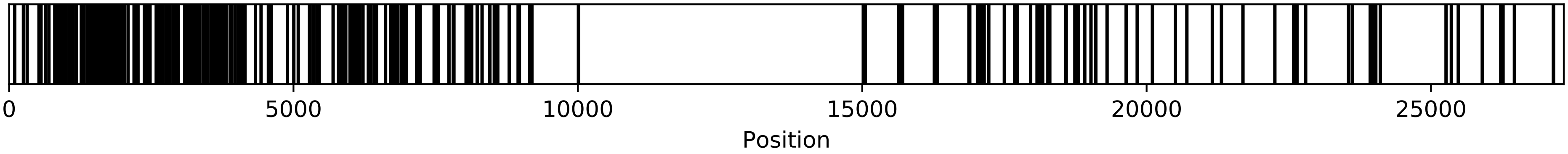


MA1 vs MM1
Number of differences: 0

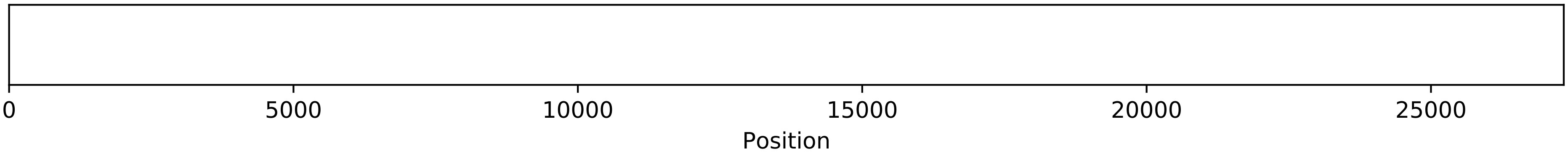


MA1 vs MM2
Number of differences: 282

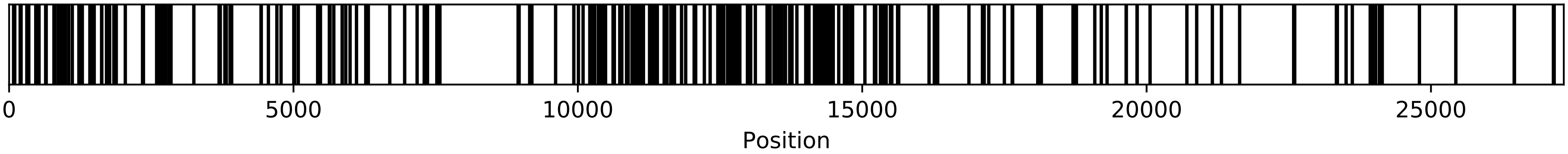


MA2 vs MM1
Number of differences: 303

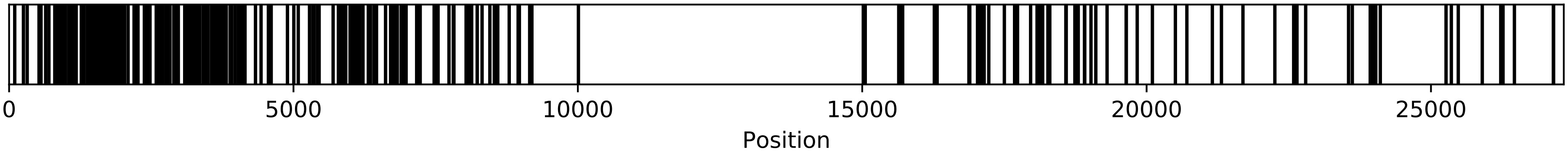


MA2 vs MM2
Number of differences: 442

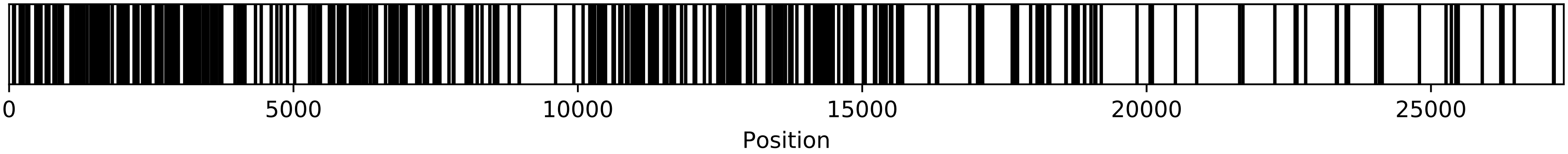


MM1 vs MM2
Number of differences: 282

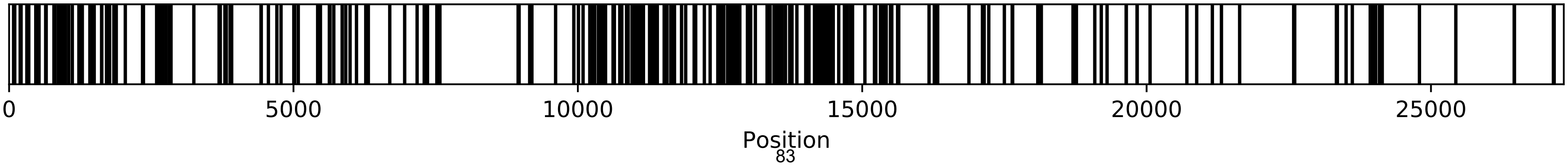

### Region: 297



MA1 vs MA2
Number of differences: 254

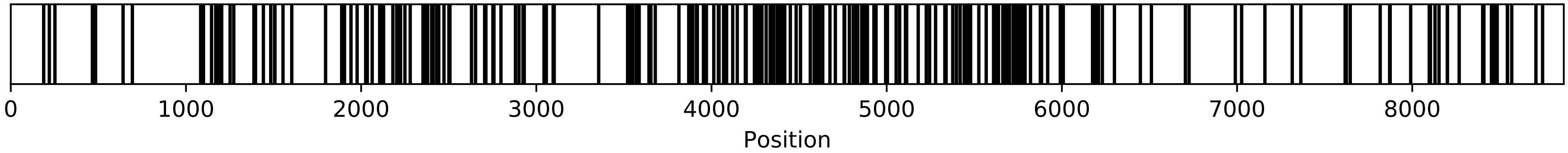


MA1 vs MM1
Number of differences: 180

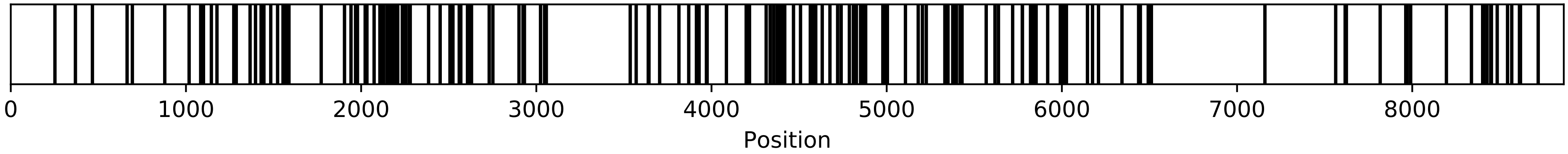


MA1 vs MM2
Number of differences: 0

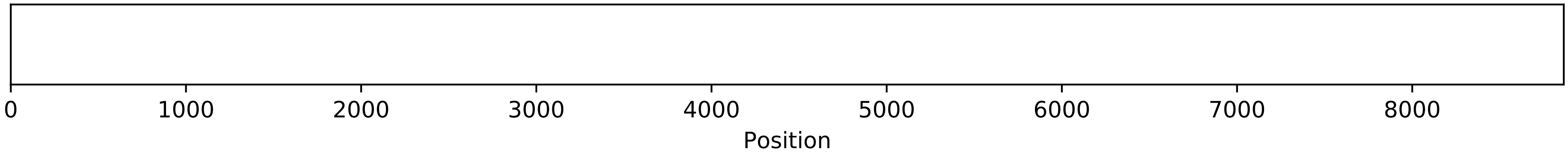


MA2 vs MM1
Number of differences: 256

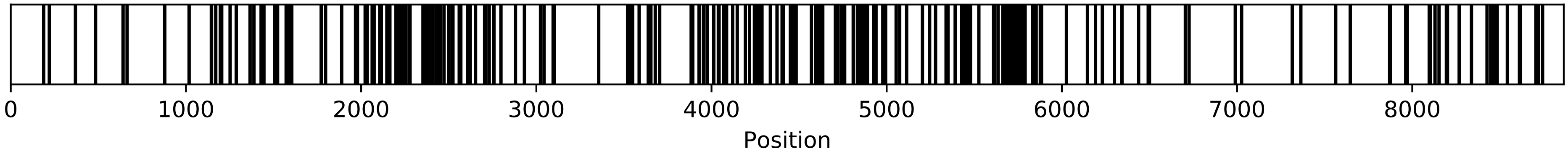


MA2 vs MM2
Number of differences: 254

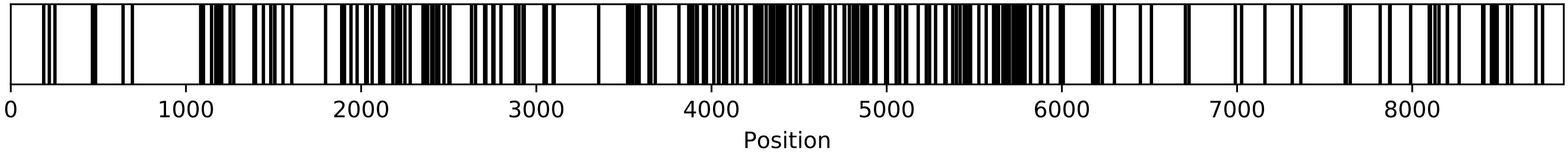


MM1 vs MM2
Number of differences: 180

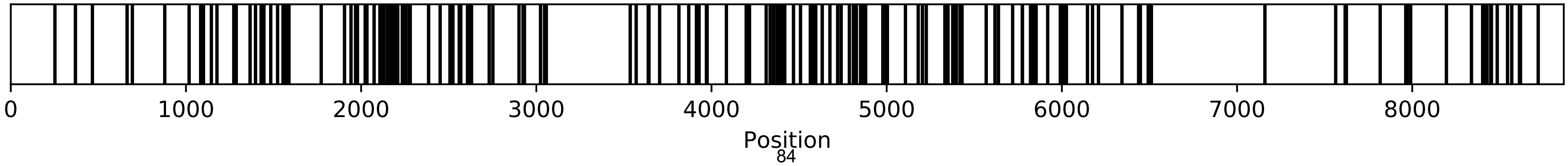

### Region: 299

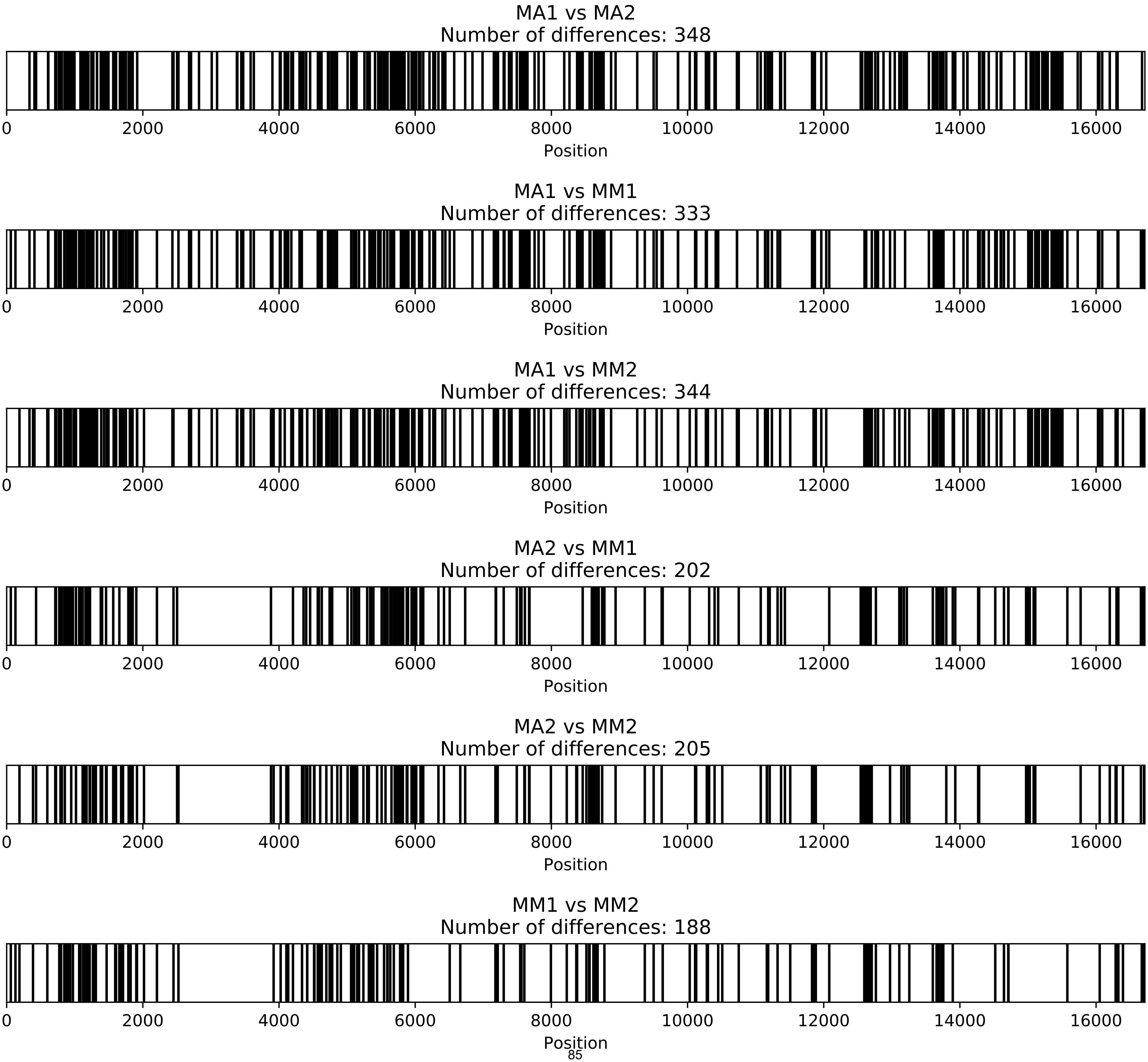
MA1 vs MA2
Number of differences: 348
MA1 vs MM1
Number of differences: 333
MA1 vs MM2
Number of differences: 344
MA2 vs MM1
Number of differences: 202
MA2 vs MM2
Number of differences: 205
MM1 vs MM2
Number of differences: 188
0
2000
4000
6000
8000
10000
12000
14000
16000
Position

### Region: 301



MA1 vs MA2
Number of differences: 183

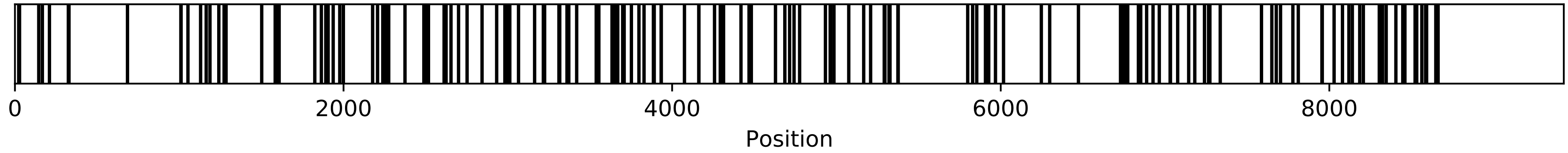


MA1 vs MM1
Number of differences: 190

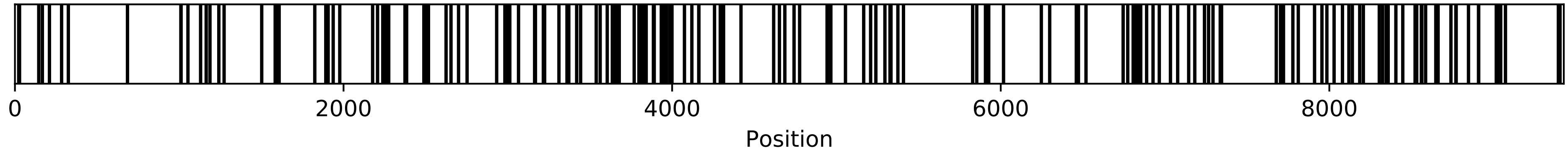


MA1 vs MM2
Number of differences: 192

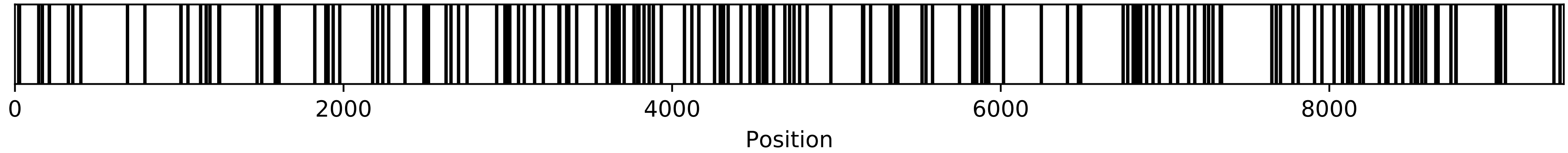


MA2 vs MM1
Number of differences: 93

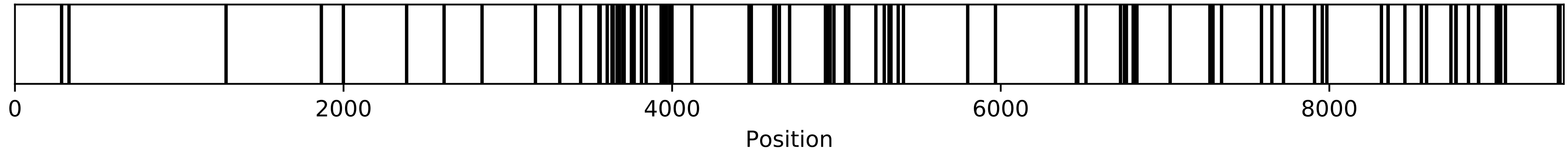


MA2 vs MM2
Number of differences: 104

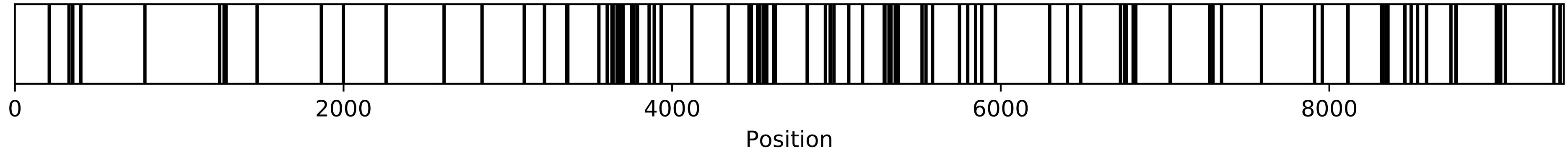


MM1 vs MM2
Number of differences: 79

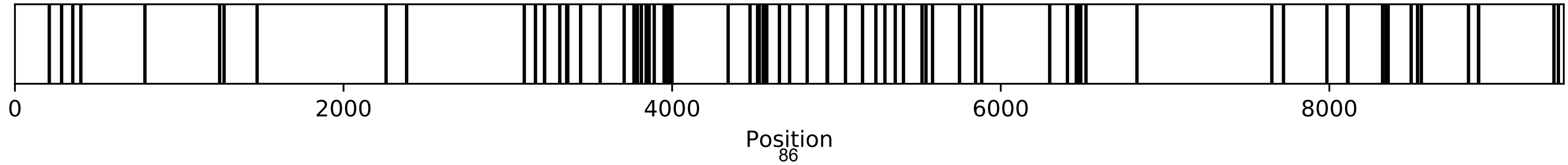

### Region: 304



MA1 vs MA2
Number of differences: 603

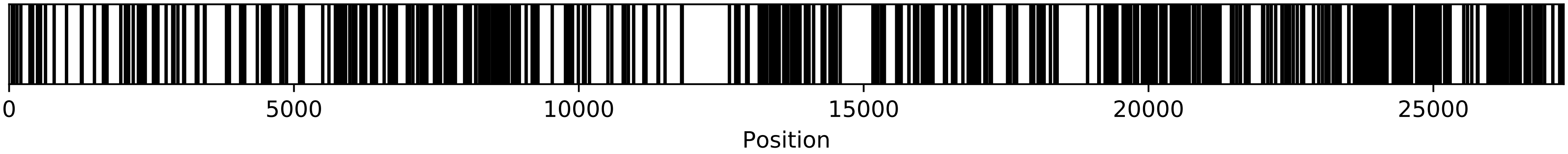


MA1 vs MM1
Number of differences: 617

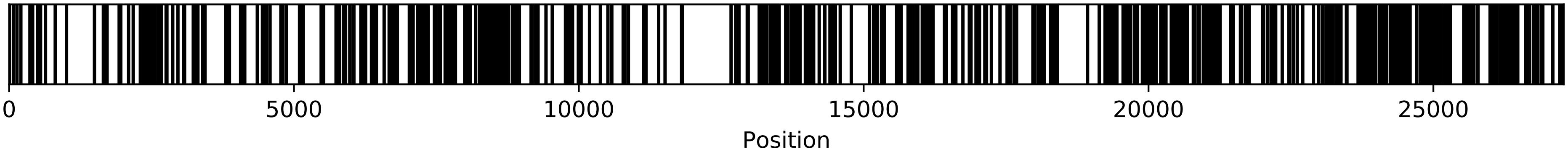


MA1 vs MM2
Number of differences: 603

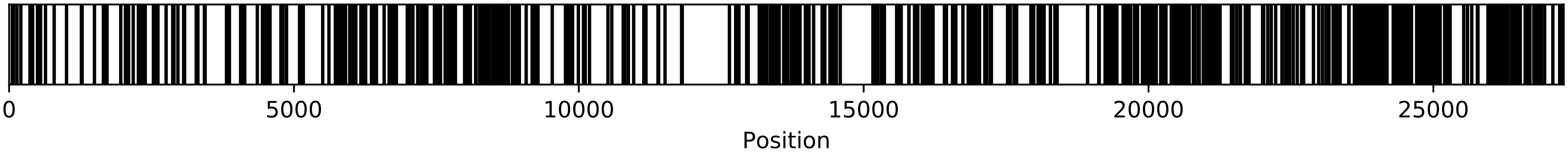


MA2 vs MM1
Number of differences: 302

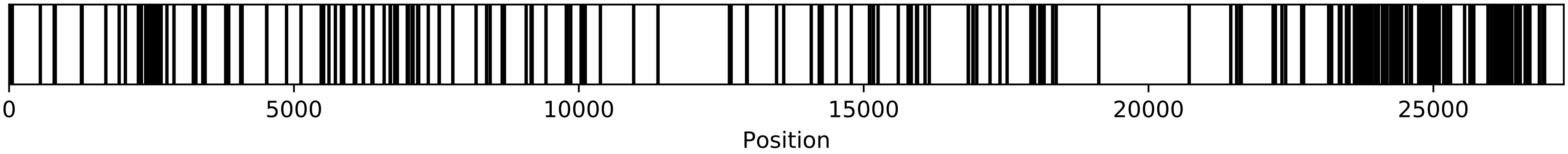


MA2 vs MM2
Number of differences: 0

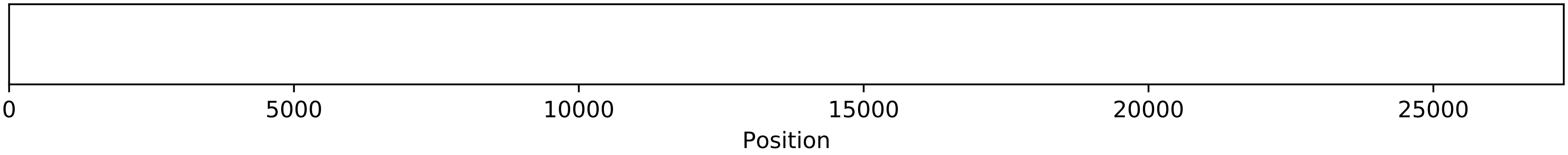


MM1 vs MM2
Number of differences: 302

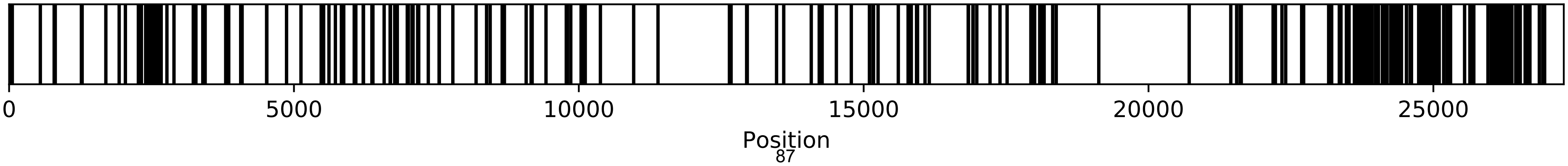

### Region: 305



MA1 vs MA2
Number of differences: 502

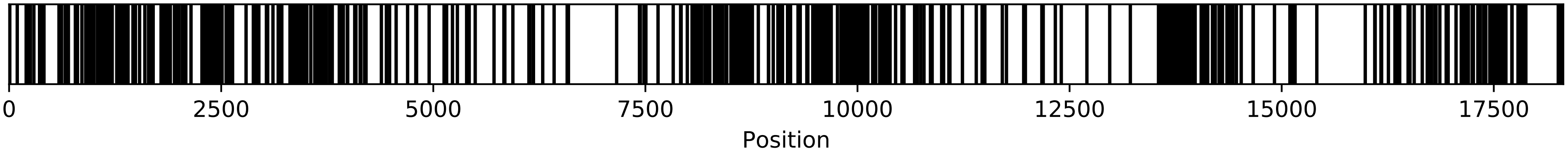


MA1 vs MM1
Number of differences: 514

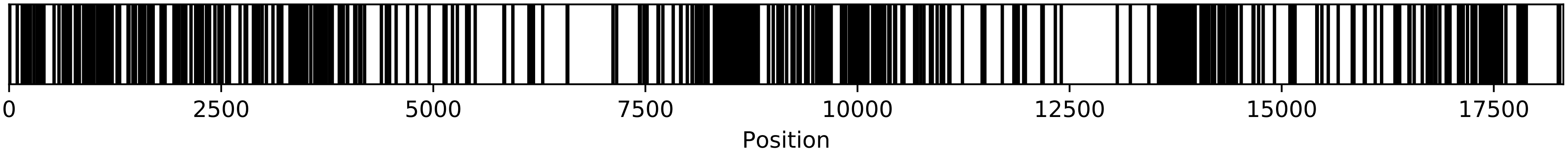


MA1 vs MM2
Number of differences: 500

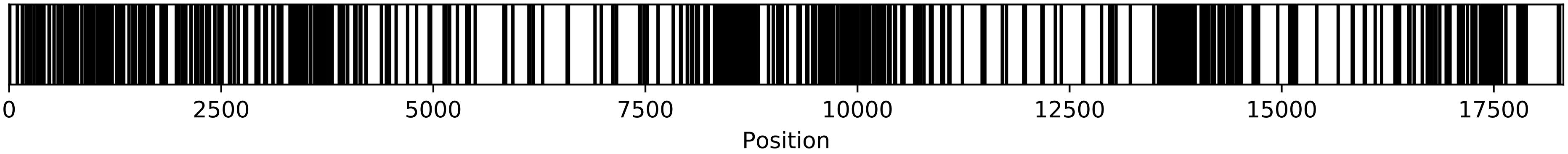


MA2 vs MM1
Number of differences: 240

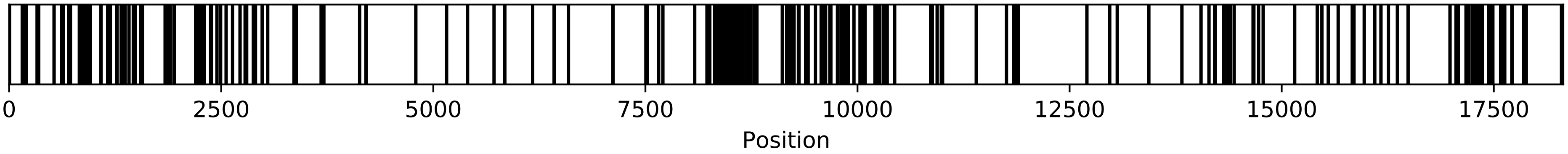


MA2 vs MM2
Number of differences: 218

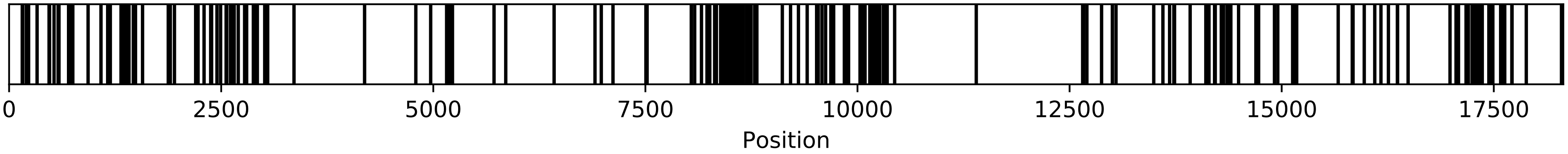


MM1 vs MM2
Number of differences: 171

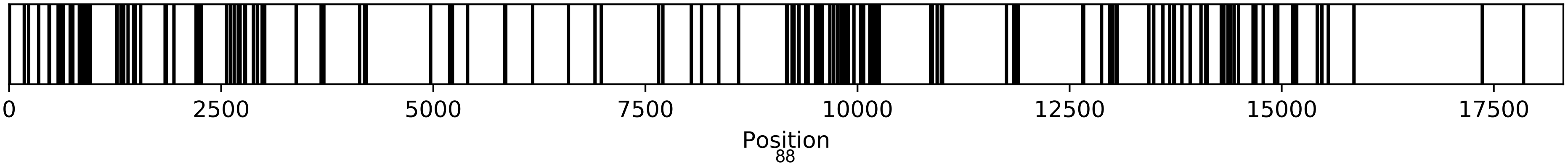

### Region: 313



MA1 vs MA2
Number of differences: 253

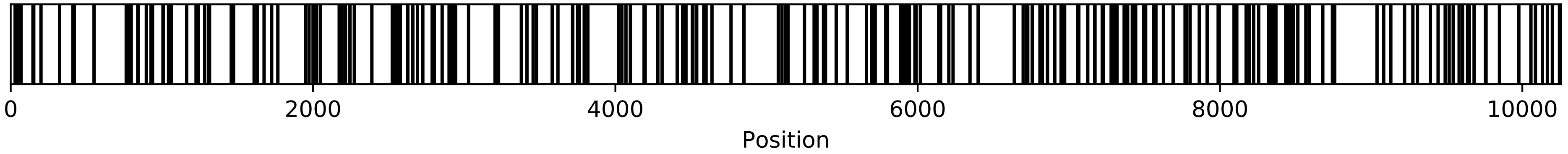


MA1 vs MM1
Number of differences: 0

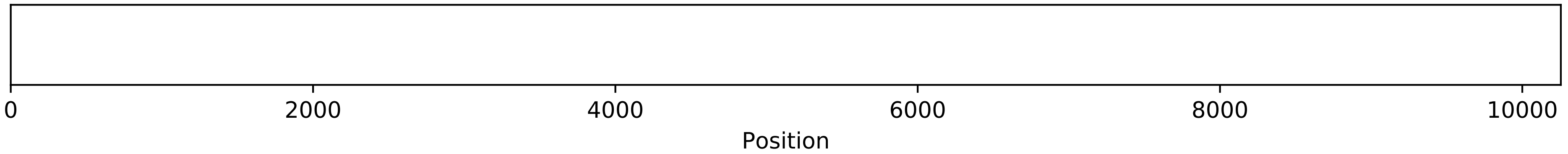


MA1 vs MM2
Number of differences: 99

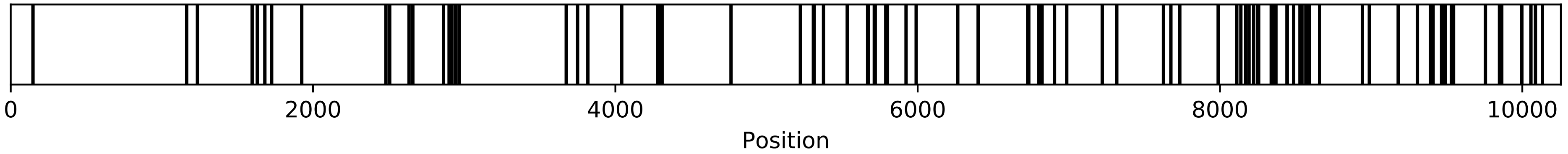


MA2 vs MM1
Number of differences: 253

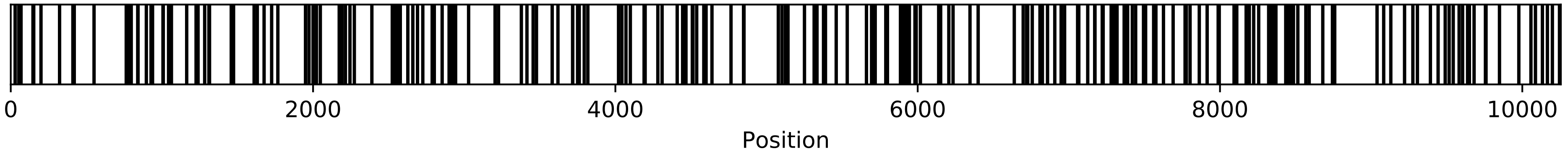


MA2 vs MM2
Number of differences: 265

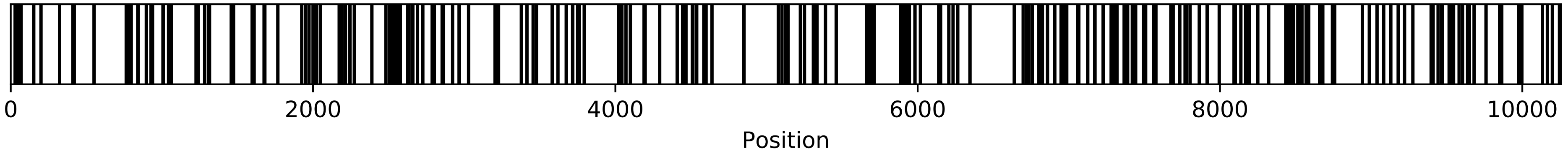


MM1 vs MM2
Number of differences: 99

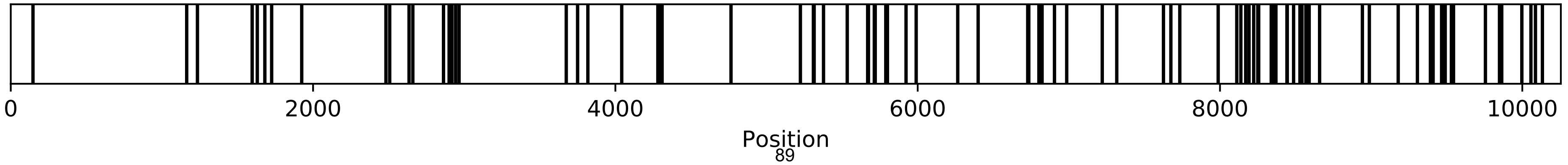

### Region: 314



MA1 vs MA2
Number of differences: 168

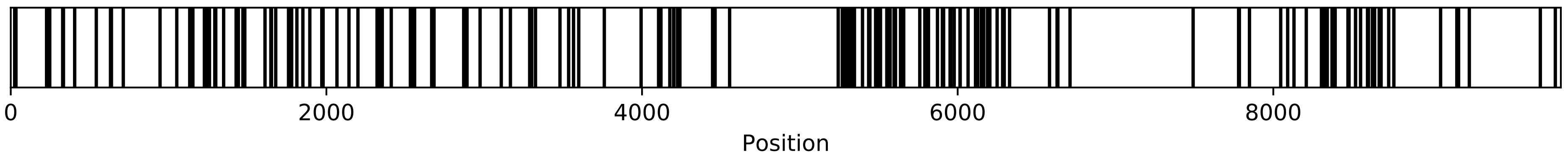


MA1 vs MM1
Number of differences: 0

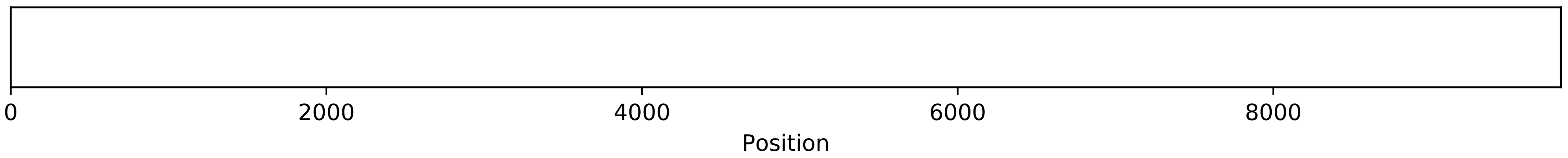


MA1 vs MM2
Number of differences: 115

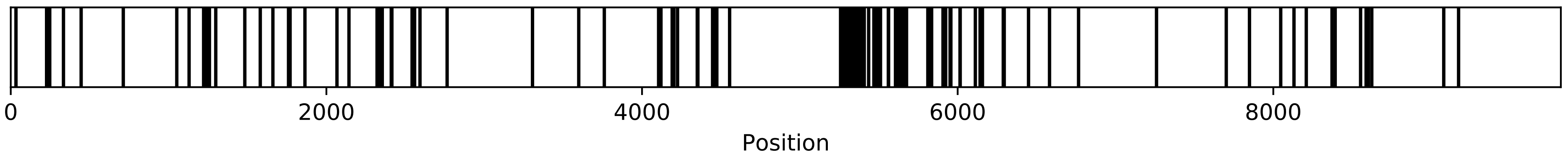


MA2 vs MM1
Number of differences: 168

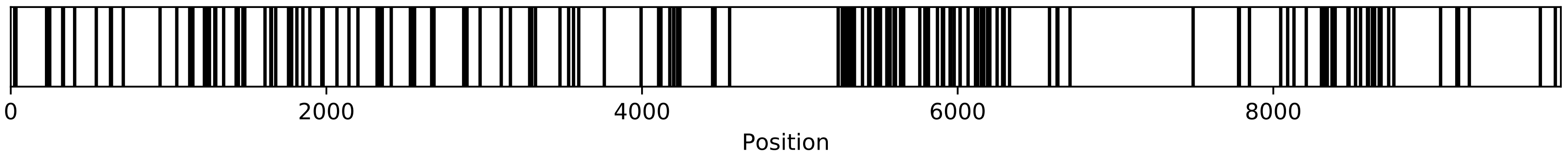


MA2 vs MM2
Number of differences: 153

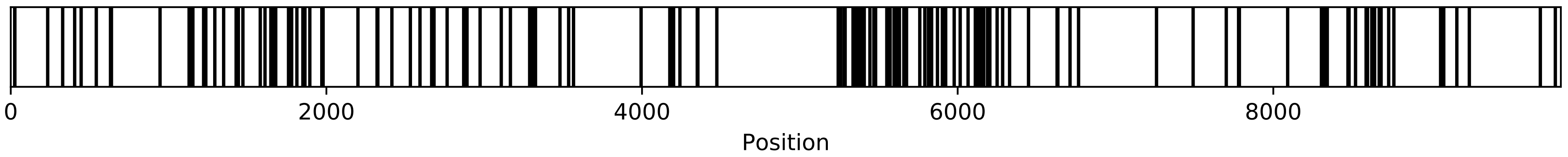


MM1 vs MM2
Number of differences: 115

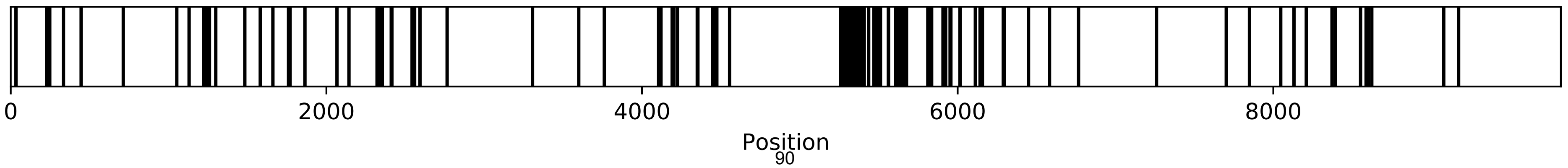

### Region: 315



MA1 vs MA2
Number of differences: 108

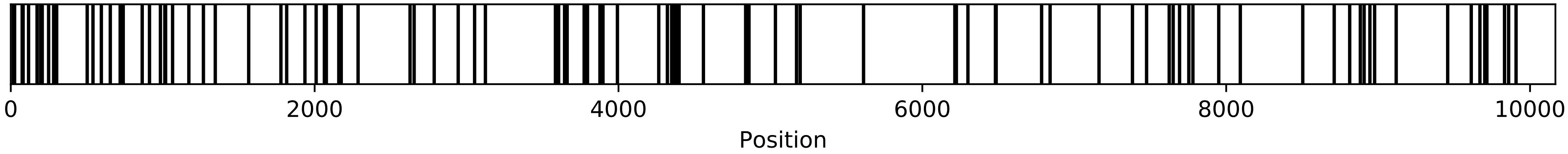


MA1 vs MM1
Number of differences: 45

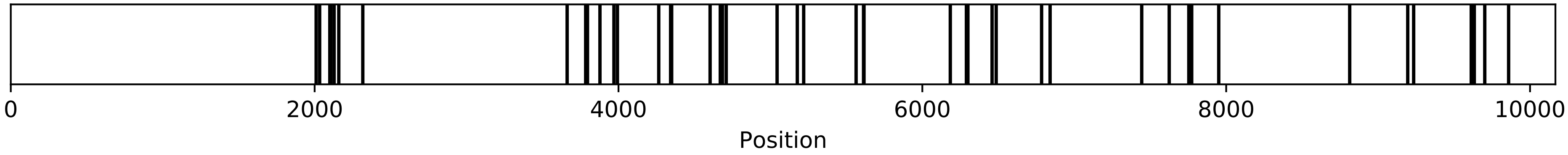


MA1 vs MM2
Number of differences: 65

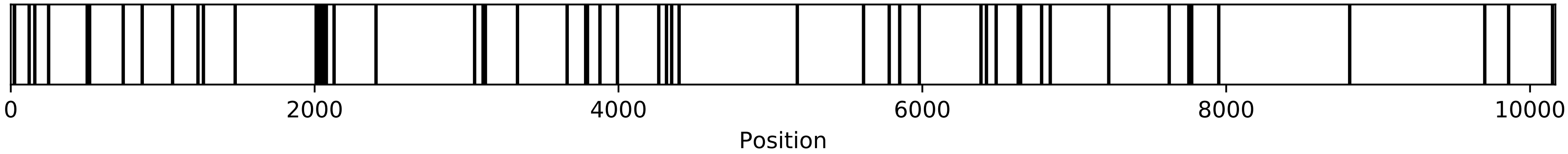


MA2 vs MM1
Number of differences: 113

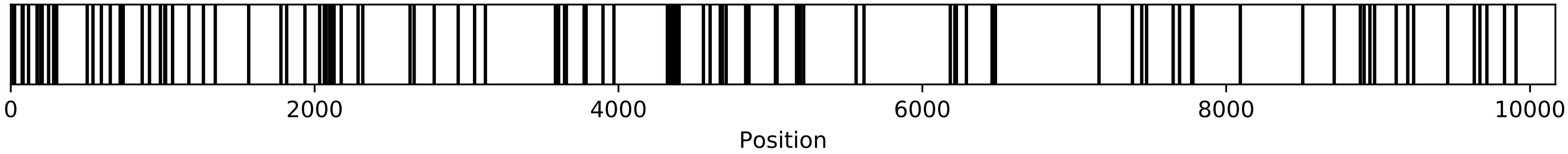


MA2 vs MM2
Number of differences: 101

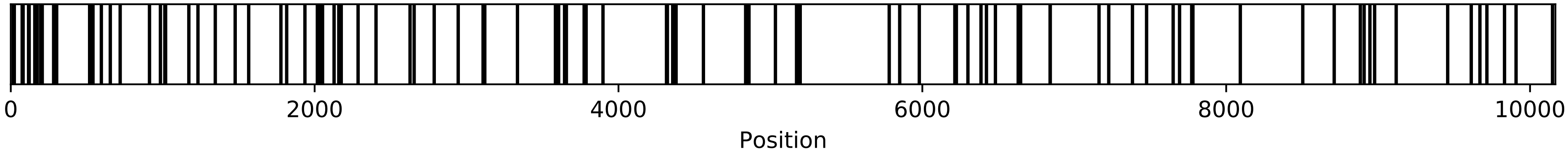


MM1 vs MM2
Number of differences: 68

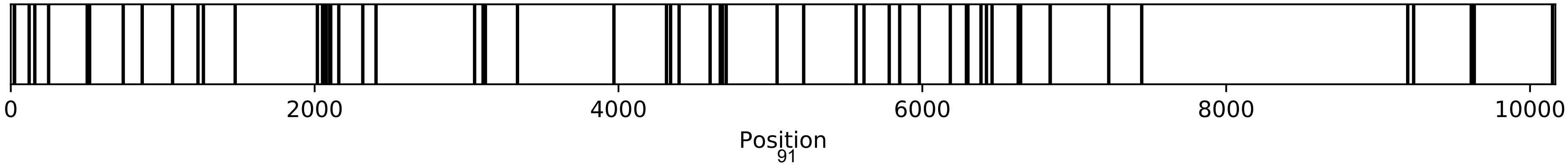

### Region: 318



MA1 vs MA2
Number of differences: 204

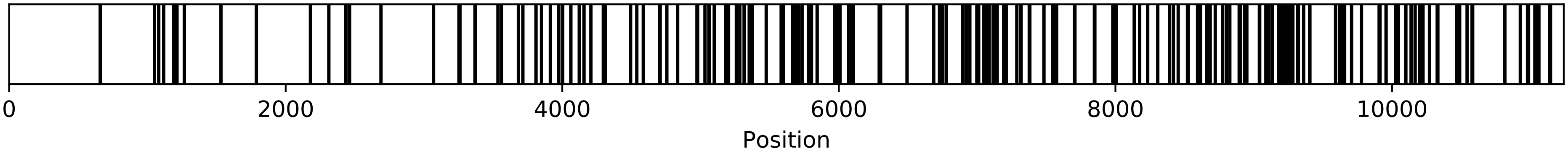


MA1 vs MM1
Number of differences: 207

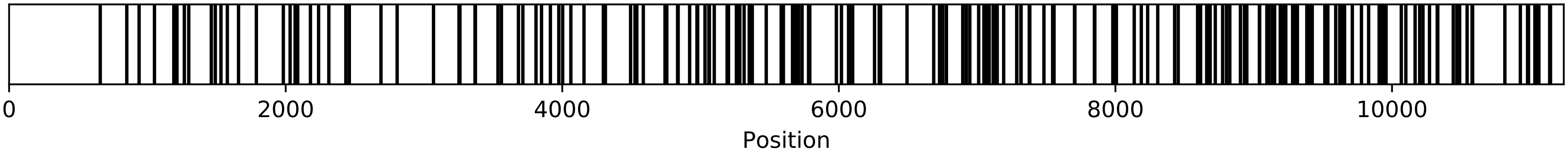


MA1 vs MM2
Number of differences: 204

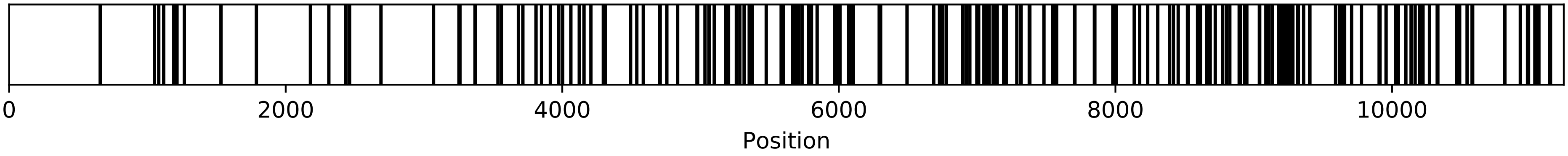


MA2 vs MM1
Number of differences: 88

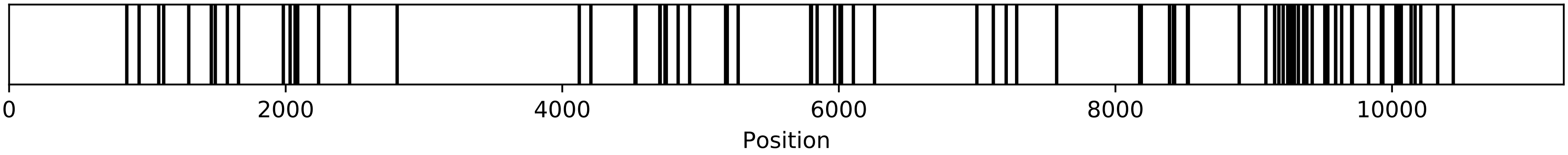


MA2 vs MM2
Number of differences: 0

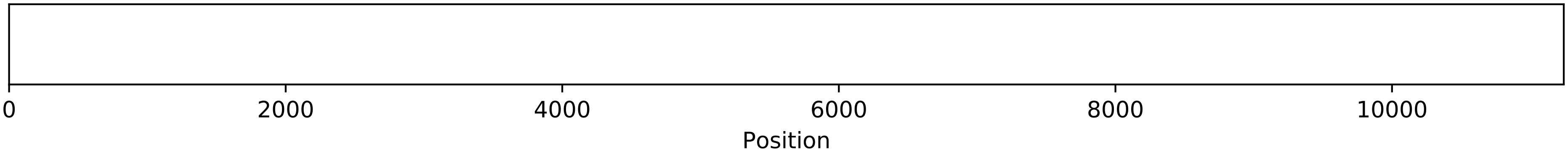


MM1 vs MM2
Number of differences: 88

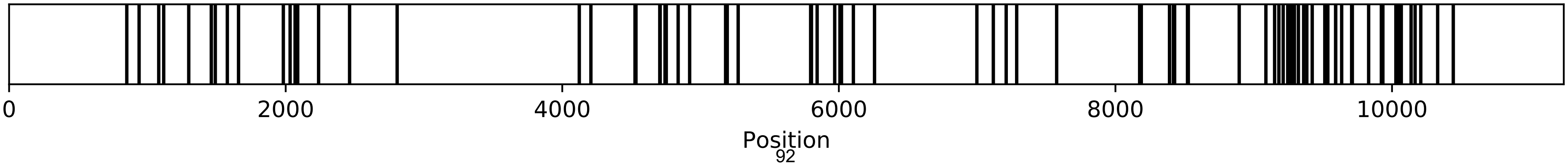

### Region: 322



MA1 vs MA2
Number of differences: 112

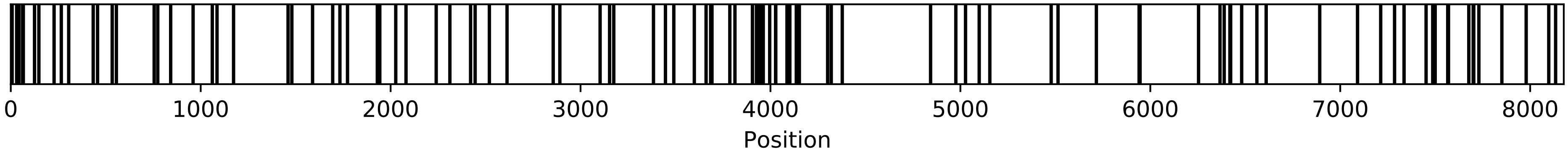


MA1 vs MM1
Number of differences: 108

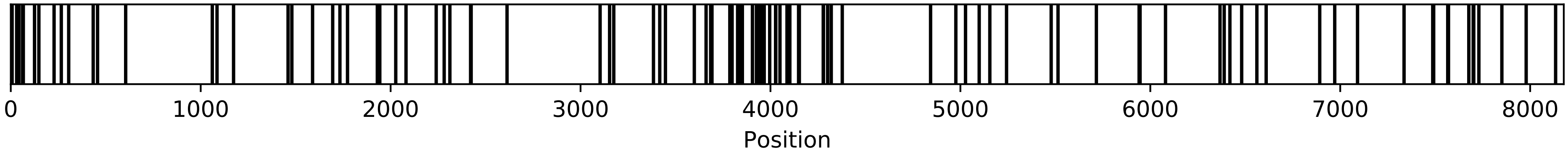


MA1 vs MM2
Number of differences: 118

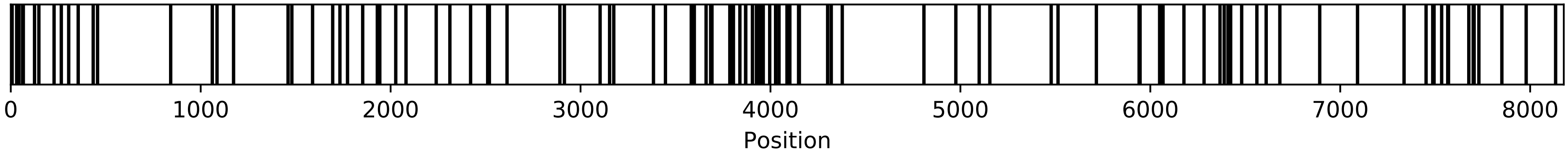


MA2 vs MM1
Number of differences: 37

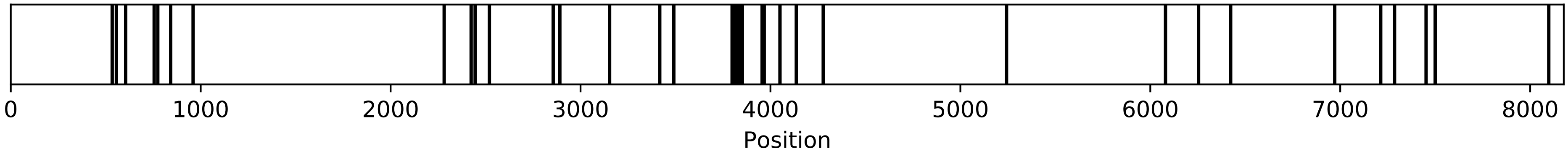


MA2 vs MM2
Number of differences: 40

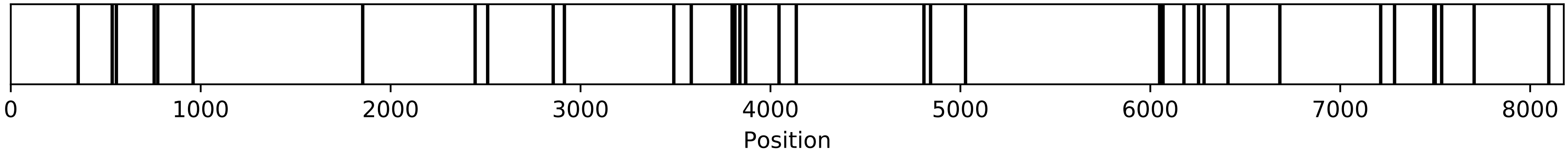


MM1 vs MM2
Number of differences: 43

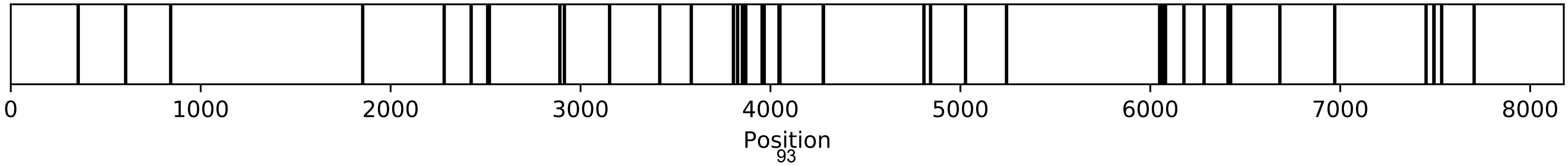

### Region: 324



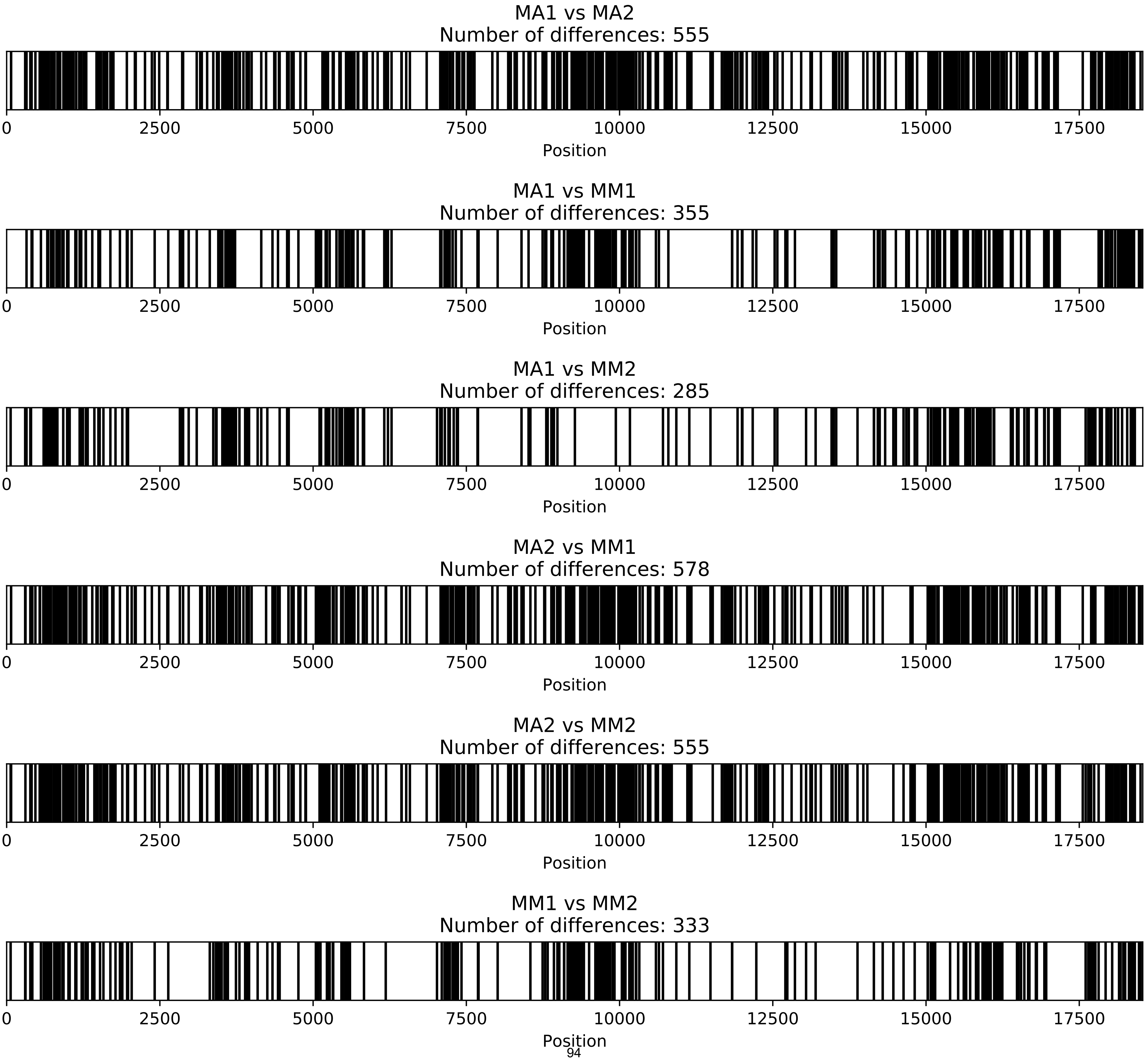
MA1 vs MA2
Number of differences: 555
MA1 vs MM1
Number of differences: 355
MA1 vs MM2
Number of differences: 285
MA2 vs MM1
Number of differences: 578
MA2 vs MM2
Number of differences: 555
MM1 vs MM2
Number of differences: 333
0
2500
5000
7500
10000
12500
15000
17500
Position

### Region: 325

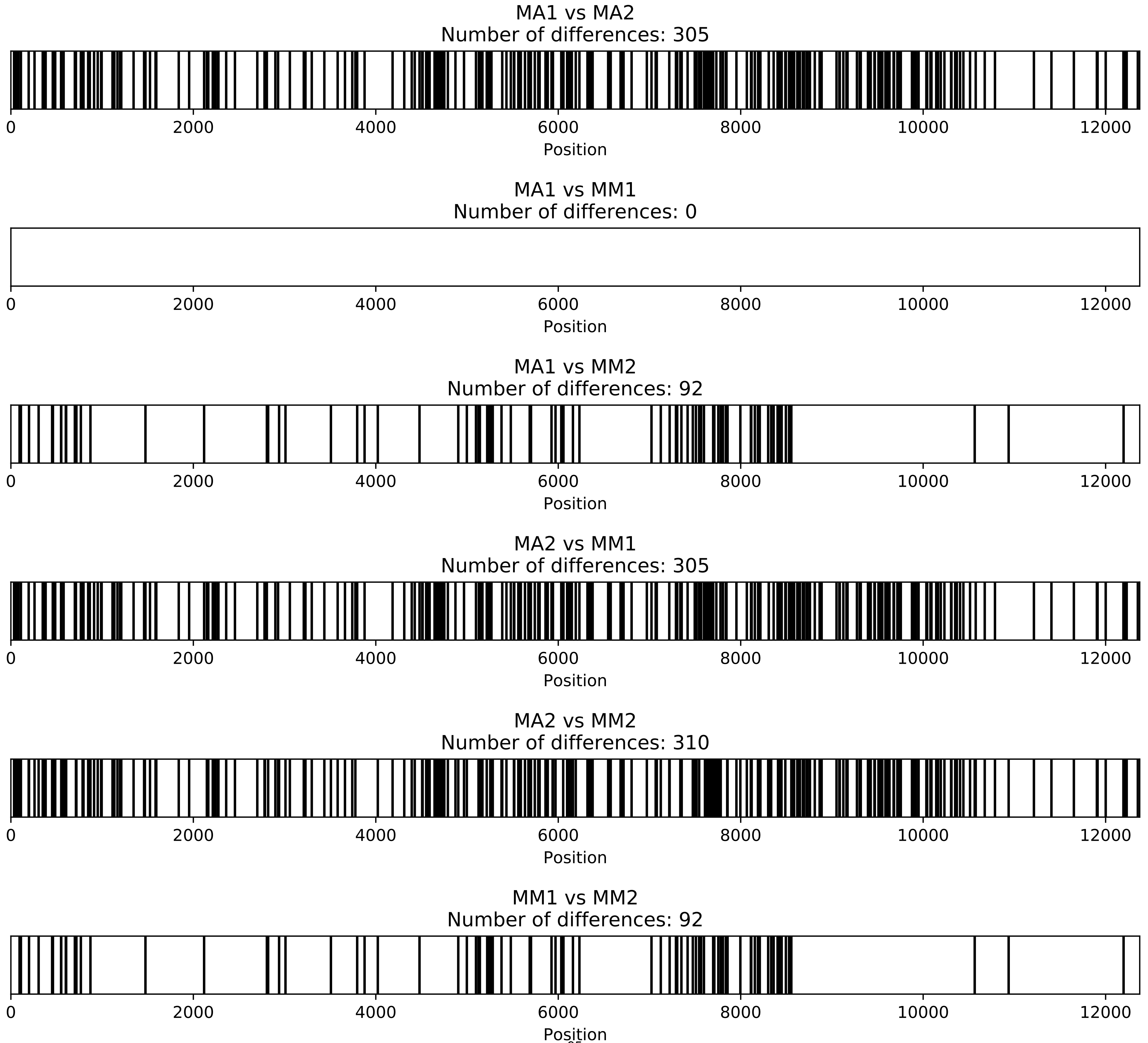
MA1 vs MA2
Number of differences: 305
MA1 vs MM1
Number of differences: 0
MA1 vs MM2
Number of differences: 92
MA2 vs MM1
Number of differences: 305
MA2 vs MM2
Number of differences: 310
MM1 vs MM2
Number of differences: 92
0
2000
4000
6000
8000
10000
12000
Position

### Region: 331



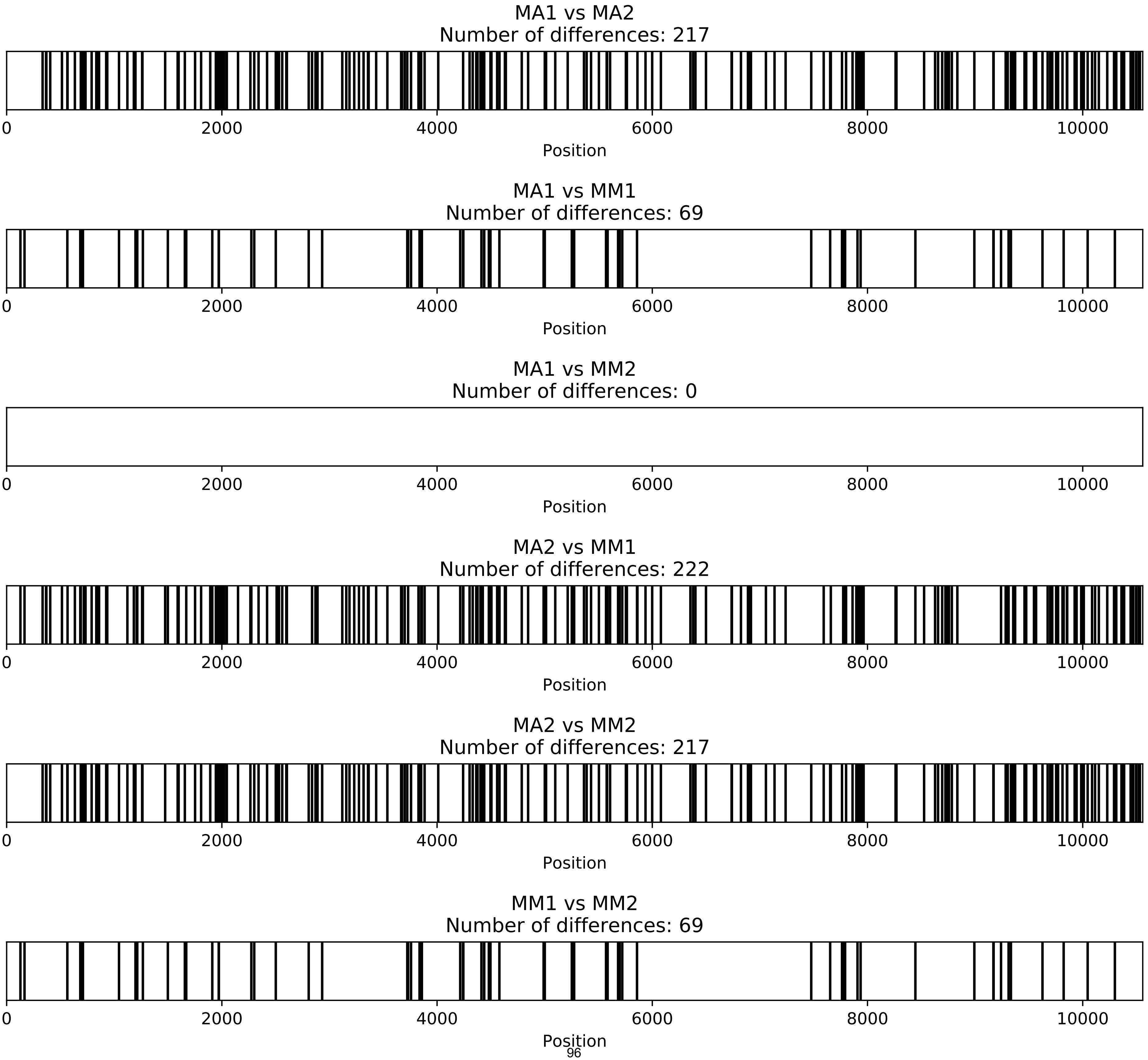
MA1 vs MA2
Number of differences: 217
0
2000
4000
6000
8000
10000
Position
MA1 vs MM1
Number of differences: 69
0
2000
4000
6000
8000
10000
Position
MA1 vs MM2
Number of differences: 0
0
2000
4000
6000
8000
10000
Position
MA2 vs MM1
Number of differences: 222
0
2000
4000
6000
8000
10000
Position
MA2 vs MM2
Number of differences: 217
0
2000
4000
6000
8000
10000
Position
MM1 vs MM2
Number of differences: 69
0
2000
4000
6000
8000
10000
Position

### Region: 333



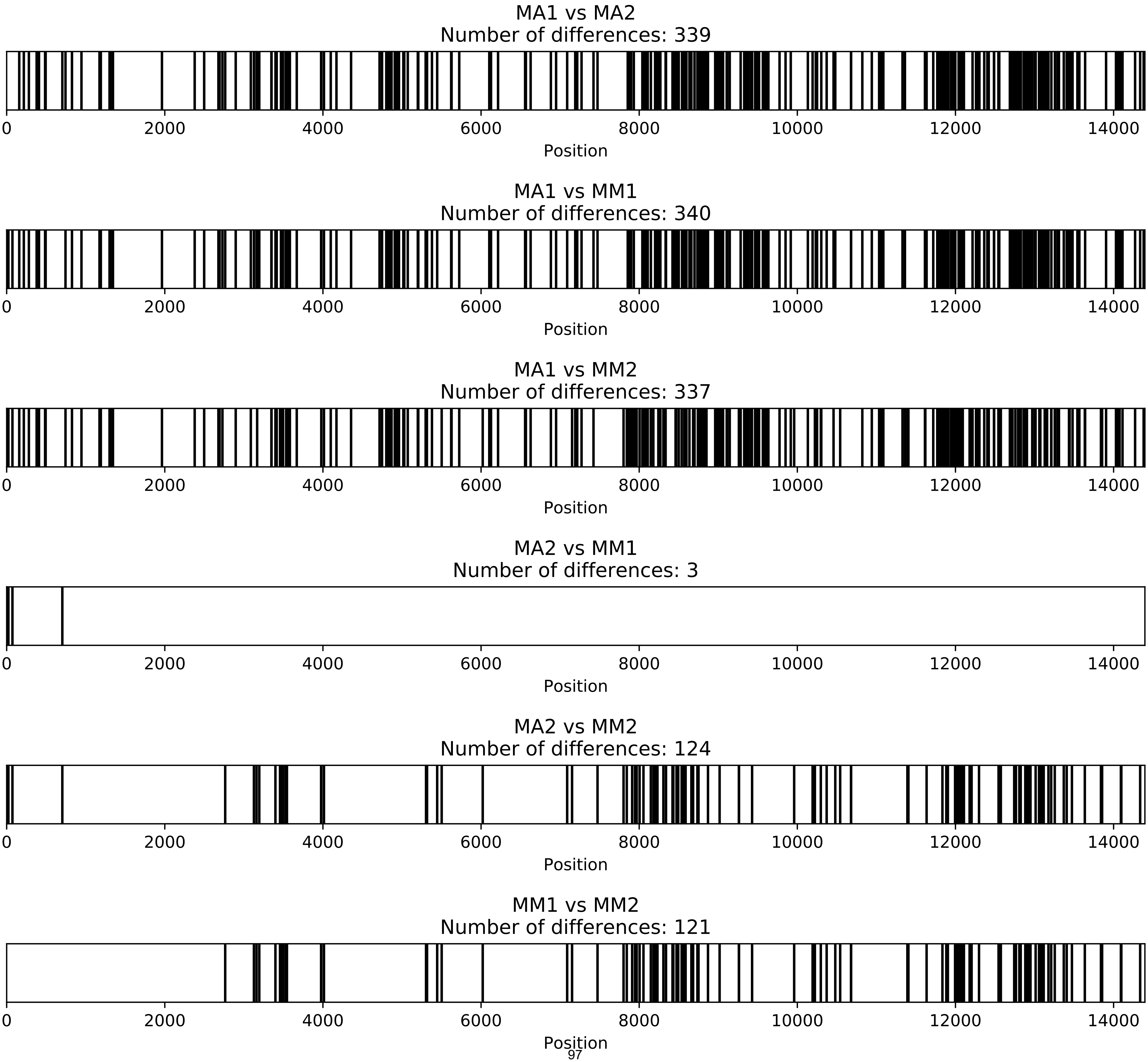
MA1 vs MA2
Number of differences: 339
MA1 vs MM1
Number of differences: 340
MA1 vs MM2
Number of differences: 337
MA2 vs MM1
Number of differences: 3
MA2 vs MM2
Number of differences: 124
MM1 vs MM2
Number of differences: 121
0
2000
4000
6000
8000
10000
12000
14000
Position

### Region: 334

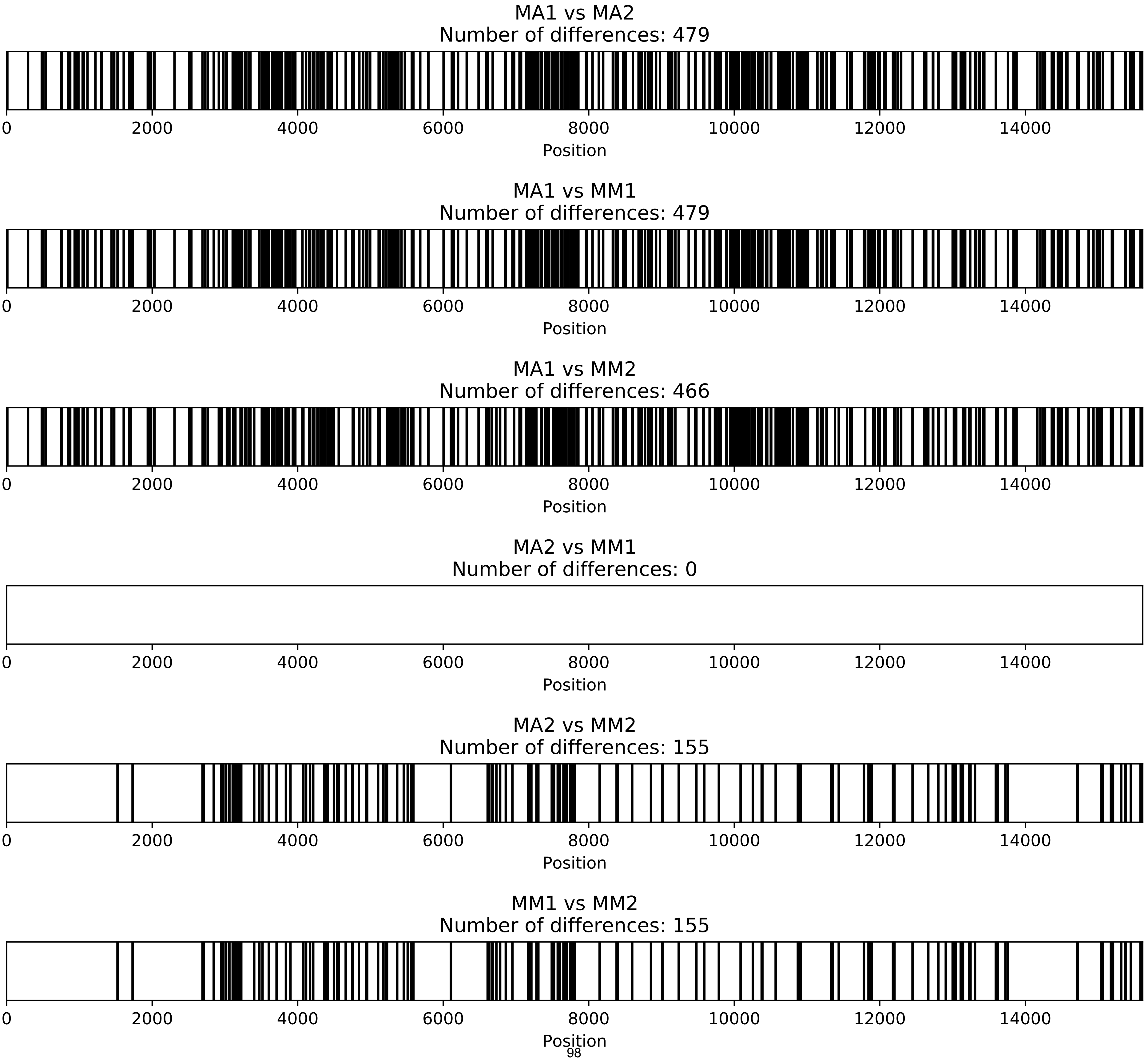
MA1 vs MA2
Number of differences: 479
MA1 vs MM1
Number of differences: 479
MA1 vs MM2
Number of differences: 466
MA2 vs MM1
Number of differences: 0
MA2 vs MM2
Number of differences: 155
MM1 vs MM2
Number of differences: 155
0
2000
4000
6000
8000
10000
12000
14000
Position

### Region: 341



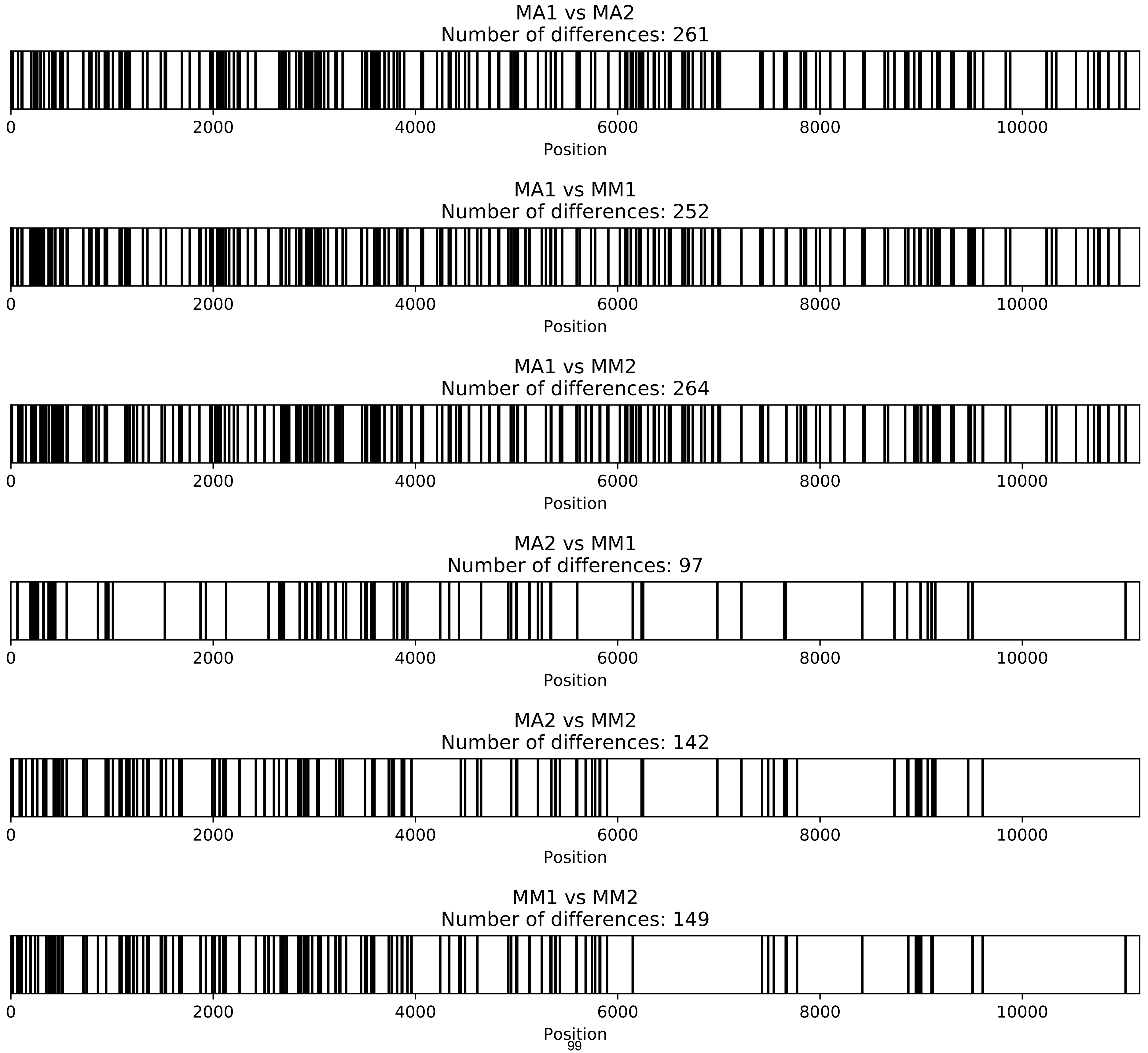
MA1 vs MA2
Number of differences: 261
MA1 vs MM1
Number of differences: 252
MA1 vs MM2
Number of differences: 264
MA2 vs MM1
Number of differences: 97
MA2 vs MM2
Number of differences: 142
MM1 vs MM2
Number of differences: 149
0
2000
4000
6000
8000
10000
Position

### Region: 343



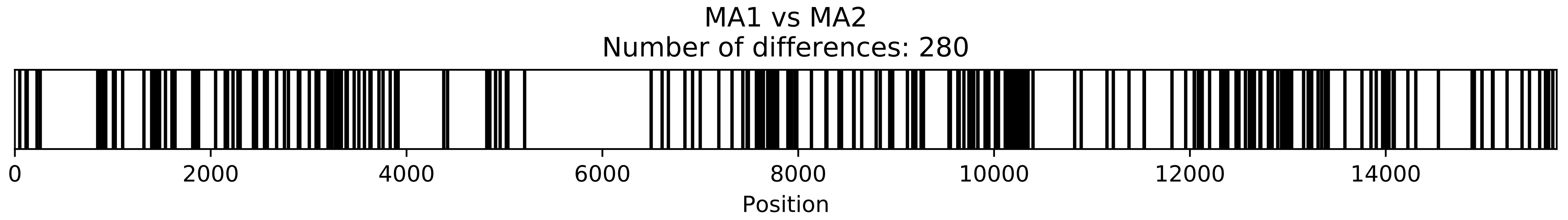
MA1 vs MA2
Number of differences: 280
0
2000
4000
6000
8000
10000
12000
14000
Position

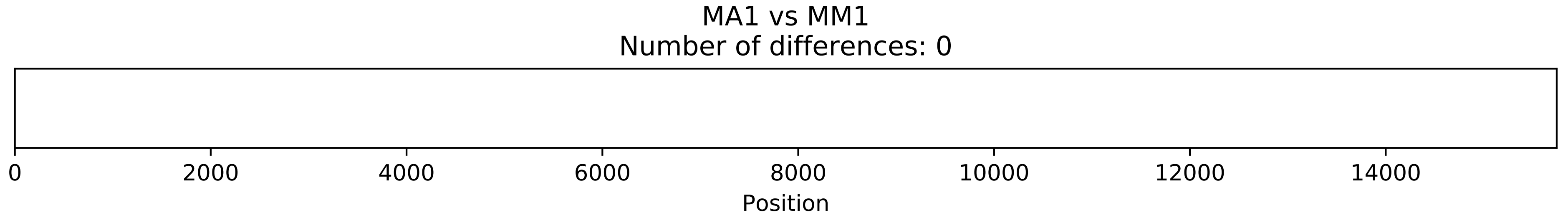
MA1 vs MM1
Number of differences: 0
0
2000
4000
6000
8000
10000
12000
14000
Position

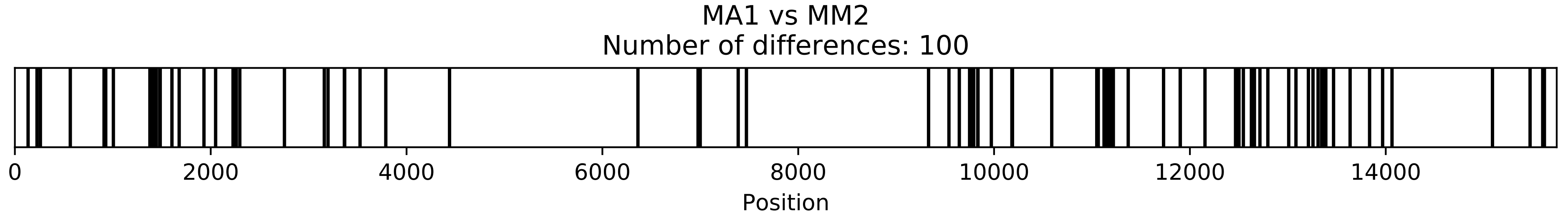
MA1 vs MM2
Number of differences: 100
0
2000
4000
6000
8000
10000
12000
14000
Position

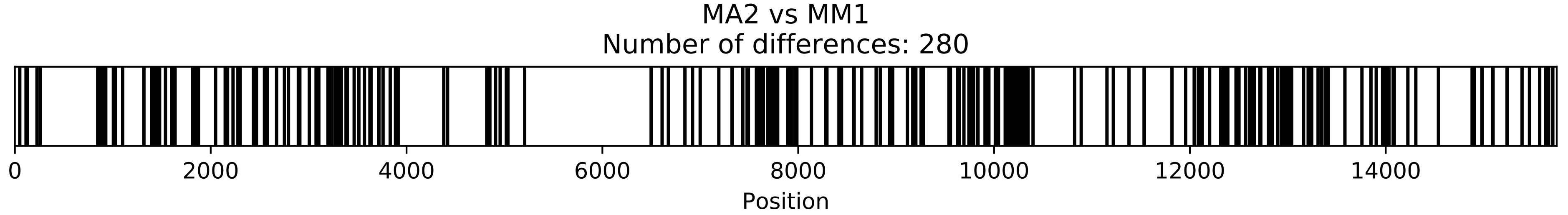
MA2 vs MM1
Number of differences: 280
0
2000
4000
6000
8000
10000
12000
14000
Position

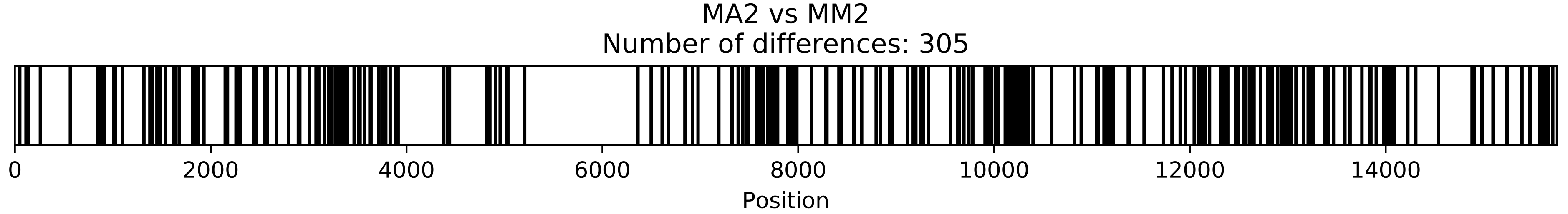
MA2 vs MM2
Number of differences: 305
0
2000
4000
6000
8000
10000
12000
14000
Position

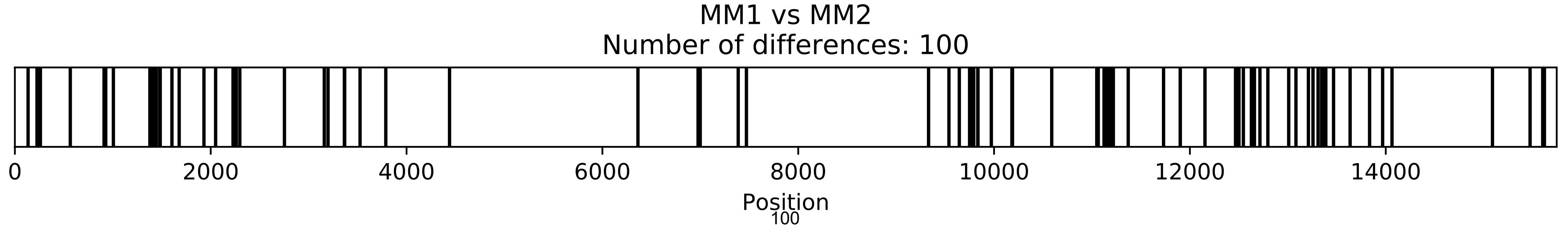
MM1 vs MM2
Number of differences: 100
0
2000
4000
6000
8000
10000
12000
14000
Position

### Region: 352



MA1 vs MA2
Number of differences: 0

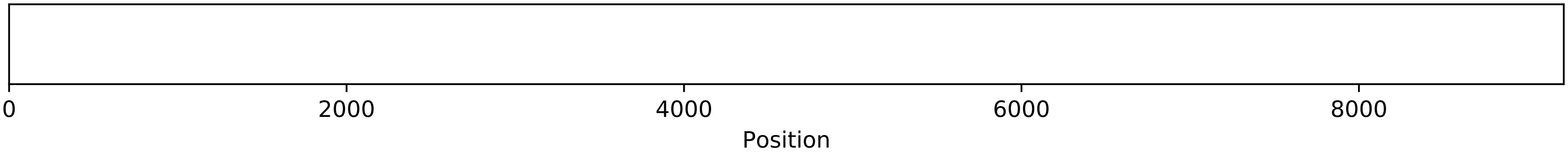


MA1 vs MM1
Number of differences: 288

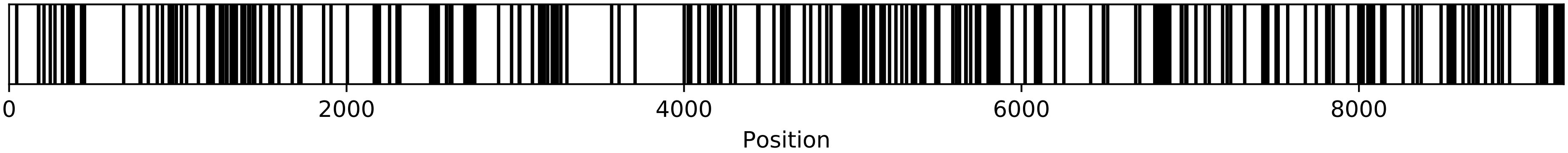


MA1 vs MM2
Number of differences: 220

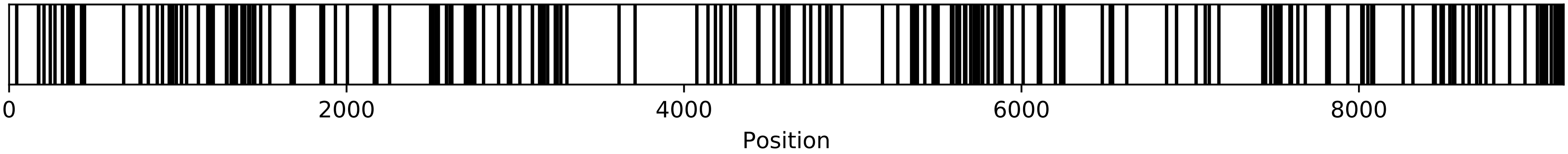


MA2 vs MM1
Number of differences: 288

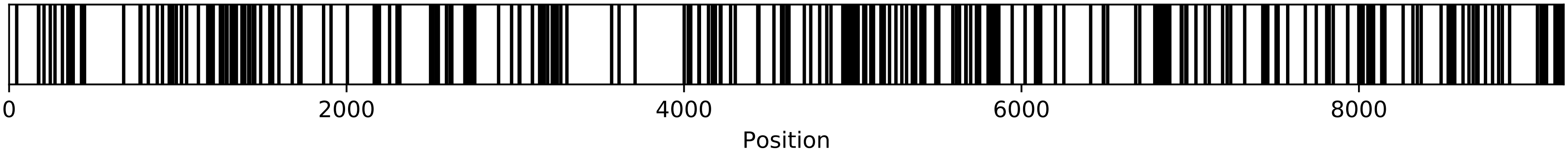


MA2 vs MM2
Number of differences: 220

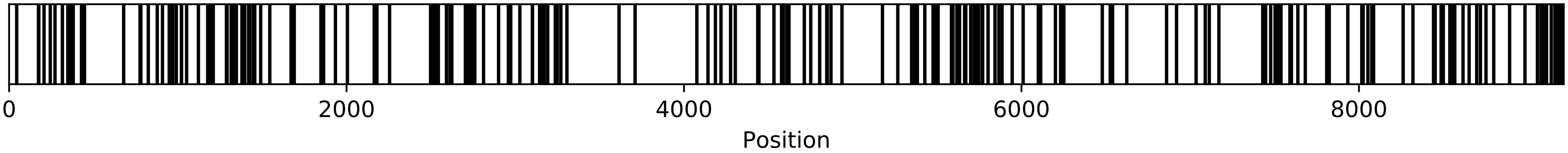


MM1 vs MM2
Number of differences: 189

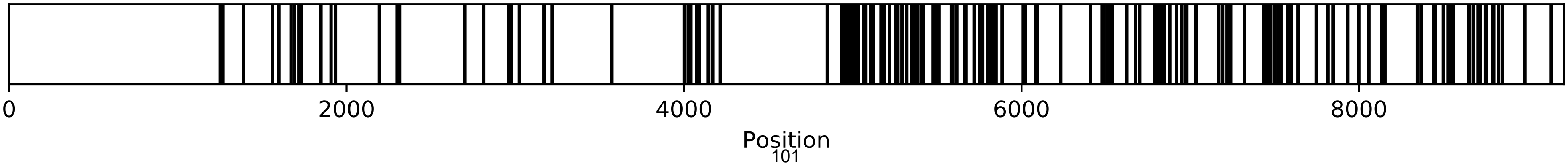

### Region: 353



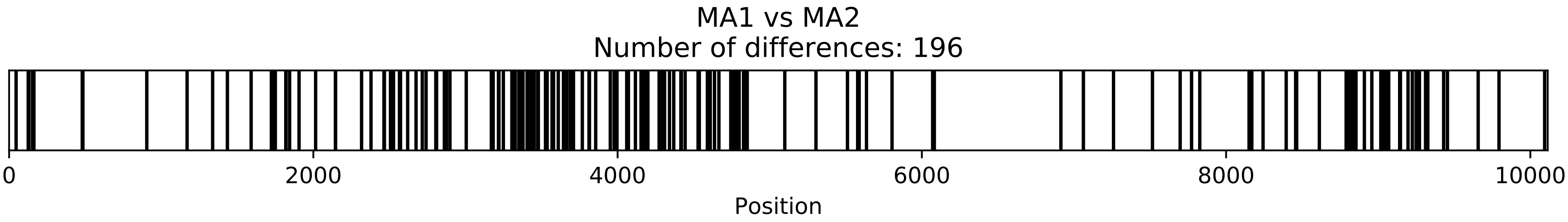
MA1 vs MA2
Number of differences: 196
0
2000
4000
6000
8000
10000
Position

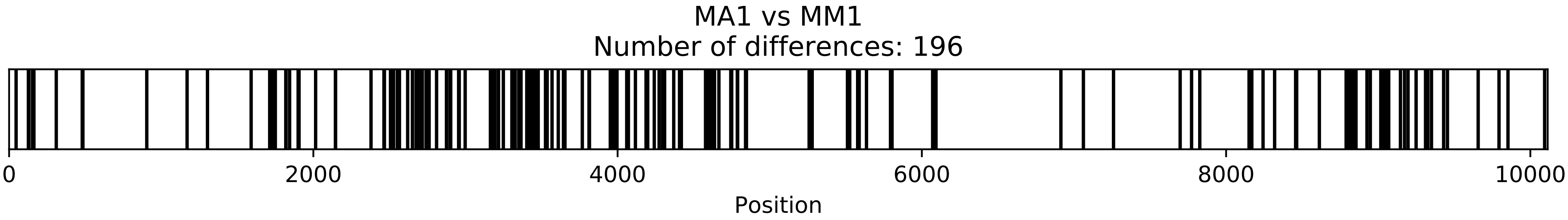
MA1 vs MM1
Number of differences: 196
0
2000
4000
6000
8000
10000
Position

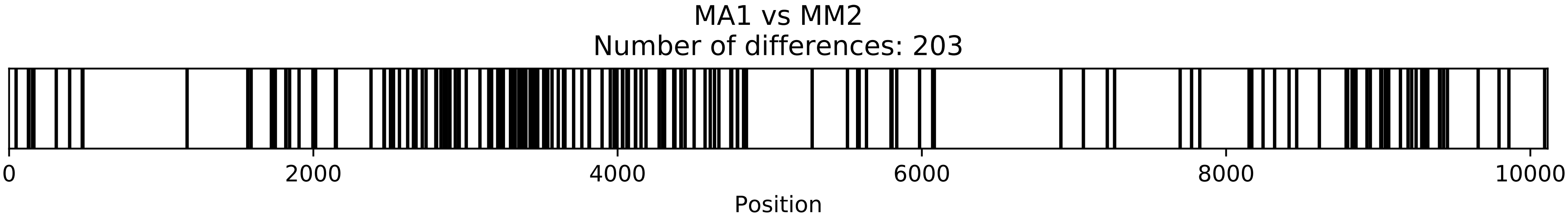
MA1 vs MM2
Number of differences: 203
0
2000
4000
6000
8000
10000
Position

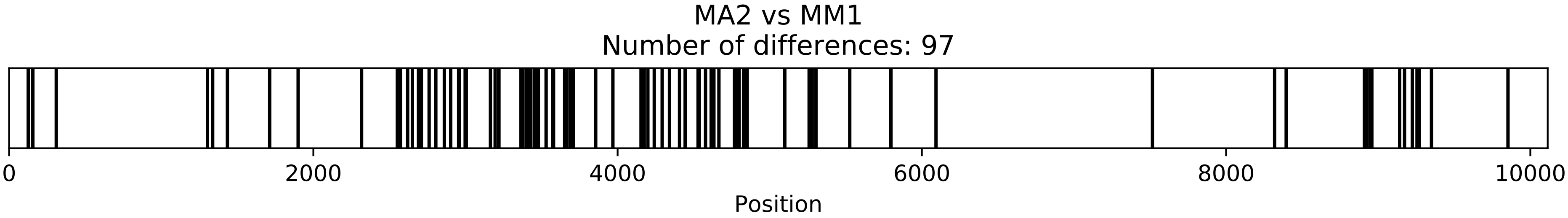
MA2 vs MM1
Number of differences: 97
0
2000
4000
6000
8000
10000
Position

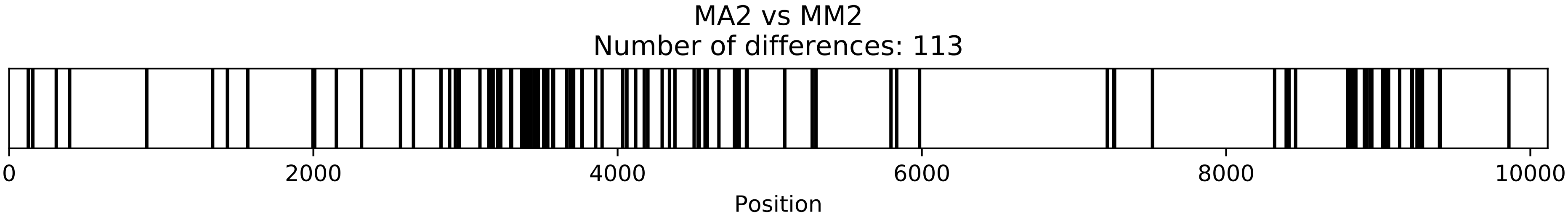
MA2 vs MM2
Number of differences: 113
0
2000
4000
6000
8000
10000
Position

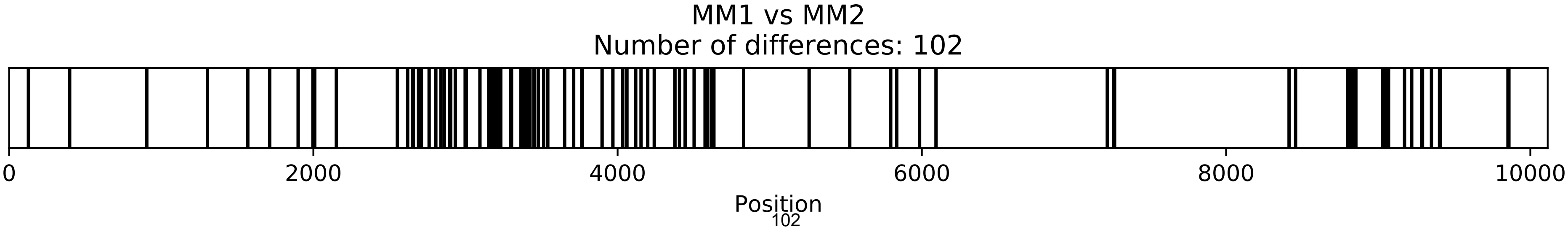
MM1 vs MM2
Number of differences: 102
0
2000
4000
6000
8000
10000
Position

### Region: 356



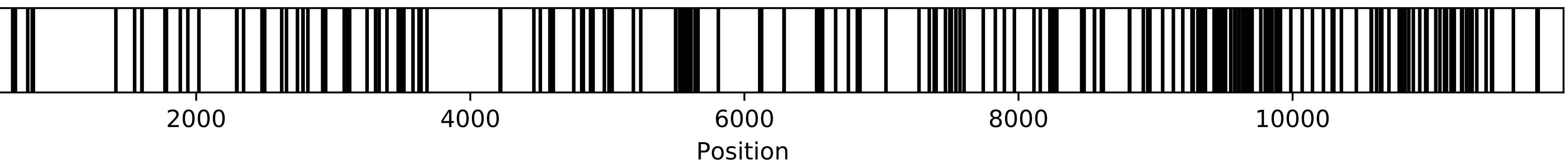
MA1 vs MA2
Number of differences: 229
0
2000
4000
6000
8000
10000
Position

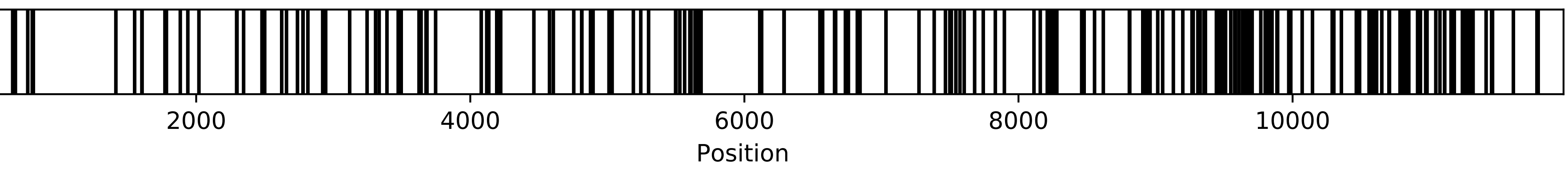
MA1 vs MM1
Number of differences: 230
0
2000
4000
6000
8000
10000
Position

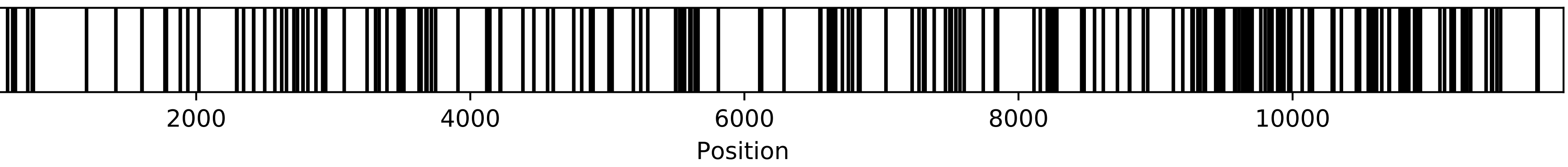
MA1 vs MM2
Number of differences: 245
0
2000
4000
6000
8000
10000
Position

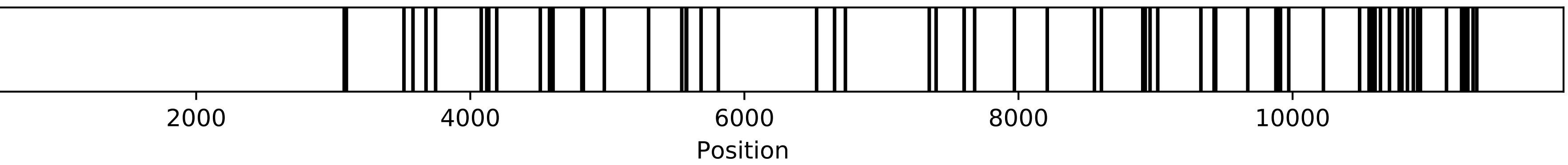
MA2 vs MM1
Number of differences: 79
0
2000
4000
6000
8000
10000
Position

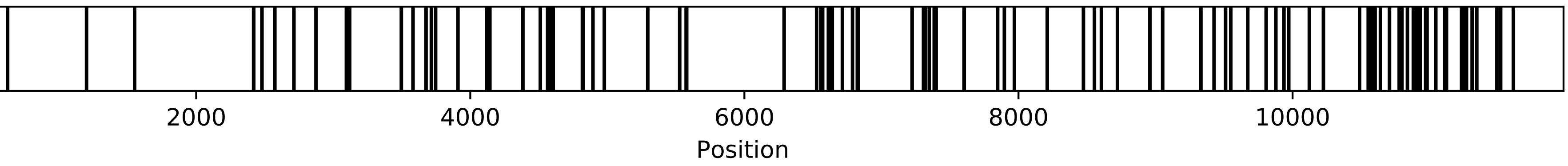
MA2 vs MM2
Number of differences: 123
0
2000
4000
6000
8000
10000
Position

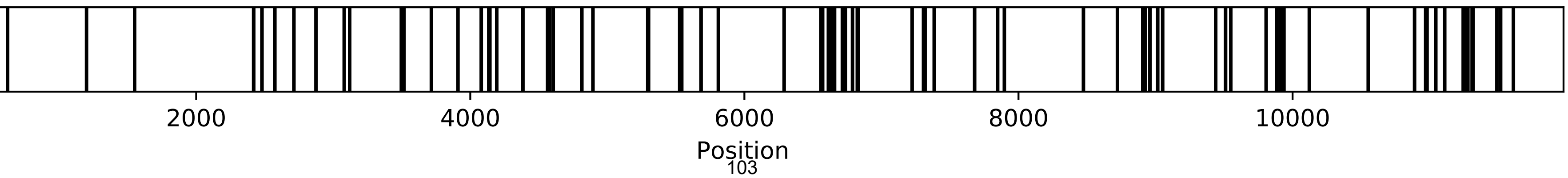
MM1 vs MM2
Number of differences: 90
0
2000
4000
6000
8000
10000
Position

### Region: 357



MA1 vs MA2
Number of differences: 214

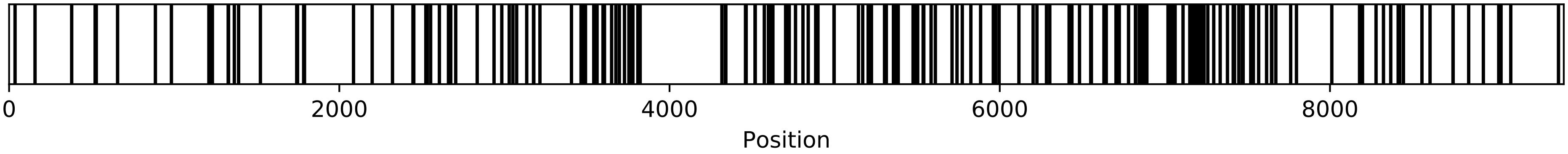


MA1 vs MM1
Number of differences: 120

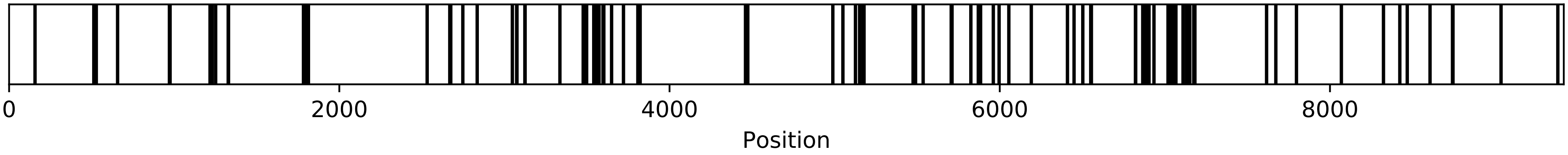


MA1 vs MM2
Number of differences: 120

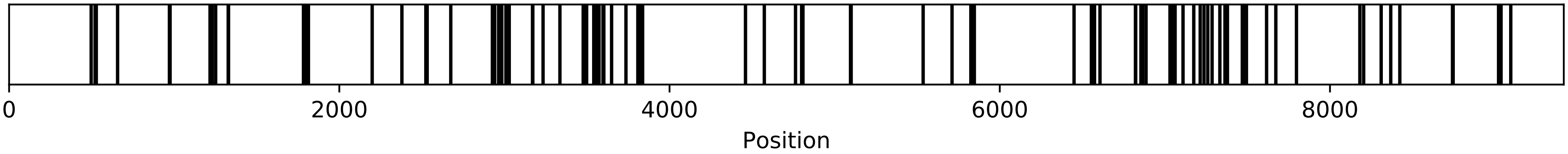


MA2 vs MM1
Number of differences: 216

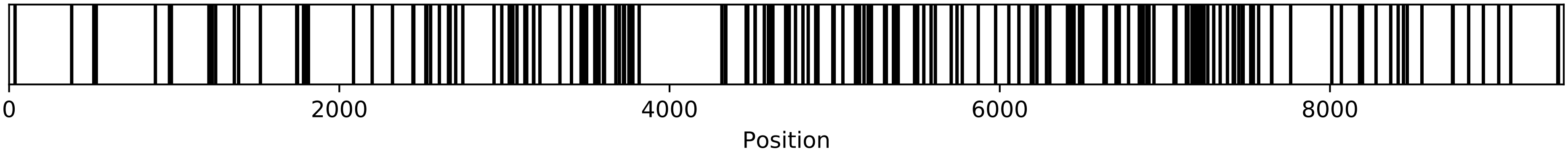


MA2 vs MM2
Number of differences: 217

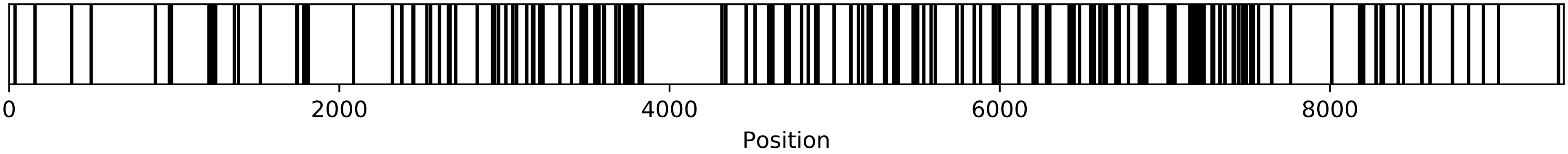


MM1 vs MM2
Number of differences: 99

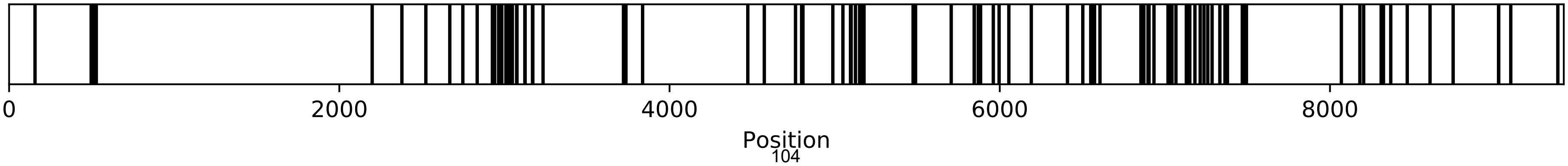

### Region: 363



MA1 vs MA2
Number of differences: 0

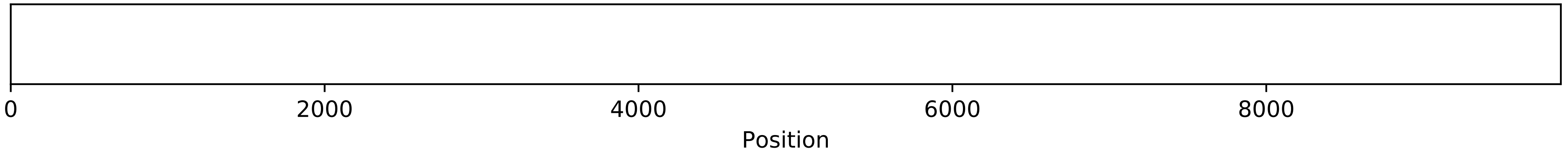


MA1 vs MM1
Number of differences: 208

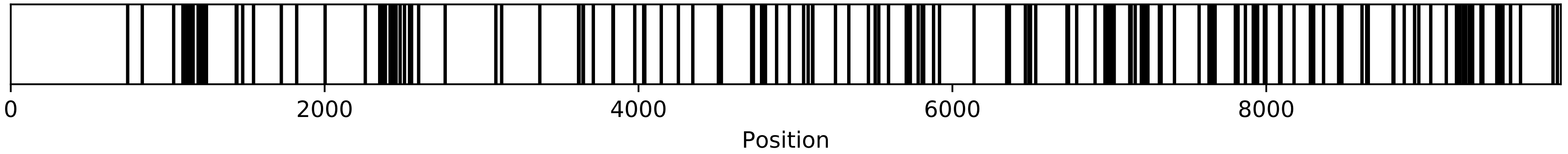


MA1 vs MM2
Number of differences: 180

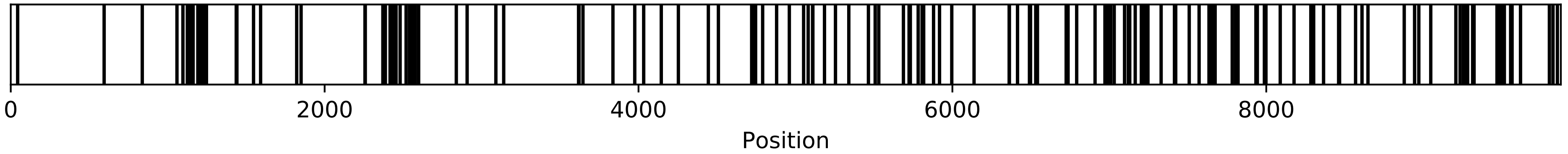


MA2 vs MM1
Number of differences: 208

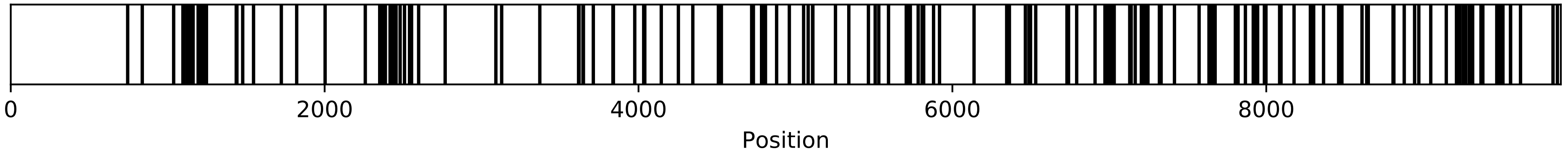


MA2 vs MM2
Number of differences: 180

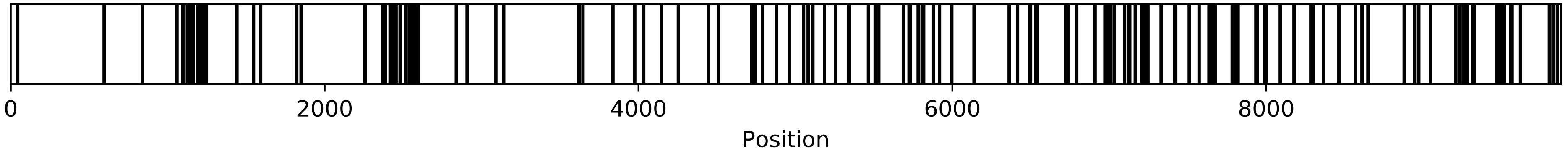


MM1 vs MM2
Number of differences: 126

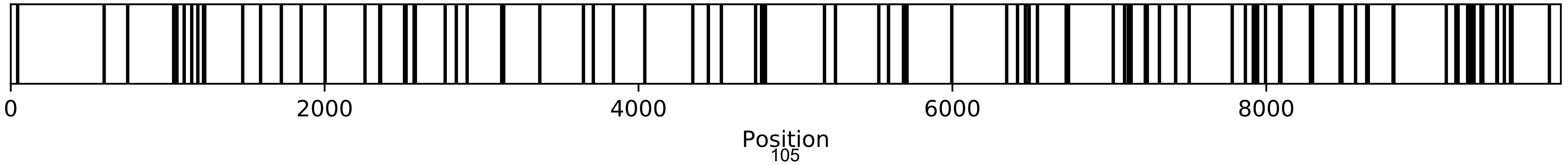

### Region: 365



MA1 vs MA2
Number of differences: 281

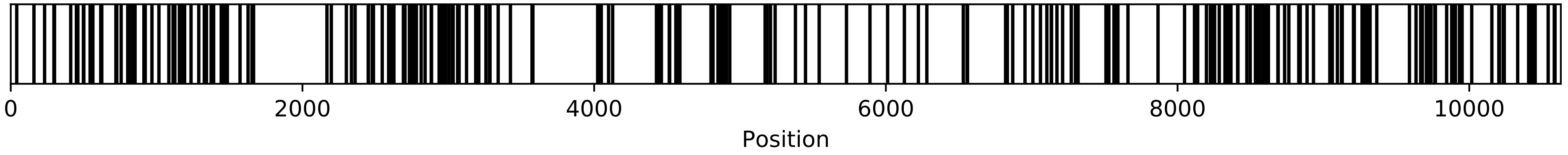


MA1 vs MM1
Number of differences: 113

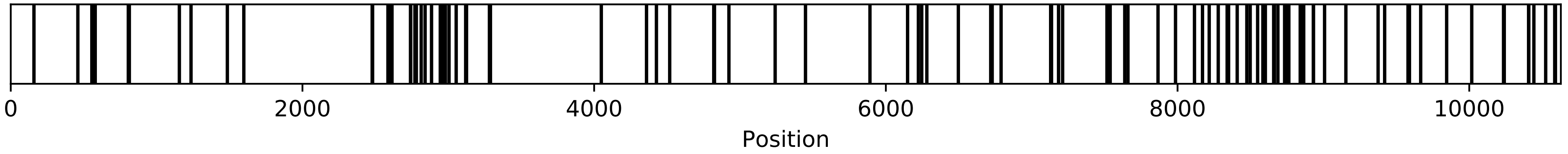


MA1 vs MM2
Number of differences: 0

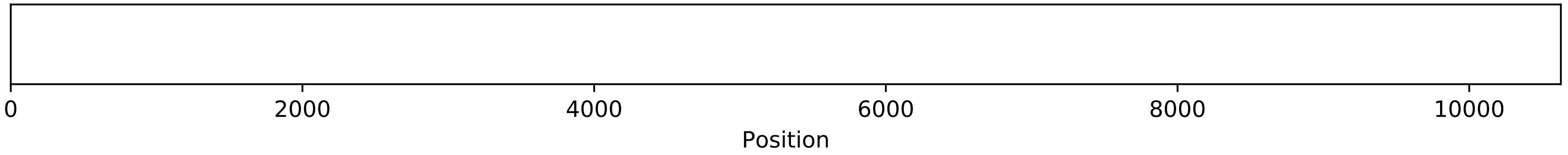


MA2 vs MM1
Number of differences: 269

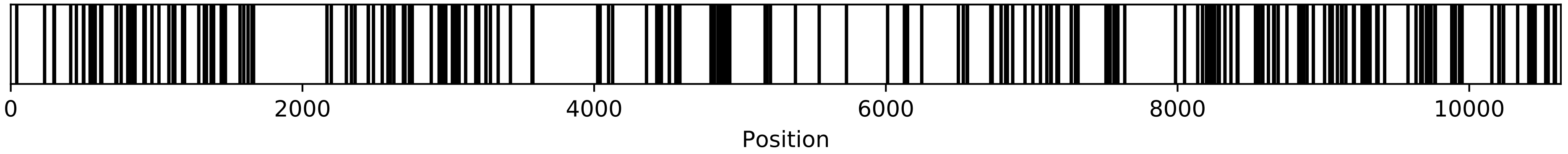


MA2 vs MM2
Number of differences: 281

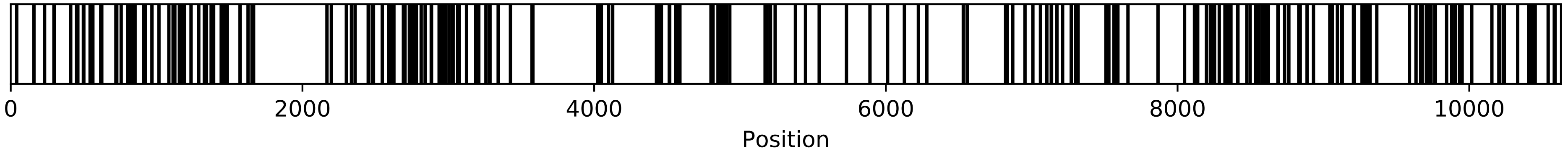


MM1 vs MM2
Number of differences: 113

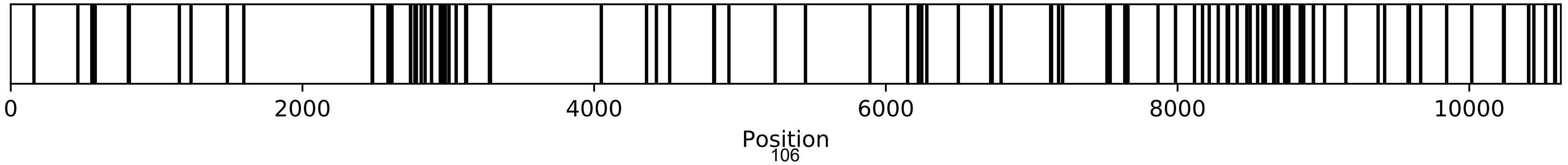

### Region: 367



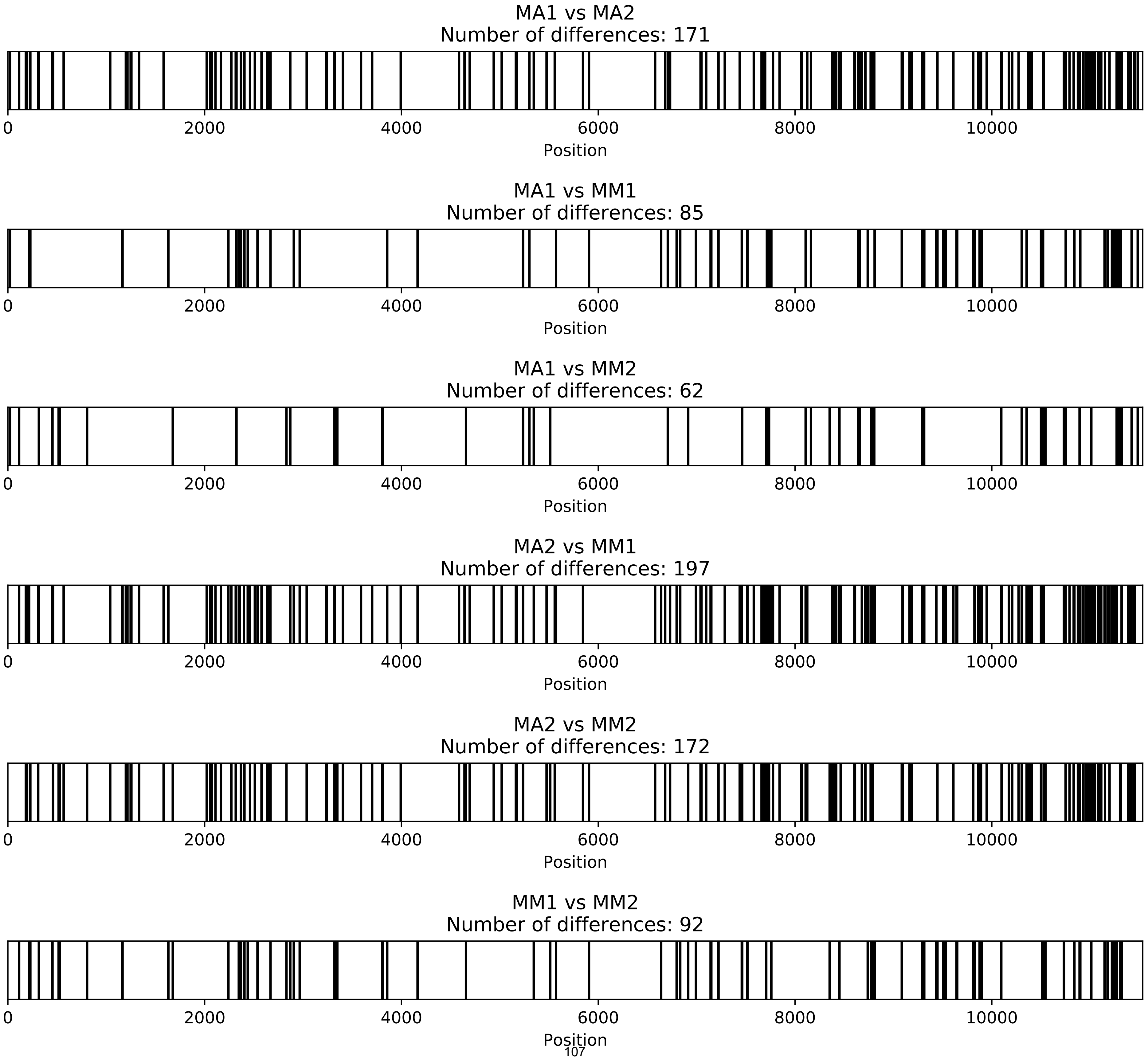
MA1 vs MA2
Number of differences: 171
MA1 vs MM1
Number of differences: 85
MA1 vs MM2
Number of differences: 62
MA2 vs MM1
Number of differences: 197
MA2 vs MM2
Number of differences: 172
MM1 vs MM2
Number of differences: 92
0
2000
4000
6000
8000
10000
Position

### Region: 368



MA1 vs MA2
Number of differences: 318

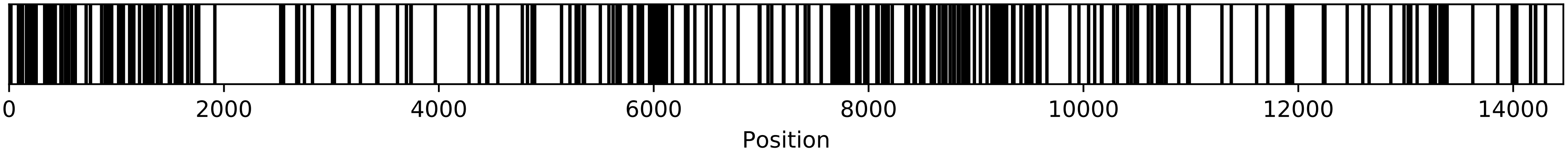


MA1 vs MM1
Number of differences: 130

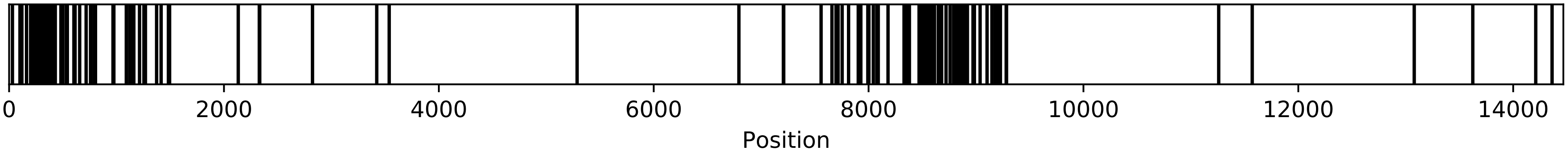


MA1 vs MM2
Number of differences: 0

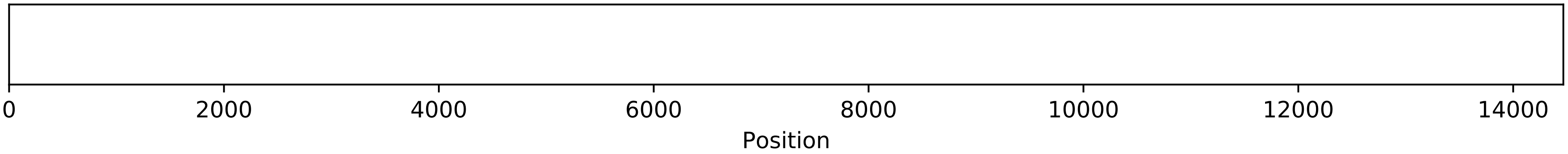


MA2 vs MM1
Number of differences: 323

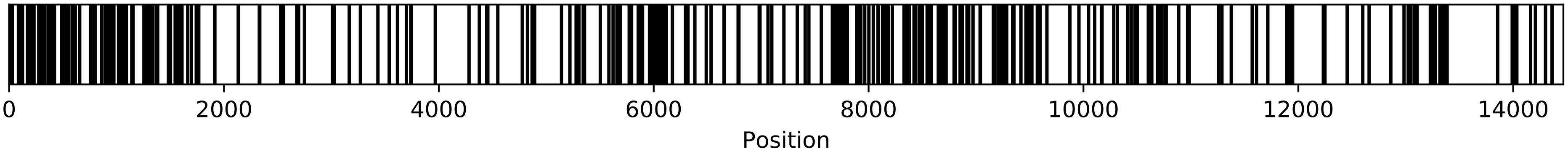


MA2 vs MM2
Number of differences: 318

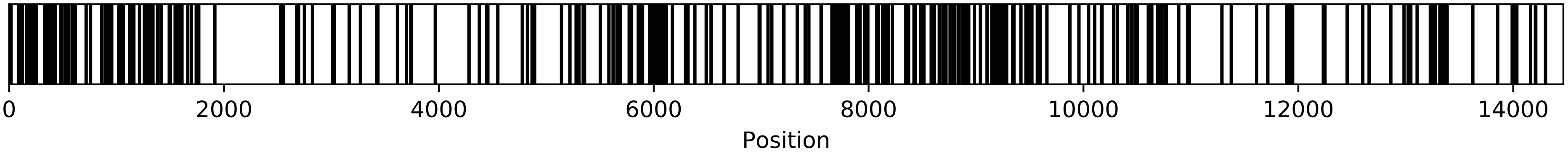


MM1 vs MM2
Number of differences: 130

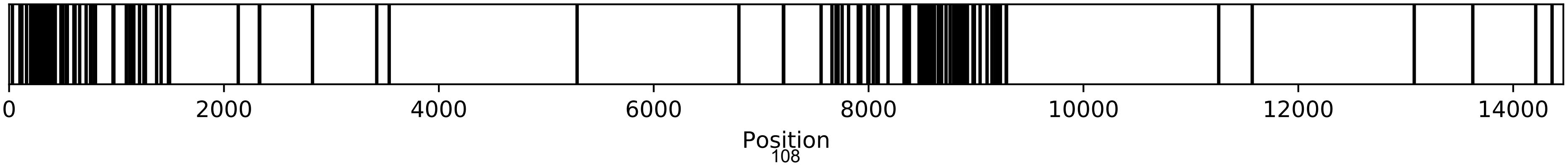

### Region: 372



MA1 vs MA2
Number of differences: 0

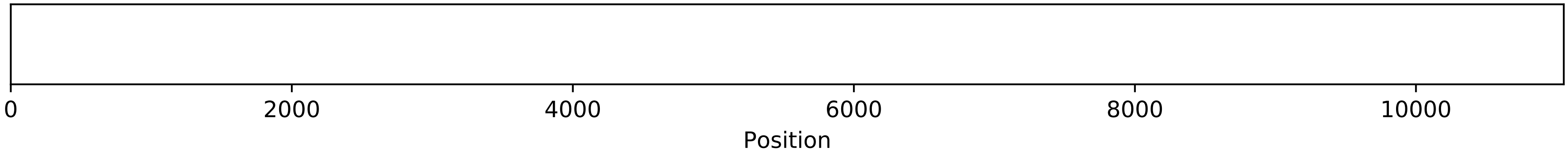


MA1 vs MM1
Number of differences: 129

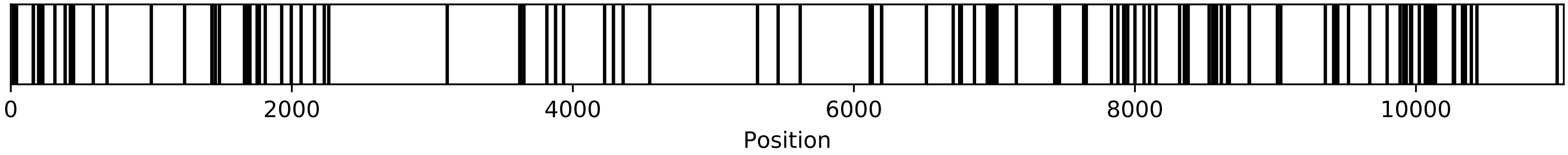


MA1 vs MM2
Number of differences: 0

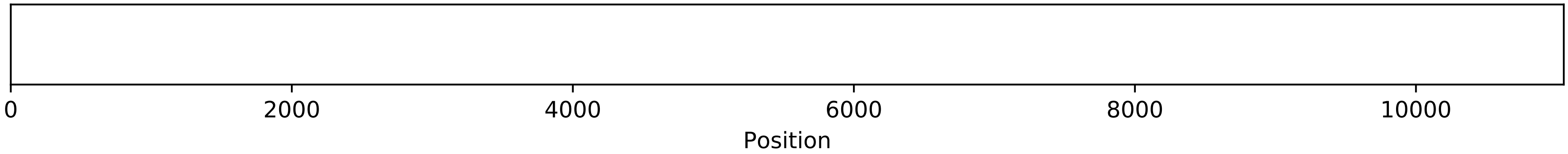


MA2 vs MM1
Number of differences: 129

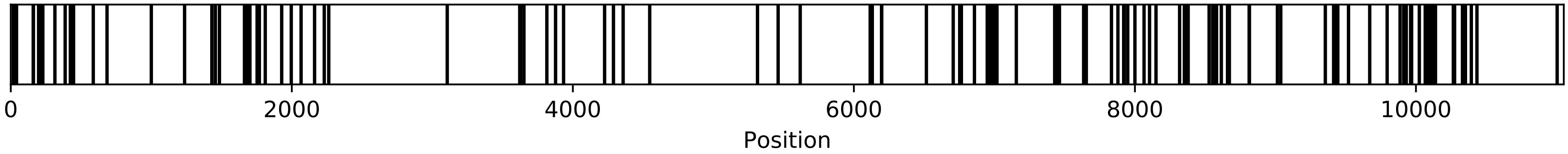


MA2 vs MM2
Number of differences: 0

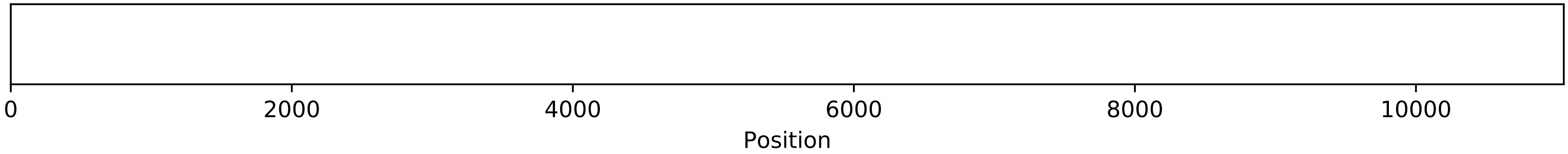


MM1 vs MM2
Number of differences: 129

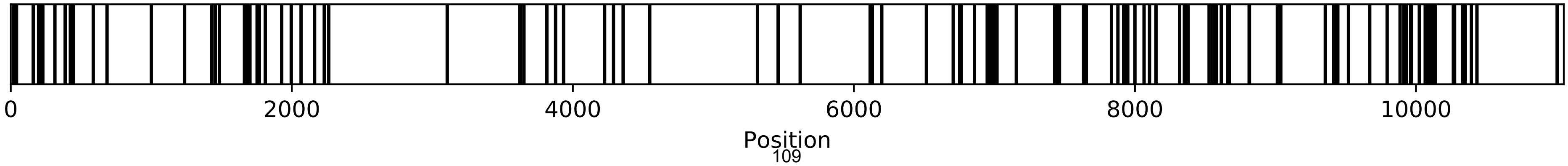

### Region: 373



MA1 vs MA2
Number of differences: 192

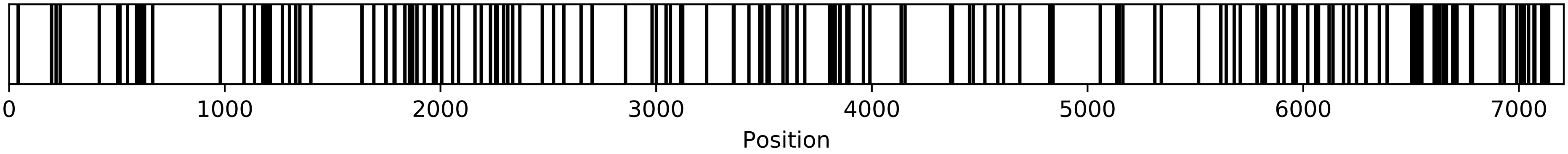


MA1 vs MM1
Number of differences: 67

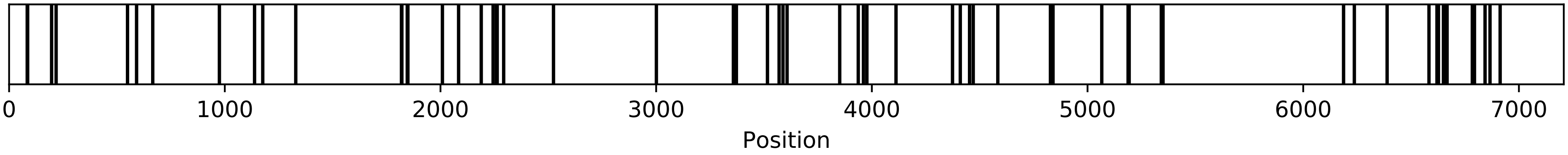


MA1 vs MM2
Number of differences: 115

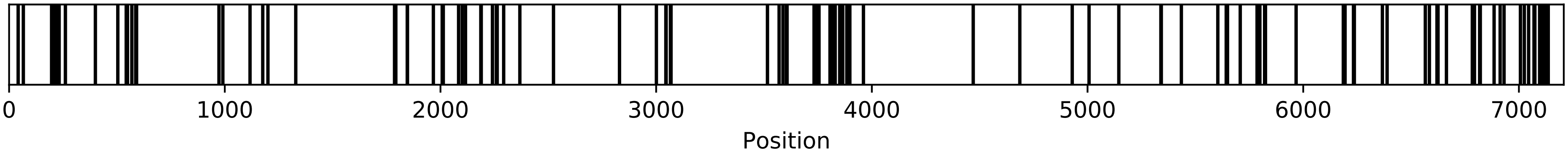


MA2 vs MM1
Number of differences: 186

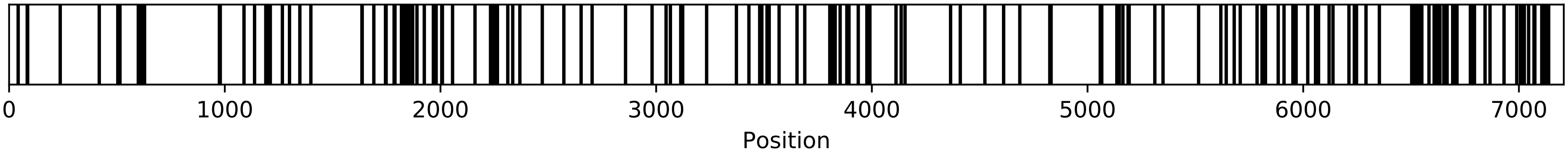


MA2 vs MM2
Number of differences: 201

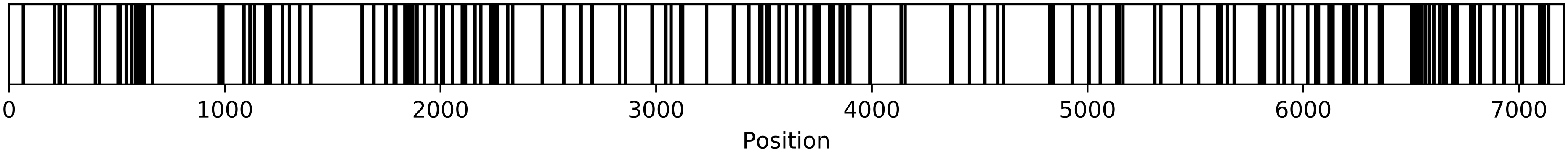


MM1 vs MM2
Number of differences: 118

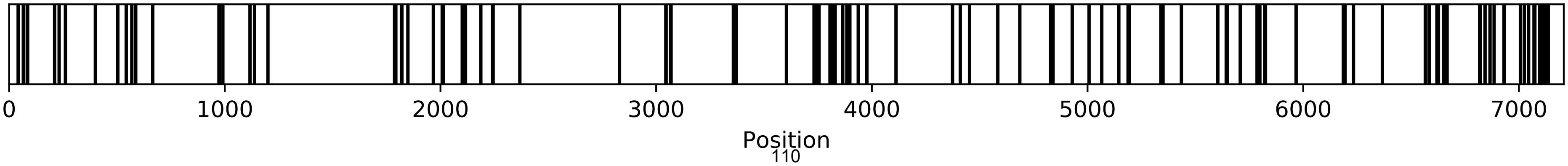

### Region: 374



MA1 vs MA2
Number of differences: 343

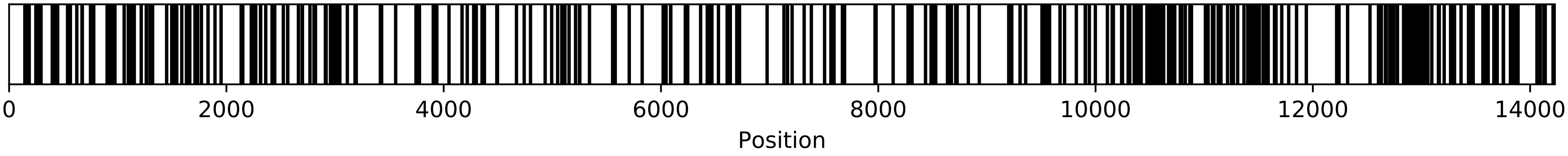


MA1 vs MM1
Number of differences: 167

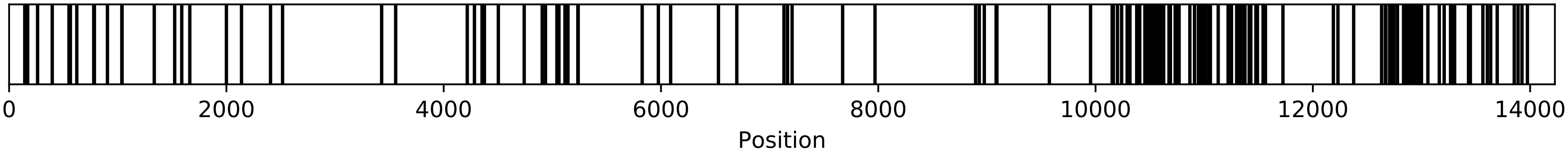


MA1 vs MM2
Number of differences: 146

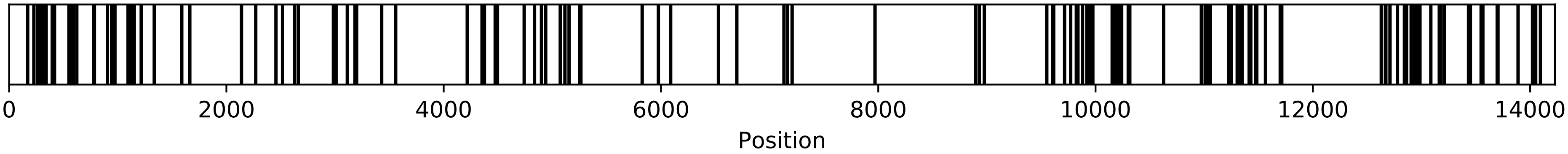


MA2 vs MM1
Number of differences: 344

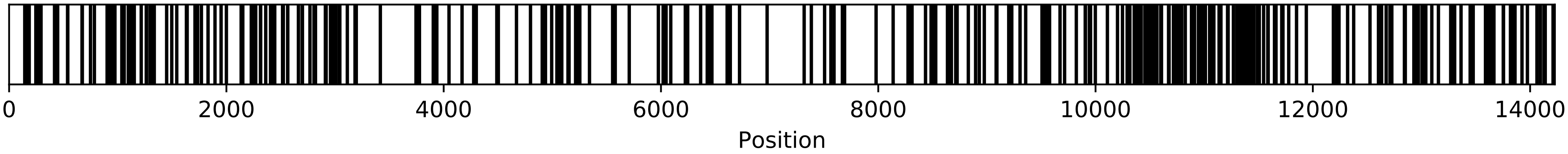


MA2 vs MM2
Number of differences: 322

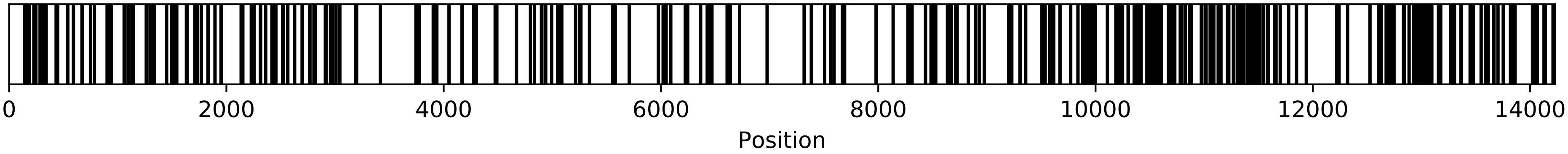


MM1 vs MM2
Number of differences: 182

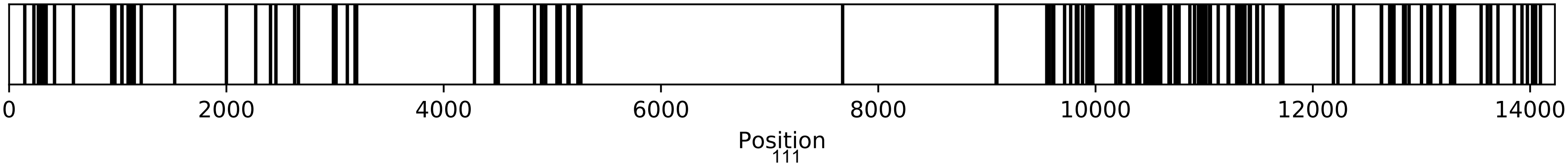

### Region: 379



MA1 vs MA2
Number of differences: 393

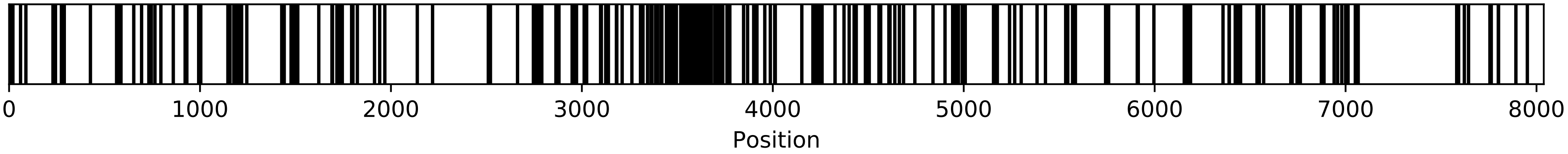


MA1 vs MM1
Number of differences: 387

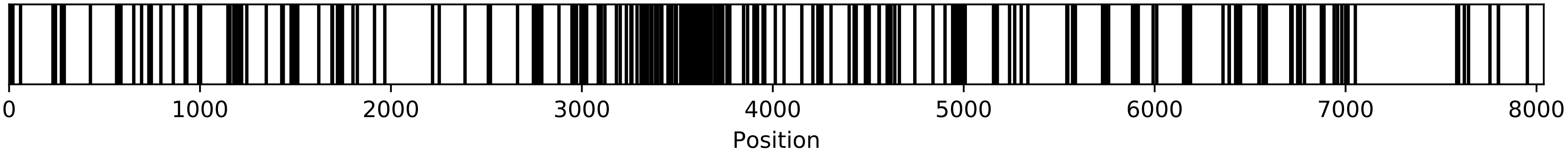


MA1 vs MM2
Number of differences: 393

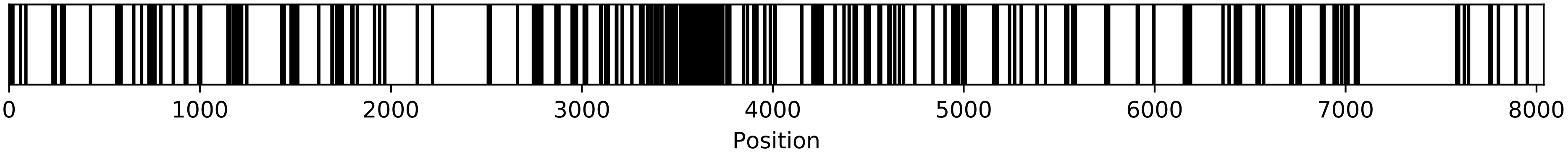


MA2 vs MM1
Number of differences: 110

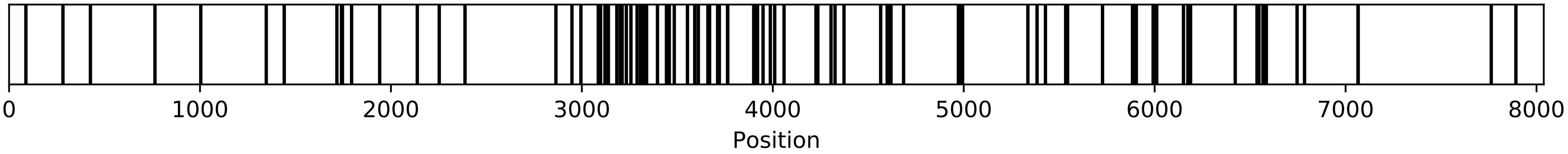


MA2 vs MM2
Number of differences: 0

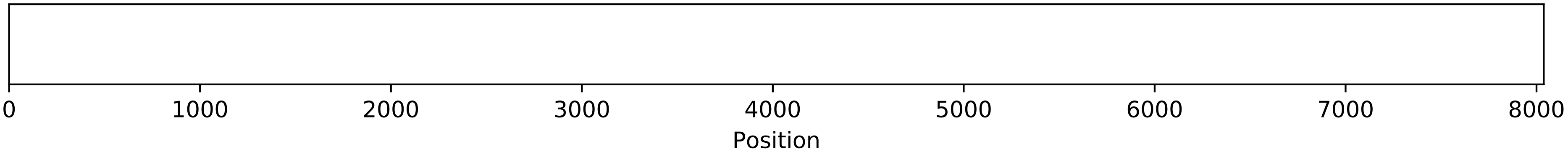


MM1 vs MM2
Number of differences: 110

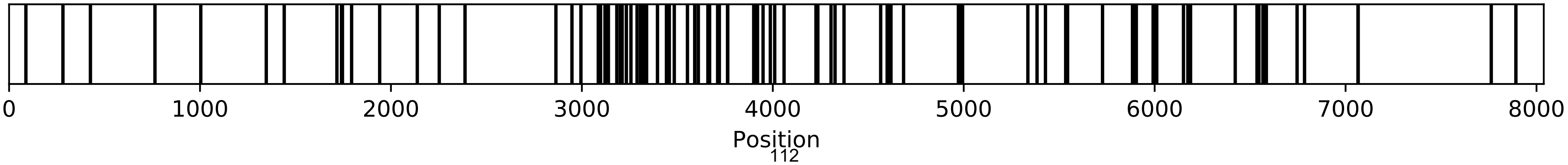

### Region: 382



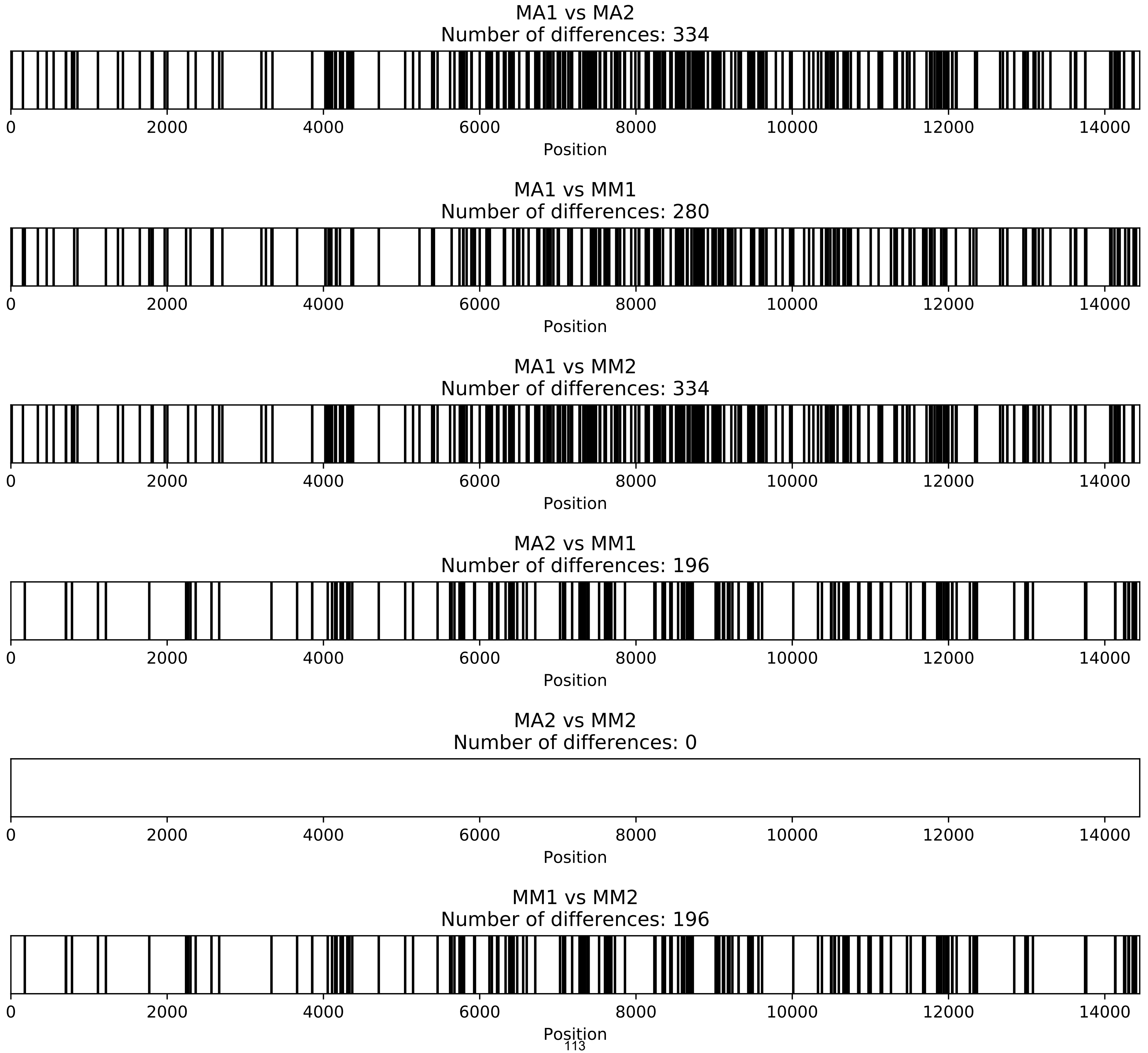
MA1 vs MA2
Number of differences: 334
MA1 vs MM1
Number of differences: 280
MA1 vs MM2
Number of differences: 334
MA2 vs MM1
Number of differences: 196
MA2 vs MM2
Number of differences: 0
MM1 vs MM2
Number of differences: 196
0
2000
4000
6000
8000
10000
12000
14000
Position

### Region: 383



MA1 vs MA2
Number of differences: 249

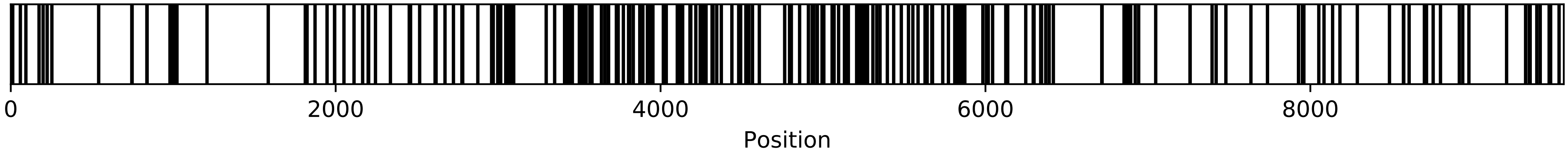


MA1 vs MM1
Number of differences: 95

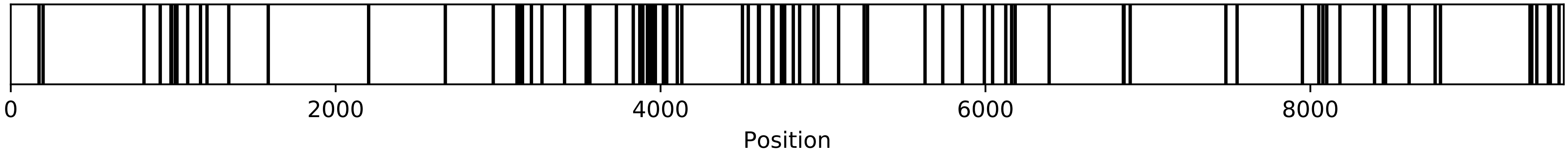


MA1 vs MM2
Number of differences: 0

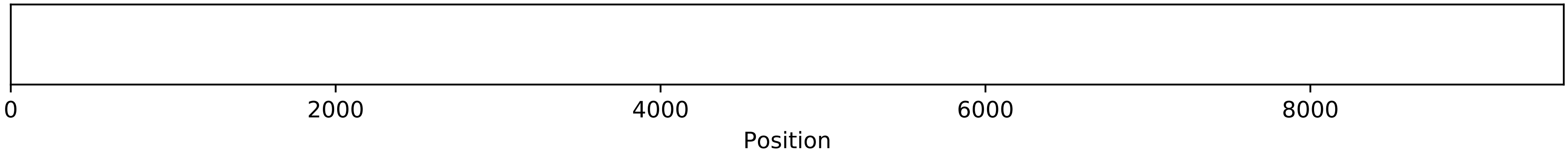


MA2 vs MM1
Number of differences: 249

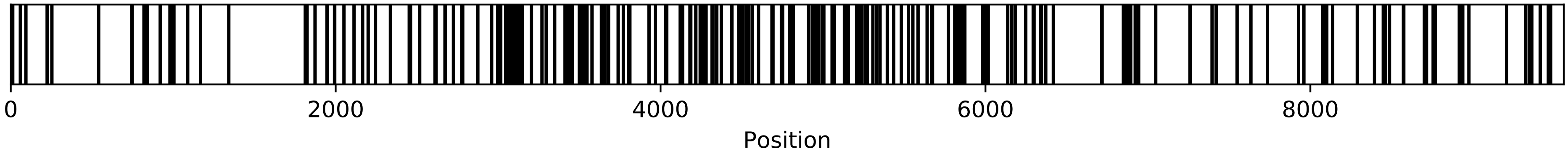


MA2 vs MM2
Number of differences: 249

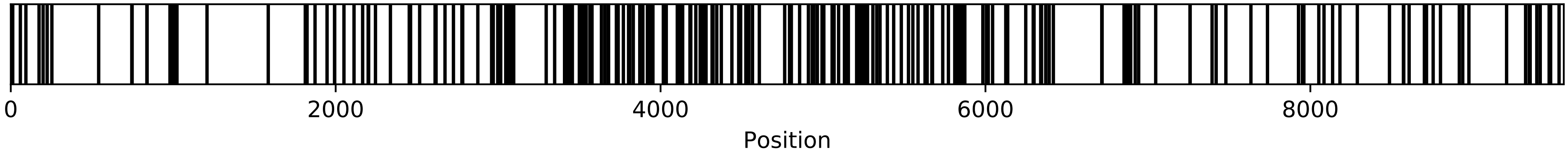


MM1 vs MM2
Number of differences: 95

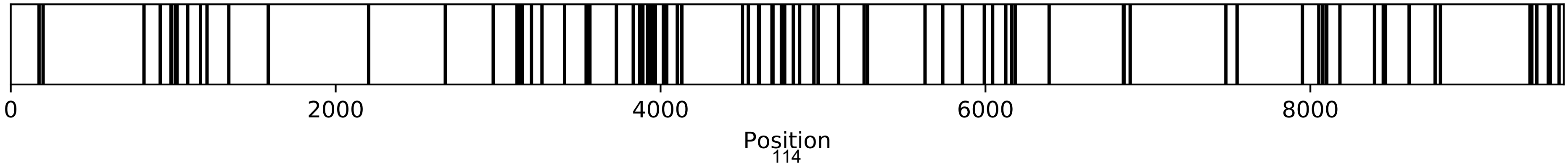

### Region: 386



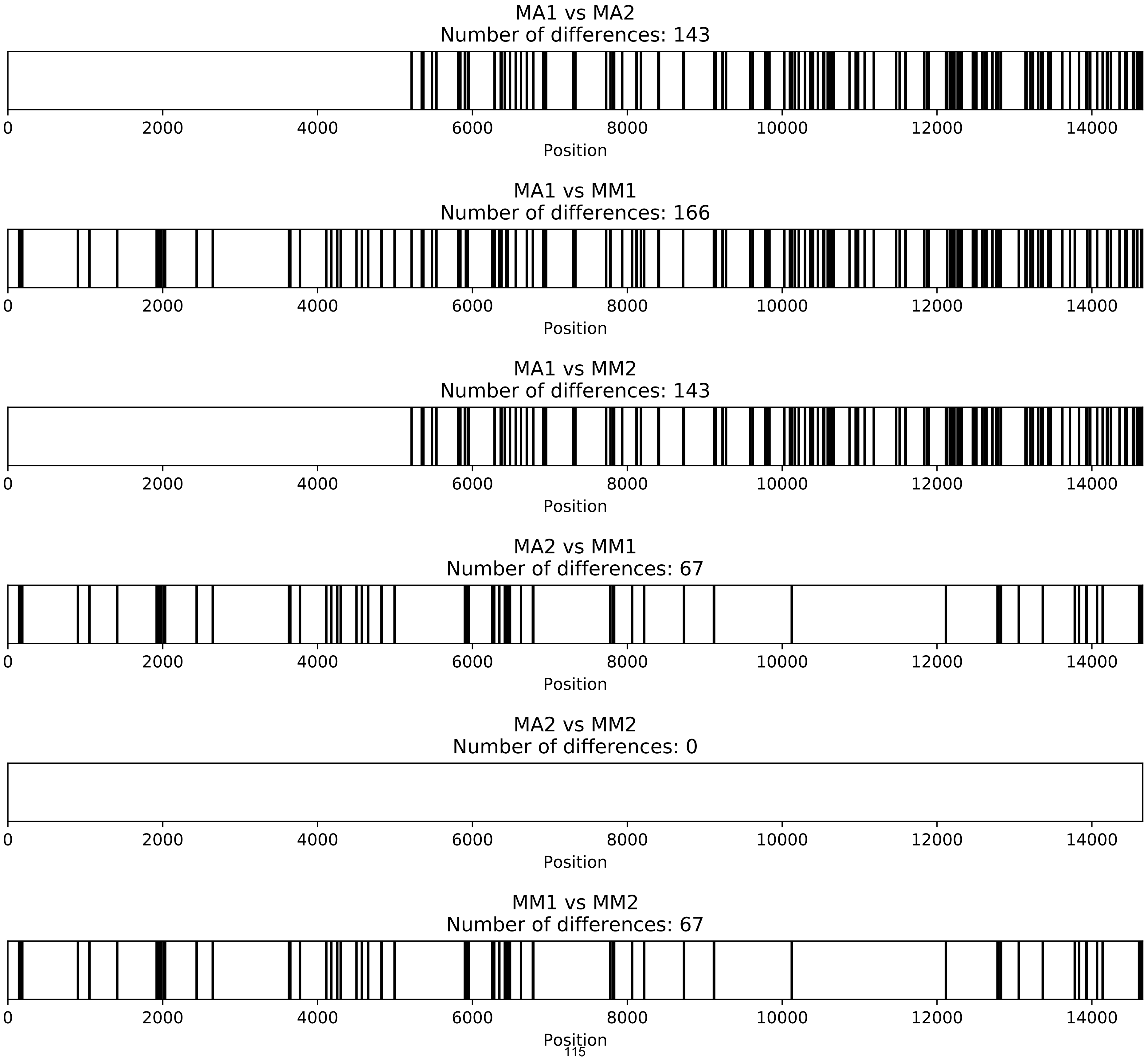
MA1 vs MA2
Number of differences: 143
0
2000
4000
6000
8000
10000
12000
14000
Position
MA1 vs MM1
Number of differences: 166
Position
MA1 vs MM2
Number of differences: 143
Position
MA2 vs MM1
Number of differences: 67
Position
MA2 vs MM2
Number of differences: 0
Position
MM1 vs MM2
Number of differences: 67
Position

### Region: 388



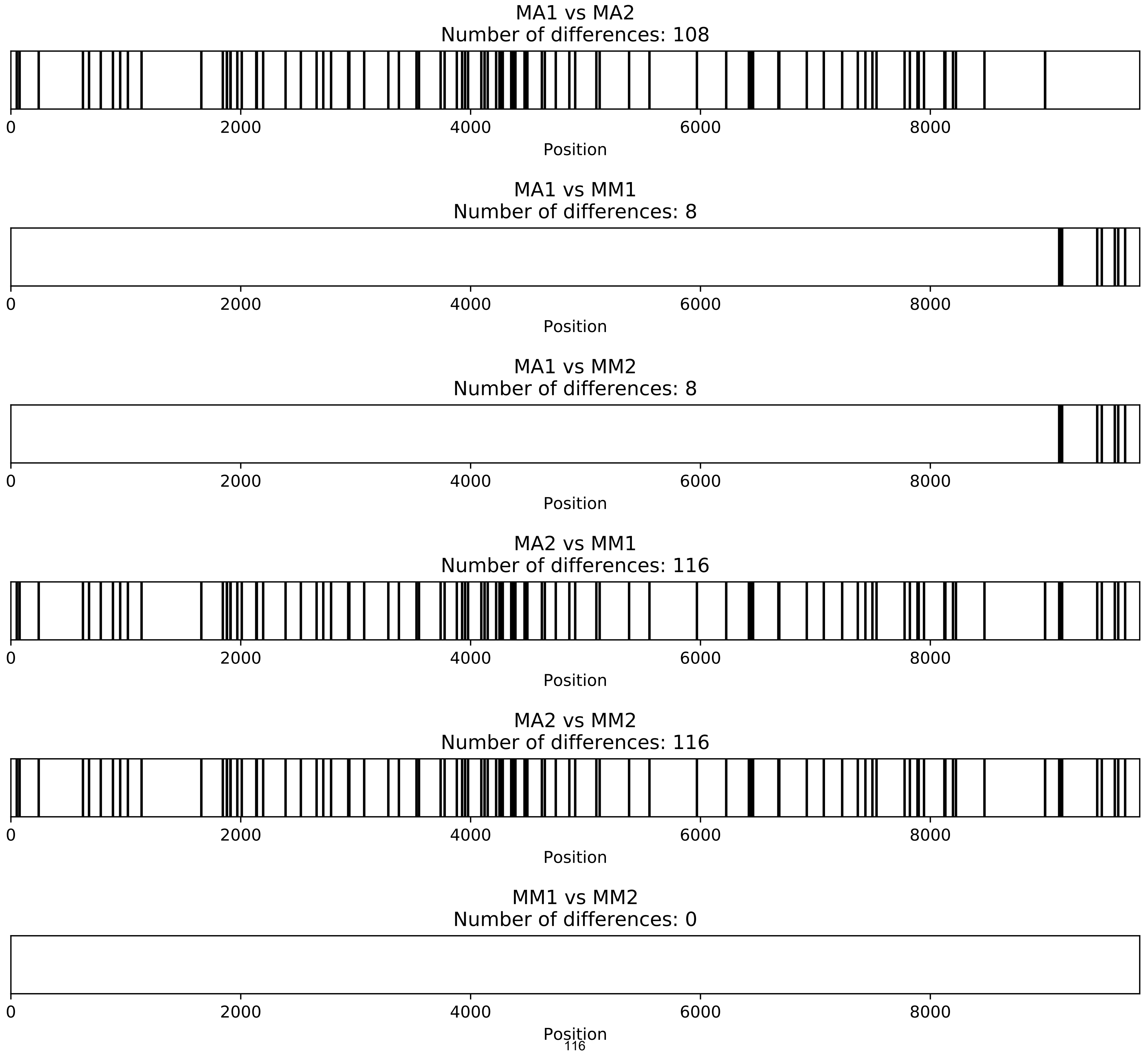
MA1 vs MA2
Number of differences: 108
0
2000
4000
6000
8000
Position
MA1 vs MM1
Number of differences: 8
0
2000
4000
6000
8000
Position
MA1 vs MM2
Number of differences: 8
0
2000
4000
6000
8000
Position
MA2 vs MM1
Number of differences: 116
0
2000
4000
6000
8000
Position
MA2 vs MM2
Number of differences: 116
0
2000
4000
6000
8000
Position
MM1 vs MM2
Number of differences: 0
0
2000
4000
6000
8000
Position

### Region: 389



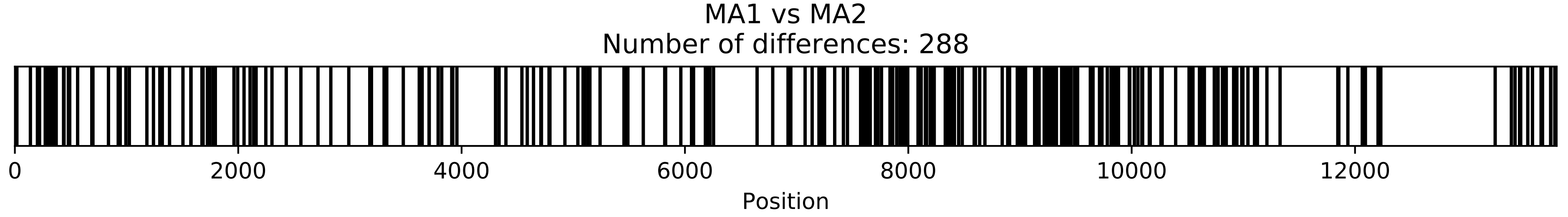
MA1 vs MA2
Number of differences: 288
0
2000
4000
6000
8000
10000
12000
Position

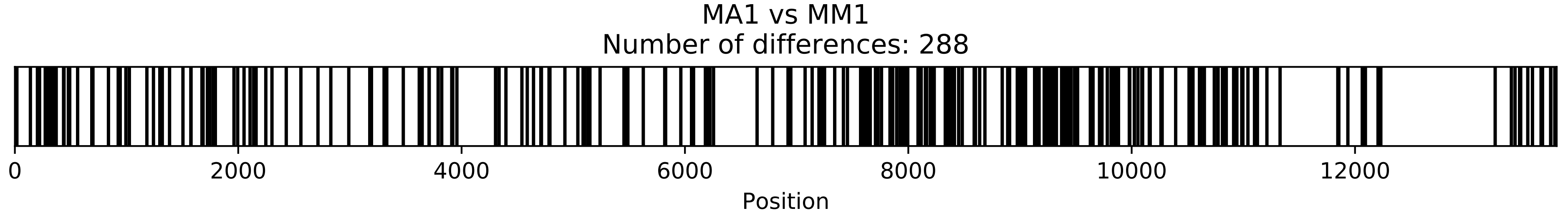
MA1 vs MM1
Number of differences: 288
0
2000
4000
6000
8000
10000
12000
Position

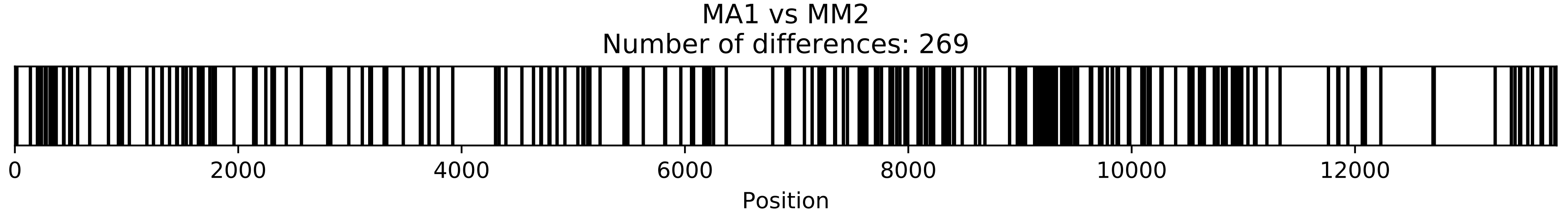
MA1 vs MM2
Number of differences: 269
0
2000
4000
6000
8000
10000
12000
Position

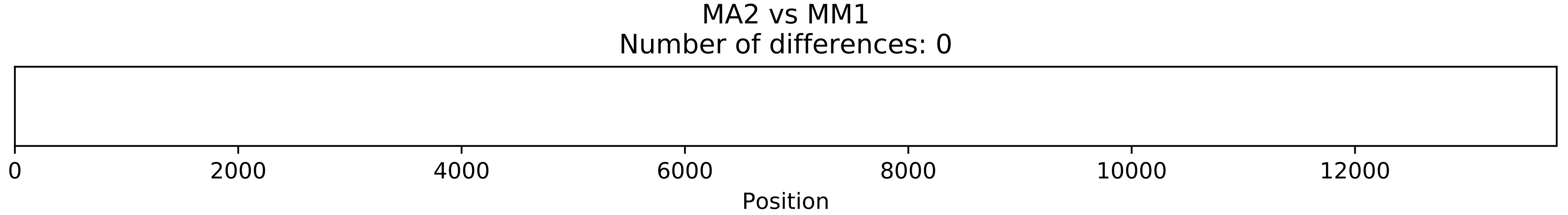
MA2 vs MM1
Number of differences: 0
0
2000
4000
6000
8000
10000
12000
Position

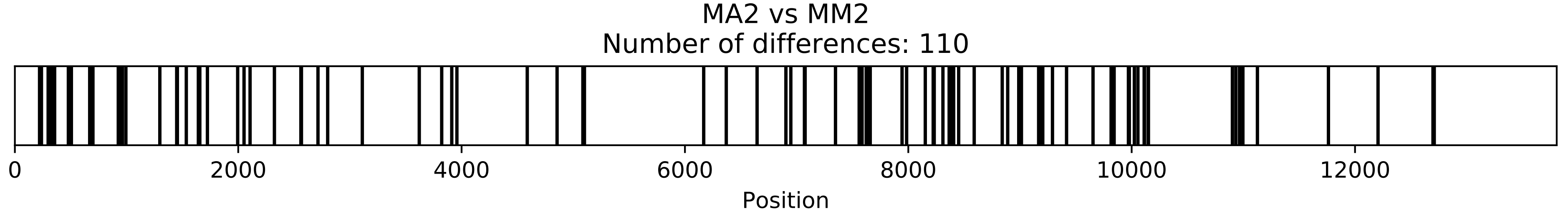
MA2 vs MM2
Number of differences: 110
0
2000
4000
6000
8000
10000
12000
Position

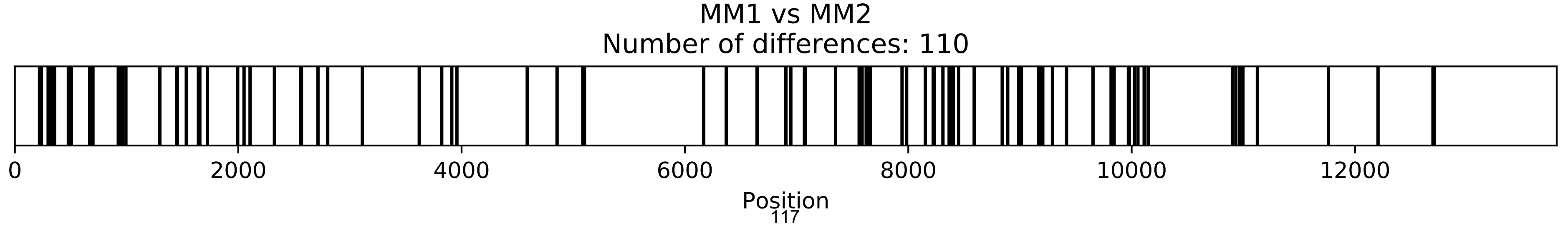
MM1 vs MM2
Number of differences: 110
0
2000
4000
6000
8000
10000
12000
Position

### Region: 392



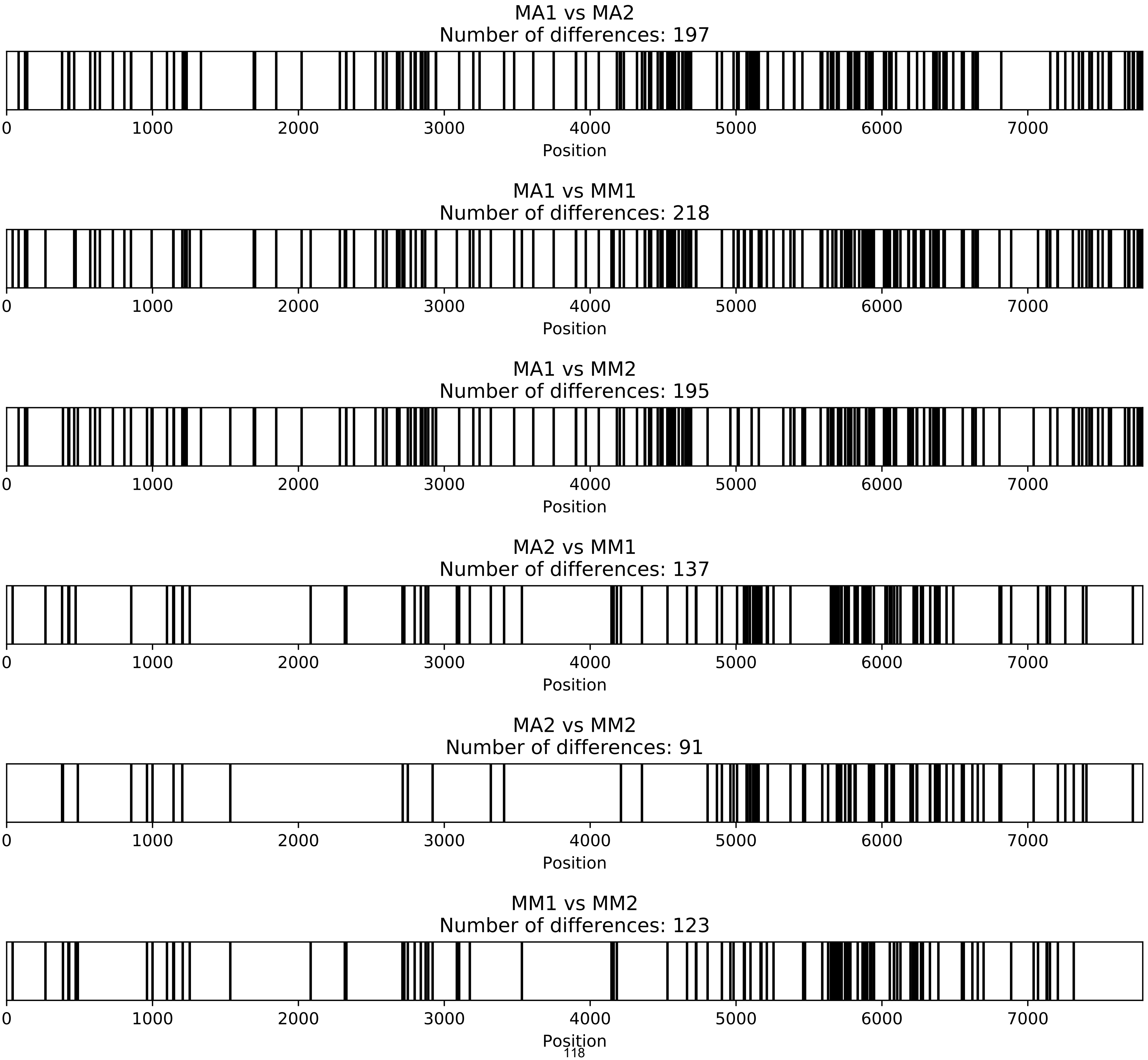
MA1 vs MA2
Number of differences: 197
MA1 vs MM1
Number of differences: 218
MA1 vs MM2
Number of differences: 195
MA2 vs MM1
Number of differences: 137
MA2 vs MM2
Number of differences: 91
MM1 vs MM2
Number of differences: 123
Position
0
1000
2000
3000
4000
5000
6000
7000

### Region: 393



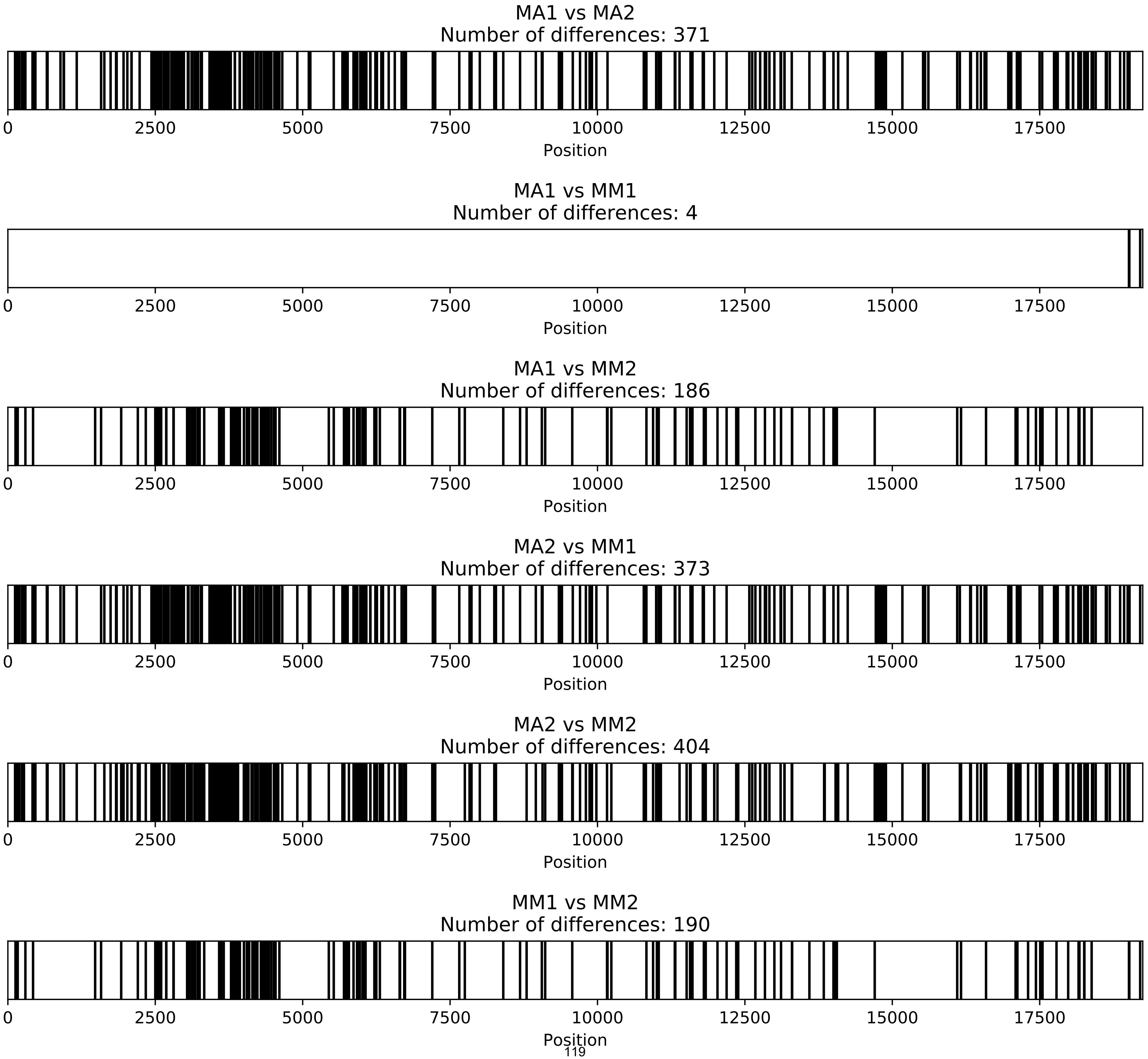
MA1 vs MA2
Number of differences: 371
MA1 vs MM1
Number of differences: 4
MA1 vs MM2
Number of differences: 186
MA2 vs MM1
Number of differences: 373
MA2 vs MM2
Number of differences: 404
MM1 vs MM2
Number of differences: 190
0
2500
5000
7500
10000
12500
15000
17500
Position

### Region: 396



MA1 vs MA2
Number of differences: 3

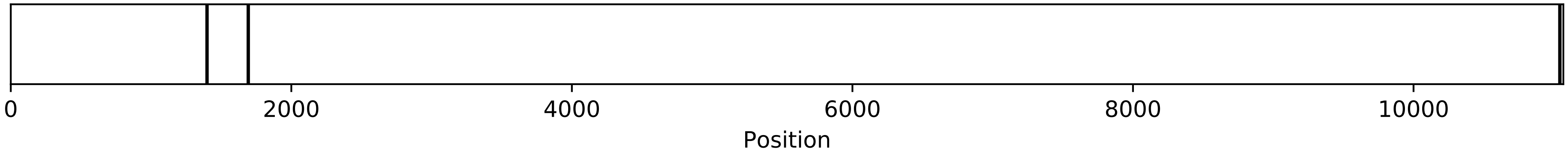


MA1 vs MM1
Number of differences: 259

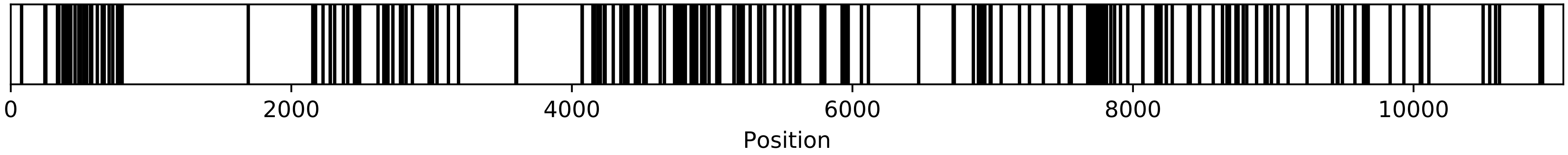


MA1 vs MM2
Number of differences: 241

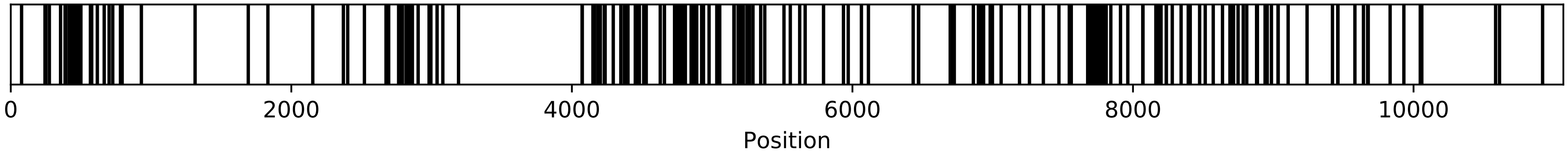


MA2 vs MM1
Number of differences: 260

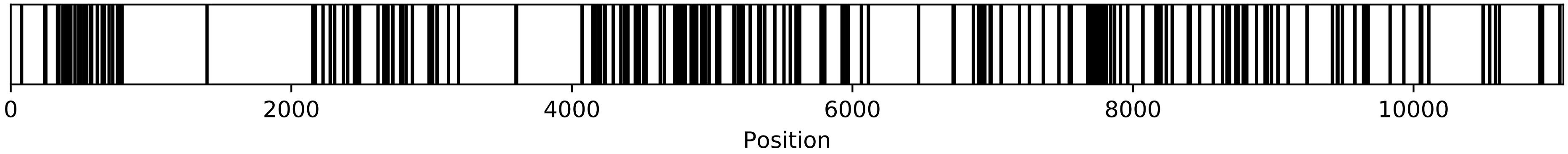


MA2 vs MM2
Number of differences: 242

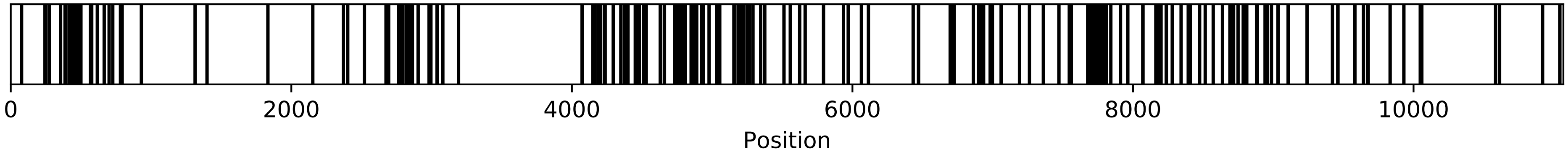


MM1 vs MM2
Number of differences: 116

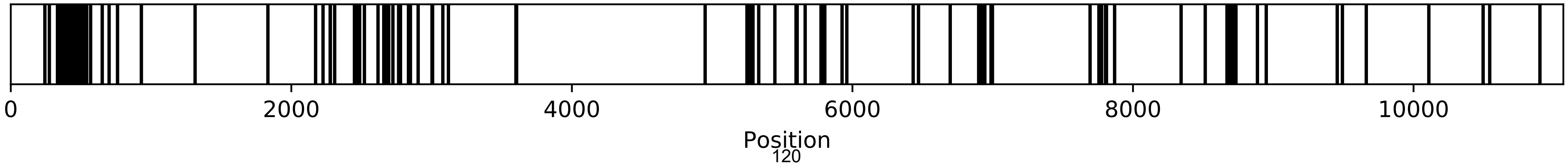

### Region: 398



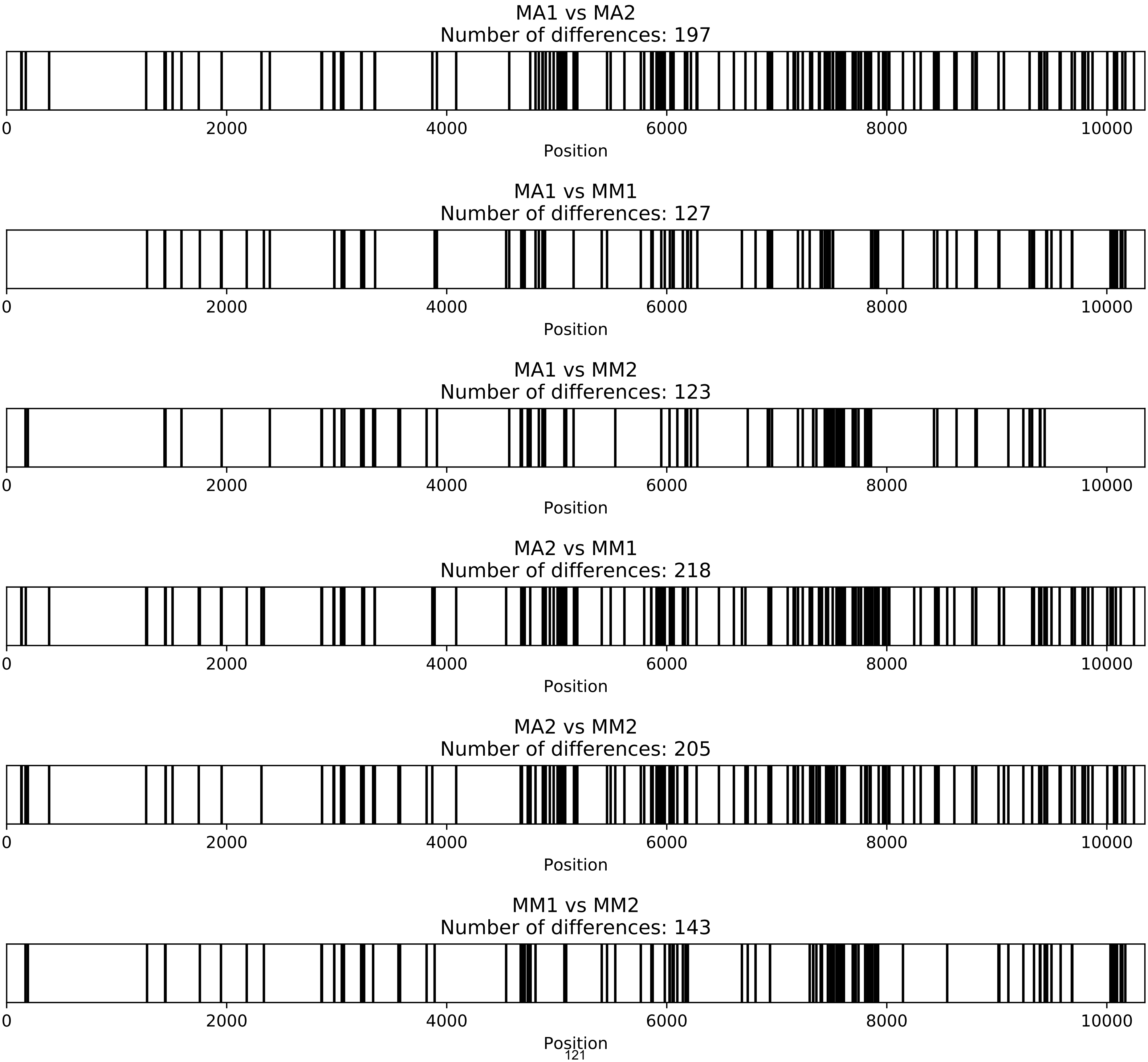
MA1 vs MA2
Number of differences: 197
MA1 vs MM1
Number of differences: 127
MA1 vs MM2
Number of differences: 123
MA2 vs MM1
Number of differences: 218
MA2 vs MM2
Number of differences: 205
MM1 vs MM2
Number of differences: 143
0
2000
4000
6000
8000
10000
Position

### Region: 404



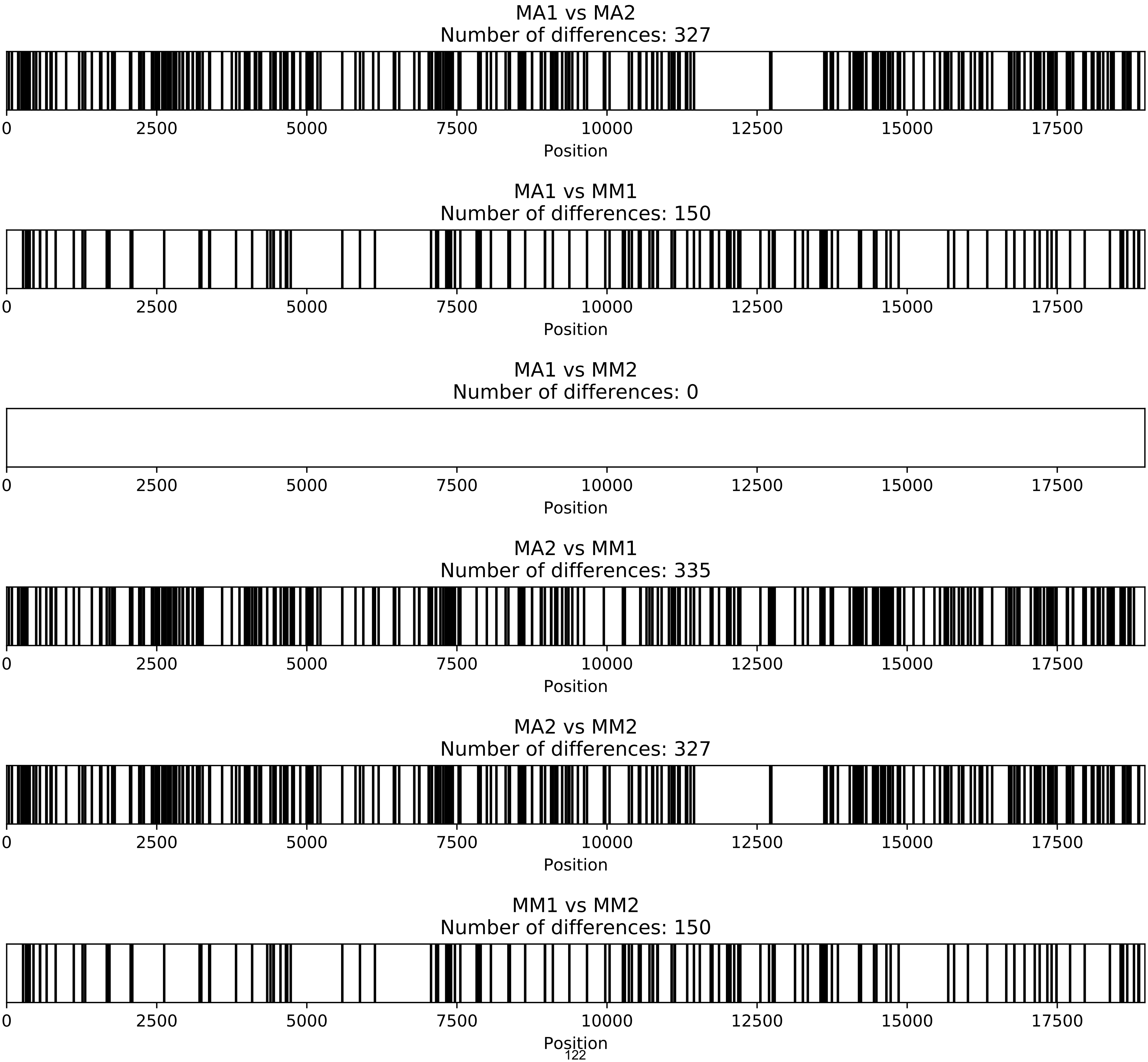
MA1 vs MA2
Number of differences: 327
MA1 vs MM1
Number of differences: 150
MA1 vs MM2
Number of differences: 0
MA2 vs MM1
Number of differences: 335
MA2 vs MM2
Number of differences: 327
MM1 vs MM2
Number of differences: 150
0
2500
5000
7500
10000
12500
15000
17500
Position

### Region: 408



MA1 vs MA2
Number of differences: 349

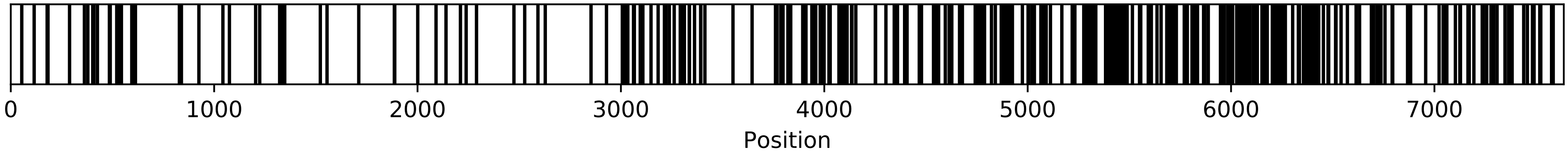


MA1 vs MM1
Number of differences: 303

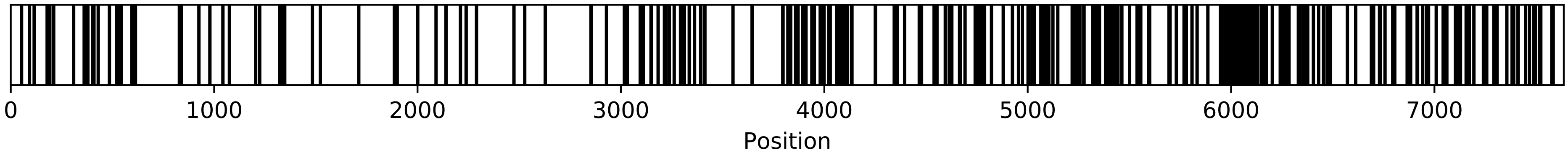


MA1 vs MM2
Number of differences: 303

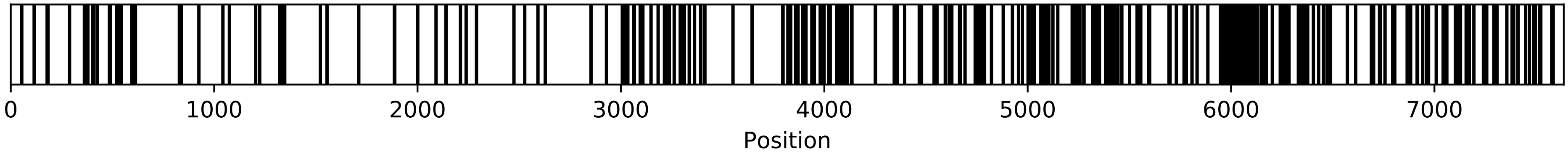


MA2 vs MM1
Number of differences: 229

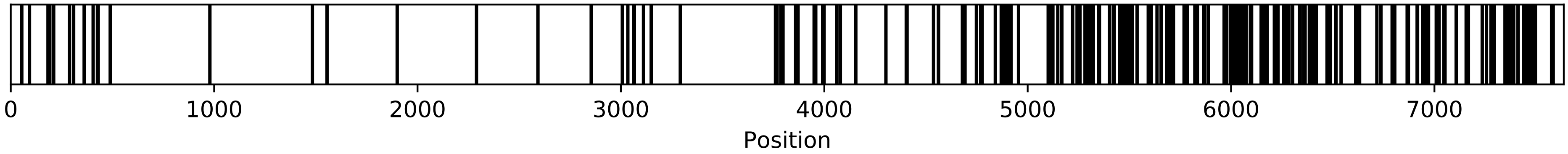


MA2 vs MM2
Number of differences: 200

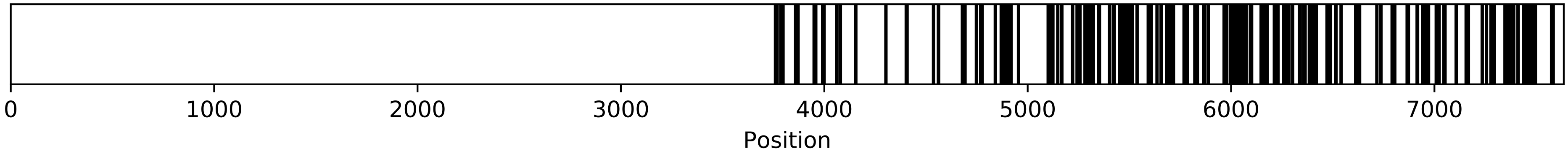


MM1 vs MM2
Number of differences: 29

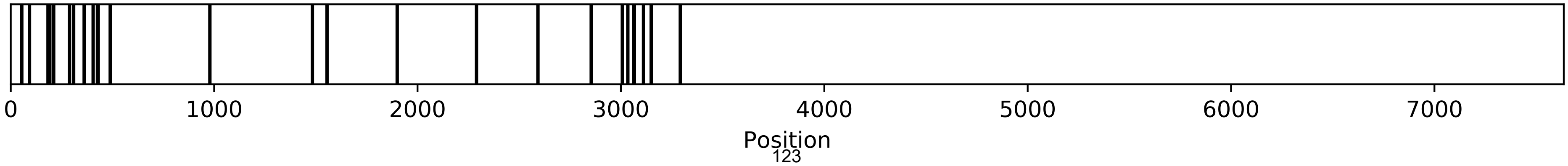

### Region: 413



MA1 vs MA2
Number of differences: 105

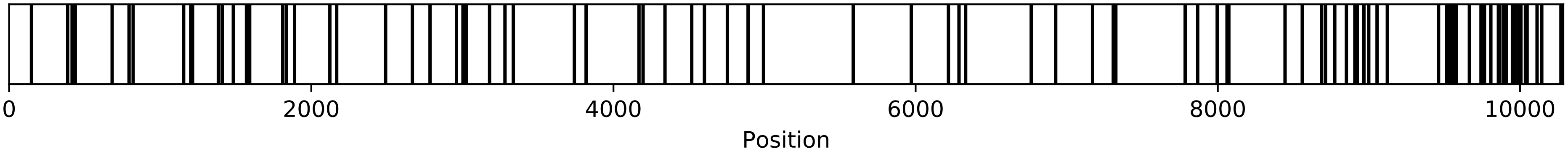


MA1 vs MM1
Number of differences: 113

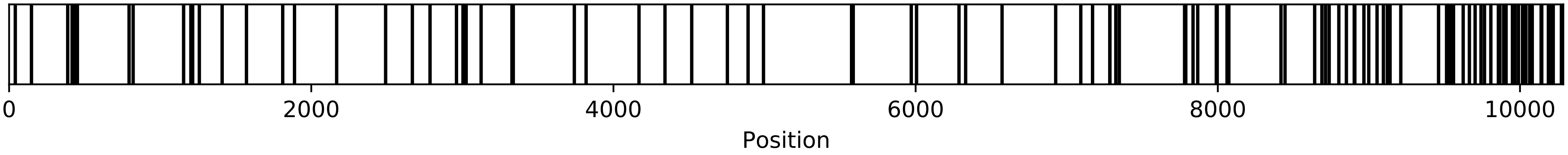


MA1 vs MM2
Number of differences: 105

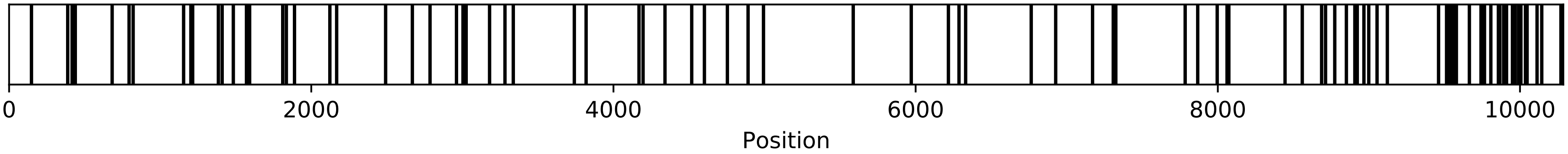


MA2 vs MM1
Number of differences: 76

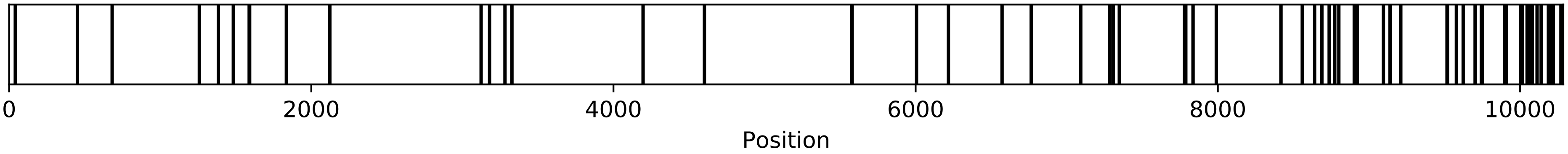


MA2 vs MM2
Number of differences: 0

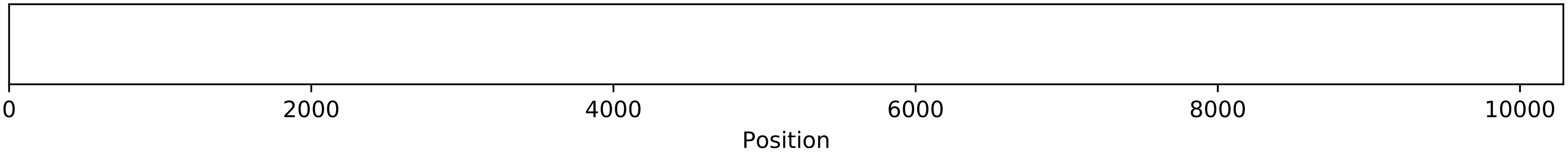


MM1 vs MM2
Number of differences: 76

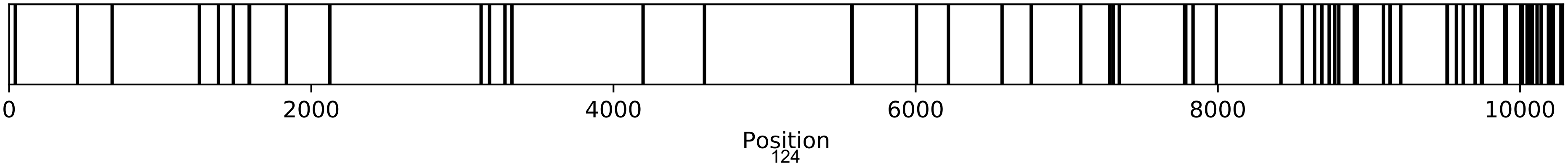

### Region: 416

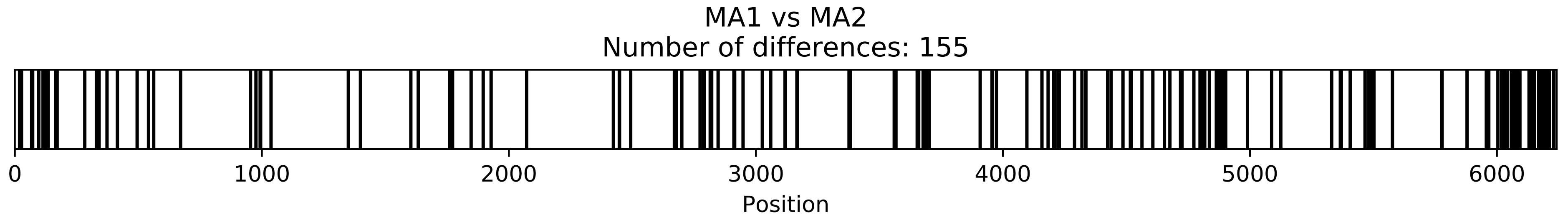
MA1 vs MA2
Number of differences: 155
0
1000
2000
3000
4000
5000
6000
Position

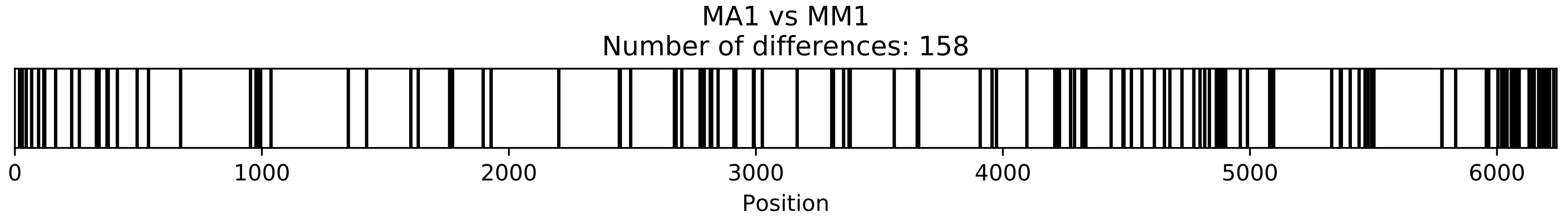
MA1 vs MM1
Number of differences: 158
0
1000
2000
3000
4000
5000
6000
Position

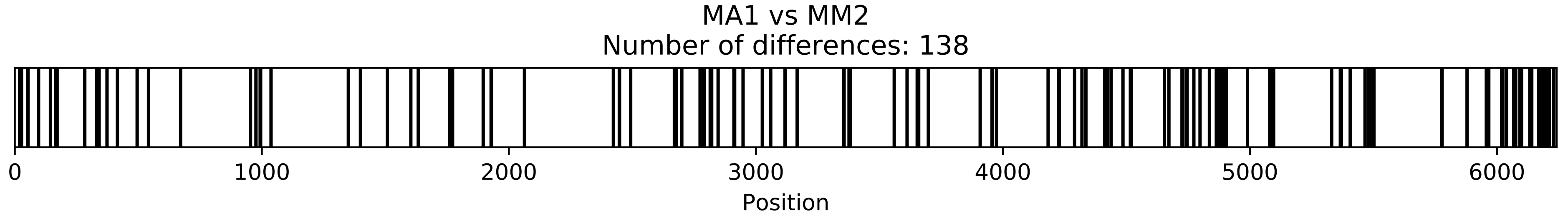
MA1 vs MM2
Number of differences: 138
0
1000
2000
3000
4000
5000
6000
Position

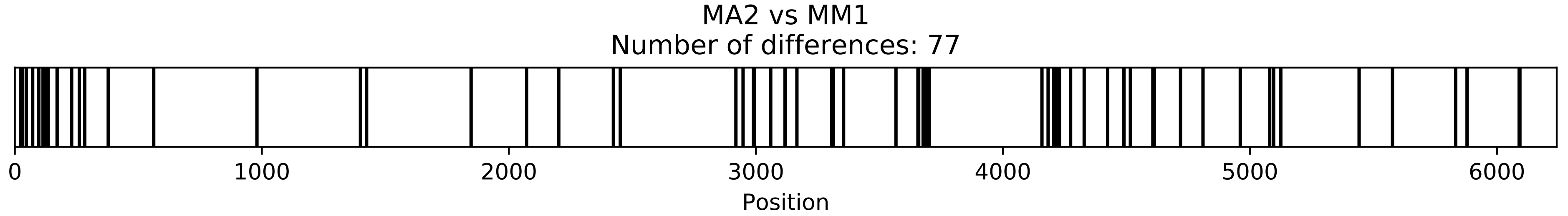
MA2 vs MM1
Number of differences: 77
0
1000
2000
3000
4000
5000
6000
Position

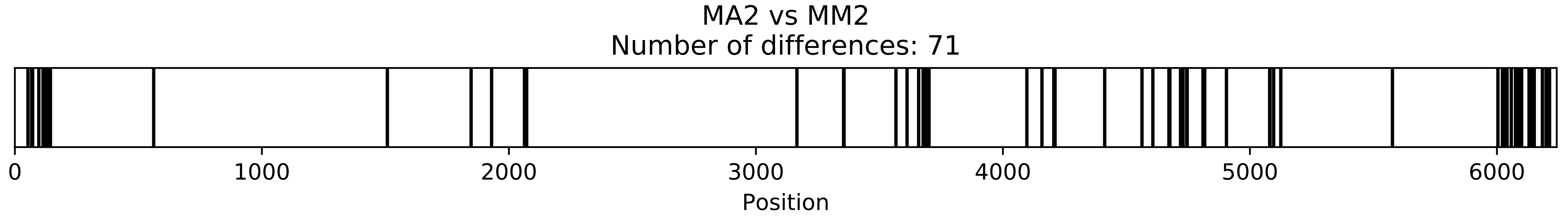
MA2 vs MM2
Number of differences: 71
0
1000
2000
3000
4000
5000
6000
Position

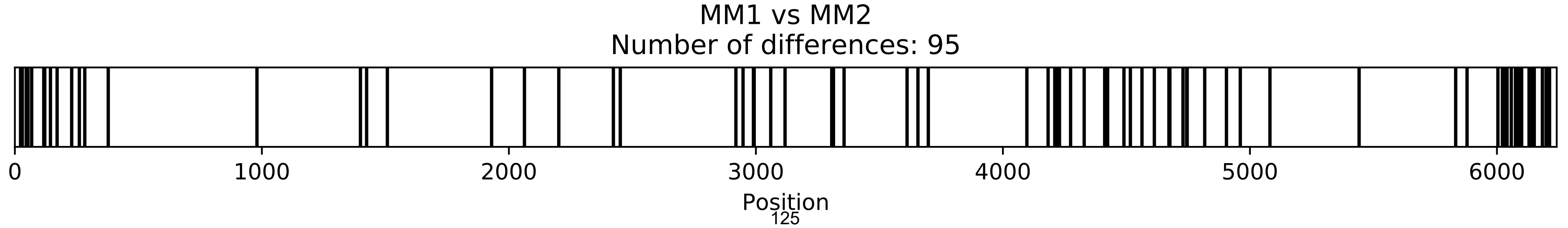
MM1 vs MM2
Number of differences: 95
0
1000
2000
3000
4000
5000
6000
Position

### Region: 420



MA1 vs MA2
Number of differences: 137

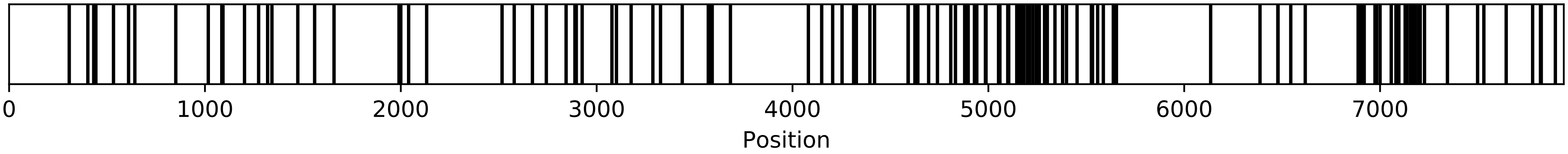


MA1 vs MM1
Number of differences: 94

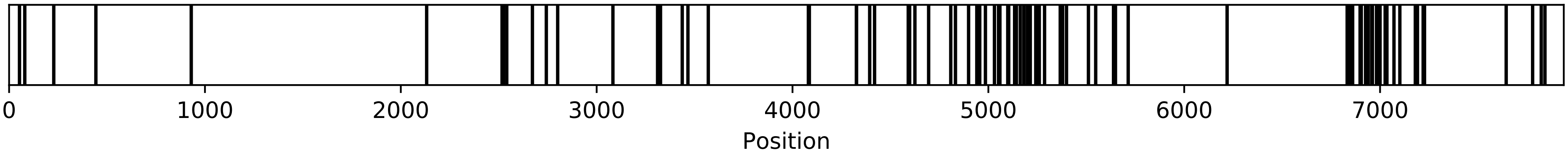


MA1 vs MM2
Number of differences: 0

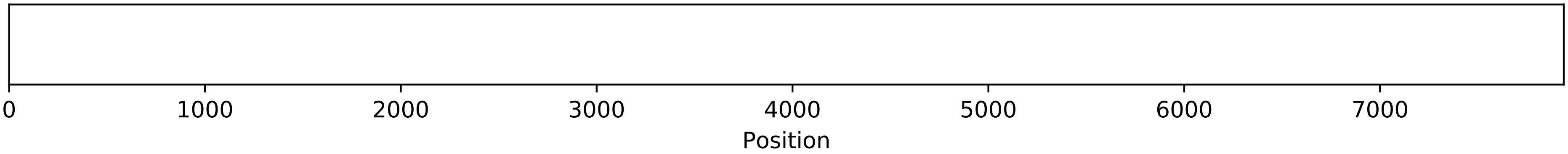


MA2 vs MM1
Number of differences: 147

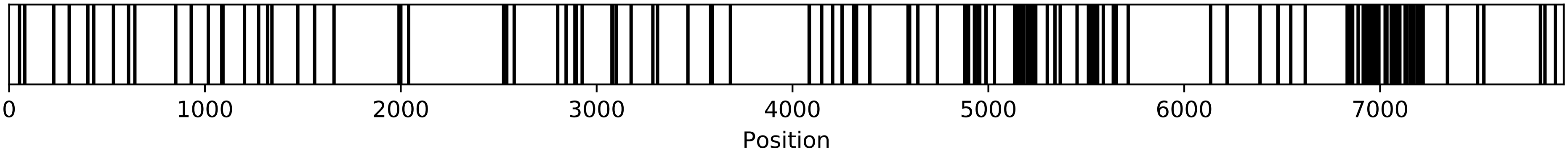


MA2 vs MM2
Number of differences: 137

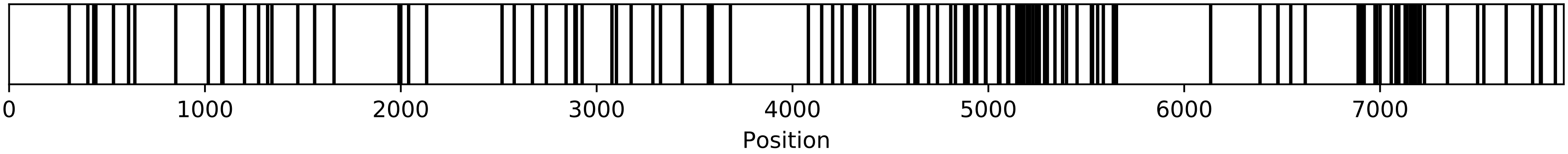


MM1 vs MM2
Number of differences: 94

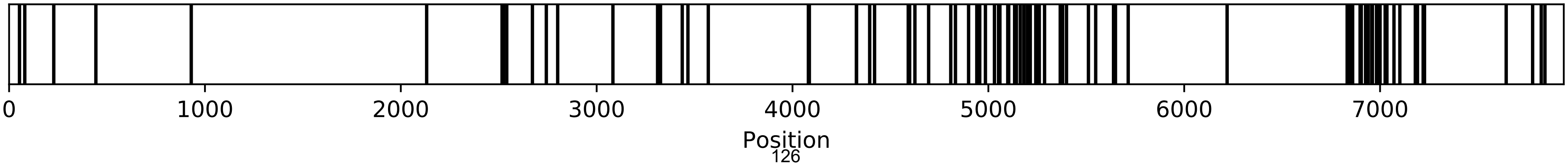

### Region: 421



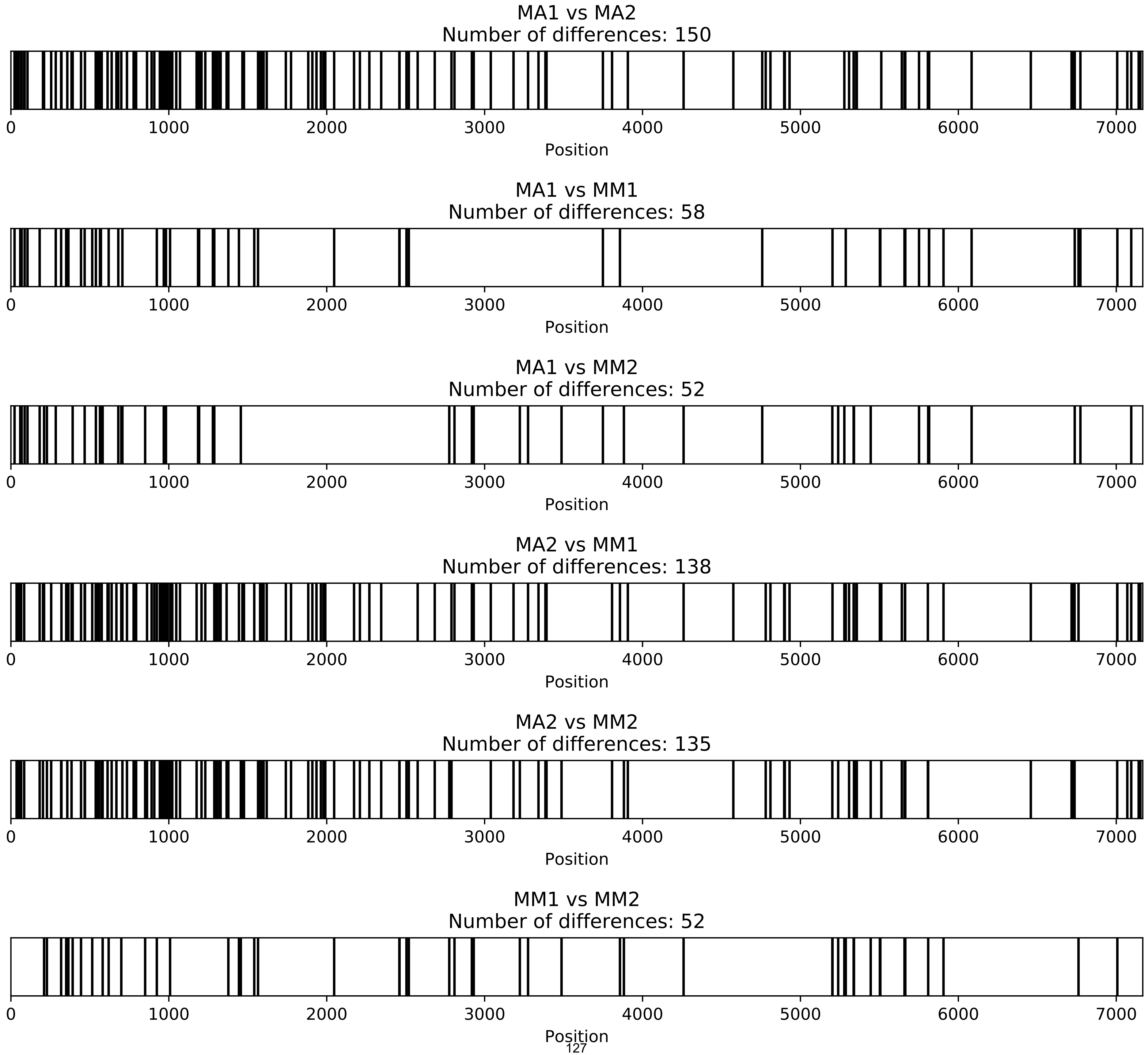
MA1 vs MA2
Number of differences: 150
MA1 vs MM1
Number of differences: 58
MA1 vs MM2
Number of differences: 52
MA2 vs MM1
Number of differences: 138
MA2 vs MM2
Number of differences: 135
MM1 vs MM2
Number of differences: 52
0
1000
2000
3000
4000
5000
6000
7000
Position

### Region: 424



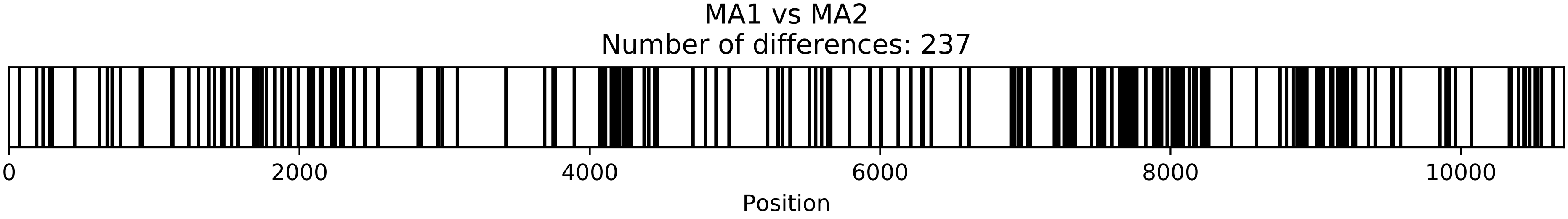
MA1 vs MA2
Number of differences: 237
0
2000
4000
6000
8000
10000
Position

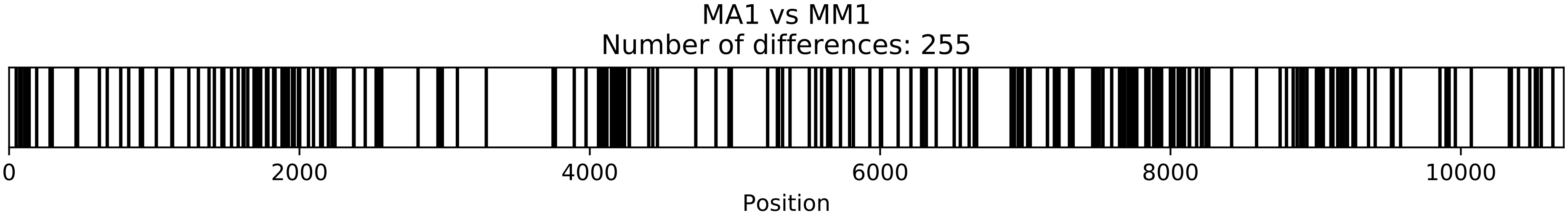
MA1 vs MM1
Number of differences: 255
0
2000
4000
6000
8000
10000
Position

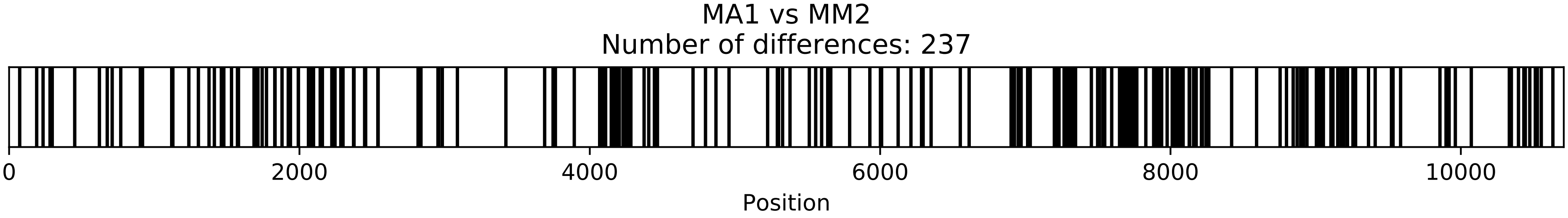
MA1 vs MM2
Number of differences: 237
0
2000
4000
6000
8000
10000
Position

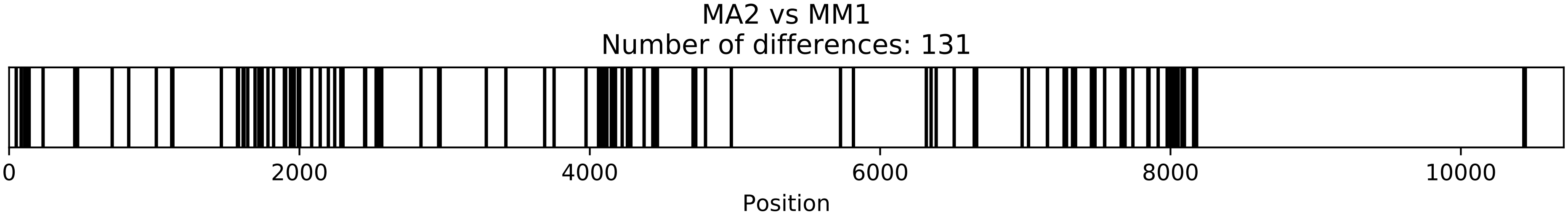
MA2 vs MM1
Number of differences: 131
0
2000
4000
6000
8000
10000
Position

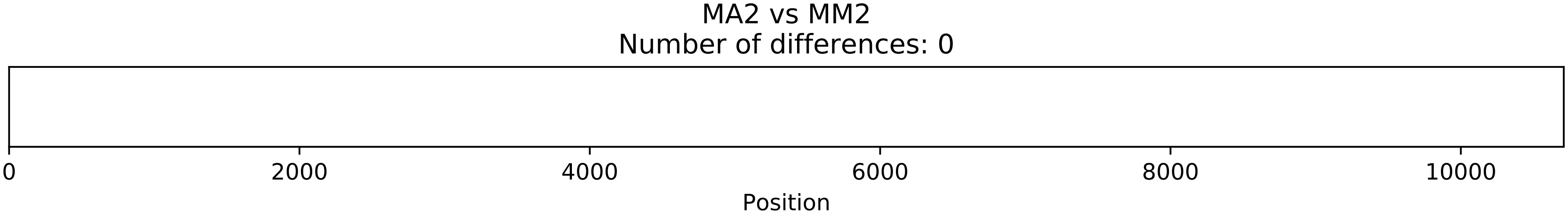
MA2 vs MM2
Number of differences: 0
0
2000
4000
6000
8000
10000
Position

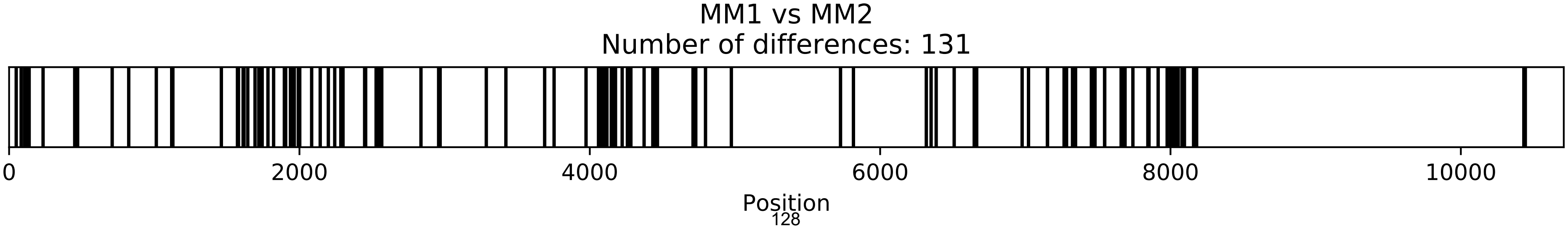
MM1 vs MM2
Number of differences: 131
0
2000
4000
6000
8000
10000
Position

### Region: 429



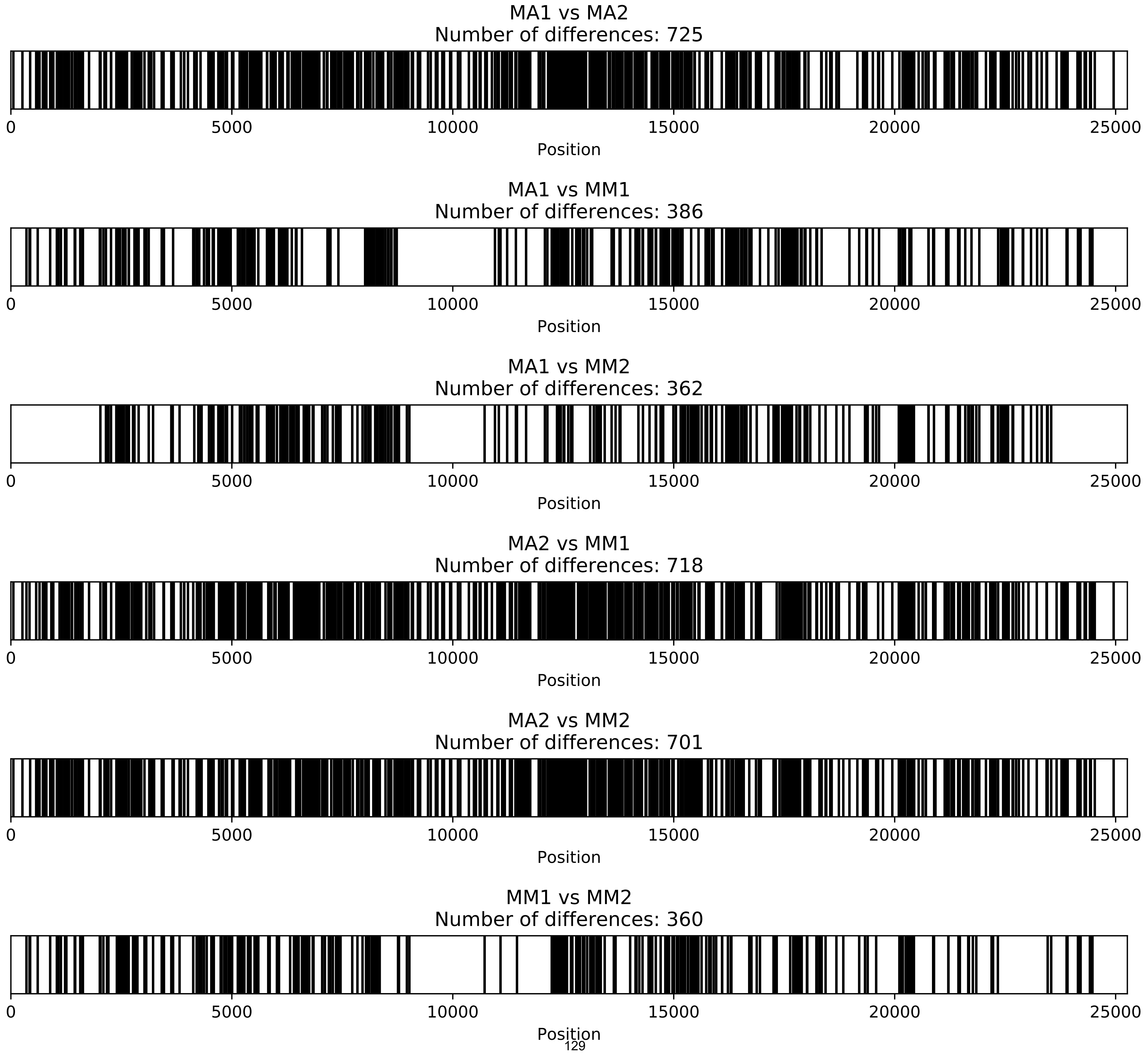
MA1 vs MA2
Number of differences: 725
MA1 vs MM1
Number of differences: 386
MA1 vs MM2
Number of differences: 362
MA2 vs MM1
Number of differences: 718
MA2 vs MM2
Number of differences: 701
MM1 vs MM2
Number of differences: 360
0
5000
10000
15000
20000
25000
Position

### Region: 430

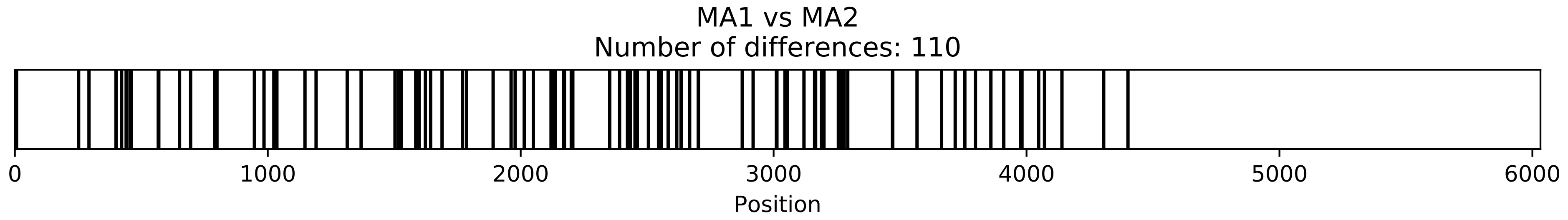
MA1 vs MA2
Number of differences: 110
0
1000
2000
3000
4000
5000
6000
Position

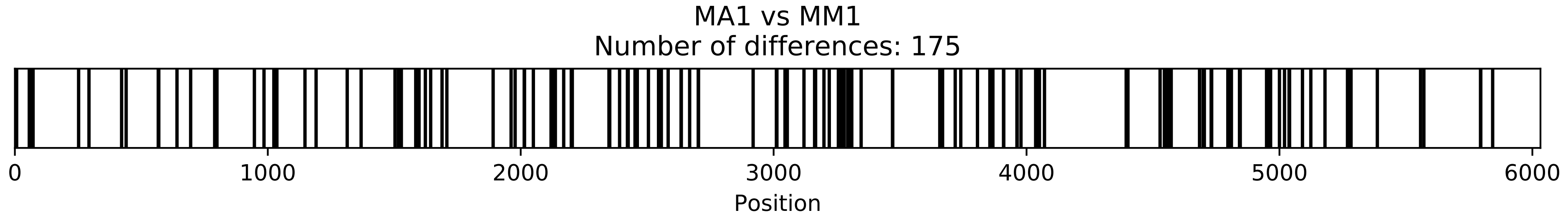
MA1 vs MM1
Number of differences: 175
0
1000
2000
3000
4000
5000
6000
Position

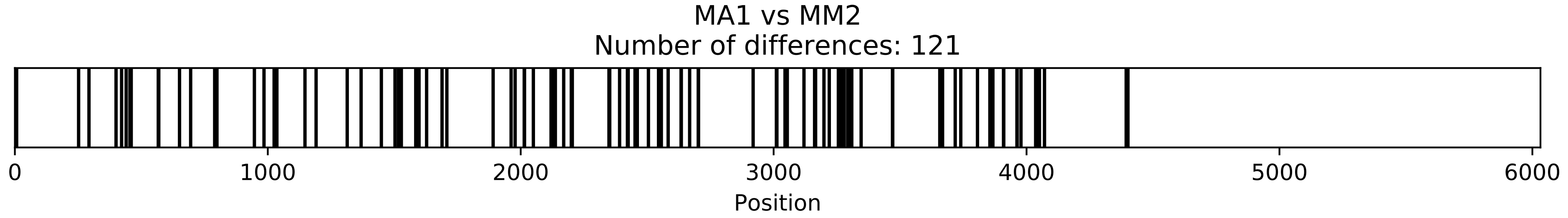
MA1 vs MM2
Number of differences: 121
0
1000
2000
3000
4000
5000
6000
Position

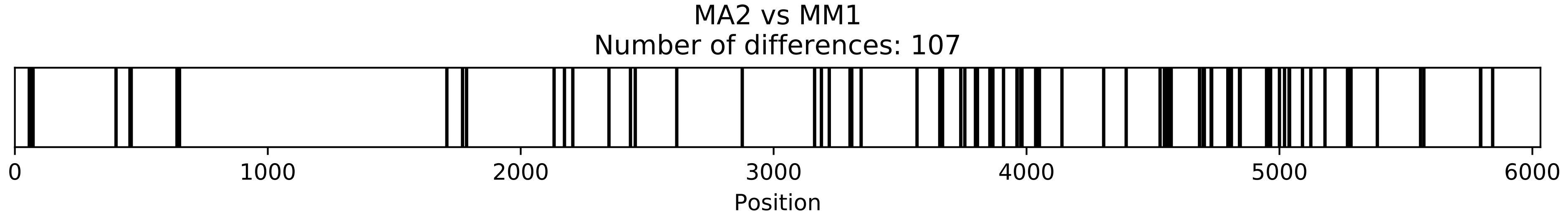
MA2 vs MM1
Number of differences: 107
0
1000
2000
3000
4000
5000
6000
Position

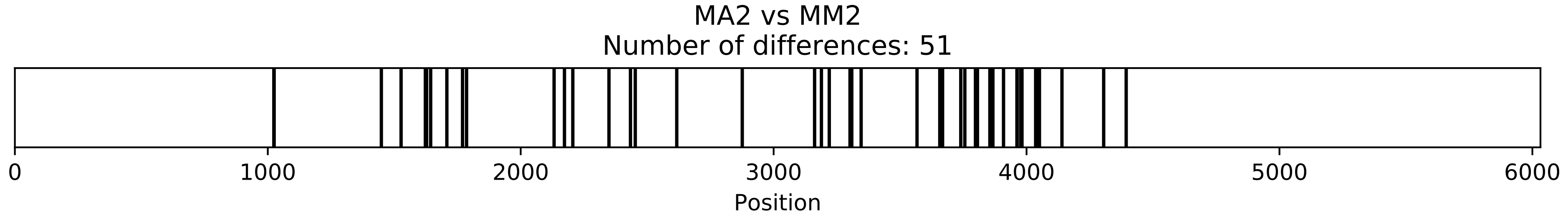
MA2 vs MM2
Number of differences: 51
0
1000
2000
3000
4000
5000
6000
Position

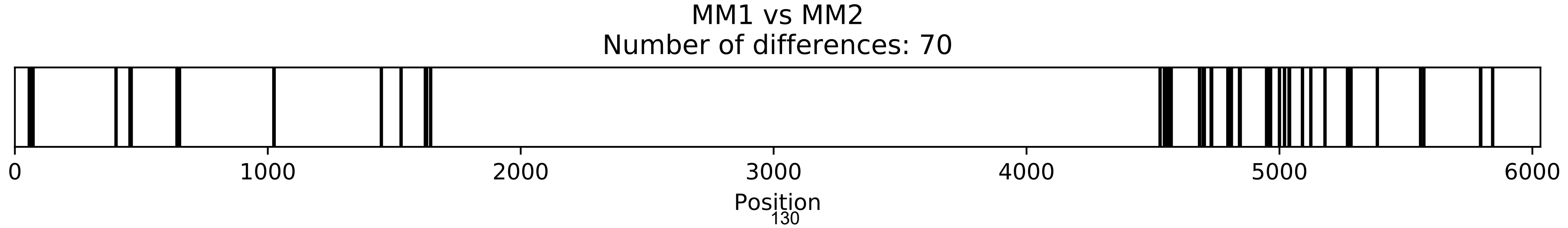
MM1 vs MM2
Number of differences: 70
0
1000
2000
3000
4000
5000
6000
Position

### Region: 433



MA1 vs MA2
Number of differences: 187

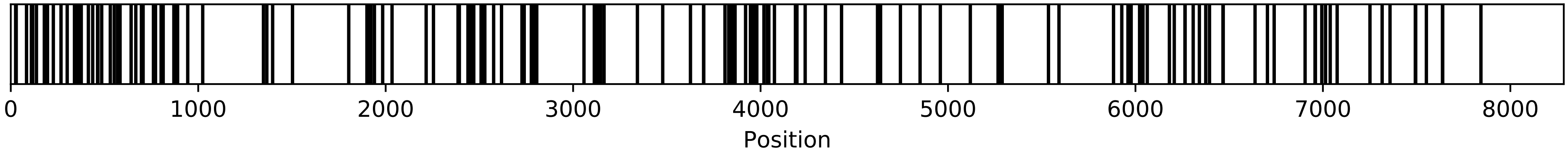


MA1 vs MM1
Number of differences: 187

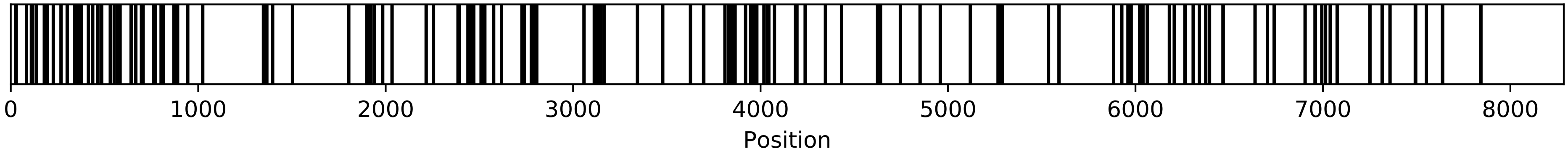


MA1 vs MM2
Number of differences: 189

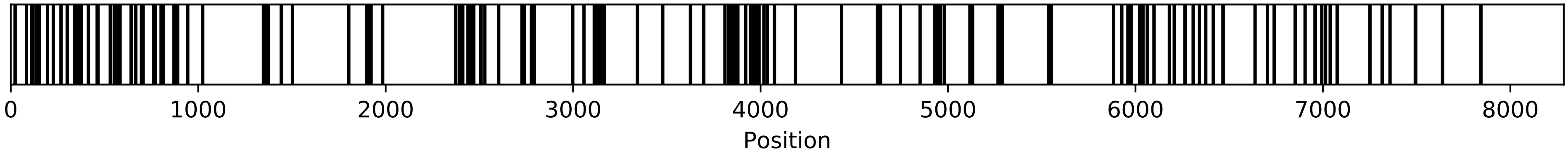


MA2 vs MM1
Number of differences: 0

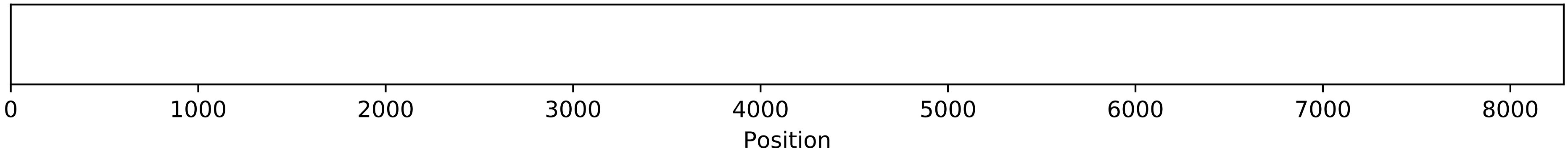


MA2 vs MM2
Number of differences: 58

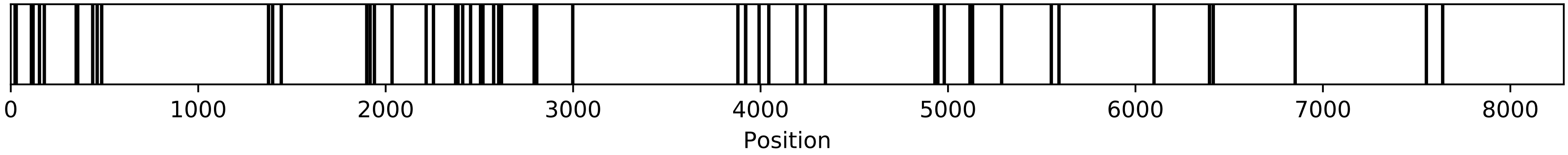


MM1 vs MM2
Number of differences: 58

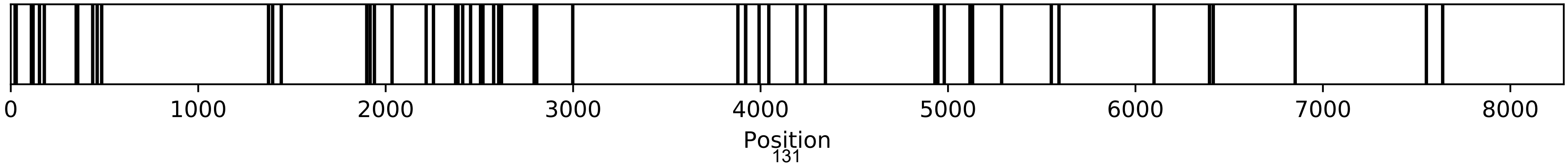

### Region: 435



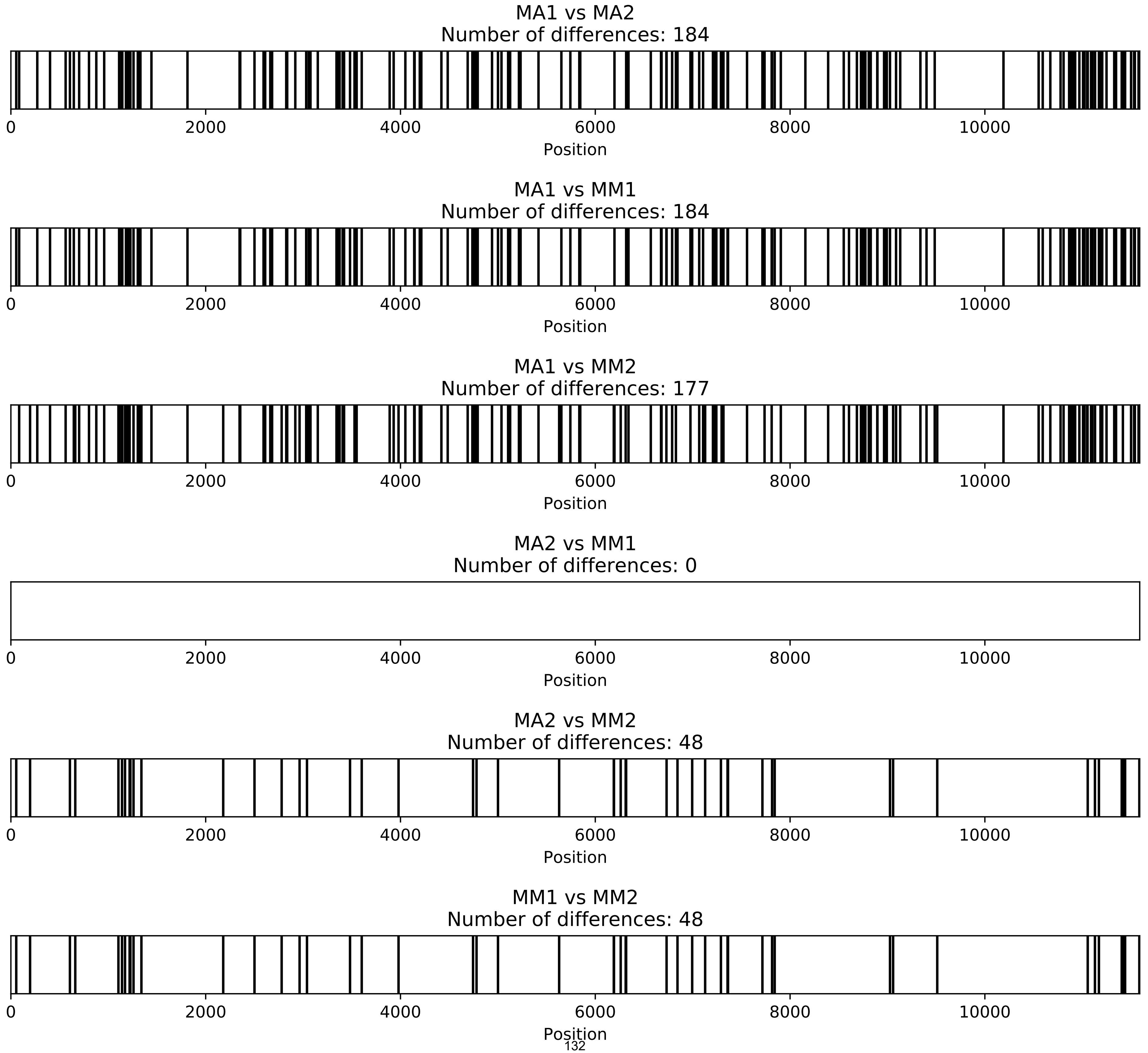
MA1 vs MA2
Number of differences: 184
MA1 vs MM1
Number of differences: 184
MA1 vs MM2
Number of differences: 177
MA2 vs MM1
Number of differences: 0
MA2 vs MM2
Number of differences: 48
MM1 vs MM2
Number of differences: 48
0
2000
4000
6000
8000
10000
Position

### Region: 436



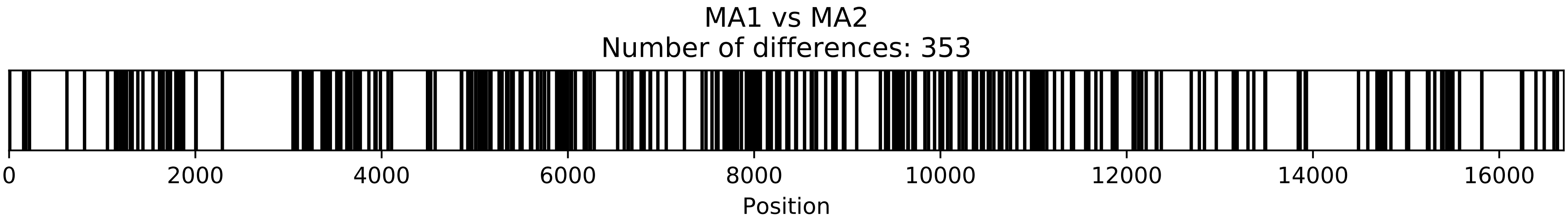
MA1 vs MA2
Number of differences: 353
0 2000 4000 6000 8000 10000 12000 14000 16000
Position

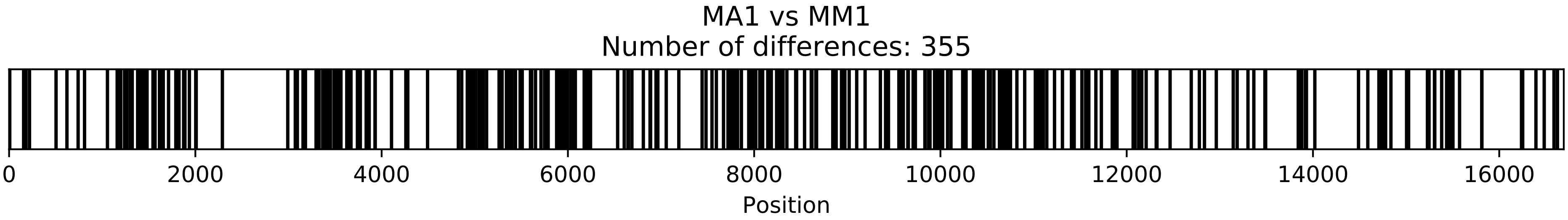
MA1 vs MM1
Number of differences: 355
0 2000 4000 6000 8000 10000 12000 14000 16000
Position

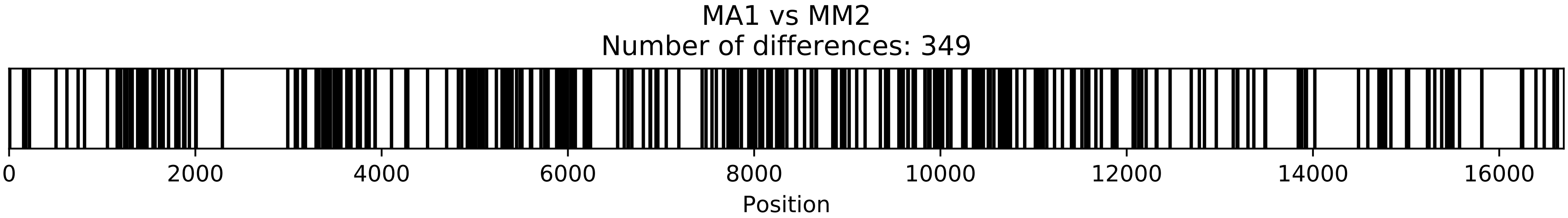
MA1 vs MM2
Number of differences: 349
0 2000 4000 6000 8000 10000 12000 14000 16000
Position

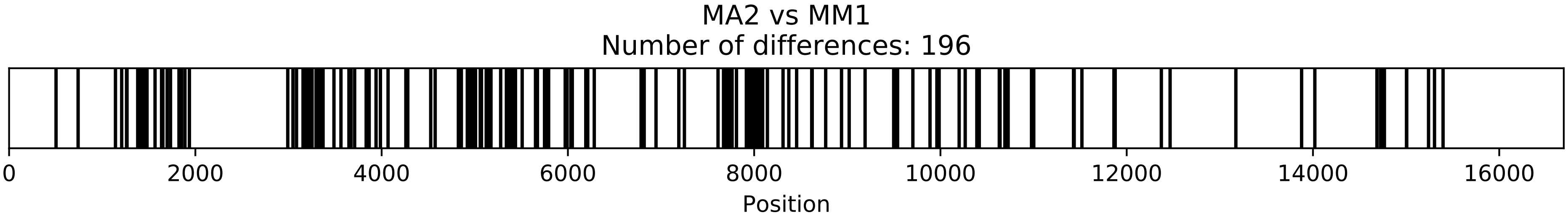
MA2 vs MM1
Number of differences: 196
0 2000 4000 6000 8000 10000 12000 14000 16000
Position

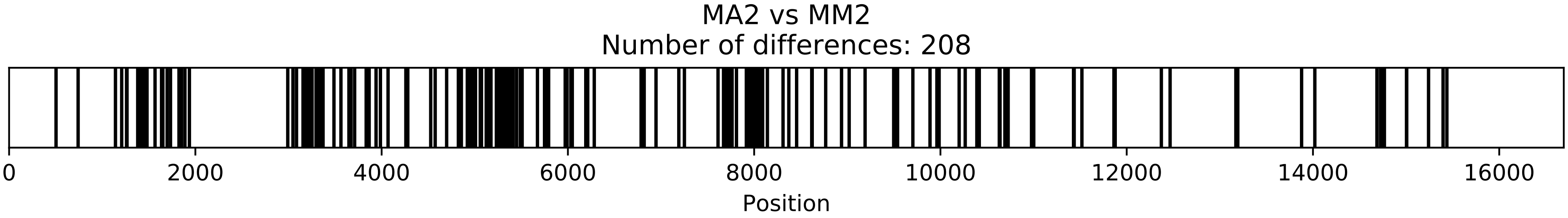
MA2 vs MM2
Number of differences: 208
0 2000 4000 6000 8000 10000 12000 14000 16000
Position

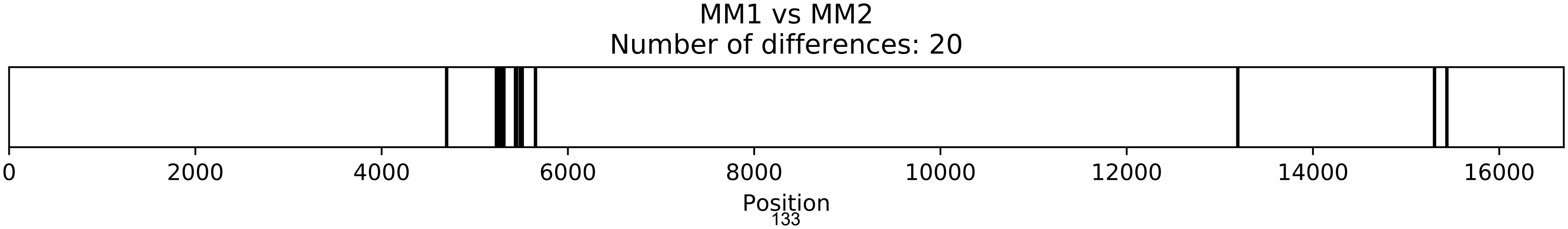
MM1 vs MM2
Number of differences: 20
0 2000 4000 6000 8000 10000 12000 14000 16000
Position

### Region: 438



MA1 vs MA2
Number of differences: 178

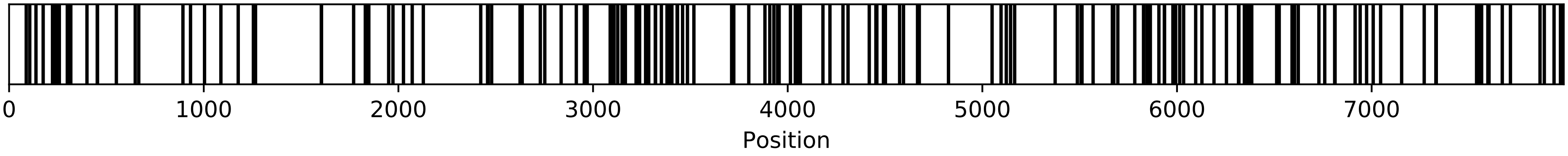


MA1 vs MM1
Number of differences: 195

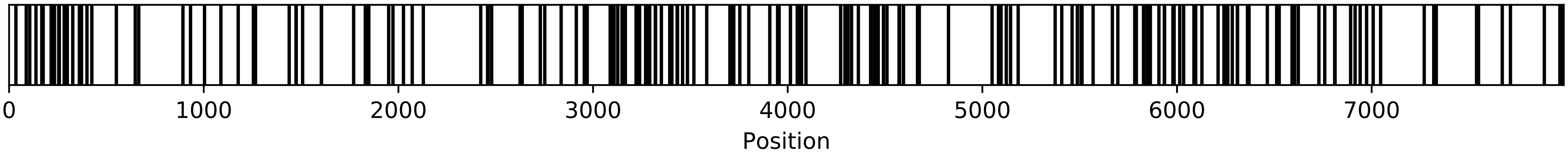


MA1 vs MM2
Number of differences: 178

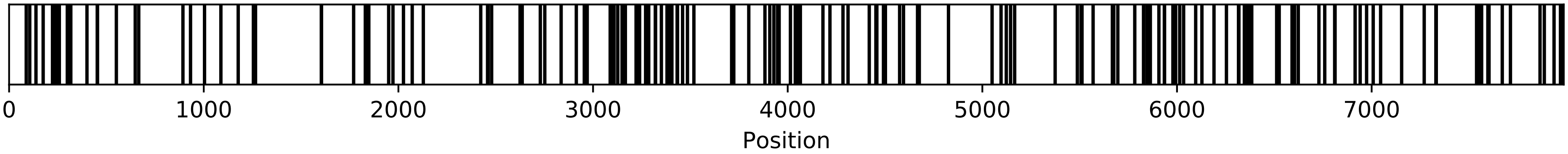


MA2 vs MM1
Number of differences: 82

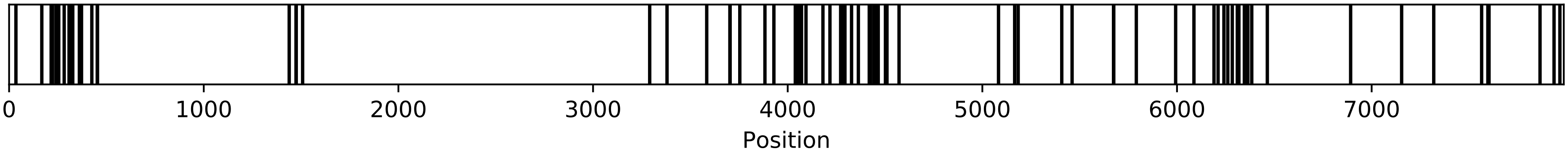


MA2 vs MM2
Number of differences: 0

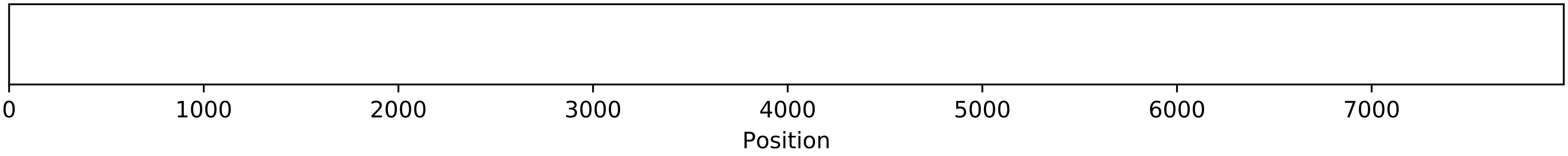


MM1 vs MM2
Number of differences: 82

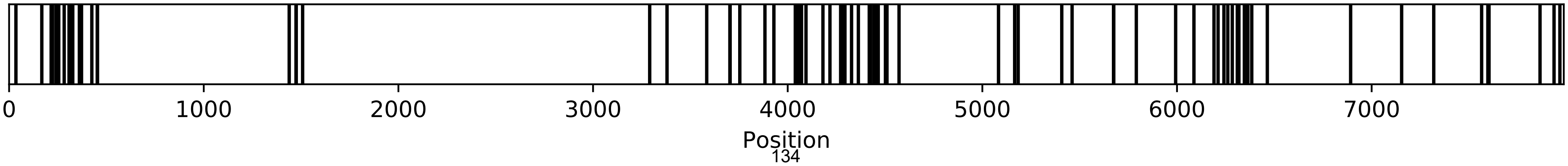

### Region: 439



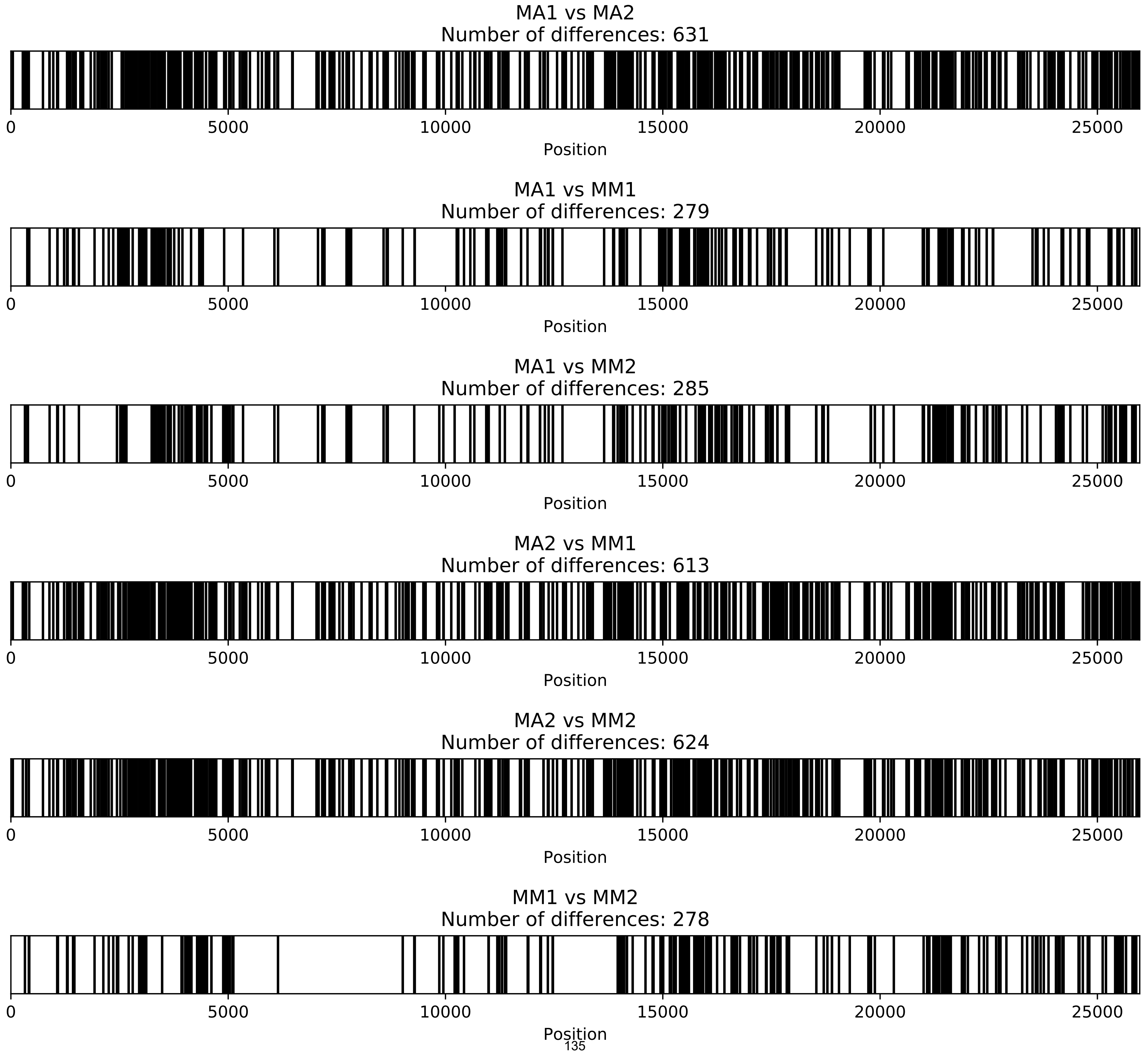
MA1 vs MA2
Number of differences: 631
MA1 vs MM1
Number of differences: 279
MA1 vs MM2
Number of differences: 285
MA2 vs MM1
Number of differences: 613
MA2 vs MM2
Number of differences: 624
MM1 vs MM2
Number of differences: 278
0
5000
10000
15000
20000
25000
Position

### Region: 441

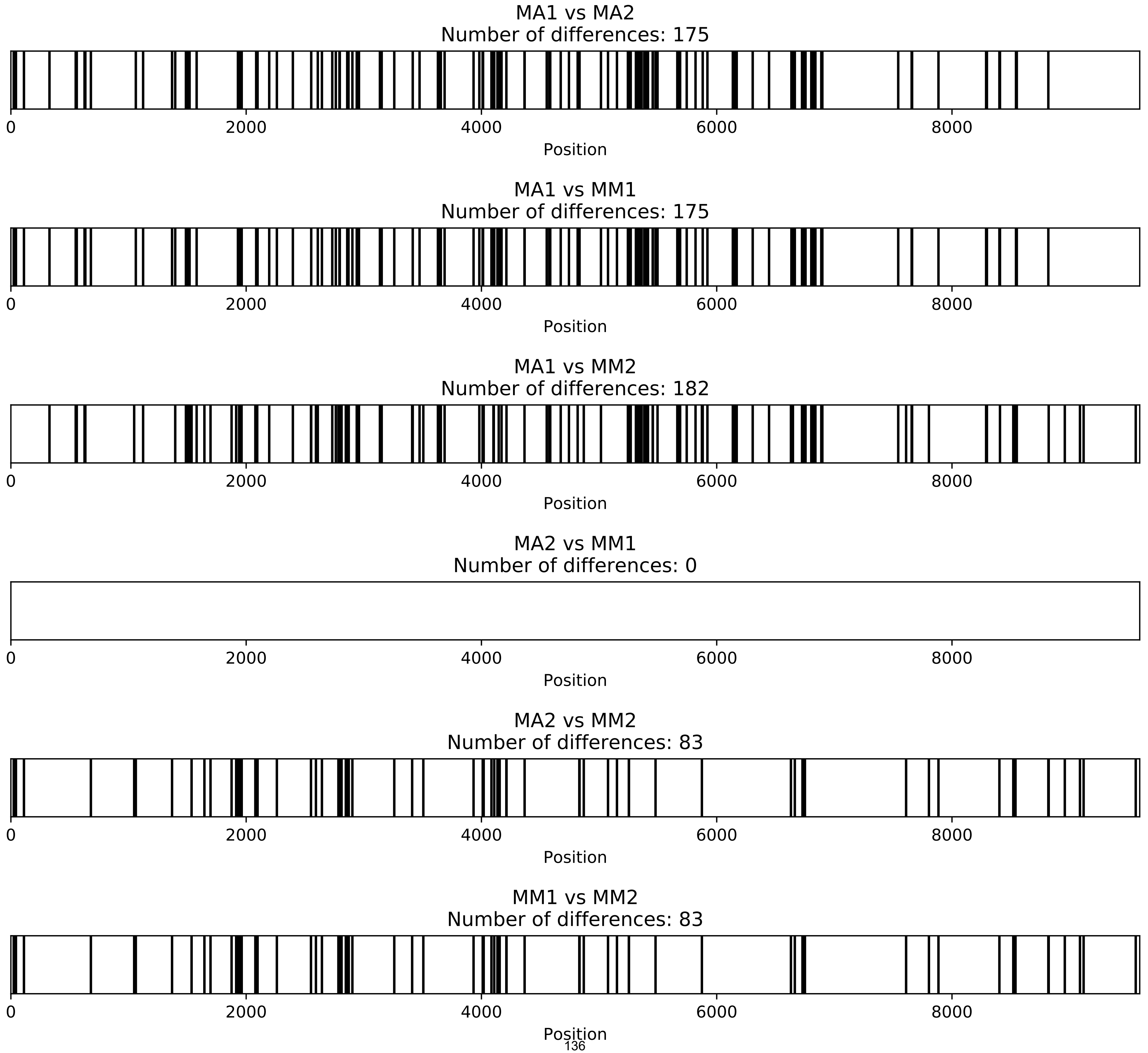
MA1 vs MA2
Number of differences: 175
0
2000
4000
6000
8000
Position
MA1 vs MM1
Number of differences: 175
0
2000
4000
6000
8000
Position
MA1 vs MM2
Number of differences: 182
0
2000
4000
6000
8000
Position
MA2 vs MM1
Number of differences: 0
0
2000
4000
6000
8000
Position
MA2 vs MM2
Number of differences: 83
0
2000
4000
6000
8000
Position
MM1 vs MM2
Number of differences: 83
0
2000
4000
6000
8000
Position

### Region: 442



MA1 vs MA2
Number of differences: 269

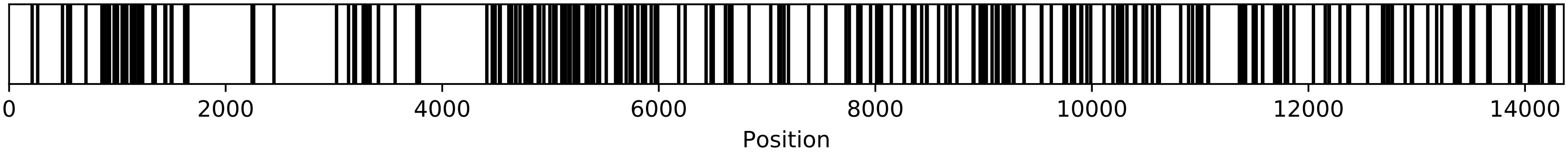


MA1 vs MM1
Number of differences: 260

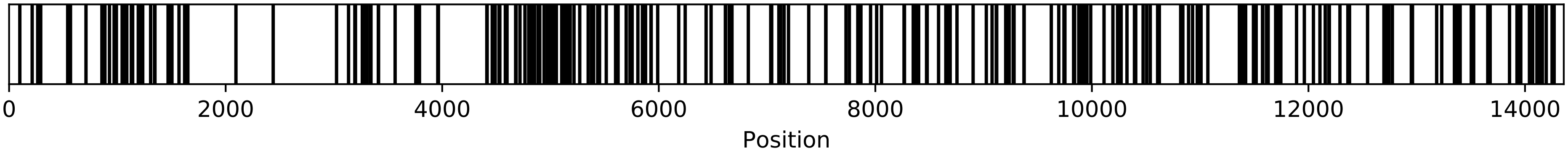


MA1 vs MM2
Number of differences: 269

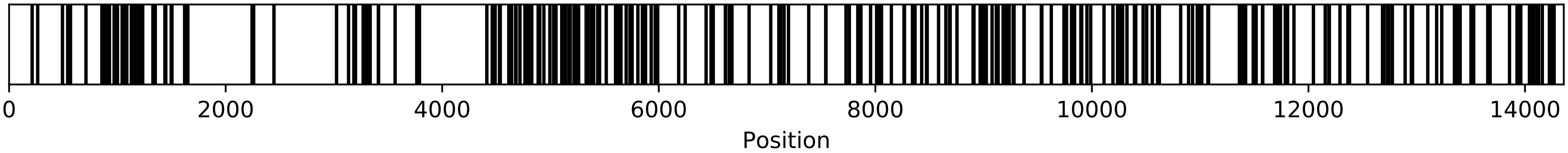


MA2 vs MM1
Number of differences: 126

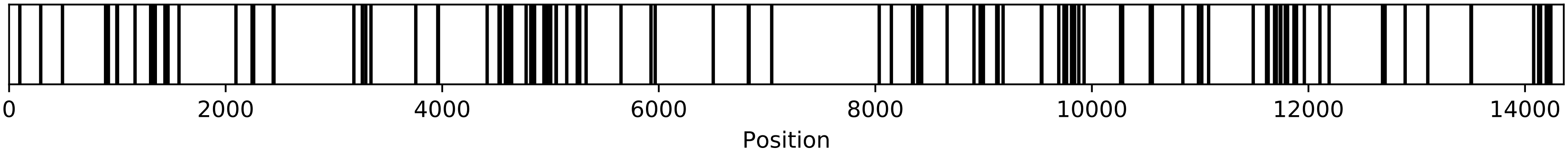


MA2 vs MM2
Number of differences: 0

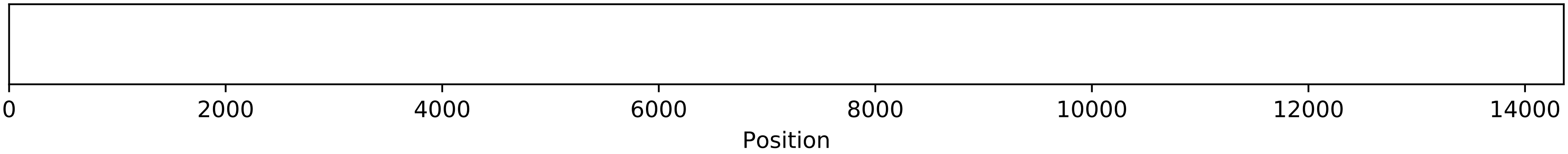


MM1 vs MM2
Number of differences: 126

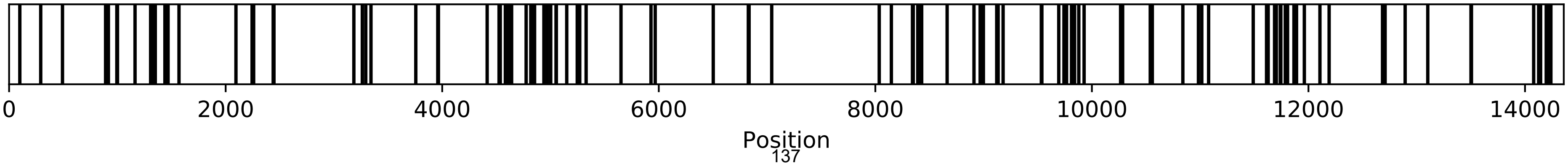

### Region: 453



## MA1 vs MA2
Number of differences: 0

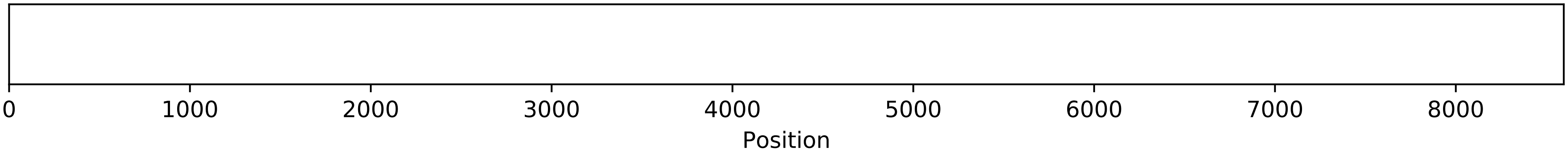


## MA1 vs MM1
Number of differences: 354

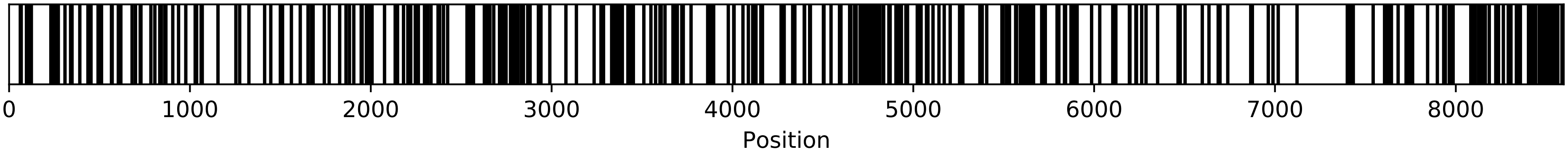


## MA1 vs MM2
Number of differences: 351

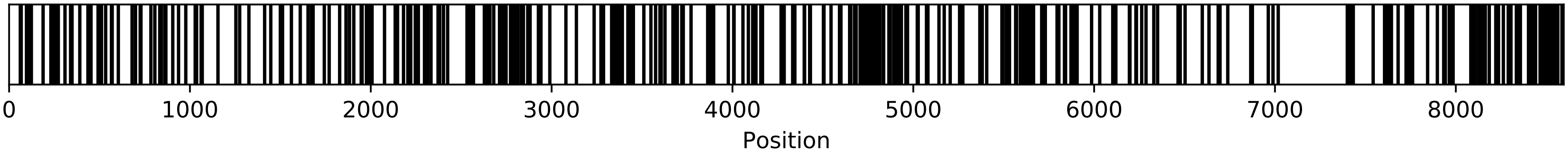


## MA2 vs MM1
Number of differences: 354

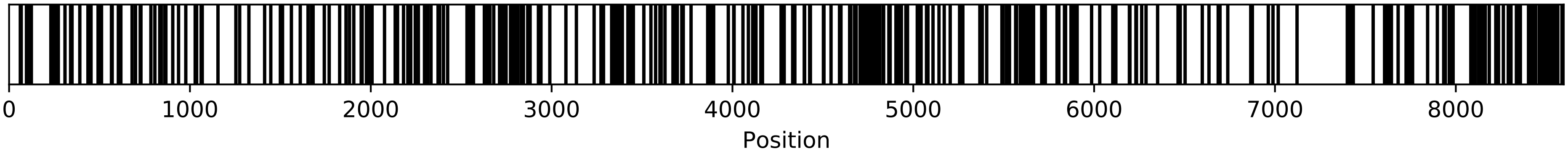


## MA2 vs MM2
Number of differences: 351

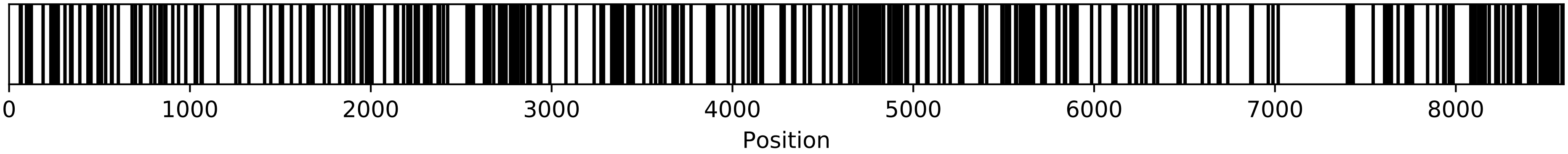


## MM1 vs MM2
Number of differences: 13

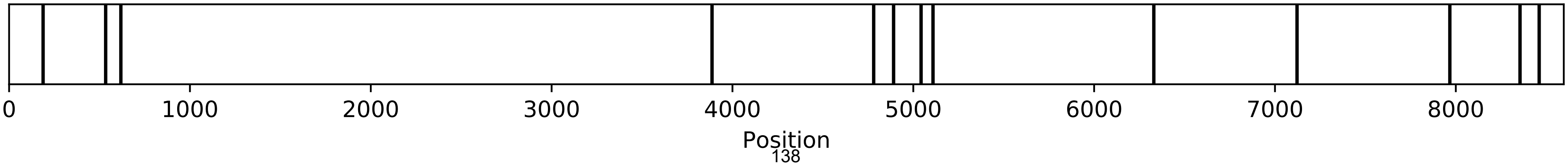

### Region: 456



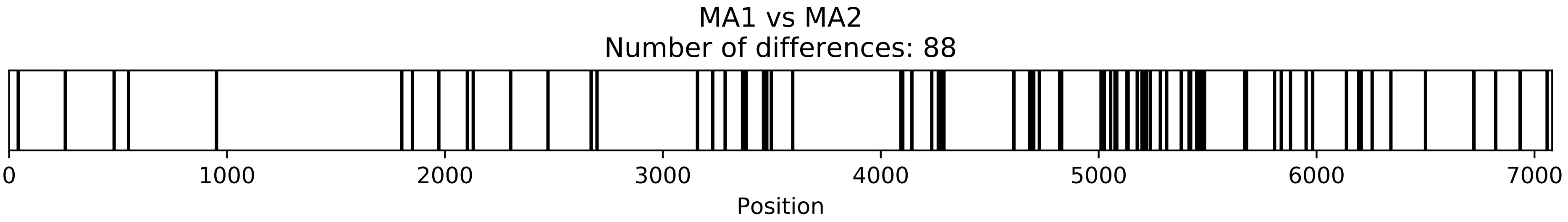
MA1 vs MA2
Number of differences: 88
0 1000 2000 3000 4000 5000 6000 7000
Position

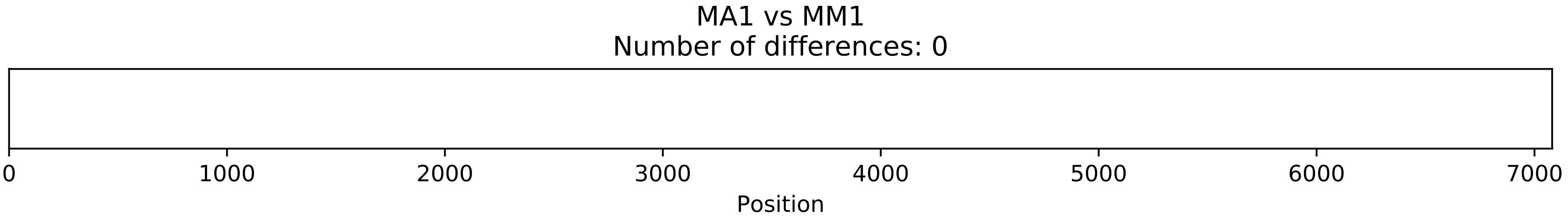
MA1 vs MM1
Number of differences: 0
0 1000 2000 3000 4000 5000 6000 7000
Position

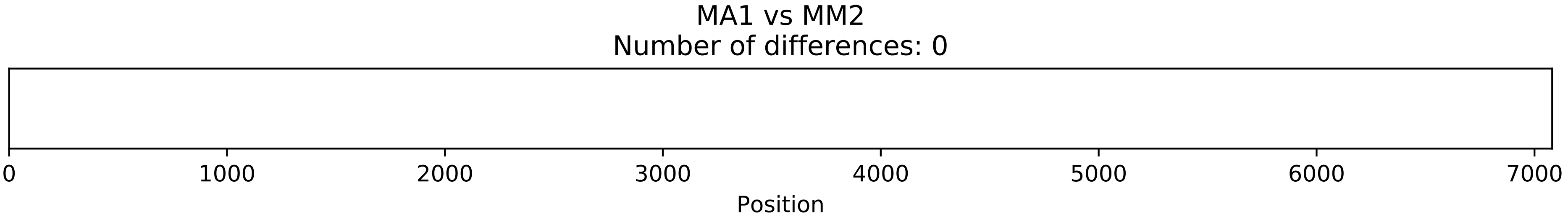
MA1 vs MM2
Number of differences: 0
0 1000 2000 3000 4000 5000 6000 7000
Position

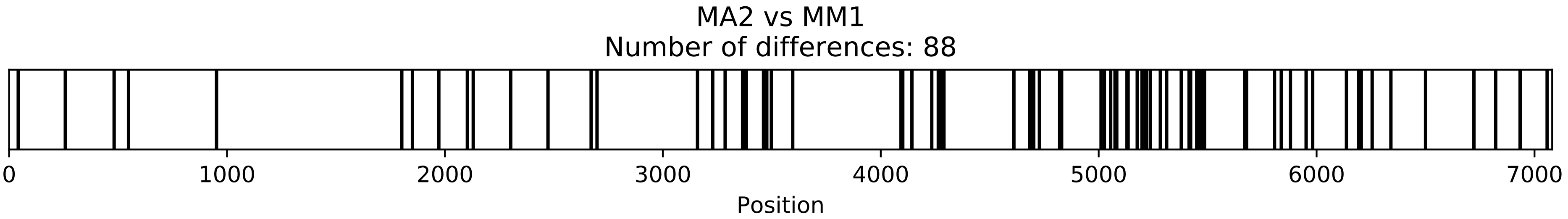
MA2 vs MM1
Number of differences: 88
0 1000 2000 3000 4000 5000 6000 7000
Position

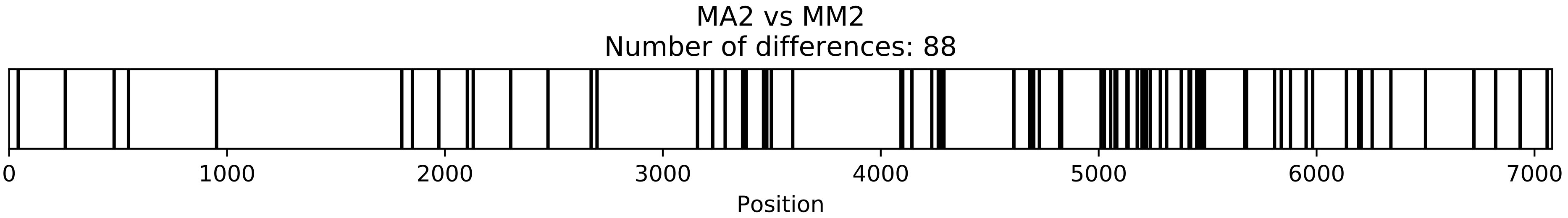
MA2 vs MM2
Number of differences: 88
0 1000 2000 3000 4000 5000 6000 7000
Position

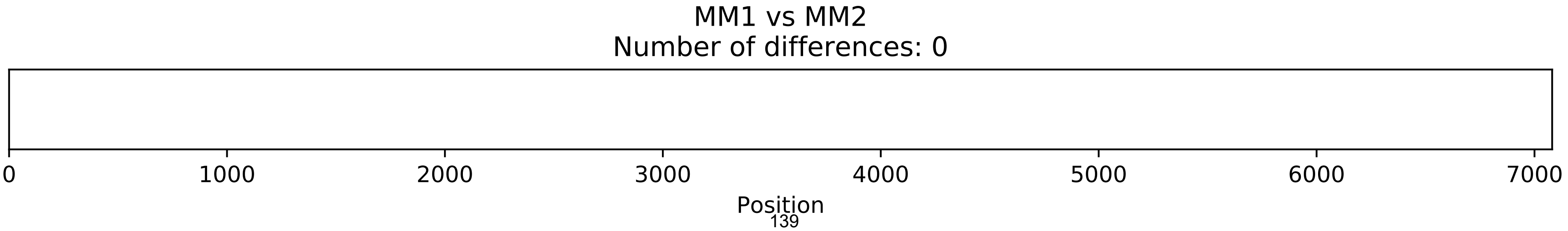
MM1 vs MM2
Number of differences: 0
0 1000 2000 3000 4000 5000 6000 7000
Position

### Region: 464



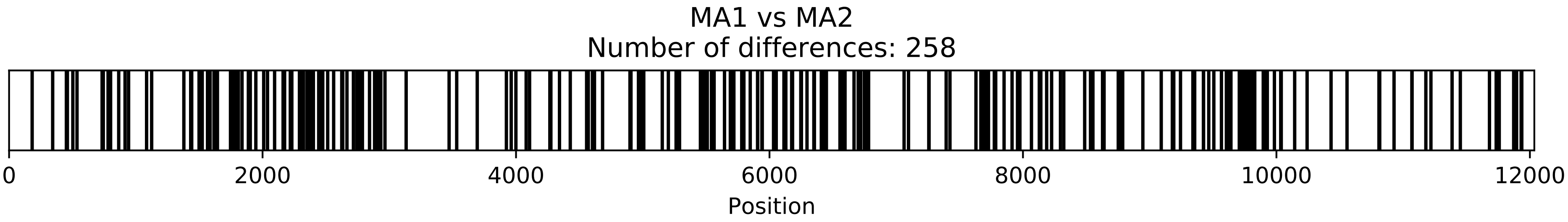
MA1 vs MA2
Number of differences: 258
0
2000
4000
6000
8000
10000
12000
Position

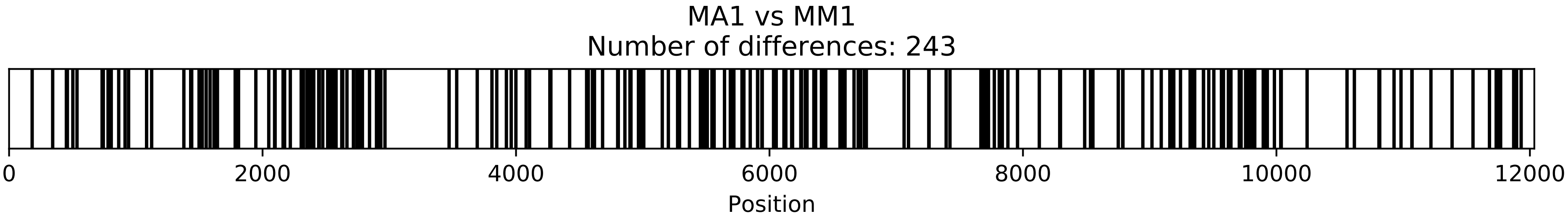
MA1 vs MM1
Number of differences: 243
0
2000
4000
6000
8000
10000
12000
Position

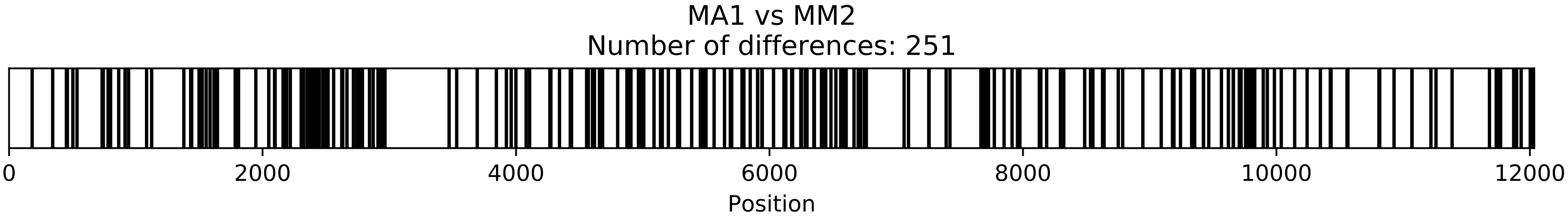
MA1 vs MM2
Number of differences: 251
0
2000
4000
6000
8000
10000
12000
Position

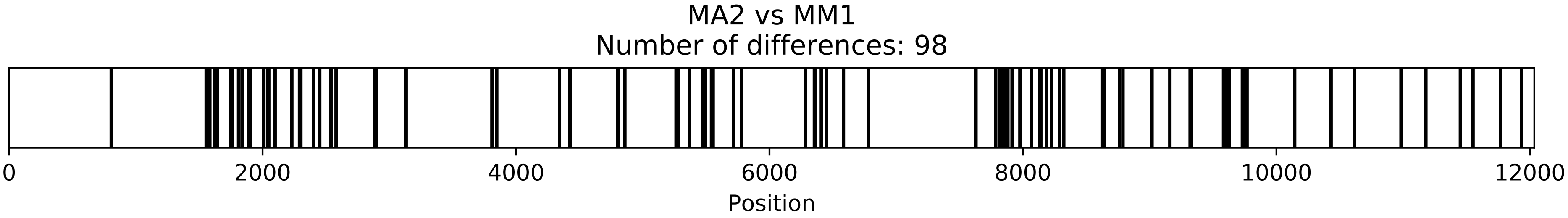
MA2 vs MM1
Number of differences: 98
0
2000
4000
6000
8000
10000
12000
Position

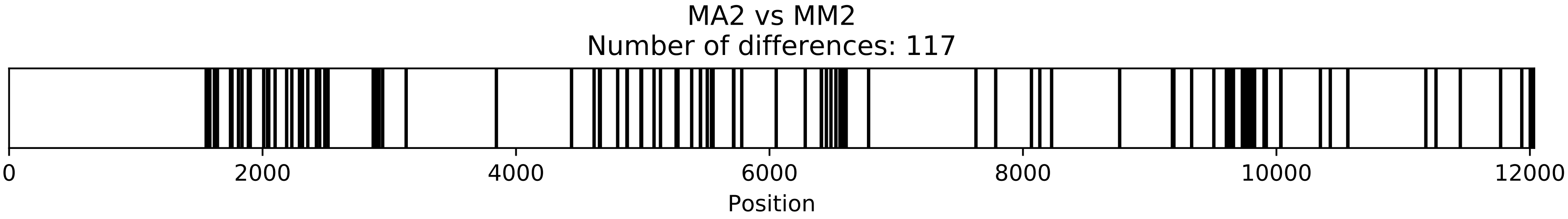
MA2 vs MM2
Number of differences: 117
0
2000
4000
6000
8000
10000
12000
Position

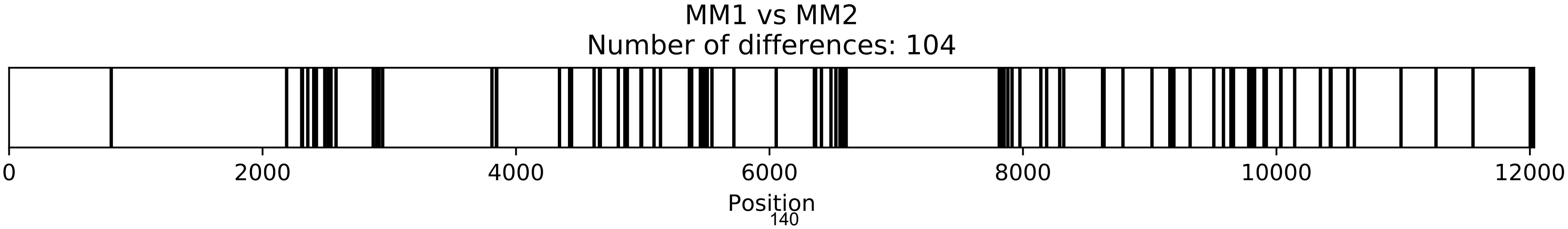
MM1 vs MM2
Number of differences: 104
0
2000
4000
6000
8000
10000
12000
Position

### Region: 468



MA1 vs MA2
Number of differences: 184

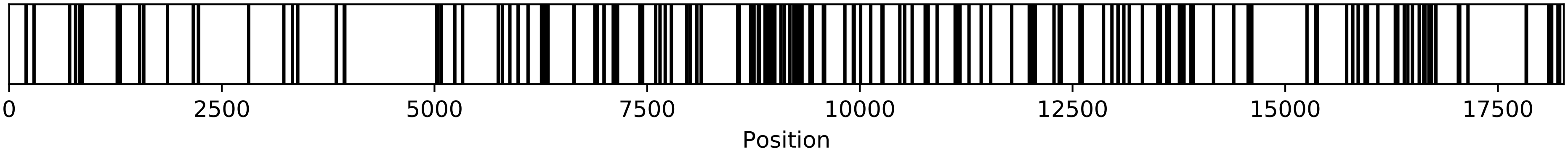


MA1 vs MM1
Number of differences: 82

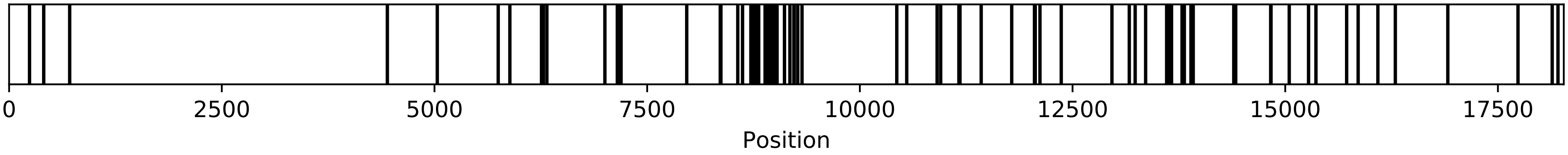


MA1 vs MM2
Number of differences: 0

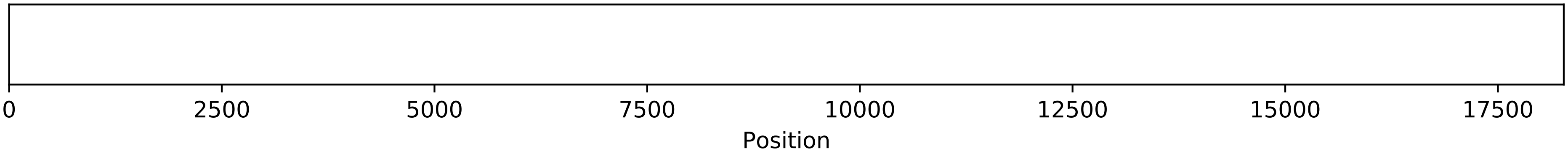


MA2 vs MM1
Number of differences: 178

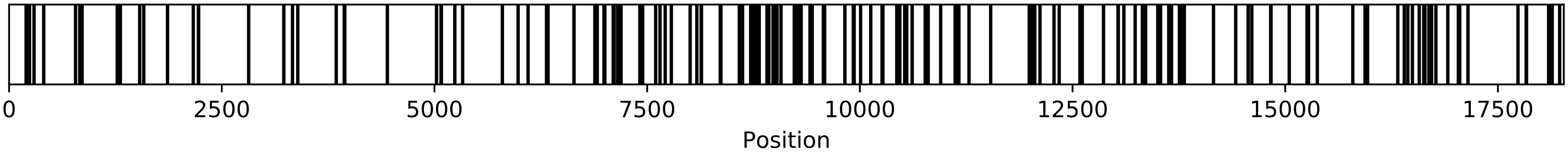


MA2 vs MM2
Number of differences: 184

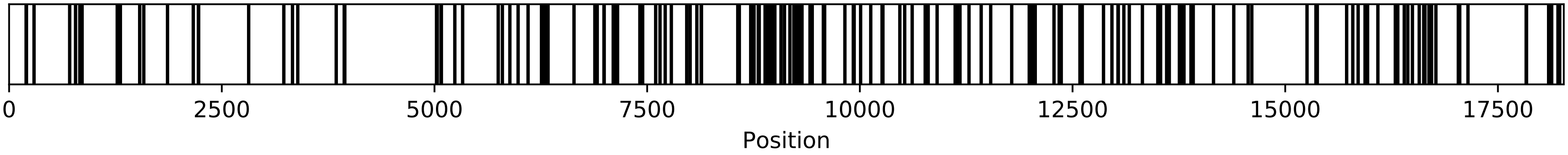


MM1 vs MM2
Number of differences: 82

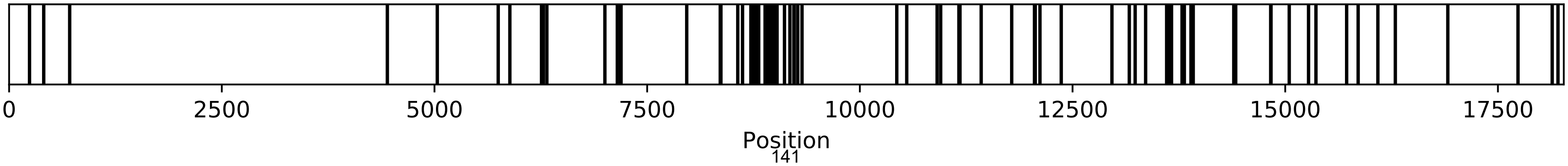

### Region: 469



MA1 vs MA2
Number of differences: 255

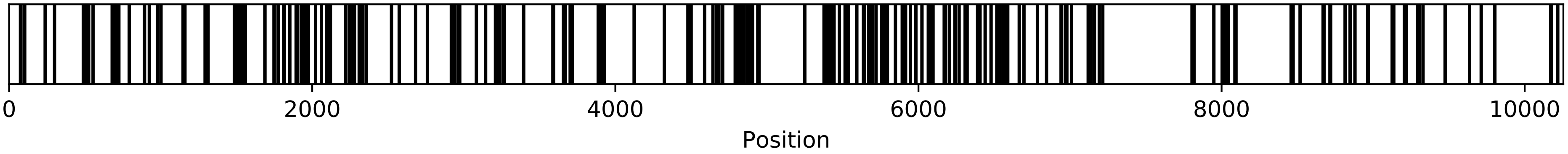


MA1 vs MM1
Number of differences: 255

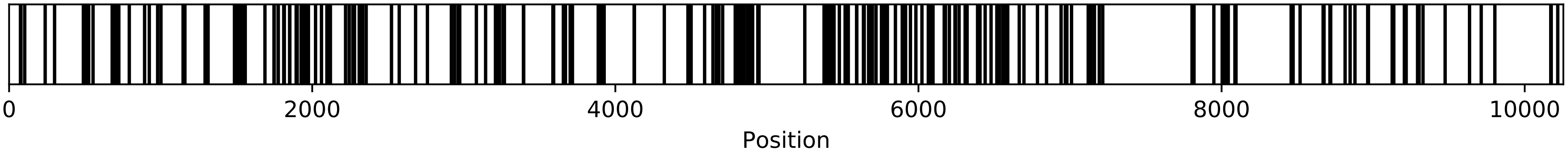


MA1 vs MM2
Number of differences: 237

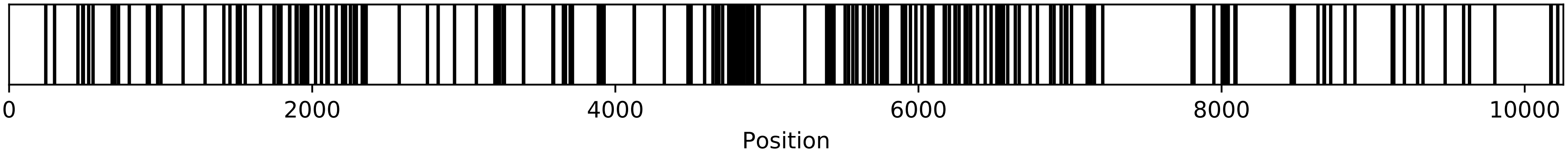


MA2 vs MM1
Number of differences: 0

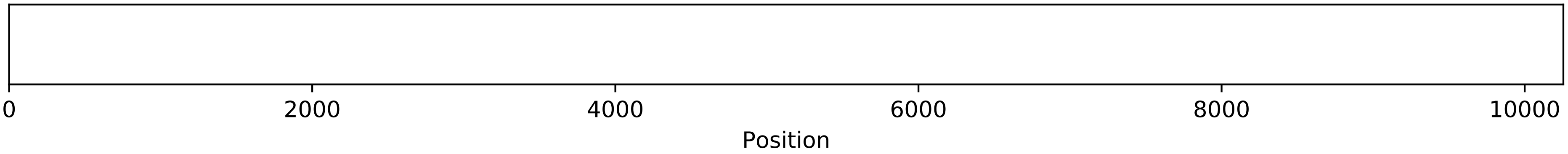


MA2 vs MM2
Number of differences: 105

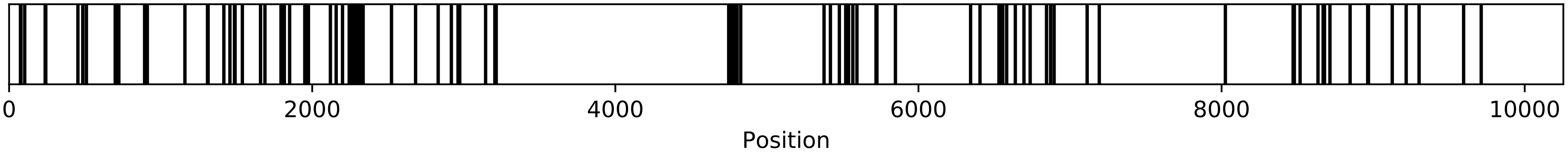


MM1 vs MM2
Number of differences: 105

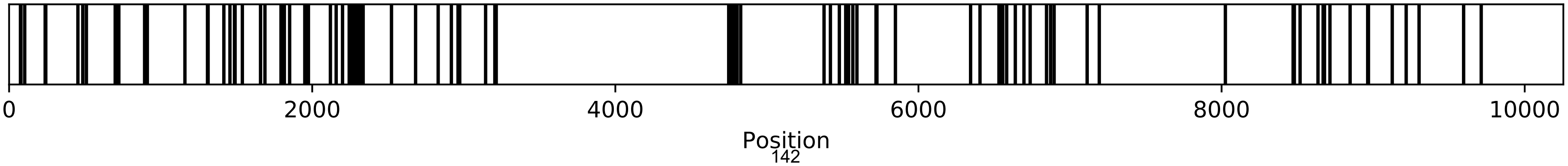

### Region: 470



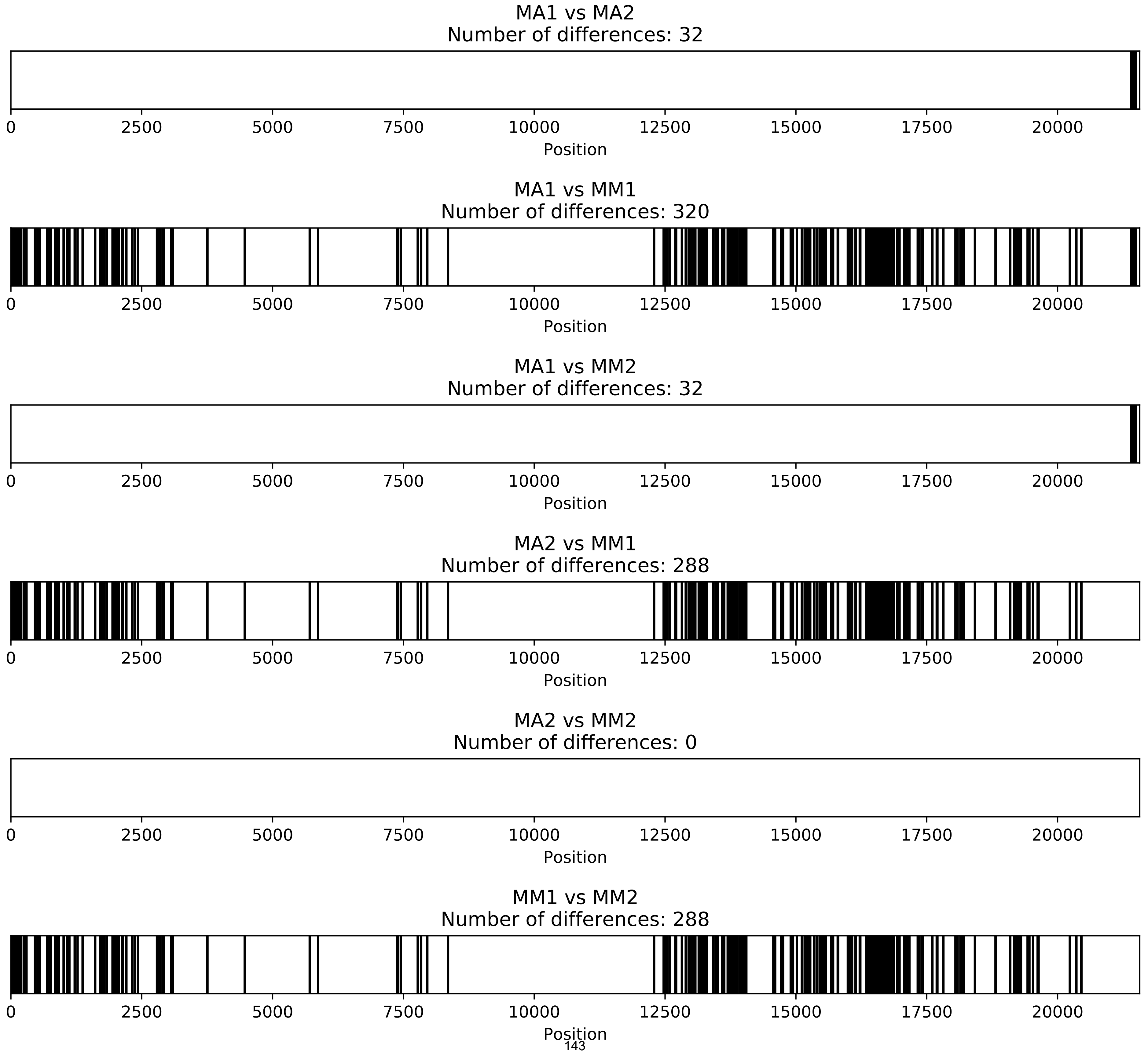
MA1 vs MA2
Number of differences: 32
MA1 vs MM1
Number of differences: 320
MA1 vs MM2
Number of differences: 32
MA2 vs MM1
Number of differences: 288
MA2 vs MM2
Number of differences: 0
MM1 vs MM2
Number of differences: 288
0
2500
5000
7500
10000
12500
15000
17500
20000
Position

### Region: 471



MA1 vs MA2
Number of differences: 148

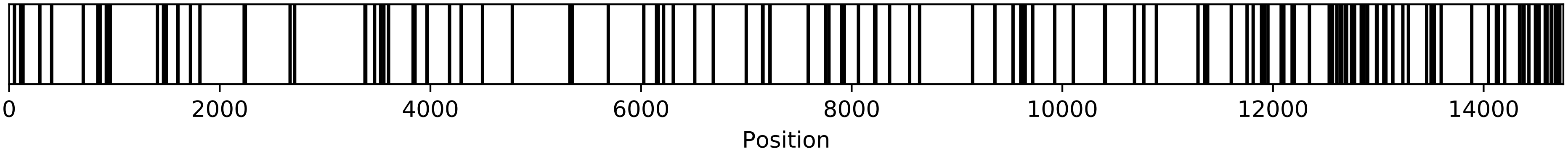


MA1 vs MM1
Number of differences: 72

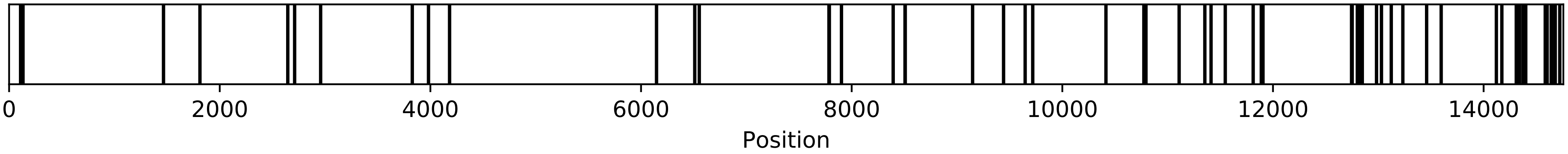


MA1 vs MM2
Number of differences: 0

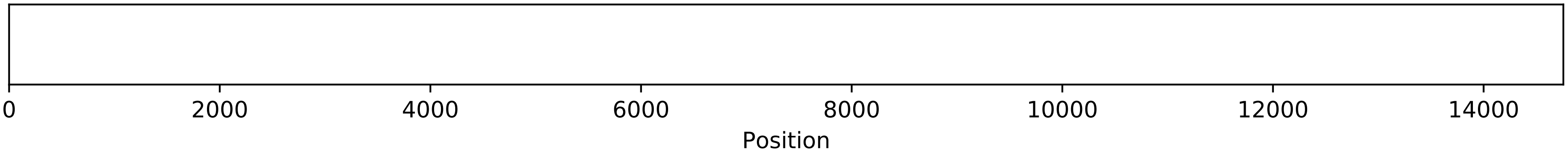


MA2 vs MM1
Number of differences: 149

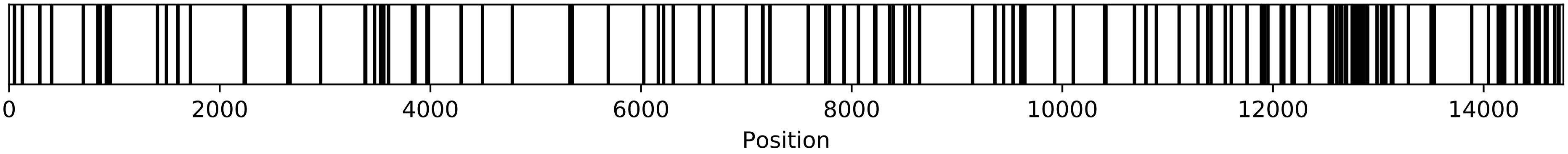


MA2 vs MM2
Number of differences: 148

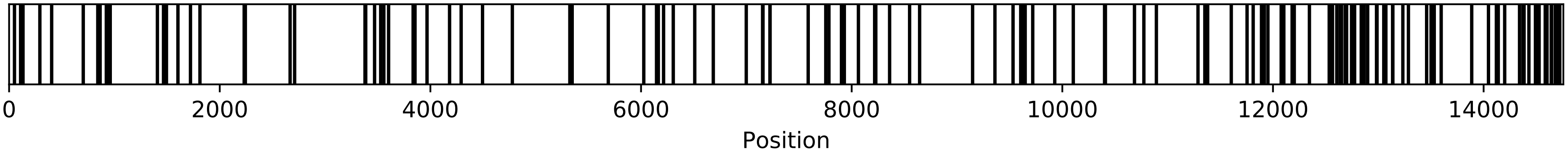


MM1 vs MM2
Number of differences: 72

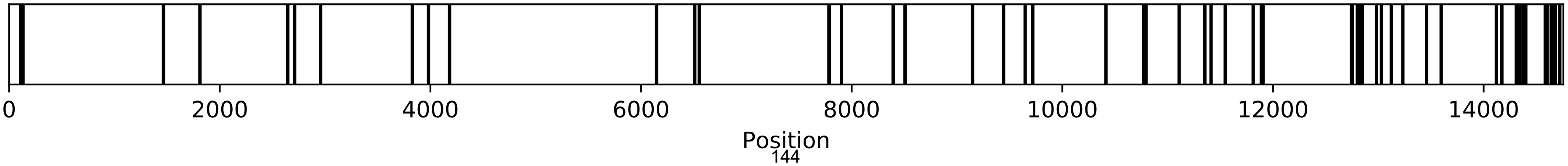

### Region: 473



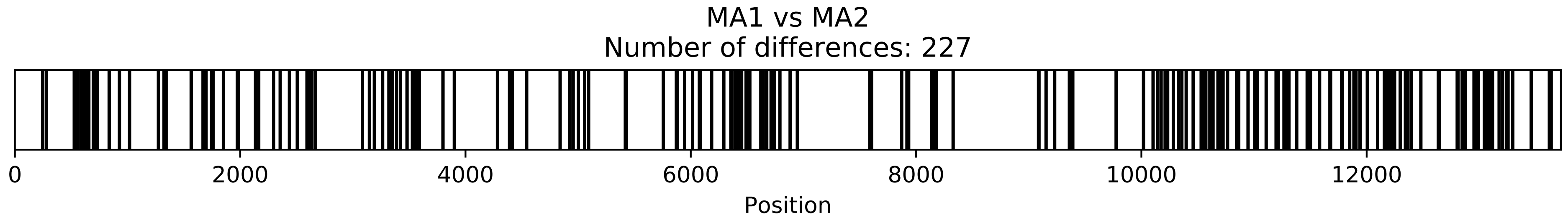
MA1 vs MA2
Number of differences: 227
0
2000
4000
6000
8000
10000
12000
Position

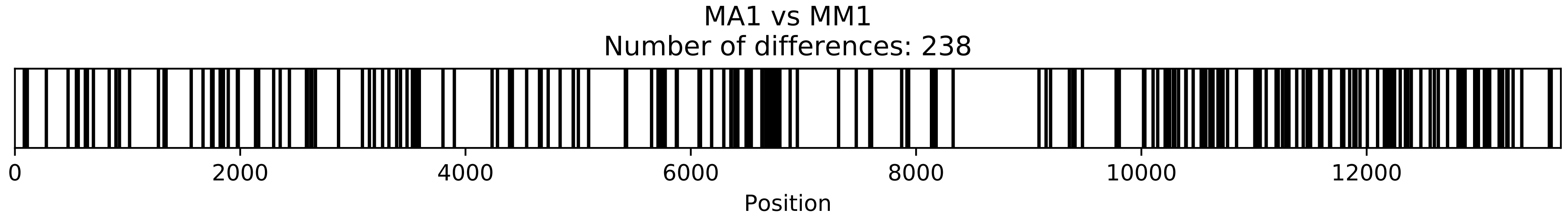
MA1 vs MM1
Number of differences: 238
0
2000
4000
6000
8000
10000
12000
Position

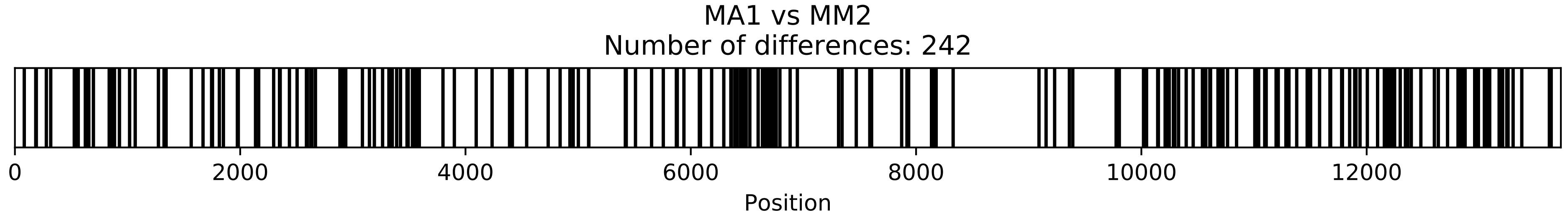
MA1 vs MM2
Number of differences: 242
0
2000
4000
6000
8000
10000
12000
Position

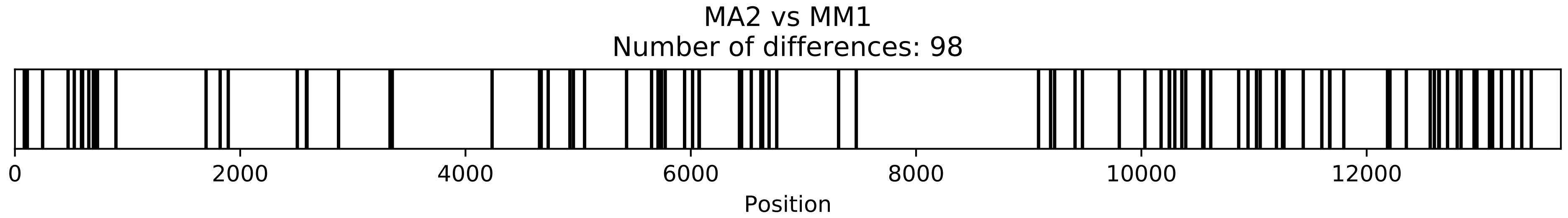
MA2 vs MM1
Number of differences: 98
0
2000
4000
6000
8000
10000
12000
Position

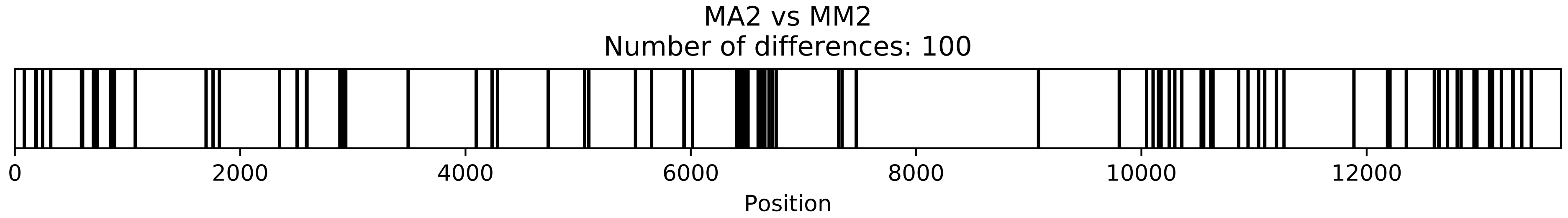
MA2 vs MM2
Number of differences: 100
0
2000
4000
6000
8000
10000
12000
Position

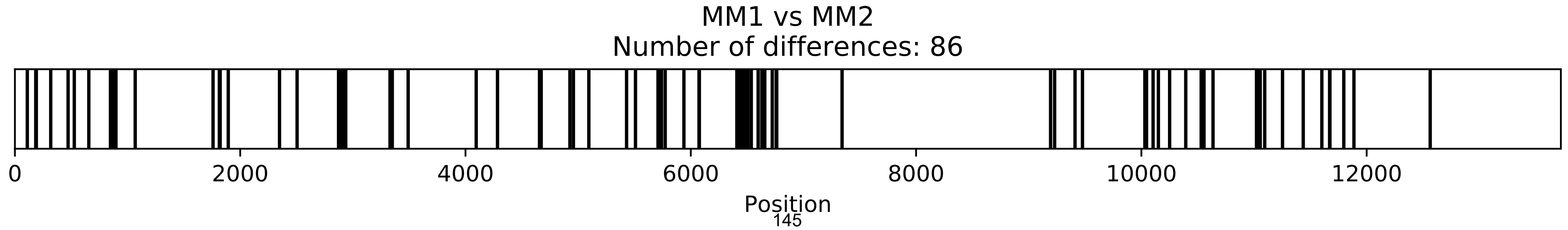
MM1 vs MM2
Number of differences: 86
0
2000
4000
6000
8000
10000
12000
Position

### Region: 480



MA1 vs MA2
Number of differences: 359

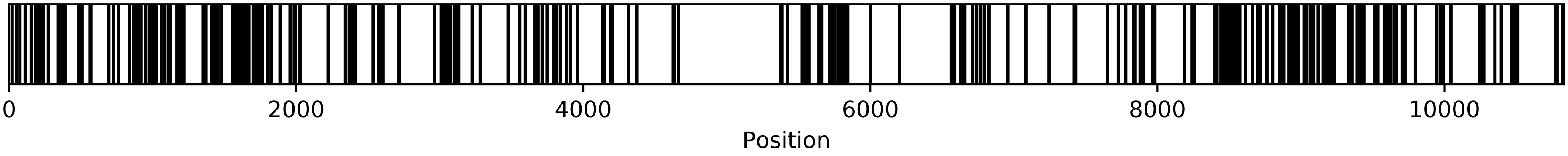


MA1 vs MM1
Number of differences: 165

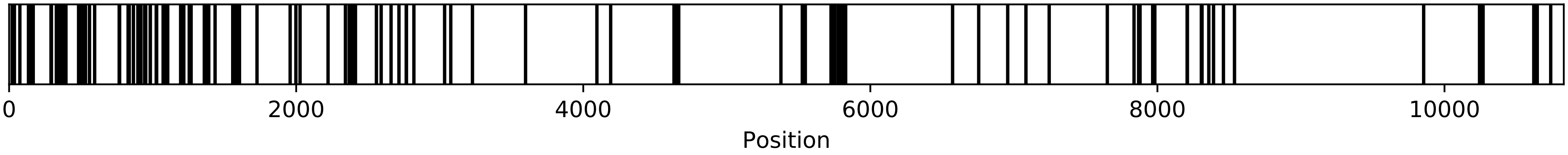


MA1 vs MM2
Number of differences: 164

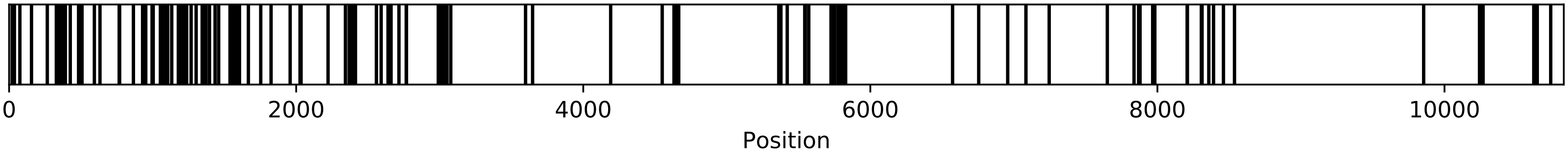


MA2 vs MM1
Number of differences: 354

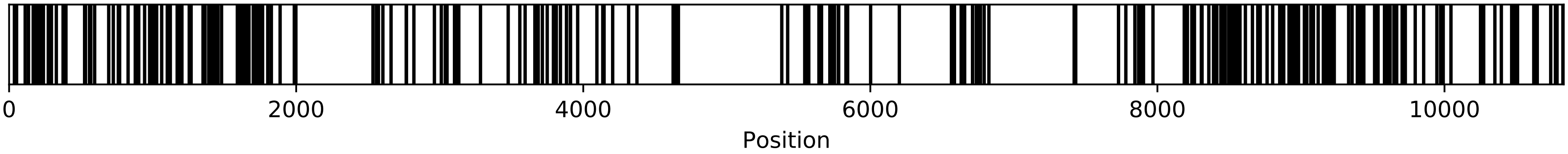


MA2 vs MM2
Number of differences: 350

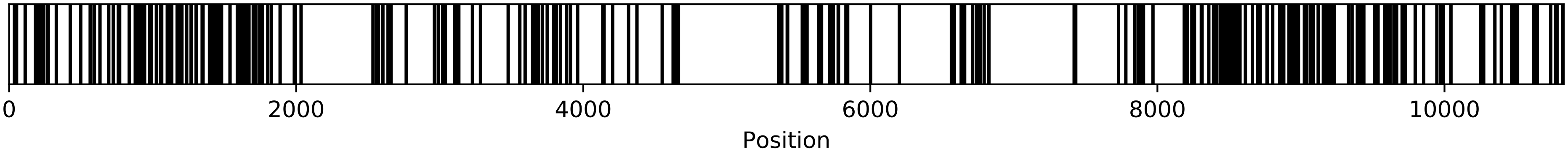


MM1 vs MM2
Number of differences: 64

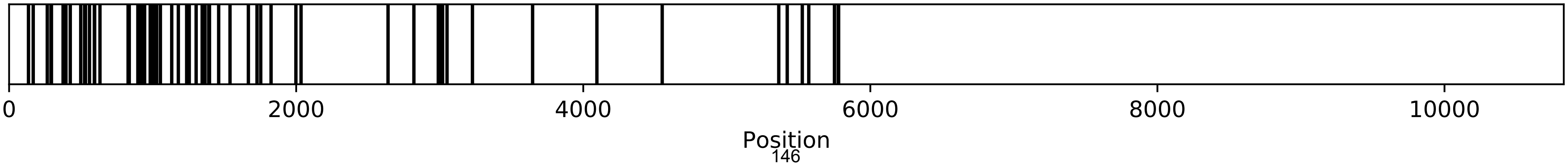

### Region: 483



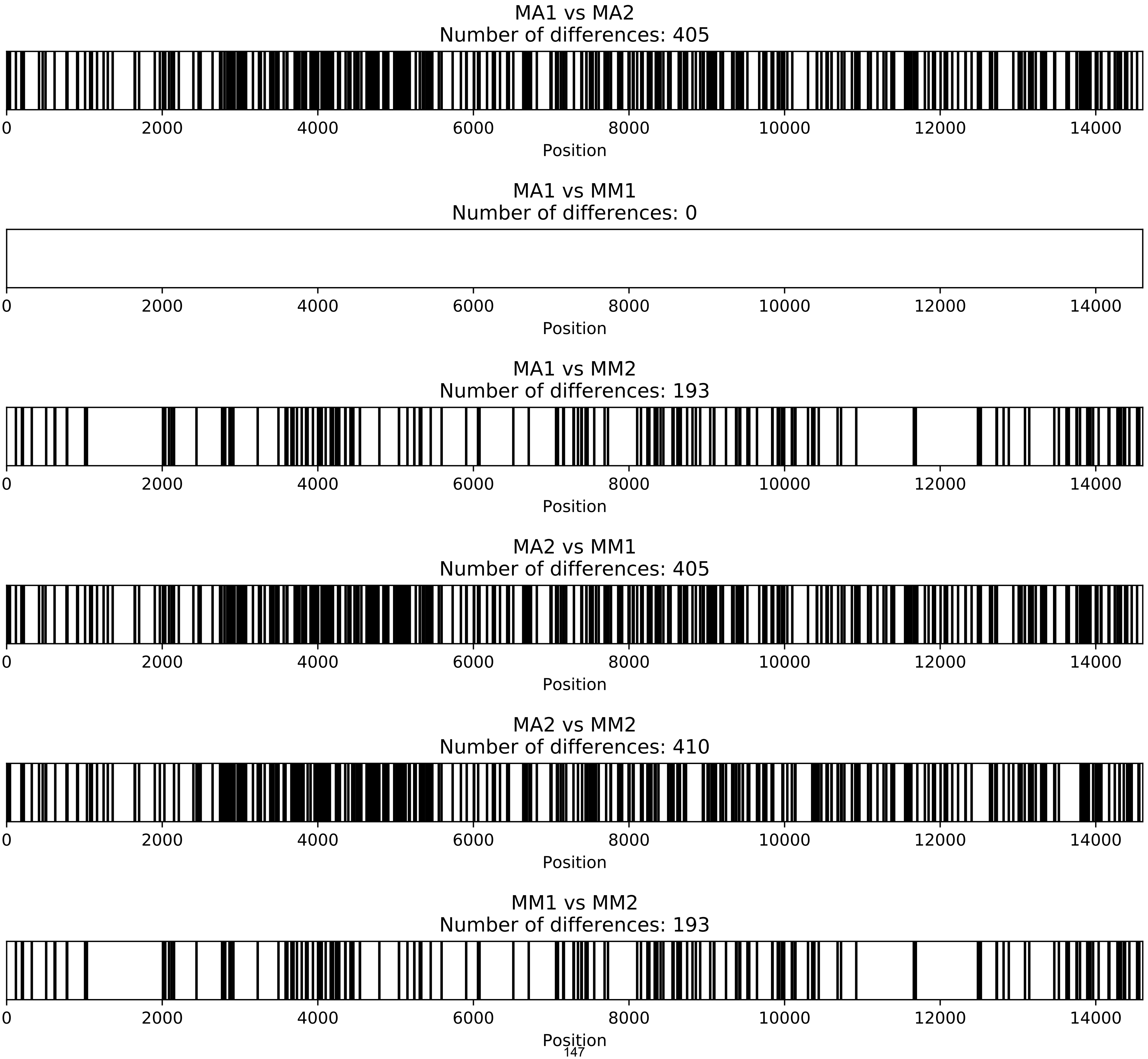
MA1 vs MA2
Number of differences: 405
0
2000
4000
6000
8000
10000
12000
14000
Position
MA1 vs MM1
Number of differences: 0
Position
MA1 vs MM2
Number of differences: 193
Position
MA2 vs MM1
Number of differences: 405
Position
MA2 vs MM2
Number of differences: 410
Position
MM1 vs MM2
Number of differences: 193
Position

### Region: 485



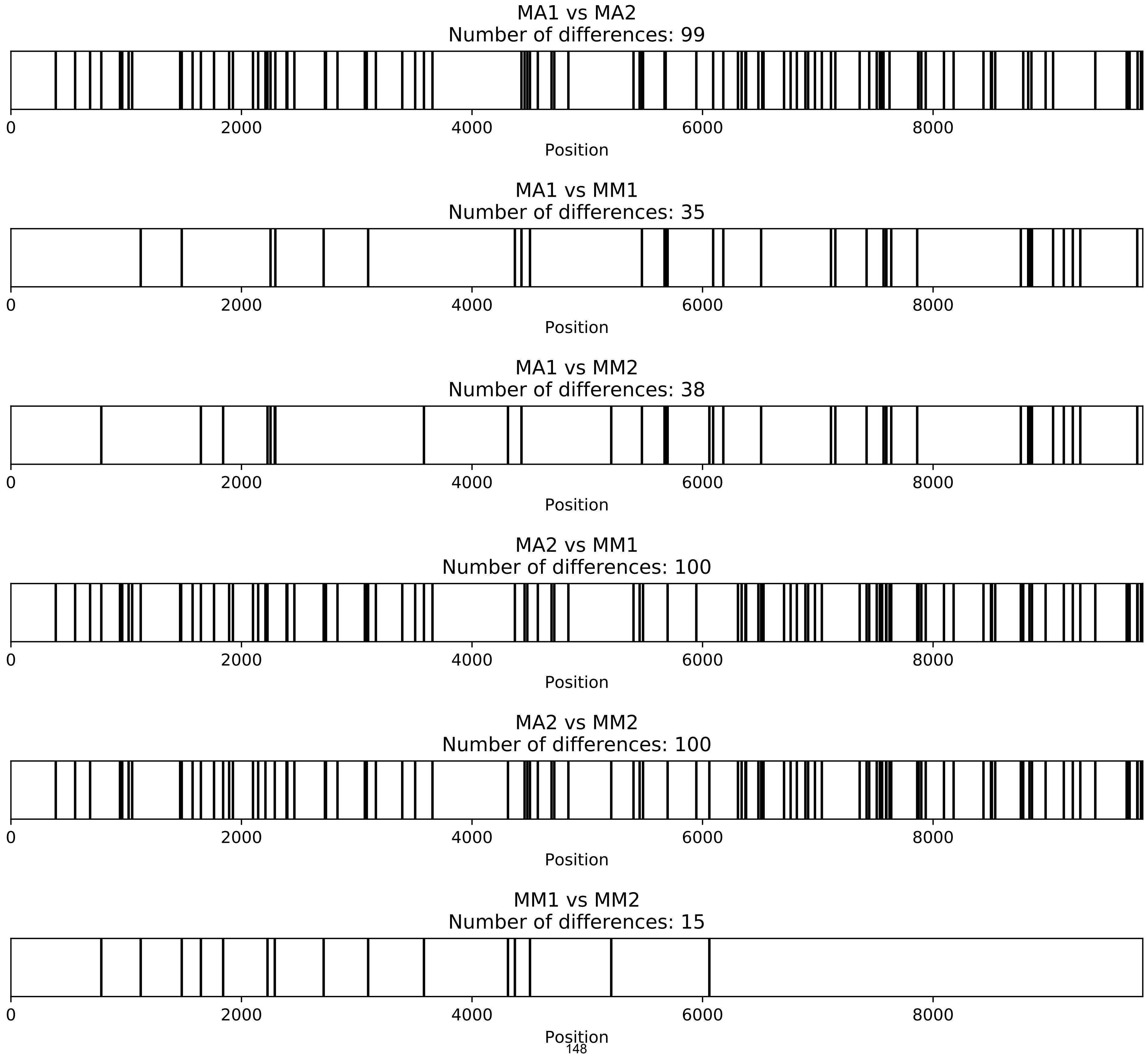
MA1 vs MA2
Number of differences: 99
MA1 vs MM1
Number of differences: 35
MA1 vs MM2
Number of differences: 38
MA2 vs MM1
Number of differences: 100
MA2 vs MM2
Number of differences: 100
MM1 vs MM2
Number of differences: 15
0
2000
4000
6000
8000
Position

### Region: 488



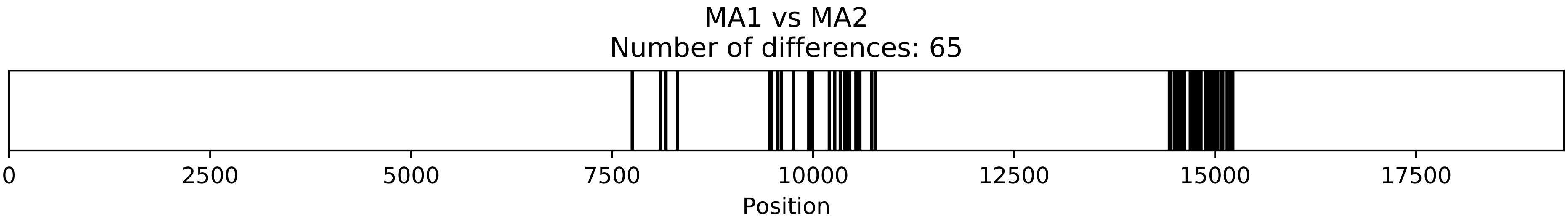
MA1 vs MA2
Number of differences: 65
0
2500
5000
7500
10000
12500
15000
17500
Position

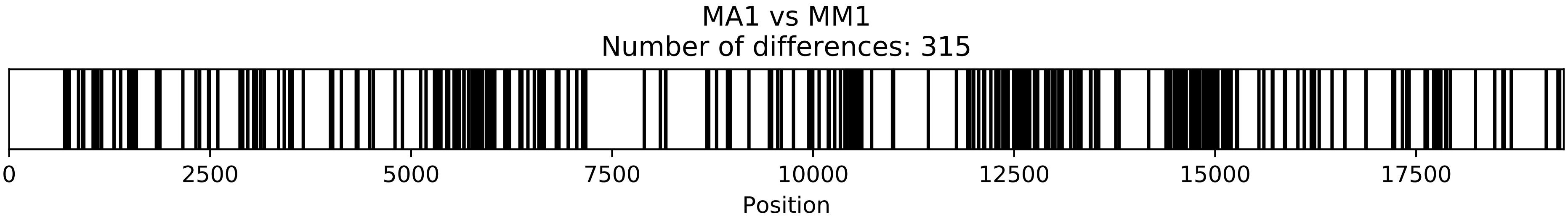
MA1 vs MM1
Number of differences: 315
0
2500
5000
7500
10000
12500
15000
17500
Position

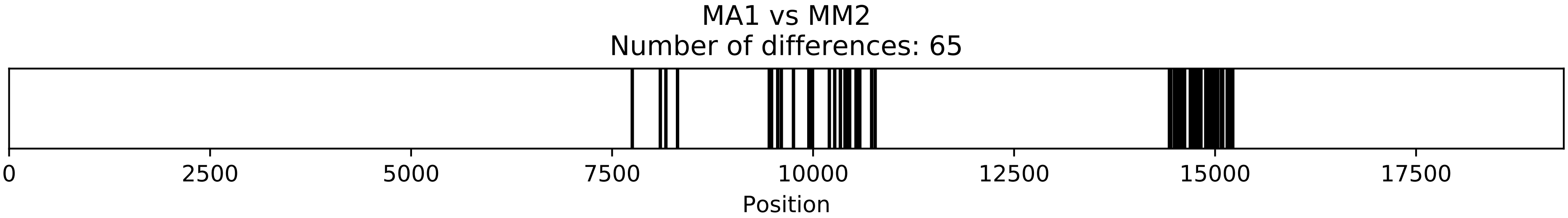
MA1 vs MM2
Number of differences: 65
0
2500
5000
7500
10000
12500
15000
17500
Position

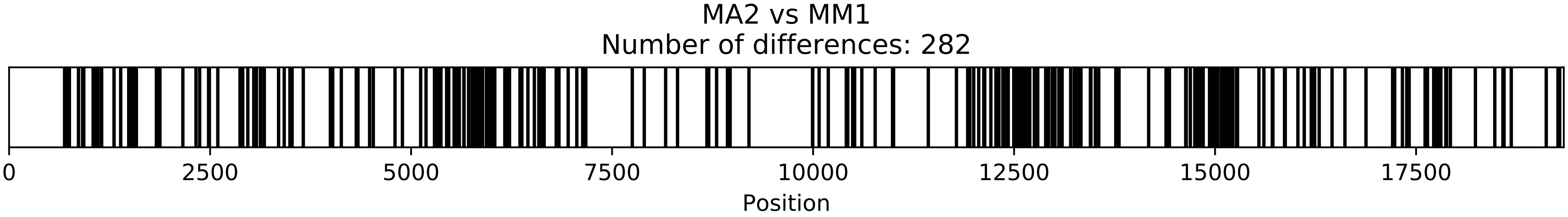
MA2 vs MM1
Number of differences: 282
0
2500
5000
7500
10000
12500
15000
17500
Position

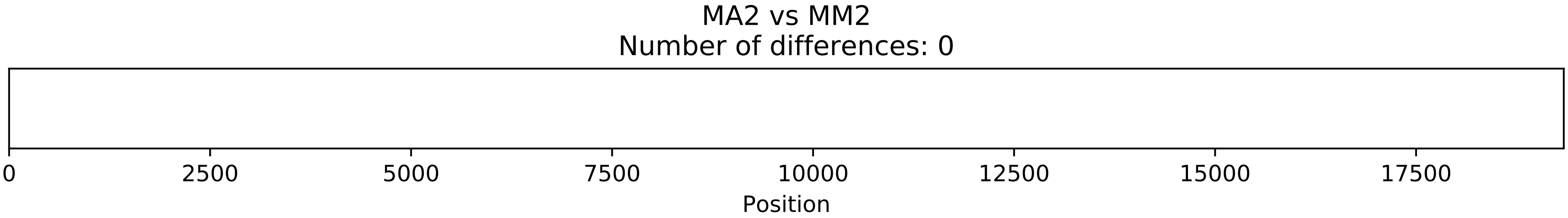
MA2 vs MM2
Number of differences: 0
0
2500
5000
7500
10000
12500
15000
17500
Position

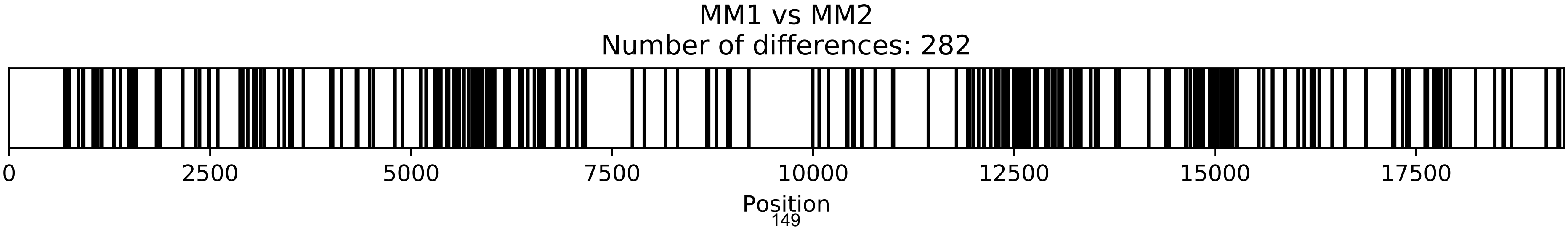
MM1 vs MM2
Number of differences: 282
0
2500
5000
7500
10000
12500
15000
17500
Position

### Region: 489



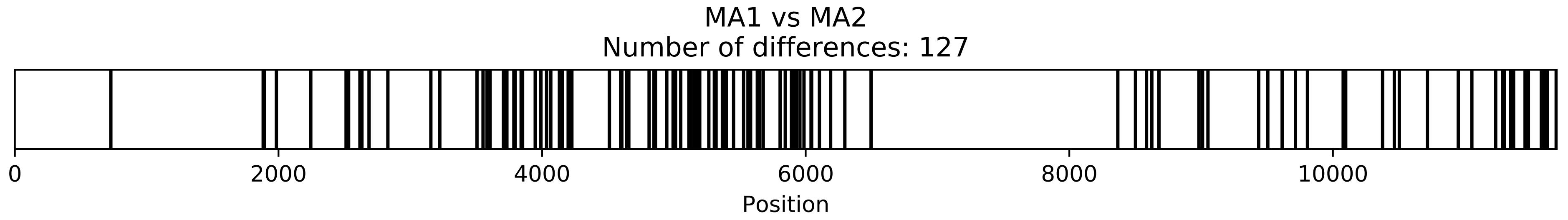
MA1 vs MA2
Number of differences: 127
0
2000
4000
6000
8000
10000
Position

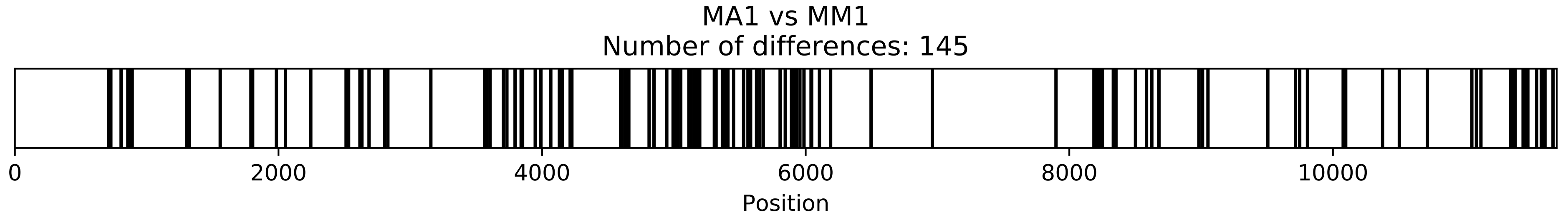
MA1 vs MM1
Number of differences: 145
0
2000
4000
6000
8000
10000
Position

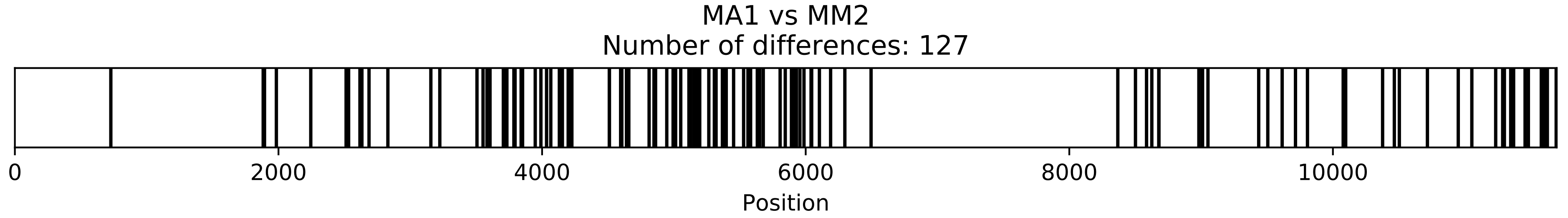
MA1 vs MM2
Number of differences: 127
0
2000
4000
6000
8000
10000
Position

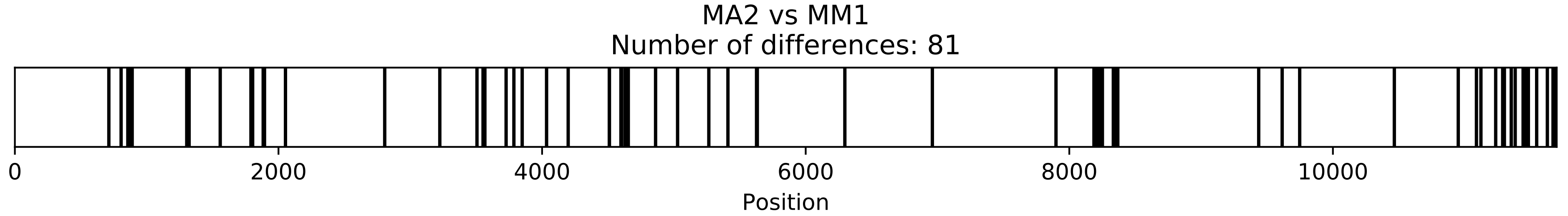
MA2 vs MM1
Number of differences: 81
0
2000
4000
6000
8000
10000
Position

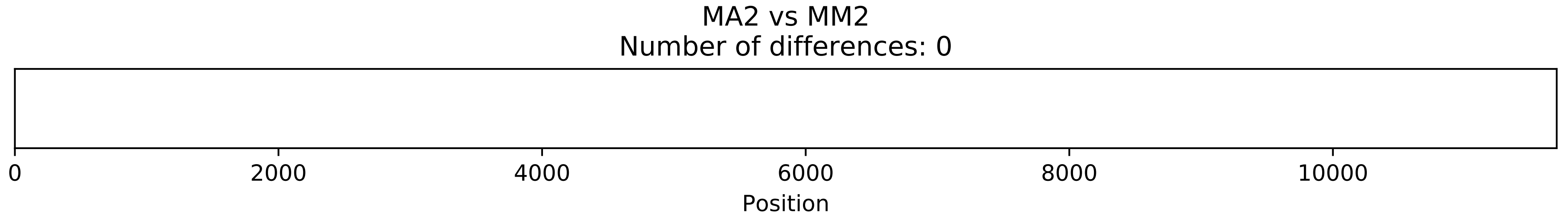
MA2 vs MM2
Number of differences: 0
0
2000
4000
6000
8000
10000
Position

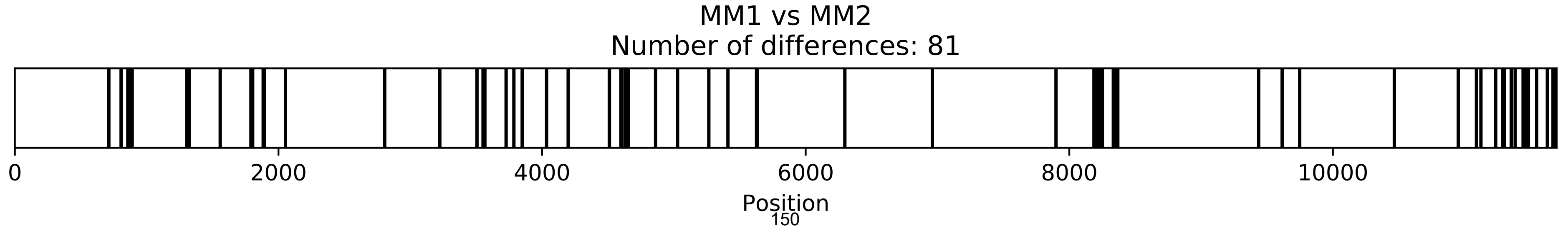
MM1 vs MM2
Number of differences: 81
0
2000
4000
6000
8000
10000
Position

### Region: 491



## MA1 vs MA2
Number of differences: 273

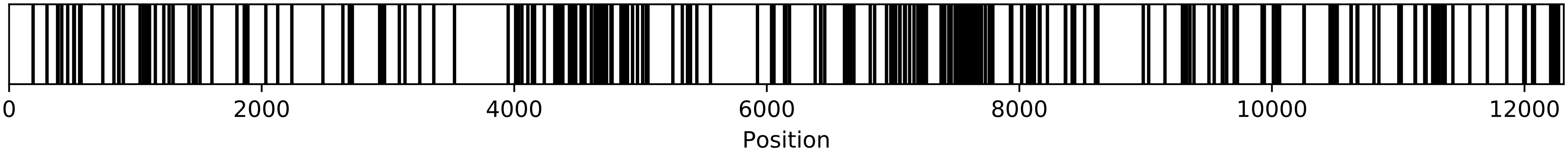


## MA1 vs MM1
Number of differences: 109

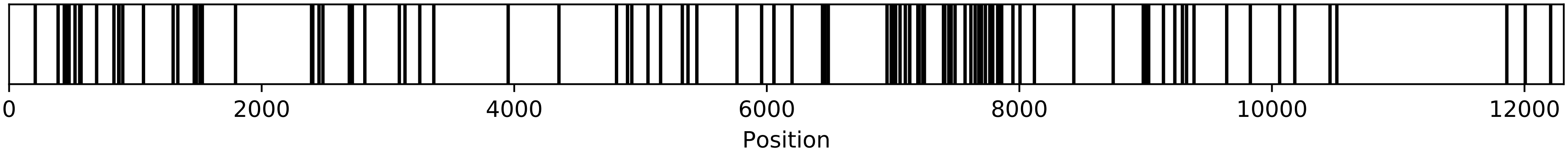


## MA1 vs MM2
Number of differences: 102

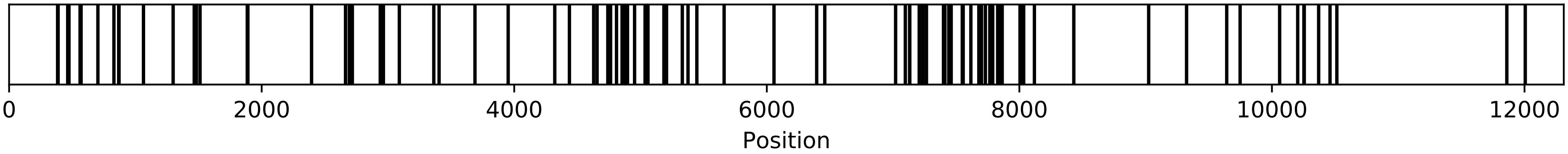


## MA2 vs MM1
Number of differences: 260

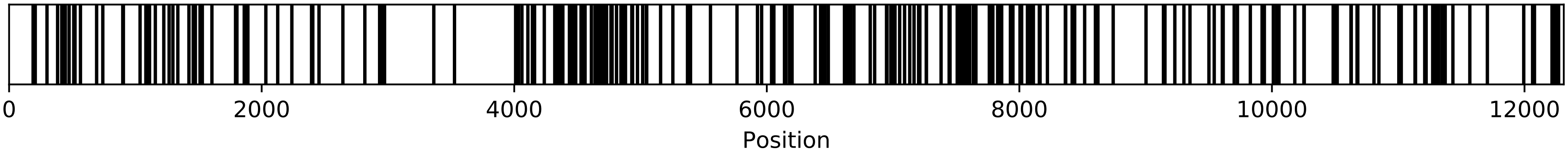


## MA2 vs MM2
Number of differences: 258

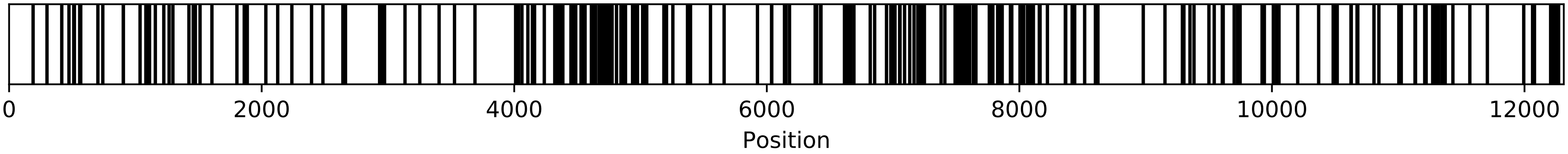


## MM1 vs MM2
Number of differences: 96

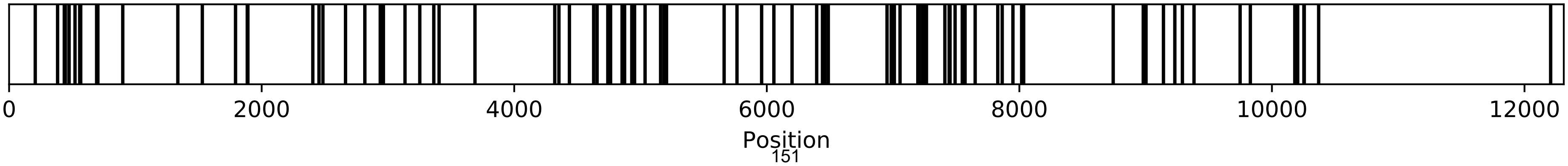

### Region: 493



MA1 vs MA2
Number of differences: 200

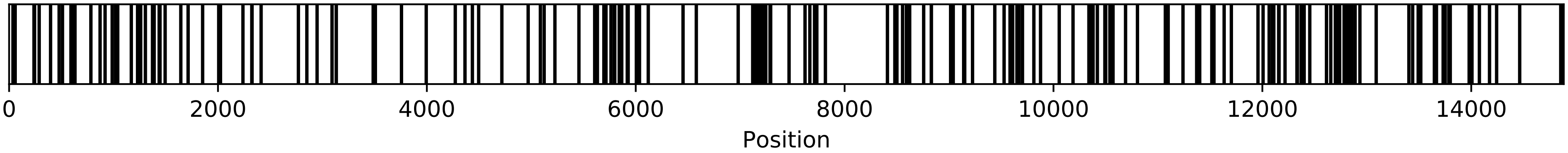


MA1 vs MM1
Number of differences: 103

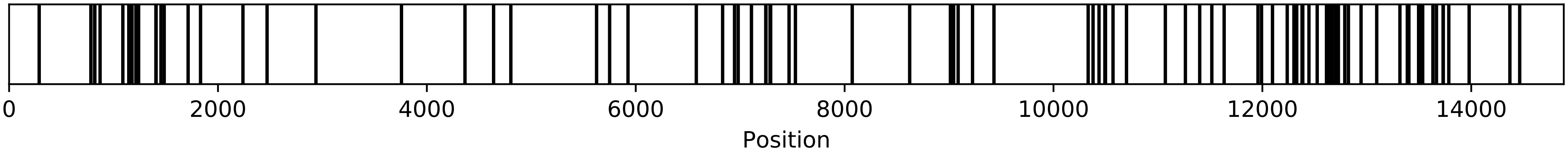


MA1 vs MM2
Number of differences: 89

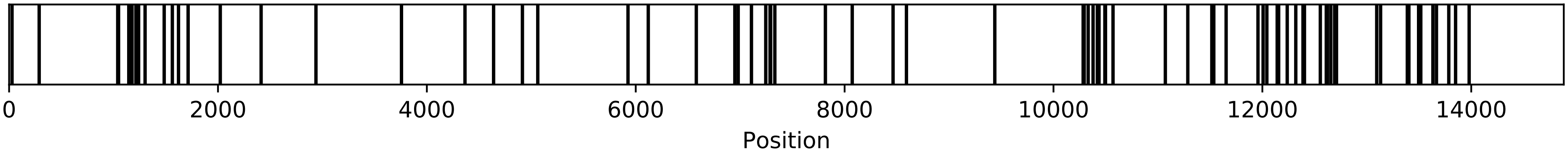


MA2 vs MM1
Number of differences: 217

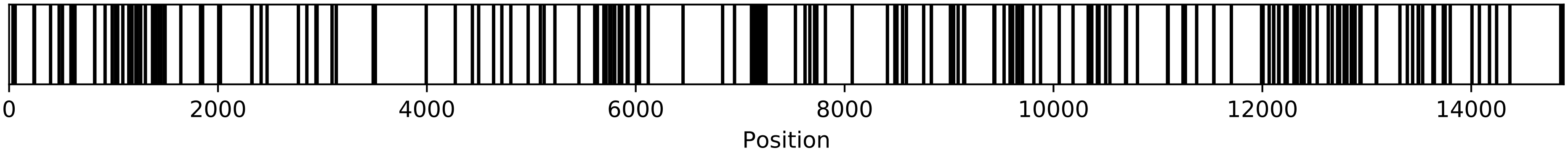


MA2 vs MM2
Number of differences: 212

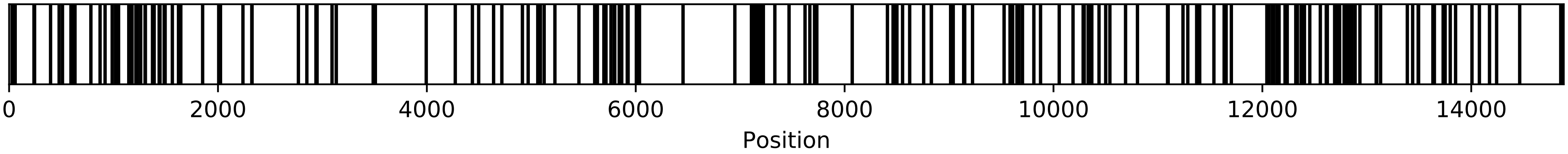


MM1 vs MM2
Number of differences: 91

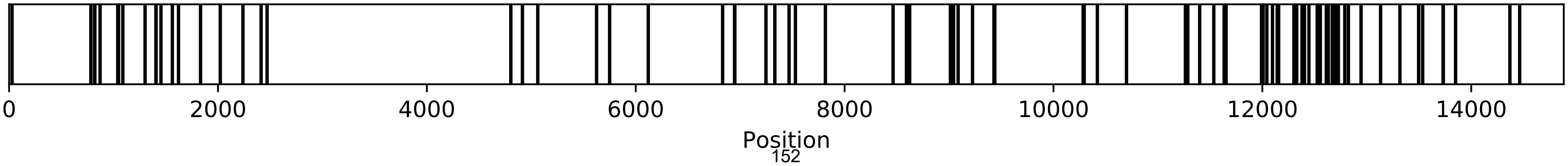

### Region: 495



MA1 vs MA2
Number of differences: 308

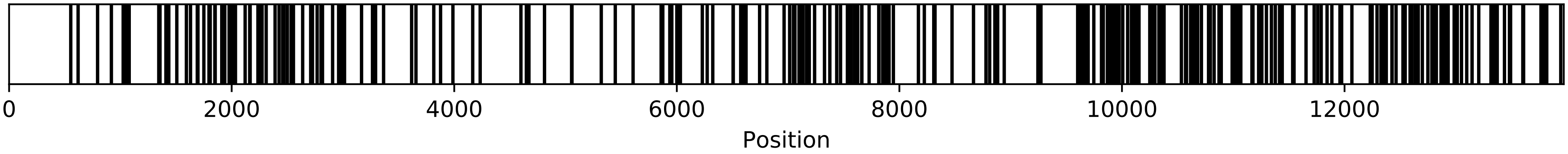


MA1 vs MM1
Number of differences: 296

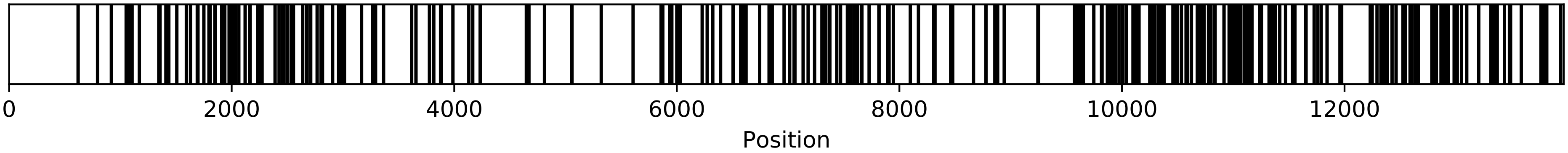


MA1 vs MM2
Number of differences: 308

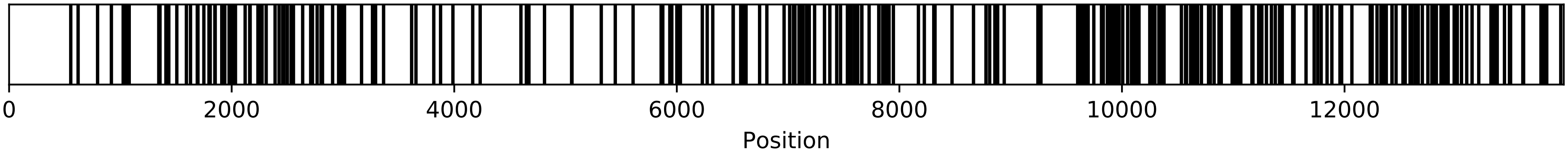


MA2 vs MM1
Number of differences: 141

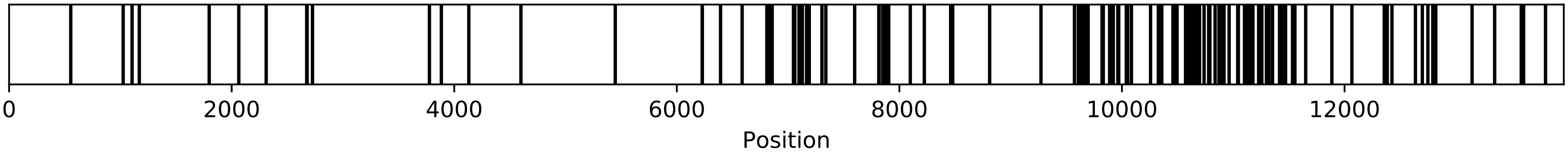


MA2 vs MM2
Number of differences: 0

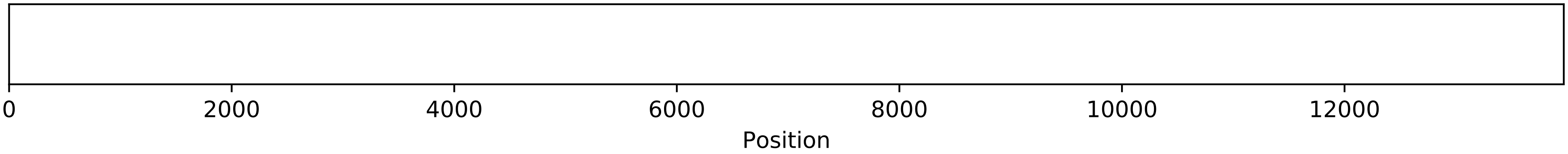


MM1 vs MM2
Number of differences: 141

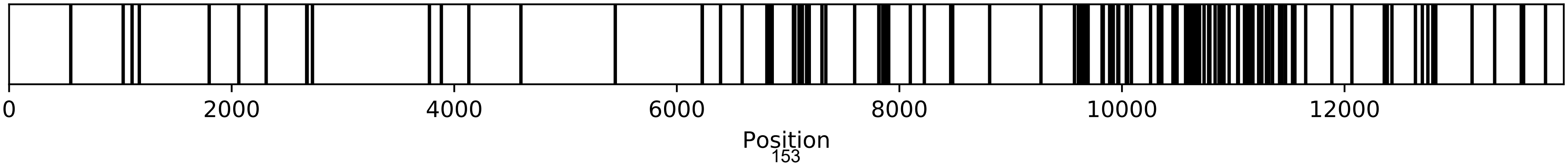

### Region: 497



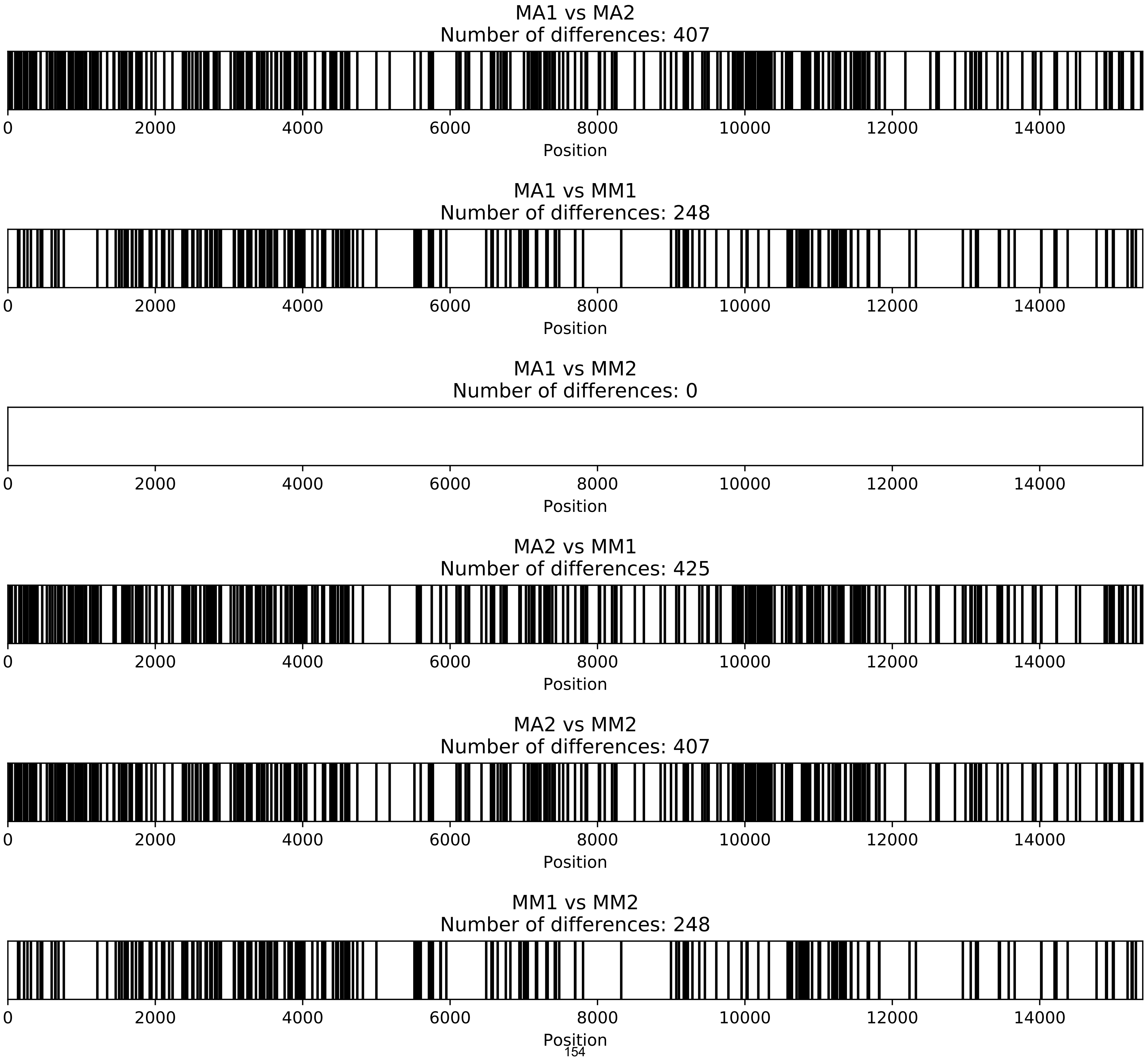
MA1 vs MA2
Number of differences: 407
MA1 vs MM1
Number of differences: 248
MA1 vs MM2
Number of differences: 0
MA2 vs MM1
Number of differences: 425
MA2 vs MM2
Number of differences: 407
MM1 vs MM2
Number of differences: 248
0
2000
4000
6000
8000
10000
12000
14000
Position

### Region: 499



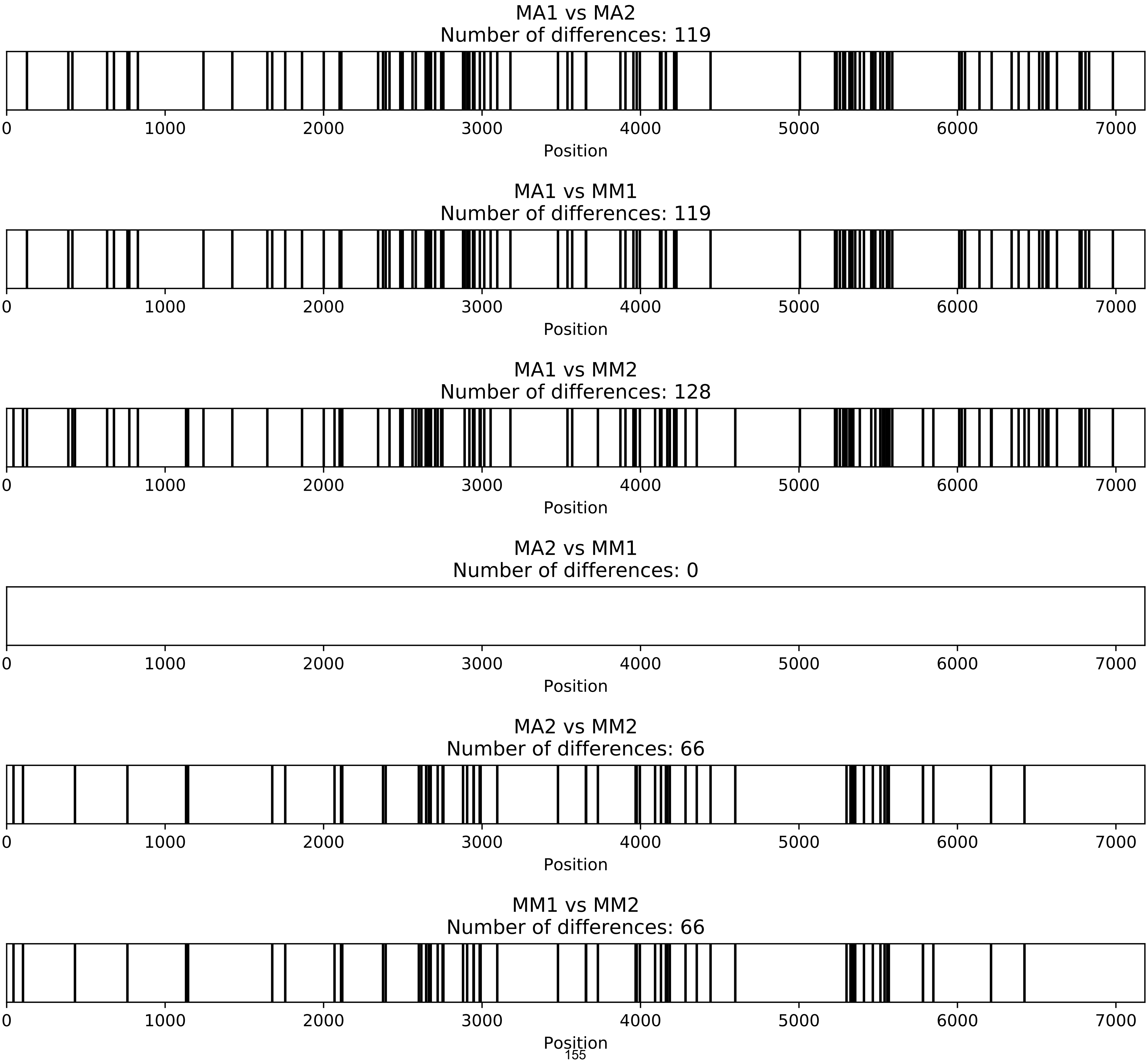
MA1 vs MA2
Number of differences: 119
MA1 vs MM1
Number of differences: 119
MA1 vs MM2
Number of differences: 128
MA2 vs MM1
Number of differences: 0
MA2 vs MM2
Number of differences: 66
MM1 vs MM2
Number of differences: 66
0 1000 2000 3000 4000 5000 6000 7000
Position

### Region: 500



MA1 vs MA2
Number of differences: 384

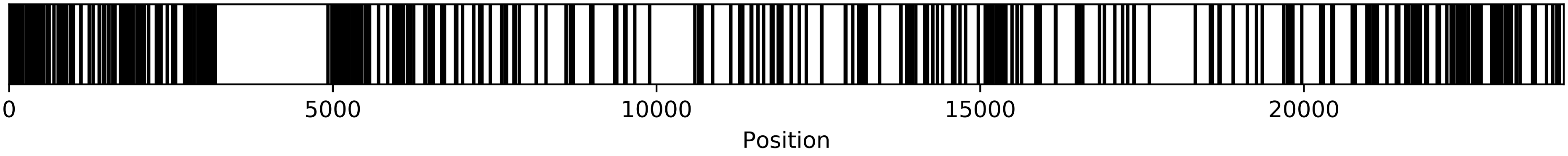


MA1 vs MM1
Number of differences: 280

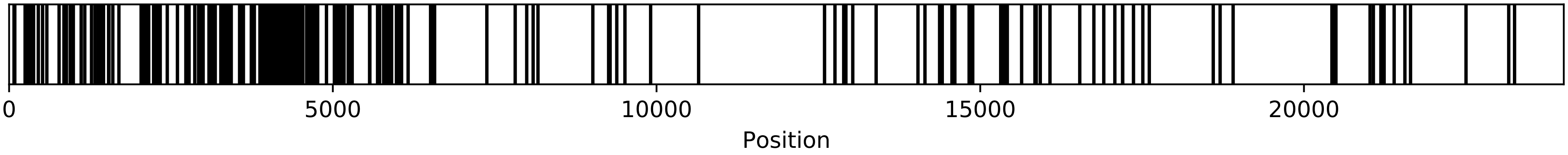


MA1 vs MM2
Number of differences: 0

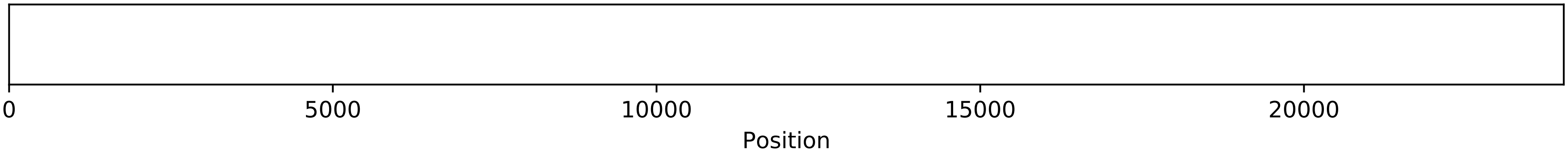


MA2 vs MM1
Number of differences: 505

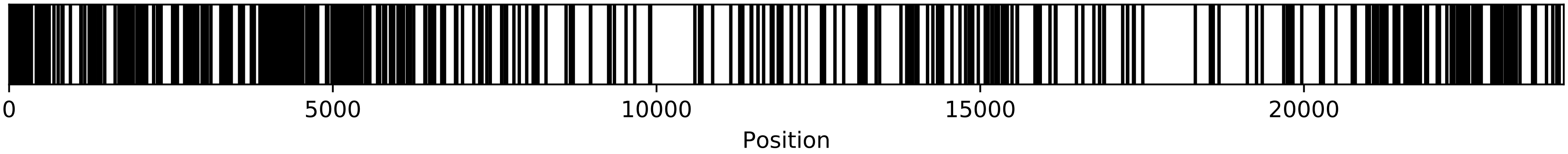


MA2 vs MM2
Number of differences: 384

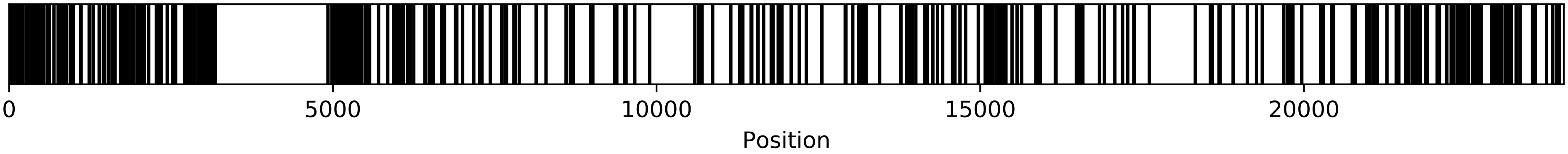


MM1 vs MM2
Number of differences: 280

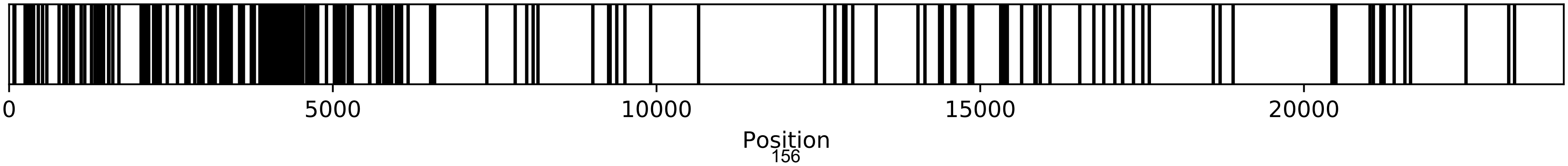

### Region: 501



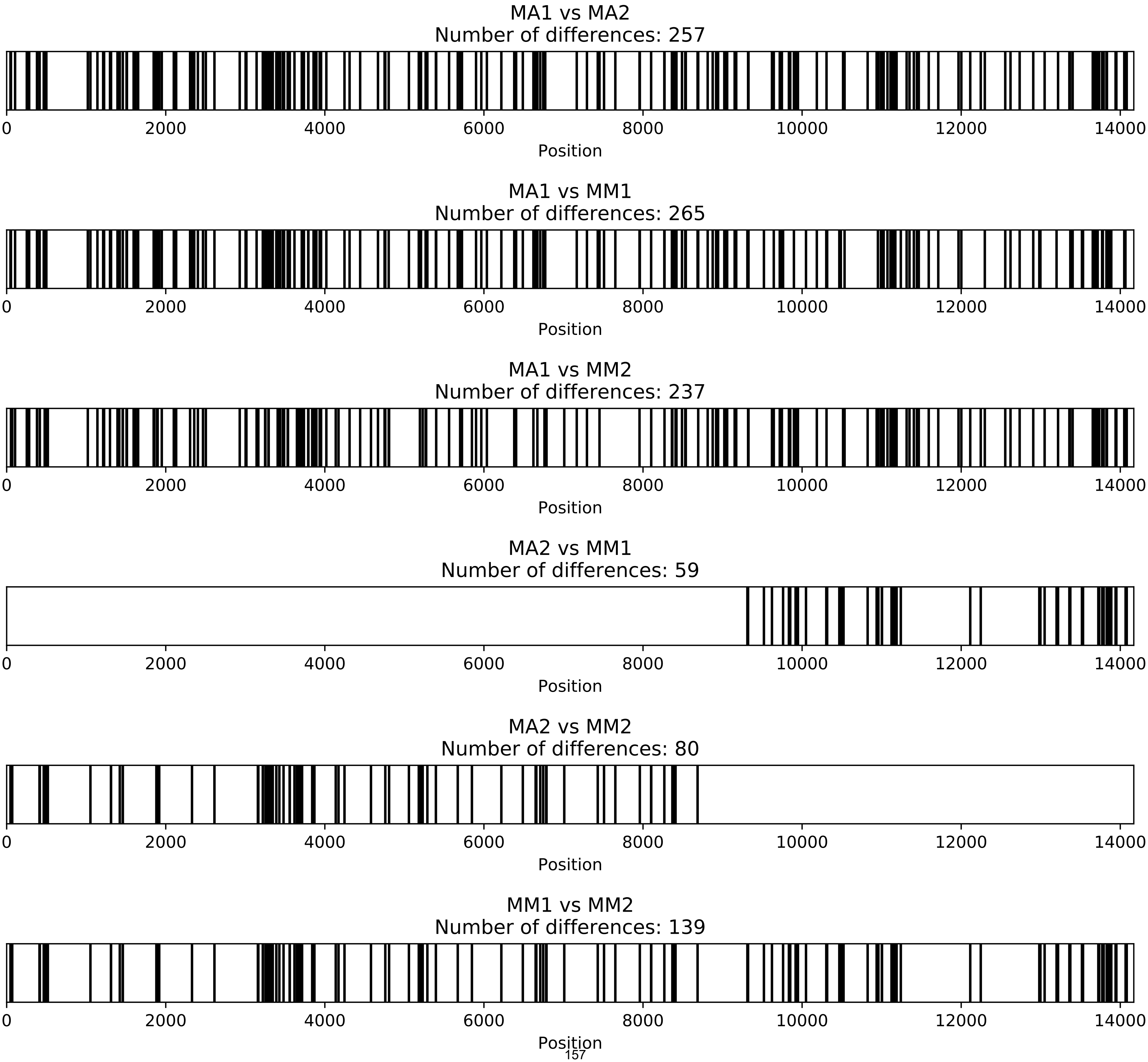
MA1 vs MA2
Number of differences: 257
Position
MA1 vs MM1
Number of differences: 265
Position
MA1 vs MM2
Number of differences: 237
Position
MA2 vs MM1
Number of differences: 59
Position
MA2 vs MM2
Number of differences: 80
Position
MM1 vs MM2
Number of differences: 139
Position
0
2000
4000
6000
8000
10000
12000
14000

### Region: 503



MA1 vs MA2
Number of differences: 277

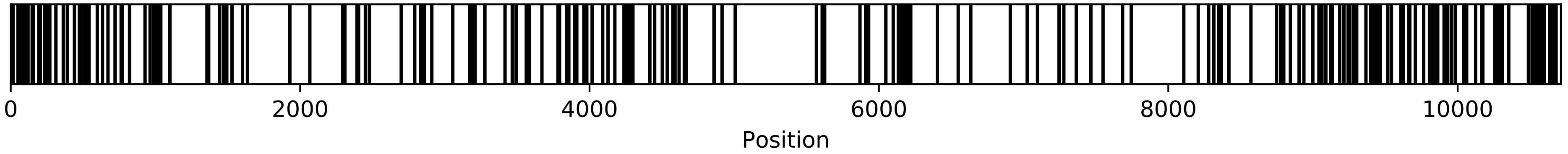


MA1 vs MM1
Number of differences: 0

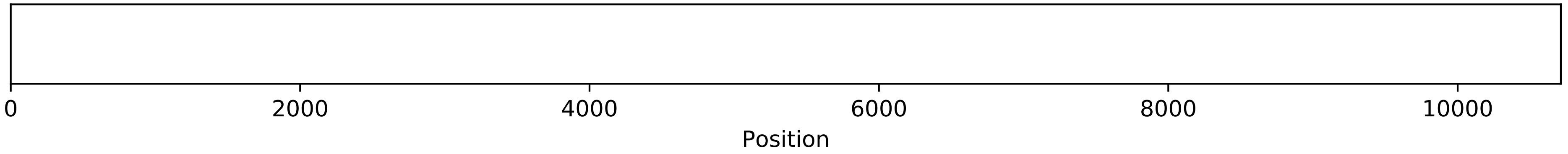


MA1 vs MM2
Number of differences: 147

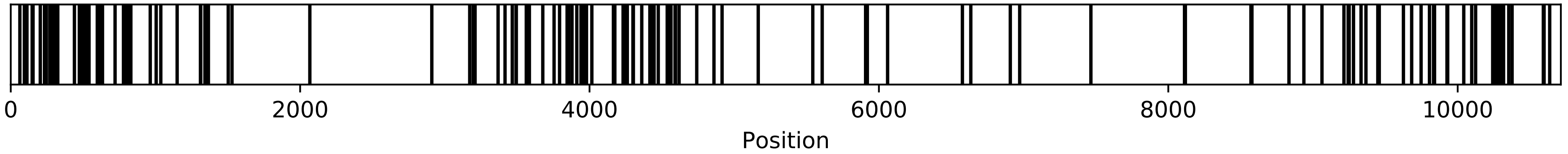


MA2 vs MM1
Number of differences: 277

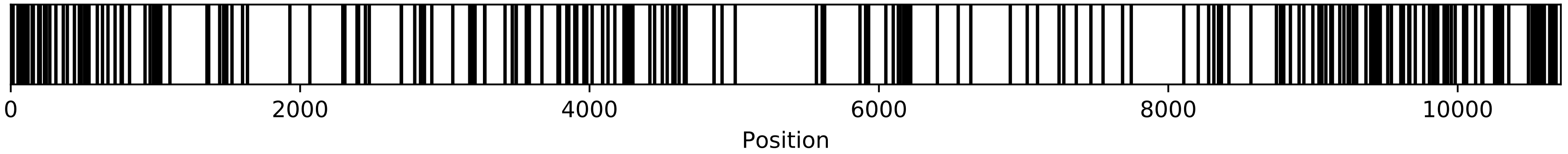


MA2 vs MM2
Number of differences: 272

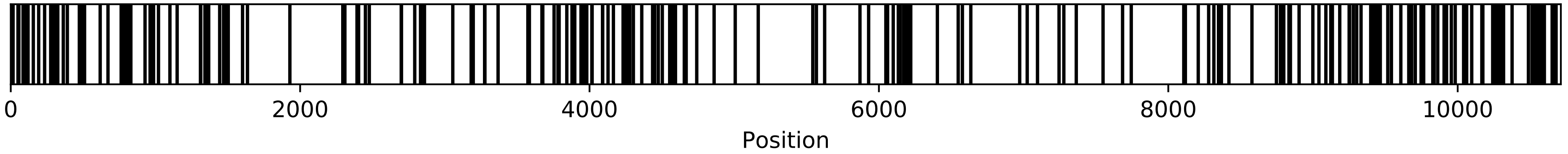


MM1 vs MM2
Number of differences: 147

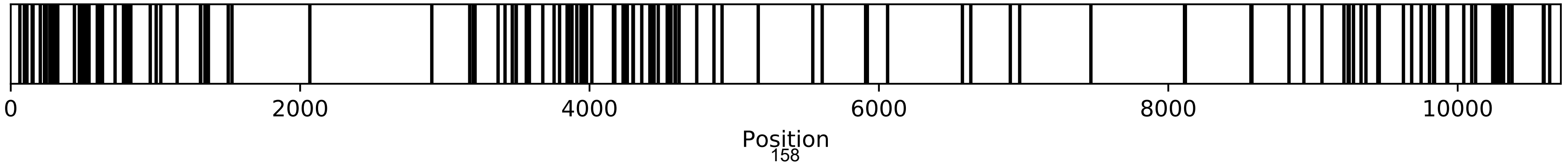

### Region: 508



MA1 vs MA2
Number of differences: 328

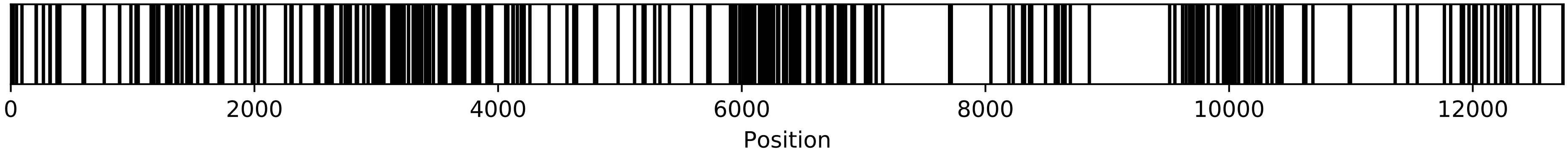


MA1 vs MM1
Number of differences: 324

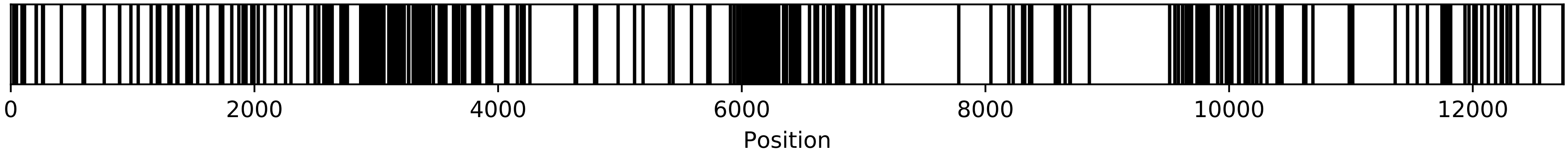


MA1 vs MM2
Number of differences: 337

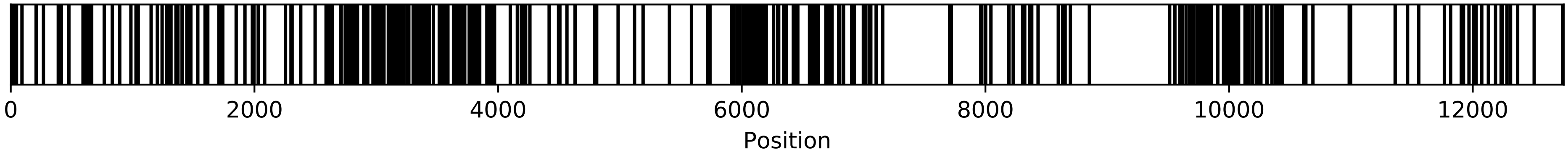


MA2 vs MM1
Number of differences: 152

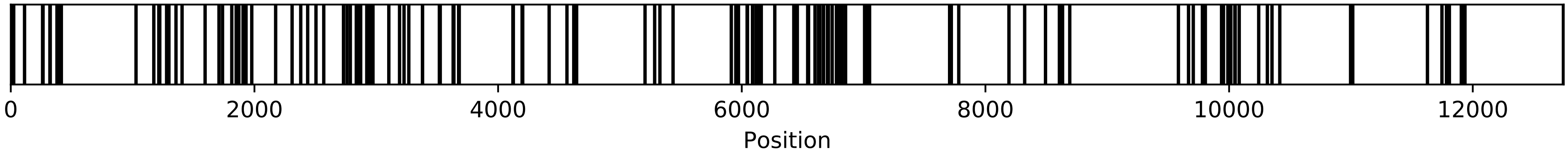


MA2 vs MM2
Number of differences: 153

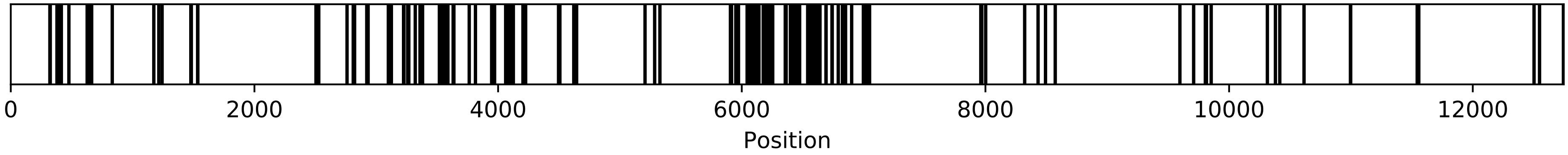


MM1 vs MM2
Number of differences: 192

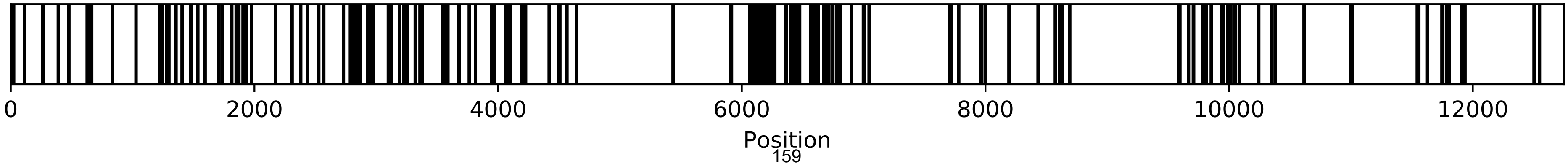

### Region: 509



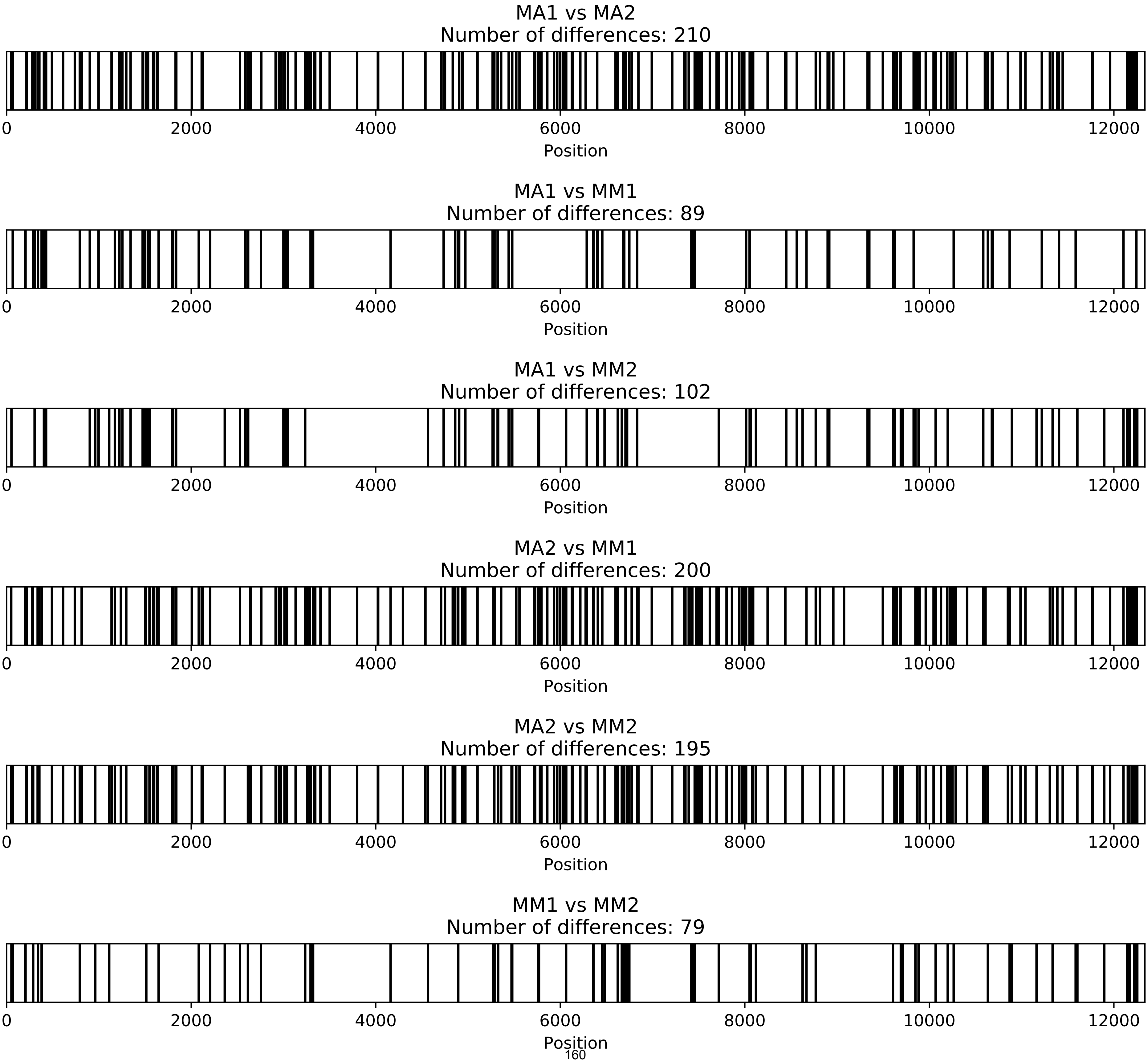
MA1 vs MA2
Number of differences: 210
MA1 vs MM1
Number of differences: 89
MA1 vs MM2
Number of differences: 102
MA2 vs MM1
Number of differences: 200
MA2 vs MM2
Number of differences: 195
MM1 vs MM2
Number of differences: 79
0
2000
4000
6000
8000
10000
12000
Position

### Region: 519



MA1 vs MA2
Number of differences: 240

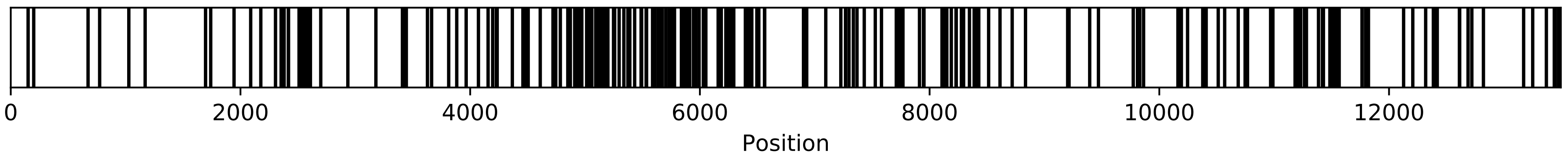


MA1 vs MM1
Number of differences: 0

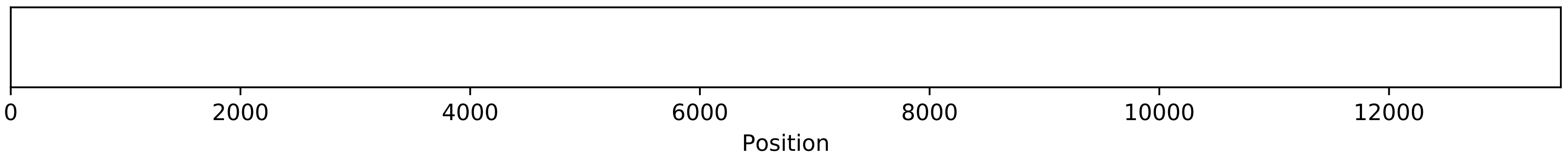


MA1 vs MM2
Number of differences: 44

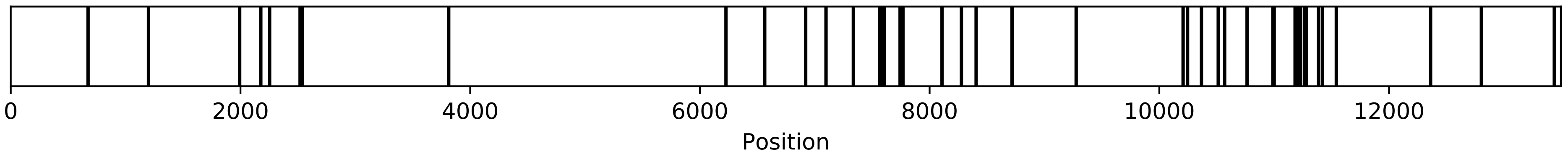


MA2 vs MM1
Number of differences: 240

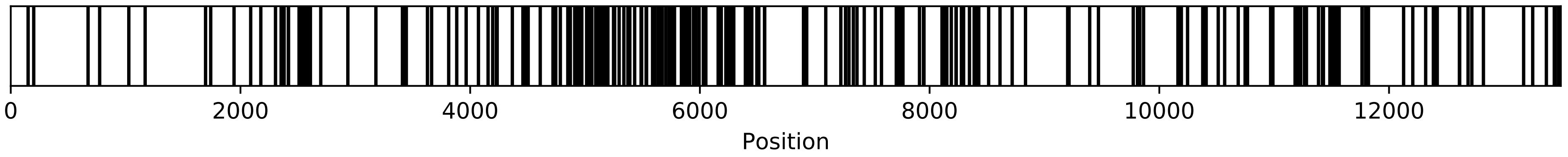


MA2 vs MM2
Number of differences: 229

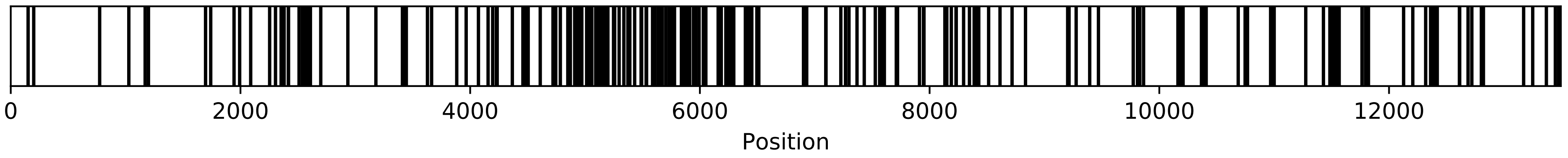


MM1 vs MM2
Number of differences: 44

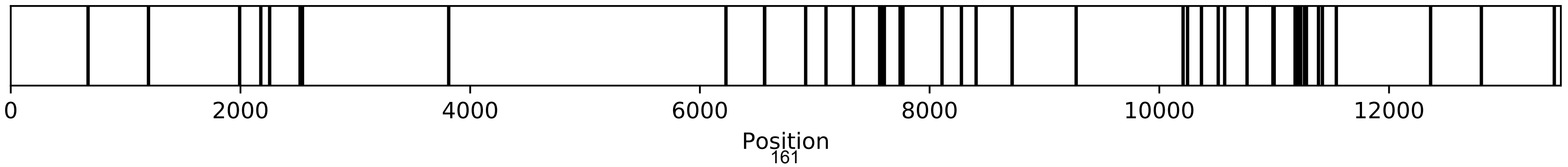

### Region: 520



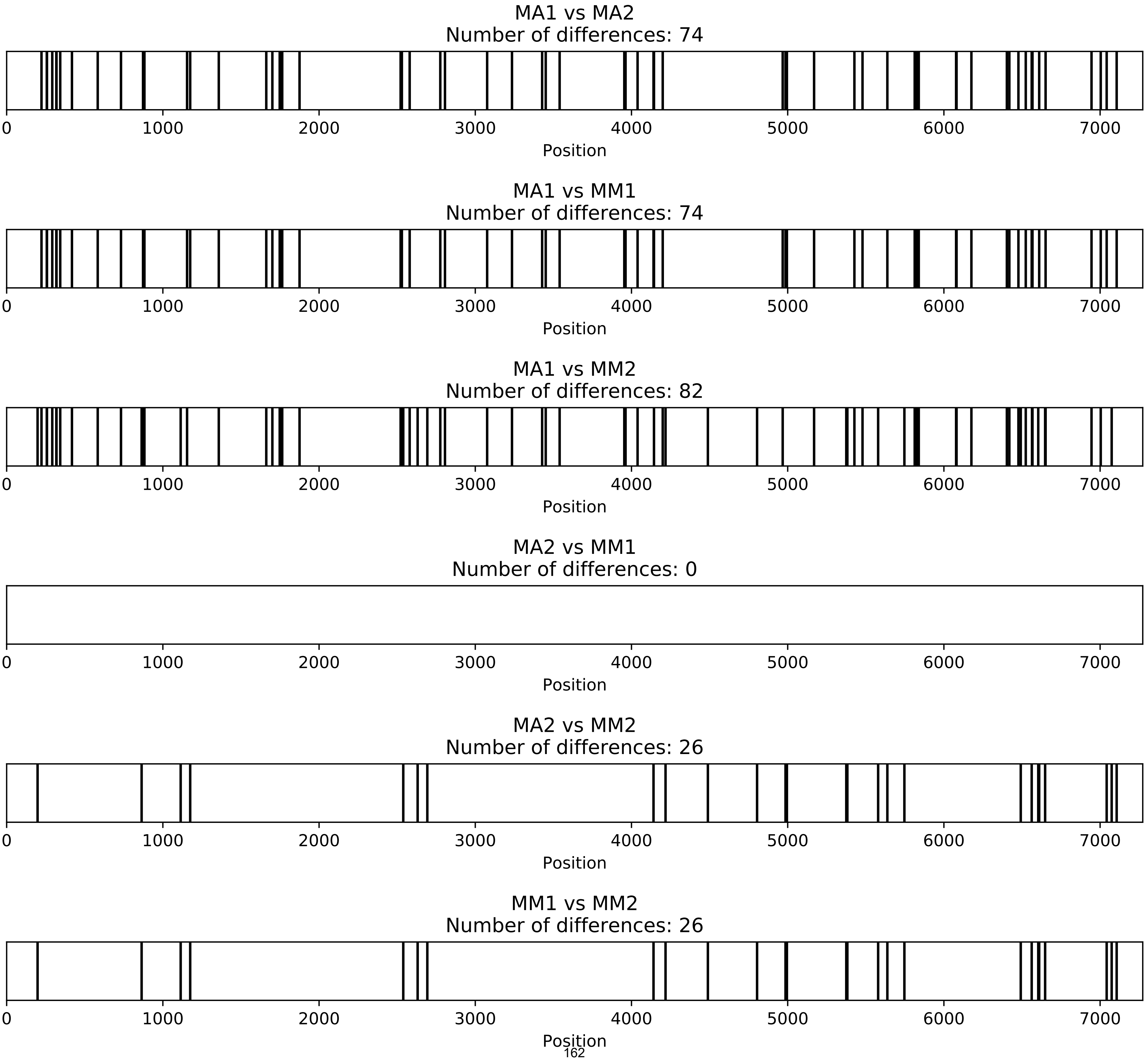
MA1 vs MA2
Number of differences: 74
MA1 vs MM1
Number of differences: 74
MA1 vs MM2
Number of differences: 82
MA2 vs MM1
Number of differences: 0
MA2 vs MM2
Number of differences: 26
MM1 vs MM2
Number of differences: 26
0
1000
2000
3000
4000
5000
6000
7000
Position

### Region: 521



MA1 vs MA2
Number of differences: 187

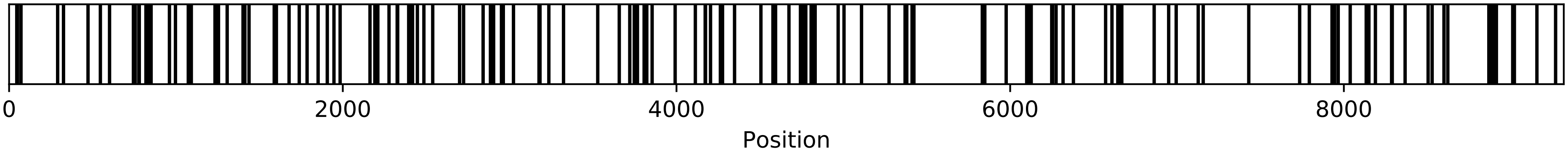


MA1 vs MM1
Number of differences: 51

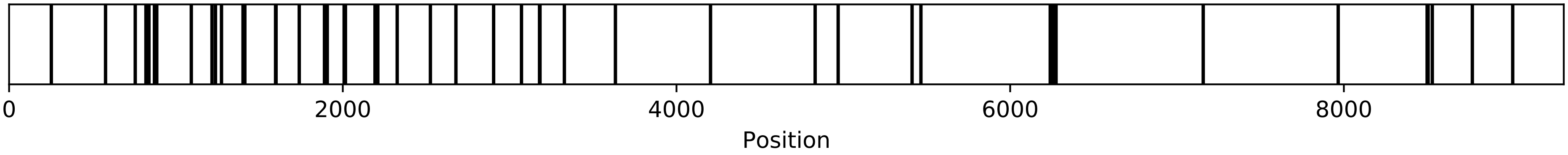


MA1 vs MM2
Number of differences: 0

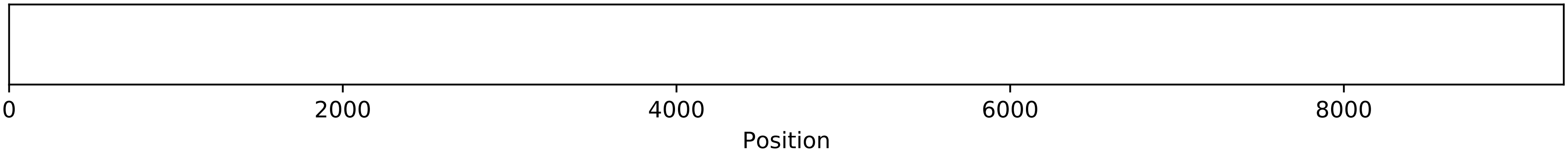


MA2 vs MM1
Number of differences: 185

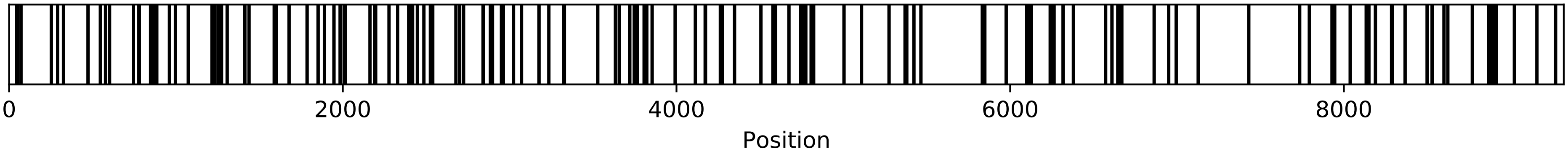


MA2 vs MM2
Number of differences: 187

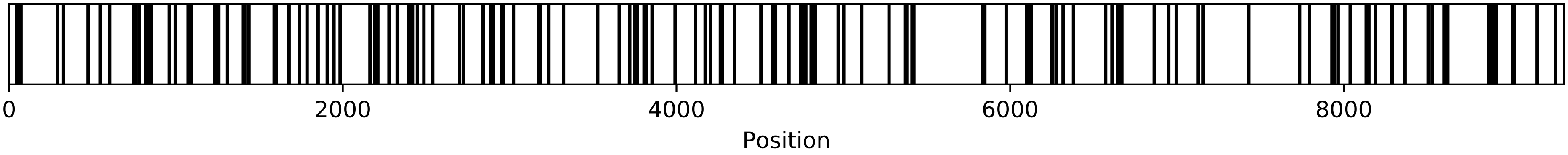


MM1 vs MM2
Number of differences: 51

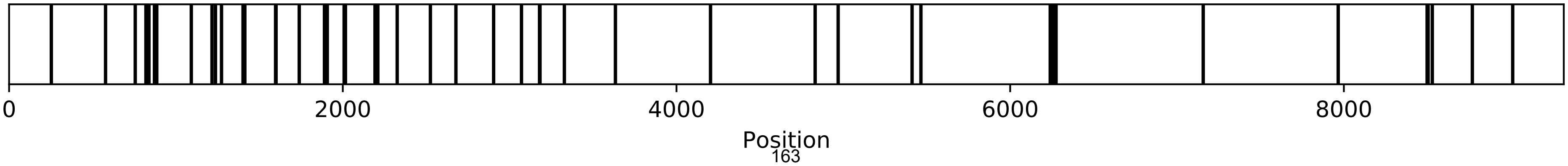

### Region: 524

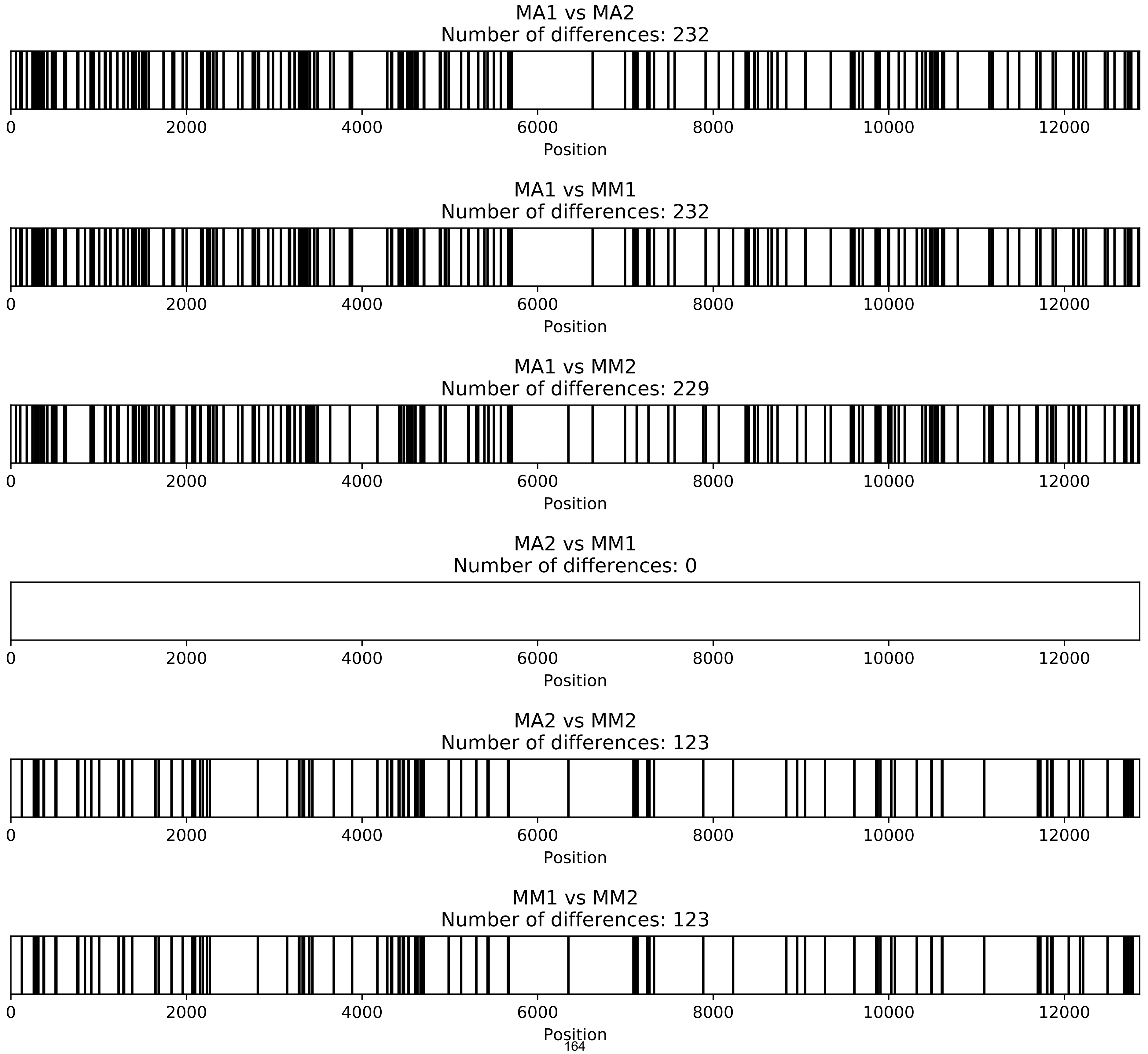
MA1 vs MA2
Number of differences: 232
Position
MA1 vs MM1
Number of differences: 232
Position
MA1 vs MM2
Number of differences: 229
Position
MA2 vs MM1
Number of differences: 0
Position
MA2 vs MM2
Number of differences: 123
Position
MM1 vs MM2
Number of differences: 123
Position
0
2000
4000
6000
8000
10000
12000

### Region: 530



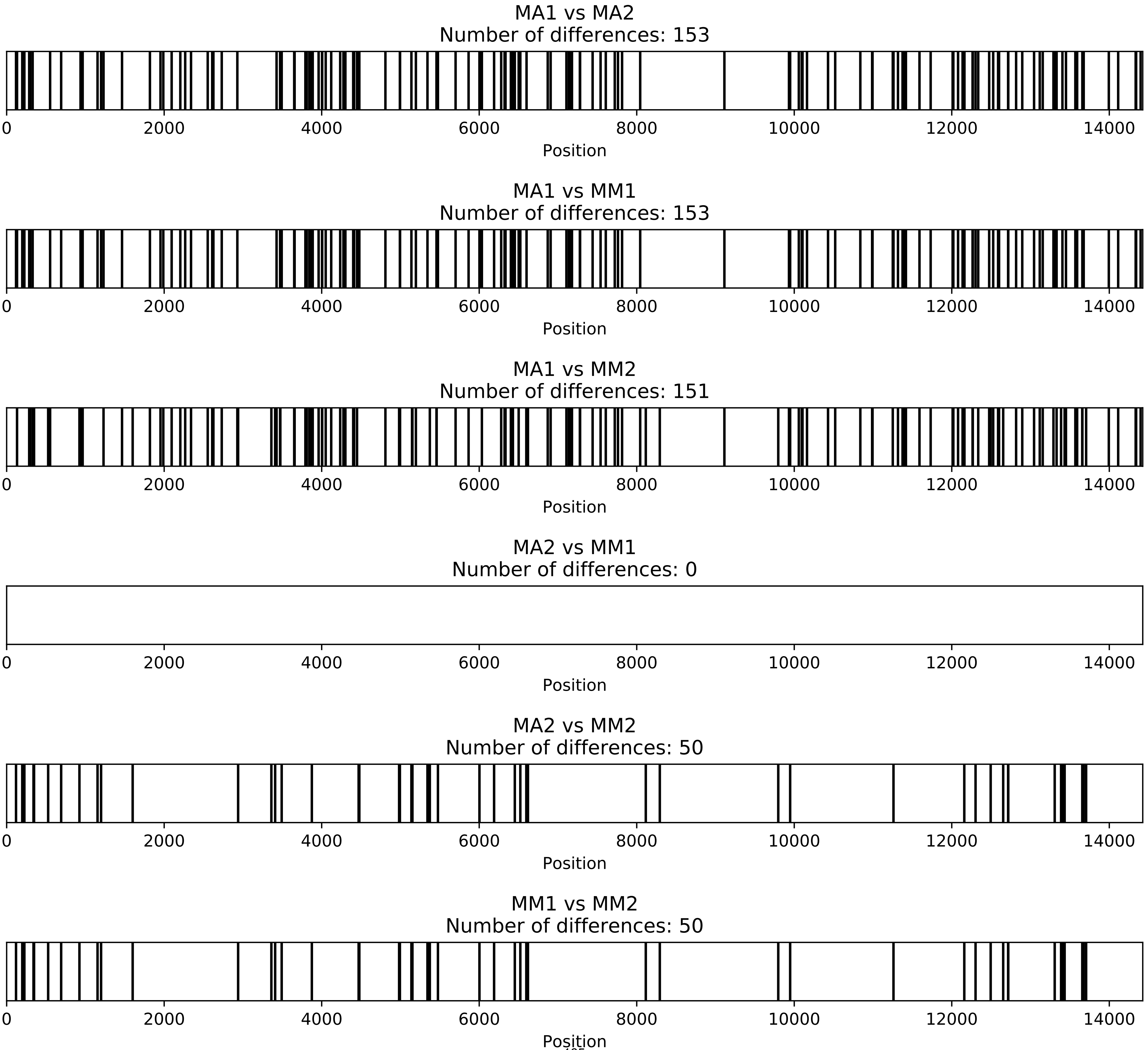
MA1 vs MA2
Number of differences: 153
MA1 vs MM1
Number of differences: 153
MA1 vs MM2
Number of differences: 151
MA2 vs MM1
Number of differences: 0
MA2 vs MM2
Number of differences: 50
MM1 vs MM2
Number of differences: 50
0
2000
4000
6000
8000
10000
12000
14000
Position

### Region: 533



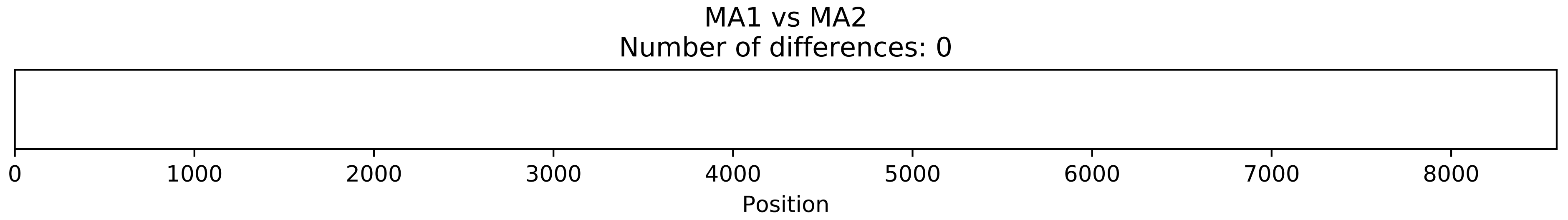
MA1 vs MA2
Number of differences: 0
0
1000
2000
3000
4000
5000
6000
7000
8000
Position

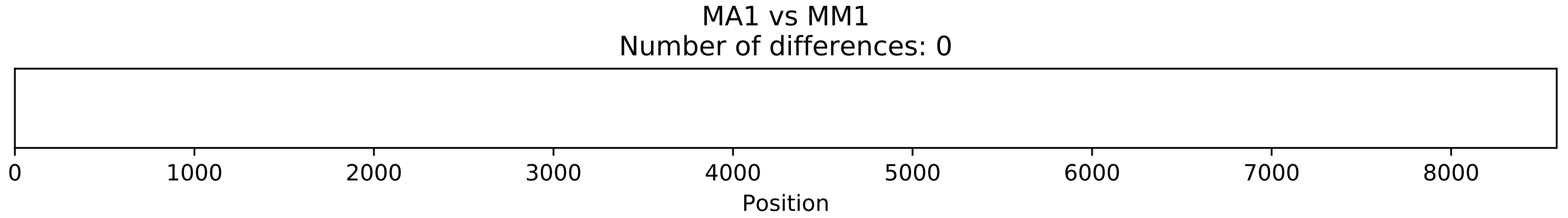
MA1 vs MM1
Number of differences: 0
Position

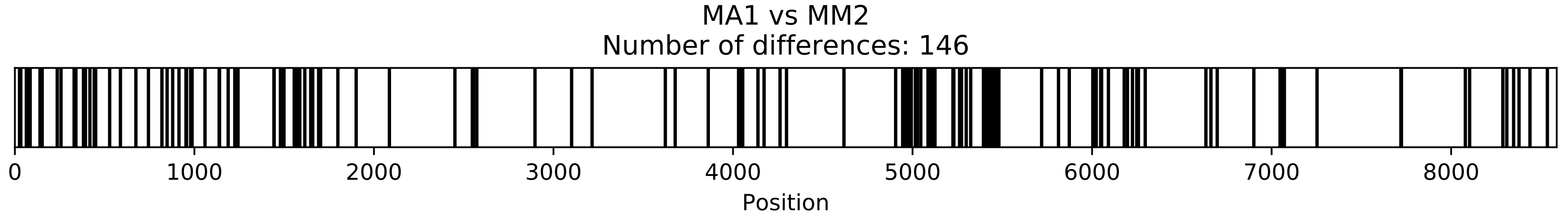
MA1 vs MM2
Number of differences: 146
Position

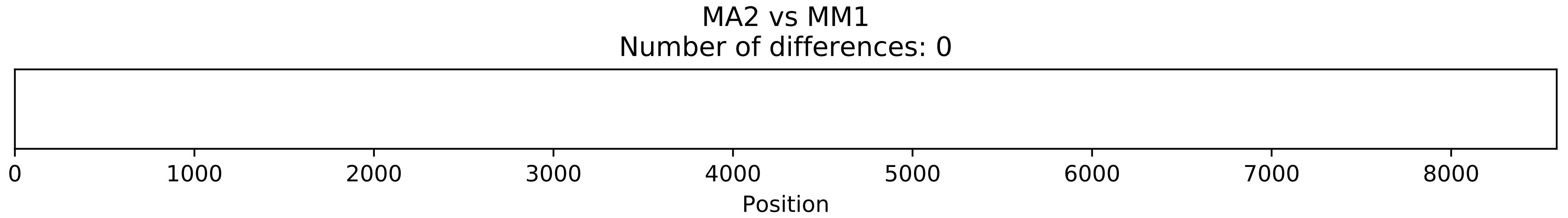
MA2 vs MM1
Number of differences: 0
Position

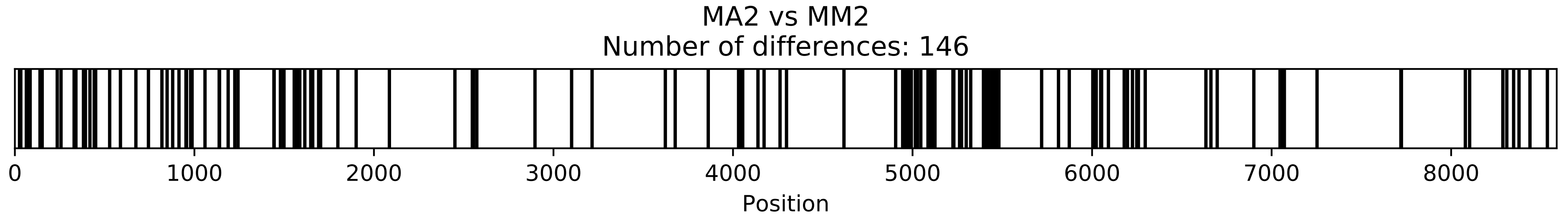
MA2 vs MM2
Number of differences: 146
Position

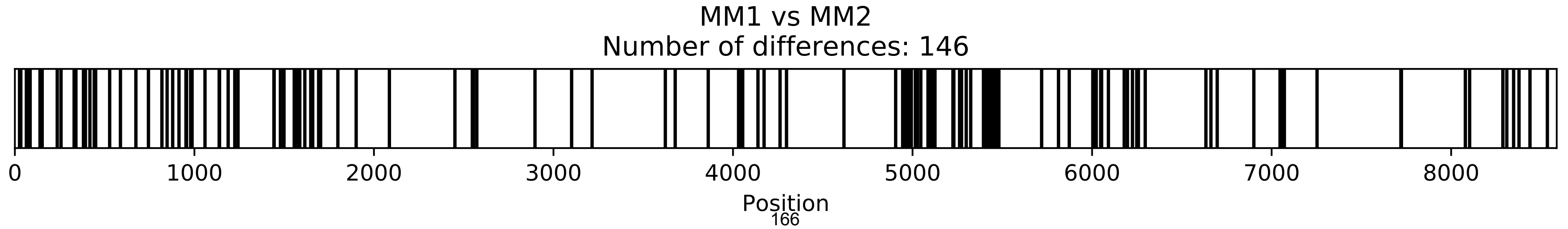
MM1 vs MM2
Number of differences: 146
Position

### Region: 535



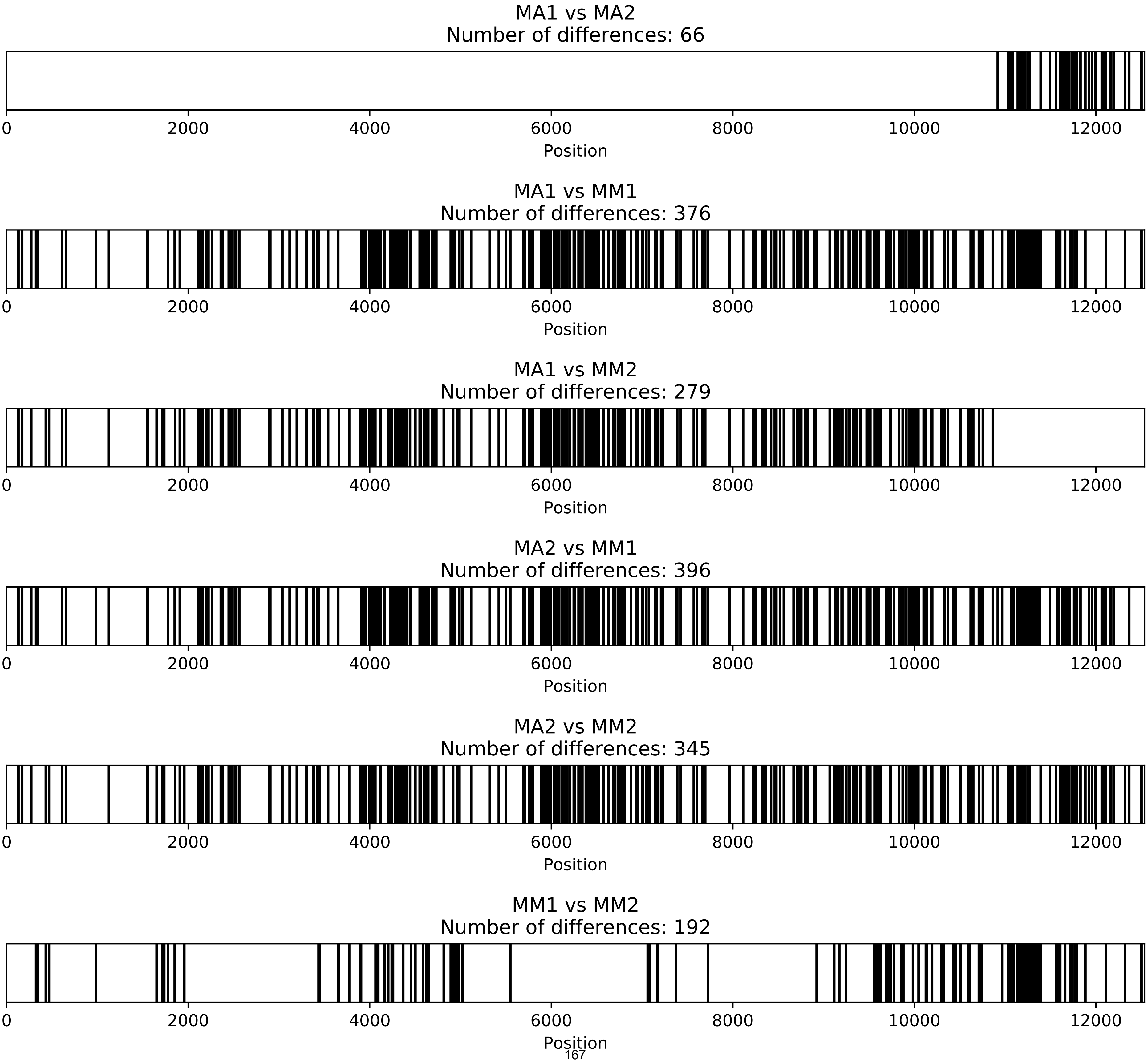
MA1 vs MA2
Number of differences: 66
MA1 vs MM1
Number of differences: 376
MA1 vs MM2
Number of differences: 279
MA2 vs MM1
Number of differences: 396
MA2 vs MM2
Number of differences: 345
MM1 vs MM2
Number of differences: 192
0
2000
4000
6000
8000
10000
12000
Position

### Region: 537



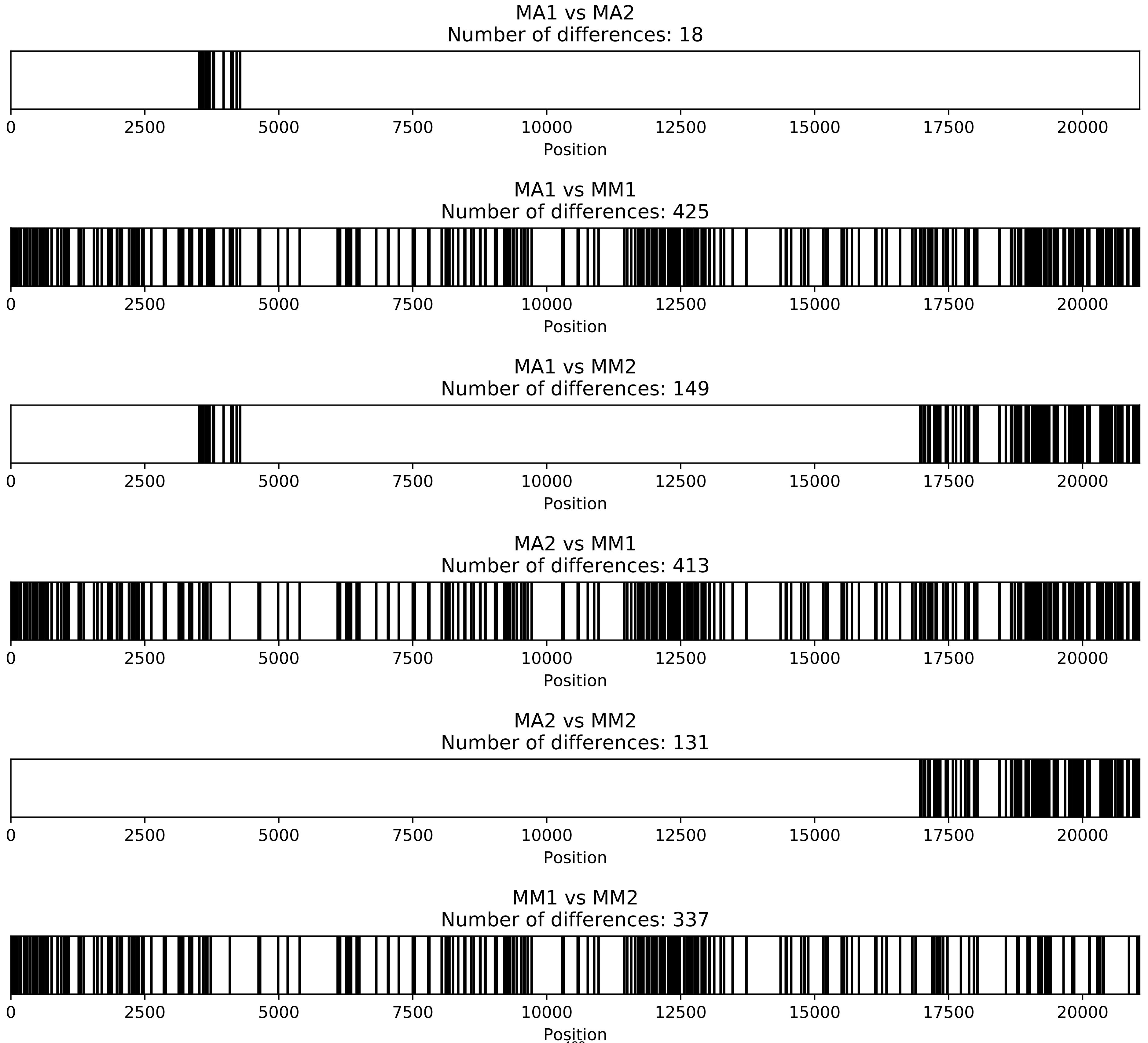
MA1 vs MA2
Number of differences: 18
MA1 vs MM1
Number of differences: 425
MA1 vs MM2
Number of differences: 149
MA2 vs MM1
Number of differences: 413
MA2 vs MM2
Number of differences: 131
MM1 vs MM2
Number of differences: 337
0
2500
5000
7500
10000
12500
15000
17500
20000
Position

### Region: 541



MA1 vs MA2
Number of differences: 334

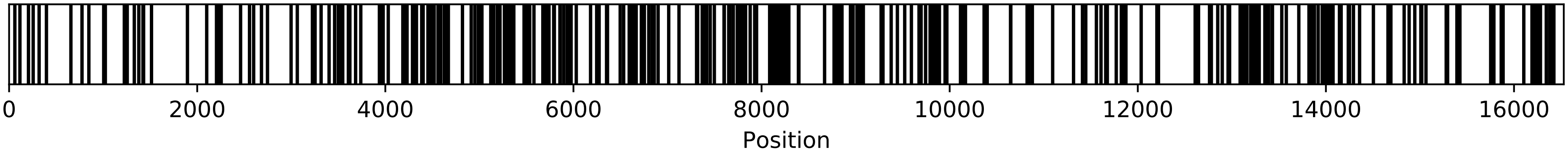


MA1 vs MM1
Number of differences: 315

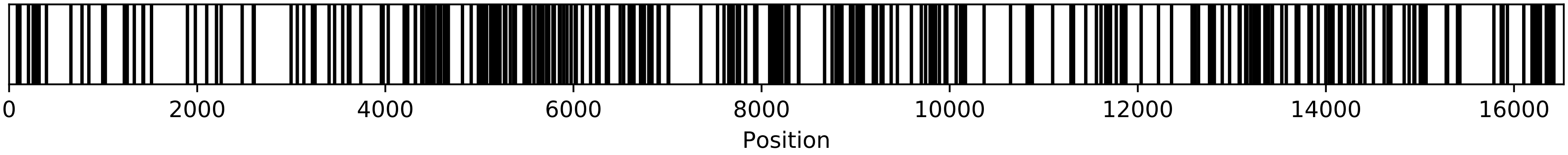


MA1 vs MM2
Number of differences: 334

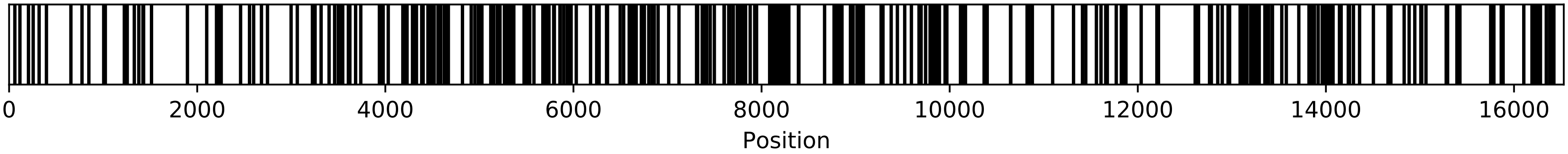


MA2 vs MM1
Number of differences: 168

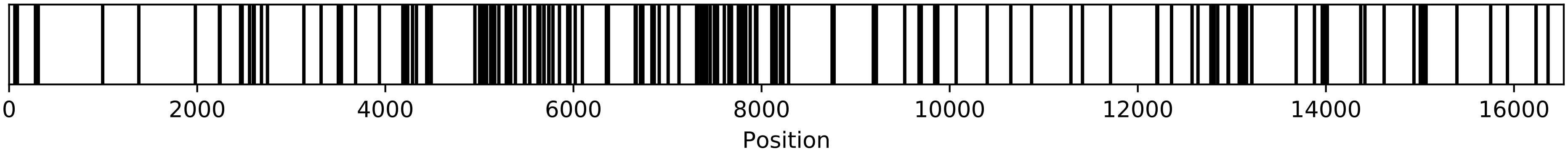


MA2 vs MM2
Number of differences: 0

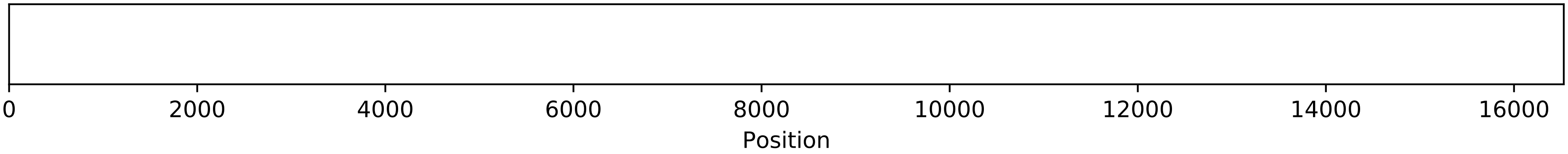


MM1 vs MM2
Number of differences: 168

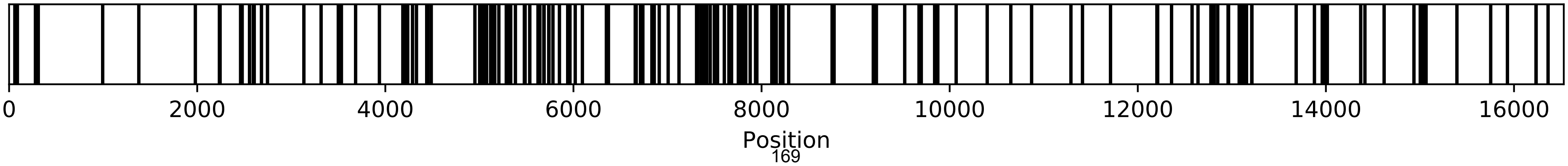

### Region: 542



MA1 vs MA2
Number of differences: 569

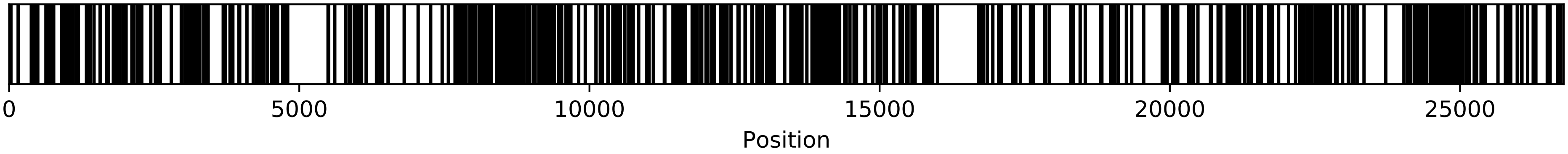


MA1 vs MM1
Number of differences: 569

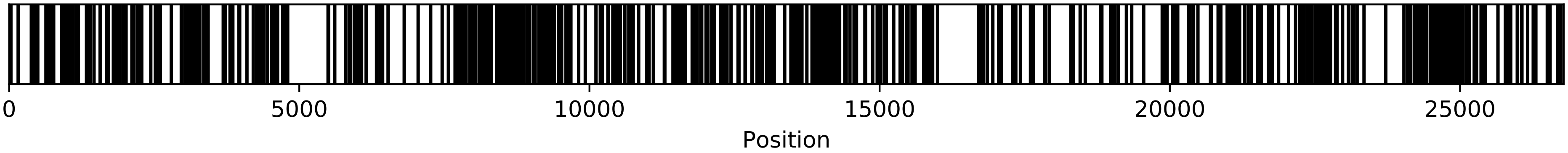


MA1 vs MM2
Number of differences: 593

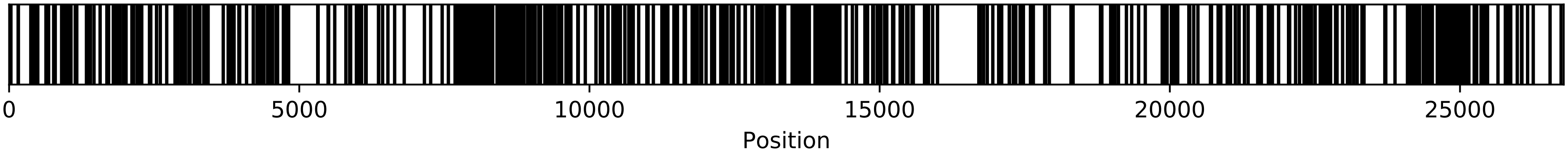


MA2 vs MM1
Number of differences: 0

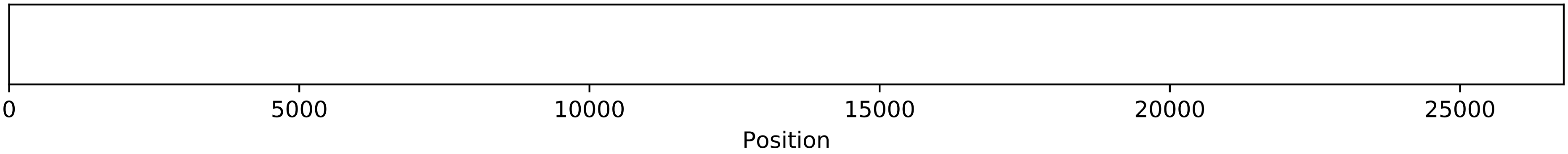


MA2 vs MM2
Number of differences: 289

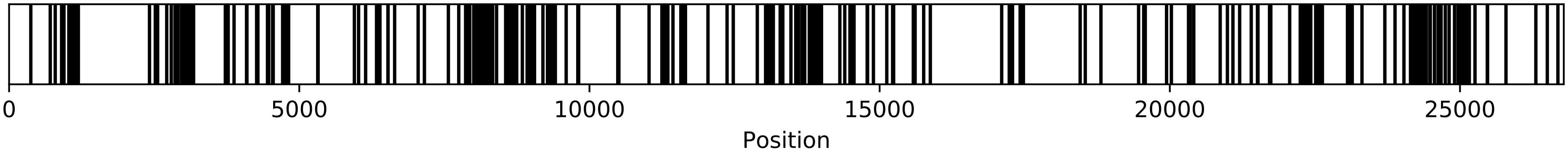


MM1 vs MM2
Number of differences: 289

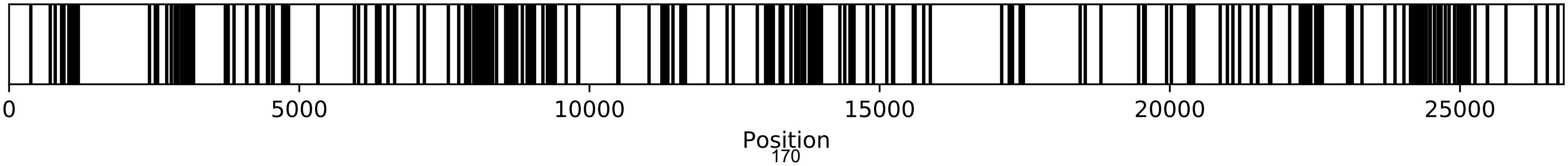

### Region: 543



MA1 vs MA2
Number of differences: 302

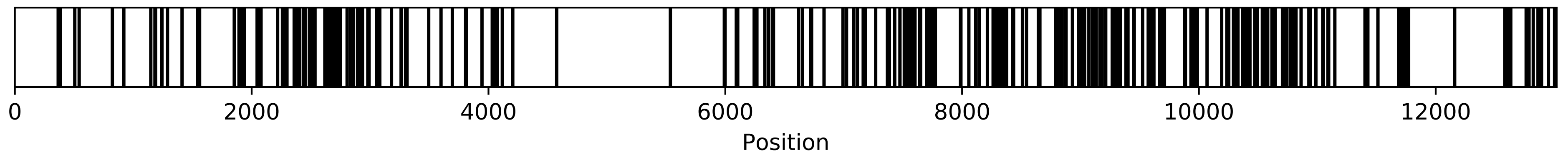

MA1 vs MM1
Number of differences: 137

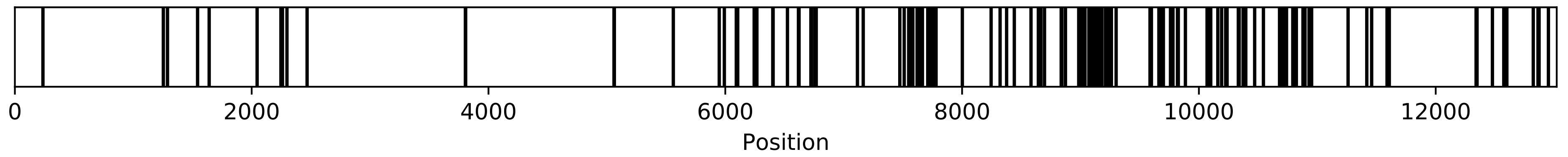

MA1 vs MM2
Number of differences: 0

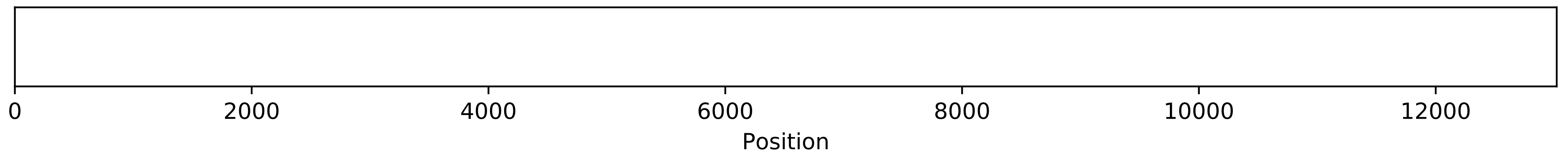

MA2 vs MM1
Number of differences: 319

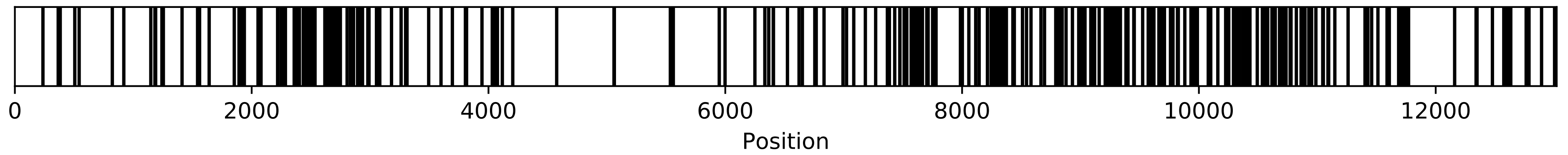

MA2 vs MM2
Number of differences: 302

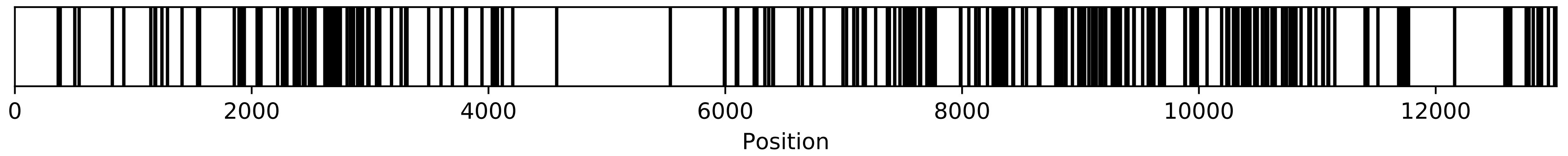

MM1 vs MM2
Number of differences: 137

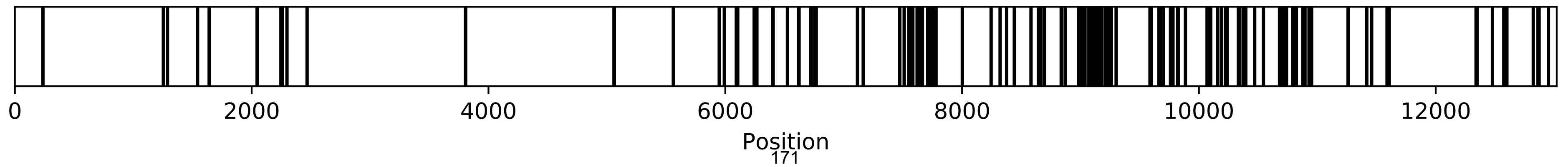

### Region: 547



## MA1 vs MA2
Number of differences: 409

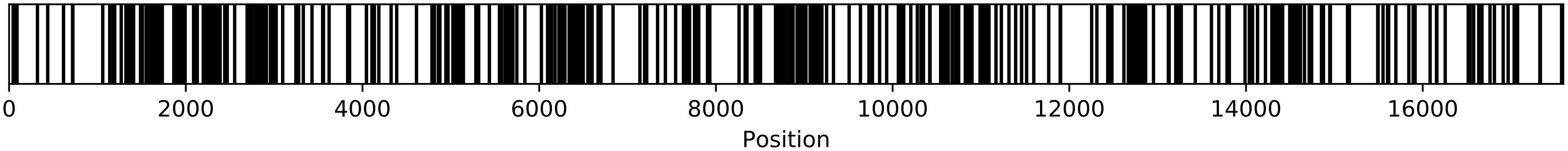


## MA1 vs MM1
Number of differences: 378

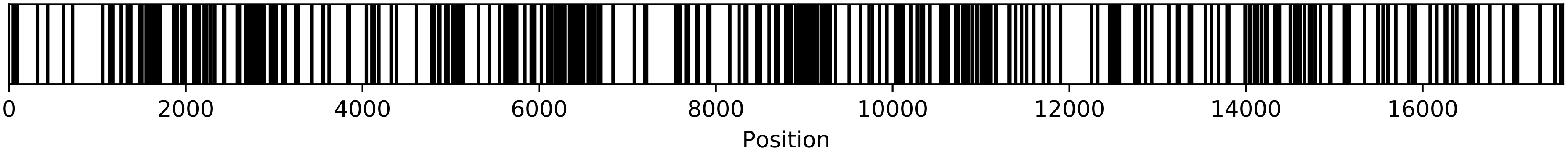


## MA1 vs MM2
Number of differences: 390

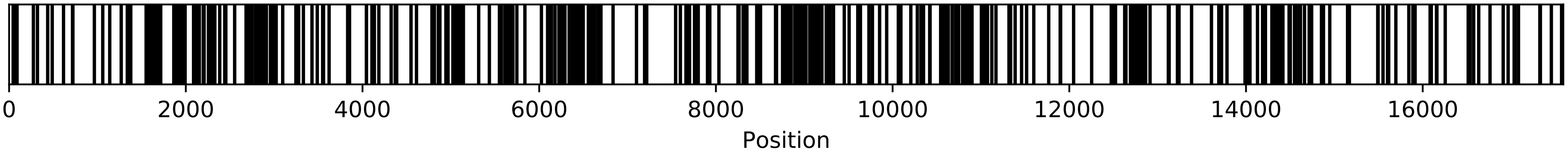


## MA2 vs MM1
Number of differences: 211

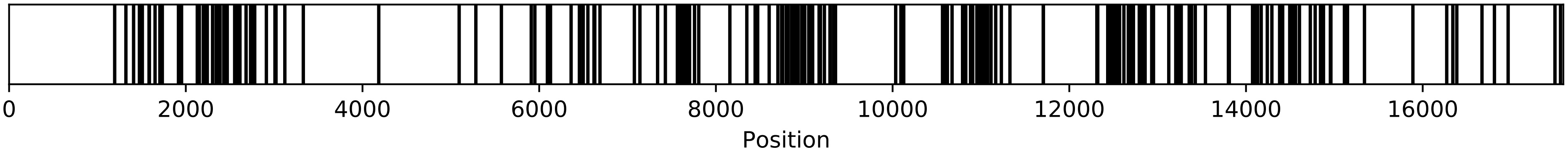


## MA2 vs MM2
Number of differences: 192

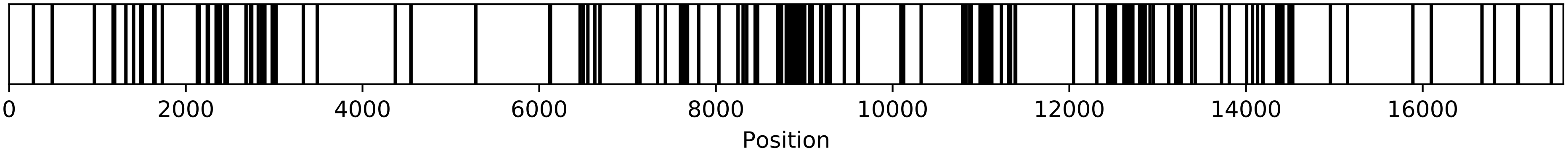


## MM1 vs MM2
Number of differences: 187

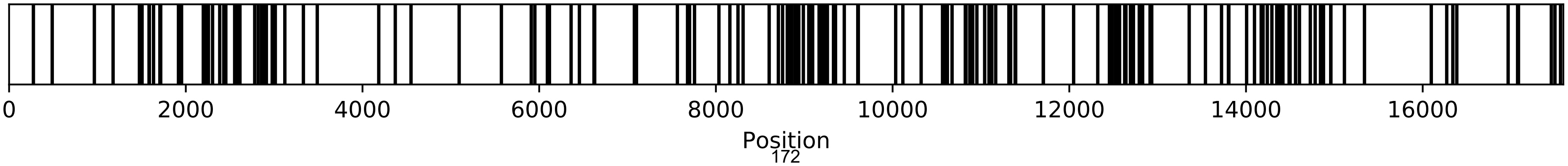

### Region: 548



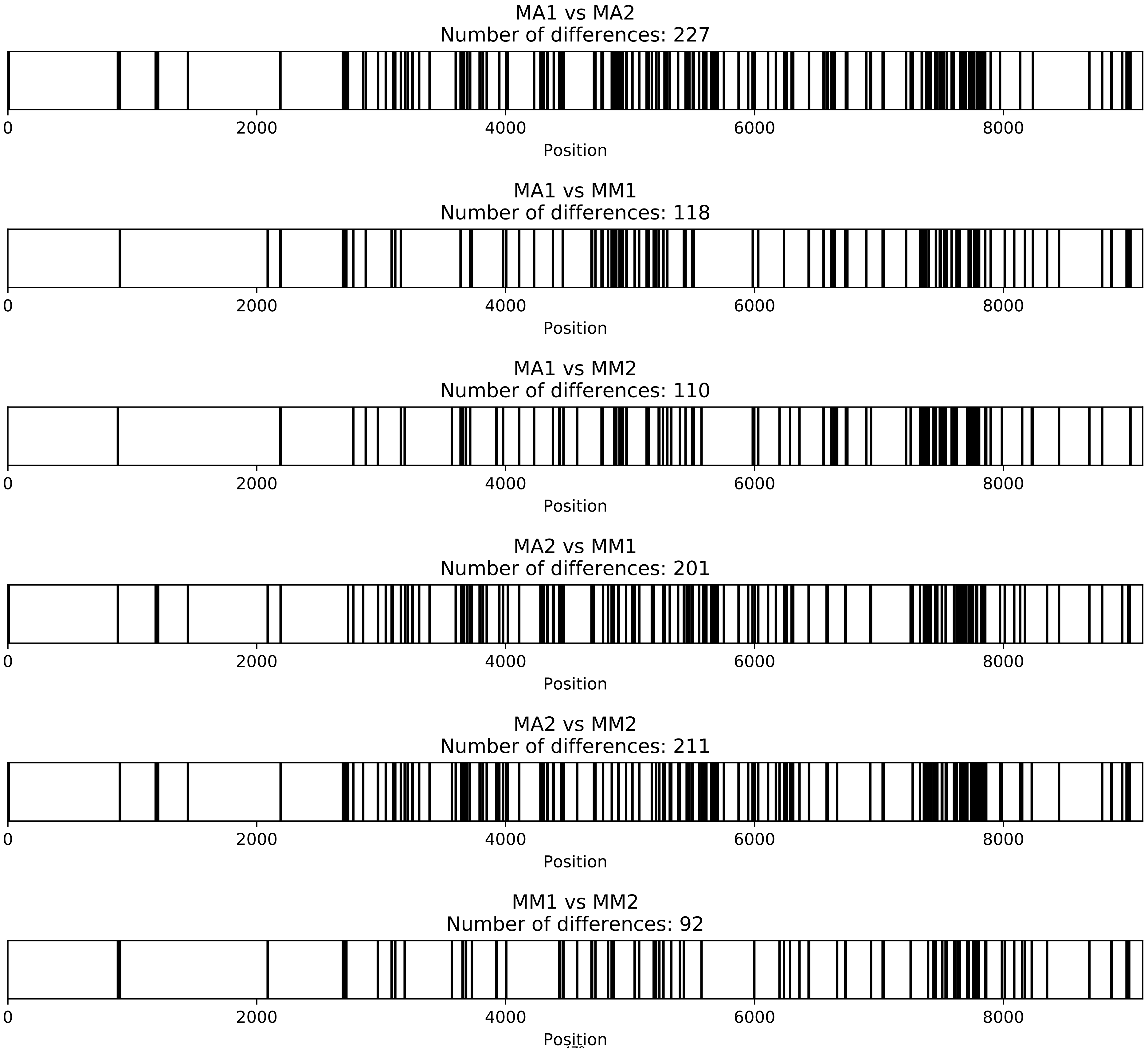
MA1 vs MA2
Number of differences: 227
MA1 vs MM1
Number of differences: 118
MA1 vs MM2
Number of differences: 110
MA2 vs MM1
Number of differences: 201
MA2 vs MM2
Number of differences: 211
MM1 vs MM2
Number of differences: 92
0
2000
4000
6000
8000
Position

### Region: 552



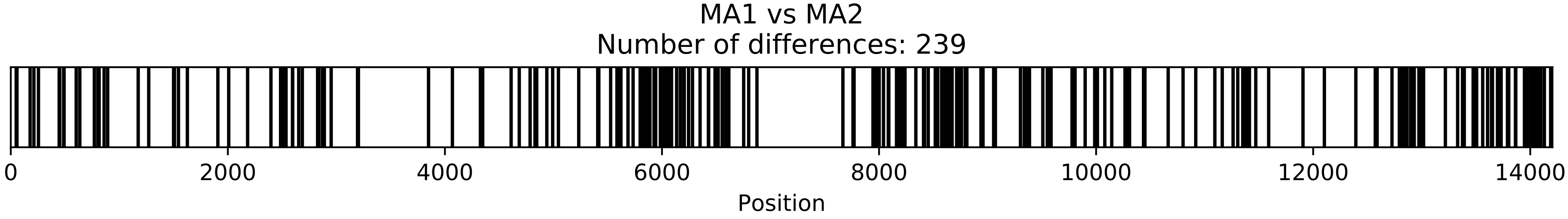
MA1 vs MA2
Number of differences: 239
0
2000
4000
6000
8000
10000
12000
14000
Position

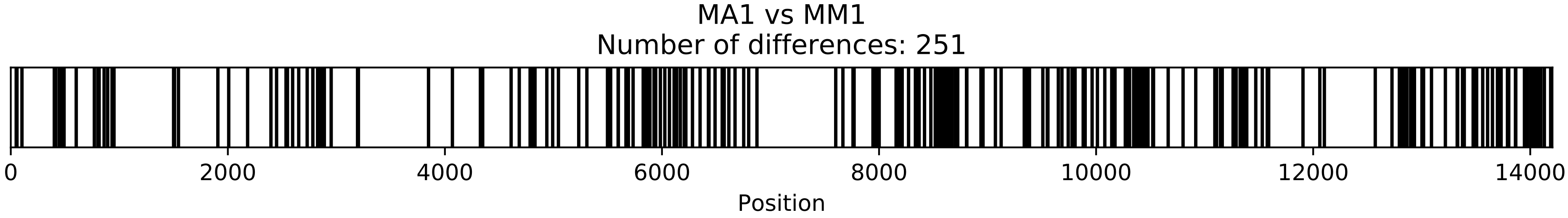
MA1 vs MM1
Number of differences: 251
0
2000
4000
6000
8000
10000
12000
14000
Position

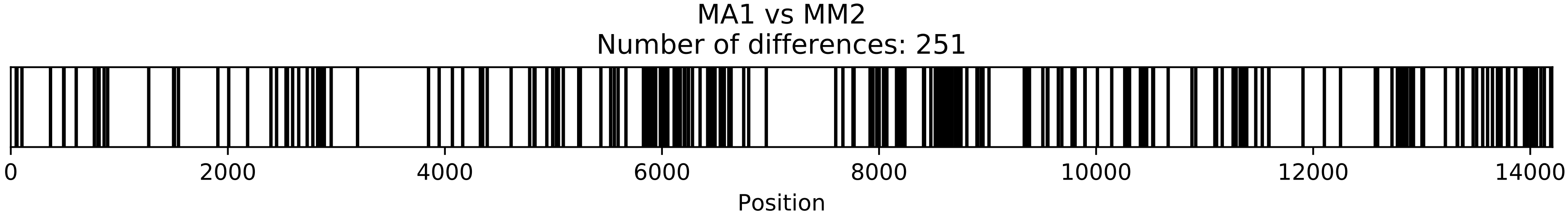
MA1 vs MM2
Number of differences: 251
0
2000
4000
6000
8000
10000
12000
14000
Position

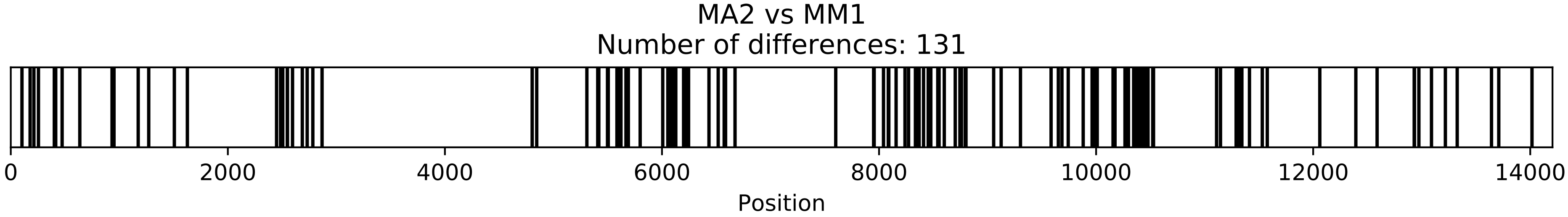
MA2 vs MM1
Number of differences: 131
0
2000
4000
6000
8000
10000
12000
14000
Position

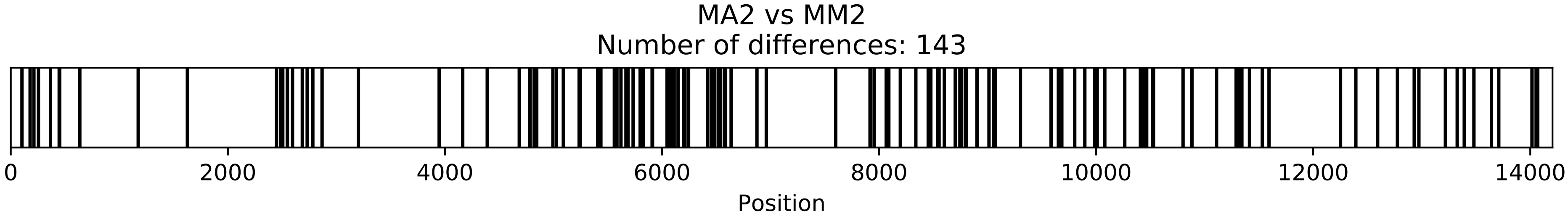
MA2 vs MM2
Number of differences: 143
0
2000
4000
6000
8000
10000
12000
14000
Position

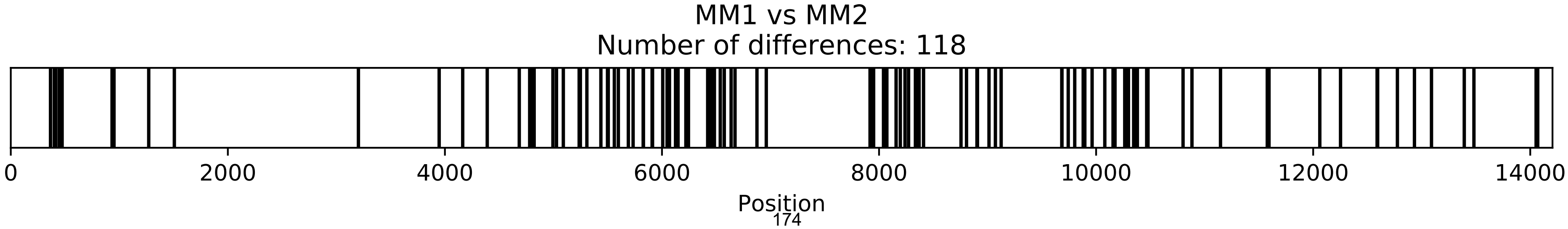
MM1 vs MM2
Number of differences: 118
0
2000
4000
6000
8000
10000
12000
14000
Position

### Region: 556



MA1 vs MA2
Number of differences: 22

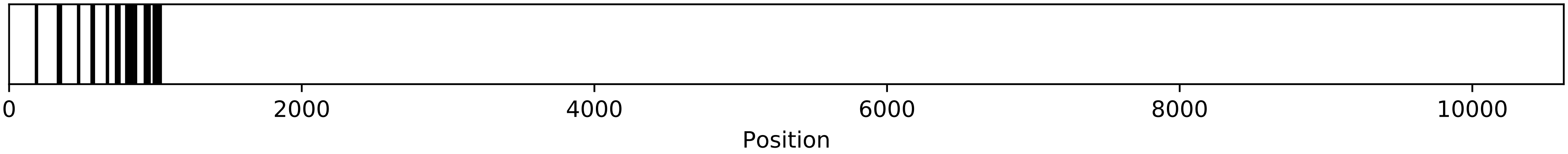


MA1 vs MM1
Number of differences: 138

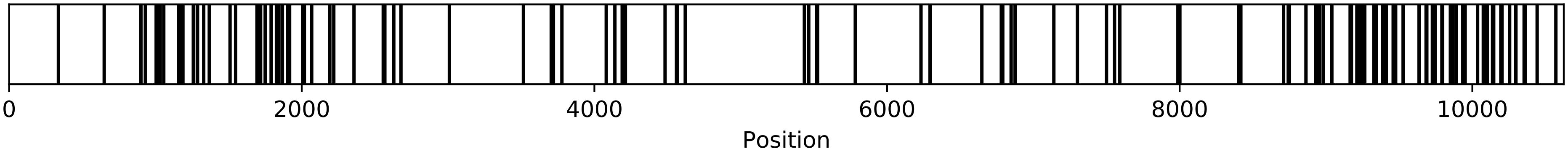


MA1 vs MM2
Number of differences: 0

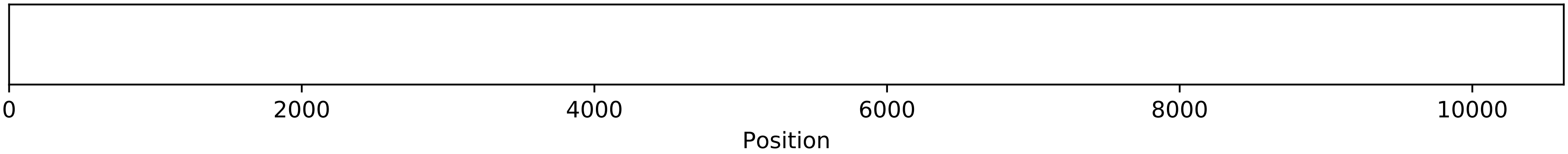


MA2 vs MM1
Number of differences: 154

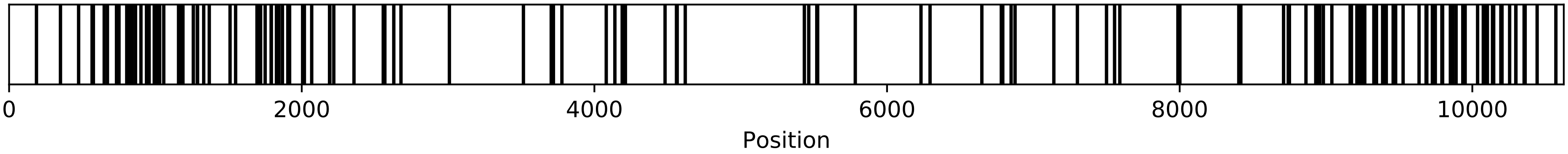


MA2 vs MM2
Number of differences: 22

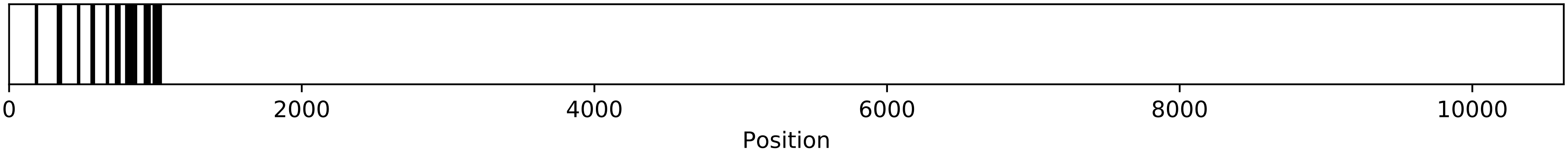


MM1 vs MM2
Number of differences: 138

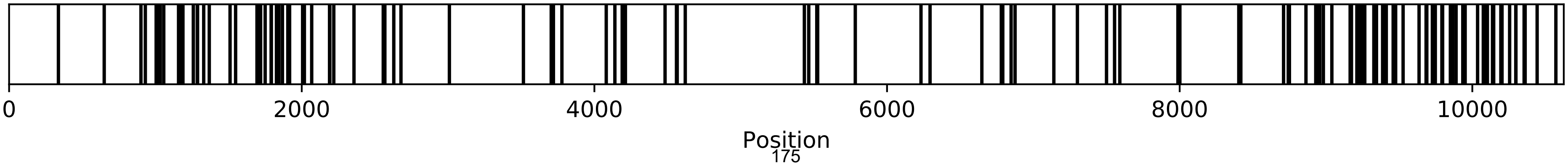

### Region: 557



MA1 vs MA2
Number of differences: 296

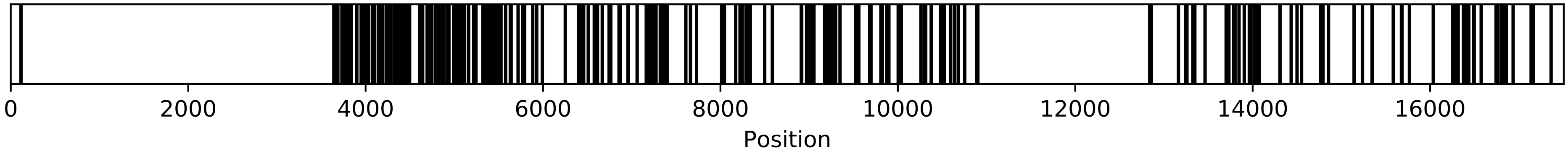


MA1 vs MM1
Number of differences: 256

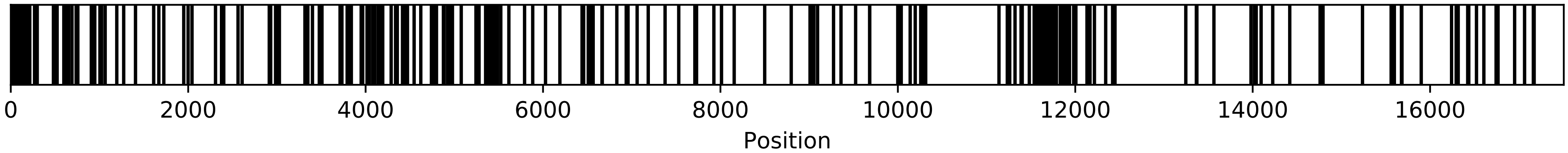


MA1 vs MM2
Number of differences: 246

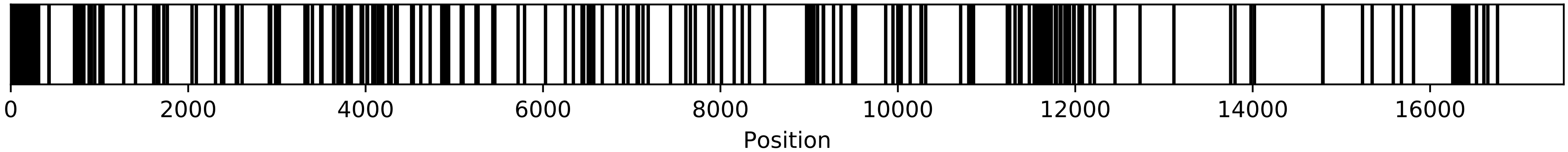


MA2 vs MM1
Number of differences: 413

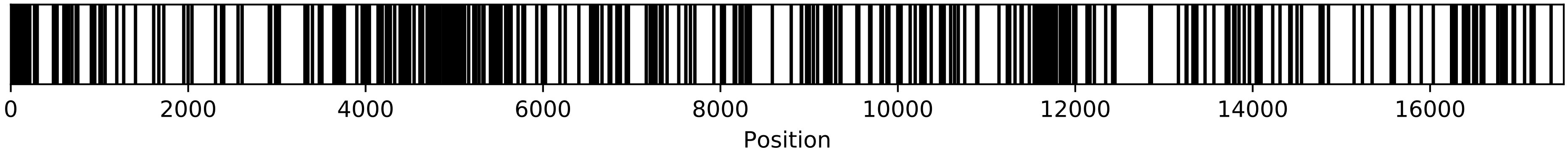


MA2 vs MM2
Number of differences: 394

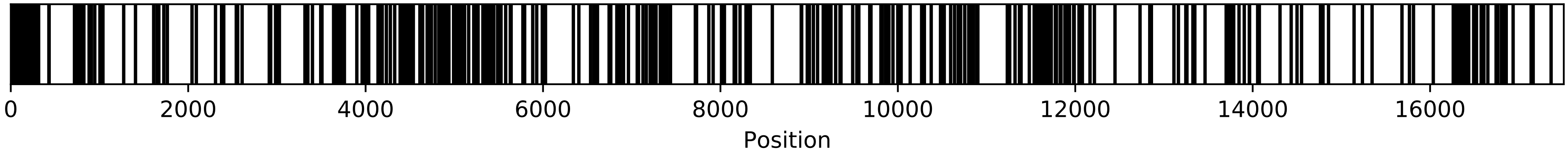


MM1 vs MM2
Number of differences: 206

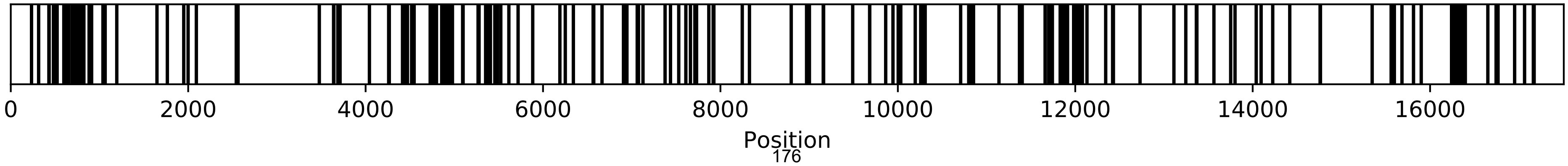

### Region: 562



MA1 vs MA2
Number of differences: 176

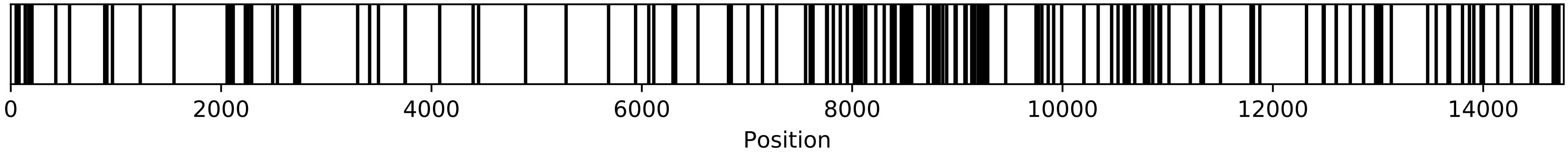


MA1 vs MM1
Number of differences: 0

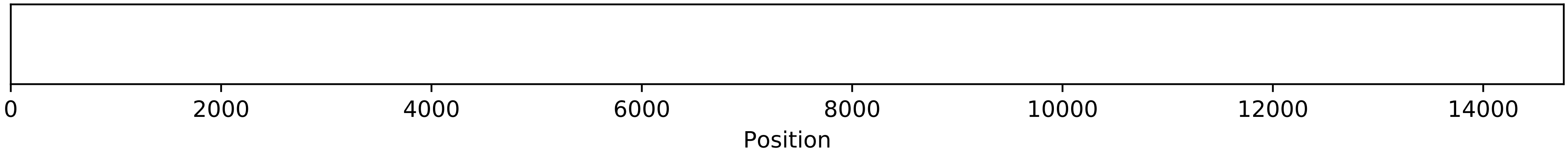


MA1 vs MM2
Number of differences: 97

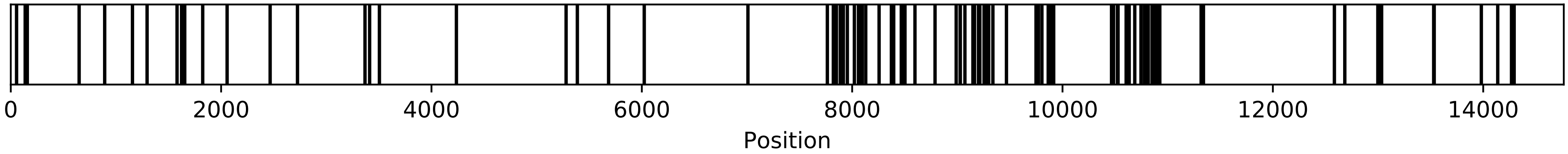


MA2 vs MM1
Number of differences: 176

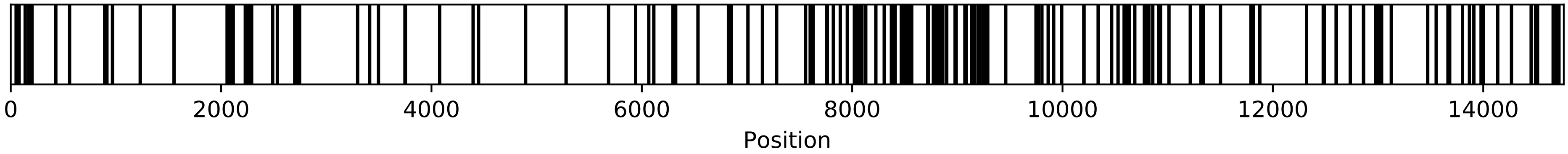


MA2 vs MM2
Number of differences: 174

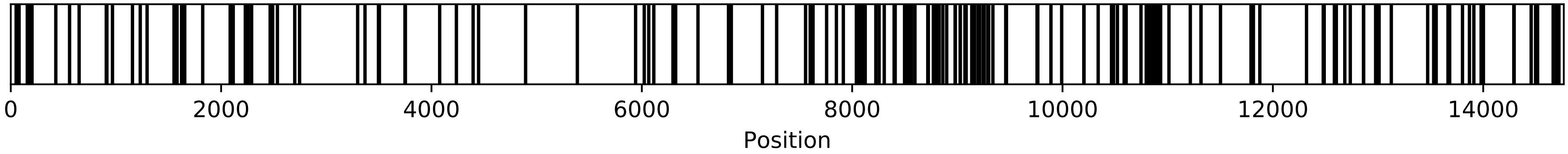


MM1 vs MM2
Number of differences: 97

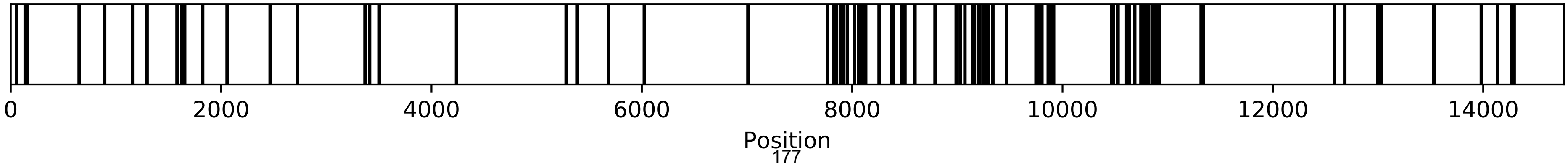

### Region: 566



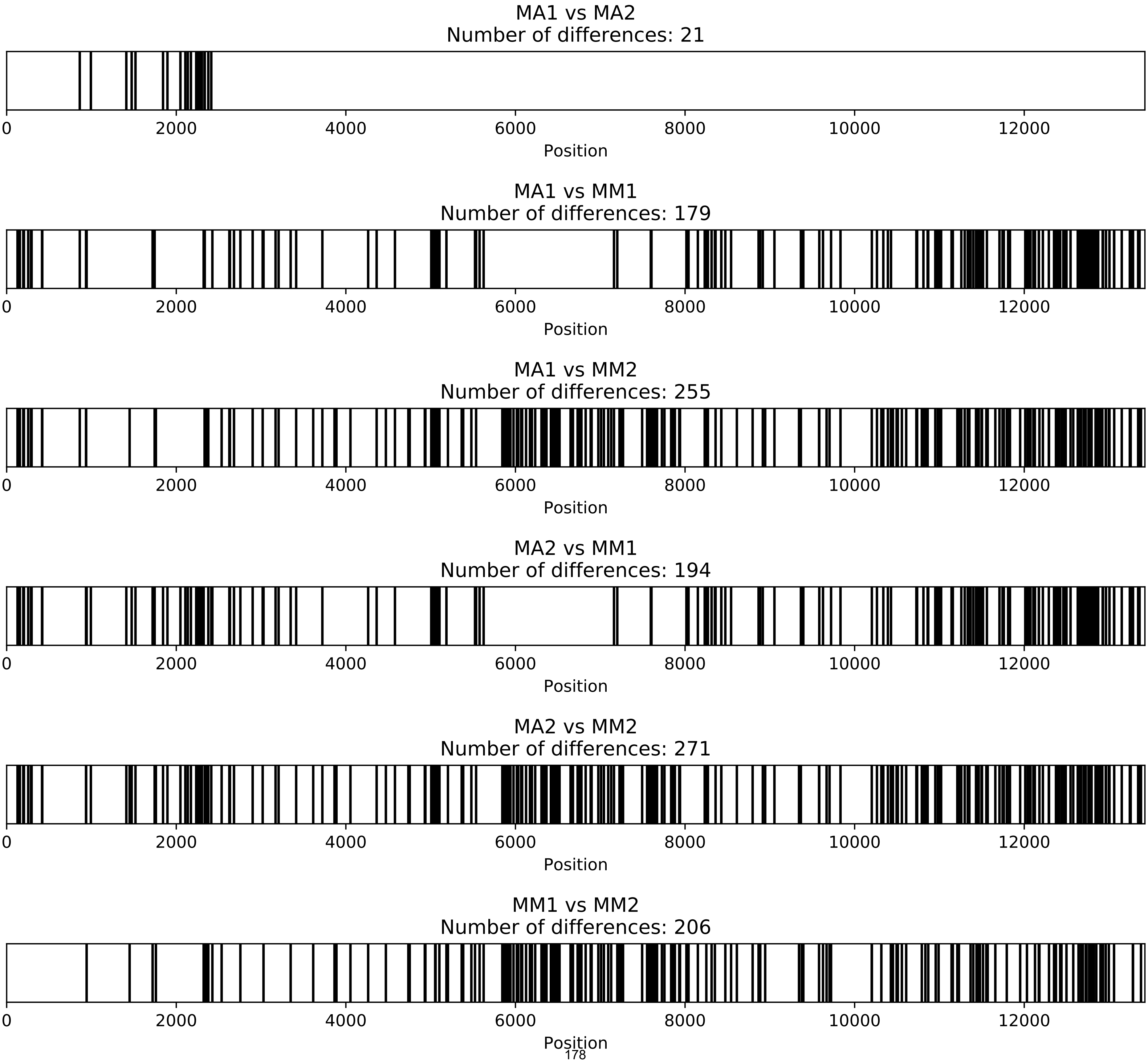
MA1 vs MA2
Number of differences: 21
MA1 vs MM1
Number of differences: 179
MA1 vs MM2
Number of differences: 255
MA2 vs MM1
Number of differences: 194
MA2 vs MM2
Number of differences: 271
MM1 vs MM2
Number of differences: 206
0
2000
4000
6000
8000
10000
12000
Position

### Region: 569



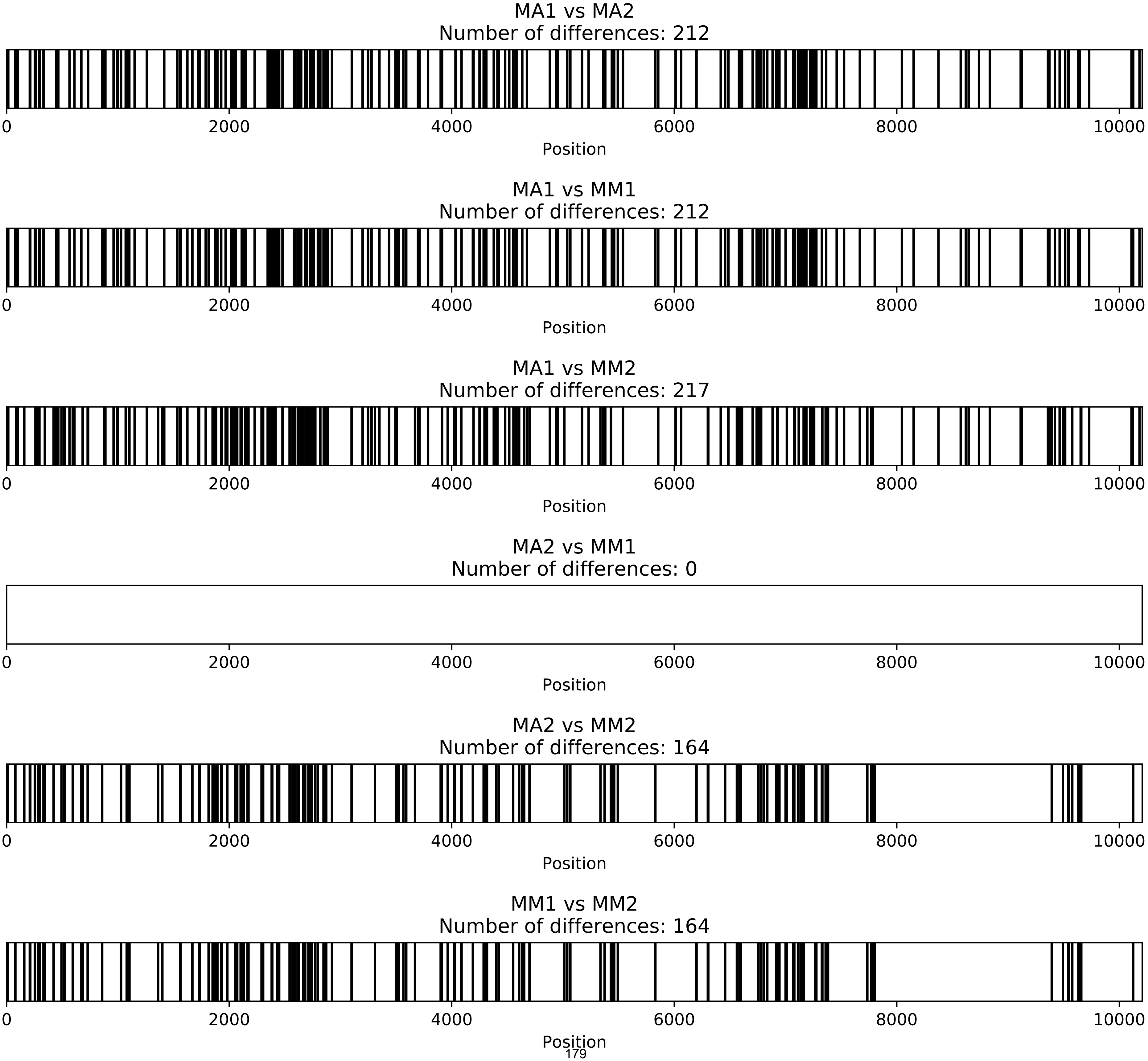
MA1 vs MA2
Number of differences: 212
Position
MA1 vs MM1
Number of differences: 212
Position
MA1 vs MM2
Number of differences: 217
Position
MA2 vs MM1
Number of differences: 0
Position
MA2 vs MM2
Number of differences: 164
Position
MM1 vs MM2
Number of differences: 164
Position
0
2000
4000
6000
8000
10000

### Region: 571



## MA1 vs MA2
Number of differences: 201

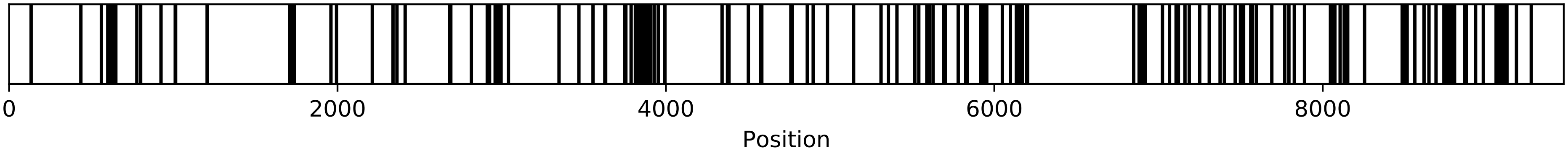


## MA1 vs MM1
Number of differences: 205

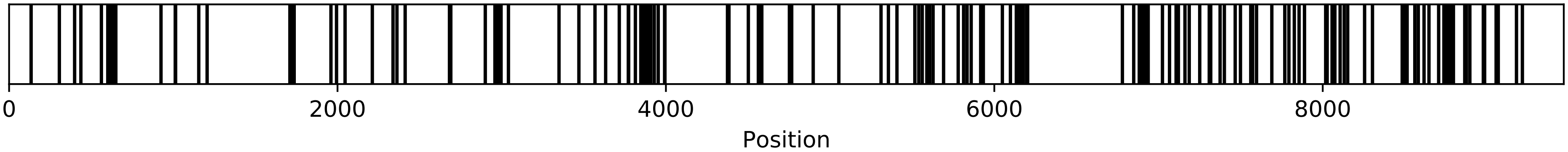


## MA1 vs MM2
Number of differences: 201

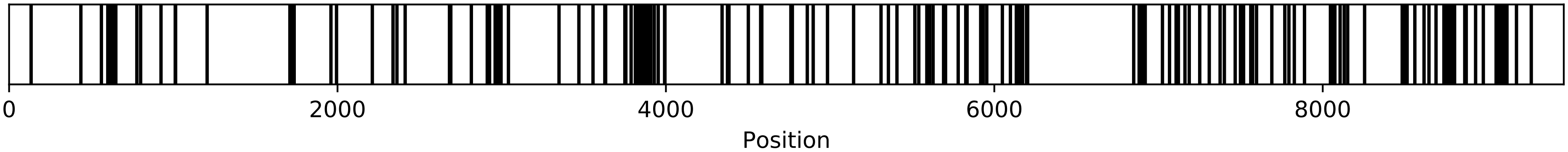


## MA2 vs MM1
Number of differences: 94

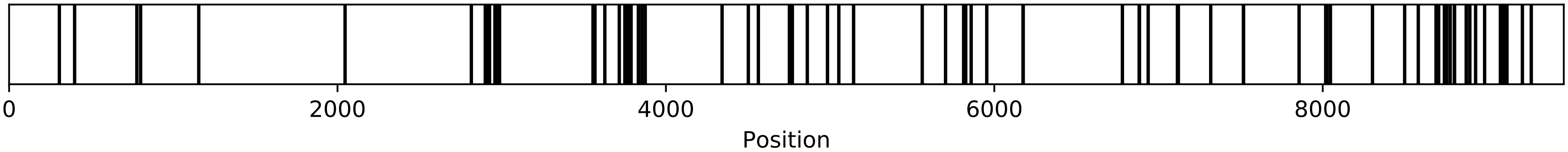


## MA2 vs MM2
Number of differences: 0

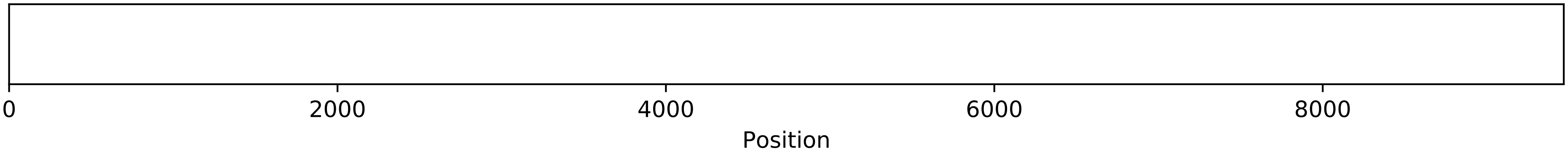


## MM1 vs MM2
Number of differences: 94

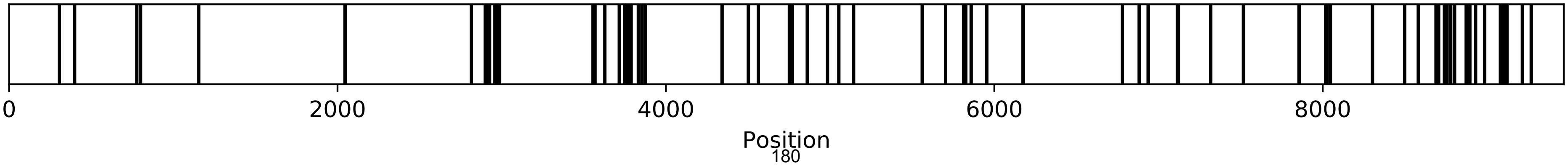

### Region: 577



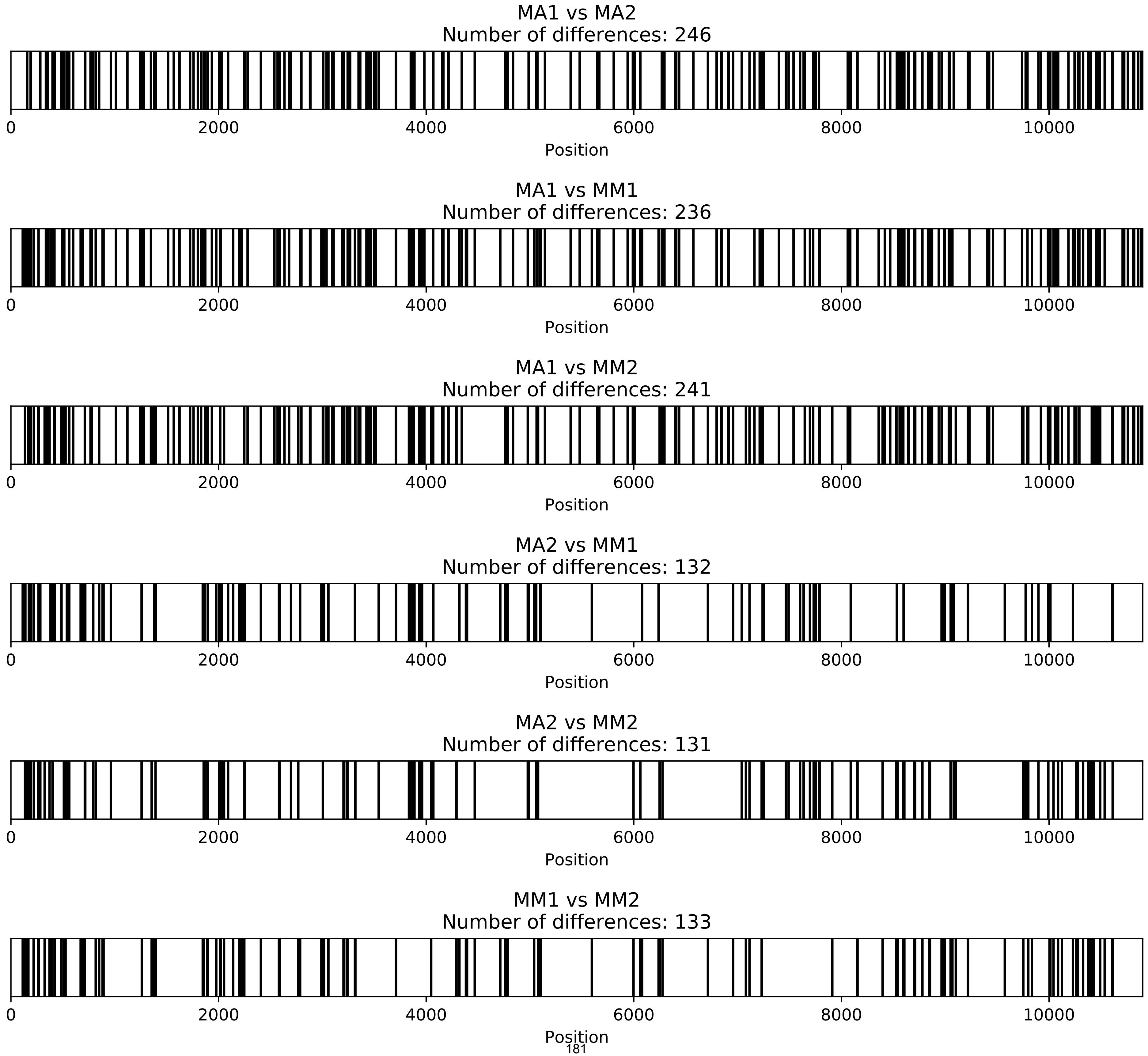
MA1 vs MA2
Number of differences: 246
0
2000
4000
6000
8000
10000
Position
MA1 vs MM1
Number of differences: 236
Position
MA1 vs MM2
Number of differences: 241
Position
MA2 vs MM1
Number of differences: 132
Position
MA2 vs MM2
Number of differences: 131
Position
MM1 vs MM2
Number of differences: 133
Position

### Region: 578



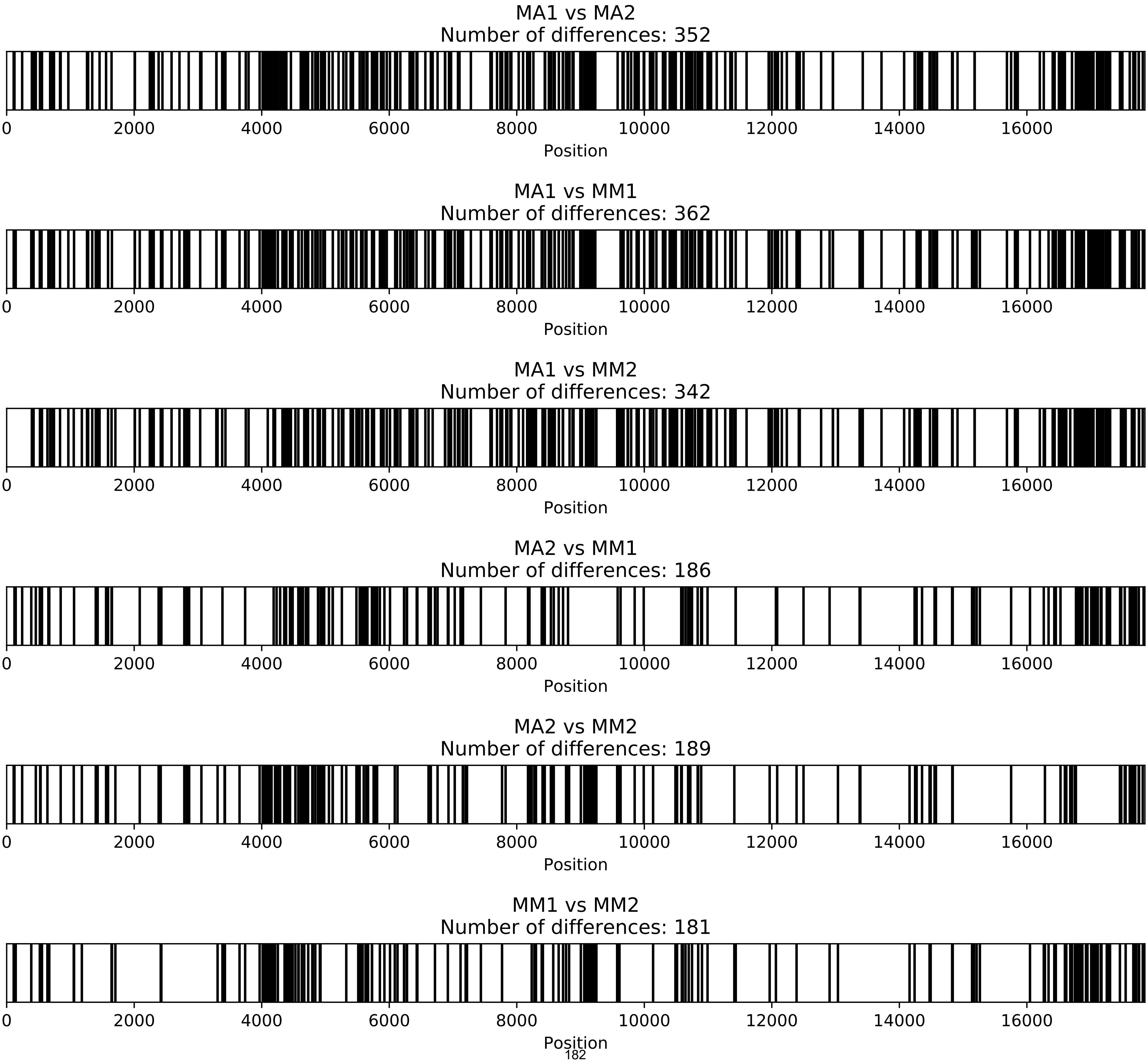
MA1 vs MA2
Number of differences: 352
MA1 vs MM1
Number of differences: 362
MA1 vs MM2
Number of differences: 342
MA2 vs MM1
Number of differences: 186
MA2 vs MM2
Number of differences: 189
MM1 vs MM2
Number of differences: 181
0
2000
4000
6000
8000
10000
12000
14000
16000
Position

### Region: 584



MA1 vs MA2
Number of differences: 375

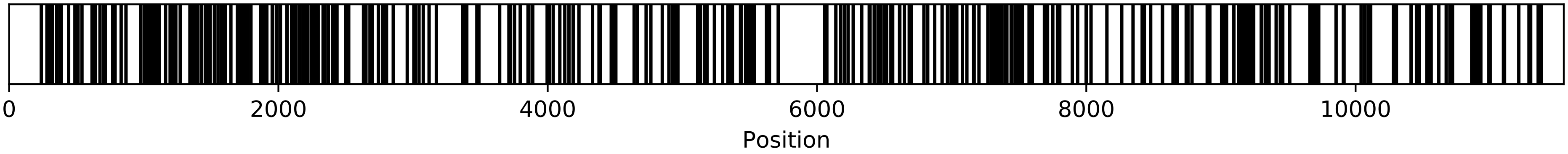


MA1 vs MM1
Number of differences: 0

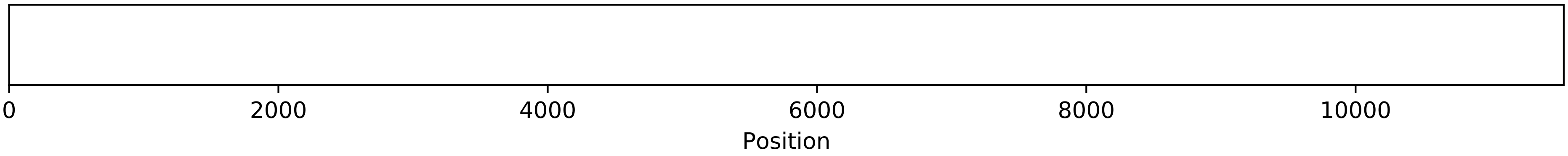


MA1 vs MM2
Number of differences: 178

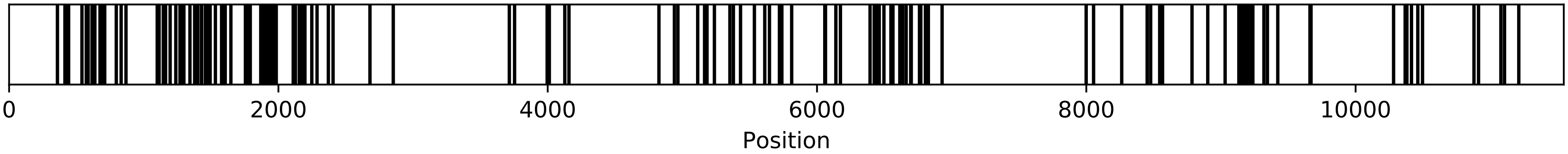


MA2 vs MM1
Number of differences: 375

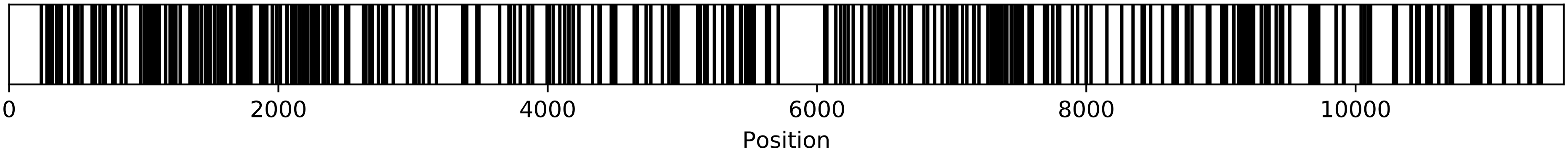


MA2 vs MM2
Number of differences: 350

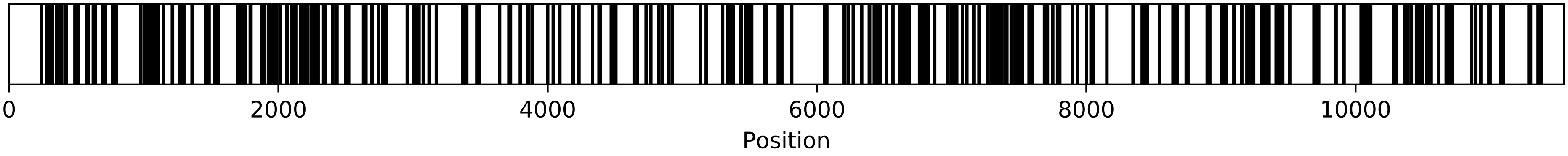


MM1 vs MM2
Number of differences: 178

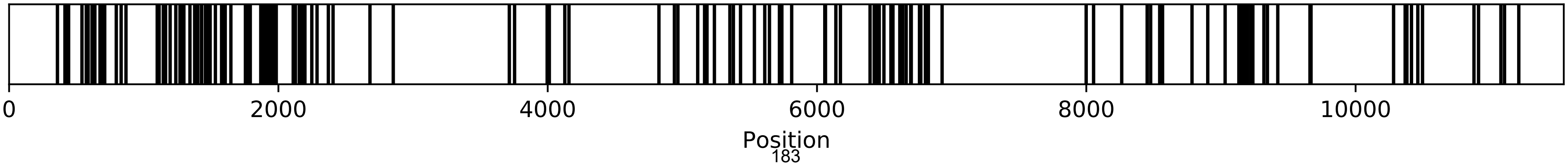

### Region: 585



MA1 vs MA2
Number of differences: 246

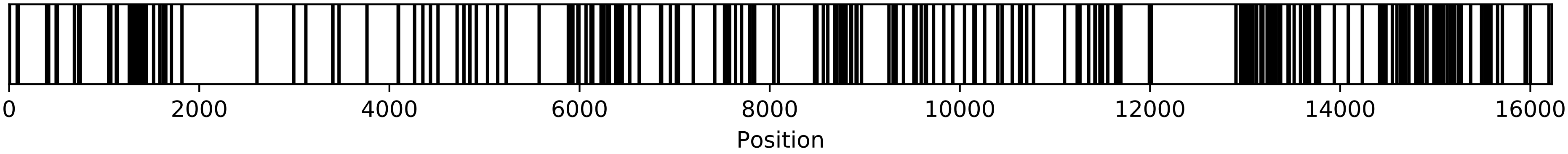


MA1 vs MM1
Number of differences: 106

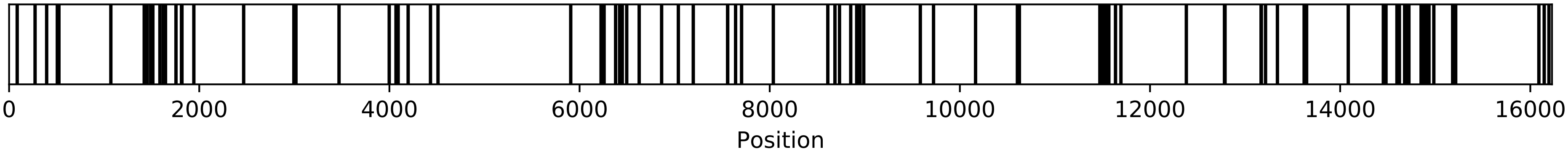


MA1 vs MM2
Number of differences: 103

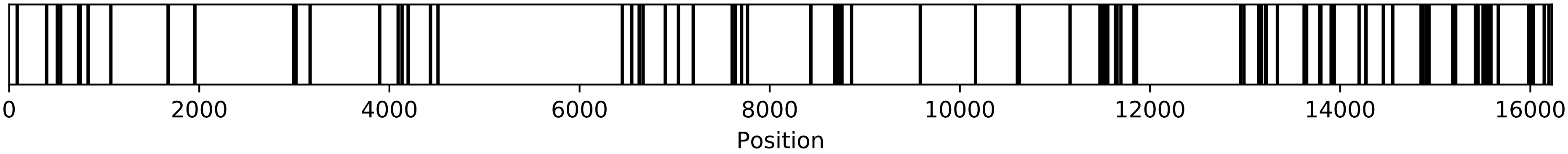


MA2 vs MM1
Number of differences: 249

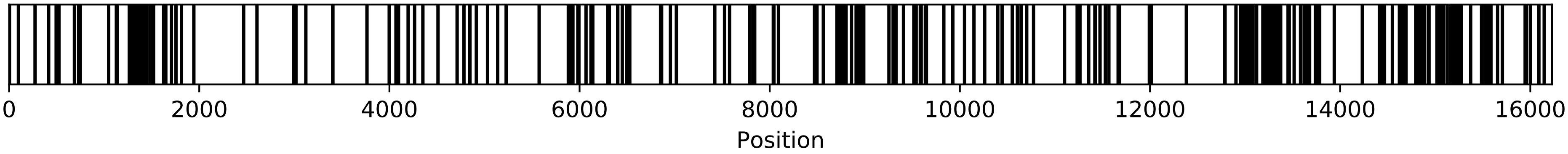


MA2 vs MM2
Number of differences: 273

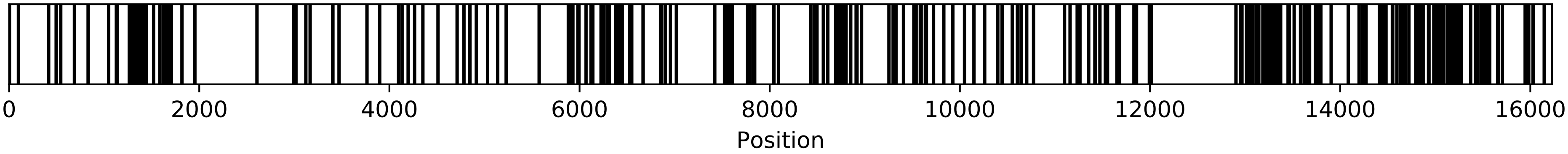


MM1 vs MM2
Number of differences: 114

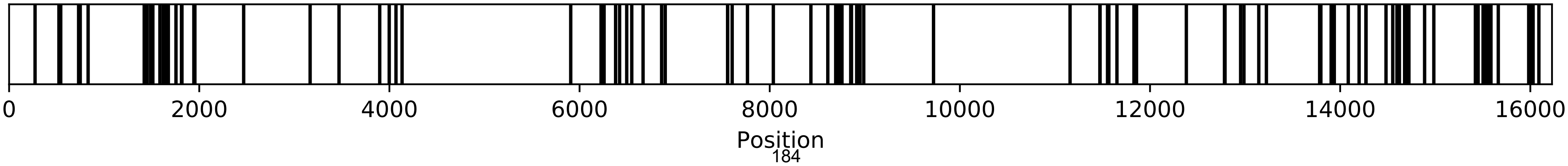

### Region: 586



MA1 vs MA2
Number of differences: 235

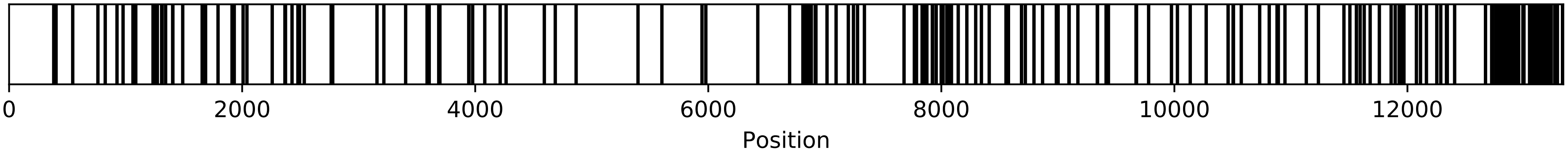


MA1 vs MM1
Number of differences: 235

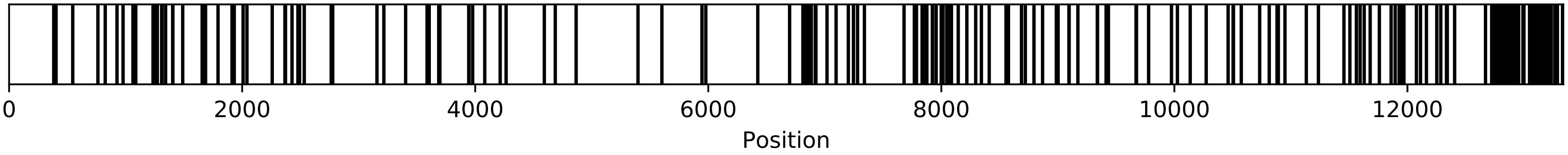


MA1 vs MM2
Number of differences: 232

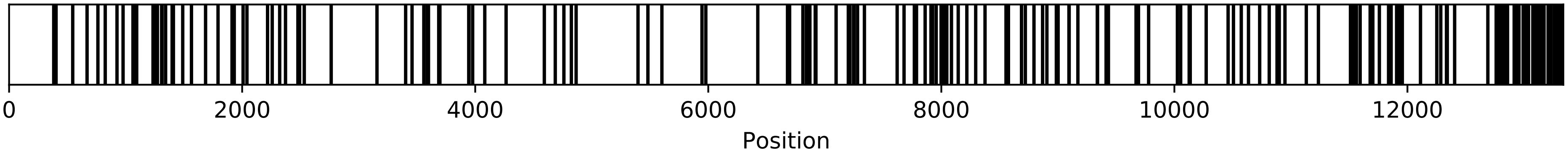


MA2 vs MM1
Number of differences: 0

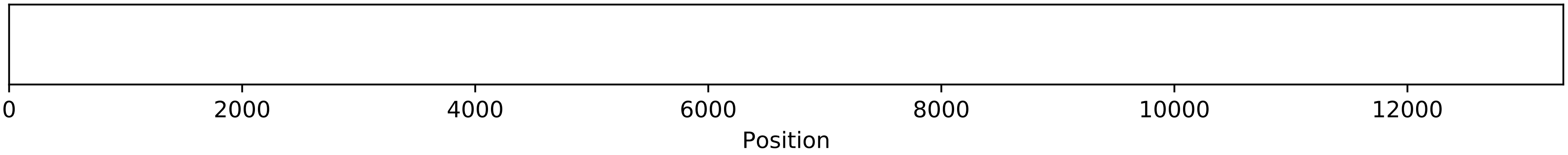


MA2 vs MM2
Number of differences: 112

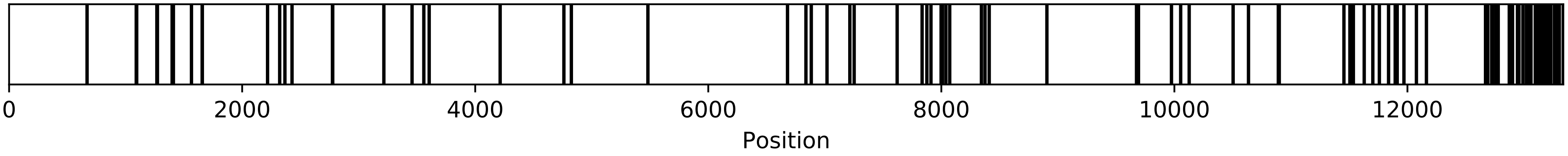


MM1 vs MM2
Number of differences: 112

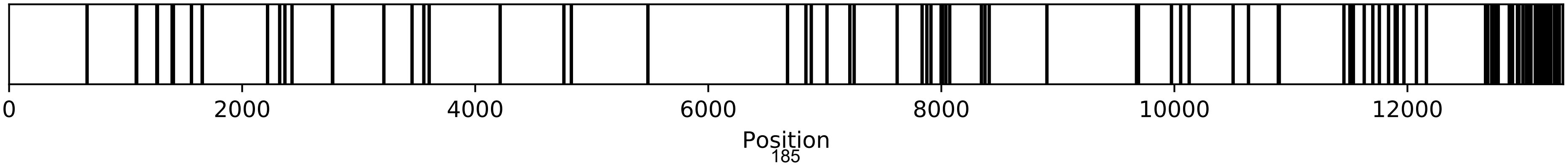

### Region: 588



MA1 vs MA2
Number of differences: 337

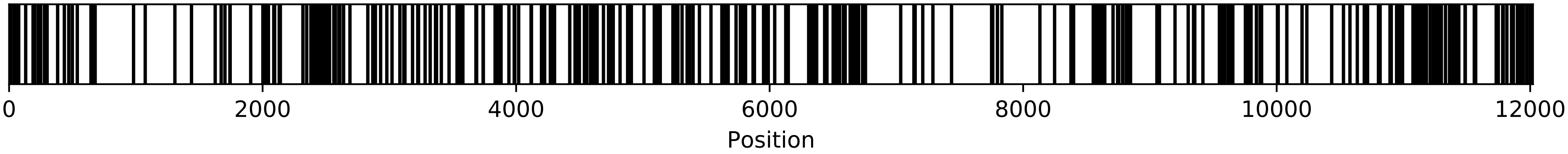


MA1 vs MM1
Number of differences: 0

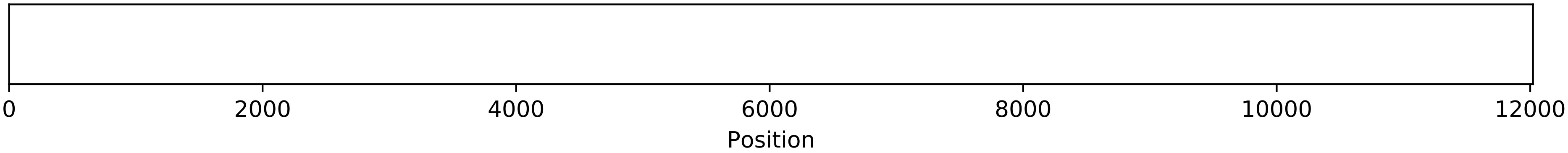


MA1 vs MM2
Number of differences: 194

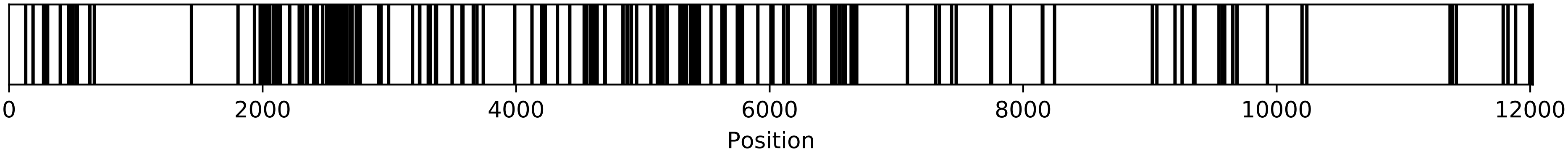


MA2 vs MM1
Number of differences: 337

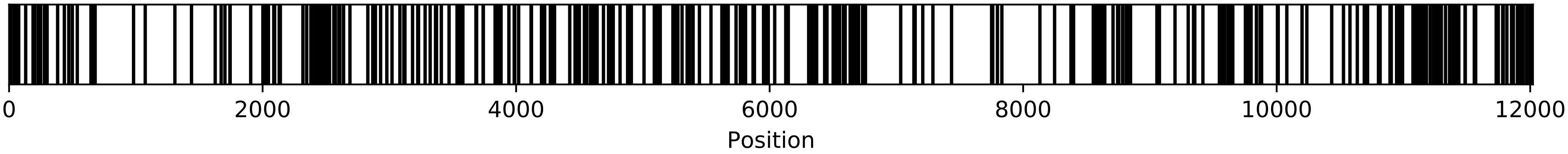


MA2 vs MM2
Number of differences: 358

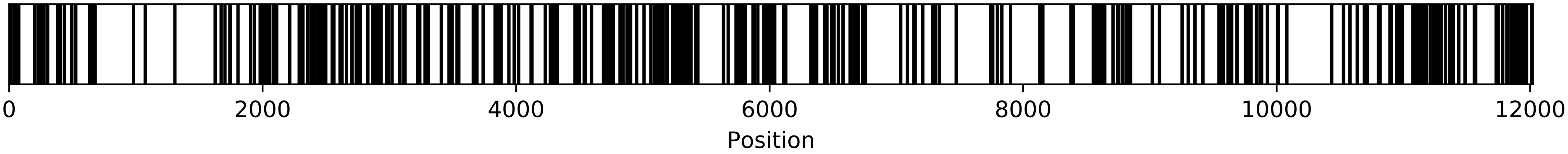


MM1 vs MM2
Number of differences: 194

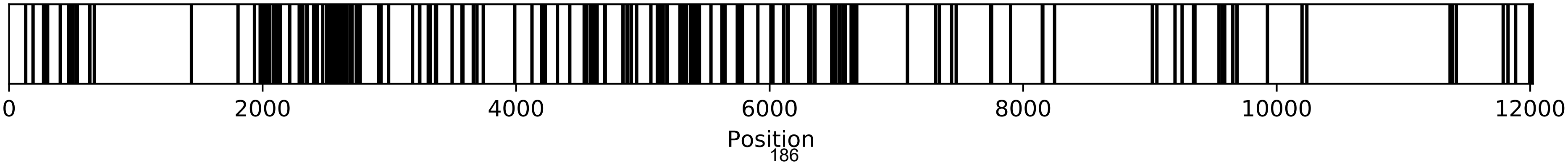

### Region: 589



MA1 vs MA2
Number of differences: 223

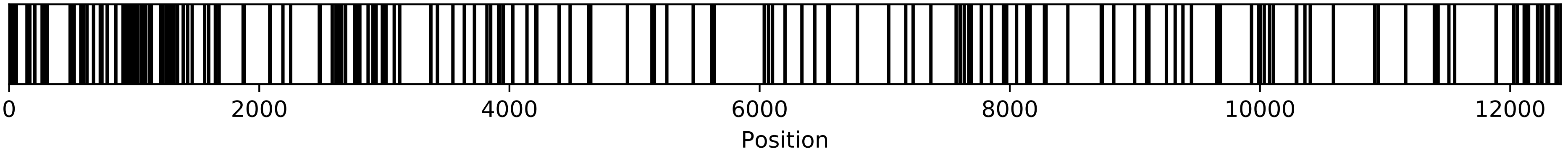


MA1 vs MM1
Number of differences: 60

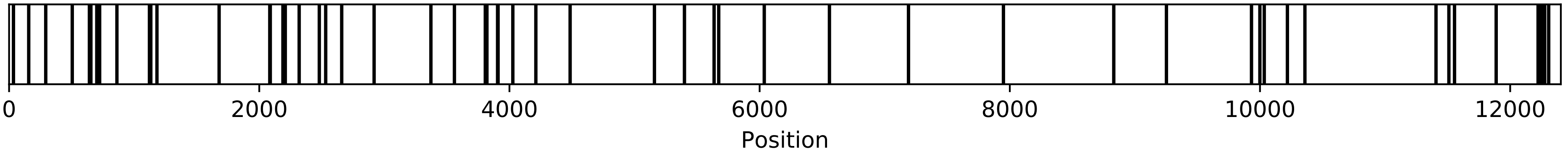


MA1 vs MM2
Number of differences: 73

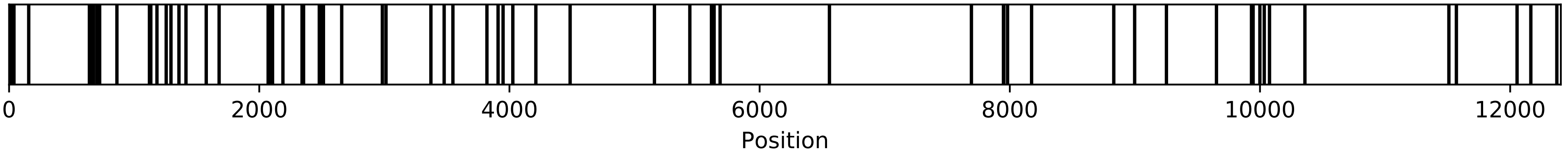


MA2 vs MM1
Number of differences: 214

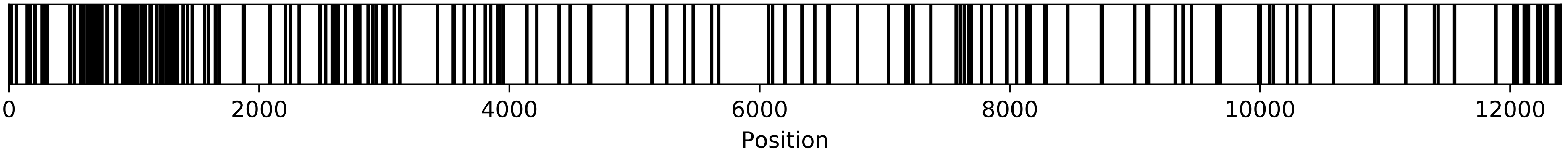


MA2 vs MM2
Number of differences: 219

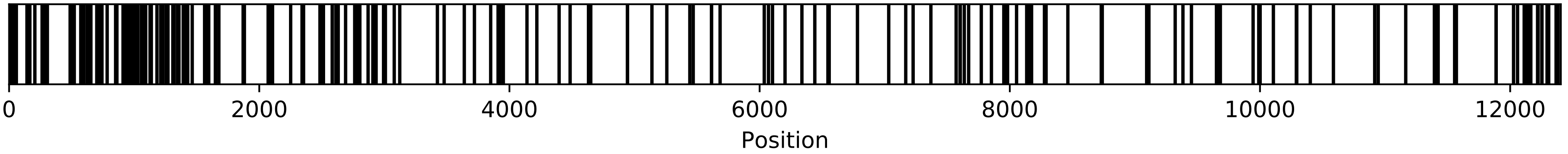


MM1 vs MM2
Number of differences: 66

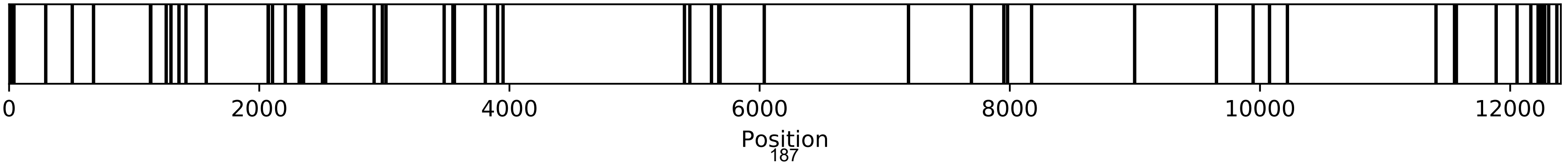

### Region: 592



## MA1 vs MA2
Number of differences: 148

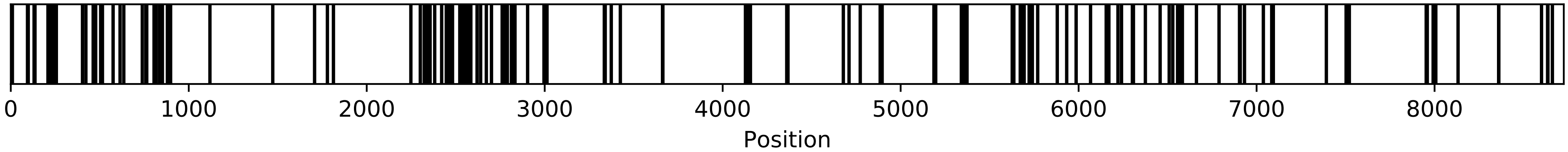


## MA1 vs MM1
Number of differences: 151

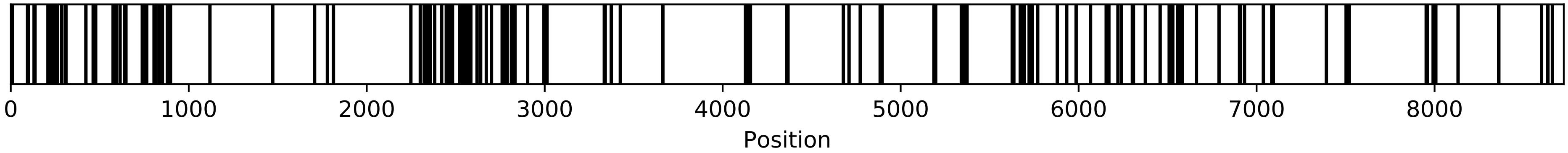


## MA1 vs MM2
Number of differences: 158

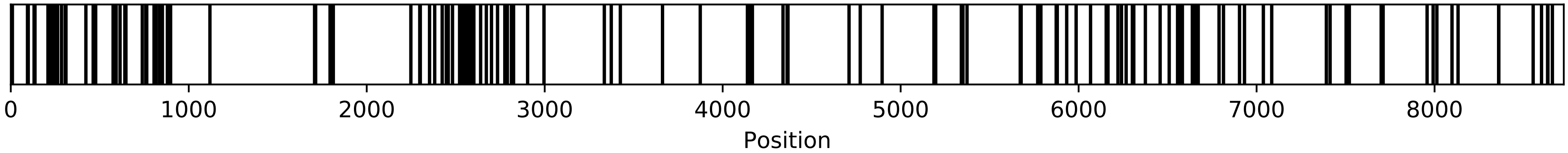


## MA2 vs MM1
Number of differences: 19

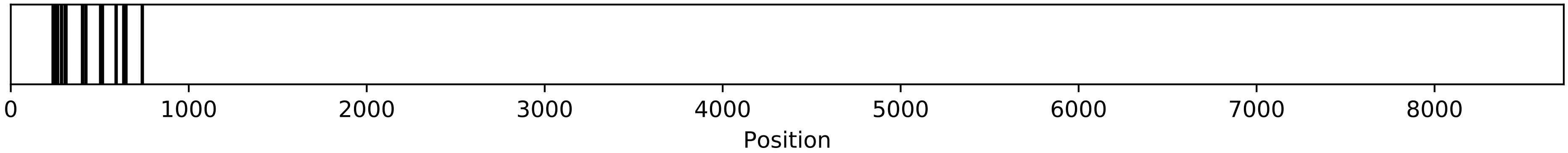


## MA2 vs MM2
Number of differences: 99

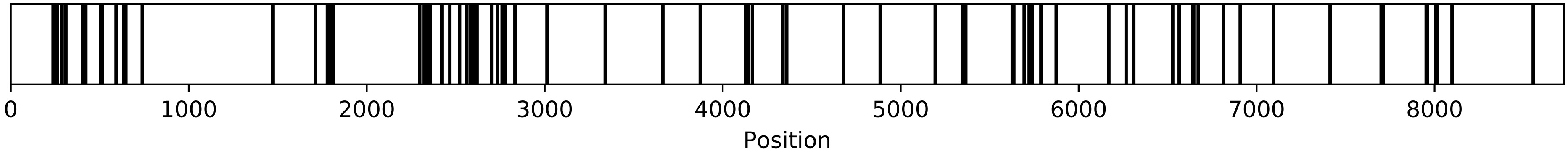


## MM1 vs MM2
Number of differences: 80

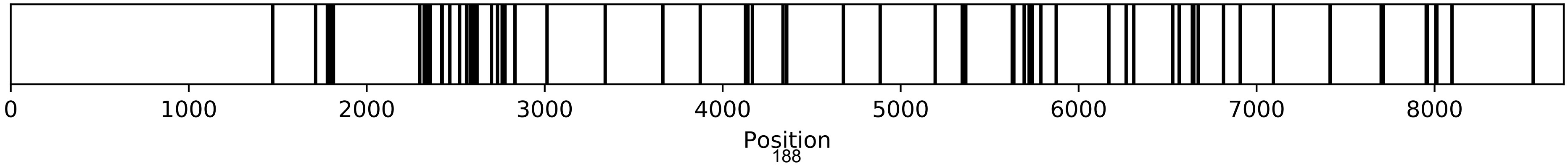

### Region: 596

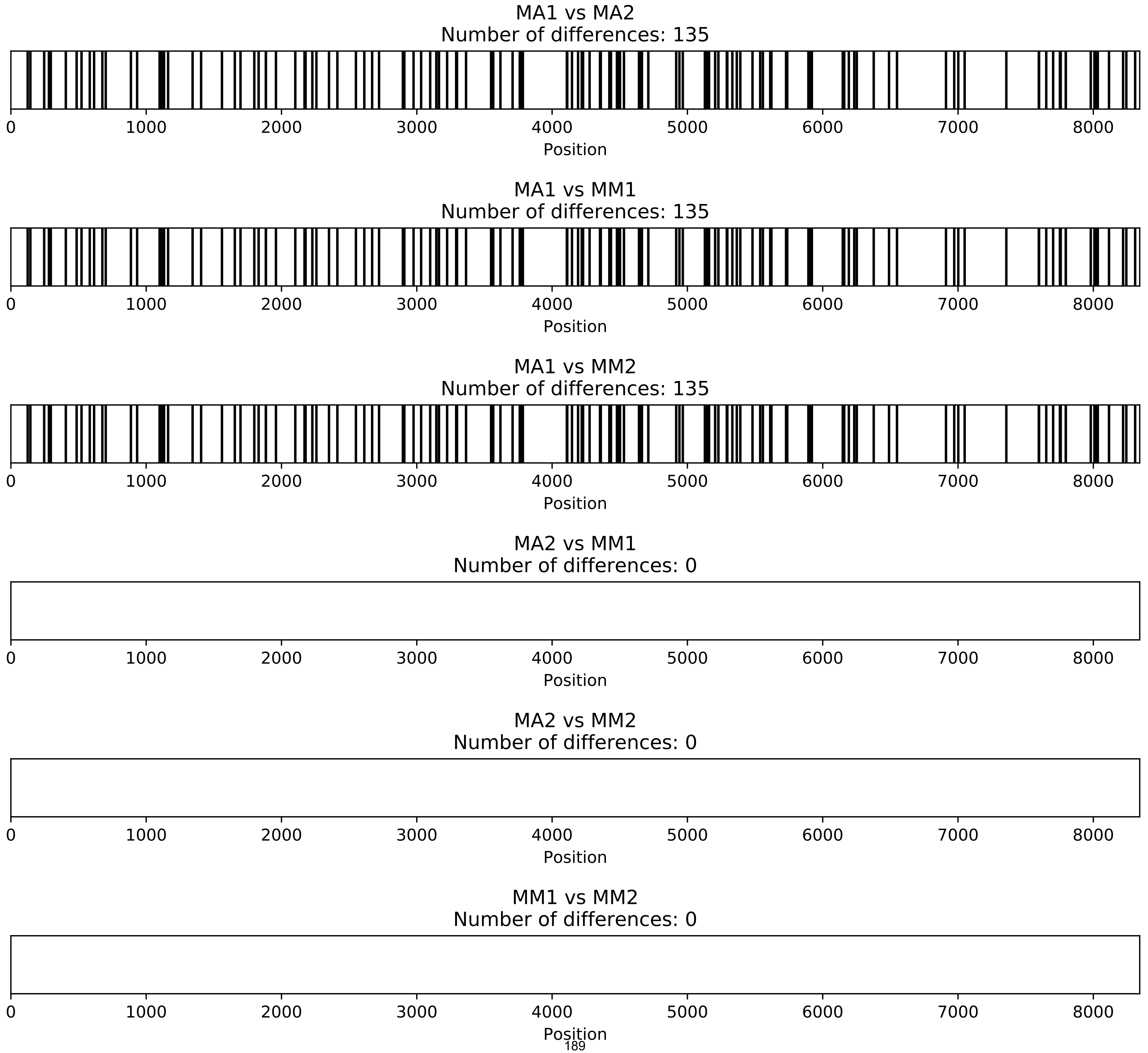
MA1 vs MA2
Number of differences: 135
MA1 vs MM1
Number of differences: 135
MA1 vs MM2
Number of differences: 135
MA2 vs MM1
Number of differences: 0
MA2 vs MM2
Number of differences: 0
MM1 vs MM2
Number of differences: 0
0
1000
2000
3000
4000
5000
6000
7000
8000
Position

### Region: 599



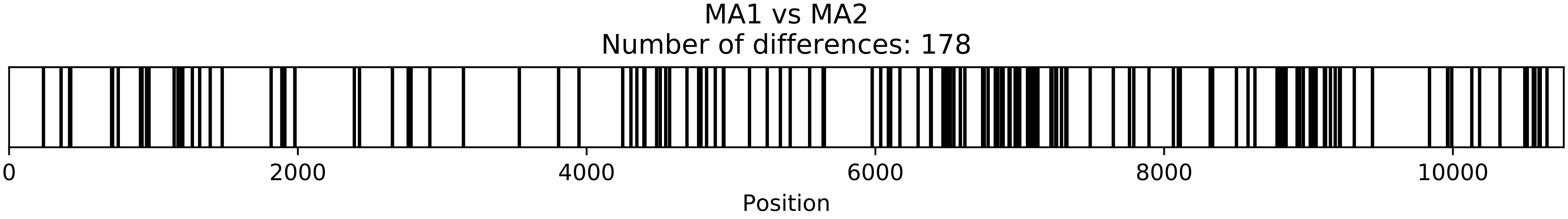
MA1 vs MA2
Number of differences: 178
0
2000
4000
6000
8000
10000
Position

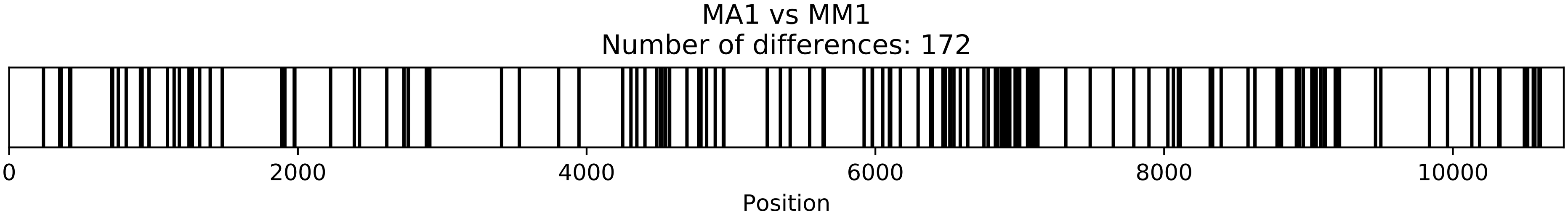
MA1 vs MM1
Number of differences: 172
0
2000
4000
6000
8000
10000
Position

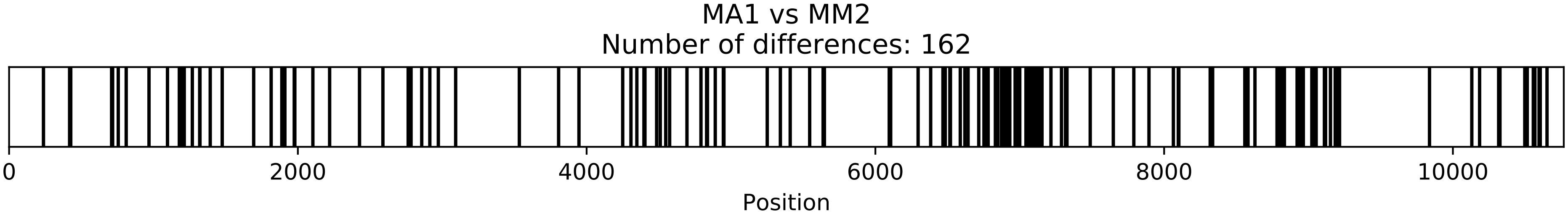
MA1 vs MM2
Number of differences: 162
0
2000
4000
6000
8000
10000
Position

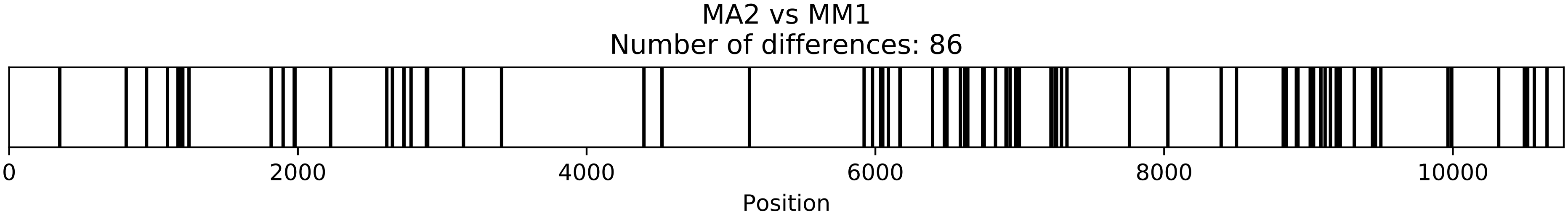
MA2 vs MM1
Number of differences: 86
0
2000
4000
6000
8000
10000
Position

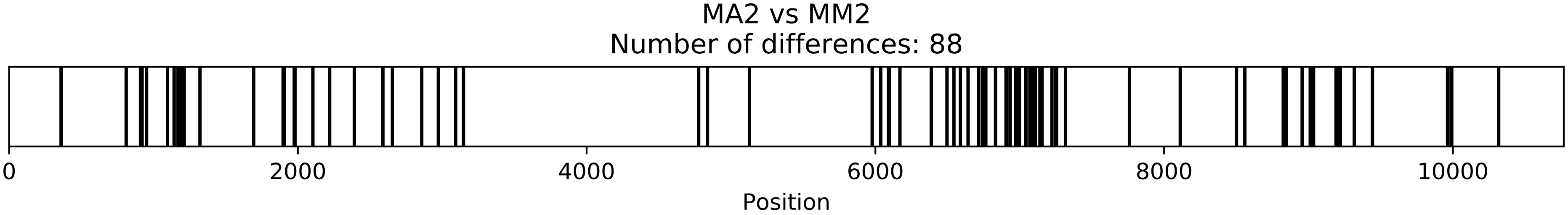
MA2 vs MM2
Number of differences: 88
0
2000
4000
6000
8000
10000
Position

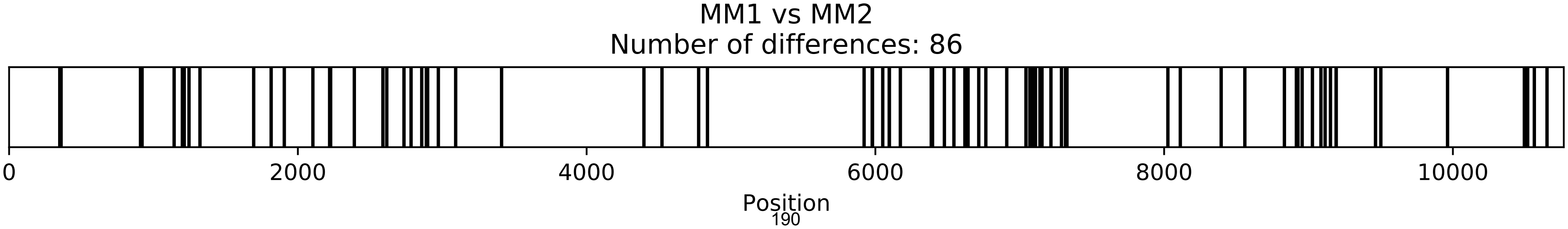
MM1 vs MM2
Number of differences: 86
0
2000
4000
6000
8000
10000
Position

### Region: 605



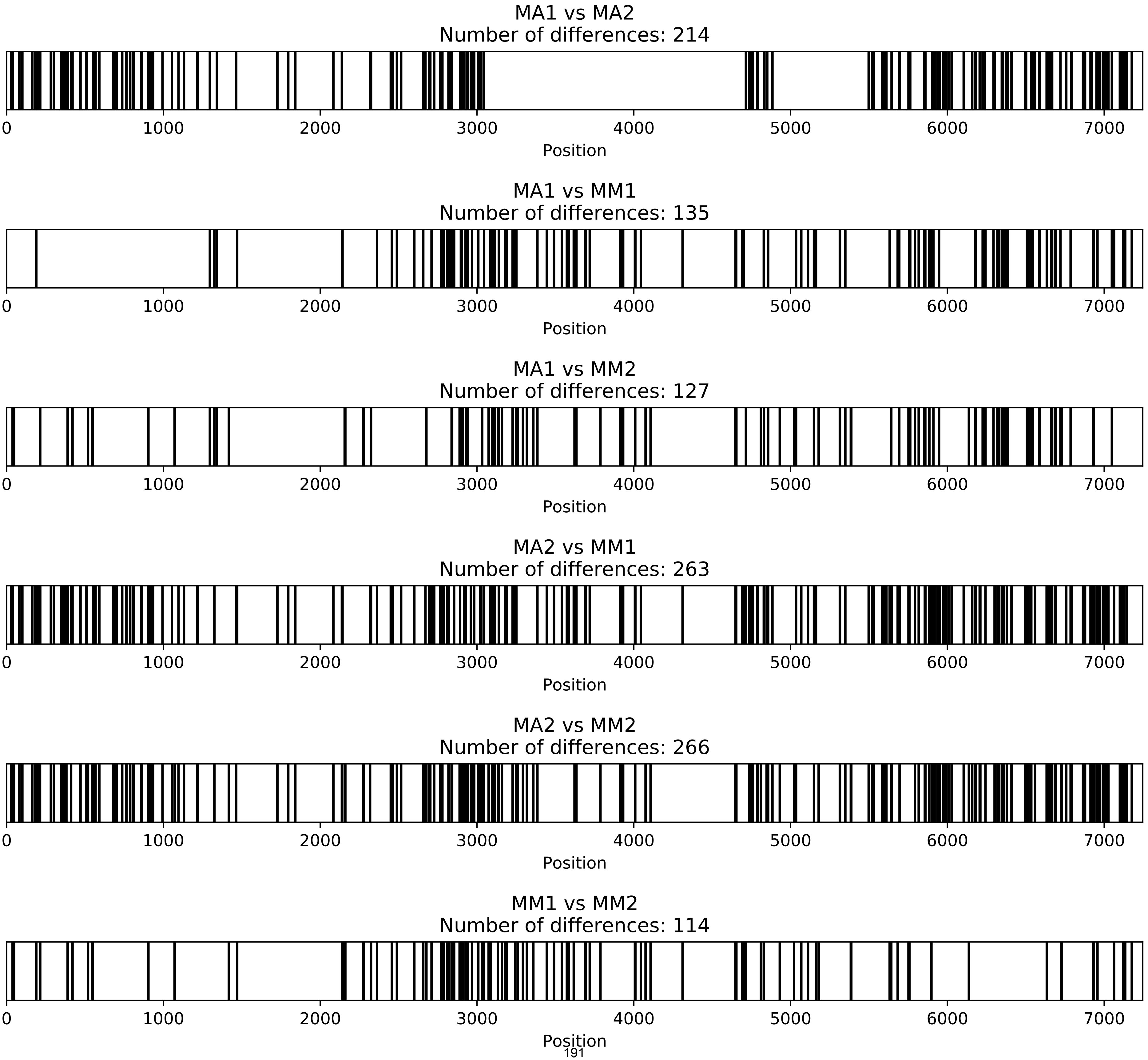
MA1 vs MA2
Number of differences: 214
MA1 vs MM1
Number of differences: 135
MA1 vs MM2
Number of differences: 127
MA2 vs MM1
Number of differences: 263
MA2 vs MM2
Number of differences: 266
MM1 vs MM2
Number of differences: 114
0
1000
2000
3000
4000
5000
6000
7000
Position

### Region: 608



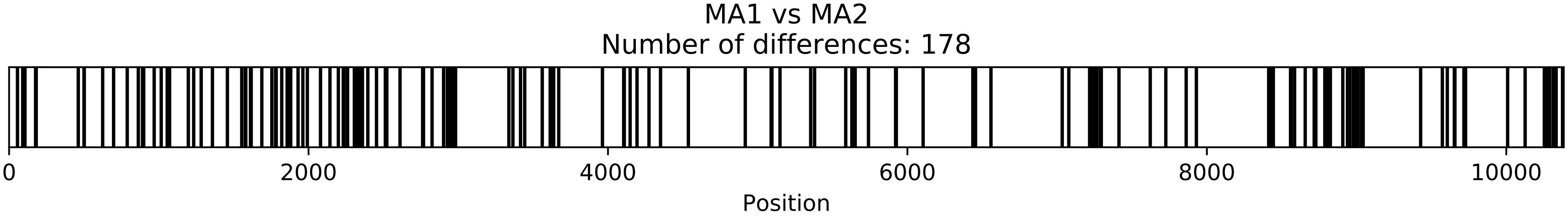
MA1 vs MA2
Number of differences: 178
0
2000
4000
6000
8000
10000
Position

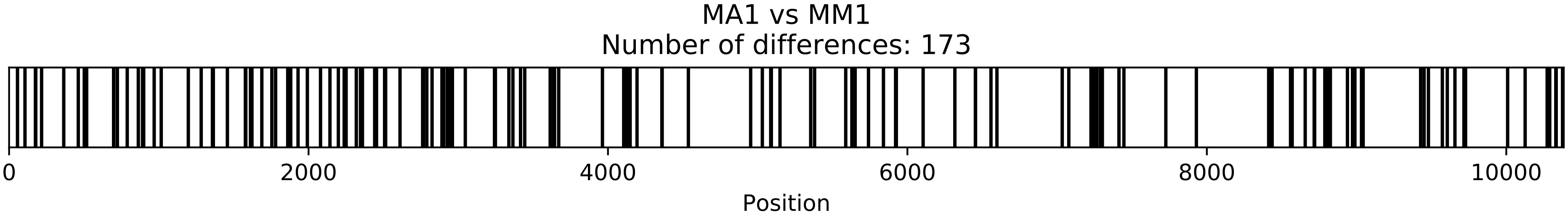
MA1 vs MM1
Number of differences: 173
0
2000
4000
6000
8000
10000
Position

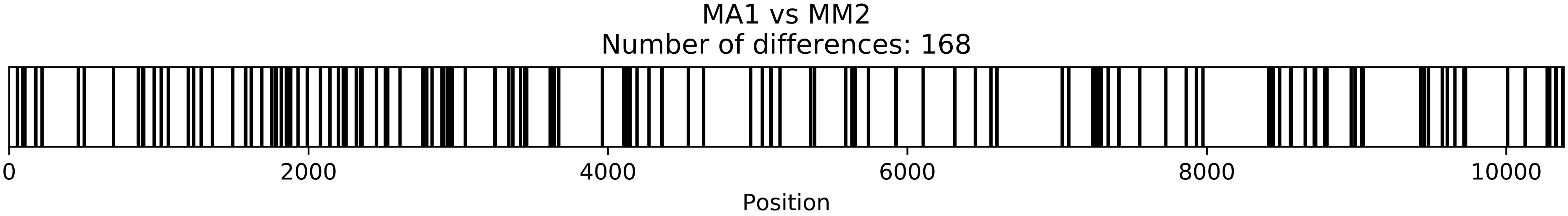
MA1 vs MM2
Number of differences: 168
0
2000
4000
6000
8000
10000
Position

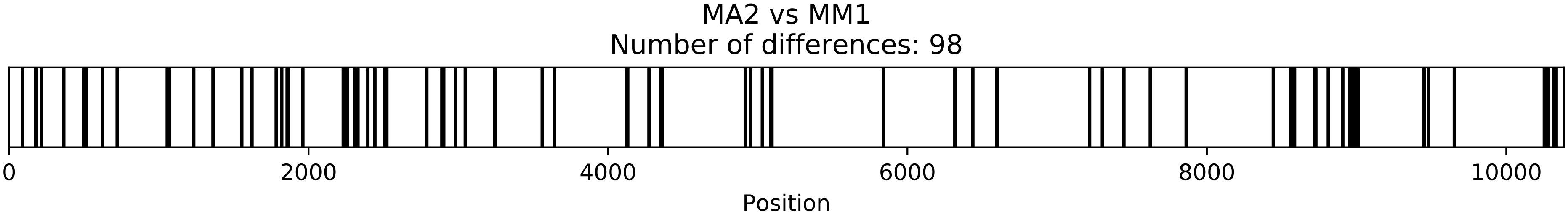
MA2 vs MM1
Number of differences: 98
0
2000
4000
6000
8000
10000
Position

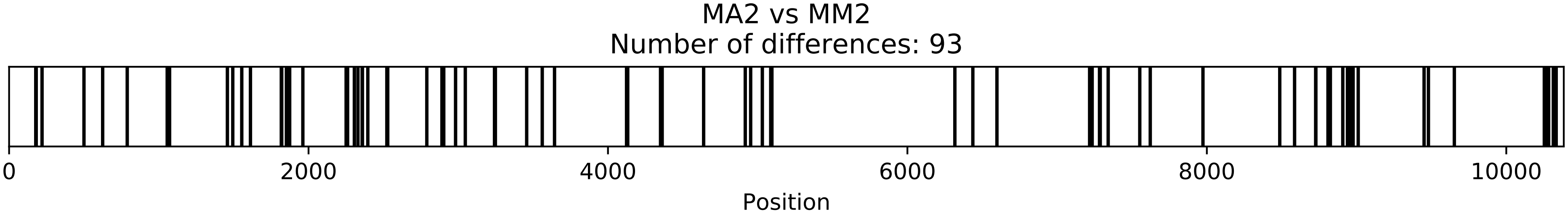
MA2 vs MM2
Number of differences: 93
0
2000
4000
6000
8000
10000
Position

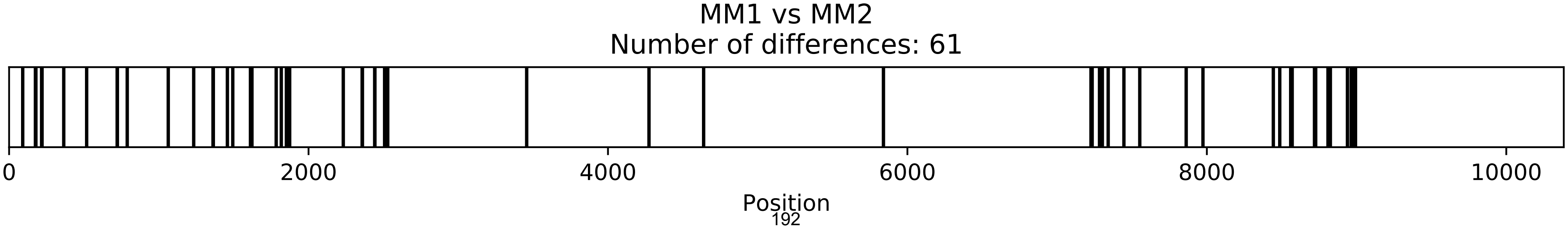
MM1 vs MM2
Number of differences: 61
0
2000
4000
6000
8000
10000
Position

### Region: 610



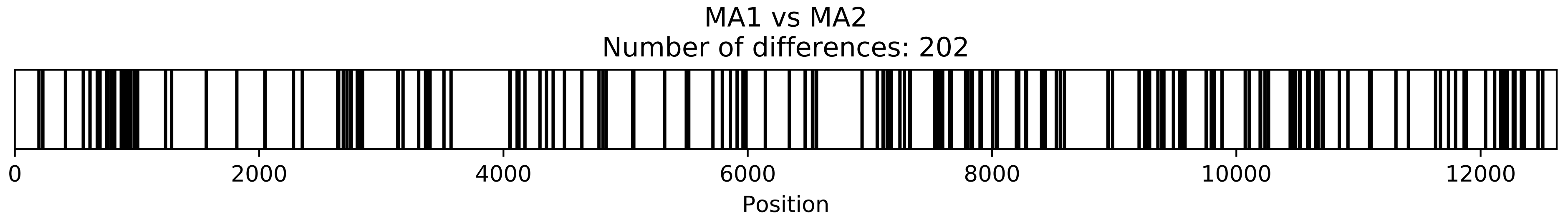
MA1 vs MA2
Number of differences: 202
0
2000
4000
6000
8000
10000
12000
Position

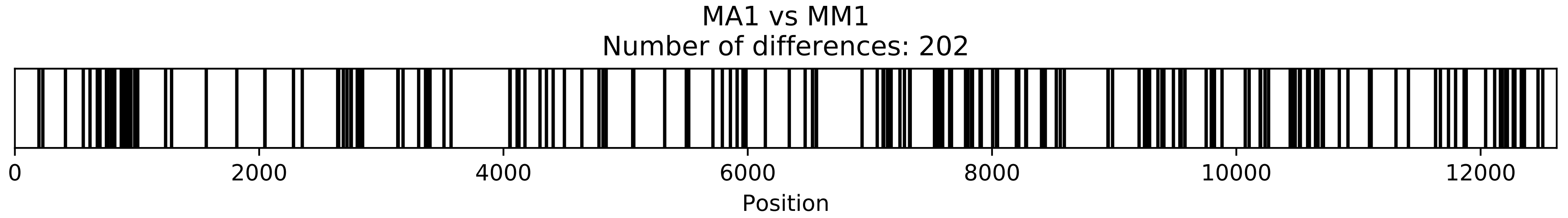
MA1 vs MM1
Number of differences: 202
0
2000
4000
6000
8000
10000
12000
Position

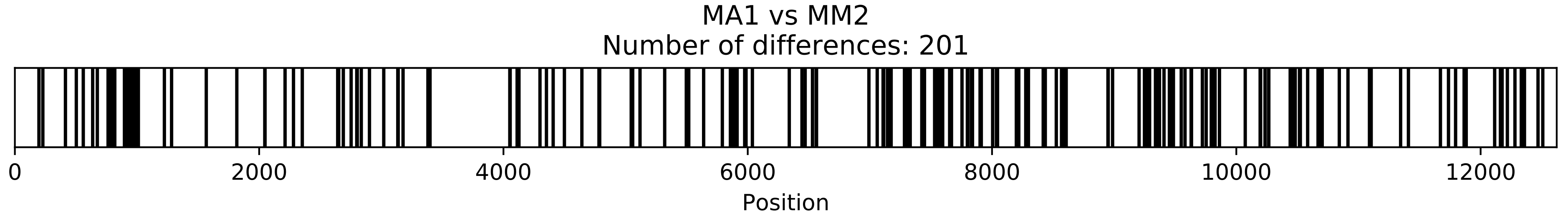
MA1 vs MM2
Number of differences: 201
0
2000
4000
6000
8000
10000
12000
Position

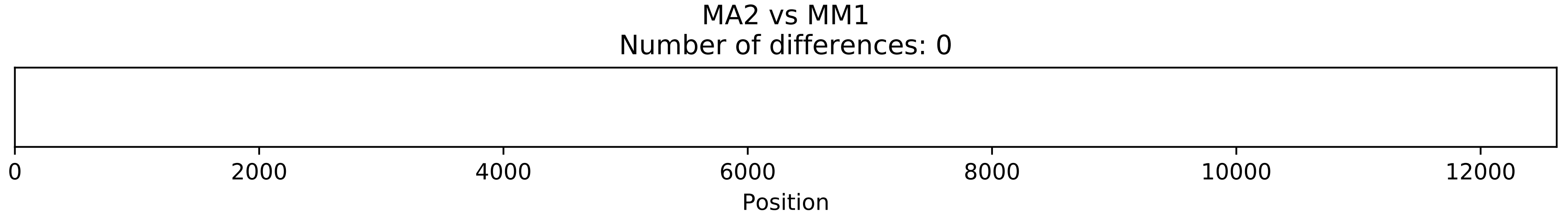
MA2 vs MM1
Number of differences: 0
0
2000
4000
6000
8000
10000
12000
Position

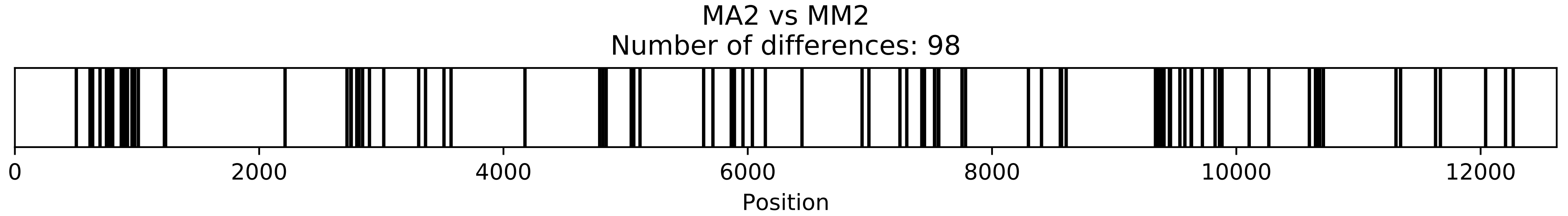
MA2 vs MM2
Number of differences: 98
0
2000
4000
6000
8000
10000
12000
Position

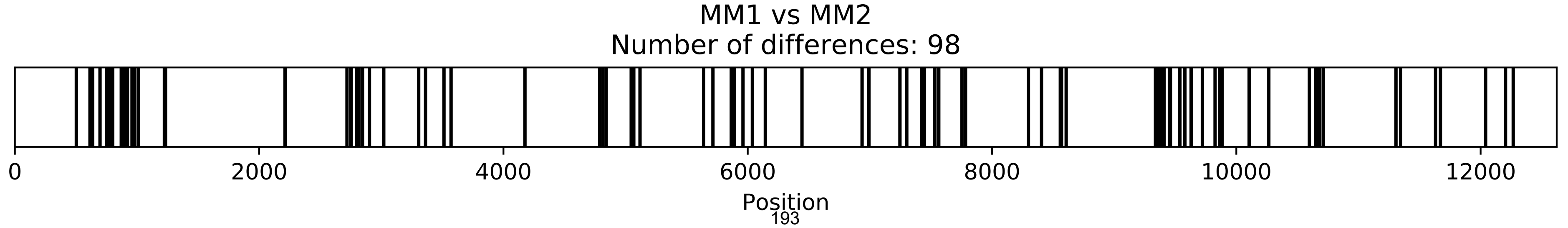
MM1 vs MM2
Number of differences: 98
0
2000
4000
6000
8000
10000
12000
Position

### Region: 613



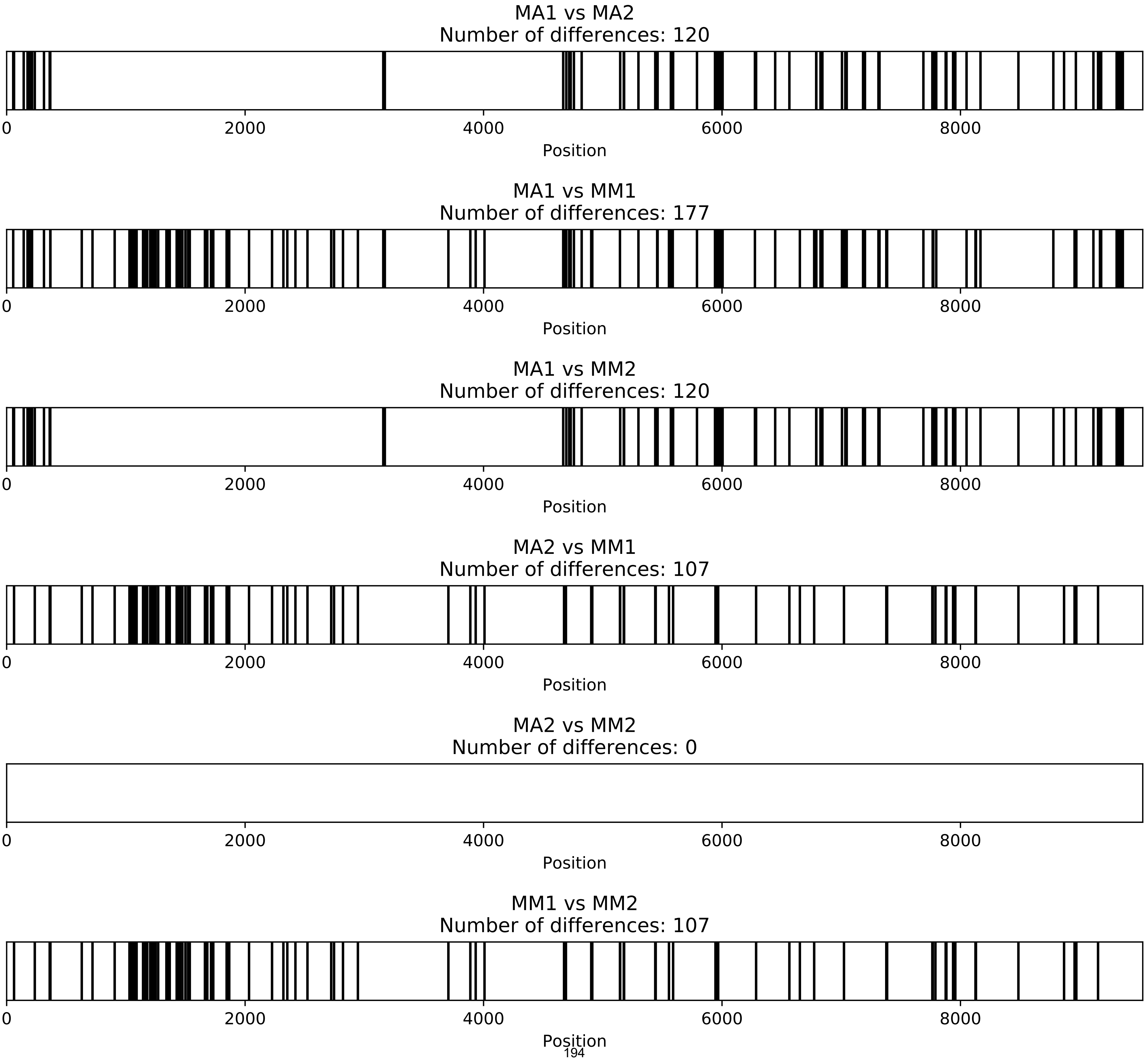
MA1 vs MA2
Number of differences: 120
0
2000
4000
6000
8000
Position
MA1 vs MM1
Number of differences: 177
0
2000
4000
6000
8000
Position
MA1 vs MM2
Number of differences: 120
0
2000
4000
6000
8000
Position
MA2 vs MM1
Number of differences: 107
0
2000
4000
6000
8000
Position
MA2 vs MM2
Number of differences: 0
0
2000
4000
6000
8000
Position
MM1 vs MM2
Number of differences: 107
0
2000
4000
6000
8000
Position

### Region: 622



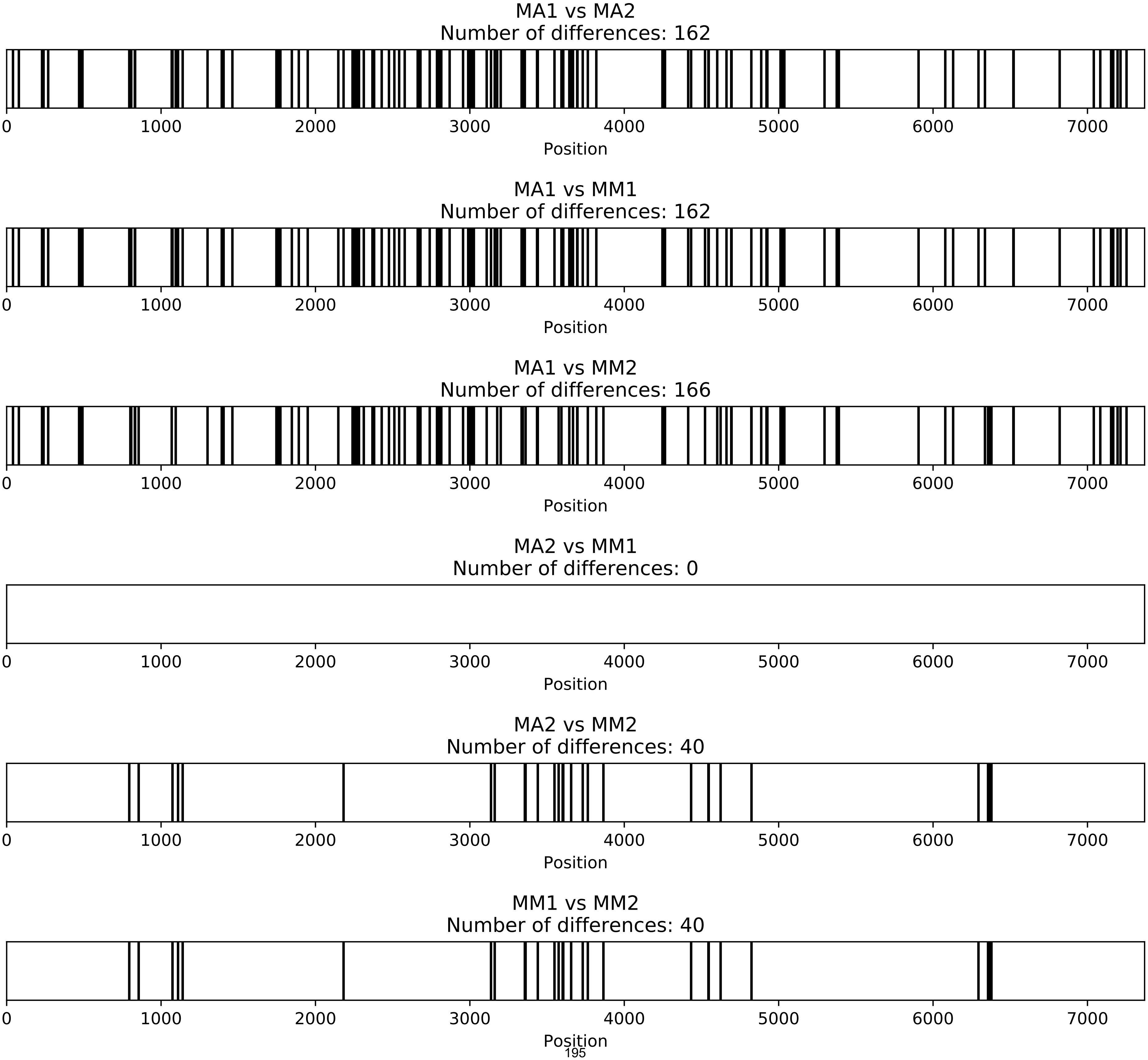
MA1 vs MA2
Number of differences: 162
MA1 vs MM1
Number of differences: 162
MA1 vs MM2
Number of differences: 166
MA2 vs MM1
Number of differences: 0
MA2 vs MM2
Number of differences: 40
MM1 vs MM2
Number of differences: 40
0
1000
2000
3000
4000
5000
6000
7000
Position

### Region: 626



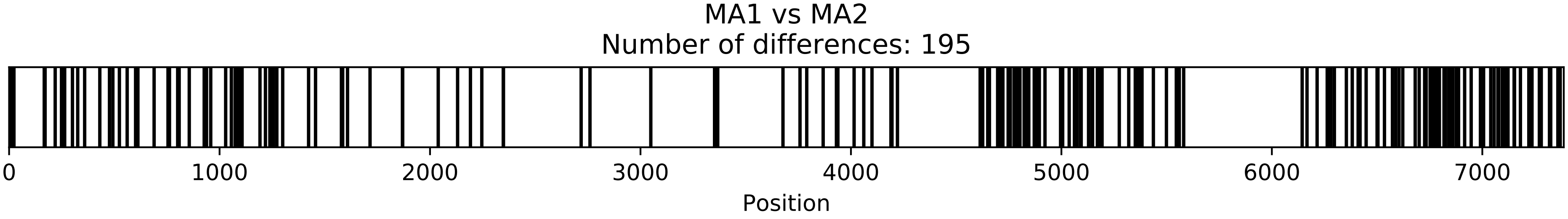
MA1 vs MA2
Number of differences: 195
0
1000
2000
3000
4000
5000
6000
7000
Position

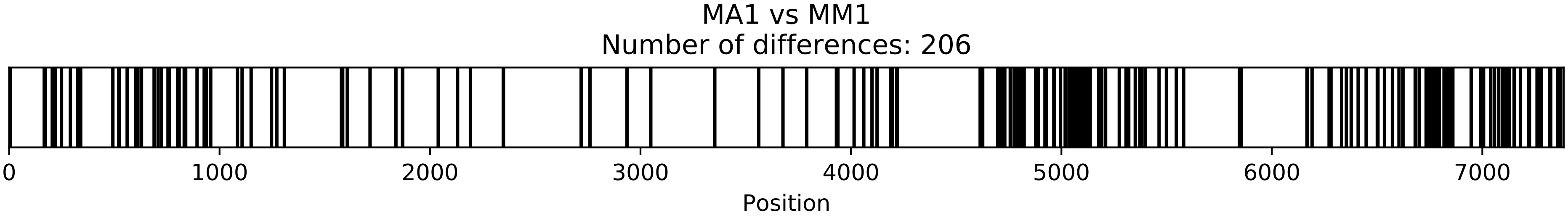
MA1 vs MM1
Number of differences: 206
0
1000
2000
3000
4000
5000
6000
7000
Position

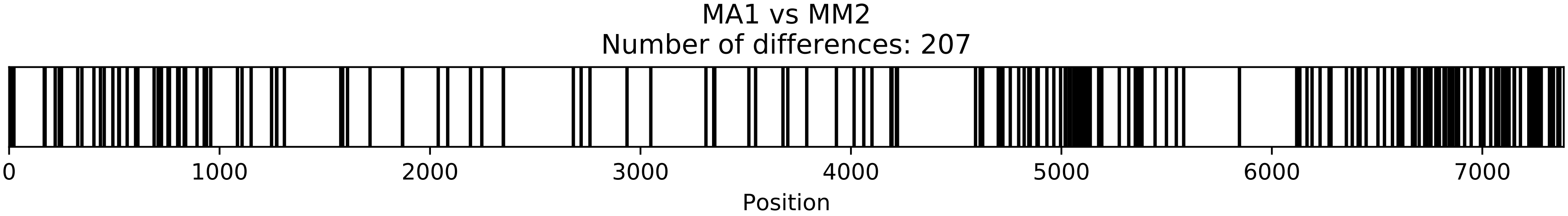
MA1 vs MM2
Number of differences: 207
0
1000
2000
3000
4000
5000
6000
7000
Position

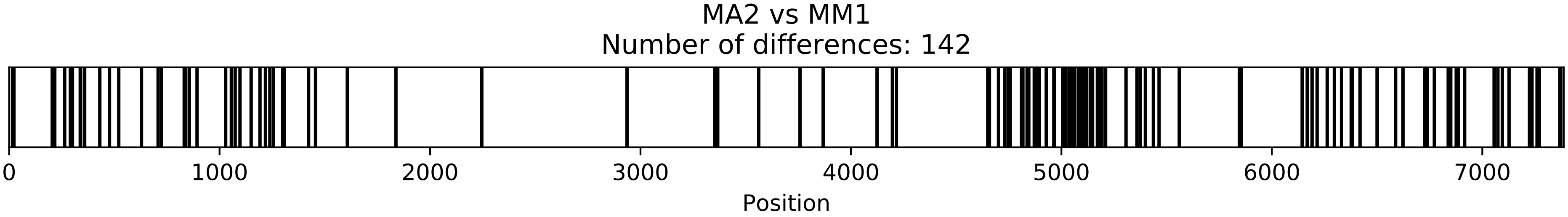
MA2 vs MM1
Number of differences: 142
0
1000
2000
3000
4000
5000
6000
7000
Position

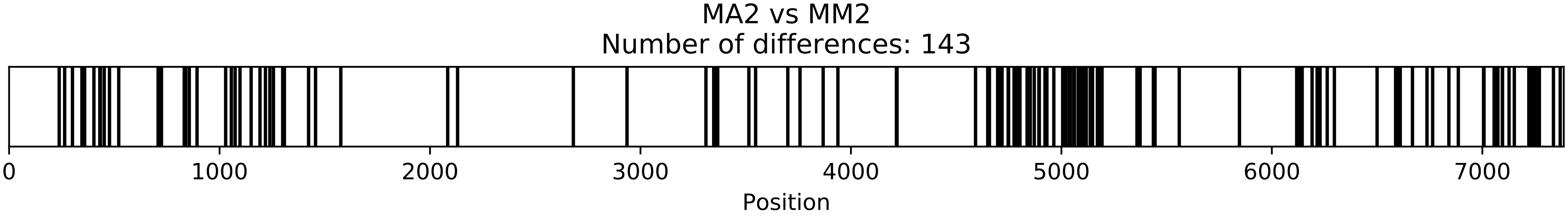
MA2 vs MM2
Number of differences: 143
0
1000
2000
3000
4000
5000
6000
7000
Position

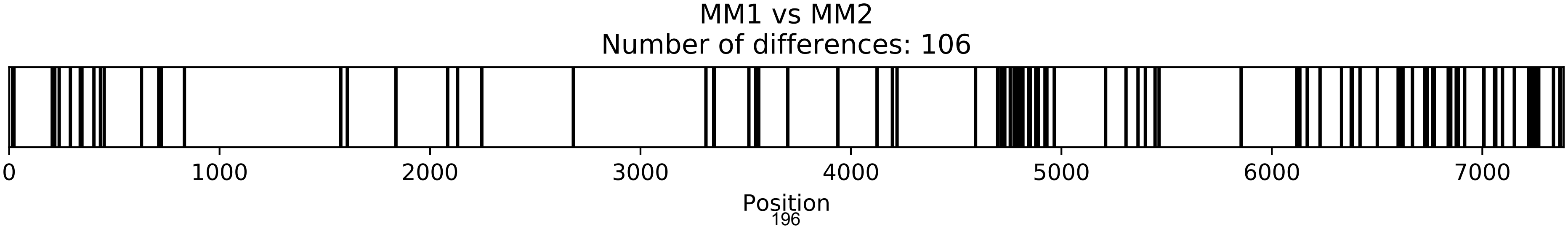
MM1 vs MM2
Number of differences: 106
0
1000
2000
3000
4000
5000
6000
7000
Position

### Region: 635



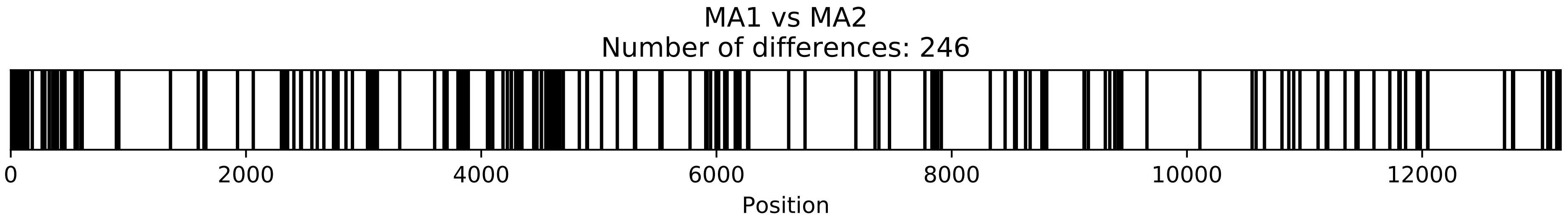
MA1 vs MA2
Number of differences: 246
0
2000
4000
6000
8000
10000
12000
Position

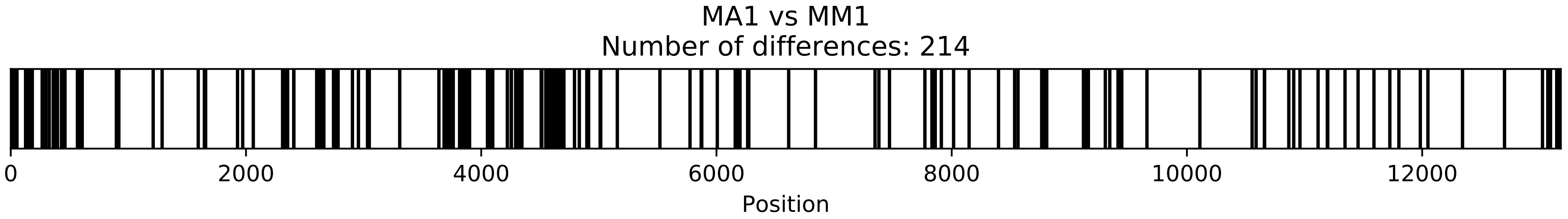
MA1 vs MM1
Number of differences: 214
0
2000
4000
6000
8000
10000
12000
Position

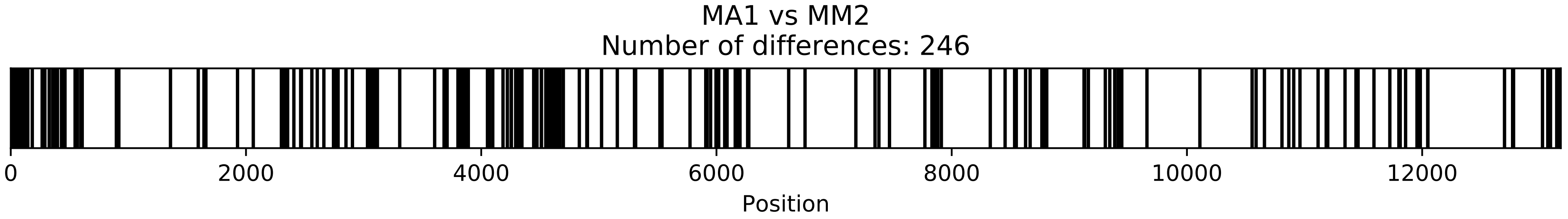
MA1 vs MM2
Number of differences: 246
0
2000
4000
6000
8000
10000
12000
Position

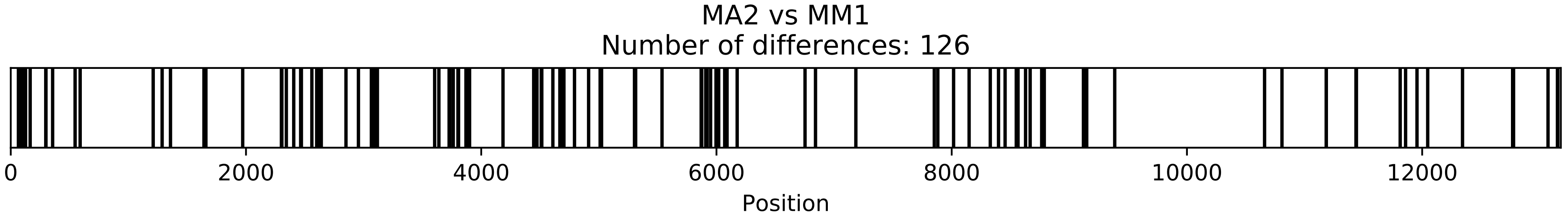
MA2 vs MM1
Number of differences: 126
0
2000
4000
6000
8000
10000
12000
Position

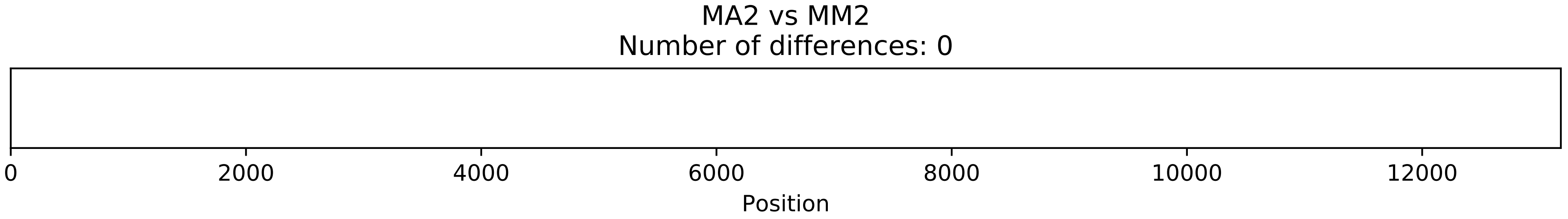
MA2 vs MM2
Number of differences: 0
0
2000
4000
6000
8000
10000
12000
Position

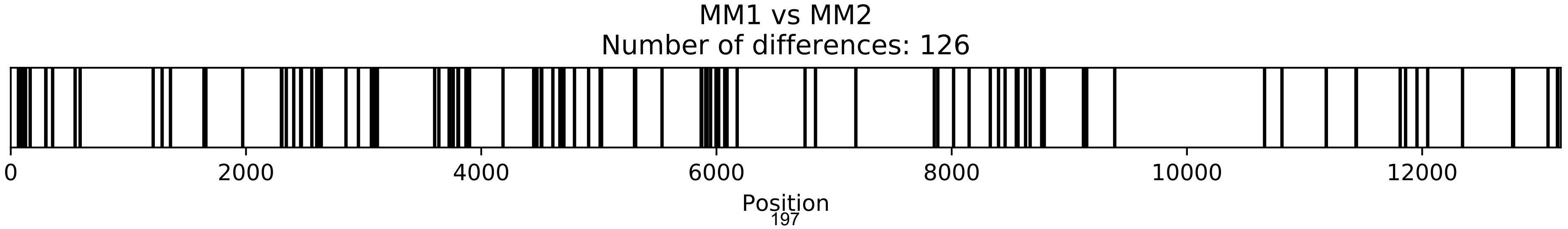
MM1 vs MM2
Number of differences: 126
0
2000
4000
6000
8000
10000
12000
Position

### Region: 638



MA1 vs MA2
Number of differences: 267

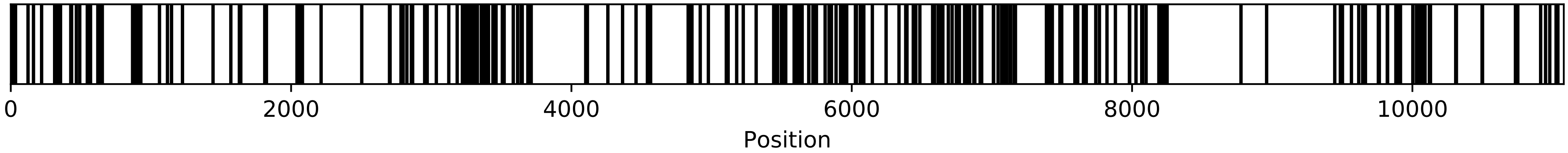


MA1 vs MM1
Number of differences: 268

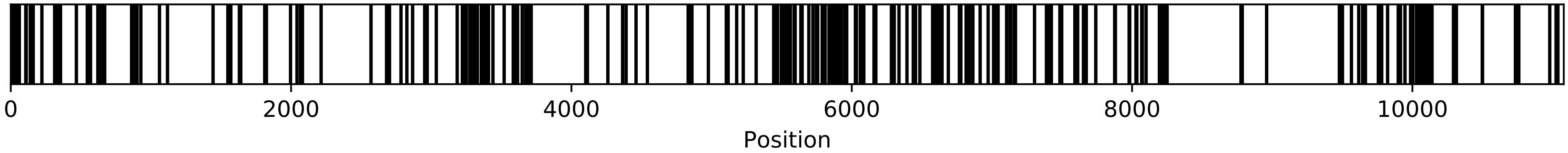


MA1 vs MM2
Number of differences: 272

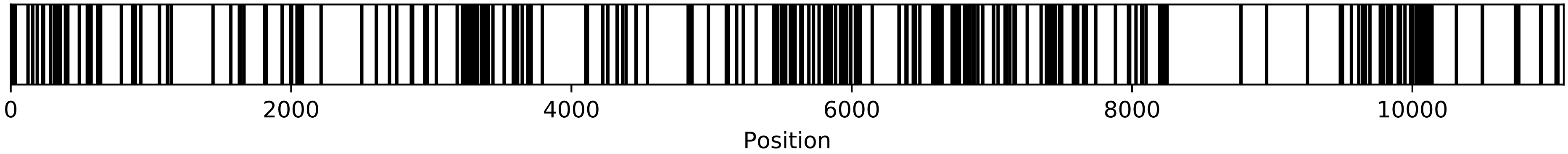


MA2 vs MM1
Number of differences: 144

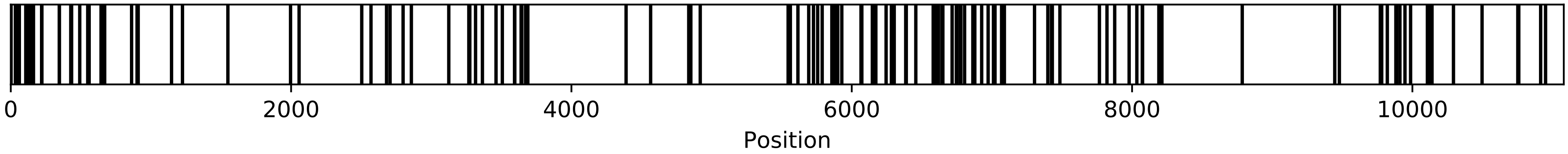


MA2 vs MM2
Number of differences: 152

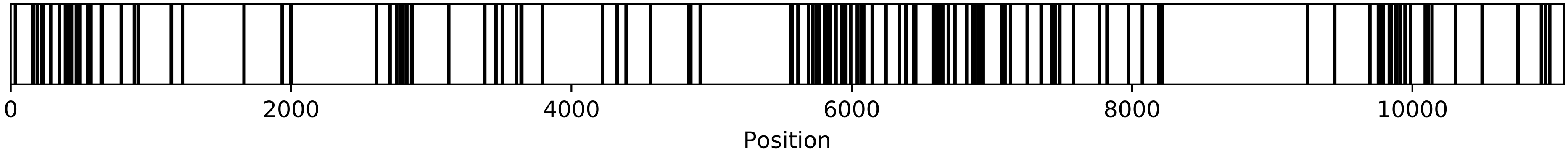


MM1 vs MM2
Number of differences: 148

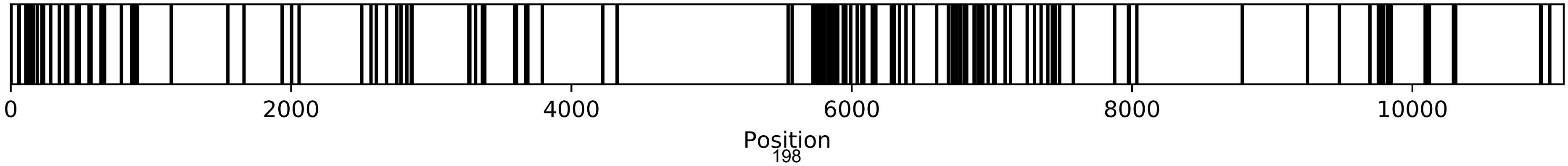

### Region: 639



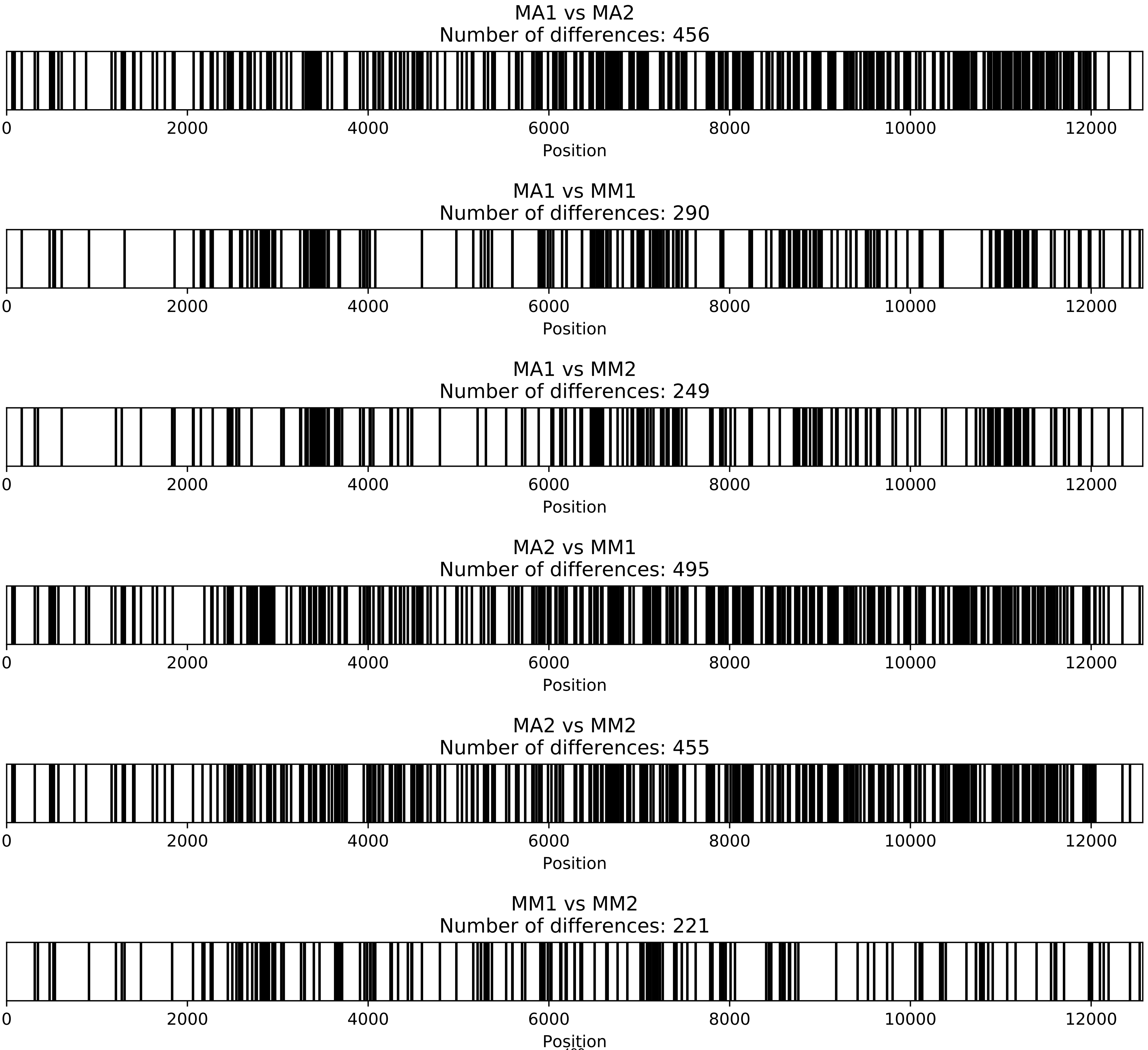
MA1 vs MA2
Number of differences: 456
MA1 vs MM1
Number of differences: 290
MA1 vs MM2
Number of differences: 249
MA2 vs MM1
Number of differences: 495
MA2 vs MM2
Number of differences: 455
MM1 vs MM2
Number of differences: 221
0
2000
4000
6000
8000
10000
12000
Position

### Region: 649



## MA1 vs MA2
Number of differences: 300

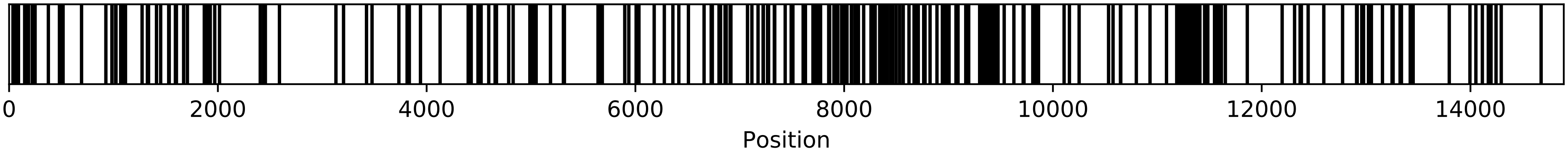


## MA1 vs MM1
Number of differences: 301

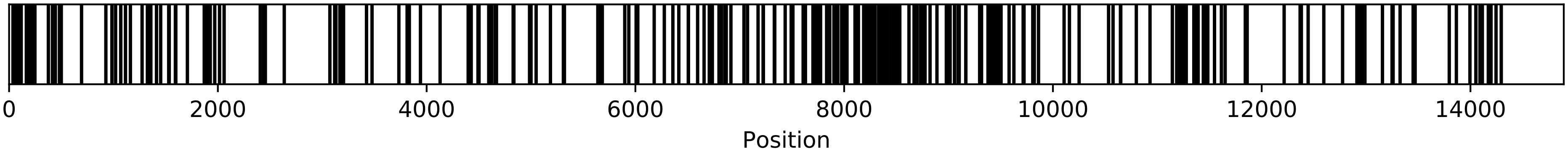


## MA1 vs MM2
Number of differences: 300

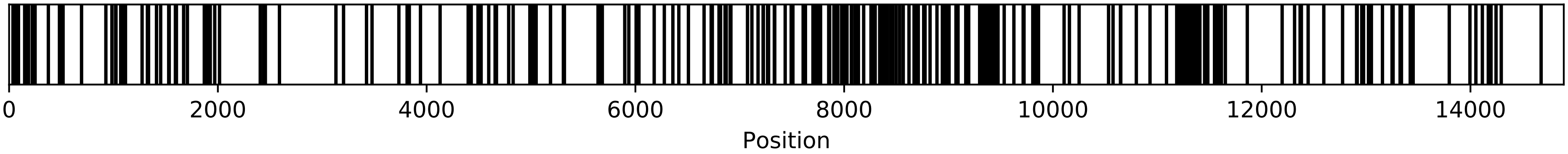


## MA2 vs MM1
Number of differences: 140

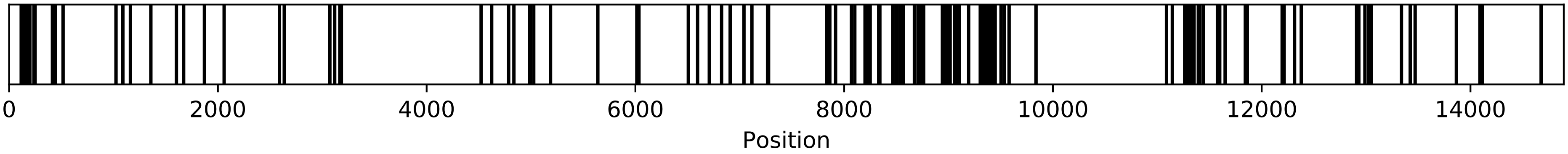


## MA2 vs MM2
Number of differences: 0

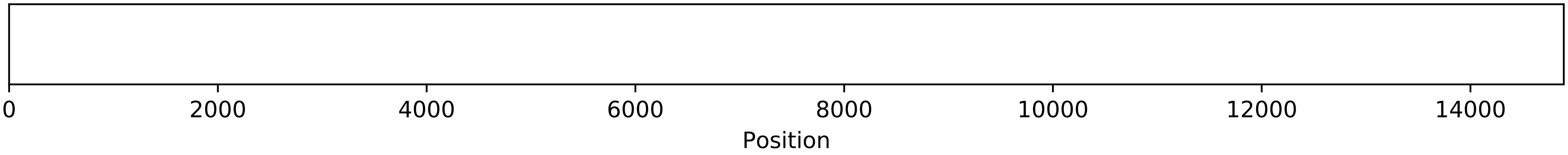


## MM1 vs MM2
Number of differences: 140

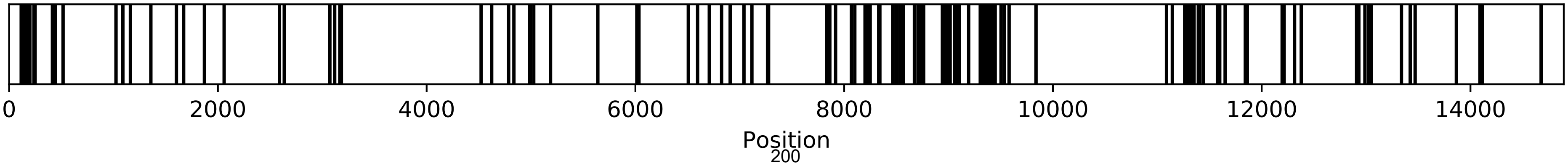

### Region: 650



MA1 vs MA2
Number of differences: 241

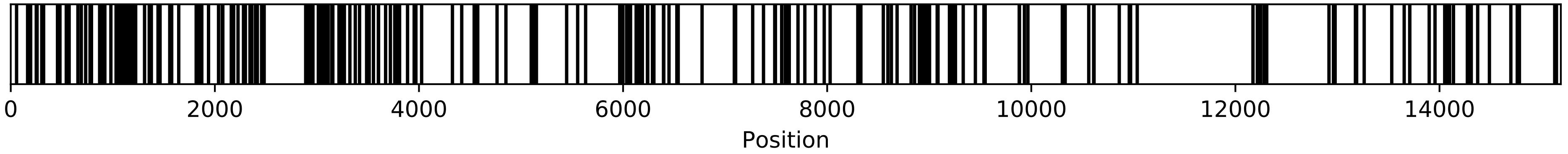


MA1 vs MM1
Number of differences: 0

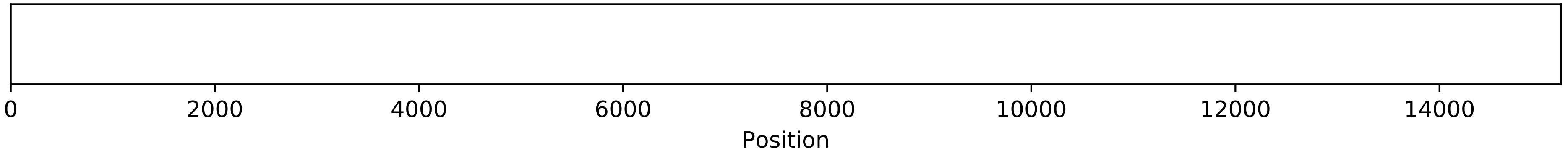


MA1 vs MM2
Number of differences: 134

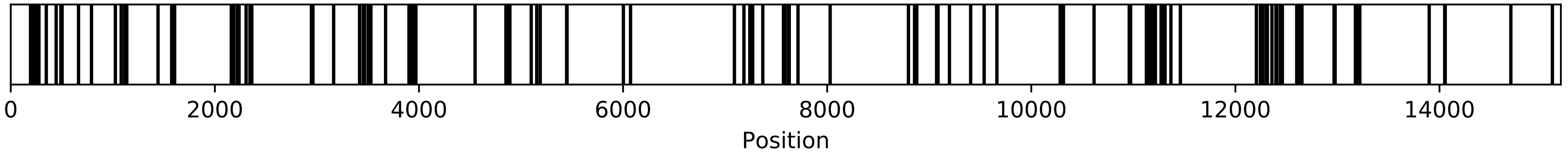


MA2 vs MM1
Number of differences: 241

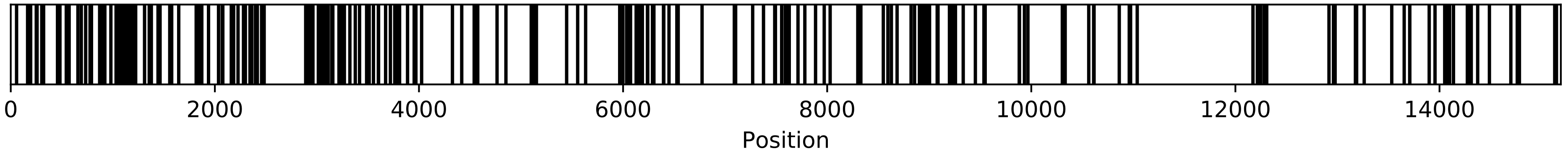


MA2 vs MM2
Number of differences: 271

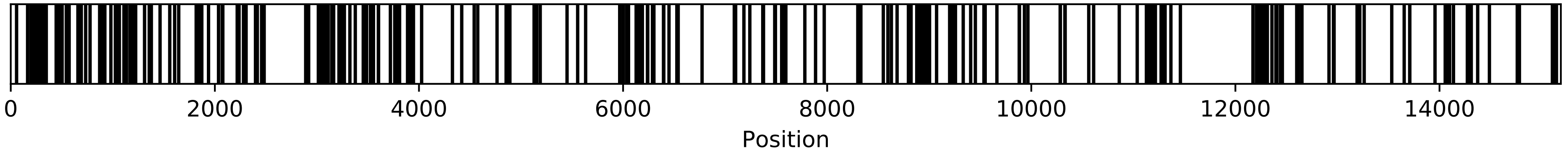


MM1 vs MM2
Number of differences: 134

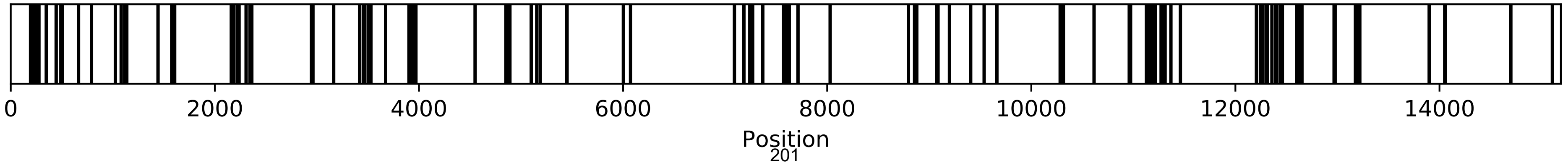

### Region: 652



MA1 vs MA2
Number of differences: 162

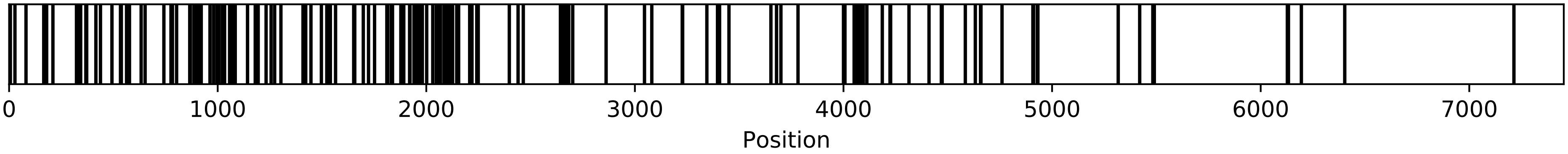


MA1 vs MM1
Number of differences: 98

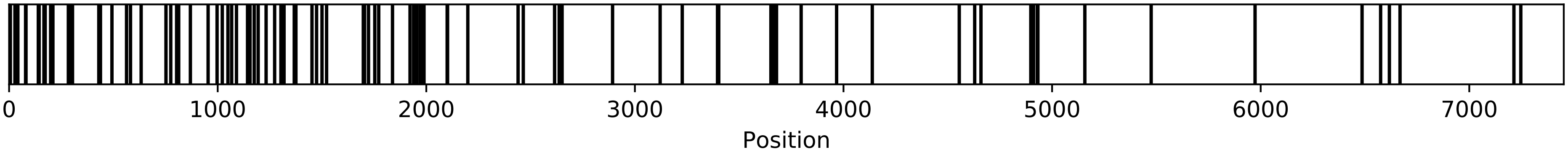


MA1 vs MM2
Number of differences: 19

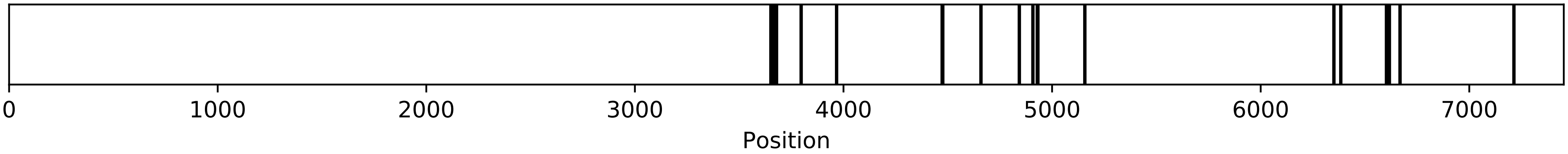


MA2 vs MM1
Number of differences: 177

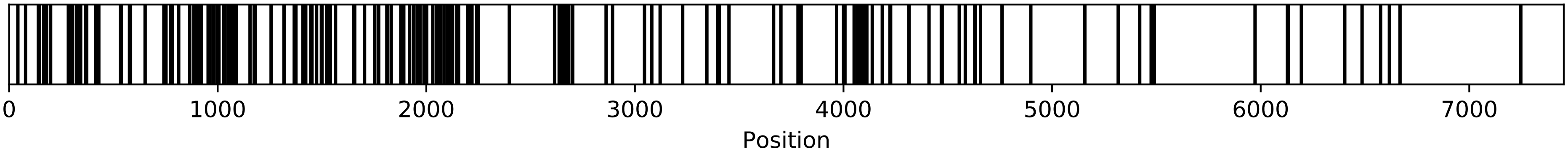


MA2 vs MM2
Number of differences: 165

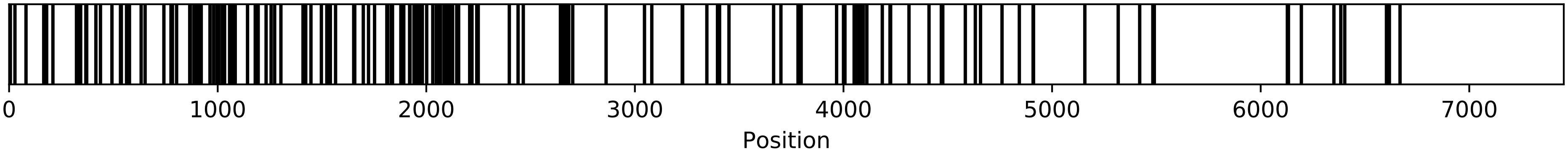


MM1 vs MM2
Number of differences: 92

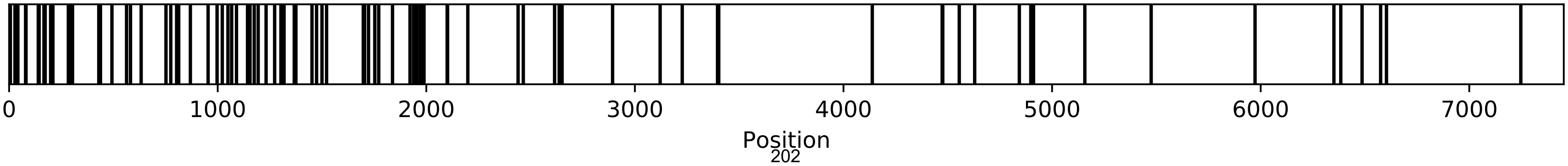

### Region: 661



MA1 vs MA2
Number of differences: 337

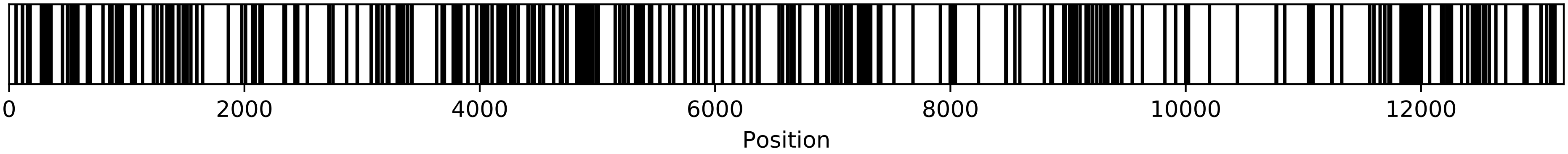


MA1 vs MM1
Number of differences: 337

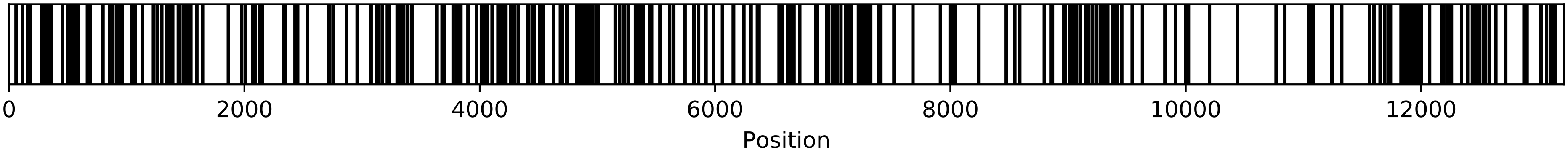


MA1 vs MM2
Number of differences: 351

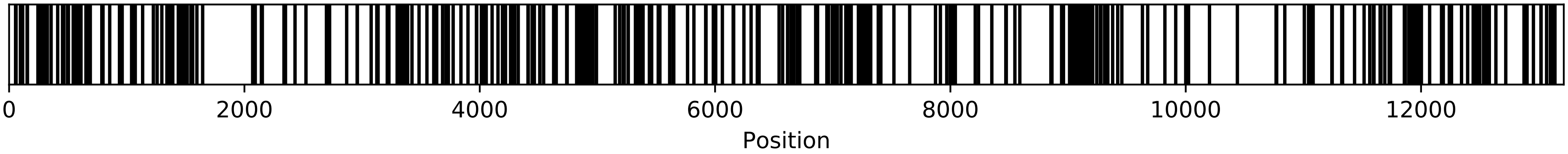


MA2 vs MM1
Number of differences: 0

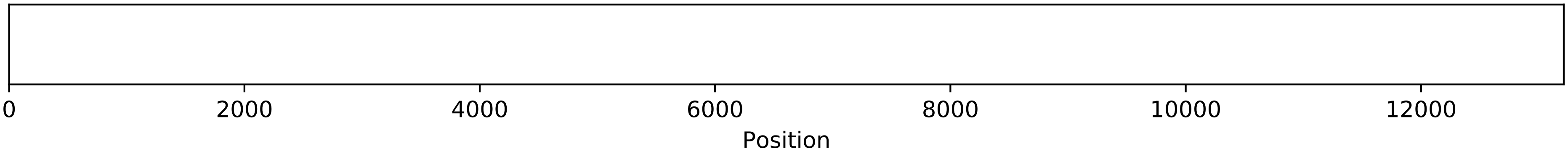


MA2 vs MM2
Number of differences: 129

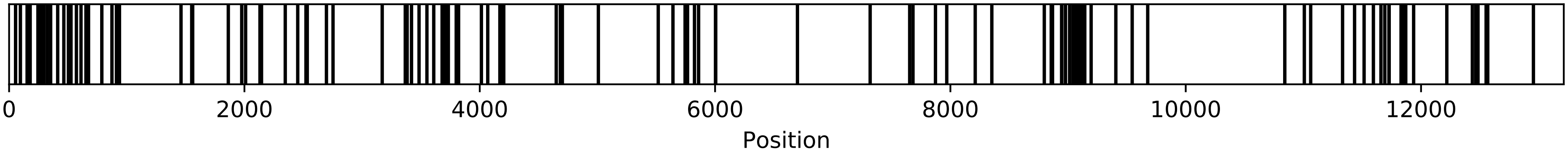


MM1 vs MM2
Number of differences: 129

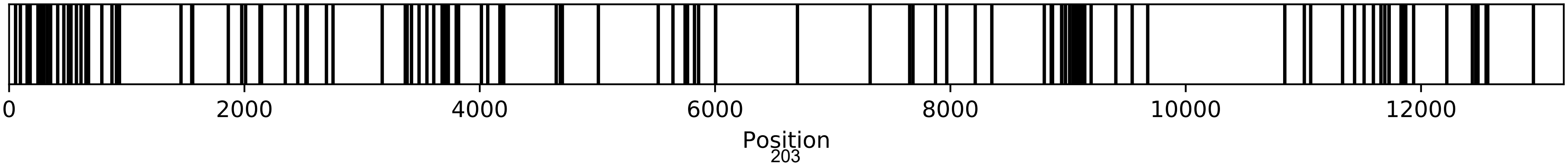

### Region: 662



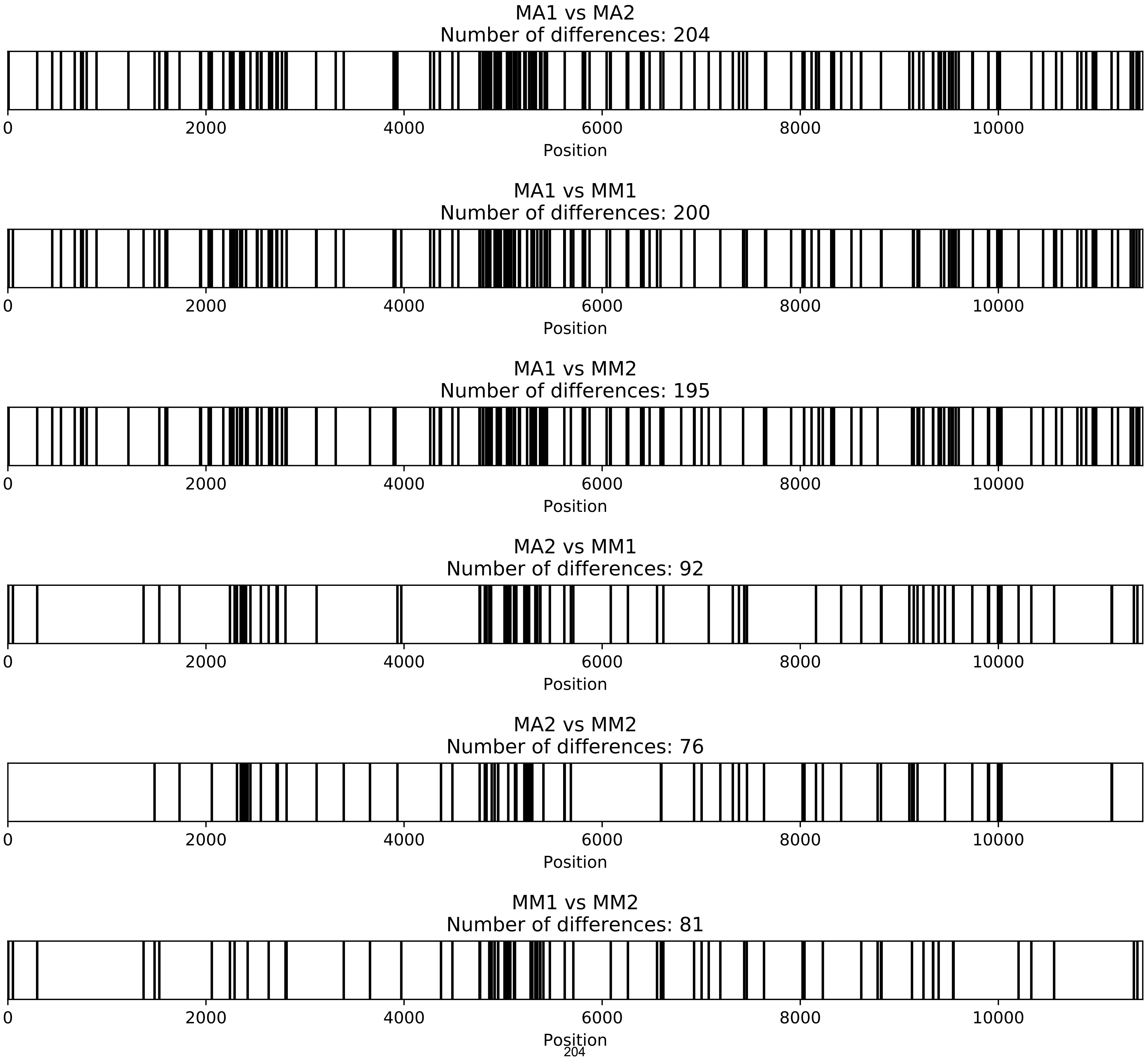
MA1 vs MA2
Number of differences: 204
MA1 vs MM1
Number of differences: 200
MA1 vs MM2
Number of differences: 195
MA2 vs MM1
Number of differences: 92
MA2 vs MM2
Number of differences: 76
MM1 vs MM2
Number of differences: 81
0
2000
4000
6000
8000
10000
Position

### Region: 665



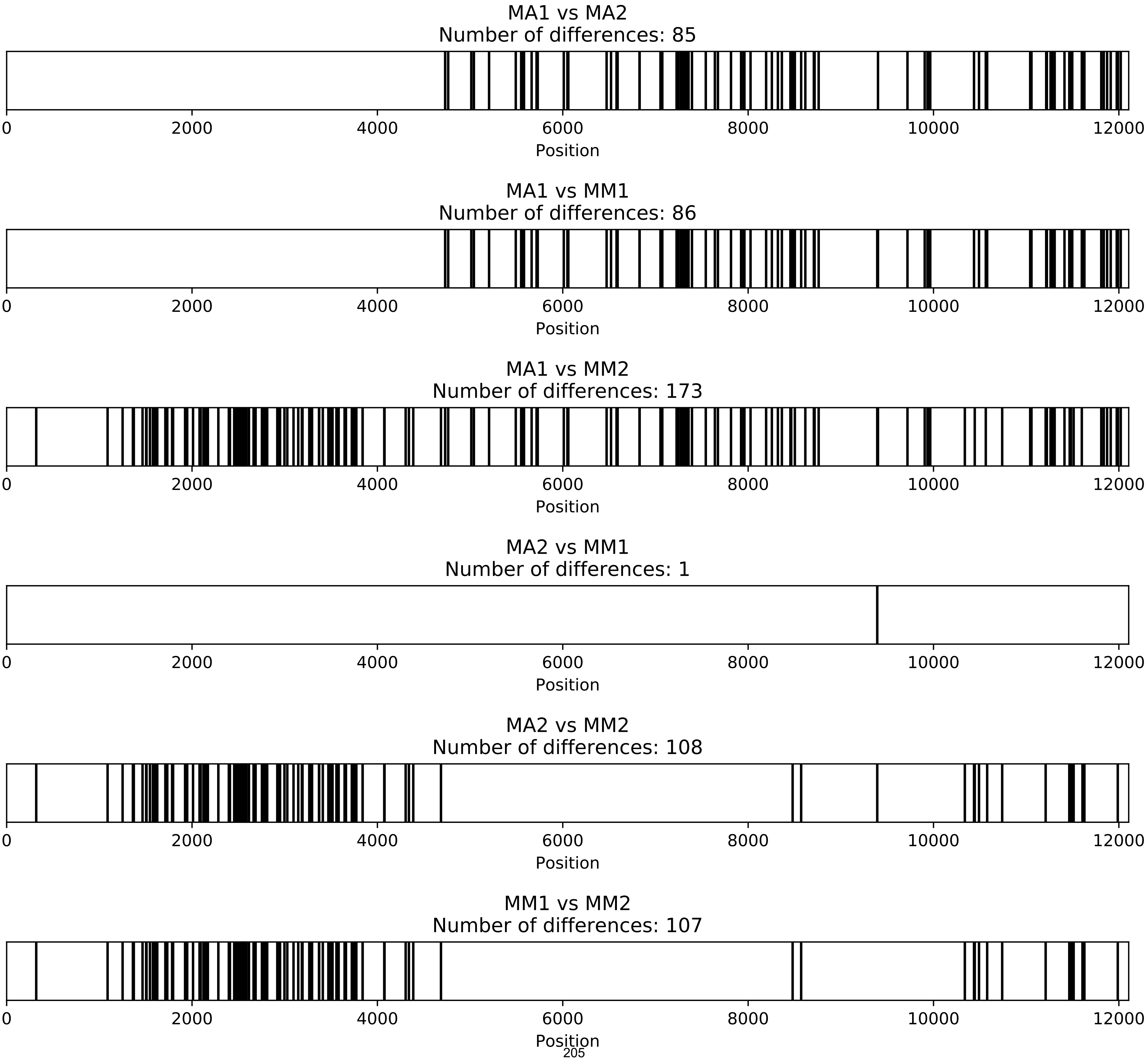
MA1 vs MA2
Number of differences: 85
MA1 vs MM1
Number of differences: 86
MA1 vs MM2
Number of differences: 173
MA2 vs MM1
Number of differences: 1
MA2 vs MM2
Number of differences: 108
MM1 vs MM2
Number of differences: 107
0
2000
4000
6000
8000
10000
12000
Position

### Region: 668



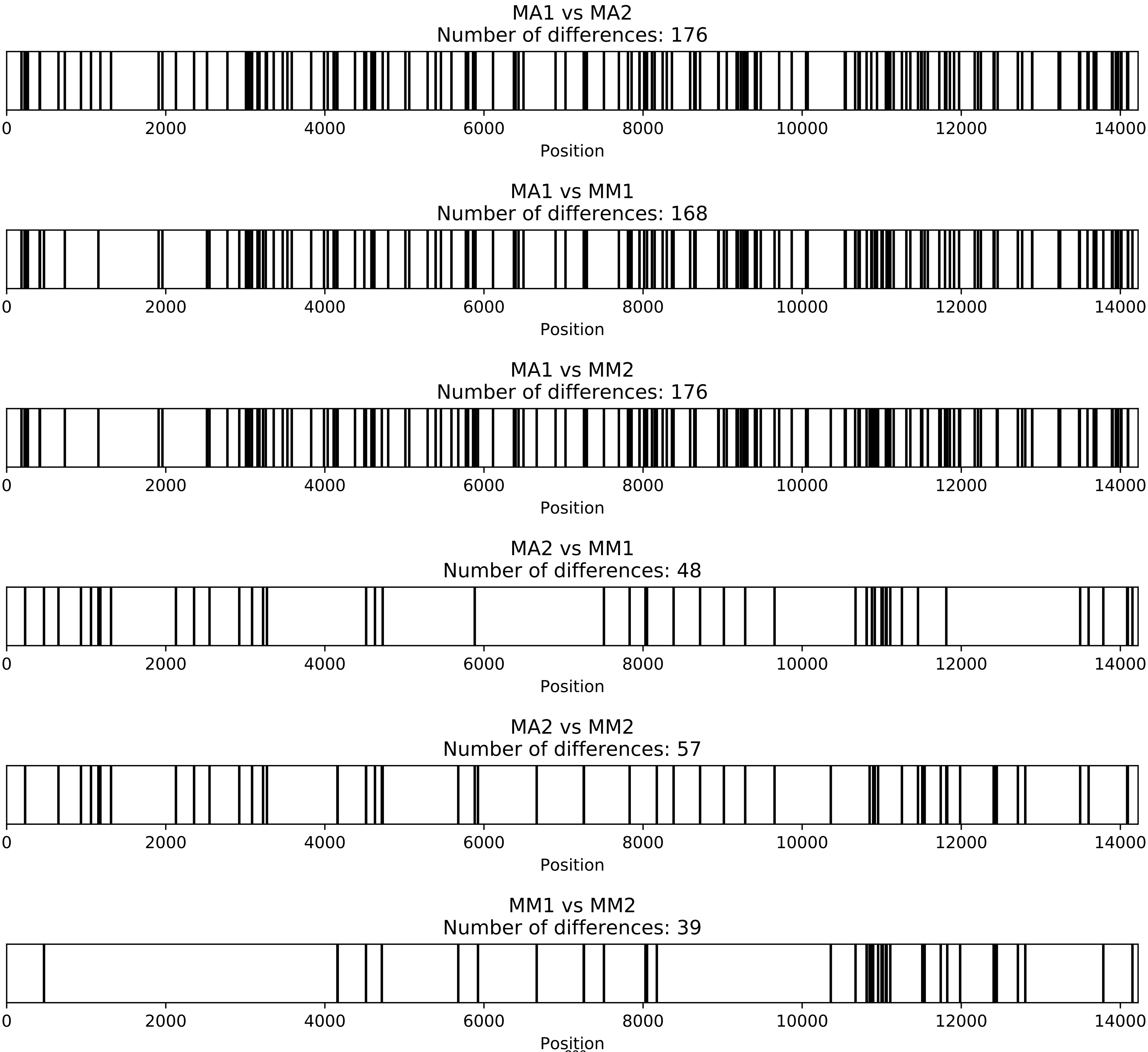
MA1 vs MA2
Number of differences: 176
MA1 vs MM1
Number of differences: 168
MA1 vs MM2
Number of differences: 176
MA2 vs MM1
Number of differences: 48
MA2 vs MM2
Number of differences: 57
MM1 vs MM2
Number of differences: 39
0
2000
4000
6000
8000
10000
12000
14000
Position

### Region: 669



## MA1 vs MA2
Number of differences: 775

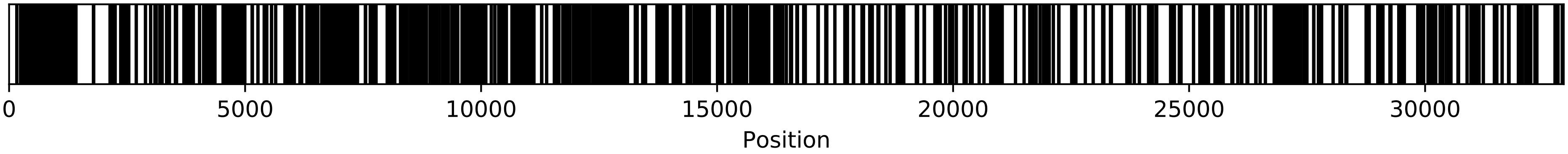


## MA1 vs MM1
Number of differences: 249

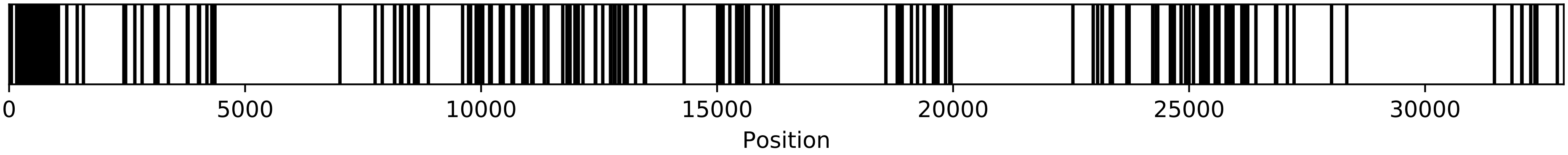


## MA1 vs MM2
Number of differences: 404

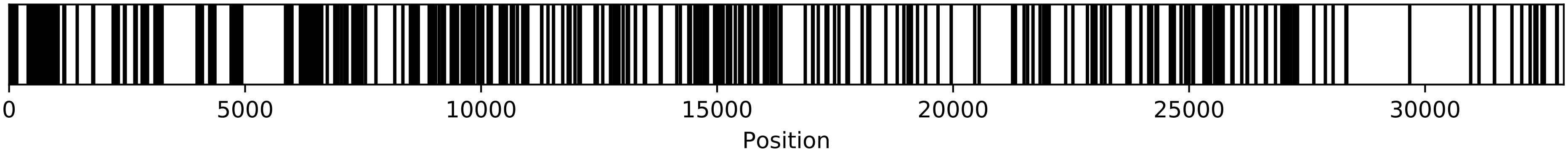


## MA2 vs MM1
Number of differences: 783

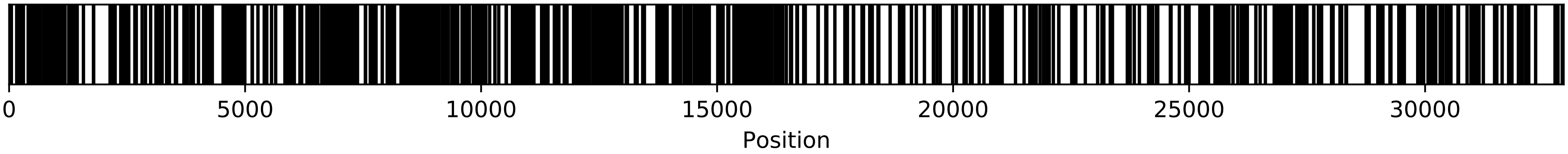


## MA2 vs MM2
Number of differences: 782

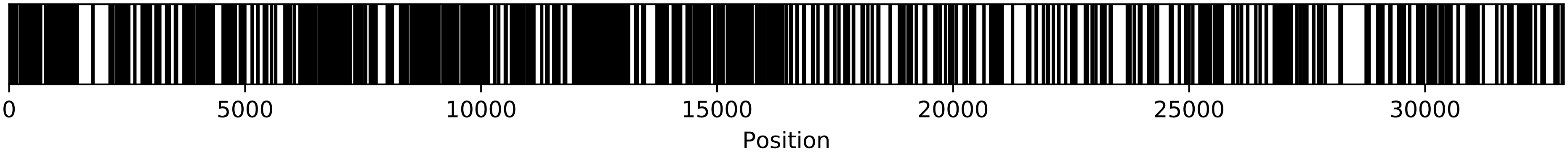


## MM1 vs MM2
Number of differences: 431

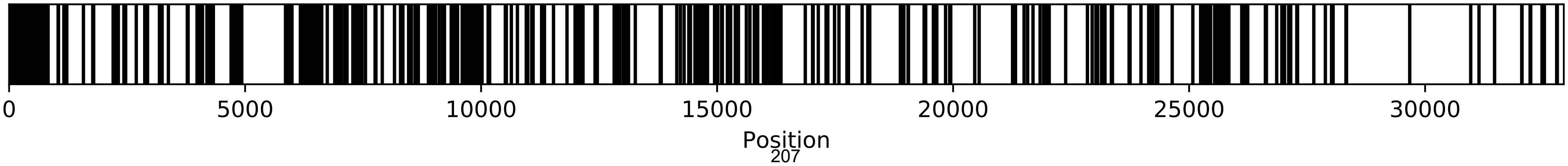

### Region: 671



MA1 vs MA2
Number of differences: 262

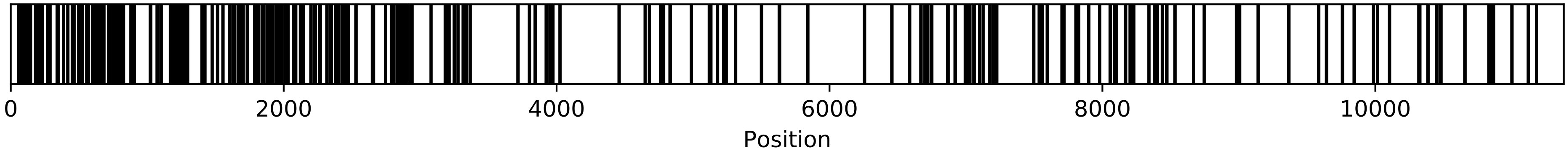


MA1 vs MM1
Number of differences: 0

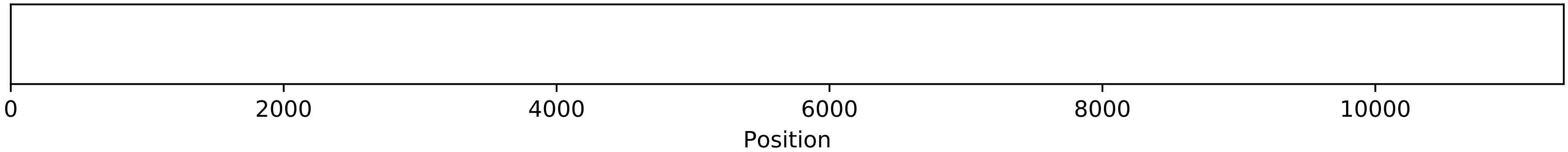


MA1 vs MM2
Number of differences: 10

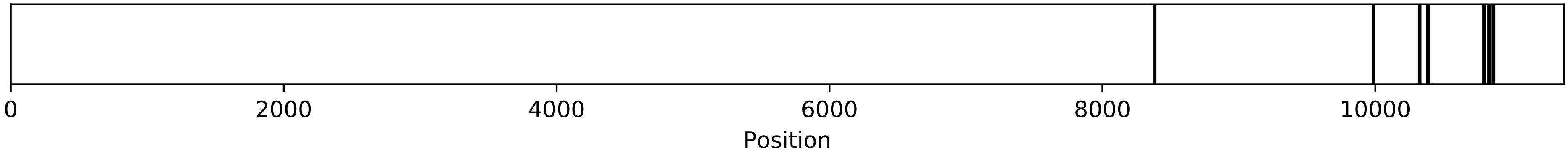


MA2 vs MM1
Number of differences: 262

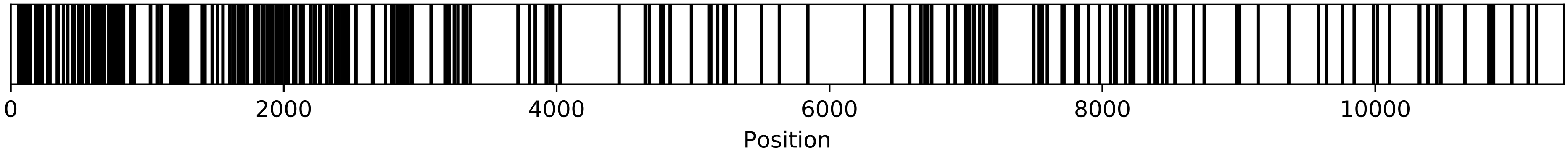


MA2 vs MM2
Number of differences: 255

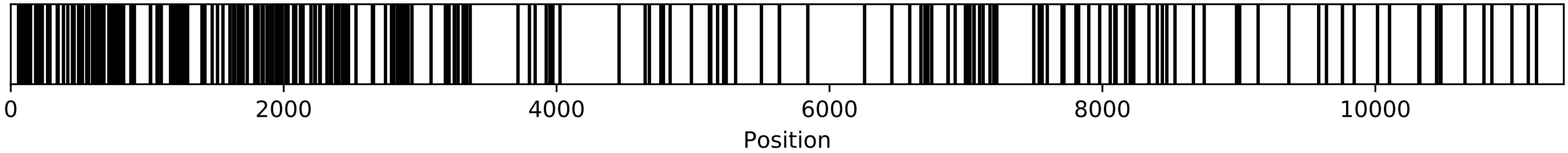


MM1 vs MM2
Number of differences: 10

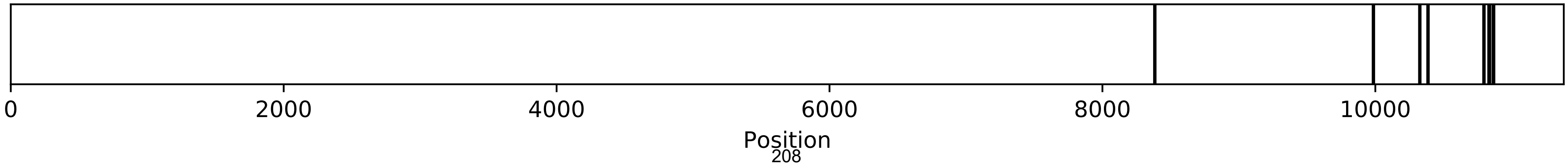

### Region: 672



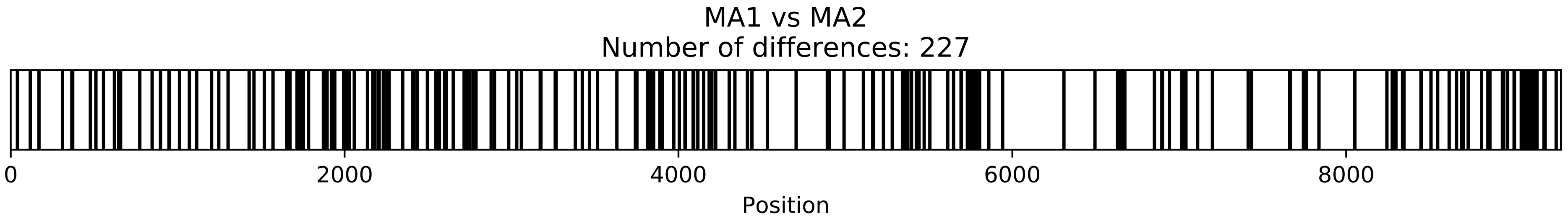
MA1 vs MA2
Number of differences: 227
0
2000
4000
6000
8000
Position

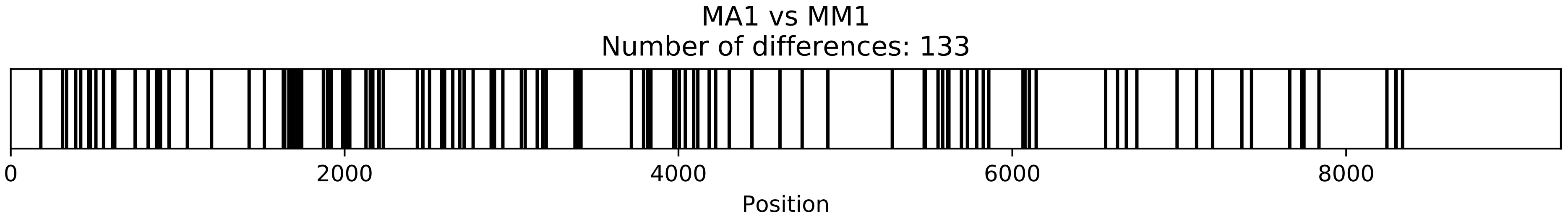
MA1 vs MM1
Number of differences: 133
0
2000
4000
6000
8000
Position

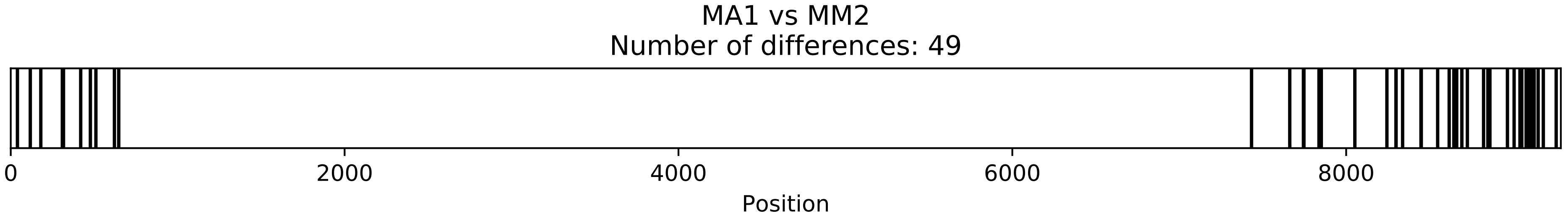
MA1 vs MM2
Number of differences: 49
0
2000
4000
6000
8000
Position

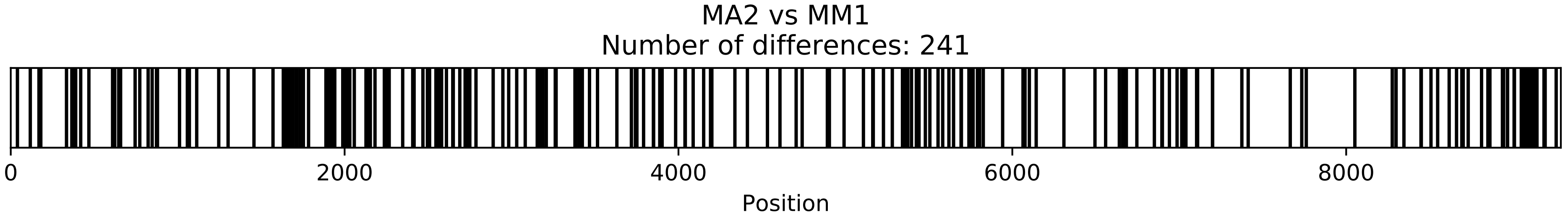
MA2 vs MM1
Number of differences: 241
0
2000
4000
6000
8000
Position

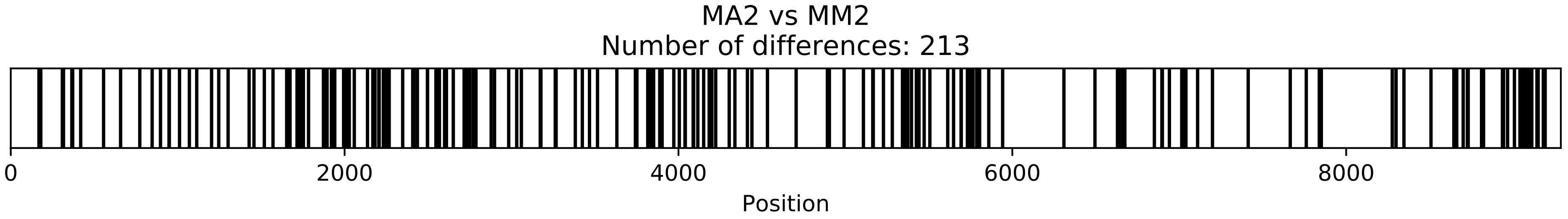
MA2 vs MM2
Number of differences: 213
0
2000
4000
6000
8000
Position

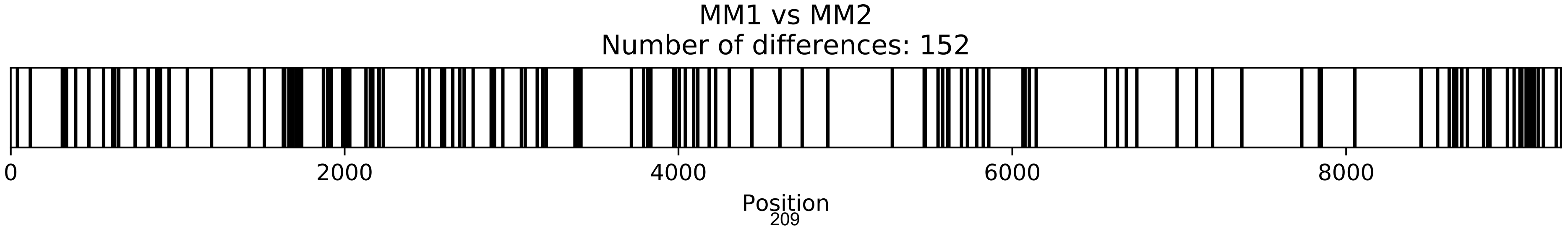
MM1 vs MM2
Number of differences: 152
0
2000
4000
6000
8000
Position

### Region: 677



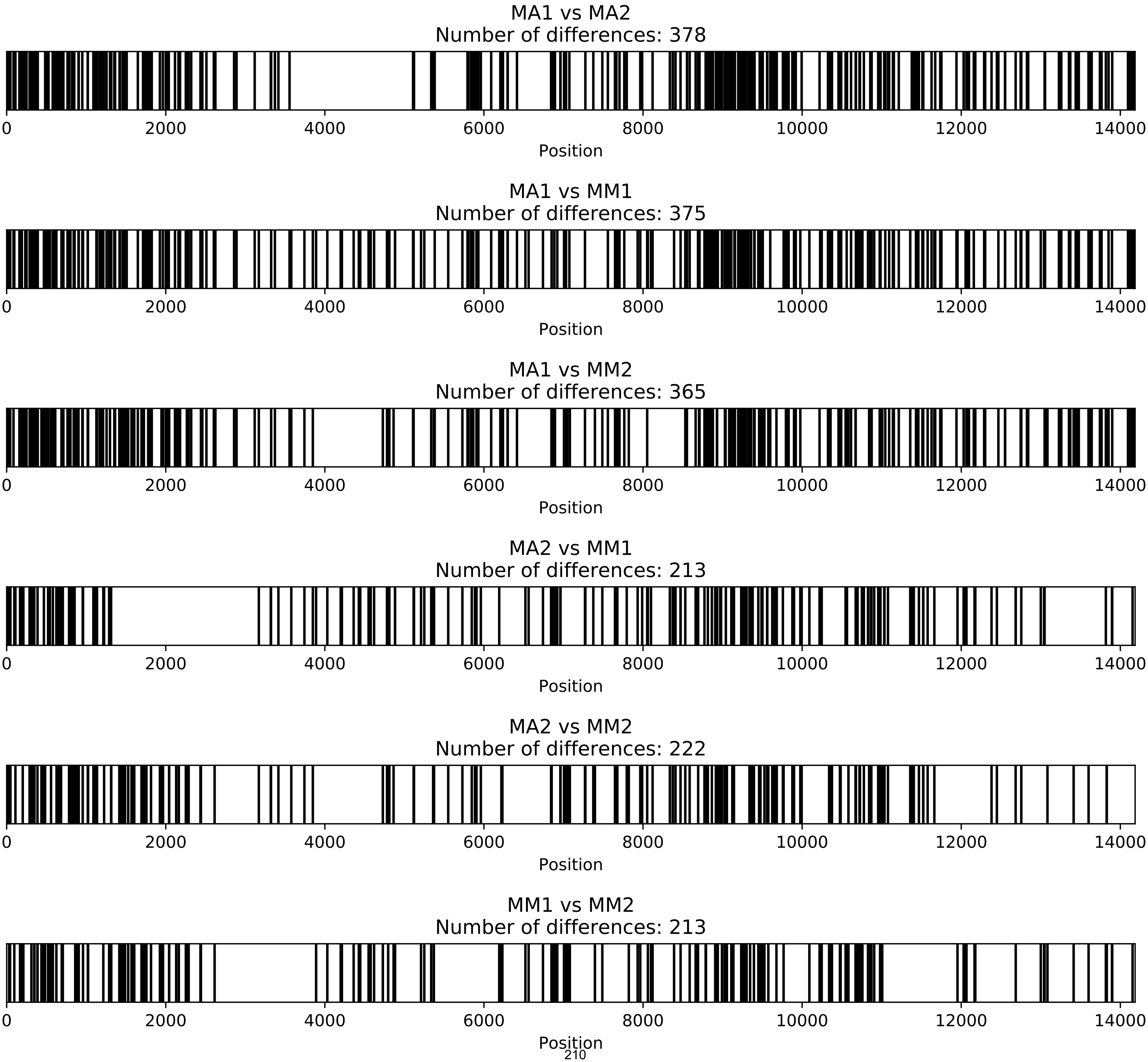
MA1 vs MA2
Number of differences: 378
MA1 vs MM1
Number of differences: 375
MA1 vs MM2
Number of differences: 365
MA2 vs MM1
Number of differences: 213
MA2 vs MM2
Number of differences: 222
MM1 vs MM2
Number of differences: 213
0
2000
4000
6000
8000
10000
12000
14000
Position

### Region: 685



MA1 vs MA2
Number of differences: 471

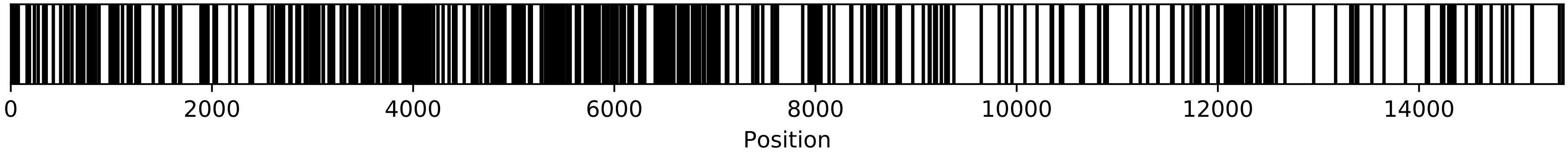


MA1 vs MM1
Number of differences: 471

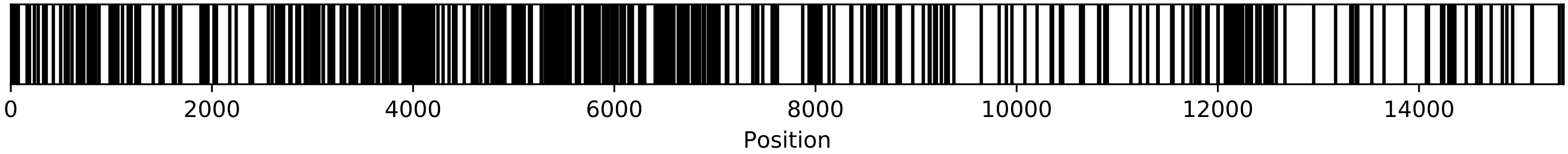


MA1 vs MM2
Number of differences: 472

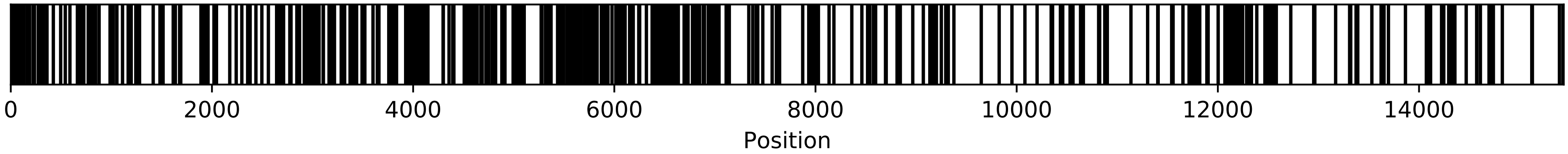


MA2 vs MM1
Number of differences: 0

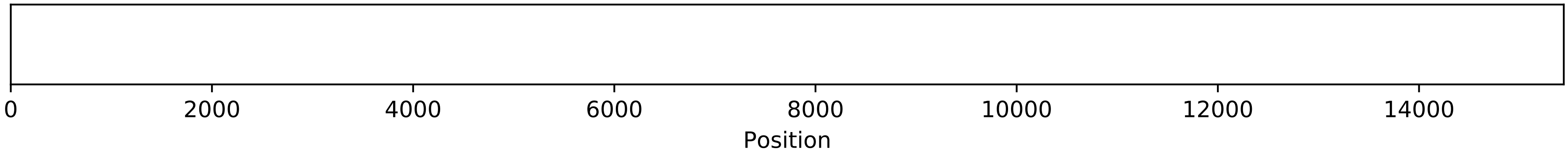


MA2 vs MM2
Number of differences: 221

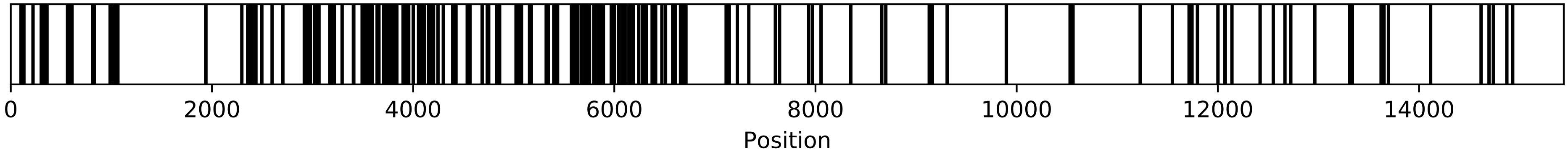


MM1 vs MM2
Number of differences: 221

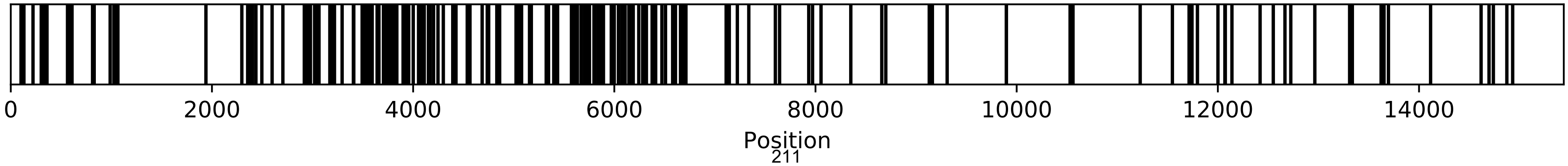

### Region: 691



MA1 vs MA2
Number of differences: 63

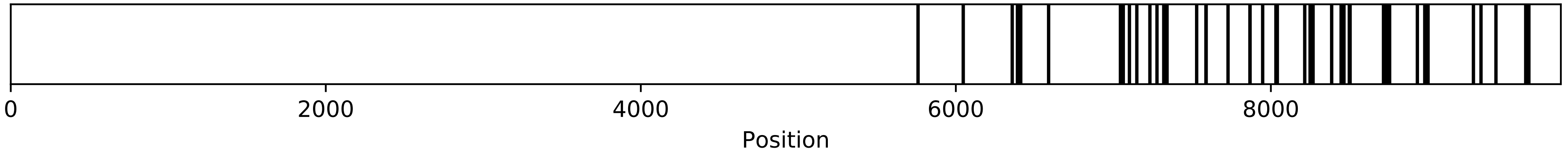


MA1 vs MM1
Number of differences: 256

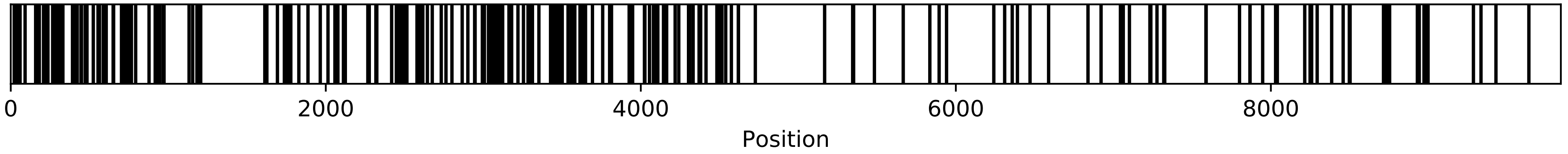


MA1 vs MM2
Number of differences: 265

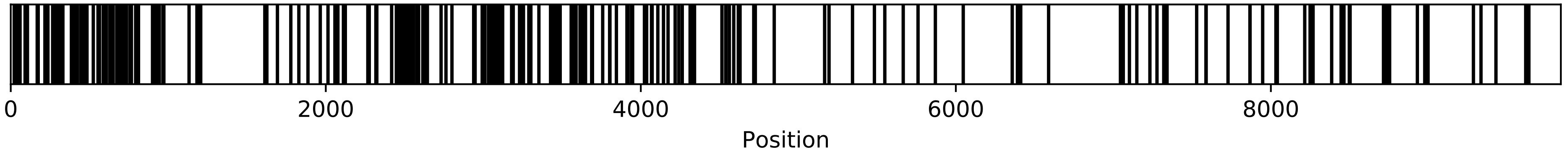


MA2 vs MM1
Number of differences: 222

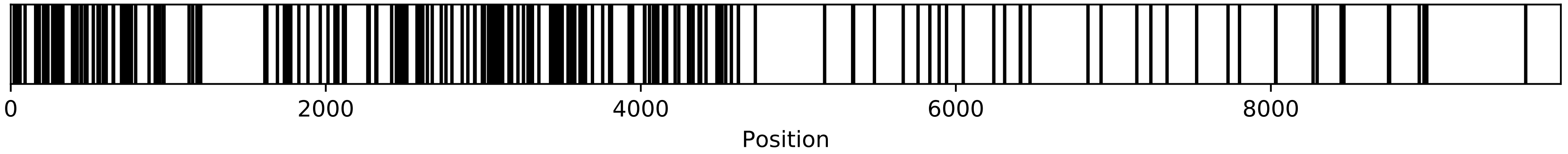


MA2 vs MM2
Number of differences: 202

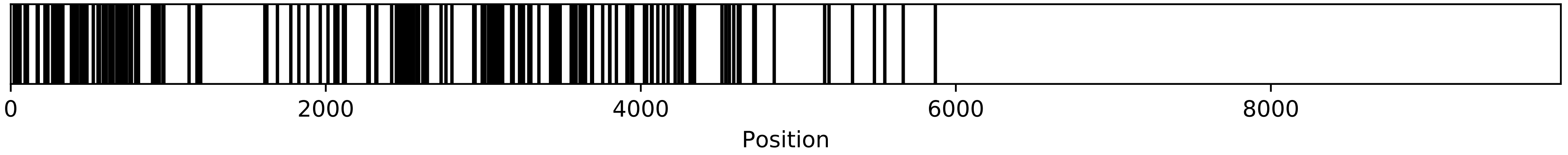


MM1 vs MM2
Number of differences: 141

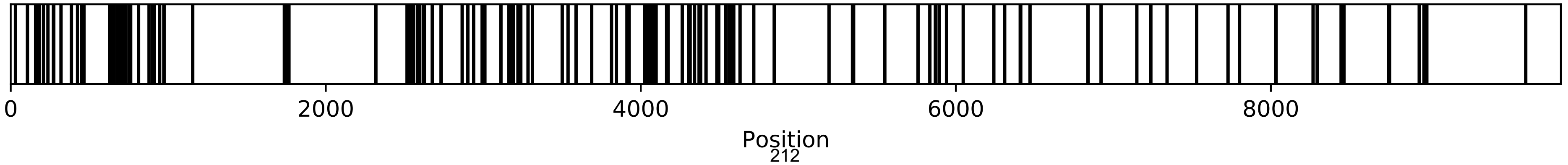

### Region: 692



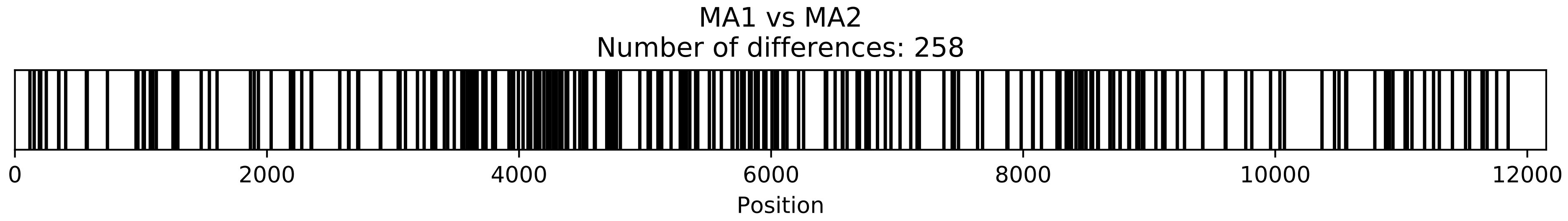
MA1 vs MA2
Number of differences: 258
0
2000
4000
6000
8000
10000
12000
Position

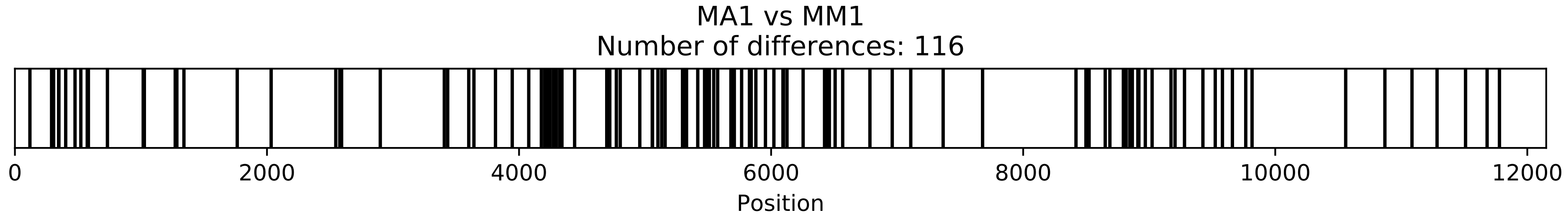
MA1 vs MM1
Number of differences: 116
0
2000
4000
6000
8000
10000
12000
Position

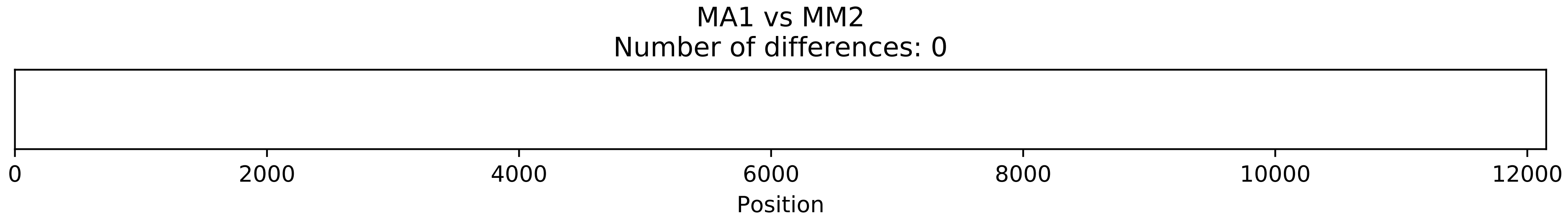
MA1 vs MM2
Number of differences: 0
0
2000
4000
6000
8000
10000
12000
Position

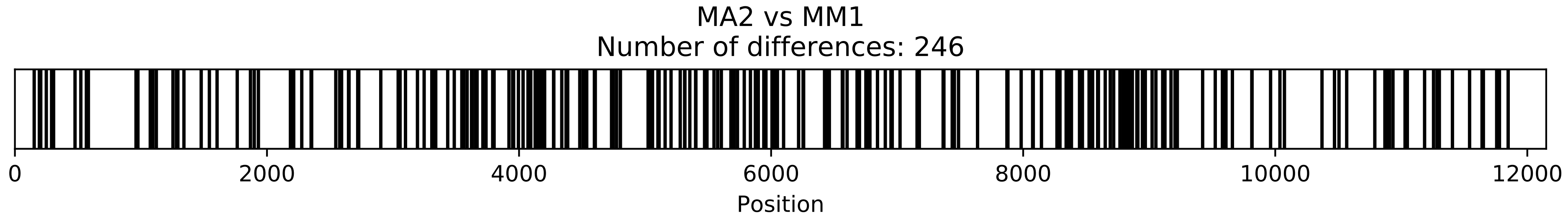
MA2 vs MM1
Number of differences: 246
0
2000
4000
6000
8000
10000
12000
Position

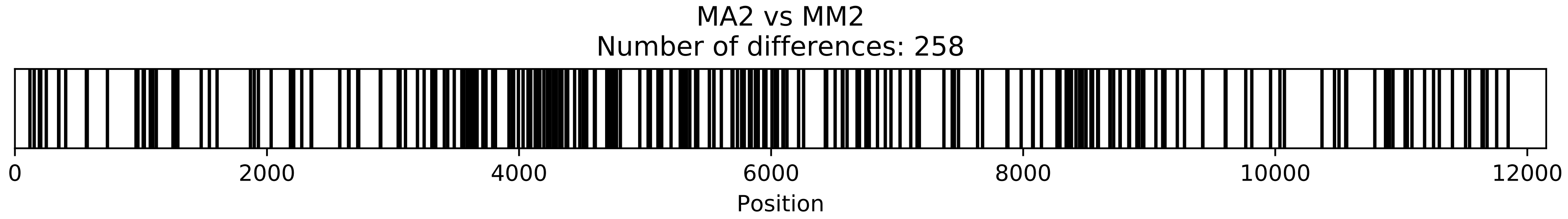
MA2 vs MM2
Number of differences: 258
0
2000
4000
6000
8000
10000
12000
Position

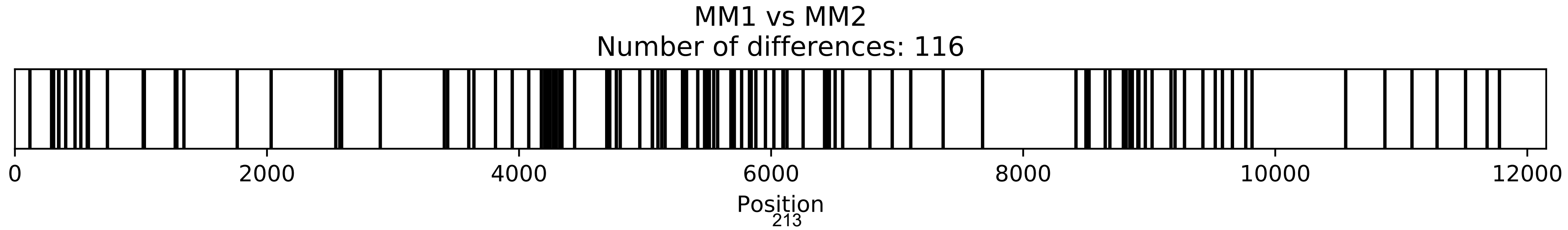
MM1 vs MM2
Number of differences: 116
0
2000
4000
6000
8000
10000
12000
Position

### Region: 693



MA1 vs MA2
Number of differences: 312

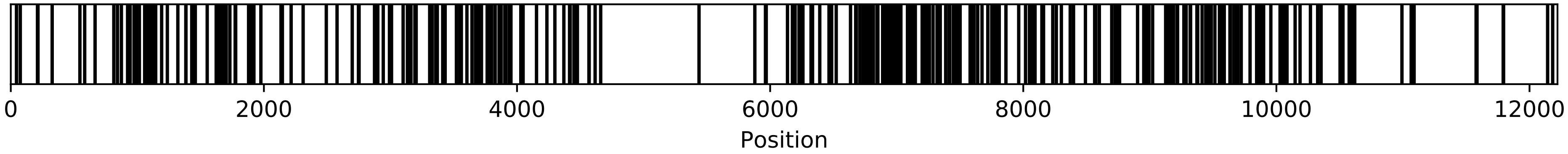


MA1 vs MM1
Number of differences: 182

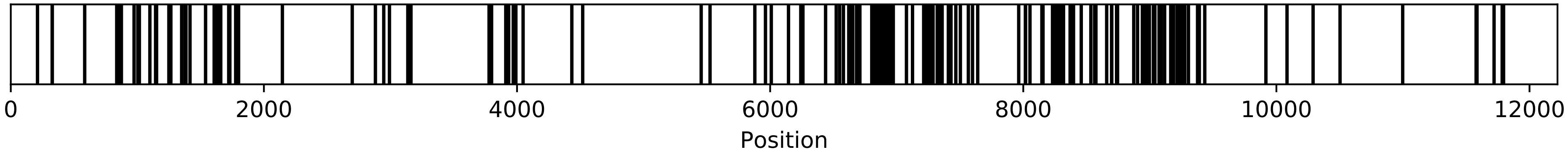


MA1 vs MM2
Number of differences: 178

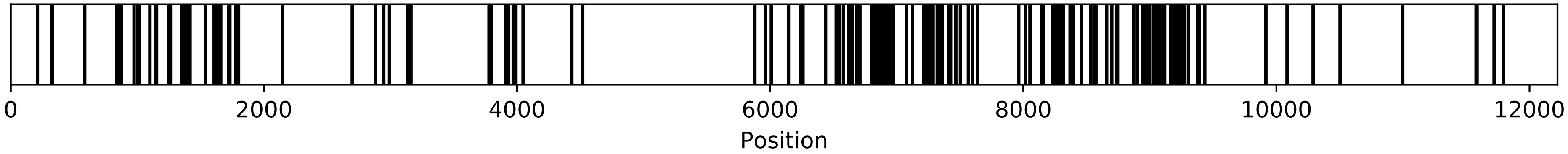


MA2 vs MM1
Number of differences: 337

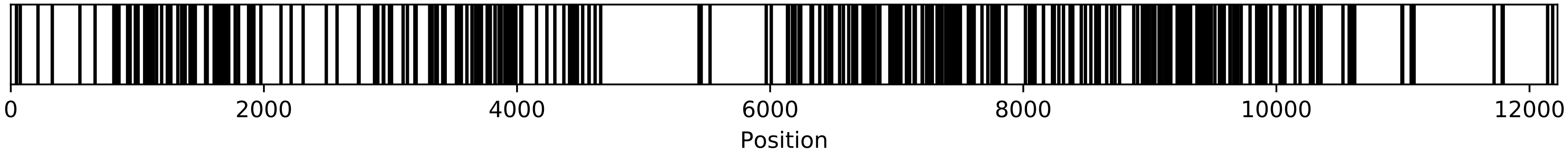


MA2 vs MM2
Number of differences: 333

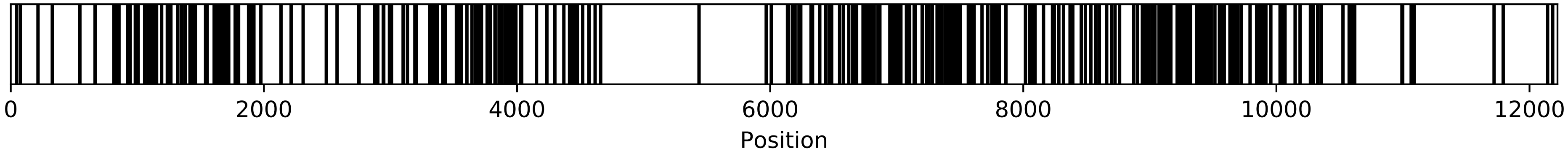


MM1 vs MM2
Number of differences: 4

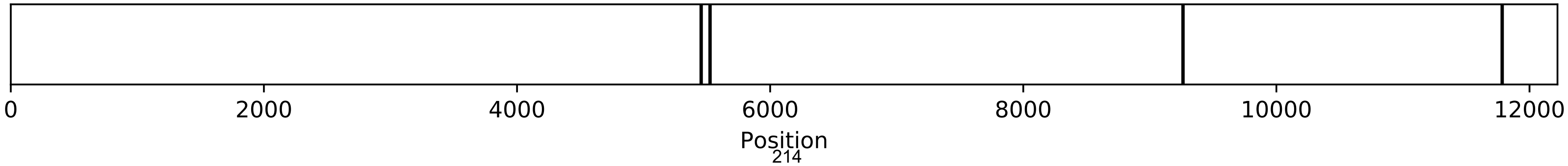

### Region: 694



MA1 vs MA2
Number of differences: 483

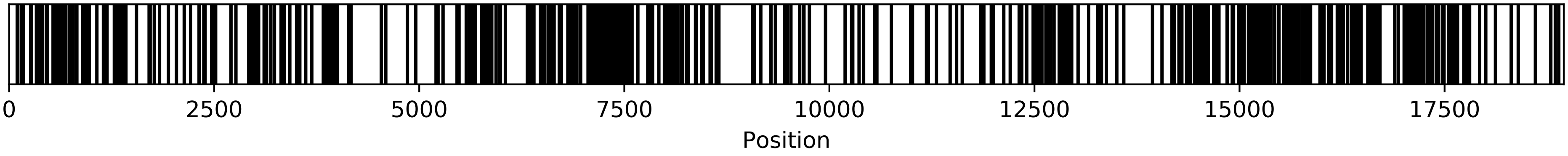


MA1 vs MM1
Number of differences: 487

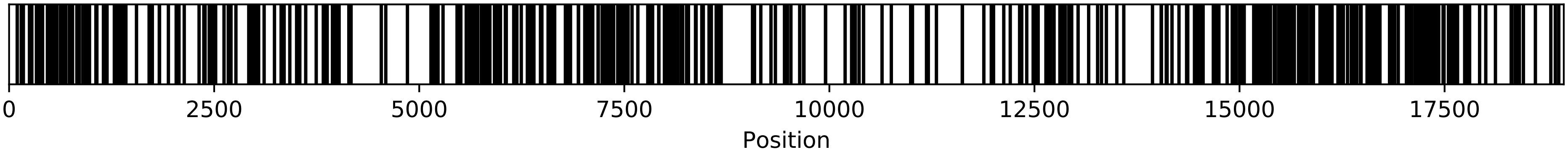


MA1 vs MM2
Number of differences: 502

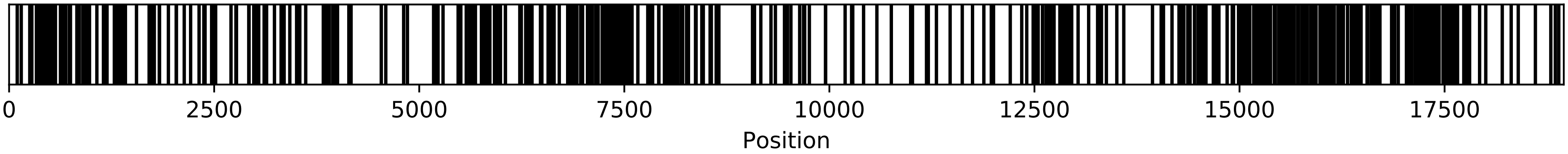


MA2 vs MM1
Number of differences: 213

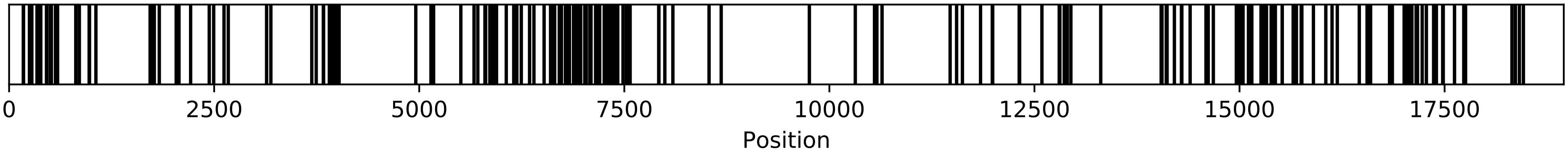


MA2 vs MM2
Number of differences: 188

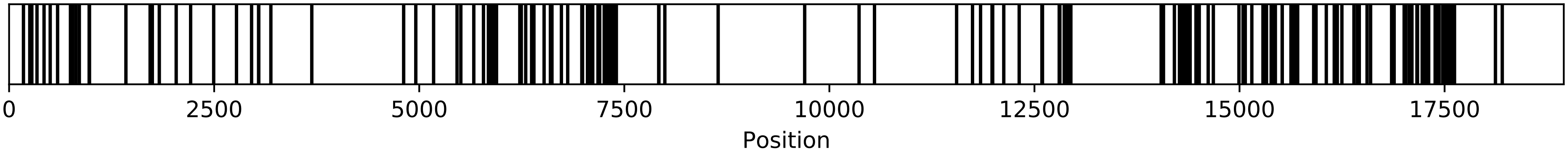


MM1 vs MM2
Number of differences: 202

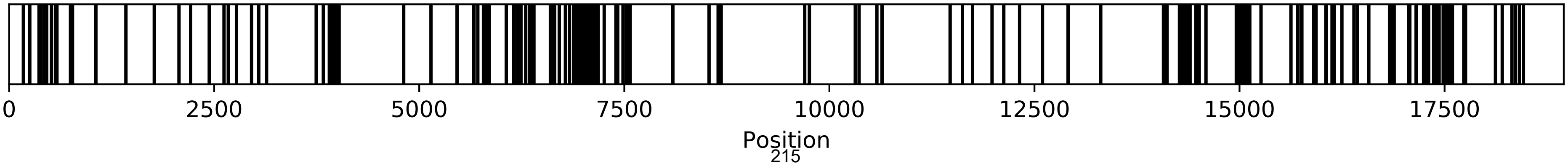

### Region: 697



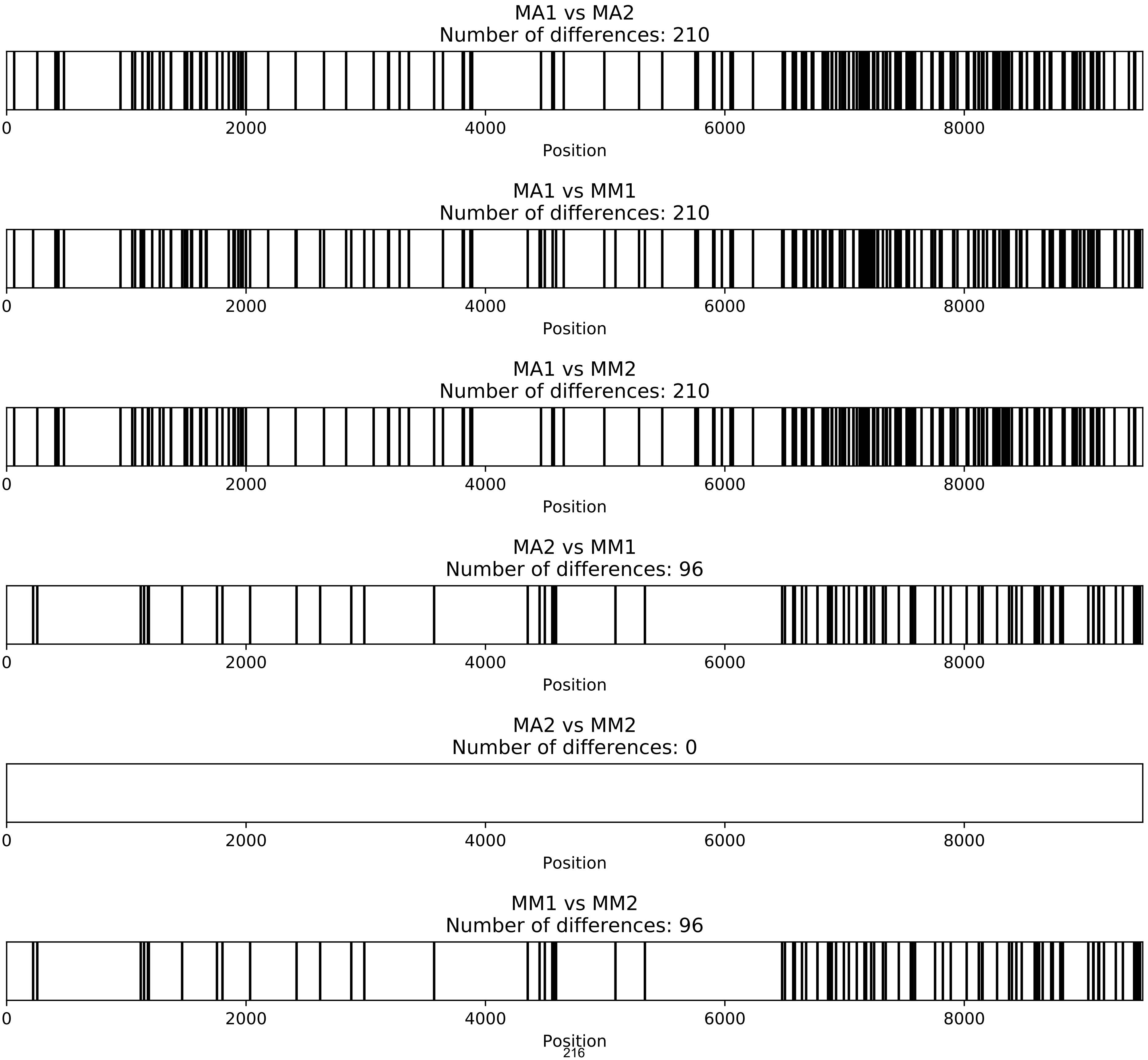
MA1 vs MA2
Number of differences: 210
MA1 vs MM1
Number of differences: 210
MA1 vs MM2
Number of differences: 210
MA2 vs MM1
Number of differences: 96
MA2 vs MM2
Number of differences: 0
MM1 vs MM2
Number of differences: 96
0
2000
4000
6000
8000
Position

### Region: 699



MA1 vs MA2
Number of differences: 407

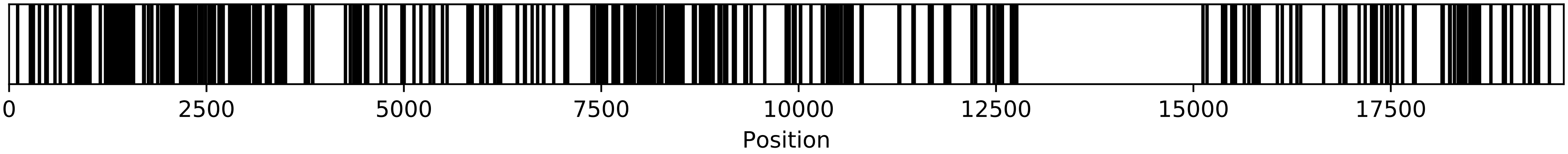


MA1 vs MM1
Number of differences: 444

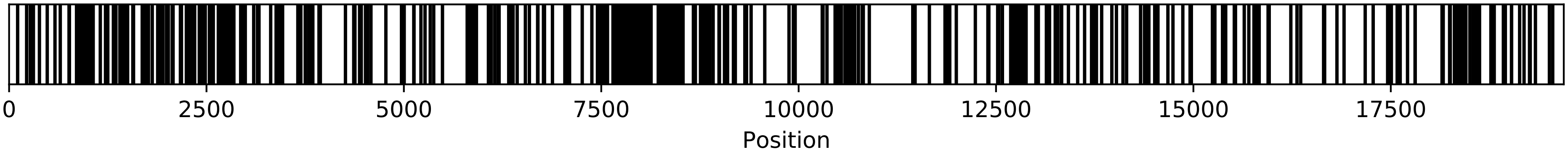


MA1 vs MM2
Number of differences: 457

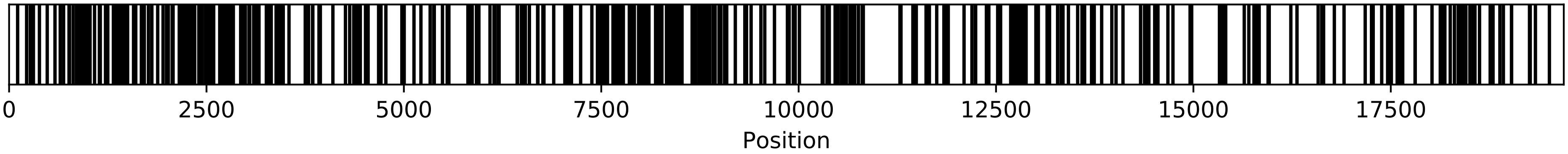


MA2 vs MM1
Number of differences: 315

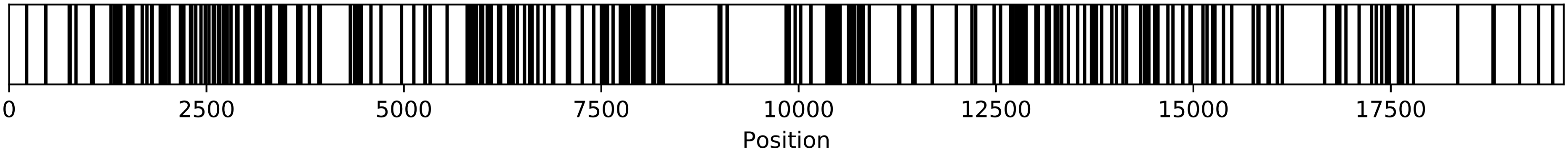


MA2 vs MM2
Number of differences: 332

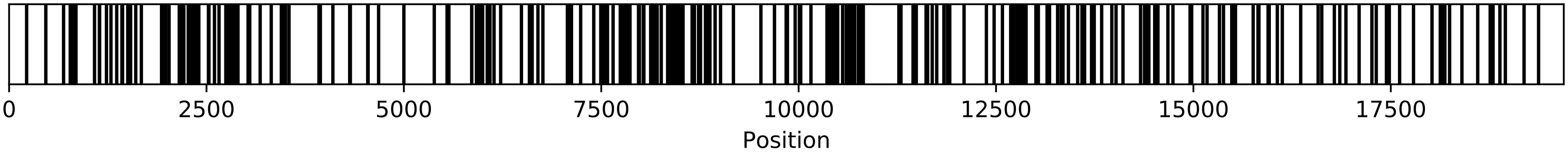


MM1 vs MM2
Number of differences: 281

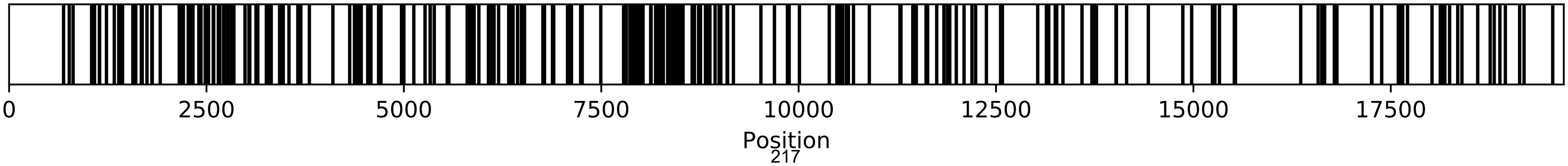

### Region: 700

MA1 vs MA2
Number of differences: 281

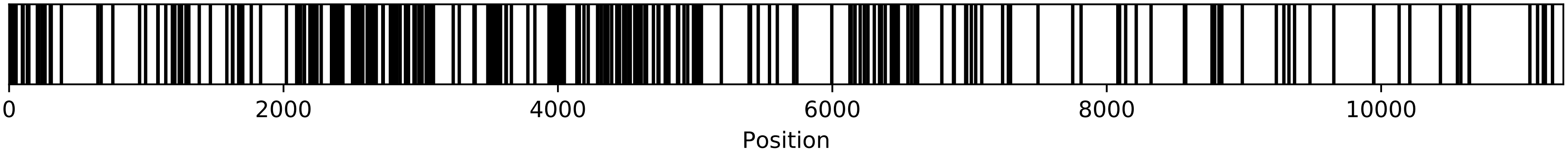


MA1 vs MM1
Number of differences: 281

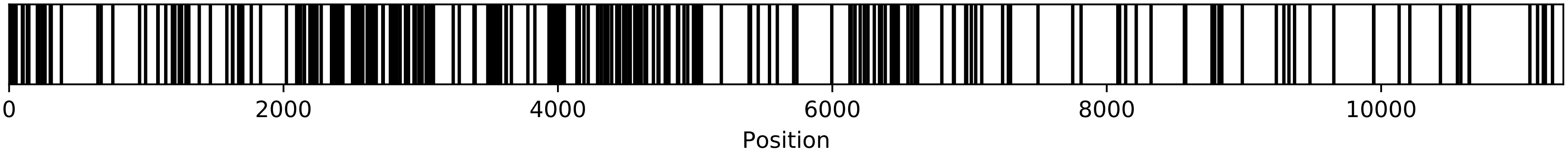


MA1 vs MM2
Number of differences: 263

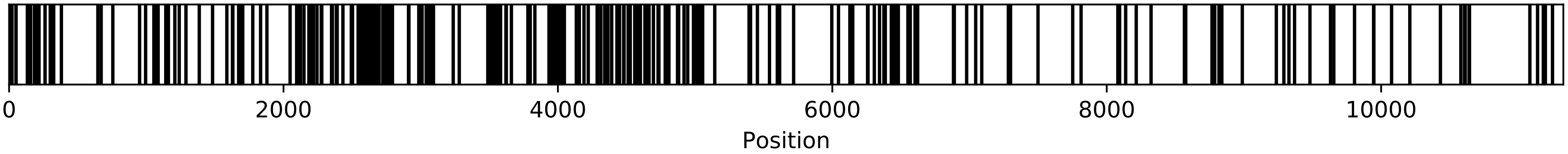


MA2 vs MM1
Number of differences: 0

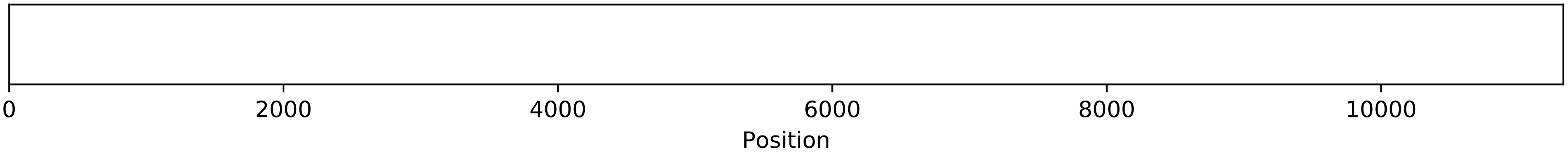


MA2 vs MM2
Number of differences: 126

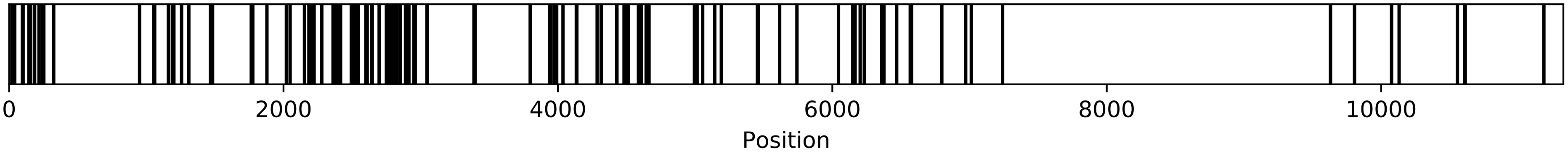


MM1 vs MM2
Number of differences: 126

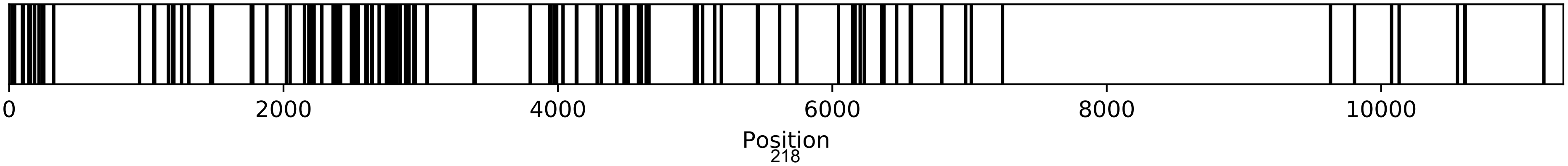

### Region: 701



MA1 vs MA2
Number of differences: 184

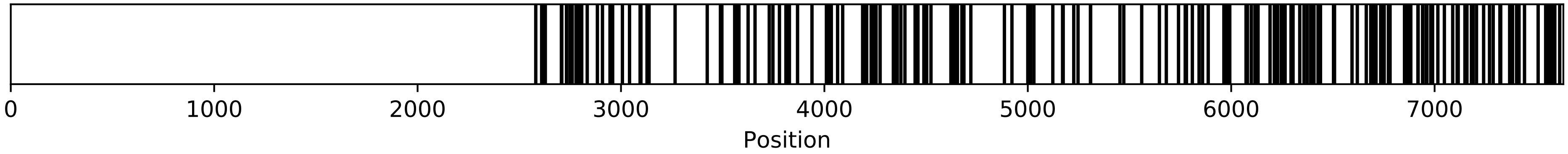


MA1 vs MM1
Number of differences: 47

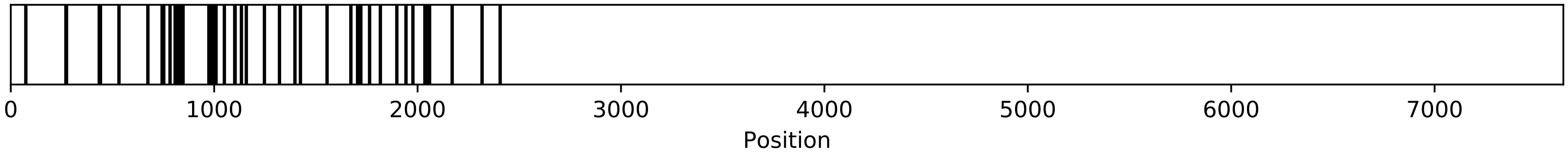


MA1 vs MM2
Number of differences: 160

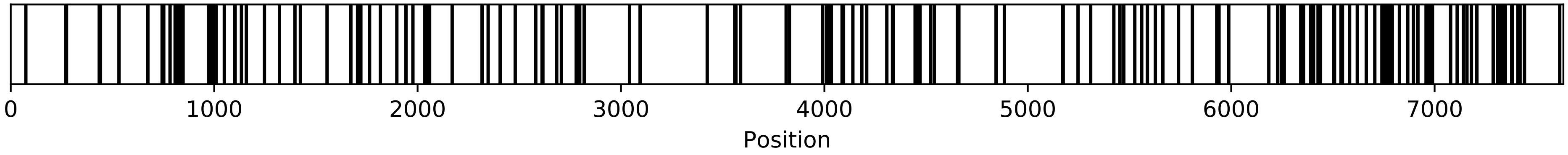


MA2 vs MM1
Number of differences: 231

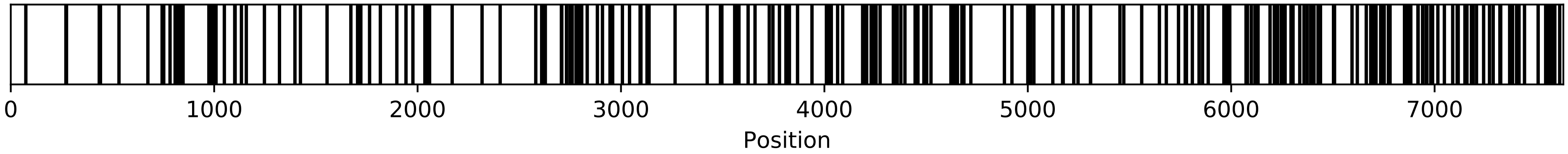


MA2 vs MM2
Number of differences: 221

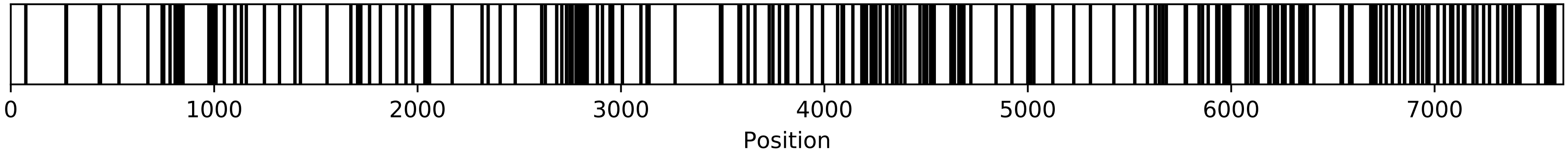


MM1 vs MM2
Number of differences: 113

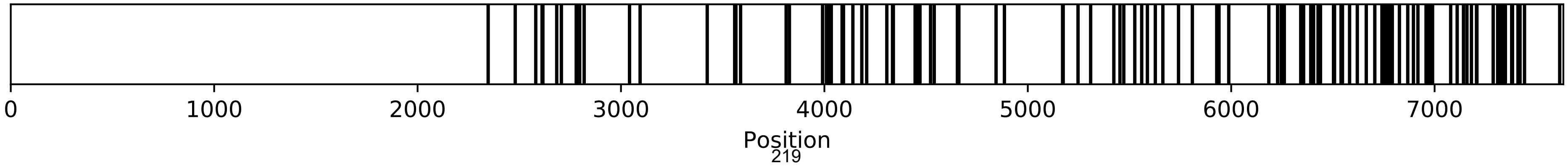

### Region: 704



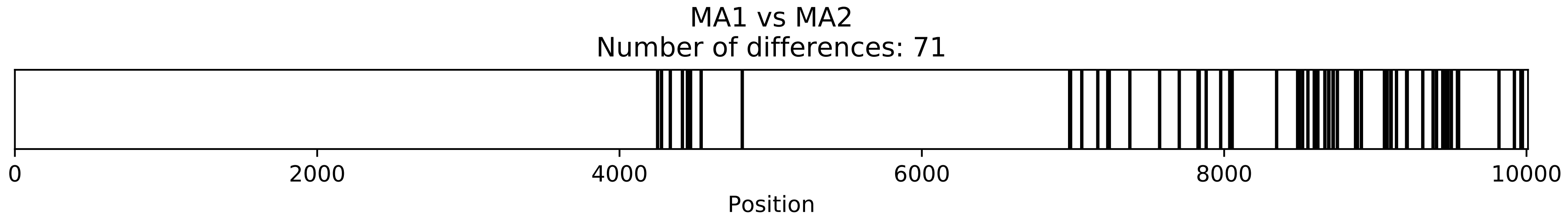
MA1 vs MA2
Number of differences: 71
0
2000
4000
6000
8000
10000
Position

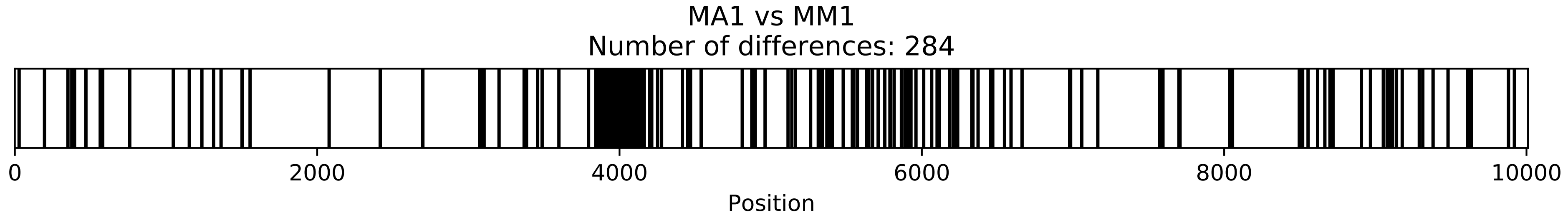
MA1 vs MM1
Number of differences: 284
0
2000
4000
6000
8000
10000
Position

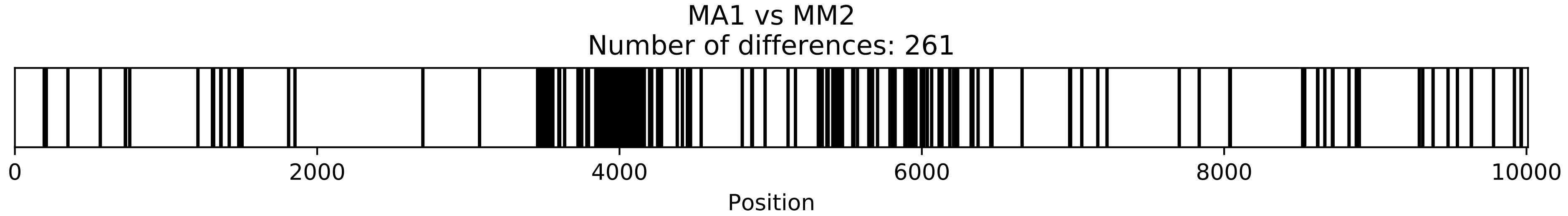
MA1 vs MM2
Number of differences: 261
0
2000
4000
6000
8000
10000
Position

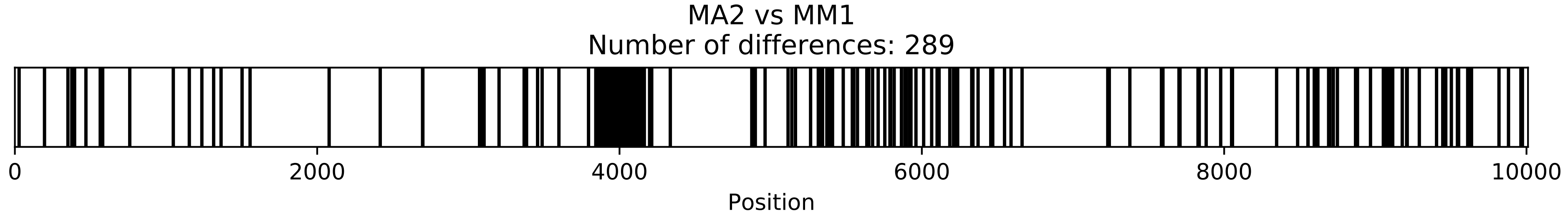
MA2 vs MM1
Number of differences: 289
0
2000
4000
6000
8000
10000
Position

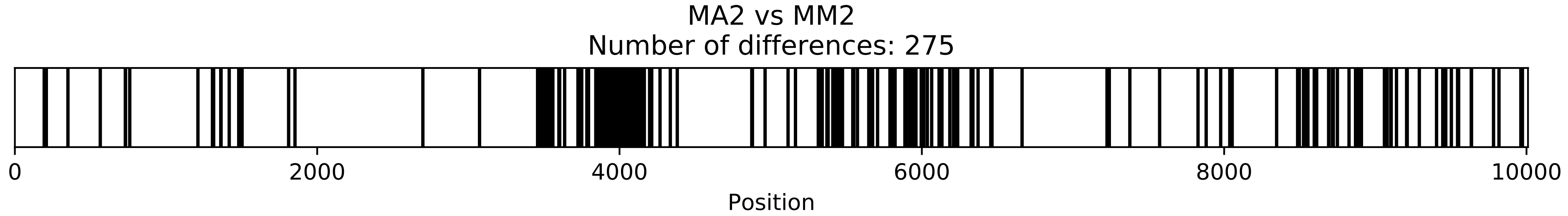
MA2 vs MM2
Number of differences: 275
0
2000
4000
6000
8000
10000
Position

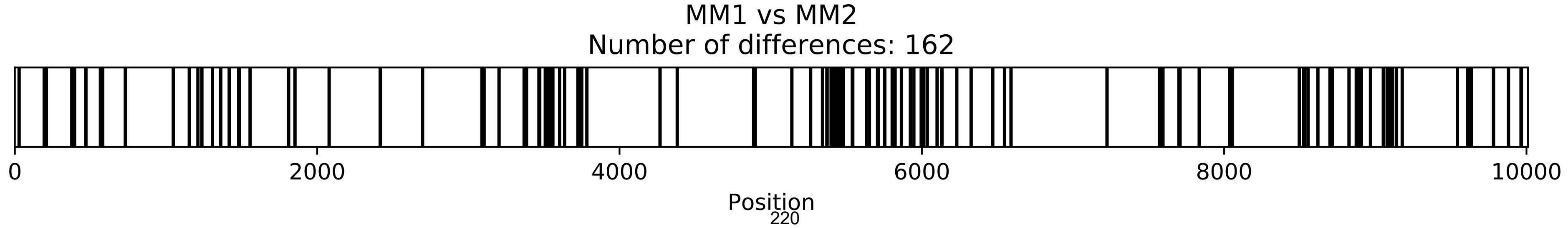
MM1 vs MM2
Number of differences: 162
0
2000
4000
6000
8000
10000
Position

### Region: 718



MA1 vs MA2
Number of differences: 334

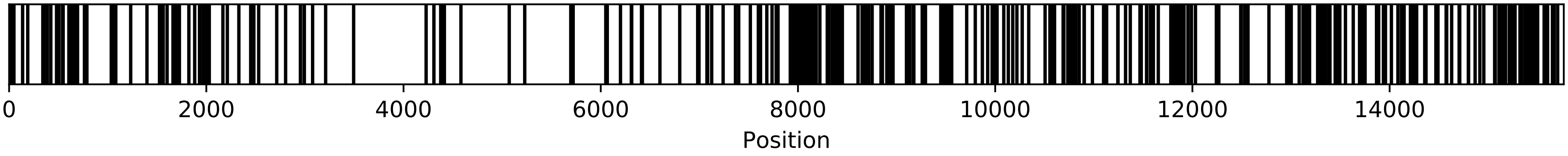


MA1 vs MM1
Number of differences: 330

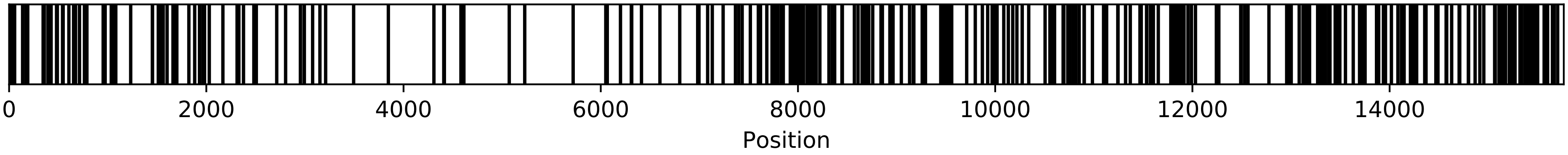


MA1 vs MM2
Number of differences: 334

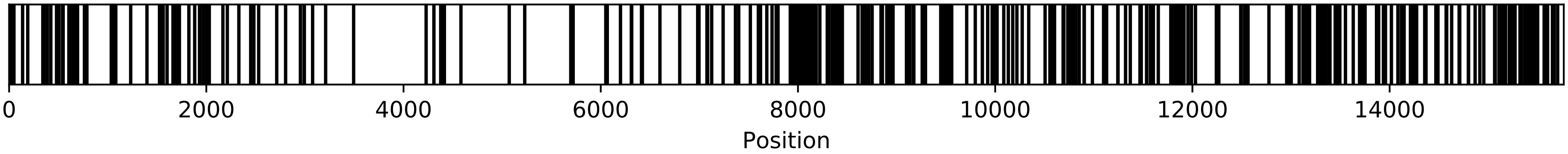


MA2 vs MM1
Number of differences: 80

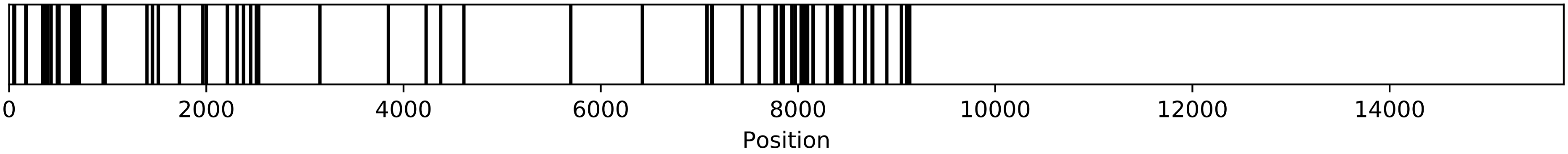


MA2 vs MM2
Number of differences: 0

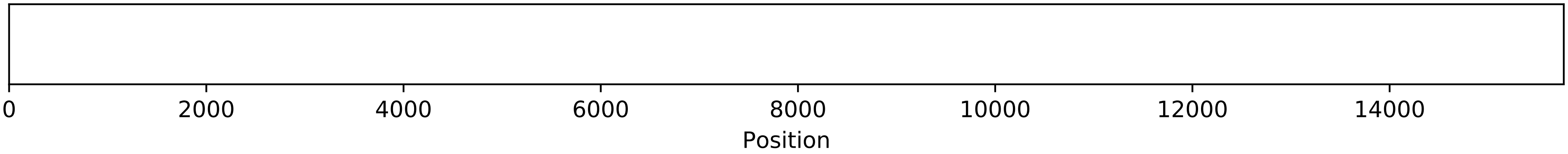


MM1 vs MM2
Number of differences: 80

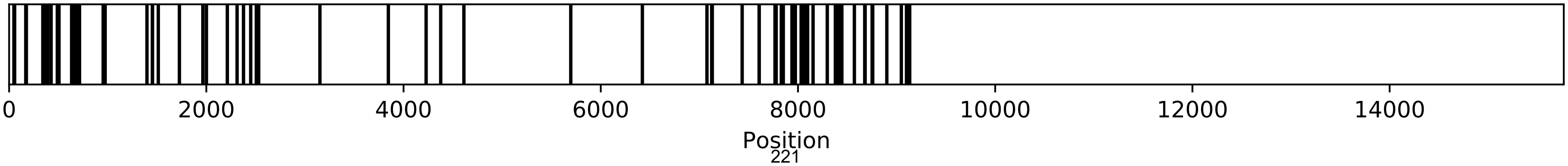

### Region: 719



MA1 vs MA2
Number of differences: 158

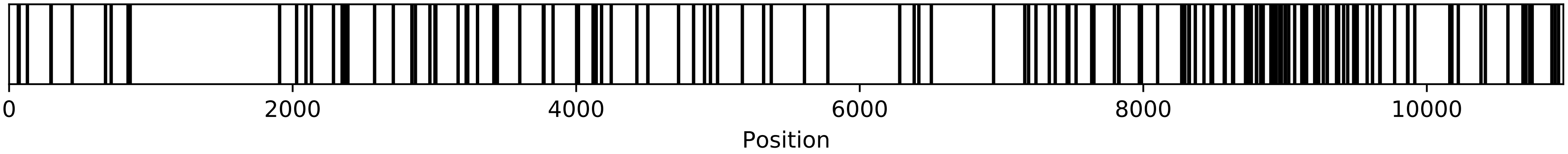


MA1 vs MM1
Number of differences: 162

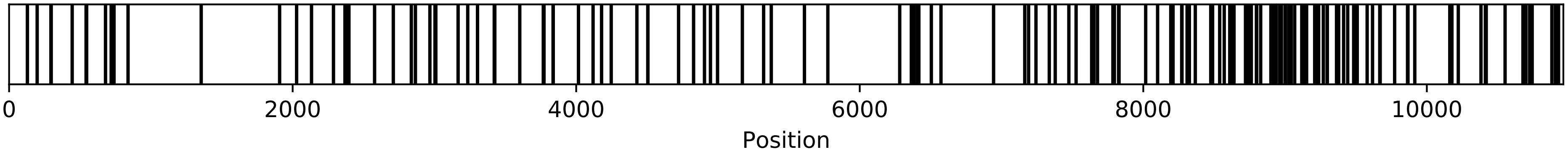


MA1 vs MM2
Number of differences: 166

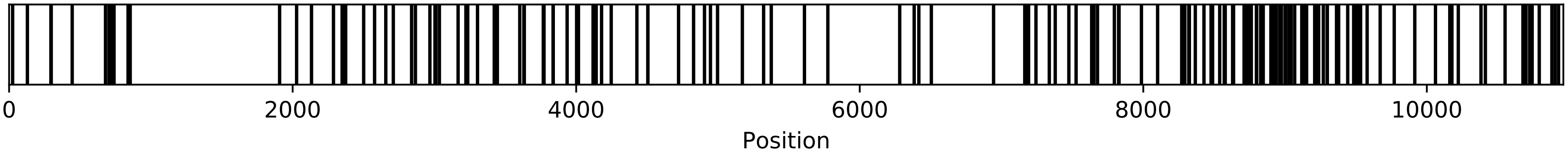


MA2 vs MM1
Number of differences: 51

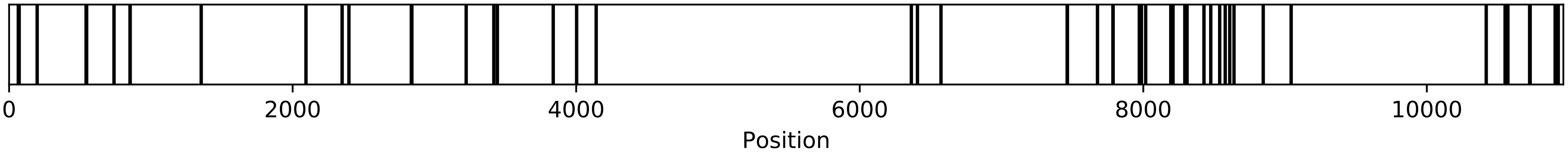


MA2 vs MM2
Number of differences: 46

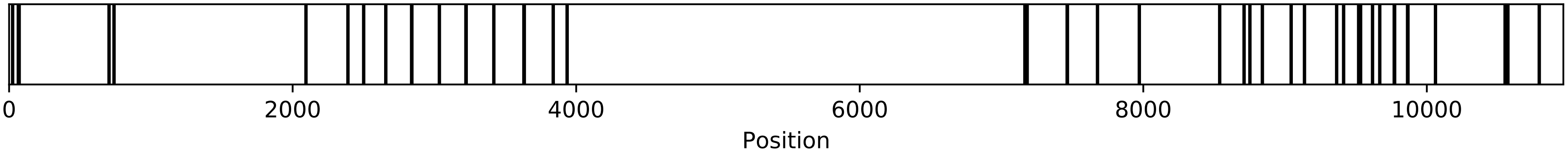


MM1 vs MM2
Number of differences: 65

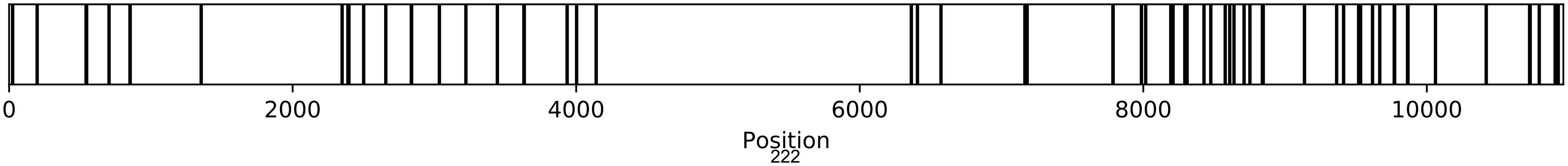

### Region: 722

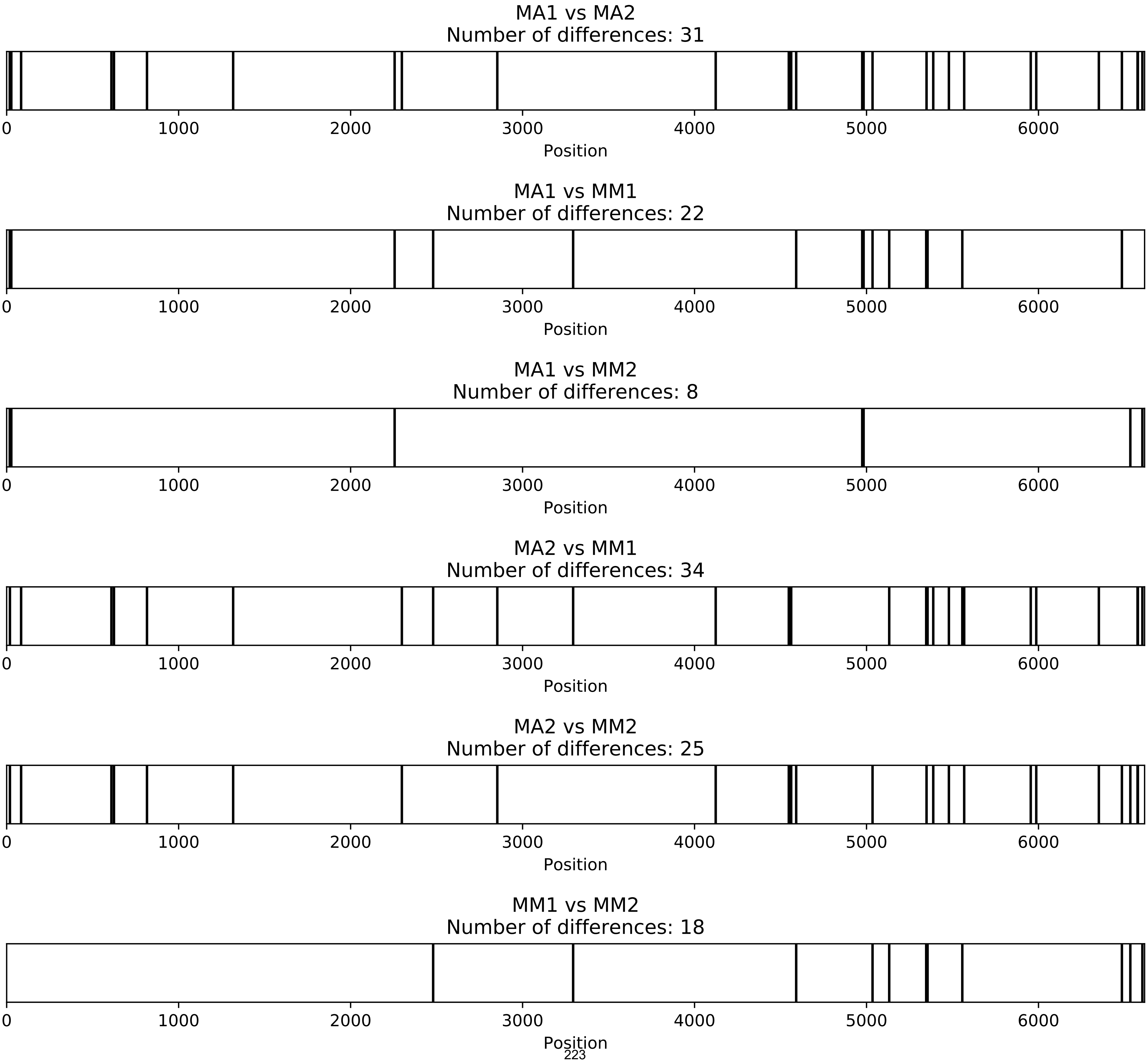
MA1 vs MA2
Number of differences: 31
0 1000 2000 3000 4000 5000 6000
Position
MA1 vs MM1
Number of differences: 22
0 1000 2000 3000 4000 5000 6000
Position
MA1 vs MM2
Number of differences: 8
0 1000 2000 3000 4000 5000 6000
Position
MA2 vs MM1
Number of differences: 34
0 1000 2000 3000 4000 5000 6000
Position
MA2 vs MM2
Number of differences: 25
0 1000 2000 3000 4000 5000 6000
Position
MM1 vs MM2
Number of differences: 18
0 1000 2000 3000 4000 5000 6000
Position

### Region: 723



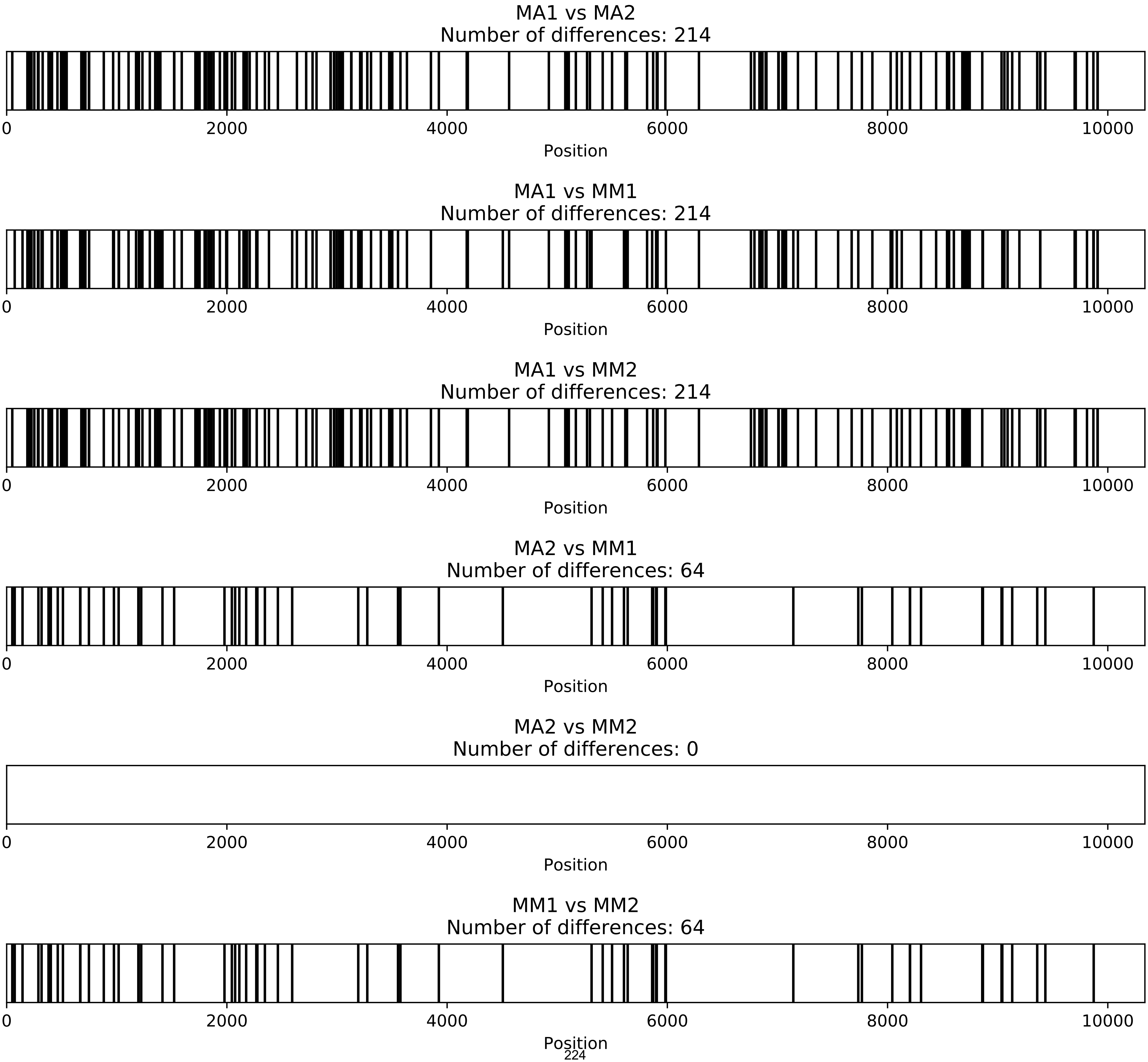
MA1 vs MA2
Number of differences: 214
0
2000
4000
6000
8000
10000
Position
MA1 vs MM1
Number of differences: 214
Position
MA1 vs MM2
Number of differences: 214
Position
MA2 vs MM1
Number of differences: 64
Position
MA2 vs MM2
Number of differences: 0
Position
MM1 vs MM2
Number of differences: 64
Position

### Region: 725



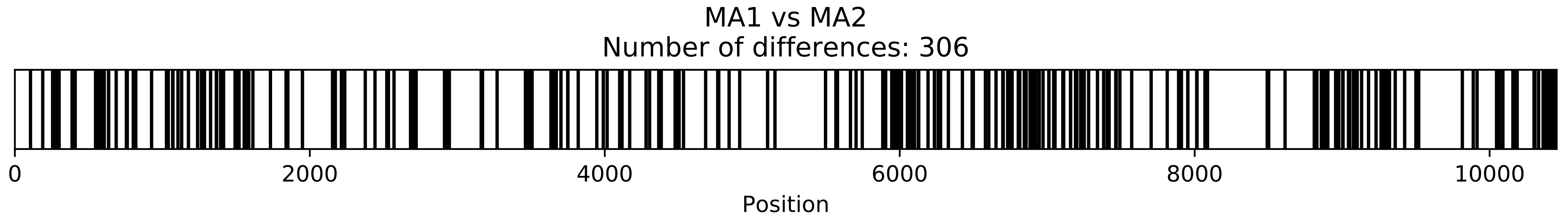
MA1 vs MA2
Number of differences: 306
0
2000
4000
6000
8000
10000
Position

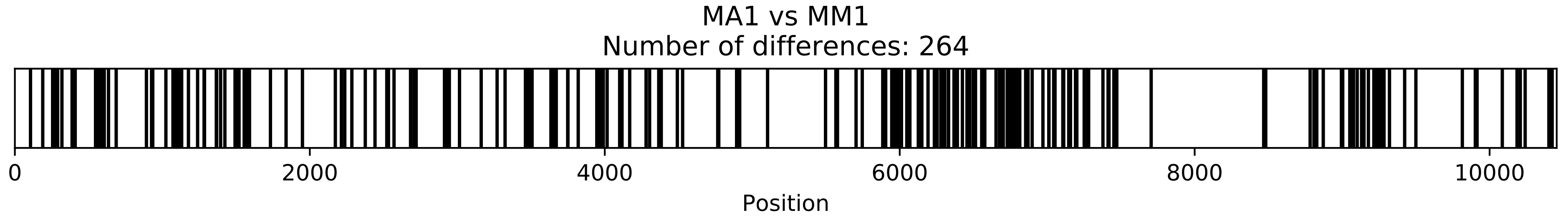
MA1 vs MM1
Number of differences: 264
0
2000
4000
6000
8000
10000
Position

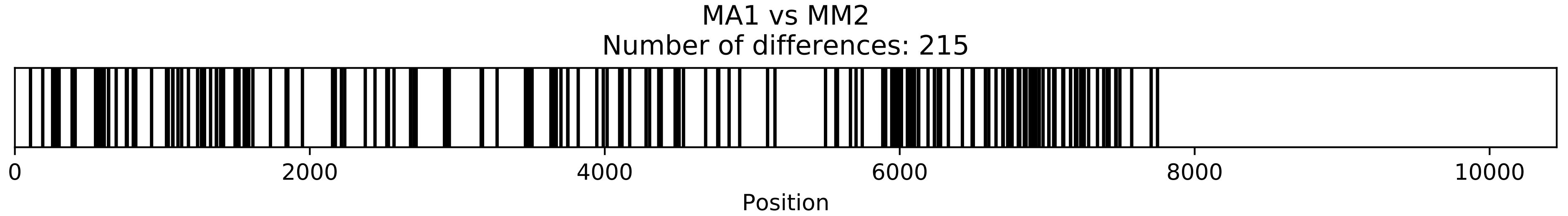
MA1 vs MM2
Number of differences: 215
0
2000
4000
6000
8000
10000
Position

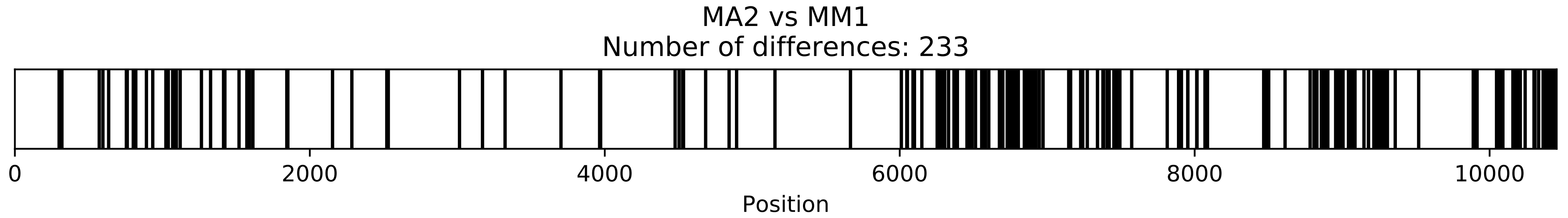
MA2 vs MM1
Number of differences: 233
0
2000
4000
6000
8000
10000
Position

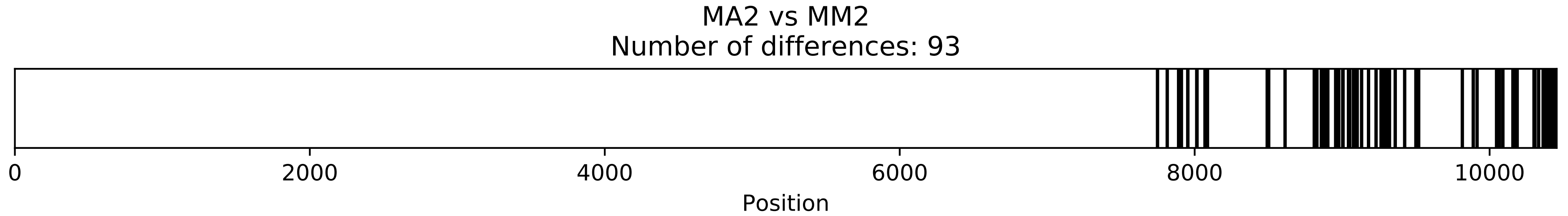
MA2 vs MM2
Number of differences: 93
0
2000
4000
6000
8000
10000
Position

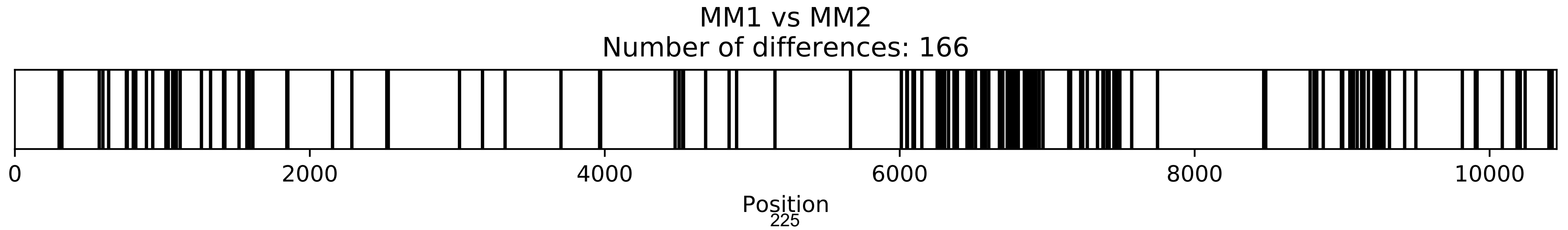
MM1 vs MM2
Number of differences: 166
0
2000
4000
6000
8000
10000
Position

### Region: 726



MA1 vs MA2
Number of differences: 249

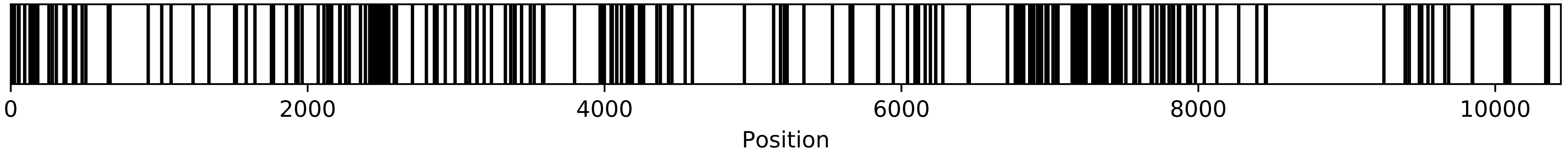


MA1 vs MM1
Number of differences: 3

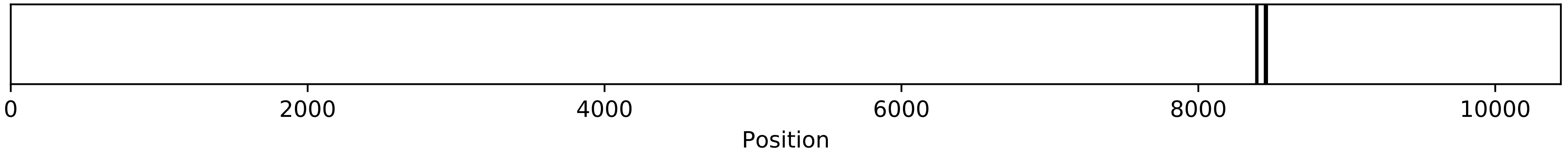


MA1 vs MM2
Number of differences: 104

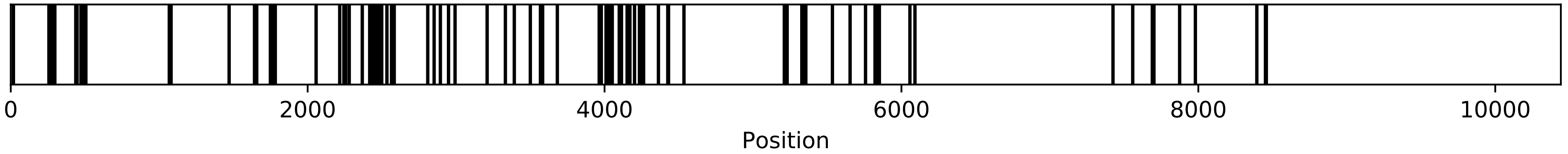


MA2 vs MM1
Number of differences: 246

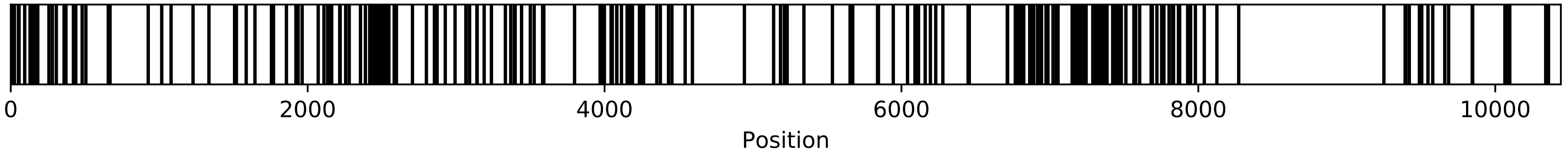


MA2 vs MM2
Number of differences: 256

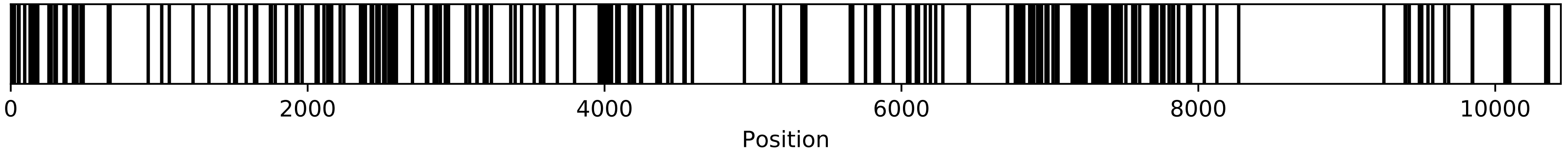


MM1 vs MM2
Number of differences: 101

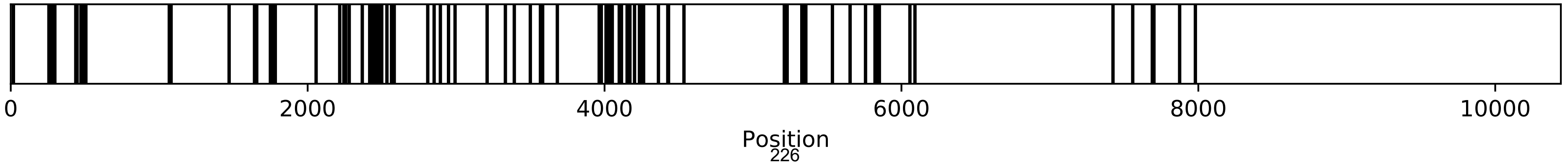

### Region: 732



MA1 vs MA2
Number of differences: 181

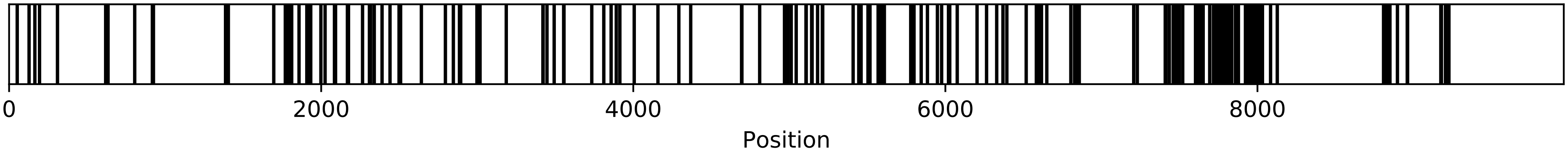


MA1 vs MM1
Number of differences: 189

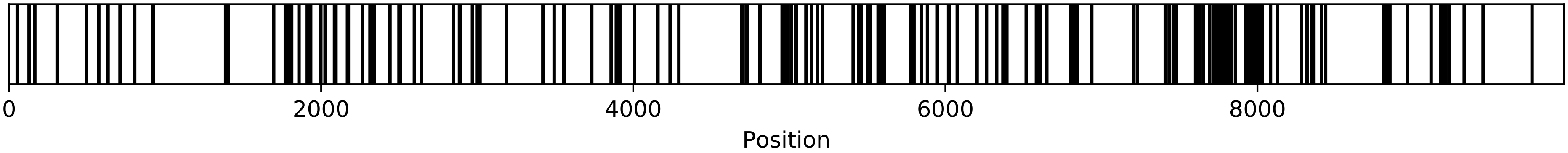


MA1 vs MM2
Number of differences: 181

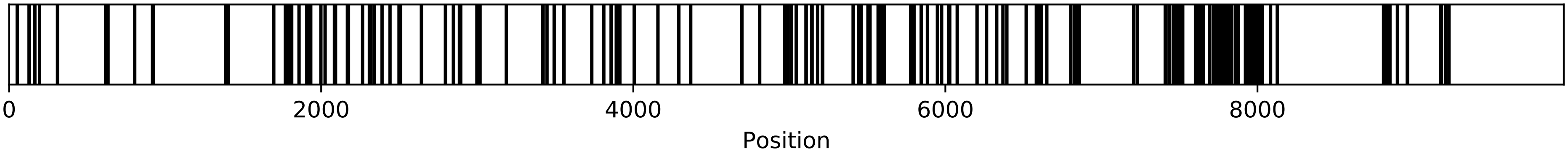


MA2 vs MM1
Number of differences: 46

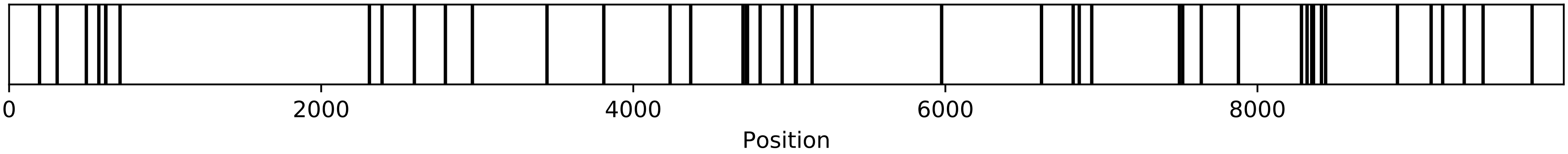


MA2 vs MM2
Number of differences: 0

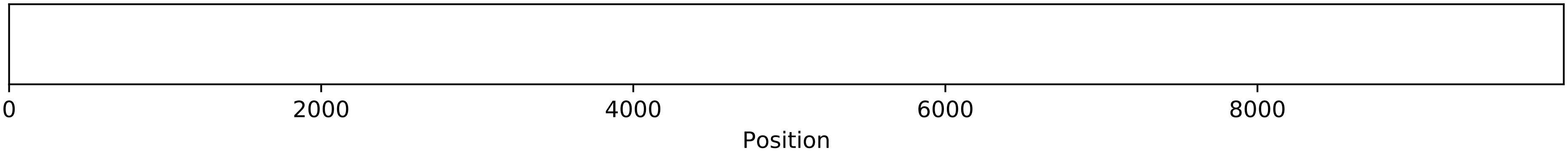


MM1 vs MM2
Number of differences: 46

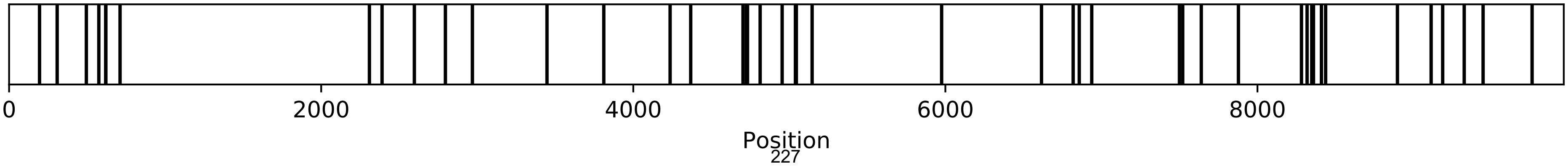

### Region: 738



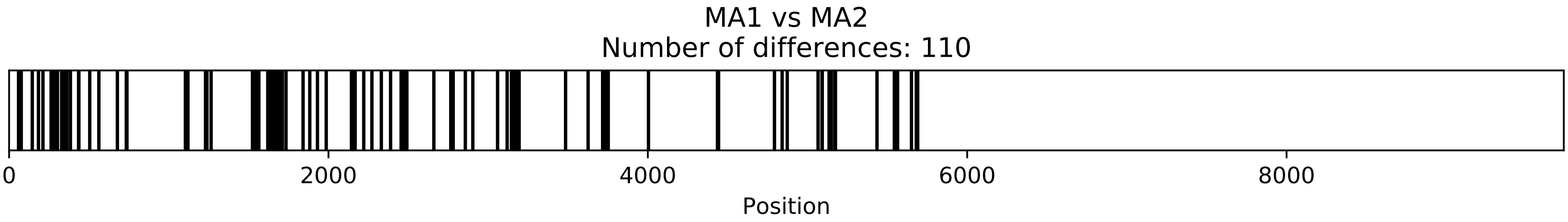
MA1 vs MA2
Number of differences: 110
0
2000
4000
6000
8000
Position

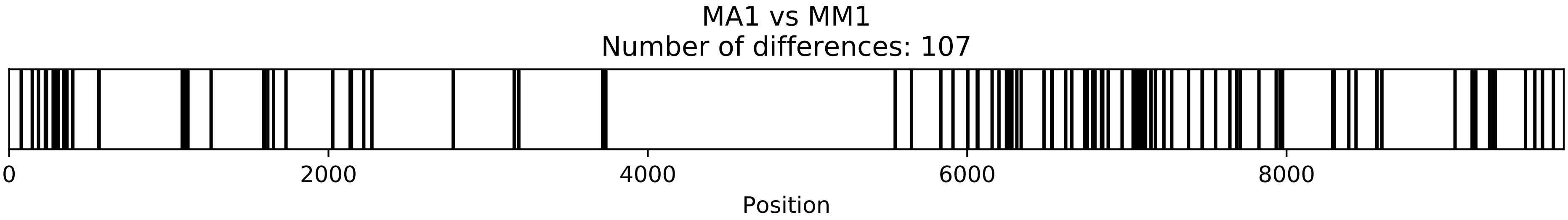
MA1 vs MM1
Number of differences: 107
0
2000
4000
6000
8000
Position

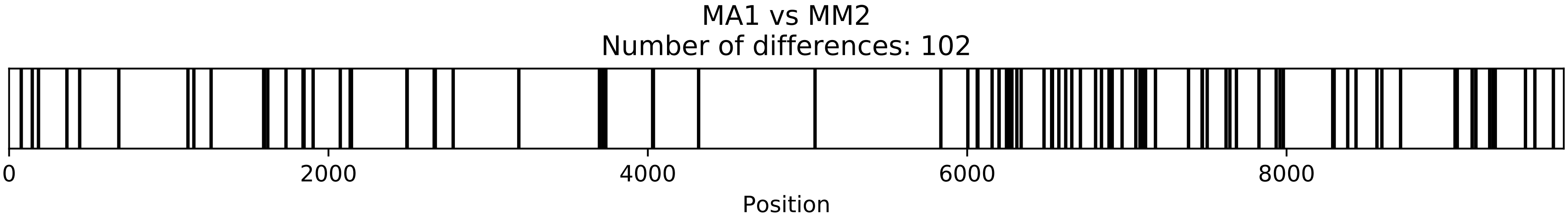
MA1 vs MM2
Number of differences: 102
0
2000
4000
6000
8000
Position

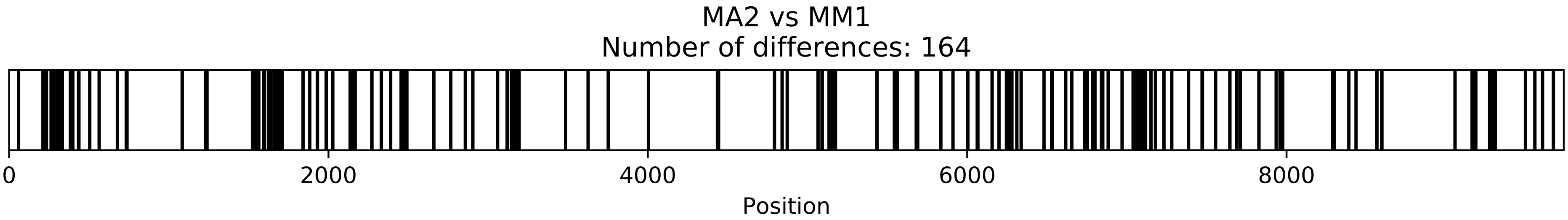
MA2 vs MM1
Number of differences: 164
0
2000
4000
6000
8000
Position

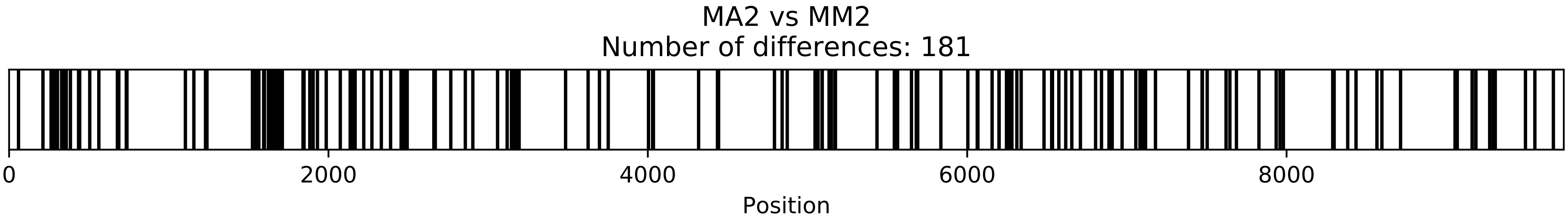
MA2 vs MM2
Number of differences: 181
0
2000
4000
6000
8000
Position

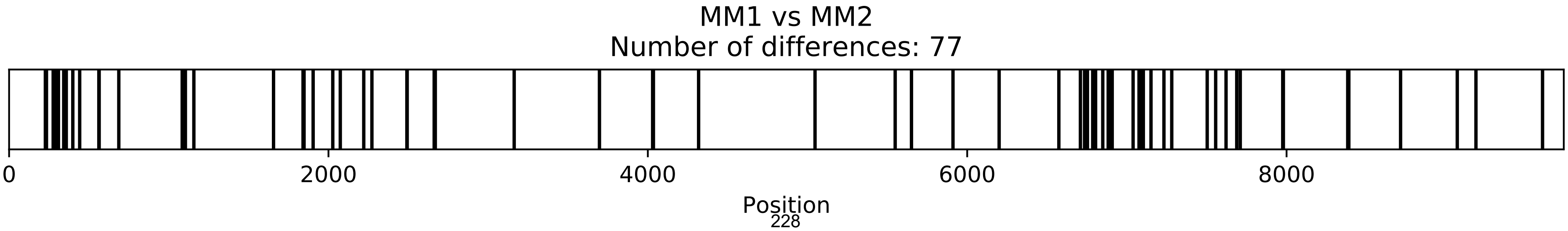
MM1 vs MM2
Number of differences: 77
0
2000
4000
6000
8000
Position

### Region: 740



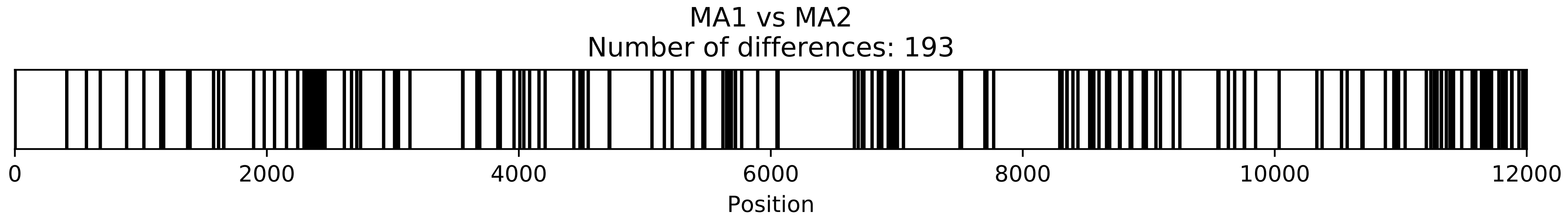
MA1 vs MA2
Number of differences: 193
0
2000
4000
6000
8000
10000
12000
Position

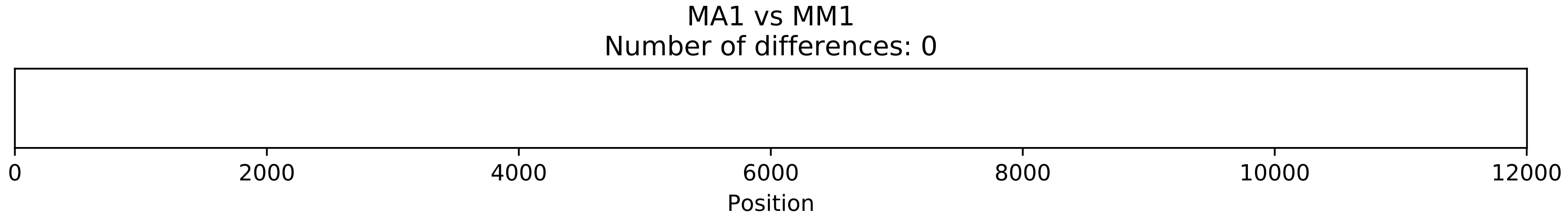
MA1 vs MM1
Number of differences: 0
0
2000
4000
6000
8000
10000
12000
Position

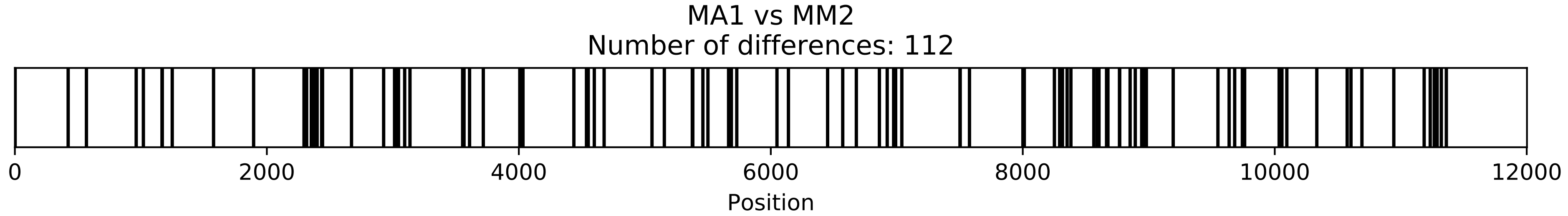
MA1 vs MM2
Number of differences: 112
0
2000
4000
6000
8000
10000
12000
Position

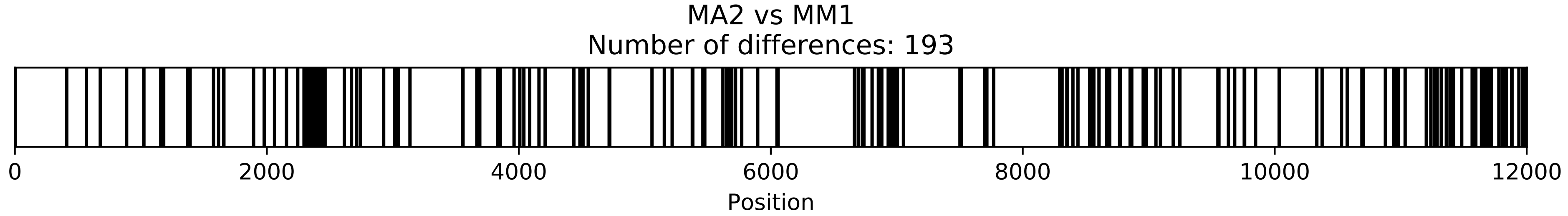
MA2 vs MM1
Number of differences: 193
0
2000
4000
6000
8000
10000
12000
Position

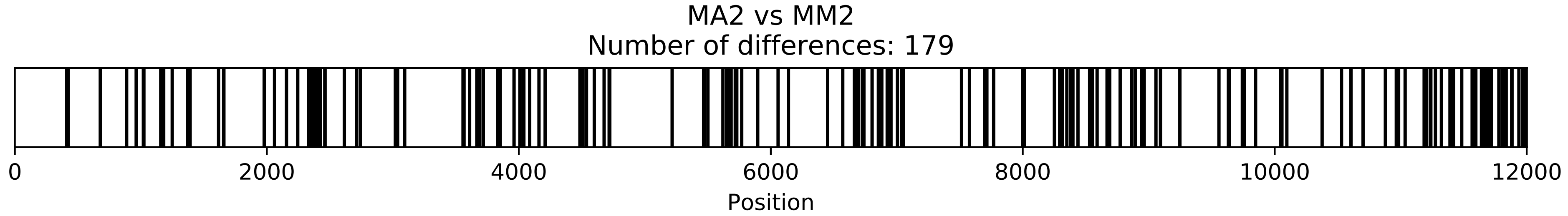
MA2 vs MM2
Number of differences: 179
0
2000
4000
6000
8000
10000
12000
Position

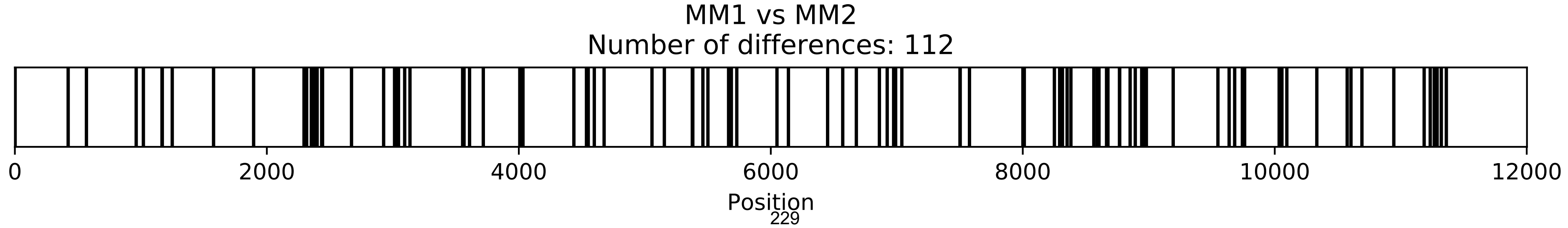
MM1 vs MM2
Number of differences: 112
0
2000
4000
6000
8000
10000
12000
Position

### Region: 743



MA1 vs MA2
Number of differences: 553

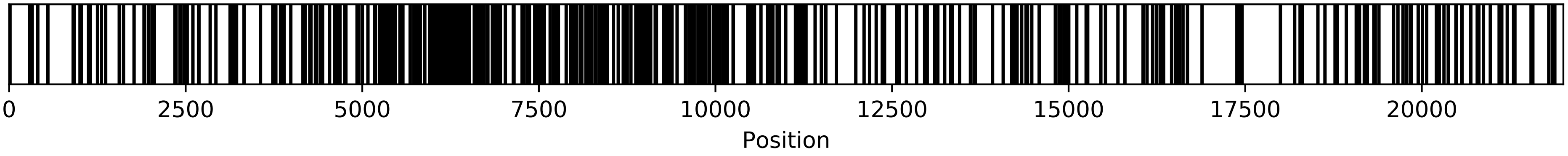


MA1 vs MM1
Number of differences: 553

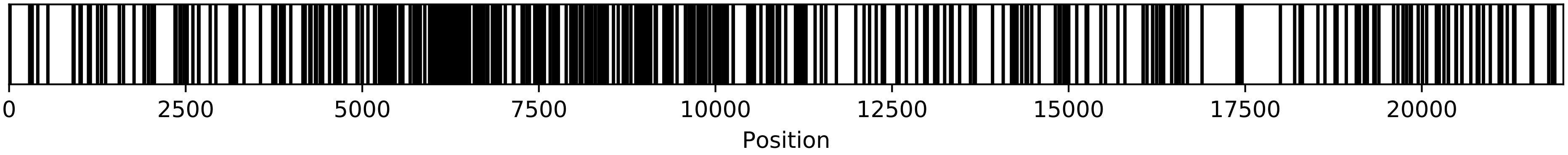


MA1 vs MM2
Number of differences: 548

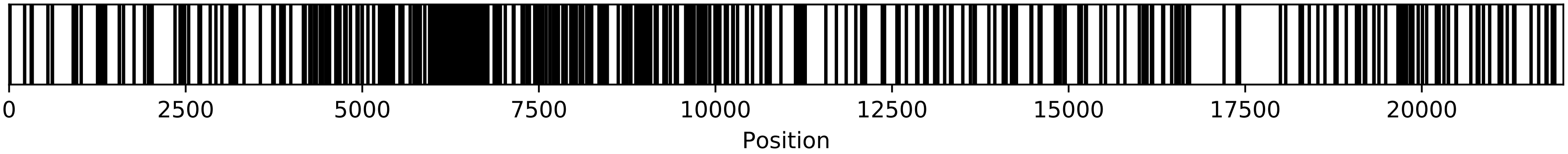


MA2 vs MM1
Number of differences: 0

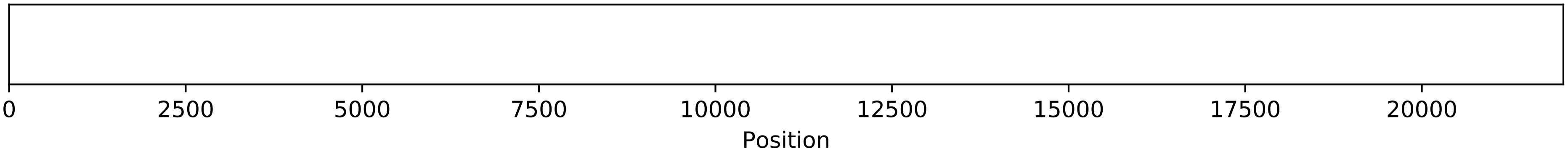


MA2 vs MM2
Number of differences: 261

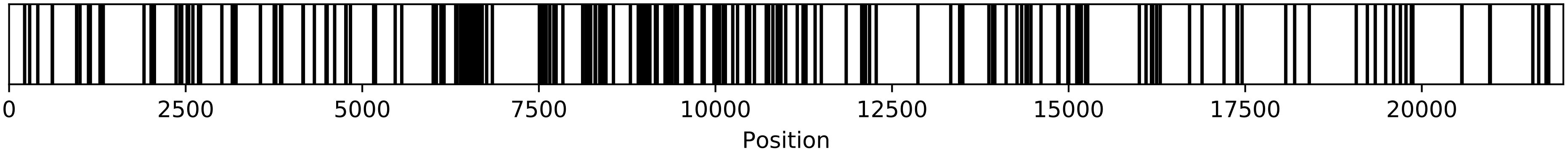


MM1 vs MM2
Number of differences: 261

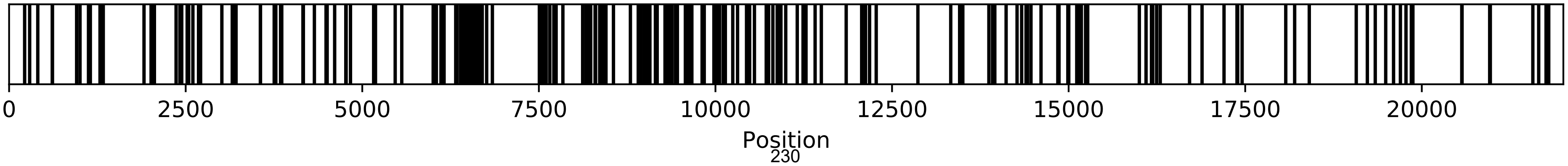

### Region: 746



MA1 vs MA2
Number of differences: 227

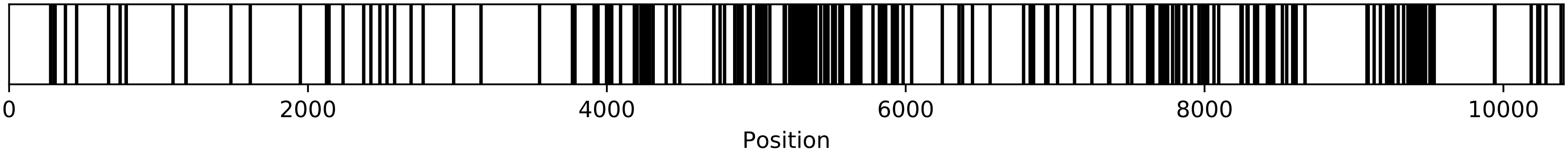


MA1 vs MM1
Number of differences: 227

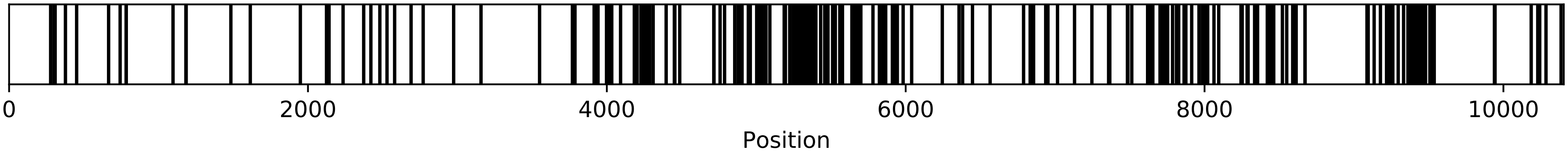


MA1 vs MM2
Number of differences: 217

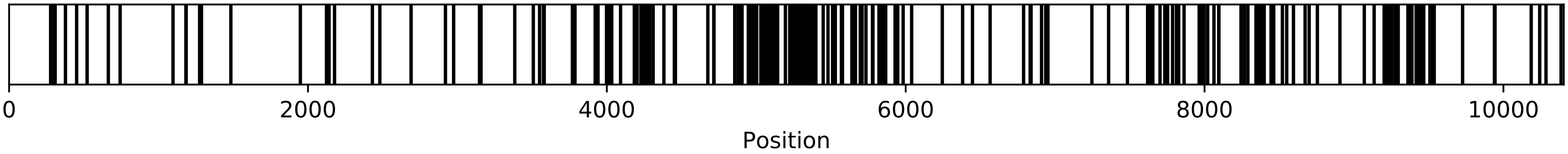


MA2 vs MM1
Number of differences: 0

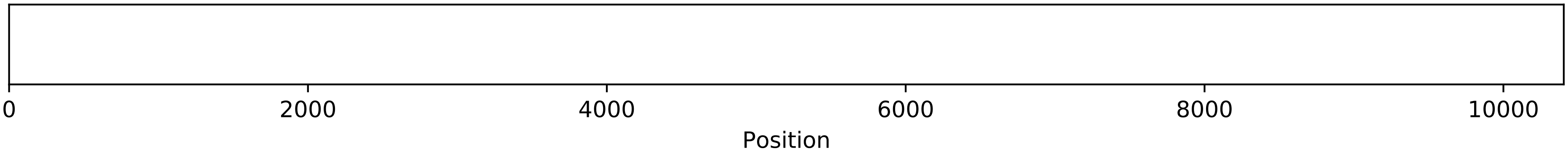


MA2 vs MM2
Number of differences: 125

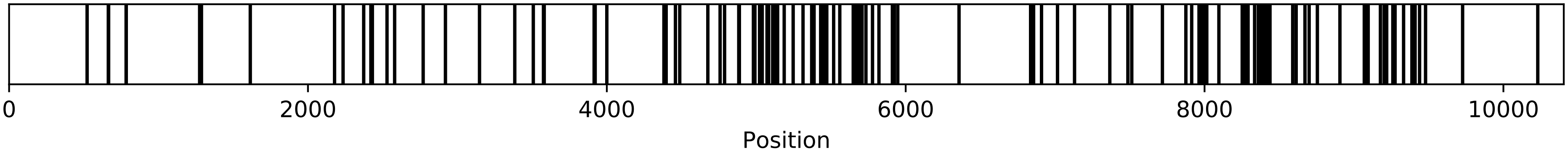


MM1 vs MM2
Number of differences: 125

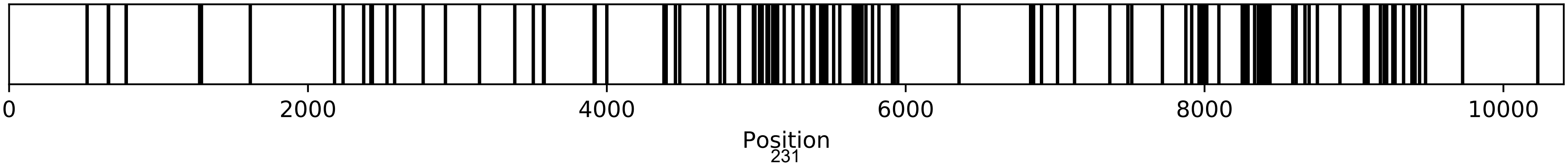

### Region: 747



MA1 vs MA2
Number of differences: 216

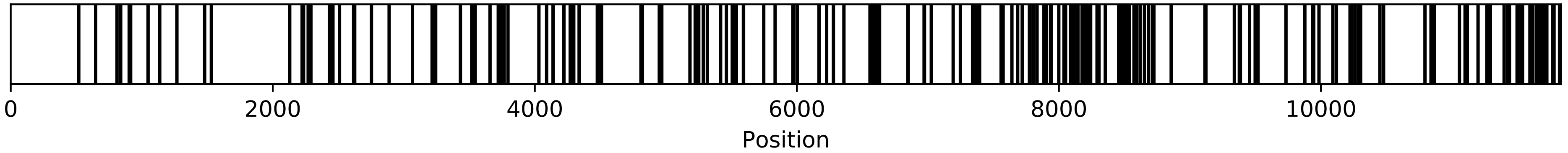


MA1 vs MM1
Number of differences: 115

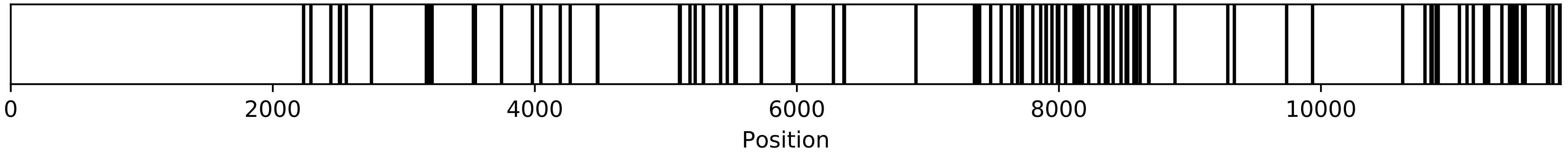


MA1 vs MM2
Number of differences: 115

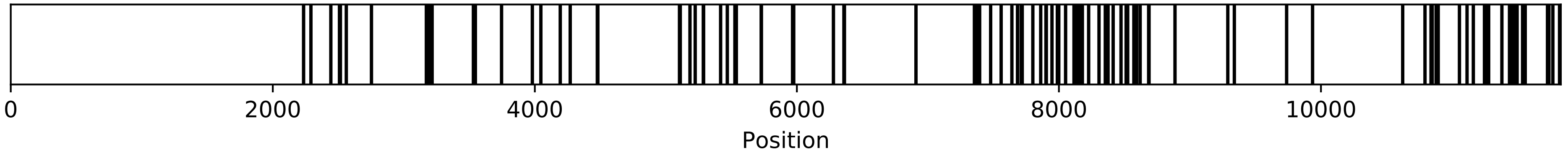


MA2 vs MM1
Number of differences: 217

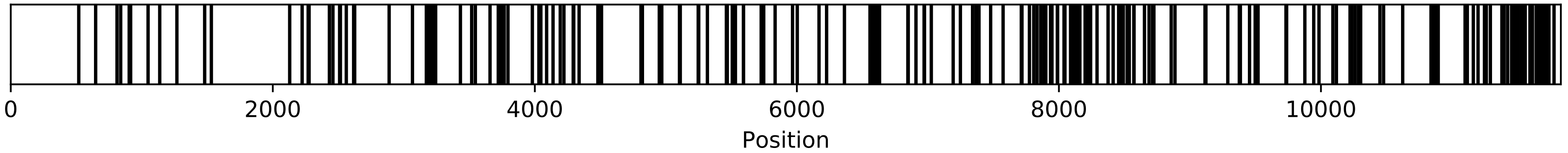


MA2 vs MM2
Number of differences: 217

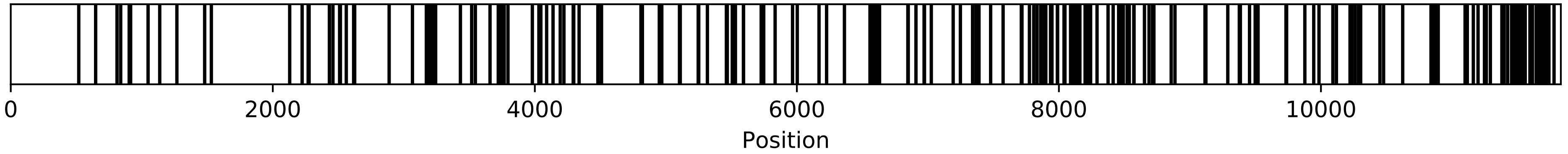


MM1 vs MM2
Number of differences: 0

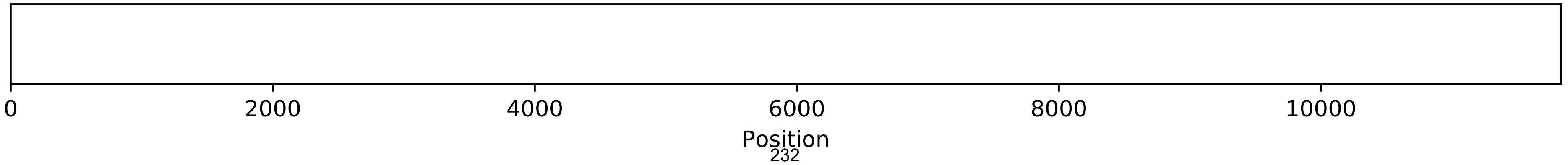

### Region: 761



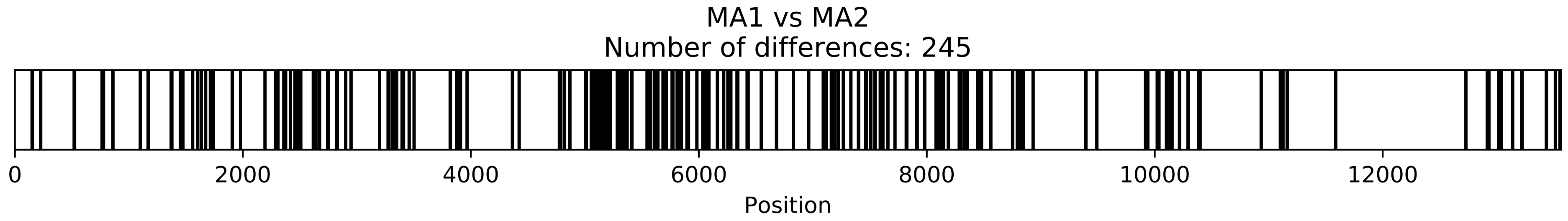
MA1 vs MA2
Number of differences: 245
0
2000
4000
6000
8000
10000
12000
Position

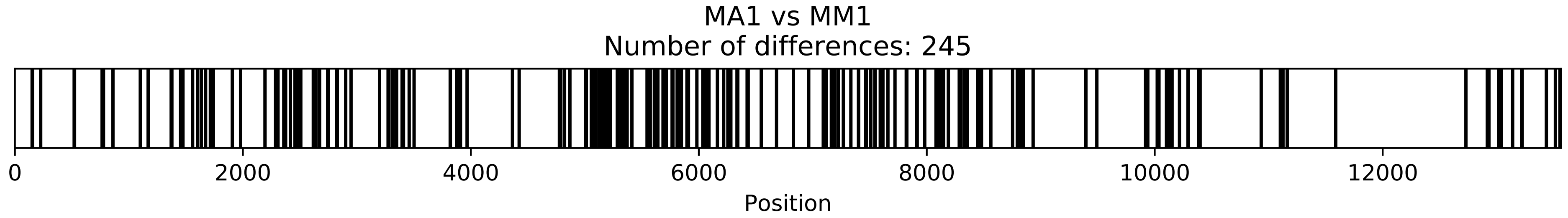
MA1 vs MM1
Number of differences: 245
0
2000
4000
6000
8000
10000
12000
Position

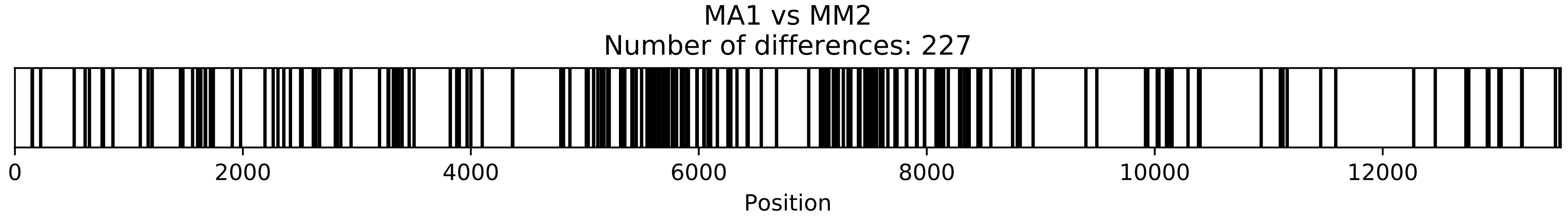
MA1 vs MM2
Number of differences: 227
0
2000
4000
6000
8000
10000
12000
Position

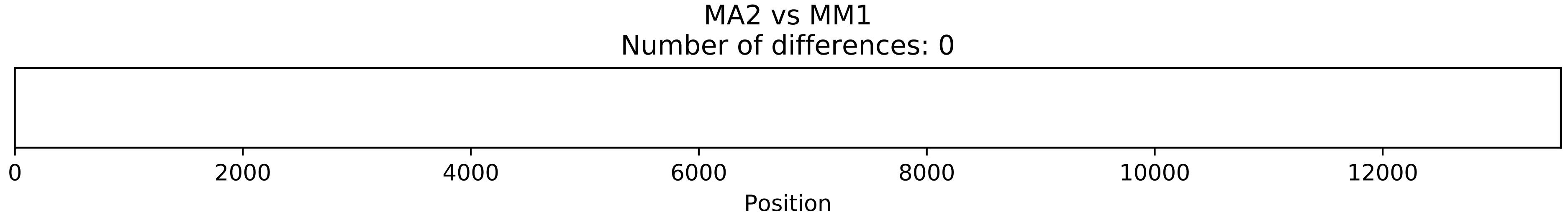
MA2 vs MM1
Number of differences: 0
0
2000
4000
6000
8000
10000
12000
Position

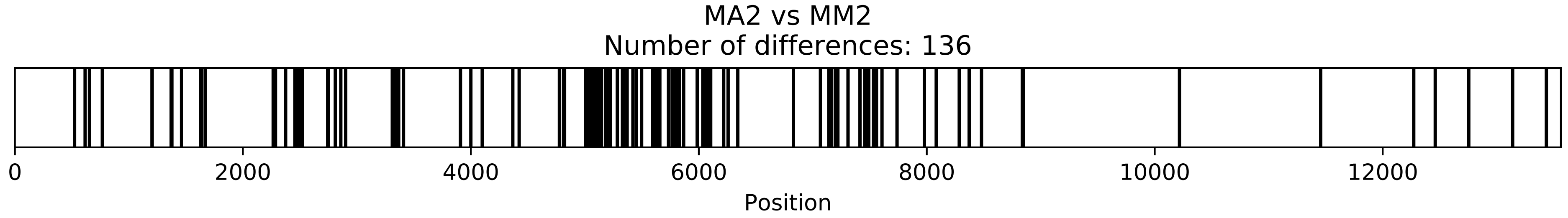
MA2 vs MM2
Number of differences: 136
0
2000
4000
6000
8000
10000
12000
Position

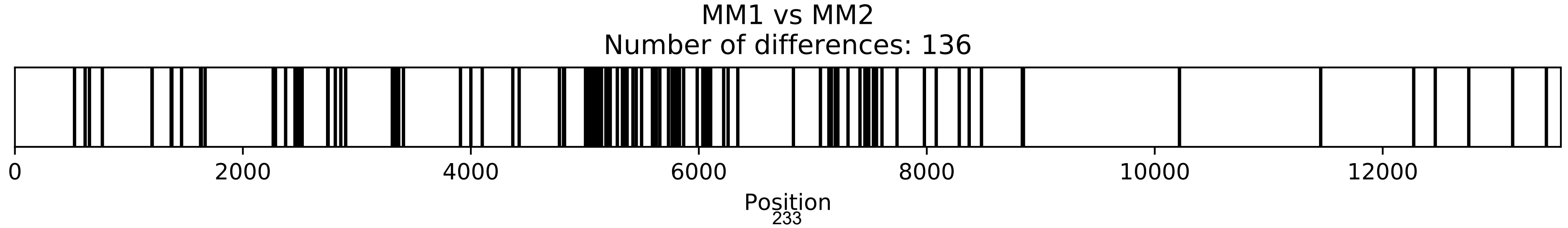
MM1 vs MM2
Number of differences: 136
0
2000
4000
6000
8000
10000
12000
Position

### Region: 762



MA1 vs MA2
Number of differences: 259

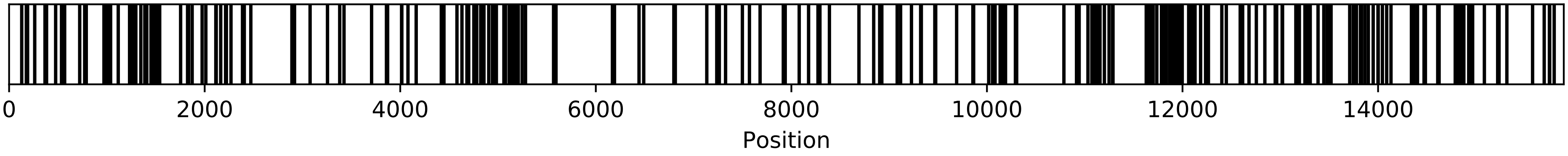


MA1 vs MM1
Number of differences: 259

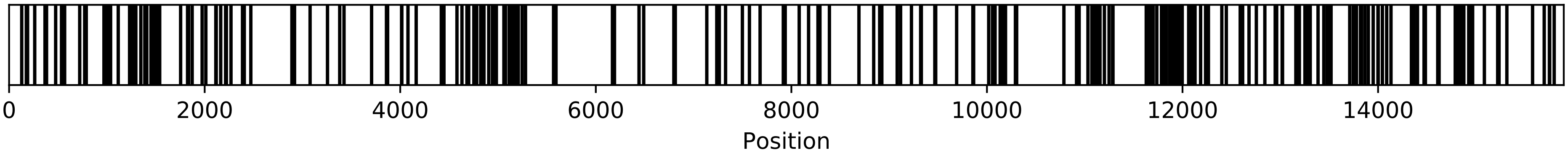


MA1 vs MM2
Number of differences: 238

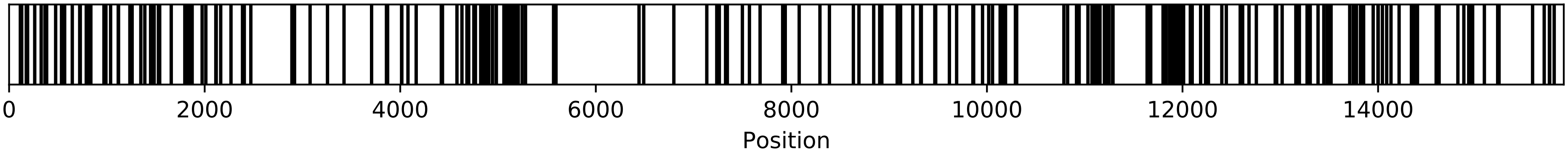


MA2 vs MM1
Number of differences: 0

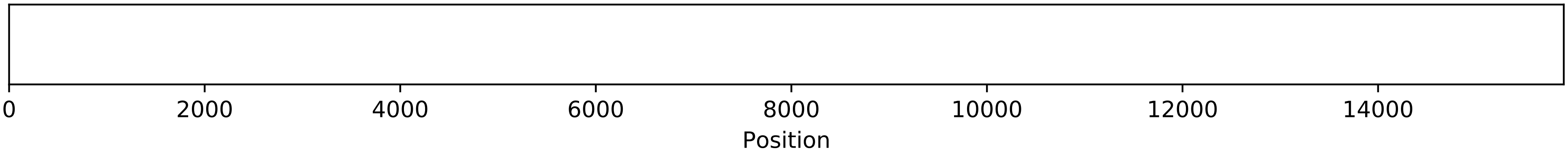


MA2 vs MM2
Number of differences: 103

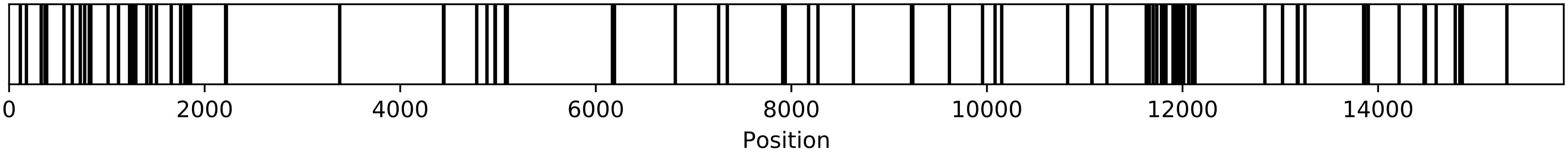


MM1 vs MM2
Number of differences: 103

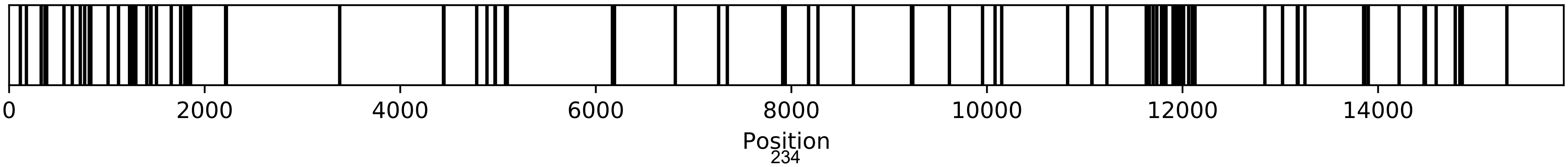

### Region: 769

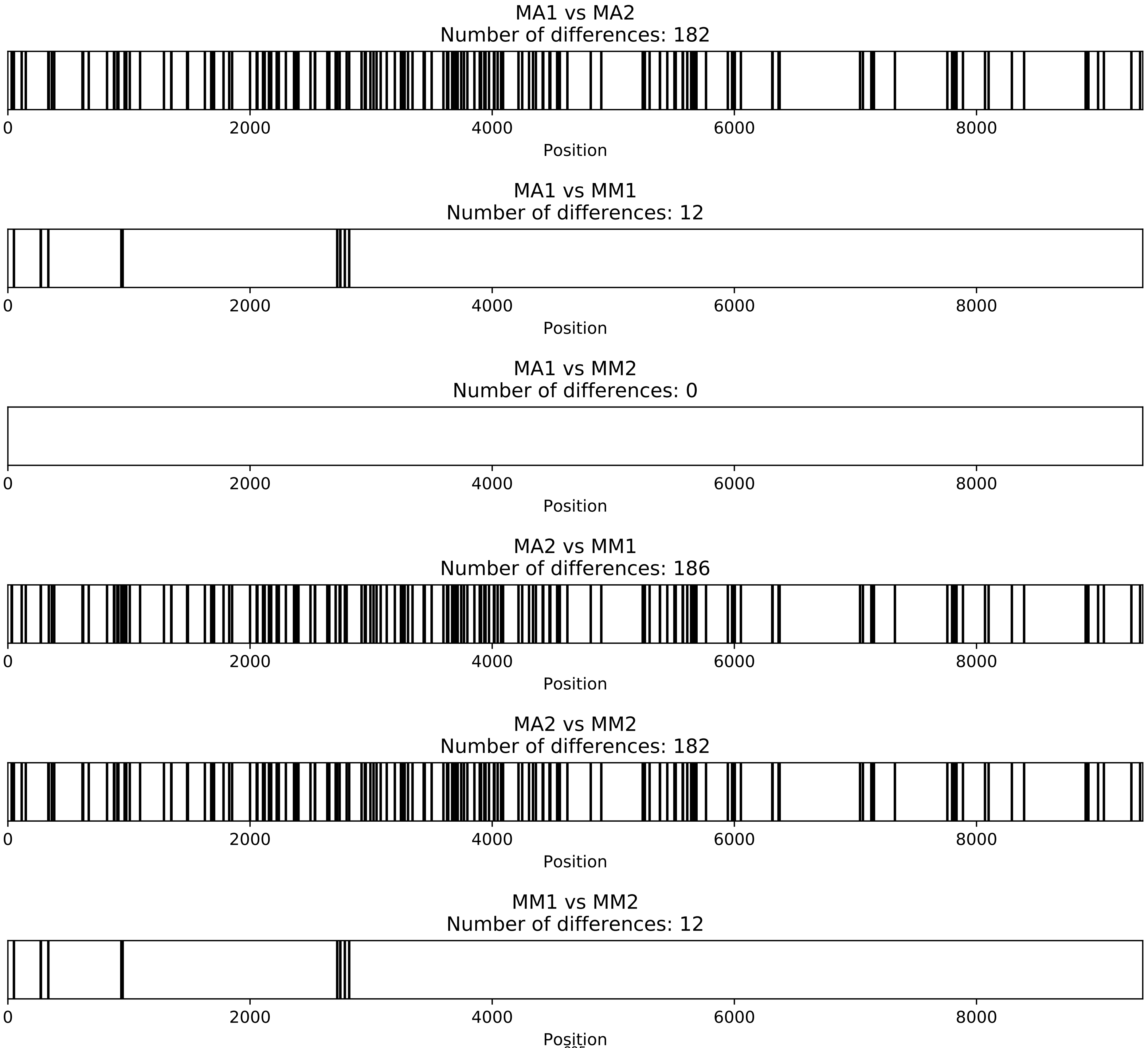
MA1 vs MA2
Number of differences: 182
0
2000
4000
6000
8000
Position
MA1 vs MM1
Number of differences: 12
0
2000
4000
6000
8000
Position
MA1 vs MM2
Number of differences: 0
0
2000
4000
6000
8000
Position
MA2 vs MM1
Number of differences: 186
0
2000
4000
6000
8000
Position
MA2 vs MM2
Number of differences: 182
0
2000
4000
6000
8000
Position
MM1 vs MM2
Number of differences: 12
0
2000
4000
6000
8000
Position

### Region: 773



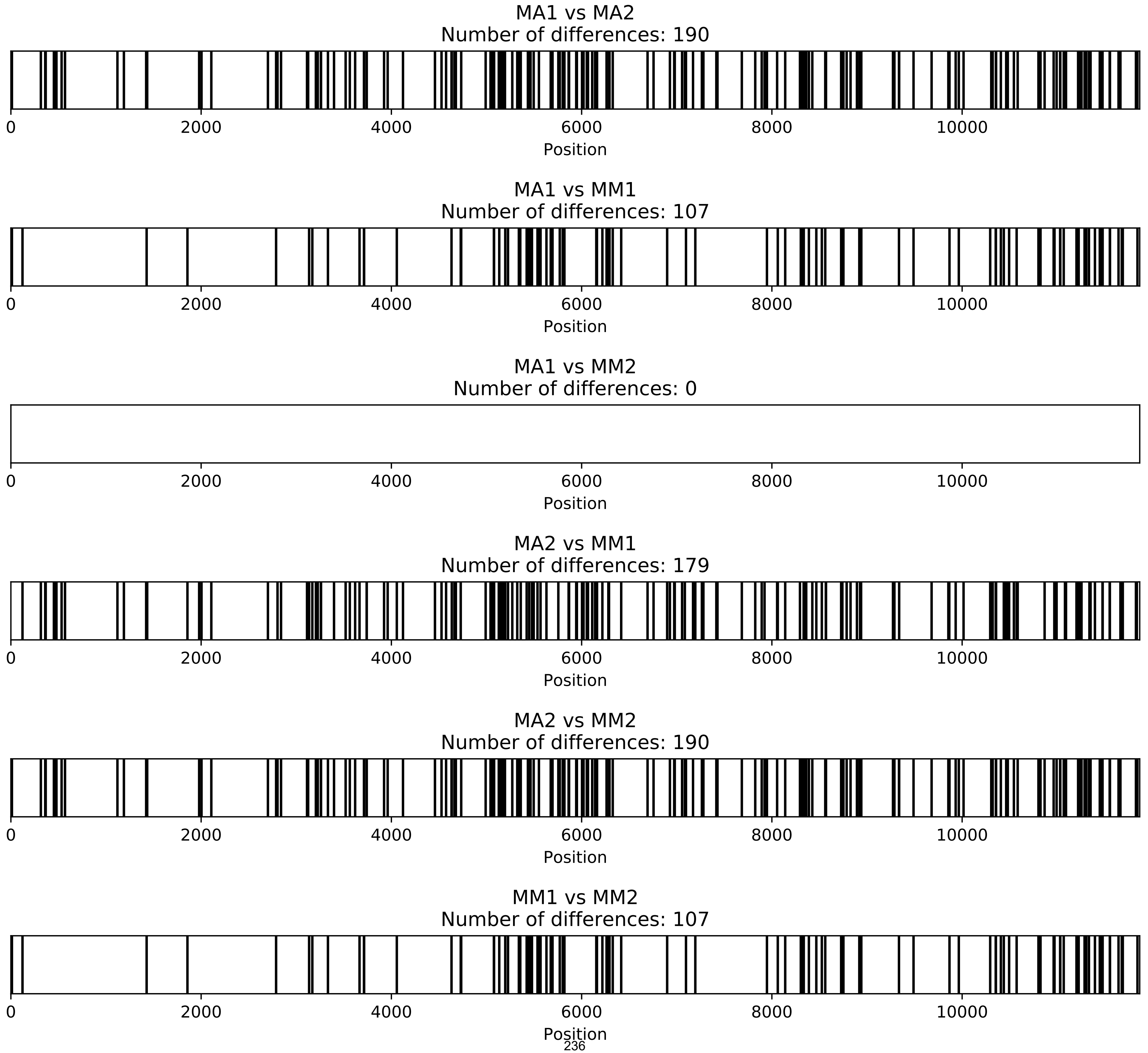
MA1 vs MA2
Number of differences: 190
MA1 vs MM1
Number of differences: 107
MA1 vs MM2
Number of differences: 0
MA2 vs MM1
Number of differences: 179
MA2 vs MM2
Number of differences: 190
MM1 vs MM2
Number of differences: 107
0
2000
4000
6000
8000
10000
Position

### Region: 776

MA1 vs MA2
Number of differences: 110

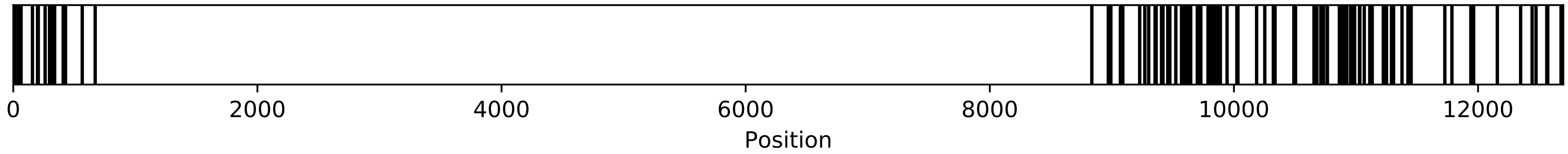


MA1 vs MM1
Number of differences: 165

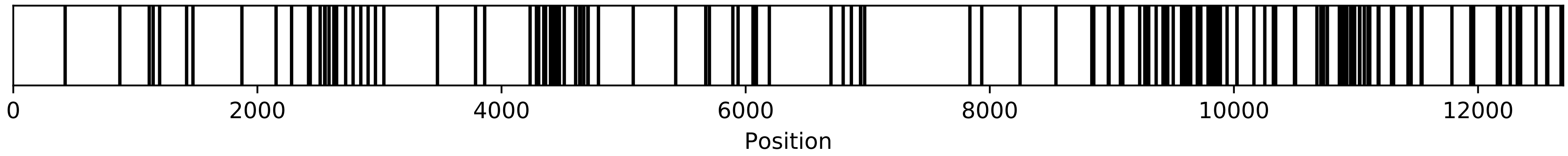


MA1 vs MM2
Number of differences: 129

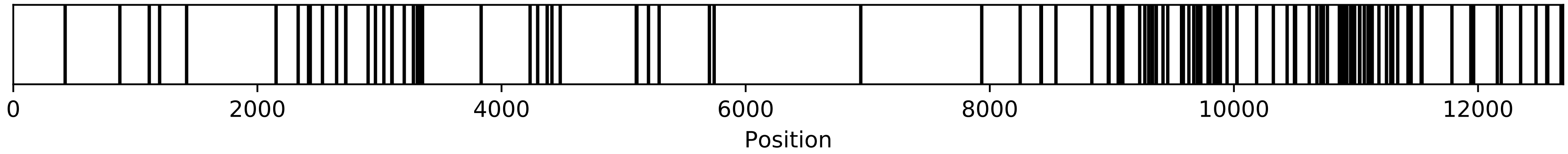


MA2 vs MM1
Number of differences: 129

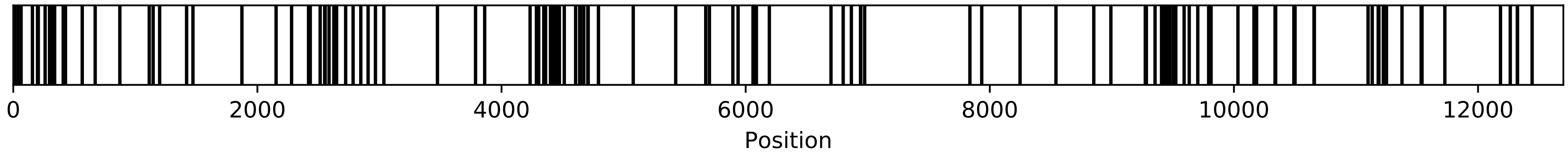


MA2 vs MM2
Number of differences: 103

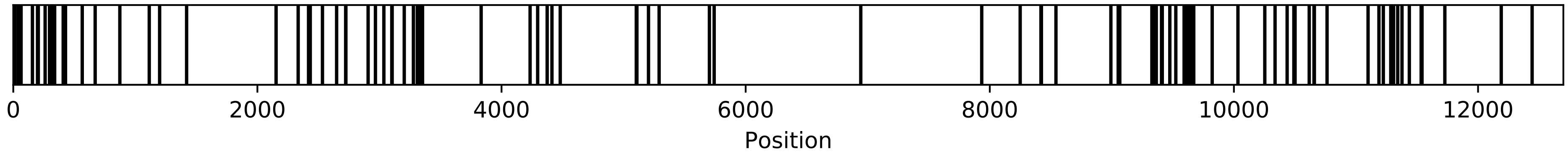


MM1 vs MM2
Number of differences: 118

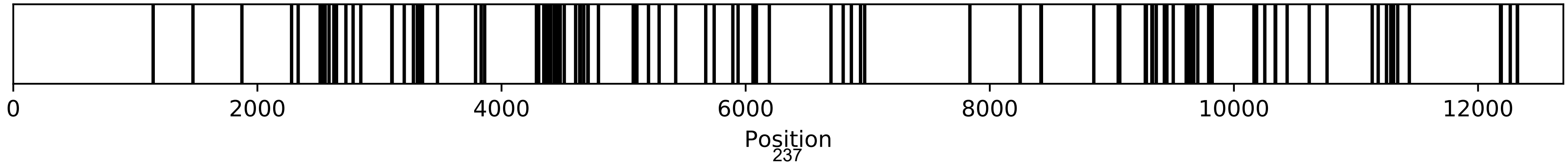

### Region: 779



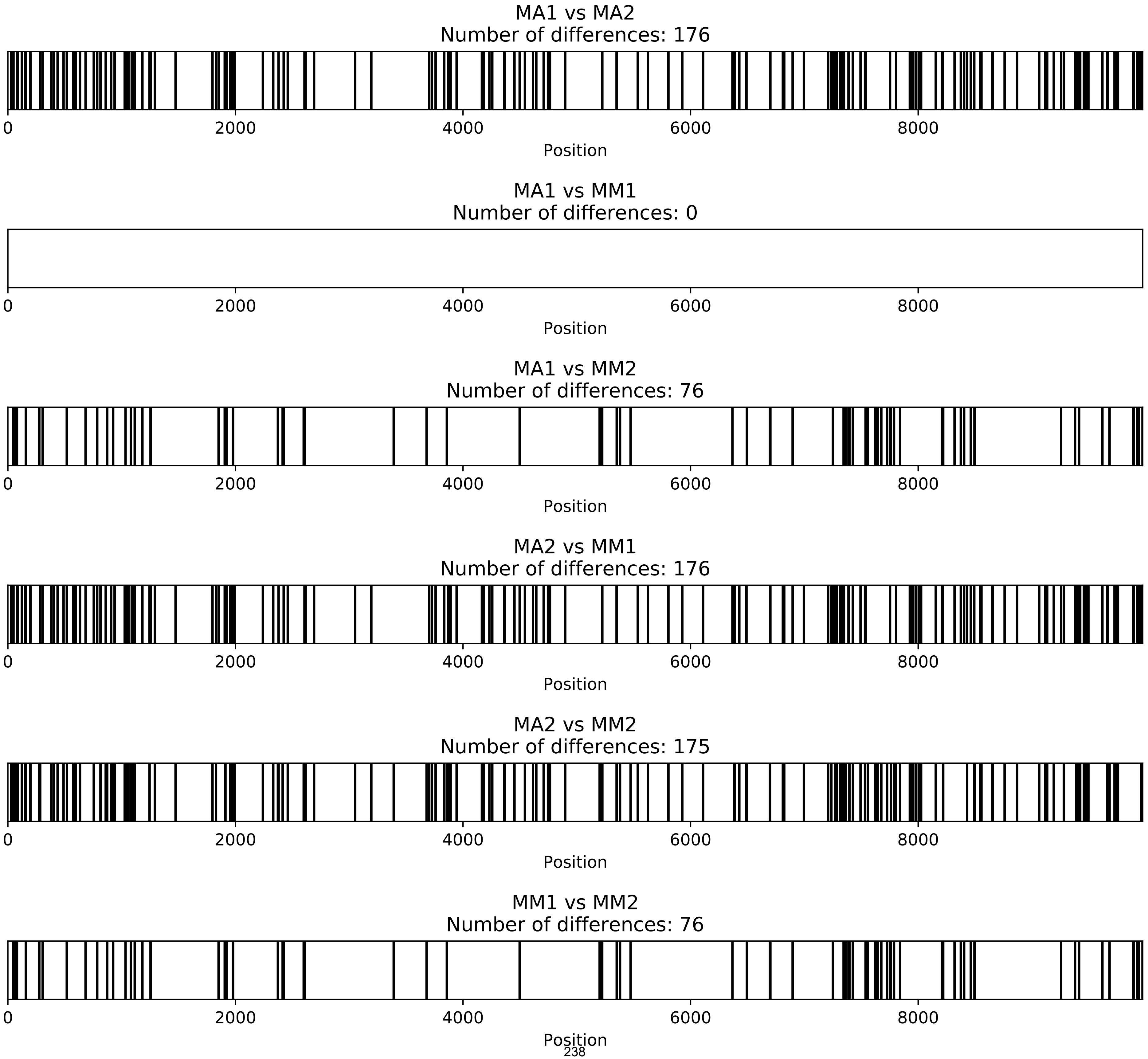
MA1 vs MA2
Number of differences: 176
0
2000
4000
6000
8000
Position
MA1 vs MM1
Number of differences: 0
0
2000
4000
6000
8000
Position
MA1 vs MM2
Number of differences: 76
0
2000
4000
6000
8000
Position
MA2 vs MM1
Number of differences: 176
0
2000
4000
6000
8000
Position
MA2 vs MM2
Number of differences: 175
0
2000
4000
6000
8000
Position
MM1 vs MM2
Number of differences: 76
0
2000
4000
6000
8000
Position

### Region: 780



MA1 vs MA2
Number of differences: 221

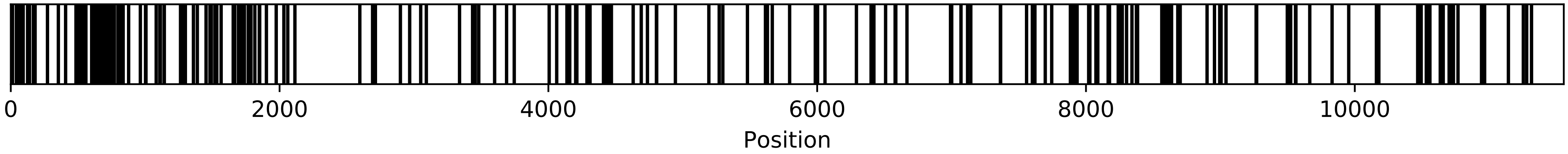


MA1 vs MM1
Number of differences: 221

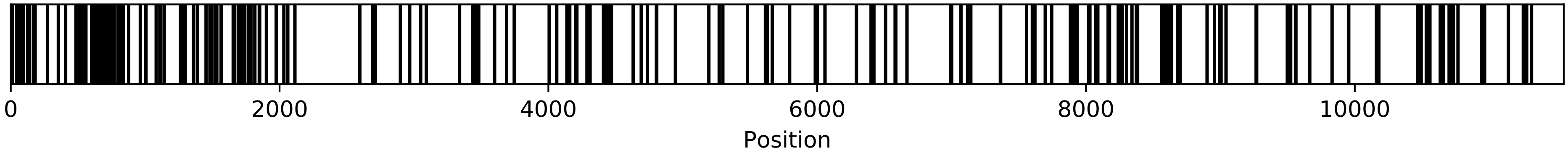


MA1 vs MM2
Number of differences: 214

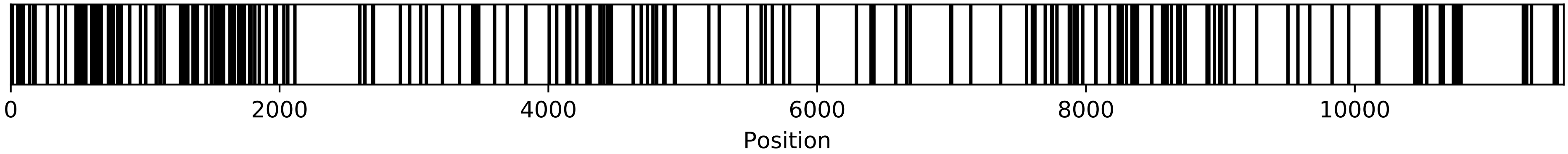


MA2 vs MM1
Number of differences: 0

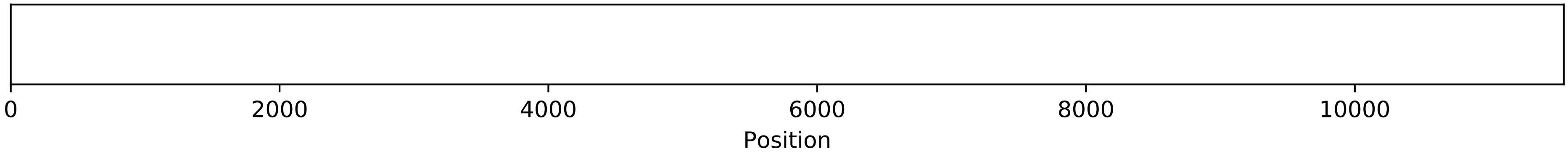


MA2 vs MM2
Number of differences: 106

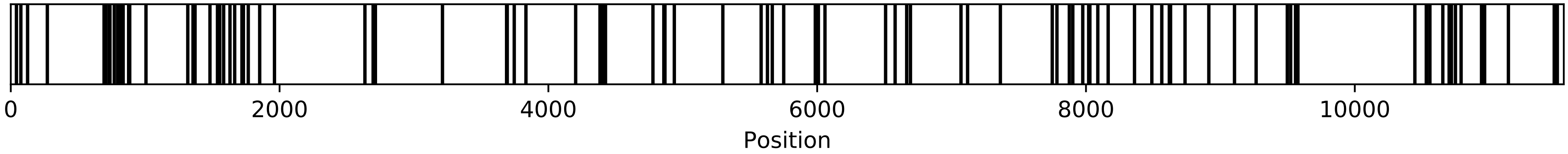


MM1 vs MM2
Number of differences: 106

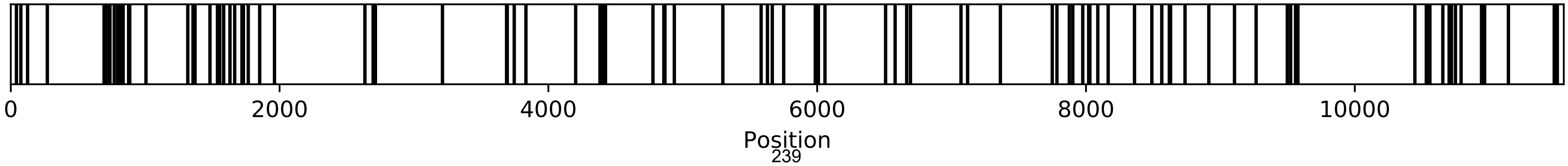

### Region: 794



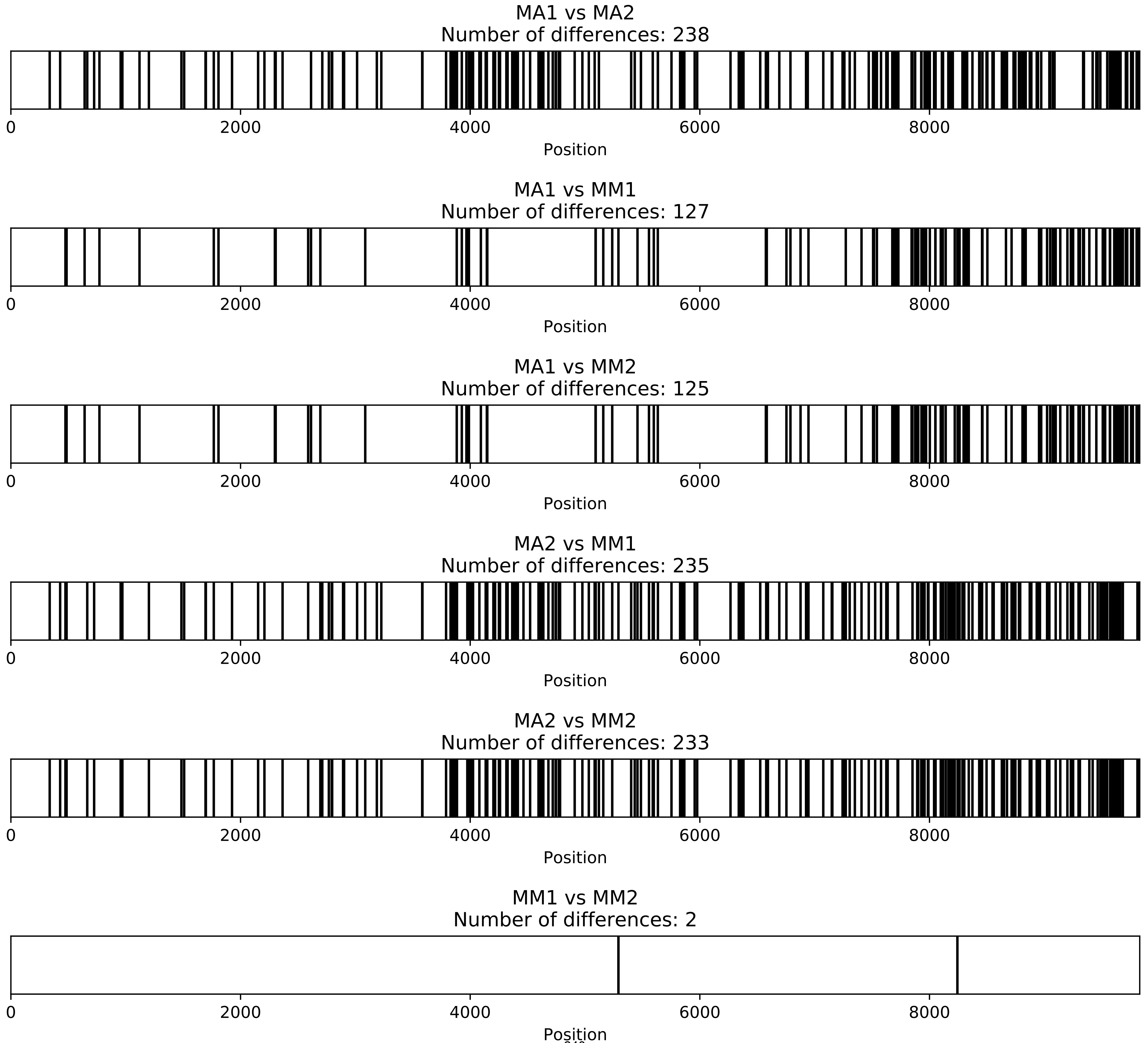
MA1 vs MA2
Number of differences: 238
MA1 vs MM1
Number of differences: 127
MA1 vs MM2
Number of differences: 125
MA2 vs MM1
Number of differences: 235
MA2 vs MM2
Number of differences: 233
MM1 vs MM2
Number of differences: 2
0
2000
4000
6000
8000
Position

### Region: 796



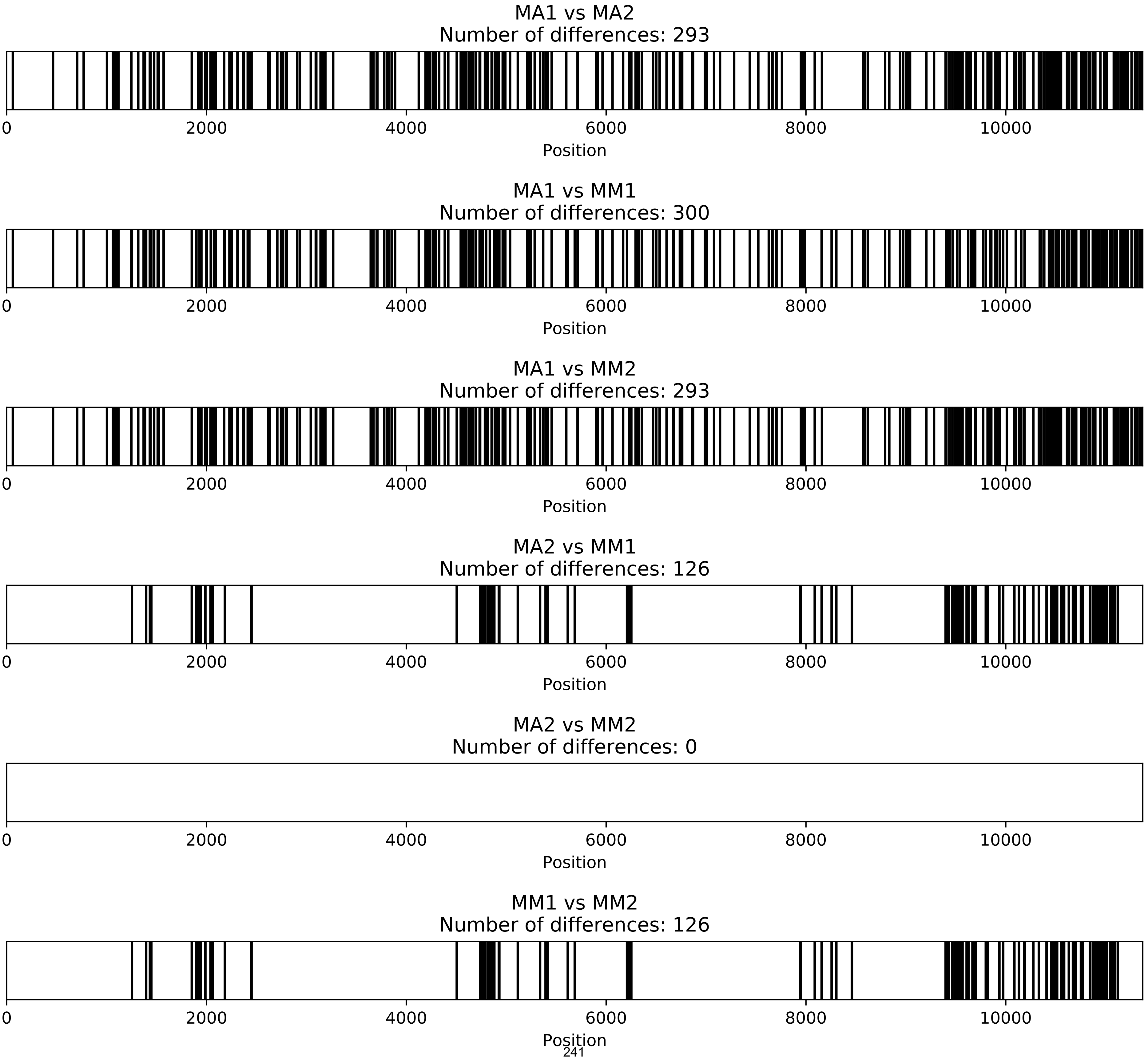
MA1 vs MA2
Number of differences: 293
MA1 vs MM1
Number of differences: 300
MA1 vs MM2
Number of differences: 293
MA2 vs MM1
Number of differences: 126
MA2 vs MM2
Number of differences: 0
MM1 vs MM2
Number of differences: 126
0
2000
4000
6000
8000
10000
Position

### Region: 797



MA1 vs MA2
Number of differences: 325

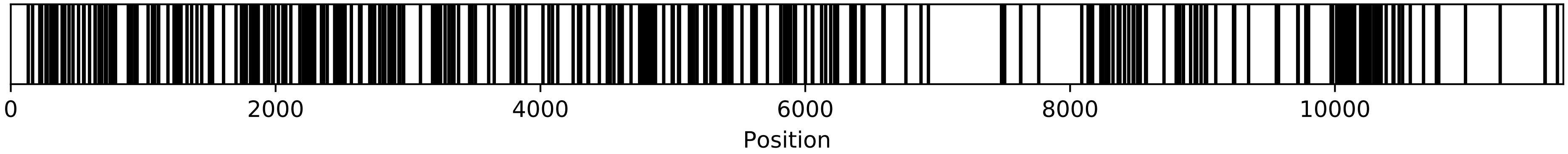


MA1 vs MM1
Number of differences: 159

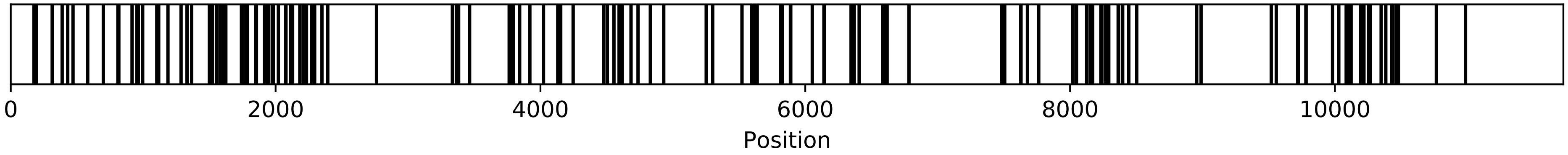


MA1 vs MM2
Number of differences: 0

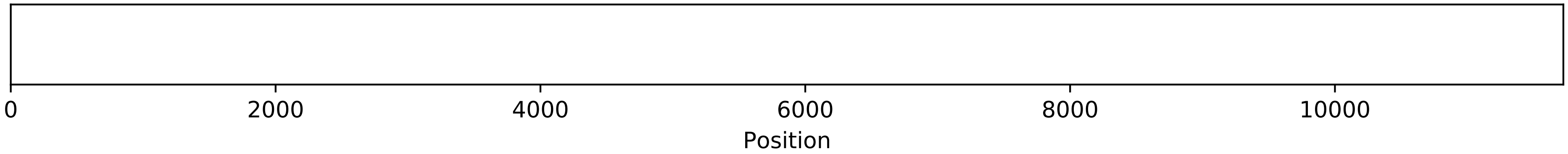


MA2 vs MM1
Number of differences: 305

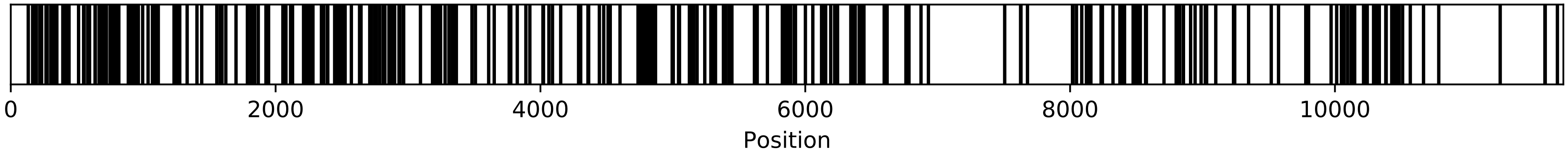


MA2 vs MM2
Number of differences: 325

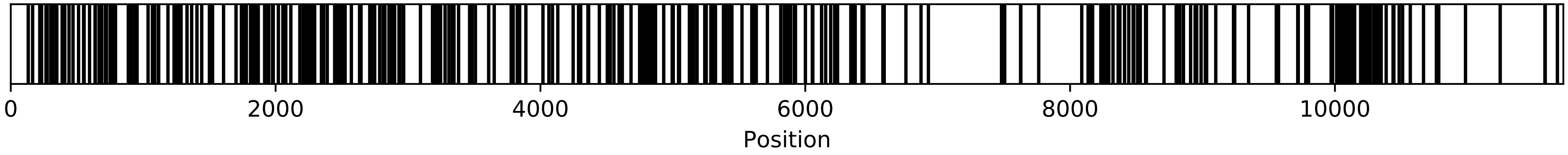


MM1 vs MM2
Number of differences: 159

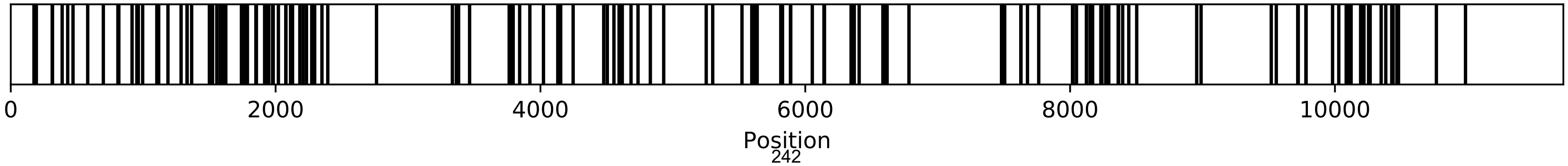

### Region: 802



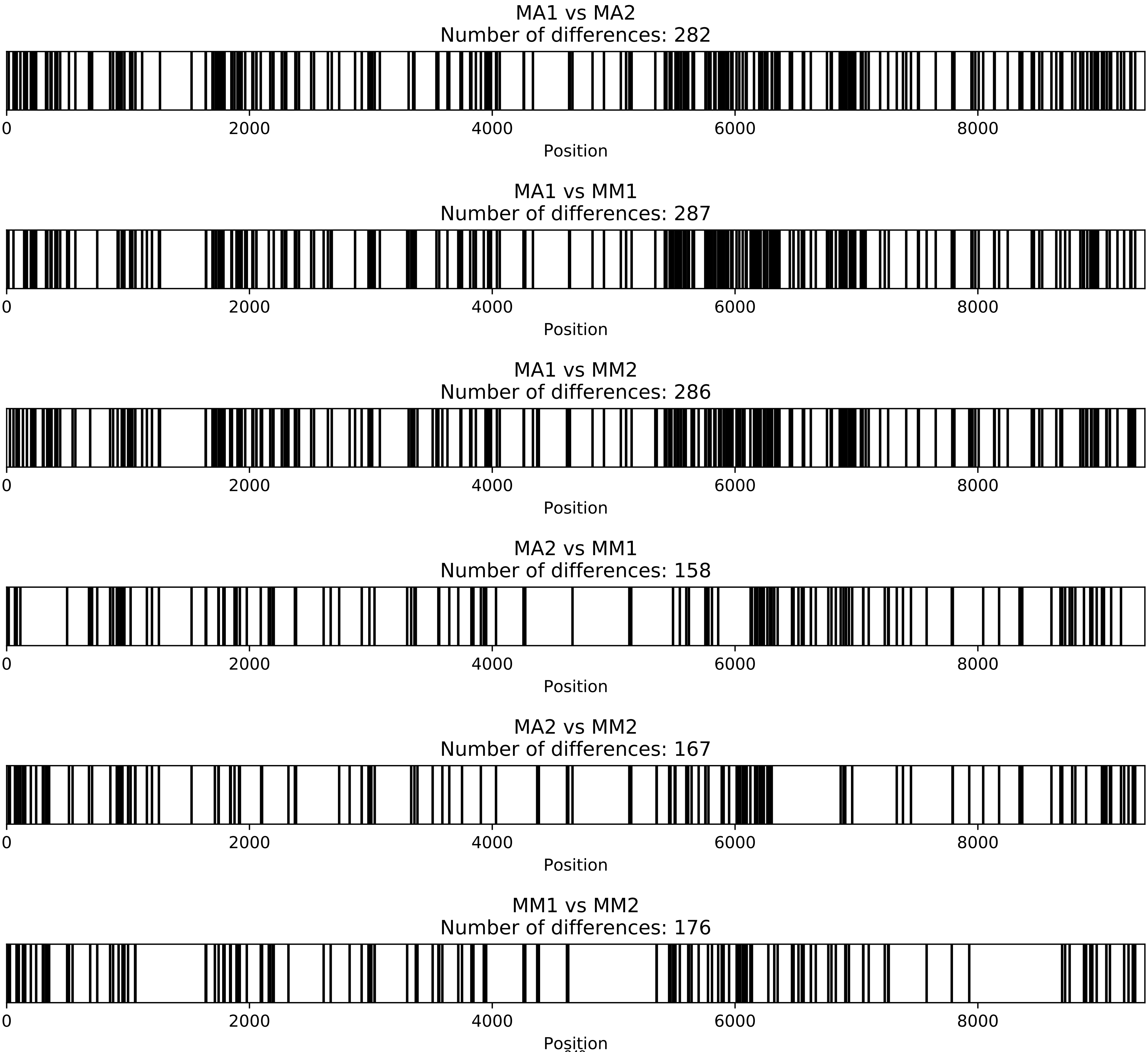
MA1 vs MA2
Number of differences: 282
Position
MA1 vs MM1
Number of differences: 287
Position
MA1 vs MM2
Number of differences: 286
Position
MA2 vs MM1
Number of differences: 158
Position
MA2 vs MM2
Number of differences: 167
Position
MM1 vs MM2
Number of differences: 176
Position
0
2000
4000
6000
8000

### Region: 808



## MA1 vs MA2
Number of differences: 142

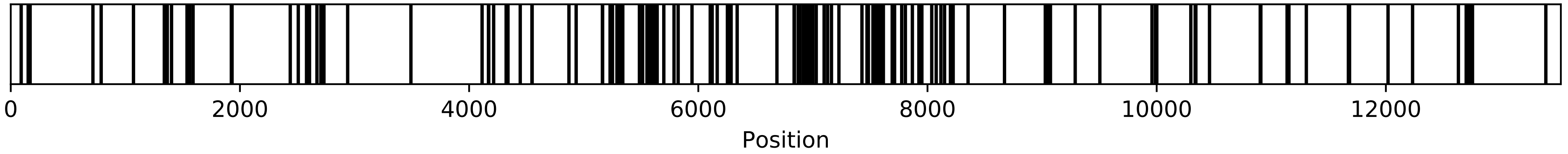


## MA1 vs MM1
Number of differences: 67

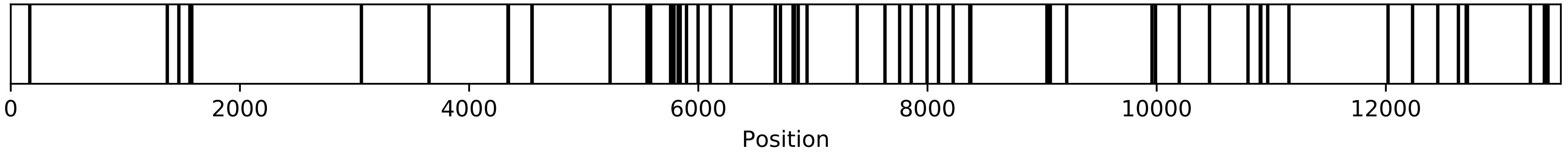


## MA1 vs MM2
Number of differences: 79

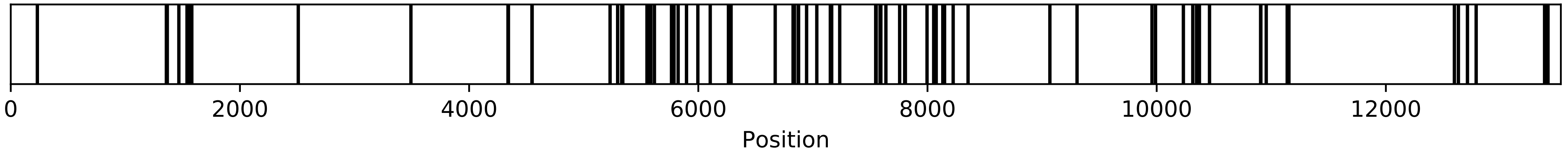


## MA2 vs MM1
Number of differences: 146

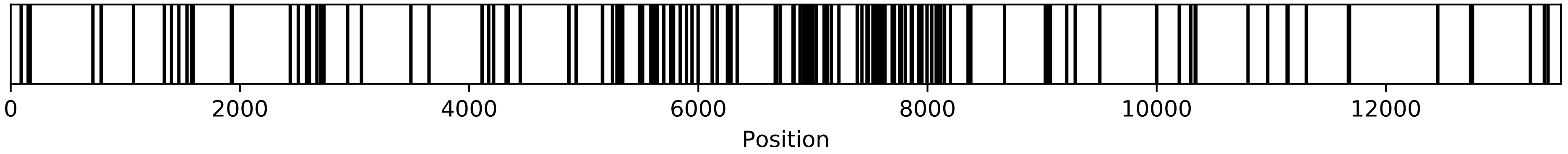


## MA2 vs MM2
Number of differences: 145

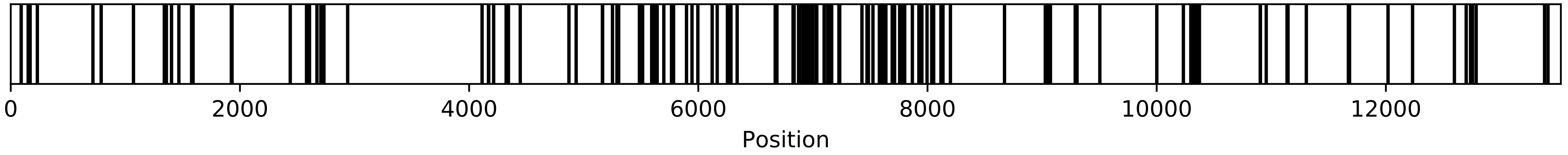


## MM1 vs MM2
Number of differences: 70

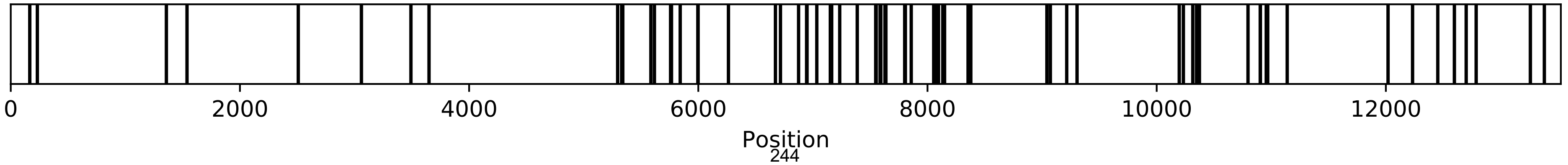

### Region: 810



MA1 vs MA2
Number of differences: 468

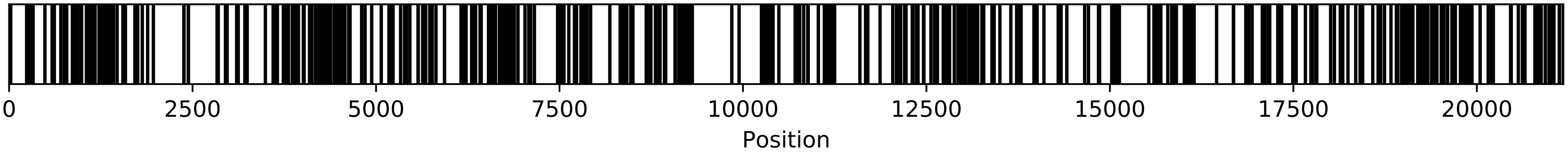


MA1 vs MM1
Number of differences: 460

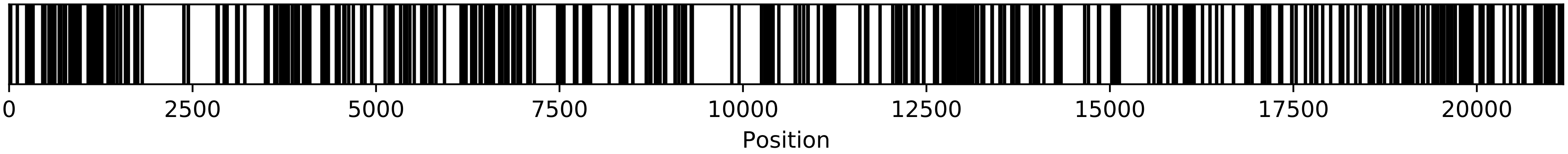


MA1 vs MM2
Number of differences: 476

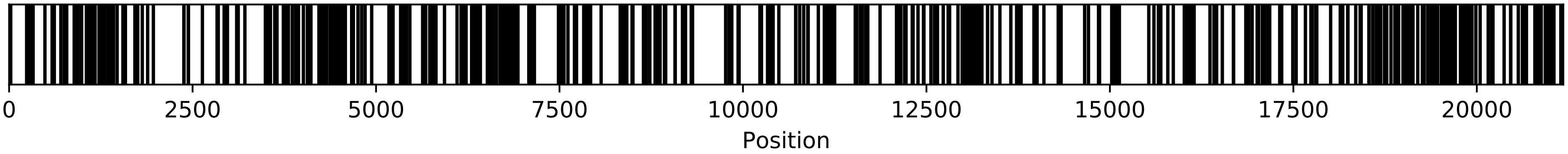


MA2 vs MM1
Number of differences: 222

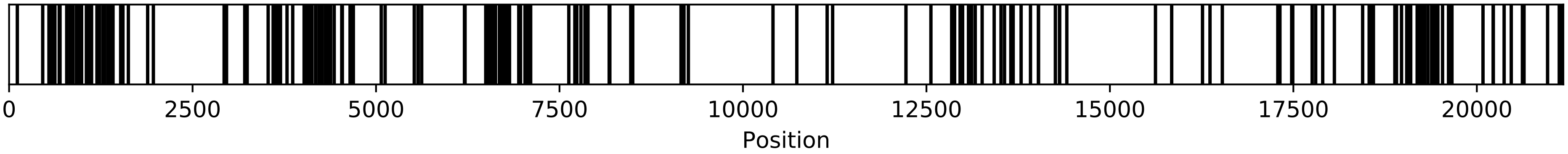


MA2 vs MM2
Number of differences: 200

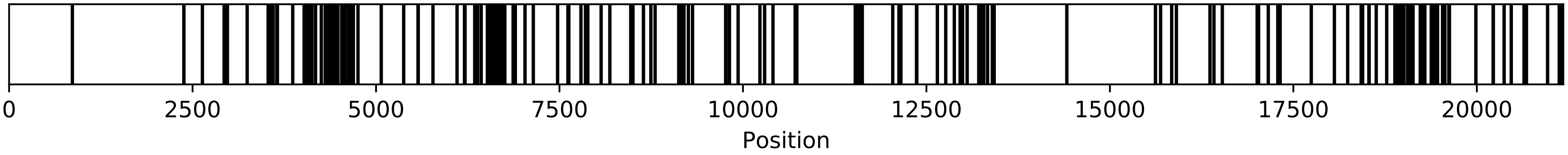


MM1 vs MM2
Number of differences: 254

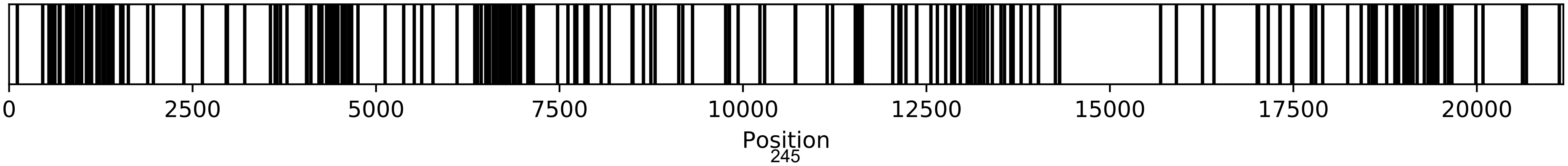

### Region: 814



MA1 vs MA2
Number of differences: 267

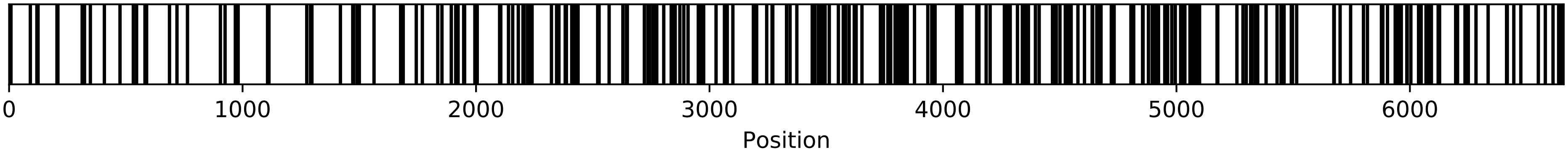


MA1 vs MM1
Number of differences: 247

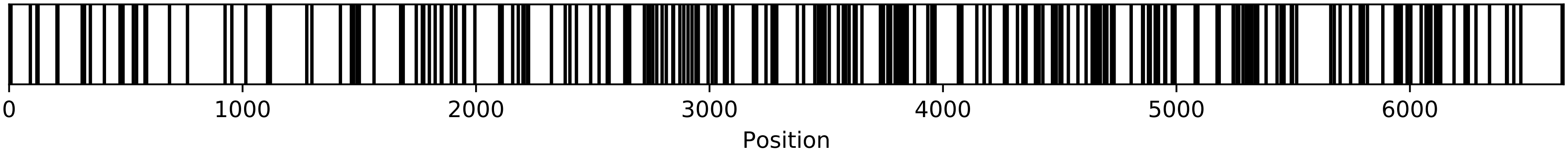


MA1 vs MM2
Number of differences: 237

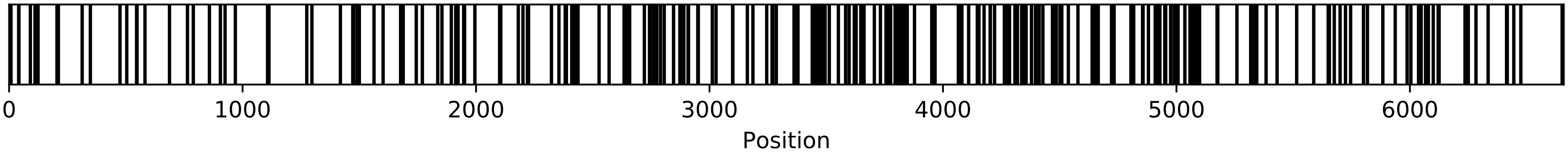


MA2 vs MM1
Number of differences: 162

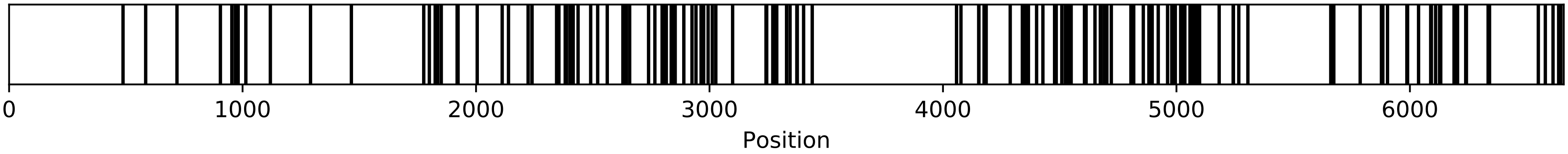


MA2 vs MM2
Number of differences: 154

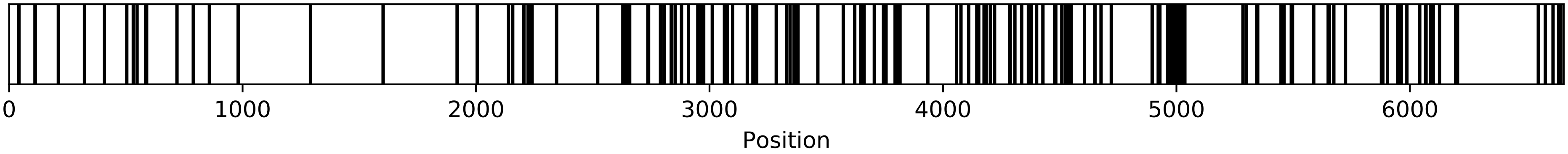


MM1 vs MM2
Number of differences: 170

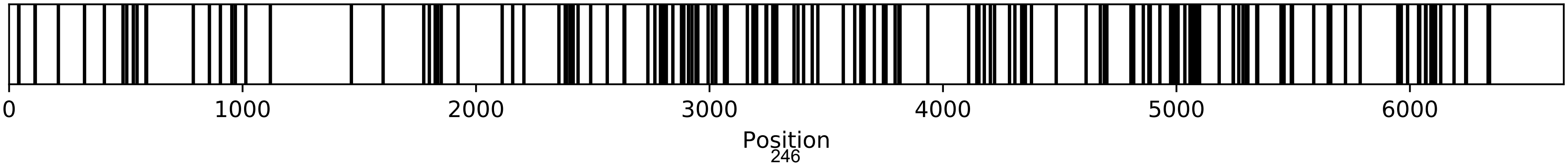

### Region: 817



MA1 vs MA2
Number of differences: 264

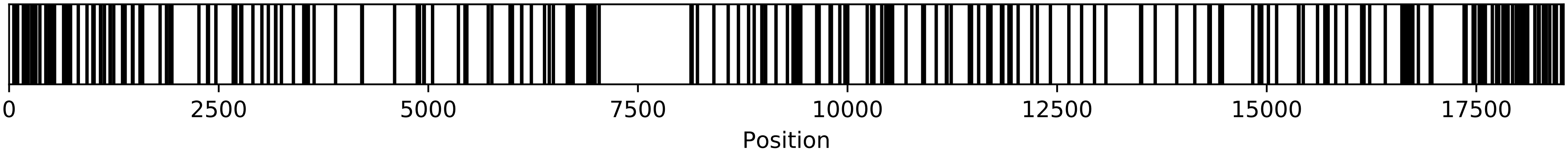


MA1 vs MM1
Number of differences: 262

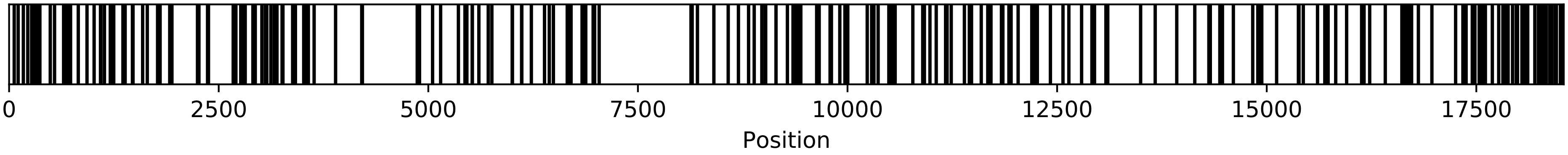


MA1 vs MM2
Number of differences: 264

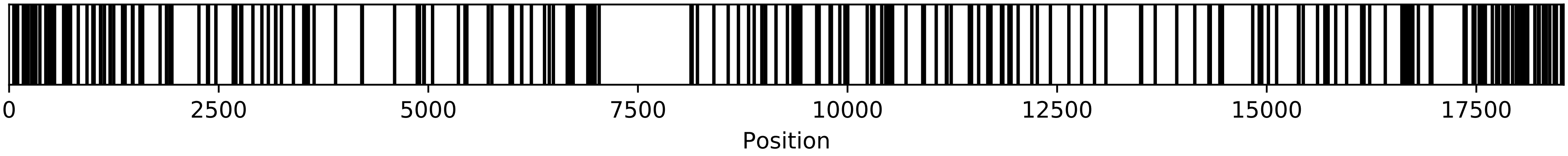


MA2 vs MM1
Number of differences: 110

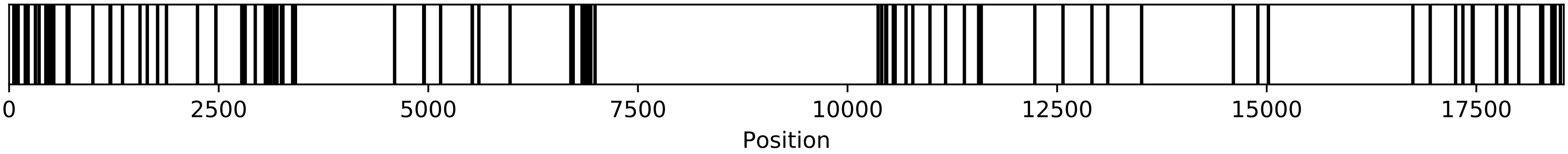


MA2 vs MM2
Number of differences: 0

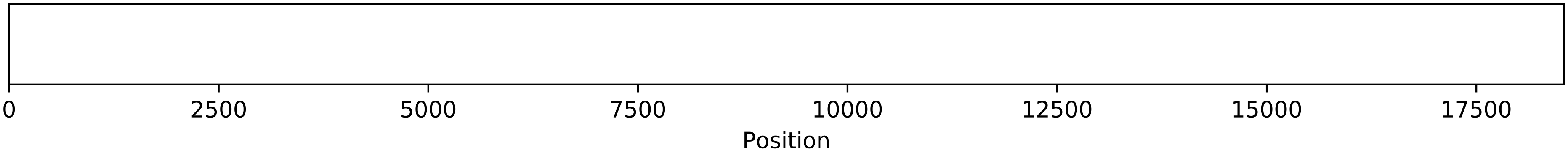


MM1 vs MM2
Number of differences: 110

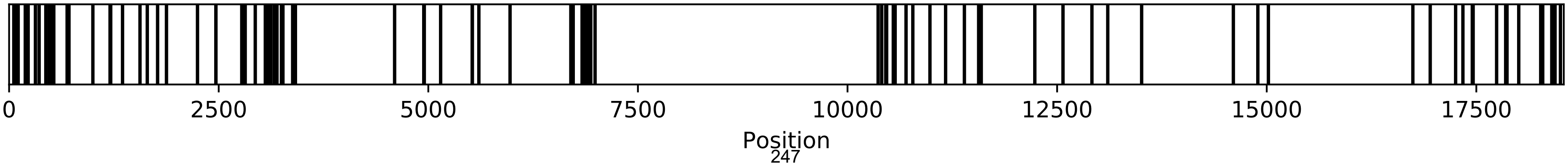

### Region: 819



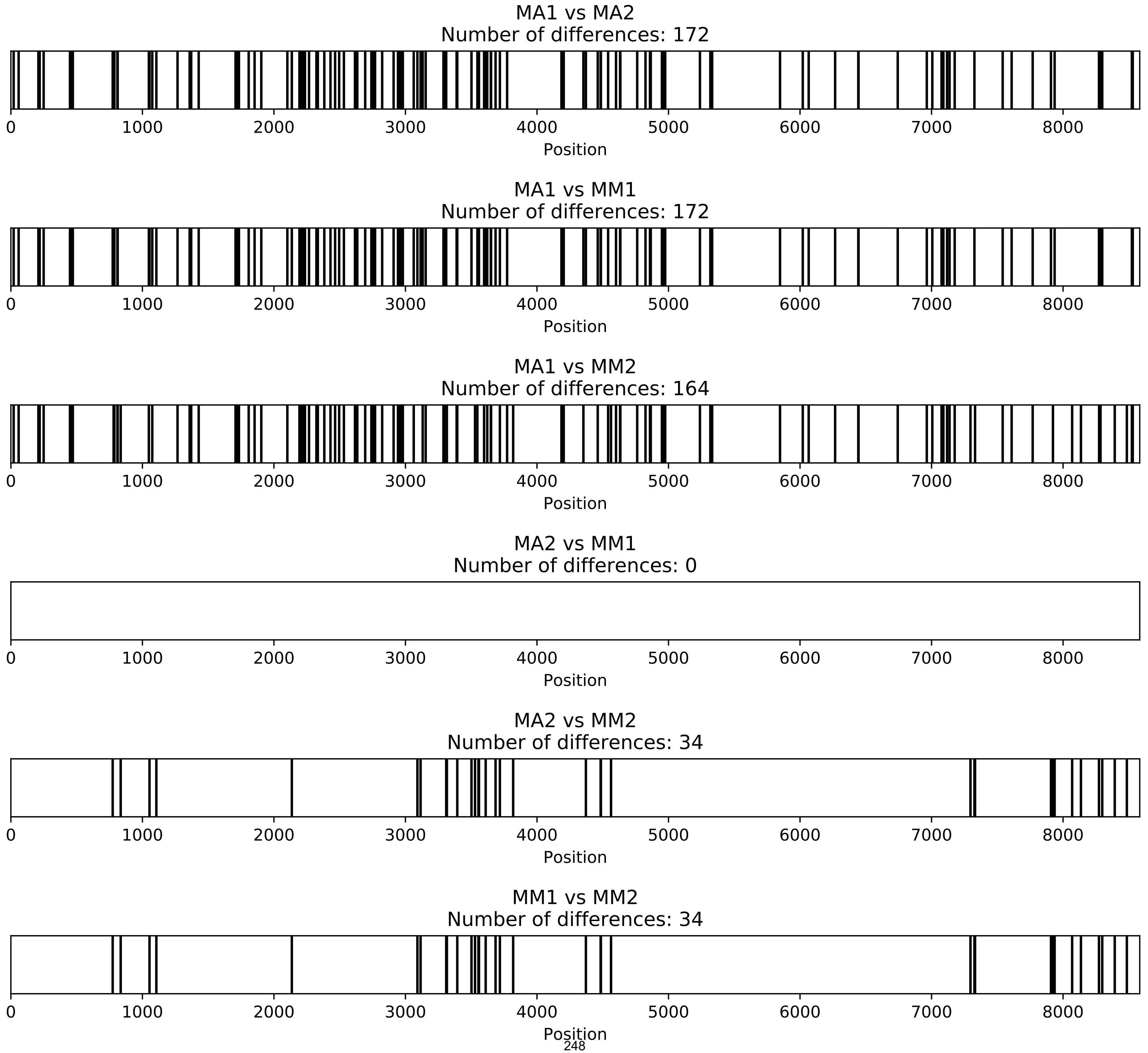
MA1 vs MA2
Number of differences: 172
Position
MA1 vs MM1
Number of differences: 172
Position
MA1 vs MM2
Number of differences: 164
Position
MA2 vs MM1
Number of differences: 0
Position
MA2 vs MM2
Number of differences: 34
Position
MM1 vs MM2
Number of differences: 34
0
1000
2000
3000
4000
5000
6000
7000
8000
Position

### Region: 820



## MA1 vs MA2
Number of differences: 267

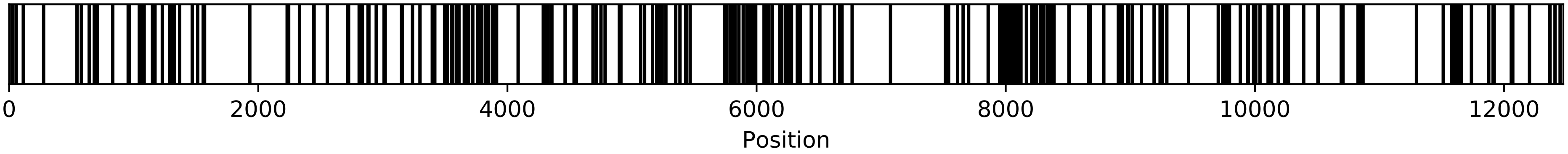


## MA1 vs MM1
Number of differences: 241

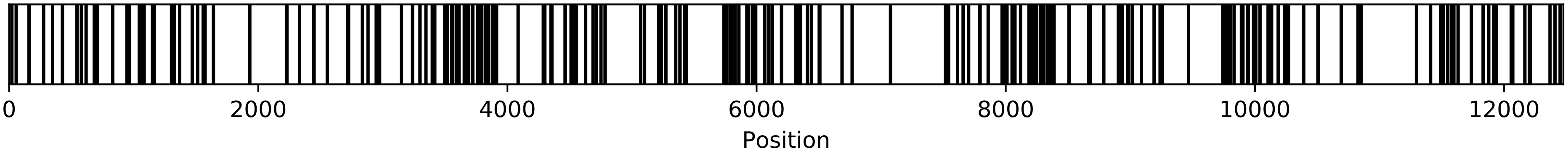


## MA1 vs MM2
Number of differences: 270

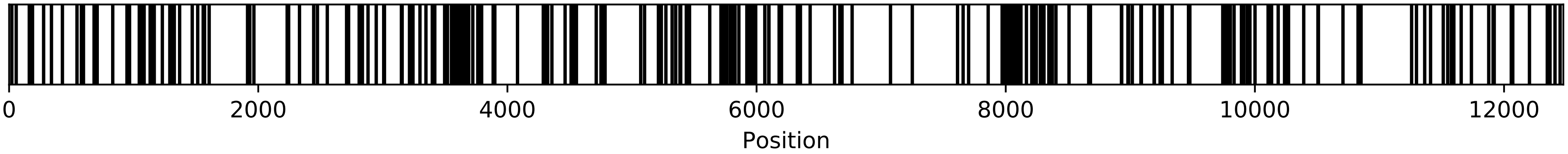


## MA2 vs MM1
Number of differences: 118

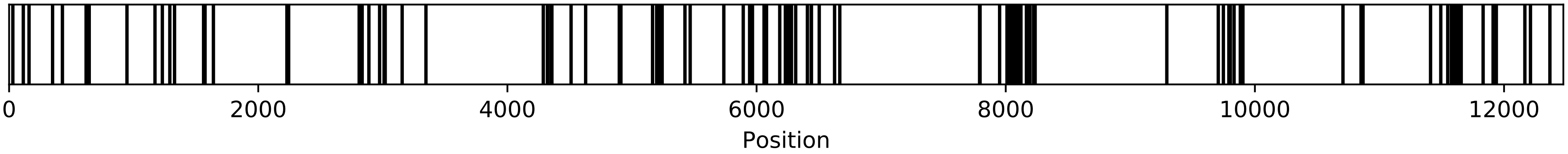


## MA2 vs MM2
Number of differences: 149

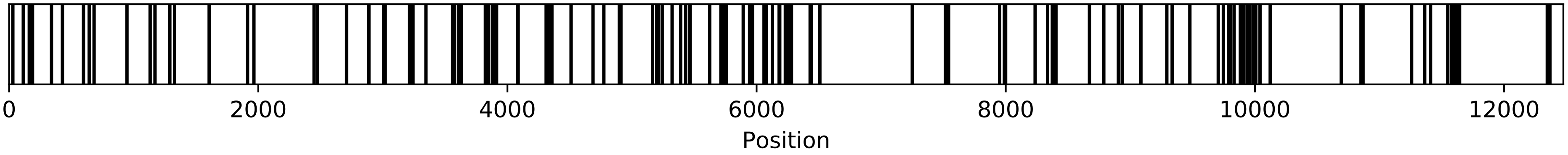


## MM1 vs MM2
Number of differences: 137

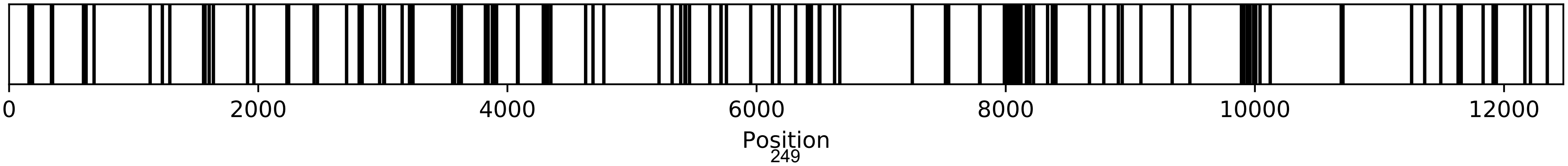

### Region: 826



MA1 vs MA2
Number of differences: 238

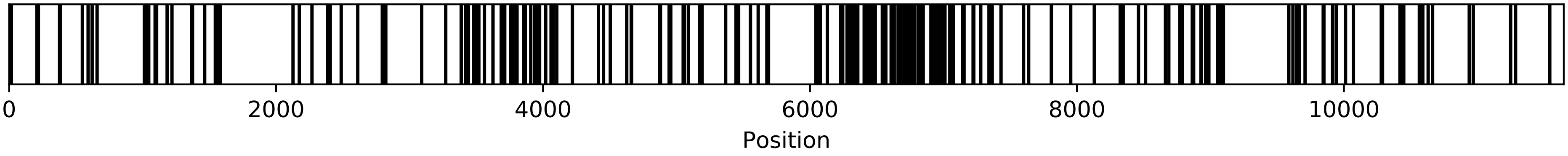


MA1 vs MM1
Number of differences: 238

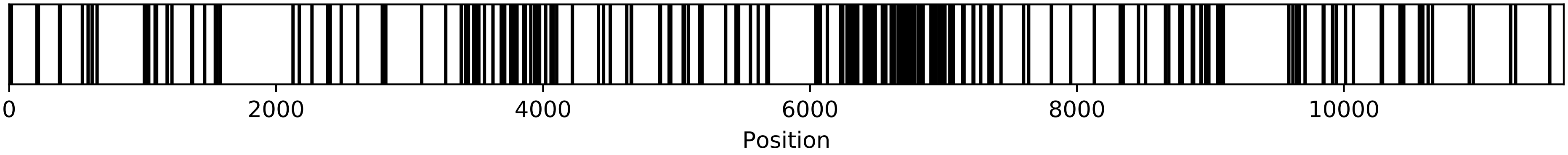


MA1 vs MM2
Number of differences: 239

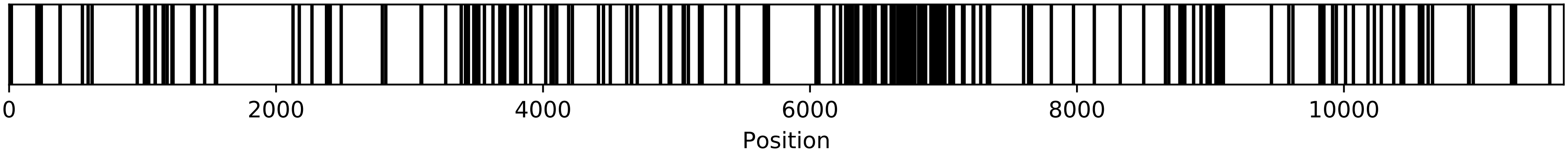


MA2 vs MM1
Number of differences: 0

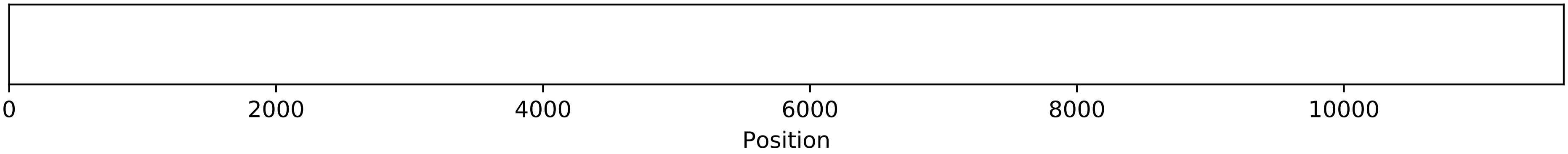


MA2 vs MM2
Number of differences: 94

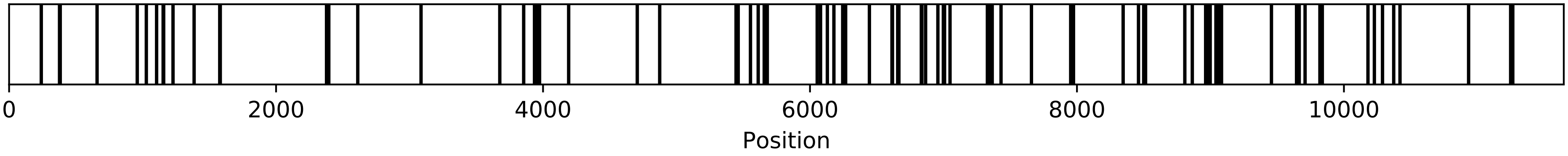


MM1 vs MM2
Number of differences: 94

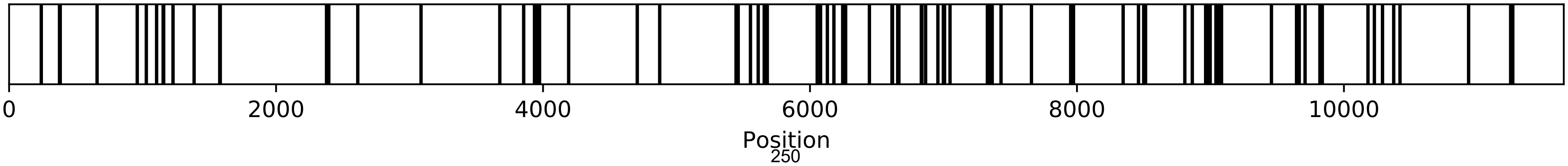

### Region: 827



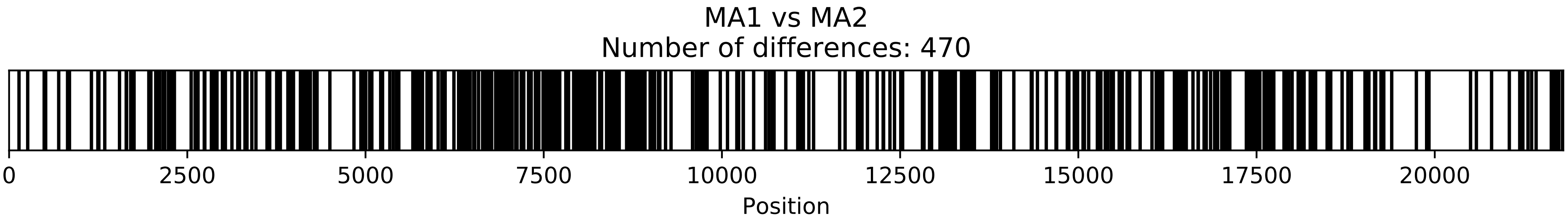
MA1 vs MA2
Number of differences: 470
0
2500
5000
7500
10000
12500
15000
17500
20000
Position

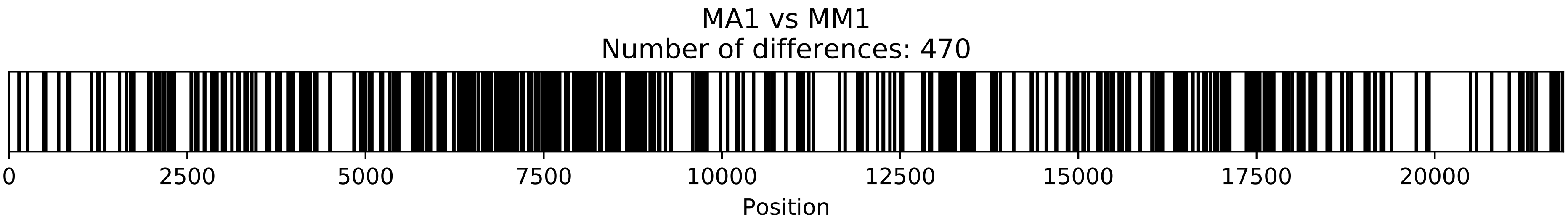
MA1 vs MM1
Number of differences: 470
0
2500
5000
7500
10000
12500
15000
17500
20000
Position

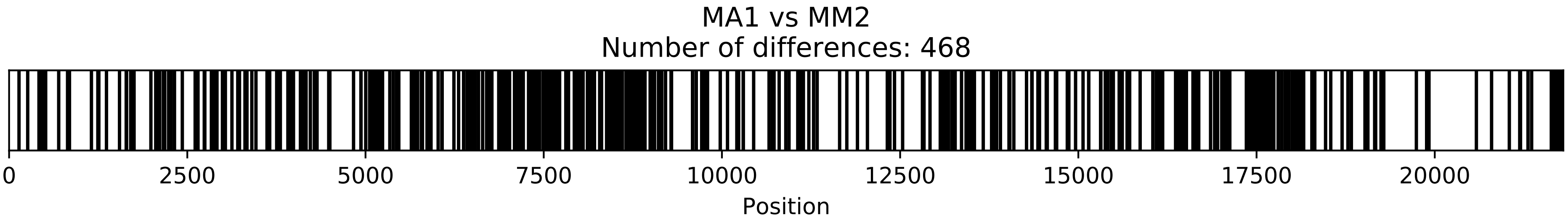
MA1 vs MM2
Number of differences: 468
0
2500
5000
7500
10000
12500
15000
17500
20000
Position

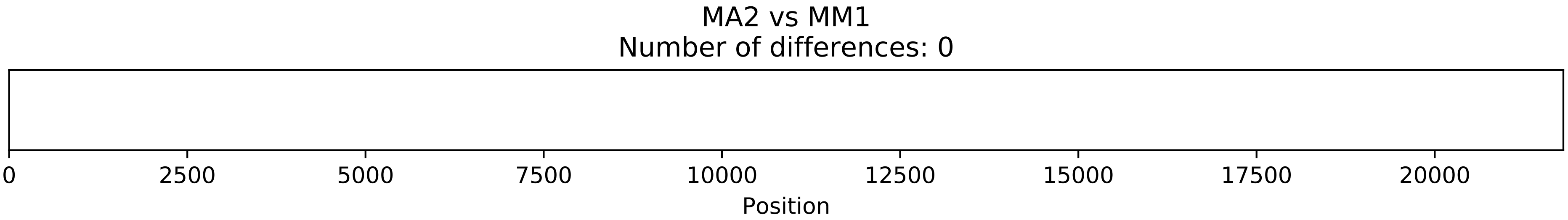
MA2 vs MM1
Number of differences: 0
0
2500
5000
7500
10000
12500
15000
17500
20000
Position

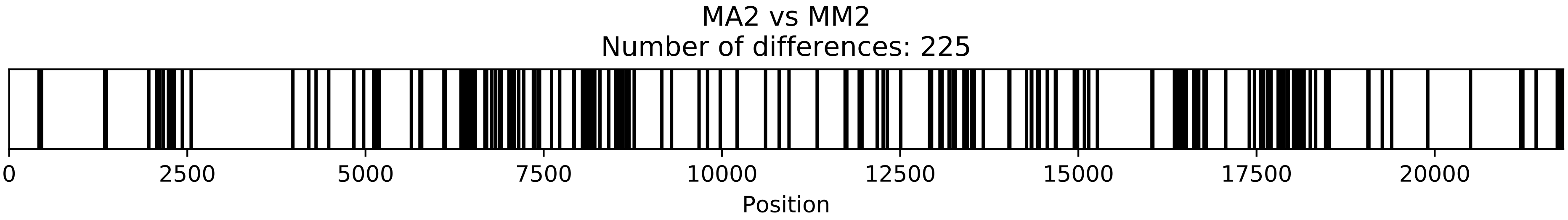
MA2 vs MM2
Number of differences: 225
0
2500
5000
7500
10000
12500
15000
17500
20000
Position

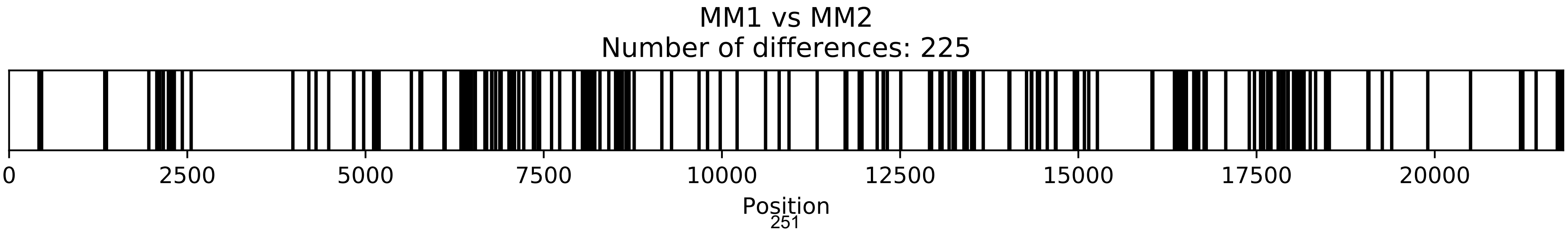
MM1 vs MM2
Number of differences: 225
0
2500
5000
7500
10000
12500
15000
17500
20000
Position

### Region: 832



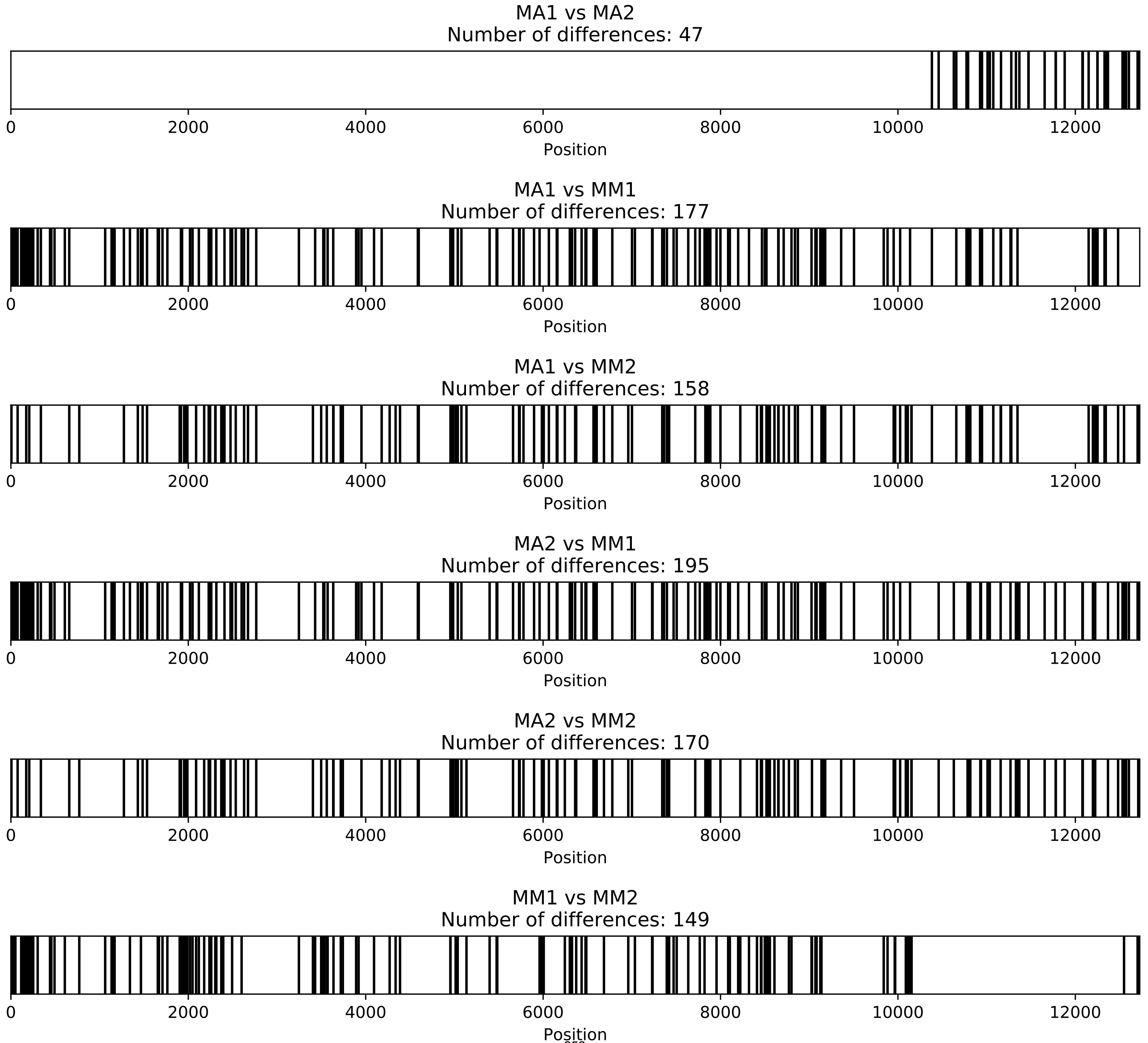
MA1 vs MA2
Number of differences: 47
MA1 vs MM1
Number of differences: 177
MA1 vs MM2
Number of differences: 158
MA2 vs MM1
Number of differences: 195
MA2 vs MM2
Number of differences: 170
MM1 vs MM2
Number of differences: 149
0
2000
4000
6000
8000
10000
12000
Position

### Region: 842



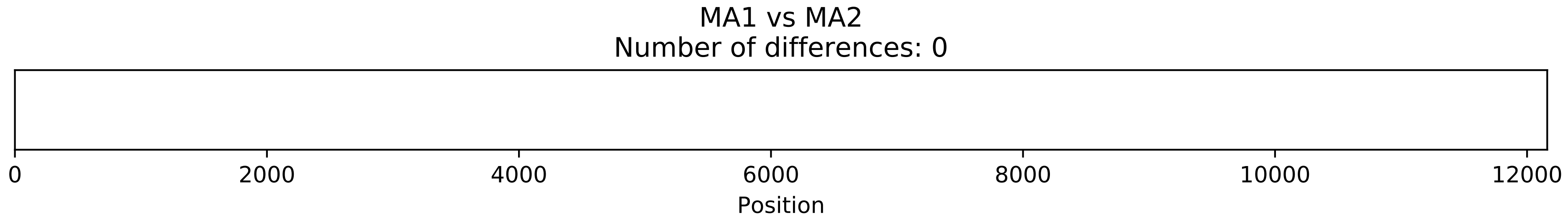
MA1 vs MA2
Number of differences: 0
0
2000
4000
6000
8000
10000
12000
Position

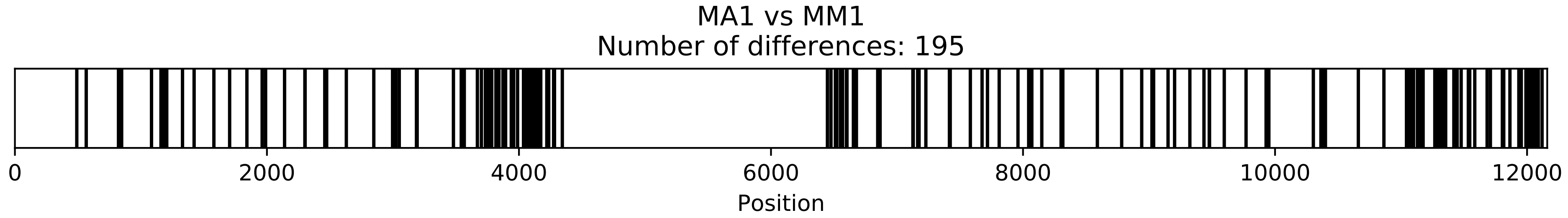
MA1 vs MM1
Number of differences: 195
0
2000
4000
6000
8000
10000
12000
Position

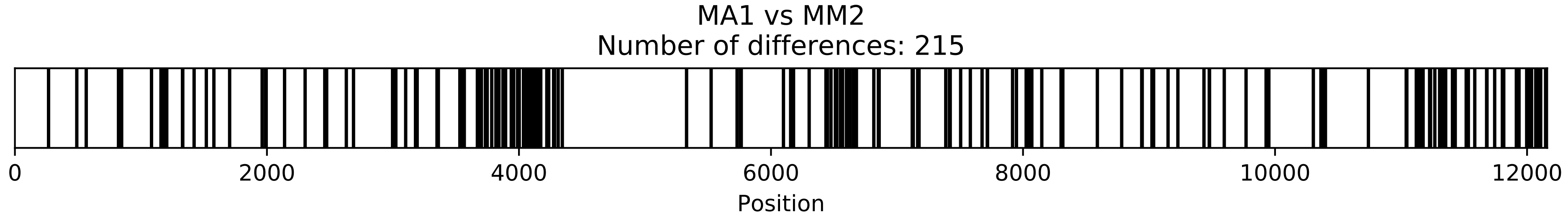
MA1 vs MM2
Number of differences: 215
0
2000
4000
6000
8000
10000
12000
Position

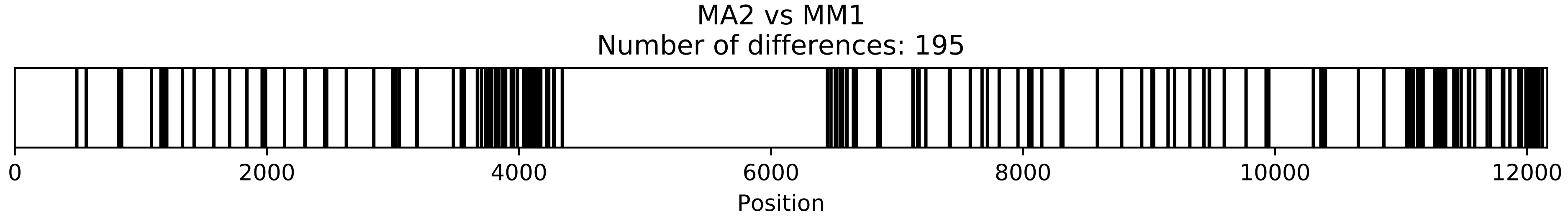
MA2 vs MM1
Number of differences: 195
0
2000
4000
6000
8000
10000
12000
Position

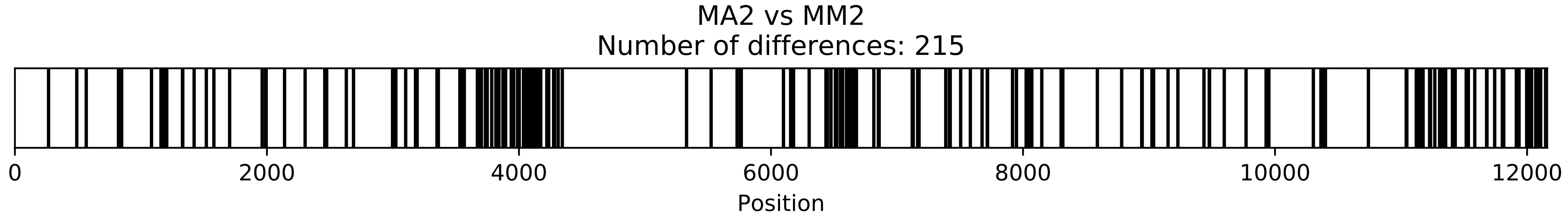
MA2 vs MM2
Number of differences: 215
0
2000
4000
6000
8000
10000
12000
Position

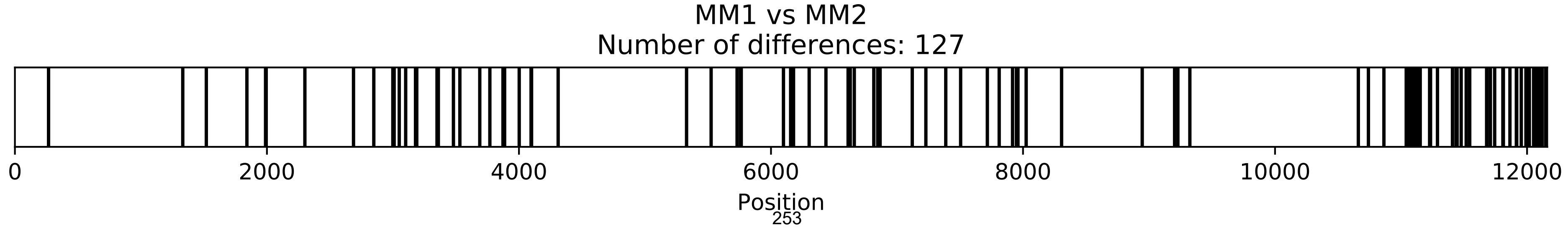
MM1 vs MM2
Number of differences: 127
0
2000
4000
6000
8000
10000
12000
Position

### Region: 845



MA1 vs MA2
Number of differences: 215

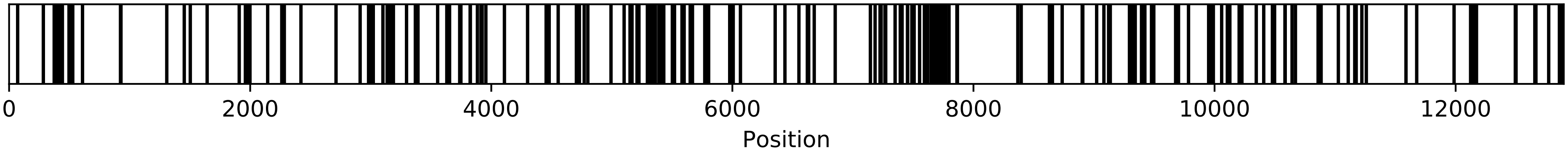


MA1 vs MM1
Number of differences: 214

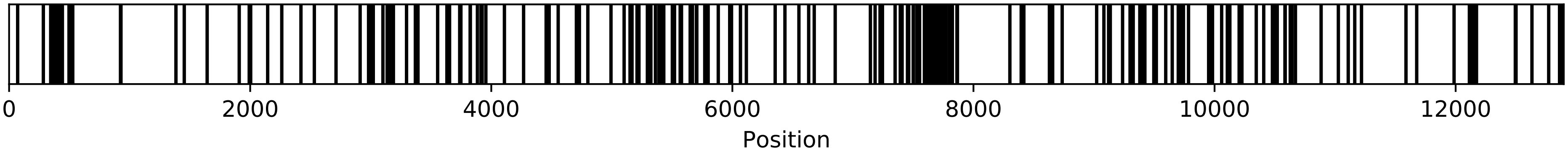


MA1 vs MM2
Number of differences: 212

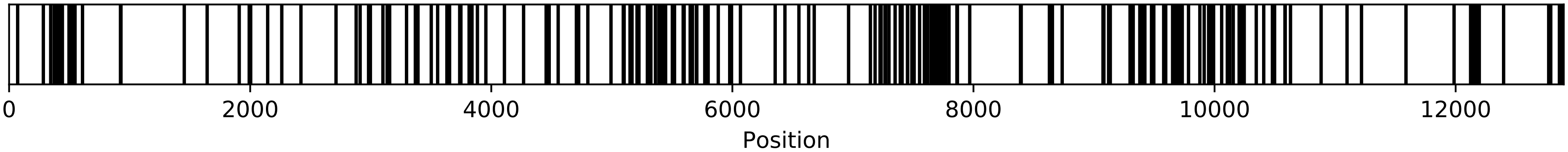


MA2 vs MM1
Number of differences: 88

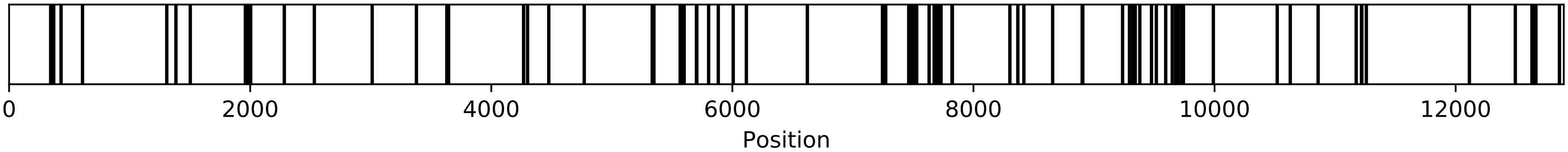


MA2 vs MM2
Number of differences: 93

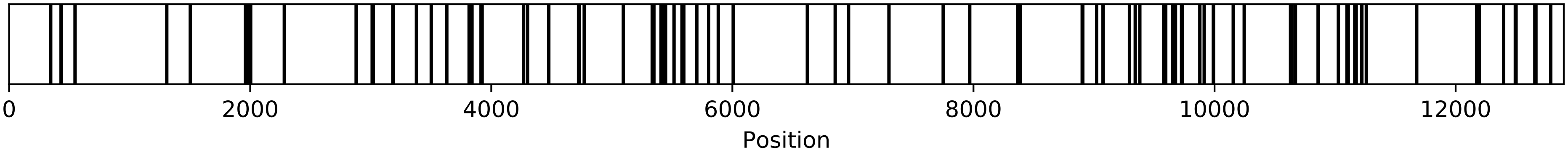


MM1 vs MM2
Number of differences: 94

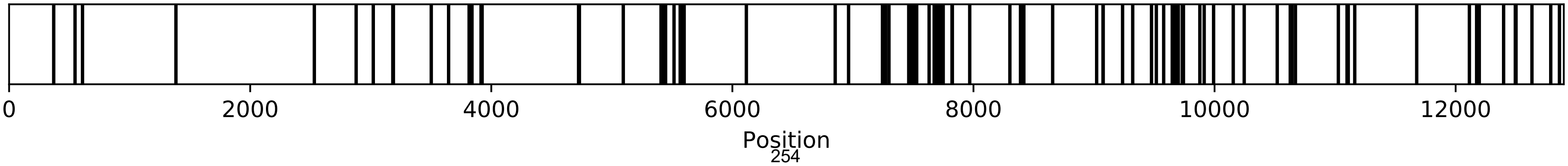

### Region: 852



MA1 vs MA2
Number of differences: 390

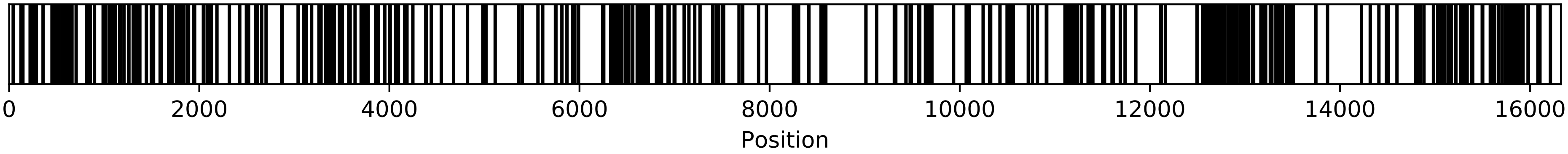


MA1 vs MM1
Number of differences: 201

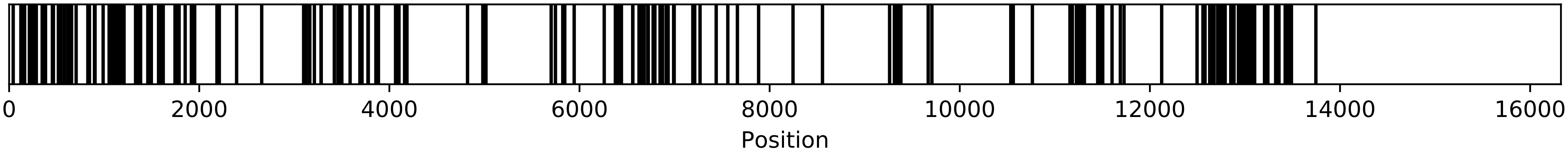


MA1 vs MM2
Number of differences: 235

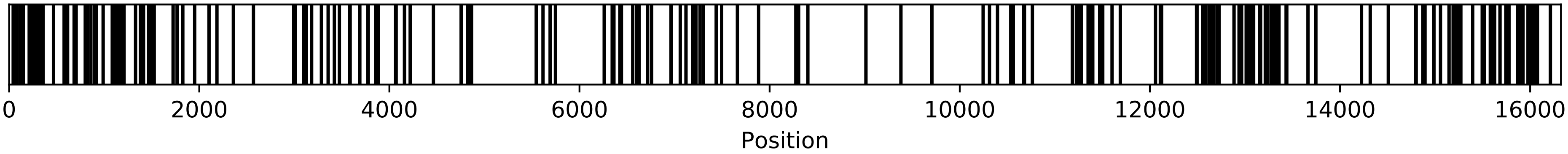


MA2 vs MM1
Number of differences: 392

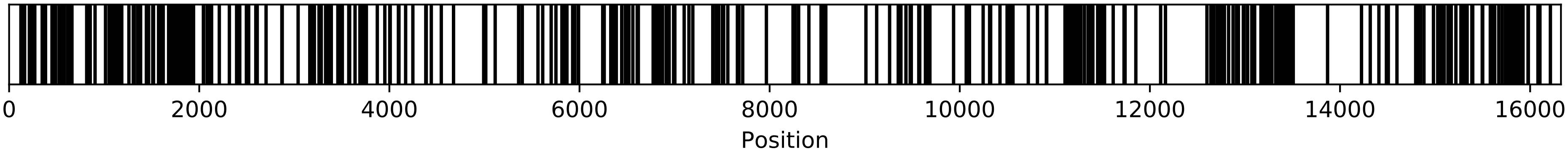


MA2 vs MM2
Number of differences: 408

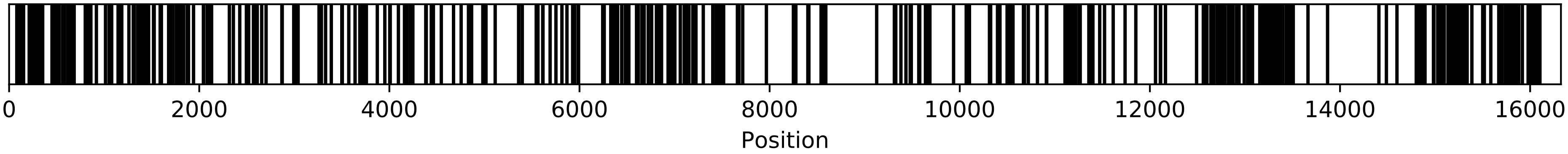


MM1 vs MM2
Number of differences: 250

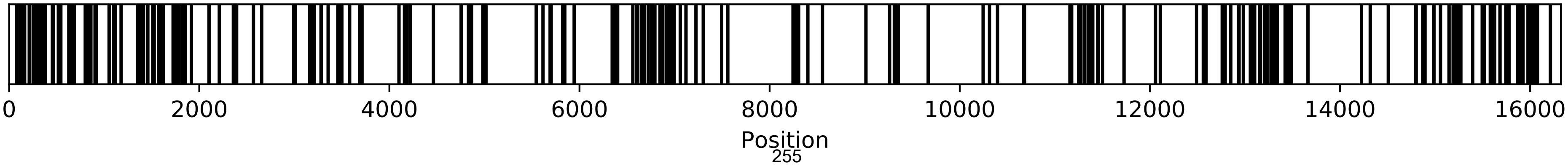

### Region: 855



MA1 vs MA2
Number of differences: 350

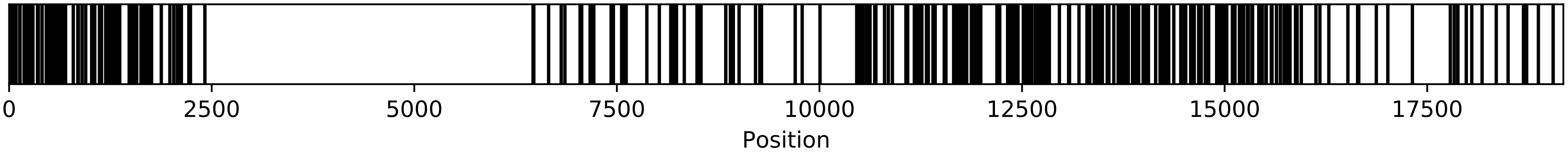


MA1 vs MM1
Number of differences: 372

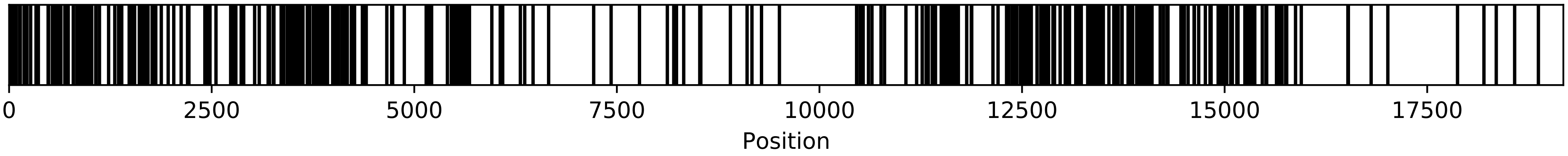


MA1 vs MM2
Number of differences: 363

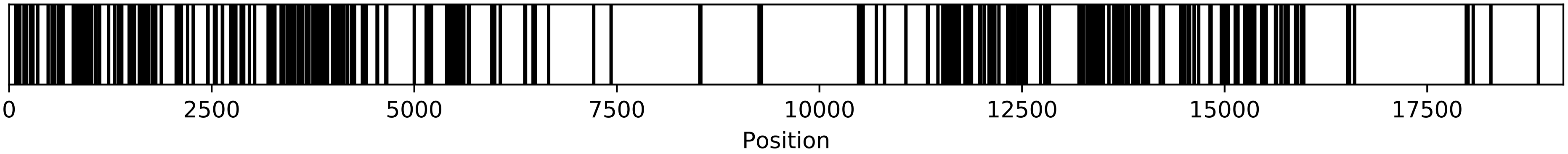


MA2 vs MM1
Number of differences: 445

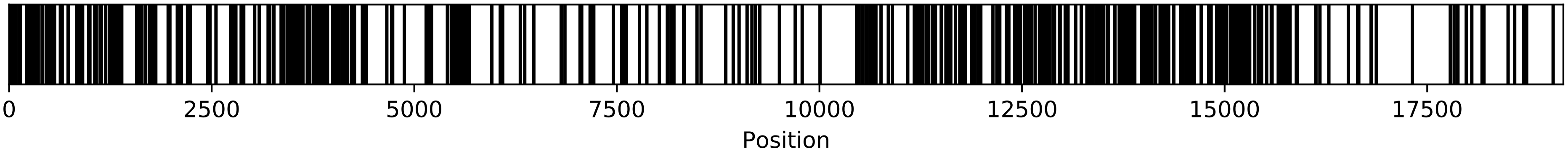


MA2 vs MM2
Number of differences: 470

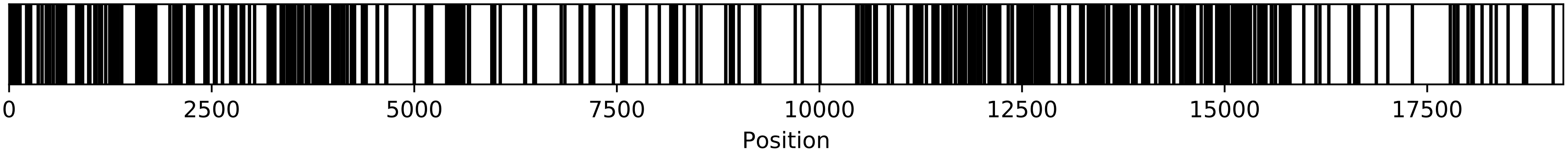


MM1 vs MM2
Number of differences: 278

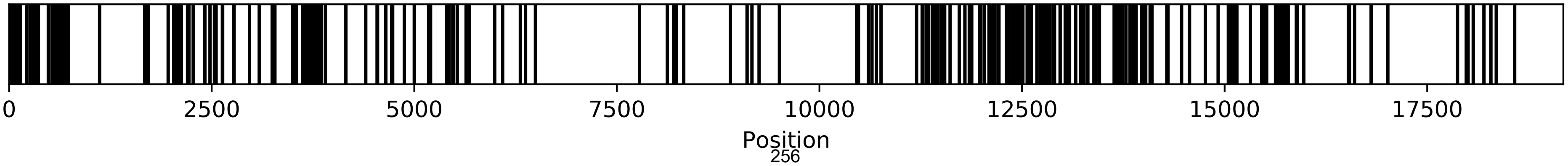

### Region: 864

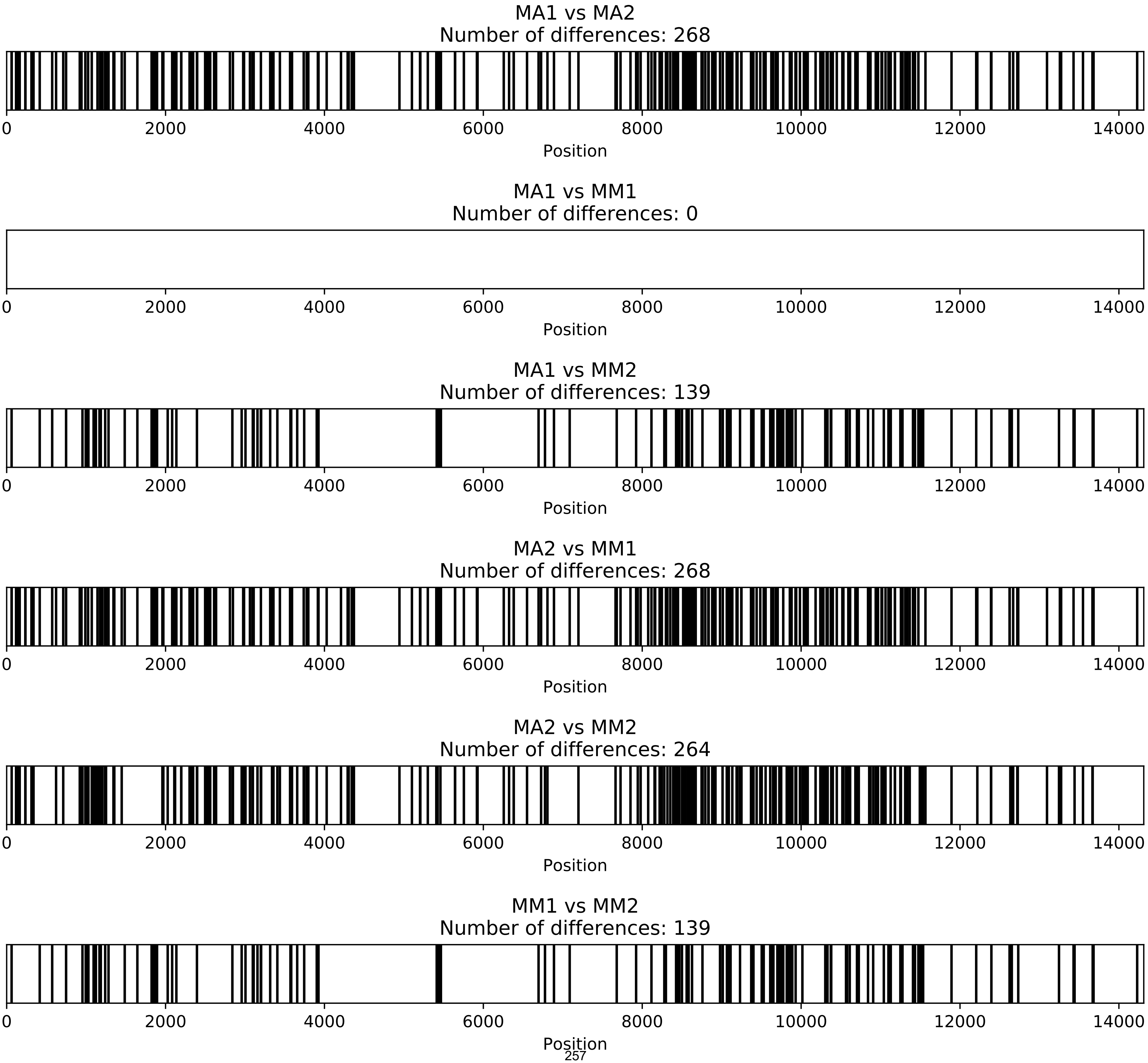
MA1 vs MA2
Number of differences: 268
MA1 vs MM1
Number of differences: 0
MA1 vs MM2
Number of differences: 139
MA2 vs MM1
Number of differences: 268
MA2 vs MM2
Number of differences: 264
MM1 vs MM2
Number of differences: 139
0
2000
4000
6000
8000
10000
12000
14000
Position

### Region: 869



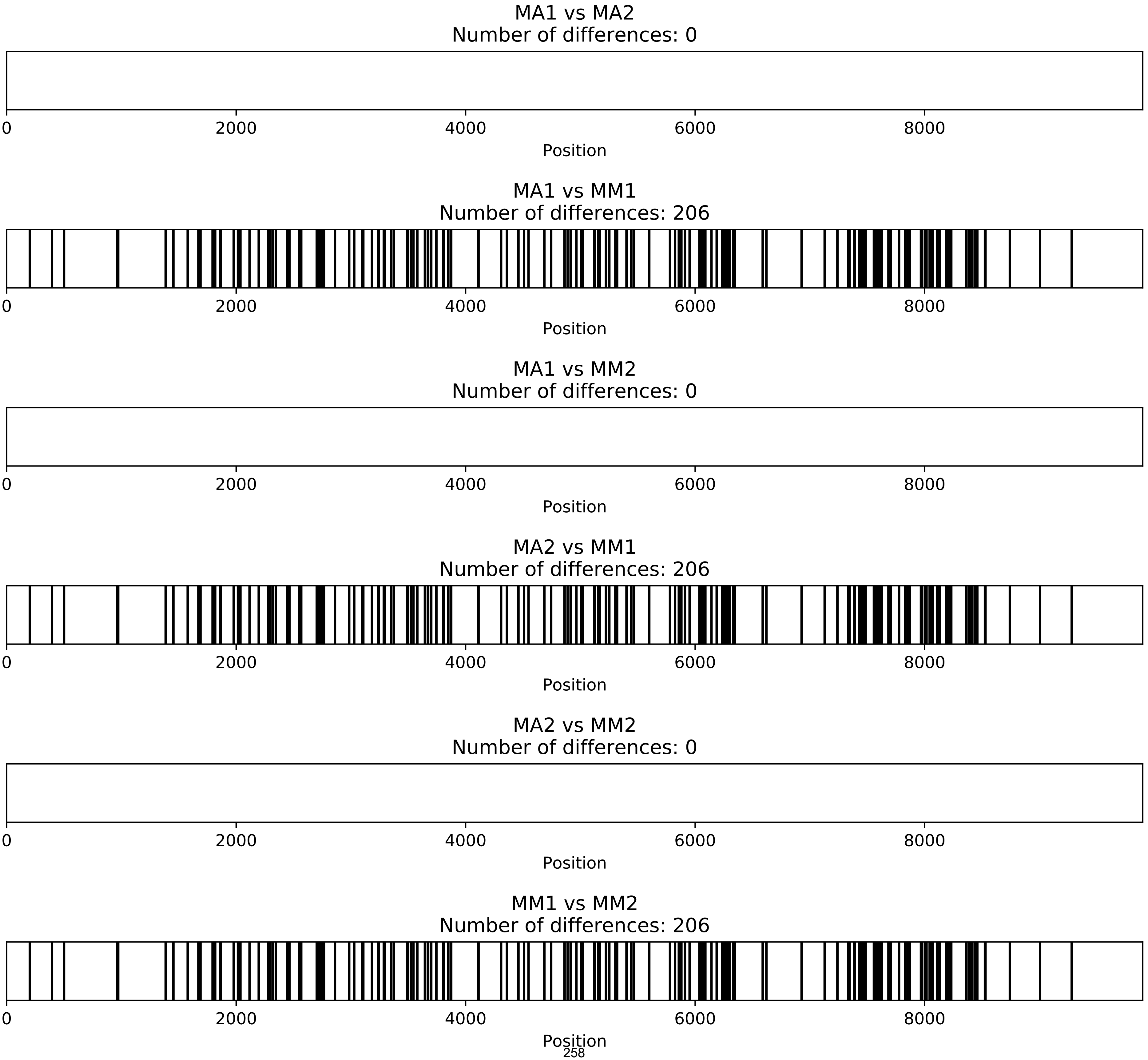
MA1 vs MA2
Number of differences: 0
MA1 vs MM1
Number of differences: 206
MA1 vs MM2
Number of differences: 0
MA2 vs MM1
Number of differences: 206
MA2 vs MM2
Number of differences: 0
MM1 vs MM2
Number of differences: 206
0
2000
4000
6000
8000
Position

### Region: 880



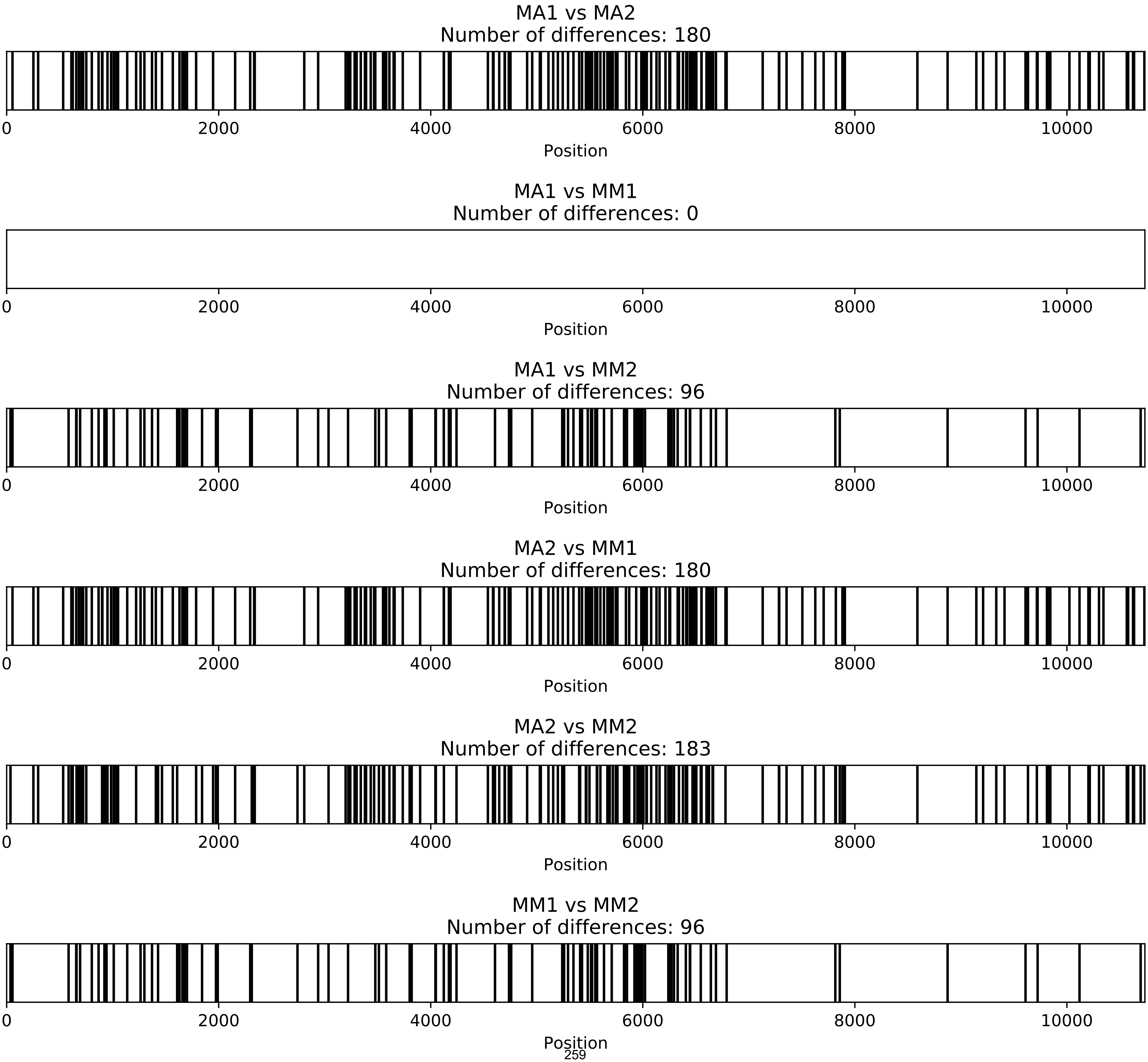
MA1 vs MA2
Number of differences: 180
0
2000
4000
6000
8000
10000
Position
MA1 vs MM1
Number of differences: 0
0
2000
4000
6000
8000
10000
Position
MA1 vs MM2
Number of differences: 96
0
2000
4000
6000
8000
10000
Position
MA2 vs MM1
Number of differences: 180
0
2000
4000
6000
8000
10000
Position
MA2 vs MM2
Number of differences: 183
0
2000
4000
6000
8000
10000
Position
MM1 vs MM2
Number of differences: 96
0
2000
4000
6000
8000
10000
Position

### Region: 894



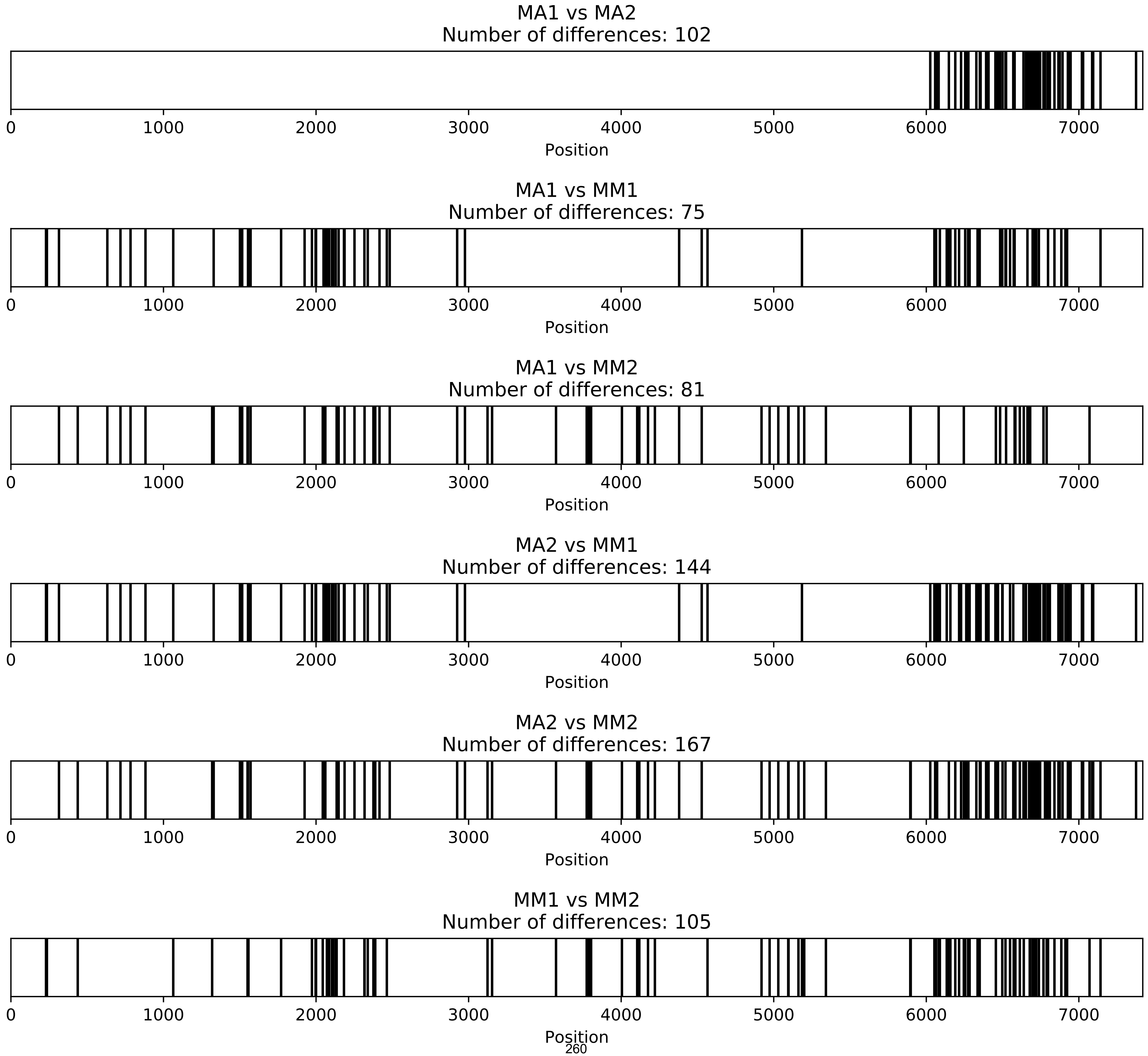
MA1 vs MA2
Number of differences: 102
Position
MA1 vs MM1
Number of differences: 75
Position
MA1 vs MM2
Number of differences: 81
Position
MA2 vs MM1
Number of differences: 144
Position
MA2 vs MM2
Number of differences: 167
Position
MM1 vs MM2
Number of differences: 105
Position
0
1000
2000
3000
4000
5000
6000
7000

### Region: 902



MA1 vs MA2
Number of differences: 614

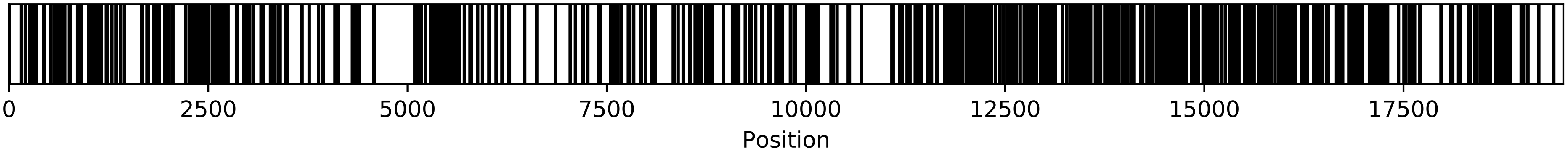


MA1 vs MM1
Number of differences: 561

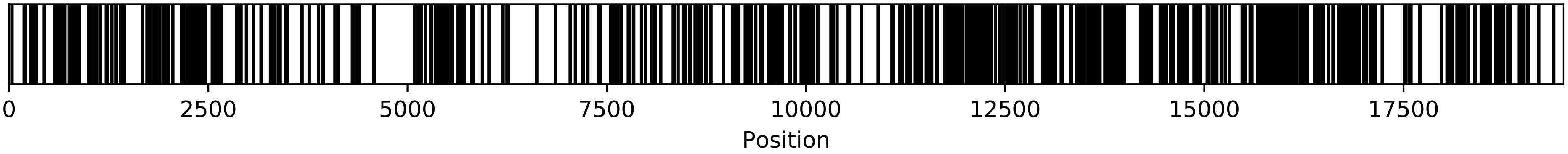


MA1 vs MM2
Number of differences: 614

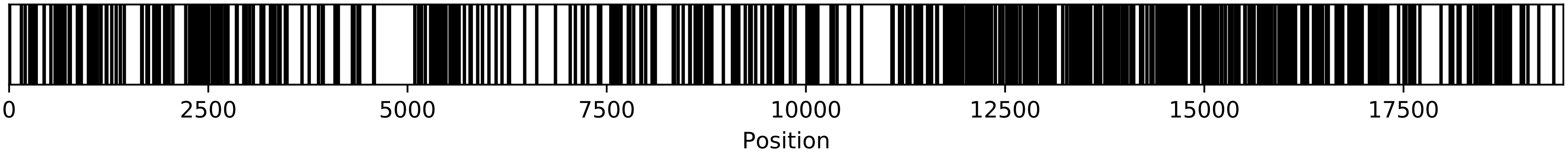


MA2 vs MM1
Number of differences: 377

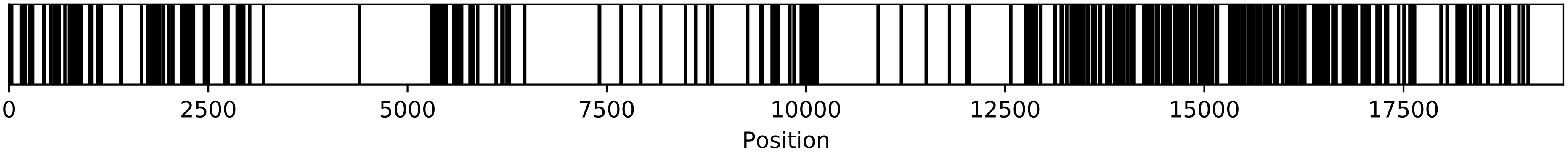


MA2 vs MM2
Number of differences: 0

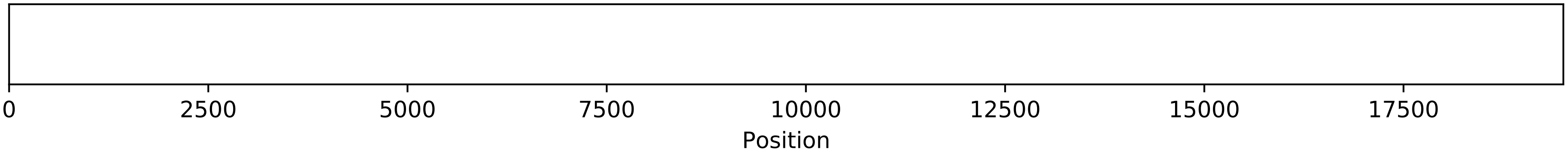


MM1 vs MM2
Number of differences: 377

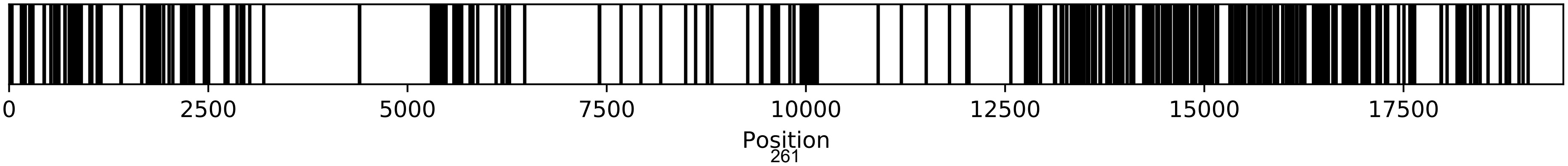

### Region: 904



MA1 vs MA2
Number of differences: 170

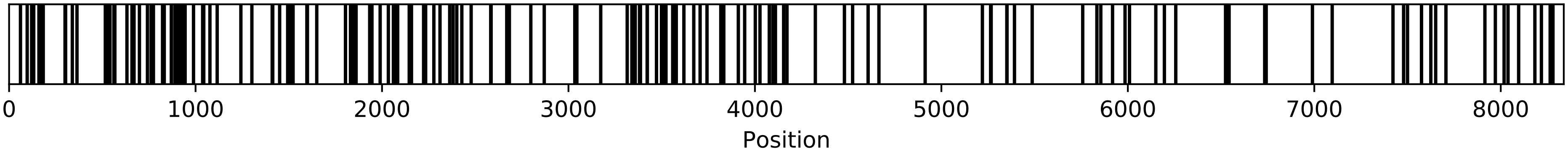


MA1 vs MM1
Number of differences: 0

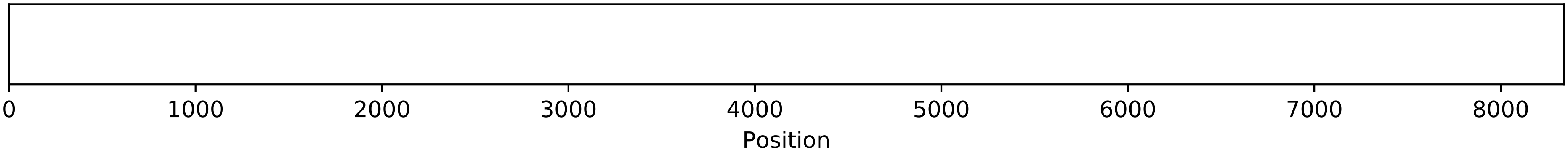


MA1 vs MM2
Number of differences: 81

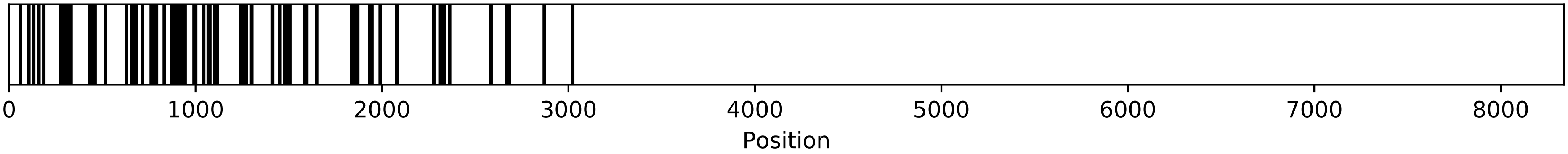


MA2 vs MM1
Number of differences: 170

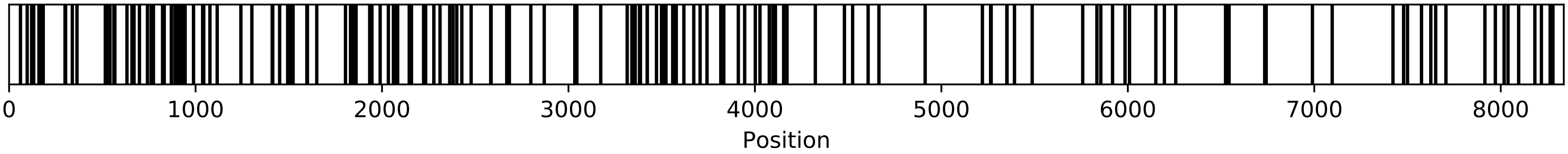


MA2 vs MM2
Number of differences: 167

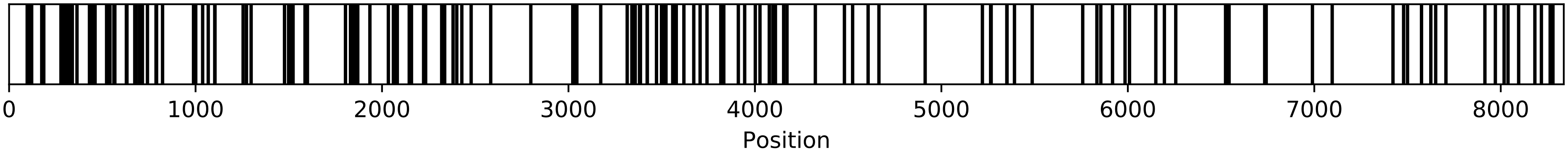


MM1 vs MM2
Number of differences: 81

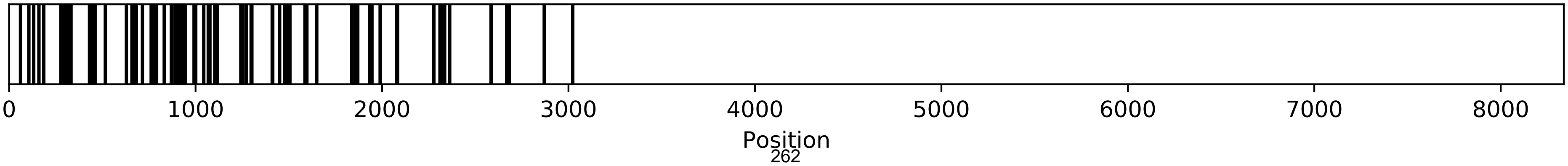

### Region: 905



MA1 vs MA2
Number of differences: 324

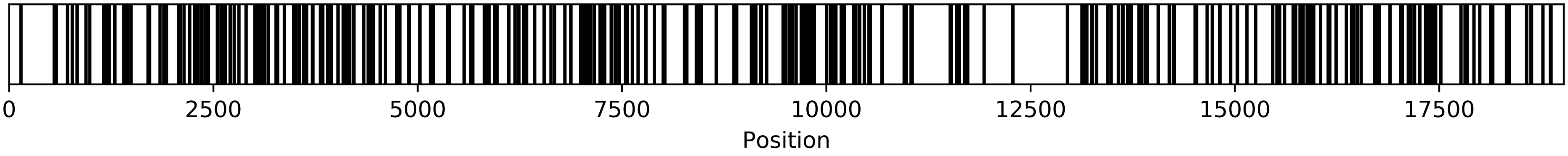


MA1 vs MM1
Number of differences: 0

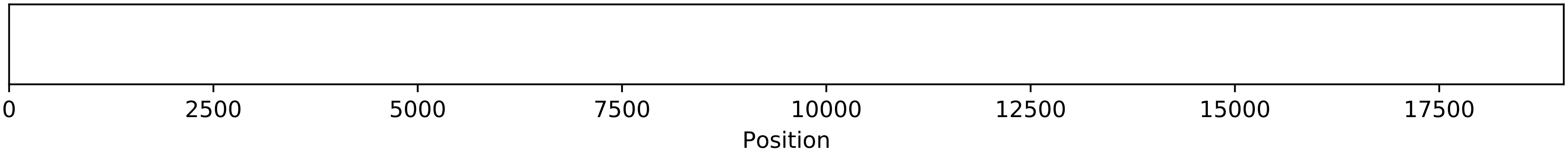


MA1 vs MM2
Number of differences: 166

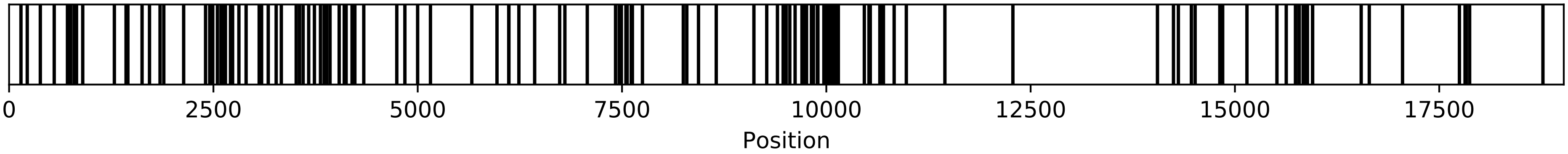


MA2 vs MM1
Number of differences: 324

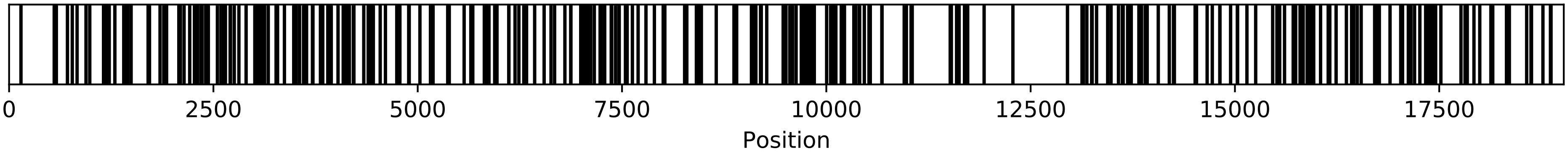


MA2 vs MM2
Number of differences: 324

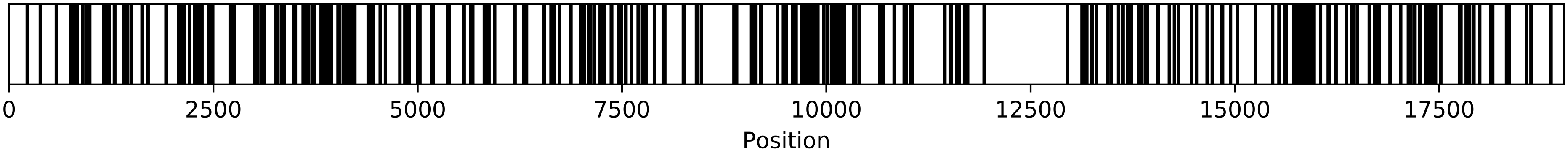


MM1 vs MM2
Number of differences: 166

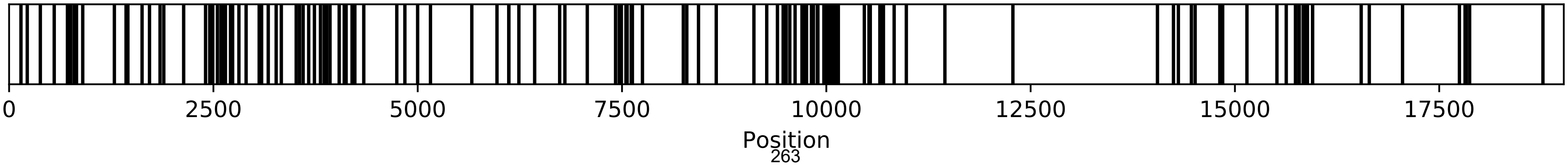

### Region: 906



MA1 vs MA2
Number of differences: 305

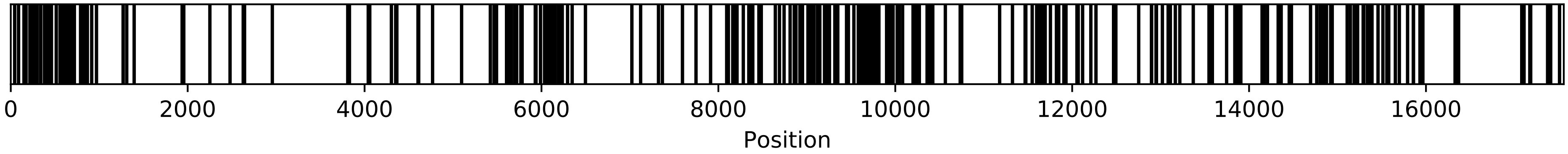


MA1 vs MM1
Number of differences: 67

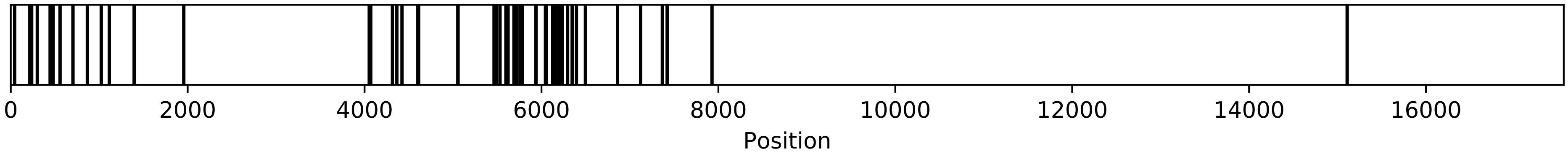


MA1 vs MM2
Number of differences: 157

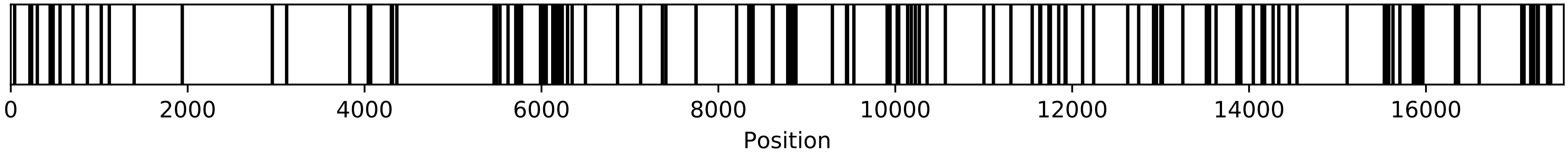


MA2 vs MM1
Number of differences: 290

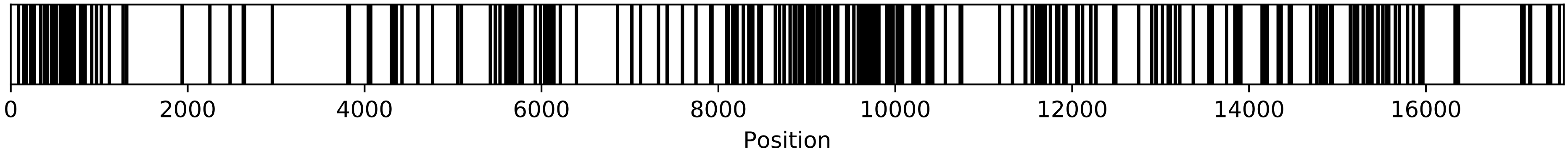


MA2 vs MM2
Number of differences: 276

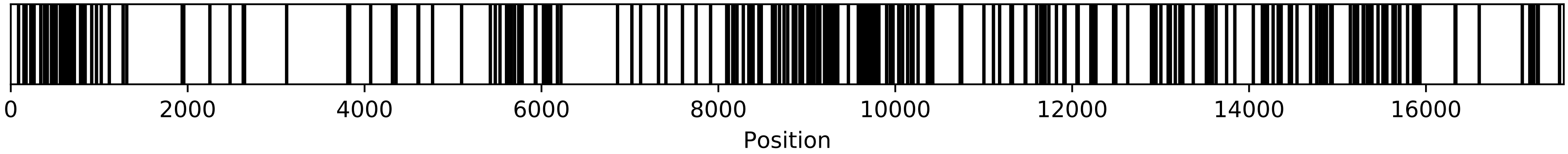


MM1 vs MM2
Number of differences: 136

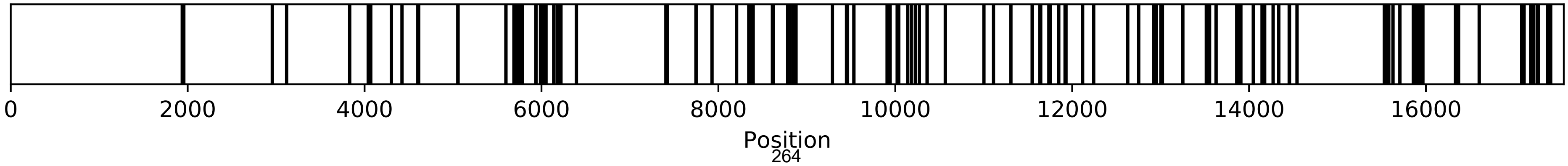

### Region: 907



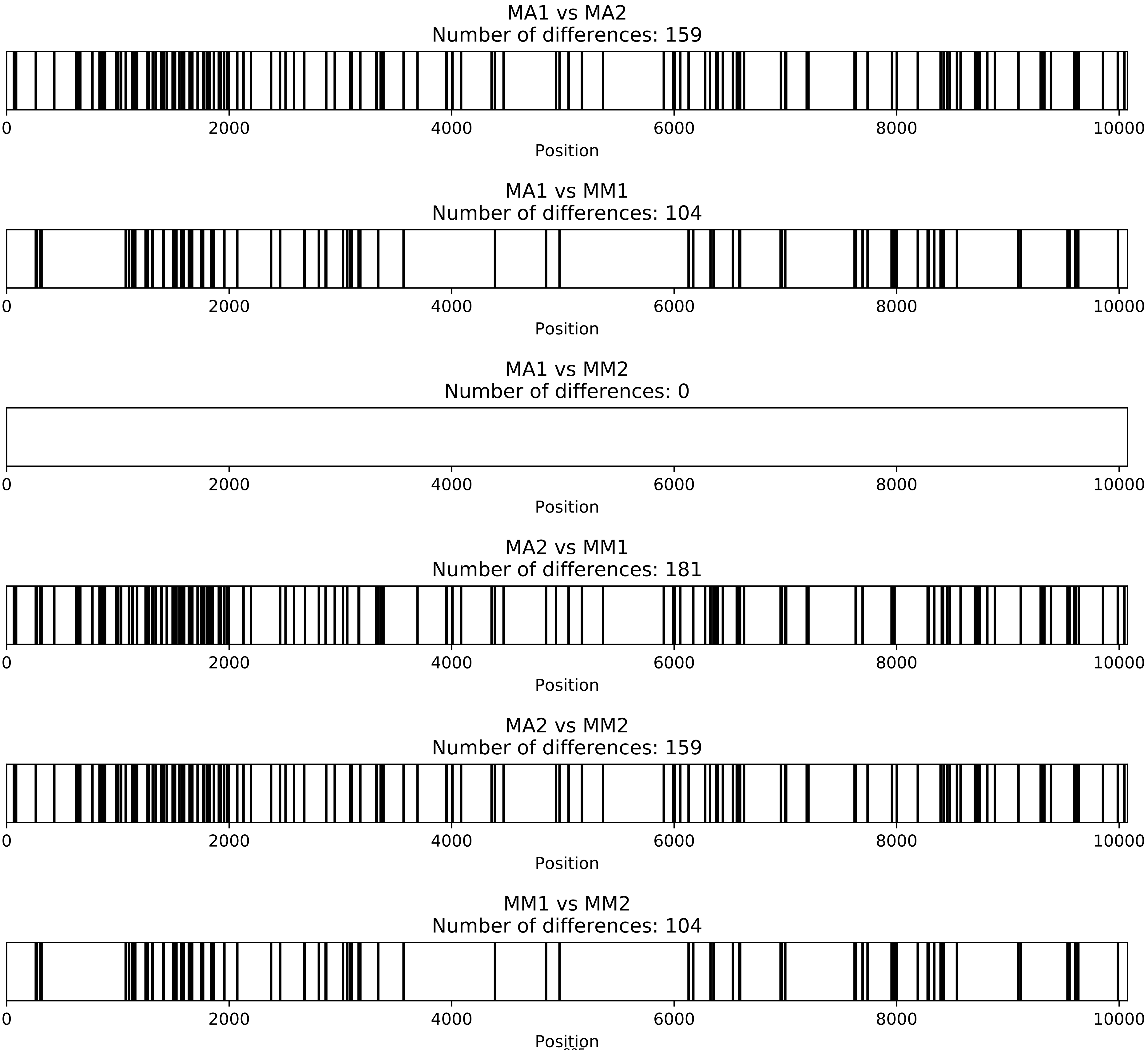
MA1 vs MA2
Number of differences: 159
MA1 vs MM1
Number of differences: 104
MA1 vs MM2
Number of differences: 0
MA2 vs MM1
Number of differences: 181
MA2 vs MM2
Number of differences: 159
MM1 vs MM2
Number of differences: 104
0
2000
4000
6000
8000
10000
Position

### Region: 908



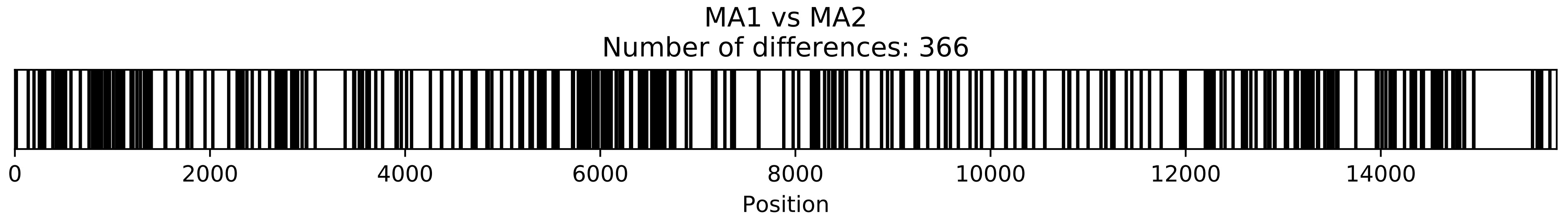
MA1 vs MA2
Number of differences: 366
0 2000 4000 6000 8000 10000 12000 14000
Position

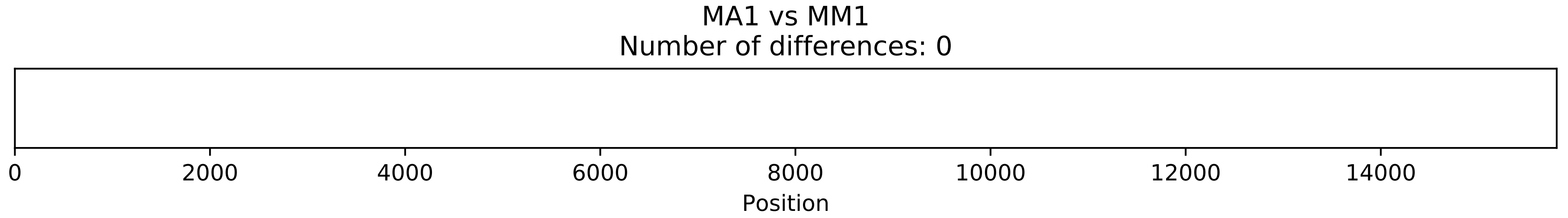
MA1 vs MM1
Number of differences: 0
0 2000 4000 6000 8000 10000 12000 14000
Position

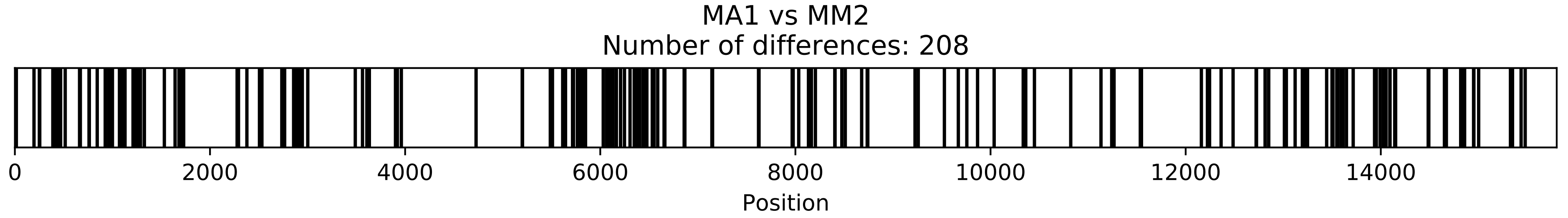
MA1 vs MM2
Number of differences: 208
0 2000 4000 6000 8000 10000 12000 14000
Position

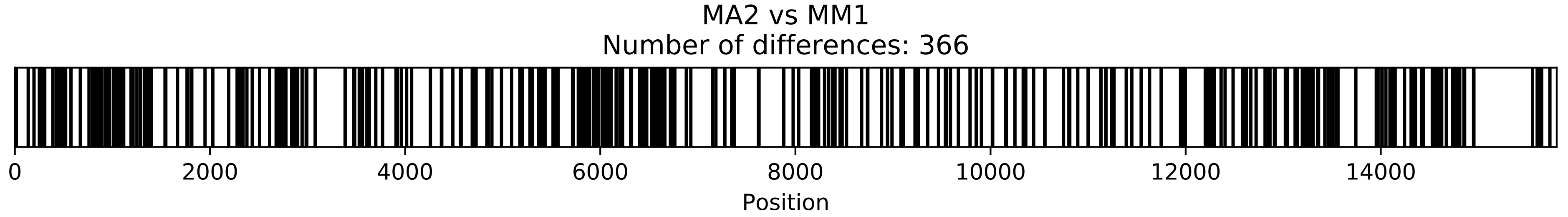
MA2 vs MM1
Number of differences: 366
0 2000 4000 6000 8000 10000 12000 14000
Position

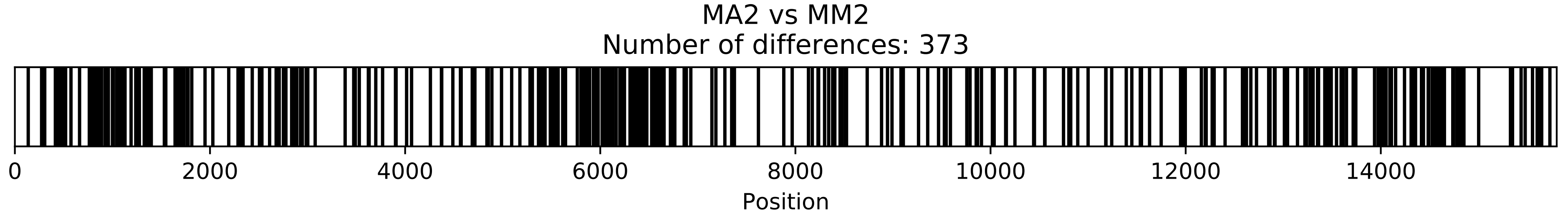
MA2 vs MM2
Number of differences: 373
0 2000 4000 6000 8000 10000 12000 14000
Position

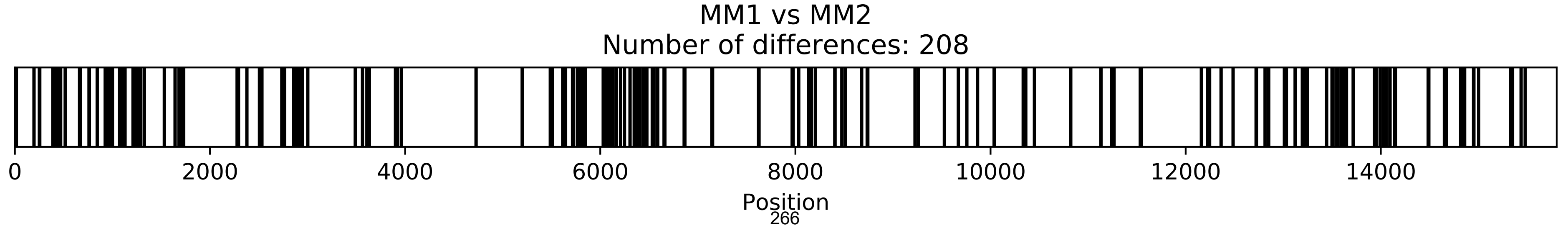
MM1 vs MM2
Number of differences: 208
0 2000 4000 6000 8000 10000 12000 14000
Position

### Region: 917



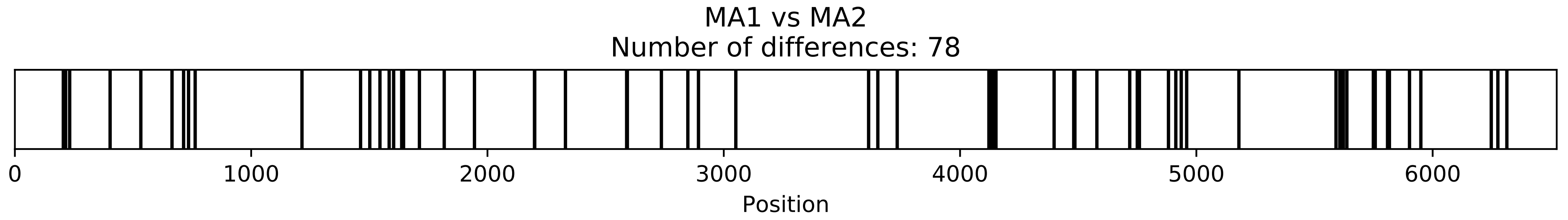
MA1 vs MA2
Number of differences: 78
0
1000
2000
3000
4000
5000
6000
Position

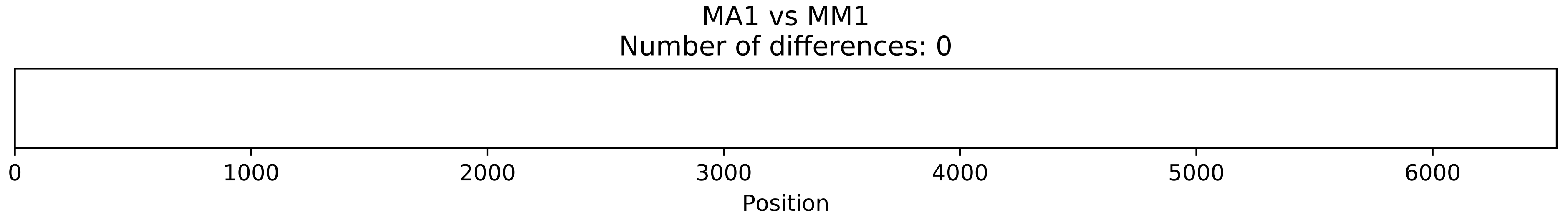
MA1 vs MM1
Number of differences: 0
0
1000
2000
3000
4000
5000
6000
Position

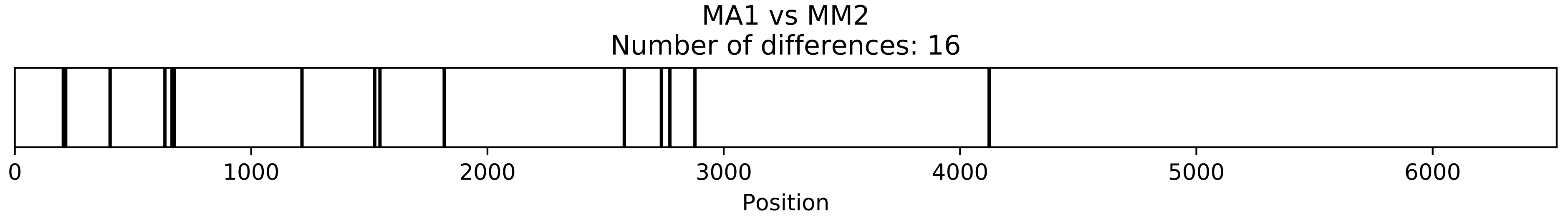
MA1 vs MM2
Number of differences: 16
0
1000
2000
3000
4000
5000
6000
Position

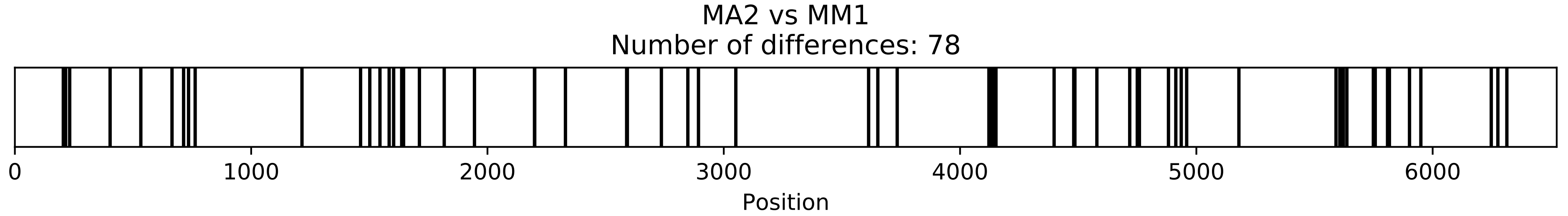
MA2 vs MM1
Number of differences: 78
0
1000
2000
3000
4000
5000
6000
Position

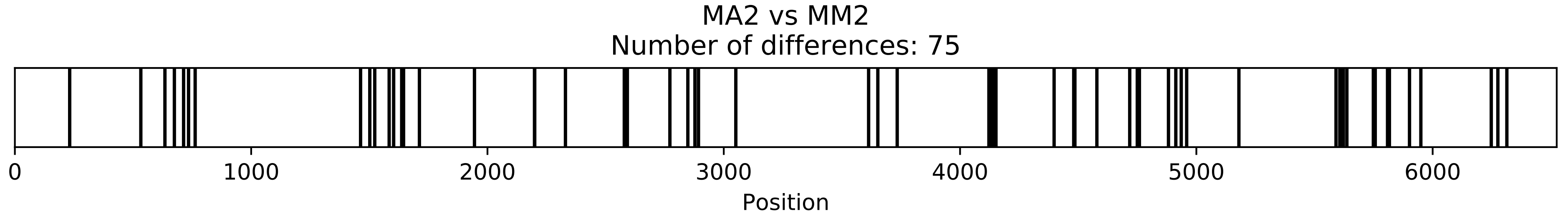
MA2 vs MM2
Number of differences: 75
0
1000
2000
3000
4000
5000
6000
Position

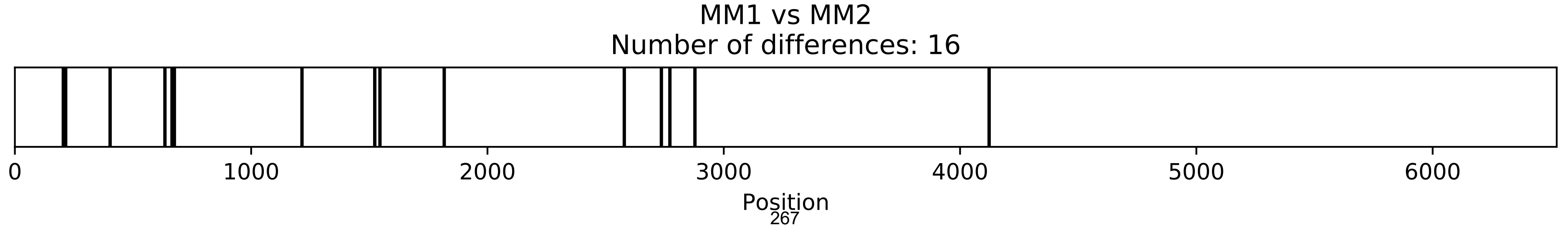
MM1 vs MM2
Number of differences: 16
0
1000
2000
3000
4000
5000
6000
Position

### Region: 919



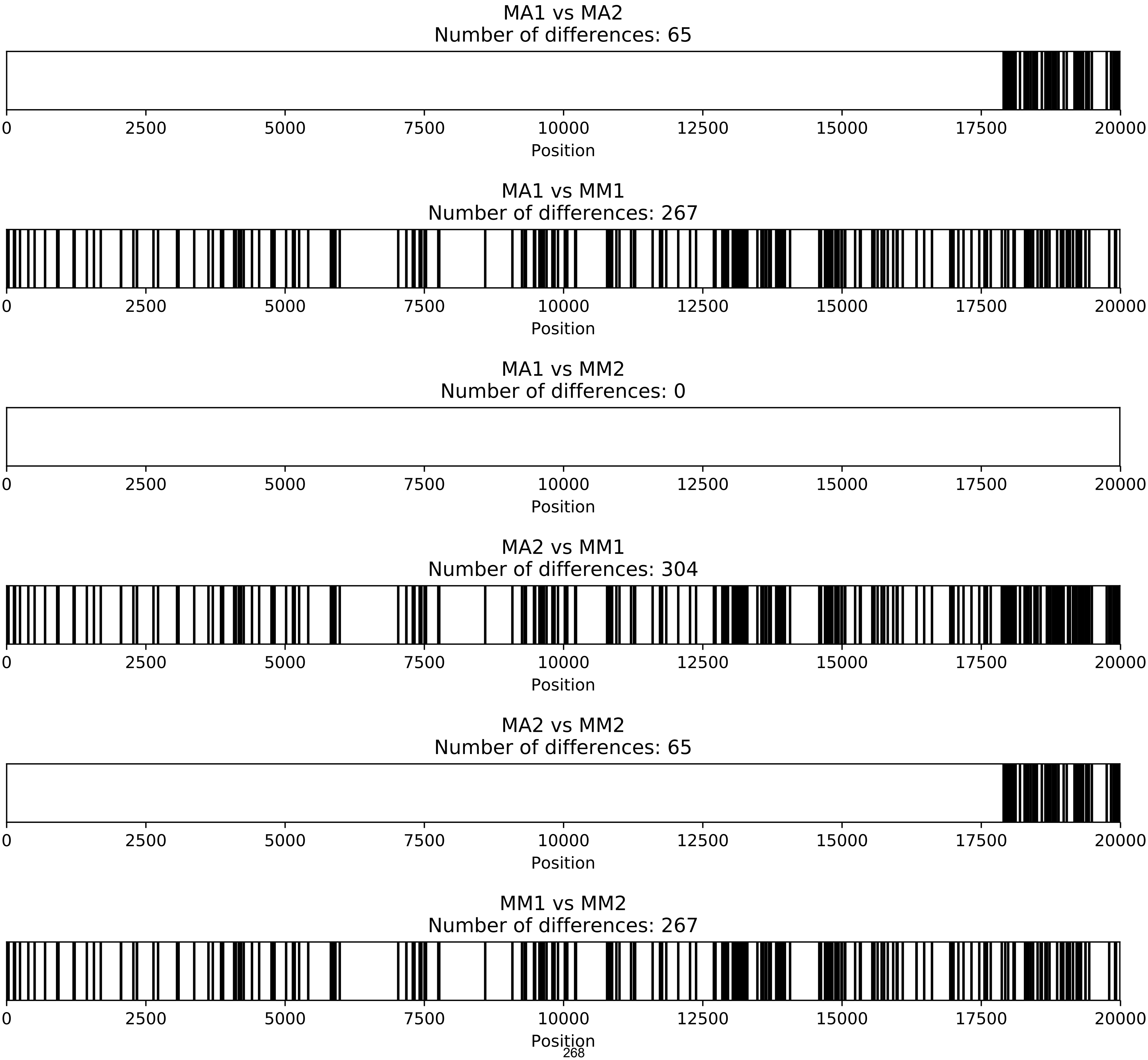
MA1 vs MA2
Number of differences: 65
MA1 vs MM1
Number of differences: 267
MA1 vs MM2
Number of differences: 0
MA2 vs MM1
Number of differences: 304
MA2 vs MM2
Number of differences: 65
MM1 vs MM2
Number of differences: 267
0
2500
5000
7500
10000
12500
15000
17500
20000
Position

### Region: 920



MA1 vs MA2
Number of differences: 88

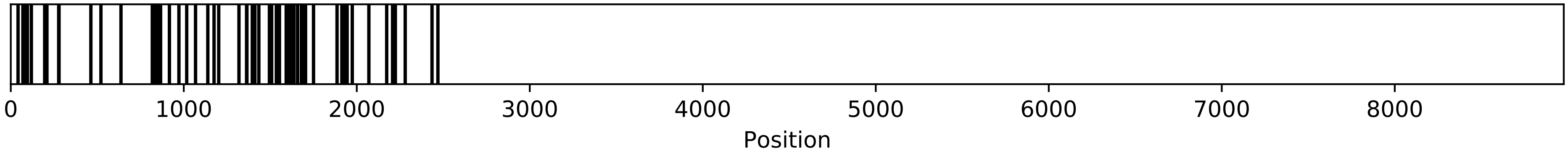


MA1 vs MM1
Number of differences: 111

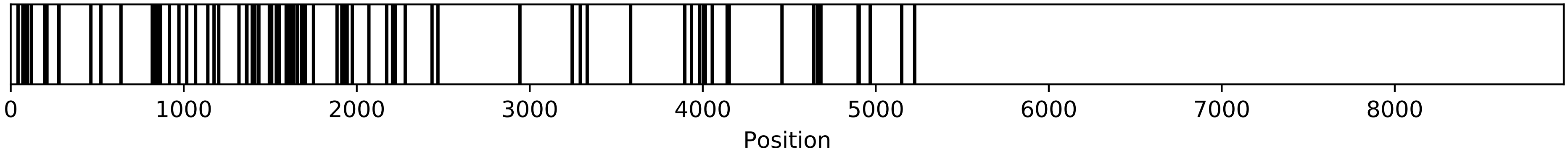


MA1 vs MM2
Number of differences: 147

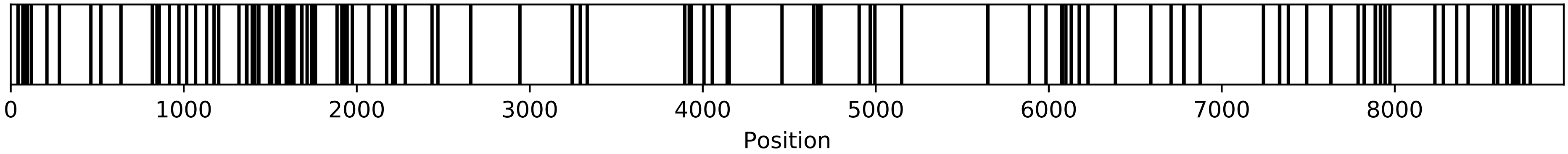


MA2 vs MM1
Number of differences: 23

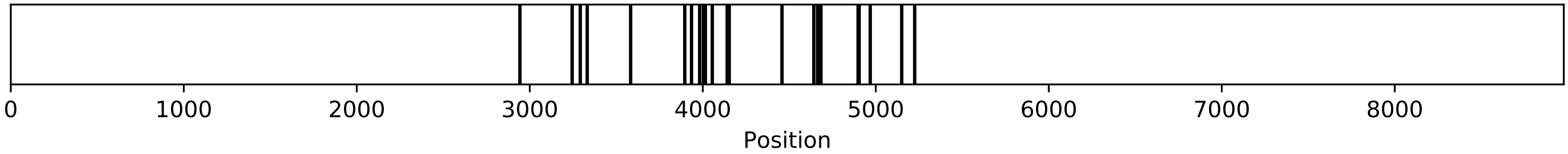


MA2 vs MM2
Number of differences: 86

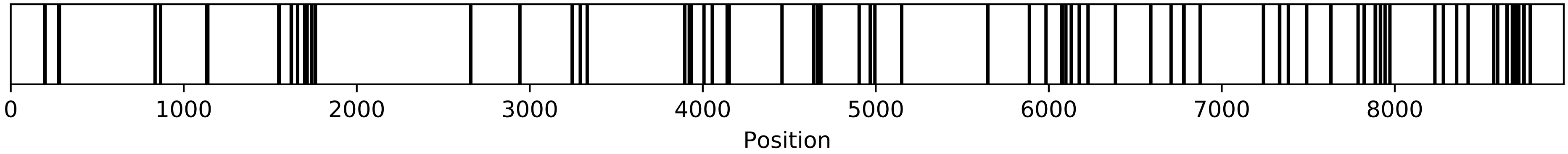


MM1 vs MM2
Number of differences: 76

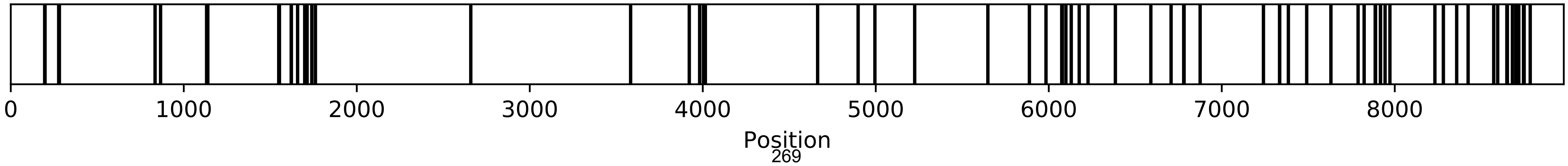

### Region: 922



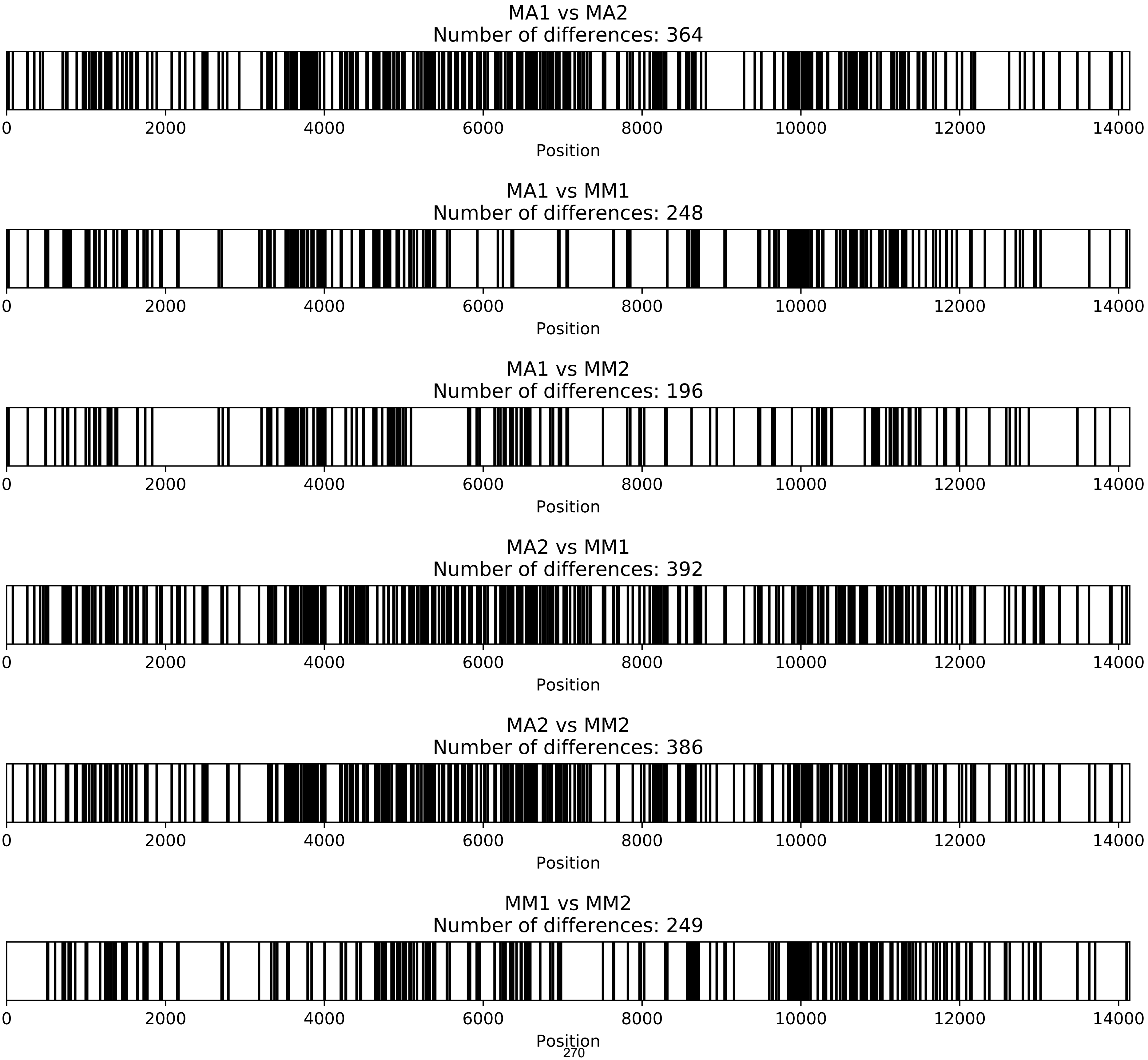
MA1 vs MA2
Number of differences: 364
MA1 vs MM1
Number of differences: 248
MA1 vs MM2
Number of differences: 196
MA2 vs MM1
Number of differences: 392
MA2 vs MM2
Number of differences: 386
MM1 vs MM2
Number of differences: 249
0
2000
4000
6000
8000
10000
12000
14000
Position

### Region: 924



## MA1 vs MA2
Number of differences: 193

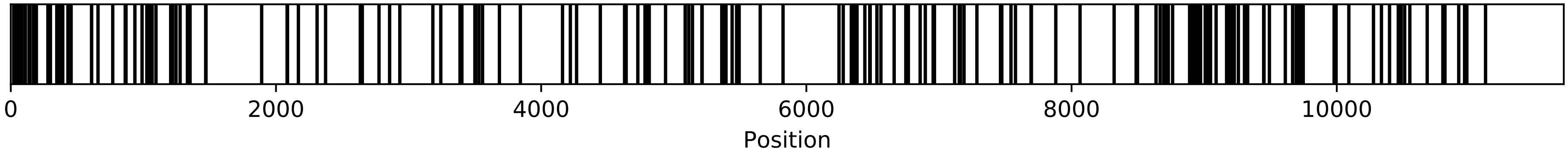


## MA1 vs MM1
Number of differences: 18

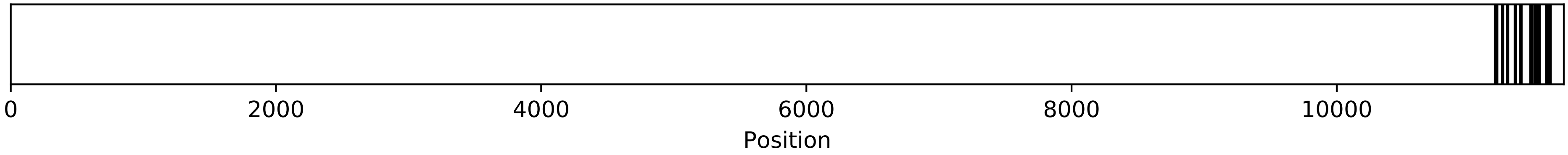


## MA1 vs MM2
Number of differences: 94

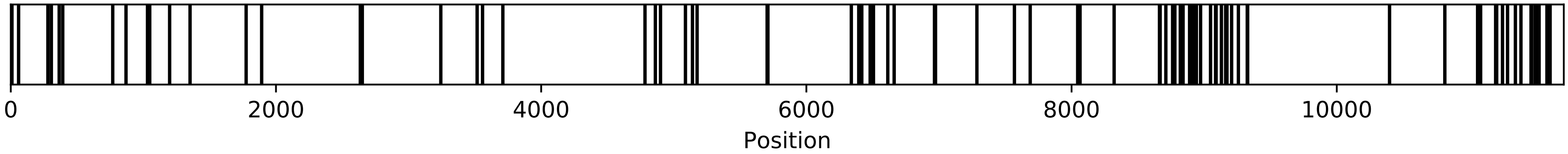


## MA2 vs MM1
Number of differences: 211

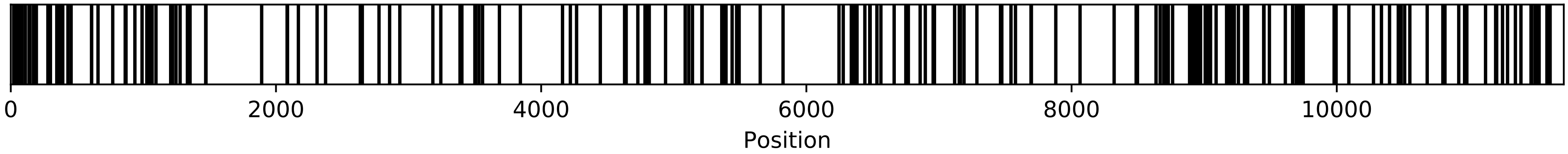


## MA2 vs MM2
Number of differences: 207

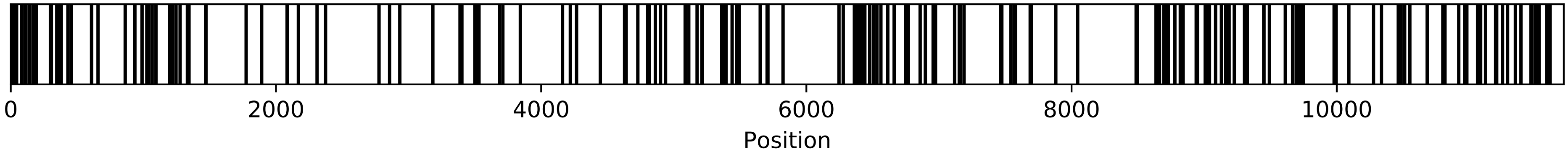


## MM1 vs MM2
Number of differences: 76

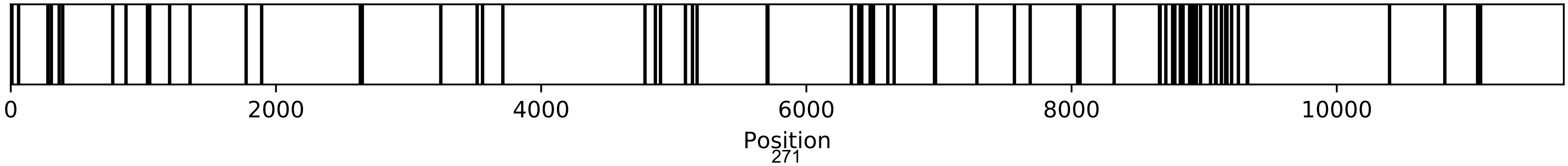

### Region: 926



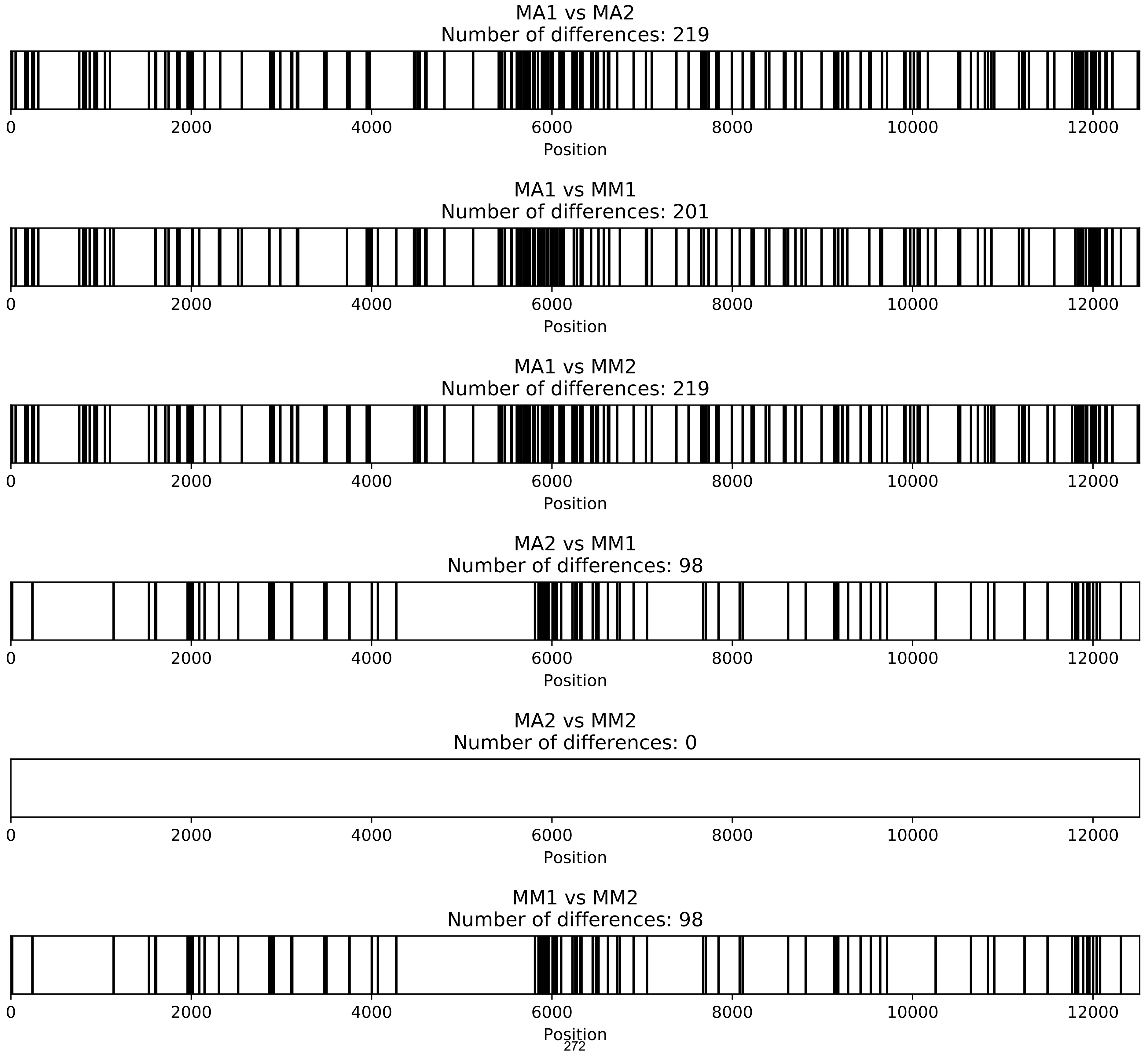
MA1 vs MA2
Number of differences: 219
MA1 vs MM1
Number of differences: 201
MA1 vs MM2
Number of differences: 219
MA2 vs MM1
Number of differences: 98
MA2 vs MM2
Number of differences: 0
MM1 vs MM2
Number of differences: 98
0
2000
4000
6000
8000
10000
12000
Position

### Region: 927



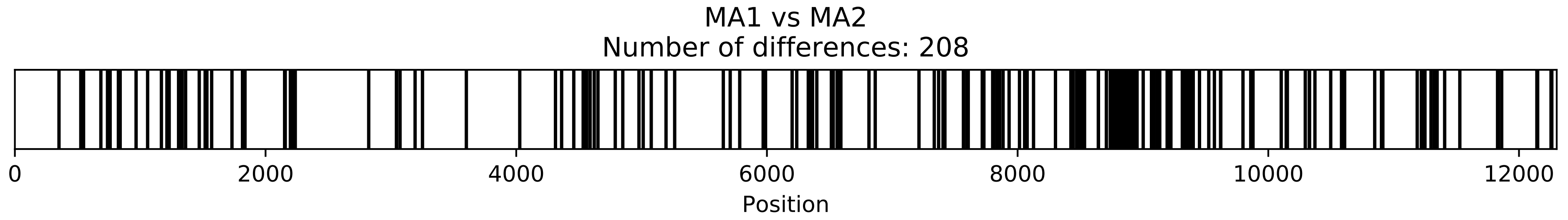
MA1 vs MA2
Number of differences: 208
0
2000
4000
6000
8000
10000
12000
Position

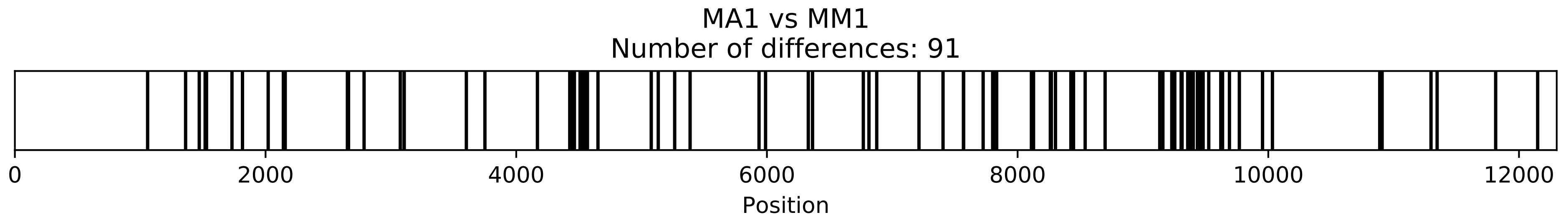
MA1 vs MM1
Number of differences: 91
0
2000
4000
6000
8000
10000
12000
Position

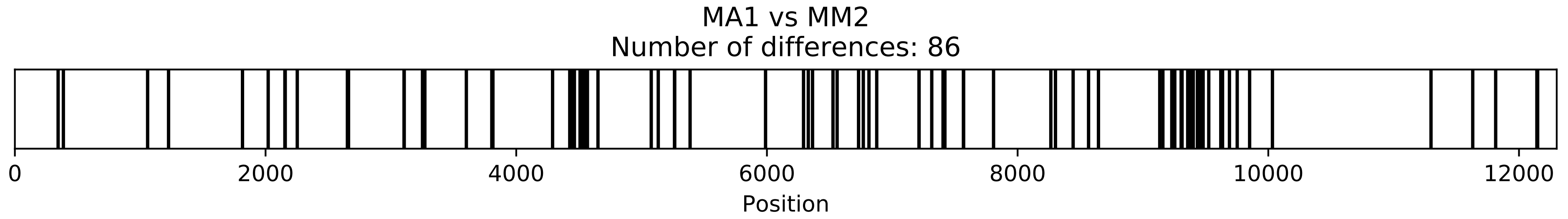
MA1 vs MM2
Number of differences: 86
0
2000
4000
6000
8000
10000
12000
Position

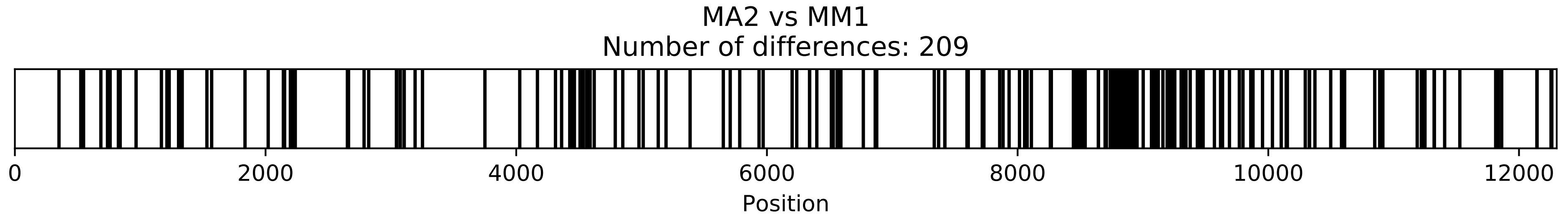
MA2 vs MM1
Number of differences: 209
0
2000
4000
6000
8000
10000
12000
Position

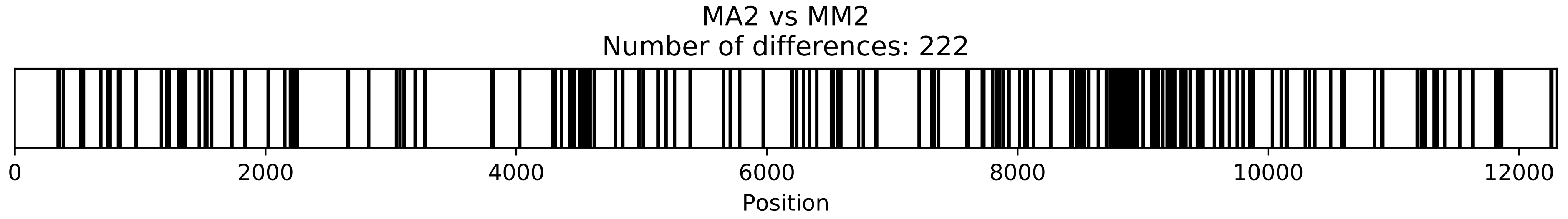
MA2 vs MM2
Number of differences: 222
0
2000
4000
6000
8000
10000
12000
Position

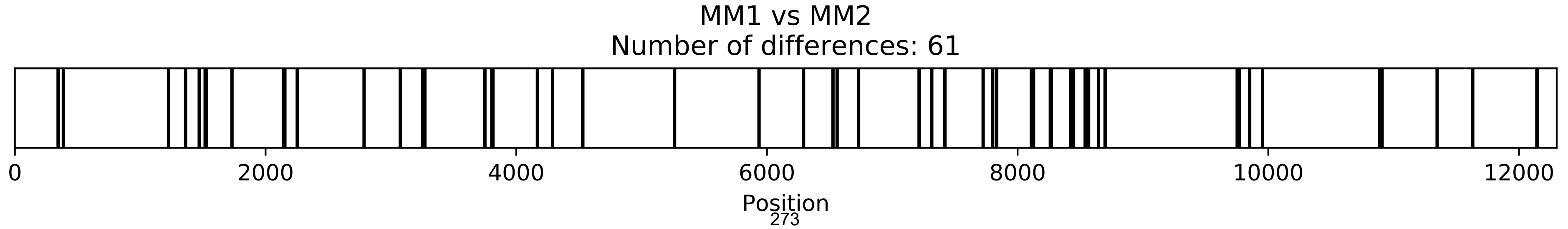
MM1 vs MM2
Number of differences: 61
0
2000
4000
6000
8000
10000
12000
Position

### Region: 928



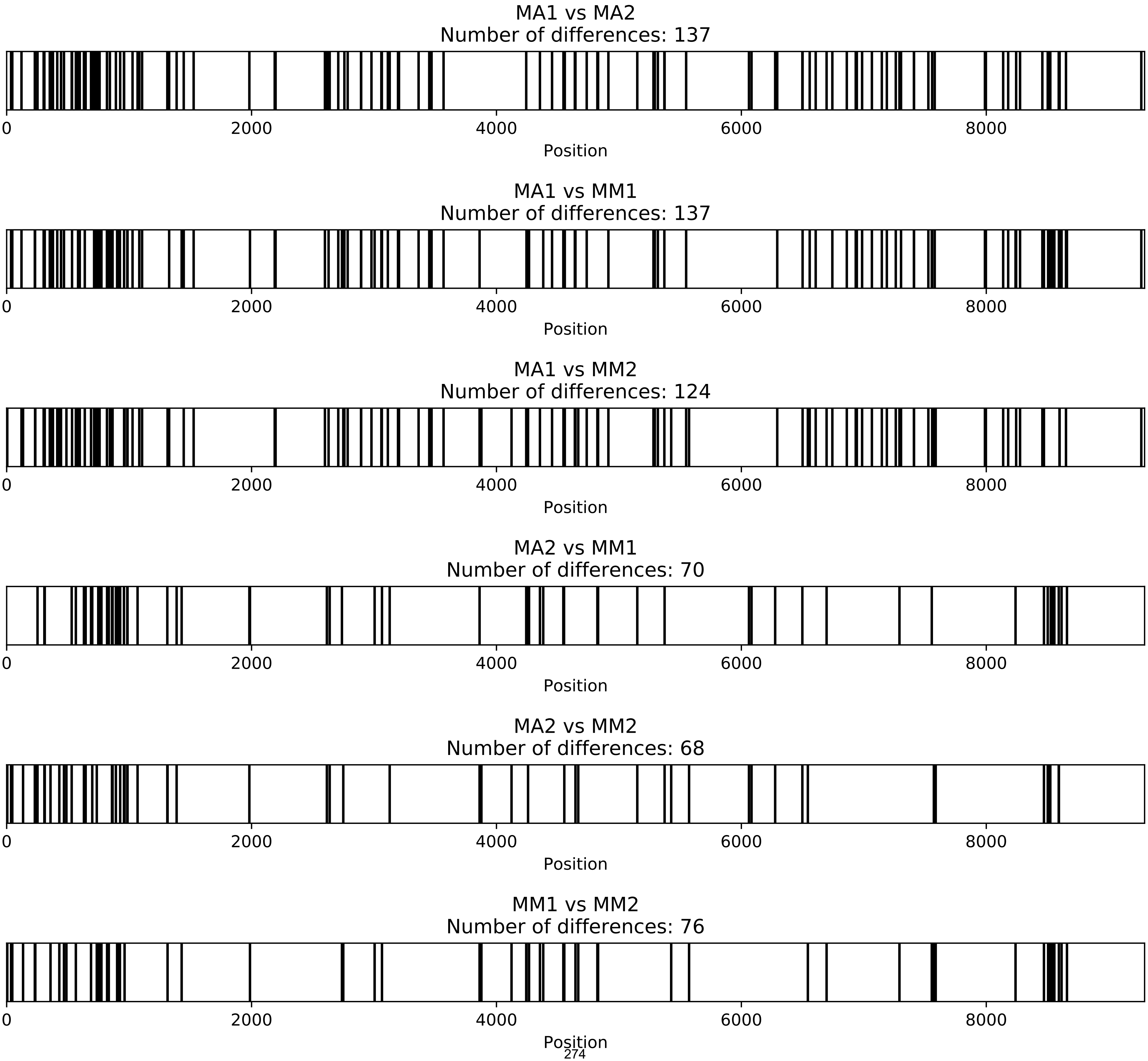
MA1 vs MA2
Number of differences: 137
0
2000
4000
6000
8000
Position
MA1 vs MM1
Number of differences: 137
0
2000
4000
6000
8000
Position
MA1 vs MM2
Number of differences: 124
0
2000
4000
6000
8000
Position
MA2 vs MM1
Number of differences: 70
0
2000
4000
6000
8000
Position
MA2 vs MM2
Number of differences: 68
0
2000
4000
6000
8000
Position
MM1 vs MM2
Number of differences: 76
0
2000
4000
6000
8000
Position

### Region: 932



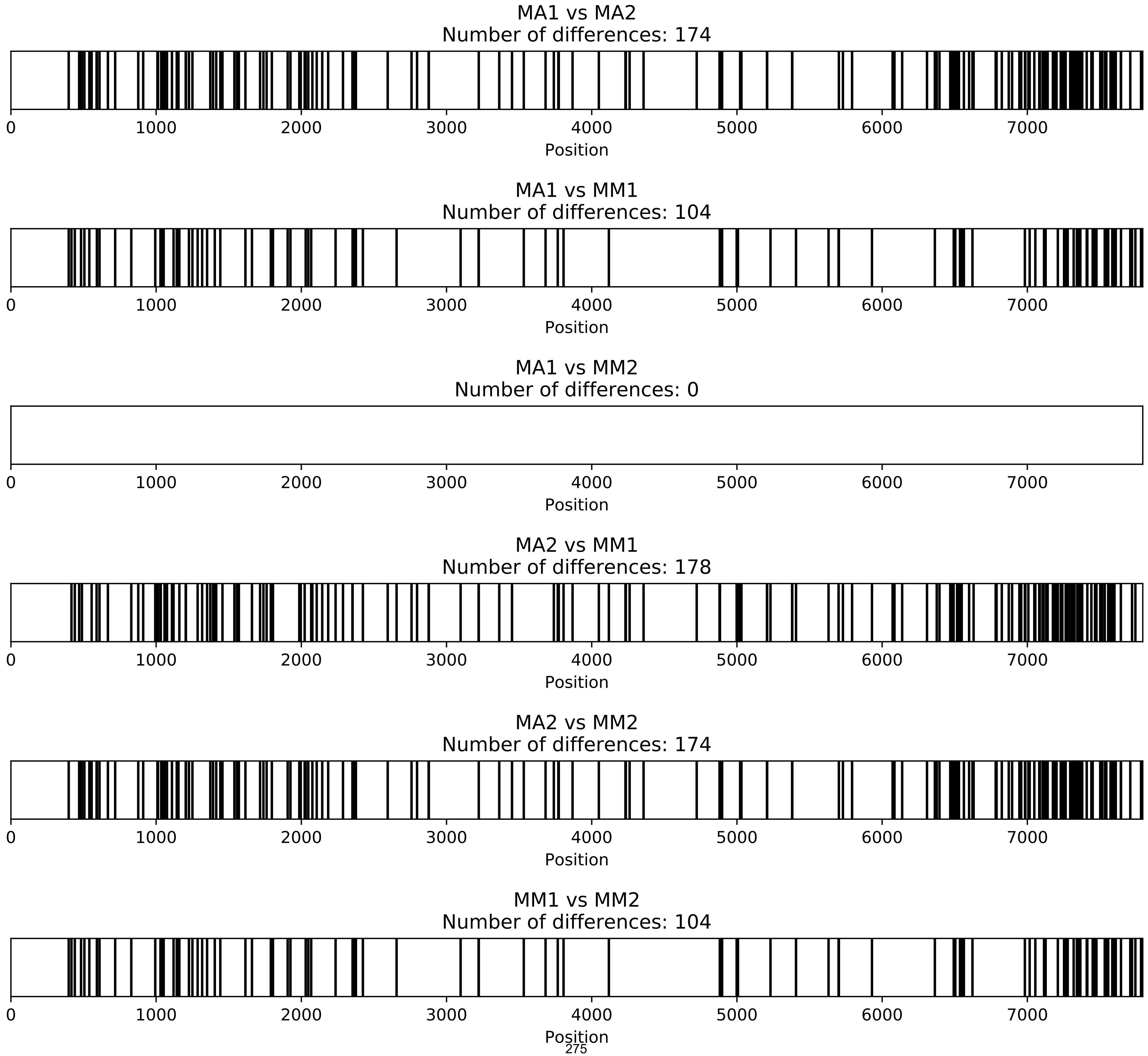
MA1 vs MA2
Number of differences: 174
Position
MA1 vs MM1
Number of differences: 104
Position
MA1 vs MM2
Number of differences: 0
Position
MA2 vs MM1
Number of differences: 178
Position
MA2 vs MM2
Number of differences: 174
Position
MM1 vs MM2
Number of differences: 104
0
1000
2000
3000
4000
5000
6000
7000
Position

### Region: 933



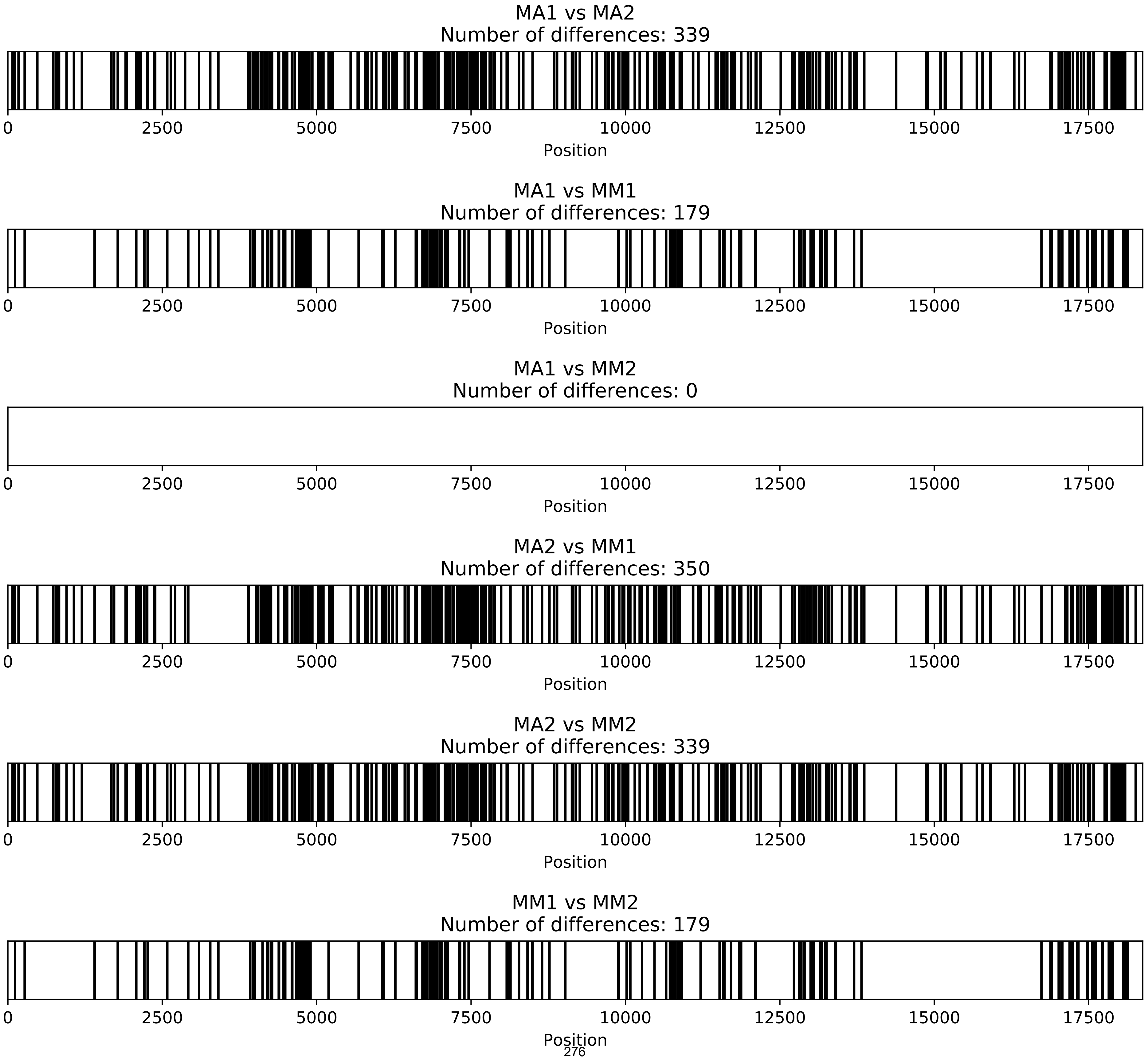
MA1 vs MA2
Number of differences: 339
MA1 vs MM1
Number of differences: 179
MA1 vs MM2
Number of differences: 0
MA2 vs MM1
Number of differences: 350
MA2 vs MM2
Number of differences: 339
MM1 vs MM2
Number of differences: 179
0
2500
5000
7500
10000
12500
15000
17500
Position

### Region: 937



MA1 vs MA2
Number of differences: 120

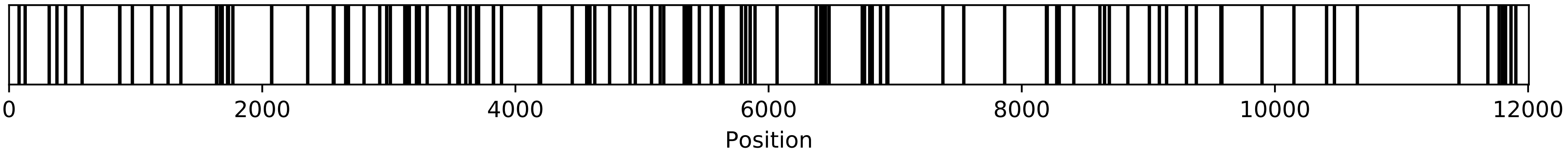


MA1 vs MM1
Number of differences: 0

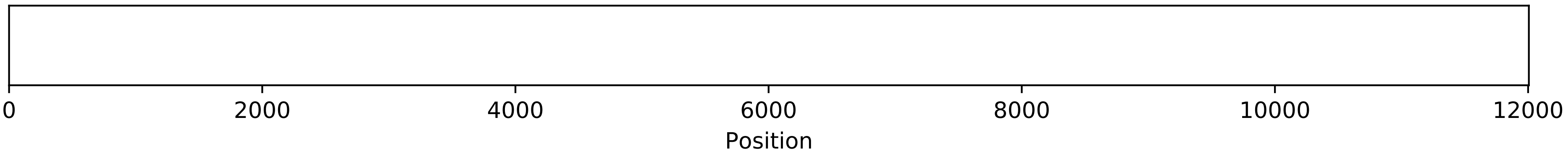


MA1 vs MM2
Number of differences: 49

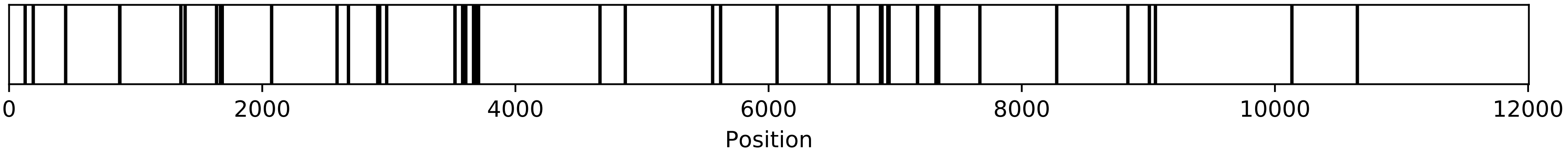


MA2 vs MM1
Number of differences: 120

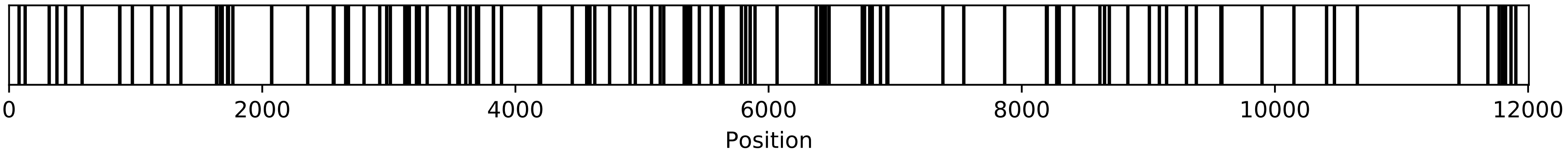


MA2 vs MM2
Number of differences: 123

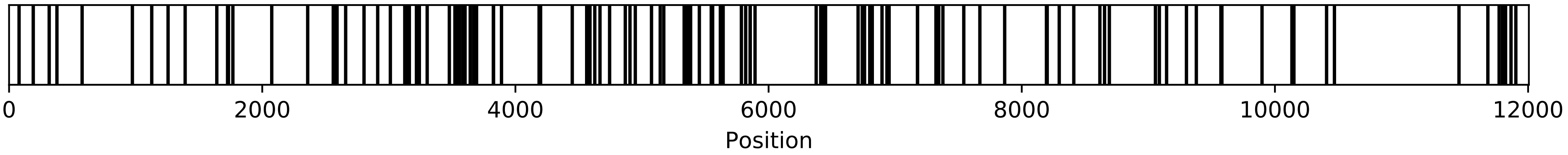


MM1 vs MM2
Number of differences: 49

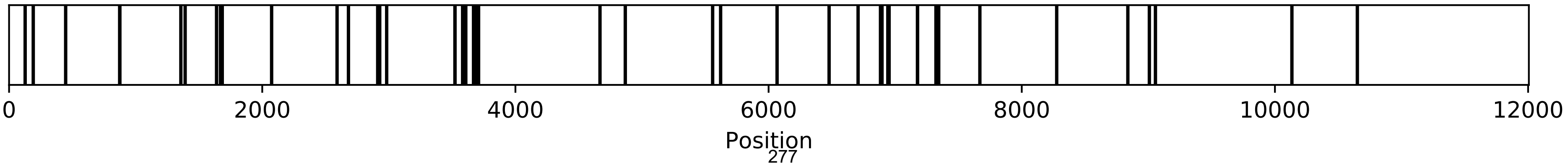

### Region: 939



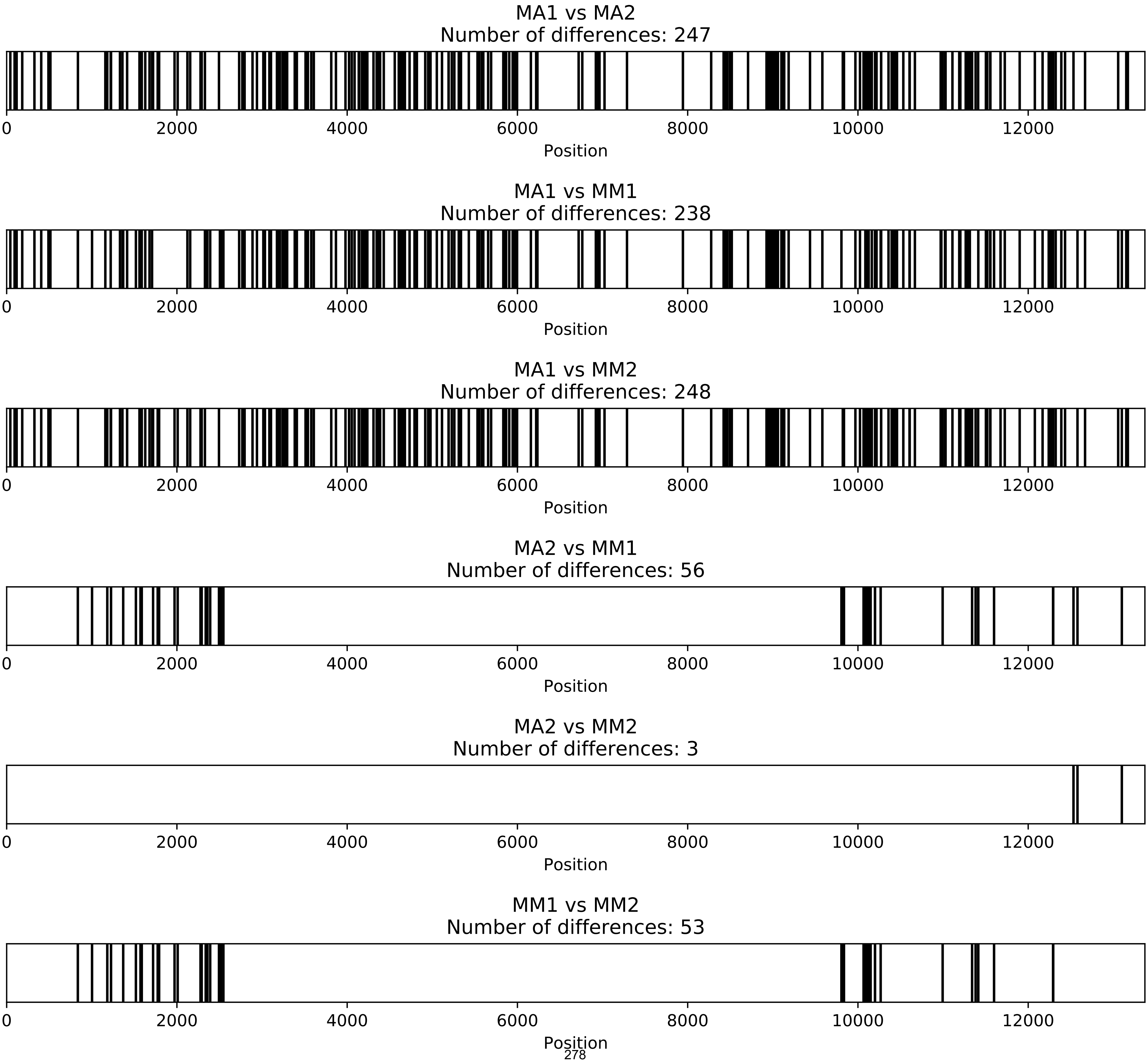
MA1 vs MA2
Number of differences: 247
MA1 vs MM1
Number of differences: 238
MA1 vs MM2
Number of differences: 248
MA2 vs MM1
Number of differences: 56
MA2 vs MM2
Number of differences: 3
MM1 vs MM2
Number of differences: 53
0
2000
4000
6000
8000
10000
12000
Position

### Region: 941



MA1 vs MA2
Number of differences: 81

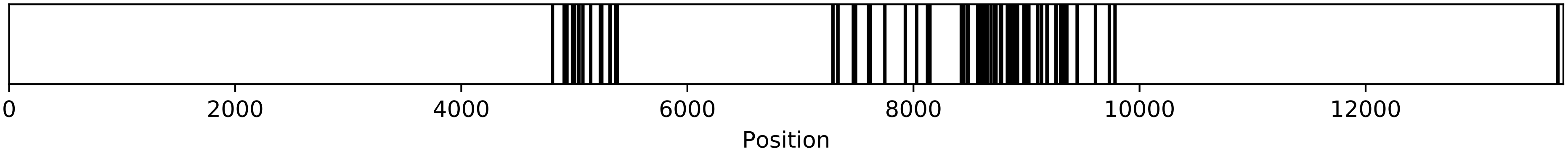


MA1 vs MM1
Number of differences: 269

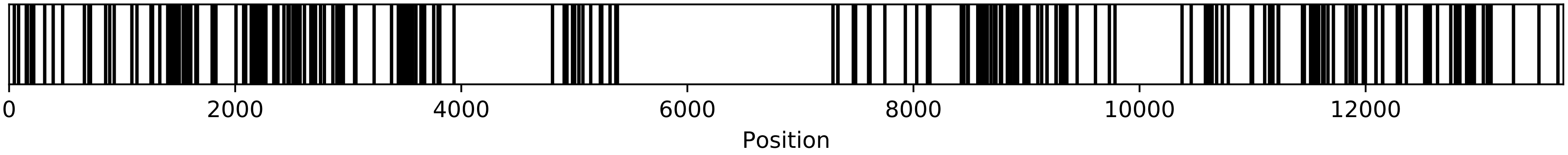


MA1 vs MM2
Number of differences: 294

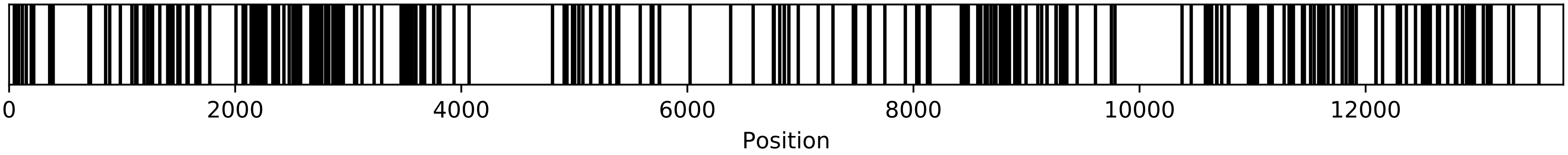


MA2 vs MM1
Number of differences: 188

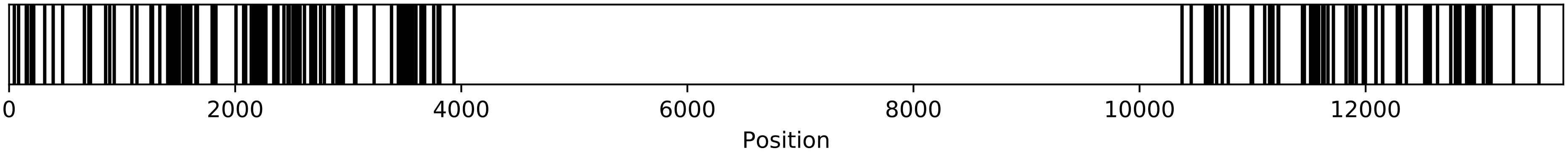


MA2 vs MM2
Number of differences: 247

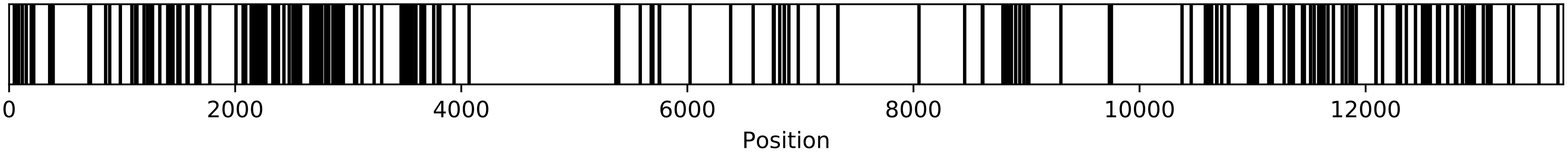


MM1 vs MM2
Number of differences: 178

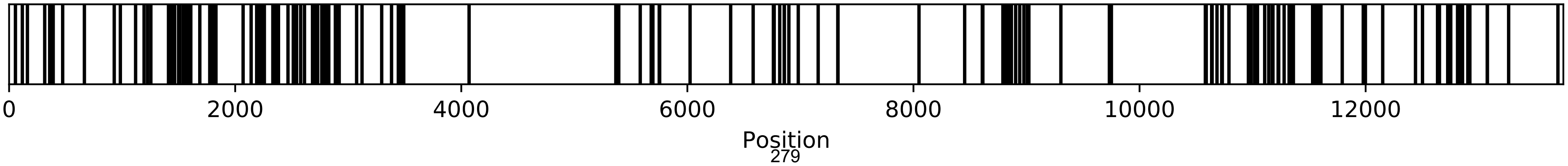

### Region: 943



MA1 vs MA2
Number of differences: 189

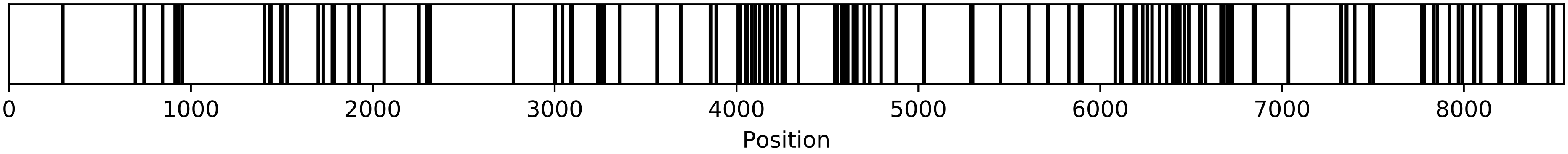


MA1 vs MM1
Number of differences: 188

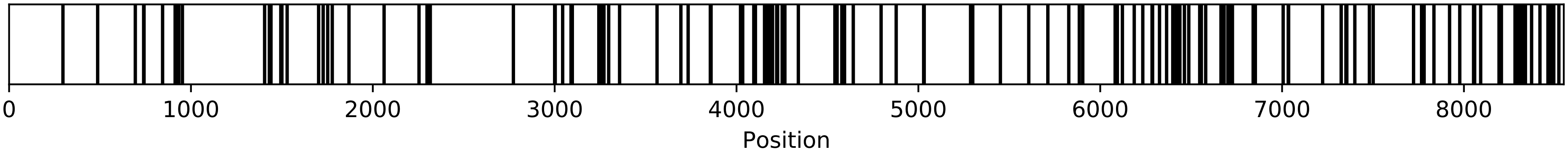


MA1 vs MM2
Number of differences: 188

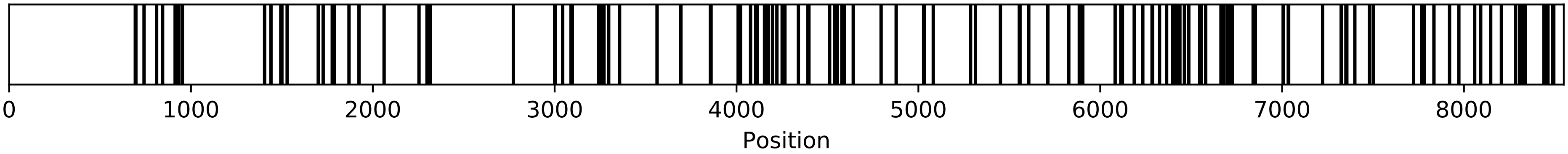


MA2 vs MM1
Number of differences: 52

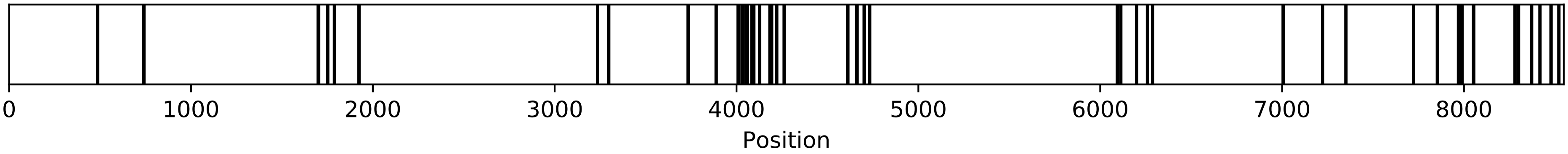


MA2 vs MM2
Number of differences: 51

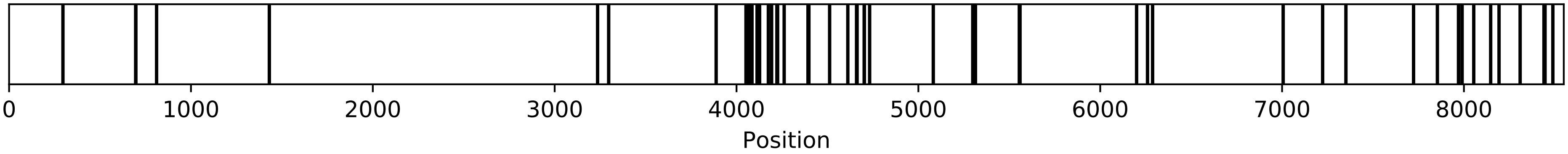


MM1 vs MM2
Number of differences: 51

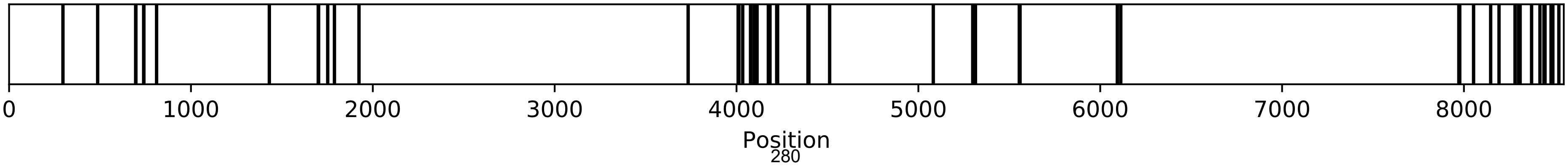

### Region: 945



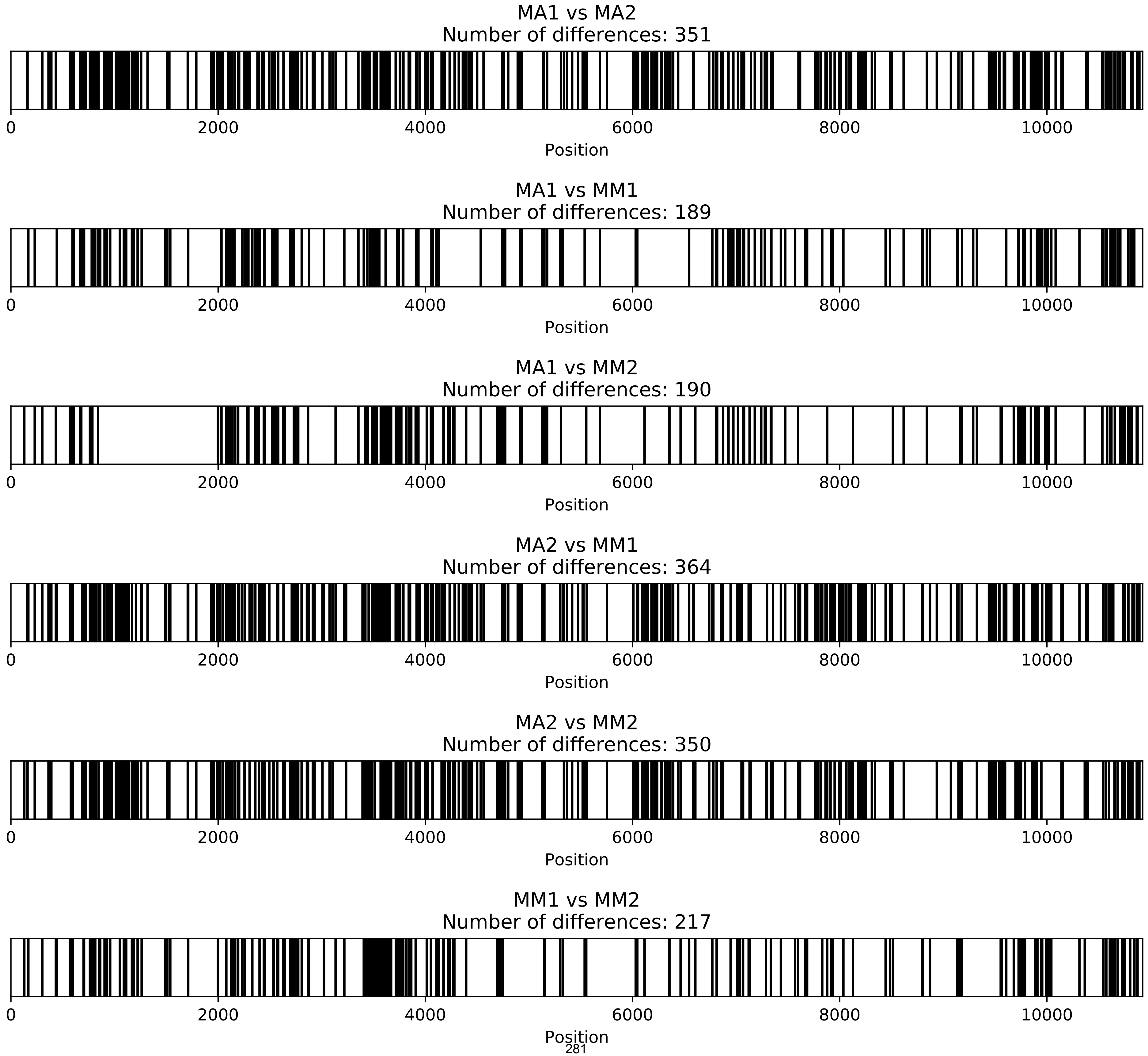
MA1 vs MA2
Number of differences: 351
0 2000 4000 6000 8000 10000
Position
MA1 vs MM1
Number of differences: 189
0 2000 4000 6000 8000 10000
Position
MA1 vs MM2
Number of differences: 190
0 2000 4000 6000 8000 10000
Position
MA2 vs MM1
Number of differences: 364
0 2000 4000 6000 8000 10000
Position
MA2 vs MM2
Number of differences: 350
0 2000 4000 6000 8000 10000
Position
MM1 vs MM2
Number of differences: 217
0 2000 4000 6000 8000 10000
Position

### Region: 953



MA1 vs MA2
Number of differences: 225

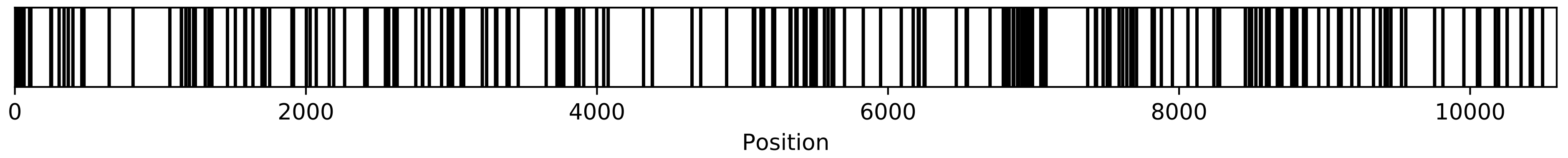

MA1 vs MM1
Number of differences: 234

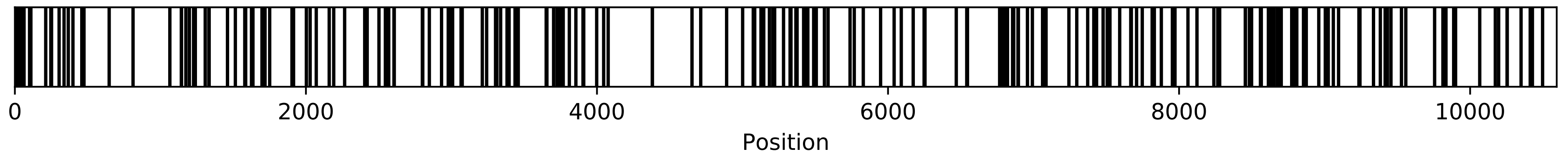

MA1 vs MM2
Number of differences: 218

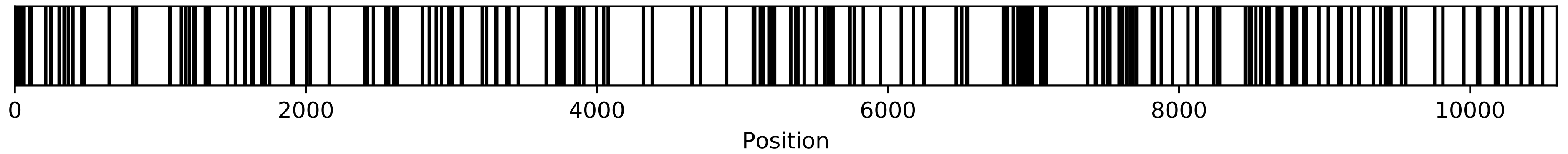

MA2 vs MM1
Number of differences: 95

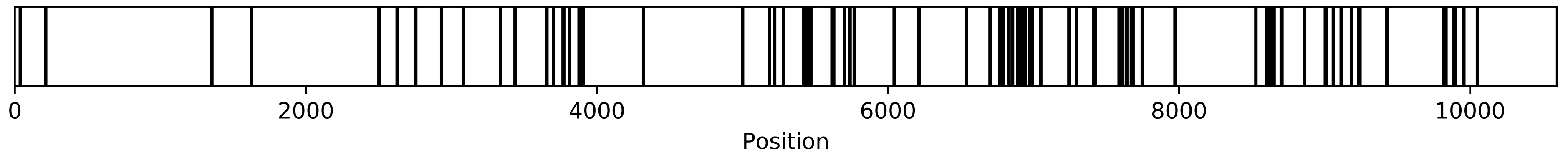

MA2 vs MM2
Number of differences: 42

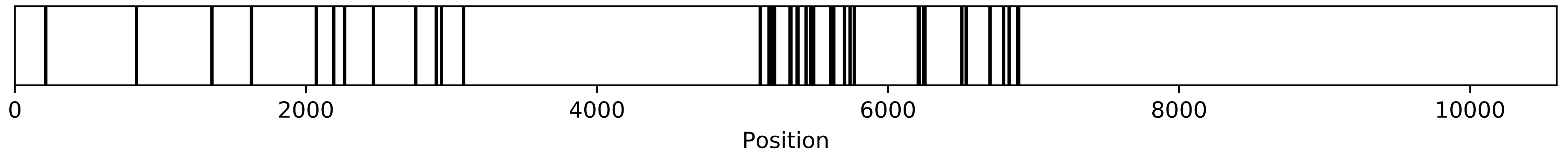

MM1 vs MM2
Number of differences: 94

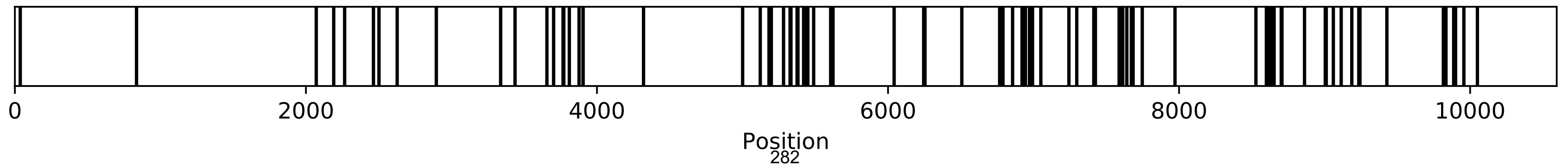

### Region: 954



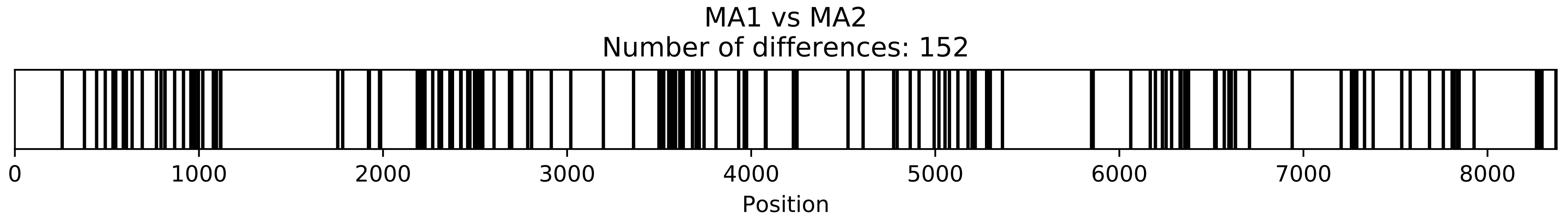
MA1 vs MA2
Number of differences: 152
0
1000
2000
3000
4000
5000
6000
7000
8000
Position

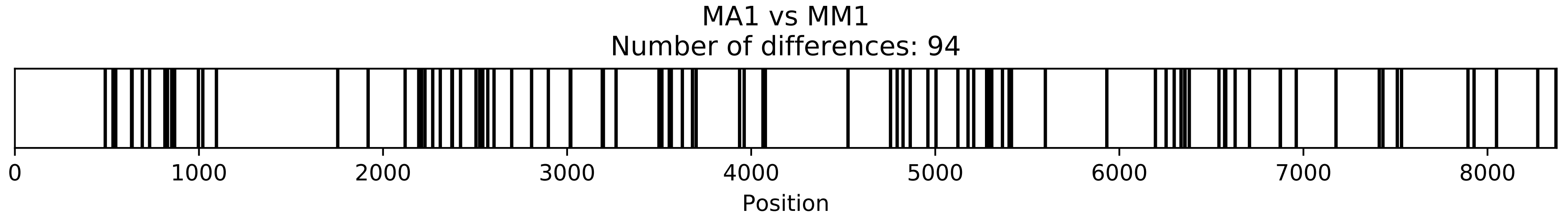
MA1 vs MM1
Number of differences: 94
0
1000
2000
3000
4000
5000
6000
7000
8000
Position

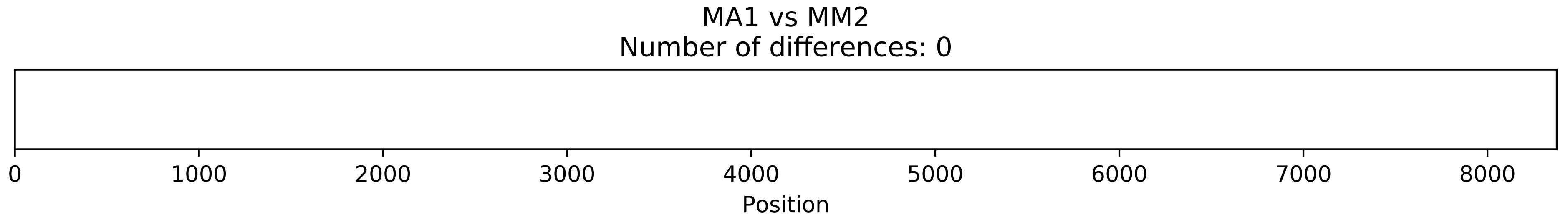
MA1 vs MM2
Number of differences: 0
0
1000
2000
3000
4000
5000
6000
7000
8000
Position

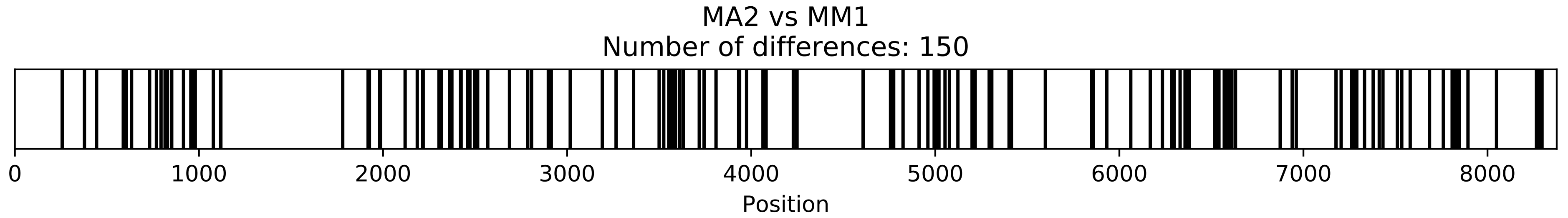
MA2 vs MM1
Number of differences: 150
0
1000
2000
3000
4000
5000
6000
7000
8000
Position

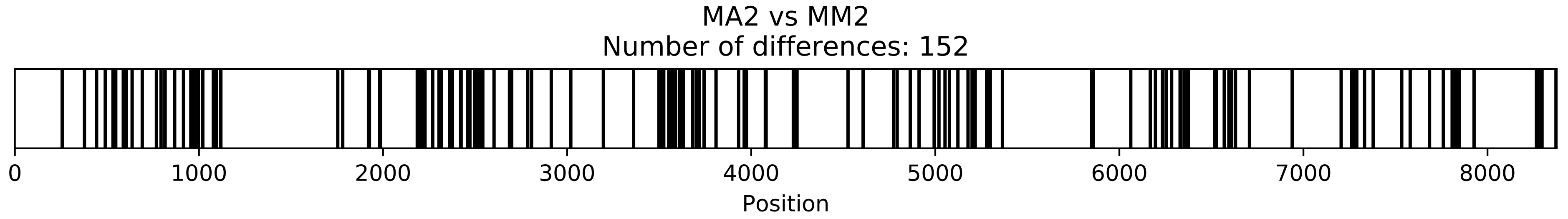
MA2 vs MM2
Number of differences: 152
0
1000
2000
3000
4000
5000
6000
7000
8000
Position

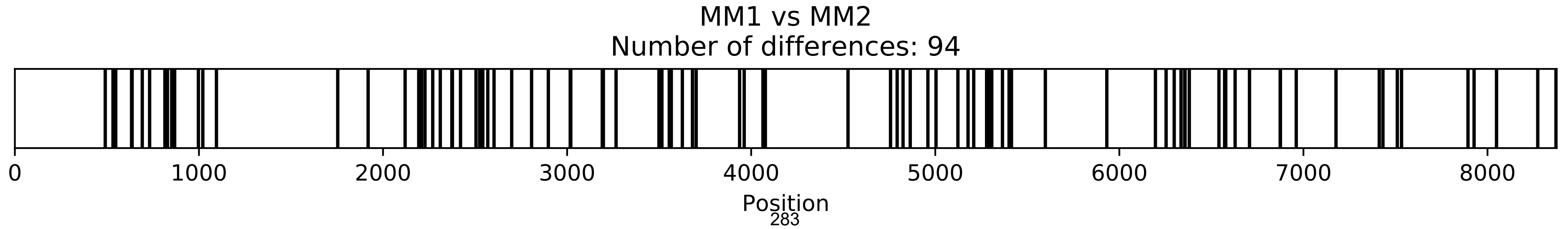
MM1 vs MM2
Number of differences: 94
0
1000
2000
3000
4000
5000
6000
7000
8000
Position

### Region: 957



MA1 vs MA2
Number of differences: 241

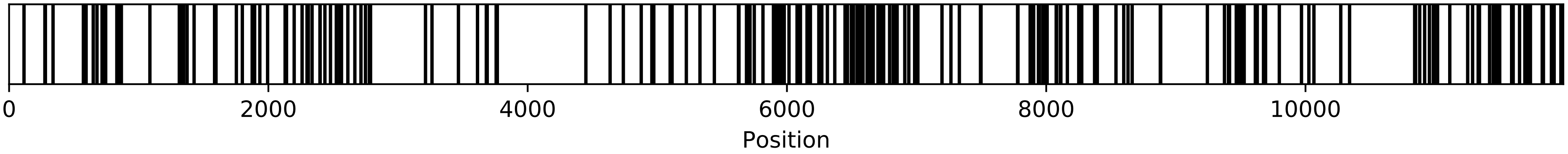


MA1 vs MM1
Number of differences: 209

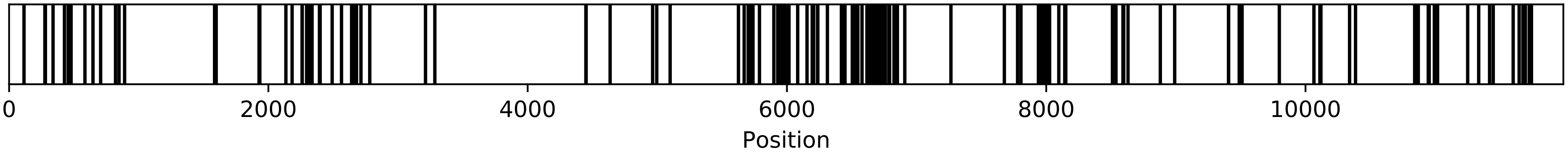


MA1 vs MM2
Number of differences: 114

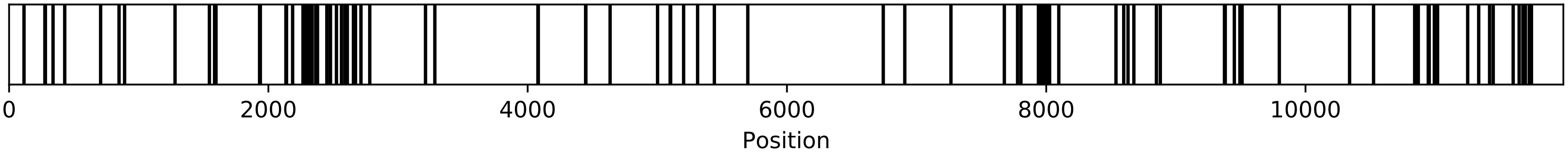


MA2 vs MM1
Number of differences: 261

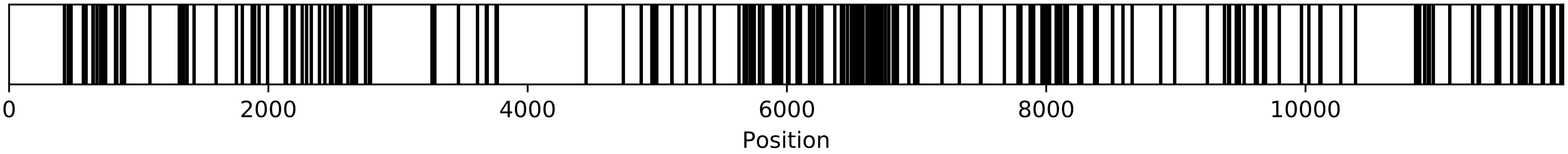


MA2 vs MM2
Number of differences: 257

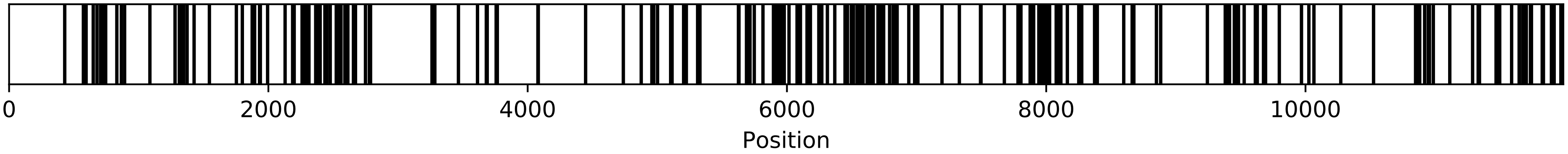


MM1 vs MM2
Number of differences: 176

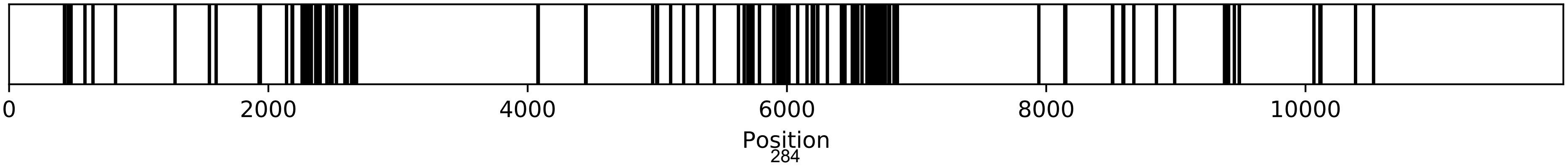

### Region: 958



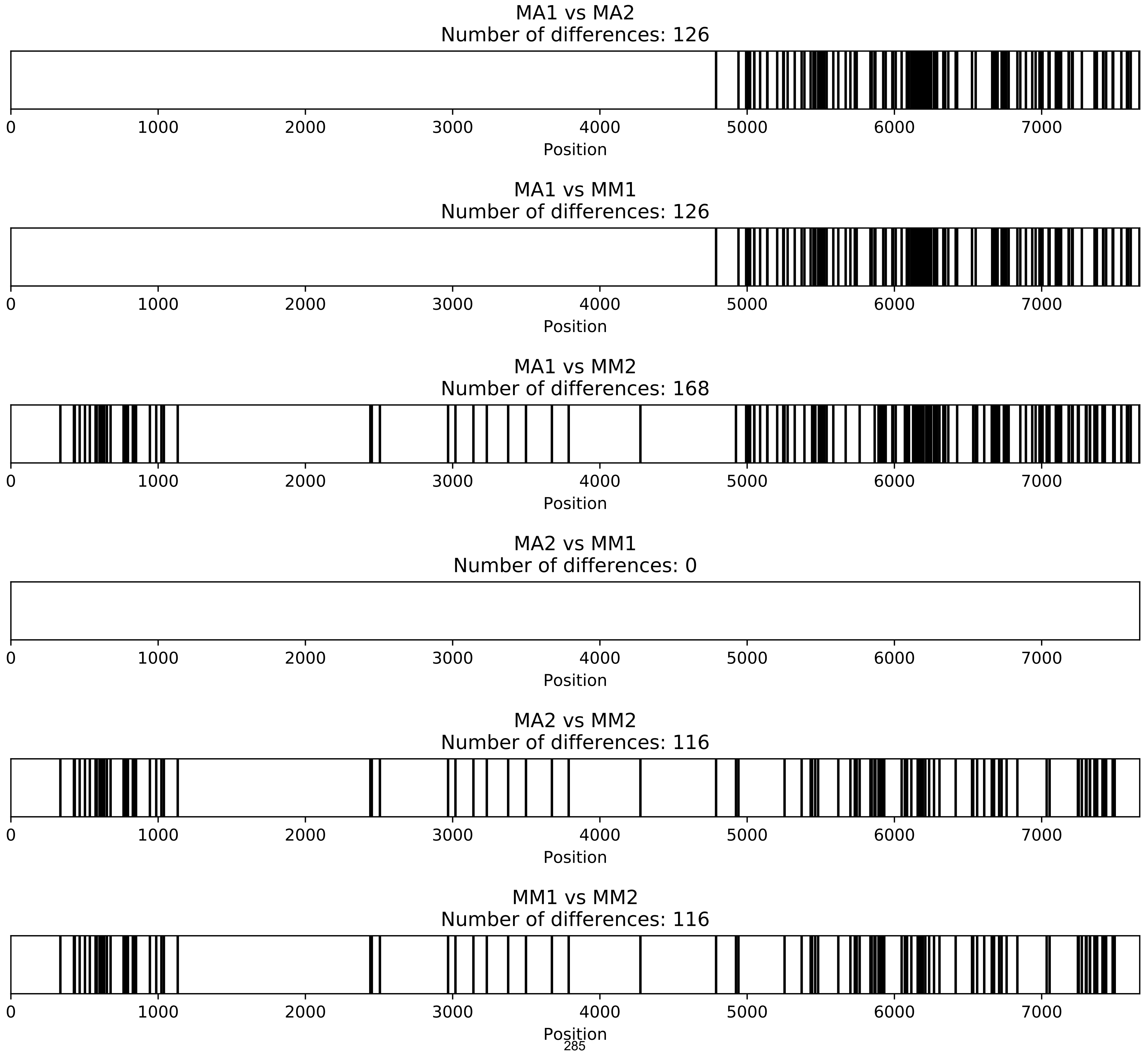
MA1 vs MA2
Number of differences: 126
0
1000
2000
3000
4000
5000
6000
7000
Position
MA1 vs MM1
Number of differences: 126
Position
MA1 vs MM2
Number of differences: 168
Position
MA2 vs MM1
Number of differences: 0
Position
MA2 vs MM2
Number of differences: 116
Position
MM1 vs MM2
Number of differences: 116
Position

### Region: 964



MA1 vs MA2
Number of differences: 325

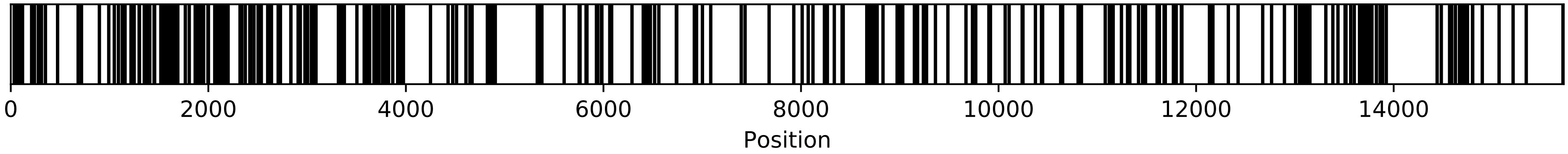


MA1 vs MM1
Number of differences: 325

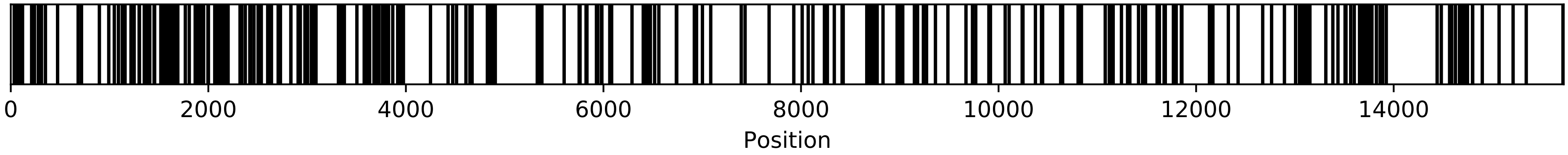


MA1 vs MM2
Number of differences: 314

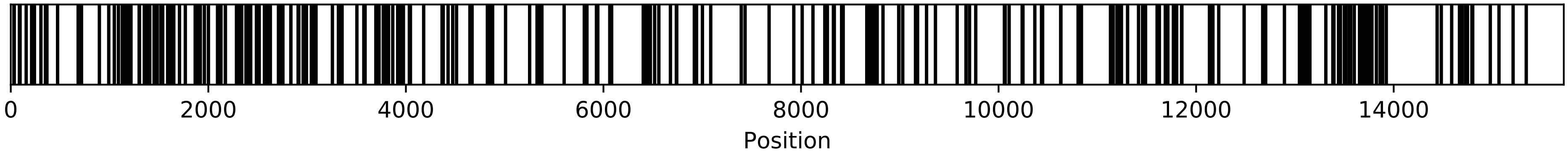


MA2 vs MM1
Number of differences: 0

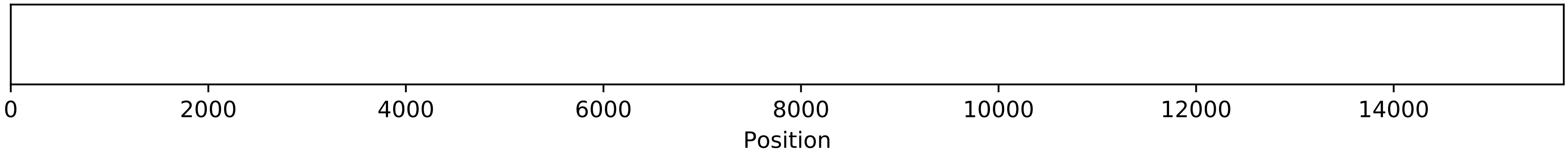


MA2 vs MM2
Number of differences: 171

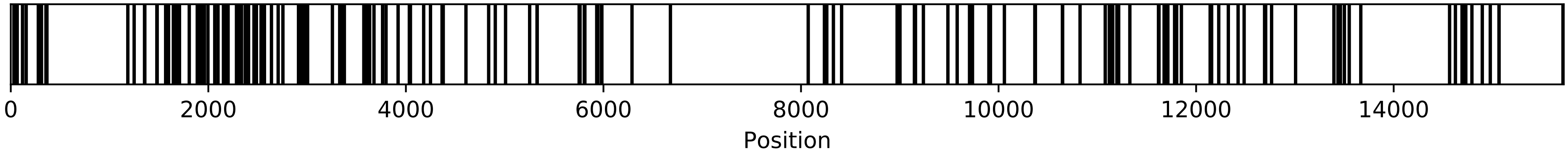


MM1 vs MM2
Number of differences: 171

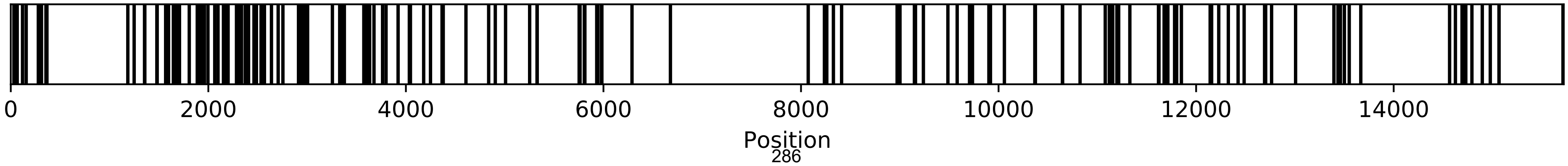

### Region: 967



MA1 vs MA2
Number of differences: 206

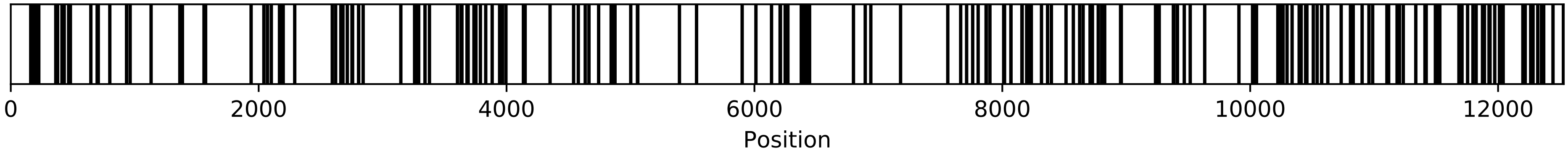


MA1 vs MM1
Number of differences: 197

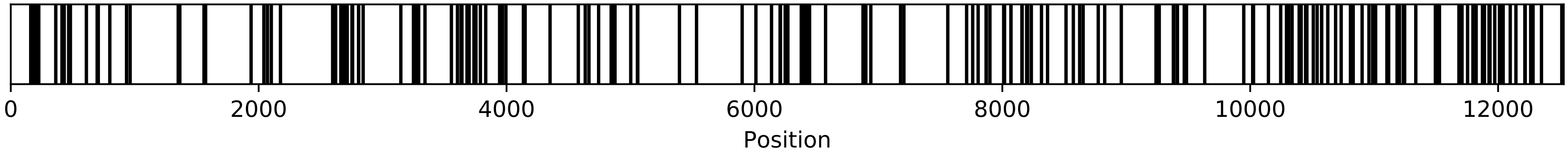


MA1 vs MM2
Number of differences: 206

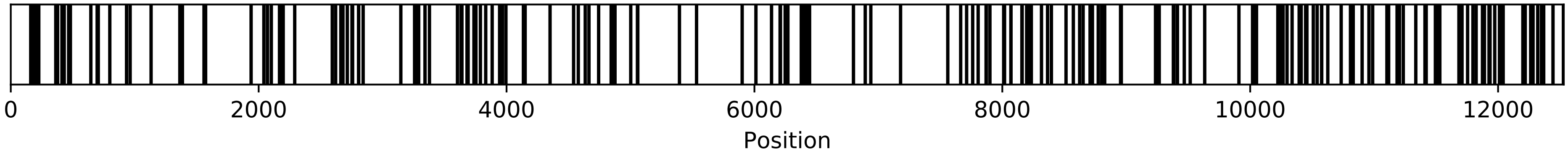


MA2 vs MM1
Number of differences: 74

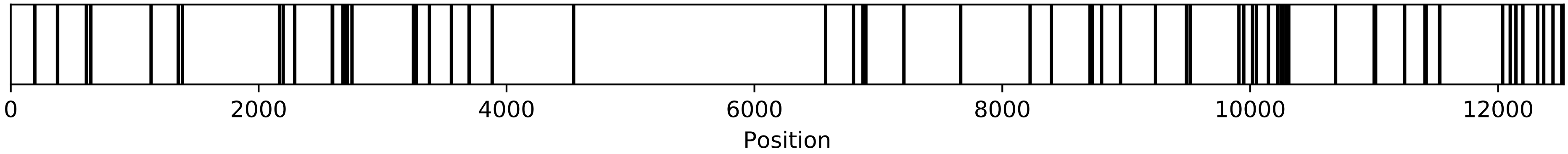


MA2 vs MM2
Number of differences: 0

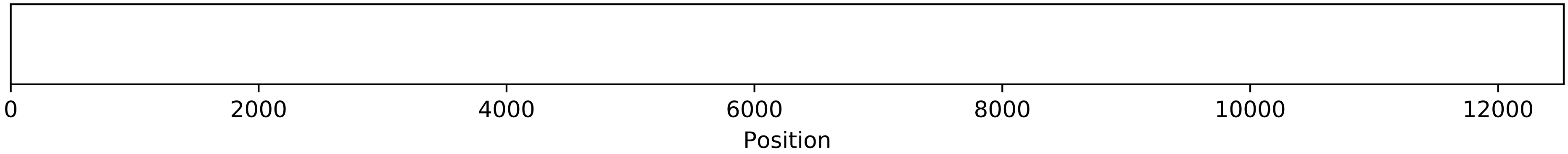


MM1 vs MM2
Number of differences: 74

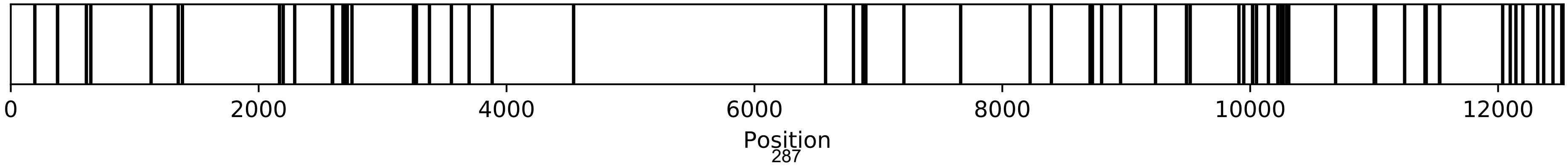

### Region: 974



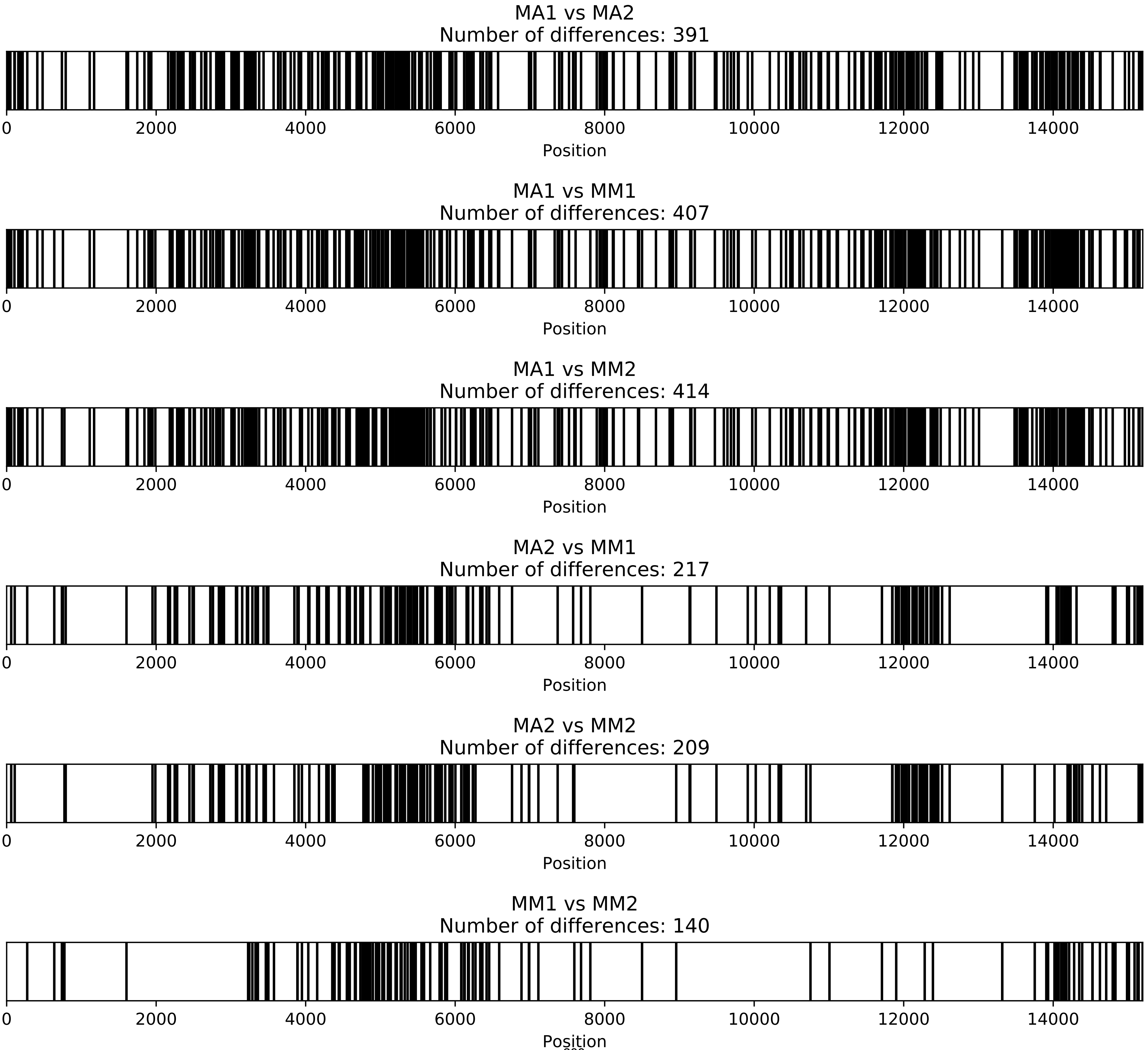
MA1 vs MA2
Number of differences: 391
MA1 vs MM1
Number of differences: 407
MA1 vs MM2
Number of differences: 414
MA2 vs MM1
Number of differences: 217
MA2 vs MM2
Number of differences: 209
MM1 vs MM2
Number of differences: 140
0
2000
4000
6000
8000
10000
12000
14000
Position

### Region: 975



MA1 vs MA2
Number of differences: 149

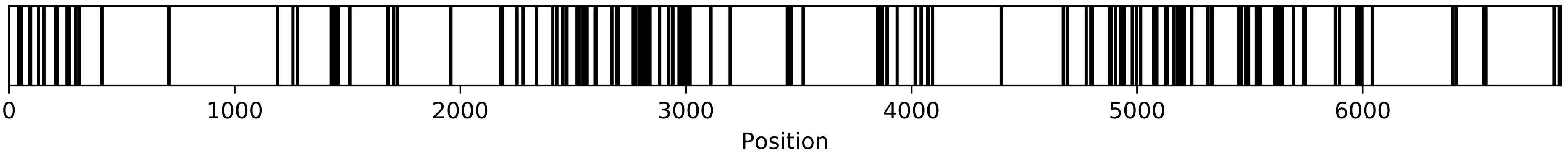


MA1 vs MM1
Number of differences: 0

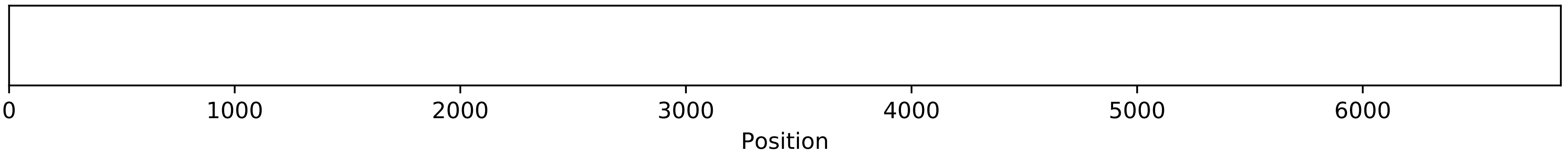


MA1 vs MM2
Number of differences: 66

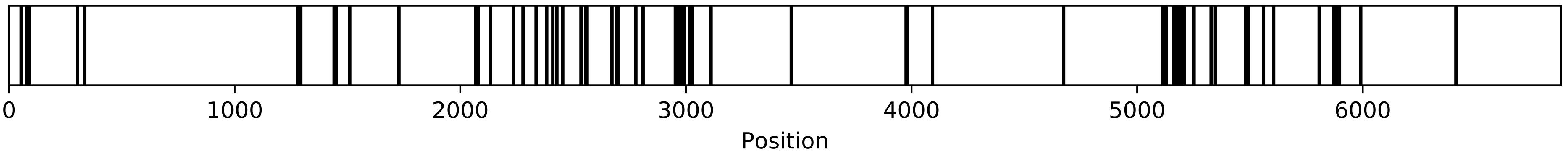


MA2 vs MM1
Number of differences: 149

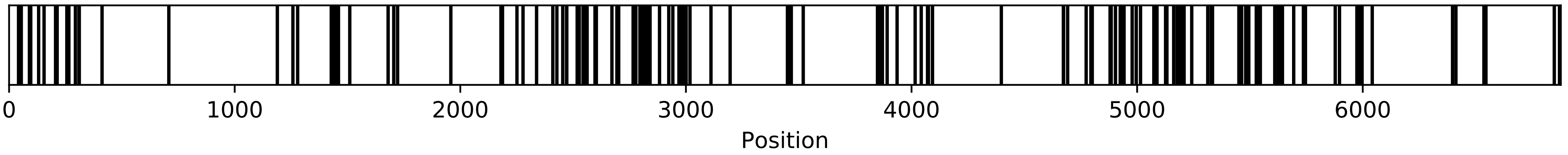


MA2 vs MM2
Number of differences: 139

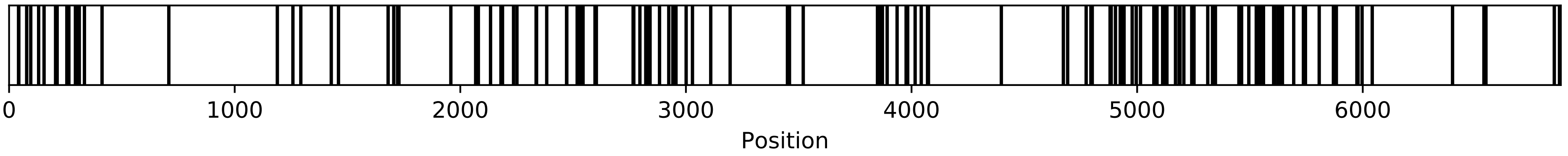


MM1 vs MM2
Number of differences: 66

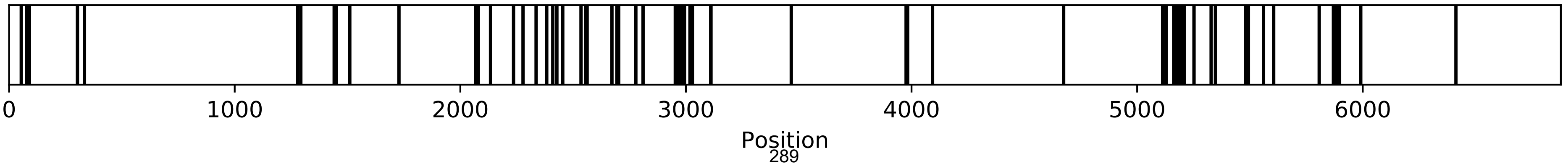

### Region: 980



MA1 vs MA2
Number of differences: 183

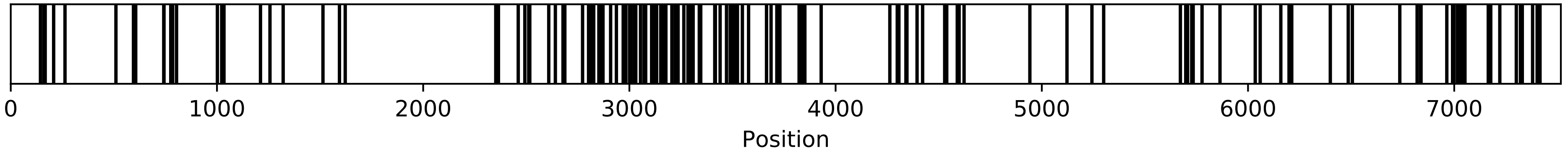

MA1 vs MM1
Number of differences: 211

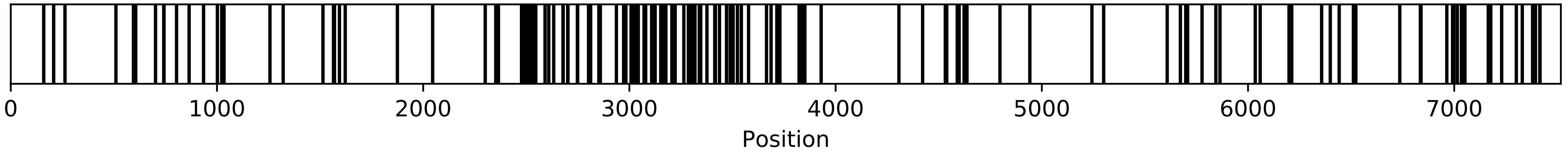

MA1 vs MM2
Number of differences: 216

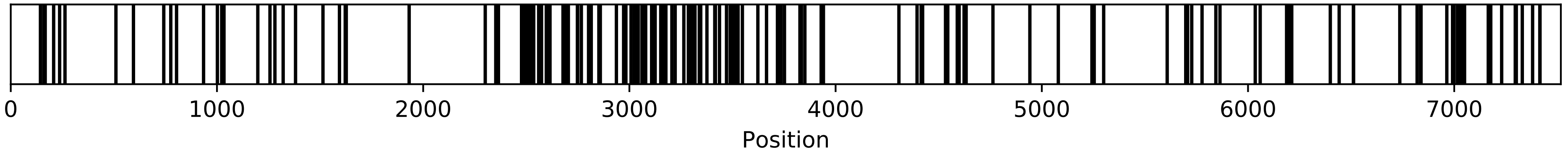

MA2 vs MM1
Number of differences: 137

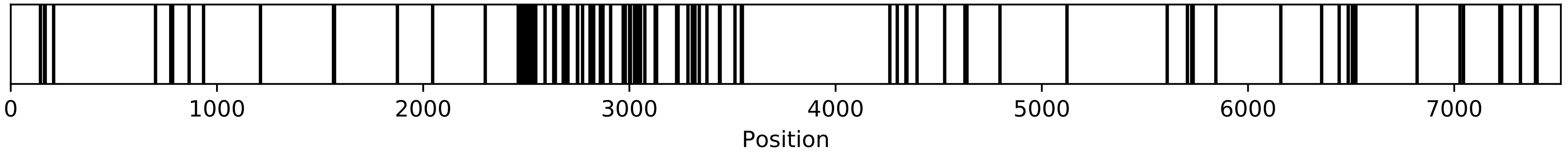

MA2 vs MM2
Number of differences: 145

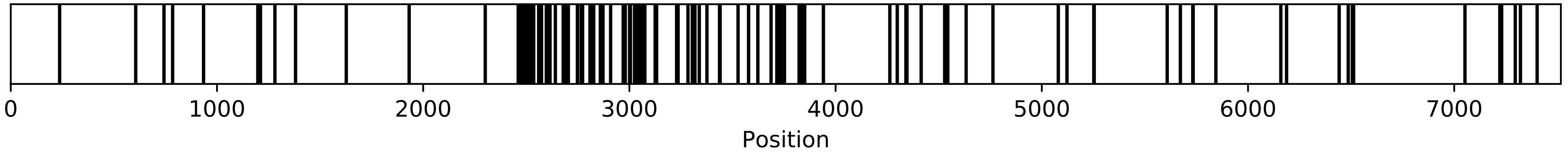

MM1 vs MM2
Number of differences: 84

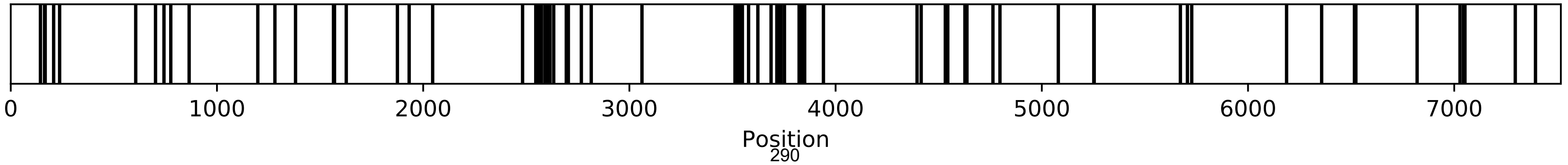

### Region: 988



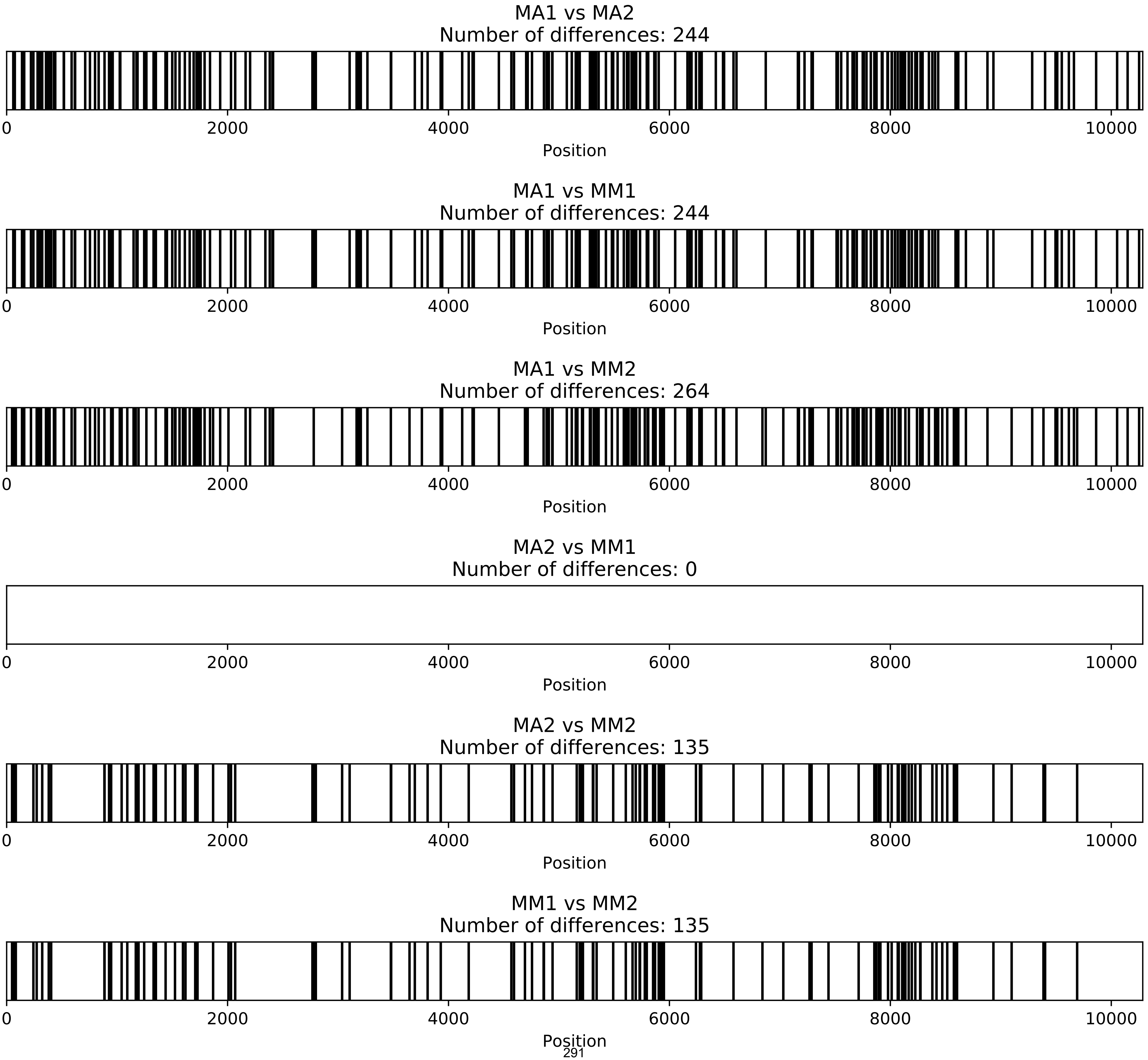
MA1 vs MA2
Number of differences: 244
MA1 vs MM1
Number of differences: 244
MA1 vs MM2
Number of differences: 264
MA2 vs MM1
Number of differences: 0
MA2 vs MM2
Number of differences: 135
MM1 vs MM2
Number of differences: 135
0
2000
4000
6000
8000
10000
Position

### Region: 8



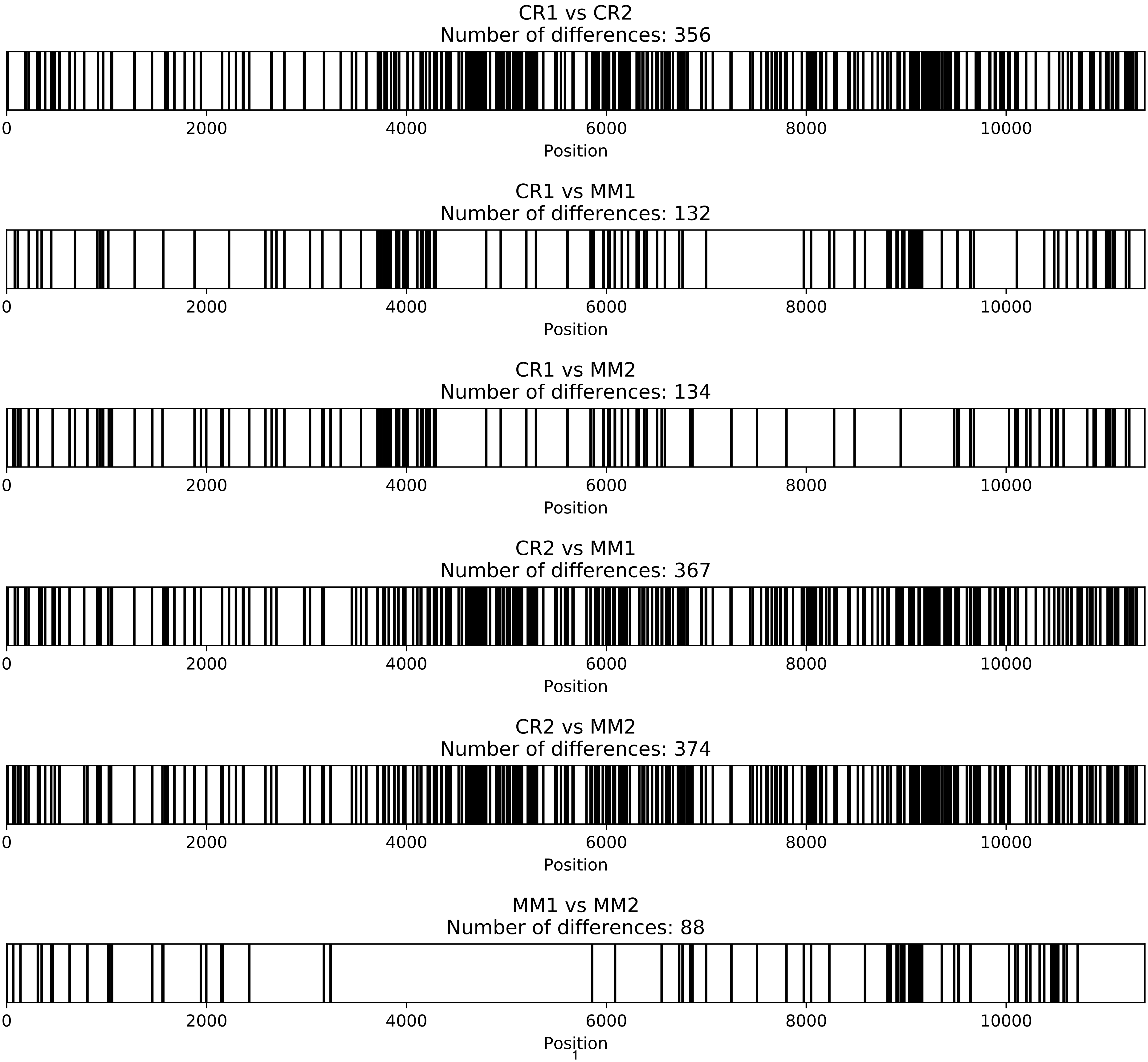
CR1 vs CR2
Number of differences: 356
CR1 vs MM1
Number of differences: 132
CR1 vs MM2
Number of differences: 134
CR2 vs MM1
Number of differences: 367
CR2 vs MM2
Number of differences: 374
MM1 vs MM2
Number of differences: 88
0
2000
4000
6000
8000
10000
Position

### Region: 16



## CR1 vs CR2
Number of differences: 333

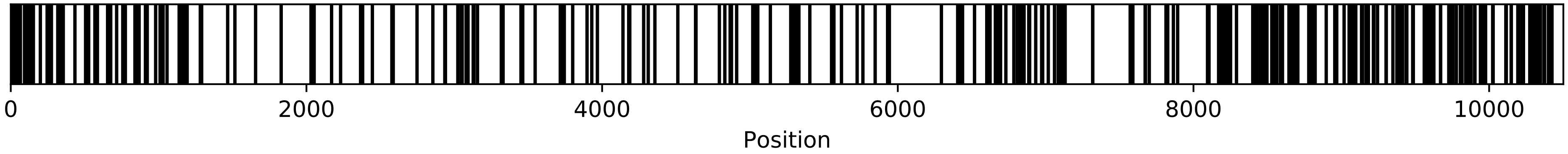


## CR1 vs MM1
Number of differences: 129

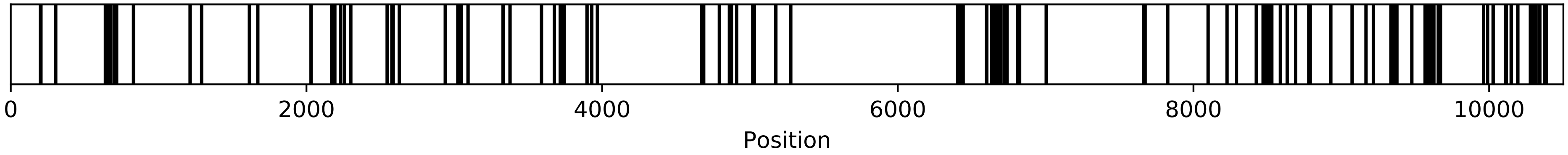


## CR1 vs MM2
Number of differences: 159

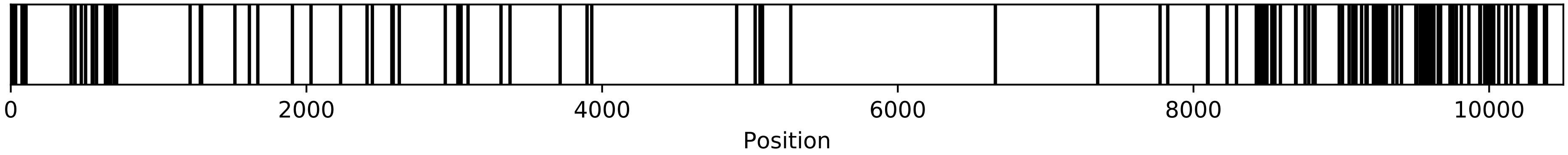


## CR2 vs MM1
Number of differences: 319

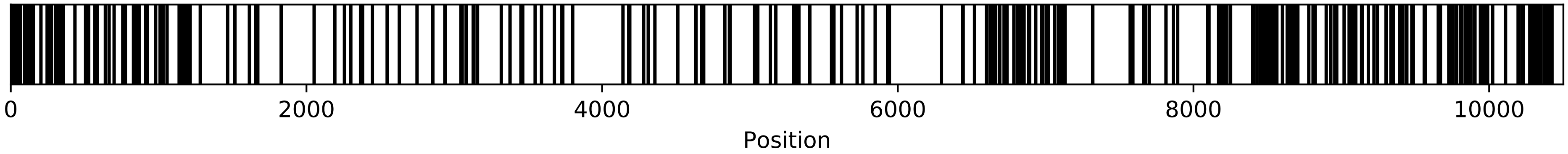


## CR2 vs MM2
Number of differences: 322

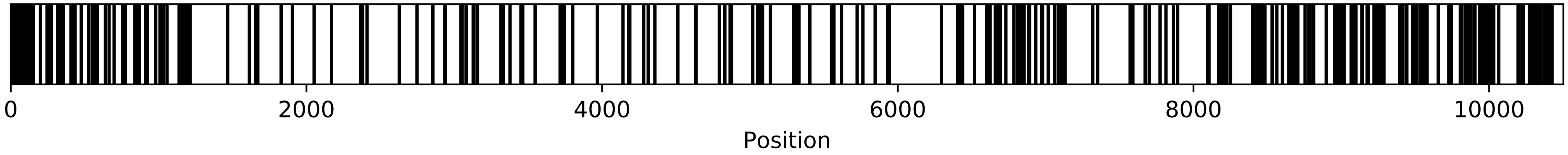


## MM1 vs MM2
Number of differences: 153

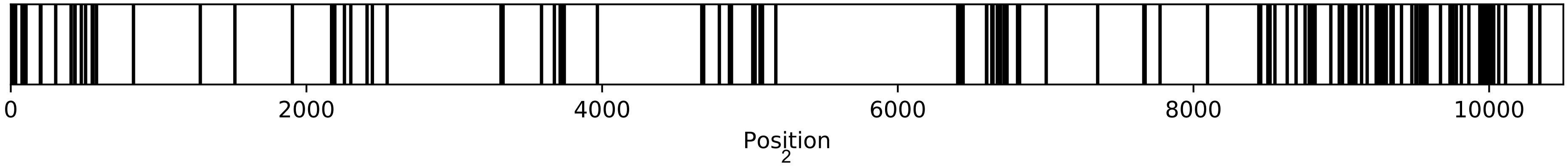

### Region: 18



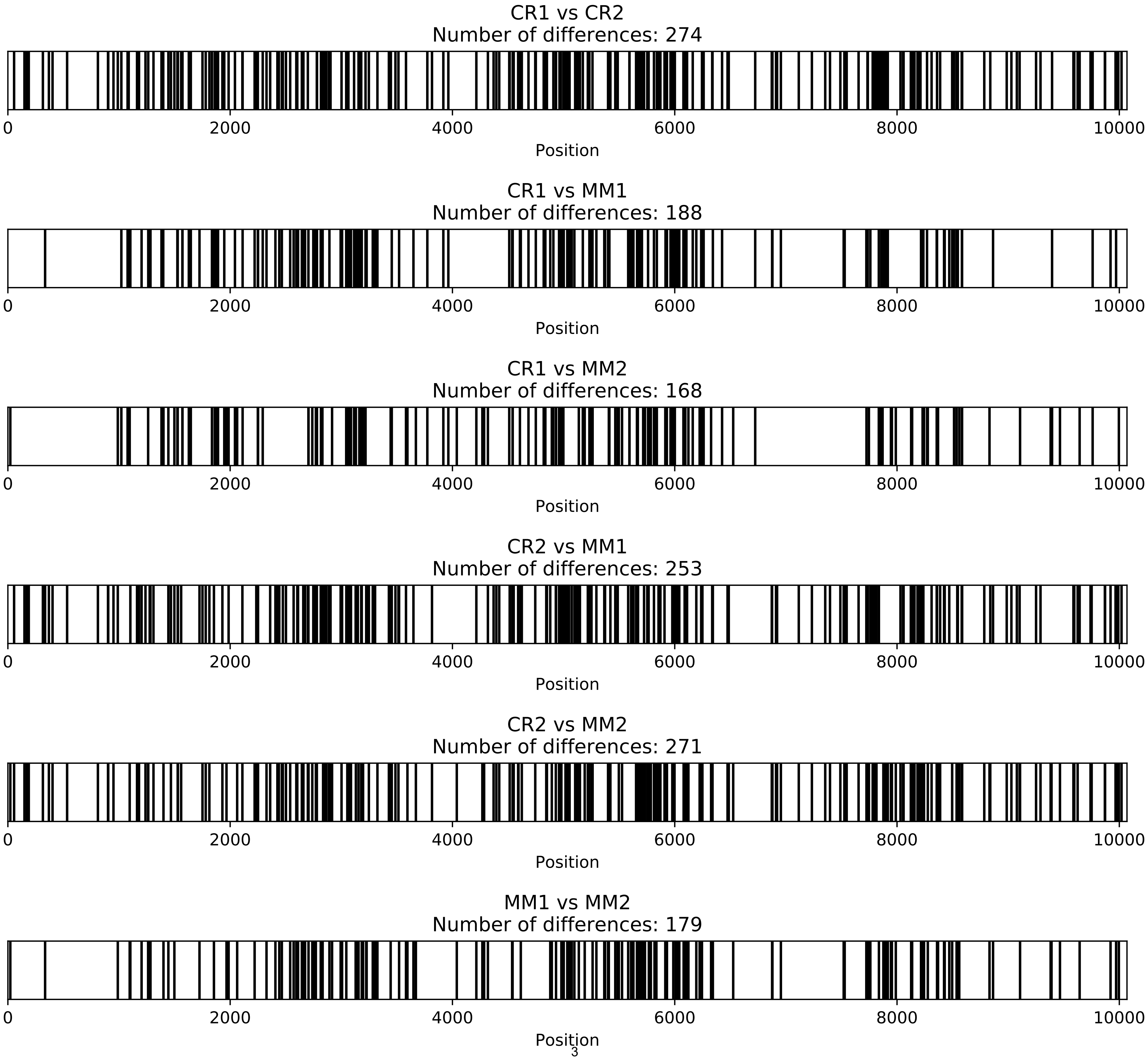
CR1 vs CR2
Number of differences: 274
CR1 vs MM1
Number of differences: 188
CR1 vs MM2
Number of differences: 168
CR2 vs MM1
Number of differences: 253
CR2 vs MM2
Number of differences: 271
MM1 vs MM2
Number of differences: 179
0
2000
4000
6000
8000
10000
Position

### Region: 20



## CR1 vs CR2
Number of differences: 143

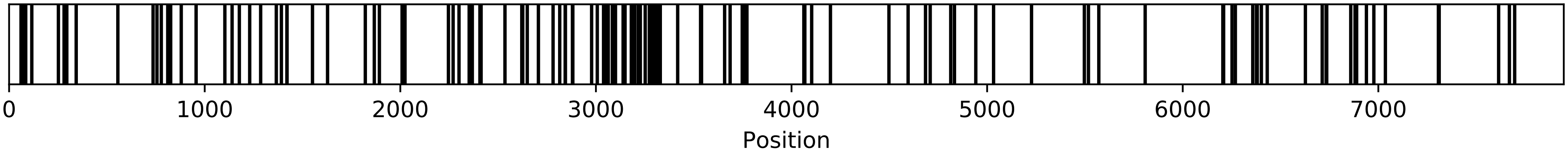


## CR1 vs MM1
Number of differences: 49

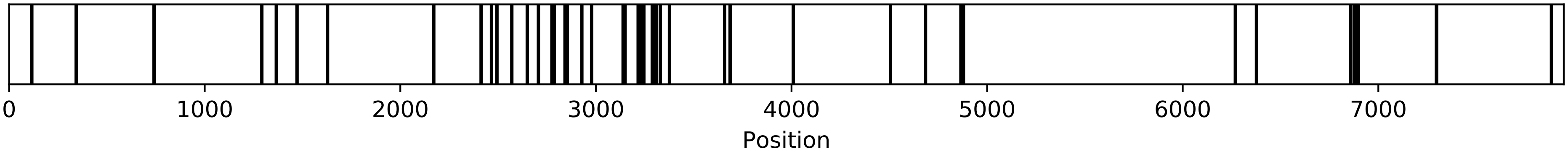


## CR1 vs MM2
Number of differences: 66

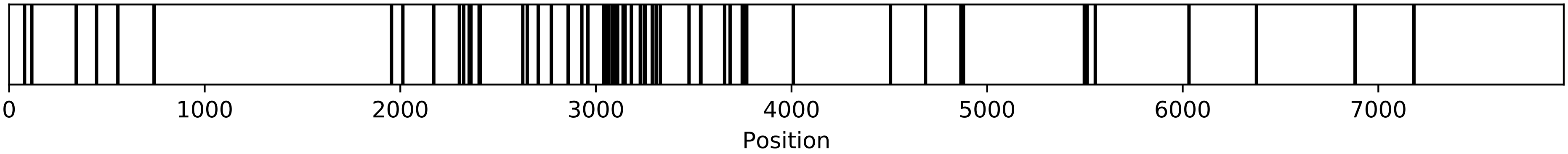


## CR2 vs MM1
Number of differences: 140

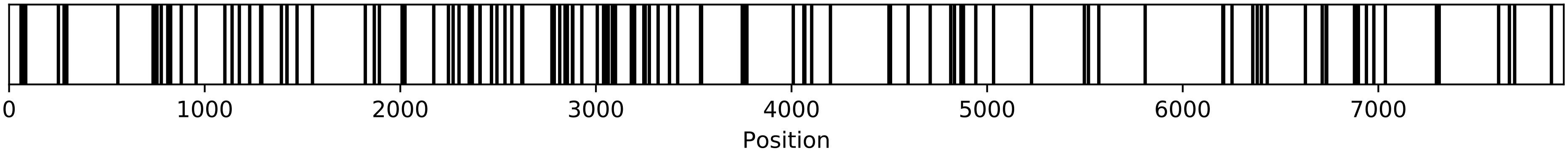


## CR2 vs MM2
Number of differences: 141

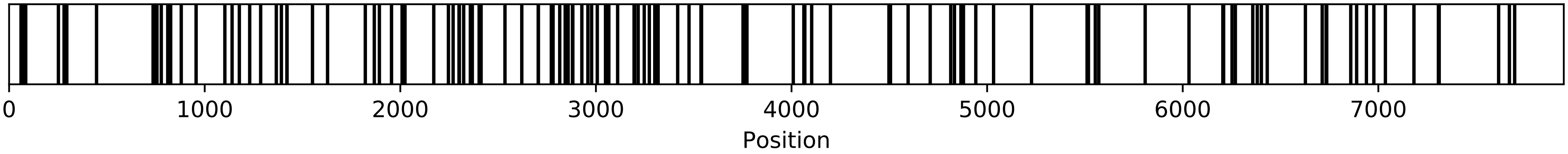


## MM1 vs MM2
Number of differences: 71

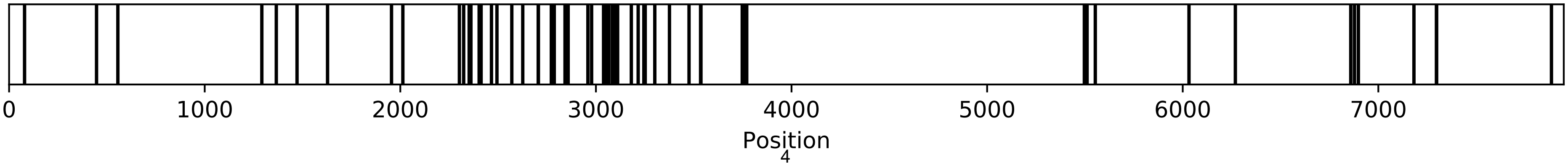

### Region: 22



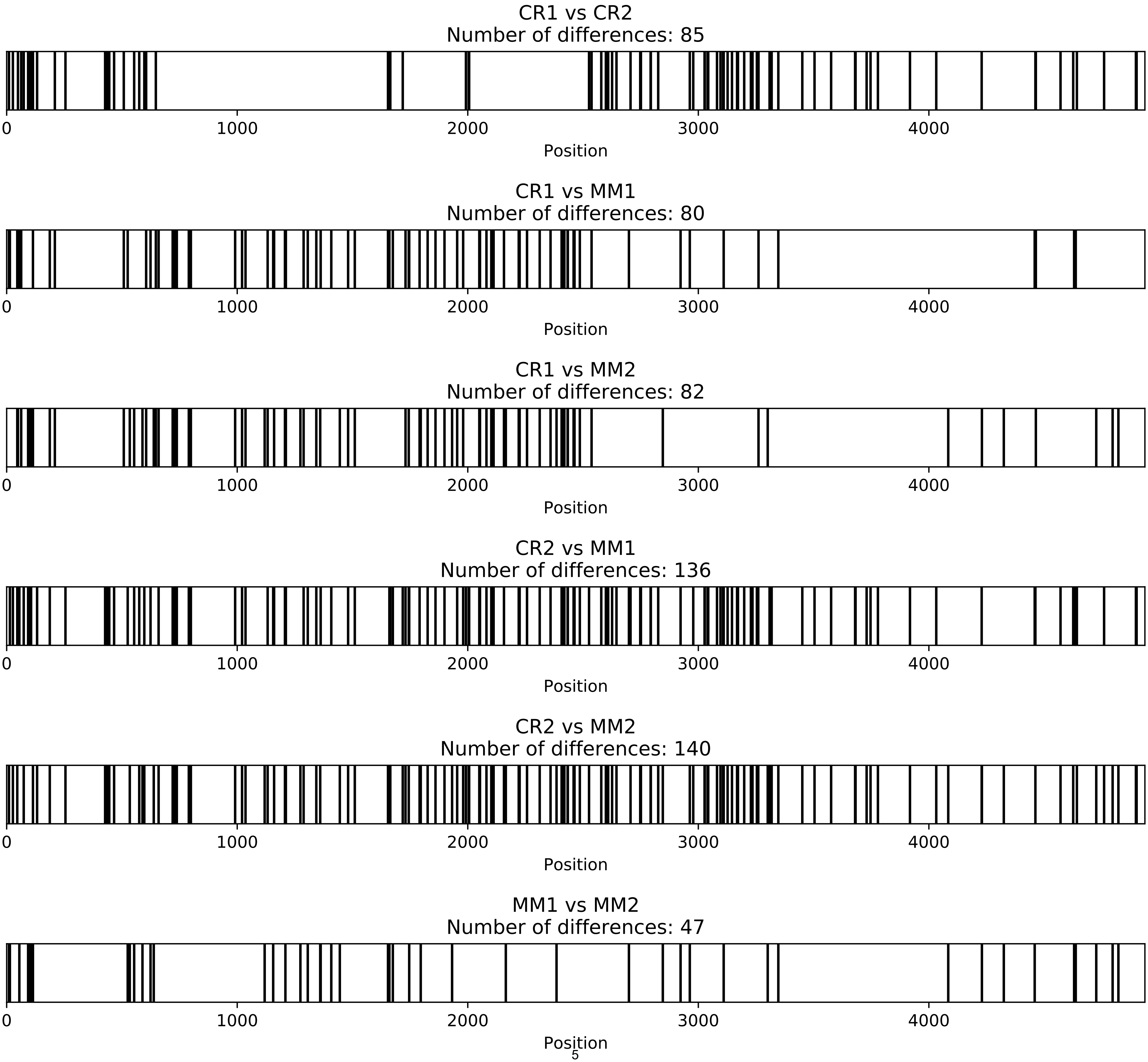
CR1 vs CR2
Number of differences: 85
CR1 vs MM1
Number of differences: 80
CR1 vs MM2
Number of differences: 82
CR2 vs MM1
Number of differences: 136
CR2 vs MM2
Number of differences: 140
MM1 vs MM2
Number of differences: 47
0
1000
2000
3000
4000
Position

### Region: 26



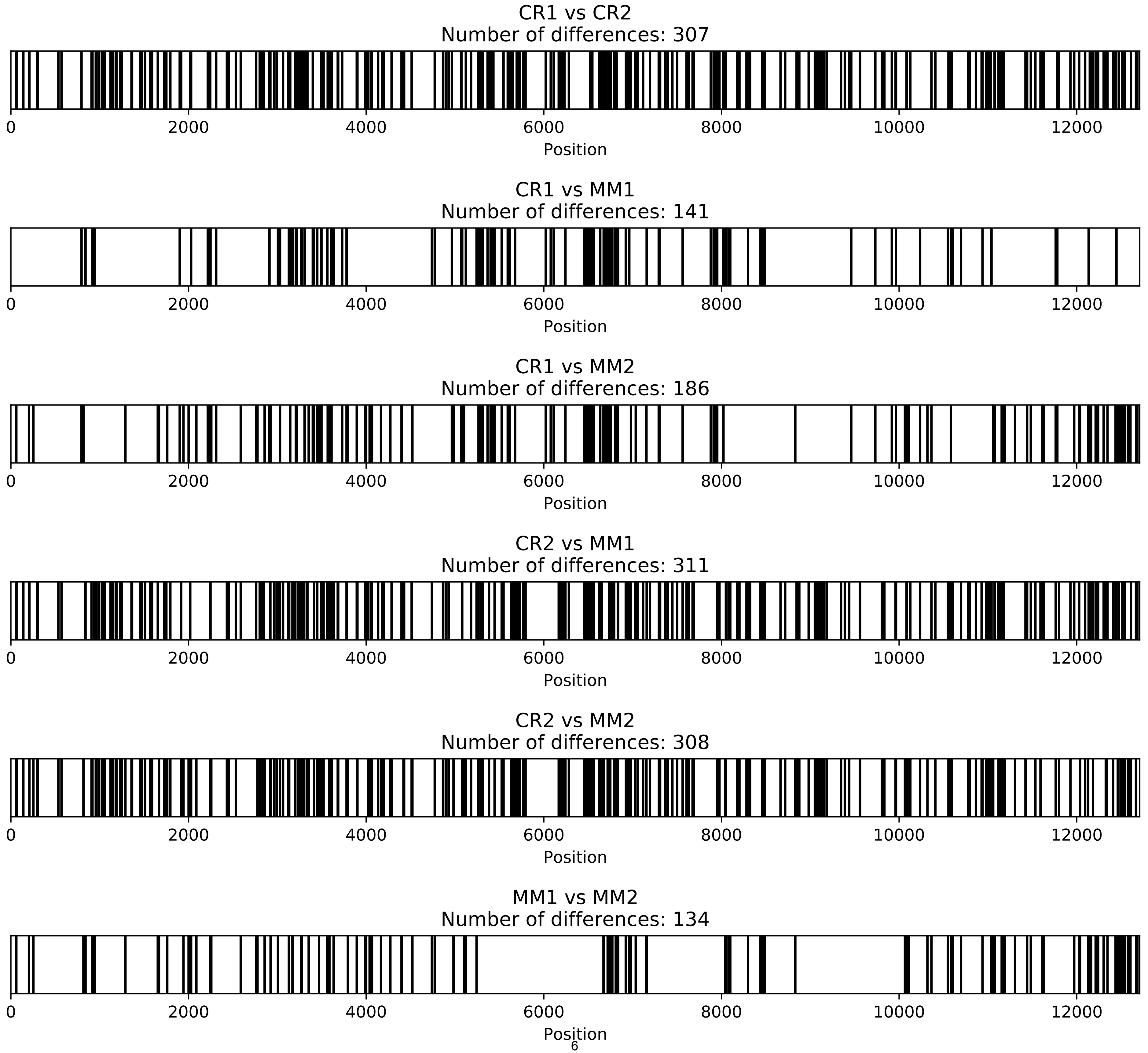
CR1 vs CR2
Number of differences: 307
CR1 vs MM1
Number of differences: 141
CR1 vs MM2
Number of differences: 186
CR2 vs MM1
Number of differences: 311
CR2 vs MM2
Number of differences: 308
MM1 vs MM2
Number of differences: 134
0
2000
4000
6000
8000
10000
12000
Position

### Region: 28



## CR1 vs CR2
Number of differences: 15

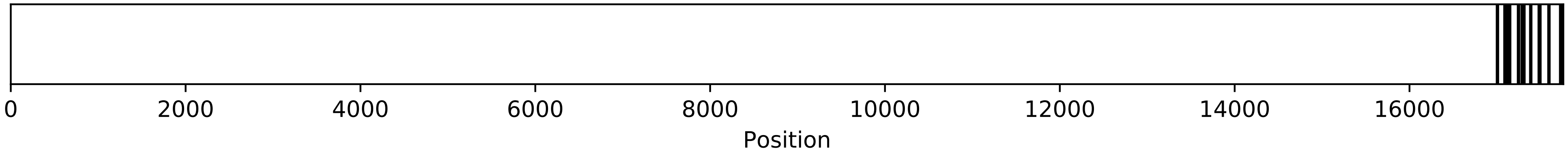


## CR1 vs MM1
Number of differences: 129

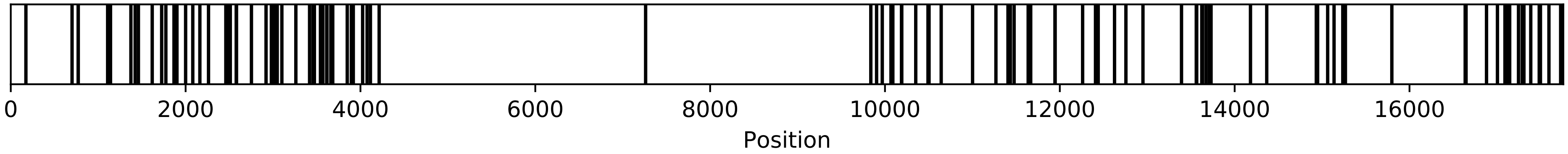


## CR1 vs MM2
Number of differences: 168

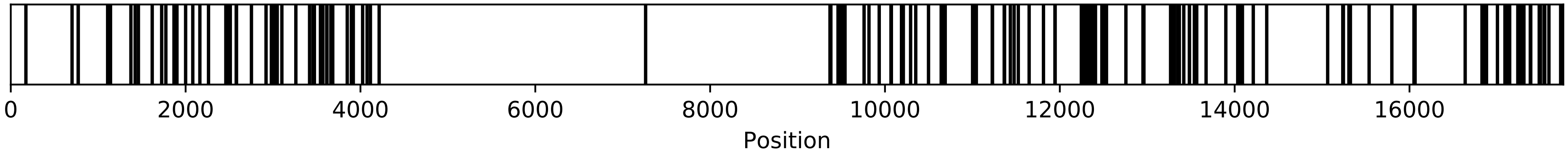


## CR2 vs MM1
Number of differences: 116

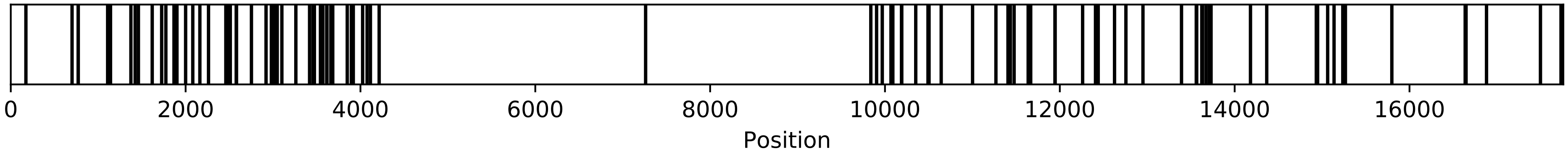


## CR2 vs MM2
Number of differences: 155

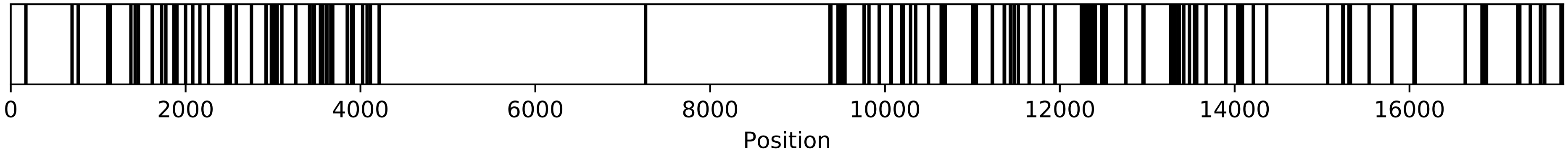


## MM1 vs MM2
Number of differences: 105

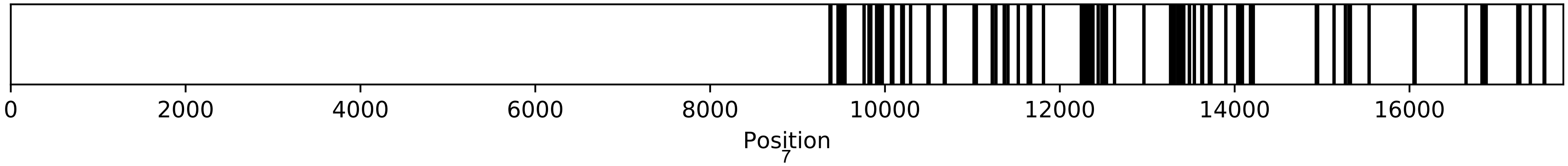

### Region: 29



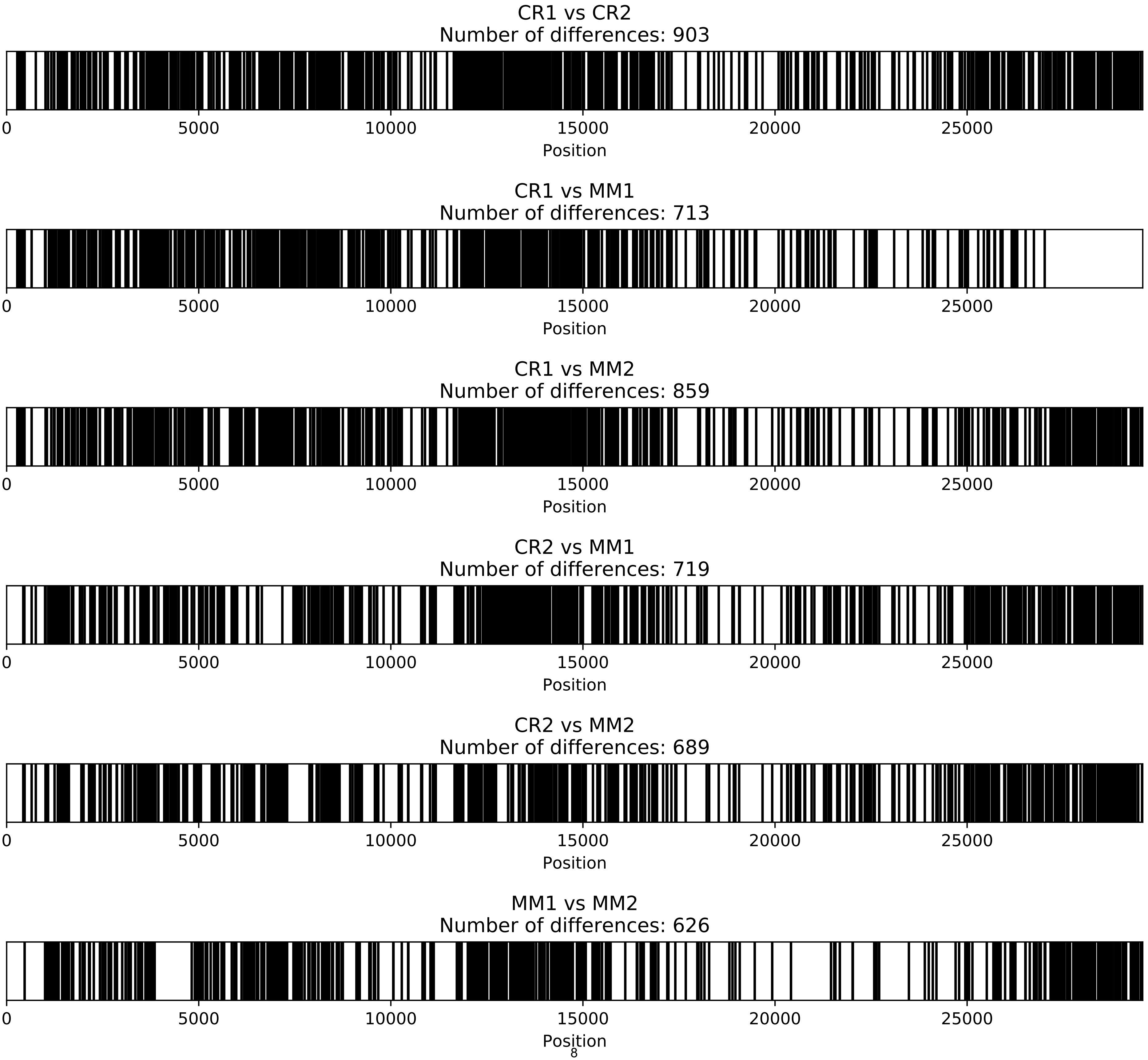
CR1 vs CR2
Number of differences: 903
0 5000 10000 15000 20000 25000
Position
CR1 vs MM1
Number of differences: 713
0 5000 10000 15000 20000 25000
Position
CR1 vs MM2
Number of differences: 859
0 5000 10000 15000 20000 25000
Position
CR2 vs MM1
Number of differences: 719
0 5000 10000 15000 20000 25000
Position
CR2 vs MM2
Number of differences: 689
0 5000 10000 15000 20000 25000
Position
MM1 vs MM2
Number of differences: 626
0 5000 10000 15000 20000 25000
Position

### Region: 32



CR1 vs CR2
Number of differences: 167

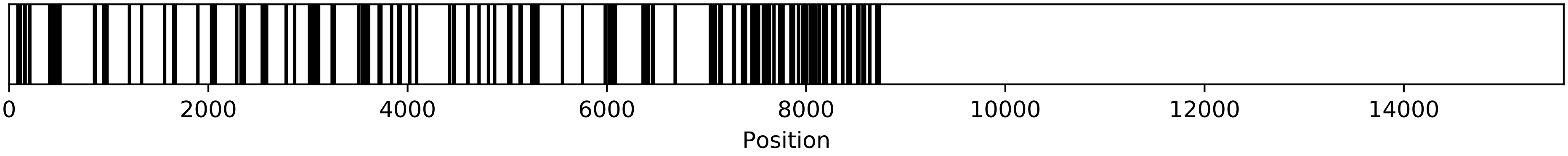


CR1 vs MM1
Number of differences: 187

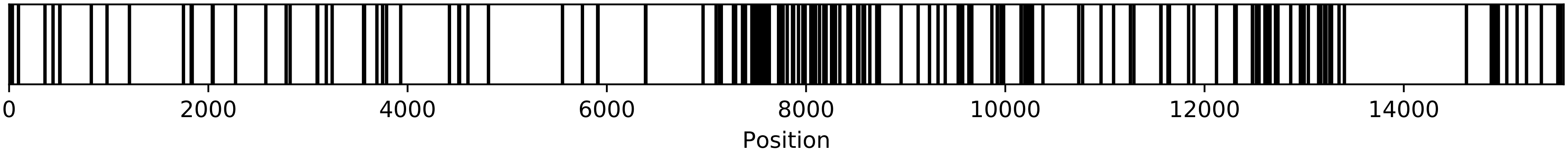


CR1 vs MM2
Number of differences: 197

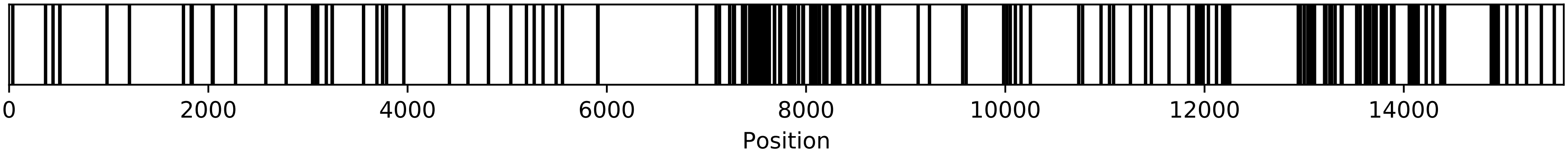


CR2 vs MM1
Number of differences: 222

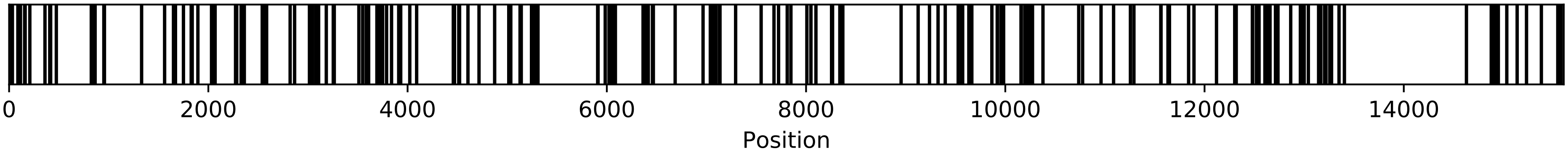


CR2 vs MM2
Number of differences: 235

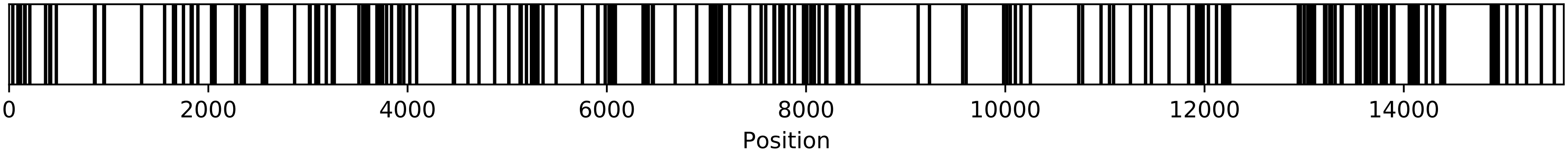


MM1 vs MM2
Number of differences: 185

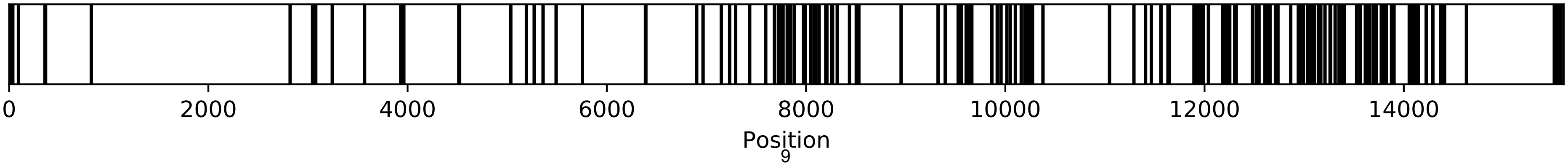

### Region: 33



## CR1 vs CR2
Number of differences: 456

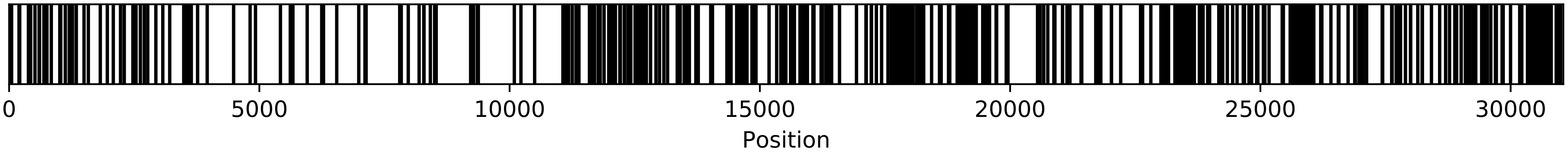


## CR1 vs MM1
Number of differences: 201

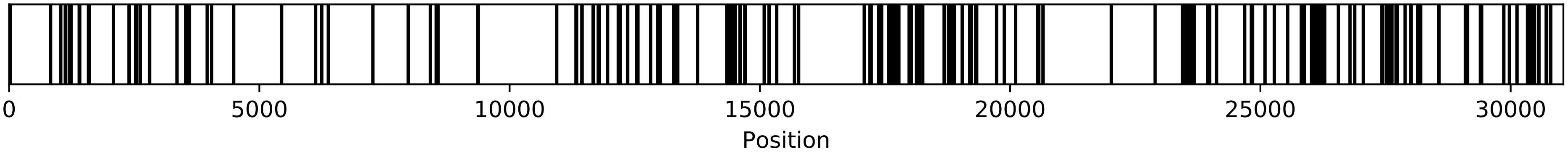


## CR1 vs MM2
Number of differences: 195

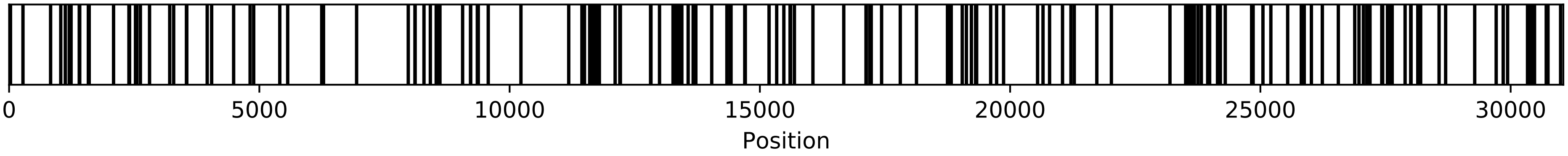


## CR2 vs MM1
Number of differences: 489

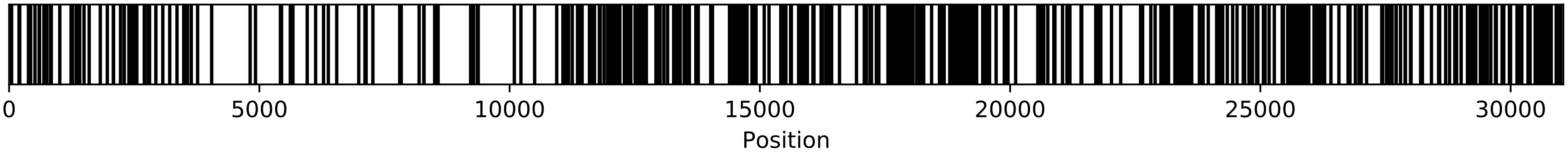


## CR2 vs MM2
Number of differences: 465

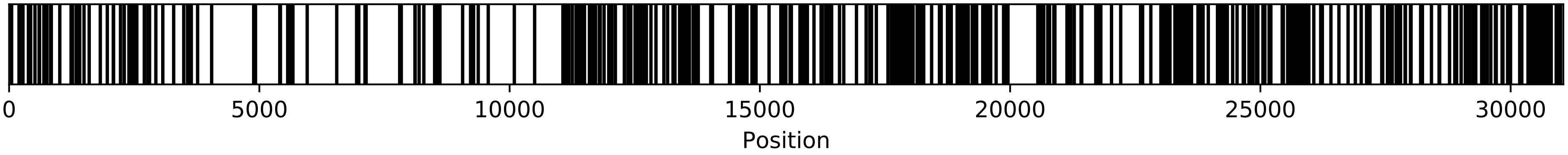


## MM1 vs MM2
Number of differences: 187

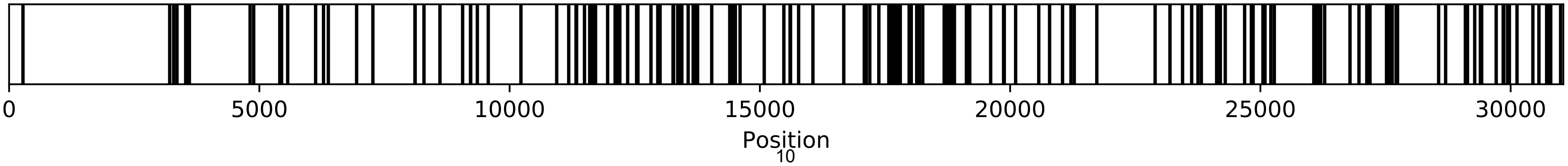

### Region: 34



CR1 vs CR2
Number of differences: 357

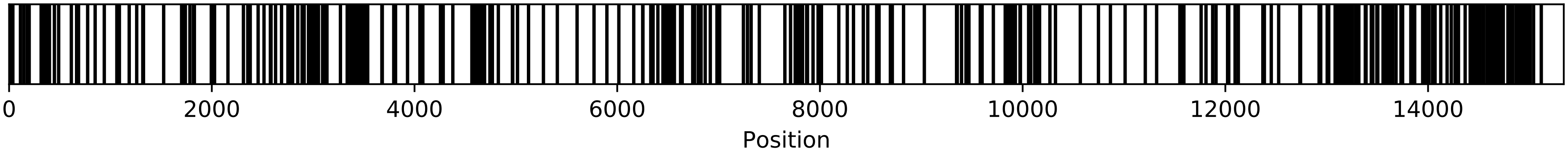


CR1 vs MM1
Number of differences: 228

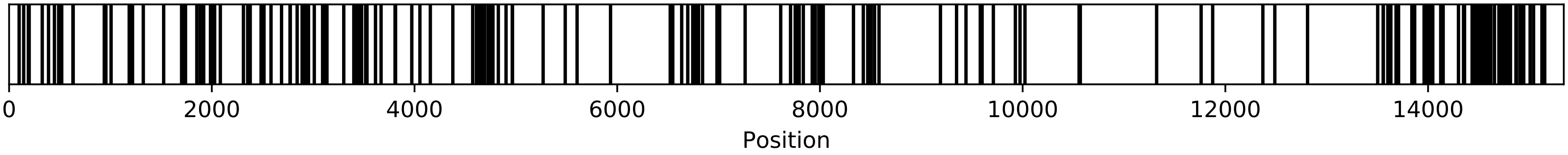


CR1 vs MM2
Number of differences: 238

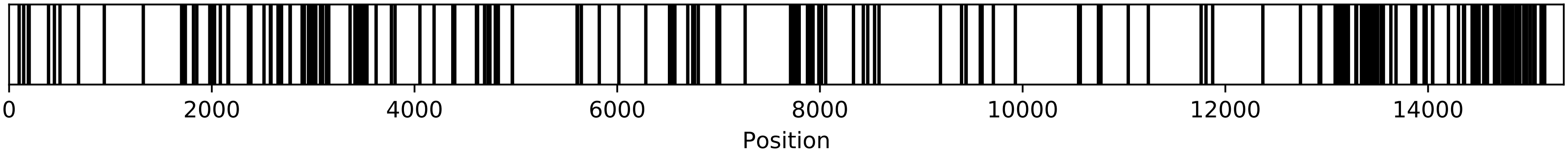


CR2 vs MM1
Number of differences: 340

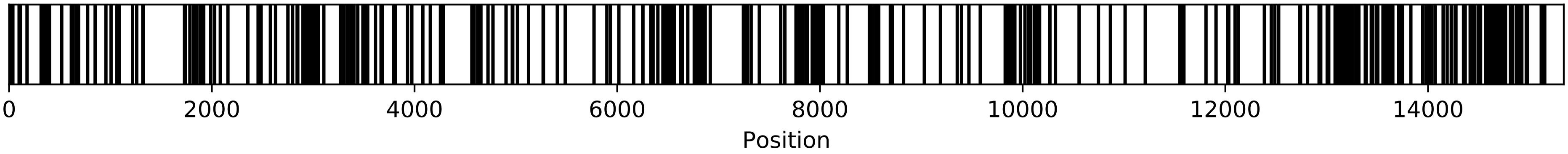


CR2 vs MM2
Number of differences: 326

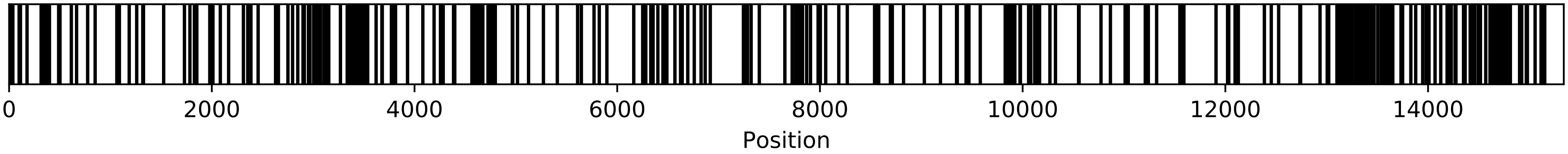


MM1 vs MM2
Number of differences: 187

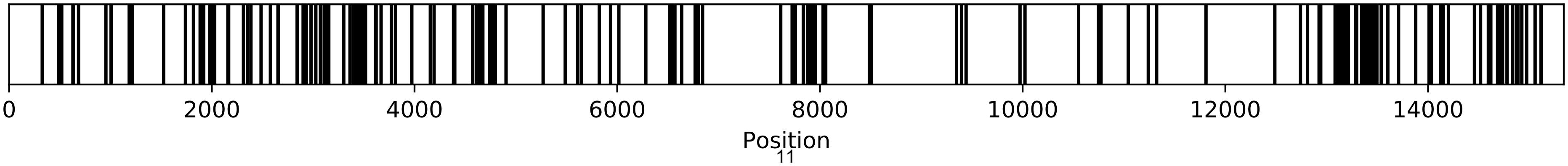

### Region: 35



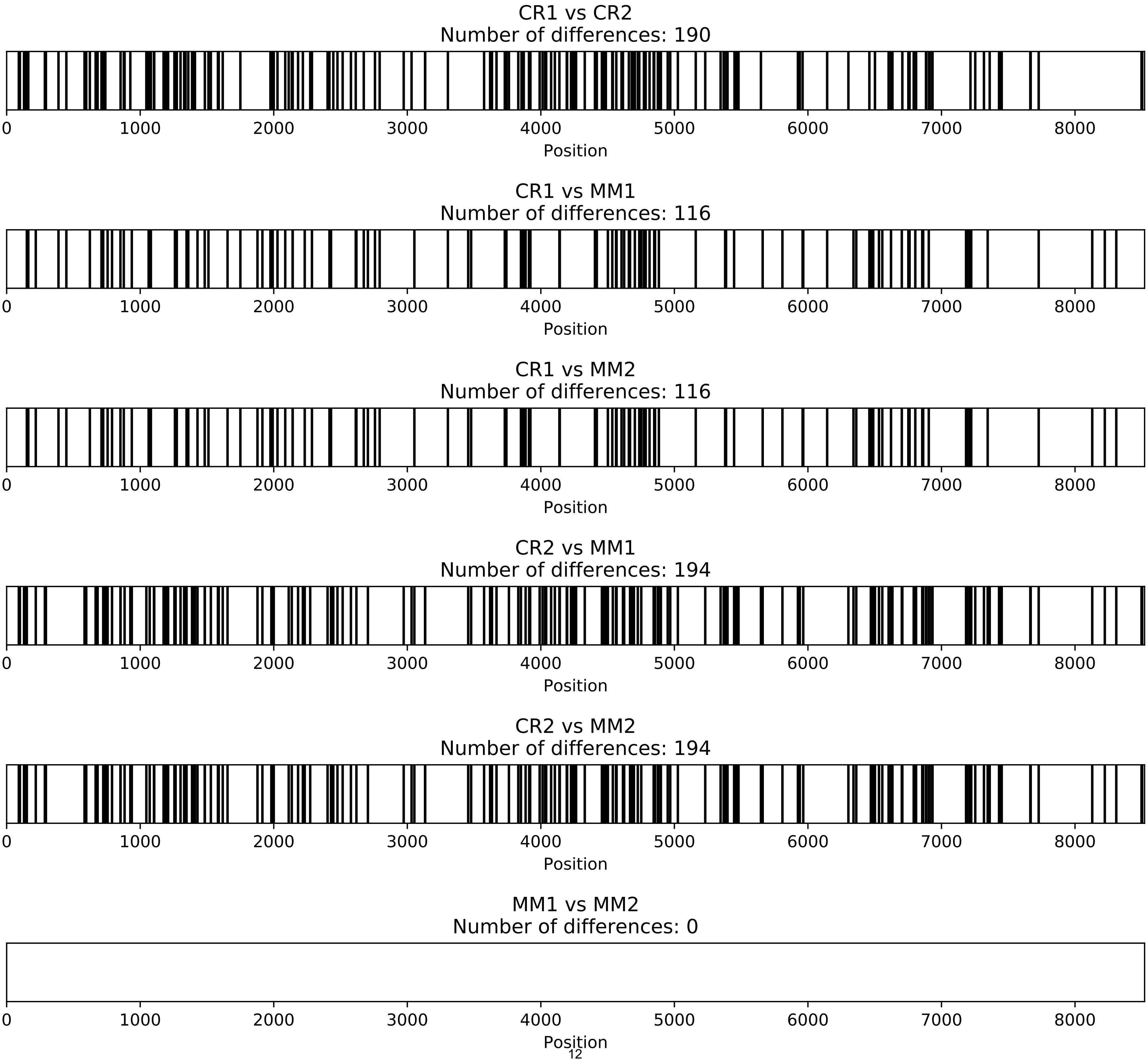
CR1 vs CR2
Number of differences: 190
CR1 vs MM1
Number of differences: 116
CR1 vs MM2
Number of differences: 116
CR2 vs MM1
Number of differences: 194
CR2 vs MM2
Number of differences: 194
MM1 vs MM2
Number of differences: 0
0 1000 2000 3000 4000 5000 6000 7000 8000
Position

### Region: 36



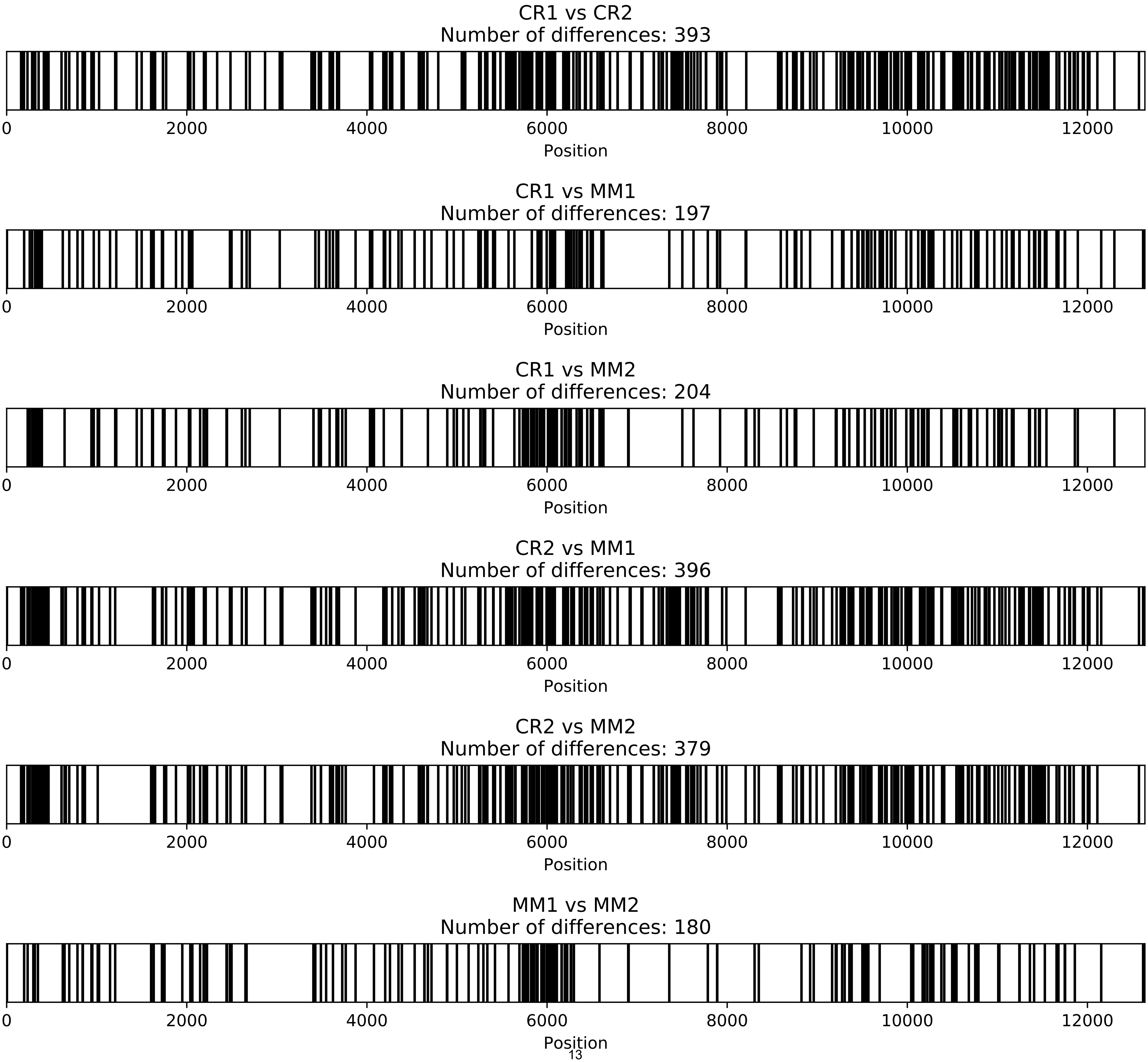
CR1 vs CR2
Number of differences: 393
CR1 vs MM1
Number of differences: 197
CR1 vs MM2
Number of differences: 204
CR2 vs MM1
Number of differences: 396
CR2 vs MM2
Number of differences: 379
MM1 vs MM2
Number of differences: 180
0
2000
4000
6000
8000
10000
12000
Position

### Region: 41



CR1 vs CR2
Number of differences: 276

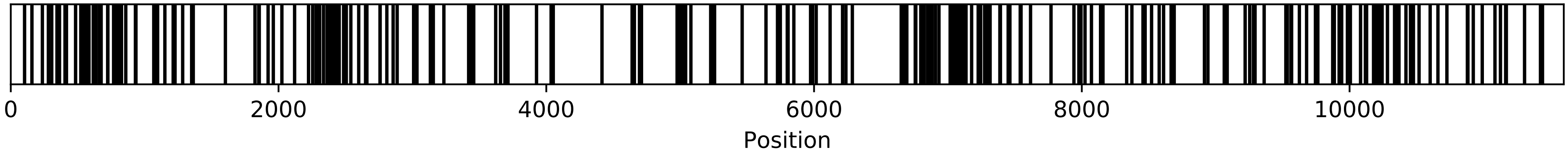


CR1 vs MM1
Number of differences: 164

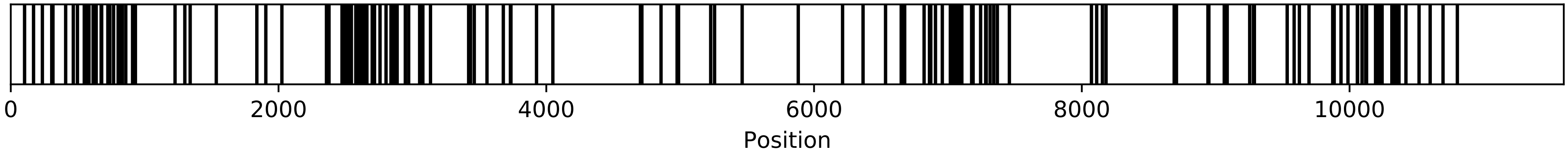


CR1 vs MM2
Number of differences: 165

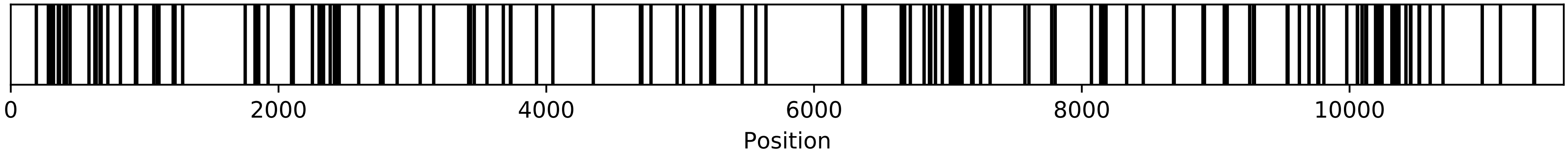


CR2 vs MM1
Number of differences: 274

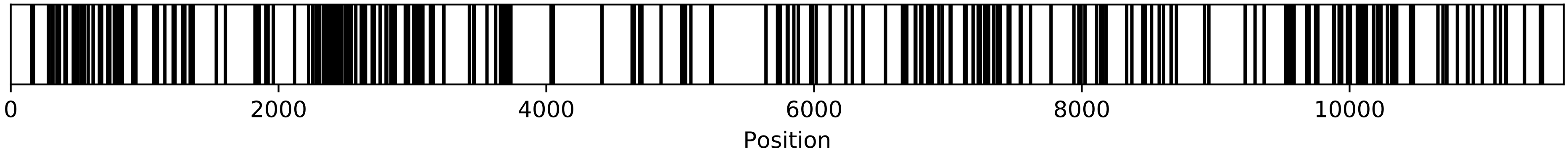


CR2 vs MM2
Number of differences: 261

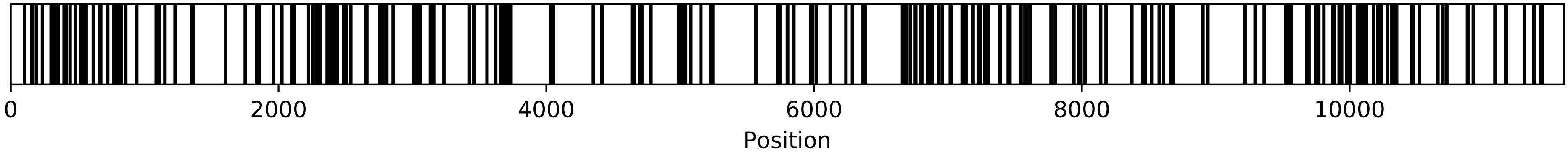


MM1 vs MM2
Number of differences: 156

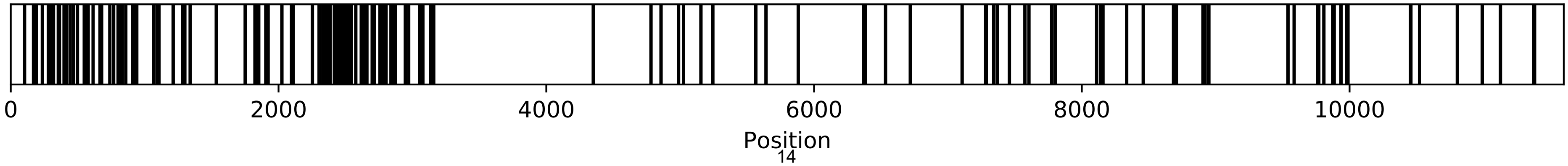

### Region: 46

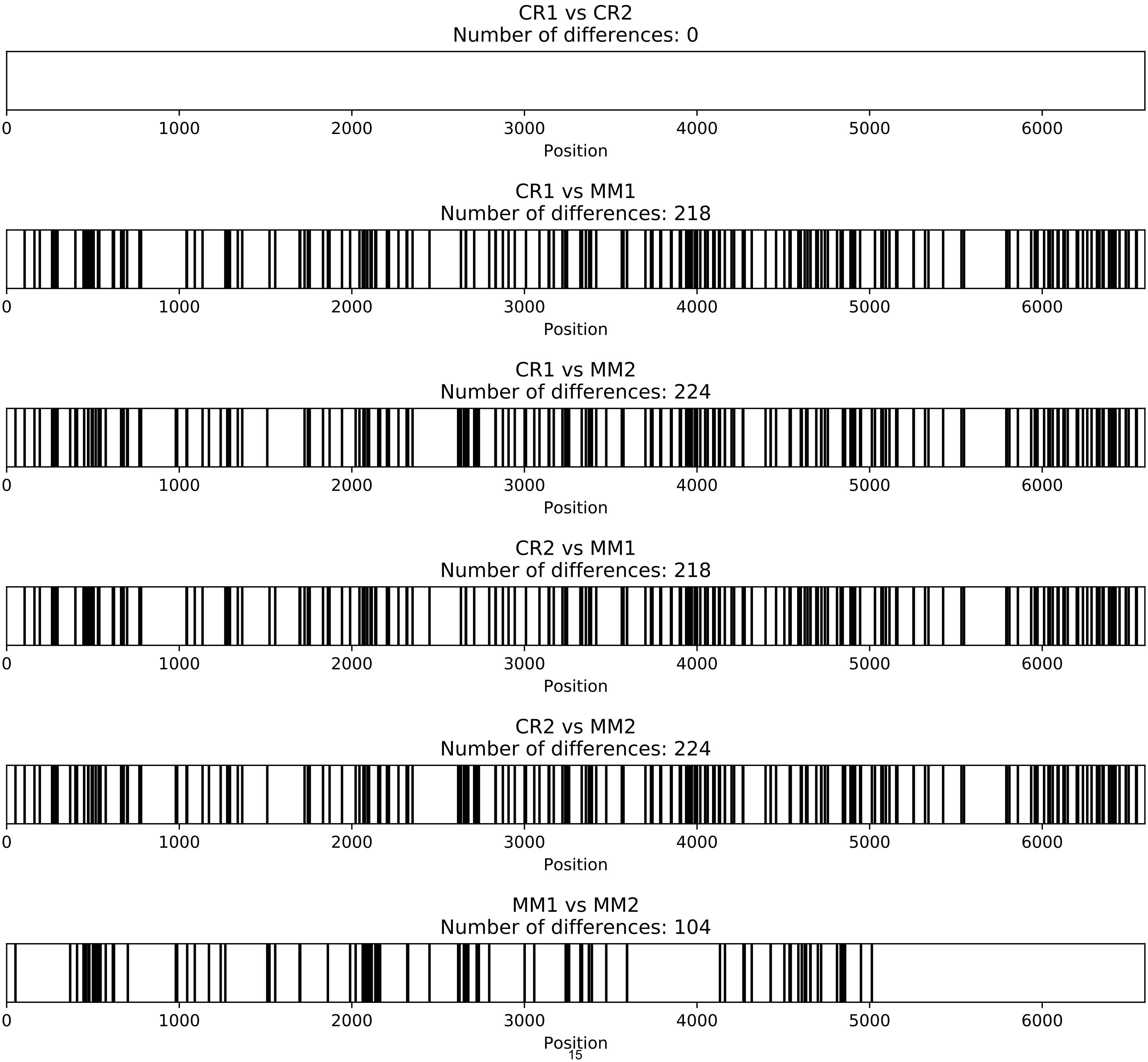
CR1 vs CR2
Number of differences: 0
CR1 vs MM1
Number of differences: 218
CR1 vs MM2
Number of differences: 224
CR2 vs MM1
Number of differences: 218
CR2 vs MM2
Number of differences: 224
MM1 vs MM2
Number of differences: 104
0
1000
2000
3000
4000
5000
6000
Position

### Region: 48



CR1 vs CR2
Number of differences: 142

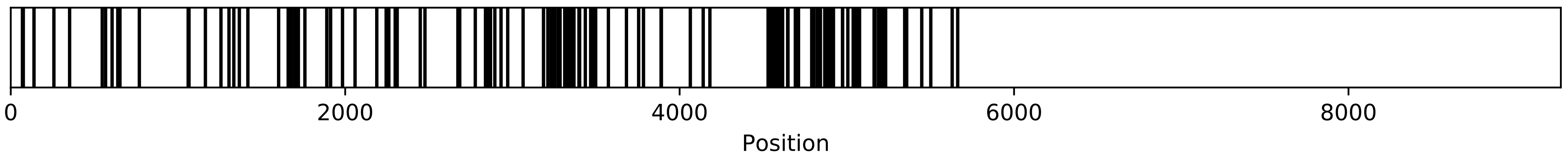


CR1 vs MM1
Number of differences: 223

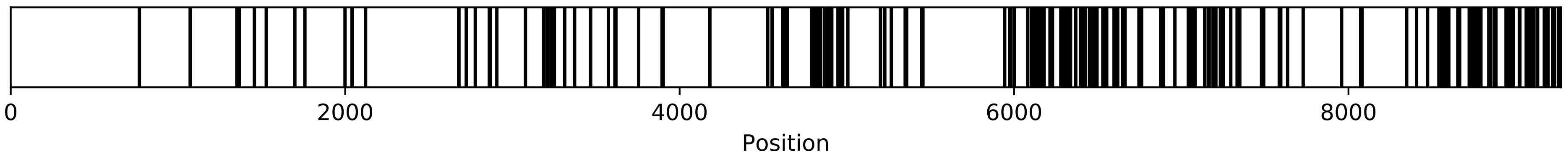


CR1 vs MM2
Number of differences: 232

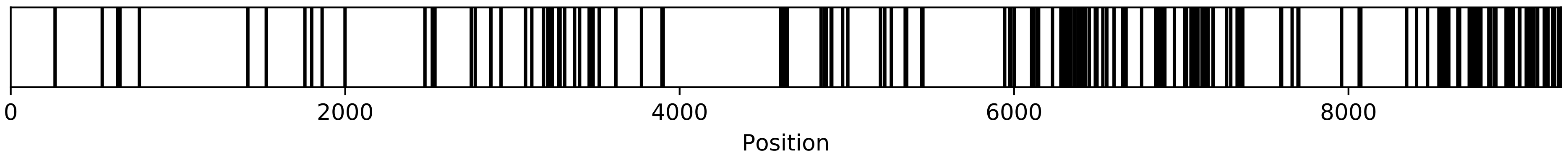


CR2 vs MM1
Number of differences: 297

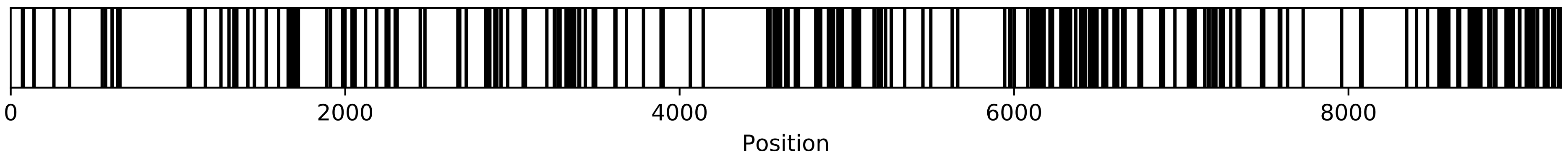


CR2 vs MM2
Number of differences: 308

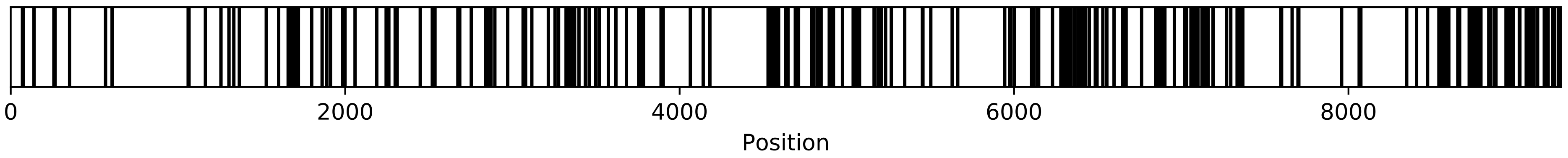


MM1 vs MM2
Number of differences: 177

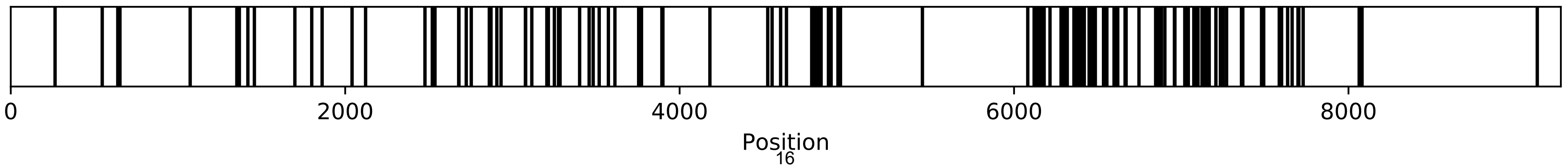

### Region: 49



CR1 vs CR2
Number of differences: 275

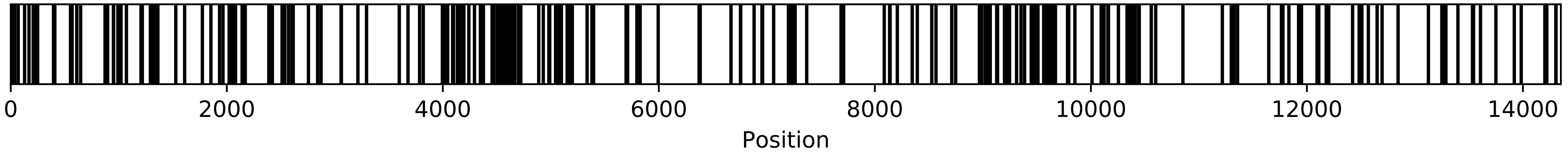


CR1 vs MM1
Number of differences: 121

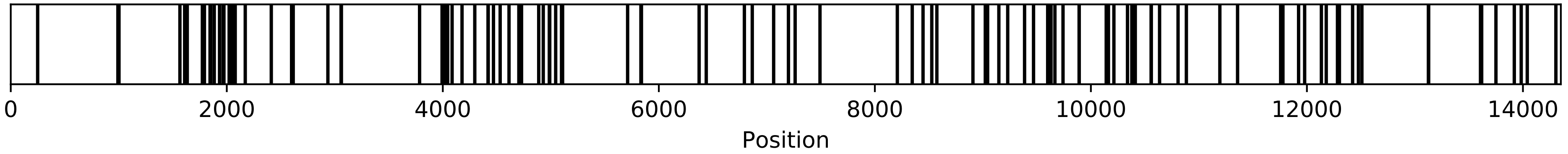


CR1 vs MM2
Number of differences: 135

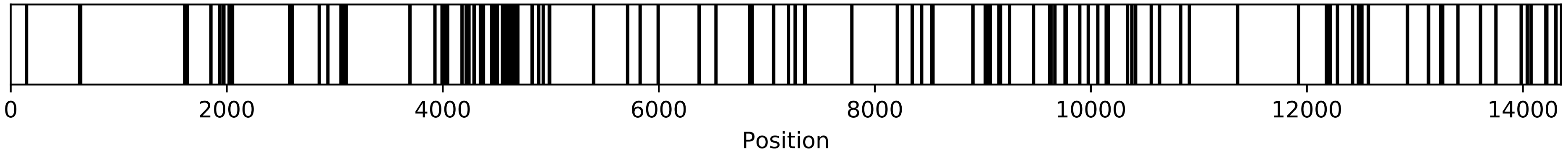


CR2 vs MM1
Number of differences: 272

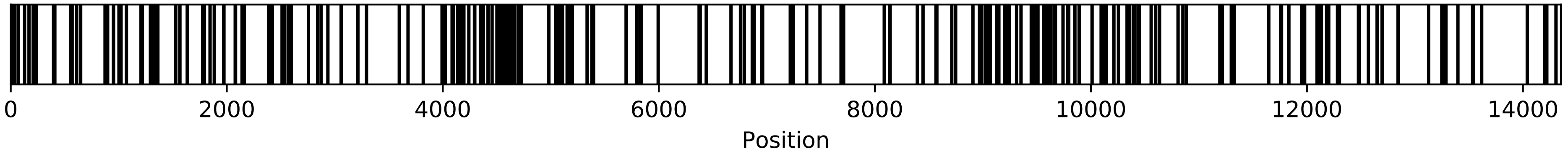


CR2 vs MM2
Number of differences: 264

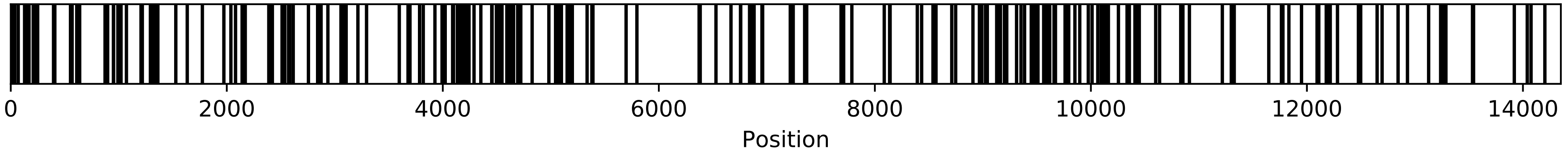


MM1 vs MM2
Number of differences: 134

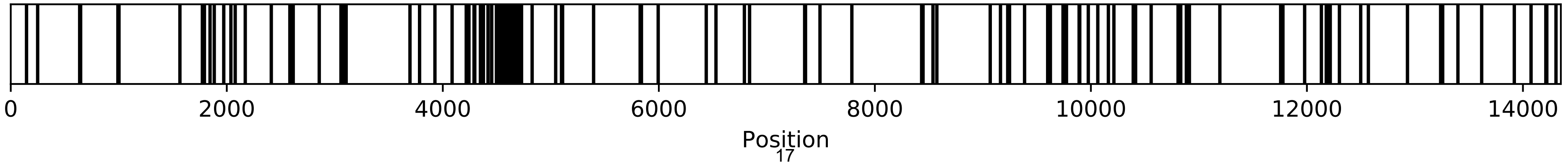

### Region: 52



CR1 vs CR2
Number of differences: 66

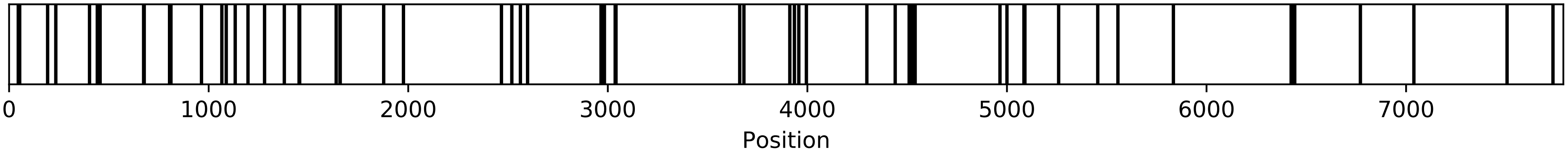


CR1 vs MM1
Number of differences: 35

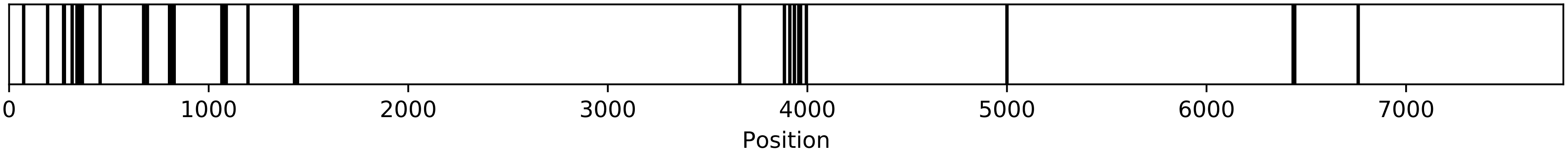


CR1 vs MM2
Number of differences: 34

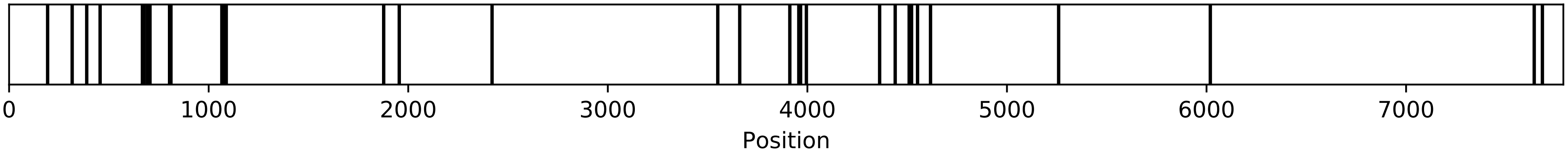


CR2 vs MM1
Number of differences: 65

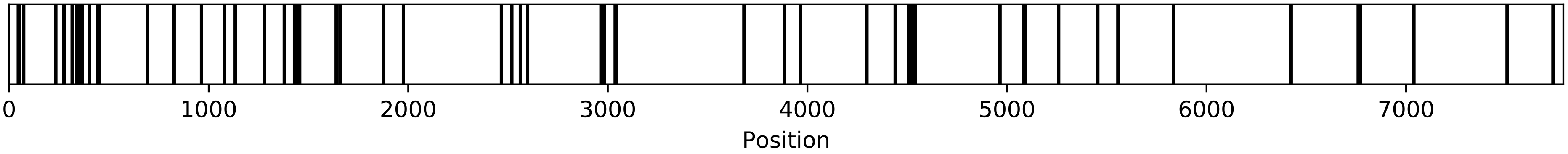


CR2 vs MM2
Number of differences: 66

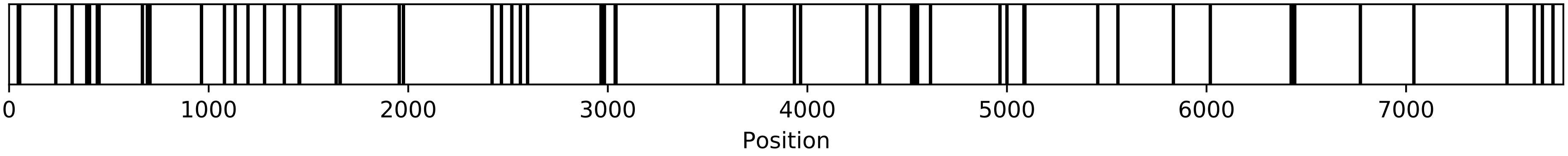


MM1 vs MM2
Number of differences: 35

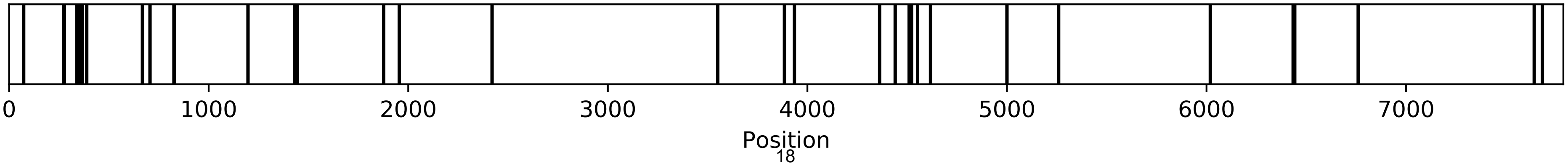

### Region: 58



CR1 vs CR2
Number of differences: 204

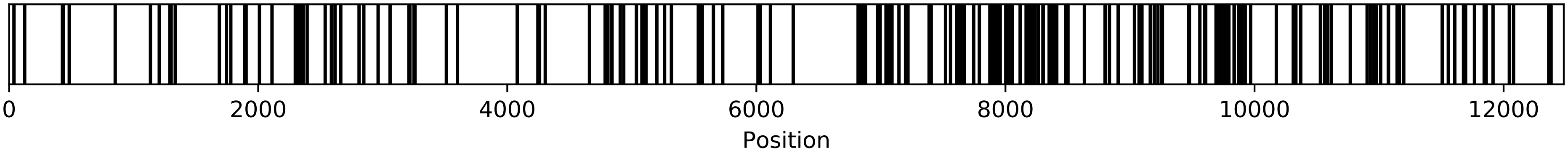


CR1 vs MM1
Number of differences: 114

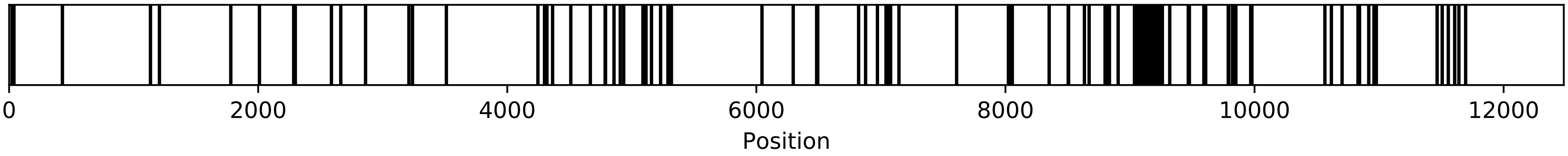


CR1 vs MM2
Number of differences: 119

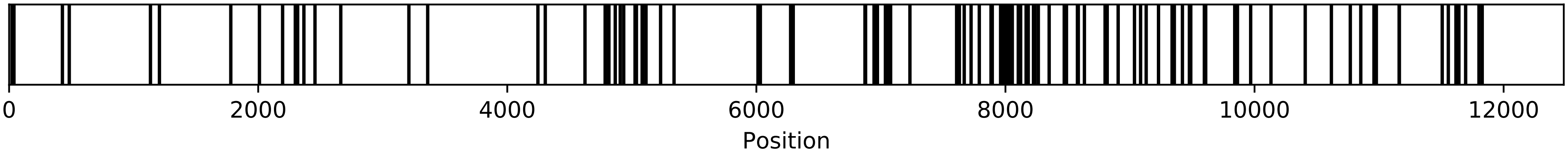


CR2 vs MM1
Number of differences: 204

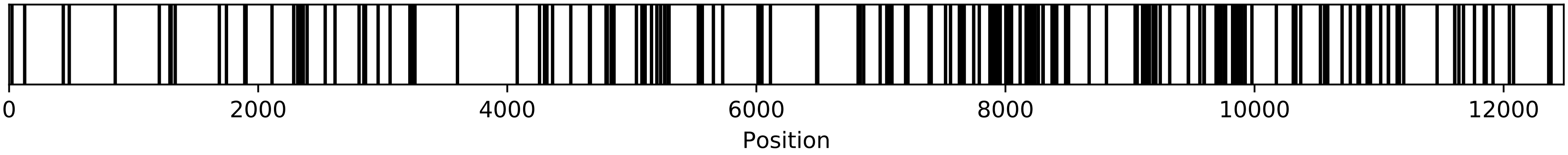


CR2 vs MM2
Number of differences: 192

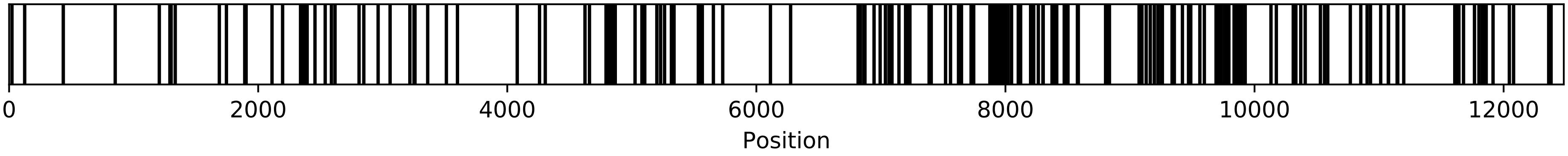


MM1 vs MM2
Number of differences: 136

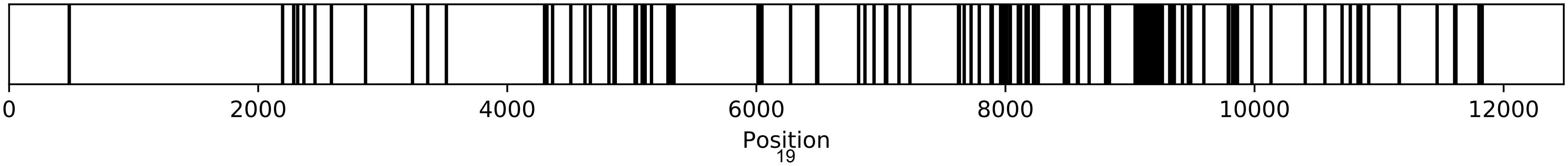

### Region: 59



CR1 vs CR2
Number of differences: 273

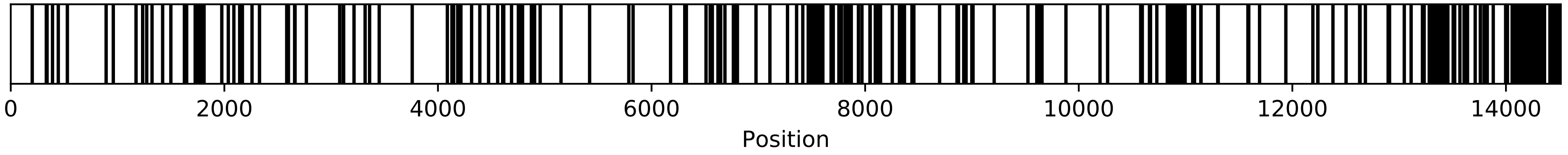


CR1 vs MM1
Number of differences: 110

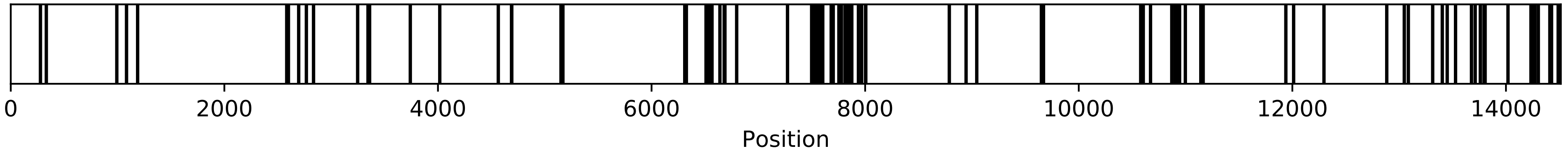


CR1 vs MM2
Number of differences: 140

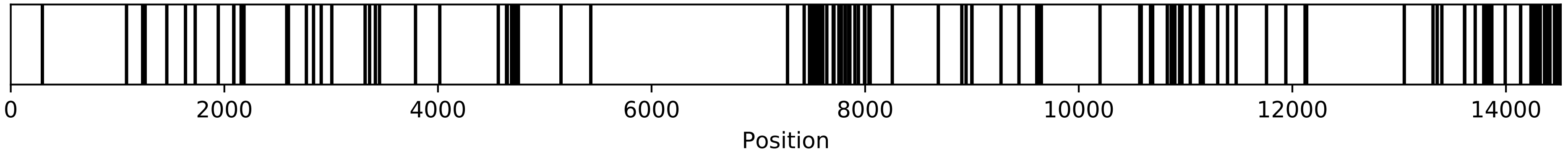


CR2 vs MM1
Number of differences: 258

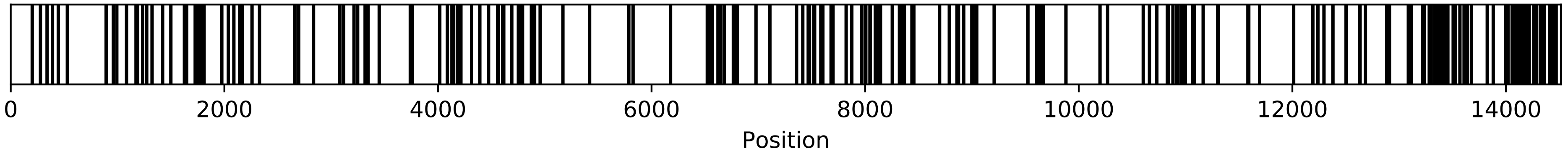


CR2 vs MM2
Number of differences: 286

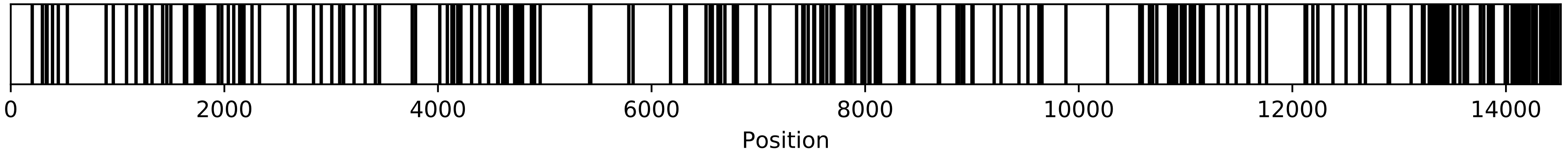


MM1 vs MM2
Number of differences: 150

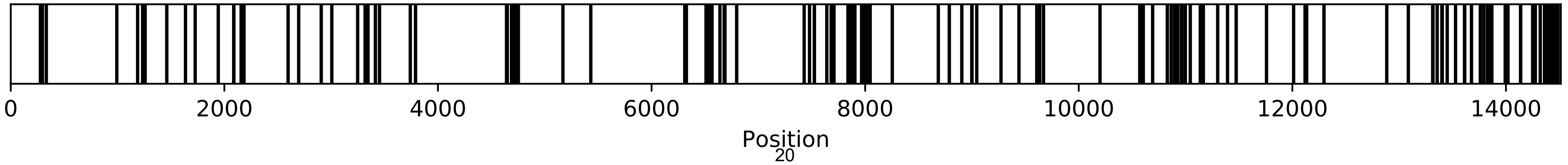

### Region: 60



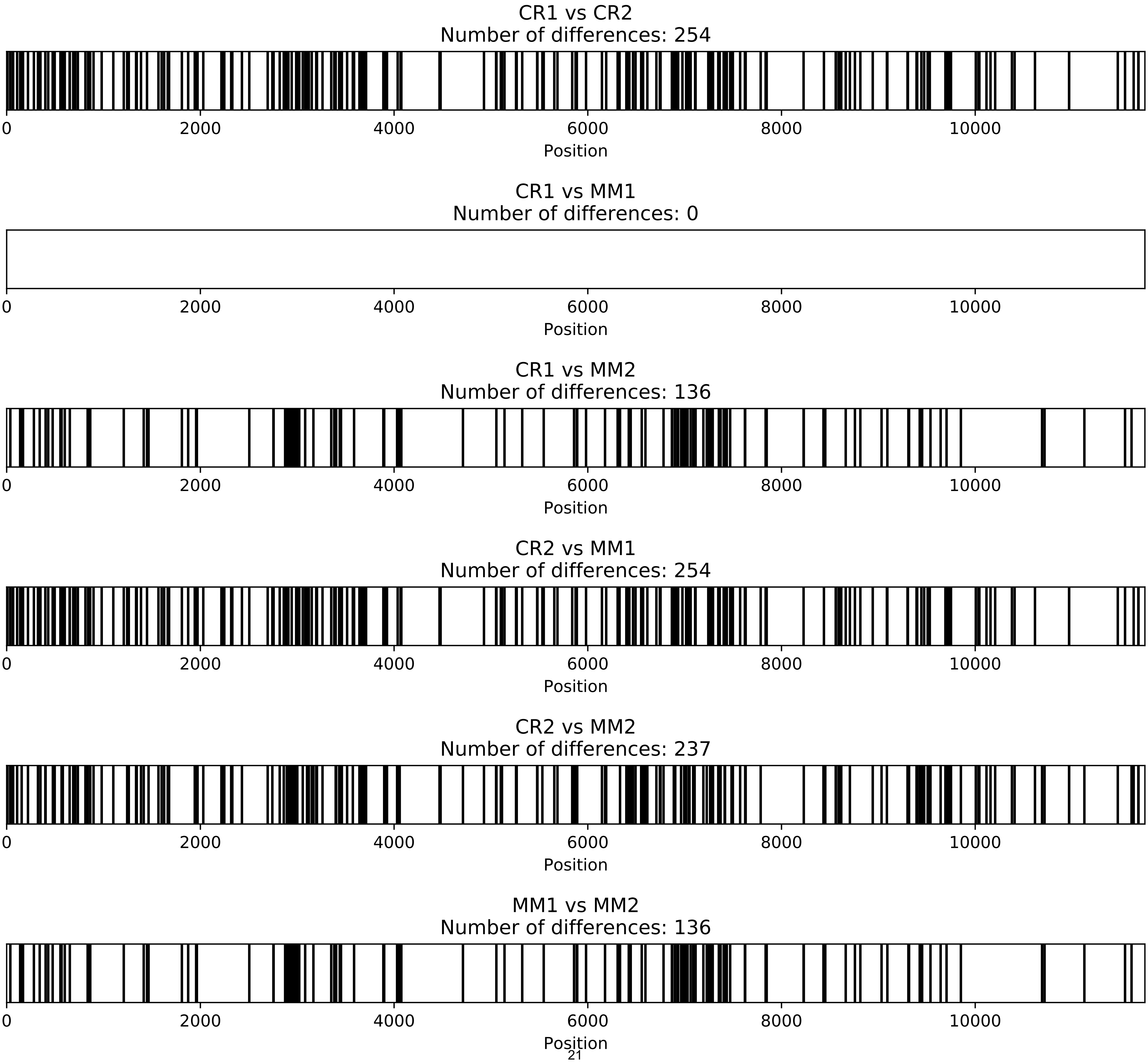
CR1 vs CR2
Number of differences: 254
0 2000 4000 6000 8000 10000
Position
CR1 vs MM1
Number of differences: 0
0 2000 4000 6000 8000 10000
Position
CR1 vs MM2
Number of differences: 136
0 2000 4000 6000 8000 10000
Position
CR2 vs MM1
Number of differences: 254
0 2000 4000 6000 8000 10000
Position
CR2 vs MM2
Number of differences: 237
0 2000 4000 6000 8000 10000
Position
MM1 vs MM2
Number of differences: 136
0 2000 4000 6000 8000 10000
Position

### Region: 62



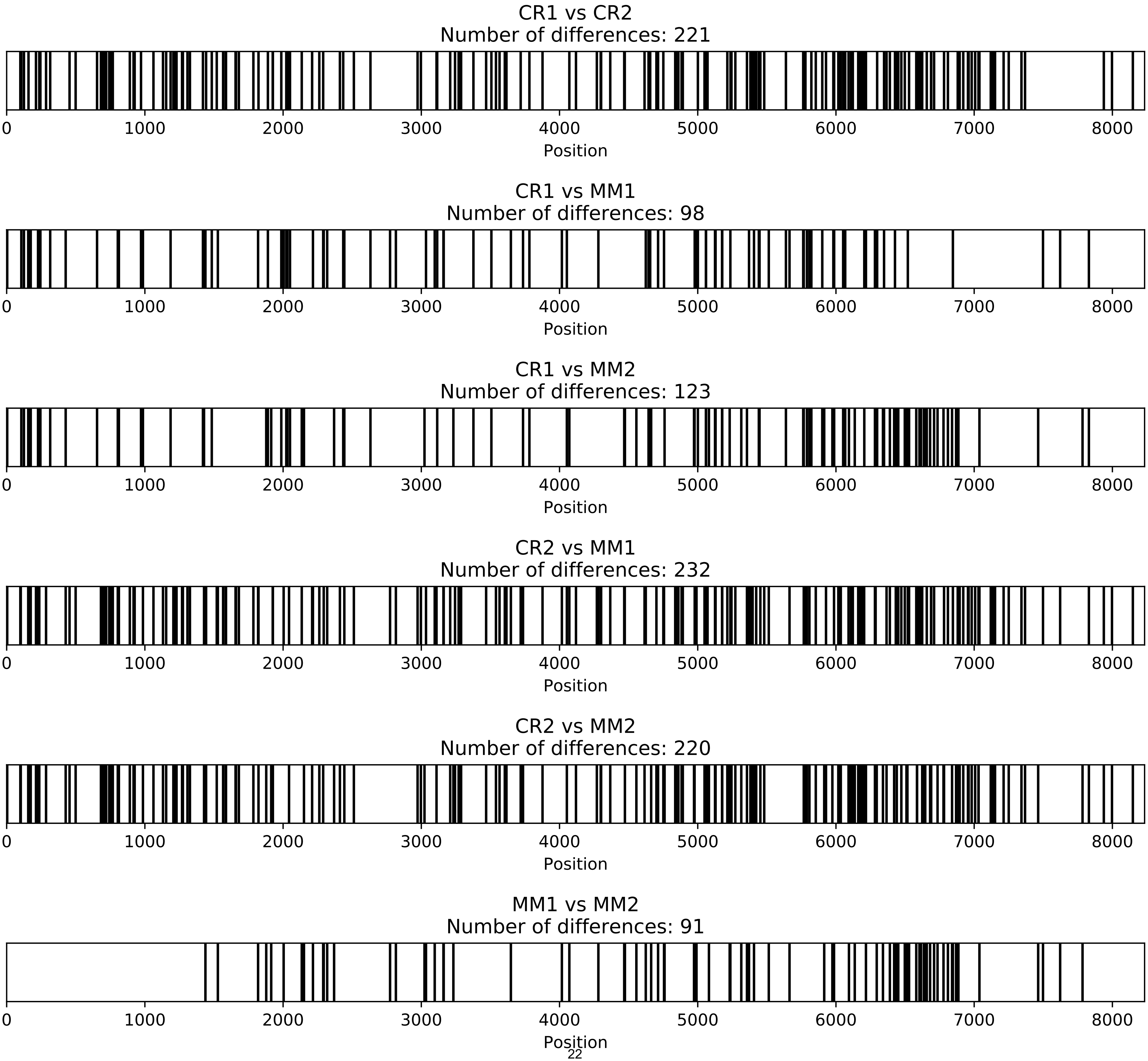
CR1 vs CR2
Number of differences: 221
CR1 vs MM1
Number of differences: 98
CR1 vs MM2
Number of differences: 123
CR2 vs MM1
Number of differences: 232
CR2 vs MM2
Number of differences: 220
MM1 vs MM2
Number of differences: 91
0
1000
2000
3000
4000
5000
6000
7000
8000
Position

### Region: 63



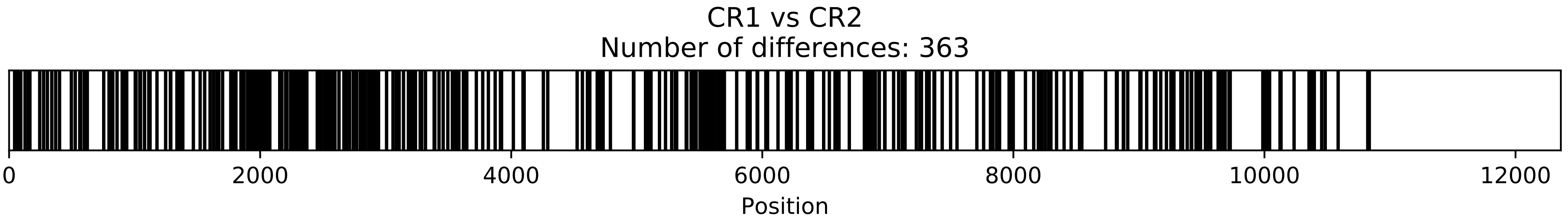
CR1 vs CR2
Number of differences: 363
0
2000
4000
6000
8000
10000
12000
Position

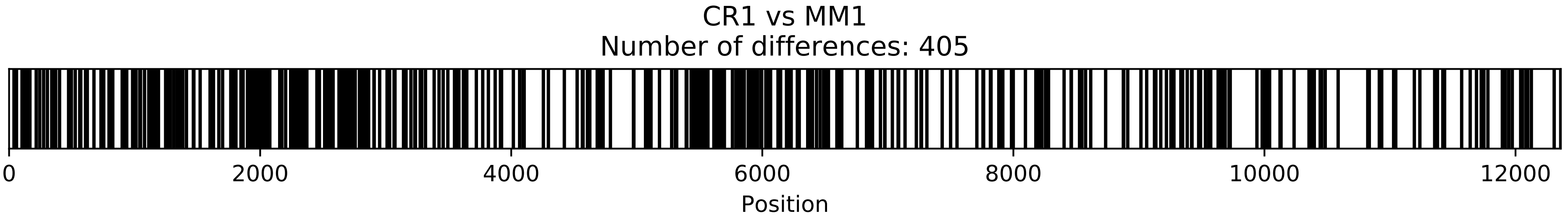
CR1 vs MM1
Number of differences: 405
0
2000
4000
6000
8000
10000
12000
Position

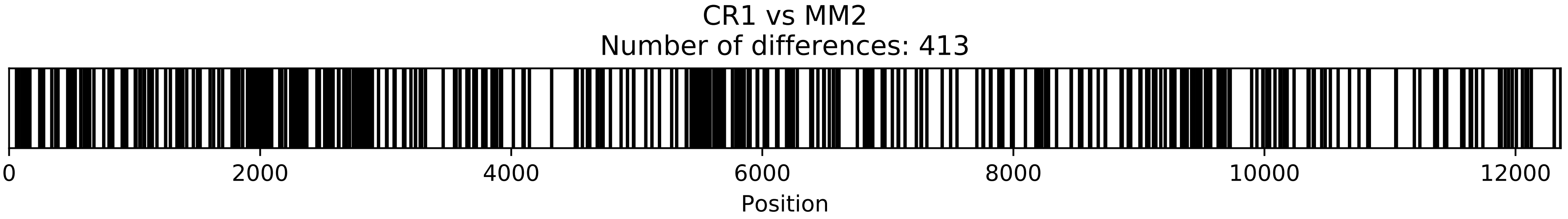
CR1 vs MM2
Number of differences: 413
0
2000
4000
6000
8000
10000
12000
Position

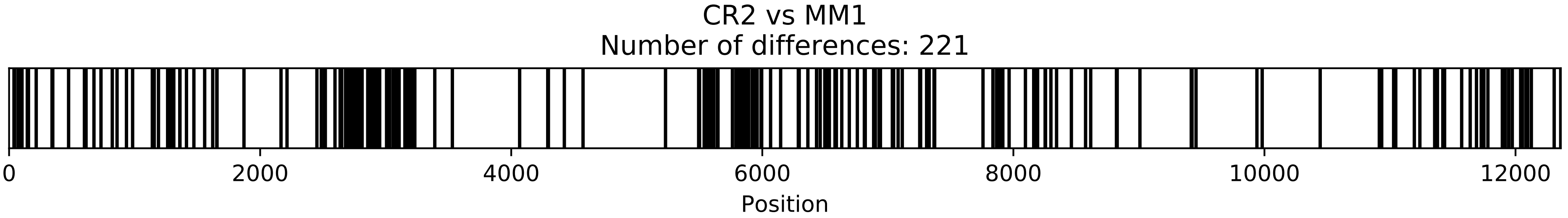
CR2 vs MM1
Number of differences: 221
0
2000
4000
6000
8000
10000
12000
Position

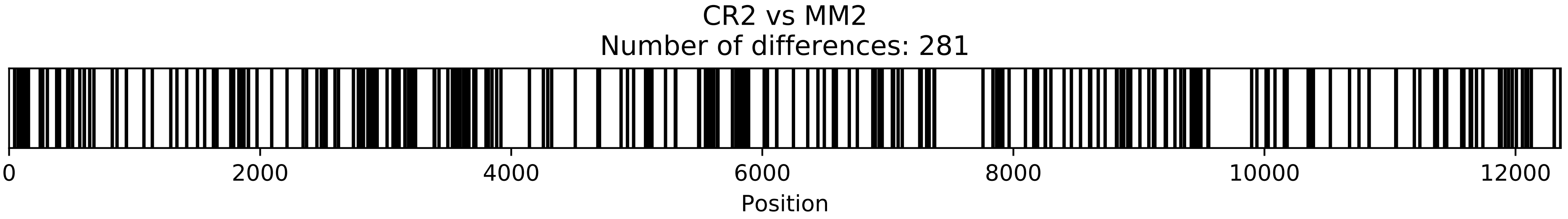
CR2 vs MM2
Number of differences: 281
0
2000
4000
6000
8000
10000
12000
Position

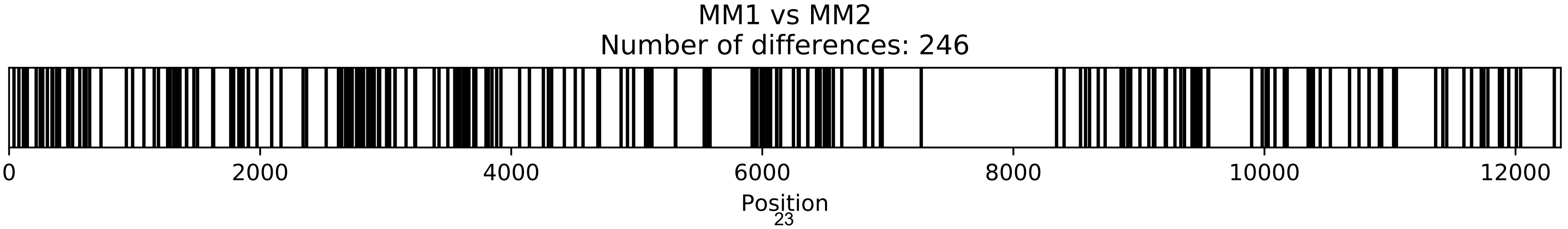
MM1 vs MM2
Number of differences: 246
0
2000
4000
6000
8000
10000
12000
Position

### Region: 64



## CR1 vs CR2
Number of differences: 227

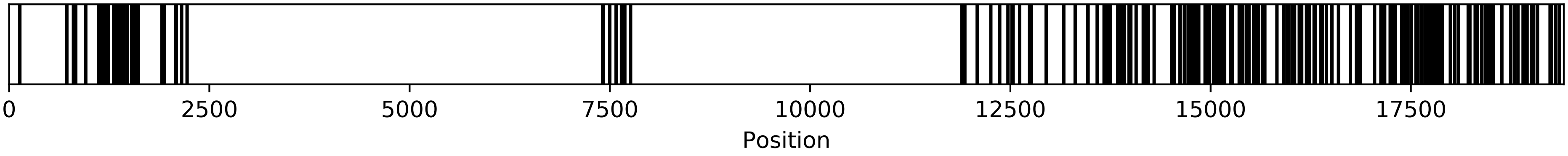


## CR1 vs MM1
Number of differences: 343

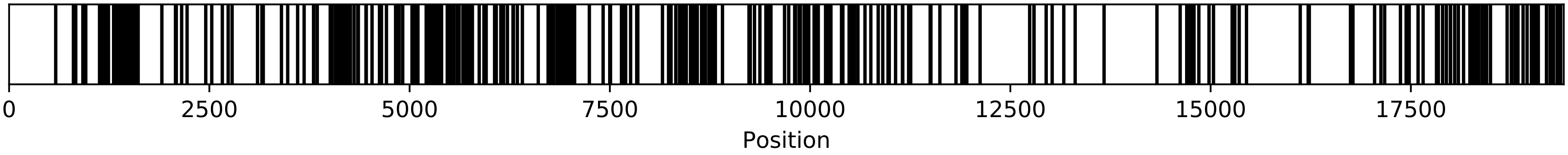


## CR1 vs MM2
Number of differences: 345

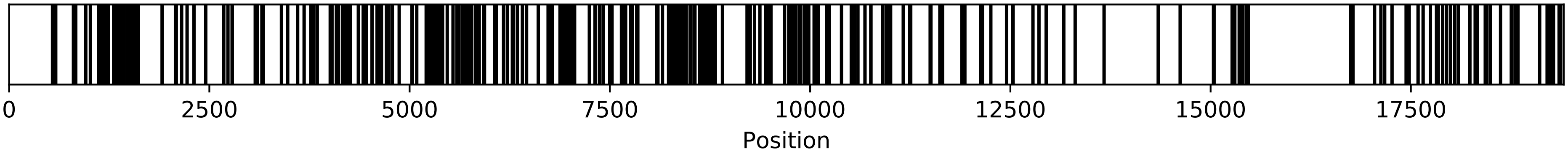


## CR2 vs MM1
Number of differences: 411

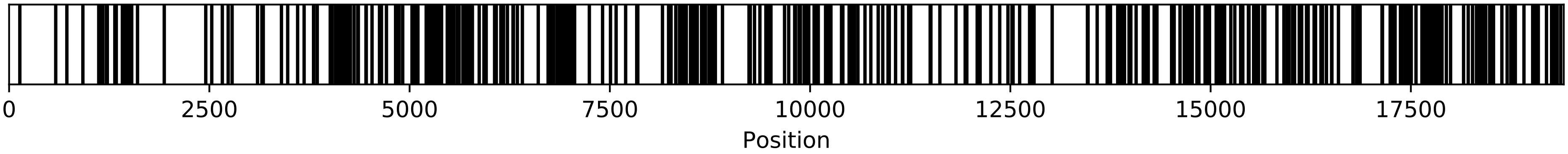


## CR2 vs MM2
Number of differences: 418

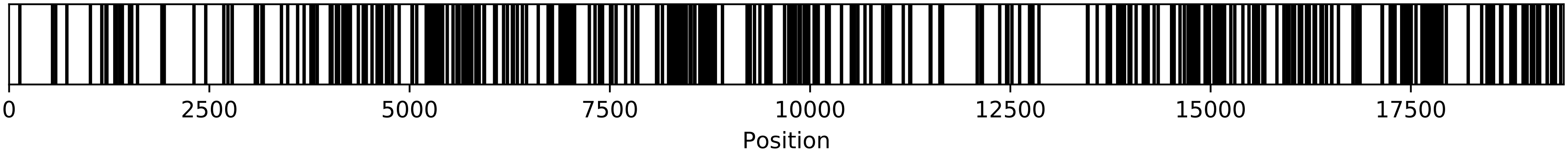


## MM1 vs MM2
Number of differences: 228

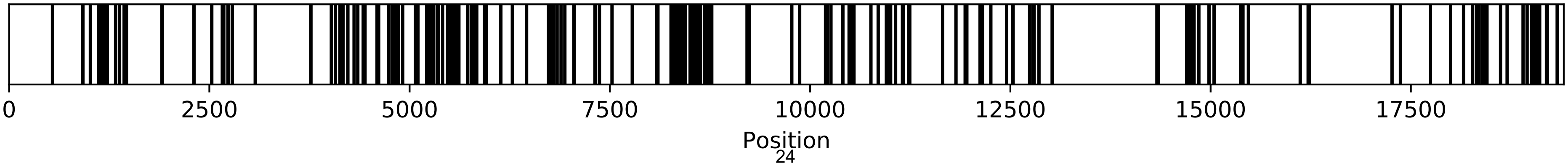

### Region: 66



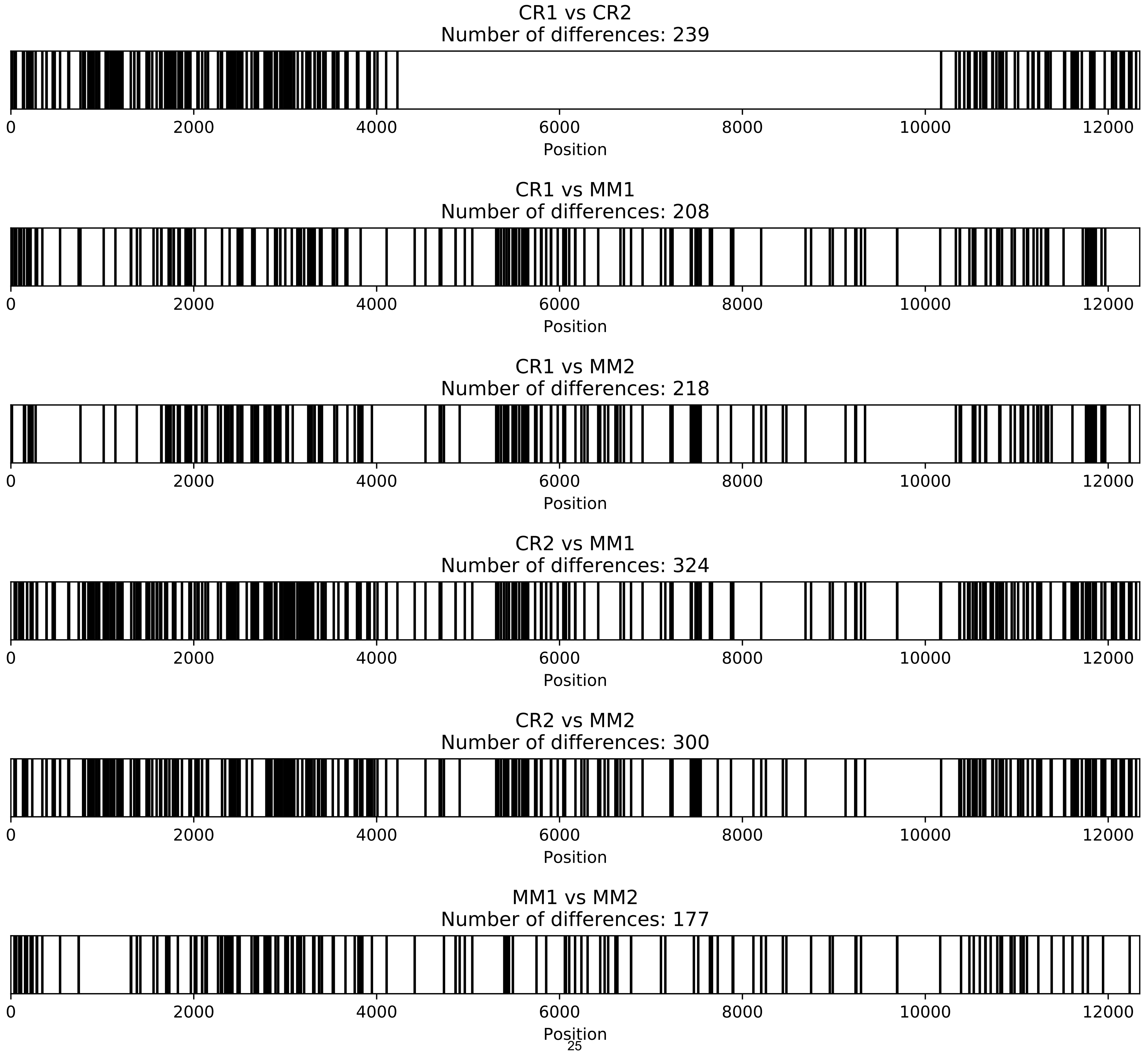
CR1 vs CR2
Number of differences: 239
CR1 vs MM1
Number of differences: 208
CR1 vs MM2
Number of differences: 218
CR2 vs MM1
Number of differences: 324
CR2 vs MM2
Number of differences: 300
MM1 vs MM2
Number of differences: 177
0
2000
4000
6000
8000
10000
12000
Position

### Region: 70



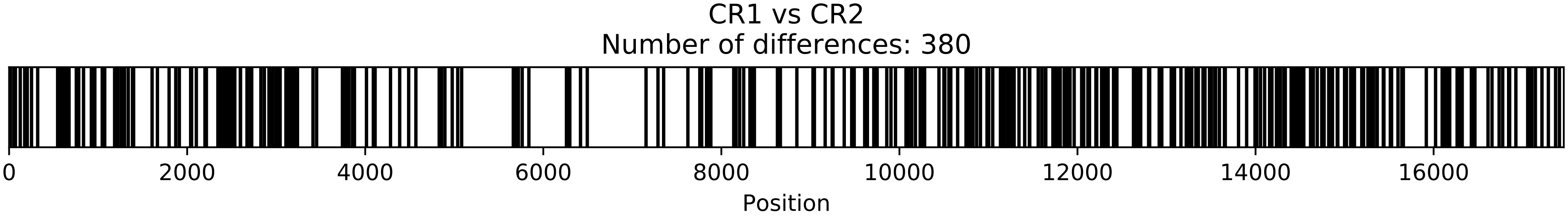
CR1 vs CR2
Number of differences: 380
0
2000
4000
6000
8000
10000
12000
14000
16000
Position

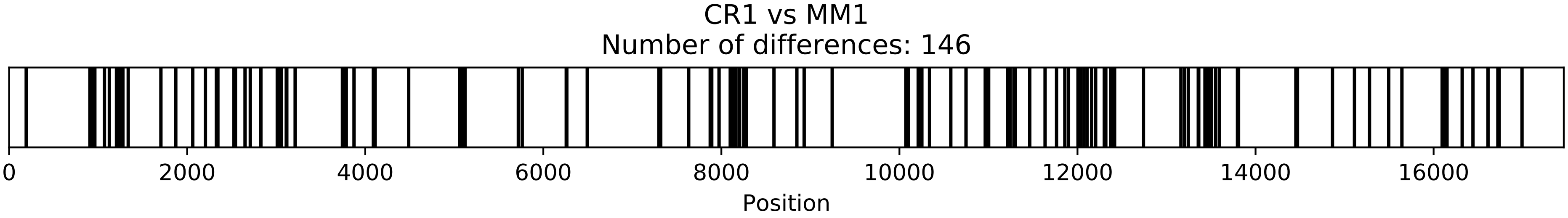
CR1 vs MM1
Number of differences: 146
0
2000
4000
6000
8000
10000
12000
14000
16000
Position

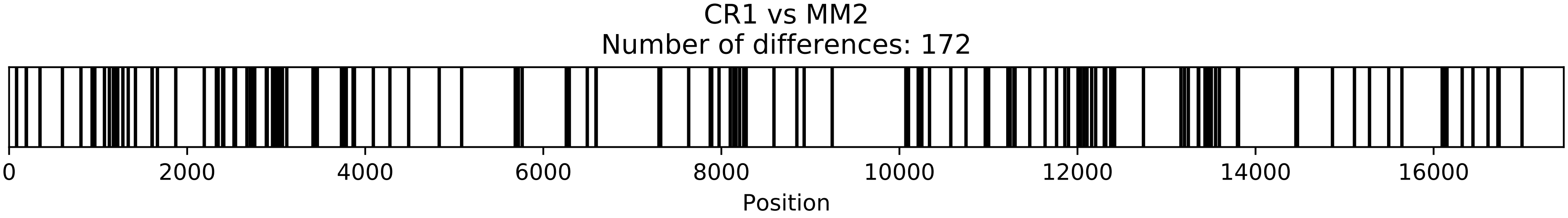
CR1 vs MM2
Number of differences: 172
0
2000
4000
6000
8000
10000
12000
14000
16000
Position

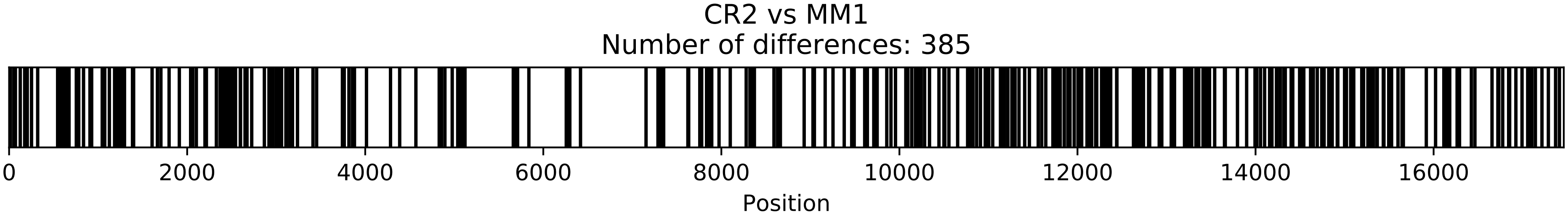
CR2 vs MM1
Number of differences: 385
0
2000
4000
6000
8000
10000
12000
14000
16000
Position

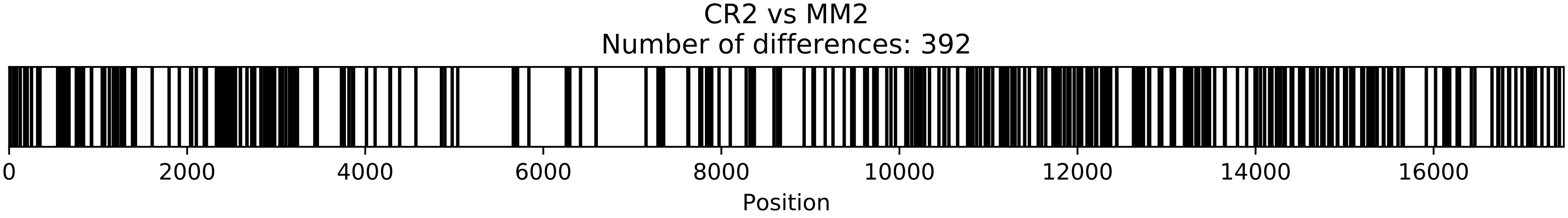
CR2 vs MM2
Number of differences: 392
0
2000
4000
6000
8000
10000
12000
14000
16000
Position

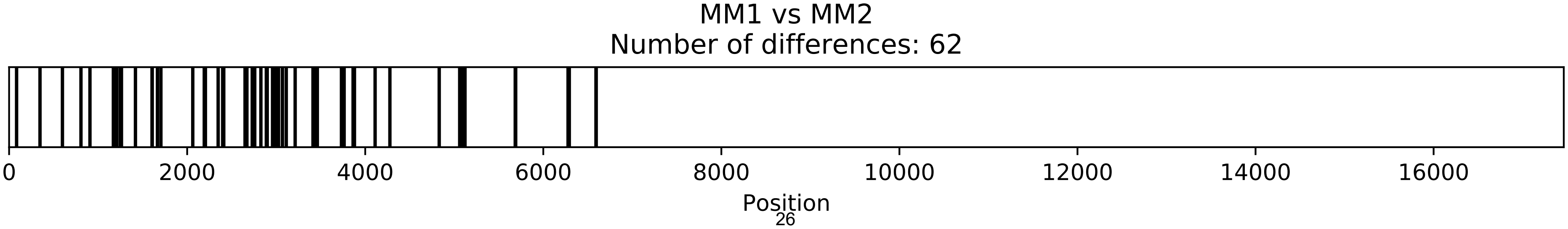
MM1 vs MM2
Number of differences: 62
0
2000
4000
6000
8000
10000
12000
14000
16000
Position

### Region: 71



## CR1 vs CR2
Number of differences: 230

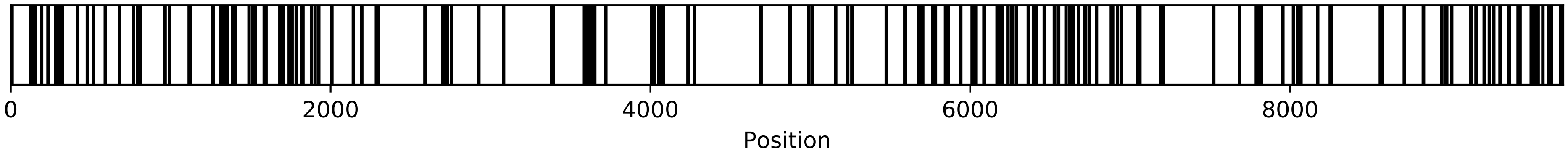


## CR1 vs MM1
Number of differences: 94

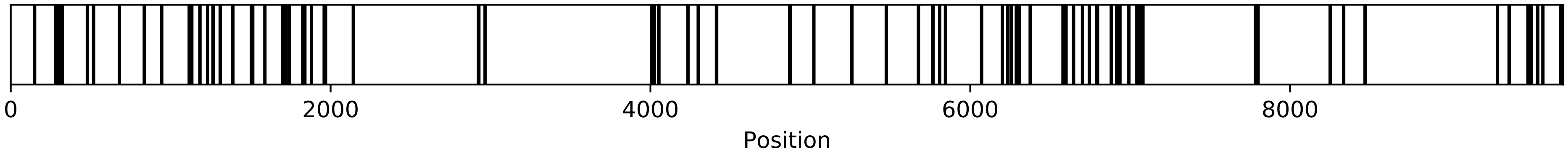


## CR1 vs MM2
Number of differences: 123

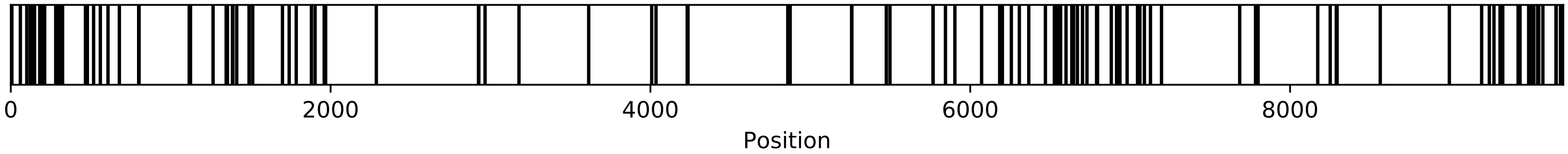


## CR2 vs MM1
Number of differences: 221

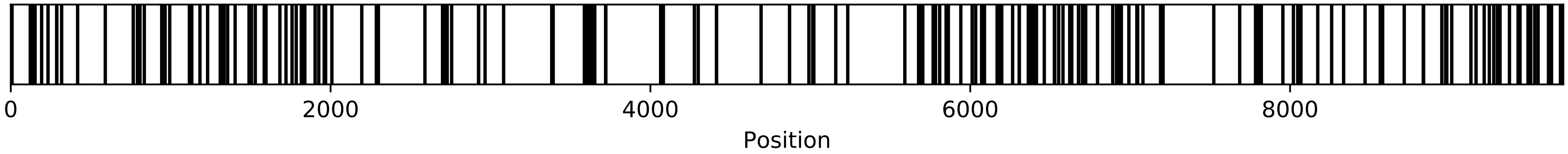


## CR2 vs MM2
Number of differences: 224

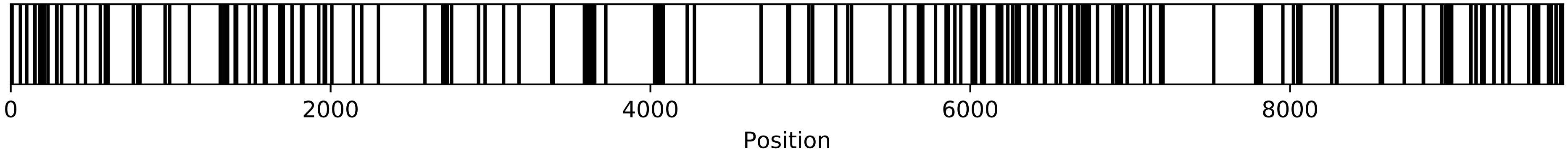


## MM1 vs MM2
Number of differences: 109

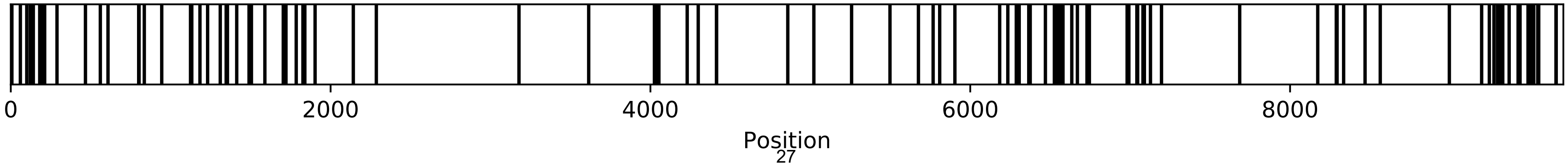

### Region: 72



CR1 vs CR2
Number of differences: 318

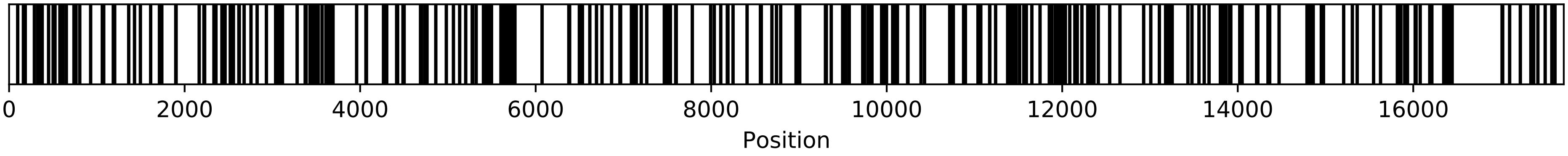


CR1 vs MM1
Number of differences: 187

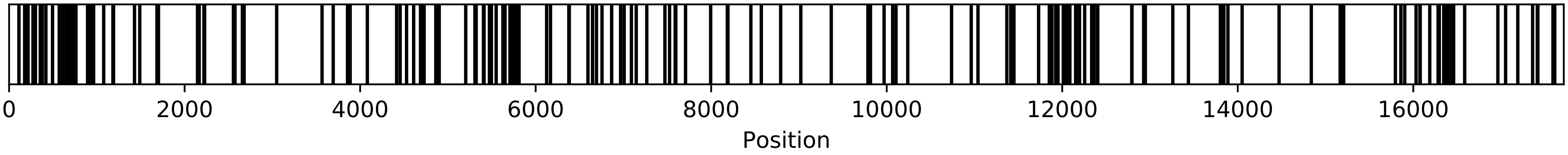


CR1 vs MM2
Number of differences: 200

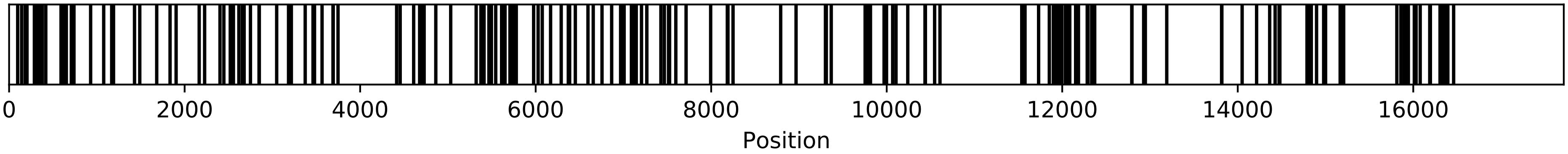


CR2 vs MM1
Number of differences: 340

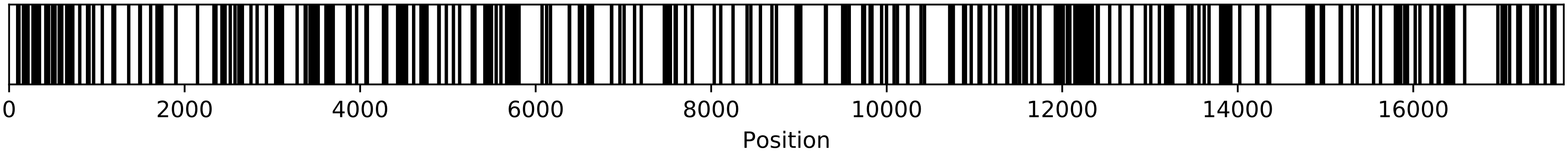


CR2 vs MM2
Number of differences: 329

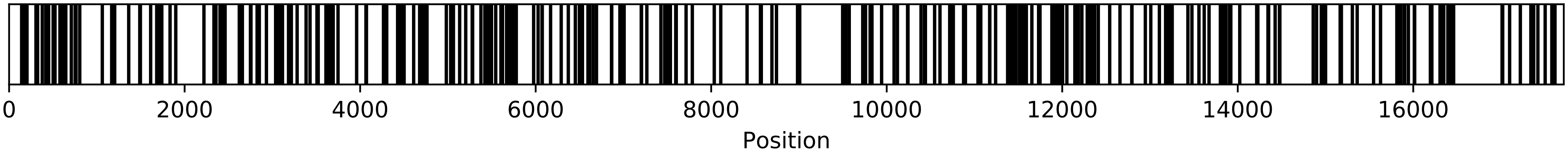


MM1 vs MM2
Number of differences: 206

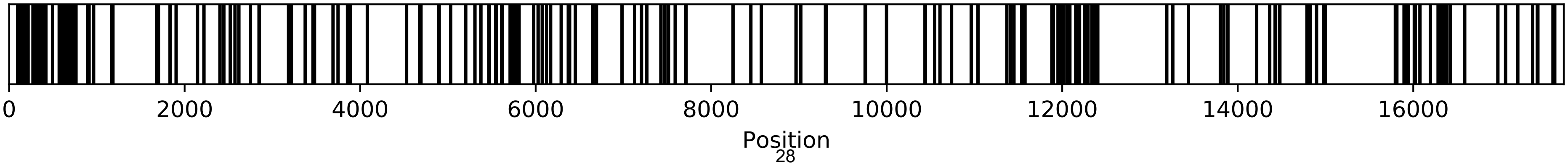

### Region: 73



CR1 vs CR2
Number of differences: 225

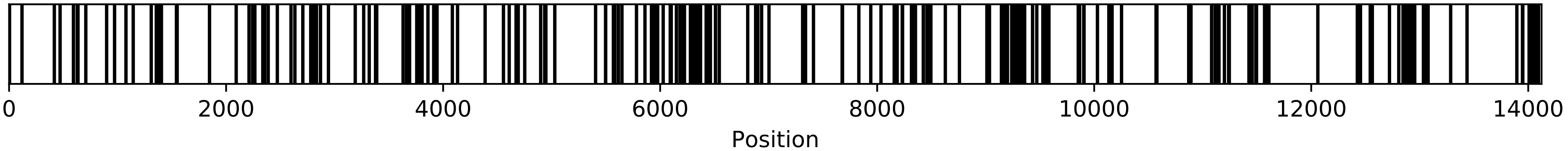


CR1 vs MM1
Number of differences: 221

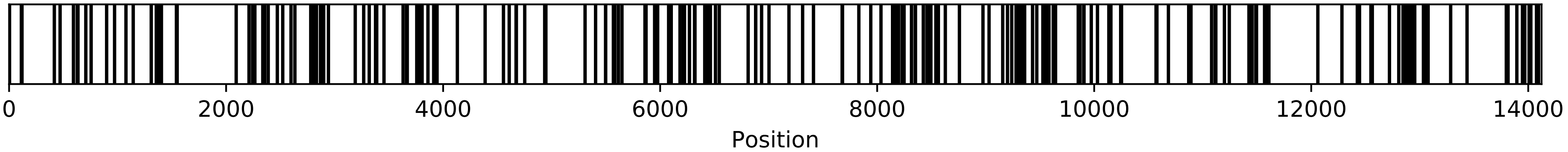


CR1 vs MM2
Number of differences: 224

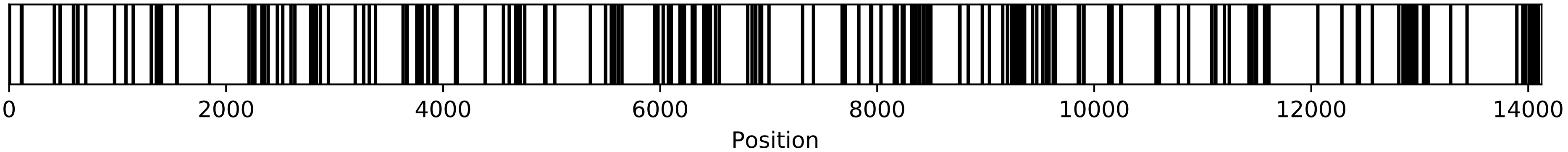


CR2 vs MM1
Number of differences: 88

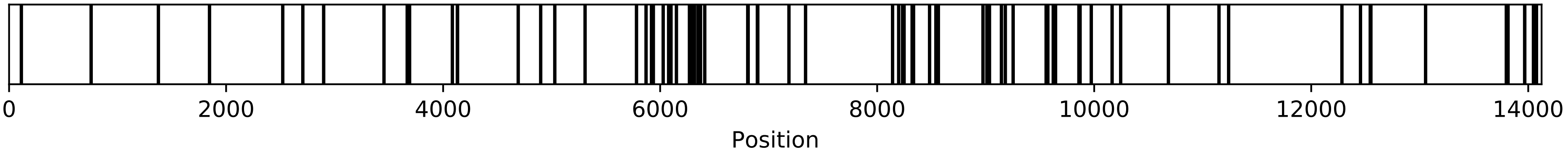


CR2 vs MM2
Number of differences: 96

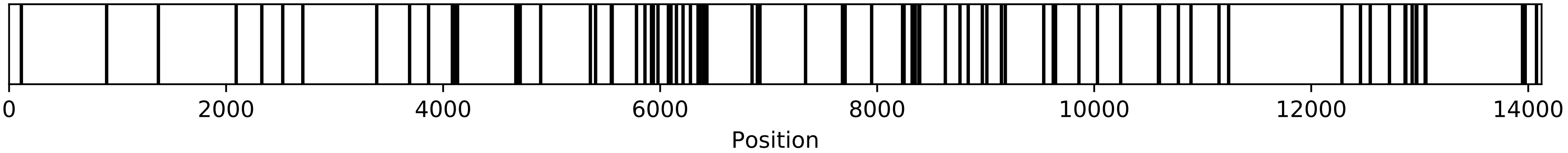


MM1 vs MM2
Number of differences: 93

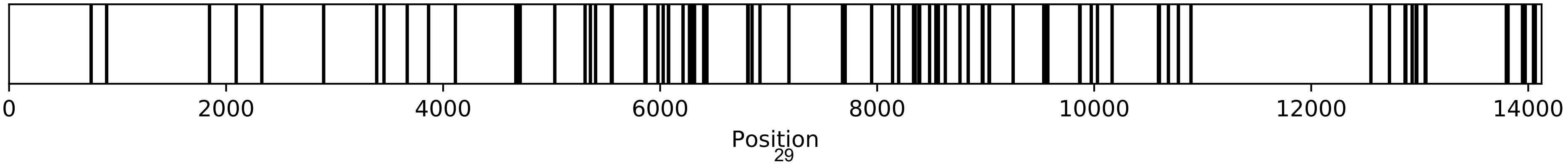

### Region: 76



CR1 vs CR2
Number of differences: 185

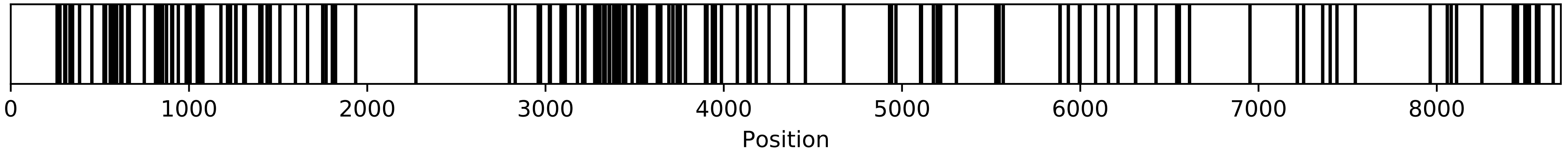


CR1 vs MM1
Number of differences: 204

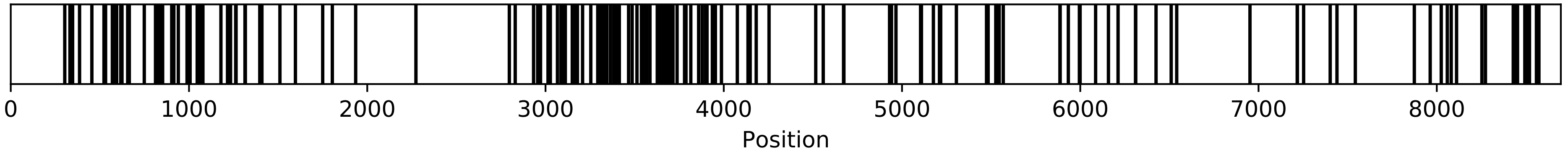


CR1 vs MM2
Number of differences: 211

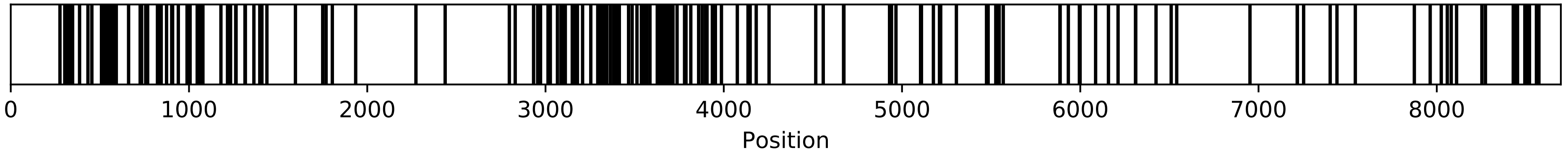


CR2 vs MM1
Number of differences: 95

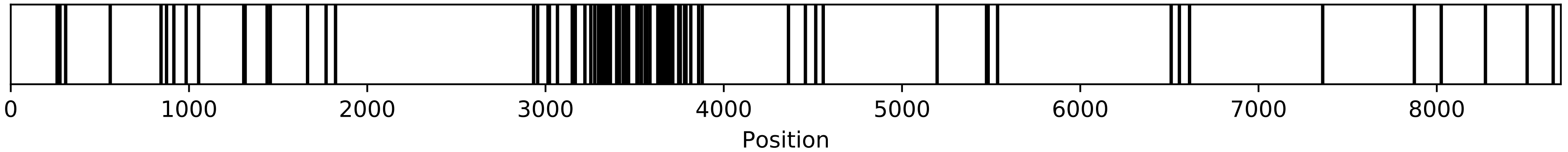


CR2 vs MM2
Number of differences: 123

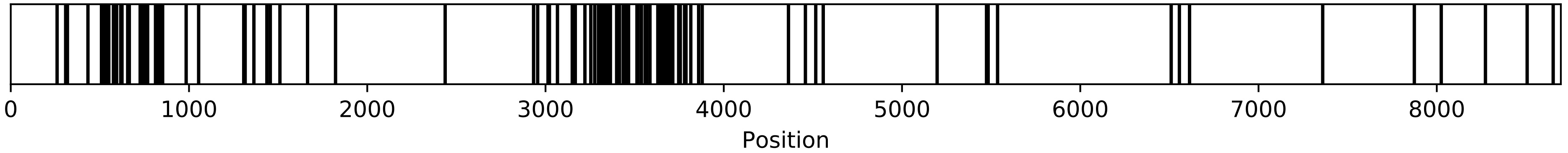


MM1 vs MM2
Number of differences: 39

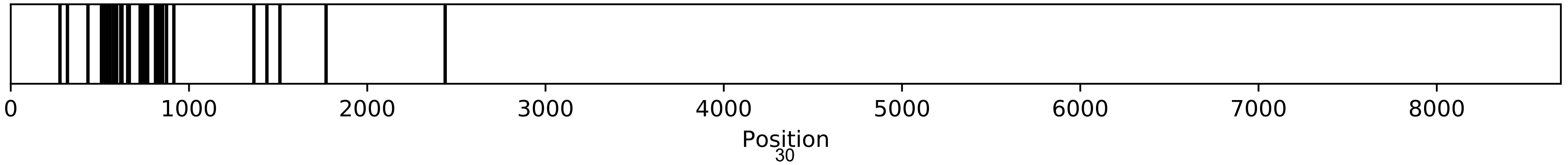

### Region: 78



CR1 vs CR2
Number of differences: 380

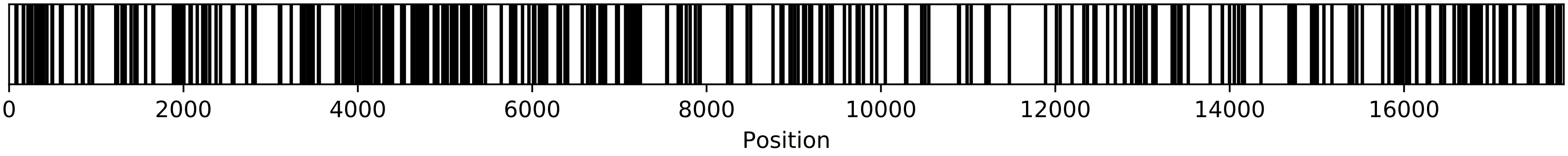


CR1 vs MM1
Number of differences: 388

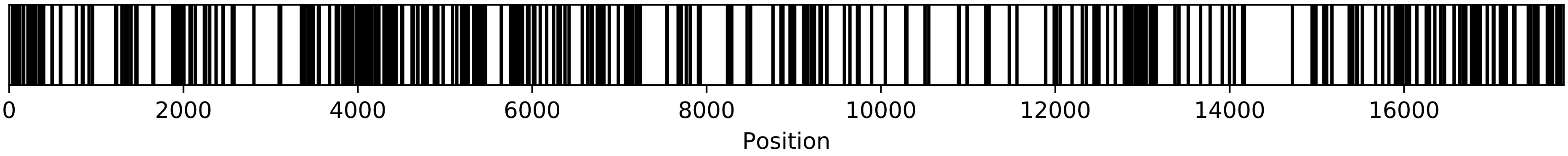


CR1 vs MM2
Number of differences: 400

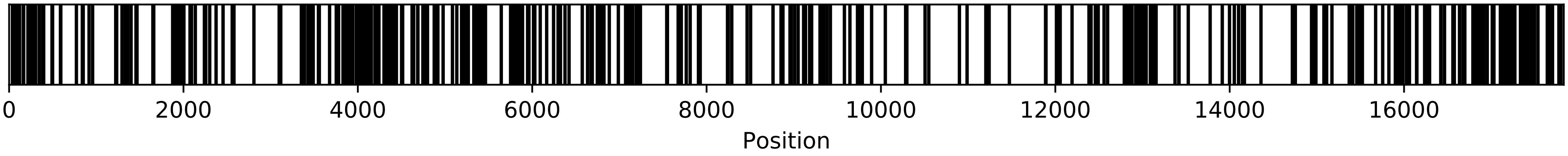


CR2 vs MM1
Number of differences: 178

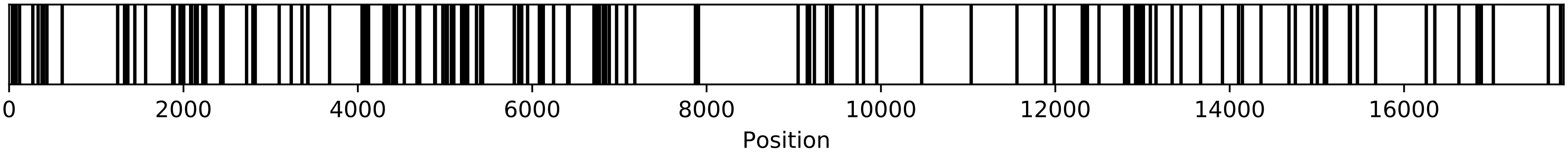


CR2 vs MM2
Number of differences: 203

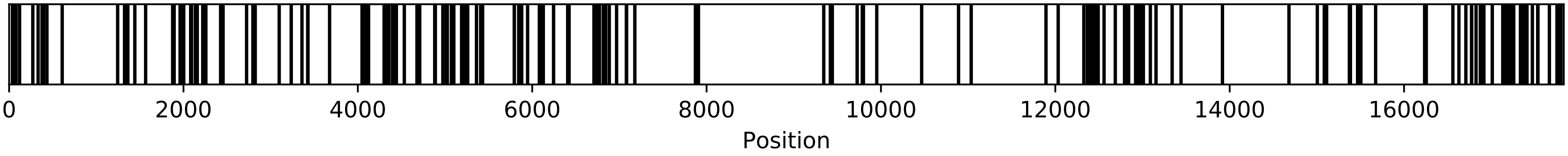


MM1 vs MM2
Number of differences: 73

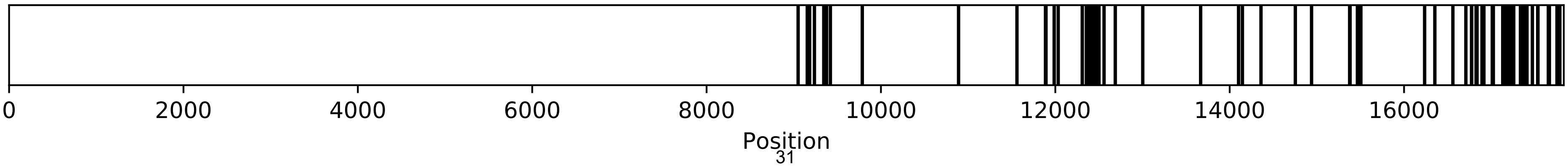

### Region: 80



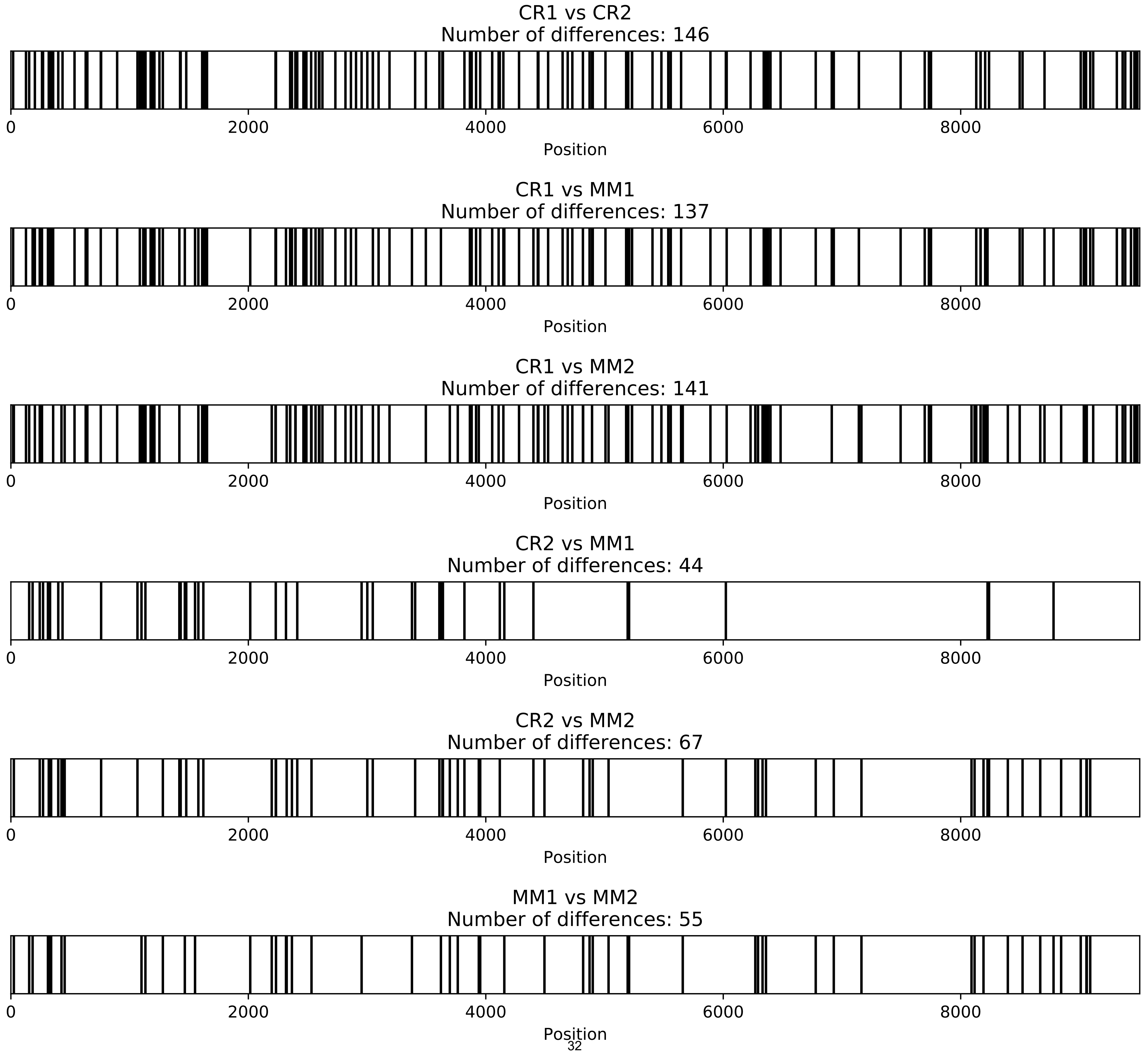
CR1 vs CR2
Number of differences: 146
0
2000
4000
6000
8000
Position
CR1 vs MM1
Number of differences: 137
0
2000
4000
6000
8000
Position
CR1 vs MM2
Number of differences: 141
0
2000
4000
6000
8000
Position
CR2 vs MM1
Number of differences: 44
0
2000
4000
6000
8000
Position
CR2 vs MM2
Number of differences: 67
0
2000
4000
6000
8000
Position
MM1 vs MM2
Number of differences: 55
0
2000
4000
6000
8000
Position

### Region: 84



CR1 vs CR2
Number of differences: 412

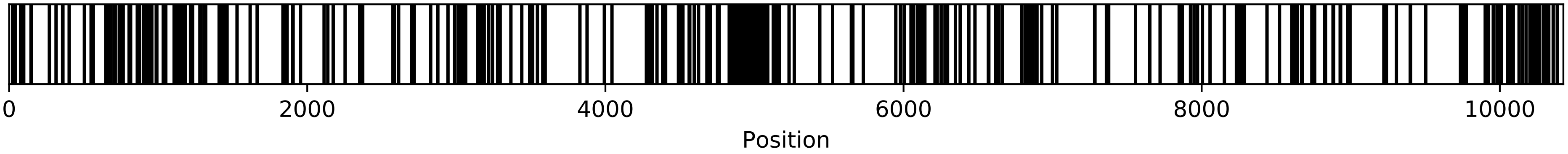


CR1 vs MM1
Number of differences: 174

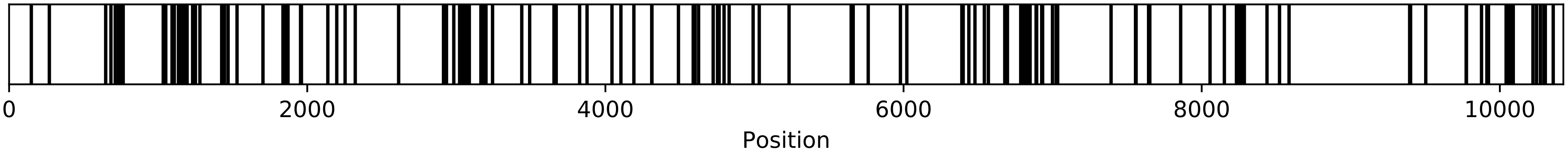


CR1 vs MM2
Number of differences: 193

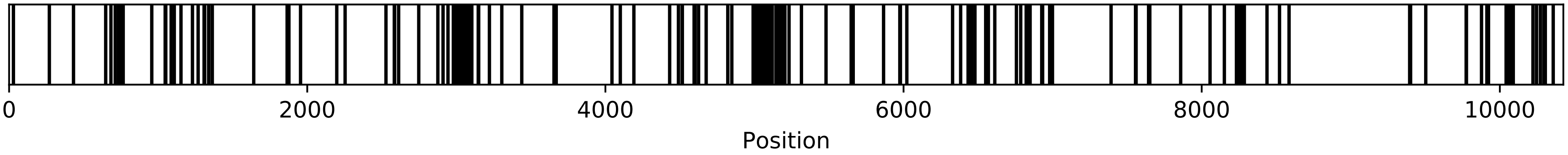


CR2 vs MM1
Number of differences: 416

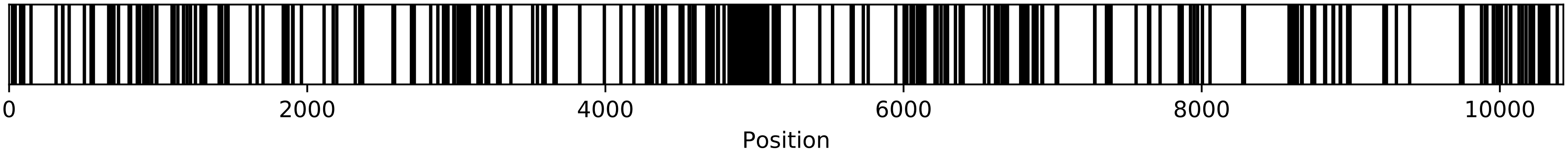


CR2 vs MM2
Number of differences: 425

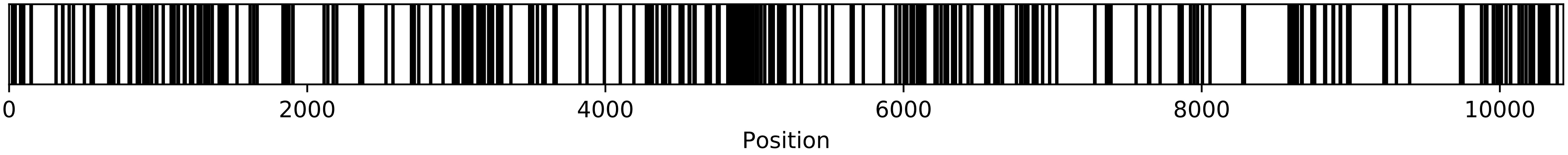


MM1 vs MM2
Number of differences: 150

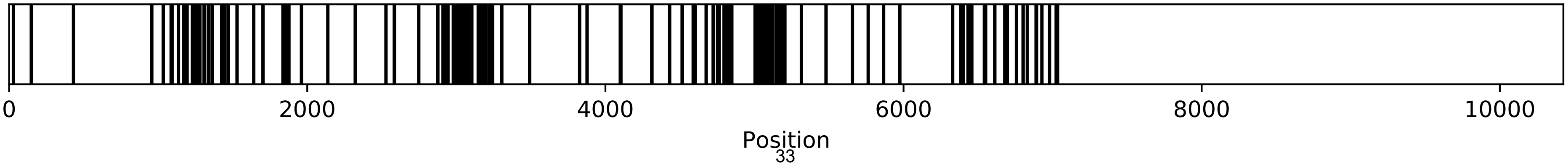

### Region: 86



CR1 vs CR2
Number of differences: 330

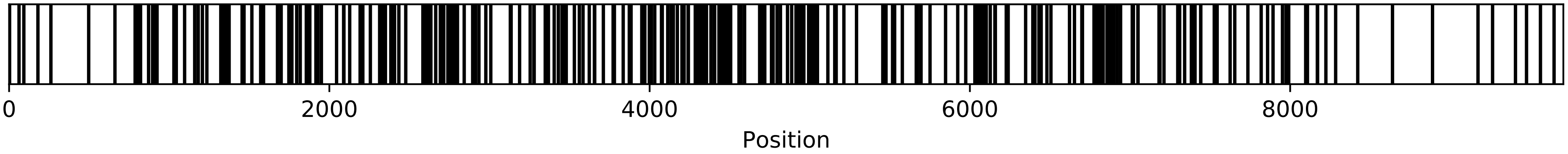


CR1 vs MM1
Number of differences: 217

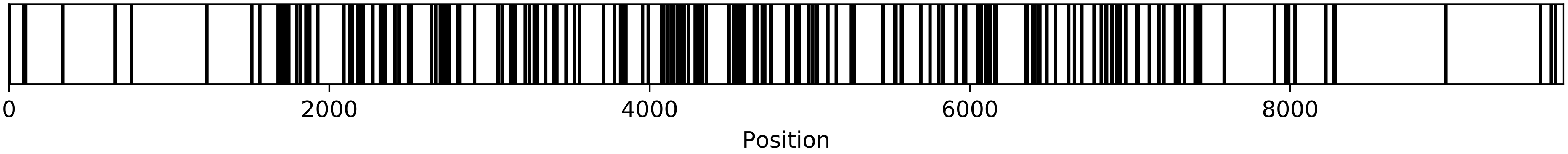


CR1 vs MM2
Number of differences: 217

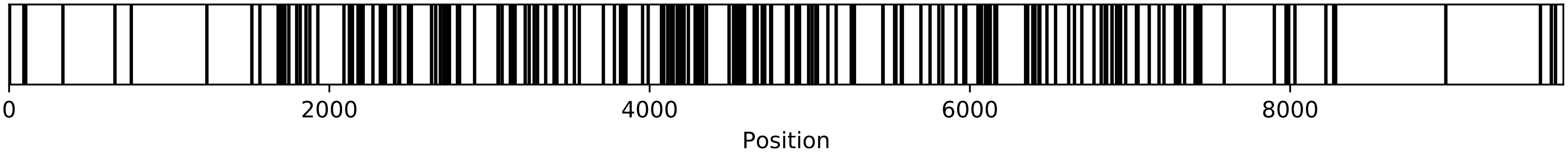


CR2 vs MM1
Number of differences: 307

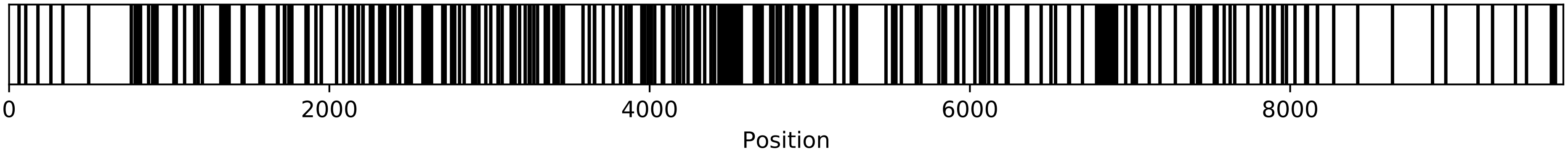


CR2 vs MM2
Number of differences: 307

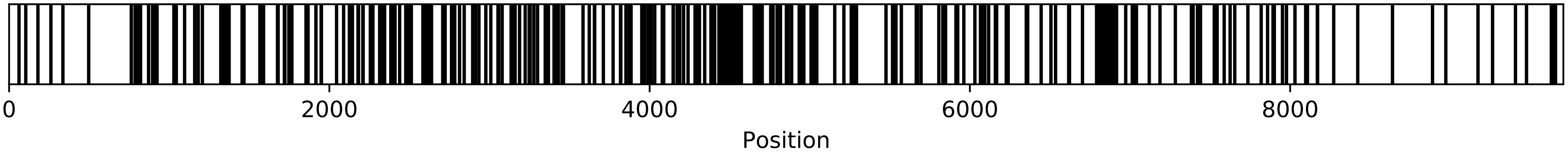


MM1 vs MM2
Number of differences: 0

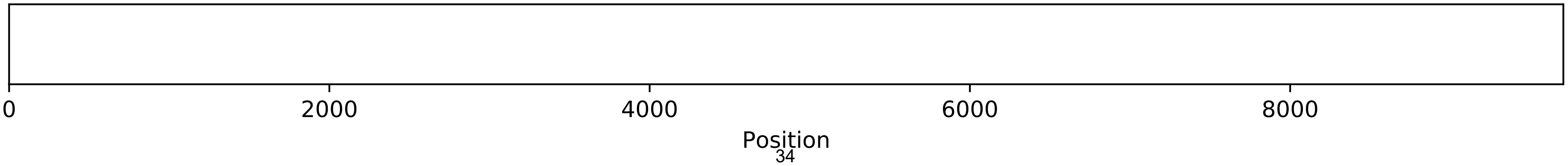

### Region: 90



## CR1 vs CR2
Number of differences: 294

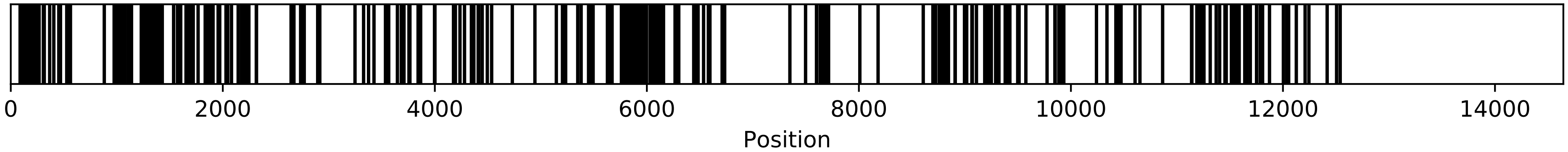


## CR1 vs MM1
Number of differences: 144

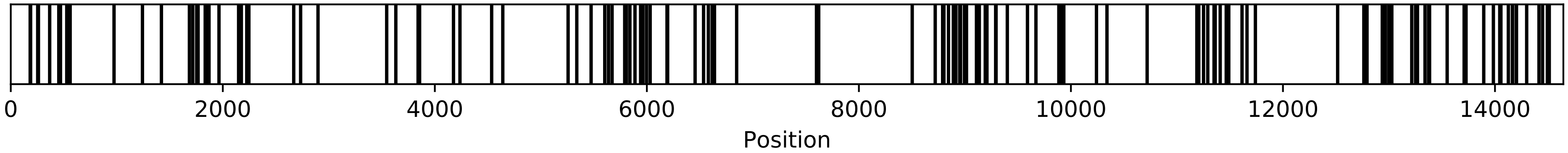


## CR1 vs MM2
Number of differences: 139

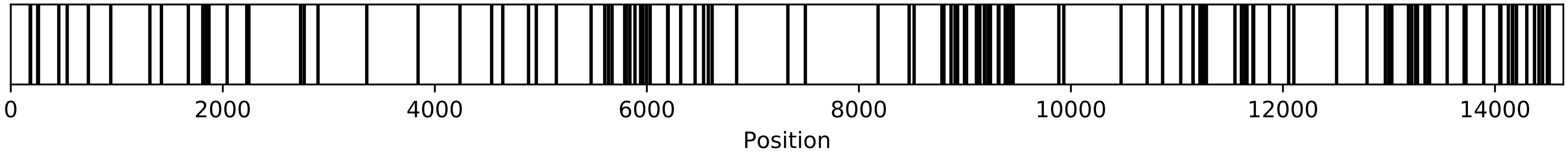


## CR2 vs MM1
Number of differences: 326

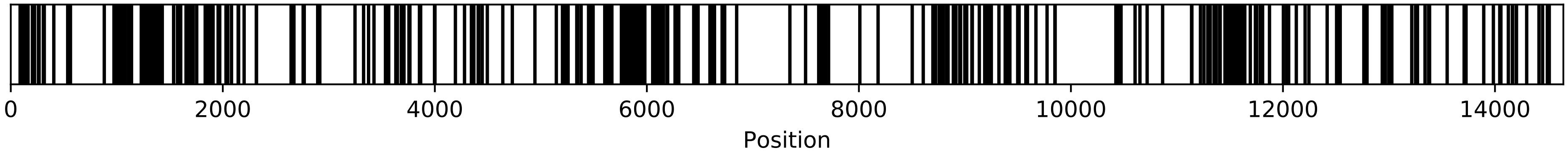


## CR2 vs MM2
Number of differences: 332

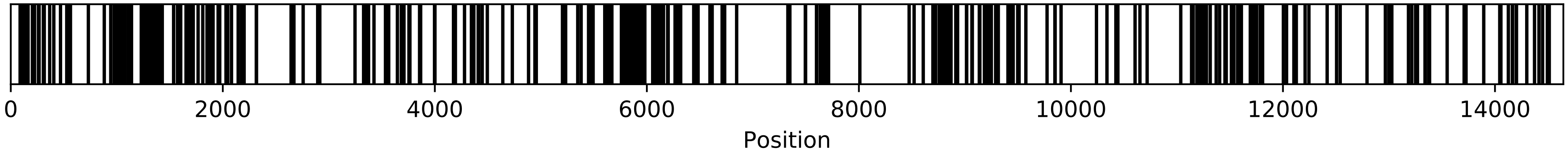


## MM1 vs MM2
Number of differences: 133

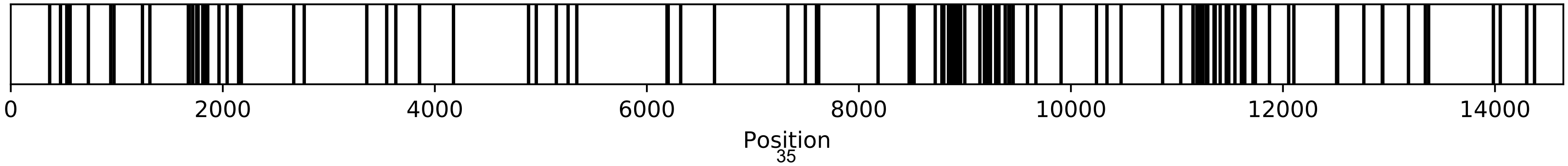

### Region: 91



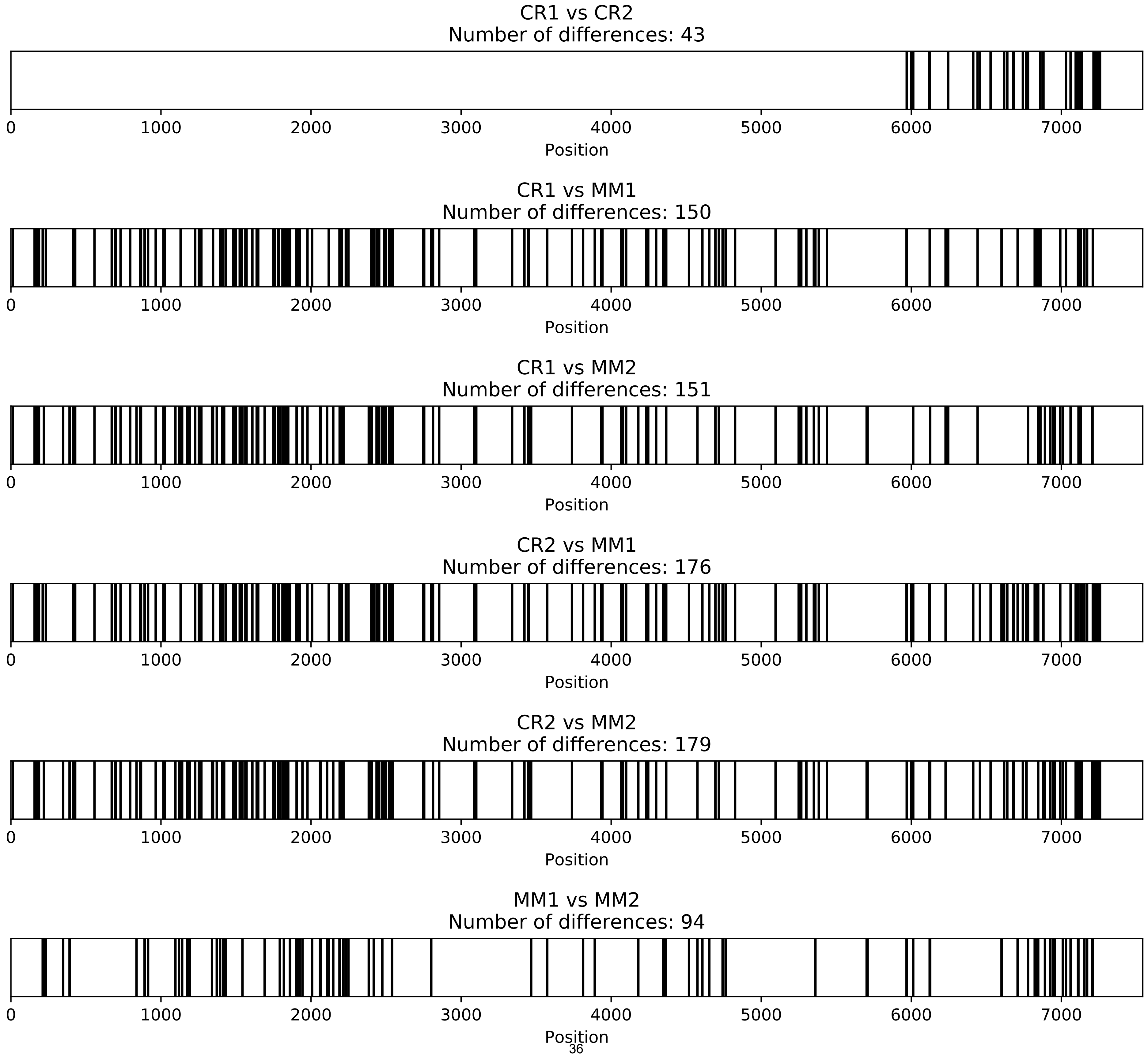
CR1 vs CR2
Number of differences: 43
CR1 vs MM1
Number of differences: 150
CR1 vs MM2
Number of differences: 151
CR2 vs MM1
Number of differences: 176
CR2 vs MM2
Number of differences: 179
MM1 vs MM2
Number of differences: 94
0 1000 2000 3000 4000 5000 6000 7000
Position

### Region: 92



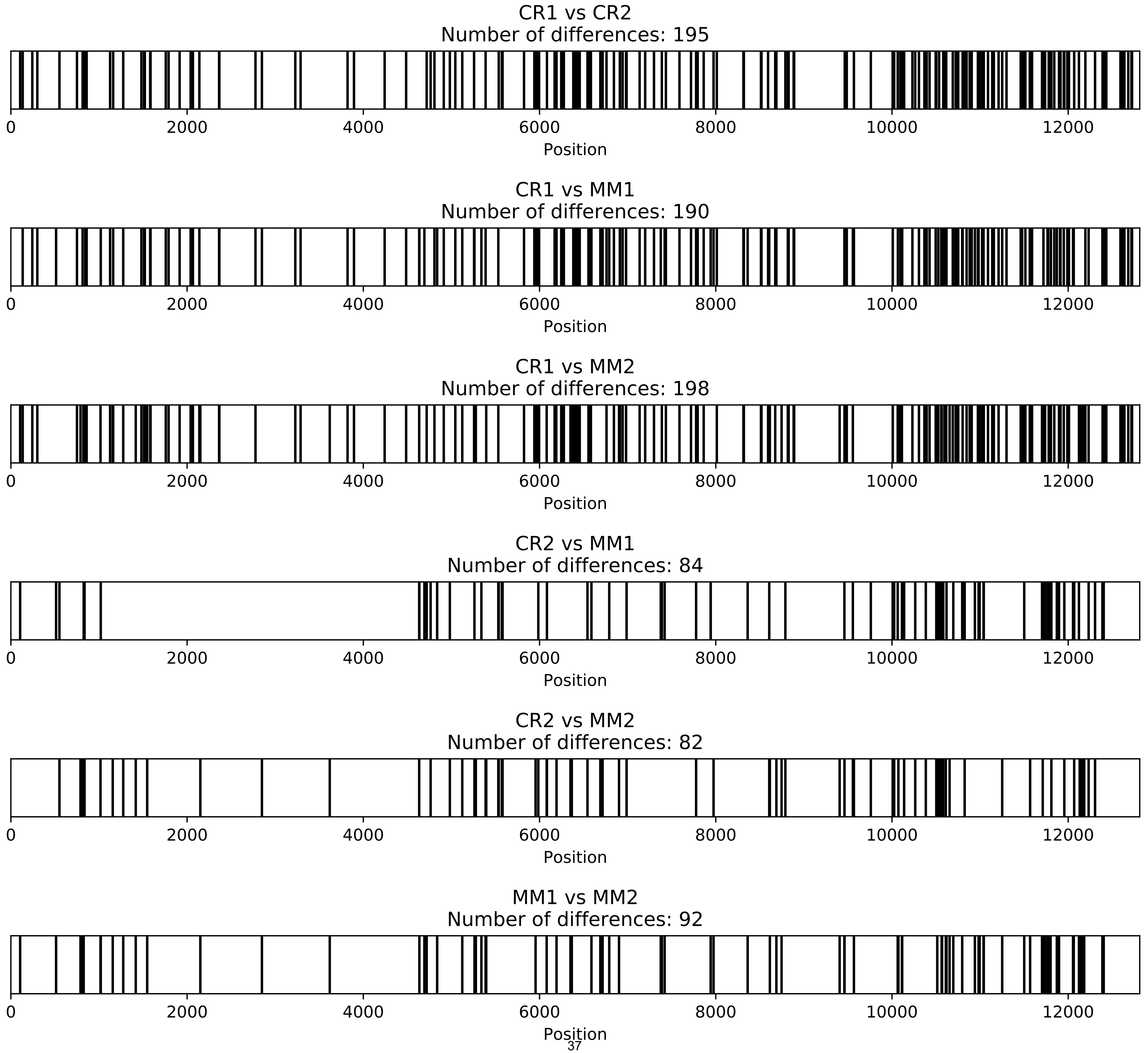
CR1 vs CR2
Number of differences: 195
CR1 vs MM1
Number of differences: 190
CR1 vs MM2
Number of differences: 198
CR2 vs MM1
Number of differences: 84
CR2 vs MM2
Number of differences: 82
MM1 vs MM2
Number of differences: 92
0
2000
4000
6000
8000
10000
12000
Position

### Region: 93



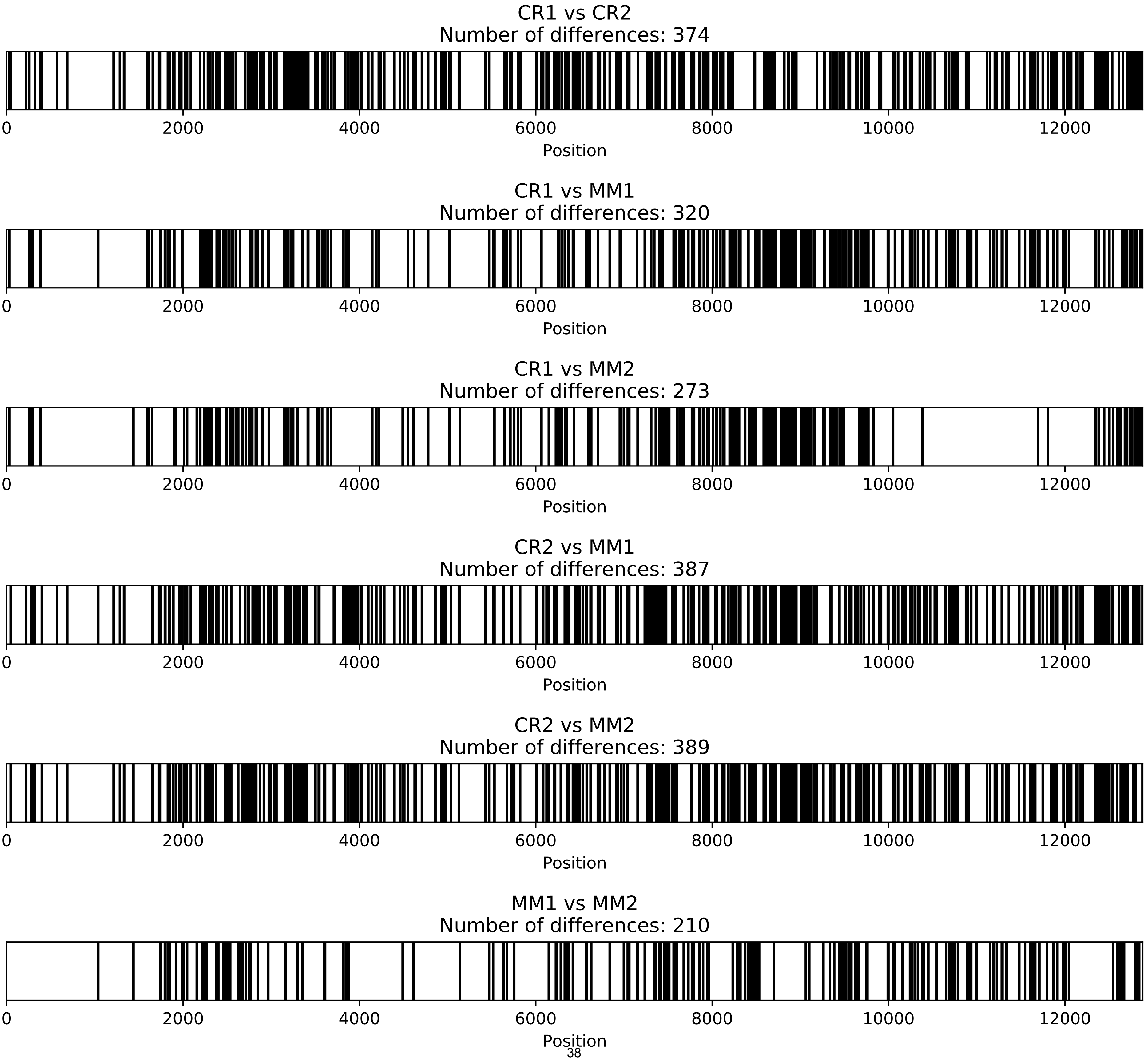
CR1 vs CR2
Number of differences: 374
0
2000
4000
6000
8000
10000
12000
Position
CR1 vs MM1
Number of differences: 320
0
2000
4000
6000
8000
10000
12000
Position
CR1 vs MM2
Number of differences: 273
0
2000
4000
6000
8000
10000
12000
Position
CR2 vs MM1
Number of differences: 387
0
2000
4000
6000
8000
10000
12000
Position
CR2 vs MM2
Number of differences: 389
0
2000
4000
6000
8000
10000
12000
Position
MM1 vs MM2
Number of differences: 210
0
2000
4000
6000
8000
10000
12000
Position

### Region: 94



CR1 vs CR2
Number of differences: 378

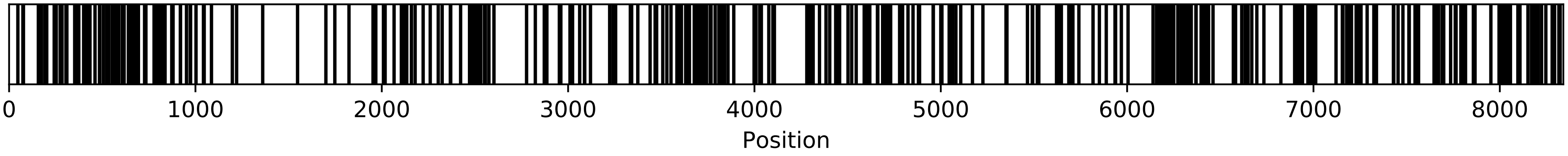


CR1 vs MM1
Number of differences: 203

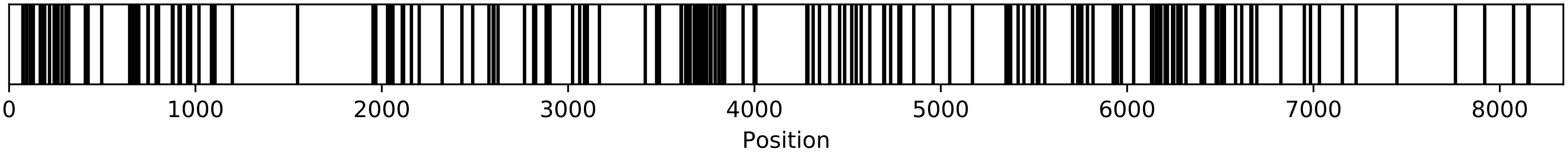


CR1 vs MM2
Number of differences: 174

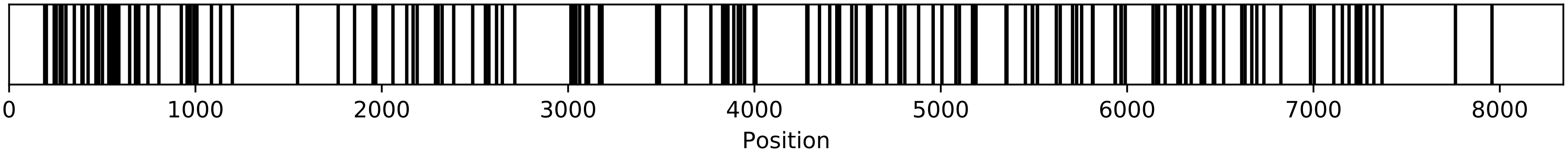


CR2 vs MM1
Number of differences: 352

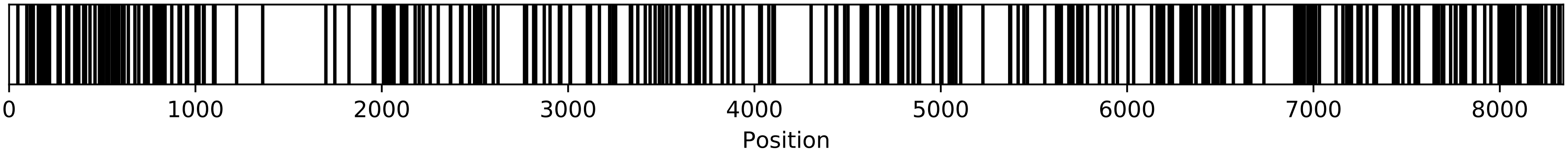


CR2 vs MM2
Number of differences: 353

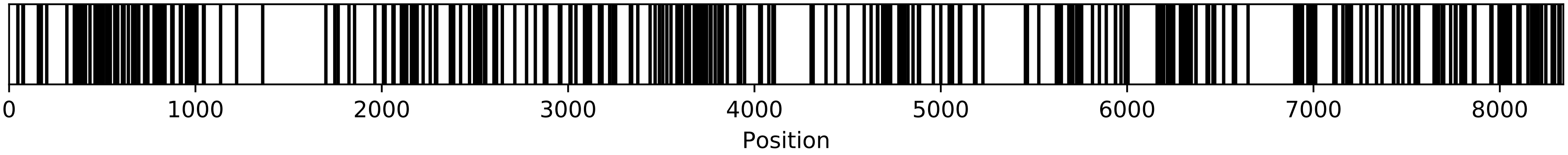


MM1 vs MM2
Number of differences: 216

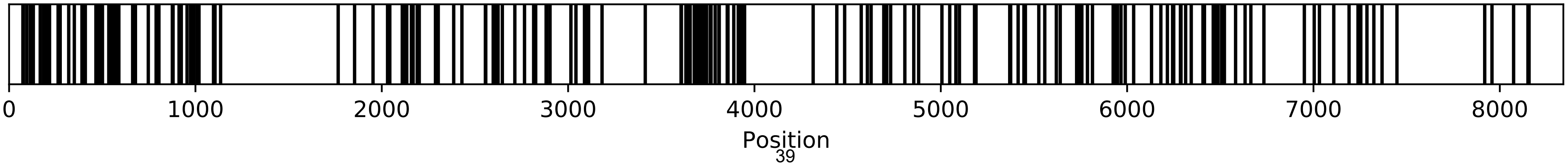

### Region: 97



CR1 vs CR2
Number of differences: 114

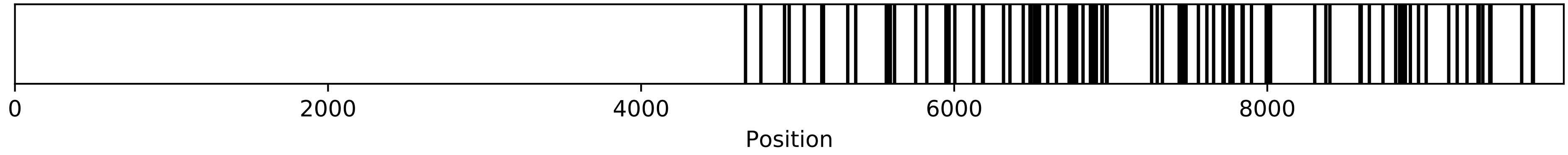


CR1 vs MM1
Number of differences: 187

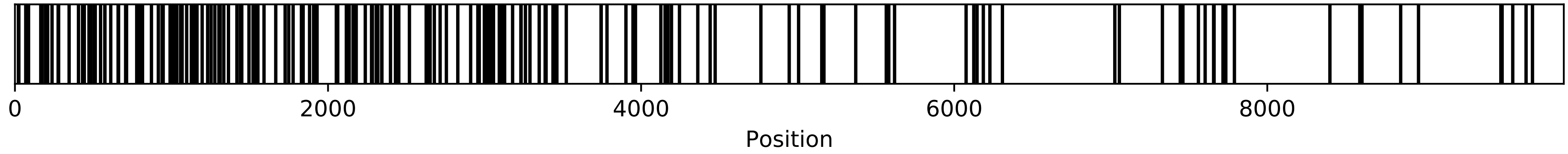


CR1 vs MM2
Number of differences: 236

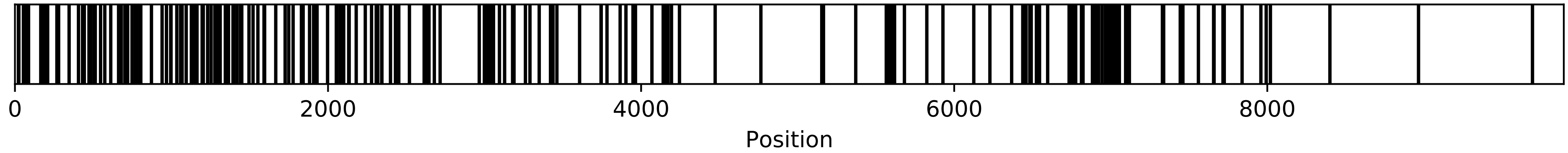


CR2 vs MM1
Number of differences: 261

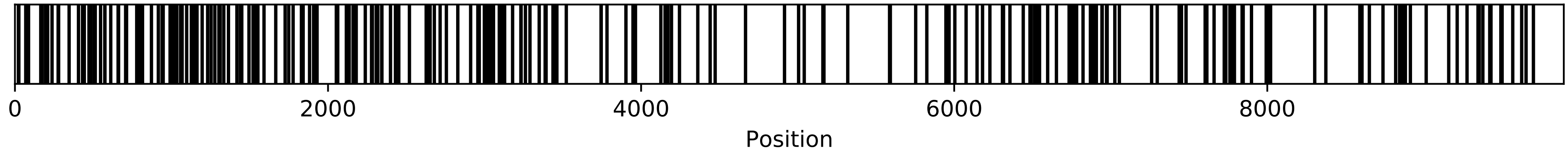


CR2 vs MM2
Number of differences: 282

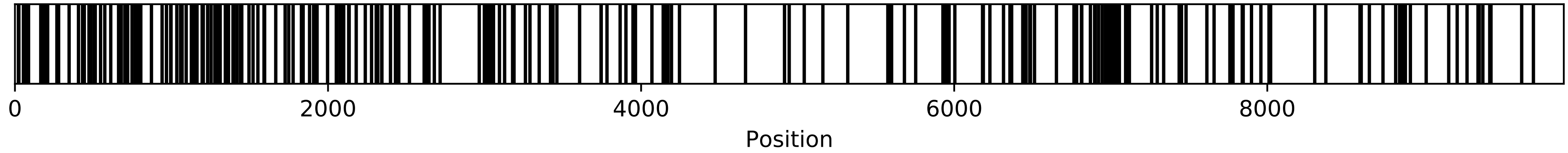


MM1 vs MM2
Number of differences: 155

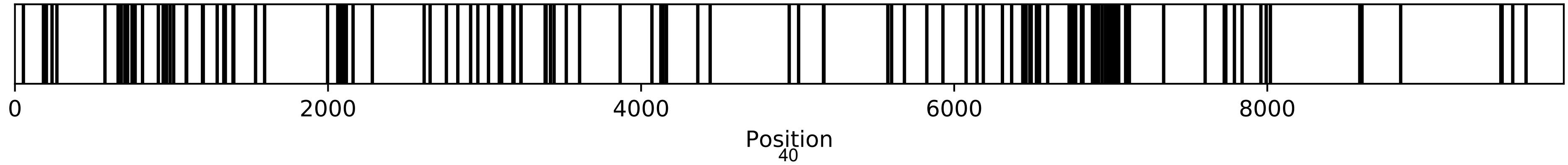

### Region: 98



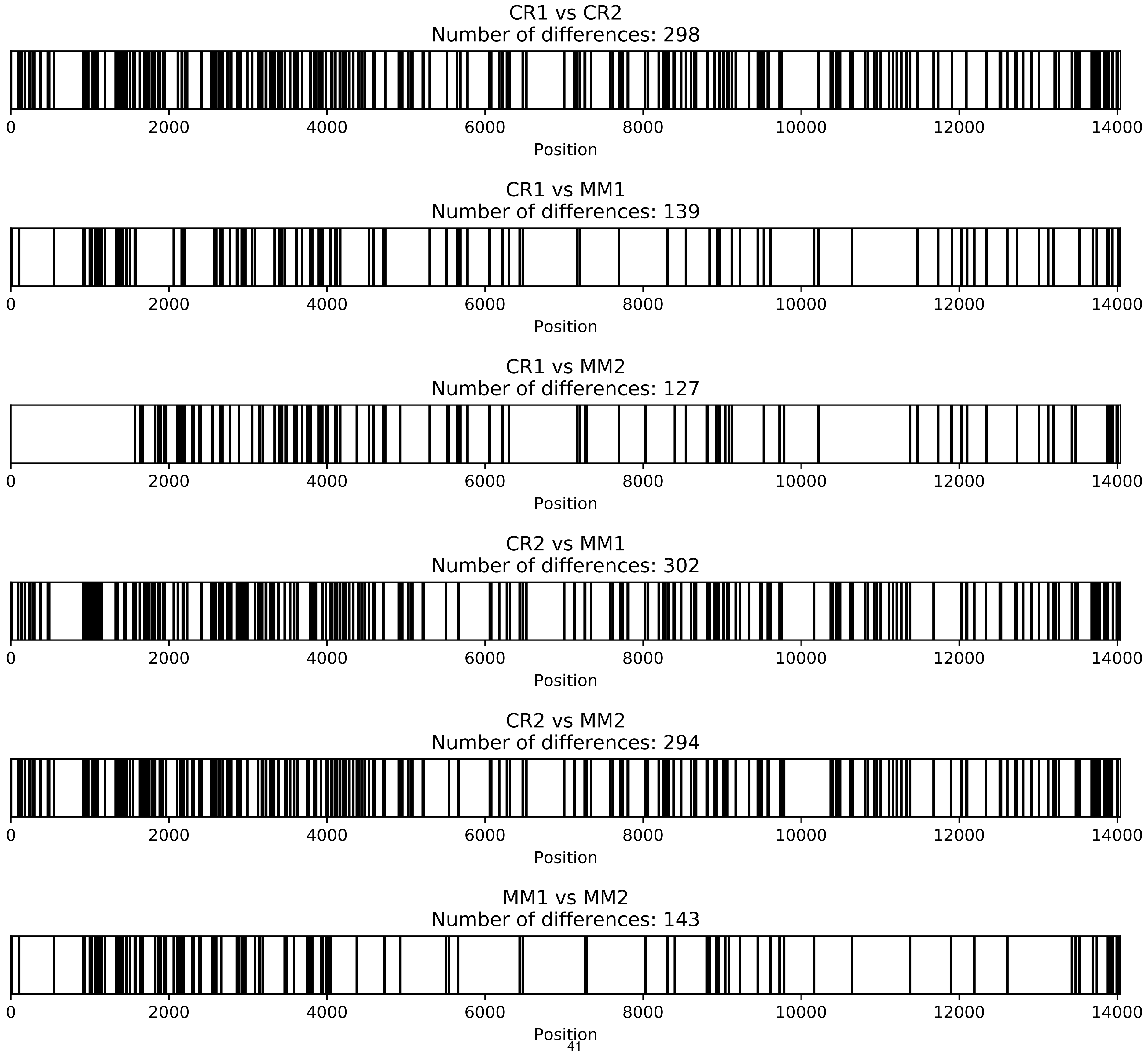
CR1 vs CR2
Number of differences: 298
0
2000
4000
6000
8000
10000
12000
14000
Position
CR1 vs MM1
Number of differences: 139
Position
CR1 vs MM2
Number of differences: 127
Position
CR2 vs MM1
Number of differences: 302
Position
CR2 vs MM2
Number of differences: 294
Position
MM1 vs MM2
Number of differences: 143
Position

### Region: 101



CR1 vs CR2
Number of differences: 359

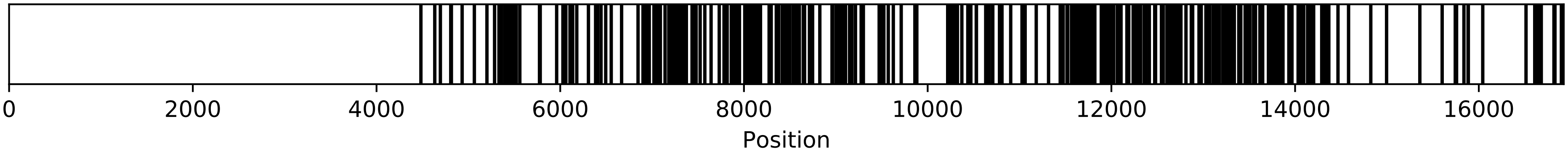


CR1 vs MM1
Number of differences: 314

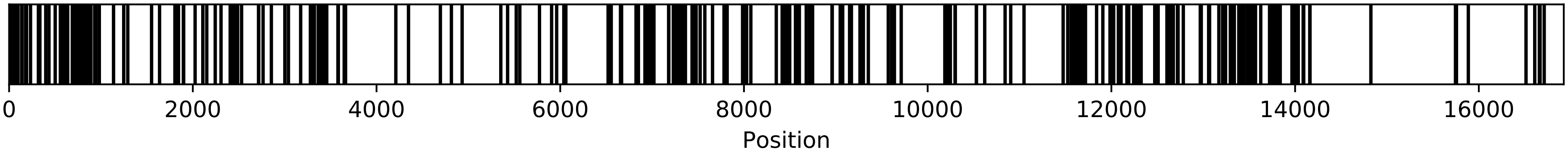


CR1 vs MM2
Number of differences: 325

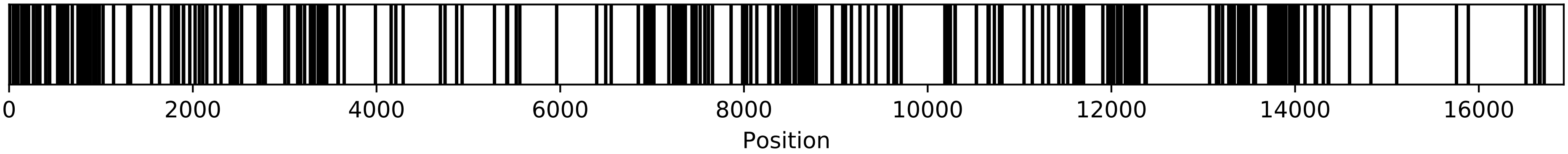


CR2 vs MM1
Number of differences: 397

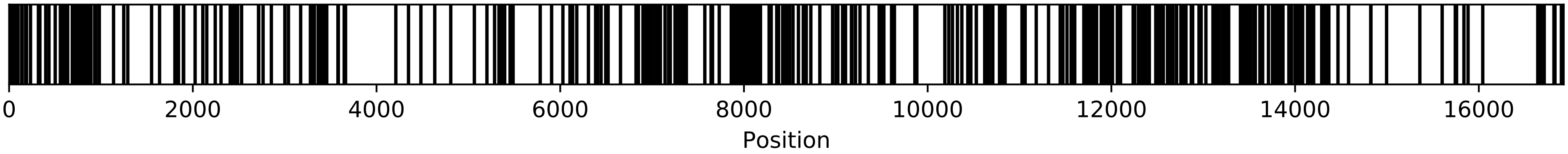


CR2 vs MM2
Number of differences: 462

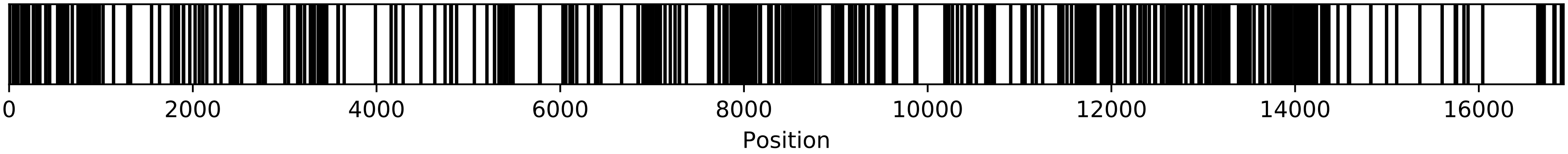


MM1 vs MM2
Number of differences: 243

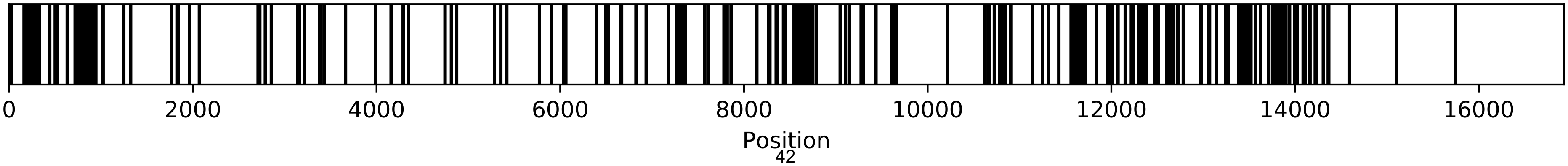

### Region: 107



CR1 vs CR2
Number of differences: 362

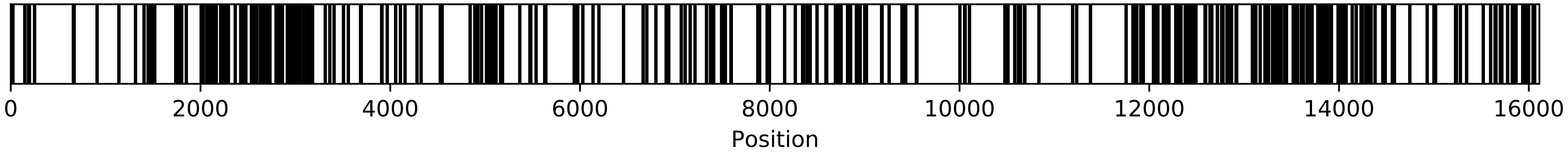


CR1 vs MM1
Number of differences: 183

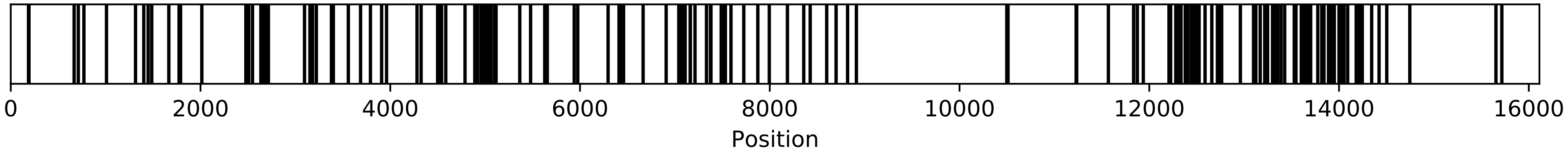


CR1 vs MM2
Number of differences: 146

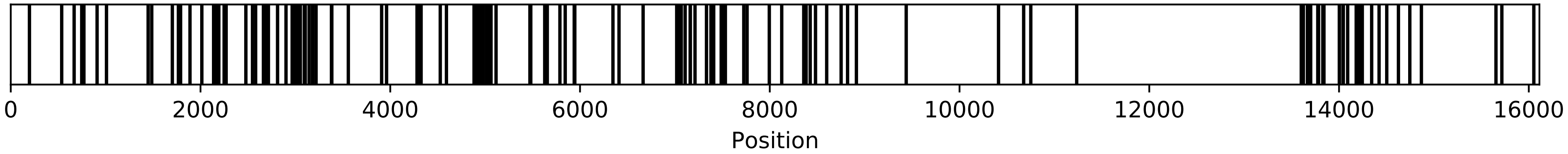


CR2 vs MM1
Number of differences: 347

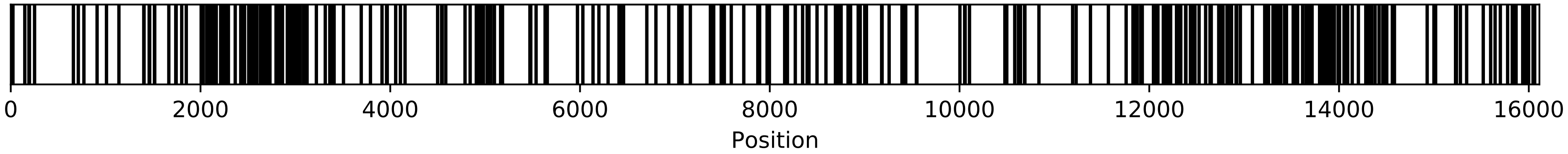


CR2 vs MM2
Number of differences: 334

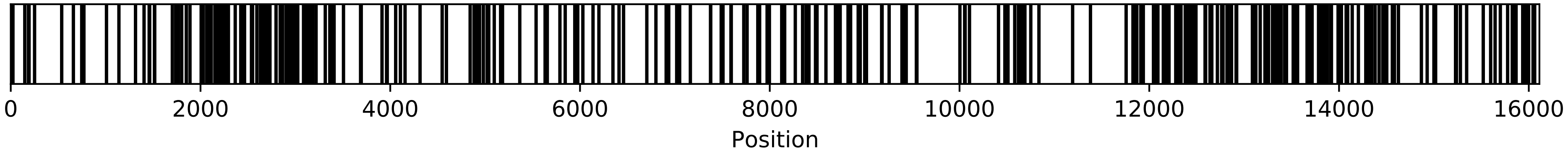


MM1 vs MM2
Number of differences: 163

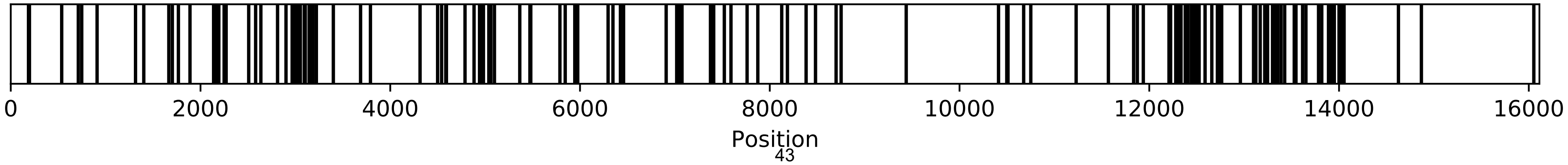

### Region: 109



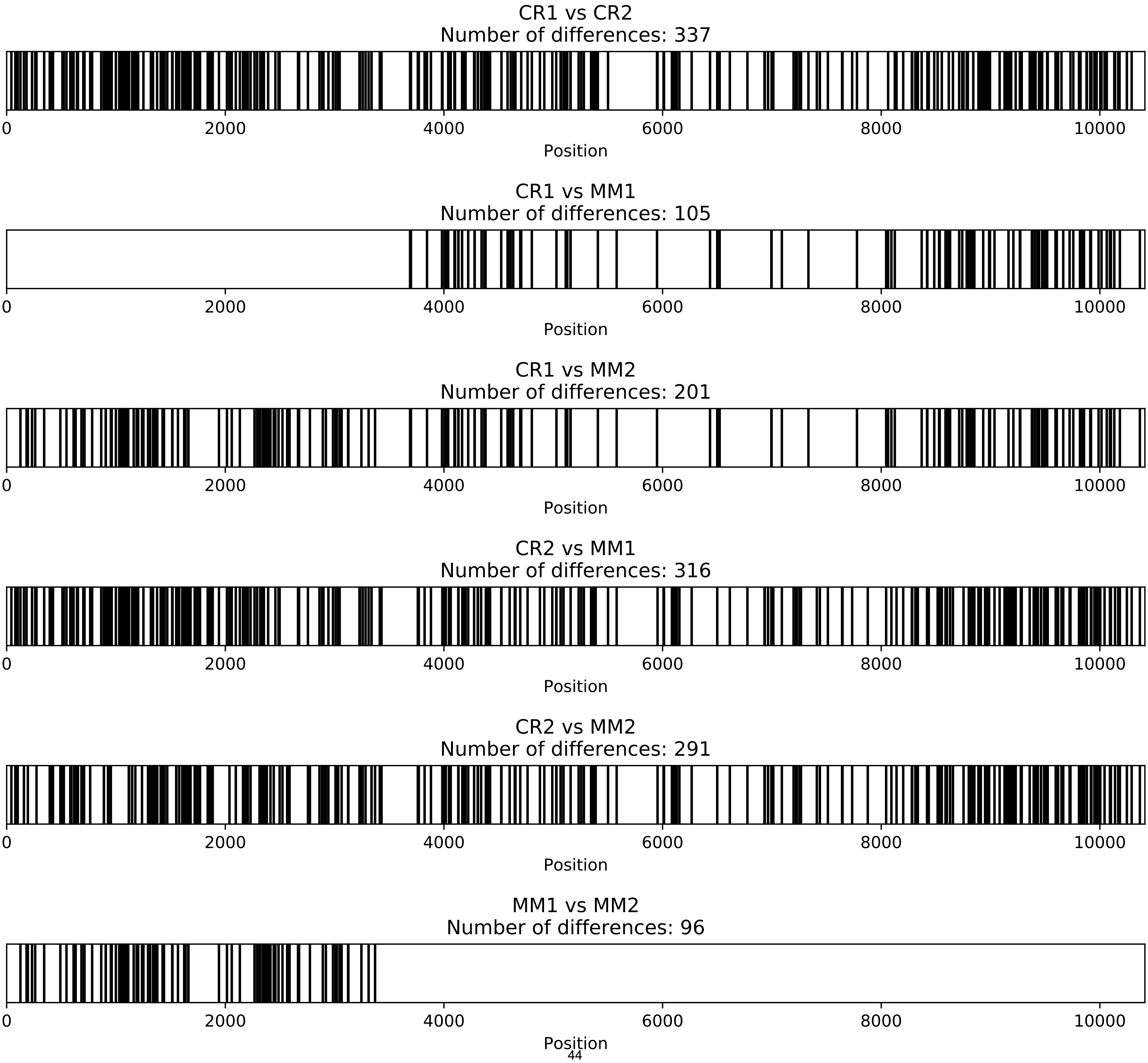
CR1 vs CR2
Number of differences: 337
Position
CR1 vs MM1
Number of differences: 105
Position
CR1 vs MM2
Number of differences: 201
Position
CR2 vs MM1
Number of differences: 316
Position
CR2 vs MM2
Number of differences: 291
Position
MM1 vs MM2
Number of differences: 96
Position
0
2000
4000
6000
8000
10000

### Region: 110



## CR1 vs CR2
Number of differences: 203

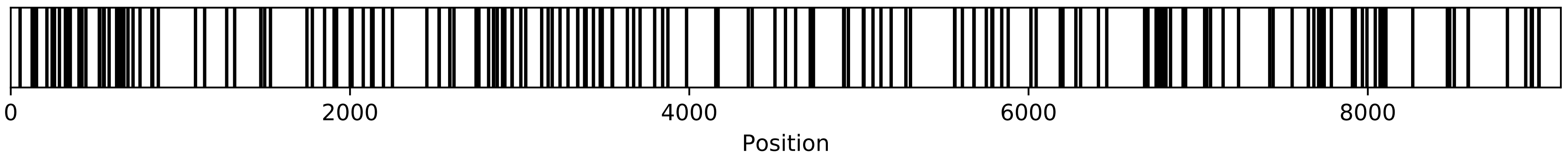


## CR1 vs MM1
Number of differences: 114

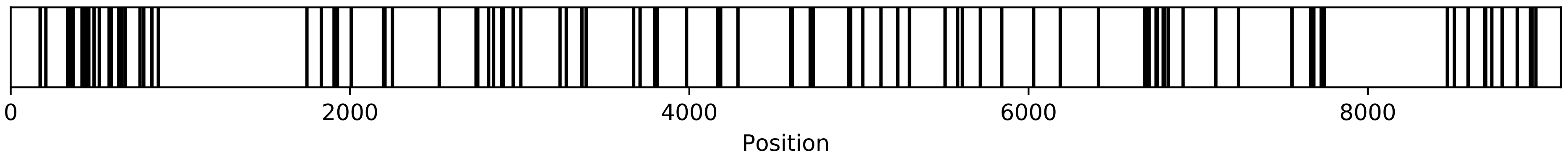


## CR1 vs MM2
Number of differences: 97

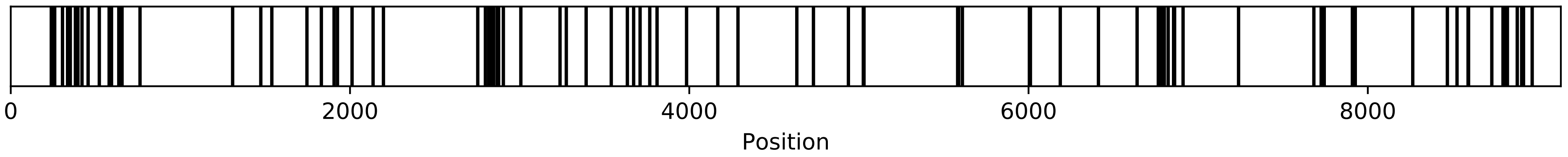


## CR2 vs MM1
Number of differences: 209

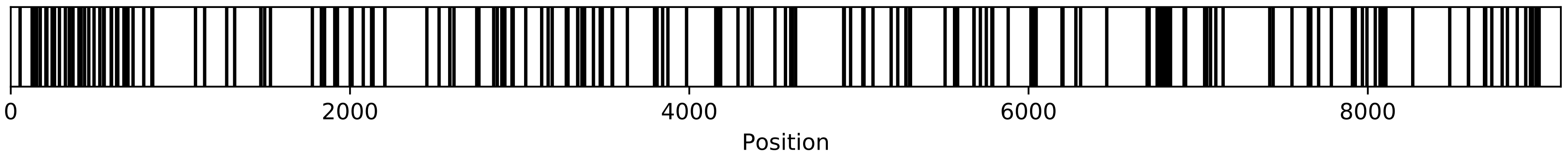


## CR2 vs MM2
Number of differences: 193

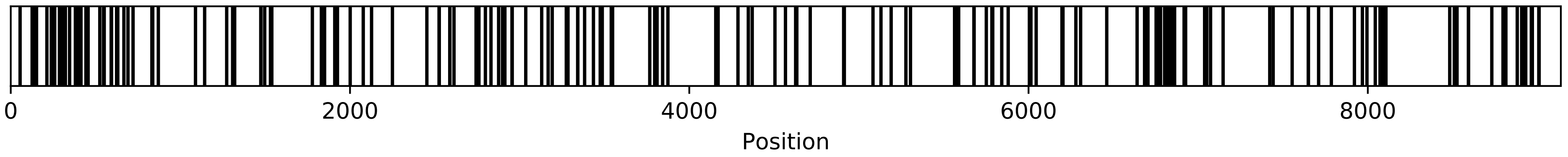


## MM1 vs MM2
Number of differences: 113

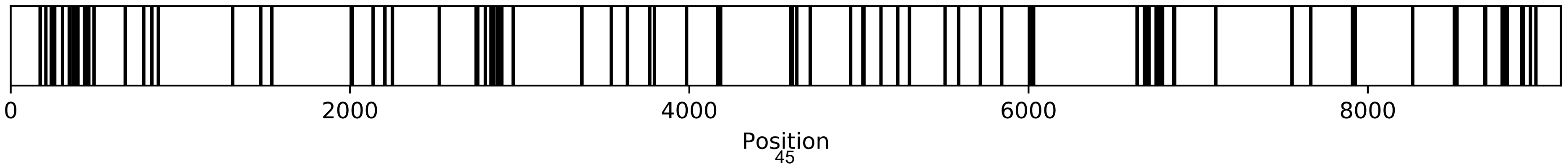

### Region: 113



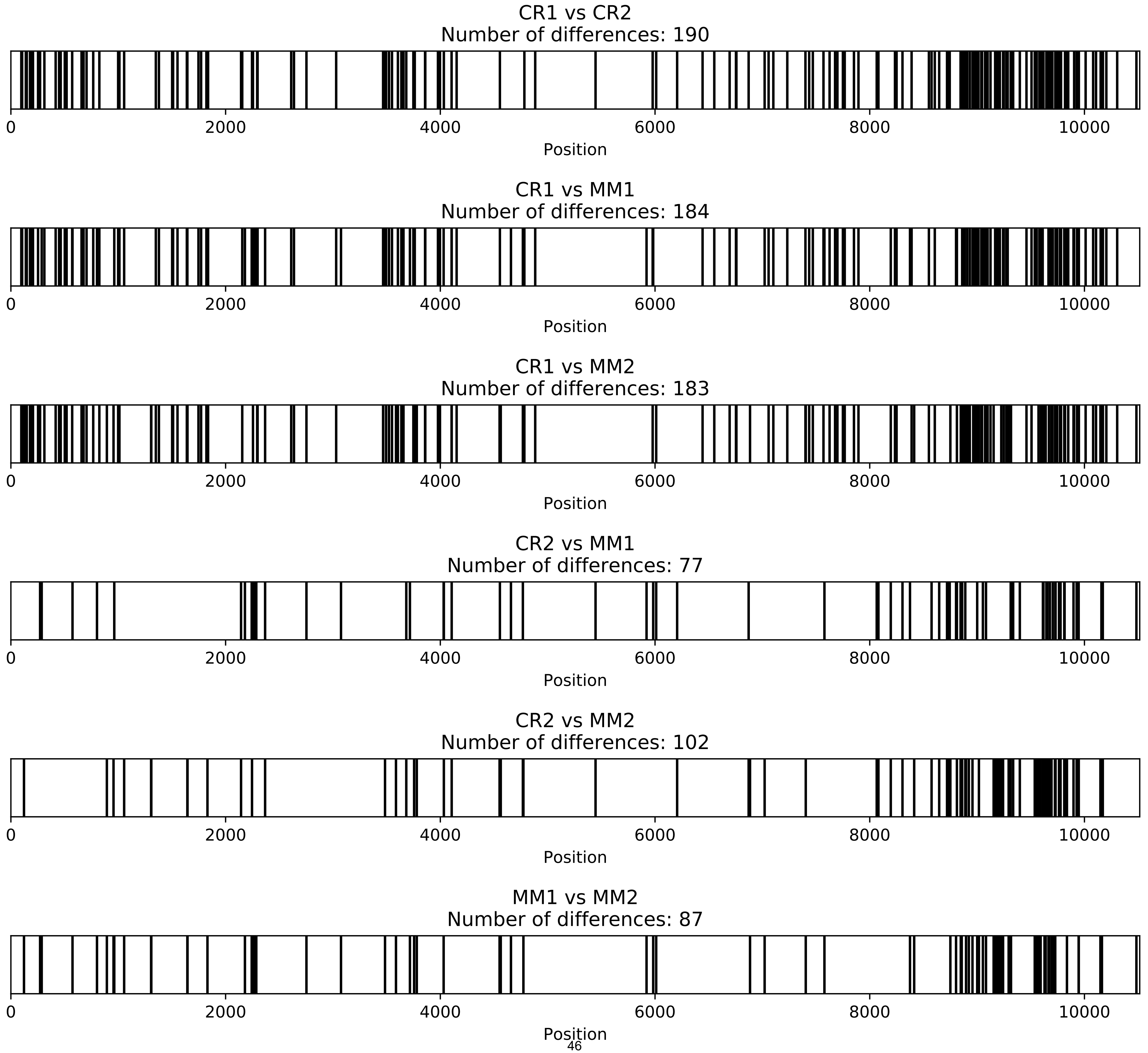
CR1 vs CR2
Number of differences: 190
CR1 vs MM1
Number of differences: 184
CR1 vs MM2
Number of differences: 183
CR2 vs MM1
Number of differences: 77
CR2 vs MM2
Number of differences: 102
MM1 vs MM2
Number of differences: 87
0
2000
4000
6000
8000
10000
Position

### Region: 114



## CR1 vs CR2
Number of differences: 242

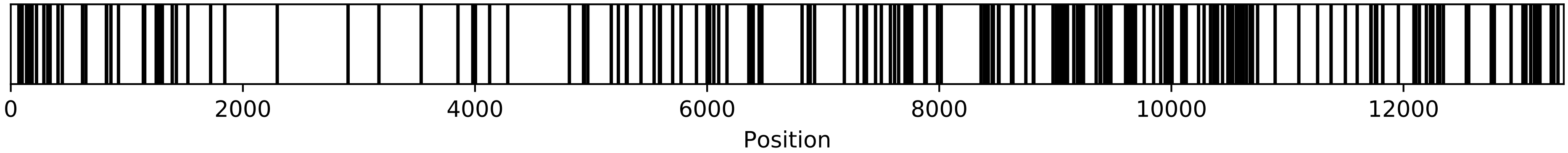


## CR1 vs MM1
Number of differences: 141

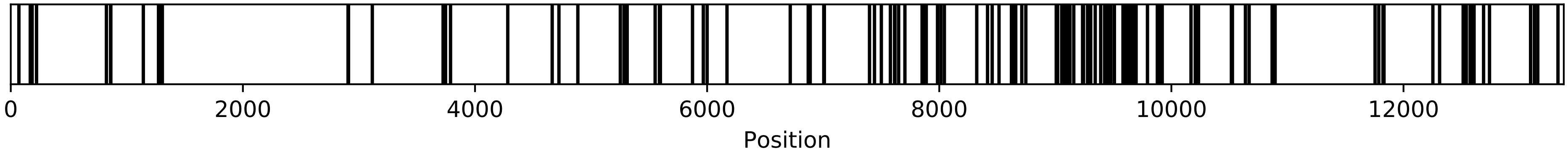


## CR1 vs MM2
Number of differences: 149

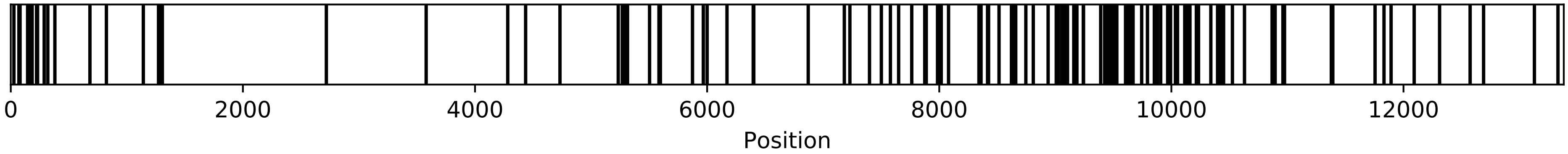


## CR2 vs MM1
Number of differences: 242

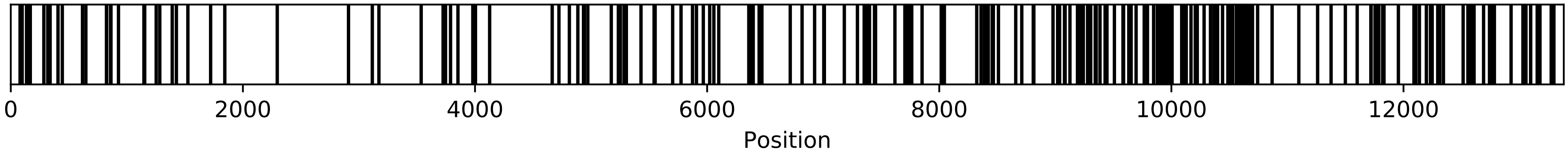


## CR2 vs MM2
Number of differences: 231

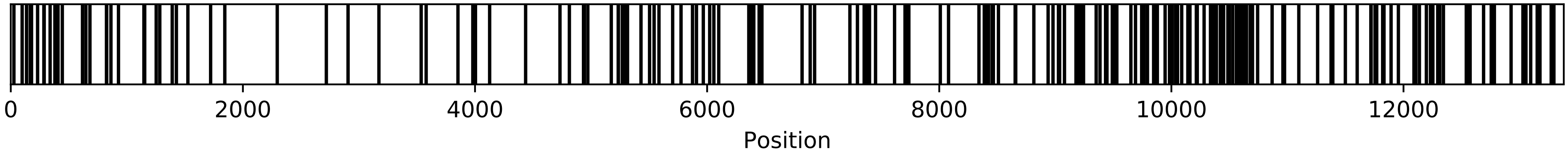


## MM1 vs MM2
Number of differences: 134

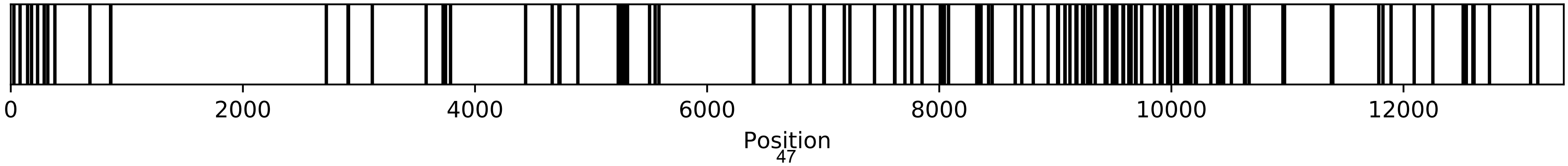

### Region: 115



CR1 vs CR2
Number of differences: 199

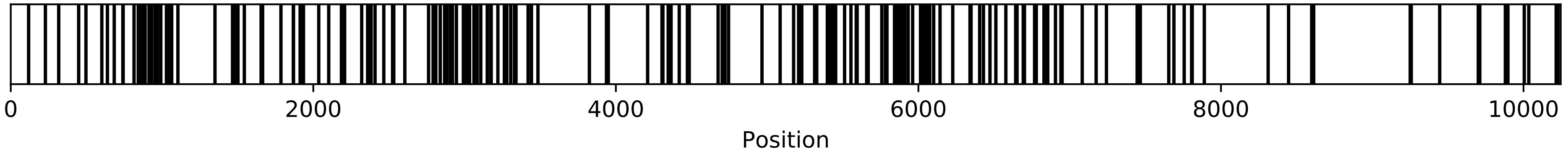


CR1 vs MM1
Number of differences: 97

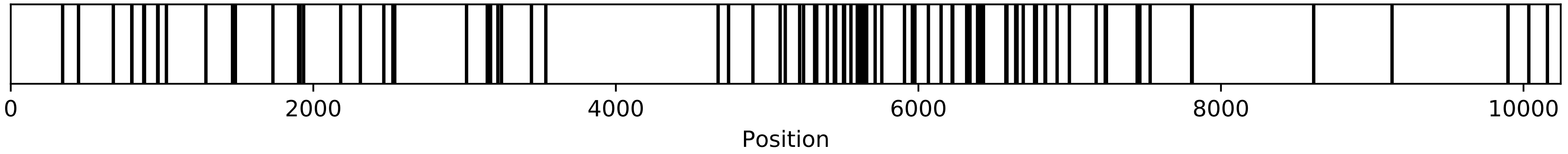


CR1 vs MM2
Number of differences: 85

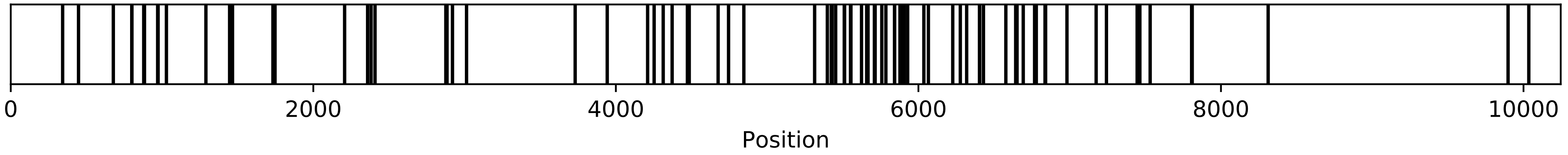


CR2 vs MM1
Number of differences: 205

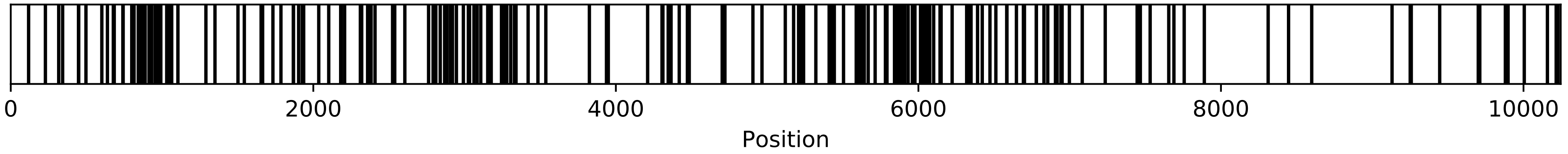


CR2 vs MM2
Number of differences: 192

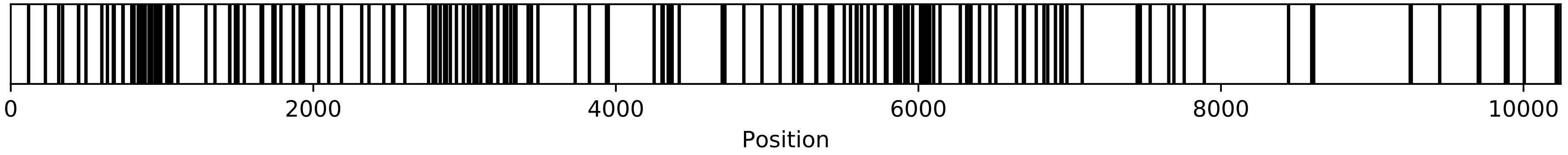


MM1 vs MM2
Number of differences: 87

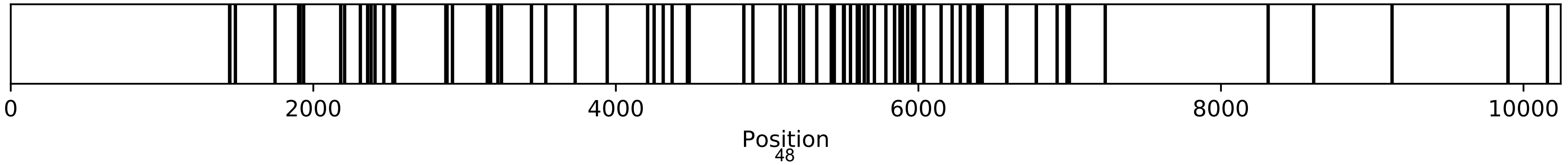

### Region: 117



CR1 vs CR2
Number of differences: 10

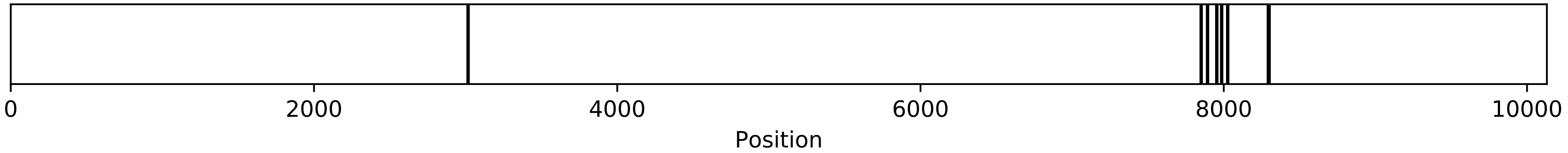


CR1 vs MM1
Number of differences: 236

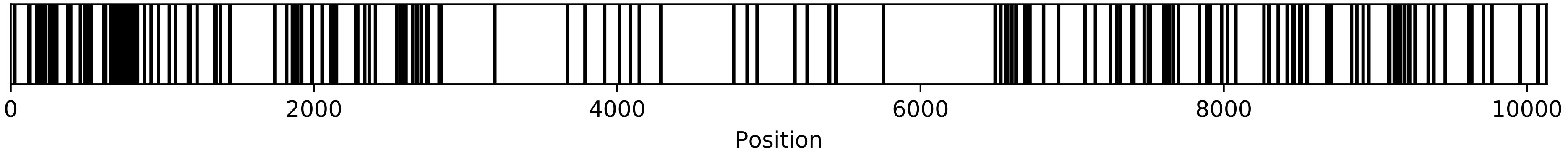


CR1 vs MM2
Number of differences: 211

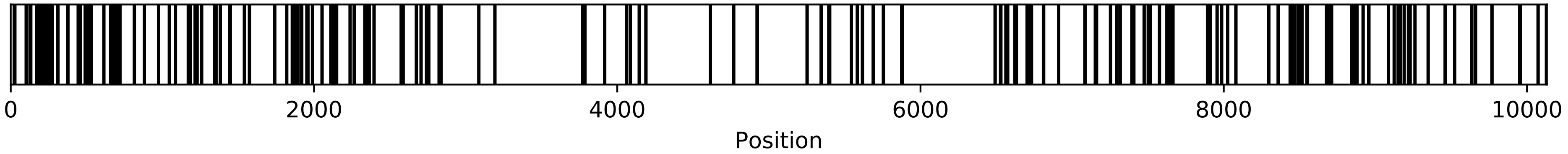


CR2 vs MM1
Number of differences: 237

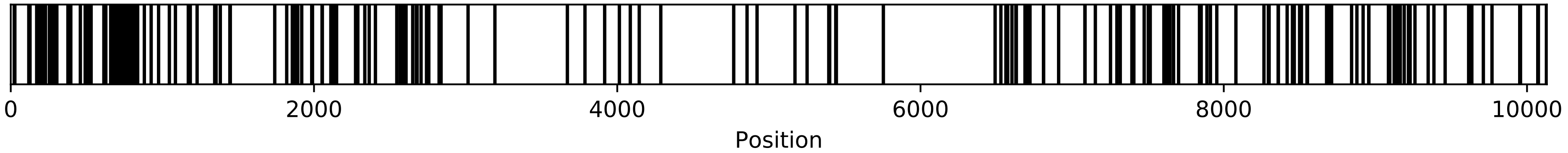


CR2 vs MM2
Number of differences: 212

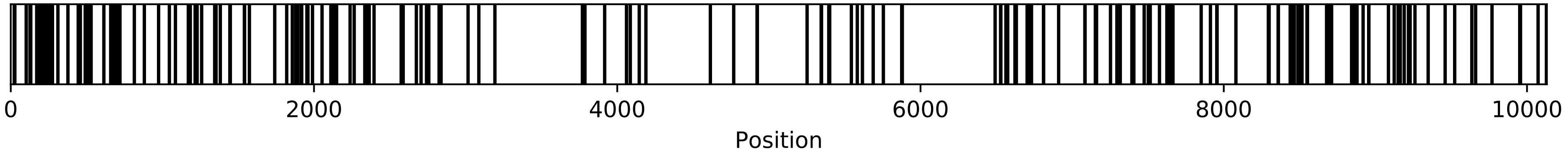


MM1 vs MM2
Number of differences: 147

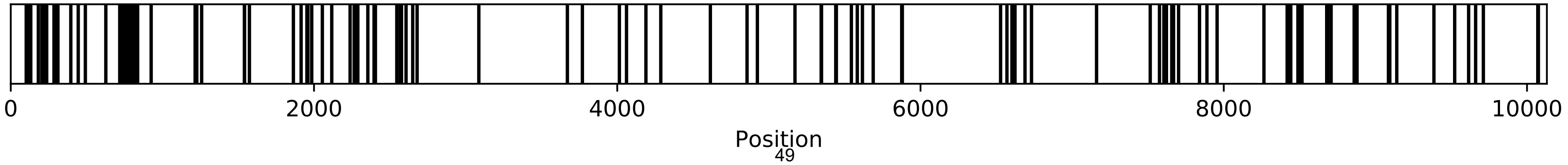

### Region: 120



CR1 vs CR2
Number of differences: 240

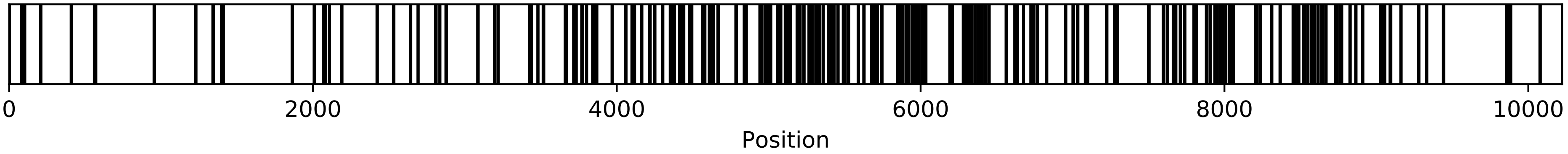


CR1 vs MM1
Number of differences: 182

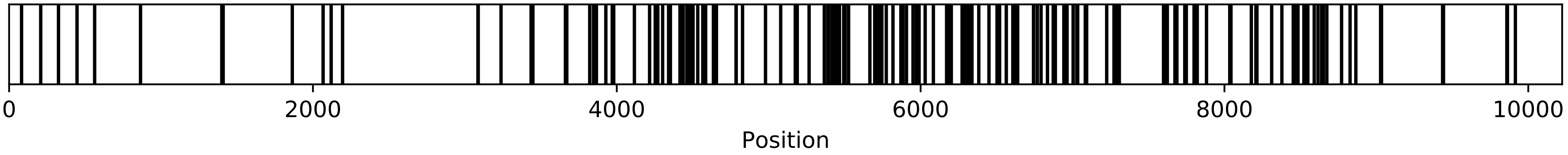


CR1 vs MM2
Number of differences: 152

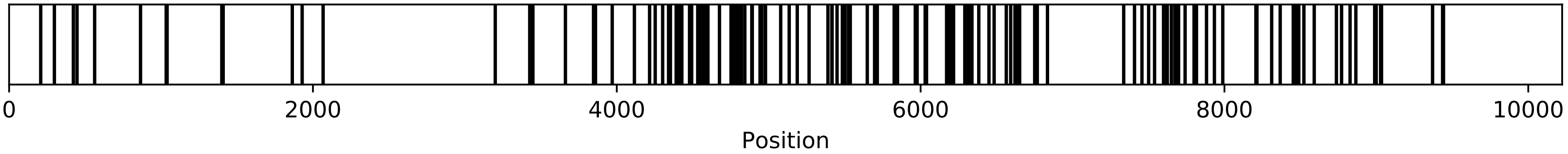


CR2 vs MM1
Number of differences: 229

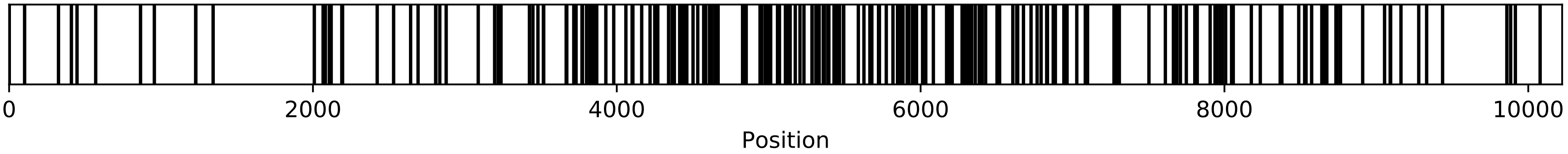


CR2 vs MM2
Number of differences: 236

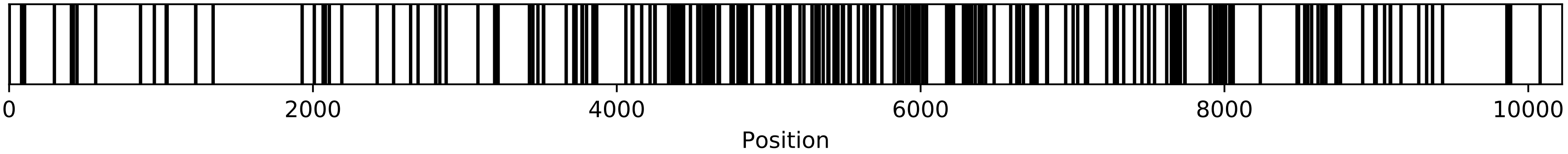


MM1 vs MM2
Number of differences: 155

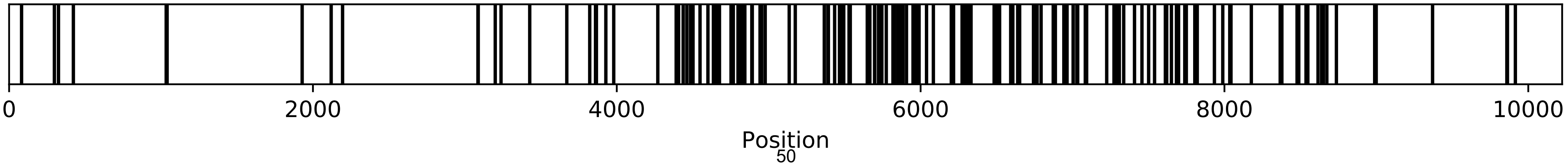

### Region: 121



CR1 vs CR2
Number of differences: 332

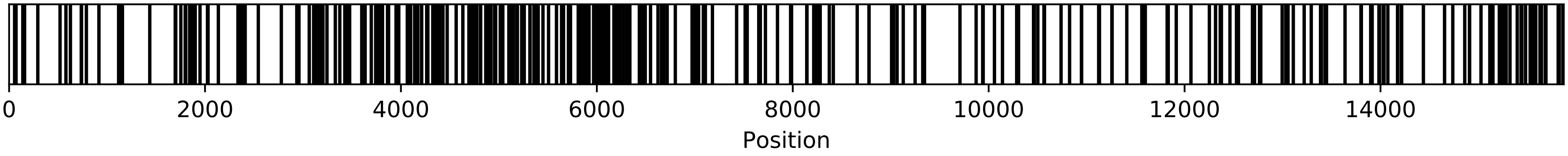


CR1 vs MM1
Number of differences: 172

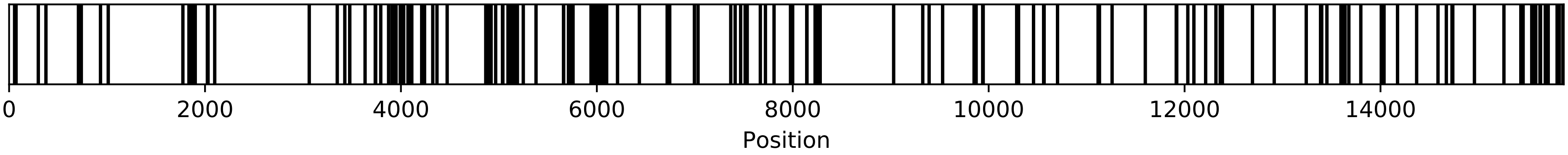


CR1 vs MM2
Number of differences: 175

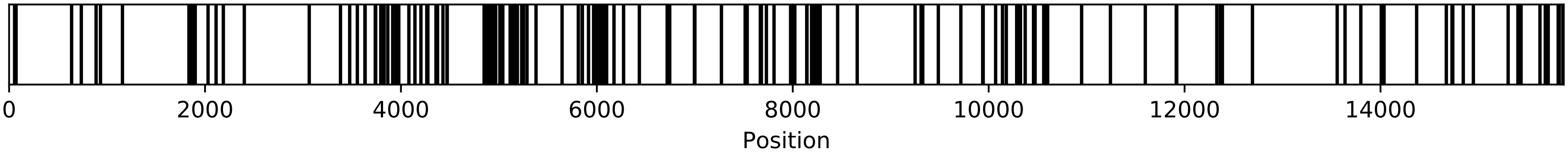


CR2 vs MM1
Number of differences: 341

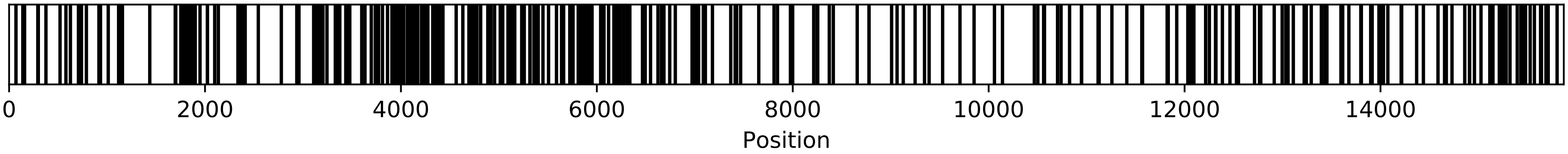


CR2 vs MM2
Number of differences: 335

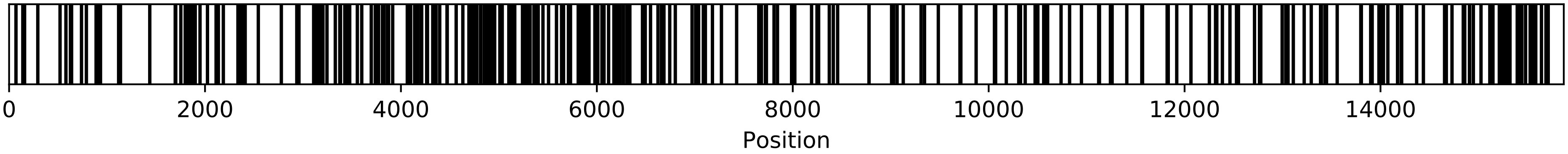


MM1 vs MM2
Number of differences: 171

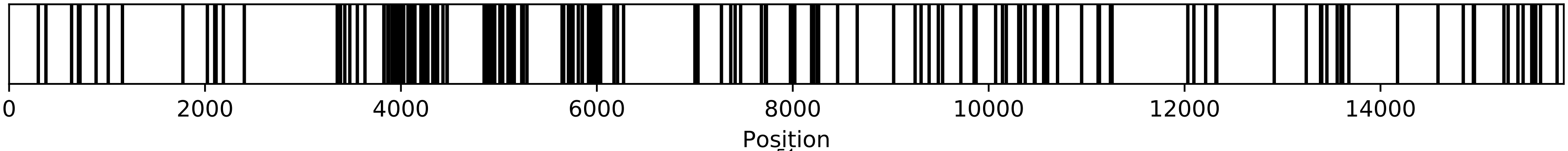

### Region: 124



CR1 vs CR2
Number of differences: 183

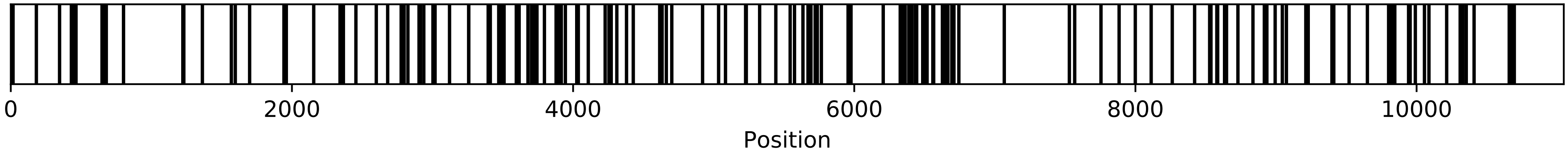


CR1 vs MM1
Number of differences: 86

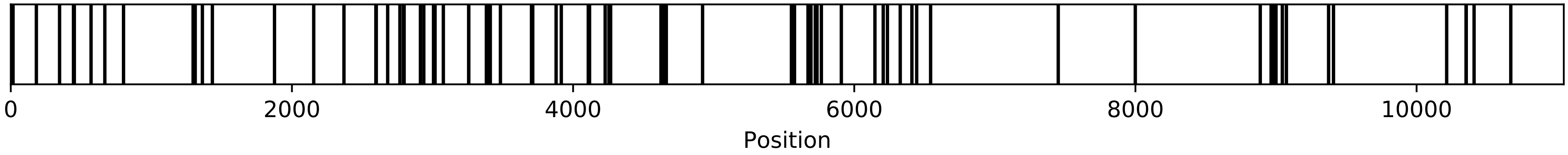


CR1 vs MM2
Number of differences: 128

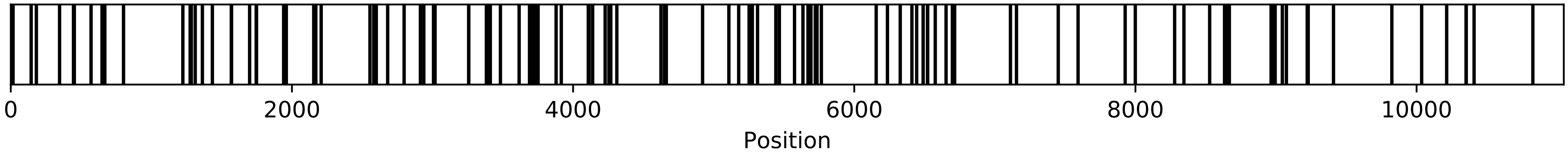


CR2 vs MM1
Number of differences: 166

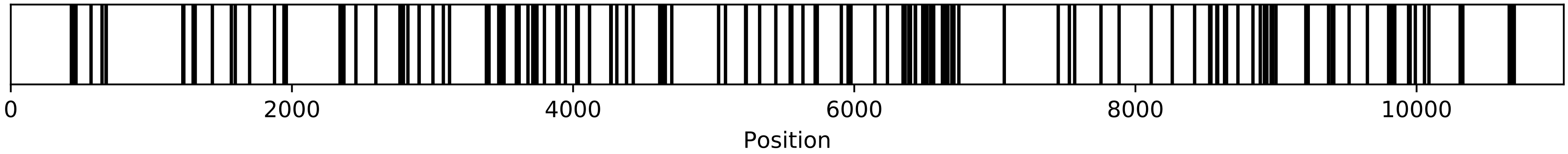


CR2 vs MM2
Number of differences: 174

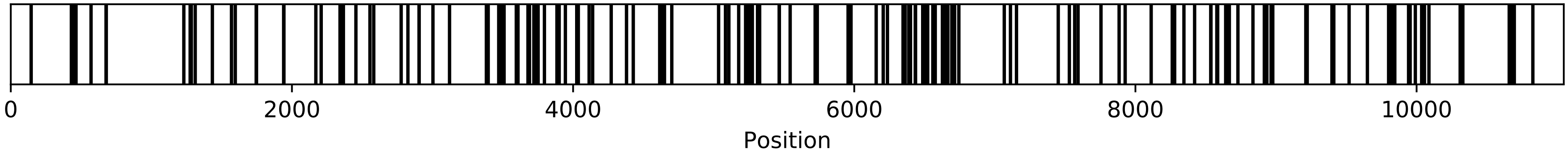


MM1 vs MM2
Number of differences: 86

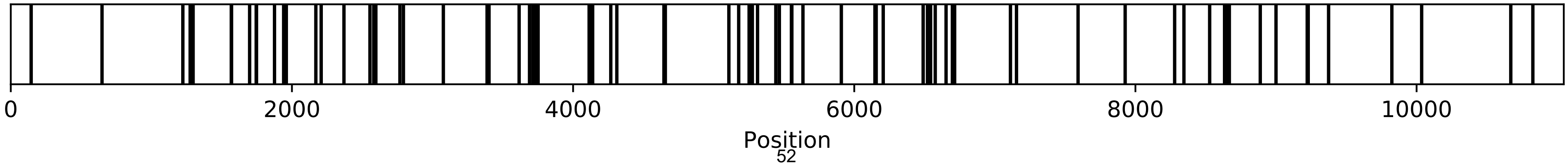

### Region: 125



CR1 vs CR2
Number of differences: 215

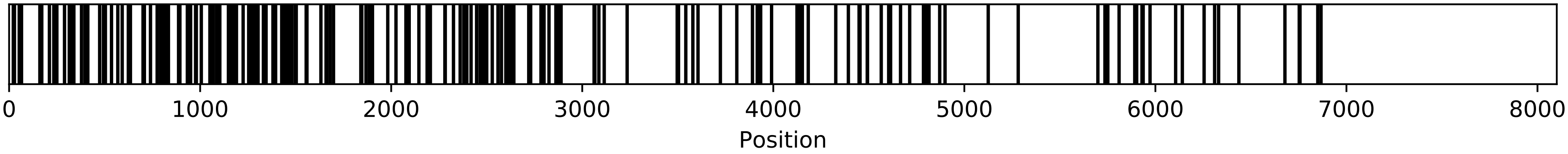


CR1 vs MM1
Number of differences: 135

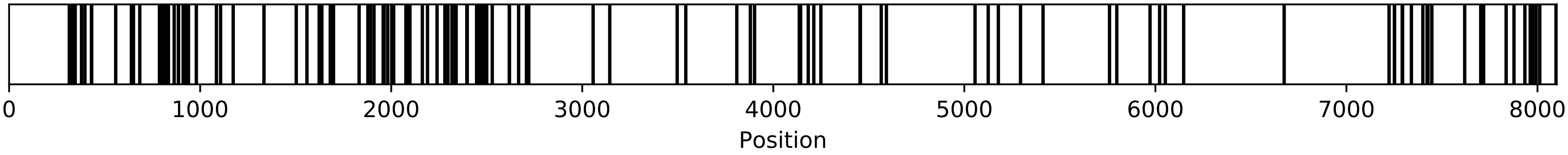


CR1 vs MM2
Number of differences: 110

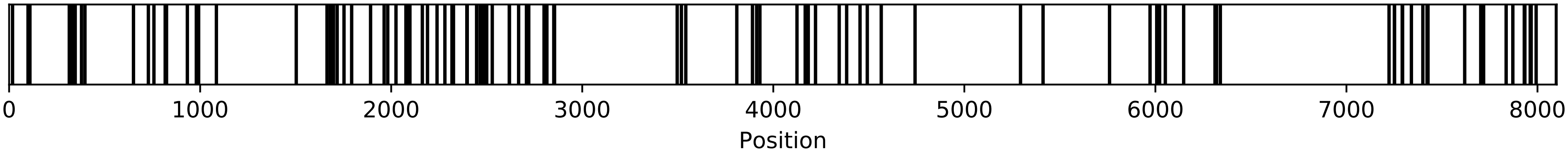


CR2 vs MM1
Number of differences: 258

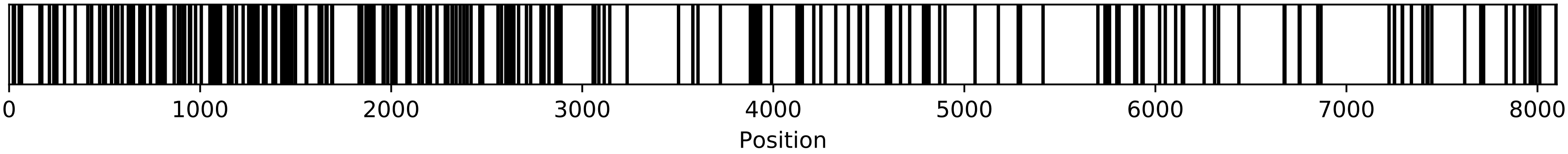


CR2 vs MM2
Number of differences: 239

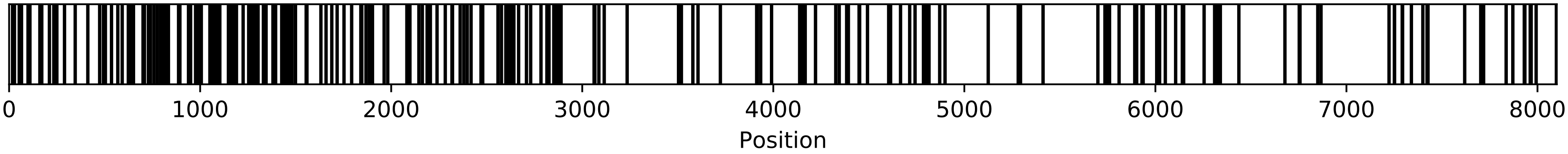


MM1 vs MM2
Number of differences: 93

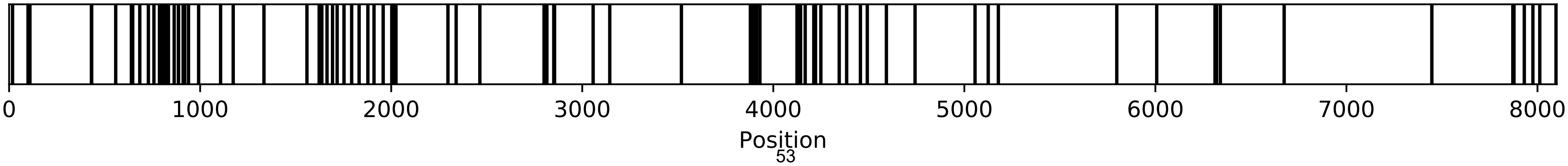

### Region: 135



CR1 vs CR2
Number of differences: 0

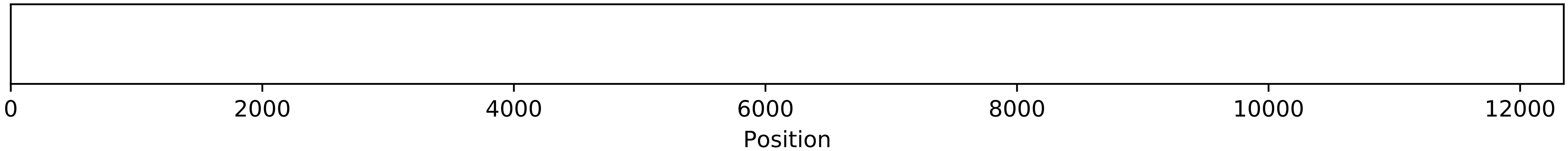


CR1 vs MM1
Number of differences: 307

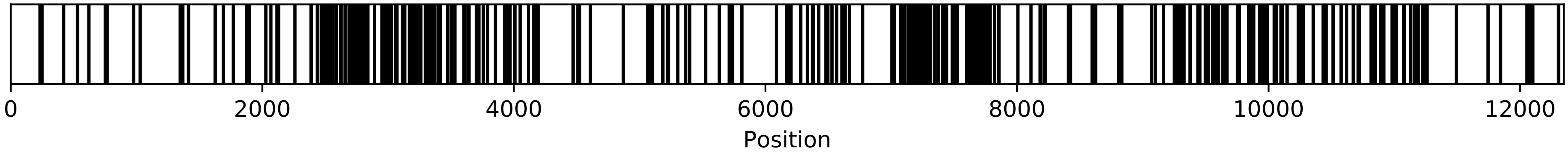


CR1 vs MM2
Number of differences: 320

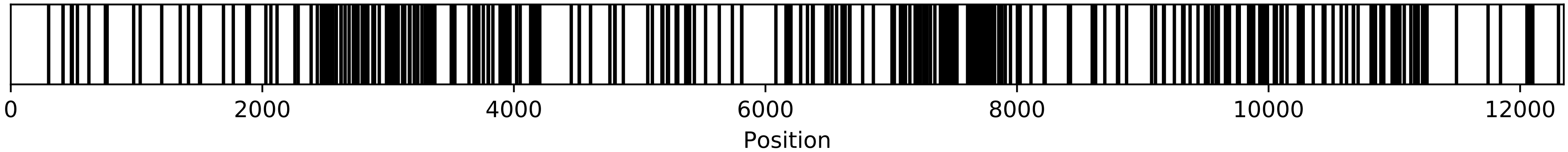


CR2 vs MM1
Number of differences: 307

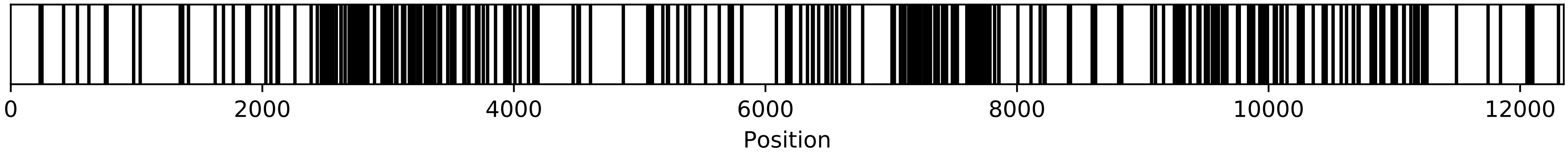


CR2 vs MM2
Number of differences: 320

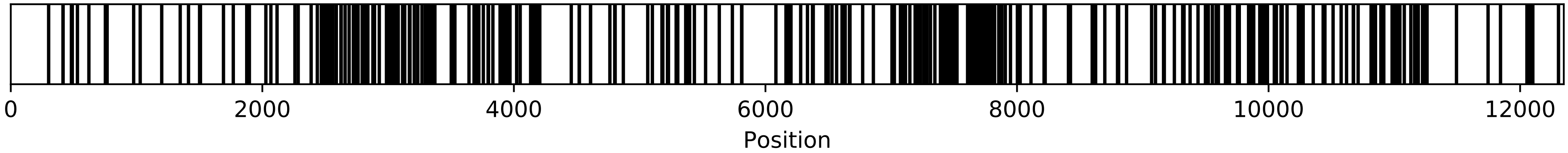


MM1 vs MM2
Number of differences: 165

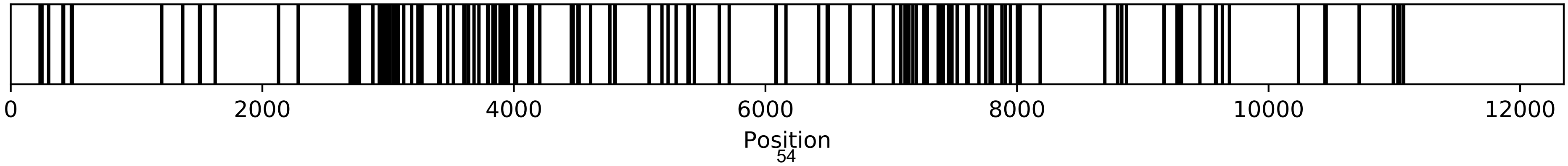

### Region: 146

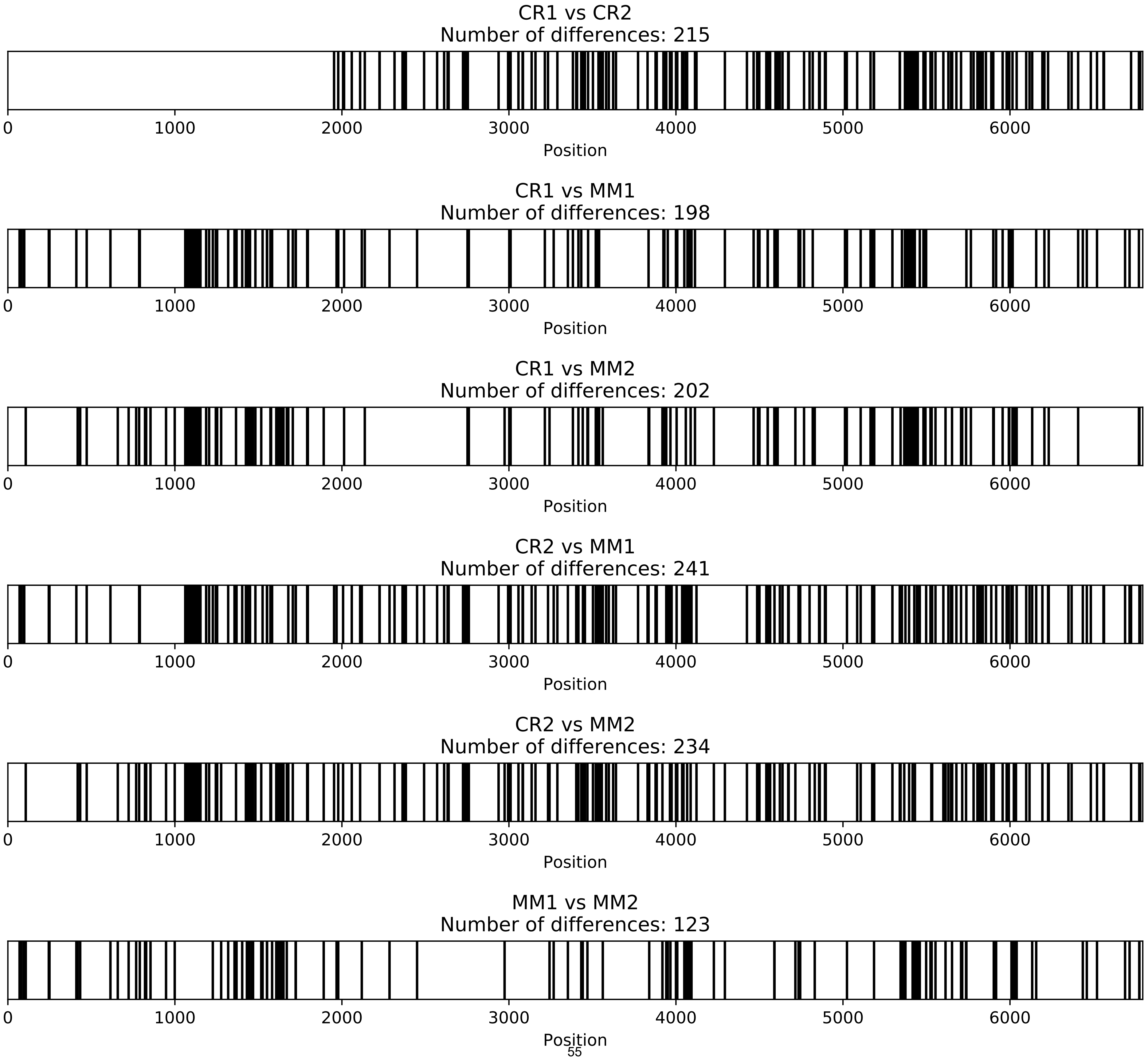
CR1 vs CR2
Number of differences: 215
CR1 vs MM1
Number of differences: 198
CR1 vs MM2
Number of differences: 202
CR2 vs MM1
Number of differences: 241
CR2 vs MM2
Number of differences: 234
MM1 vs MM2
Number of differences: 123
0
1000
2000
3000
4000
5000
6000
Position

### Region: 150



CR1 vs CR2
Number of differences: 264

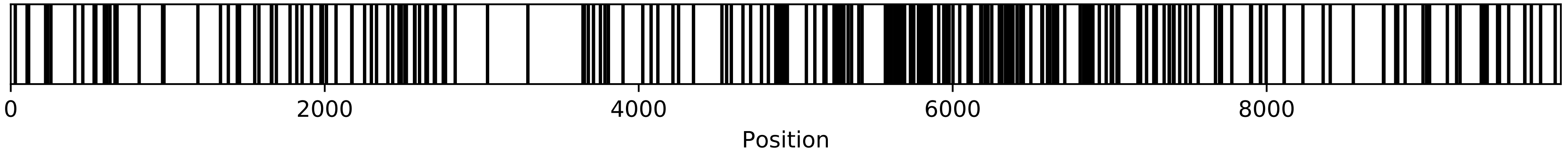


CR1 vs MM1
Number of differences: 249

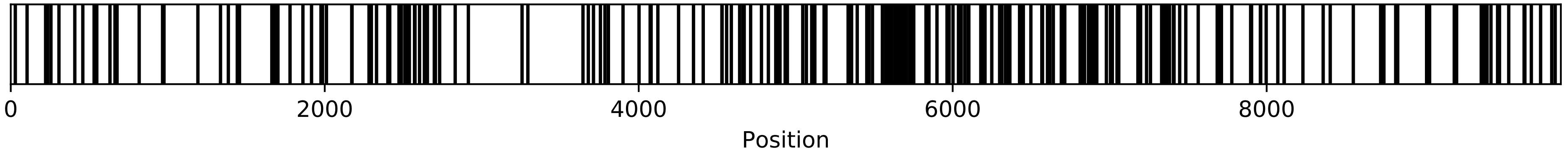


CR1 vs MM2
Number of differences: 263

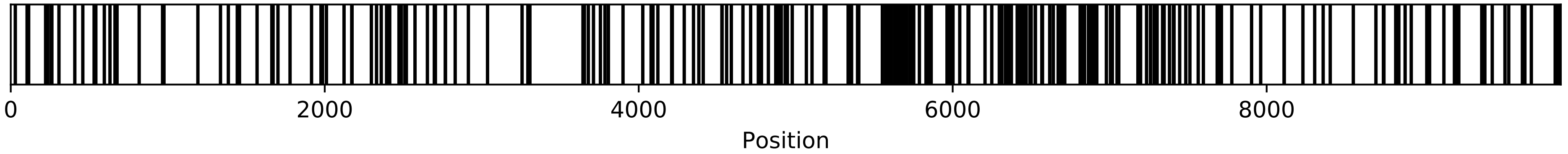


CR2 vs MM1
Number of differences: 143

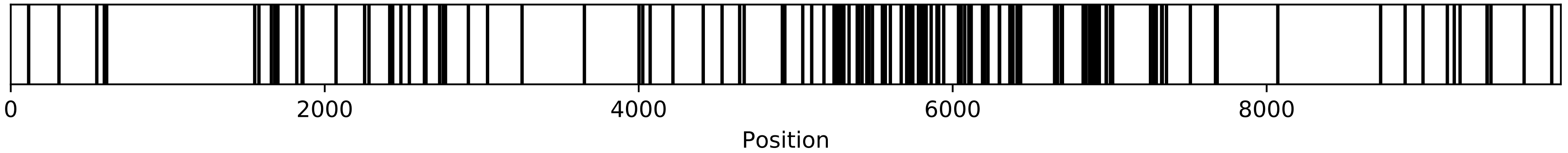


CR2 vs MM2
Number of differences: 164

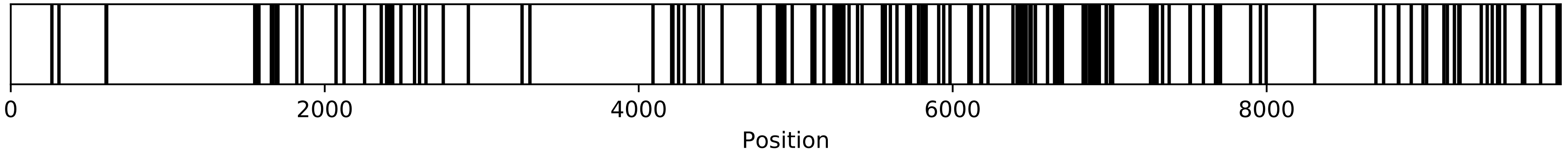


MM1 vs MM2
Number of differences: 131

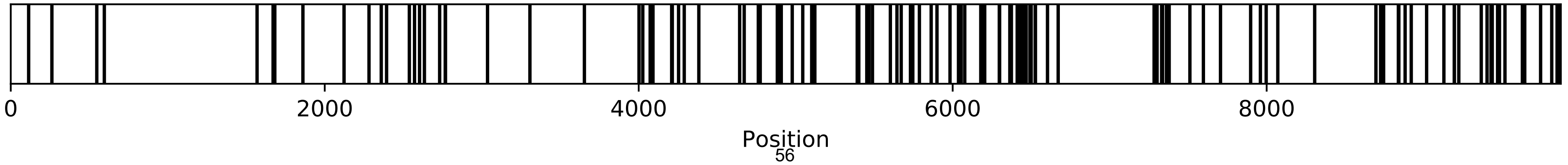

### Region: 151



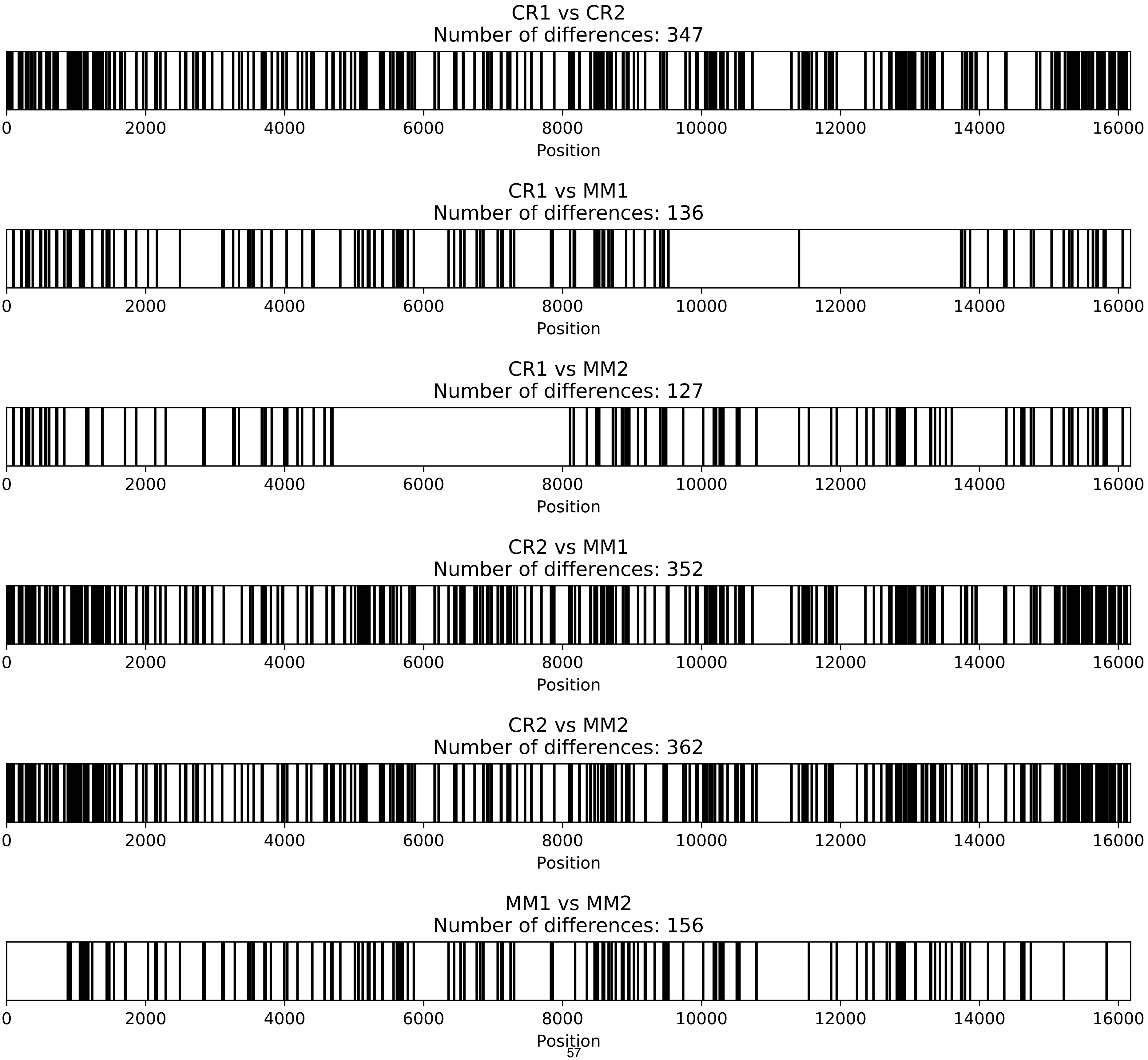
CR1 vs CR2
Number of differences: 347
CR1 vs MM1
Number of differences: 136
CR1 vs MM2
Number of differences: 127
CR2 vs MM1
Number of differences: 352
CR2 vs MM2
Number of differences: 362
MM1 vs MM2
Number of differences: 156
0 2000 4000 6000 8000 10000 12000 14000 16000
Position

### Region: 152



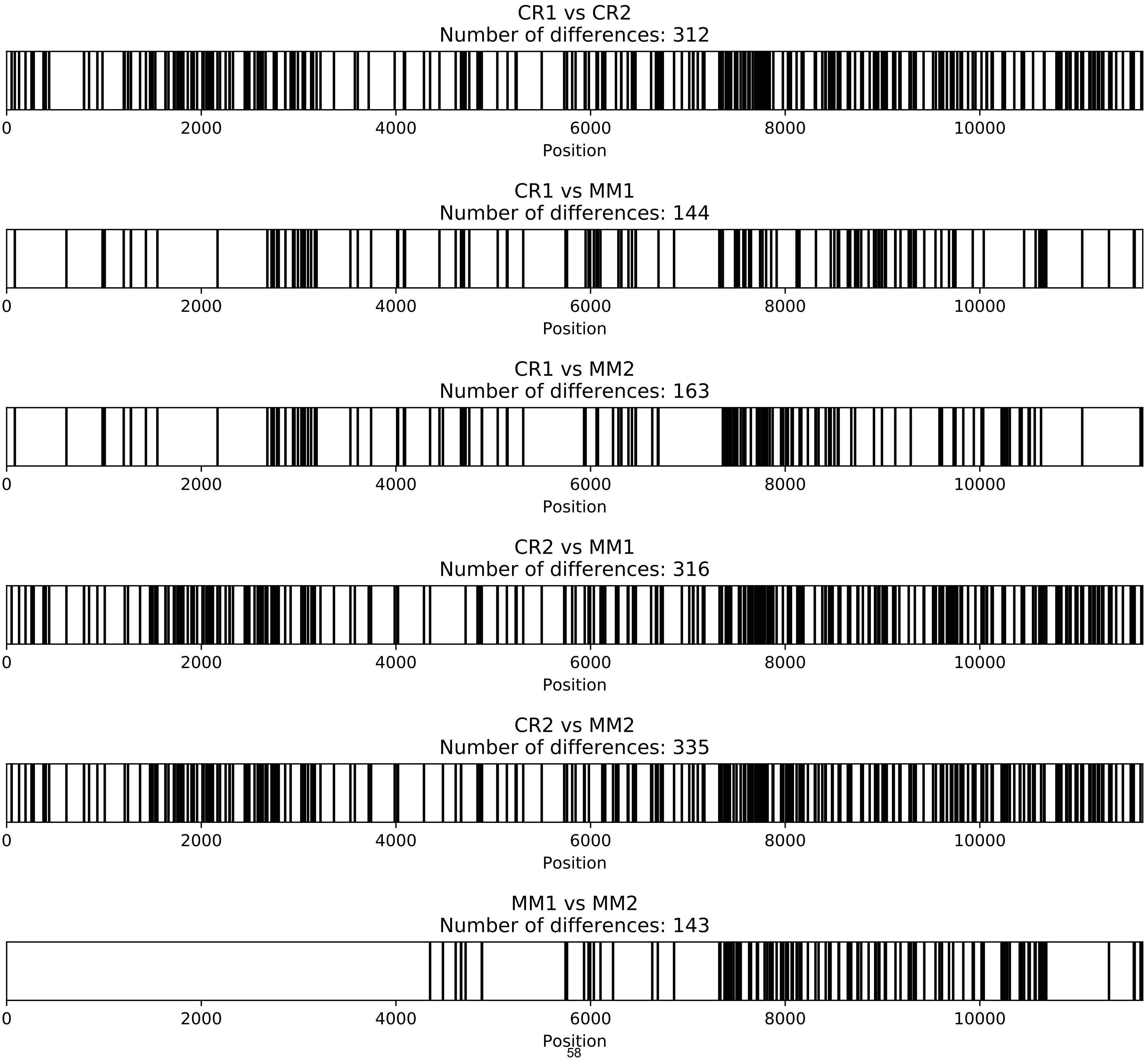
CR1 vs CR2
Number of differences: 312
0
2000
4000
6000
8000
10000
Position
CR1 vs MM1
Number of differences: 144
0
2000
4000
6000
8000
10000
Position
CR1 vs MM2
Number of differences: 163
0
2000
4000
6000
8000
10000
Position
CR2 vs MM1
Number of differences: 316
0
2000
4000
6000
8000
10000
Position
CR2 vs MM2
Number of differences: 335
0
2000
4000
6000
8000
10000
Position
MM1 vs MM2
Number of differences: 143
0
2000
4000
6000
8000
10000
Position

### Region: 156



CR1 vs CR2
Number of differences: 363

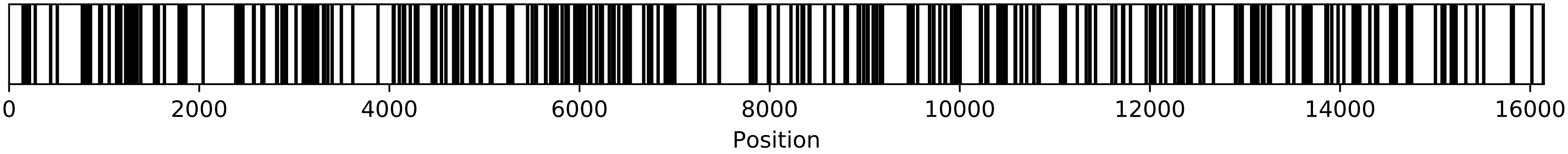


CR1 vs MM1
Number of differences: 372

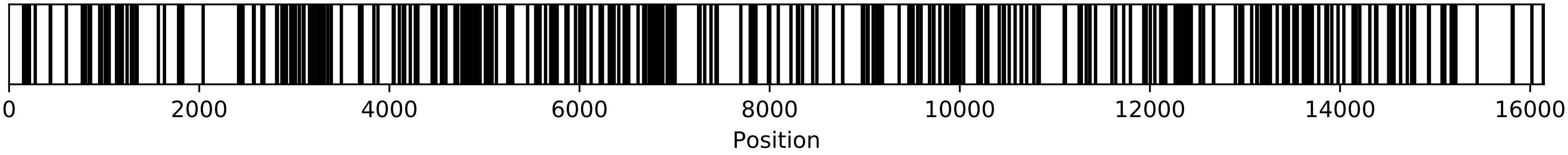


CR1 vs MM2
Number of differences: 339

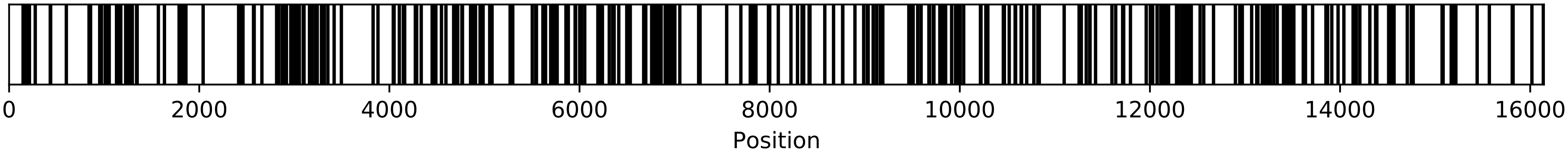


CR2 vs MM1
Number of differences: 245

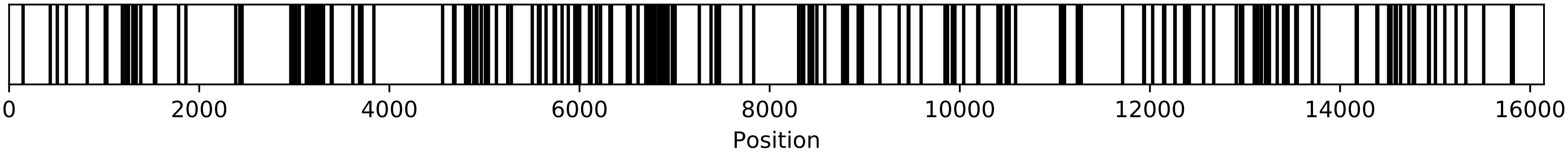


CR2 vs MM2
Number of differences: 206

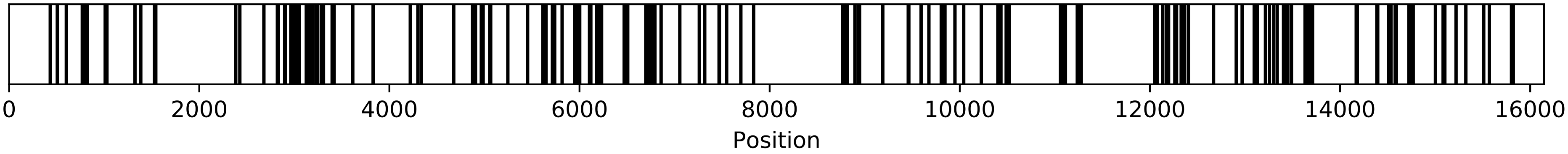


MM1 vs MM2
Number of differences: 189

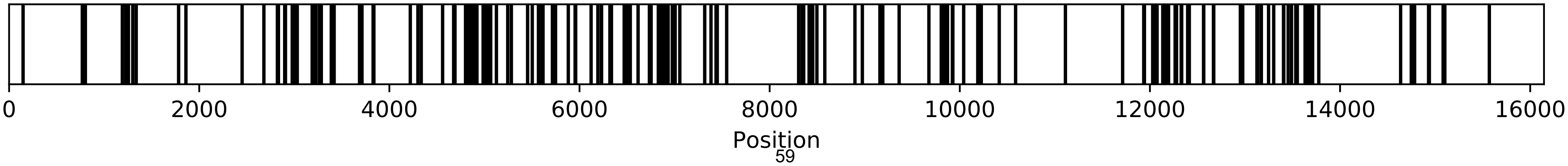

### Region: 158



CR1 vs CR2
Number of differences: 232

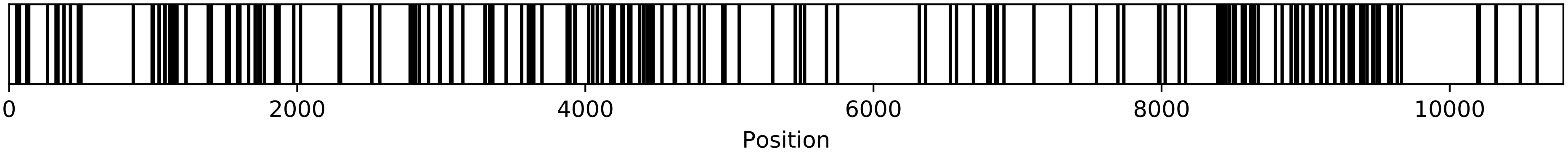


CR1 vs MM1
Number of differences: 74

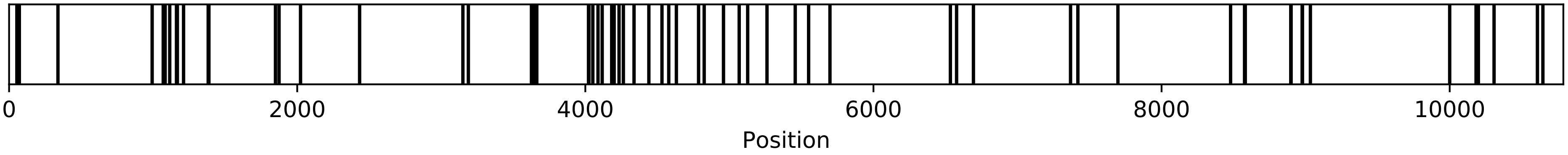


CR1 vs MM2
Number of differences: 14

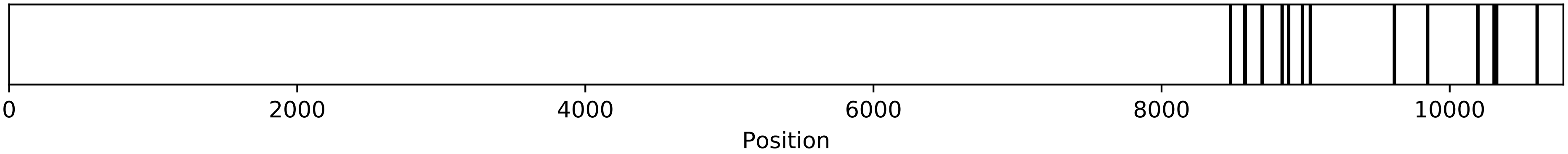


CR2 vs MM1
Number of differences: 244

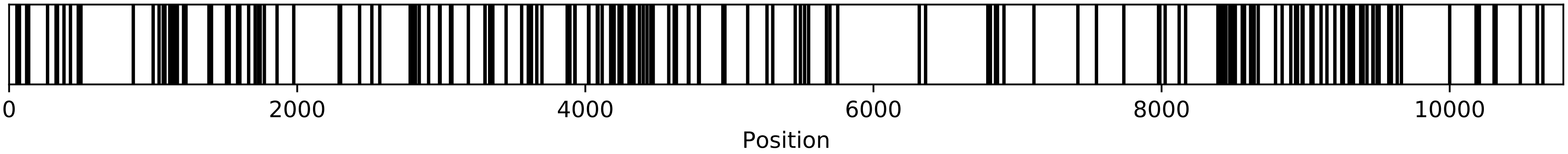


CR2 vs MM2
Number of differences: 236

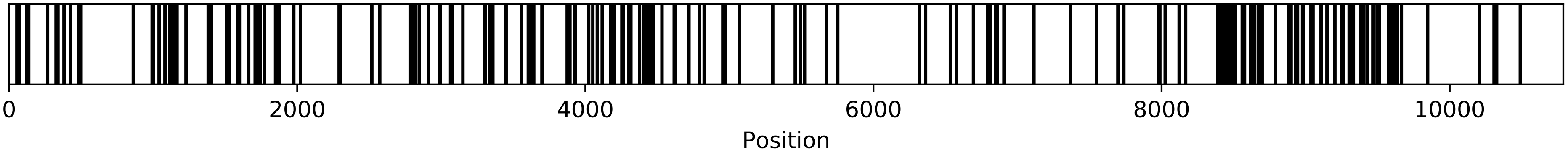


MM1 vs MM2
Number of differences: 75

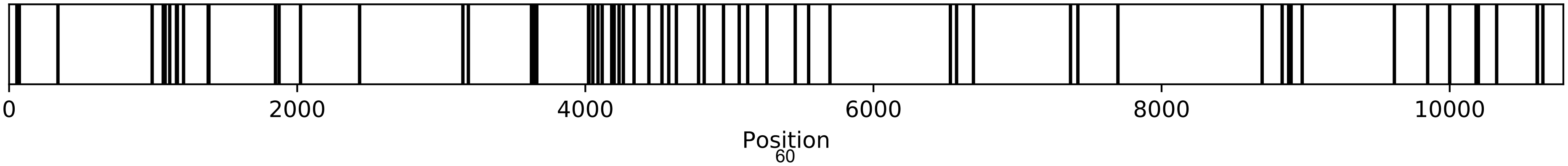

### Region: 160



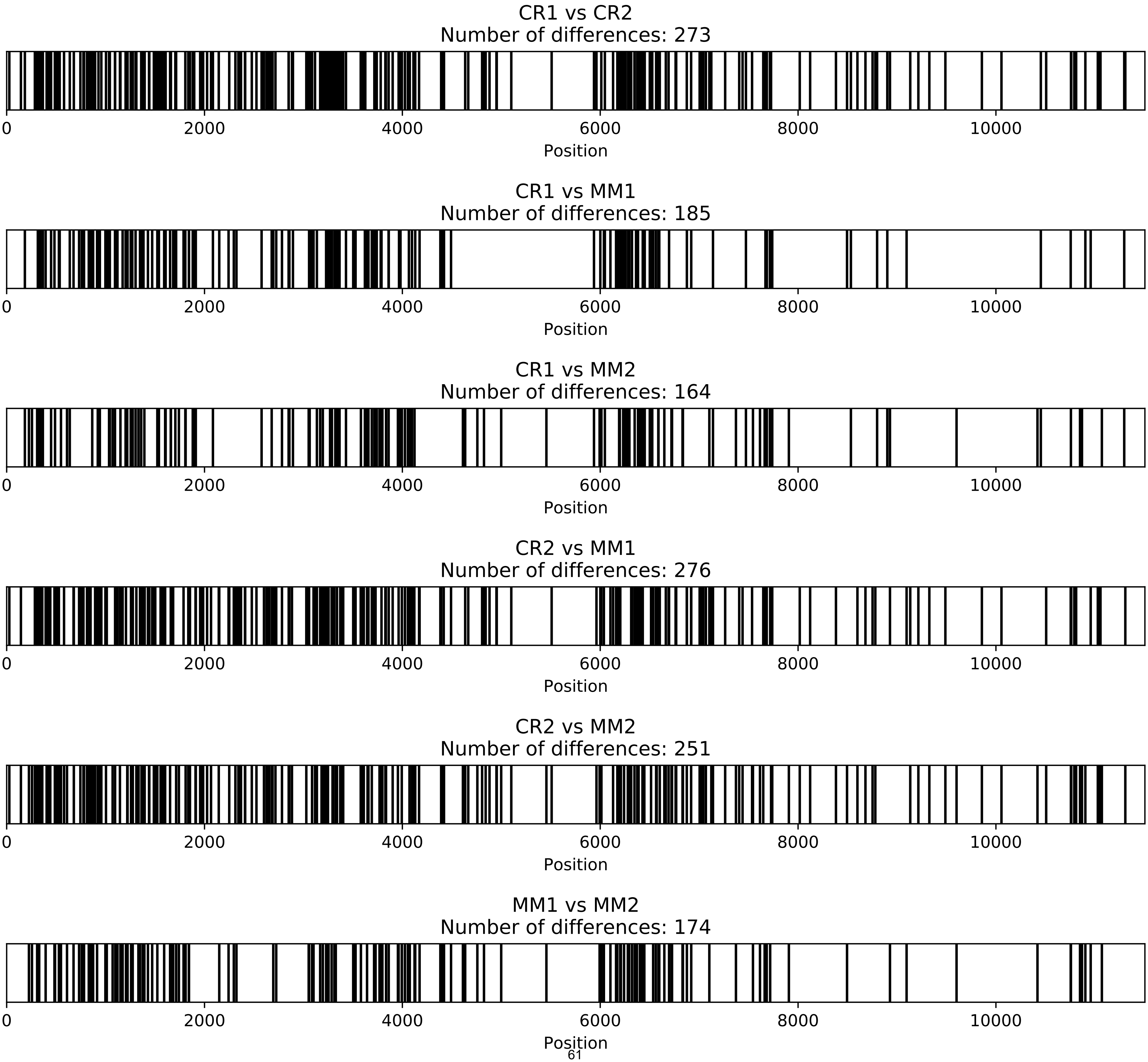
CR1 vs CR2
Number of differences: 273
CR1 vs MM1
Number of differences: 185
CR1 vs MM2
Number of differences: 164
CR2 vs MM1
Number of differences: 276
CR2 vs MM2
Number of differences: 251
MM1 vs MM2
Number of differences: 174
0
2000
4000
6000
8000
10000
Position

### Region: 164



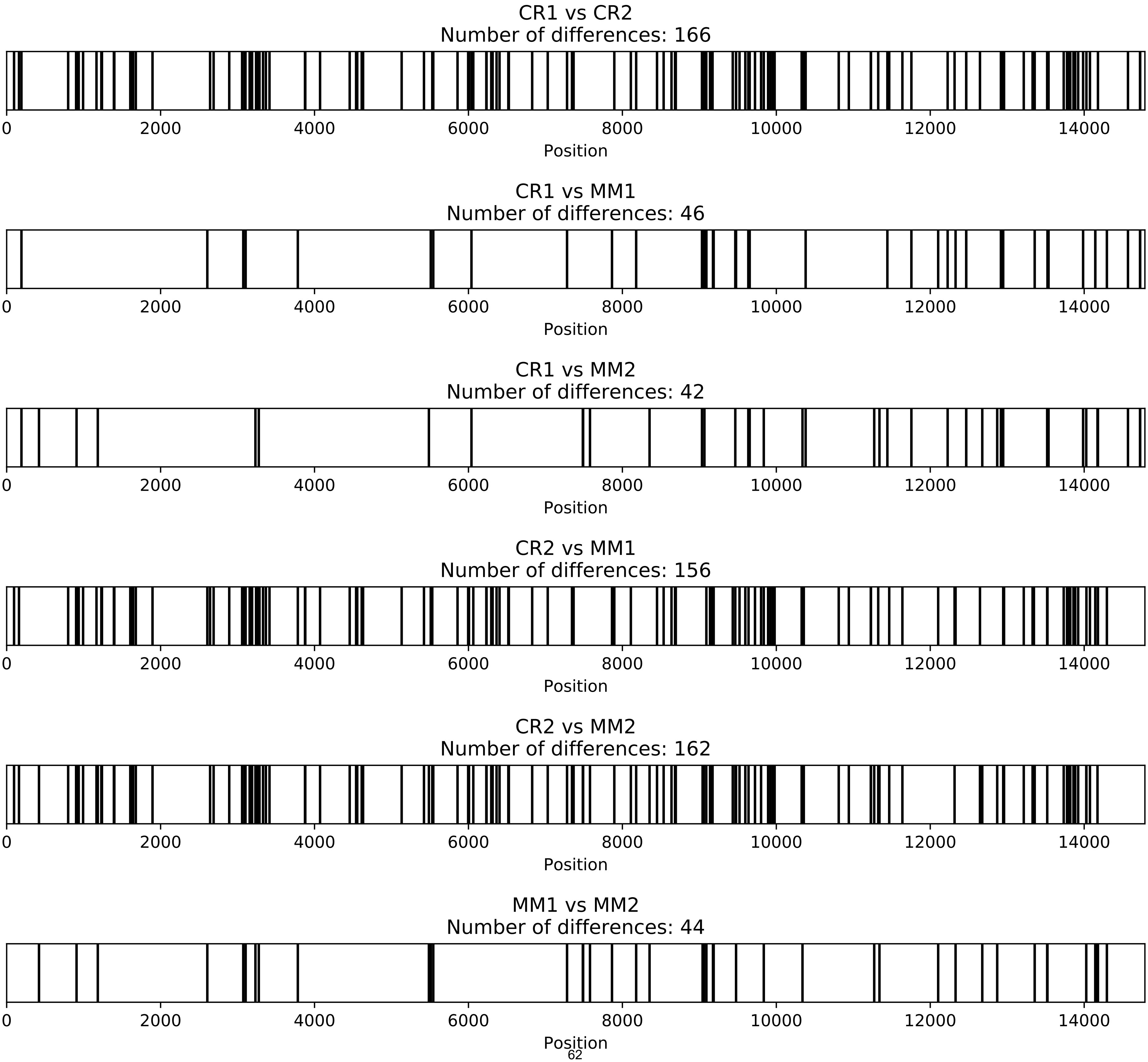
CR1 vs CR2
Number of differences: 166
CR1 vs MM1
Number of differences: 46
CR1 vs MM2
Number of differences: 42
CR2 vs MM1
Number of differences: 156
CR2 vs MM2
Number of differences: 162
MM1 vs MM2
Number of differences: 44
0
2000
4000
6000
8000
10000
12000
14000
Position

### Region: 165



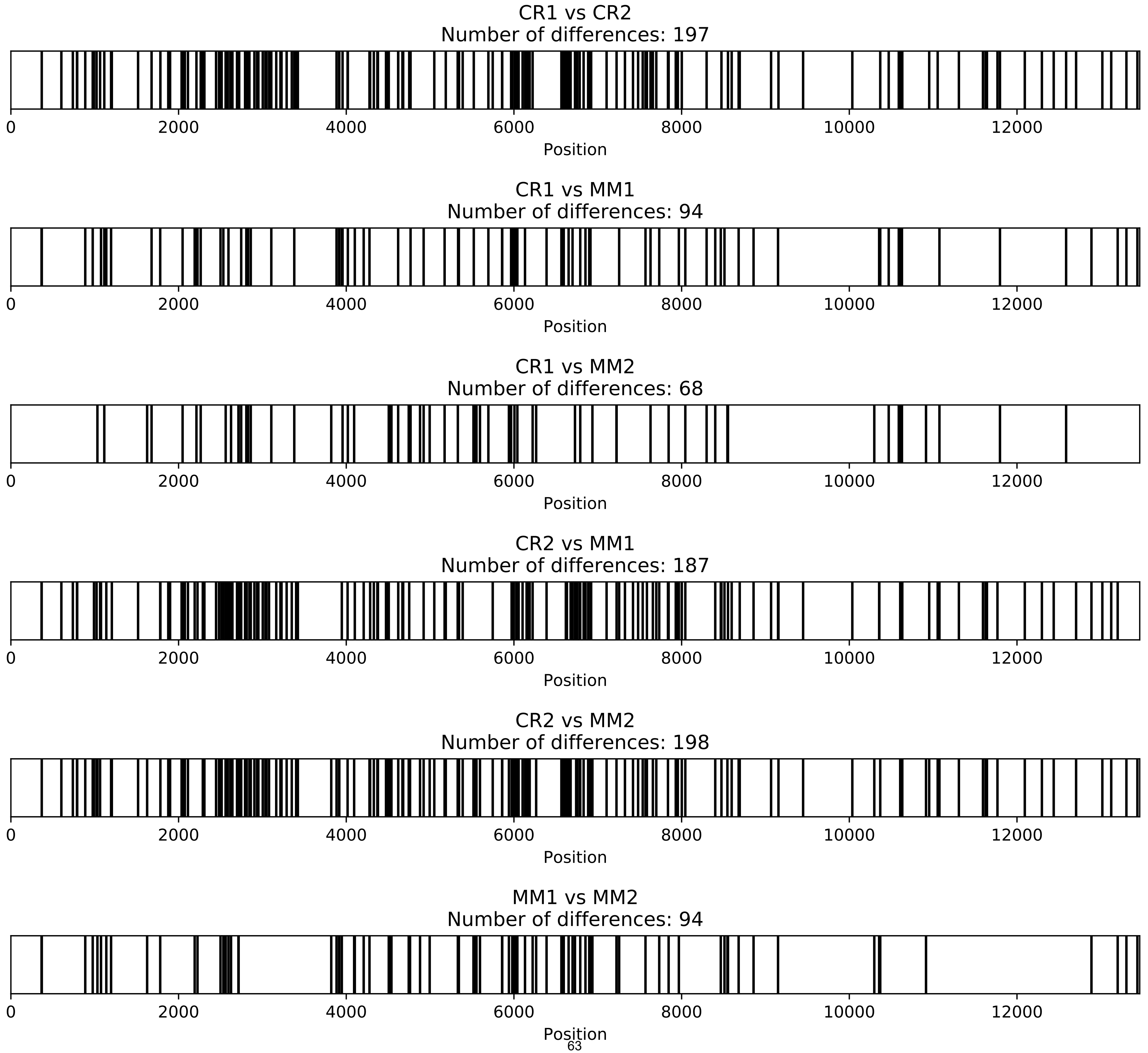
CR1 vs CR2
Number of differences: 197
0
2000
4000
6000
8000
10000
12000
Position
CR1 vs MM1
Number of differences: 94
Position
CR1 vs MM2
Number of differences: 68
Position
CR2 vs MM1
Number of differences: 187
Position
CR2 vs MM2
Number of differences: 198
Position
MM1 vs MM2
Number of differences: 94
Position

### Region: 166



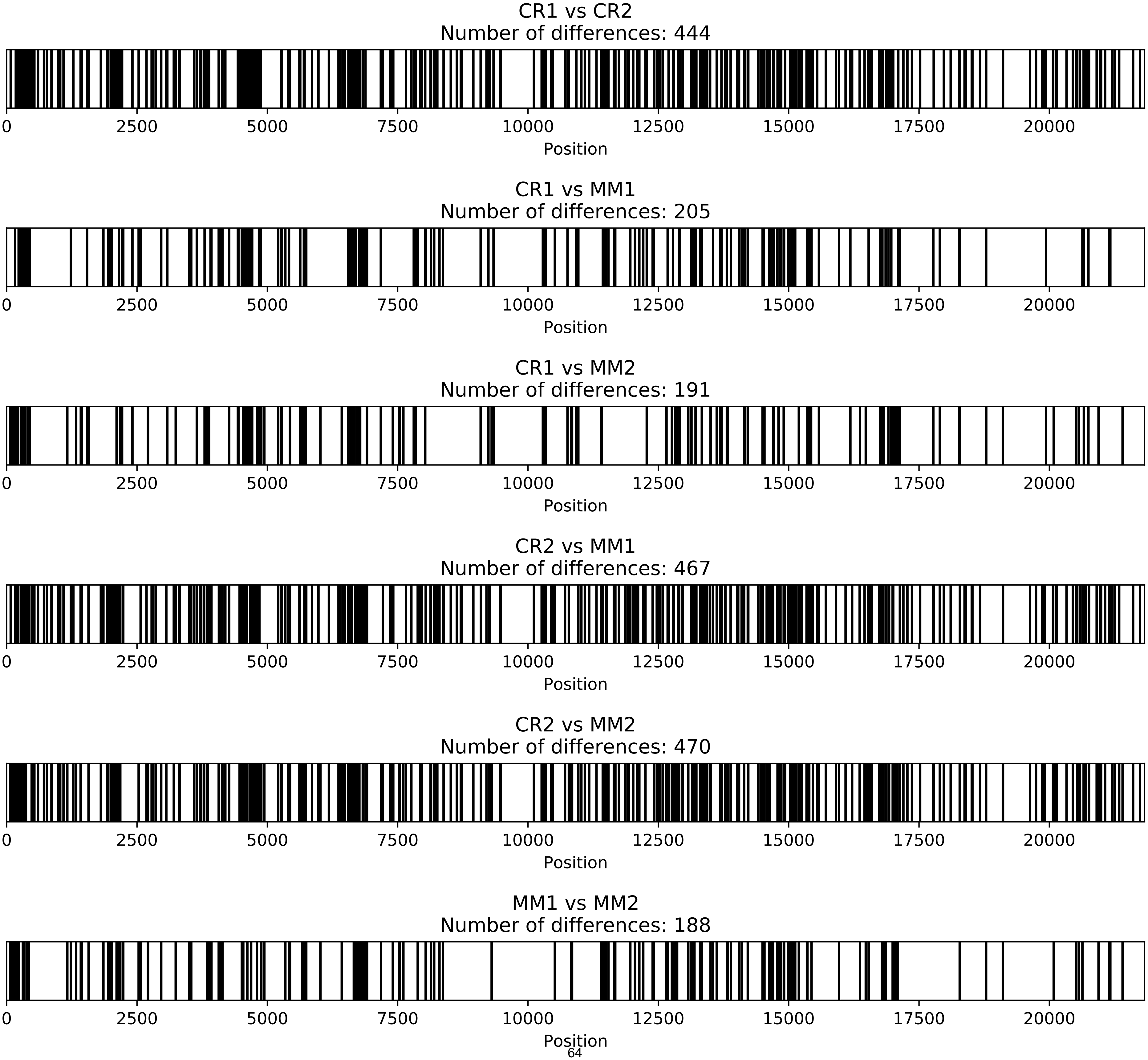
CR1 vs CR2
Number of differences: 444
CR1 vs MM1
Number of differences: 205
CR1 vs MM2
Number of differences: 191
CR2 vs MM1
Number of differences: 467
CR2 vs MM2
Number of differences: 470
MM1 vs MM2
Number of differences: 188
0
2500
5000
7500
10000
12500
15000
17500
20000
Position

### Region: 171



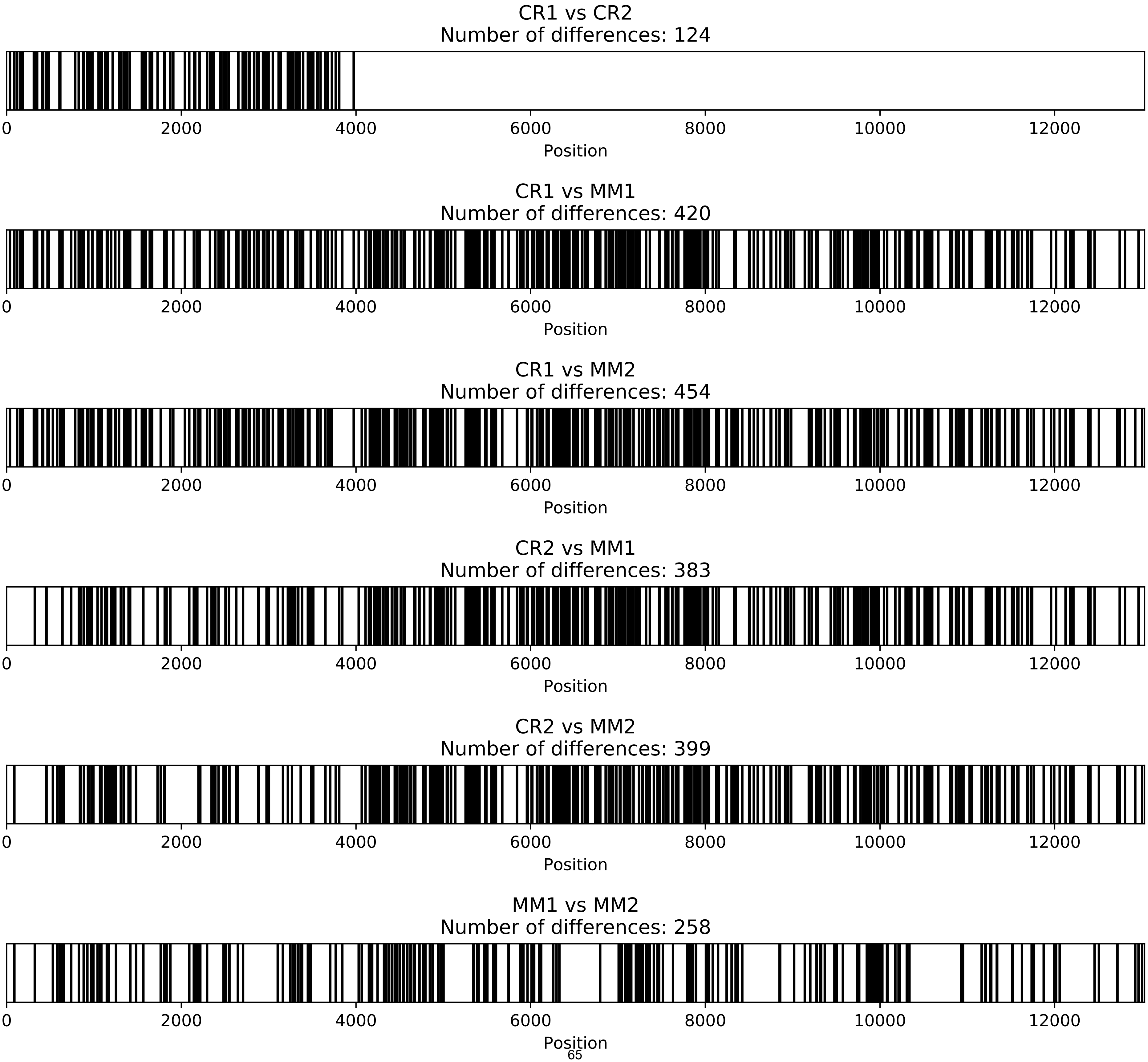
CR1 vs CR2
Number of differences: 124
CR1 vs MM1
Number of differences: 420
CR1 vs MM2
Number of differences: 454
CR2 vs MM1
Number of differences: 383
CR2 vs MM2
Number of differences: 399
MM1 vs MM2
Number of differences: 258
0
2000
4000
6000
8000
10000
12000
Position

### Region: 174



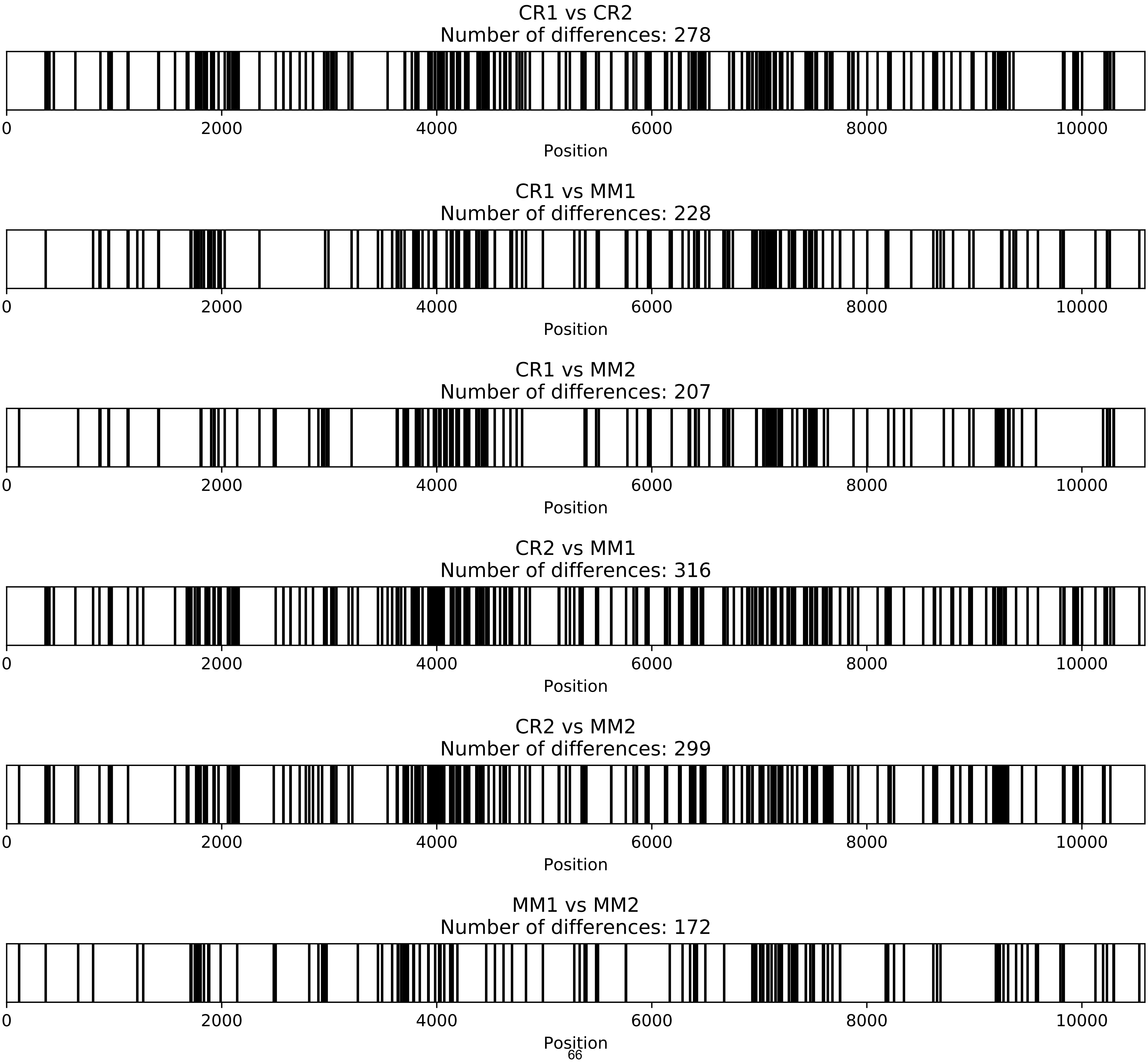
CR1 vs CR2
Number of differences: 278
CR1 vs MM1
Number of differences: 228
CR1 vs MM2
Number of differences: 207
CR2 vs MM1
Number of differences: 316
CR2 vs MM2
Number of differences: 299
MM1 vs MM2
Number of differences: 172
0
2000
4000
6000
8000
10000
Position

### Region: 175



CR1 vs CR2
Number of differences: 325

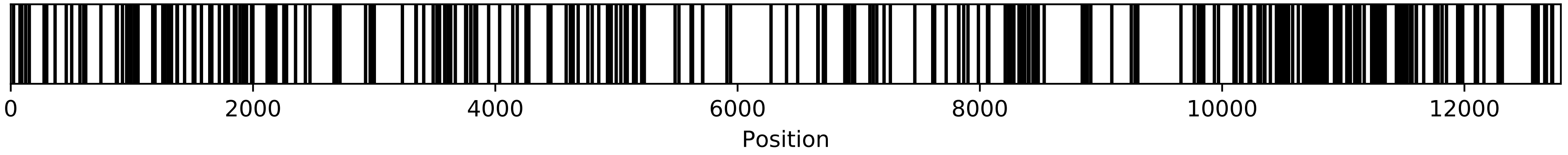


CR1 vs MM1
Number of differences: 161

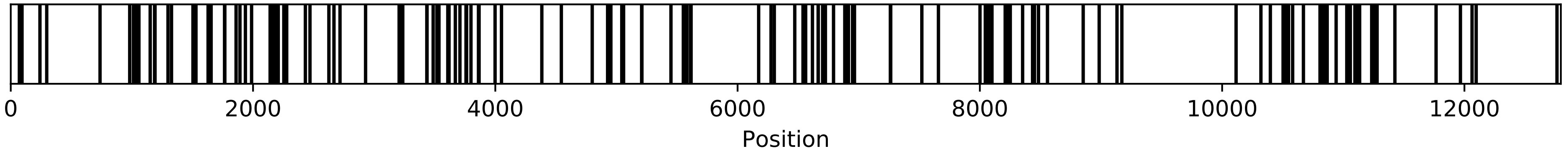


CR1 vs MM2
Number of differences: 159

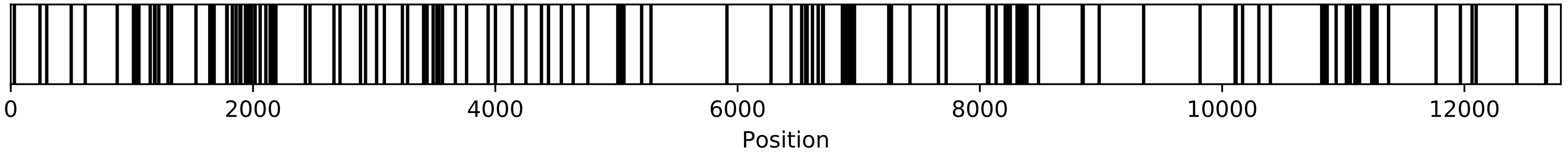


CR2 vs MM1
Number of differences: 320

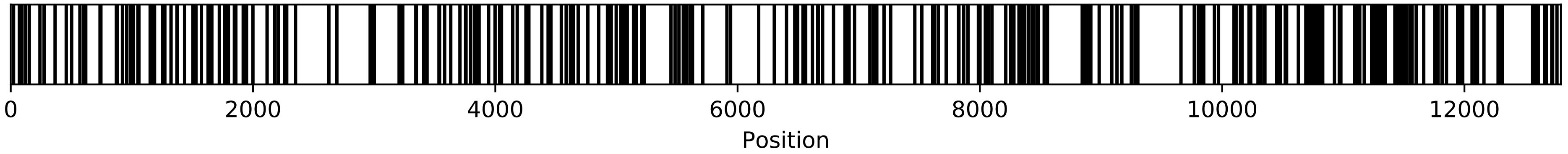


CR2 vs MM2
Number of differences: 328

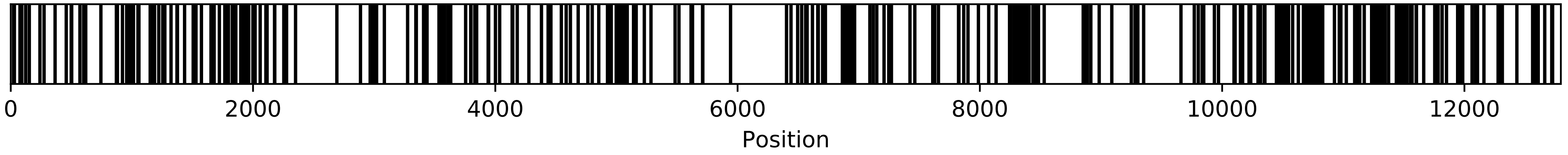


MM1 vs MM2
Number of differences: 133

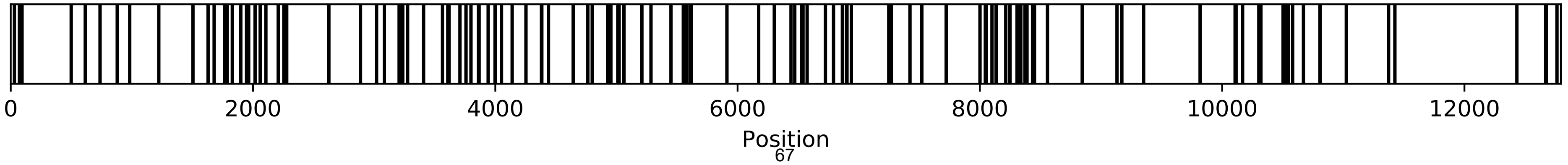

### Region: 179



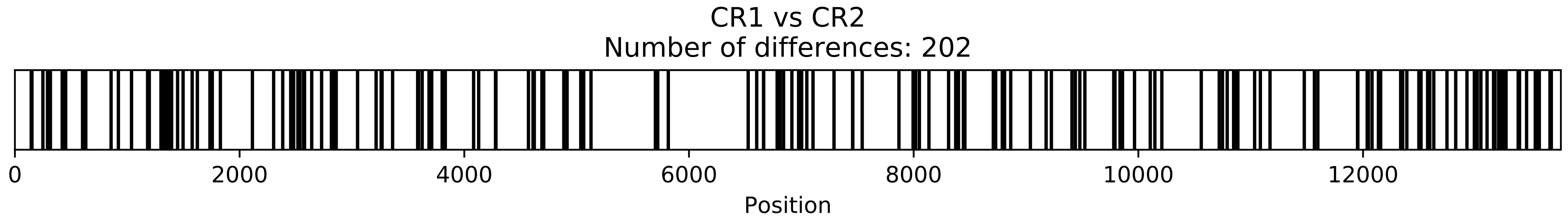
CR1 vs CR2
Number of differences: 202
0
2000
4000
6000
8000
10000
12000
Position

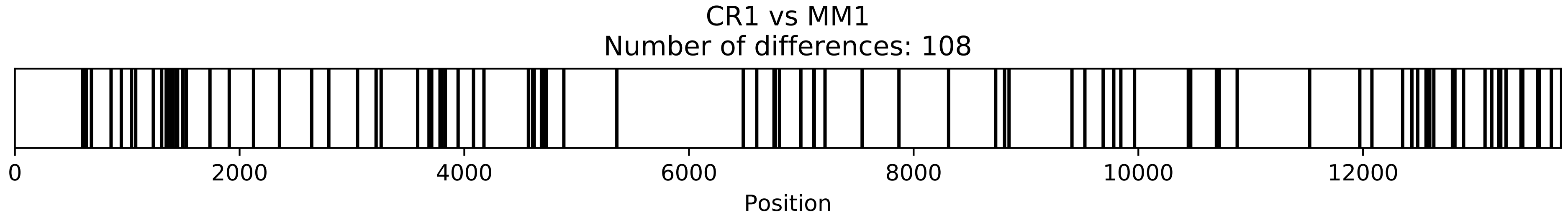
CR1 vs MM1
Number of differences: 108
0
2000
4000
6000
8000
10000
12000
Position

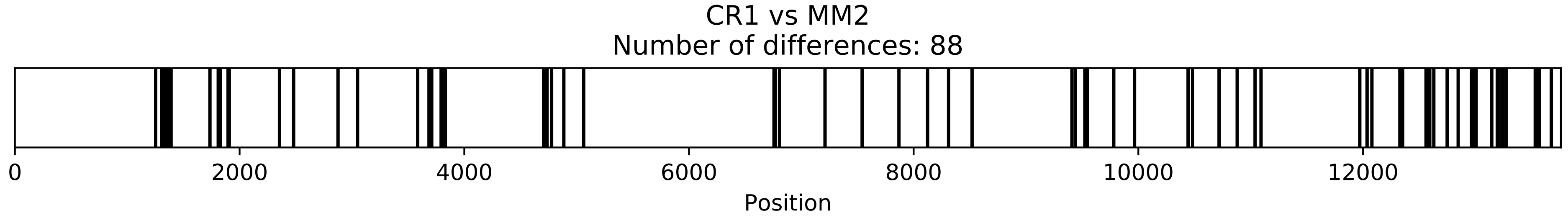
CR1 vs MM2
Number of differences: 88
0
2000
4000
6000
8000
10000
12000
Position

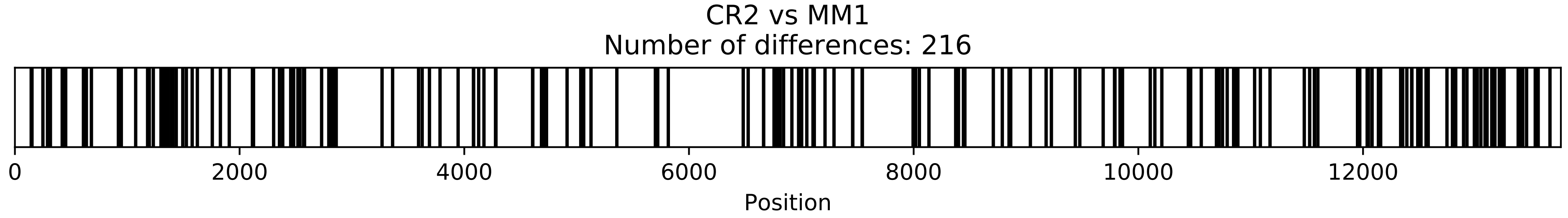
CR2 vs MM1
Number of differences: 216
0
2000
4000
6000
8000
10000
12000
Position

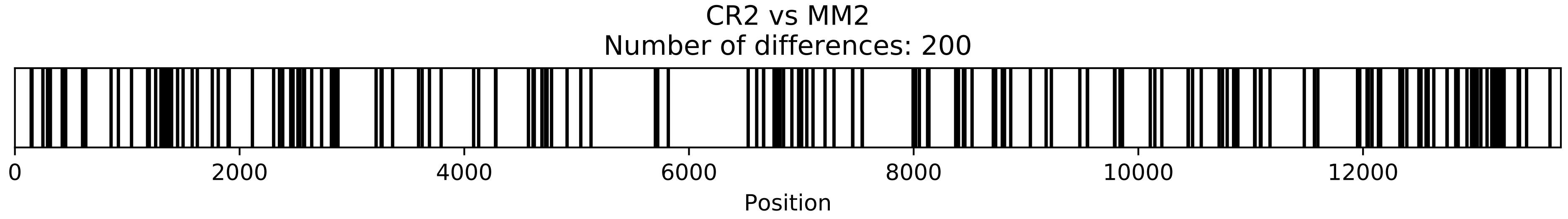
CR2 vs MM2
Number of differences: 200
0
2000
4000
6000
8000
10000
12000
Position

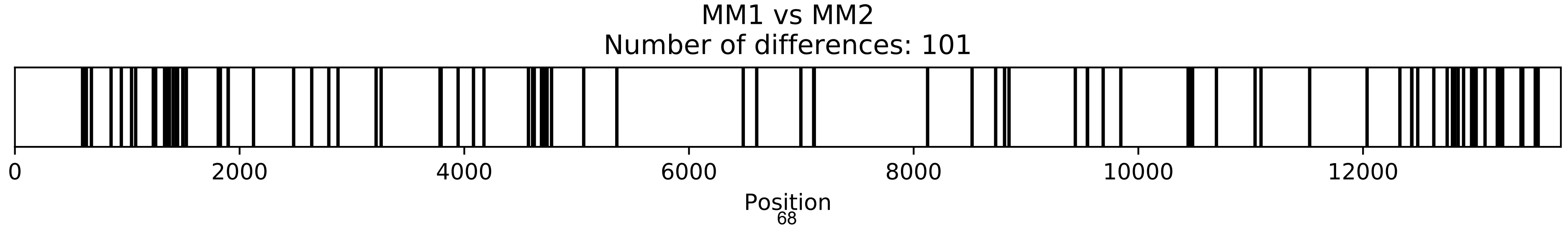
MM1 vs MM2
Number of differences: 101
0
2000
4000
6000
8000
10000
12000
Position

### Region: 183



CR1 vs CR2
Number of differences: 373

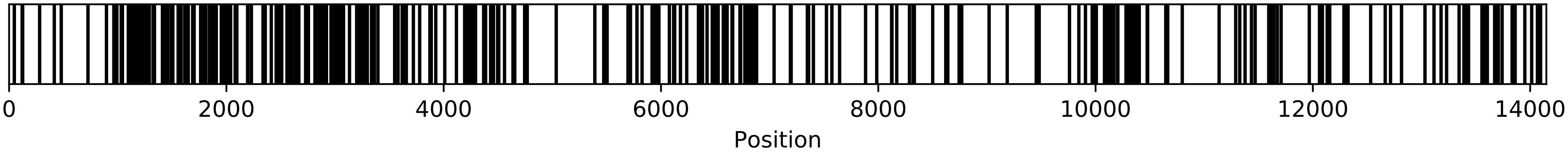


CR1 vs MM1
Number of differences: 166

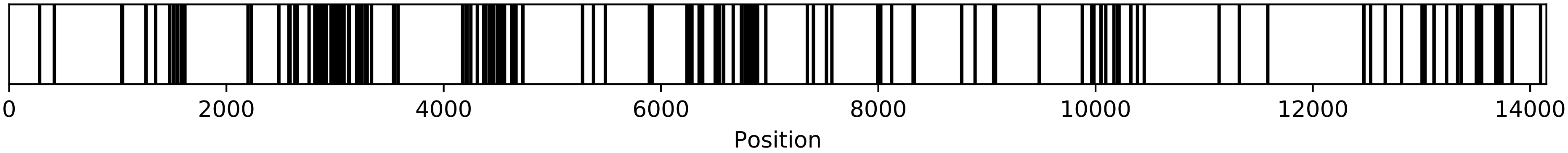


CR1 vs MM2
Number of differences: 44

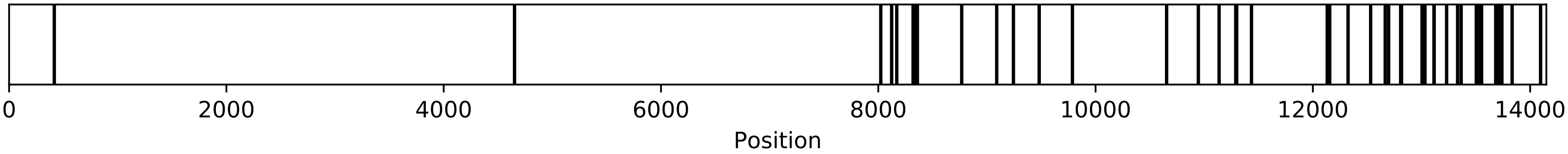


CR2 vs MM1
Number of differences: 380

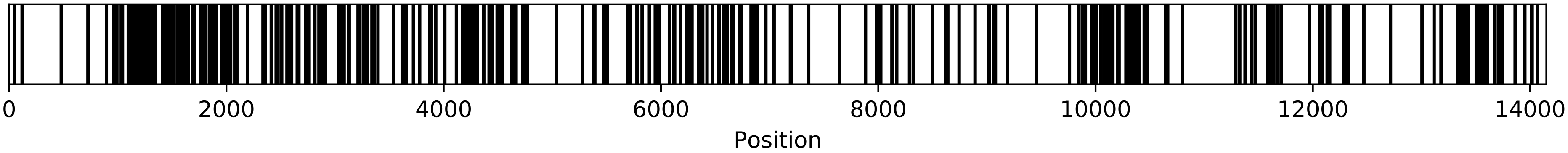


CR2 vs MM2
Number of differences: 372

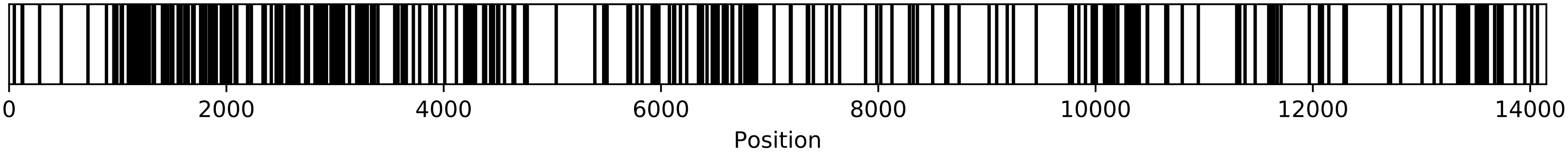


MM1 vs MM2
Number of differences: 154

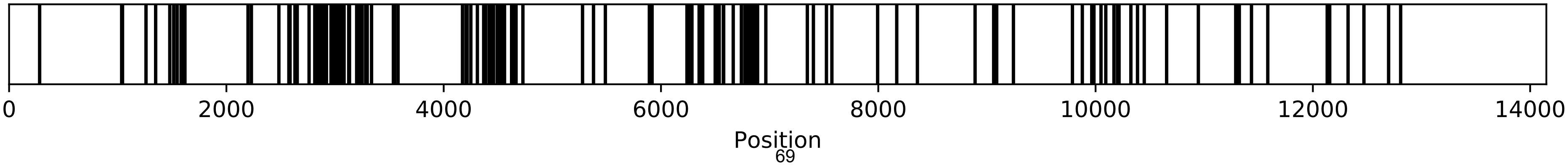

### Region: 185



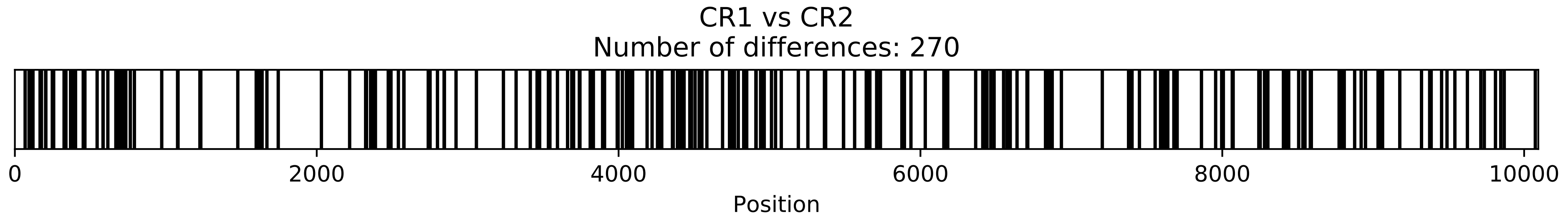
CR1 vs CR2
Number of differences: 270
0
2000
4000
6000
8000
10000
Position

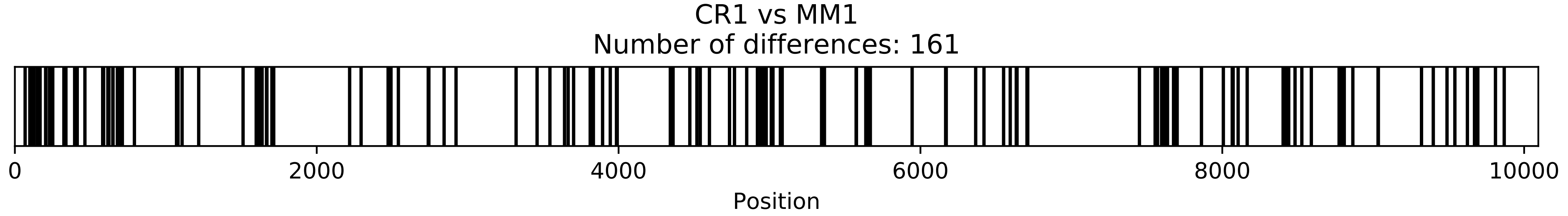
CR1 vs MM1
Number of differences: 161
0
2000
4000
6000
8000
10000
Position

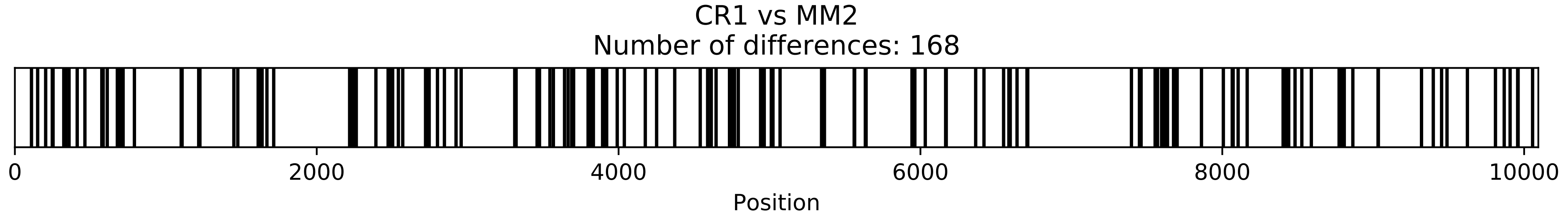
CR1 vs MM2
Number of differences: 168
0
2000
4000
6000
8000
10000
Position

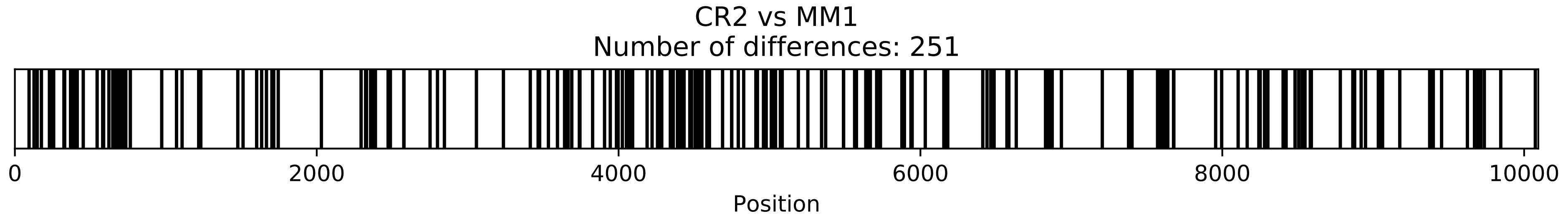
CR2 vs MM1
Number of differences: 251
0
2000
4000
6000
8000
10000
Position

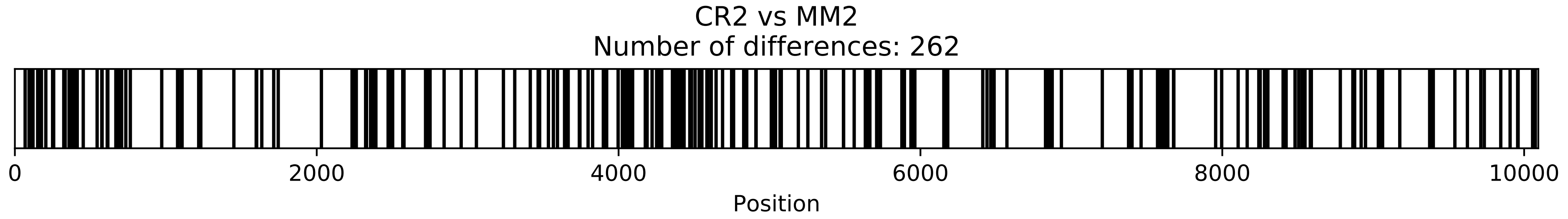
CR2 vs MM2
Number of differences: 262
0
2000
4000
6000
8000
10000
Position

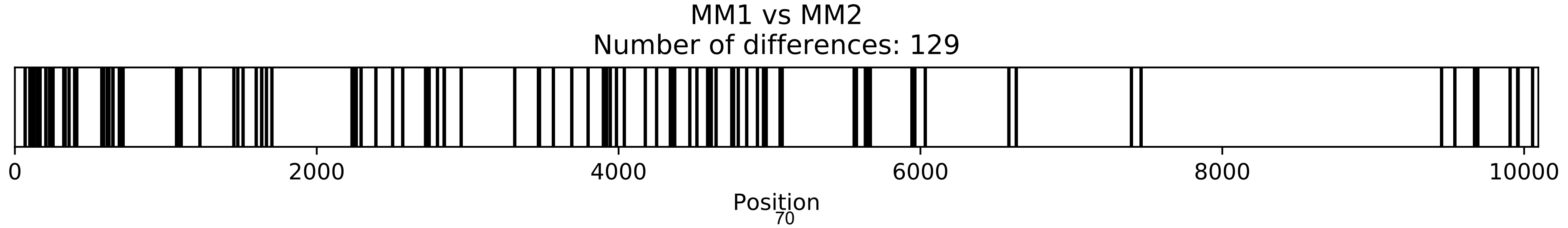
MM1 vs MM2
Number of differences: 129
0
2000
4000
6000
8000
10000
Position

### Region: 195



CR1 vs CR2
Number of differences: 271

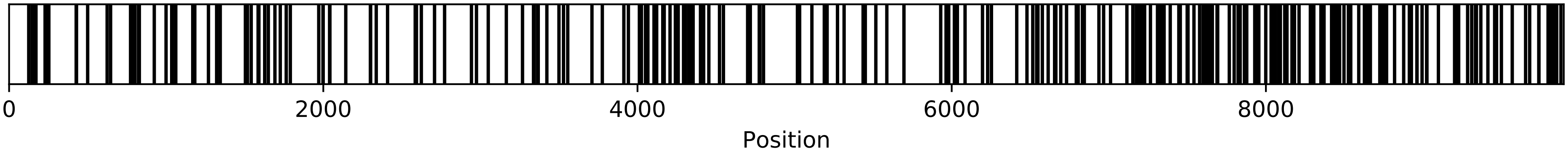


CR1 vs MM1
Number of differences: 140

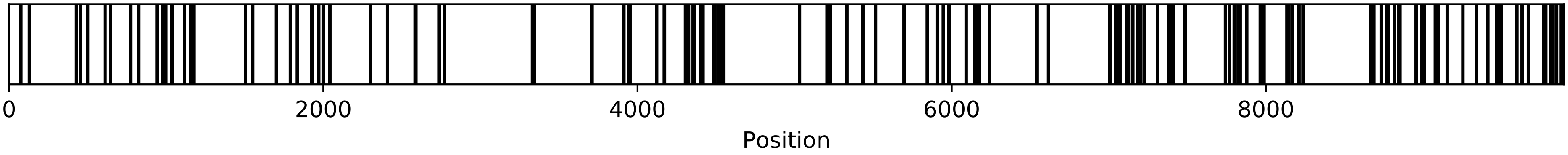


CR1 vs MM2
Number of differences: 111

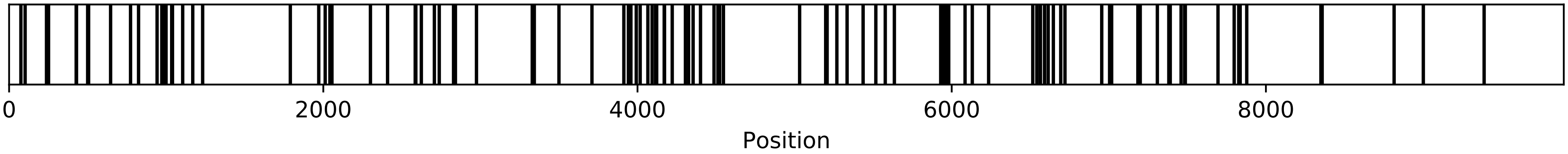


CR2 vs MM1
Number of differences: 288

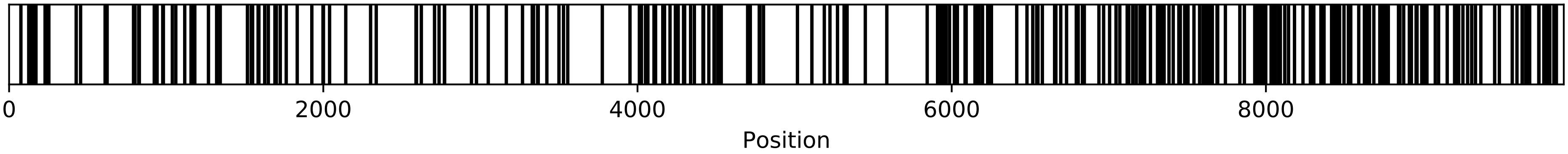


CR2 vs MM2
Number of differences: 281

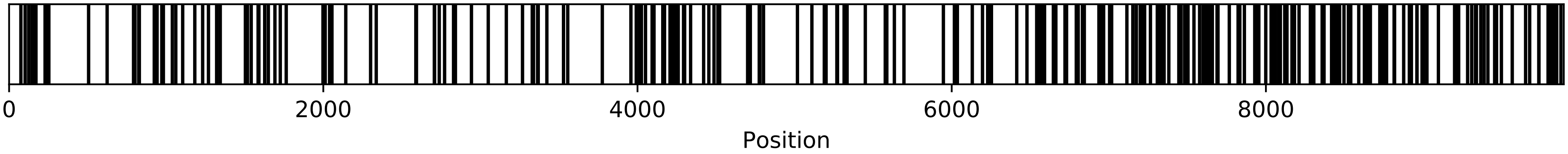


MM1 vs MM2
Number of differences: 151

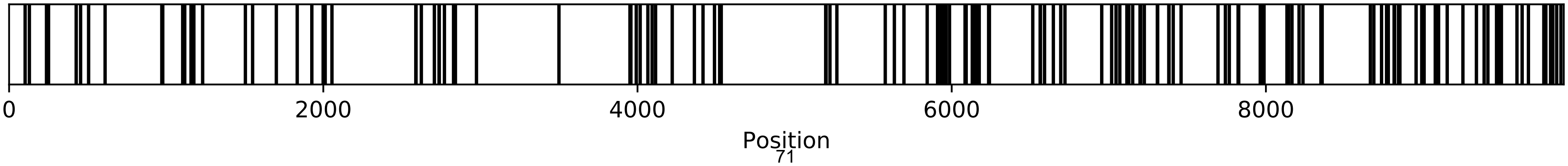

### Region: 196



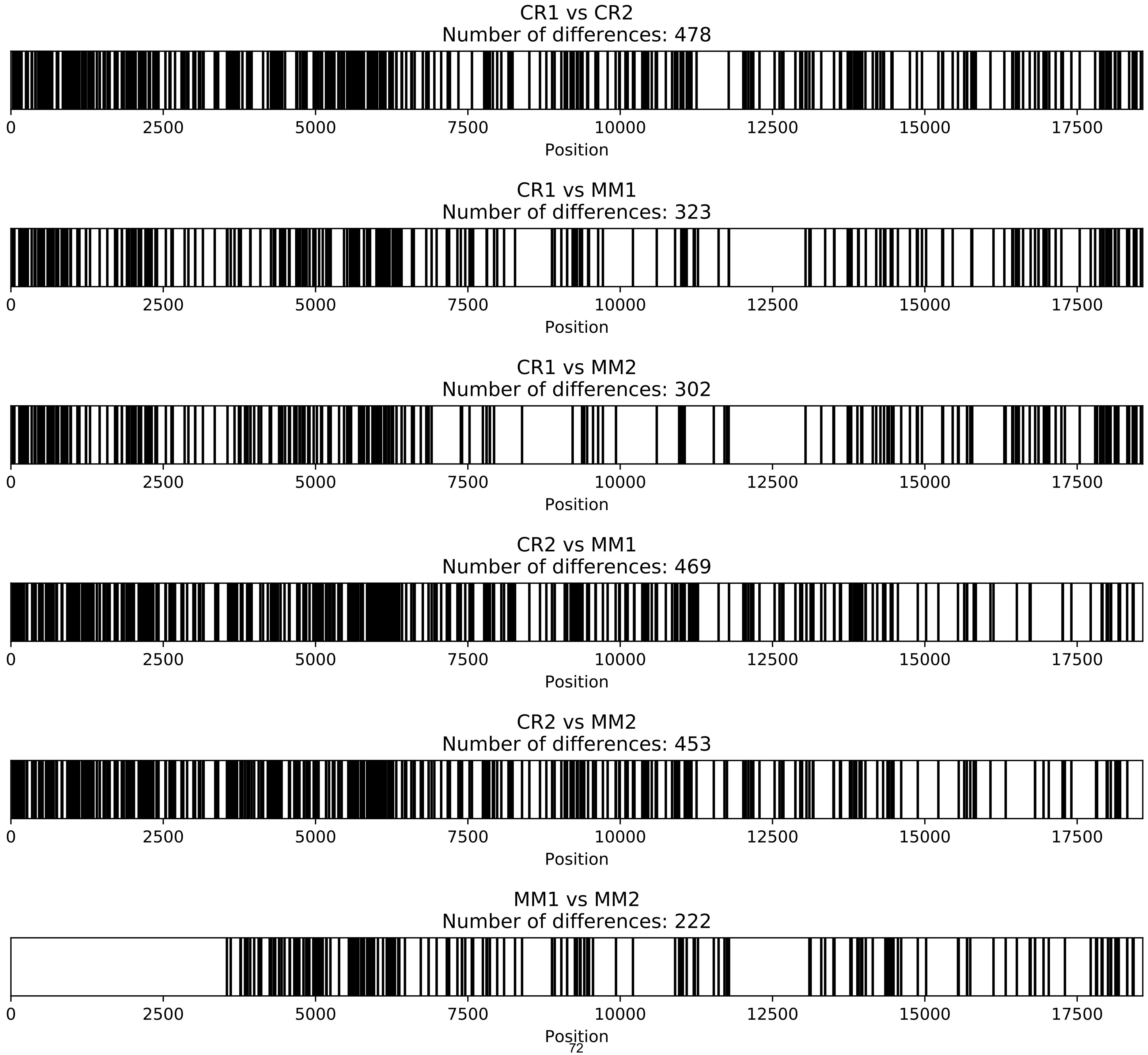
CR1 vs CR2
Number of differences: 478
CR1 vs MM1
Number of differences: 323
CR1 vs MM2
Number of differences: 302
CR2 vs MM1
Number of differences: 469
CR2 vs MM2
Number of differences: 453
MM1 vs MM2
Number of differences: 222
0
2500
5000
7500
10000
12500
15000
17500
Position

### Region: 198

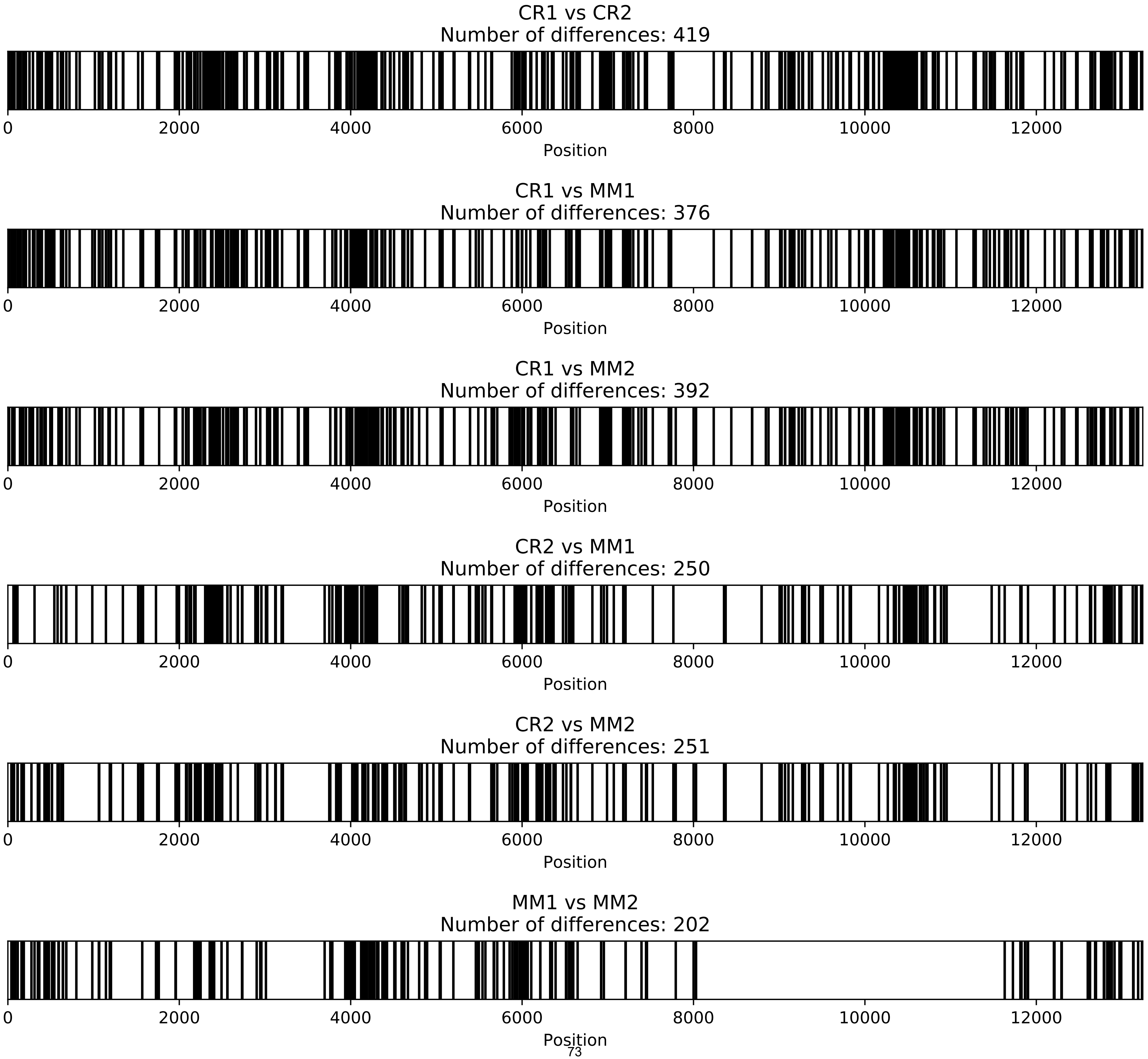
CR1 vs CR2
Number of differences: 419
0
2000
4000
6000
8000
10000
12000
Position
CR1 vs MM1
Number of differences: 376
0
2000
4000
6000
8000
10000
12000
Position
CR1 vs MM2
Number of differences: 392
0
2000
4000
6000
8000
10000
12000
Position
CR2 vs MM1
Number of differences: 250
0
2000
4000
6000
8000
10000
12000
Position
CR2 vs MM2
Number of differences: 251
0
2000
4000
6000
8000
10000
12000
Position
MM1 vs MM2
Number of differences: 202
0
2000
4000
6000
8000
10000
12000
Position

### Region: 209

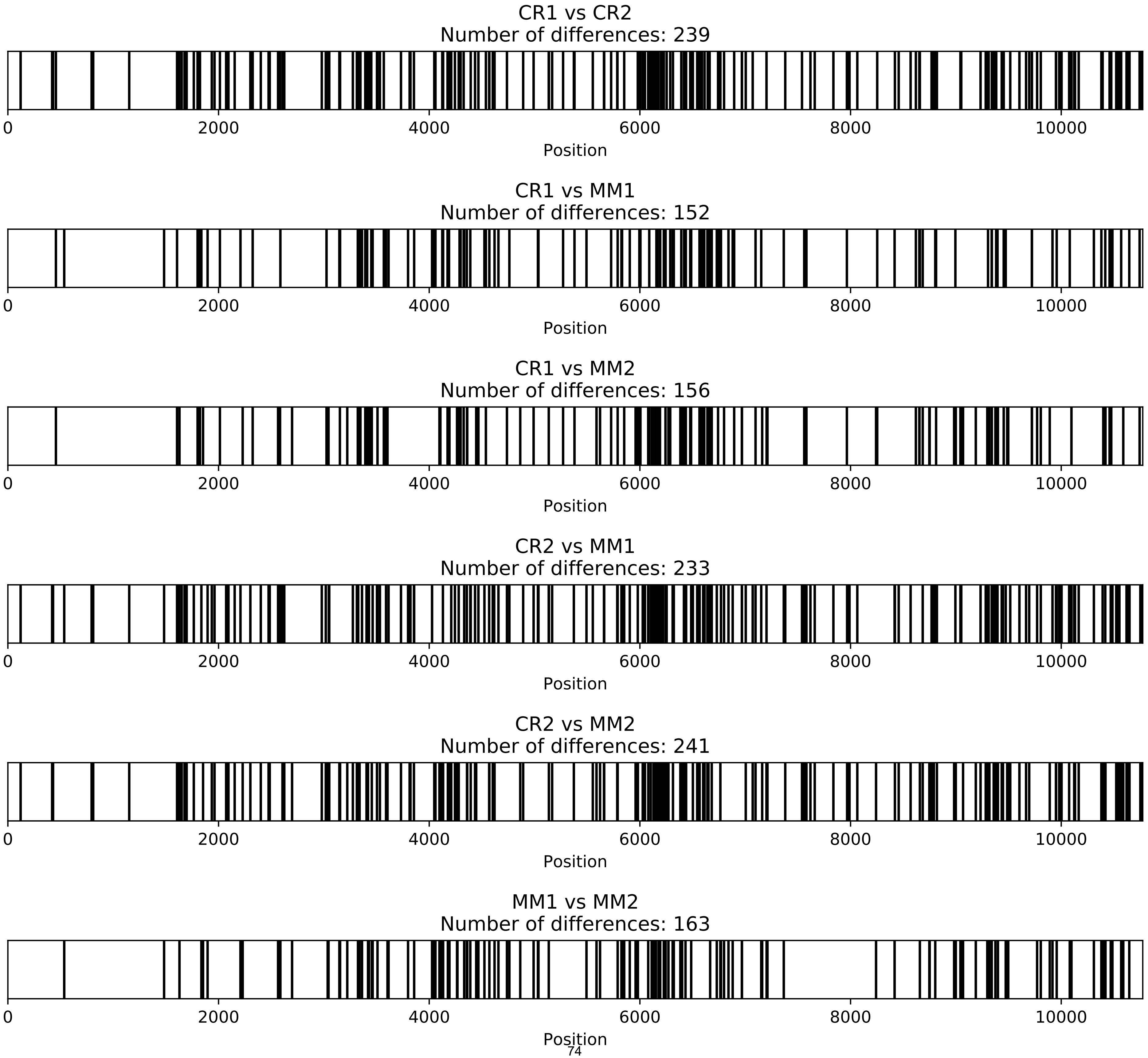
CR1 vs CR2
Number of differences: 239
0
2000
4000
6000
8000
10000
Position
CR1 vs MM1
Number of differences: 152
0
2000
4000
6000
8000
10000
Position
CR1 vs MM2
Number of differences: 156
0
2000
4000
6000
8000
10000
Position
CR2 vs MM1
Number of differences: 233
0
2000
4000
6000
8000
10000
Position
CR2 vs MM2
Number of differences: 241
0
2000
4000
6000
8000
10000
Position
MM1 vs MM2
Number of differences: 163
0
2000
4000
6000
8000
10000
Position

### Region: 213



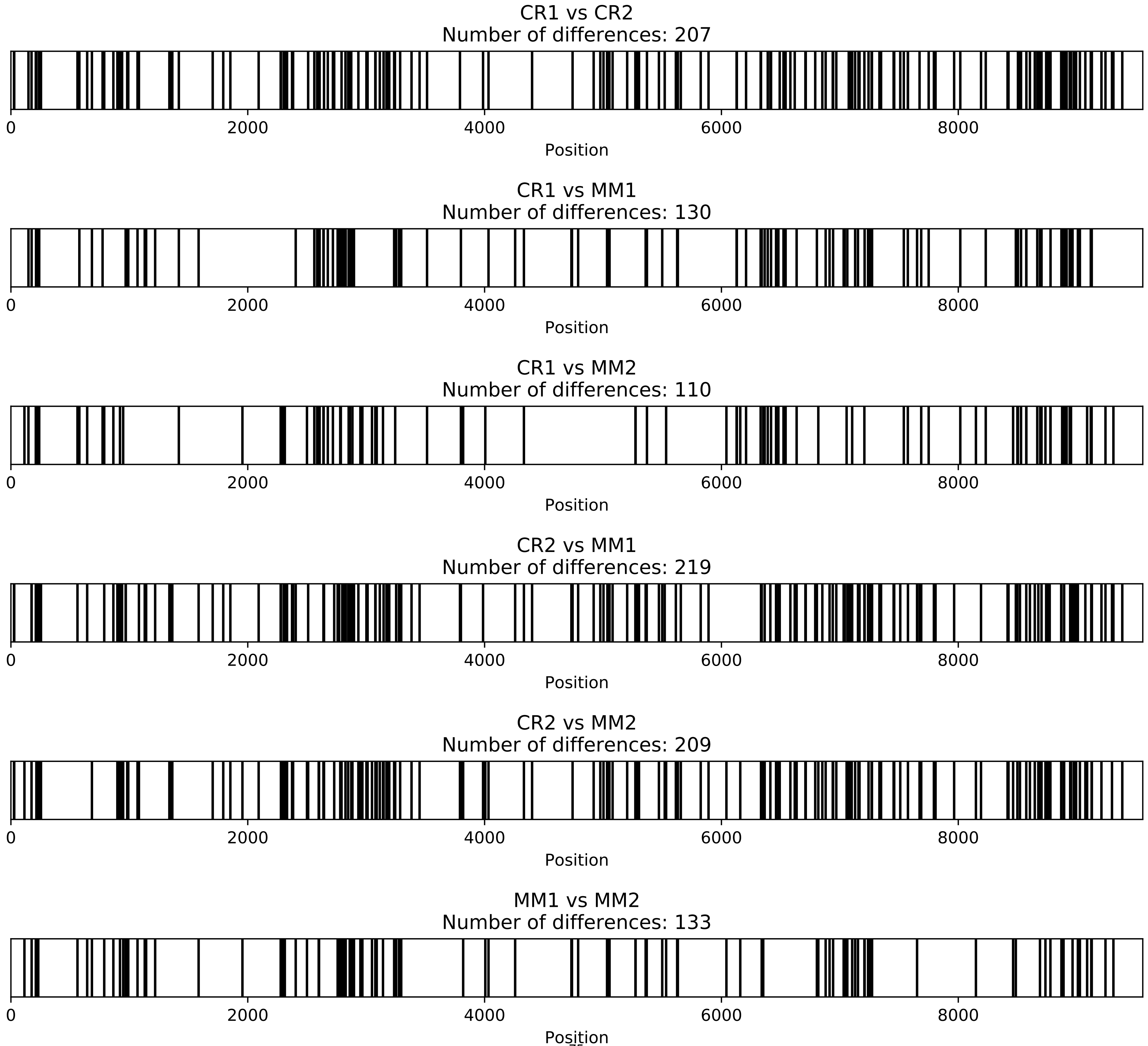
CR1 vs CR2
Number of differences: 207
0
2000
4000
6000
8000
Position
CR1 vs MM1
Number of differences: 130
0
2000
4000
6000
8000
Position
CR1 vs MM2
Number of differences: 110
0
2000
4000
6000
8000
Position
CR2 vs MM1
Number of differences: 219
0
2000
4000
6000
8000
Position
CR2 vs MM2
Number of differences: 209
0
2000
4000
6000
8000
Position
MM1 vs MM2
Number of differences: 133
0
2000
4000
6000
8000
Position

### Region: 214



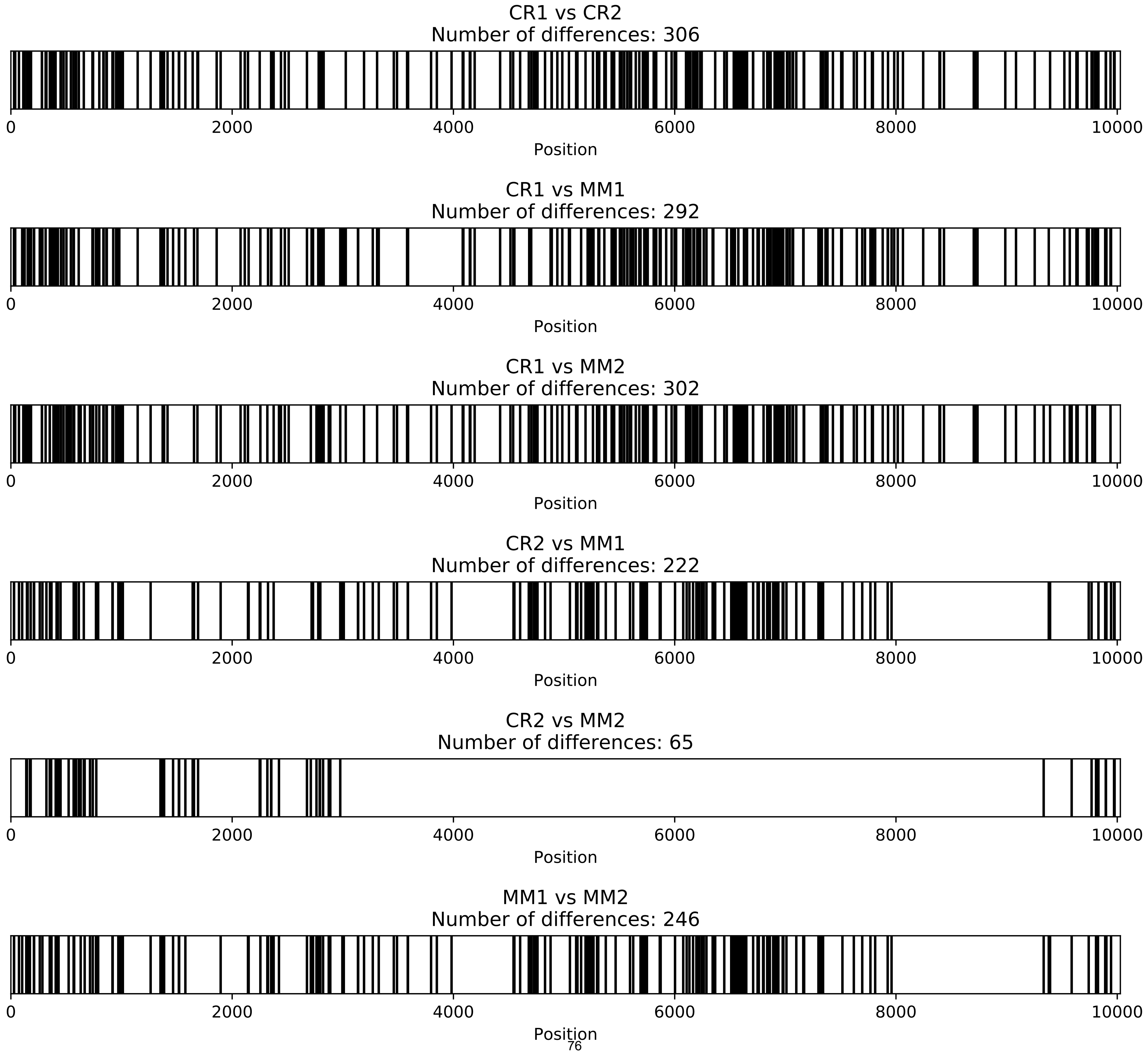
CR1 vs CR2
Number of differences: 306
CR1 vs MM1
Number of differences: 292
CR1 vs MM2
Number of differences: 302
CR2 vs MM1
Number of differences: 222
CR2 vs MM2
Number of differences: 65
MM1 vs MM2
Number of differences: 246
0
2000
4000
6000
8000
10000
Position

### Region: 215



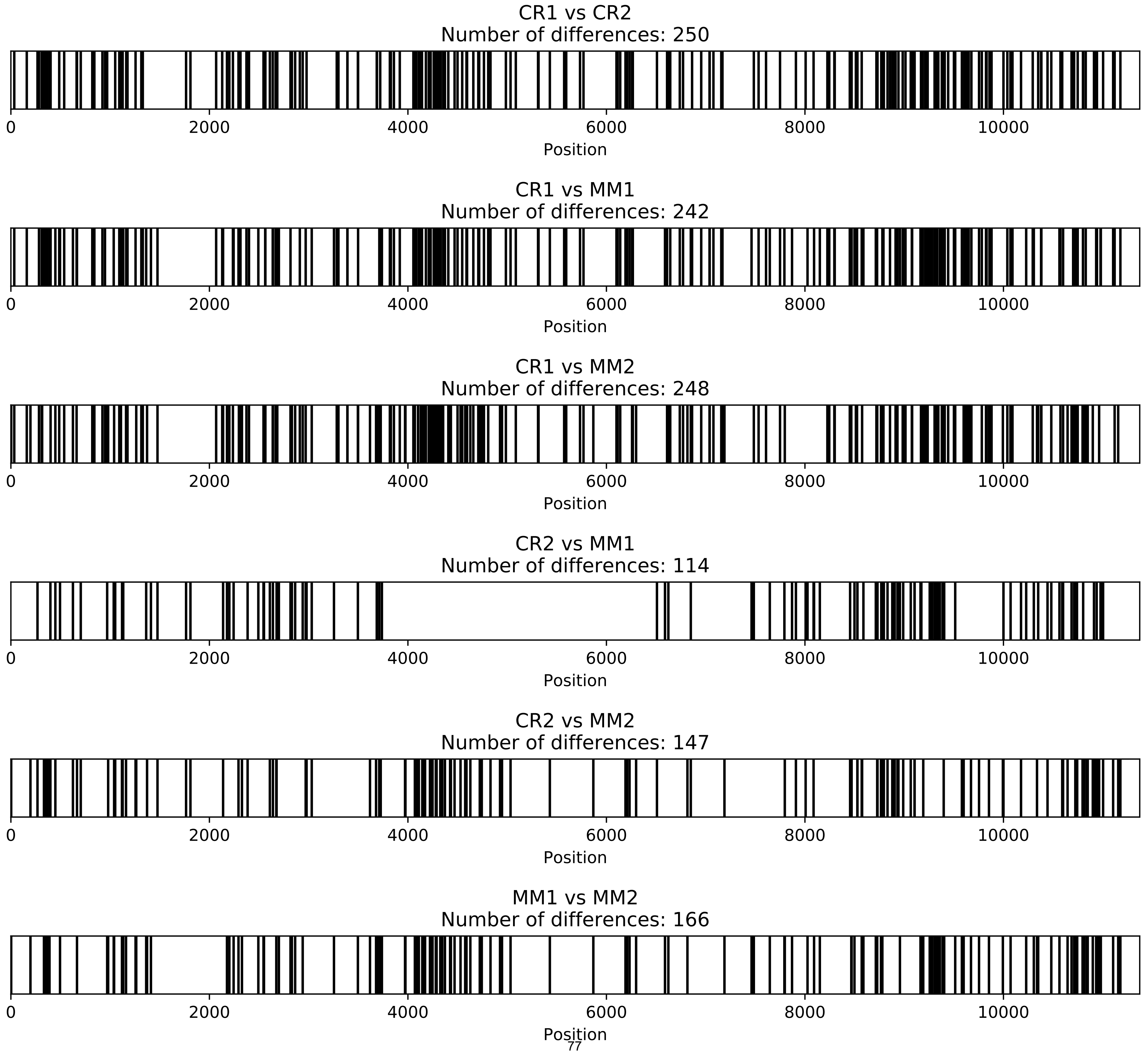
CR1 vs CR2
Number of differences: 250
CR1 vs MM1
Number of differences: 242
CR1 vs MM2
Number of differences: 248
CR2 vs MM1
Number of differences: 114
CR2 vs MM2
Number of differences: 147
MM1 vs MM2
Number of differences: 166
0
2000
4000
6000
8000
10000
Position

### Region: 219



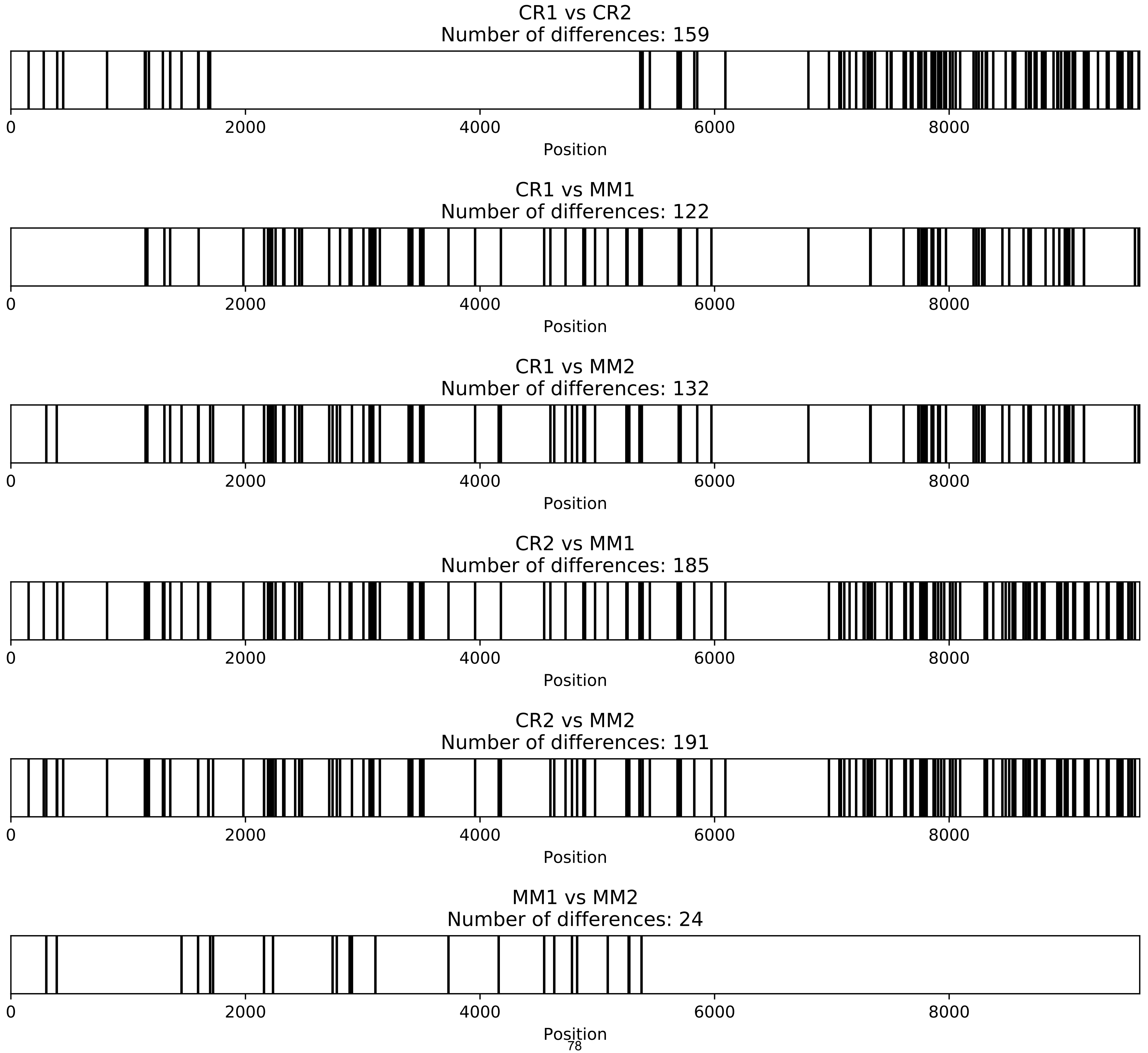
CR1 vs CR2
Number of differences: 159
0
2000
4000
6000
8000
Position
CR1 vs MM1
Number of differences: 122
0
2000
4000
6000
8000
Position
CR1 vs MM2
Number of differences: 132
0
2000
4000
6000
8000
Position
CR2 vs MM1
Number of differences: 185
0
2000
4000
6000
8000
Position
CR2 vs MM2
Number of differences: 191
0
2000
4000
6000
8000
Position
MM1 vs MM2
Number of differences: 24
0
2000
4000
6000
8000
Position

### Region: 222



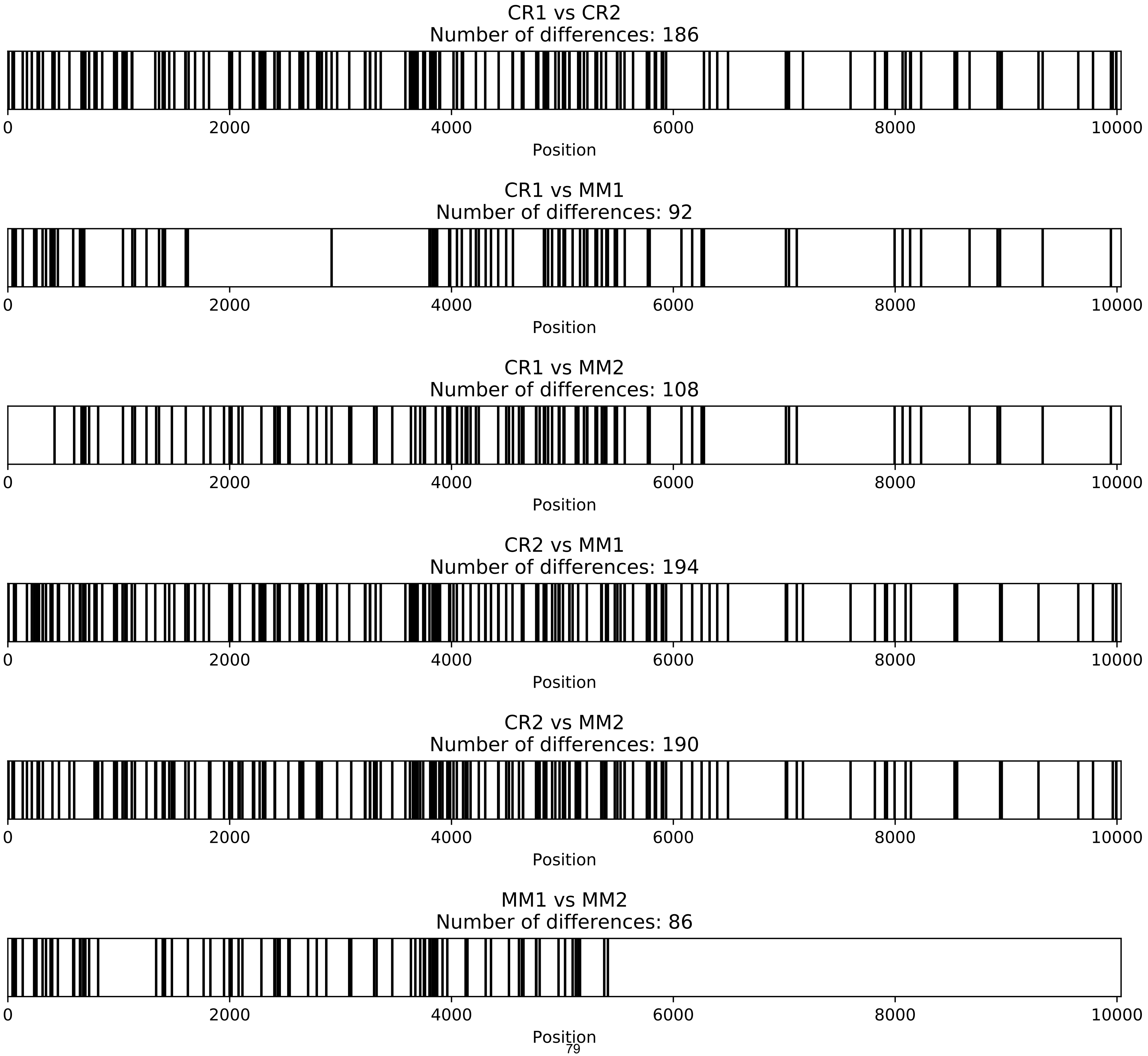
CR1 vs CR2
Number of differences: 186
0
2000
4000
6000
8000
10000
Position
CR1 vs MM1
Number of differences: 92
0
2000
4000
6000
8000
10000
Position
CR1 vs MM2
Number of differences: 108
0
2000
4000
6000
8000
10000
Position
CR2 vs MM1
Number of differences: 194
0
2000
4000
6000
8000
10000
Position
CR2 vs MM2
Number of differences: 190
0
2000
4000
6000
8000
10000
Position
MM1 vs MM2
Number of differences: 86
0
2000
4000
6000
8000
10000
Position

### Region: 223



CR1 vs CR2
Number of differences: 218

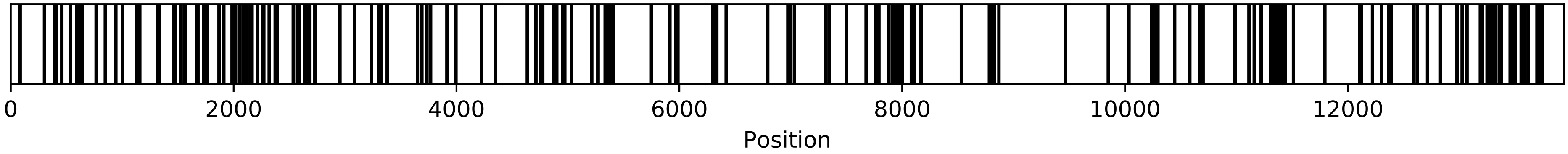


CR1 vs MM1
Number of differences: 231

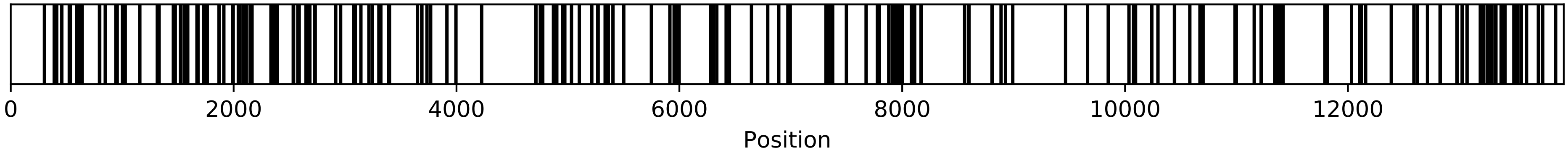


CR1 vs MM2
Number of differences: 222

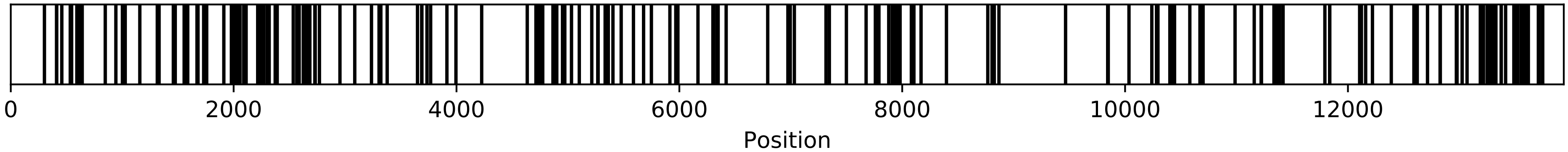


CR2 vs MM1
Number of differences: 118

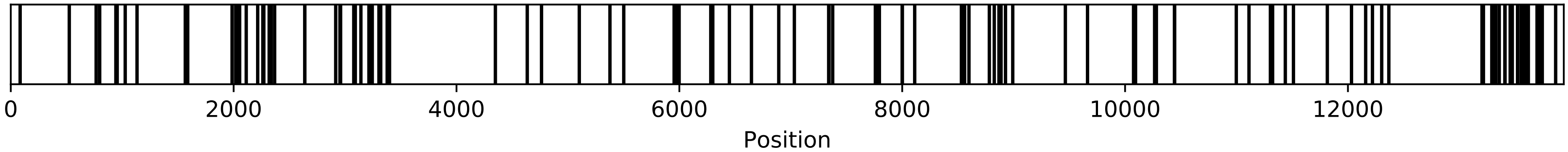


CR2 vs MM2
Number of differences: 109

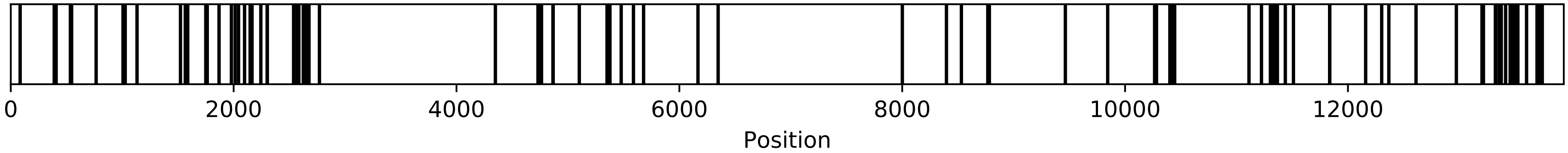


MM1 vs MM2
Number of differences: 143

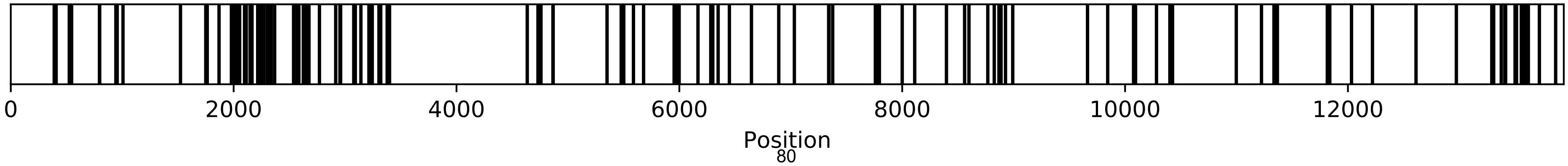

### Region: 229



CR1 vs CR2
Number of differences: 202

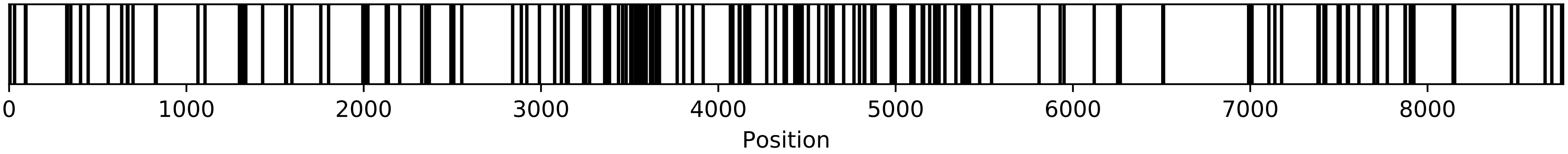


CR1 vs MM1
Number of differences: 206

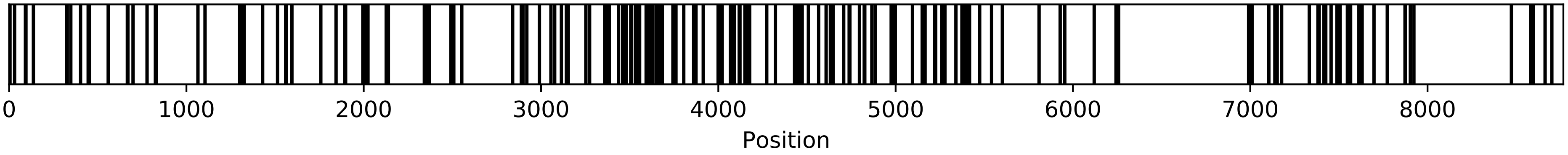


CR1 vs MM2
Number of differences: 206

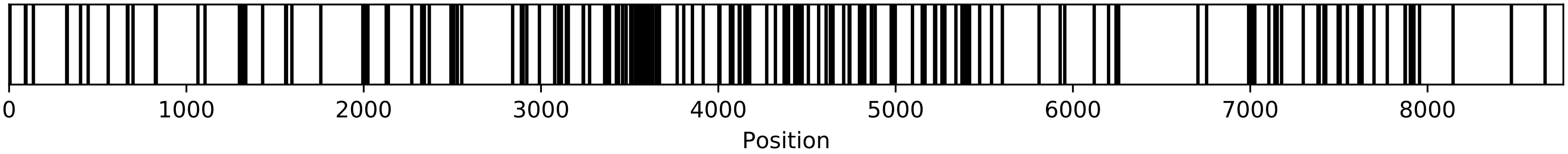


CR2 vs MM1
Number of differences: 89

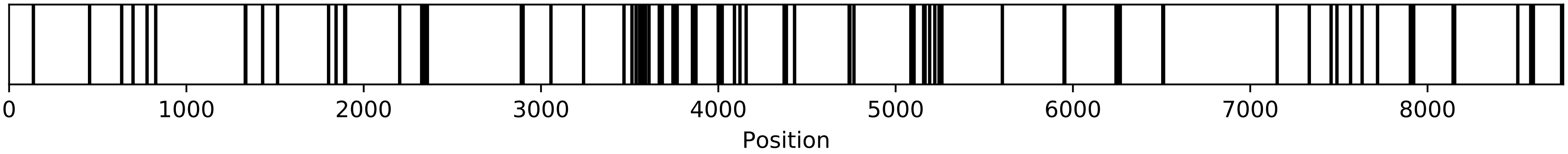


CR2 vs MM2
Number of differences: 70

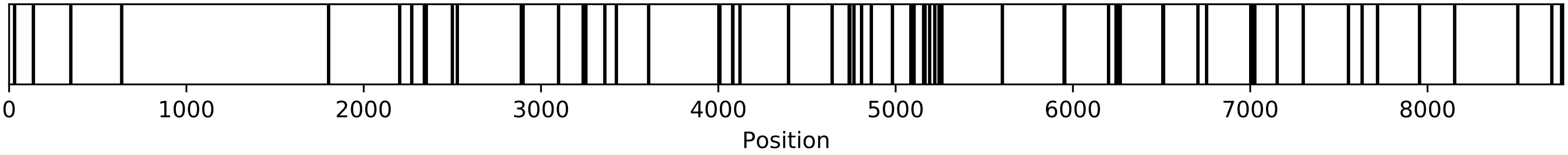


MM1 vs MM2
Number of differences: 91

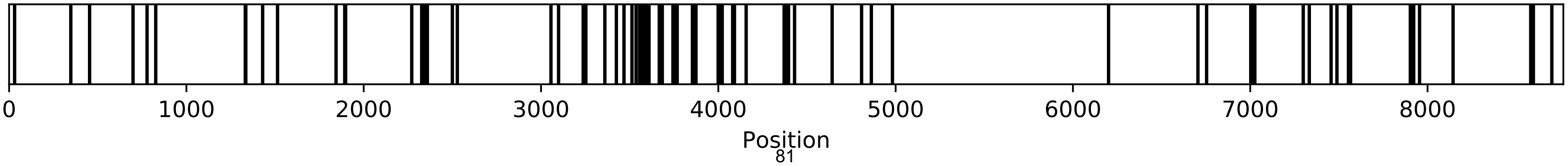

### Region: 231



## CR1 vs CR2
Number of differences: 130

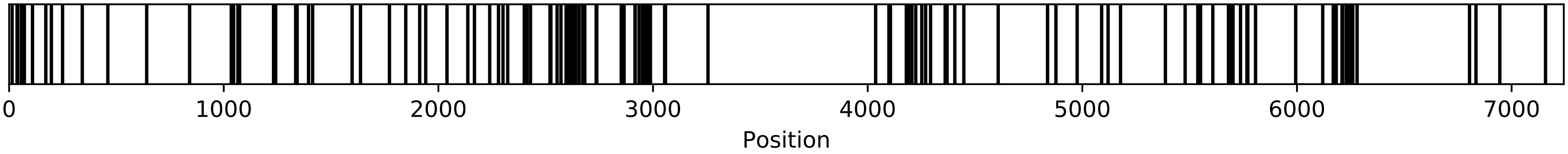


## CR1 vs MM1
Number of differences: 113

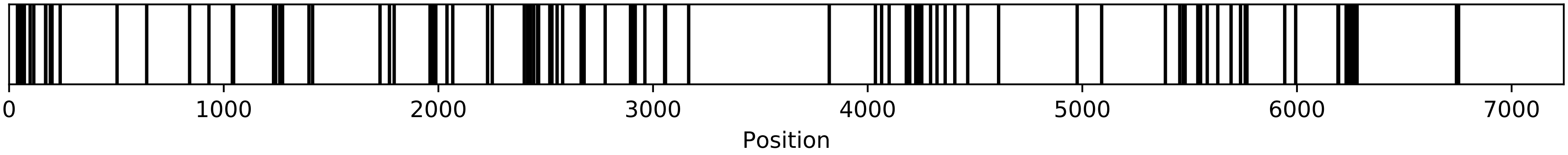


## CR1 vs MM2
Number of differences: 105

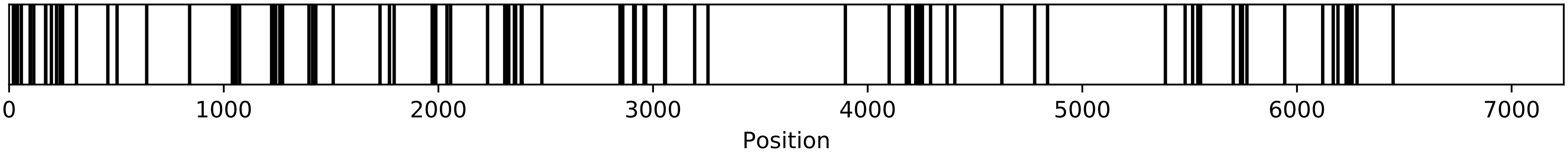


## CR2 vs MM1
Number of differences: 147

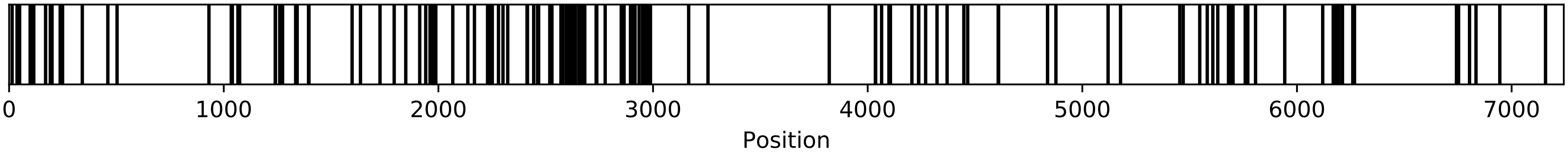


## CR2 vs MM2
Number of differences: 145

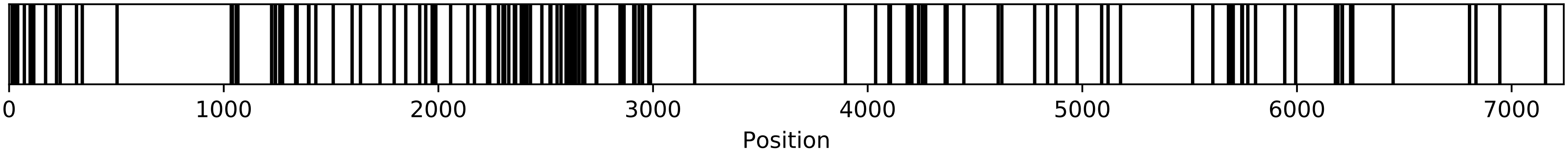


## MM1 vs MM2
Number of differences: 94

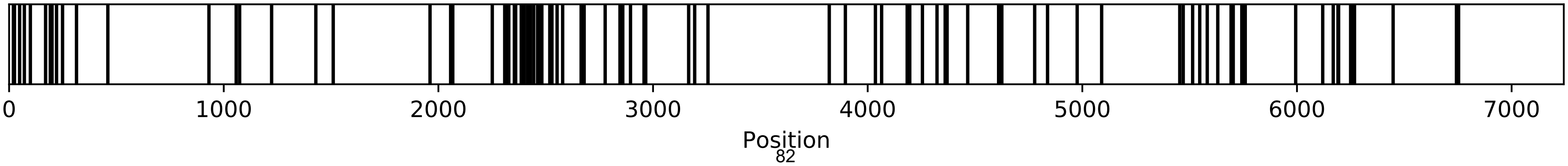

### Region: 234



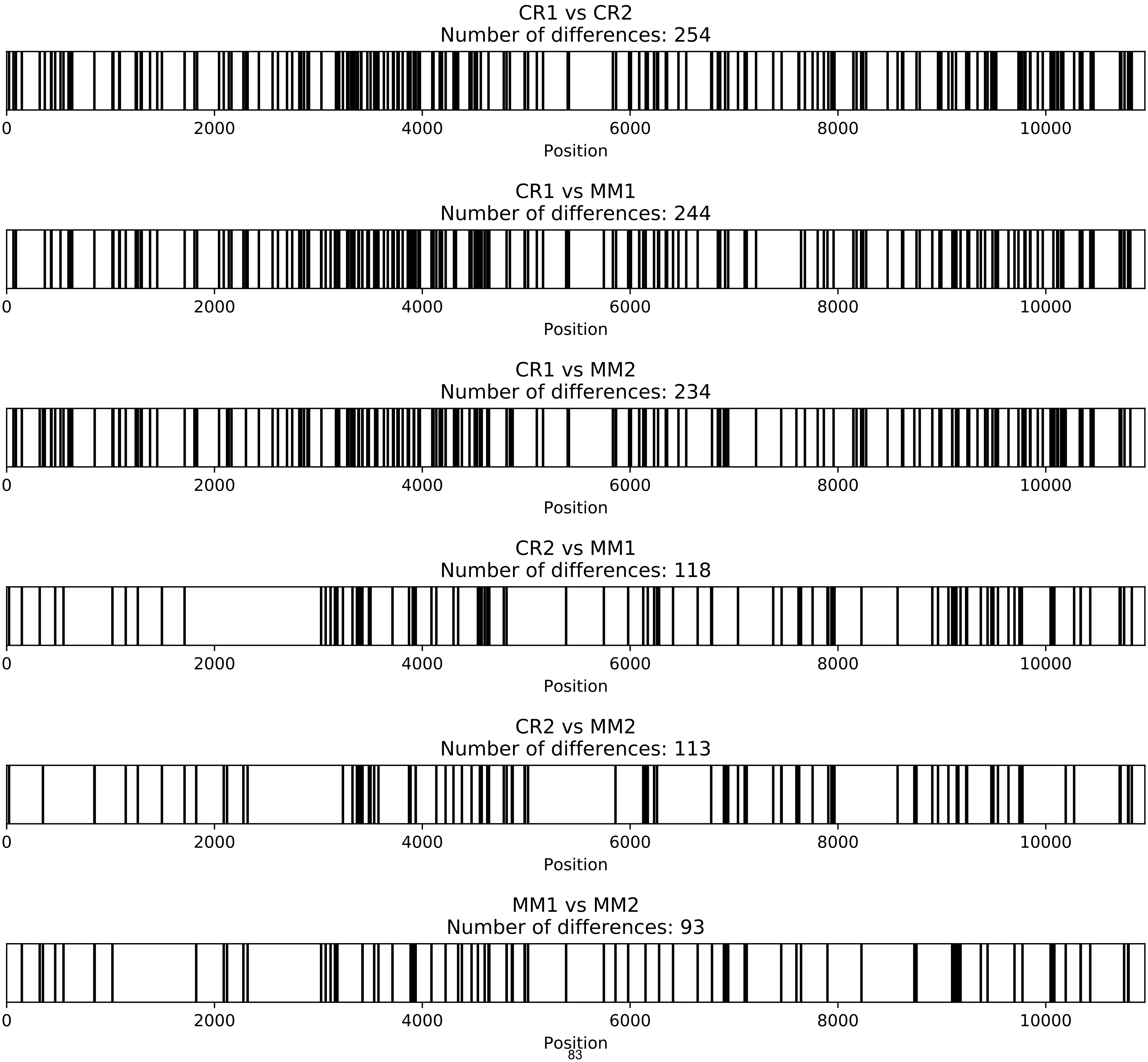
CR1 vs CR2
Number of differences: 254
CR1 vs MM1
Number of differences: 244
CR1 vs MM2
Number of differences: 234
CR2 vs MM1
Number of differences: 118
CR2 vs MM2
Number of differences: 113
MM1 vs MM2
Number of differences: 93
0
2000
4000
6000
8000
10000
Position

### Region: 239



CR1 vs CR2
Number of differences: 235

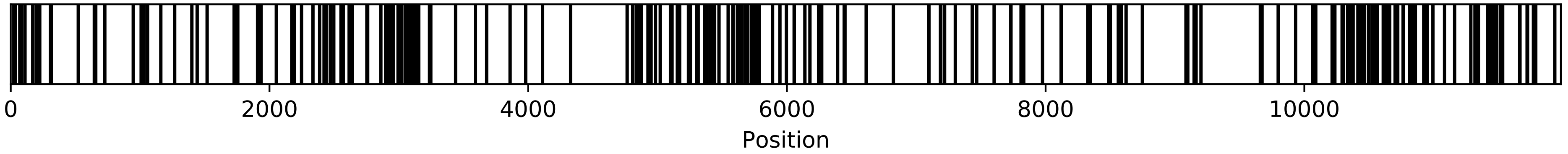


CR1 vs MM1
Number of differences: 239

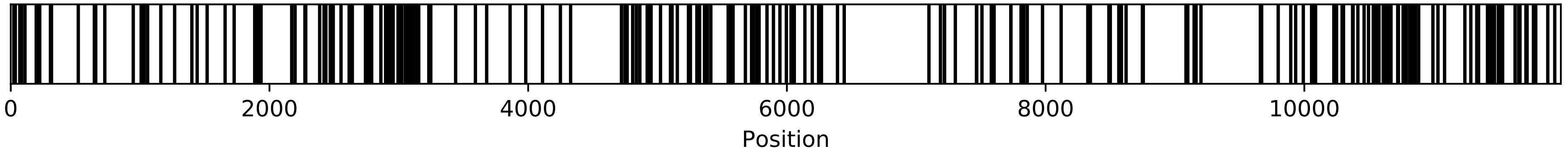


CR1 vs MM2
Number of differences: 243

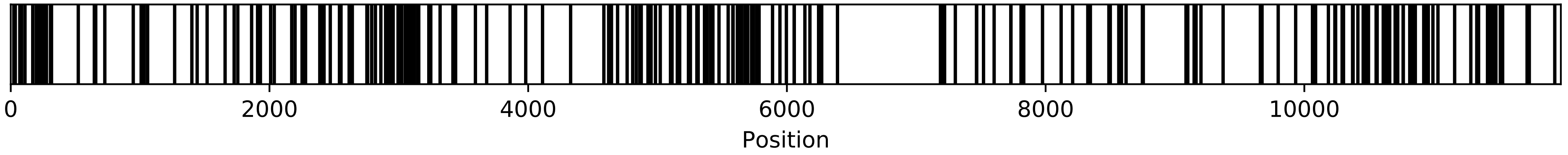


CR2 vs MM1
Number of differences: 129

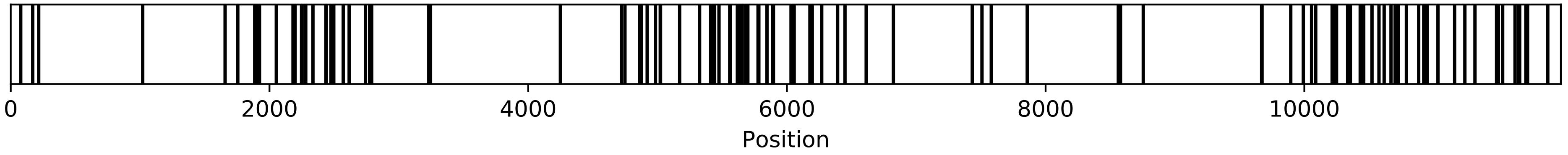


CR2 vs MM2
Number of differences: 95

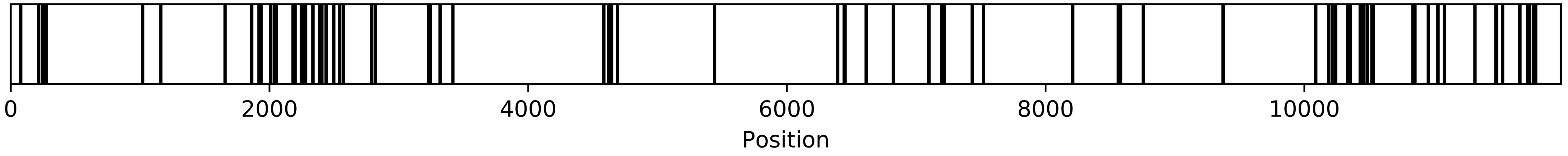


MM1 vs MM2
Number of differences: 122

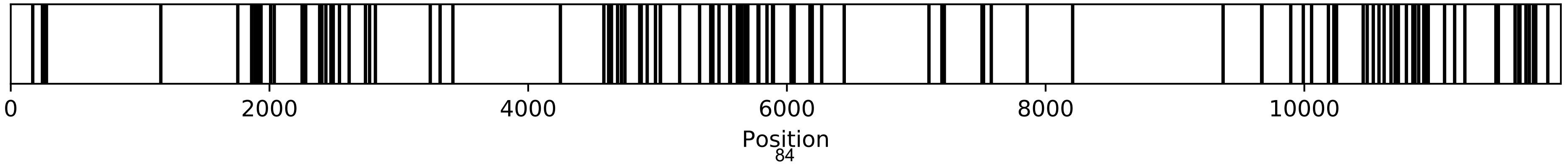

### Region: 244



CR1 vs CR2
Number of differences: 11

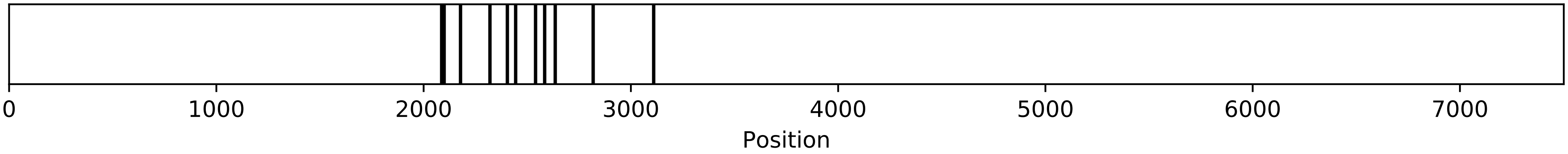


CR1 vs MM1
Number of differences: 166

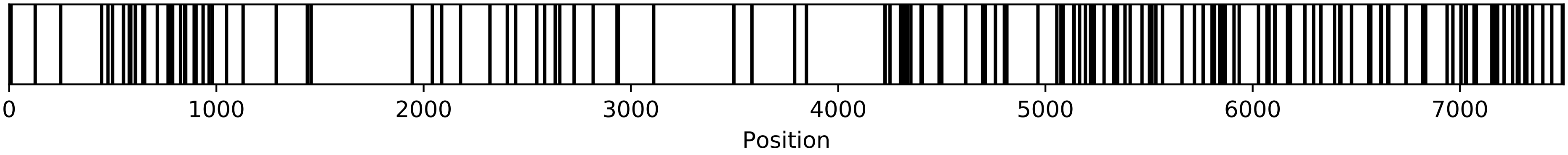


CR1 vs MM2
Number of differences: 167

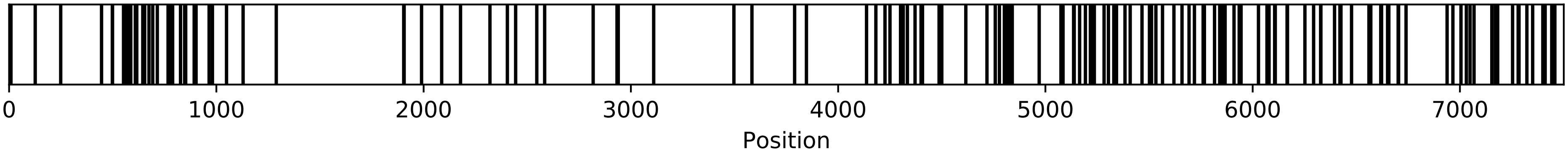


CR2 vs MM1
Number of differences: 159

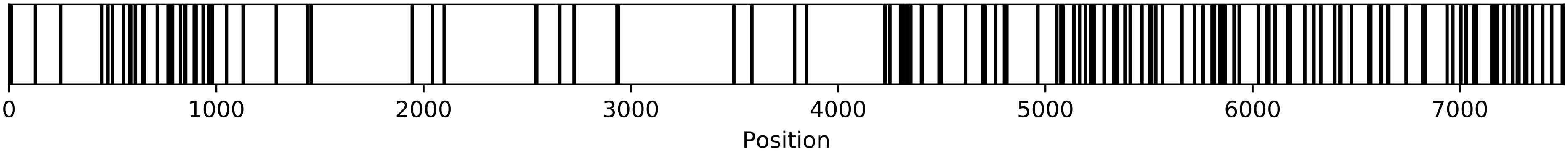


CR2 vs MM2
Number of differences: 162

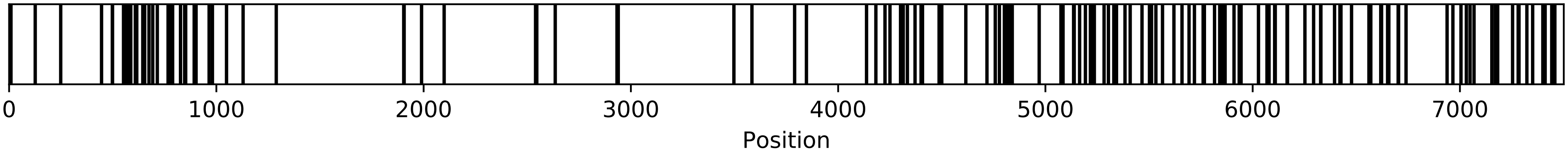


MM1 vs MM2
Number of differences: 69

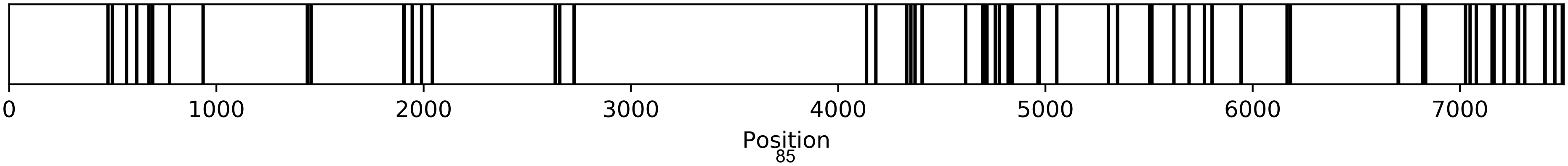

### Region: 247



CR1 vs CR2
Number of differences: 228

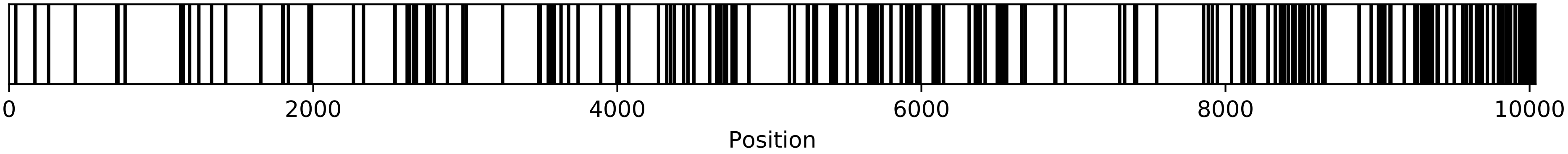


CR1 vs MM1
Number of differences: 214

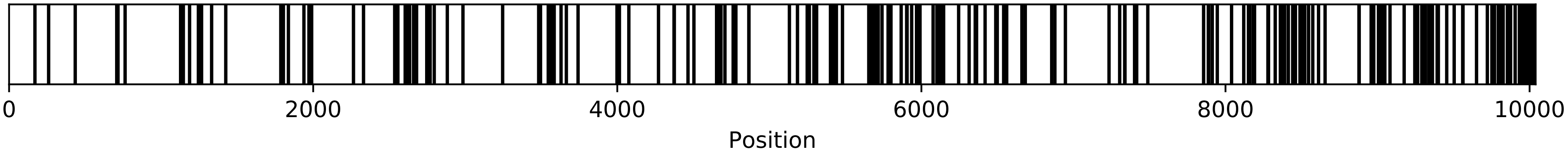


CR1 vs MM2
Number of differences: 217

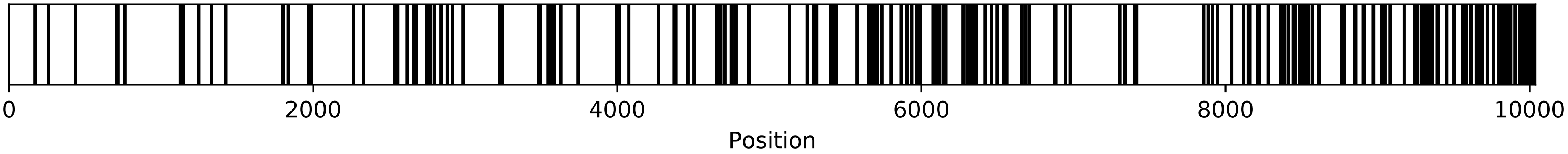


CR2 vs MM1
Number of differences: 84

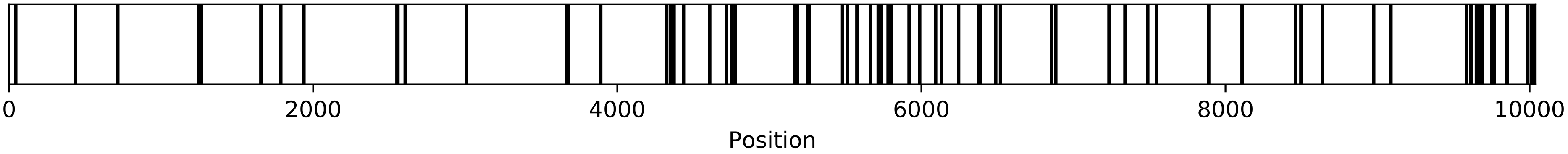


CR2 vs MM2
Number of differences: 89

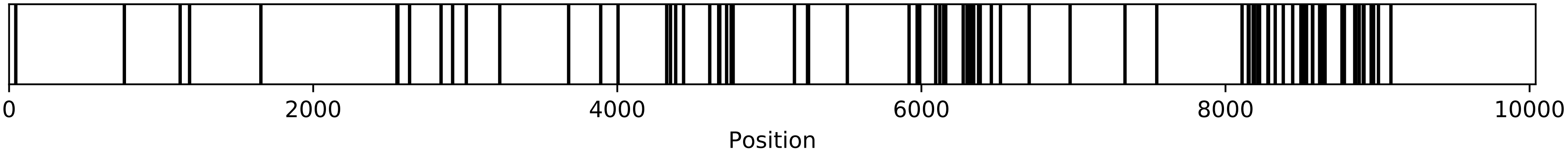


MM1 vs MM2
Number of differences: 113

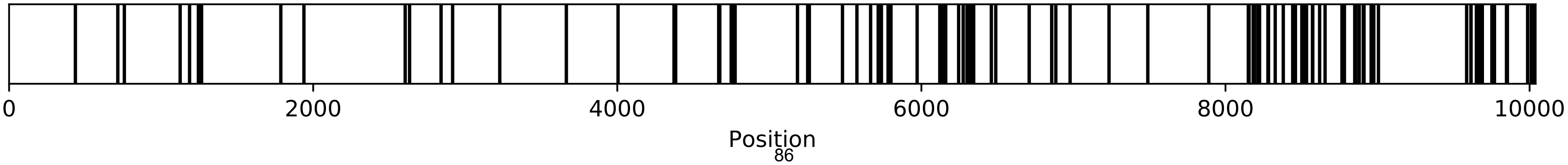

### Region: 253



CR1 vs CR2
Number of differences: 162

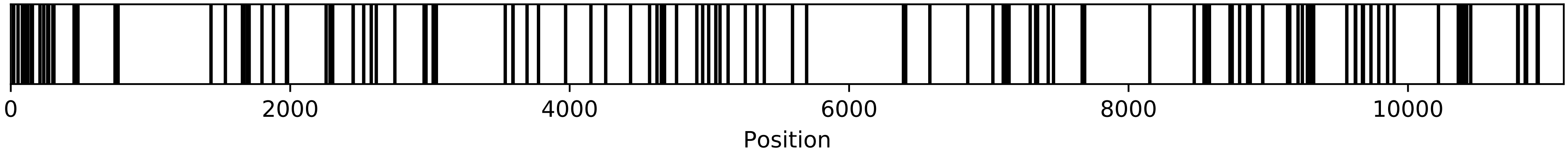


CR1 vs MM1
Number of differences: 161

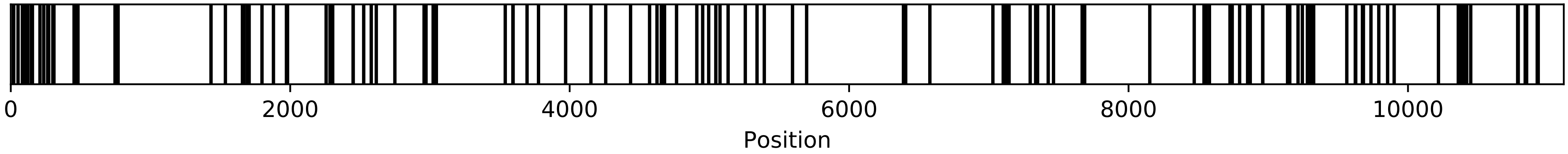


CR1 vs MM2
Number of differences: 166

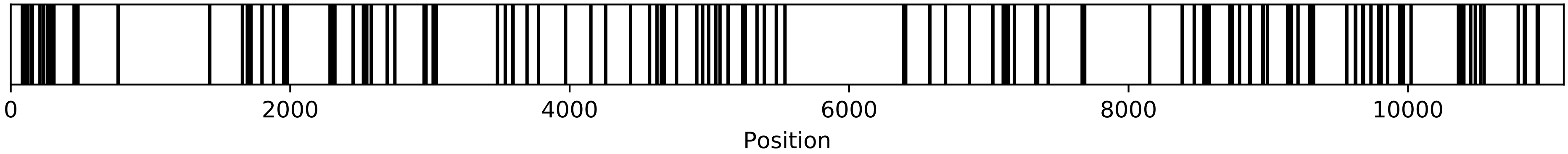


CR2 vs MM1
Number of differences: 1

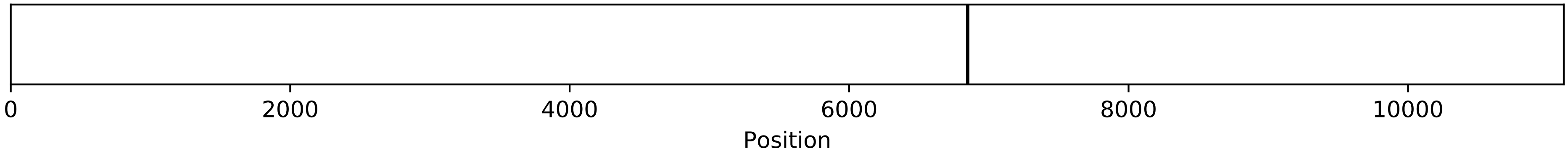


CR2 vs MM2
Number of differences: 80

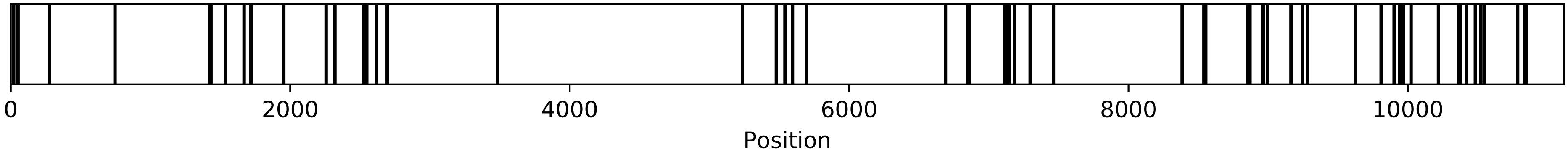


MM1 vs MM2
Number of differences: 79

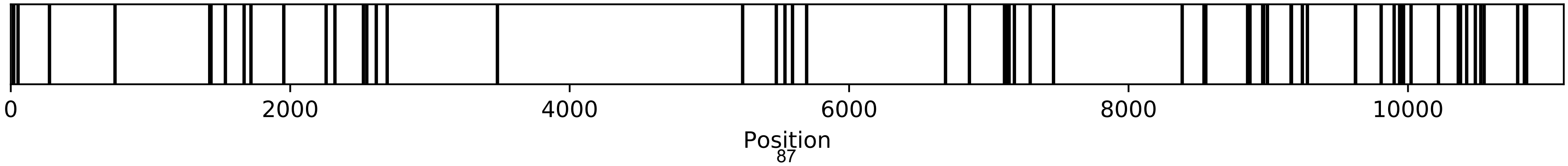

### Region: 254



CR1 vs CR2
Number of differences: 672

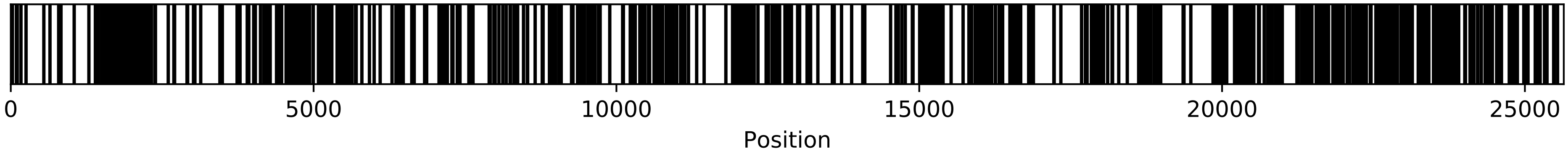


CR1 vs MM1
Number of differences: 226

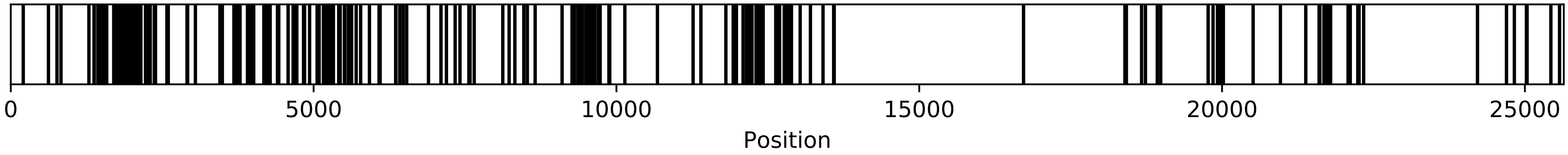


CR1 vs MM2
Number of differences: 226

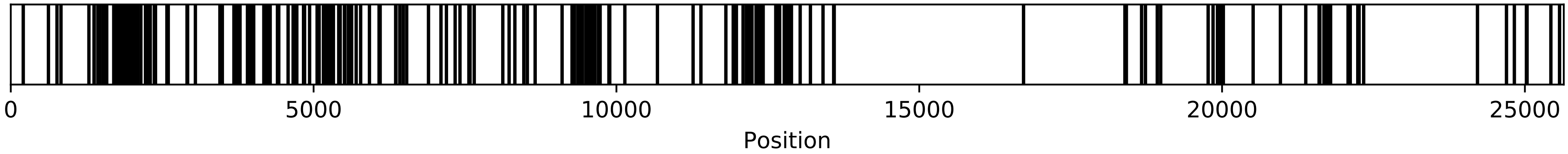


CR2 vs MM1
Number of differences: 693

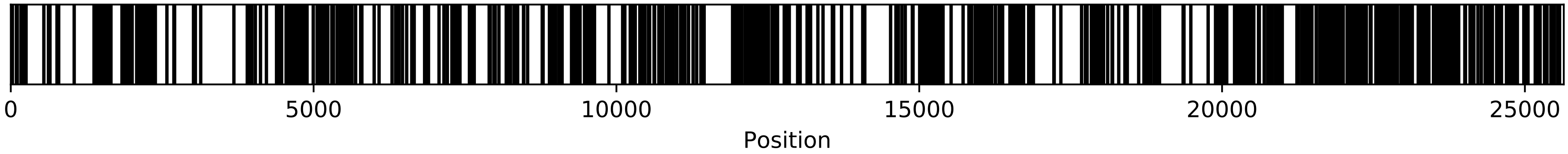


CR2 vs MM2
Number of differences: 693

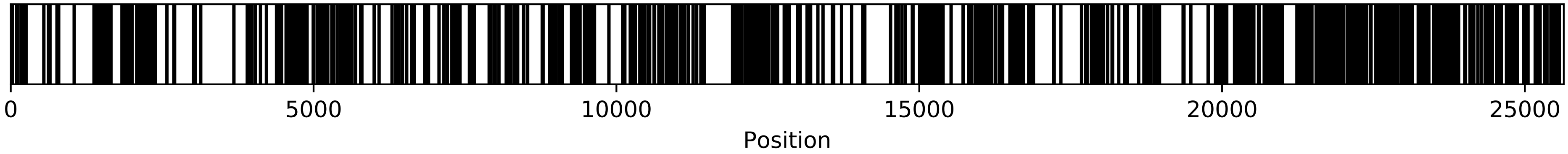


MM1 vs MM2
Number of differences: 0

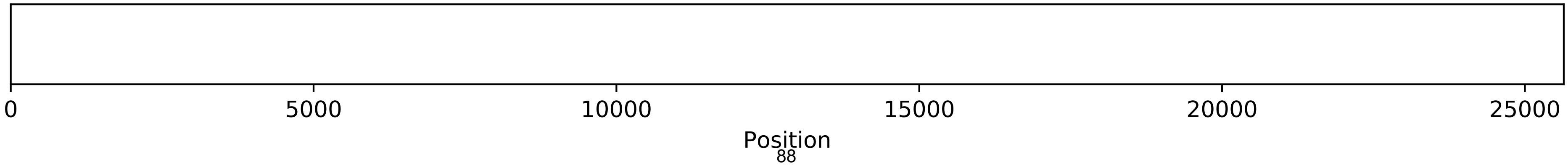

### Region: 256



CR1 vs CR2
Number of differences: 300

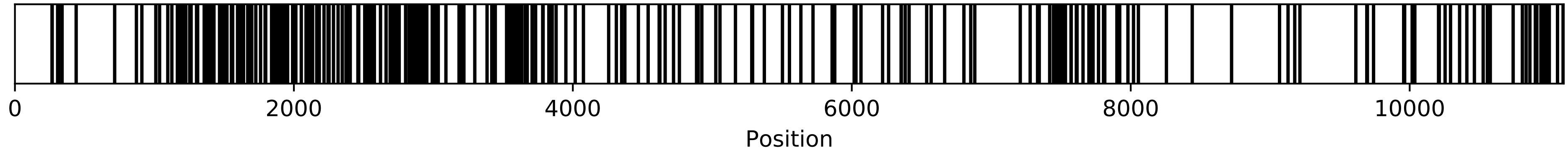


CR1 vs MM1
Number of differences: 167

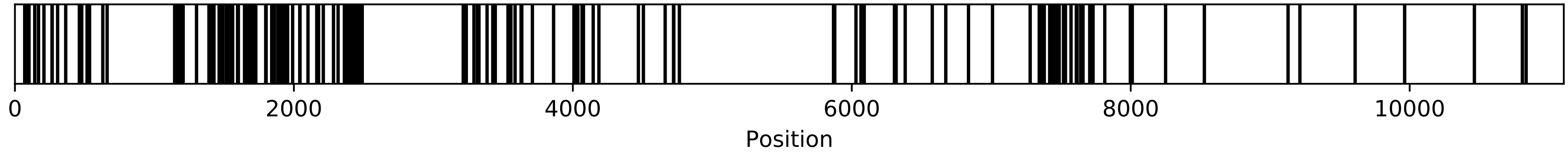


CR1 vs MM2
Number of differences: 175

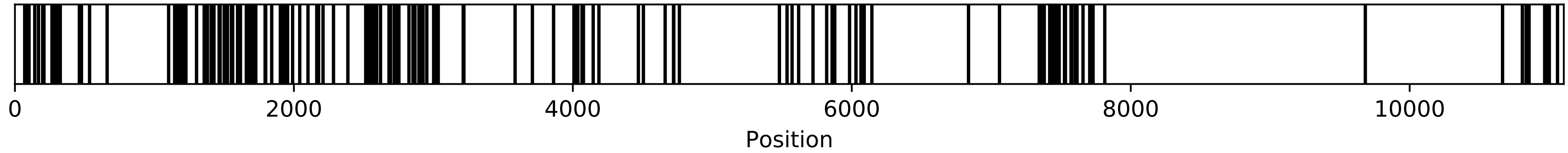


CR2 vs MM1
Number of differences: 302

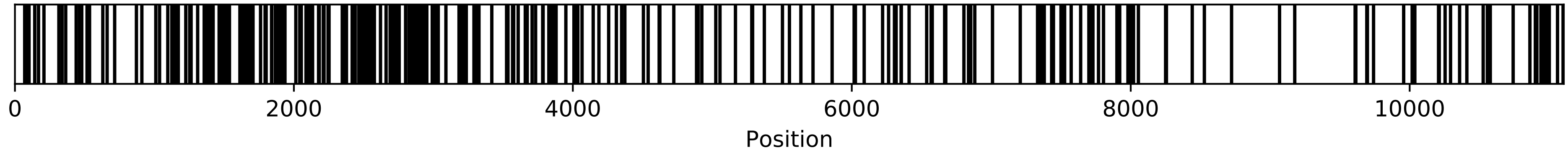


CR2 vs MM2
Number of differences: 283

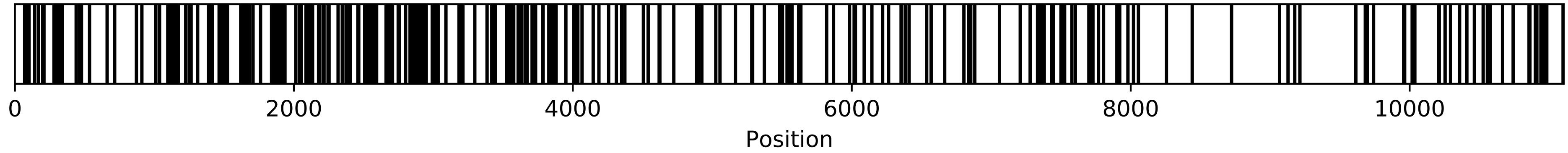


MM1 vs MM2
Number of differences: 154

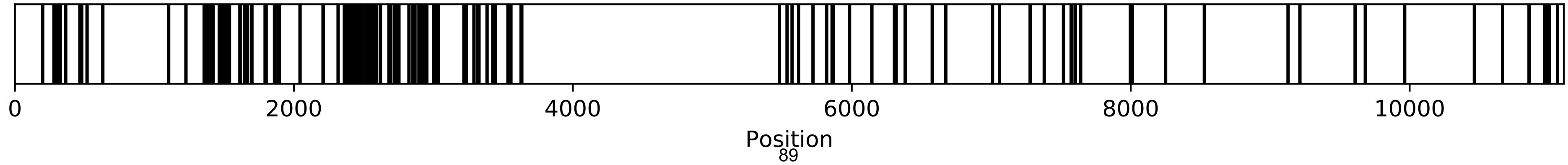

### Region: 262

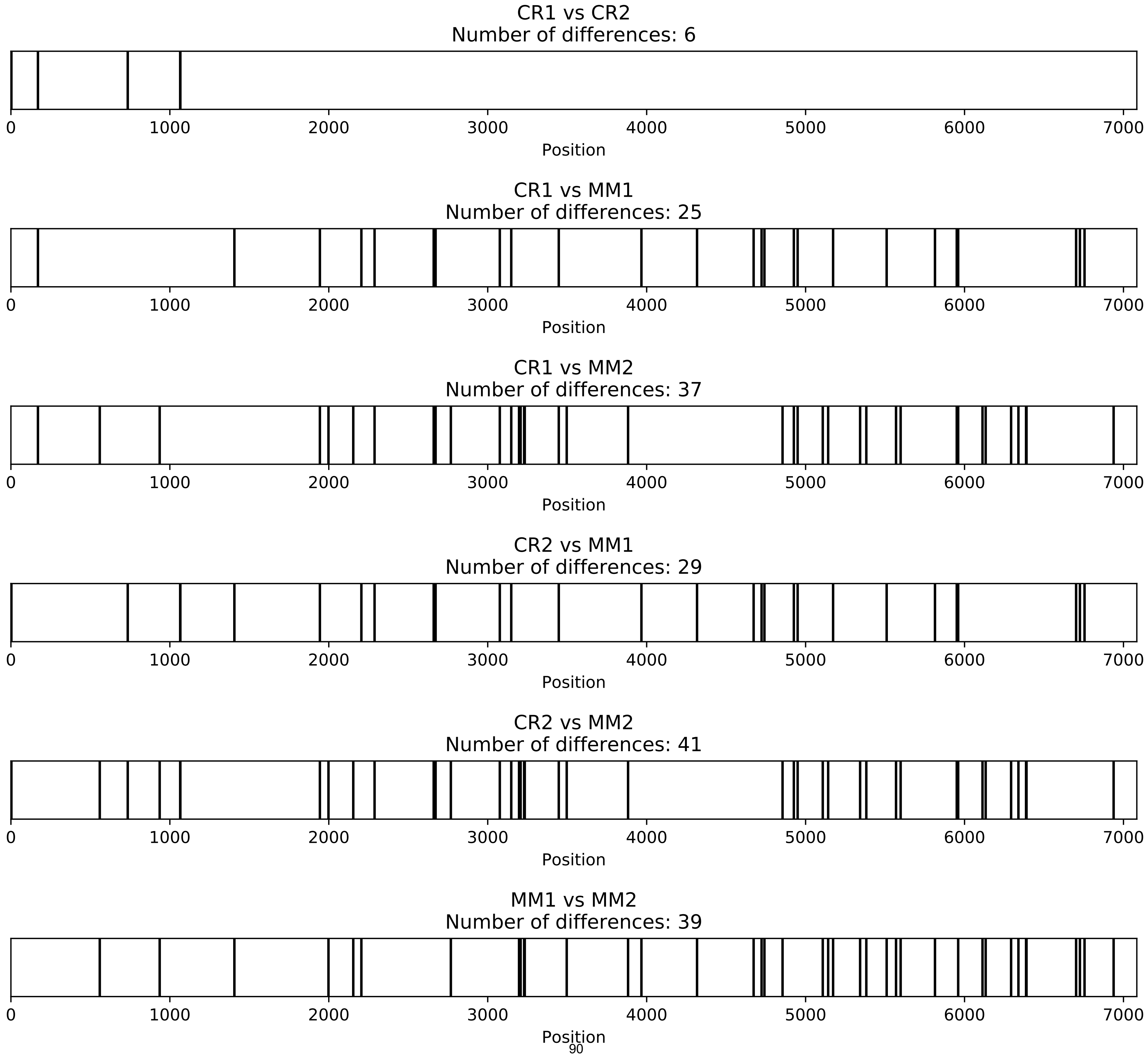
CR1 vs CR2
Number of differences: 6
CR1 vs MM1
Number of differences: 25
CR1 vs MM2
Number of differences: 37
CR2 vs MM1
Number of differences: 29
CR2 vs MM2
Number of differences: 41
MM1 vs MM2
Number of differences: 39
0
1000
2000
3000
4000
5000
6000
7000
Position

### Region: 265



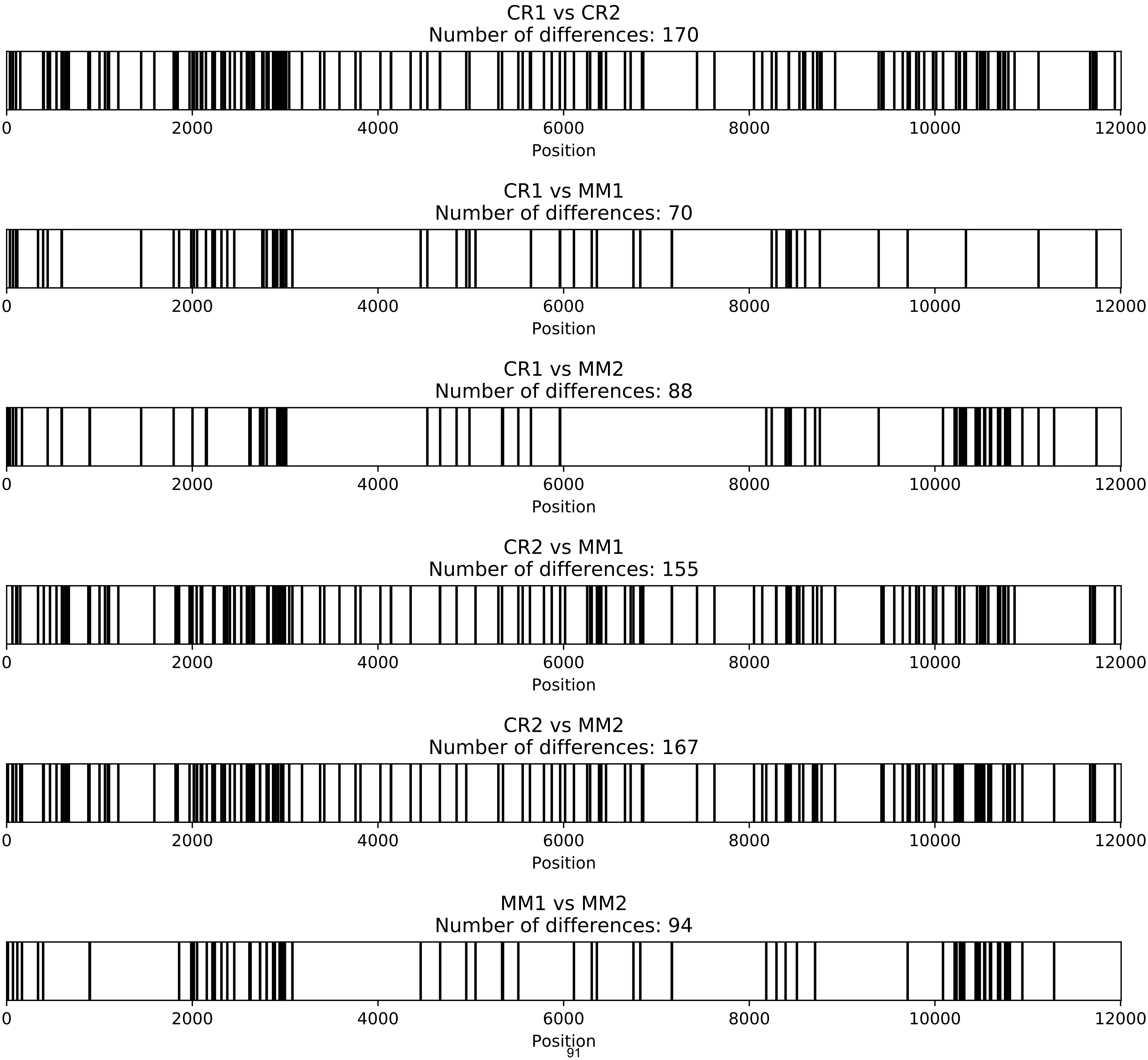
CR1 vs CR2
Number of differences: 170
CR1 vs MM1
Number of differences: 70
CR1 vs MM2
Number of differences: 88
CR2 vs MM1
Number of differences: 155
CR2 vs MM2
Number of differences: 167
MM1 vs MM2
Number of differences: 94
0
2000
4000
6000
8000
10000
12000
Position

### Region: 270

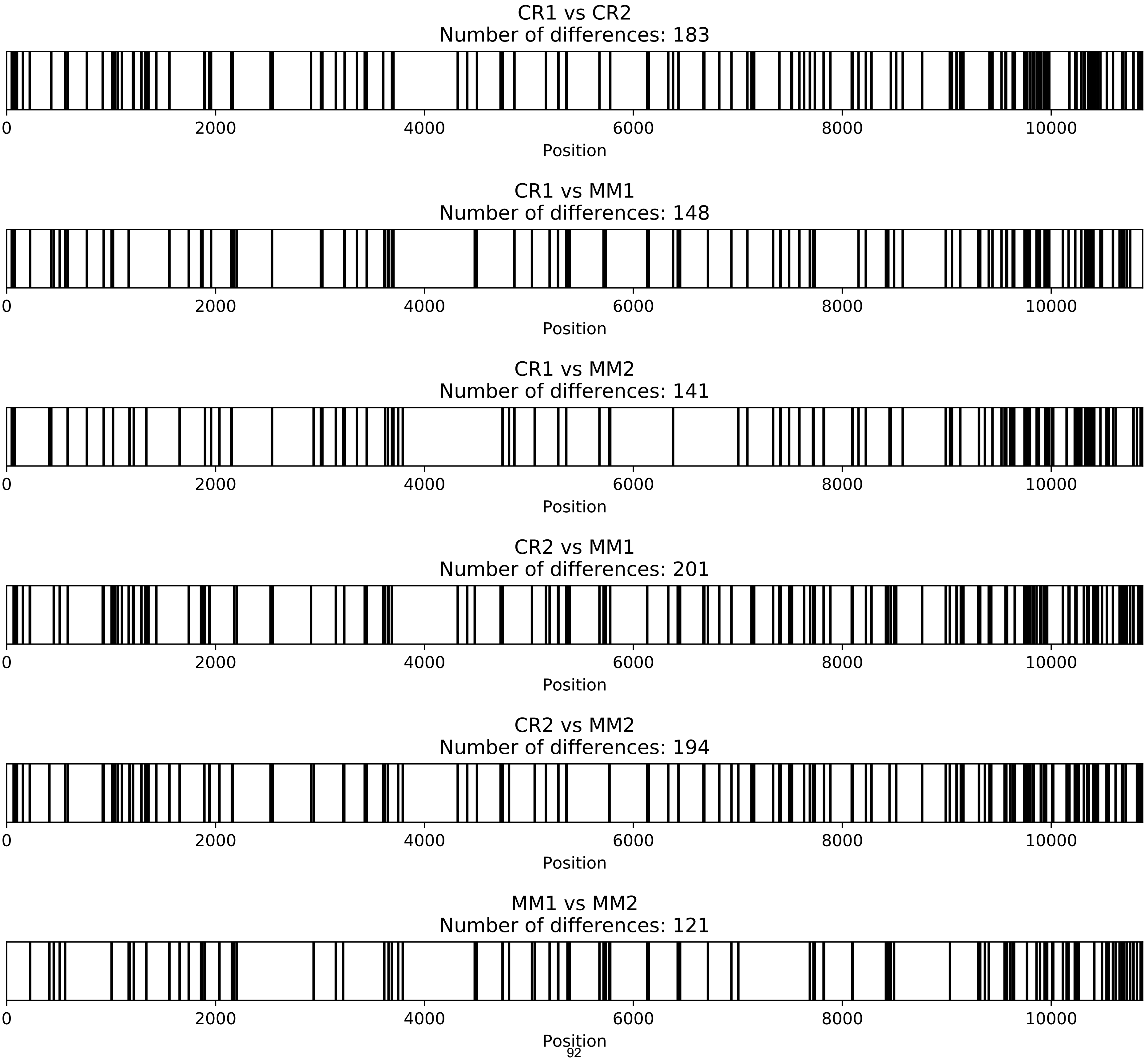
CR1 vs CR2
Number of differences: 183
Position
CR1 vs MM1
Number of differences: 148
Position
CR1 vs MM2
Number of differences: 141
Position
CR2 vs MM1
Number of differences: 201
Position
CR2 vs MM2
Number of differences: 194
Position
MM1 vs MM2
Number of differences: 121
Position
0
2000
4000
6000
8000
10000

### Region: 272



CR1 vs CR2
Number of differences: 207

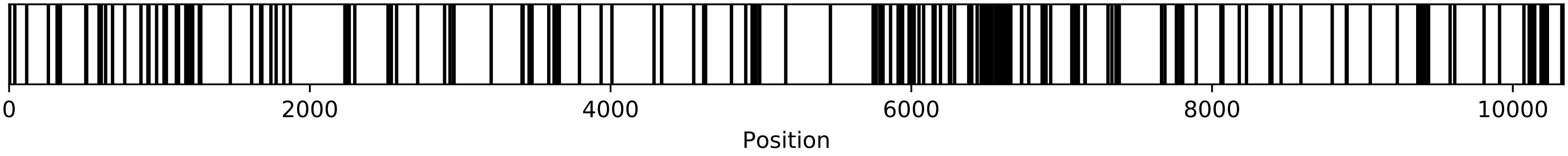


CR1 vs MM1
Number of differences: 217

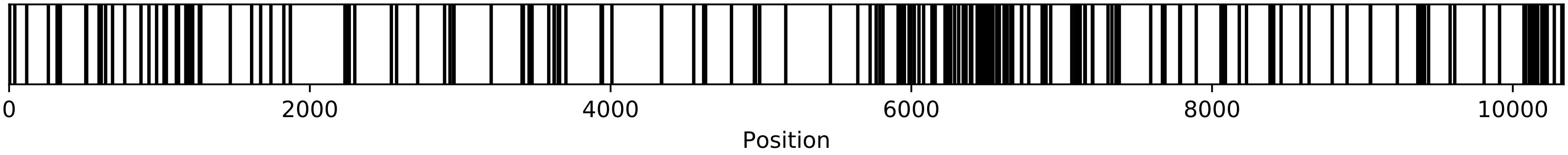


CR1 vs MM2
Number of differences: 220

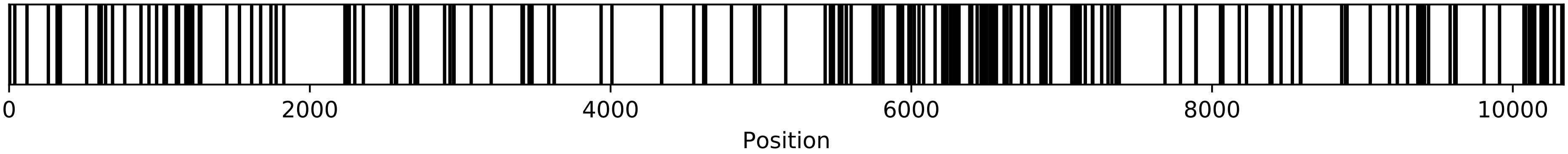


CR2 vs MM1
Number of differences: 89

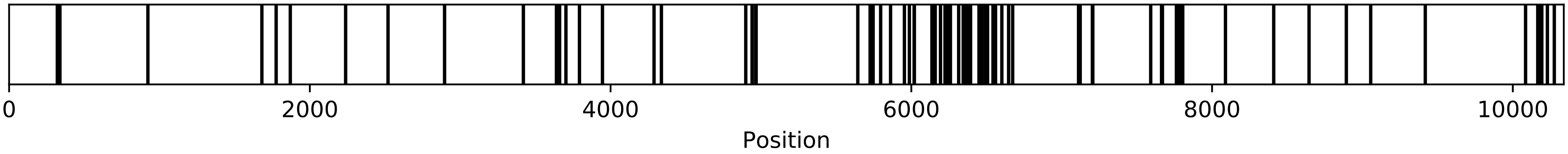


CR2 vs MM2
Number of differences: 113

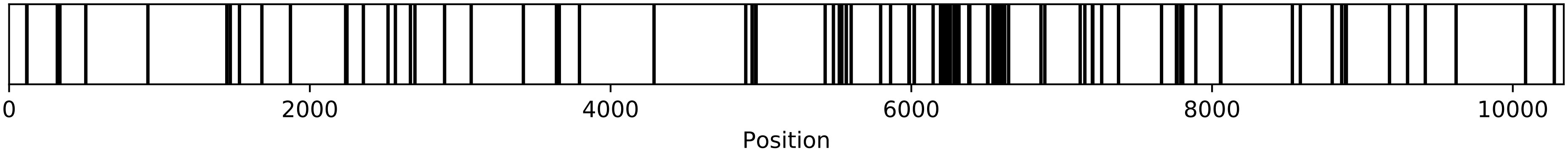


MM1 vs MM2
Number of differences: 105

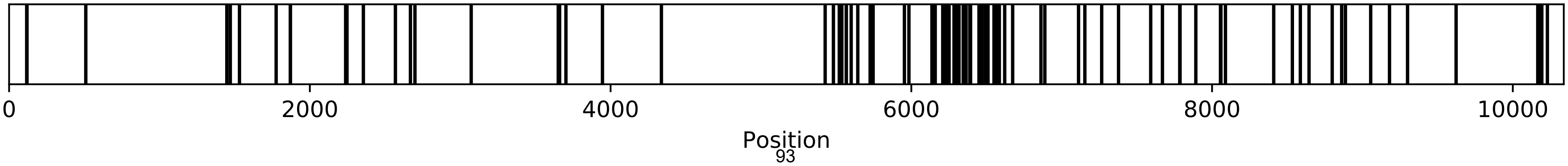

### Region: 275



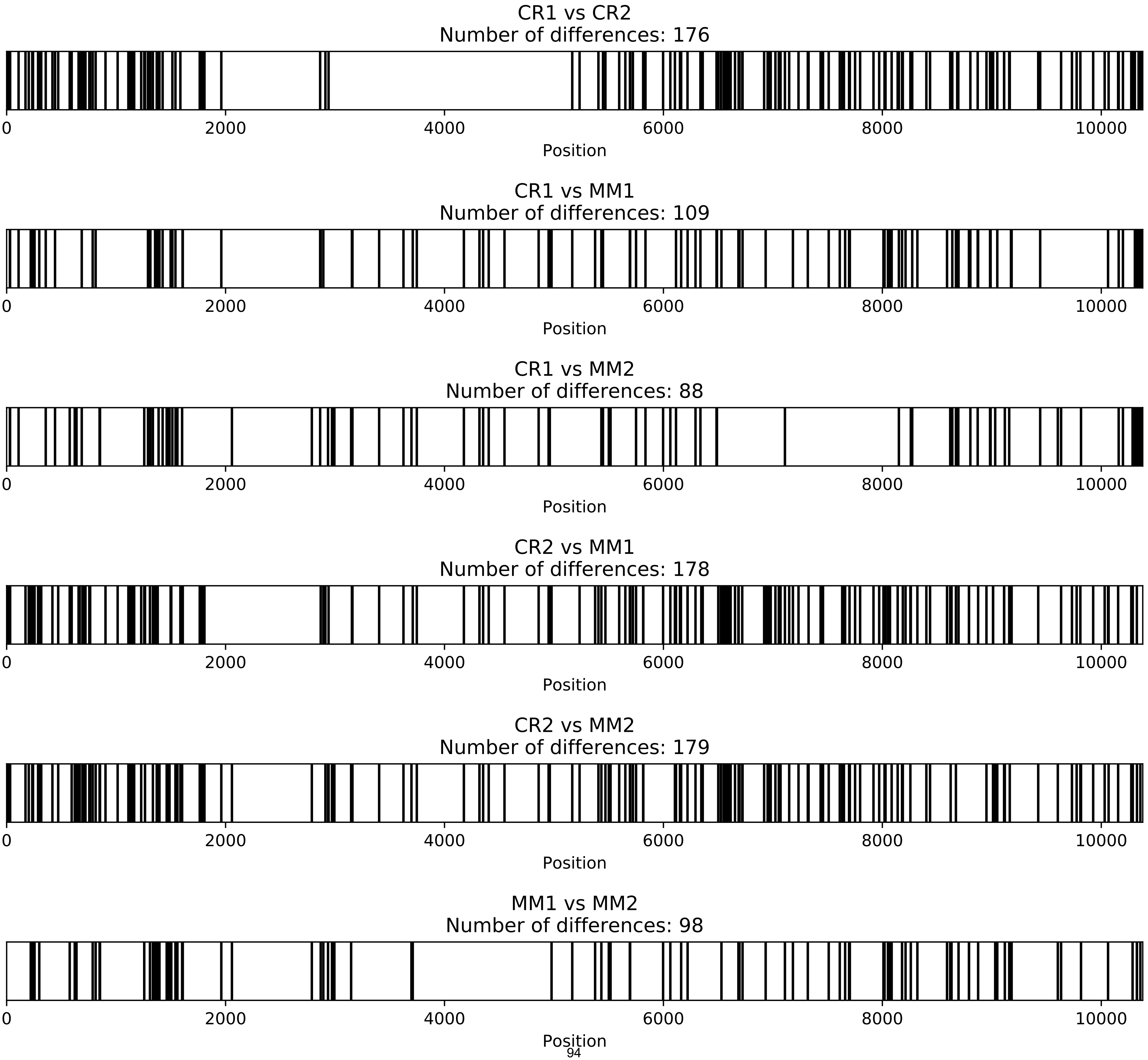
CR1 vs CR2
Number of differences: 176
CR1 vs MM1
Number of differences: 109
CR1 vs MM2
Number of differences: 88
CR2 vs MM1
Number of differences: 178
CR2 vs MM2
Number of differences: 179
MM1 vs MM2
Number of differences: 98
0
2000
4000
6000
8000
10000
Position

### Region: 277



CR1 vs CR2
Number of differences: 461

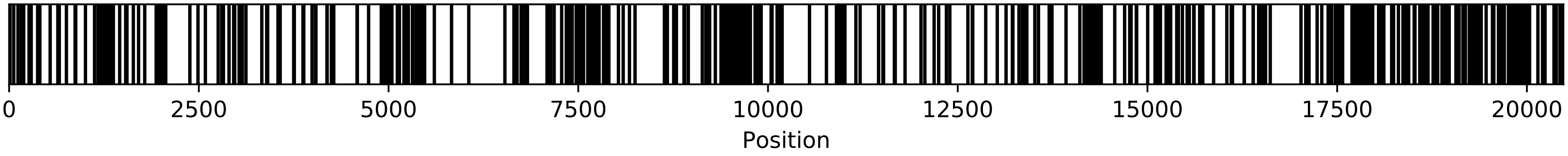


CR1 vs MM1
Number of differences: 70

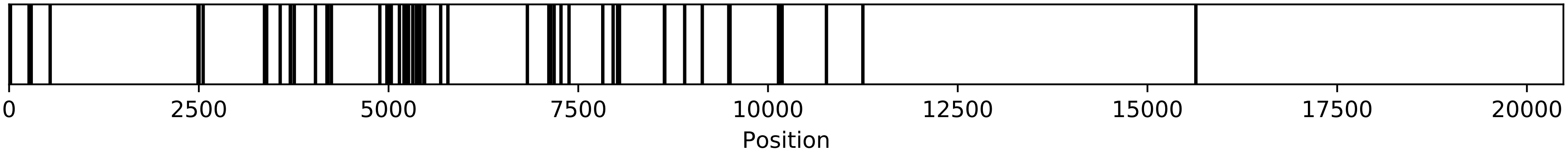


CR1 vs MM2
Number of differences: 100

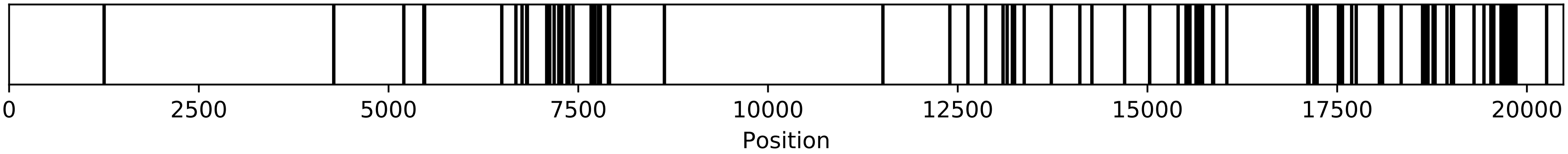


CR2 vs MM1
Number of differences: 453

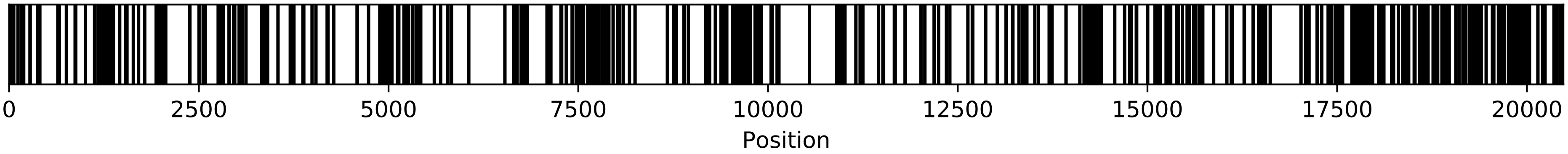


CR2 vs MM2
Number of differences: 451

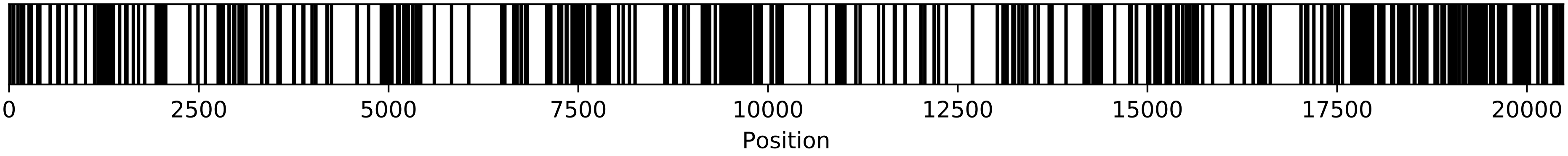


MM1 vs MM2
Number of differences: 152

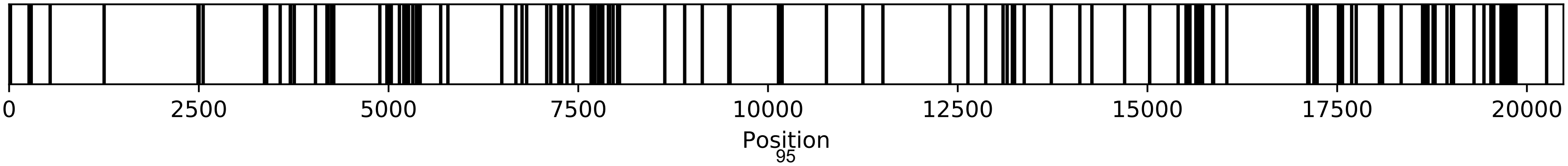

### Region: 281



CR1 vs CR2
Number of differences: 346

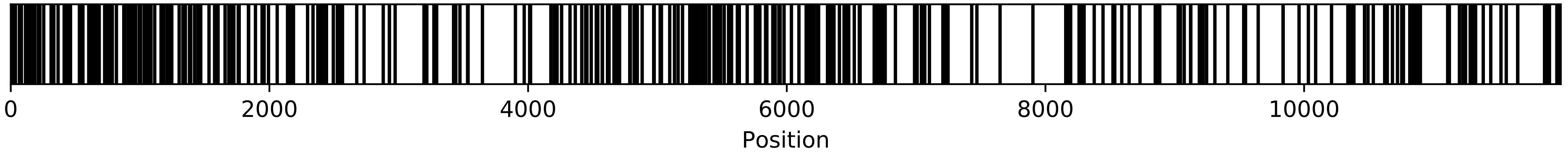


CR1 vs MM1
Number of differences: 173

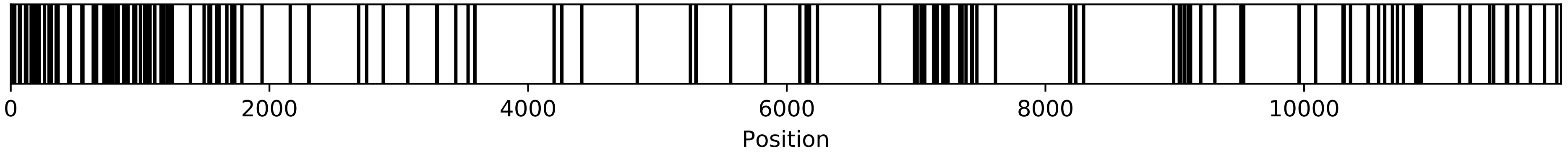


CR1 vs MM2
Number of differences: 173

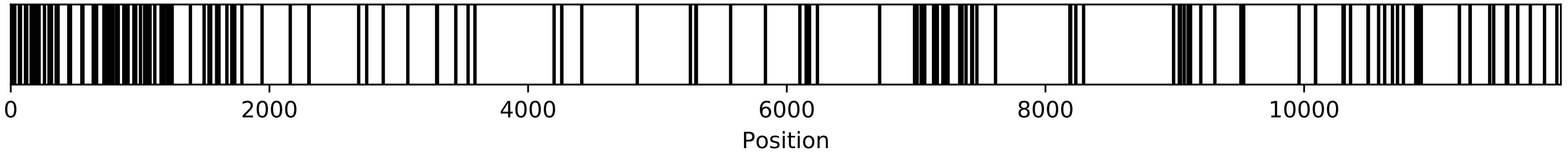


CR2 vs MM1
Number of differences: 330

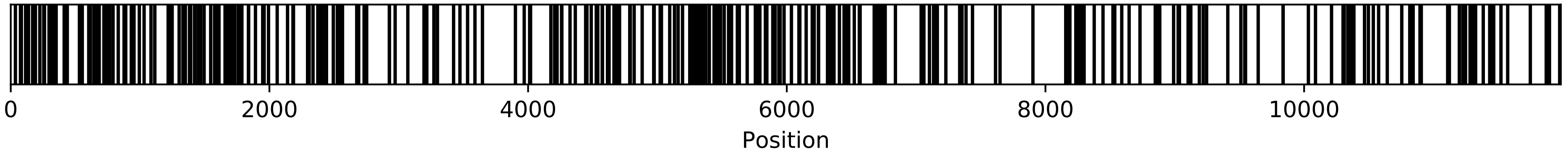


CR2 vs MM2
Number of differences: 330

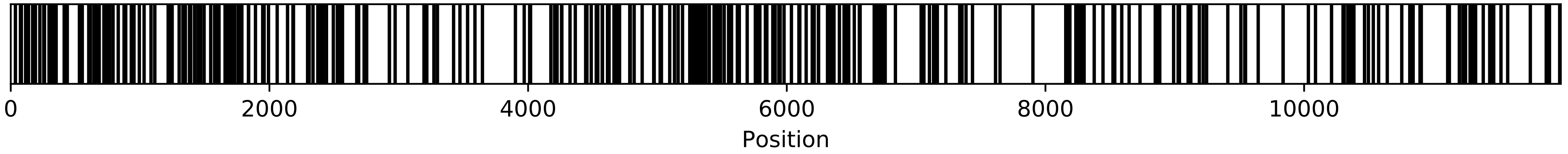


MM1 vs MM2
Number of differences: 0

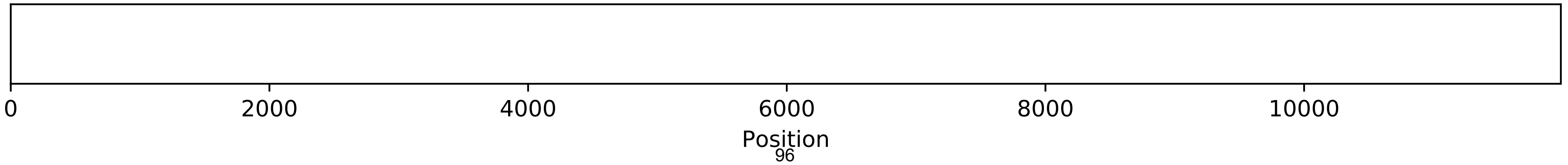

### Region: 282



CR1 vs CR2
Number of differences: 209

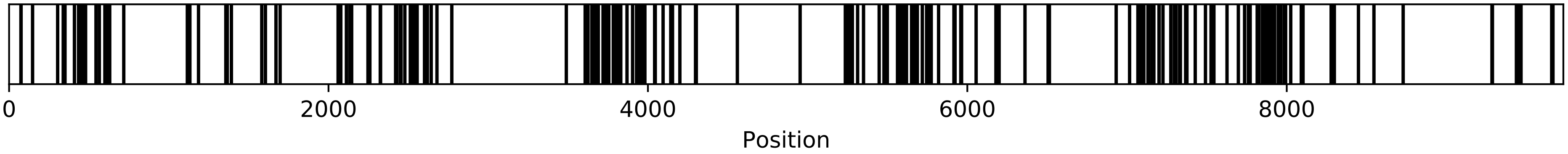


CR1 vs MM1
Number of differences: 197

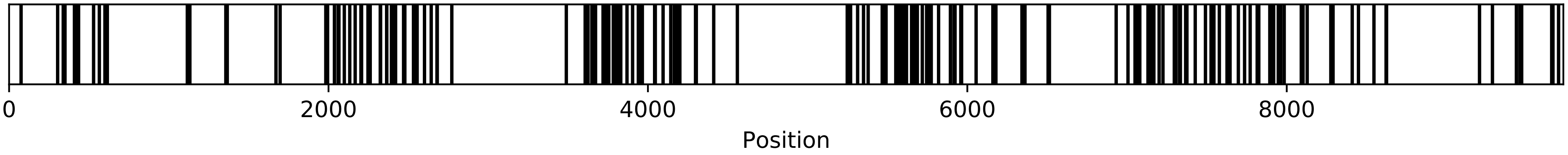


CR1 vs MM2
Number of differences: 245

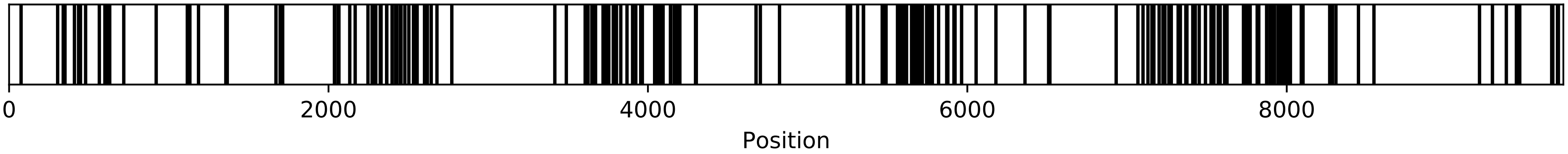


CR2 vs MM1
Number of differences: 115

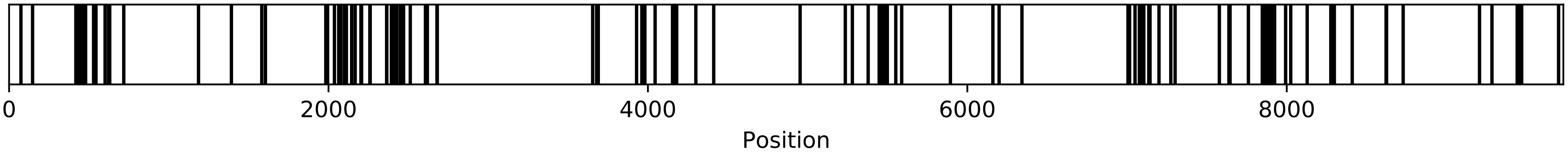


CR2 vs MM2
Number of differences: 185

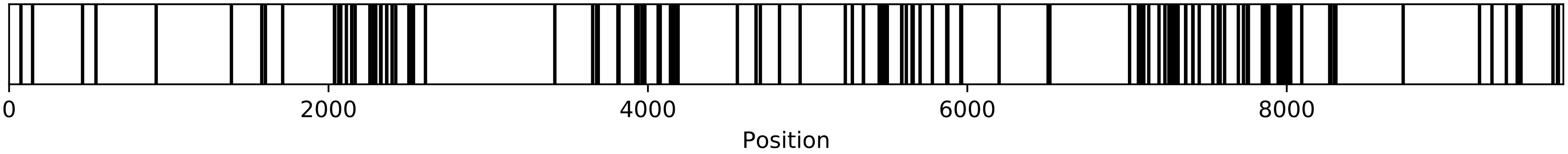


MM1 vs MM2
Number of differences: 178

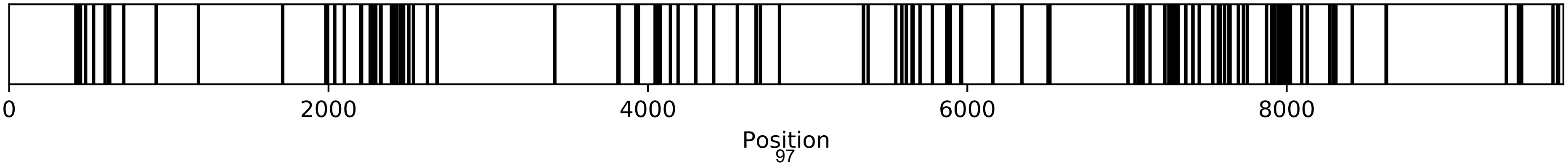

### Region: 285



CR1 vs CR2
Number of differences: 22

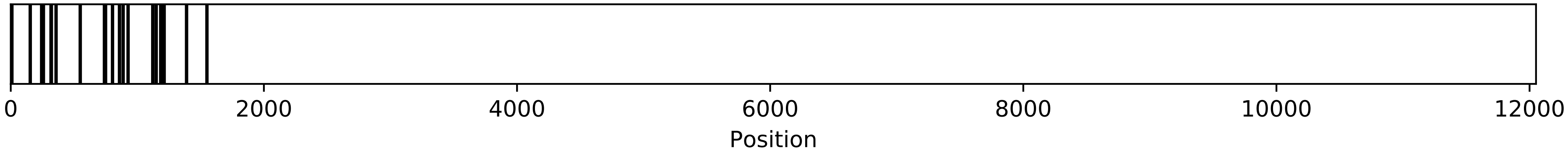


CR1 vs MM1
Number of differences: 95

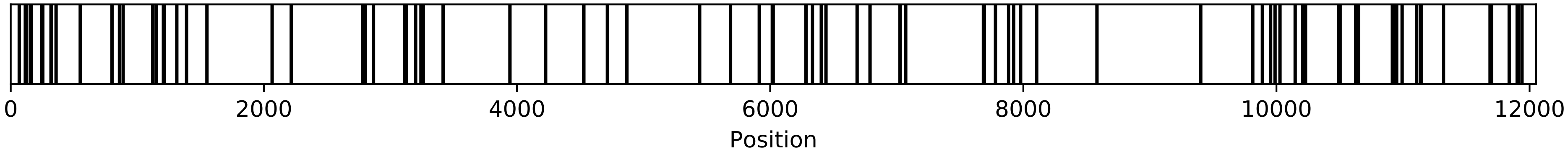


CR1 vs MM2
Number of differences: 108

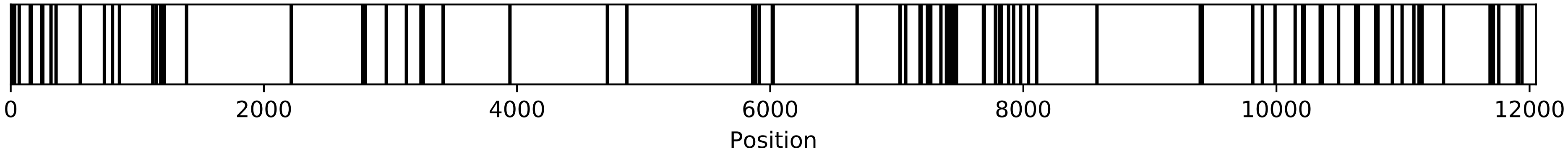


CR2 vs MM1
Number of differences: 87

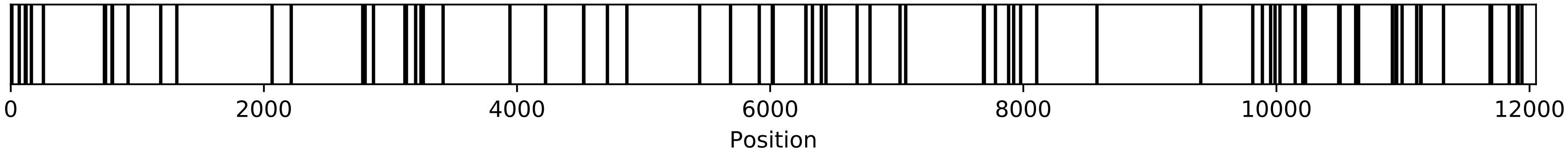


CR2 vs MM2
Number of differences: 98

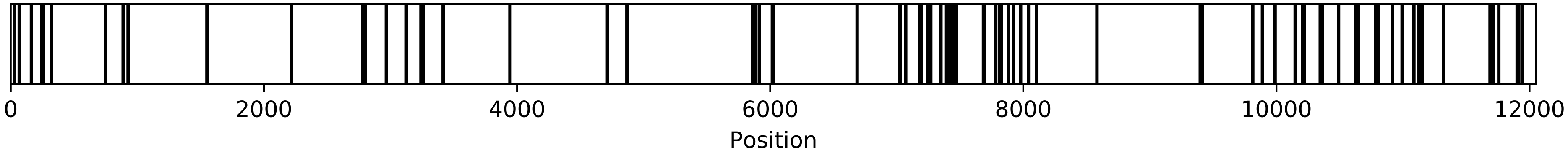


MM1 vs MM2
Number of differences: 87

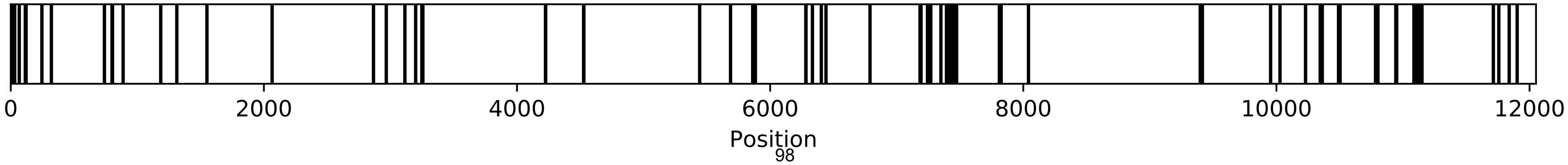

### Region: 287



## CR1 vs CR2
Number of differences: 331

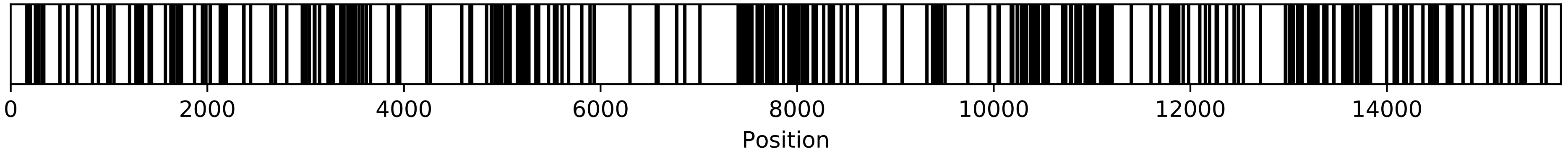


## CR1 vs MM1
Number of differences: 212

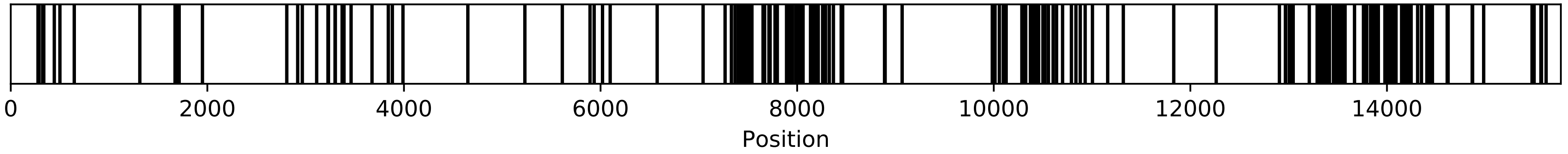


## CR1 vs MM2
Number of differences: 189

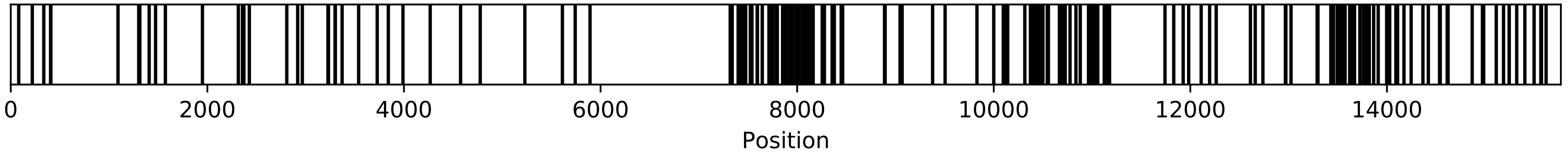


## CR2 vs MM1
Number of differences: 360

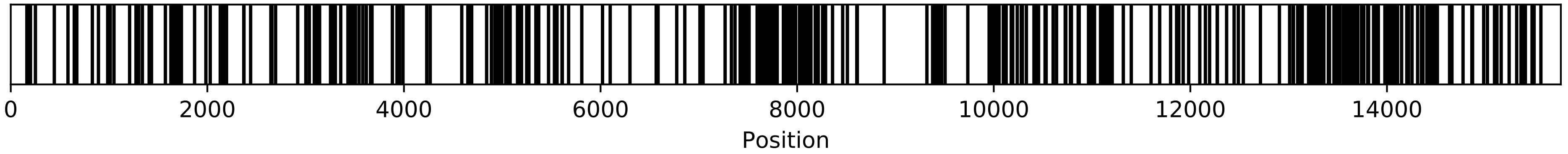


## CR2 vs MM2
Number of differences: 326

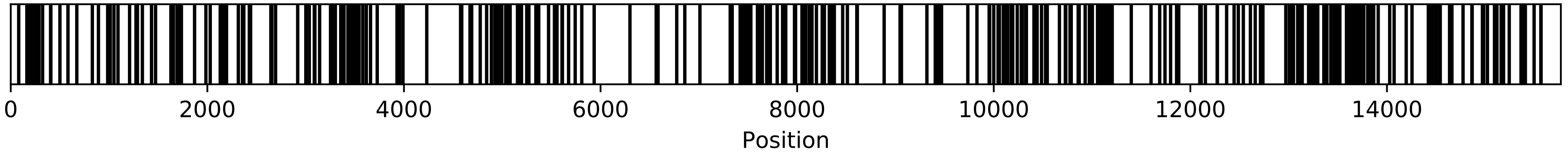


## MM1 vs MM2
Number of differences: 219

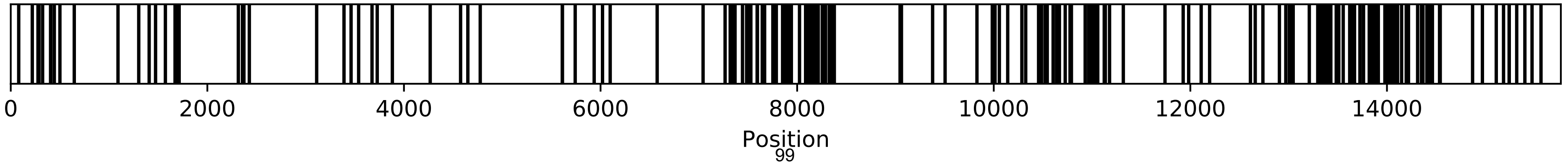

### Region: 288



CR1 vs CR2
Number of differences: 228

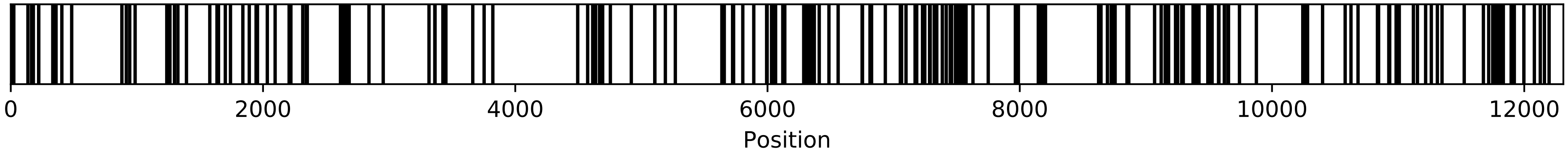


CR1 vs MM1
Number of differences: 135

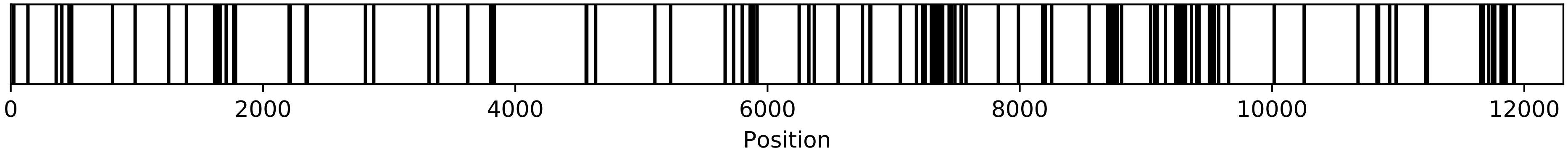


CR1 vs MM2
Number of differences: 142

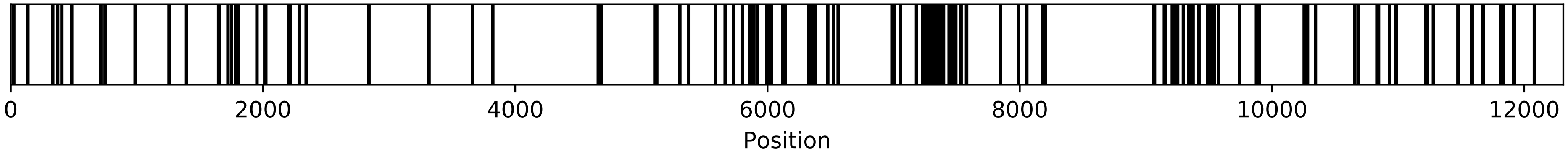


CR2 vs MM1
Number of differences: 222

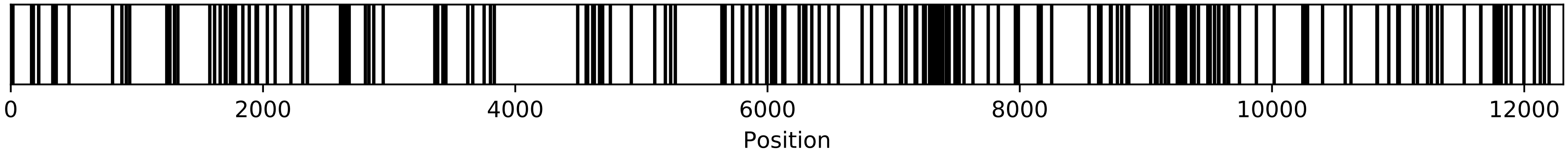


CR2 vs MM2
Number of differences: 225

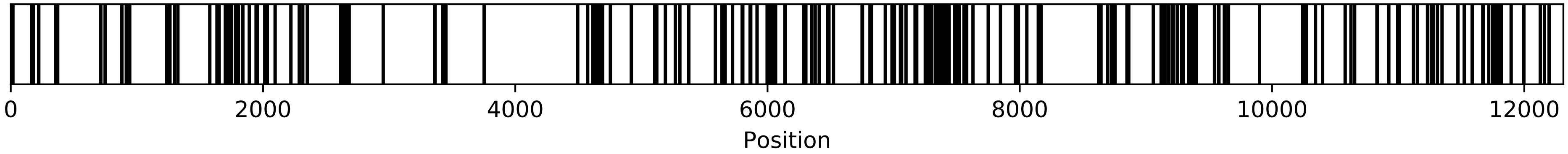


MM1 vs MM2
Number of differences: 129

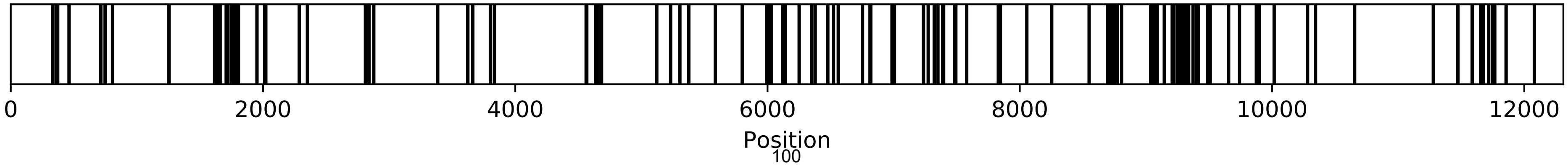

### Region: 294



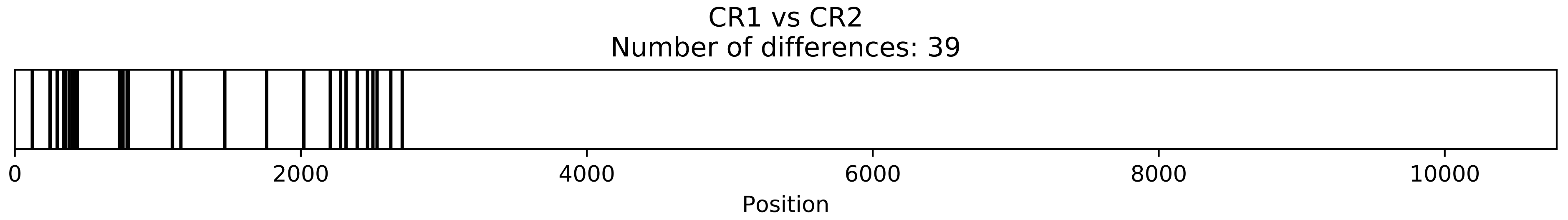
CR1 vs CR2
Number of differences: 39
0
2000
4000
6000
8000
10000
Position

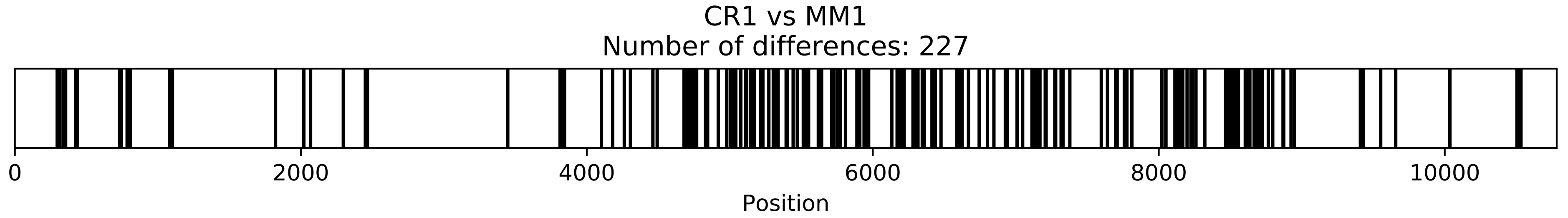
CR1 vs MM1
Number of differences: 227
0
2000
4000
6000
8000
10000
Position

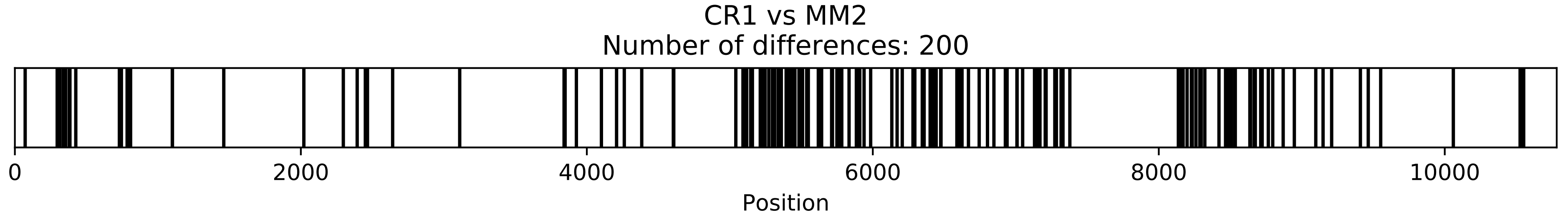
CR1 vs MM2
Number of differences: 200
0
2000
4000
6000
8000
10000
Position

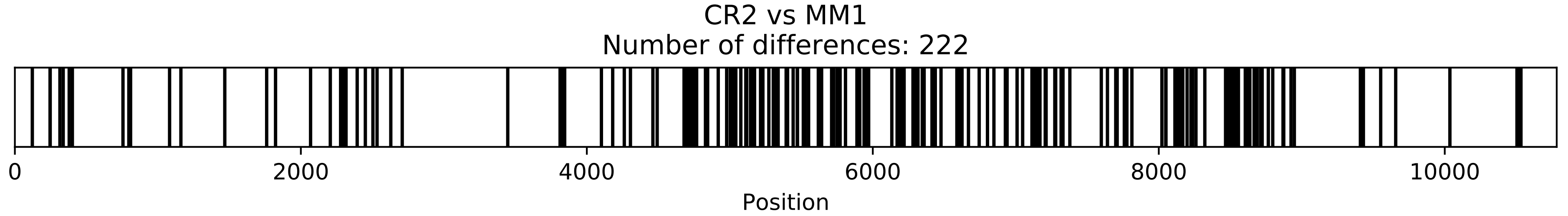
CR2 vs MM1
Number of differences: 222
0
2000
4000
6000
8000
10000
Position

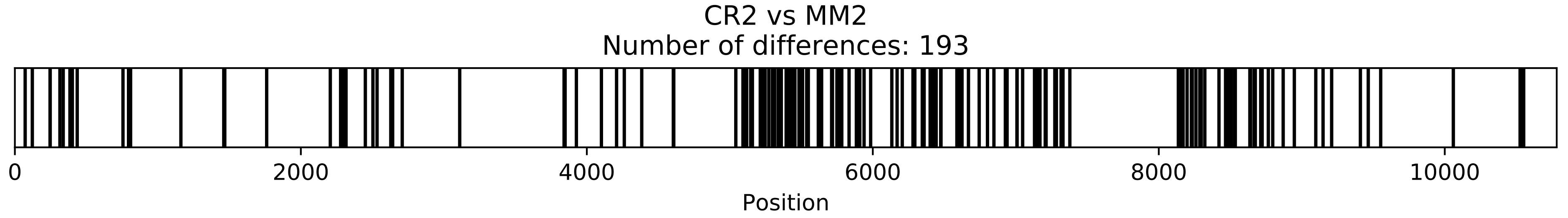
CR2 vs MM2
Number of differences: 193
0
2000
4000
6000
8000
10000
Position

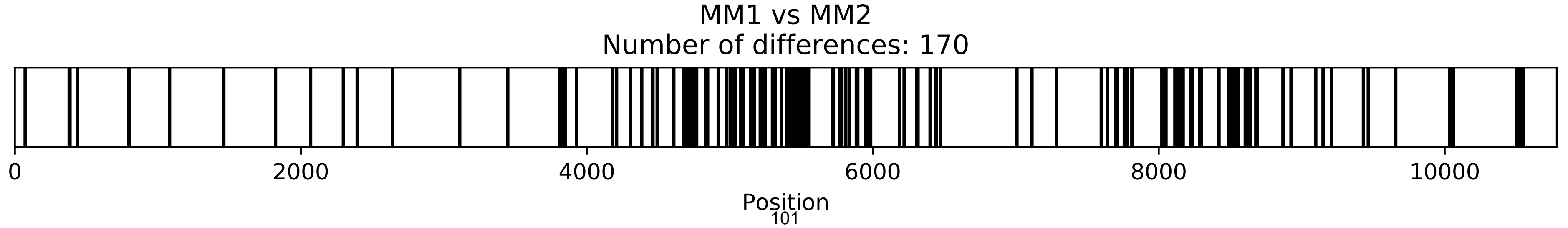
MM1 vs MM2
Number of differences: 170
0
2000
4000
6000
8000
10000
Position

### Region: 295



CR1 vs CR2
Number of differences: 97

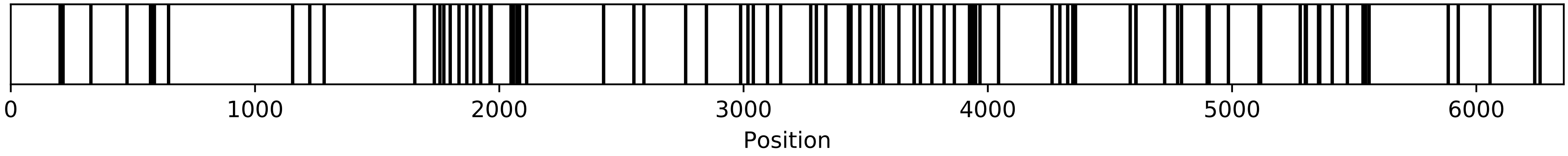


CR1 vs MM1
Number of differences: 44

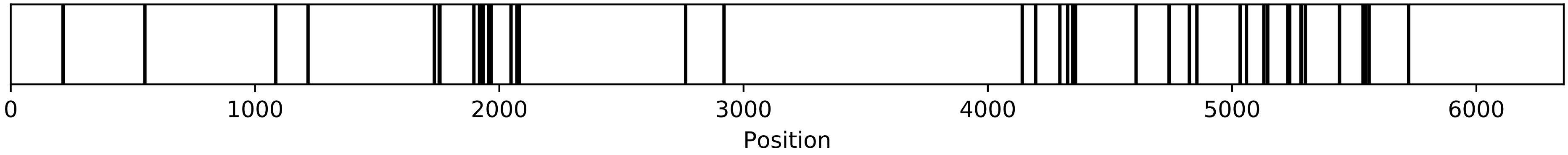


CR1 vs MM2
Number of differences: 0

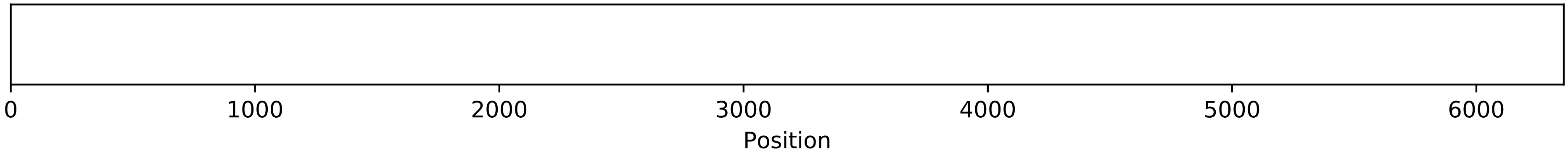


CR2 vs MM1
Number of differences: 101

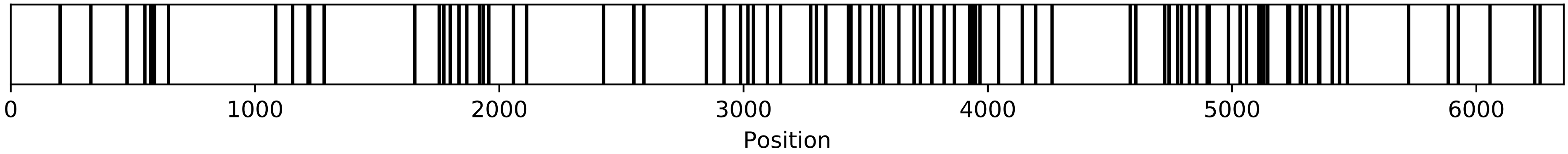


CR2 vs MM2
Number of differences: 97

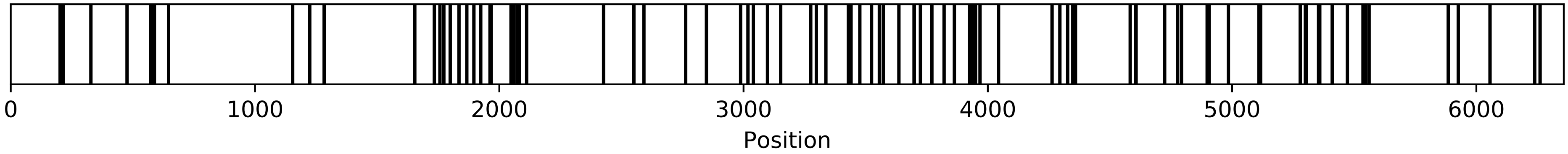


MM1 vs MM2
Number of differences: 44

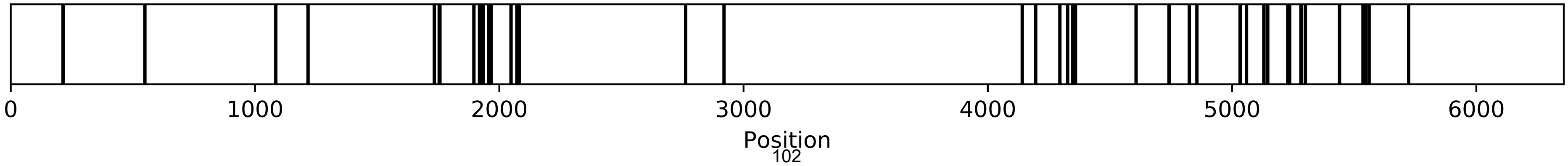

### Region: 296



CR1 vs CR2
Number of differences: 170

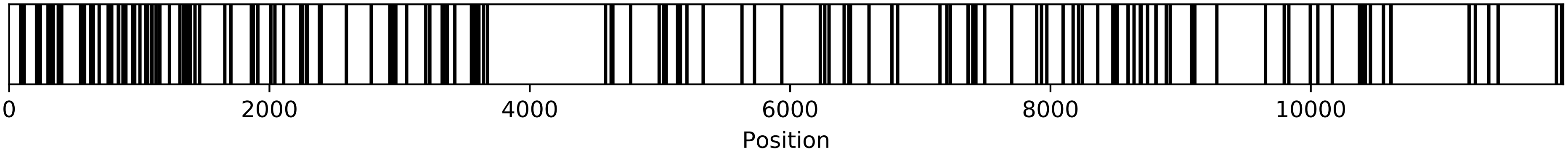


CR1 vs MM1
Number of differences: 76

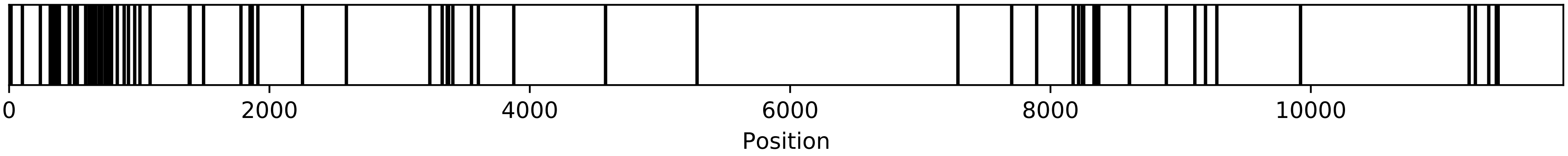


CR1 vs MM2
Number of differences: 101

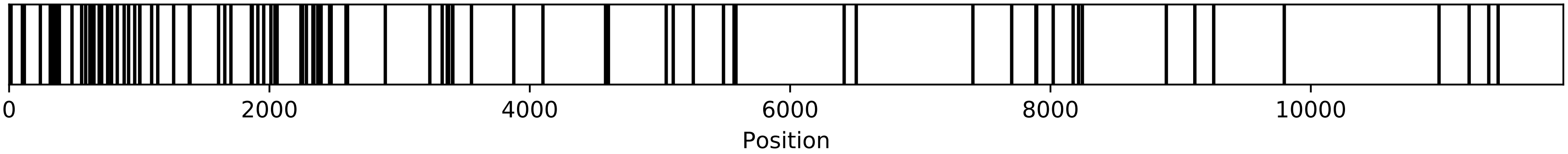


CR2 vs MM1
Number of differences: 176

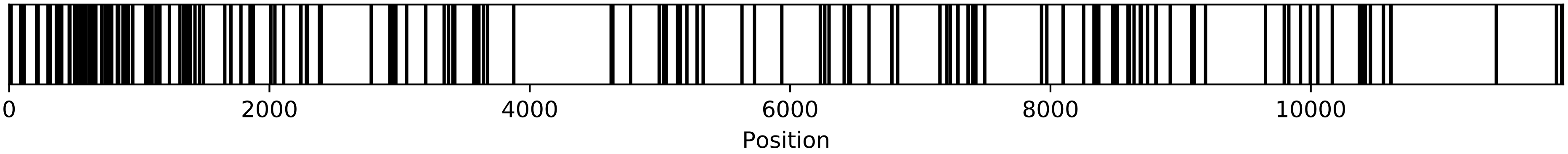


CR2 vs MM2
Number of differences: 182

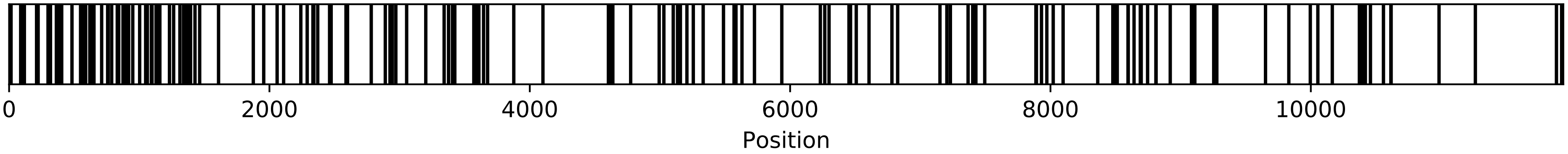


MM1 vs MM2
Number of differences: 82

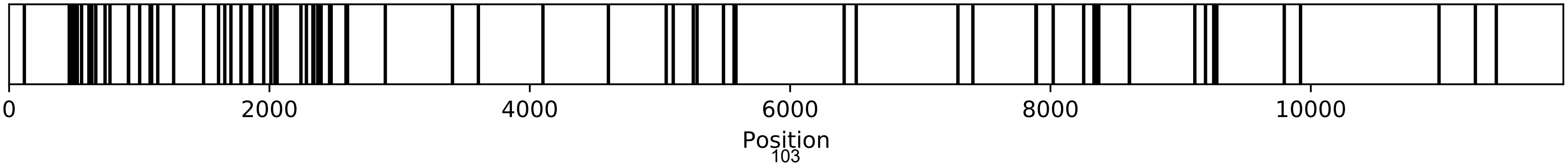

### Region: 301



CR1 vs CR2
Number of differences: 1

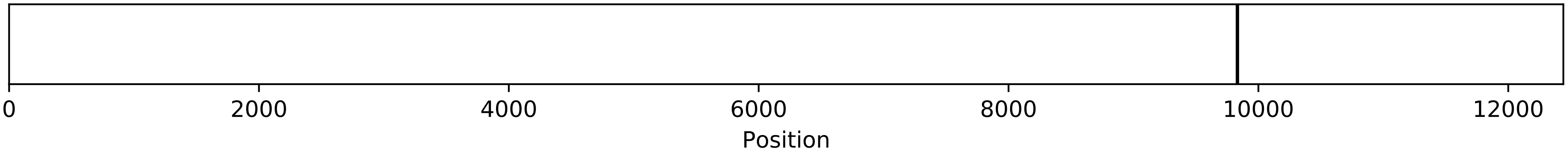


CR1 vs MM1
Number of differences: 203

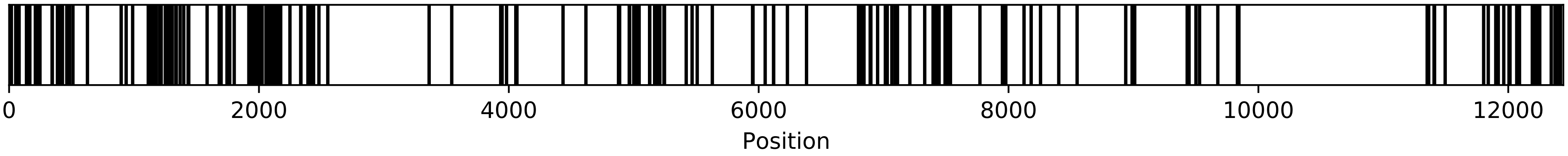


CR1 vs MM2
Number of differences: 167

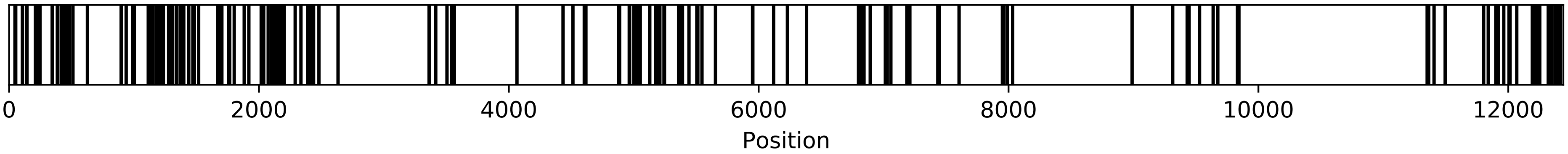


CR2 vs MM1
Number of differences: 202

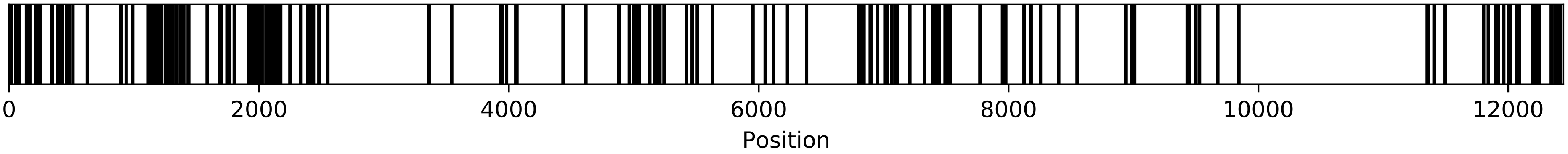


CR2 vs MM2
Number of differences: 166

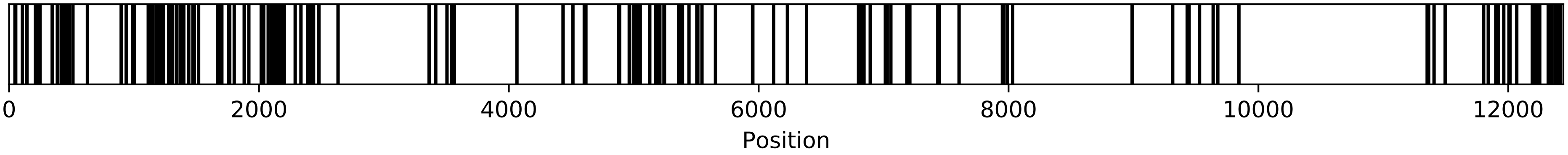


MM1 vs MM2
Number of differences: 143

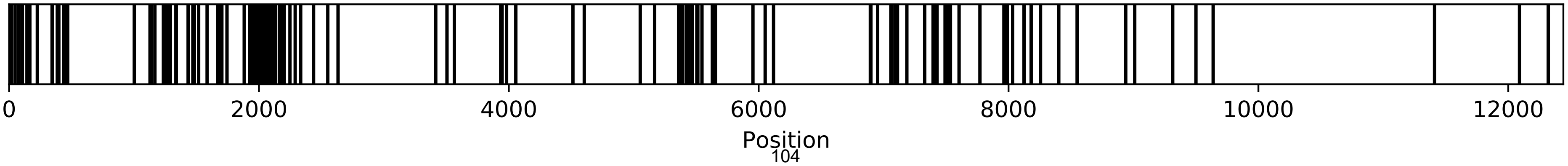

### Region: 302



CR1 vs CR2
Number of differences: 65

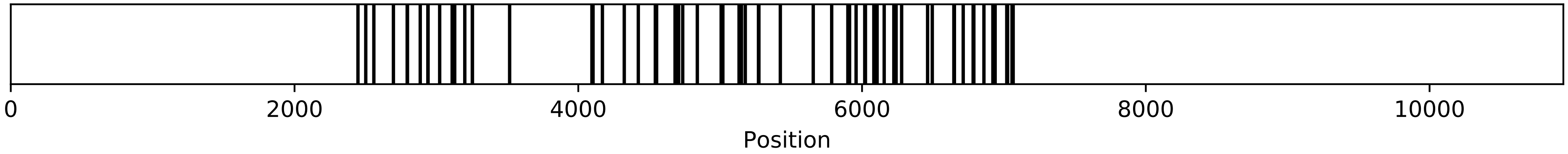


CR1 vs MM1
Number of differences: 334

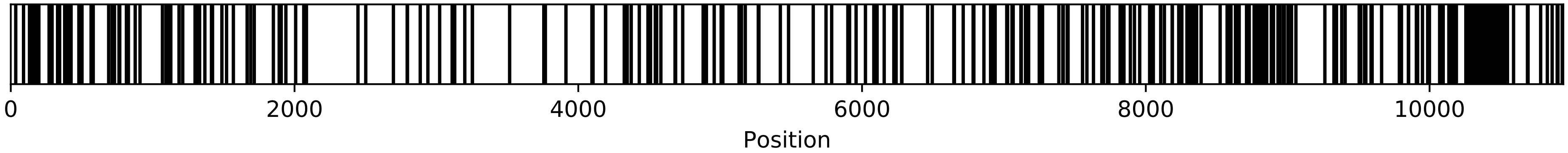


CR1 vs MM2
Number of differences: 329

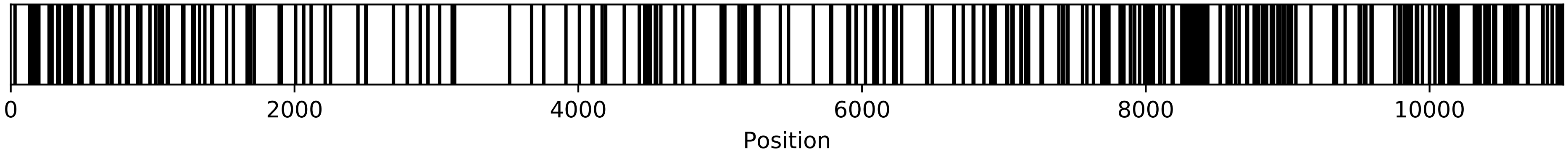


CR2 vs MM1
Number of differences: 284

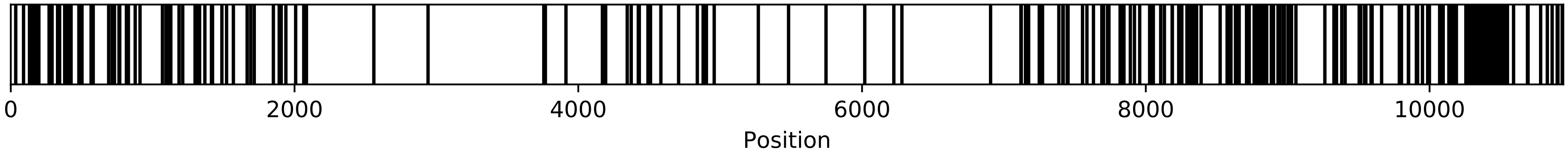


CR2 vs MM2
Number of differences: 284

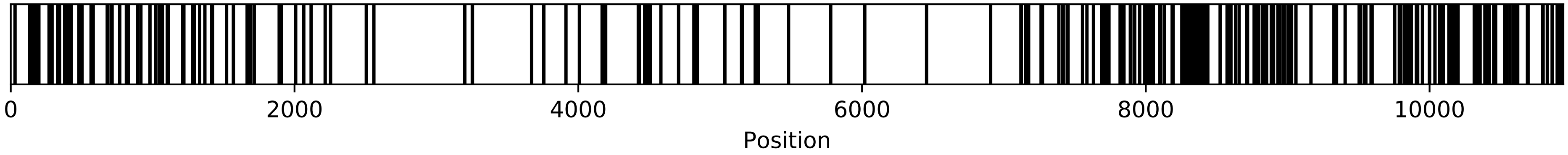


MM1 vs MM2
Number of differences: 190

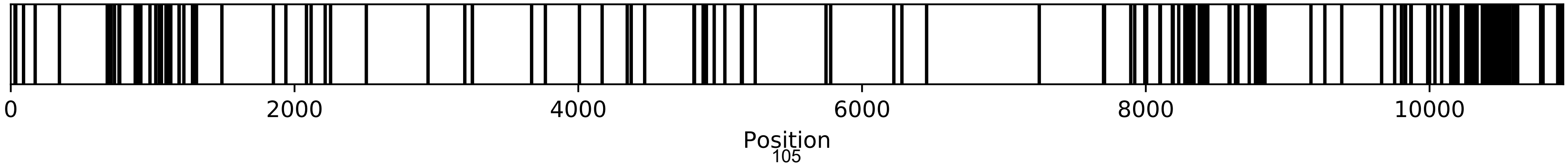

### Region: 304



CR1 vs CR2
Number of differences: 52

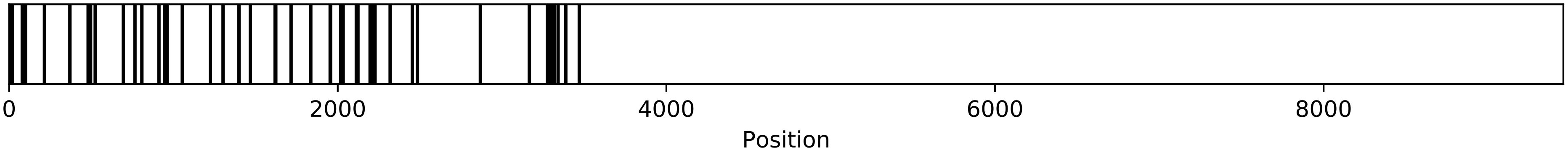


CR1 vs MM1
Number of differences: 235

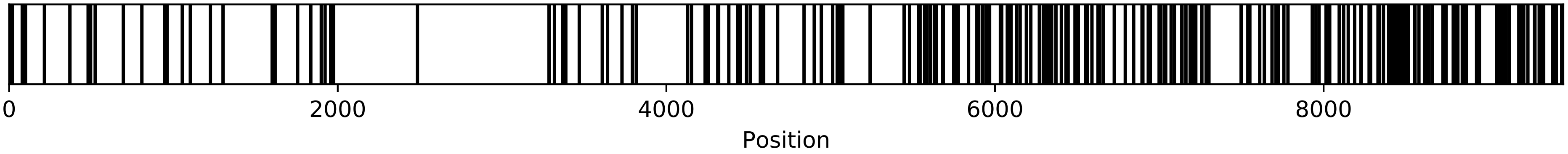


CR1 vs MM2
Number of differences: 230

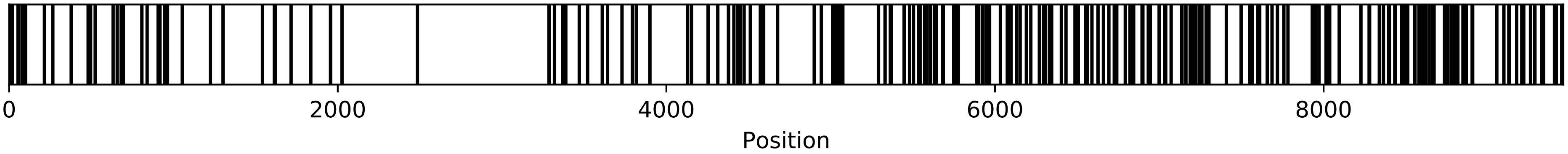


CR2 vs MM1
Number of differences: 233

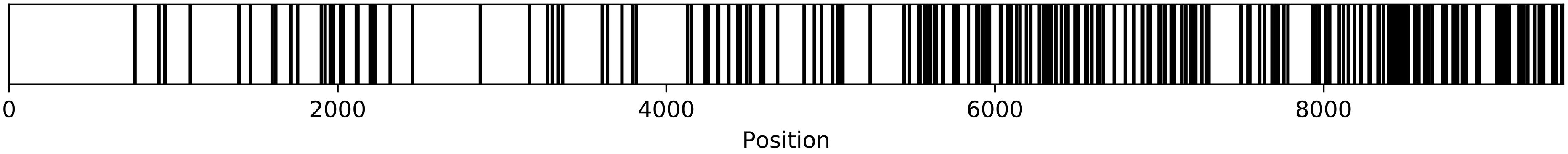


CR2 vs MM2
Number of differences: 226

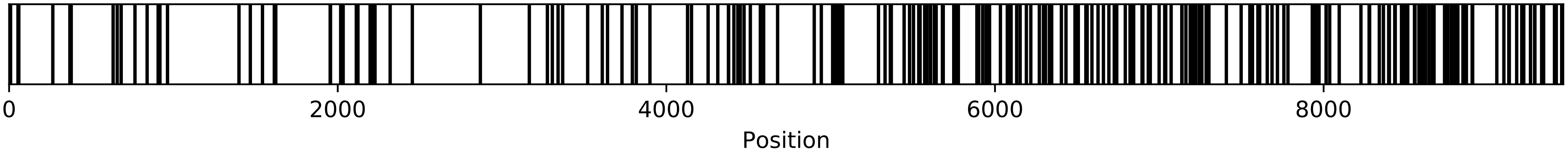


MM1 vs MM2
Number of differences: 126

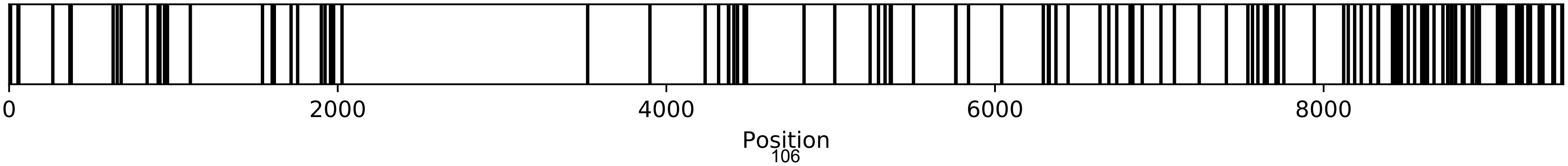

### Region: 309



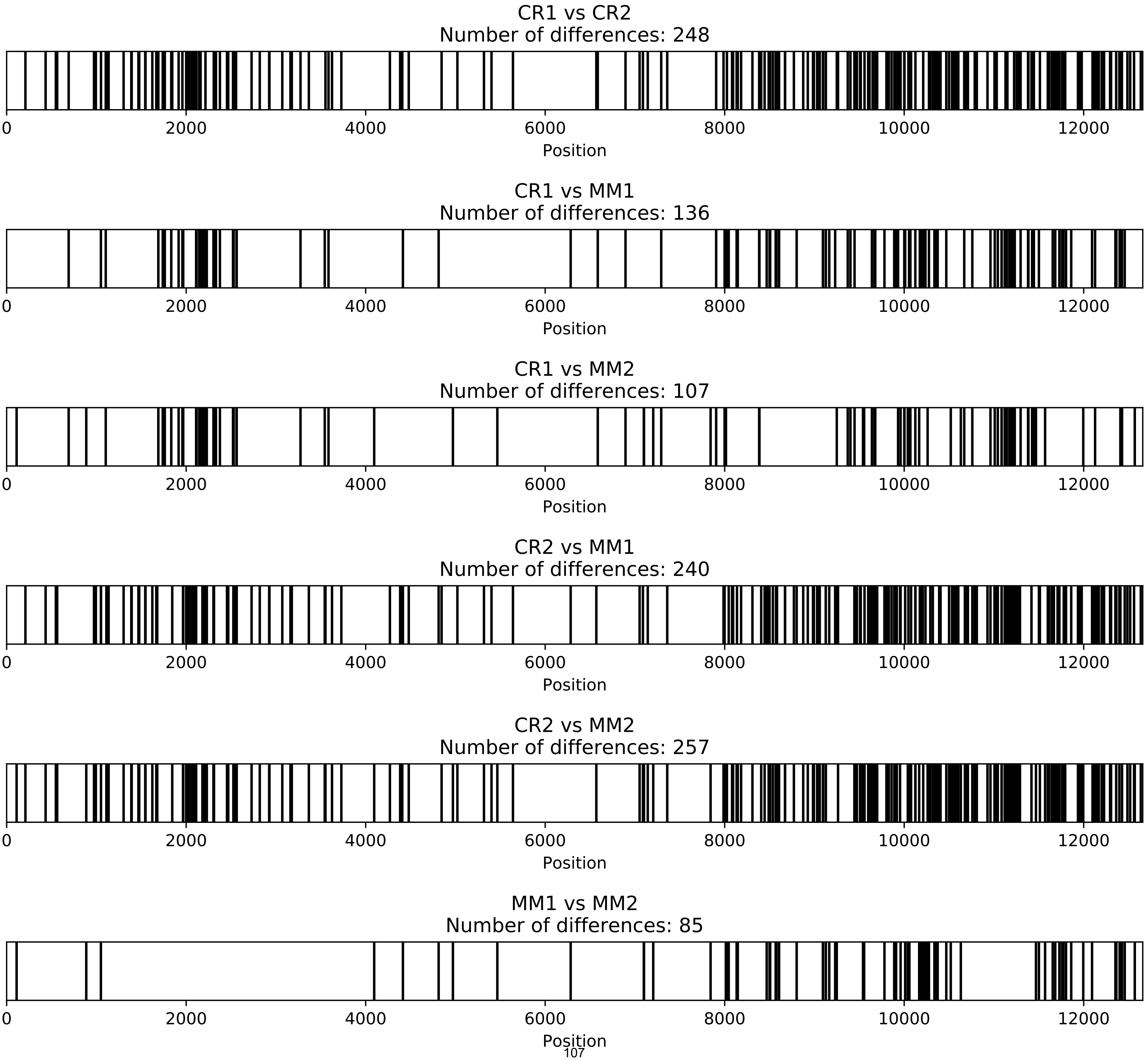
CR1 vs CR2
Number of differences: 248
0 2000 4000 6000 8000 10000 12000
Position
CR1 vs MM1
Number of differences: 136
0 2000 4000 6000 8000 10000 12000
Position
CR1 vs MM2
Number of differences: 107
0 2000 4000 6000 8000 10000 12000
Position
CR2 vs MM1
Number of differences: 240
0 2000 4000 6000 8000 10000 12000
Position
CR2 vs MM2
Number of differences: 257
0 2000 4000 6000 8000 10000 12000
Position
MM1 vs MM2
Number of differences: 85
0 2000 4000 6000 8000 10000 12000
Position

### Region: 322



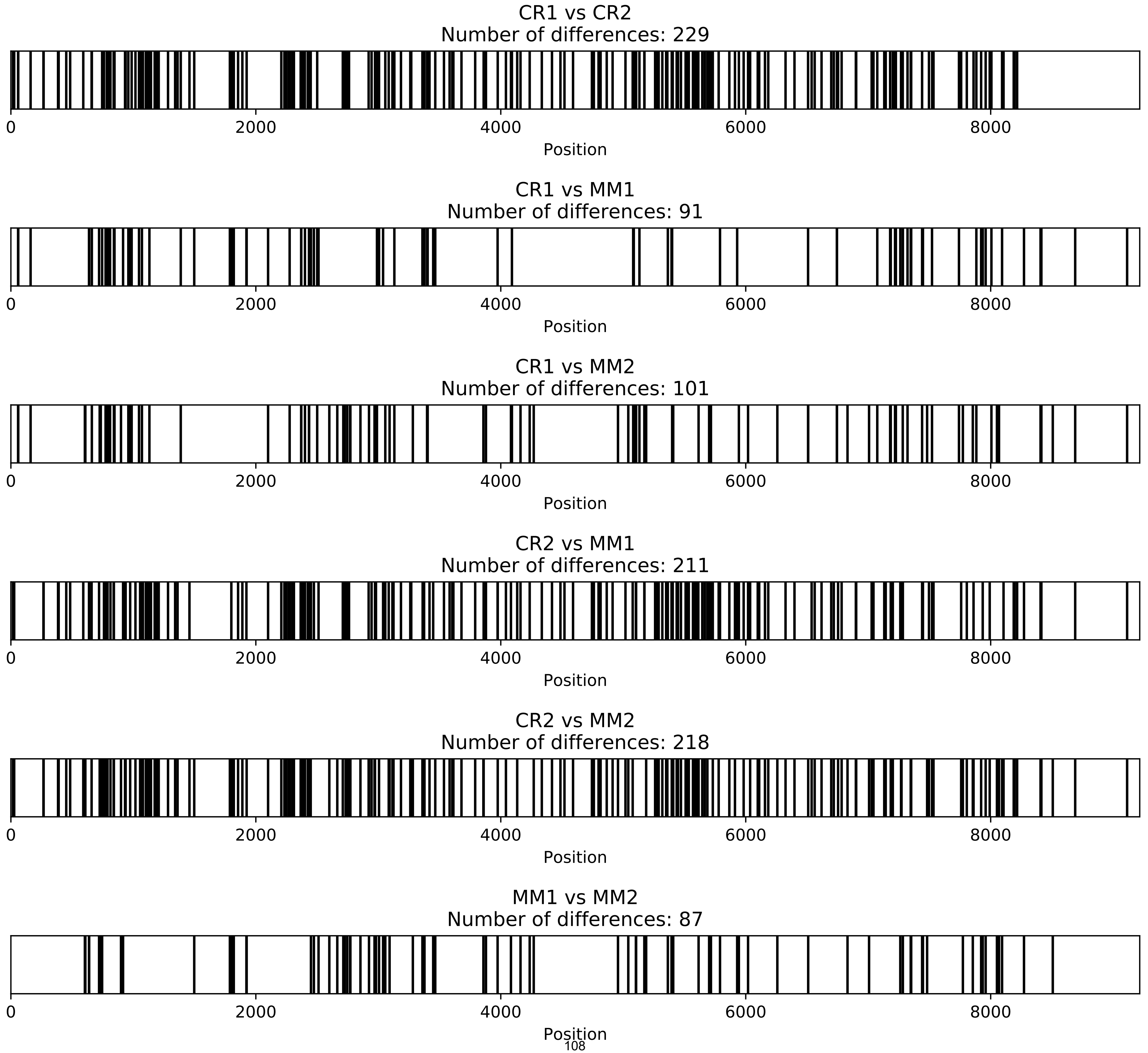
CR1 vs CR2
Number of differences: 229
0
2000
4000
6000
8000
Position
CR1 vs MM1
Number of differences: 91
0
2000
4000
6000
8000
Position
CR1 vs MM2
Number of differences: 101
0
2000
4000
6000
8000
Position
CR2 vs MM1
Number of differences: 211
0
2000
4000
6000
8000
Position
CR2 vs MM2
Number of differences: 218
0
2000
4000
6000
8000
Position
MM1 vs MM2
Number of differences: 87
0
2000
4000
6000
8000
Position

### Region: 326



## CR1 vs CR2
Number of differences: 167

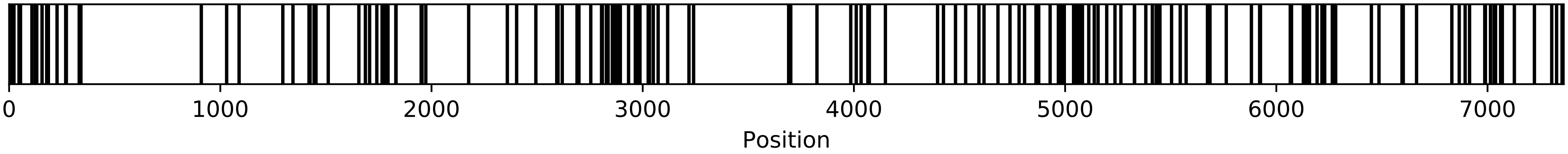


## CR1 vs MM1
Number of differences: 69

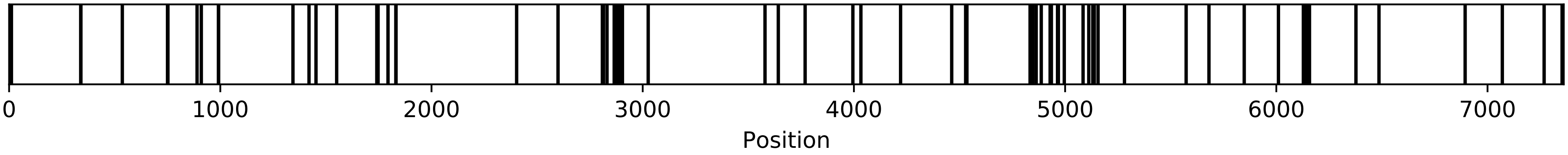


## CR1 vs MM2
Number of differences: 71

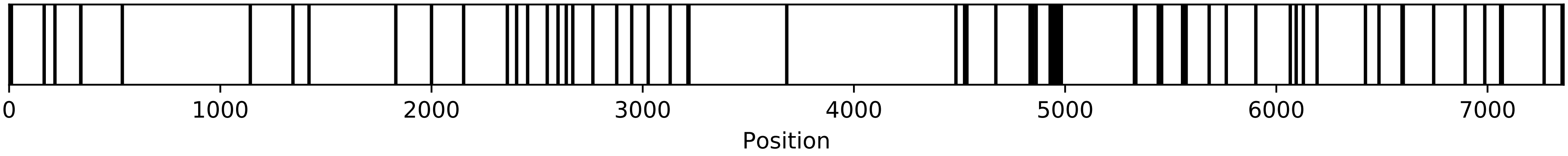


## CR2 vs MM1
Number of differences: 171

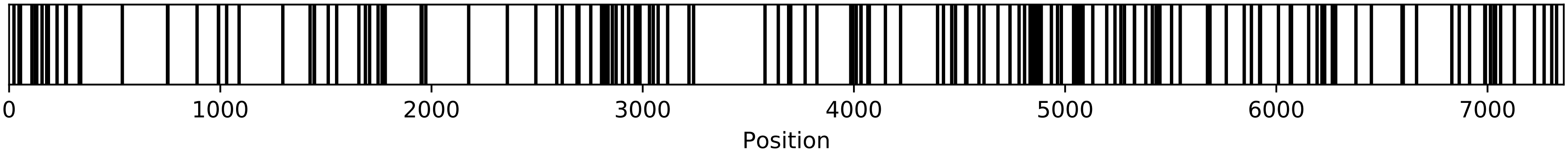


## CR2 vs MM2
Number of differences: 179

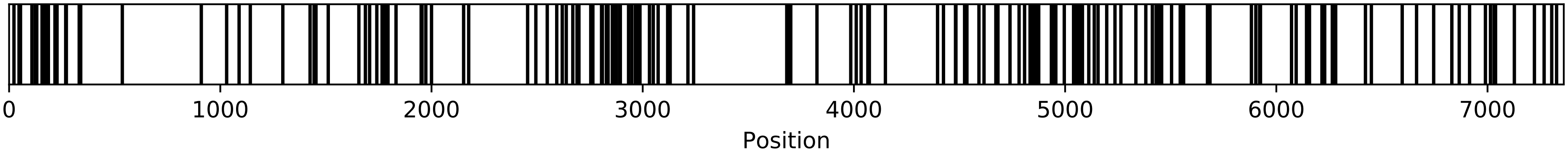


## MM1 vs MM2
Number of differences: 84

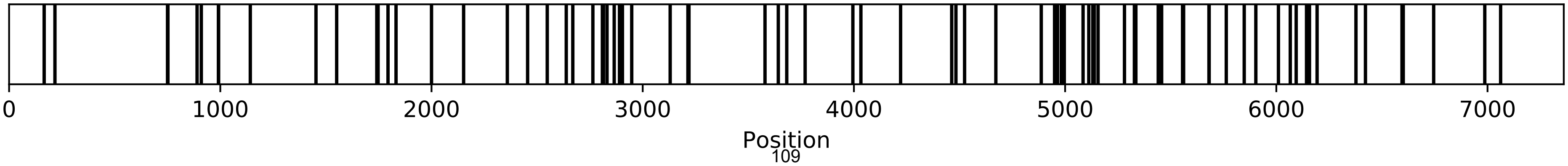

### Region: 328



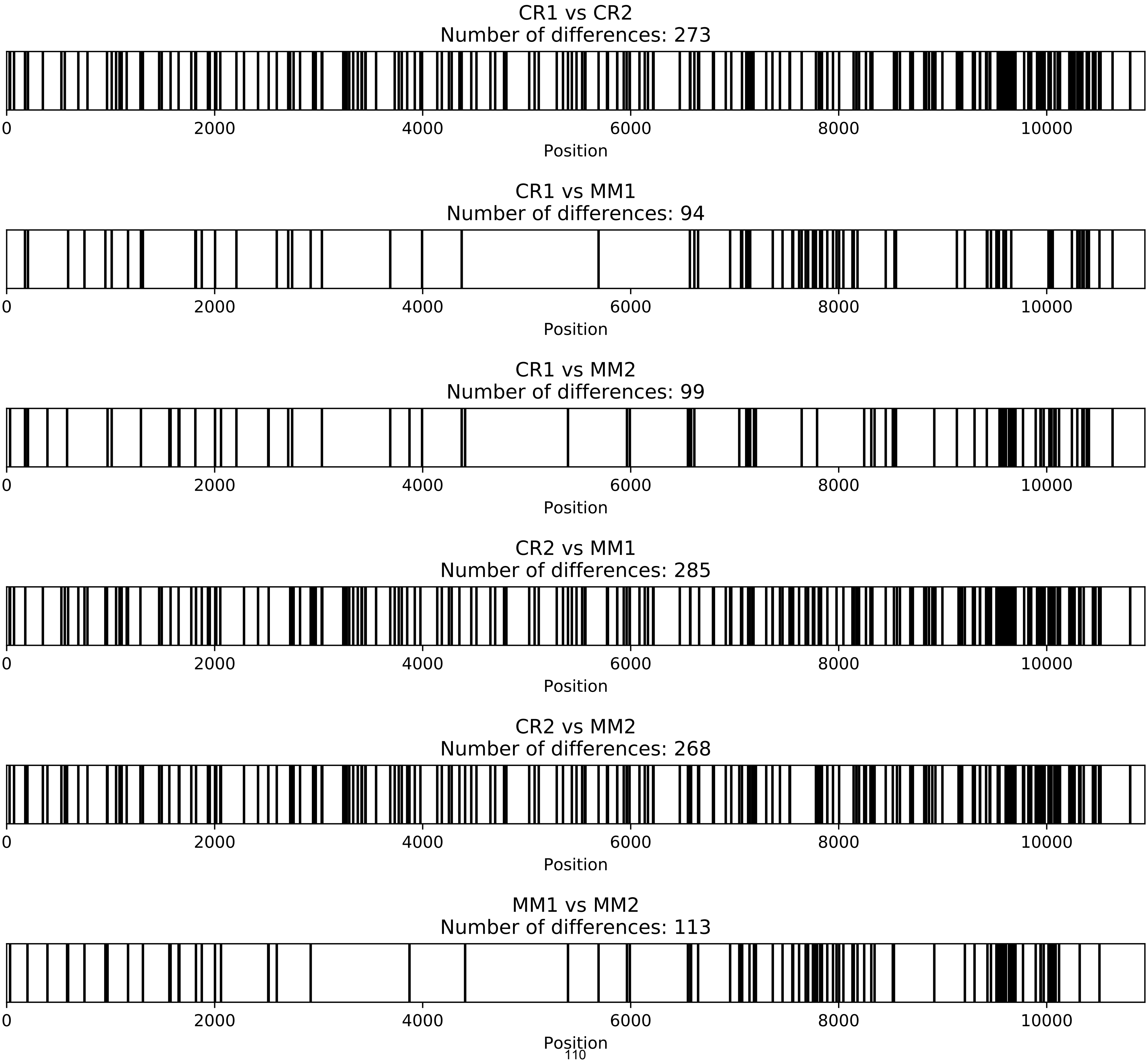
CR1 vs CR2
Number of differences: 273
CR1 vs MM1
Number of differences: 94
CR1 vs MM2
Number of differences: 99
CR2 vs MM1
Number of differences: 285
CR2 vs MM2
Number of differences: 268
MM1 vs MM2
Number of differences: 113
0
2000
4000
6000
8000
10000
Position

### Region: 329



## CR1 vs CR2
Number of differences: 229

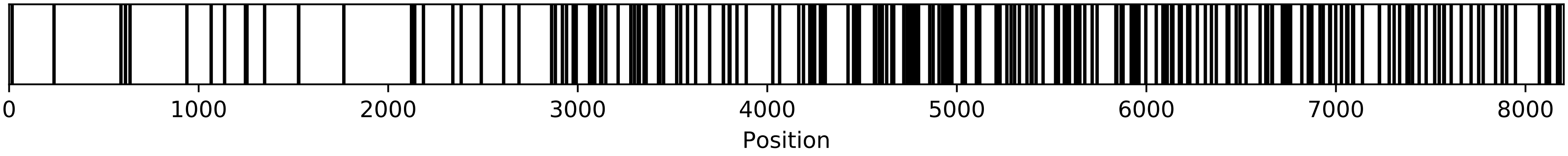


## CR1 vs MM1
Number of differences: 70

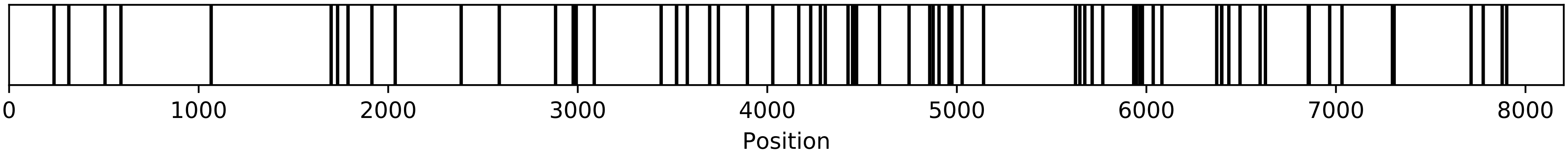


## CR1 vs MM2
Number of differences: 65

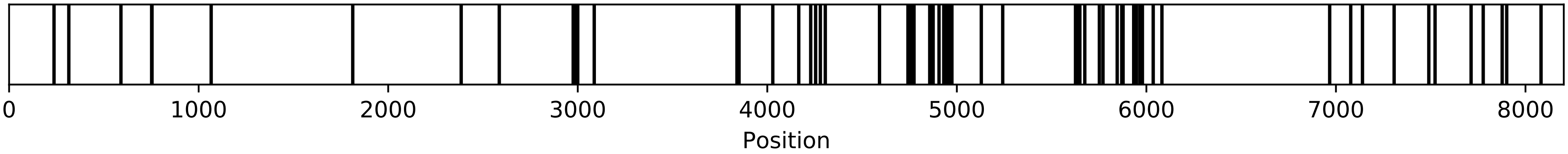


## CR2 vs MM1
Number of differences: 217

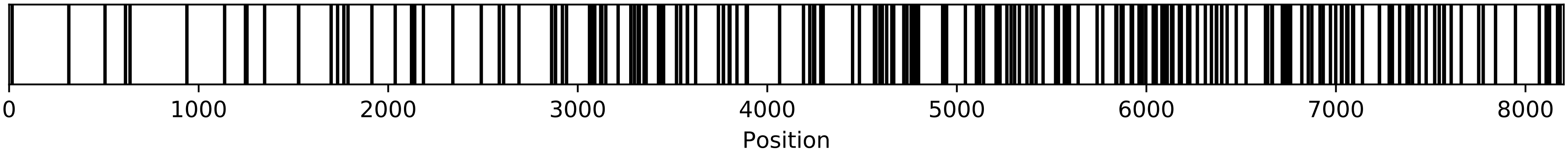


## CR2 vs MM2
Number of differences: 218

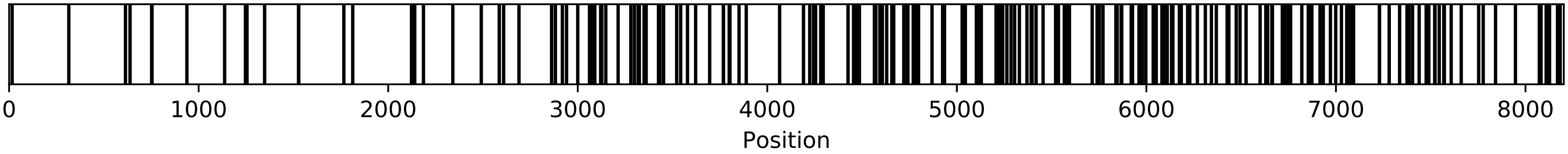


## MM1 vs MM2
Number of differences: 62

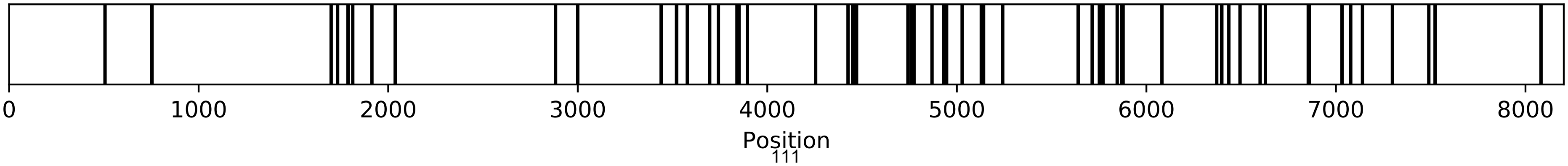

### Region: 330



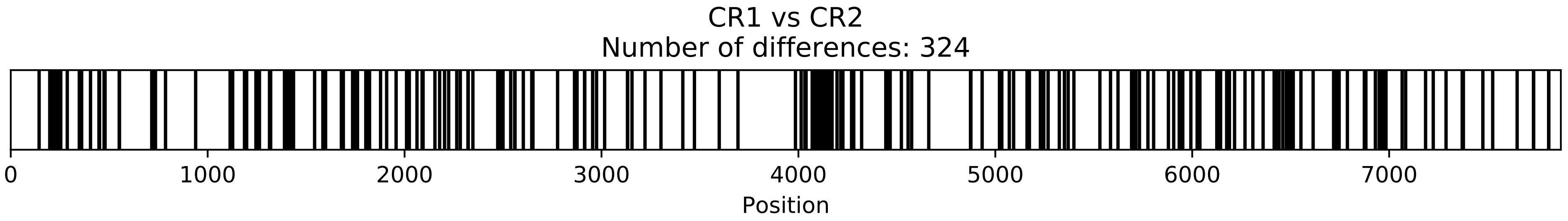
CR1 vs CR2
Number of differences: 324
0
1000
2000
3000
4000
5000
6000
7000
Position

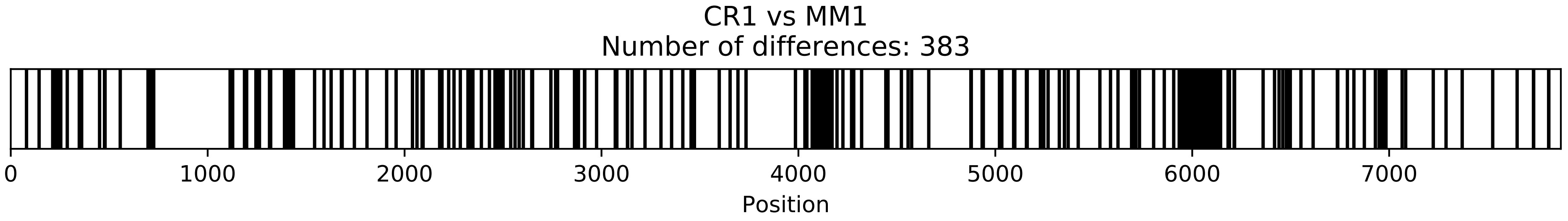
CR1 vs MM1
Number of differences: 383
0
1000
2000
3000
4000
5000
6000
7000
Position

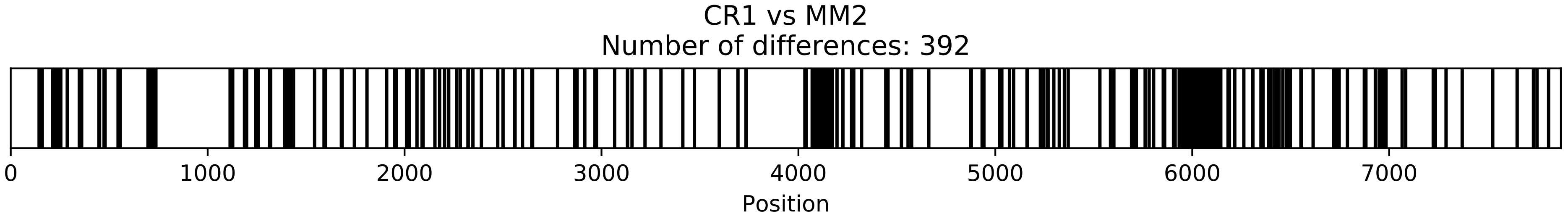
CR1 vs MM2
Number of differences: 392
0
1000
2000
3000
4000
5000
6000
7000
Position

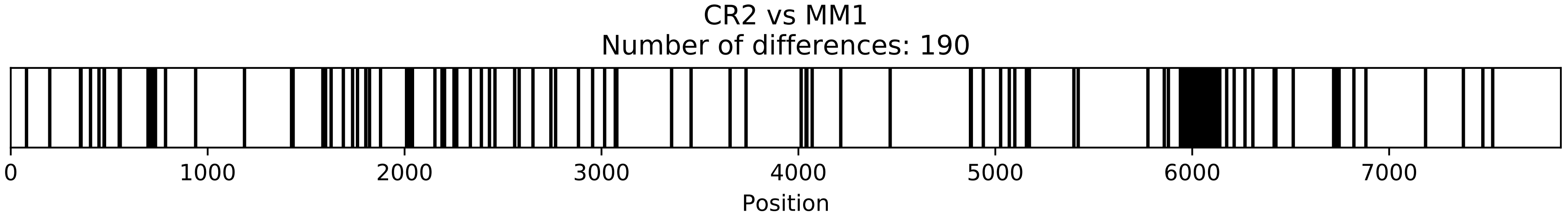
CR2 vs MM1
Number of differences: 190
0
1000
2000
3000
4000
5000
6000
7000
Position

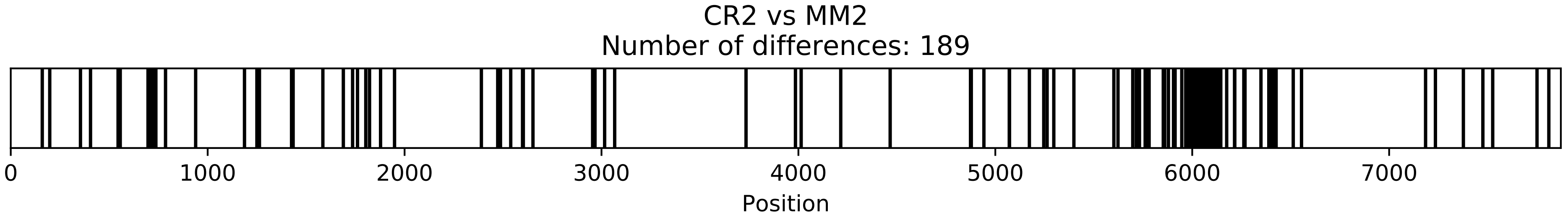
CR2 vs MM2
Number of differences: 189
0
1000
2000
3000
4000
5000
6000
7000
Position

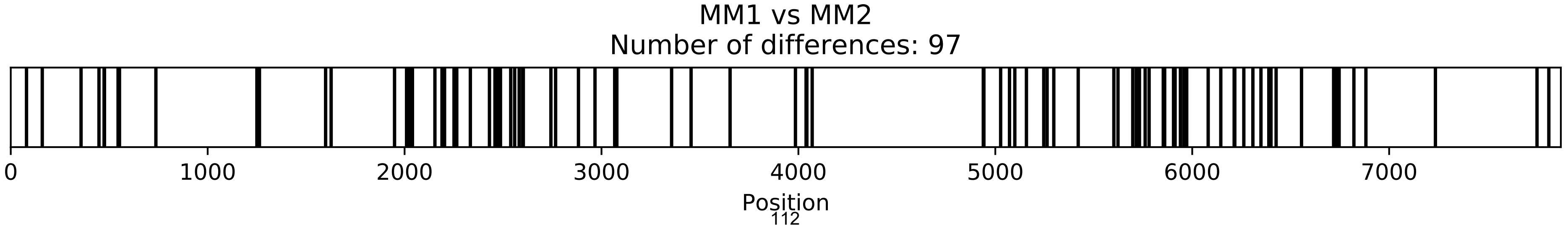
MM1 vs MM2
Number of differences: 97
0
1000
2000
3000
4000
5000
6000
7000
Position

### Region: 331



CR1 vs CR2
Number of differences: 322

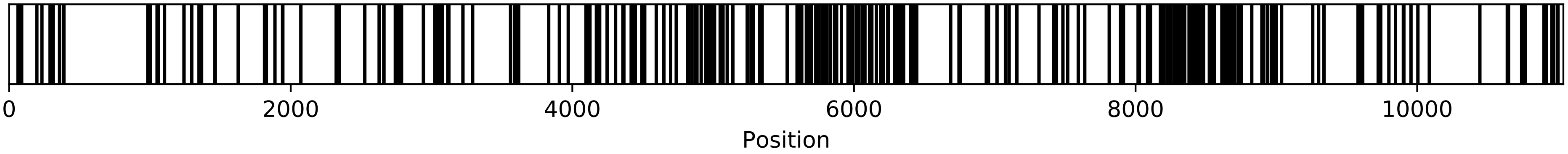


CR1 vs MM1
Number of differences: 138

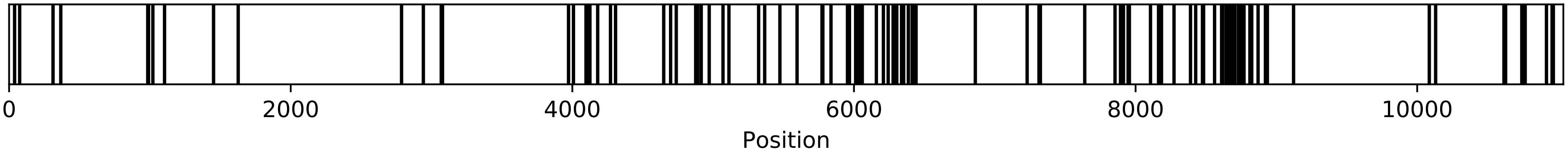


CR1 vs MM2
Number of differences: 137

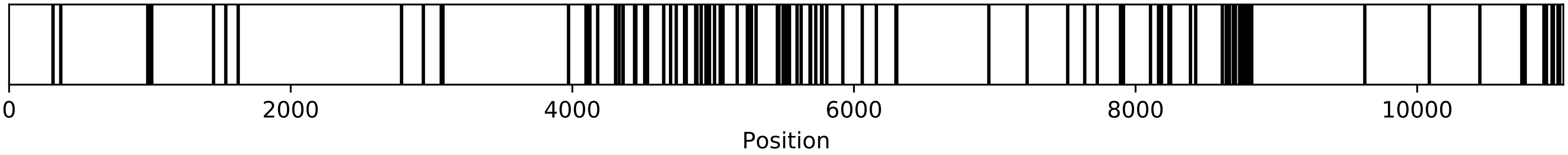


CR2 vs MM1
Number of differences: 300

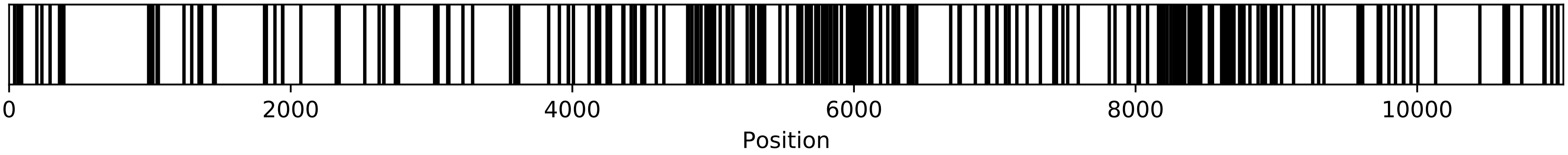


CR2 vs MM2
Number of differences: 310

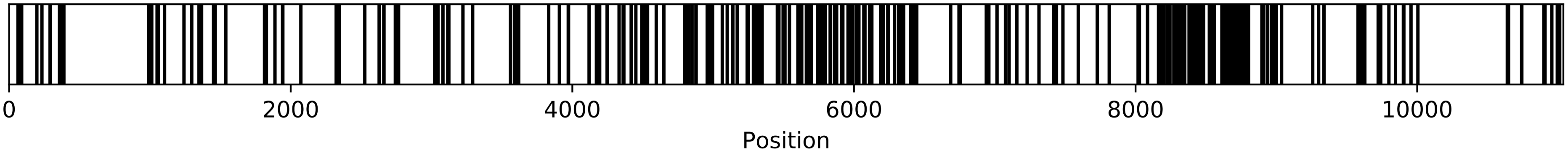


MM1 vs MM2
Number of differences: 138

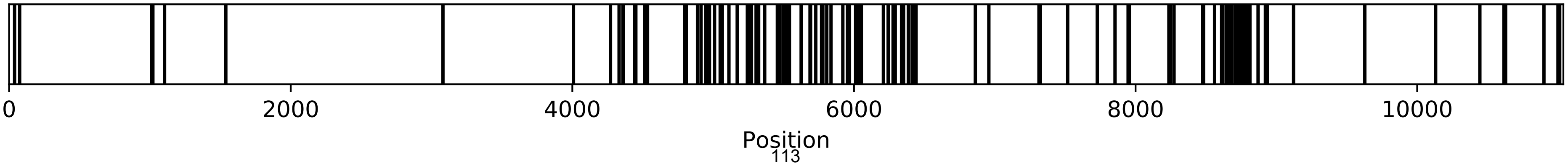

### Region: 333



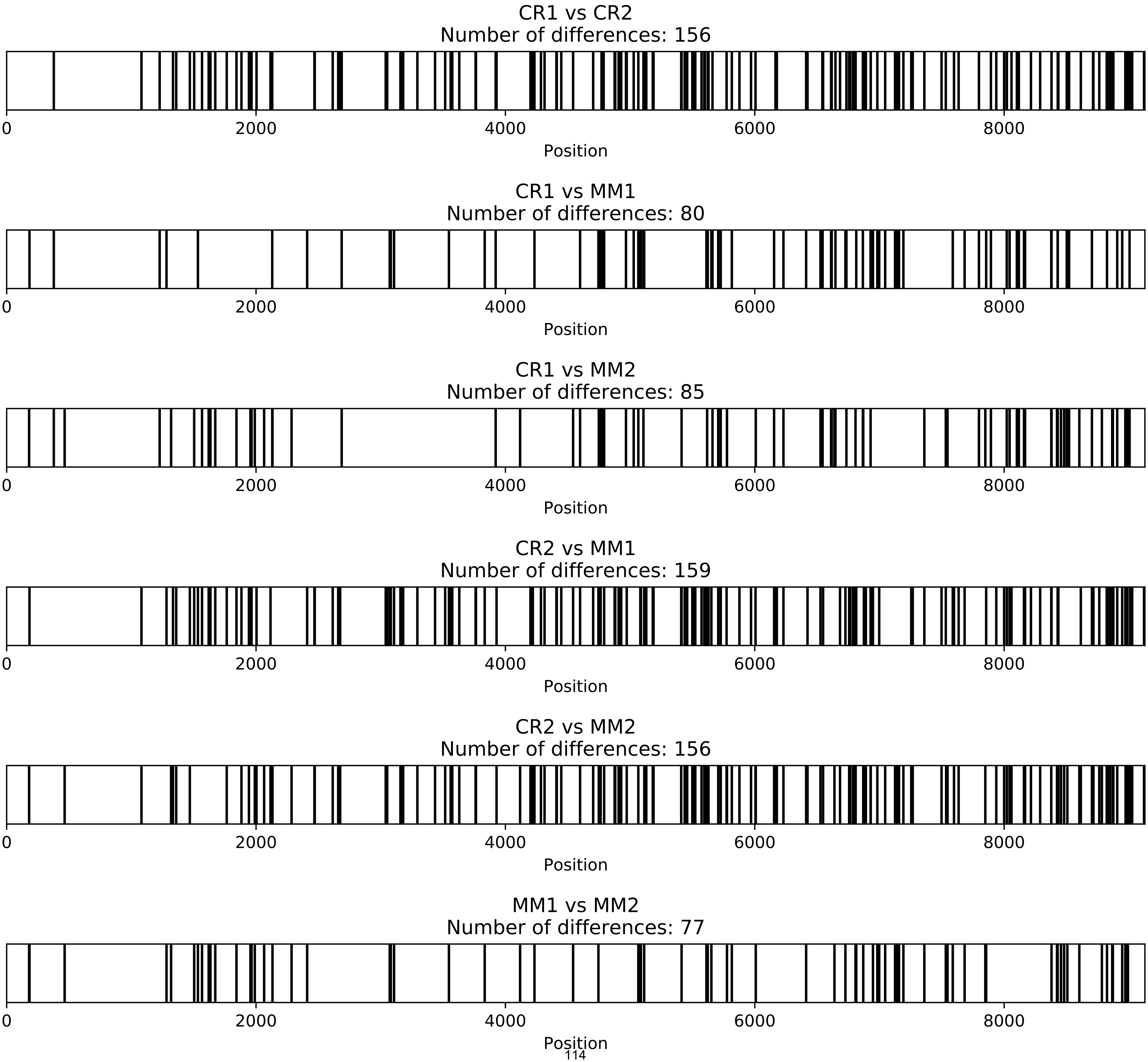
CR1 vs CR2
Number of differences: 156
0
2000
4000
6000
8000
Position
CR1 vs MM1
Number of differences: 80
0
2000
4000
6000
8000
Position
CR1 vs MM2
Number of differences: 85
0
2000
4000
6000
8000
Position
CR2 vs MM1
Number of differences: 159
0
2000
4000
6000
8000
Position
CR2 vs MM2
Number of differences: 156
0
2000
4000
6000
8000
Position
MM1 vs MM2
Number of differences: 77
0
2000
4000
6000
8000
Position

### Region: 334



CR1 vs CR2
Number of differences: 289

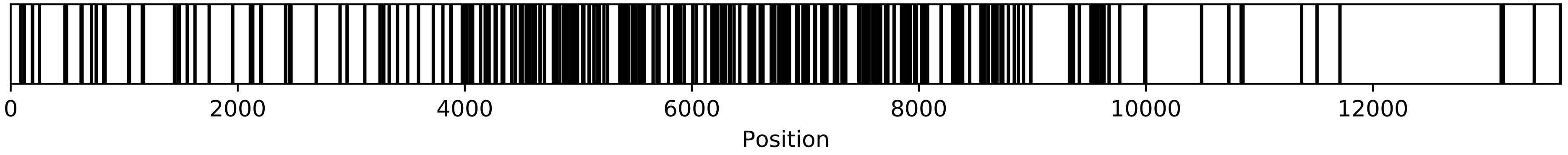

CR1 vs MM1
Number of differences: 117

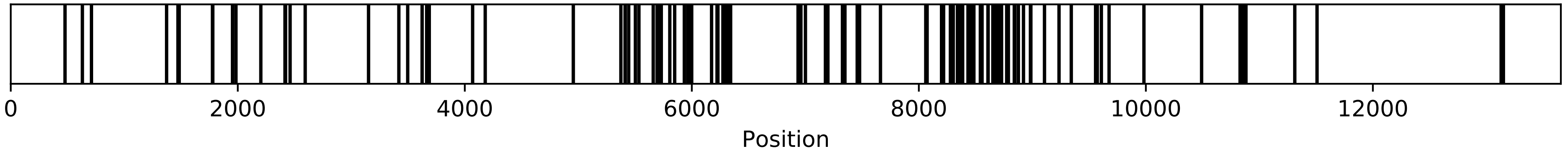

CR1 vs MM2
Number of differences: 185

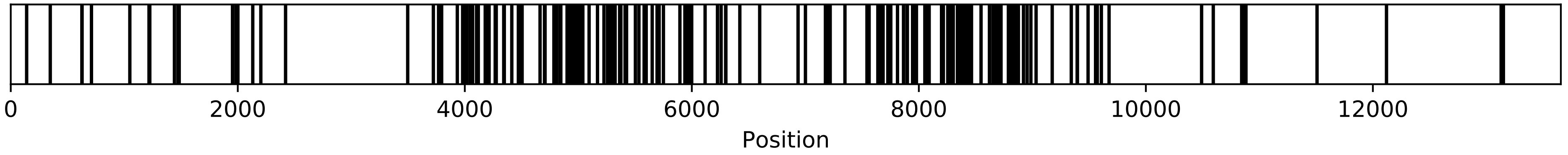

CR2 vs MM1
Number of differences: 296

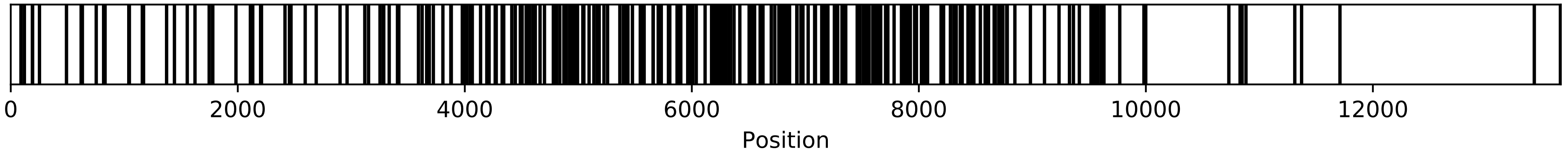

CR2 vs MM2
Number of differences: 289

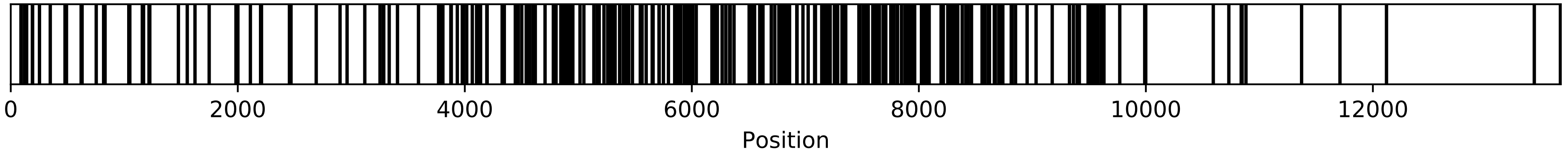

MM1 vs MM2
Number of differences: 172

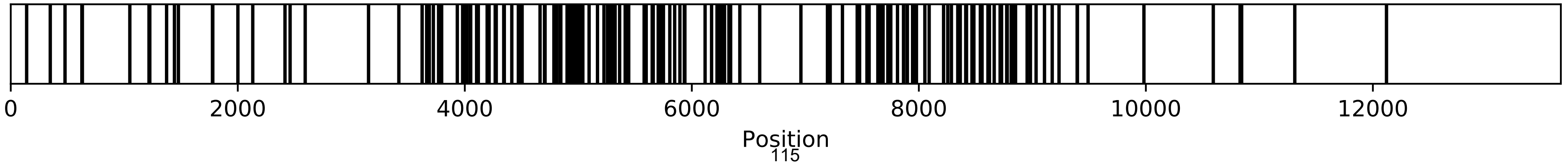

### Region: 338



CR1 vs CR2
Number of differences: 269

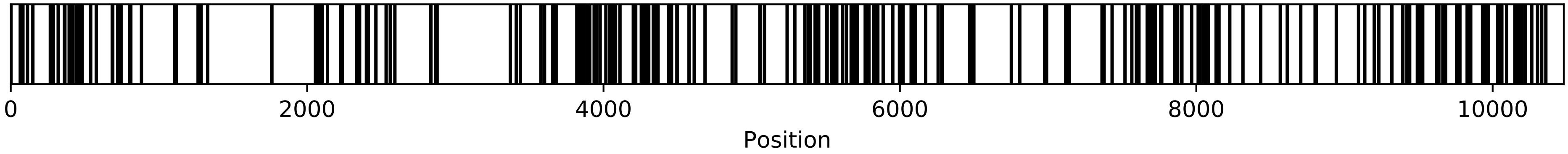


CR1 vs MM1
Number of differences: 174

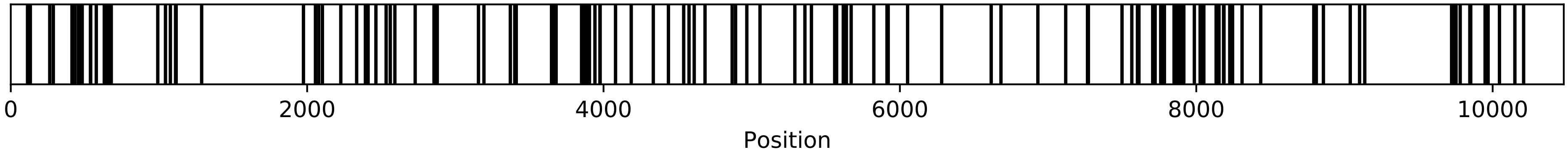


CR1 vs MM2
Number of differences: 155

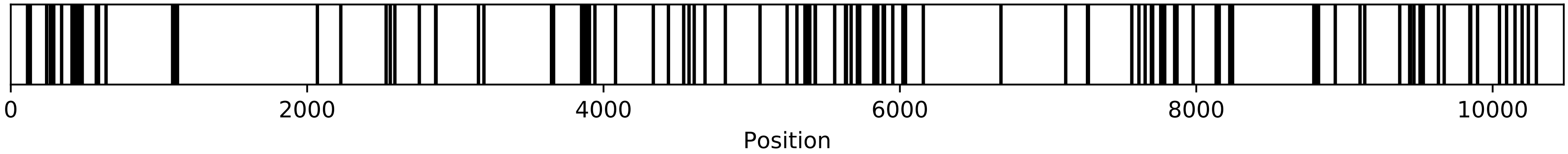


CR2 vs MM1
Number of differences: 269

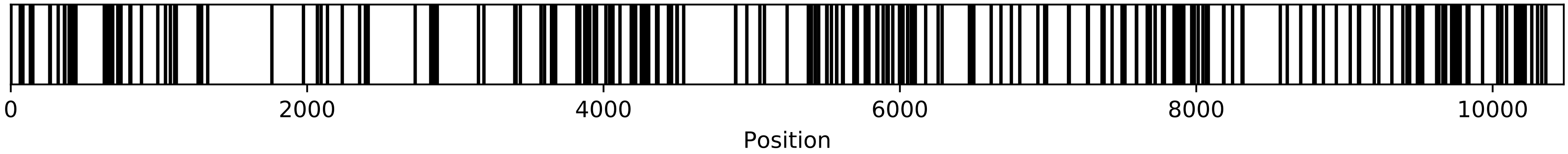


CR2 vs MM2
Number of differences: 260

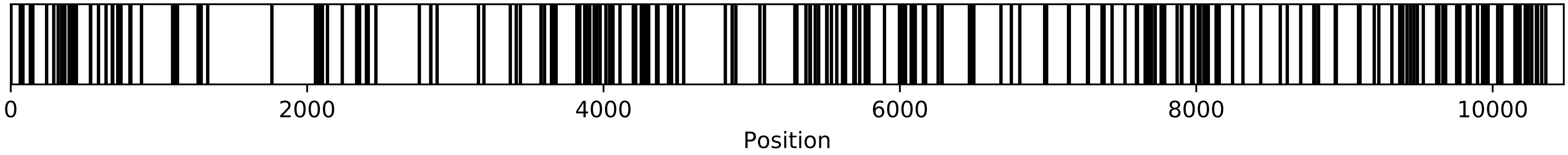


MM1 vs MM2
Number of differences: 149

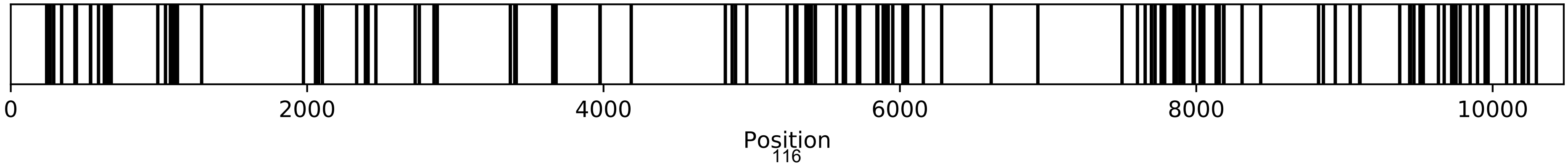

### Region: 343



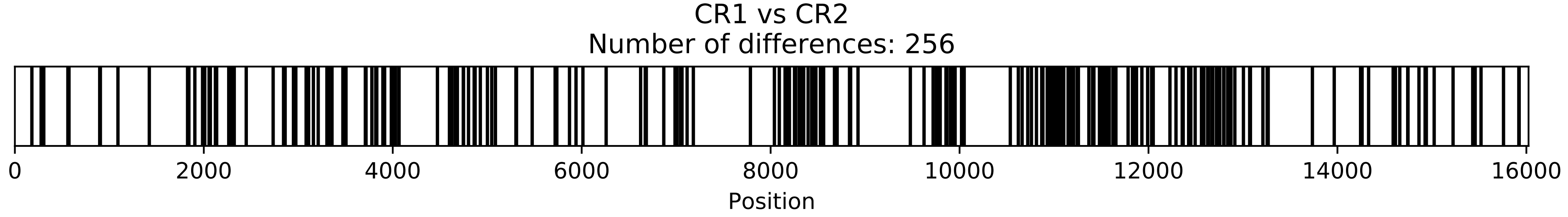
CR1 vs CR2
Number of differences: 256
0
2000
4000
6000
8000
10000
12000
14000
16000
Position

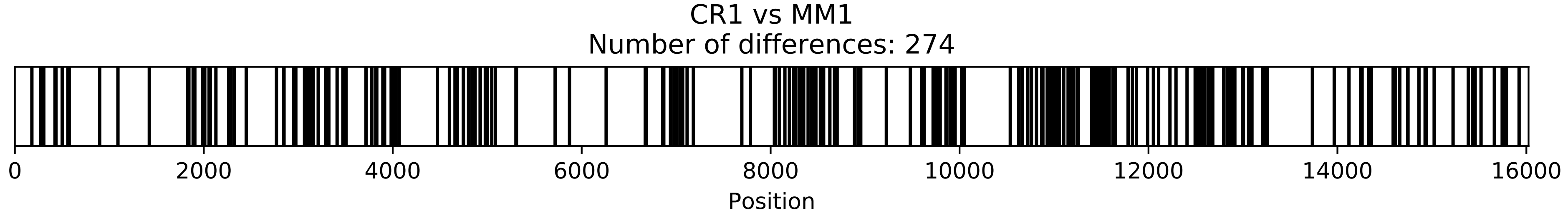
CR1 vs MM1
Number of differences: 274
0
2000
4000
6000
8000
10000
12000
14000
16000
Position

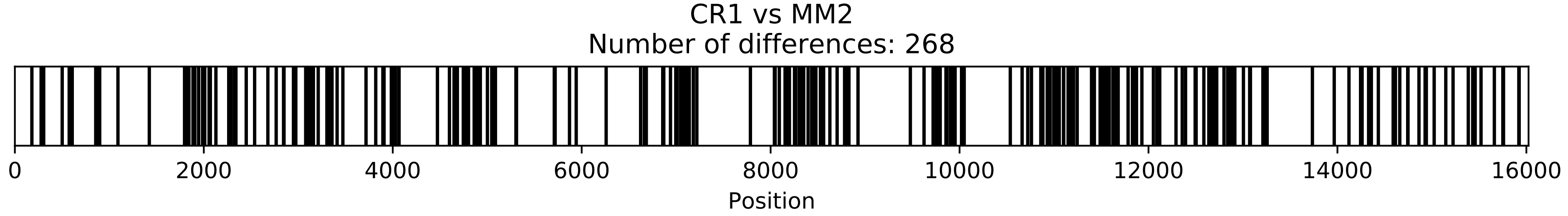
CR1 vs MM2
Number of differences: 268
0
2000
4000
6000
8000
10000
12000
14000
16000
Position

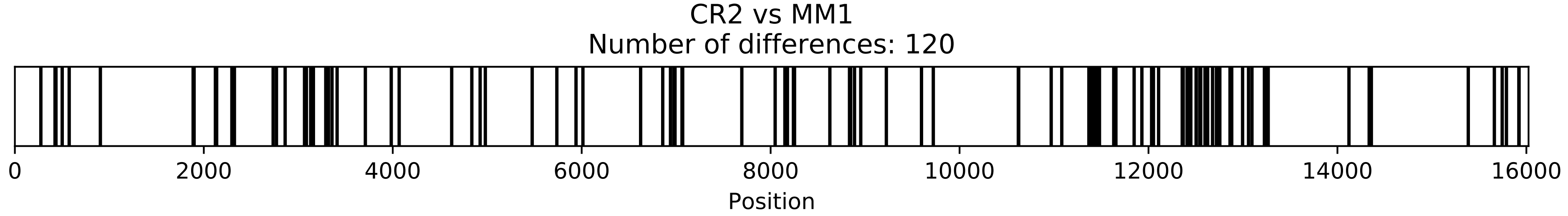
CR2 vs MM1
Number of differences: 120
0
2000
4000
6000
8000
10000
12000
14000
16000
Position

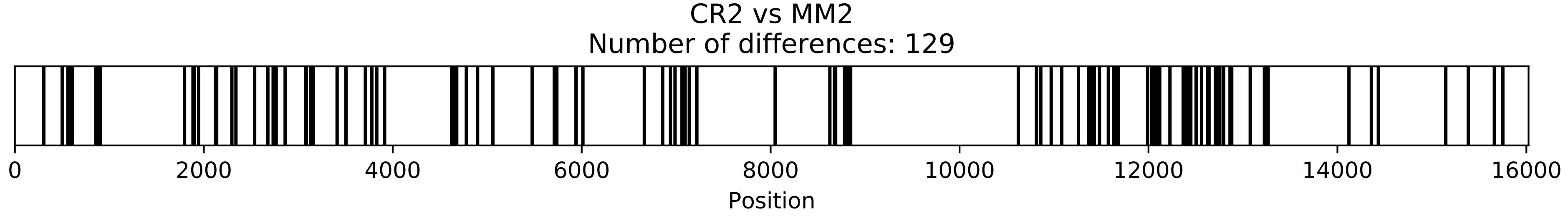
CR2 vs MM2
Number of differences: 129
0
2000
4000
6000
8000
10000
12000
14000
16000
Position

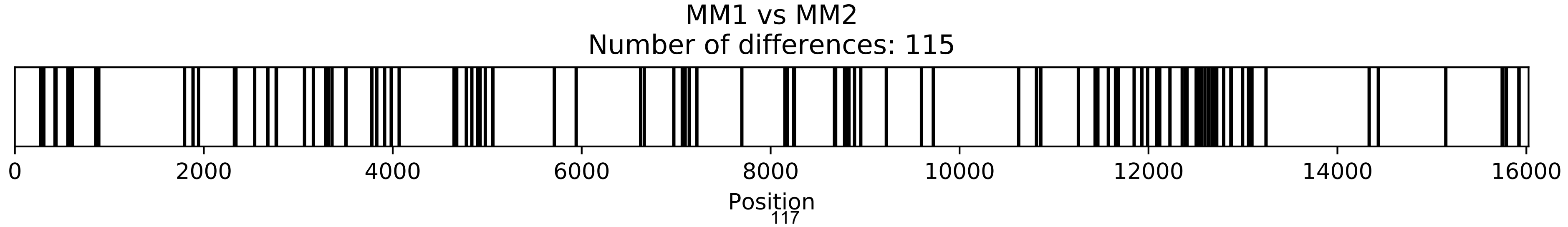
MM1 vs MM2
Number of differences: 115
0
2000
4000
6000
8000
10000
12000
14000
16000
Position

### Region: 345

CR1 vs CR2
Number of differences: 271

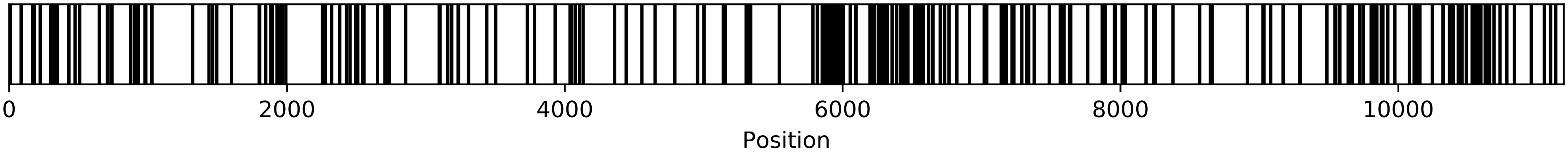


CR1 vs MM1
Number of differences: 104

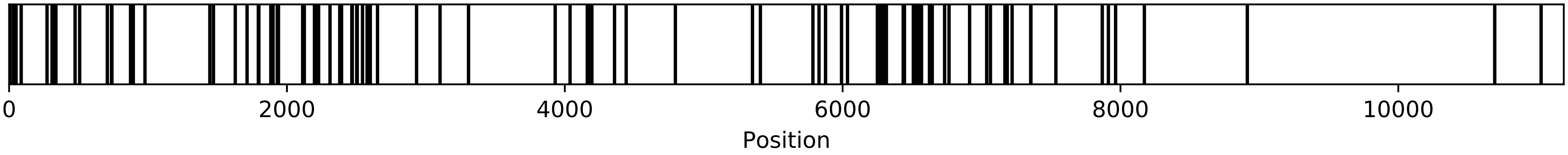


CR1 vs MM2
Number of differences: 127

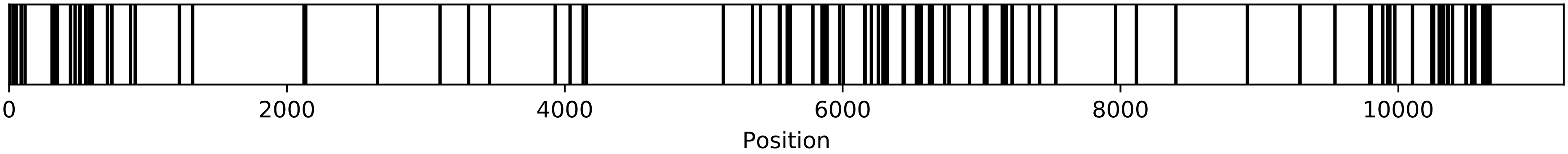


CR2 vs MM1
Number of differences: 280

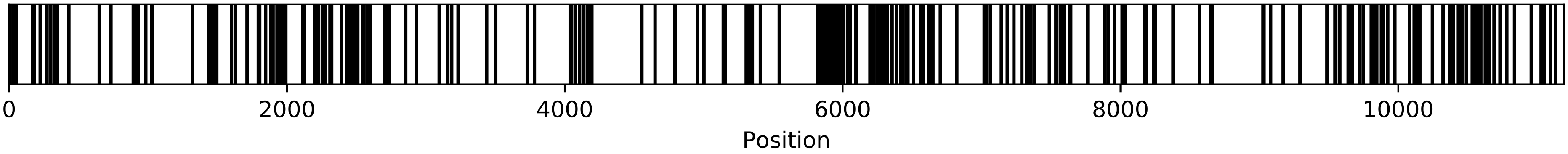


CR2 vs MM2
Number of differences: 274

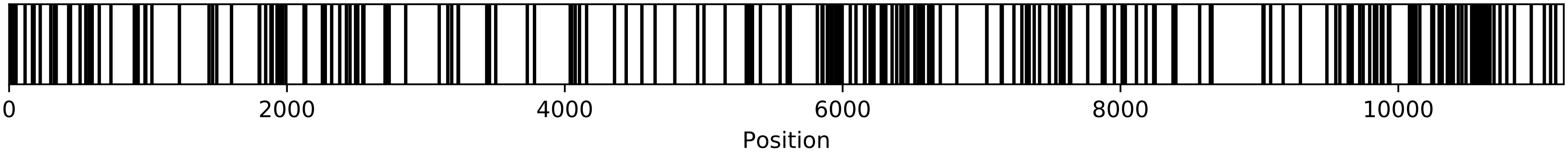


MM1 vs MM2
Number of differences: 137

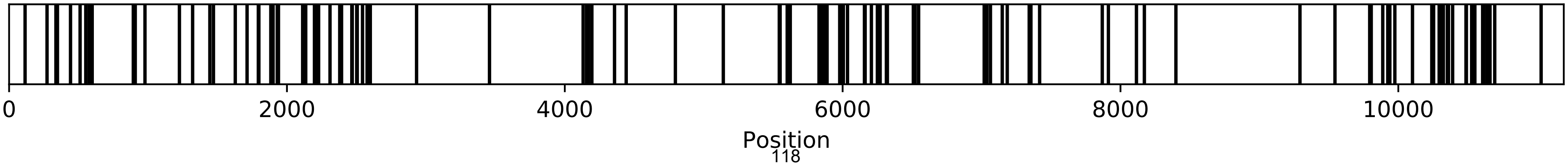

### Region: 346



CR1 vs CR2
Number of differences: 278

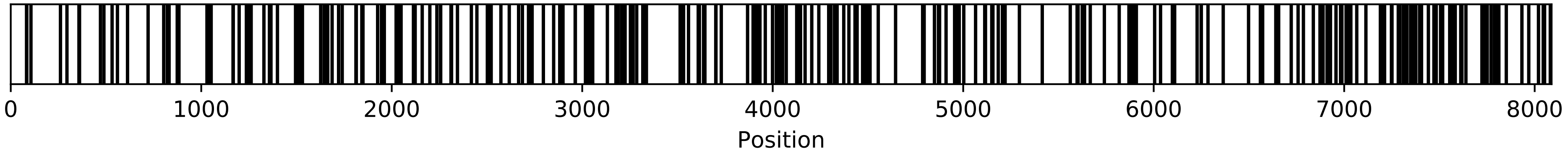


CR1 vs MM1
Number of differences: 302

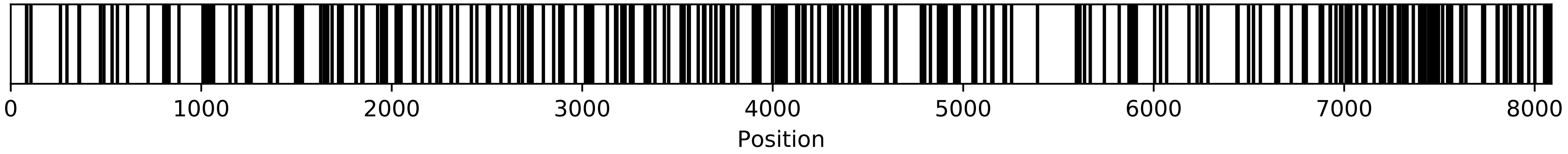


CR1 vs MM2
Number of differences: 268

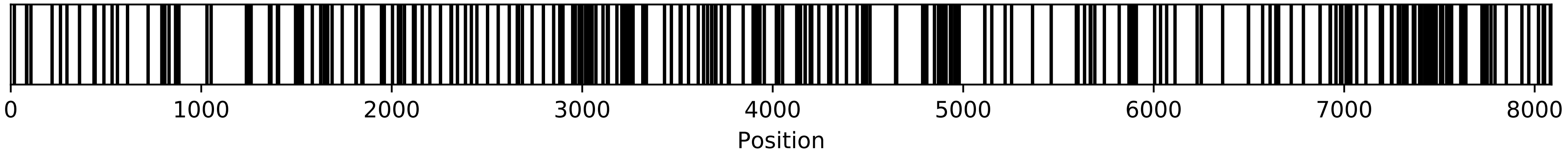


CR2 vs MM1
Number of differences: 149

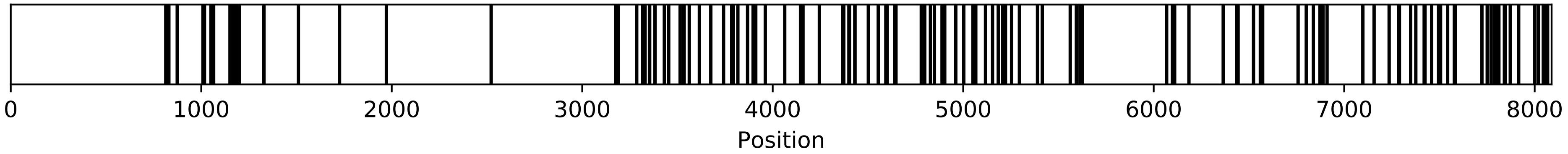


CR2 vs MM2
Number of differences: 148

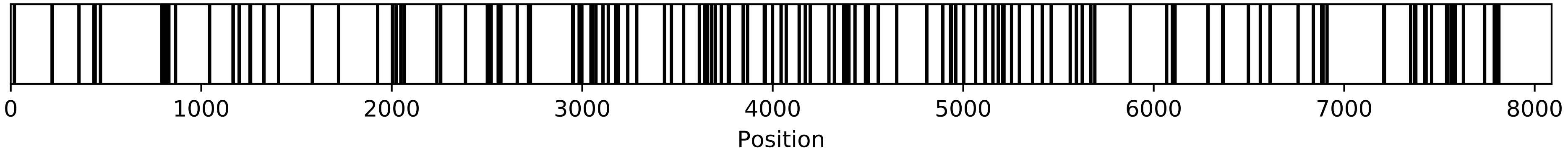


MM1 vs MM2
Number of differences: 193

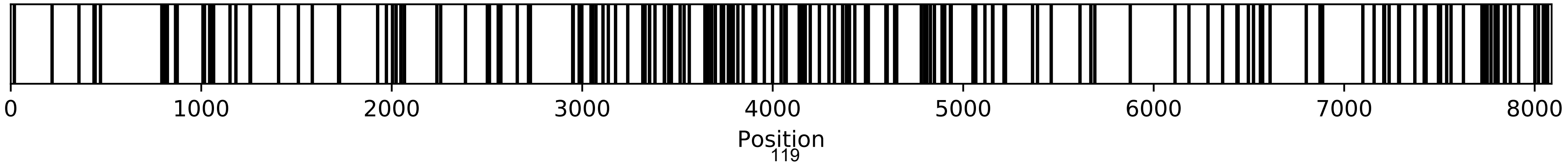

### Region: 348



## CR1 vs CR2
Number of differences: 354

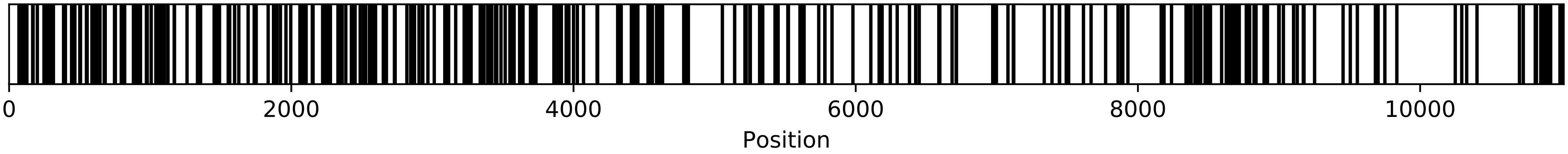


## CR1 vs MM1
Number of differences: 364

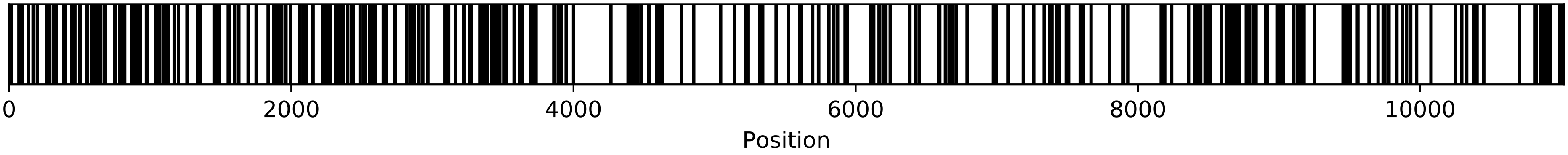


## CR1 vs MM2
Number of differences: 364

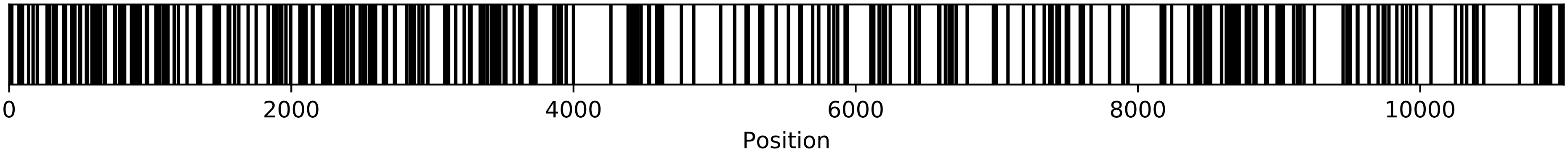


## CR2 vs MM1
Number of differences: 172

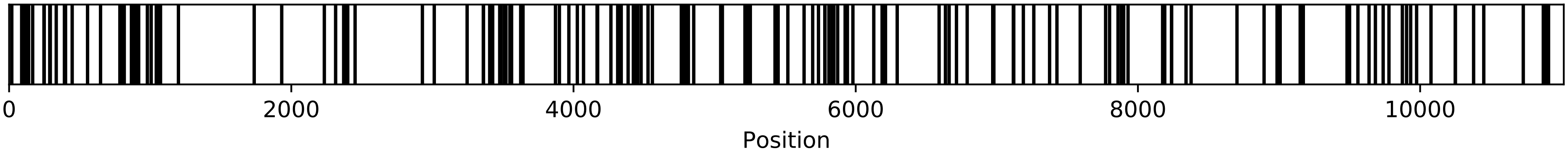


## CR2 vs MM2
Number of differences: 172

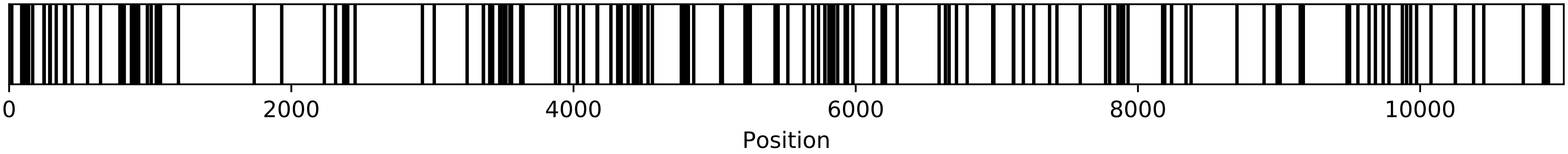


## MM1 vs MM2
Number of differences: 0

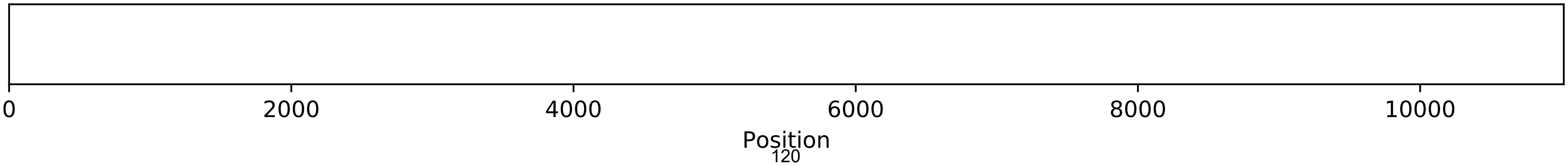

### Region: 352



## CR1 vs CR2
Number of differences: 16

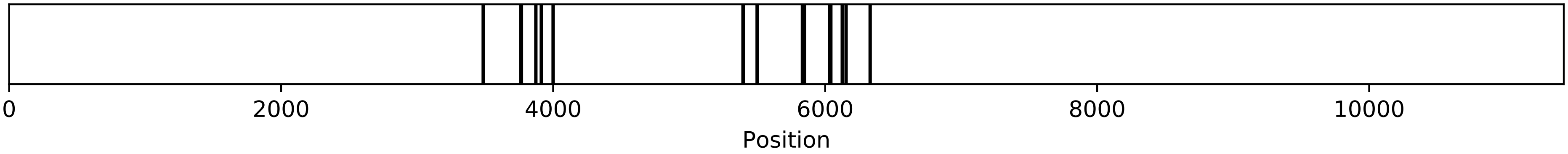


## CR1 vs MM1
Number of differences: 131

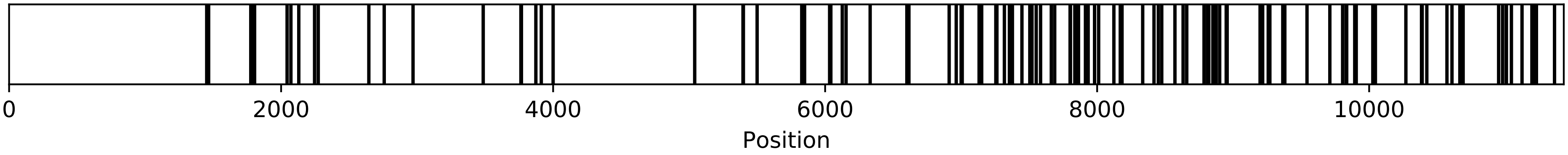


## CR1 vs MM2
Number of differences: 152

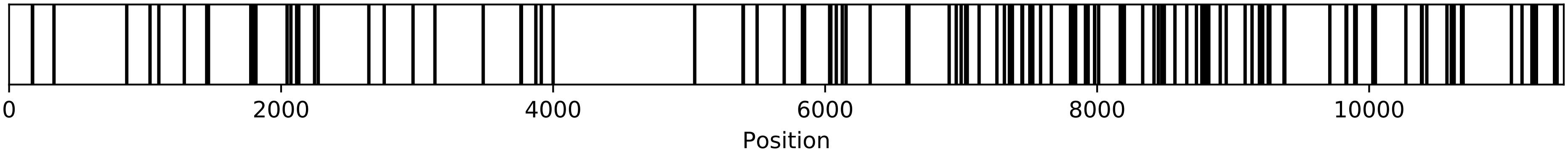


## CR2 vs MM1
Number of differences: 115

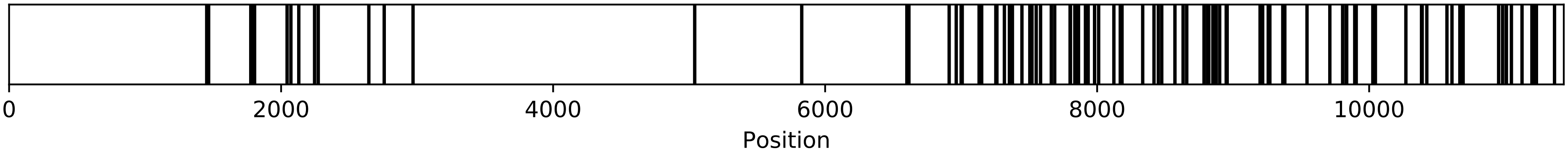


## CR2 vs MM2
Number of differences: 136

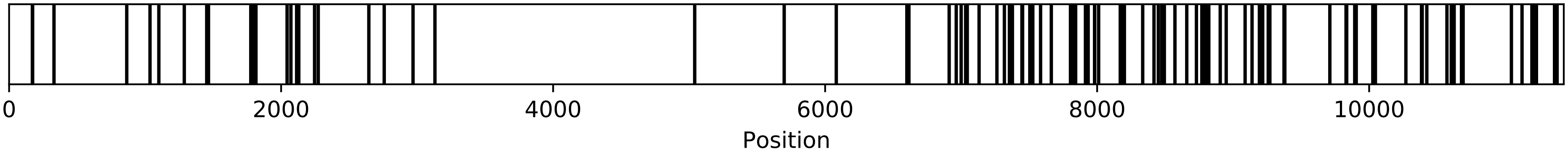


## MM1 vs MM2
Number of differences: 73

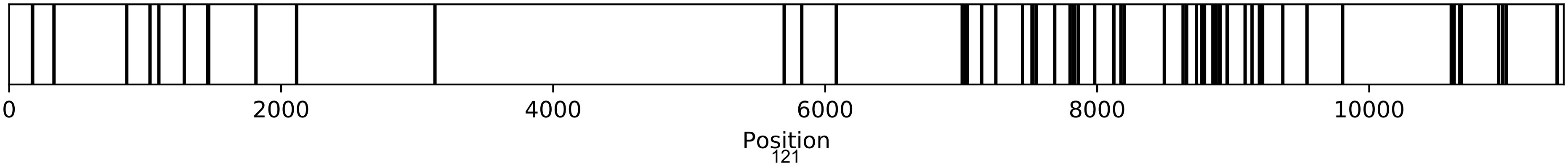

### Region: 353



CR1 vs CR2
Number of differences: 0

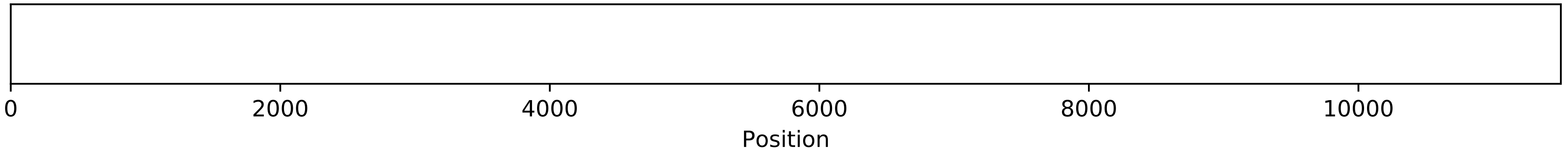


CR1 vs MM1
Number of differences: 347

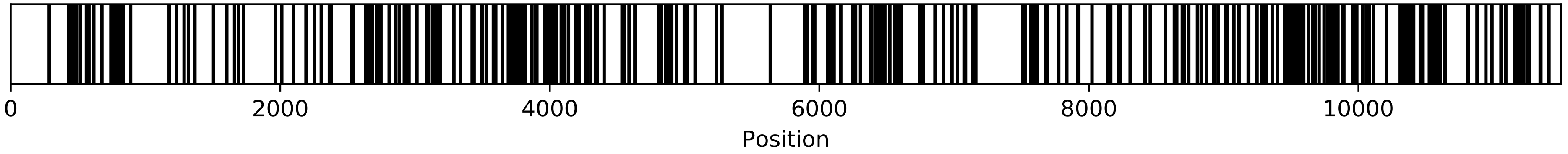


CR1 vs MM2
Number of differences: 347

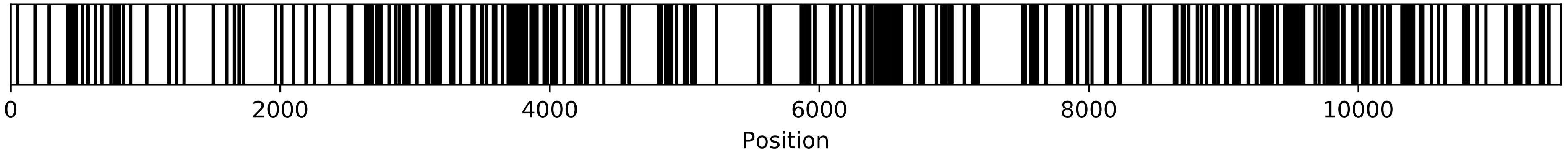


CR2 vs MM1
Number of differences: 347

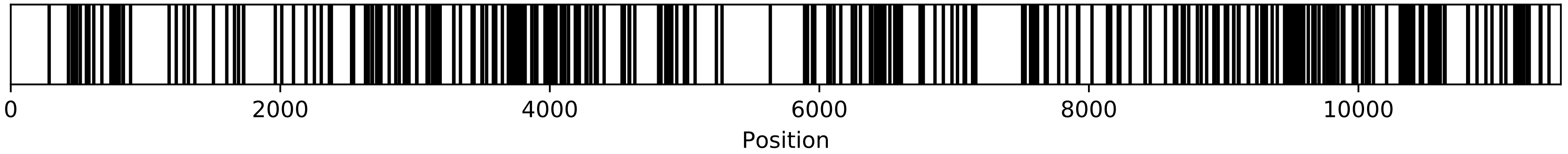


CR2 vs MM2
Number of differences: 347

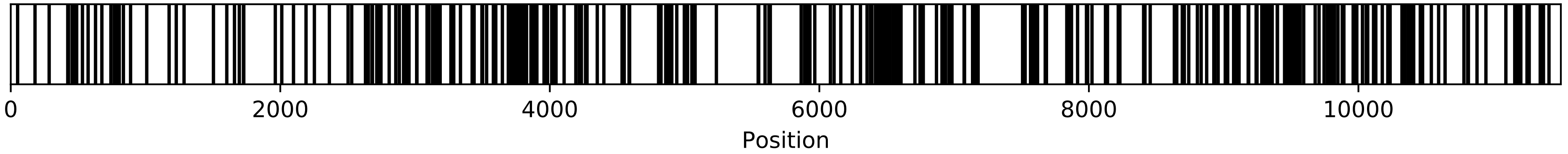


MM1 vs MM2
Number of differences: 153

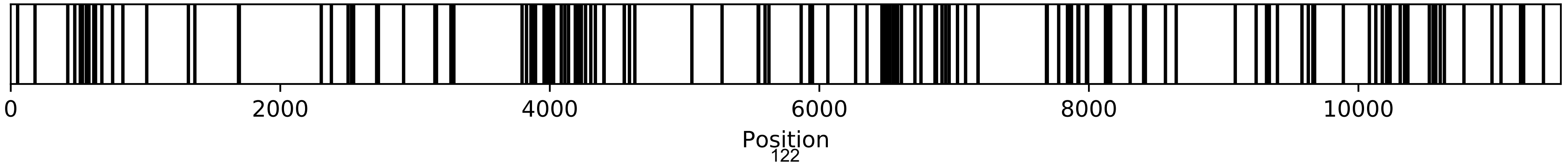

### Region: 354



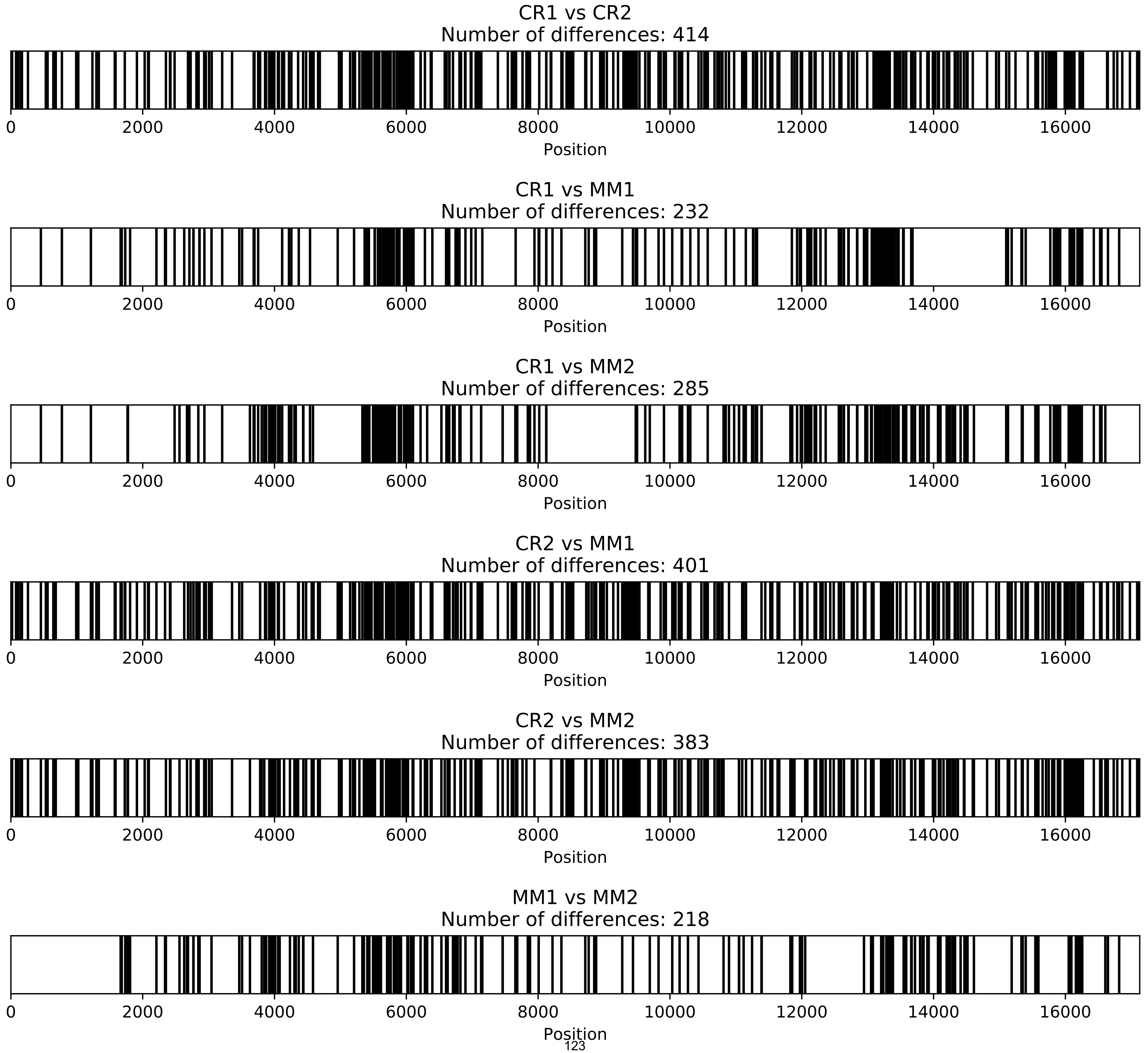
CR1 vs CR2
Number of differences: 414
CR1 vs MM1
Number of differences: 232
CR1 vs MM2
Number of differences: 285
CR2 vs MM1
Number of differences: 401
CR2 vs MM2
Number of differences: 383
MM1 vs MM2
Number of differences: 218
0
2000
4000
6000
8000
10000
12000
14000
16000
Position

### Region: 355



CR1 vs CR2
Number of differences: 407

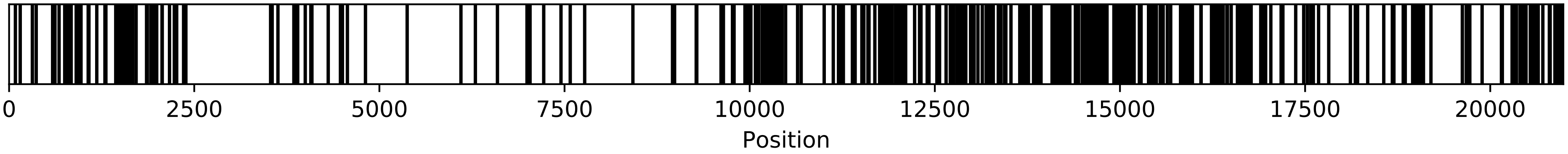


CR1 vs MM1
Number of differences: 188

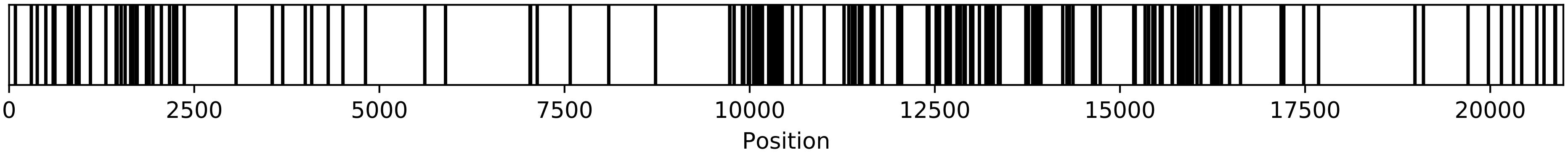


CR1 vs MM2
Number of differences: 205

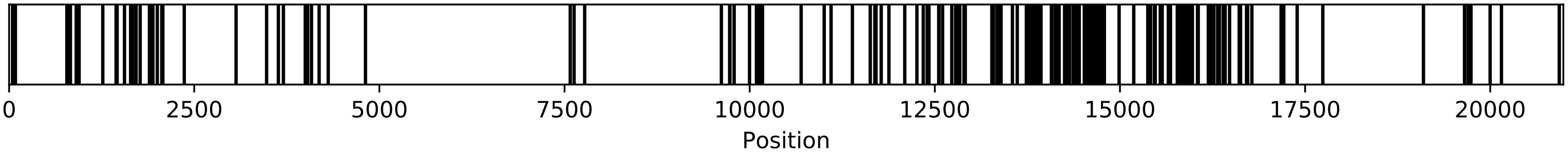


CR2 vs MM1
Number of differences: 379

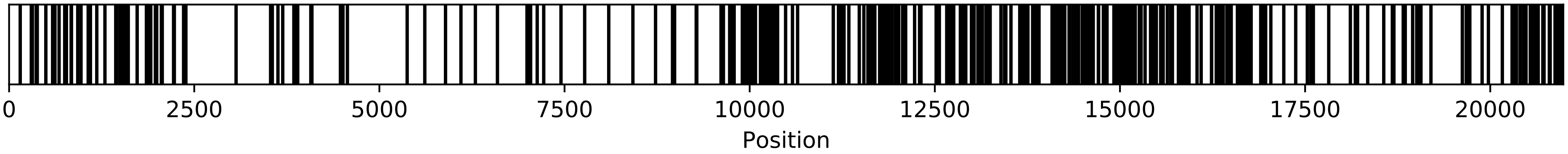


CR2 vs MM2
Number of differences: 419

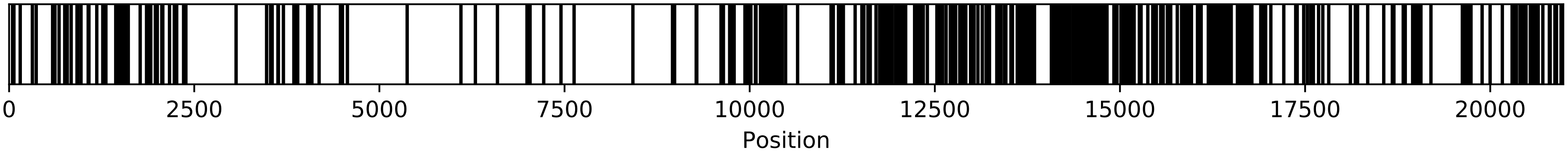


MM1 vs MM2
Number of differences: 240

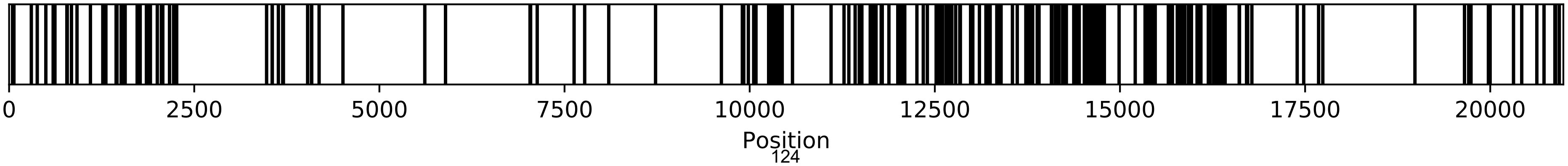

### Region: 356



CR1 vs CR2
Number of differences: 117

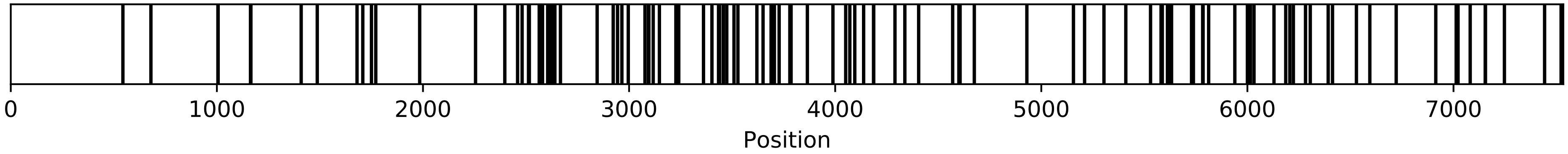


CR1 vs MM1
Number of differences: 25

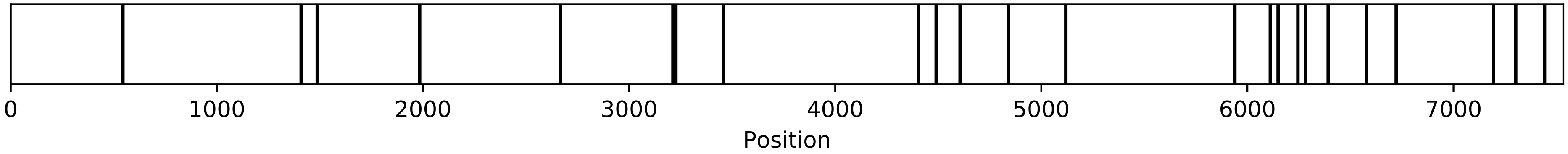


CR1 vs MM2
Number of differences: 37

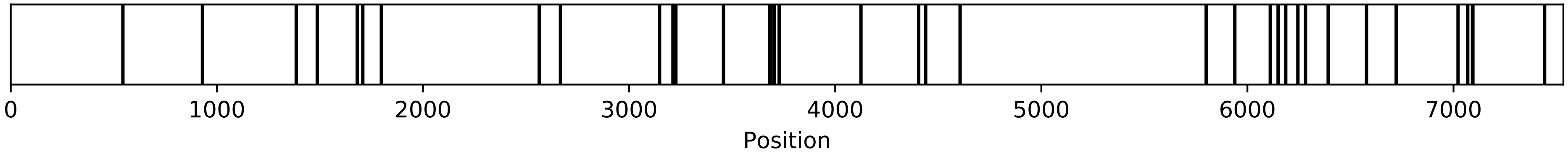


CR2 vs MM1
Number of differences: 114

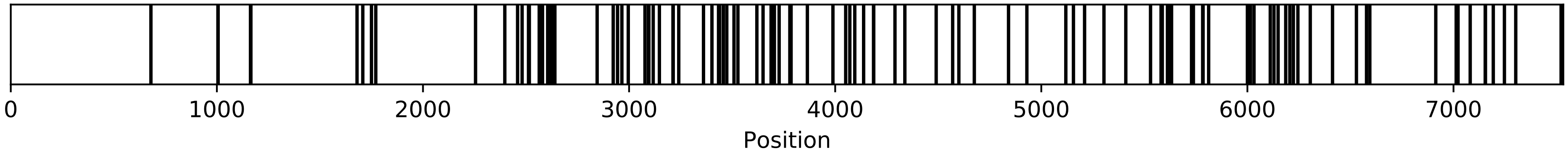


CR2 vs MM2
Number of differences: 117

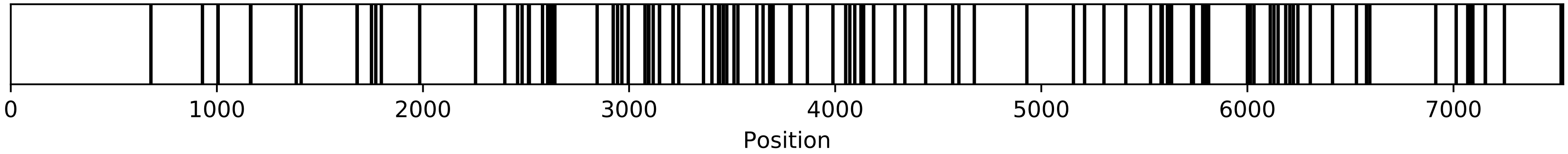


MM1 vs MM2
Number of differences: 26

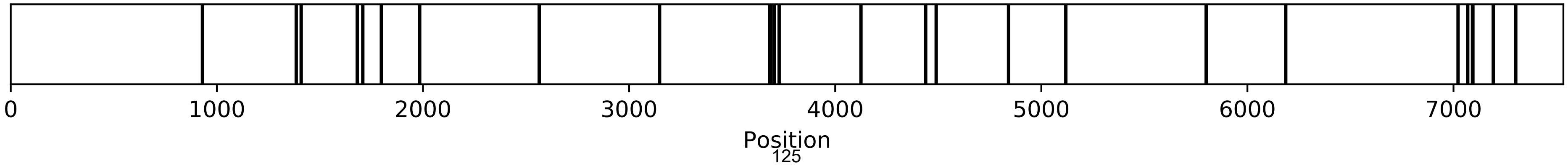

### Region: 358



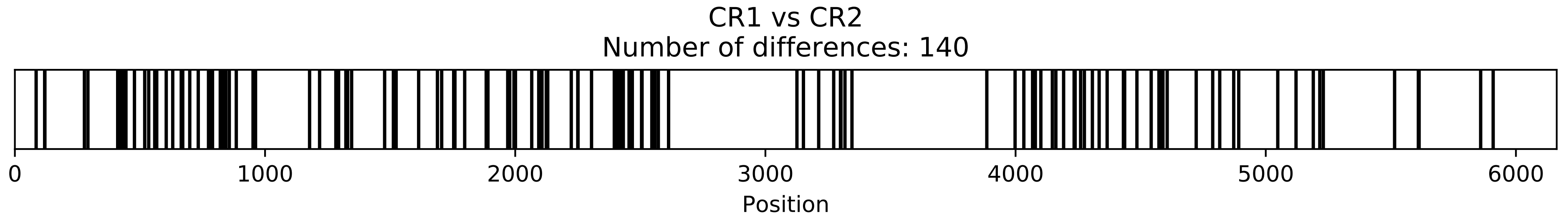
CR1 vs CR2
Number of differences: 140
Position

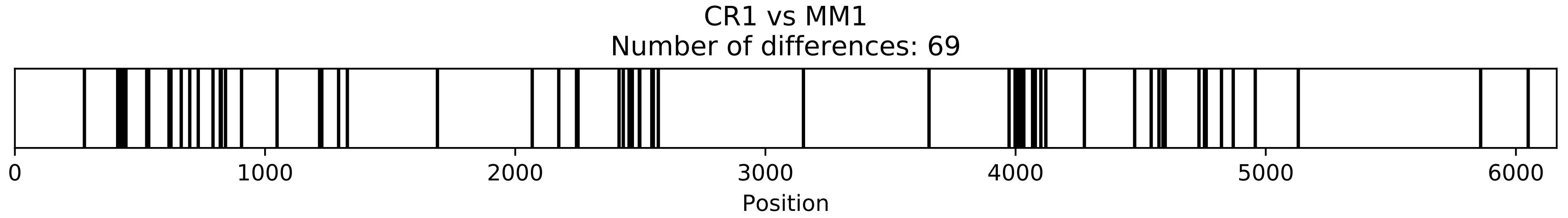
CR1 vs MM1
Number of differences: 69
Position

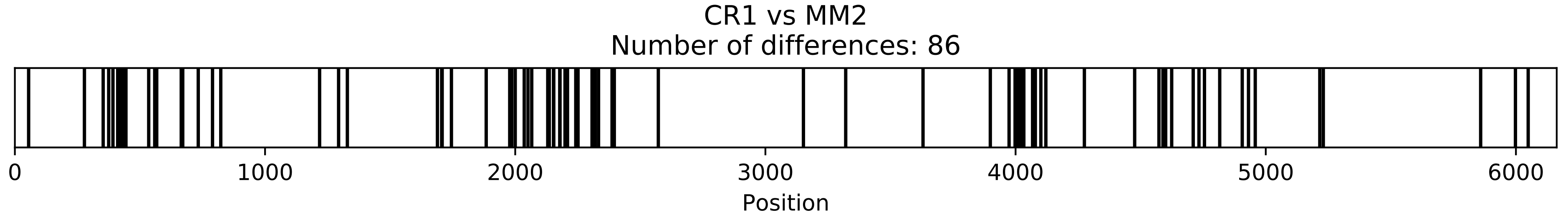
CR1 vs MM2
Number of differences: 86
Position

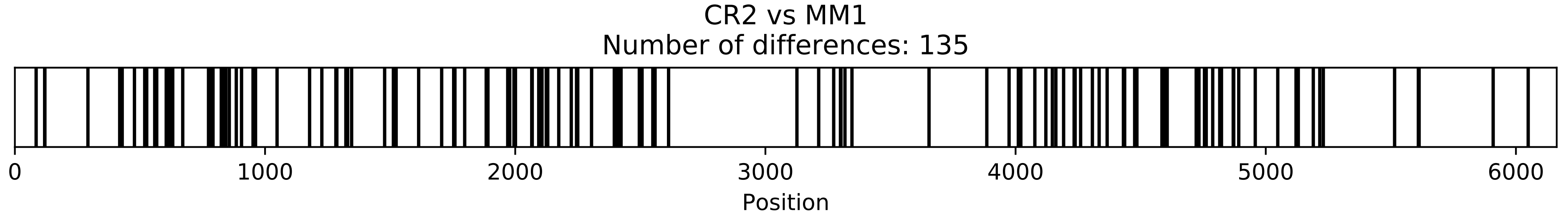
CR2 vs MM1
Number of differences: 135
Position

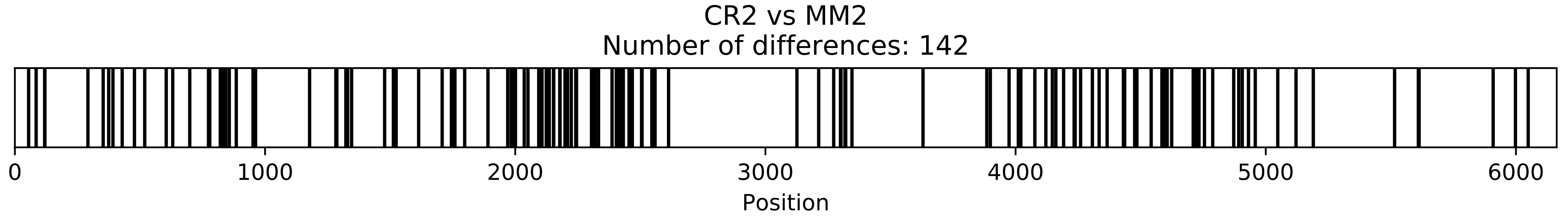
CR2 vs MM2
Number of differences: 142
Position

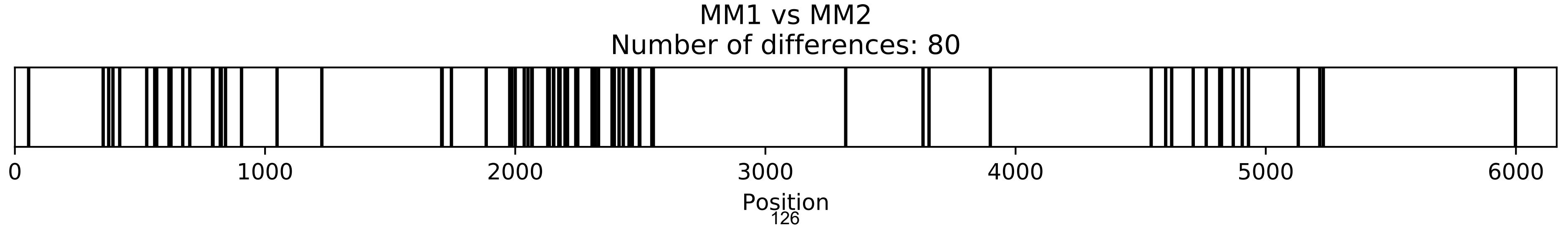
MM1 vs MM2
Number of differences: 80
Position

### Region: 359



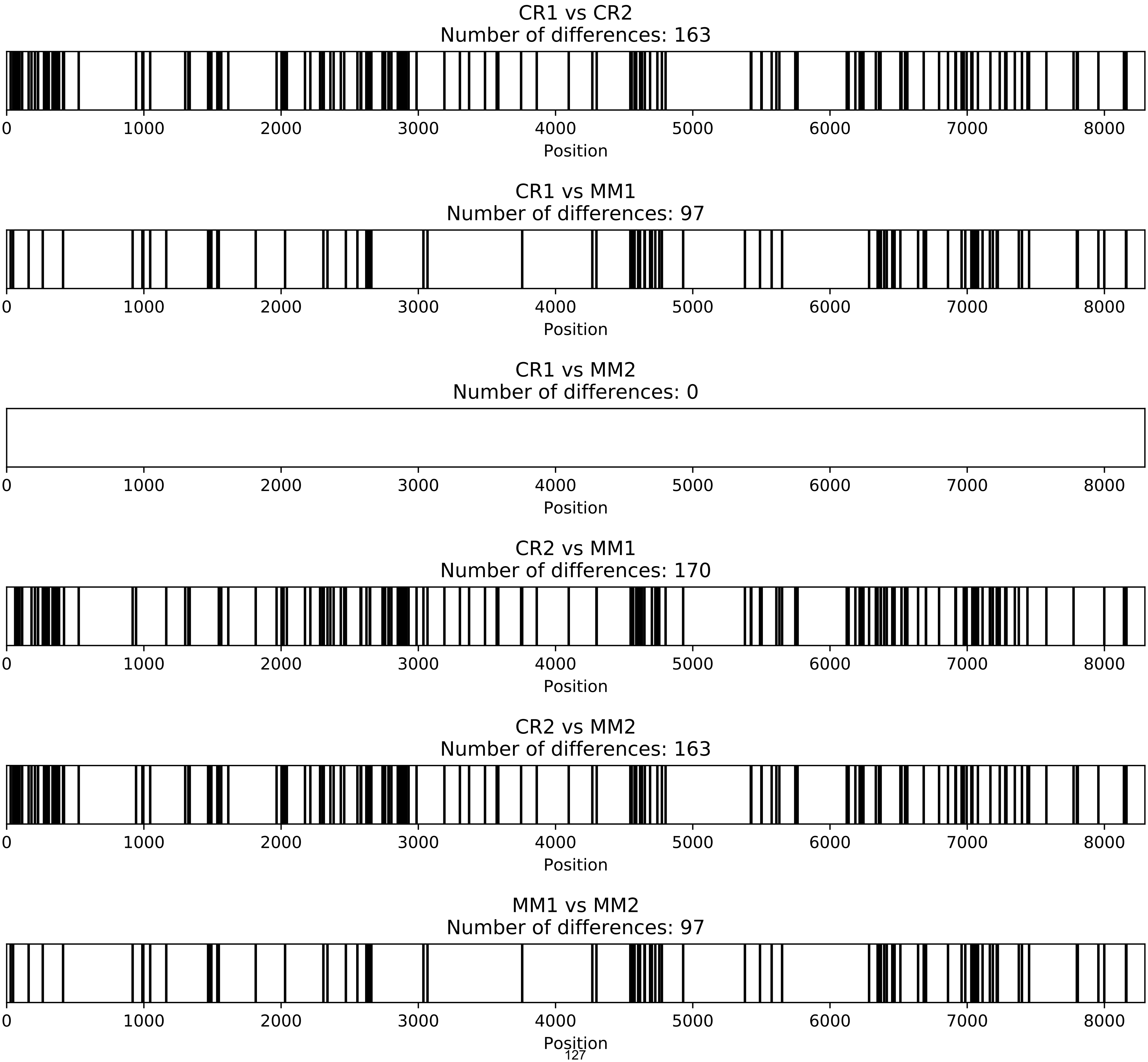
CR1 vs CR2
Number of differences: 163
CR1 vs MM1
Number of differences: 97
CR1 vs MM2
Number of differences: 0
CR2 vs MM1
Number of differences: 170
CR2 vs MM2
Number of differences: 163
MM1 vs MM2
Number of differences: 97
0
1000
2000
3000
4000
5000
6000
7000
8000
Position

### Region: 368



CR1 vs CR2
Number of differences: 255

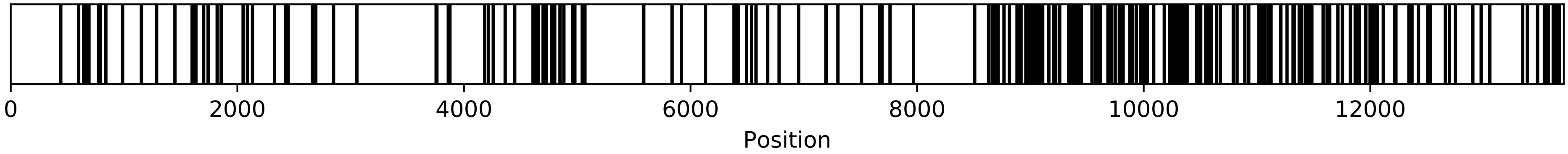


CR1 vs MM1
Number of differences: 273

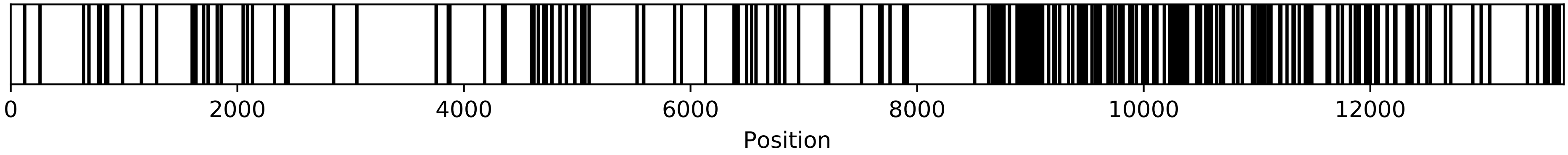


CR1 vs MM2
Number of differences: 250

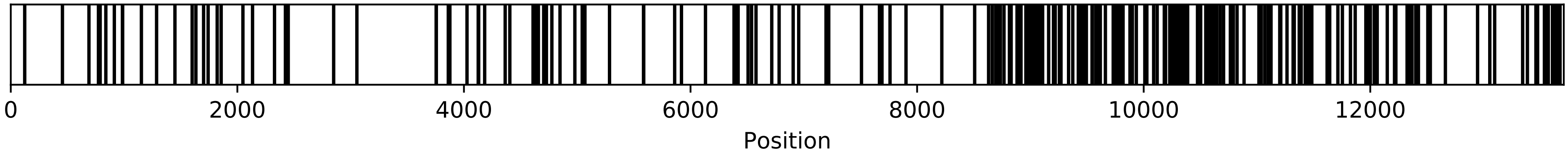


CR2 vs MM1
Number of differences: 115

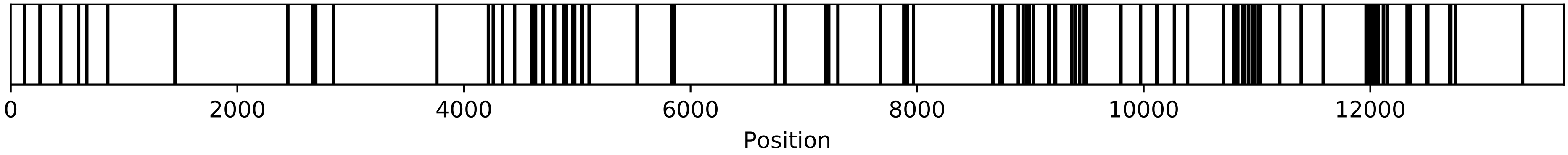


CR2 vs MM2
Number of differences: 130

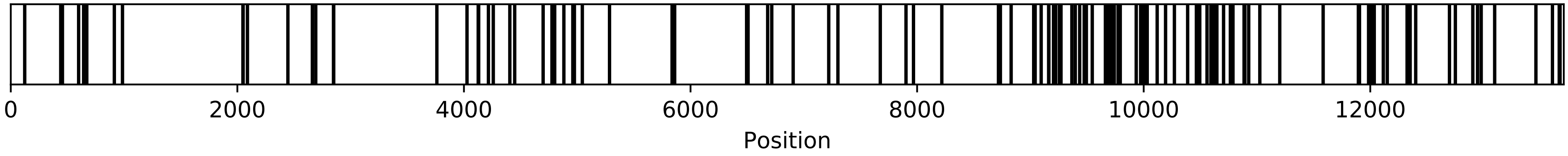


MM1 vs MM2
Number of differences: 134

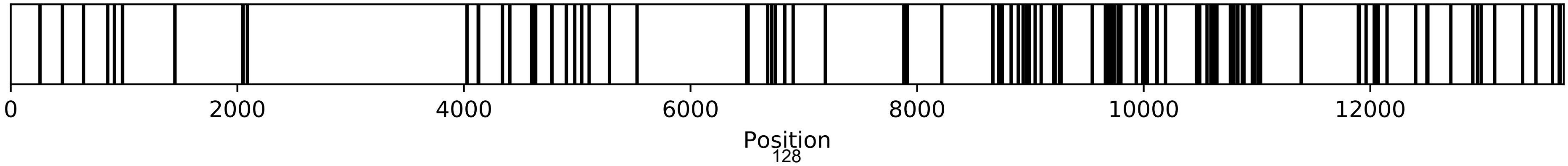

### Region: 383



CR1 vs CR2
Number of differences: 229

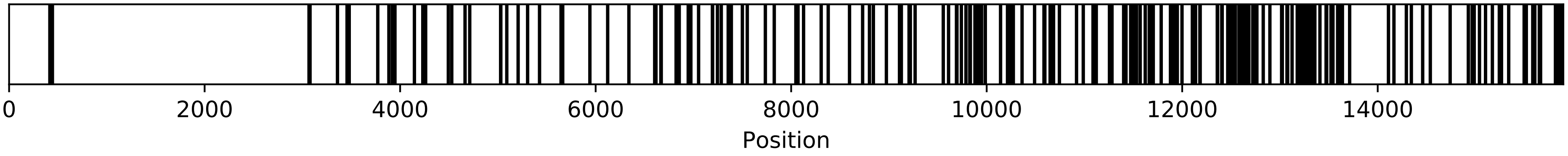


CR1 vs MM1
Number of differences: 276

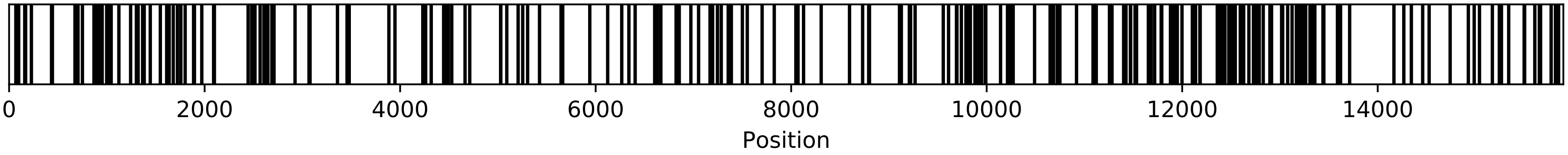


CR1 vs MM2
Number of differences: 274

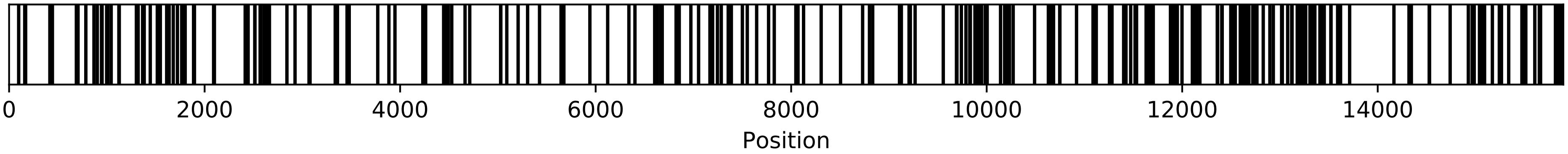


CR2 vs MM1
Number of differences: 168

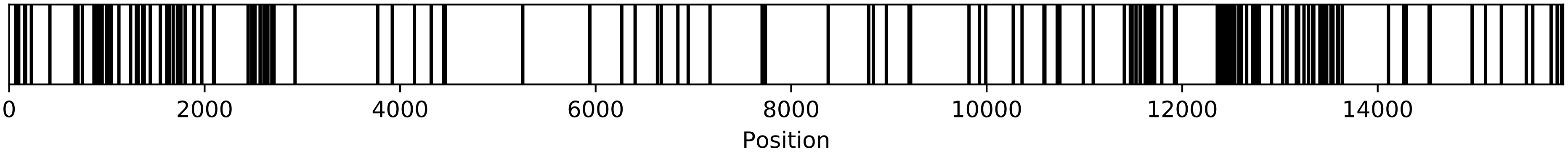


CR2 vs MM2
Number of differences: 136

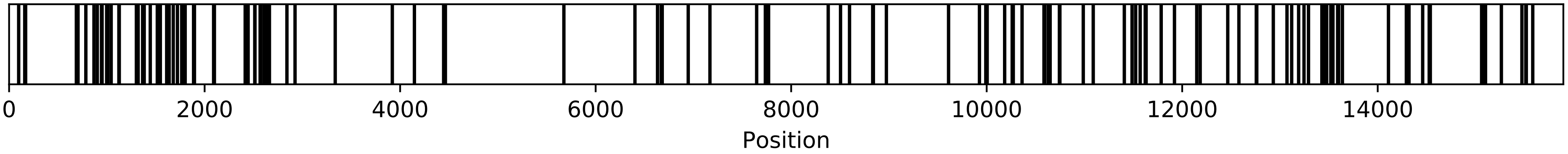


MM1 vs MM2
Number of differences: 138

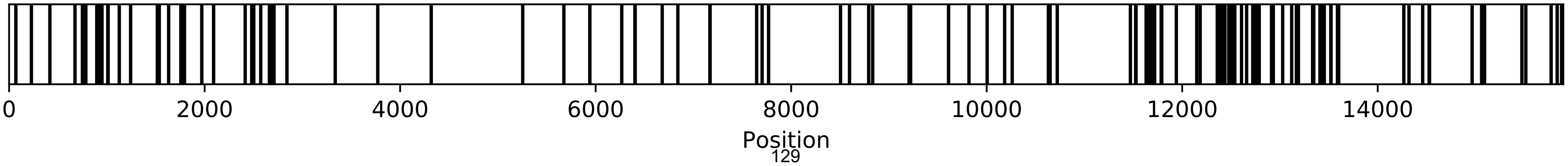

### Region: 387



CR1 vs CR2
Number of differences: 423

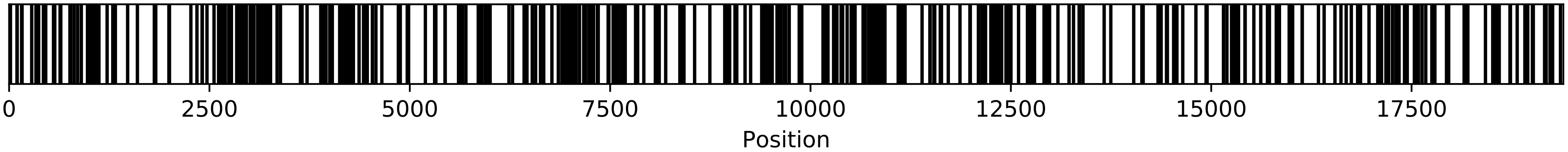


CR1 vs MM1
Number of differences: 413

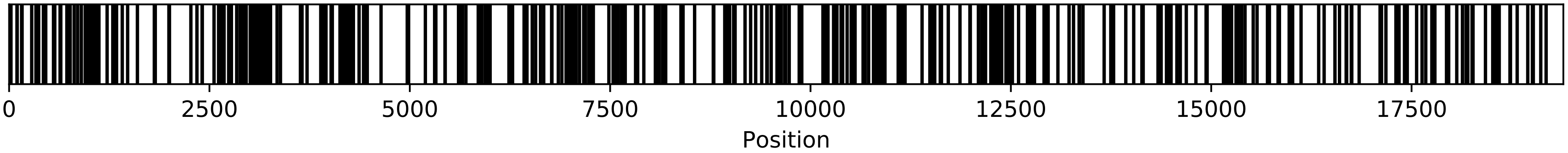


CR1 vs MM2
Number of differences: 395

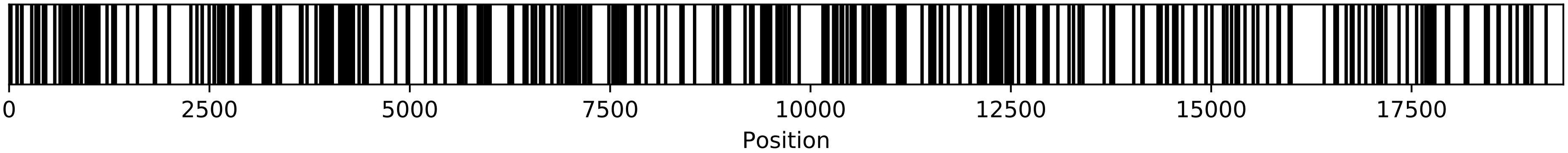


CR2 vs MM1
Number of differences: 135

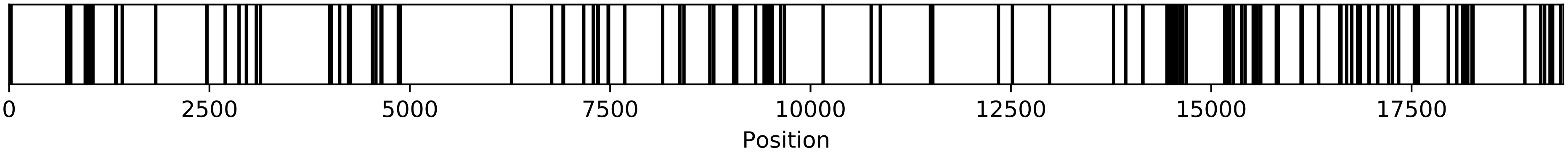


CR2 vs MM2
Number of differences: 171

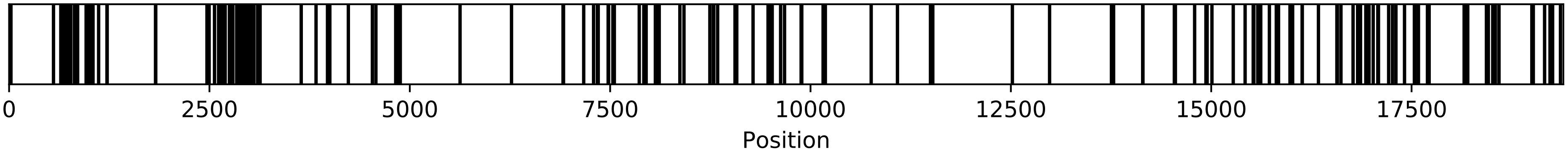


MM1 vs MM2
Number of differences: 155

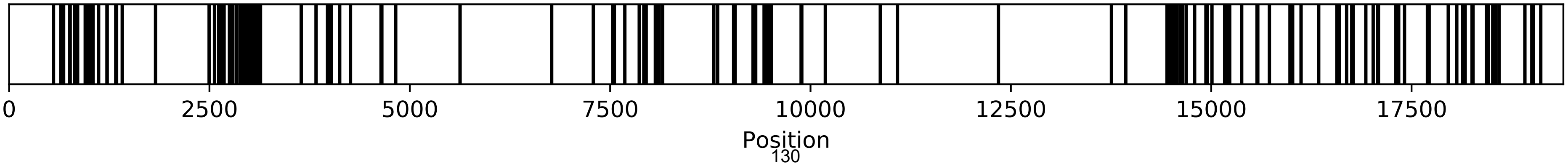

### Region: 389



CR1 vs CR2
Number of differences: 280

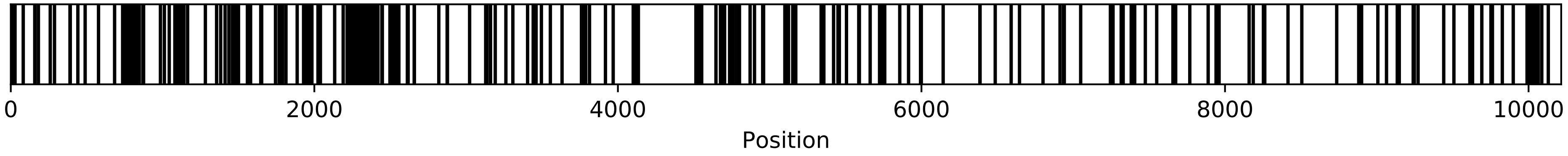


CR1 vs MM1
Number of differences: 271

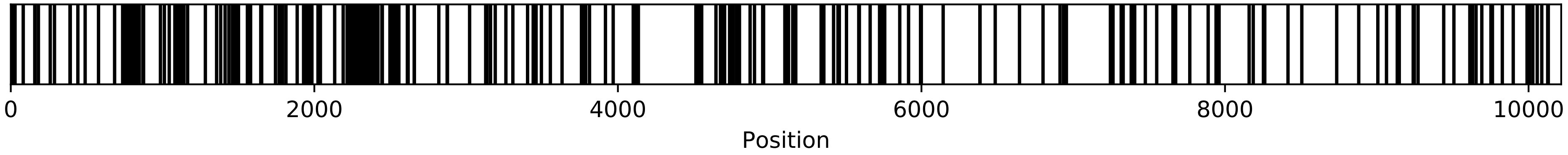


CR1 vs MM2
Number of differences: 277

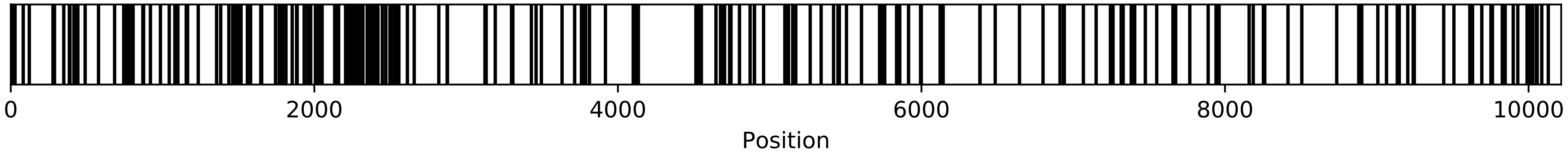


CR2 vs MM1
Number of differences: 15

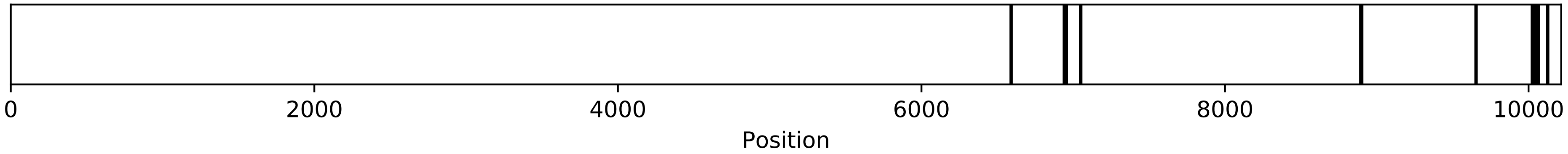


CR2 vs MM2
Number of differences: 126

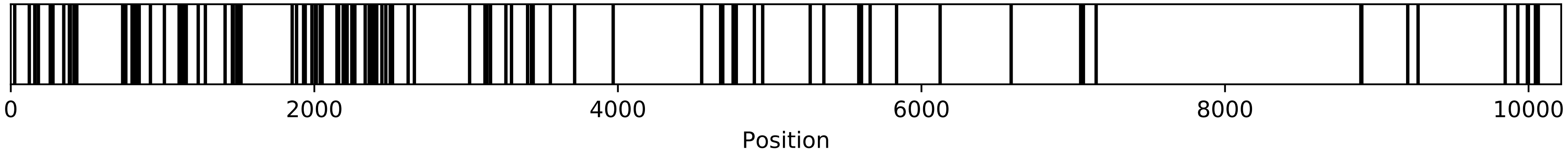


MM1 vs MM2
Number of differences: 123

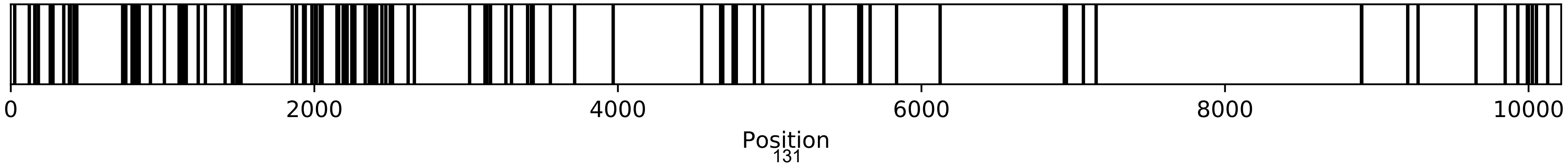

### Region: 390



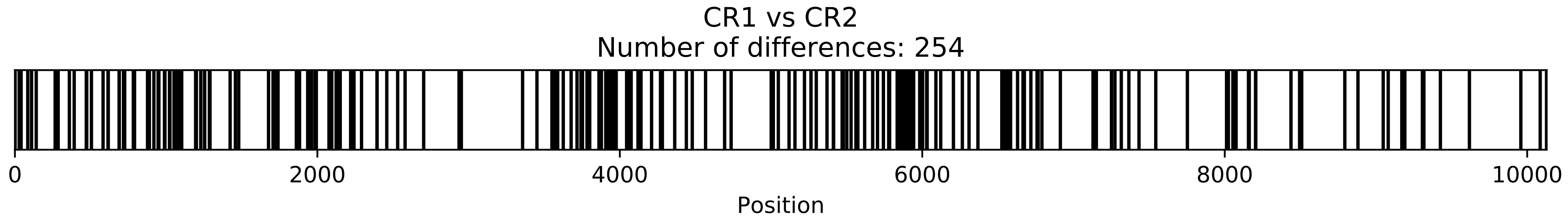
CR1 vs CR2
Number of differences: 254
0
2000
4000
6000
8000
10000
Position

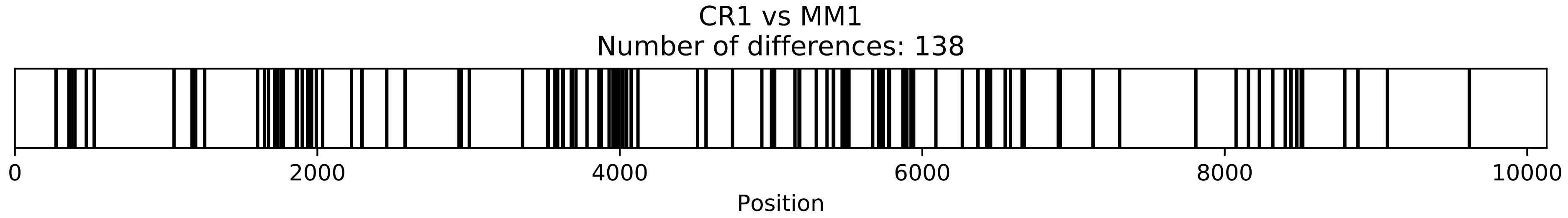
CR1 vs MM1
Number of differences: 138
0
2000
4000
6000
8000
10000
Position

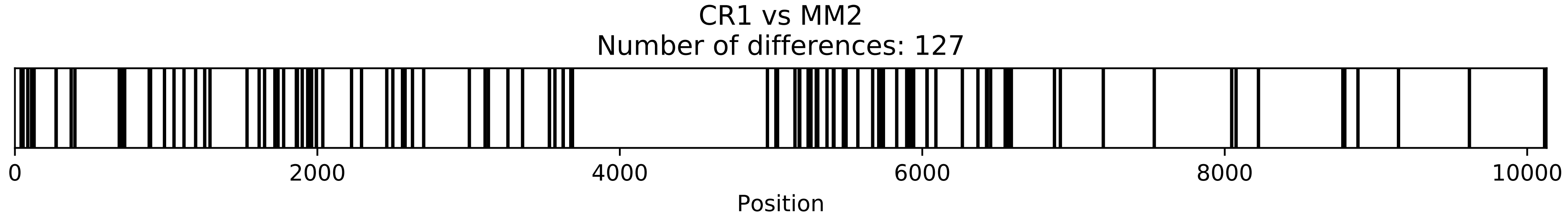
CR1 vs MM2
Number of differences: 127
0
2000
4000
6000
8000
10000
Position

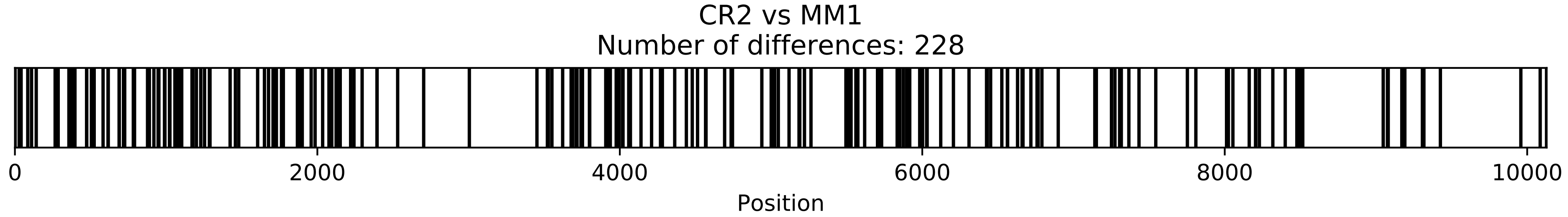
CR2 vs MM1
Number of differences: 228
0
2000
4000
6000
8000
10000
Position

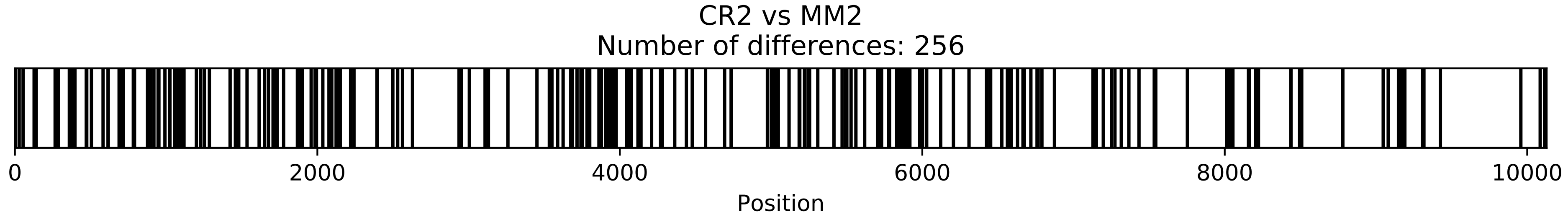
CR2 vs MM2
Number of differences: 256
0
2000
4000
6000
8000
10000
Position

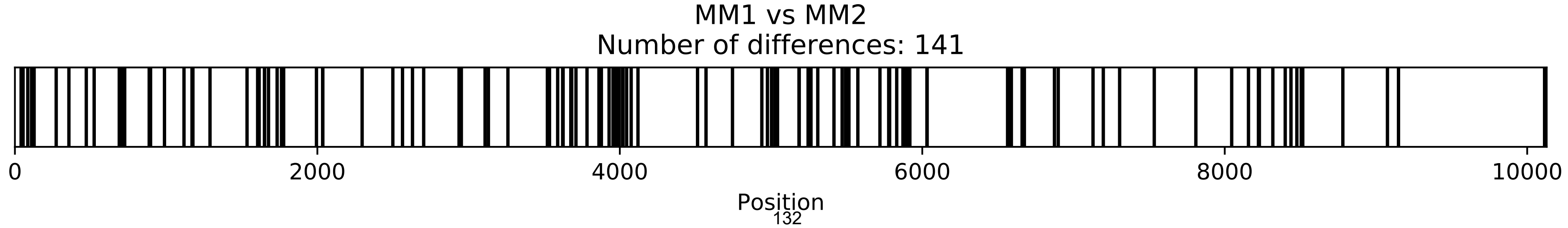
MM1 vs MM2
Number of differences: 141
0
2000
4000
6000
8000
10000
Position

### Region: 400

## CR1 vs CR2
Number of differences: 227

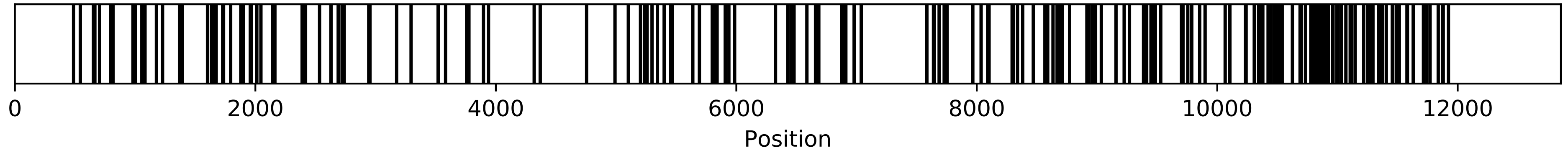


## CR1 vs MM1
Number of differences: 231

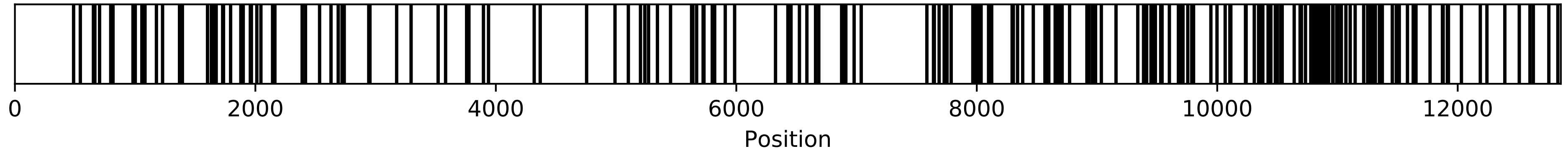


## CR1 vs MM2
Number of differences: 248

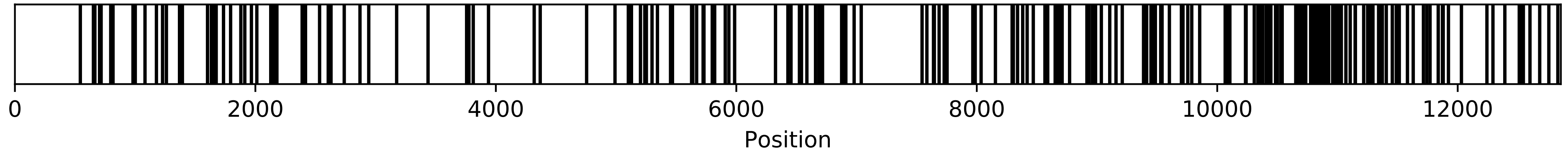


## CR2 vs MM1
Number of differences: 72

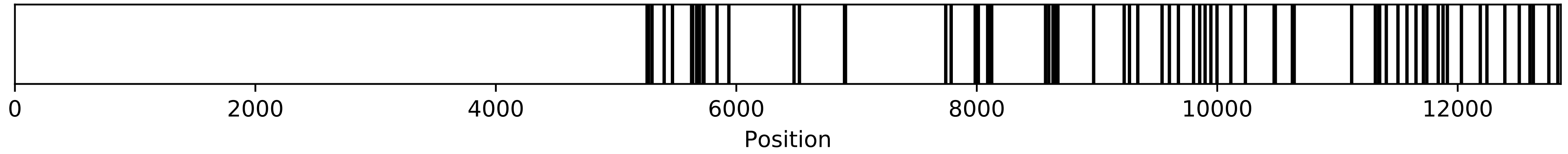


## CR2 vs MM2
Number of differences: 118

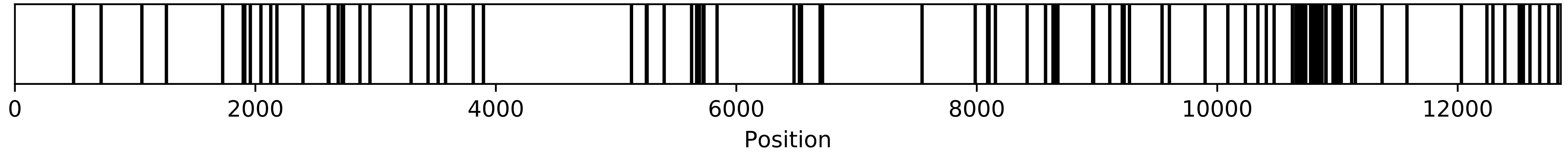


## MM1 vs MM2
Number of differences: 116

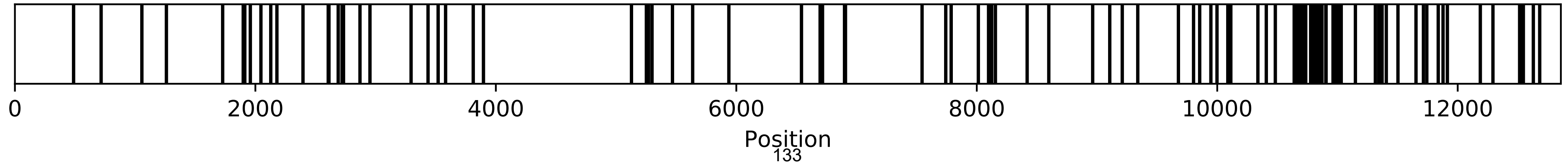

### Region: 401



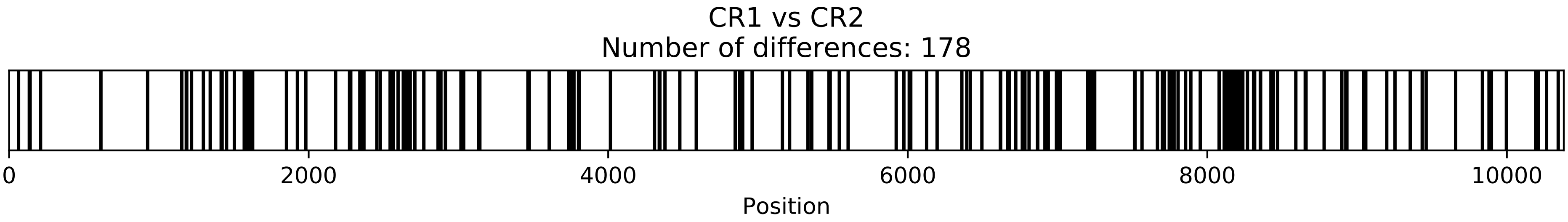
CR1 vs CR2
Number of differences: 178
0
2000
4000
6000
8000
10000
Position

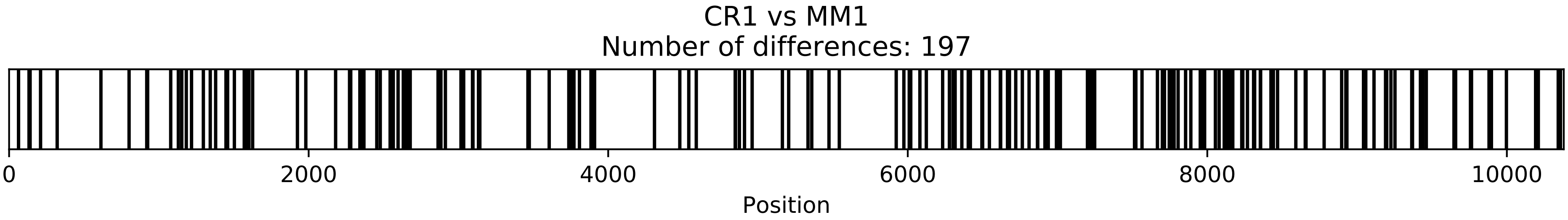
CR1 vs MM1
Number of differences: 197
0
2000
4000
6000
8000
10000
Position

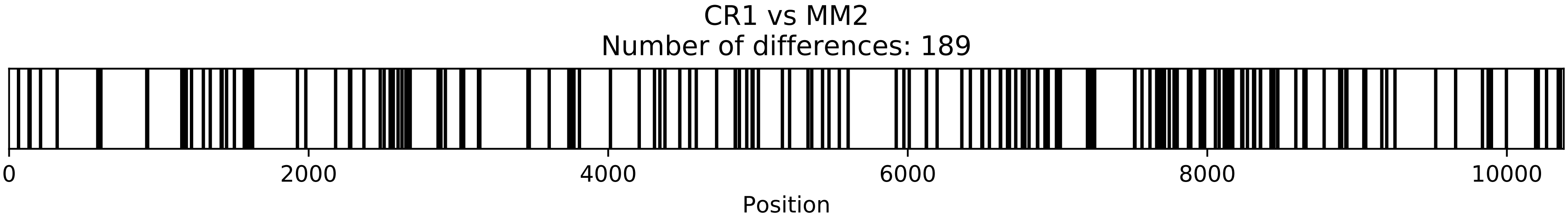
CR1 vs MM2
Number of differences: 189
0
2000
4000
6000
8000
10000
Position

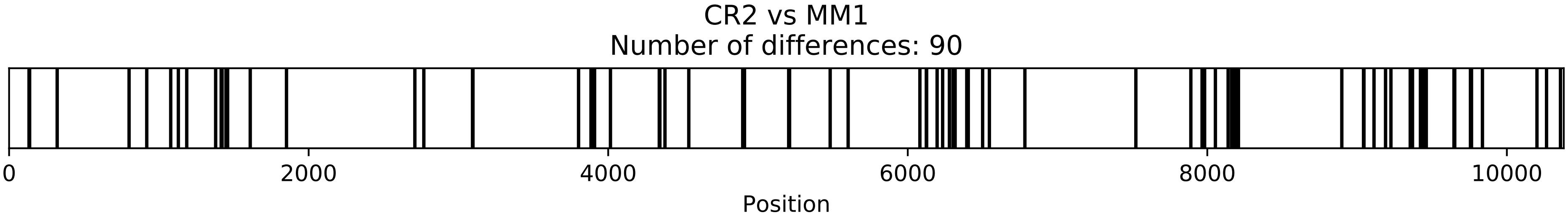
CR2 vs MM1
Number of differences: 90
0
2000
4000
6000
8000
10000
Position

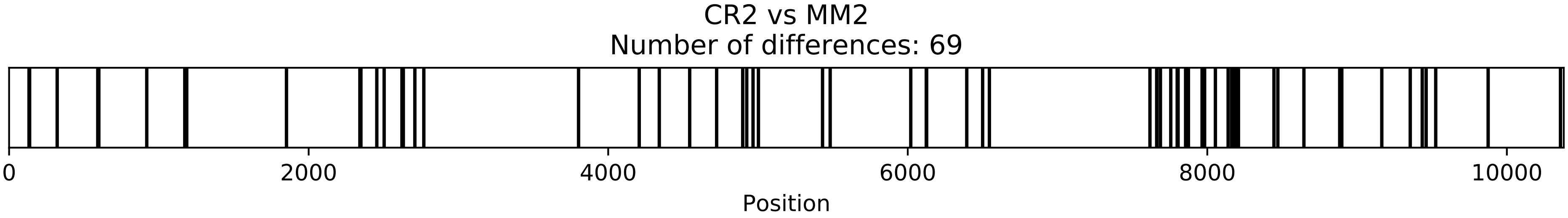
CR2 vs MM2
Number of differences: 69
0
2000
4000
6000
8000
10000
Position

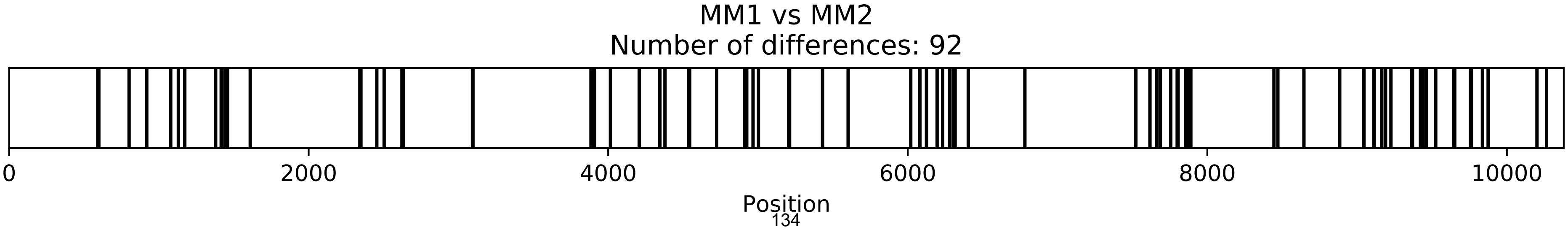
MM1 vs MM2
Number of differences: 92
0
2000
4000
6000
8000
10000
Position

### Region: 402

## CR1 vs CR2
Number of differences: 401

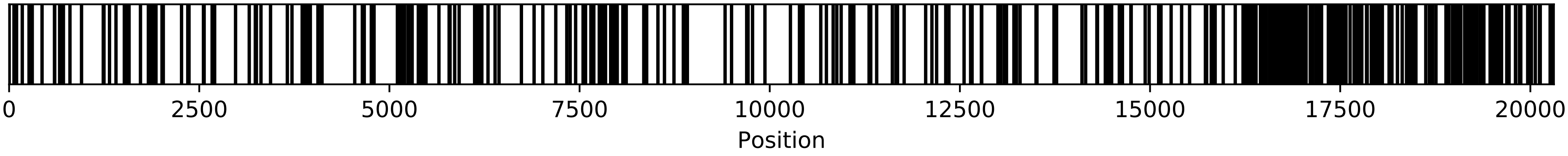


## CR1 vs MM1
Number of differences: 247

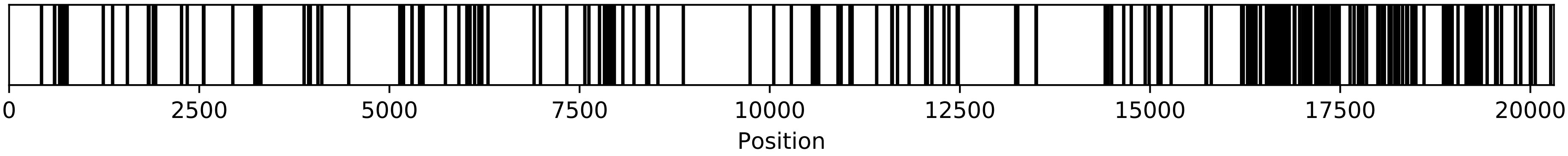


## CR1 vs MM2
Number of differences: 211

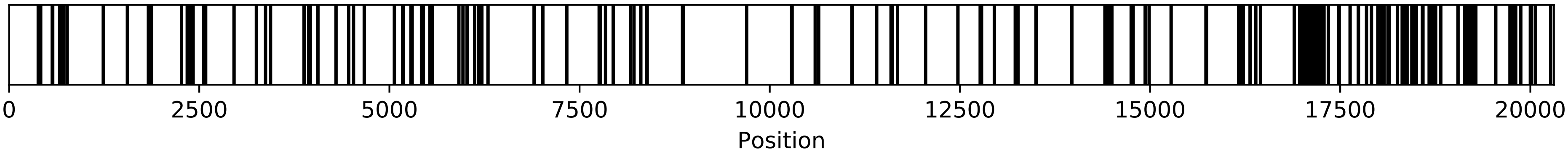


## CR2 vs MM1
Number of differences: 374

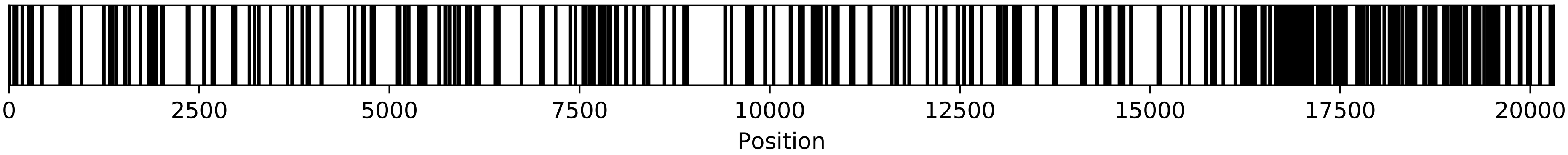


## CR2 vs MM2
Number of differences: 384

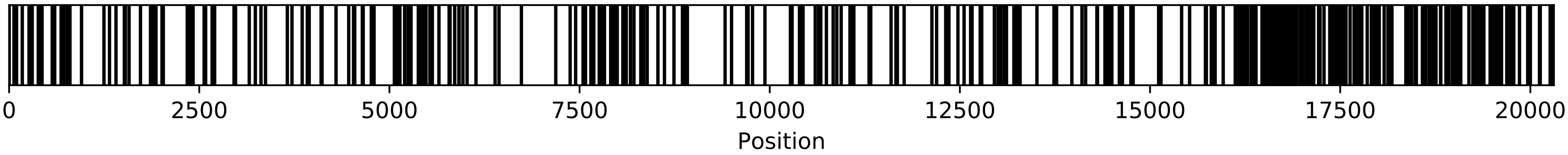


## MM1 vs MM2
Number of differences: 234

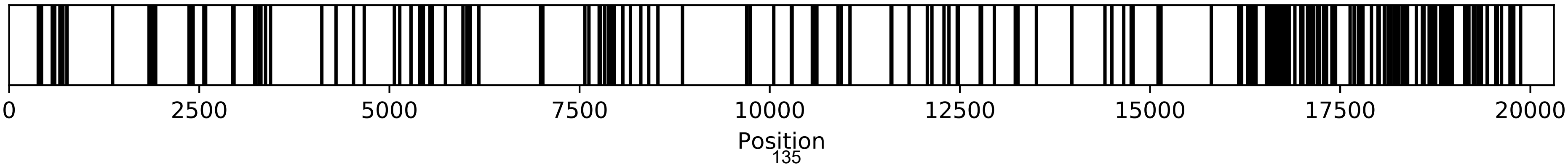

### Region: 403



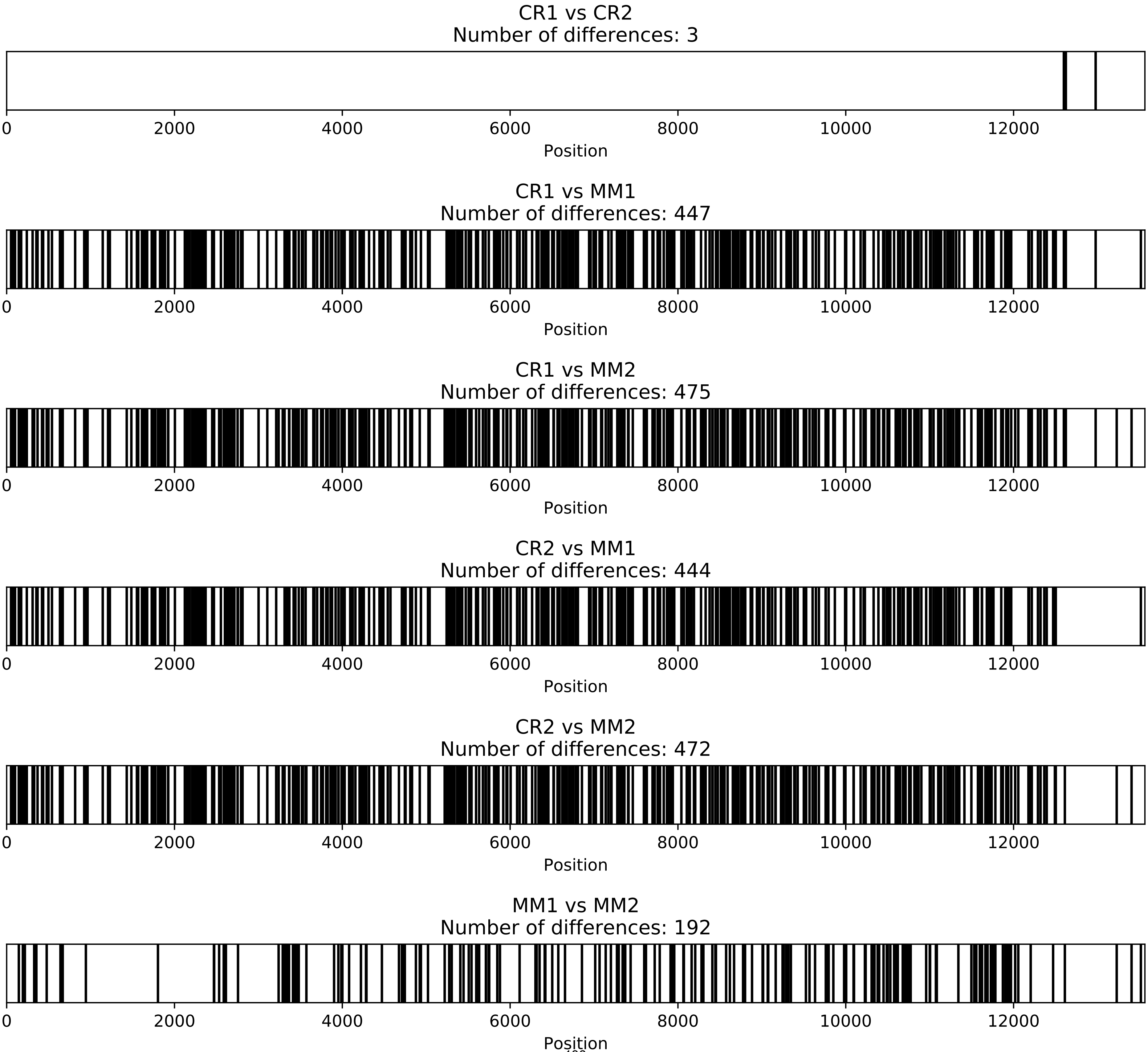
CR1 vs CR2
Number of differences: 3
CR1 vs MM1
Number of differences: 447
CR1 vs MM2
Number of differences: 475
CR2 vs MM1
Number of differences: 444
CR2 vs MM2
Number of differences: 472
MM1 vs MM2
Number of differences: 192
0
2000
4000
6000
8000
10000
12000
Position

### Region: 408



## CR1 vs CR2
Number of differences: 321

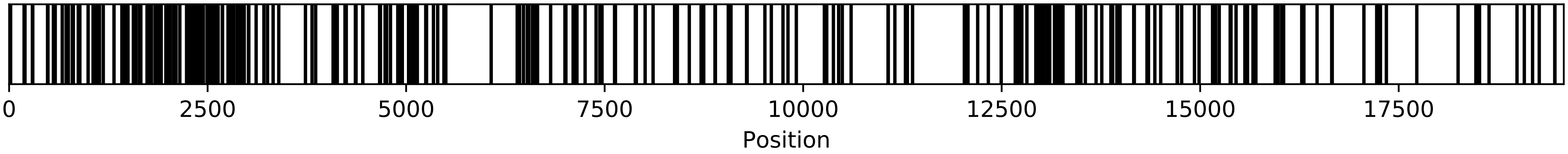


## CR1 vs MM1
Number of differences: 179

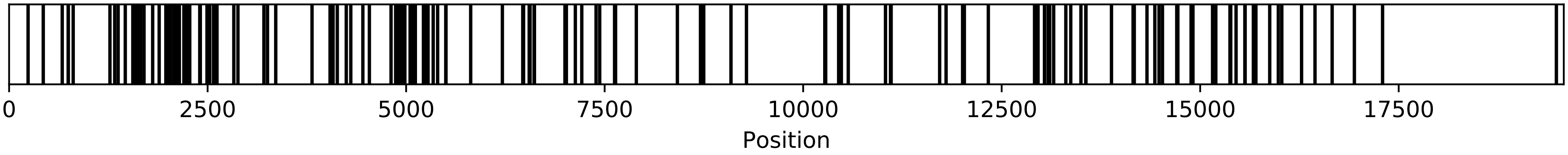


## CR1 vs MM2
Number of differences: 190

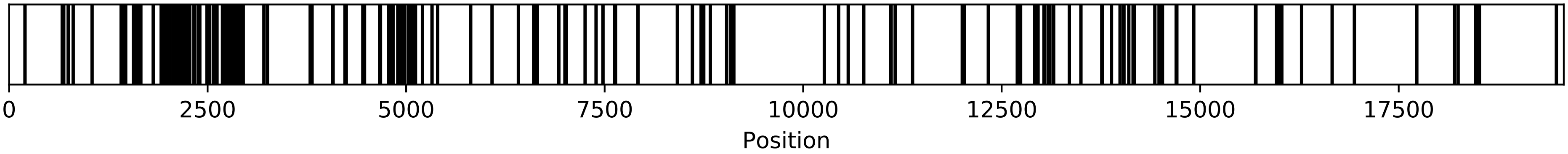


## CR2 vs MM1
Number of differences: 362

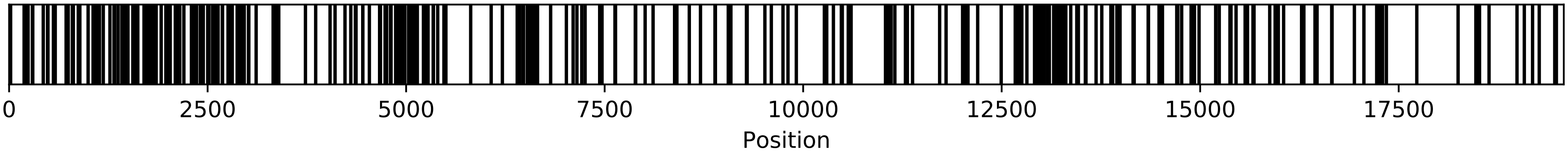


## CR2 vs MM2
Number of differences: 339

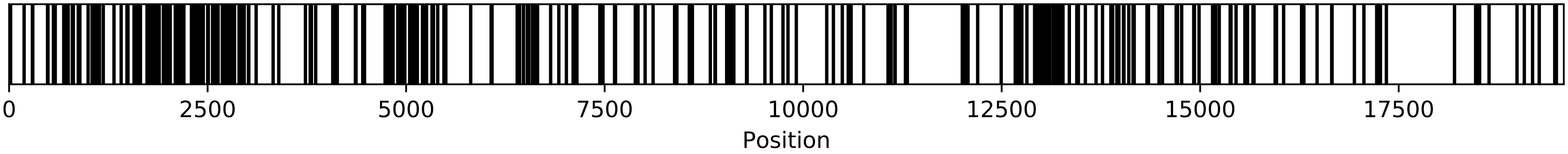


## MM1 vs MM2
Number of differences: 194

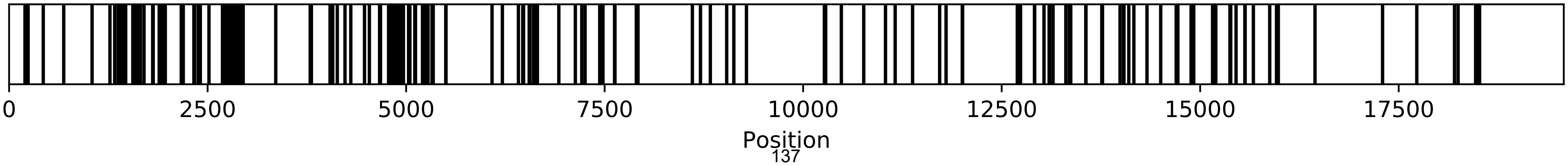

### Region: 411



CR1 vs CR2
Number of differences: 37

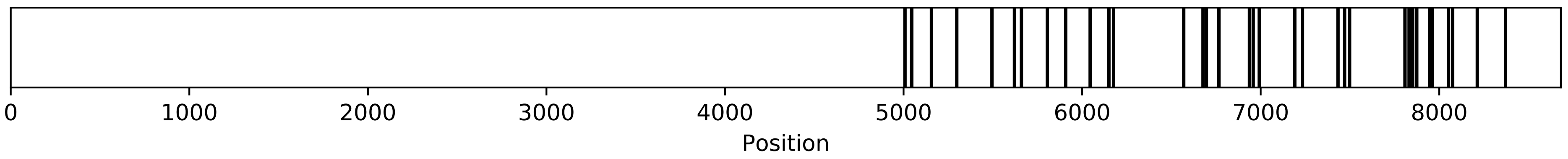


CR1 vs MM1
Number of differences: 191

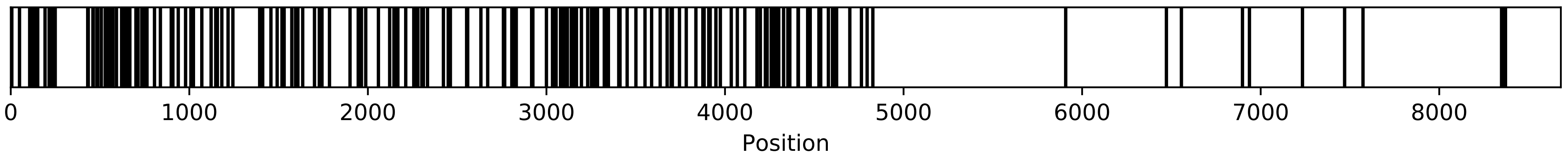


CR1 vs MM2
Number of differences: 194

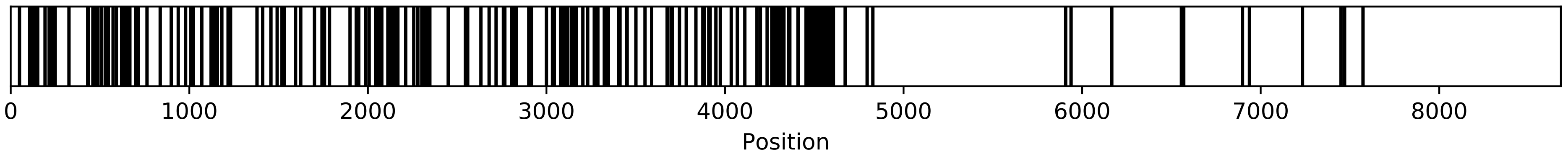


CR2 vs MM1
Number of differences: 218

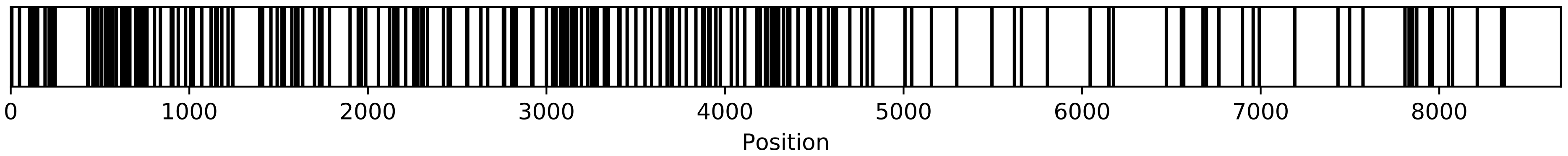


CR2 vs MM2
Number of differences: 221

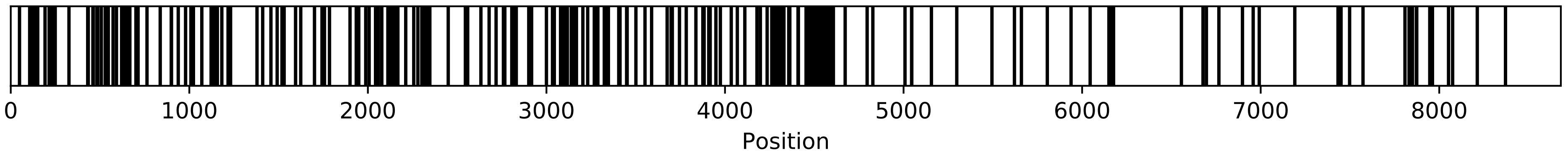


MM1 vs MM2
Number of differences: 94

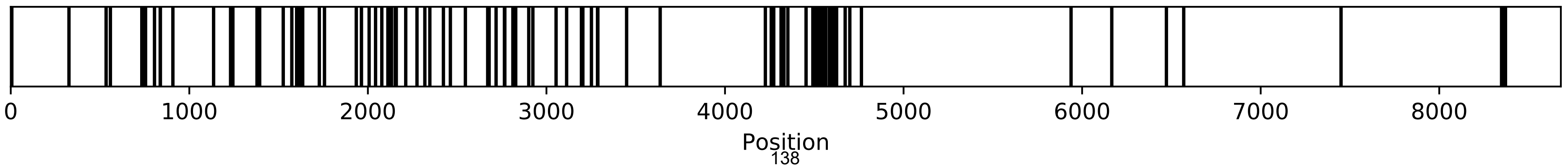

### Region: 414



CR1 vs CR2
Number of differences: 237

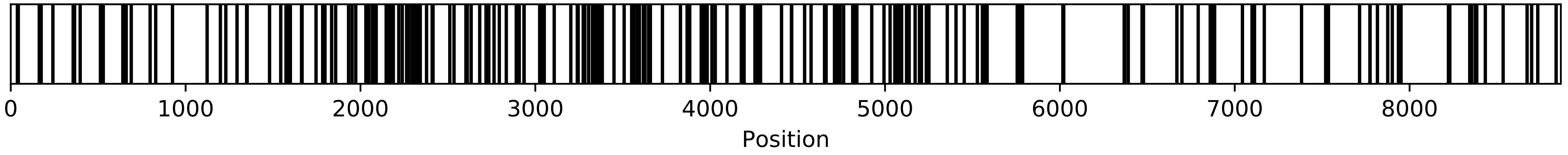


CR1 vs MM1
Number of differences: 128

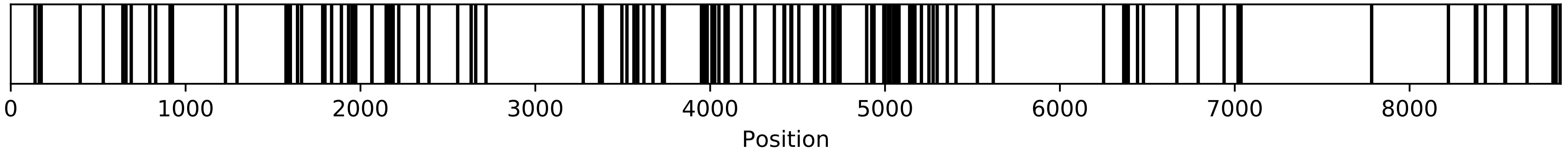


CR1 vs MM2
Number of differences: 109

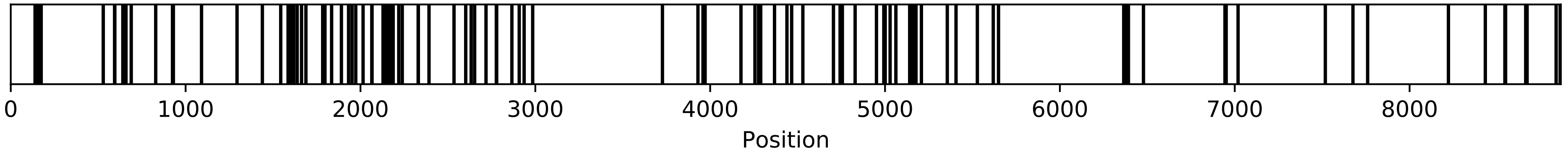


CR2 vs MM1
Number of differences: 235

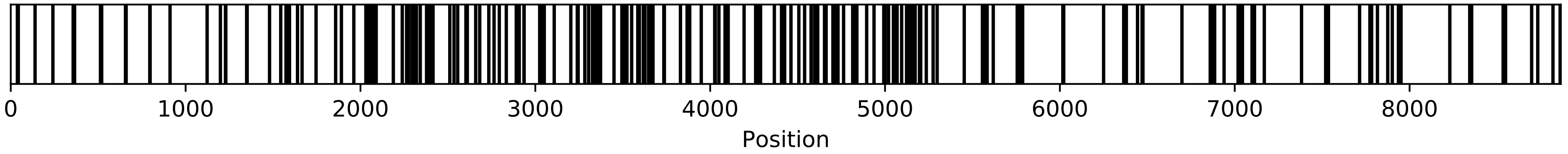


CR2 vs MM2
Number of differences: 245

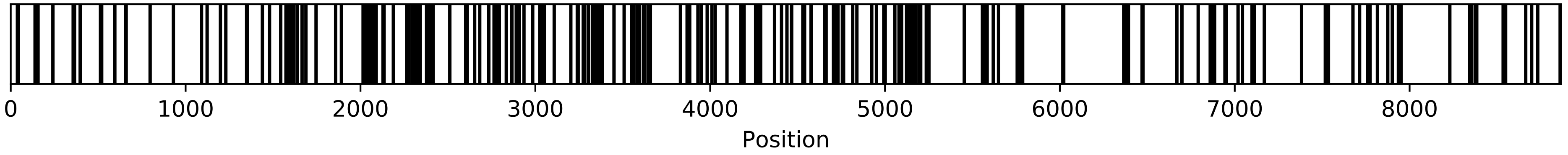


MM1 vs MM2
Number of differences: 110

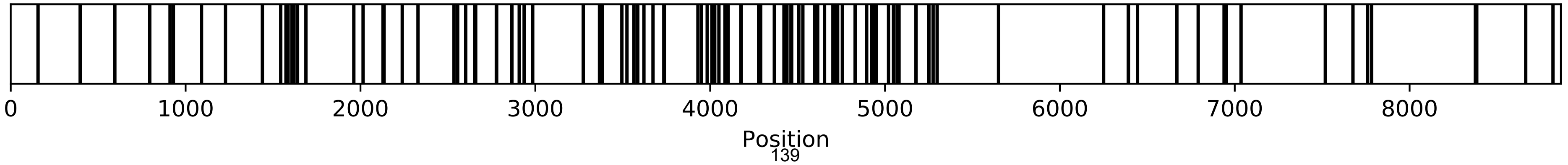

### Region: 418



CR1 vs CR2
Number of differences: 93

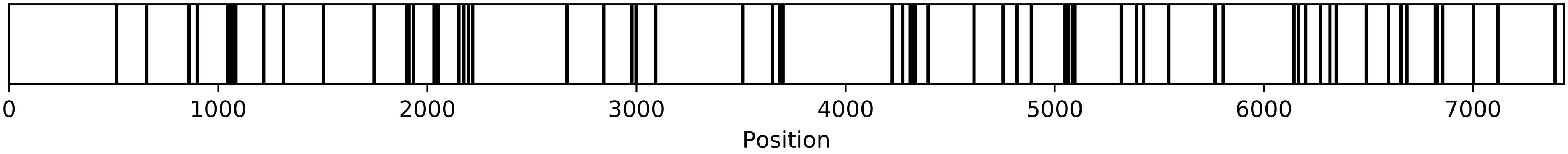


CR1 vs MM1
Number of differences: 36

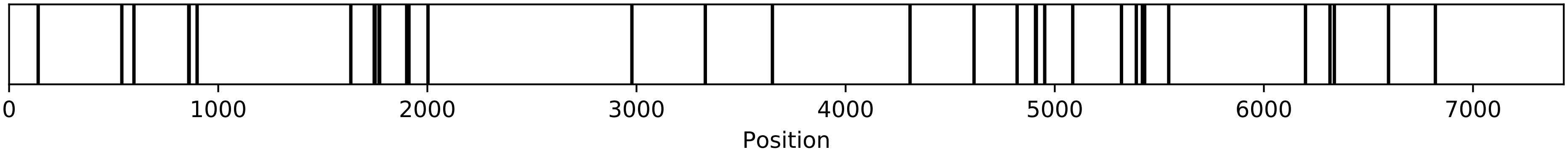


CR1 vs MM2
Number of differences: 37

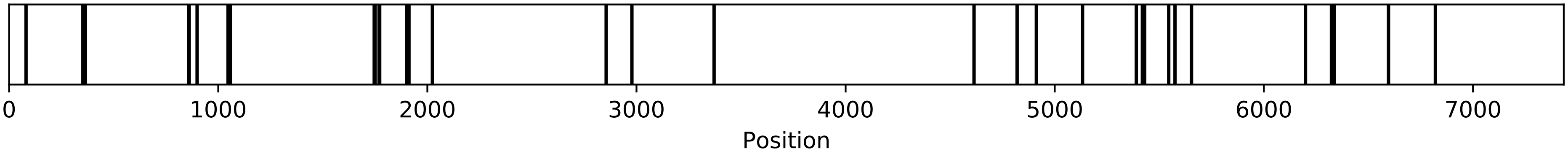


CR2 vs MM1
Number of differences: 95

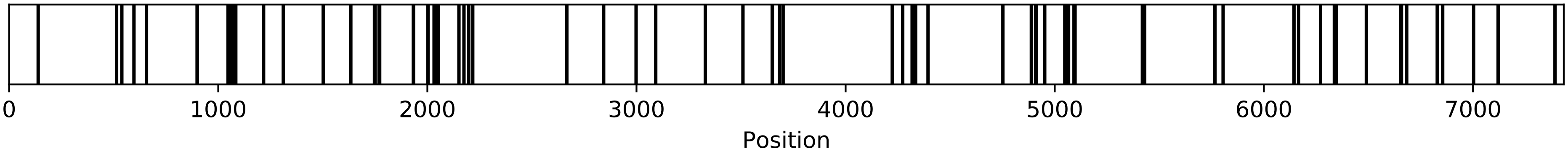


CR2 vs MM2
Number of differences: 98

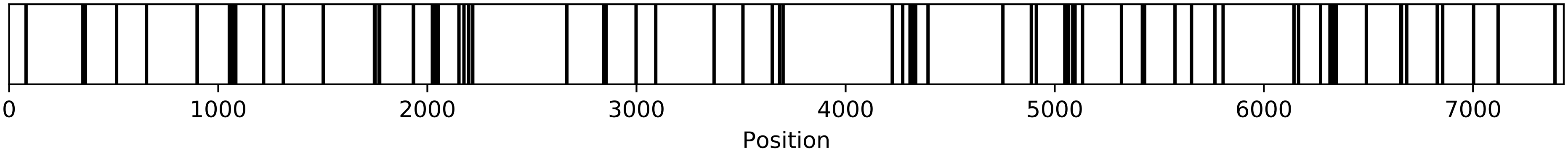


MM1 vs MM2
Number of differences: 28

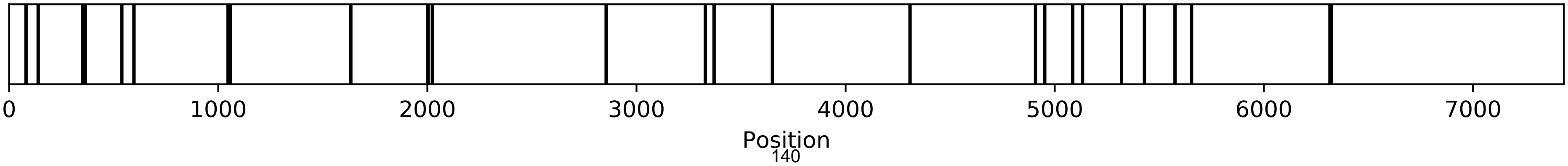

### Region: 419



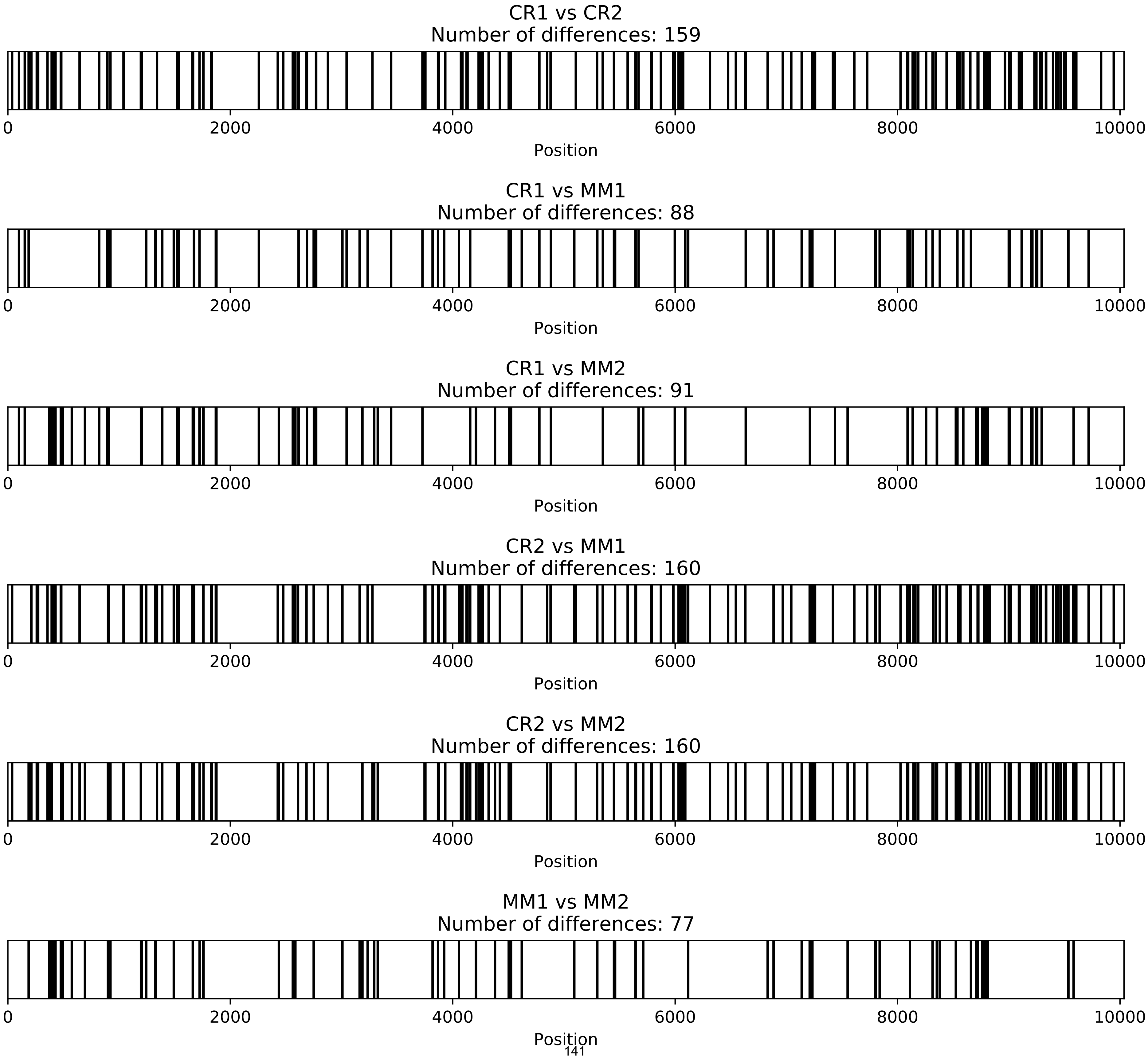
CR1 vs CR2
Number of differences: 159
0
2000
4000
6000
8000
10000
Position
CR1 vs MM1
Number of differences: 88
Position
CR1 vs MM2
Number of differences: 91
Position
CR2 vs MM1
Number of differences: 160
Position
CR2 vs MM2
Number of differences: 160
Position
MM1 vs MM2
Number of differences: 77
Position

### Region: 420



## CR1 vs CR2
Number of differences: 385

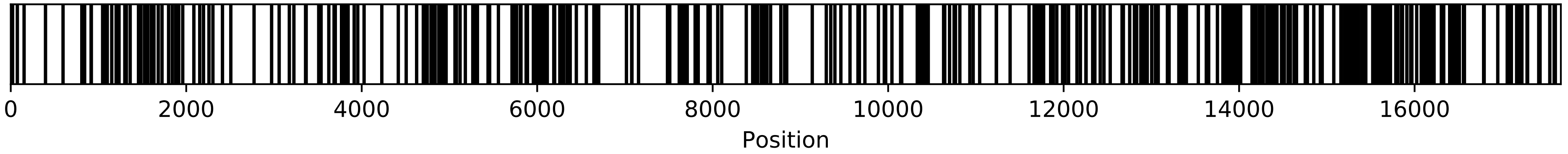


## CR1 vs MM1
Number of differences: 134

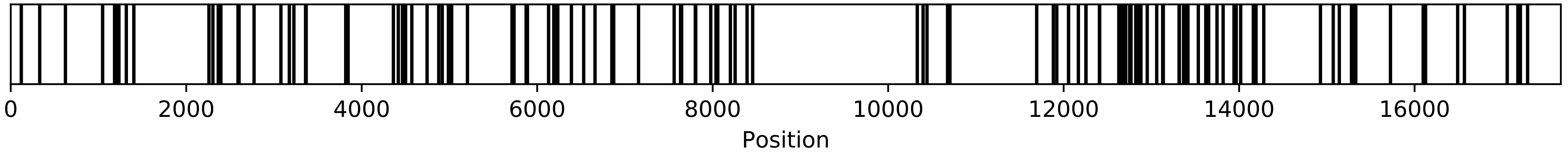


## CR1 vs MM2
Number of differences: 152

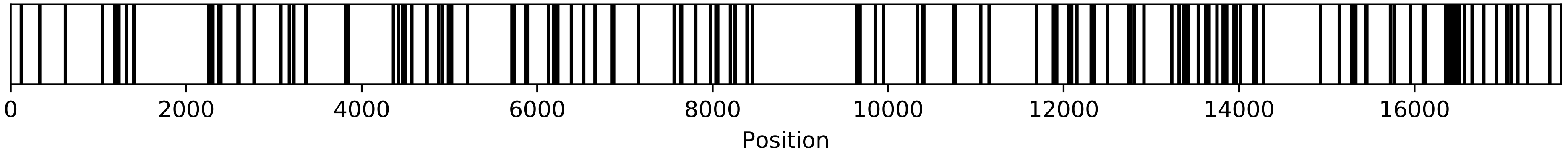


## CR2 vs MM1
Number of differences: 393

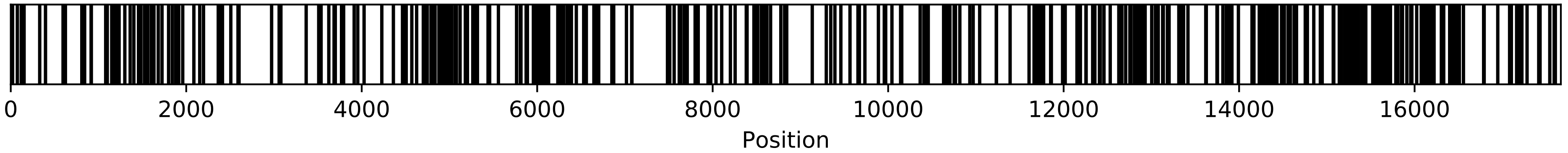


## CR2 vs MM2
Number of differences: 409

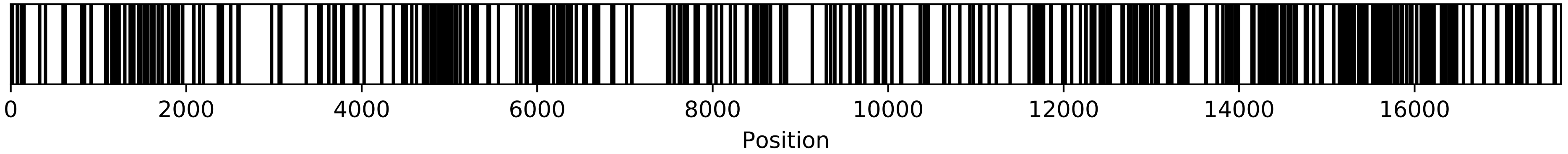


## MM1 vs MM2
Number of differences: 73

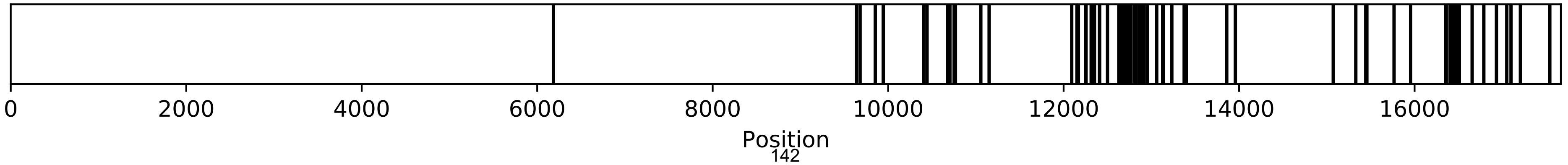

### Region: 423



CR1 vs CR2
Number of differences: 292

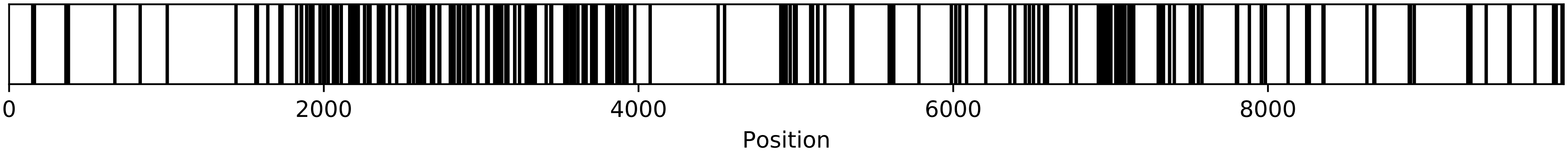


CR1 vs MM1
Number of differences: 288

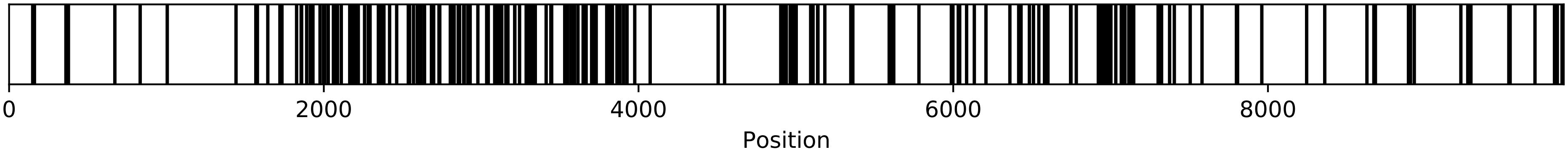


CR1 vs MM2
Number of differences: 289

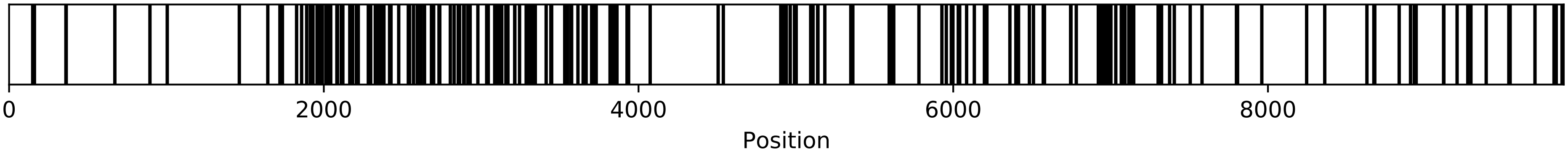


CR2 vs MM1
Number of differences: 52

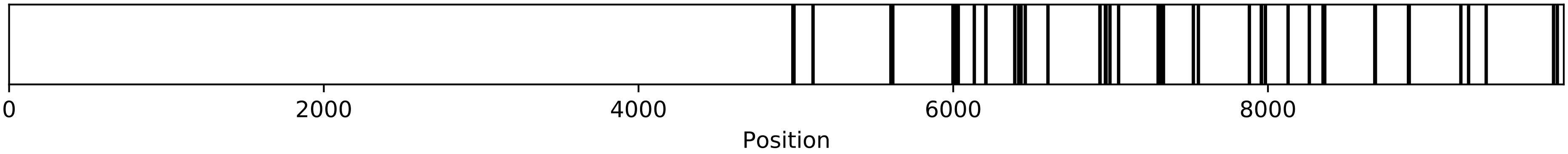


CR2 vs MM2
Number of differences: 123

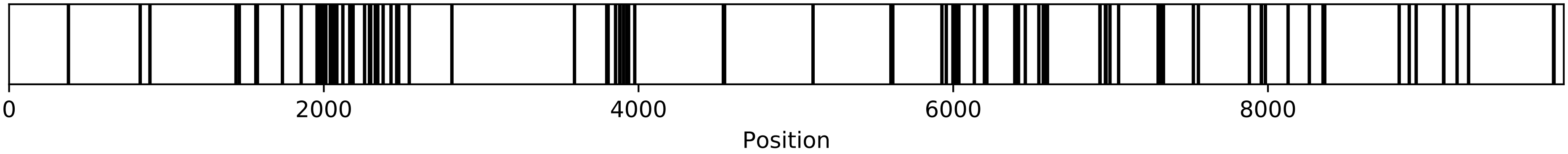


MM1 vs MM2
Number of differences: 97

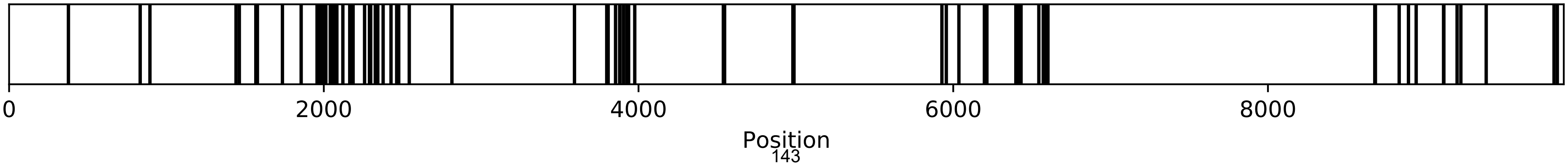

### Region: 427



CR1 vs CR2
Number of differences: 253

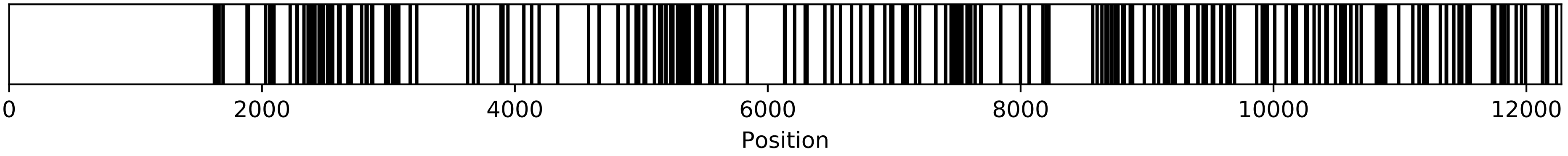


CR1 vs MM1
Number of differences: 165

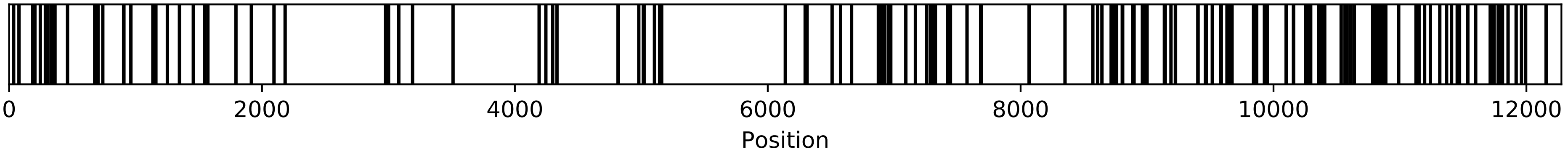


CR1 vs MM2
Number of differences: 167

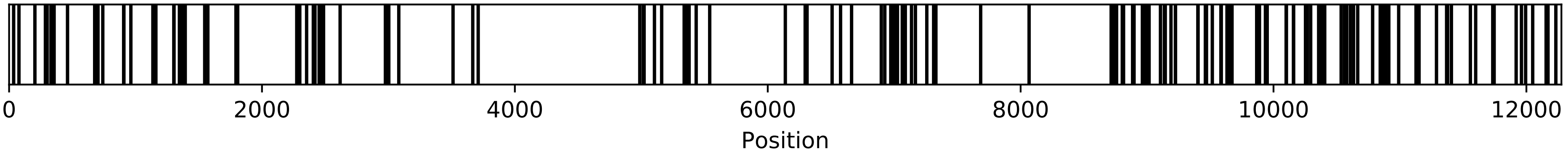


CR2 vs MM1
Number of differences: 284

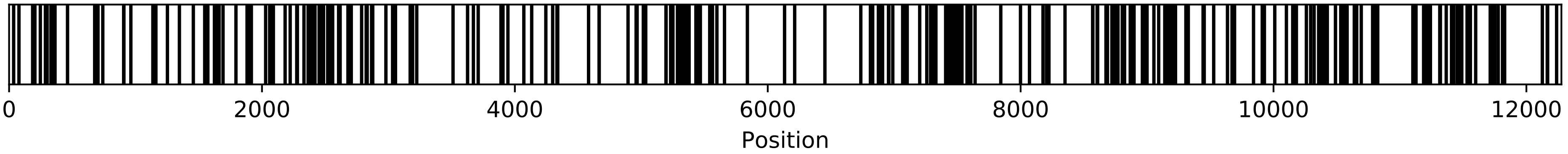


CR2 vs MM2
Number of differences: 279

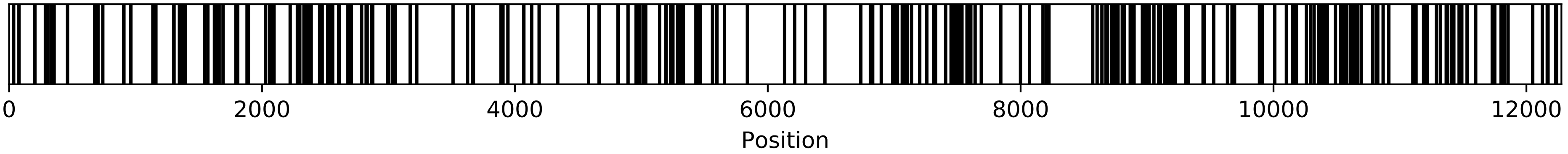


MM1 vs MM2
Number of differences: 108

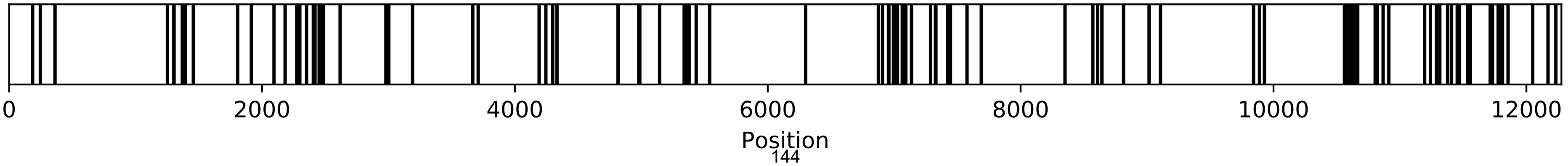

### Region: 430



CR1 vs CR2
Number of differences: 236

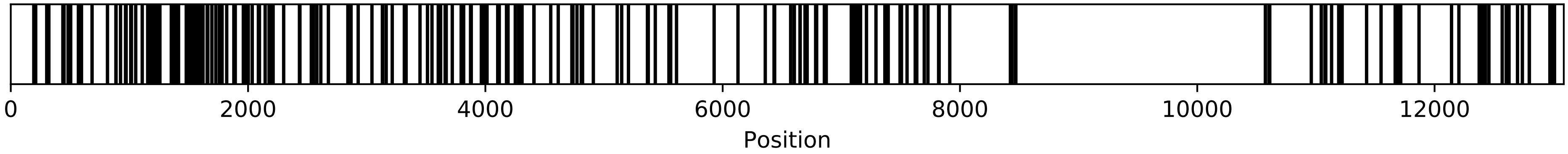


CR1 vs MM1
Number of differences: 165

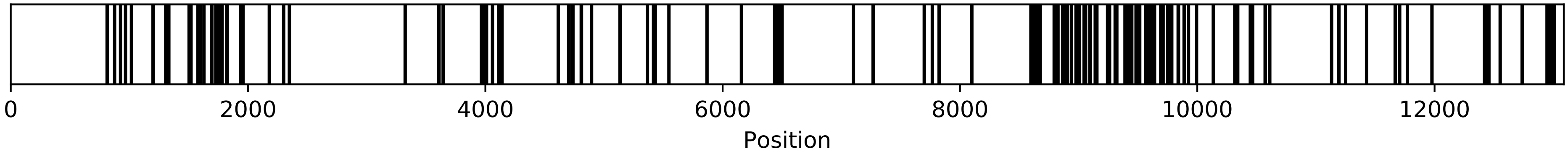


CR1 vs MM2
Number of differences: 196

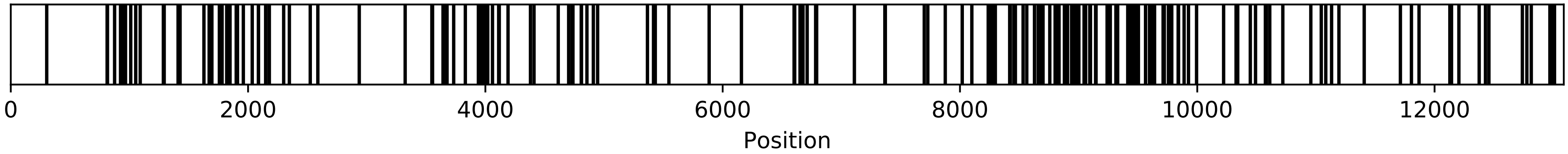


CR2 vs MM1
Number of differences: 318

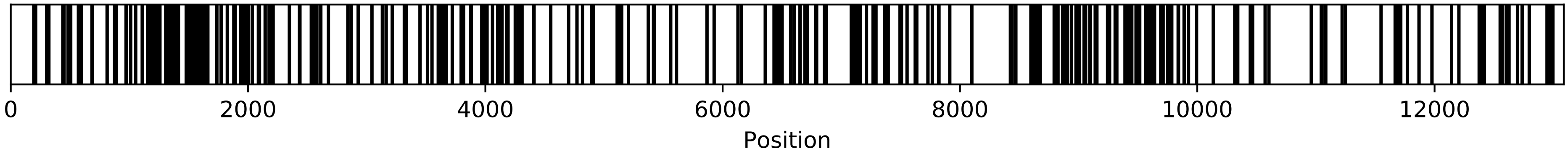


CR2 vs MM2
Number of differences: 307

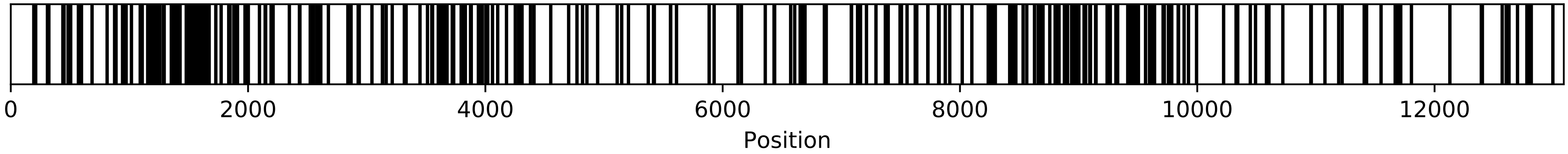


MM1 vs MM2
Number of differences: 190

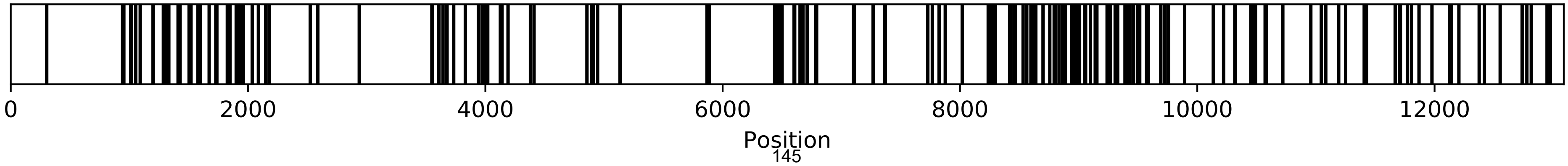

### Region: 431



CR1 vs CR2
Number of differences: 229

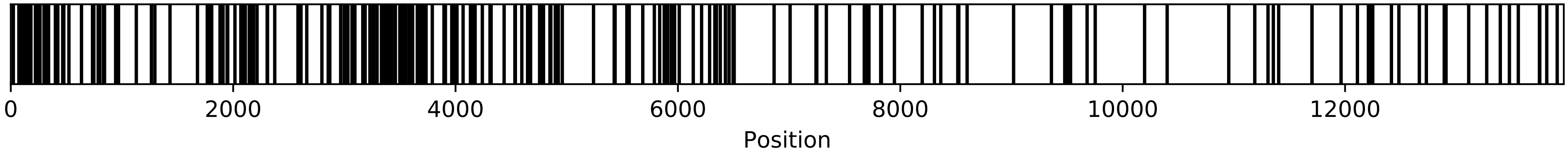


CR1 vs MM1
Number of differences: 214

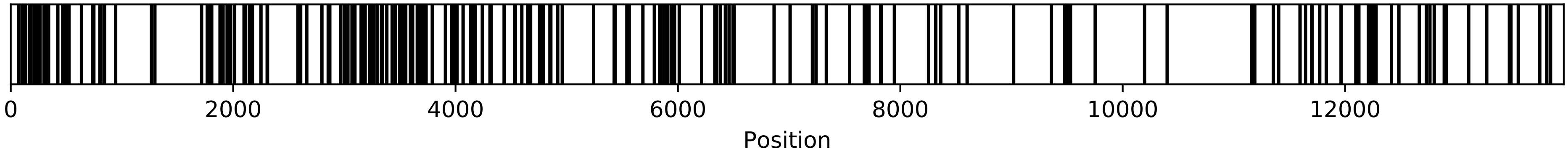


CR1 vs MM2
Number of differences: 225

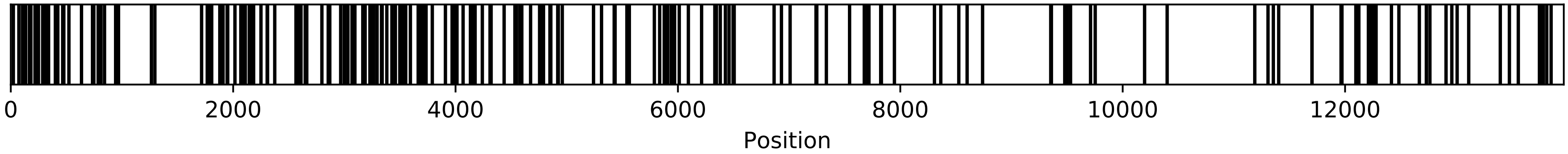


CR2 vs MM1
Number of differences: 84

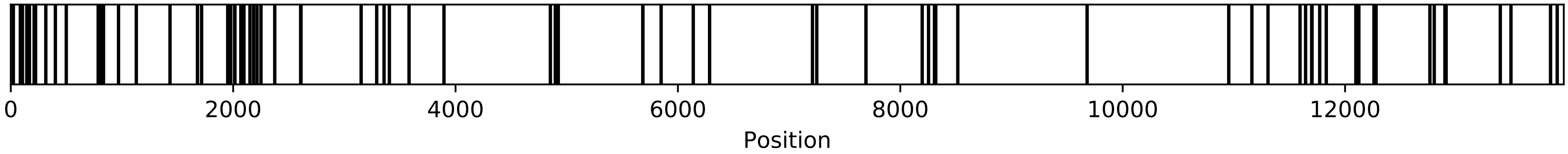


CR2 vs MM2
Number of differences: 80

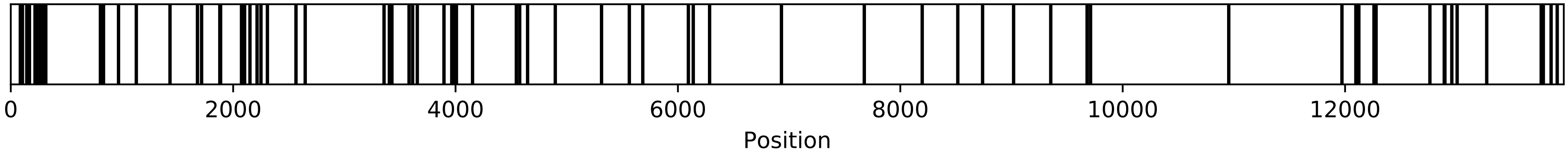


MM1 vs MM2
Number of differences: 83

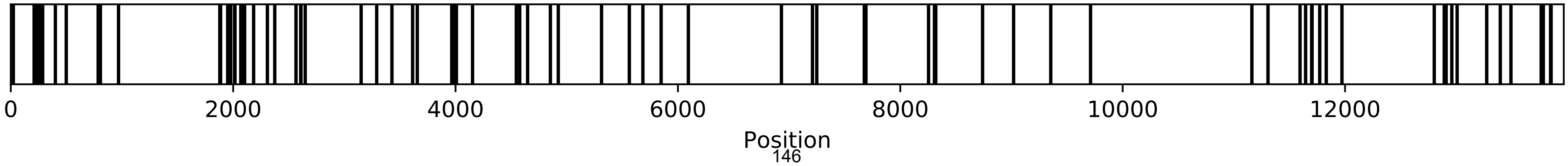

### Region: 442



CR1 vs CR2
Number of differences: 290

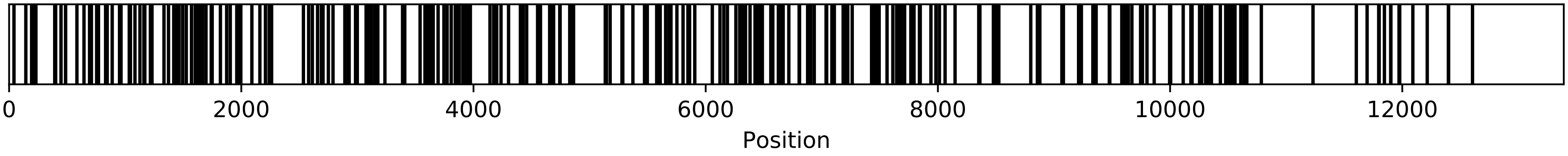


CR1 vs MM1
Number of differences: 136

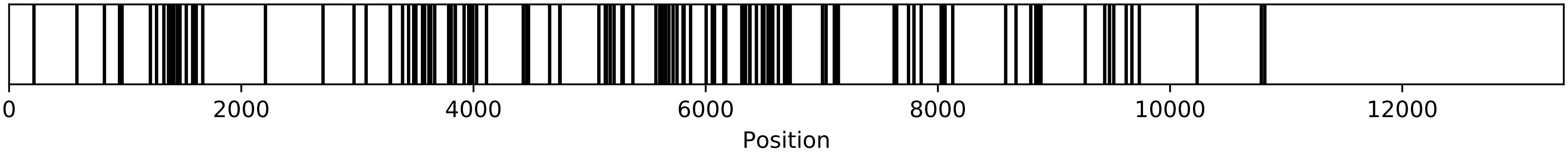


CR1 vs MM2
Number of differences: 155

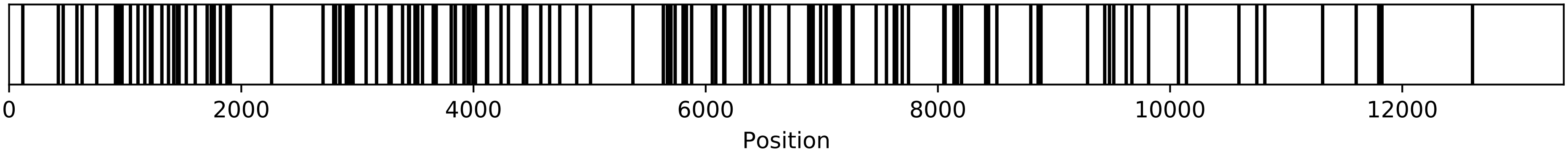


CR2 vs MM1
Number of differences: 307

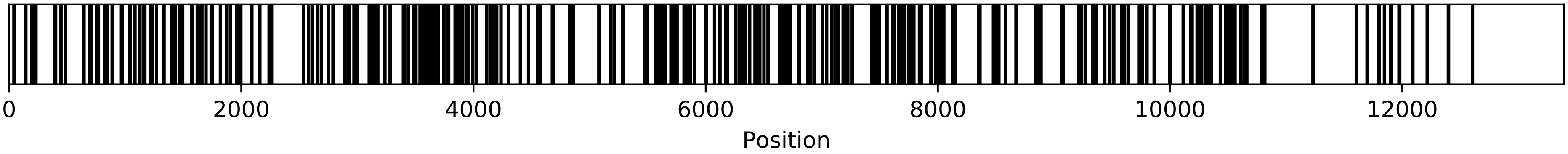


CR2 vs MM2
Number of differences: 316

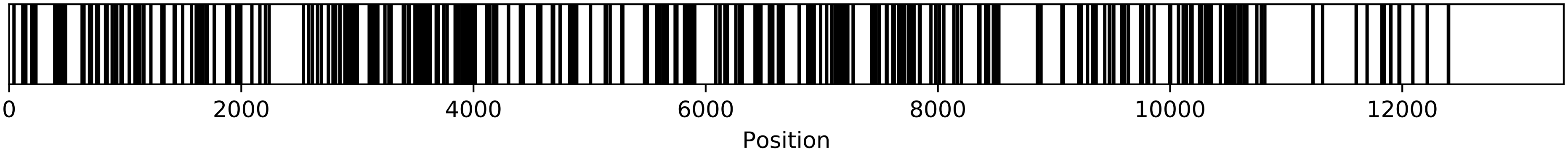


MM1 vs MM2
Number of differences: 174

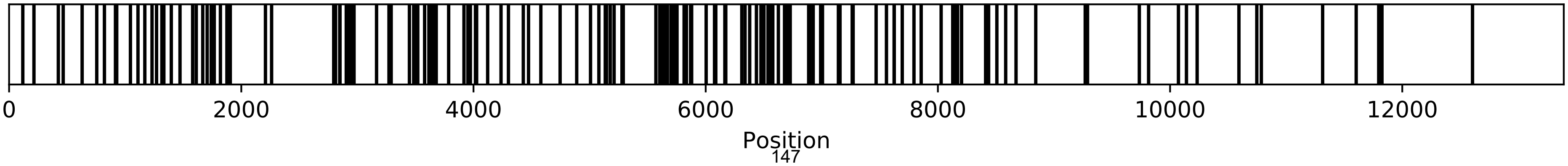

### Region: 443



CR1 vs CR2
Number of differences: 51

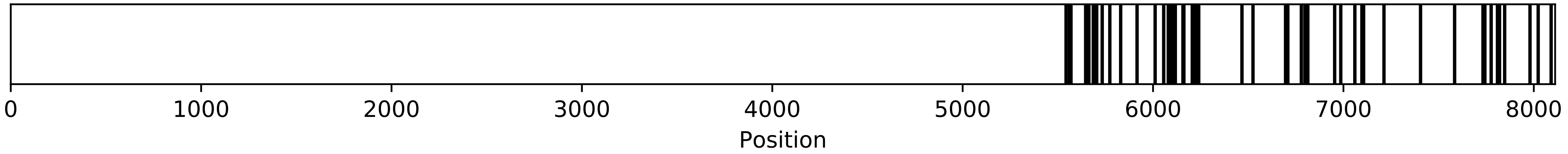


CR1 vs MM1
Number of differences: 253

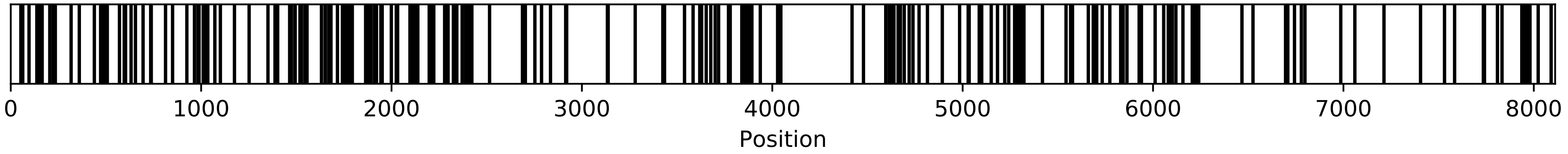


CR1 vs MM2
Number of differences: 241

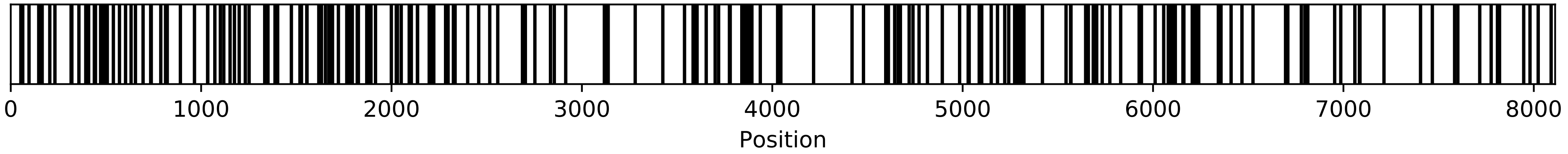


CR2 vs MM1
Number of differences: 232

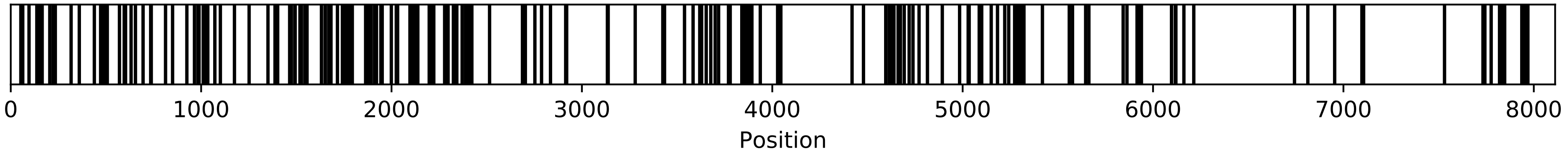


CR2 vs MM2
Number of differences: 206

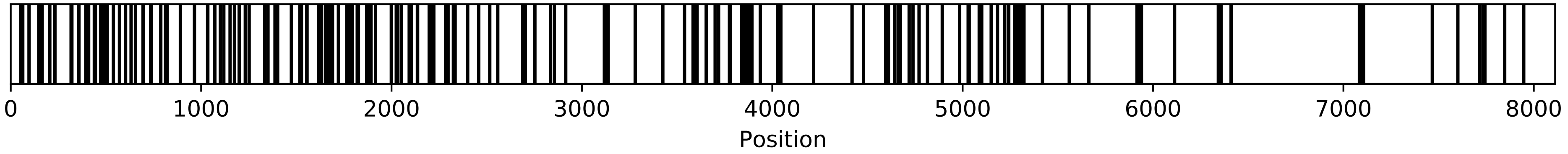


MM1 vs MM2
Number of differences: 133

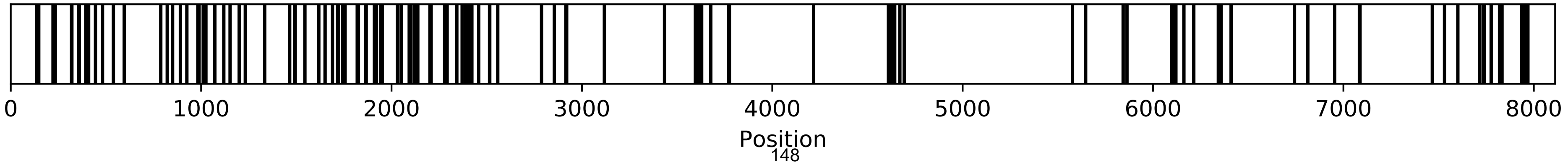

### Region: 448



CR1 vs CR2
Number of differences: 195

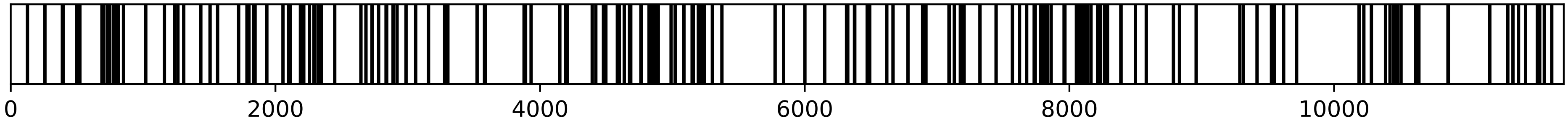

Position

CR1 vs MM1
Number of differences: 84

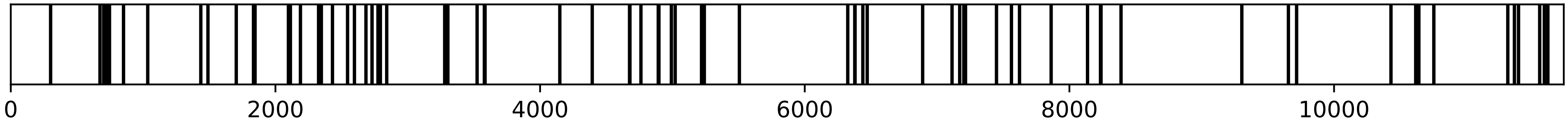

Position

CR1 vs MM2
Number of differences: 91

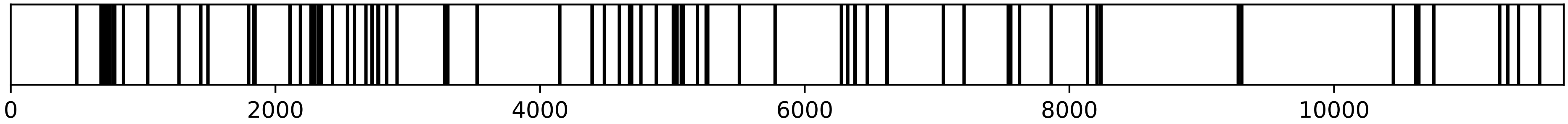

Position

CR2 vs MM1
Number of differences: 180

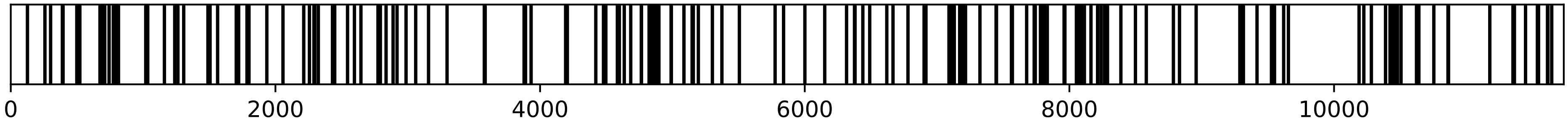

Position

CR2 vs MM2
Number of differences: 192

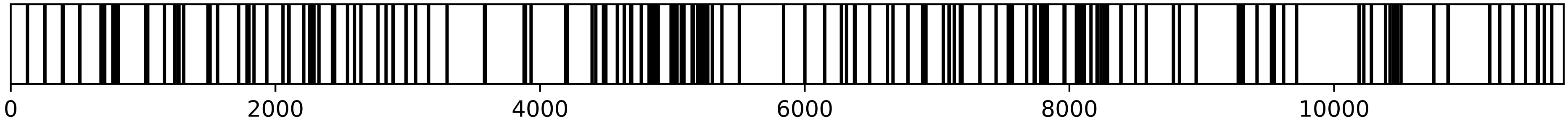

Position

MM1 vs MM2
Number of differences: 70

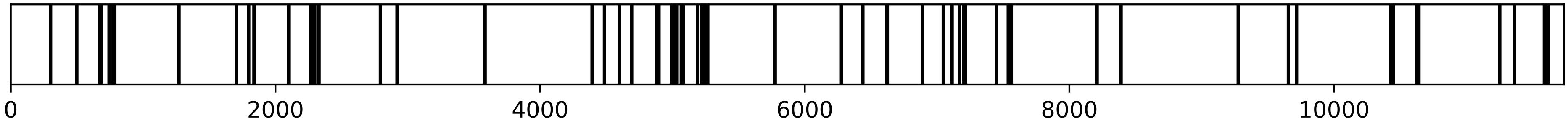

Position

### Region: 450



CR1 vs CR2
Number of differences: 111

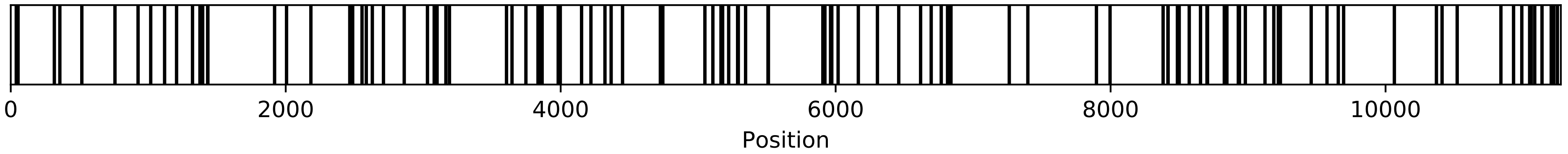


CR1 vs MM1
Number of differences: 42

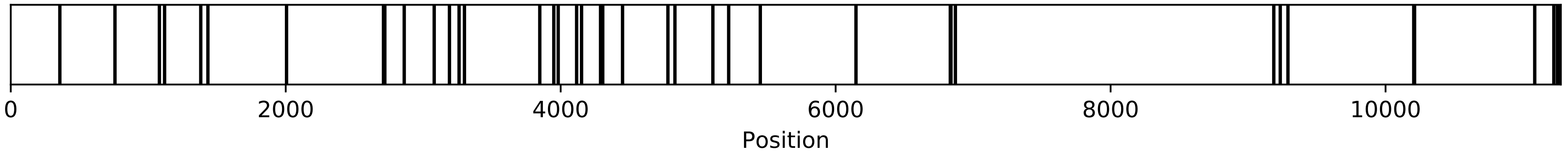


CR1 vs MM2
Number of differences: 48

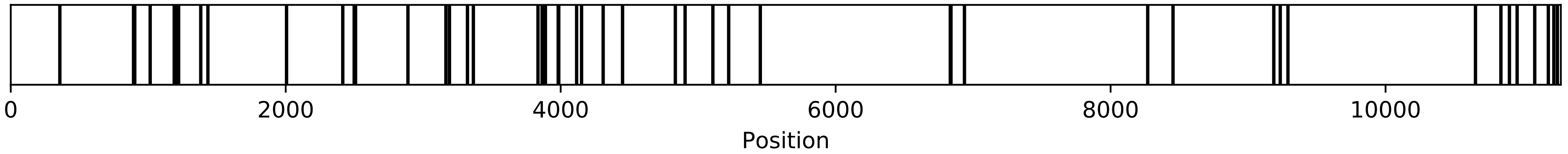


CR2 vs MM1
Number of differences: 113

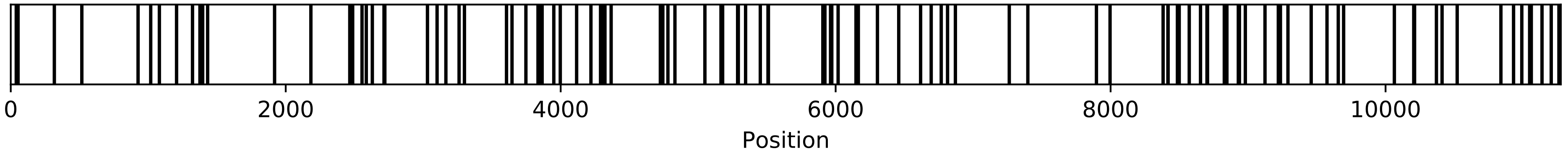


CR2 vs MM2
Number of differences: 119

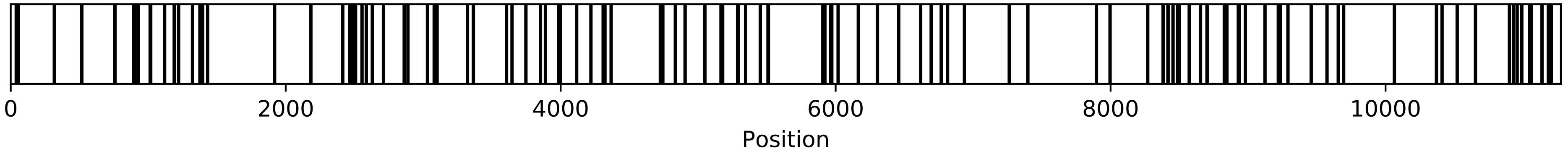


MM1 vs MM2
Number of differences: 50

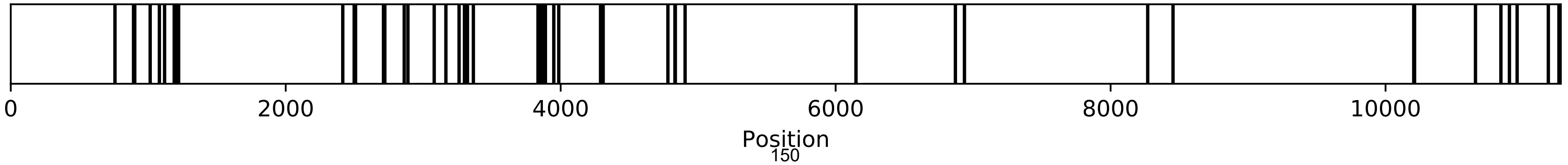

### Region: 452



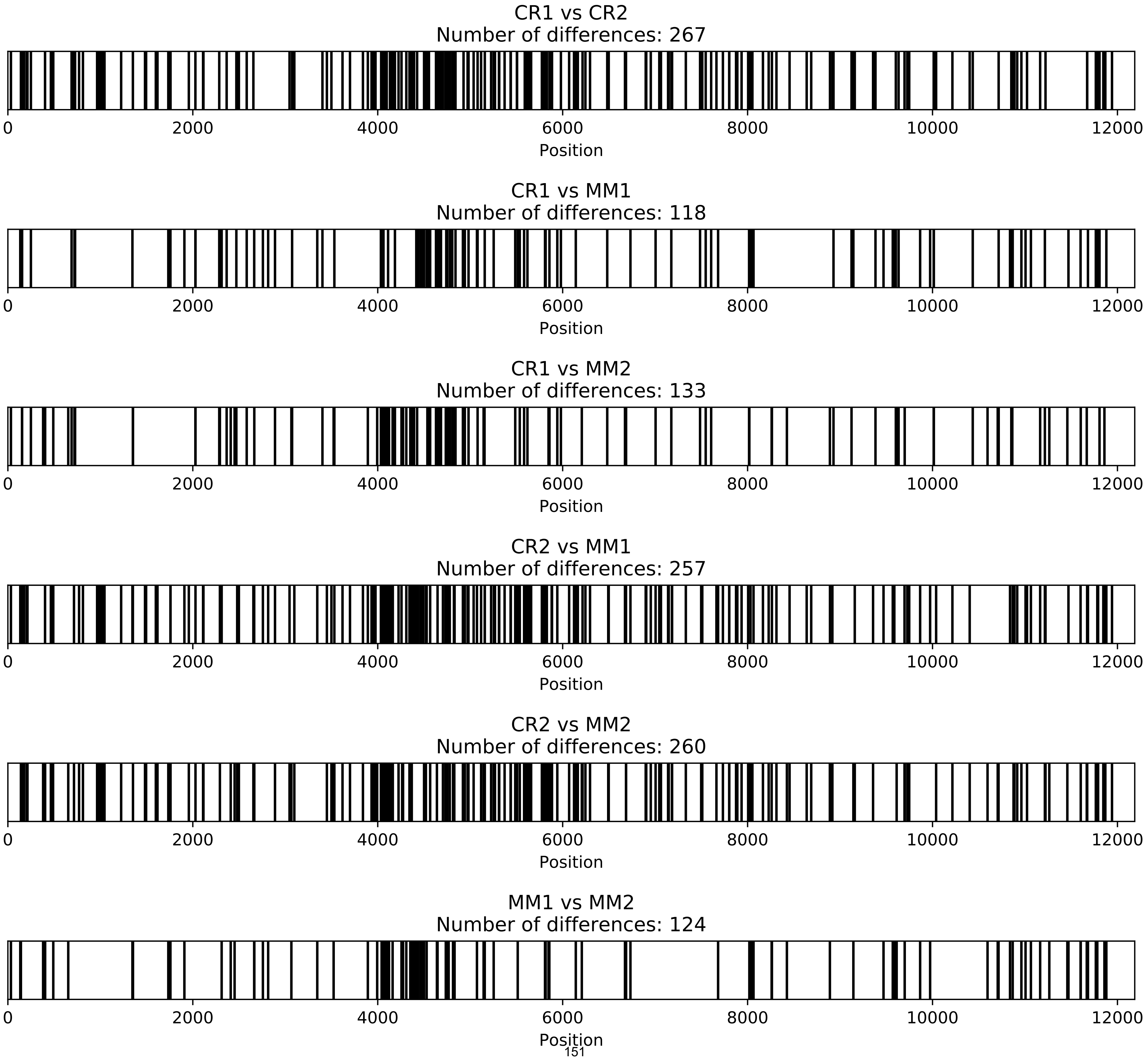
CR1 vs CR2
Number of differences: 267
CR1 vs MM1
Number of differences: 118
CR1 vs MM2
Number of differences: 133
CR2 vs MM1
Number of differences: 257
CR2 vs MM2
Number of differences: 260
MM1 vs MM2
Number of differences: 124
0 2000 4000 6000 8000 10000 12000
Position

### Region: 454



CR1 vs CR2
Number of differences: 158

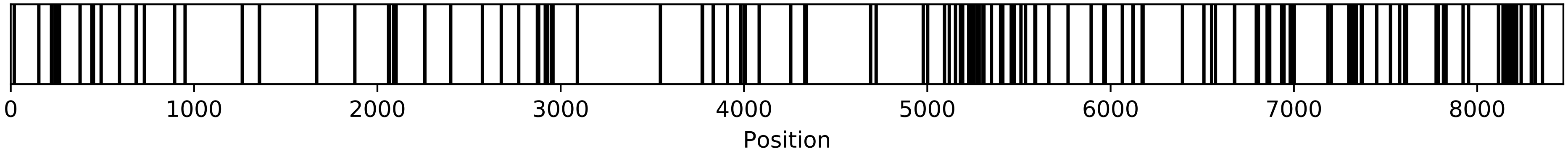


CR1 vs MM1
Number of differences: 59

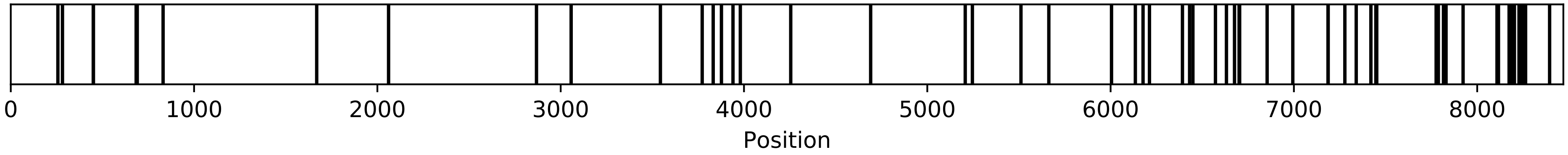


CR1 vs MM2
Number of differences: 56

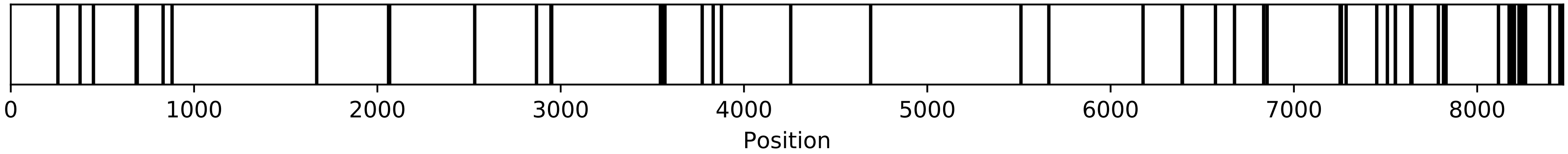


CR2 vs MM1
Number of differences: 154

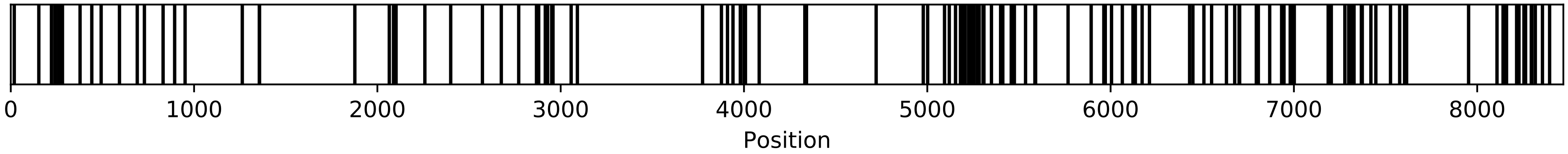


CR2 vs MM2
Number of differences: 158

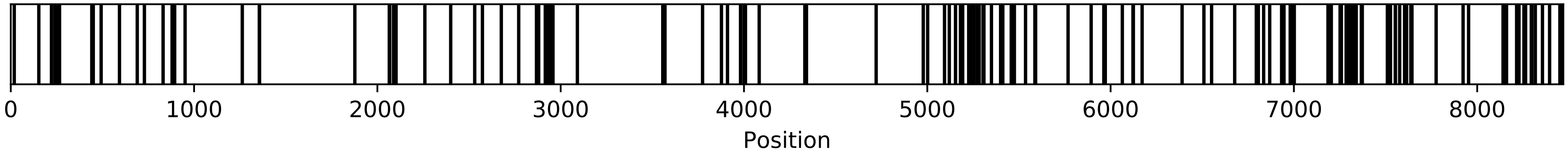


MM1 vs MM2
Number of differences: 47

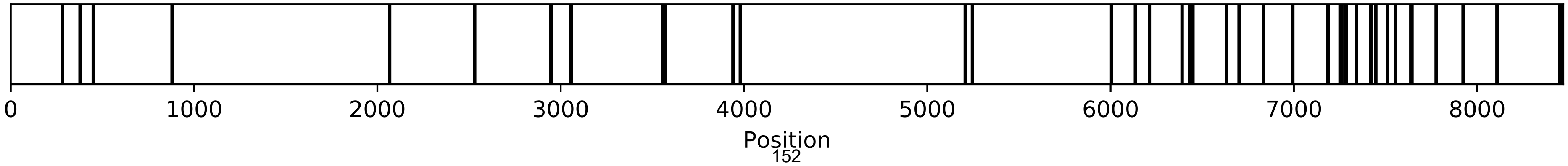

### Region: 455



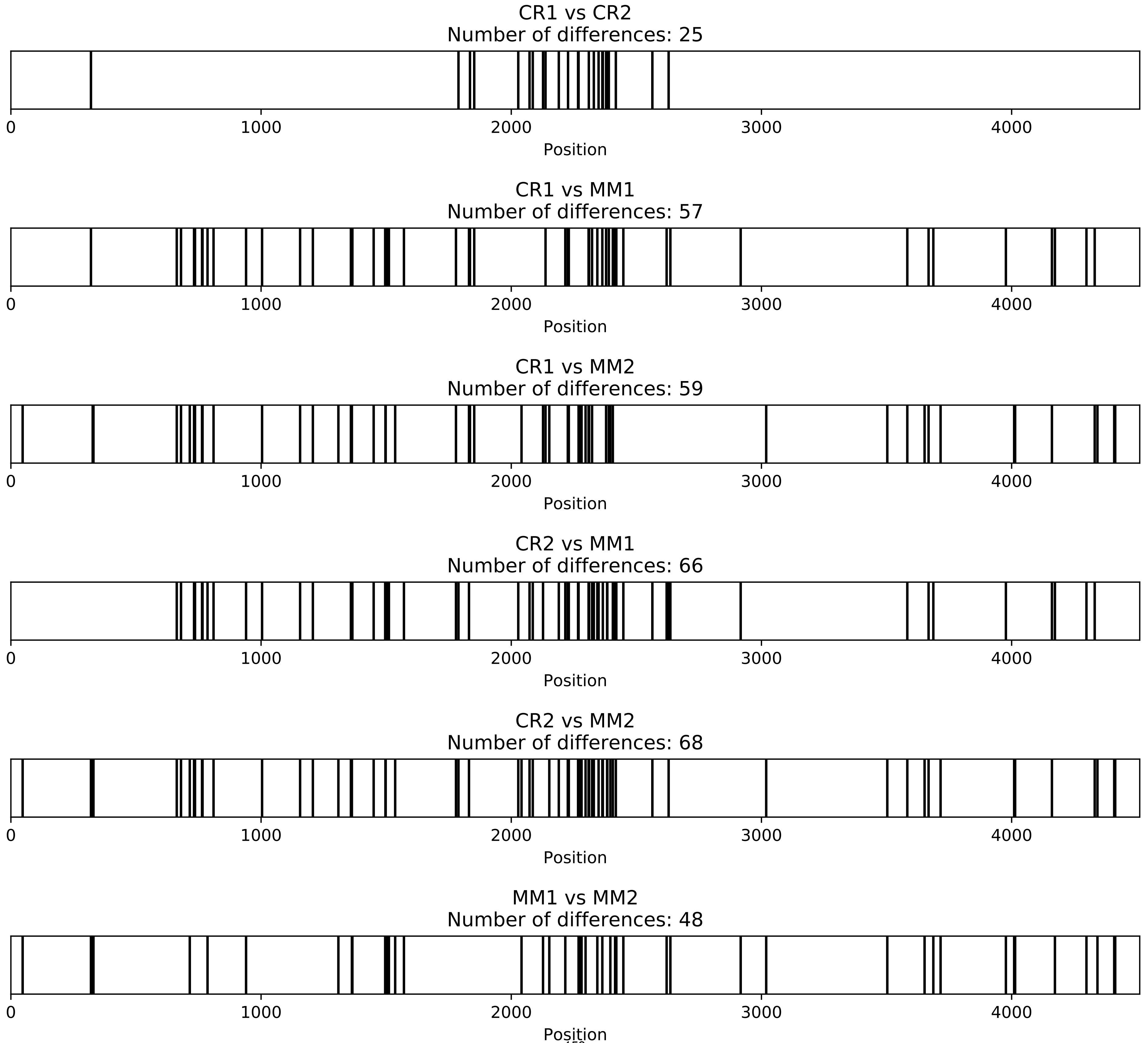
CR1 vs CR2
Number of differences: 25
0
1000
2000
3000
4000
Position
CR1 vs MM1
Number of differences: 57
0
1000
2000
3000
4000
Position
CR1 vs MM2
Number of differences: 59
0
1000
2000
3000
4000
Position
CR2 vs MM1
Number of differences: 66
0
1000
2000
3000
4000
Position
CR2 vs MM2
Number of differences: 68
0
1000
2000
3000
4000
Position
MM1 vs MM2
Number of differences: 48
0
1000
2000
3000
4000
Position

### Region: 460



CR1 vs CR2
Number of differences: 149

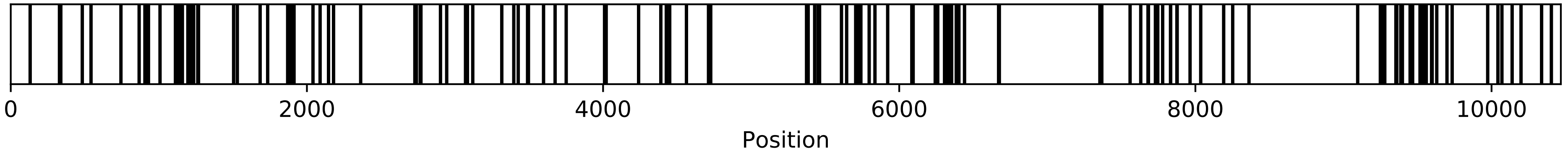


CR1 vs MM1
Number of differences: 30

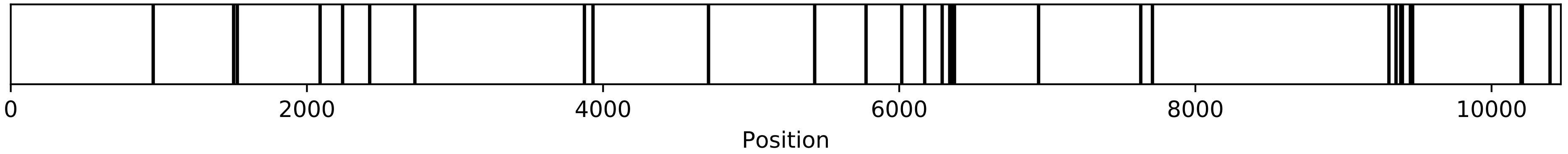


CR1 vs MM2
Number of differences: 15

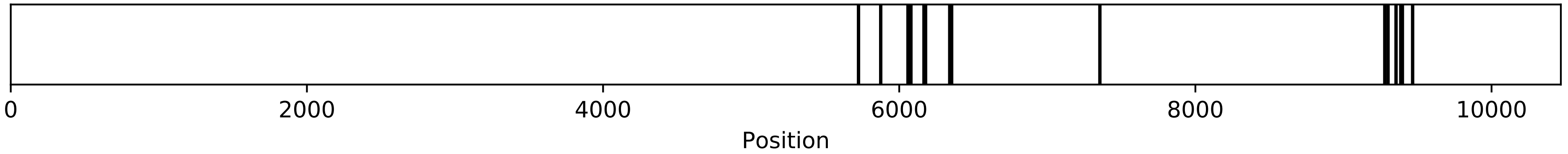


CR2 vs MM1
Number of differences: 153

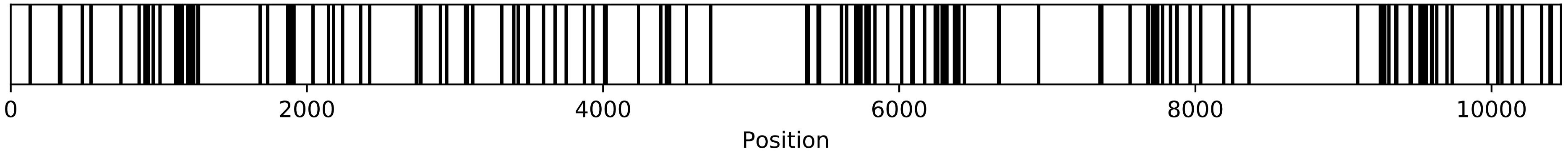


CR2 vs MM2
Number of differences: 149

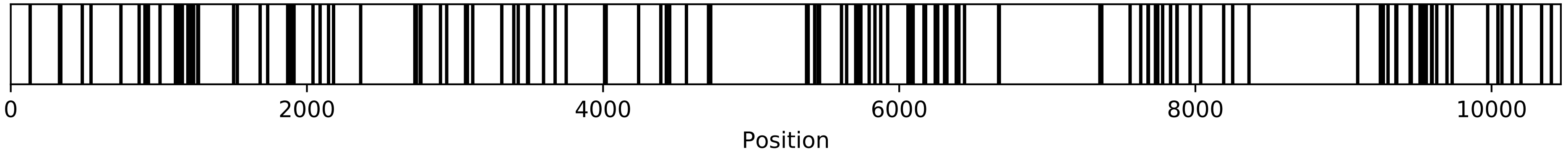


MM1 vs MM2
Number of differences: 33

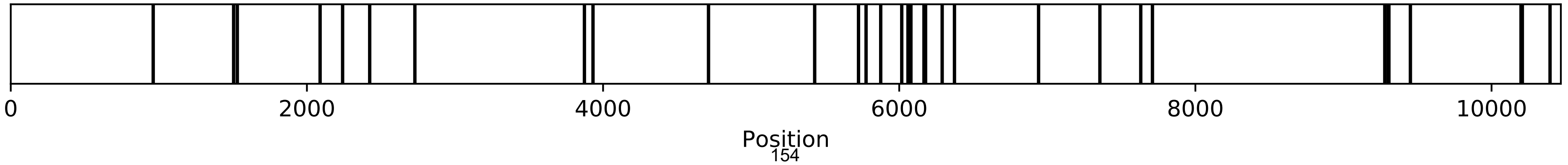

### Region: 462



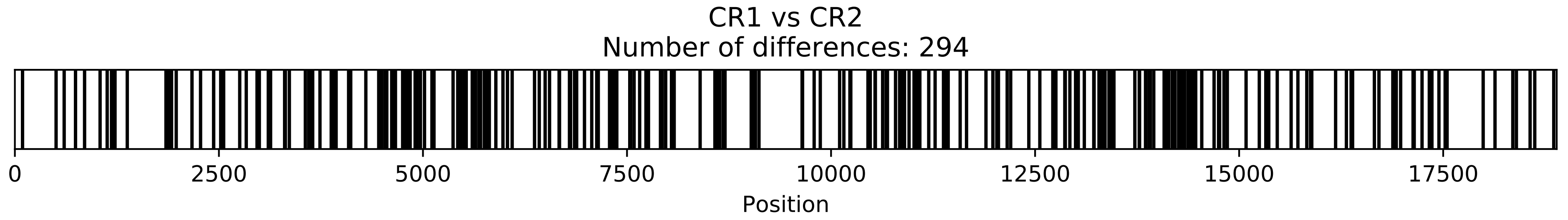
CR1 vs CR2
Number of differences: 294
0
2500
5000
7500
10000
12500
15000
17500
Position

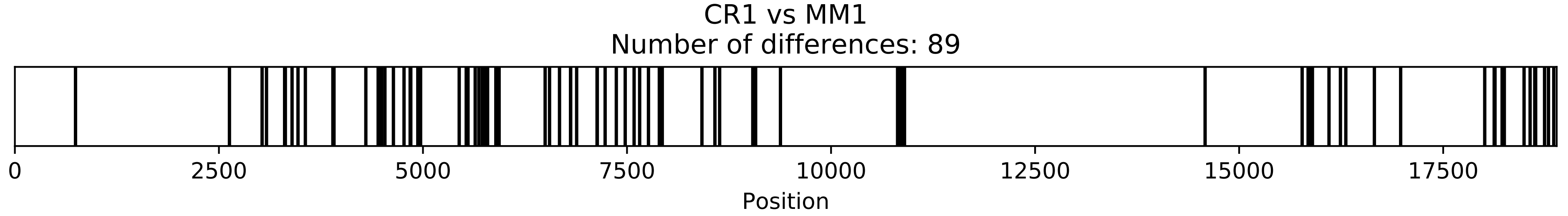
CR1 vs MM1
Number of differences: 89
0
2500
5000
7500
10000
12500
15000
17500
Position

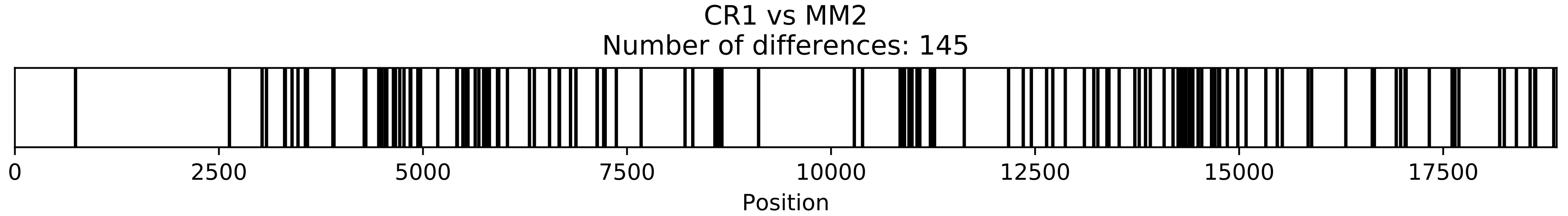
CR1 vs MM2
Number of differences: 145
0
2500
5000
7500
10000
12500
15000
17500
Position

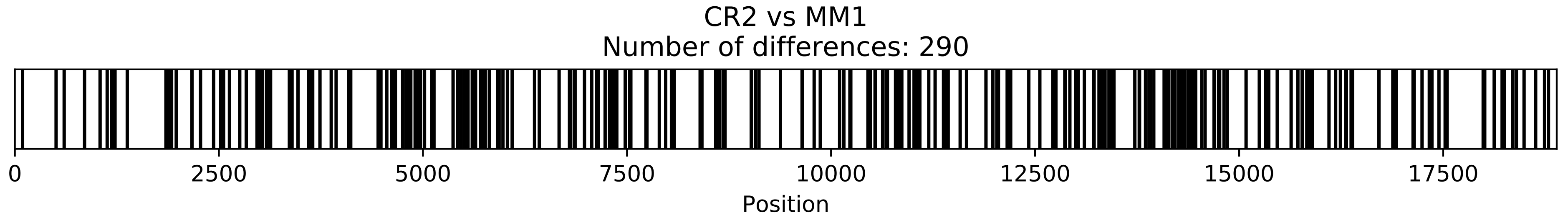
CR2 vs MM1
Number of differences: 290
0
2500
5000
7500
10000
12500
15000
17500
Position

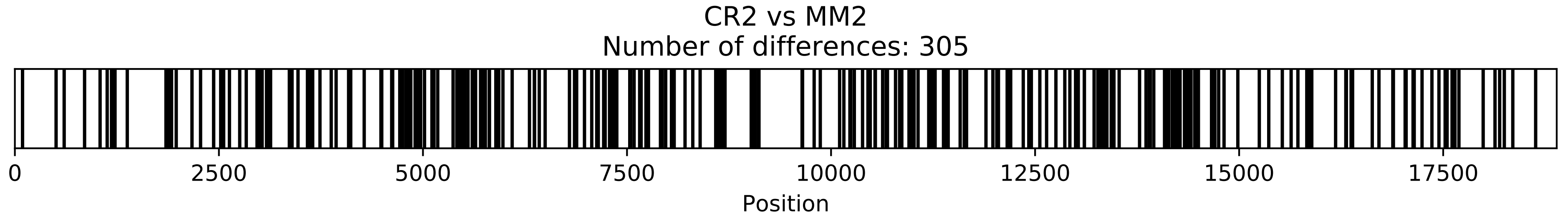
CR2 vs MM2
Number of differences: 305
0
2500
5000
7500
10000
12500
15000
17500
Position

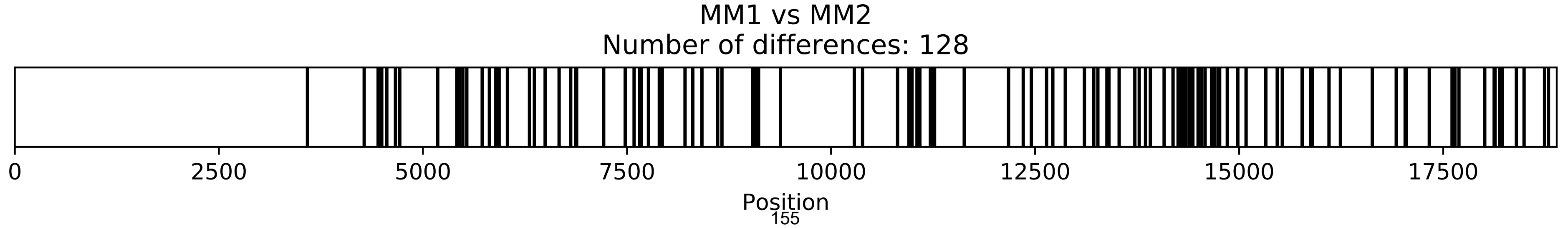
MM1 vs MM2
Number of differences: 128
0
2500
5000
7500
10000
12500
15000
17500
Position

### Region: 465



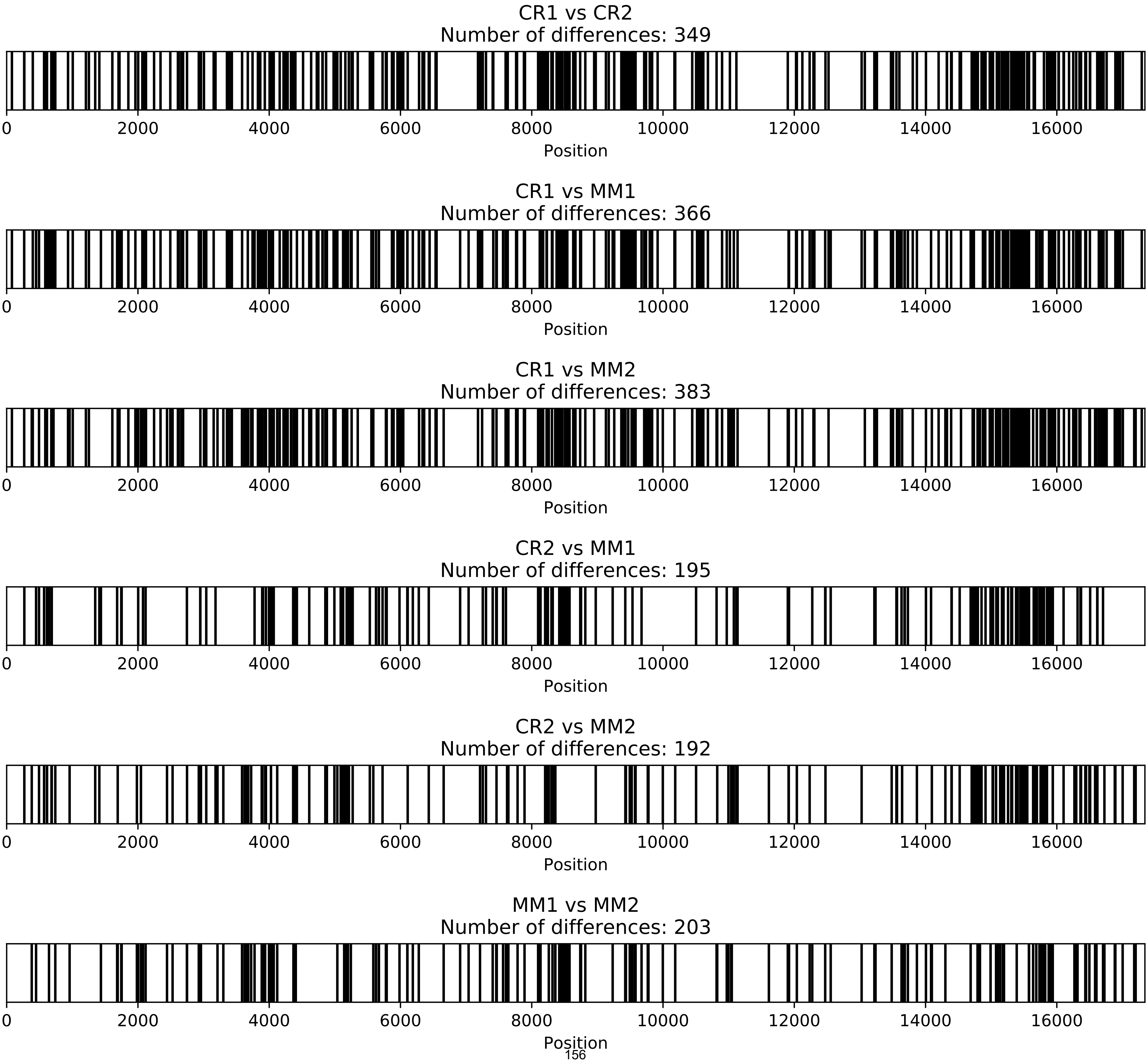
CR1 vs CR2
Number of differences: 349
Position
CR1 vs MM1
Number of differences: 366
Position
CR1 vs MM2
Number of differences: 383
Position
CR2 vs MM1
Number of differences: 195
Position
CR2 vs MM2
Number of differences: 192
Position
MM1 vs MM2
Number of differences: 203
0
2000
4000
6000
8000
10000
12000
14000
16000
Position

### Region: 467



## CR1 vs CR2
Number of differences: 334

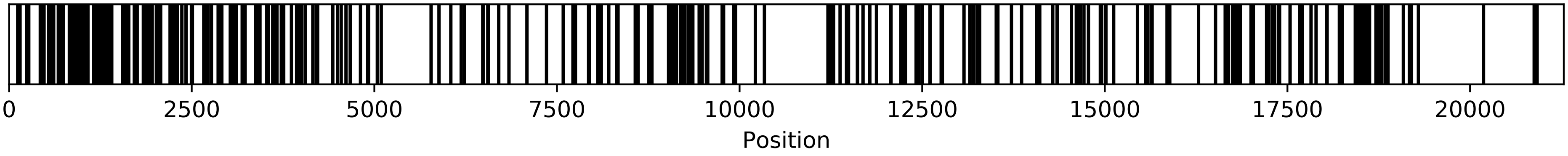


## CR1 vs MM1
Number of differences: 163

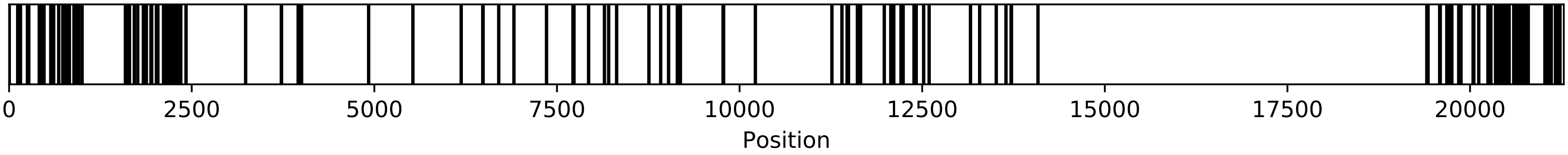


## CR1 vs MM2
Number of differences: 193

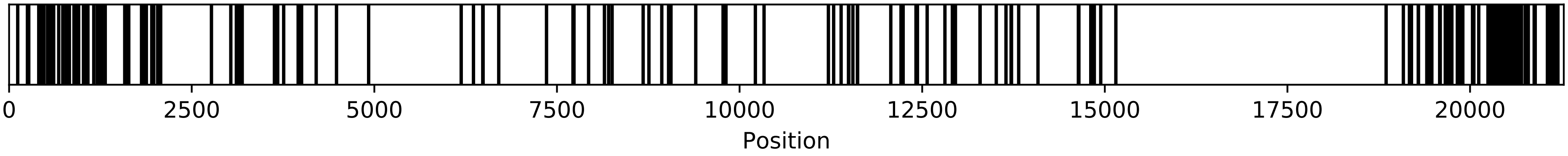


## CR2 vs MM1
Number of differences: 374

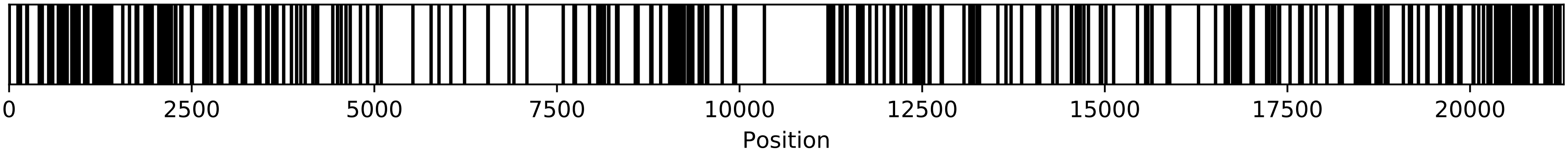


## CR2 vs MM2
Number of differences: 382

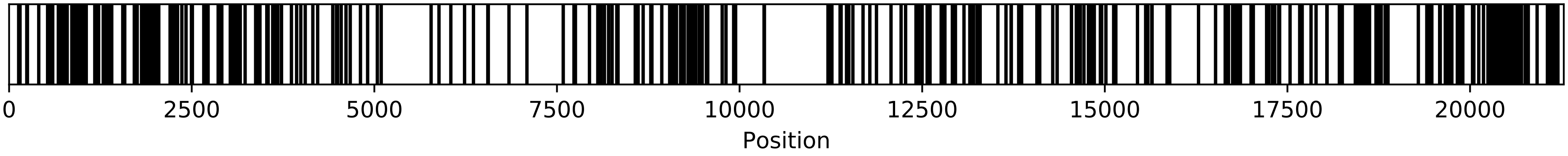


## MM1 vs MM2
Number of differences: 161

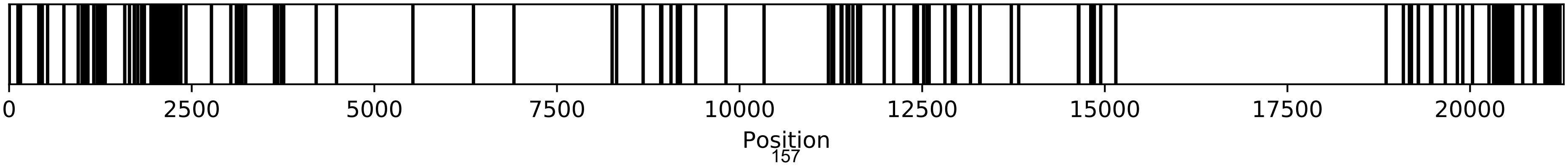

### Region: 471



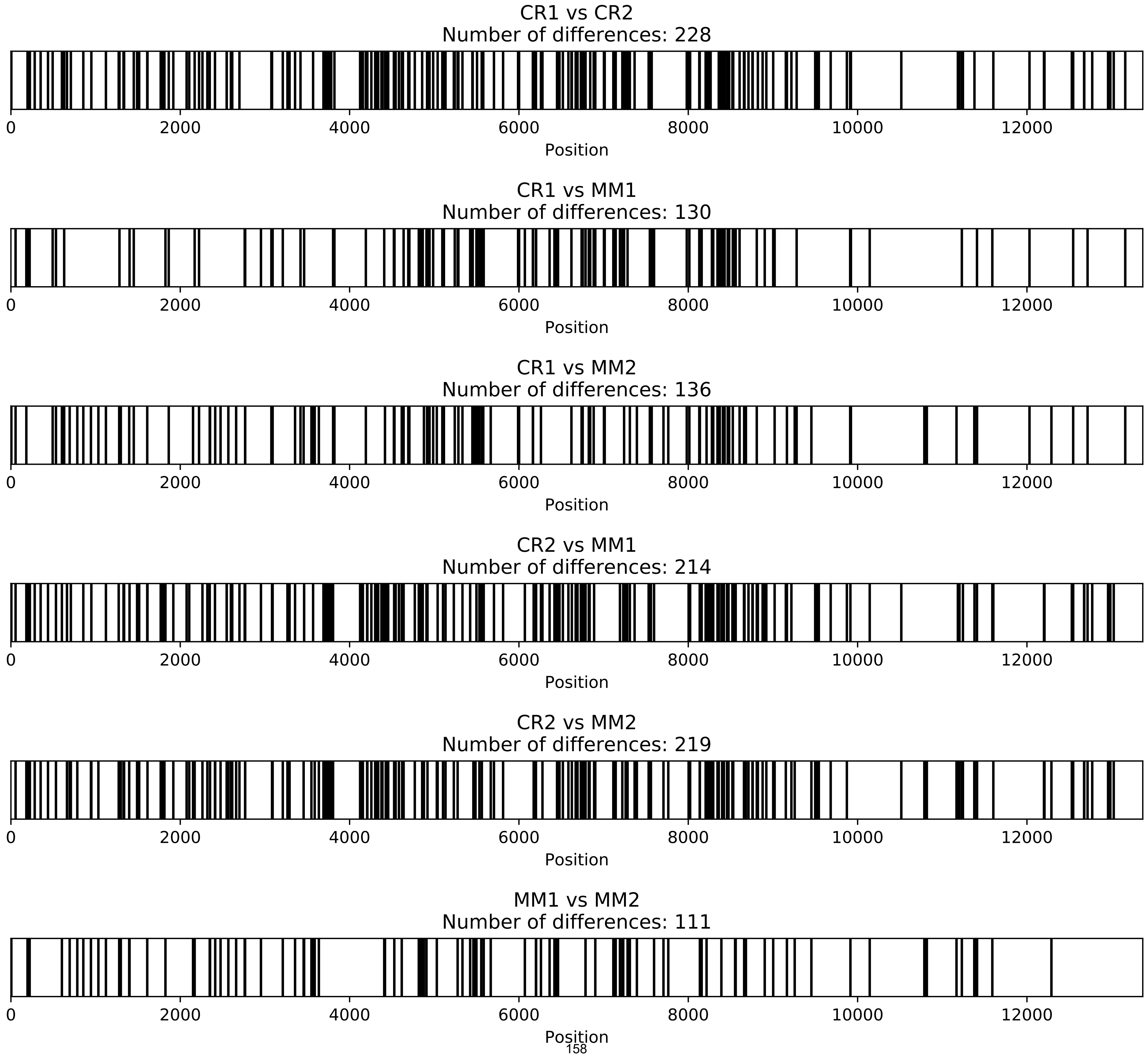
CR1 vs CR2
Number of differences: 228
CR1 vs MM1
Number of differences: 130
CR1 vs MM2
Number of differences: 136
CR2 vs MM1
Number of differences: 214
CR2 vs MM2
Number of differences: 219
MM1 vs MM2
Number of differences: 111
0
2000
4000
6000
8000
10000
12000
Position

### Region: 473



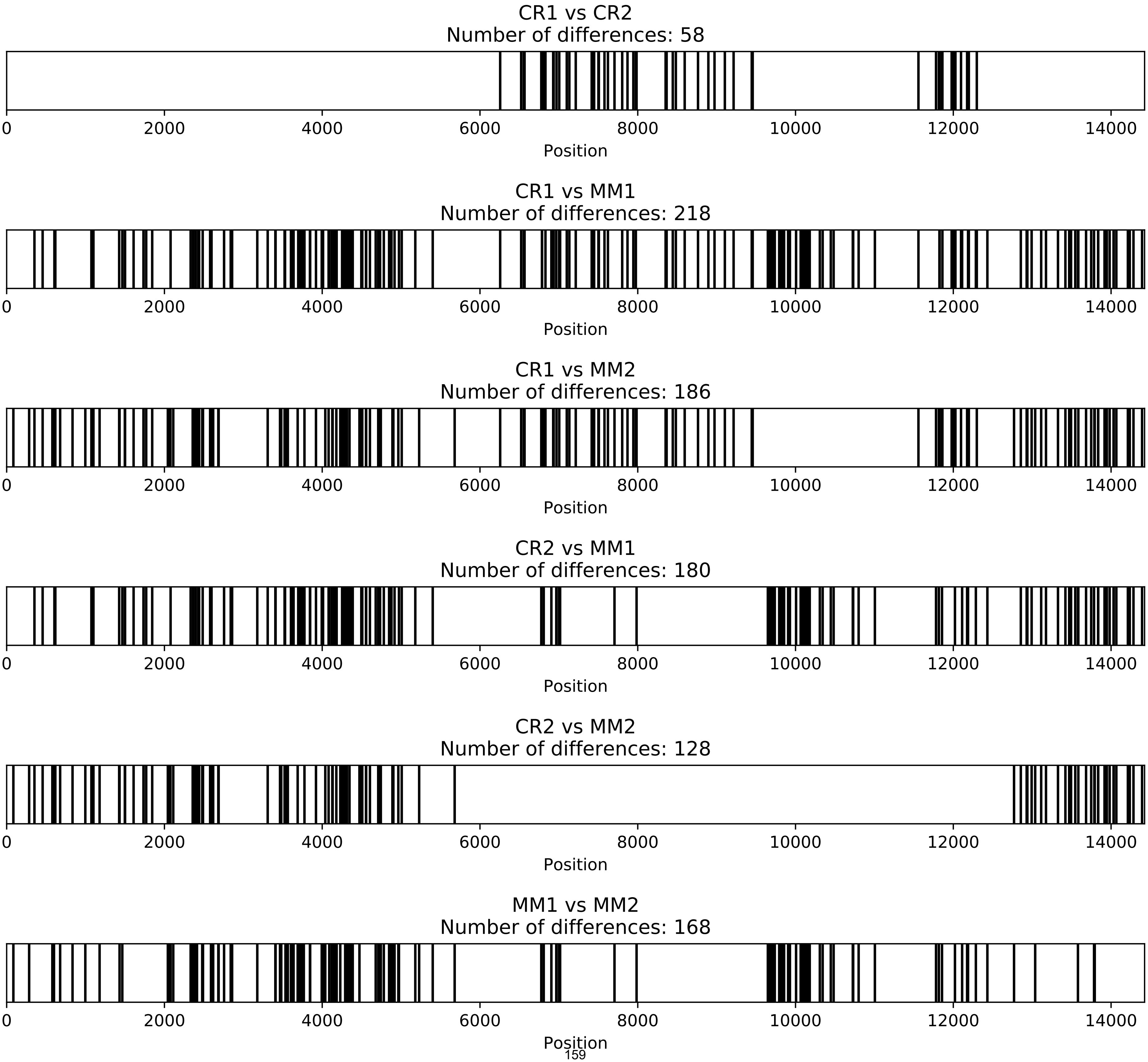
CR1 vs CR2
Number of differences: 58
CR1 vs MM1
Number of differences: 218
CR1 vs MM2
Number of differences: 186
CR2 vs MM1
Number of differences: 180
CR2 vs MM2
Number of differences: 128
MM1 vs MM2
Number of differences: 168
0
2000
4000
6000
8000
10000
12000
14000
Position

### Region: 475



CR1 vs CR2
Number of differences: 279

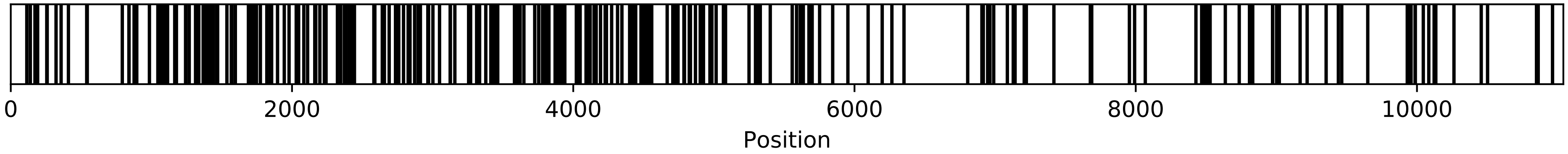


CR1 vs MM1
Number of differences: 156

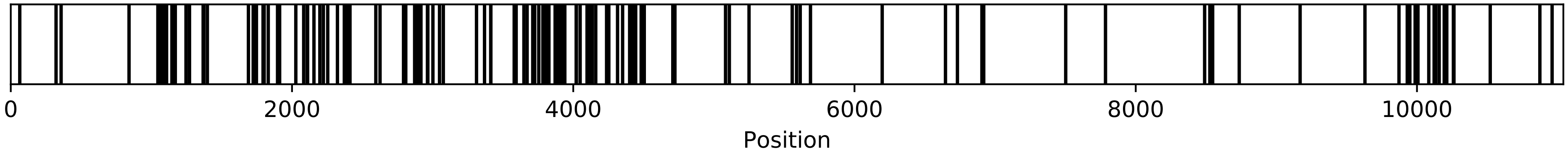


CR1 vs MM2
Number of differences: 156

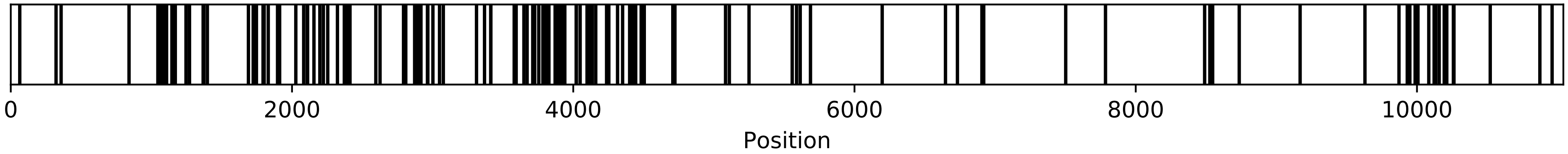


CR2 vs MM1
Number of differences: 255

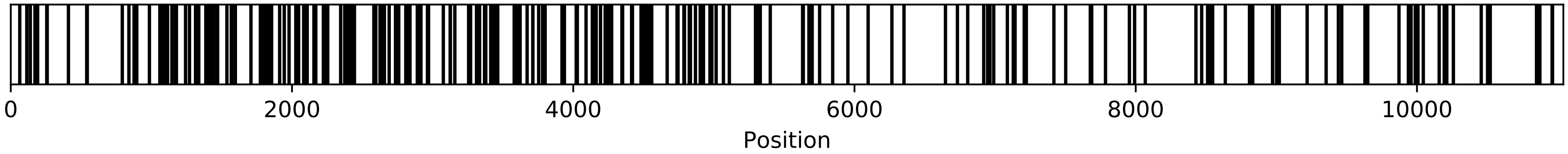


CR2 vs MM2
Number of differences: 255

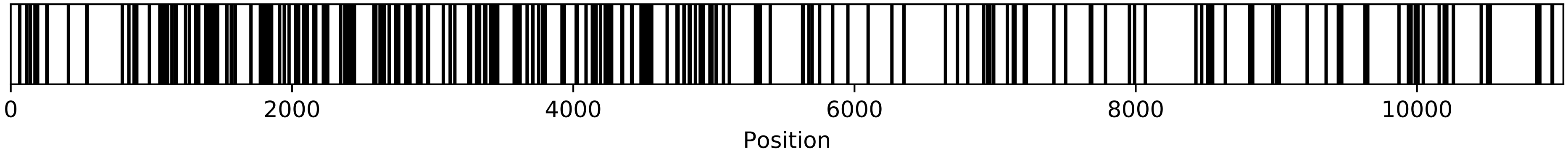


MM1 vs MM2
Number of differences: 0

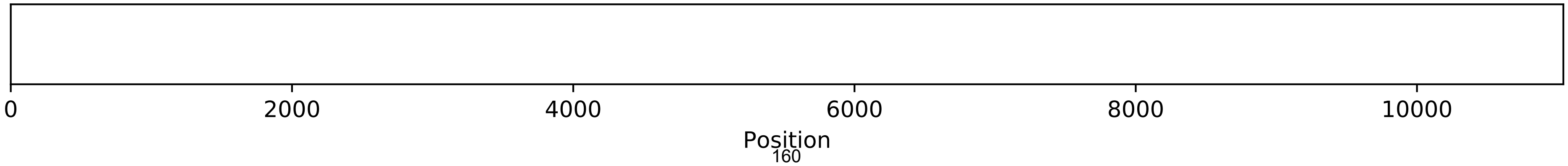

### Region: 480



CR1 vs CR2
Number of differences: 152

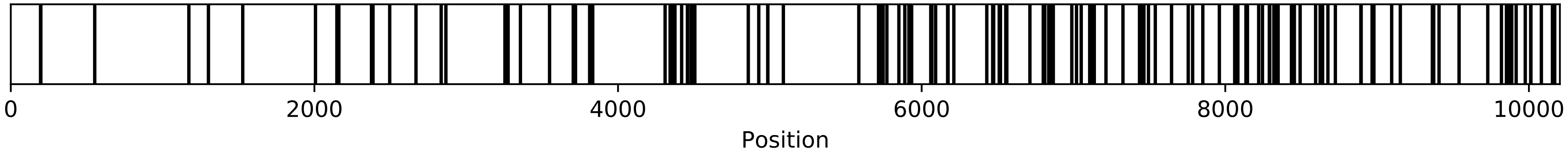


CR1 vs MM1
Number of differences: 150

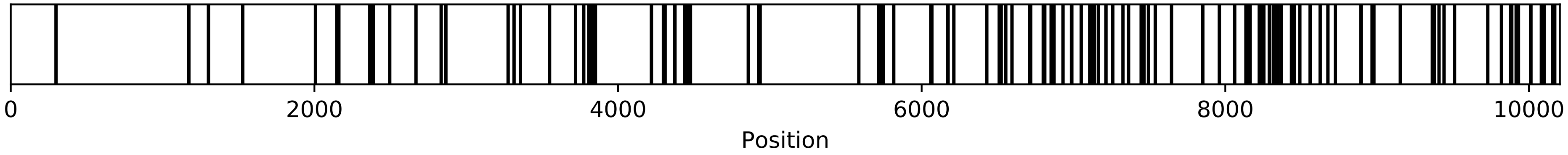


CR1 vs MM2
Number of differences: 150

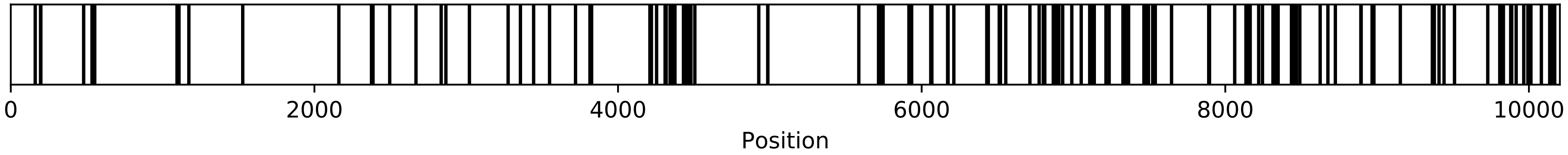


CR2 vs MM1
Number of differences: 108

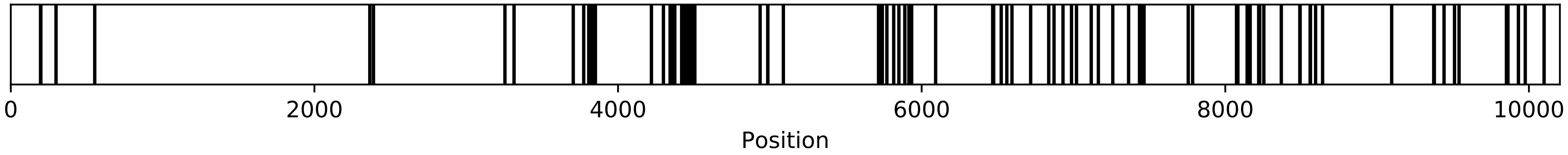


CR2 vs MM2
Number of differences: 102

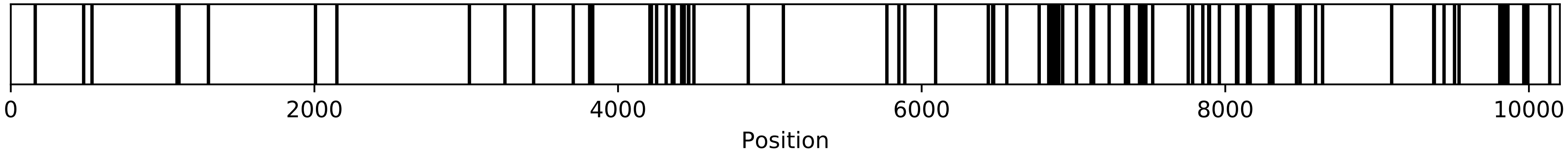


MM1 vs MM2
Number of differences: 111

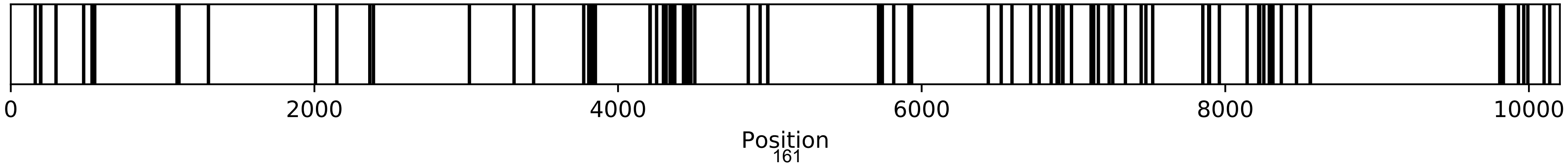

### Region: 481



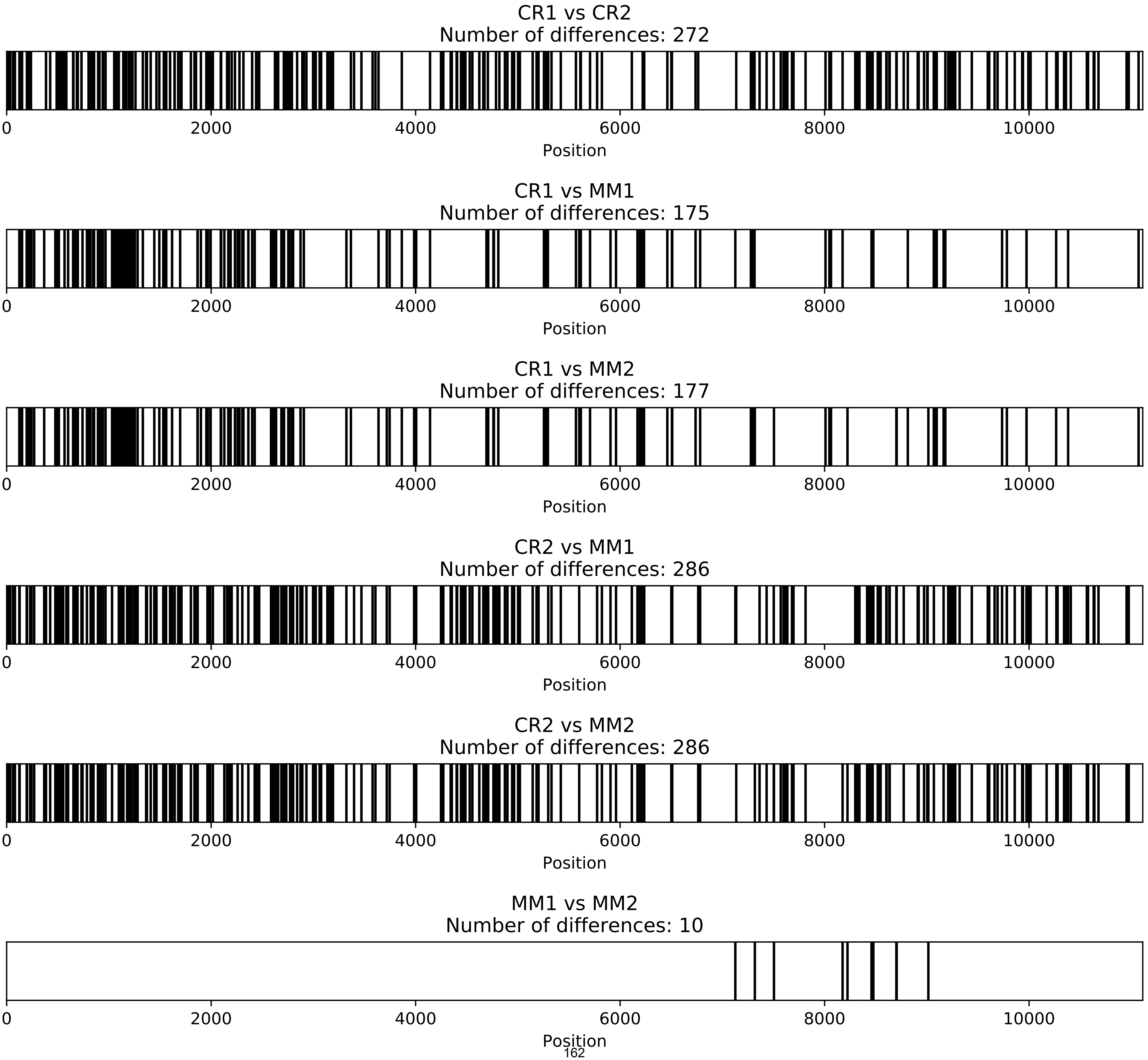
CR1 vs CR2
Number of differences: 272
CR1 vs MM1
Number of differences: 175
CR1 vs MM2
Number of differences: 177
CR2 vs MM1
Number of differences: 286
CR2 vs MM2
Number of differences: 286
MM1 vs MM2
Number of differences: 10
0
2000
4000
6000
8000
10000
Position

### Region: 484



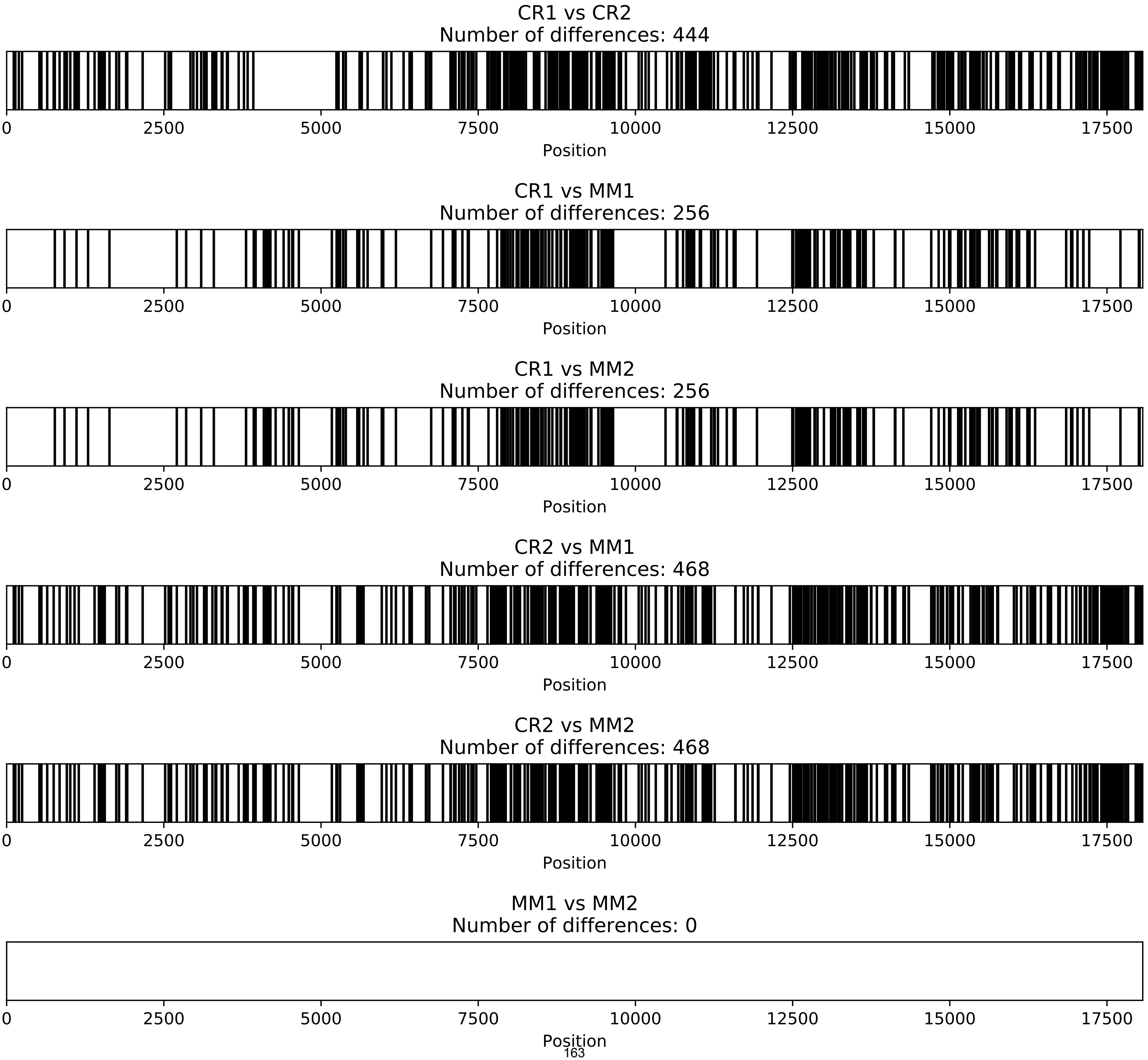
CR1 vs CR2
Number of differences: 444
CR1 vs MM1
Number of differences: 256
CR1 vs MM2
Number of differences: 256
CR2 vs MM1
Number of differences: 468
CR2 vs MM2
Number of differences: 468
MM1 vs MM2
Number of differences: 0
0
2500
5000
7500
10000
12500
15000
17500
Position

### Region: 485



## CR1 vs CR2
Number of differences: 244

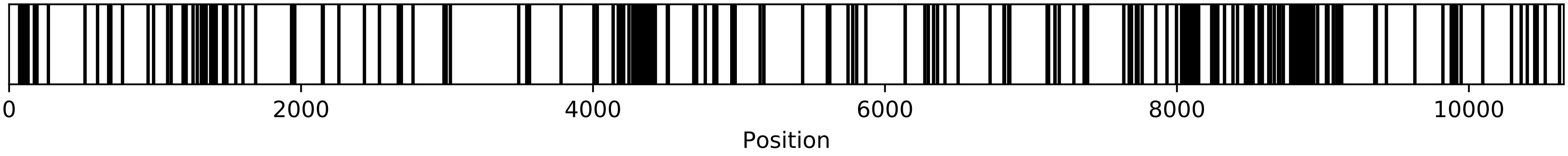


## CR1 vs MM1
Number of differences: 255

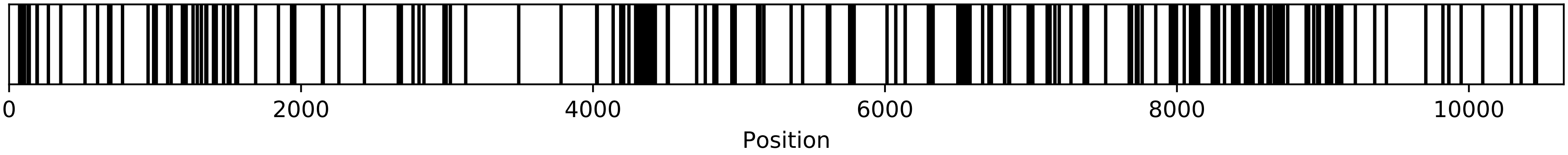


## CR1 vs MM2
Number of differences: 252

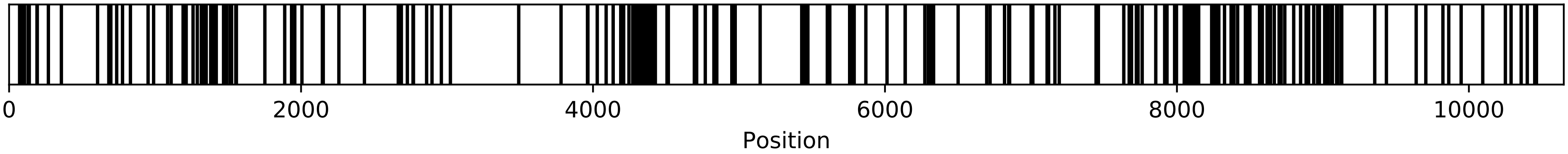


## CR2 vs MM1
Number of differences: 141

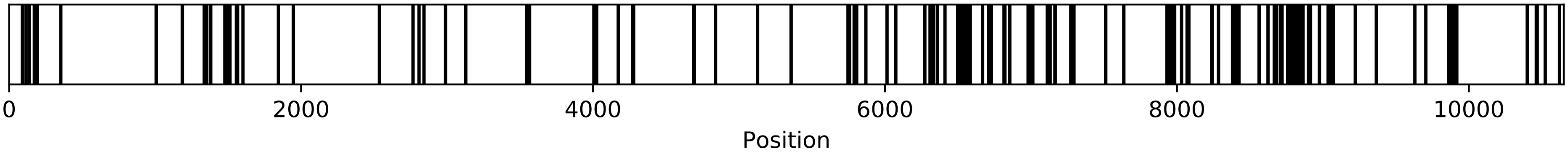


## CR2 vs MM2
Number of differences: 139

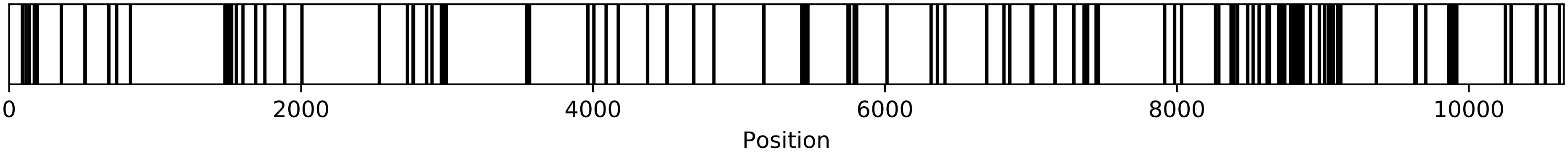


## MM1 vs MM2
Number of differences: 159

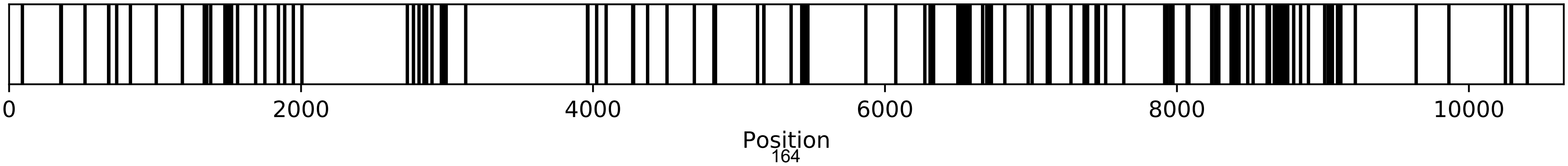

### Region: 488



CR1 vs CR2
Number of differences: 403

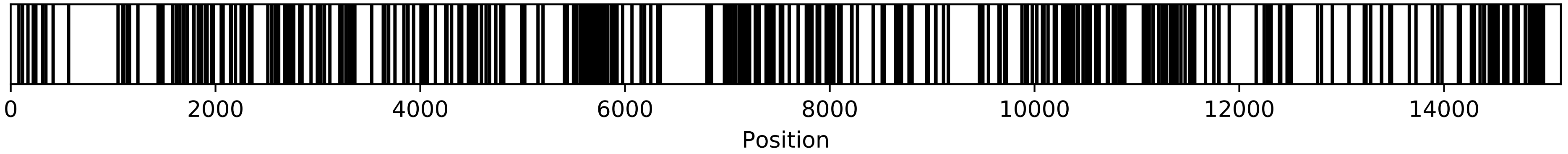


CR1 vs MM1
Number of differences: 174

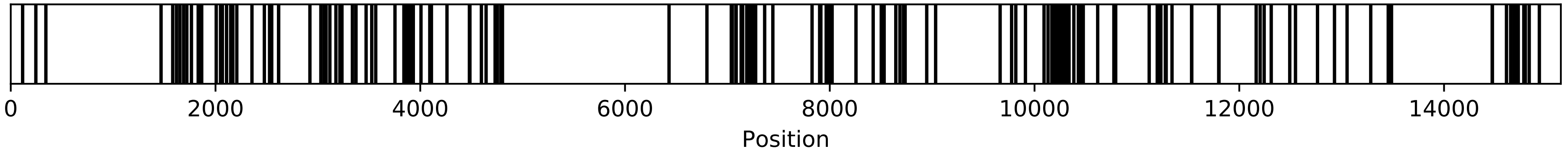


CR1 vs MM2
Number of differences: 174

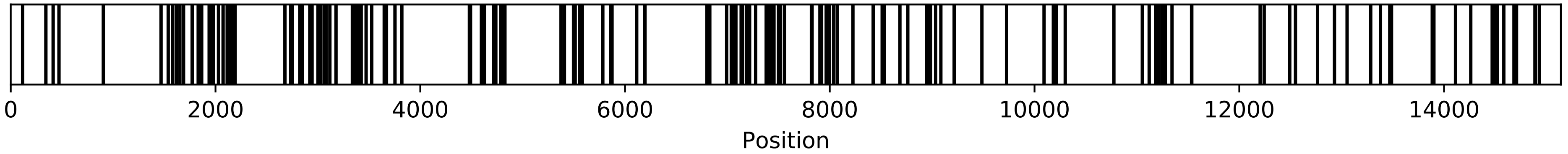


CR2 vs MM1
Number of differences: 380

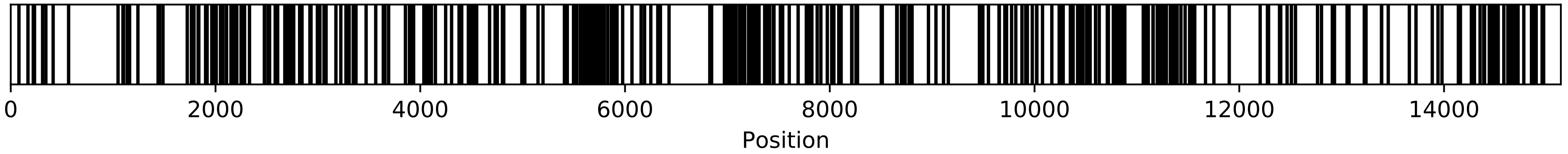


CR2 vs MM2
Number of differences: 397

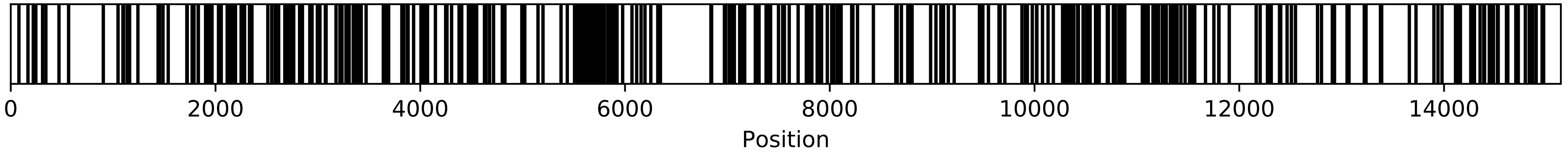


MM1 vs MM2
Number of differences: 193

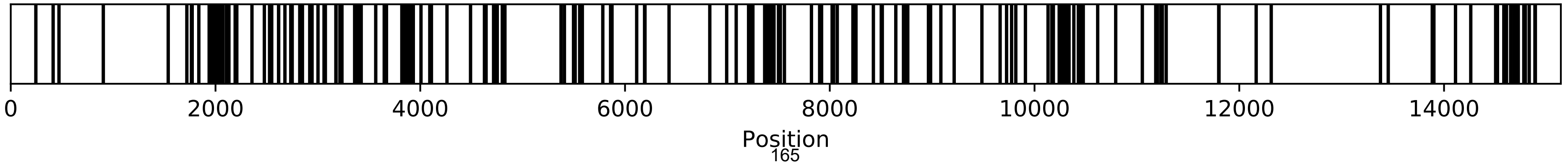

### Region: 489



CR1 vs CR2
Number of differences: 255

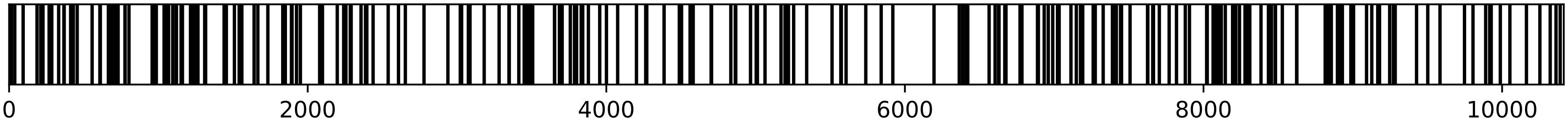


CR1 vs MM1
Number of differences: 267

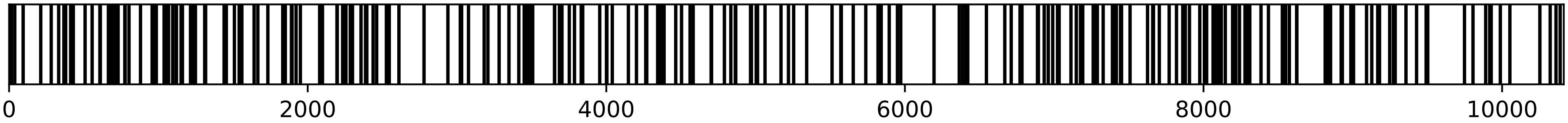


CR1 vs MM2
Number of differences: 254

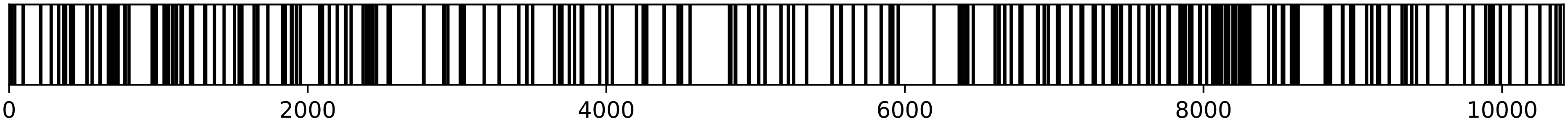


CR2 vs MM1
Number of differences: 96

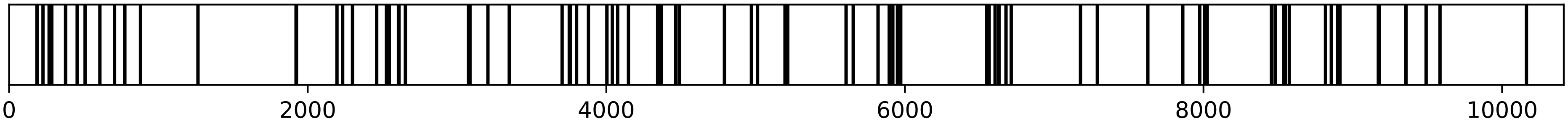


CR2 vs MM2
Number of differences: 127

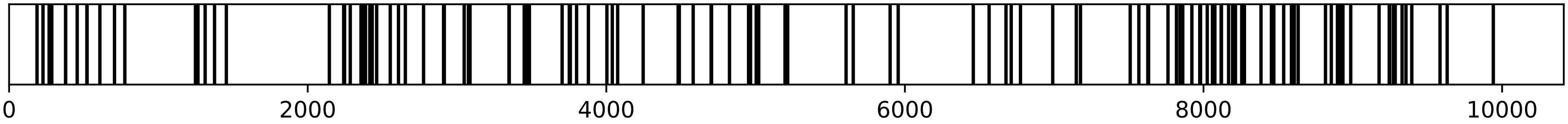


MM1 vs MM2
Number of differences: 130

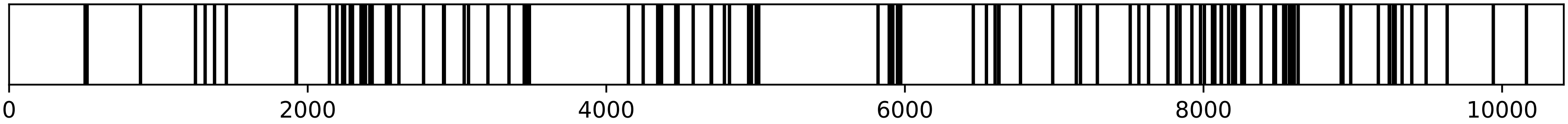

### Region: 490



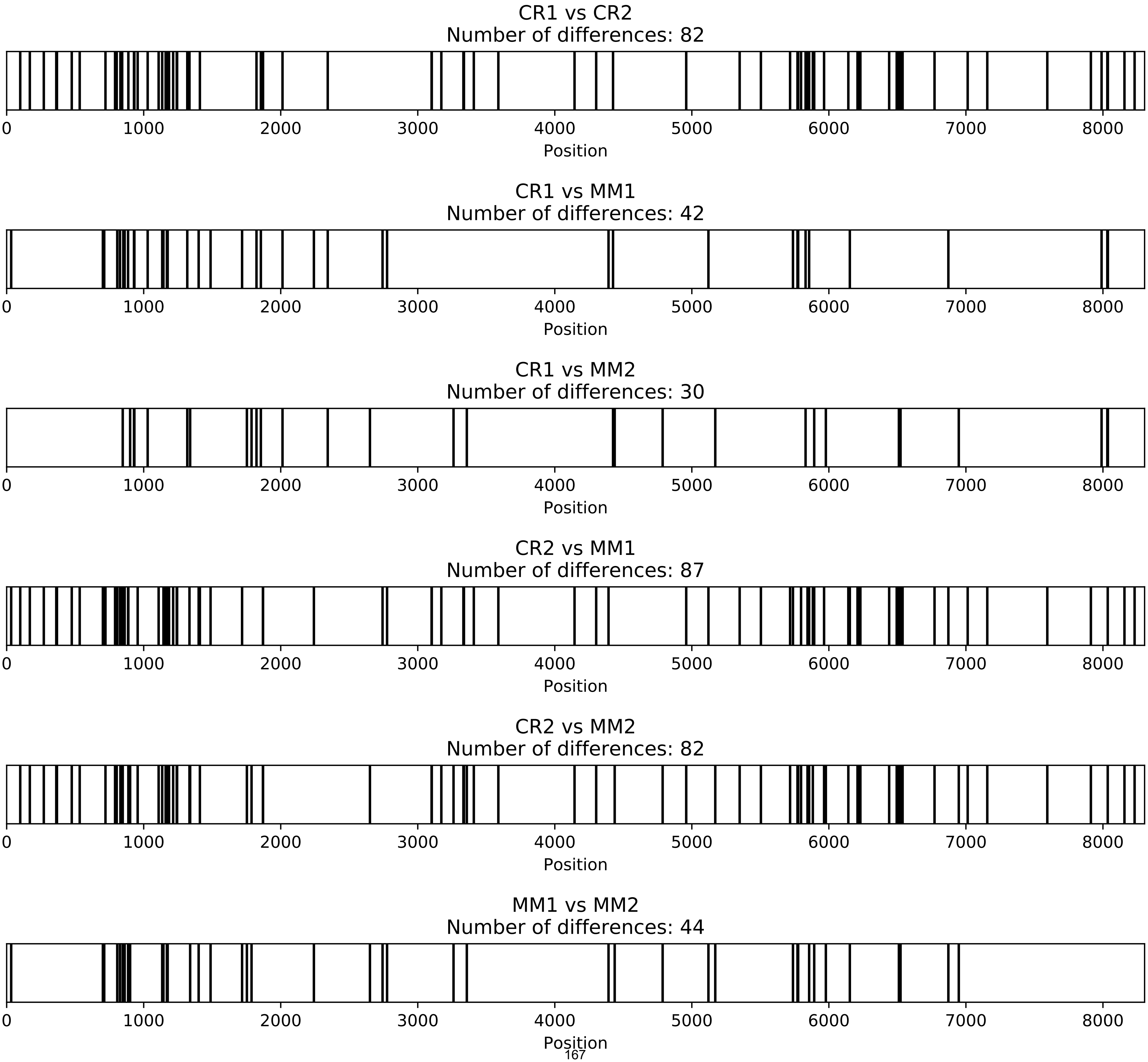
CR1 vs CR2
Number of differences: 82
CR1 vs MM1
Number of differences: 42
CR1 vs MM2
Number of differences: 30
CR2 vs MM1
Number of differences: 87
CR2 vs MM2
Number of differences: 82
MM1 vs MM2
Number of differences: 44
0
1000
2000
3000
4000
5000
6000
7000
8000
Position

### Region: 491



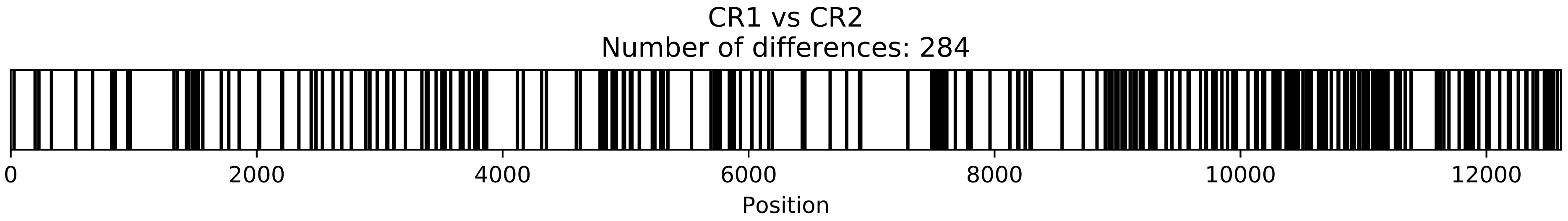
CR1 vs CR2
Number of differences: 284
0
2000
4000
6000
8000
10000
12000
Position

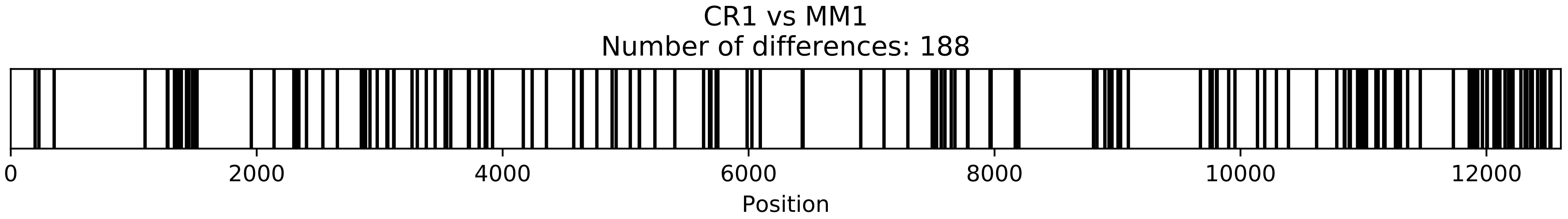
CR1 vs MM1
Number of differences: 188
0
2000
4000
6000
8000
10000
12000
Position

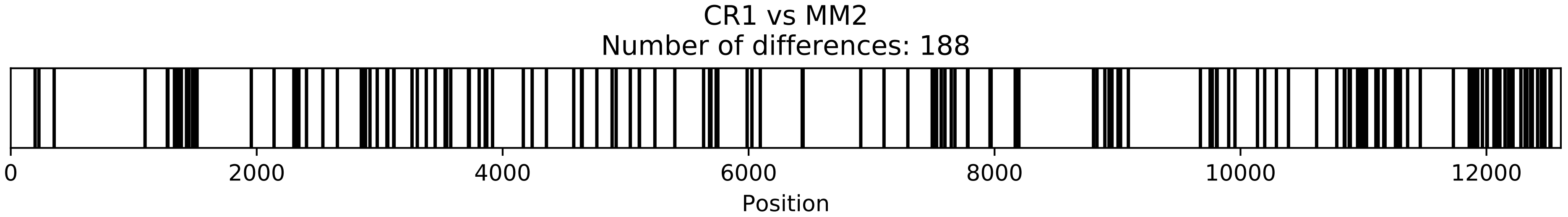
CR1 vs MM2
Number of differences: 188
0
2000
4000
6000
8000
10000
12000
Position

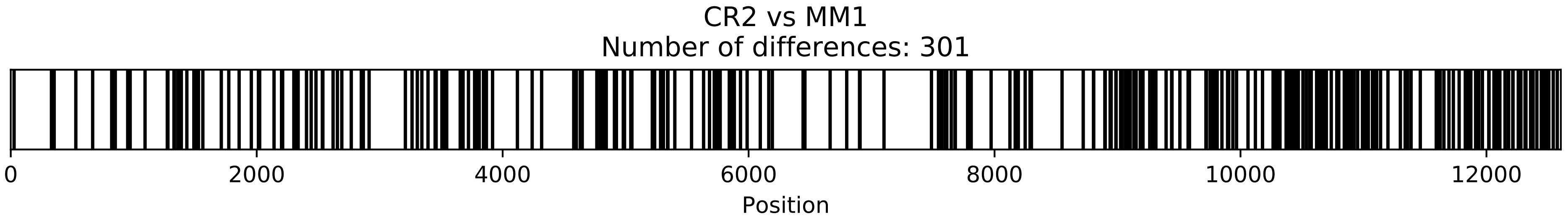
CR2 vs MM1
Number of differences: 301
0
2000
4000
6000
8000
10000
12000
Position

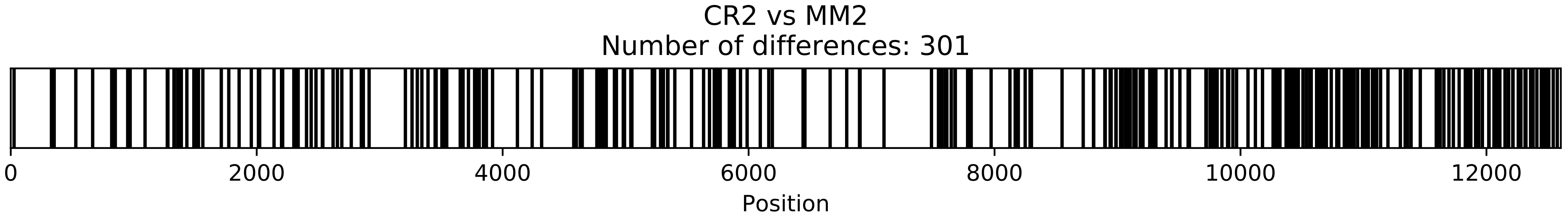
CR2 vs MM2
Number of differences: 301
0
2000
4000
6000
8000
10000
12000
Position

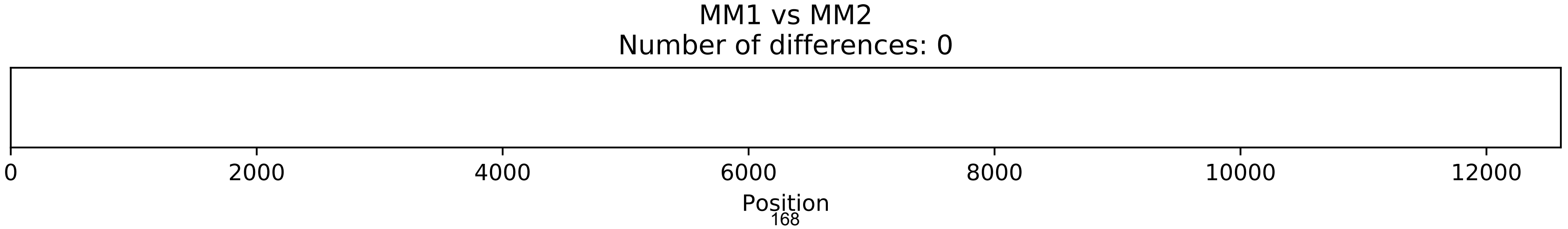
MM1 vs MM2
Number of differences: 0
0
2000
4000
6000
8000
10000
12000
Position

### Region: 494



CR1 vs CR2
Number of differences: 373

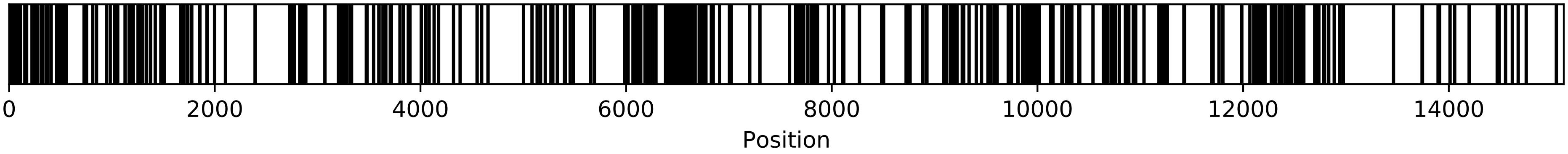


CR1 vs MM1
Number of differences: 236

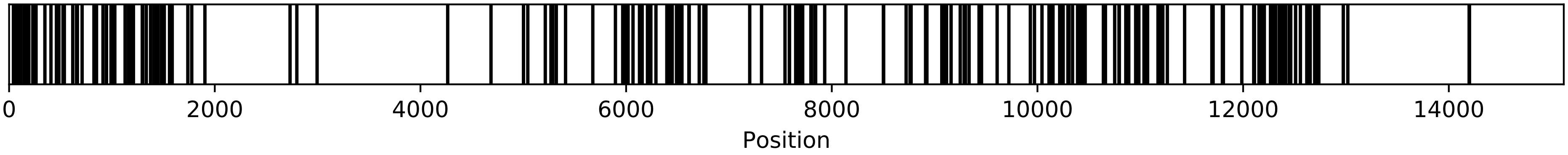


CR1 vs MM2
Number of differences: 233

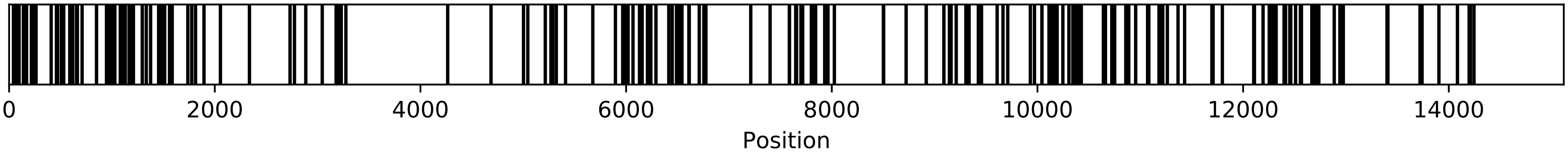


CR2 vs MM1
Number of differences: 378

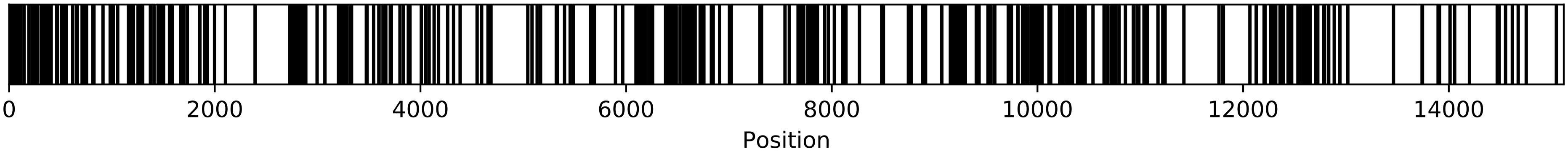


CR2 vs MM2
Number of differences: 381

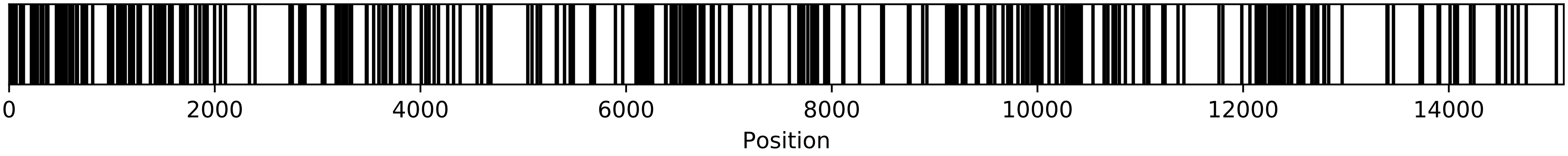


MM1 vs MM2
Number of differences: 178

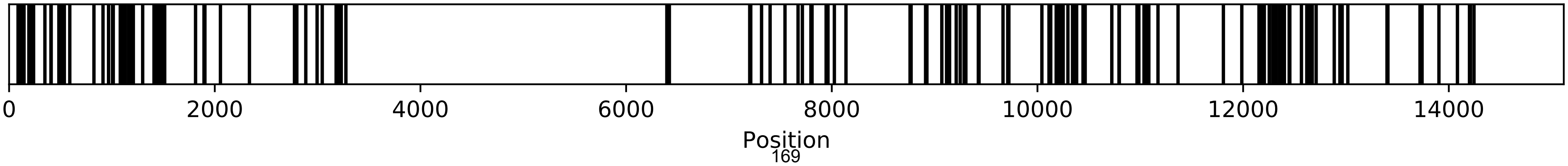

### Region: 497



## CR1 vs CR2
Number of differences: 297

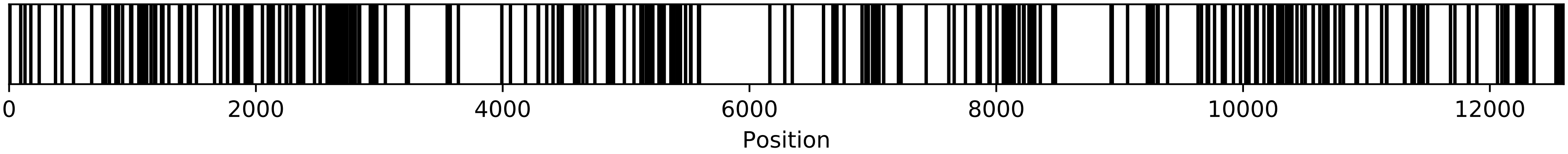


## CR1 vs MM1
Number of differences: 150

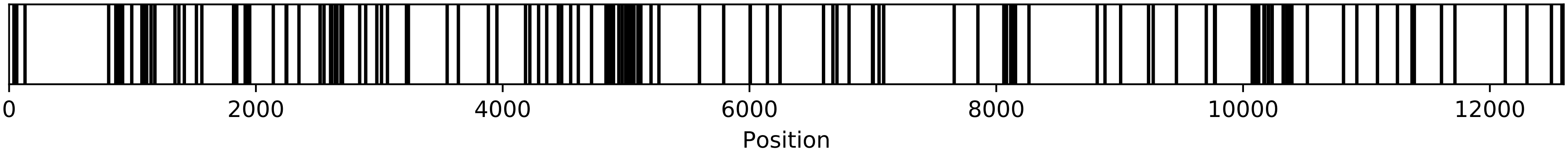


## CR1 vs MM2
Number of differences: 154

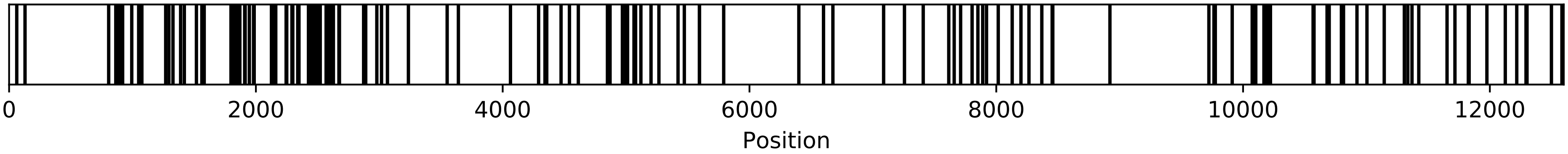


## CR2 vs MM1
Number of differences: 292

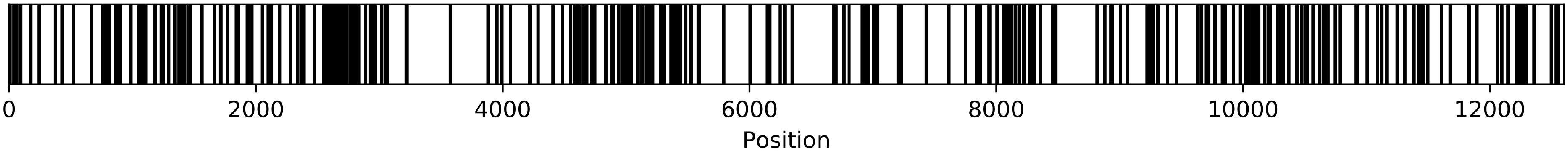


## CR2 vs MM2
Number of differences: 316

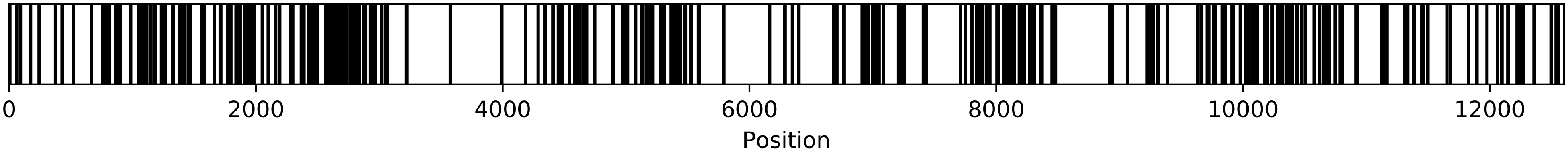


## MM1 vs MM2
Number of differences: 160

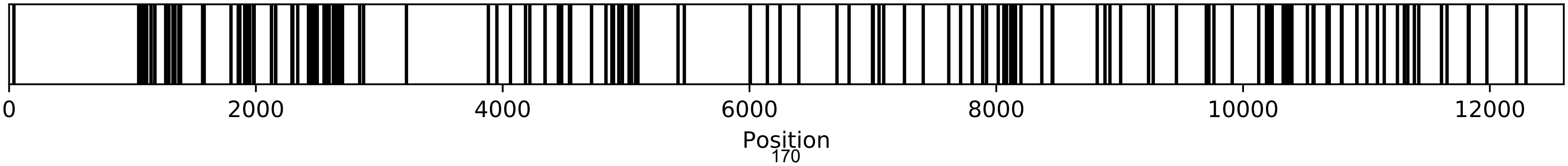

### Region: 498



CR1 vs CR2
Number of differences: 300

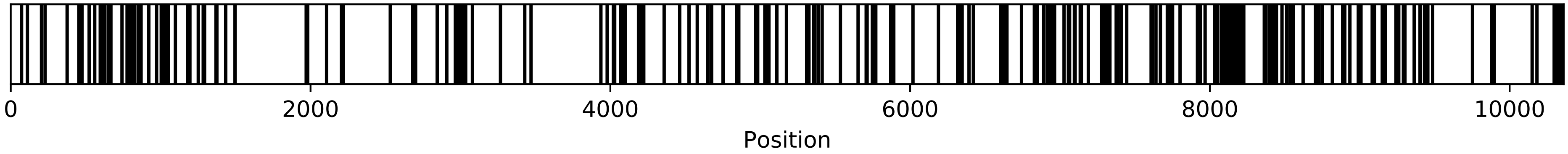


CR1 vs MM1
Number of differences: 129

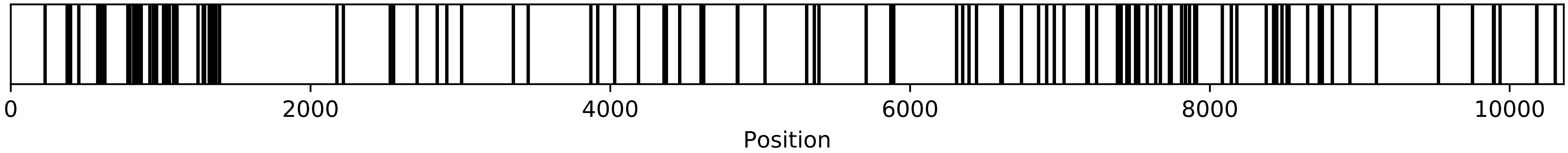


CR1 vs MM2
Number of differences: 166

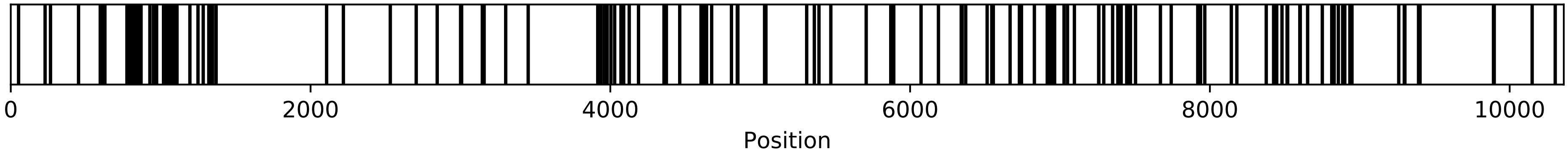


CR2 vs MM1
Number of differences: 308

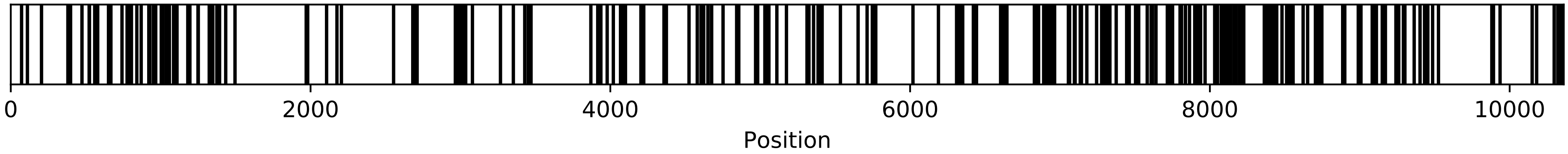


CR2 vs MM2
Number of differences: 302

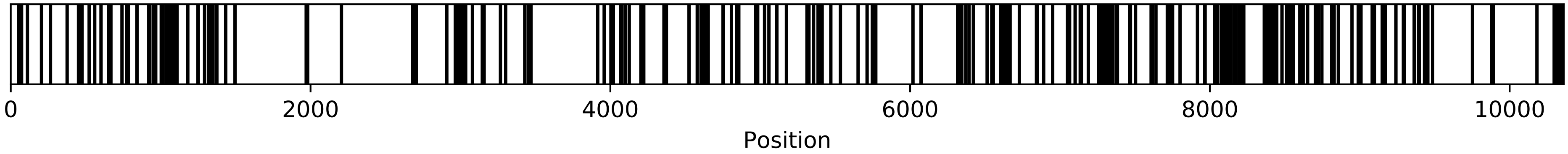


MM1 vs MM2
Number of differences: 139

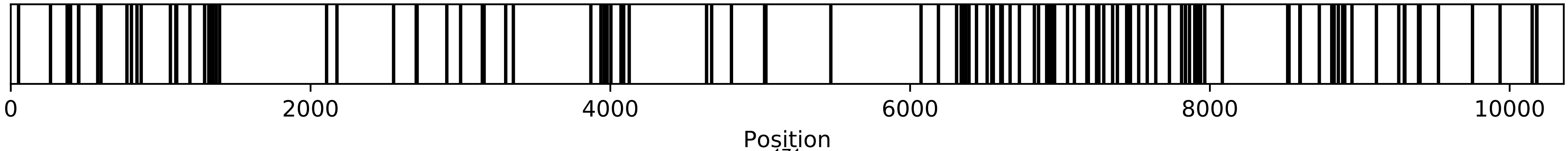

### Region: 499



## CR1 vs CR2
Number of differences: 164

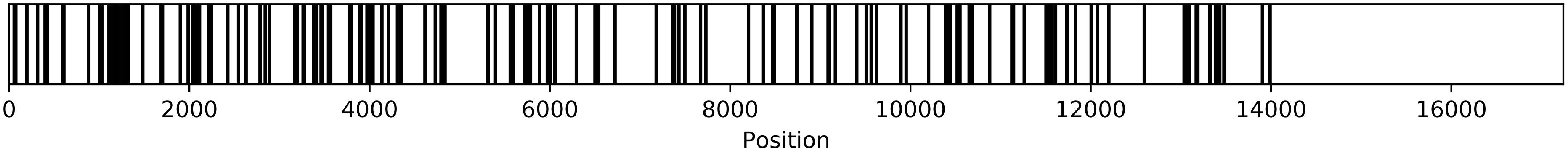


## CR1 vs MM1
Number of differences: 138

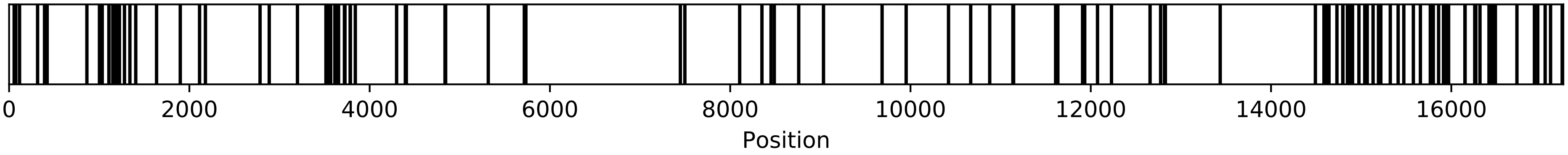


## CR1 vs MM2
Number of differences: 123

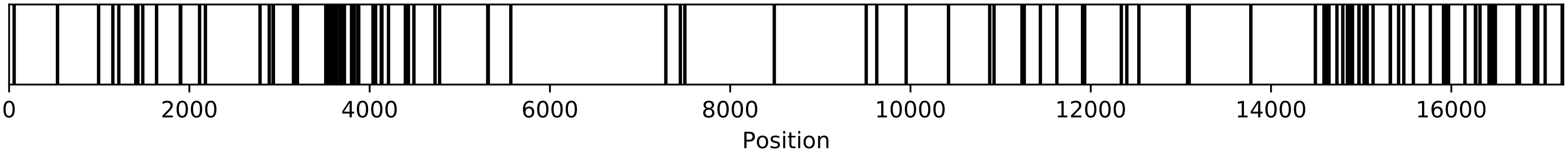


## CR2 vs MM1
Number of differences: 234

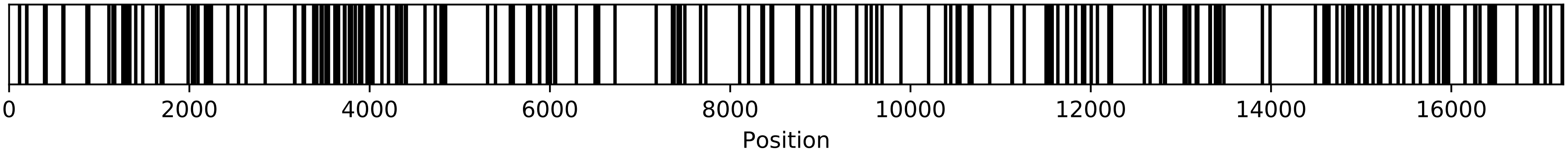


## CR2 vs MM2
Number of differences: 240

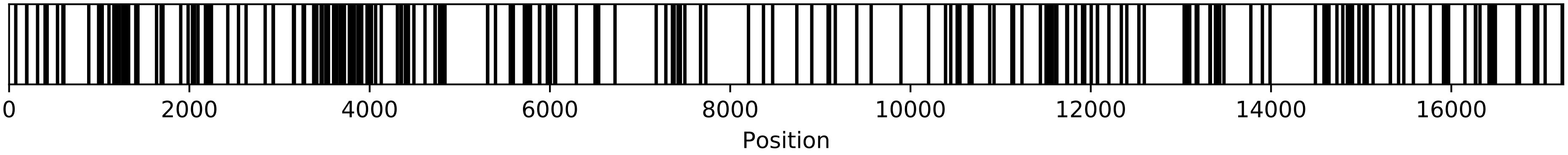


## MM1 vs MM2
Number of differences: 134

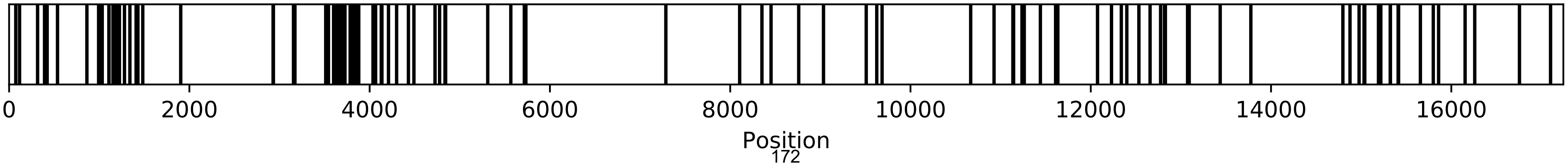

### Region: 501



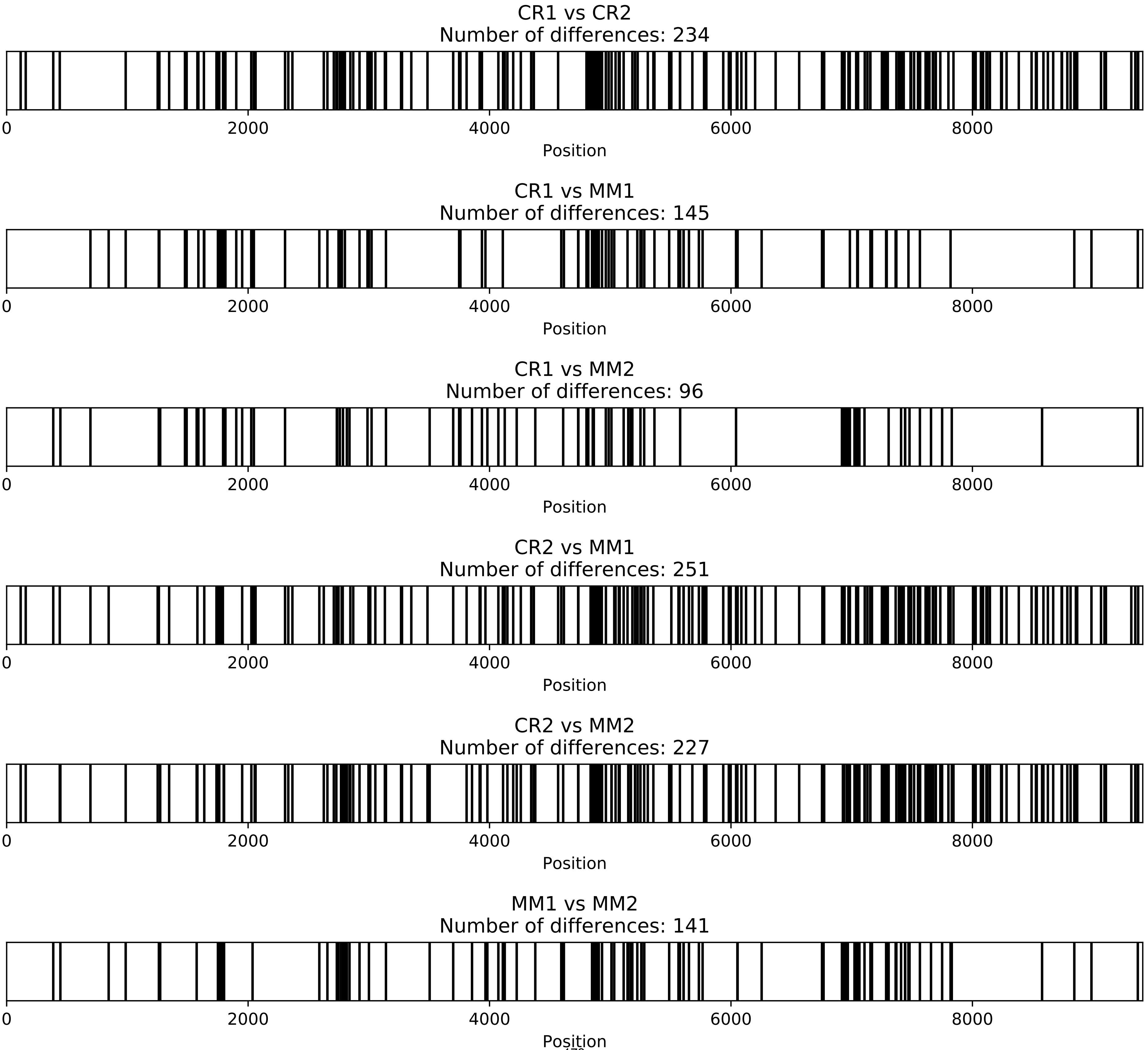
CR1 vs CR2
Number of differences: 234
CR1 vs MM1
Number of differences: 145
CR1 vs MM2
Number of differences: 96
CR2 vs MM1
Number of differences: 251
CR2 vs MM2
Number of differences: 227
MM1 vs MM2
Number of differences: 141
0
2000
4000
6000
8000
Position

### Region: 502



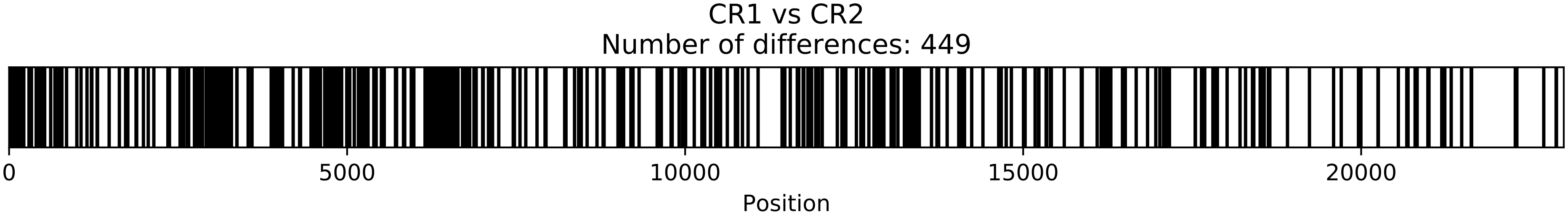
CR1 vs CR2
Number of differences: 449
0
5000
10000
15000
20000
Position

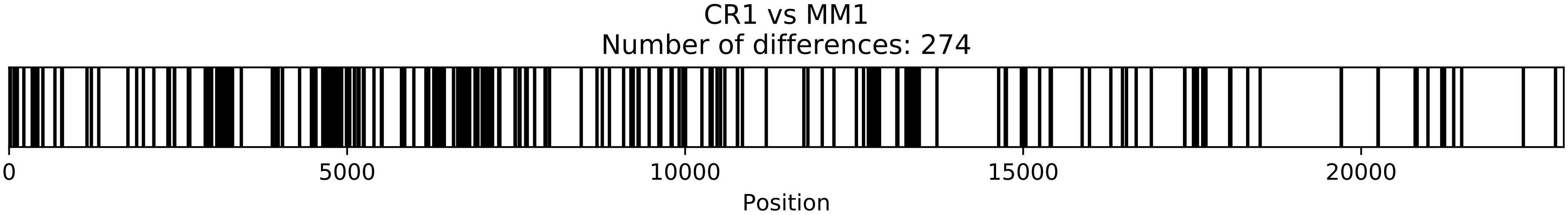
CR1 vs MM1
Number of differences: 274
0
5000
10000
15000
20000
Position

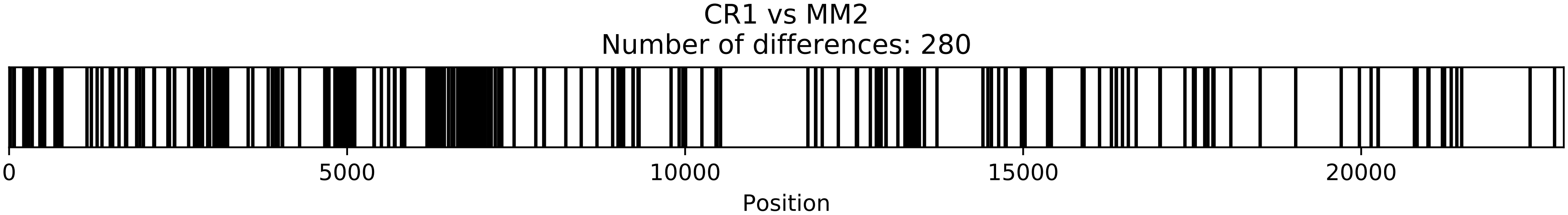
CR1 vs MM2
Number of differences: 280
0
5000
10000
15000
20000
Position

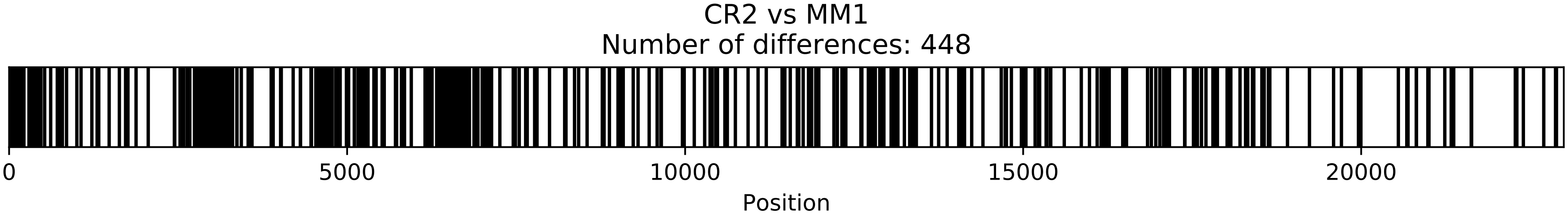
CR2 vs MM1
Number of differences: 448
0
5000
10000
15000
20000
Position

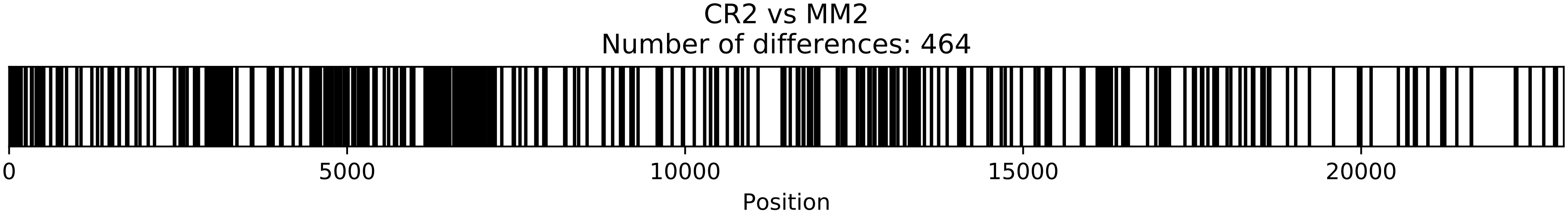
CR2 vs MM2
Number of differences: 464
0
5000
10000
15000
20000
Position

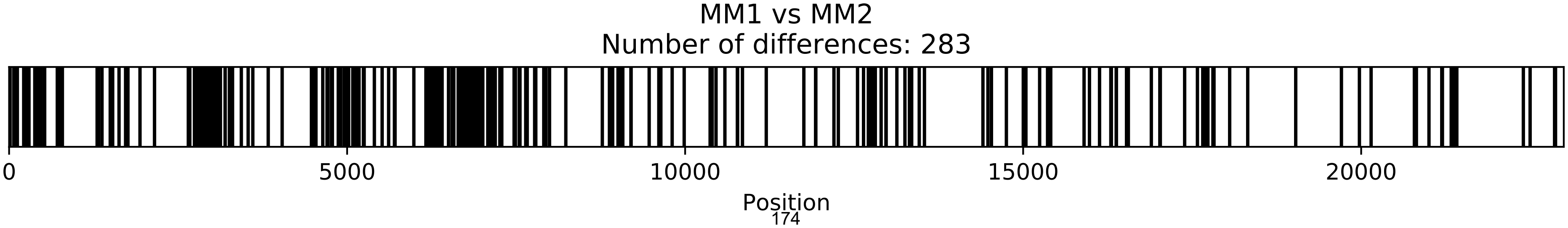
MM1 vs MM2
Number of differences: 283
0
5000
10000
15000
20000
Position

### Region: 504



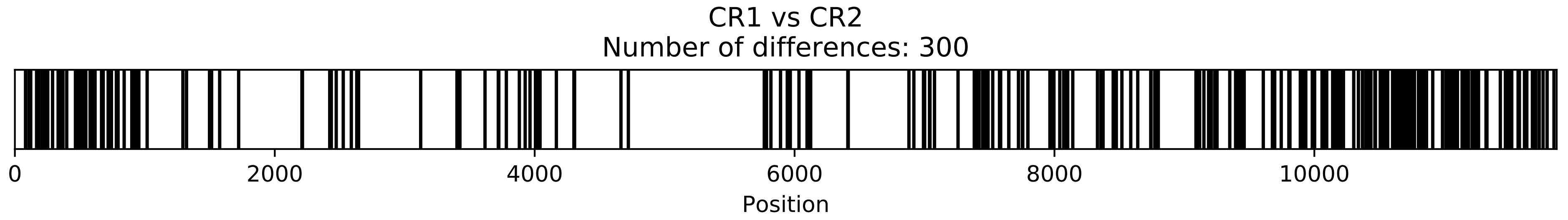
CR1 vs CR2
Number of differences: 300
0
2000
4000
6000
8000
10000
Position

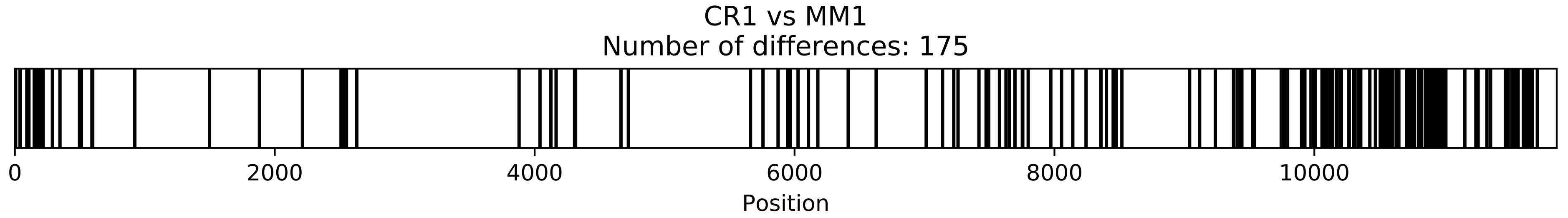
CR1 vs MM1
Number of differences: 175
0
2000
4000
6000
8000
10000
Position

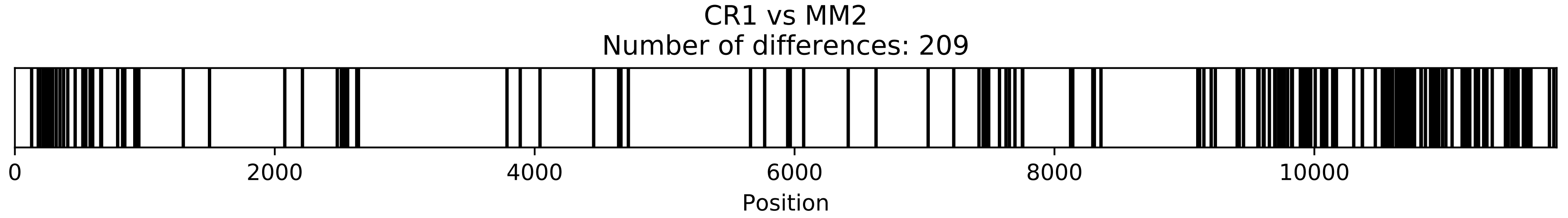
CR1 vs MM2
Number of differences: 209
0
2000
4000
6000
8000
10000
Position

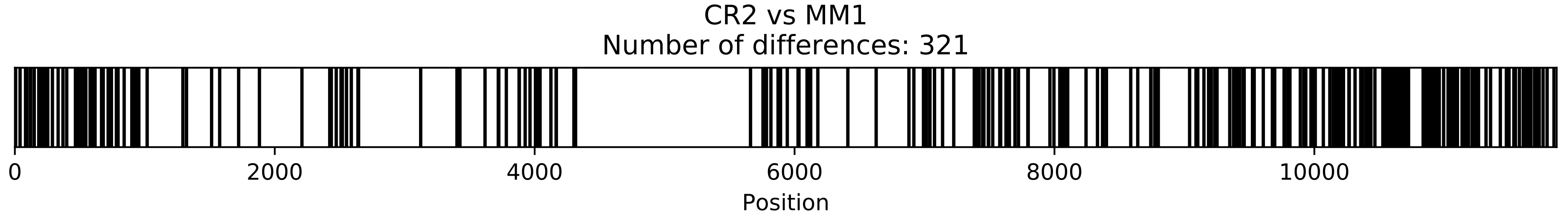
CR2 vs MM1
Number of differences: 321
0
2000
4000
6000
8000
10000
Position

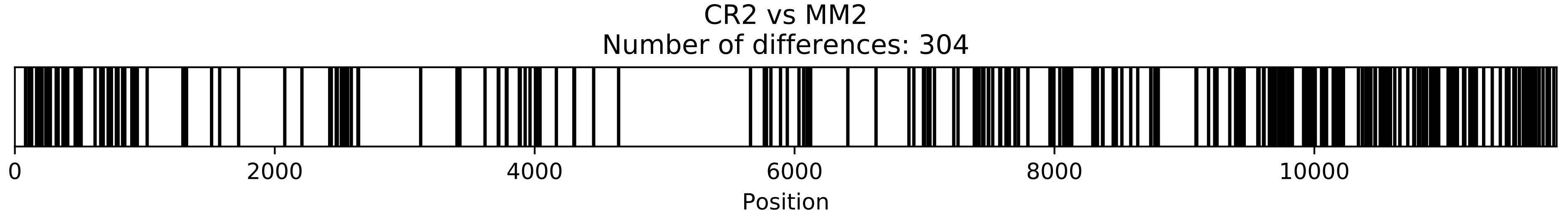
CR2 vs MM2
Number of differences: 304
0
2000
4000
6000
8000
10000
Position

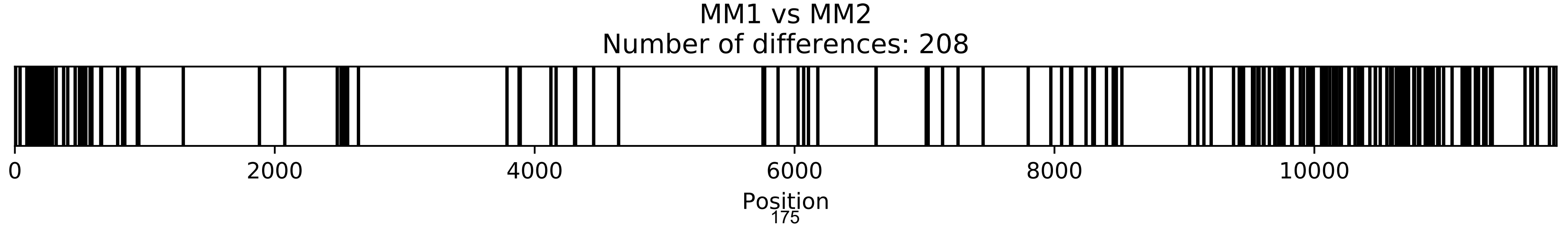
MM1 vs MM2
Number of differences: 208
0
2000
4000
6000
8000
10000
Position

### Region: 505



CR1 vs CR2
Number of differences: 209

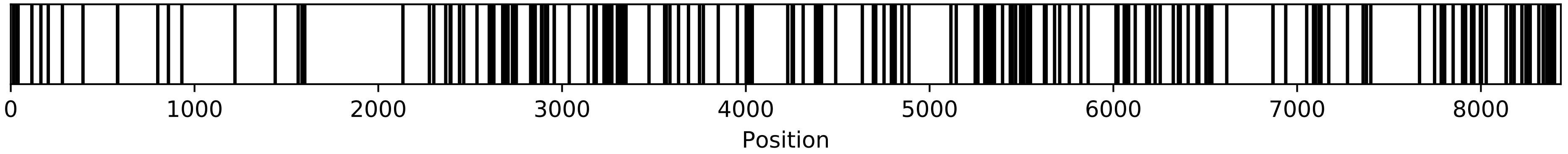


CR1 vs MM1
Number of differences: 202

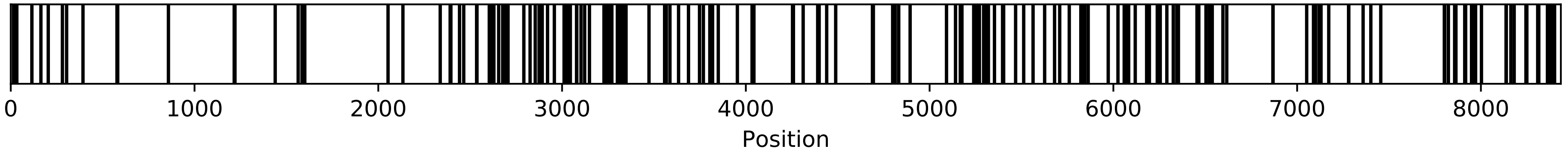


CR1 vs MM2
Number of differences: 203

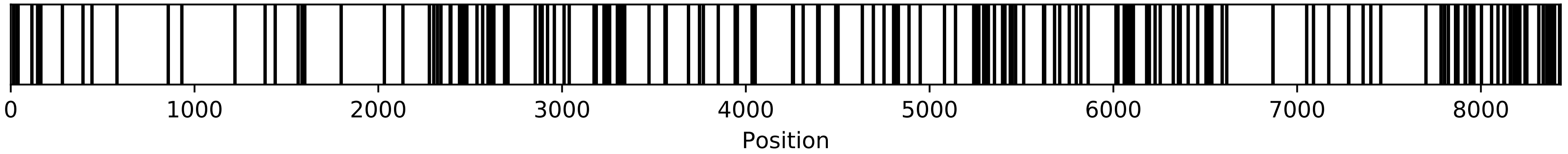


CR2 vs MM1
Number of differences: 135

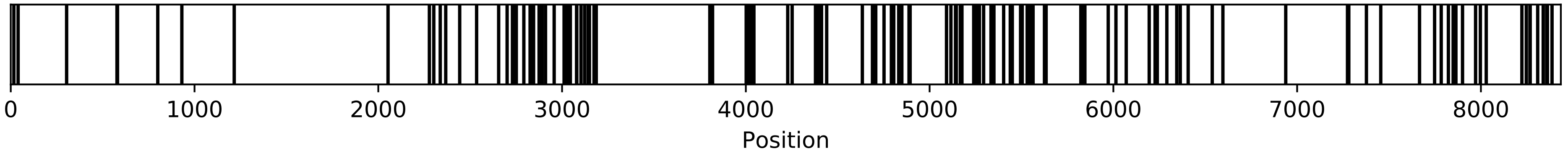


CR2 vs MM2
Number of differences: 146

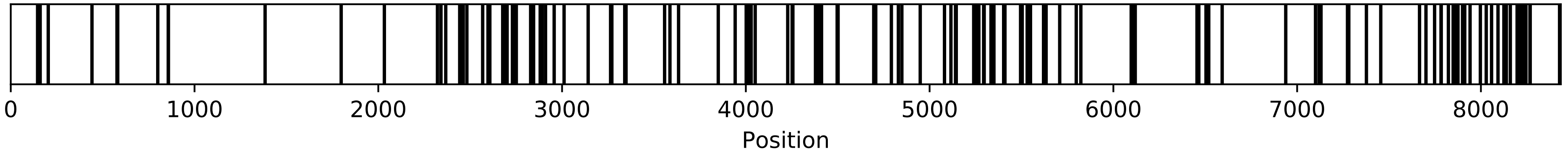


MM1 vs MM2
Number of differences: 145

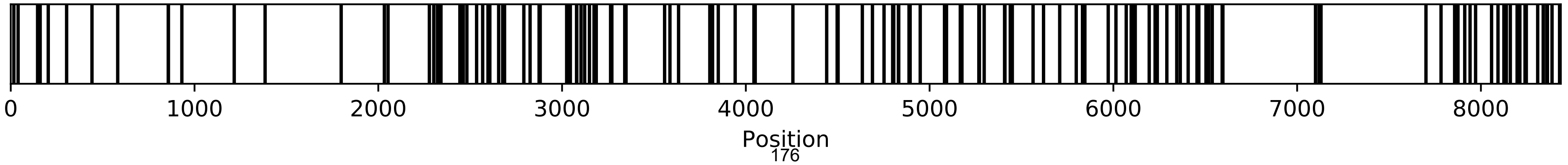

### Region: 509



CR1 vs CR2
Number of differences: 156

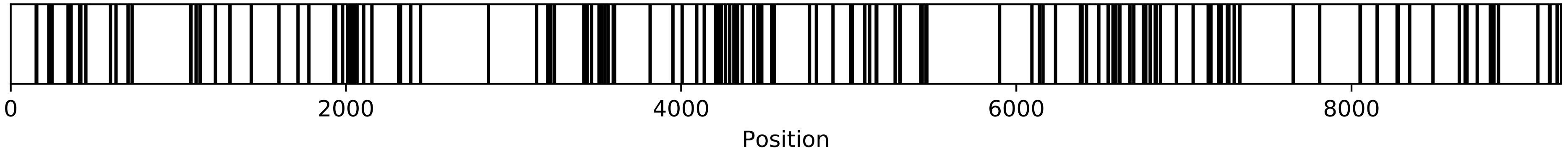


CR1 vs MM1
Number of differences: 84

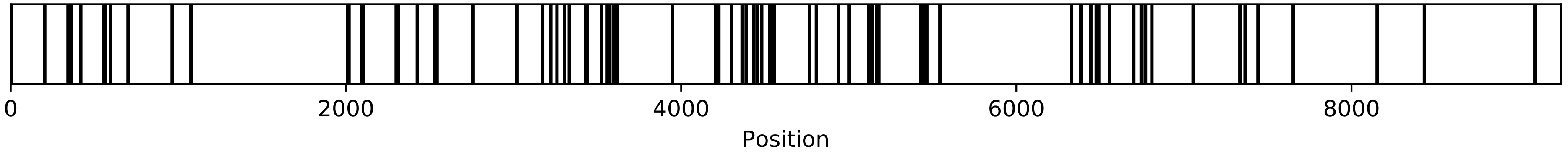


CR1 vs MM2
Number of differences: 58

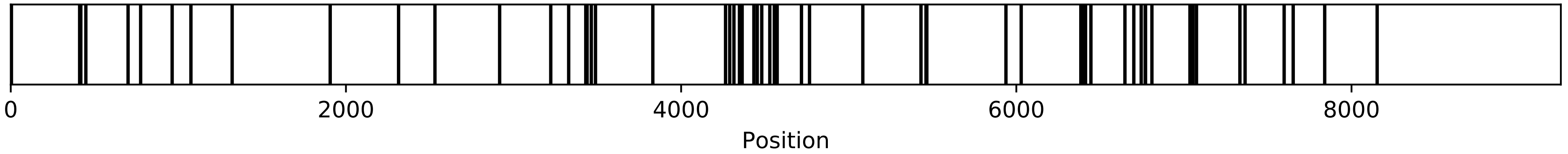


CR2 vs MM1
Number of differences: 167

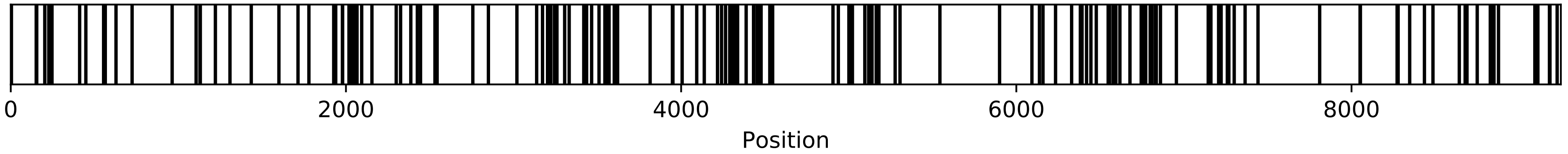


CR2 vs MM2
Number of differences: 162

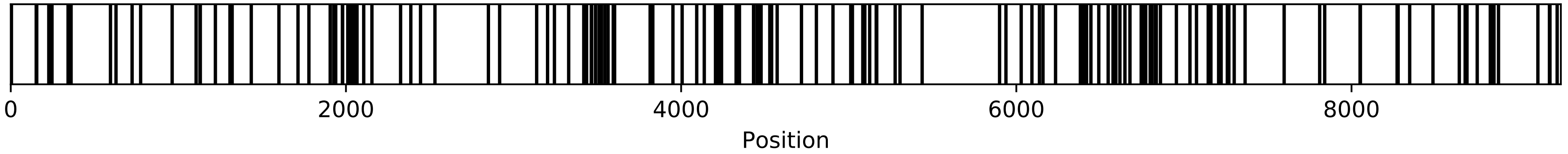


MM1 vs MM2
Number of differences: 79

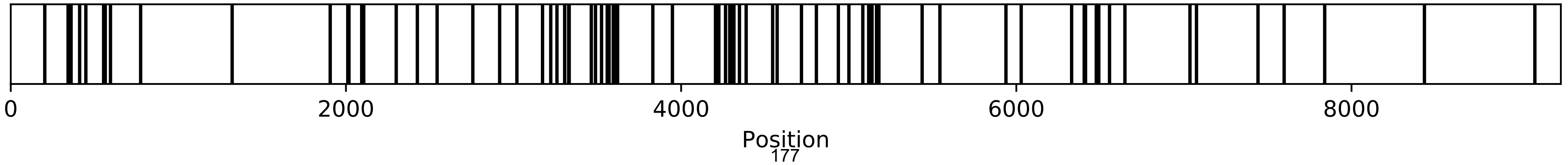

### Region: 511



CR1 vs CR2
Number of differences: 280

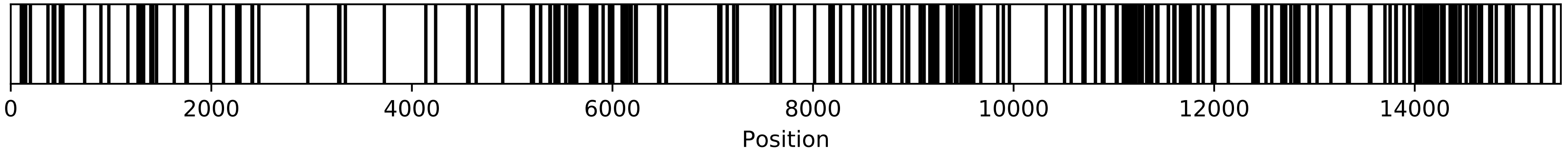


CR1 vs MM1
Number of differences: 141

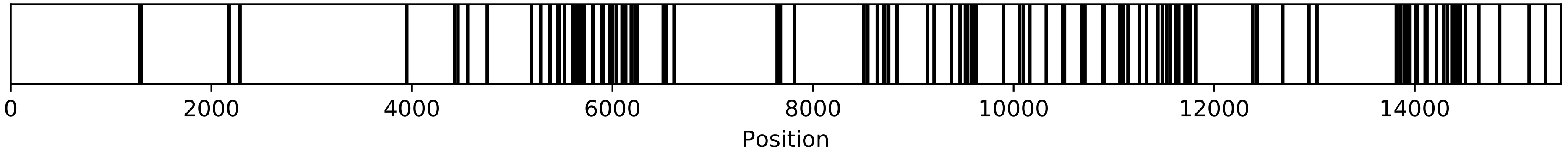


CR1 vs MM2
Number of differences: 158

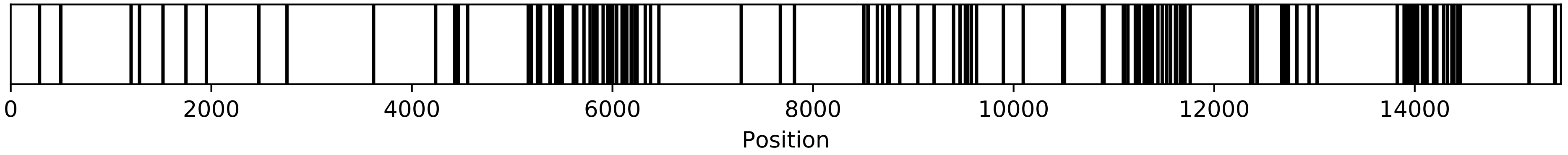


CR2 vs MM1
Number of differences: 283

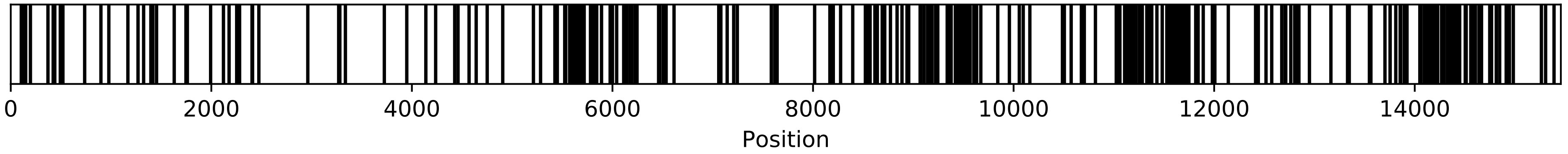


CR2 vs MM2
Number of differences: 290

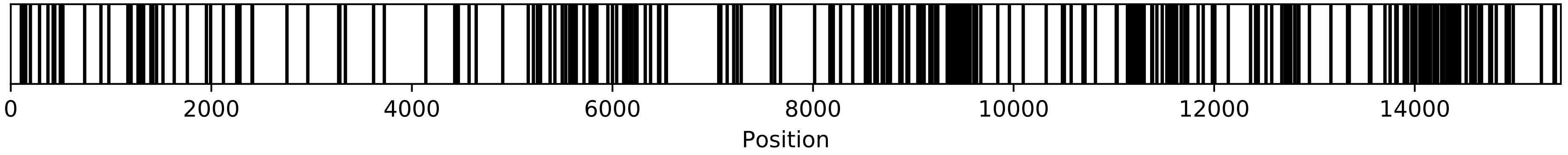


MM1 vs MM2
Number of differences: 139

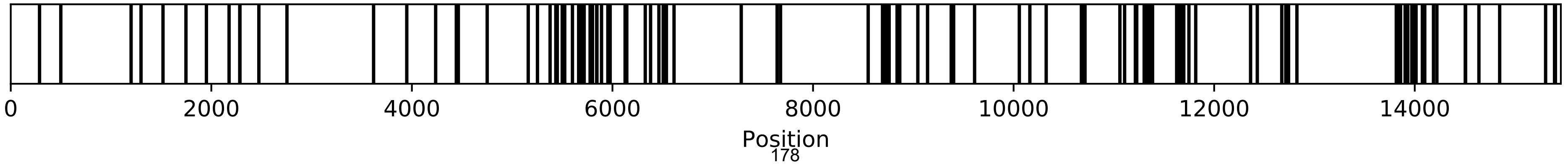

### Region: 512



## CR1 vs CR2
Number of differences: 264

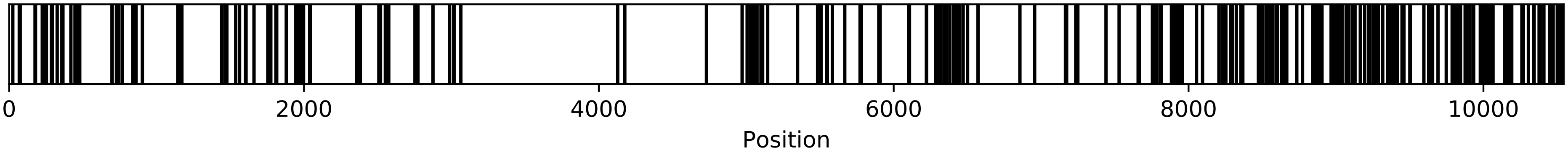


## CR1 vs MM1
Number of differences: 142

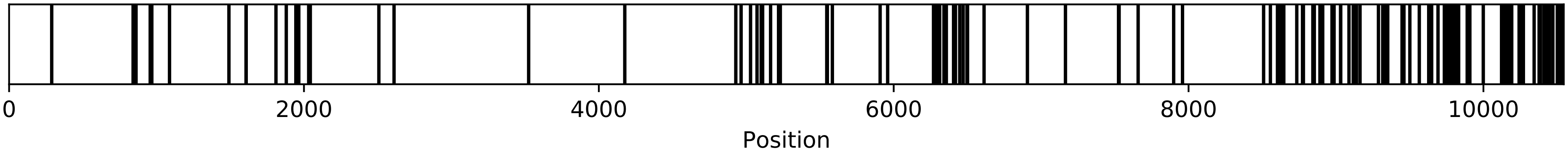


## CR1 vs MM2
Number of differences: 142

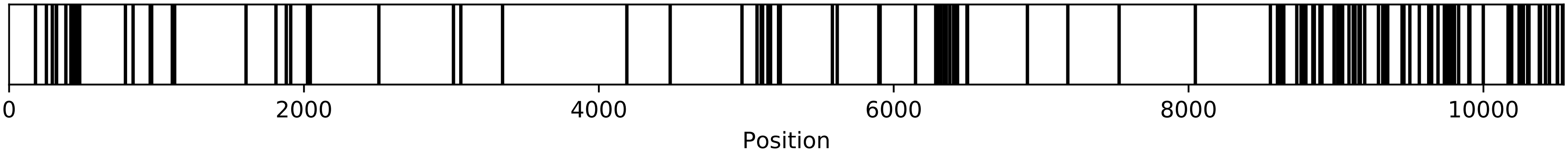


## CR2 vs MM1
Number of differences: 230

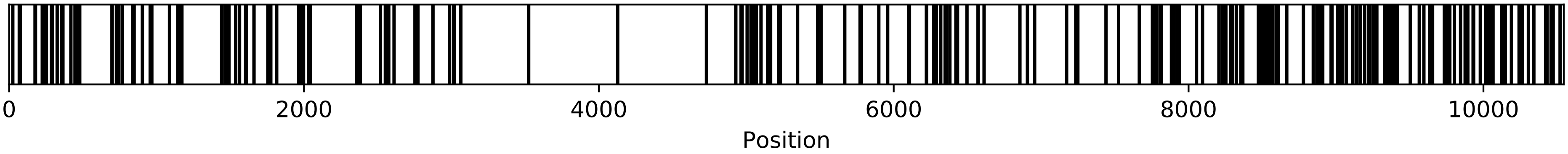


## CR2 vs MM2
Number of differences: 237

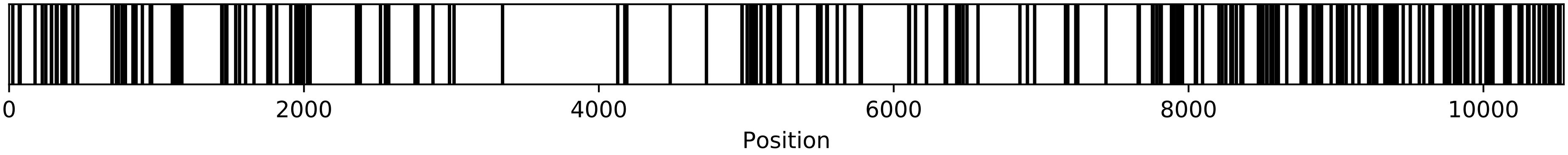


## MM1 vs MM2
Number of differences: 110

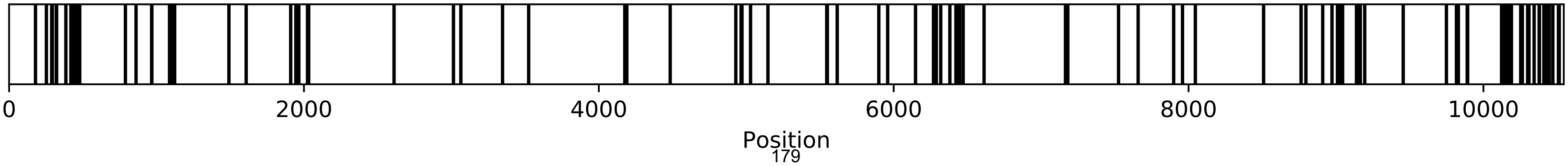

### Region: 517



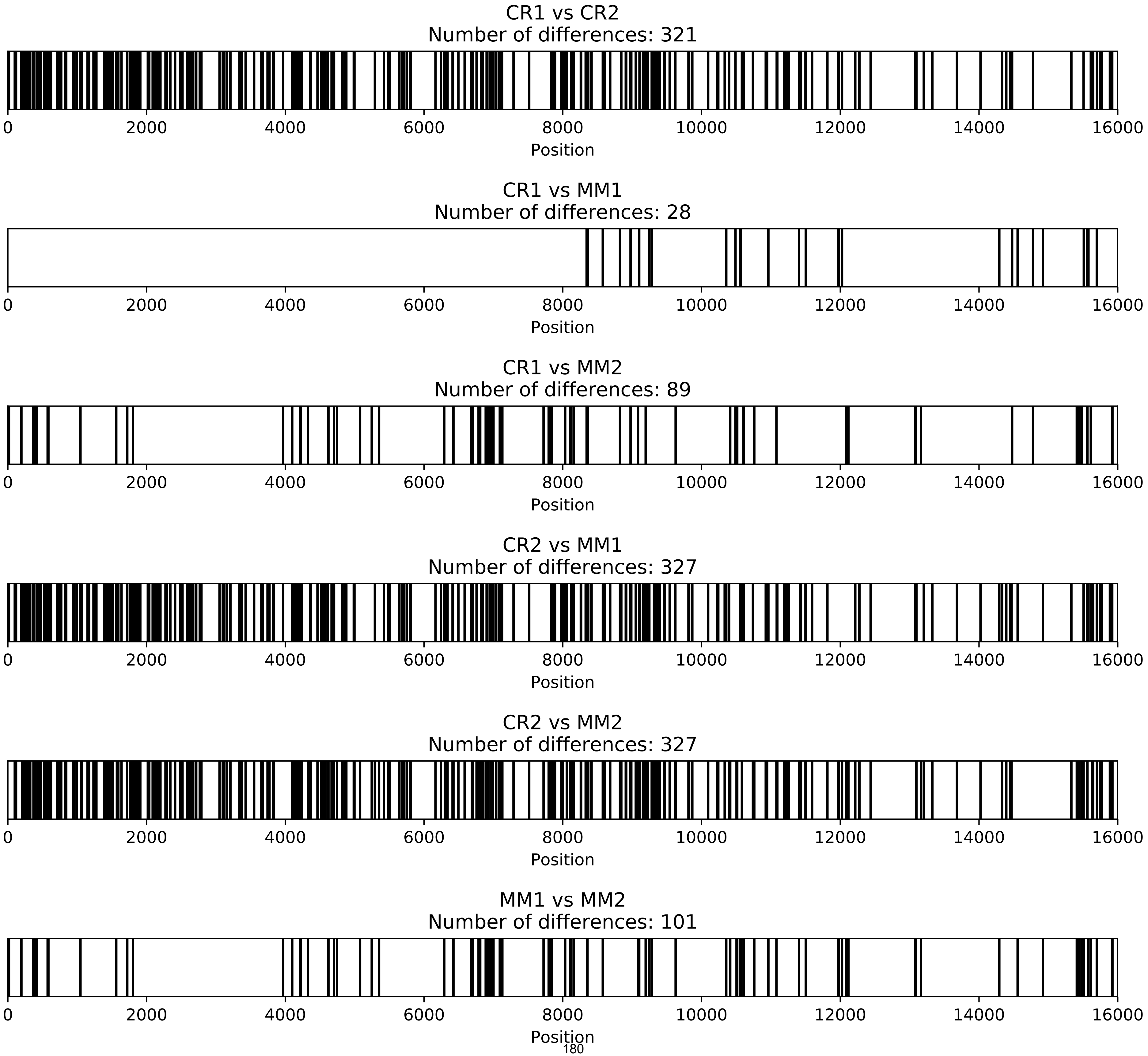
CR1 vs CR2
Number of differences: 321
CR1 vs MM1
Number of differences: 28
CR1 vs MM2
Number of differences: 89
CR2 vs MM1
Number of differences: 327
CR2 vs MM2
Number of differences: 327
MM1 vs MM2
Number of differences: 101
0 2000 4000 6000 8000 10000 12000 14000 16000
Position

### Region: 518



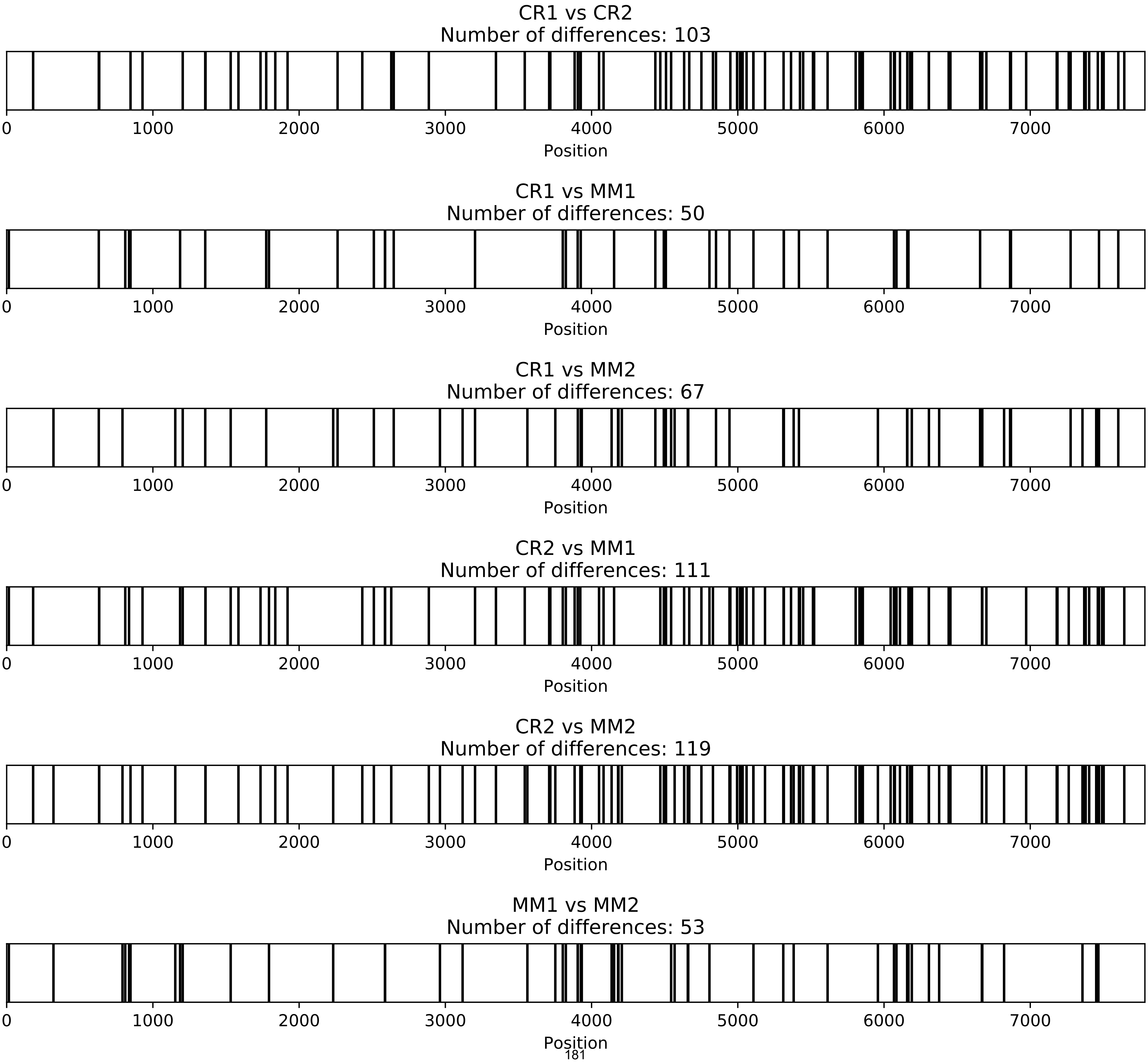
CR1 vs CR2
Number of differences: 103
Position
CR1 vs MM1
Number of differences: 50
Position
CR1 vs MM2
Number of differences: 67
Position
CR2 vs MM1
Number of differences: 111
Position
CR2 vs MM2
Number of differences: 119
Position
MM1 vs MM2
Number of differences: 53
Position
0
1000
2000
3000
4000
5000
6000
7000

### Region: 524



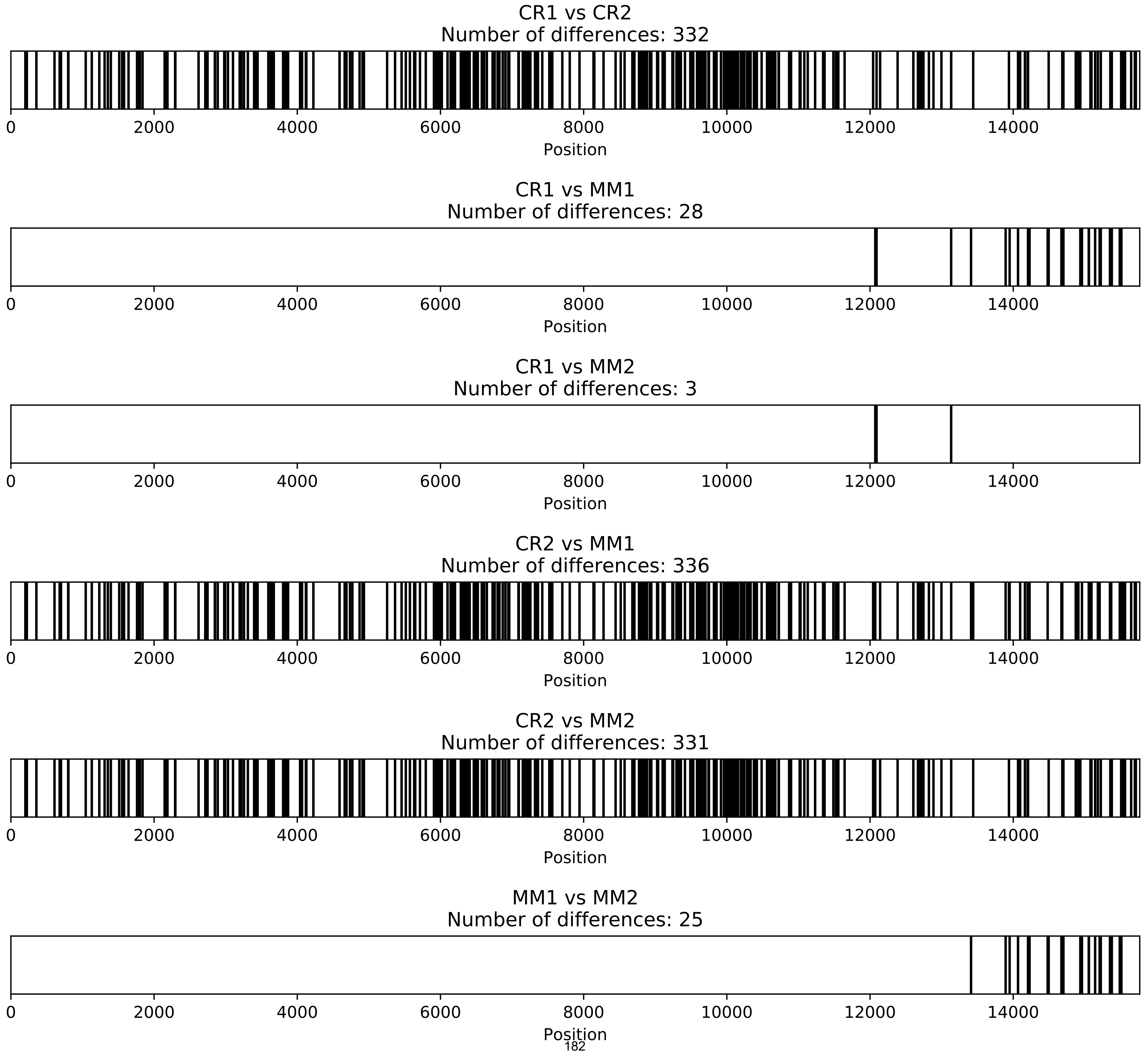
CR1 vs CR2
Number of differences: 332
CR1 vs MM1
Number of differences: 28
CR1 vs MM2
Number of differences: 3
CR2 vs MM1
Number of differences: 336
CR2 vs MM2
Number of differences: 331
MM1 vs MM2
Number of differences: 25
0
2000
4000
6000
8000
10000
12000
14000
Position

### Region: 526



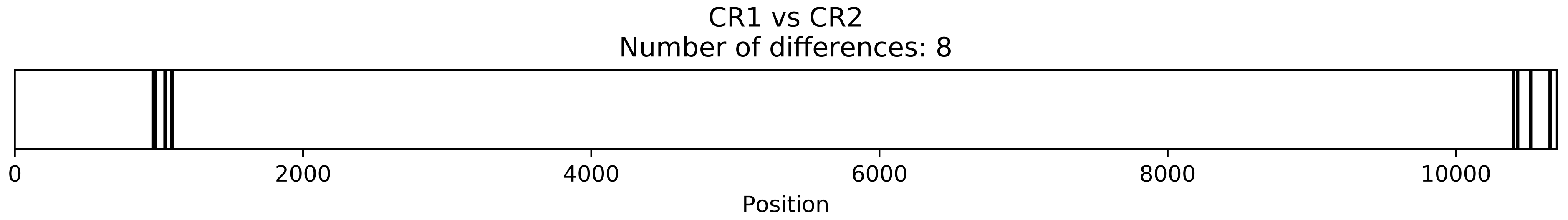
CR1 vs CR2
Number of differences: 8
0
2000
4000
6000
8000
10000
Position

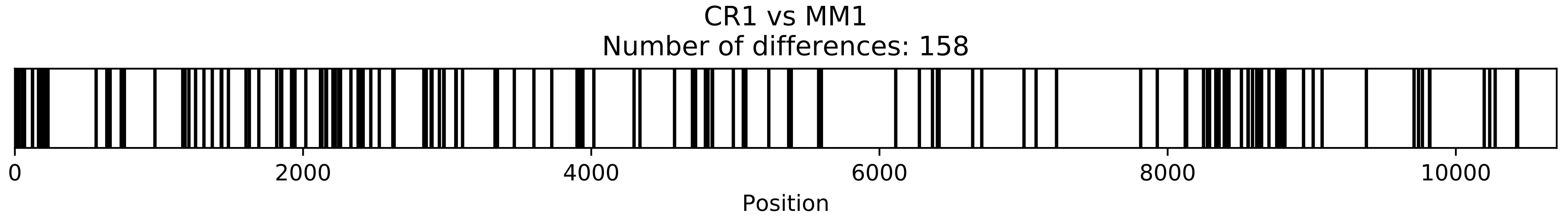
CR1 vs MM1
Number of differences: 158
0
2000
4000
6000
8000
10000
Position

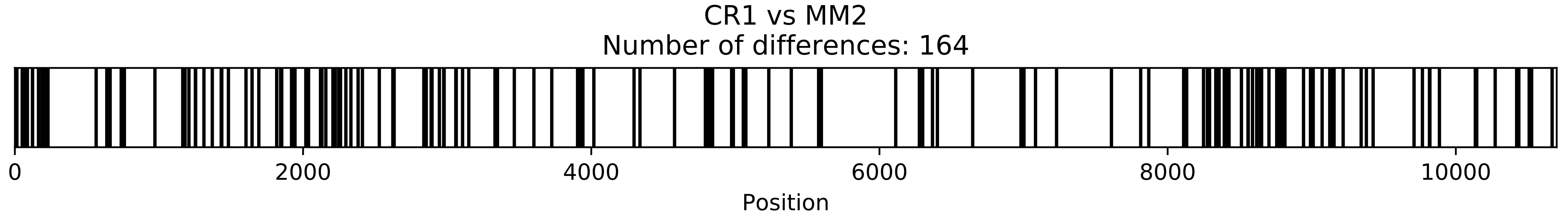
CR1 vs MM2
Number of differences: 164
0
2000
4000
6000
8000
10000
Position

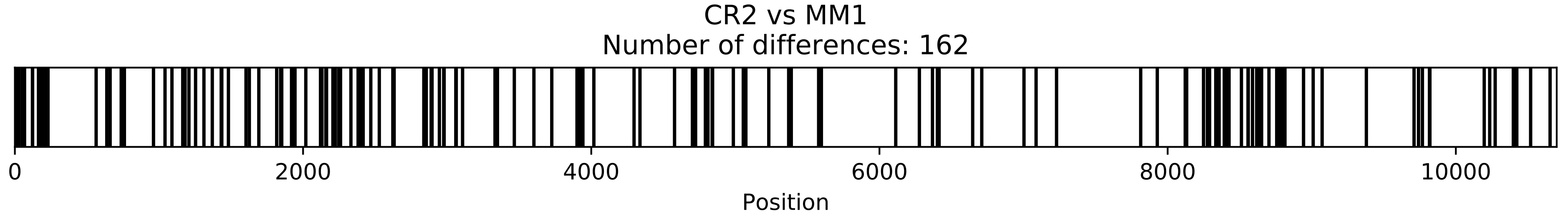
CR2 vs MM1
Number of differences: 162
0
2000
4000
6000
8000
10000
Position

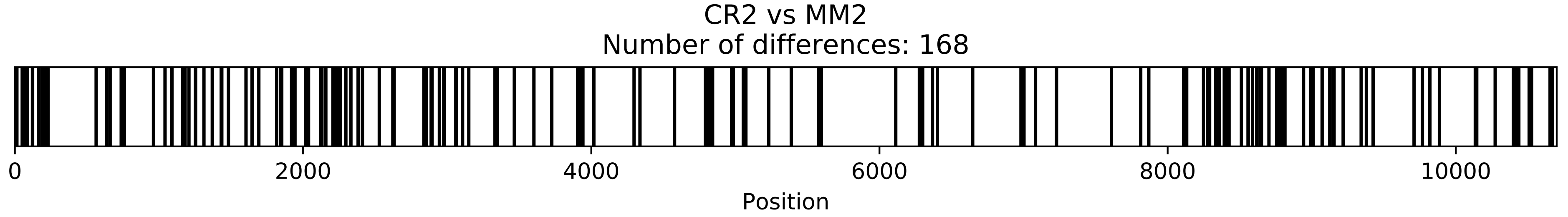
CR2 vs MM2
Number of differences: 168
0
2000
4000
6000
8000
10000
Position

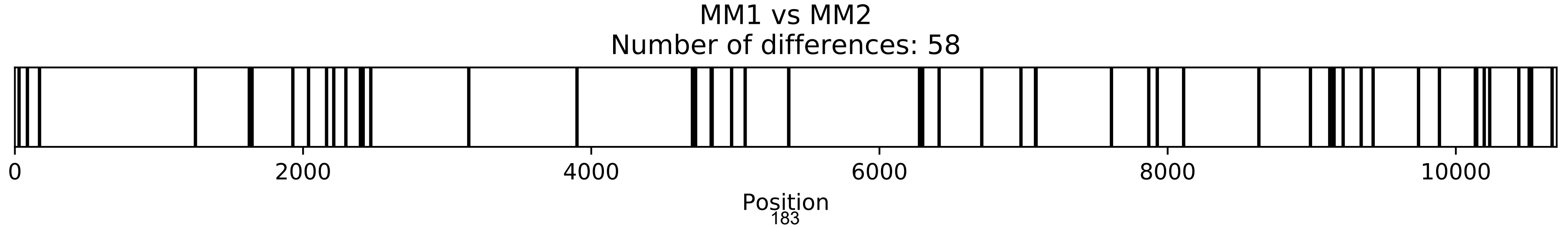
MM1 vs MM2
Number of differences: 58
0
2000
4000
6000
8000
10000
Position

### Region: 527



## CR1 vs CR2
Number of differences: 282

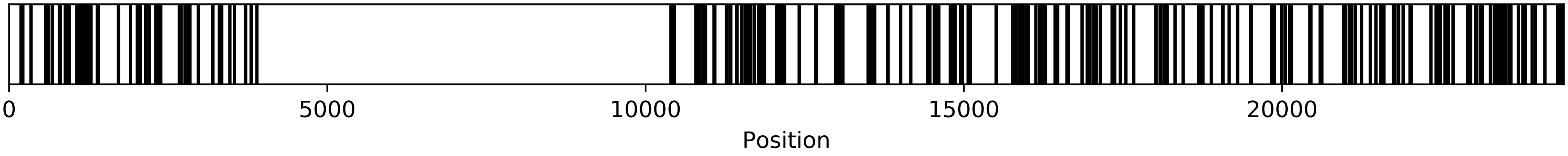


## CR1 vs MM1
Number of differences: 241

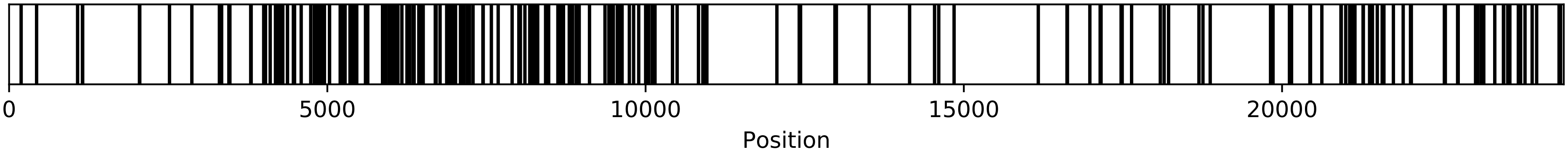


## CR1 vs MM2
Number of differences: 224

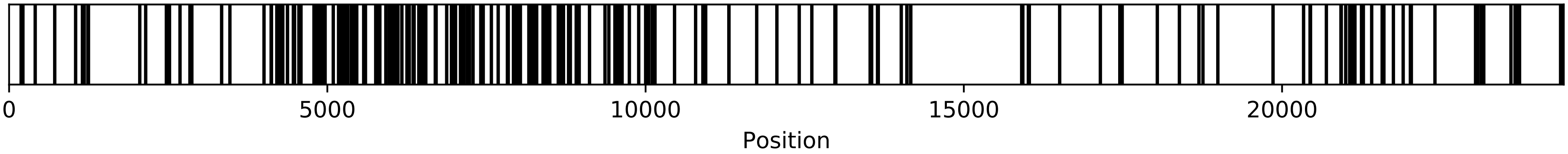


## CR2 vs MM1
Number of differences: 420

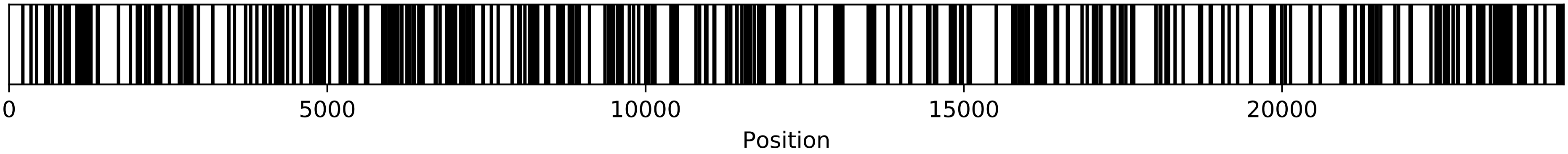


## CR2 vs MM2
Number of differences: 412

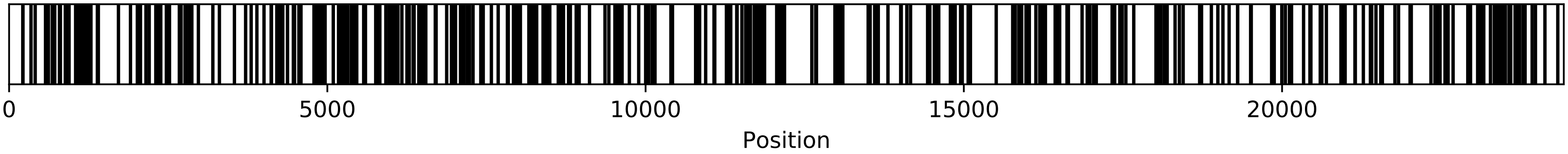


## MM1 vs MM2
Number of differences: 194

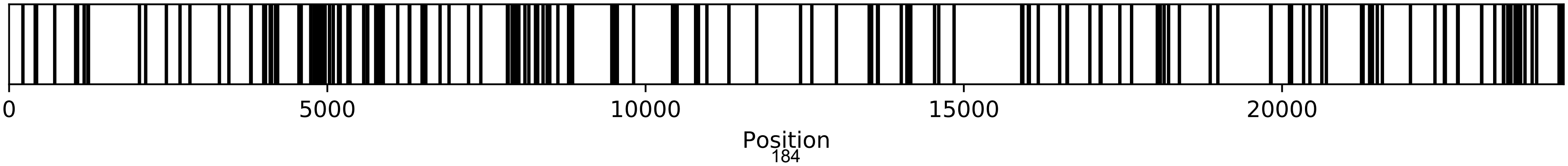

### Region: 528



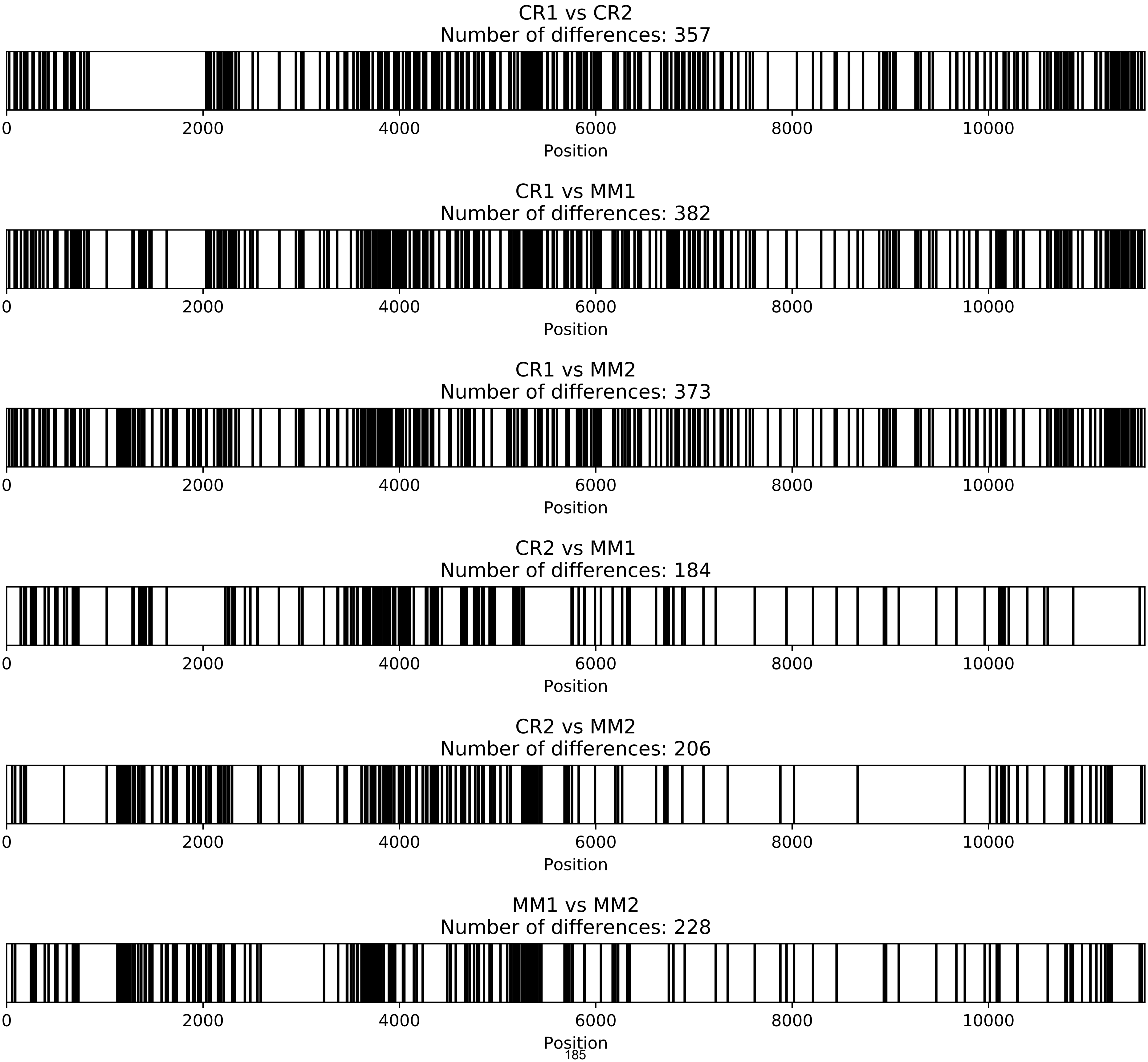
CR1 vs CR2
Number of differences: 357
CR1 vs MM1
Number of differences: 382
CR1 vs MM2
Number of differences: 373
CR2 vs MM1
Number of differences: 184
CR2 vs MM2
Number of differences: 206
MM1 vs MM2
Number of differences: 228
0
2000
4000
6000
8000
10000
Position

### Region: 529



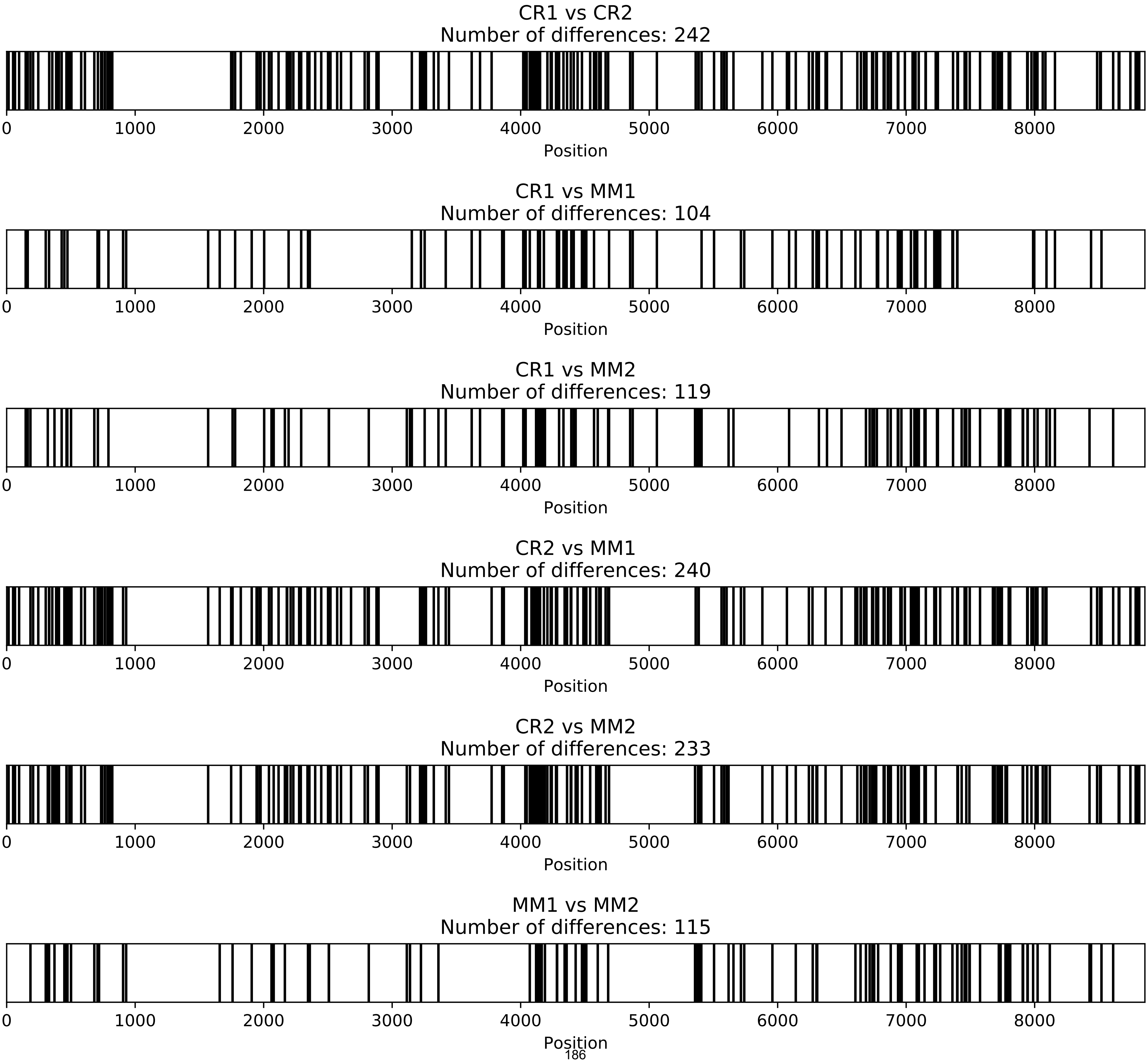
CR1 vs CR2
Number of differences: 242
0
1000
2000
3000
4000
5000
6000
7000
8000
Position
CR1 vs MM1
Number of differences: 104
Position
CR1 vs MM2
Number of differences: 119
Position
CR2 vs MM1
Number of differences: 240
Position
CR2 vs MM2
Number of differences: 233
Position
MM1 vs MM2
Number of differences: 115
Position

### Region: 531



CR1 vs CR2
Number of differences: 0

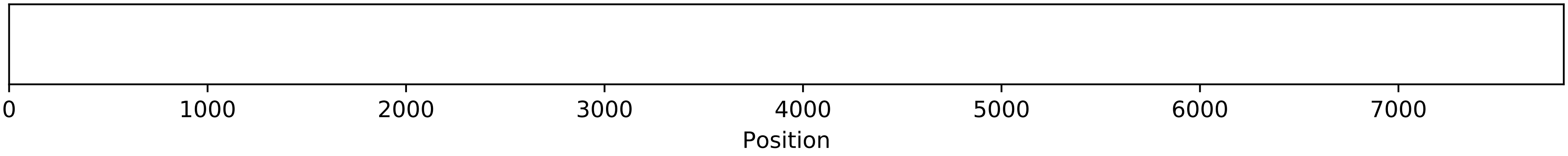


CR1 vs MM1
Number of differences: 238

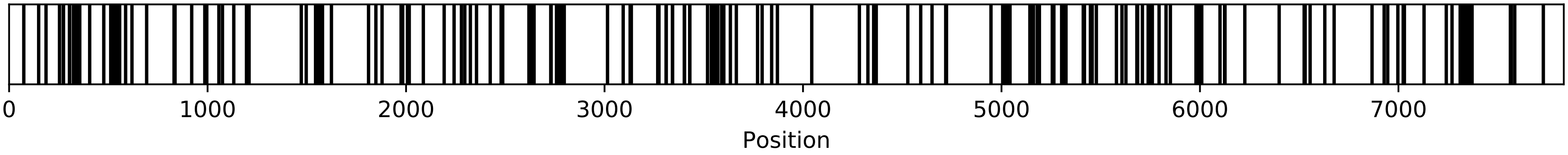


CR1 vs MM2
Number of differences: 257

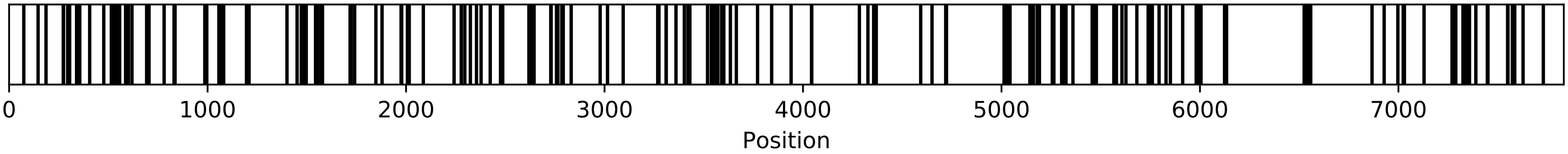


CR2 vs MM1
Number of differences: 238

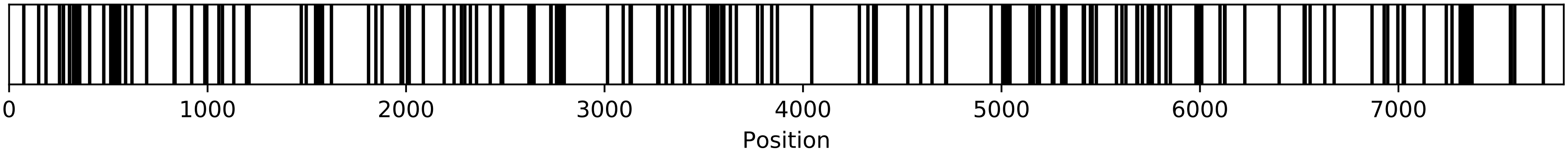


CR2 vs MM2
Number of differences: 257

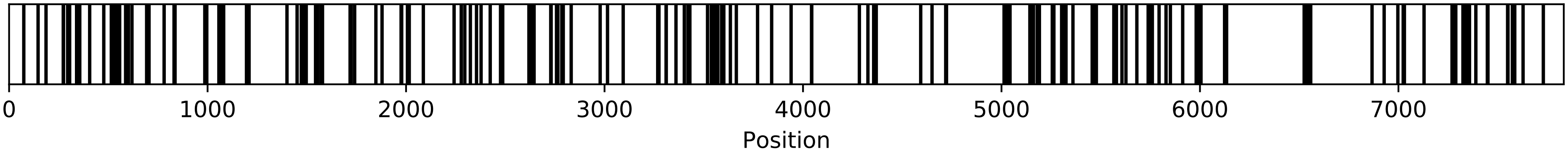


MM1 vs MM2
Number of differences: 141

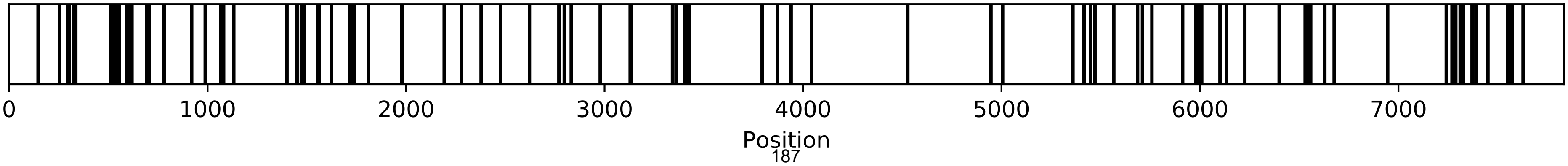

### Region: 532



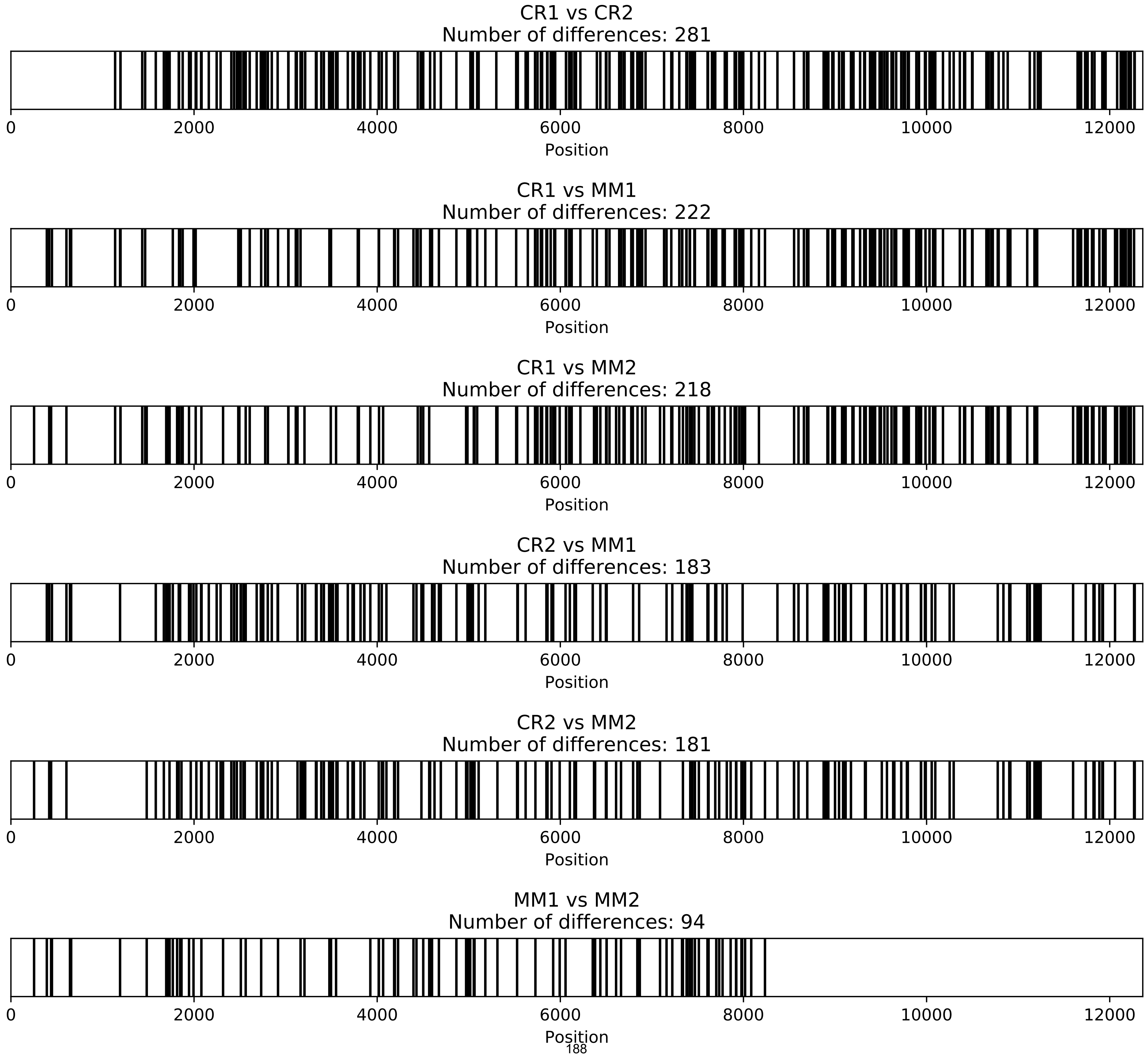
CR1 vs CR2
Number of differences: 281
CR1 vs MM1
Number of differences: 222
CR1 vs MM2
Number of differences: 218
CR2 vs MM1
Number of differences: 183
CR2 vs MM2
Number of differences: 181
MM1 vs MM2
Number of differences: 94
0
2000
4000
6000
8000
10000
12000
Position

### Region: 535



## CR1 vs CR2
Number of differences: 151

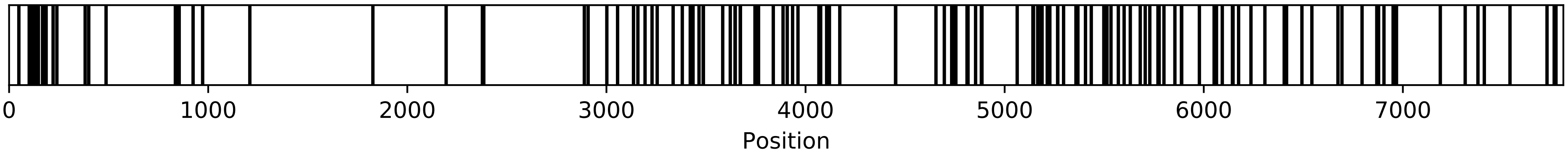


## CR1 vs MM1
Number of differences: 68

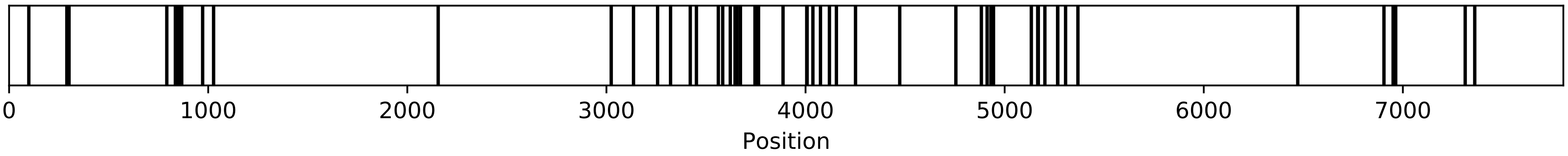


## CR1 vs MM2
Number of differences: 72

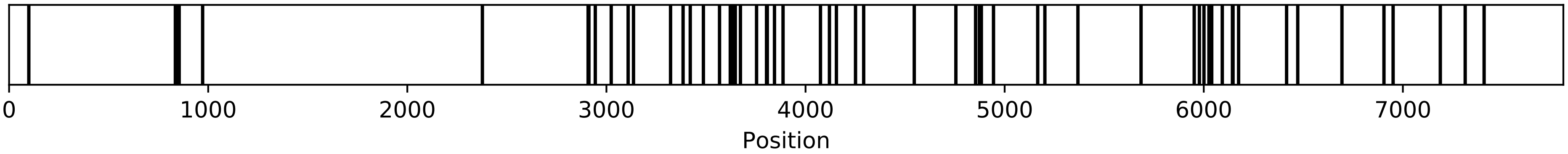


## CR2 vs MM1
Number of differences: 146

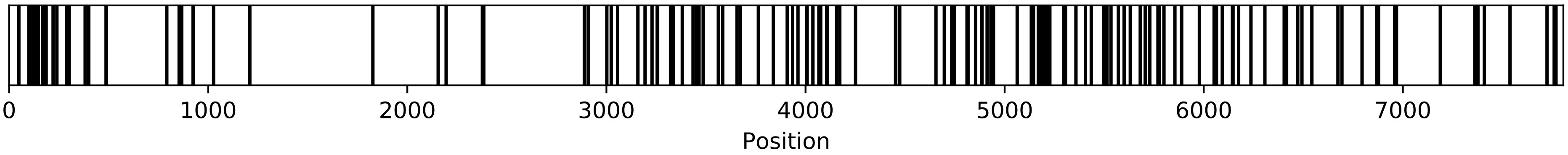


## CR2 vs MM2
Number of differences: 136

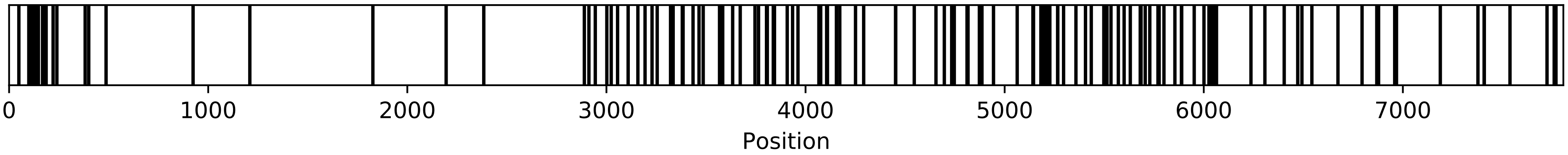


## MM1 vs MM2
Number of differences: 60

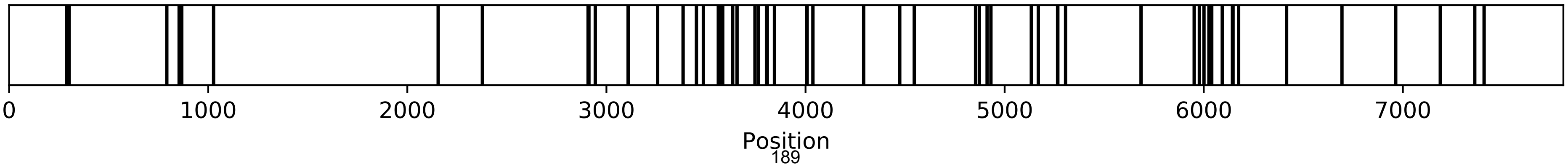

### Region: 537



CR1 vs CR2
Number of differences: 307

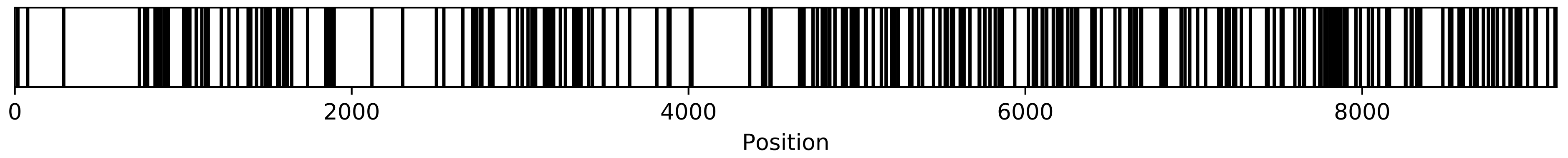


CR1 vs MM1
Number of differences: 159

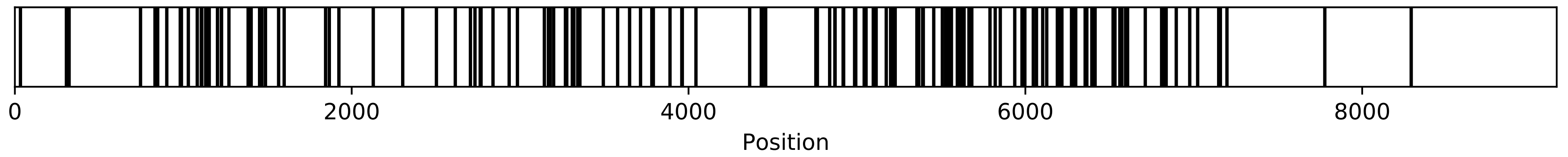


CR1 vs MM2
Number of differences: 207

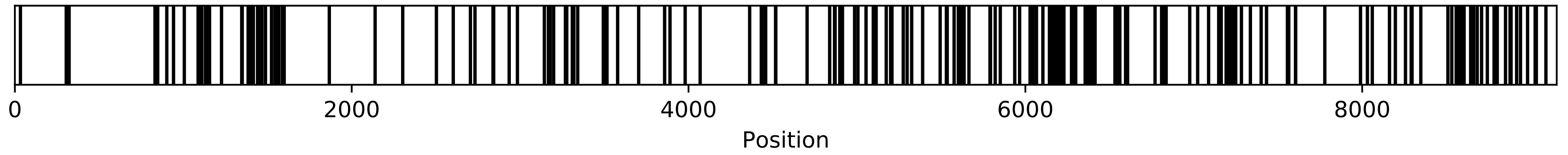


CR2 vs MM1
Number of differences: 296

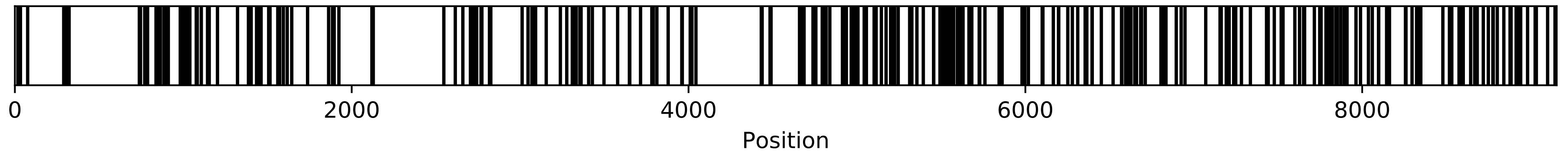


CR2 vs MM2
Number of differences: 300

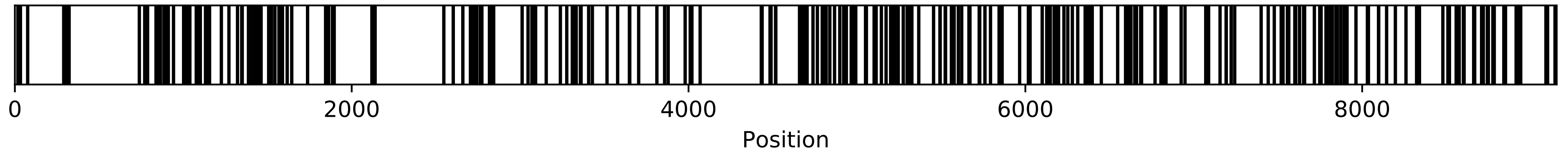


MM1 vs MM2
Number of differences: 168

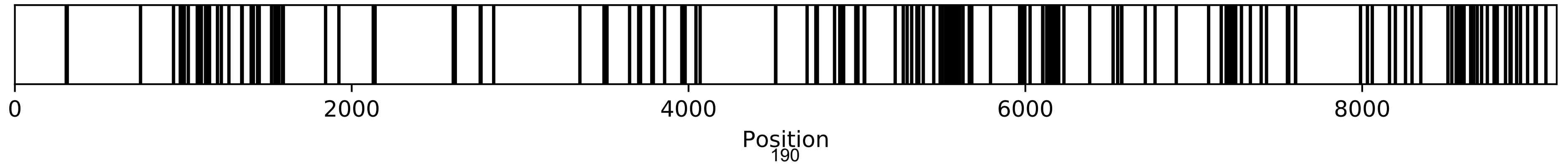

### Region: 538



CR1 vs CR2
Number of differences: 360

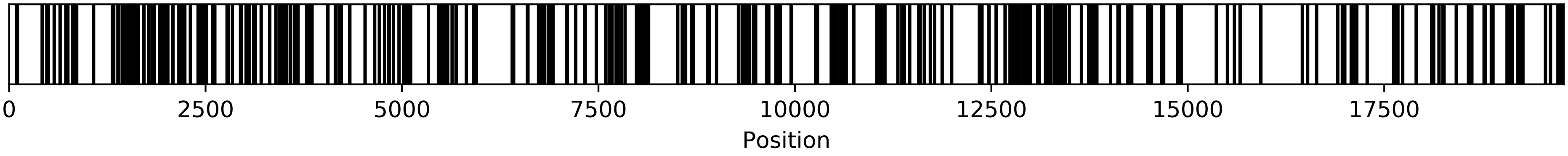


CR1 vs MM1
Number of differences: 165

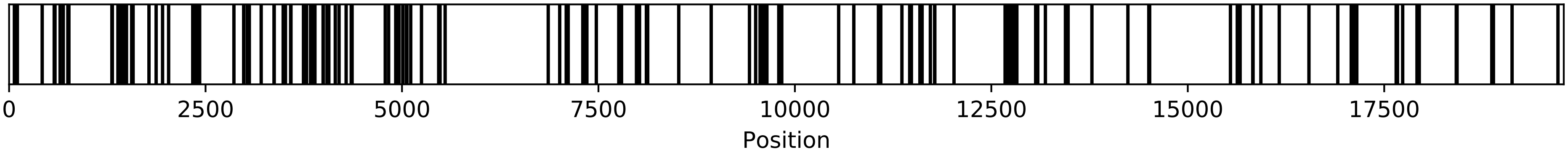


CR1 vs MM2
Number of differences: 89

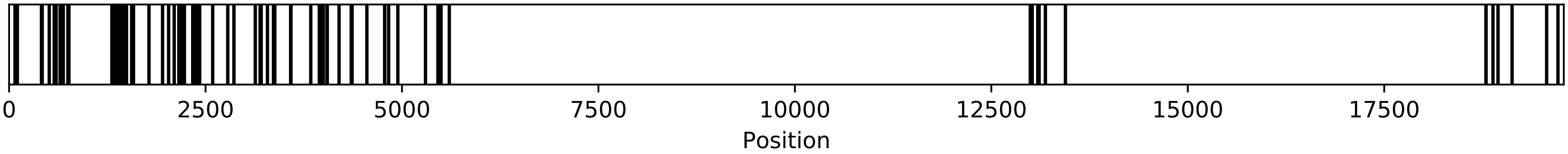


CR2 vs MM1
Number of differences: 356

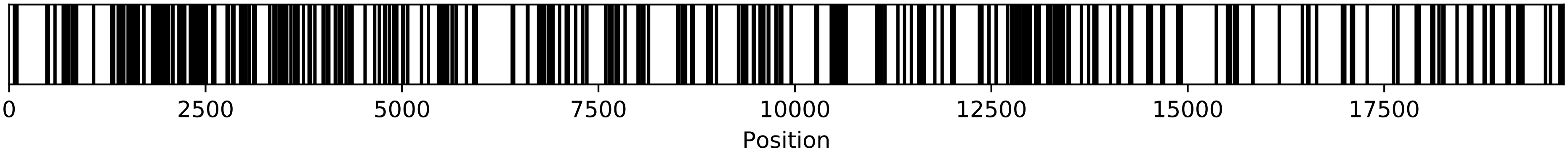


CR2 vs MM2
Number of differences: 373

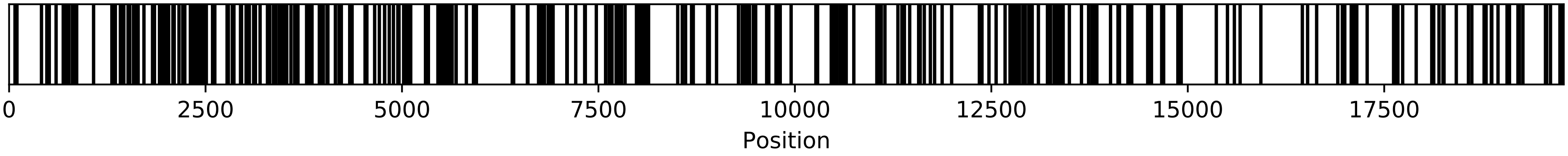


MM1 vs MM2
Number of differences: 162

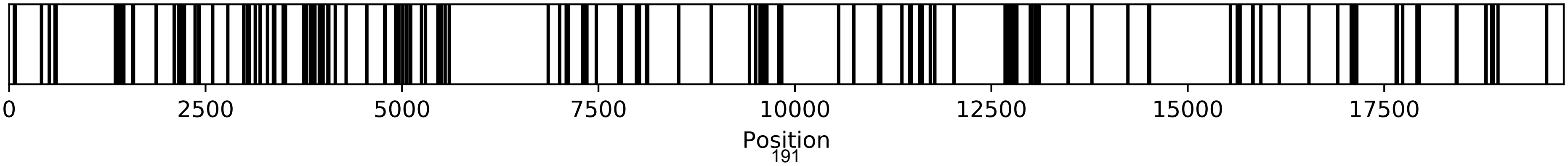

### Region: 546



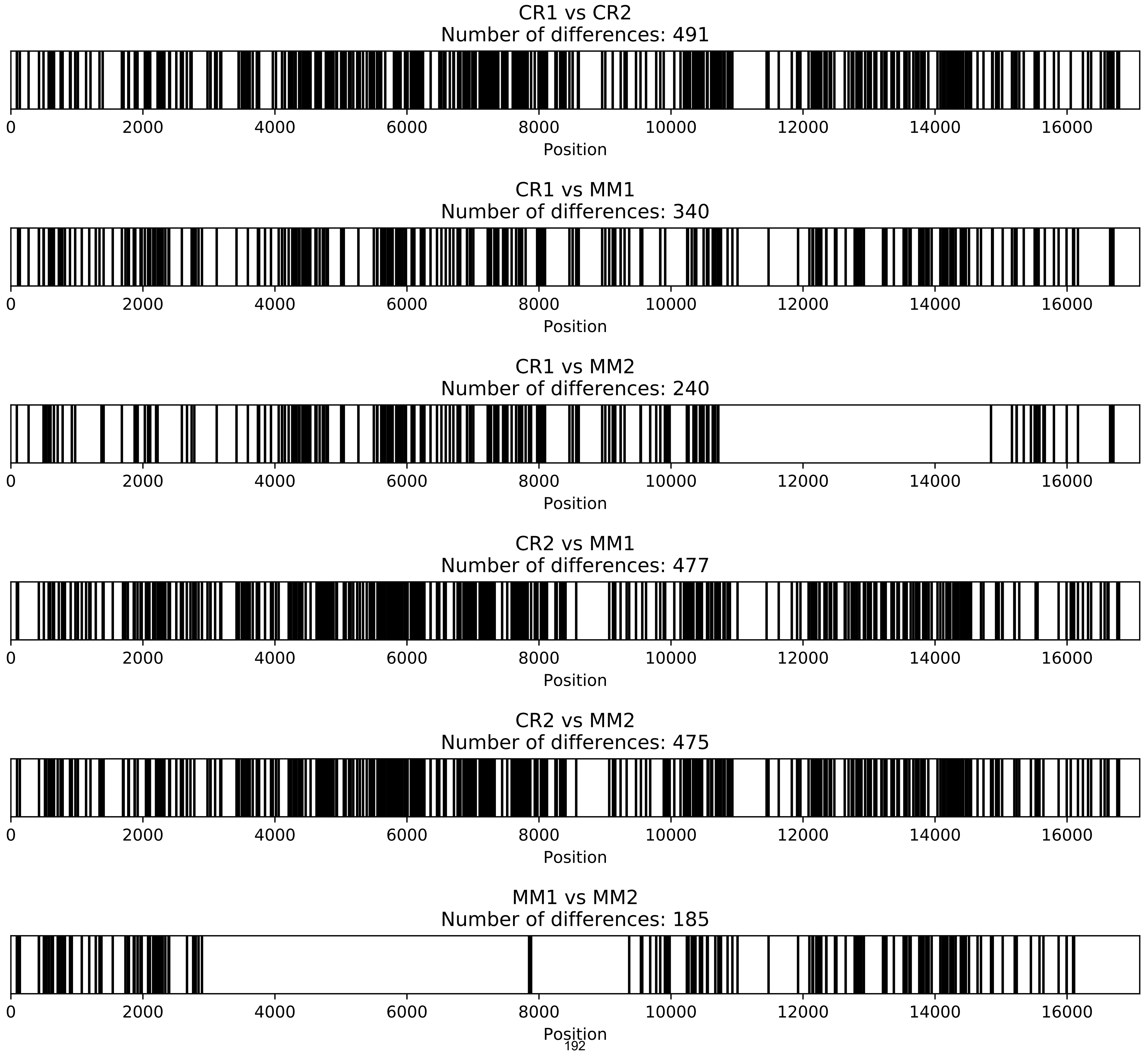
CR1 vs CR2
Number of differences: 491
0
2000
4000
6000
8000
10000
12000
14000
16000
Position
CR1 vs MM1
Number of differences: 340
Position
CR1 vs MM2
Number of differences: 240
Position
CR2 vs MM1
Number of differences: 477
Position
CR2 vs MM2
Number of differences: 475
Position
MM1 vs MM2
Number of differences: 185
Position

### Region: 551



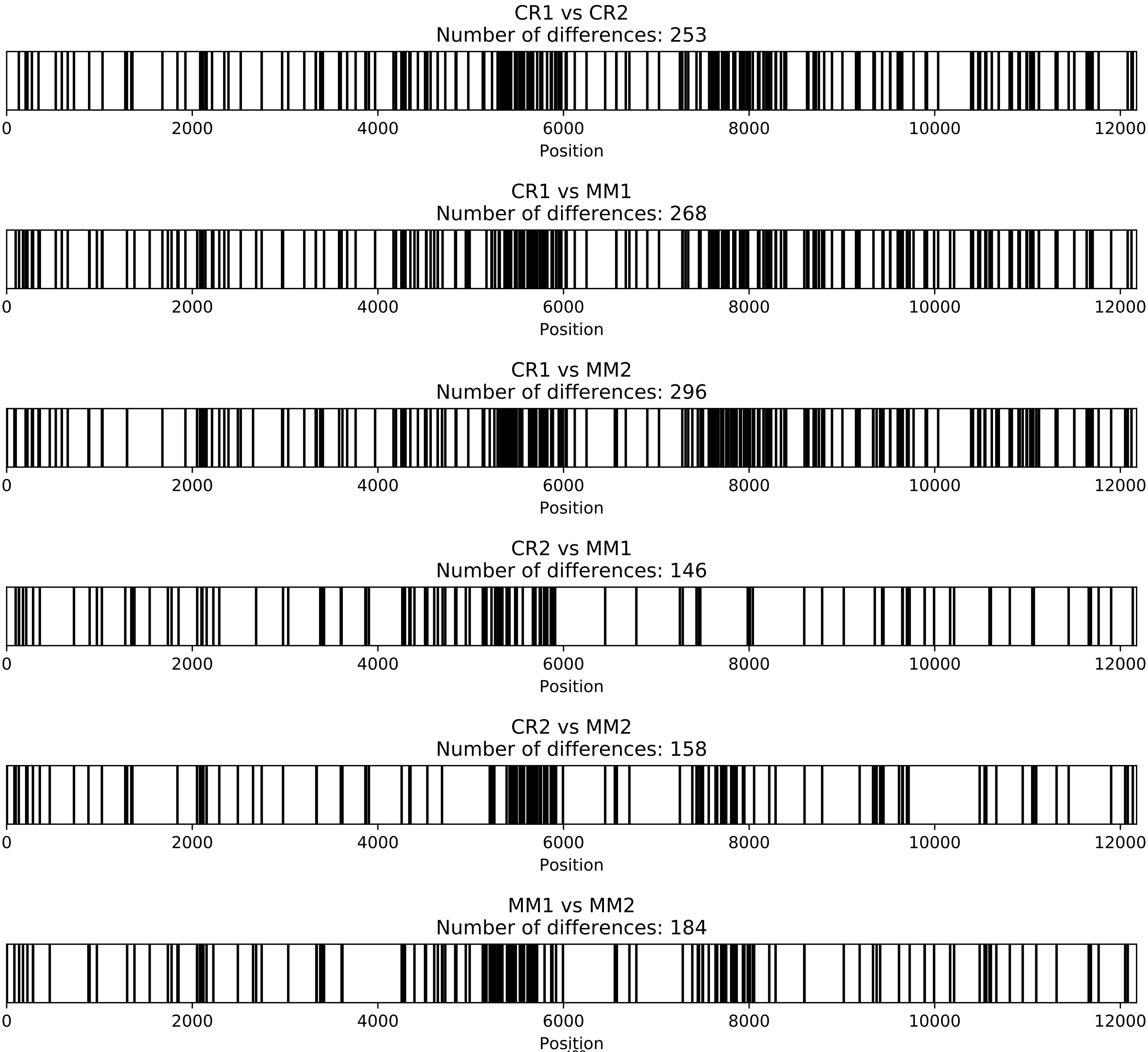
CR1 vs CR2
Number of differences: 253
CR1 vs MM1
Number of differences: 268
CR1 vs MM2
Number of differences: 296
CR2 vs MM1
Number of differences: 146
CR2 vs MM2
Number of differences: 158
MM1 vs MM2
Number of differences: 184
0
2000
4000
6000
8000
10000
12000
Position

### Region: 552



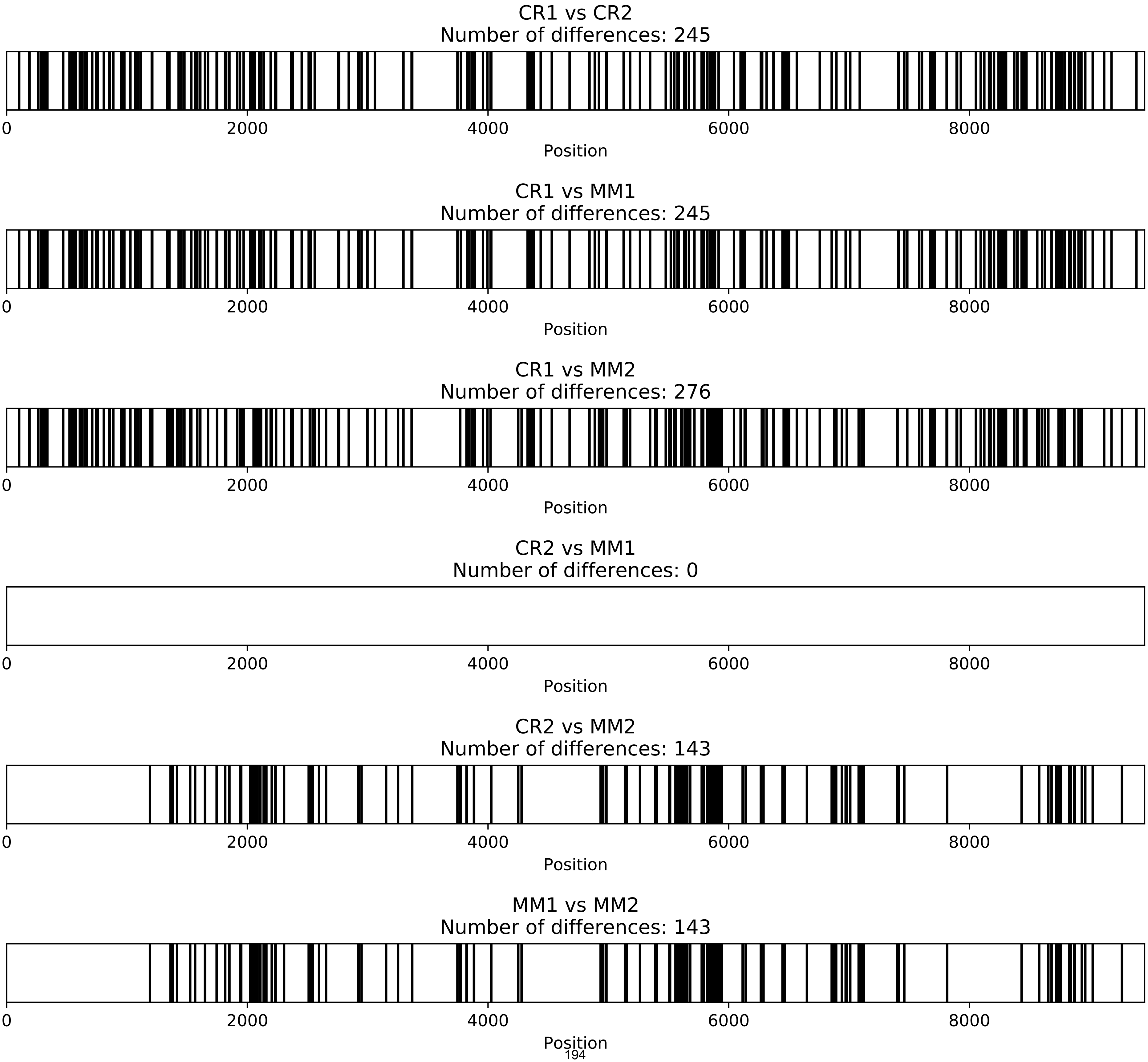
CR1 vs CR2
Number of differences: 245
CR1 vs MM1
Number of differences: 245
CR1 vs MM2
Number of differences: 276
CR2 vs MM1
Number of differences: 0
CR2 vs MM2
Number of differences: 143
MM1 vs MM2
Number of differences: 143
0
2000
4000
6000
8000
Position

### Region: 555



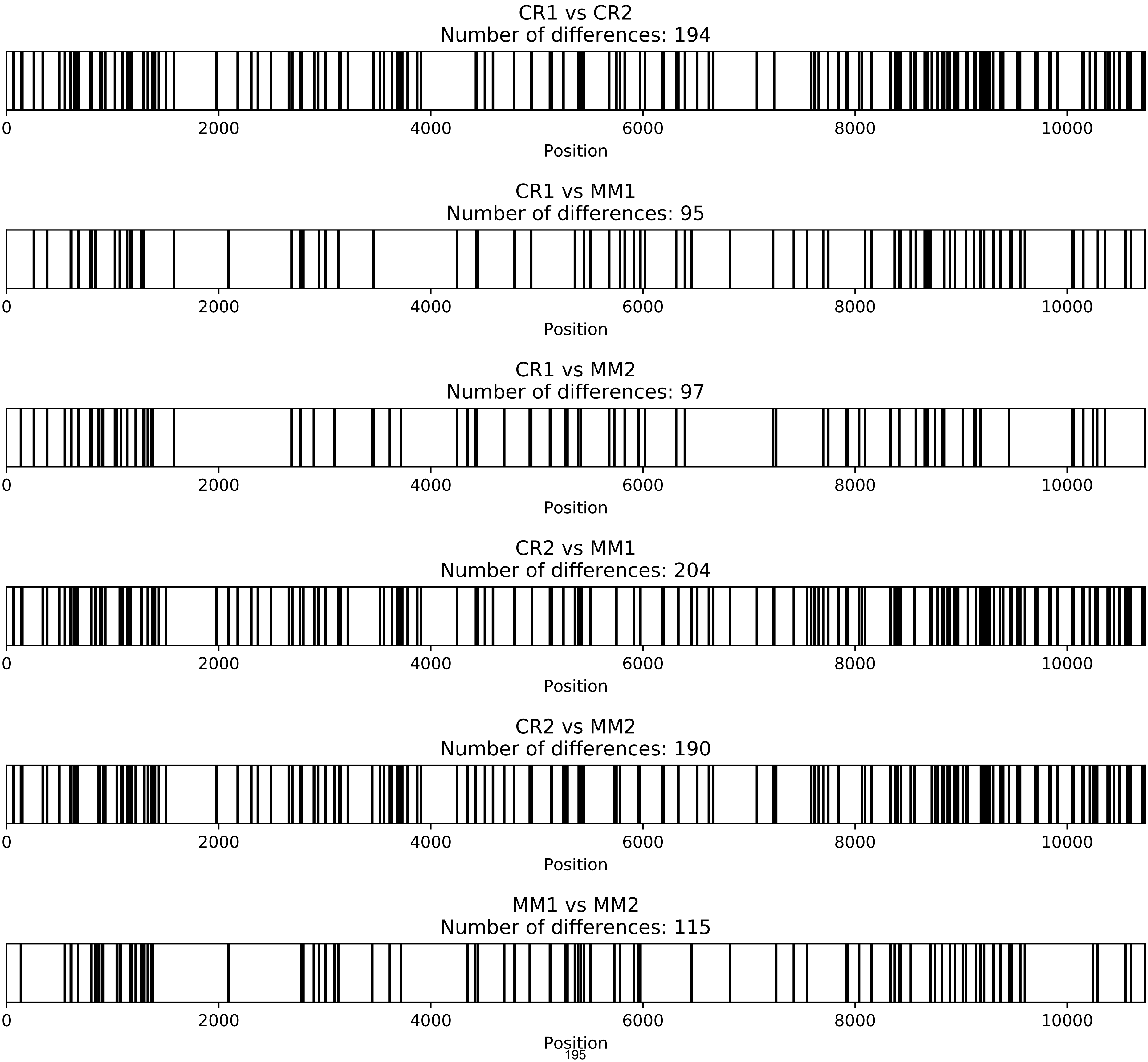
CR1 vs CR2
Number of differences: 194
CR1 vs MM1
Number of differences: 95
CR1 vs MM2
Number of differences: 97
CR2 vs MM1
Number of differences: 204
CR2 vs MM2
Number of differences: 190
MM1 vs MM2
Number of differences: 115
0
2000
4000
6000
8000
10000
Position

### Region: 556



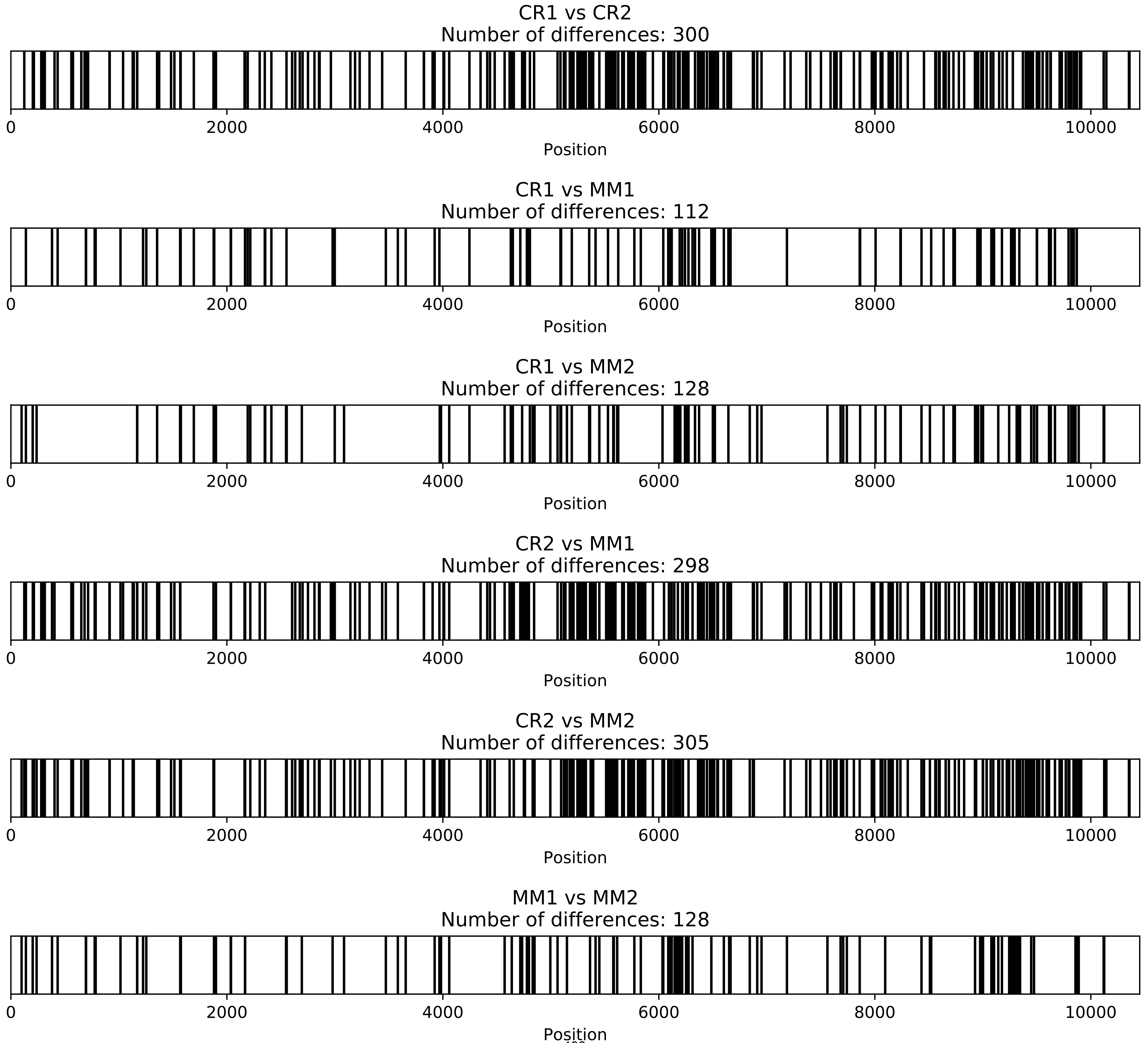
CR1 vs CR2
Number of differences: 300
CR1 vs MM1
Number of differences: 112
CR1 vs MM2
Number of differences: 128
CR2 vs MM1
Number of differences: 298
CR2 vs MM2
Number of differences: 305
MM1 vs MM2
Number of differences: 128
0
2000
4000
6000
8000
10000
Position

### Region: 561



CR1 vs CR2
Number of differences: 151

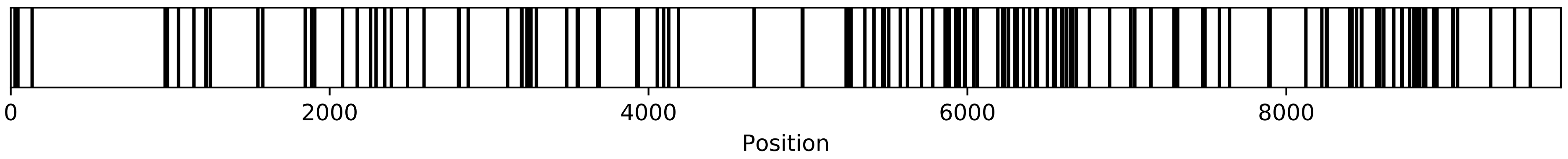


CR1 vs MM1
Number of differences: 70

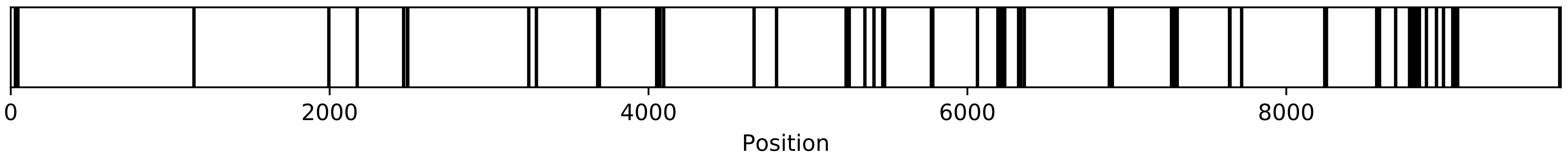


CR1 vs MM2
Number of differences: 76

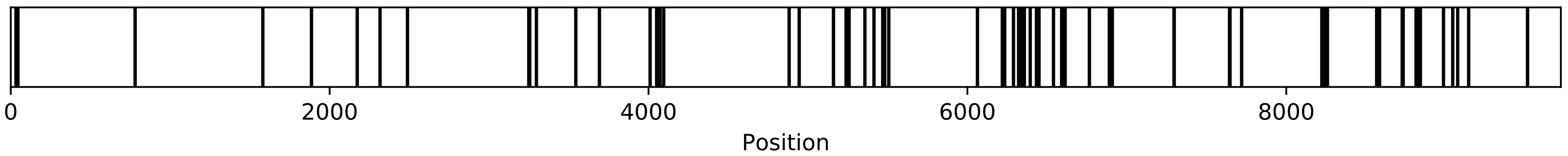


CR2 vs MM1
Number of differences: 150

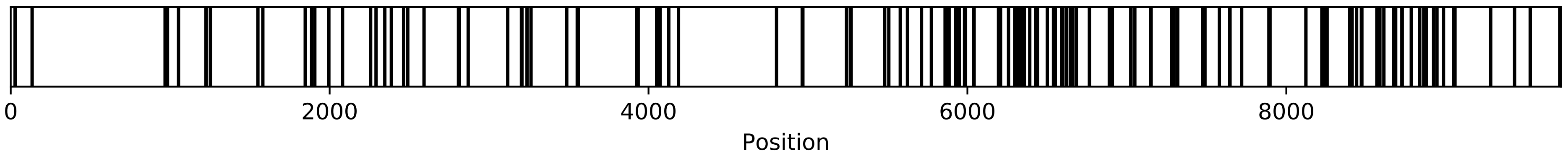


CR2 vs MM2
Number of differences: 154

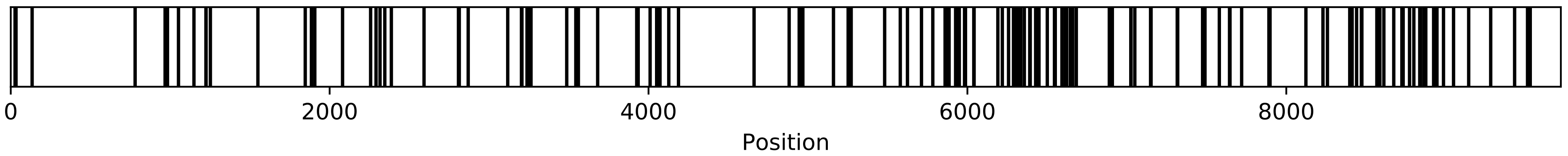


MM1 vs MM2
Number of differences: 64

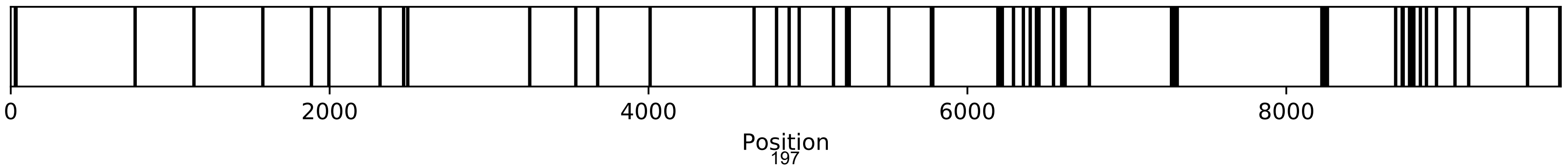

### Region: 562



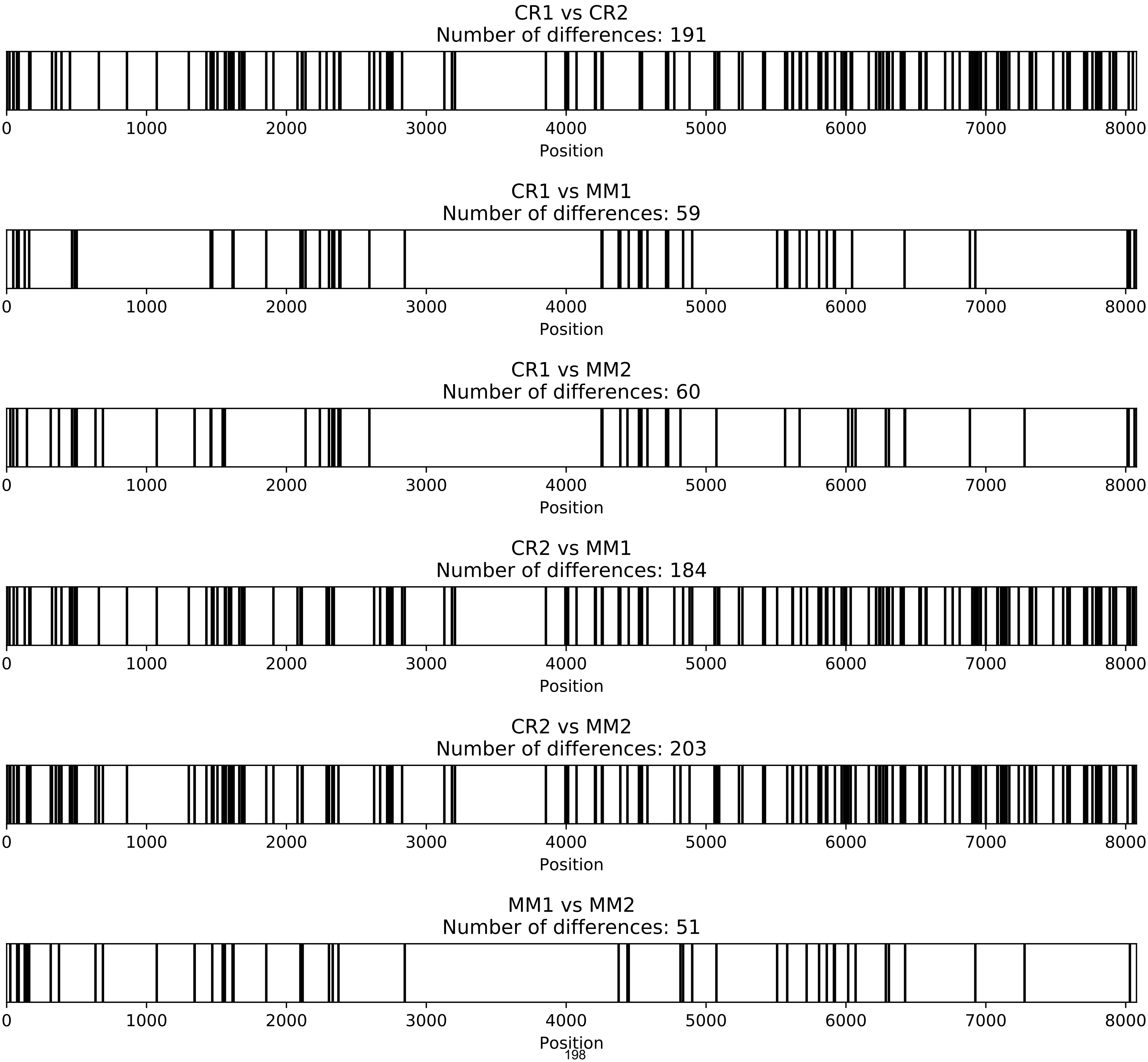
CR1 vs CR2
Number of differences: 191
CR1 vs MM1
Number of differences: 59
CR1 vs MM2
Number of differences: 60
CR2 vs MM1
Number of differences: 184
CR2 vs MM2
Number of differences: 203
MM1 vs MM2
Number of differences: 51
0
1000
2000
3000
4000
5000
6000
7000
8000
Position

### Region: 567



CR1 vs CR2
Number of differences: 0

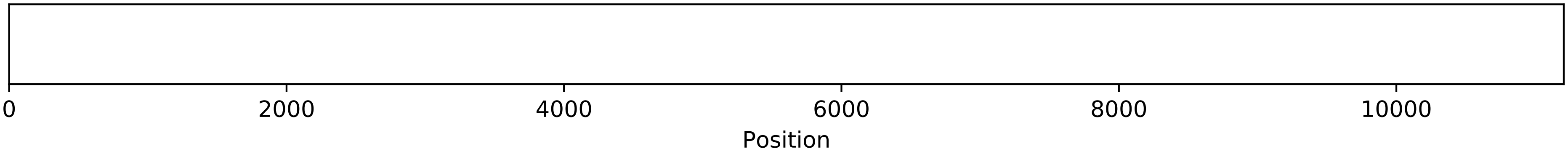


CR1 vs MM1
Number of differences: 216

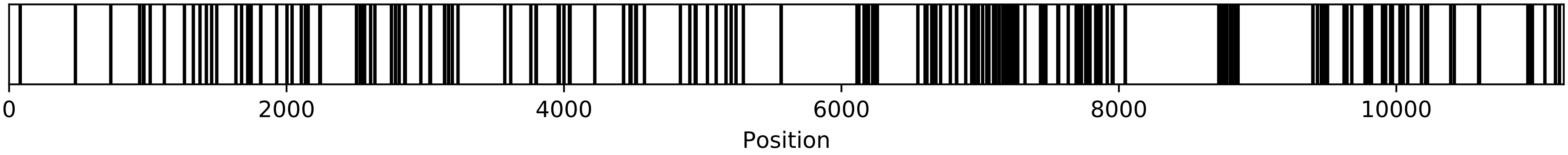


CR1 vs MM2
Number of differences: 241

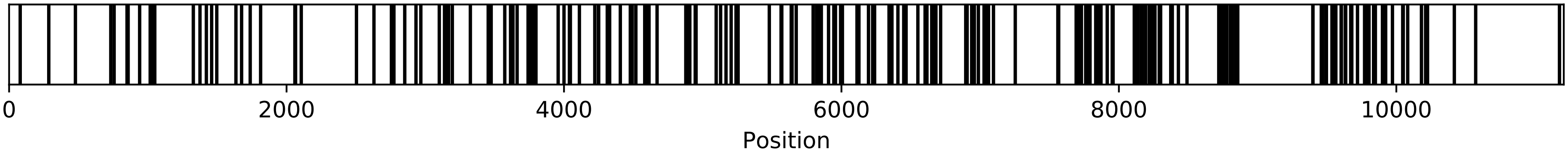


CR2 vs MM1
Number of differences: 216

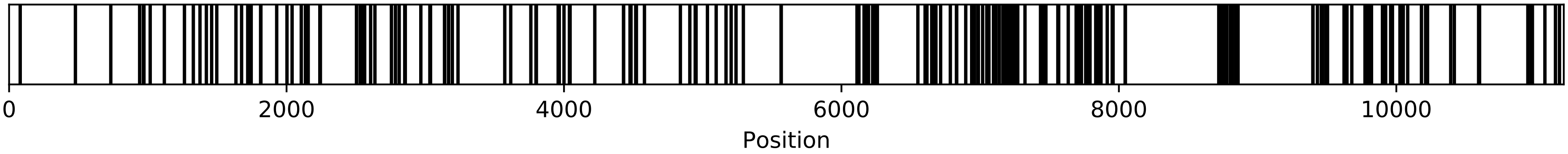


CR2 vs MM2
Number of differences: 241

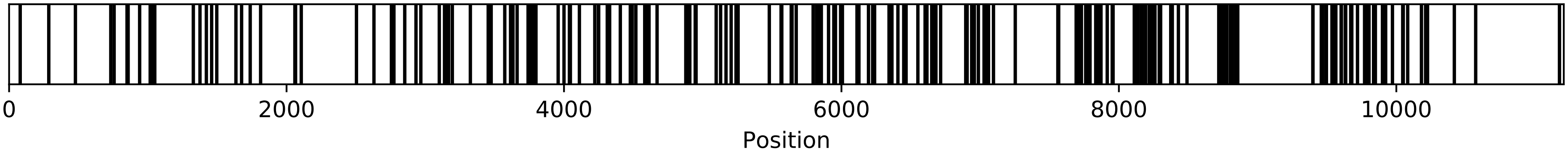


MM1 vs MM2
Number of differences: 205

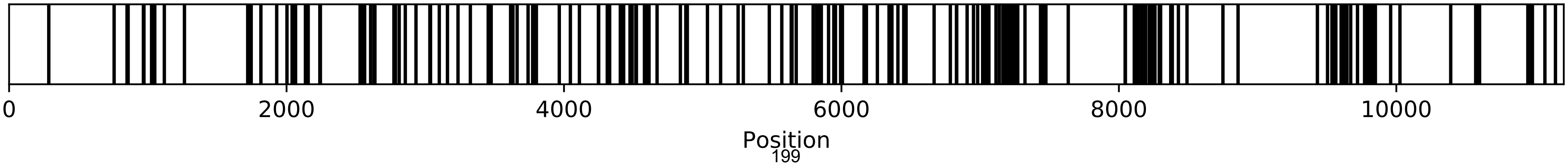

### Region: 572



CR1 vs CR2
Number of differences: 243

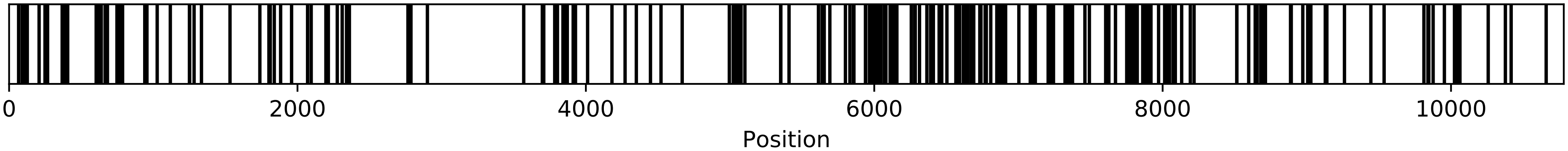


CR1 vs MM1
Number of differences: 162

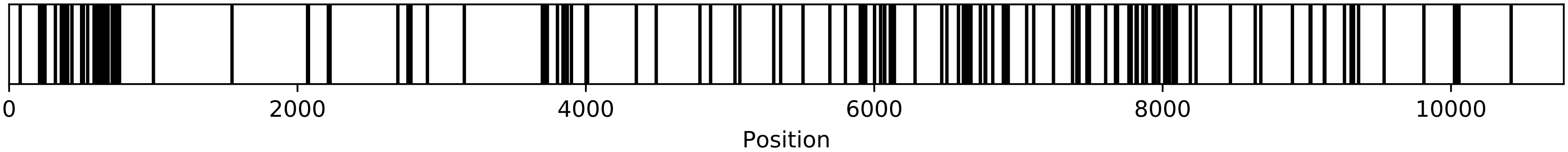


CR1 vs MM2
Number of differences: 211

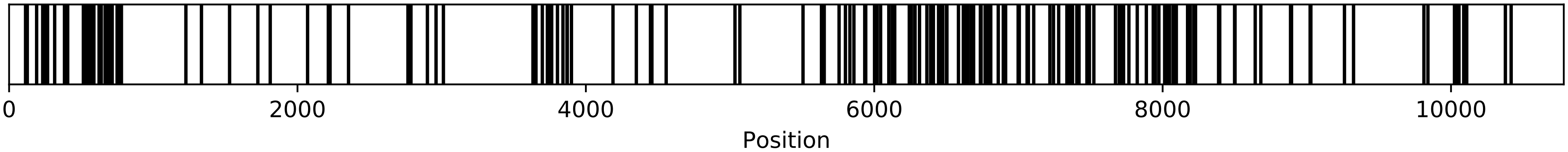


CR2 vs MM1
Number of differences: 219

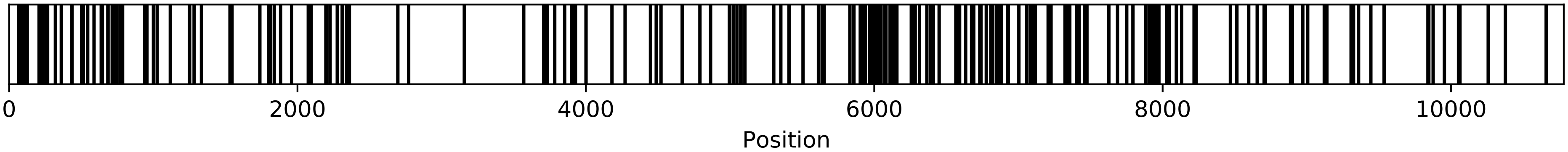


CR2 vs MM2
Number of differences: 236

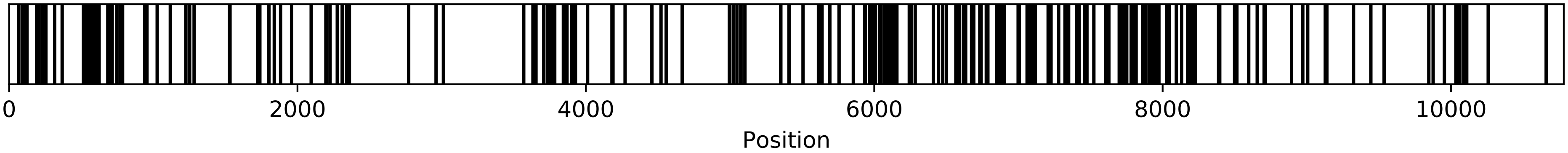


MM1 vs MM2
Number of differences: 168

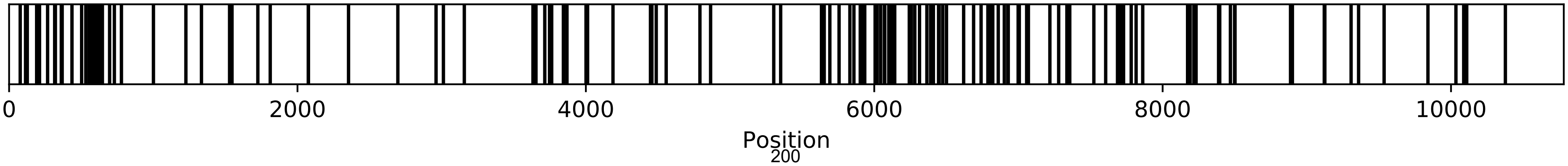

### Region: 576



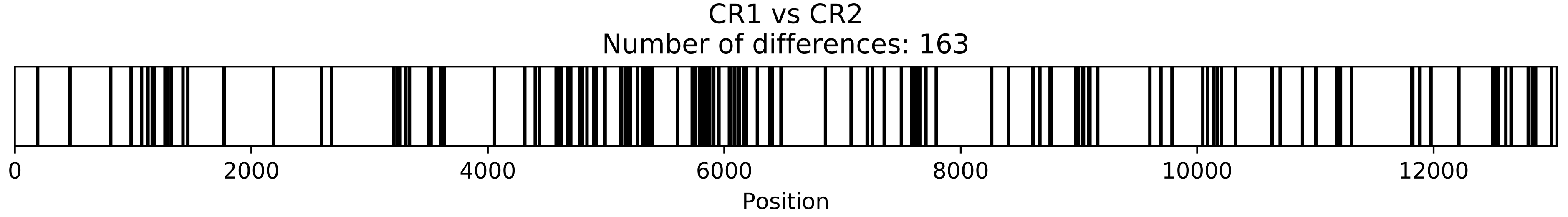
CR1 vs CR2
Number of differences: 163
0
2000
4000
6000
8000
10000
12000
Position

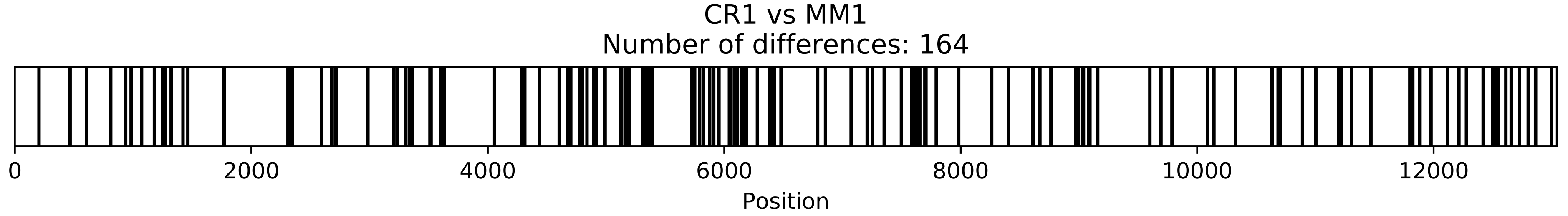
CR1 vs MM1
Number of differences: 164
0
2000
4000
6000
8000
10000
12000
Position

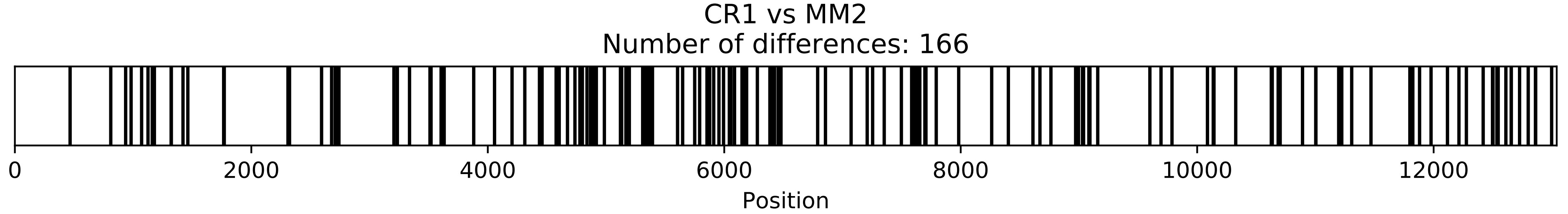
CR1 vs MM2
Number of differences: 166
0
2000
4000
6000
8000
10000
12000
Position

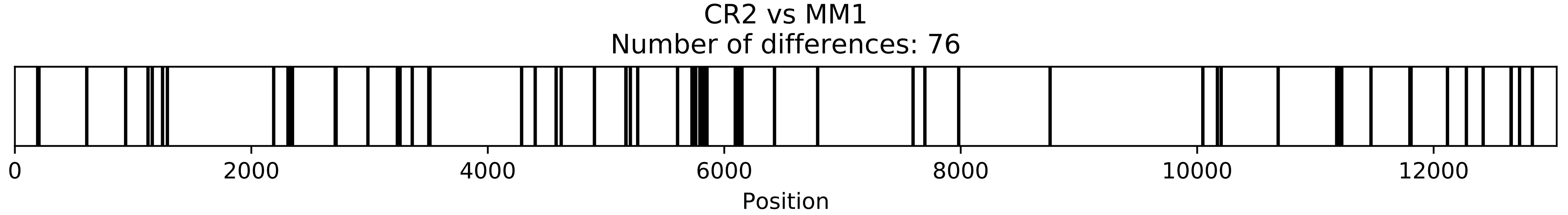
CR2 vs MM1
Number of differences: 76
0
2000
4000
6000
8000
10000
12000
Position

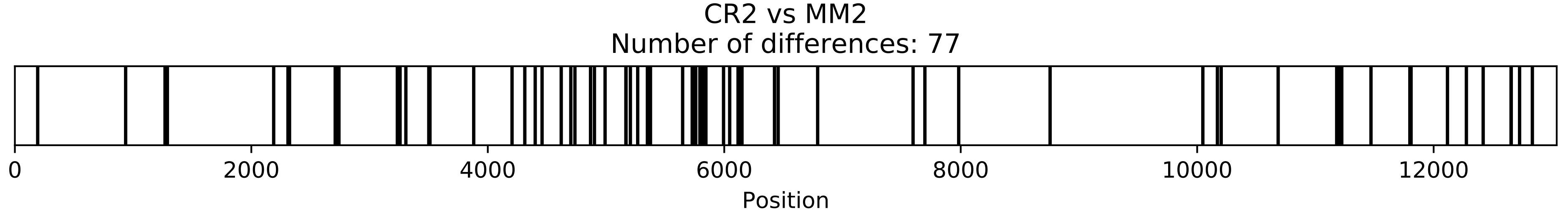
CR2 vs MM2
Number of differences: 77
0
2000
4000
6000
8000
10000
12000
Position

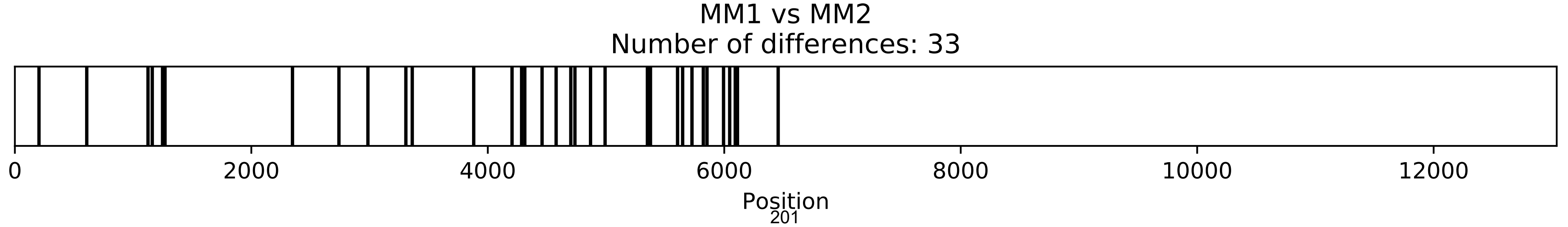
MM1 vs MM2
Number of differences: 33
0
2000
4000
6000
8000
10000
12000
Position

### Region: 577



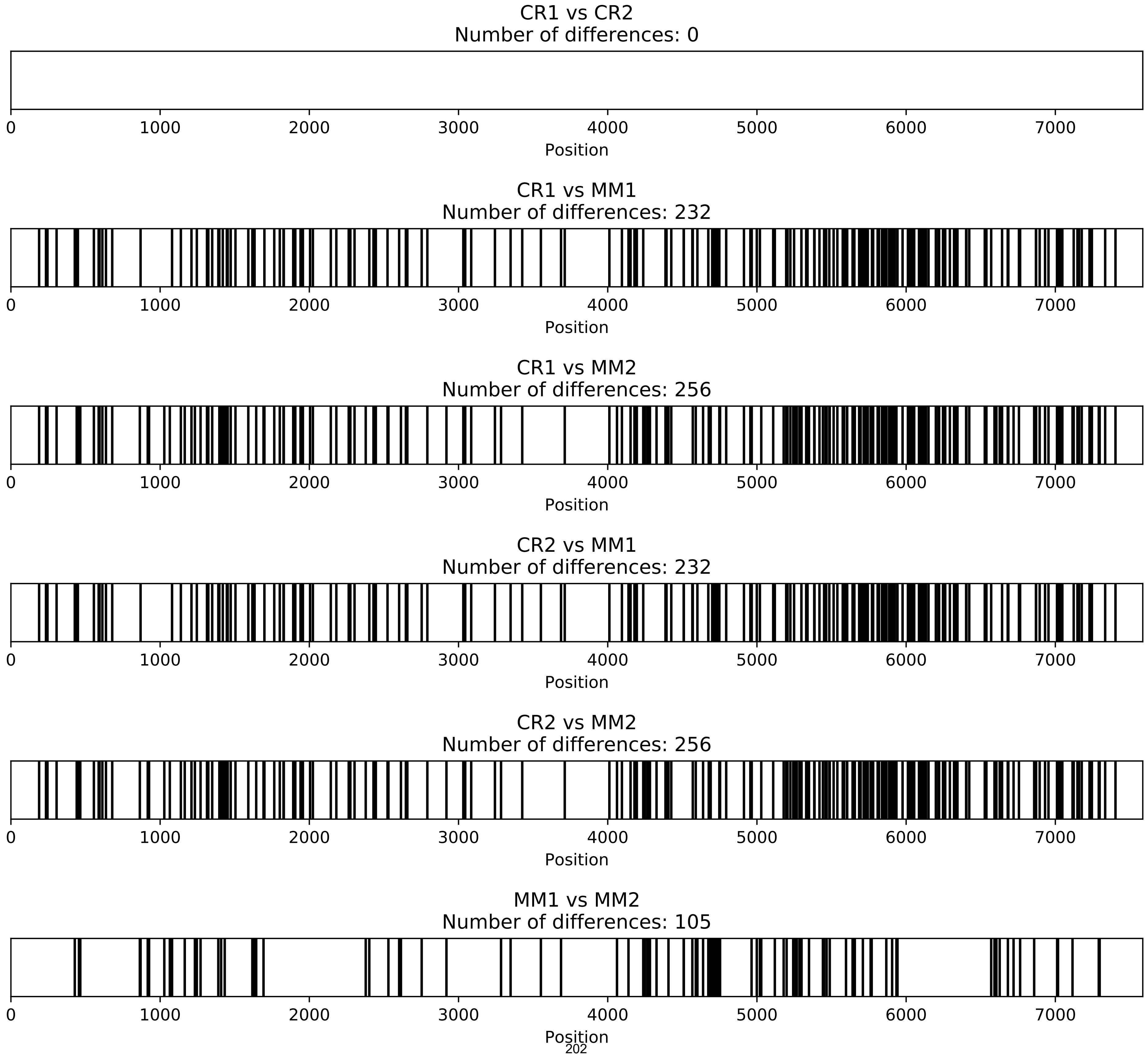
CR1 vs CR2
Number of differences: 0
CR1 vs MM1
Number of differences: 232
CR1 vs MM2
Number of differences: 256
CR2 vs MM1
Number of differences: 232
CR2 vs MM2
Number of differences: 256
MM1 vs MM2
Number of differences: 105
0 1000 2000 3000 4000 5000 6000 7000
Position

### Region: 590

CR1 vs CR2
Number of differences: 228

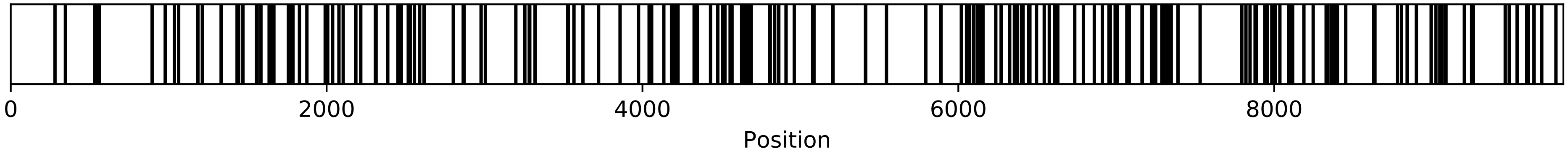


CR1 vs MM1
Number of differences: 69

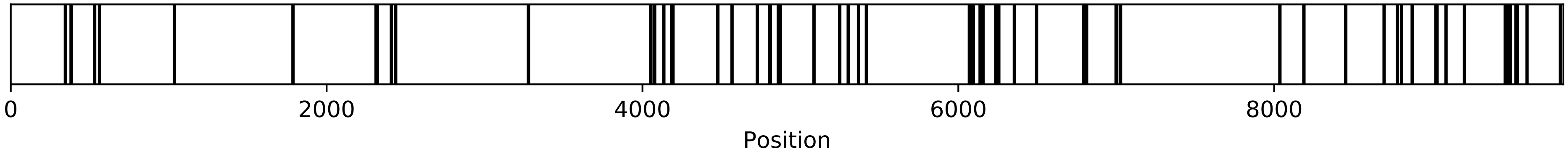


CR1 vs MM2
Number of differences: 71

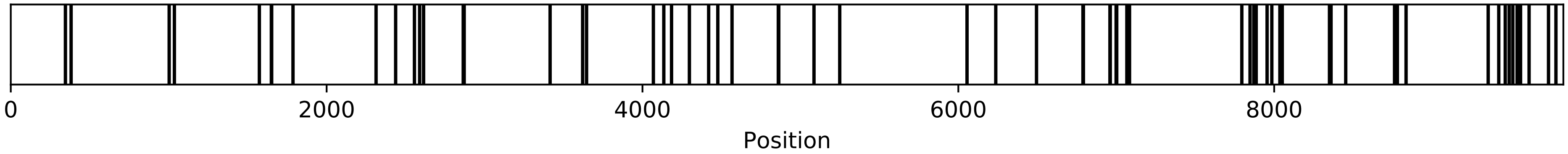


CR2 vs MM1
Number of differences: 229

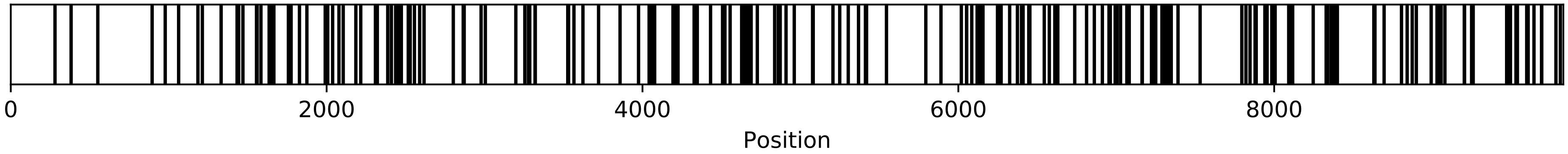


CR2 vs MM2
Number of differences: 223

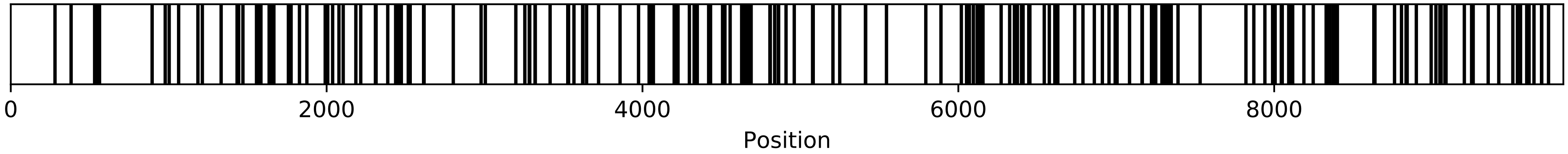


MM1 vs MM2
Number of differences: 89

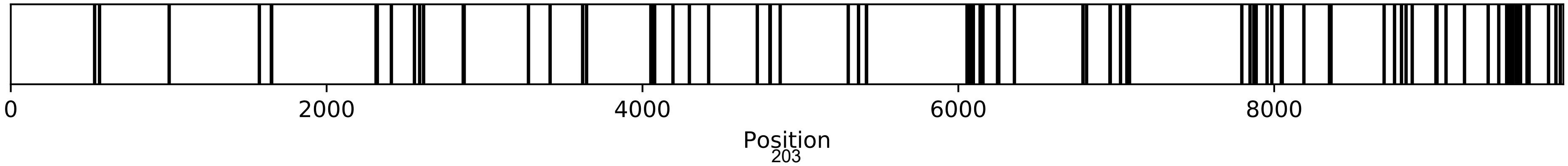

### Region: 597



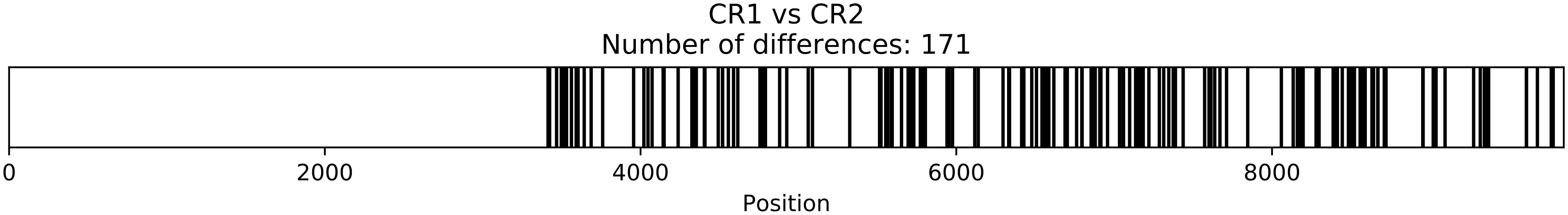
CR1 vs CR2
Number of differences: 171
0
2000
4000
6000
8000
Position

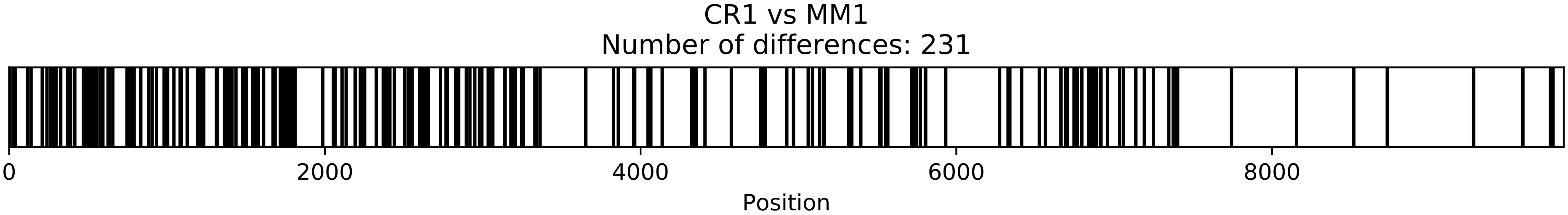
CR1 vs MM1
Number of differences: 231
0
2000
4000
6000
8000
Position

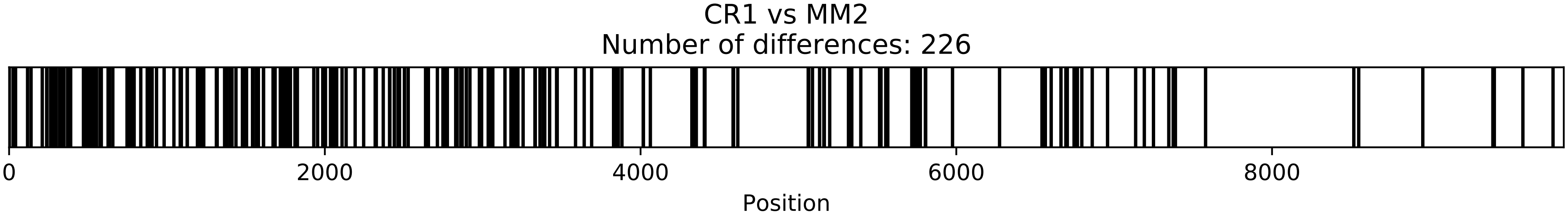
CR1 vs MM2
Number of differences: 226
0
2000
4000
6000
8000
Position

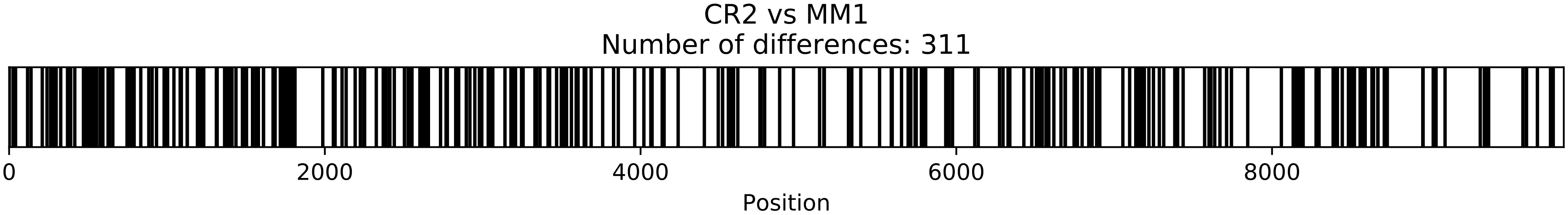
CR2 vs MM1
Number of differences: 311
0
2000
4000
6000
8000
Position

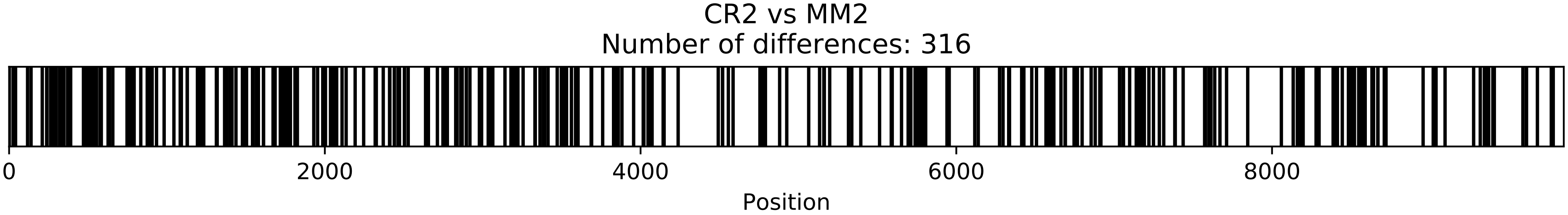
CR2 vs MM2
Number of differences: 316
0
2000
4000
6000
8000
Position

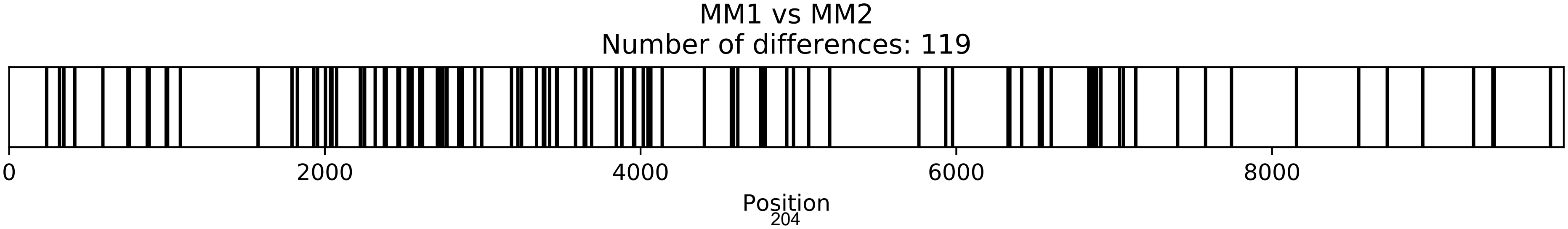
MM1 vs MM2
Number of differences: 119
0
2000
4000
6000
8000
Position

### Region: 598



## CR1 vs CR2
Number of differences: 297

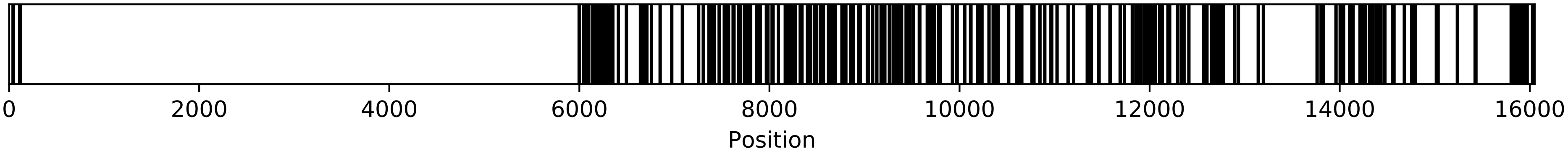


## CR1 vs MM1
Number of differences: 389

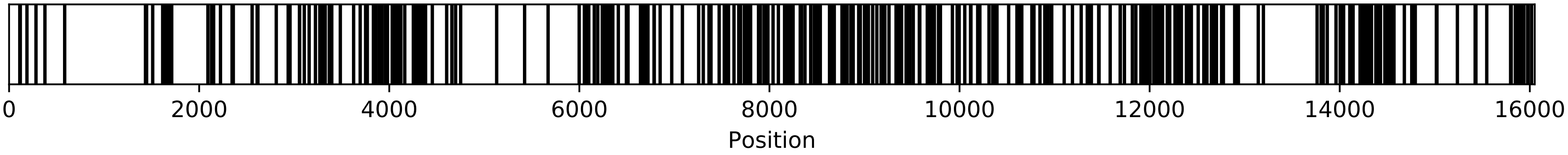


## CR1 vs MM2
Number of differences: 390

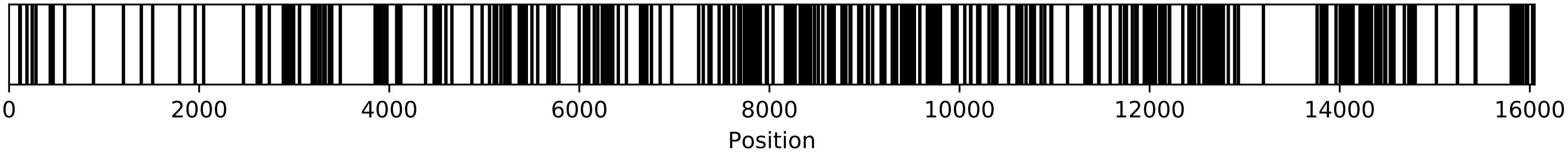


## CR2 vs MM1
Number of differences: 215

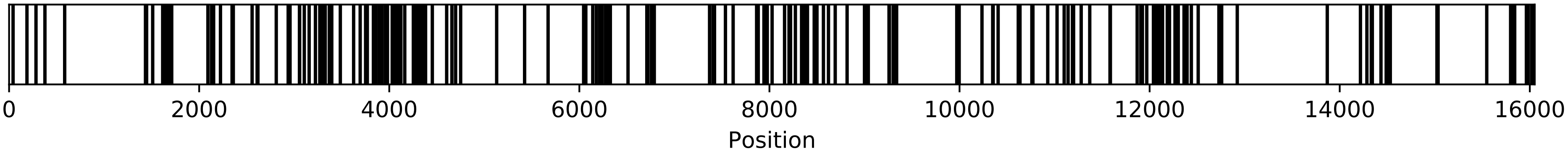


## CR2 vs MM2
Number of differences: 263

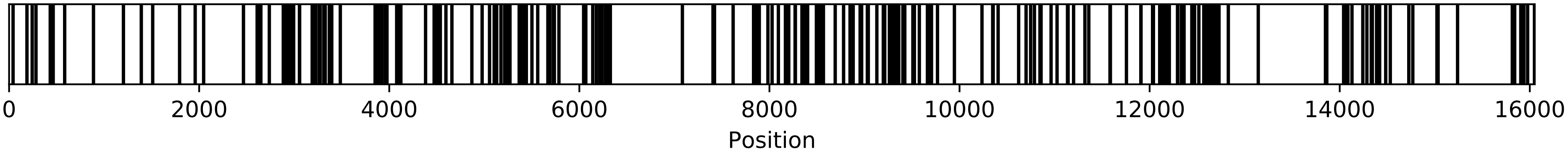


## MM1 vs MM2
Number of differences: 300

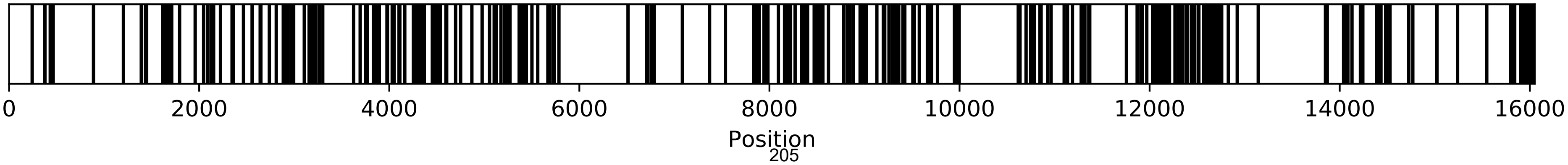

### Region: 600



## CR1 vs CR2
Number of differences: 173

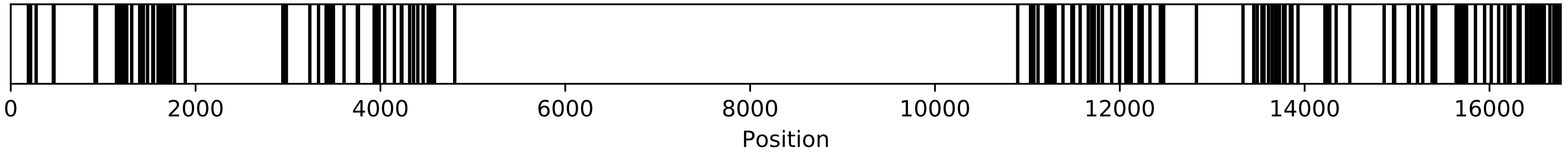


## CR1 vs MM1
Number of differences: 159

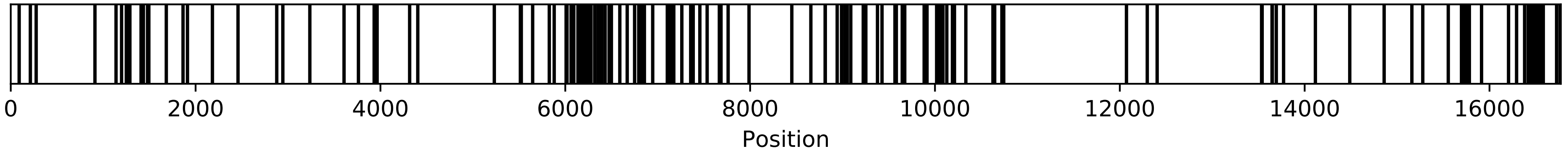


## CR1 vs MM2
Number of differences: 180

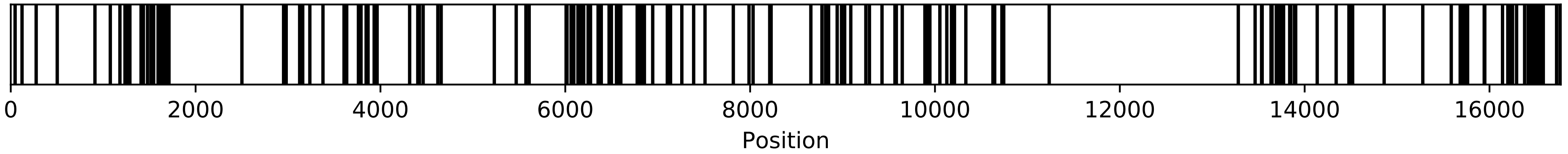


## CR2 vs MM1
Number of differences: 260

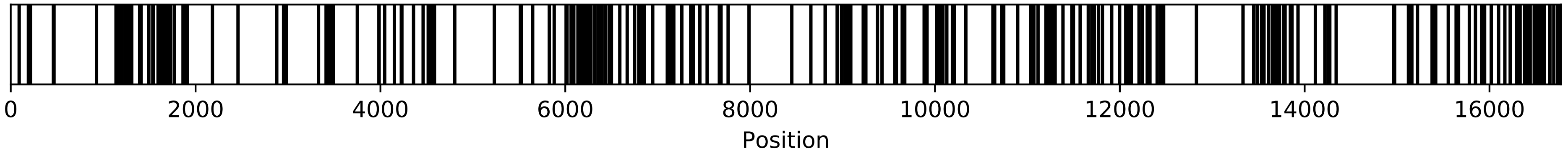


## CR2 vs MM2
Number of differences: 264

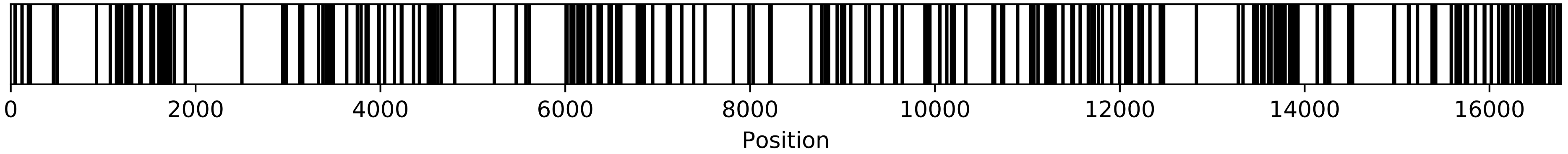


## MM1 vs MM2
Number of differences: 157

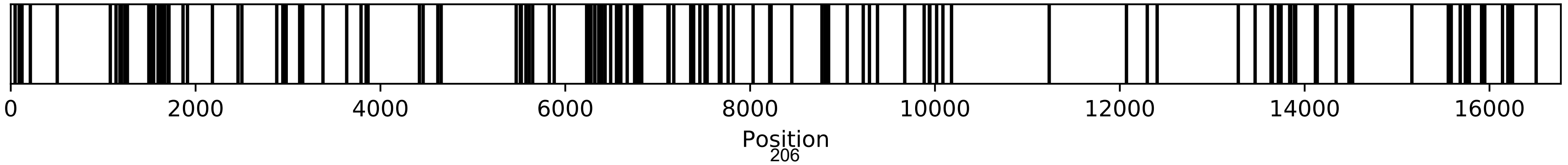

### Region: 601



CR1 vs CR2
Number of differences: 268

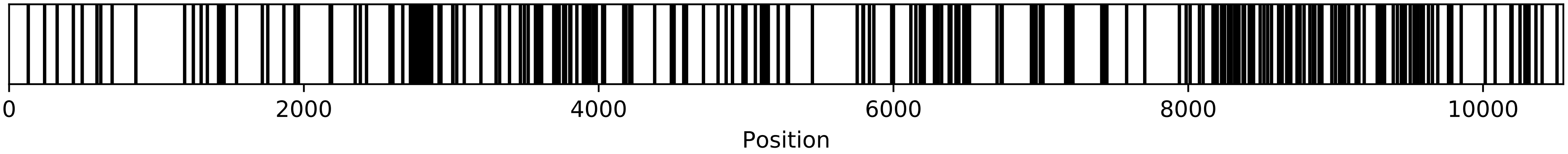


CR1 vs MM1
Number of differences: 145

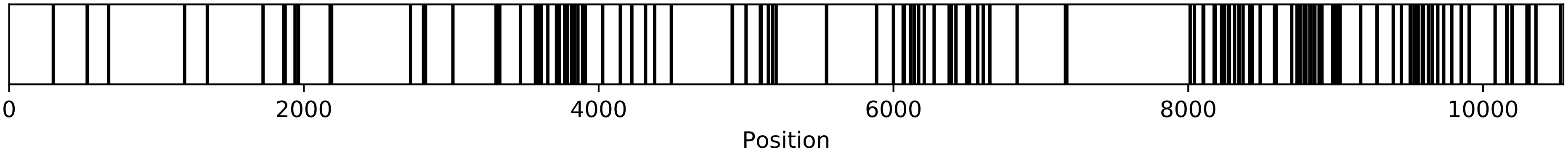


CR1 vs MM2
Number of differences: 144

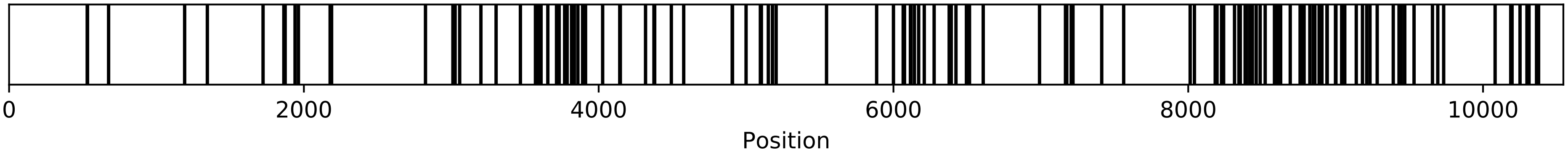


CR2 vs MM1
Number of differences: 262

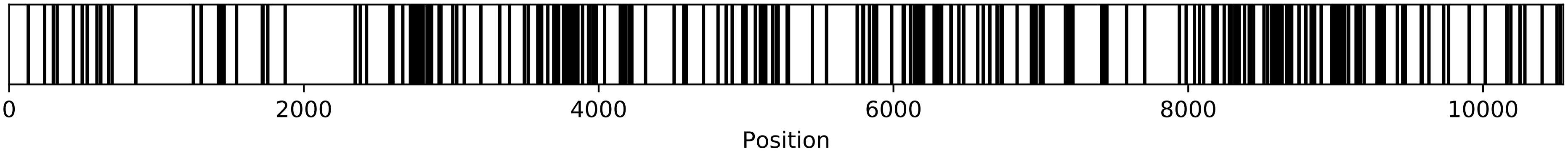


CR2 vs MM2
Number of differences: 280

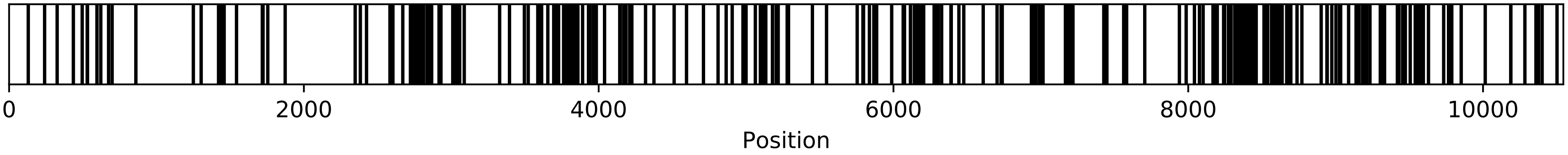


MM1 vs MM2
Number of differences: 97

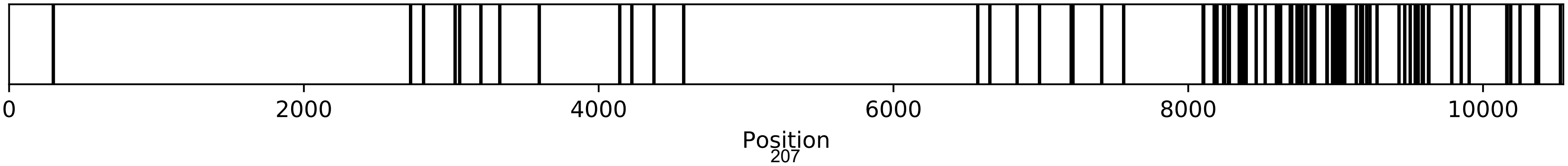

### Region: 602



## CR1 vs CR2
Number of differences: 115

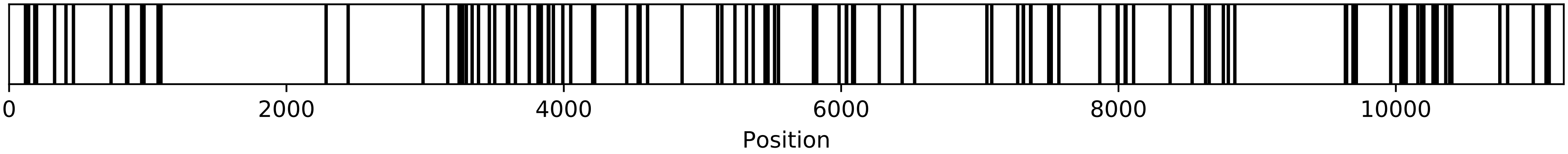


## CR1 vs MM1
Number of differences: 33

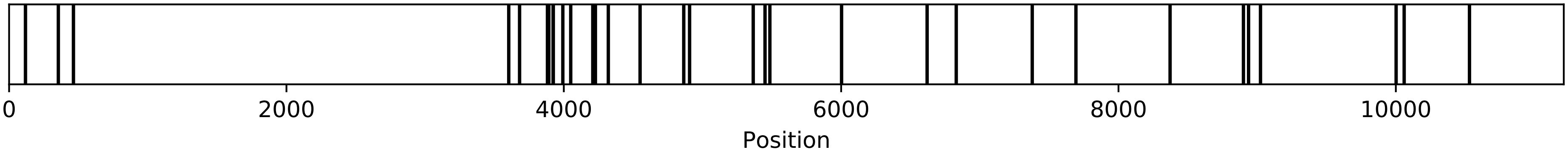


## CR1 vs MM2
Number of differences: 36

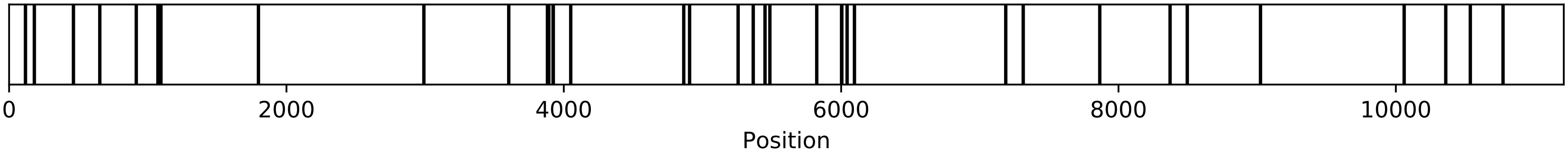


## CR2 vs MM1
Number of differences: 126

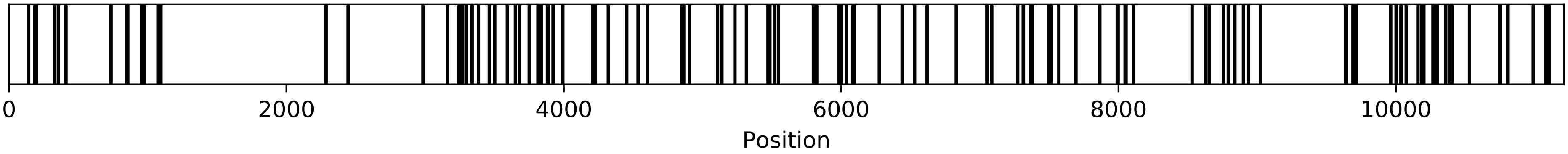


## CR2 vs MM2
Number of differences: 122

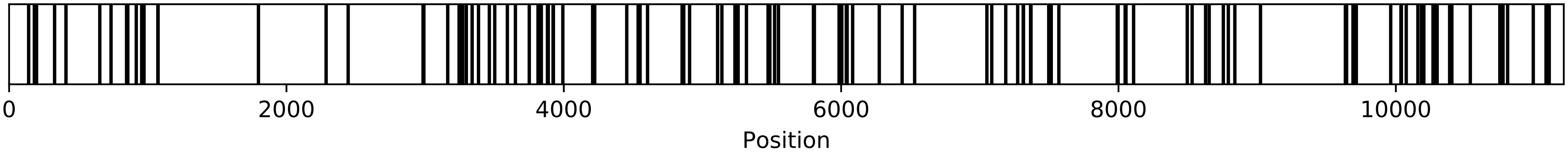


## MM1 vs MM2
Number of differences: 35

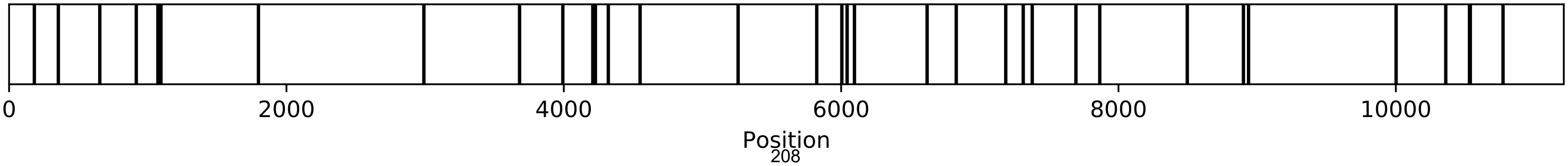

### Region: 606



CR1 vs CR2
Number of differences: 139

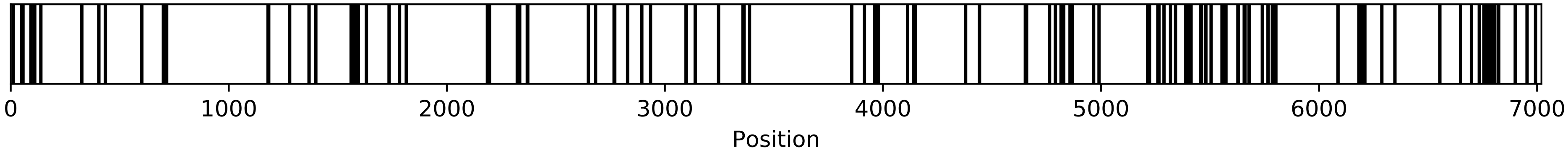


CR1 vs MM1
Number of differences: 148

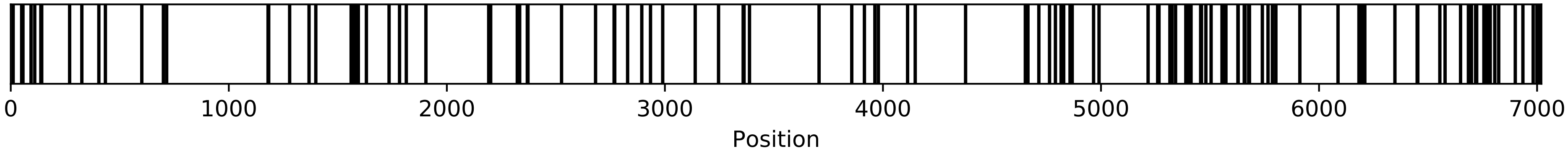


CR1 vs MM2
Number of differences: 146

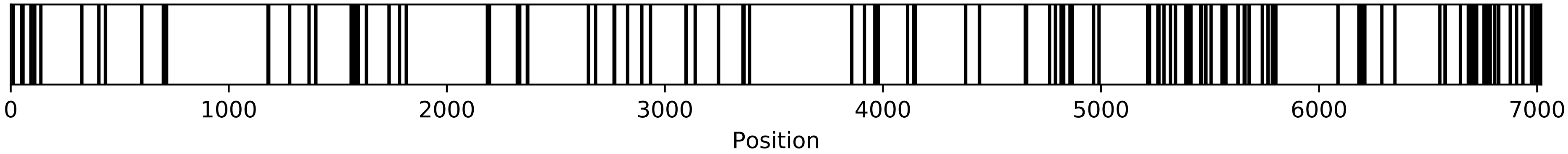


CR2 vs MM1
Number of differences: 51

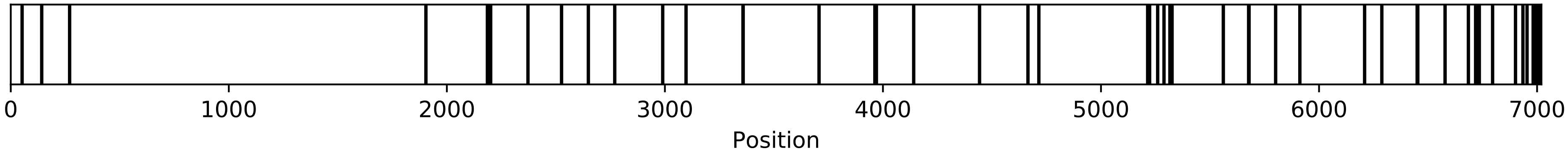


CR2 vs MM2
Number of differences: 21

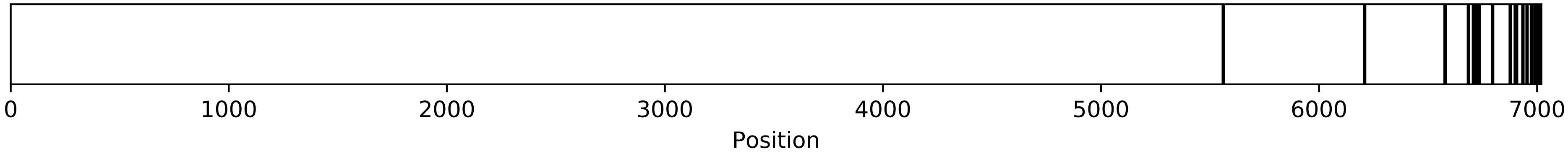


MM1 vs MM2
Number of differences: 42

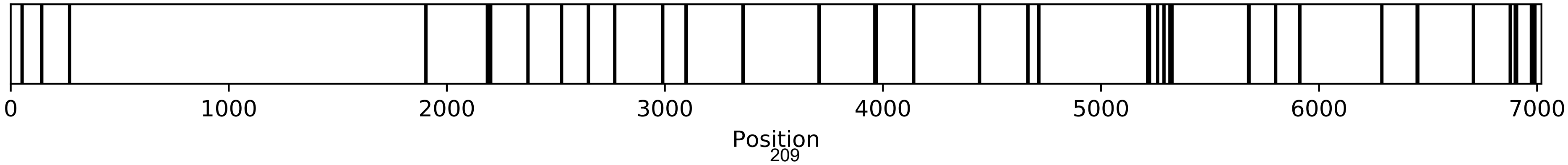

### Region: 608



CR1 vs CR2
Number of differences: 357

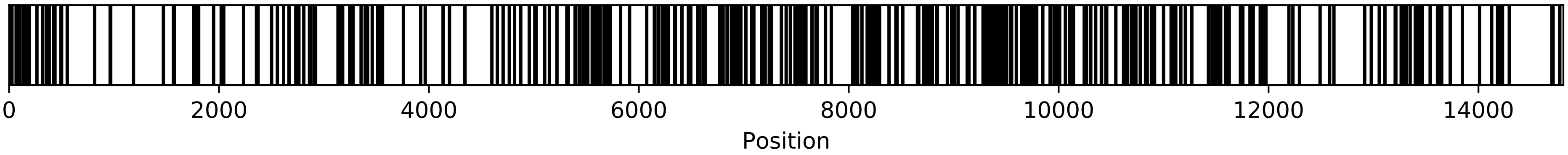


CR1 vs MM1
Number of differences: 173

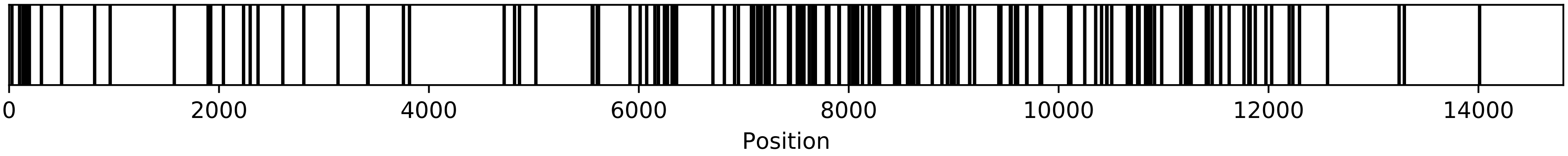


CR1 vs MM2
Number of differences: 144

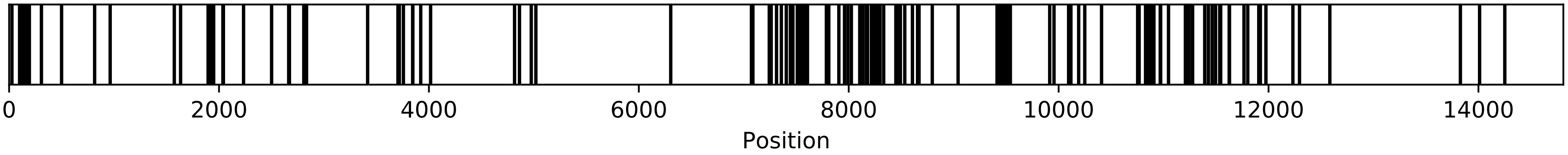


CR2 vs MM1
Number of differences: 367

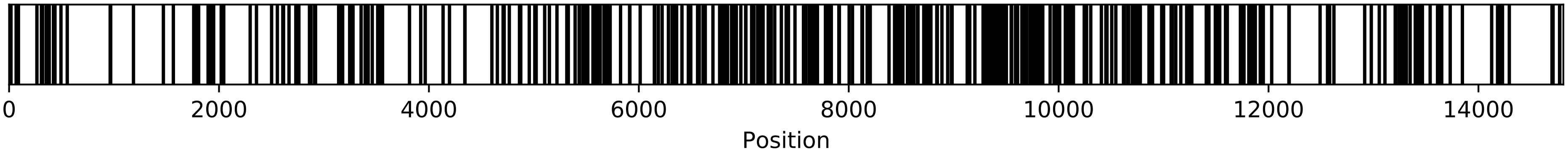


CR2 vs MM2
Number of differences: 369

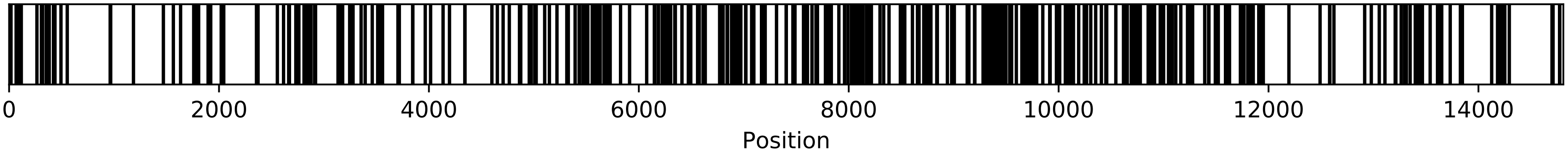


MM1 vs MM2
Number of differences: 177

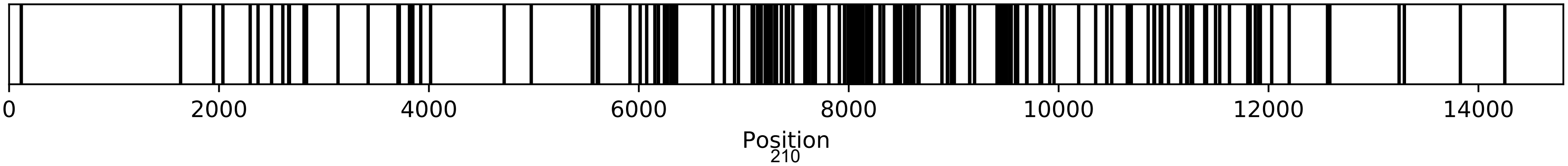

### Region: 613



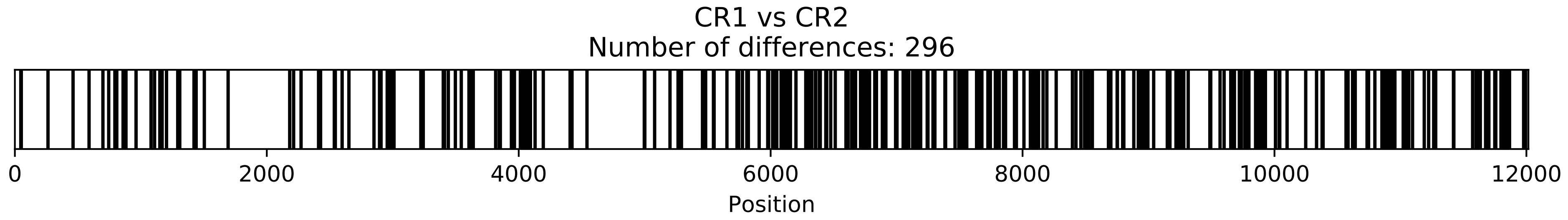
CR1 vs CR2
Number of differences: 296
0
2000
4000
6000
8000
10000
12000
Position

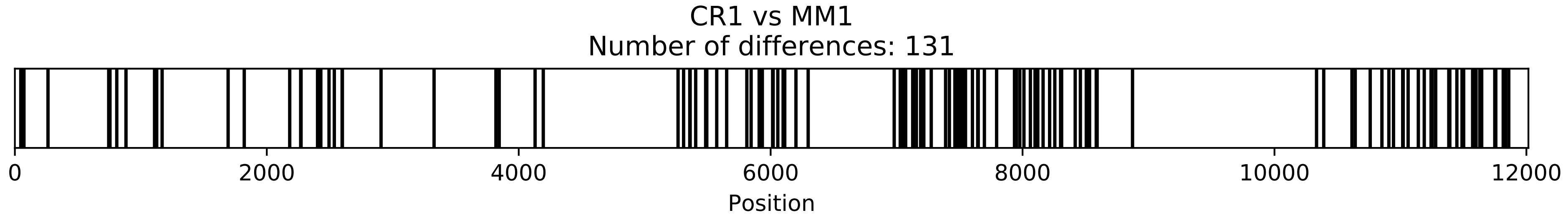
CR1 vs MM1
Number of differences: 131
0
2000
4000
6000
8000
10000
12000
Position

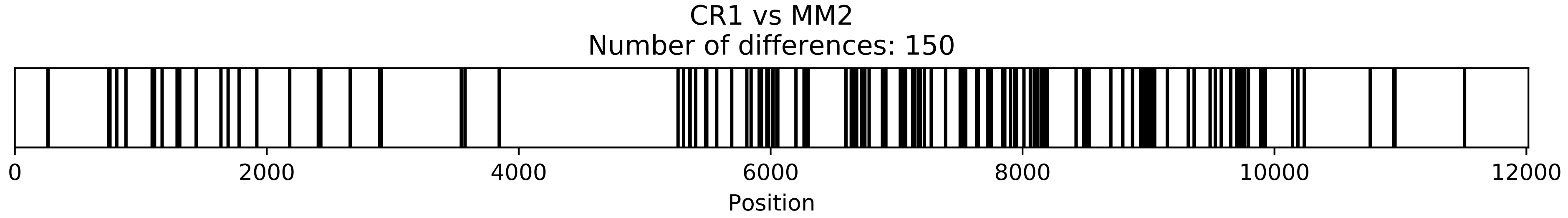
CR1 vs MM2
Number of differences: 150
0
2000
4000
6000
8000
10000
12000
Position

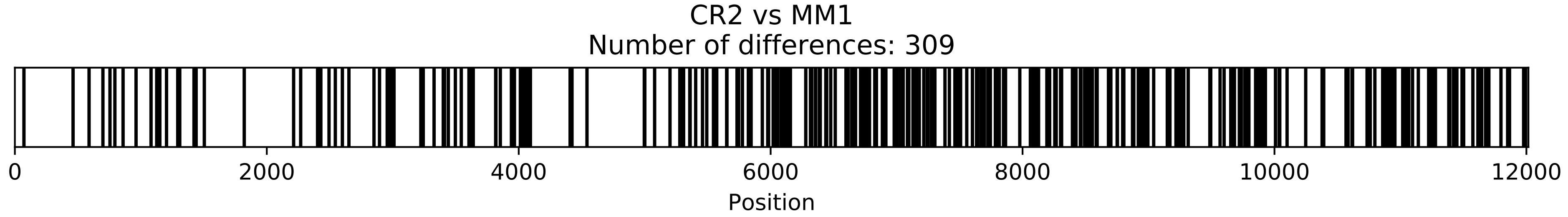
CR2 vs MM1
Number of differences: 309
0
2000
4000
6000
8000
10000
12000
Position

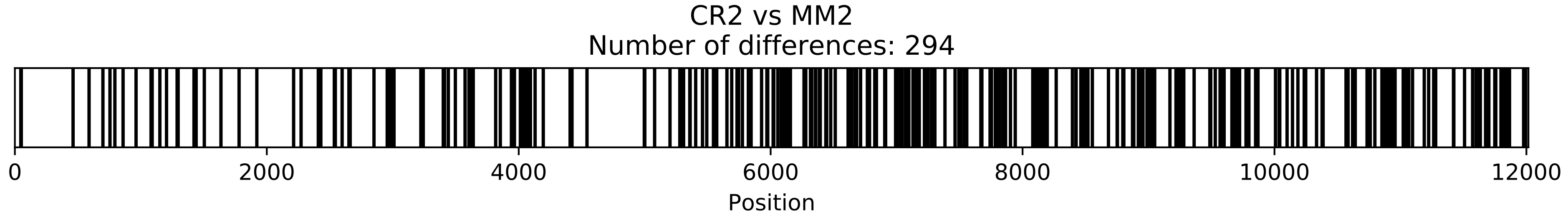
CR2 vs MM2
Number of differences: 294
0
2000
4000
6000
8000
10000
12000
Position

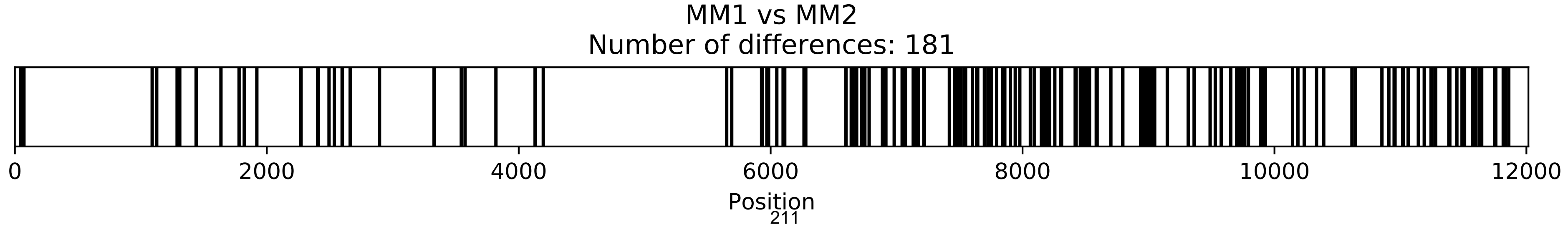
MM1 vs MM2
Number of differences: 181
0
2000
4000
6000
8000
10000
12000
Position

### Region: 614



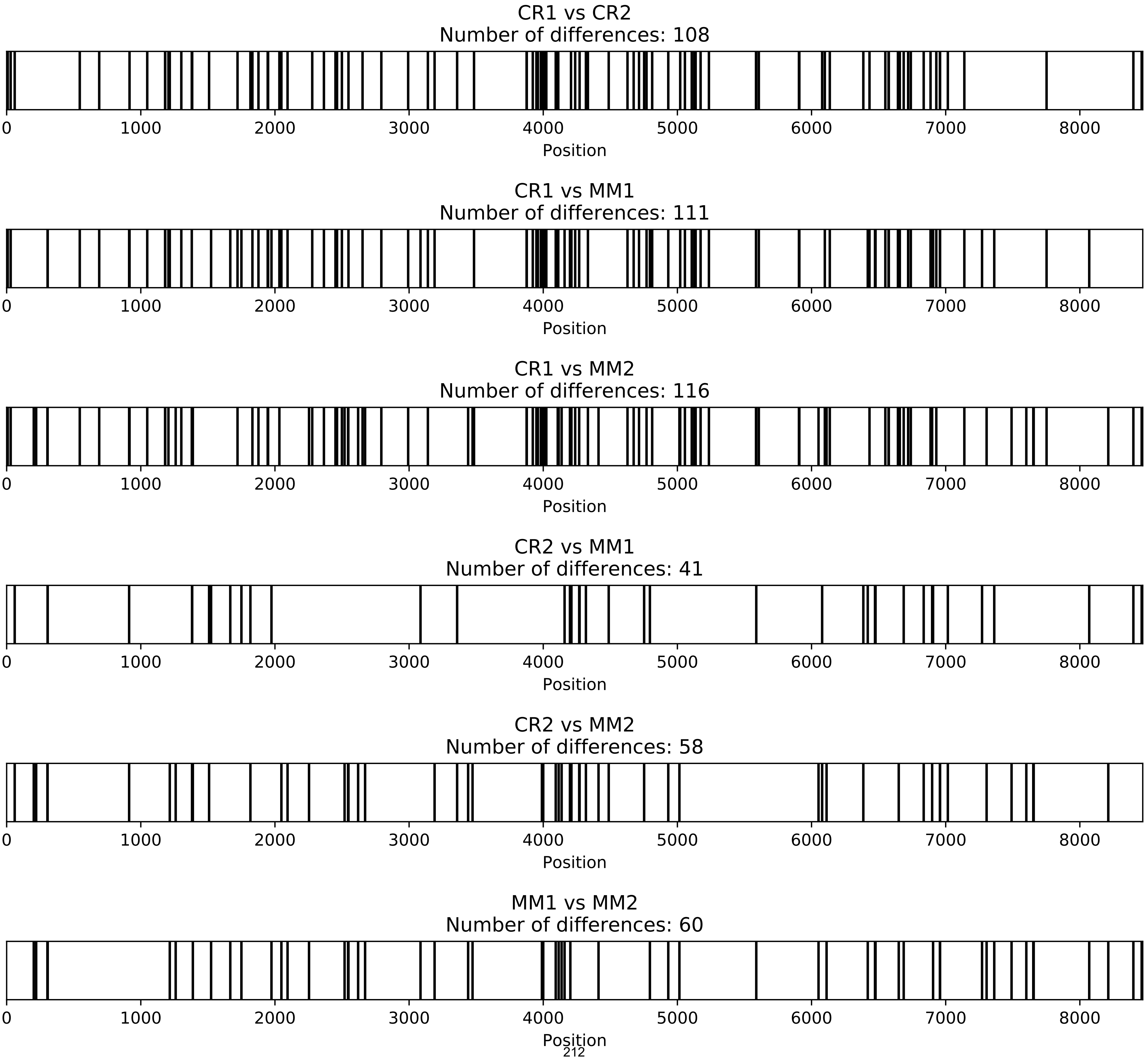
CR1 vs CR2
Number of differences: 108
0 1000 2000 3000 4000 5000 6000 7000 8000
Position
CR1 vs MM1
Number of differences: 111
0 1000 2000 3000 4000 5000 6000 7000 8000
Position
CR1 vs MM2
Number of differences: 116
0 1000 2000 3000 4000 5000 6000 7000 8000
Position
CR2 vs MM1
Number of differences: 41
0 1000 2000 3000 4000 5000 6000 7000 8000
Position
CR2 vs MM2
Number of differences: 58
0 1000 2000 3000 4000 5000 6000 7000 8000
Position
MM1 vs MM2
Number of differences: 60
0 1000 2000 3000 4000 5000 6000 7000 8000
Position

### Region: 616



## CR1 vs CR2
Number of differences: 119

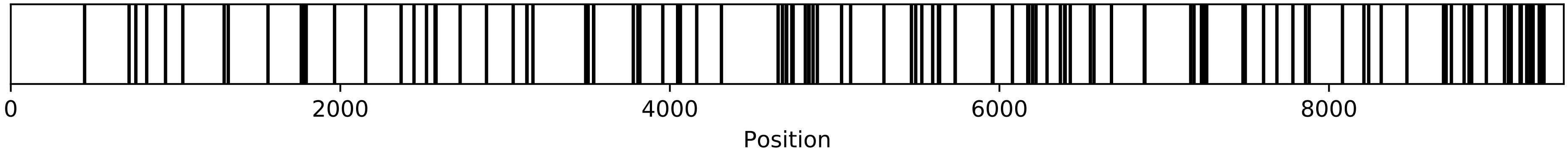


## CR1 vs MM1
Number of differences: 45

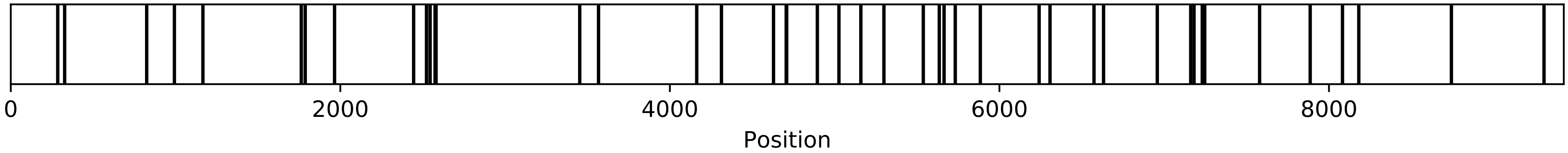


## CR1 vs MM2
Number of differences: 44

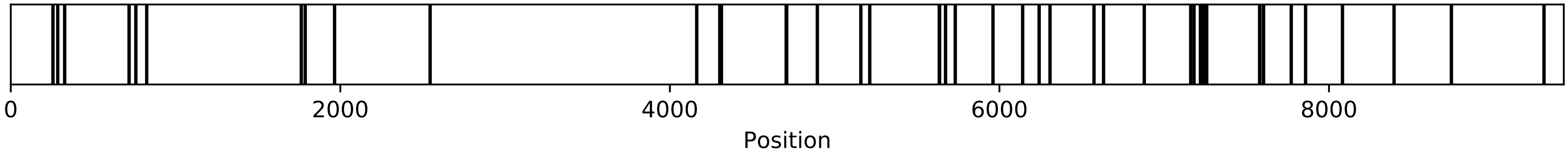


## CR2 vs MM1
Number of differences: 121

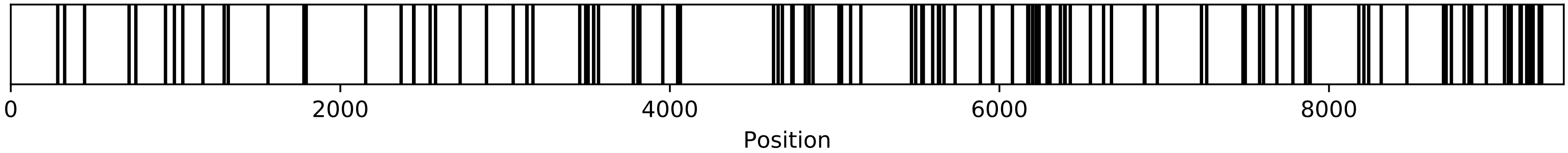


## CR2 vs MM2
Number of differences: 111

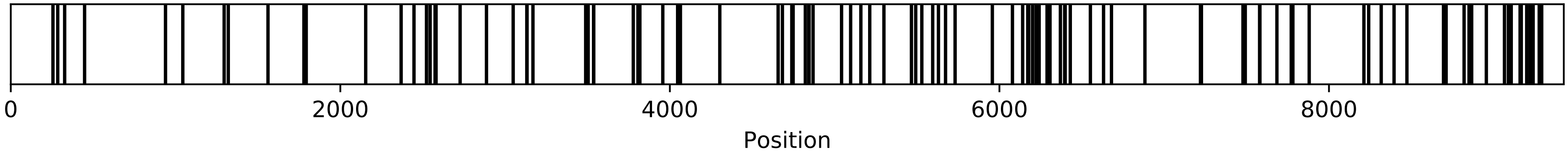


## MM1 vs MM2
Number of differences: 37

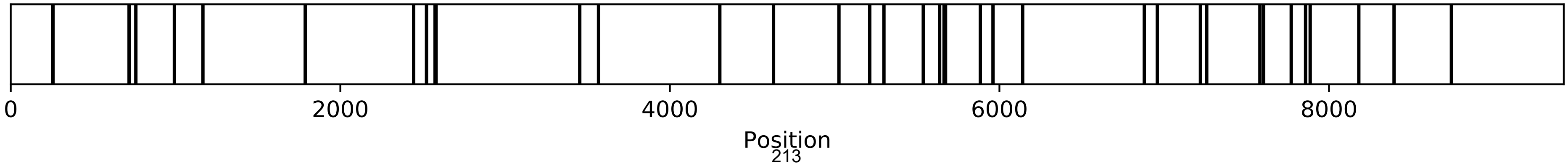

### Region: 619



CR1 vs CR2
Number of differences: 205

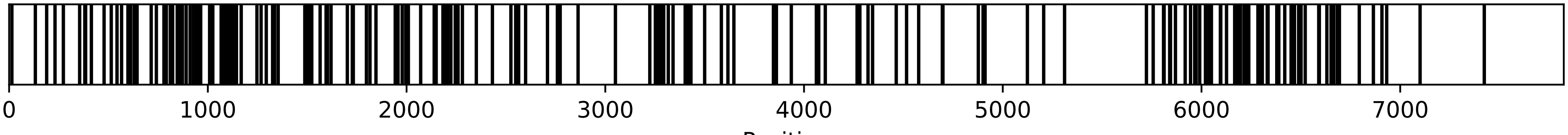


CR1 vs MM1
Number of differences: 203

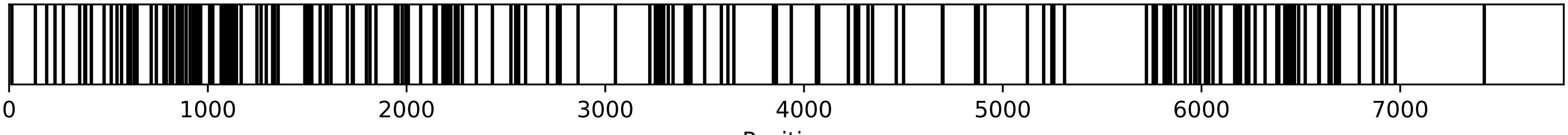


CR1 vs MM2
Number of differences: 194

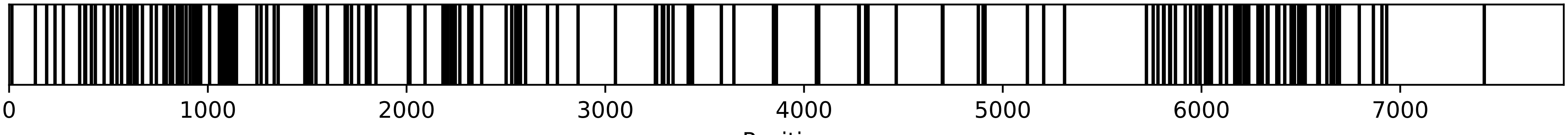


CR2 vs MM1
Number of differences: 48

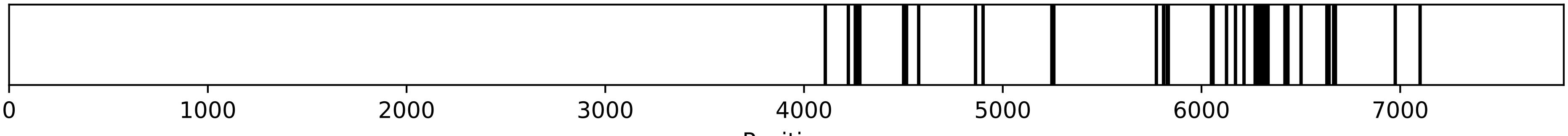


CR2 vs MM2
Number of differences: 86

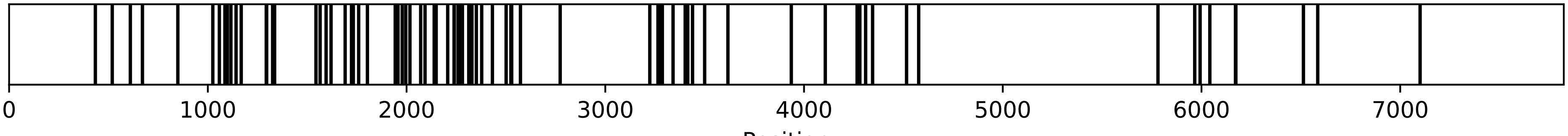


MM1 vs MM2
Number of differences: 110

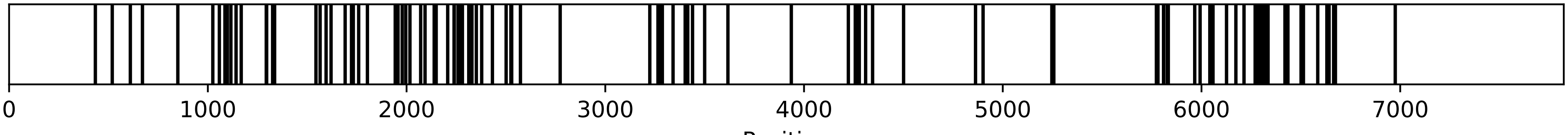

### Region: 620



CR1 vs CR2
Number of differences: 158

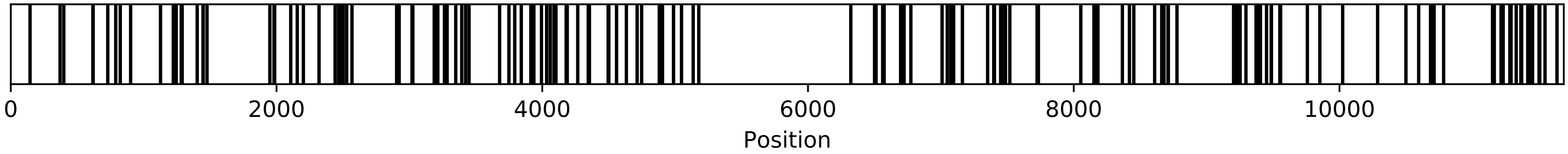


CR1 vs MM1
Number of differences: 74

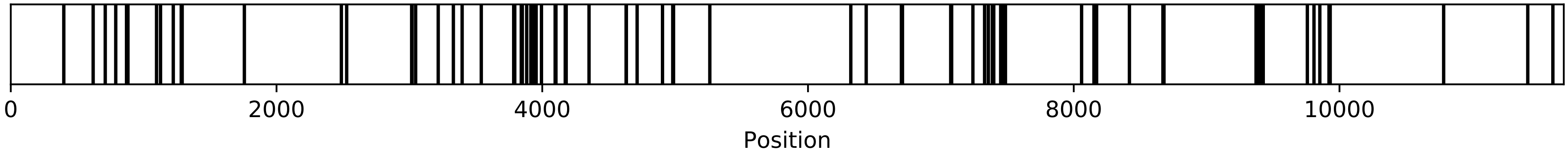


CR1 vs MM2
Number of differences: 78

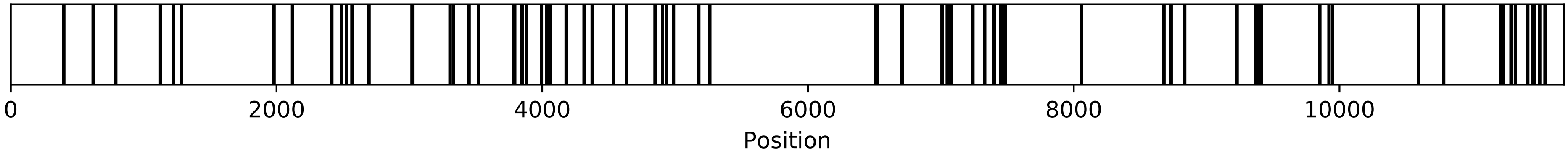


CR2 vs MM1
Number of differences: 156

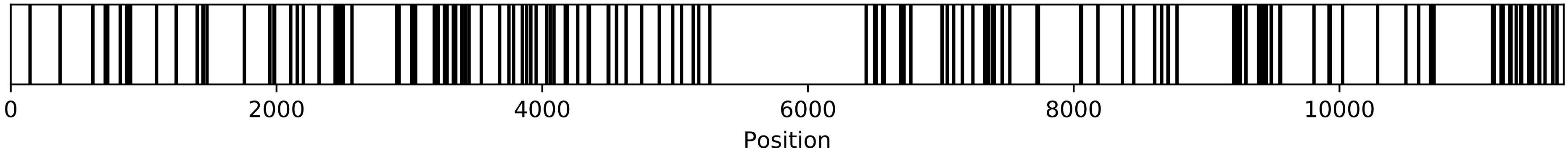


CR2 vs MM2
Number of differences: 150

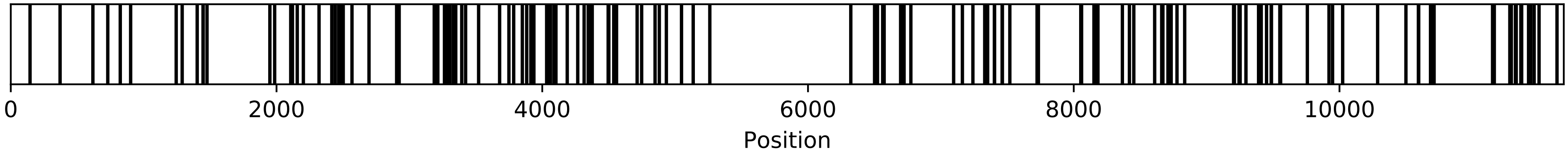


MM1 vs MM2
Number of differences: 74

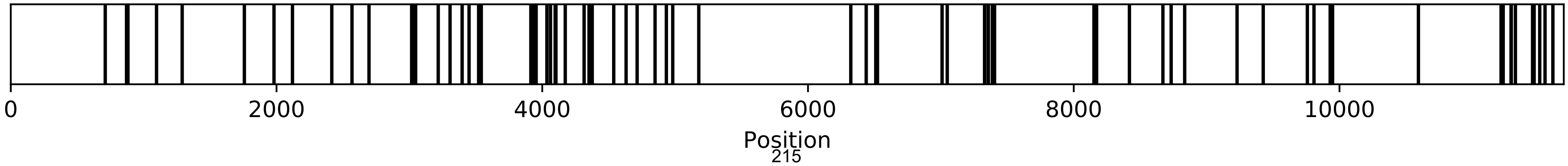

### Region: 621



CR1 vs CR2
Number of differences: 300

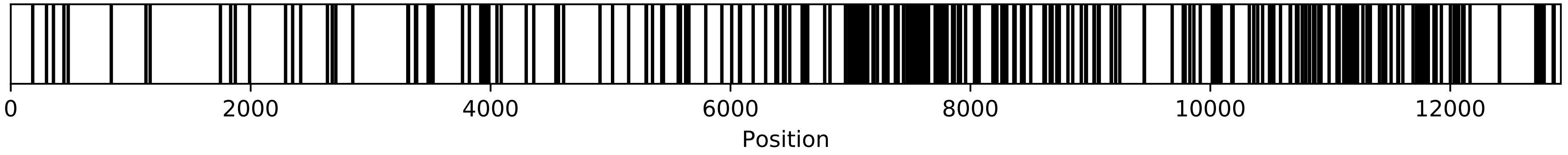


CR1 vs MM1
Number of differences: 109

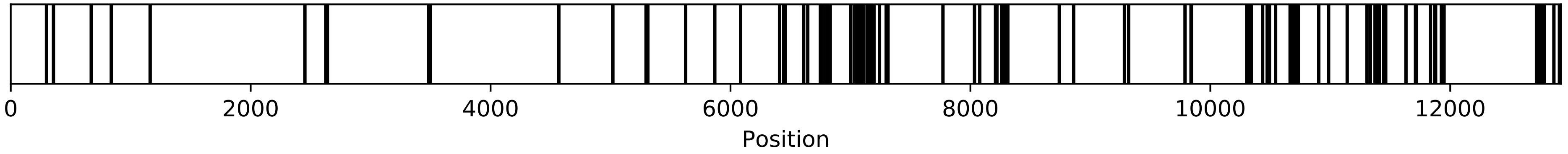


CR1 vs MM2
Number of differences: 78

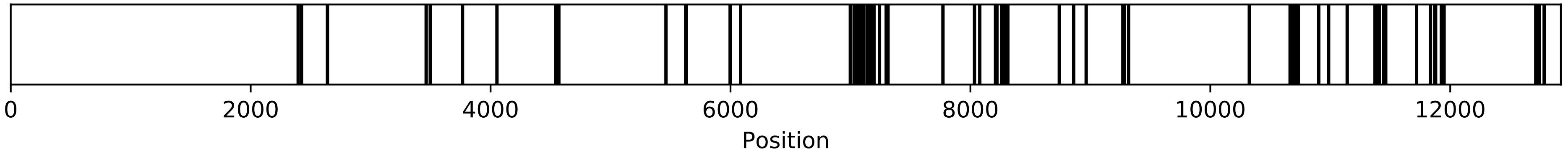


CR2 vs MM1
Number of differences: 303

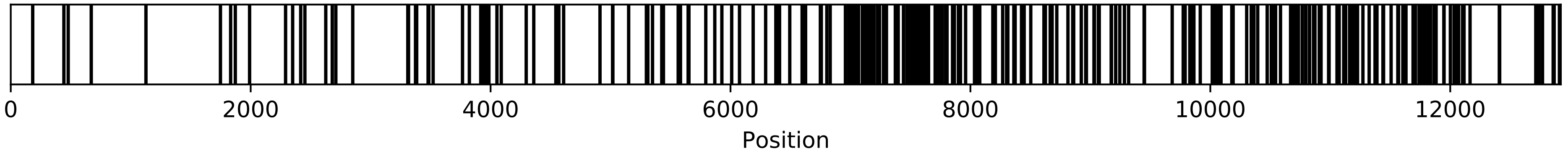


CR2 vs MM2
Number of differences: 300

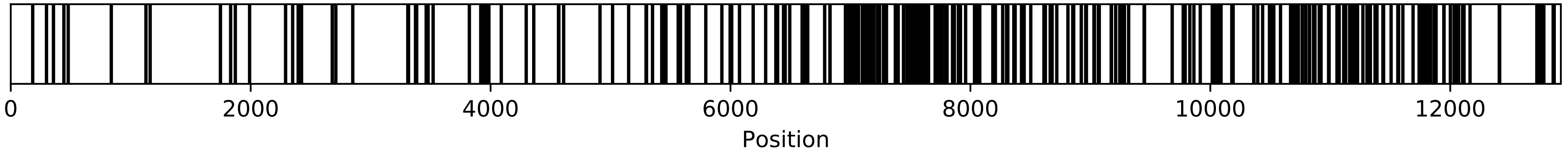


MM1 vs MM2
Number of differences: 58

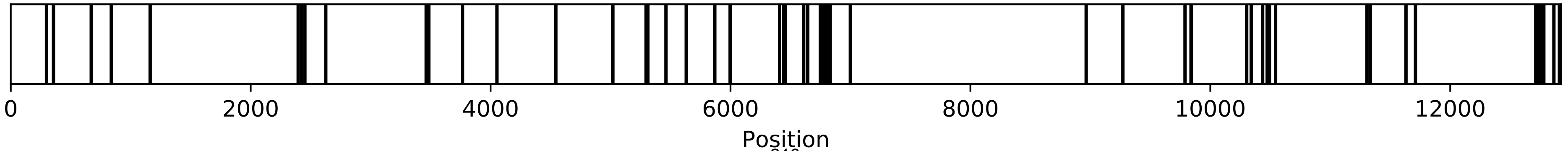

### Region: 623



CR1 vs CR2
Number of differences: 258

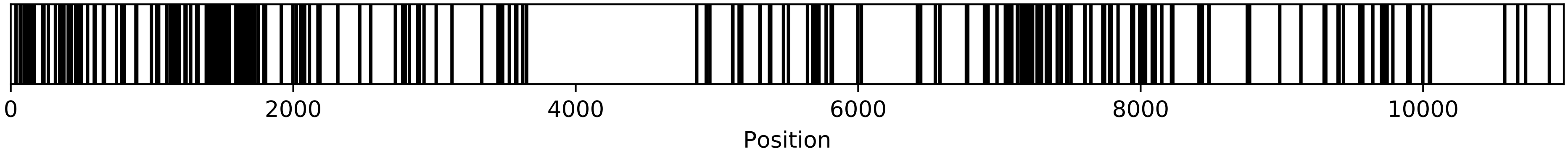


CR1 vs MM1
Number of differences: 181

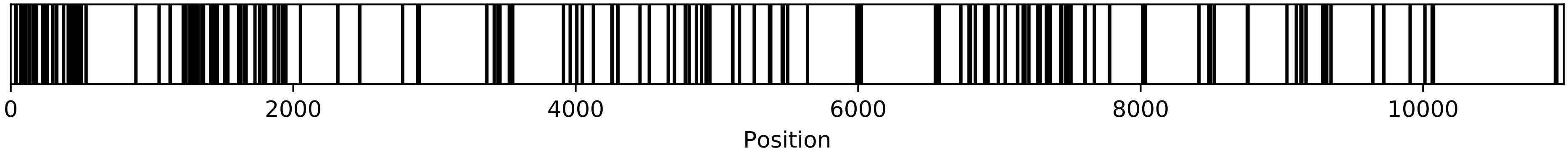


CR1 vs MM2
Number of differences: 77

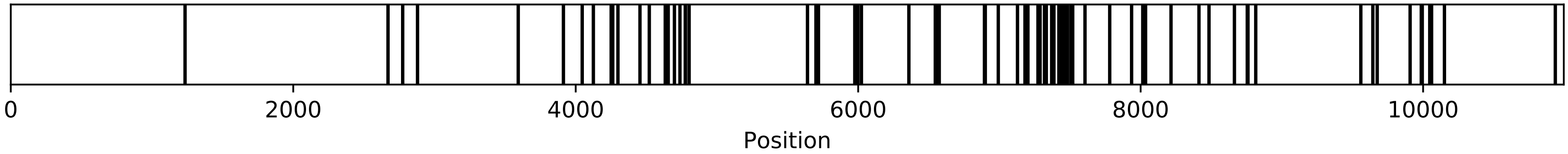


CR2 vs MM1
Number of differences: 285

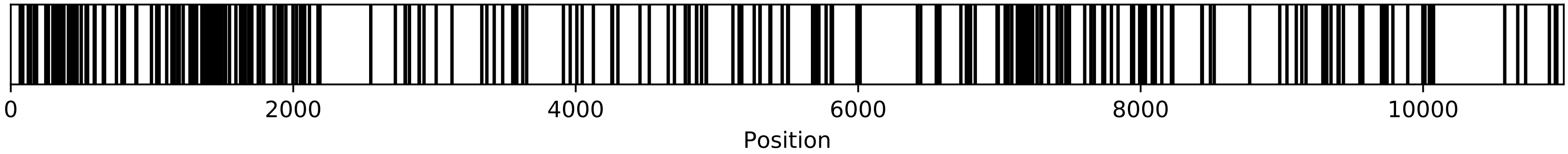


CR2 vs MM2
Number of differences: 276

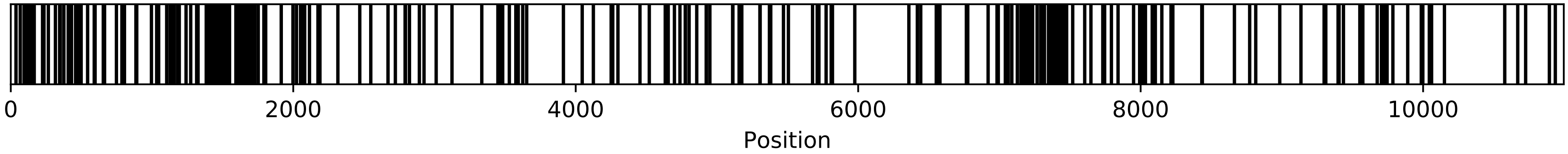


MM1 vs MM2
Number of differences: 164

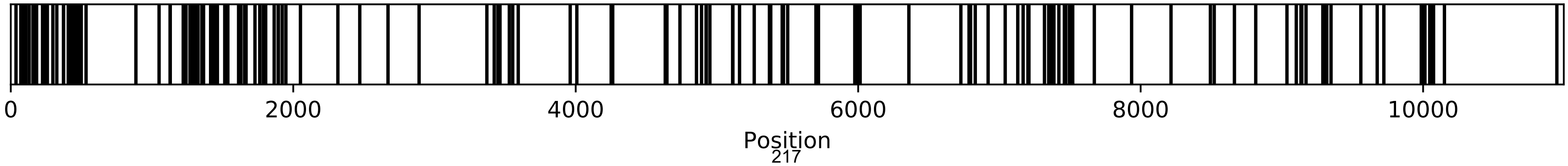

### Region: 627

CR1 vs CR2
Number of differences: 204

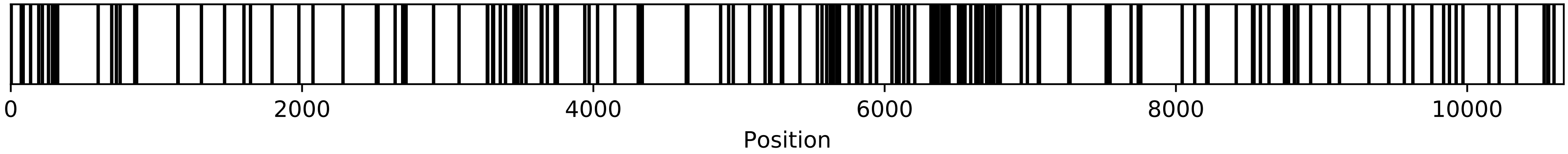


CR1 vs MM1
Number of differences: 86

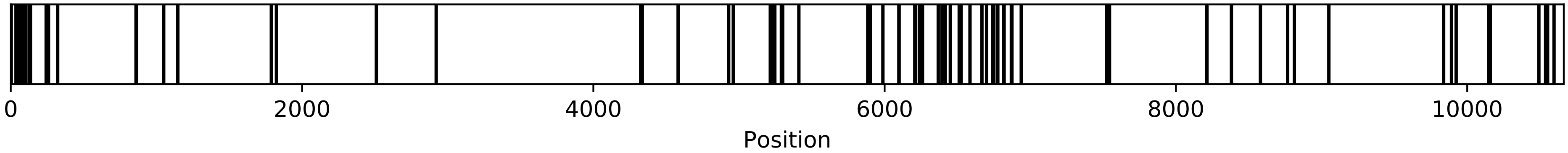


CR1 vs MM2
Number of differences: 83

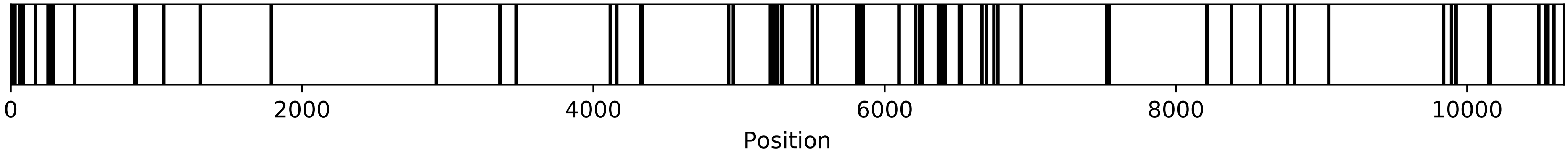


CR2 vs MM1
Number of differences: 200

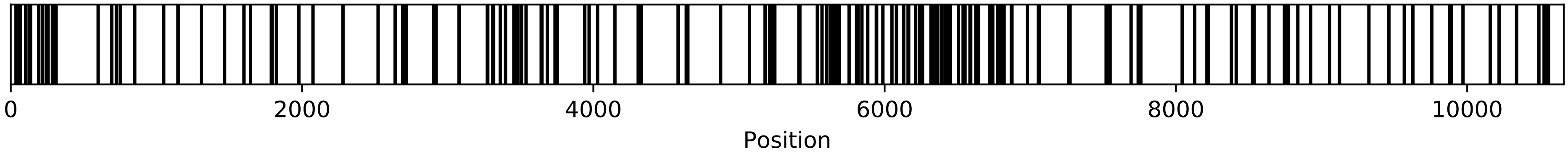


CR2 vs MM2
Number of differences: 193

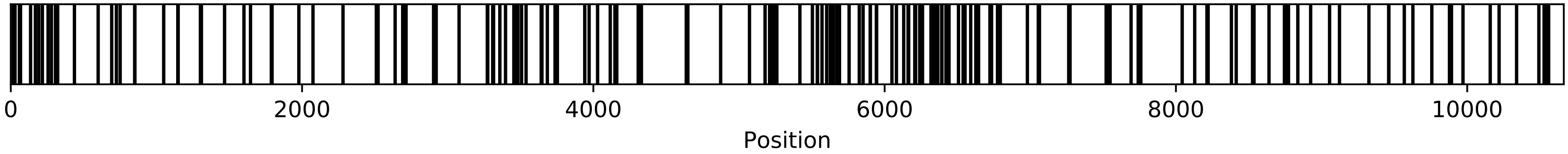


MM1 vs MM2
Number of differences: 55

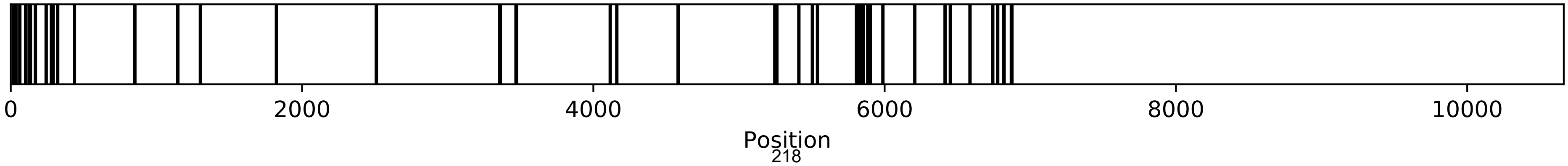

### Region: 628



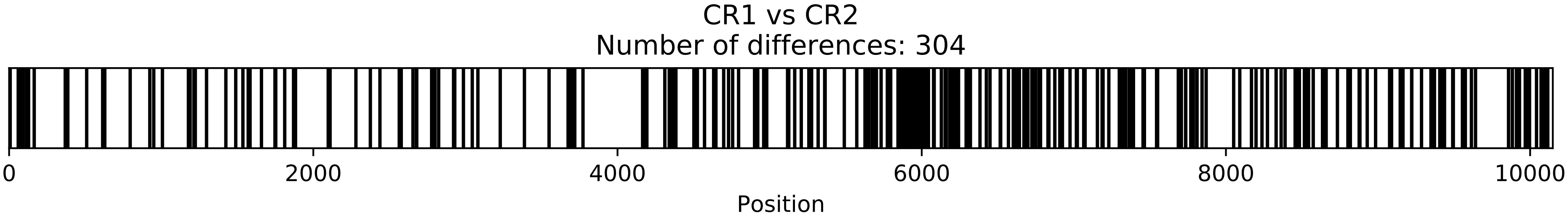
CR1 vs CR2
Number of differences: 304
0
2000
4000
6000
8000
10000
Position

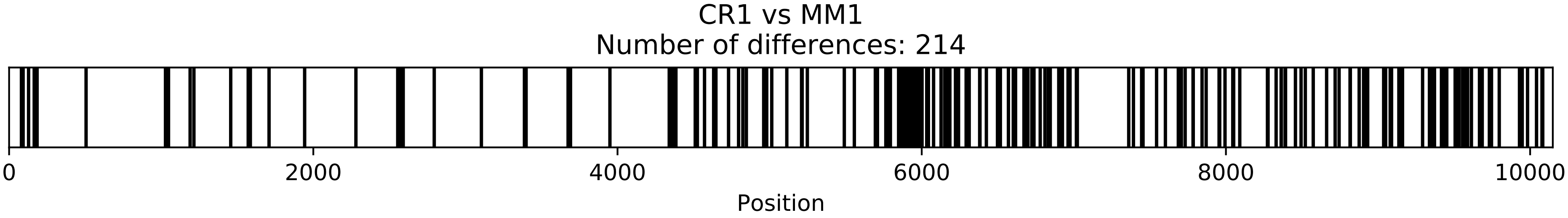
CR1 vs MM1
Number of differences: 214
0
2000
4000
6000
8000
10000
Position

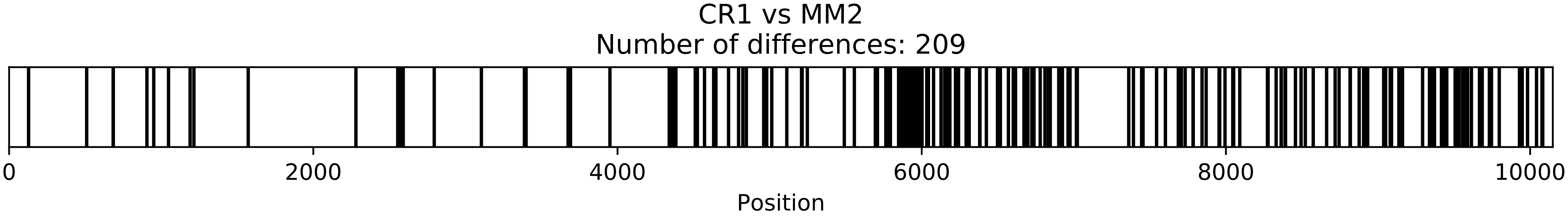
CR1 vs MM2
Number of differences: 209
0
2000
4000
6000
8000
10000
Position

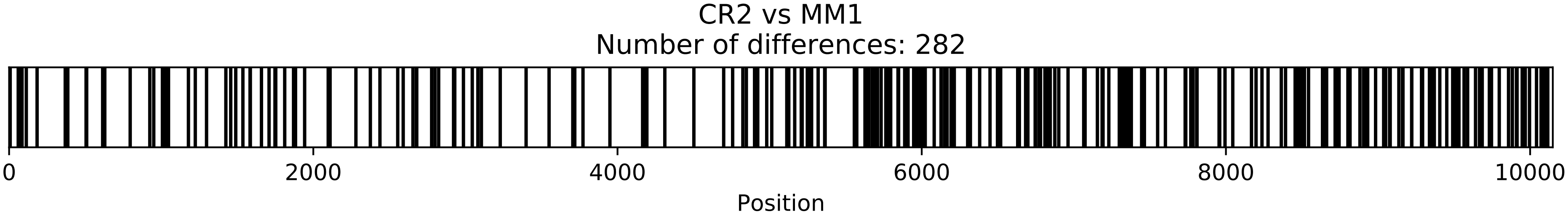
CR2 vs MM1
Number of differences: 282
0
2000
4000
6000
8000
10000
Position

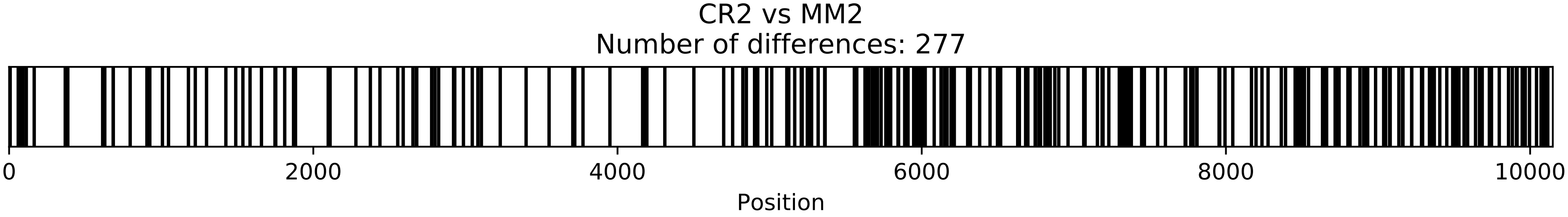
CR2 vs MM2
Number of differences: 277
0
2000
4000
6000
8000
10000
Position

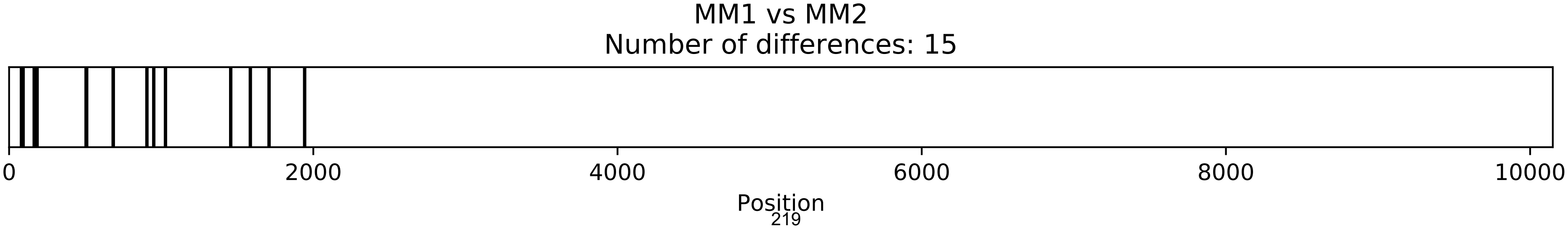
MM1 vs MM2
Number of differences: 15
0
2000
4000
6000
8000
10000
Position

### Region: 629



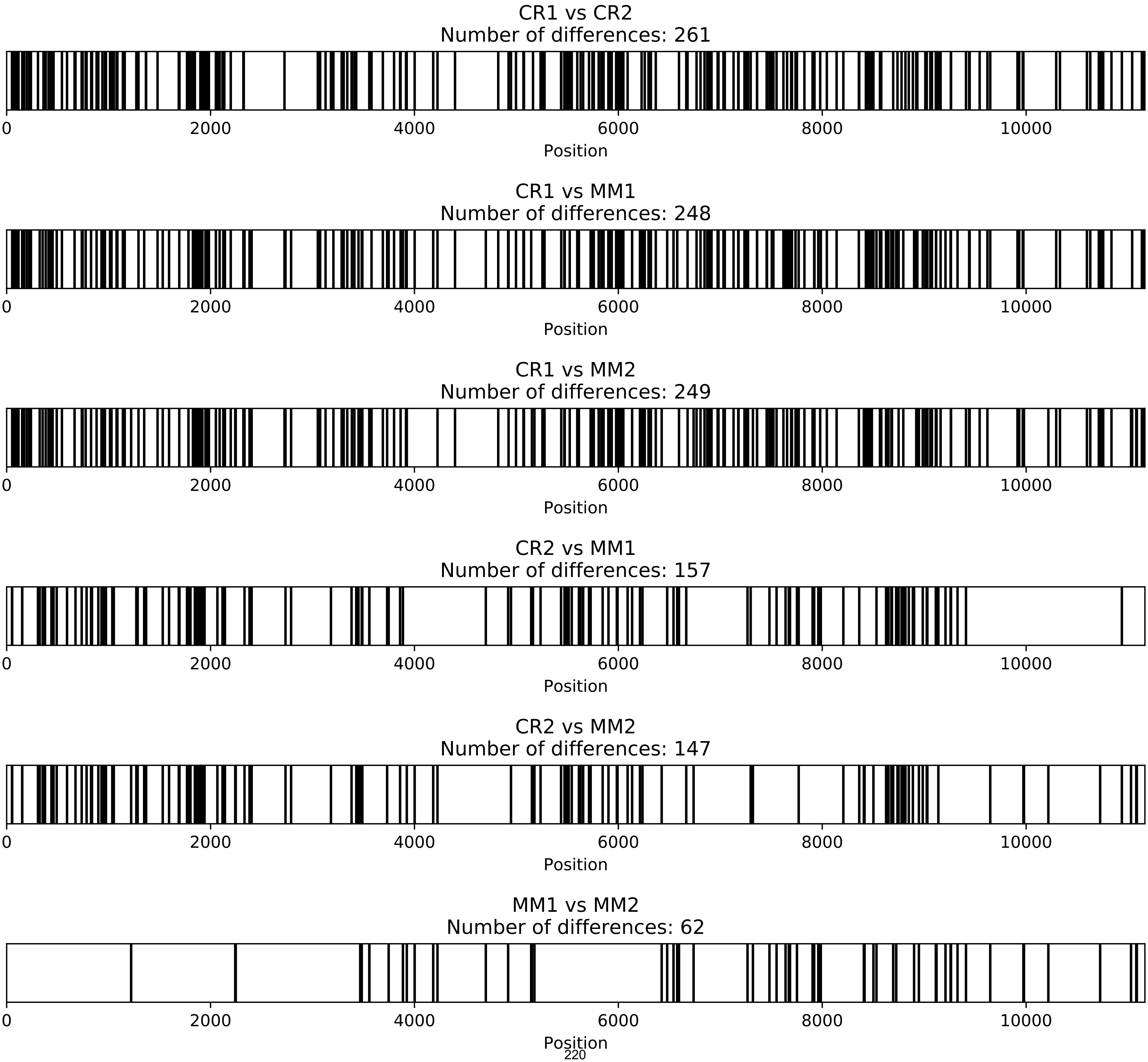
CR1 vs CR2
Number of differences: 261
CR1 vs MM1
Number of differences: 248
CR1 vs MM2
Number of differences: 249
CR2 vs MM1
Number of differences: 157
CR2 vs MM2
Number of differences: 147
MM1 vs MM2
Number of differences: 62
0
2000
4000
6000
8000
10000
Position

### Region: 630



## CR1 vs CR2
Number of differences: 284

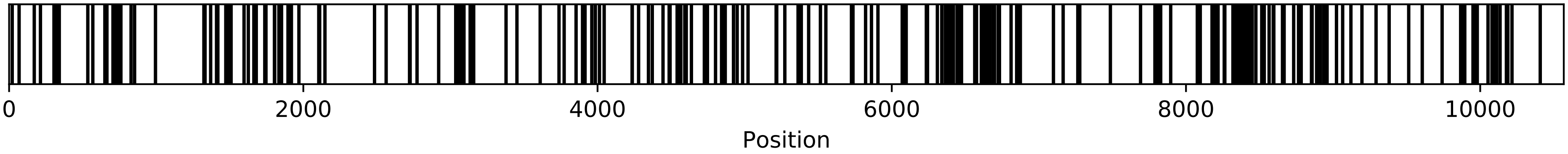


## CR1 vs MM1
Number of differences: 129

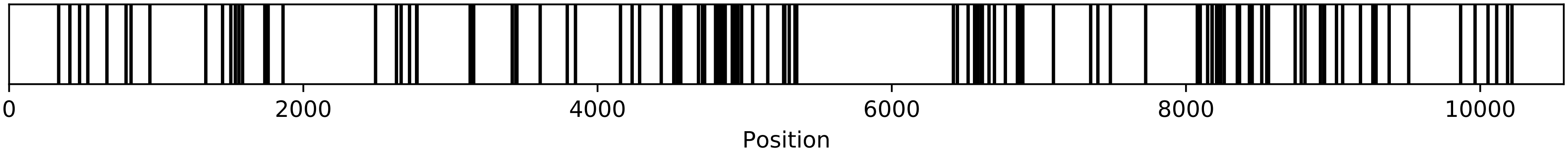


## CR1 vs MM2
Number of differences: 158

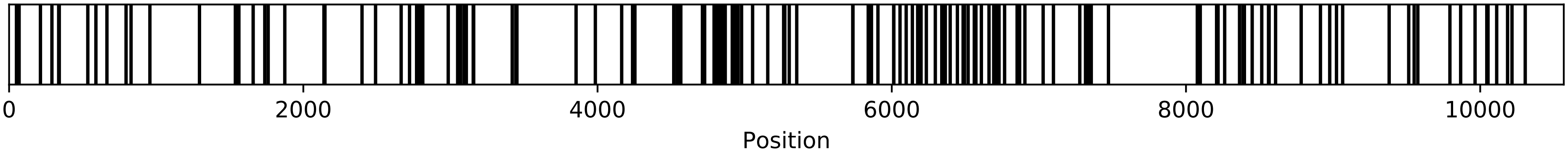


## CR2 vs MM1
Number of differences: 291

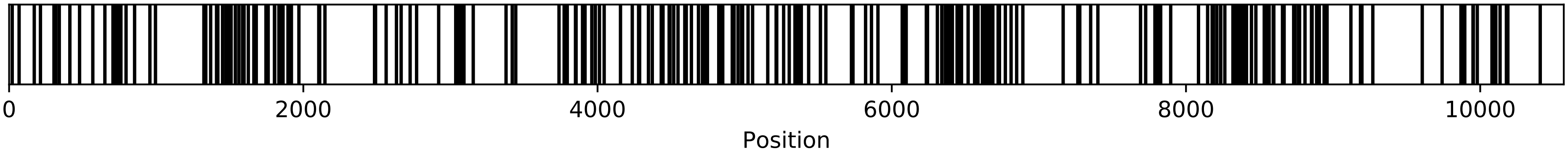


## CR2 vs MM2
Number of differences: 309

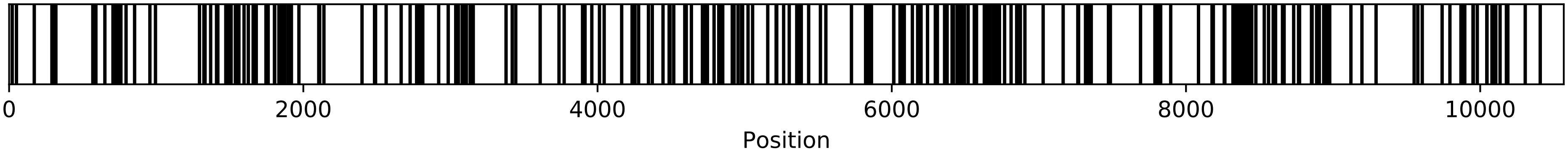


## MM1 vs MM2
Number of differences: 130

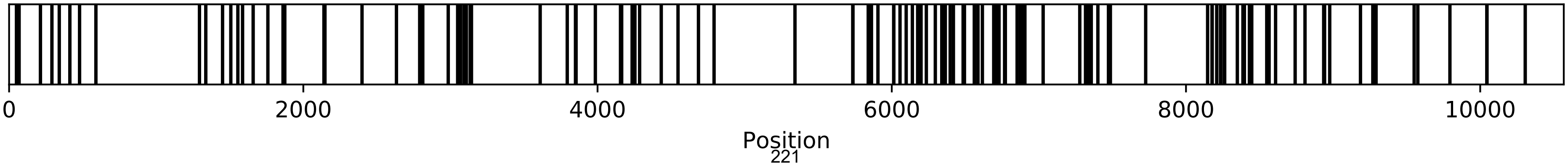

### Region: 633



CR1 vs CR2
Number of differences: 491

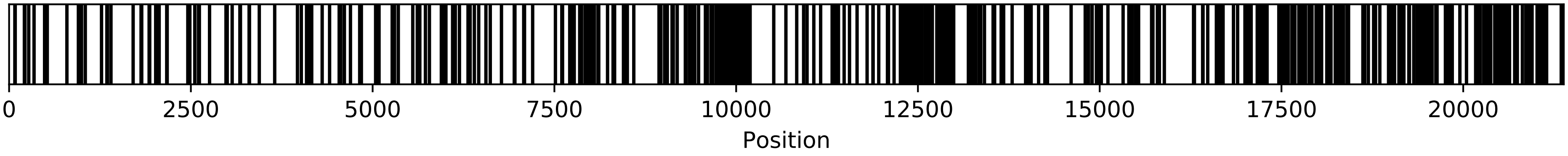


CR1 vs MM1
Number of differences: 216

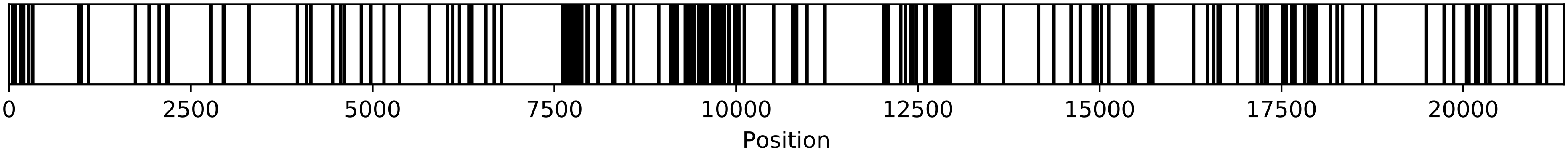


CR1 vs MM2
Number of differences: 195

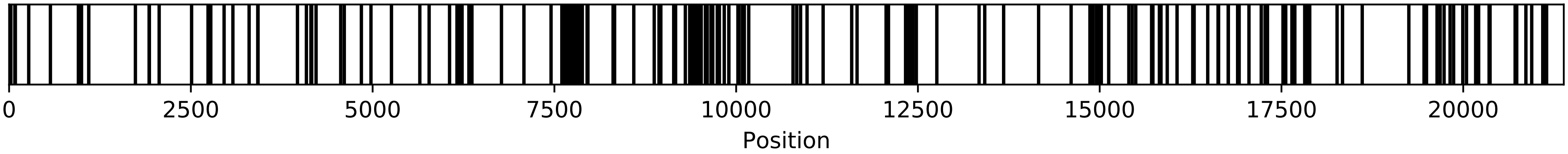


CR2 vs MM1
Number of differences: 512

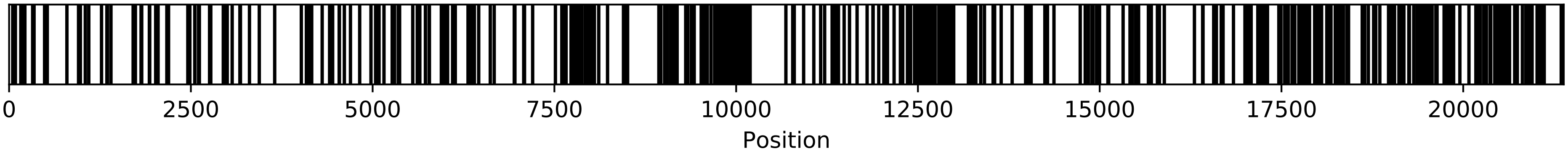


CR2 vs MM2
Number of differences: 498

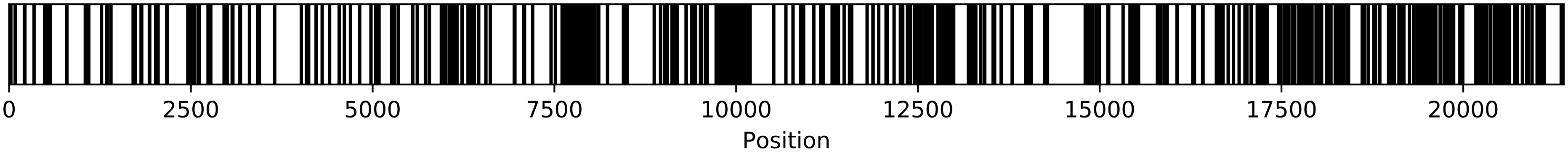


MM1 vs MM2
Number of differences: 172

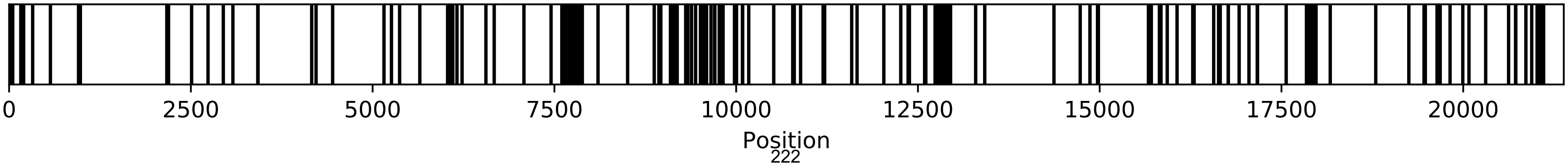

### Region: 636



CR1 vs CR2
Number of differences: 142

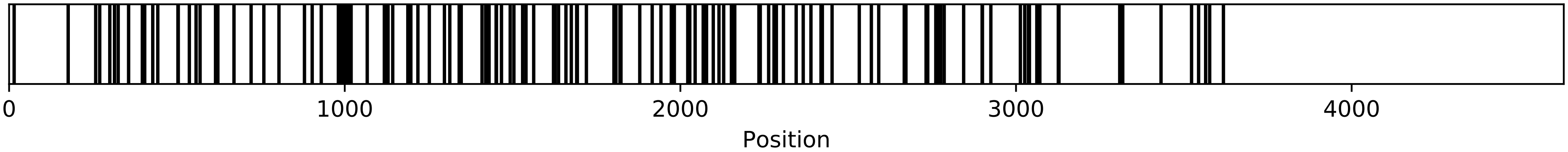


CR1 vs MM1
Number of differences: 95

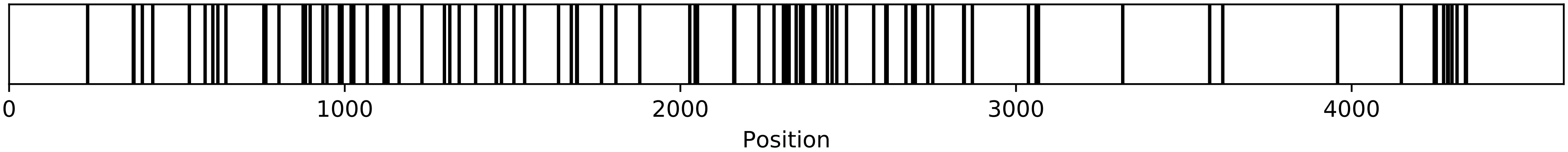


CR1 vs MM2
Number of differences: 116

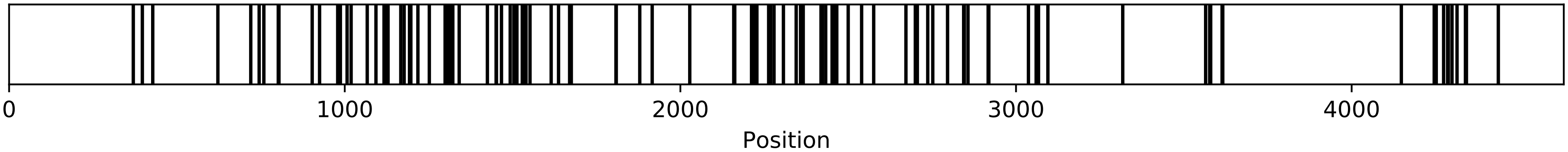


CR2 vs MM1
Number of differences: 157

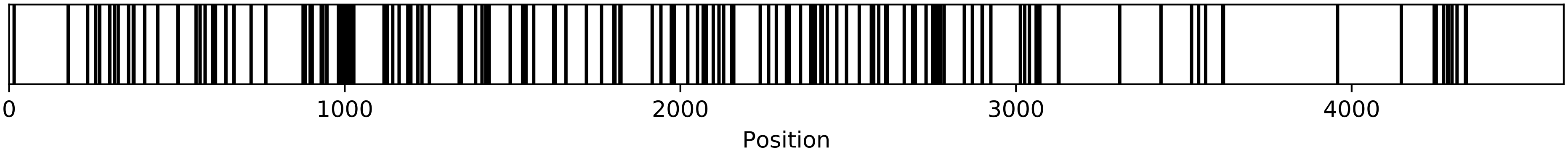


CR2 vs MM2
Number of differences: 167

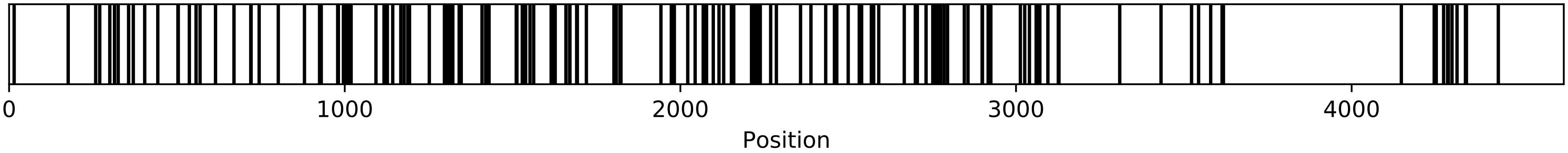


MM1 vs MM2
Number of differences: 95

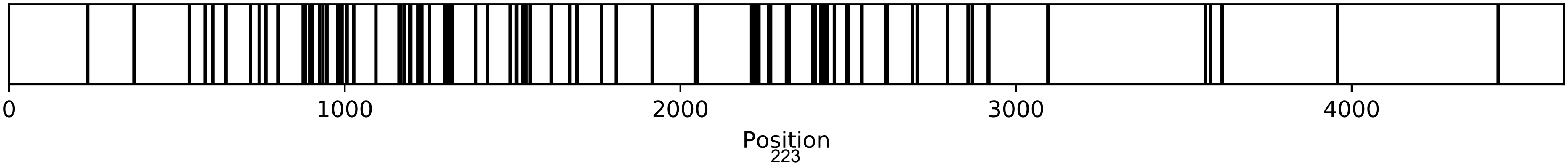

### Region: 637



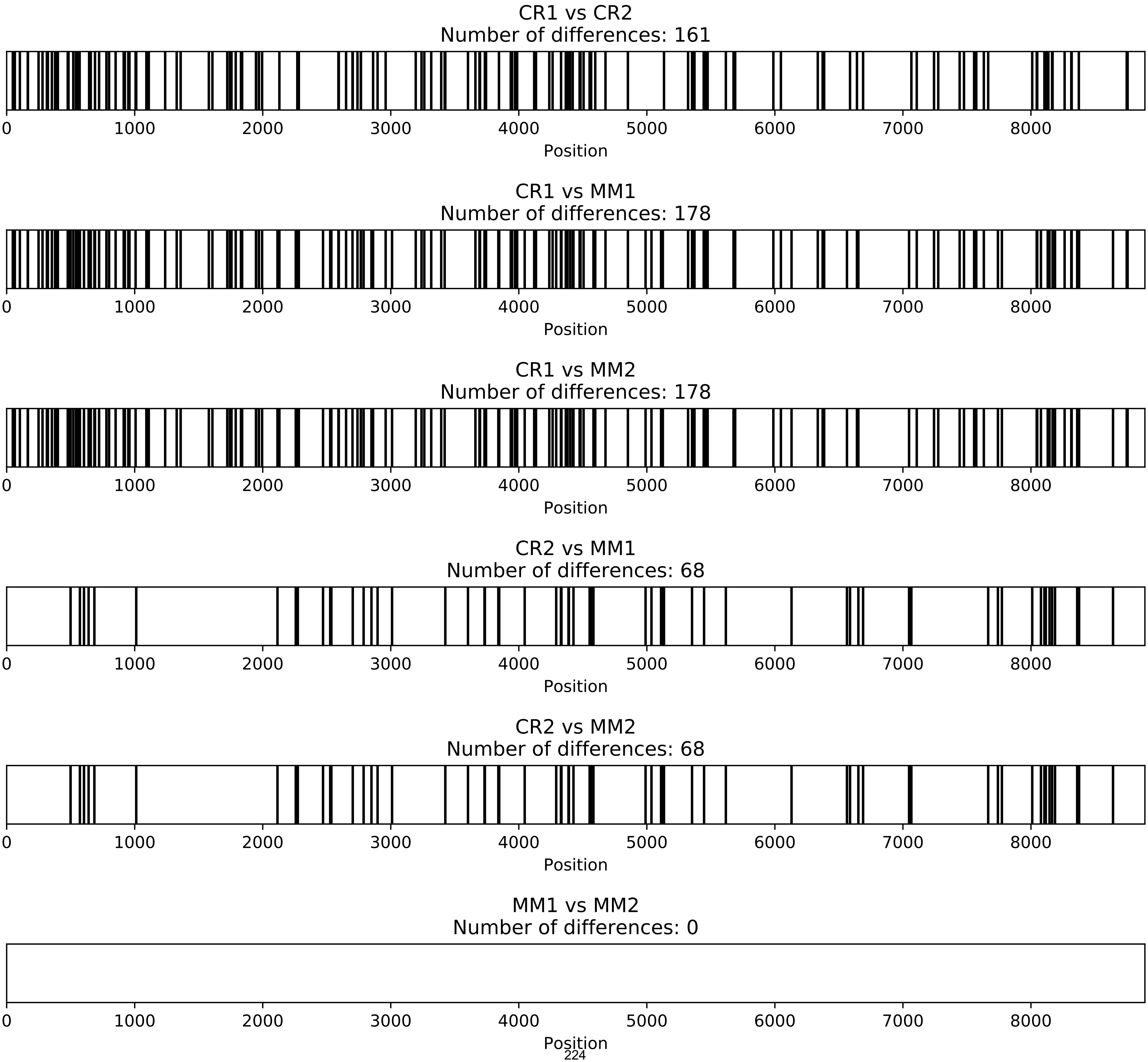
CR1 vs CR2
Number of differences: 161
CR1 vs MM1
Number of differences: 178
CR1 vs MM2
Number of differences: 178
CR2 vs MM1
Number of differences: 68
CR2 vs MM2
Number of differences: 68
MM1 vs MM2
Number of differences: 0
0
1000
2000
3000
4000
5000
6000
7000
8000
Position

### Region: 642



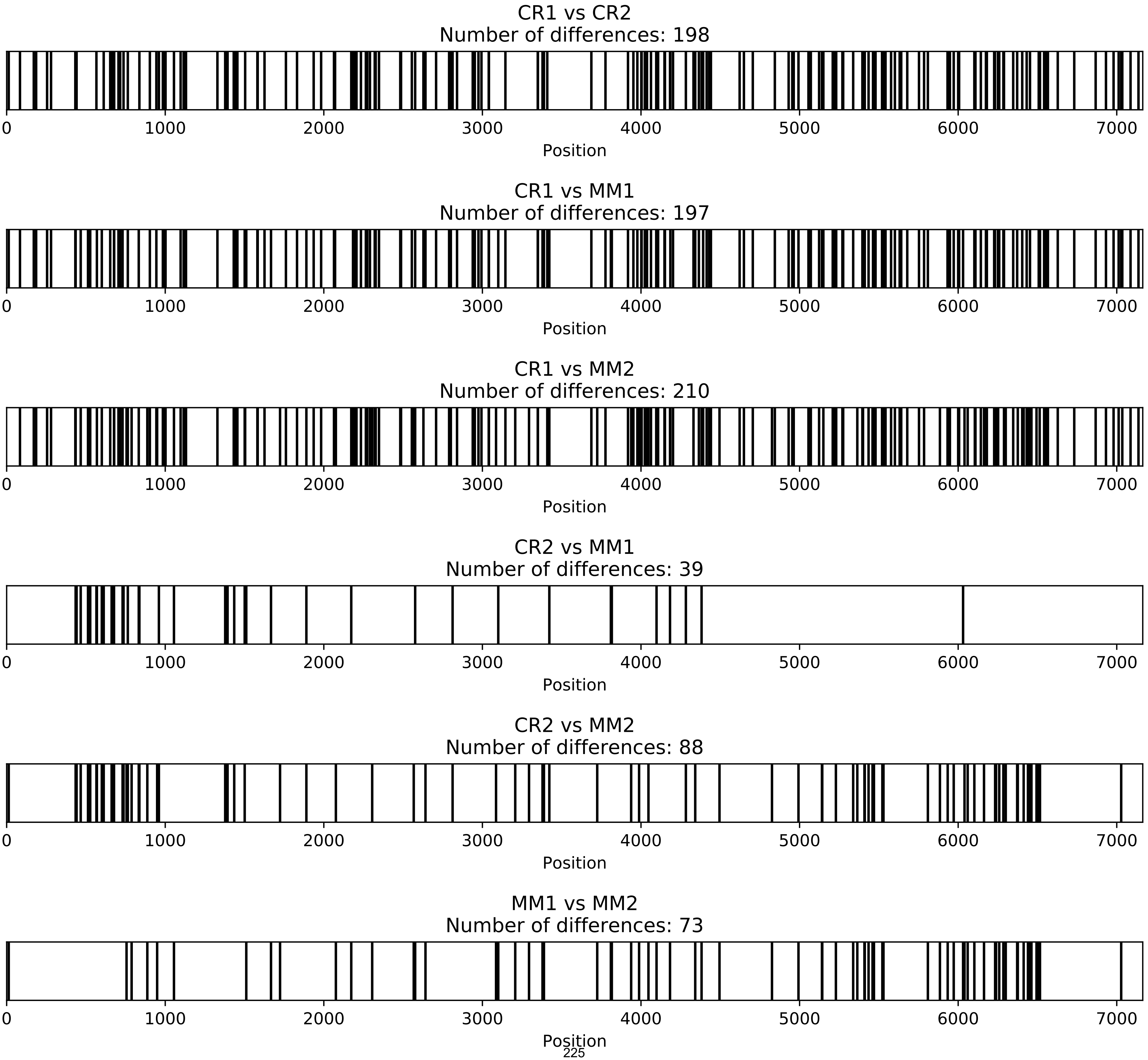
CR1 vs CR2
Number of differences: 198
CR1 vs MM1
Number of differences: 197
CR1 vs MM2
Number of differences: 210
CR2 vs MM1
Number of differences: 39
CR2 vs MM2
Number of differences: 88
MM1 vs MM2
Number of differences: 73
0 1000 2000 3000 4000 5000 6000 7000
Position

### Region: 644



CR1 vs CR2
Number of differences: 261

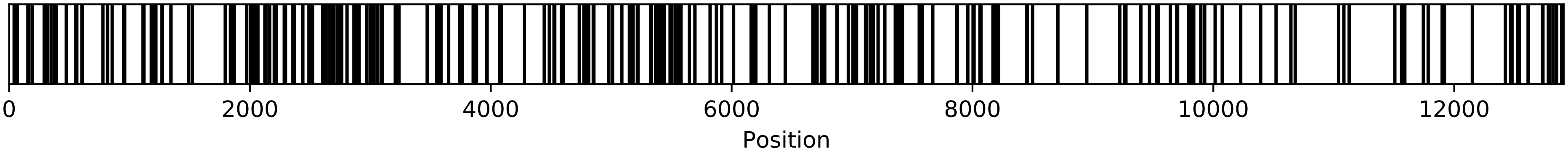


CR1 vs MM1
Number of differences: 254

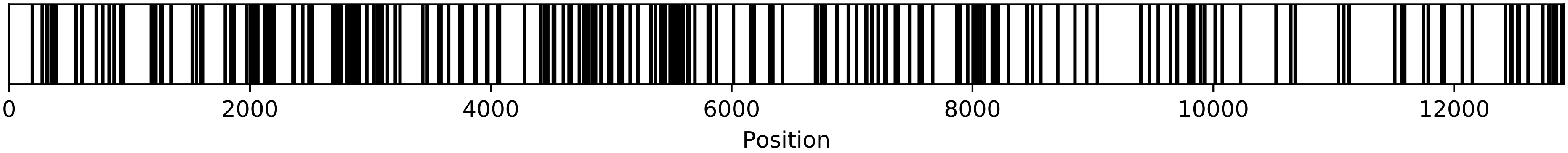


CR1 vs MM2
Number of differences: 232

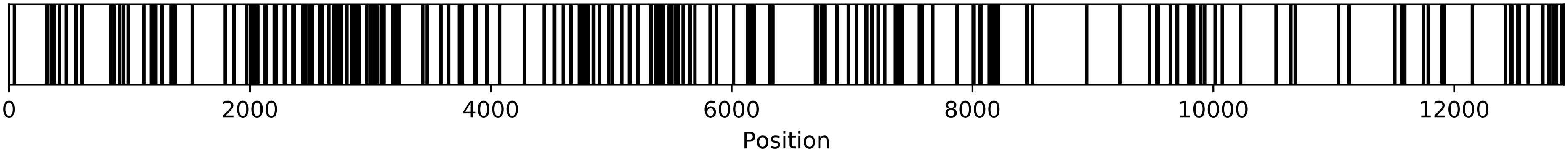


CR2 vs MM1
Number of differences: 153

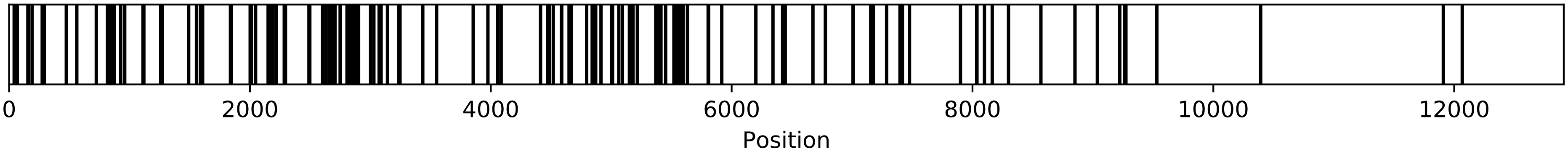


CR2 vs MM2
Number of differences: 99

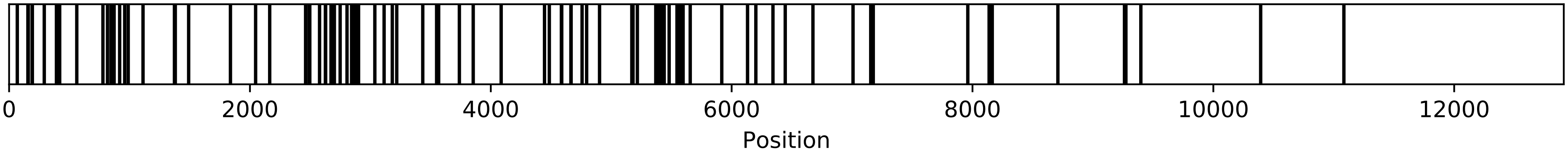


MM1 vs MM2
Number of differences: 122

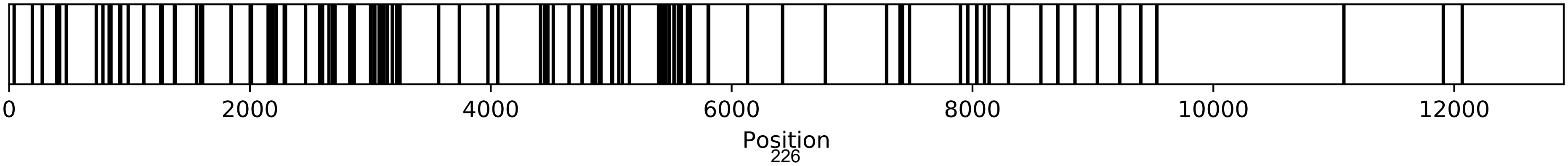

### Region: 645



CR1 vs CR2
Number of differences: 202

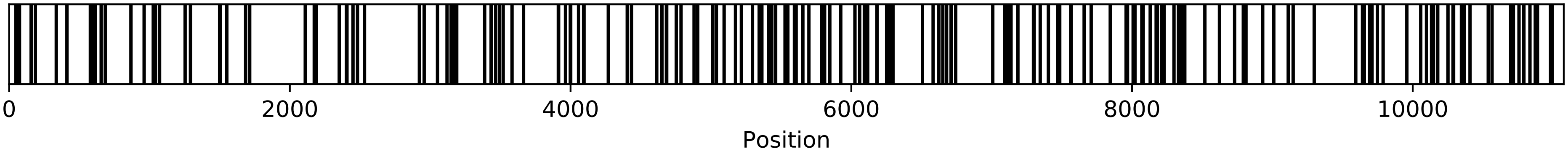


CR1 vs MM1
Number of differences: 44

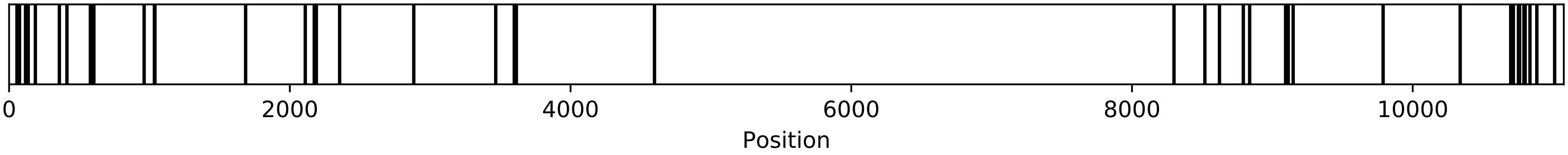


CR1 vs MM2
Number of differences: 77

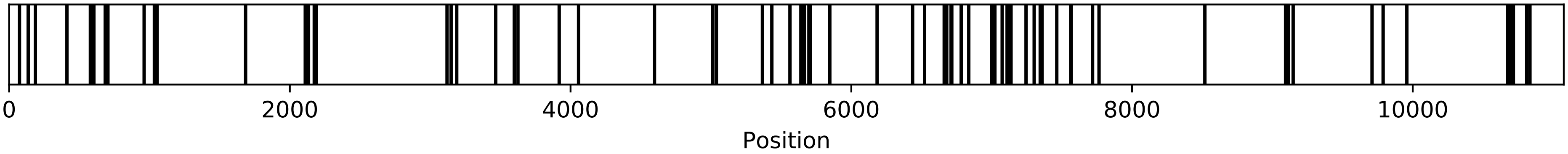


CR2 vs MM1
Number of differences: 197

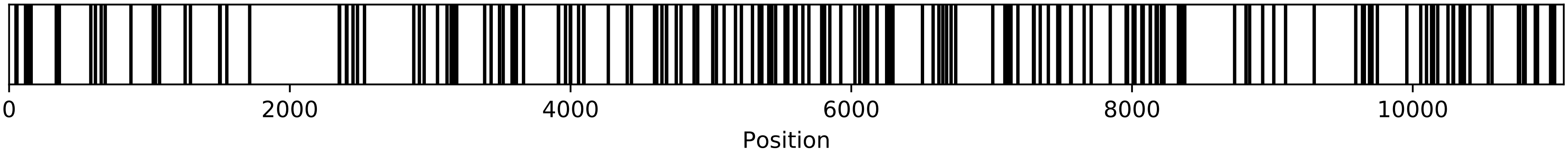


CR2 vs MM2
Number of differences: 198

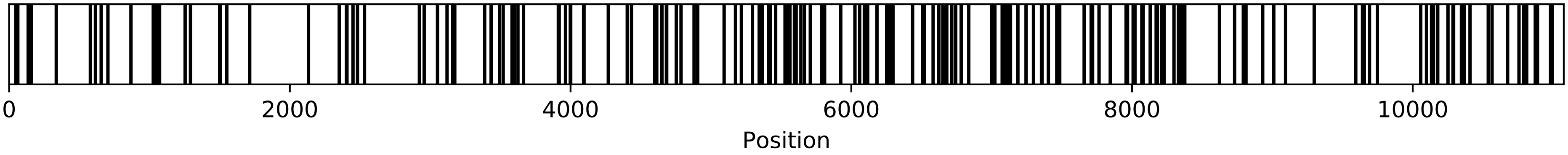


MM1 vs MM2
Number of differences: 72

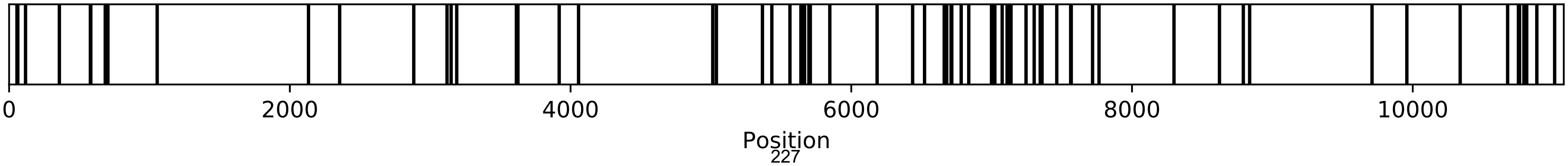

### Region: 647



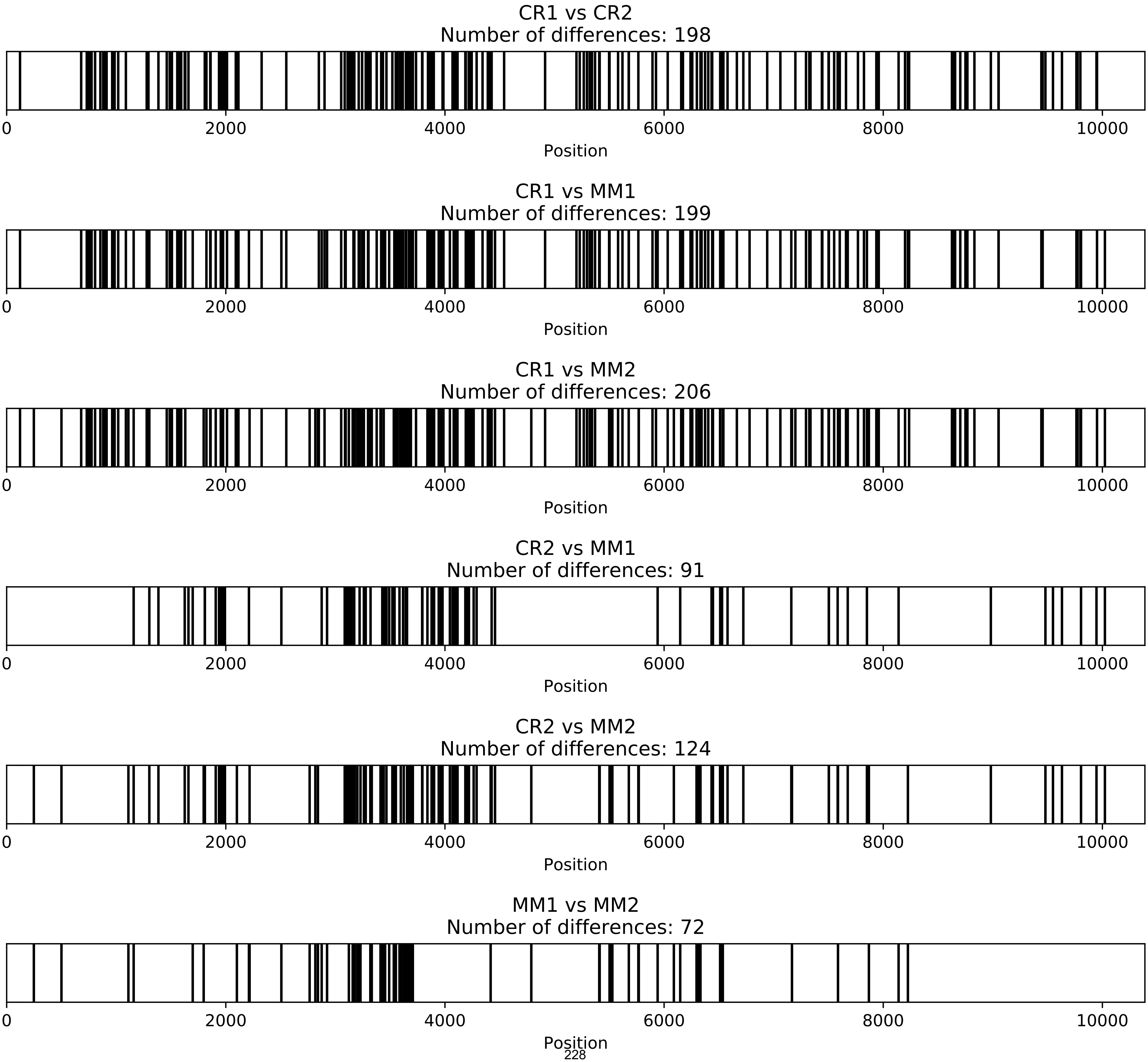
CR1 vs CR2
Number of differences: 198
CR1 vs MM1
Number of differences: 199
CR1 vs MM2
Number of differences: 206
CR2 vs MM1
Number of differences: 91
CR2 vs MM2
Number of differences: 124
MM1 vs MM2
Number of differences: 72
0
2000
4000
6000
8000
10000
Position

### Region: 651



CR1 vs CR2
Number of differences: 239

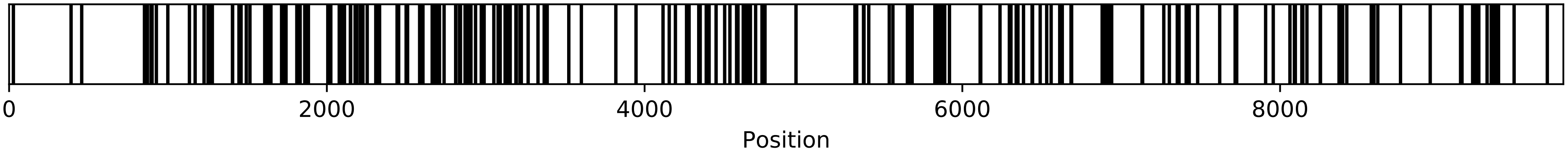


CR1 vs MM1
Number of differences: 138

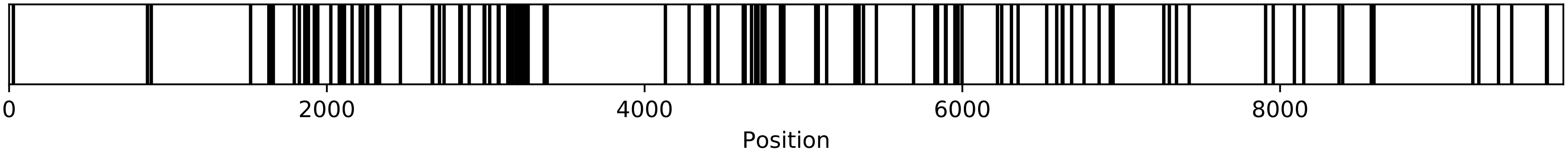


CR1 vs MM2
Number of differences: 106

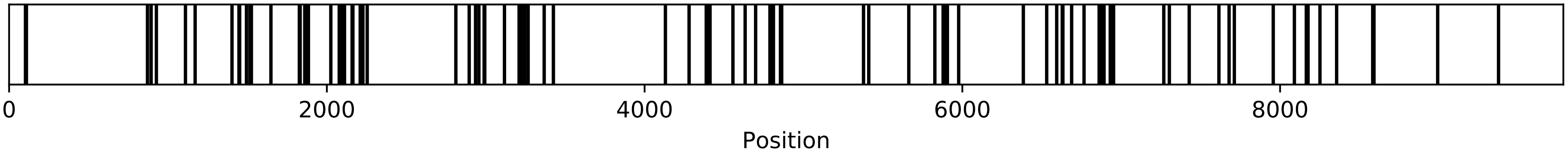


CR2 vs MM1
Number of differences: 250

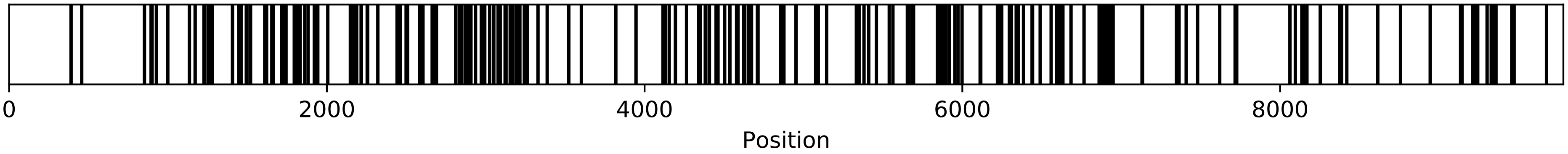


CR2 vs MM2
Number of differences: 229

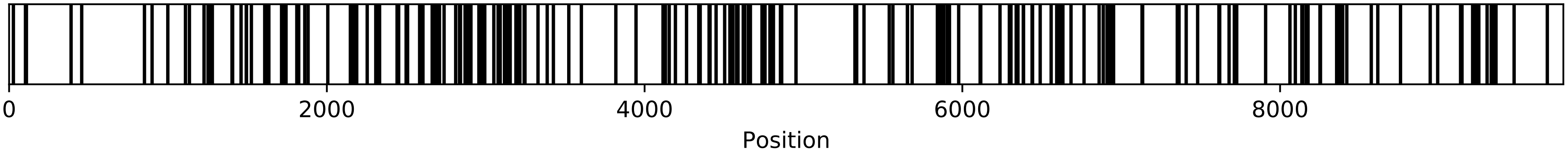


MM1 vs MM2
Number of differences: 136

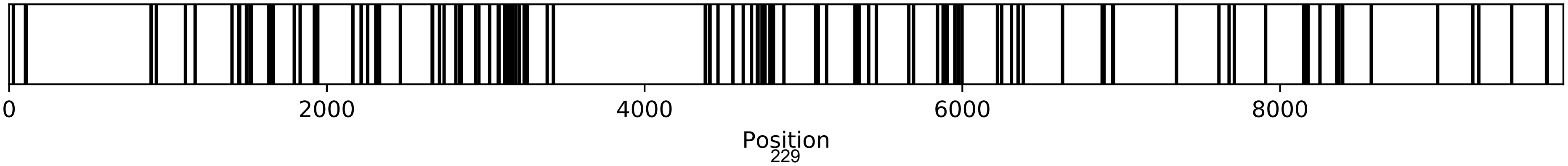

### Region: 654



CR1 vs CR2
Number of differences: 271

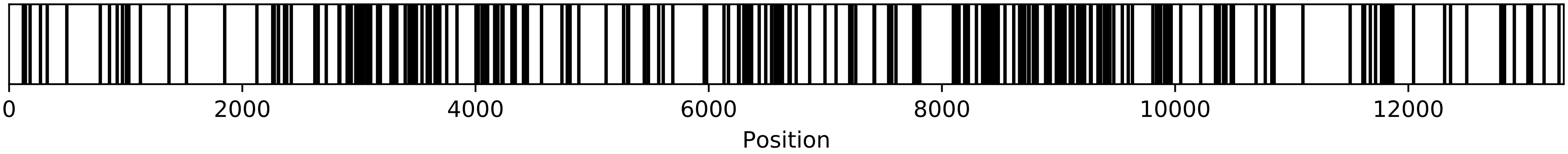


CR1 vs MM1
Number of differences: 175

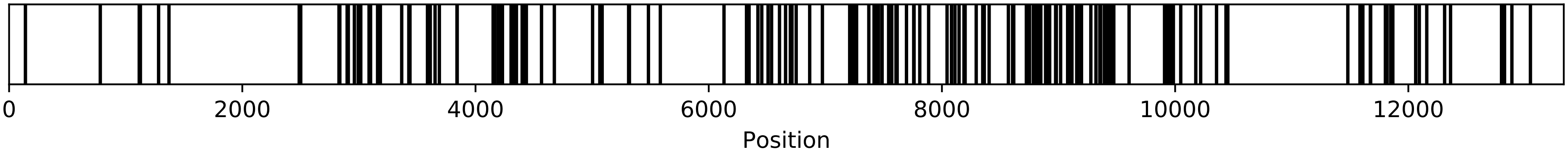


CR1 vs MM2
Number of differences: 140

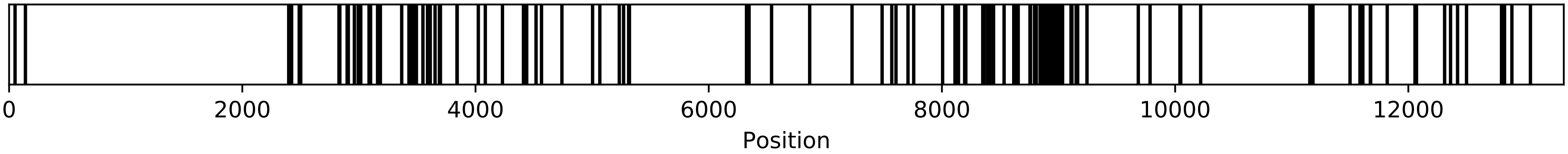


CR2 vs MM1
Number of differences: 270

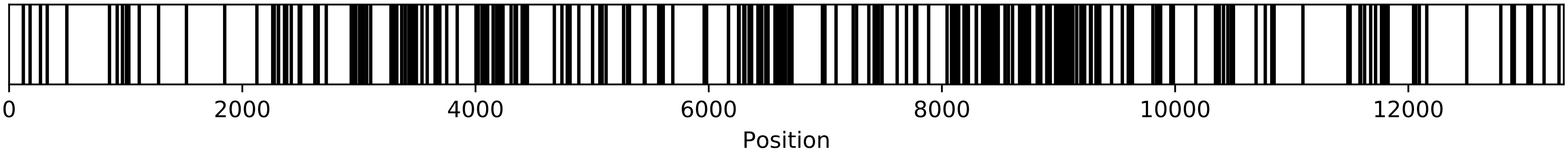


CR2 vs MM2
Number of differences: 274

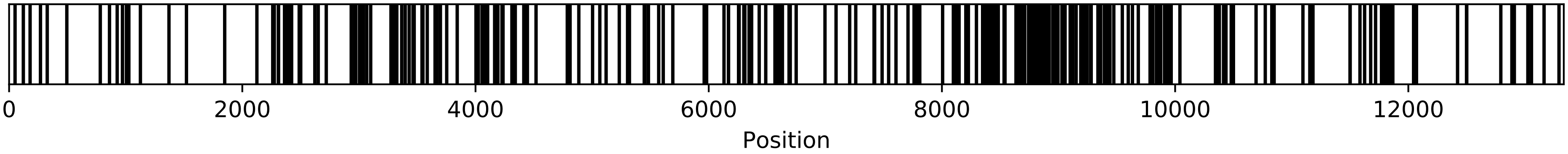


MM1 vs MM2
Number of differences: 178

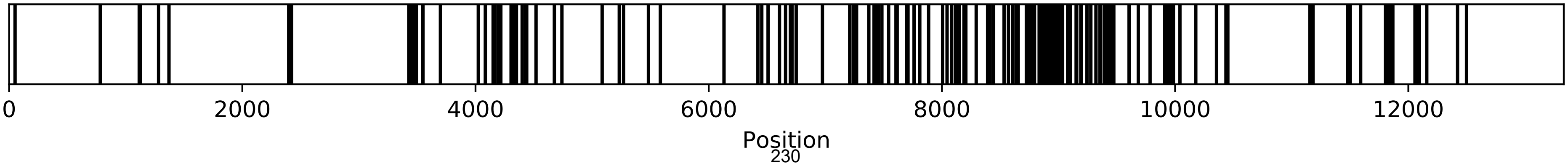

### Region: 657



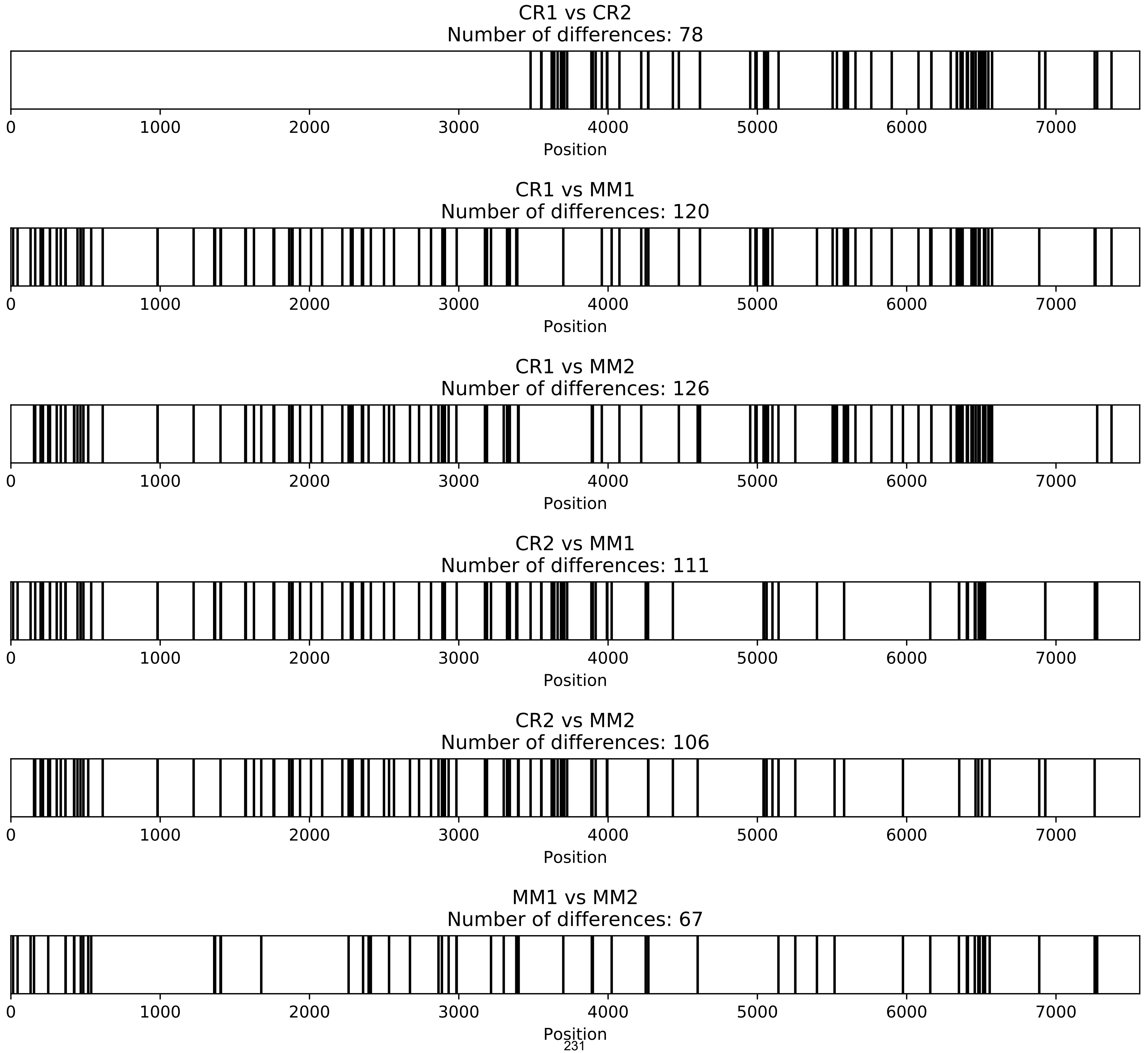
CR1 vs CR2
Number of differences: 78
CR1 vs MM1
Number of differences: 120
CR1 vs MM2
Number of differences: 126
CR2 vs MM1
Number of differences: 111
CR2 vs MM2
Number of differences: 106
MM1 vs MM2
Number of differences: 67
0
1000
2000
3000
4000
5000
6000
7000
Position

### Region: 658



CR1 vs CR2
Number of differences: 377

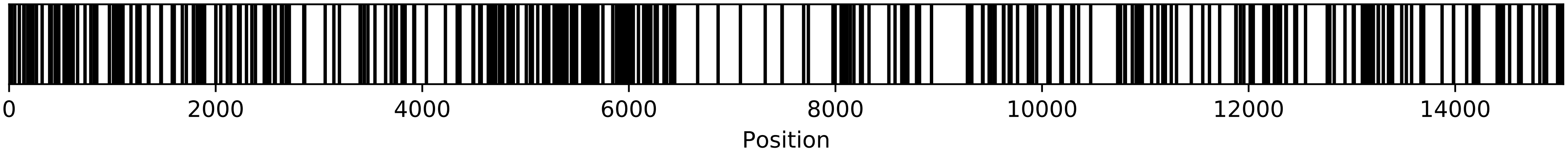


CR1 vs MM1
Number of differences: 146

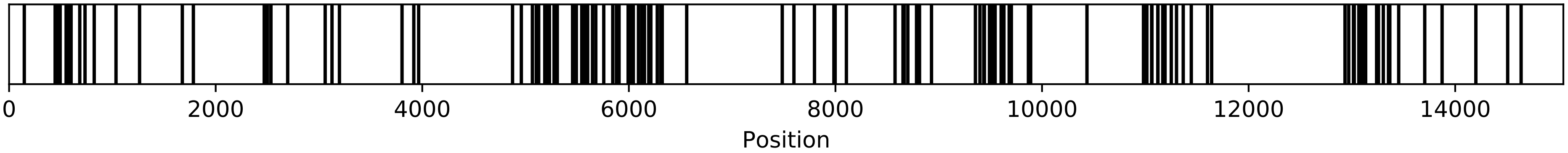


CR1 vs MM2
Number of differences: 134

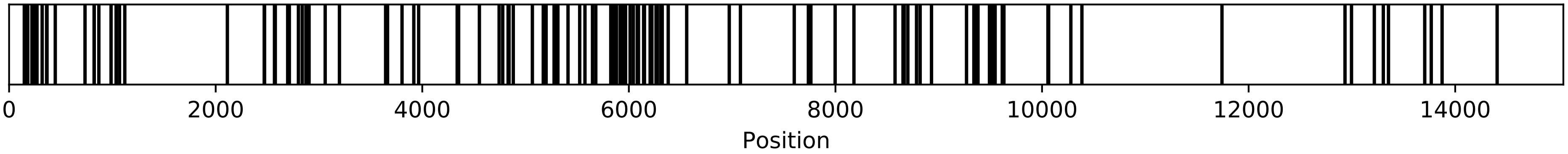


CR2 vs MM1
Number of differences: 361

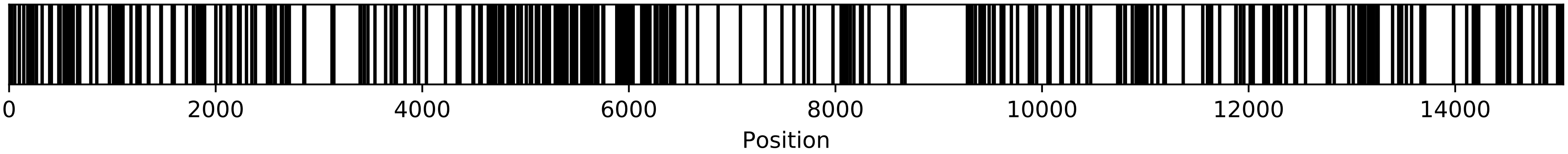


CR2 vs MM2
Number of differences: 384

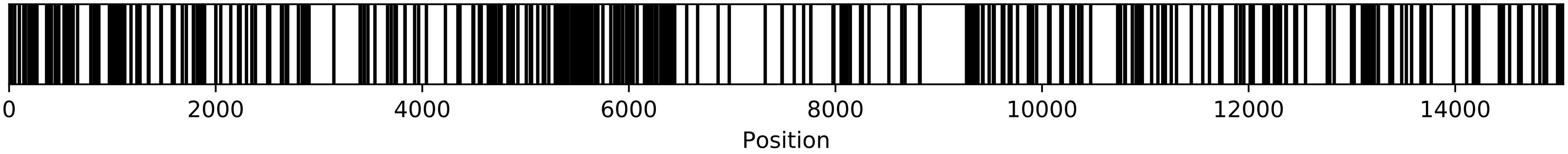


MM1 vs MM2
Number of differences: 177

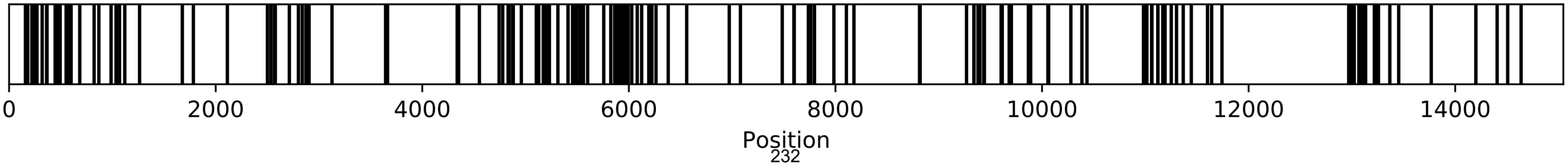

### Region: 661



CR1 vs CR2
Number of differences: 201

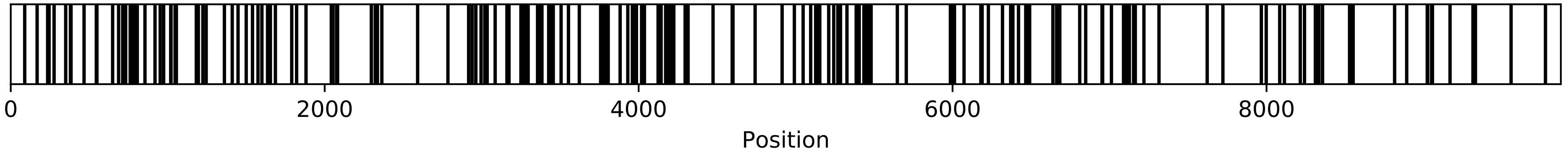


CR1 vs MM1
Number of differences: 143

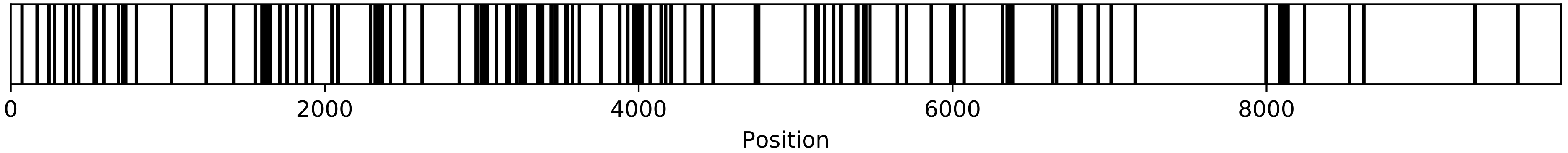


CR1 vs MM2
Number of differences: 135

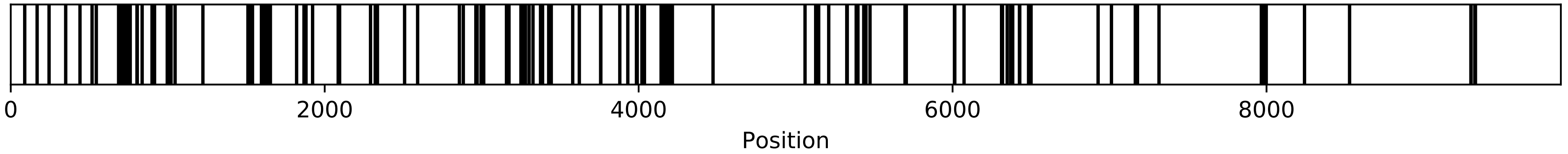


CR2 vs MM1
Number of differences: 198

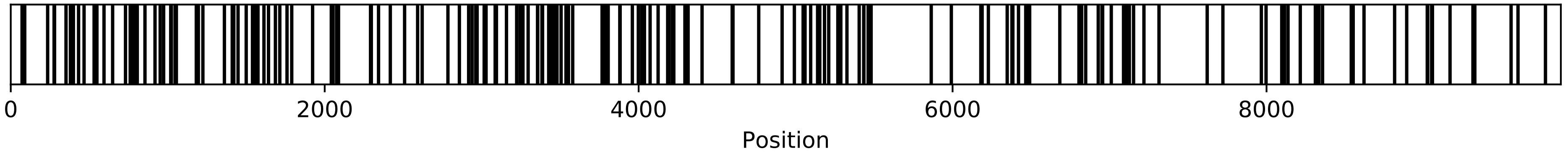


CR2 vs MM2
Number of differences: 185

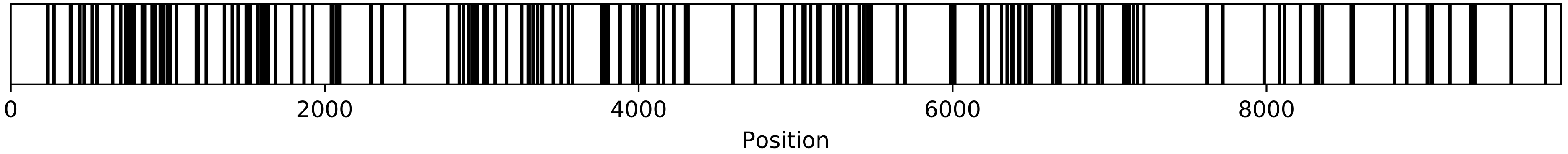


MM1 vs MM2
Number of differences: 132

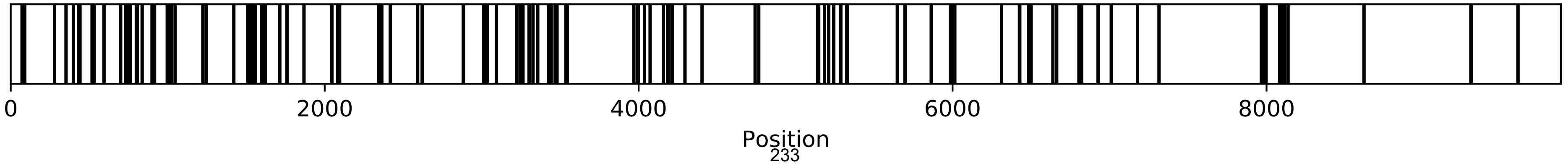

### Region: 662



## CR1 vs CR2
Number of differences: 188

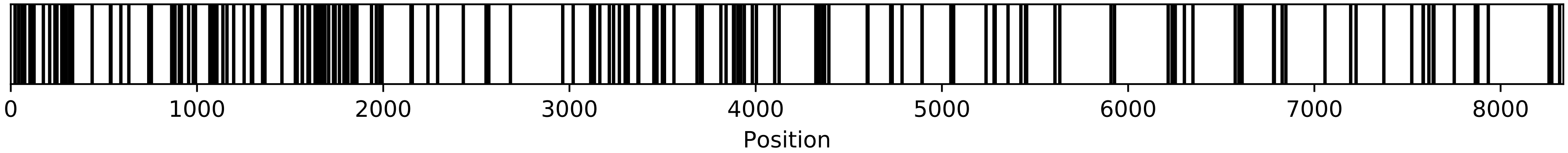


## CR1 vs MM1
Number of differences: 104

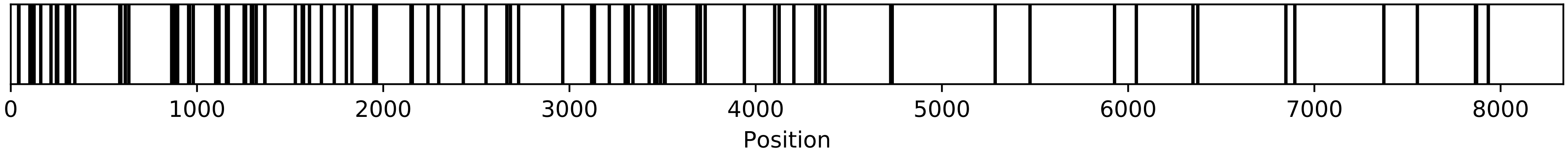


## CR1 vs MM2
Number of differences: 98

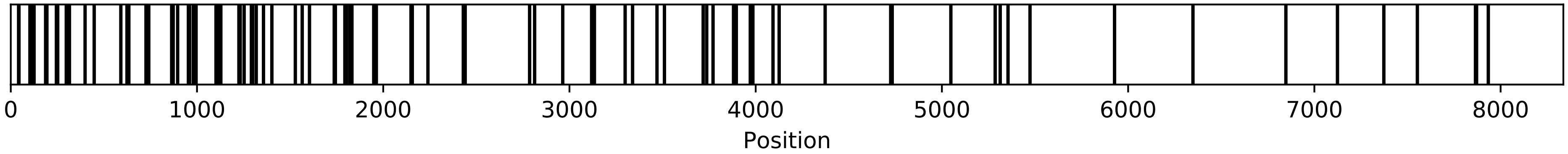


## CR2 vs MM1
Number of differences: 169

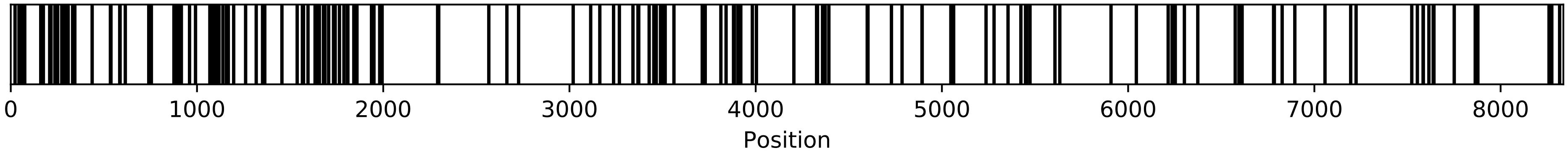


## CR2 vs MM2
Number of differences: 181

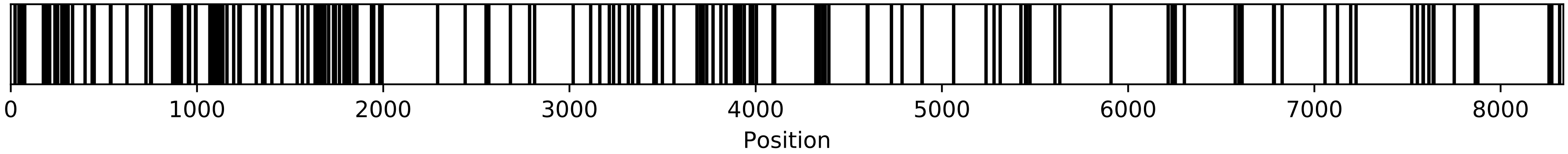


## MM1 vs MM2
Number of differences: 80

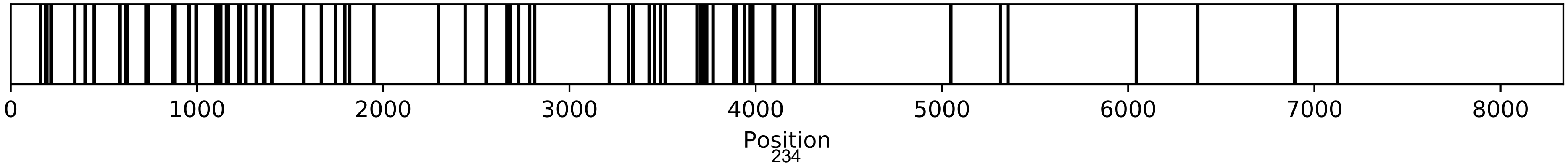

### Region: 663



CR1 vs CR2
Number of differences: 192

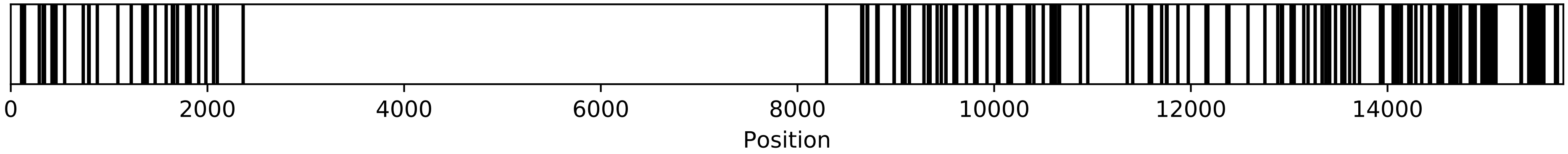


CR1 vs MM1
Number of differences: 283

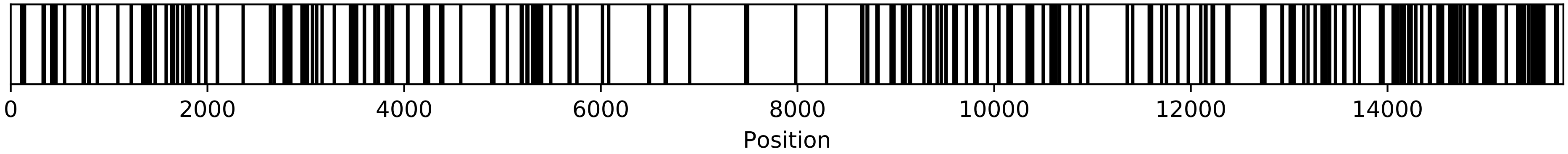


CR1 vs MM2
Number of differences: 268

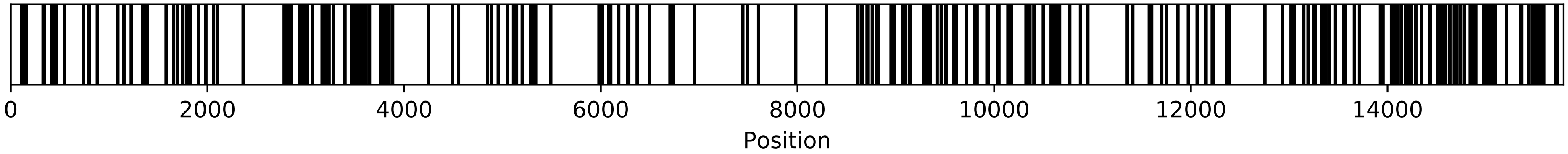


CR2 vs MM1
Number of differences: 143

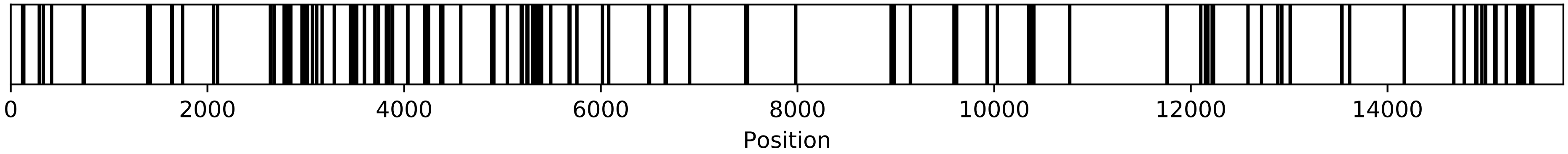


CR2 vs MM2
Number of differences: 136

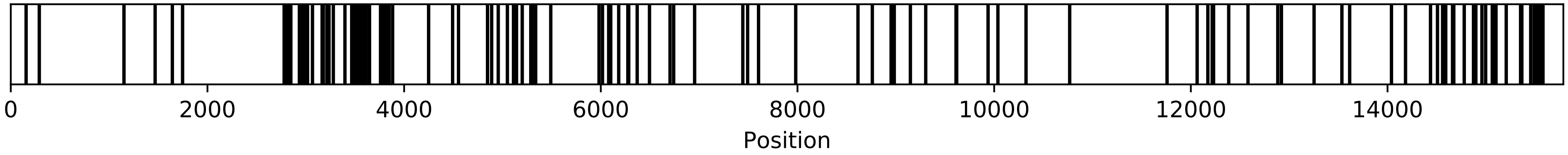


MM1 vs MM2
Number of differences: 156

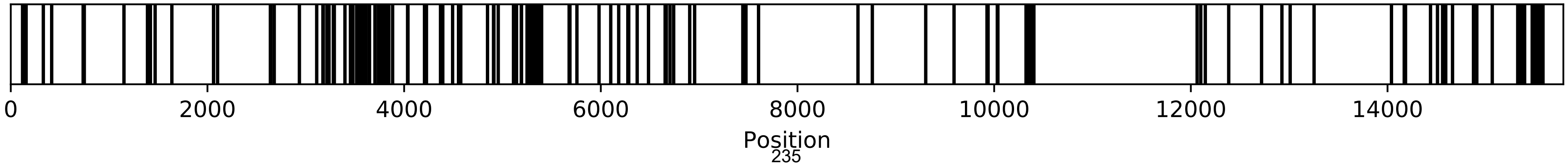

### Region: 664

## CR1 vs CR2
Number of differences: 402

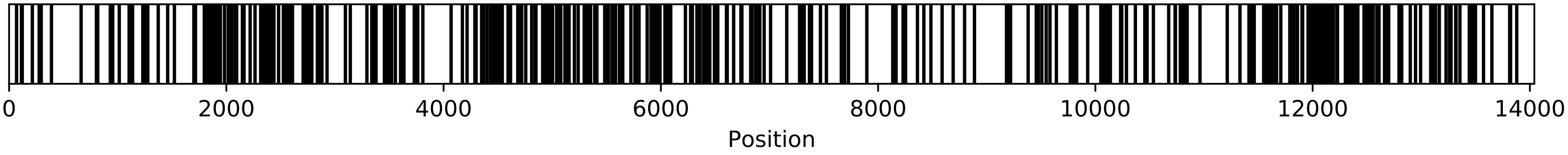


## CR1 vs MM1
Number of differences: 174

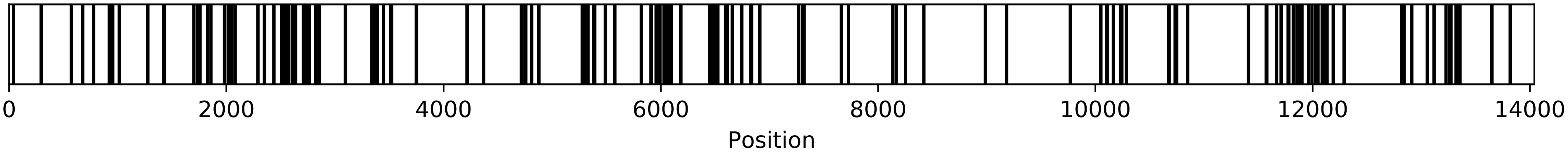


## CR1 vs MM2
Number of differences: 186

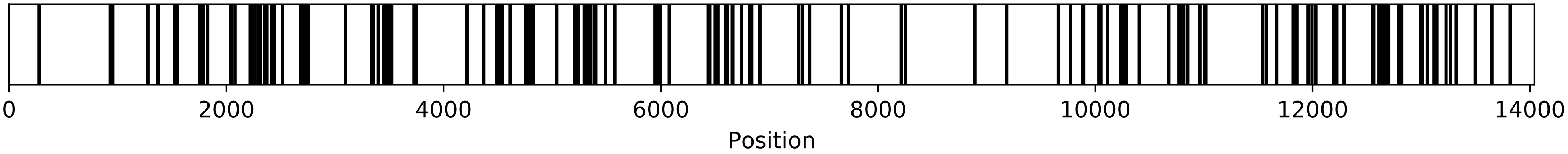


## CR2 vs MM1
Number of differences: 382

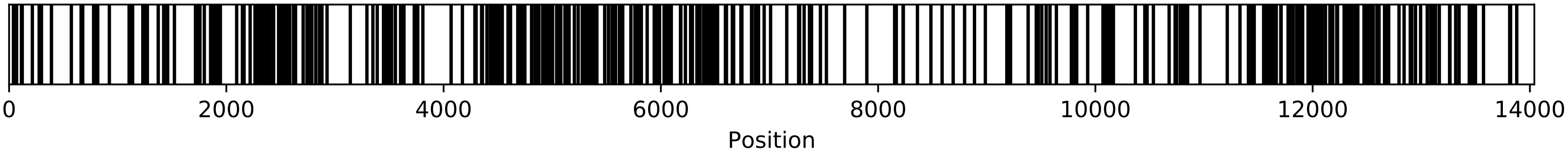


## CR2 vs MM2
Number of differences: 405

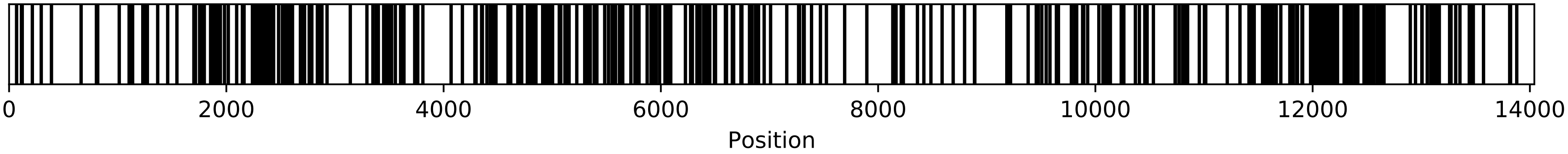


## MM1 vs MM2
Number of differences: 211

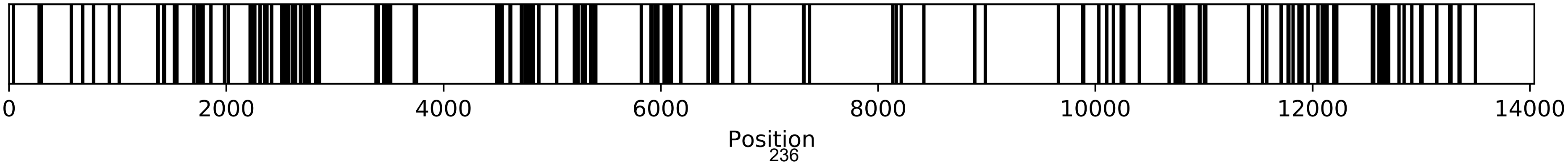

### Region: 665



CR1 vs CR2
Number of differences: 189

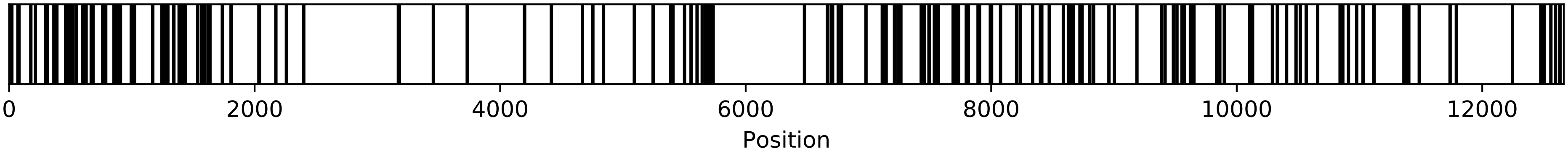


CR1 vs MM1
Number of differences: 106

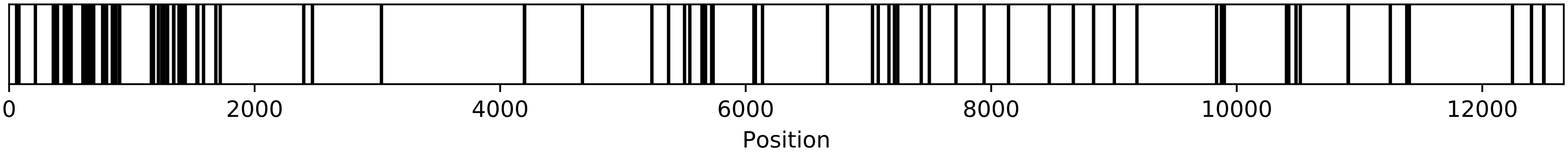


CR1 vs MM2
Number of differences: 78

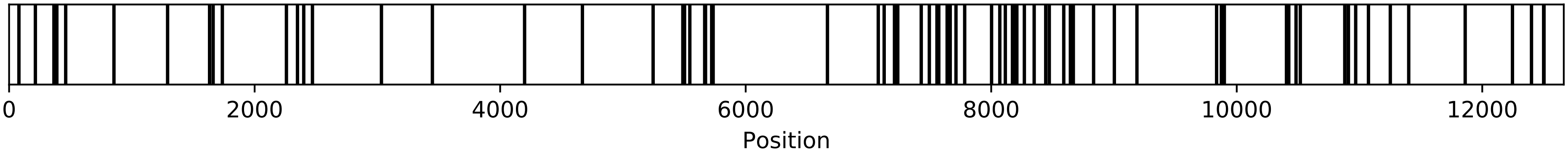


CR2 vs MM1
Number of differences: 182

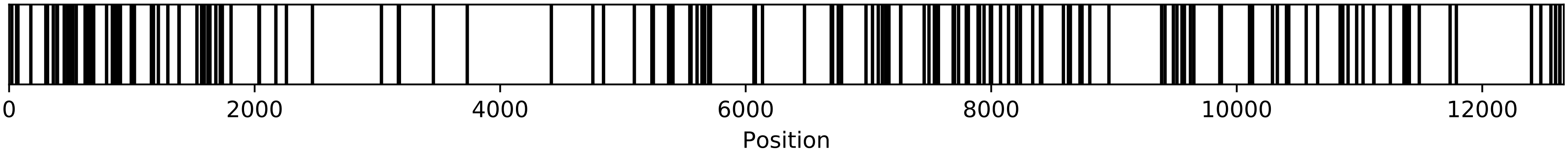


CR2 vs MM2
Number of differences: 185

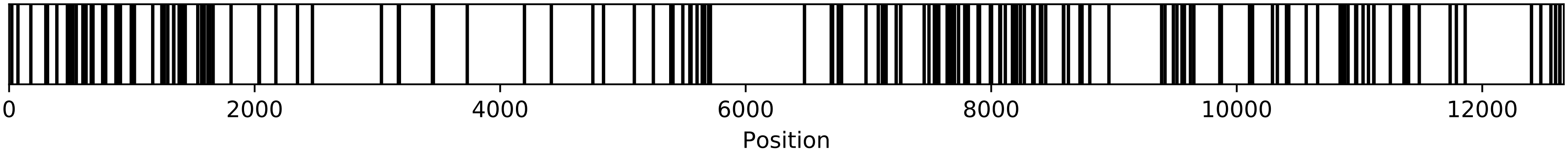


MM1 vs MM2
Number of differences: 92

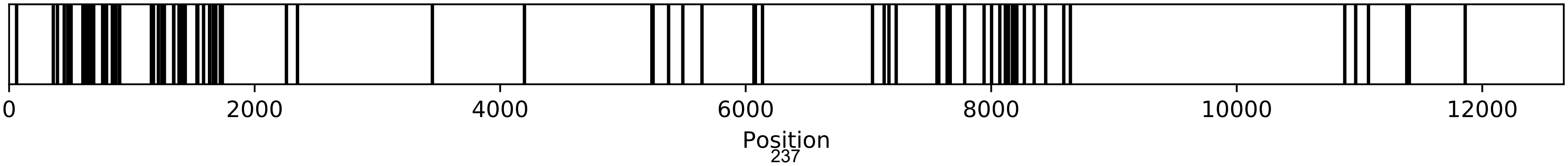

### Region: 669



## CR1 vs CR2
Number of differences: 233

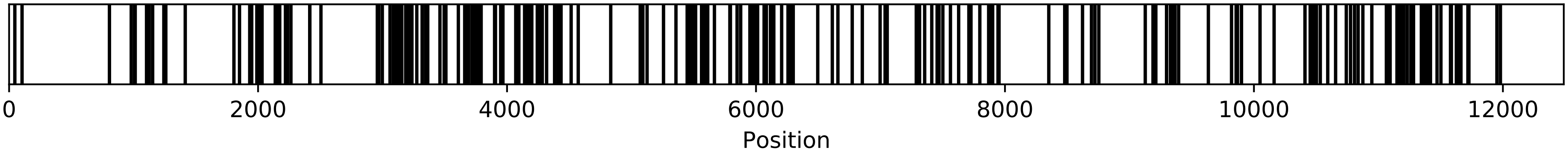


## CR1 vs MM1
Number of differences: 125

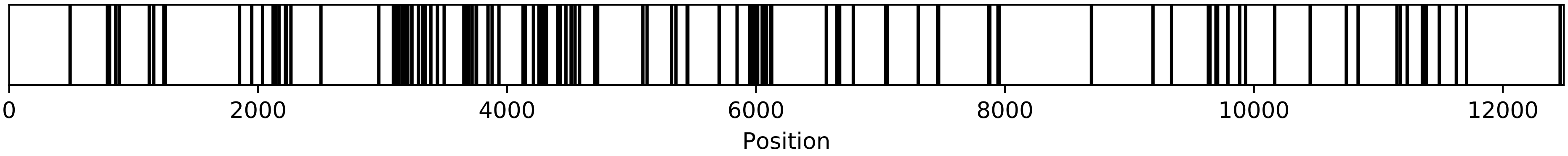


## CR1 vs MM2
Number of differences: 106

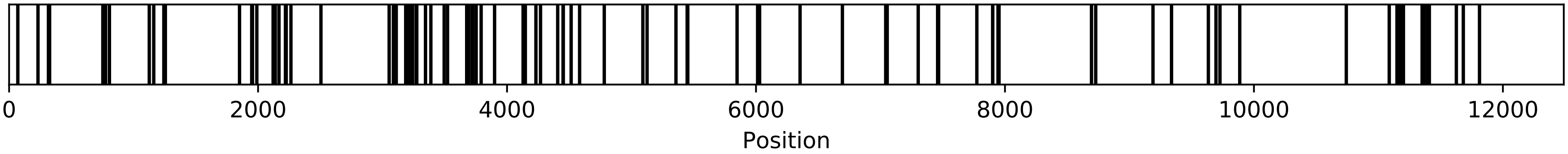


## CR2 vs MM1
Number of differences: 237

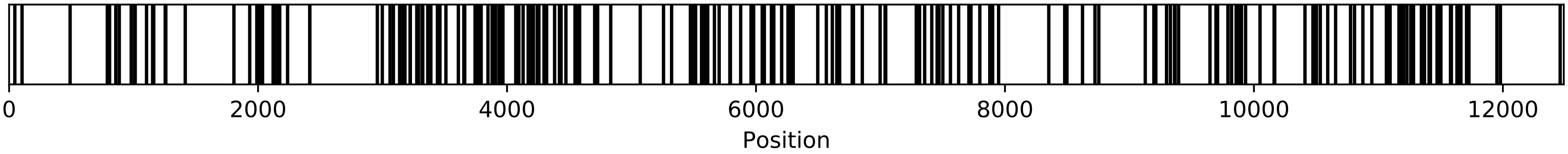


## CR2 vs MM2
Number of differences: 219

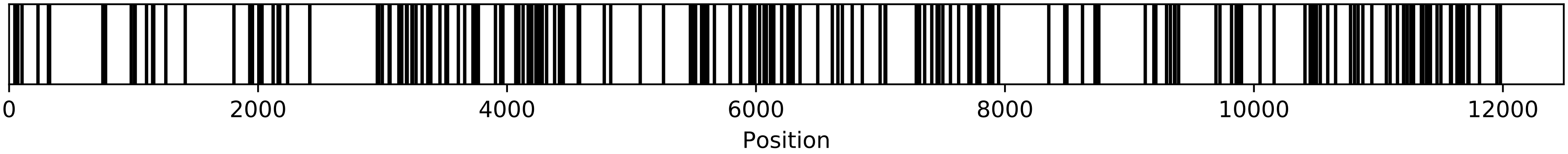


## MM1 vs MM2
Number of differences: 120

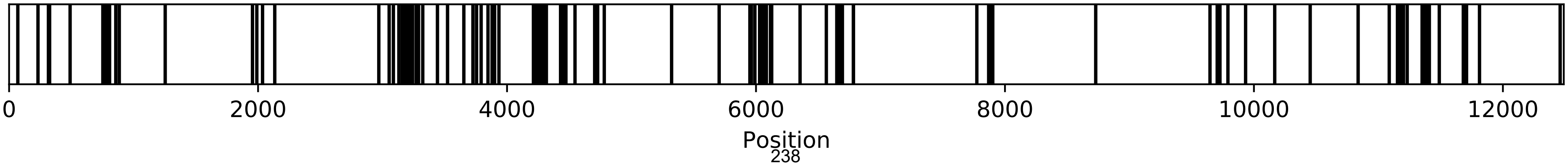

### Region: 671



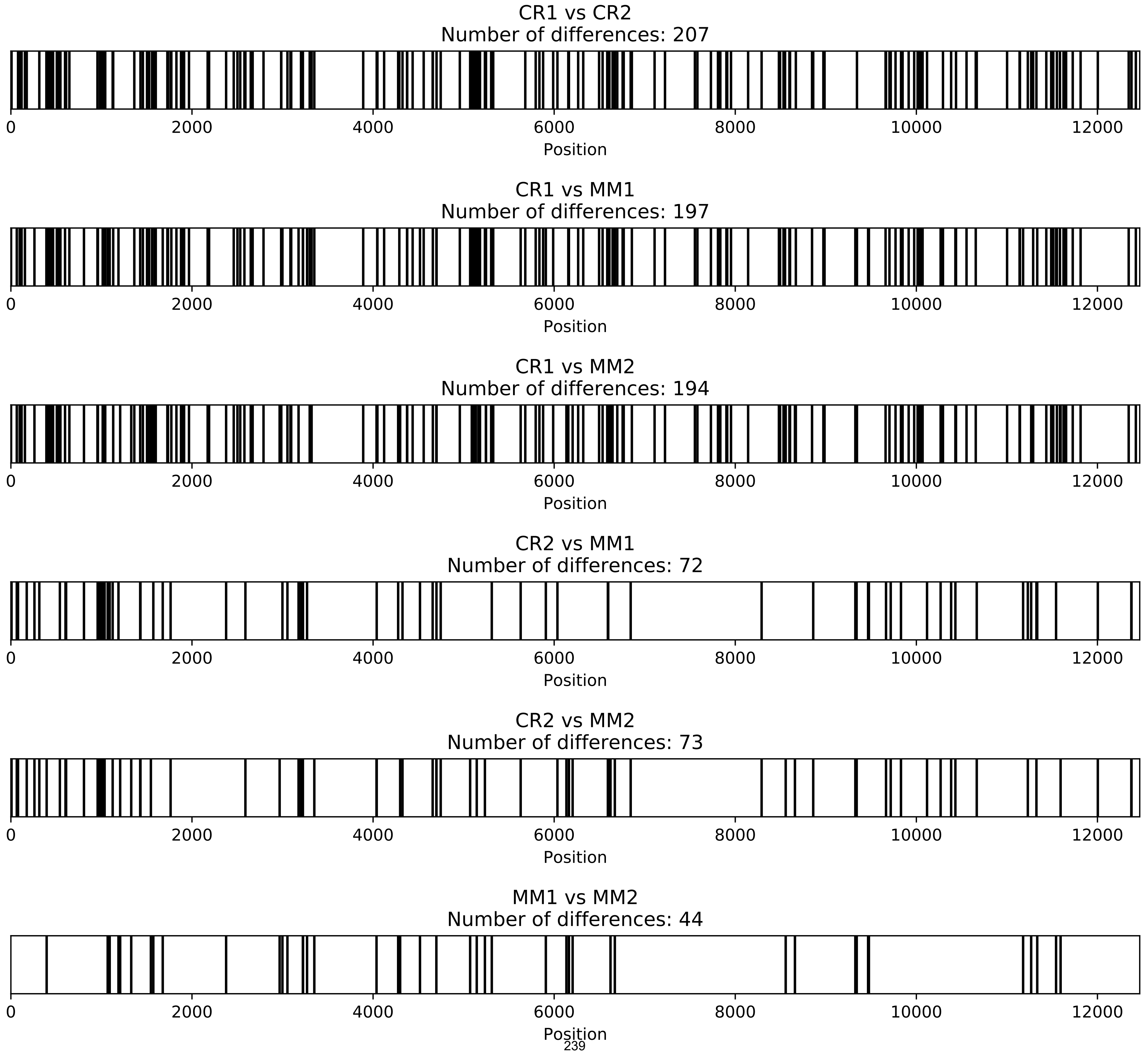
CR1 vs CR2
Number of differences: 207
CR1 vs MM1
Number of differences: 197
CR1 vs MM2
Number of differences: 194
CR2 vs MM1
Number of differences: 72
CR2 vs MM2
Number of differences: 73
MM1 vs MM2
Number of differences: 44
0
2000
4000
6000
8000
10000
12000
Position

### Region: 672



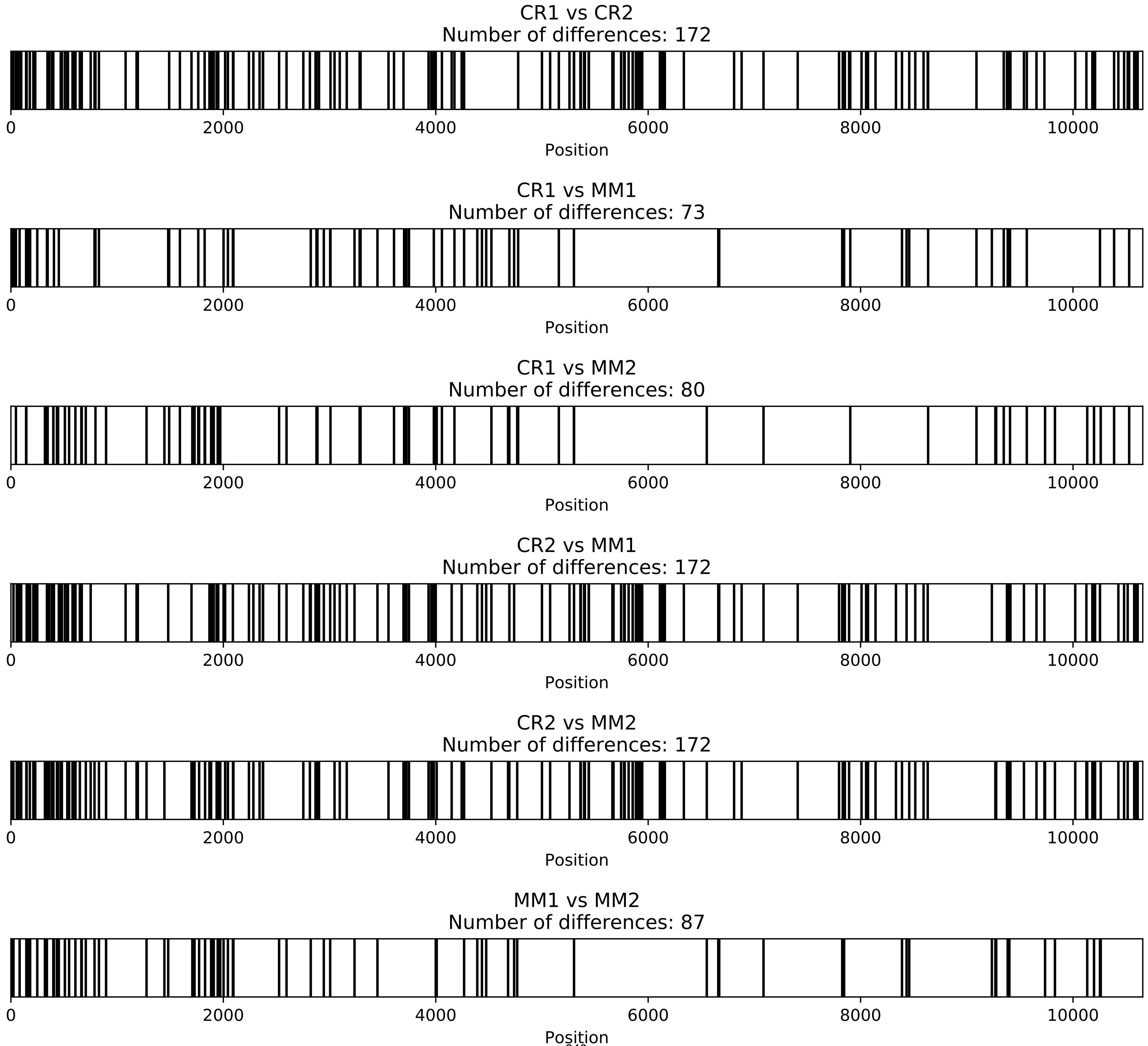
CR1 vs CR2
Number of differences: 172
0
2000
4000
6000
8000
10000
Position
CR1 vs MM1
Number of differences: 73
Position
CR1 vs MM2
Number of differences: 80
Position
CR2 vs MM1
Number of differences: 172
Position
CR2 vs MM2
Number of differences: 172
Position
MM1 vs MM2
Number of differences: 87
Position

### Region: 674



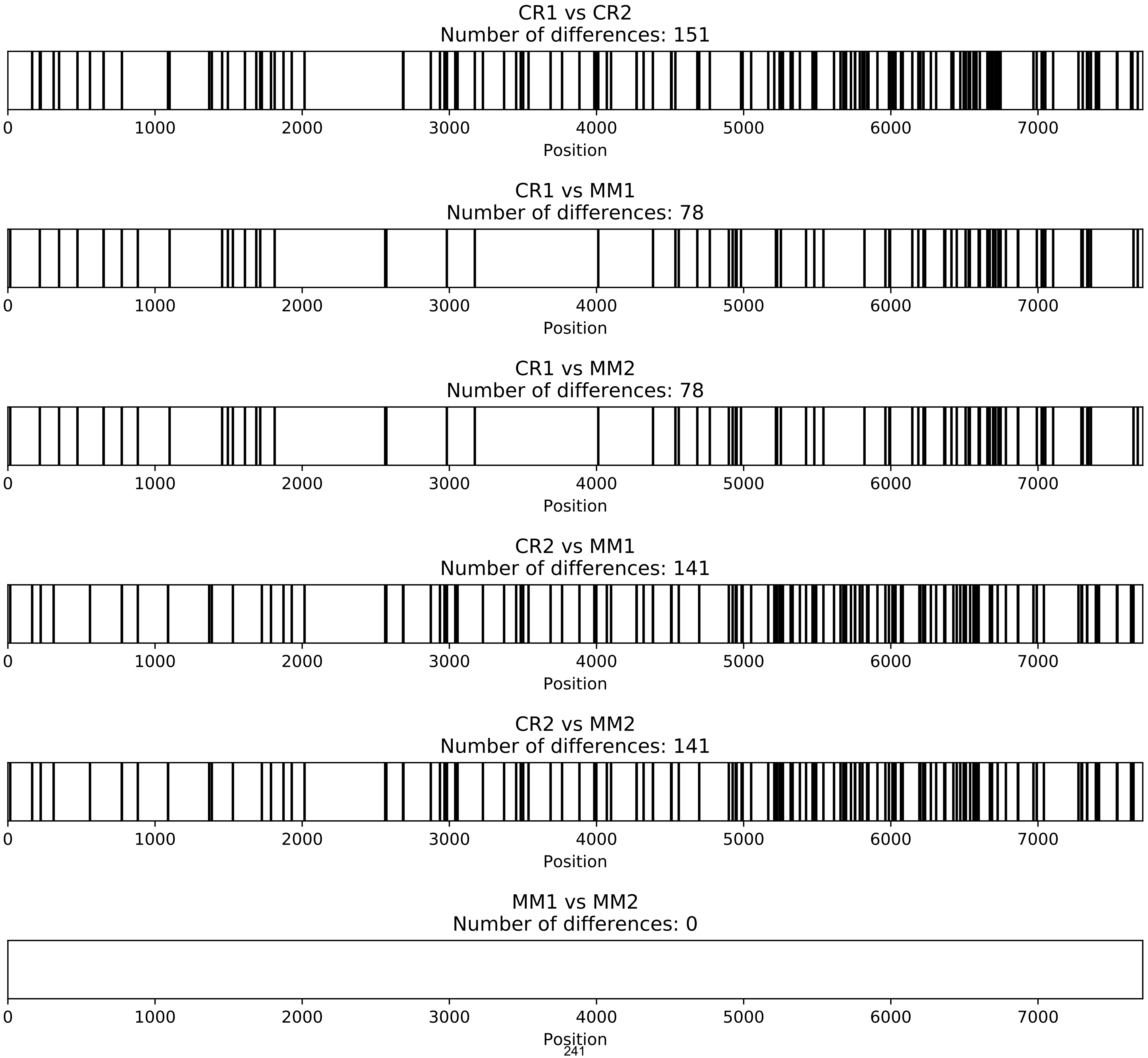
CR1 vs CR2
Number of differences: 151
CR1 vs MM1
Number of differences: 78
CR1 vs MM2
Number of differences: 78
CR2 vs MM1
Number of differences: 141
CR2 vs MM2
Number of differences: 141
MM1 vs MM2
Number of differences: 0
0 1000 2000 3000 4000 5000 6000 7000
Position

### Region: 675



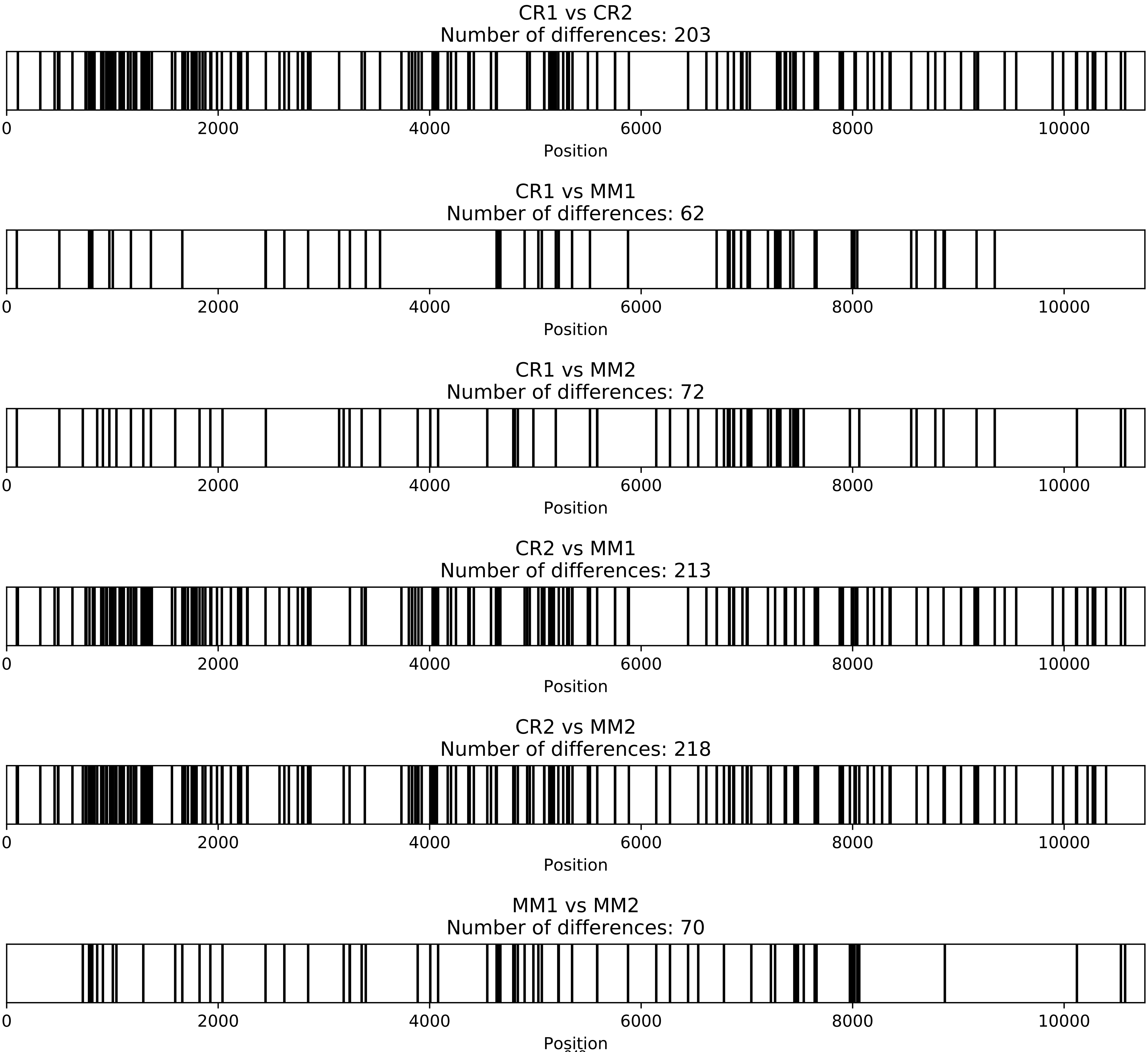
CR1 vs CR2
Number of differences: 203
CR1 vs MM1
Number of differences: 62
CR1 vs MM2
Number of differences: 72
CR2 vs MM1
Number of differences: 213
CR2 vs MM2
Number of differences: 218
MM1 vs MM2
Number of differences: 70
0
2000
4000
6000
8000
10000
Position

### Region: 676



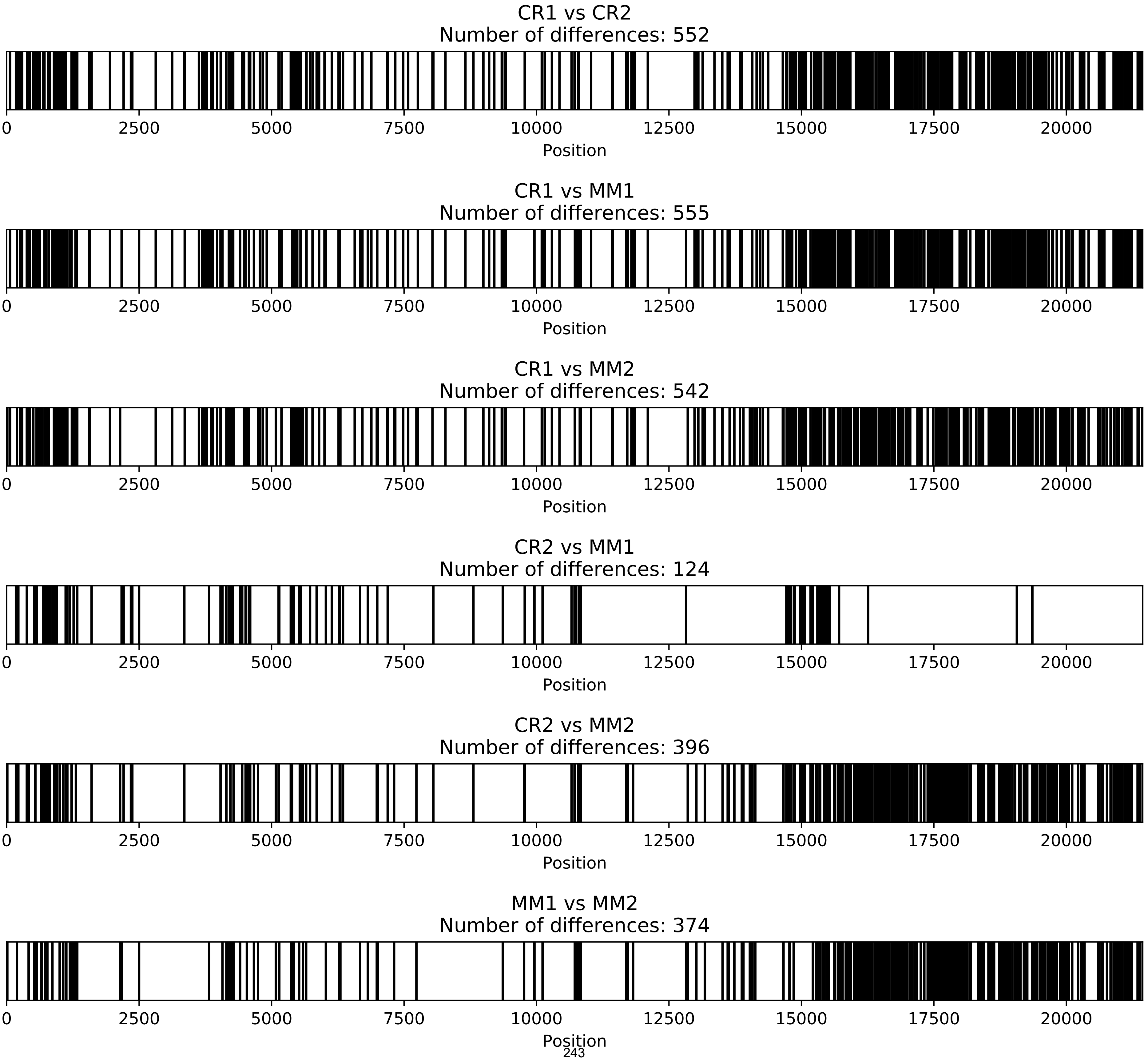
CR1 vs CR2
Number of differences: 552
CR1 vs MM1
Number of differences: 555
CR1 vs MM2
Number of differences: 542
CR2 vs MM1
Number of differences: 124
CR2 vs MM2
Number of differences: 396
MM1 vs MM2
Number of differences: 374
0
2500
5000
7500
10000
12500
15000
17500
20000
Position

### Region: 678



CR1 vs CR2
Number of differences: 275

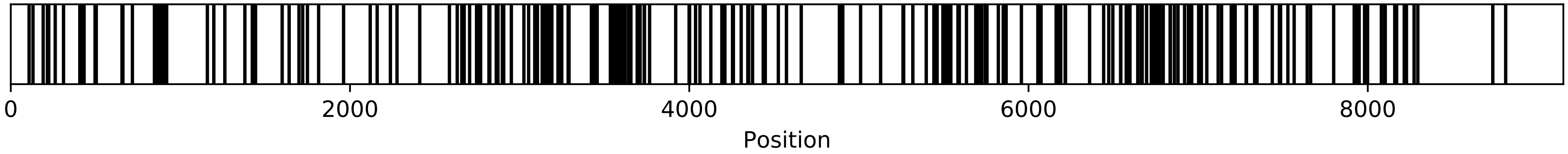


CR1 vs MM1
Number of differences: 184

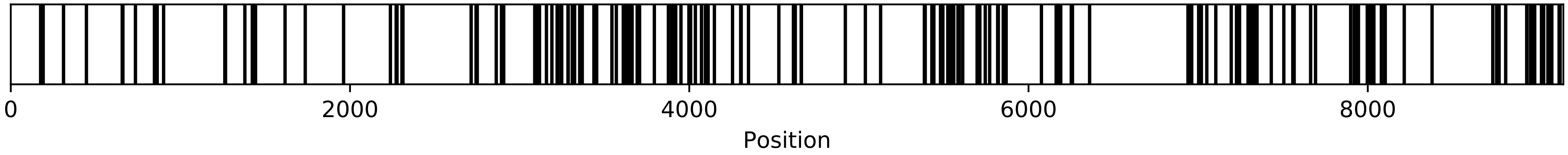


CR1 vs MM2
Number of differences: 211

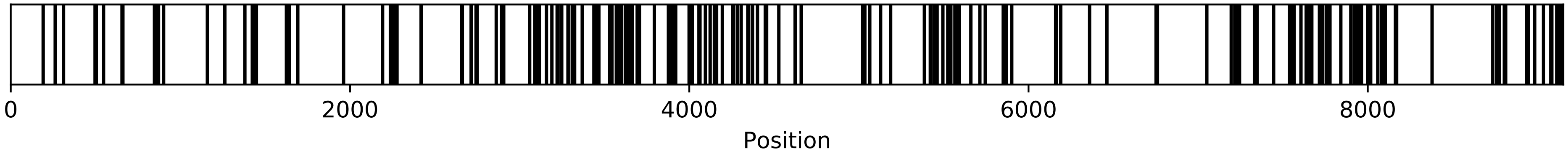


CR2 vs MM1
Number of differences: 305

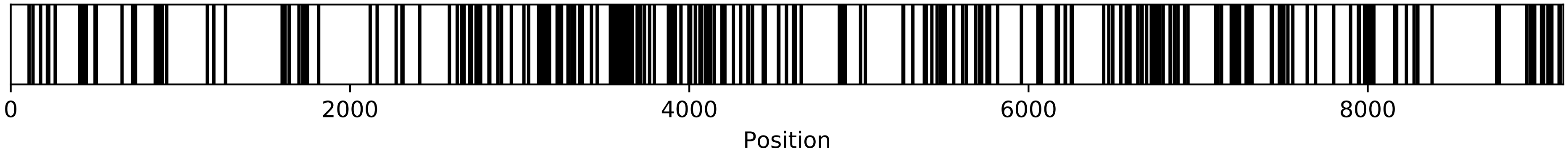


CR2 vs MM2
Number of differences: 305

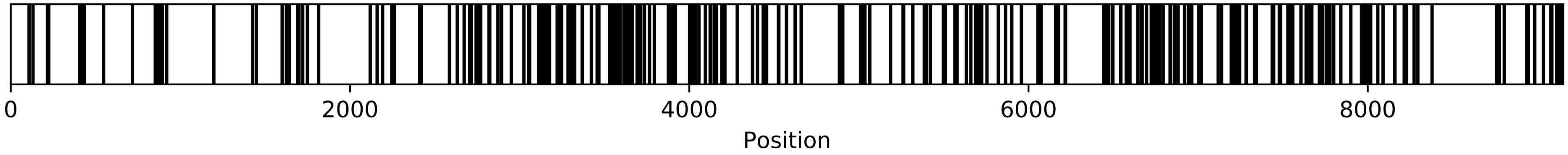


MM1 vs MM2
Number of differences: 158

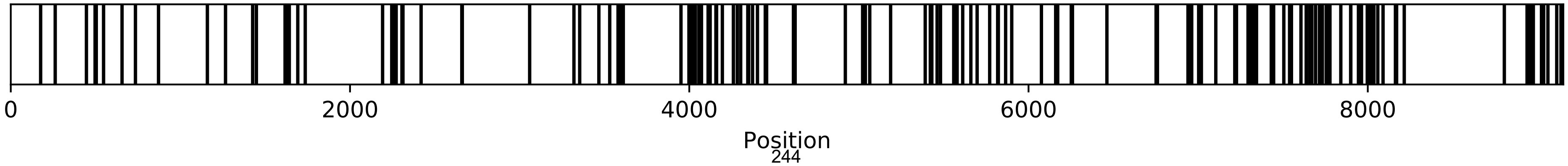

### Region: 682



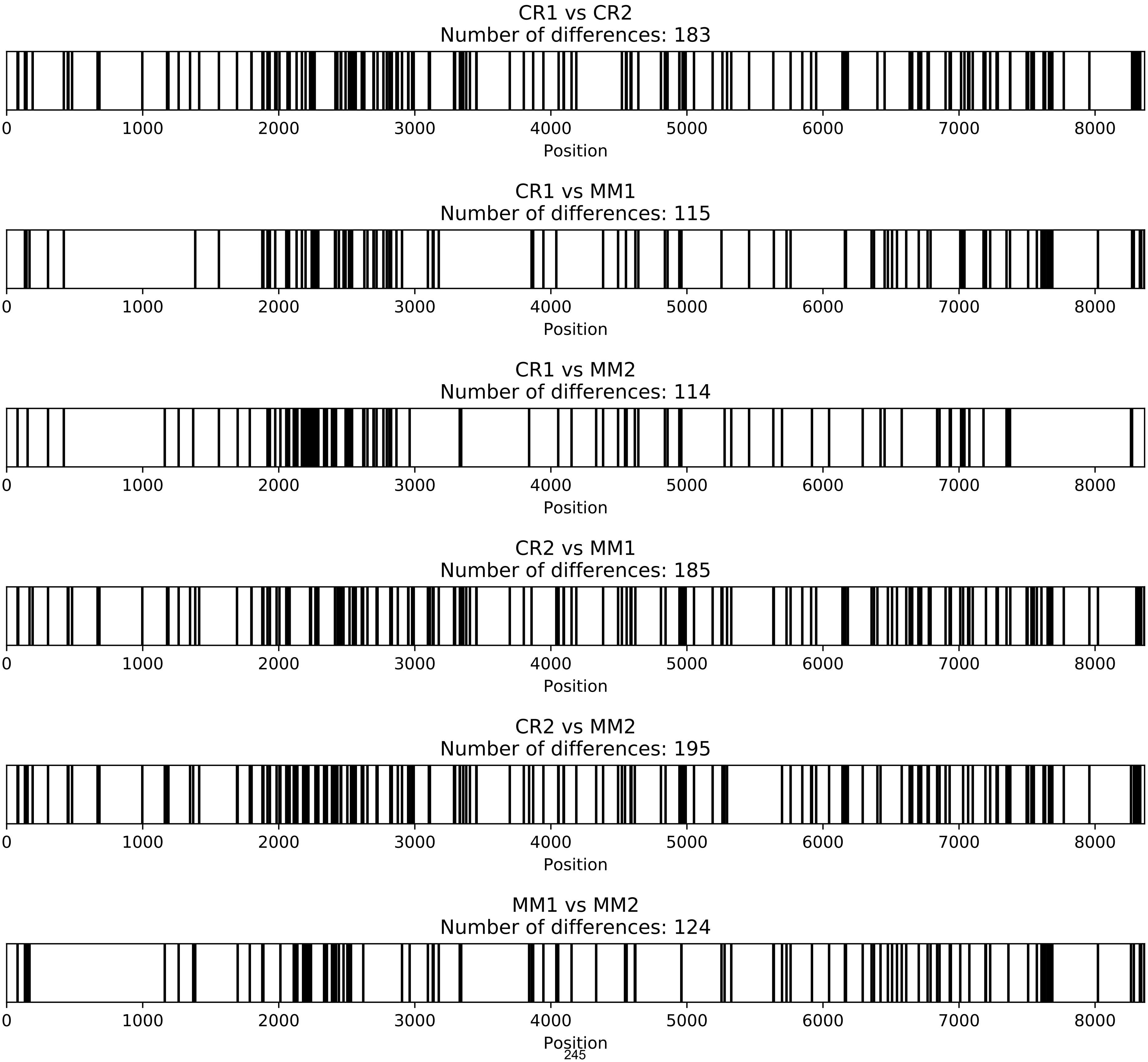
CR1 vs CR2
Number of differences: 183
CR1 vs MM1
Number of differences: 115
CR1 vs MM2
Number of differences: 114
CR2 vs MM1
Number of differences: 185
CR2 vs MM2
Number of differences: 195
MM1 vs MM2
Number of differences: 124
0 1000 2000 3000 4000 5000 6000 7000 8000
Position

### Region: 686



CR1 vs CR2
Number of differences: 104

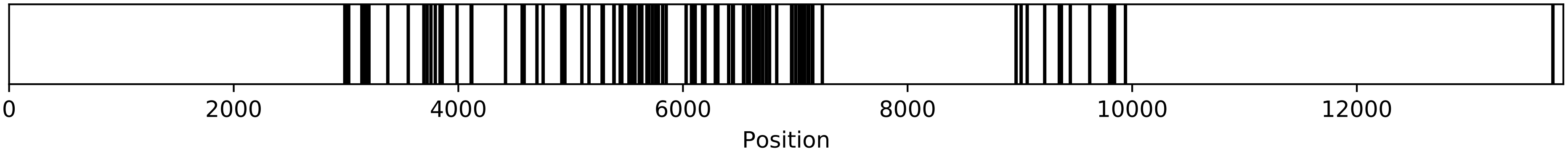


CR1 vs MM1
Number of differences: 195

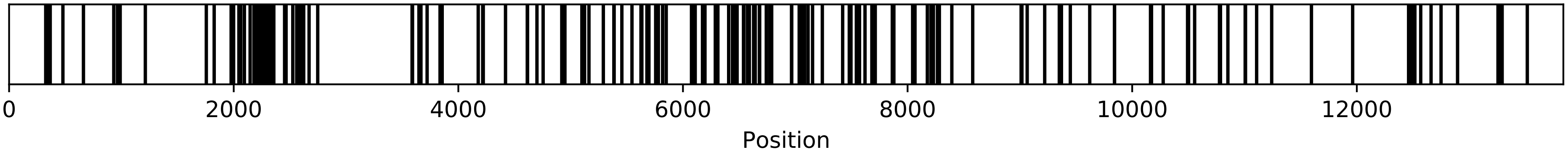


CR1 vs MM2
Number of differences: 175

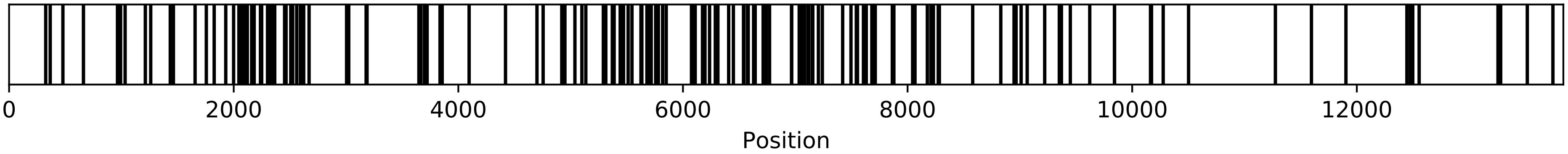


CR2 vs MM1
Number of differences: 179

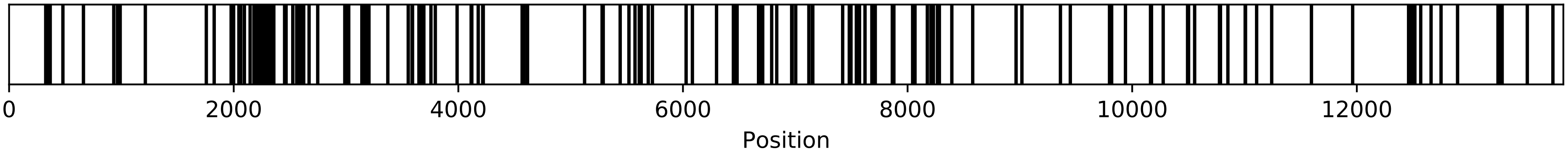


CR2 vs MM2
Number of differences: 149

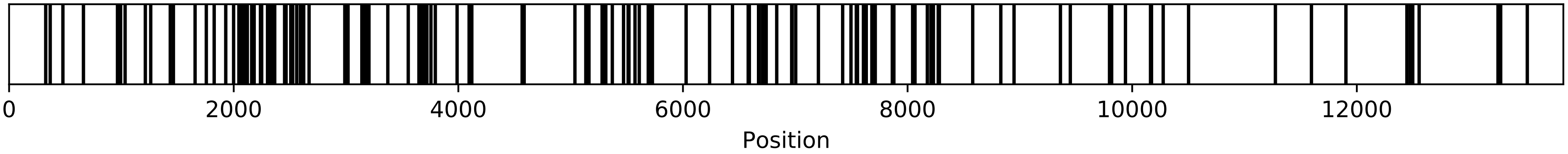


MM1 vs MM2
Number of differences: 123

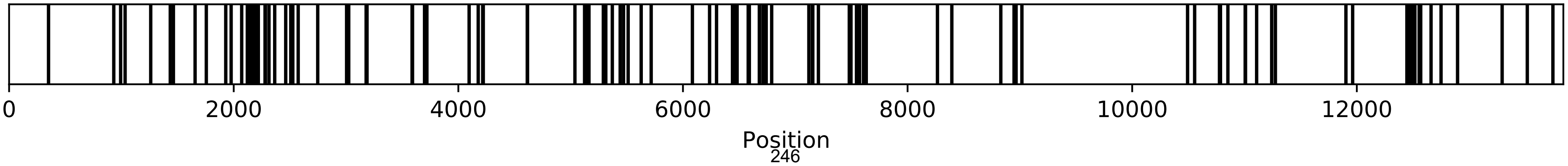

### Region: 692

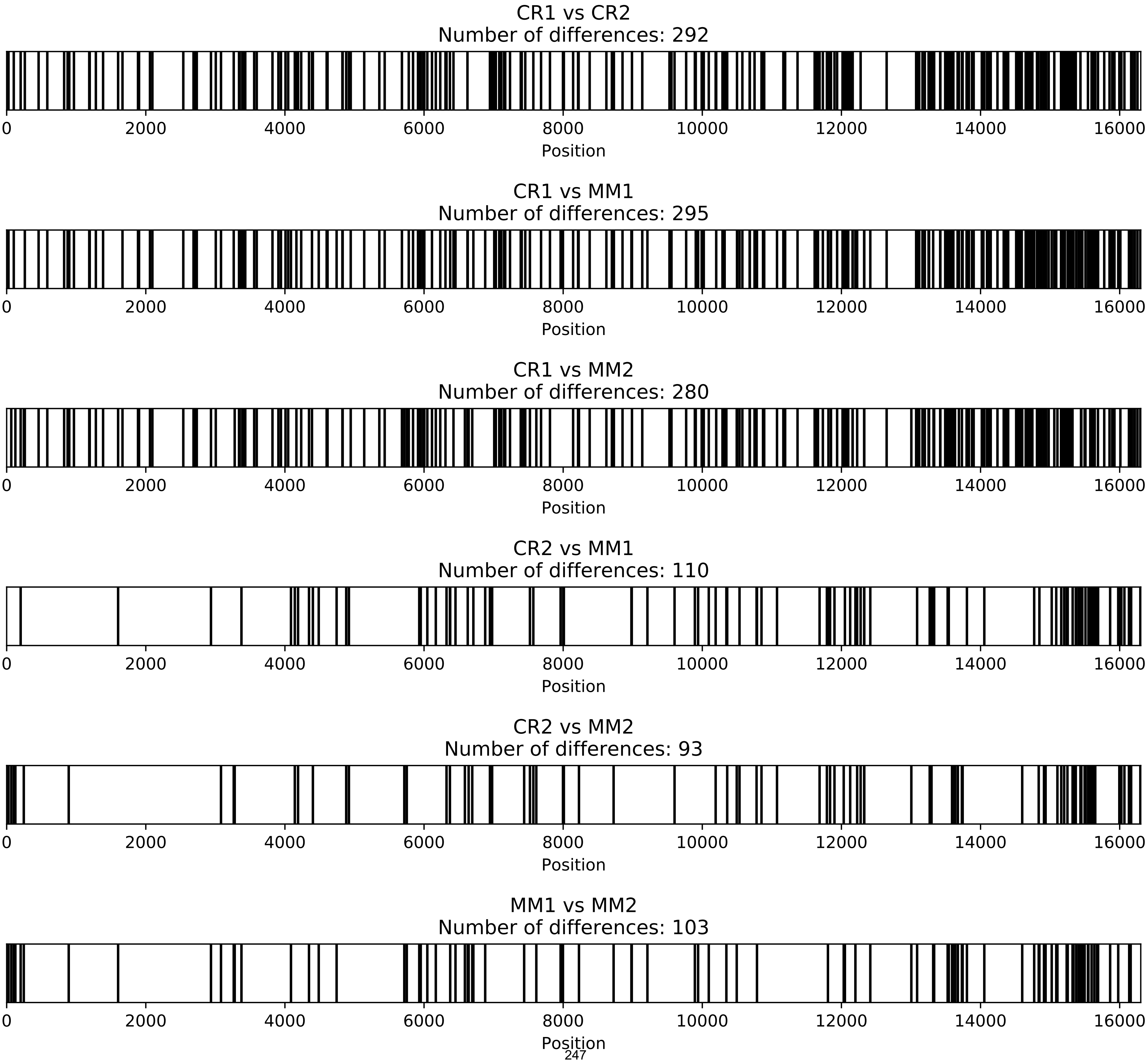
CR1 vs CR2
Number of differences: 292
CR1 vs MM1
Number of differences: 295
CR1 vs MM2
Number of differences: 280
CR2 vs MM1
Number of differences: 110
CR2 vs MM2
Number of differences: 93
MM1 vs MM2
Number of differences: 103
0
2000
4000
6000
8000
10000
12000
14000
16000
Position

### Region: 693



CR1 vs CR2
Number of differences: 324

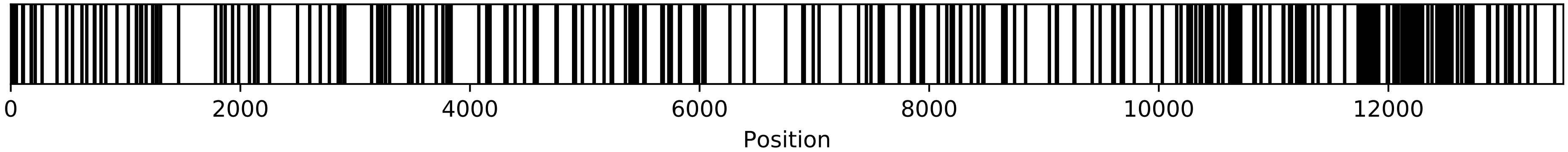


CR1 vs MM1
Number of differences: 310

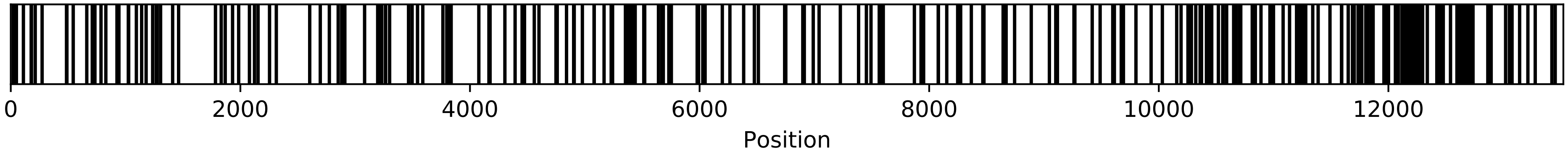


CR1 vs MM2
Number of differences: 318

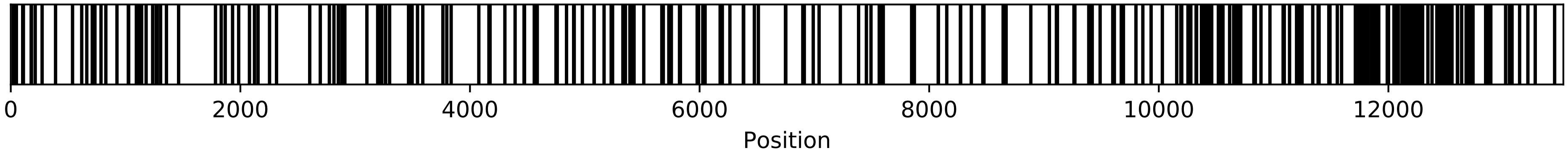


CR2 vs MM1
Number of differences: 145

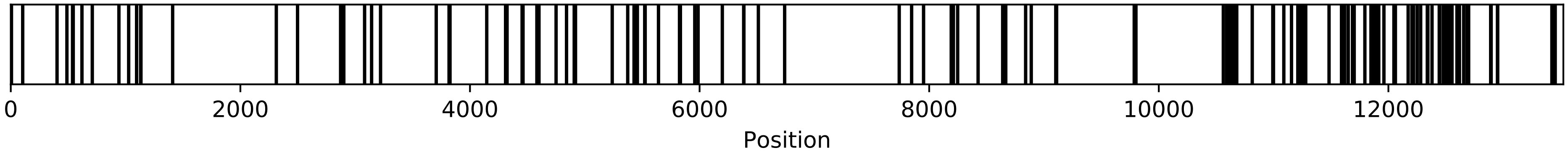


CR2 vs MM2
Number of differences: 105

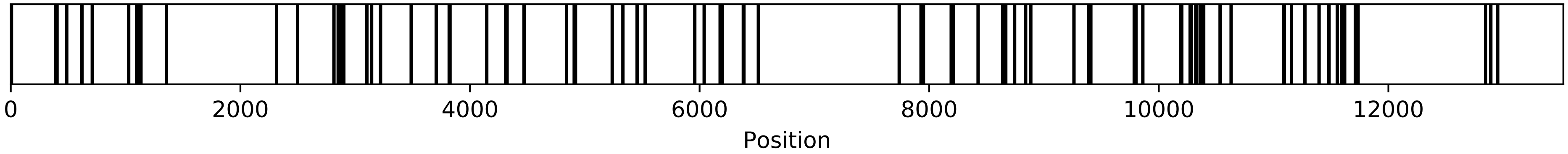


MM1 vs MM2
Number of differences: 136

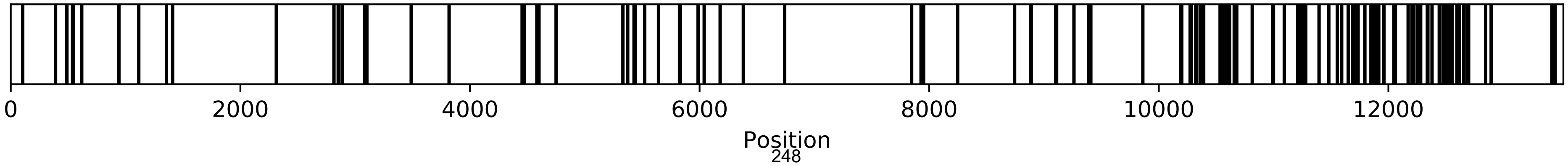

### Region: 694



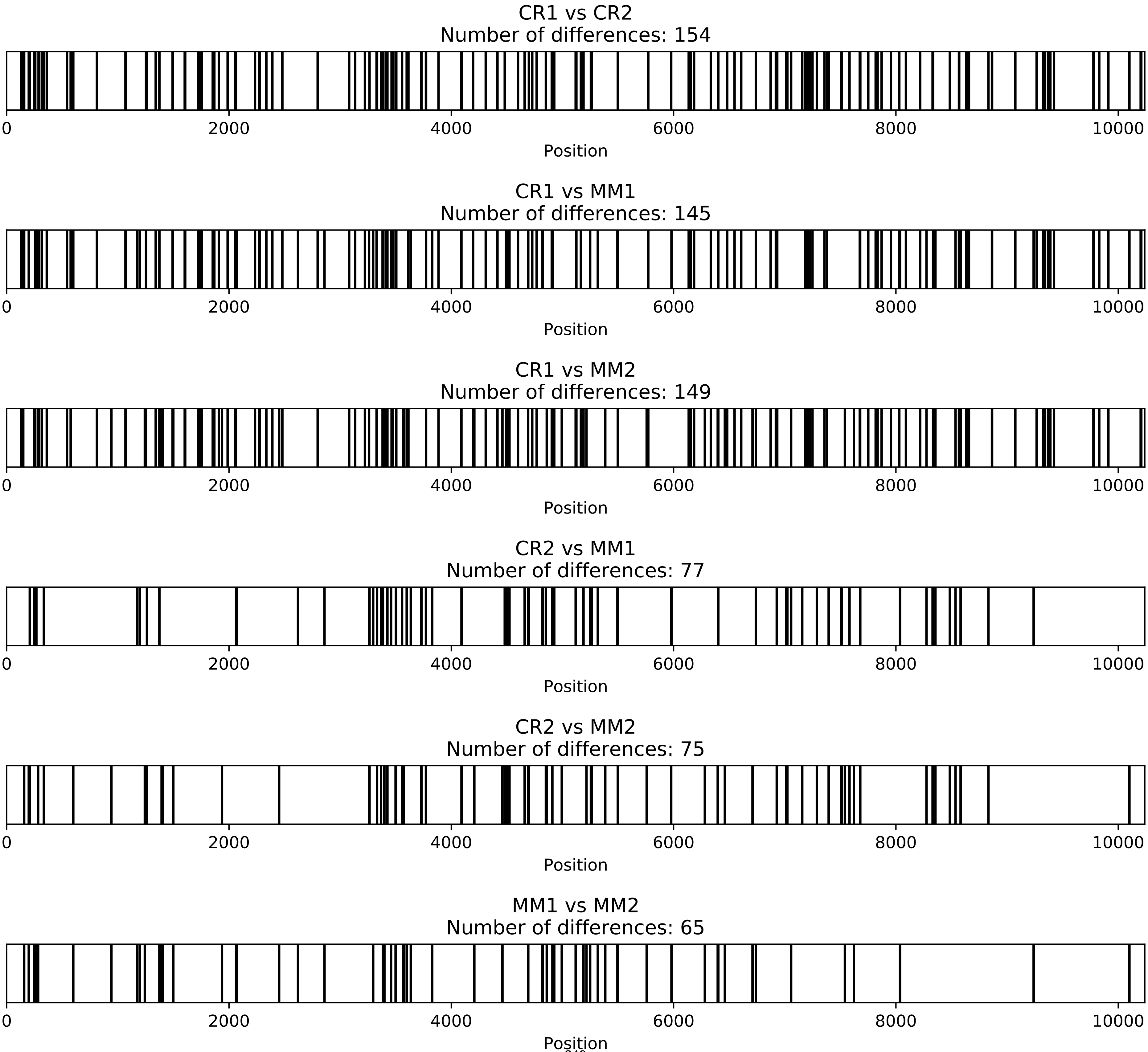
CR1 vs CR2
Number of differences: 154
CR1 vs MM1
Number of differences: 145
CR1 vs MM2
Number of differences: 149
CR2 vs MM1
Number of differences: 77
CR2 vs MM2
Number of differences: 75
MM1 vs MM2
Number of differences: 65
0
2000
4000
6000
8000
10000
Position

### Region: 695



CR1 vs CR2
Number of differences: 310

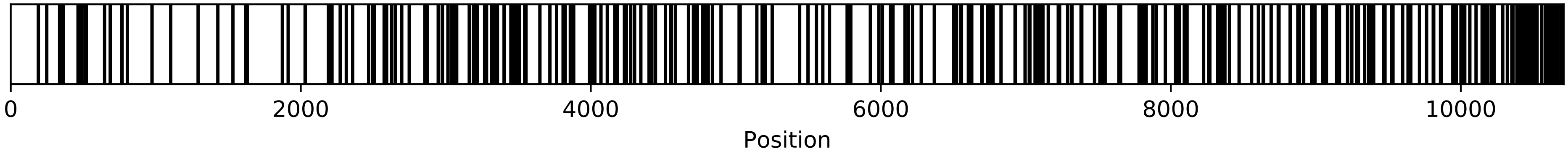


CR1 vs MM1
Number of differences: 161

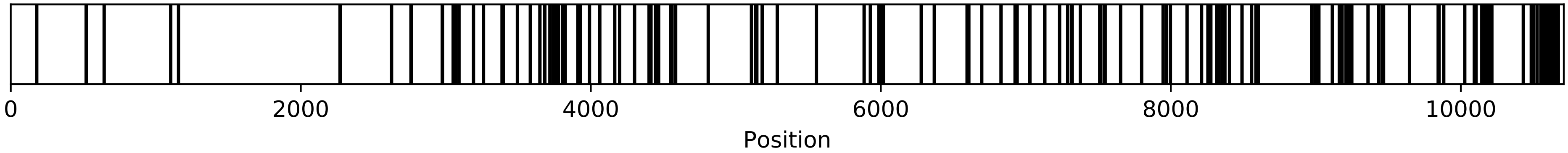


CR1 vs MM2
Number of differences: 162

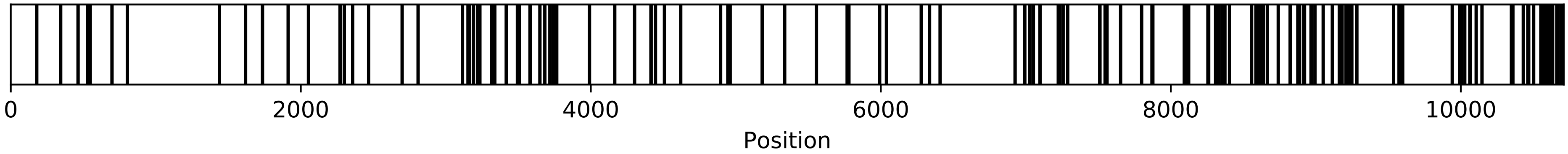


CR2 vs MM1
Number of differences: 316

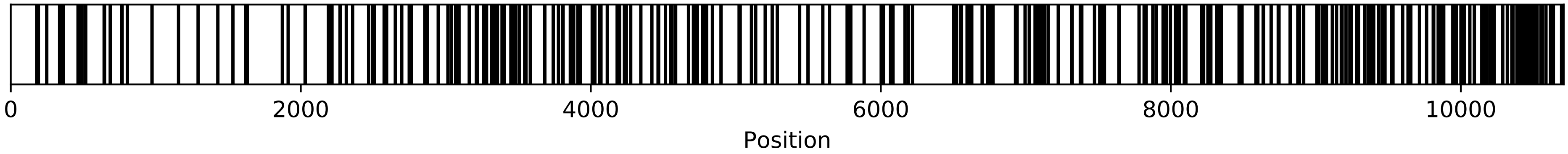


CR2 vs MM2
Number of differences: 318

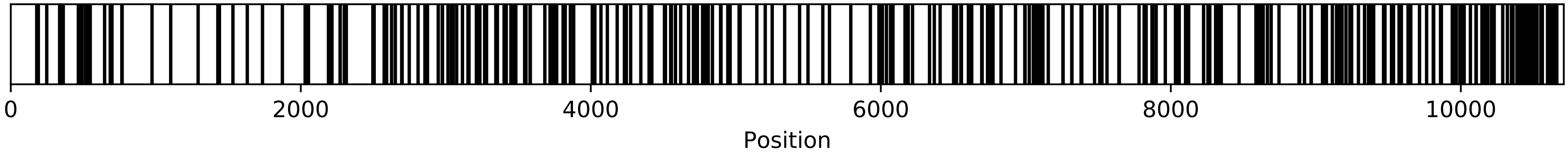


MM1 vs MM2
Number of differences: 191

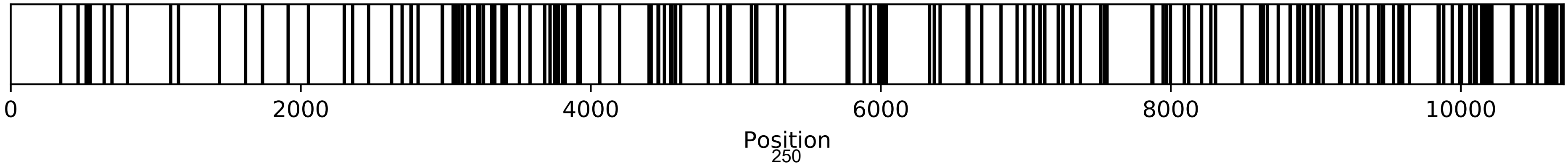

### Region: 701



CR1 vs CR2
Number of differences: 286

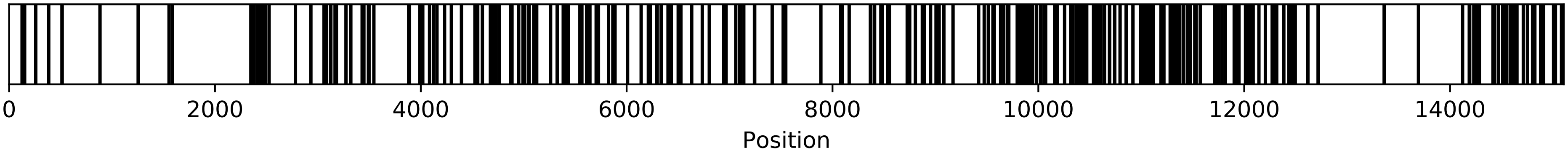


CR1 vs MM1
Number of differences: 127

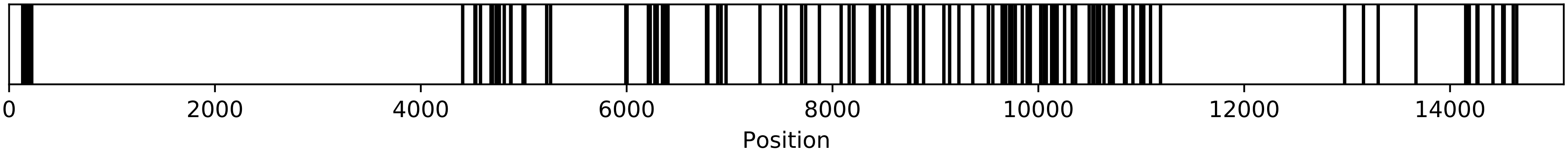


CR1 vs MM2
Number of differences: 135

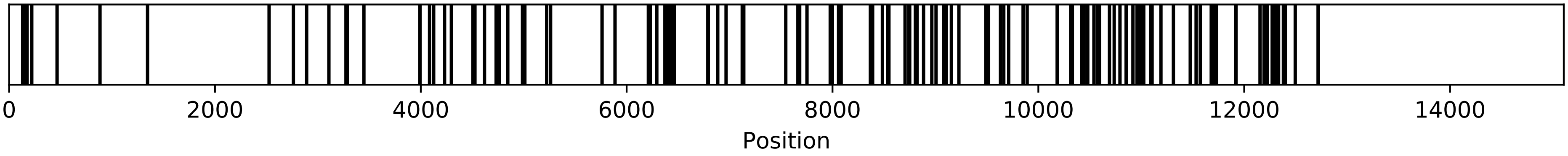


CR2 vs MM1
Number of differences: 307

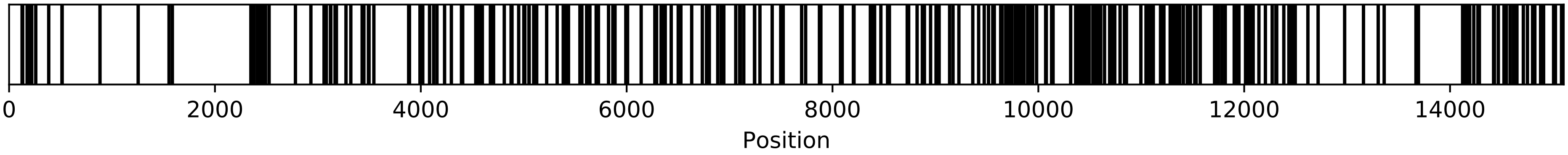


CR2 vs MM2
Number of differences: 296

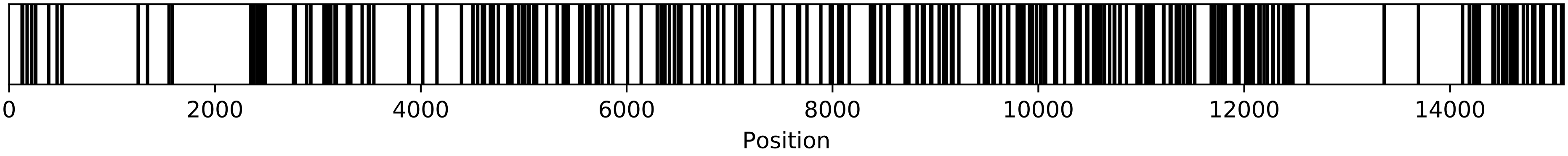


MM1 vs MM2
Number of differences: 172

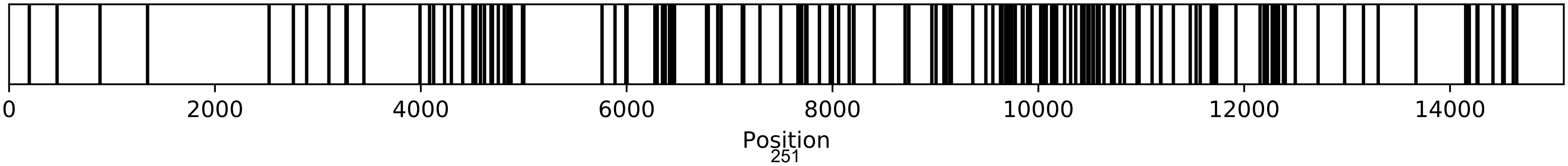

### Region: 702

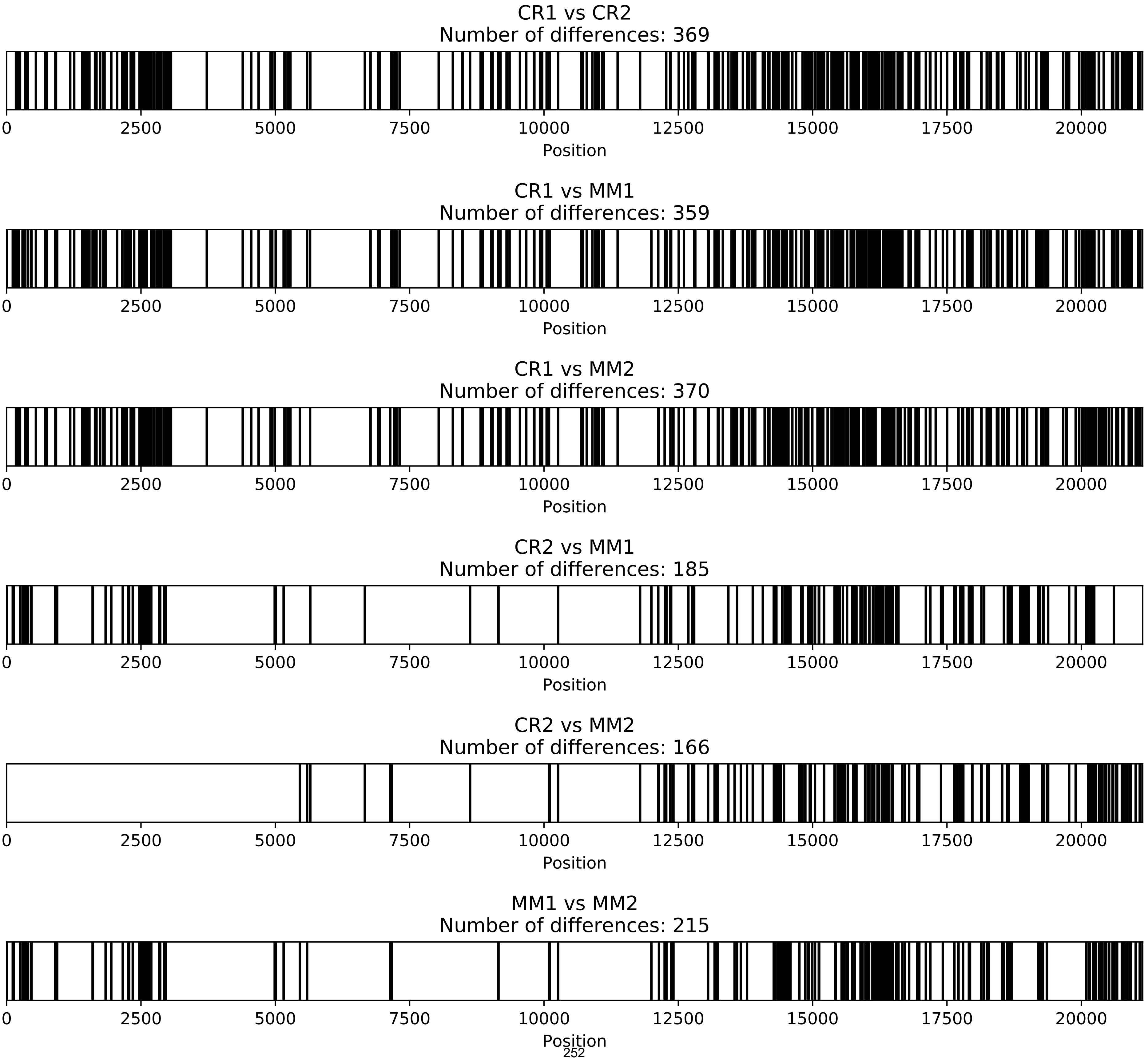
CR1 vs CR2
Number of differences: 369
CR1 vs MM1
Number of differences: 359
CR1 vs MM2
Number of differences: 370
CR2 vs MM1
Number of differences: 185
CR2 vs MM2
Number of differences: 166
MM1 vs MM2
Number of differences: 215
0
2500
5000
7500
10000
12500
15000
17500
20000
Position

### Region: 704



CR1 vs CR2
Number of differences: 260

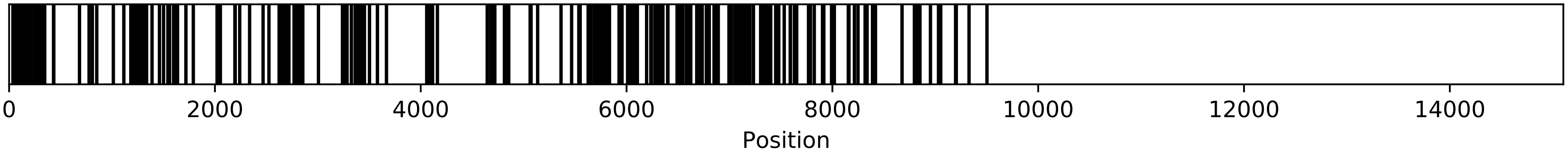


CR1 vs MM1
Number of differences: 158

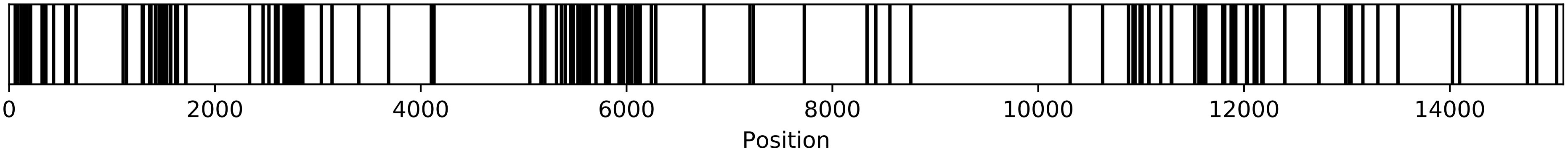


CR1 vs MM2
Number of differences: 254

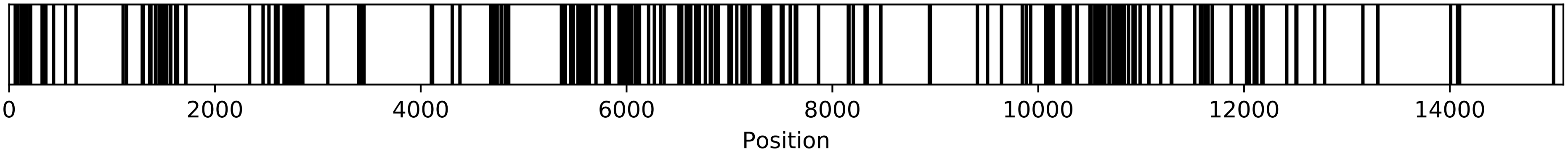


CR2 vs MM1
Number of differences: 310

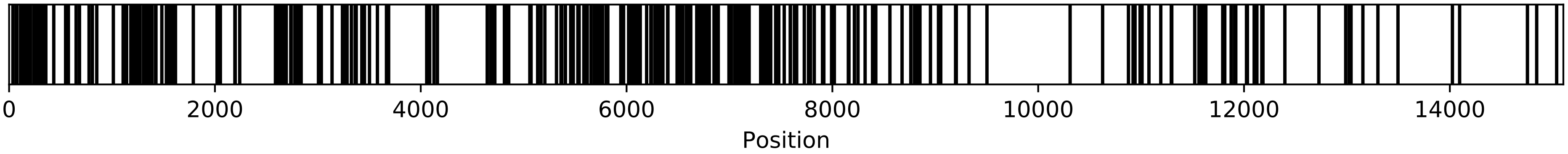


CR2 vs MM2
Number of differences: 333

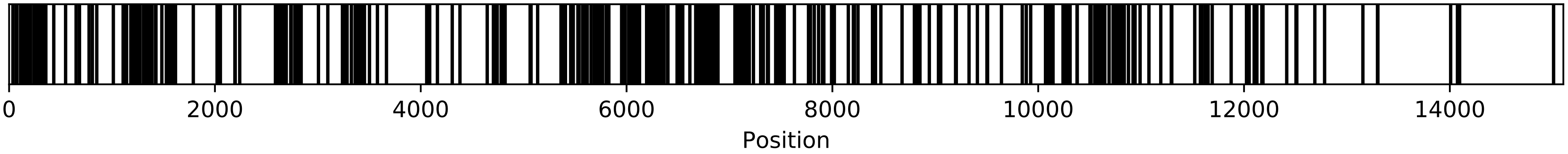


MM1 vs MM2
Number of differences: 180

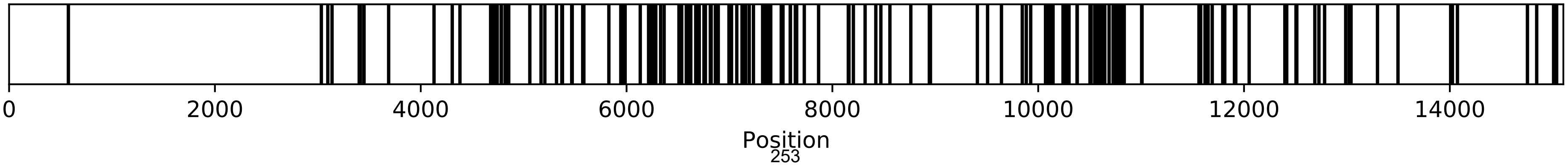

### Region: 707



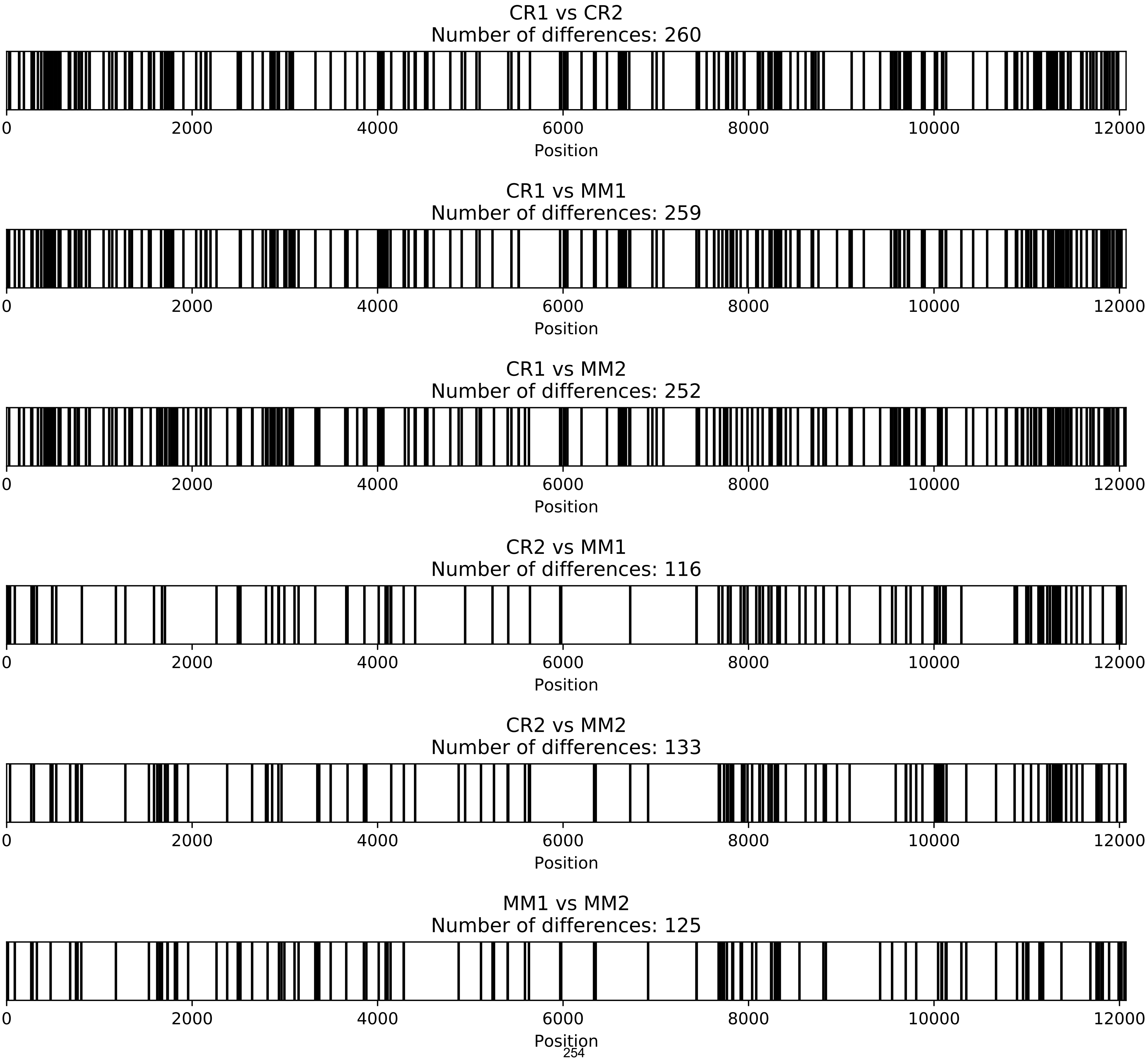
CR1 vs CR2
Number of differences: 260
CR1 vs MM1
Number of differences: 259
CR1 vs MM2
Number of differences: 252
CR2 vs MM1
Number of differences: 116
CR2 vs MM2
Number of differences: 133
MM1 vs MM2
Number of differences: 125
0
2000
4000
6000
8000
10000
12000
Position

### Region: 708

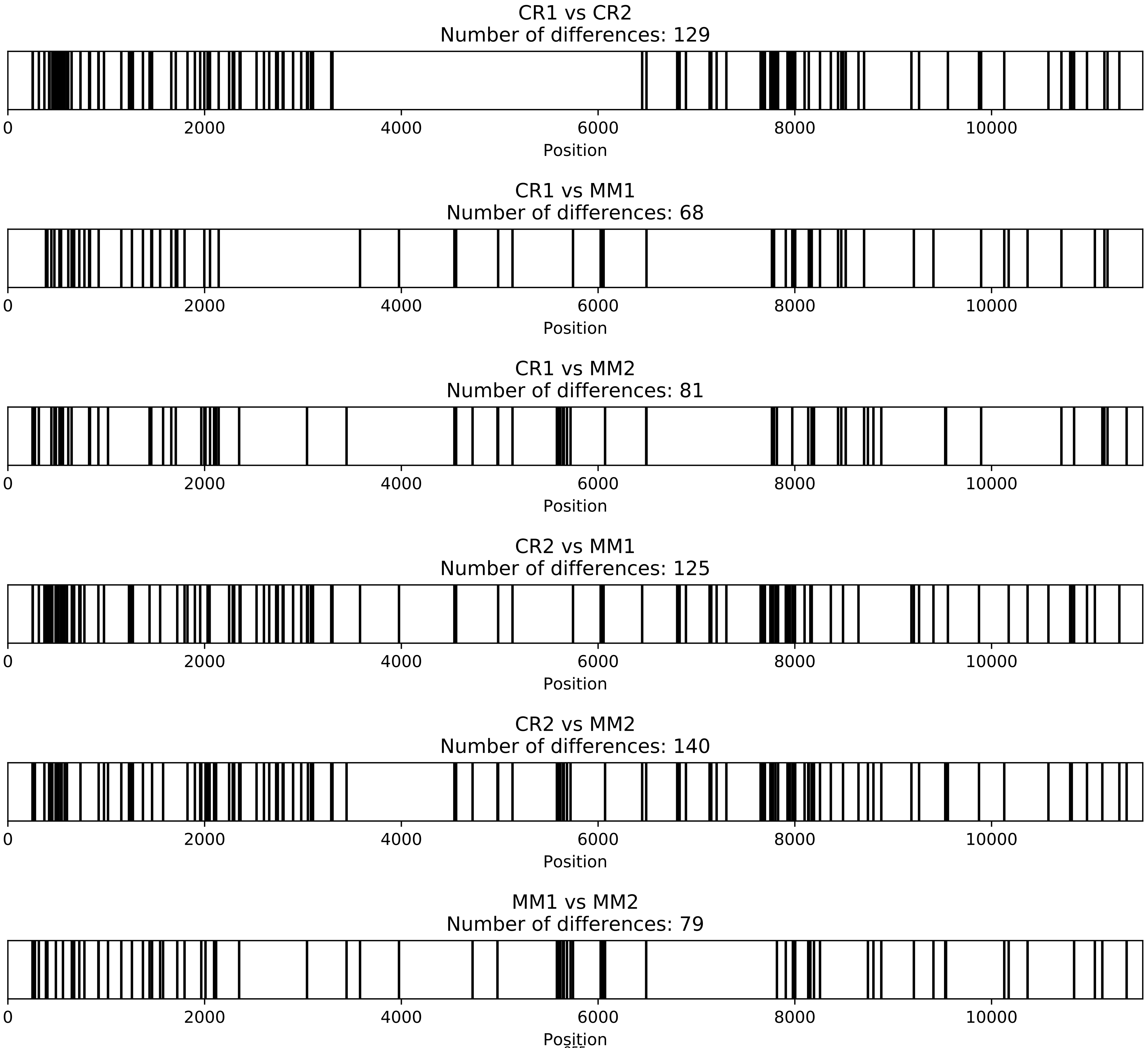
CR1 vs CR2
Number of differences: 129
CR1 vs MM1
Number of differences: 68
CR1 vs MM2
Number of differences: 81
CR2 vs MM1
Number of differences: 125
CR2 vs MM2
Number of differences: 140
MM1 vs MM2
Number of differences: 79
0
2000
4000
6000
8000
10000
Position

### Region: 712



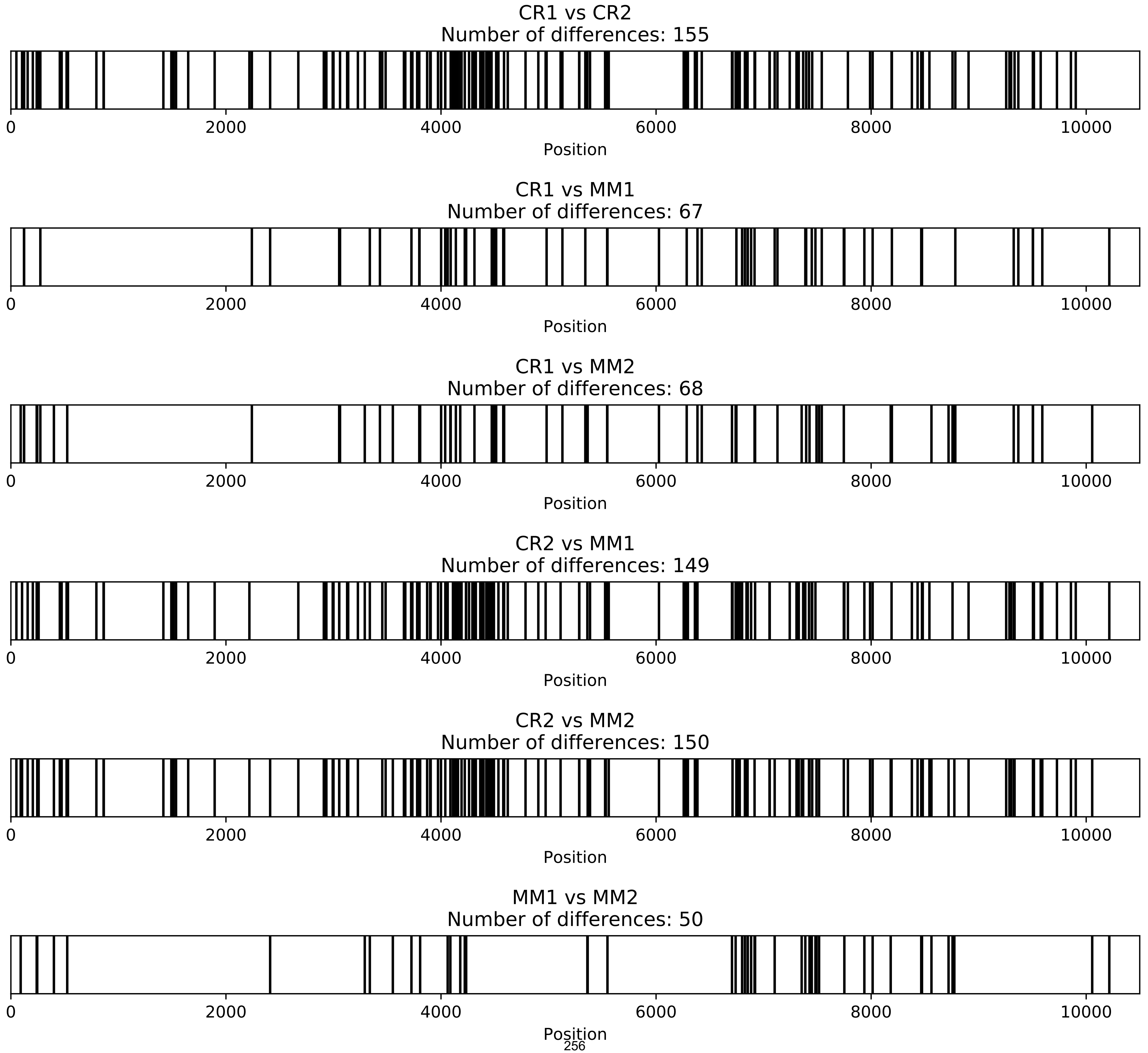
CR1 vs CR2
Number of differences: 155
CR1 vs MM1
Number of differences: 67
CR1 vs MM2
Number of differences: 68
CR2 vs MM1
Number of differences: 149
CR2 vs MM2
Number of differences: 150
MM1 vs MM2
Number of differences: 50
0
2000
4000
6000
8000
10000
Position

### Region: 713



CR1 vs CR2
Number of differences: 277

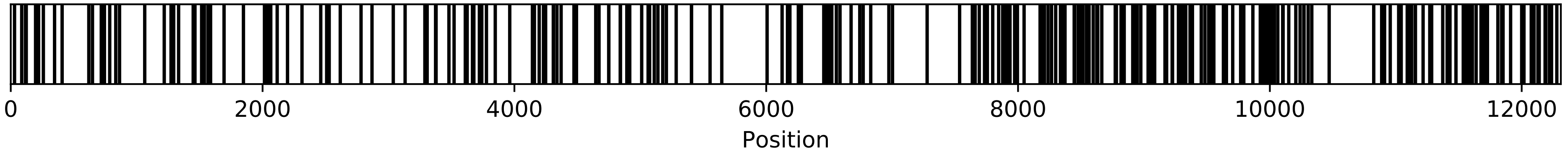


CR1 vs MM1
Number of differences: 106

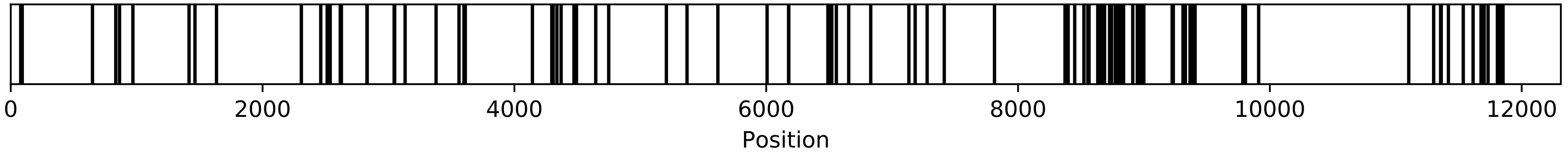


CR1 vs MM2
Number of differences: 117

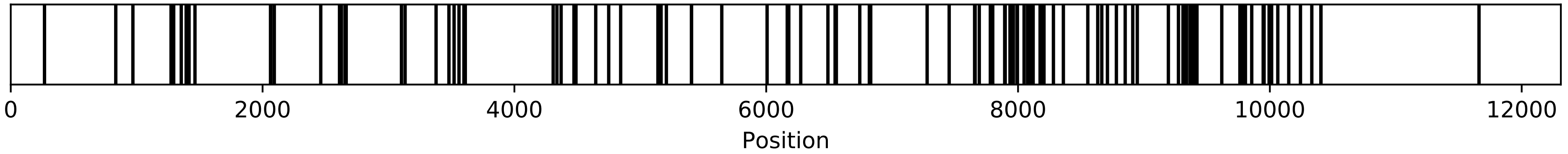


CR2 vs MM1
Number of differences: 281

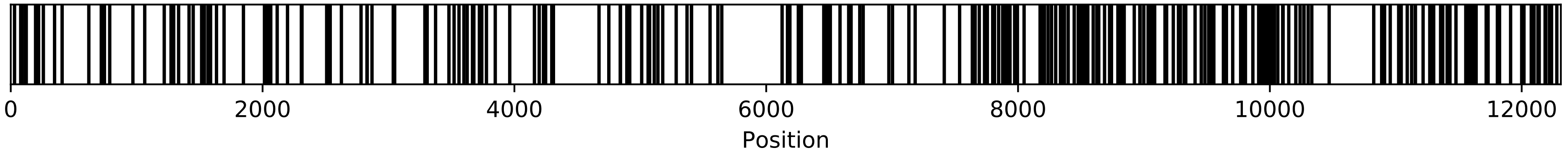


CR2 vs MM2
Number of differences: 280

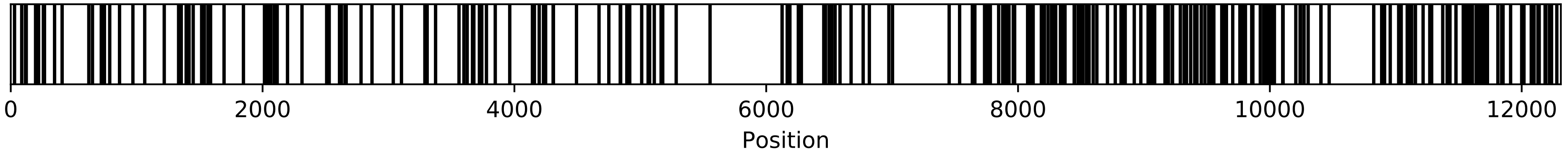


MM1 vs MM2
Number of differences: 146

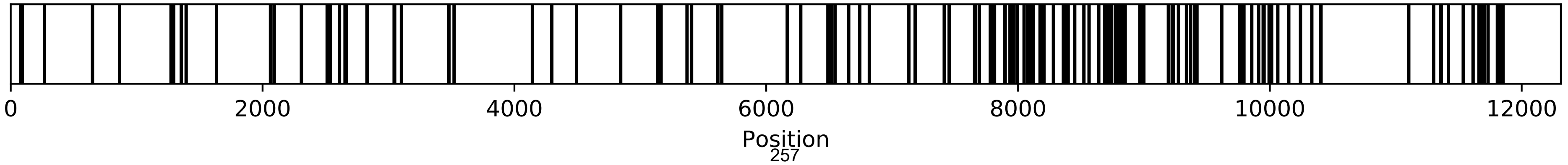

### Region: 714



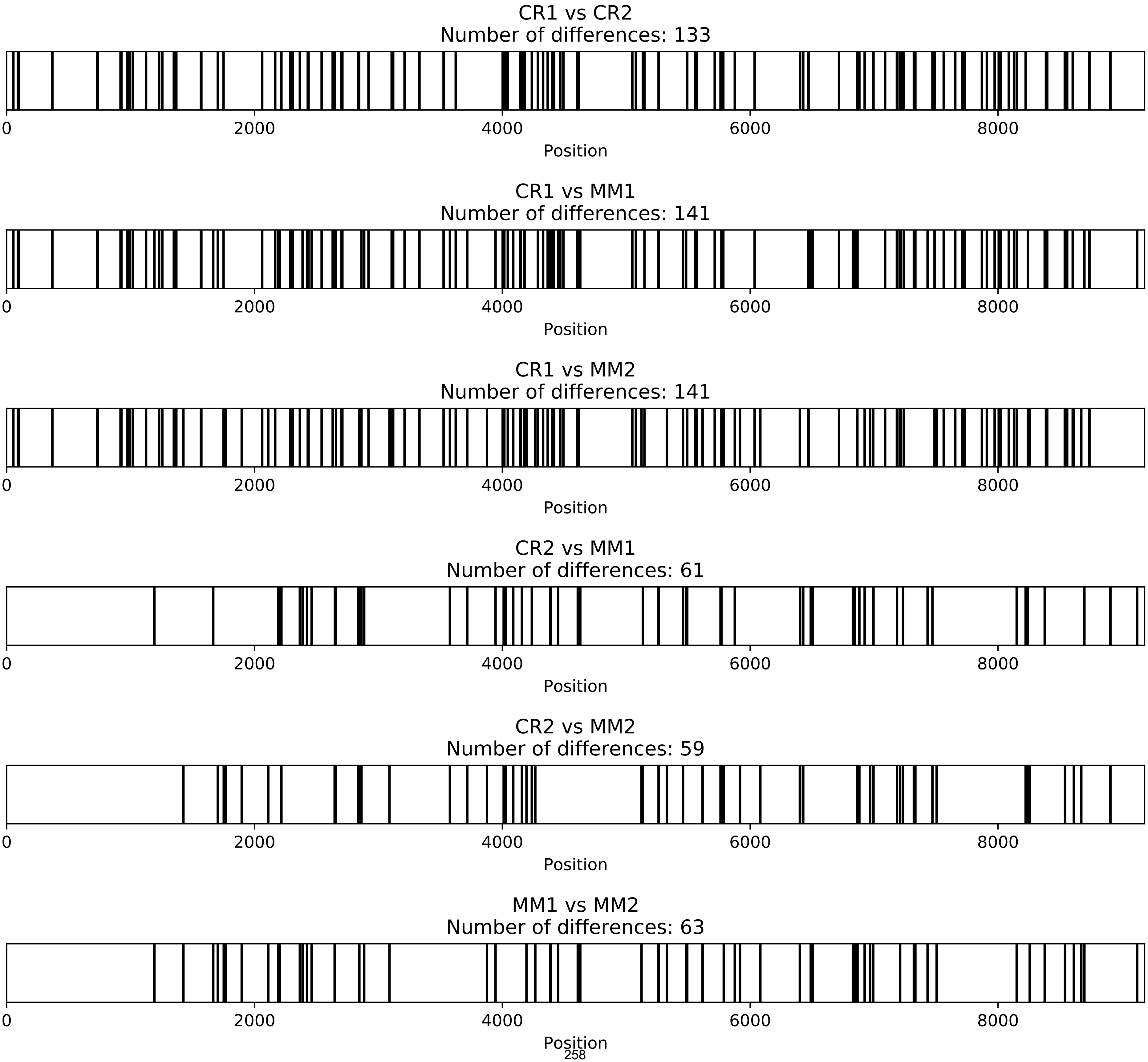
CR1 vs CR2
Number of differences: 133
CR1 vs MM1
Number of differences: 141
CR1 vs MM2
Number of differences: 141
CR2 vs MM1
Number of differences: 61
CR2 vs MM2
Number of differences: 59
MM1 vs MM2
Number of differences: 63
0
2000
4000
6000
8000
Position

### Region: 716



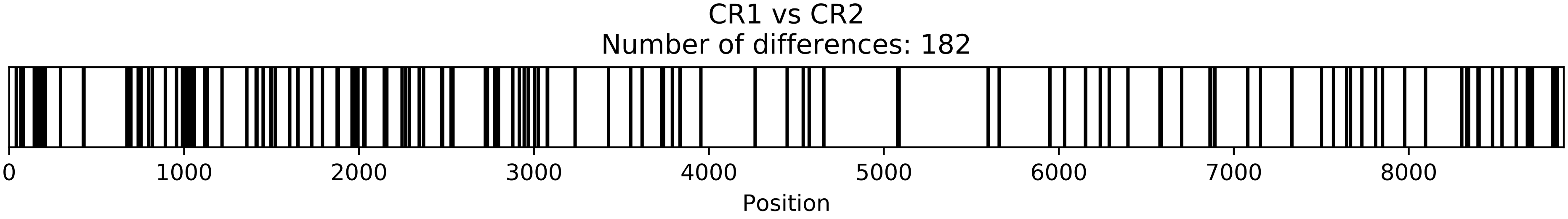
CR1 vs CR2
Number of differences: 182
0
1000
2000
3000
4000
5000
6000
7000
8000
Position

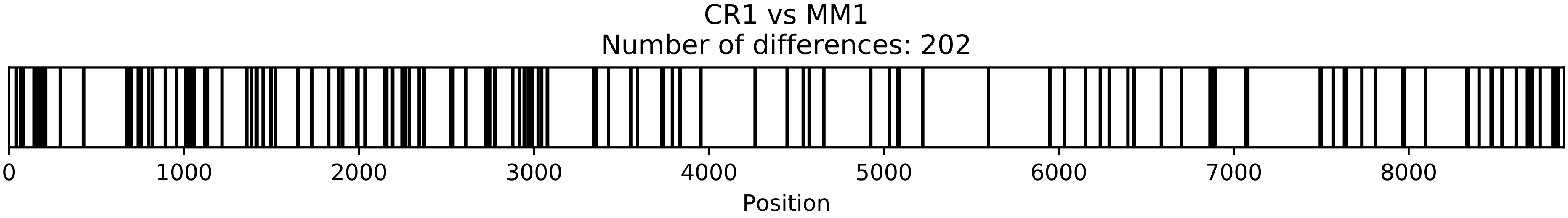
CR1 vs MM1
Number of differences: 202
0
1000
2000
3000
4000
5000
6000
7000
8000
Position

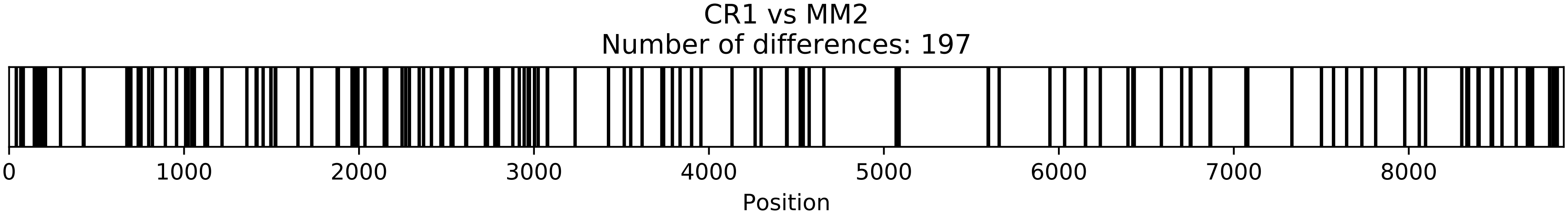
CR1 vs MM2
Number of differences: 197
0
1000
2000
3000
4000
5000
6000
7000
8000
Position

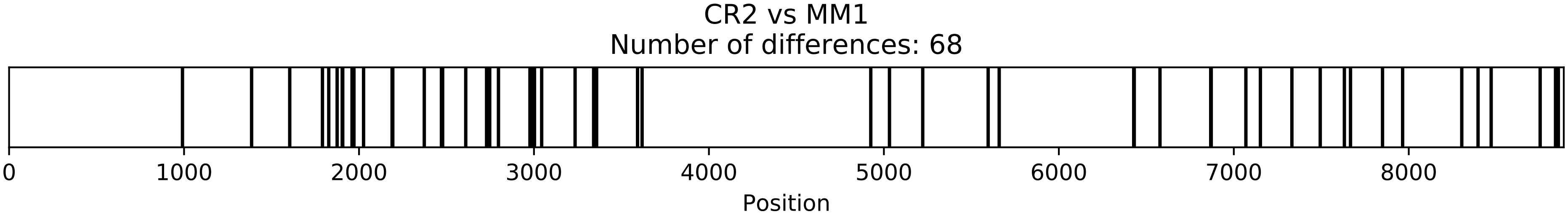
CR2 vs MM1
Number of differences: 68
0
1000
2000
3000
4000
5000
6000
7000
8000
Position

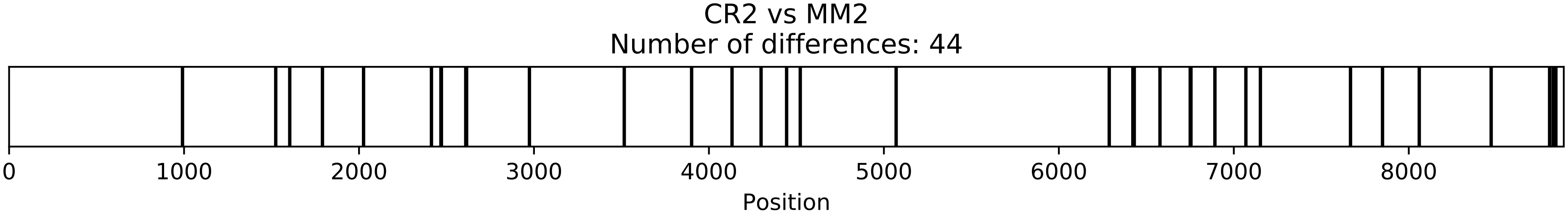
CR2 vs MM2
Number of differences: 44
0
1000
2000
3000
4000
5000
6000
7000
8000
Position

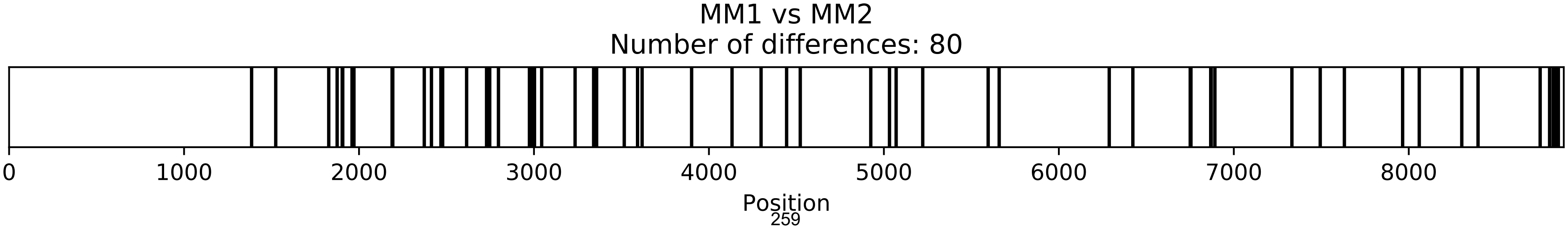
MM1 vs MM2
Number of differences: 80
0
1000
2000
3000
4000
5000
6000
7000
8000
Position

### Region: 719



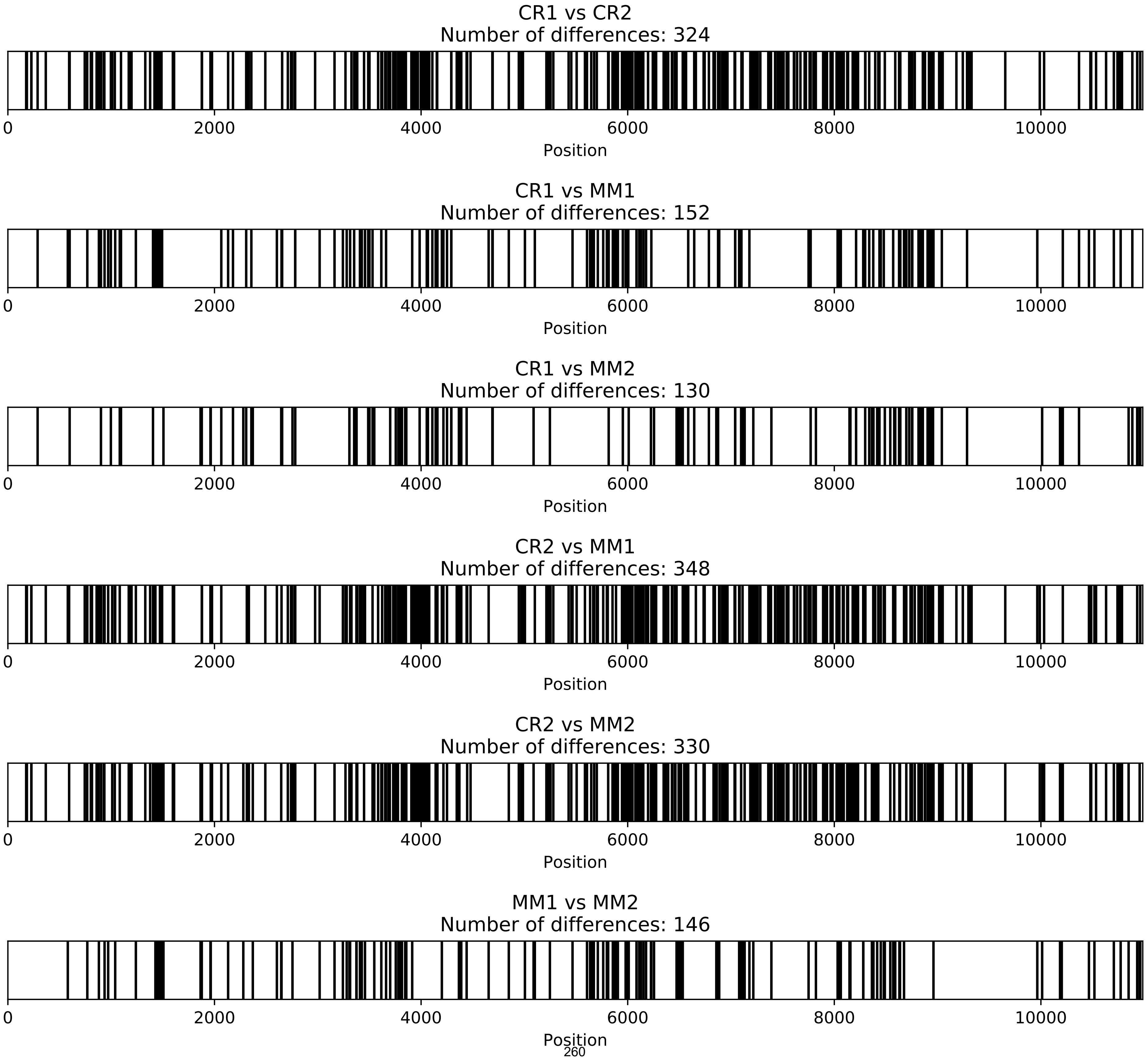
CR1 vs CR2
Number of differences: 324
Position
CR1 vs MM1
Number of differences: 152
Position
CR1 vs MM2
Number of differences: 130
Position
CR2 vs MM1
Number of differences: 348
Position
CR2 vs MM2
Number of differences: 330
Position
MM1 vs MM2
Number of differences: 146
Position
0
2000
4000
6000
8000
10000

### Region: 720



CR1 vs CR2
Number of differences: 147

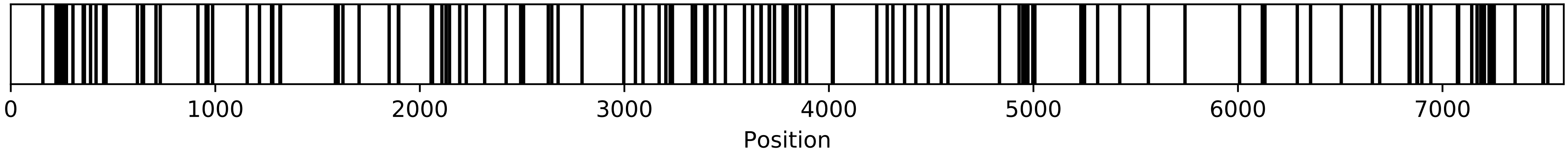


CR1 vs MM1
Number of differences: 139

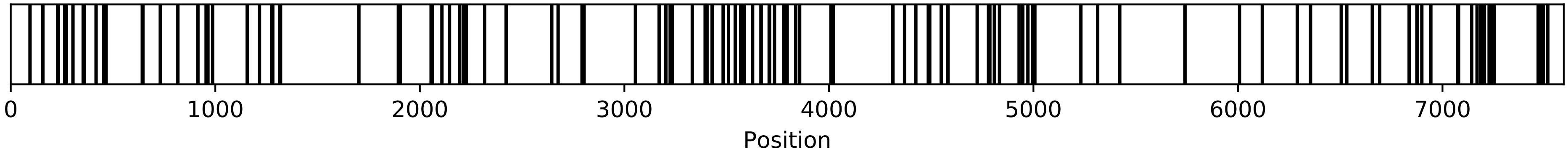


CR1 vs MM2
Number of differences: 138

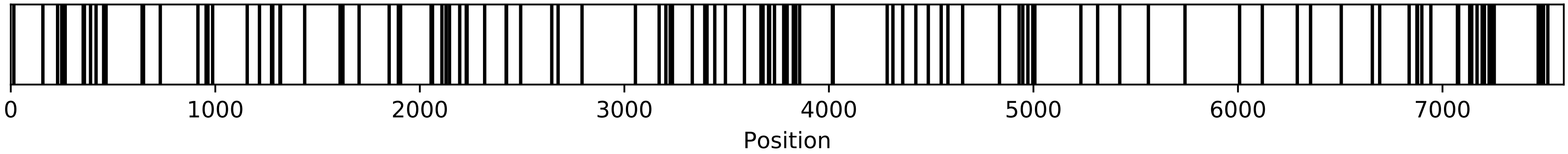


CR2 vs MM1
Number of differences: 80

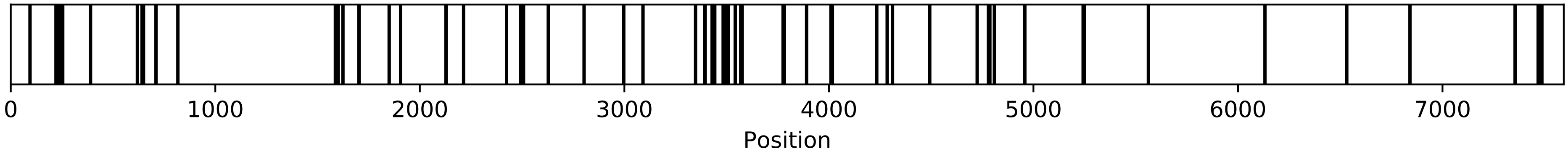


CR2 vs MM2
Number of differences: 60

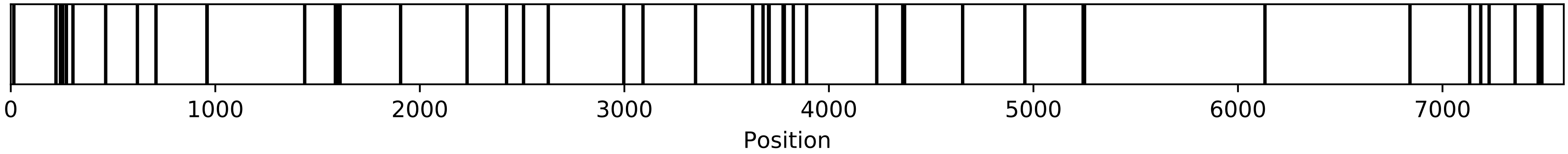


MM1 vs MM2
Number of differences: 61

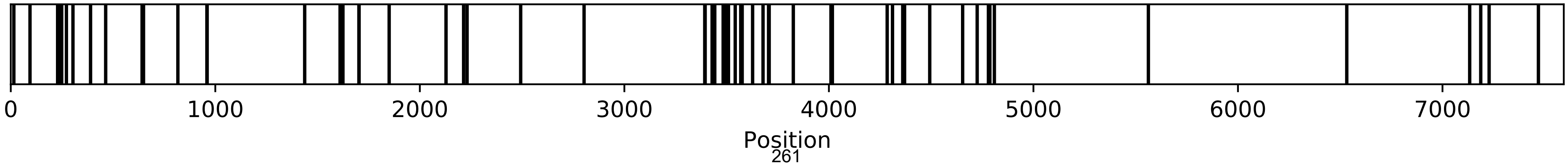

### Region: 721



CR1 vs CR2
Number of differences: 225

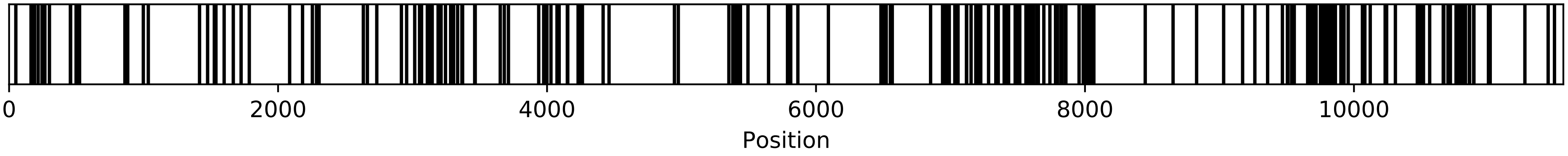


CR1 vs MM1
Number of differences: 112

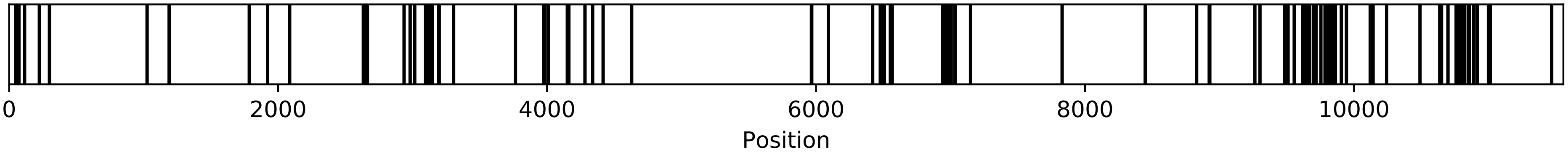


CR1 vs MM2
Number of differences: 148

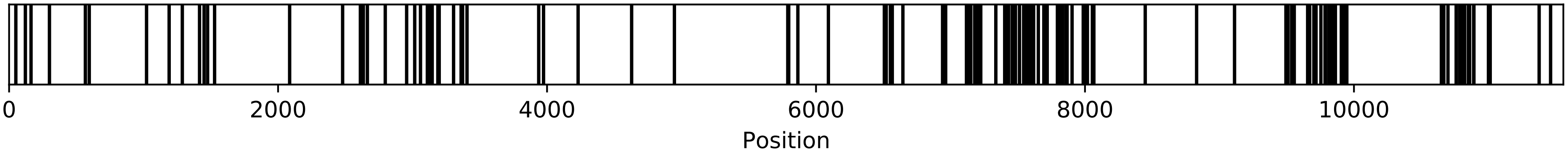


CR2 vs MM1
Number of differences: 222

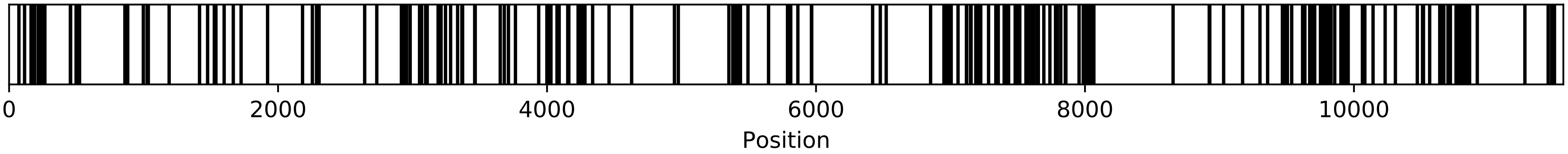


CR2 vs MM2
Number of differences: 200

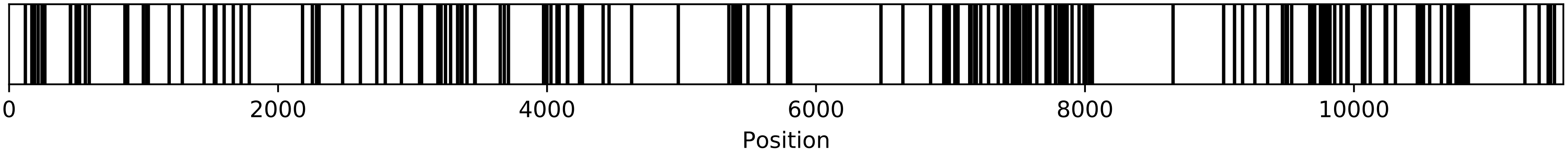


MM1 vs MM2
Number of differences: 152

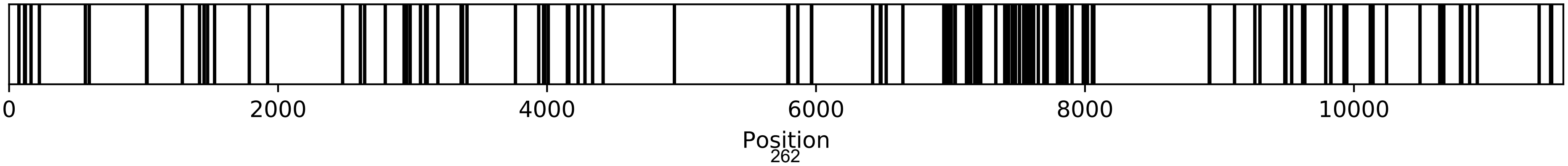

### Region: 728



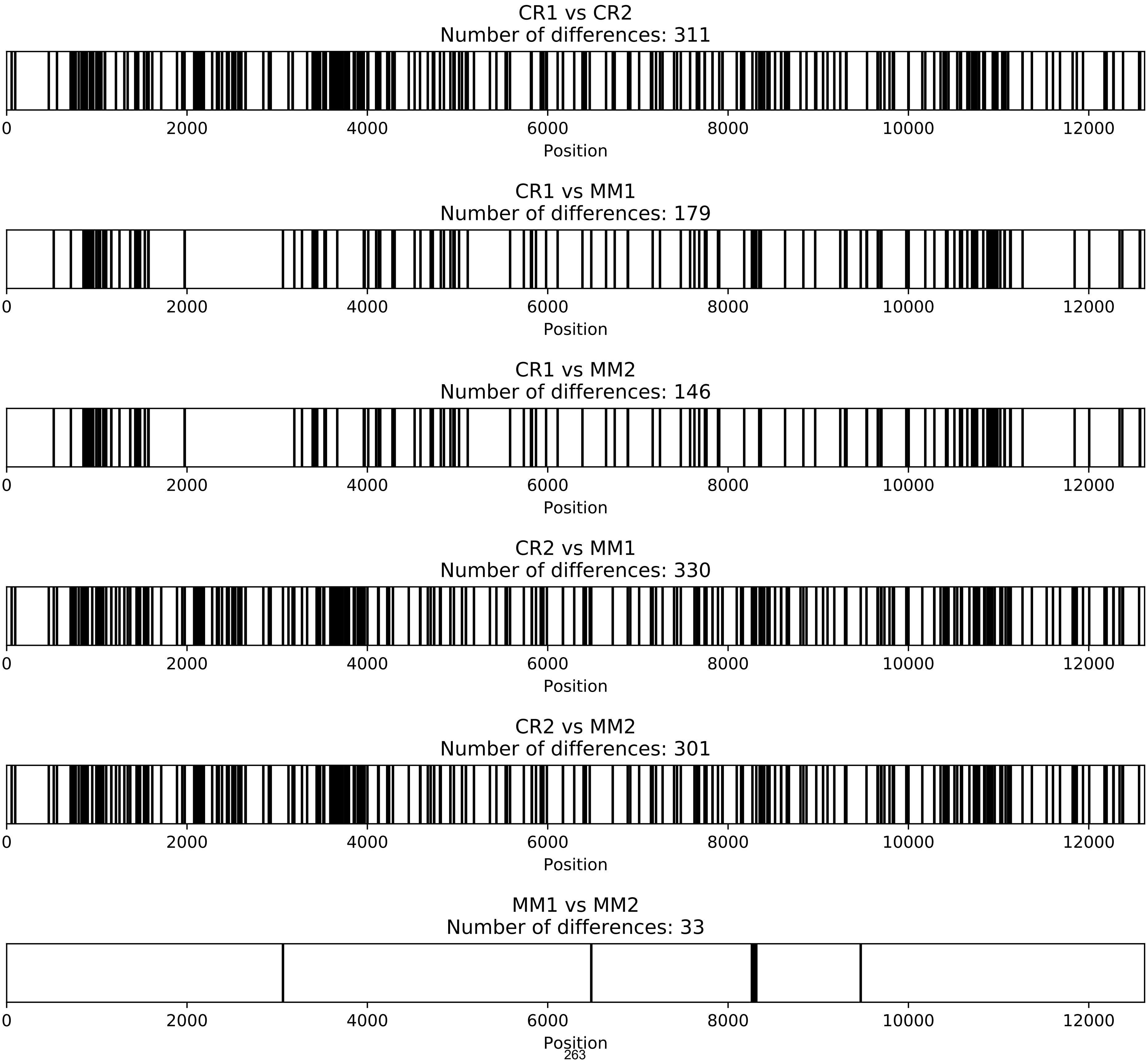
CR1 vs CR2
Number of differences: 311
0
2000
4000
6000
8000
10000
12000
Position
CR1 vs MM1
Number of differences: 179
0
2000
4000
6000
8000
10000
12000
Position
CR1 vs MM2
Number of differences: 146
0
2000
4000
6000
8000
10000
12000
Position
CR2 vs MM1
Number of differences: 330
0
2000
4000
6000
8000
10000
12000
Position
CR2 vs MM2
Number of differences: 301
0
2000
4000
6000
8000
10000
12000
Position
MM1 vs MM2
Number of differences: 33
0
2000
4000
6000
8000
10000
12000
Position

### Region: 733



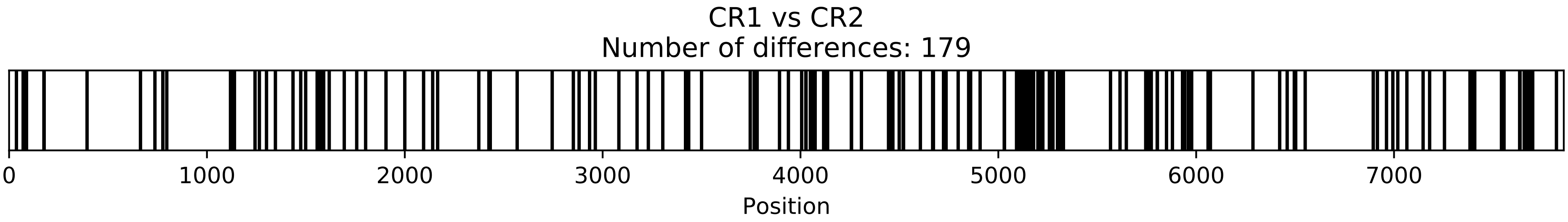
CR1 vs CR2
Number of differences: 179
0
1000
2000
3000
4000
5000
6000
7000
Position

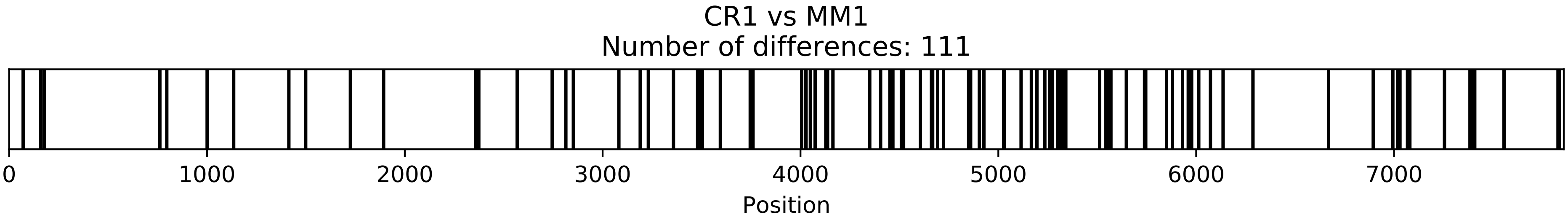
CR1 vs MM1
Number of differences: 111
0
1000
2000
3000
4000
5000
6000
7000
Position

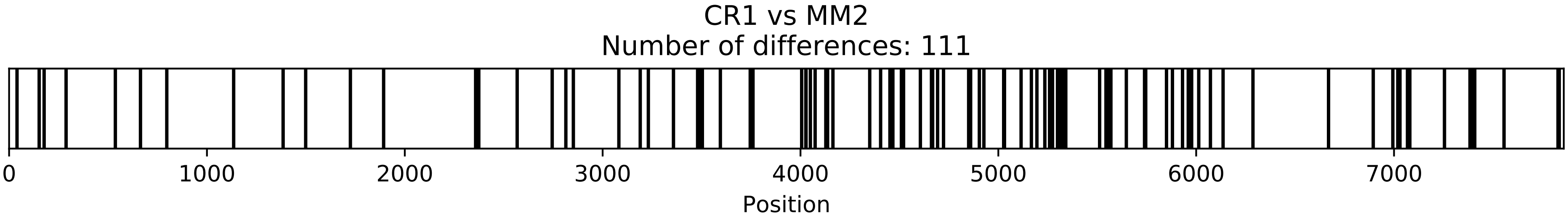
CR1 vs MM2
Number of differences: 111
0
1000
2000
3000
4000
5000
6000
7000
Position

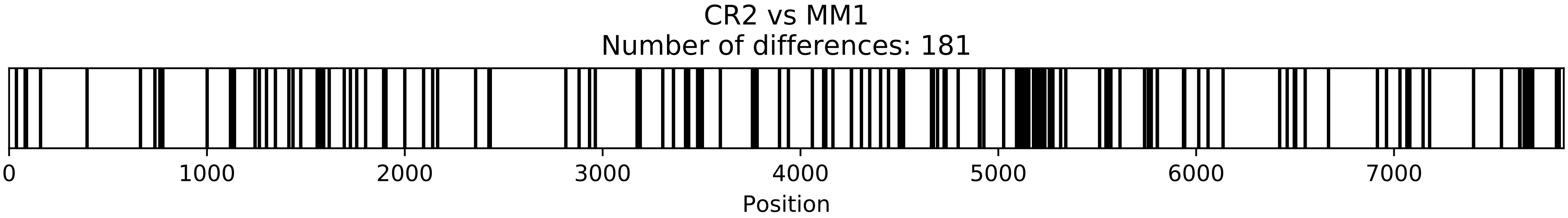
CR2 vs MM1
Number of differences: 181
0
1000
2000
3000
4000
5000
6000
7000
Position

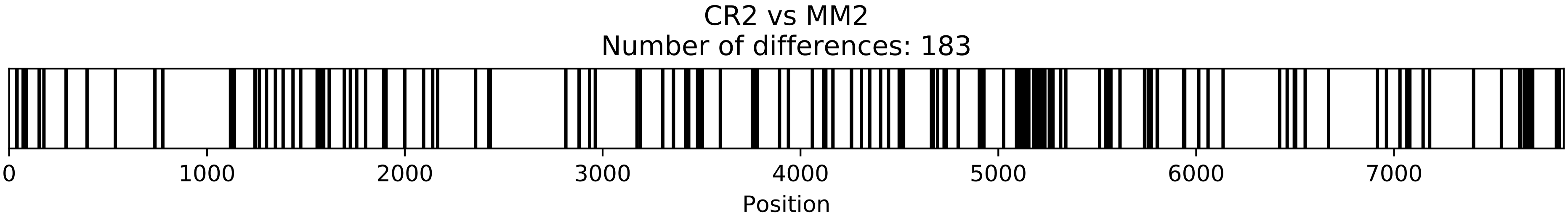
CR2 vs MM2
Number of differences: 183
0
1000
2000
3000
4000
5000
6000
7000
Position

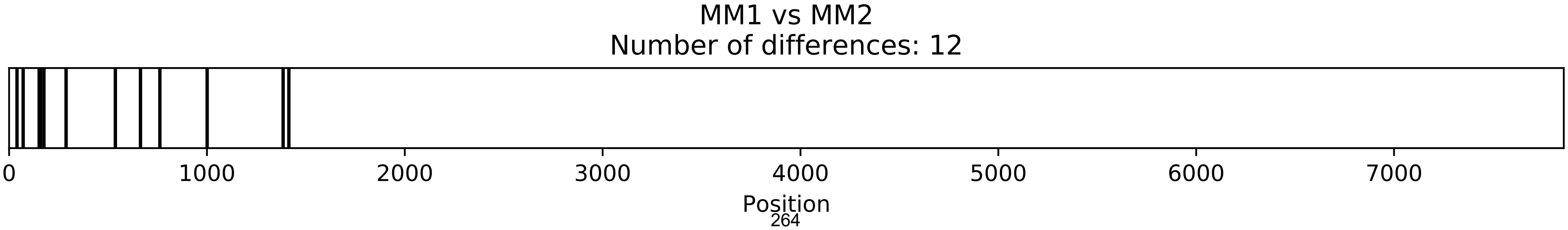
MM1 vs MM2
Number of differences: 12
0
1000
2000
3000
4000
5000
6000
7000
Position

### Region: 737



## CR1 vs CR2
Number of differences: 302

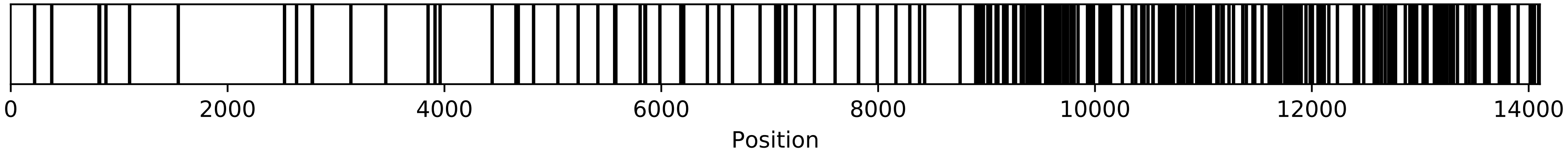


## CR1 vs MM1
Number of differences: 167

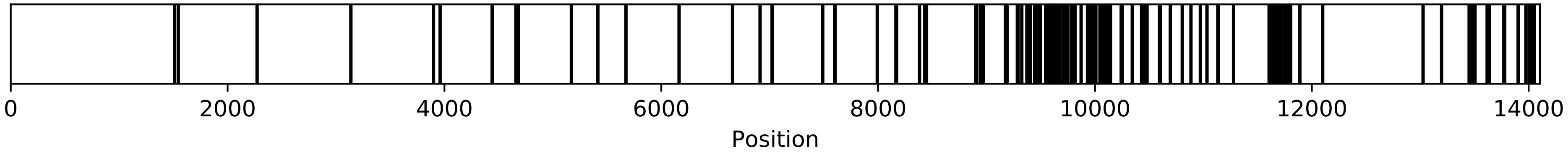


## CR1 vs MM2
Number of differences: 153

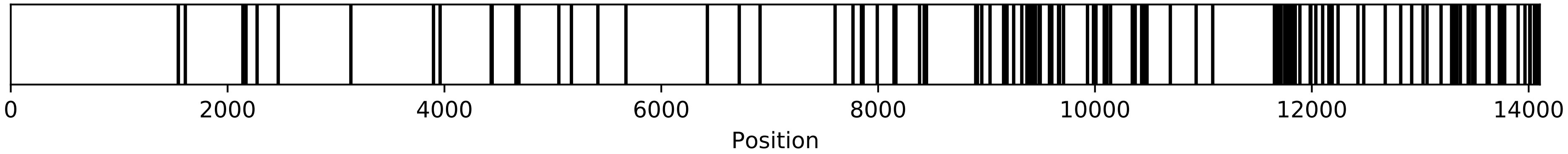


## CR2 vs MM1
Number of differences: 257

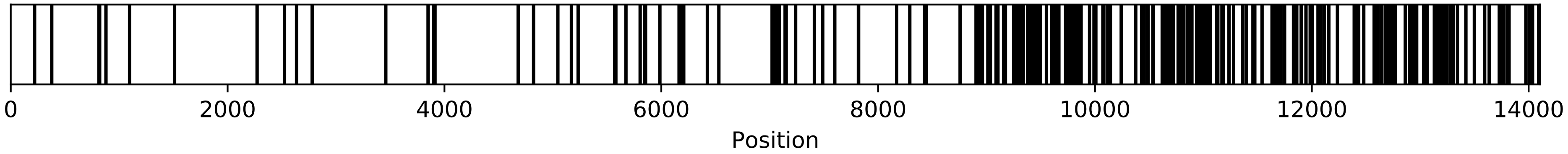


## CR2 vs MM2
Number of differences: 295

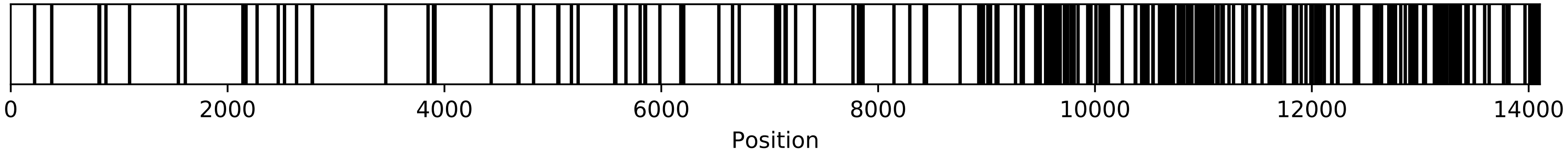


## MM1 vs MM2
Number of differences: 156

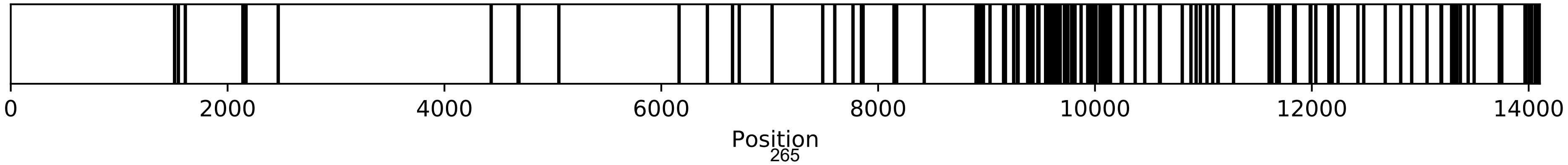

### Region: 738



CR1 vs CR2
Number of differences: 370

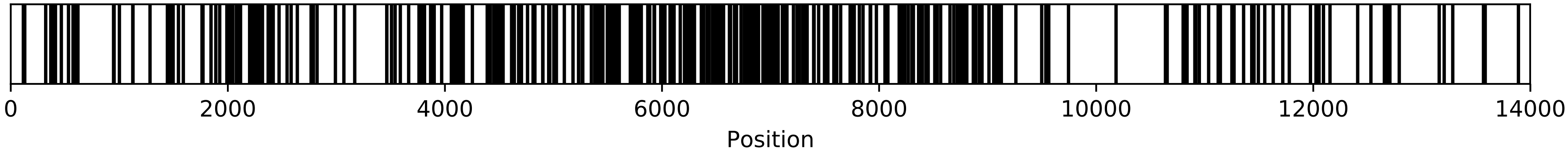


CR1 vs MM1
Number of differences: 184

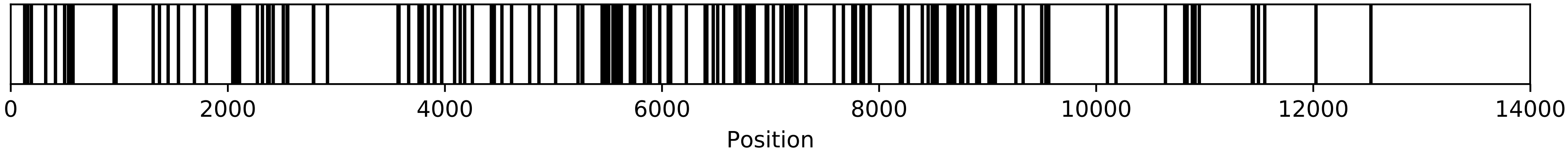


CR1 vs MM2
Number of differences: 160

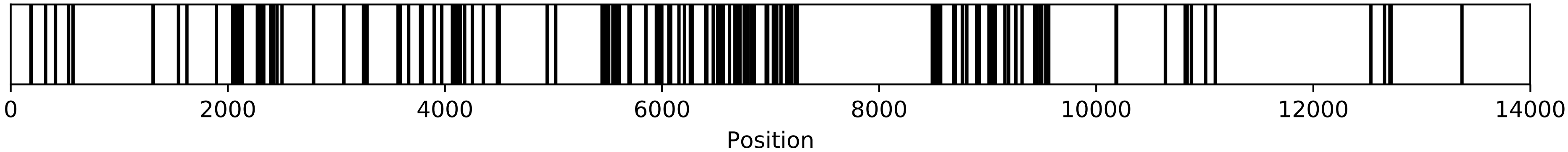


CR2 vs MM1
Number of differences: 371

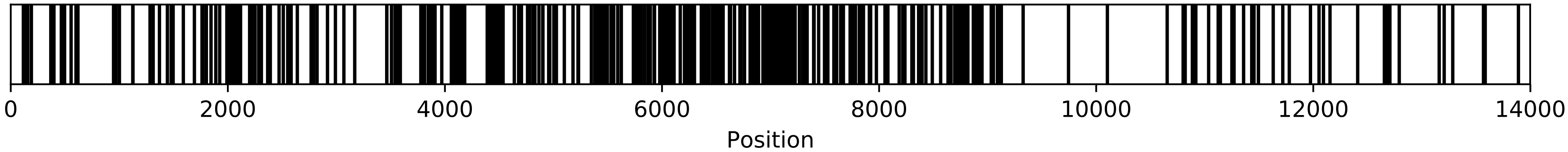


CR2 vs MM2
Number of differences: 377

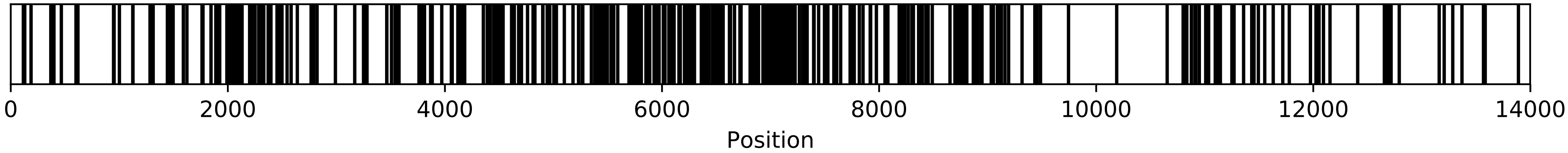


MM1 vs MM2
Number of differences: 156

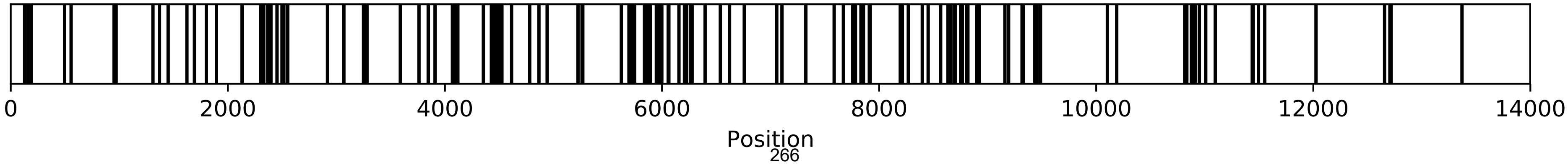

### Region: 739



CR1 vs CR2
Number of differences: 376

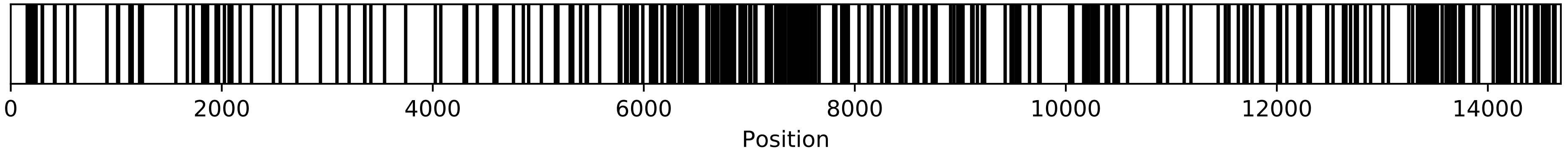


CR1 vs MM1
Number of differences: 139

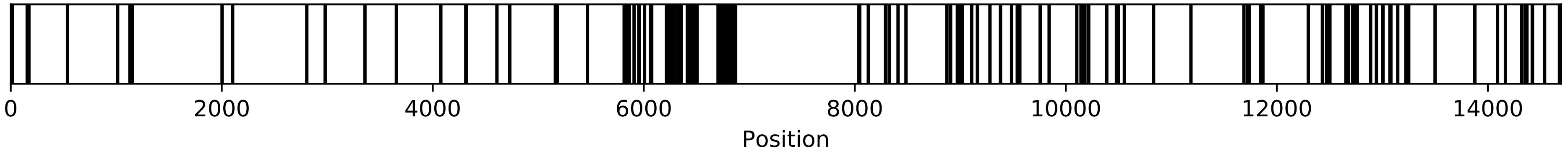


CR1 vs MM2
Number of differences: 169

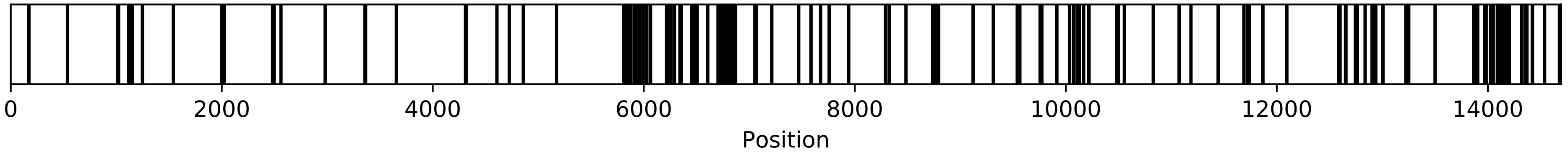


CR2 vs MM1
Number of differences: 398

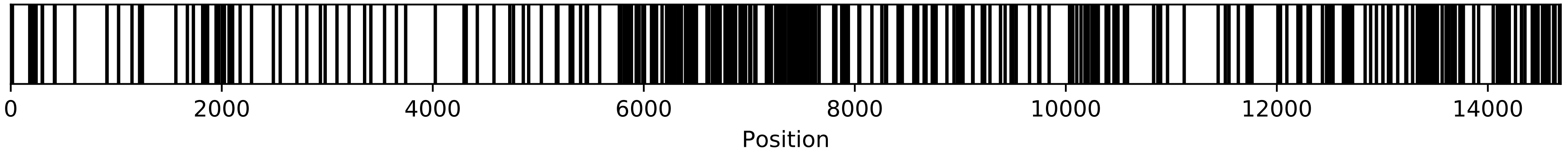


CR2 vs MM2
Number of differences: 367

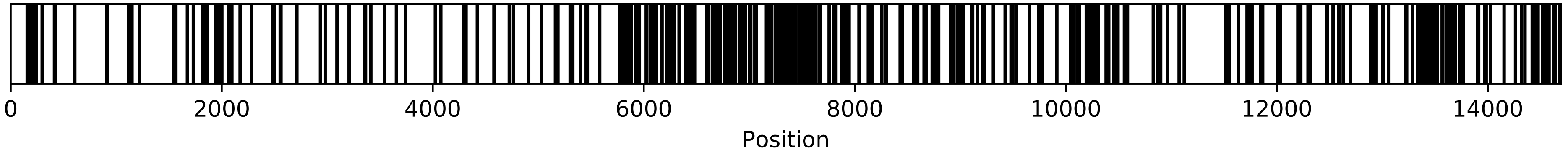


MM1 vs MM2
Number of differences: 161

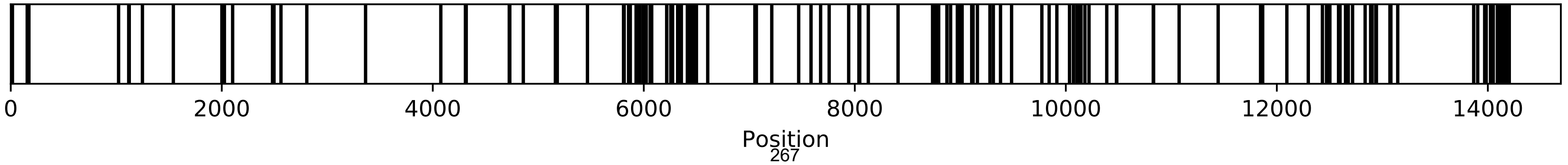

### Region: 740



CR1 vs CR2
Number of differences: 65

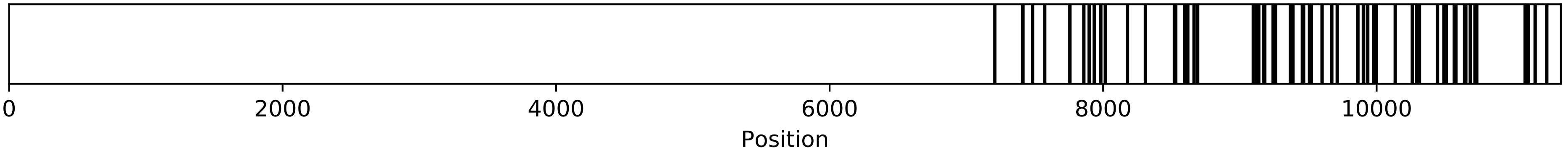


CR1 vs MM1
Number of differences: 81

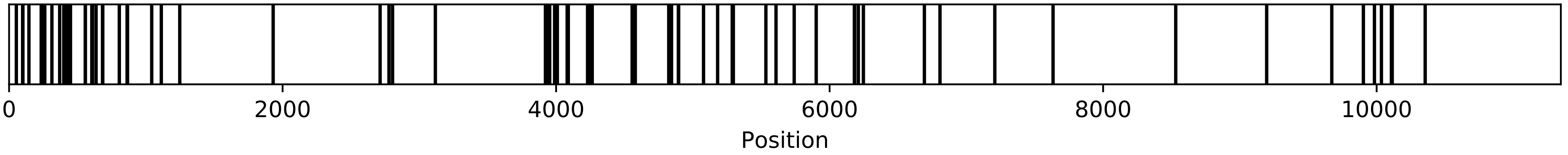


CR1 vs MM2
Number of differences: 123

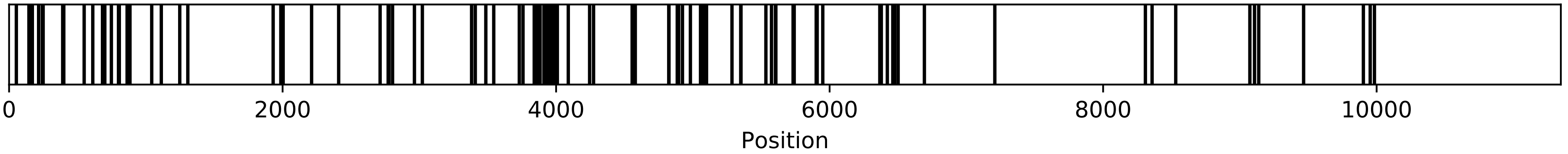


CR2 vs MM1
Number of differences: 139

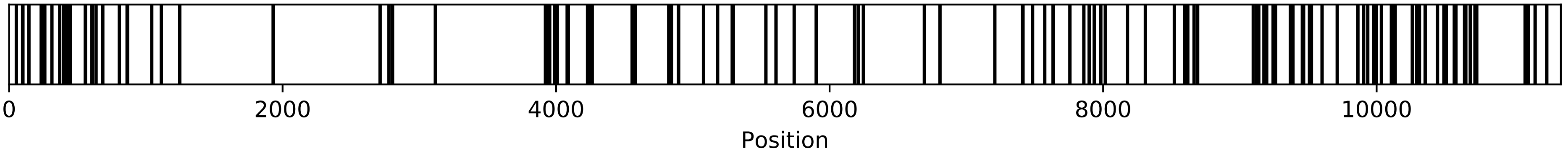


CR2 vs MM2
Number of differences: 177

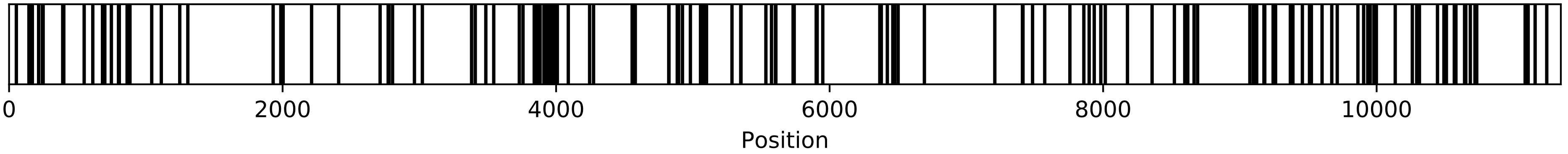


MM1 vs MM2
Number of differences: 128

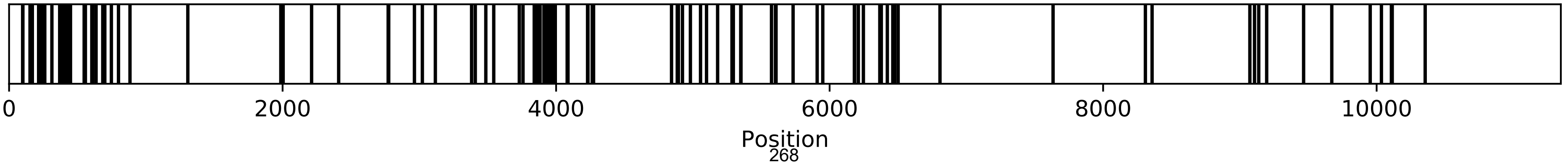

### Region: 742



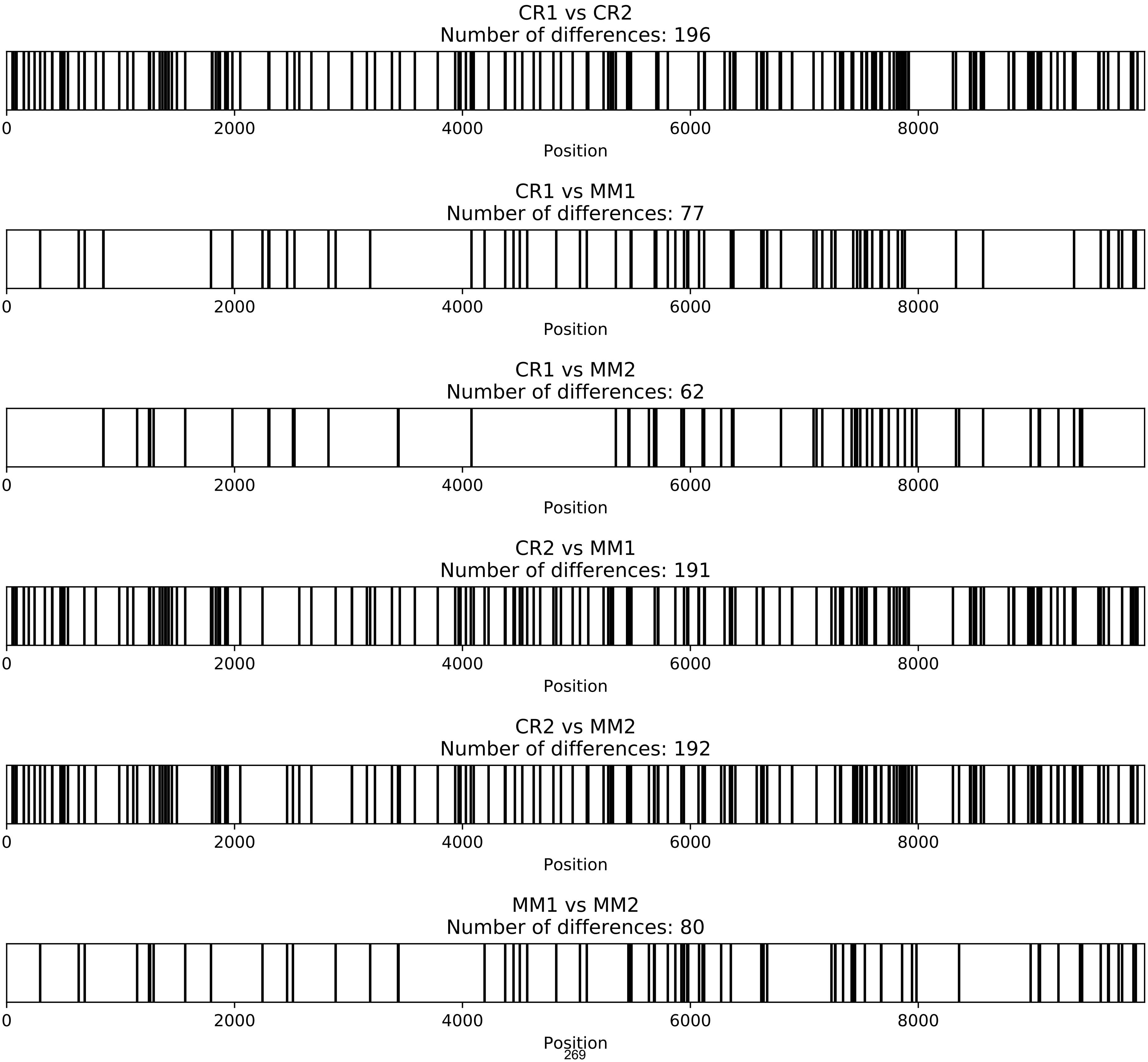
CR1 vs CR2
Number of differences: 196
CR1 vs MM1
Number of differences: 77
CR1 vs MM2
Number of differences: 62
CR2 vs MM1
Number of differences: 191
CR2 vs MM2
Number of differences: 192
MM1 vs MM2
Number of differences: 80
0
2000
4000
6000
8000
Position

### Region: 743



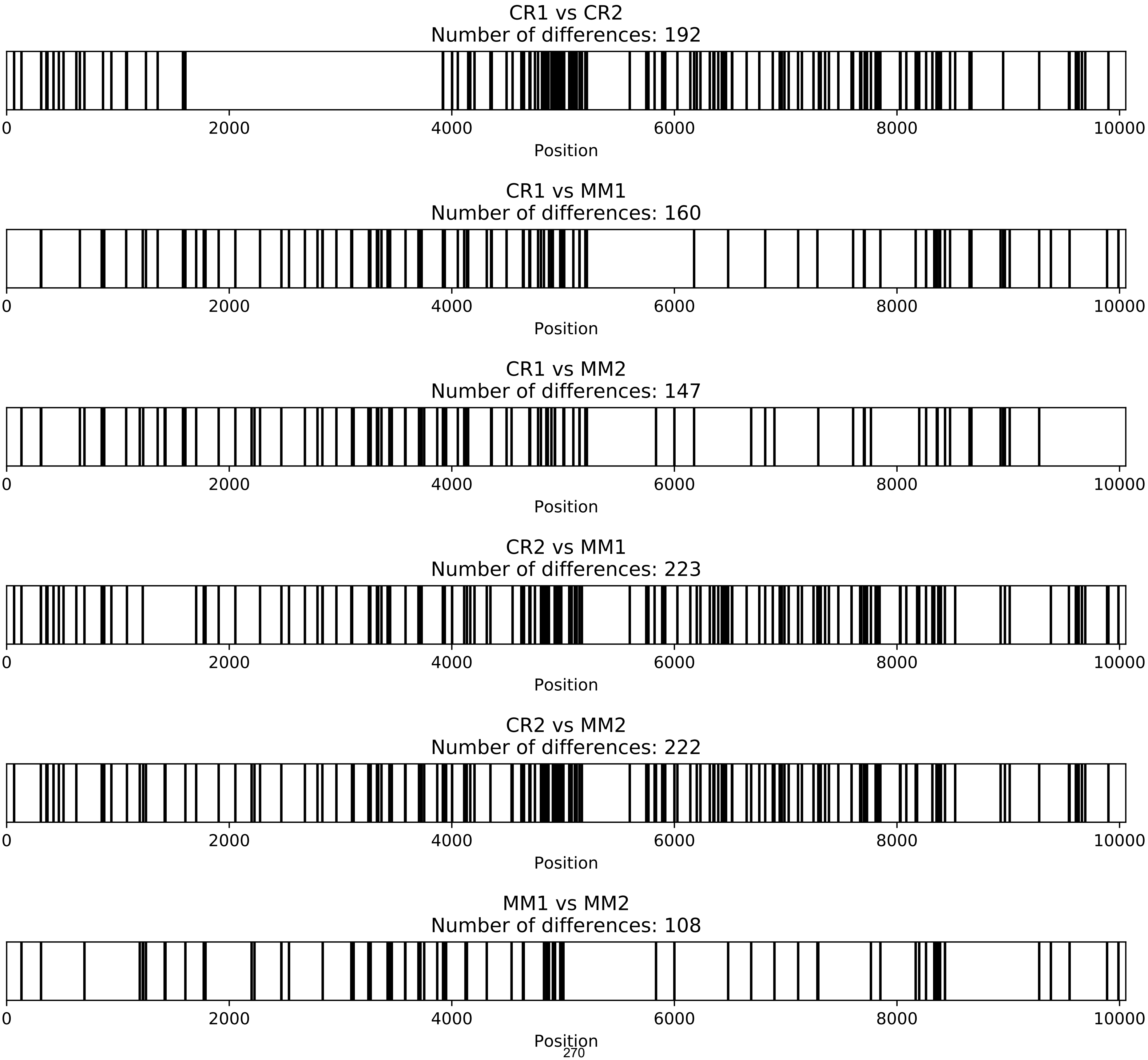
CR1 vs CR2
Number of differences: 192
CR1 vs MM1
Number of differences: 160
CR1 vs MM2
Number of differences: 147
CR2 vs MM1
Number of differences: 223
CR2 vs MM2
Number of differences: 222
MM1 vs MM2
Number of differences: 108
0
2000
4000
6000
8000
10000
Position

### Region: 745



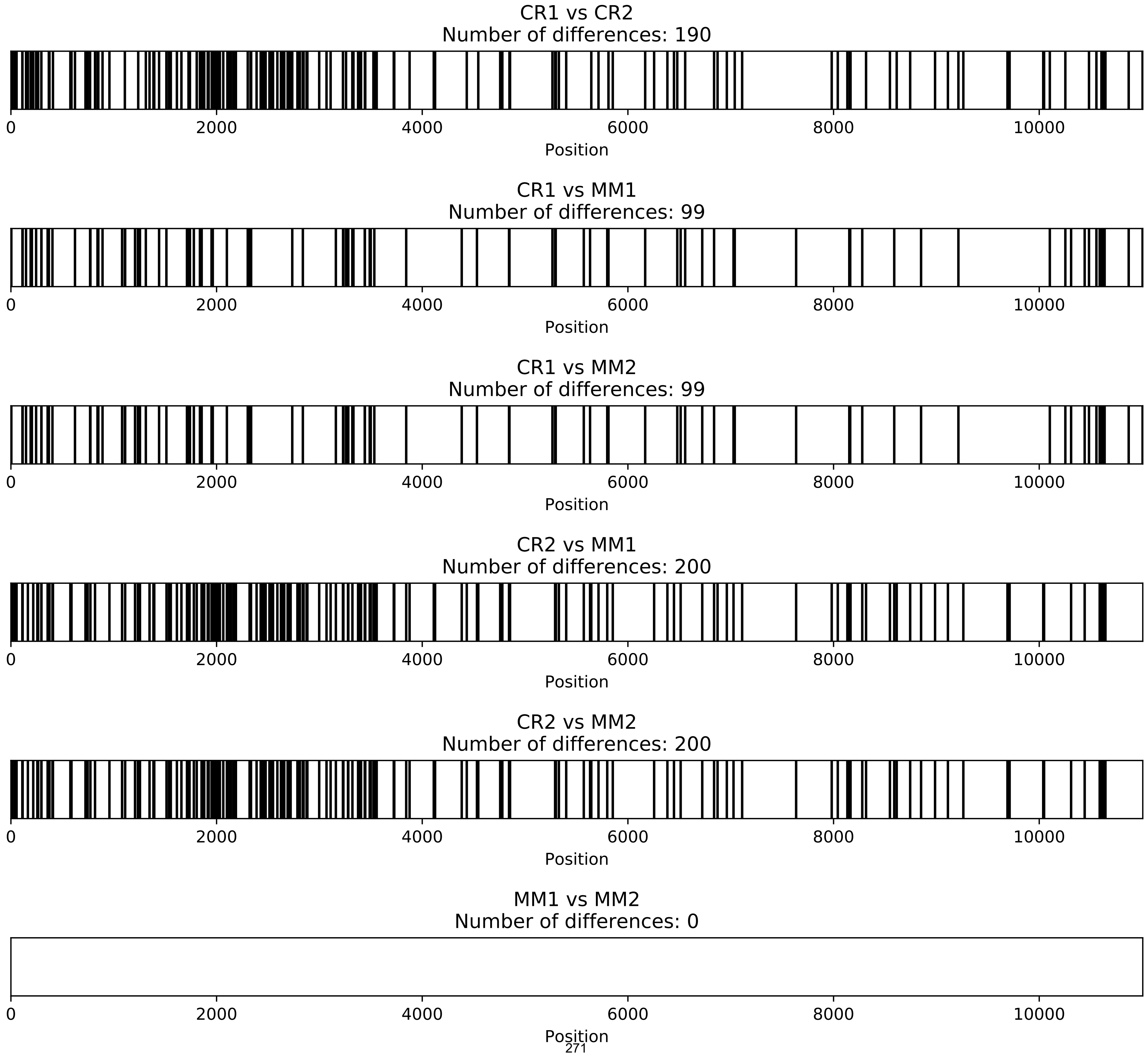
CR1 vs CR2
Number of differences: 190
CR1 vs MM1
Number of differences: 99
CR1 vs MM2
Number of differences: 99
CR2 vs MM1
Number of differences: 200
CR2 vs MM2
Number of differences: 200
MM1 vs MM2
Number of differences: 0
0
2000
4000
6000
8000
10000
Position

### Region: 746



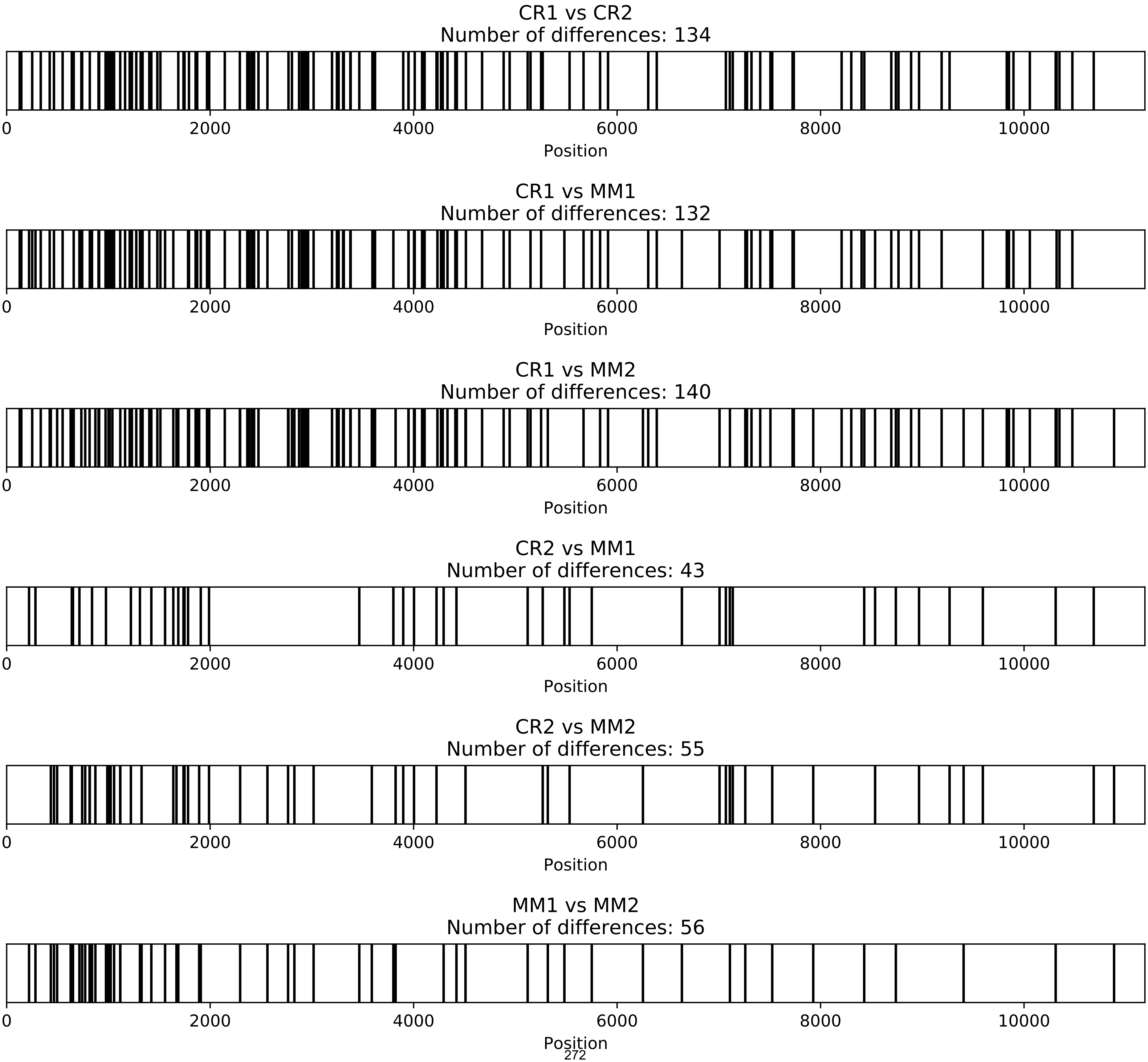
CR1 vs CR2
Number of differences: 134
CR1 vs MM1
Number of differences: 132
CR1 vs MM2
Number of differences: 140
CR2 vs MM1
Number of differences: 43
CR2 vs MM2
Number of differences: 55
MM1 vs MM2
Number of differences: 56
0
2000
4000
6000
8000
10000
Position

### Region: 747



CR1 vs CR2
Number of differences: 530

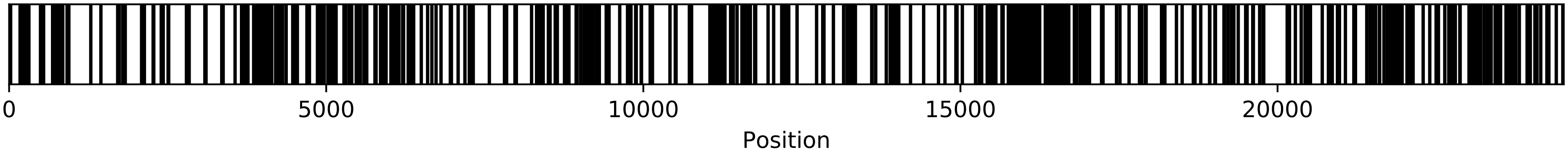


CR1 vs MM1
Number of differences: 250

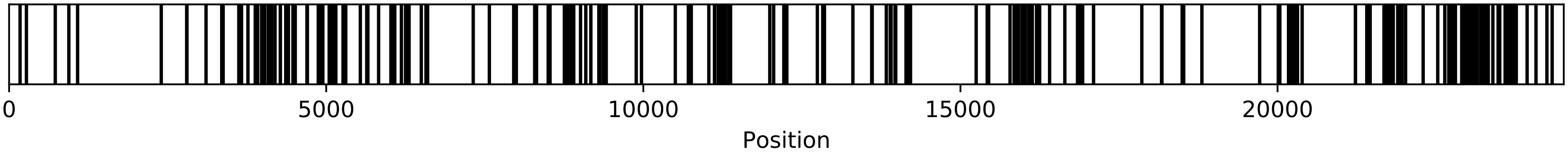


CR1 vs MM2
Number of differences: 260

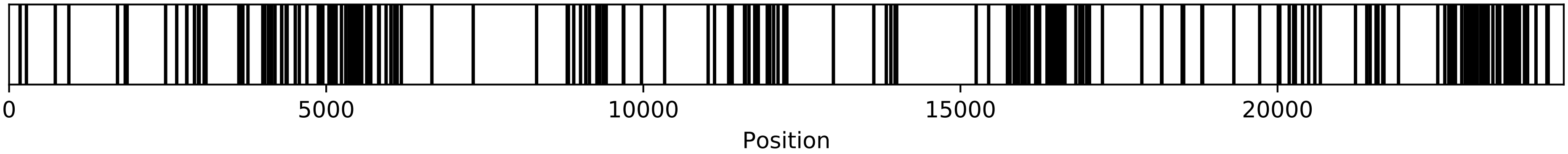


CR2 vs MM1
Number of differences: 497

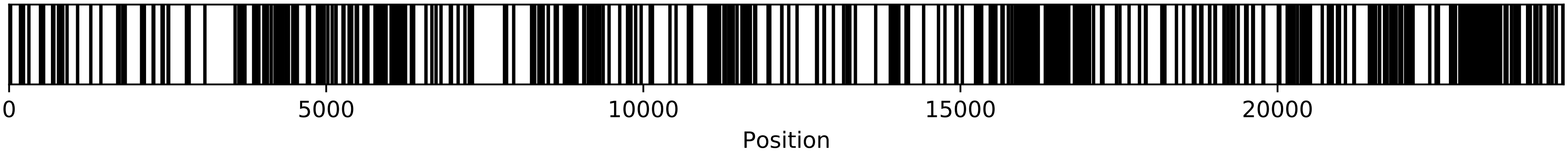


CR2 vs MM2
Number of differences: 522

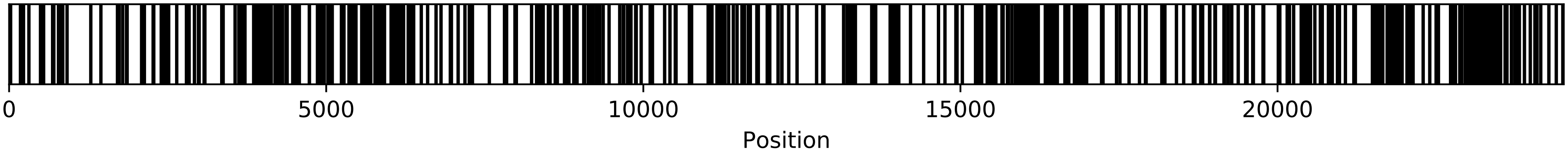


MM1 vs MM2
Number of differences: 217

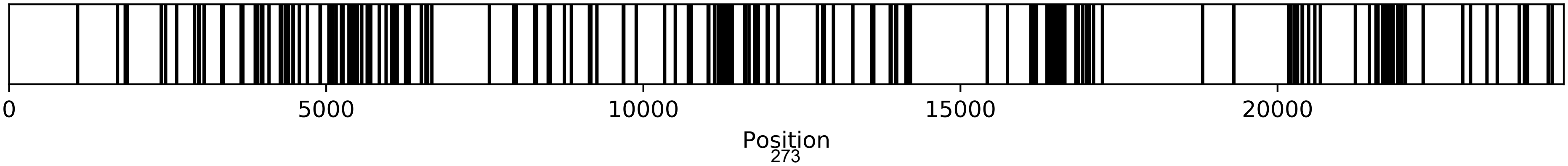

### Region: 750



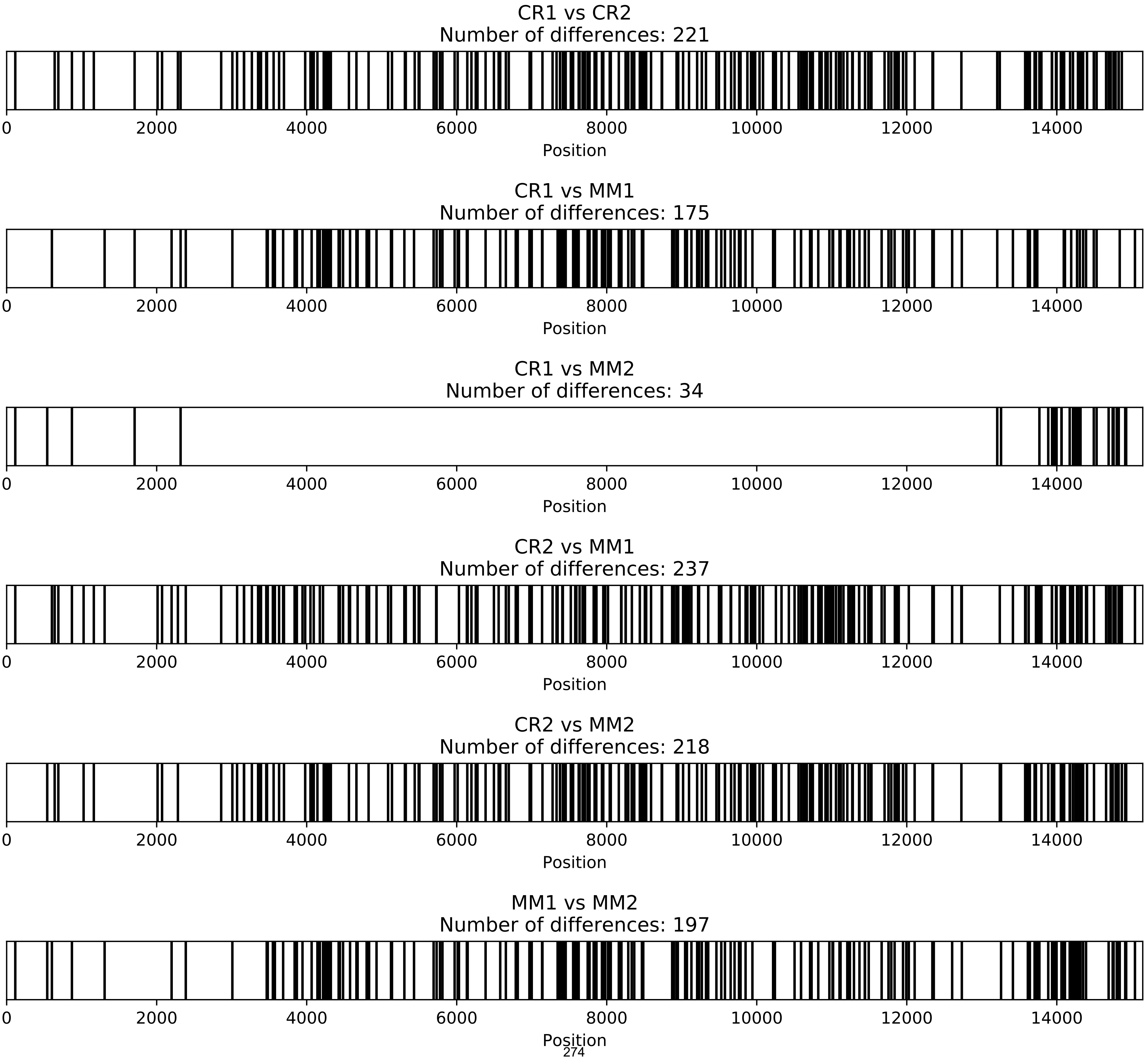
CR1 vs CR2
Number of differences: 221
0 2000 4000 6000 8000 10000 12000 14000
Position
CR1 vs MM1
Number of differences: 175
0 2000 4000 6000 8000 10000 12000 14000
Position
CR1 vs MM2
Number of differences: 34
0 2000 4000 6000 8000 10000 12000 14000
Position
CR2 vs MM1
Number of differences: 237
0 2000 4000 6000 8000 10000 12000 14000
Position
CR2 vs MM2
Number of differences: 218
0 2000 4000 6000 8000 10000 12000 14000
Position
MM1 vs MM2
Number of differences: 197
0 2000 4000 6000 8000 10000 12000 14000
Position

### Region: 754



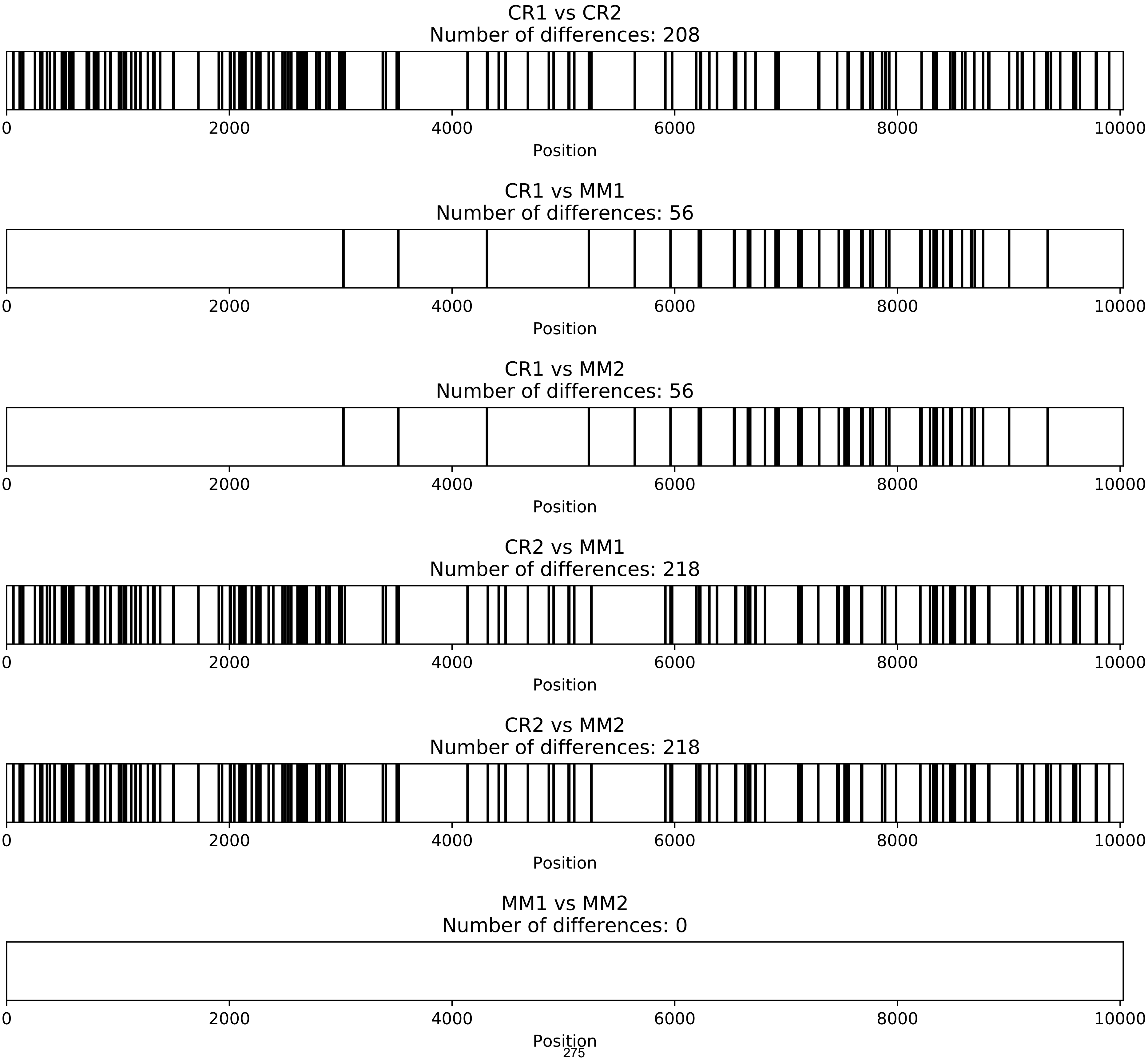
CR1 vs CR2
Number of differences: 208
0
2000
4000
6000
8000
10000
Position
CR1 vs MM1
Number of differences: 56
0
2000
4000
6000
8000
10000
Position
CR1 vs MM2
Number of differences: 56
0
2000
4000
6000
8000
10000
Position
CR2 vs MM1
Number of differences: 218
0
2000
4000
6000
8000
10000
Position
CR2 vs MM2
Number of differences: 218
0
2000
4000
6000
8000
10000
Position
MM1 vs MM2
Number of differences: 0
0
2000
4000
6000
8000
10000
Position

### Region: 755



## CR1 vs CR2
Number of differences: 167

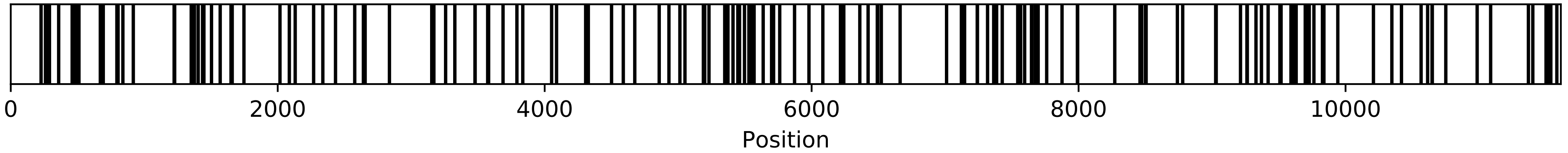


## CR1 vs MM1
Number of differences: 76

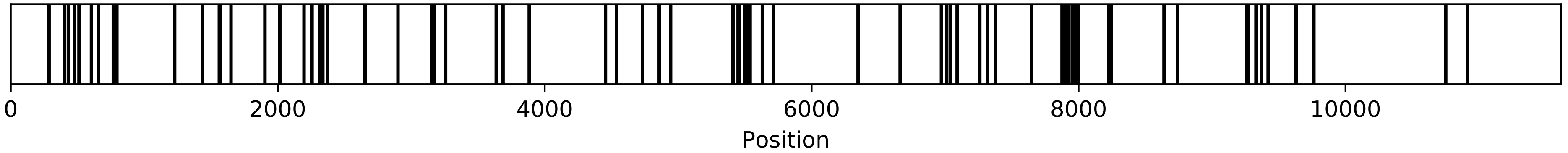


## CR1 vs MM2
Number of differences: 70

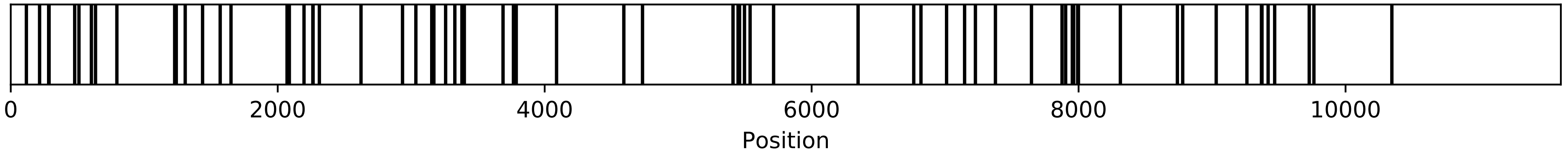


## CR2 vs MM1
Number of differences: 171

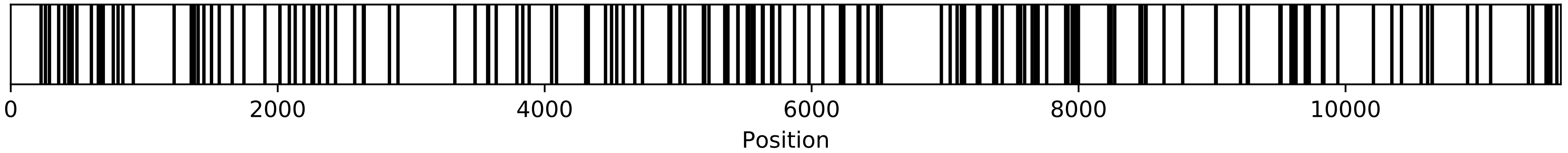


## CR2 vs MM2
Number of differences: 166

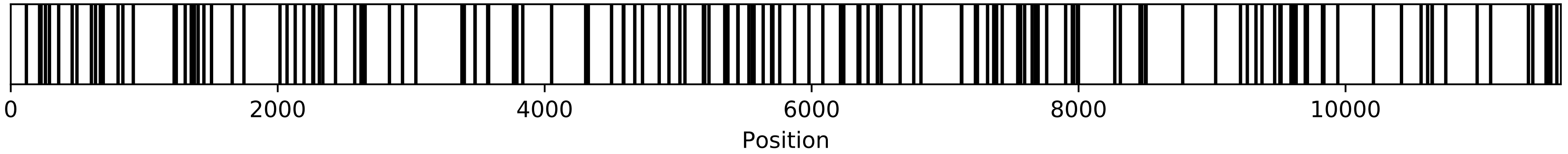


## MM1 vs MM2
Number of differences: 74

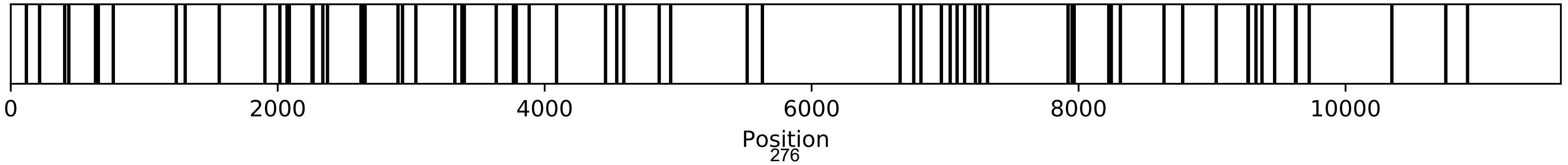

### Region: 756



CR1 vs CR2
Number of differences: 355

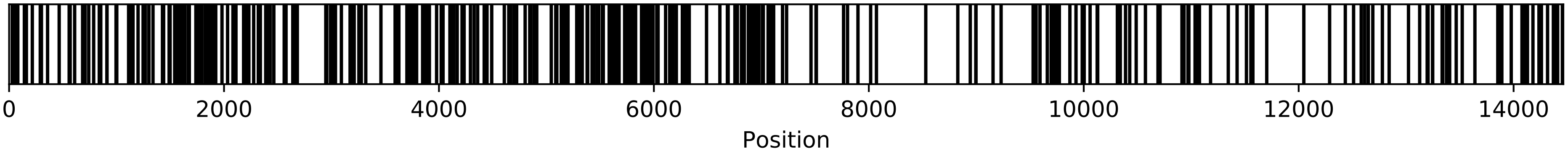


CR1 vs MM1
Number of differences: 127

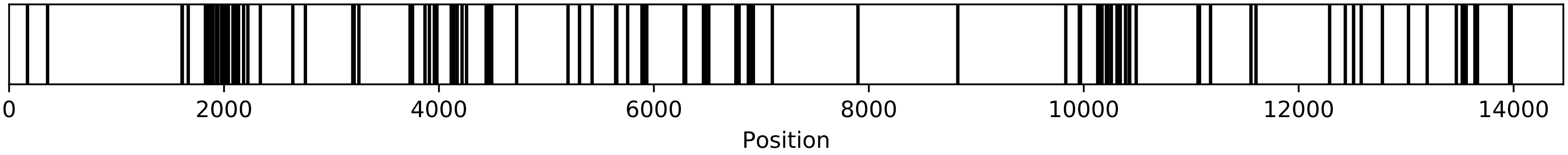


CR1 vs MM2
Number of differences: 127

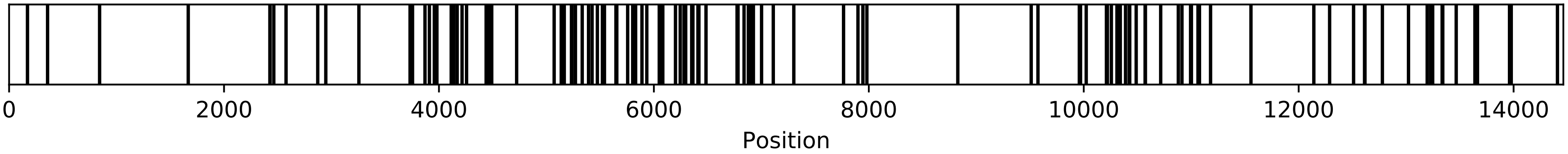


CR2 vs MM1
Number of differences: 358

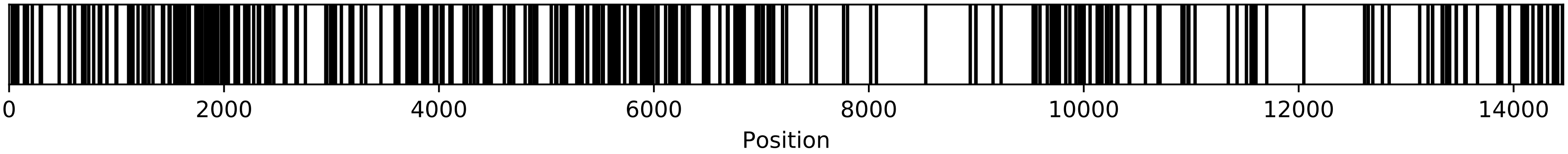


CR2 vs MM2
Number of differences: 351

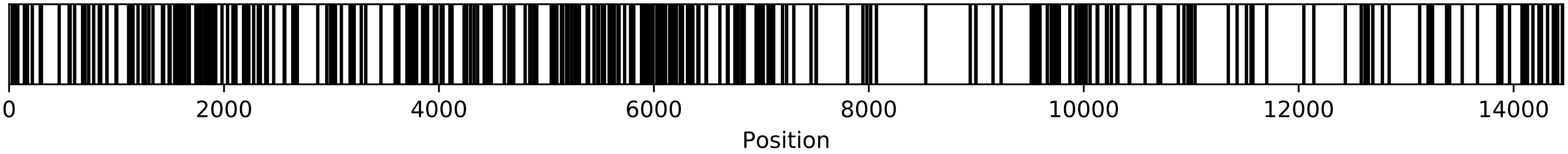


MM1 vs MM2
Number of differences: 125

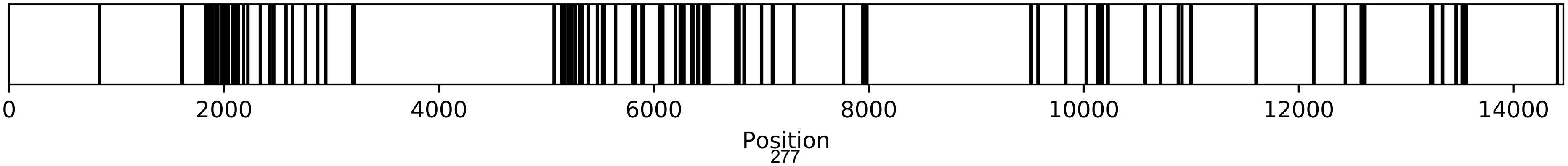

### Region: 763



CR1 vs CR2
Number of differences: 152

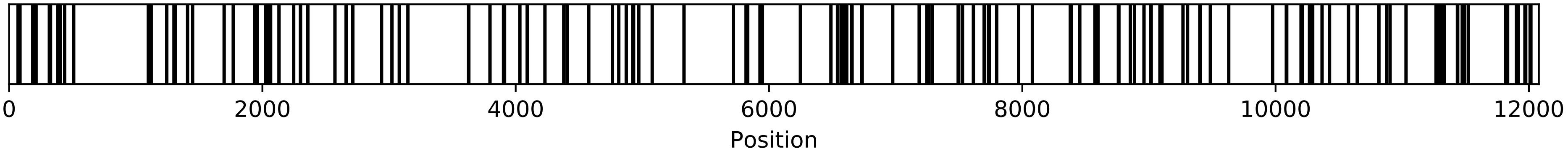


CR1 vs MM1
Number of differences: 56

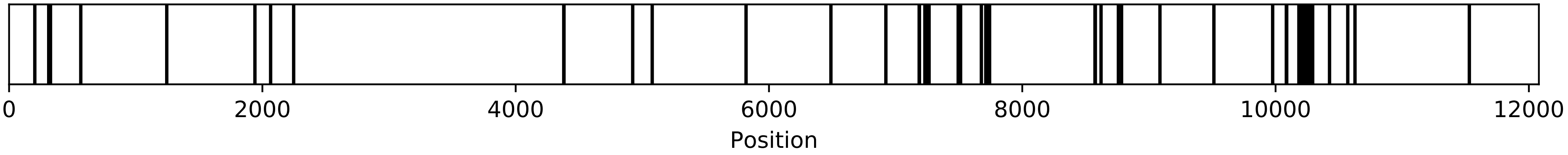


CR1 vs MM2
Number of differences: 62

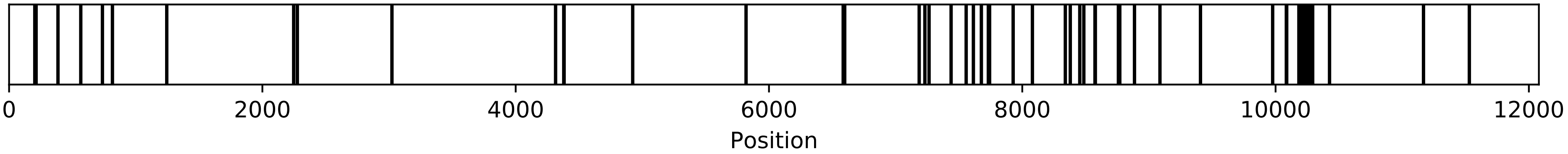


CR2 vs MM1
Number of differences: 147

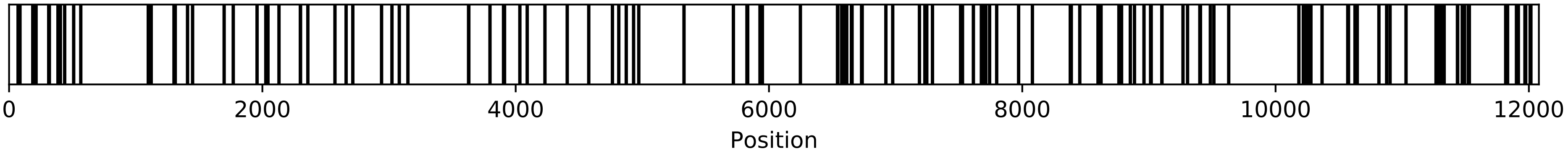


CR2 vs MM2
Number of differences: 147

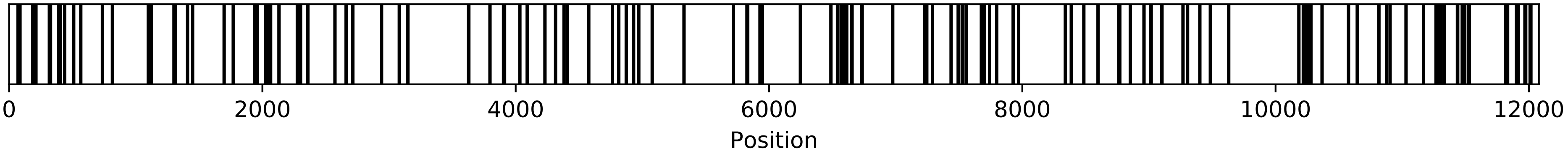


MM1 vs MM2
Number of differences: 42

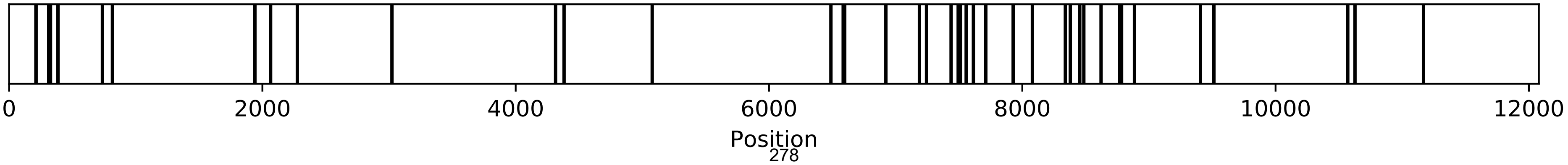

### Region: 764



CR1 vs CR2
Number of differences: 40

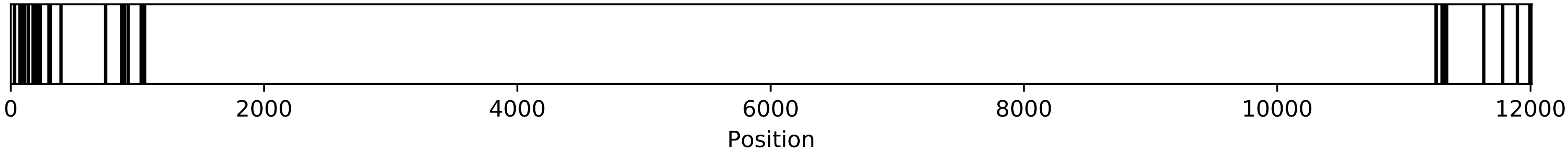


CR1 vs MM1
Number of differences: 191

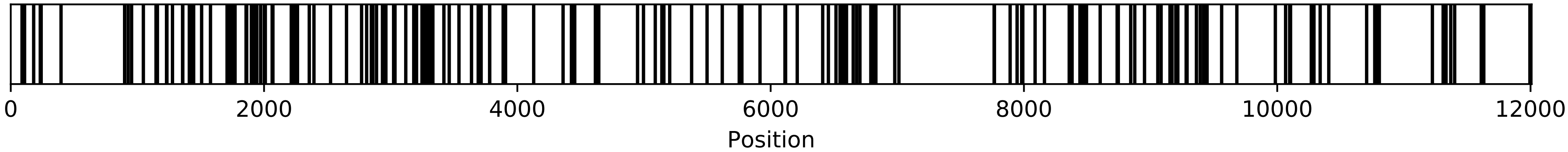


CR1 vs MM2
Number of differences: 235

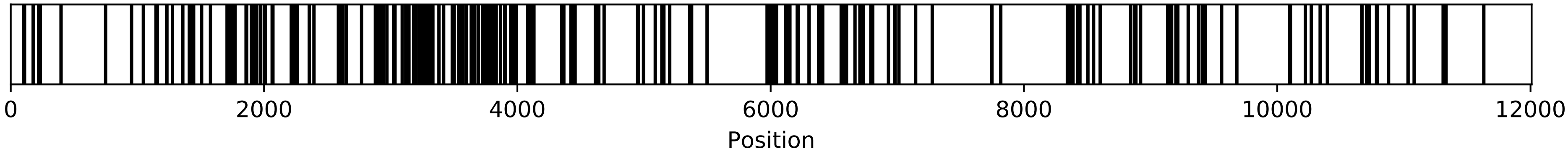


CR2 vs MM1
Number of differences: 210

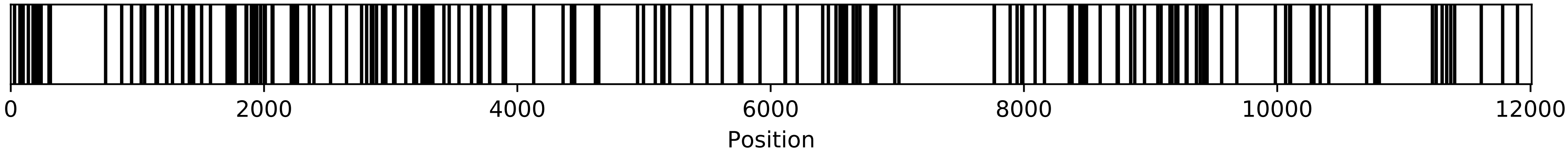


CR2 vs MM2
Number of differences: 257

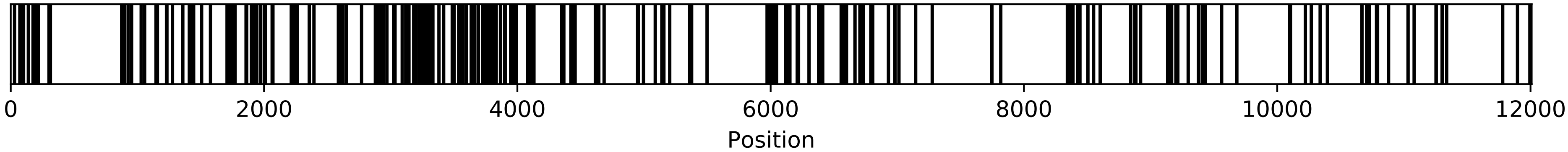


MM1 vs MM2
Number of differences: 202

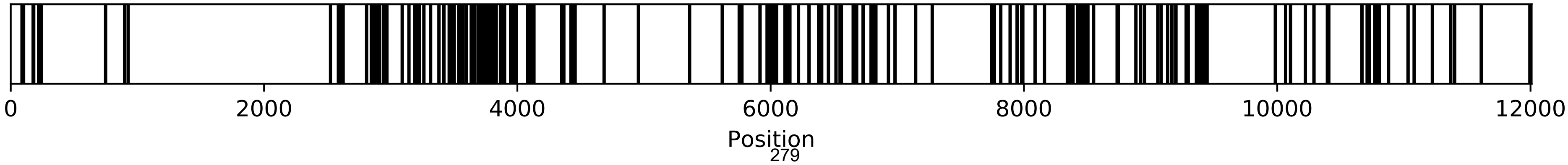

### Region: 765



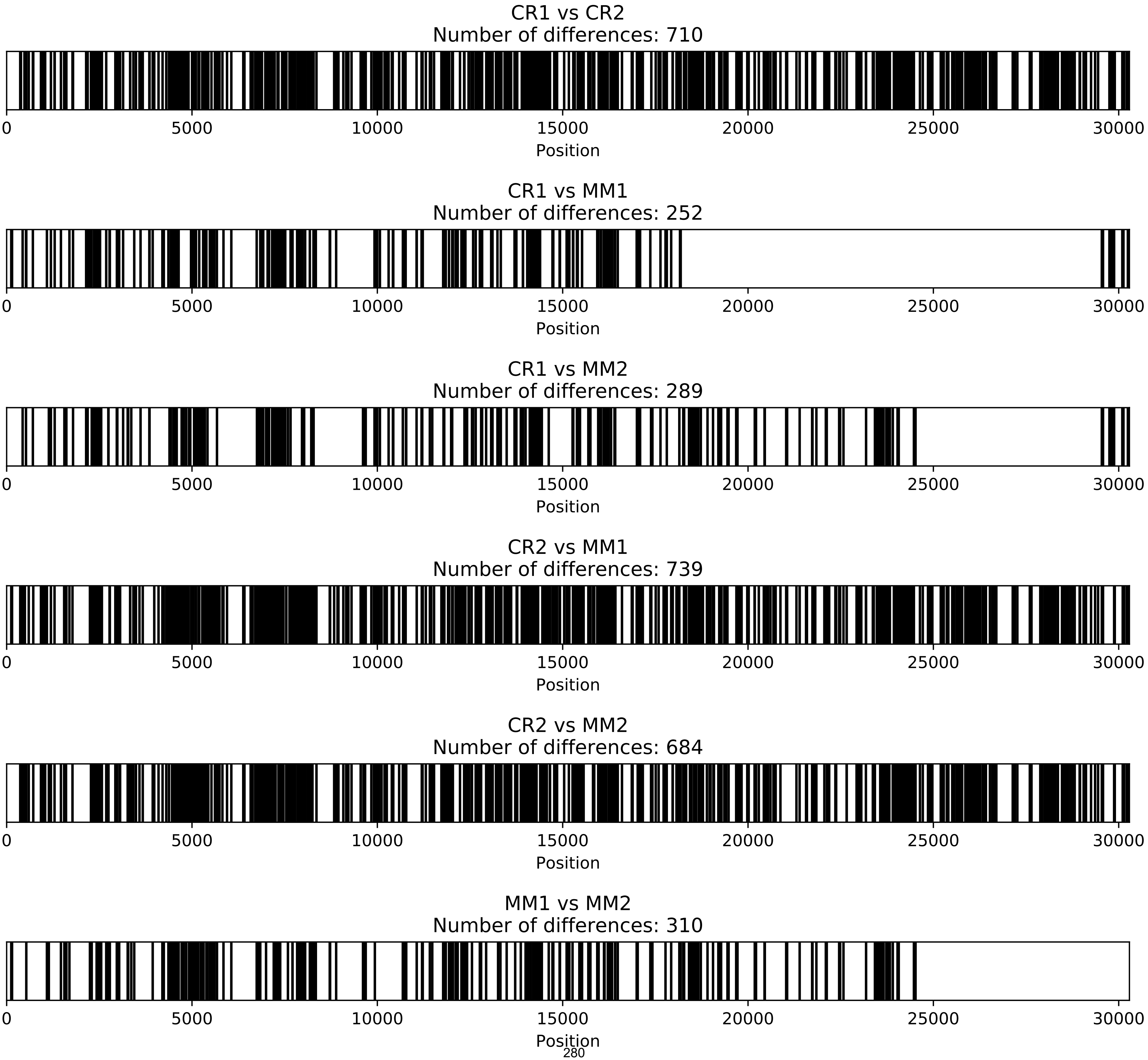
CR1 vs CR2
Number of differences: 710
CR1 vs MM1
Number of differences: 252
CR1 vs MM2
Number of differences: 289
CR2 vs MM1
Number of differences: 739
CR2 vs MM2
Number of differences: 684
MM1 vs MM2
Number of differences: 310
0
5000
10000
15000
20000
25000
30000
Position

### Region: 766



CR1 vs CR2
Number of differences: 325

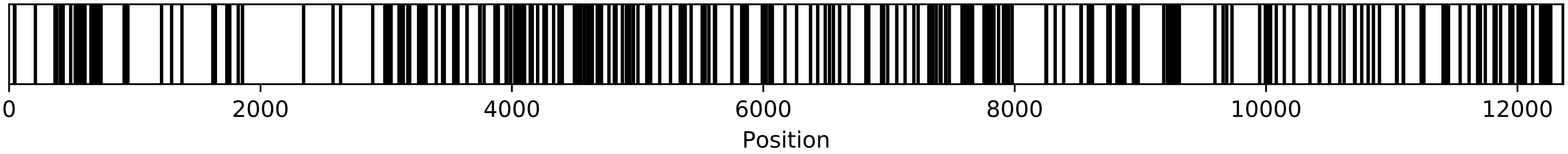


CR1 vs MM1
Number of differences: 301

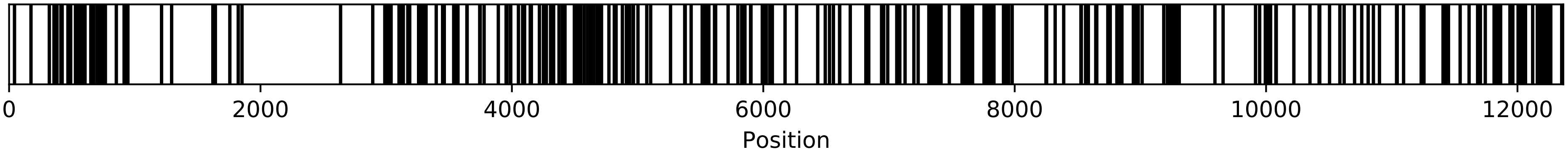


CR1 vs MM2
Number of differences: 322

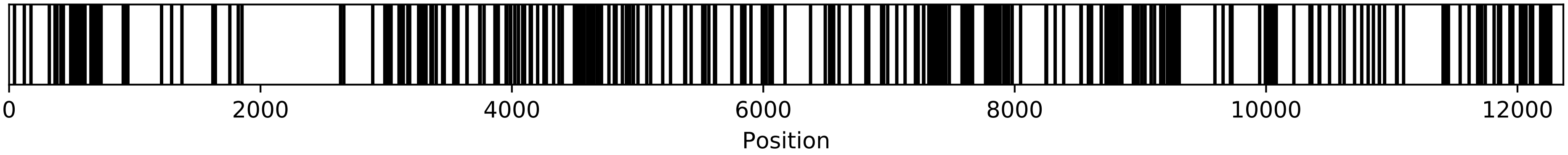


CR2 vs MM1
Number of differences: 118

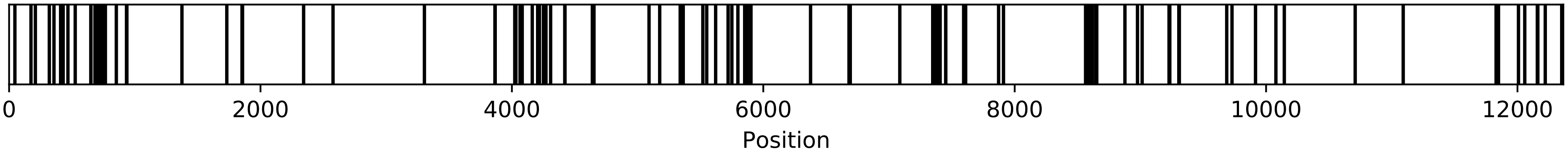


CR2 vs MM2
Number of differences: 139

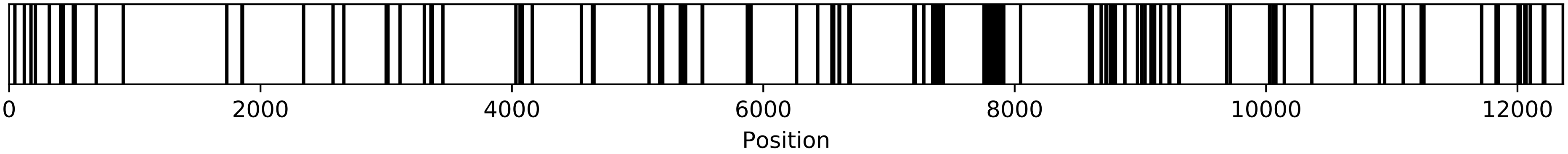


MM1 vs MM2
Number of differences: 144

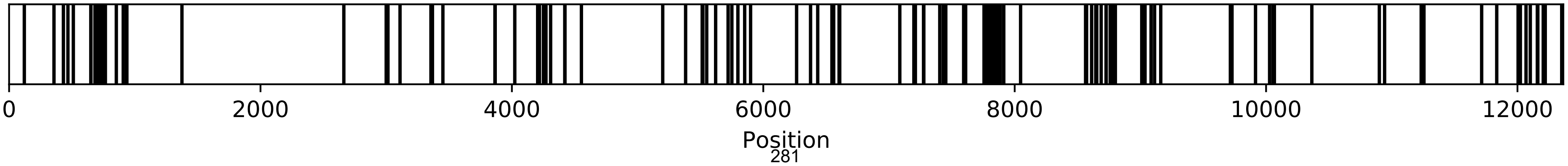

### Region: 768



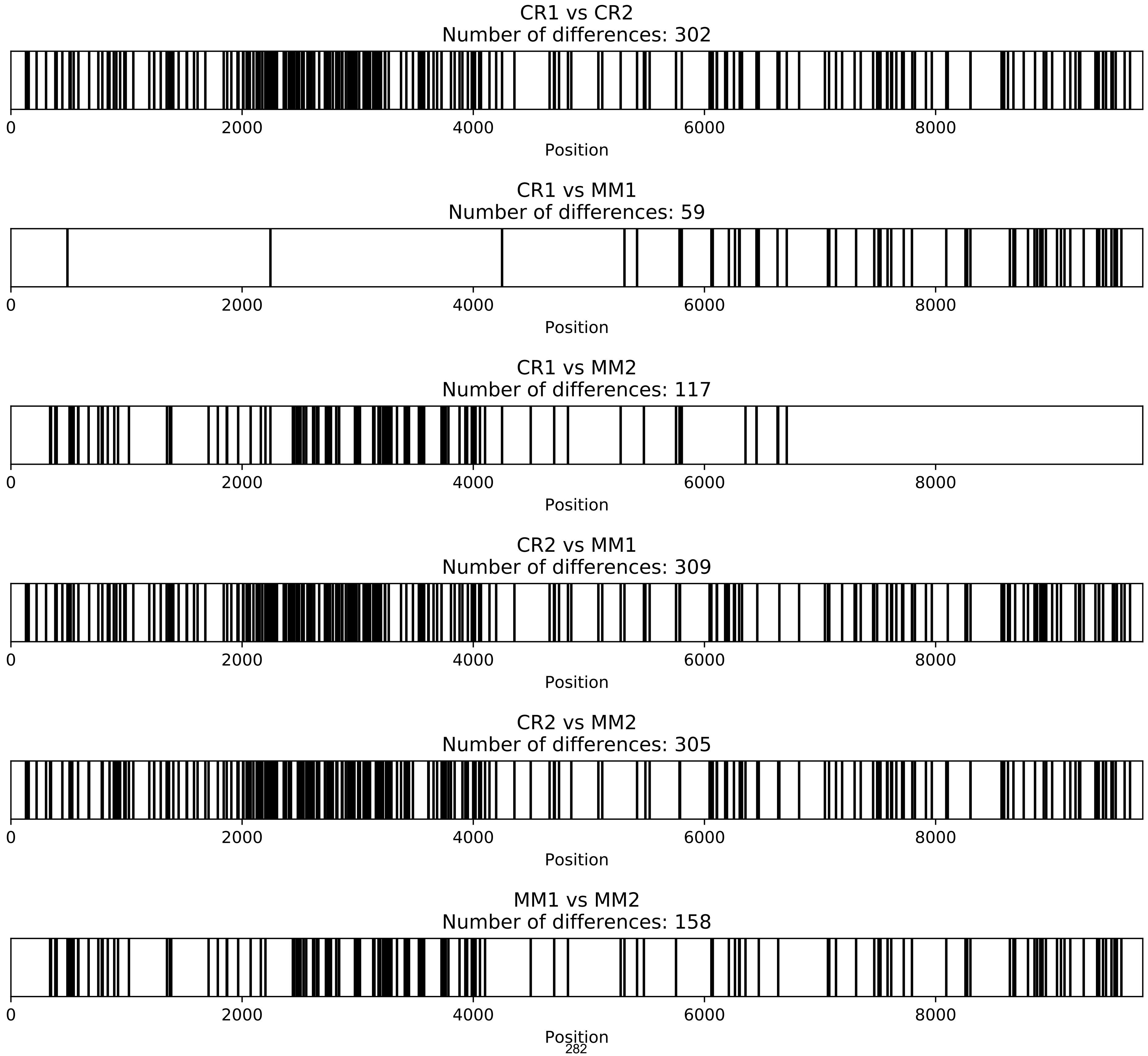
CR1 vs CR2
Number of differences: 302
0
2000
4000
6000
8000
Position
CR1 vs MM1
Number of differences: 59
Position
CR1 vs MM2
Number of differences: 117
Position
CR2 vs MM1
Number of differences: 309
Position
CR2 vs MM2
Number of differences: 305
Position
MM1 vs MM2
Number of differences: 158
Position

### Region: 774



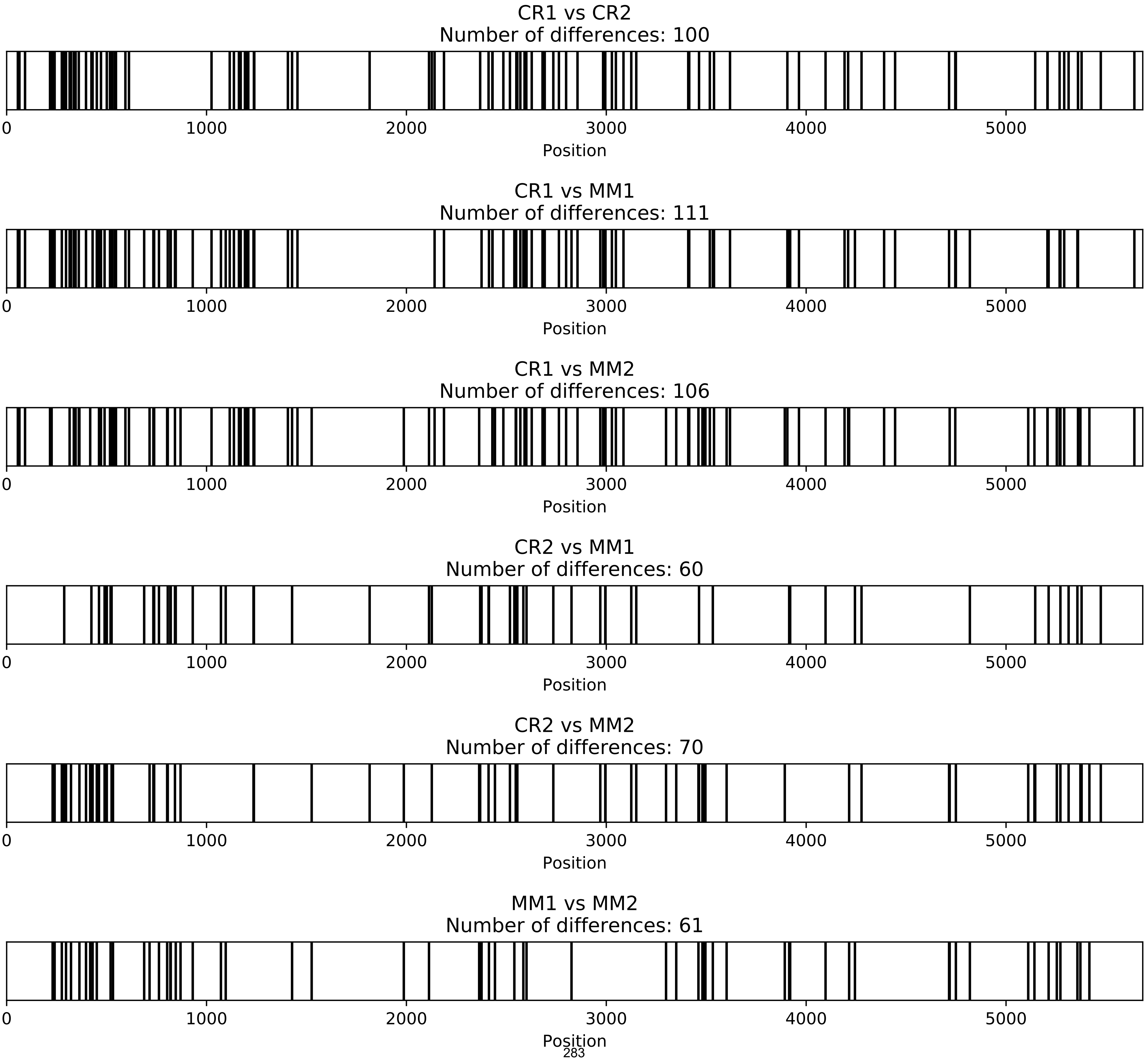
CR1 vs CR2
Number of differences: 100
CR1 vs MM1
Number of differences: 111
CR1 vs MM2
Number of differences: 106
CR2 vs MM1
Number of differences: 60
CR2 vs MM2
Number of differences: 70
MM1 vs MM2
Number of differences: 61
0
1000
2000
3000
4000
5000
Position

### Region: 775



CR1 vs CR2
Number of differences: 283

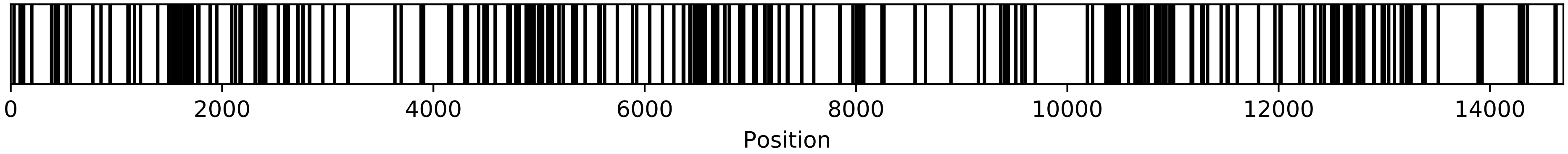


CR1 vs MM1
Number of differences: 283

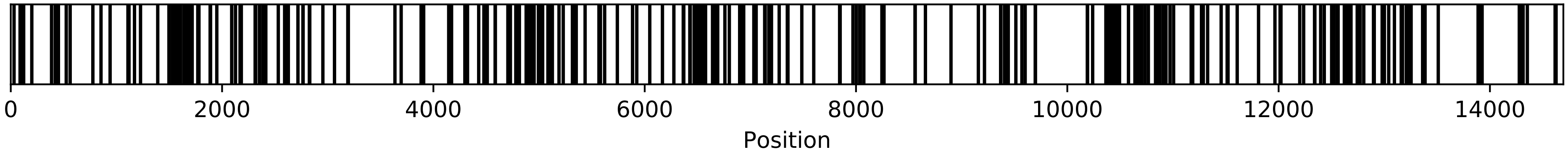


CR1 vs MM2
Number of differences: 275

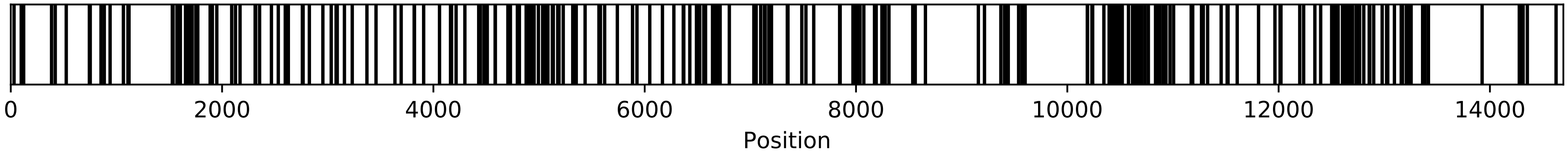


CR2 vs MM1
Number of differences: 0

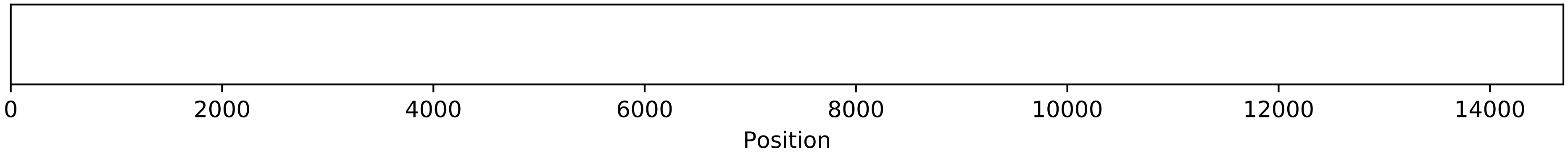


CR2 vs MM2
Number of differences: 140

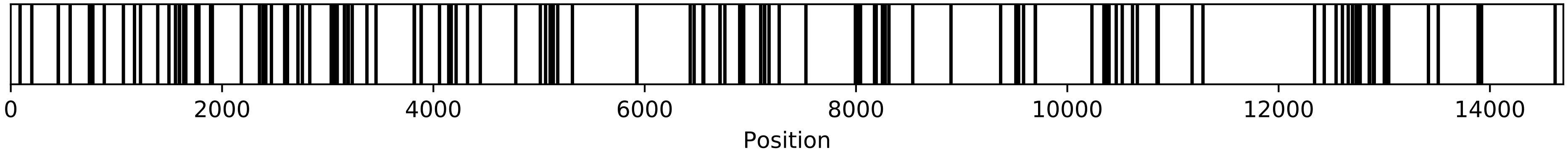


MM1 vs MM2
Number of differences: 140

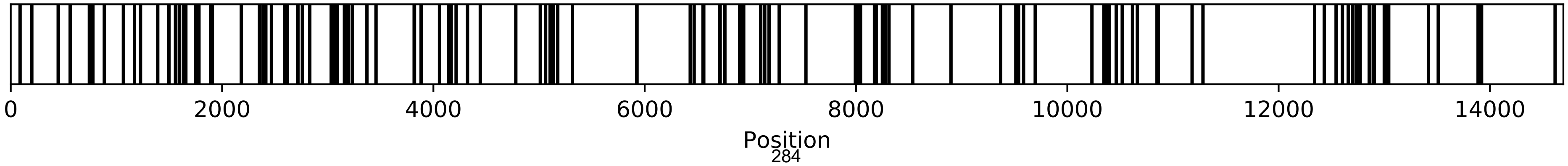

### Region: 776



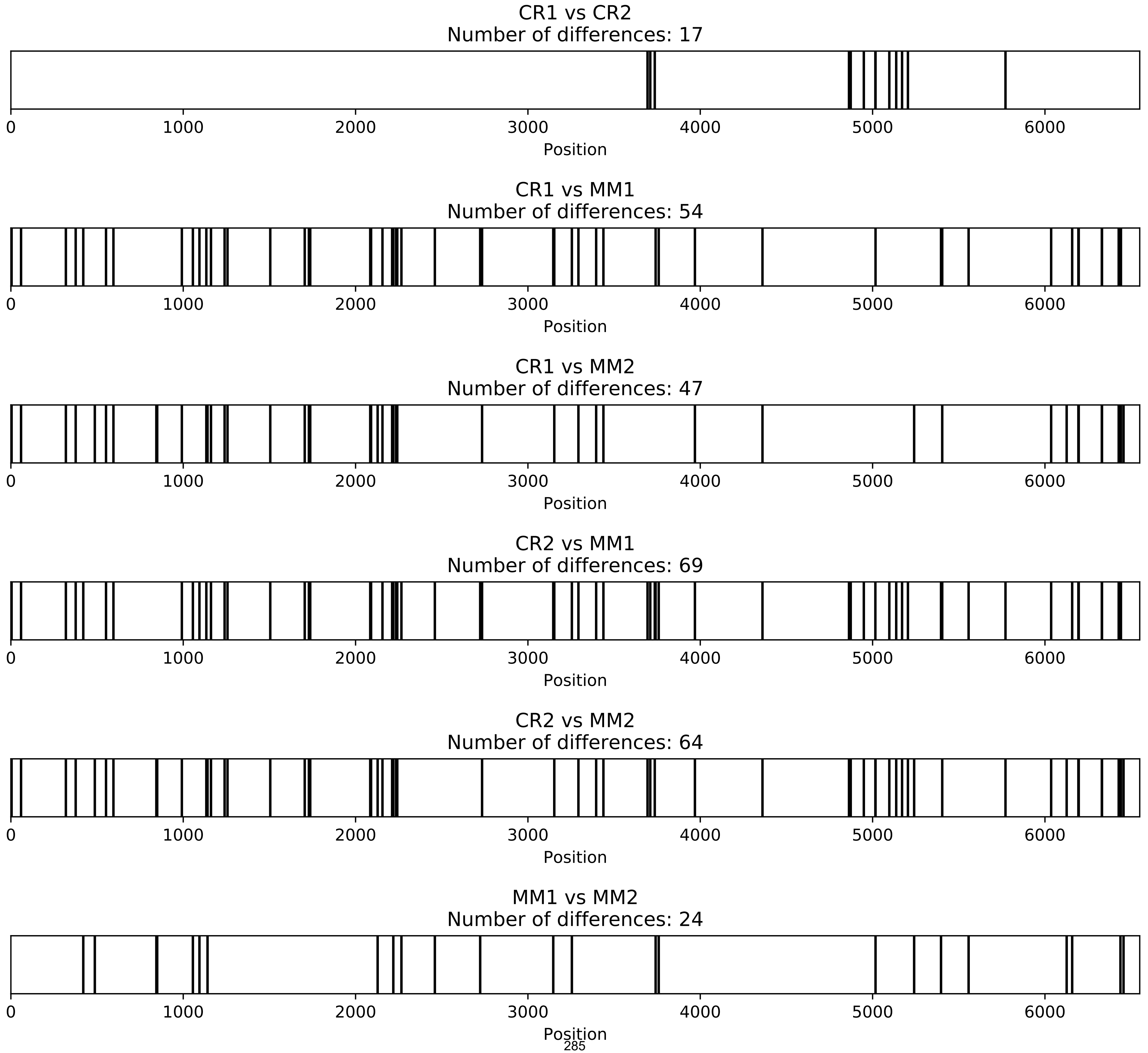
CR1 vs CR2
Number of differences: 17
CR1 vs MM1
Number of differences: 54
CR1 vs MM2
Number of differences: 47
CR2 vs MM1
Number of differences: 69
CR2 vs MM2
Number of differences: 64
MM1 vs MM2
Number of differences: 24
0
1000
2000
3000
4000
5000
6000
Position

### Region: 779



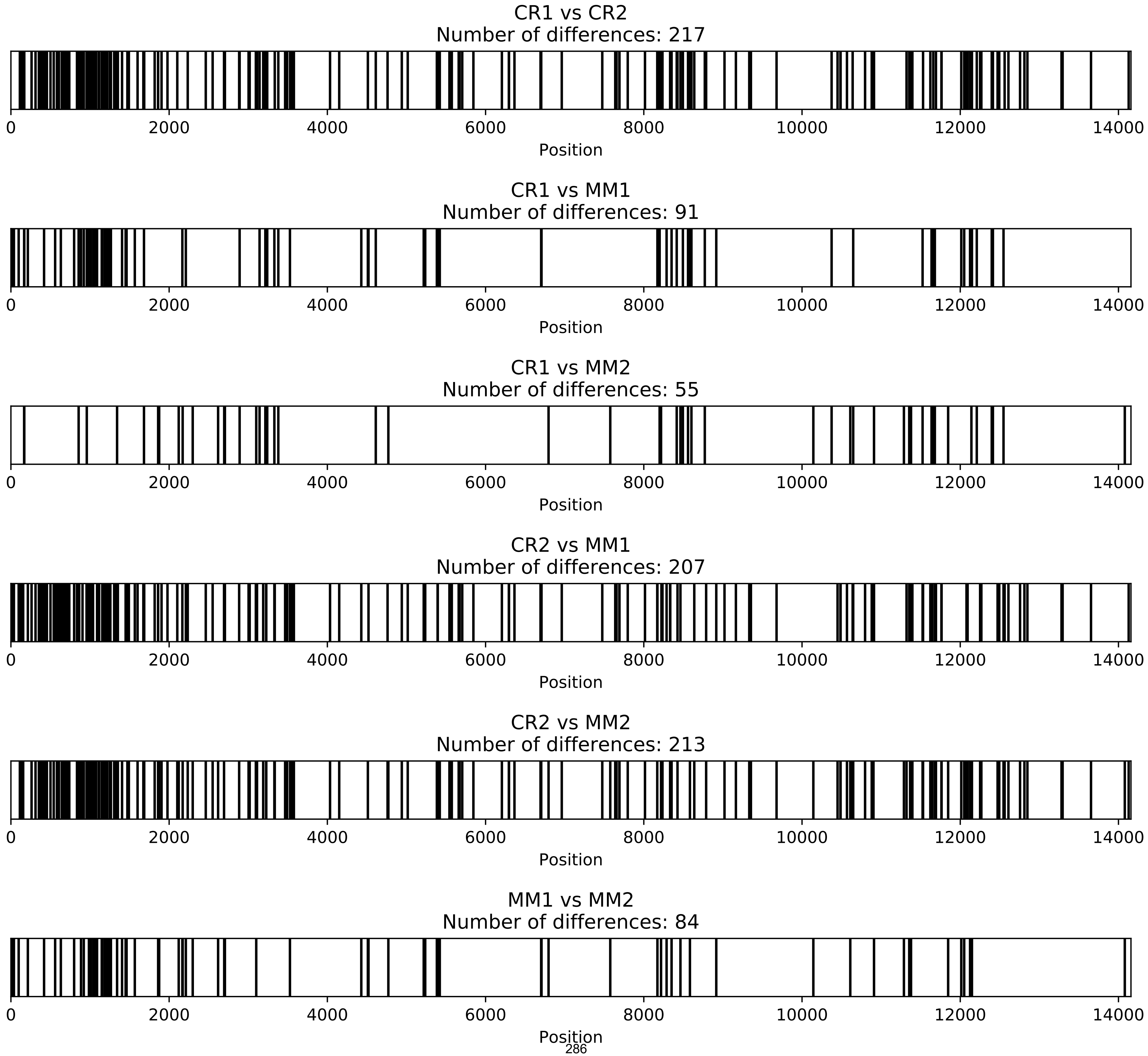
CR1 vs CR2
Number of differences: 217
0
2000
4000
6000
8000
10000
12000
14000
Position
CR1 vs MM1
Number of differences: 91
Position
CR1 vs MM2
Number of differences: 55
Position
CR2 vs MM1
Number of differences: 207
Position
CR2 vs MM2
Number of differences: 213
Position
MM1 vs MM2
Number of differences: 84
Position

### Region: 780



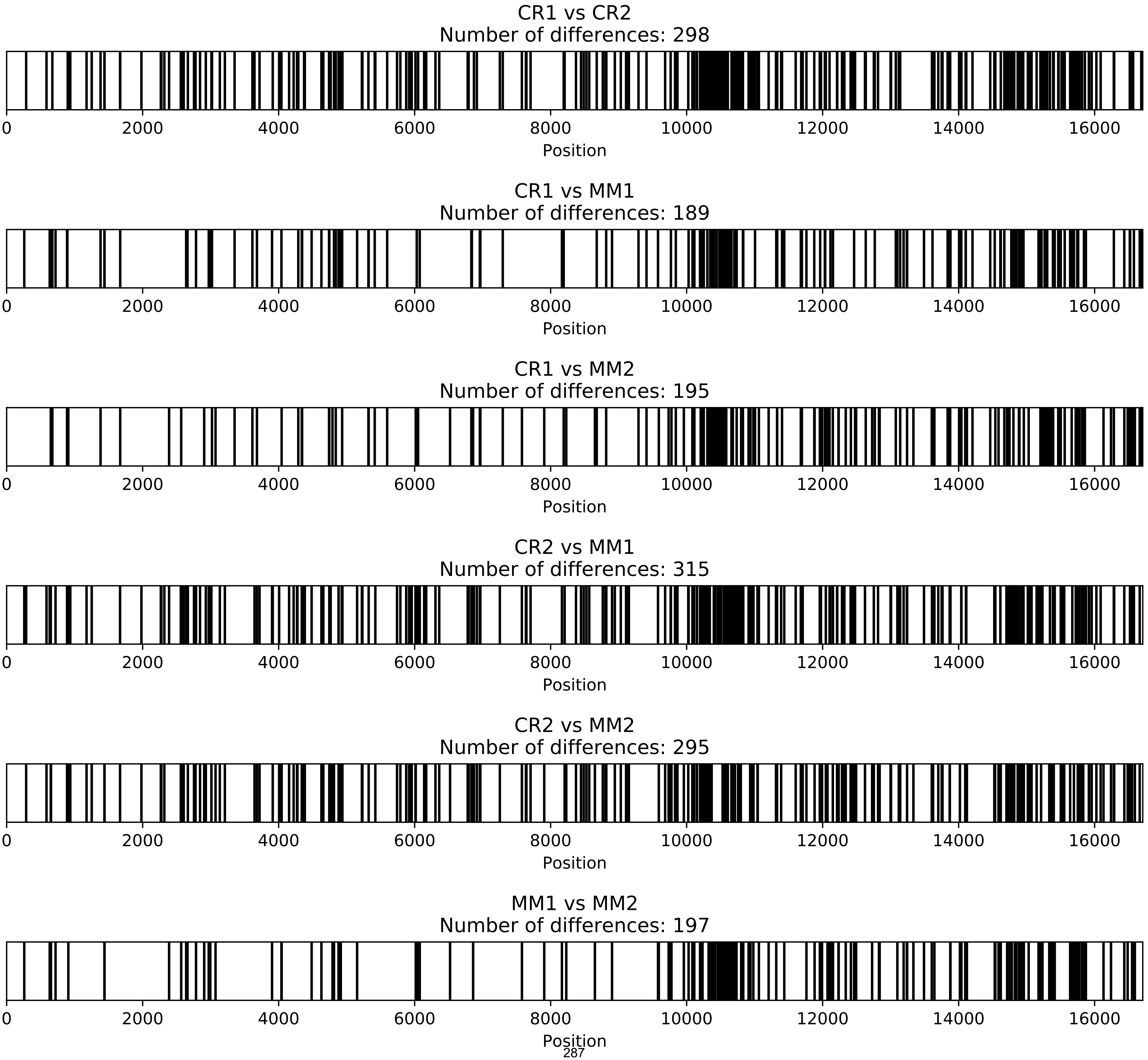
CR1 vs CR2
Number of differences: 298
Position
CR1 vs MM1
Number of differences: 189
Position
CR1 vs MM2
Number of differences: 195
Position
CR2 vs MM1
Number of differences: 315
Position
CR2 vs MM2
Number of differences: 295
Position
MM1 vs MM2
Number of differences: 197
Position
0
2000
4000
6000
8000
10000
12000
14000
16000

### Region: 781



CR1 vs CR2
Number of differences: 197

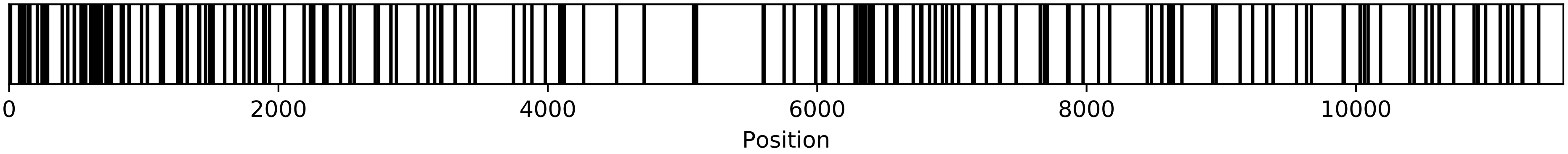


CR1 vs MM1
Number of differences: 194

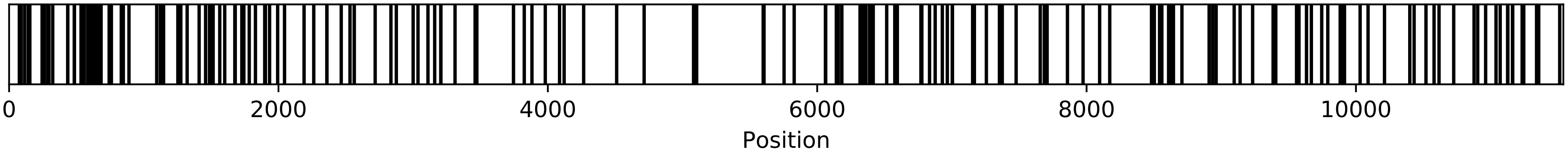


CR1 vs MM2
Number of differences: 199

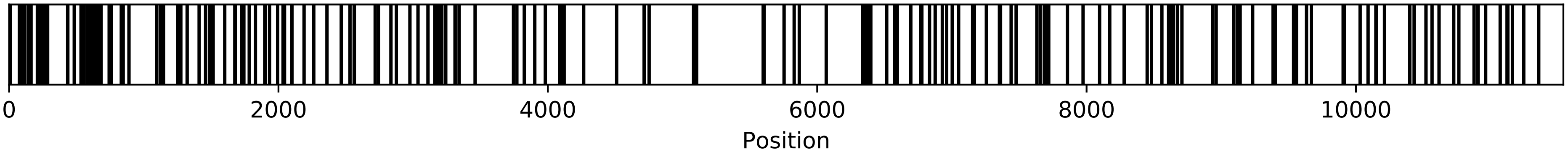


CR2 vs MM1
Number of differences: 91

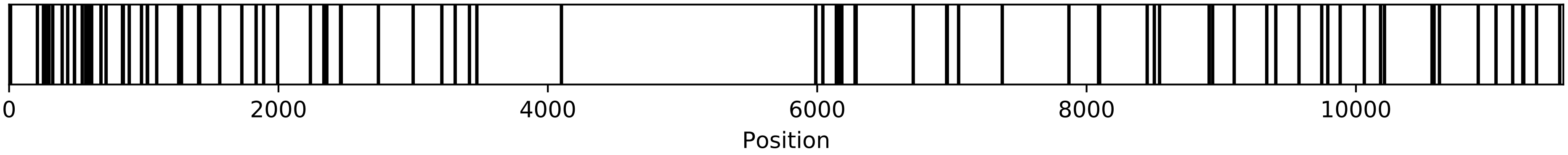


CR2 vs MM2
Number of differences: 89

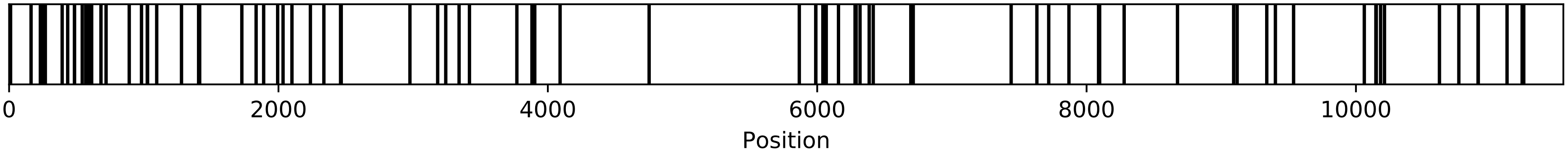


MM1 vs MM2
Number of differences: 83

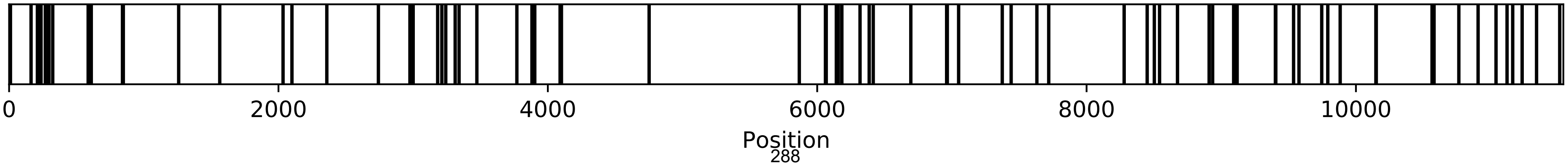

### Region: 783



CR1 vs CR2
Number of differences: 199

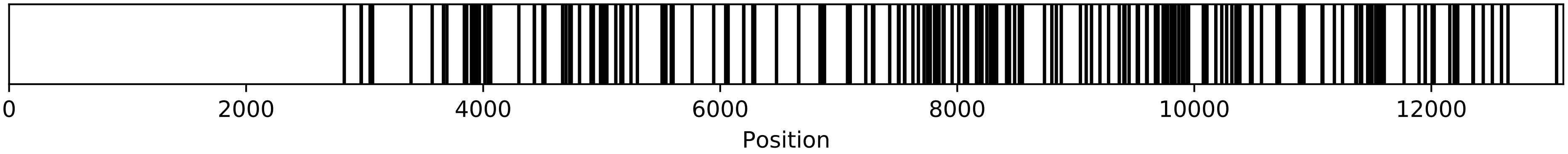


CR1 vs MM1
Number of differences: 108

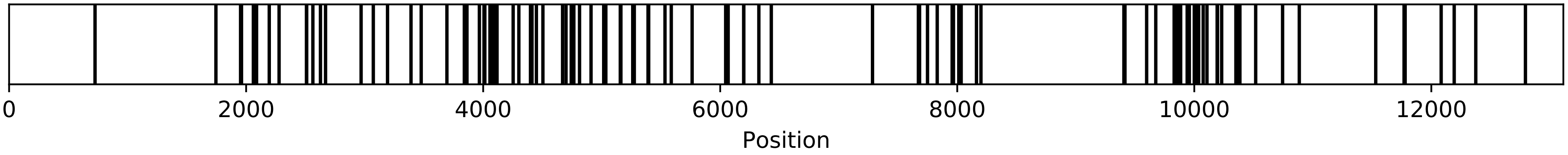


CR1 vs MM2
Number of differences: 73

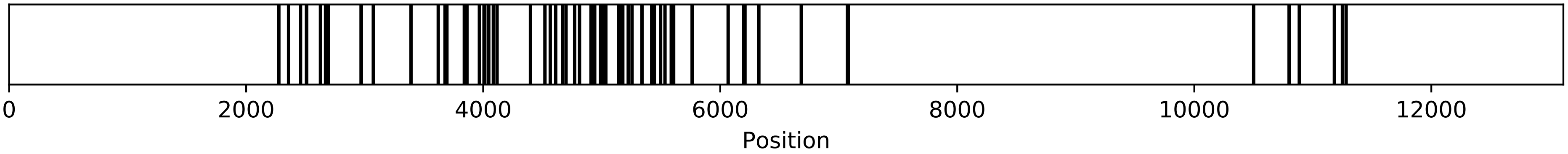


CR2 vs MM1
Number of differences: 213

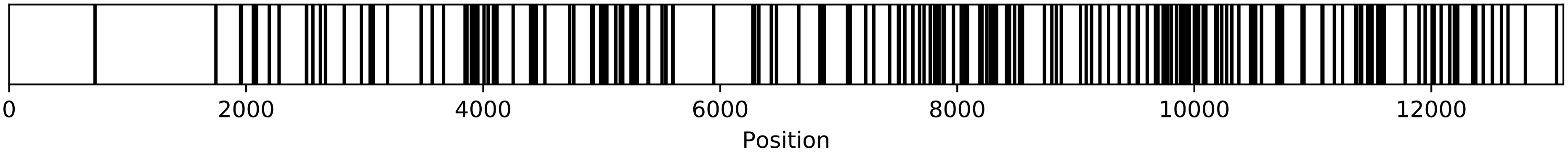


CR2 vs MM2
Number of differences: 214

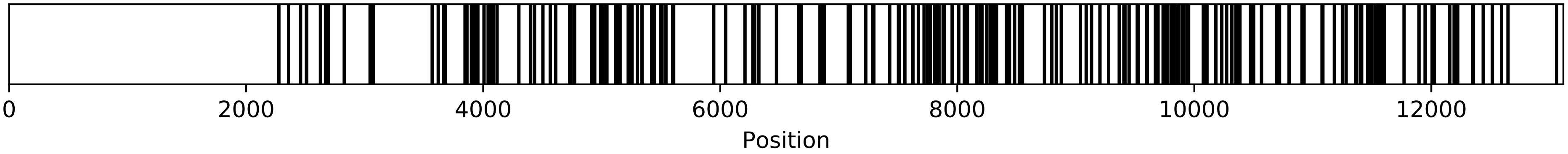


MM1 vs MM2
Number of differences: 111

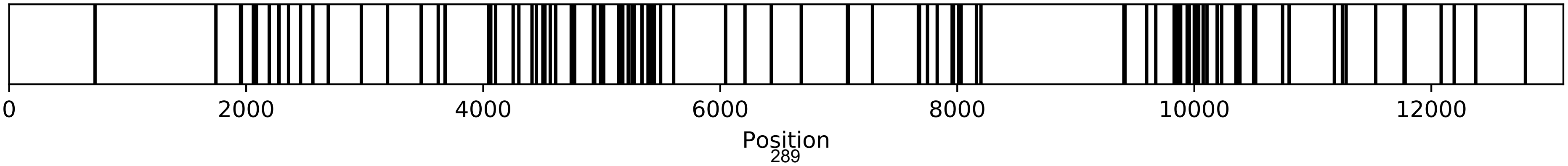

### Region: 785



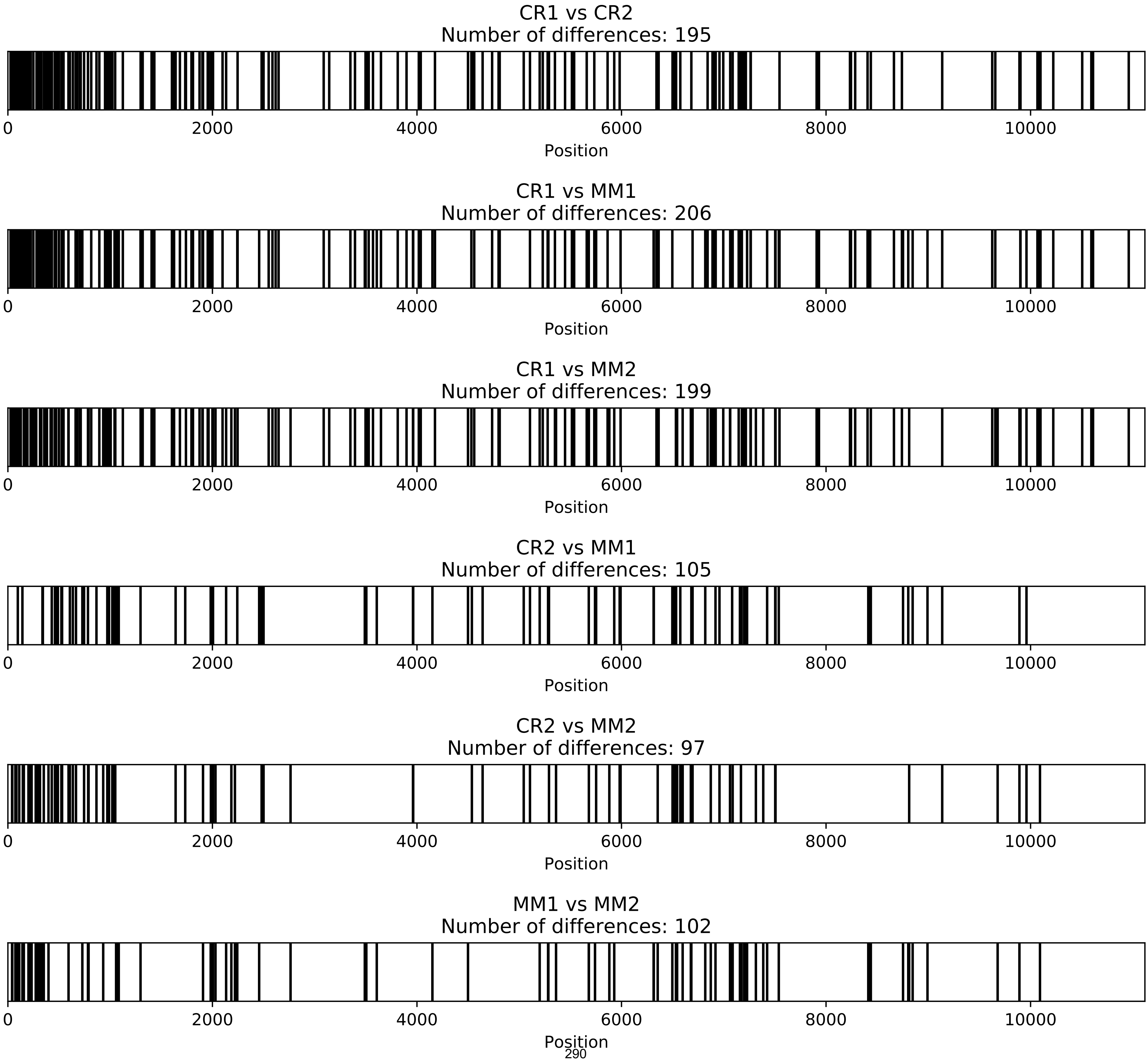
CR1 vs CR2
Number of differences: 195
CR1 vs MM1
Number of differences: 206
CR1 vs MM2
Number of differences: 199
CR2 vs MM1
Number of differences: 105
CR2 vs MM2
Number of differences: 97
MM1 vs MM2
Number of differences: 102
0
2000
4000
6000
8000
10000
Position

### Region: 786



## CR1 vs CR2
Number of differences: 286

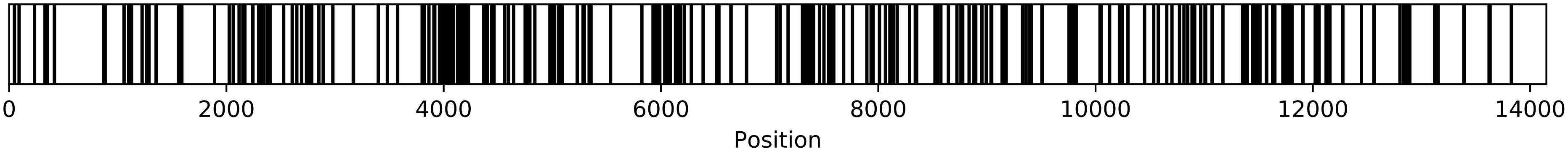


## CR1 vs MM1
Number of differences: 105

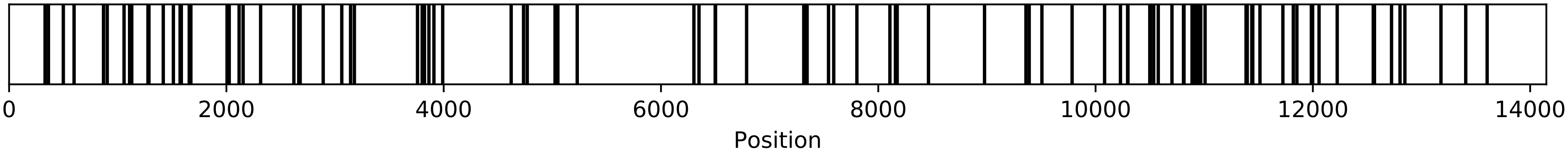


## CR1 vs MM2
Number of differences: 106

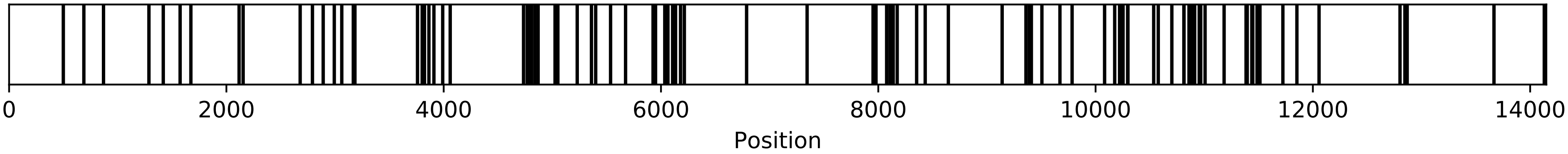


## CR2 vs MM1
Number of differences: 296

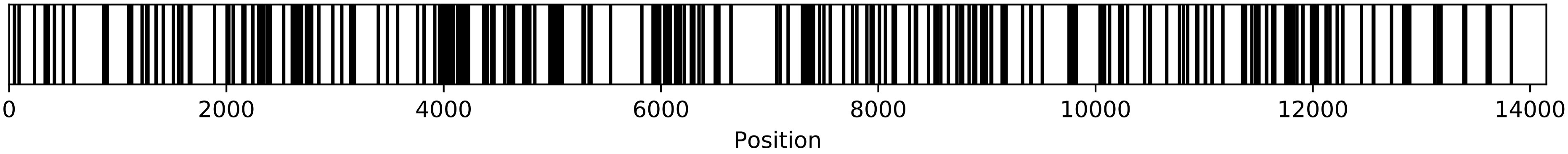


## CR2 vs MM2
Number of differences: 277

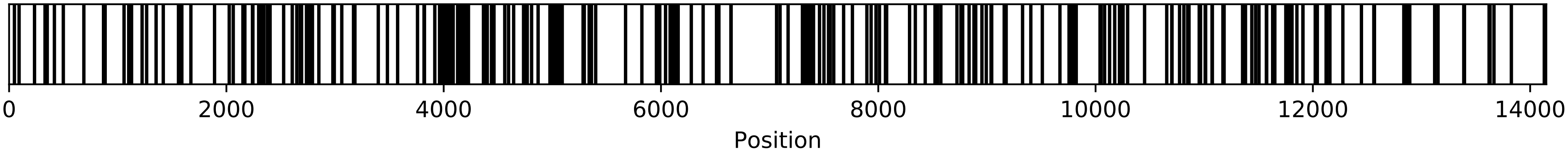


## MM1 vs MM2
Number of differences: 101

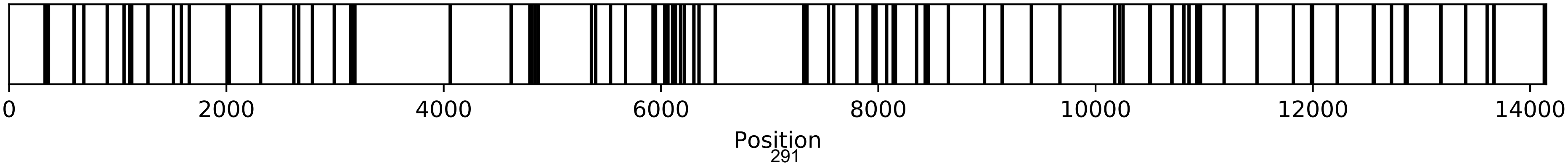

### Region: 788



CR1 vs CR2
Number of differences: 200

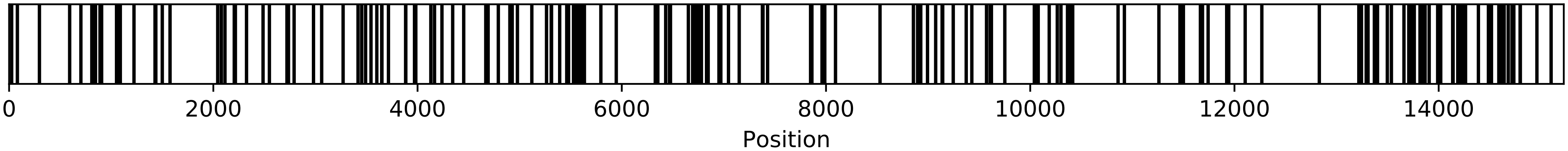


CR1 vs MM1
Number of differences: 207

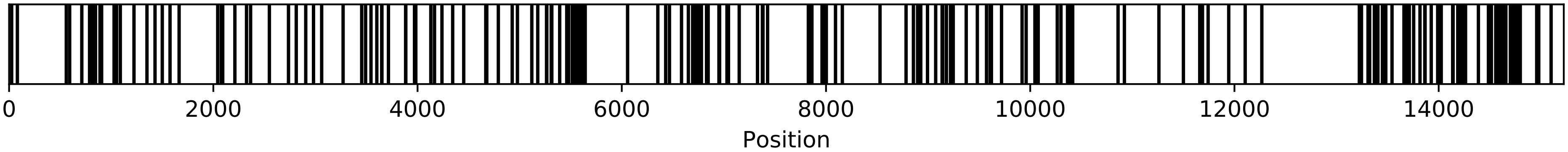


CR1 vs MM2
Number of differences: 207

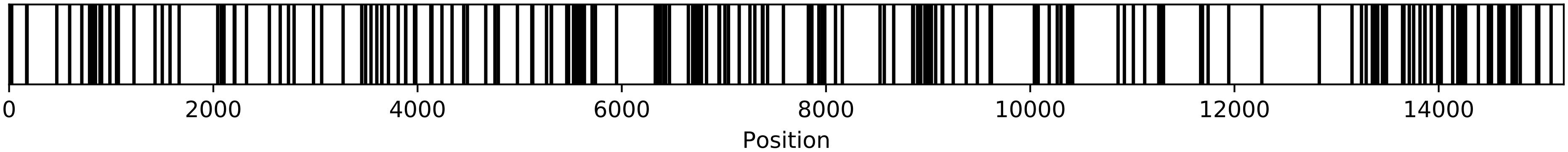


CR2 vs MM1
Number of differences: 110

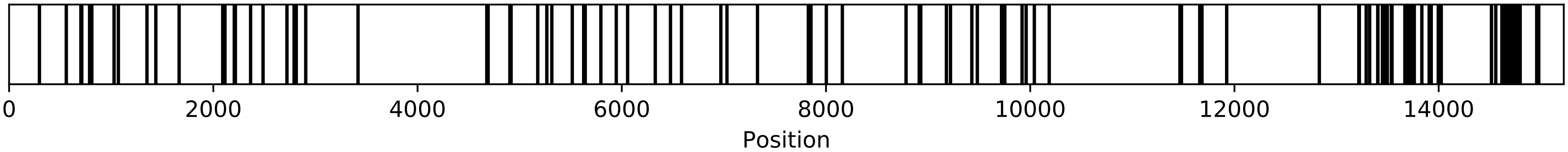


CR2 vs MM2
Number of differences: 116

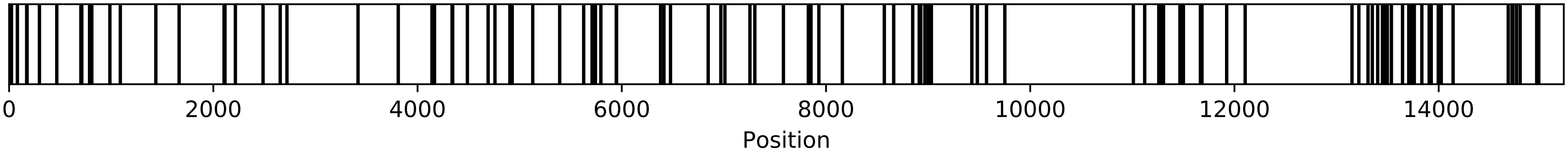


MM1 vs MM2
Number of differences: 104

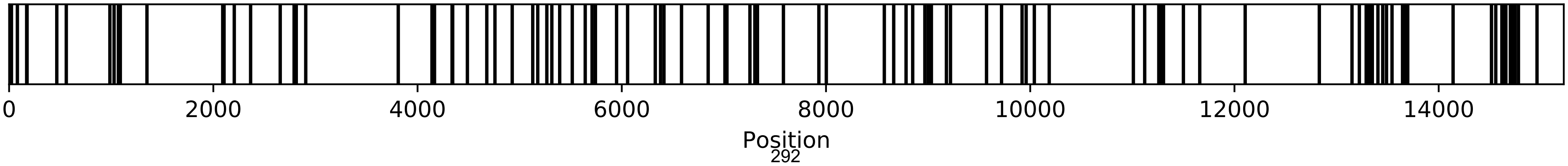

### Region: 792



CR1 vs CR2
Number of differences: 165

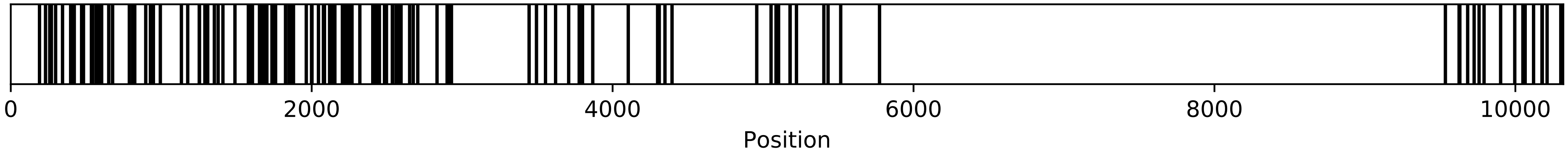


CR1 vs MM1
Number of differences: 110

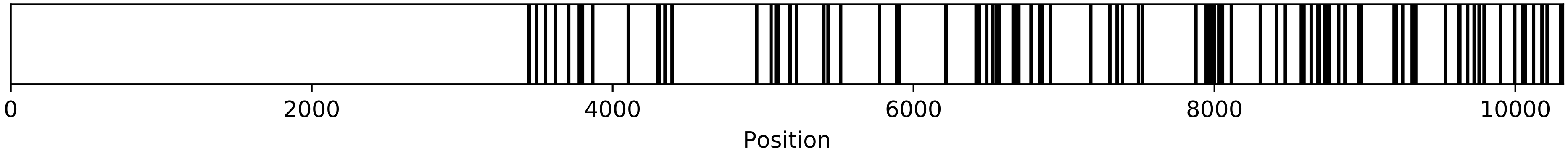


CR1 vs MM2
Number of differences: 150

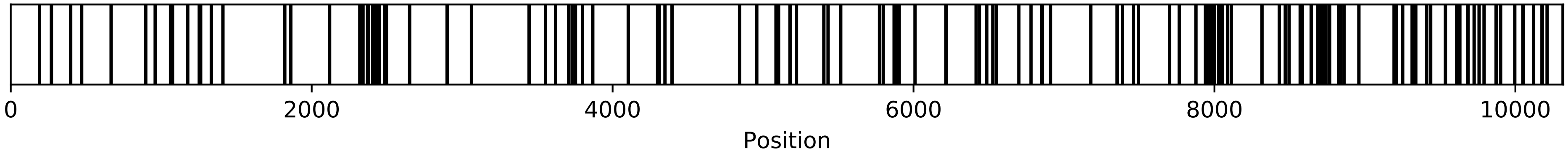


CR2 vs MM1
Number of differences: 189

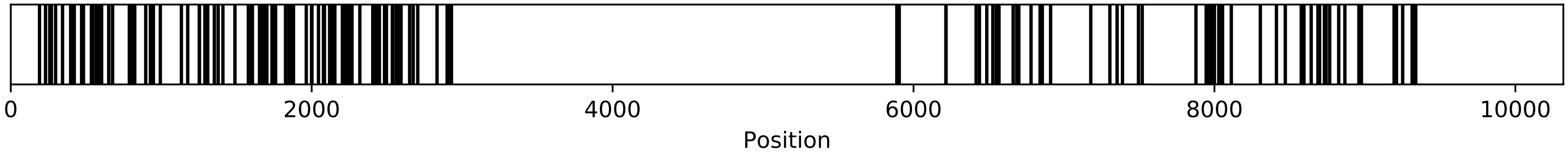


CR2 vs MM2
Number of differences: 197

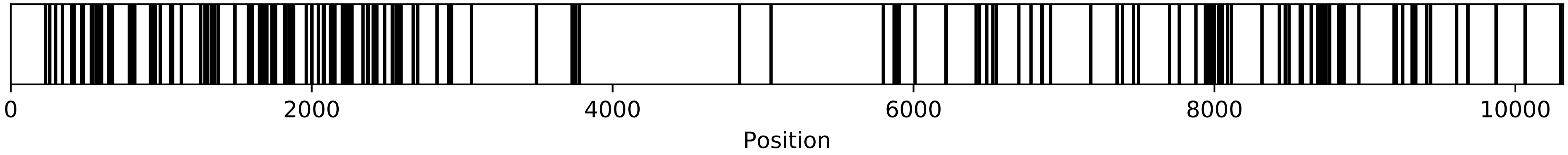


MM1 vs MM2
Number of differences: 93

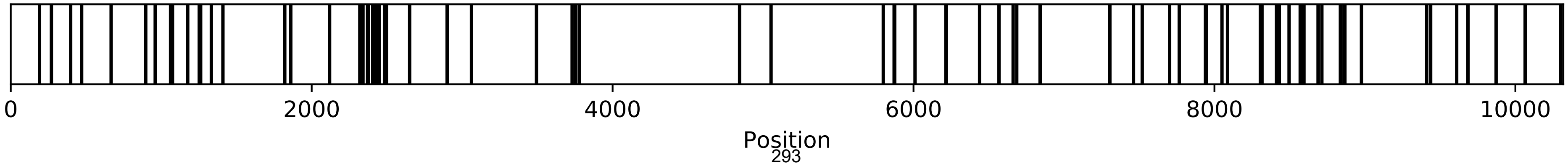

### Region: 795



## CR1 vs CR2
Number of differences: 215

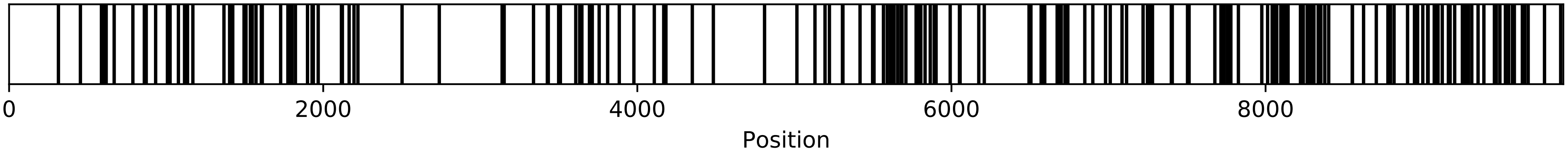


## CR1 vs MM1
Number of differences: 217

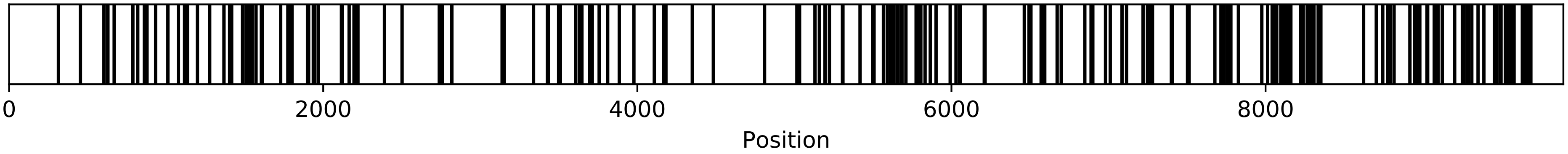


## CR1 vs MM2
Number of differences: 206

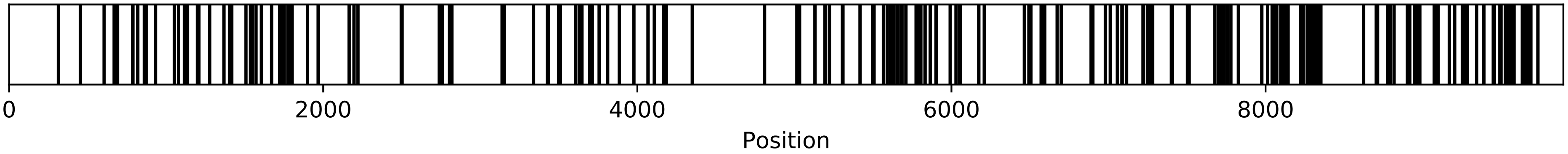


## CR2 vs MM1
Number of differences: 73

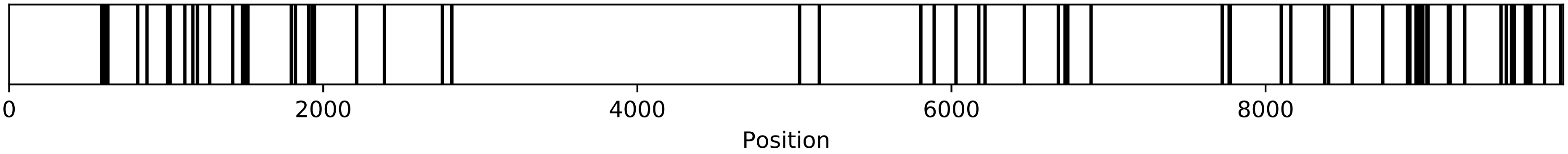


## CR2 vs MM2
Number of differences: 97

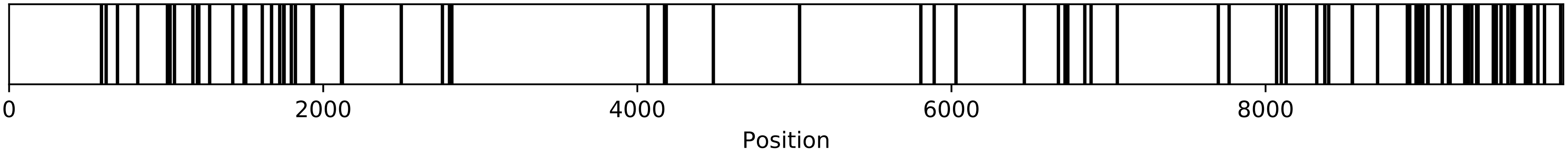


## MM1 vs MM2
Number of differences: 69

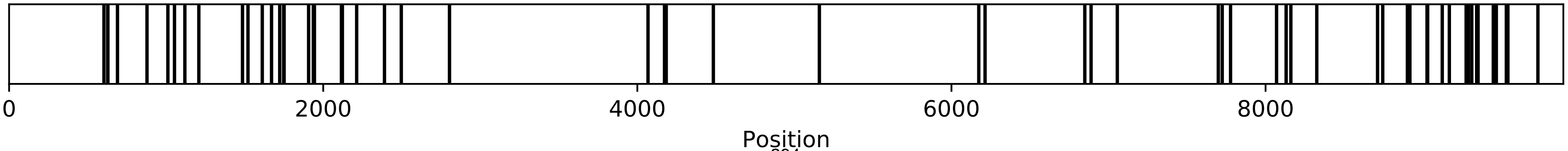

### Region: 796



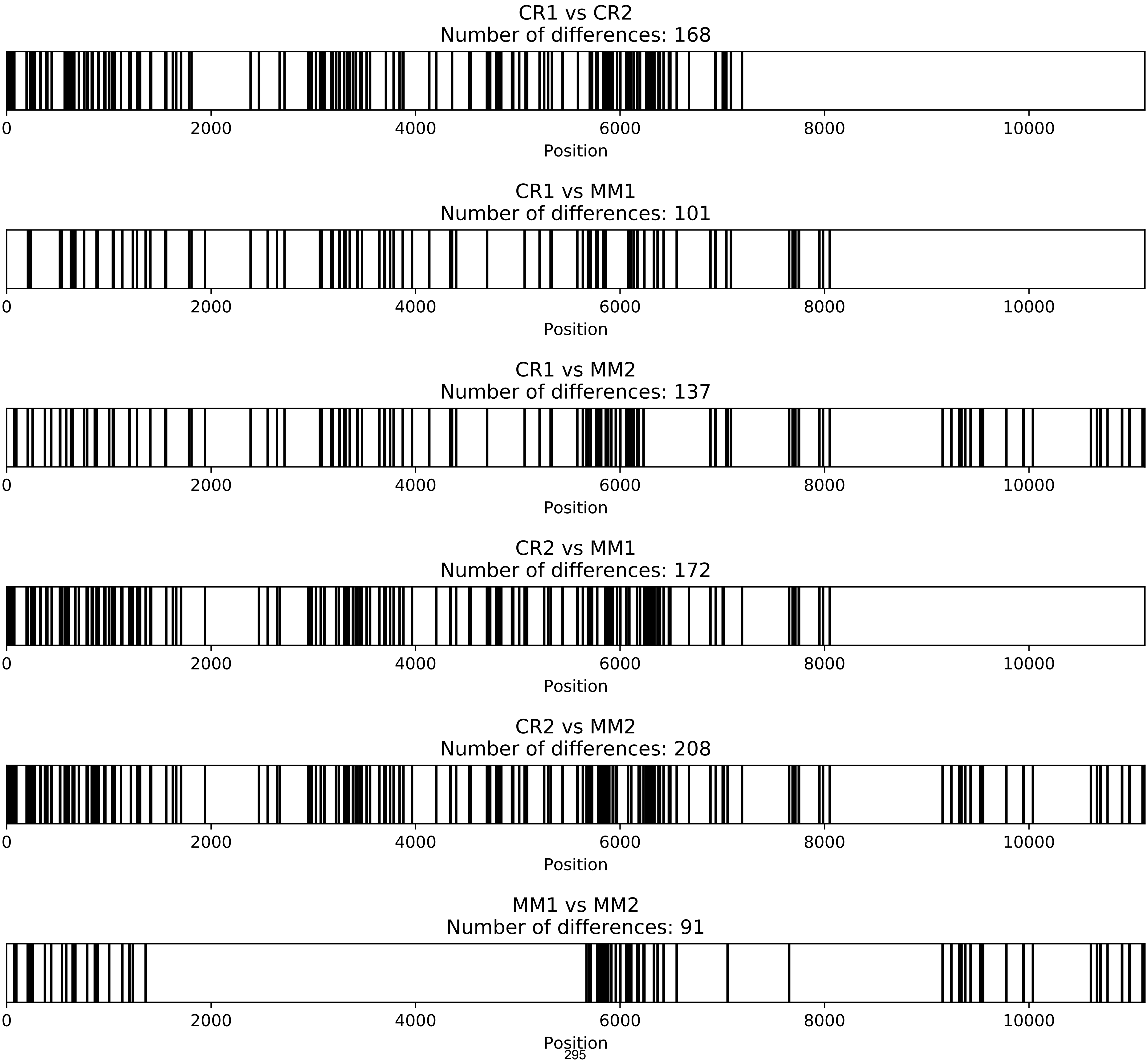
CR1 vs CR2
Number of differences: 168
CR1 vs MM1
Number of differences: 101
CR1 vs MM2
Number of differences: 137
CR2 vs MM1
Number of differences: 172
CR2 vs MM2
Number of differences: 208
MM1 vs MM2
Number of differences: 91
0
2000
4000
6000
8000
10000
Position

### Region: 801



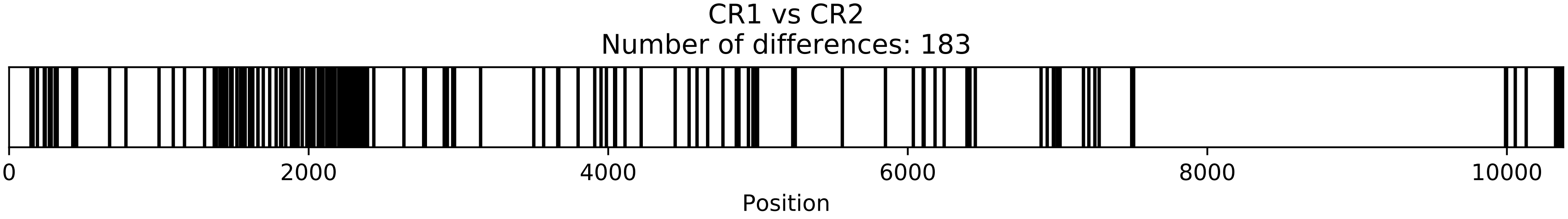
CR1 vs CR2
Number of differences: 183
0
2000
4000
6000
8000
10000
Position

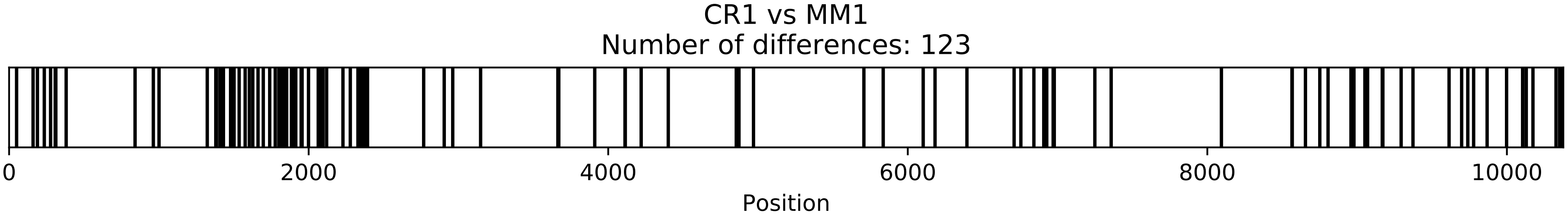
CR1 vs MM1
Number of differences: 123
0
2000
4000
6000
8000
10000
Position

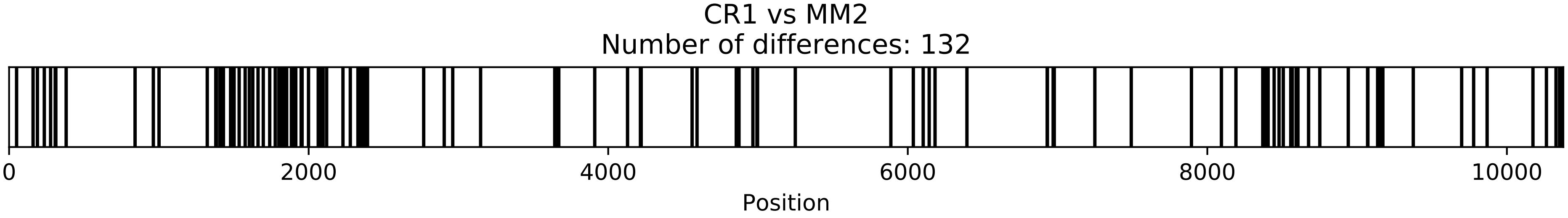
CR1 vs MM2
Number of differences: 132
0
2000
4000
6000
8000
10000
Position

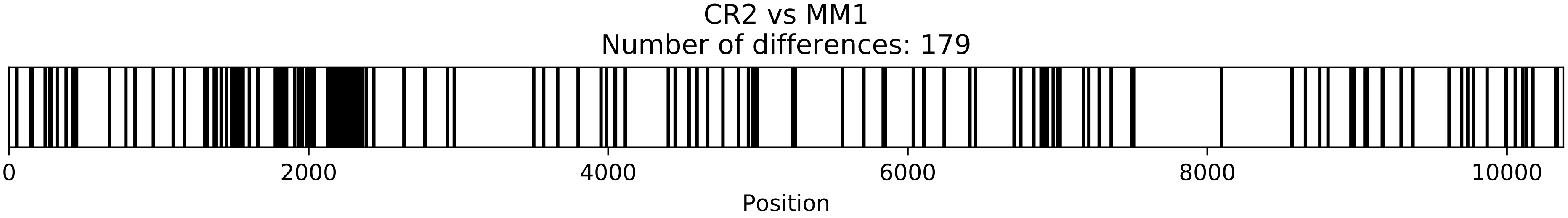
CR2 vs MM1
Number of differences: 179
0
2000
4000
6000
8000
10000
Position

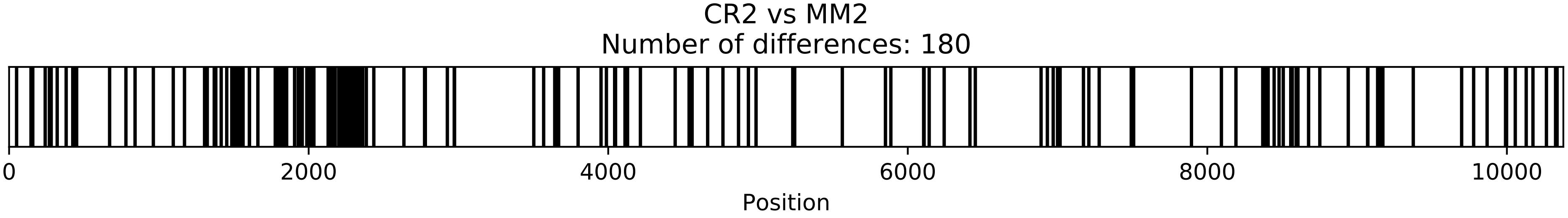
CR2 vs MM2
Number of differences: 180
0
2000
4000
6000
8000
10000
Position

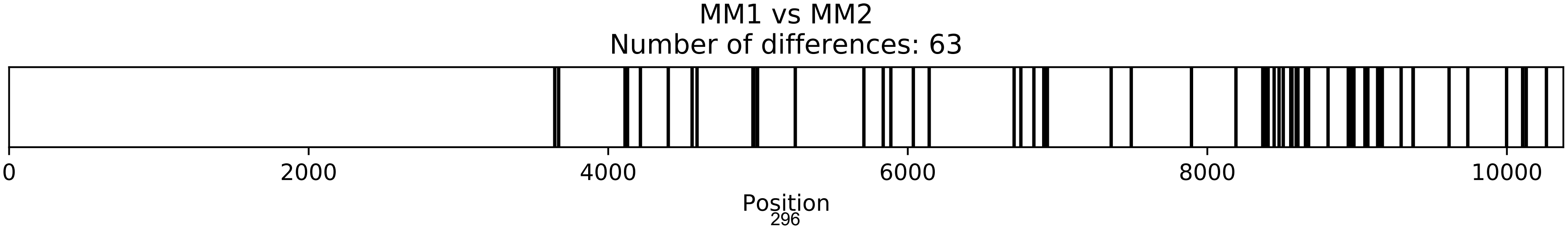
MM1 vs MM2
Number of differences: 63
0
2000
4000
6000
8000
10000
Position

### Region: 804



CR1 vs CR2
Number of differences: 251

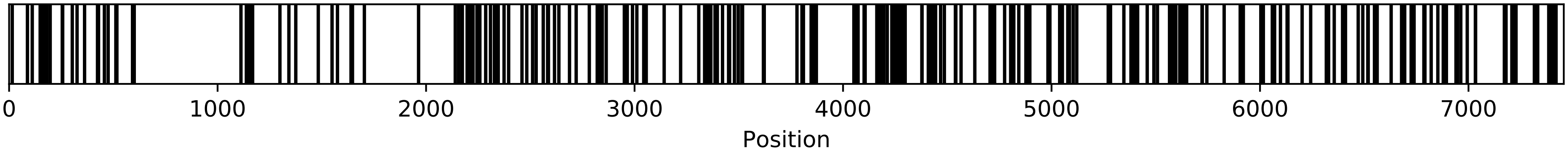


CR1 vs MM1
Number of differences: 249

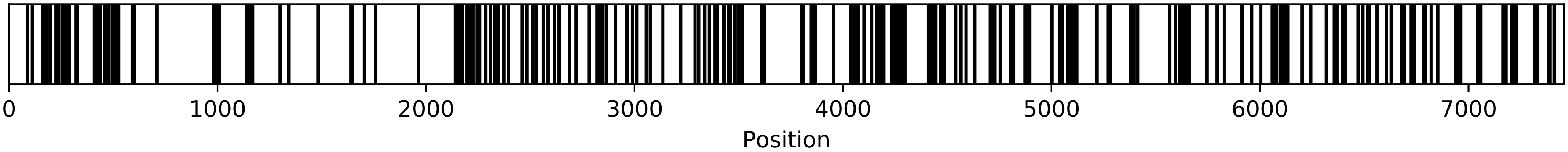


CR1 vs MM2
Number of differences: 249

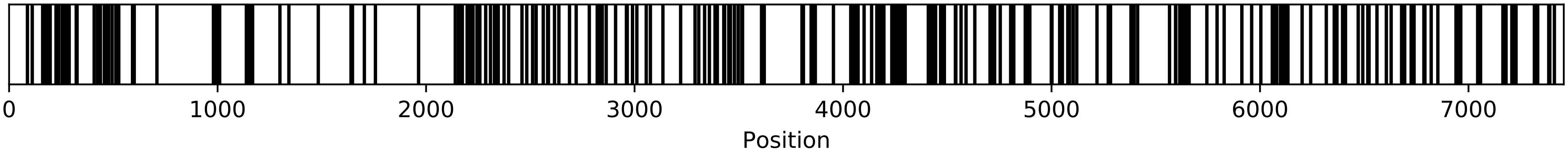


CR2 vs MM1
Number of differences: 122

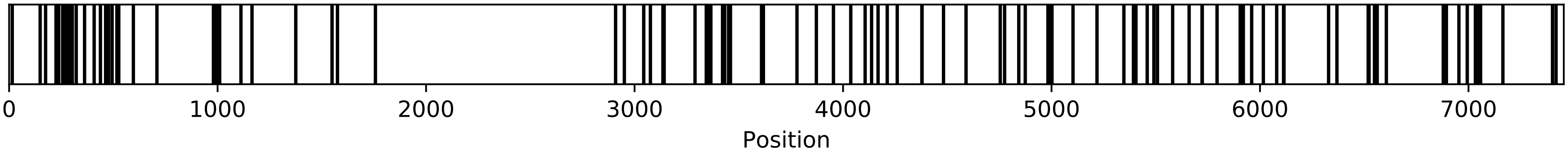


CR2 vs MM2
Number of differences: 122

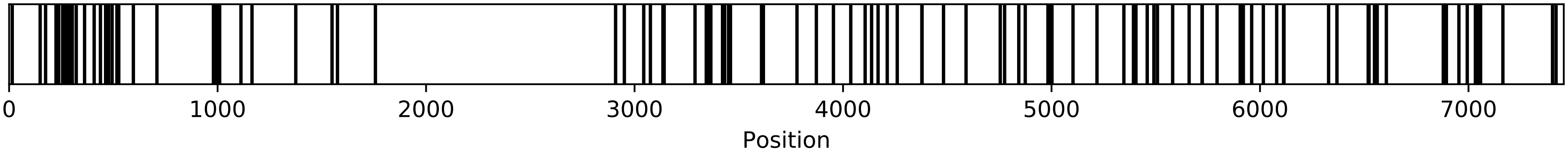


MM1 vs MM2
Number of differences: 0

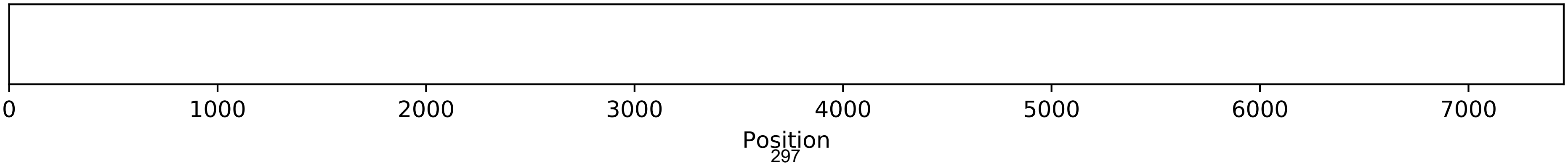

### Region: 807



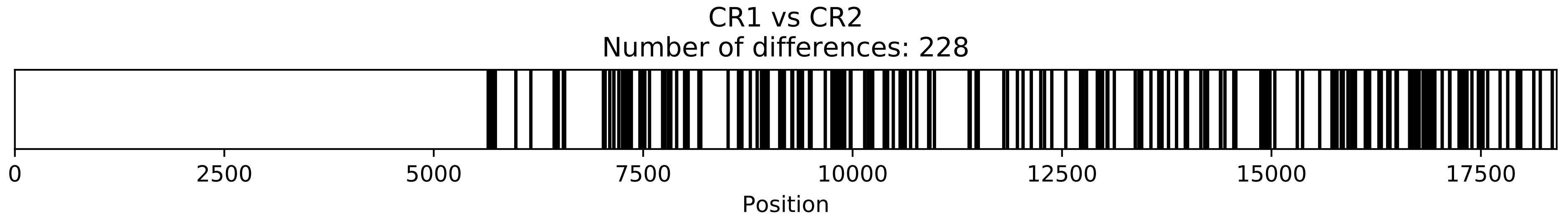
CR1 vs CR2
Number of differences: 228
0
2500
5000
7500
10000
12500
15000
17500
Position

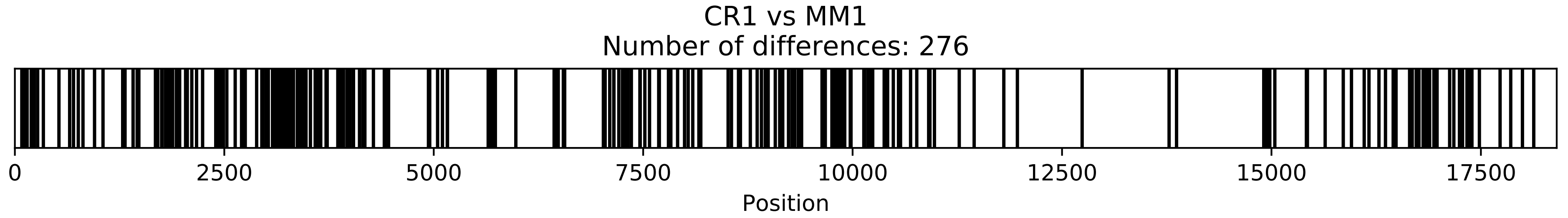
CR1 vs MM1
Number of differences: 276
0
2500
5000
7500
10000
12500
15000
17500
Position

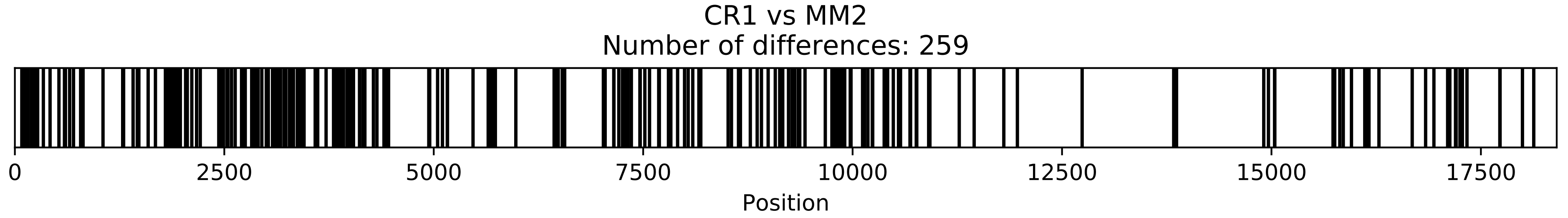
CR1 vs MM2
Number of differences: 259
0
2500
5000
7500
10000
12500
15000
17500
Position

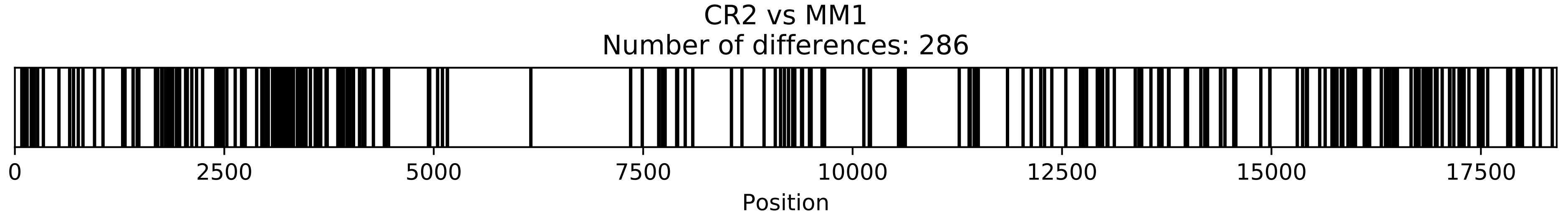
CR2 vs MM1
Number of differences: 286
0
2500
5000
7500
10000
12500
15000
17500
Position

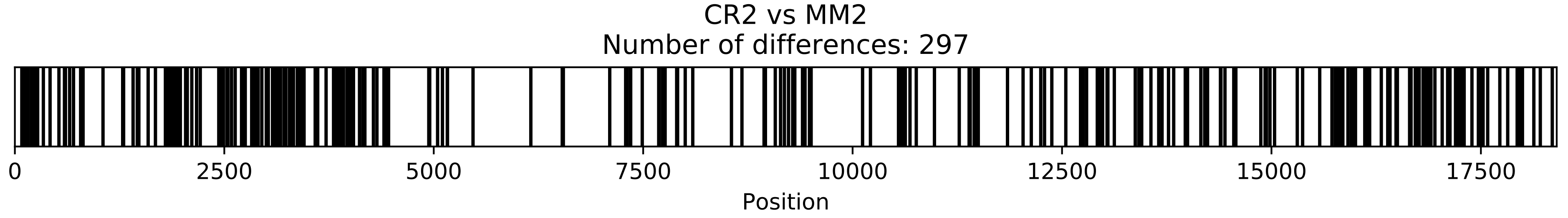
CR2 vs MM2
Number of differences: 297
0
2500
5000
7500
10000
12500
15000
17500
Position

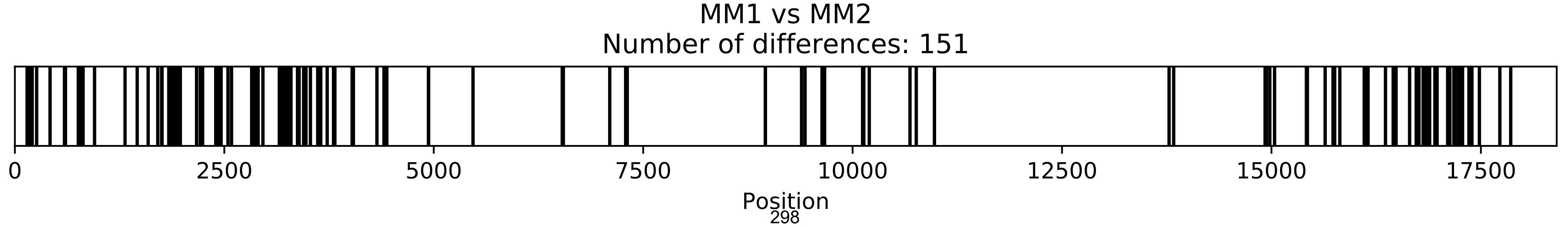
MM1 vs MM2
Number of differences: 151
0
2500
5000
7500
10000
12500
15000
17500
Position

### Region: 808



## CR1 vs CR2
Number of differences: 289

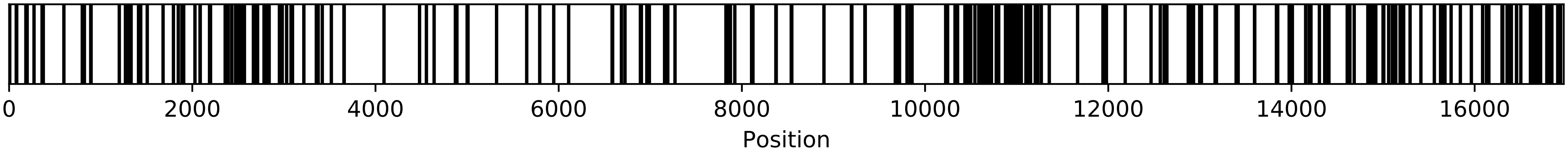


## CR1 vs MM1
Number of differences: 119

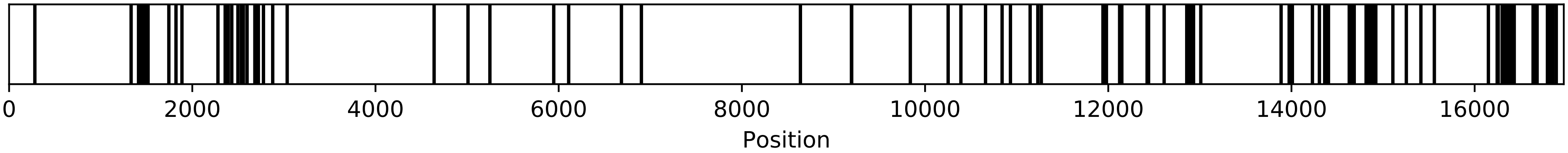


## CR1 vs MM2
Number of differences: 113

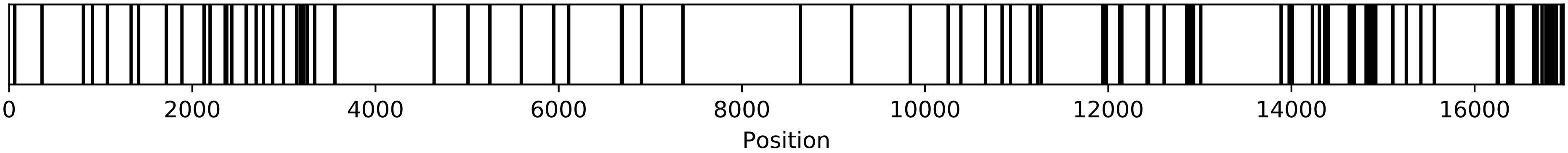


## CR2 vs MM1
Number of differences: 296

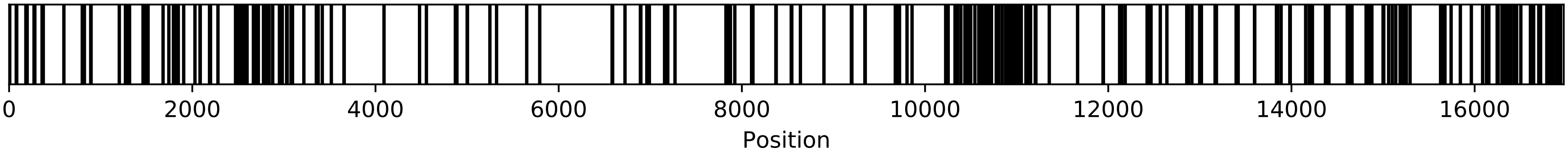


## CR2 vs MM2
Number of differences: 296

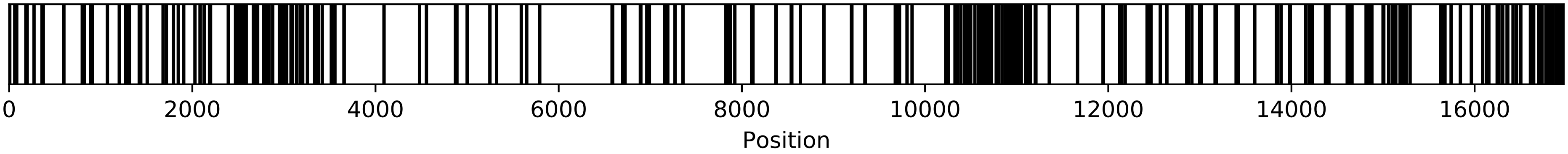


## MM1 vs MM2
Number of differences: 62

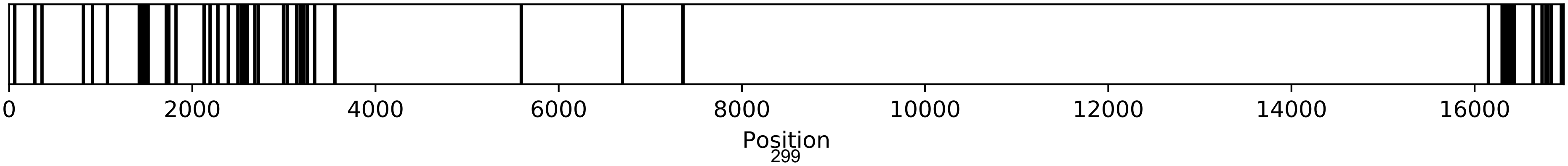

### Region: 813



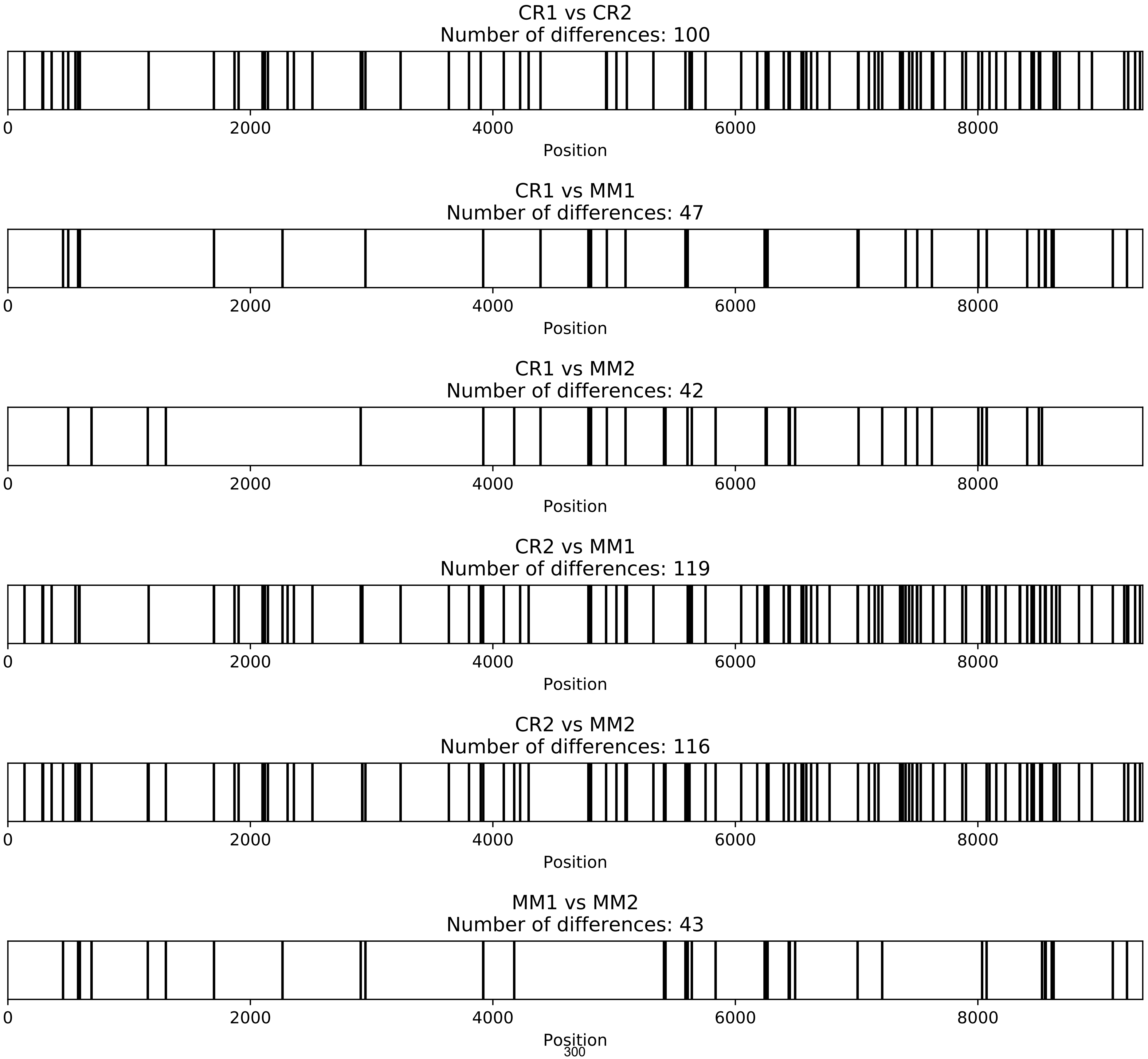
CR1 vs CR2
Number of differences: 100
CR1 vs MM1
Number of differences: 47
CR1 vs MM2
Number of differences: 42
CR2 vs MM1
Number of differences: 119
CR2 vs MM2
Number of differences: 116
MM1 vs MM2
Number of differences: 43
0
2000
4000
6000
8000
Position

### Region: 814



CR1 vs CR2
Number of differences: 216

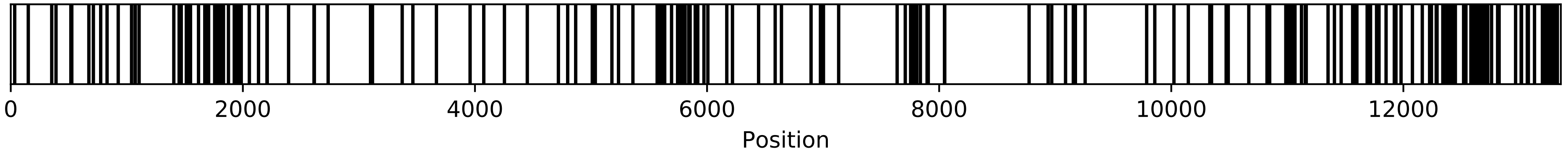


CR1 vs MM1
Number of differences: 106

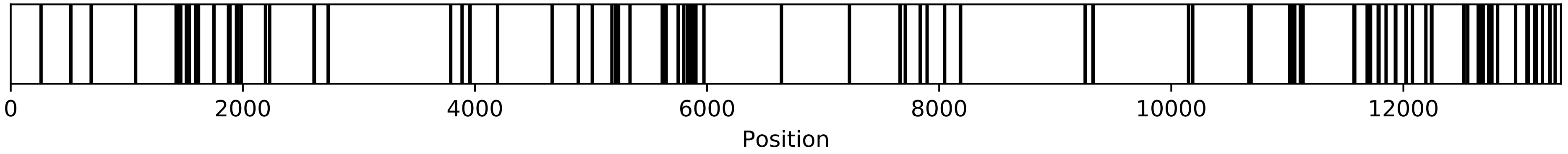


CR1 vs MM2
Number of differences: 102

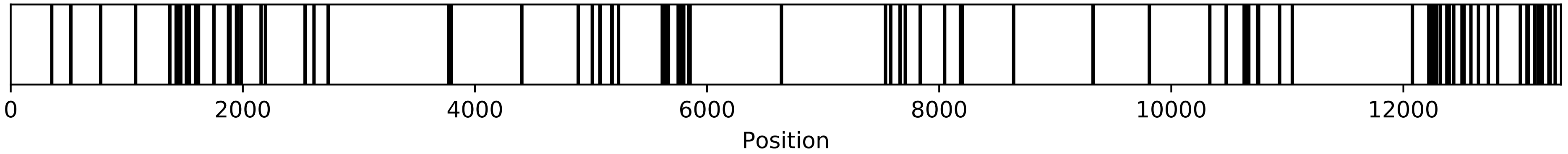


CR2 vs MM1
Number of differences: 227

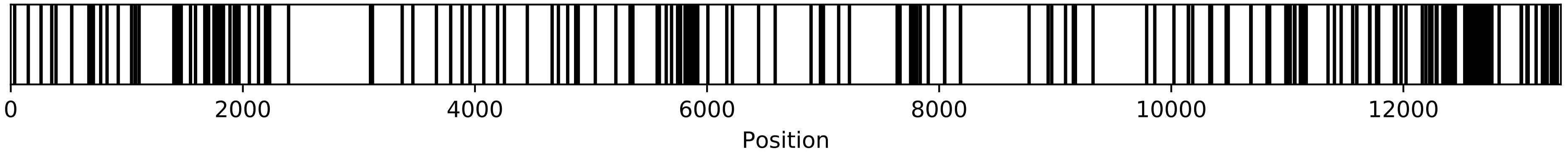


CR2 vs MM2
Number of differences: 237

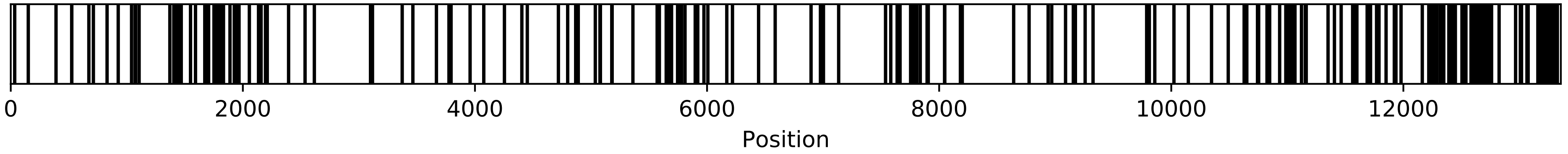


MM1 vs MM2
Number of differences: 106

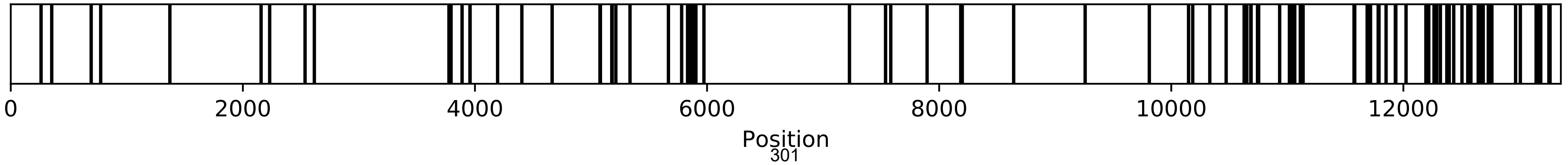

### Region: 815

CR1 vs CR2
Number of differences: 55

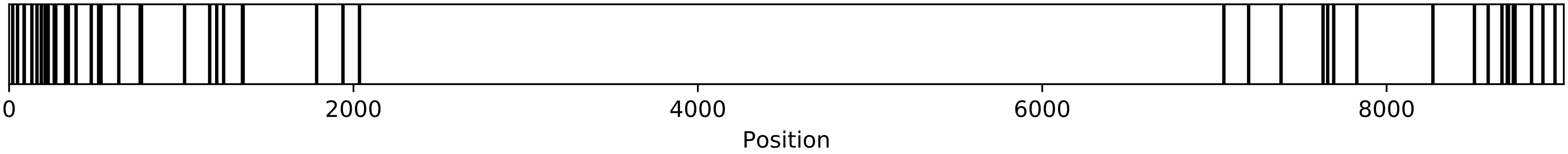


CR1 vs MM1
Number of differences: 183

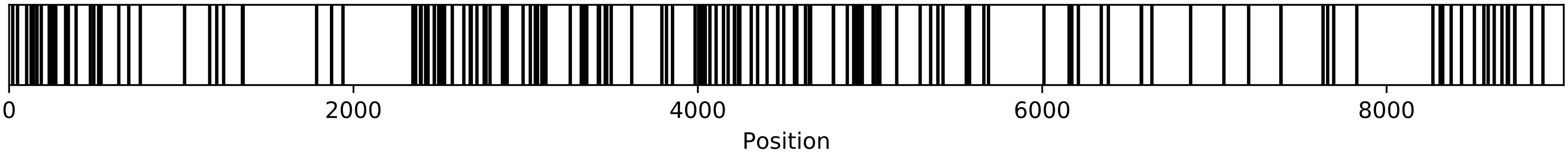


CR1 vs MM2
Number of differences: 182

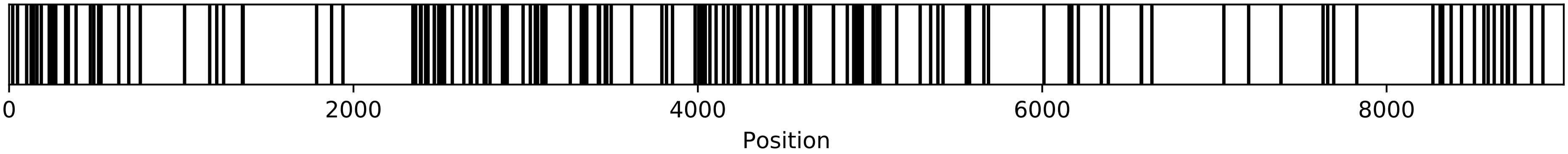


CR2 vs MM1
Number of differences: 151

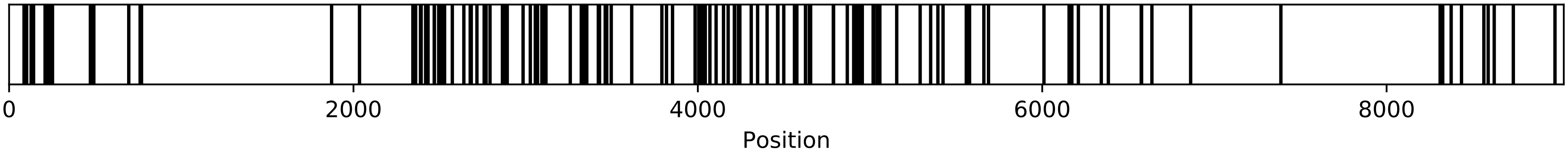


CR2 vs MM2
Number of differences: 150

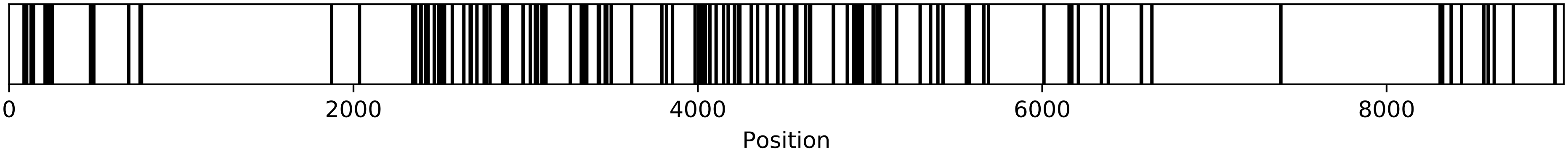


MM1 vs MM2
Number of differences: 1

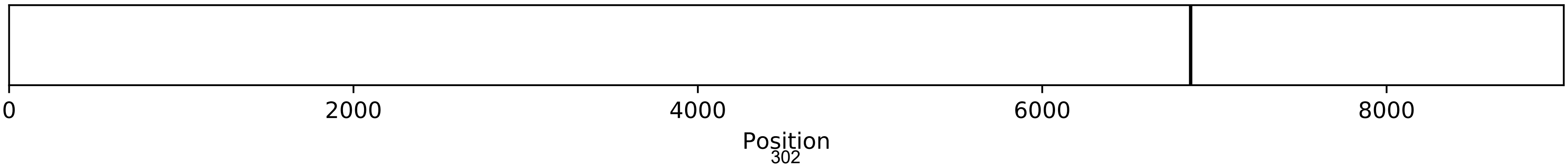

### Region: 821



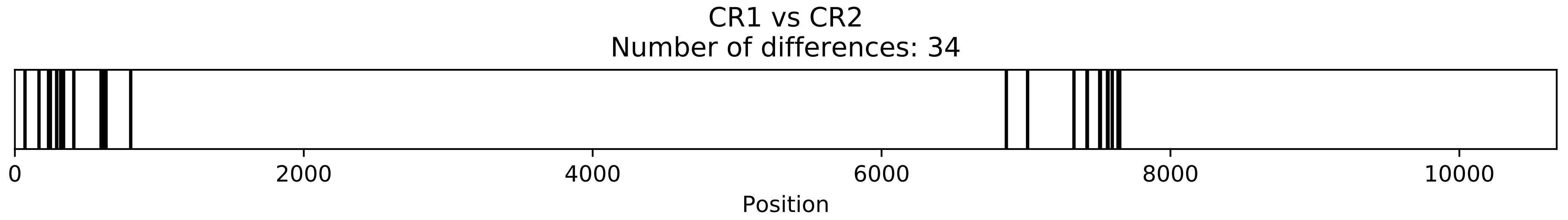
CR1 vs CR2
Number of differences: 34
0
2000
4000
6000
8000
10000
Position

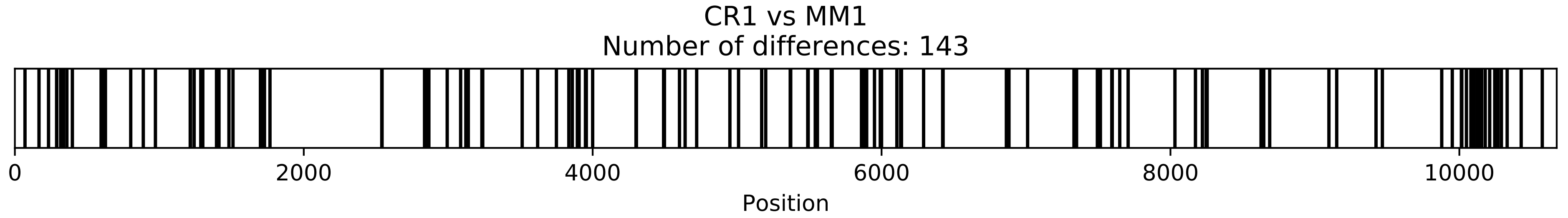
CR1 vs MM1
Number of differences: 143
0
2000
4000
6000
8000
10000
Position

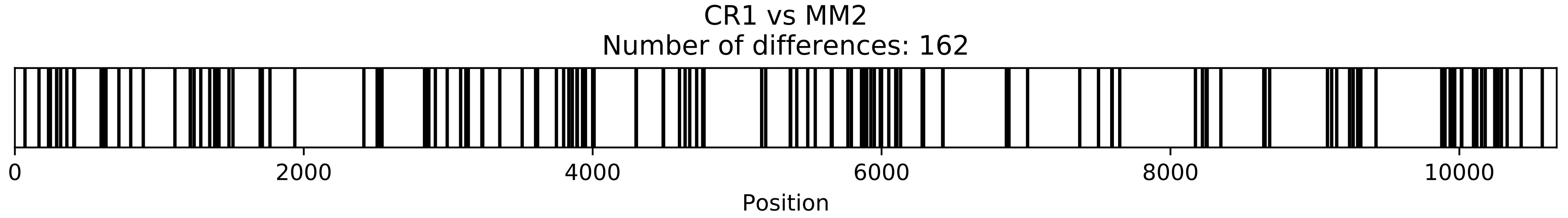
CR1 vs MM2
Number of differences: 162
0
2000
4000
6000
8000
10000
Position

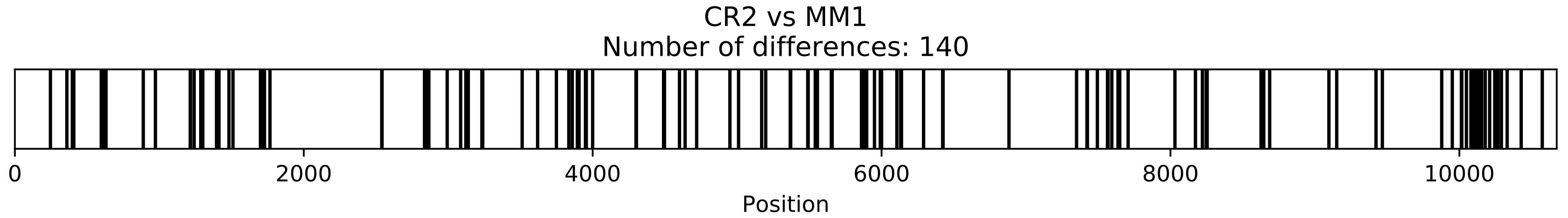
CR2 vs MM1
Number of differences: 140
0
2000
4000
6000
8000
10000
Position

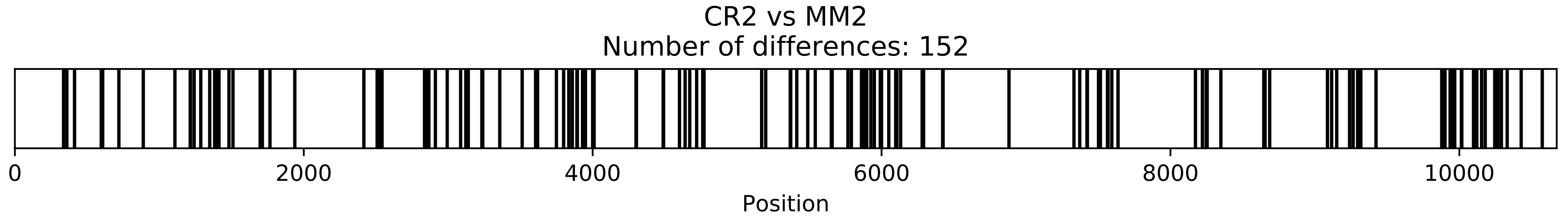
CR2 vs MM2
Number of differences: 152
0
2000
4000
6000
8000
10000
Position

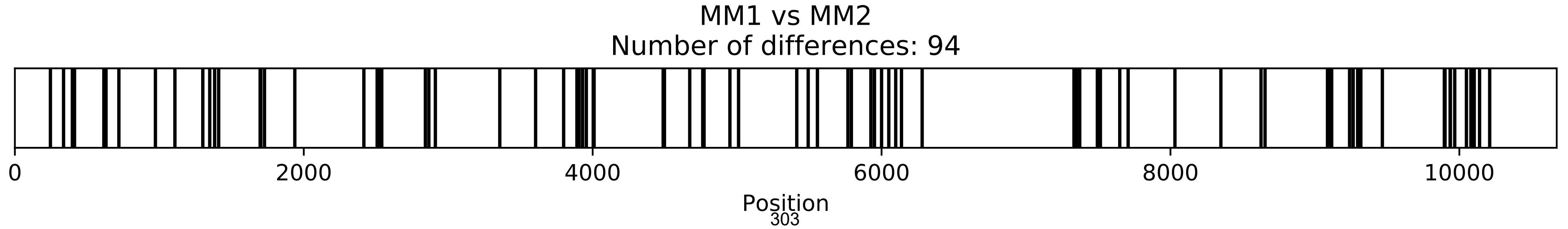
MM1 vs MM2
Number of differences: 94
0
2000
4000
6000
8000
10000
Position

### Region: 822



## CR1 vs CR2
Number of differences: 231

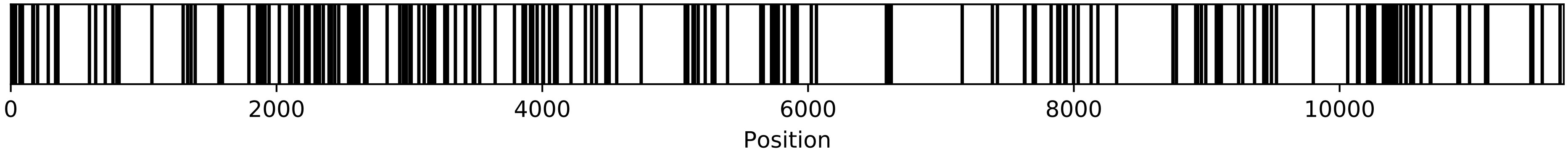


## CR1 vs MM1
Number of differences: 154

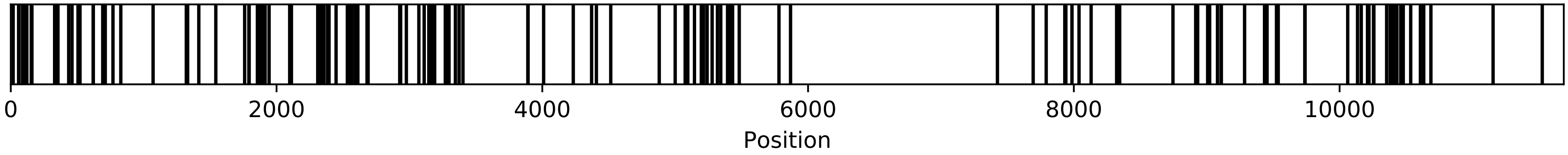


## CR1 vs MM2
Number of differences: 121

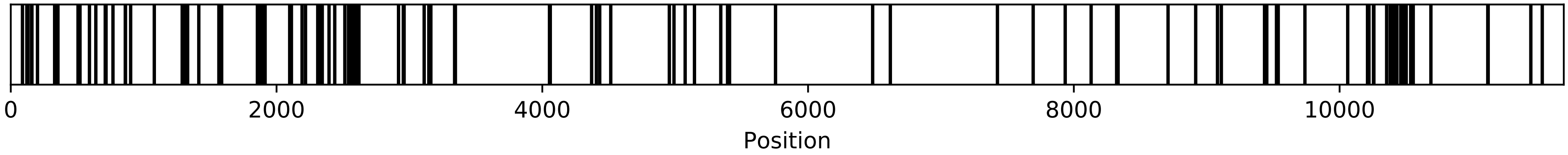


## CR2 vs MM1
Number of differences: 256

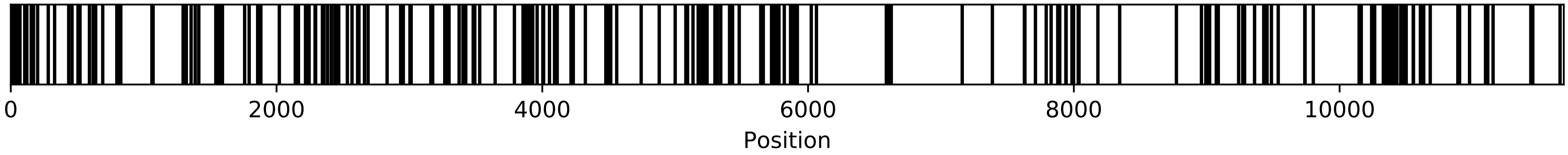


## CR2 vs MM2
Number of differences: 225

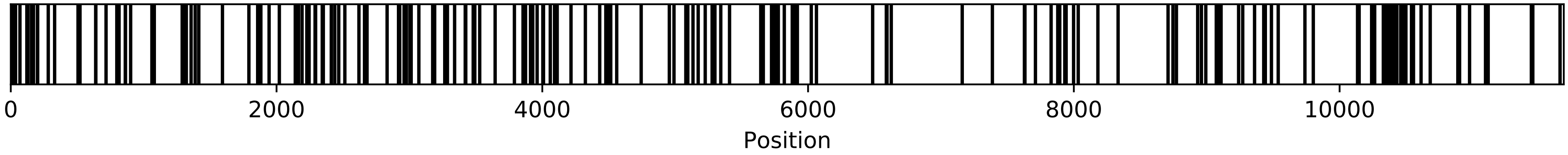


## MM1 vs MM2
Number of differences: 131

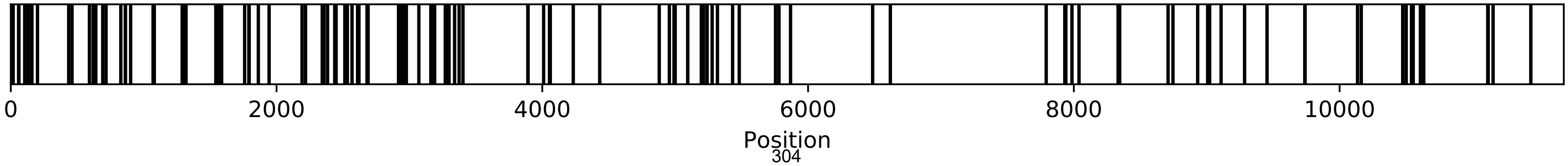

### Region: 824



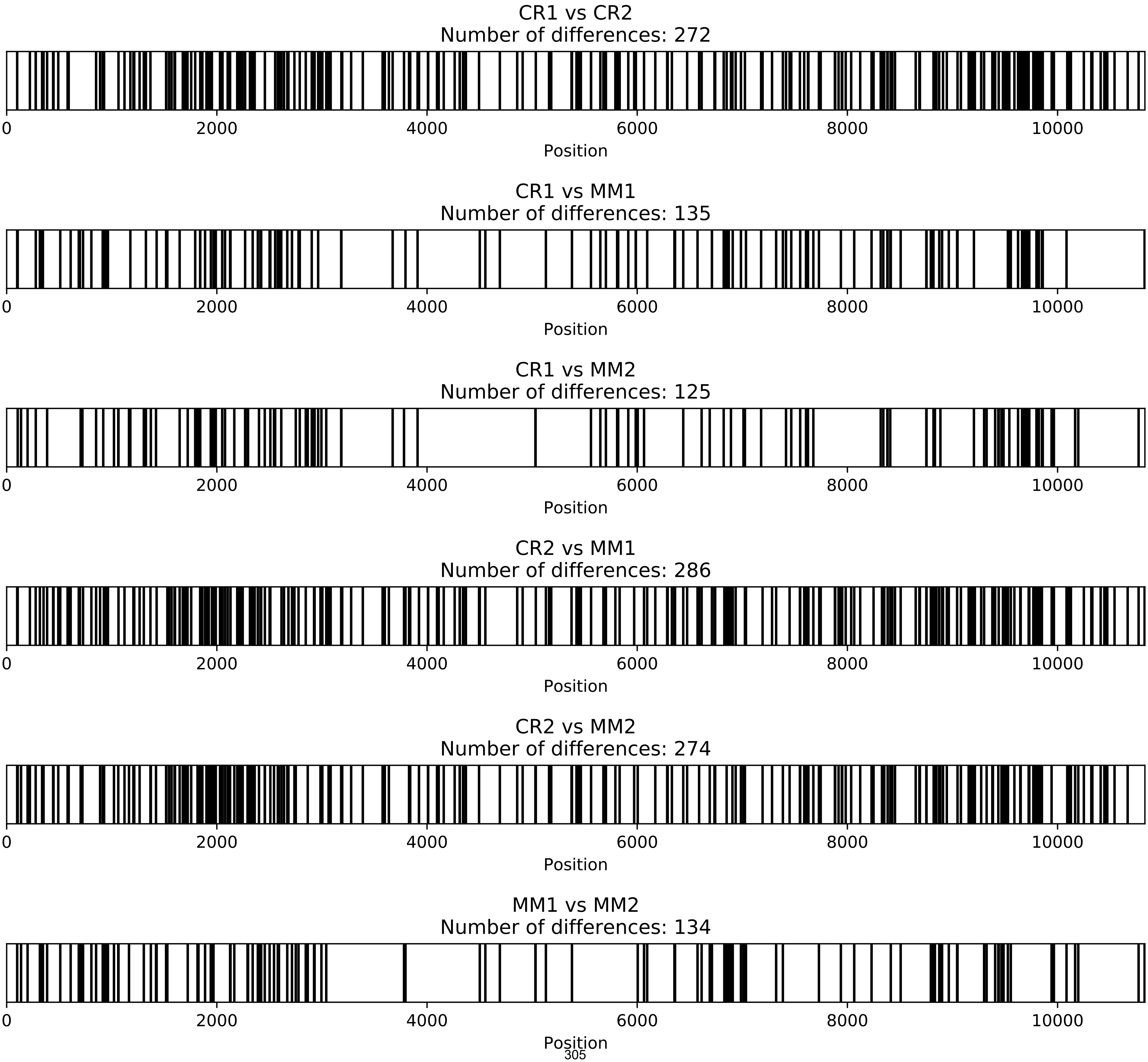
CR1 vs CR2
Number of differences: 272
0
2000
4000
6000
8000
10000
Position
CR1 vs MM1
Number of differences: 135
0
2000
4000
6000
8000
10000
Position
CR1 vs MM2
Number of differences: 125
0
2000
4000
6000
8000
10000
Position
CR2 vs MM1
Number of differences: 286
0
2000
4000
6000
8000
10000
Position
CR2 vs MM2
Number of differences: 274
0
2000
4000
6000
8000
10000
Position
MM1 vs MM2
Number of differences: 134
0
2000
4000
6000
8000
10000
Position

### Region: 826



CR1 vs CR2
Number of differences: 325

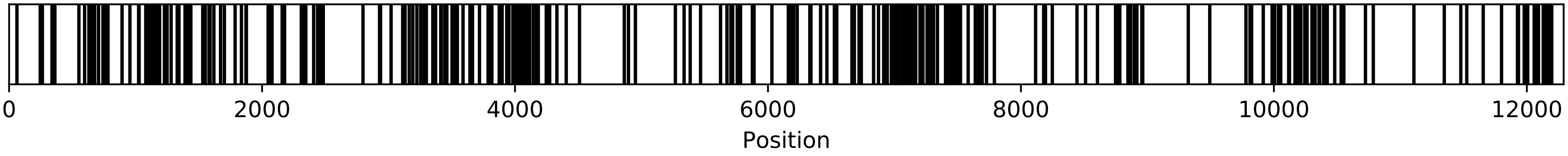


CR1 vs MM1
Number of differences: 172

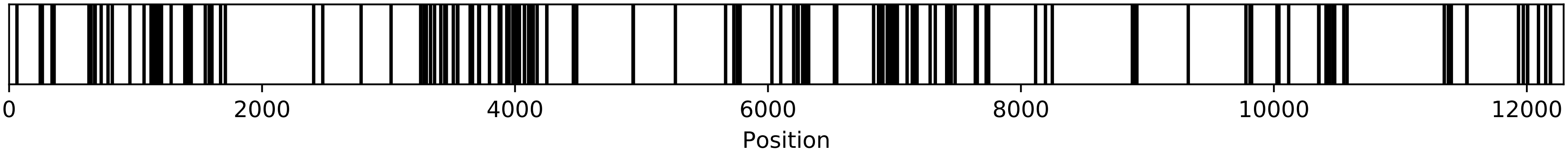


CR1 vs MM2
Number of differences: 193

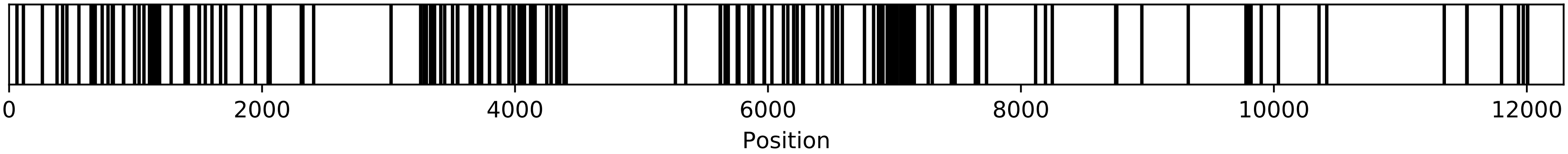


CR2 vs MM1
Number of differences: 312

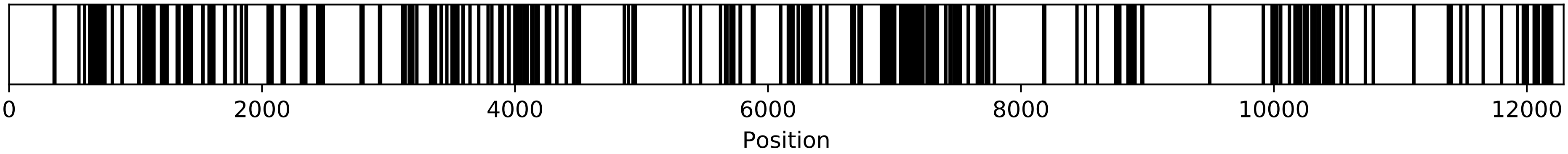


CR2 vs MM2
Number of differences: 324

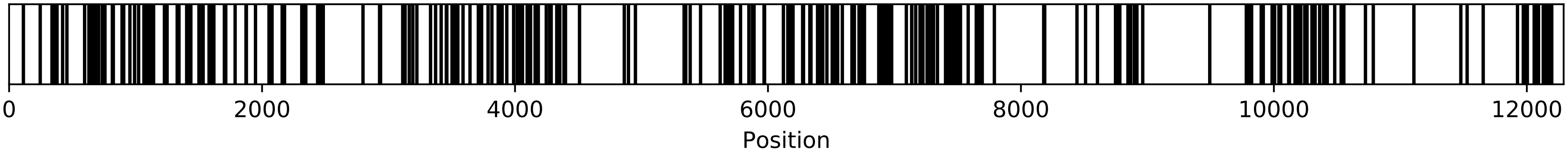


MM1 vs MM2
Number of differences: 191

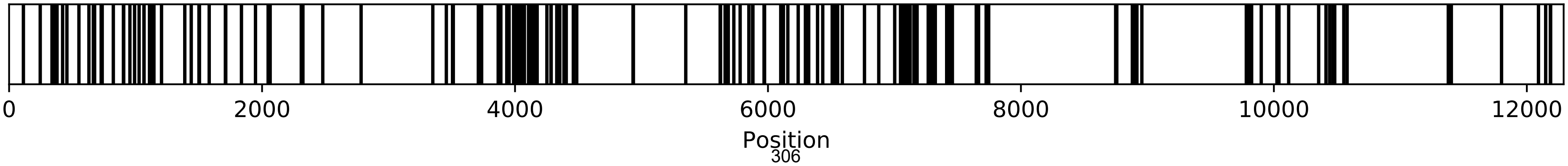

### Region: 828



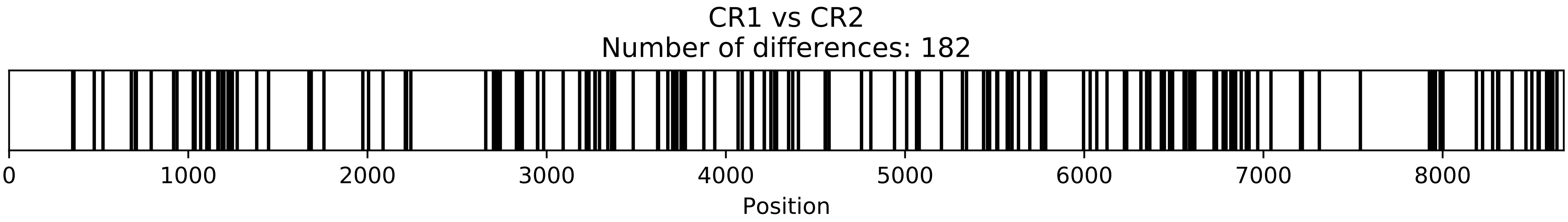
CR1 vs CR2
Number of differences: 182
0
1000
2000
3000
4000
5000
6000
7000
8000
Position

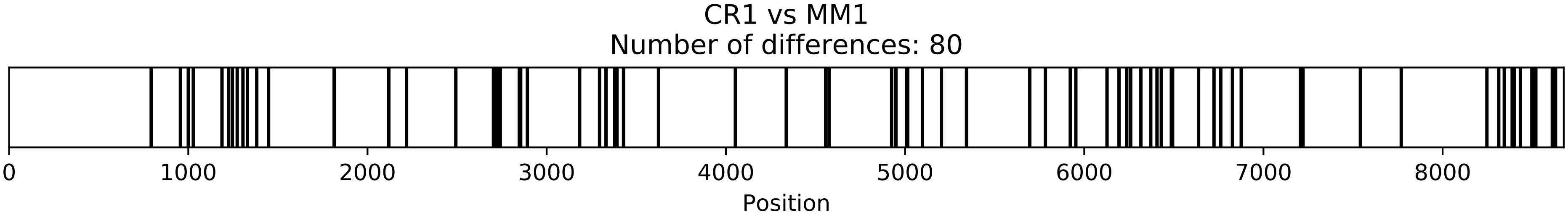
CR1 vs MM1
Number of differences: 80
0
1000
2000
3000
4000
5000
6000
7000
8000
Position

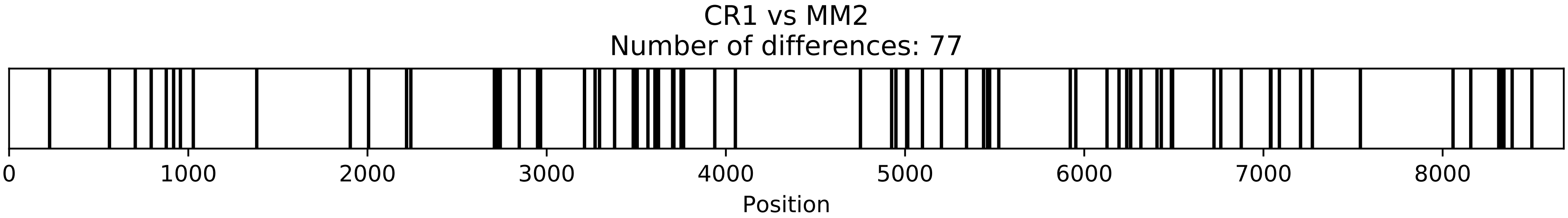
CR1 vs MM2
Number of differences: 77
0
1000
2000
3000
4000
5000
6000
7000
8000
Position

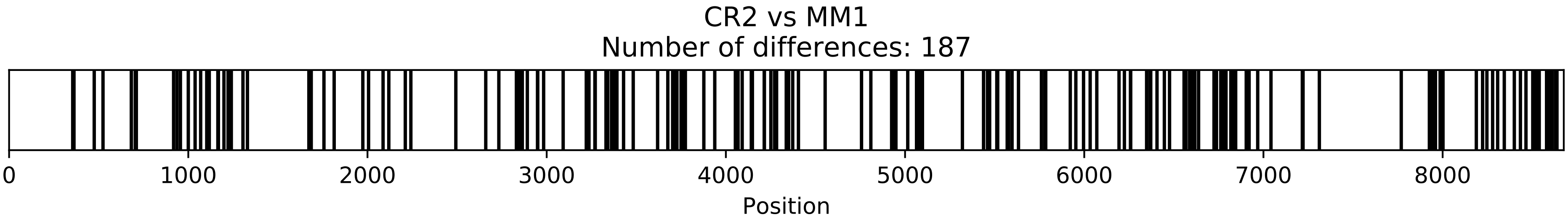
CR2 vs MM1
Number of differences: 187
0
1000
2000
3000
4000
5000
6000
7000
8000
Position

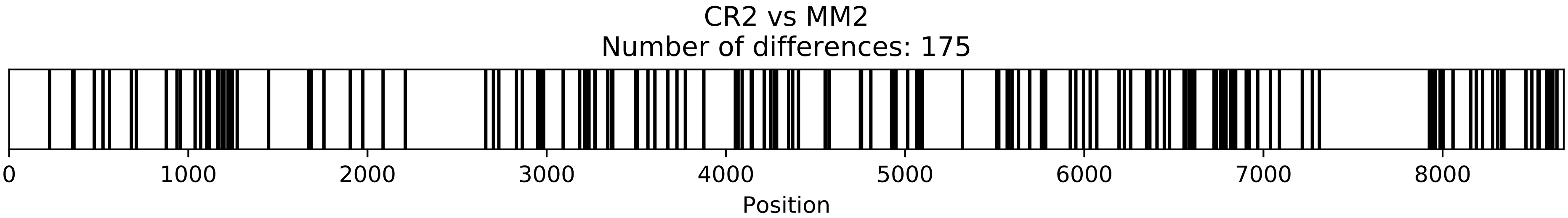
CR2 vs MM2
Number of differences: 175
0
1000
2000
3000
4000
5000
6000
7000
8000
Position

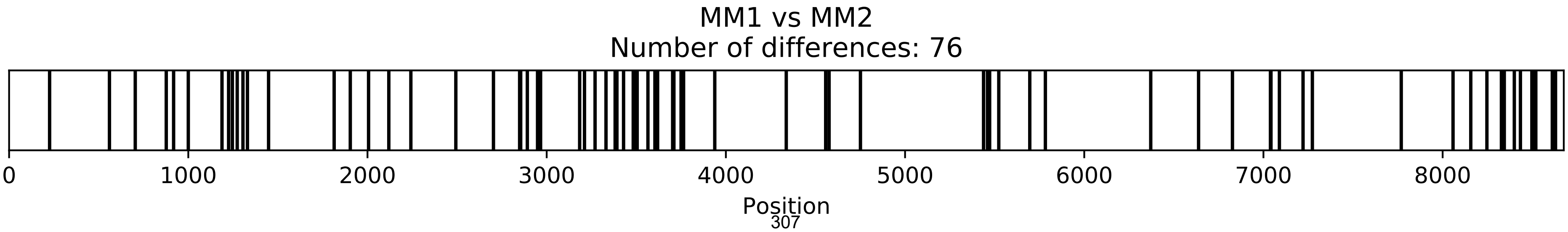
MM1 vs MM2
Number of differences: 76
0
1000
2000
3000
4000
5000
6000
7000
8000
Position

### Region: 832



## CR1 vs CR2
Number of differences: 280

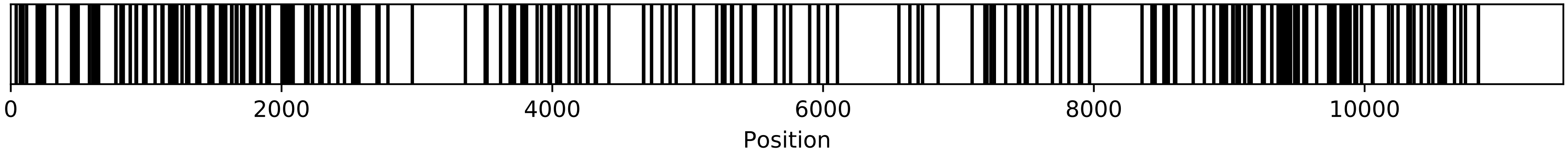


## CR1 vs MM1
Number of differences: 140

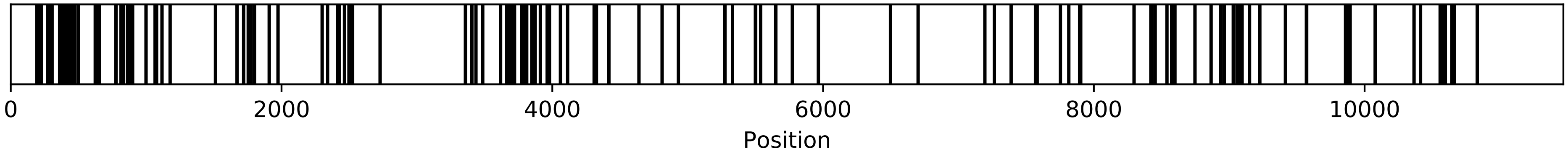


## CR1 vs MM2
Number of differences: 93

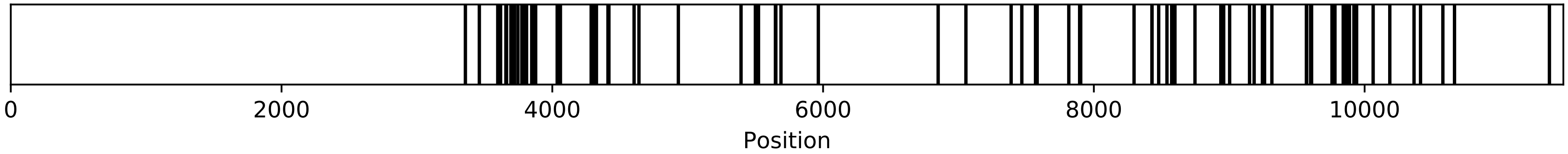


## CR2 vs MM1
Number of differences: 281

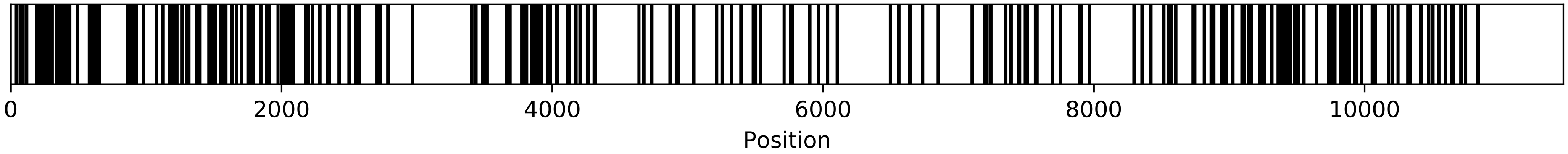


## CR2 vs MM2
Number of differences: 288

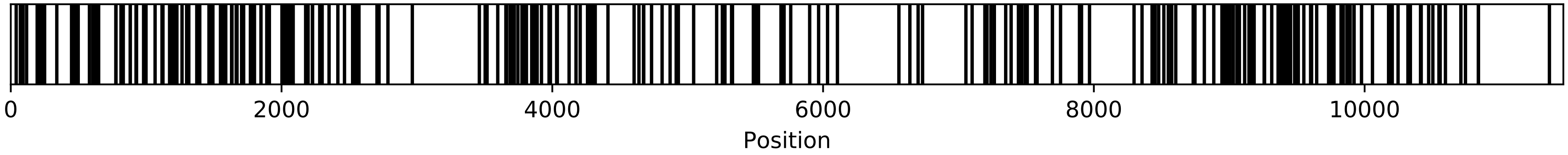


## MM1 vs MM2
Number of differences: 136

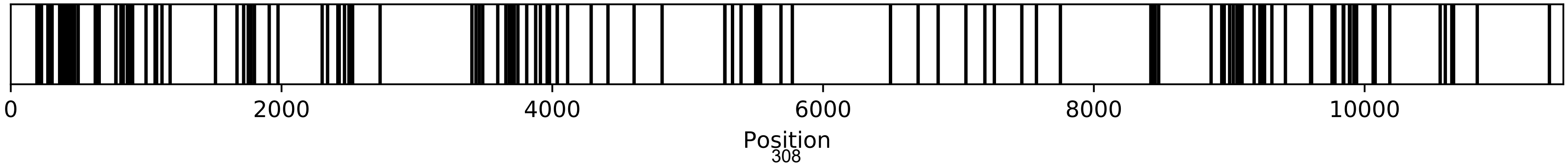

### Region: 833



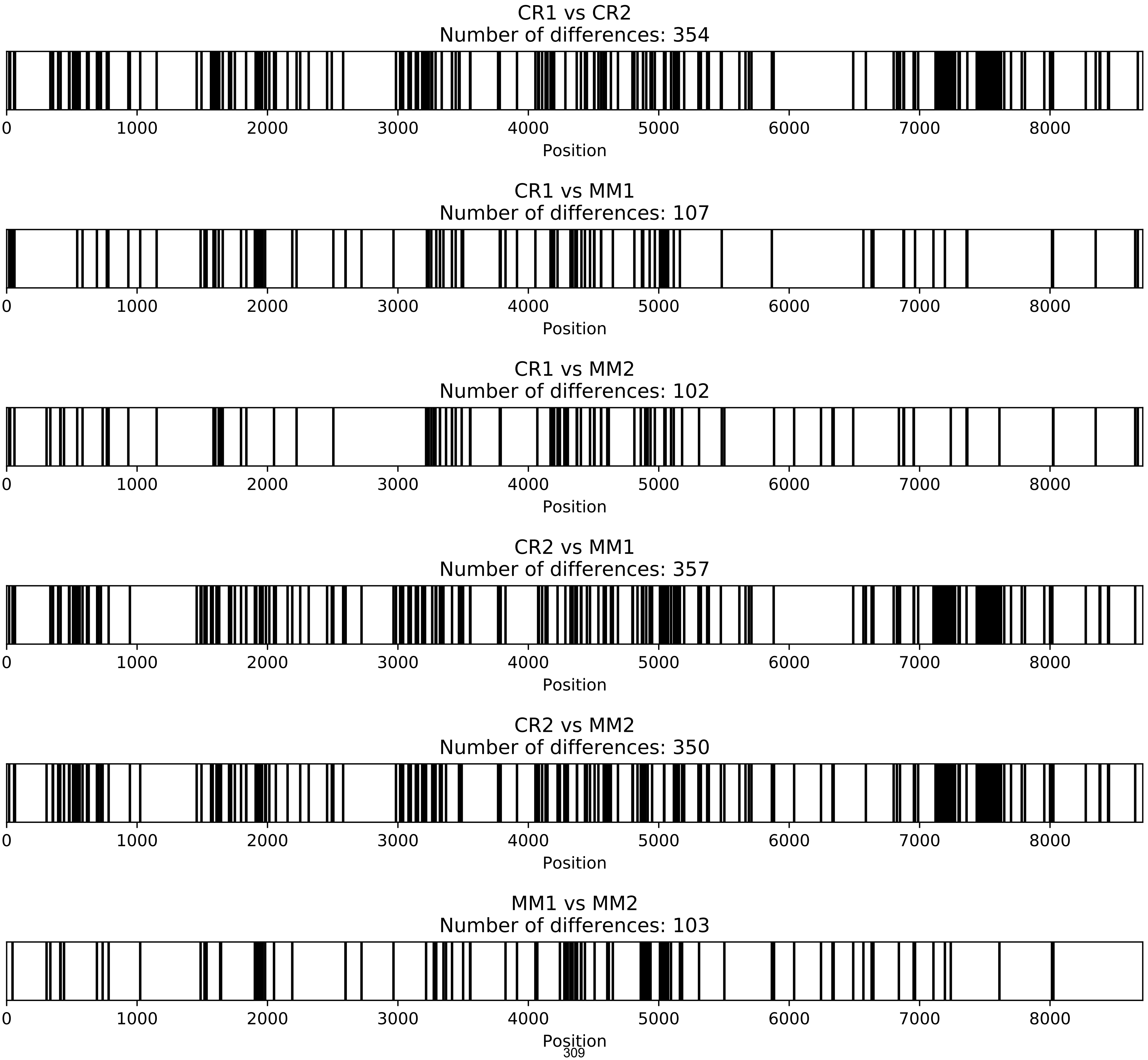
CR1 vs CR2
Number of differences: 354
CR1 vs MM1
Number of differences: 107
CR1 vs MM2
Number of differences: 102
CR2 vs MM1
Number of differences: 357
CR2 vs MM2
Number of differences: 350
MM1 vs MM2
Number of differences: 103
0
1000
2000
3000
4000
5000
6000
7000
8000
Position

### Region: 834



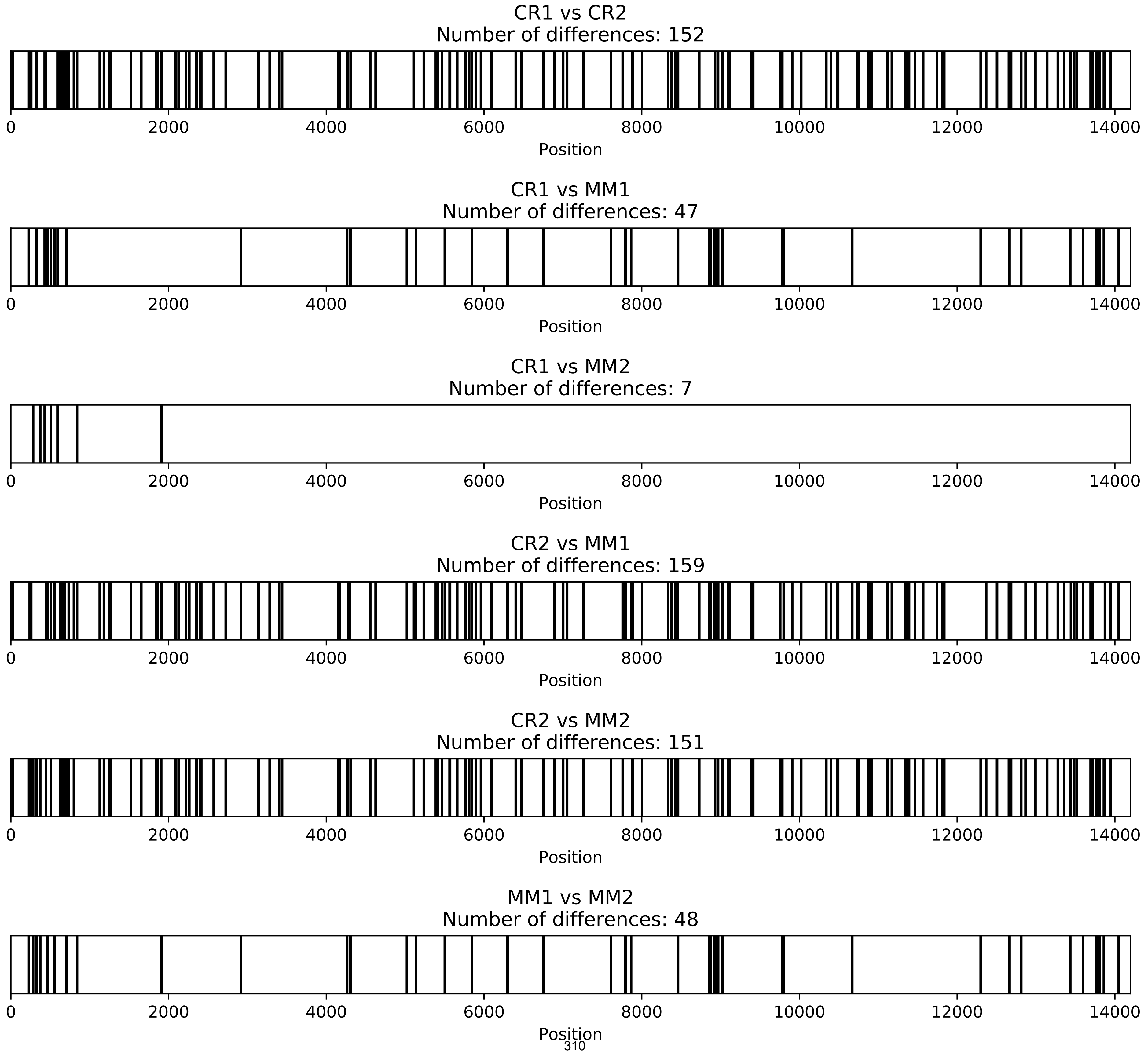
CR1 vs CR2
Number of differences: 152
0
2000
4000
6000
8000
10000
12000
14000
Position
CR1 vs MM1
Number of differences: 47
0
2000
4000
6000
8000
10000
12000
14000
Position
CR1 vs MM2
Number of differences: 7
0
2000
4000
6000
8000
10000
12000
14000
Position
CR2 vs MM1
Number of differences: 159
0
2000
4000
6000
8000
10000
12000
14000
Position
CR2 vs MM2
Number of differences: 151
0
2000
4000
6000
8000
10000
12000
14000
Position
MM1 vs MM2
Number of differences: 48
0
2000
4000
6000
8000
10000
12000
14000
Position

### Region: 835



CR1 vs CR2
Number of differences: 206

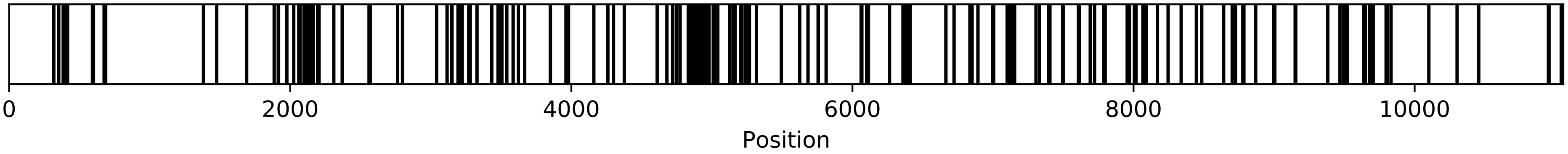


CR1 vs MM1
Number of differences: 121

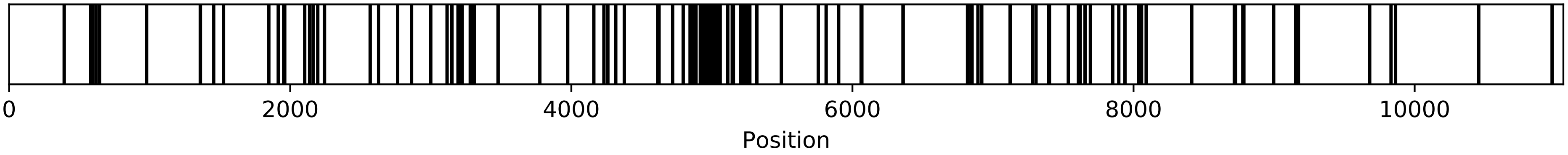


CR1 vs MM2
Number of differences: 125

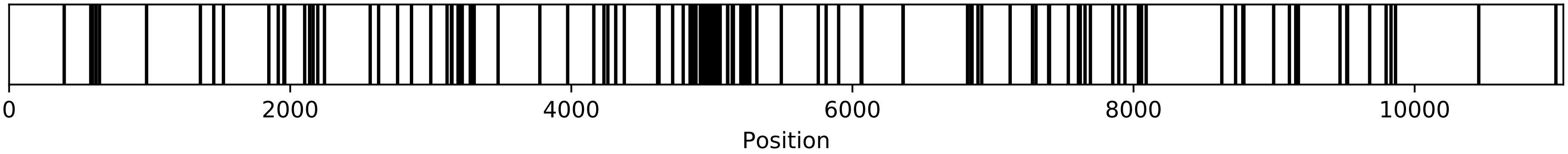


CR2 vs MM1
Number of differences: 195

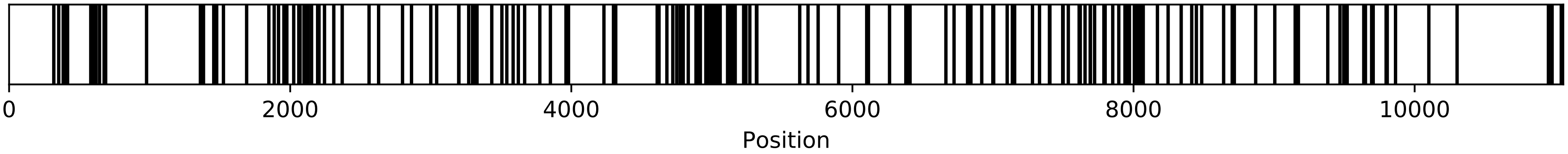


CR2 vs MM2
Number of differences: 195

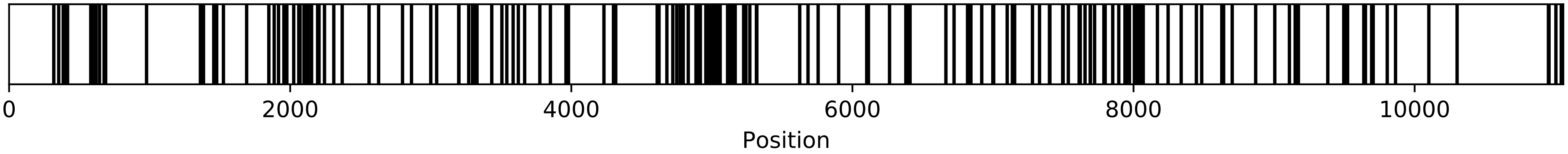


MM1 vs MM2
Number of differences: 10

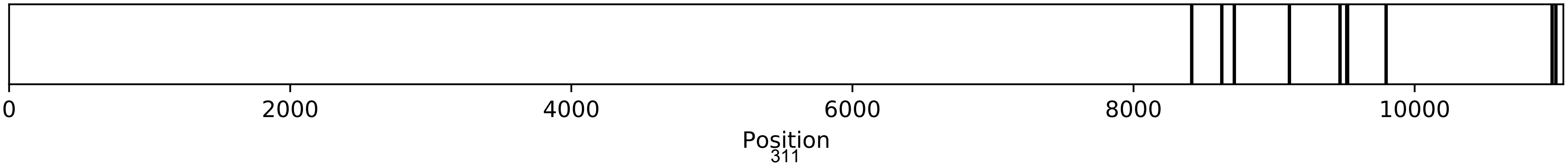

### Region: 836



CR1 vs CR2
Number of differences: 182

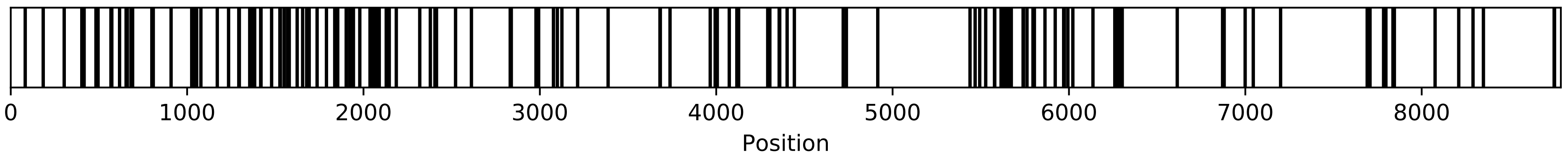


CR1 vs MM1
Number of differences: 104

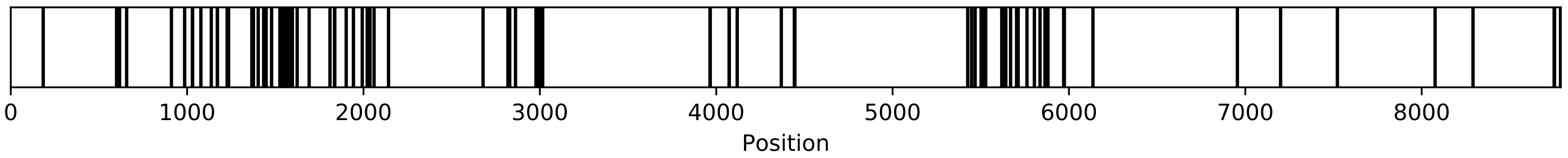


CR1 vs MM2
Number of differences: 61

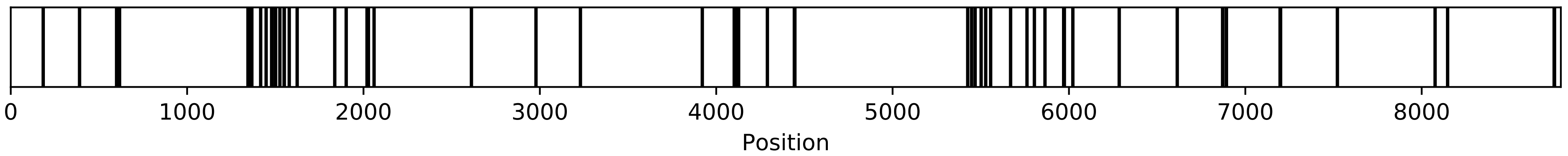


CR2 vs MM1
Number of differences: 202

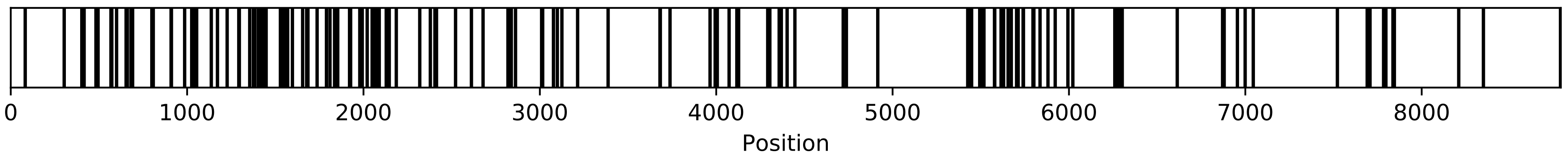


CR2 vs MM2
Number of differences: 170

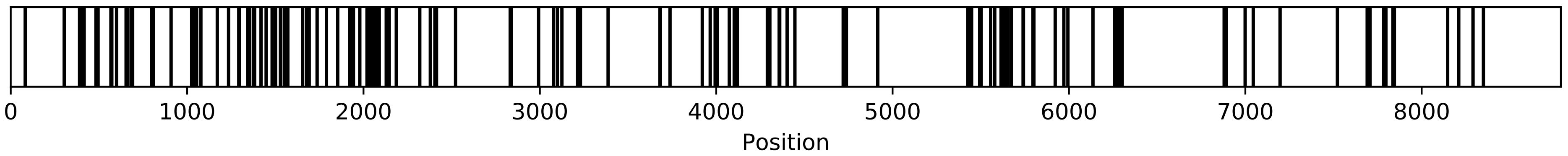


MM1 vs MM2
Number of differences: 99

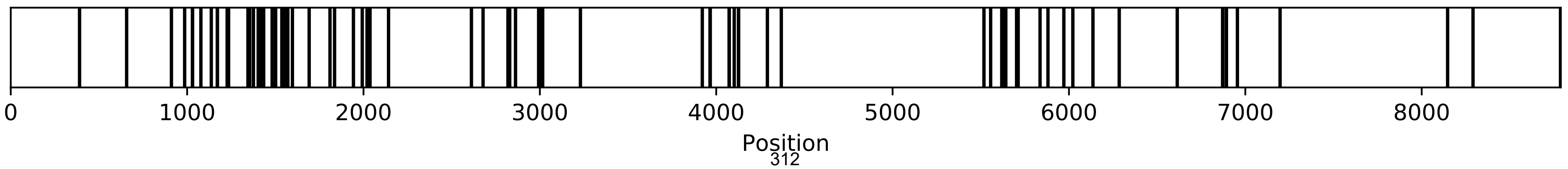

### Region: 838



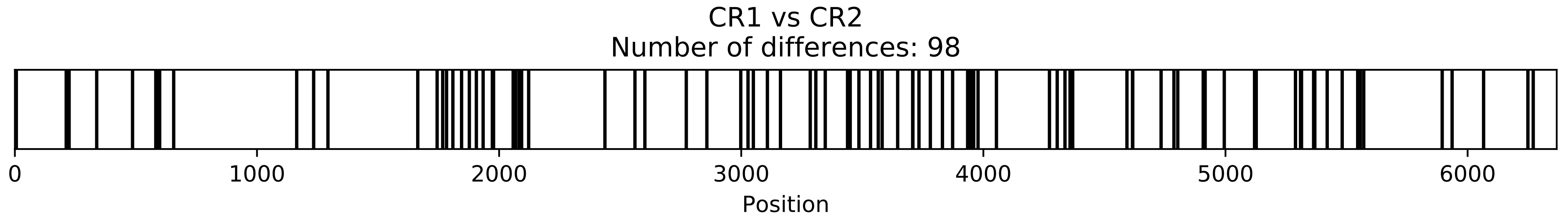
CR1 vs CR2
Number of differences: 98
0
1000
2000
3000
4000
5000
6000
Position

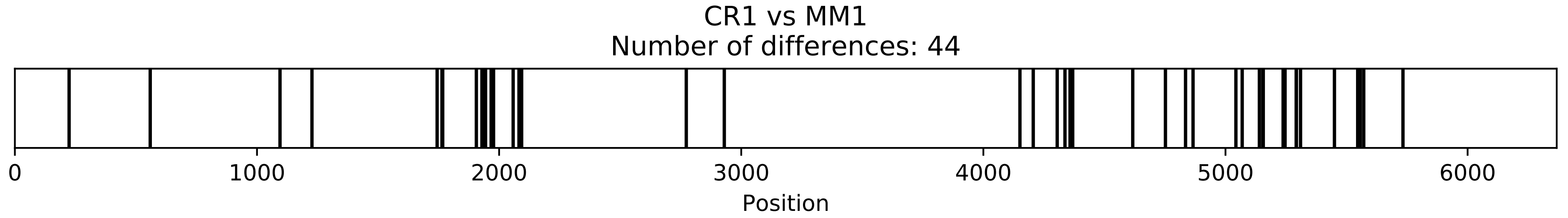
CR1 vs MM1
Number of differences: 44
0
1000
2000
3000
4000
5000
6000
Position

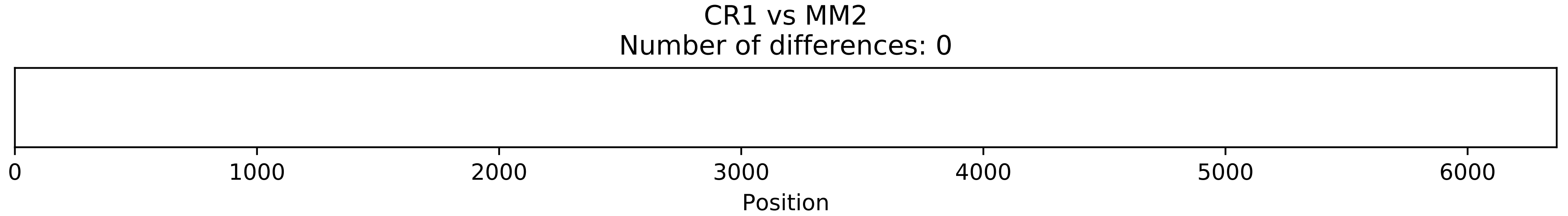
CR1 vs MM2
Number of differences: 0
0
1000
2000
3000
4000
5000
6000
Position

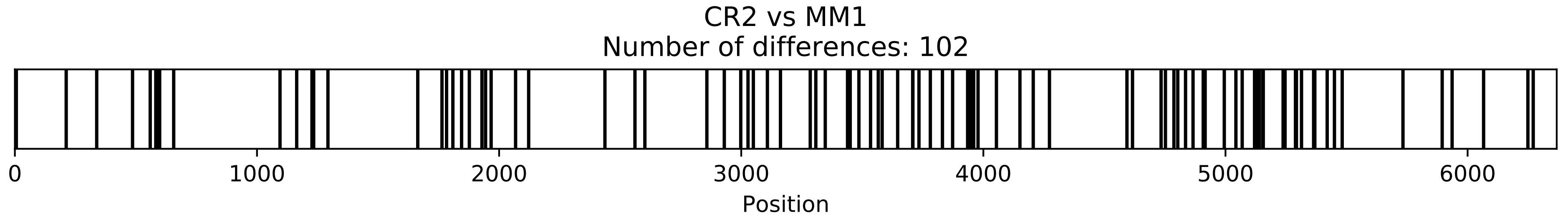
CR2 vs MM1
Number of differences: 102
0
1000
2000
3000
4000
5000
6000
Position

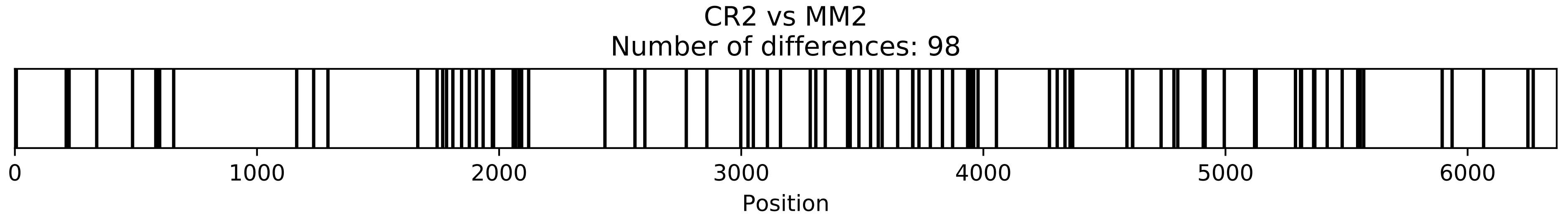
CR2 vs MM2
Number of differences: 98
0
1000
2000
3000
4000
5000
6000
Position

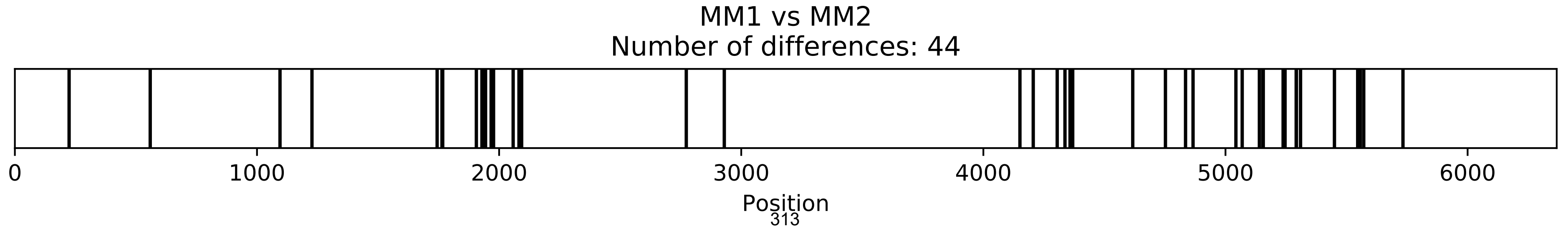
MM1 vs MM2
Number of differences: 44
0
1000
2000
3000
4000
5000
6000
Position

### Region: 839



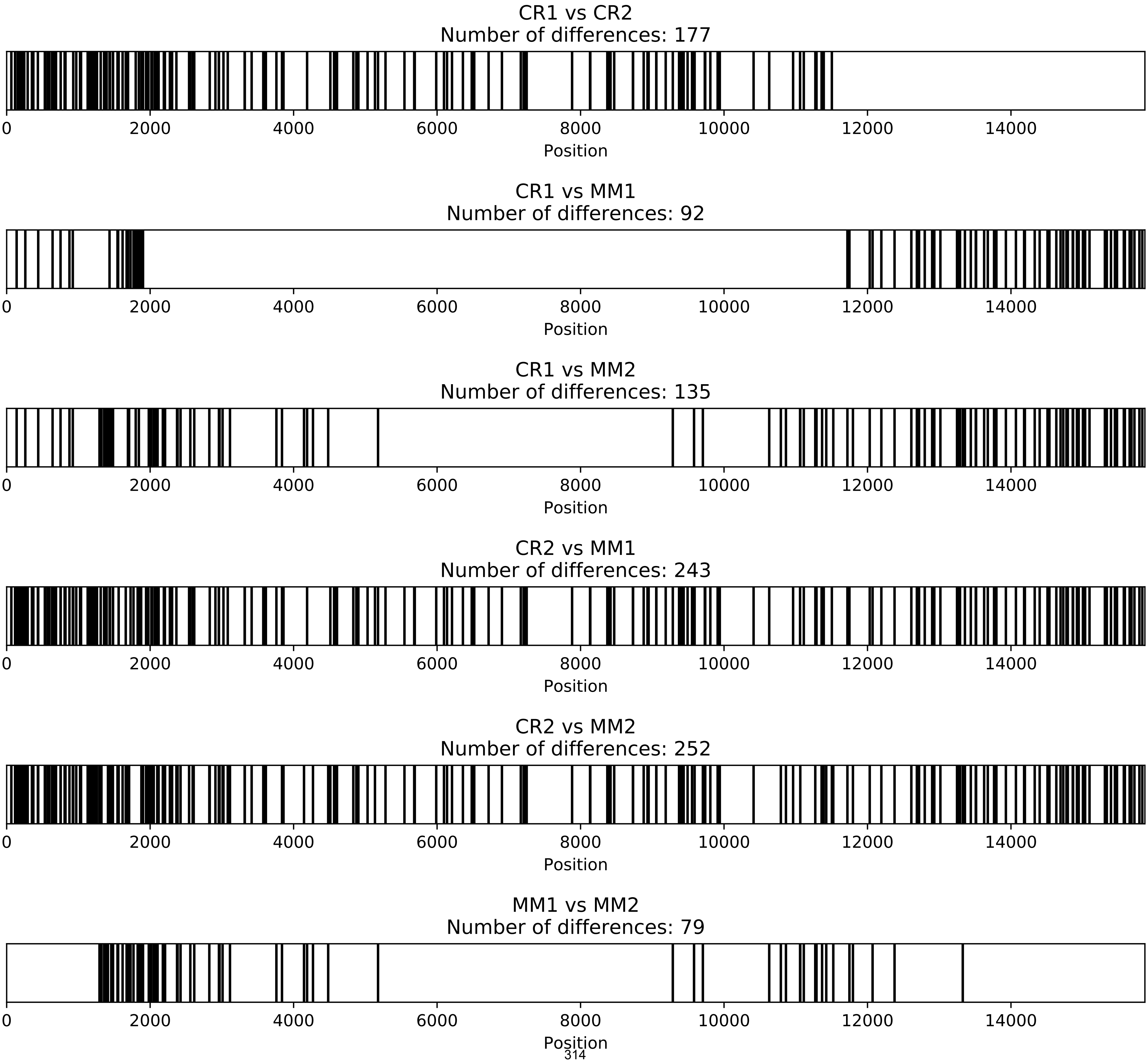
CR1 vs CR2
Number of differences: 177
CR1 vs MM1
Number of differences: 92
CR1 vs MM2
Number of differences: 135
CR2 vs MM1
Number of differences: 243
CR2 vs MM2
Number of differences: 252
MM1 vs MM2
Number of differences: 79
0
2000
4000
6000
8000
10000
12000
14000
Position

### Region: 842



CR1 vs CR2
Number of differences: 235

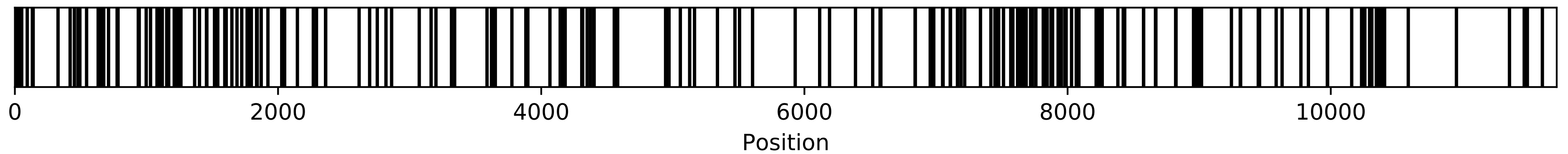


CR1 vs MM1
Number of differences: 3

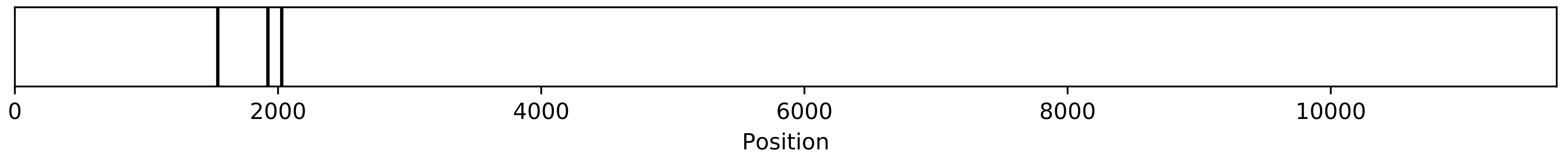


CR1 vs MM2
Number of differences: 107

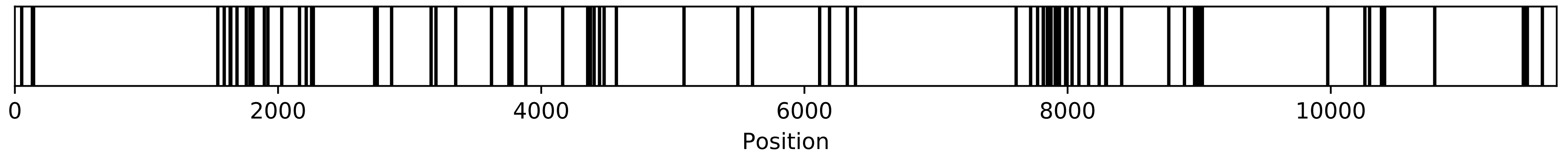


CR2 vs MM1
Number of differences: 232

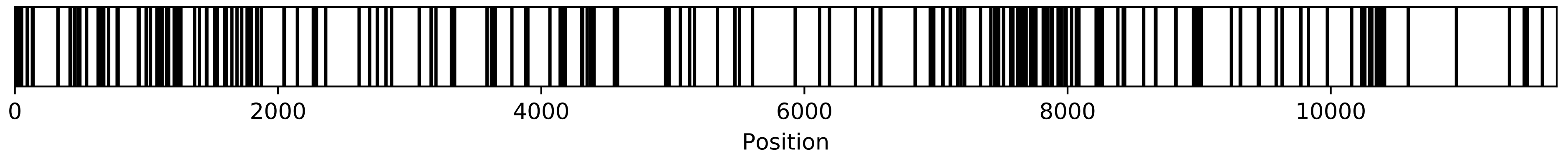


CR2 vs MM2
Number of differences: 236

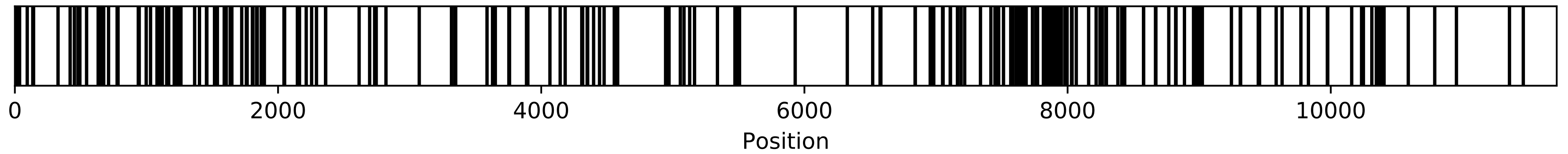


MM1 vs MM2
Number of differences: 104

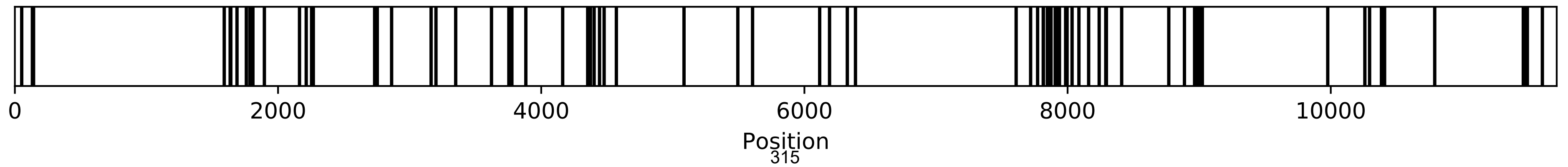

### Region: 843



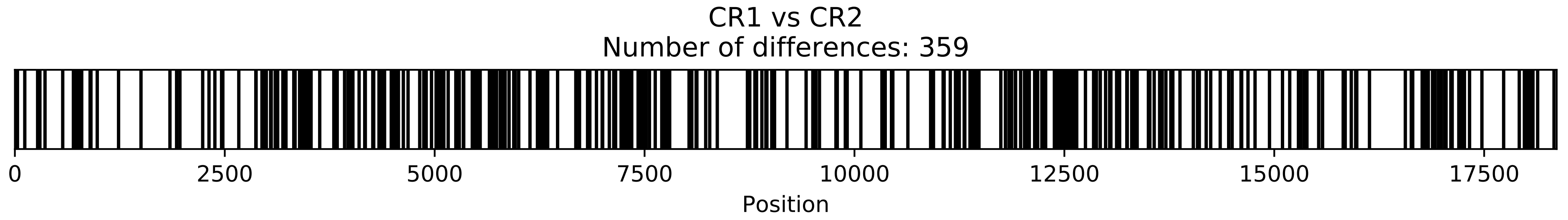
CR1 vs CR2
Number of differences: 359
0
2500
5000
7500
10000
12500
15000
17500
Position

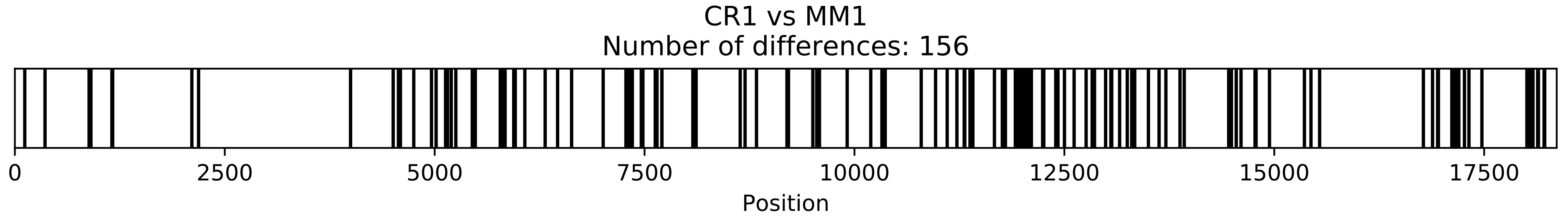
CR1 vs MM1
Number of differences: 156
0
2500
5000
7500
10000
12500
15000
17500
Position

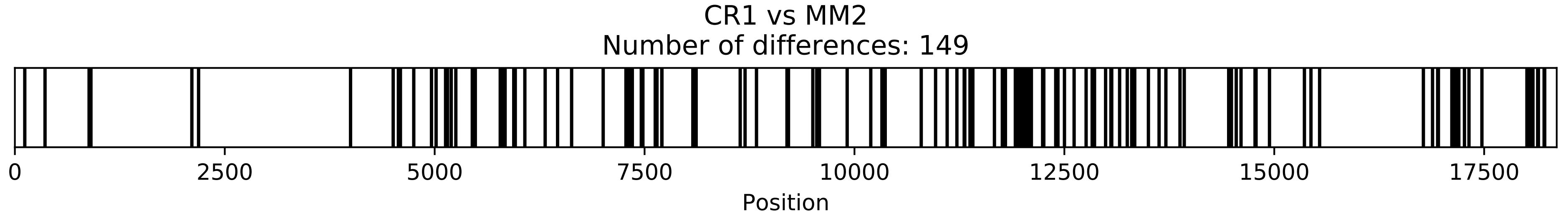
CR1 vs MM2
Number of differences: 149
0
2500
5000
7500
10000
12500
15000
17500
Position

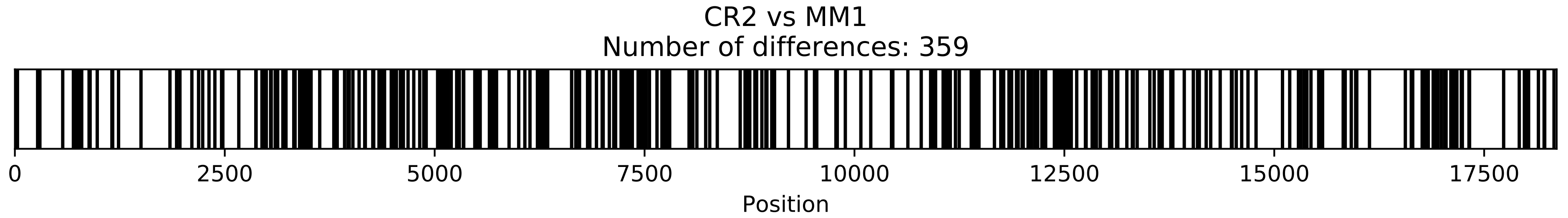
CR2 vs MM1
Number of differences: 359
0
2500
5000
7500
10000
12500
15000
17500
Position

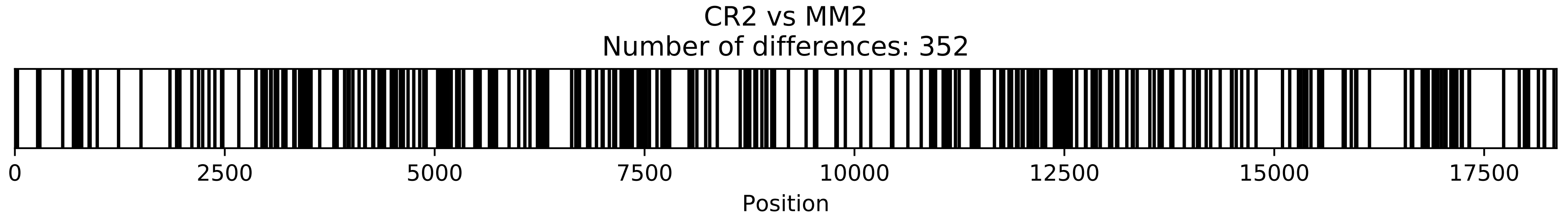
CR2 vs MM2
Number of differences: 352
0
2500
5000
7500
10000
12500
15000
17500
Position

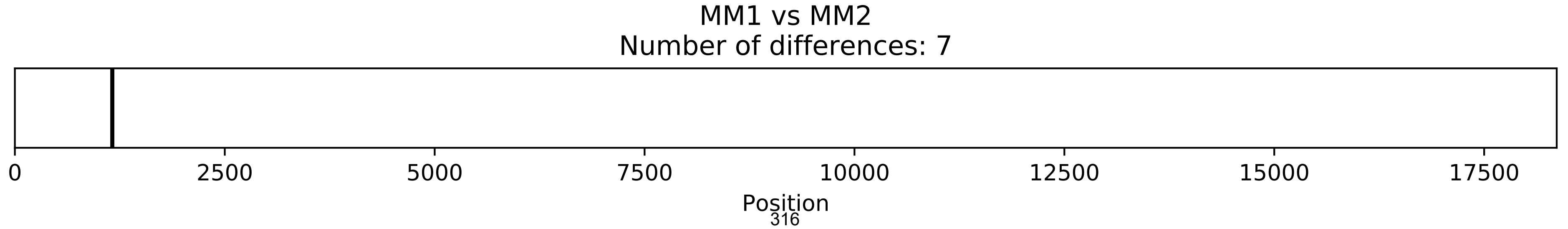
MM1 vs MM2
Number of differences: 7
0
2500
5000
7500
10000
12500
15000
17500
Position

### Region: 844



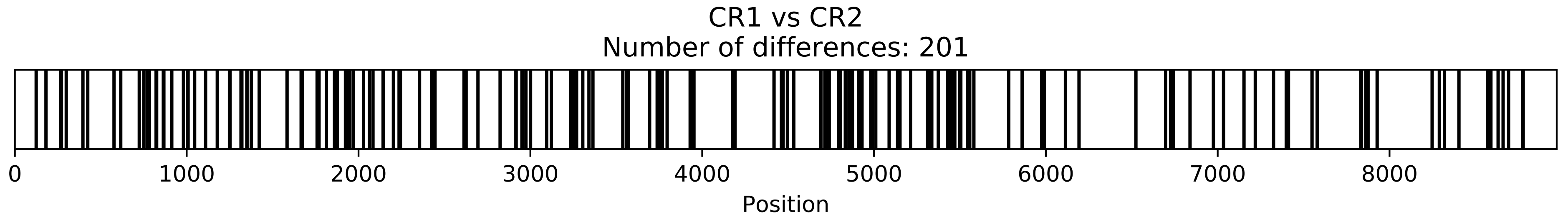
CR1 vs CR2
Number of differences: 201
0
1000
2000
3000
4000
5000
6000
7000
8000
Position

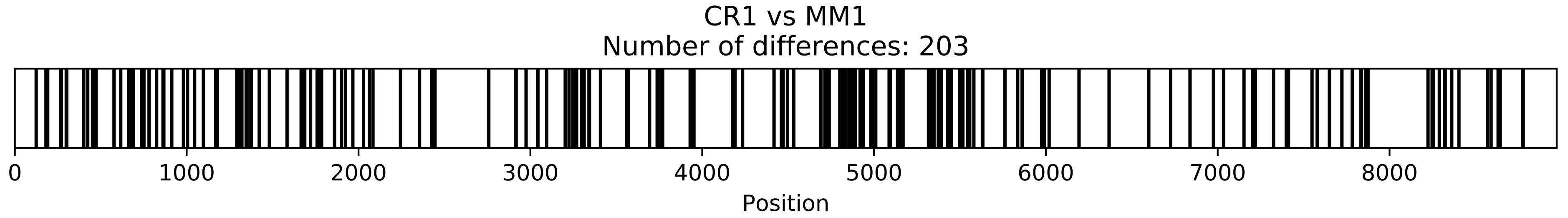
CR1 vs MM1
Number of differences: 203
0
1000
2000
3000
4000
5000
6000
7000
8000
Position

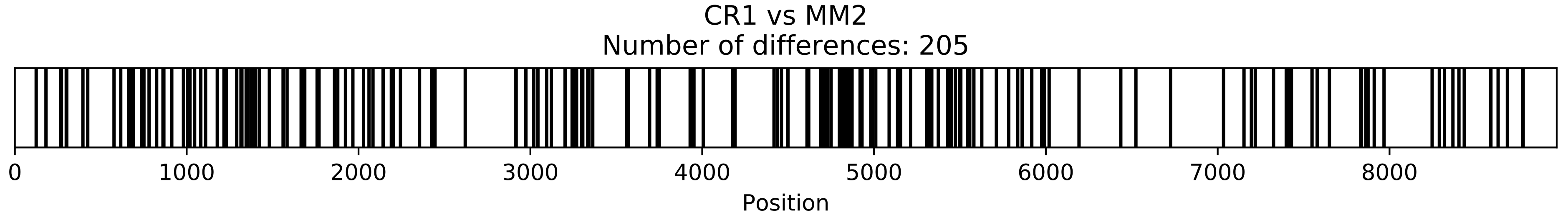
CR1 vs MM2
Number of differences: 205
0
1000
2000
3000
4000
5000
6000
7000
8000
Position

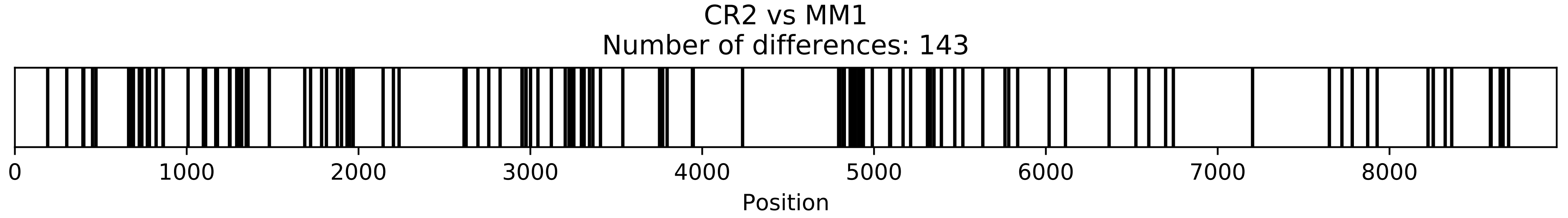
CR2 vs MM1
Number of differences: 143
0
1000
2000
3000
4000
5000
6000
7000
8000
Position

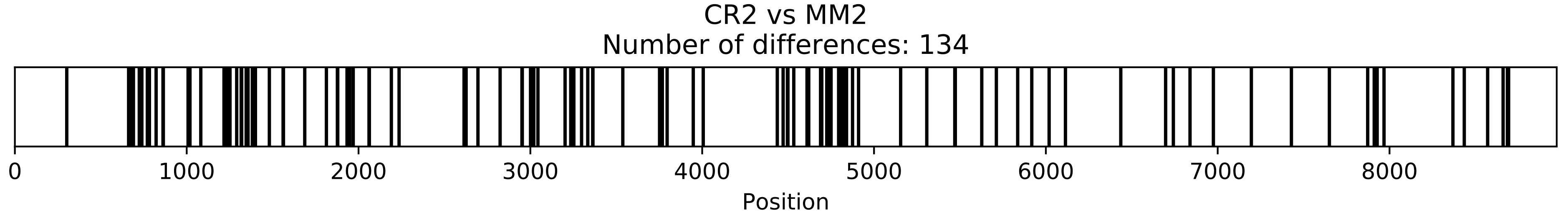
CR2 vs MM2
Number of differences: 134
0
1000
2000
3000
4000
5000
6000
7000
8000
Position

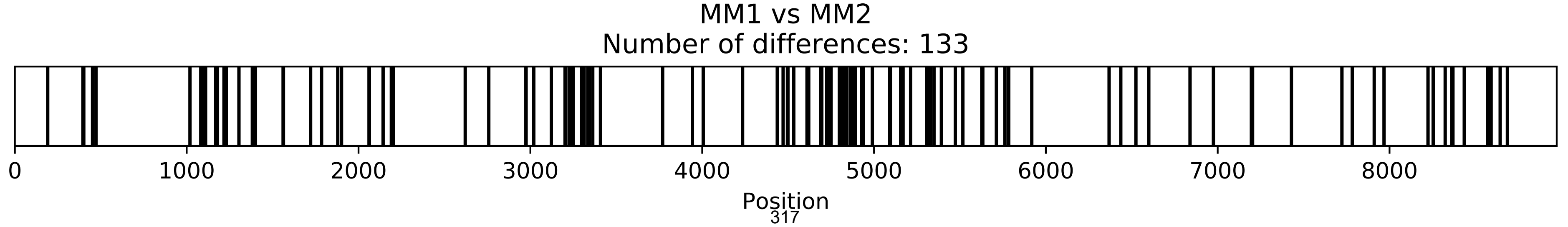
MM1 vs MM2
Number of differences: 133
0
1000
2000
3000
4000
5000
6000
7000
8000
Position

### Region: 849



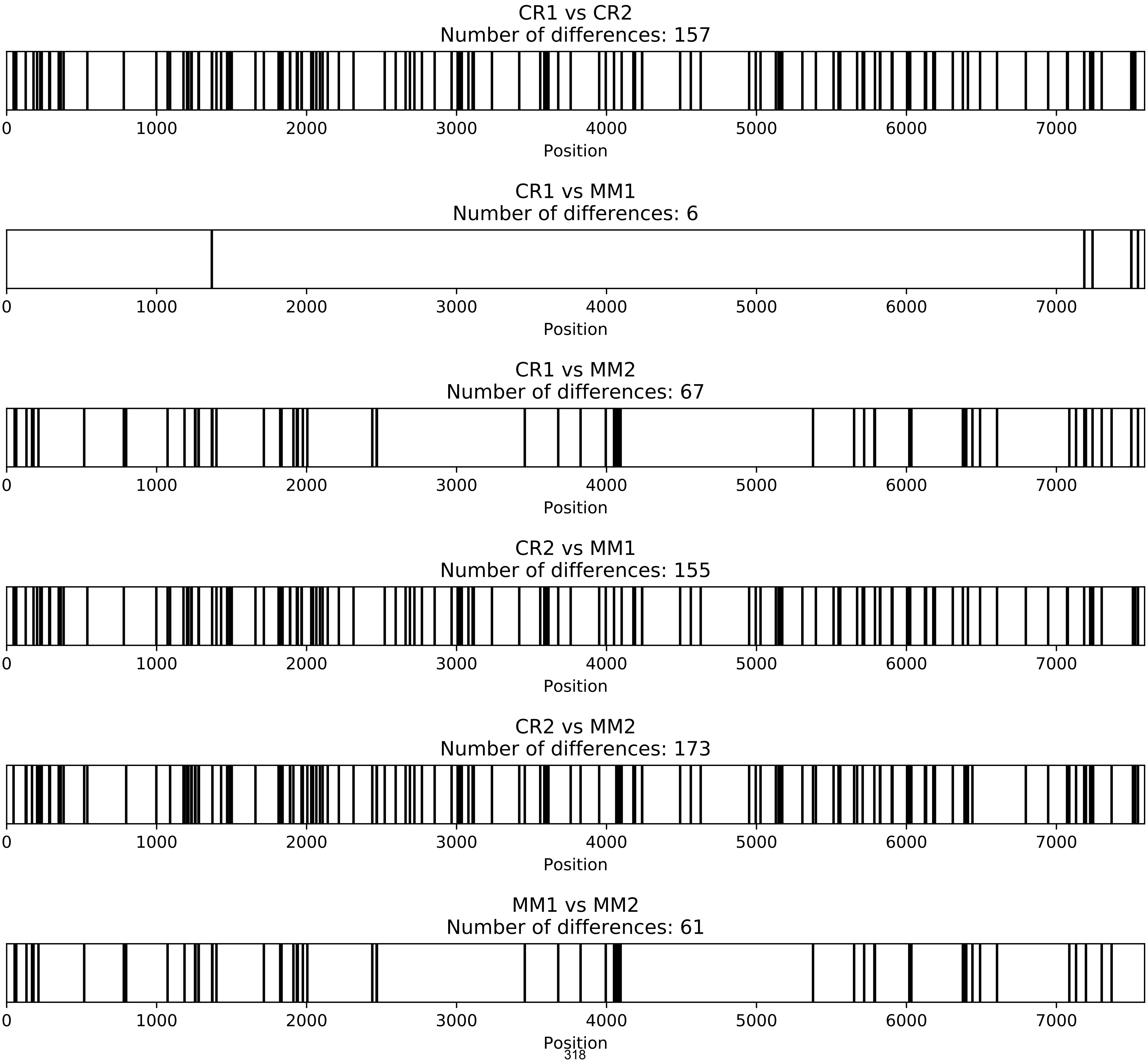
CR1 vs CR2
Number of differences: 157
CR1 vs MM1
Number of differences: 6
CR1 vs MM2
Number of differences: 67
CR2 vs MM1
Number of differences: 155
CR2 vs MM2
Number of differences: 173
MM1 vs MM2
Number of differences: 61
0
1000
2000
3000
4000
5000
6000
7000
Position

### Region: 850



## CR1 vs CR2
Number of differences: 210

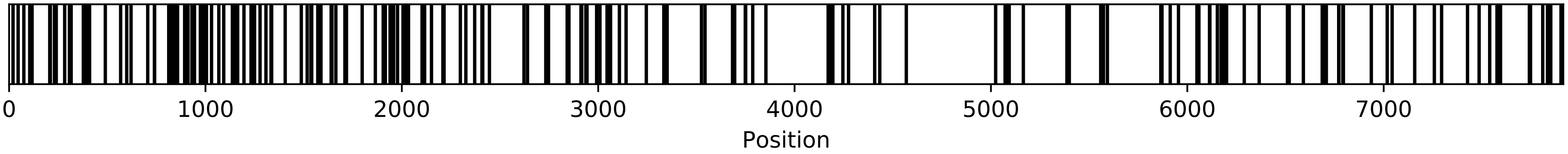


## CR1 vs MM1
Number of differences: 111

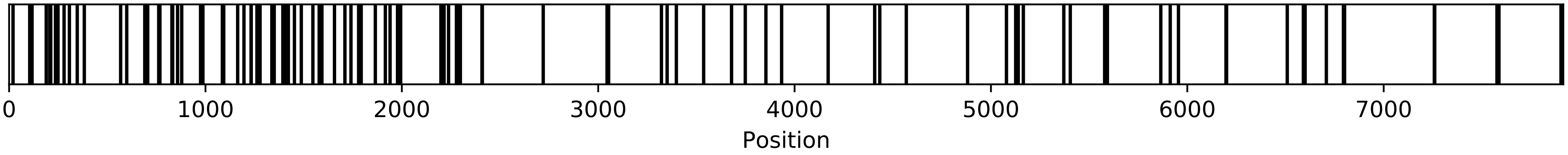


## CR1 vs MM2
Number of differences: 121

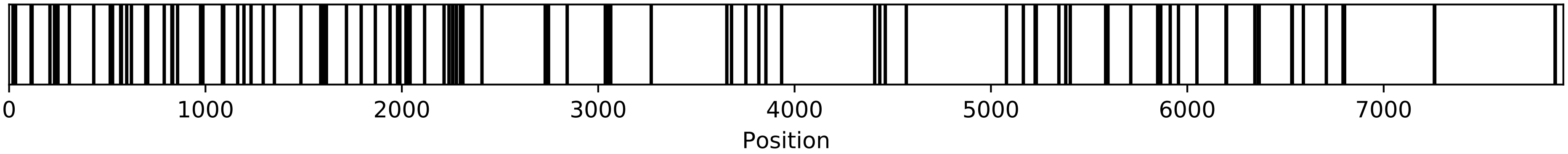


## CR2 vs MM1
Number of differences: 210

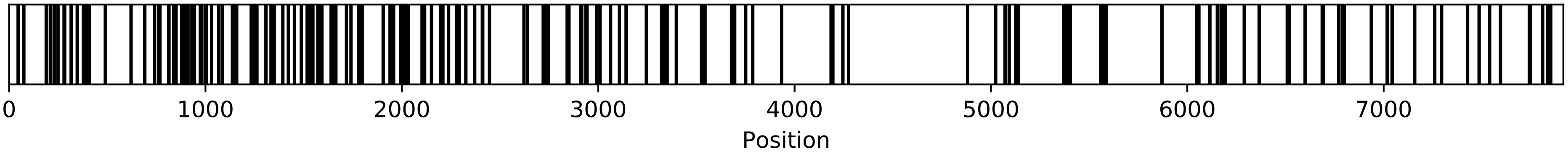


## CR2 vs MM2
Number of differences: 229

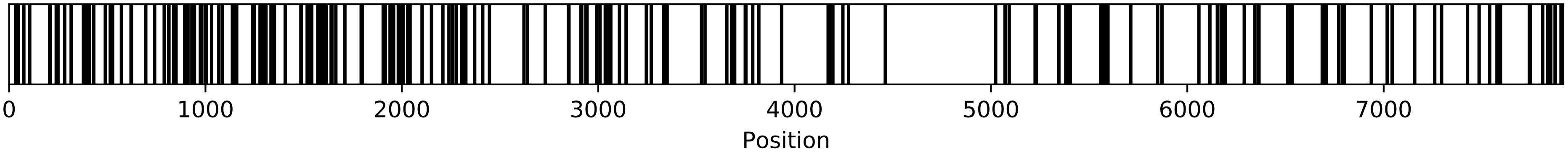


## MM1 vs MM2
Number of differences: 123

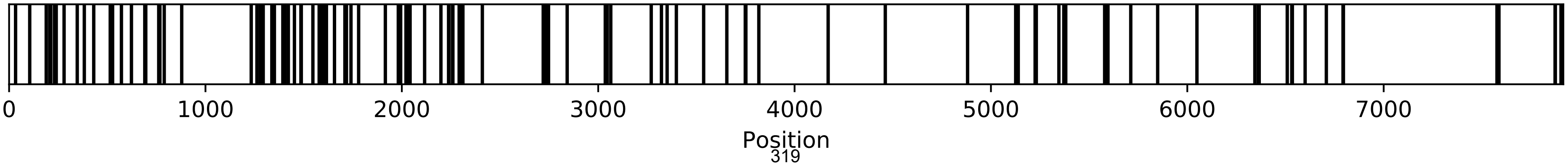

### Region: 851



CR1 vs CR2
Number of differences: 318

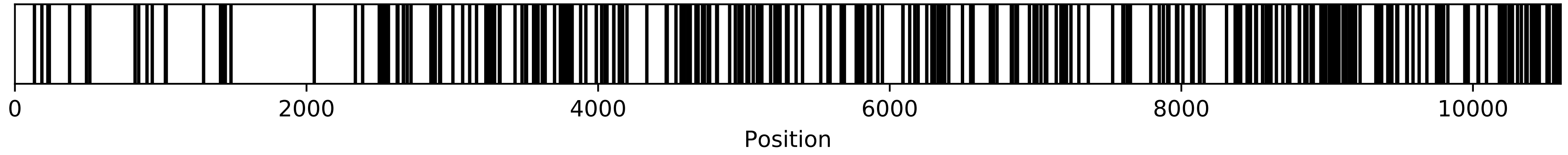


CR1 vs MM1
Number of differences: 124

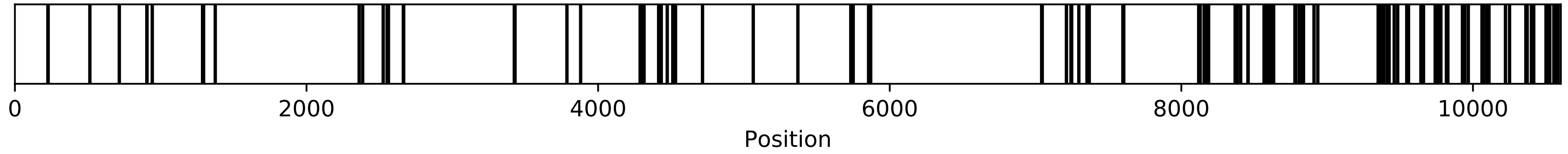


CR1 vs MM2
Number of differences: 177

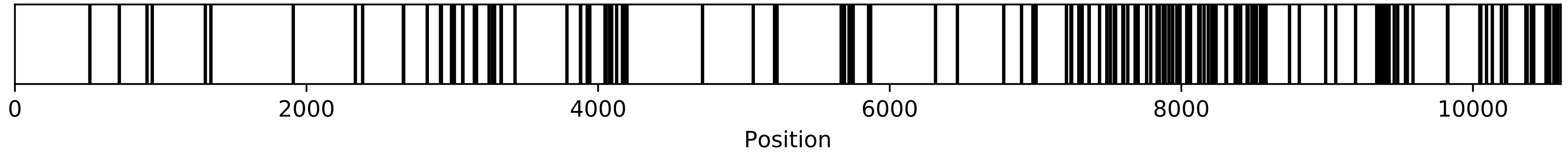


CR2 vs MM1
Number of differences: 334

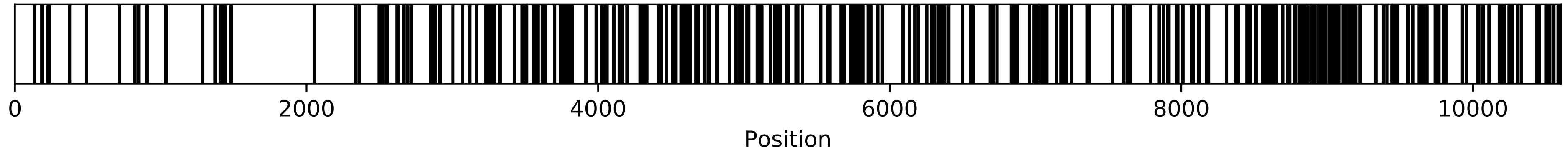


CR2 vs MM2
Number of differences: 343

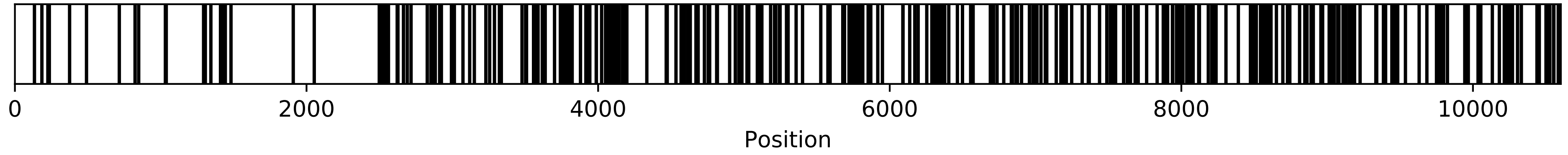


MM1 vs MM2
Number of differences: 175

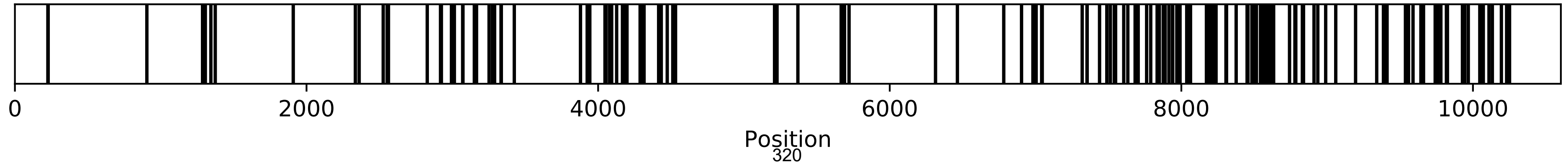

### Region: 854



CR1 vs CR2
Number of differences: 131

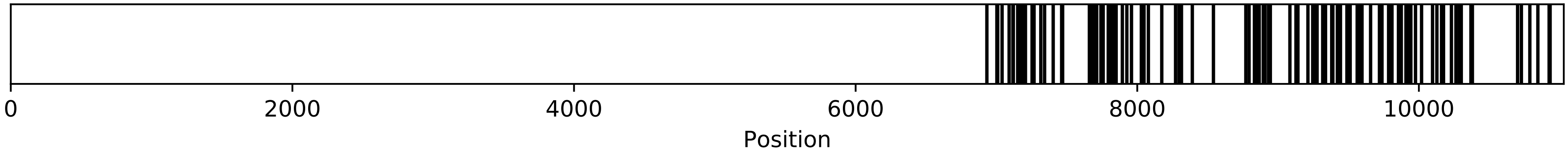


CR1 vs MM1
Number of differences: 256

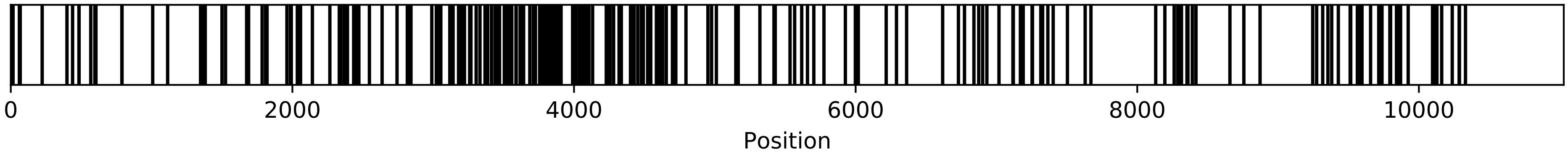


CR1 vs MM2
Number of differences: 252

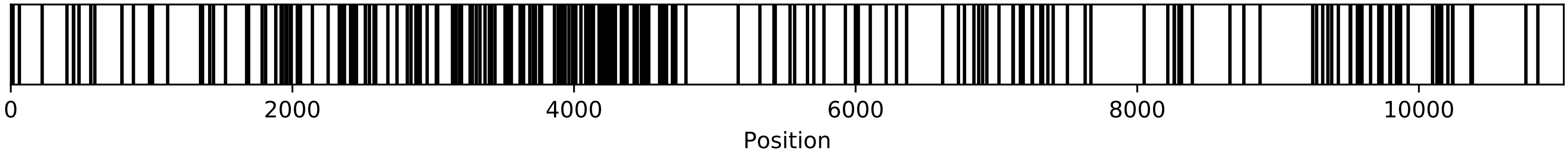


CR2 vs MM1
Number of differences: 318

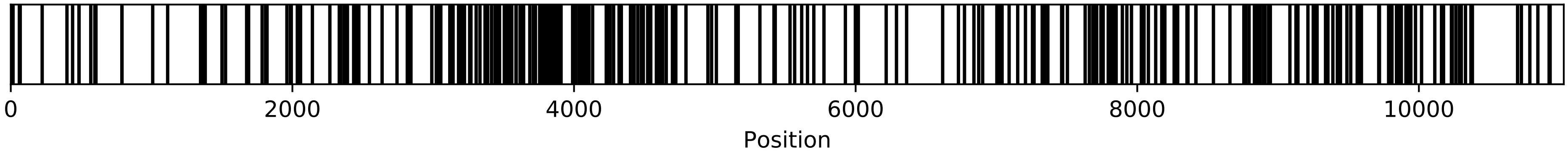


CR2 vs MM2
Number of differences: 308

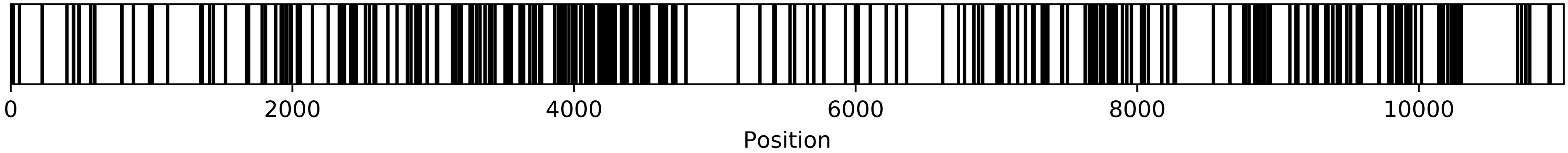


MM1 vs MM2
Number of differences: 174

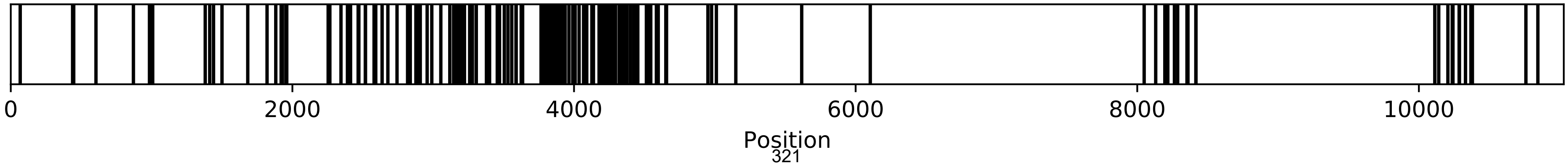

### Region: 858



CR1 vs CR2
Number of differences: 251

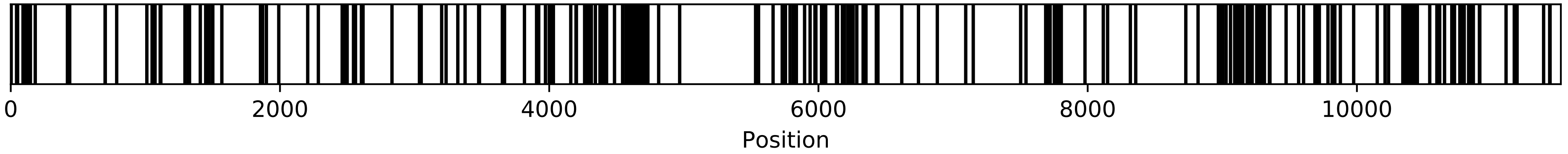


CR1 vs MM1
Number of differences: 191

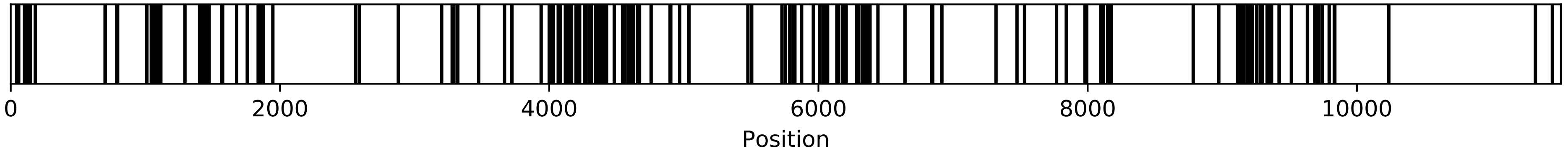


CR1 vs MM2
Number of differences: 189

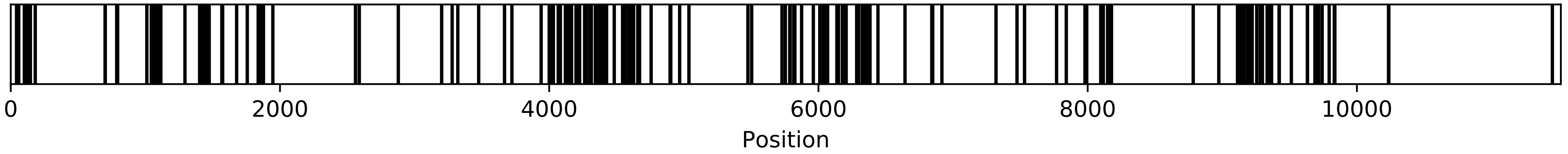


CR2 vs MM1
Number of differences: 286

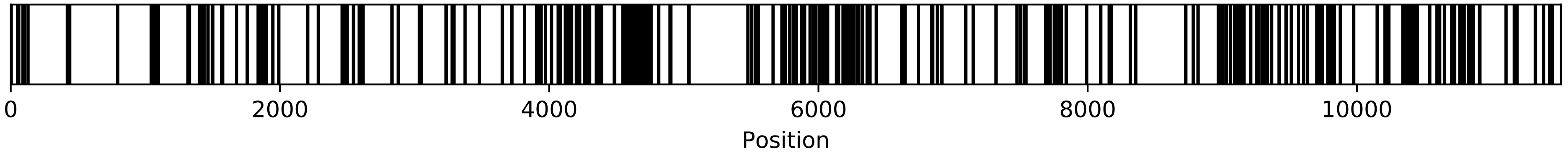


CR2 vs MM2
Number of differences: 284

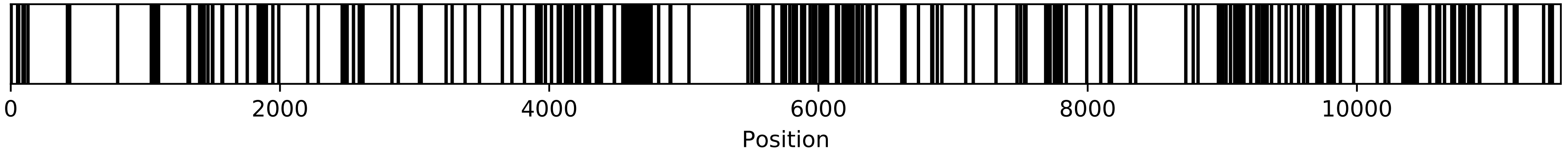


MM1 vs MM2
Number of differences: 2

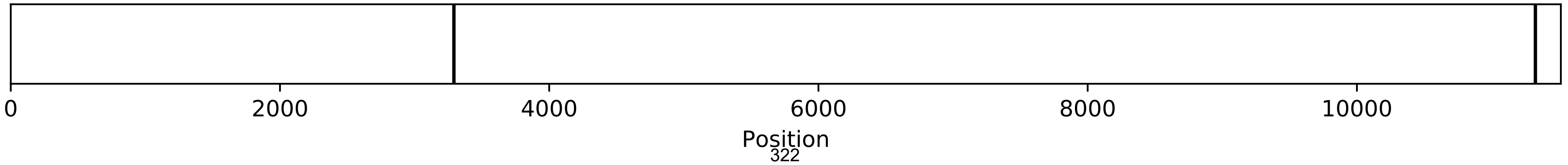

### Region: 862



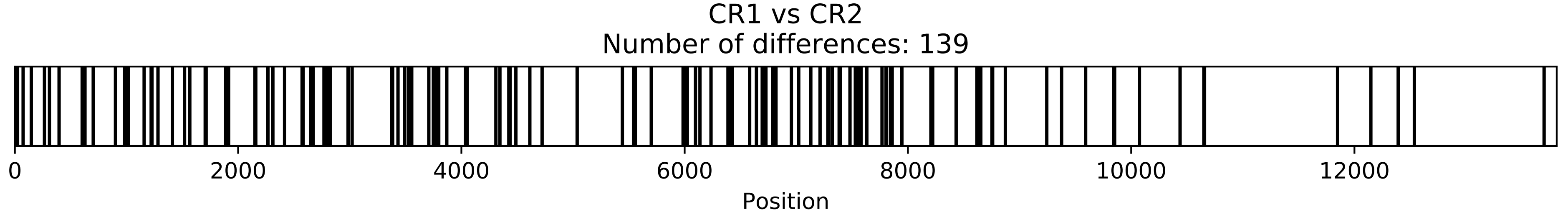
CR1 vs CR2
Number of differences: 139
Position

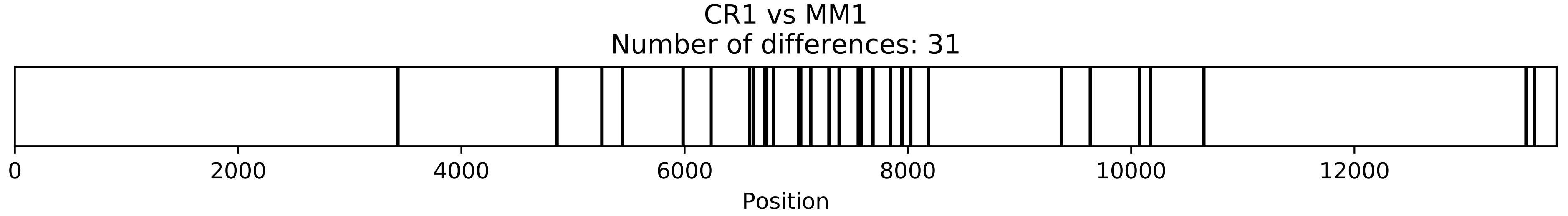
CR1 vs MM1
Number of differences: 31
Position

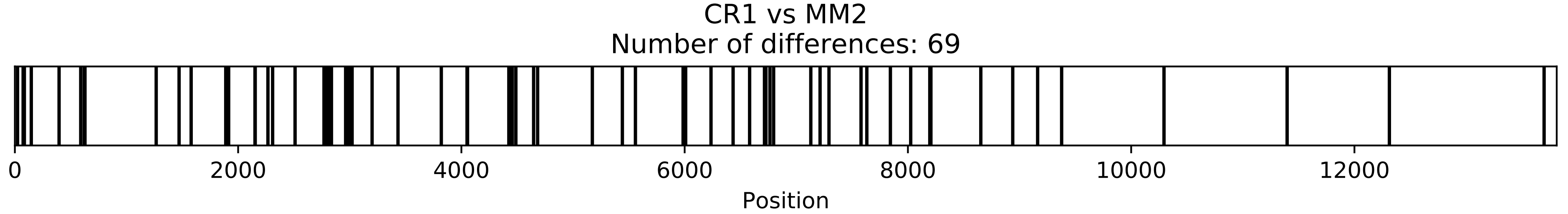
CR1 vs MM2
Number of differences: 69
Position

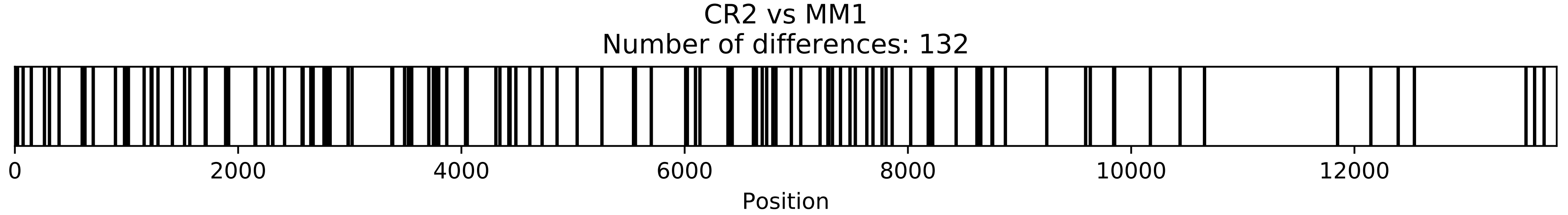
CR2 vs MM1
Number of differences: 132
Position

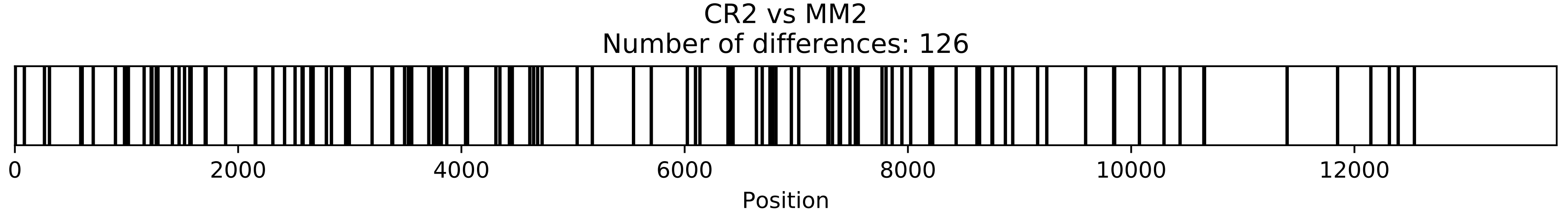
CR2 vs MM2
Number of differences: 126
Position

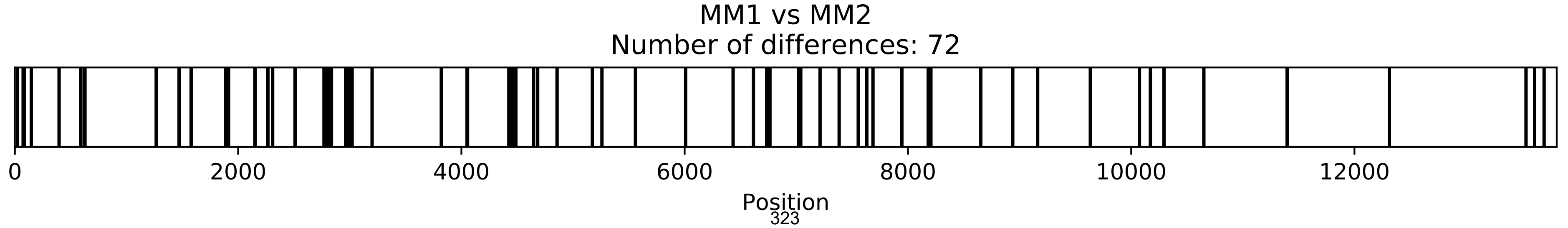
MM1 vs MM2
Number of differences: 72
Position

### Region: 863



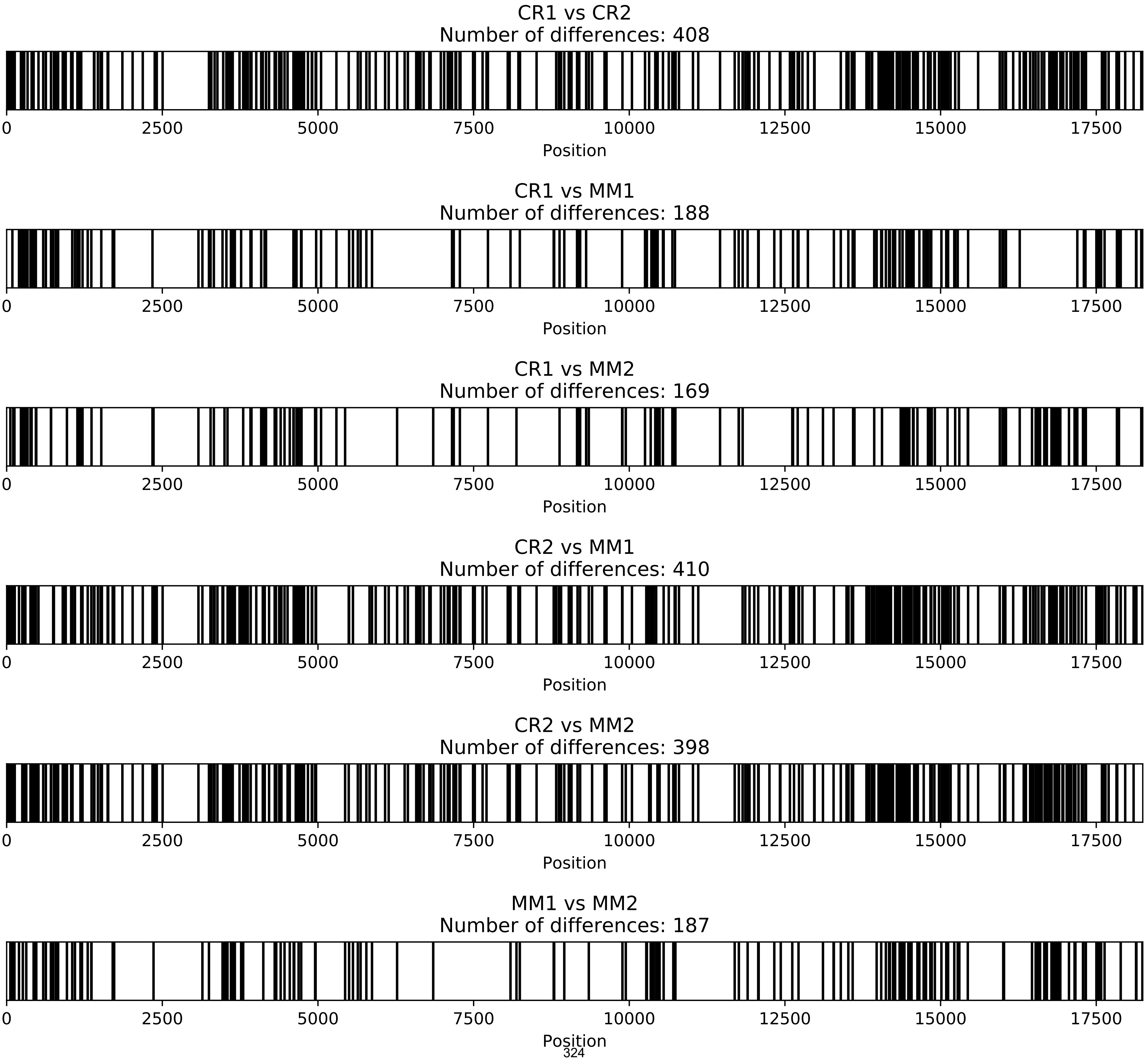
CR1 vs CR2
Number of differences: 408
CR1 vs MM1
Number of differences: 188
CR1 vs MM2
Number of differences: 169
CR2 vs MM1
Number of differences: 410
CR2 vs MM2
Number of differences: 398
MM1 vs MM2
Number of differences: 187
0
2500
5000
7500
10000
12500
15000
17500
Position

### Region: 864



CR1 vs CR2
Number of differences: 294

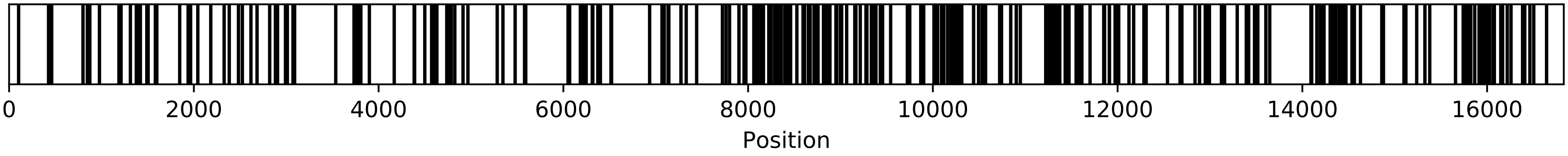


CR1 vs MM1
Number of differences: 312

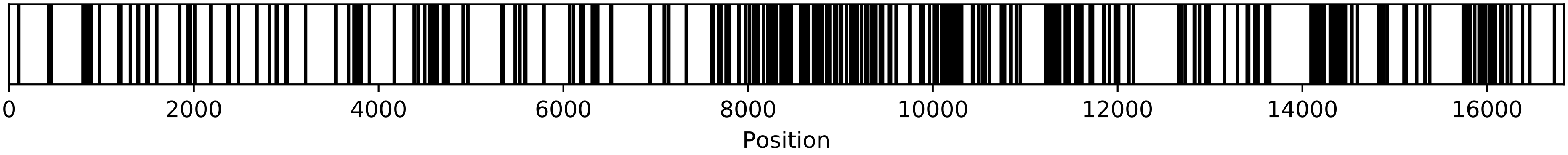


CR1 vs MM2
Number of differences: 339

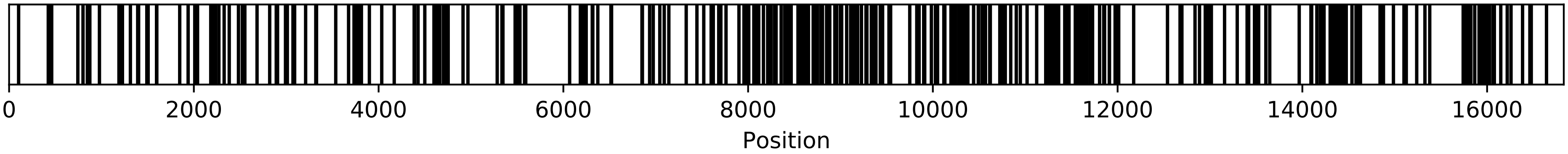


CR2 vs MM1
Number of differences: 154

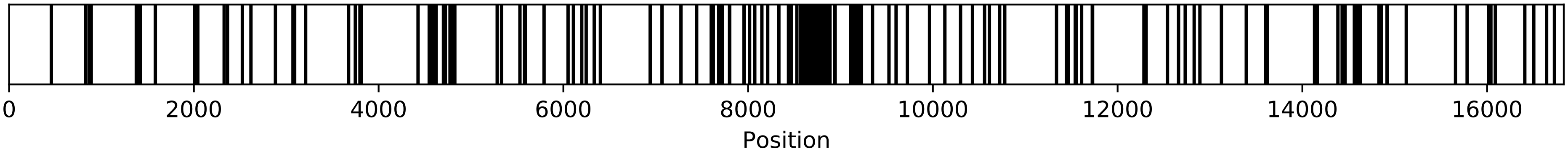


CR2 vs MM2
Number of differences: 187

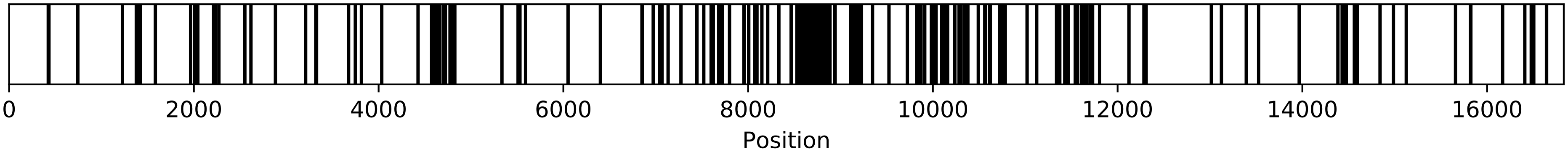


MM1 vs MM2
Number of differences: 141

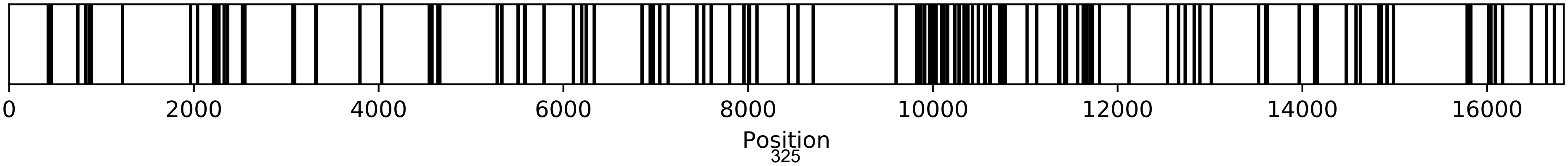

### Region: 866



CR1 vs CR2
Number of differences: 83

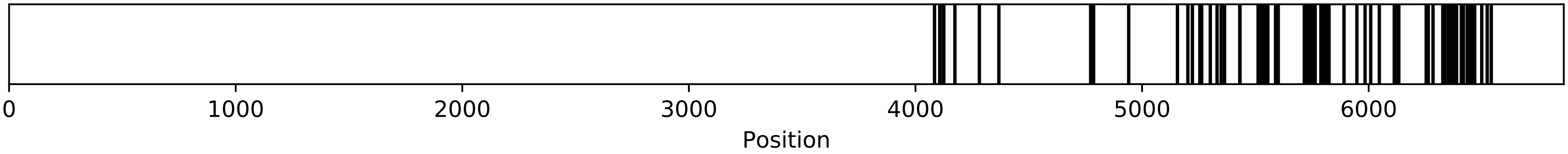


CR1 vs MM1
Number of differences: 147

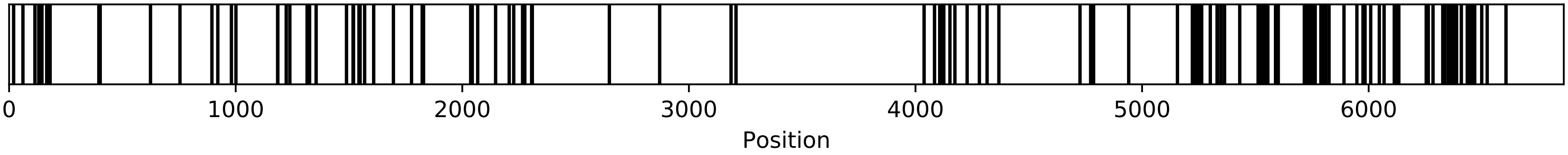


CR1 vs MM2
Number of differences: 147

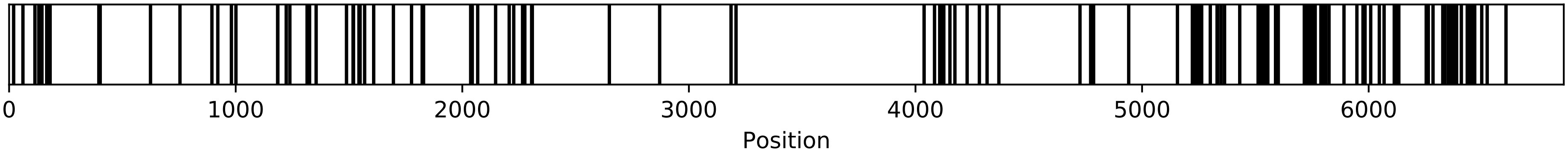


CR2 vs MM1
Number of differences: 81

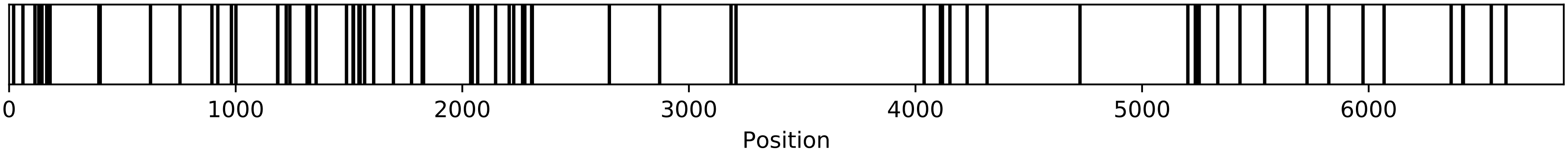


CR2 vs MM2
Number of differences: 81

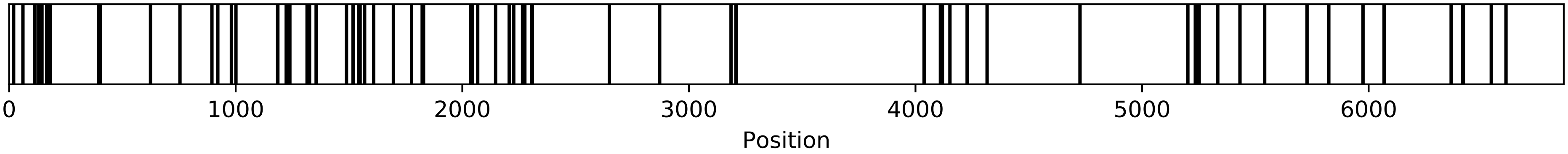


MM1 vs MM2
Number of differences: 0

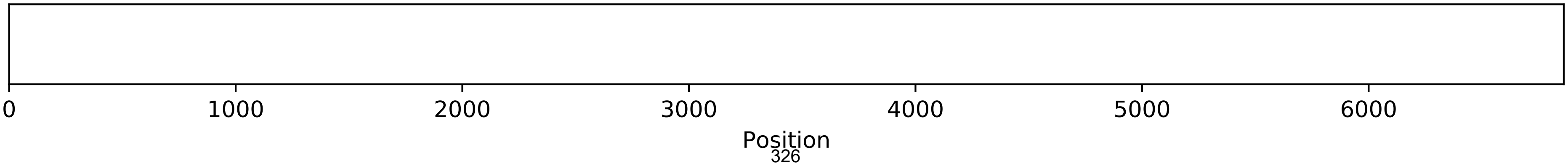

### Region: 867



CR1 vs CR2
Number of differences: 80

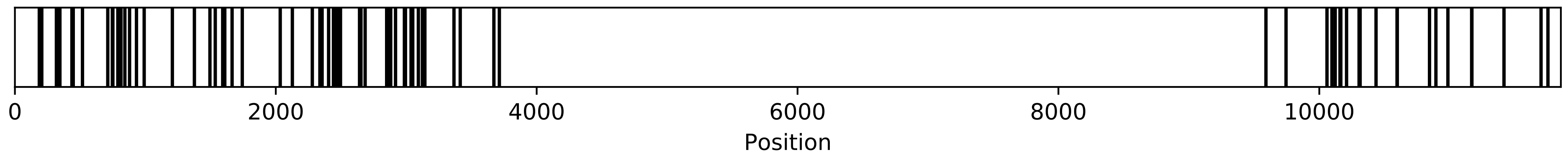


CR1 vs MM1
Number of differences: 105

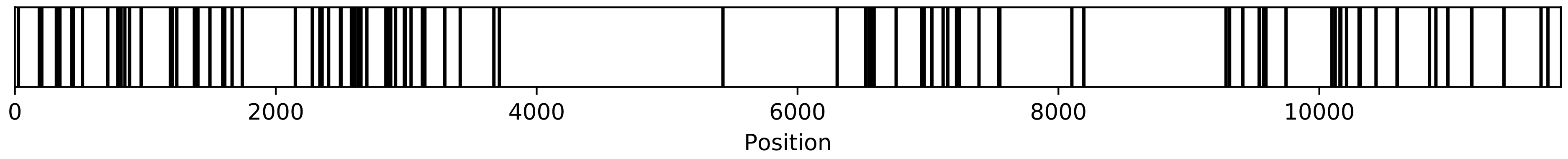


CR1 vs MM2
Number of differences: 143

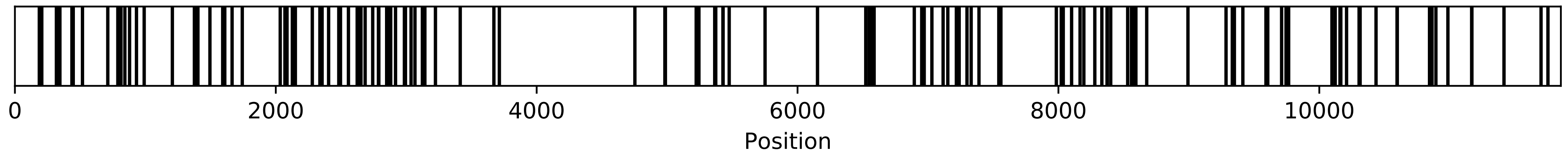


CR2 vs MM1
Number of differences: 57

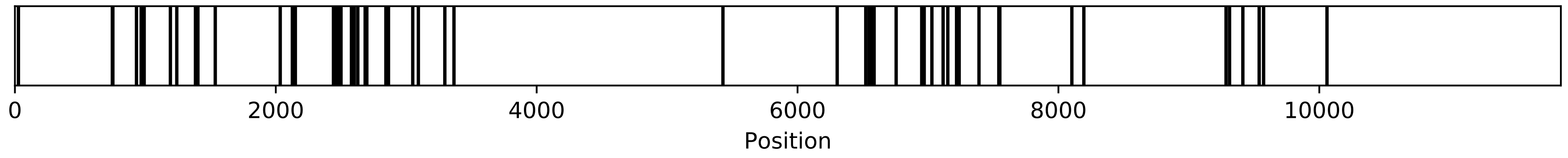


CR2 vs MM2
Number of differences: 85

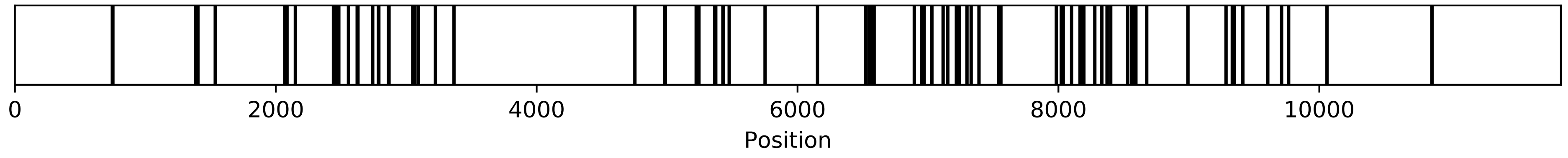


MM1 vs MM2
Number of differences: 76

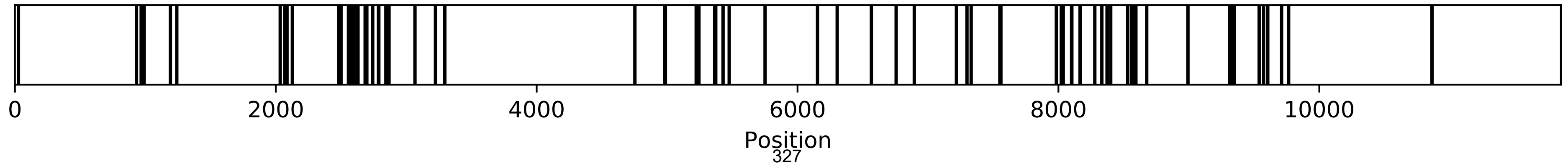

### Region: 871



CR1 vs CR2
Number of differences: 92

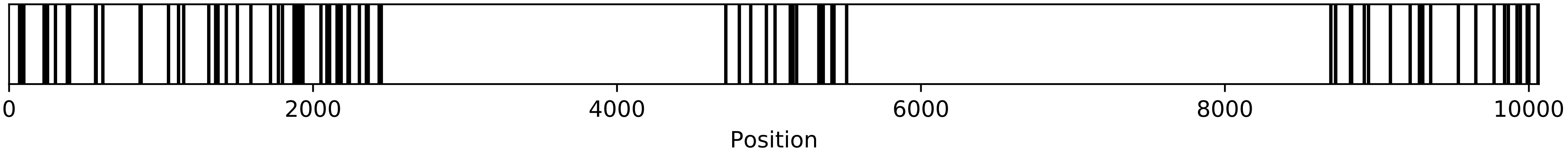


CR1 vs MM1
Number of differences: 193

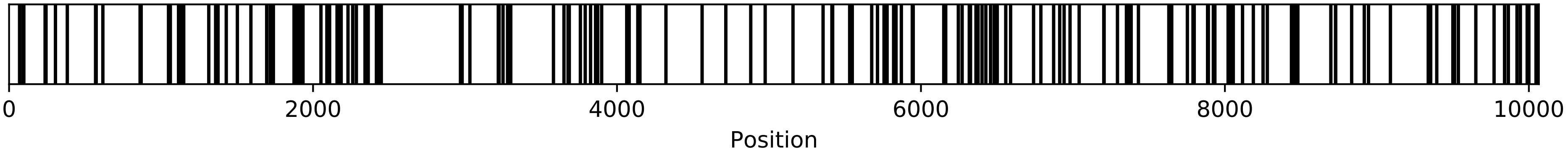


CR1 vs MM2
Number of differences: 193

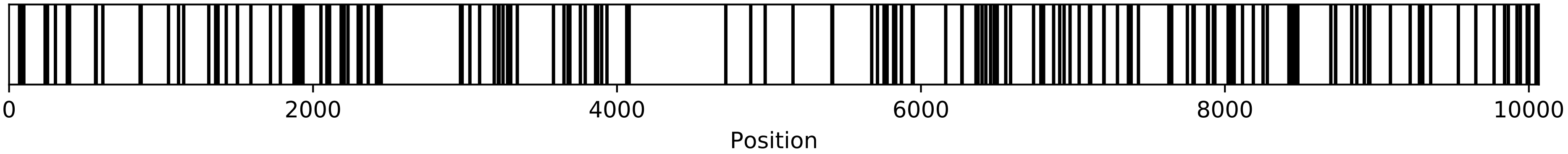


CR2 vs MM1
Number of differences: 153

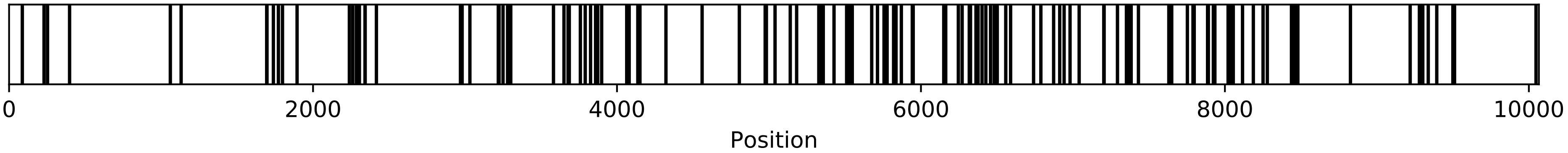


CR2 vs MM2
Number of differences: 141

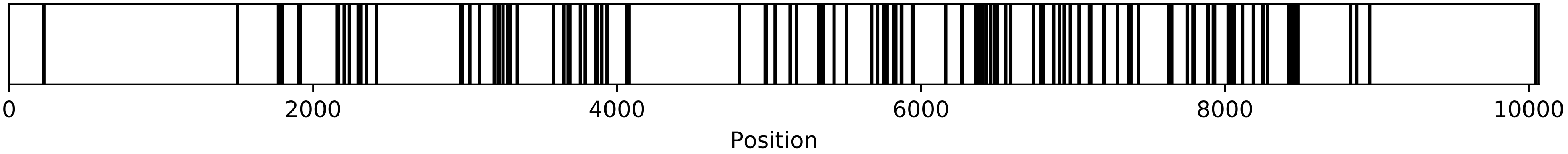


MM1 vs MM2
Number of differences: 65

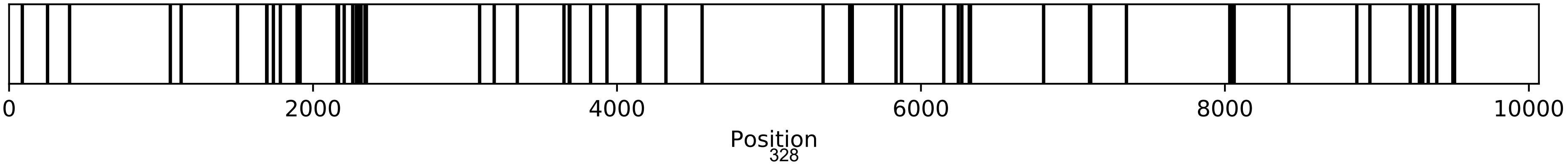

### Region: 874



CR1 vs CR2
Number of differences: 393

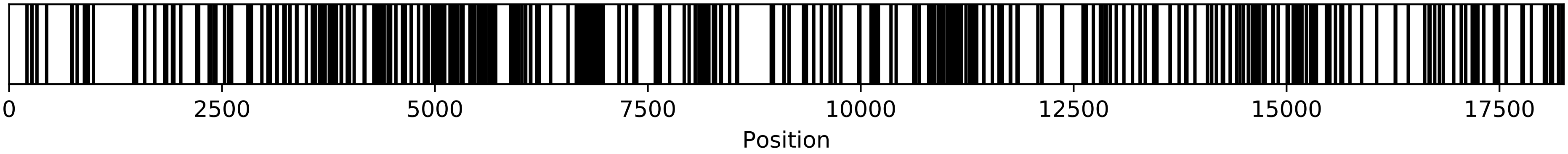


CR1 vs MM1
Number of differences: 394

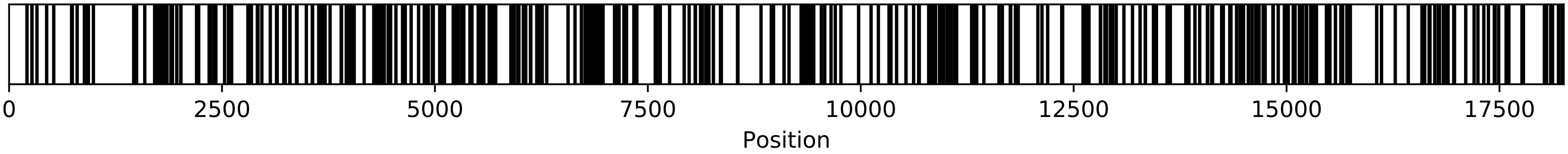


CR1 vs MM2
Number of differences: 389

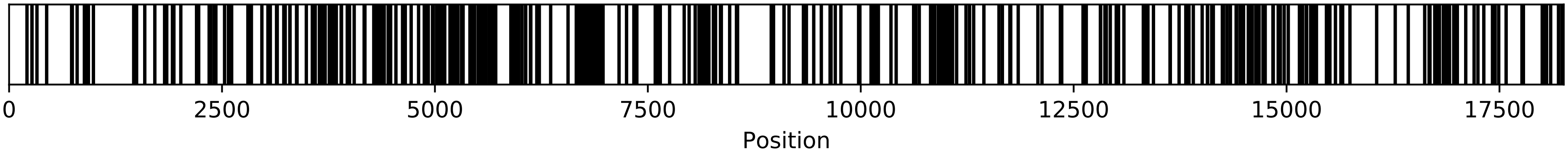


CR2 vs MM1
Number of differences: 182

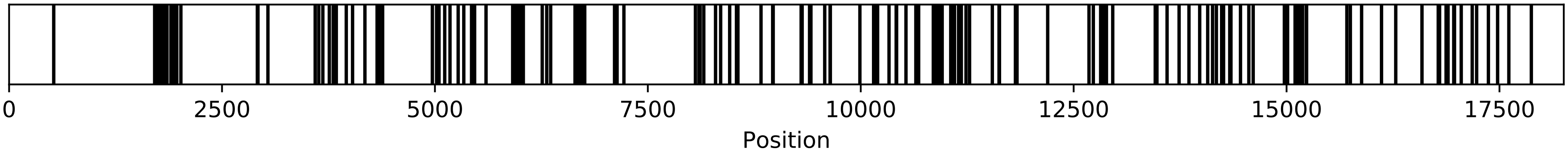


CR2 vs MM2
Number of differences: 83

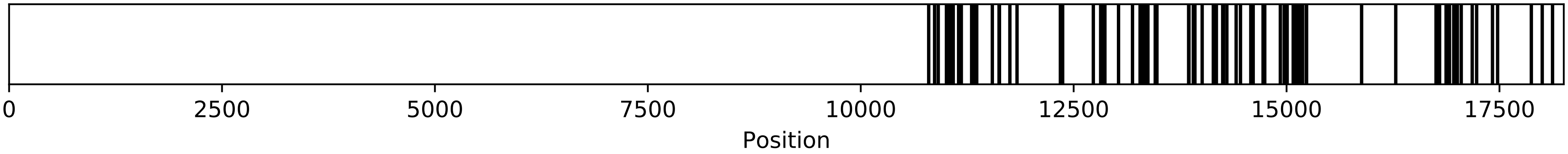


MM1 vs MM2
Number of differences: 180

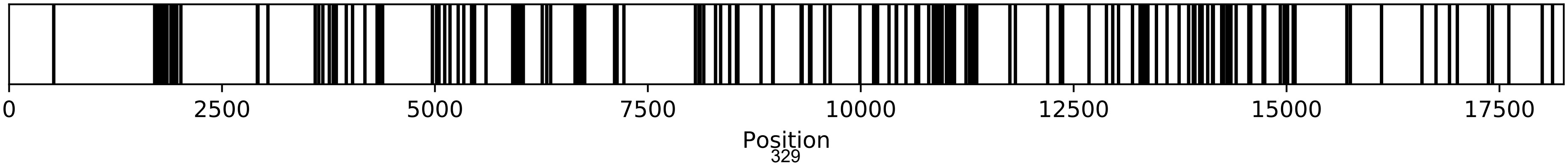

### Region: 876



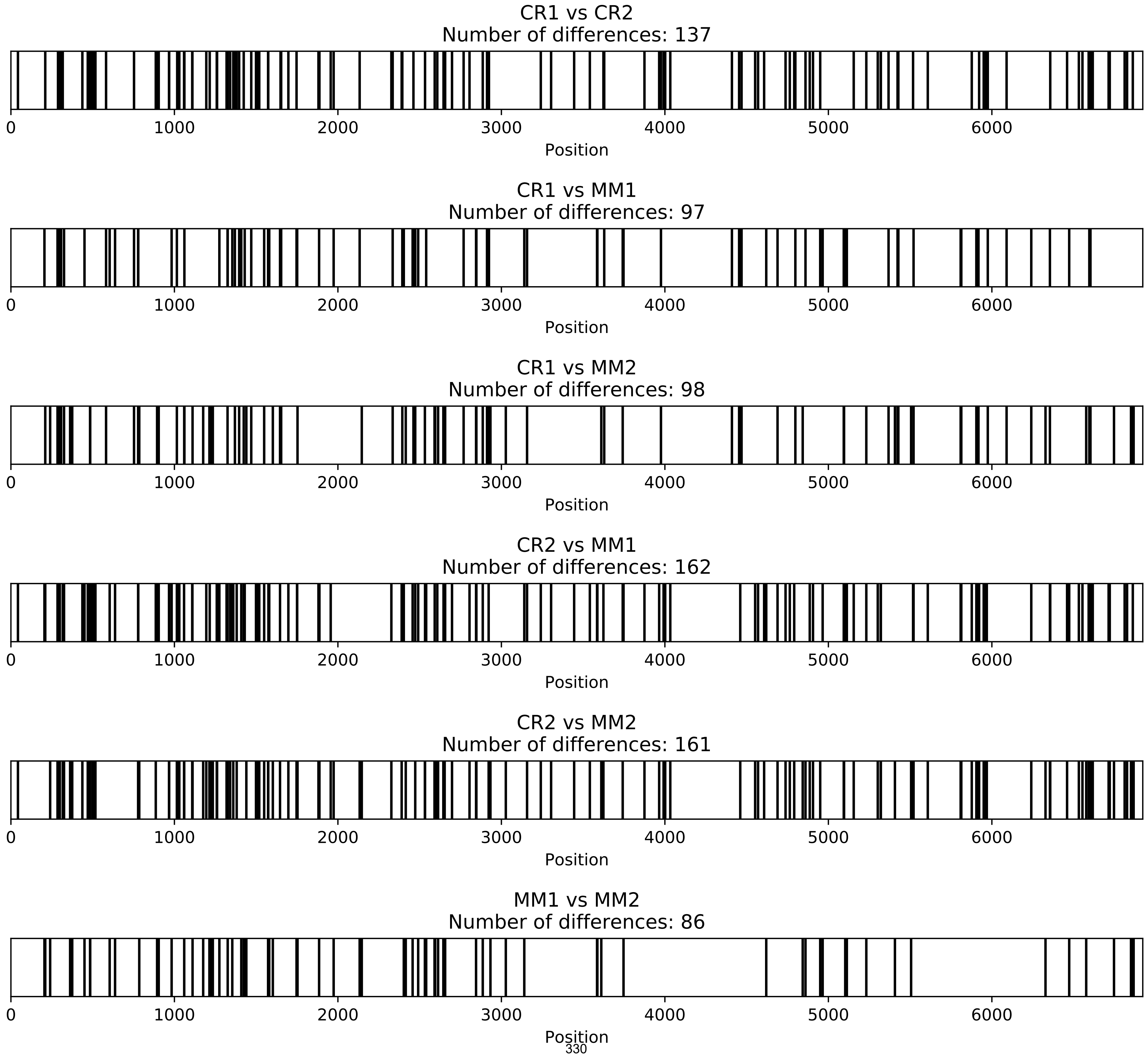
CR1 vs CR2
Number of differences: 137
CR1 vs MM1
Number of differences: 97
CR1 vs MM2
Number of differences: 98
CR2 vs MM1
Number of differences: 162
CR2 vs MM2
Number of differences: 161
MM1 vs MM2
Number of differences: 86
0
1000
2000
3000
4000
5000
6000
Position

### Region: 877



CR1 vs CR2
Number of differences: 287

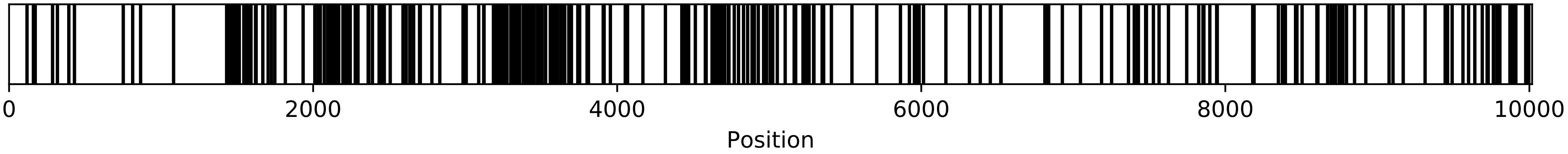


CR1 vs MM1
Number of differences: 180

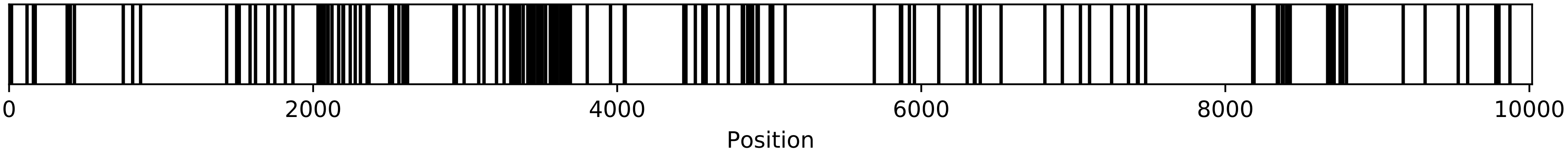


CR1 vs MM2
Number of differences: 181

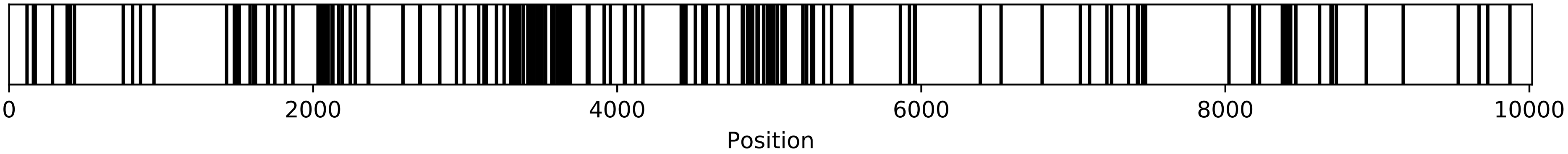


CR2 vs MM1
Number of differences: 238

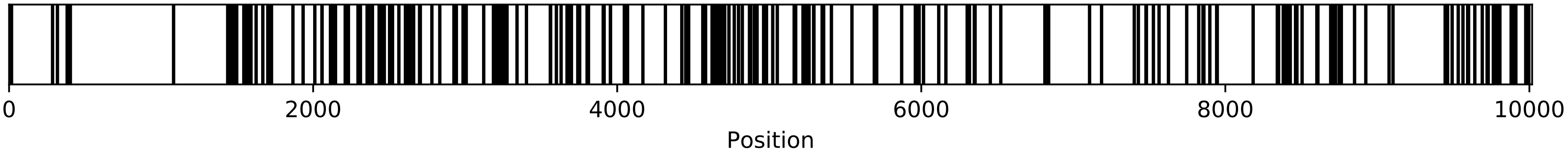


CR2 vs MM2
Number of differences: 232

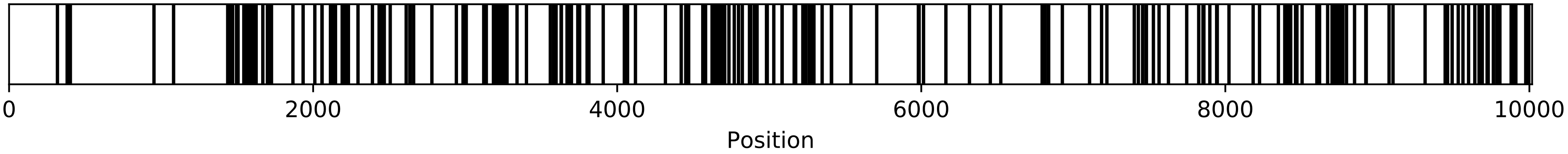


MM1 vs MM2
Number of differences: 112

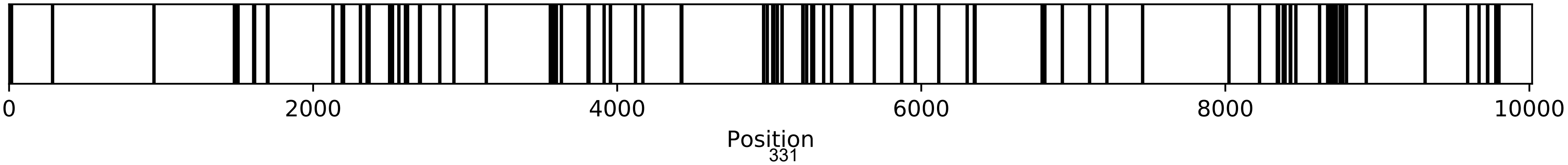

### Region: 878



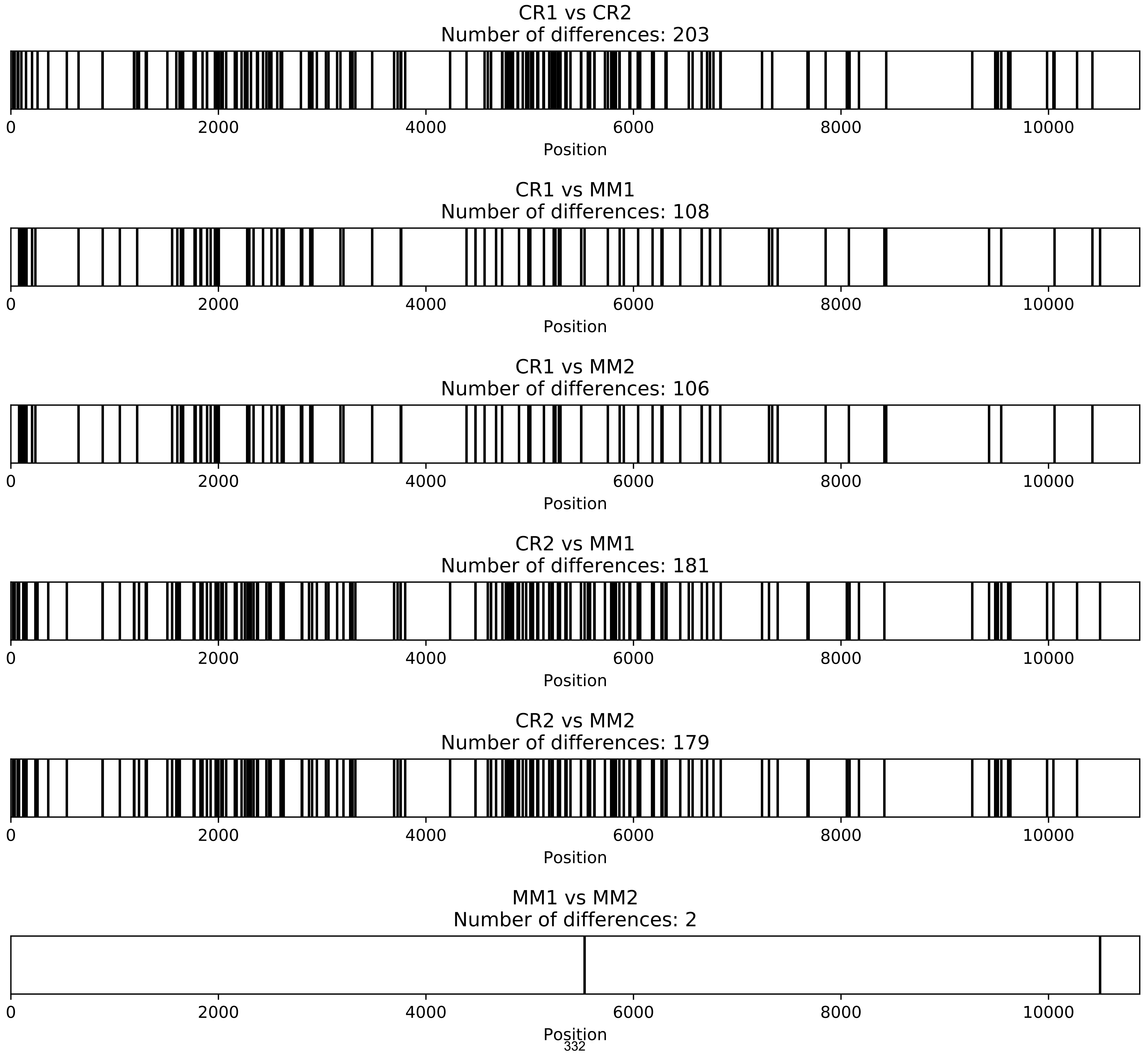
CR1 vs CR2
Number of differences: 203
CR1 vs MM1
Number of differences: 108
CR1 vs MM2
Number of differences: 106
CR2 vs MM1
Number of differences: 181
CR2 vs MM2
Number of differences: 179
MM1 vs MM2
Number of differences: 2
0
2000
4000
6000
8000
10000
Position

### Region: 880



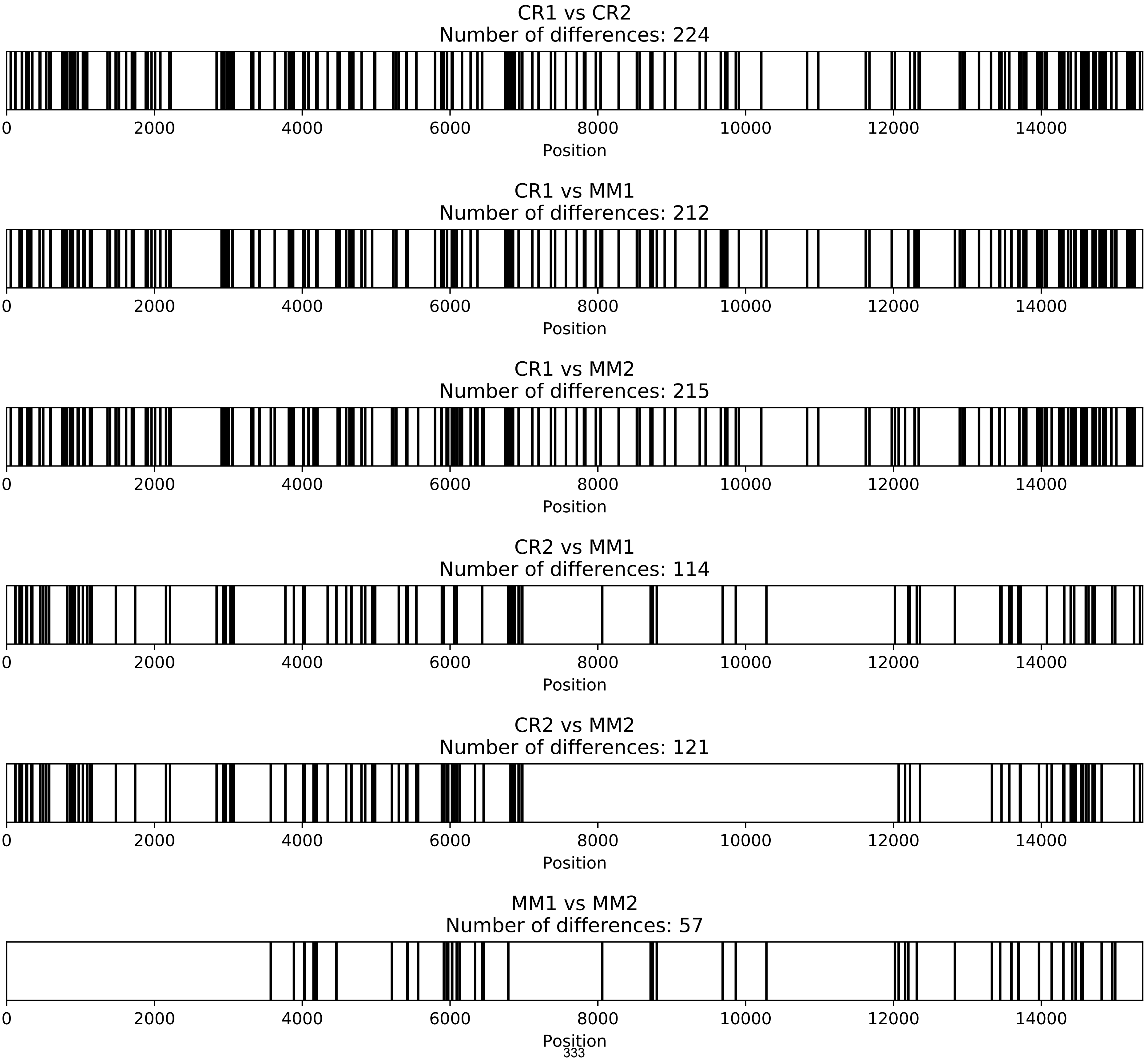
CR1 vs CR2
Number of differences: 224
CR1 vs MM1
Number of differences: 212
CR1 vs MM2
Number of differences: 215
CR2 vs MM1
Number of differences: 114
CR2 vs MM2
Number of differences: 121
MM1 vs MM2
Number of differences: 57
0
2000
4000
6000
8000
10000
12000
14000
Position

### Region: 882



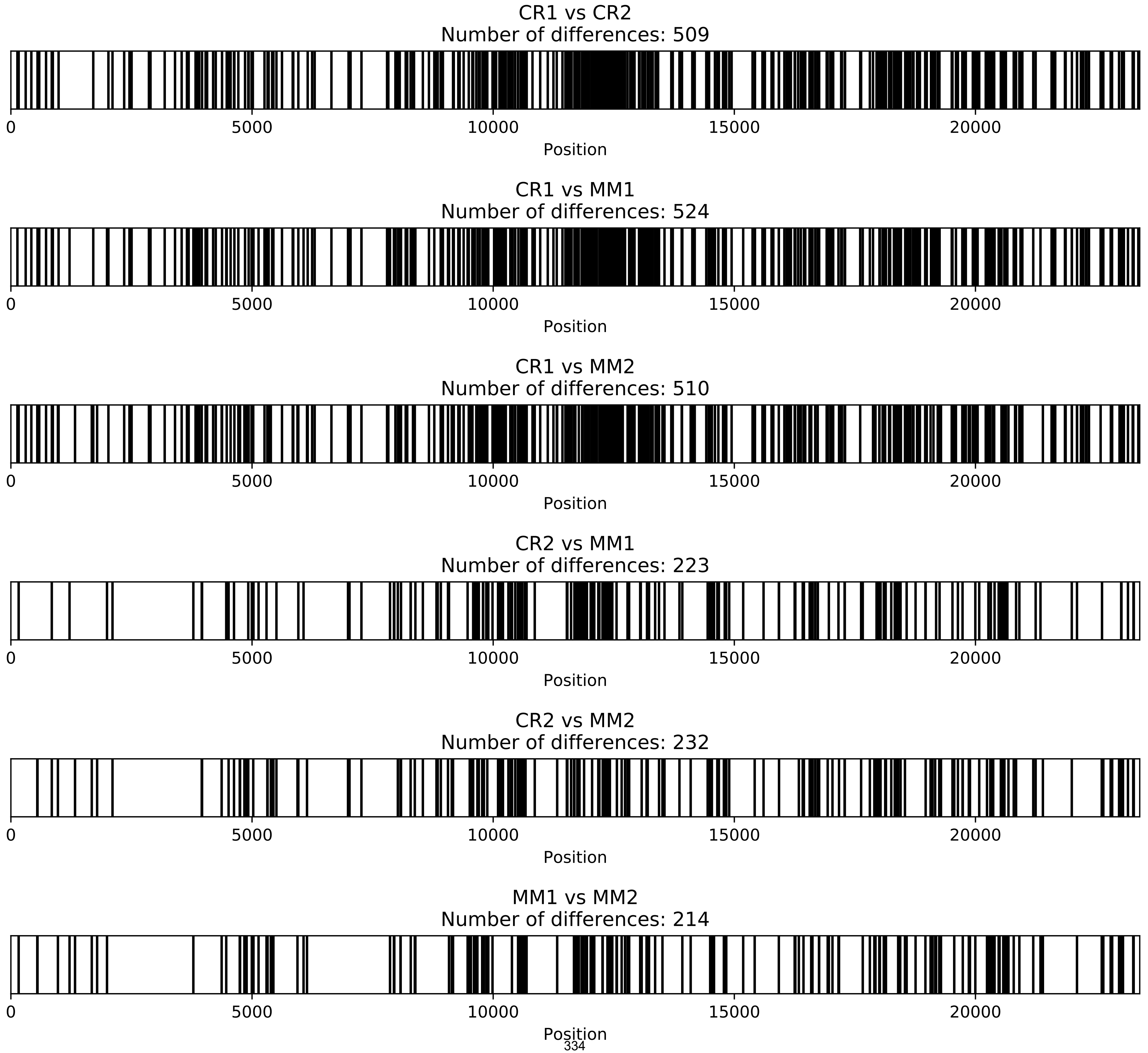
CR1 vs CR2
Number of differences: 509
0
5000
10000
15000
20000
Position
CR1 vs MM1
Number of differences: 524
0
5000
10000
15000
20000
Position
CR1 vs MM2
Number of differences: 510
0
5000
10000
15000
20000
Position
CR2 vs MM1
Number of differences: 223
0
5000
10000
15000
20000
Position
CR2 vs MM2
Number of differences: 232
0
5000
10000
15000
20000
Position
MM1 vs MM2
Number of differences: 214
0
5000
10000
15000
20000
Position

### Region: 884



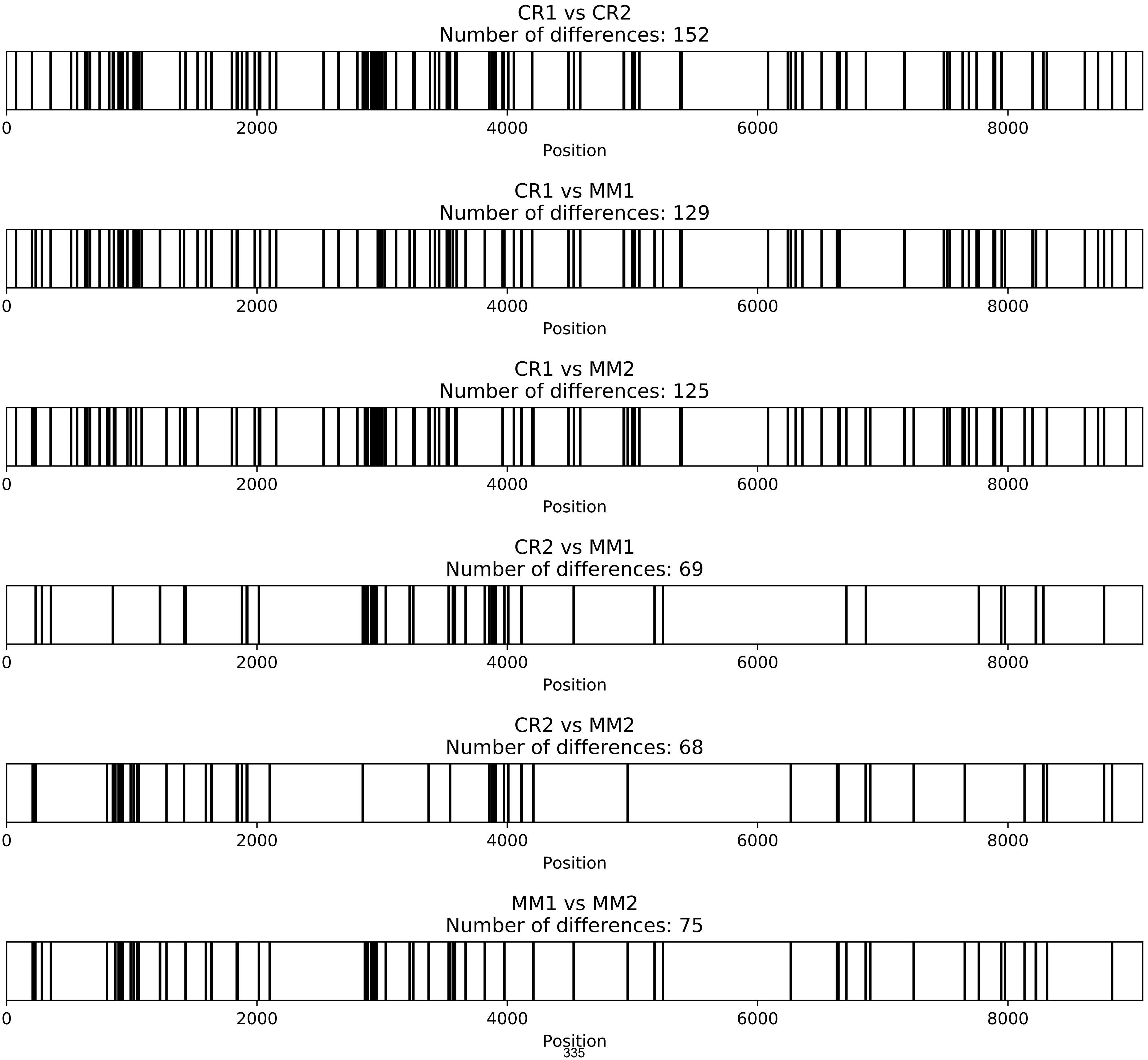
CR1 vs CR2
Number of differences: 152
0
2000
4000
6000
8000
Position
CR1 vs MM1
Number of differences: 129
0
2000
4000
6000
8000
Position
CR1 vs MM2
Number of differences: 125
0
2000
4000
6000
8000
Position
CR2 vs MM1
Number of differences: 69
0
2000
4000
6000
8000
Position
CR2 vs MM2
Number of differences: 68
0
2000
4000
6000
8000
Position
MM1 vs MM2
Number of differences: 75
0
2000
4000
6000
8000
Position

### Region: 885



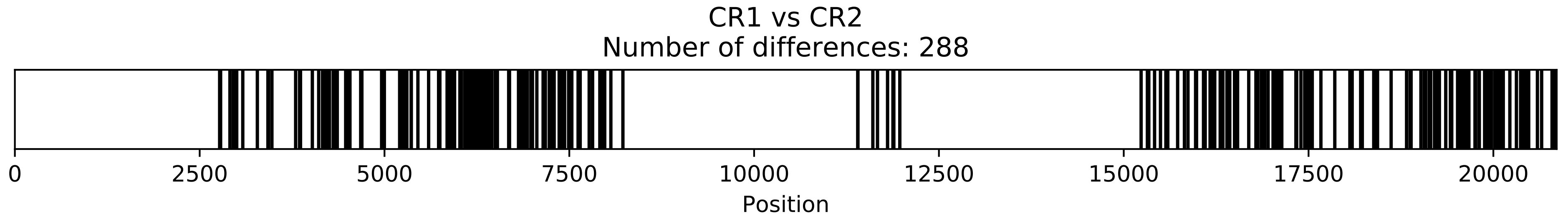
CR1 vs CR2
Number of differences: 288
0
2500
5000
7500
10000
12500
15000
17500
20000
Position

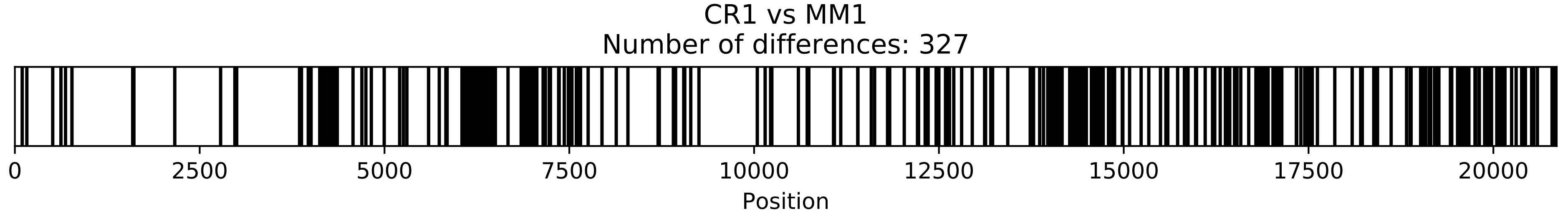
CR1 vs MM1
Number of differences: 327
0
2500
5000
7500
10000
12500
15000
17500
20000
Position

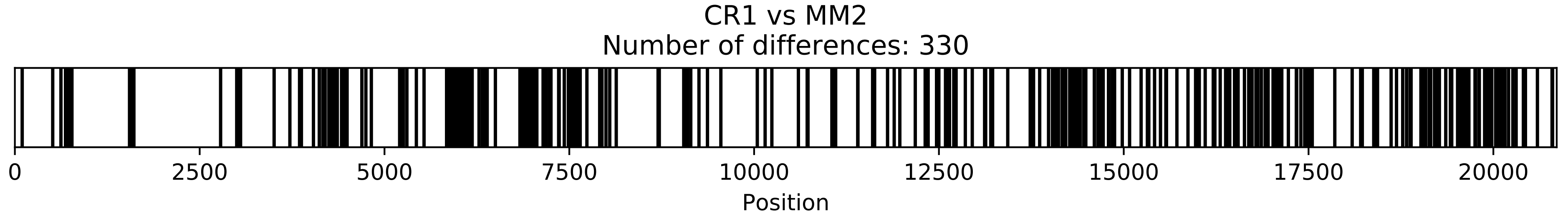
CR1 vs MM2
Number of differences: 330
0
2500
5000
7500
10000
12500
15000
17500
20000
Position

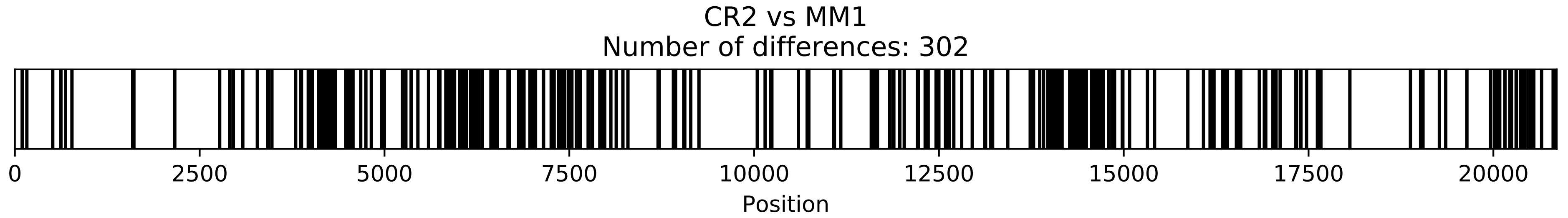
CR2 vs MM1
Number of differences: 302
0
2500
5000
7500
10000
12500
15000
17500
20000
Position

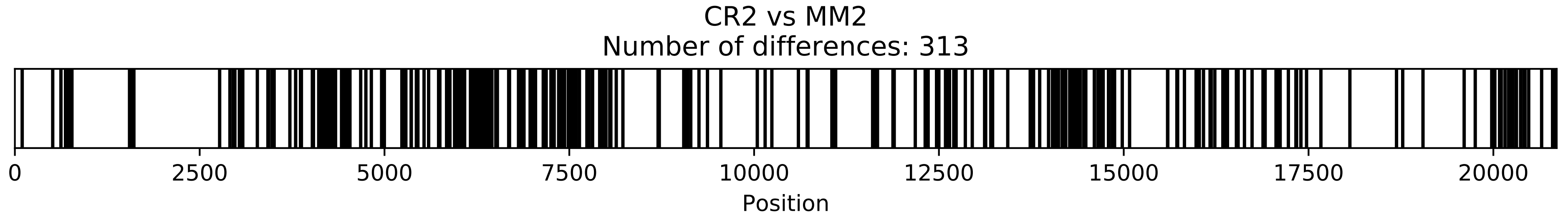
CR2 vs MM2
Number of differences: 313
0
2500
5000
7500
10000
12500
15000
17500
20000
Position

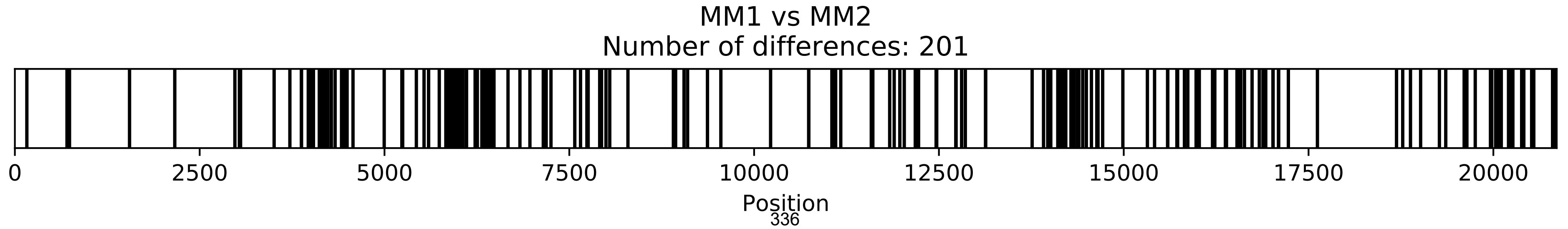
MM1 vs MM2
Number of differences: 201
0
2500
5000
7500
10000
12500
15000
17500
20000
Position

### Region: 891



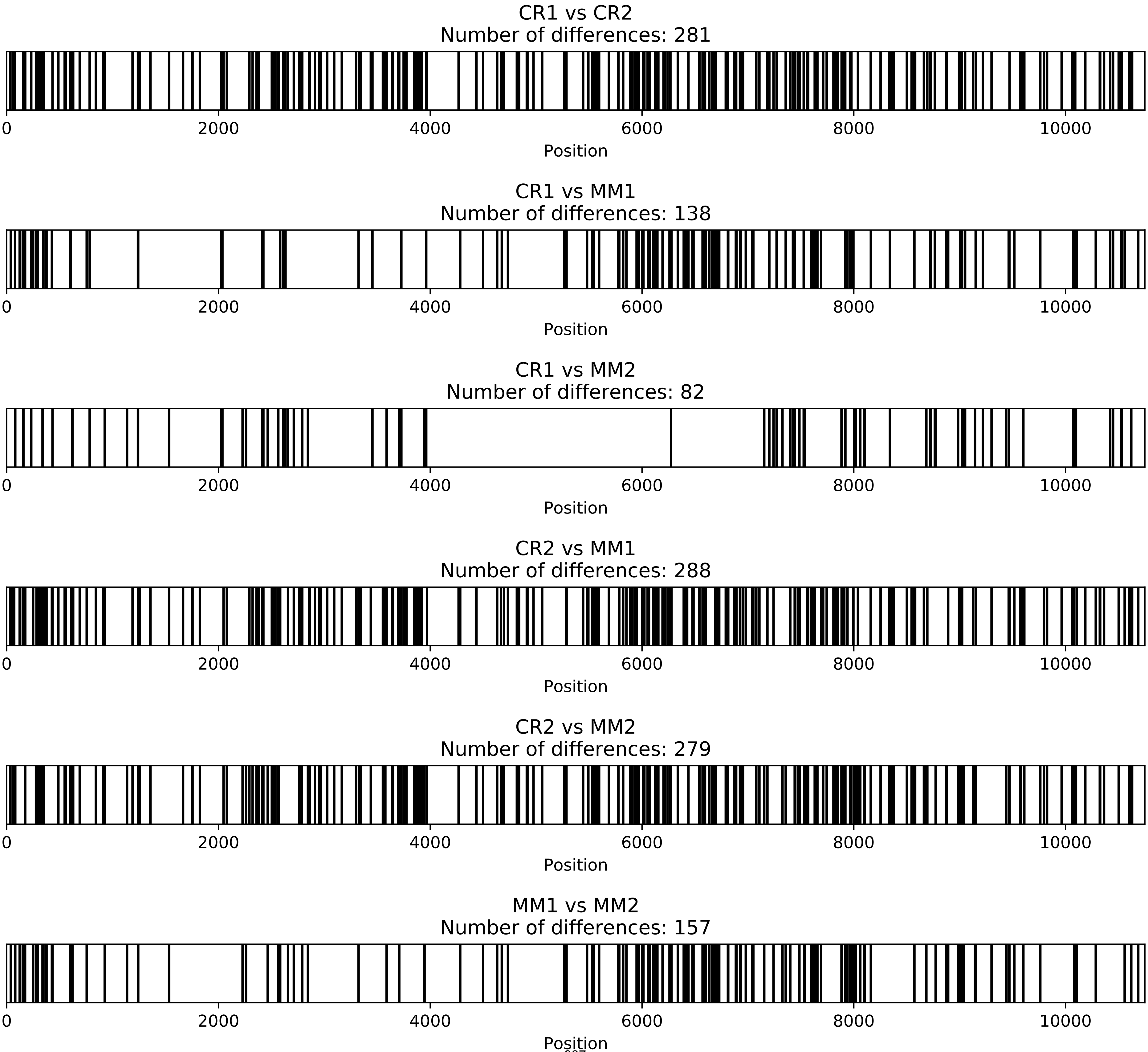
CR1 vs CR2
Number of differences: 281
0
2000
4000
6000
8000
10000
Position
CR1 vs MM1
Number of differences: 138
Position
CR1 vs MM2
Number of differences: 82
Position
CR2 vs MM1
Number of differences: 288
Position
CR2 vs MM2
Number of differences: 279
Position
MM1 vs MM2
Number of differences: 157
Position

### Region: 892



CR1 vs CR2
Number of differences: 184

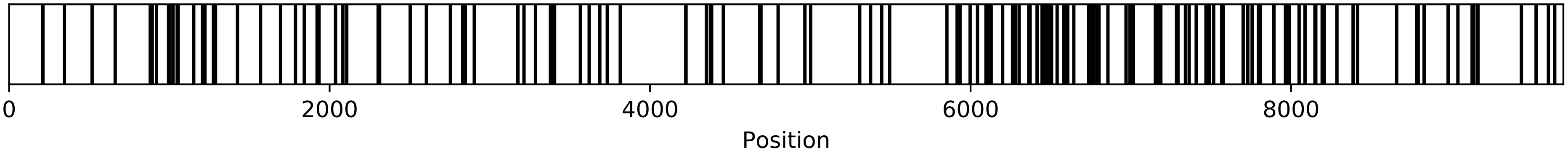


CR1 vs MM1
Number of differences: 43

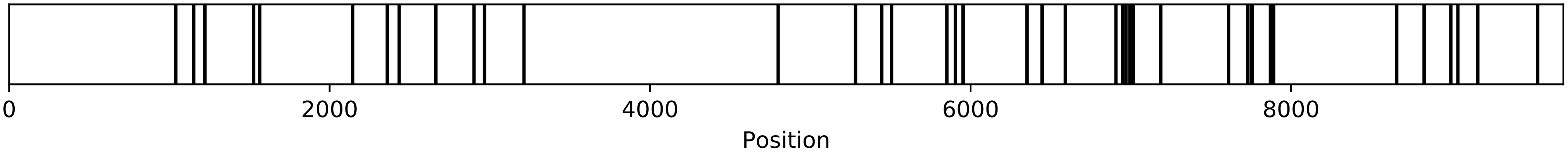


CR1 vs MM2
Number of differences: 43

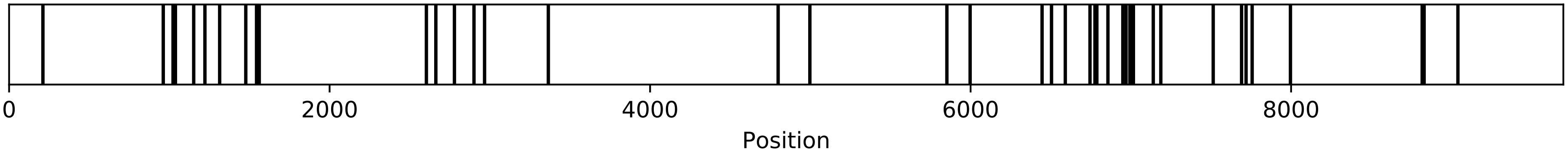


CR2 vs MM1
Number of differences: 185

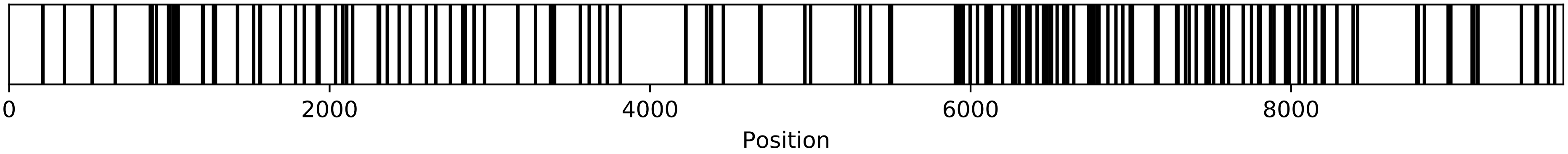


CR2 vs MM2
Number of differences: 183

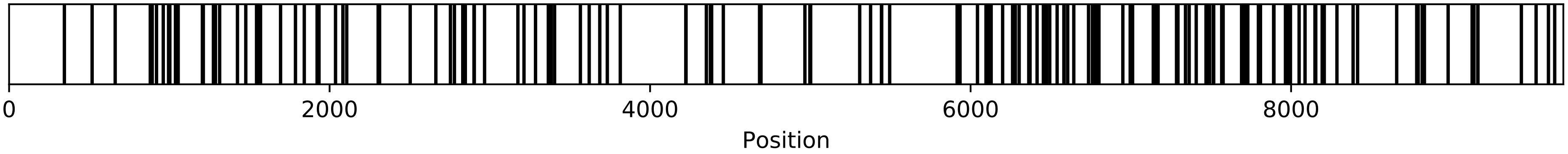


MM1 vs MM2
Number of differences: 48

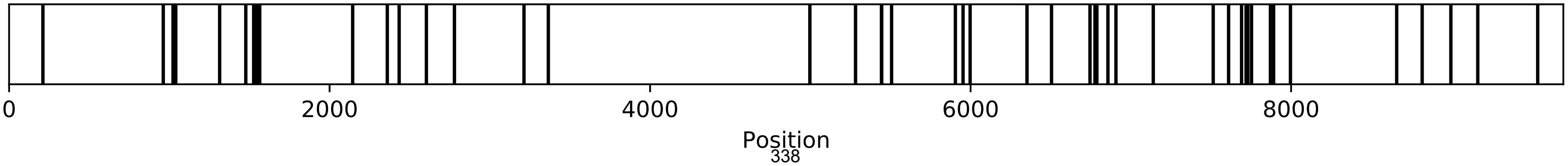

### Region: 893



CR1 vs CR2
Number of differences: 313

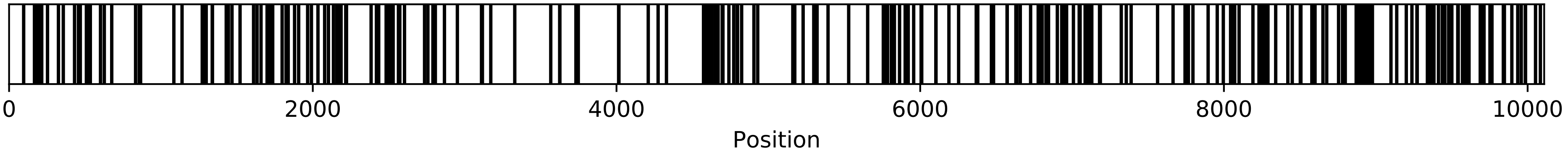


CR1 vs MM1
Number of differences: 132

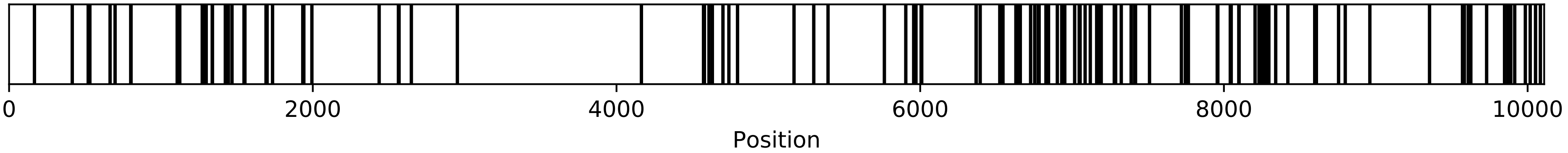


CR1 vs MM2
Number of differences: 142

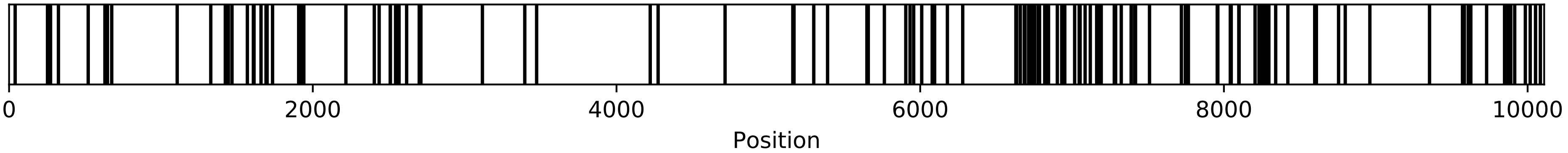


CR2 vs MM1
Number of differences: 304

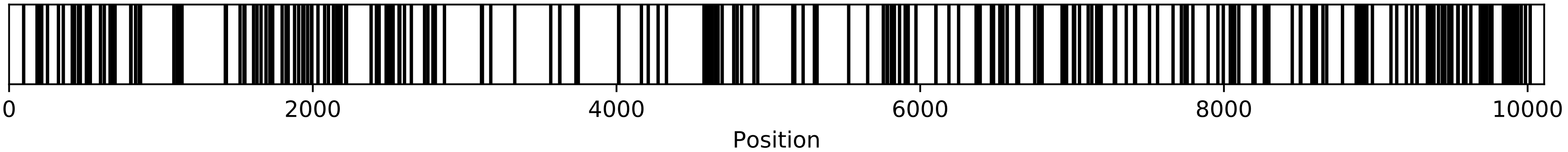


CR2 vs MM2
Number of differences: 323

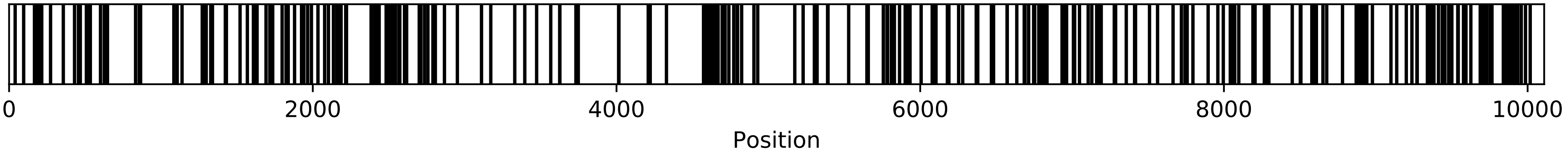


MM1 vs MM2
Number of differences: 85

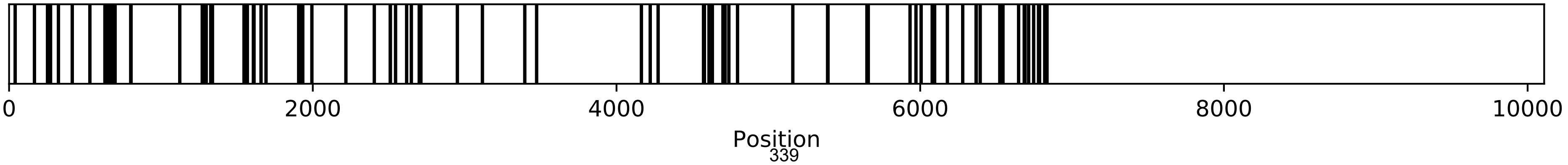

### Region: 894



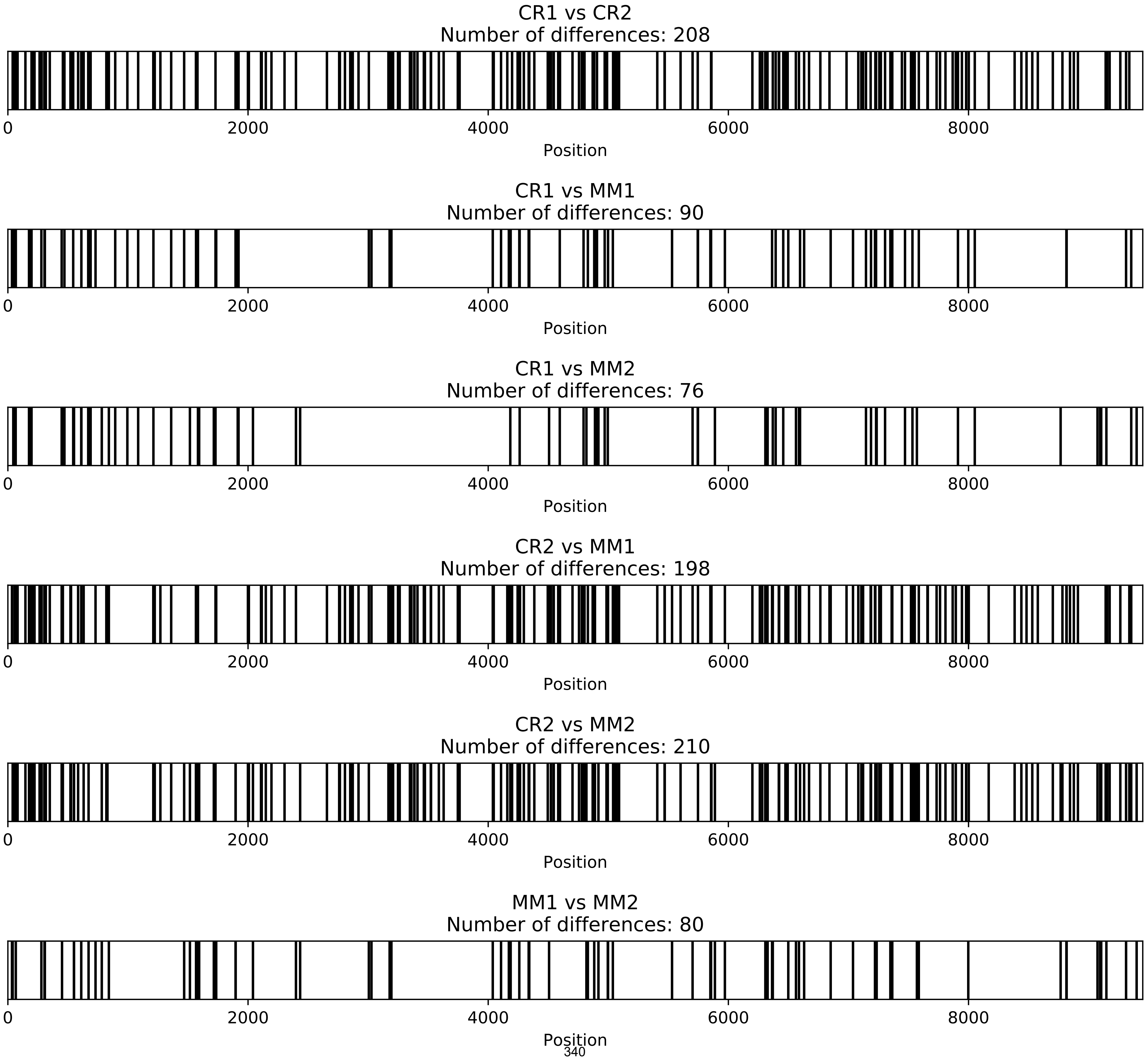
CR1 vs CR2
Number of differences: 208
CR1 vs MM1
Number of differences: 90
CR1 vs MM2
Number of differences: 76
CR2 vs MM1
Number of differences: 198
CR2 vs MM2
Number of differences: 210
MM1 vs MM2
Number of differences: 80
0
2000
4000
6000
8000
Position

### Region: 895



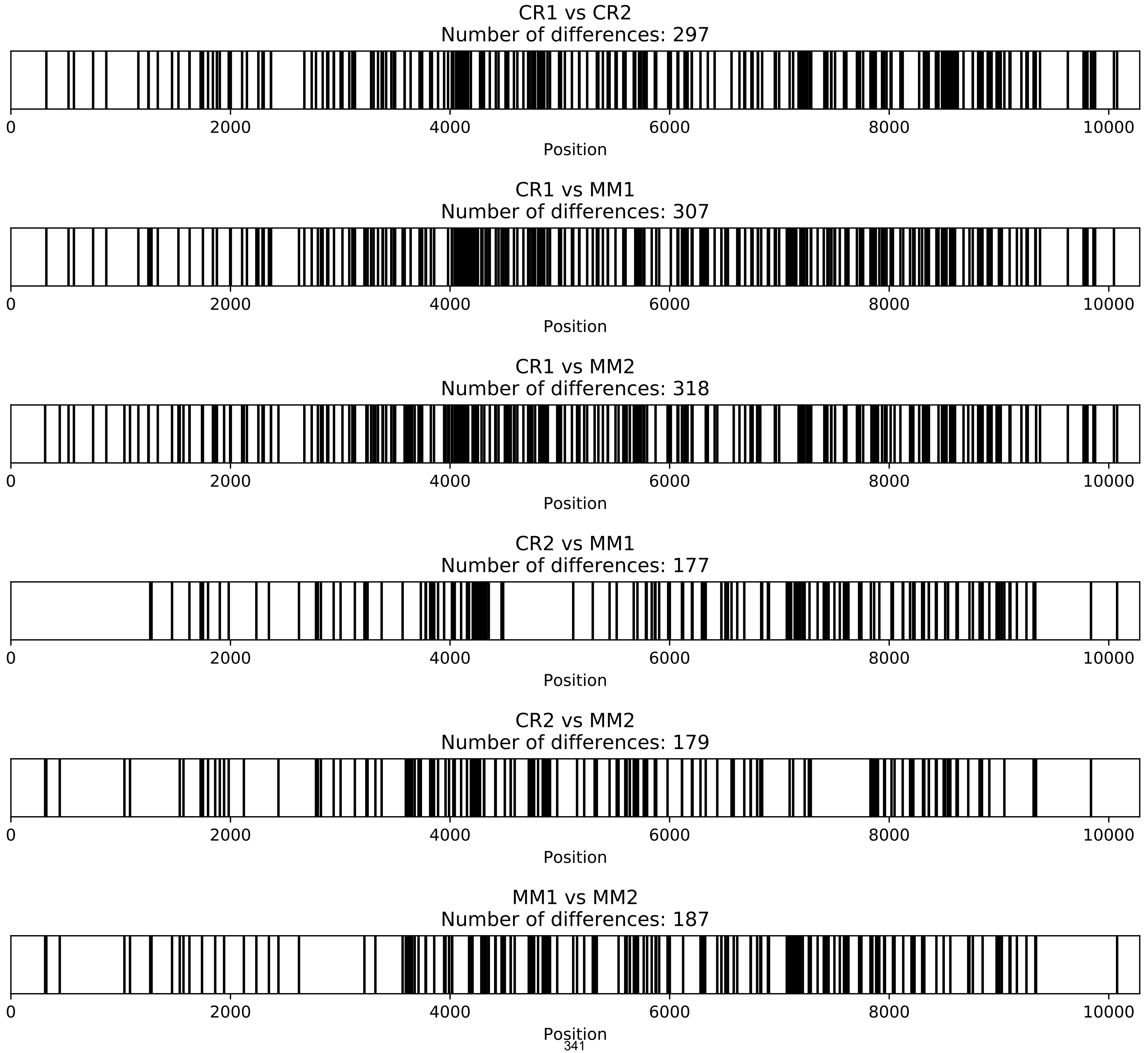
CR1 vs CR2
Number of differences: 297
CR1 vs MM1
Number of differences: 307
CR1 vs MM2
Number of differences: 318
CR2 vs MM1
Number of differences: 177
CR2 vs MM2
Number of differences: 179
MM1 vs MM2
Number of differences: 187
Position
0
2000
4000
6000
8000
10000

### Region: 896



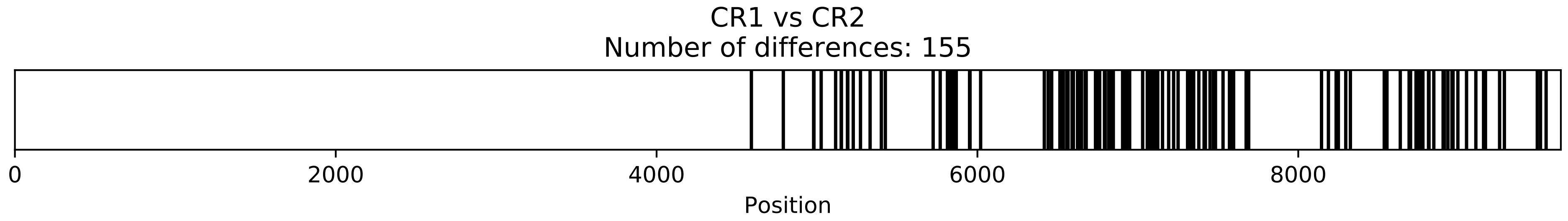
CR1 vs CR2
Number of differences: 155
0
2000
4000
6000
8000
Position

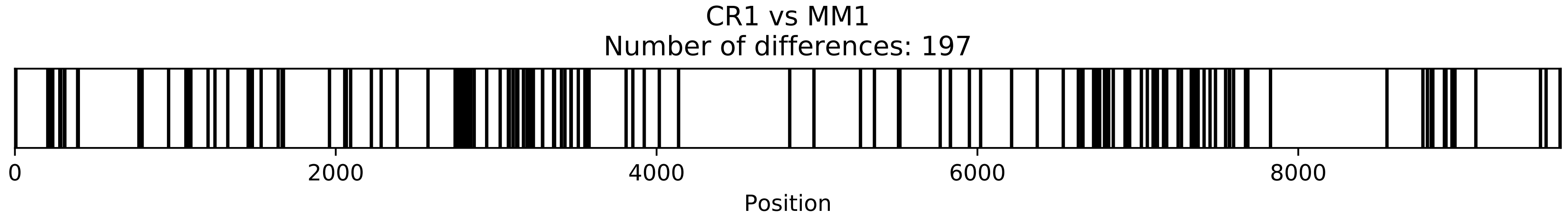
CR1 vs MM1
Number of differences: 197
0
2000
4000
6000
8000
Position

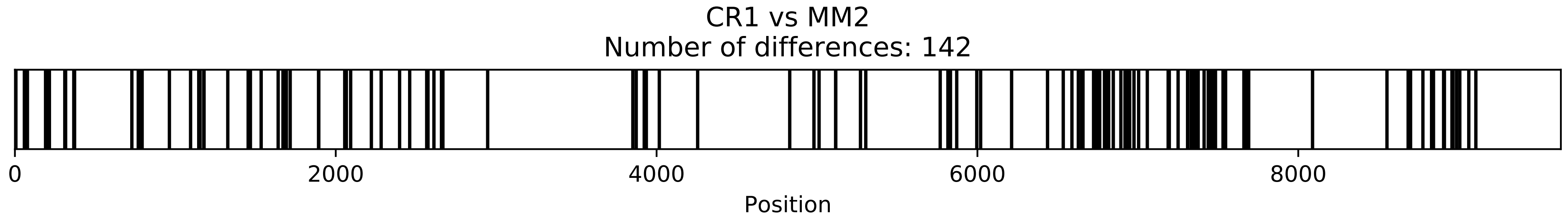
CR1 vs MM2
Number of differences: 142
0
2000
4000
6000
8000
Position

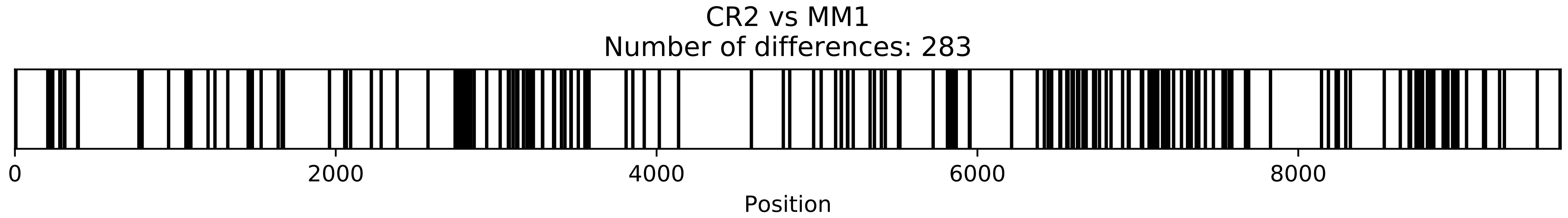
CR2 vs MM1
Number of differences: 283
0
2000
4000
6000
8000
Position

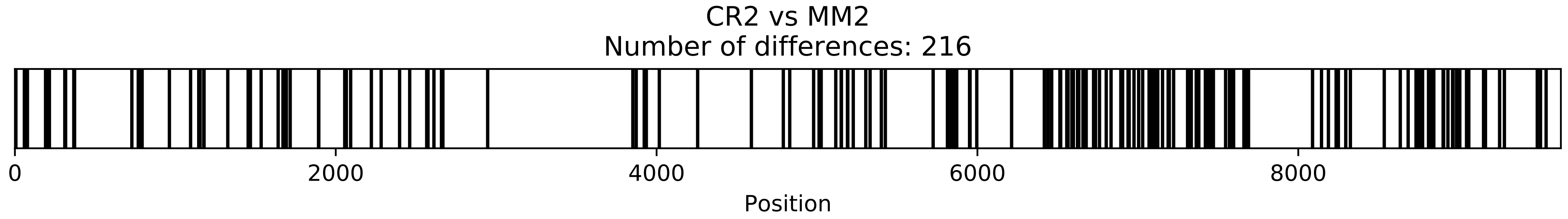
CR2 vs MM2
Number of differences: 216
0
2000
4000
6000
8000
Position

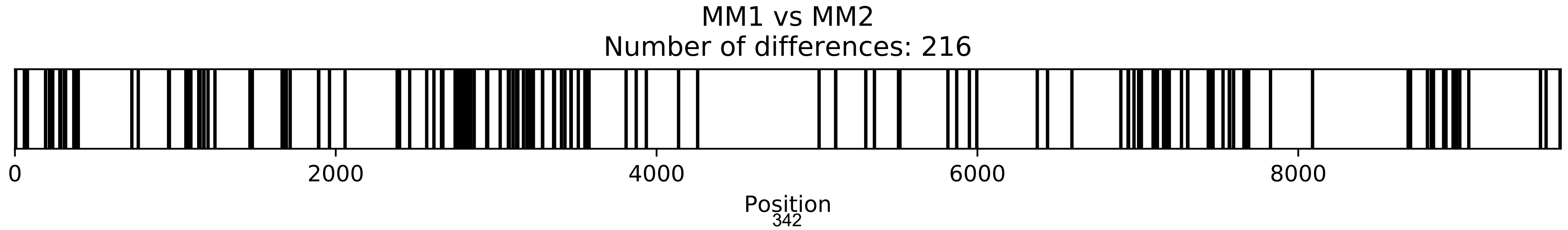
MM1 vs MM2
Number of differences: 216
0
2000
4000
6000
8000
Position

### Region: 899



CR1 vs CR2
Number of differences: 348

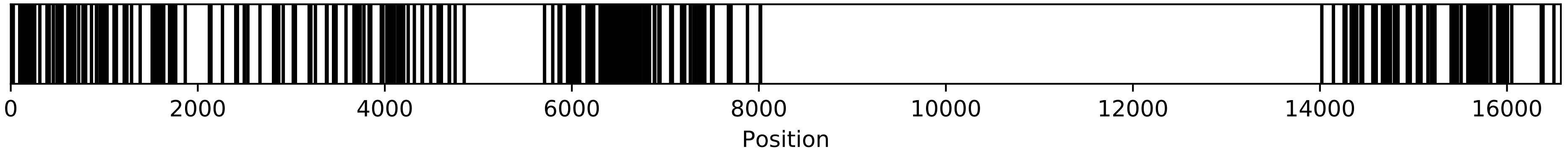


CR1 vs MM1
Number of differences: 306

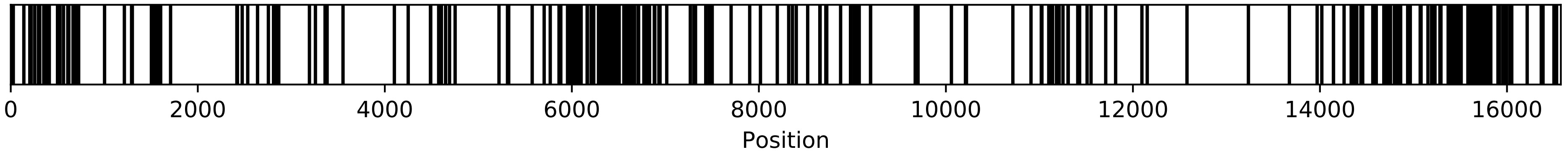


CR1 vs MM2
Number of differences: 333

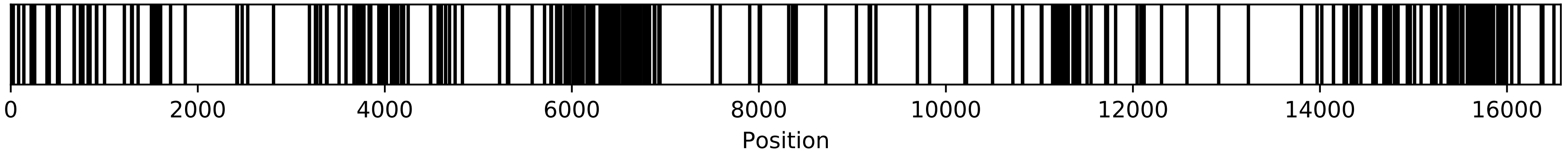


CR2 vs MM1
Number of differences: 281

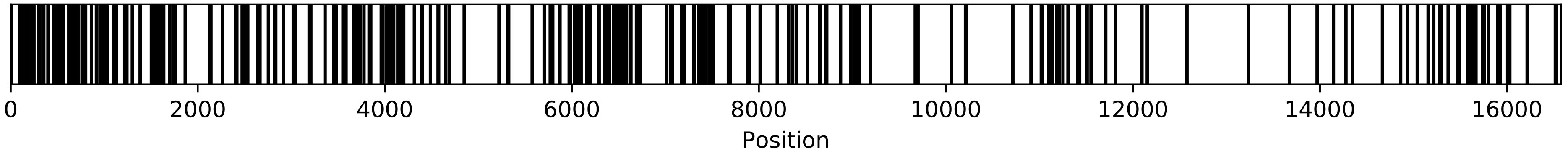


CR2 vs MM2
Number of differences: 288

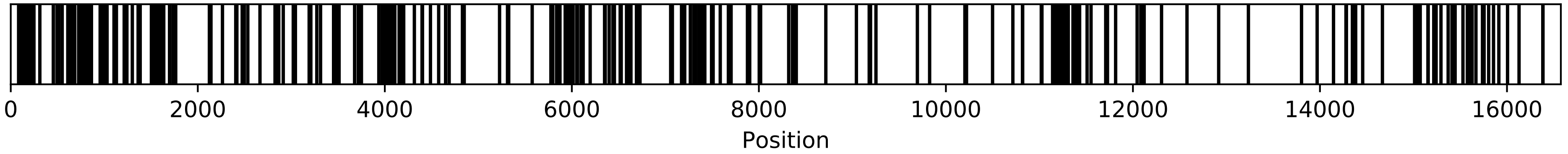


MM1 vs MM2
Number of differences: 204

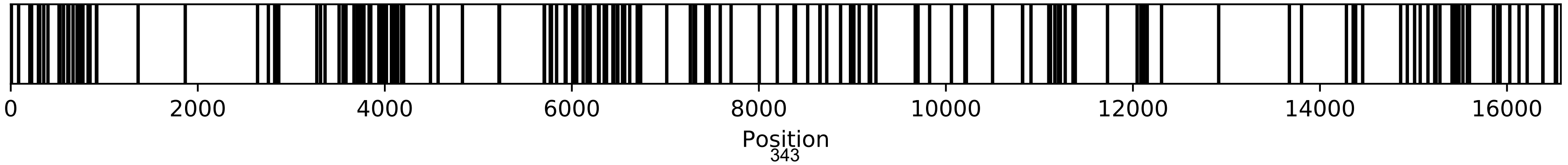

### Region: 900



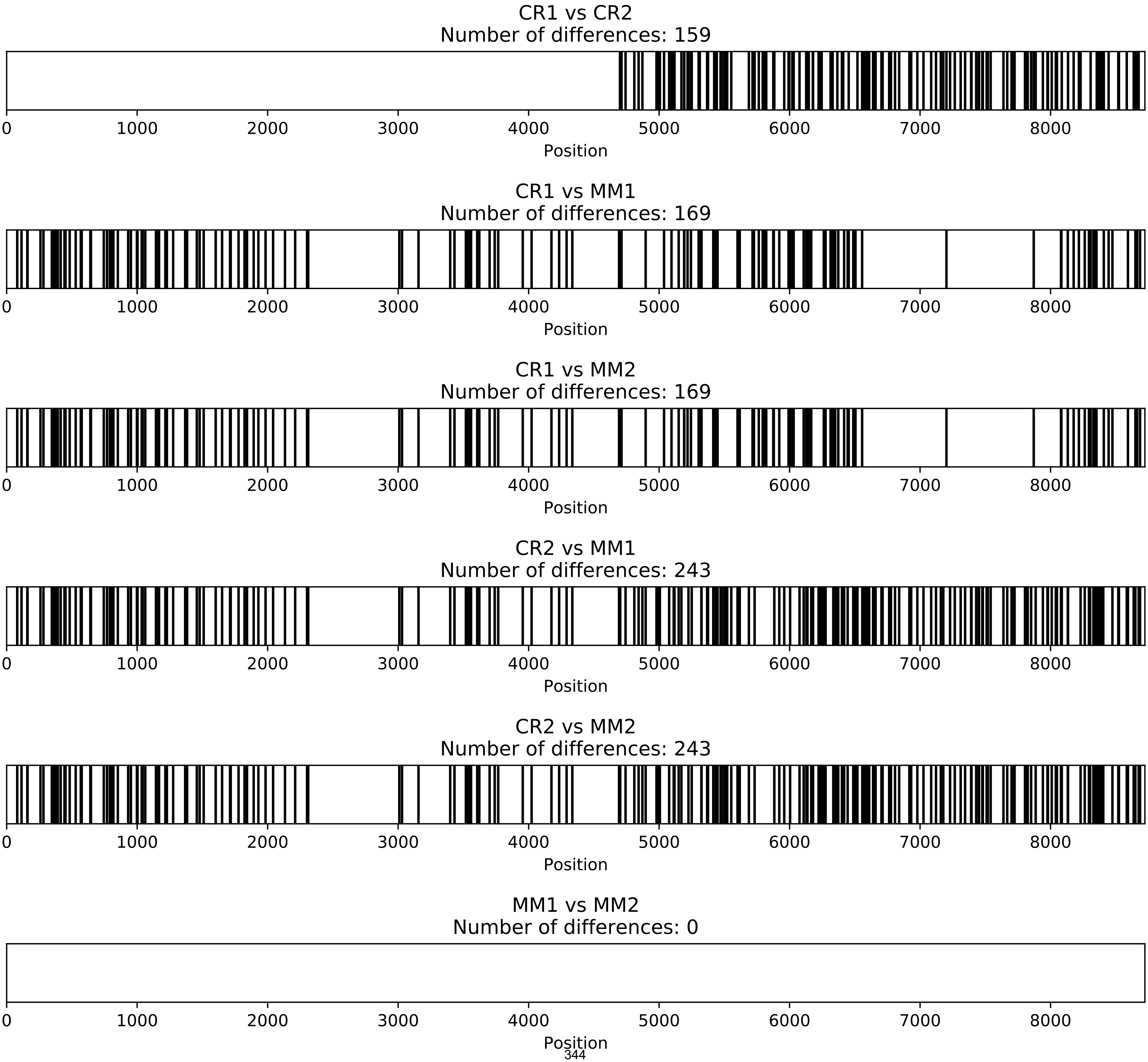
CR1 vs CR2
Number of differences: 159
CR1 vs MM1
Number of differences: 169
CR1 vs MM2
Number of differences: 169
CR2 vs MM1
Number of differences: 243
CR2 vs MM2
Number of differences: 243
MM1 vs MM2
Number of differences: 0
0
1000
2000
3000
4000
5000
6000
7000
8000
Position

### Region: 901



CR1 vs CR2
Number of differences: 220

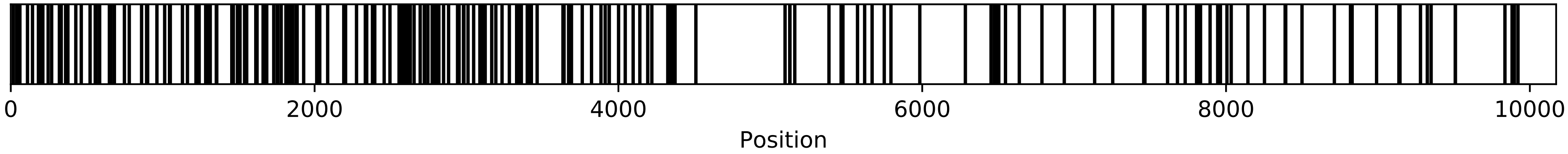


CR1 vs MM1
Number of differences: 62

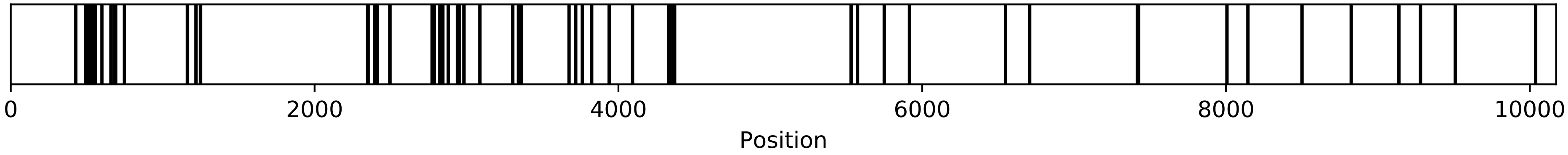


CR1 vs MM2
Number of differences: 113

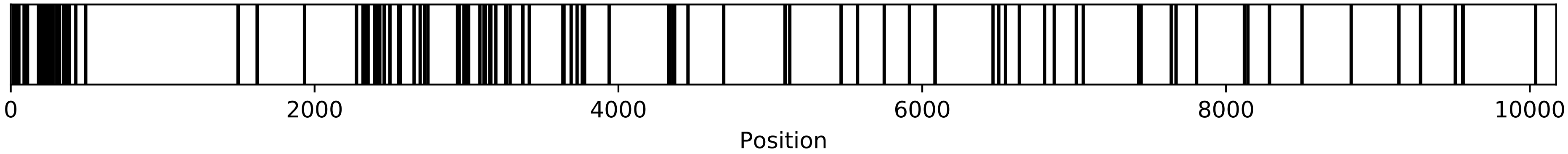


CR2 vs MM1
Number of differences: 216

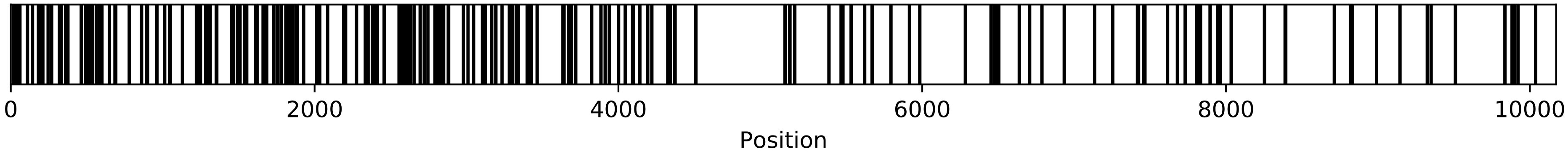


CR2 vs MM2
Number of differences: 240

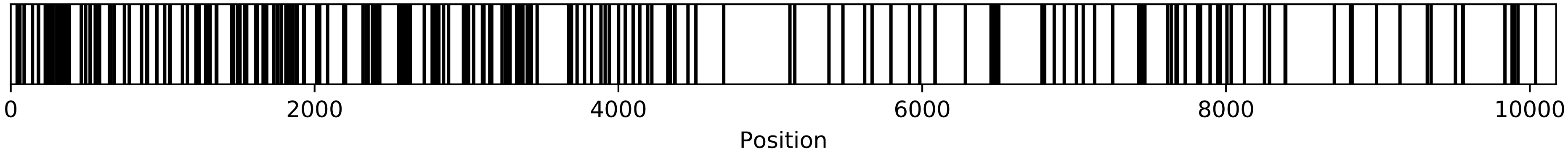


MM1 vs MM2
Number of differences: 117

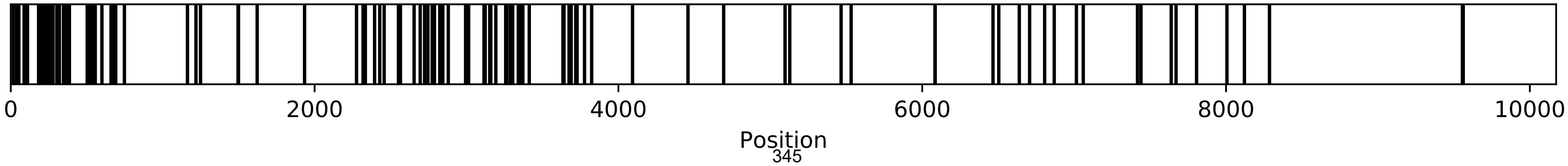

### Region: 909



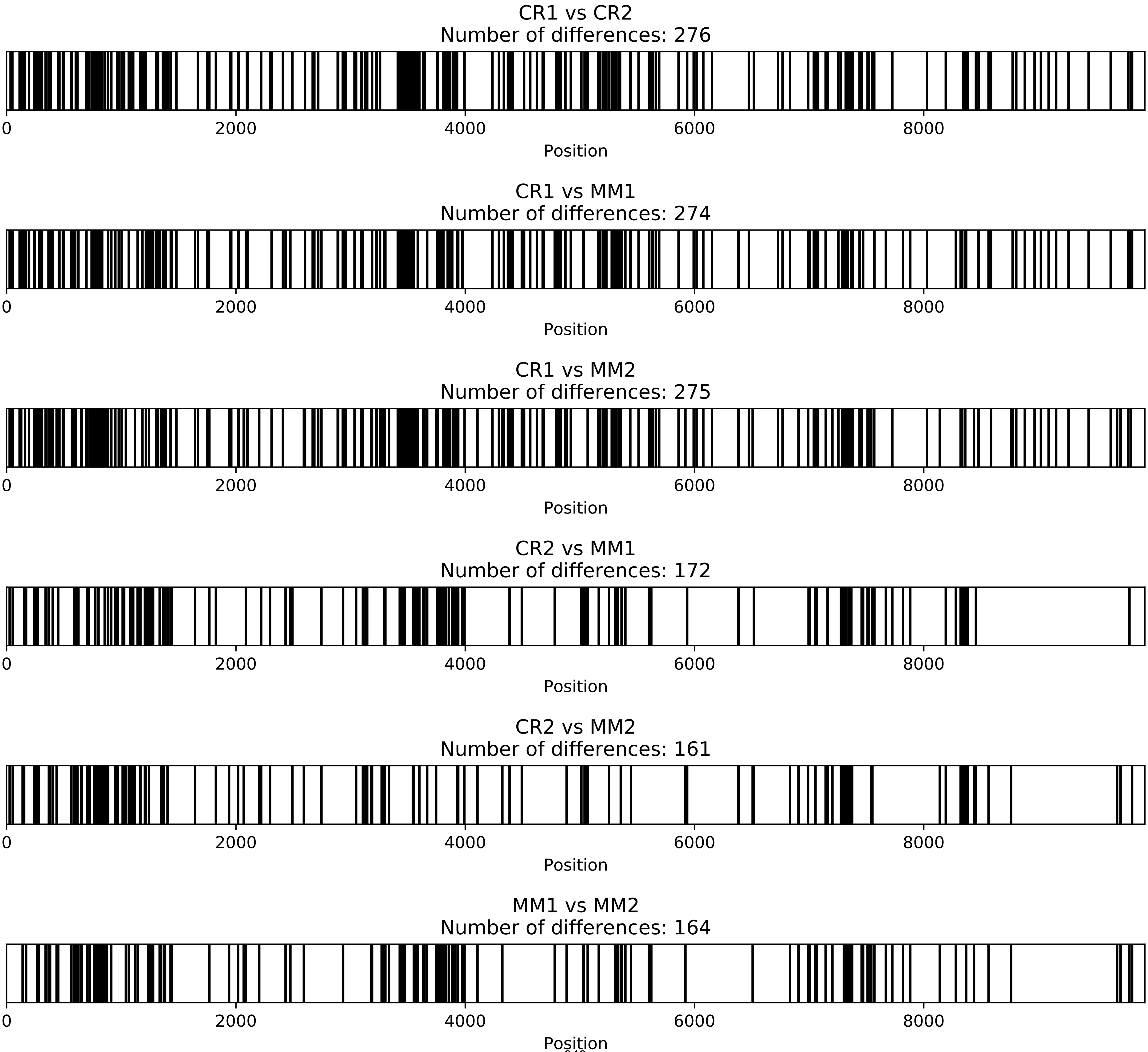
CR1 vs CR2
Number of differences: 276
0
2000
4000
6000
8000
Position
CR1 vs MM1
Number of differences: 274
0
2000
4000
6000
8000
Position
CR1 vs MM2
Number of differences: 275
0
2000
4000
6000
8000
Position
CR2 vs MM1
Number of differences: 172
0
2000
4000
6000
8000
Position
CR2 vs MM2
Number of differences: 161
0
2000
4000
6000
8000
Position
MM1 vs MM2
Number of differences: 164
0
2000
4000
6000
8000
Position

### Region: 912



CR1 vs CR2
Number of differences: 316

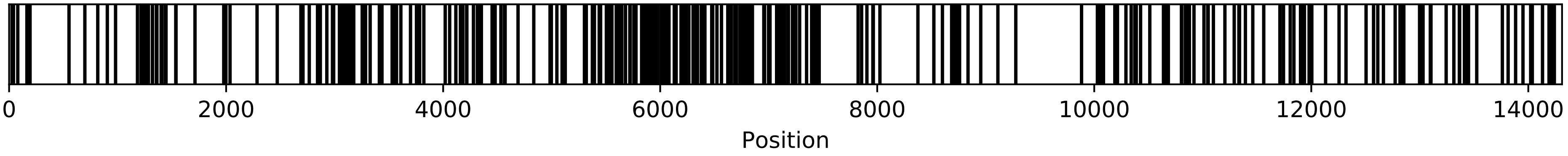


CR1 vs MM1
Number of differences: 151

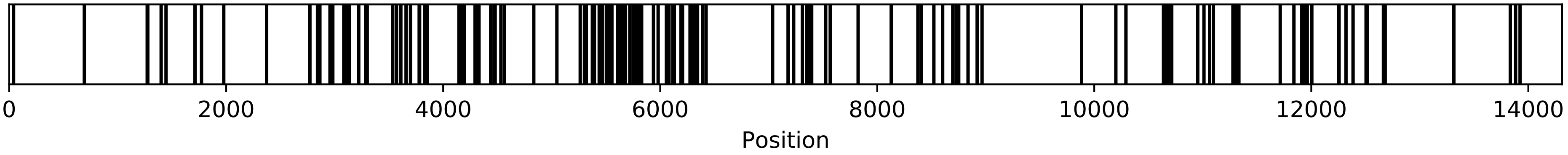


CR1 vs MM2
Number of differences: 168

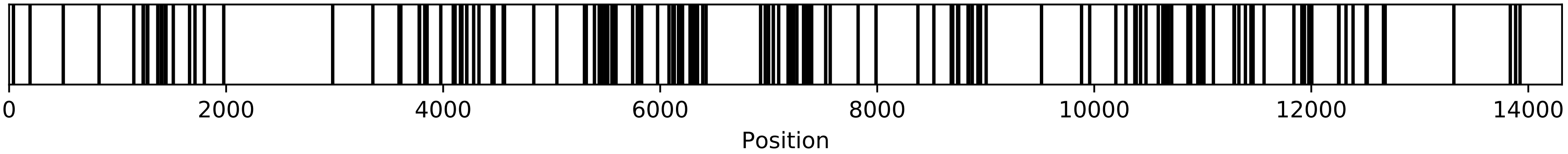


CR2 vs MM1
Number of differences: 287

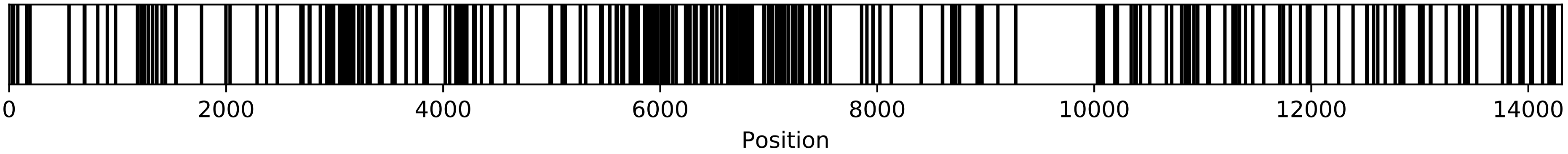


CR2 vs MM2
Number of differences: 312

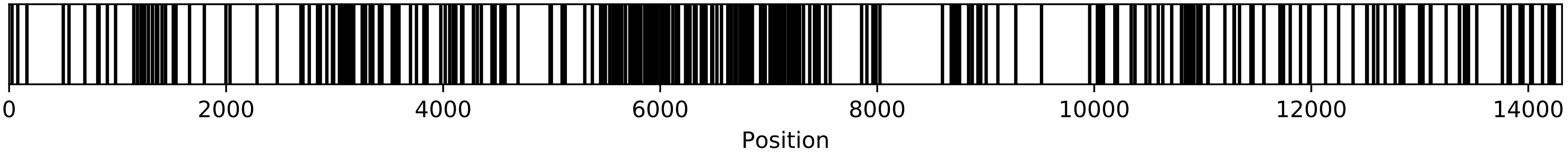


MM1 vs MM2
Number of differences: 137

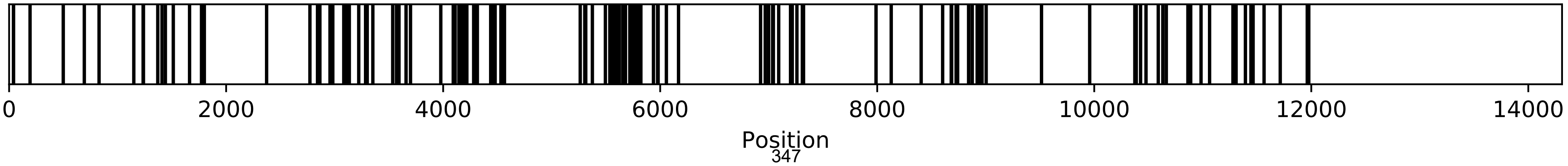

### Region: 913



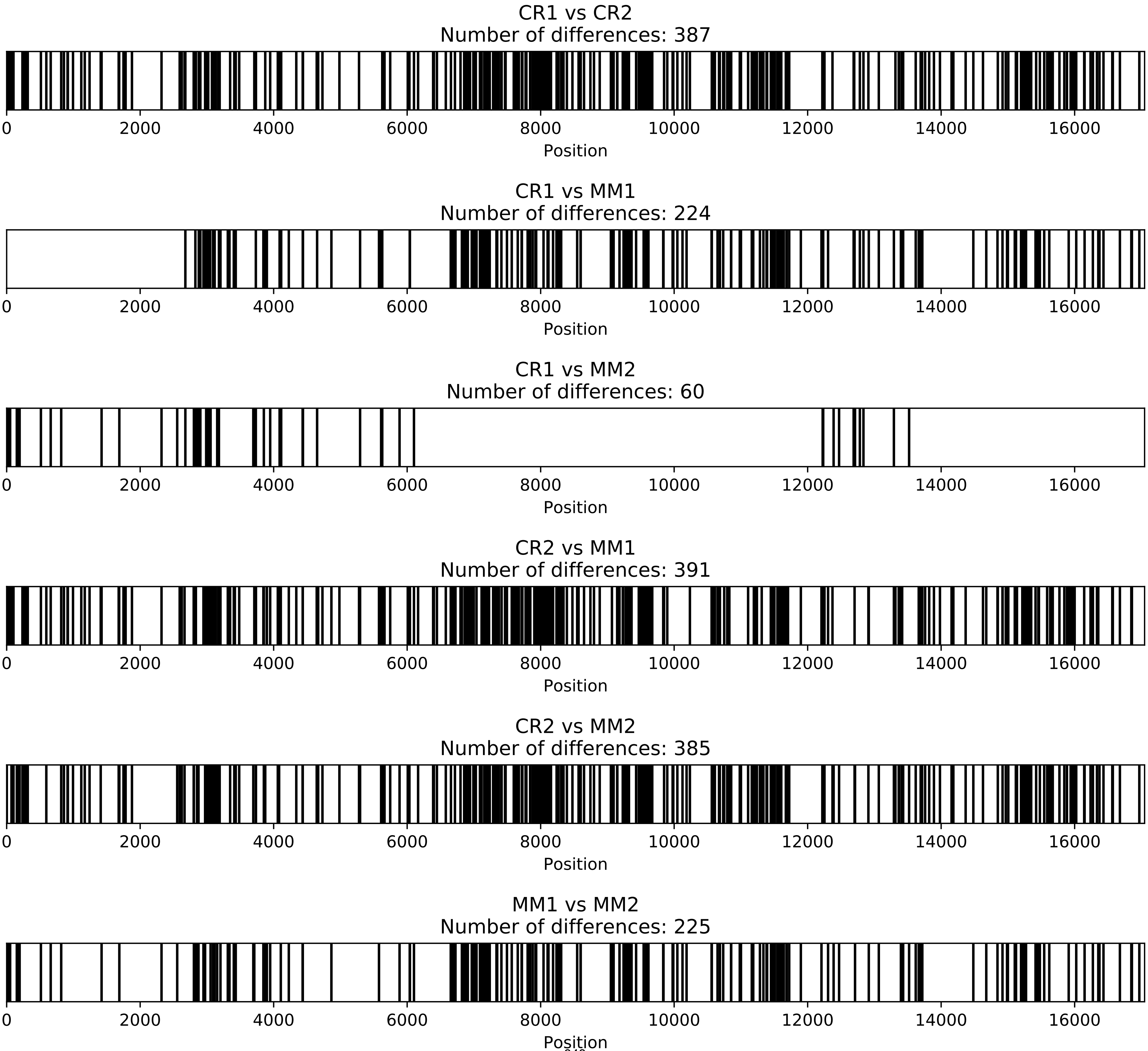
CR1 vs CR2
Number of differences: 387
CR1 vs MM1
Number of differences: 224
CR1 vs MM2
Number of differences: 60
CR2 vs MM1
Number of differences: 391
CR2 vs MM2
Number of differences: 385
MM1 vs MM2
Number of differences: 225
0
2000
4000
6000
8000
10000
12000
14000
16000
Position

### Region: 921



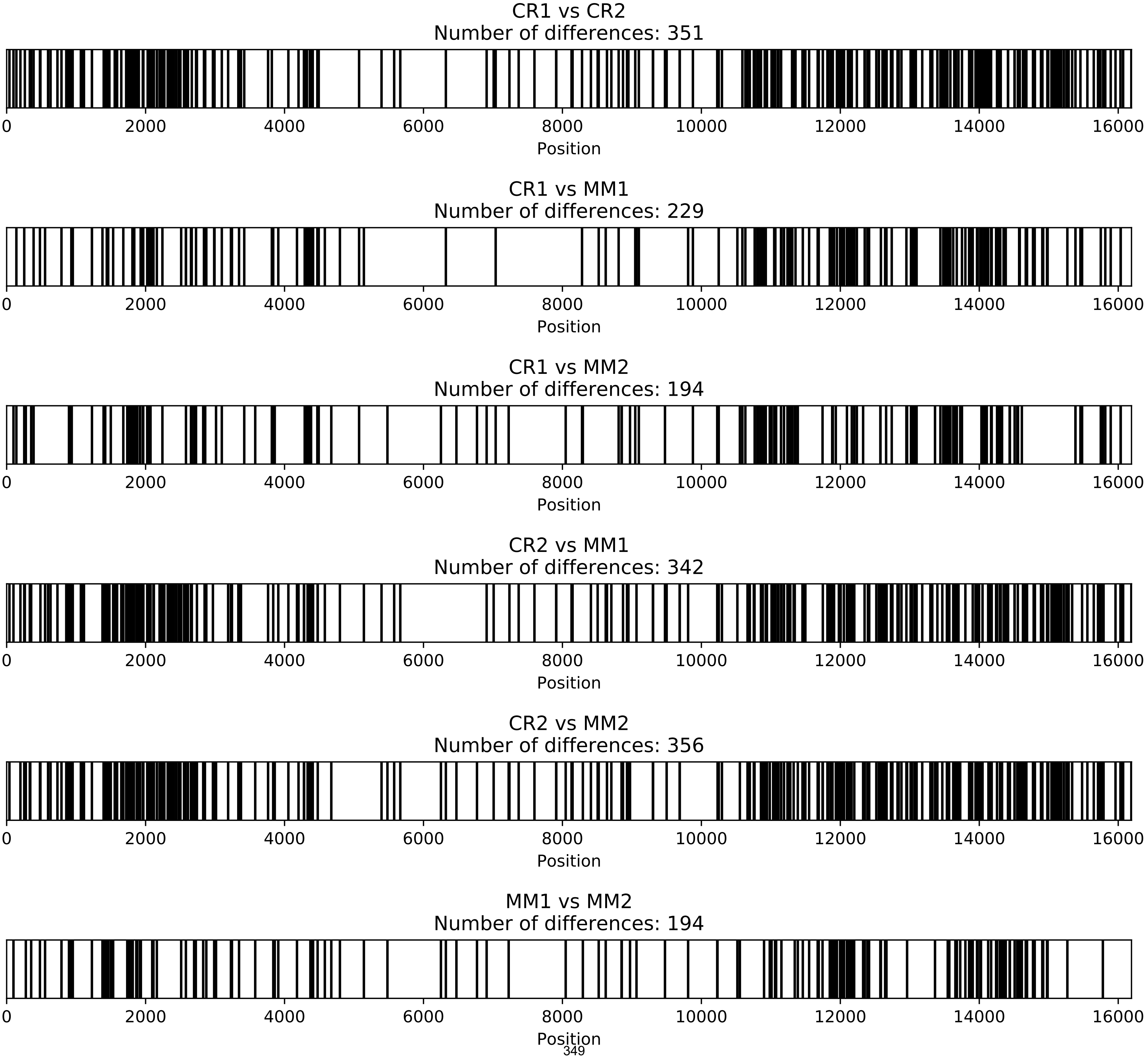
CR1 vs CR2
Number of differences: 351
0
2000
4000
6000
8000
10000
12000
14000
16000
Position
CR1 vs MM1
Number of differences: 229
0
2000
4000
6000
8000
10000
12000
14000
16000
Position
CR1 vs MM2
Number of differences: 194
0
2000
4000
6000
8000
10000
12000
14000
16000
Position
CR2 vs MM1
Number of differences: 342
0
2000
4000
6000
8000
10000
12000
14000
16000
Position
CR2 vs MM2
Number of differences: 356
0
2000
4000
6000
8000
10000
12000
14000
16000
Position
MM1 vs MM2
Number of differences: 194
0
2000
4000
6000
8000
10000
12000
14000
16000
Position

### Region: 924



CR1 vs CR2
Number of differences: 418

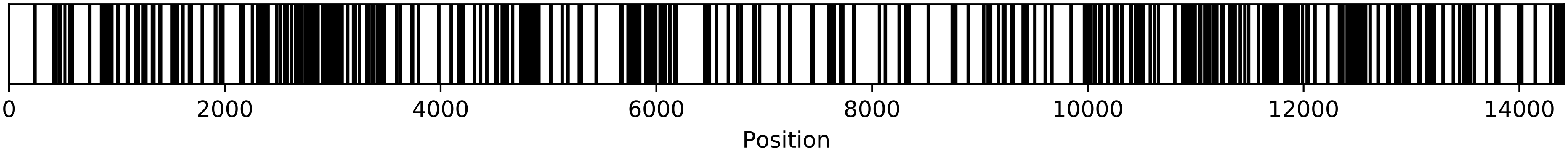


CR1 vs MM1
Number of differences: 148

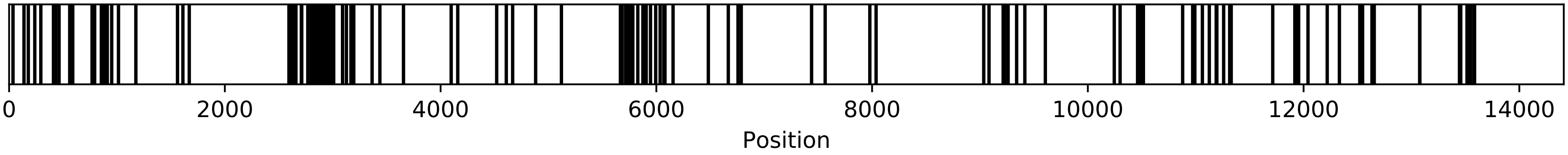


CR1 vs MM2
Number of differences: 164

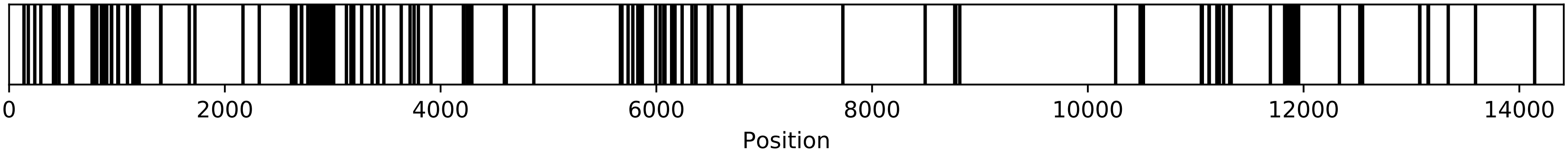


CR2 vs MM1
Number of differences: 401

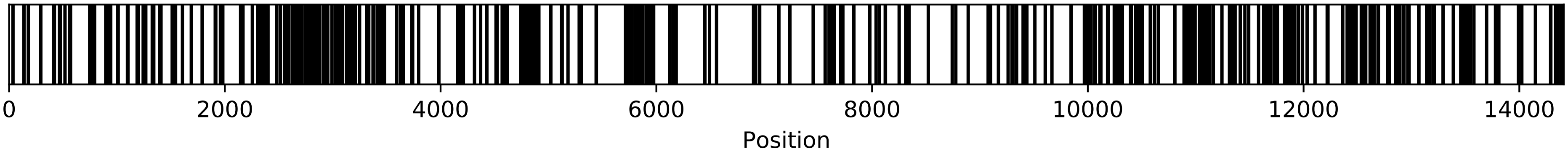


CR2 vs MM2
Number of differences: 424

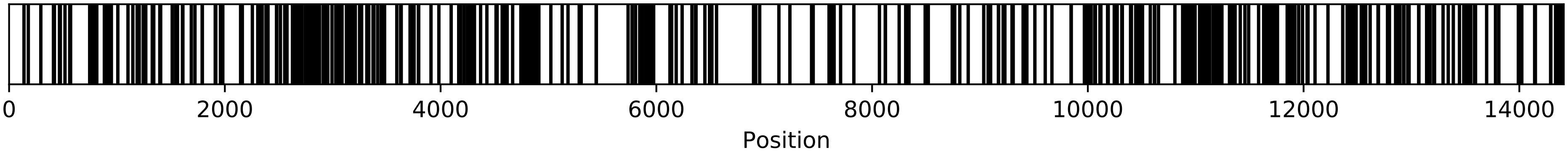


MM1 vs MM2
Number of differences: 132

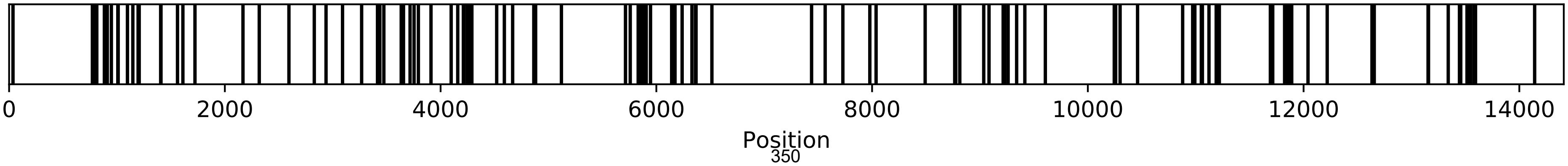

### Region: 926



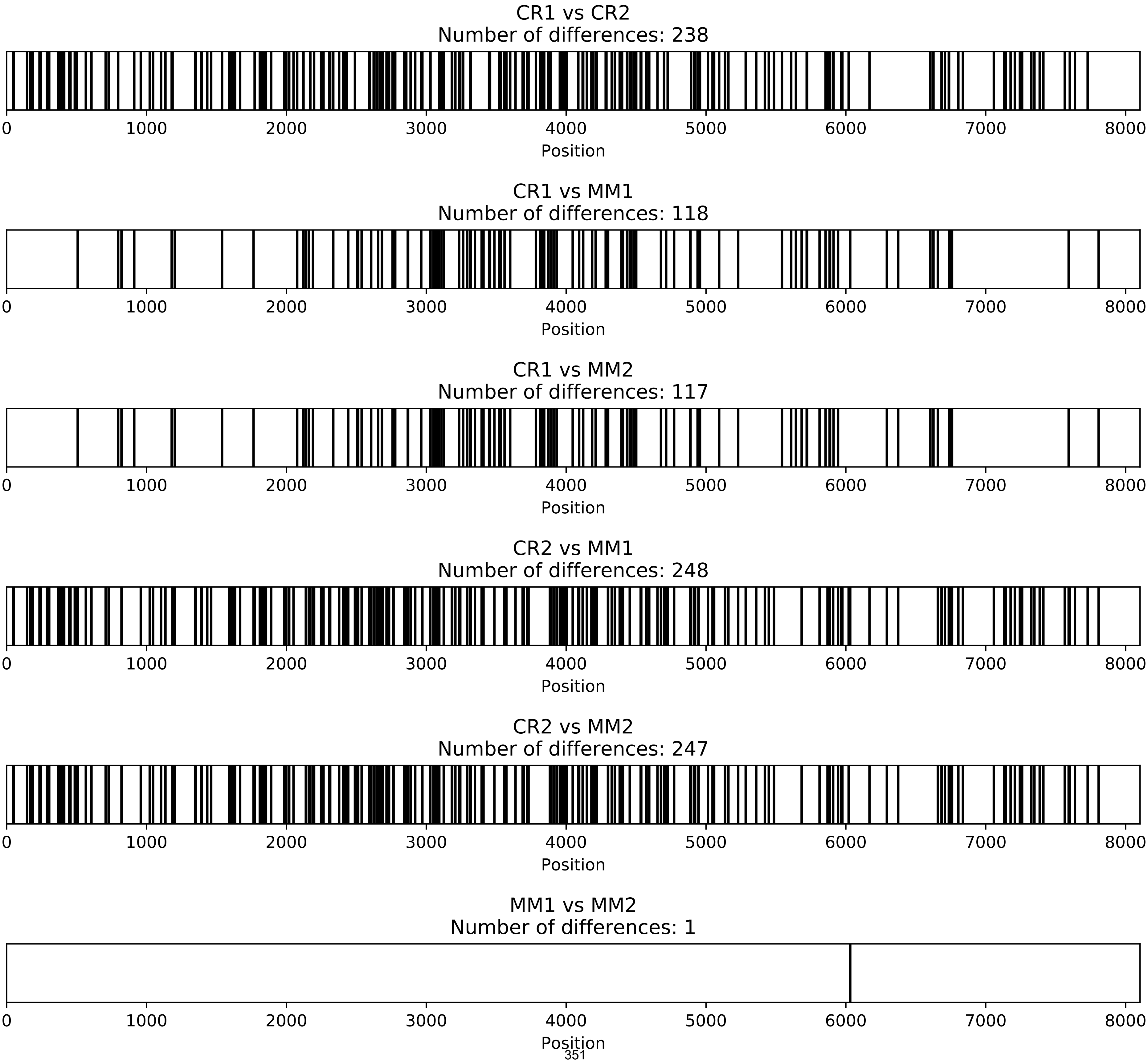
CR1 vs CR2
Number of differences: 238
CR1 vs MM1
Number of differences: 118
CR1 vs MM2
Number of differences: 117
CR2 vs MM1
Number of differences: 248
CR2 vs MM2
Number of differences: 247
MM1 vs MM2
Number of differences: 1
0
1000
2000
3000
4000
5000
6000
7000
8000
Position

### Region: 933



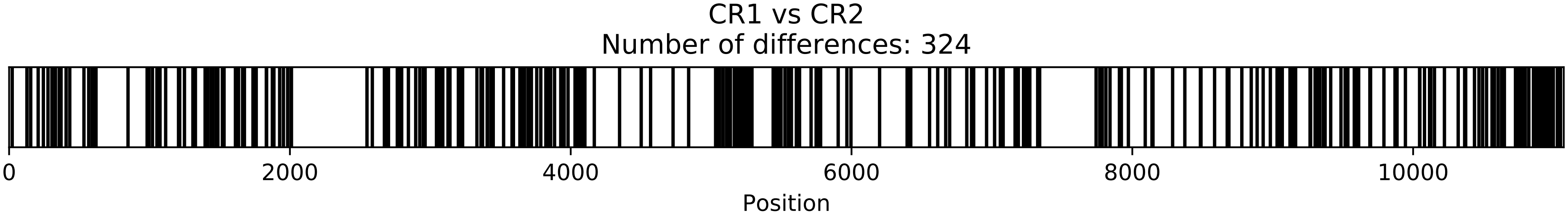
CR1 vs CR2
Number of differences: 324
0
2000
4000
6000
8000
10000
Position

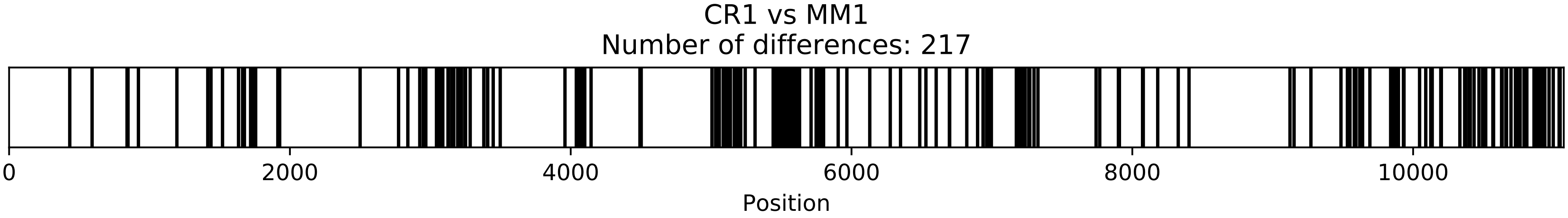
CR1 vs MM1
Number of differences: 217
0
2000
4000
6000
8000
10000
Position

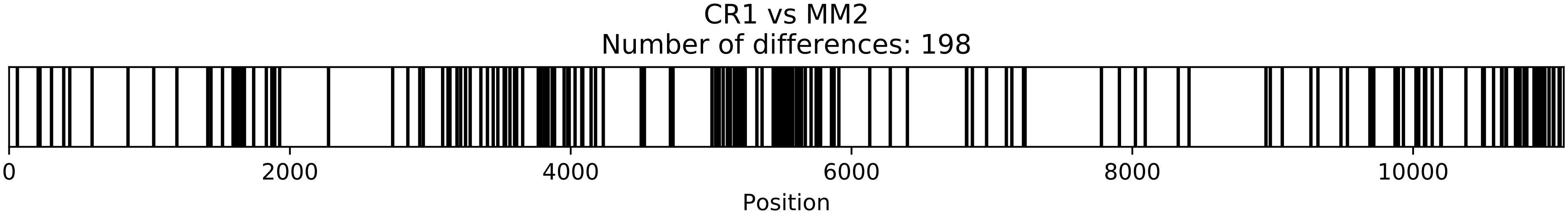
CR1 vs MM2
Number of differences: 198
0
2000
4000
6000
8000
10000
Position

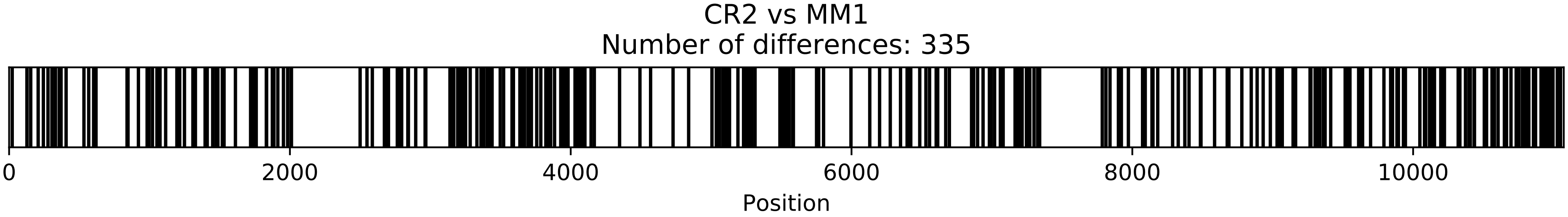
CR2 vs MM1
Number of differences: 335
0
2000
4000
6000
8000
10000
Position

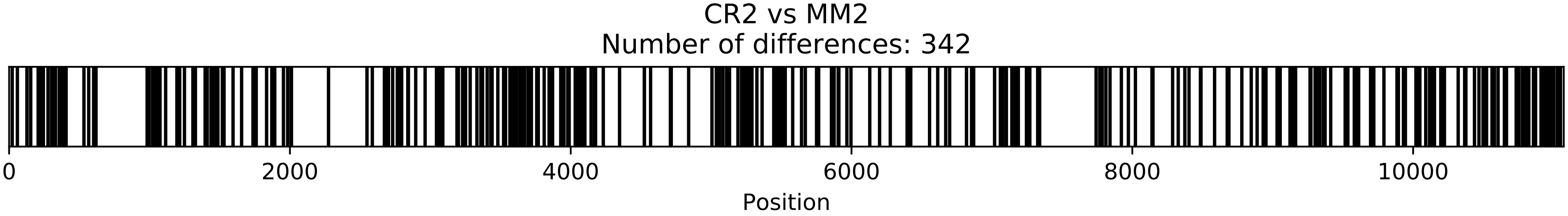
CR2 vs MM2
Number of differences: 342
0
2000
4000
6000
8000
10000
Position

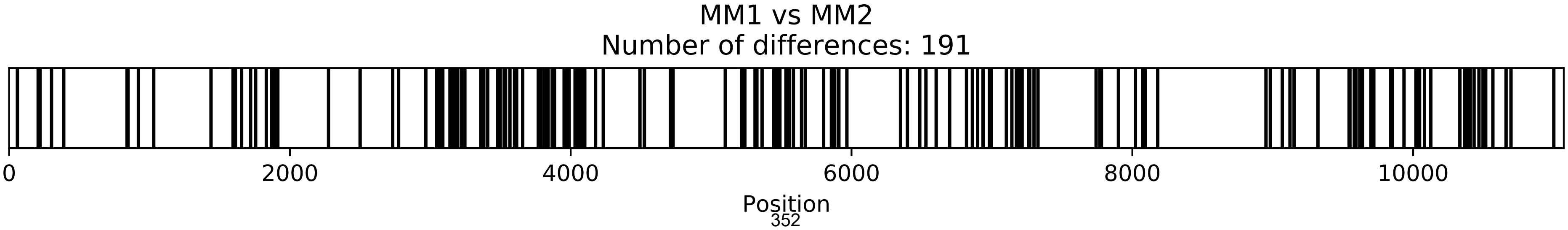
MM1 vs MM2
Number of differences: 191
0
2000
4000
6000
8000
10000
Position

### Region: 934



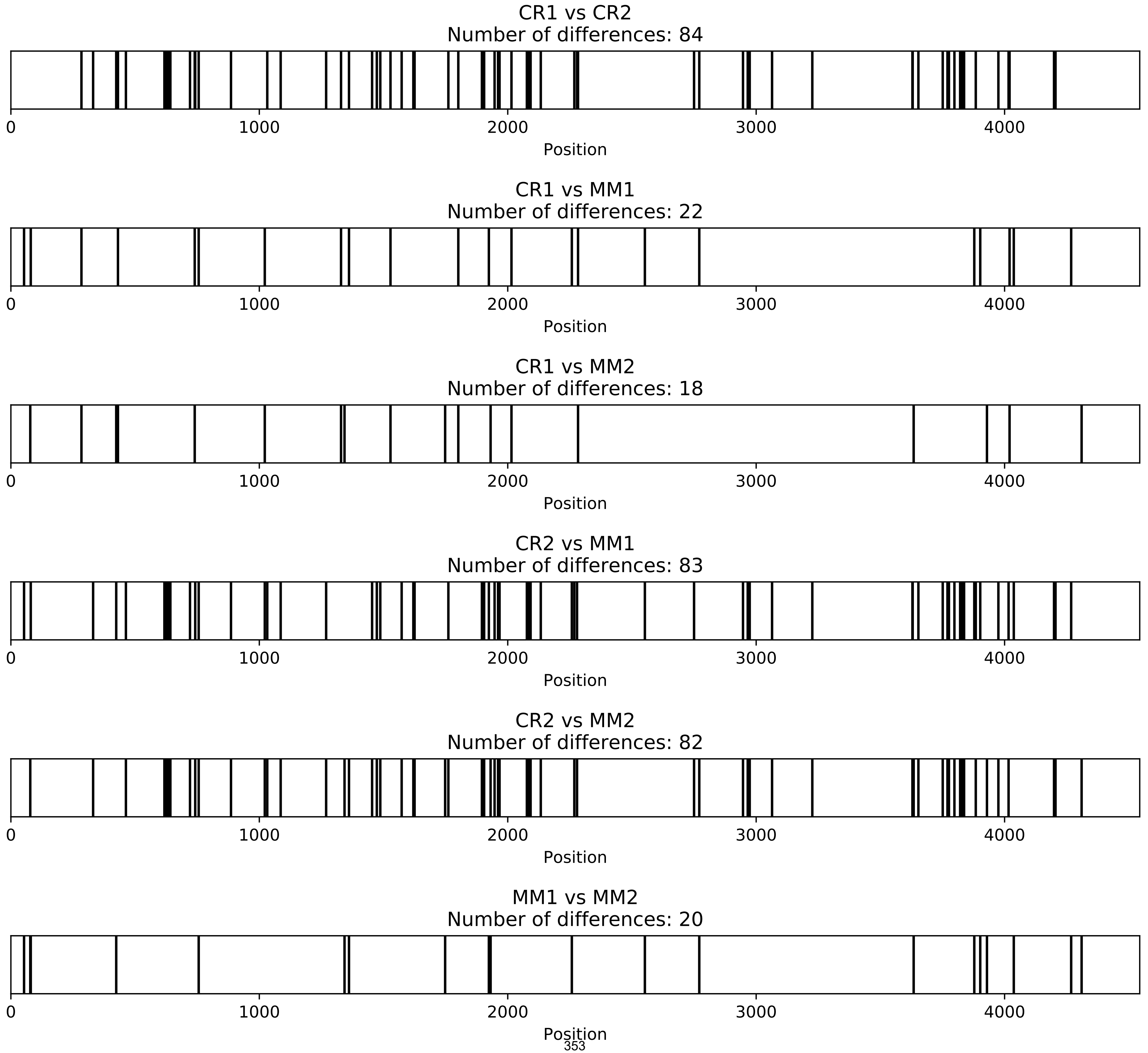
CR1 vs CR2
Number of differences: 84
0
1000
2000
3000
4000
Position
CR1 vs MM1
Number of differences: 22
0
1000
2000
3000
4000
Position
CR1 vs MM2
Number of differences: 18
0
1000
2000
3000
4000
Position
CR2 vs MM1
Number of differences: 83
0
1000
2000
3000
4000
Position
CR2 vs MM2
Number of differences: 82
0
1000
2000
3000
4000
Position
MM1 vs MM2
Number of differences: 20
0
1000
2000
3000
4000
Position

### Region: 939



CR1 vs CR2
Number of differences: 17

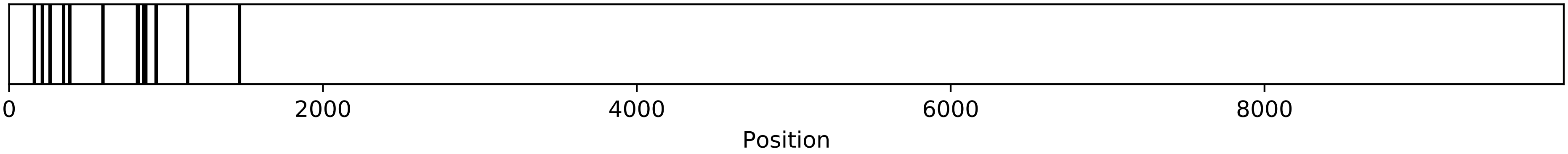


CR1 vs MM1
Number of differences: 0

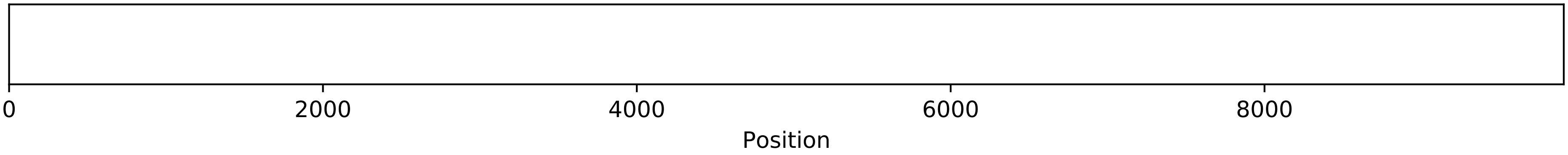


CR1 vs MM2
Number of differences: 71

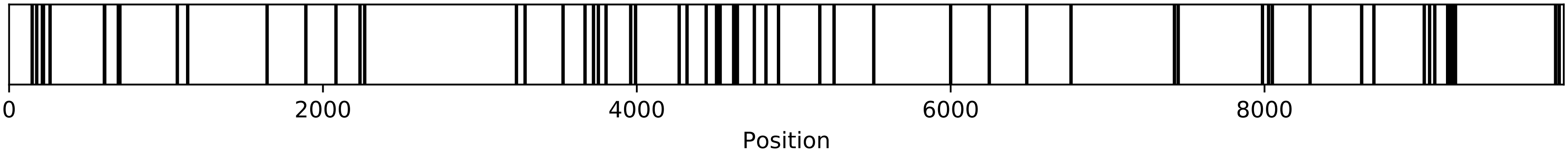


CR2 vs MM1
Number of differences: 17

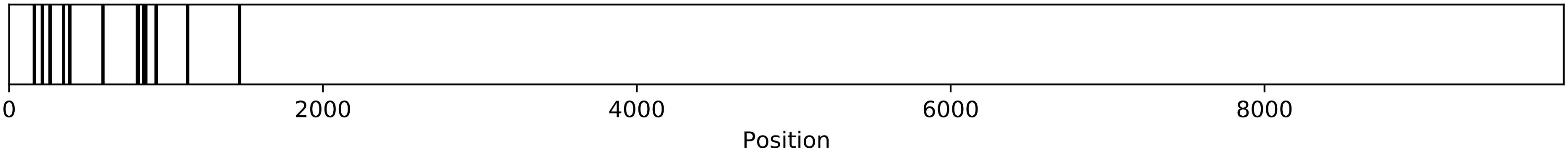


CR2 vs MM2
Number of differences: 82

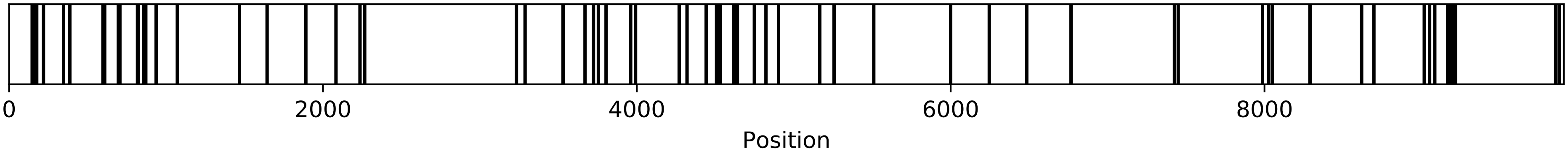


MM1 vs MM2
Number of differences: 71

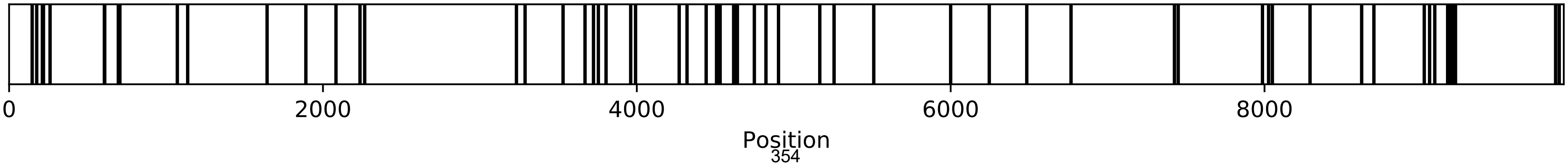

### Region: 941



CR1 vs CR2
Number of differences: 347

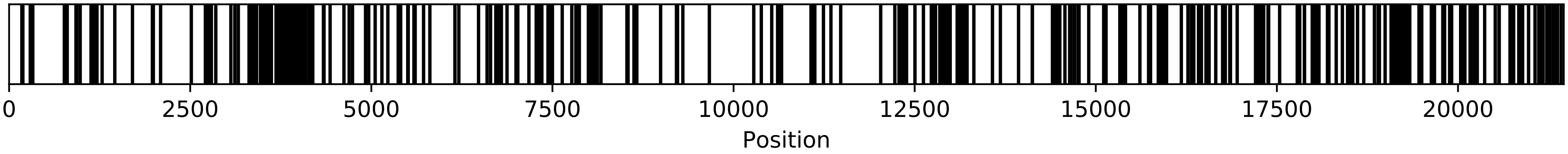


CR1 vs MM1
Number of differences: 181

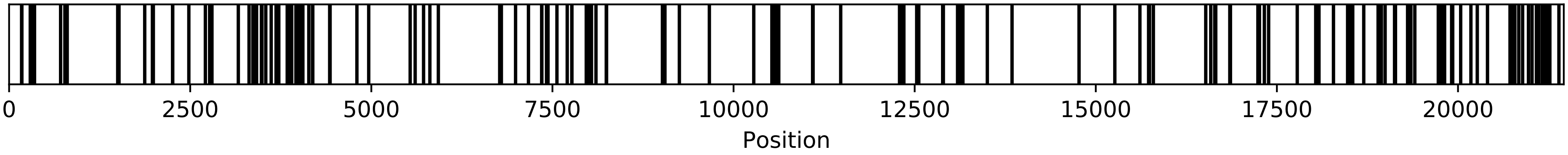


CR1 vs MM2
Number of differences: 173

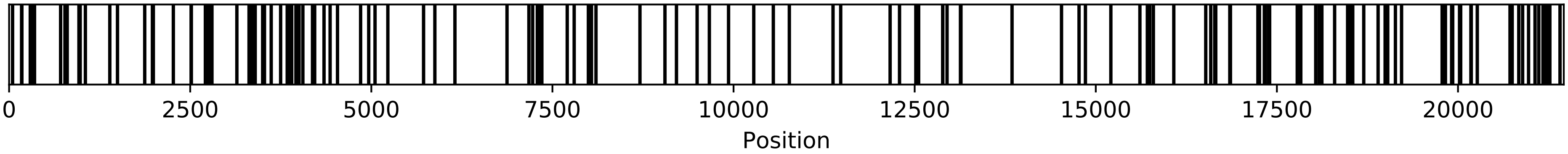


CR2 vs MM1
Number of differences: 330

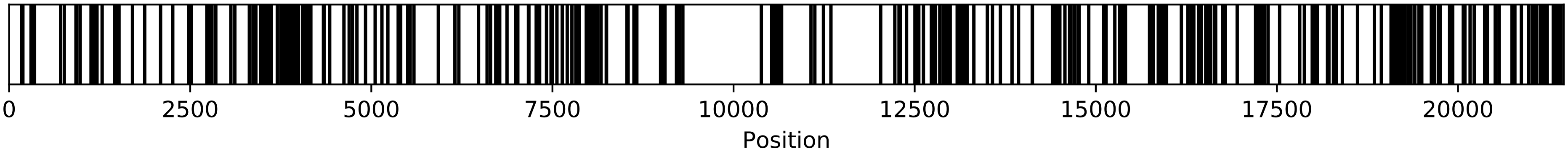


CR2 vs MM2
Number of differences: 354

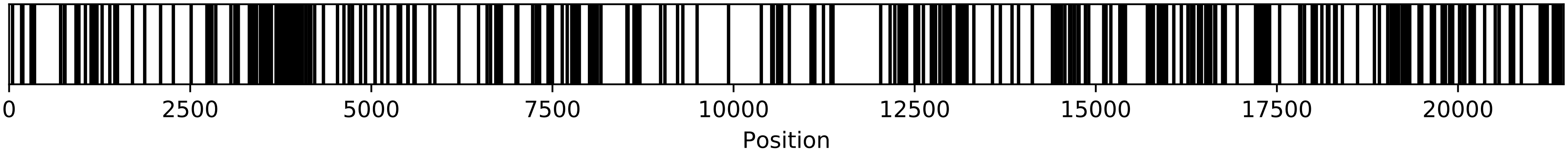


MM1 vs MM2
Number of differences: 160

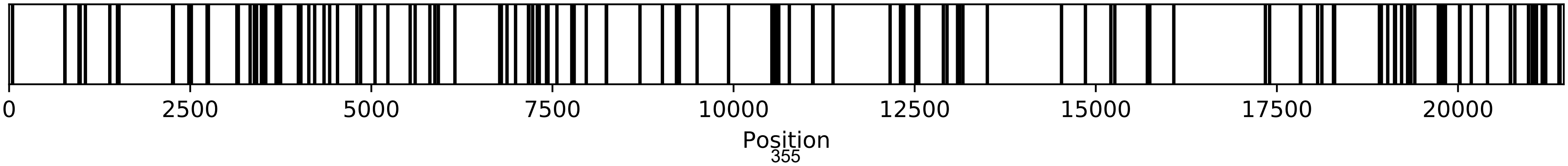

### Region: 943



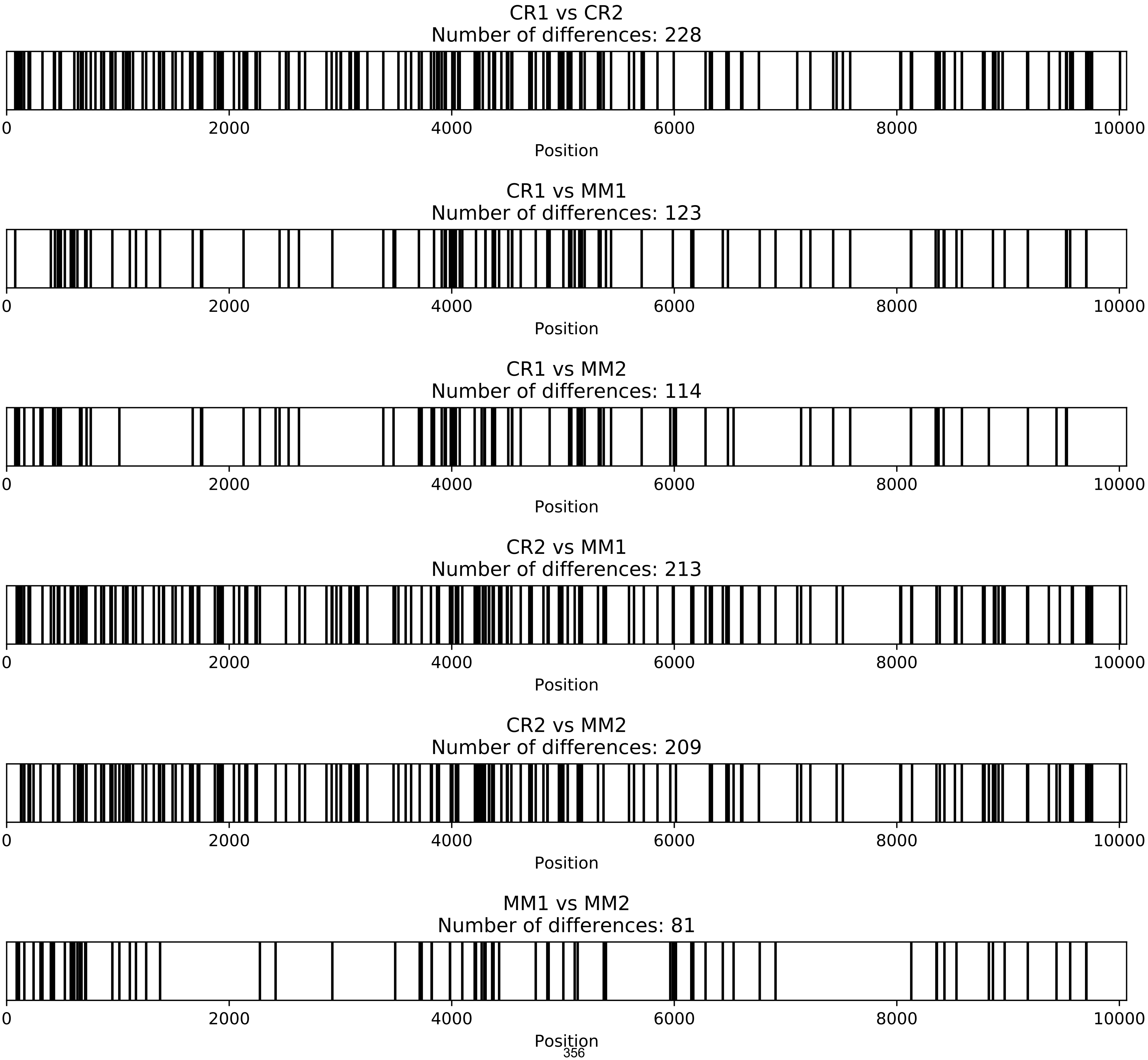
CR1 vs CR2
Number of differences: 228
CR1 vs MM1
Number of differences: 123
CR1 vs MM2
Number of differences: 114
CR2 vs MM1
Number of differences: 213
CR2 vs MM2
Number of differences: 209
MM1 vs MM2
Number of differences: 81
0
2000
4000
6000
8000
10000
Position

### Region: 944



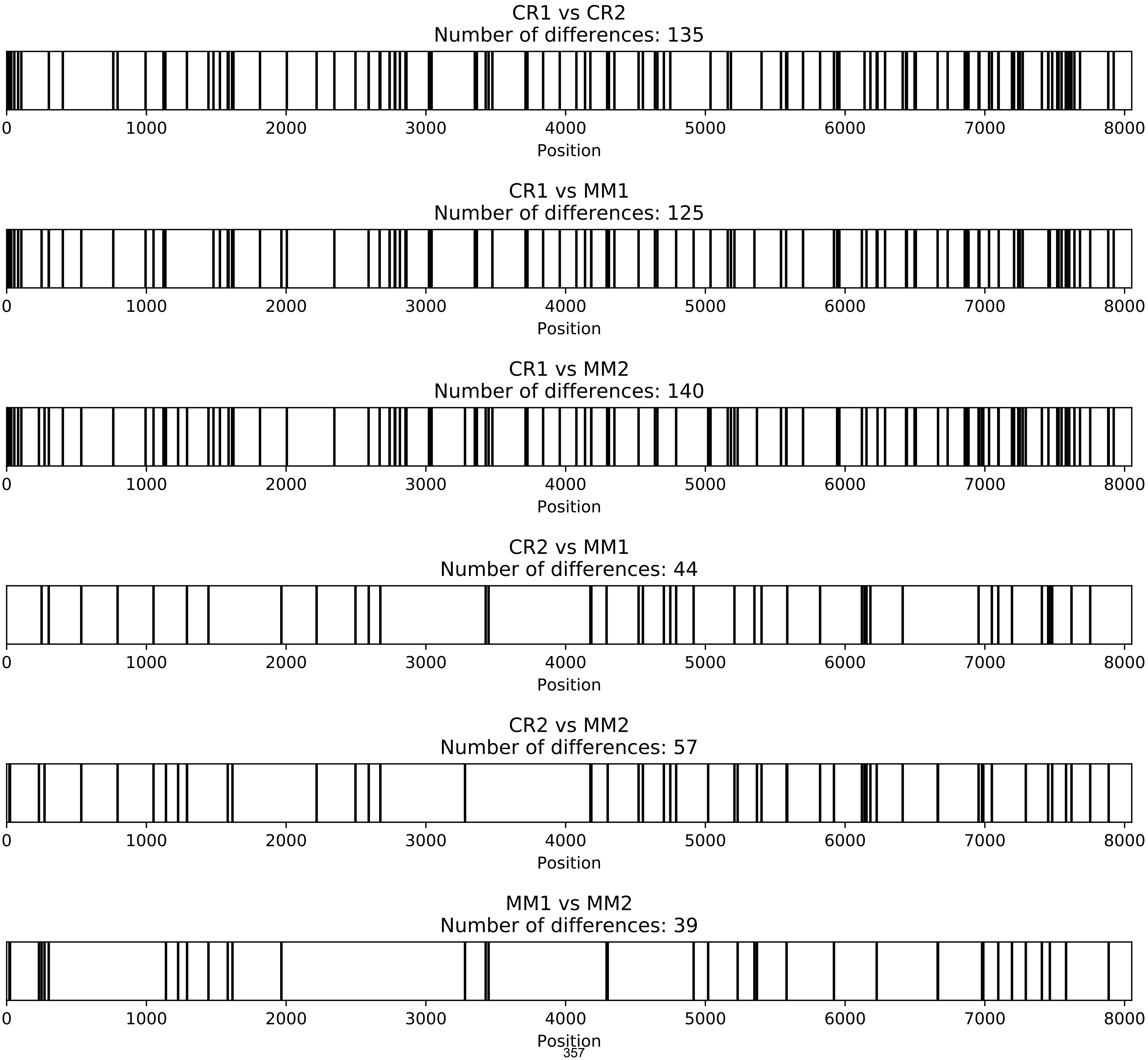
CR1 vs CR2
Number of differences: 135
CR1 vs MM1
Number of differences: 125
CR1 vs MM2
Number of differences: 140
CR2 vs MM1
Number of differences: 44
CR2 vs MM2
Number of differences: 57
MM1 vs MM2
Number of differences: 39
0
1000
2000
3000
4000
5000
6000
7000
8000
Position

### Region: 946



CR1 vs CR2
Number of differences: 288

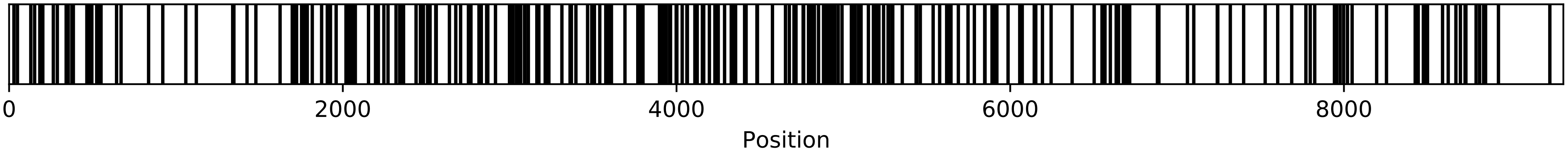


CR1 vs MM1
Number of differences: 118

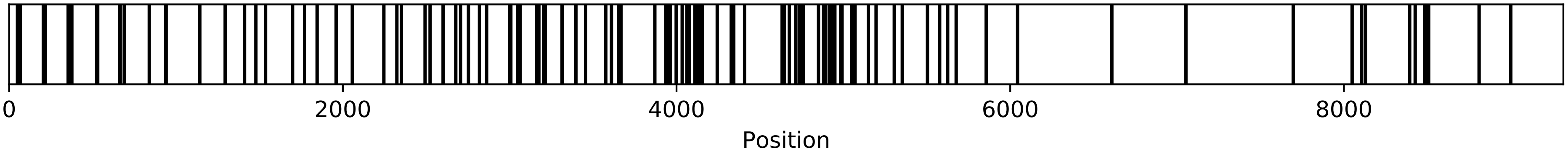


CR1 vs MM2
Number of differences: 125

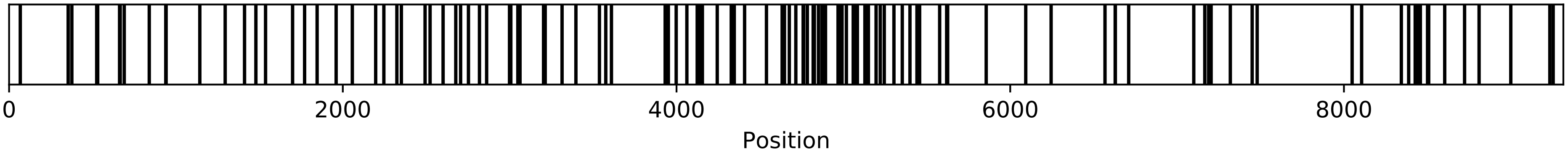


CR2 vs MM1
Number of differences: 285

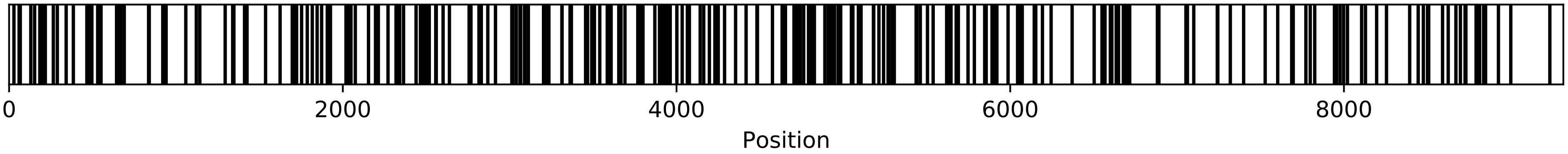


CR2 vs MM2
Number of differences: 294

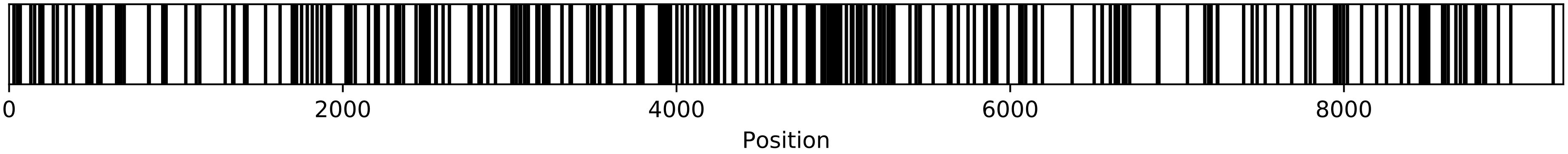


MM1 vs MM2
Number of differences: 87

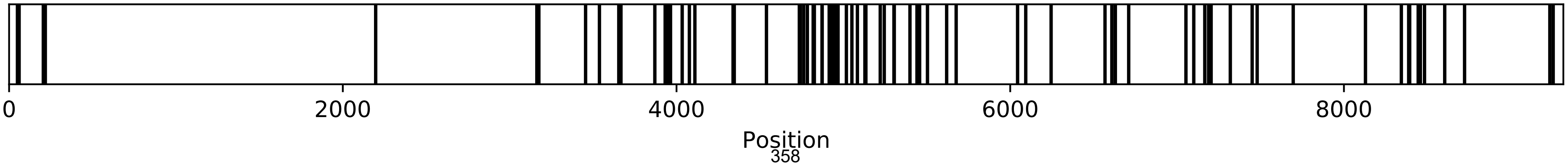

### Region: 947



CR1 vs CR2
Number of differences: 331

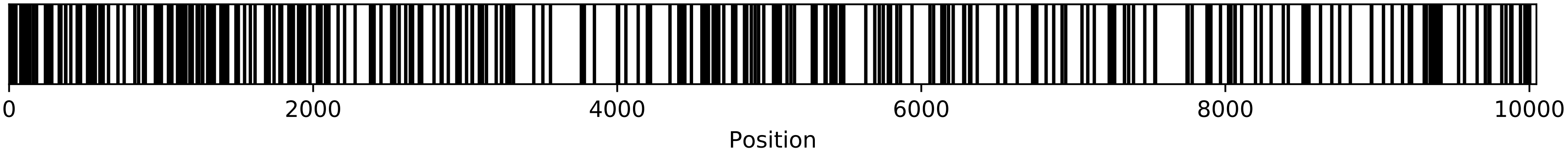


CR1 vs MM1
Number of differences: 162

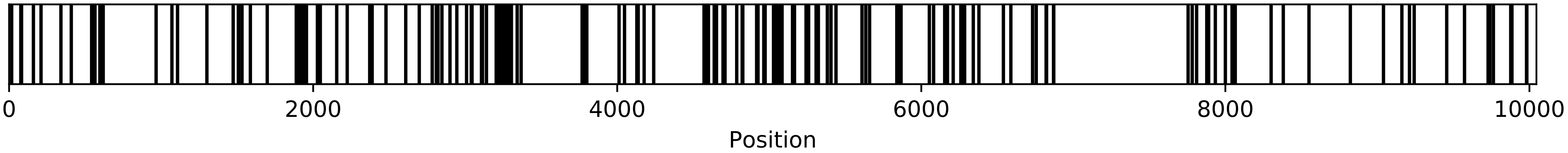


CR1 vs MM2
Number of differences: 145

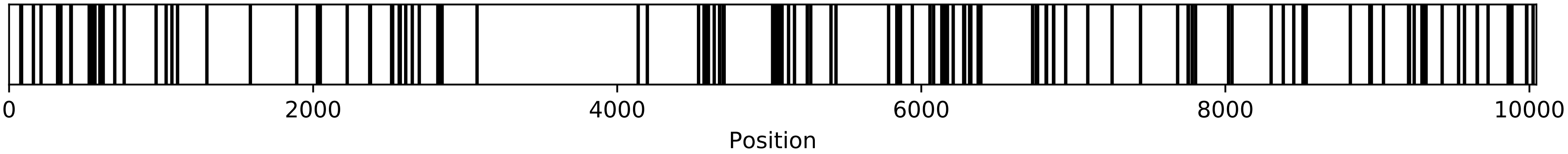


CR2 vs MM1
Number of differences: 332

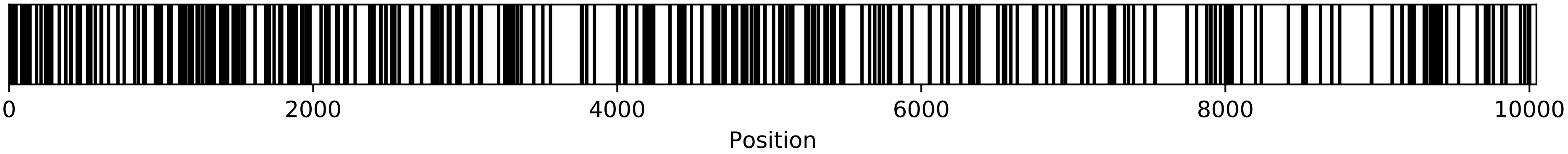


CR2 vs MM2
Number of differences: 301

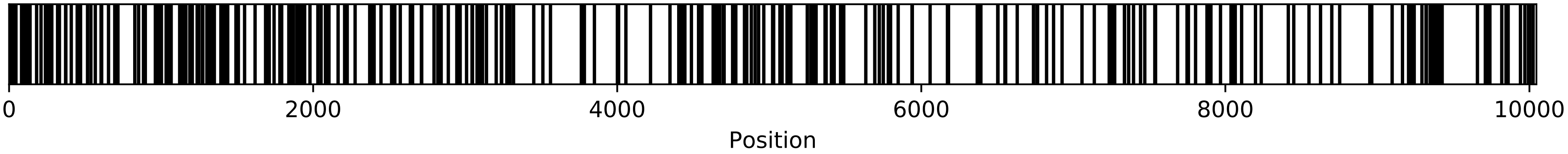


MM1 vs MM2
Number of differences: 161

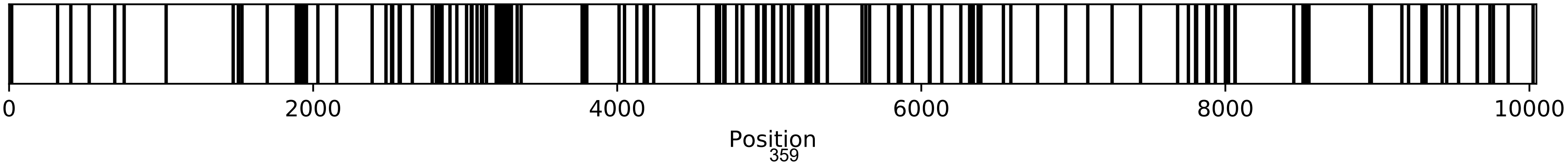

### Region: 952



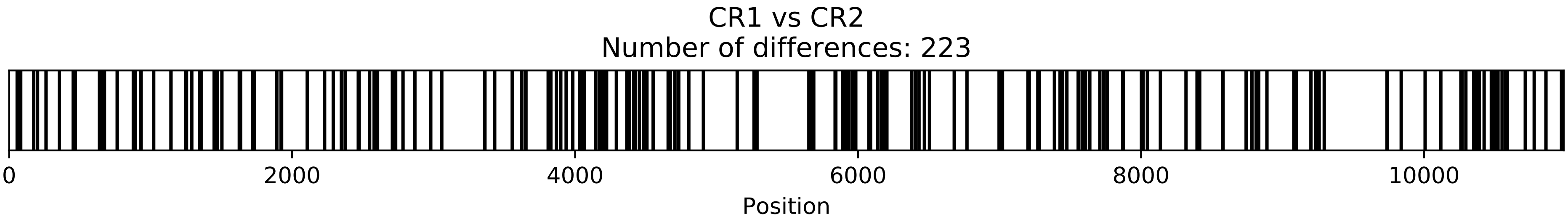
CR1 vs CR2
Number of differences: 223
0
2000
4000
6000
8000
10000
Position

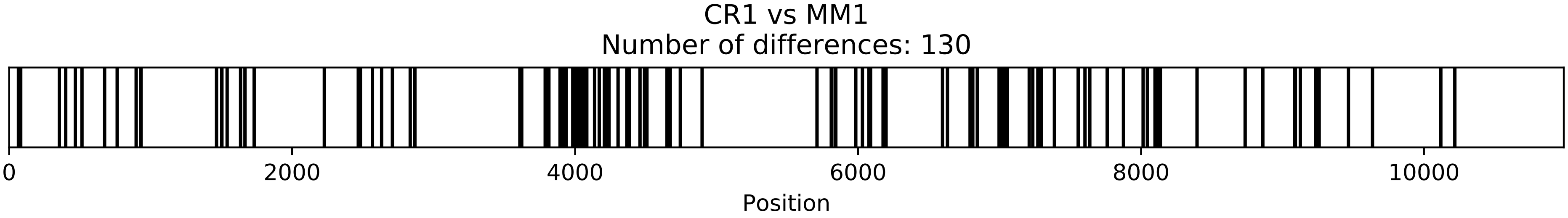
CR1 vs MM1
Number of differences: 130
0
2000
4000
6000
8000
10000
Position

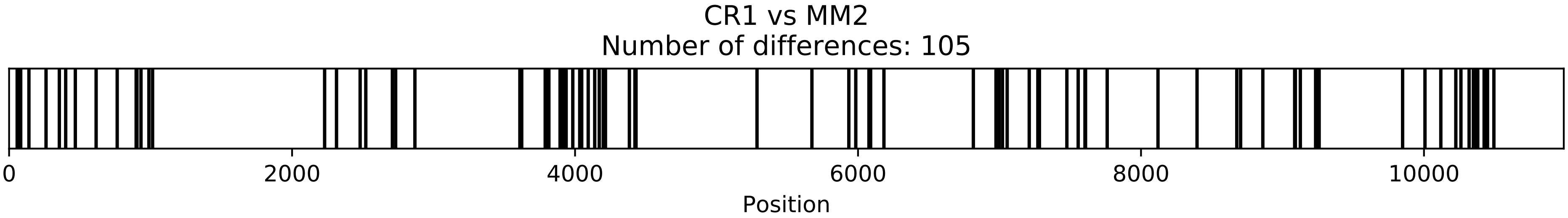
CR1 vs MM2
Number of differences: 105
0
2000
4000
6000
8000
10000
Position

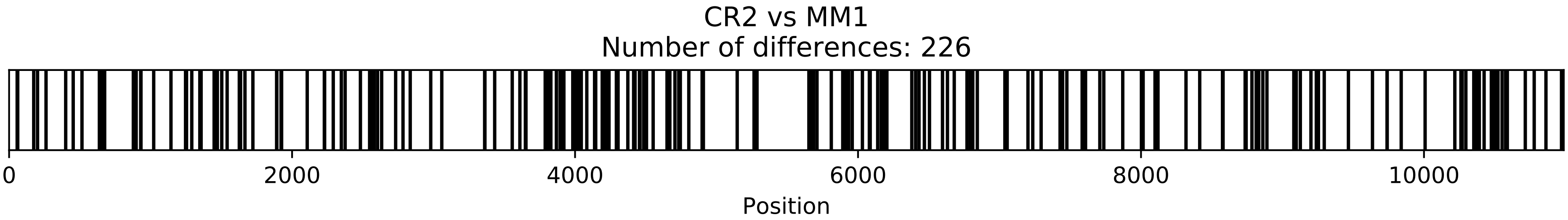
CR2 vs MM1
Number of differences: 226
0
2000
4000
6000
8000
10000
Position

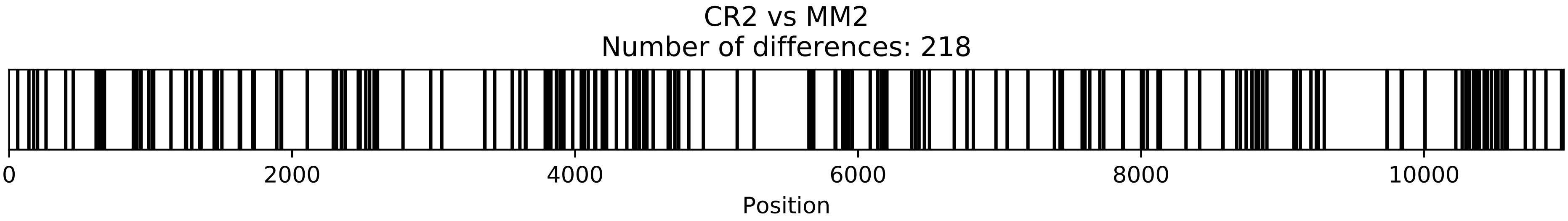
CR2 vs MM2
Number of differences: 218
0
2000
4000
6000
8000
10000
Position

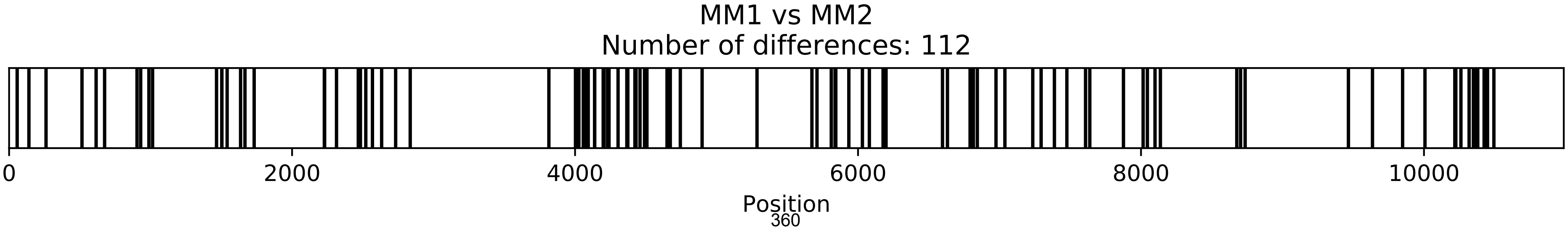
MM1 vs MM2
Number of differences: 112
0
2000
4000
6000
8000
10000
Position

### Region: 953



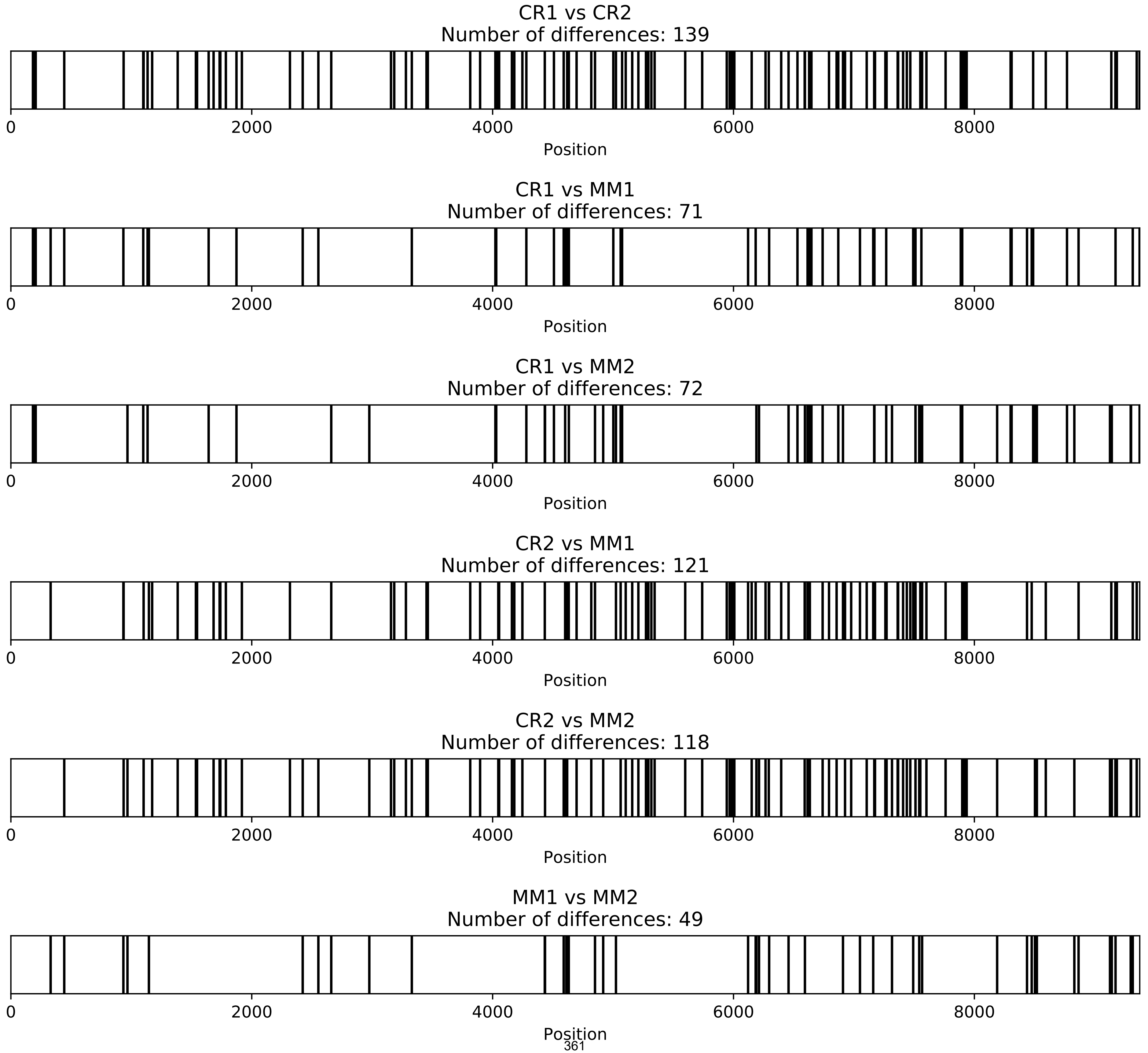
CR1 vs CR2
Number of differences: 139
CR1 vs MM1
Number of differences: 71
CR1 vs MM2
Number of differences: 72
CR2 vs MM1
Number of differences: 121
CR2 vs MM2
Number of differences: 118
MM1 vs MM2
Number of differences: 49
0
2000
4000
6000
8000
Position

### Region: 954



CR1 vs CR2
Number of differences: 218

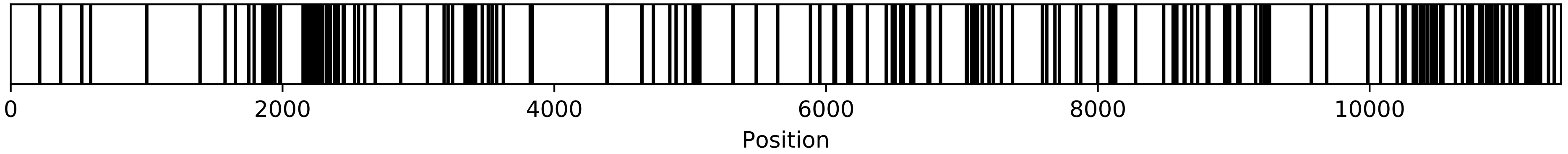


CR1 vs MM1
Number of differences: 110

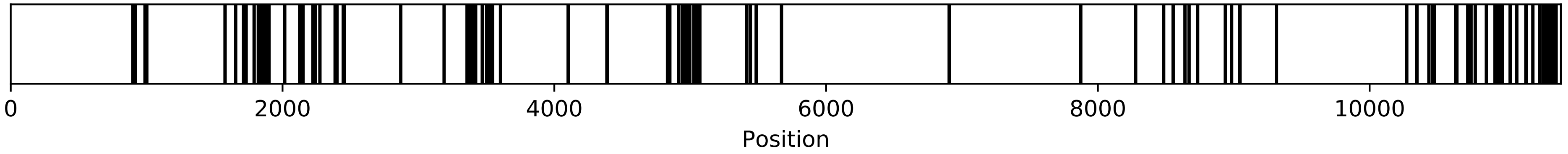


CR1 vs MM2
Number of differences: 90

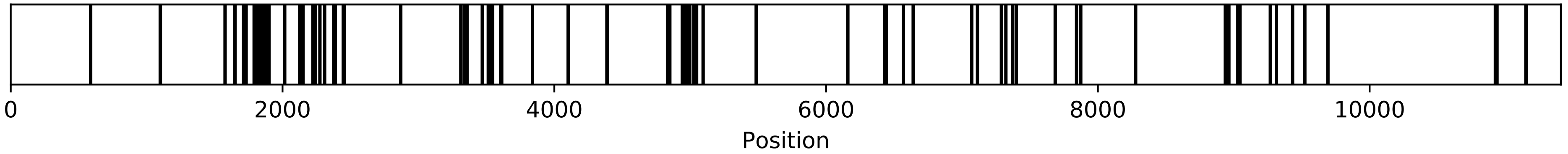


CR2 vs MM1
Number of differences: 218

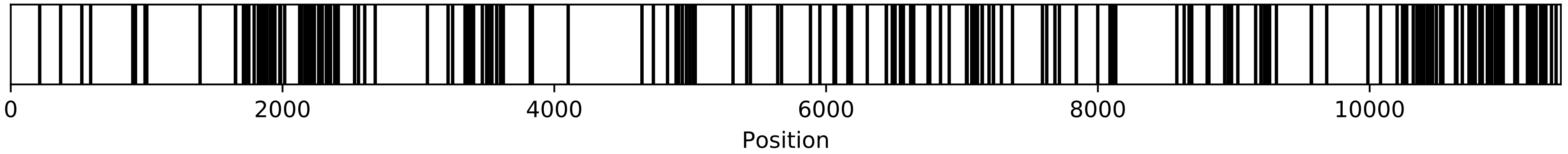


CR2 vs MM2
Number of differences: 224

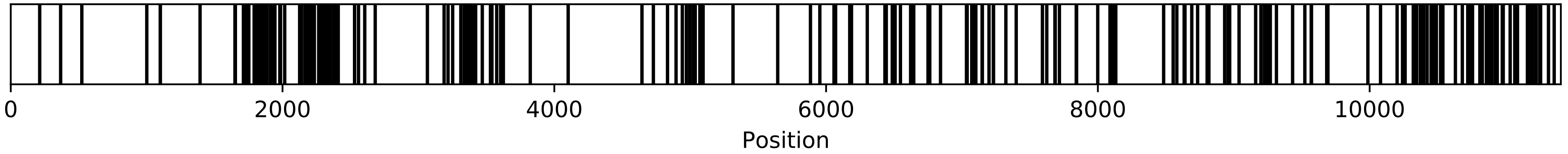


MM1 vs MM2
Number of differences: 96

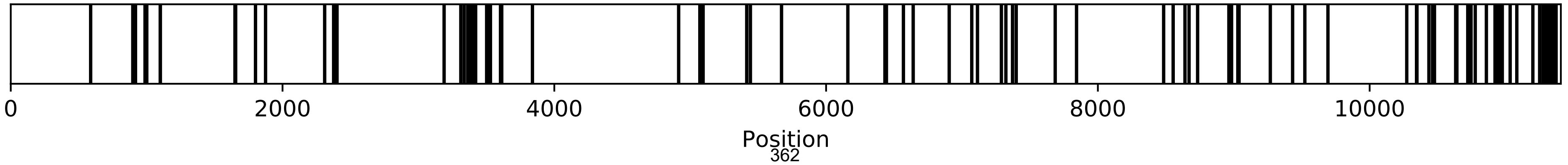

### Region: 956



## CR1 vs CR2
Number of differences: 293

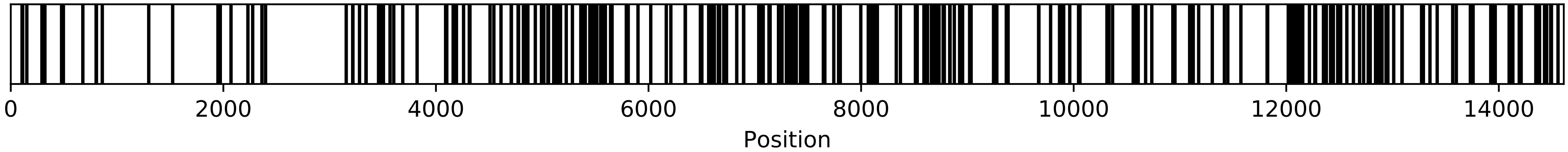


## CR1 vs MM1
Number of differences: 309

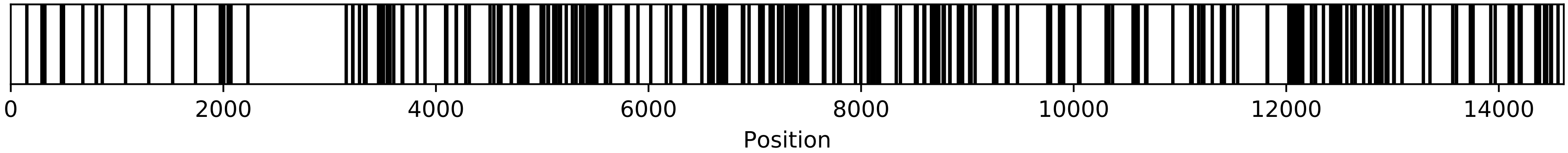


## CR1 vs MM2
Number of differences: 295

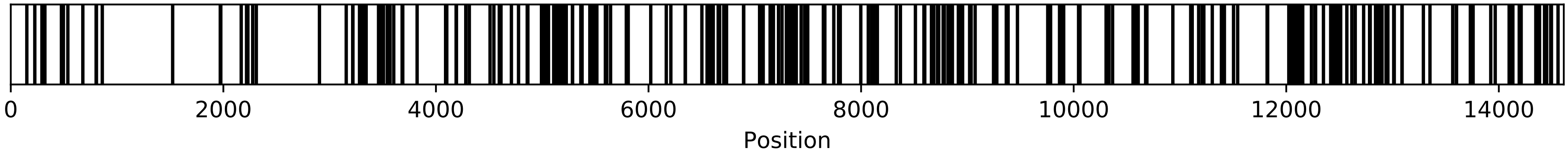


## CR2 vs MM1
Number of differences: 142

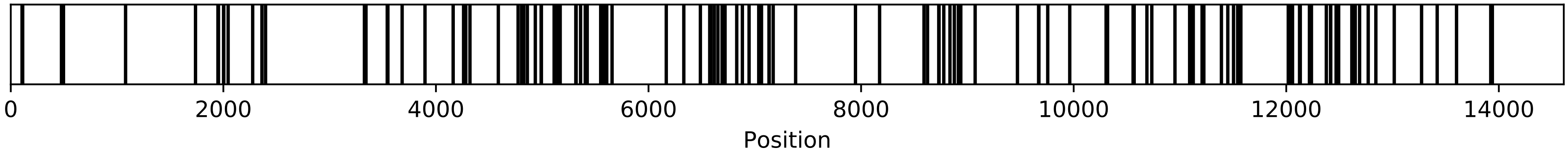


## CR2 vs MM2
Number of differences: 145

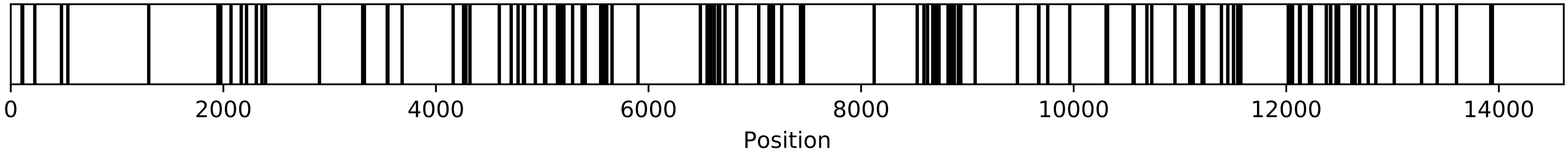


## MM1 vs MM2
Number of differences: 84

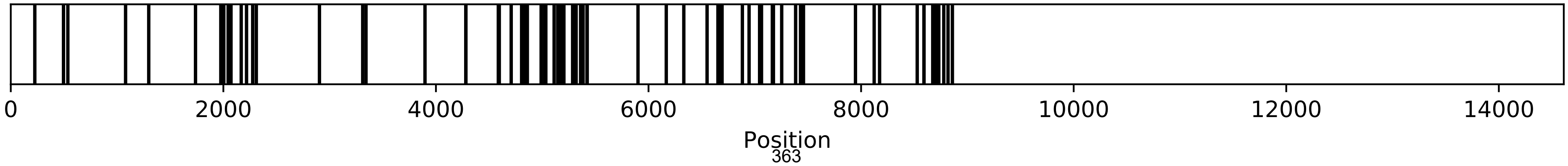

### Region: 961



## CR1 vs CR2
Number of differences: 185

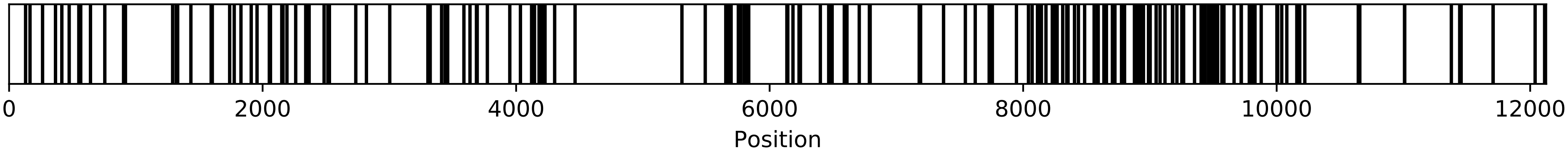


## CR1 vs MM1
Number of differences: 66

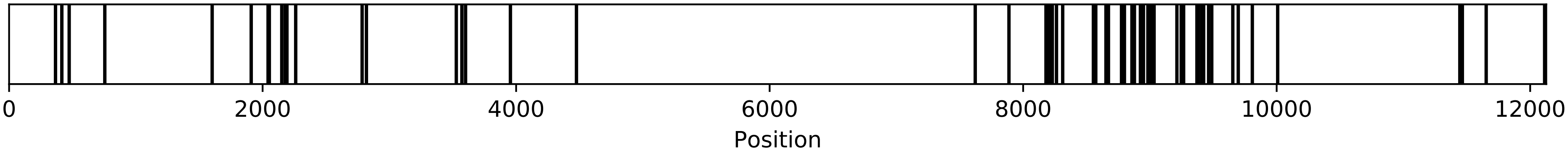


## CR1 vs MM2
Number of differences: 95

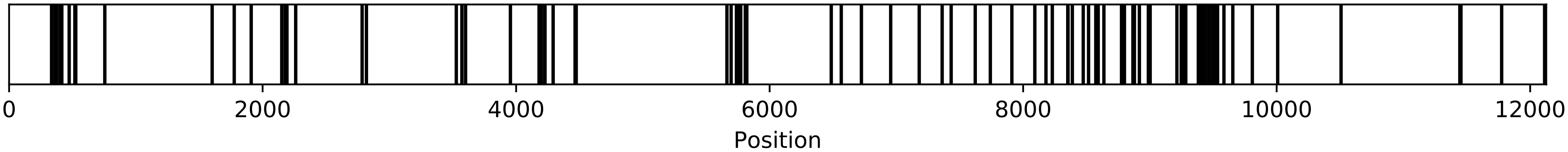


## CR2 vs MM1
Number of differences: 185

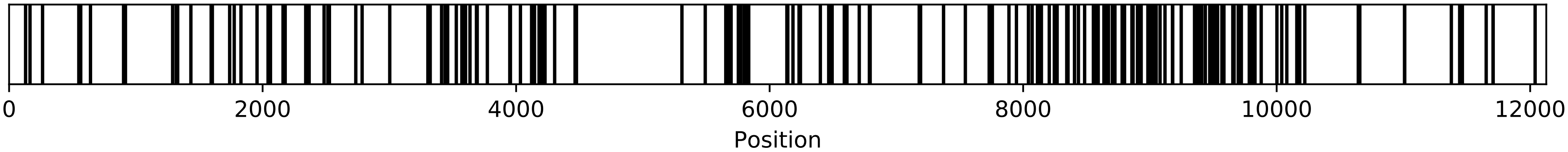


## CR2 vs MM2
Number of differences: 186

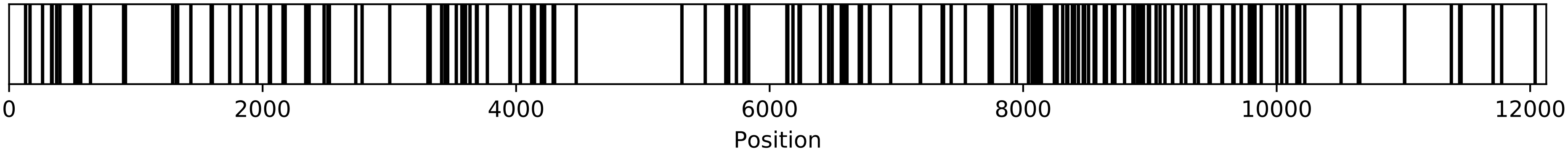


## MM1 vs MM2
Number of differences: 83

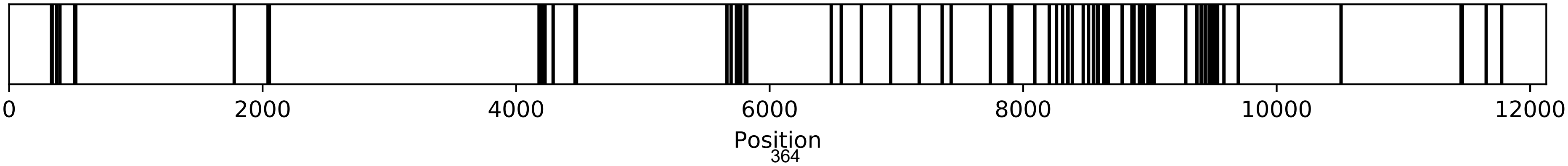

### Region: 964



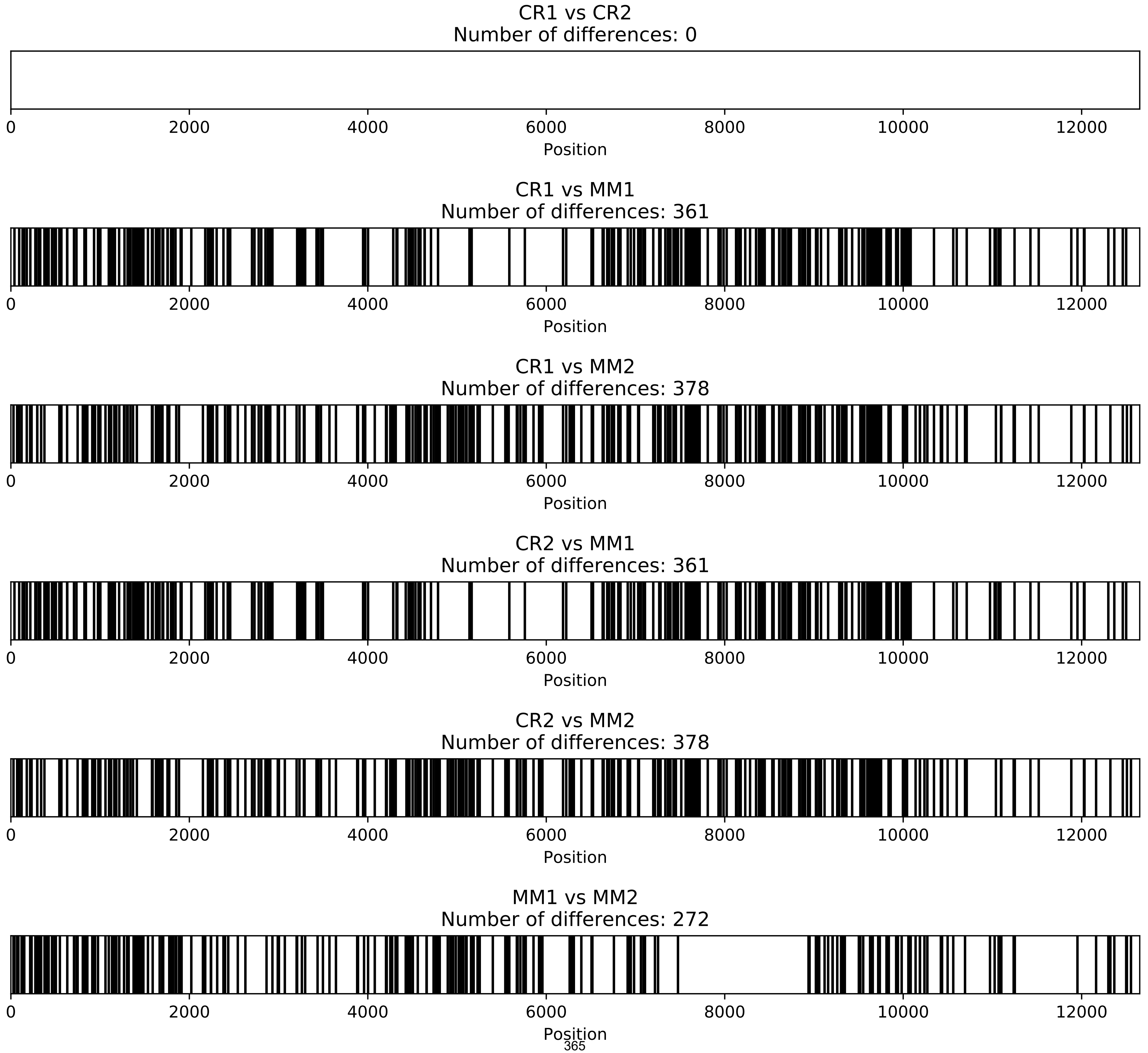
CR1 vs CR2
Number of differences: 0
CR1 vs MM1
Number of differences: 361
CR1 vs MM2
Number of differences: 378
CR2 vs MM1
Number of differences: 361
CR2 vs MM2
Number of differences: 378
MM1 vs MM2
Number of differences: 272
0
2000
4000
6000
8000
10000
12000
Position

### Region: 965



CR1 vs CR2
Number of differences: 117

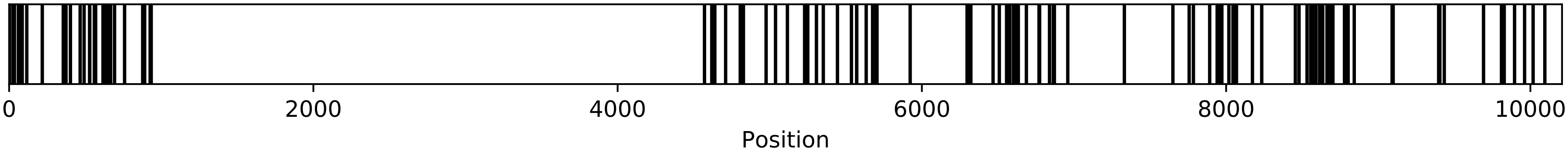


CR1 vs MM1
Number of differences: 201

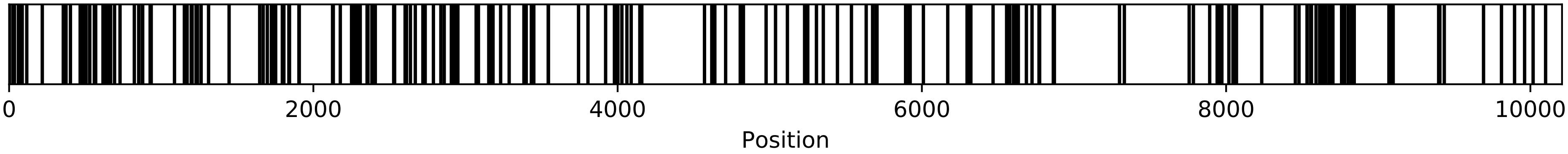


CR1 vs MM2
Number of differences: 221

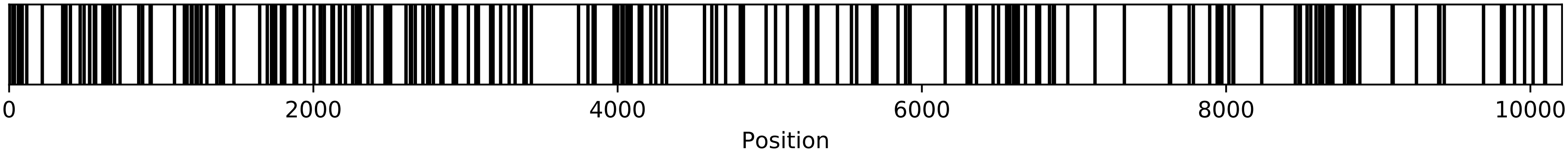


CR2 vs MM1
Number of differences: 121

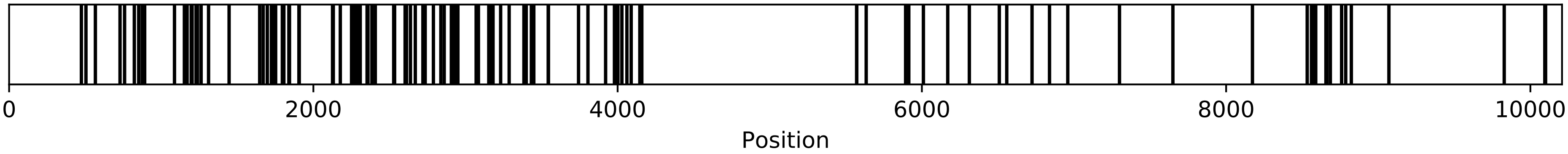


CR2 vs MM2
Number of differences: 140

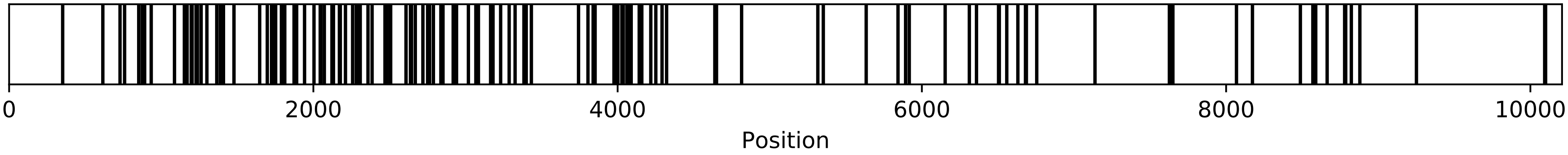


MM1 vs MM2
Number of differences: 127

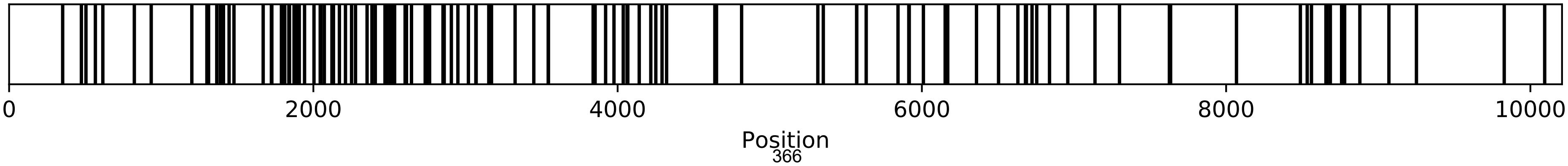

### Region: 966



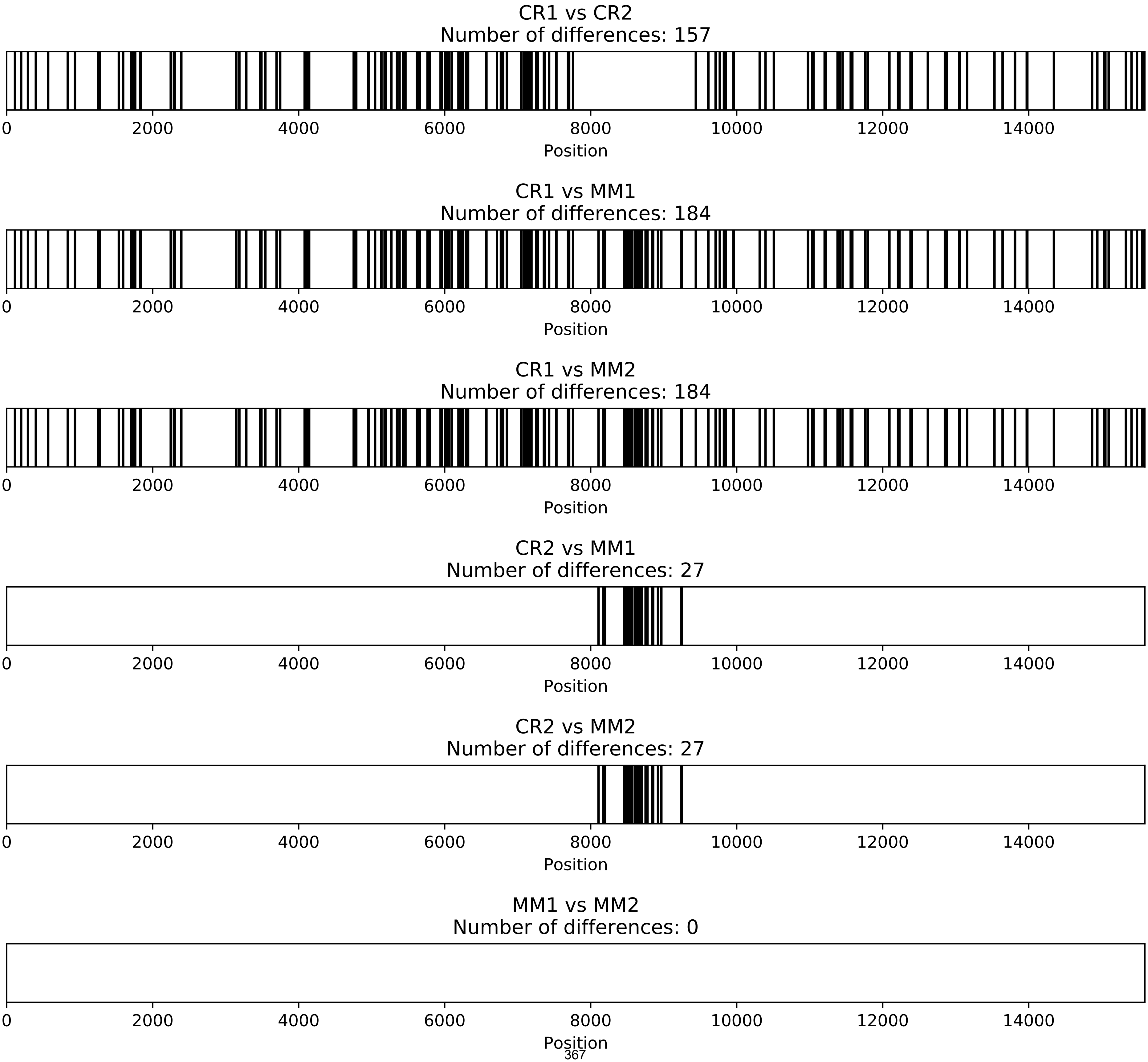
CR1 vs CR2
Number of differences: 157
CR1 vs MM1
Number of differences: 184
CR1 vs MM2
Number of differences: 184
CR2 vs MM1
Number of differences: 27
CR2 vs MM2
Number of differences: 27
MM1 vs MM2
Number of differences: 0
0
2000
4000
6000
8000
10000
12000
14000
Position

### Region: 970



## CR1 vs CR2
Number of differences: 266

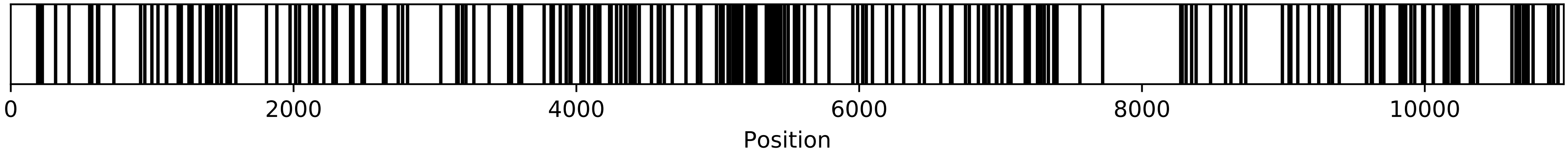


## CR1 vs MM1
Number of differences: 124

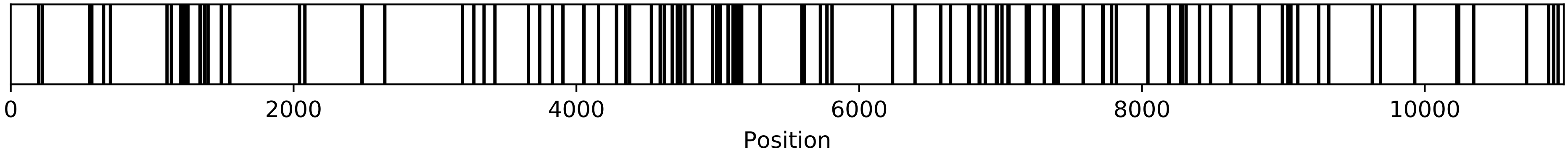


## CR1 vs MM2
Number of differences: 129

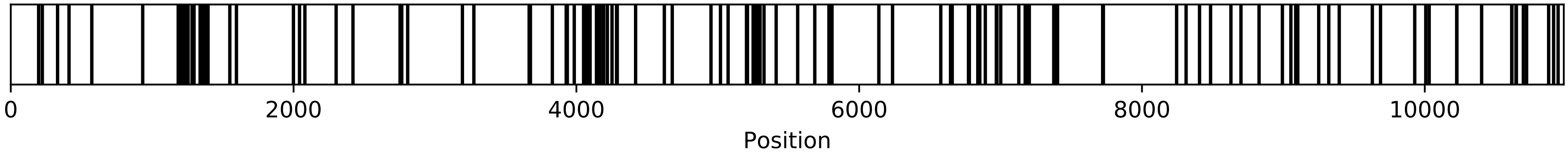


## CR2 vs MM1
Number of differences: 251

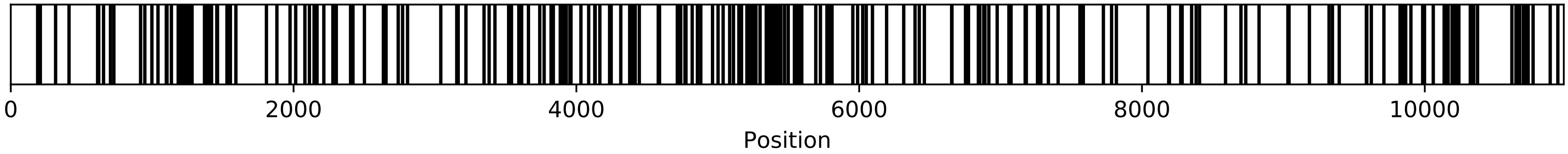


## CR2 vs MM2
Number of differences: 264

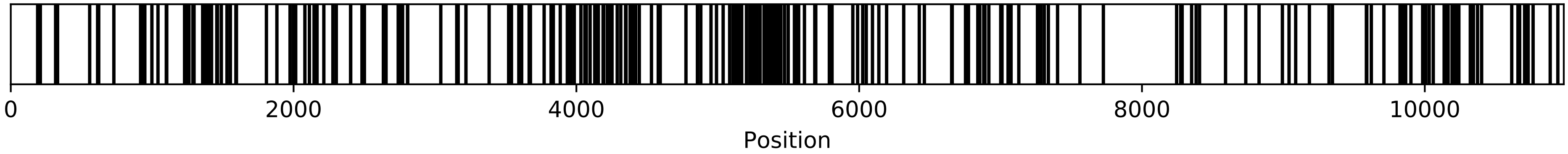


## MM1 vs MM2
Number of differences: 136

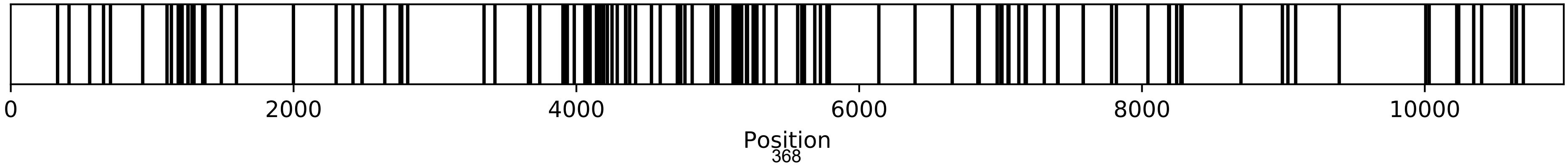

### Region: 973



CR1 vs CR2
Number of differences: 275

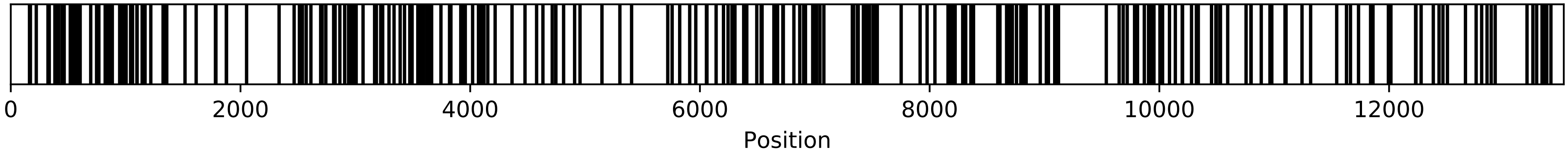


CR1 vs MM1
Number of differences: 47

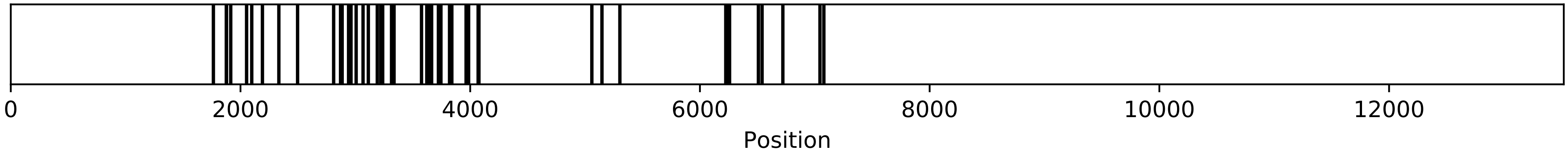


CR1 vs MM2
Number of differences: 126

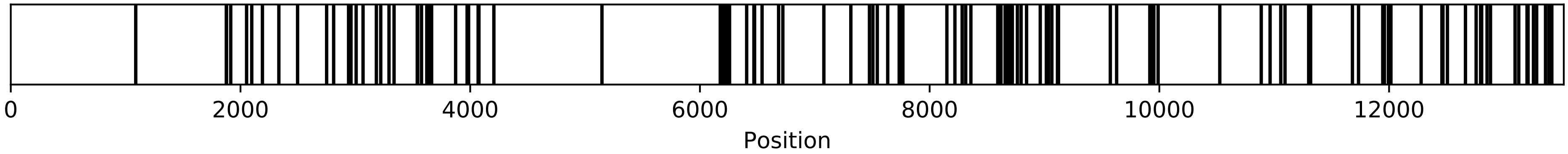


CR2 vs MM1
Number of differences: 277

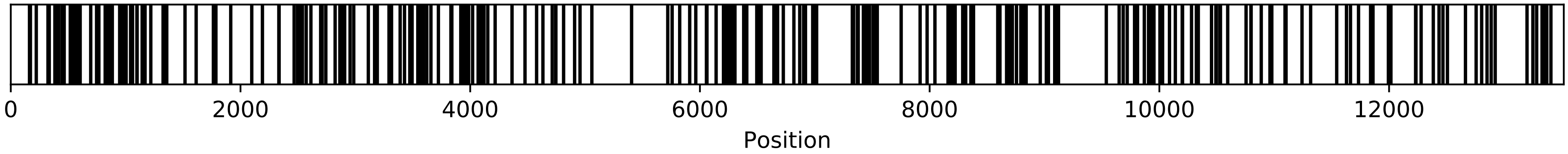


CR2 vs MM2
Number of differences: 275

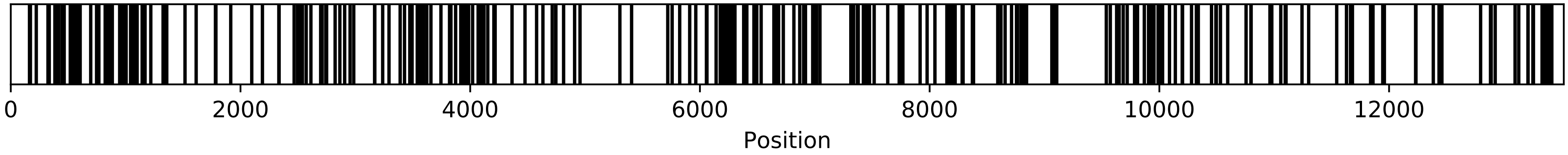


MM1 vs MM2
Number of differences: 111

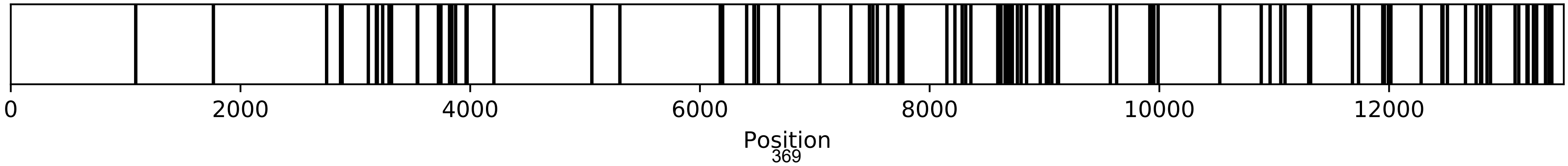

### Region: 977



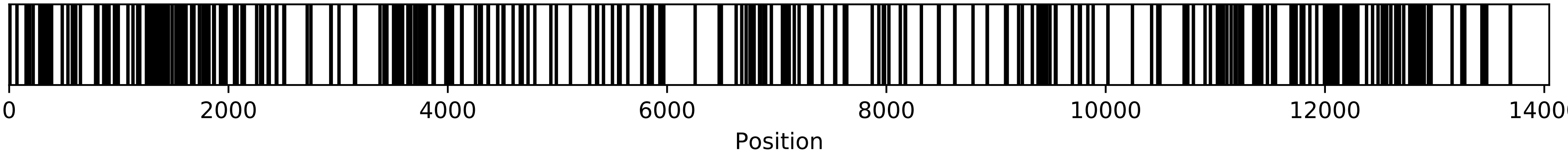
CR1 vs CR2
Number of differences: 362
0
2000
4000
6000
8000
10000
12000
14000
Position

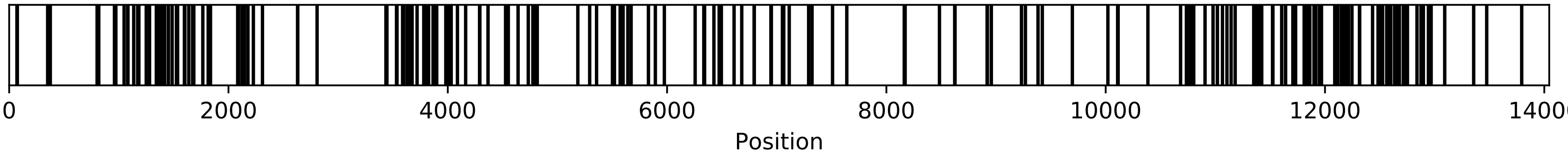
CR1 vs MM1
Number of differences: 238
0
2000
4000
6000
8000
10000
12000
14000
Position

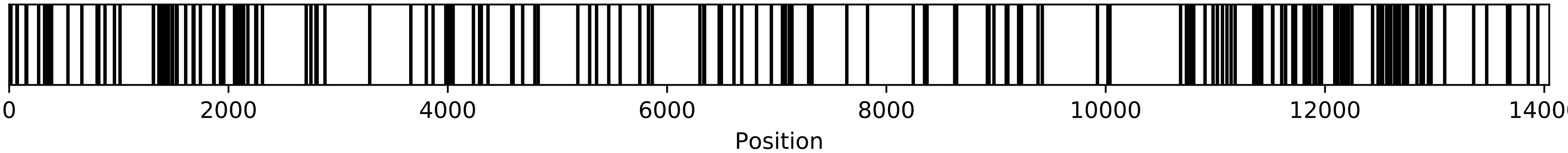
CR1 vs MM2
Number of differences: 227
0
2000
4000
6000
8000
10000
12000
14000
Position

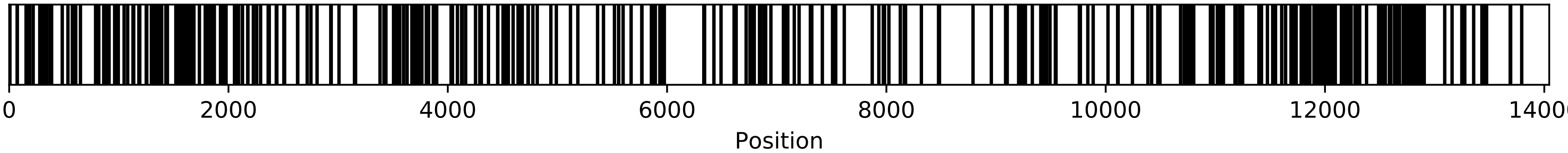
CR2 vs MM1
Number of differences: 375
0
2000
4000
6000
8000
10000
12000
14000
Position

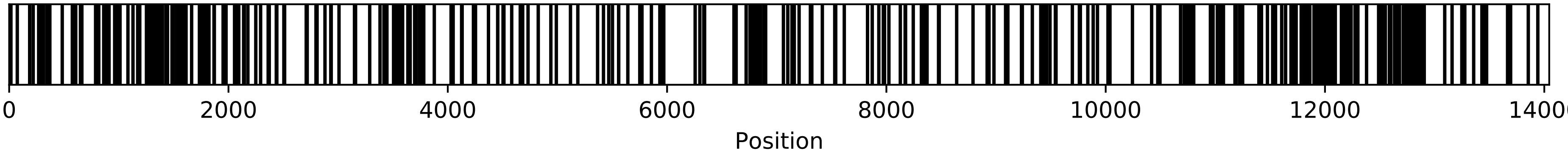
CR2 vs MM2
Number of differences: 369
0
2000
4000
6000
8000
10000
12000
14000
Position

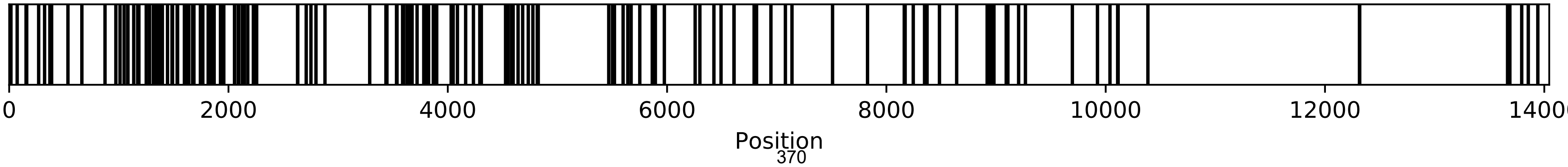
MM1 vs MM2
Number of differences: 169
0
2000
4000
6000
8000
10000
12000
14000
Position

### Region: 978



CR1 vs CR2
Number of differences: 223

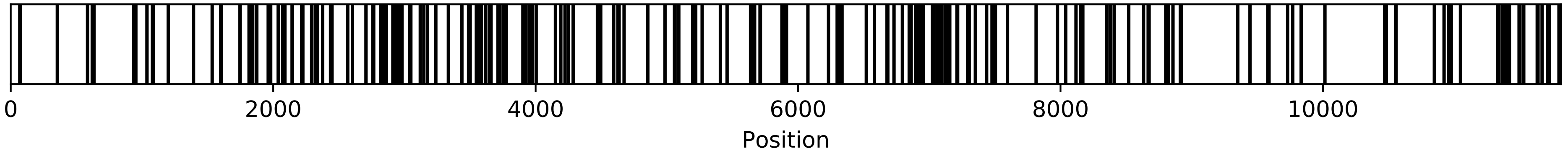


CR1 vs MM1
Number of differences: 100

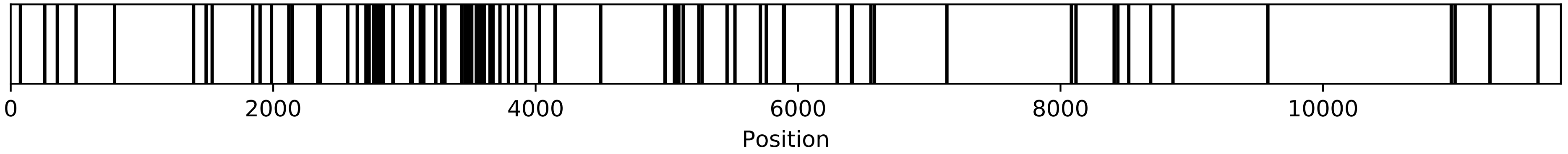


CR1 vs MM2
Number of differences: 92

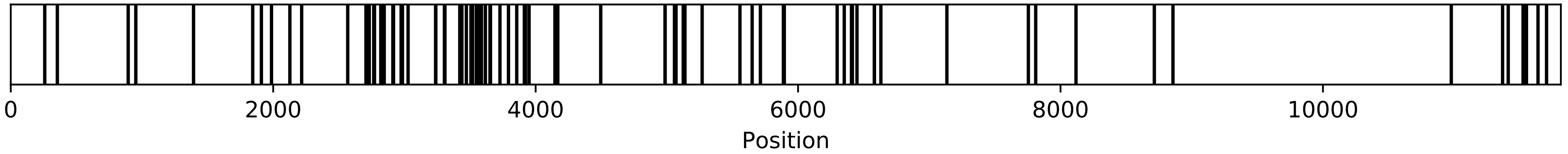


CR2 vs MM1
Number of differences: 237

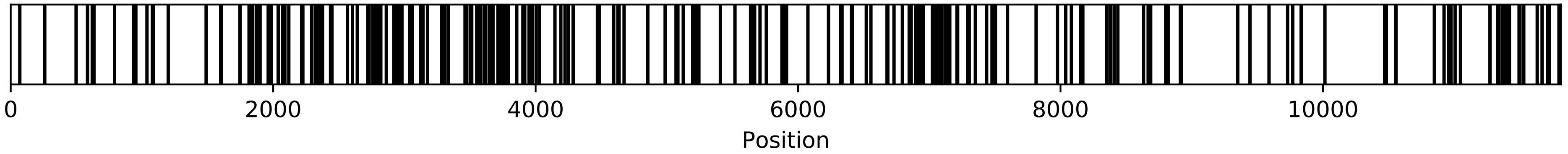


CR2 vs MM2
Number of differences: 230

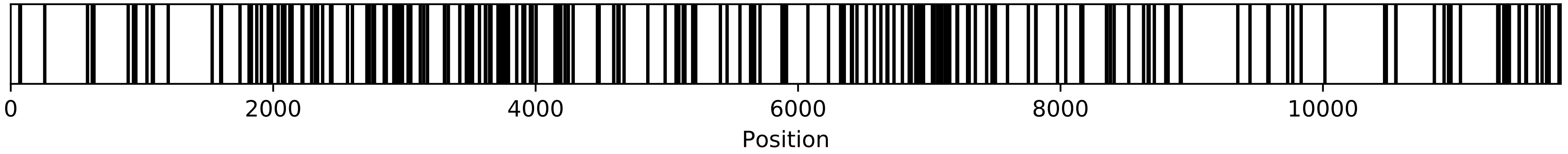


MM1 vs MM2
Number of differences: 90

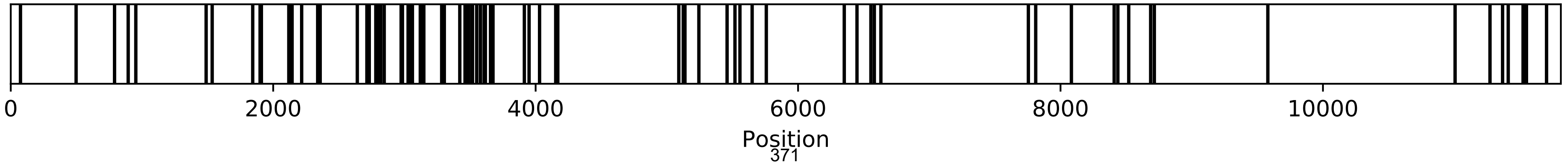

### Region: 982



CR1 vs CR2
Number of differences: 196

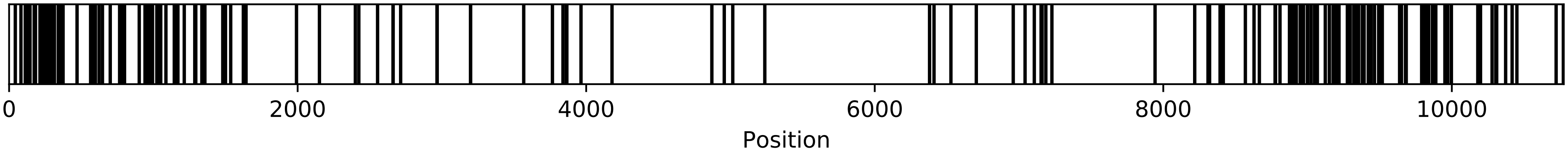


CR1 vs MM1
Number of differences: 189

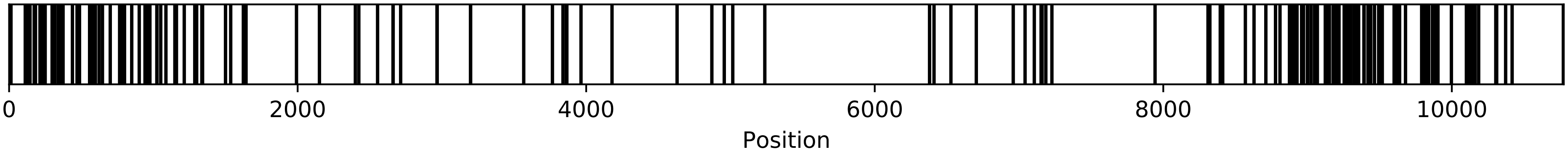


CR1 vs MM2
Number of differences: 199

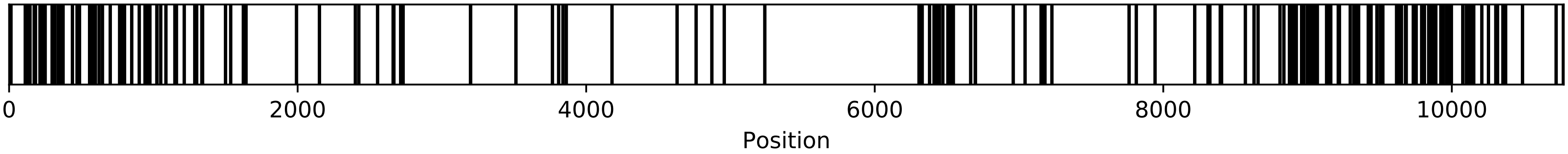


CR2 vs MM1
Number of differences: 85

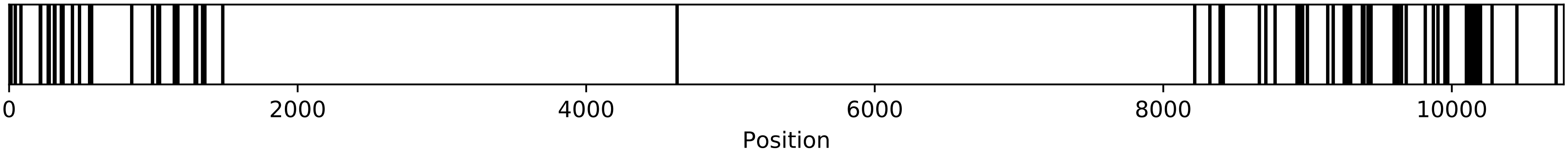


CR2 vs MM2
Number of differences: 135

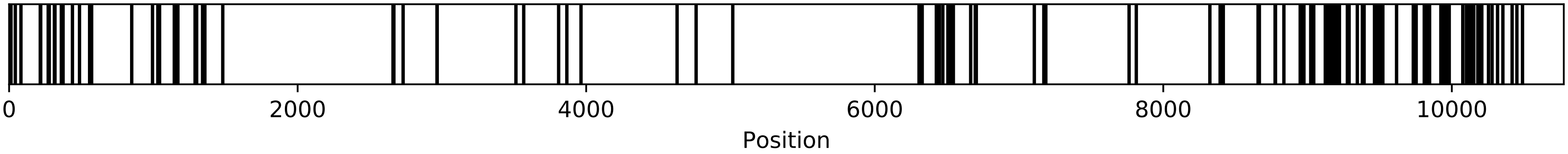


MM1 vs MM2
Number of differences: 112

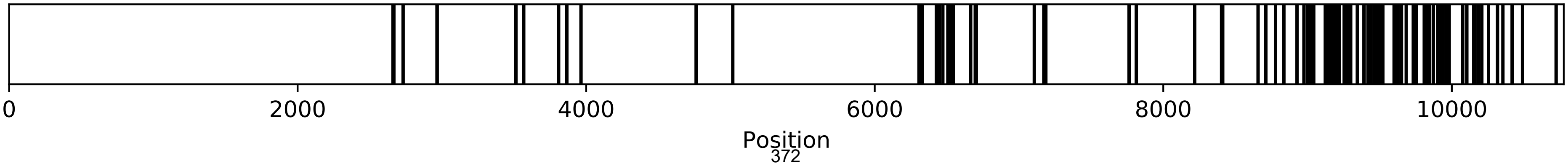

### Region: 983



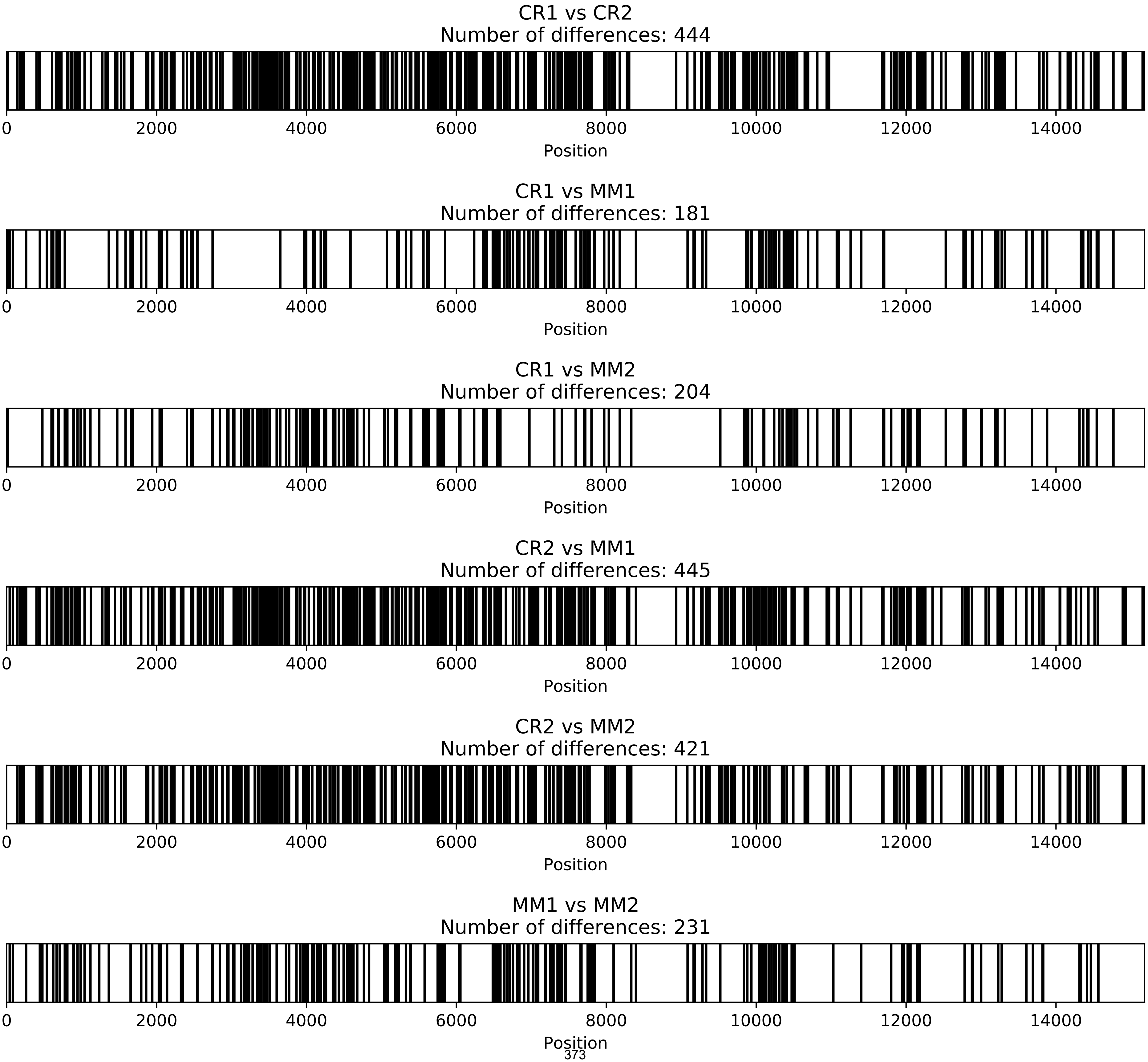
CR1 vs CR2
Number of differences: 444
CR1 vs MM1
Number of differences: 181
CR1 vs MM2
Number of differences: 204
CR2 vs MM1
Number of differences: 445
CR2 vs MM2
Number of differences: 421
MM1 vs MM2
Number of differences: 231
0
2000
4000
6000
8000
10000
12000
14000
Position

### Region: 985



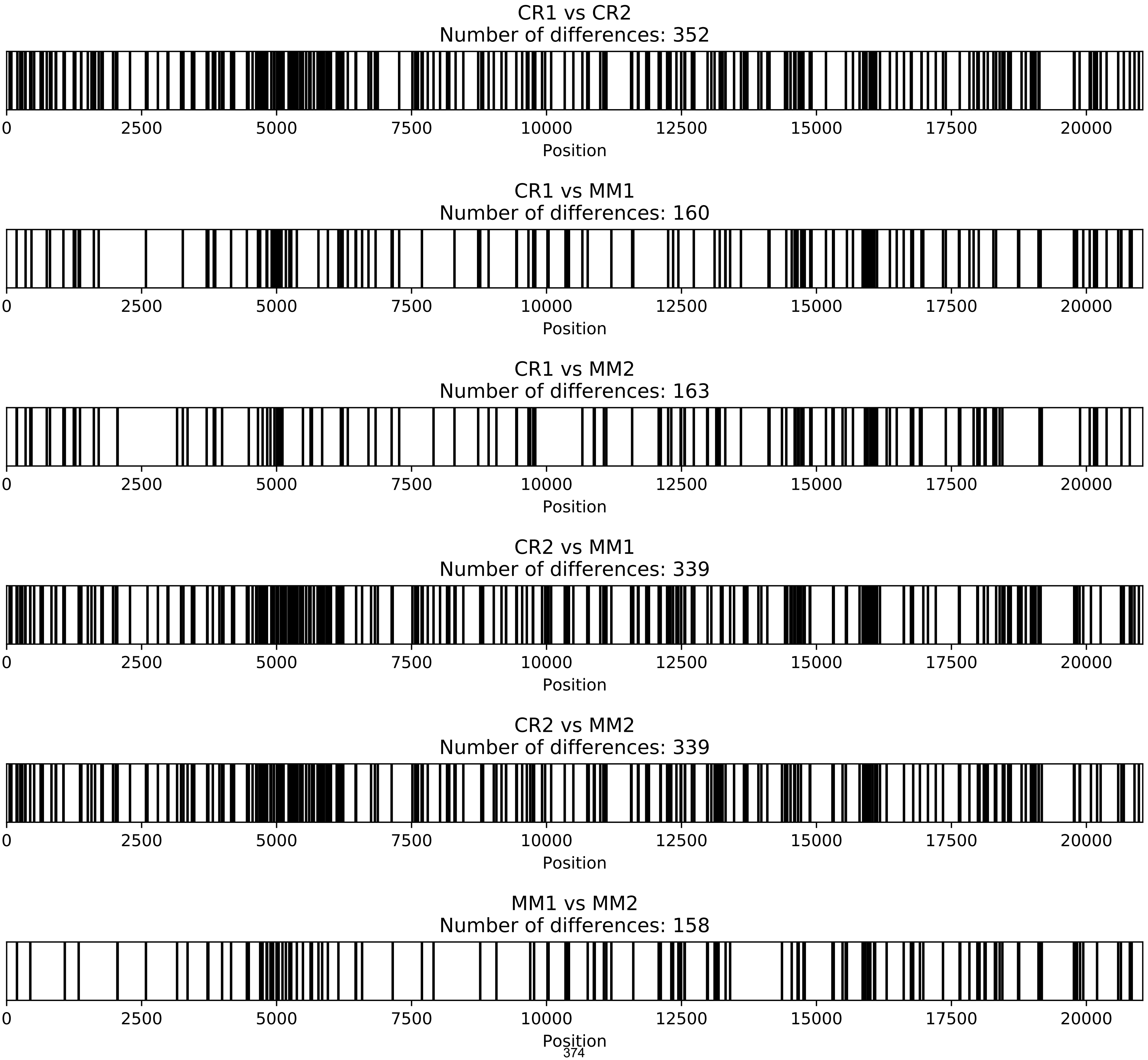
CR1 vs CR2
Number of differences: 352
CR1 vs MM1
Number of differences: 160
CR1 vs MM2
Number of differences: 163
CR2 vs MM1
Number of differences: 339
CR2 vs MM2
Number of differences: 339
MM1 vs MM2
Number of differences: 158
0
2500
5000
7500
10000
12500
15000
17500
20000
Position

### Region: 986



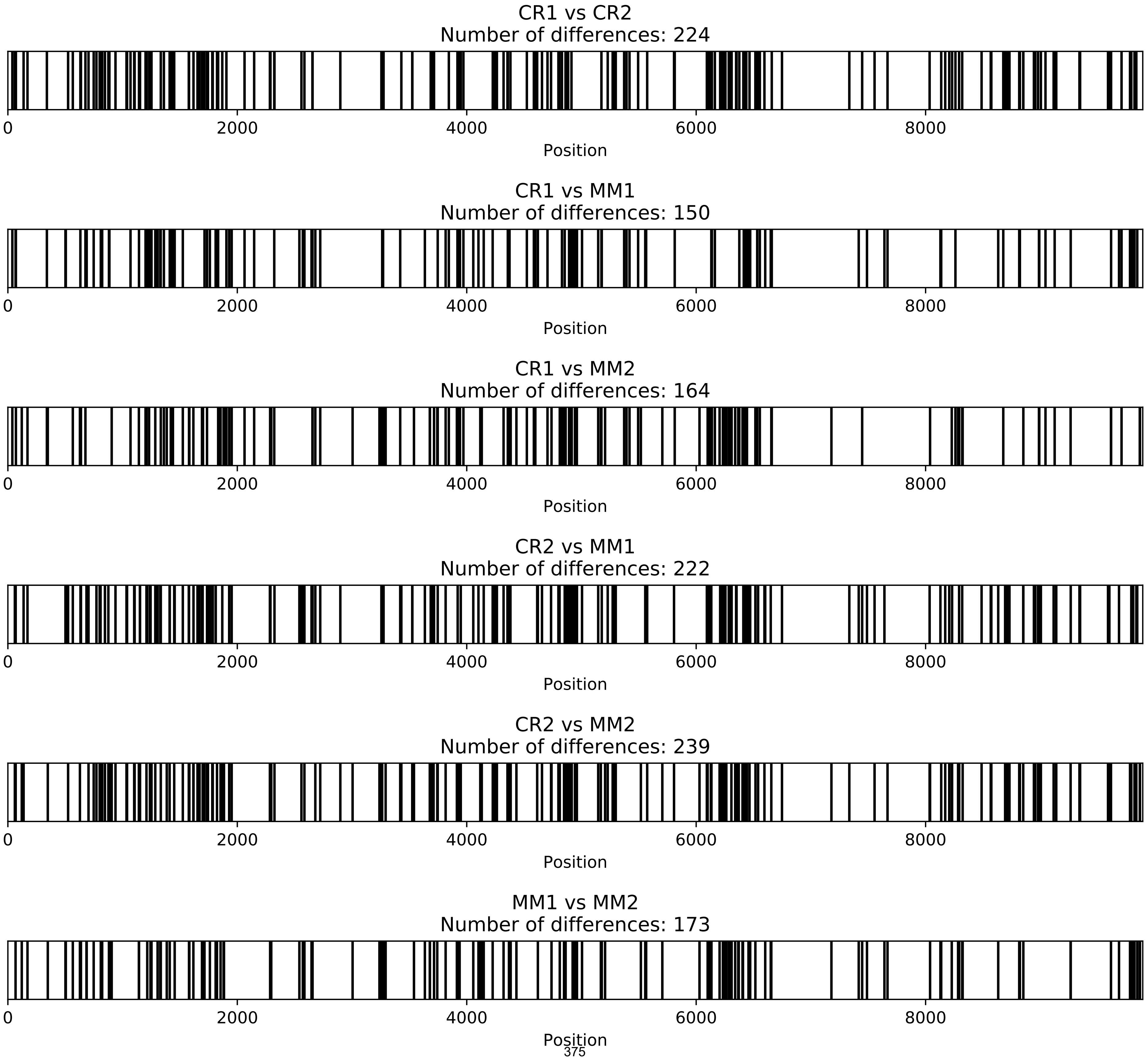
CR1 vs CR2
Number of differences: 224
CR1 vs MM1
Number of differences: 150
CR1 vs MM2
Number of differences: 164
CR2 vs MM1
Number of differences: 222
CR2 vs MM2
Number of differences: 239
MM1 vs MM2
Number of differences: 173
0
2000
4000
6000
8000
Position

### Region: 987



CR1 vs CR2
Number of differences: 337

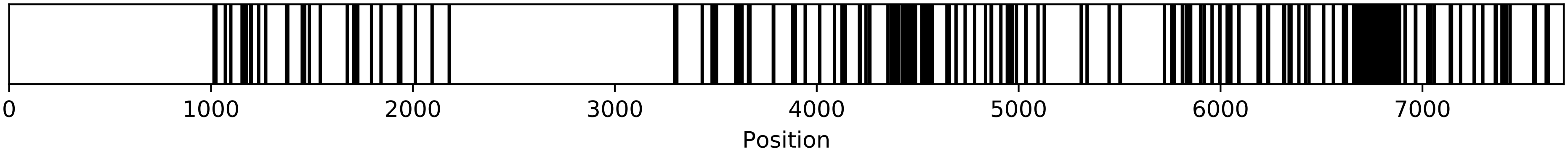


CR1 vs MM1
Number of differences: 123

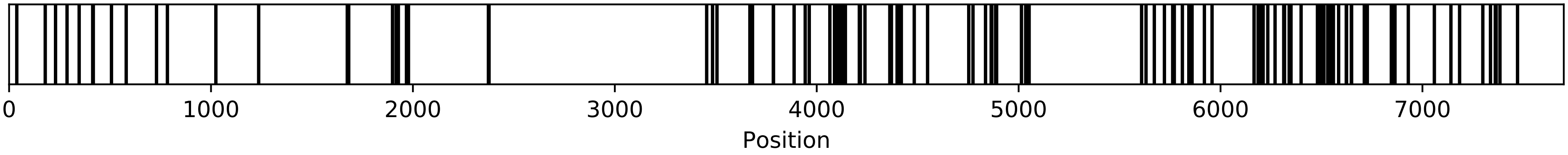


CR1 vs MM2
Number of differences: 117

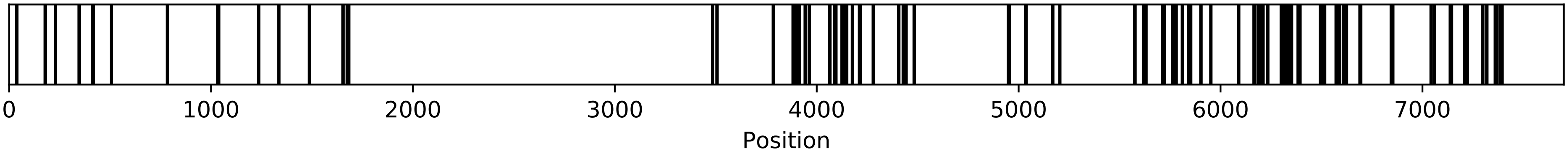


CR2 vs MM1
Number of differences: 366

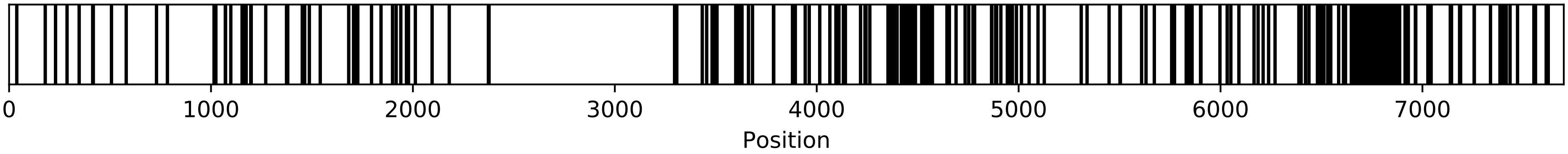


CR2 vs MM2
Number of differences: 360

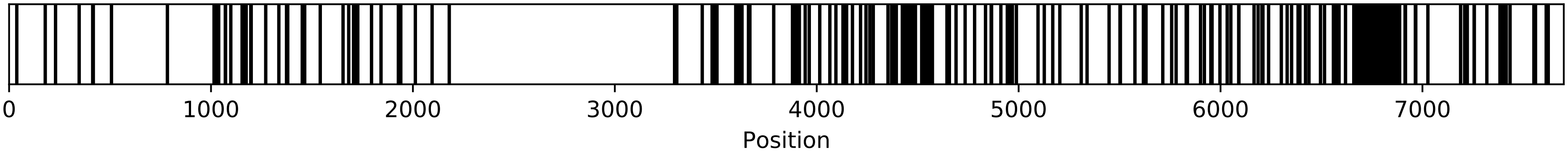


MM1 vs MM2
Number of differences: 124

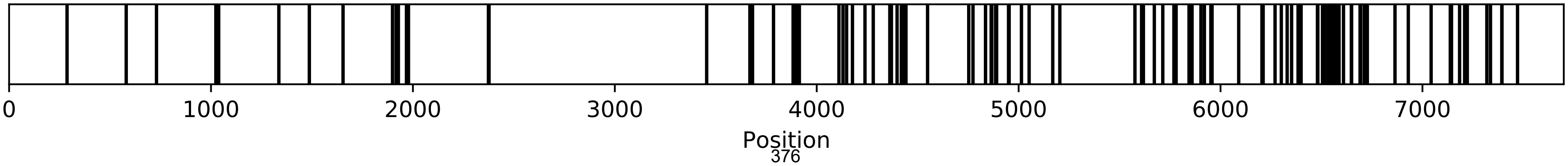

### Region: 990



CR1 vs CR2
Number of differences: 231

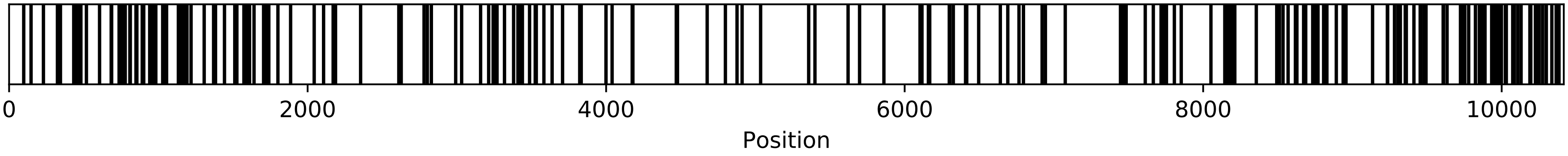


CR1 vs MM1
Number of differences: 134

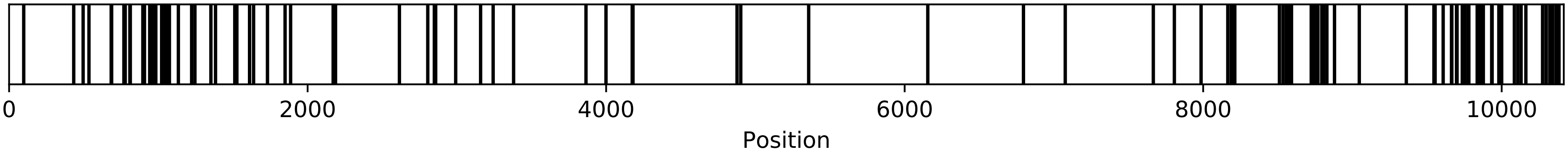


CR1 vs MM2
Number of differences: 124

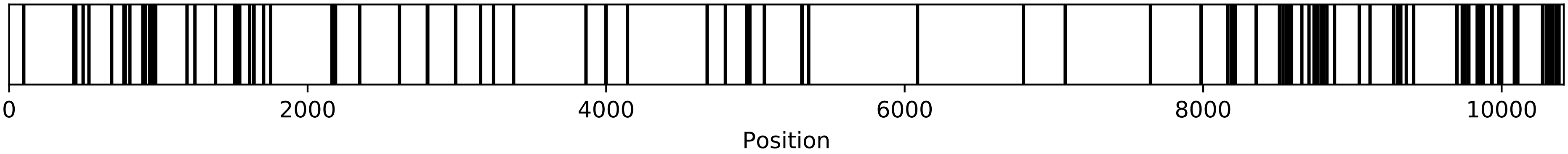


CR2 vs MM1
Number of differences: 217

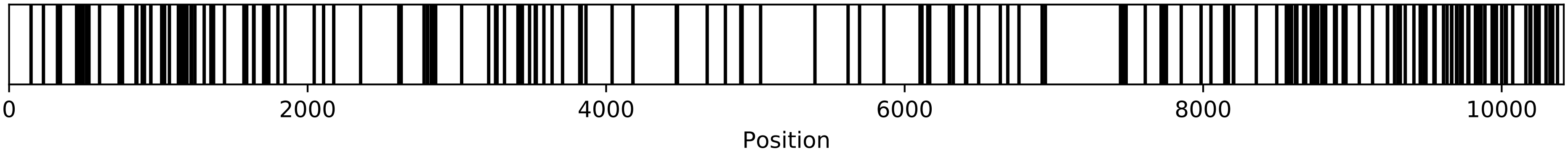


CR2 vs MM2
Number of differences: 230

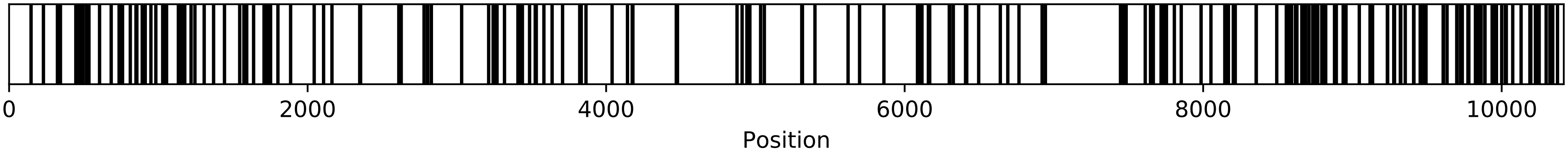


MM1 vs MM2
Number of differences: 76

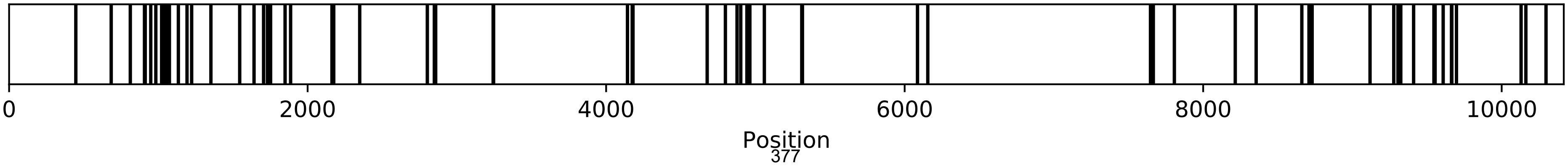

### Region: 991



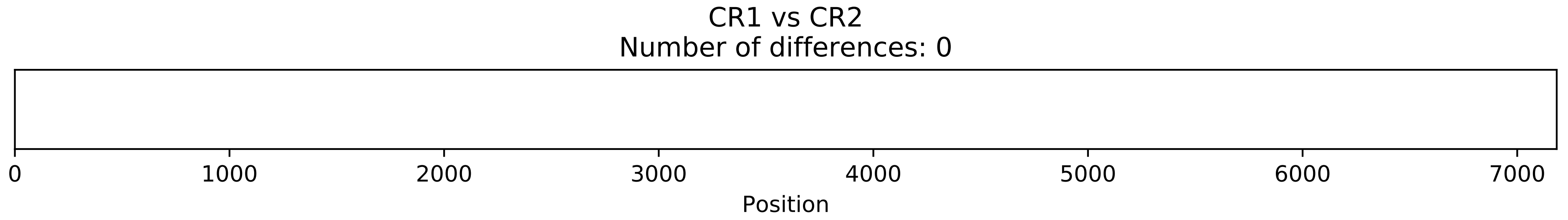
CR1 vs CR2
Number of differences: 0
0
1000
2000
3000
4000
5000
6000
7000
Position

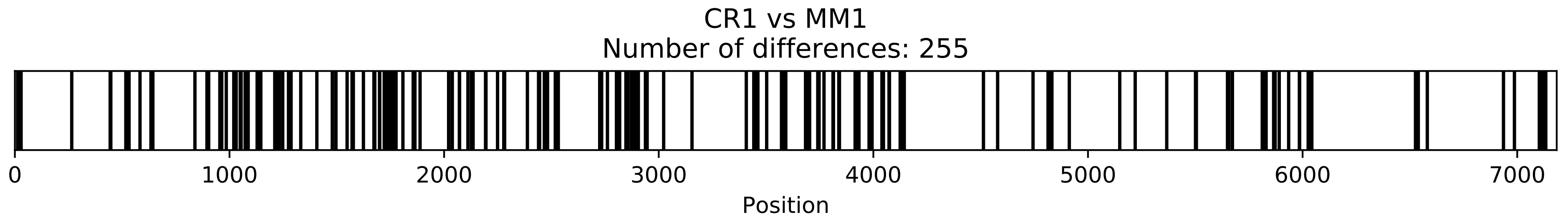
CR1 vs MM1
Number of differences: 255
0
1000
2000
3000
4000
5000
6000
7000
Position

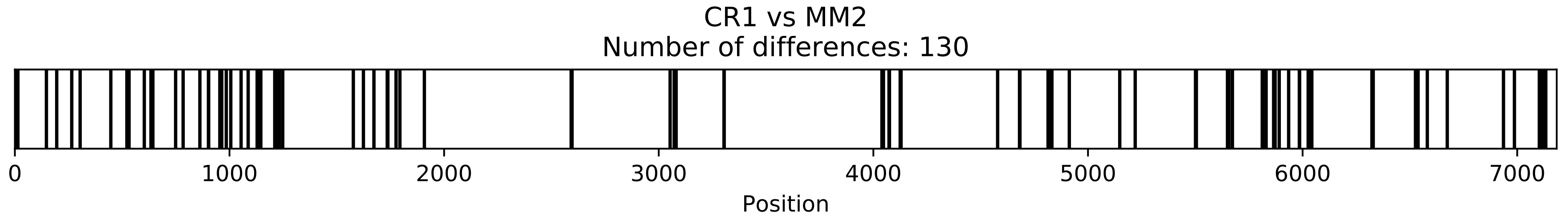
CR1 vs MM2
Number of differences: 130
0
1000
2000
3000
4000
5000
6000
7000
Position

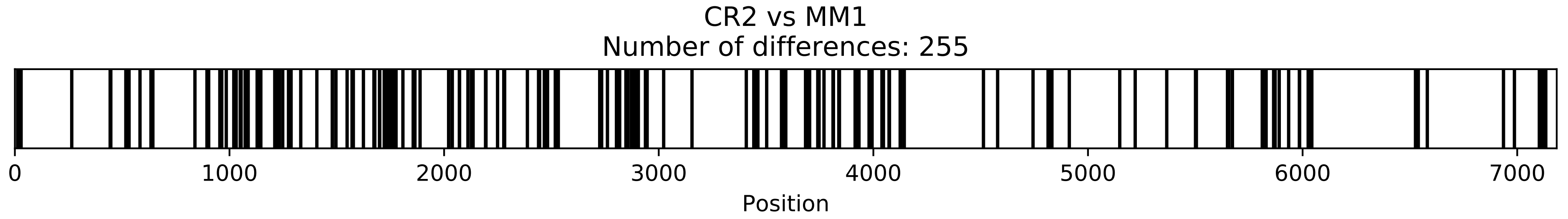
CR2 vs MM1
Number of differences: 255
0
1000
2000
3000
4000
5000
6000
7000
Position

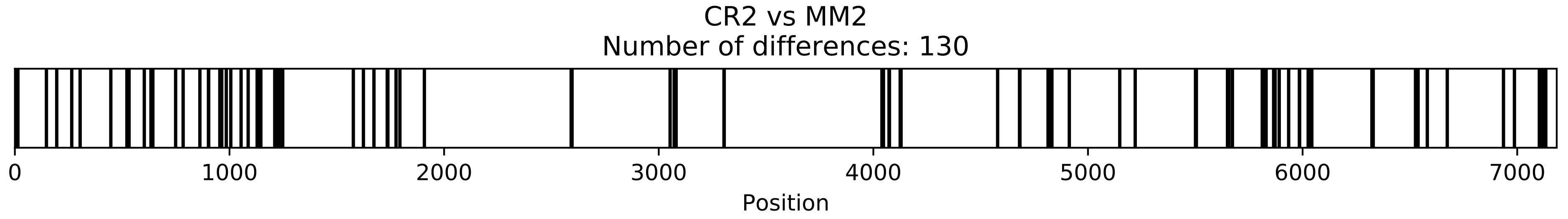
CR2 vs MM2
Number of differences: 130
0
1000
2000
3000
4000
5000
6000
7000
Position

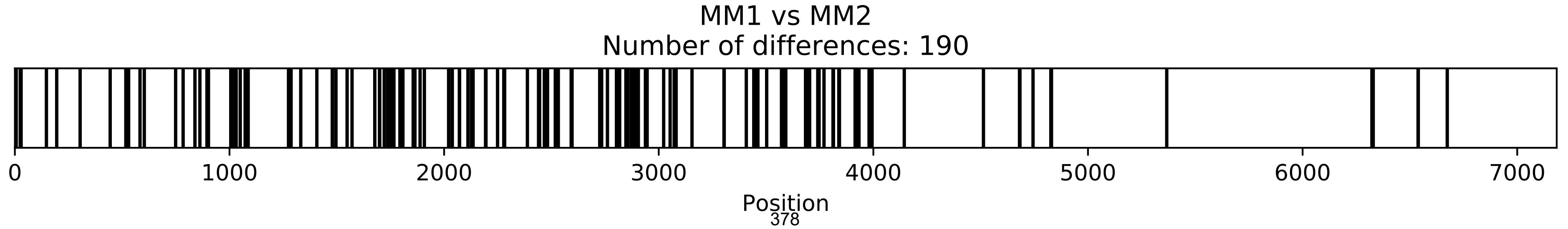
MM1 vs MM2
Number of differences: 190
0
1000
2000
3000
4000
5000
6000
7000
Position

### Region: 992



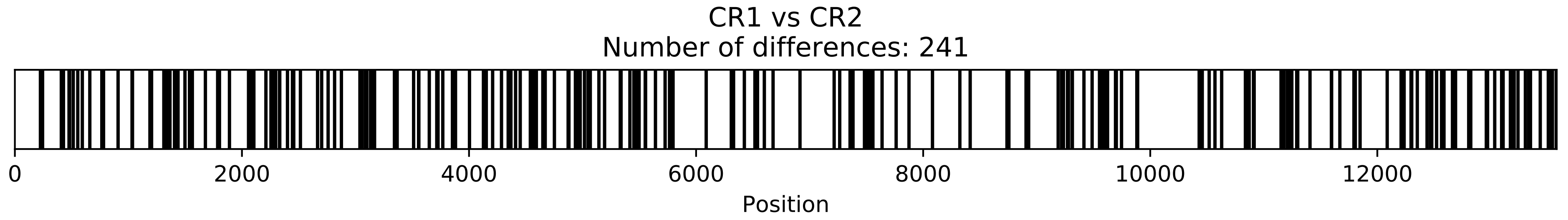
CR1 vs CR2
Number of differences: 241
0
2000
4000
6000
8000
10000
12000
Position

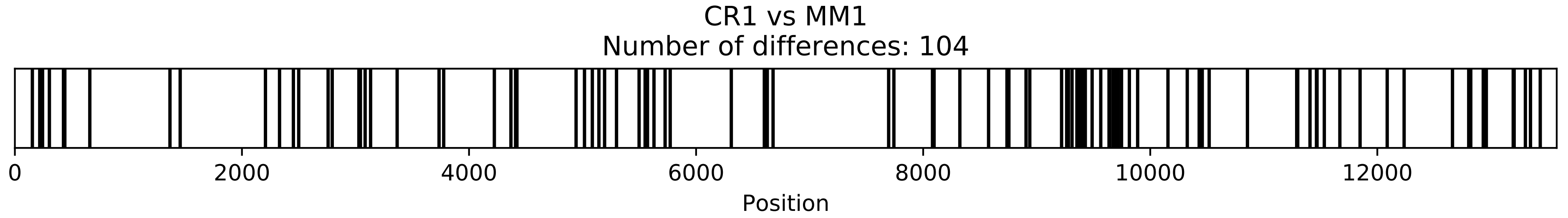
CR1 vs MM1
Number of differences: 104
0
2000
4000
6000
8000
10000
12000
Position

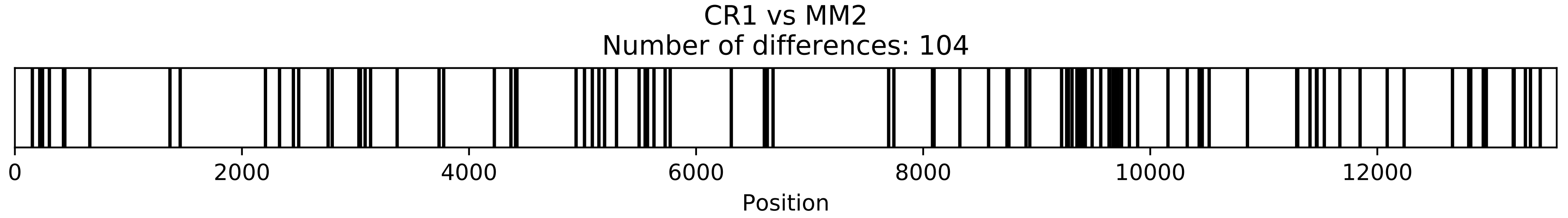
CR1 vs MM2
Number of differences: 104
0
2000
4000
6000
8000
10000
12000
Position

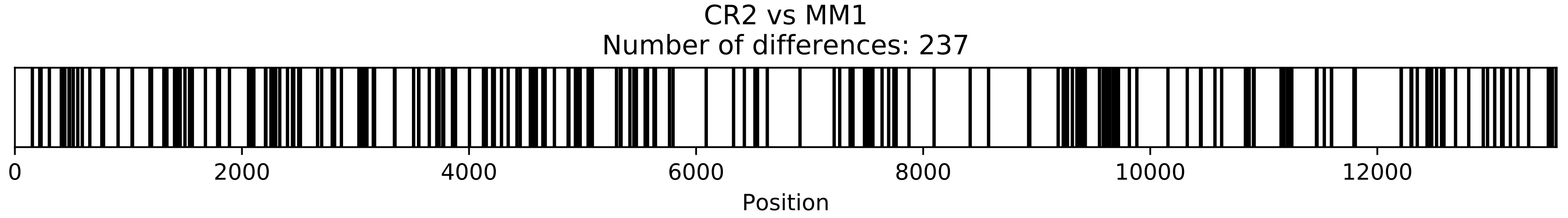
CR2 vs MM1
Number of differences: 237
0
2000
4000
6000
8000
10000
12000
Position

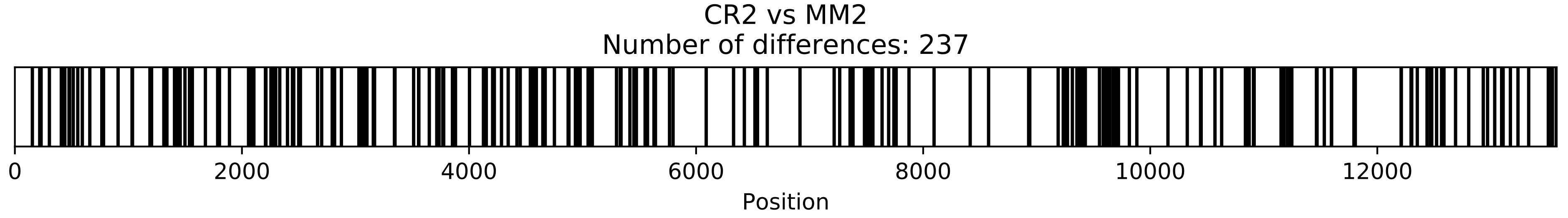
CR2 vs MM2
Number of differences: 237
0
2000
4000
6000
8000
10000
12000
Position

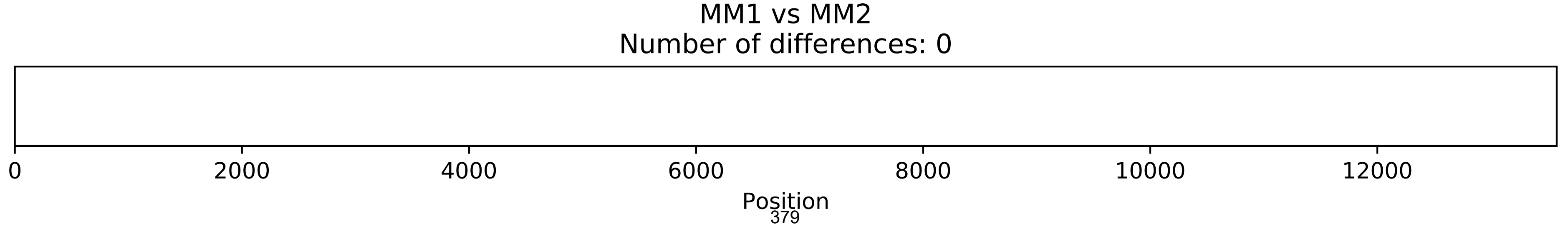
MM1 vs MM2
Number of differences: 0
0
2000
4000
6000
8000
10000
12000
Position

### Region: 996



CR1 vs CR2
Number of differences: 465

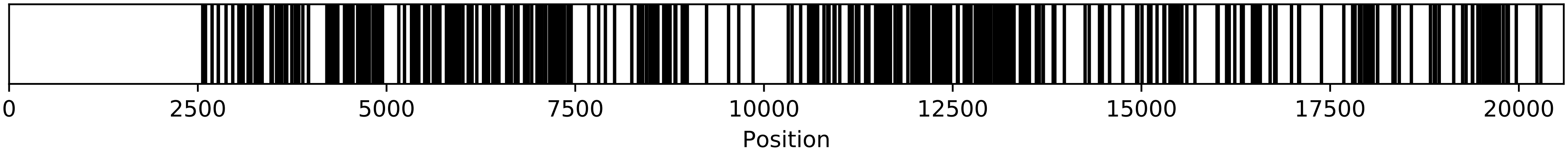


CR1 vs MM1
Number of differences: 274

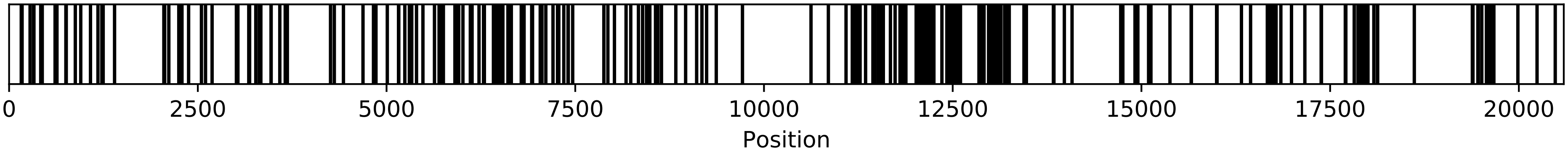


CR1 vs MM2
Number of differences: 320

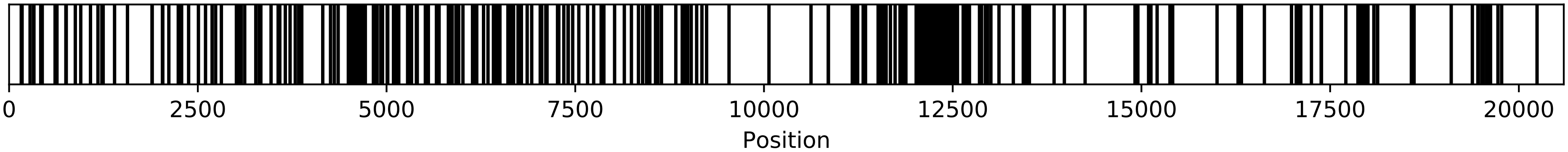


CR2 vs MM1
Number of differences: 501

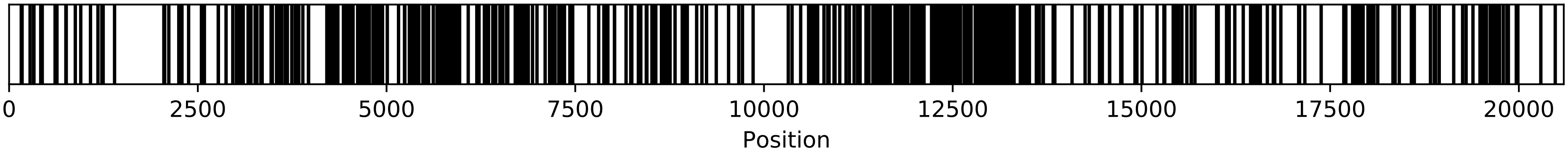


CR2 vs MM2
Number of differences: 495

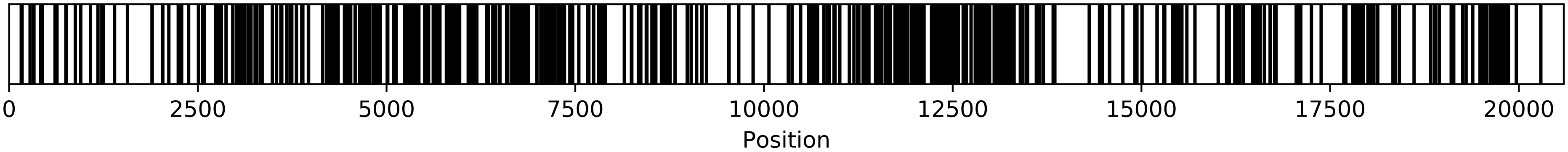


MM1 vs MM2
Number of differences: 269

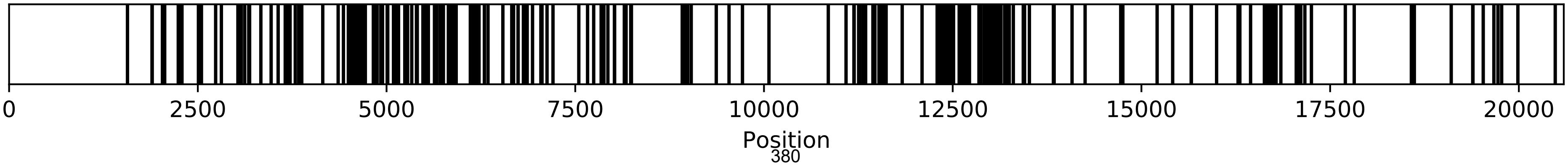

### Region: 997

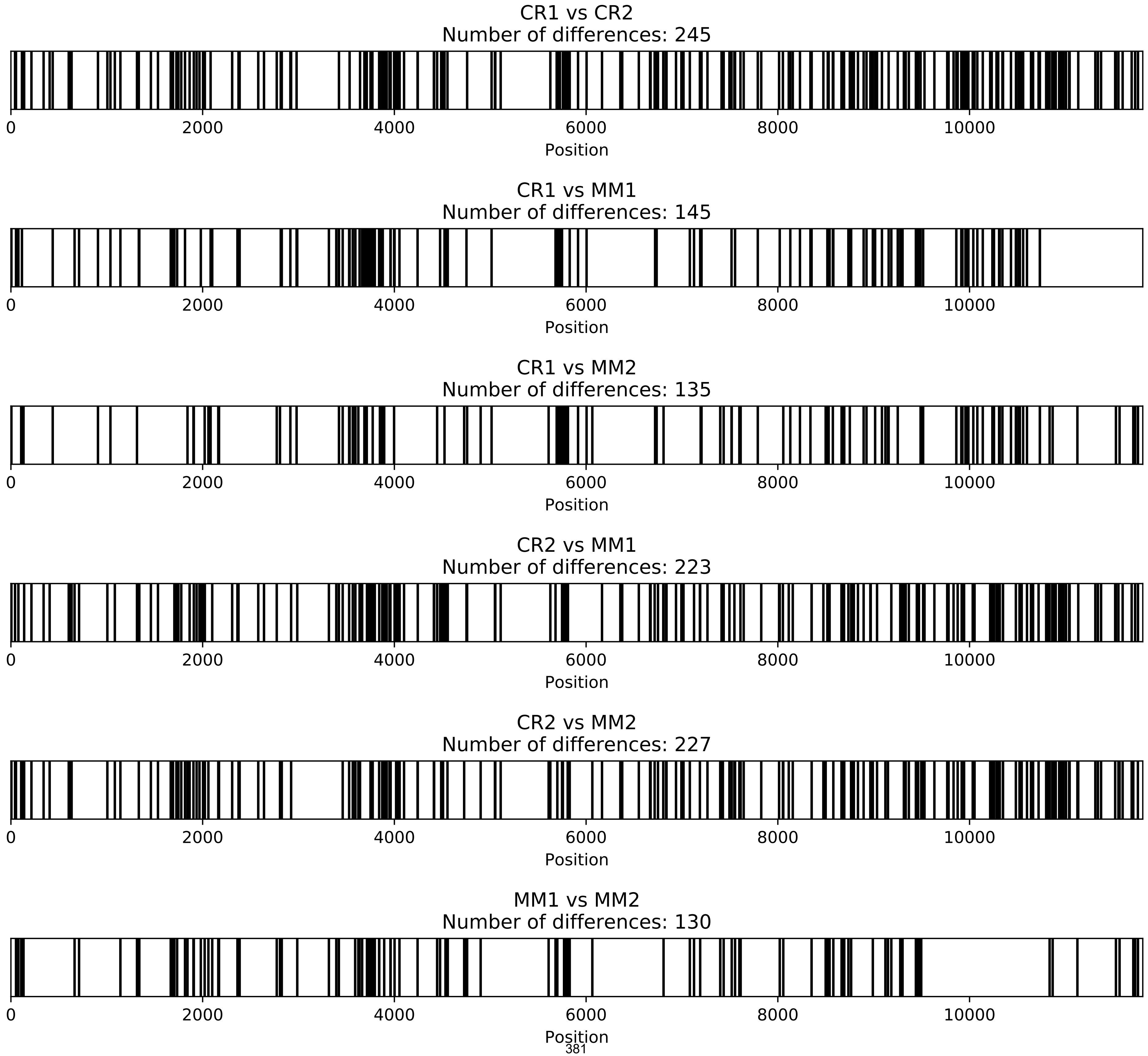
CR1 vs CR2
Number of differences: 245
CR1 vs MM1
Number of differences: 145
CR1 vs MM2
Number of differences: 135
CR2 vs MM1
Number of differences: 223
CR2 vs MM2
Number of differences: 227
MM1 vs MM2
Number of differences: 130
0
2000
4000
6000
8000
10000
Position

### Region: 1003



## CR1 vs CR2
Number of differences: 28

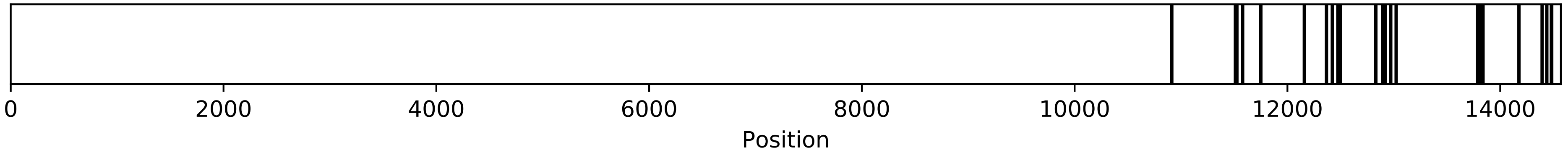


## CR1 vs MM1
Number of differences: 197

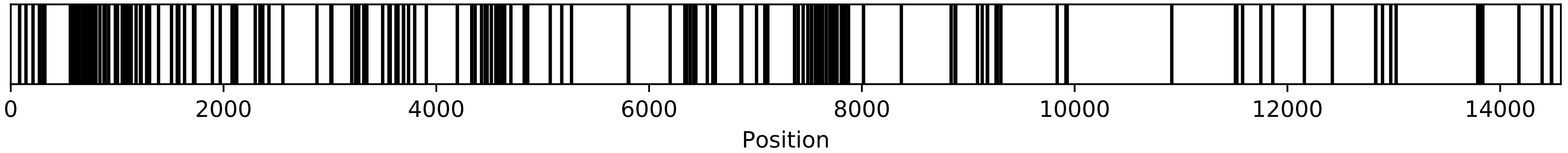


## CR1 vs MM2
Number of differences: 214

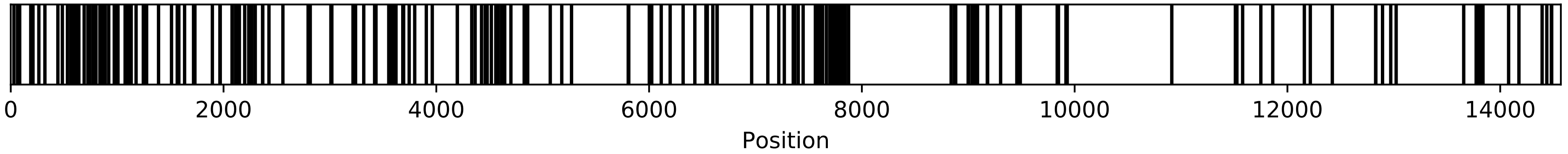


## CR2 vs MM1
Number of differences: 183

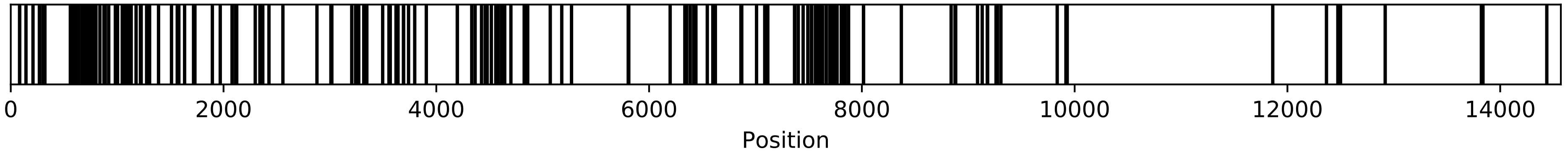


## CR2 vs MM2
Number of differences: 194

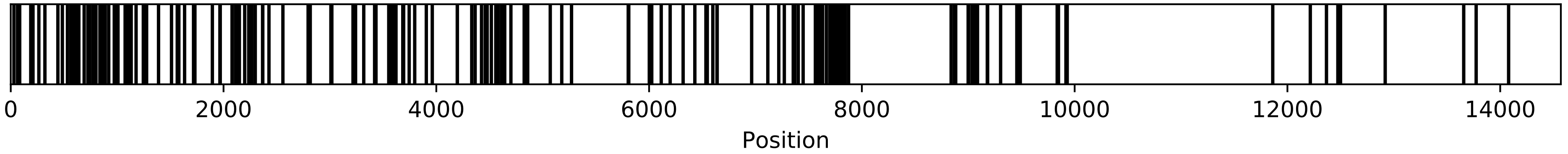


## MM1 vs MM2
Number of differences: 133

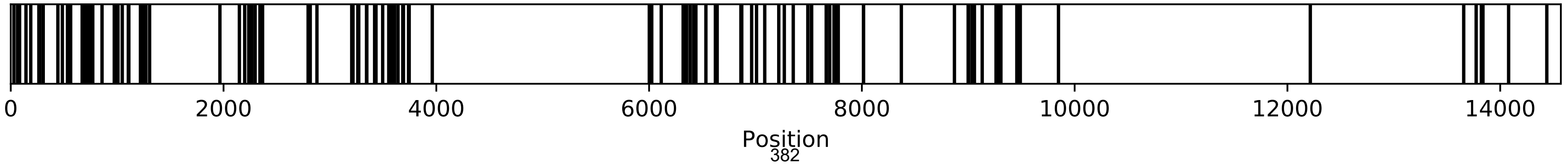

### Region: 1005



CR1 vs CR2
Number of differences: 162

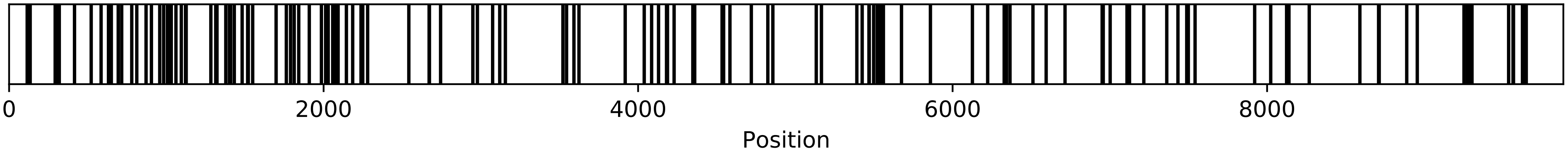


CR1 vs MM1
Number of differences: 81

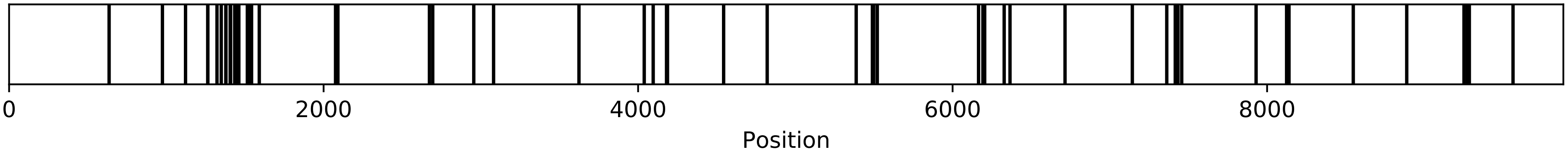


CR1 vs MM2
Number of differences: 61

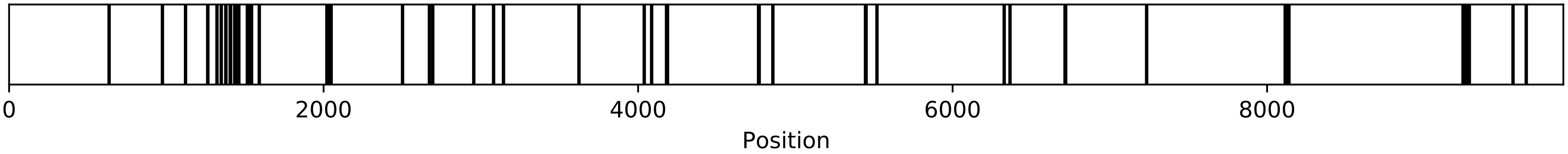


CR2 vs MM1
Number of differences: 163

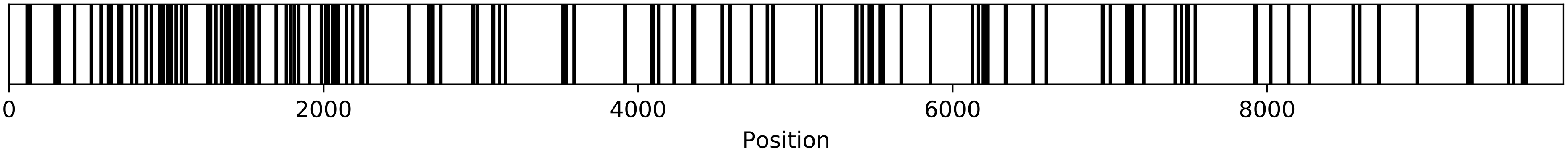


CR2 vs MM2
Number of differences: 159

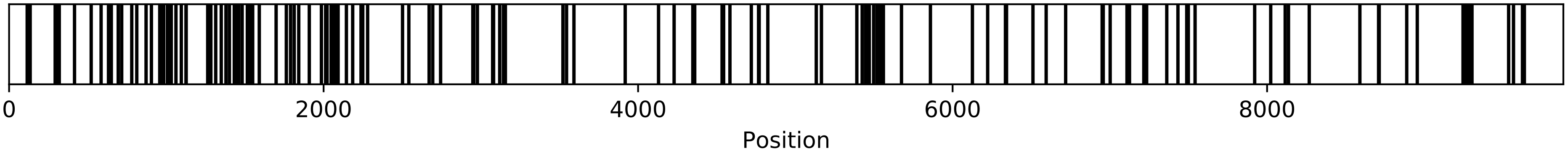


MM1 vs MM2
Number of differences: 56

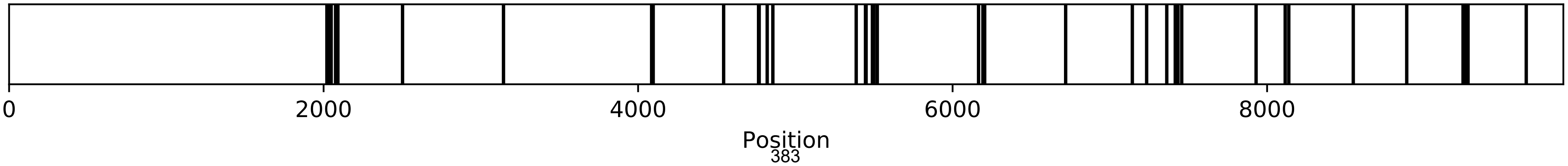

### Region: 1010



CR1 vs CR2
Number of differences: 618

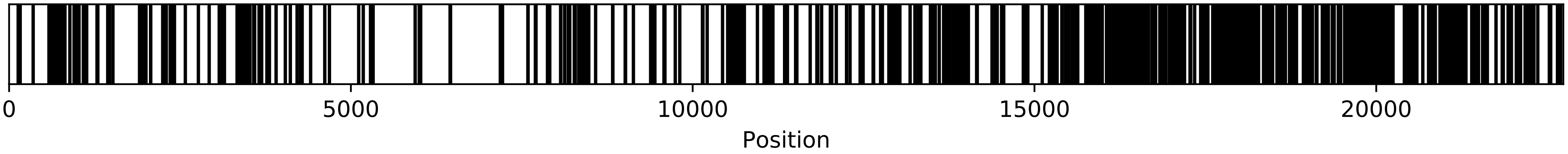


CR1 vs MM1
Number of differences: 260

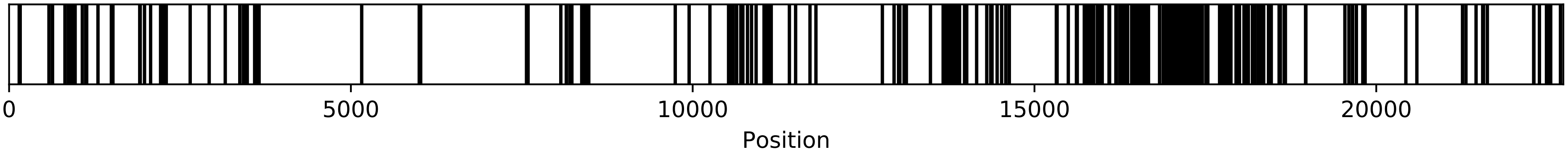


CR1 vs MM2
Number of differences: 211

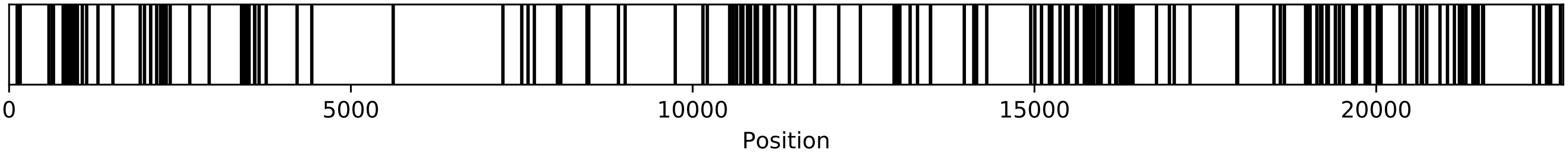


CR2 vs MM1
Number of differences: 571

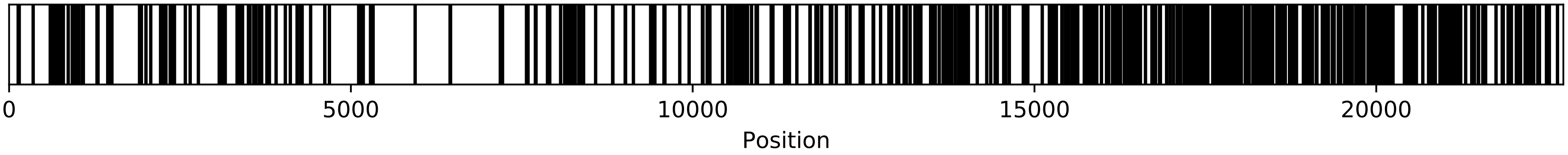


CR2 vs MM2
Number of differences: 612

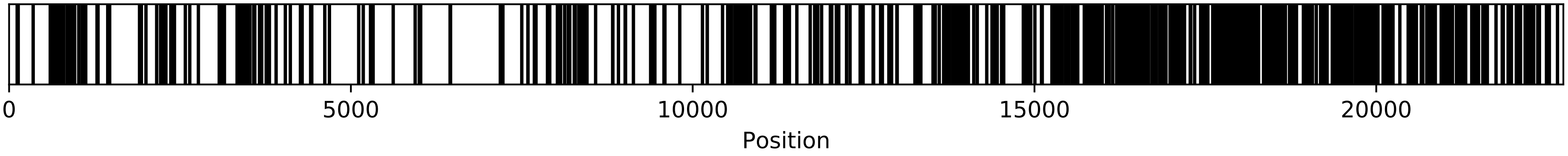


MM1 vs MM2
Number of differences: 284

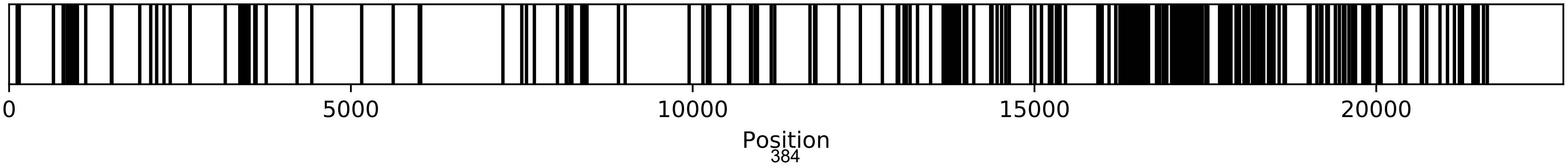

### Region: 1011



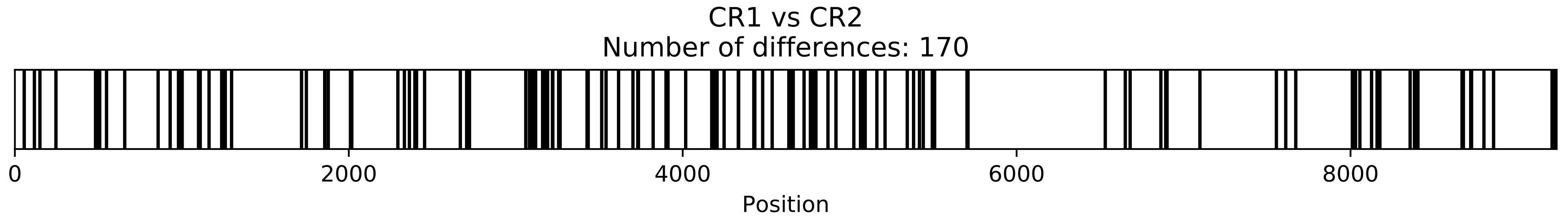
CR1 vs CR2
Number of differences: 170
0
2000
4000
6000
8000
Position

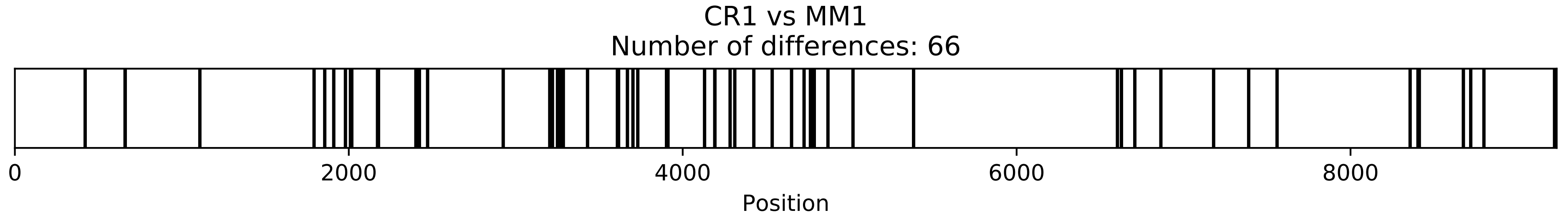
CR1 vs MM1
Number of differences: 66
0
2000
4000
6000
8000
Position

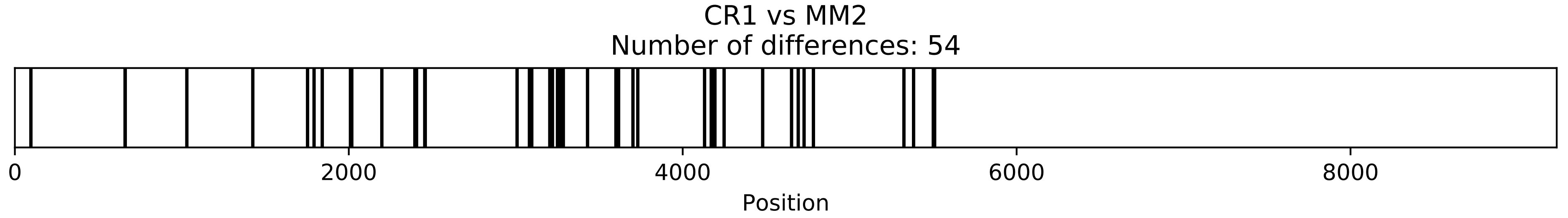
CR1 vs MM2
Number of differences: 54
0
2000
4000
6000
8000
Position

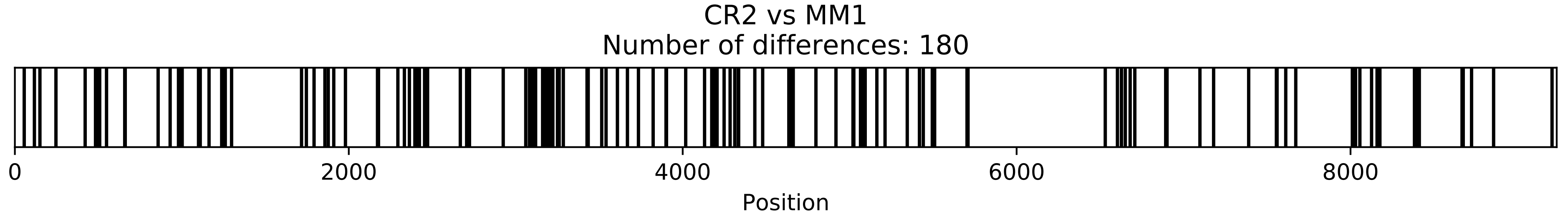
CR2 vs MM1
Number of differences: 180
0
2000
4000
6000
8000
Position

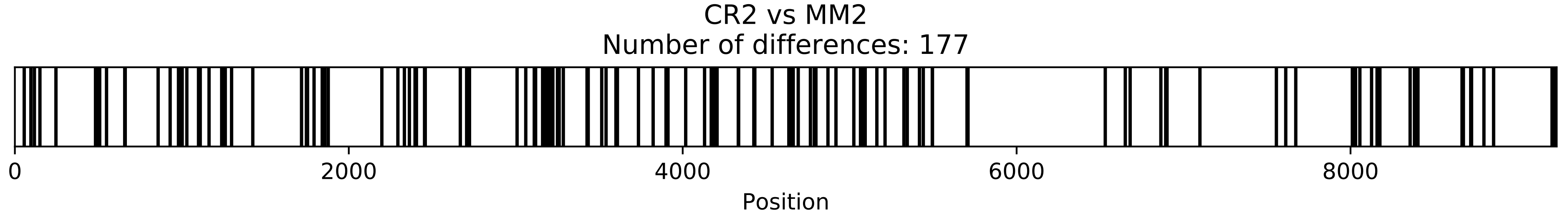
CR2 vs MM2
Number of differences: 177
0
2000
4000
6000
8000
Position

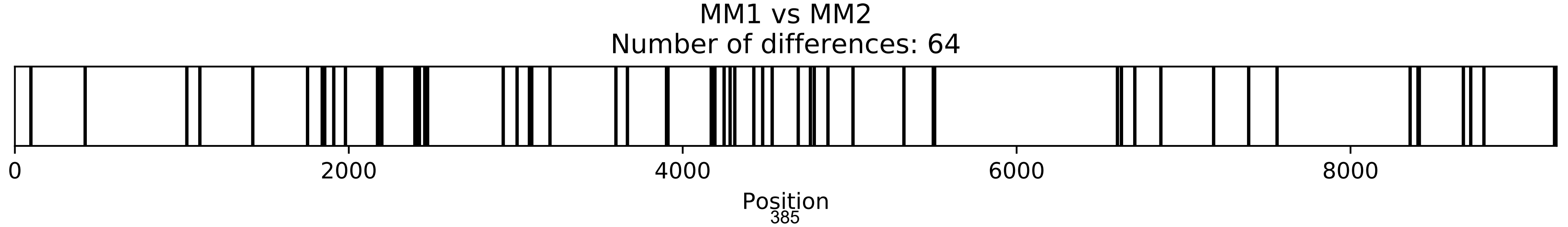
MM1 vs MM2
Number of differences: 64
0
2000
4000
6000
8000
Position

### Region: 1015



CR1 vs CR2
Number of differences: 408

CR1 vs MM1
Number of differences: 132

CR1 vs MM2
Number of differences: 155

CR2 vs MM1
Number of differences: 408

CR2 vs MM2
Number of differences: 422

MM1 vs MM2
Number of differences: 163

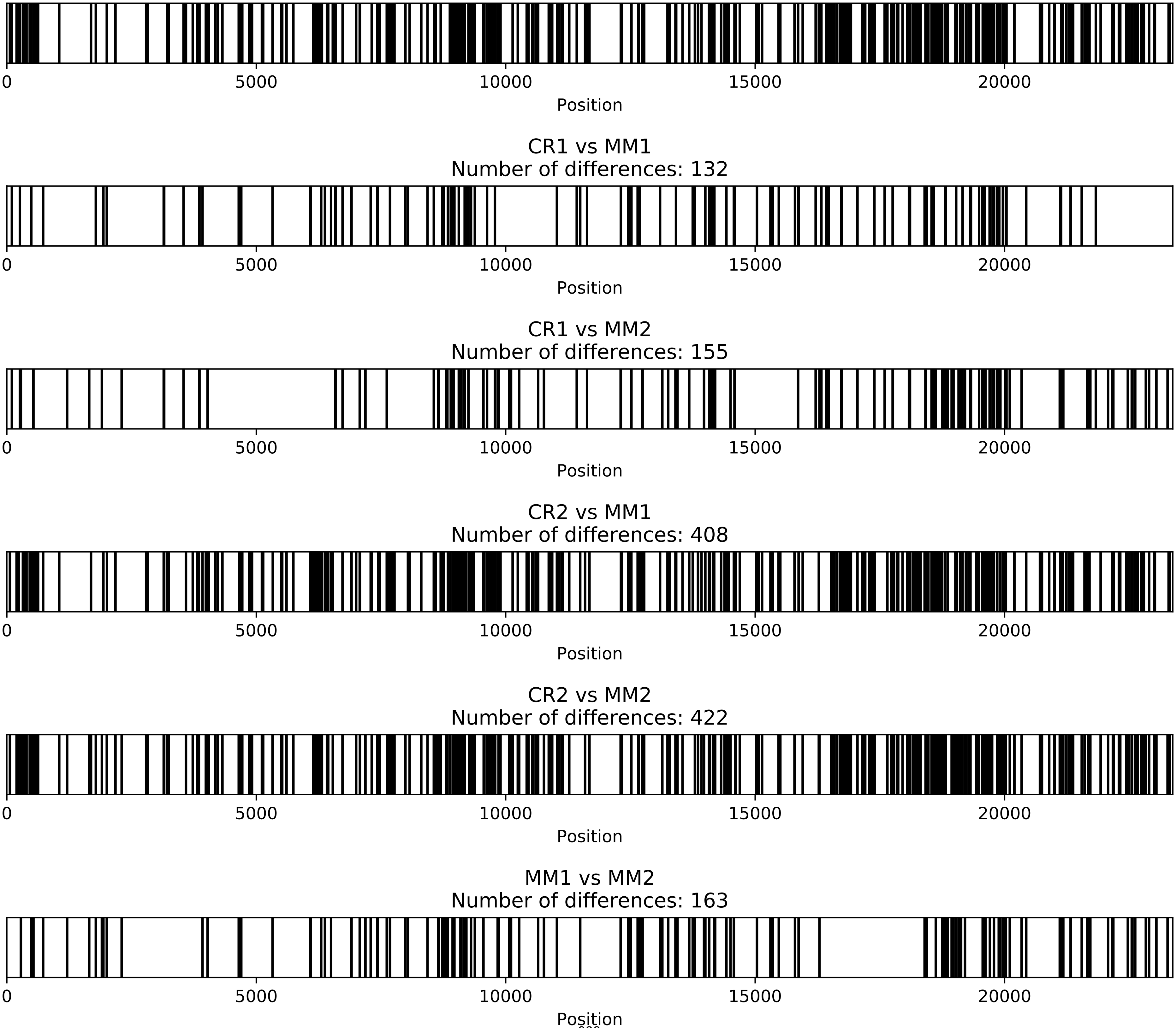

### Region: 1022



CR1 vs CR2
Number of differences: 150

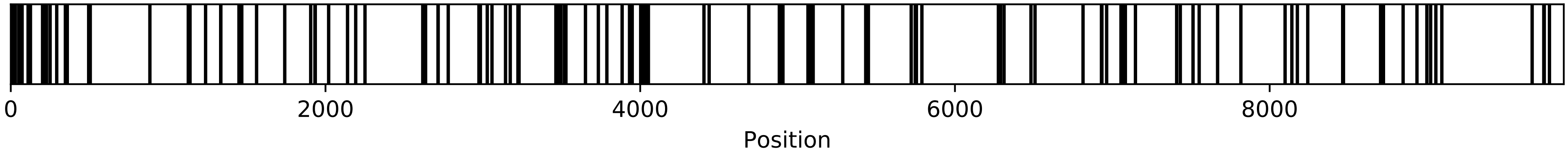


CR1 vs MM1
Number of differences: 160

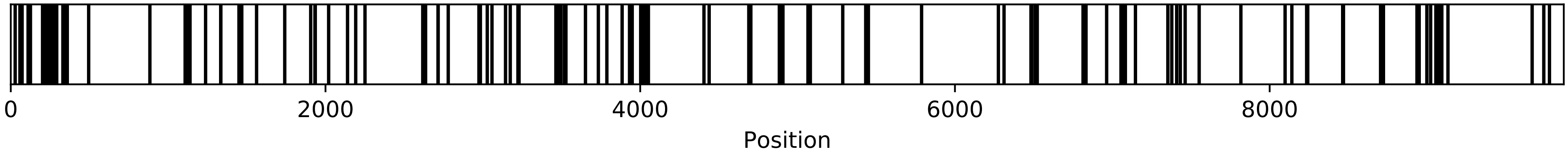


CR1 vs MM2
Number of differences: 148

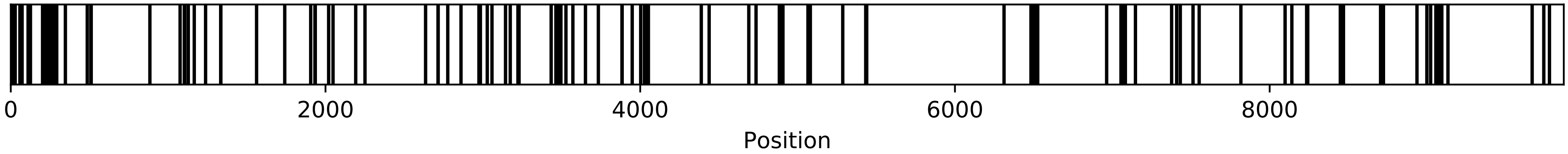


CR2 vs MM1
Number of differences: 55

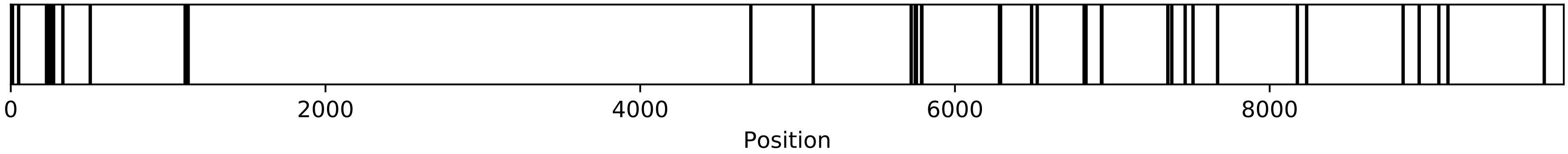


CR2 vs MM2
Number of differences: 71

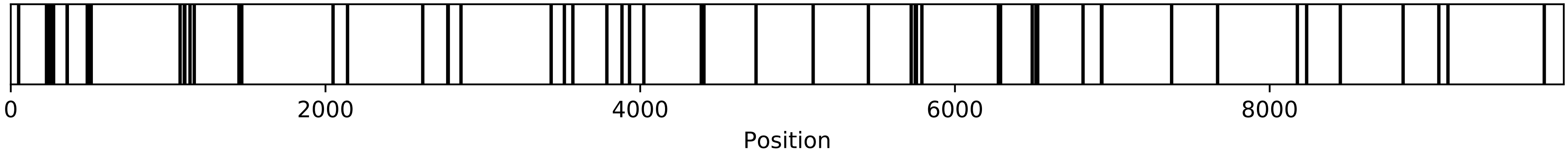


MM1 vs MM2
Number of differences: 69

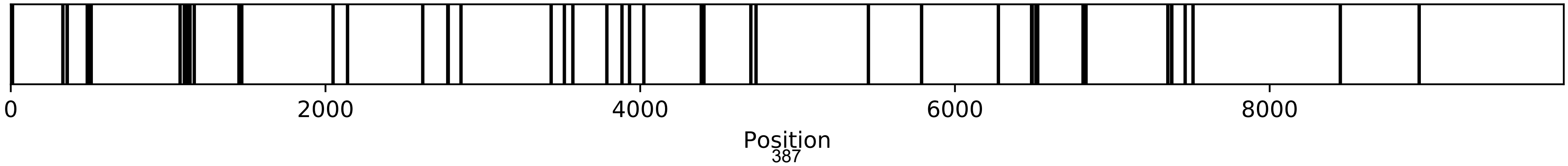

### Region: 1024



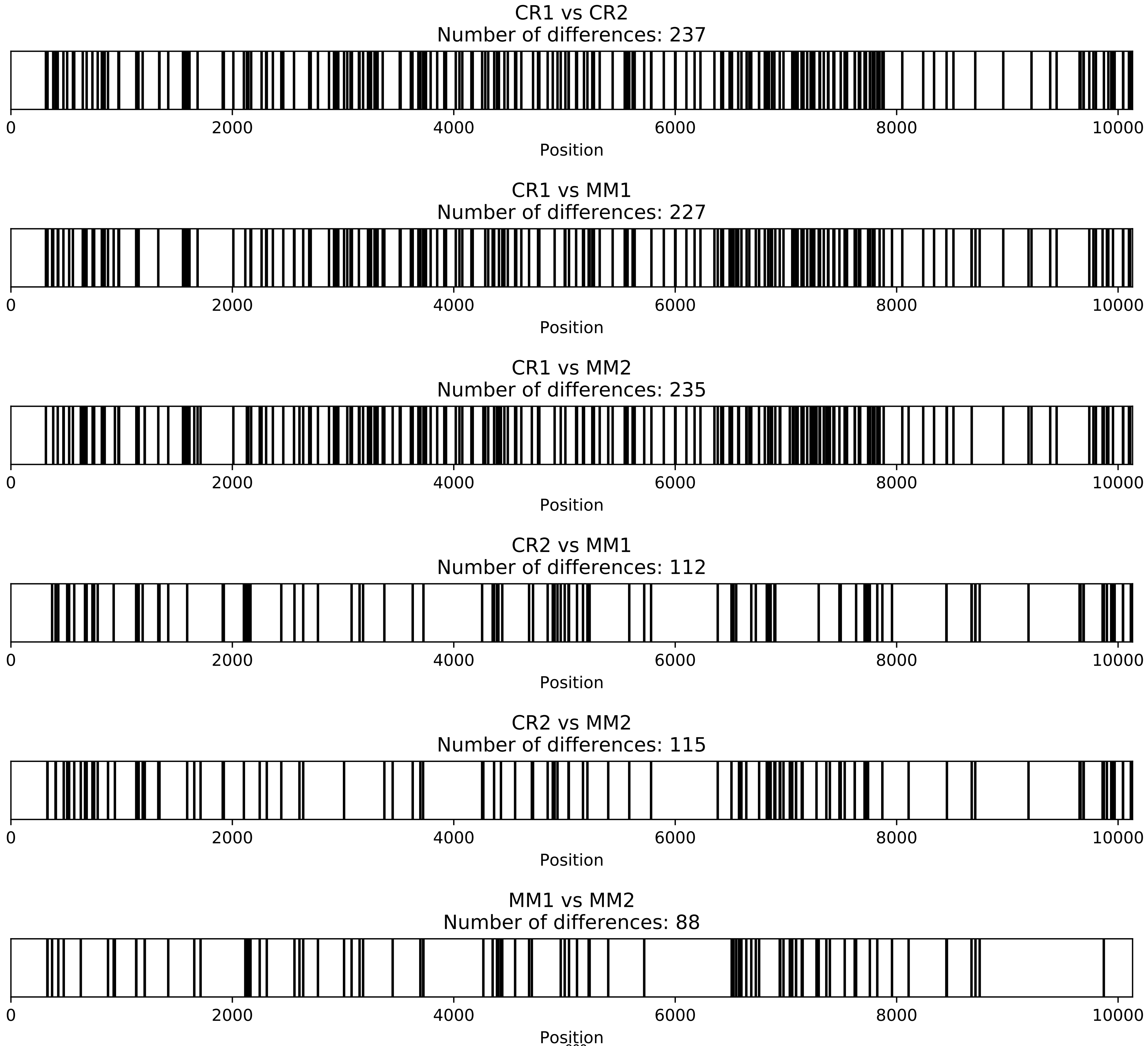
CR1 vs CR2
Number of differences: 237
CR1 vs MM1
Number of differences: 227
CR1 vs MM2
Number of differences: 235
CR2 vs MM1
Number of differences: 112
CR2 vs MM2
Number of differences: 115
MM1 vs MM2
Number of differences: 88
0
2000
4000
6000
8000
10000
Position

### Region: 1028



CR1 vs CR2
Number of differences: 50

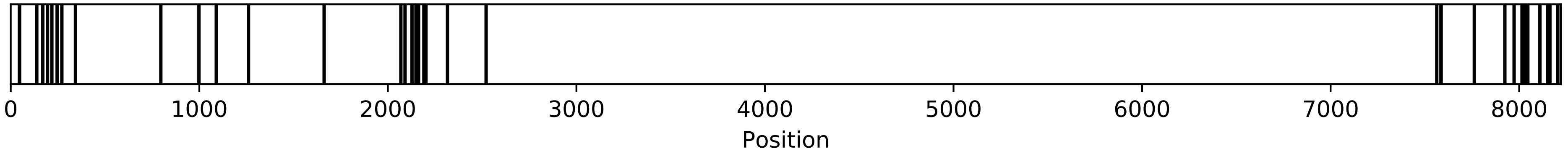


CR1 vs MM1
Number of differences: 97

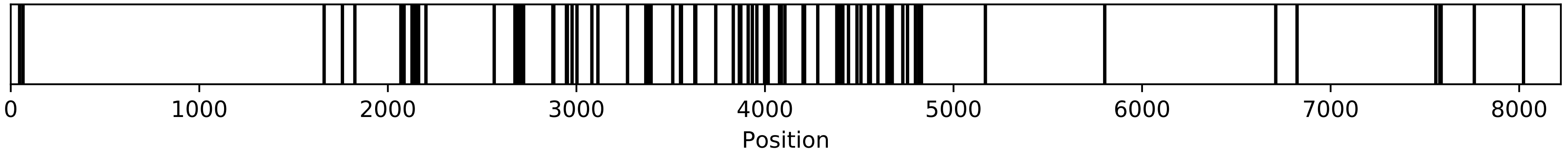


CR1 vs MM2
Number of differences: 92

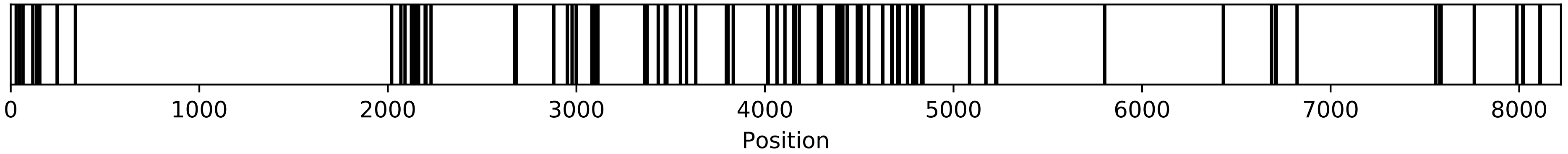


CR2 vs MM1
Number of differences: 128

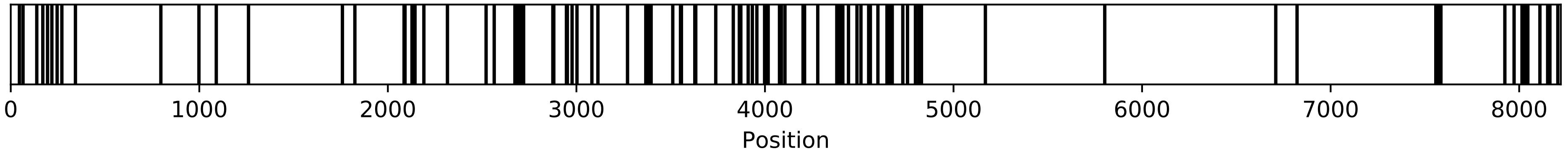


CR2 vs MM2
Number of differences: 114

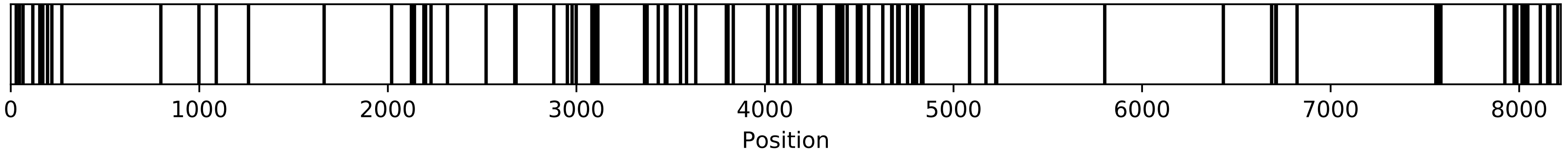


MM1 vs MM2
Number of differences: 108

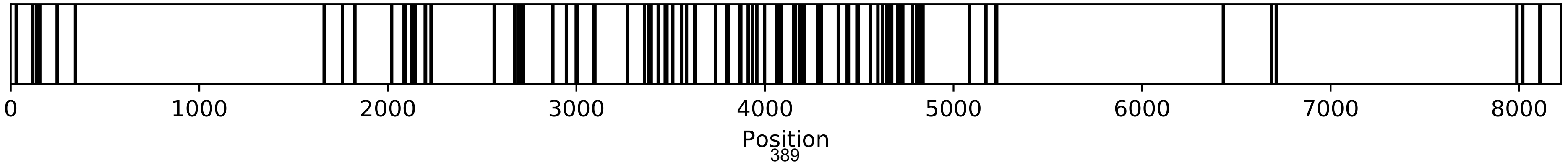

### Region: 1029



CR1 vs CR2
Number of differences: 231

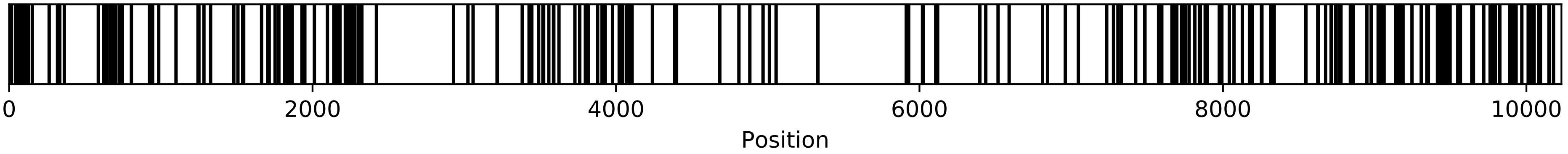


CR1 vs MM1
Number of differences: 113

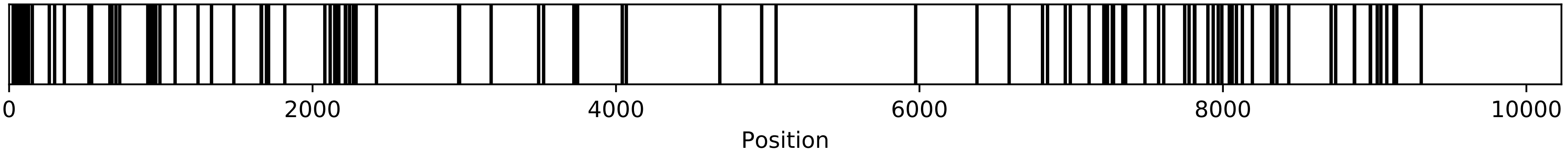


CR1 vs MM2
Number of differences: 127

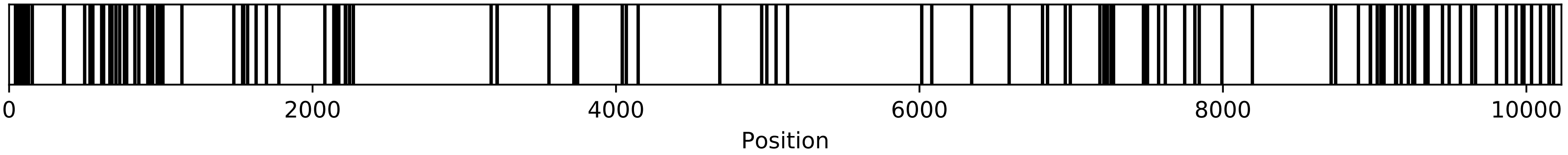


CR2 vs MM1
Number of differences: 234

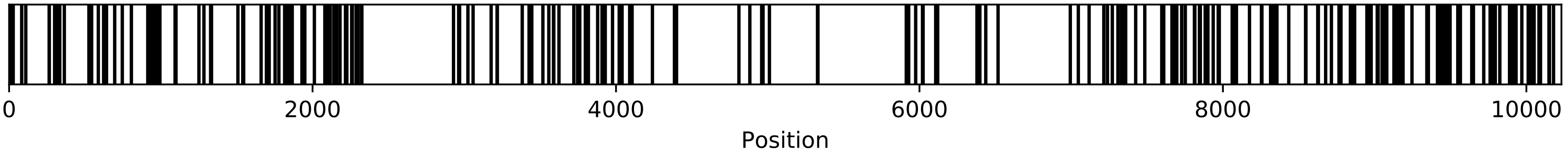


CR2 vs MM2
Number of differences: 235

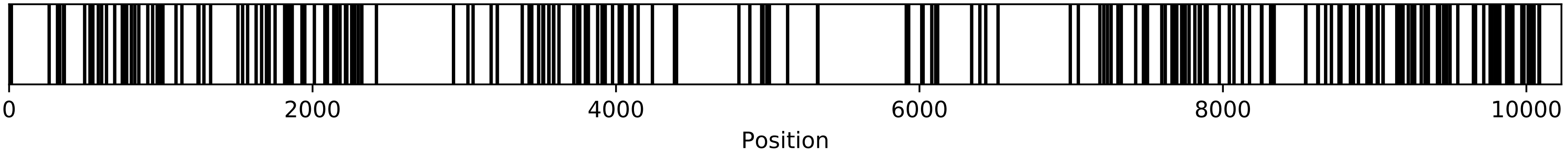


MM1 vs MM2
Number of differences: 127

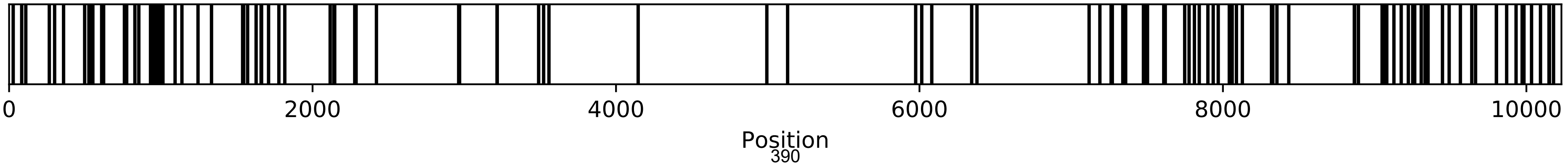

### Region: 1030



CR1 vs CR2
Number of differences: 224

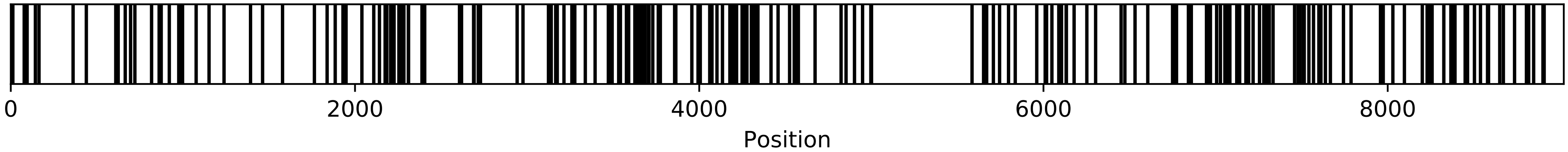


CR1 vs MM1
Number of differences: 54

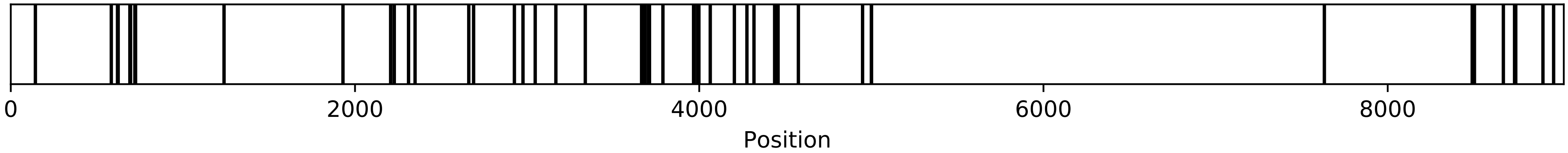


CR1 vs MM2
Number of differences: 101

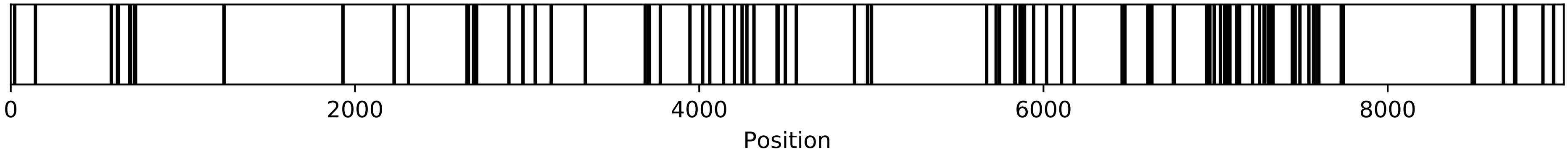


CR2 vs MM1
Number of differences: 218

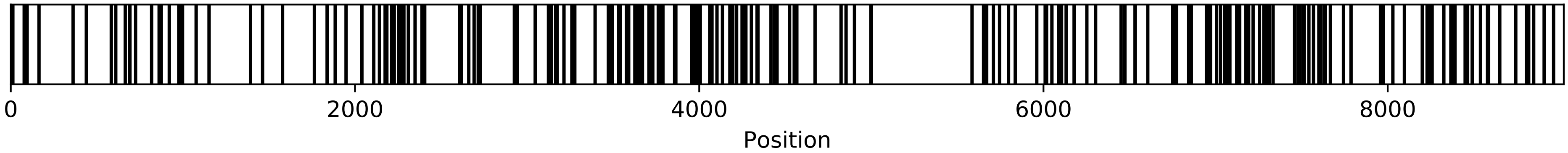


CR2 vs MM2
Number of differences: 207

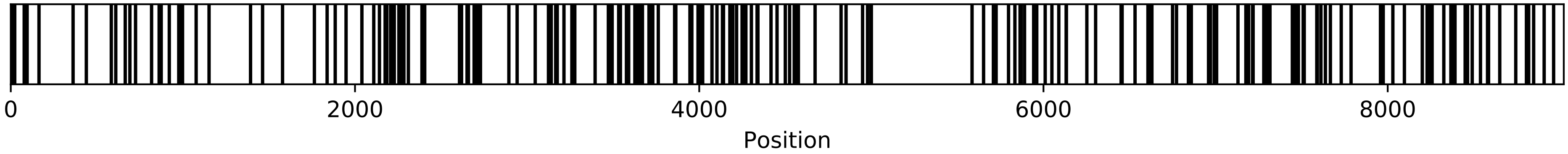


MM1 vs MM2
Number of differences: 85

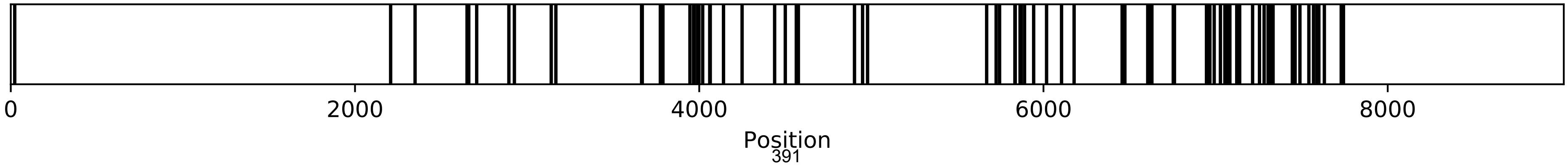

### Region: 1031



CR1 vs CR2
Number of differences: 473

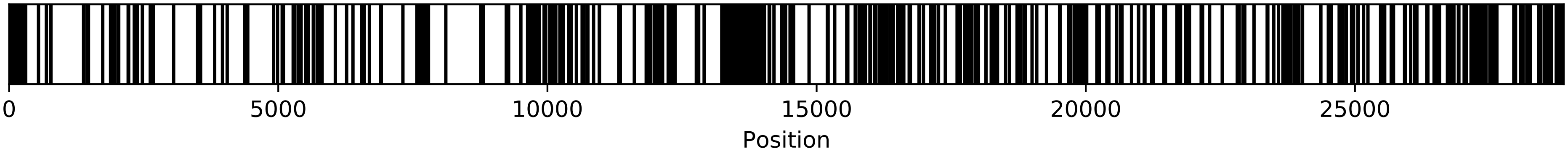


CR1 vs MM1
Number of differences: 275

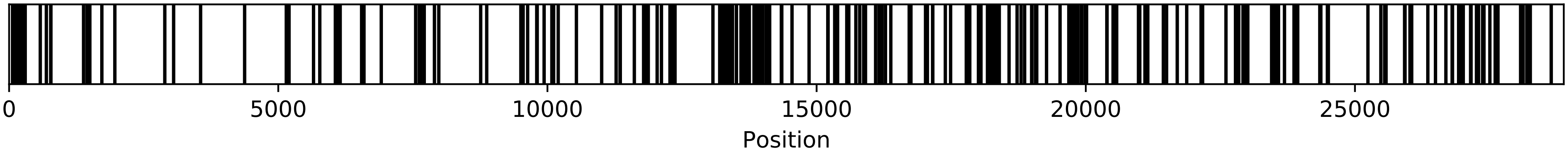


CR1 vs MM2
Number of differences: 281

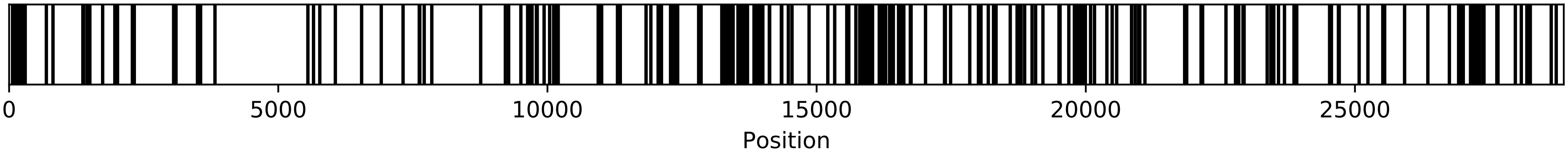


CR2 vs MM1
Number of differences: 462

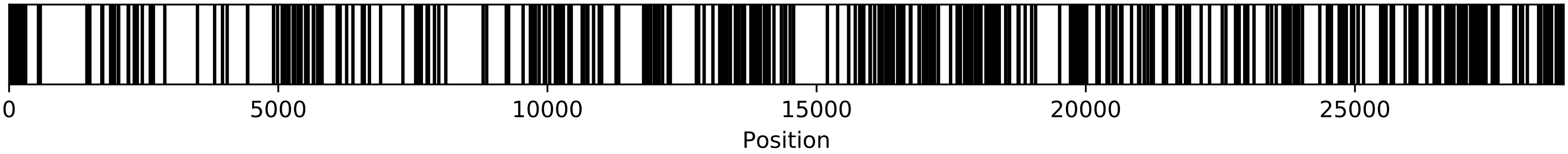


CR2 vs MM2
Number of differences: 452

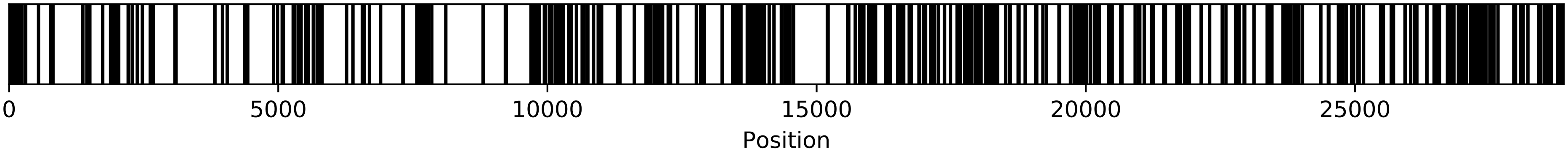


MM1 vs MM2
Number of differences: 253

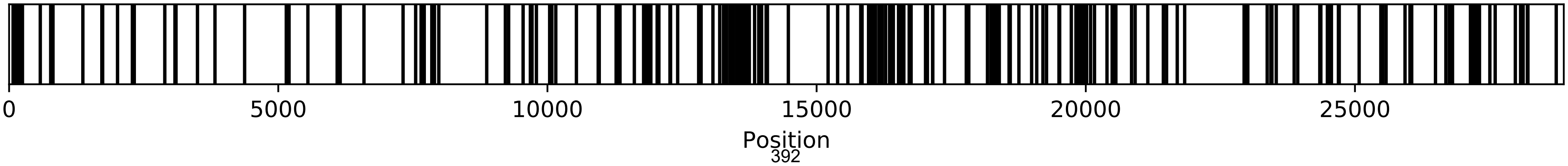

### Region: 1036



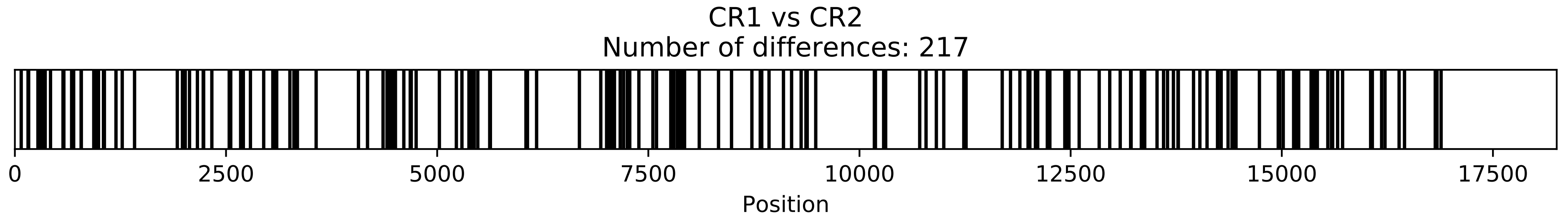
CR1 vs CR2
Number of differences: 217
0
2500
5000
7500
10000
12500
15000
17500
Position

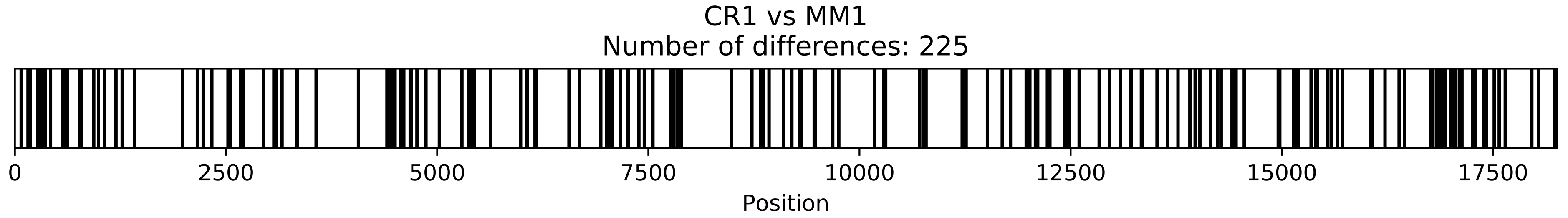
CR1 vs MM1
Number of differences: 225
0
2500
5000
7500
10000
12500
15000
17500
Position

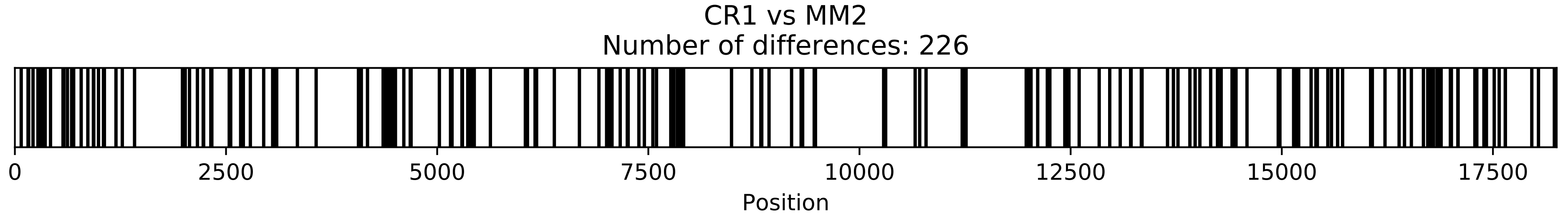
CR1 vs MM2
Number of differences: 226
0
2500
5000
7500
10000
12500
15000
17500
Position

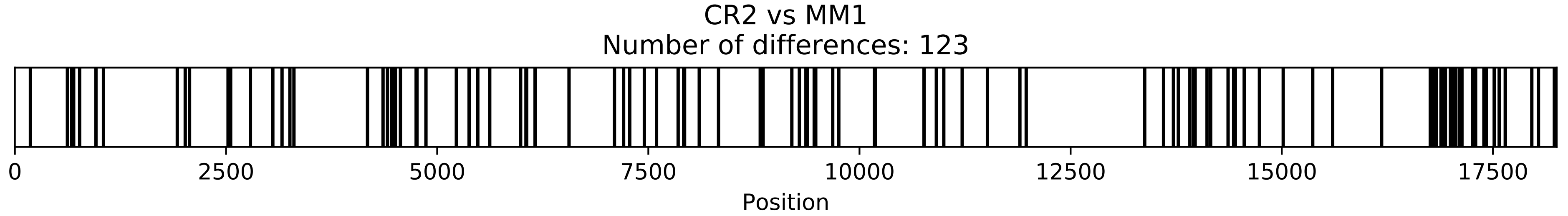
CR2 vs MM1
Number of differences: 123
0
2500
5000
7500
10000
12500
15000
17500
Position

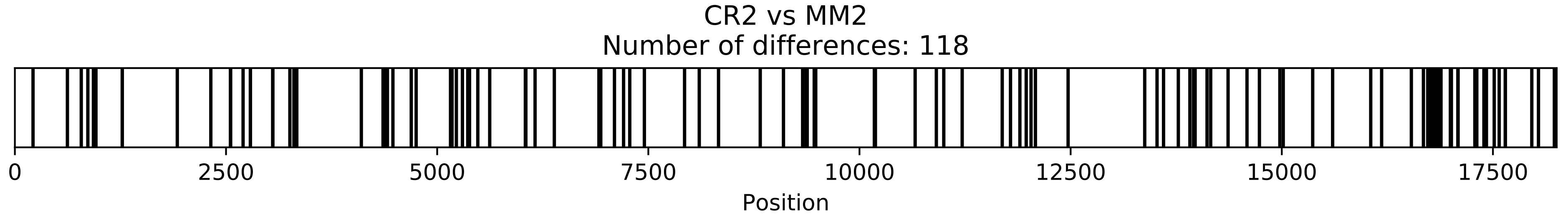
CR2 vs MM2
Number of differences: 118
0
2500
5000
7500
10000
12500
15000
17500
Position

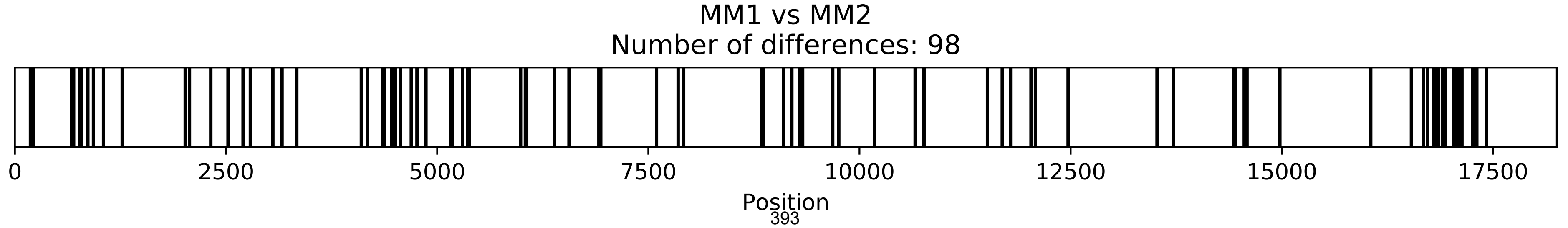
MM1 vs MM2
Number of differences: 98
0
2500
5000
7500
10000
12500
15000
17500
Position

### Region: 1037



## CR1 vs CR2
Number of differences: 540

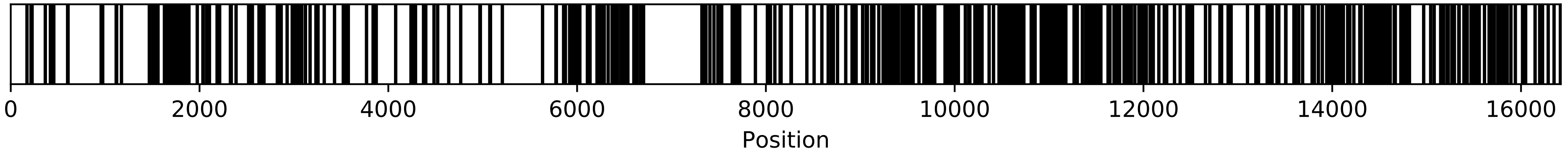


## CR1 vs MM1
Number of differences: 559

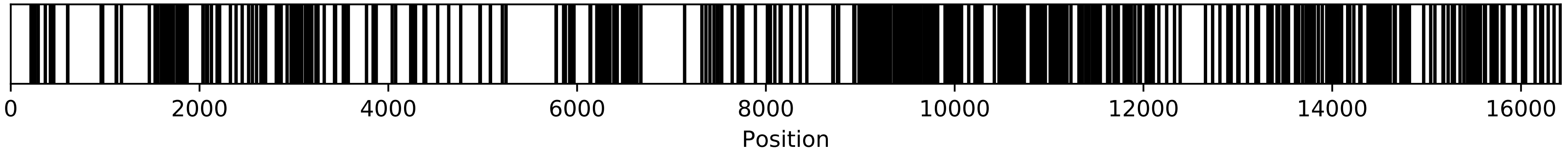


## CR1 vs MM2
Number of differences: 526

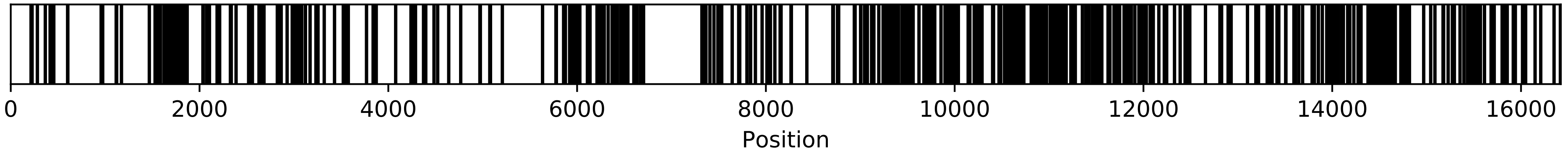


## CR2 vs MM1
Number of differences: 368

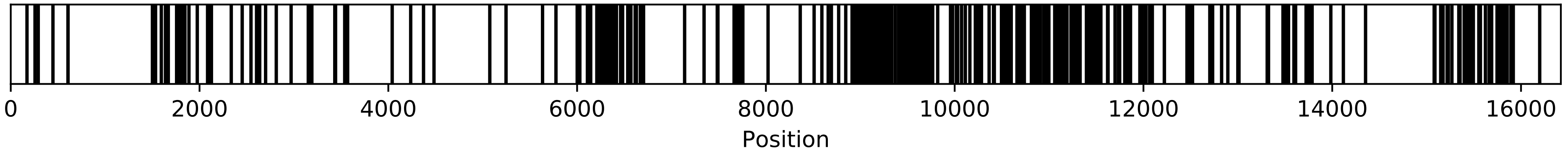


## CR2 vs MM2
Number of differences: 143

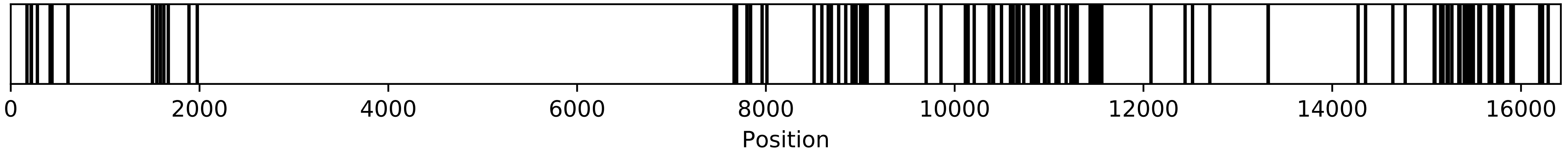


## MM1 vs MM2
Number of differences: 326

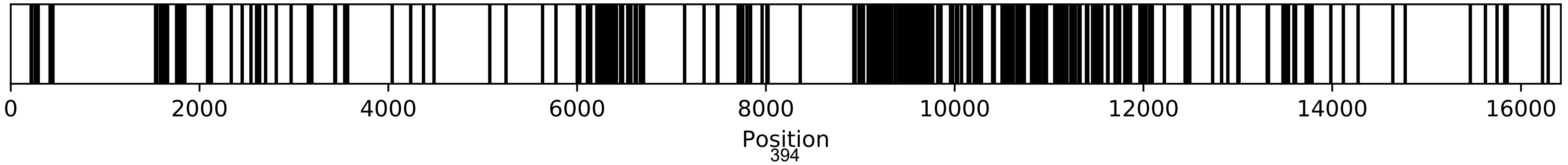

### Region: 1045



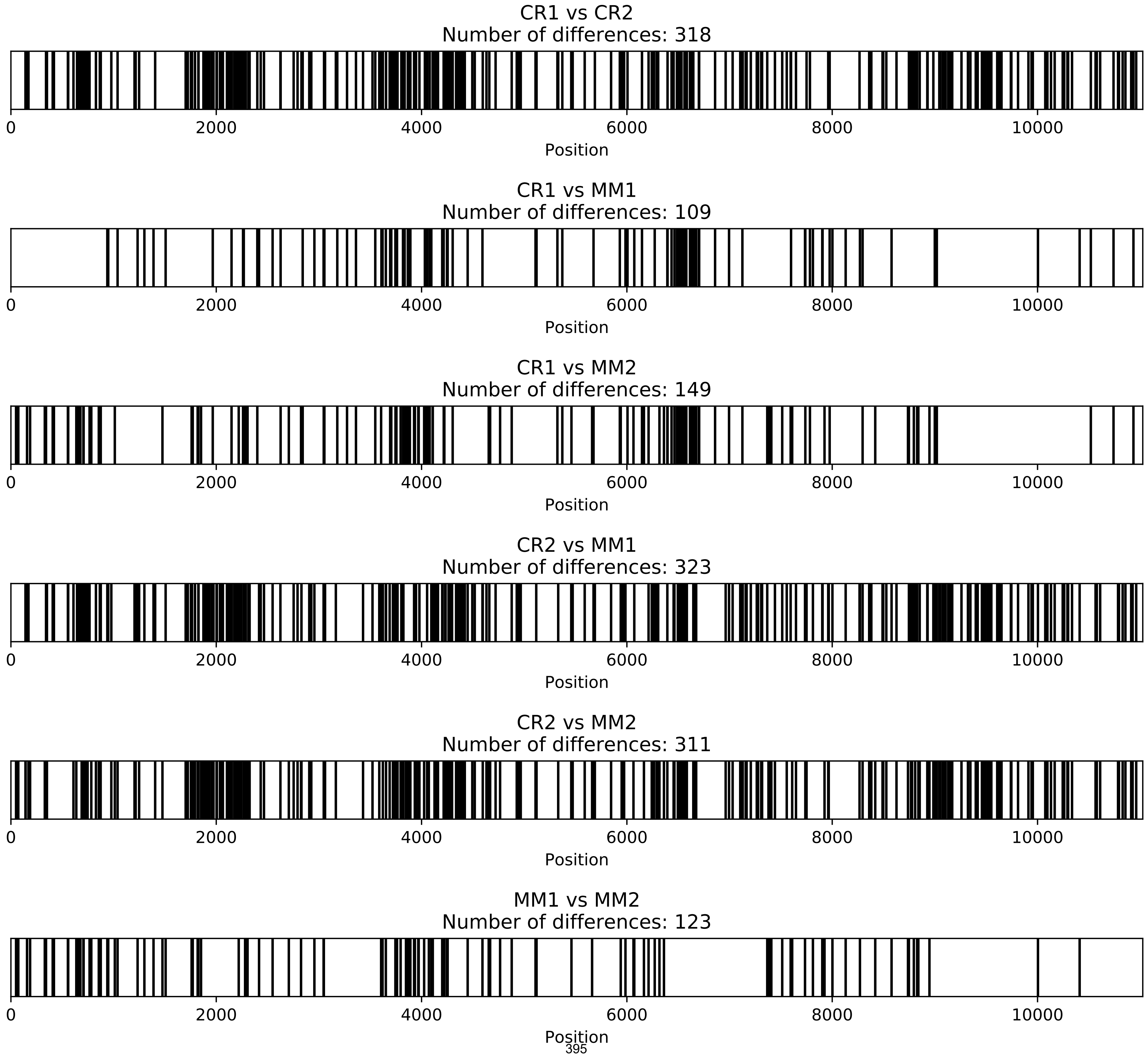
CR1 vs CR2
Number of differences: 318
CR1 vs MM1
Number of differences: 109
CR1 vs MM2
Number of differences: 149
CR2 vs MM1
Number of differences: 323
CR2 vs MM2
Number of differences: 311
MM1 vs MM2
Number of differences: 123
0
2000
4000
6000
8000
10000
Position

### Region: 1047



## CR1 vs CR2
Number of differences: 356

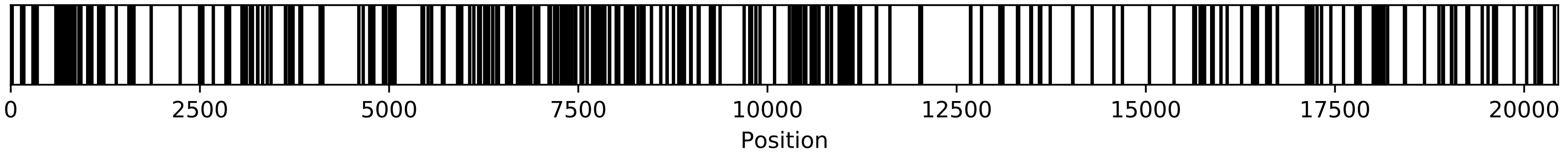


## CR1 vs MM1
Number of differences: 165

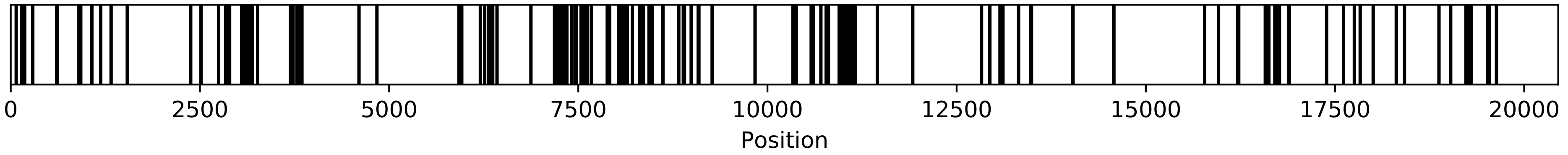


## CR1 vs MM2
Number of differences: 236

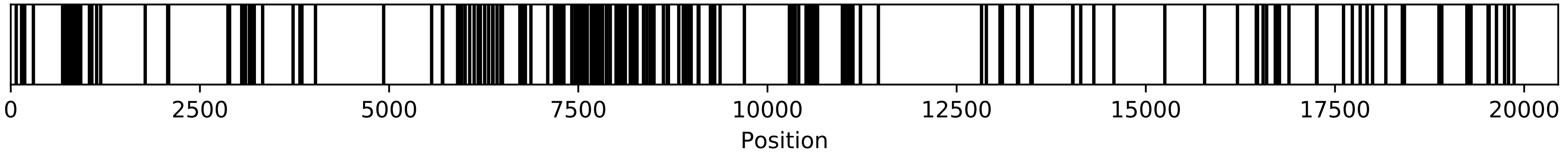


## CR2 vs MM1
Number of differences: 351

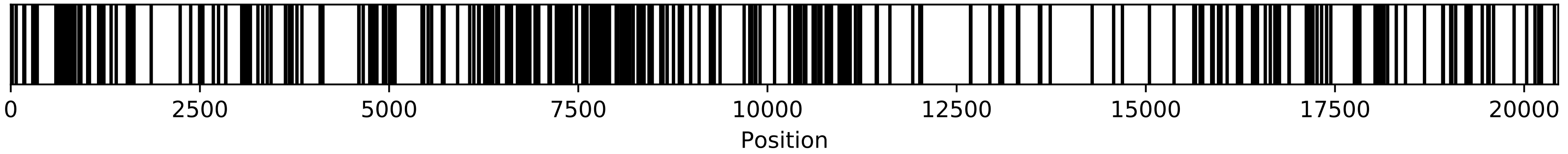


## CR2 vs MM2
Number of differences: 376

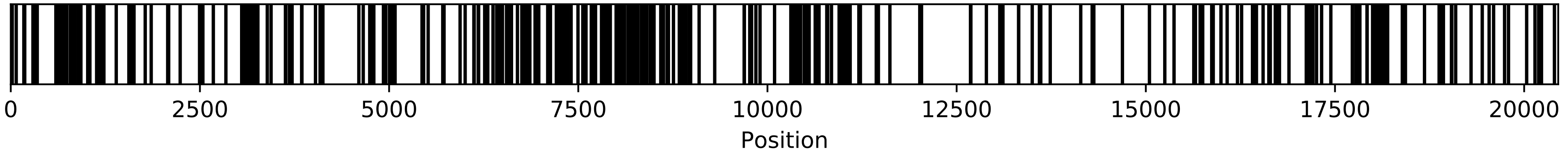


## MM1 vs MM2
Number of differences: 214

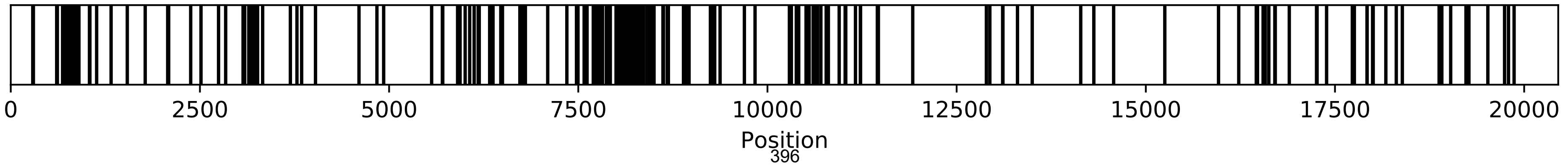

### Region: 1050



CR1 vs CR2
Number of differences: 192

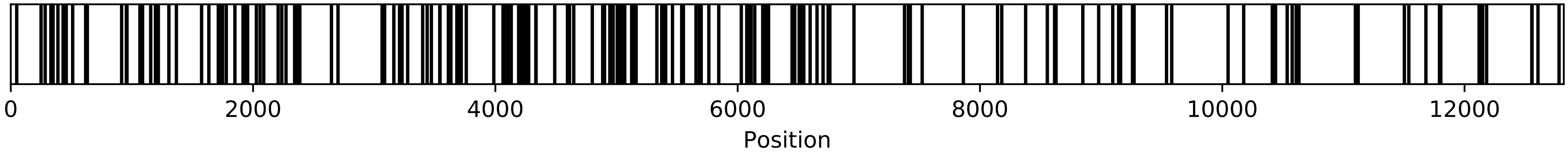


CR1 vs MM1
Number of differences: 88

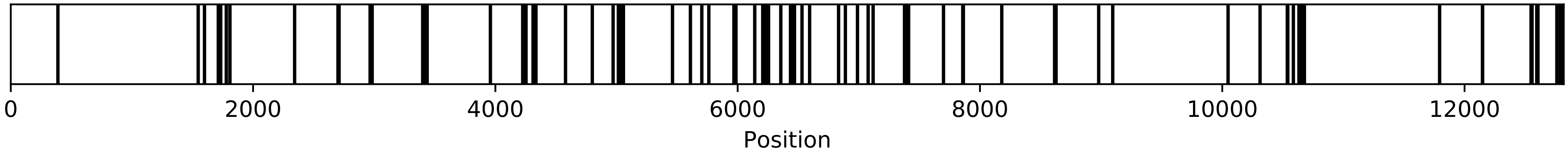


CR1 vs MM2
Number of differences: 91

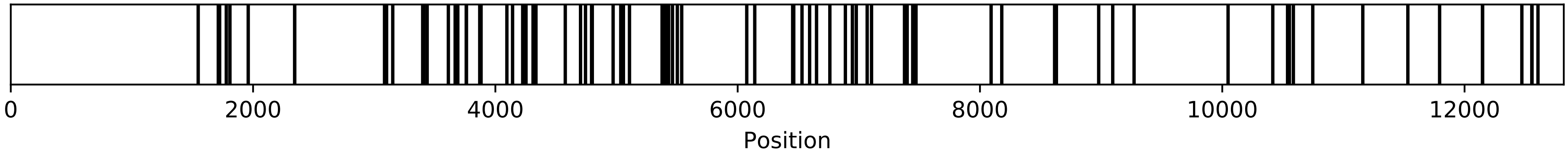


CR2 vs MM1
Number of differences: 210

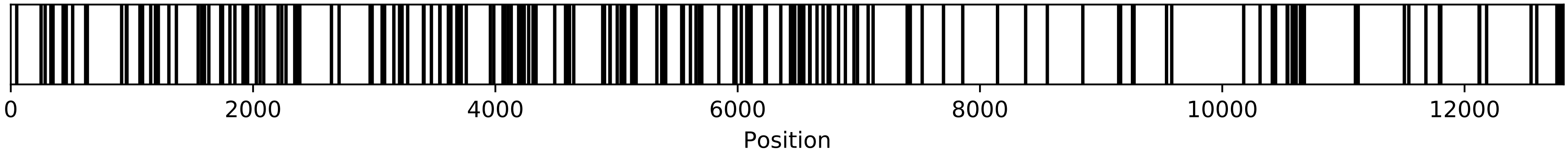


CR2 vs MM2
Number of differences: 206

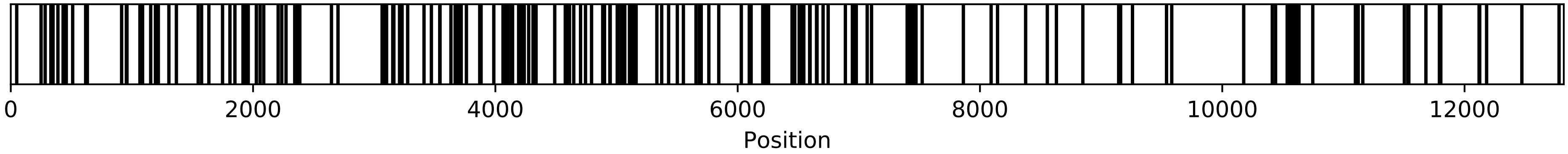


MM1 vs MM2
Number of differences: 93

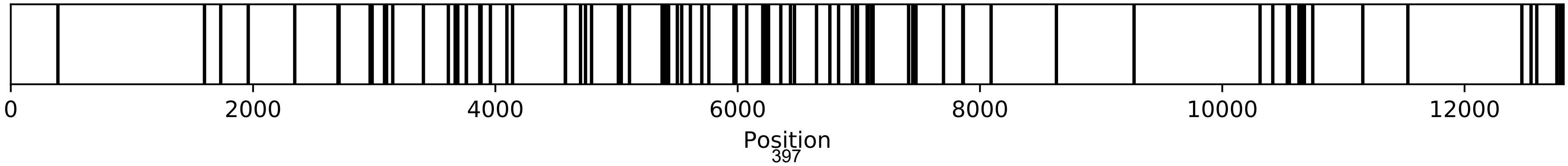

### Region: 1051



CR1 vs CR2
Number of differences: 294

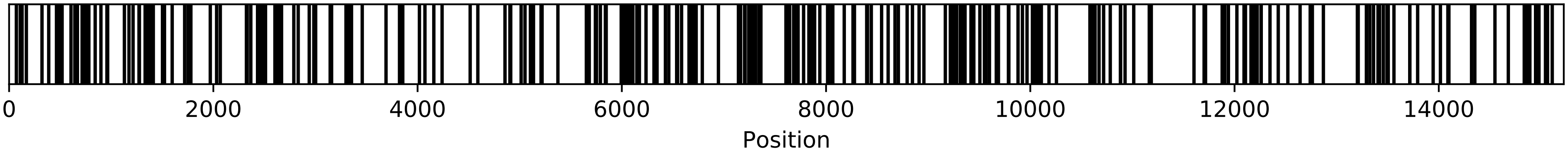


CR1 vs MM1
Number of differences: 135

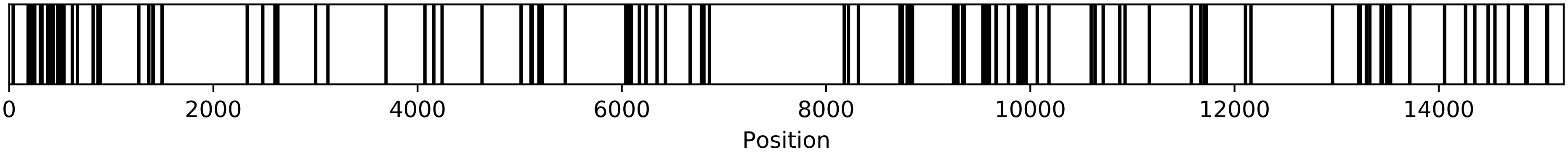


CR1 vs MM2
Number of differences: 151

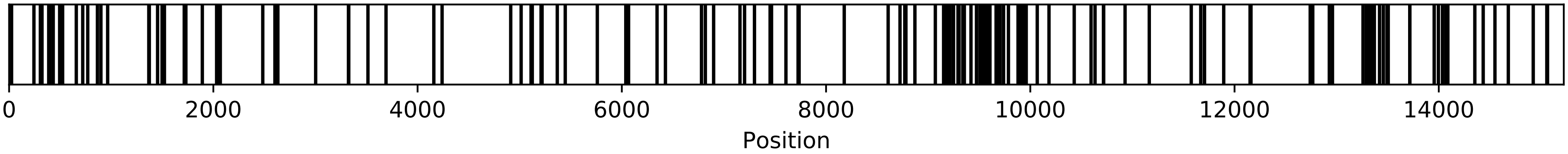


CR2 vs MM1
Number of differences: 308

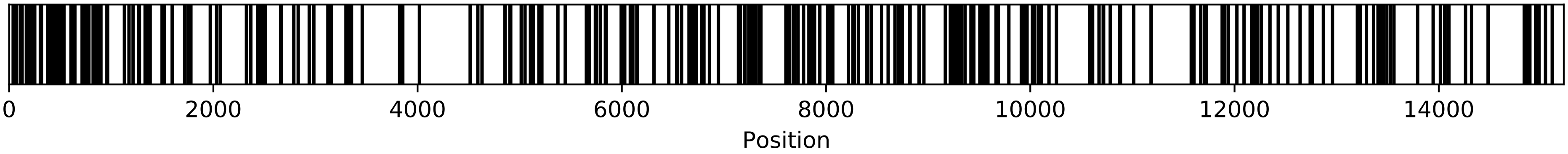


CR2 vs MM2
Number of differences: 301

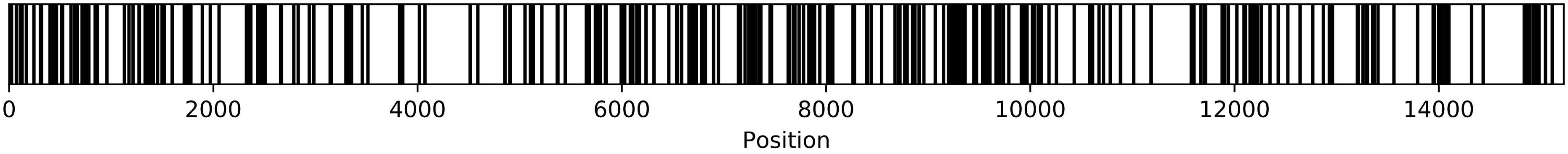


MM1 vs MM2
Number of differences: 156

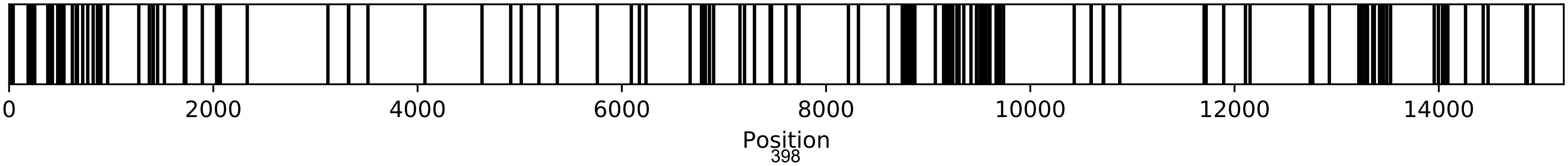

### Region: 1062



CR1 vs CR2
Number of differences: 331

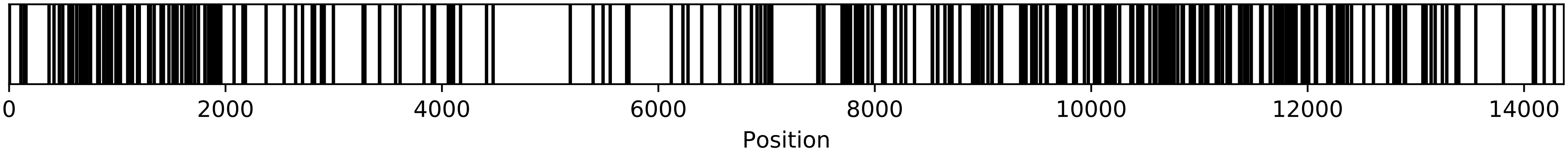


CR1 vs MM1
Number of differences: 206

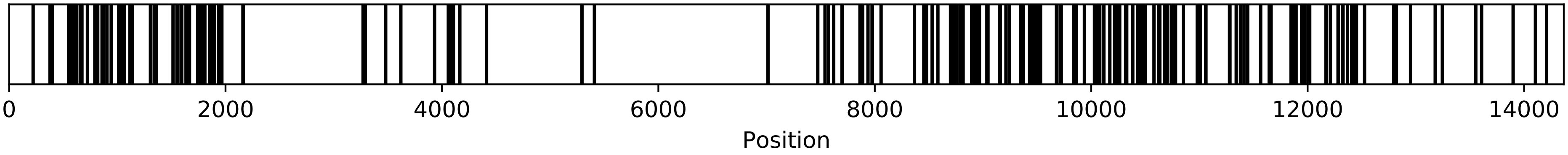


CR1 vs MM2
Number of differences: 195

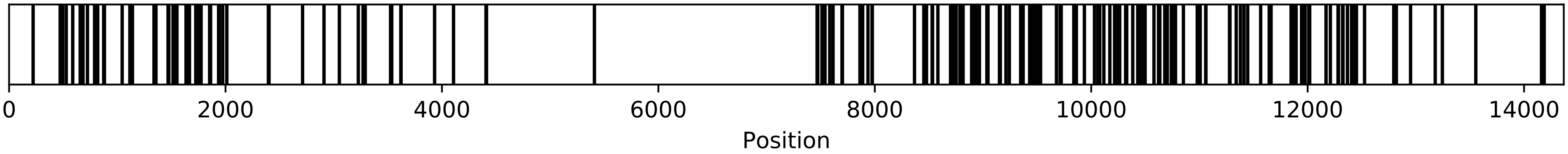


CR2 vs MM1
Number of differences: 334

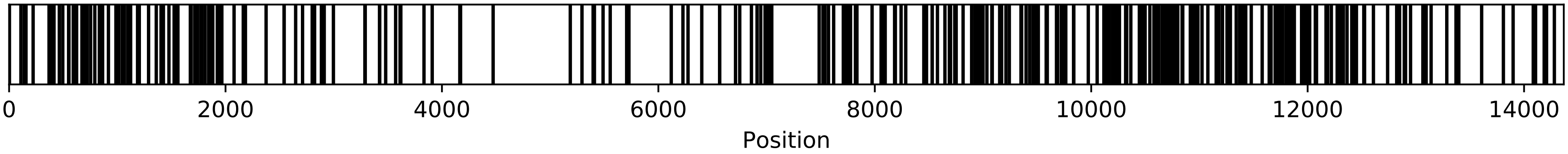


CR2 vs MM2
Number of differences: 345

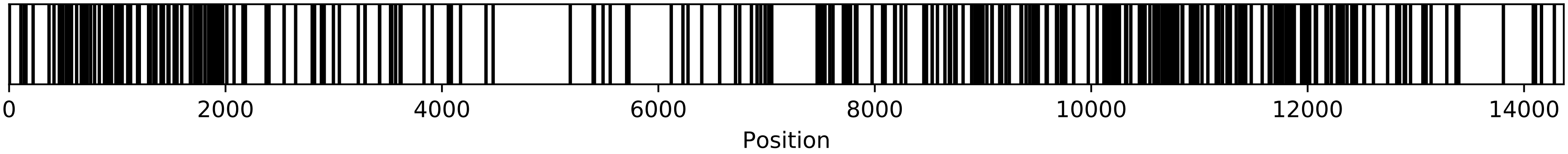


MM1 vs MM2
Number of differences: 81

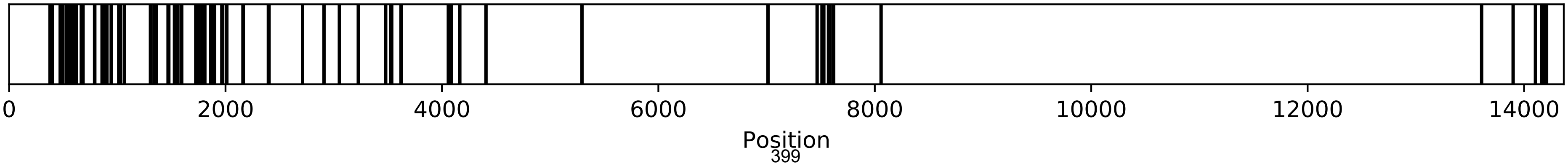

### Region: 1067



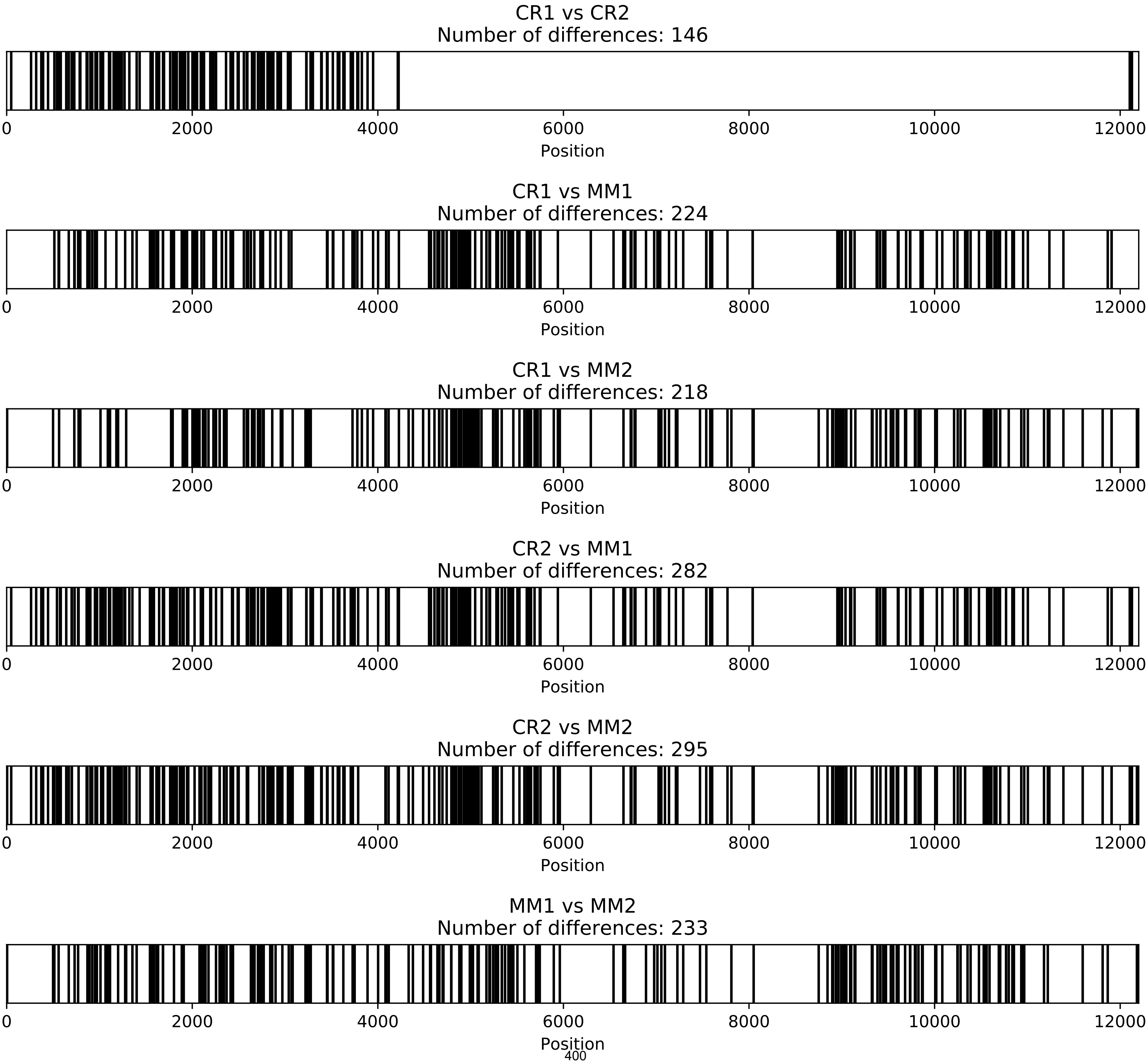
CR1 vs CR2
Number of differences: 146
CR1 vs MM1
Number of differences: 224
CR1 vs MM2
Number of differences: 218
CR2 vs MM1
Number of differences: 282
CR2 vs MM2
Number of differences: 295
MM1 vs MM2
Number of differences: 233
0
2000
4000
6000
8000
10000
12000
Position

### Region: 1068

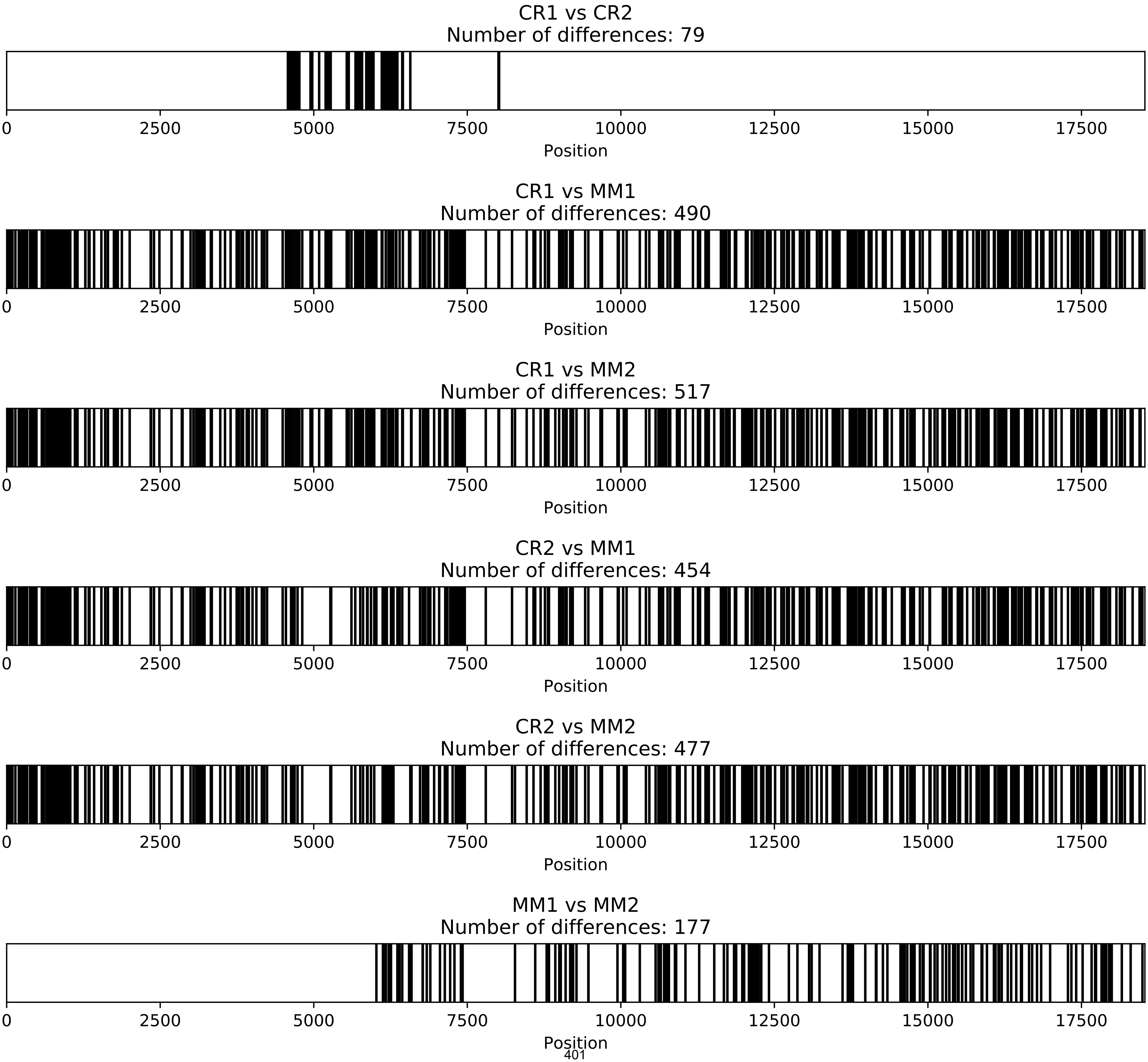
CR1 vs CR2
Number of differences: 79
CR1 vs MM1
Number of differences: 490
CR1 vs MM2
Number of differences: 517
CR2 vs MM1
Number of differences: 454
CR2 vs MM2
Number of differences: 477
MM1 vs MM2
Number of differences: 177
0
2500
5000
7500
10000
12500
15000
17500
Position

### Region: 1070



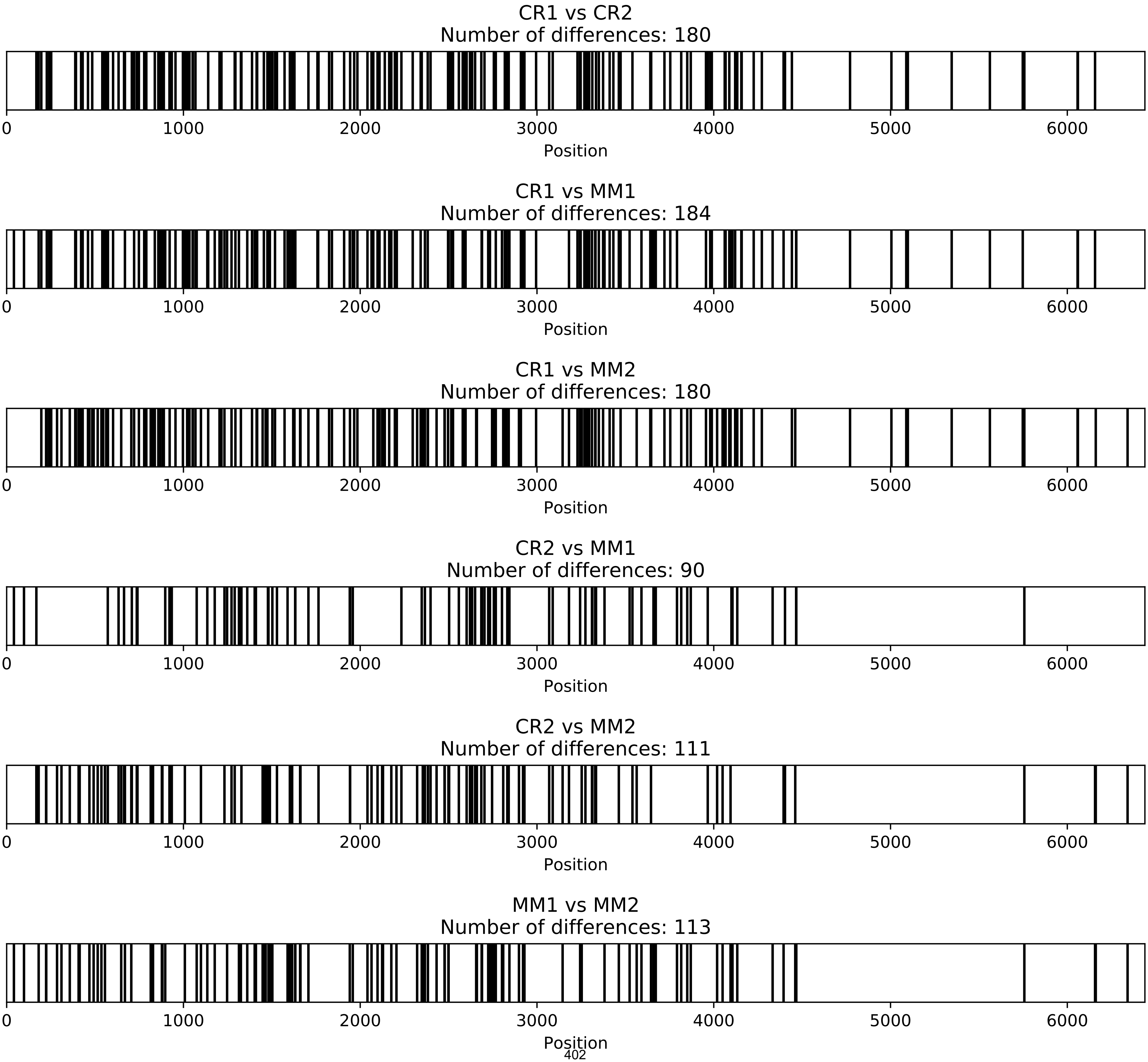
CR1 vs CR2
Number of differences: 180
CR1 vs MM1
Number of differences: 184
CR1 vs MM2
Number of differences: 180
CR2 vs MM1
Number of differences: 90
CR2 vs MM2
Number of differences: 111
MM1 vs MM2
Number of differences: 113
0
1000
2000
3000
4000
5000
6000
Position

### Region: 1075



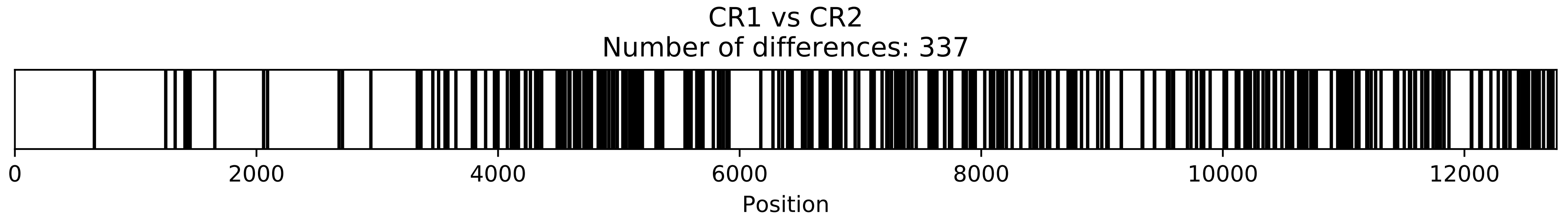
CR1 vs CR2
Number of differences: 337
0
2000
4000
6000
8000
10000
12000
Position

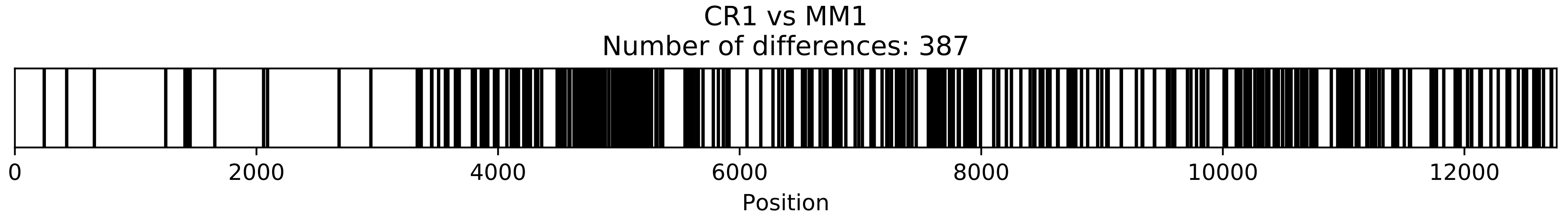
CR1 vs MM1
Number of differences: 387
0
2000
4000
6000
8000
10000
12000
Position

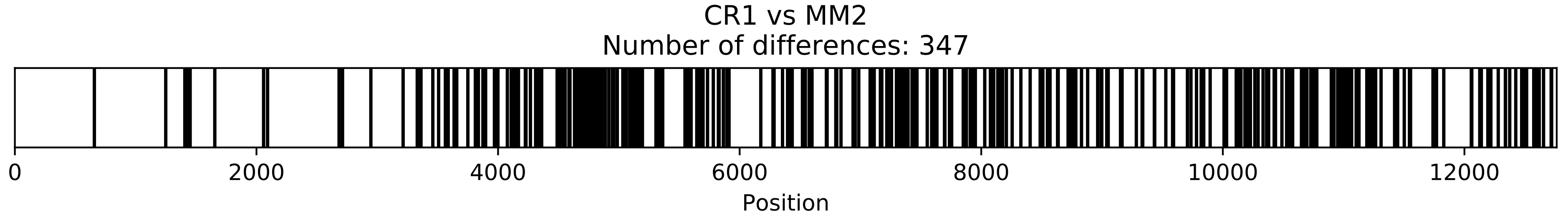
CR1 vs MM2
Number of differences: 347
0
2000
4000
6000
8000
10000
12000
Position

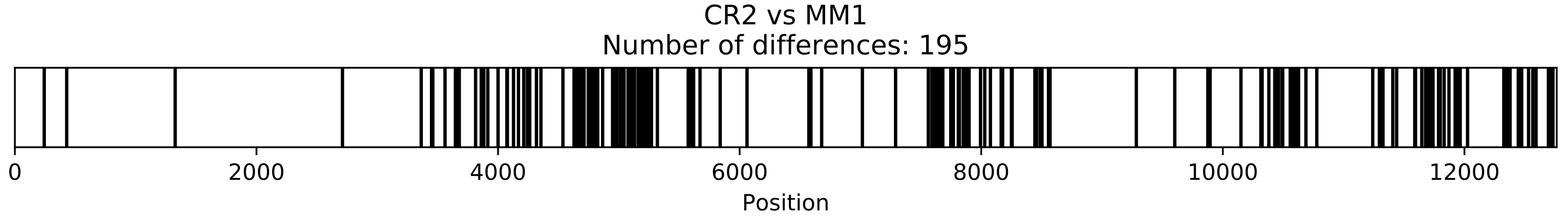
CR2 vs MM1
Number of differences: 195
0
2000
4000
6000
8000
10000
12000
Position

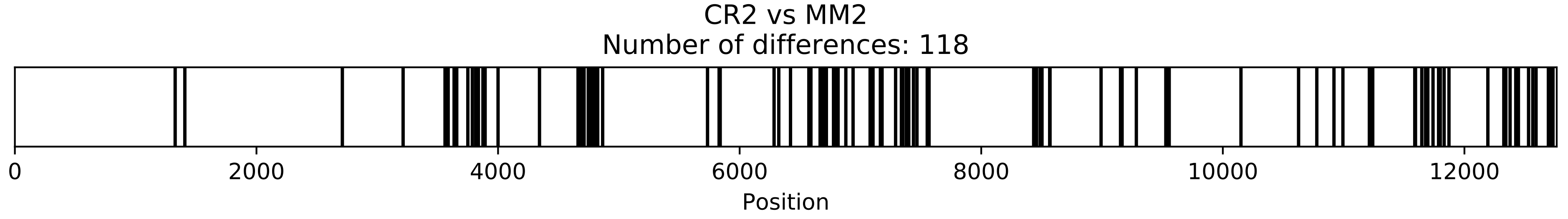
CR2 vs MM2
Number of differences: 118
0
2000
4000
6000
8000
10000
12000
Position

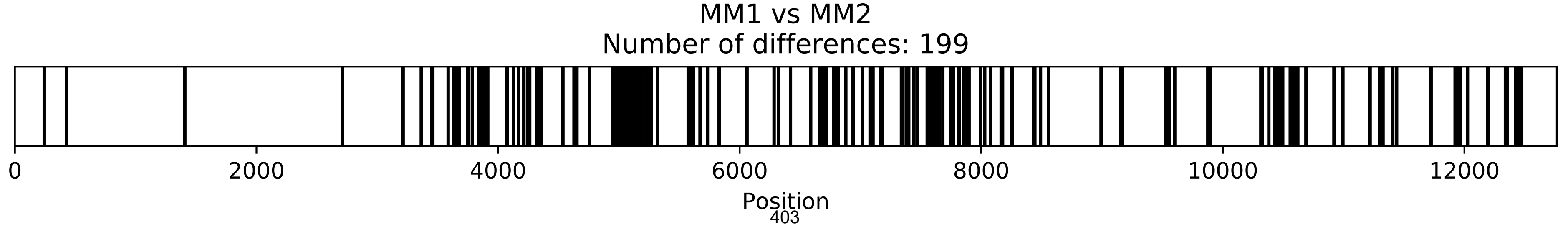
MM1 vs MM2
Number of differences: 199
0
2000
4000
6000
8000
10000
12000
Position

### Region: 1077



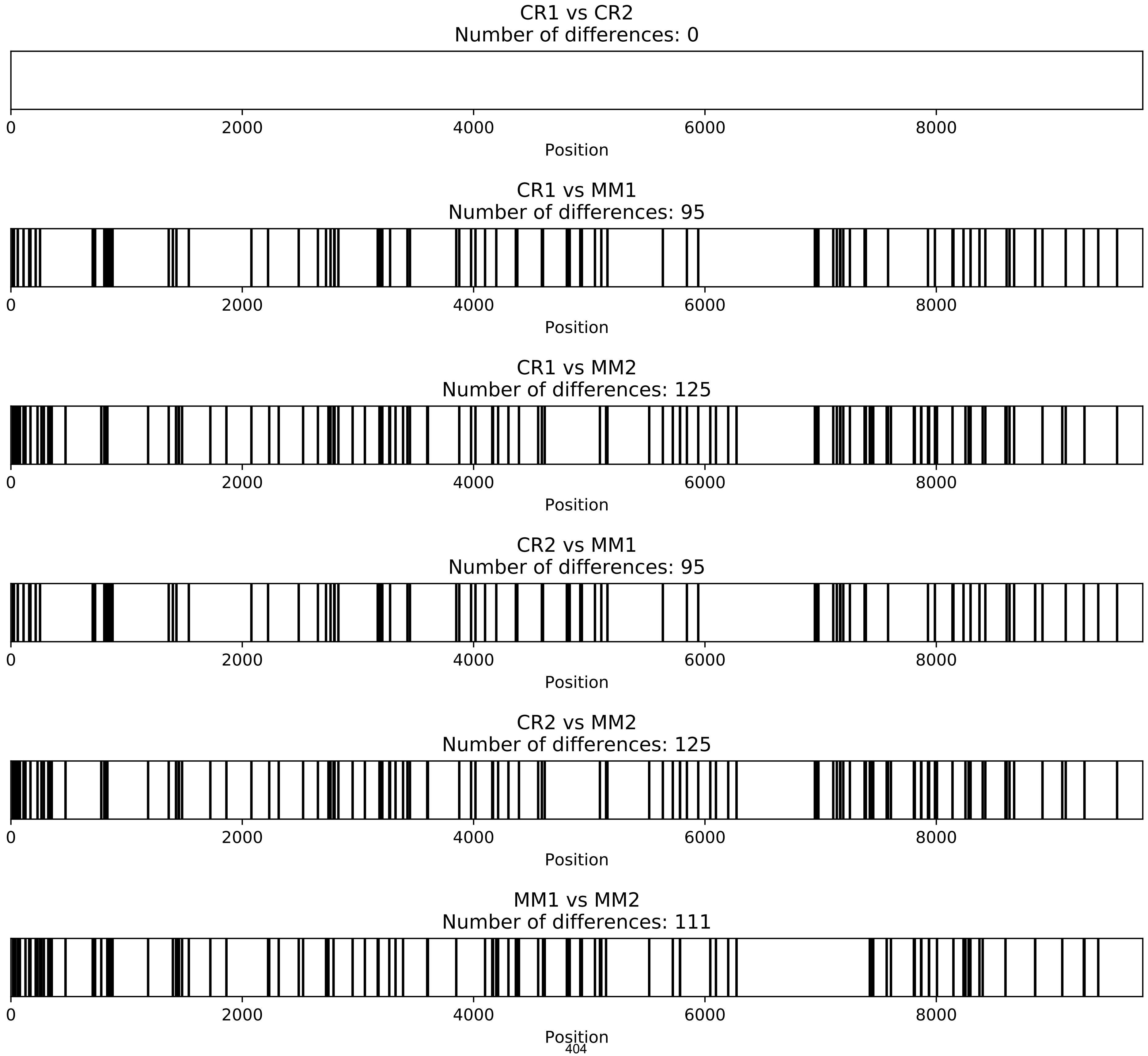
CR1 vs CR2
Number of differences: 0
0
2000
4000
6000
8000
Position
CR1 vs MM1
Number of differences: 95
0
2000
4000
6000
8000
Position
CR1 vs MM2
Number of differences: 125
0
2000
4000
6000
8000
Position
CR2 vs MM1
Number of differences: 95
0
2000
4000
6000
8000
Position
CR2 vs MM2
Number of differences: 125
0
2000
4000
6000
8000
Position
MM1 vs MM2
Number of differences: 111
0
2000
4000
6000
8000
Position

### Region: 1078



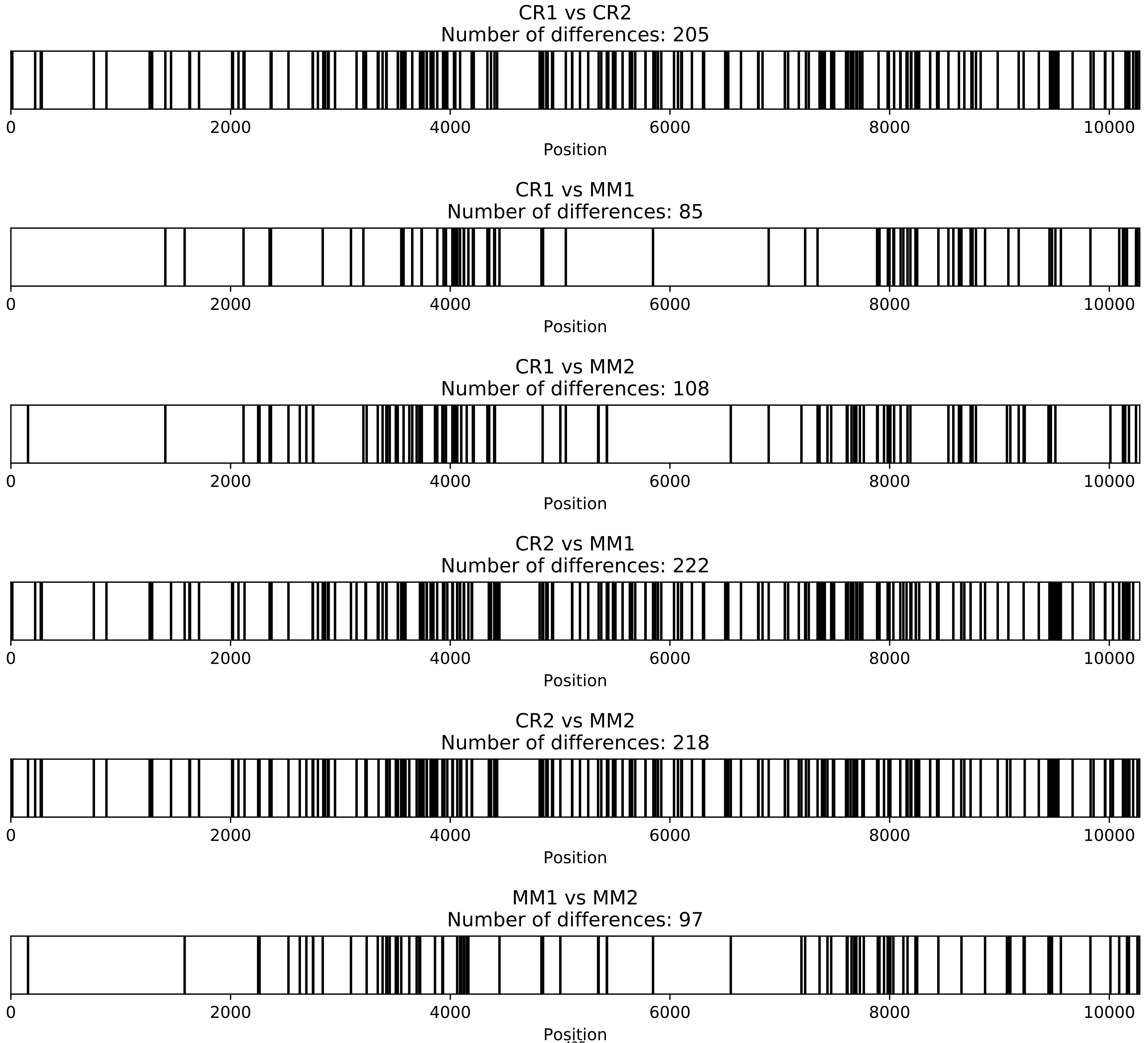
CR1 vs CR2
Number of differences: 205
Position
CR1 vs MM1
Number of differences: 85
Position
CR1 vs MM2
Number of differences: 108
Position
CR2 vs MM1
Number of differences: 222
Position
CR2 vs MM2
Number of differences: 218
Position
MM1 vs MM2
Number of differences: 97
Position
0
2000
4000
6000
8000
10000

### Region: 1080



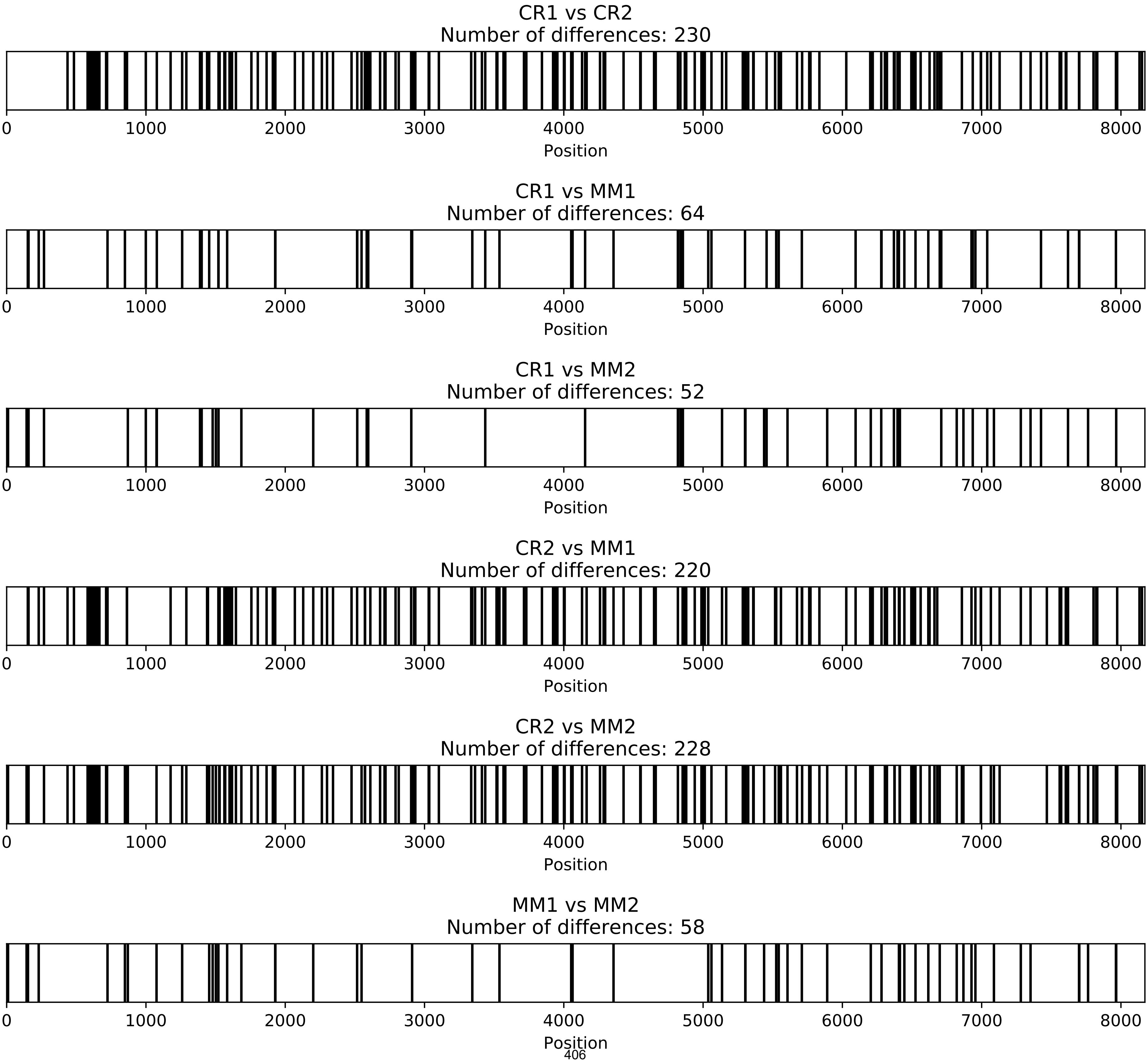
CR1 vs CR2
Number of differences: 230
0
1000
2000
3000
4000
5000
6000
7000
8000
Position
CR1 vs MM1
Number of differences: 64
Position
CR1 vs MM2
Number of differences: 52
Position
CR2 vs MM1
Number of differences: 220
Position
CR2 vs MM2
Number of differences: 228
Position
MM1 vs MM2
Number of differences: 58
Position

### Region: 1083



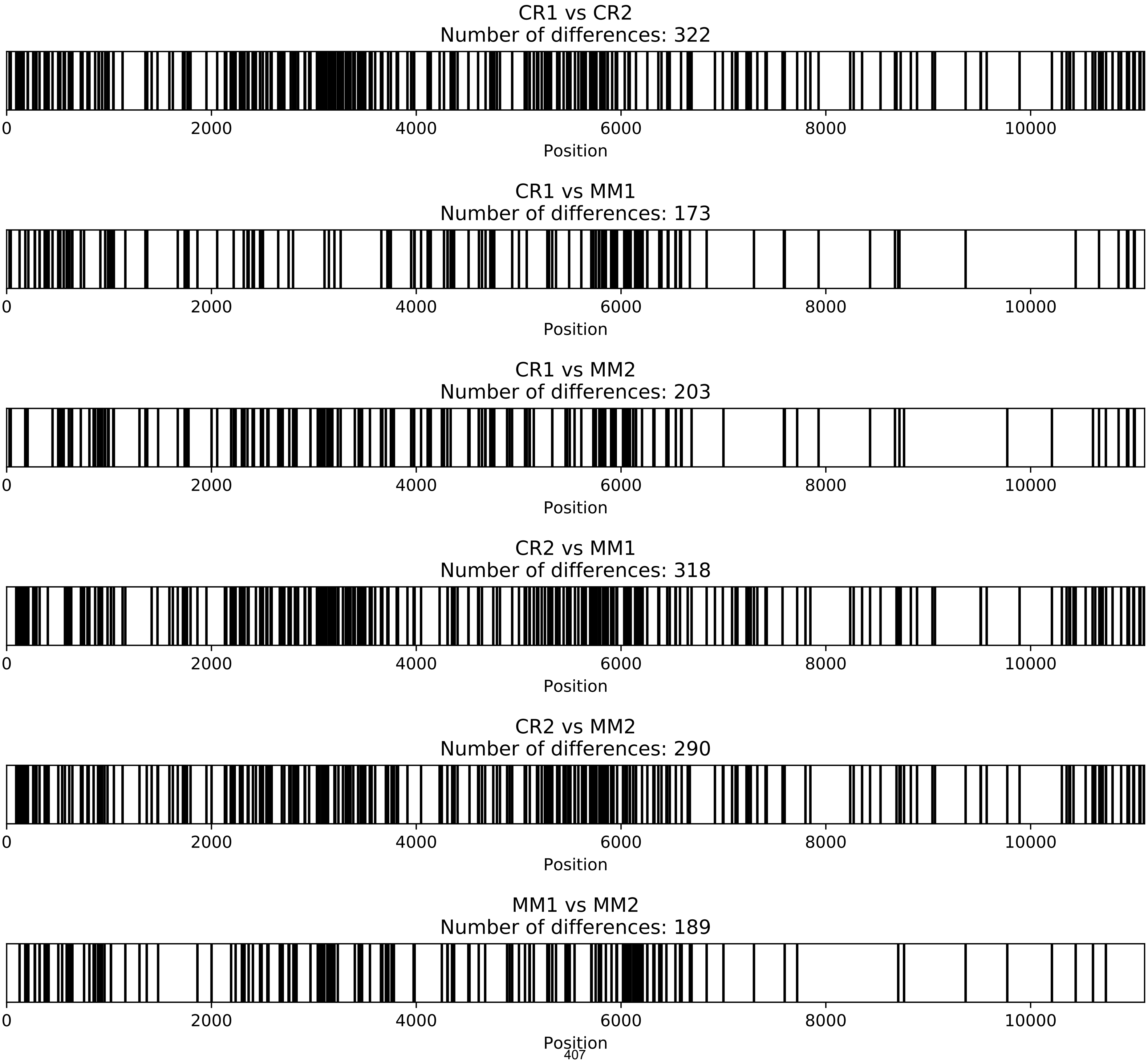
CR1 vs CR2
Number of differences: 322
CR1 vs MM1
Number of differences: 173
CR1 vs MM2
Number of differences: 203
CR2 vs MM1
Number of differences: 318
CR2 vs MM2
Number of differences: 290
MM1 vs MM2
Number of differences: 189
0
2000
4000
6000
8000
10000
Position

### Region: 1085



CR1 vs CR2
Number of differences: 0

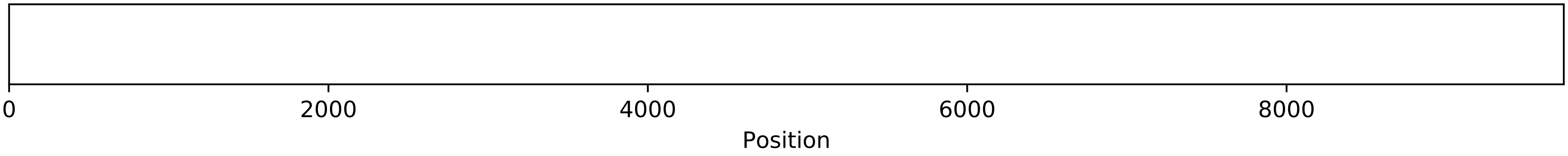


CR1 vs MM1
Number of differences: 91

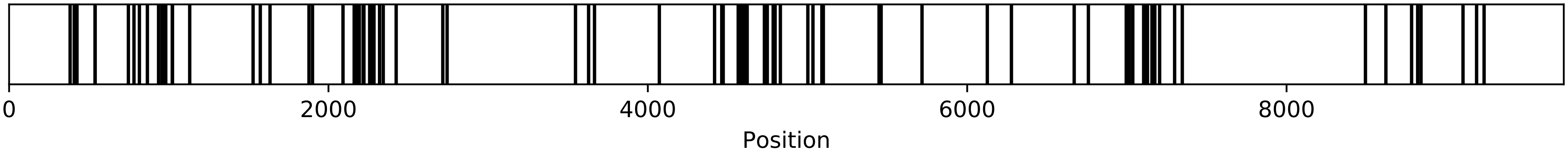


CR1 vs MM2
Number of differences: 136

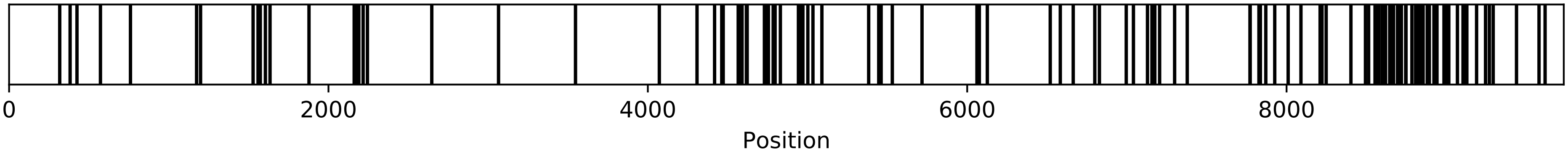


CR2 vs MM1
Number of differences: 91

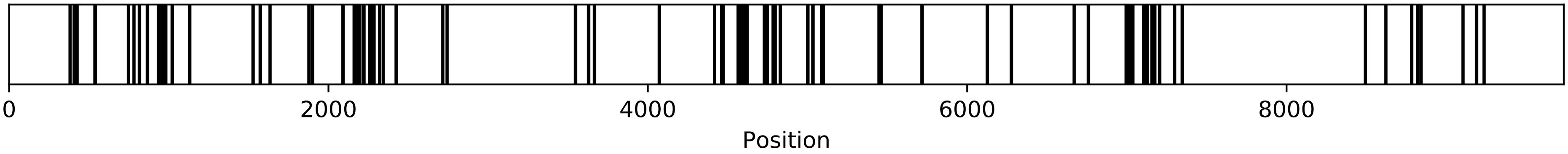


CR2 vs MM2
Number of differences: 136

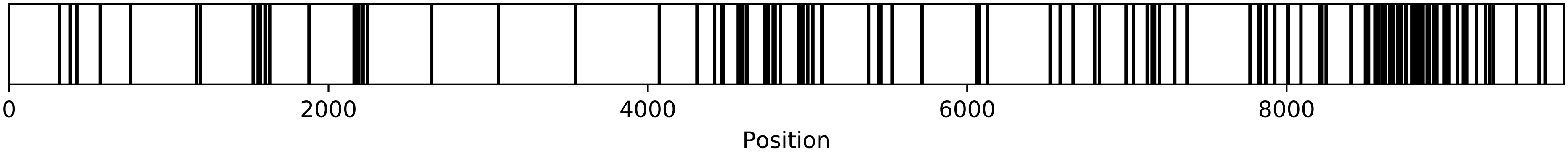


MM1 vs MM2
Number of differences: 129

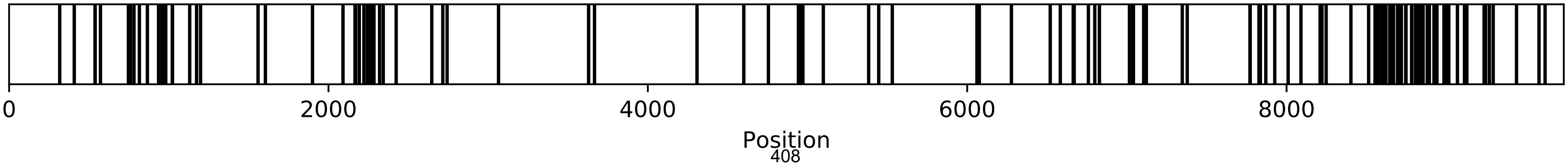

### Region: 1086



CR1 vs CR2
Number of differences: 219

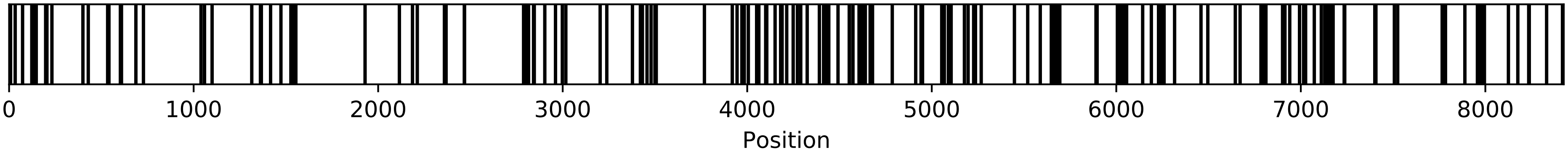


CR1 vs MM1
Number of differences: 93

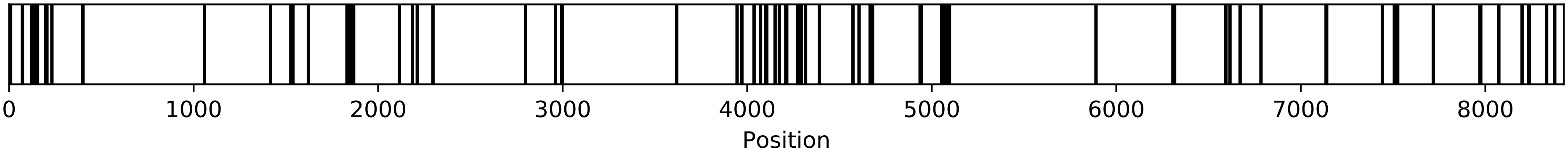


CR1 vs MM2
Number of differences: 76

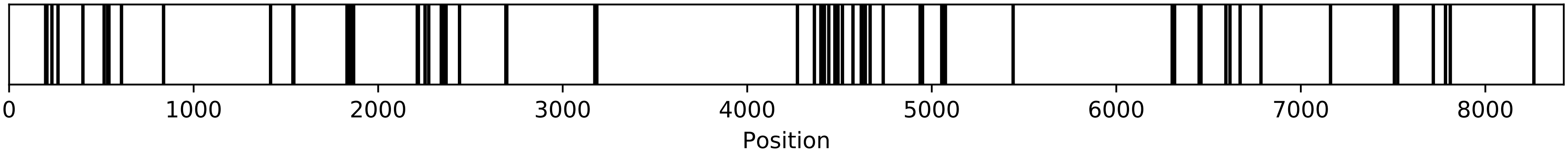


CR2 vs MM1
Number of differences: 210

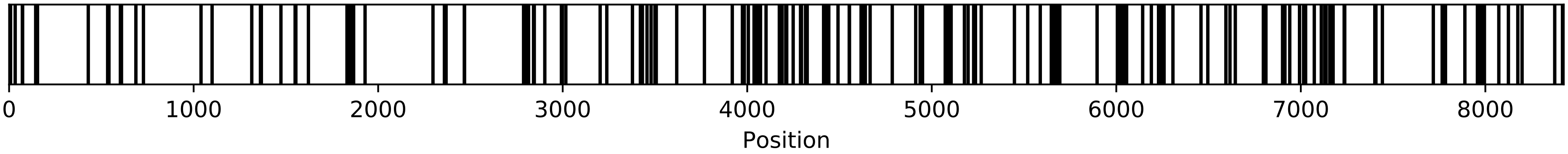


CR2 vs MM2
Number of differences: 231

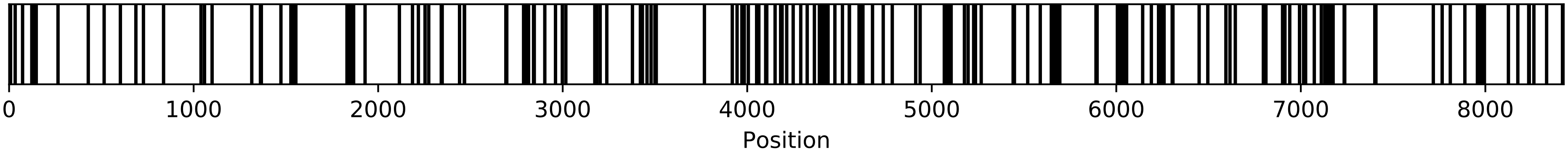


MM1 vs MM2
Number of differences: 112

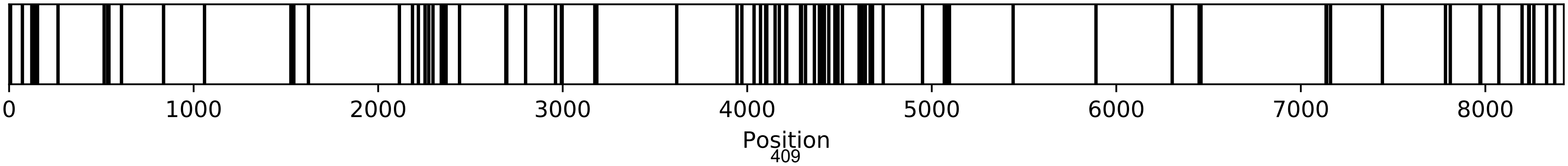

### Region: 1087



CR1 vs CR2
Number of differences: 135

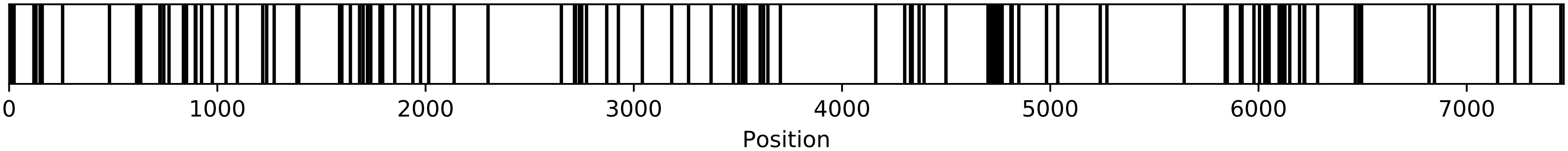


CR1 vs MM1
Number of differences: 43

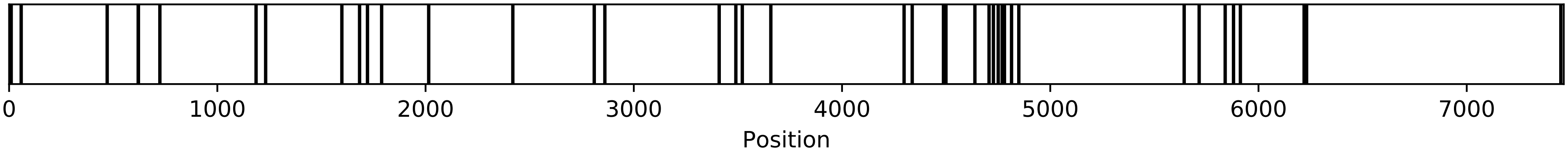


CR1 vs MM2
Number of differences: 59

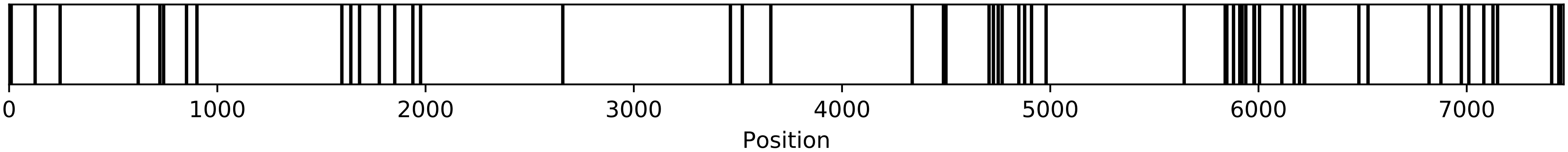


CR2 vs MM1
Number of differences: 134

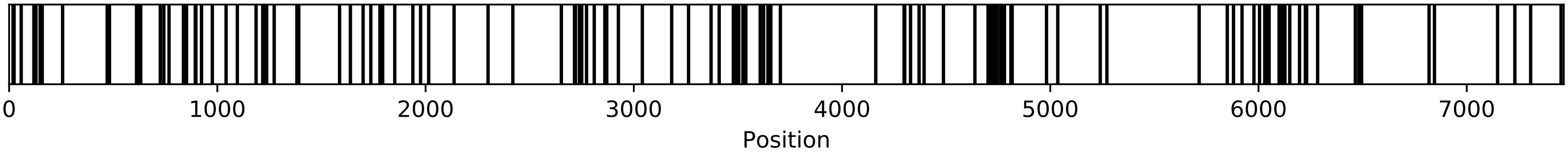


CR2 vs MM2
Number of differences: 135

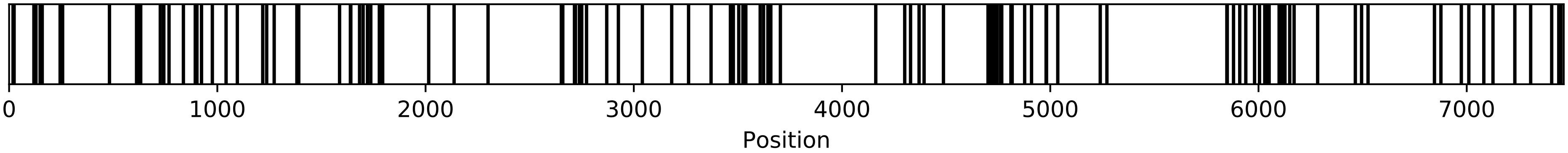


MM1 vs MM2
Number of differences: 57

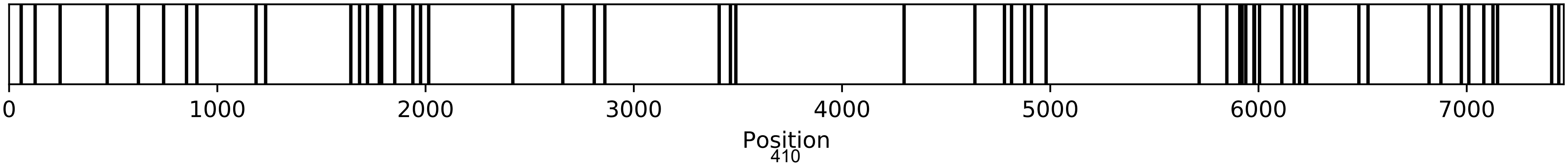

### Region: 1088



CR1 vs CR2
Number of differences: 365

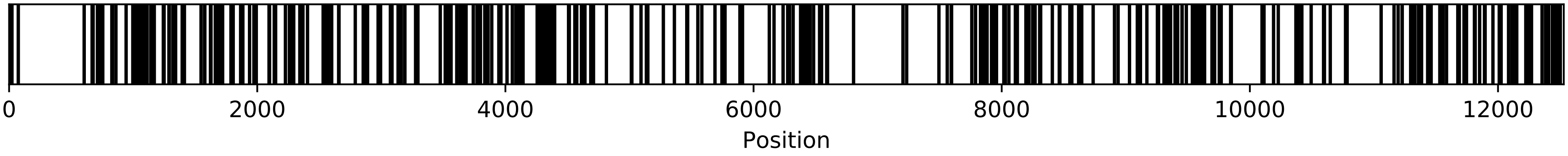


CR1 vs MM1
Number of differences: 169

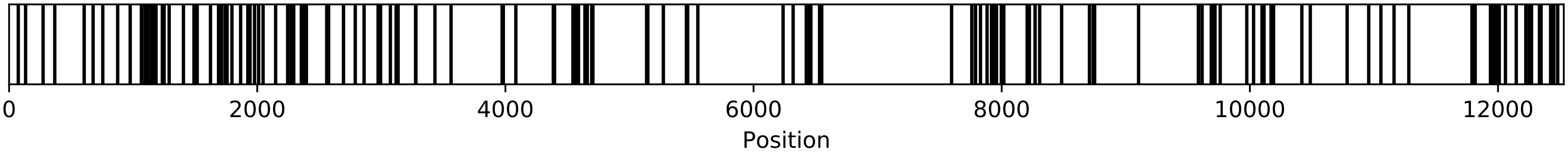


CR1 vs MM2
Number of differences: 154

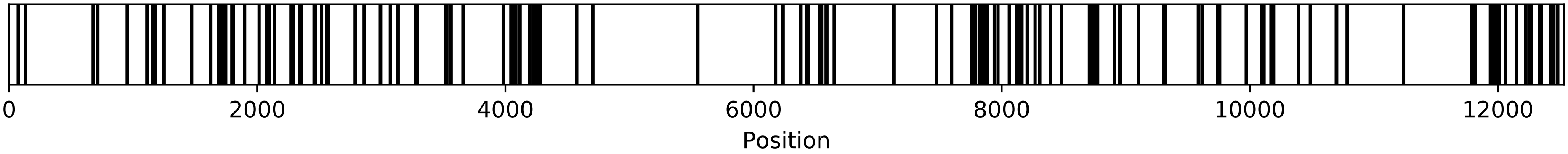


CR2 vs MM1
Number of differences: 361

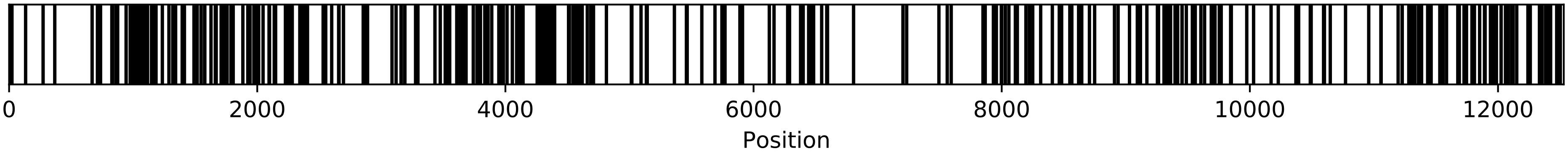


CR2 vs MM2
Number of differences: 366

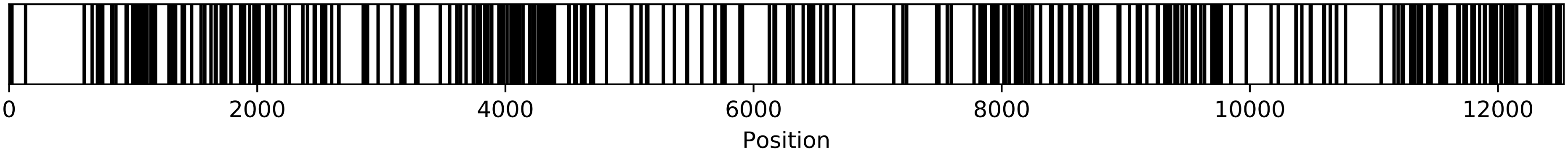


MM1 vs MM2
Number of differences: 159

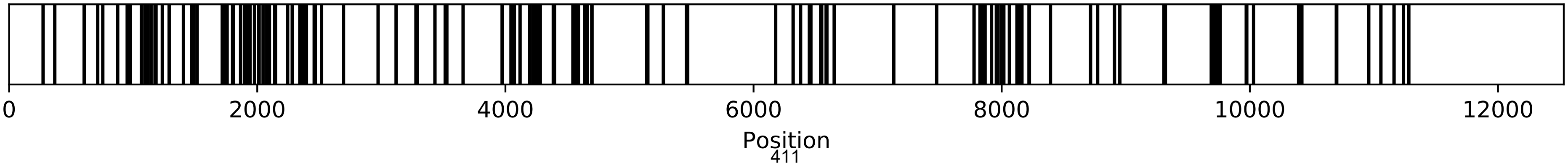

### Region: 1089



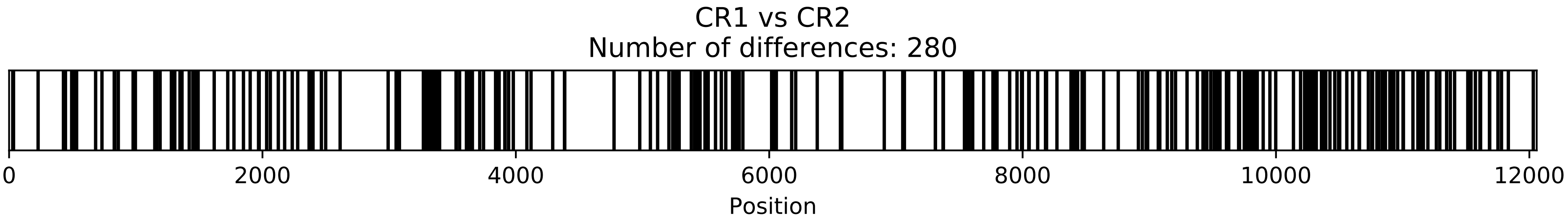
CR1 vs CR2
Number of differences: 280
0
2000
4000
6000
8000
10000
12000
Position

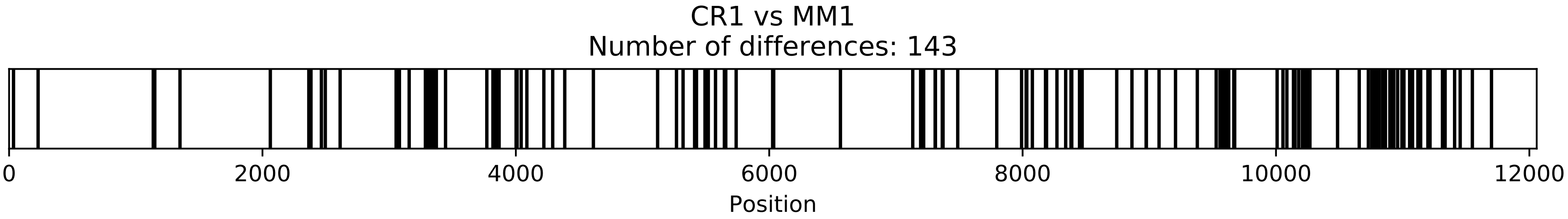
CR1 vs MM1
Number of differences: 143
0
2000
4000
6000
8000
10000
12000
Position

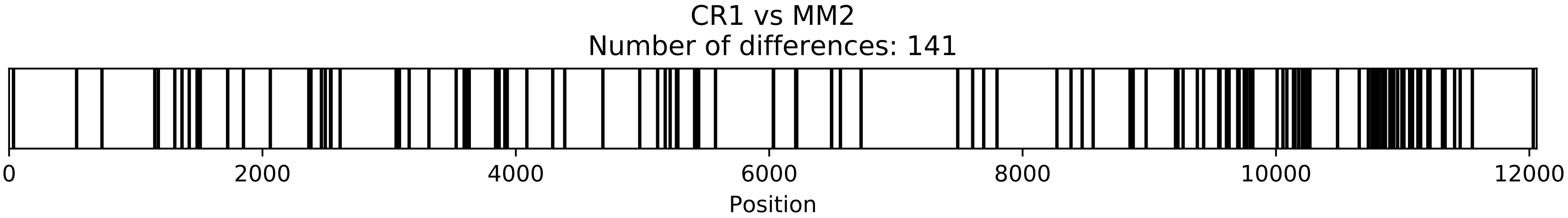
CR1 vs MM2
Number of differences: 141
0
2000
4000
6000
8000
10000
12000
Position

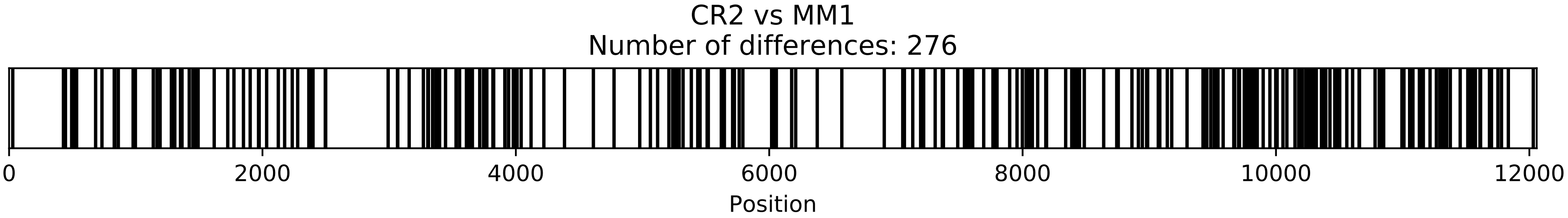
CR2 vs MM1
Number of differences: 276
0
2000
4000
6000
8000
10000
12000
Position

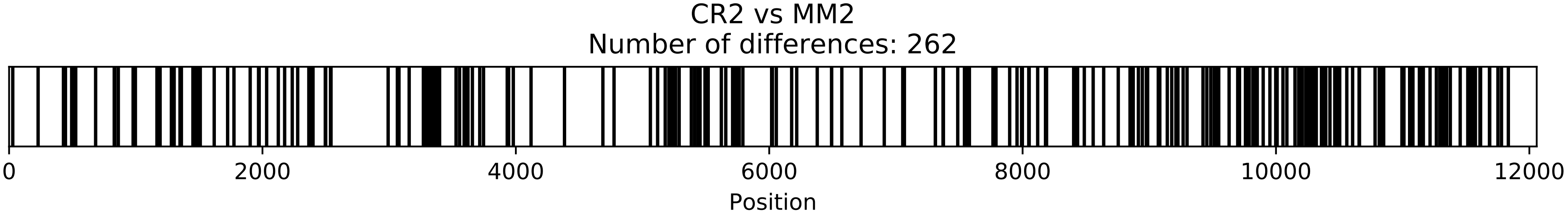
CR2 vs MM2
Number of differences: 262
0
2000
4000
6000
8000
10000
12000
Position

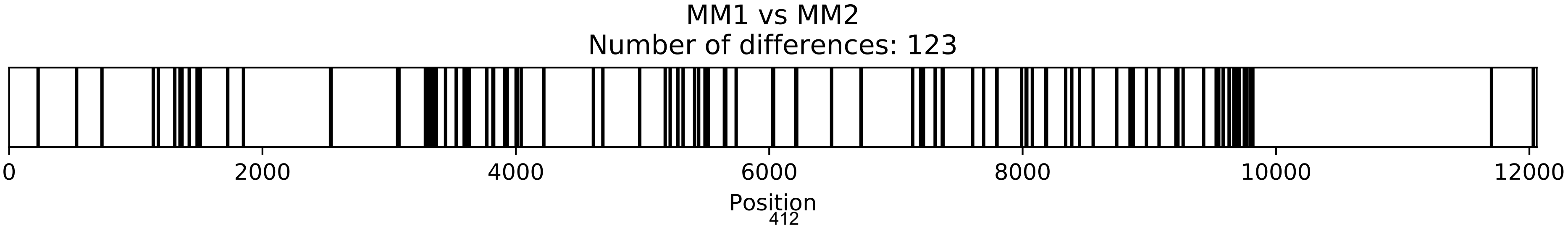
MM1 vs MM2
Number of differences: 123
0
2000
4000
6000
8000
10000
12000
Position

### Region: 1093



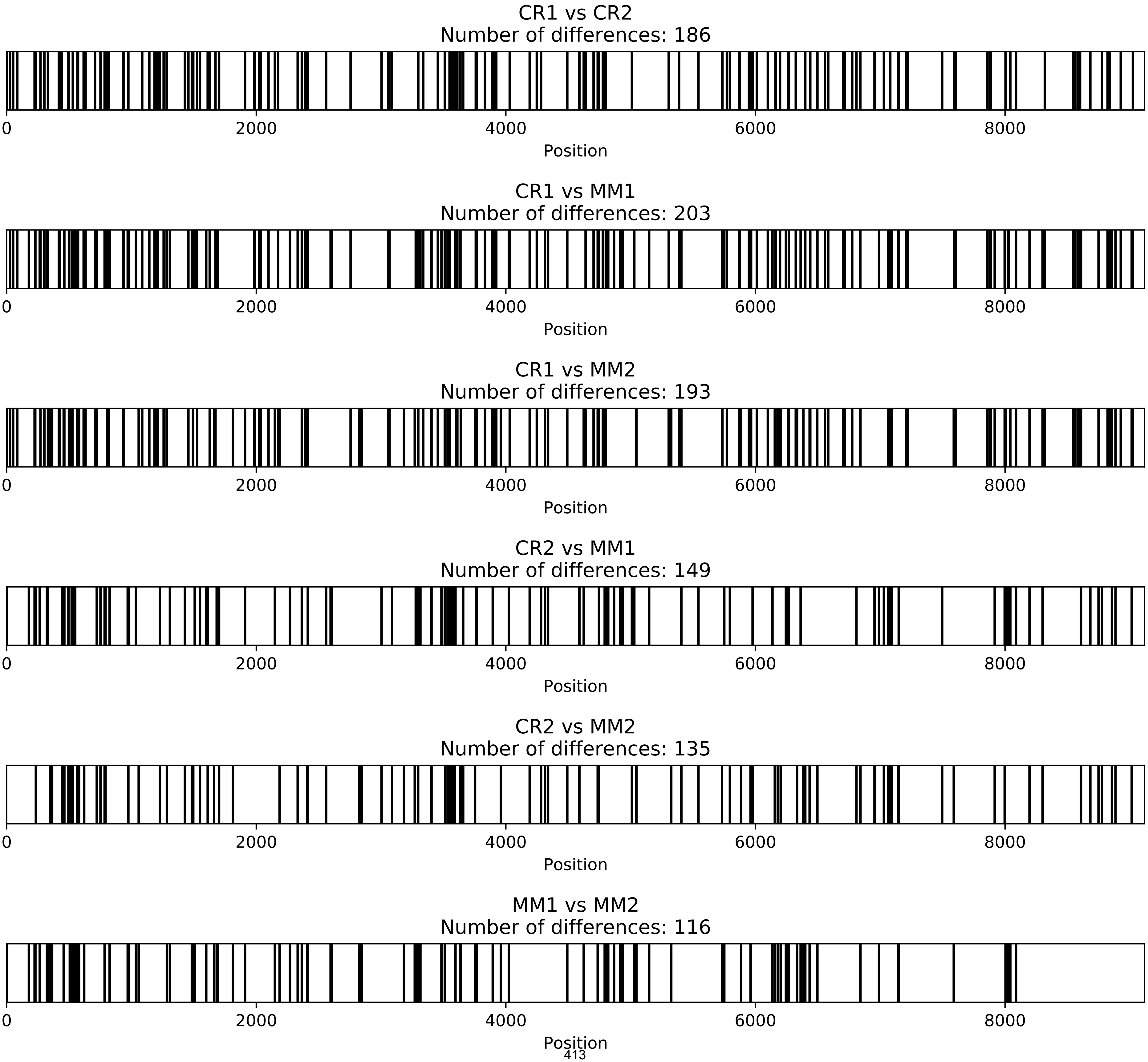
CR1 vs CR2
Number of differences: 186
0
2000
4000
6000
8000
Position
CR1 vs MM1
Number of differences: 203
0
2000
4000
6000
8000
Position
CR1 vs MM2
Number of differences: 193
0
2000
4000
6000
8000
Position
CR2 vs MM1
Number of differences: 149
0
2000
4000
6000
8000
Position
CR2 vs MM2
Number of differences: 135
0
2000
4000
6000
8000
Position
MM1 vs MM2
Number of differences: 116
0
2000
4000
6000
8000
Position

### Region: 1097



CR1 vs CR2
Number of differences: 324

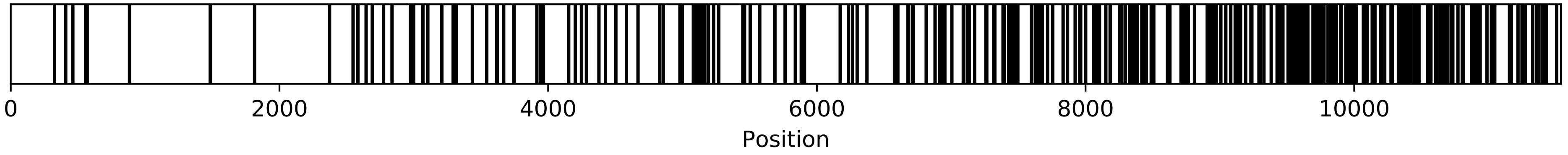


CR1 vs MM1
Number of differences: 178

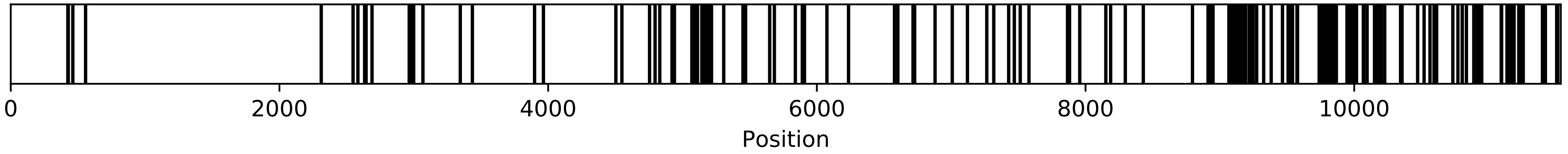


CR1 vs MM2
Number of differences: 151

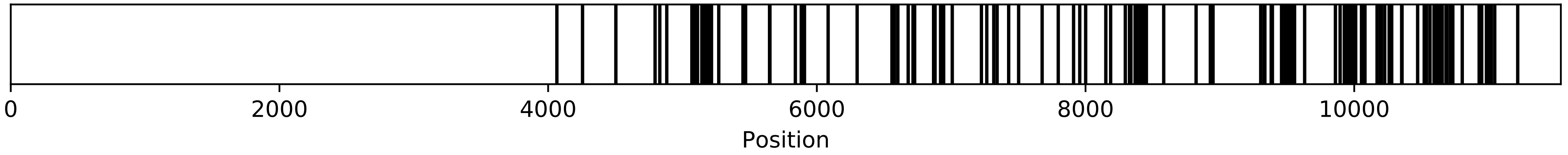


CR2 vs MM1
Number of differences: 323

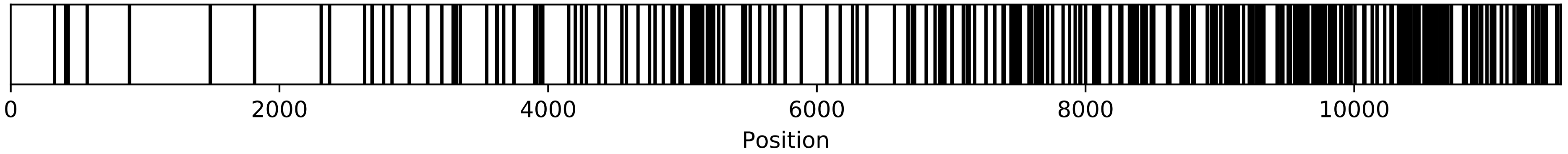


CR2 vs MM2
Number of differences: 334

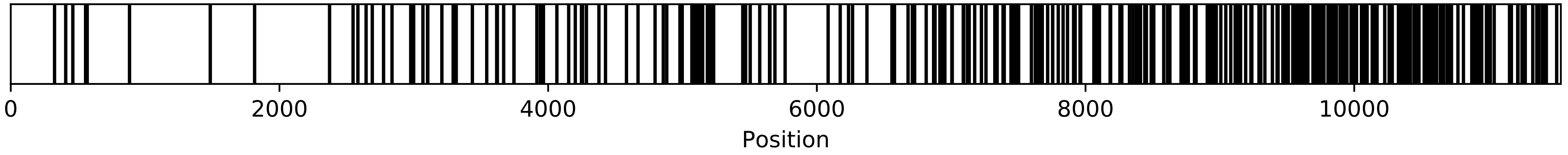


MM1 vs MM2
Number of differences: 187

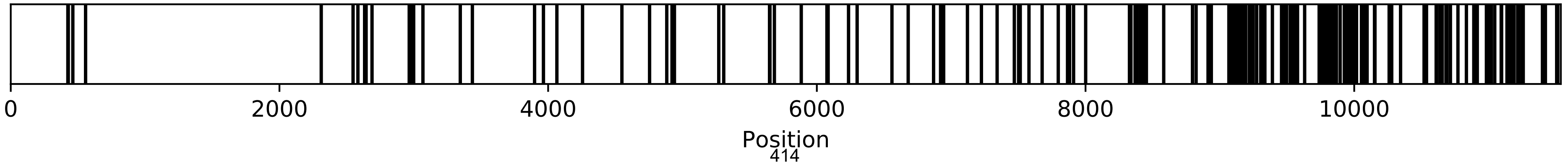

### Region: 1103



CR1 vs CR2
Number of differences: 239

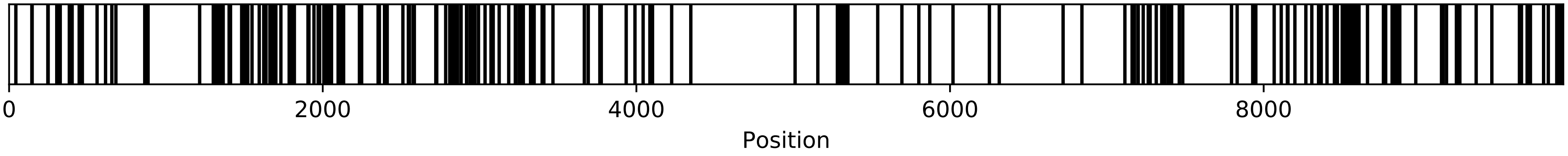


CR1 vs MM1
Number of differences: 123

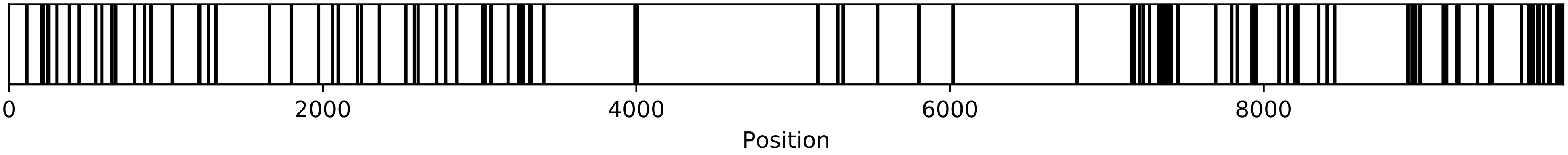


CR1 vs MM2
Number of differences: 73

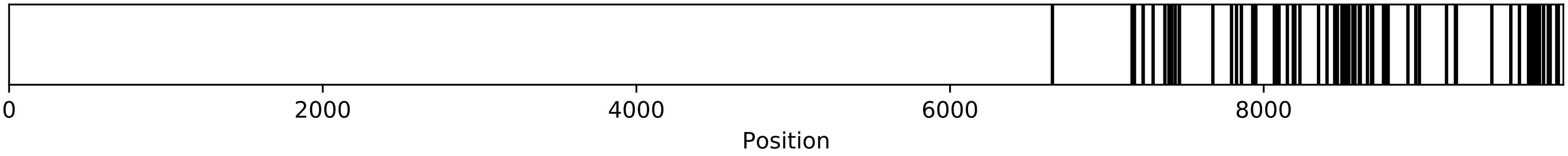


CR2 vs MM1
Number of differences: 244

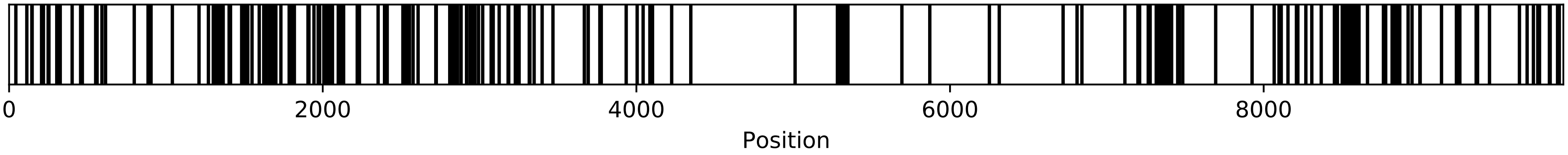


CR2 vs MM2
Number of differences: 235

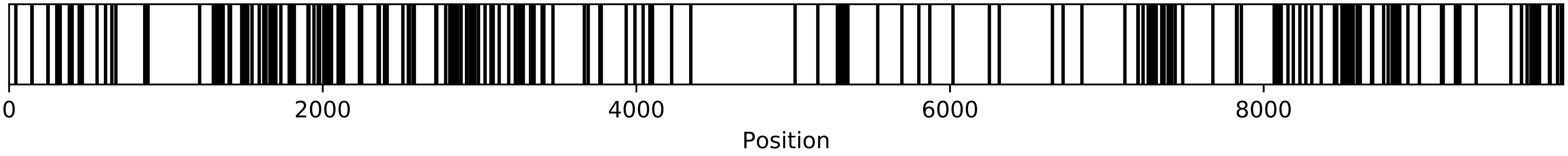


MM1 vs MM2
Number of differences: 138

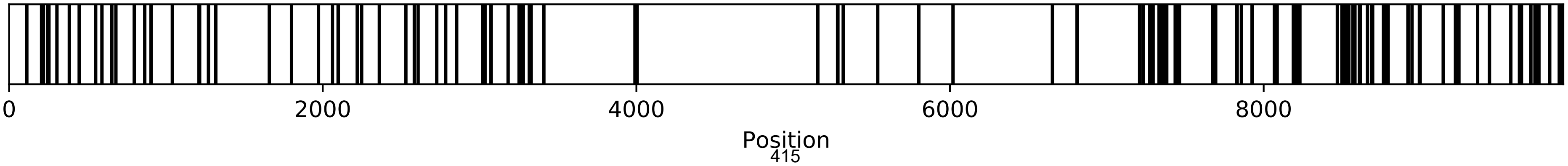

### Region: 1104



CR1 vs CR2
Number of differences: 439

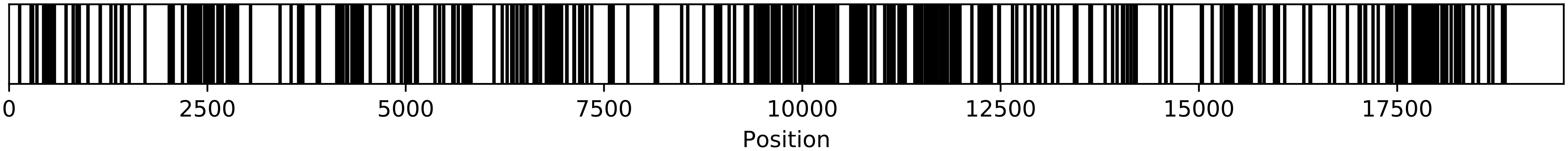


CR1 vs MM1
Number of differences: 221

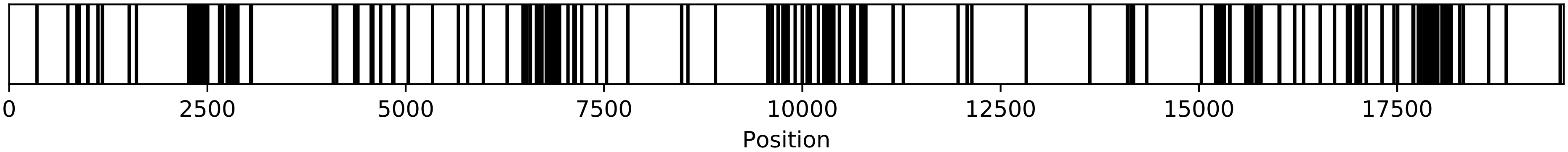


CR1 vs MM2
Number of differences: 222

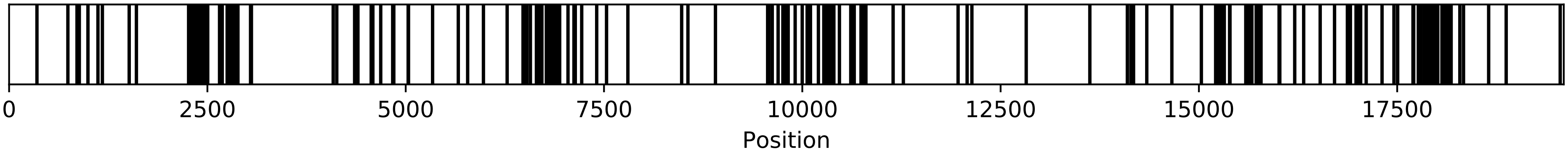


CR2 vs MM1
Number of differences: 421

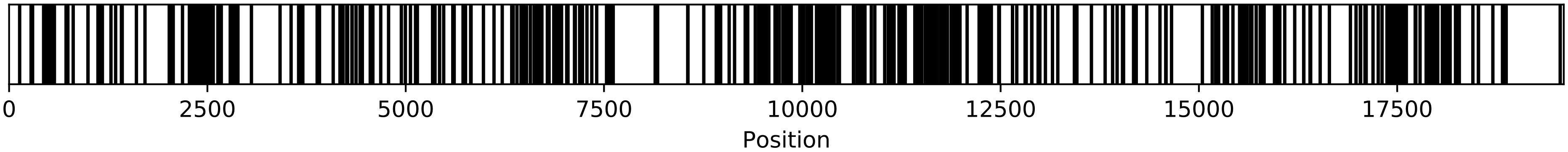


CR2 vs MM2
Number of differences: 422

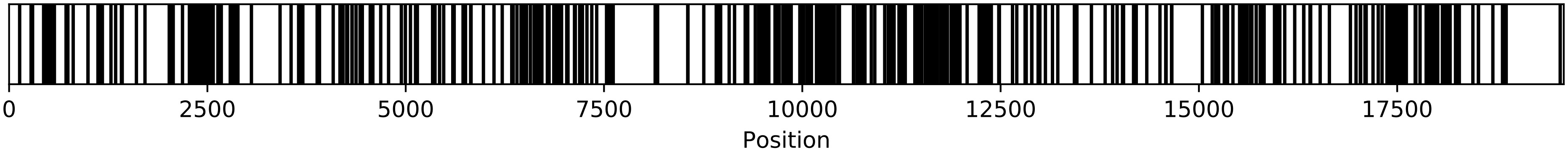


MM1 vs MM2
Number of differences: 1

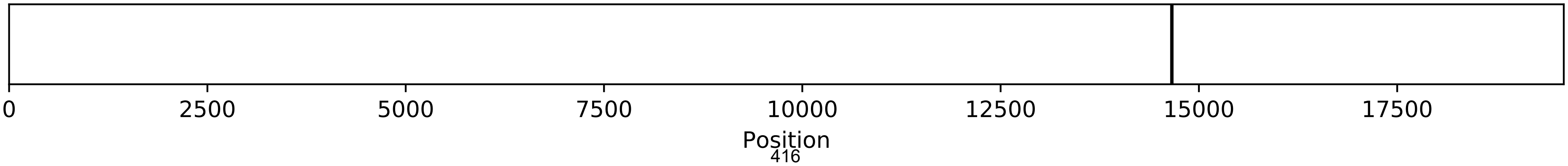

### Region: 1105



CR1 vs CR2
Number of differences: 450

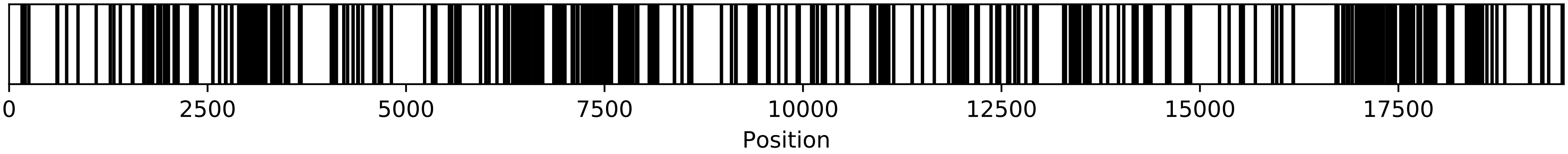


CR1 vs MM1
Number of differences: 441

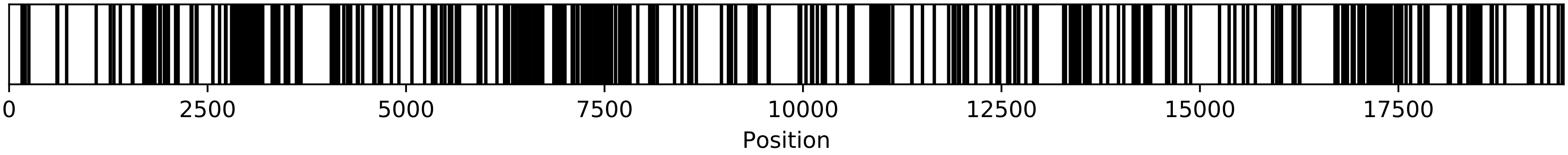


CR1 vs MM2
Number of differences: 424

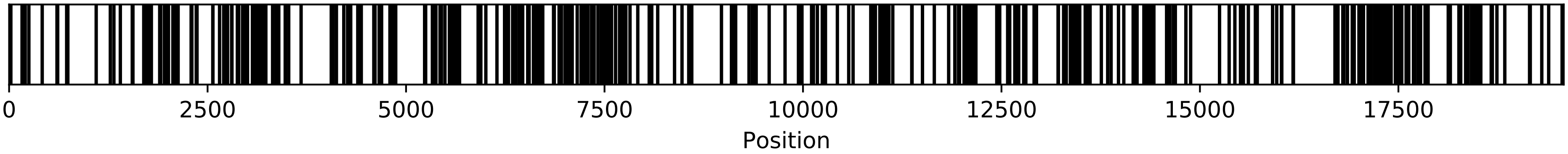


CR2 vs MM1
Number of differences: 217

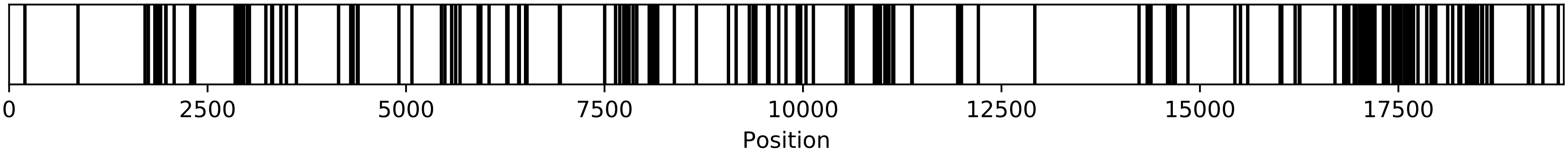


CR2 vs MM2
Number of differences: 292

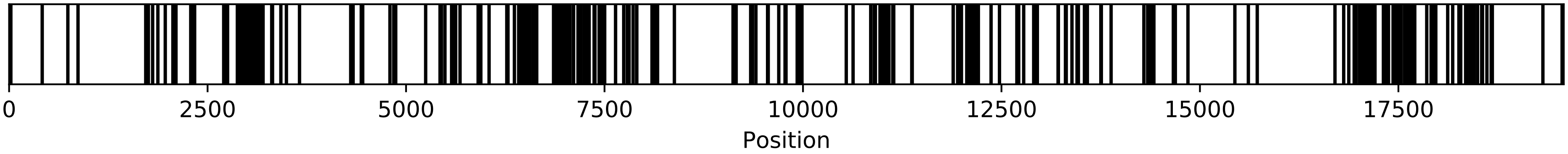


MM1 vs MM2
Number of differences: 235

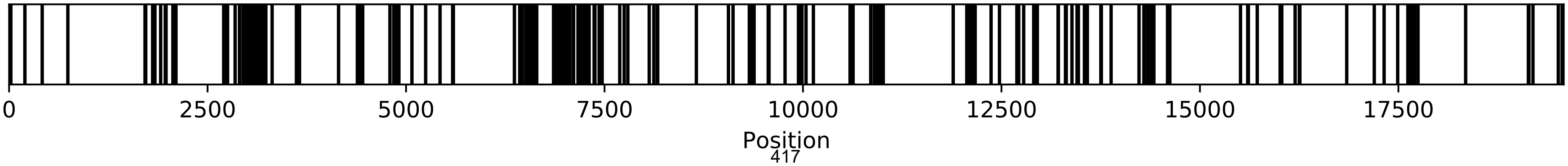

### Region: 1112



CR1 vs CR2
Number of differences: 203

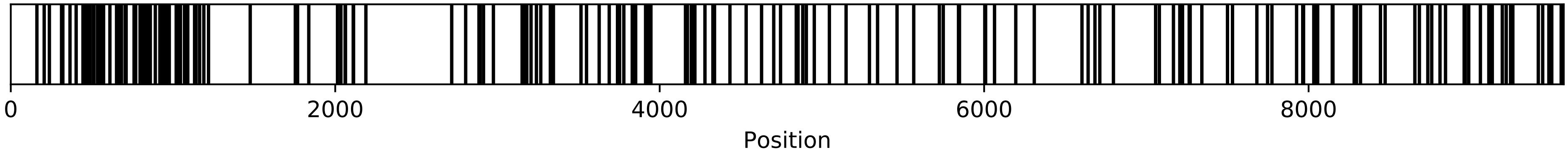


CR1 vs MM1
Number of differences: 108

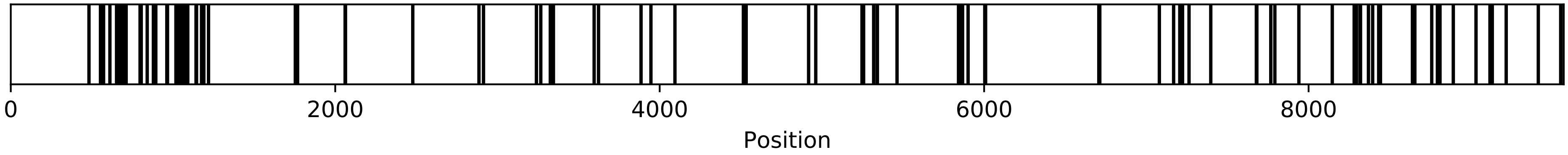


CR1 vs MM2
Number of differences: 88

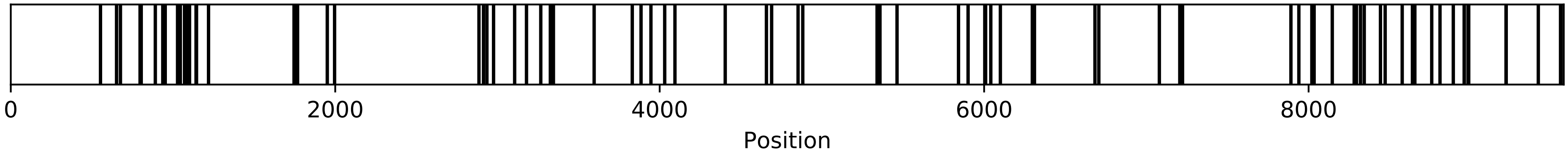


CR2 vs MM1
Number of differences: 178

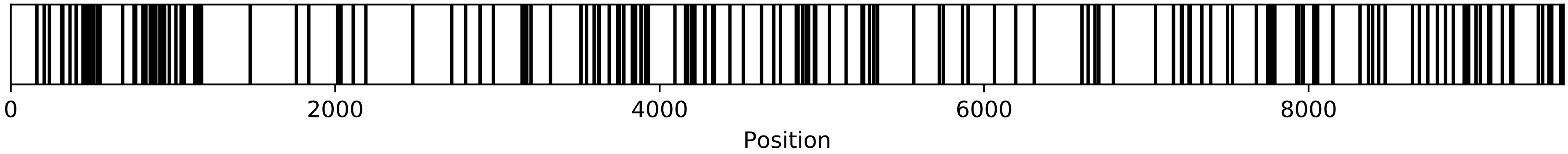


CR2 vs MM2
Number of differences: 194

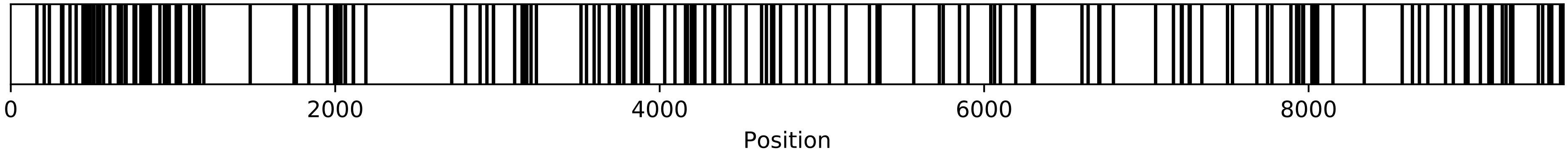


MM1 vs MM2
Number of differences: 100

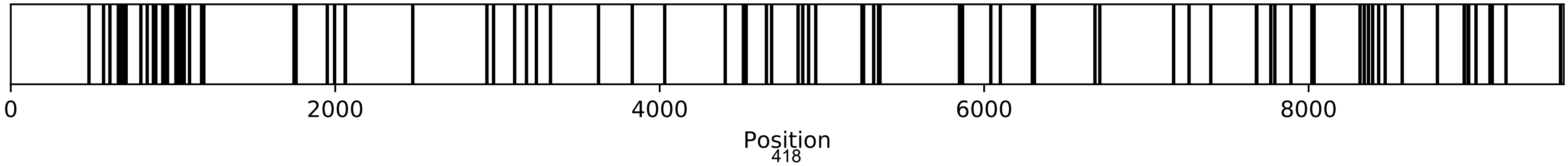

### Region: 1116



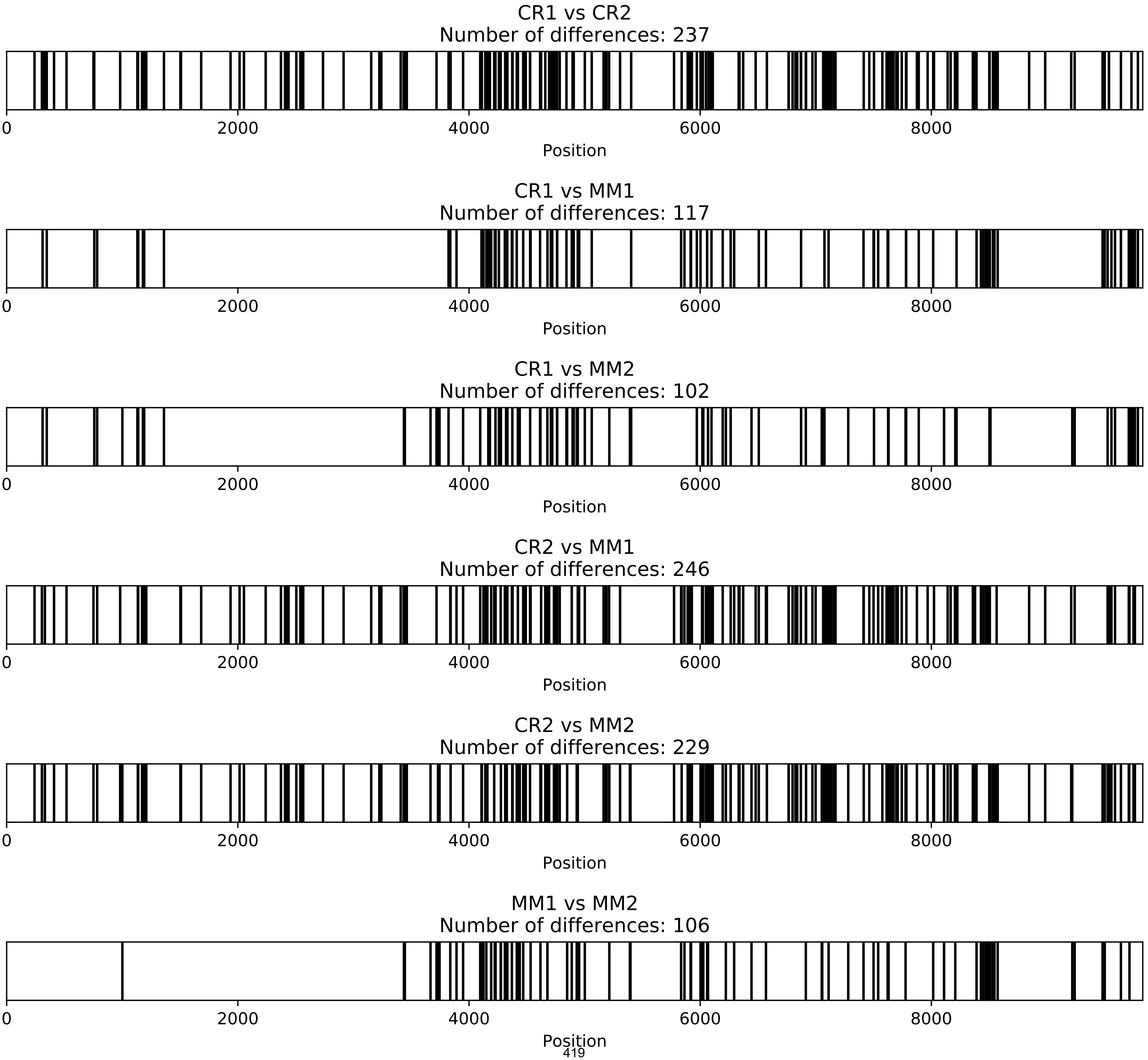
CR1 vs CR2
Number of differences: 237
0
2000
4000
6000
8000
Position
CR1 vs MM1
Number of differences: 117
0
2000
4000
6000
8000
Position
CR1 vs MM2
Number of differences: 102
0
2000
4000
6000
8000
Position
CR2 vs MM1
Number of differences: 246
0
2000
4000
6000
8000
Position
CR2 vs MM2
Number of differences: 229
0
2000
4000
6000
8000
Position
MM1 vs MM2
Number of differences: 106
0
2000
4000
6000
8000
Position

### Region: 1117



CR1 vs CR2
Number of differences: 367

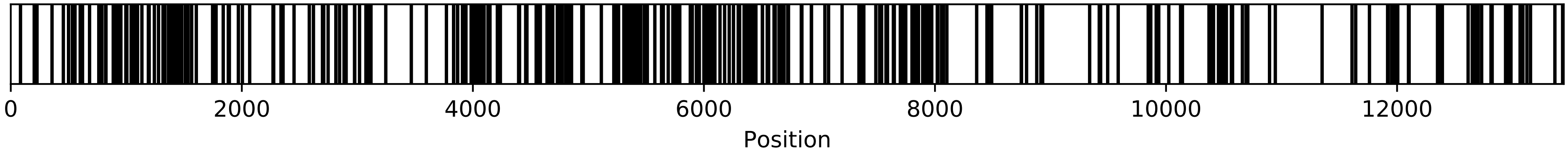


CR1 vs MM1
Number of differences: 193

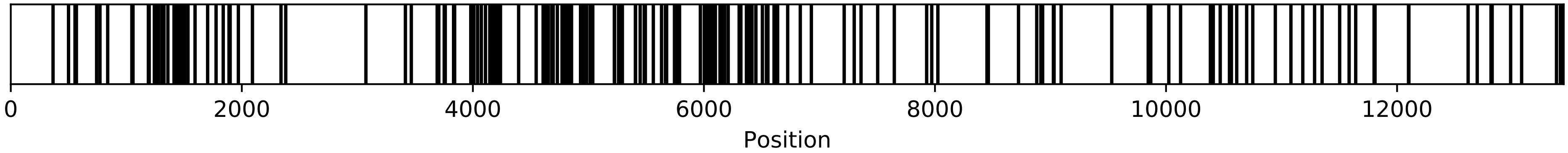


CR1 vs MM2
Number of differences: 180

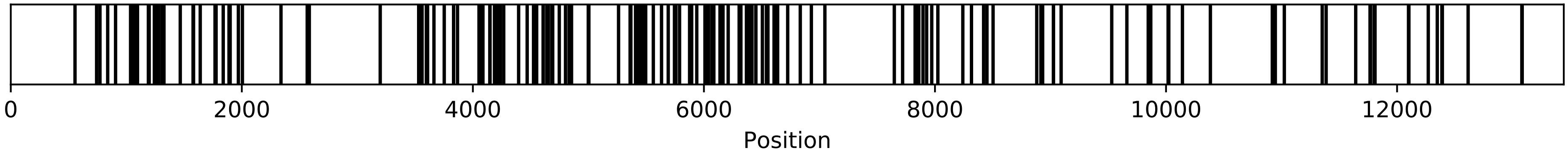


CR2 vs MM1
Number of differences: 386

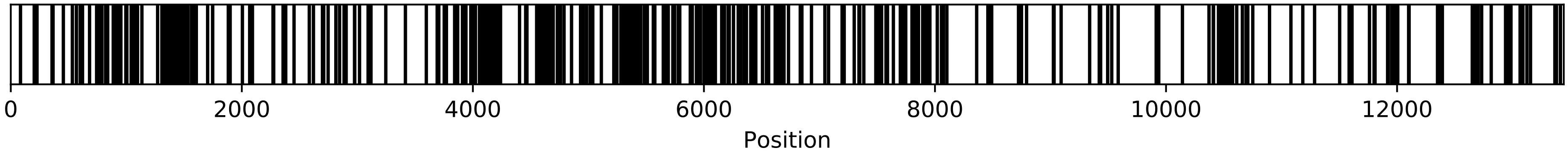


CR2 vs MM2
Number of differences: 365

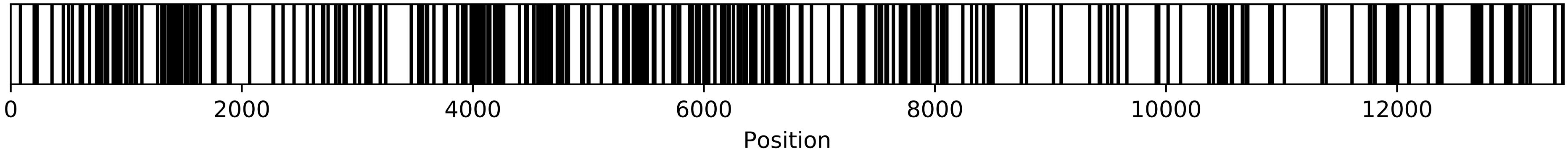


MM1 vs MM2
Number of differences: 178

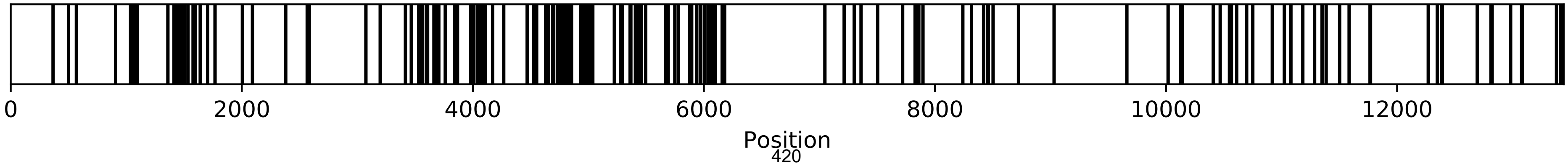

### Region: 1121



CR1 vs CR2
Number of differences: 286

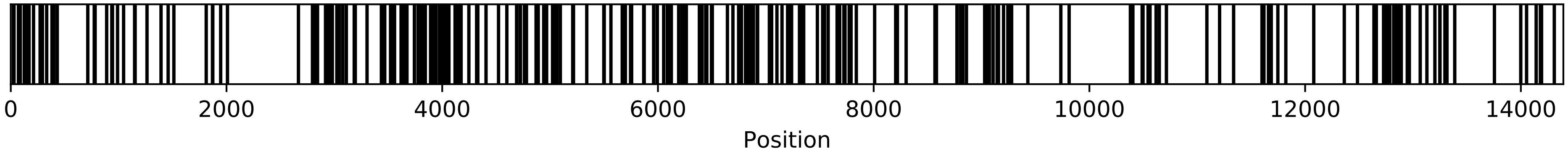


CR1 vs MM1
Number of differences: 287

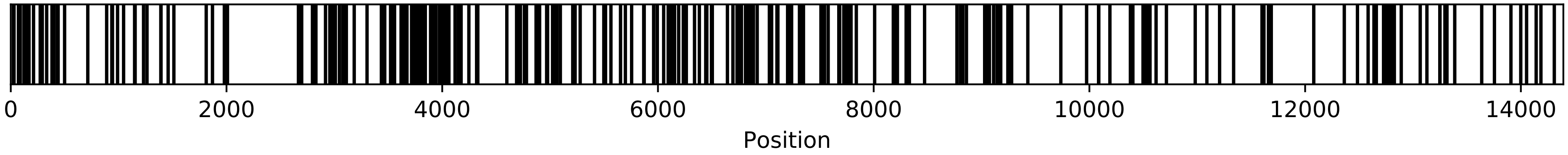


CR1 vs MM2
Number of differences: 278

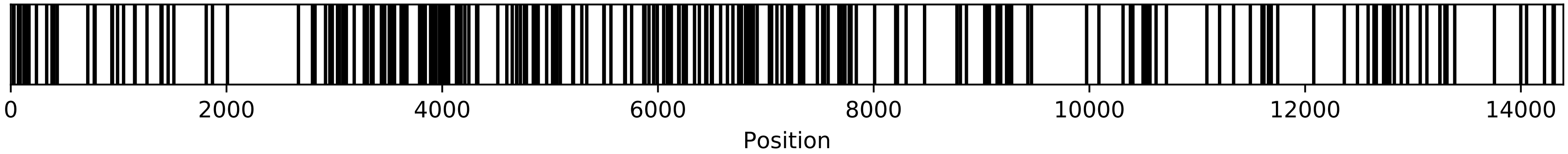


CR2 vs MM1
Number of differences: 115

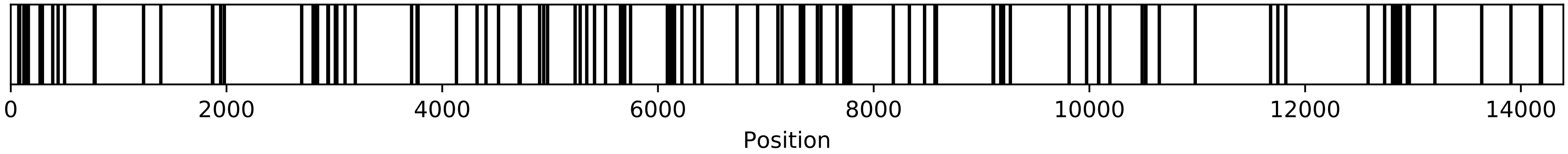


CR2 vs MM2
Number of differences: 116

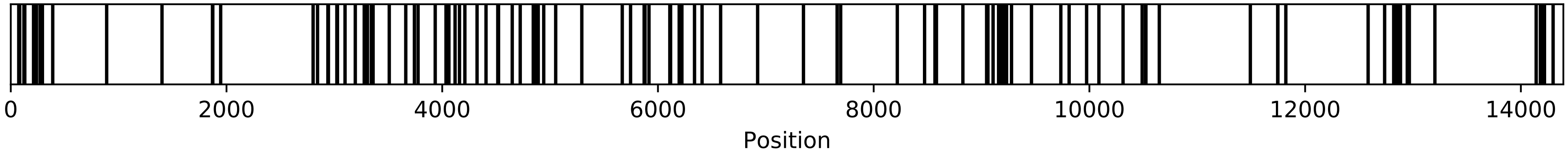


MM1 vs MM2
Number of differences: 113

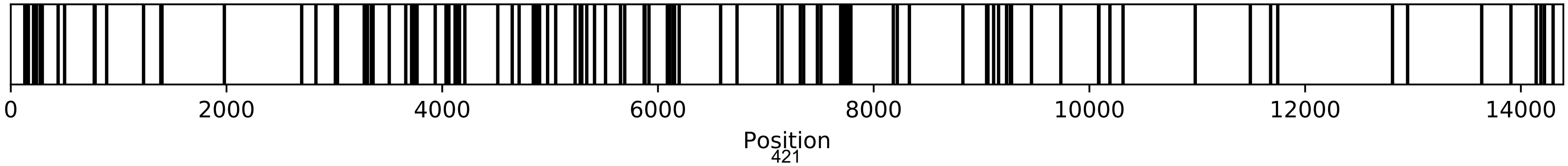

### Region: 1123



CR1 vs CR2
Number of differences: 32

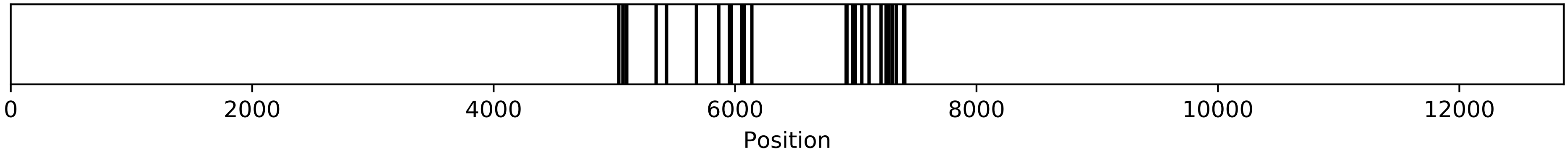


CR1 vs MM1
Number of differences: 202

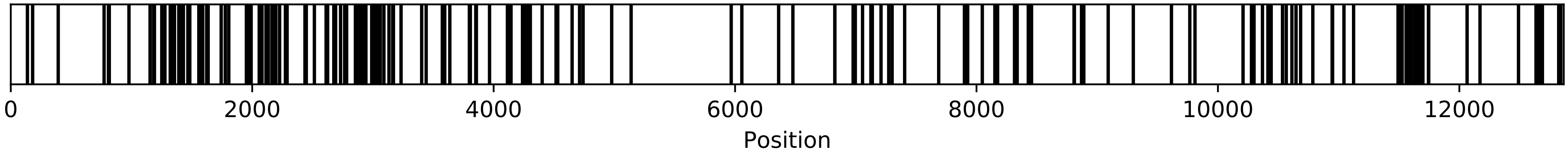


CR1 vs MM2
Number of differences: 184

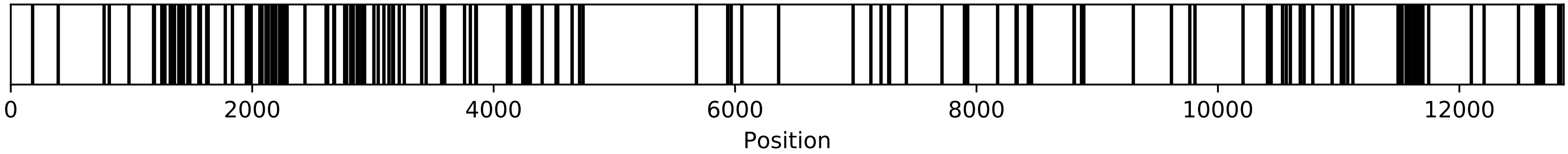


CR2 vs MM1
Number of differences: 219

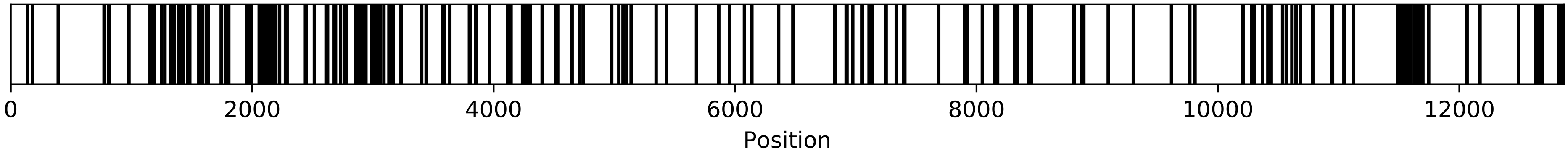


CR2 vs MM2
Number of differences: 204

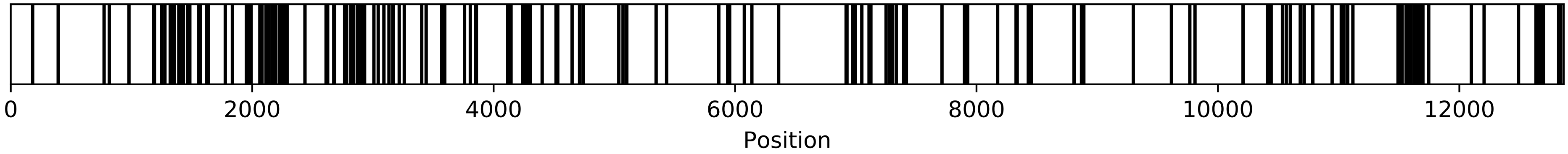


MM1 vs MM2
Number of differences: 103

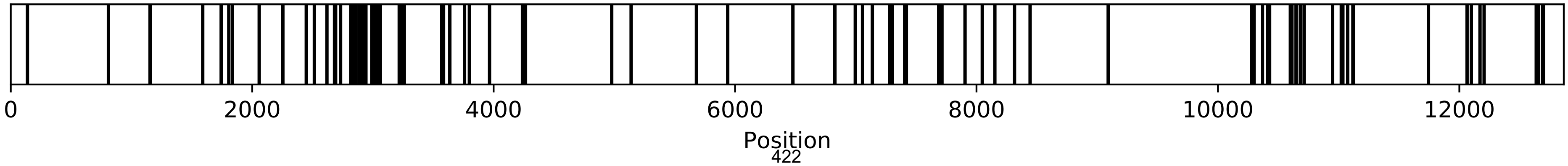

### Region: 1125



## CR1 vs CR2
Number of differences: 256

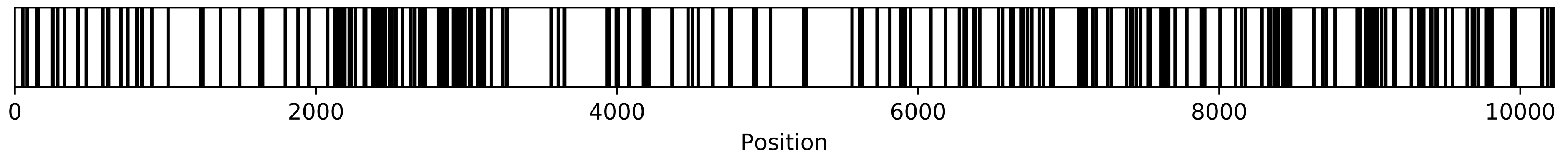


## CR1 vs MM1
Number of differences: 3

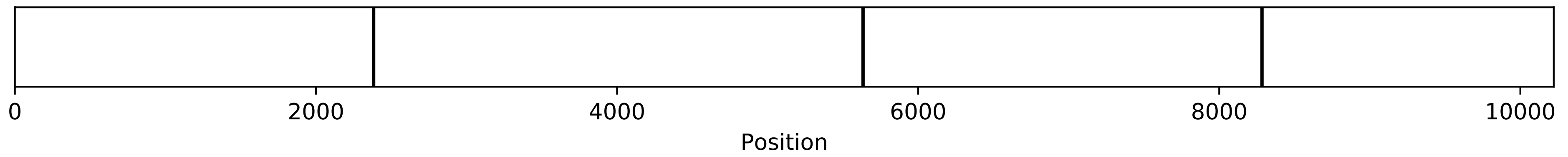


## CR1 vs MM2
Number of differences: 131

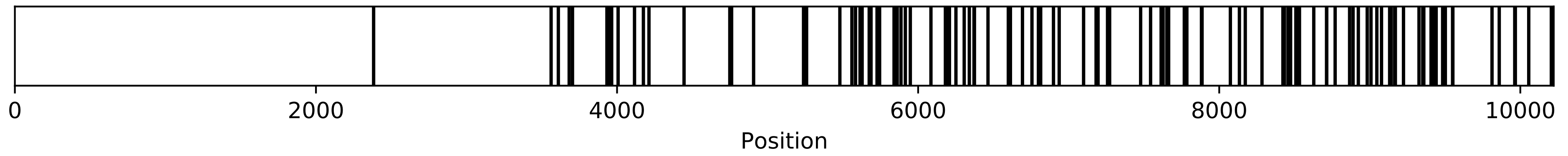


## CR2 vs MM1
Number of differences: 257

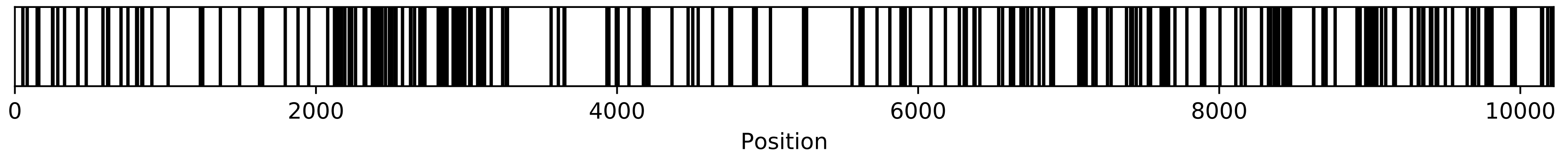


## CR2 vs MM2
Number of differences: 251

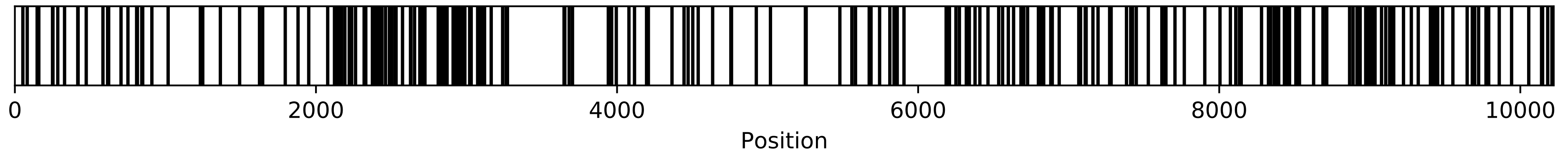


## MM1 vs MM2
Number of differences: 130

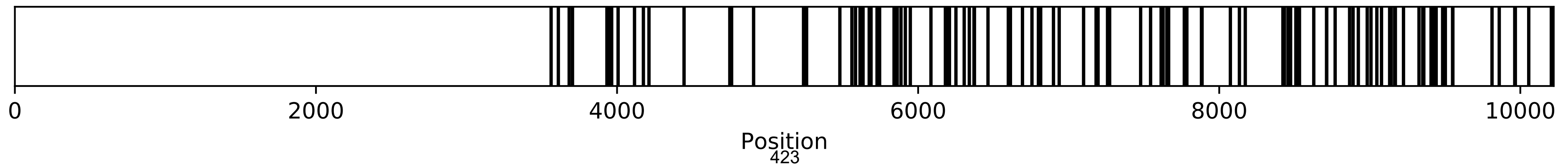

### Region: 1130



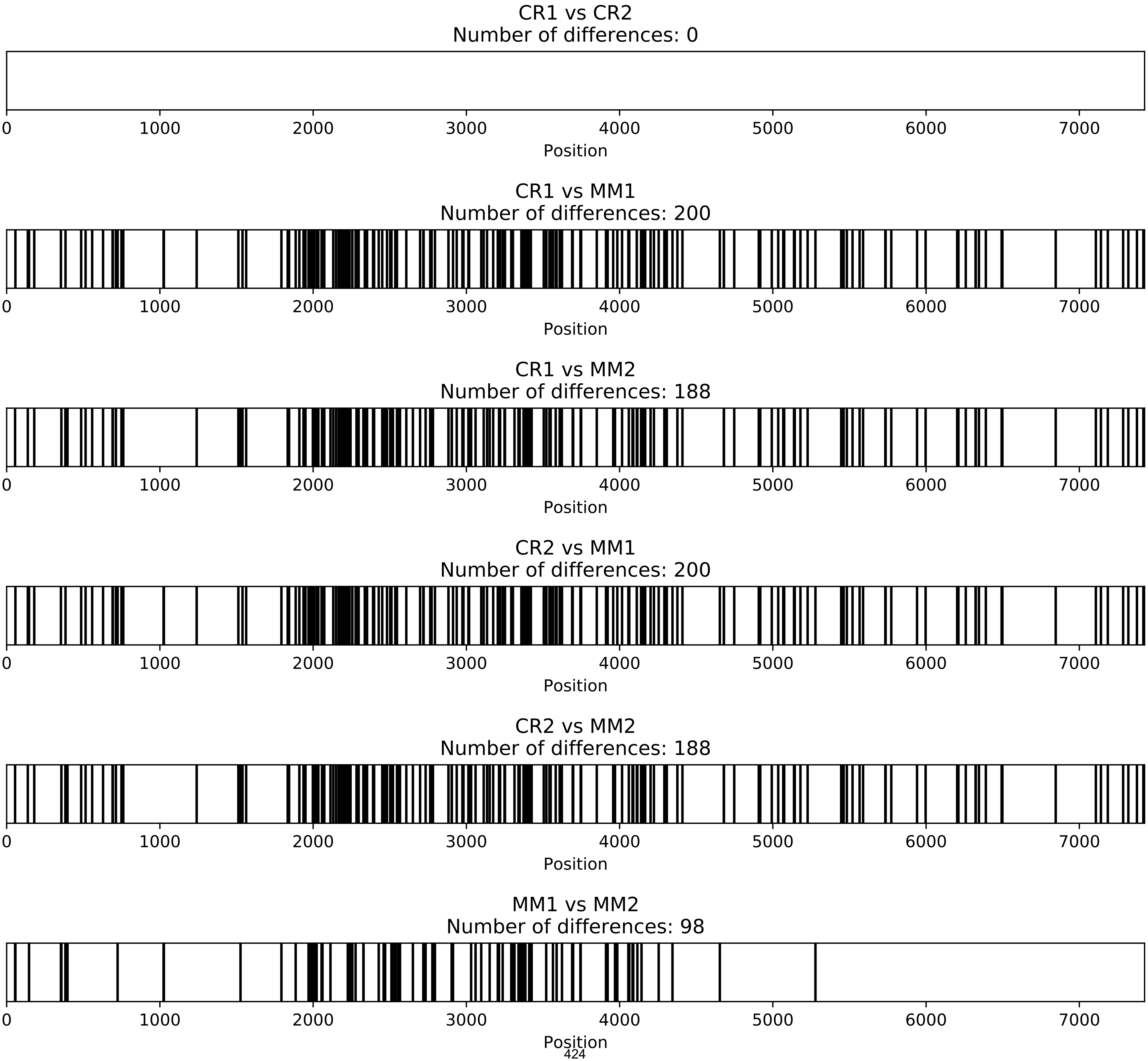
CR1 vs CR2
Number of differences: 0
CR1 vs MM1
Number of differences: 200
CR1 vs MM2
Number of differences: 188
CR2 vs MM1
Number of differences: 200
CR2 vs MM2
Number of differences: 188
MM1 vs MM2
Number of differences: 98
0 1000 2000 3000 4000 5000 6000 7000
Position

### Region: 1131



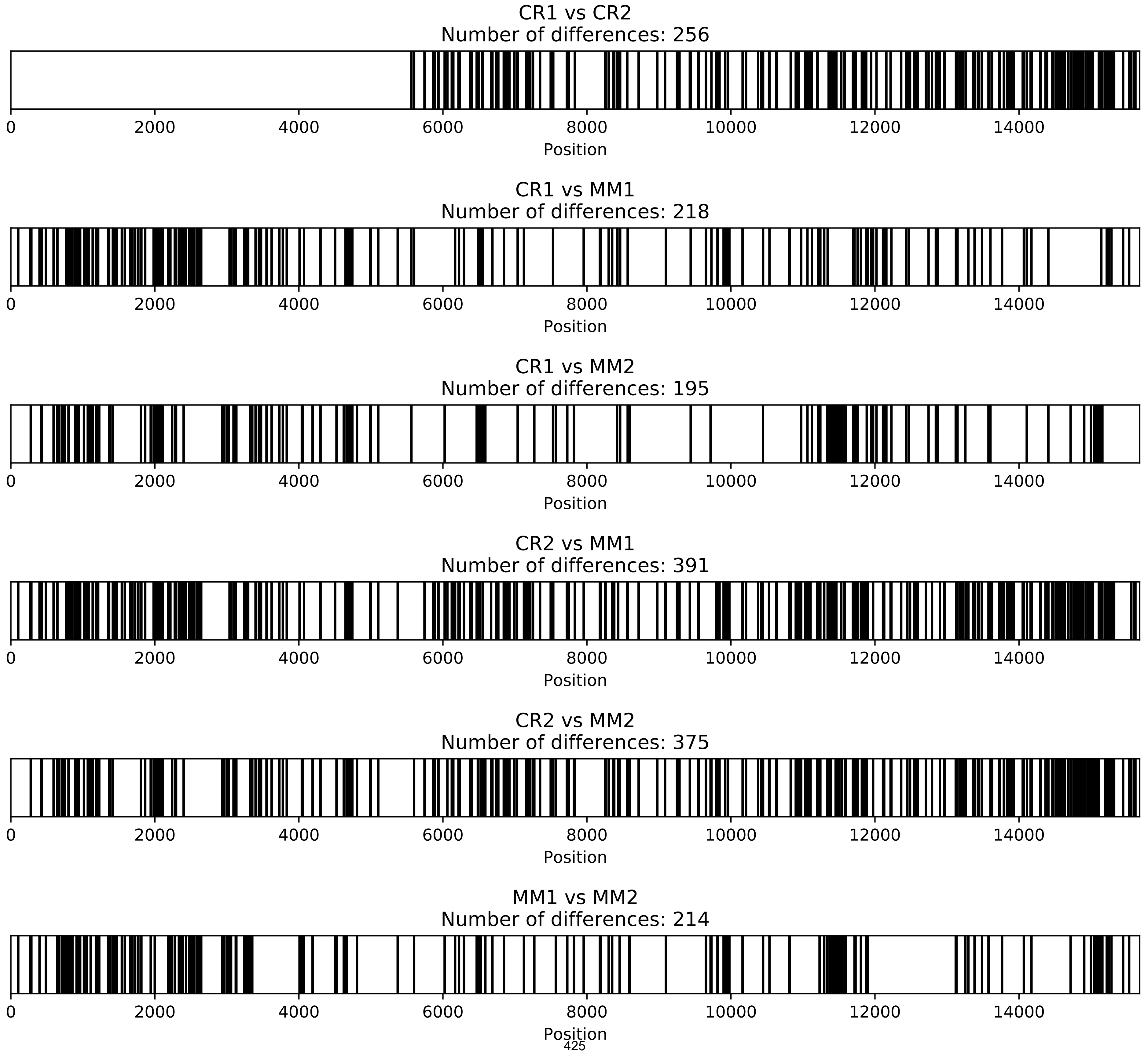
CR1 vs CR2
Number of differences: 256
0
2000
4000
6000
8000
10000
12000
14000
Position
CR1 vs MM1
Number of differences: 218
Position
CR1 vs MM2
Number of differences: 195
Position
CR2 vs MM1
Number of differences: 391
Position
CR2 vs MM2
Number of differences: 375
Position
MM1 vs MM2
Number of differences: 214
Position

### Region: 1132



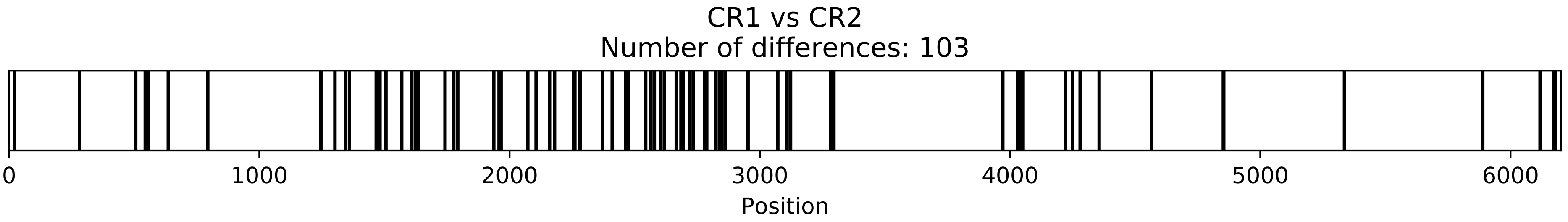
CR1 vs CR2
Number of differences: 103
0
1000
2000
3000
4000
5000
6000
Position

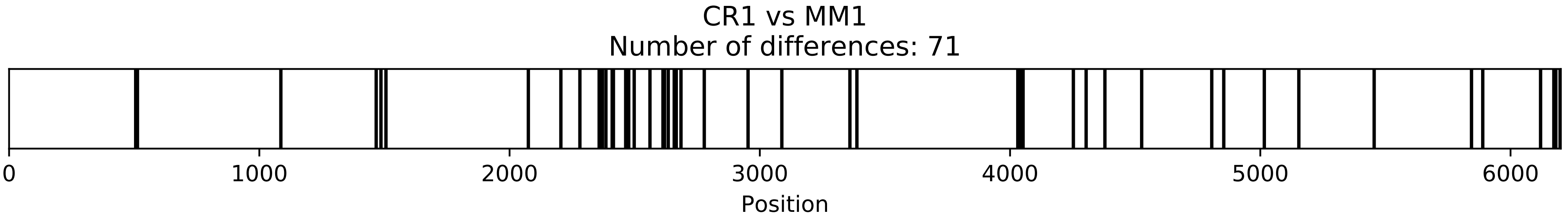
CR1 vs MM1
Number of differences: 71
0
1000
2000
3000
4000
5000
6000
Position

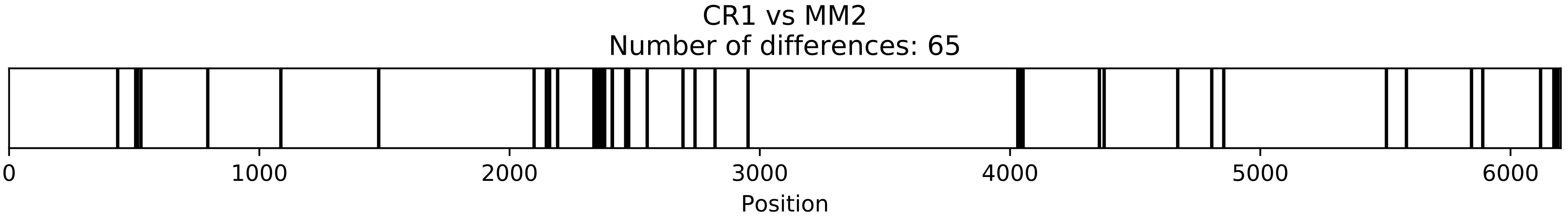
CR1 vs MM2
Number of differences: 65
0
1000
2000
3000
4000
5000
6000
Position

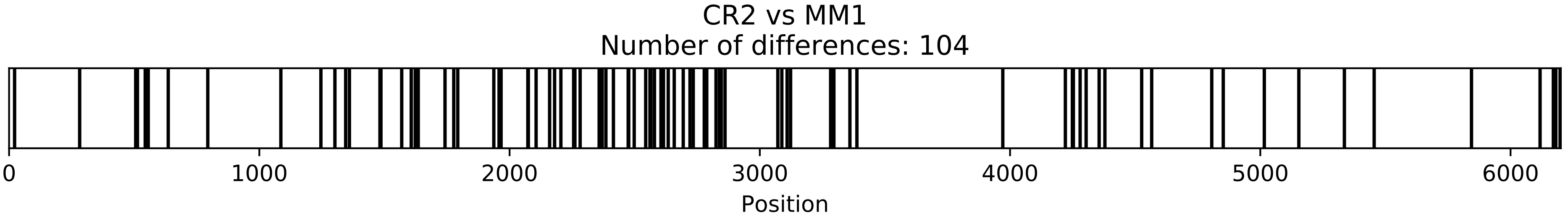
CR2 vs MM1
Number of differences: 104
0
1000
2000
3000
4000
5000
6000
Position

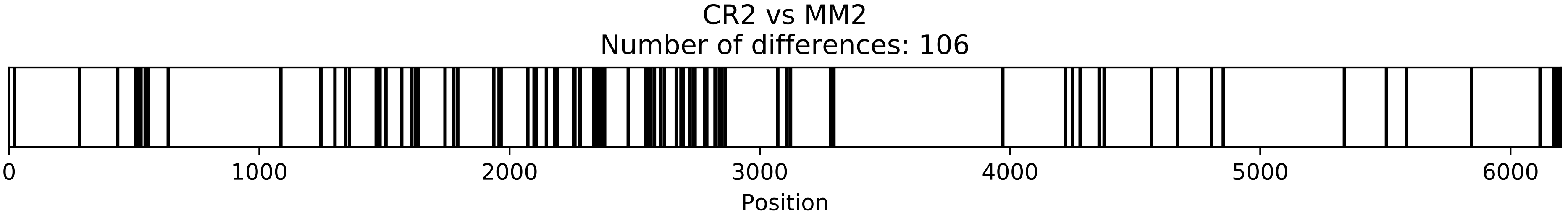
CR2 vs MM2
Number of differences: 106
0
1000
2000
3000
4000
5000
6000
Position

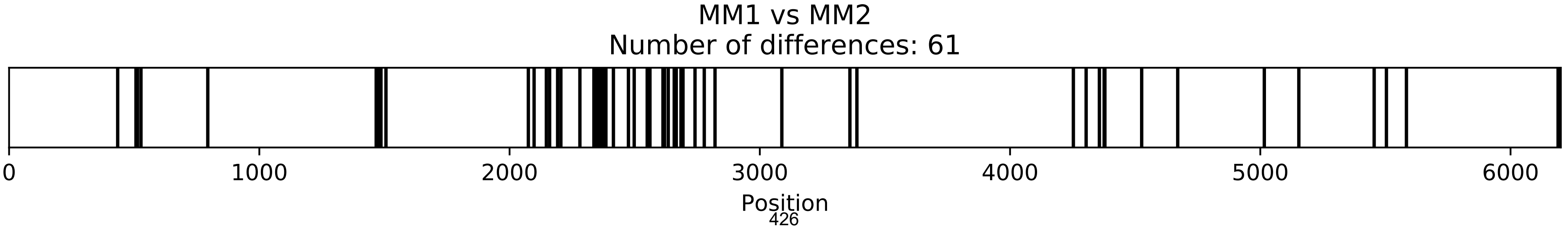
MM1 vs MM2
Number of differences: 61
0
1000
2000
3000
4000
5000
6000
Position

### Region: 1140

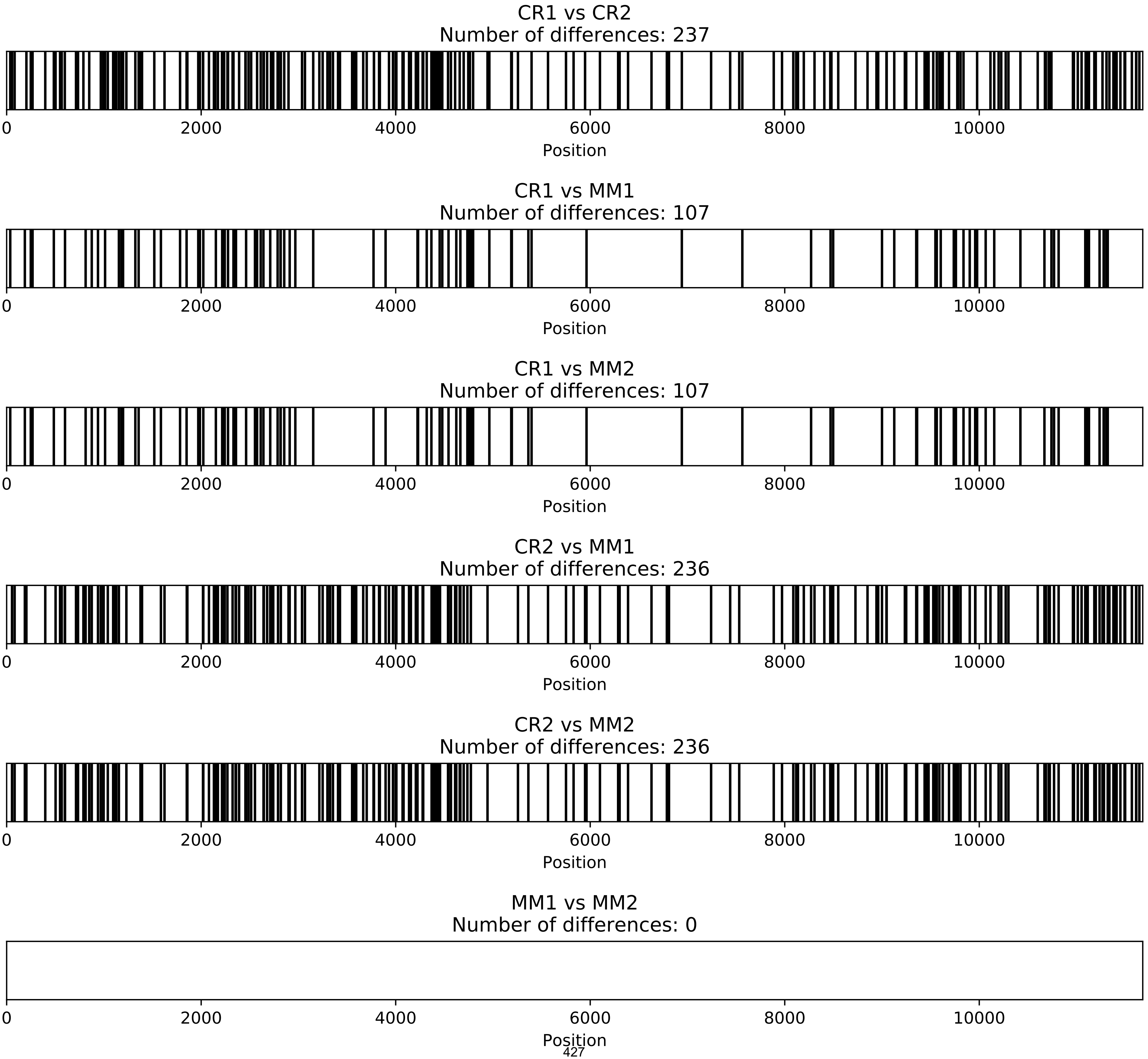
CR1 vs CR2
Number of differences: 237
CR1 vs MM1
Number of differences: 107
CR1 vs MM2
Number of differences: 107
CR2 vs MM1
Number of differences: 236
CR2 vs MM2
Number of differences: 236
MM1 vs MM2
Number of differences: 0
0
2000
4000
6000
8000
10000
Position

### Region: 1141



CR1 vs CR2
Number of differences: 3

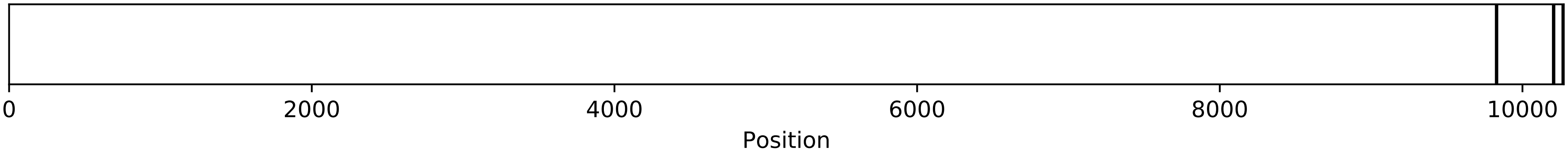


CR1 vs MM1
Number of differences: 179

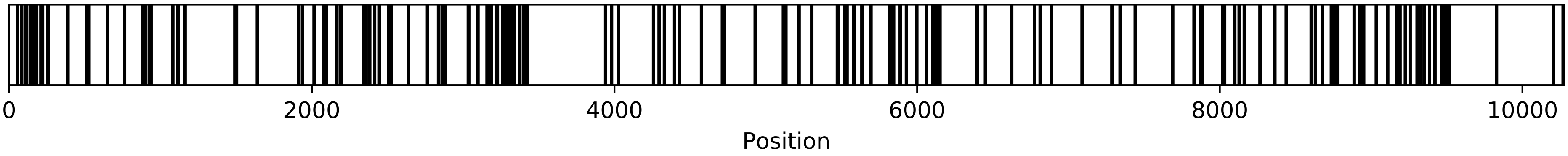


CR1 vs MM2
Number of differences: 197

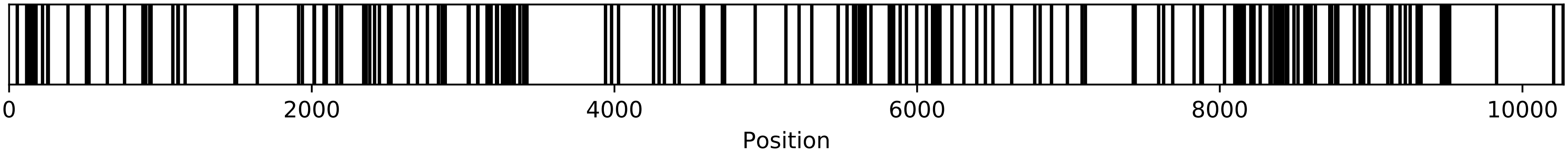


CR2 vs MM1
Number of differences: 176

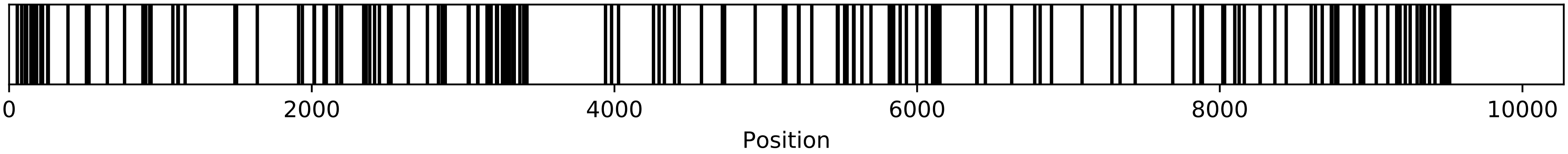


CR2 vs MM2
Number of differences: 194

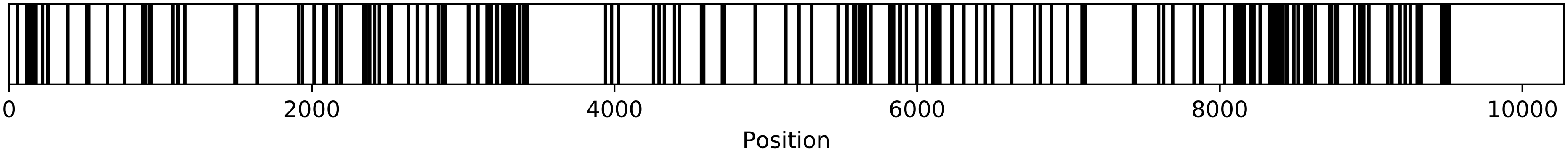


MM1 vs MM2
Number of differences: 76

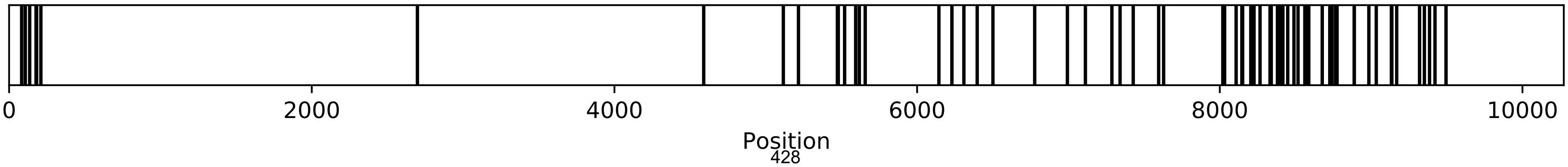

### Region: 1143



CR1 vs CR2
Number of differences: 190

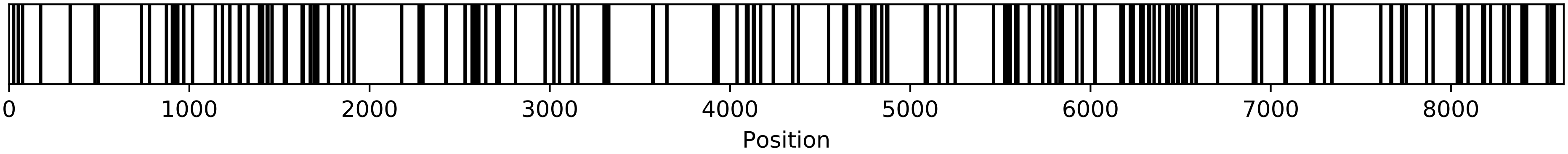


CR1 vs MM1
Number of differences: 101

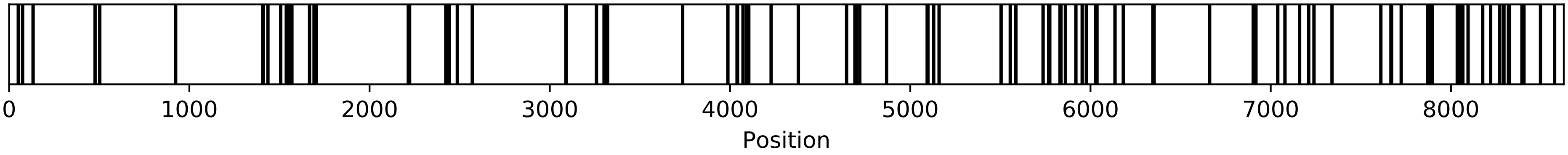


CR1 vs MM2
Number of differences: 96

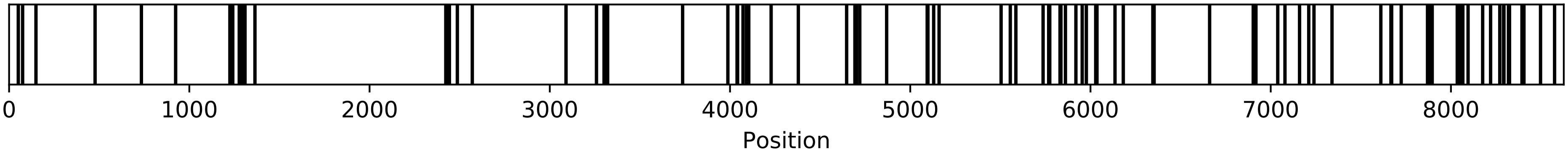


CR2 vs MM1
Number of differences: 183

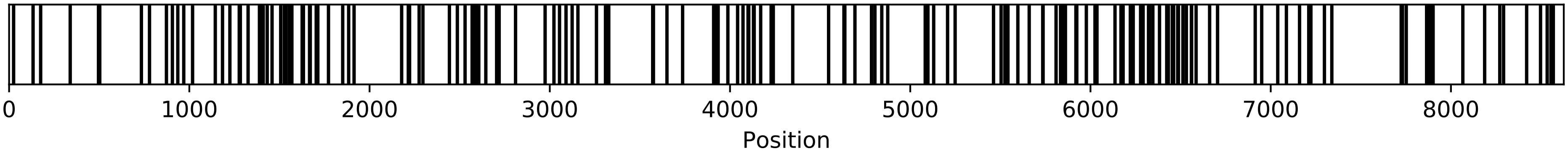


CR2 vs MM2
Number of differences: 181

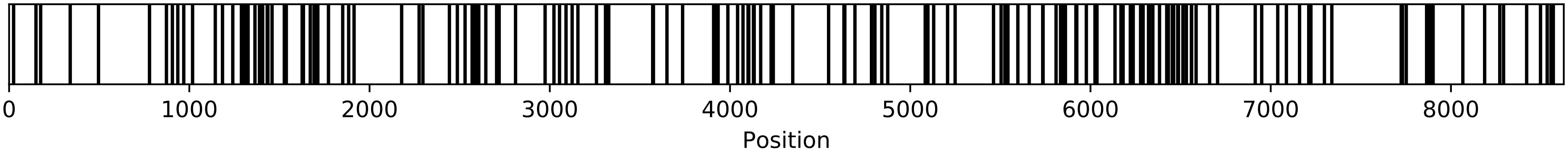


MM1 vs MM2
Number of differences: 25

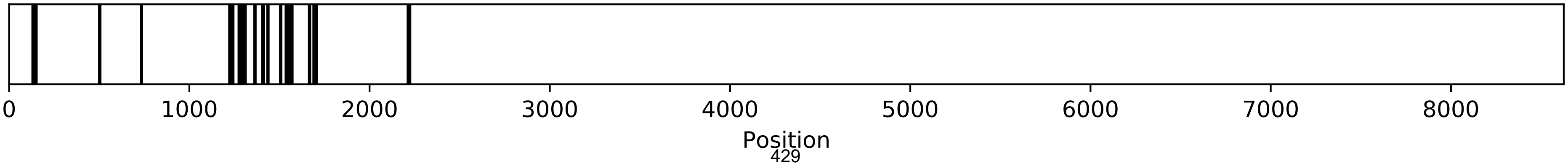

### Region: 1147

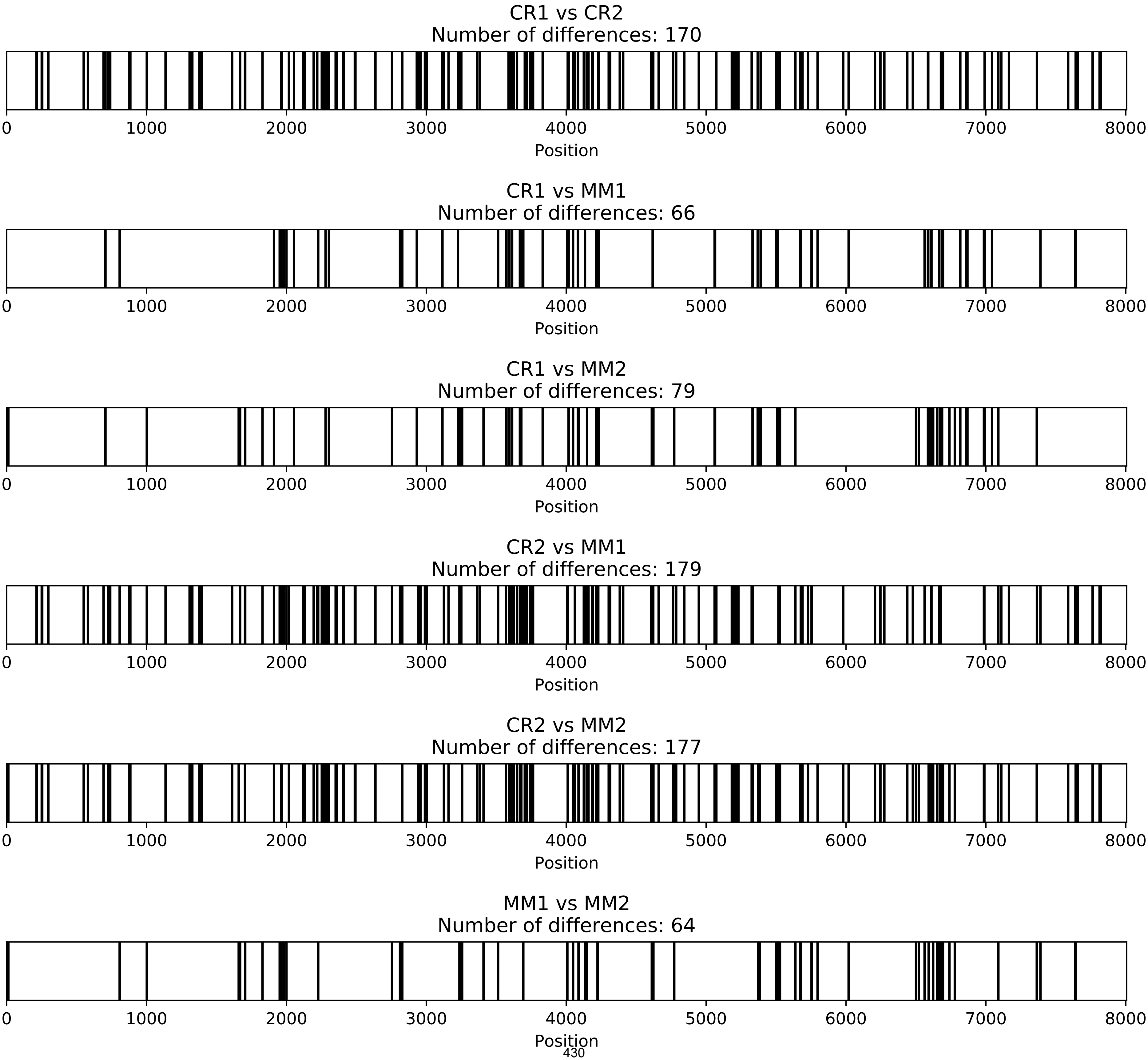
CR1 vs CR2
Number of differences: 170
CR1 vs MM1
Number of differences: 66
CR1 vs MM2
Number of differences: 79
CR2 vs MM1
Number of differences: 179
CR2 vs MM2
Number of differences: 177
MM1 vs MM2
Number of differences: 64
0
1000
2000
3000
4000
5000
6000
7000
8000
Position

### Region: 1148



CR1 vs CR2
Number of differences: 382

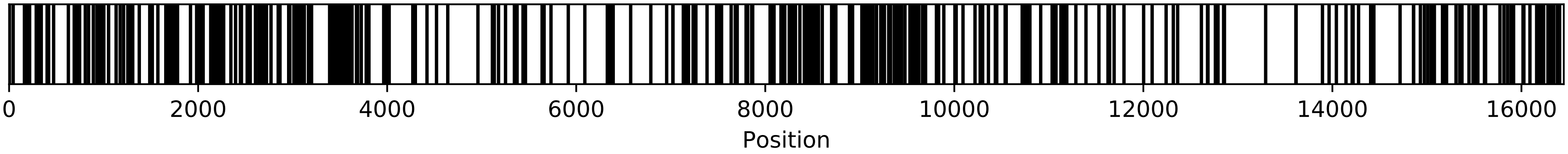


CR1 vs MM1
Number of differences: 164

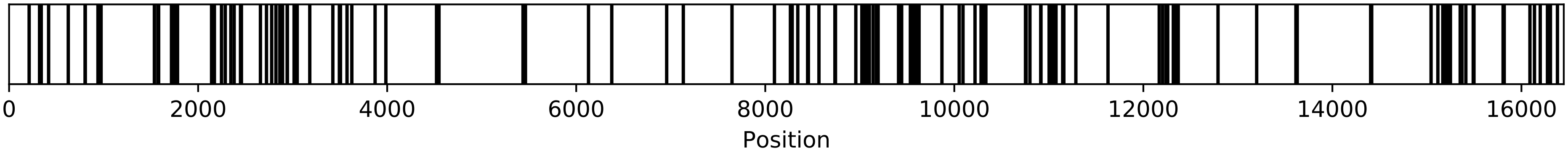


CR1 vs MM2
Number of differences: 147

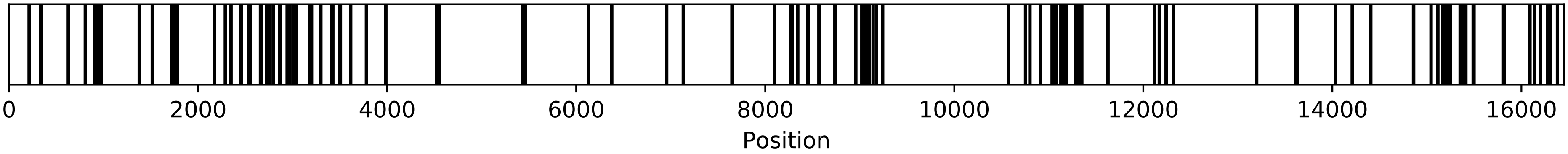


CR2 vs MM1
Number of differences: 369

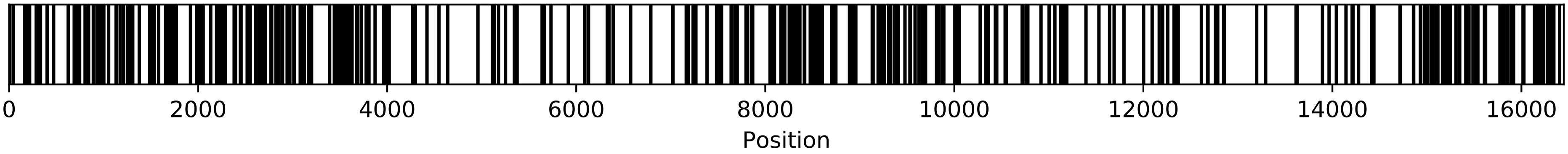


CR2 vs MM2
Number of differences: 364

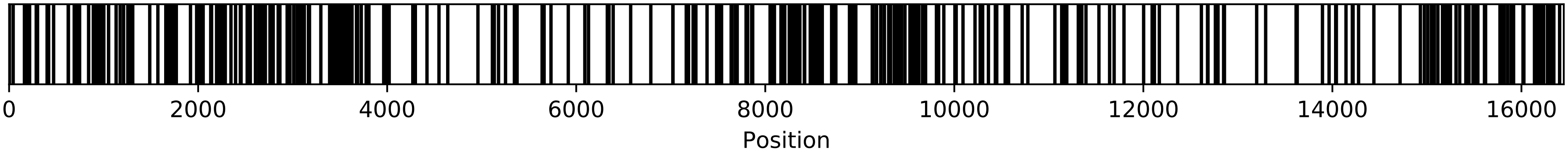


MM1 vs MM2
Number of differences: 86

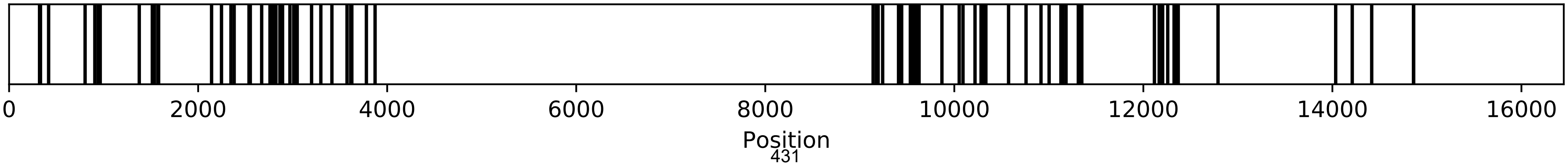

### Region: 1149



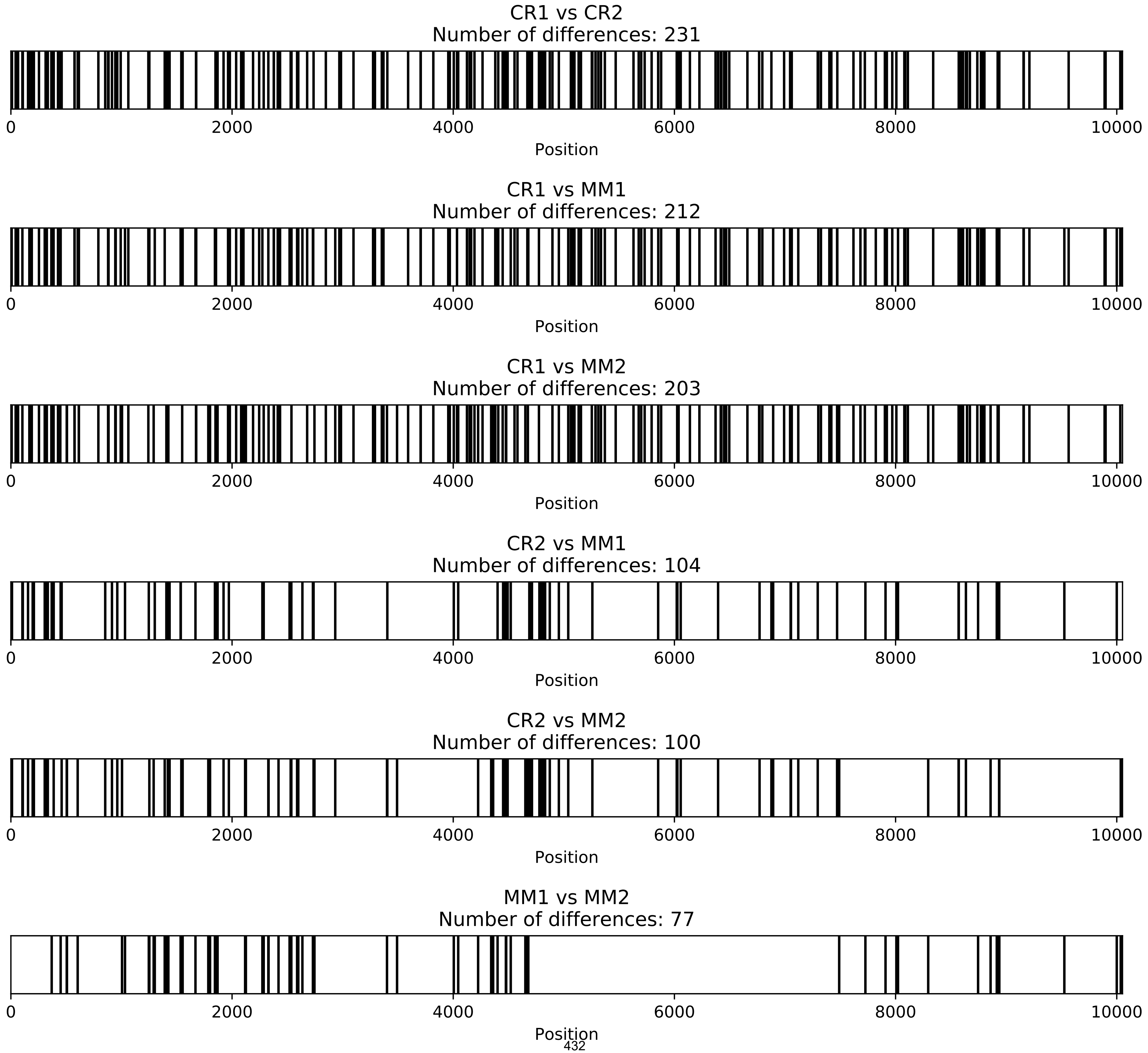
CR1 vs CR2
Number of differences: 231
CR1 vs MM1
Number of differences: 212
CR1 vs MM2
Number of differences: 203
CR2 vs MM1
Number of differences: 104
CR2 vs MM2
Number of differences: 100
MM1 vs MM2
Number of differences: 77
0
2000
4000
6000
8000
10000
Position

### Region: 1150



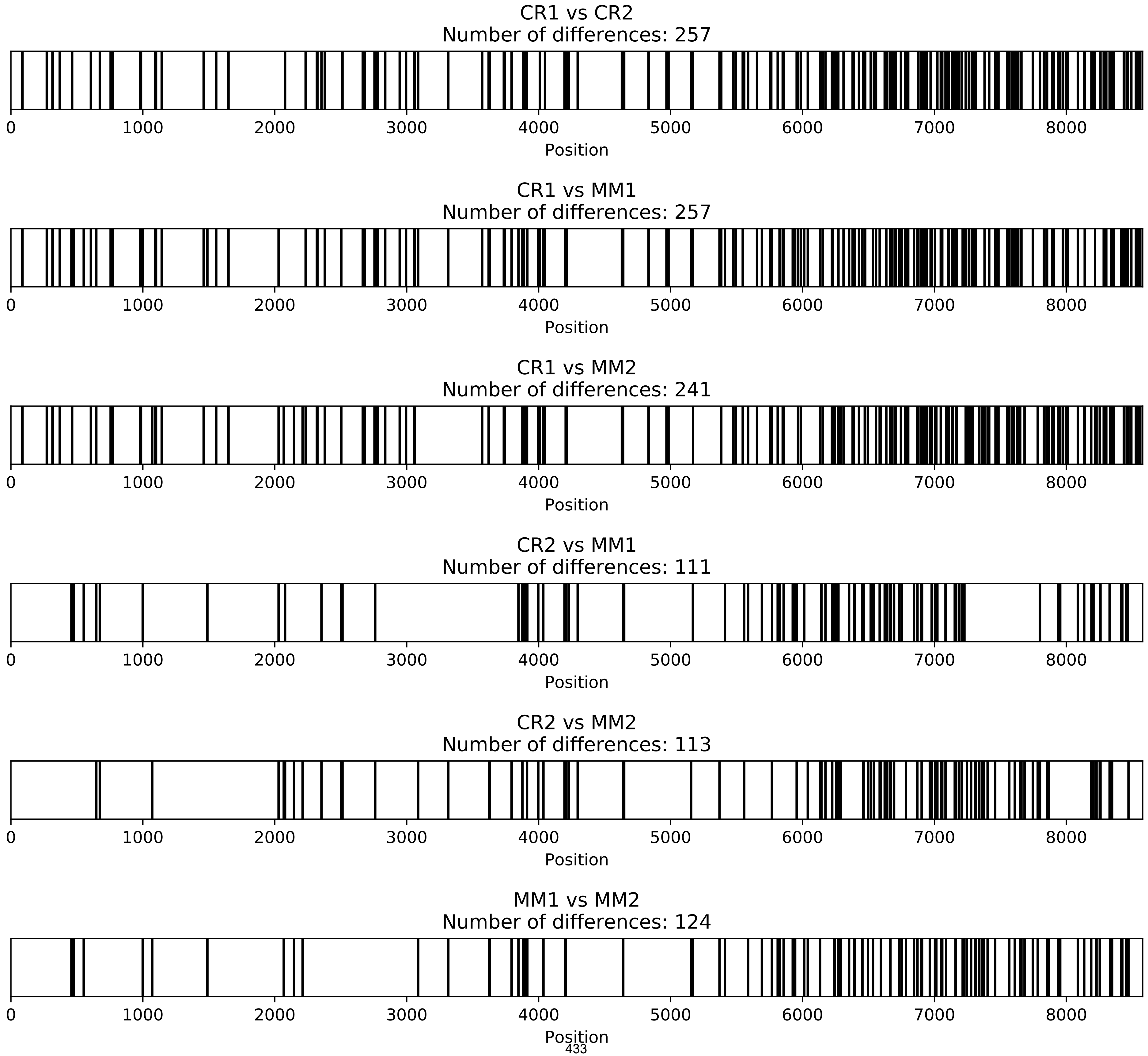
CR1 vs CR2
Number of differences: 257
Position
CR1 vs MM1
Number of differences: 257
Position
CR1 vs MM2
Number of differences: 241
Position
CR2 vs MM1
Number of differences: 111
Position
CR2 vs MM2
Number of differences: 113
Position
MM1 vs MM2
Number of differences: 124
Position
0
1000
2000
3000
4000
5000
6000
7000
8000

### Region: 1151



CR1 vs CR2
Number of differences: 247

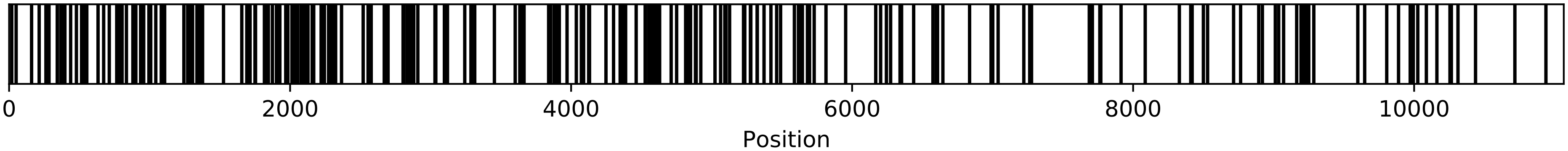


CR1 vs MM1
Number of differences: 111

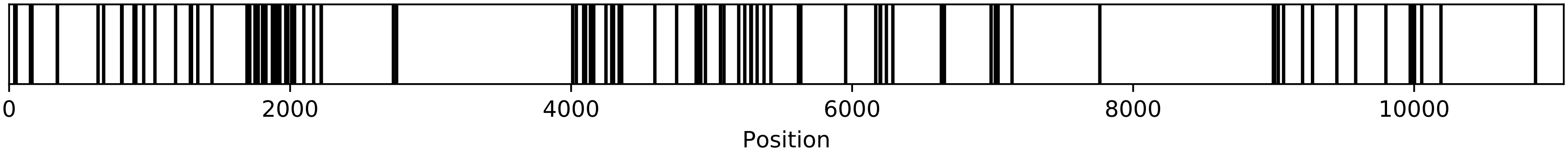


CR1 vs MM2
Number of differences: 131

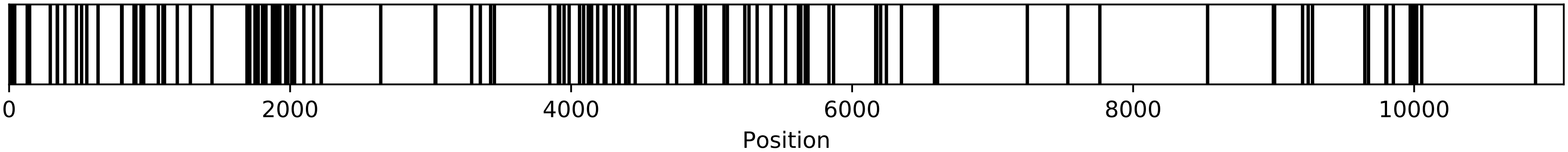


CR2 vs MM1
Number of differences: 244

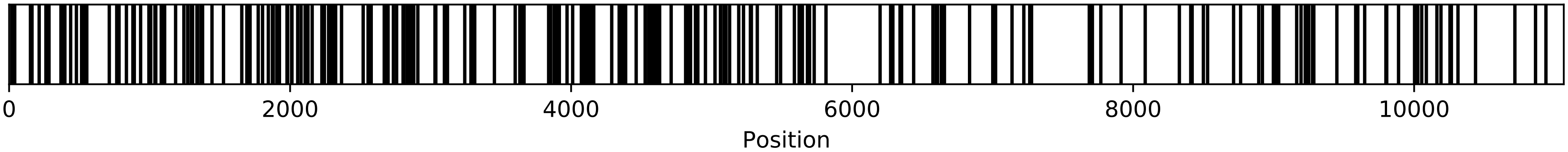


CR2 vs MM2
Number of differences: 253

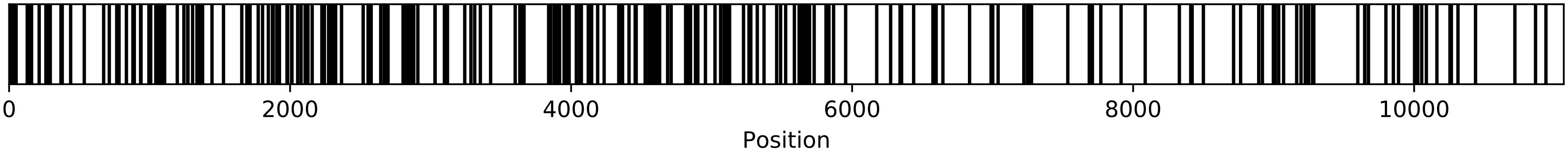


MM1 vs MM2
Number of differences: 103

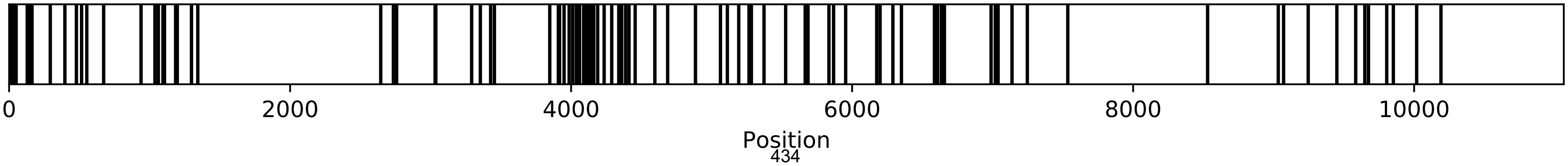

### Region: 1156



CR1 vs CR2
Number of differences: 251

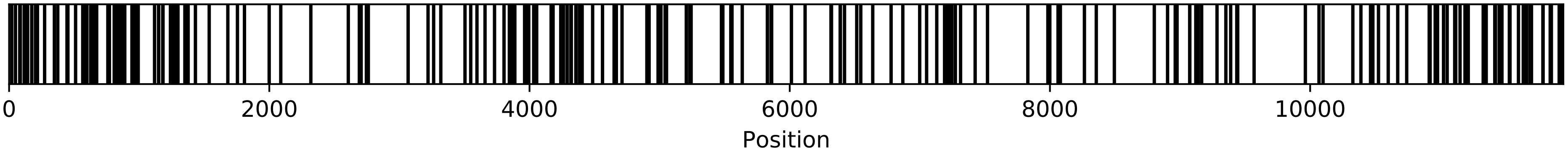


CR1 vs MM1
Number of differences: 120

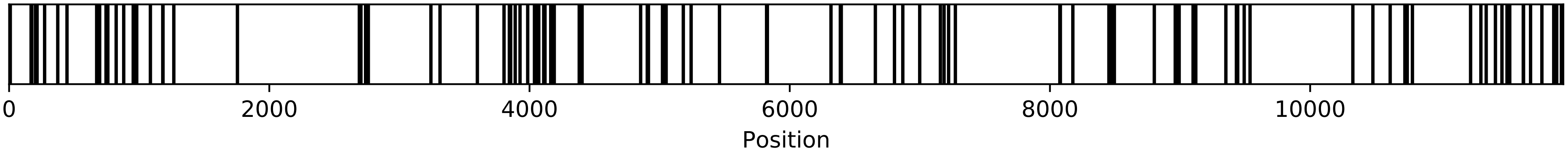


CR1 vs MM2
Number of differences: 121

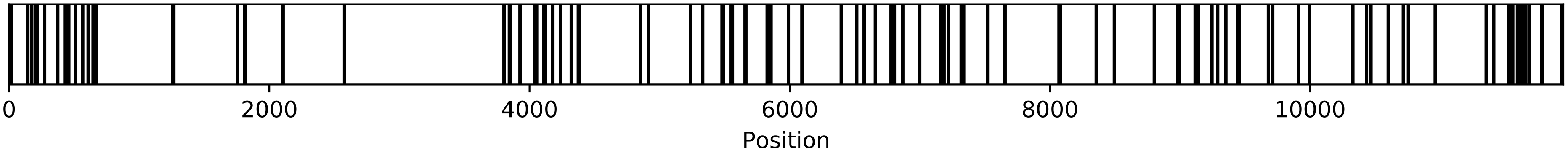


CR2 vs MM1
Number of differences: 263

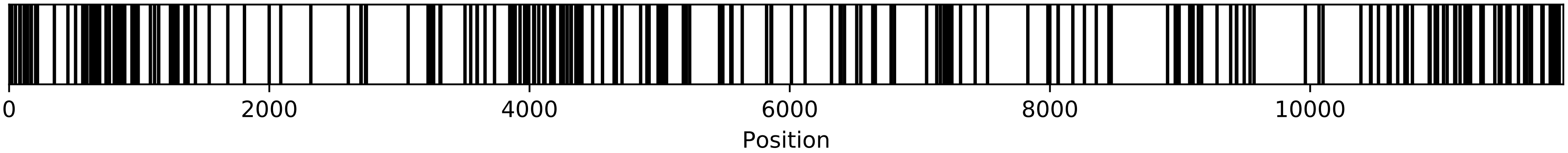


CR2 vs MM2
Number of differences: 264

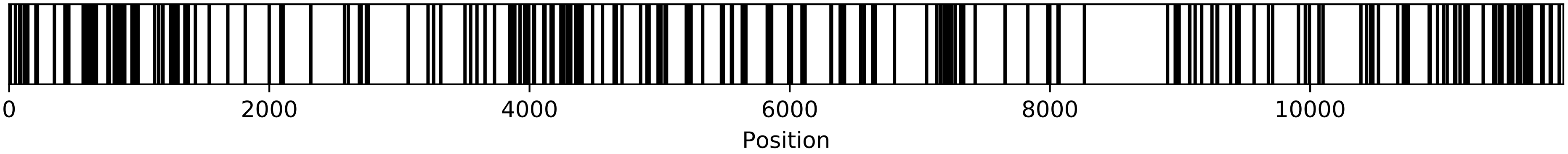


MM1 vs MM2
Number of differences: 139

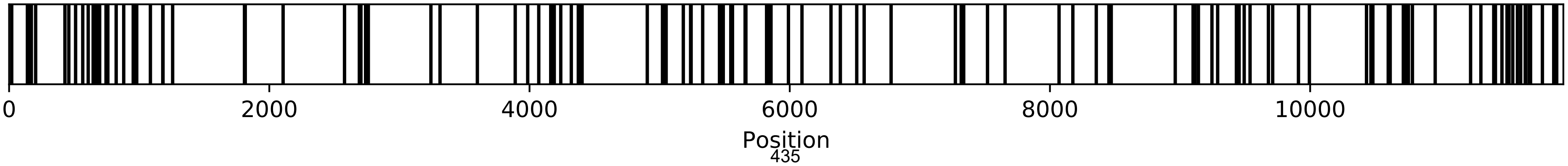

### Region: 1168



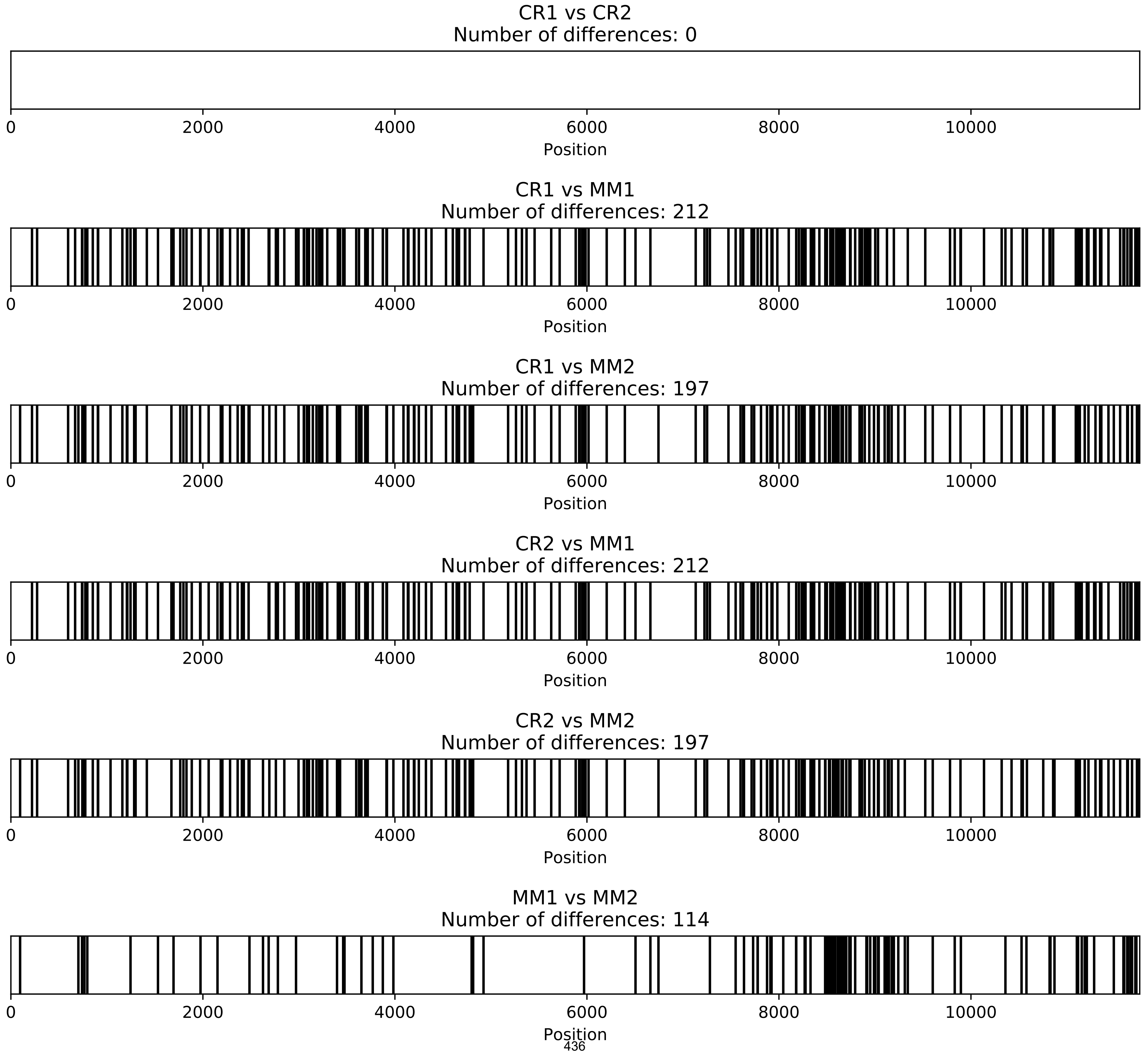
CR1 vs CR2
Number of differences: 0
CR1 vs MM1
Number of differences: 212
CR1 vs MM2
Number of differences: 197
CR2 vs MM1
Number of differences: 212
CR2 vs MM2
Number of differences: 197
MM1 vs MM2
Number of differences: 114
0
2000
4000
6000
8000
10000
Position

### Region: 1171



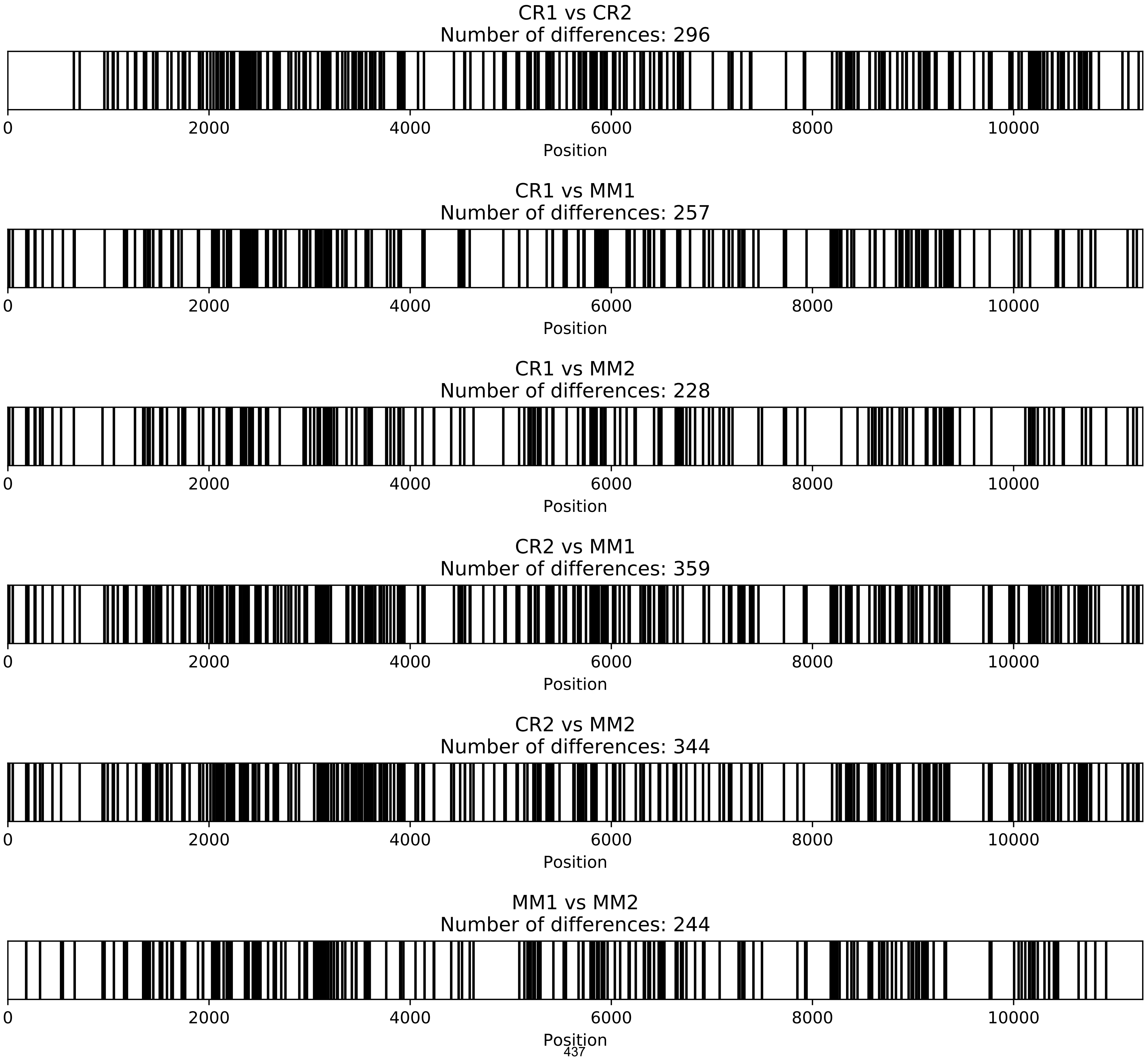
CR1 vs CR2
Number of differences: 296
0
2000
4000
6000
8000
10000
Position
CR1 vs MM1
Number of differences: 257
0
2000
4000
6000
8000
10000
Position
CR1 vs MM2
Number of differences: 228
0
2000
4000
6000
8000
10000
Position
CR2 vs MM1
Number of differences: 359
0
2000
4000
6000
8000
10000
Position
CR2 vs MM2
Number of differences: 344
0
2000
4000
6000
8000
10000
Position
MM1 vs MM2
Number of differences: 244
0
2000
4000
6000
8000
10000
Position

### Region: 1173



CR1 vs CR2
Number of differences: 143

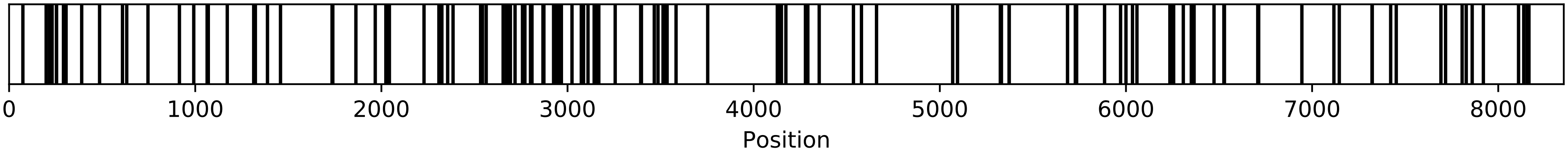


CR1 vs MM1
Number of differences: 80

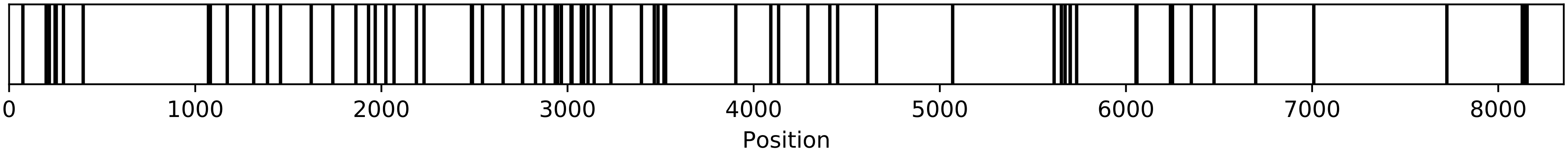


CR1 vs MM2
Number of differences: 79

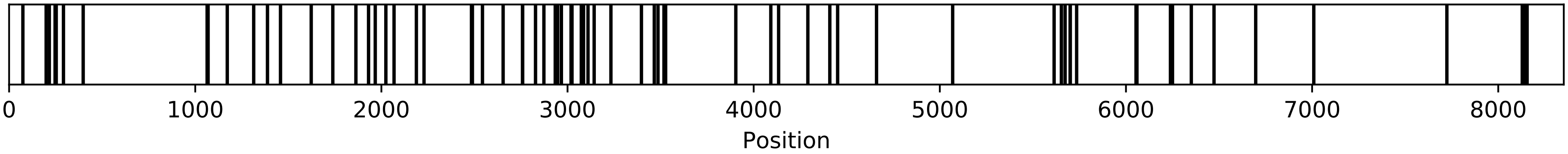


CR2 vs MM1
Number of differences: 142

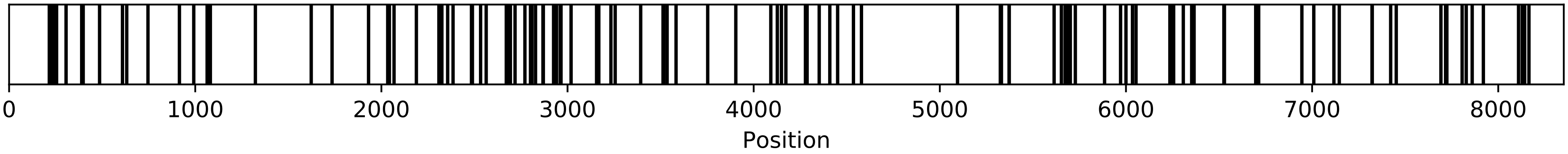


CR2 vs MM2
Number of differences: 135

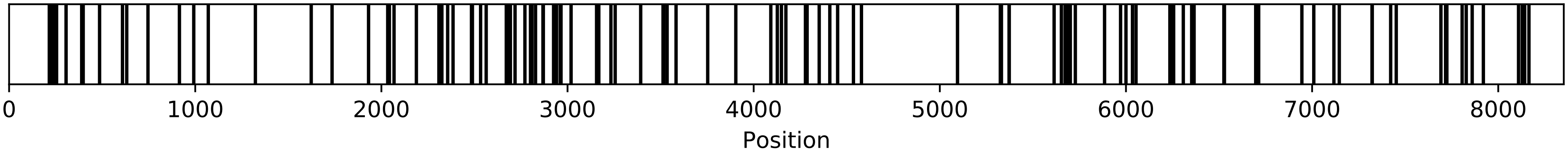


MM1 vs MM2
Number of differences: 7

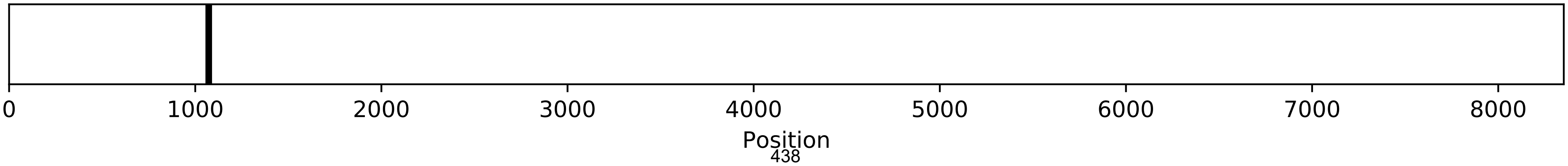

### Region: 1174



CR1 vs CR2
Number of differences: 149

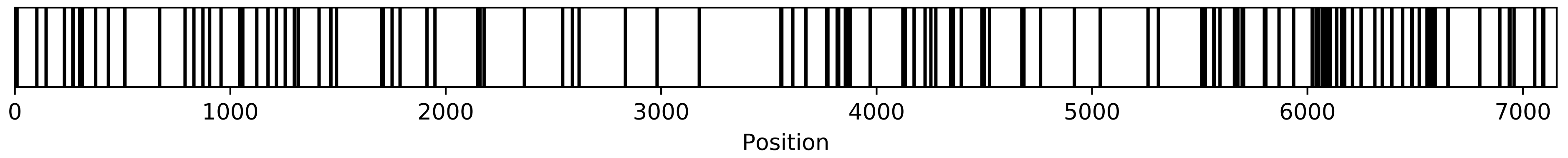


CR1 vs MM1
Number of differences: 169

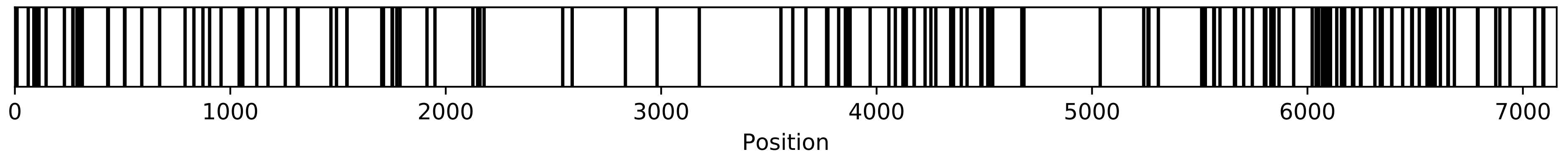


CR1 vs MM2
Number of differences: 150

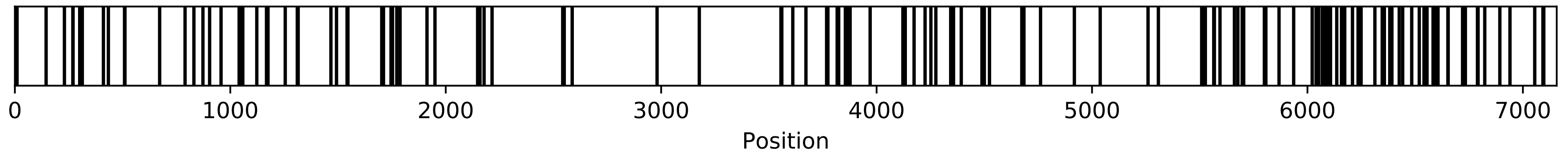


CR2 vs MM1
Number of differences: 64

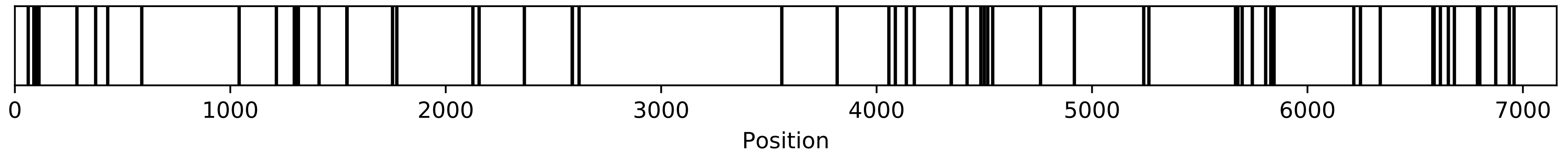


CR2 vs MM2
Number of differences: 49

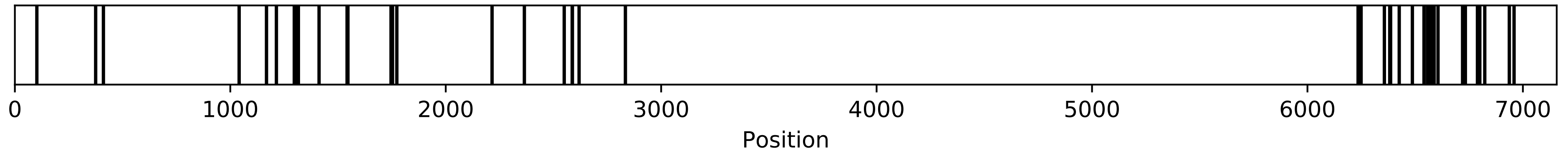


MM1 vs MM2
Number of differences: 69

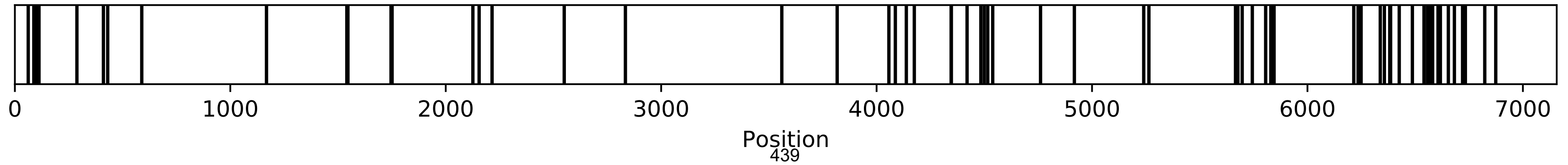

### Region: 1179



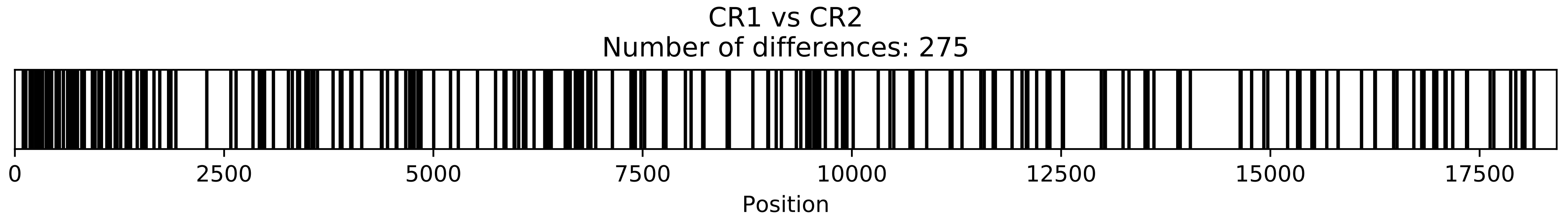
CR1 vs CR2
Number of differences: 275
0
2500
5000
7500
10000
12500
15000
17500
Position

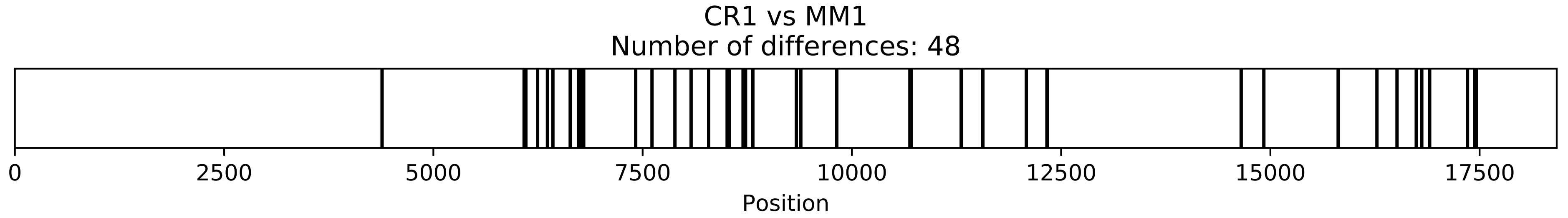
CR1 vs MM1
Number of differences: 48
0
2500
5000
7500
10000
12500
15000
17500
Position

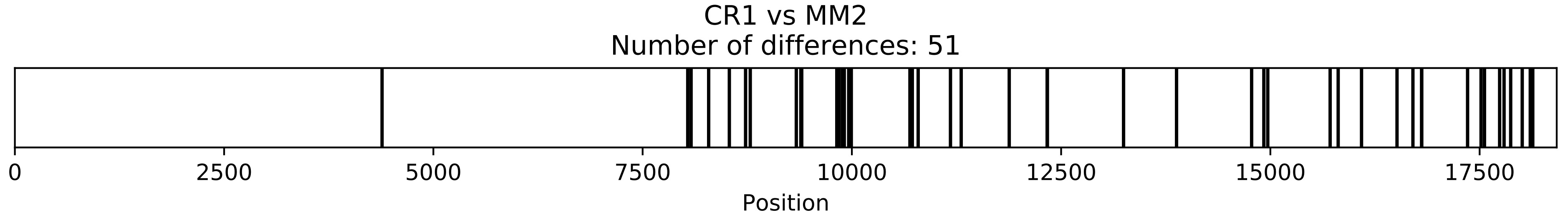
CR1 vs MM2
Number of differences: 51
0
2500
5000
7500
10000
12500
15000
17500
Position

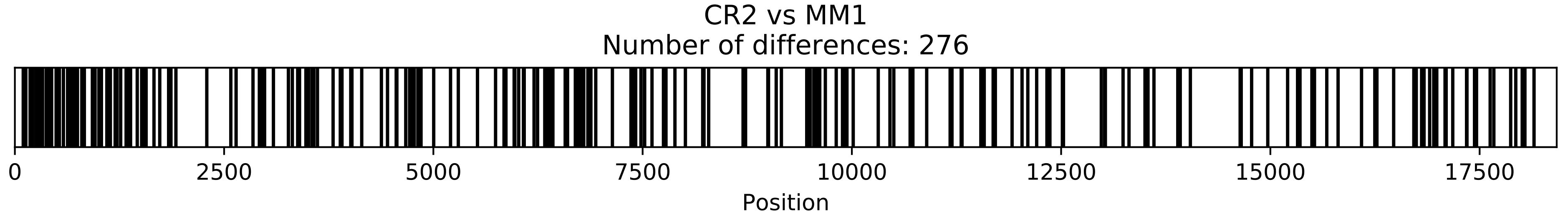
CR2 vs MM1
Number of differences: 276
0
2500
5000
7500
10000
12500
15000
17500
Position

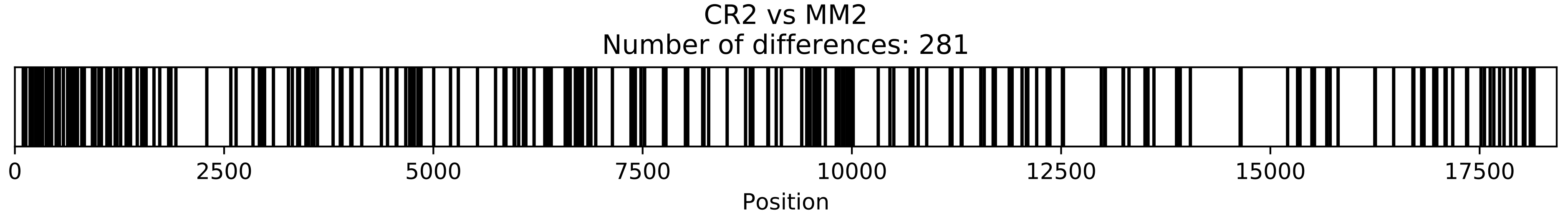
CR2 vs MM2
Number of differences: 281
0
2500
5000
7500
10000
12500
15000
17500
Position

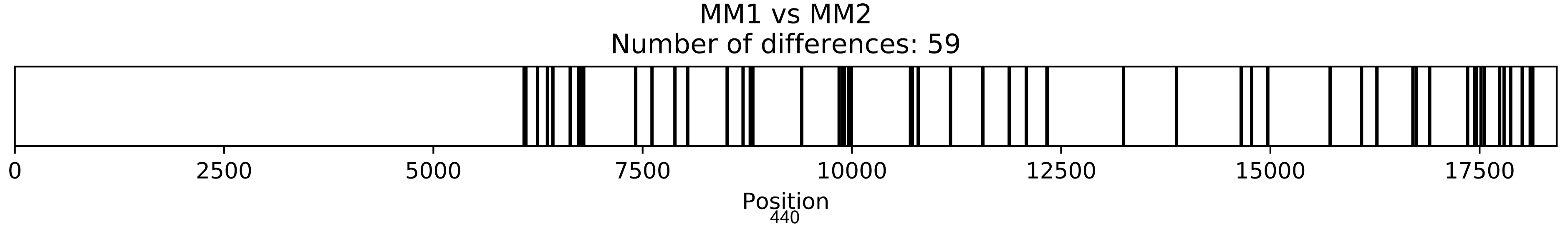
MM1 vs MM2
Number of differences: 59
0
2500
5000
7500
10000
12500
15000
17500
Position

### Region: 1180



CR1 vs CR2
Number of differences: 228

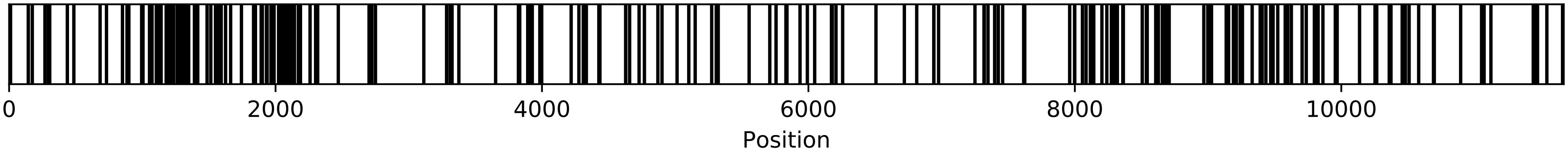


CR1 vs MM1
Number of differences: 206

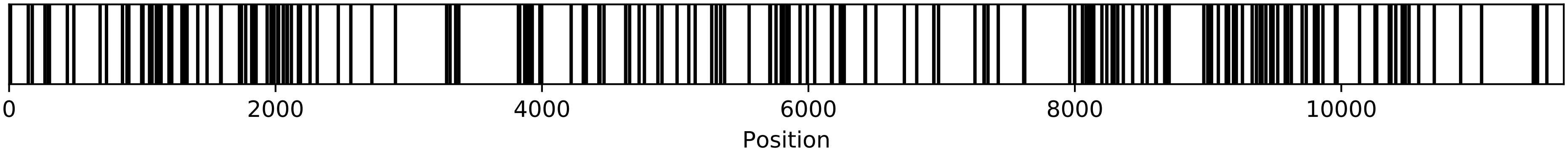


CR1 vs MM2
Number of differences: 221

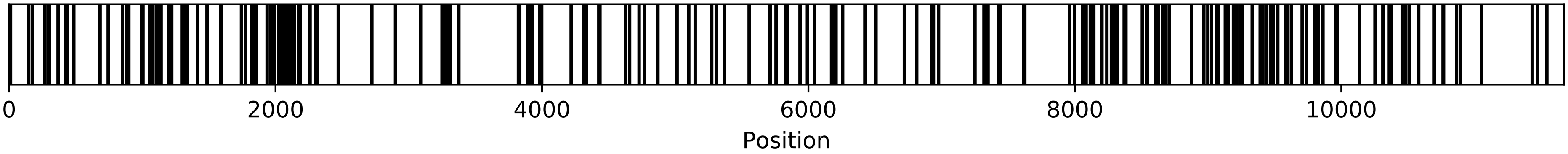


CR2 vs MM1
Number of differences: 91

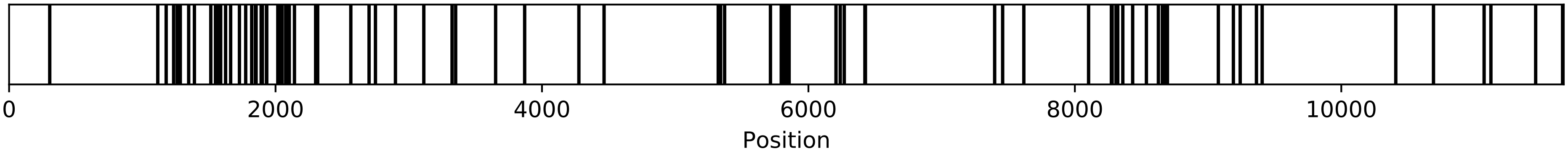


CR2 vs MM2
Number of differences: 77

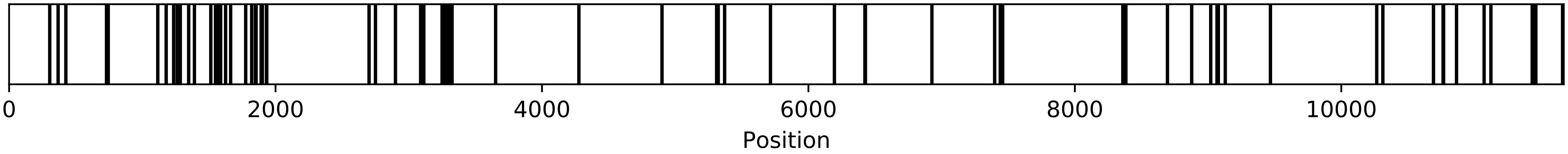


MM1 vs MM2
Number of differences: 80

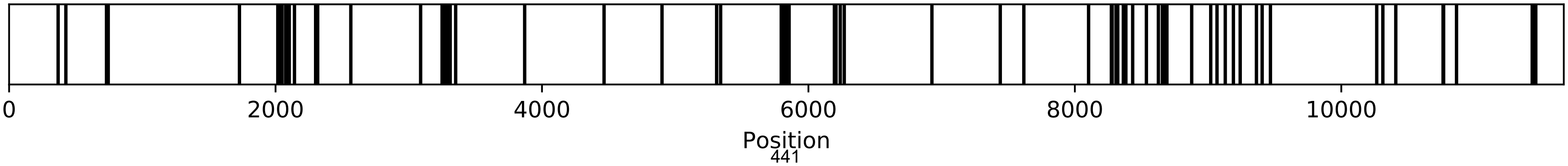

### Region: 1185



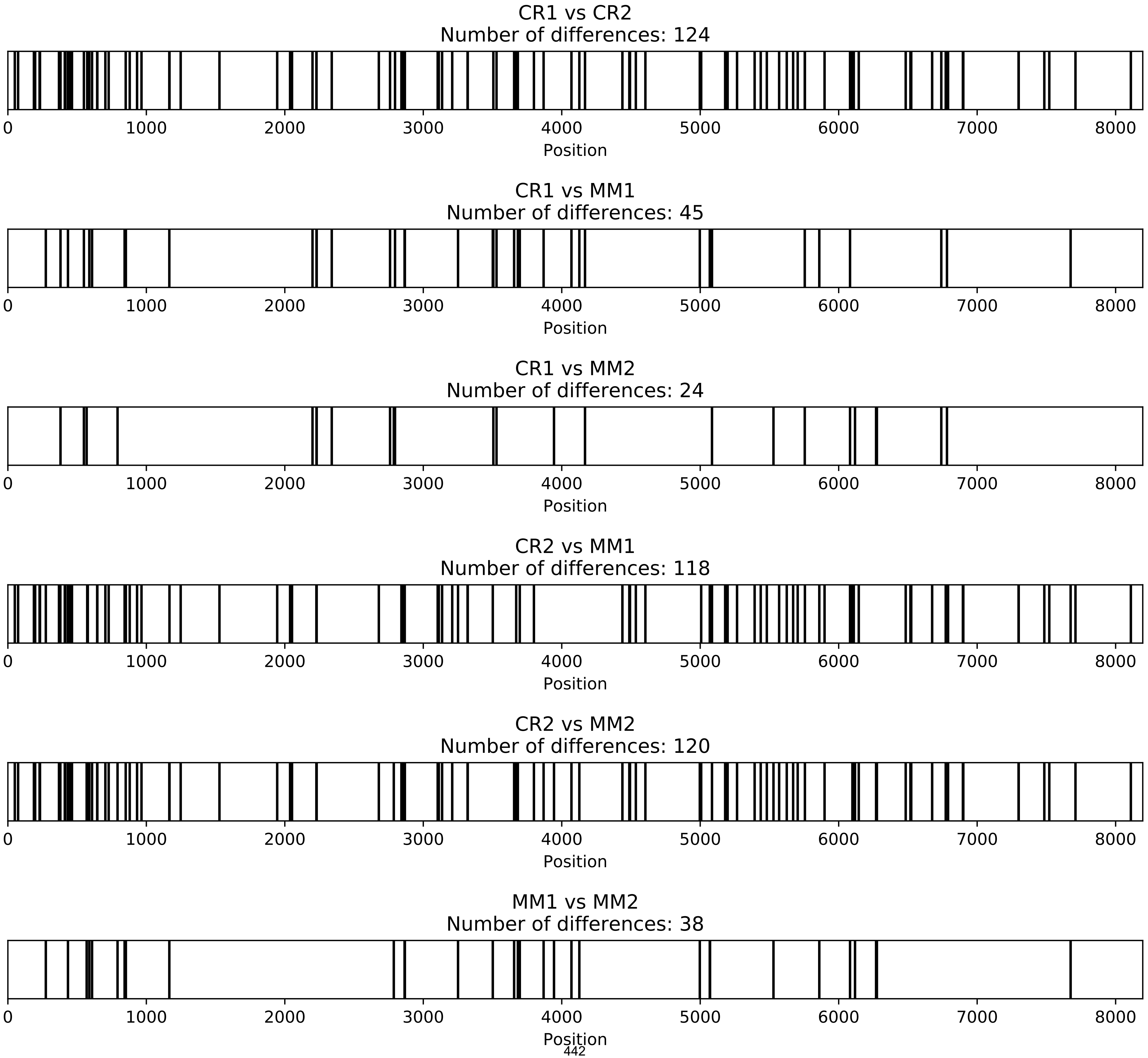
CR1 vs CR2
Number of differences: 124
0 1000 2000 3000 4000 5000 6000 7000 8000
Position
CR1 vs MM1
Number of differences: 45
0 1000 2000 3000 4000 5000 6000 7000 8000
Position
CR1 vs MM2
Number of differences: 24
0 1000 2000 3000 4000 5000 6000 7000 8000
Position
CR2 vs MM1
Number of differences: 118
0 1000 2000 3000 4000 5000 6000 7000 8000
Position
CR2 vs MM2
Number of differences: 120
0 1000 2000 3000 4000 5000 6000 7000 8000
Position
MM1 vs MM2
Number of differences: 38
0 1000 2000 3000 4000 5000 6000 7000 8000
Position

### Region: 1186



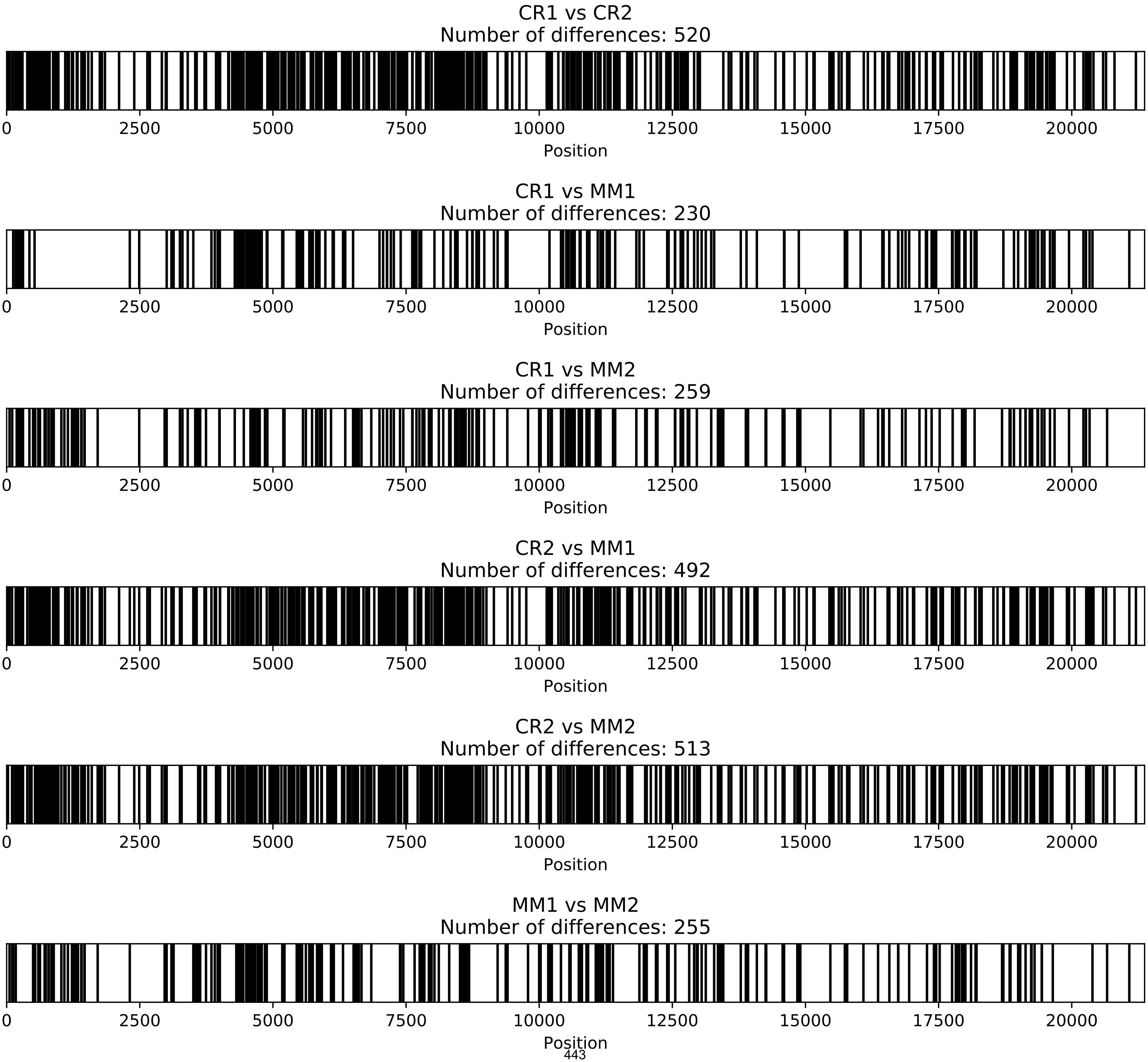
CR1 vs CR2
Number of differences: 520
0 2500 5000 7500 10000 12500 15000 17500 20000
Position
CR1 vs MM1
Number of differences: 230
0 2500 5000 7500 10000 12500 15000 17500 20000
Position
CR1 vs MM2
Number of differences: 259
0 2500 5000 7500 10000 12500 15000 17500 20000
Position
CR2 vs MM1
Number of differences: 492
0 2500 5000 7500 10000 12500 15000 17500 20000
Position
CR2 vs MM2
Number of differences: 513
0 2500 5000 7500 10000 12500 15000 17500 20000
Position
MM1 vs MM2
Number of differences: 255
0 2500 5000 7500 10000 12500 15000 17500 20000
Position

### Region: 1187



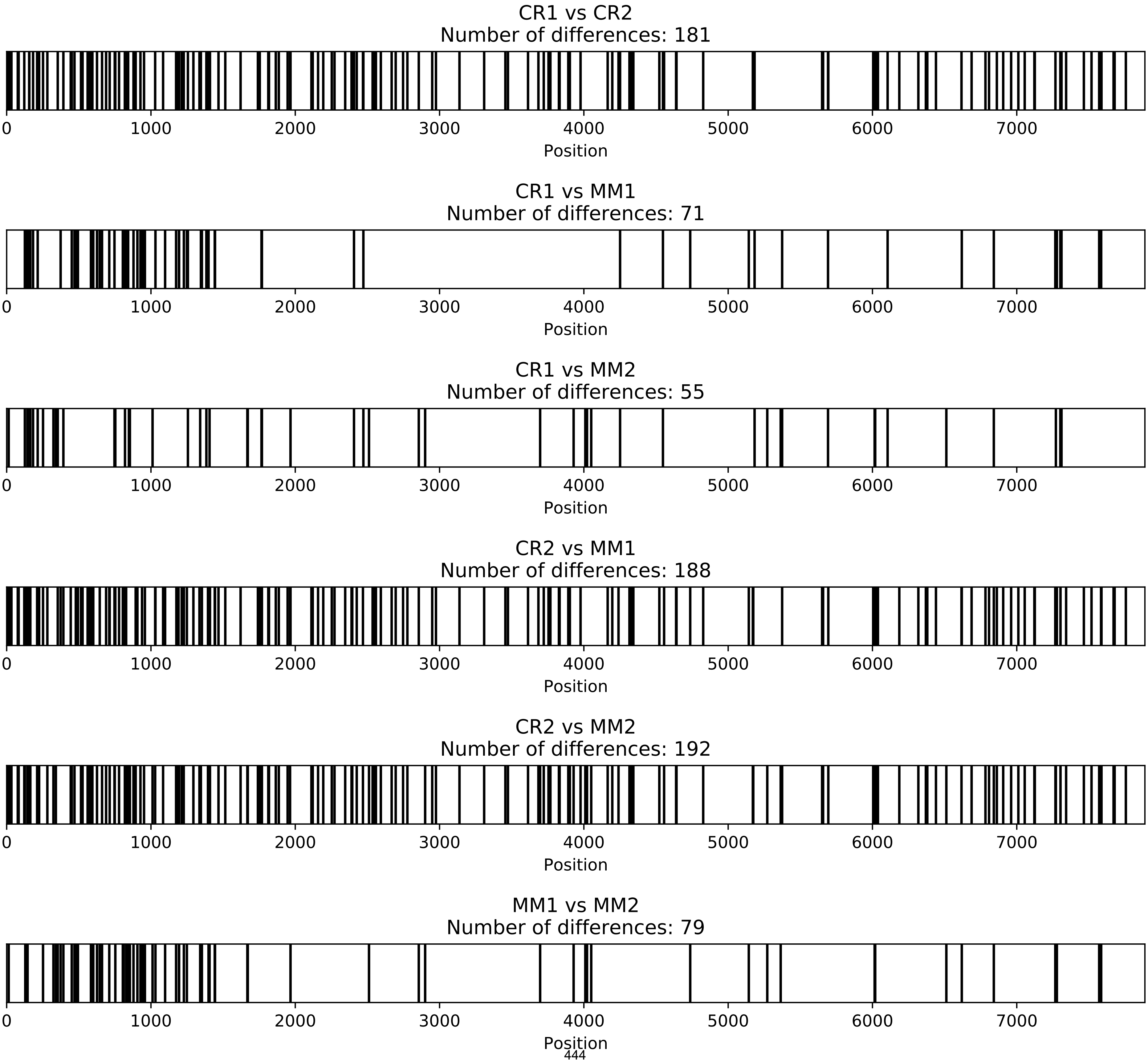
CR1 vs CR2
Number of differences: 181
0 1000 2000 3000 4000 5000 6000 7000
Position
CR1 vs MM1
Number of differences: 71
0 1000 2000 3000 4000 5000 6000 7000
Position
CR1 vs MM2
Number of differences: 55
0 1000 2000 3000 4000 5000 6000 7000
Position
CR2 vs MM1
Number of differences: 188
0 1000 2000 3000 4000 5000 6000 7000
Position
CR2 vs MM2
Number of differences: 192
0 1000 2000 3000 4000 5000 6000 7000
Position
MM1 vs MM2
Number of differences: 79
0 1000 2000 3000 4000 5000 6000 7000
Position

### Region: 1191



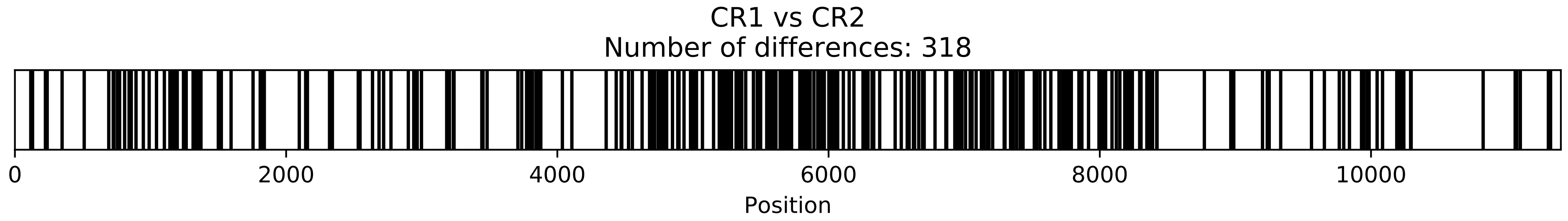
CR1 vs CR2
Number of differences: 318
0
2000
4000
6000
8000
10000
Position

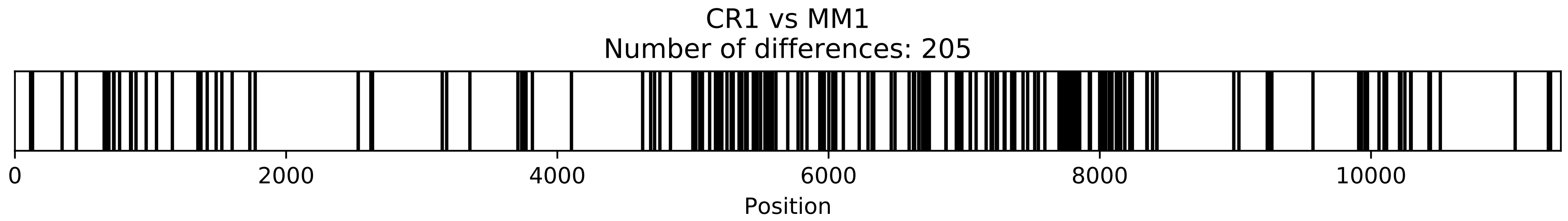
CR1 vs MM1
Number of differences: 205
0
2000
4000
6000
8000
10000
Position

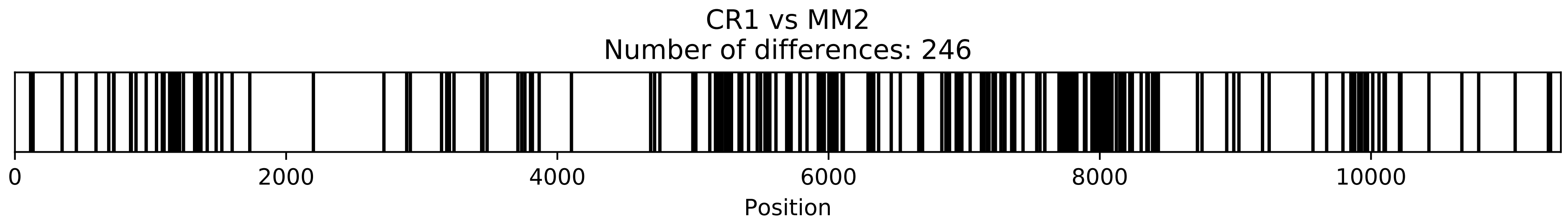
CR1 vs MM2
Number of differences: 246
0
2000
4000
6000
8000
10000
Position

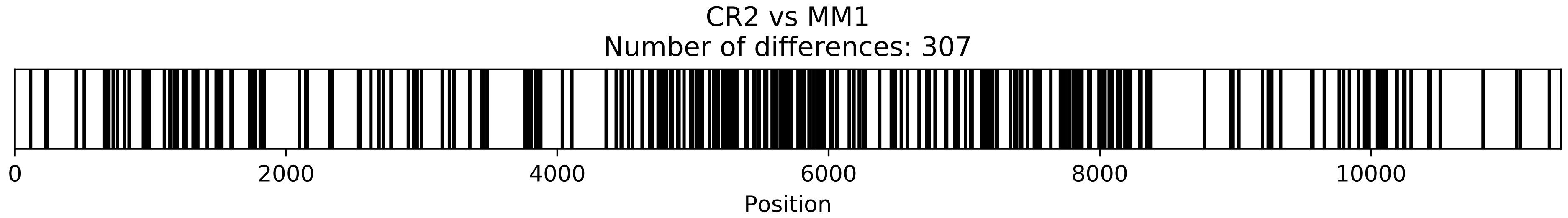
CR2 vs MM1
Number of differences: 307
0
2000
4000
6000
8000
10000
Position

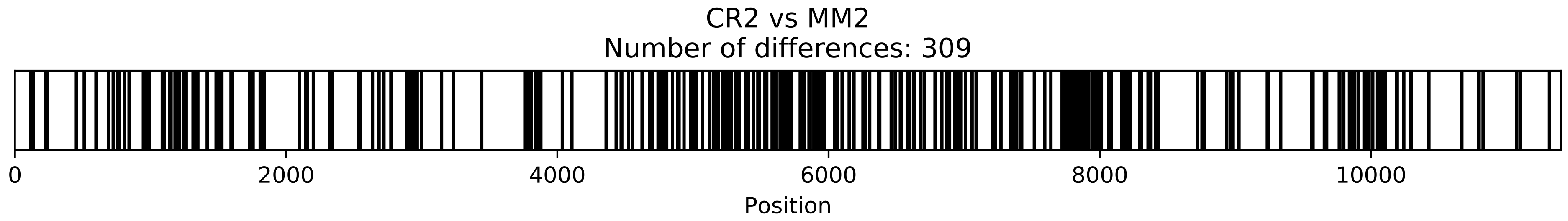
CR2 vs MM2
Number of differences: 309
0
2000
4000
6000
8000
10000
Position

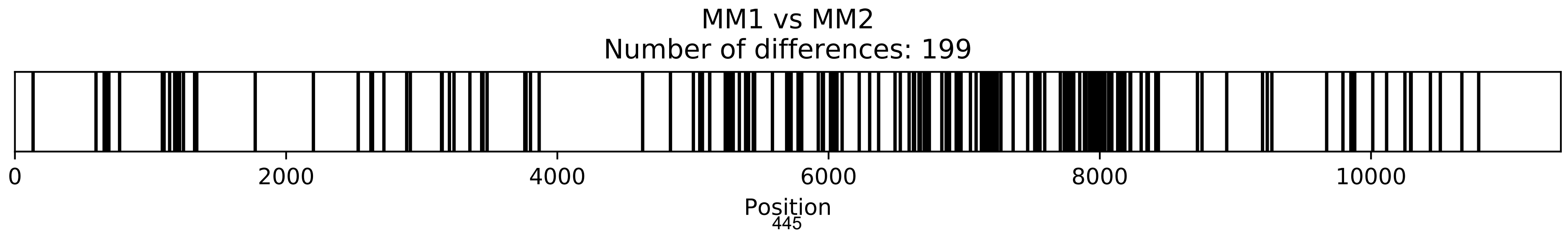
MM1 vs MM2
Number of differences: 199
0
2000
4000
6000
8000
10000
Position

### Region: 6

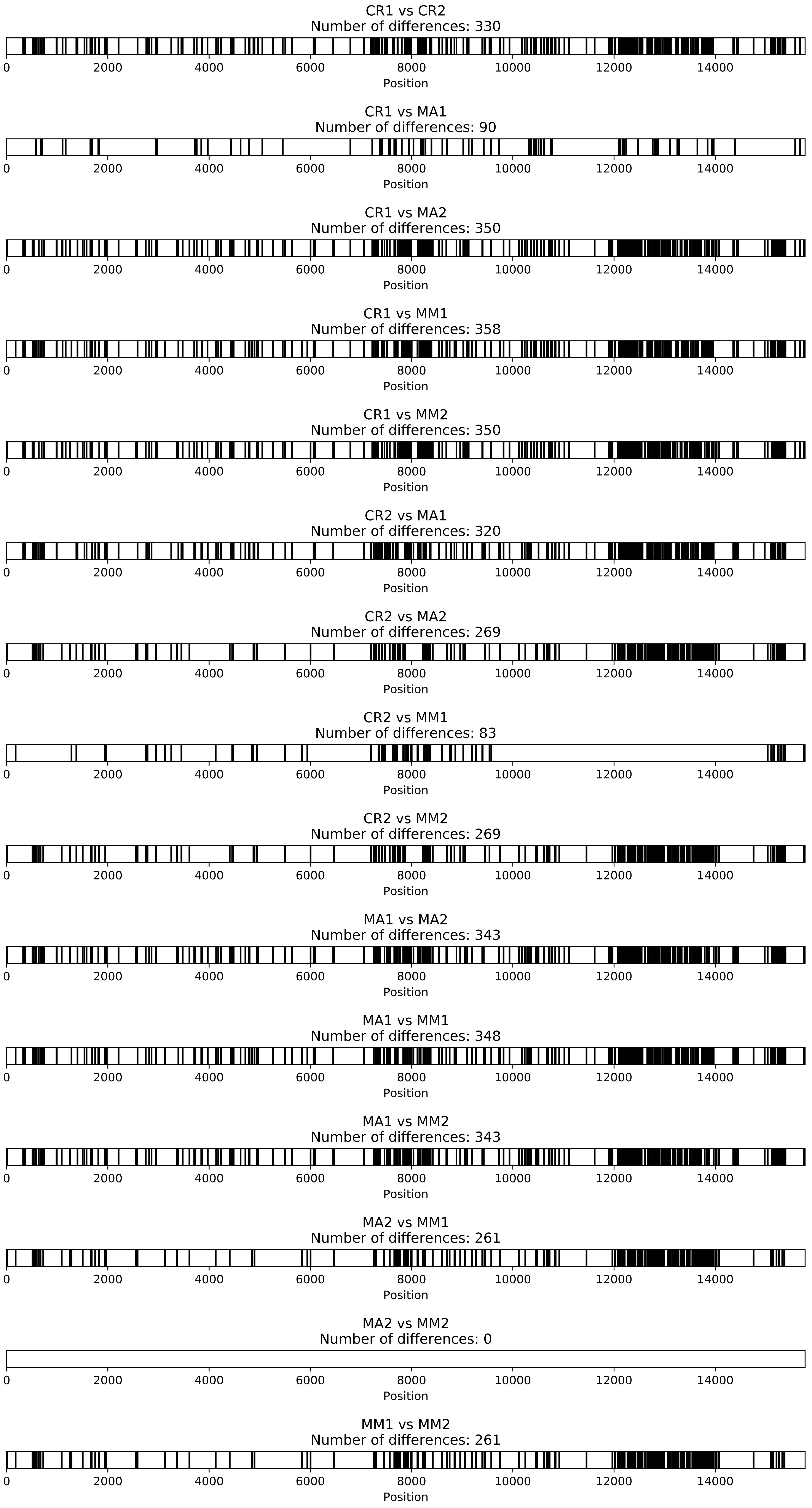
CR1 vs CR2
Number of differences: 330
CR1 vs MA1
Number of differences: 90
CR1 vs MA2
Number of differences: 350
CR1 vs MM1
Number of differences: 358
CR1 vs MM2
Number of differences: 350
CR2 vs MA1
Number of differences: 320
CR2 vs MA2
Number of differences: 269
CR2 vs MM1
Number of differences: 83
CR2 vs MM2
Number of differences: 269
MA1 vs MA2
Number of differences: 343
MA1 vs MM1
Number of differences: 348
MA1 vs MM2
Number of differences: 343
MA2 vs MM1
Number of differences: 261
MA2 vs MM2
Number of differences: 0
MM1 vs MM2
Number of differences: 261
0
2000
4000
6000
8000
10000
12000
14000
Position

### Region: 7



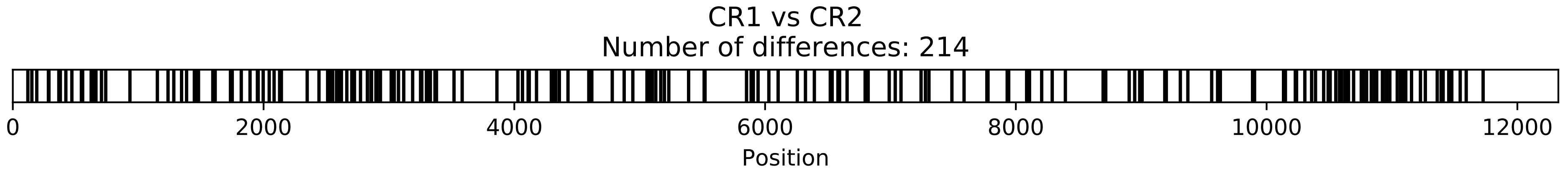
CR1 vs CR2
Number of differences: 214
0 2000 4000 6000 8000 10000 12000
Position

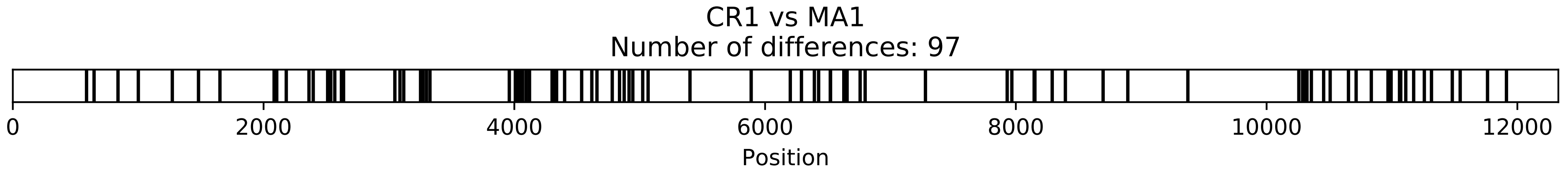
CR1 vs MA1
Number of differences: 97
0 2000 4000 6000 8000 10000 12000
Position

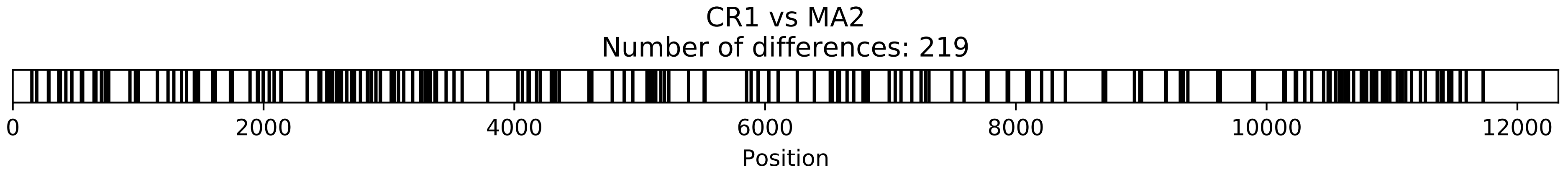
CR1 vs MA2
Number of differences: 219
0 2000 4000 6000 8000 10000 12000
Position

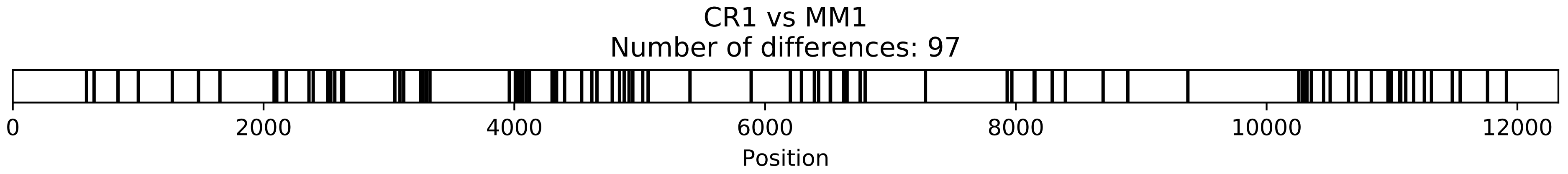
CR1 vs MM1
Number of differences: 97
0 2000 4000 6000 8000 10000 12000
Position

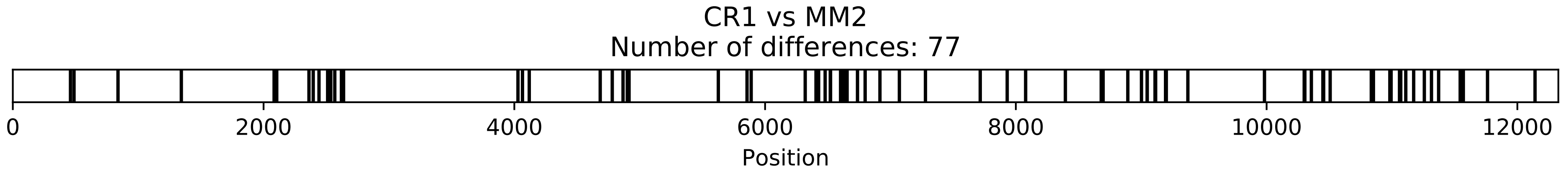
CR1 vs MM2
Number of differences: 77
0 2000 4000 6000 8000 10000 12000
Position

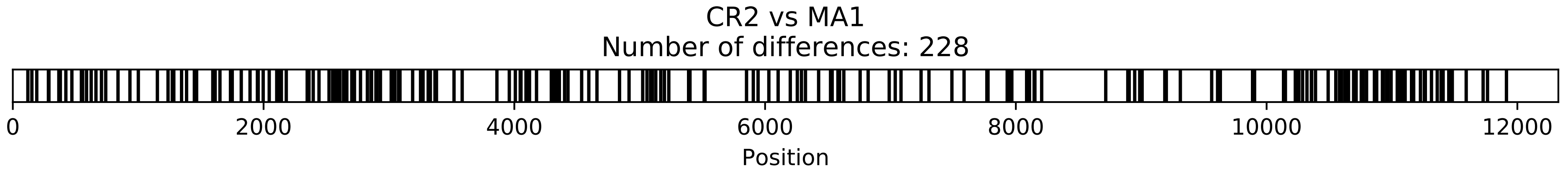
CR2 vs MA1
Number of differences: 228
0 2000 4000 6000 8000 10000 12000
Position

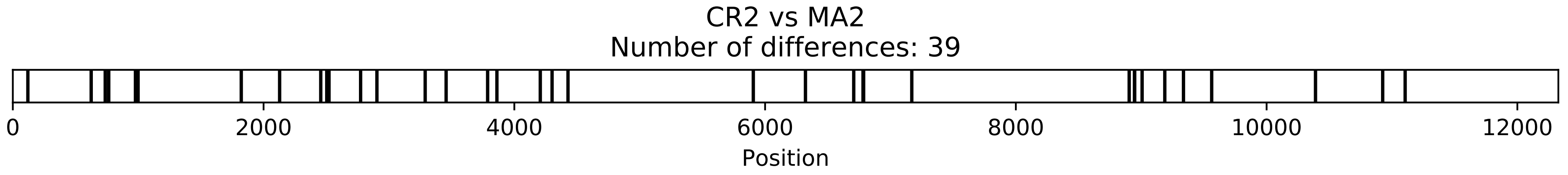
CR2 vs MA2
Number of differences: 39
0 2000 4000 6000 8000 10000 12000
Position

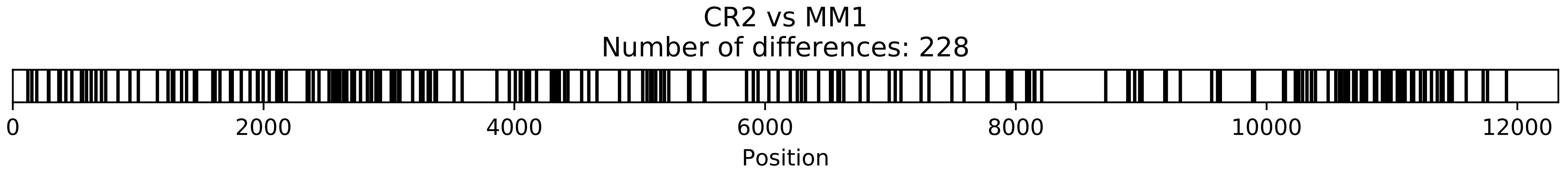
CR2 vs MM1
Number of differences: 228
0 2000 4000 6000 8000 10000 12000
Position

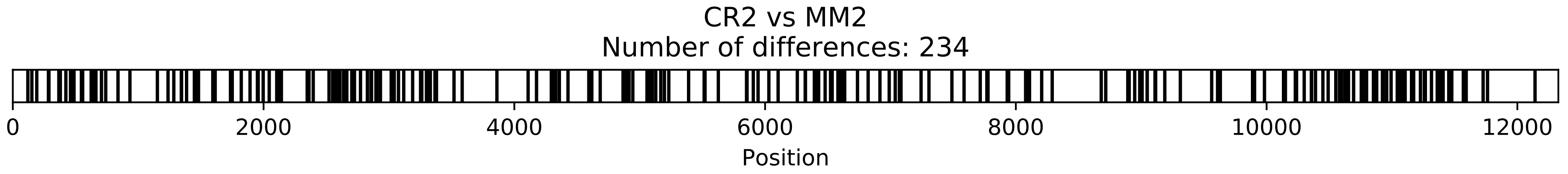
CR2 vs MM2
Number of differences: 234
0 2000 4000 6000 8000 10000 12000
Position

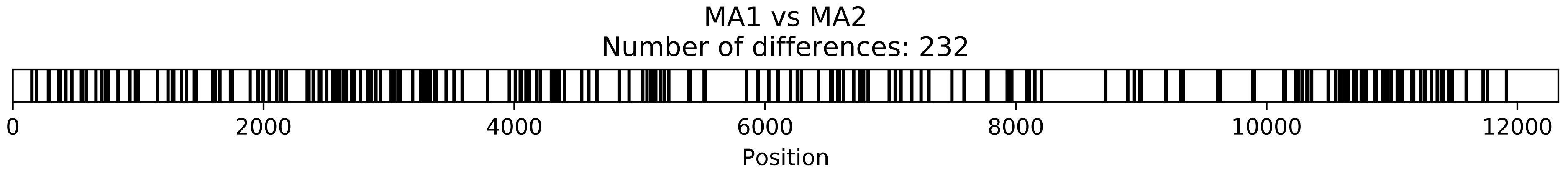
MA1 vs MA2
Number of differences: 232
0 2000 4000 6000 8000 10000 12000
Position

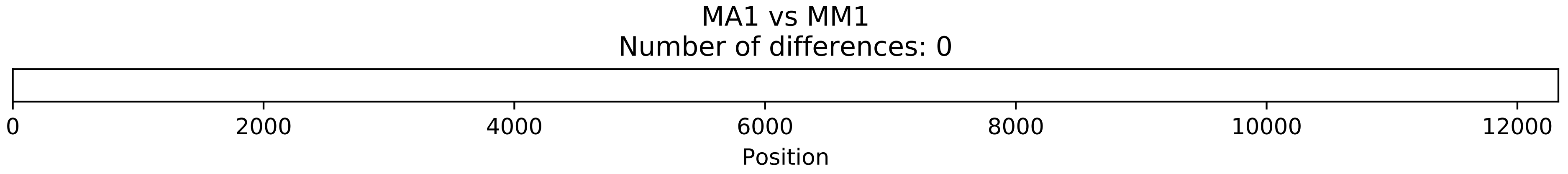
MA1 vs MM1
Number of differences: 0
0 2000 4000 6000 8000 10000 12000
Position

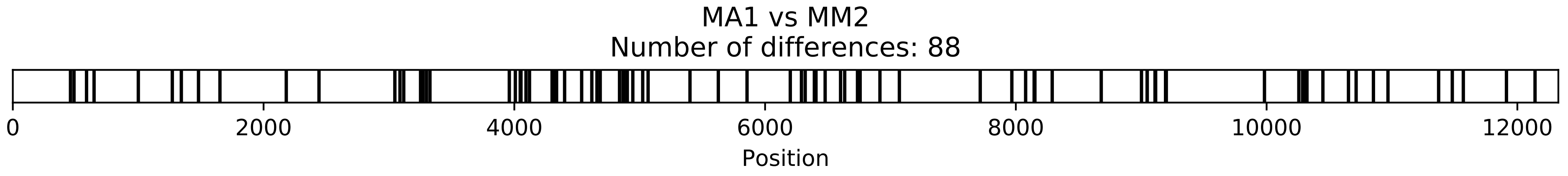
MA1 vs MM2
Number of differences: 88
0 2000 4000 6000 8000 10000 12000
Position

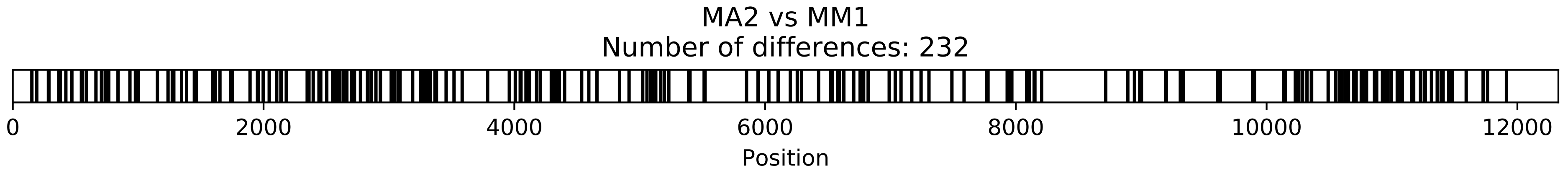
MA2 vs MM1
Number of differences: 232
0 2000 4000 6000 8000 10000 12000
Position

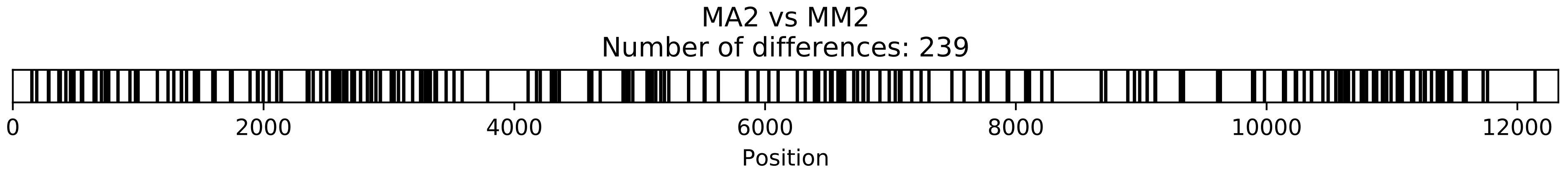
MA2 vs MM2
Number of differences: 239
0 2000 4000 6000 8000 10000 12000
Position

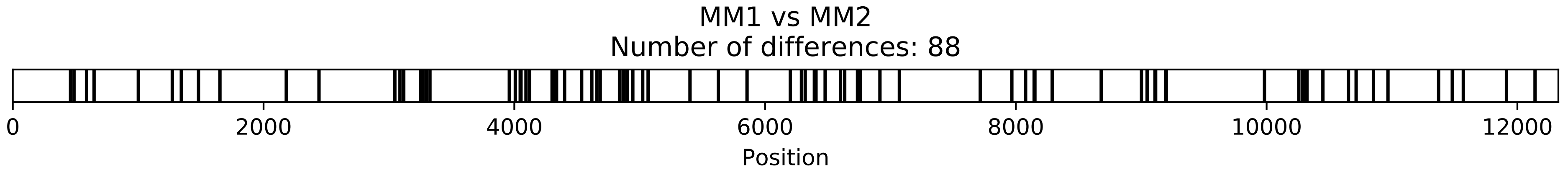
MM1 vs MM2
Number of differences: 88
0 2000 4000 6000 8000 10000 12000
Position

### Region: 8

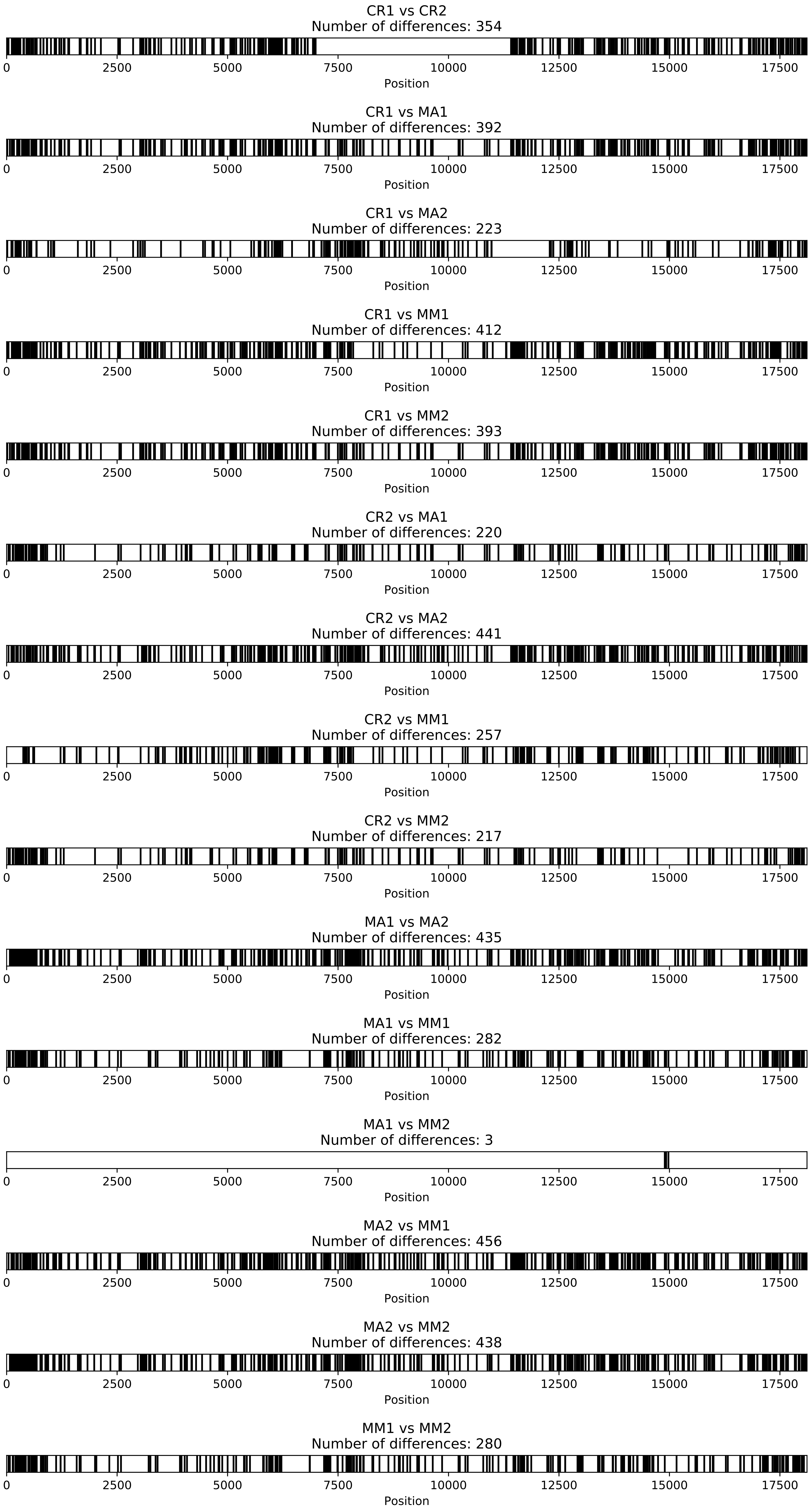
CR1 vs CR2
Number of differences: 354
CR1 vs MA1
Number of differences: 392
CR1 vs MA2
Number of differences: 223
CR1 vs MM1
Number of differences: 412
CR1 vs MM2
Number of differences: 393
CR2 vs MA1
Number of differences: 220
CR2 vs MA2
Number of differences: 441
CR2 vs MM1
Number of differences: 257
CR2 vs MM2
Number of differences: 217
MA1 vs MA2
Number of differences: 435
MA1 vs MM1
Number of differences: 282
MA1 vs MM2
Number of differences: 3
MA2 vs MM1
Number of differences: 456
MA2 vs MM2
Number of differences: 438
MM1 vs MM2
Number of differences: 280
0 2500 5000 7500 10000 12500 15000 17500
Position

### Region: 10



CR1 vs CR2
Number of differences: 201

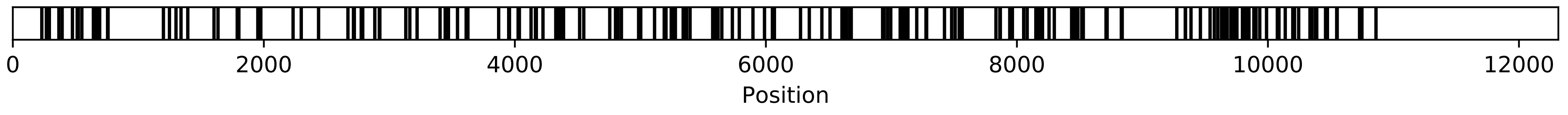


CR1 vs MA1
Number of differences: 83

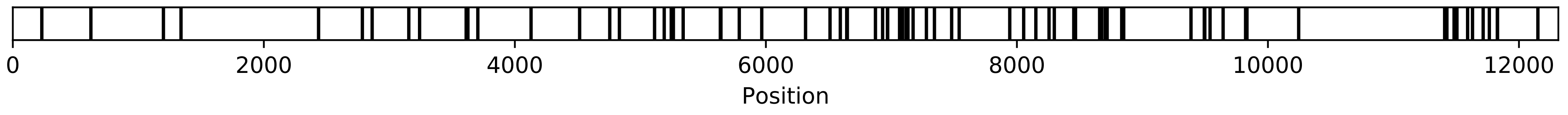


CR1 vs MA2
Number of differences: 222

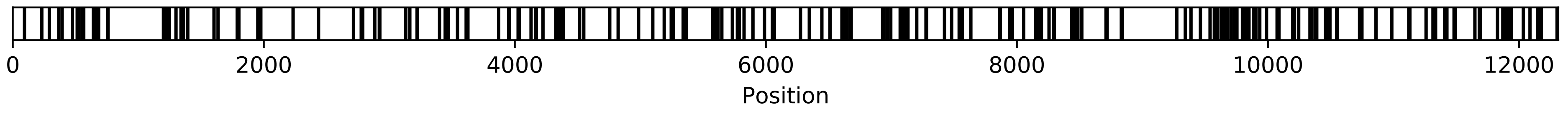


CR1 vs MM1
Number of differences: 83

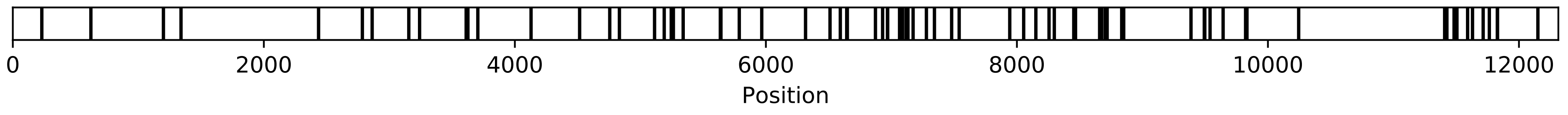


CR1 vs MM2
Number of differences: 76

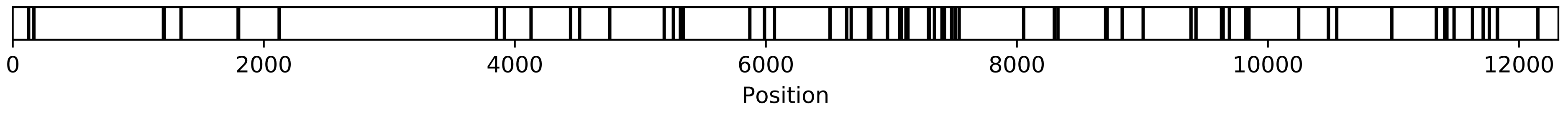


CR2 vs MA1
Number of differences: 196

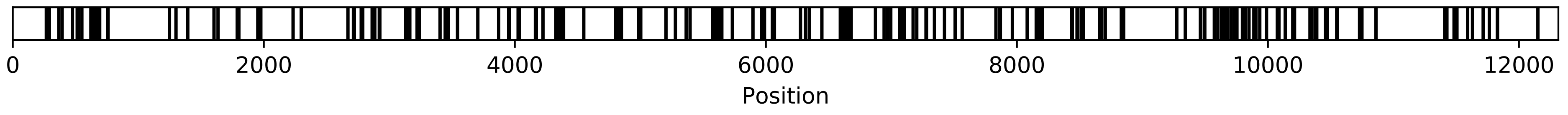


CR2 vs MA2
Number of differences: 61

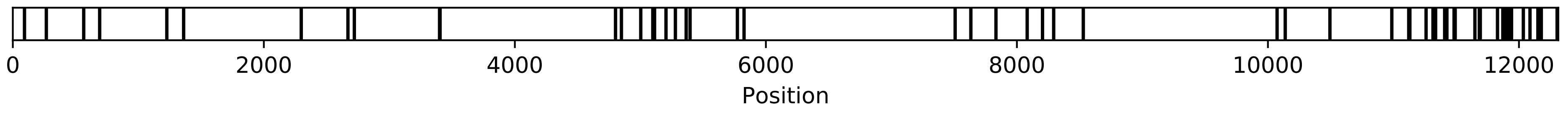


CR2 vs MM1
Number of differences: 196

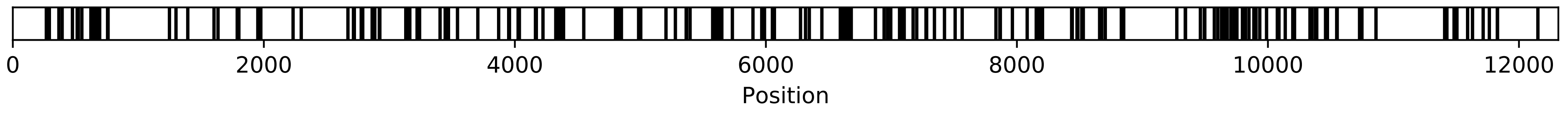


CR2 vs MM2
Number of differences: 198

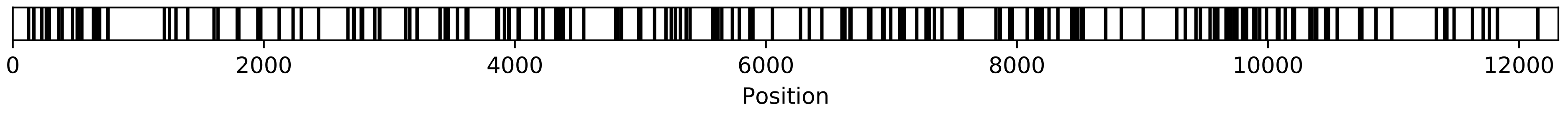


MA1 vs MA2
Number of differences: 210

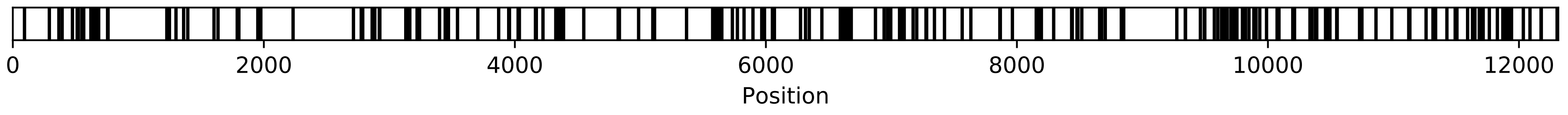


MA1 vs MM1
Number of differences: 0

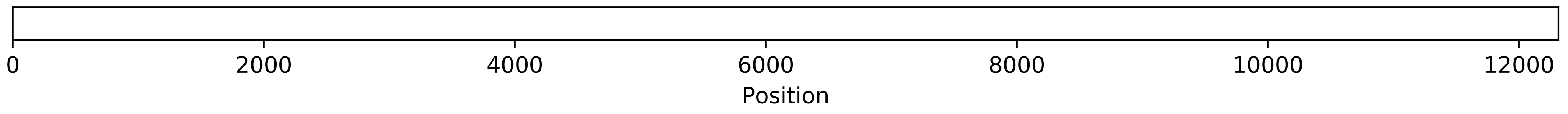


MA1 vs MM2
Number of differences: 83

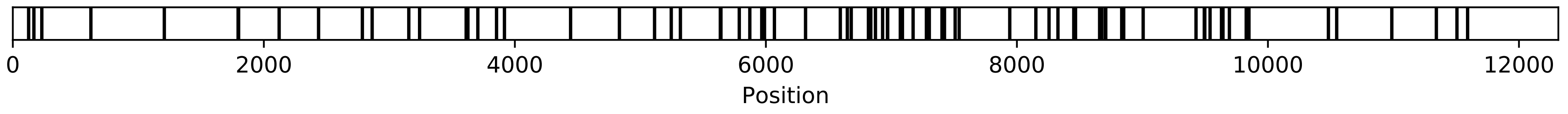


MA2 vs MM1
Number of differences: 210

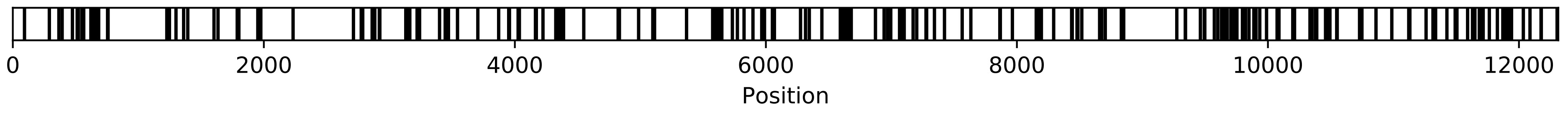


MA2 vs MM2
Number of differences: 210

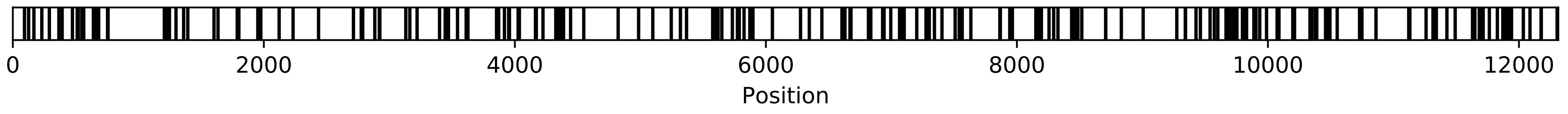


MM1 vs MM2
Number of differences: 83

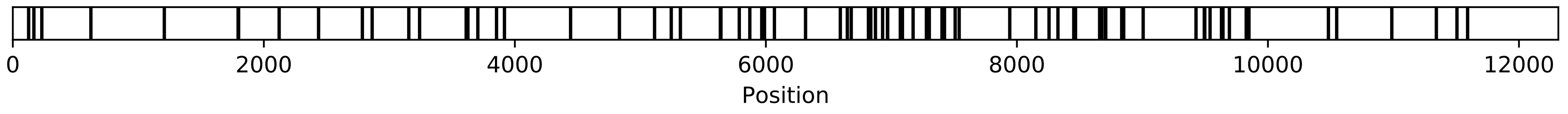

### Region: 12



CR1 vs CR2
Number of differences: 315

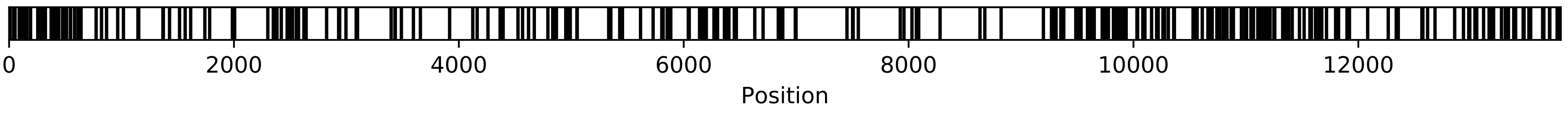


CR1 vs MA1
Number of differences: 318

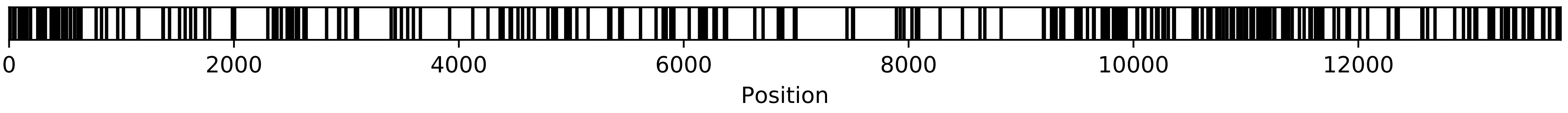


CR1 vs MA2
Number of differences: 168

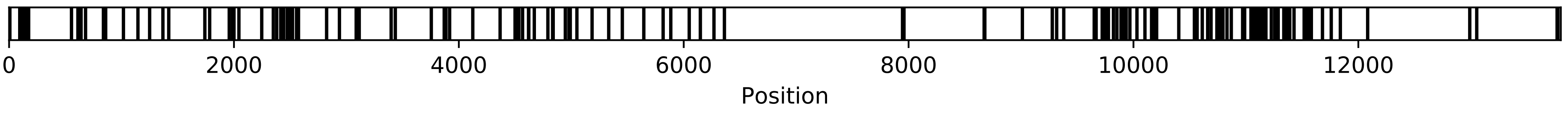


CR1 vs MM1
Number of differences: 168

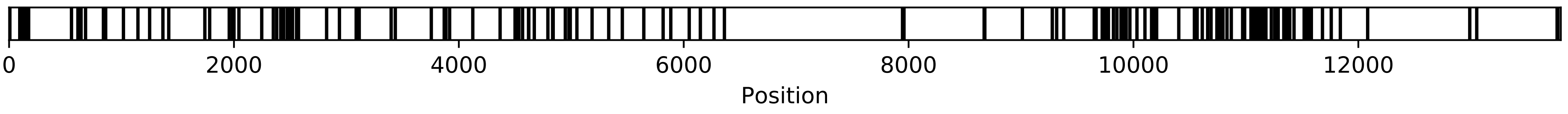


CR1 vs MM2
Number of differences: 121

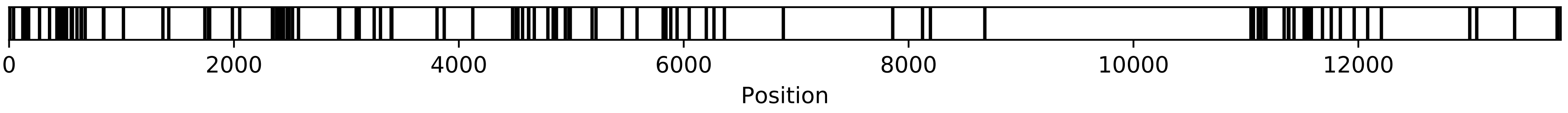


CR2 vs MA1
Number of differences: 53

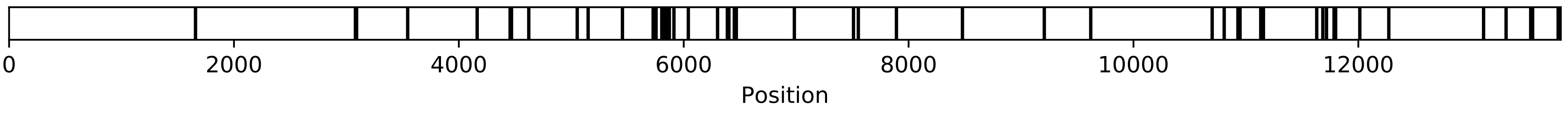


CR2 vs MA2
Number of differences: 303

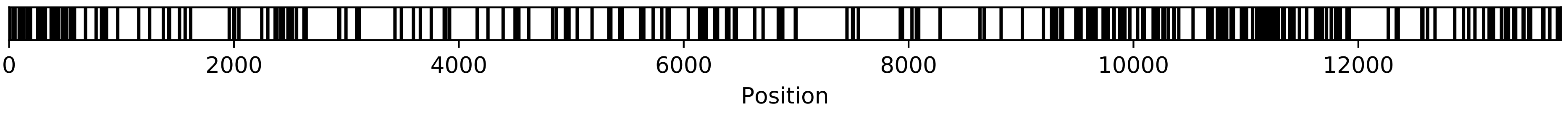


CR2 vs MM1
Number of differences: 303

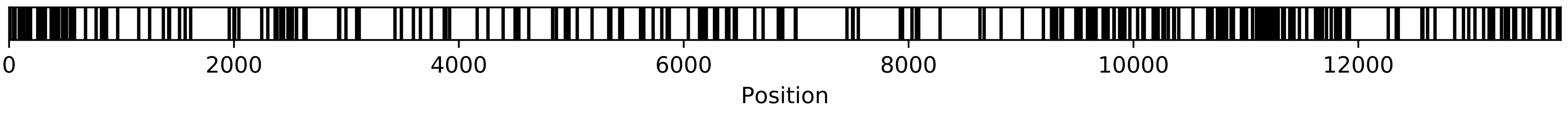


CR2 vs MM2
Number of differences: 292

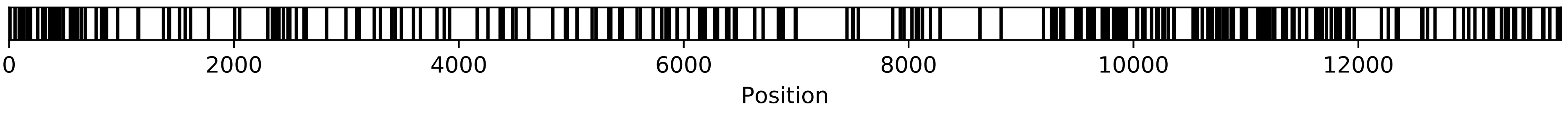


MA1 vs MA2
Number of differences: 304

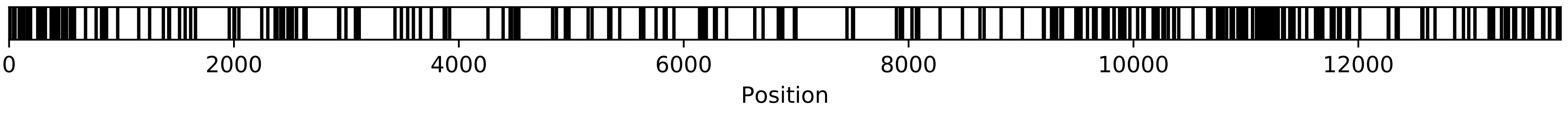


MA1 vs MM1
Number of differences: 304

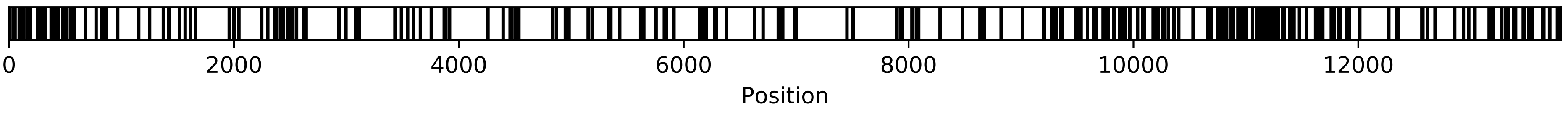


MA1 vs MM2
Number of differences: 293

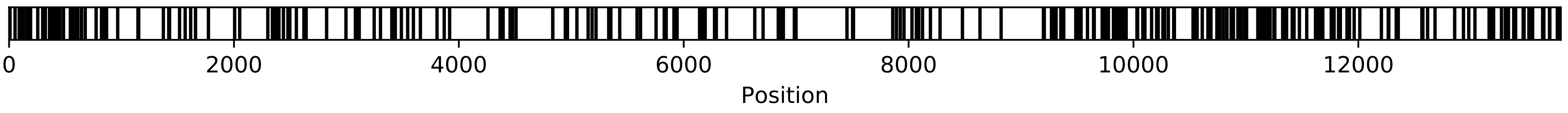


MA2 vs MM1
Number of differences: 0

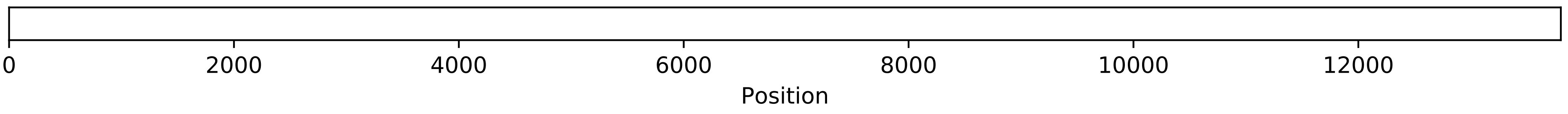


MA2 vs MM2
Number of differences: 139

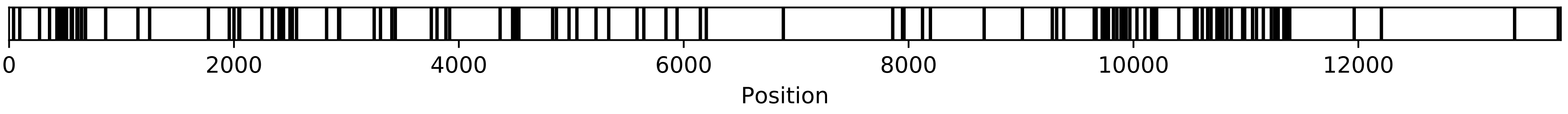


MM1 vs MM2
Number of differences: 139

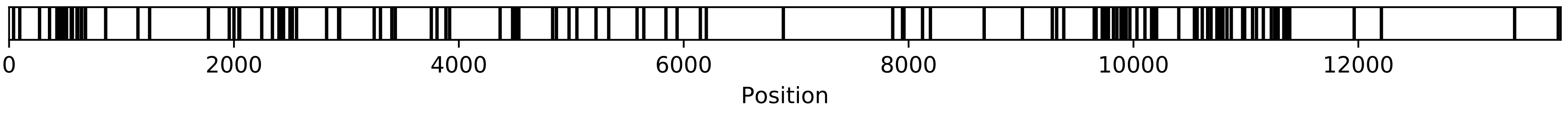

### Region: 13



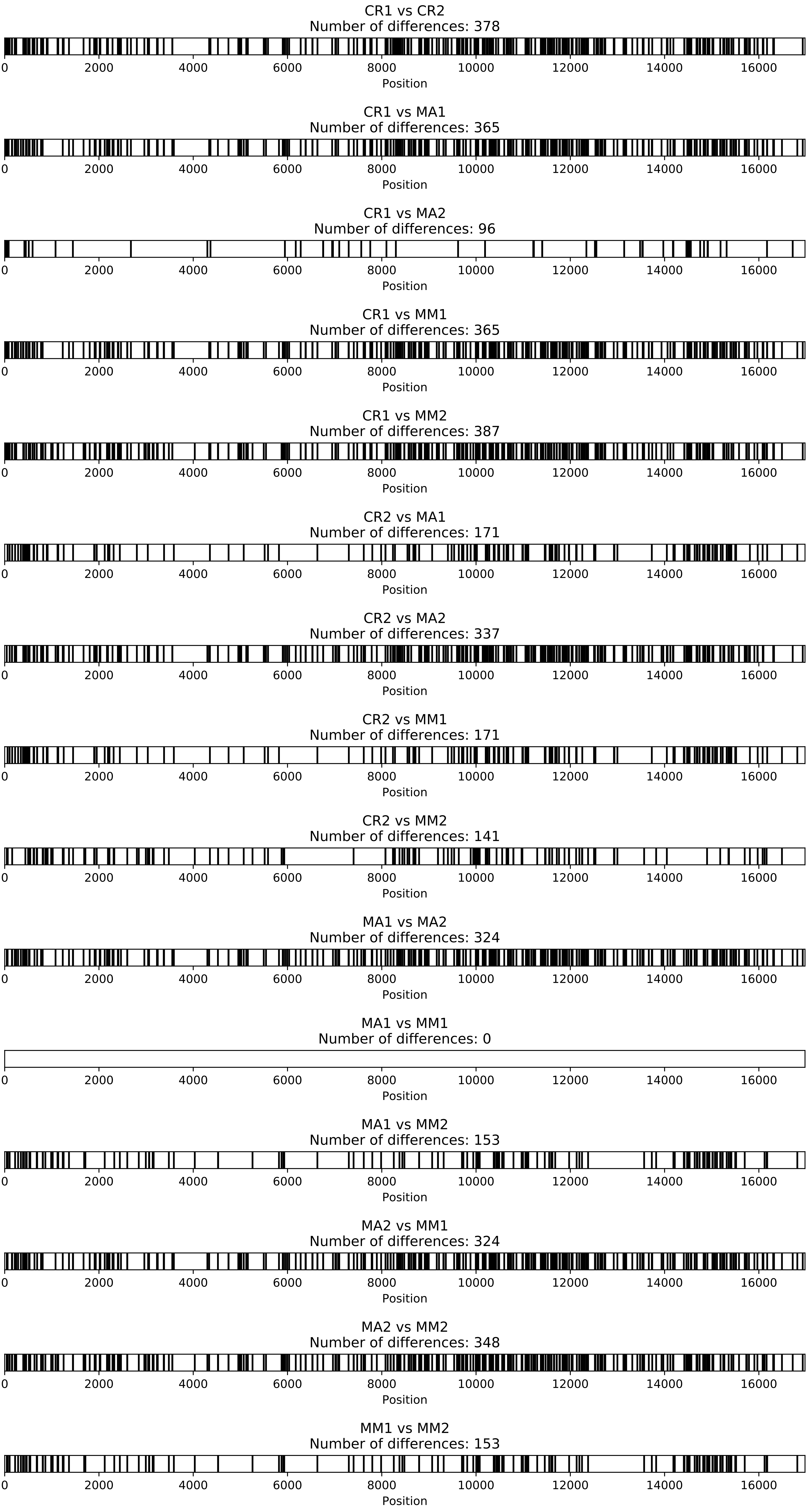
CR1 vs CR2
Number of differences: 378
CR1 vs MA1
Number of differences: 365
CR1 vs MA2
Number of differences: 96
CR1 vs MM1
Number of differences: 365
CR1 vs MM2
Number of differences: 387
CR2 vs MA1
Number of differences: 171
CR2 vs MA2
Number of differences: 337
CR2 vs MM1
Number of differences: 171
CR2 vs MM2
Number of differences: 141
MA1 vs MA2
Number of differences: 324
MA1 vs MM1
Number of differences: 0
MA1 vs MM2
Number of differences: 153
MA2 vs MM1
Number of differences: 324
MA2 vs MM2
Number of differences: 348
MM1 vs MM2
Number of differences: 153
0
2000
4000
6000
8000
10000
12000
14000
16000
Position

### Region: 14

CR1 vs CR2
Number of differences: 283

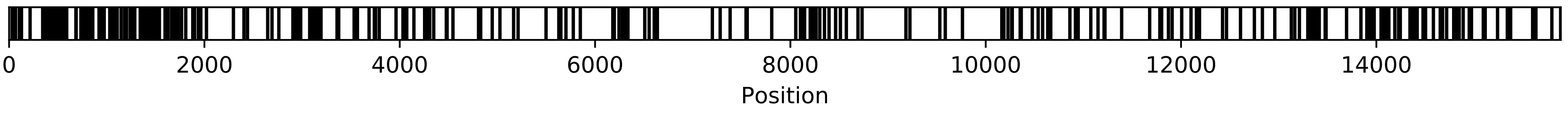


CR1 vs MA1
Number of differences: 102

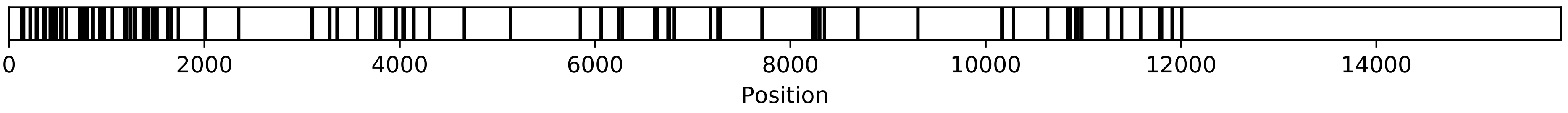


CR1 vs MA2
Number of differences: 280

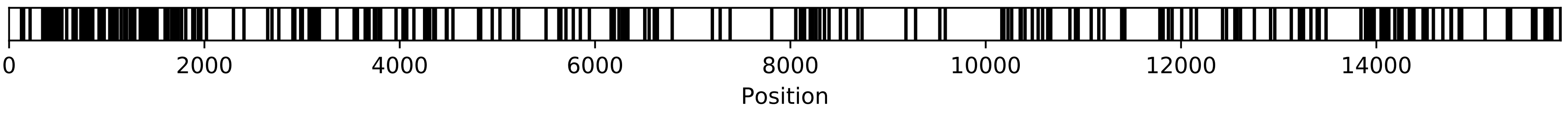


CR1 vs MM1
Number of differences: 102

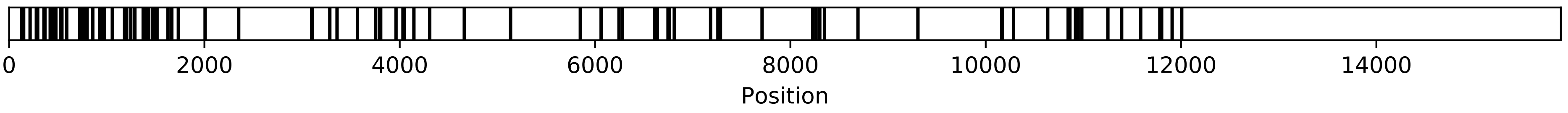


CR1 vs MM2
Number of differences: 158

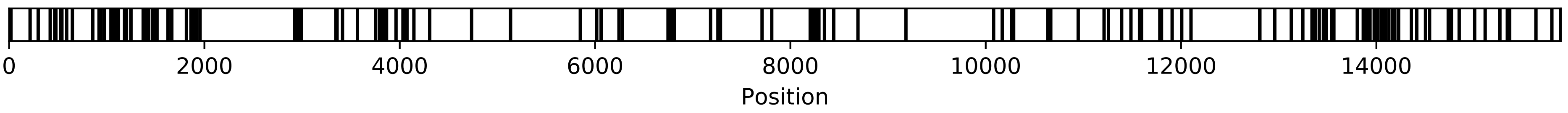


CR2 vs MA1
Number of differences: 265

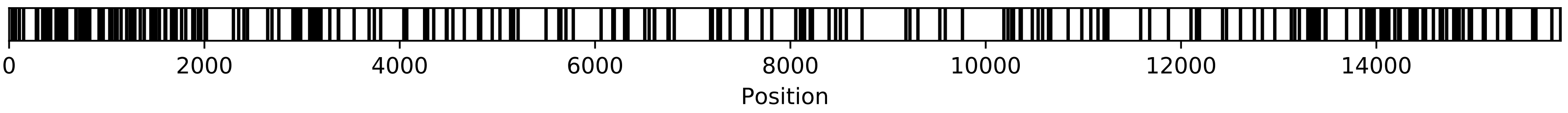


CR2 vs MA2
Number of differences: 94

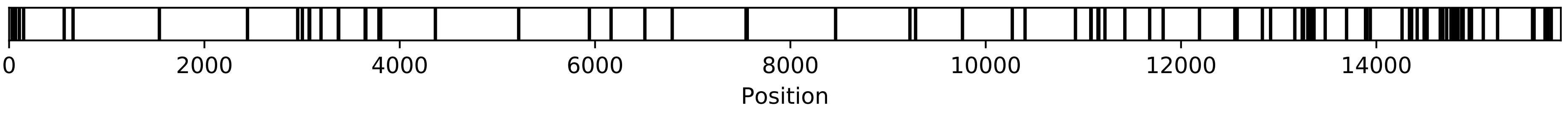


CR2 vs MM1
Number of differences: 265

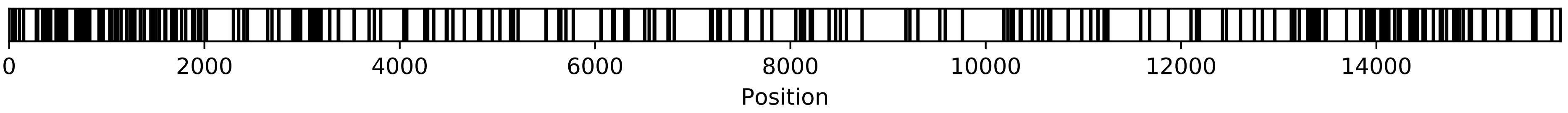


CR2 vs MM2
Number of differences: 279

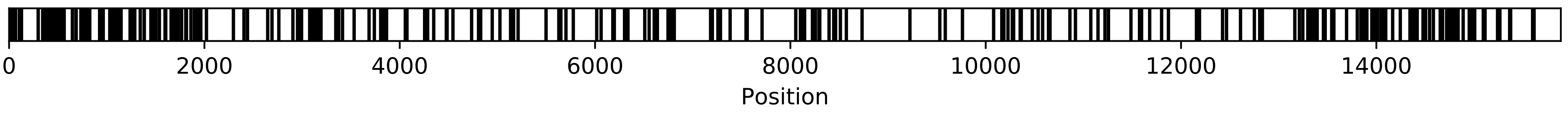


MA1 vs MA2
Number of differences: 259

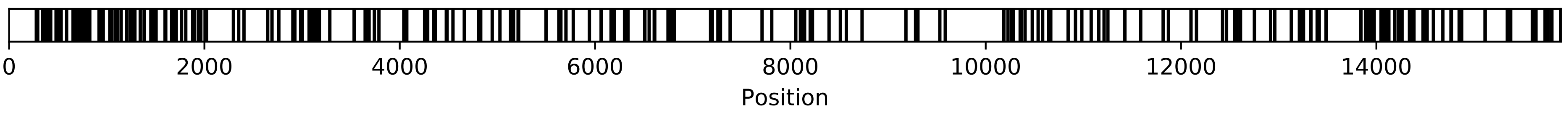


MA1 vs MM1
Number of differences: 0

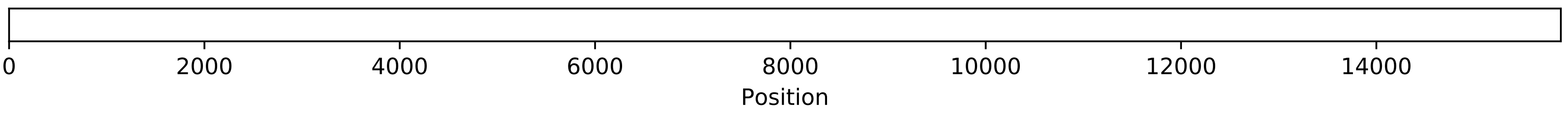


MA1 vs MM2
Number of differences: 142

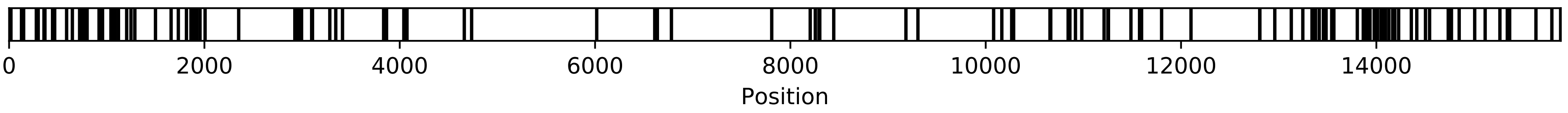


MA2 vs MM1
Number of differences: 259

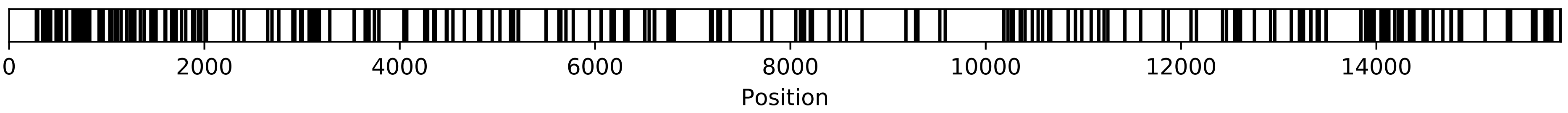


MA2 vs MM2
Number of differences: 270

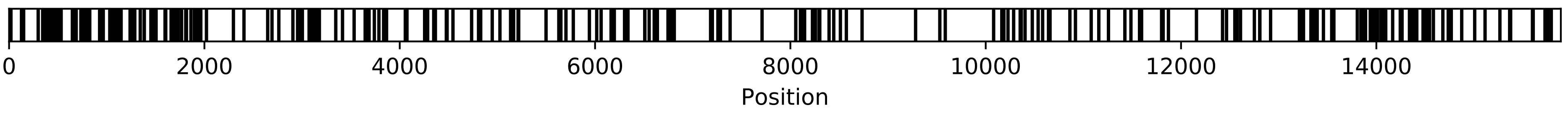


MM1 vs MM2
Number of differences: 142

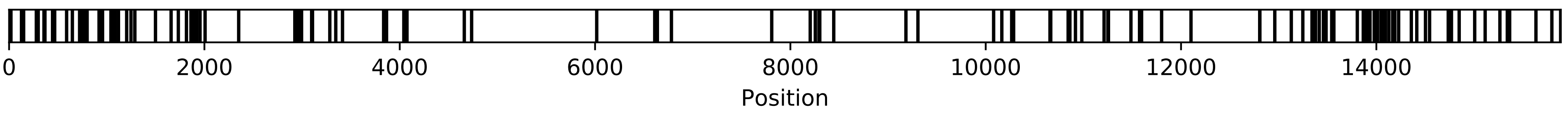

### Region: 15



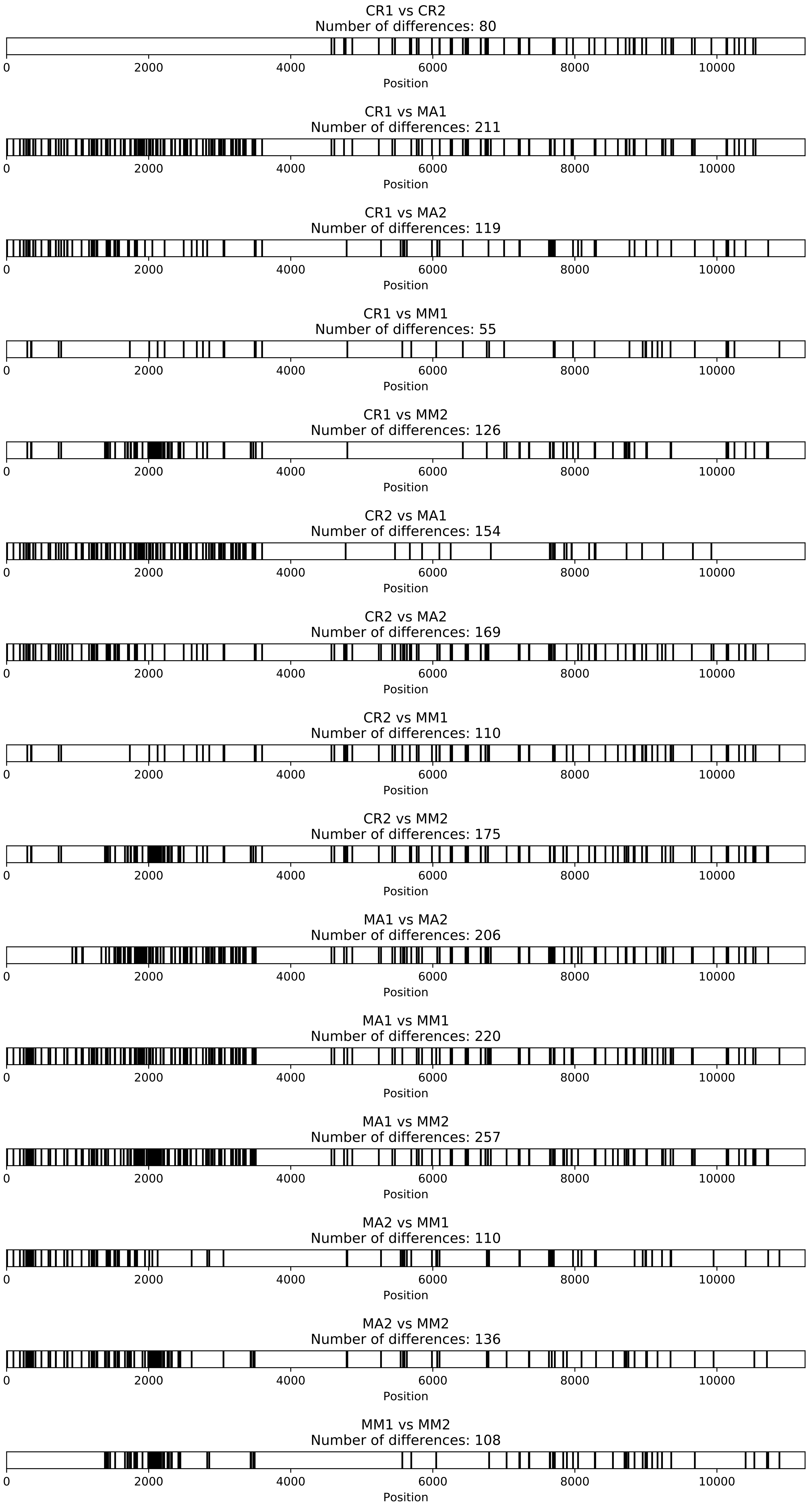
CR1 vs CR2
Number of differences: 80
CR1 vs MA1
Number of differences: 211
CR1 vs MA2
Number of differences: 119
CR1 vs MM1
Number of differences: 55
CR1 vs MM2
Number of differences: 126
CR2 vs MA1
Number of differences: 154
CR2 vs MA2
Number of differences: 169
CR2 vs MM1
Number of differences: 110
CR2 vs MM2
Number of differences: 175
MA1 vs MA2
Number of differences: 206
MA1 vs MM1
Number of differences: 220
MA1 vs MM2
Number of differences: 257
MA2 vs MM1
Number of differences: 110
MA2 vs MM2
Number of differences: 136
MM1 vs MM2
Number of differences: 108
0
2000
4000
6000
8000
10000
Position

### Region: 18



CR1 vs CR2
Number of differences: 228

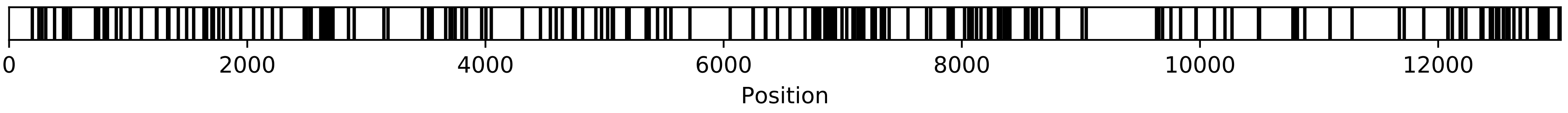


CR1 vs MA1
Number of differences: 87

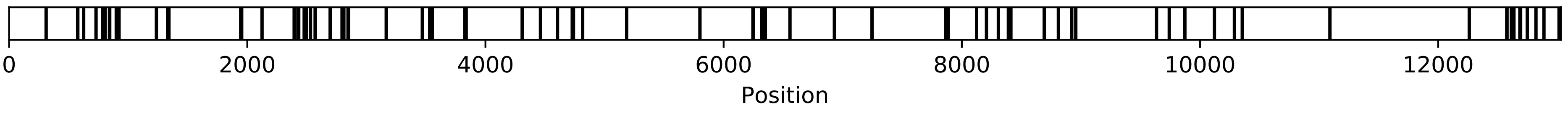


CR1 vs MA2
Number of differences: 233

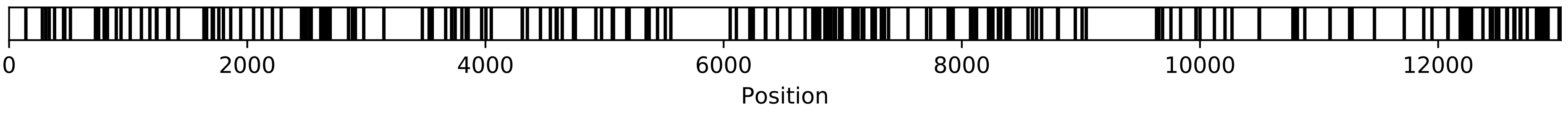


CR1 vs MM1
Number of differences: 96

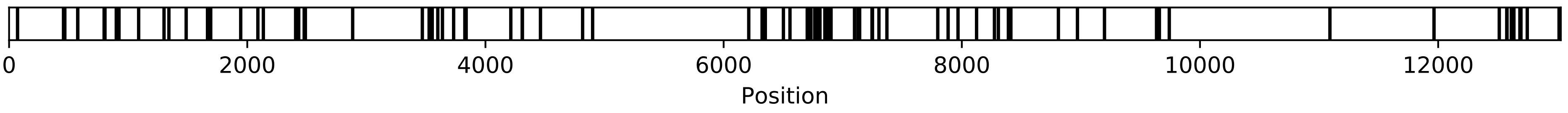


CR1 vs MM2
Number of differences: 87

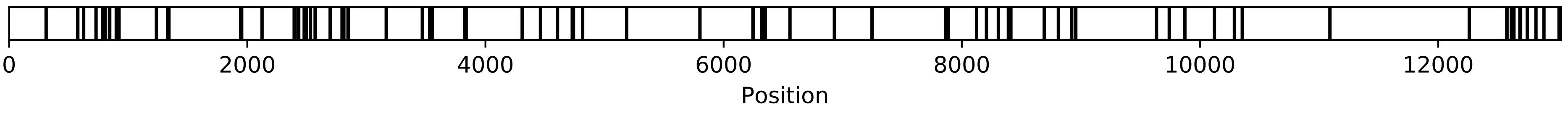


CR2 vs MA1
Number of differences: 236

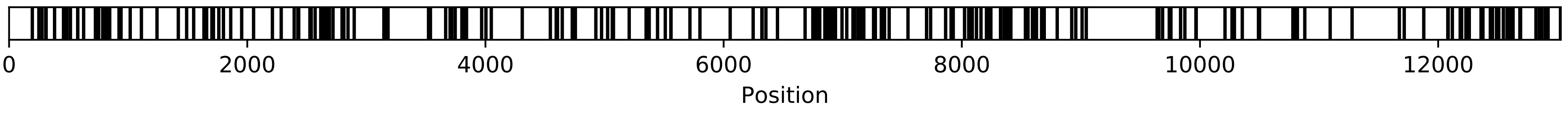


CR2 vs MA2
Number of differences: 85

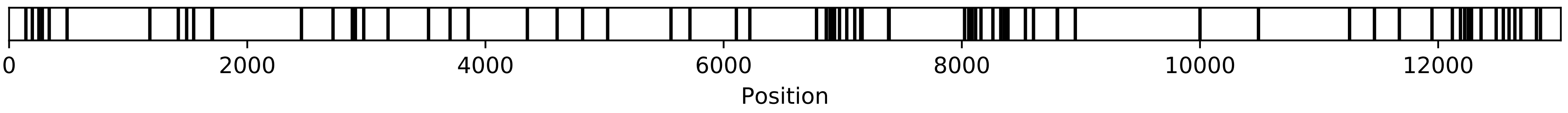


CR2 vs MM1
Number of differences: 243

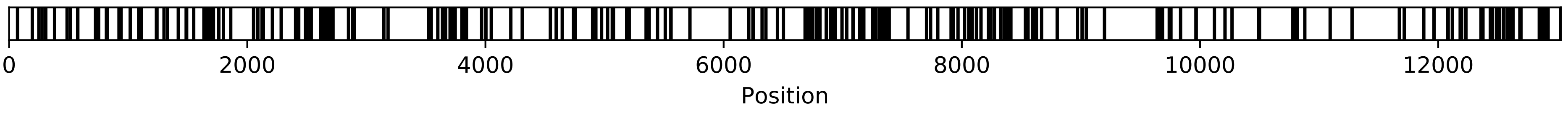


CR2 vs MM2
Number of differences: 236

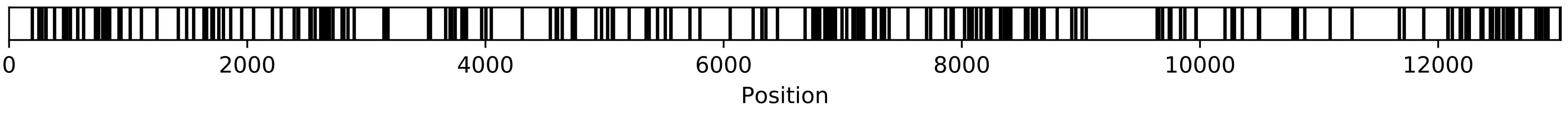


MA1 vs MA2
Number of differences: 242

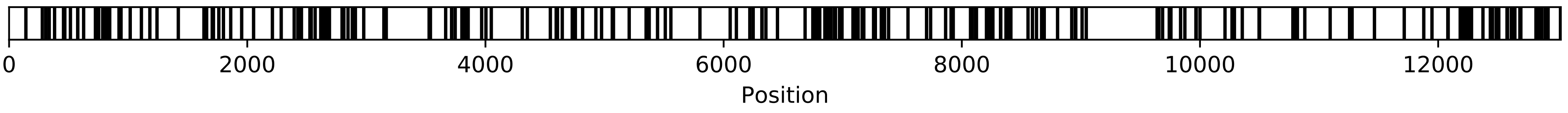


MA1 vs MM1
Number of differences: 97

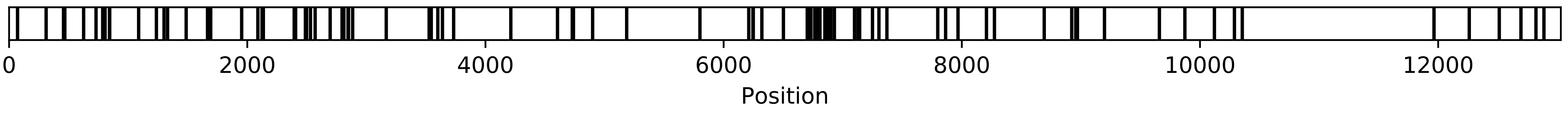


MA1 vs MM2
Number of differences: 0

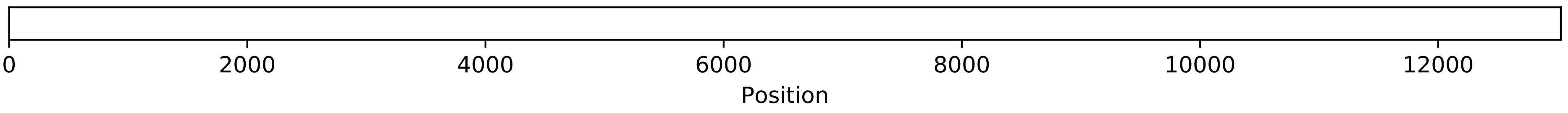


MA2 vs MM1
Number of differences: 245

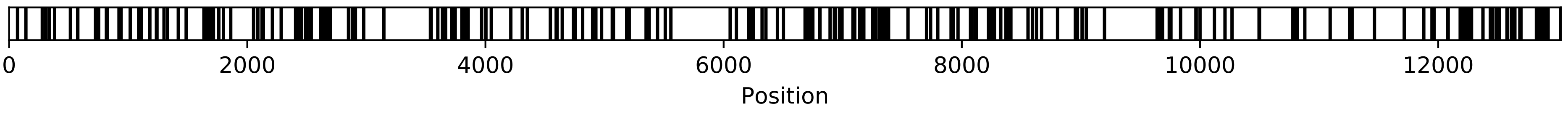


MA2 vs MM2
Number of differences: 242

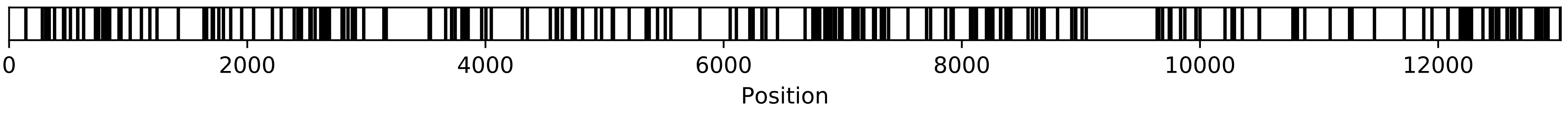


MM1 vs MM2
Number of differences: 97

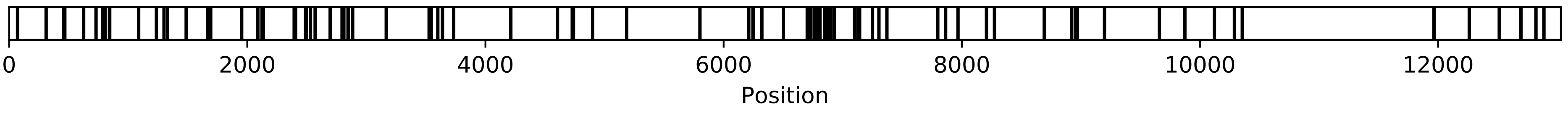

### Region: 20



CR1 vs CR2
Number of differences: 191

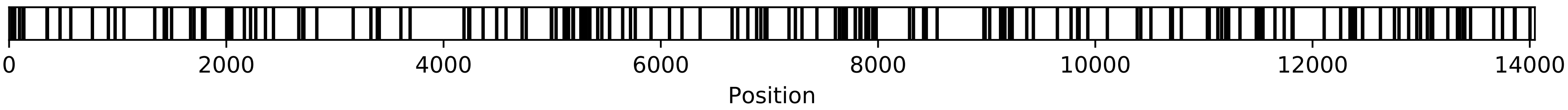

CR1 vs MA1
Number of differences: 91

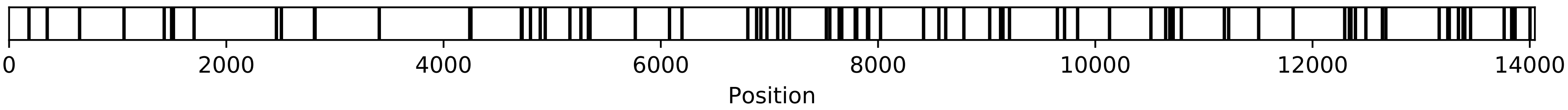

CR1 vs MA2
Number of differences: 188

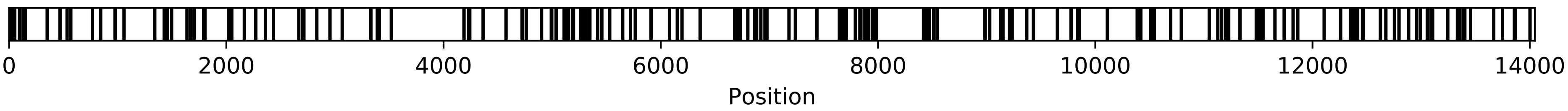

CR1 vs MM1
Number of differences: 91

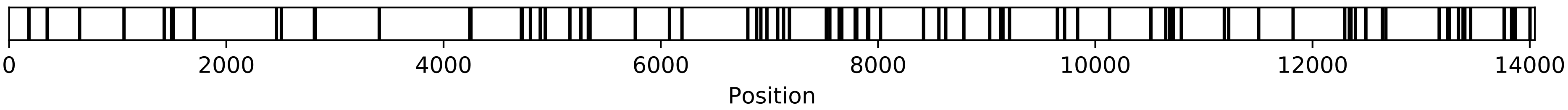

CR1 vs MM2
Number of differences: 112

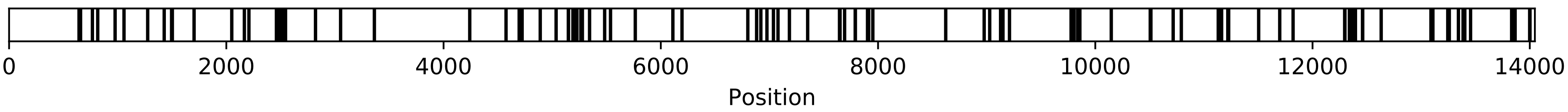

CR2 vs MA1
Number of differences: 187

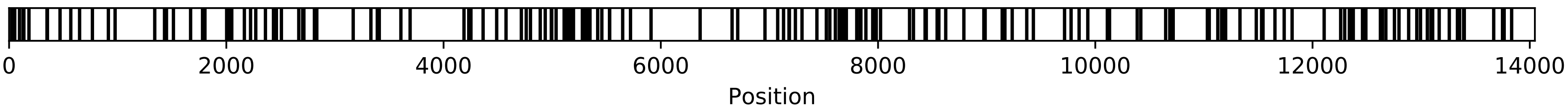

CR2 vs MA2
Number of differences: 58

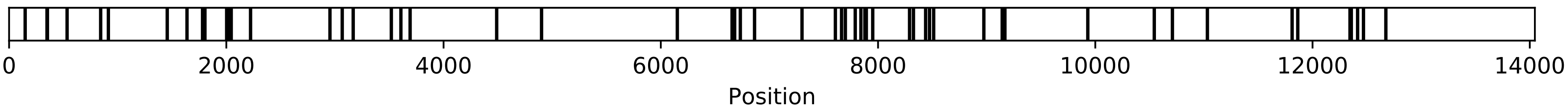

CR2 vs MM1
Number of differences: 187

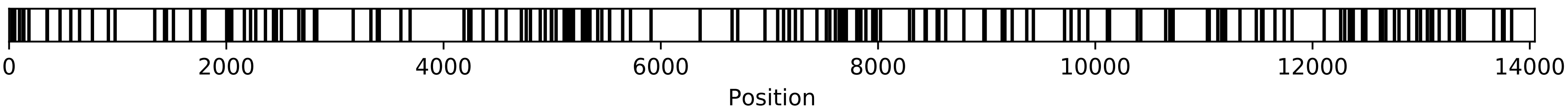

CR2 vs MM2
Number of differences: 185

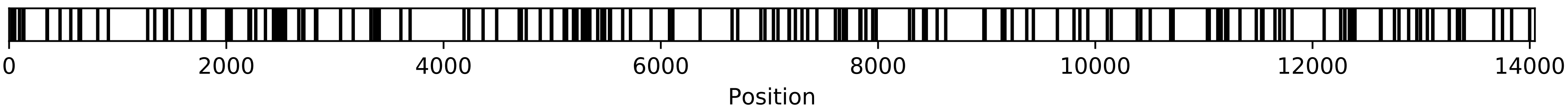

MA1 vs MA2
Number of differences: 184

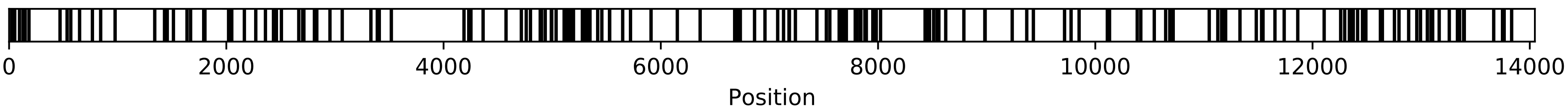

MA1 vs MM1
Number of differences: 0

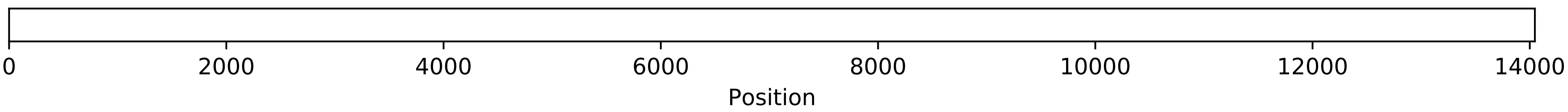

MA1 vs MM2
Number of differences: 100

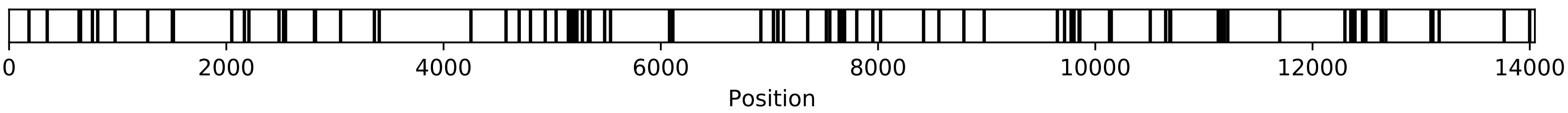

MA2 vs MM1
Number of differences: 184

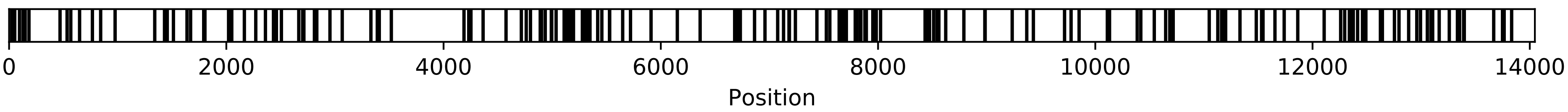

MA2 vs MM2
Number of differences: 184

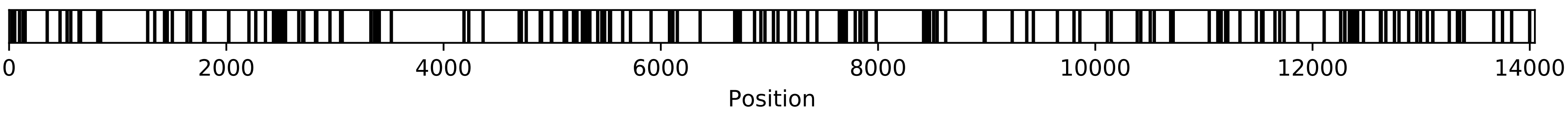

MM1 vs MM2
Number of differences: 100

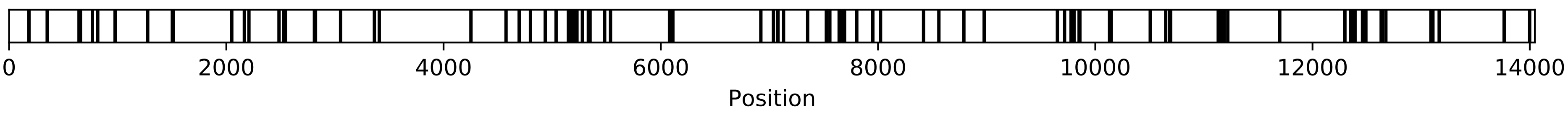

### Region: 23

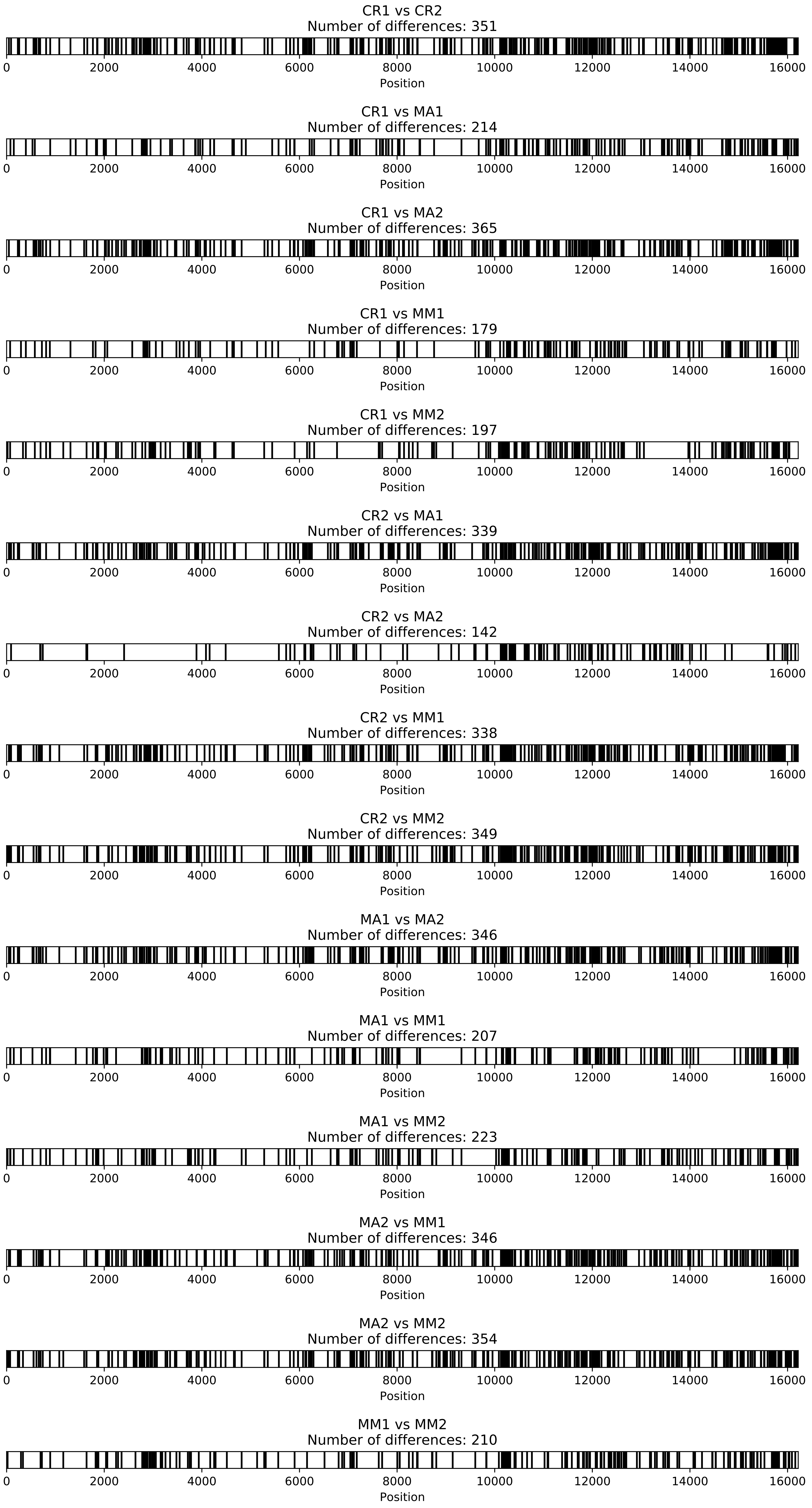
CR1 vs CR2
Number of differences: 351
CR1 vs MA1
Number of differences: 214
CR1 vs MA2
Number of differences: 365
CR1 vs MM1
Number of differences: 179
CR1 vs MM2
Number of differences: 197
CR2 vs MA1
Number of differences: 339
CR2 vs MA2
Number of differences: 142
CR2 vs MM1
Number of differences: 338
CR2 vs MM2
Number of differences: 349
MA1 vs MA2
Number of differences: 346
MA1 vs MM1
Number of differences: 207
MA1 vs MM2
Number of differences: 223
MA2 vs MM1
Number of differences: 346
MA2 vs MM2
Number of differences: 354
MM1 vs MM2
Number of differences: 210
Position

### Region: 25

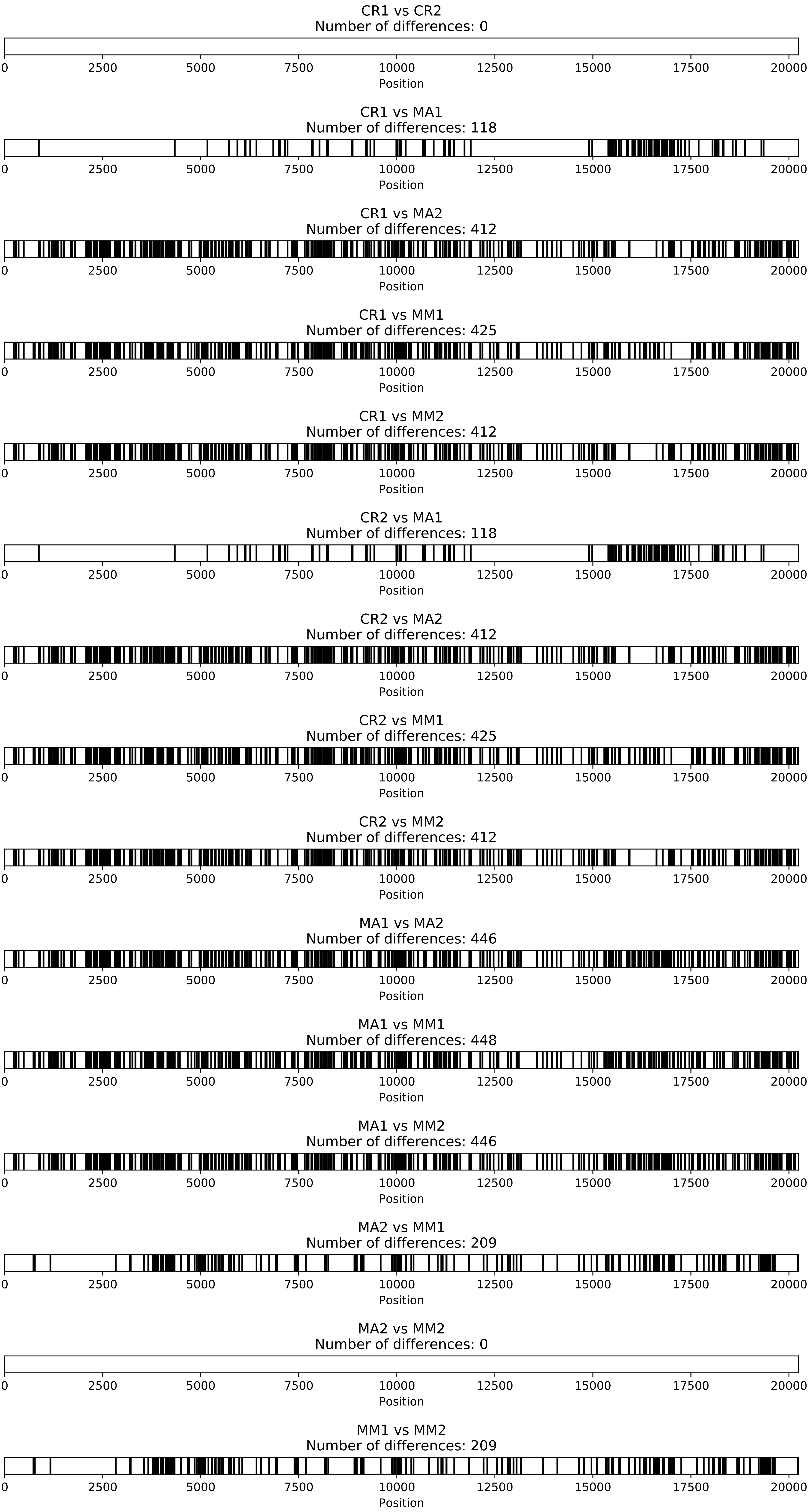
CR1 vs CR2
Number of differences: 0
CR1 vs MA1
Number of differences: 118
CR1 vs MA2
Number of differences: 412
CR1 vs MM1
Number of differences: 425
CR1 vs MM2
Number of differences: 412
CR2 vs MA1
Number of differences: 118
CR2 vs MA2
Number of differences: 412
CR2 vs MM1
Number of differences: 425
CR2 vs MM2
Number of differences: 412
MA1 vs MA2
Number of differences: 446
MA1 vs MM1
Number of differences: 448
MA1 vs MM2
Number of differences: 446
MA2 vs MM1
Number of differences: 209
MA2 vs MM2
Number of differences: 0
MM1 vs MM2
Number of differences: 209
0
2500
5000
7500
10000
12500
15000
17500
20000
Position

### Region: 27



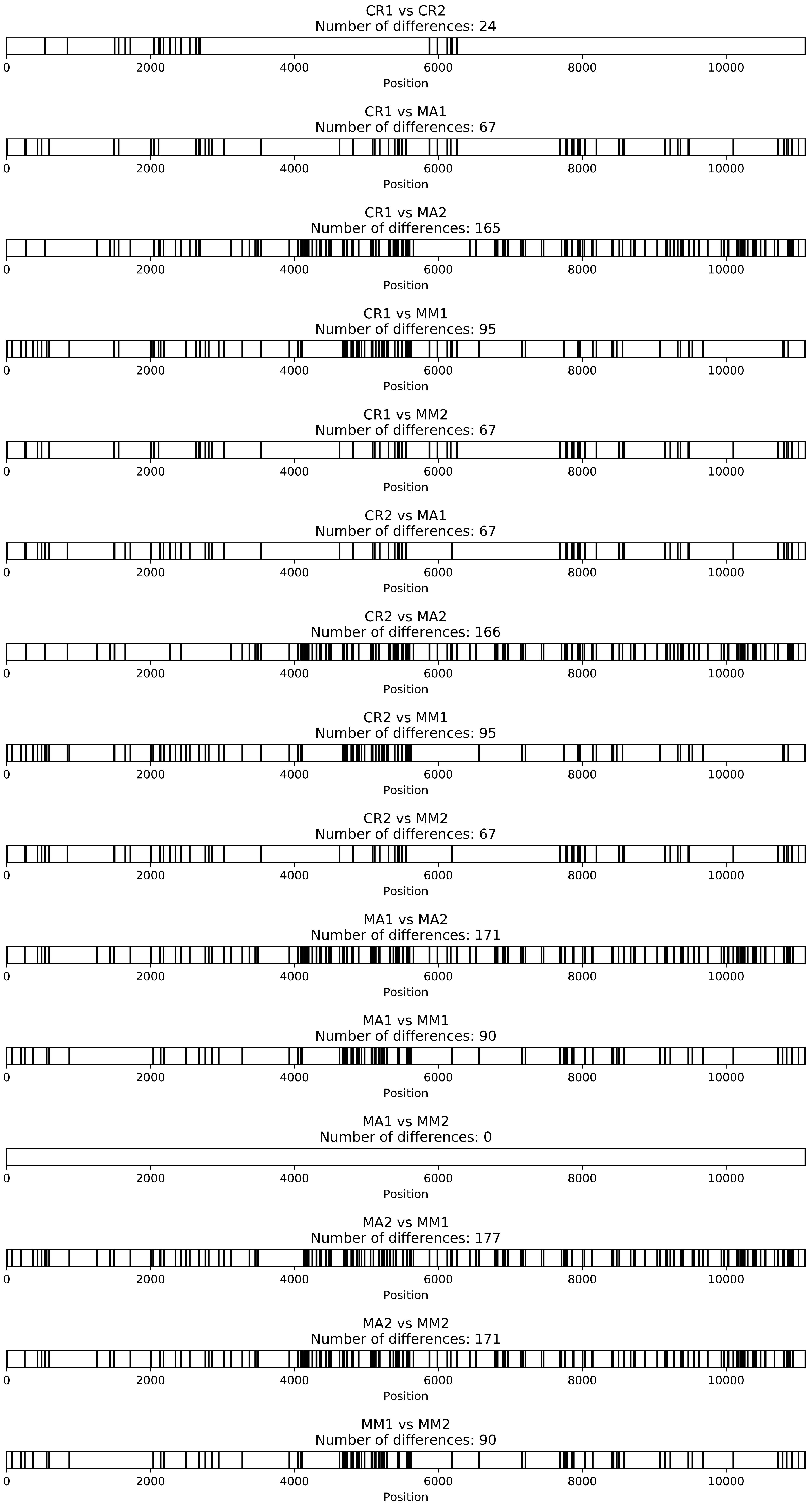
CR1 vs CR2
Number of differences: 24
CR1 vs MA1
Number of differences: 67
CR1 vs MA2
Number of differences: 165
CR1 vs MM1
Number of differences: 95
CR1 vs MM2
Number of differences: 67
CR2 vs MA1
Number of differences: 67
CR2 vs MA2
Number of differences: 166
CR2 vs MM1
Number of differences: 95
CR2 vs MM2
Number of differences: 67
MA1 vs MA2
Number of differences: 171
MA1 vs MM1
Number of differences: 90
MA1 vs MM2
Number of differences: 0
MA2 vs MM1
Number of differences: 177
MA2 vs MM2
Number of differences: 171
MM1 vs MM2
Number of differences: 90
0
2000
4000
6000
8000
10000
Position

### Region: 28



CR1 vs CR2
Number of differences: 256

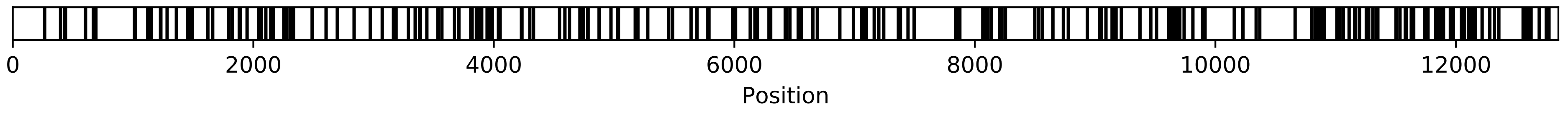


CR1 vs MA1
Number of differences: 253

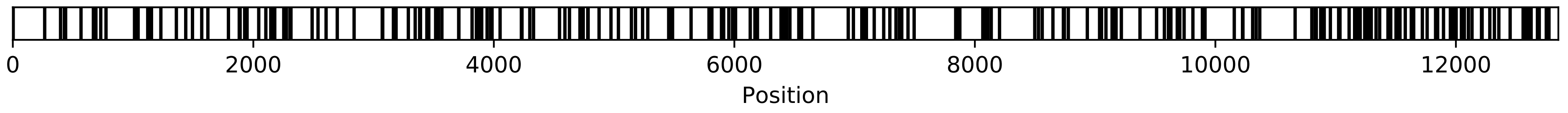


CR1 vs MA2
Number of differences: 137

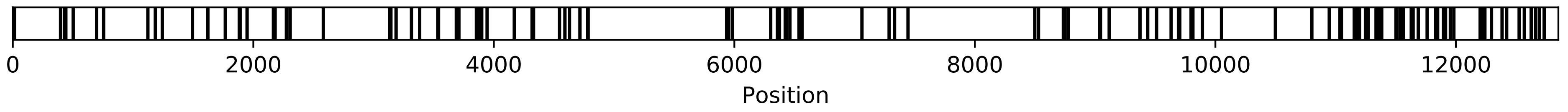


CR1 vs MM1
Number of differences: 129

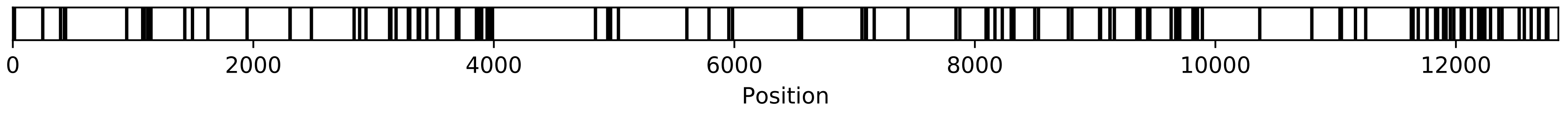


CR1 vs MM2
Number of differences: 41

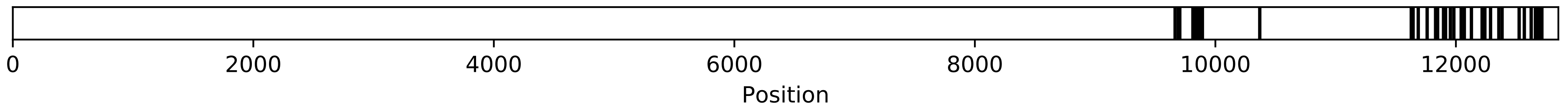


CR2 vs MA1
Number of differences: 105

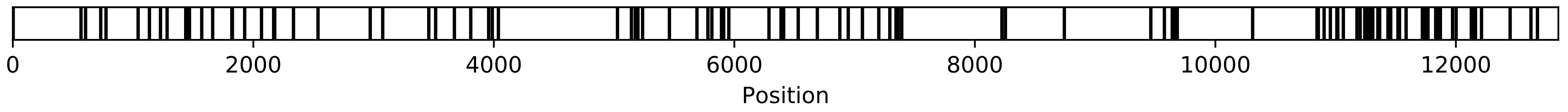


CR2 vs MA2
Number of differences: 259

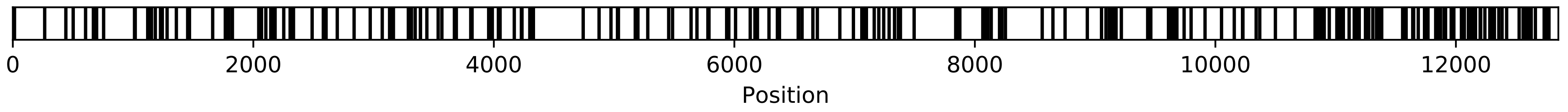


CR2 vs MM1
Number of differences: 258

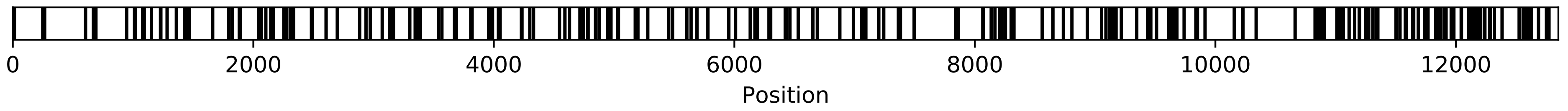


CR2 vs MM2
Number of differences: 267

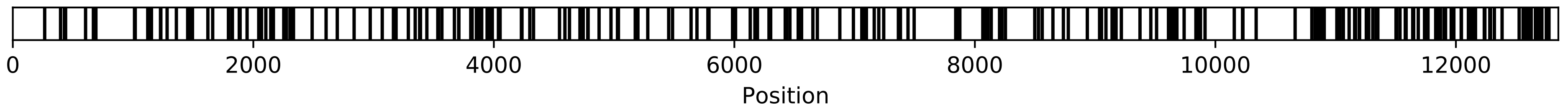


MA1 vs MA2
Number of differences: 252

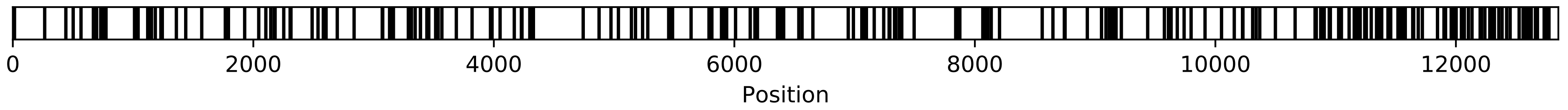


MA1 vs MM1
Number of differences: 253

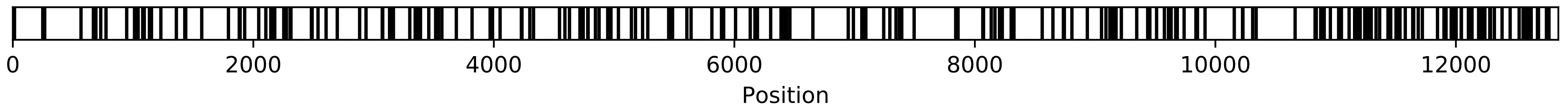


MA1 vs MM2
Number of differences: 264

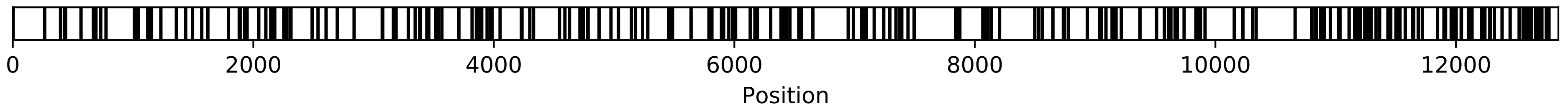


MA2 vs MM1
Number of differences: 132

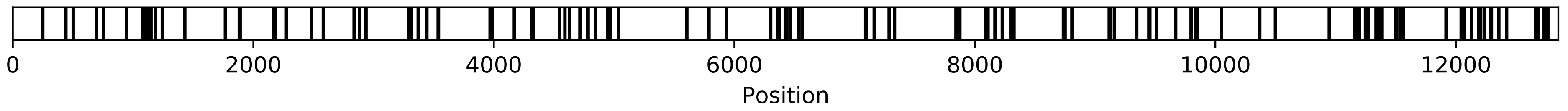


MA2 vs MM2
Number of differences: 124

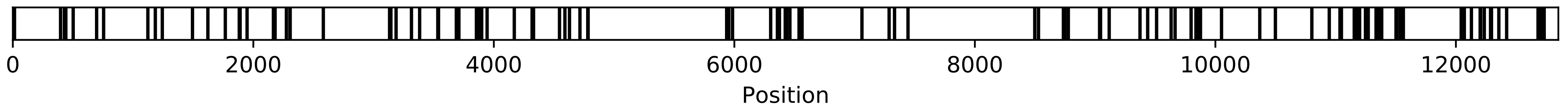


MM1 vs MM2
Number of differences: 100

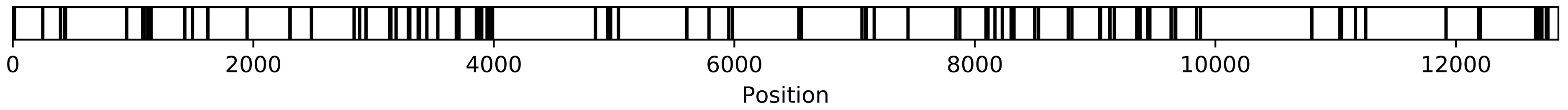

### Region: 30



CR1 vs CR2
Number of differences: 250

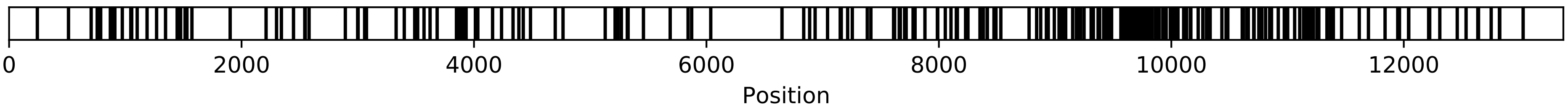


CR1 vs MA1
Number of differences: 255

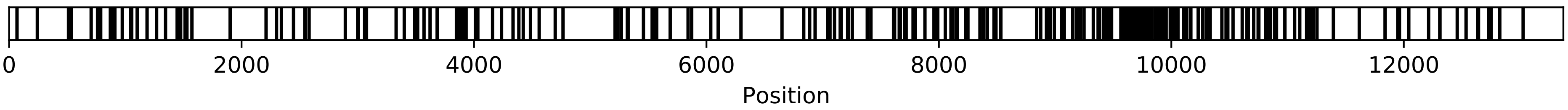


CR1 vs MA2
Number of differences: 106

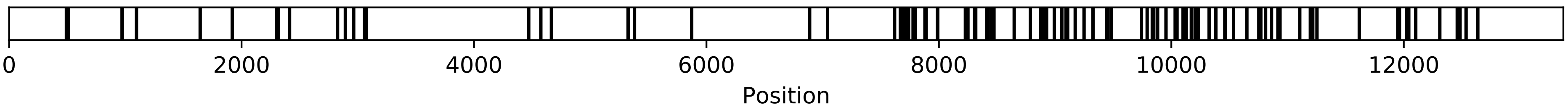


CR1 vs MM1
Number of differences: 112

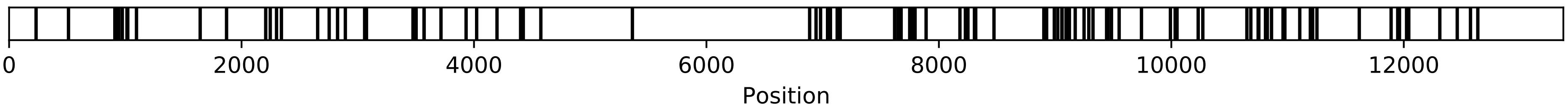


CR1 vs MM2
Number of differences: 106

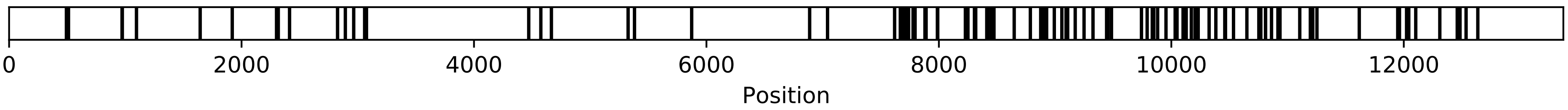


CR2 vs MA1
Number of differences: 51

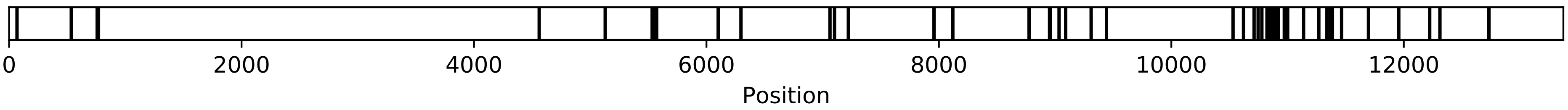


CR2 vs MA2
Number of differences: 251

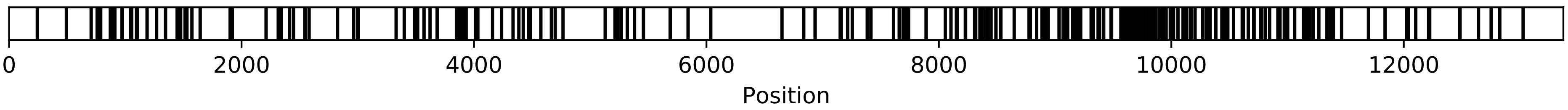


CR2 vs MM1
Number of differences: 264

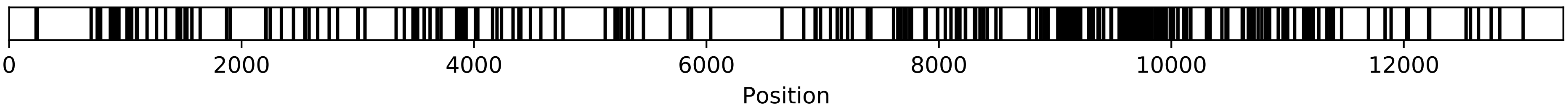


CR2 vs MM2
Number of differences: 251

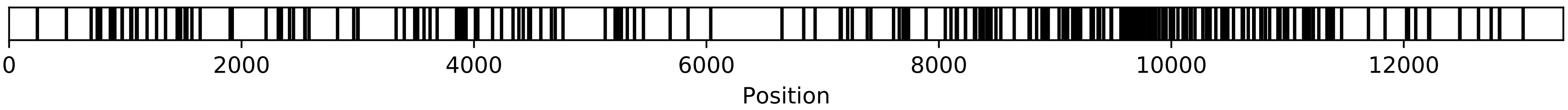


MA1 vs MA2
Number of differences: 258

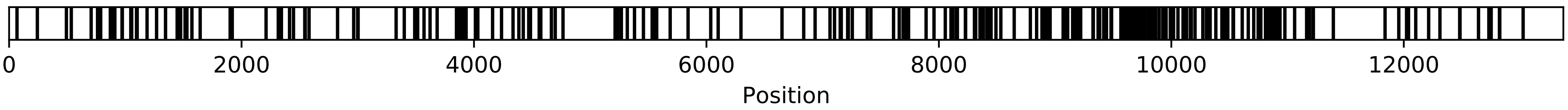


MA1 vs MM1
Number of differences: 271

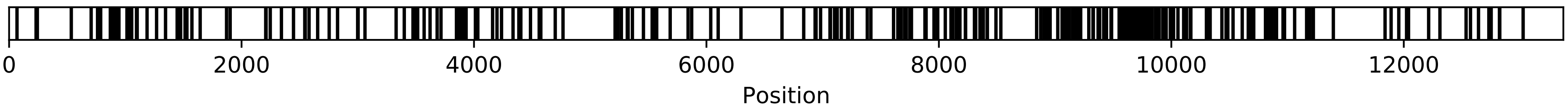


MA1 vs MM2
Number of differences: 258

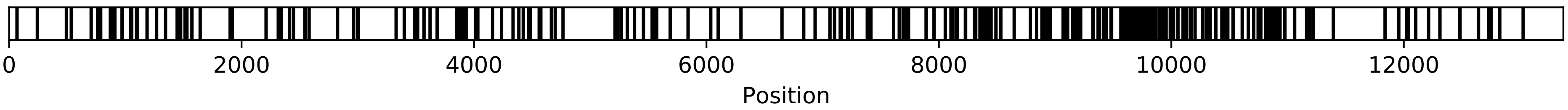


MA2 vs MM1
Number of differences: 106

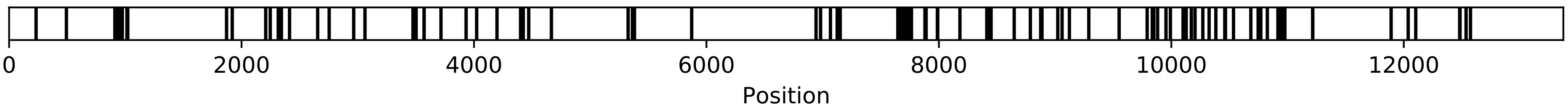


MA2 vs MM2
Number of differences: 0

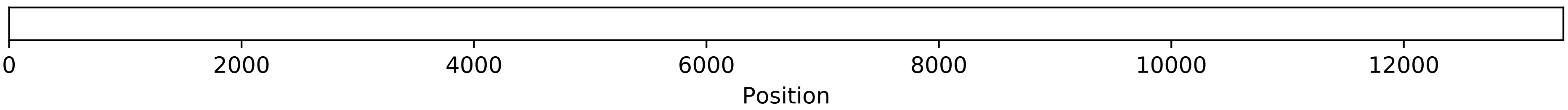


MM1 vs MM2
Number of differences: 106

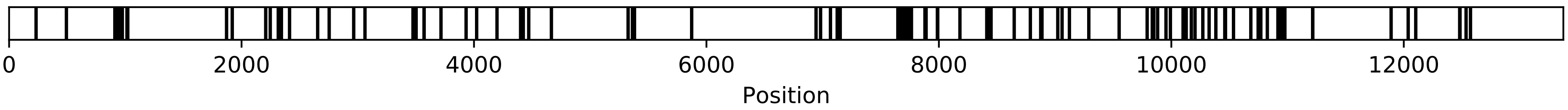

### Region: 31



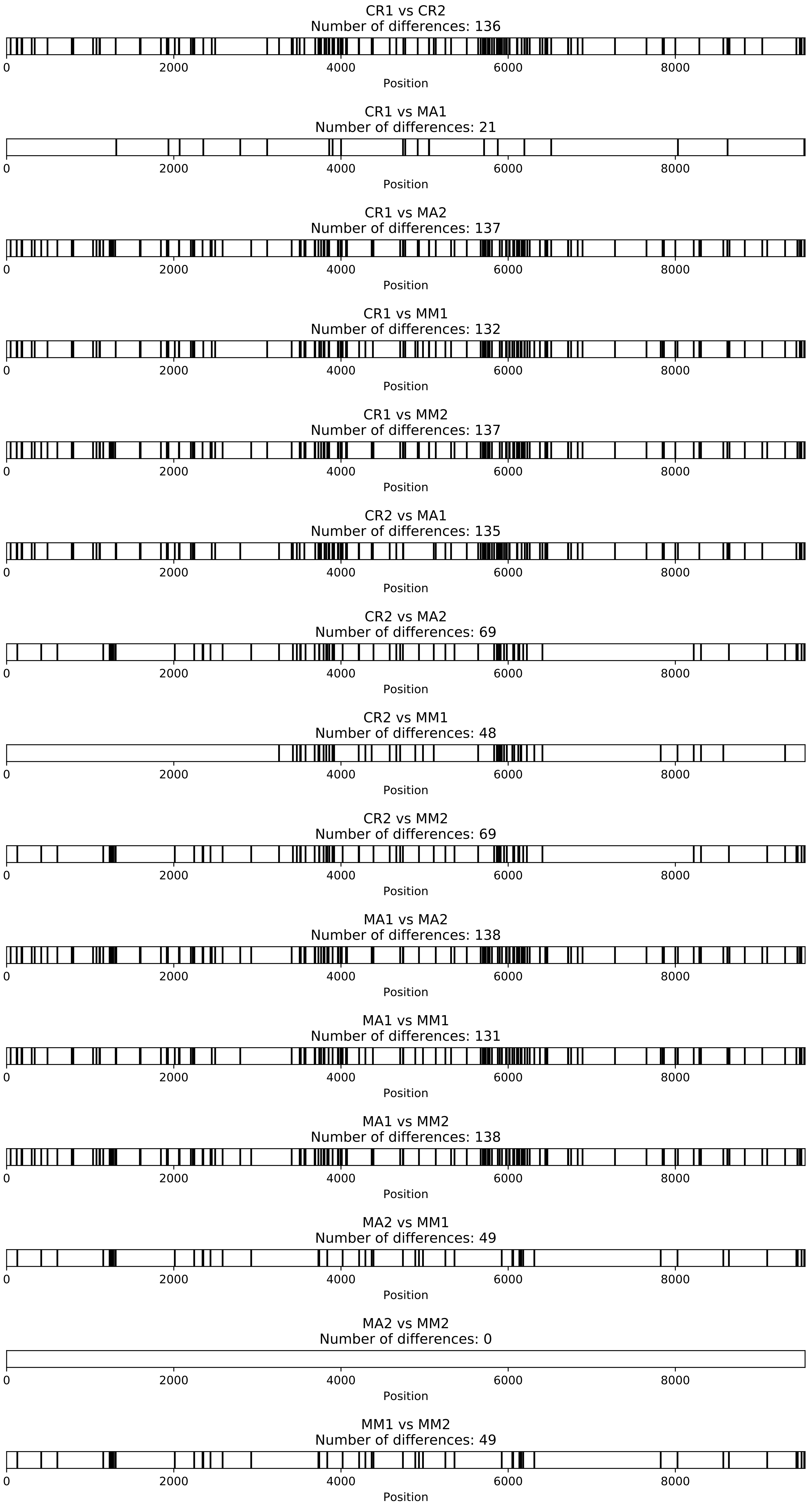
CR1 vs CR2
Number of differences: 136
CR1 vs MA1
Number of differences: 21
CR1 vs MA2
Number of differences: 137
CR1 vs MM1
Number of differences: 132
CR1 vs MM2
Number of differences: 137
CR2 vs MA1
Number of differences: 135
CR2 vs MA2
Number of differences: 69
CR2 vs MM1
Number of differences: 48
CR2 vs MM2
Number of differences: 69
MA1 vs MA2
Number of differences: 138
MA1 vs MM1
Number of differences: 131
MA1 vs MM2
Number of differences: 138
MA2 vs MM1
Number of differences: 49
MA2 vs MM2
Number of differences: 0
MM1 vs MM2
Number of differences: 49
0
2000
4000
6000
8000
Position

### Region: 33

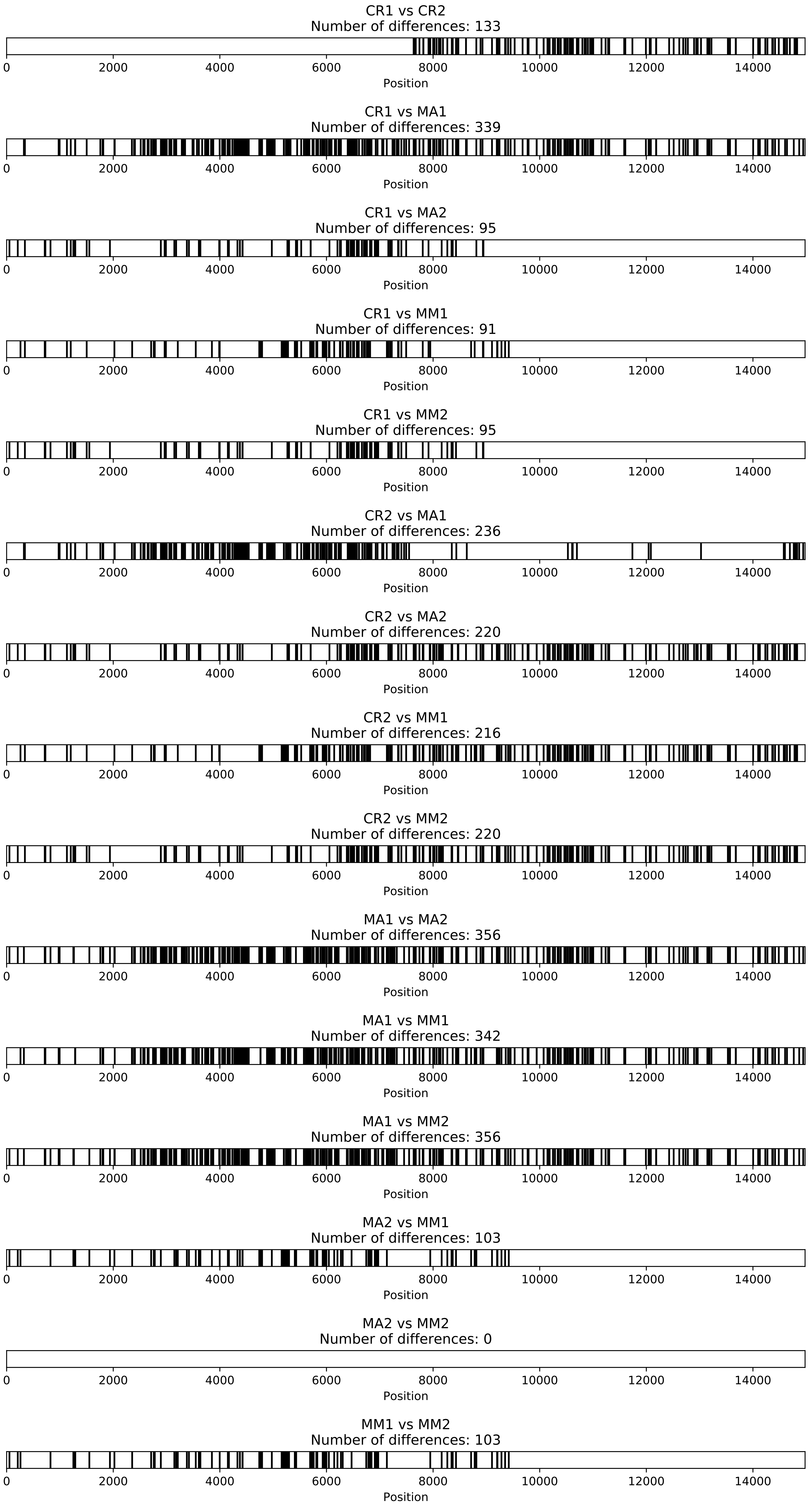
CR1 vs CR2
Number of differences: 133
CR1 vs MA1
Number of differences: 339
CR1 vs MA2
Number of differences: 95
CR1 vs MM1
Number of differences: 91
CR1 vs MM2
Number of differences: 95
CR2 vs MA1
Number of differences: 236
CR2 vs MA2
Number of differences: 220
CR2 vs MM1
Number of differences: 216
CR2 vs MM2
Number of differences: 220
MA1 vs MA2
Number of differences: 356
MA1 vs MM1
Number of differences: 342
MA1 vs MM2
Number of differences: 356
MA2 vs MM1
Number of differences: 103
MA2 vs MM2
Number of differences: 0
MM1 vs MM2
Number of differences: 103
0
2000
4000
6000
8000
10000
12000
14000
Position

### Region: 34



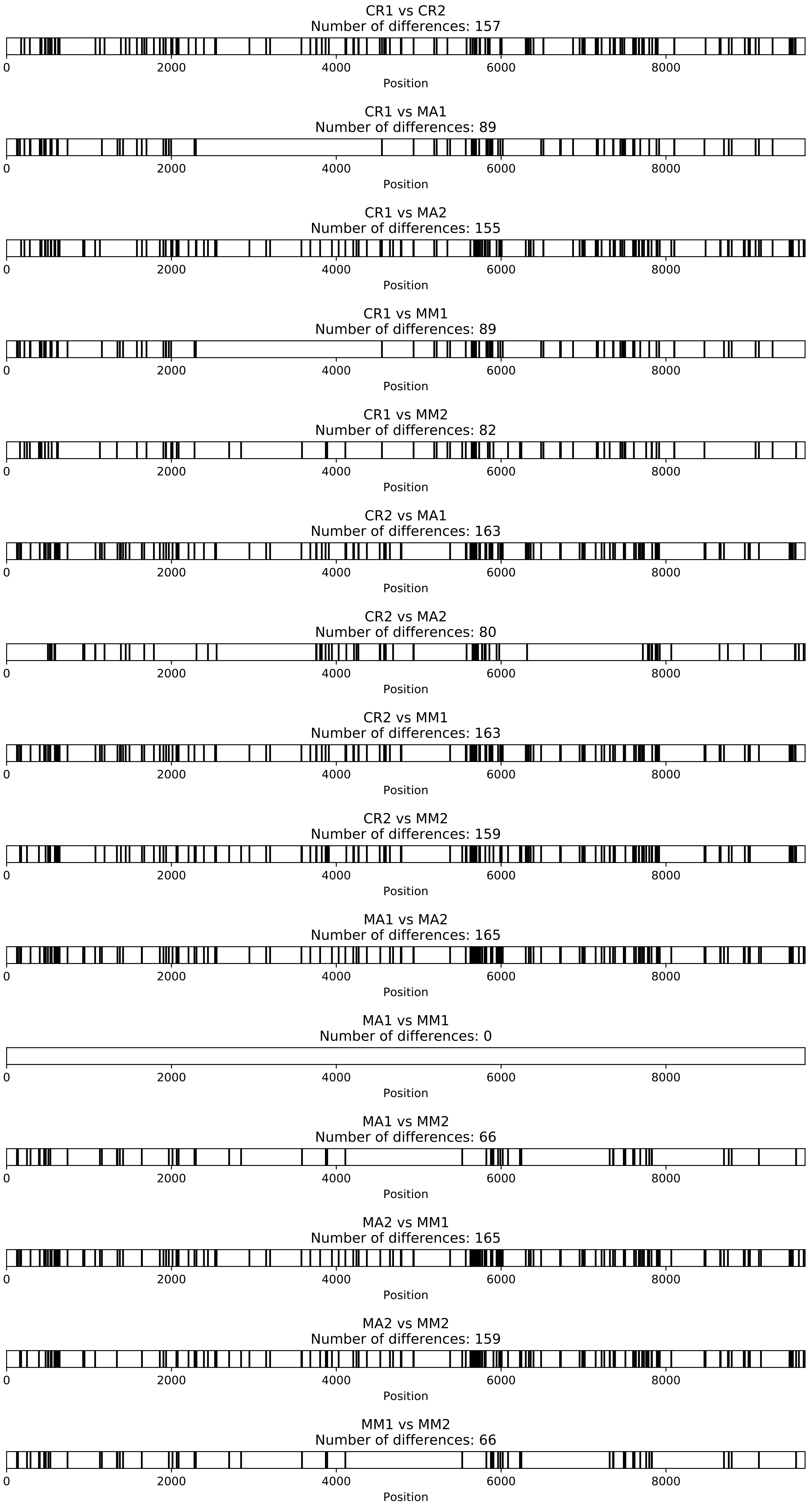
CR1 vs CR2
Number of differences: 157
CR1 vs MA1
Number of differences: 89
CR1 vs MA2
Number of differences: 155
CR1 vs MM1
Number of differences: 89
CR1 vs MM2
Number of differences: 82
CR2 vs MA1
Number of differences: 163
CR2 vs MA2
Number of differences: 80
CR2 vs MM1
Number of differences: 163
CR2 vs MM2
Number of differences: 159
MA1 vs MA2
Number of differences: 165
MA1 vs MM1
Number of differences: 0
MA1 vs MM2
Number of differences: 66
MA2 vs MM1
Number of differences: 165
MA2 vs MM2
Number of differences: 159
MM1 vs MM2
Number of differences: 66
0
2000
4000
6000
8000
Position

### Region: 36



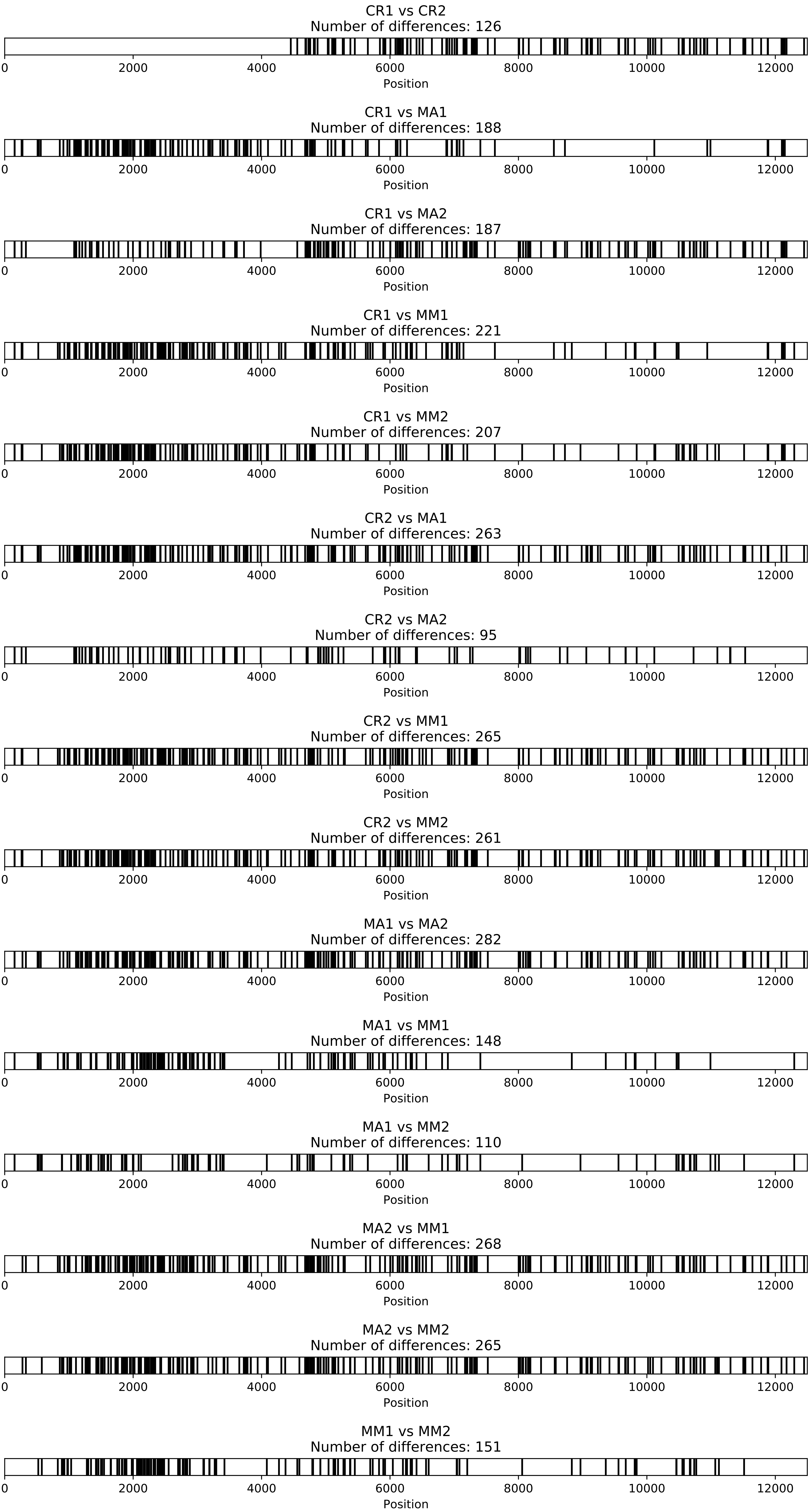
CR1 vs CR2
Number of differences: 126
CR1 vs MA1
Number of differences: 188
CR1 vs MA2
Number of differences: 187
CR1 vs MM1
Number of differences: 221
CR1 vs MM2
Number of differences: 207
CR2 vs MA1
Number of differences: 263
CR2 vs MA2
Number of differences: 95
CR2 vs MM1
Number of differences: 265
CR2 vs MM2
Number of differences: 261
MA1 vs MA2
Number of differences: 282
MA1 vs MM1
Number of differences: 148
MA1 vs MM2
Number of differences: 110
MA2 vs MM1
Number of differences: 268
MA2 vs MM2
Number of differences: 265
MM1 vs MM2
Number of differences: 151
0
2000
4000
6000
8000
10000
12000
Position

### Region: 37

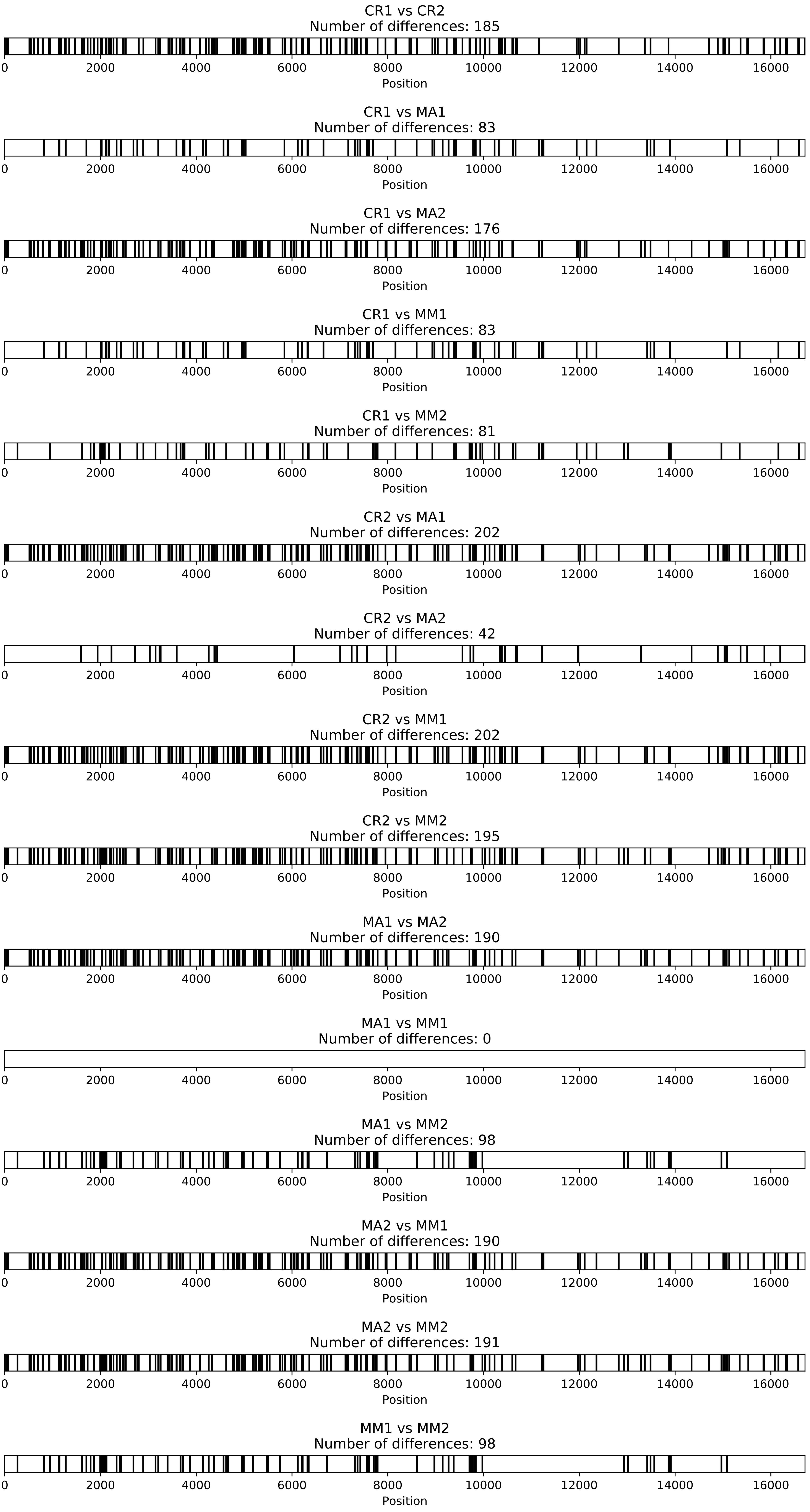
CR1 vs CR2
Number of differences: 185
CR1 vs MA1
Number of differences: 83
CR1 vs MA2
Number of differences: 176
CR1 vs MM1
Number of differences: 83
CR1 vs MM2
Number of differences: 81
CR2 vs MA1
Number of differences: 202
CR2 vs MA2
Number of differences: 42
CR2 vs MM1
Number of differences: 202
CR2 vs MM2
Number of differences: 195
MA1 vs MA2
Number of differences: 190
MA1 vs MM1
Number of differences: 0
MA1 vs MM2
Number of differences: 98
MA2 vs MM1
Number of differences: 190
MA2 vs MM2
Number of differences: 191
MM1 vs MM2
Number of differences: 98
0
2000
4000
6000
8000
10000
12000
14000
16000
Position

### Region: 38

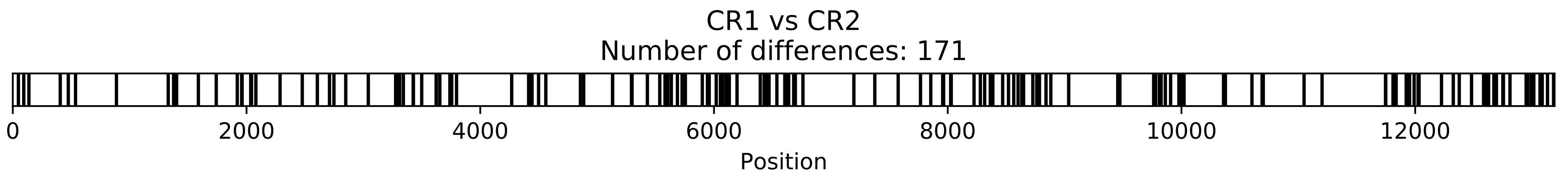
CR1 vs CR2
Number of differences: 171
0 2000 4000 6000 8000 10000 12000
Position

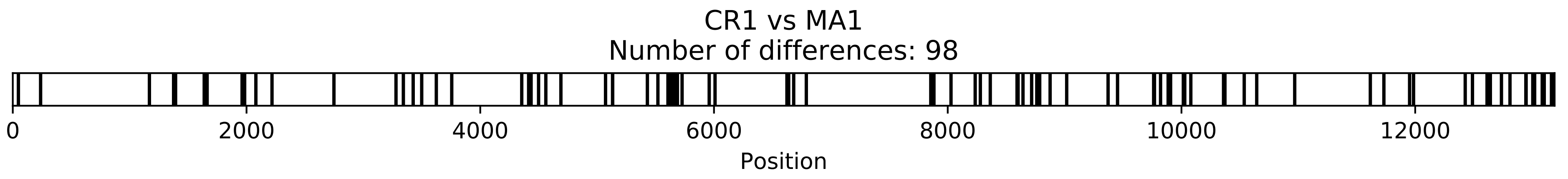
CR1 vs MA1
Number of differences: 98
0 2000 4000 6000 8000 10000 12000
Position

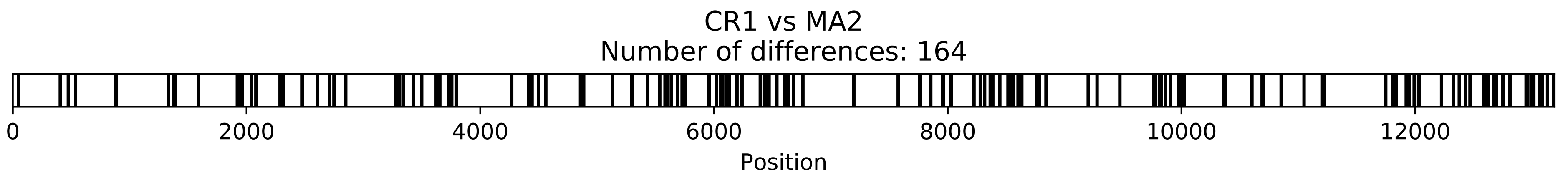
CR1 vs MA2
Number of differences: 164
0 2000 4000 6000 8000 10000 12000
Position

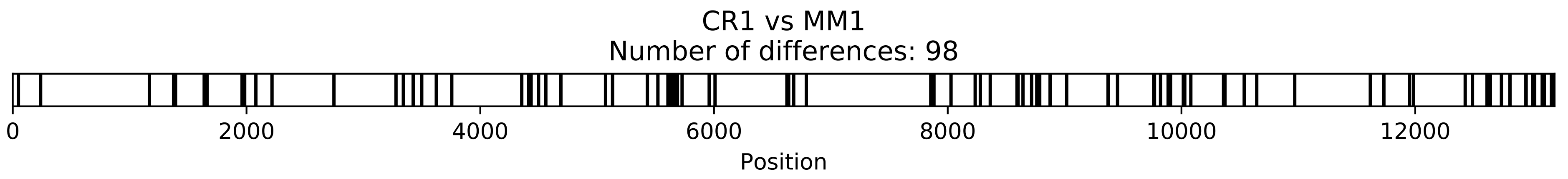
CR1 vs MM1
Number of differences: 98
0 2000 4000 6000 8000 10000 12000
Position

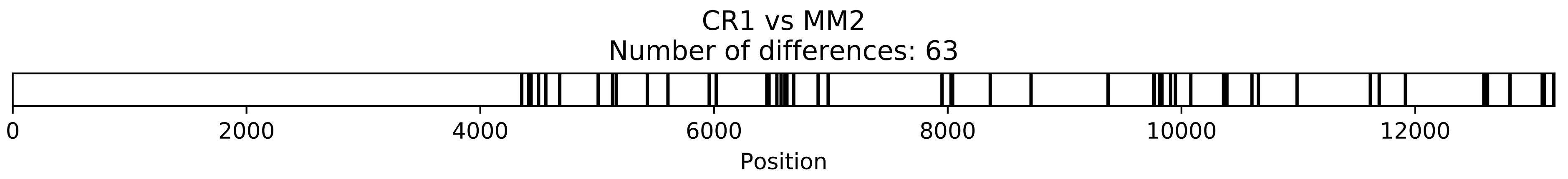
CR1 vs MM2
Number of differences: 63
0 2000 4000 6000 8000 10000 12000
Position

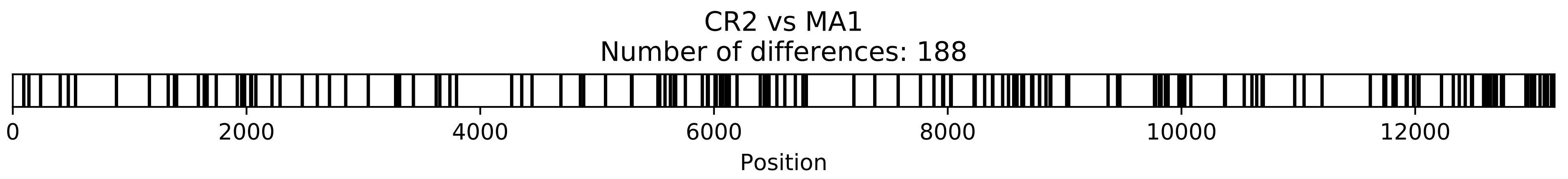
CR2 vs MA1
Number of differences: 188
0 2000 4000 6000 8000 10000 12000
Position

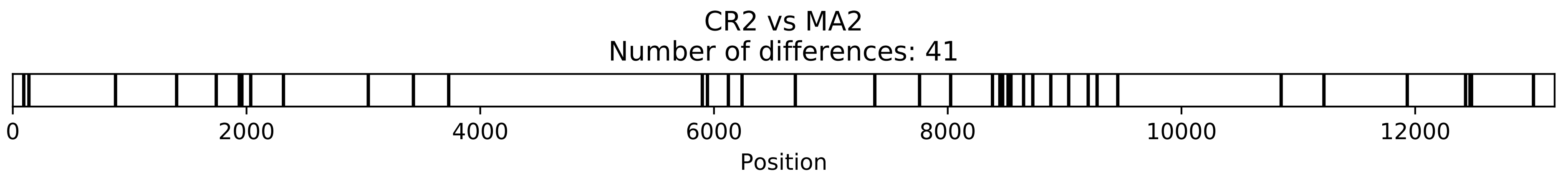
CR2 vs MA2
Number of differences: 41
0 2000 4000 6000 8000 10000 12000
Position

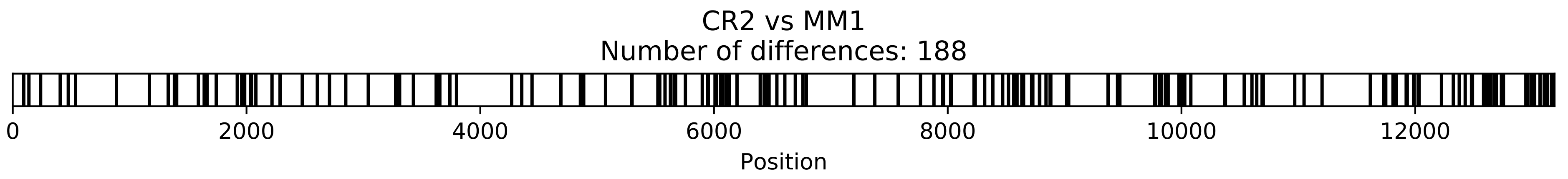
CR2 vs MM1
Number of differences: 188
0 2000 4000 6000 8000 10000 12000
Position

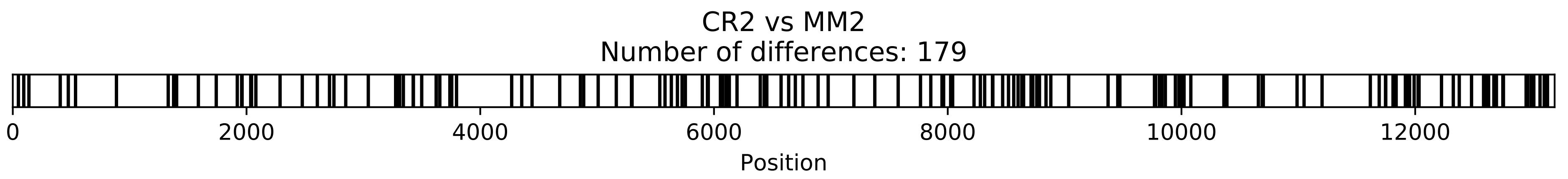
CR2 vs MM2
Number of differences: 179
0 2000 4000 6000 8000 10000 12000
Position

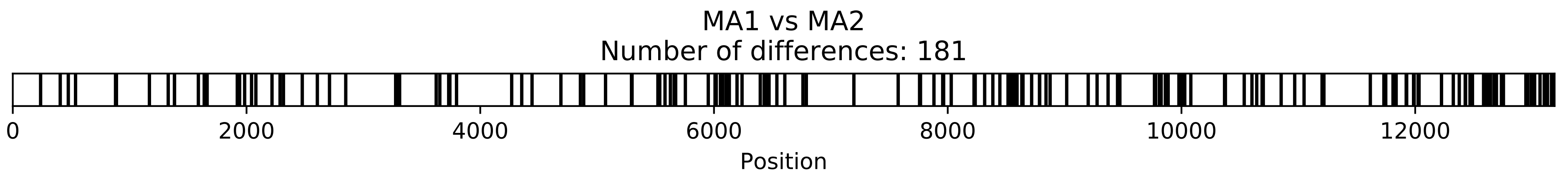
MA1 vs MA2
Number of differences: 181
0 2000 4000 6000 8000 10000 12000
Position

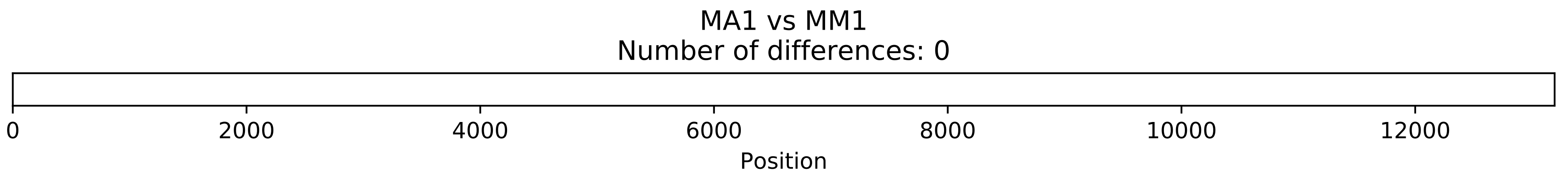
MA1 vs MM1
Number of differences: 0
0 2000 4000 6000 8000 10000 12000
Position

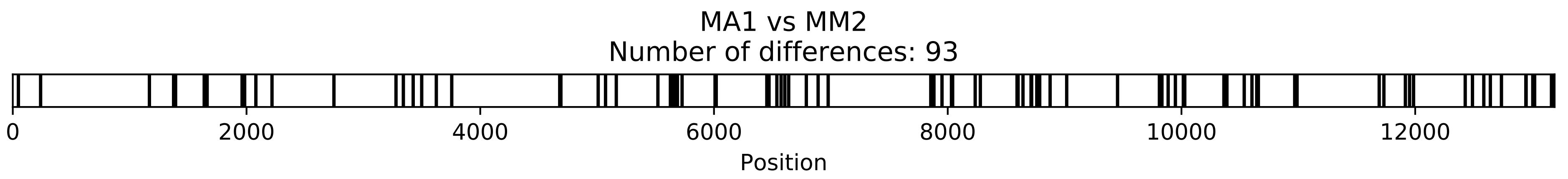
MA1 vs MM2
Number of differences: 93
0 2000 4000 6000 8000 10000 12000
Position

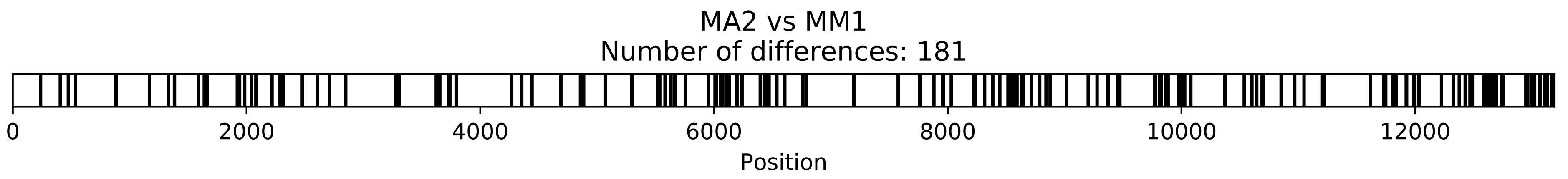
MA2 vs MM1
Number of differences: 181
0 2000 4000 6000 8000 10000 12000
Position

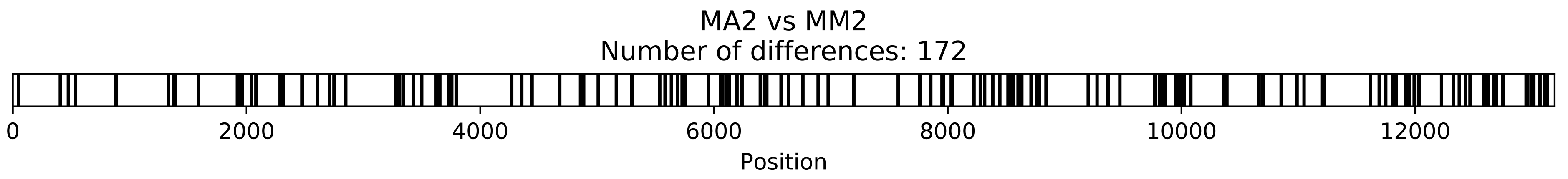
MA2 vs MM2
Number of differences: 172
0 2000 4000 6000 8000 10000 12000
Position

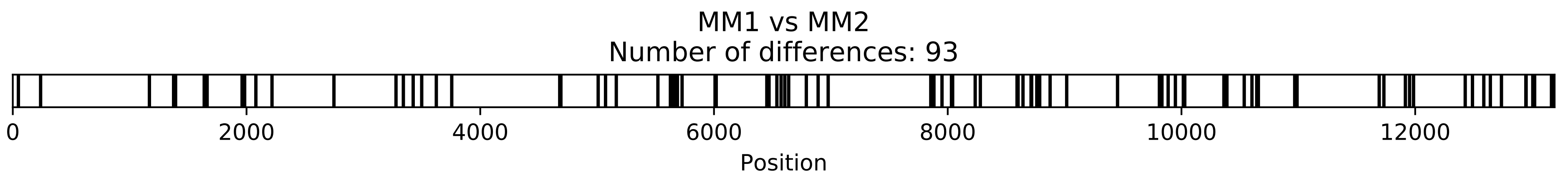
MM1 vs MM2
Number of differences: 93
0 2000 4000 6000 8000 10000 12000
Position

### Region: 39



CR1 vs CR2
Number of differences: 258

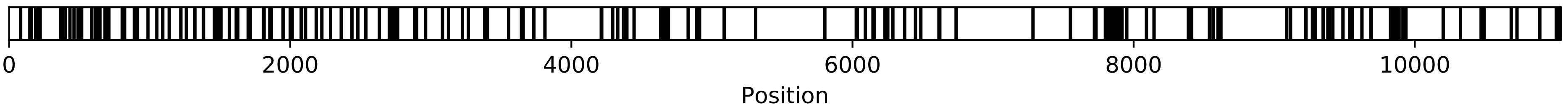


CR1 vs MA1
Number of differences: 90

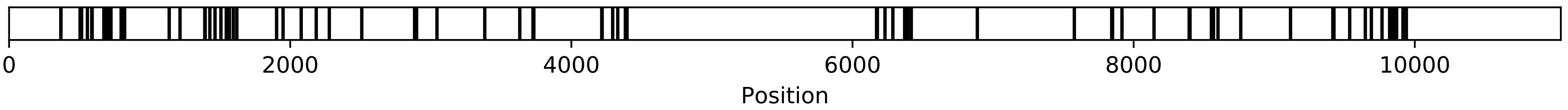


CR1 vs MA2
Number of differences: 252

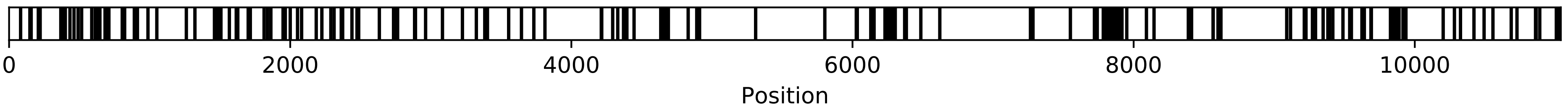


CR1 vs MM1
Number of differences: 90

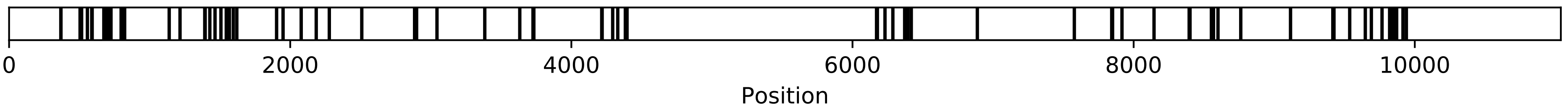


CR1 vs MM2
Number of differences: 108

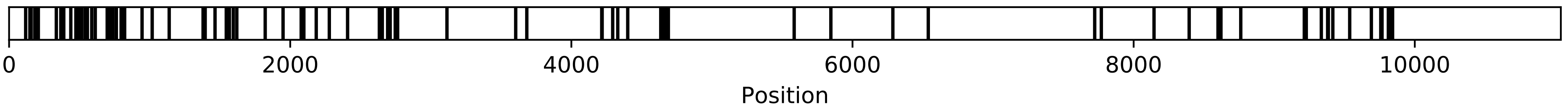


CR2 vs MA1
Number of differences: 258

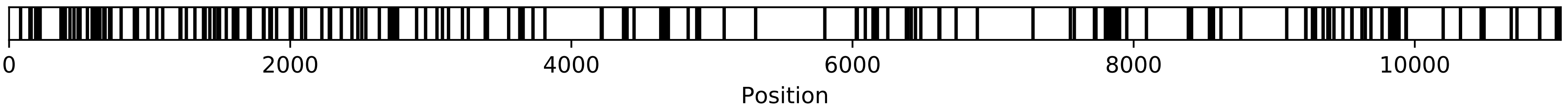


CR2 vs MA2
Number of differences: 67

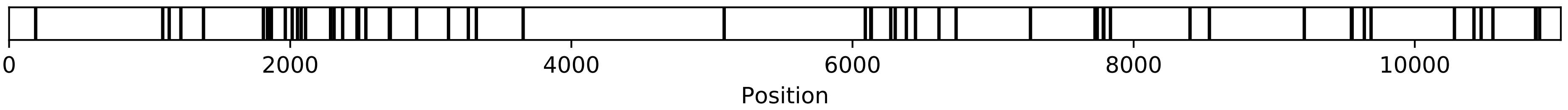


CR2 vs MM1
Number of differences: 258

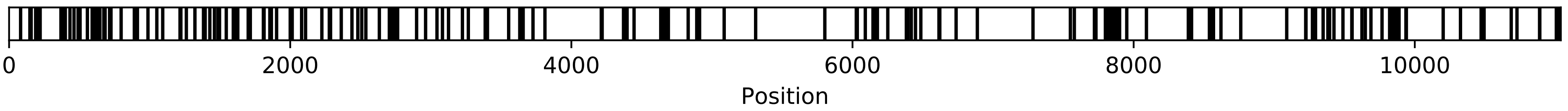


CR2 vs MM2
Number of differences: 263

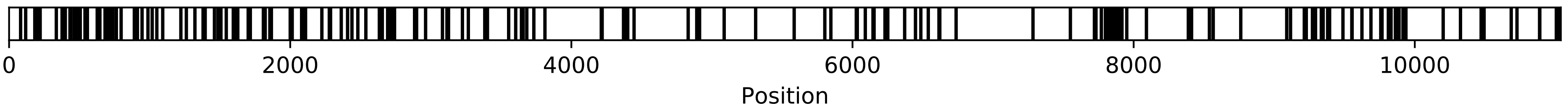


MA1 vs MA2
Number of differences: 257

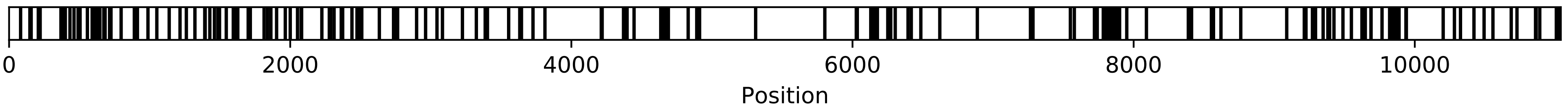


MA1 vs MM1
Number of differences: 0

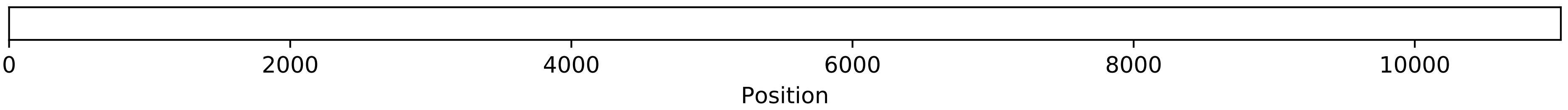


MA1 vs MM2
Number of differences: 126

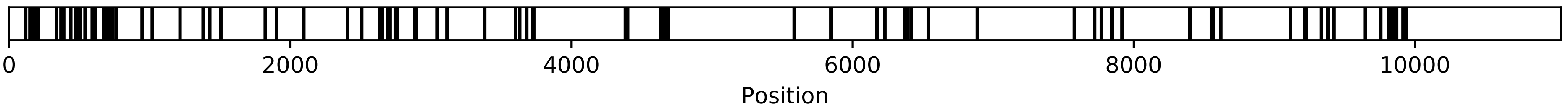


MA2 vs MM1
Number of differences: 257

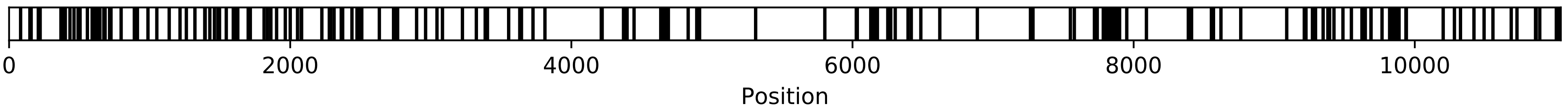


MA2 vs MM2
Number of differences: 260

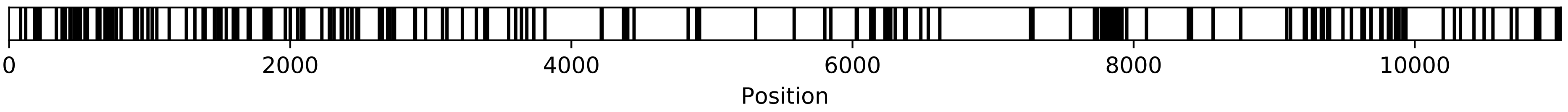


MM1 vs MM2
Number of differences: 126

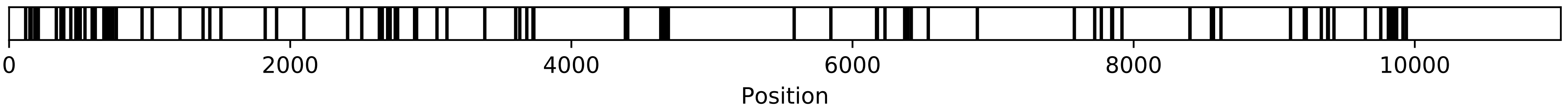

### Region: 40



CR1 vs CR2
Number of differences: 164

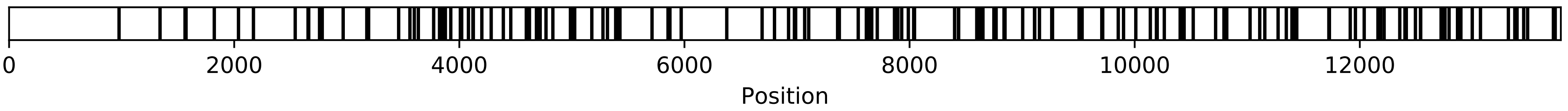


CR1 vs MA1
Number of differences: 133

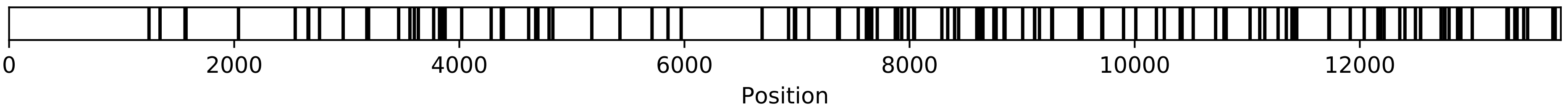


CR1 vs MA2
Number of differences: 121

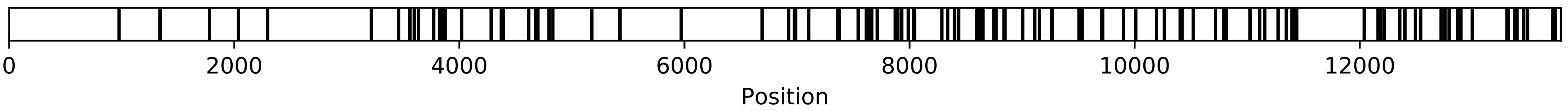


CR1 vs MM1
Number of differences: 28

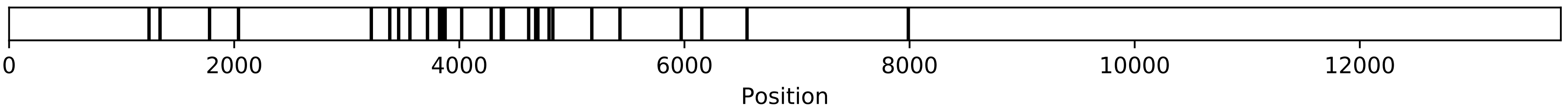


CR1 vs MM2
Number of differences: 79

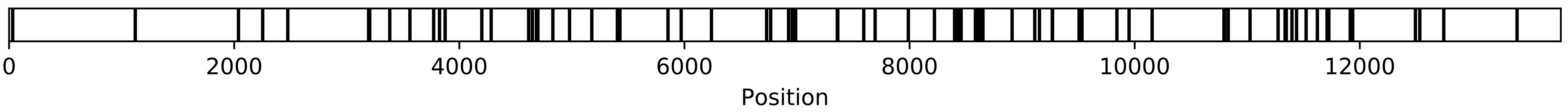


CR2 vs MA1
Number of differences: 55

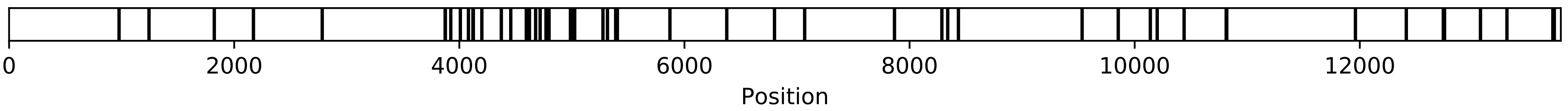


CR2 vs MA2
Number of differences: 71

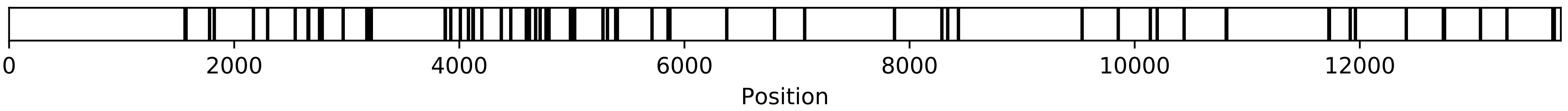


CR2 vs MM1
Number of differences: 158

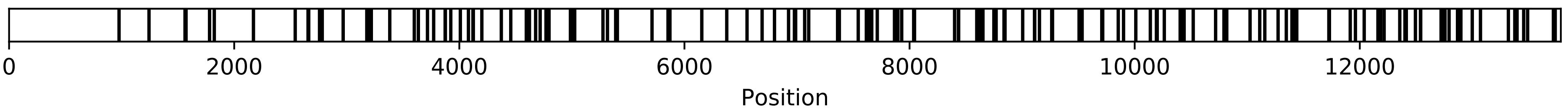


CR2 vs MM2
Number of differences: 157

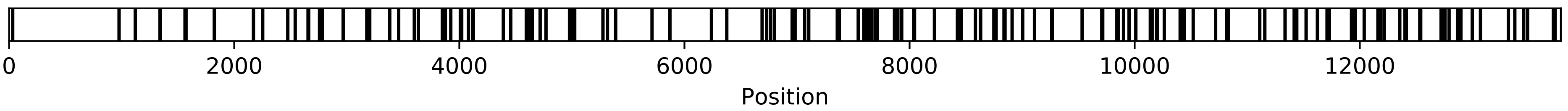


MA1 vs MA2
Number of differences: 20

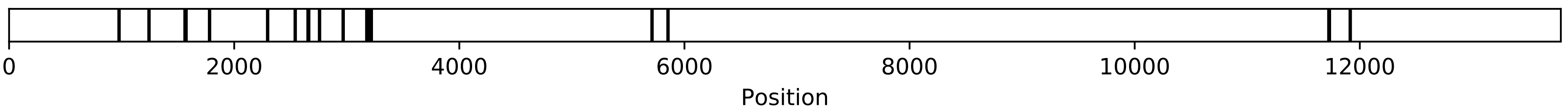


MA1 vs MM1
Number of differences: 117

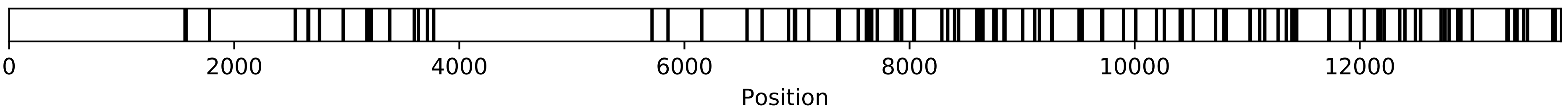


MA1 vs MM2
Number of differences: 128

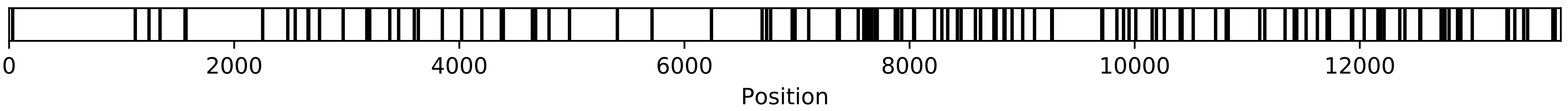


MA2 vs MM1
Number of differences: 103

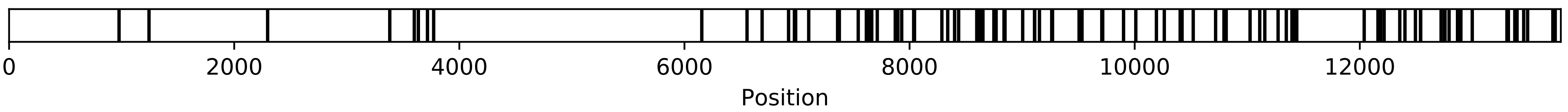


MA2 vs MM2
Number of differences: 124

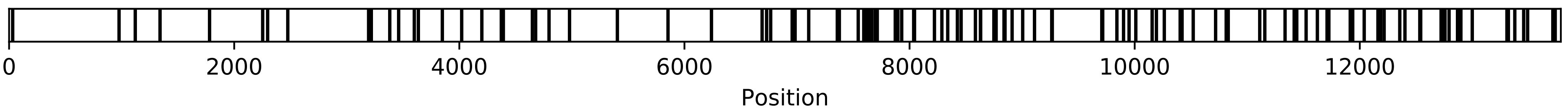


MM1 vs MM2
Number of differences: 79

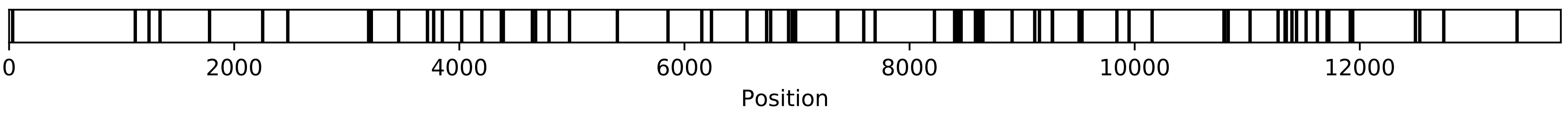

### Region: 42



CR1 vs CR2
Number of differences: 239

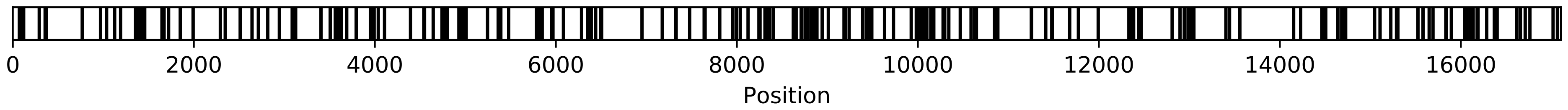


CR1 vs MA1
Number of differences: 167

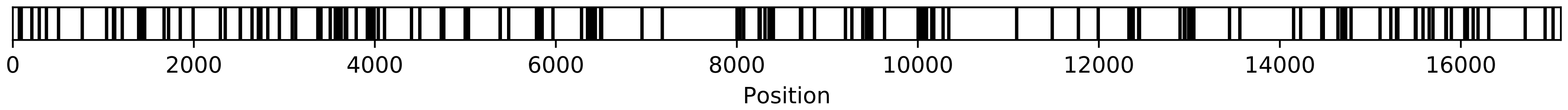


CR1 vs MA2
Number of differences: 173

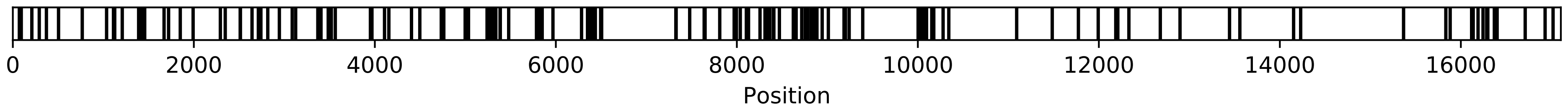


CR1 vs MM1
Number of differences: 95

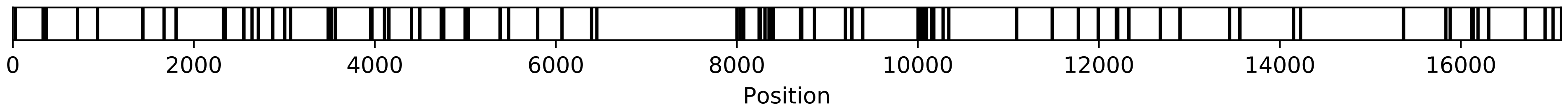


CR1 vs MM2
Number of differences: 98

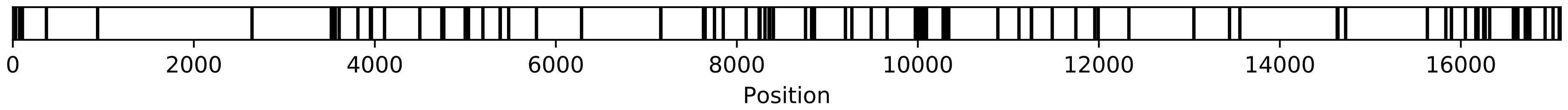


CR2 vs MA1
Number of differences: 150

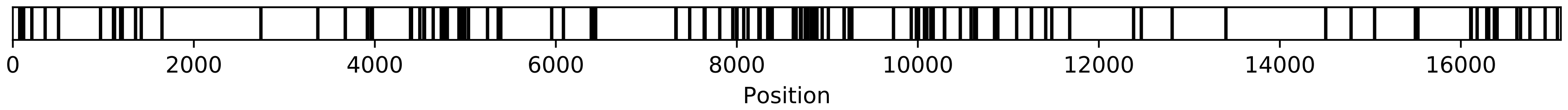


CR2 vs MA2
Number of differences: 194

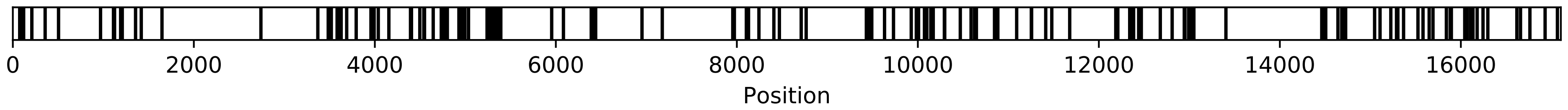


CR2 vs MM1
Number of differences: 234

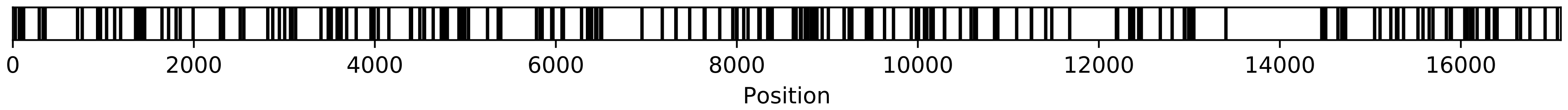


CR2 vs MM2
Number of differences: 243

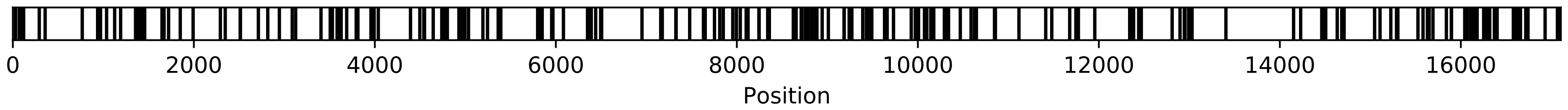


MA1 vs MA2
Number of differences: 136

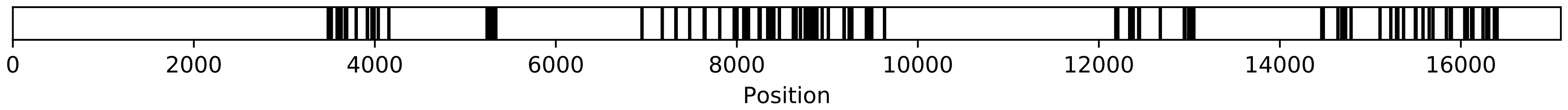


MA1 vs MM1
Number of differences: 126

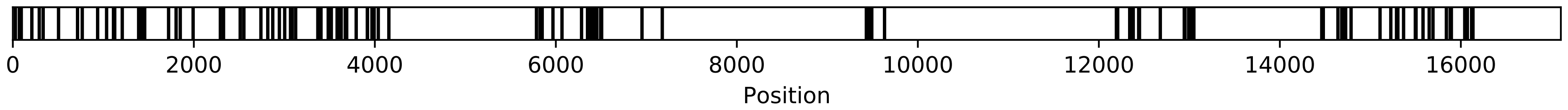


MA1 vs MM2
Number of differences: 169

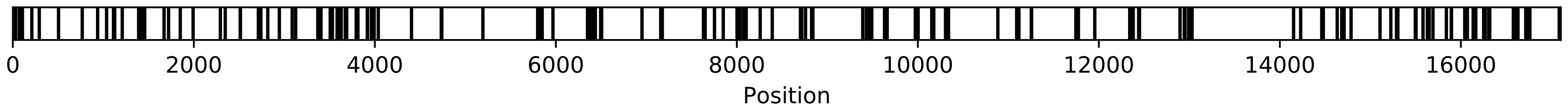


MA2 vs MM1
Number of differences: 124

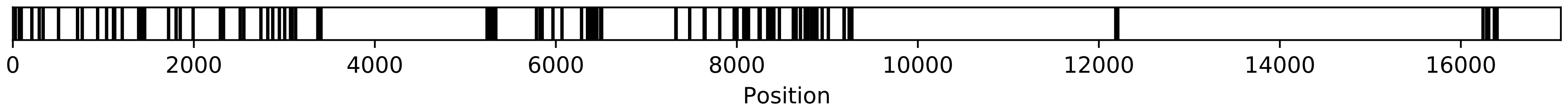


MA2 vs MM2
Number of differences: 184

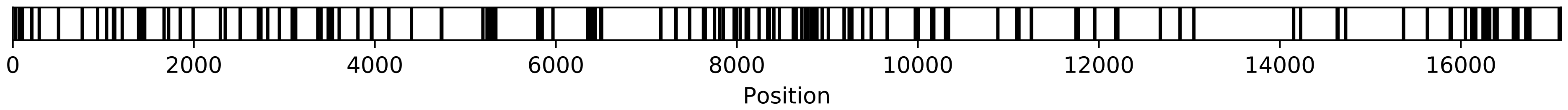


MM1 vs MM2
Number of differences: 106

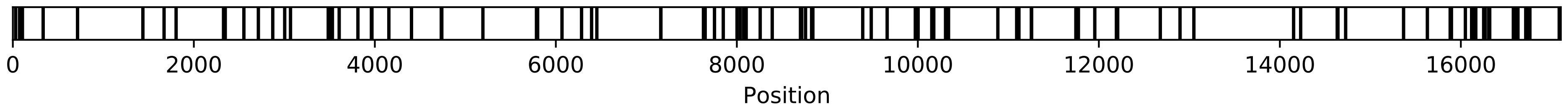

### Region: 44



CR1 vs CR2
Number of differences: 192

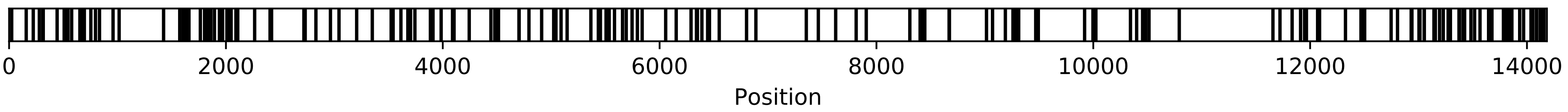


CR1 vs MA1
Number of differences: 87

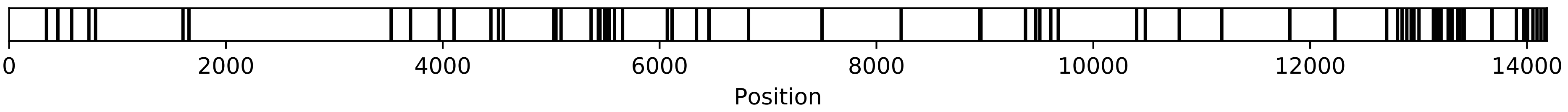


CR1 vs MA2
Number of differences: 196

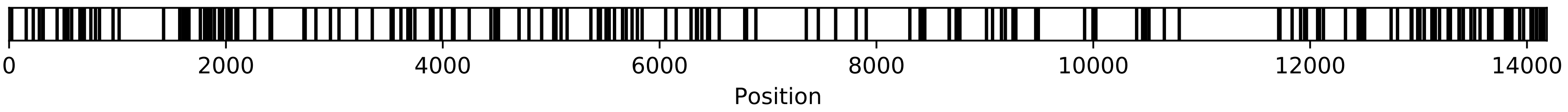


CR1 vs MM1
Number of differences: 117

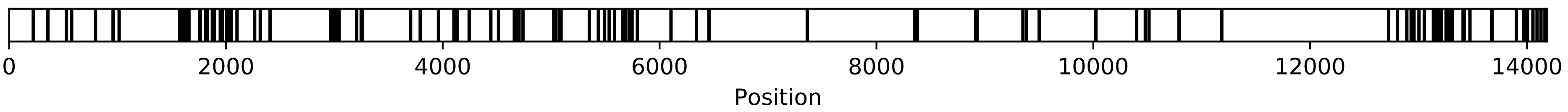


CR1 vs MM2
Number of differences: 87

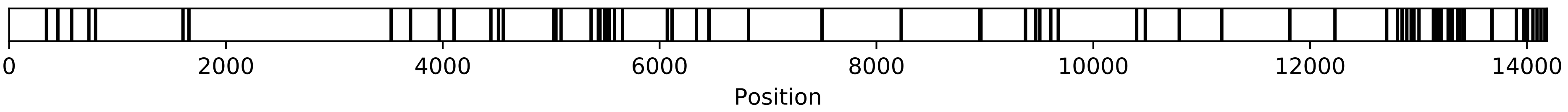


CR2 vs MA1
Number of differences: 198

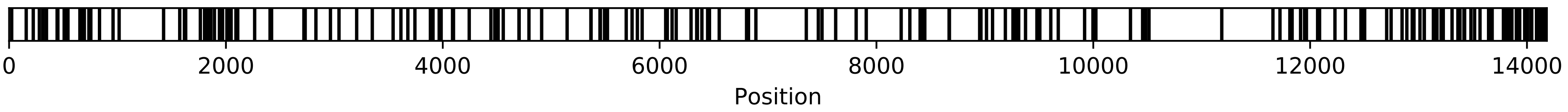


CR2 vs MA2
Number of differences: 20

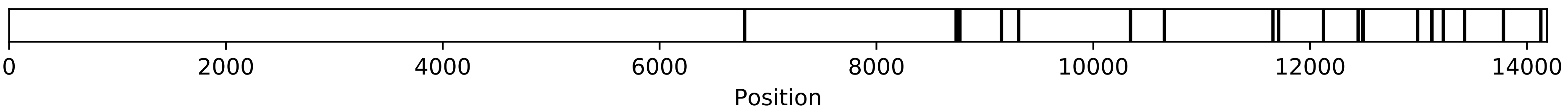


CR2 vs MM1
Number of differences: 197

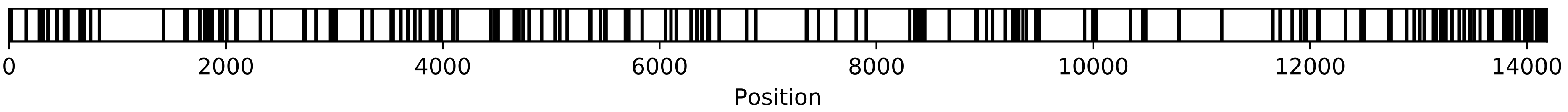


CR2 vs MM2
Number of differences: 198

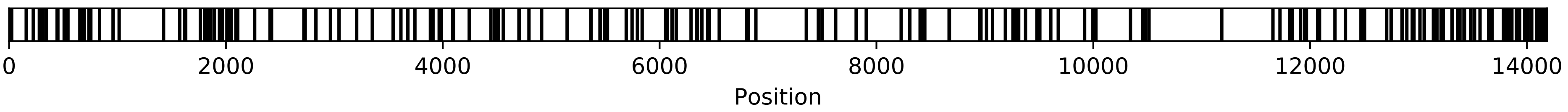


MA1 vs MA2
Number of differences: 202

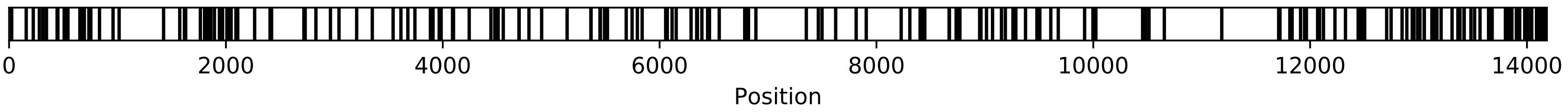


MA1 vs MM1
Number of differences: 98

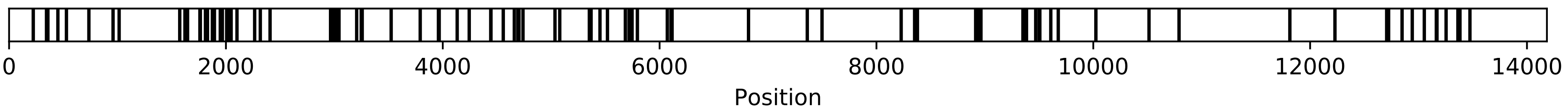


MA1 vs MM2
Number of differences: 0

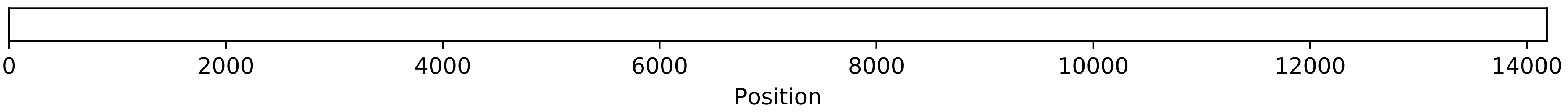


MA2 vs MM1
Number of differences: 201

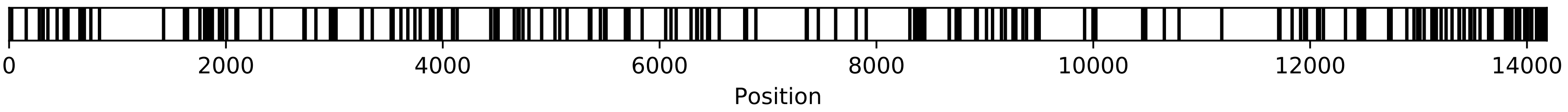


MA2 vs MM2
Number of differences: 202

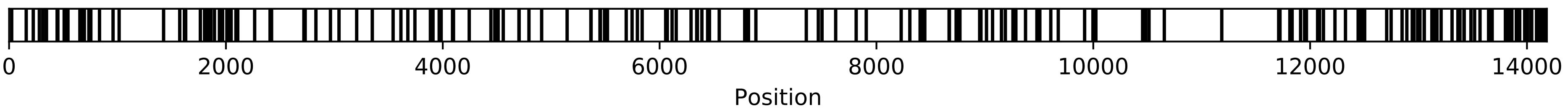


MM1 vs MM2
Number of differences: 98

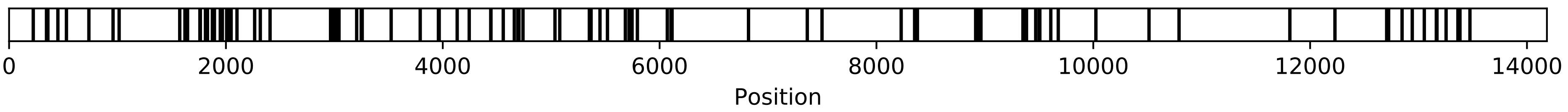

### Region: 46



CR1 vs CR2
Number of differences: 204

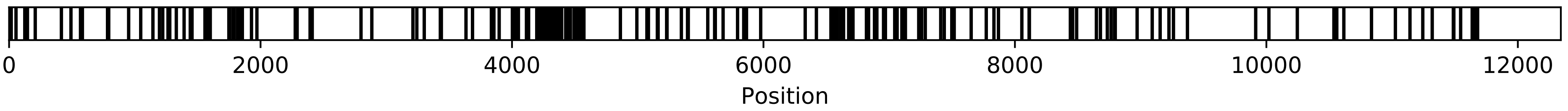


CR1 vs MA1
Number of differences: 60

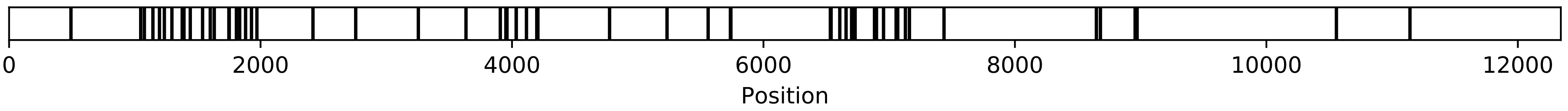


CR1 vs MA2
Number of differences: 202

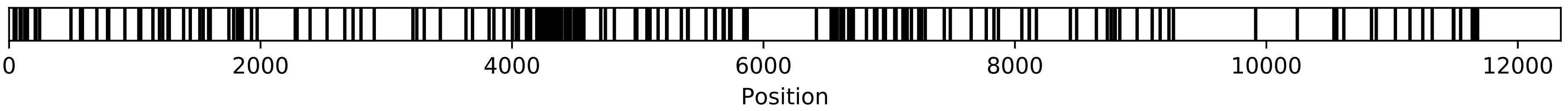


CR1 vs MM1
Number of differences: 211

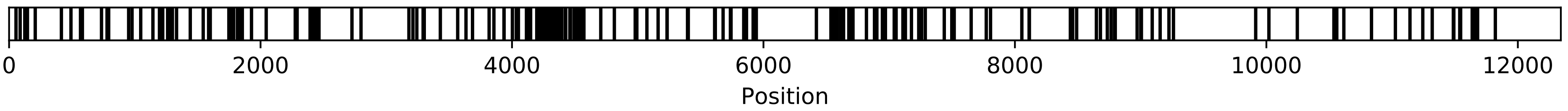


CR1 vs MM2
Number of differences: 202

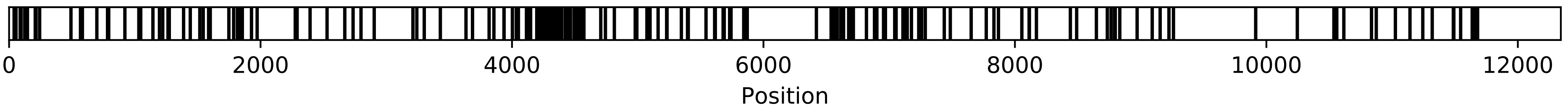


CR2 vs MA1
Number of differences: 214

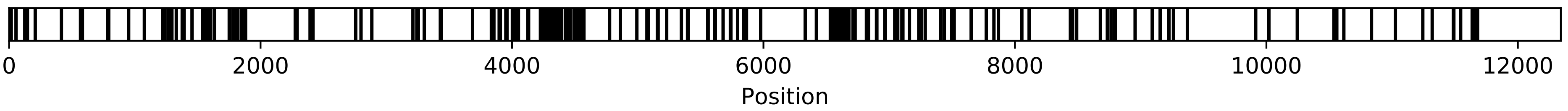


CR2 vs MA2
Number of differences: 104

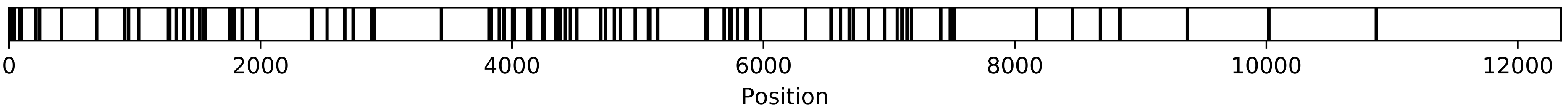


CR2 vs MM1
Number of differences: 101

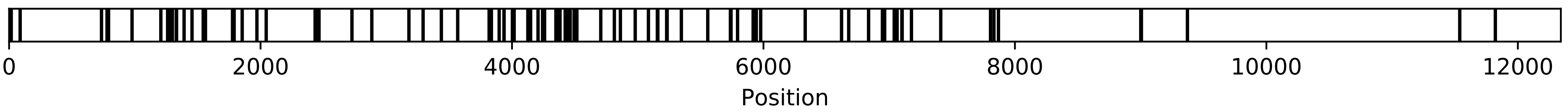


CR2 vs MM2
Number of differences: 104

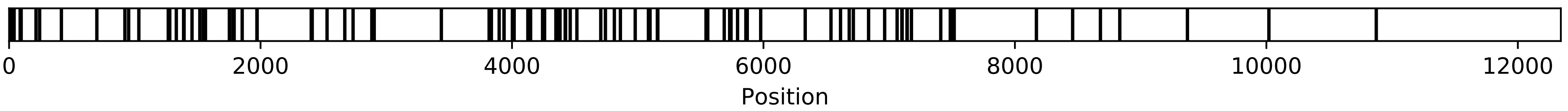


MA1 vs MA2
Number of differences: 213

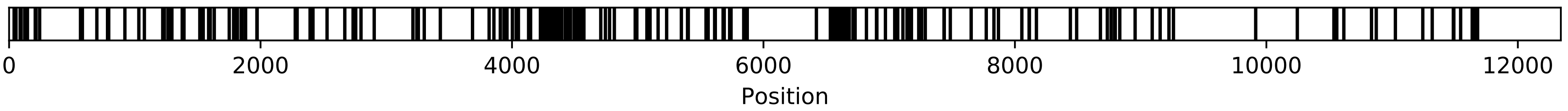


MA1 vs MM1
Number of differences: 221

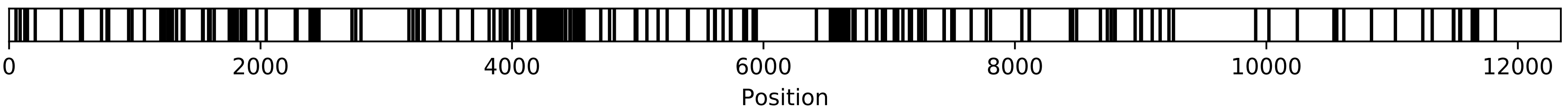


MA1 vs MM2
Number of differences: 213

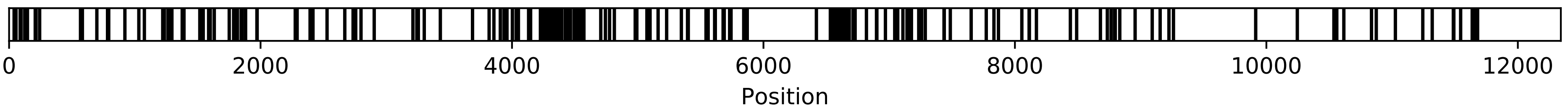


MA2 vs MM1
Number of differences: 83

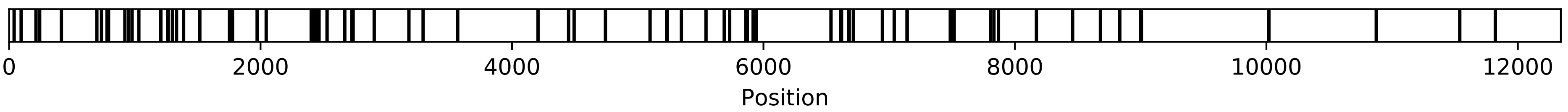


MA2 vs MM2
Number of differences: 0

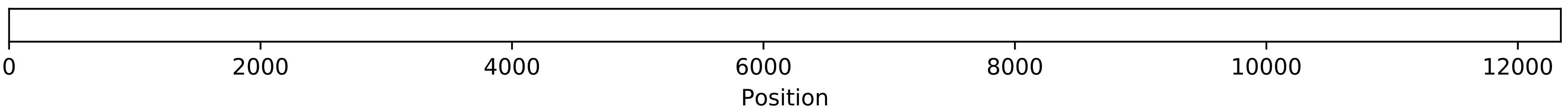


MM1 vs MM2
Number of differences: 83

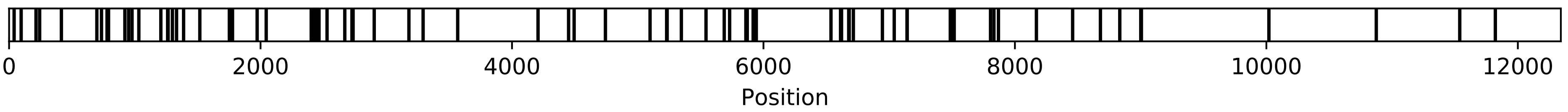

### Region: 47



CR1 vs CR2
Number of differences: 266

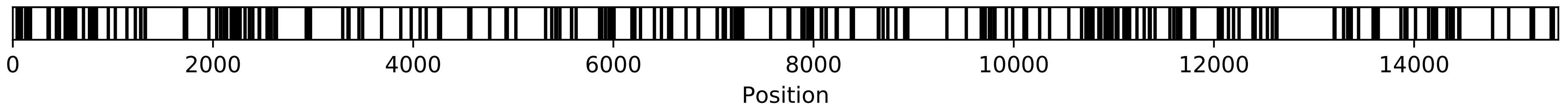


CR1 vs MA1
Number of differences: 275

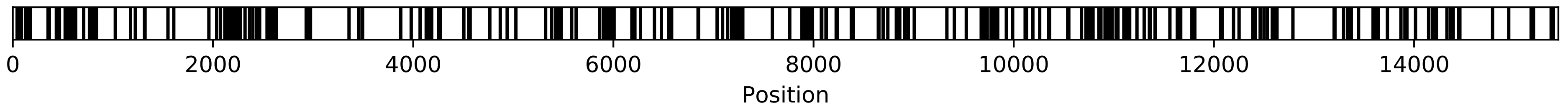


CR1 vs MA2
Number of differences: 151

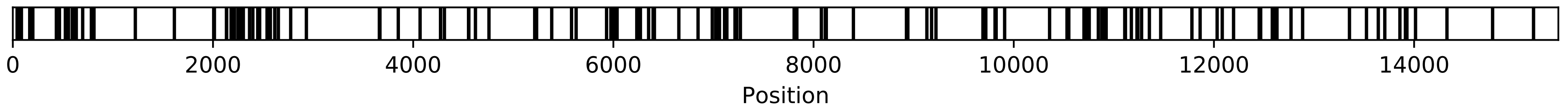


CR1 vs MM1
Number of differences: 80

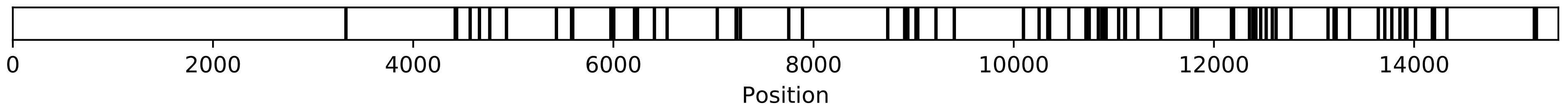


CR1 vs MM2
Number of differences: 151

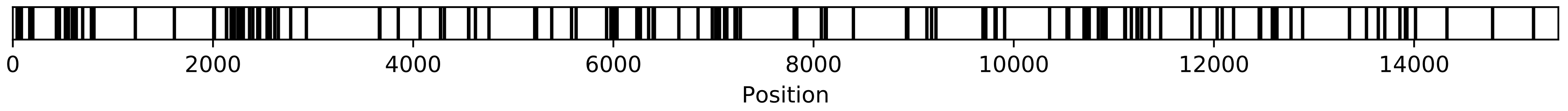


CR2 vs MA1
Number of differences: 82

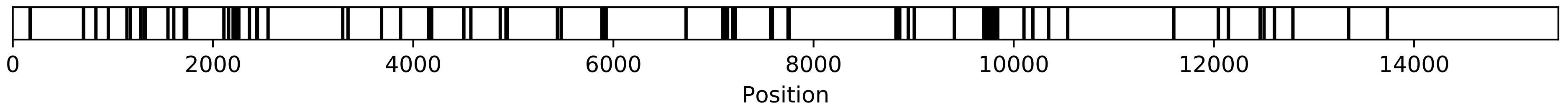


CR2 vs MA2
Number of differences: 290

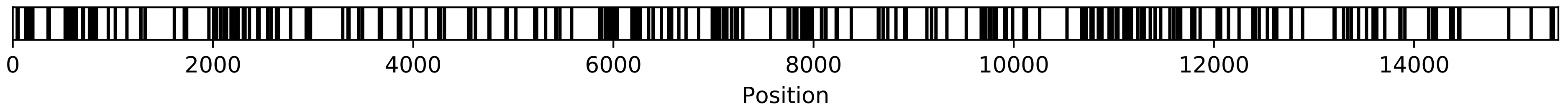


CR2 vs MM1
Number of differences: 272

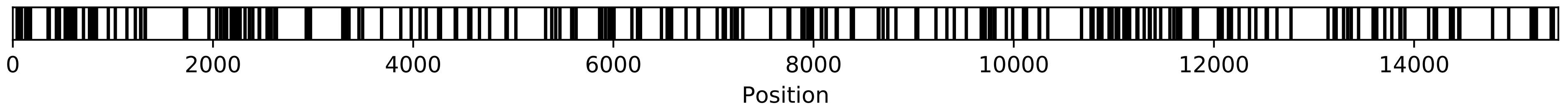


CR2 vs MM2
Number of differences: 290

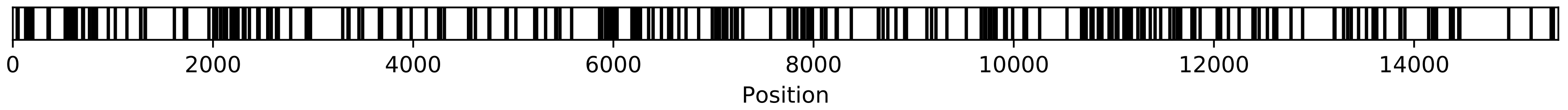


MA1 vs MA2
Number of differences: 298

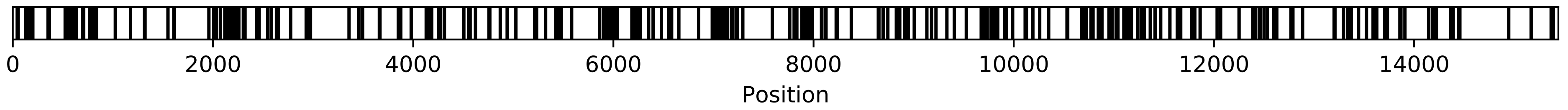


MA1 vs MM1
Number of differences: 279

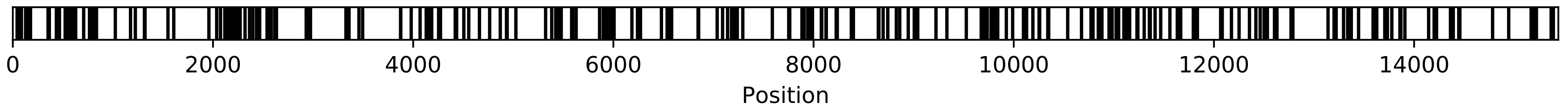


MA1 vs MM2
Number of differences: 298

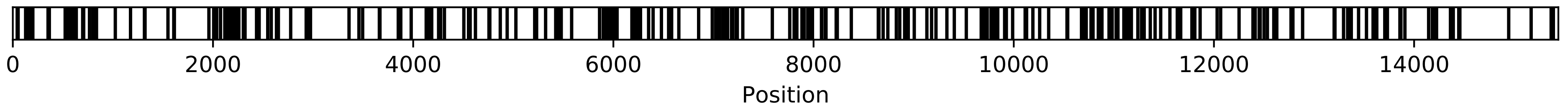


MA2 vs MM1
Number of differences: 154

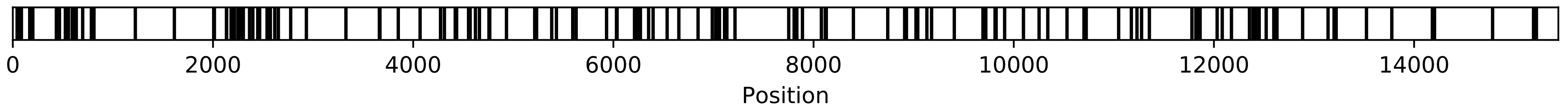


MA2 vs MM2
Number of differences: 0

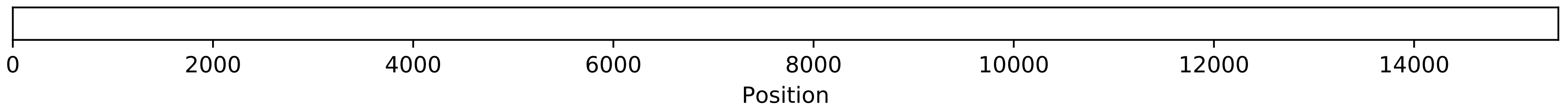


MM1 vs MM2
Number of differences: 154

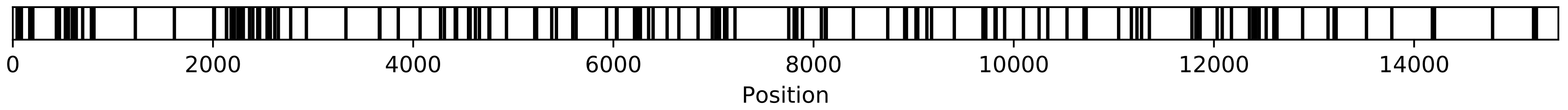

### Region: 49

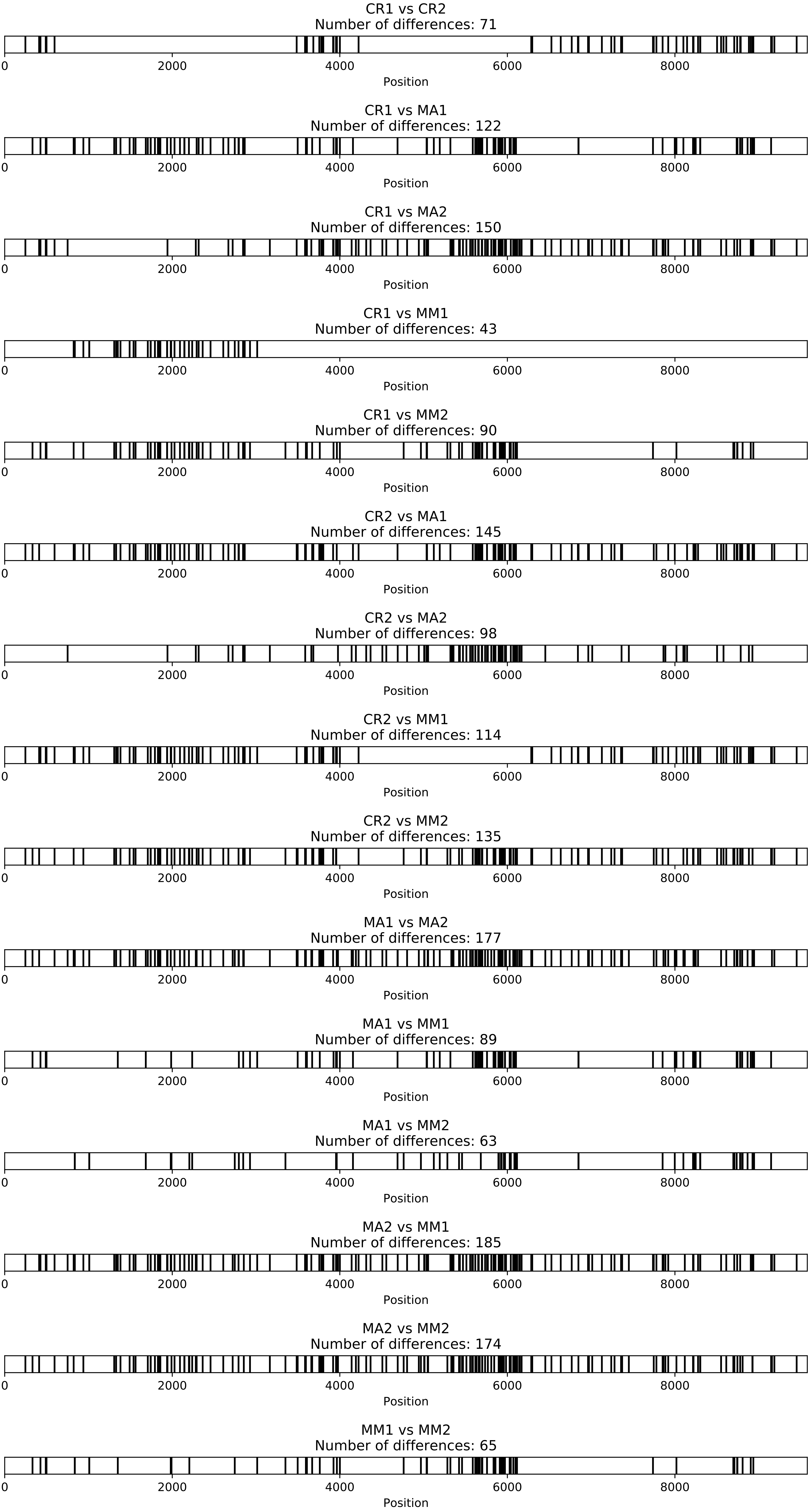
CR1 vs CR2
Number of differences: 71
CR1 vs MA1
Number of differences: 122
CR1 vs MA2
Number of differences: 150
CR1 vs MM1
Number of differences: 43
CR1 vs MM2
Number of differences: 90
CR2 vs MA1
Number of differences: 145
CR2 vs MA2
Number of differences: 98
CR2 vs MM1
Number of differences: 114
CR2 vs MM2
Number of differences: 135
MA1 vs MA2
Number of differences: 177
MA1 vs MM1
Number of differences: 89
MA1 vs MM2
Number of differences: 63
MA2 vs MM1
Number of differences: 185
MA2 vs MM2
Number of differences: 174
MM1 vs MM2
Number of differences: 65
0
2000
4000
6000
8000
Position

### Region: 50

CR1 vs CR2
Number of differences: 607

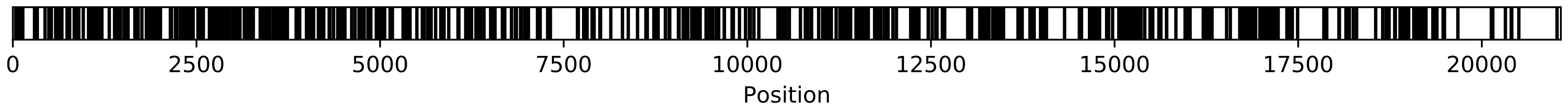


CR1 vs MA1
Number of differences: 627

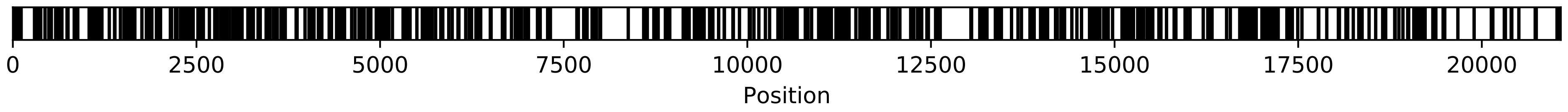


CR1 vs MA2
Number of differences: 300

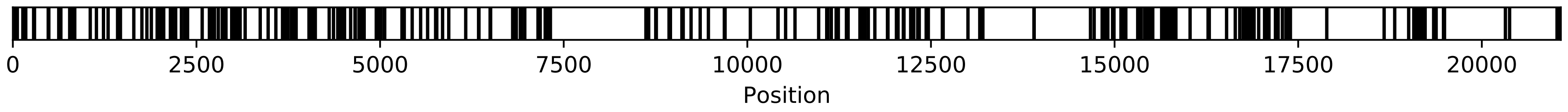


CR1 vs MM1
Number of differences: 658

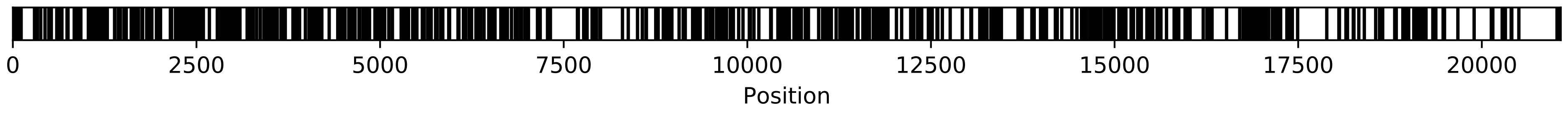


CR1 vs MM2
Number of differences: 628

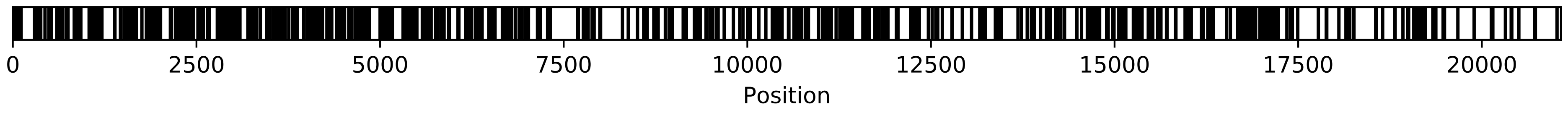


CR2 vs MA1
Number of differences: 366

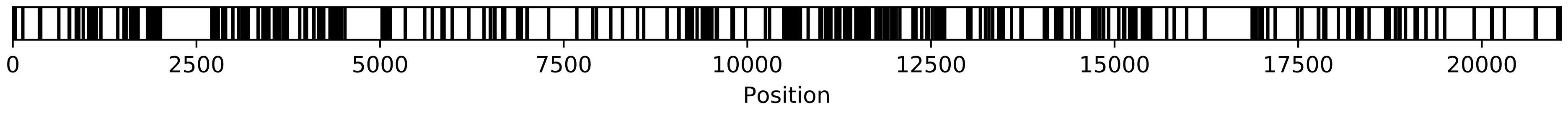


CR2 vs MA2
Number of differences: 589

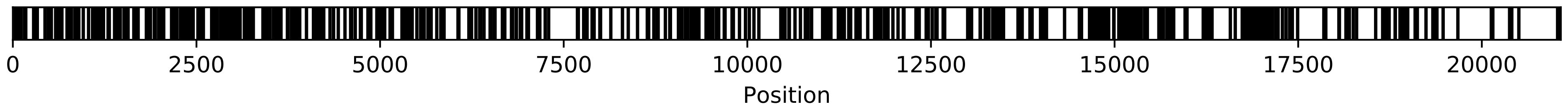


CR2 vs MM1
Number of differences: 393

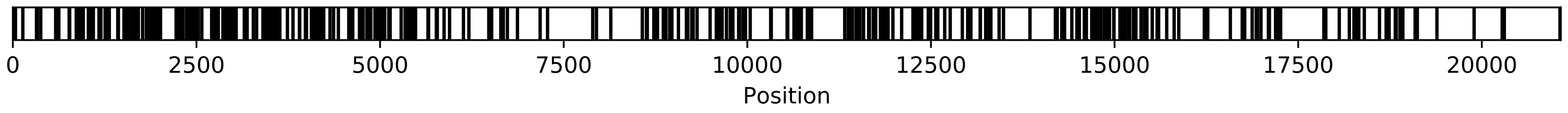


CR2 vs MM2
Number of differences: 382

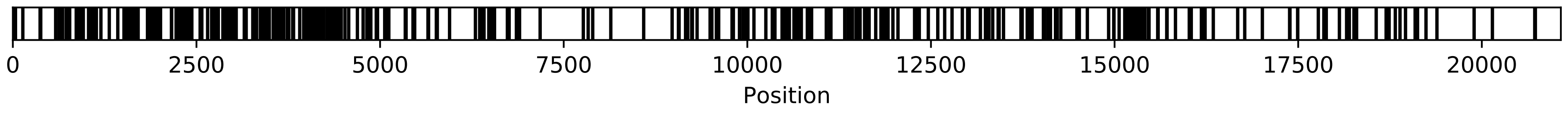


MA1 vs MA2
Number of differences: 572

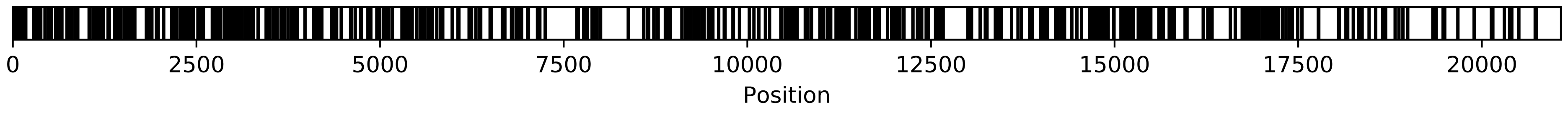


MA1 vs MM1
Number of differences: 452

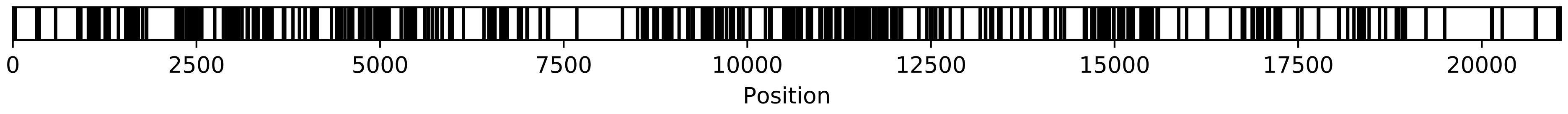


MA1 vs MM2
Number of differences: 392

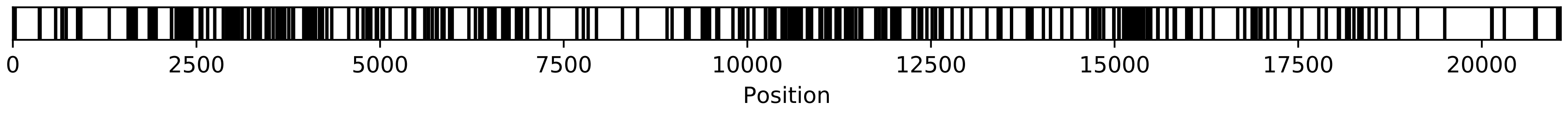


MA2 vs MM1
Number of differences: 640

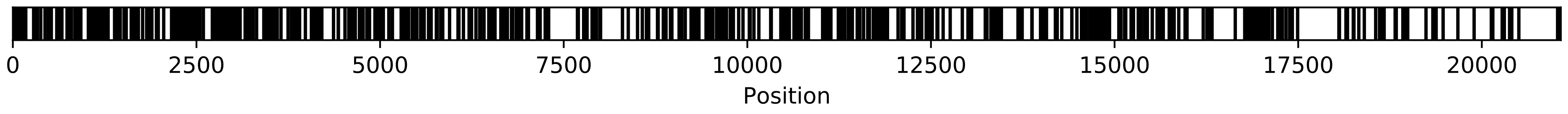


MA2 vs MM2
Number of differences: 619

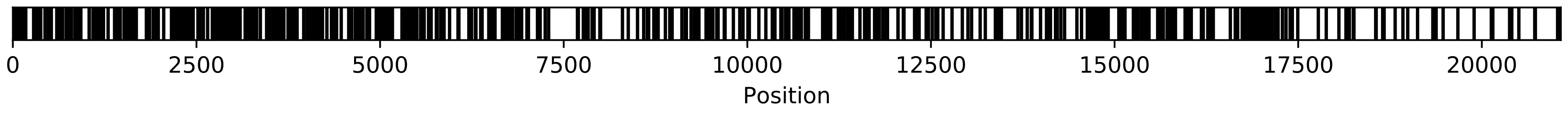


MM1 vs MM2
Number of differences: 380

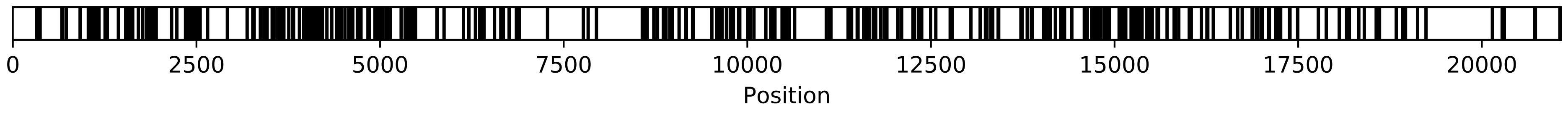

### Region: 51



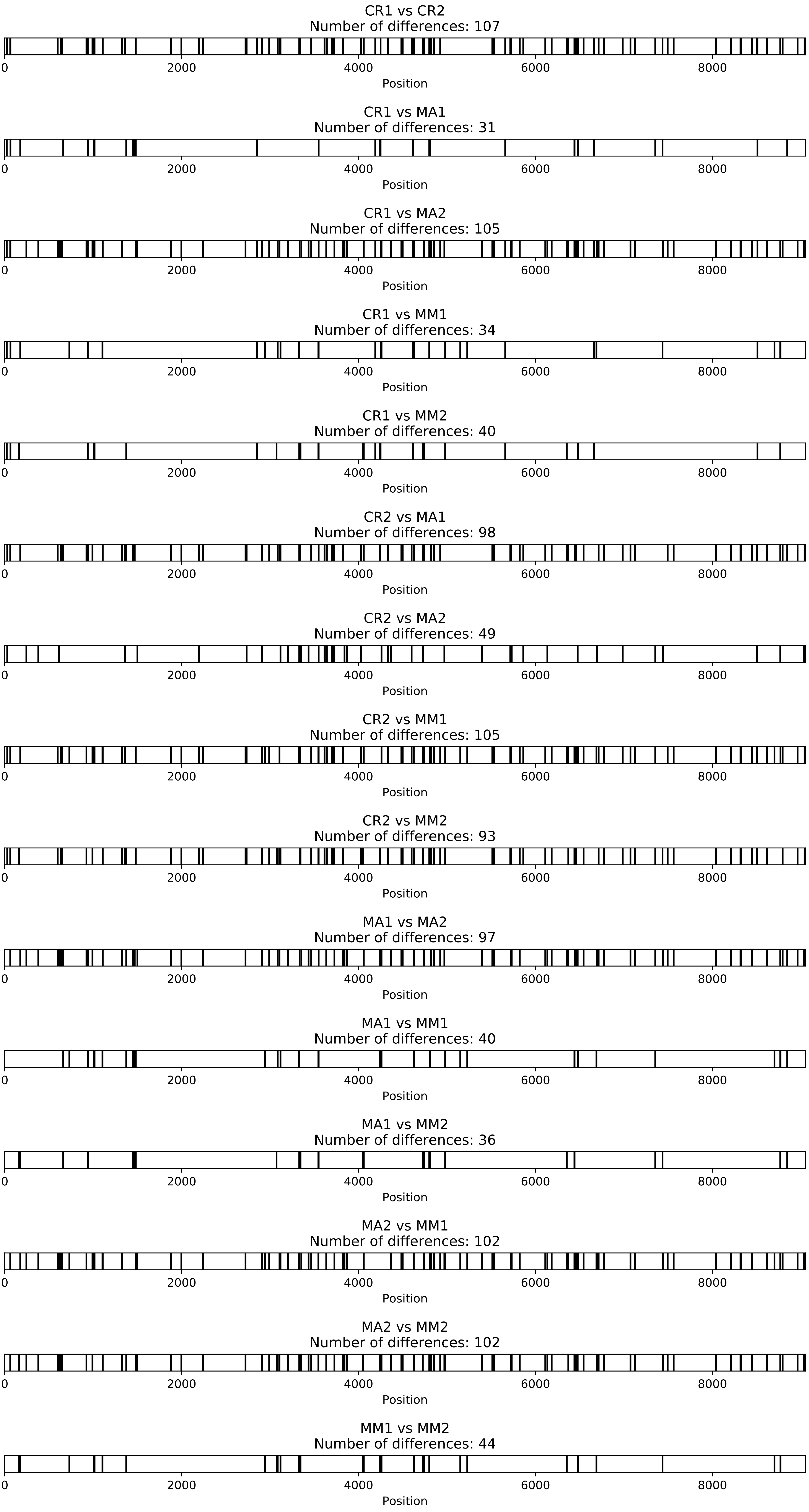
CR1 vs CR2
Number of differences: 107
CR1 vs MA1
Number of differences: 31
CR1 vs MA2
Number of differences: 105
CR1 vs MM1
Number of differences: 34
CR1 vs MM2
Number of differences: 40
CR2 vs MA1
Number of differences: 98
CR2 vs MA2
Number of differences: 49
CR2 vs MM1
Number of differences: 105
CR2 vs MM2
Number of differences: 93
MA1 vs MA2
Number of differences: 97
MA1 vs MM1
Number of differences: 40
MA1 vs MM2
Number of differences: 36
MA2 vs MM1
Number of differences: 102
MA2 vs MM2
Number of differences: 102
MM1 vs MM2
Number of differences: 44
0
2000
4000
6000
8000
Position

### Region: 52



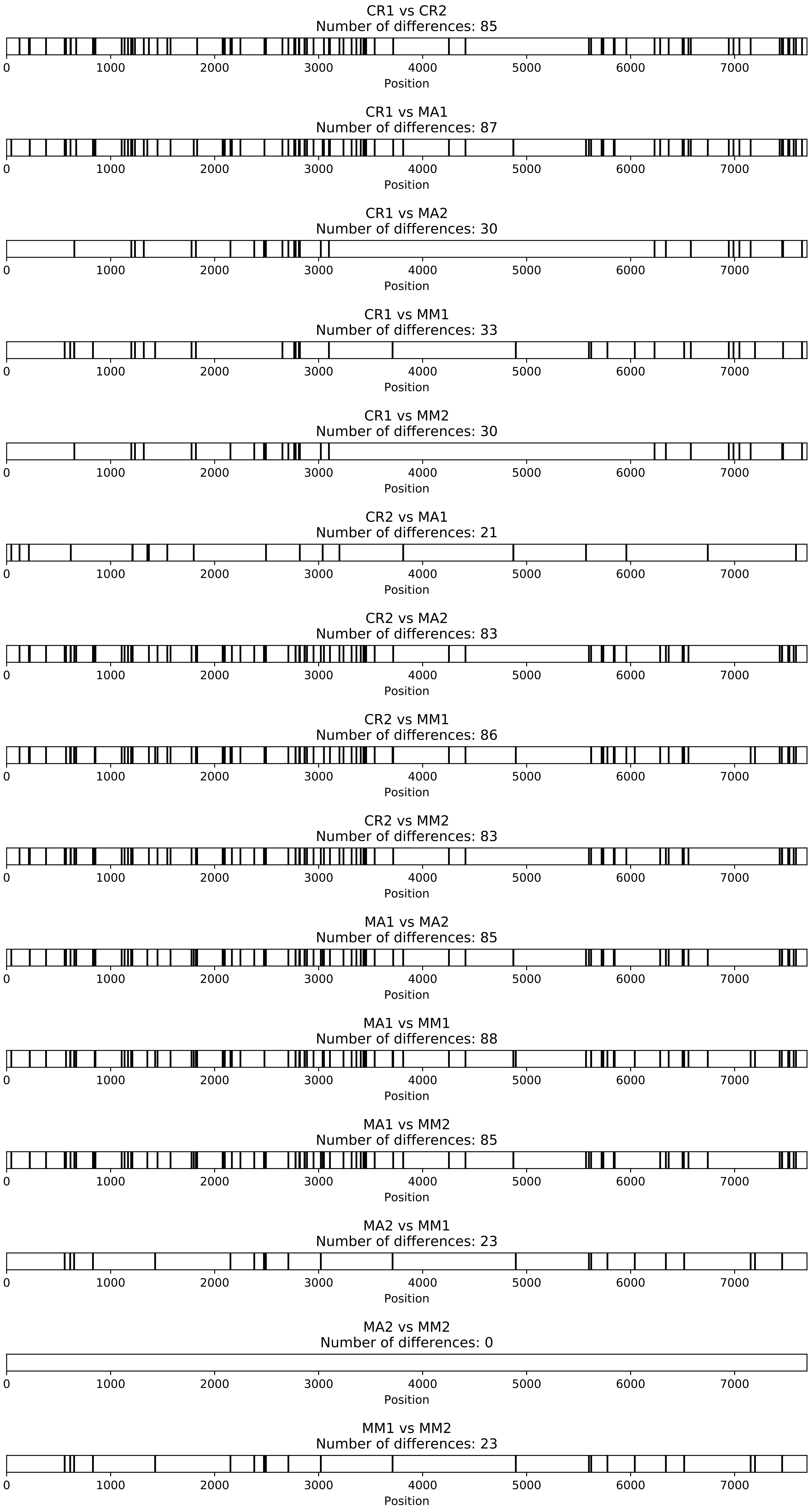
CR1 vs CR2
Number of differences: 85
CR1 vs MA1
Number of differences: 87
CR1 vs MA2
Number of differences: 30
CR1 vs MM1
Number of differences: 33
CR1 vs MM2
Number of differences: 30
CR2 vs MA1
Number of differences: 21
CR2 vs MA2
Number of differences: 83
CR2 vs MM1
Number of differences: 86
CR2 vs MM2
Number of differences: 83
MA1 vs MA2
Number of differences: 85
MA1 vs MM1
Number of differences: 88
MA1 vs MM2
Number of differences: 85
MA2 vs MM1
Number of differences: 23
MA2 vs MM2
Number of differences: 0
MM1 vs MM2
Number of differences: 23
0
1000
2000
3000
4000
5000
6000
7000
Position

### Region: 53



CR1 vs CR2
Number of differences: 368

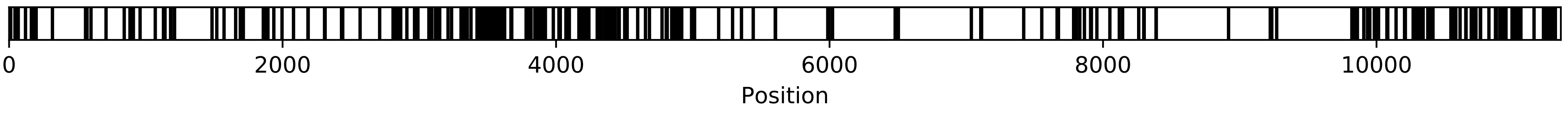


CR1 vs MA1
Number of differences: 174

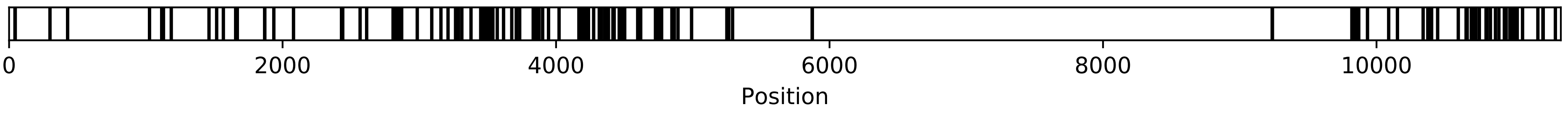


CR1 vs MA2
Number of differences: 384

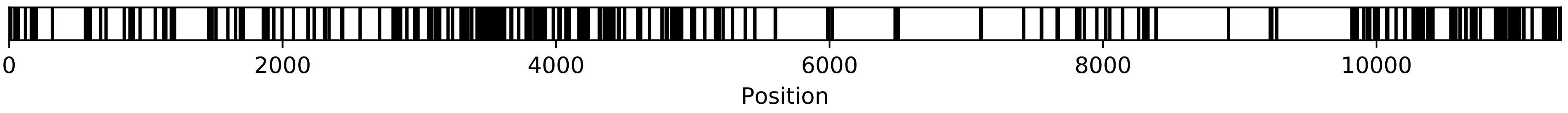


CR1 vs MM1
Number of differences: 173

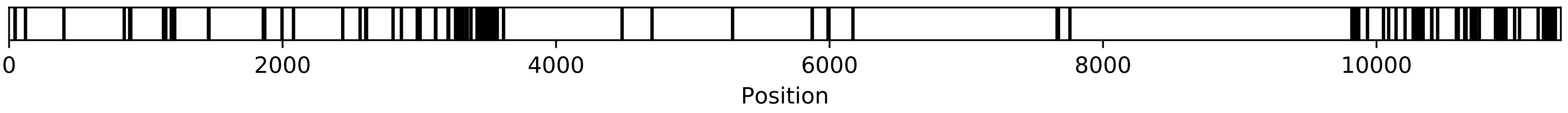


CR1 vs MM2
Number of differences: 136

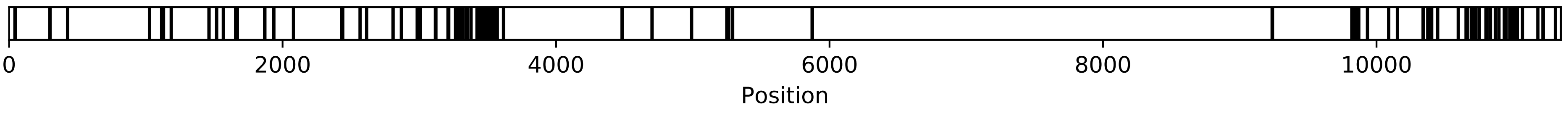


CR2 vs MA1
Number of differences: 372

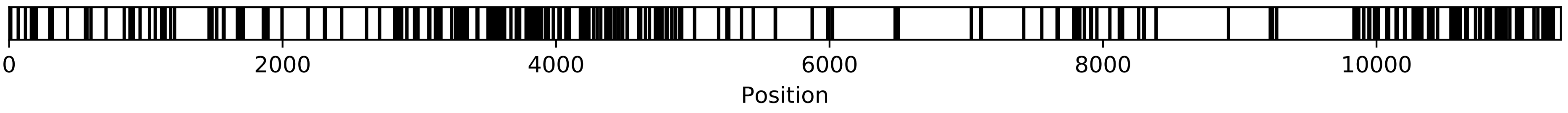


CR2 vs MA2
Number of differences: 81

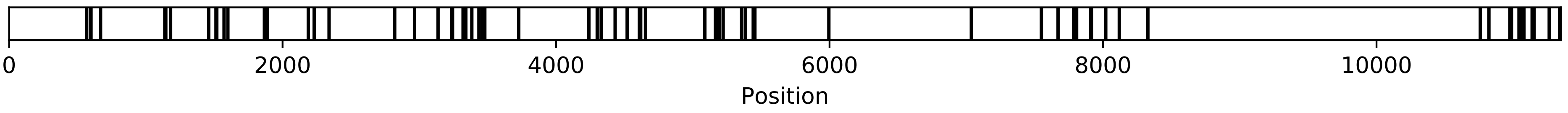


CR2 vs MM1
Number of differences: 333

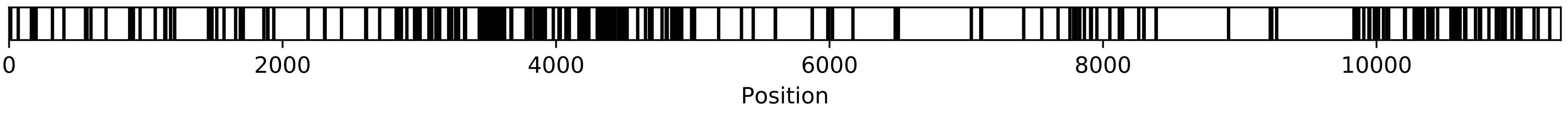


CR2 vs MM2
Number of differences: 365

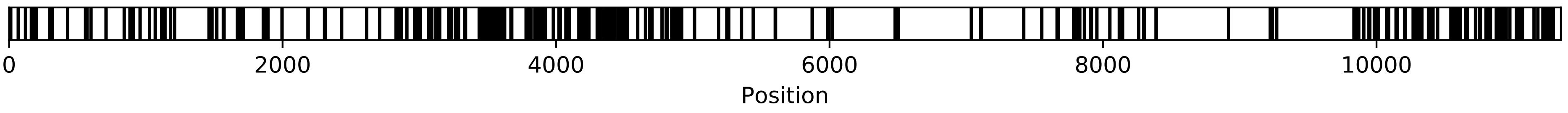


MA1 vs MA2
Number of differences: 385

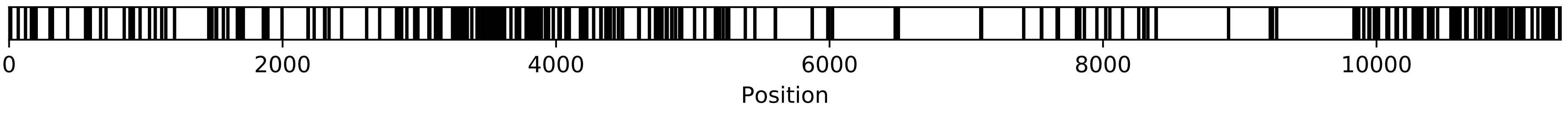


MA1 vs MM1
Number of differences: 196

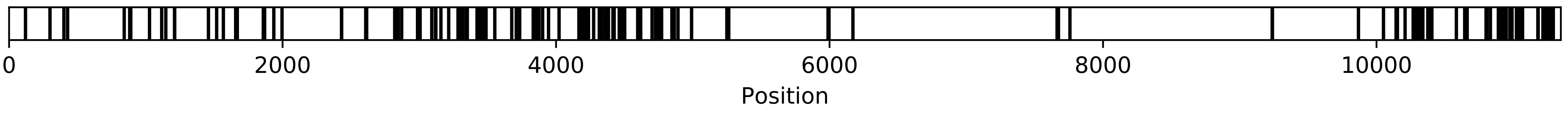


MA1 vs MM2
Number of differences: 83

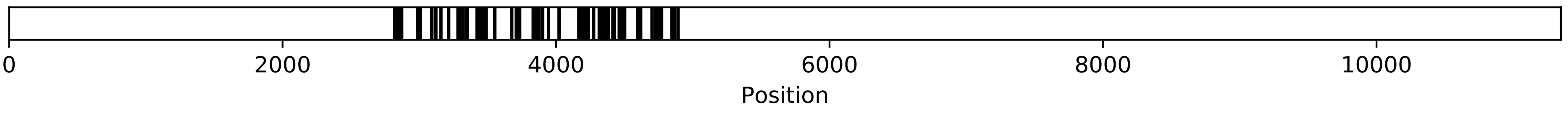


MA2 vs MM1
Number of differences: 348

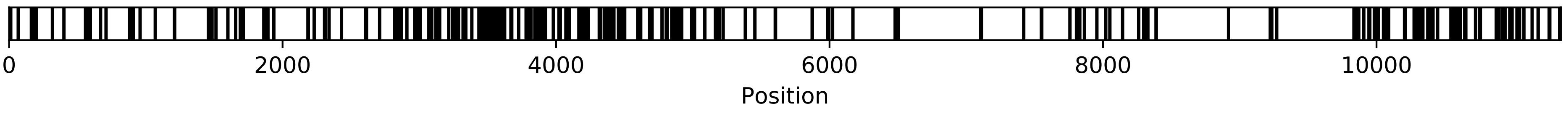


MA2 vs MM2
Number of differences: 380

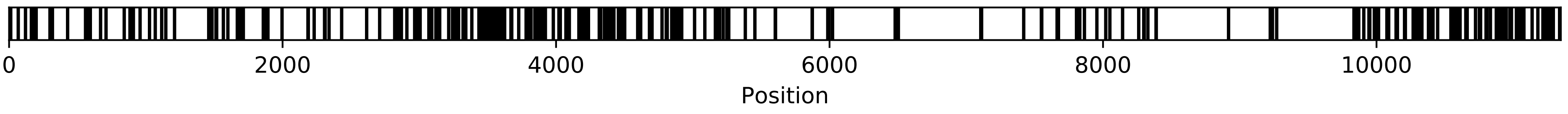


MM1 vs MM2
Number of differences: 113

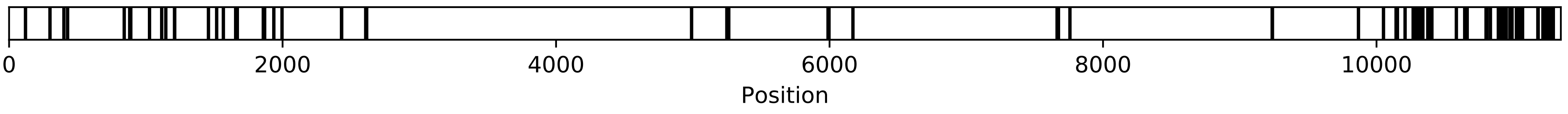

### Region: 54



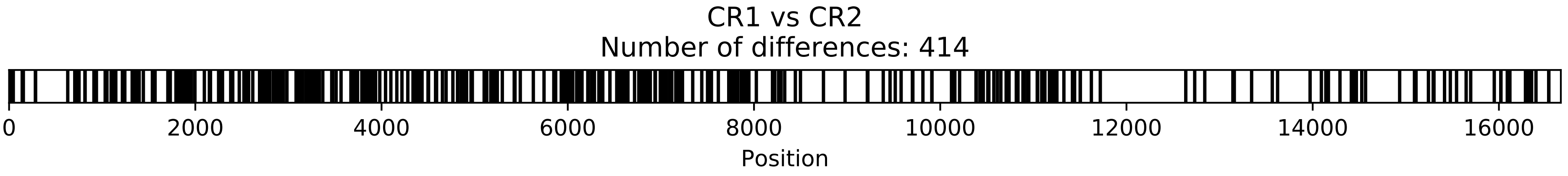
CR1 vs CR2
Number of differences: 414
0 2000 4000 6000 8000 10000 12000 14000 16000
Position

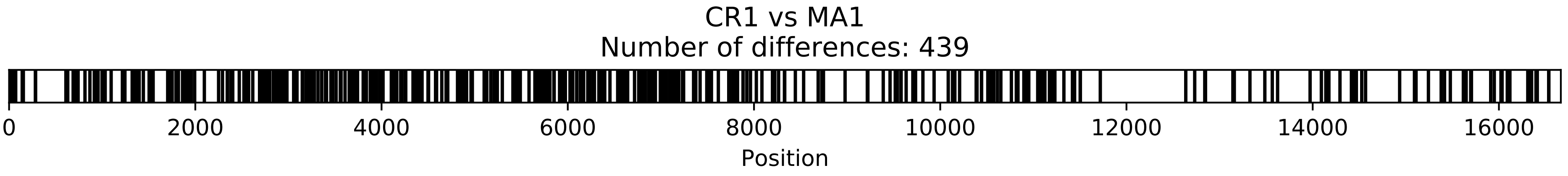
CR1 vs MA1
Number of differences: 439
0 2000 4000 6000 8000 10000 12000 14000 16000
Position

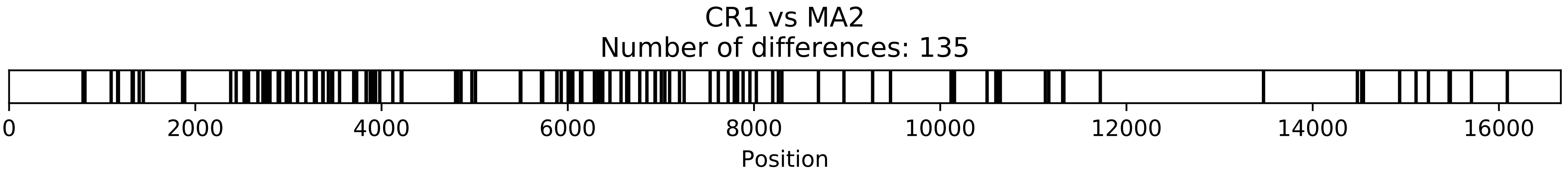
CR1 vs MA2
Number of differences: 135
0 2000 4000 6000 8000 10000 12000 14000 16000
Position

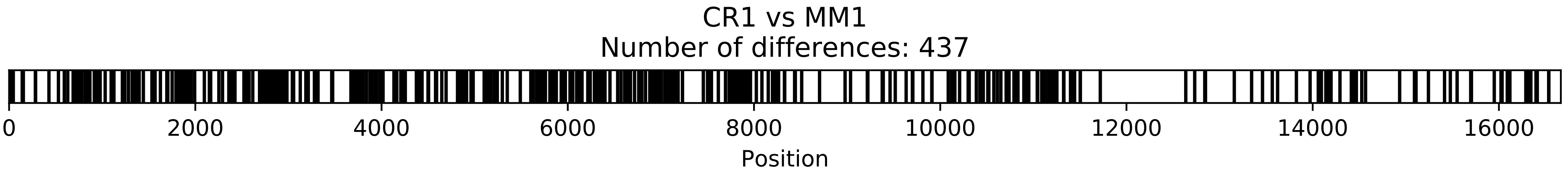
CR1 vs MM1
Number of differences: 437
0 2000 4000 6000 8000 10000 12000 14000 16000
Position

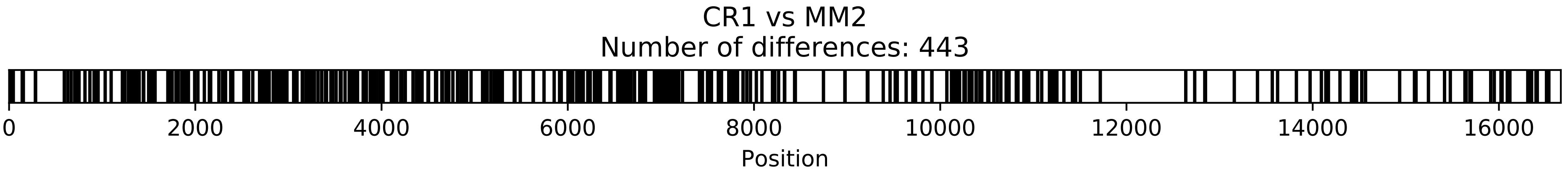
CR1 vs MM2
Number of differences: 443
0 2000 4000 6000 8000 10000 12000 14000 16000
Position

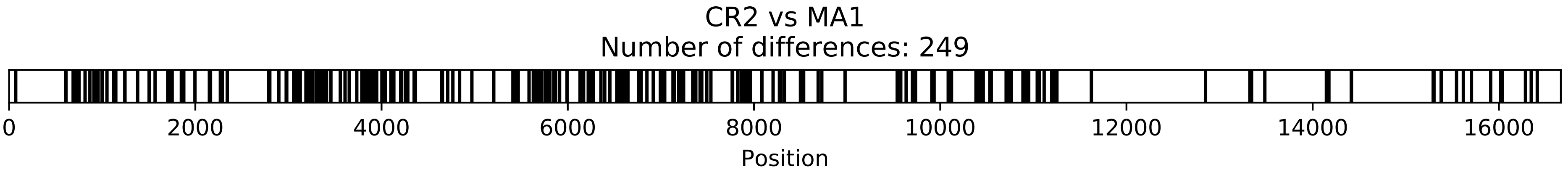
CR2 vs MA1
Number of differences: 249
0 2000 4000 6000 8000 10000 12000 14000 16000
Position

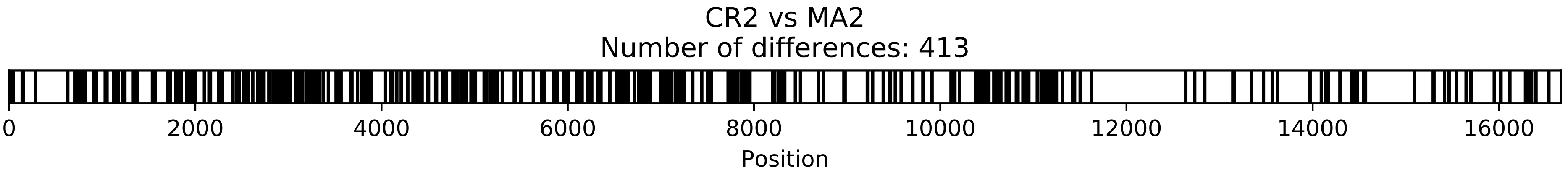
CR2 vs MA2
Number of differences: 413
0 2000 4000 6000 8000 10000 12000 14000 16000
Position

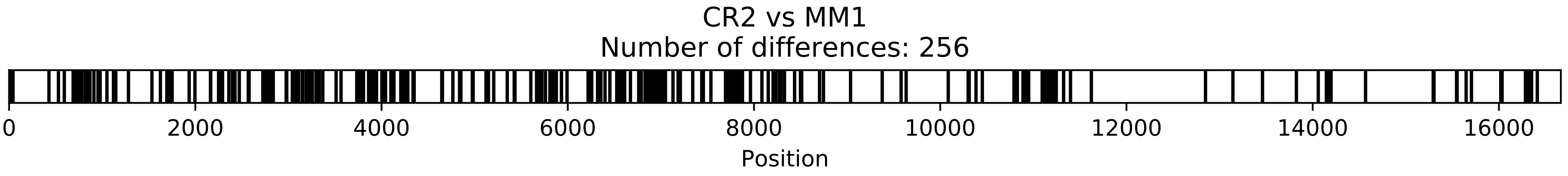
CR2 vs MM1
Number of differences: 256
0 2000 4000 6000 8000 10000 12000 14000 16000
Position

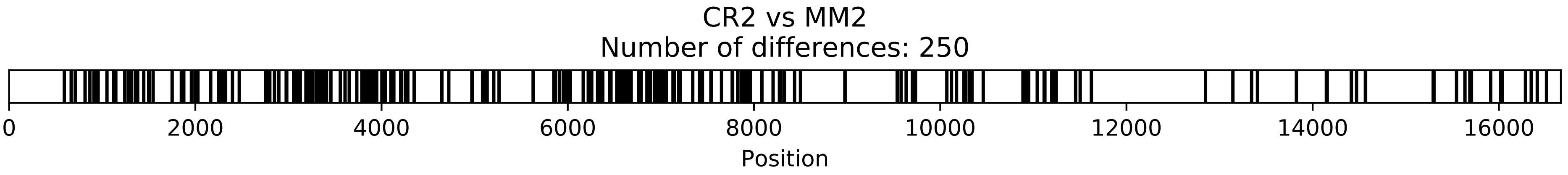
CR2 vs MM2
Number of differences: 250
0 2000 4000 6000 8000 10000 12000 14000 16000
Position

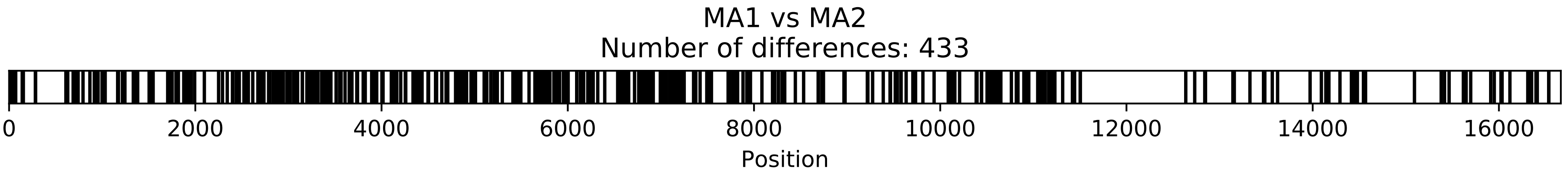
MA1 vs MA2
Number of differences: 433
0 2000 4000 6000 8000 10000 12000 14000 16000
Position

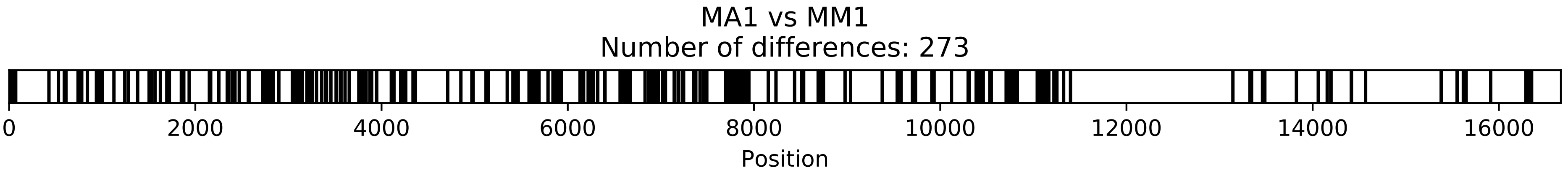
MA1 vs MM1
Number of differences: 273
0 2000 4000 6000 8000 10000 12000 14000 16000
Position

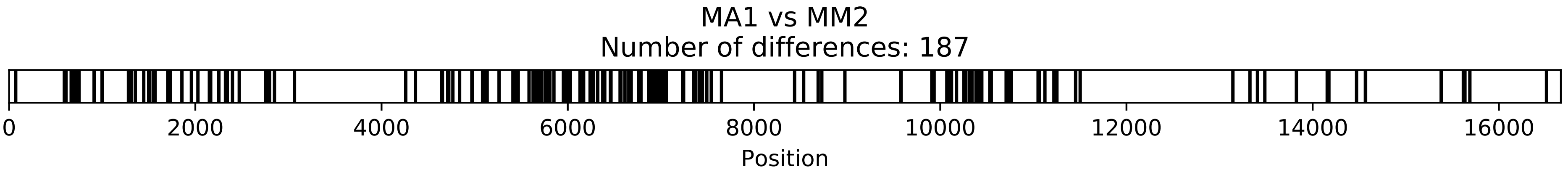
MA1 vs MM2
Number of differences: 187
0 2000 4000 6000 8000 10000 12000 14000 16000
Position

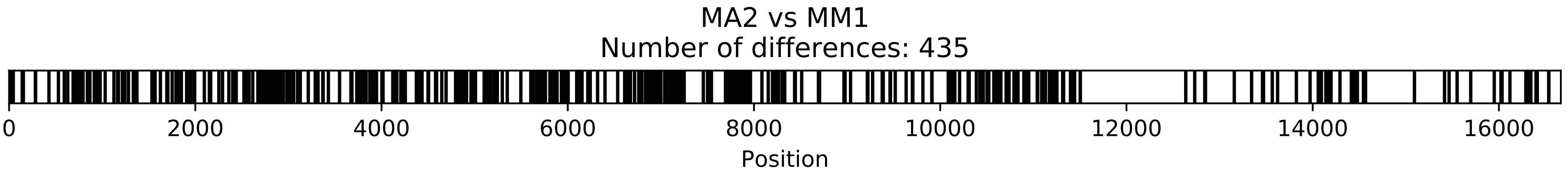
MA2 vs MM1
Number of differences: 435
0 2000 4000 6000 8000 10000 12000 14000 16000
Position

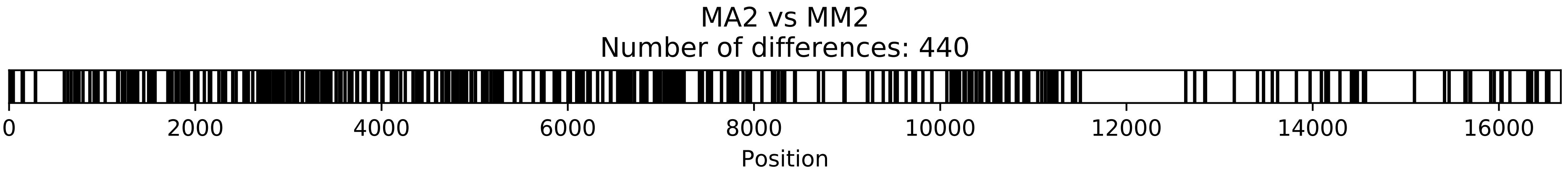
MA2 vs MM2
Number of differences: 440
0 2000 4000 6000 8000 10000 12000 14000 16000
Position

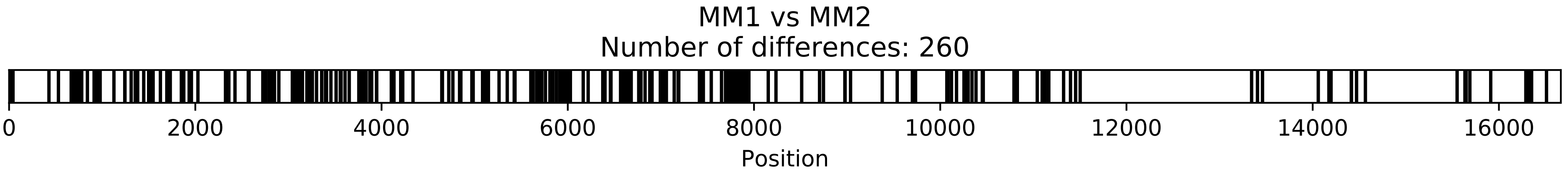
MM1 vs MM2
Number of differences: 260
0 2000 4000 6000 8000 10000 12000 14000 16000
Position

### Region: 55



CR1 vs CR2
Number of differences: 174

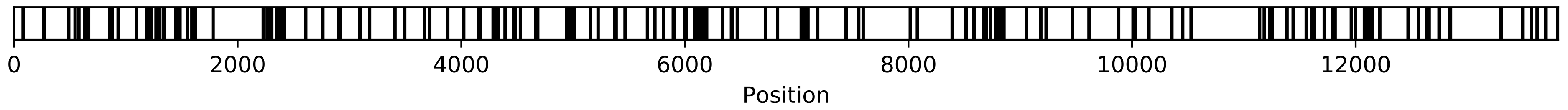


CR1 vs MA1
Number of differences: 75

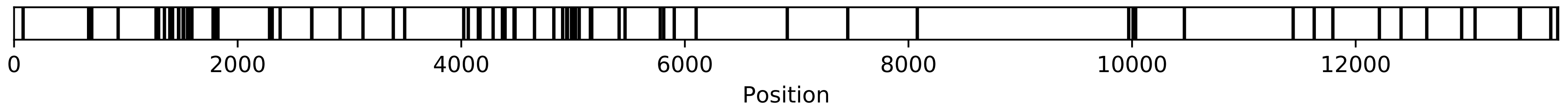


CR1 vs MA2
Number of differences: 167

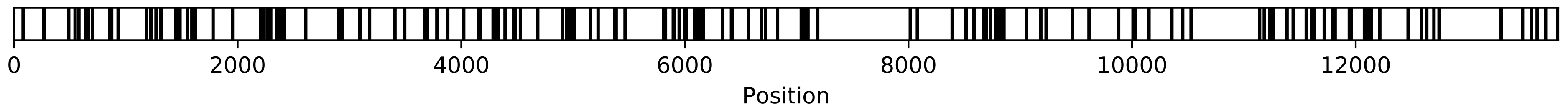


CR1 vs MM1
Number of differences: 62

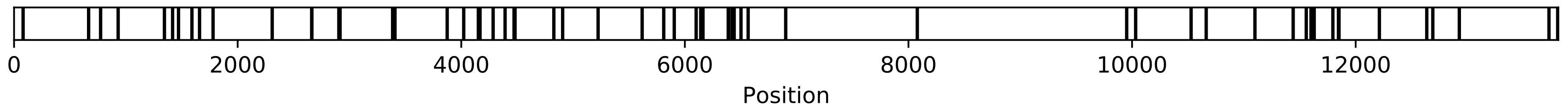


CR1 vs MM2
Number of differences: 55

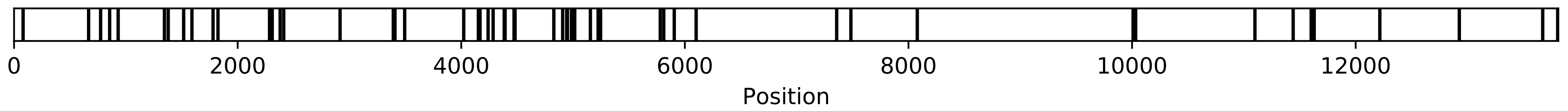


CR2 vs MA1
Number of differences: 177

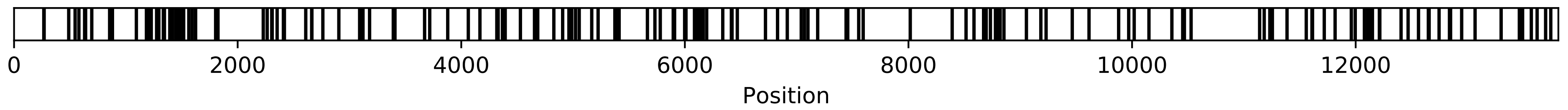


CR2 vs MA2
Number of differences: 53

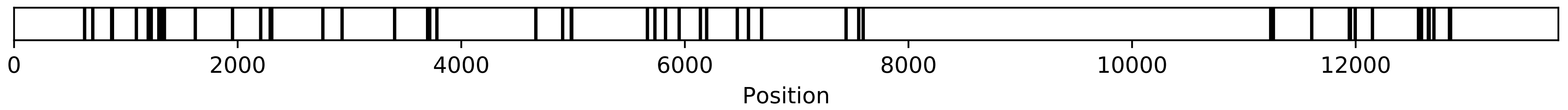


CR2 vs MM1
Number of differences: 175

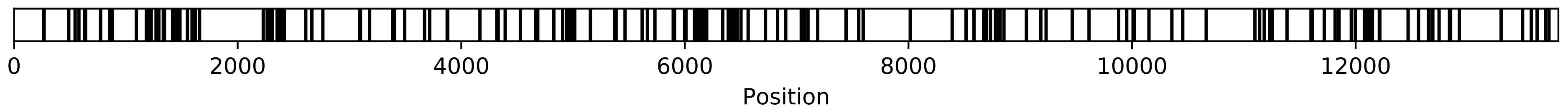


CR2 vs MM2
Number of differences: 160

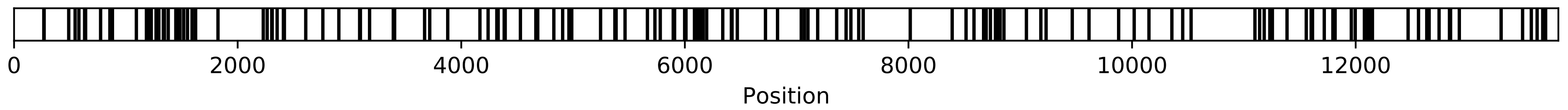


MA1 vs MA2
Number of differences: 170

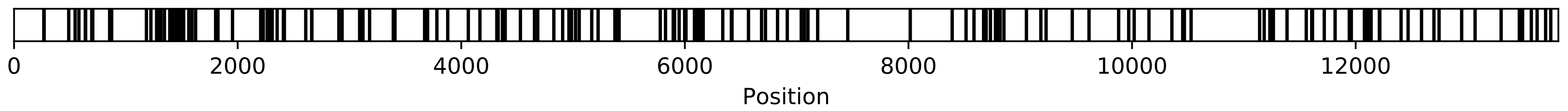


MA1 vs MM1
Number of differences: 72

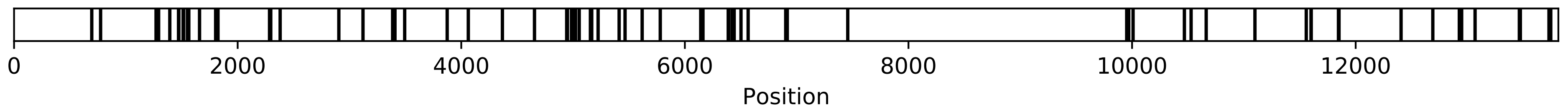


MA1 vs MM2
Number of differences: 52

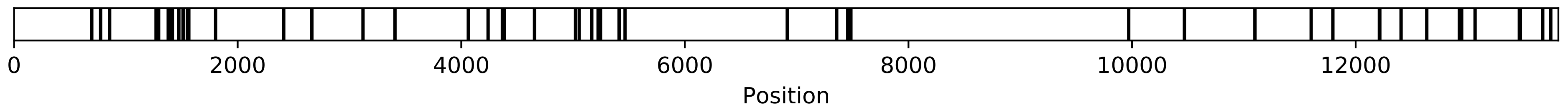


MA2 vs MM1
Number of differences: 168

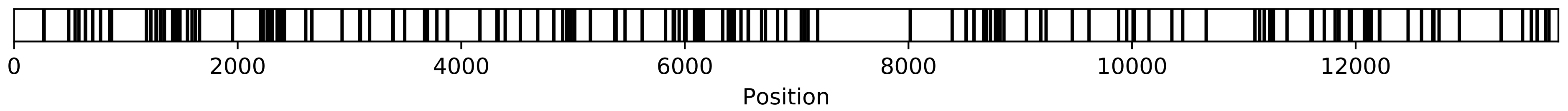


MA2 vs MM2
Number of differences: 153

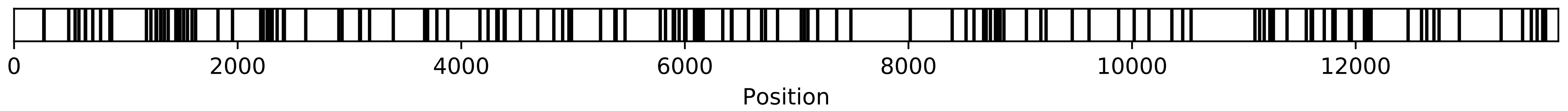


MM1 vs MM2
Number of differences: 53

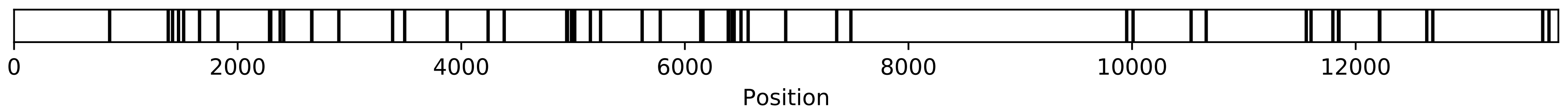

### Region: 57

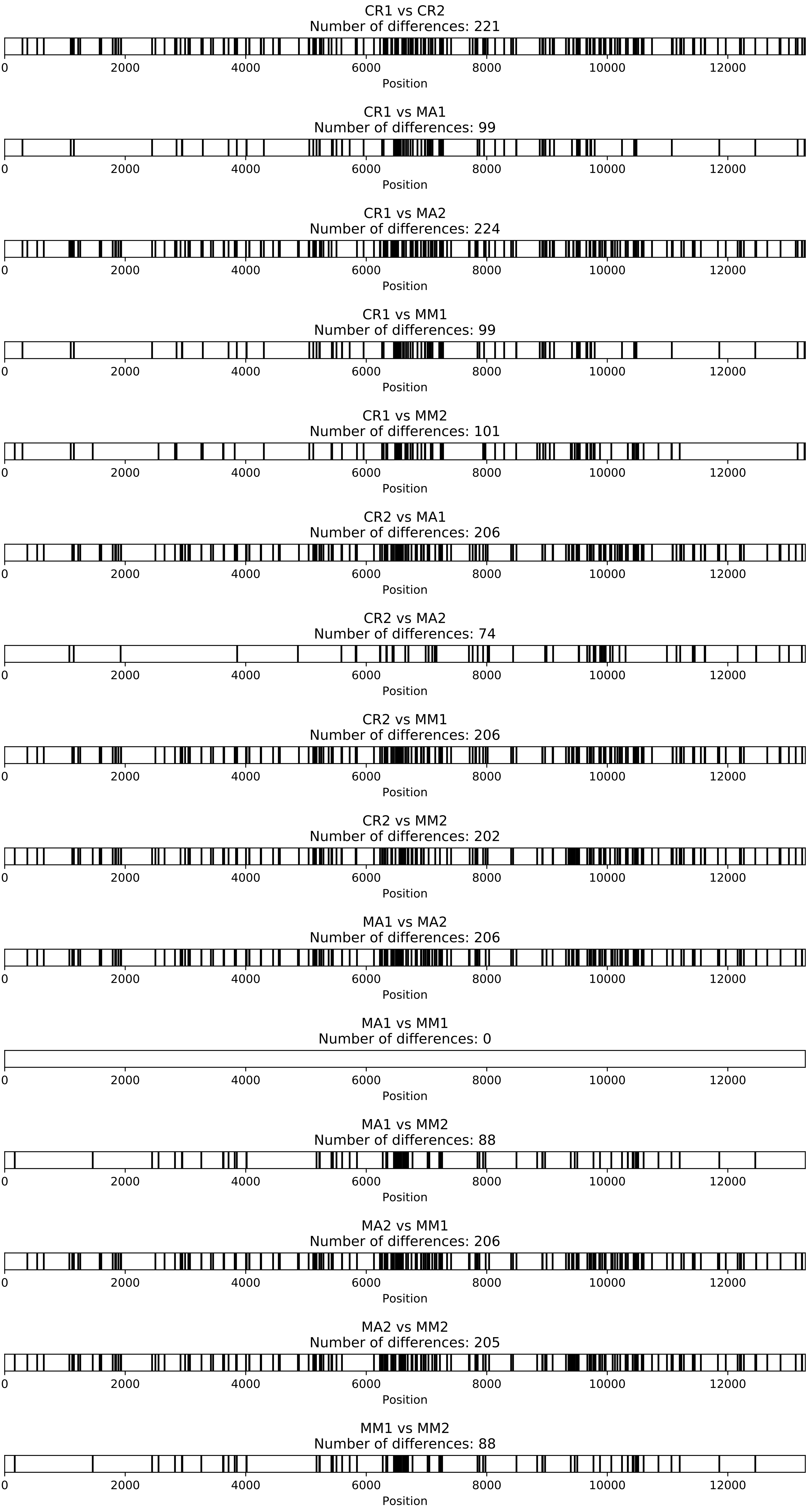
CR1 vs CR2
Number of differences: 221
CR1 vs MA1
Number of differences: 99
CR1 vs MA2
Number of differences: 224
CR1 vs MM1
Number of differences: 99
CR1 vs MM2
Number of differences: 101
CR2 vs MA1
Number of differences: 206
CR2 vs MA2
Number of differences: 74
CR2 vs MM1
Number of differences: 206
CR2 vs MM2
Number of differences: 202
MA1 vs MA2
Number of differences: 206
MA1 vs MM1
Number of differences: 0
MA1 vs MM2
Number of differences: 88
MA2 vs MM1
Number of differences: 206
MA2 vs MM2
Number of differences: 205
MM1 vs MM2
Number of differences: 88
0
2000
4000
6000
8000
10000
12000
Position

### Region: 64



CR1 vs CR2
Number of differences: 5

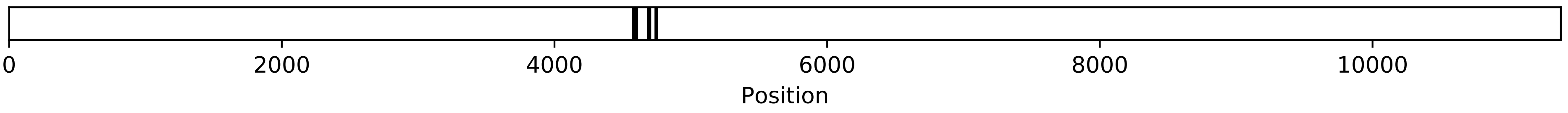


CR1 vs MA1
Number of differences: 207

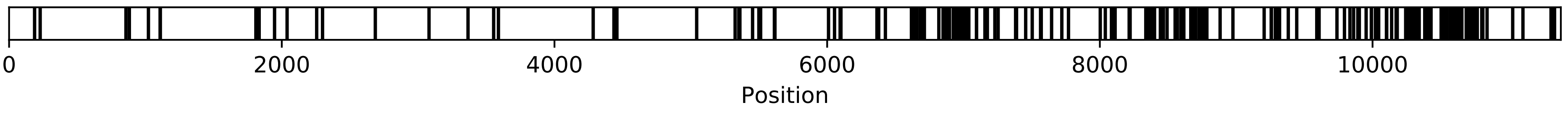


CR1 vs MA2
Number of differences: 151

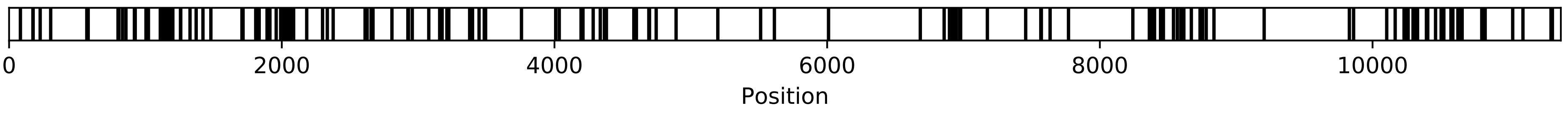


CR1 vs MM1
Number of differences: 220

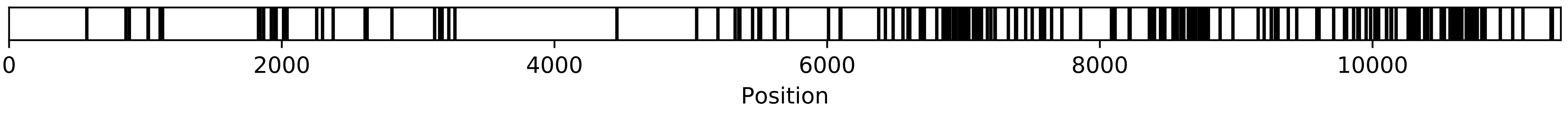


CR1 vs MM2
Number of differences: 230

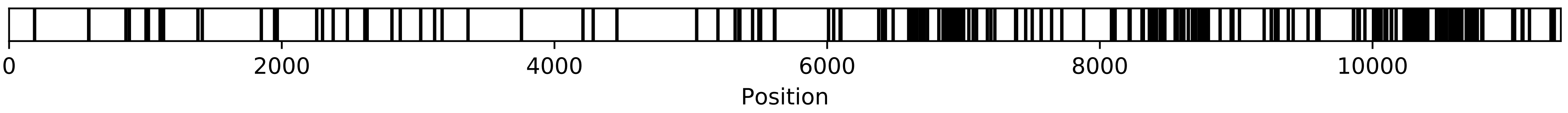


CR2 vs MA1
Number of differences: 212

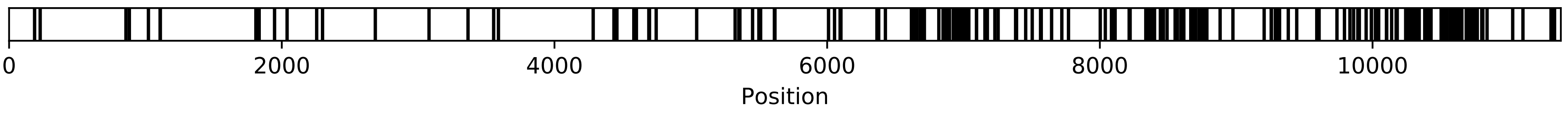


CR2 vs MA2
Number of differences: 146

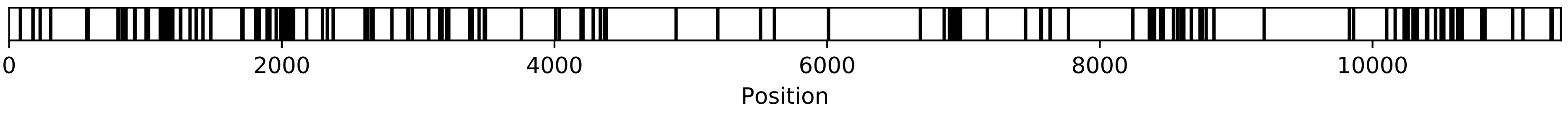


CR2 vs MM1
Number of differences: 225

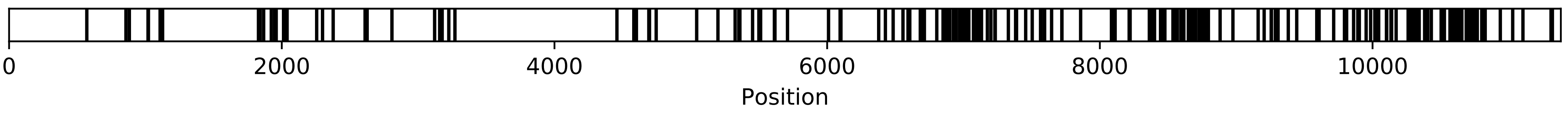


CR2 vs MM2
Number of differences: 235

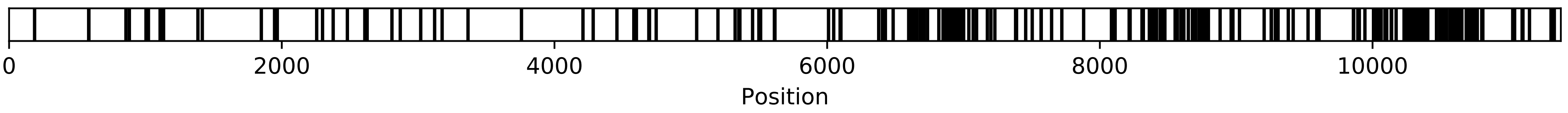


MA1 vs MA2
Number of differences: 248

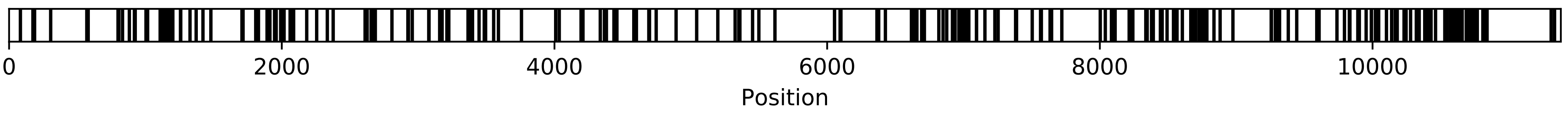


MA1 vs MM1
Number of differences: 136

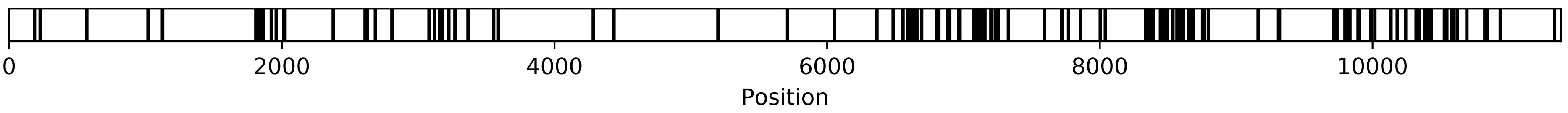


MA1 vs MM2
Number of differences: 139

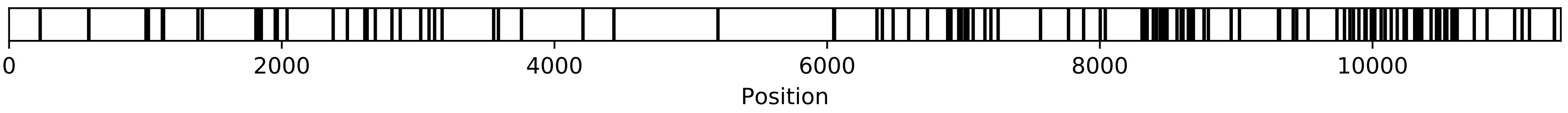


MA2 vs MM1
Number of differences: 251

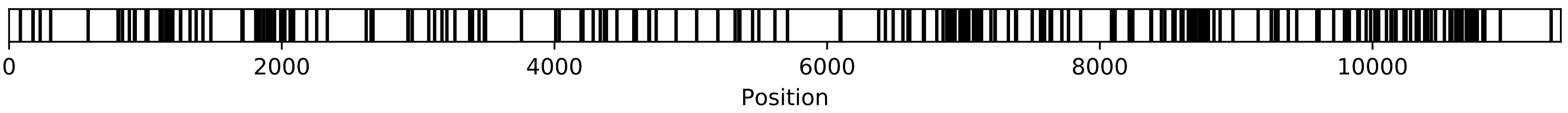


MA2 vs MM2
Number of differences: 264

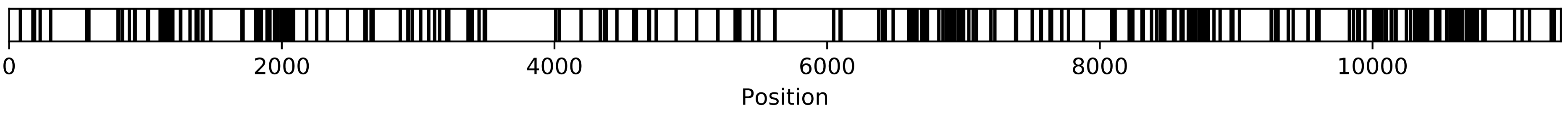


MM1 vs MM2
Number of differences: 128

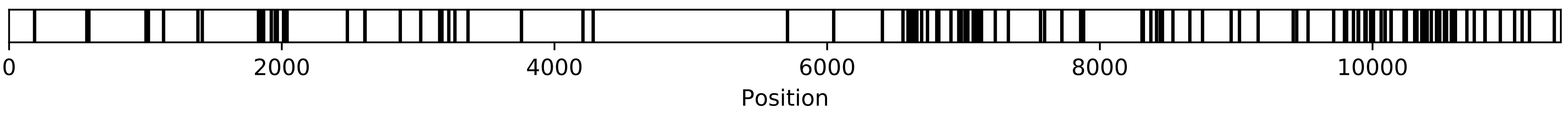

### Region: 65

CR1 vs CR2
Number of differences: 319

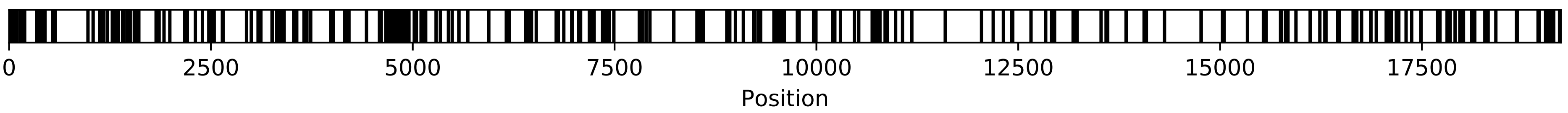


CR1 vs MA1
Number of differences: 228

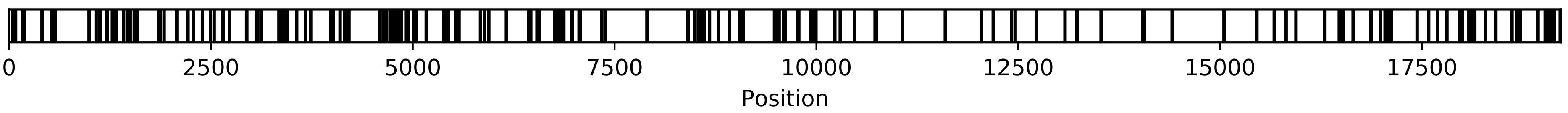


CR1 vs MA2
Number of differences: 328

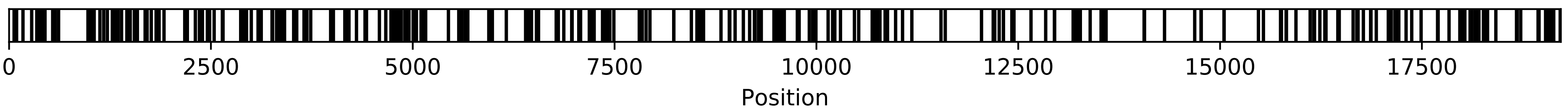


CR1 vs MM1
Number of differences: 186

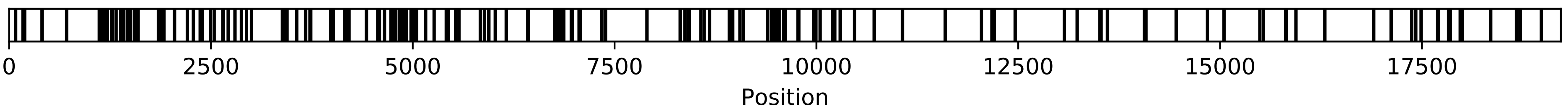


CR1 vs MM2
Number of differences: 197

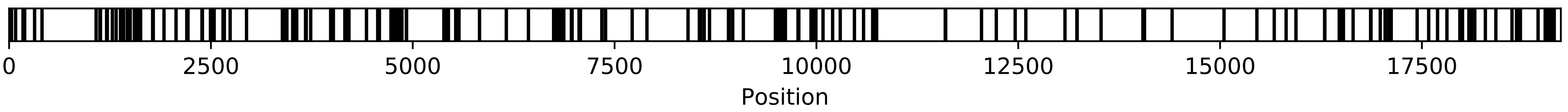


CR2 vs MA1
Number of differences: 316

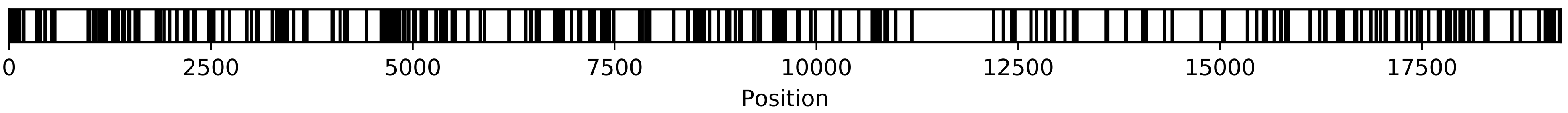


CR2 vs MA2
Number of differences: 140

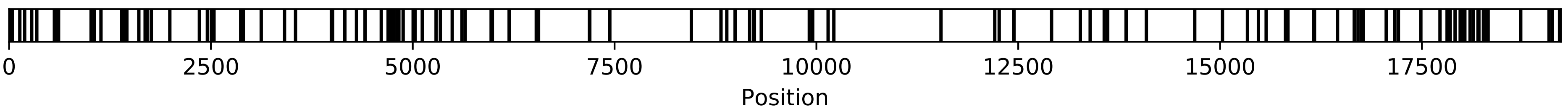


CR2 vs MM1
Number of differences: 318

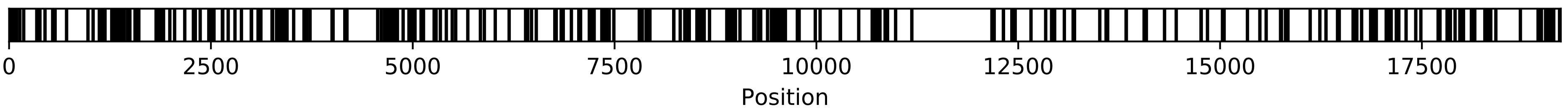


CR2 vs MM2
Number of differences: 321

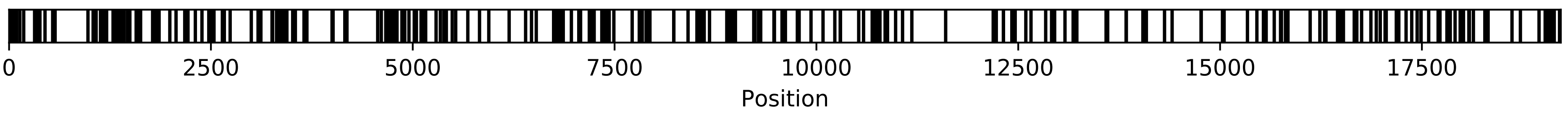


MA1 vs MA2
Number of differences: 323

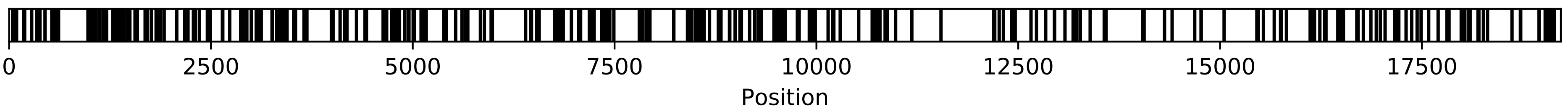


MA1 vs MM1
Number of differences: 176

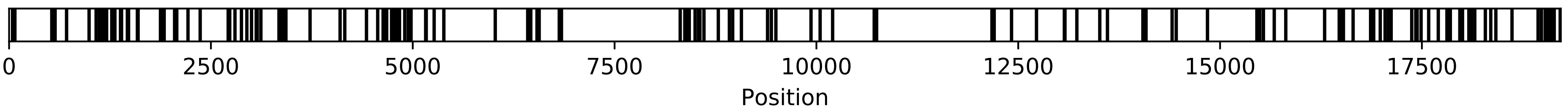


MA1 vs MM2
Number of differences: 135

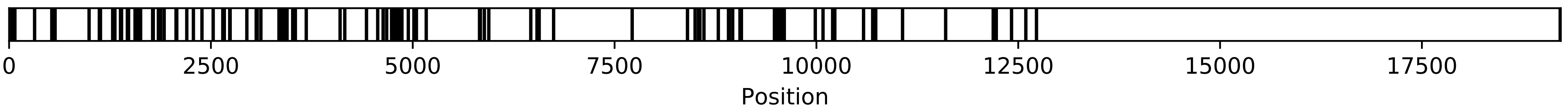


MA2 vs MM1
Number of differences: 325

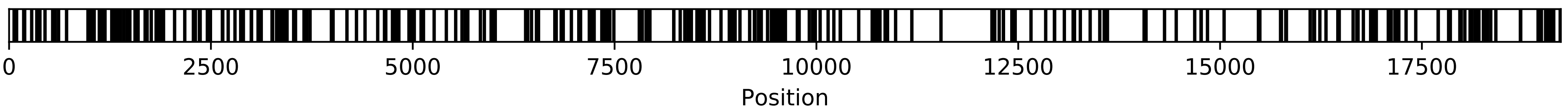


MA2 vs MM2
Number of differences: 330

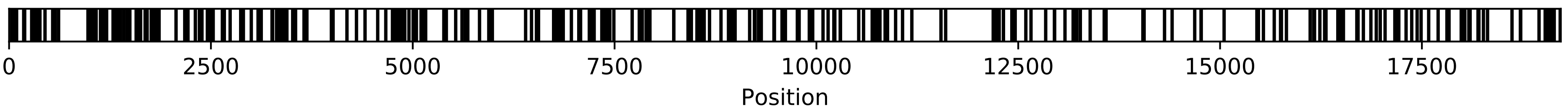


MM1 vs MM2
Number of differences: 162

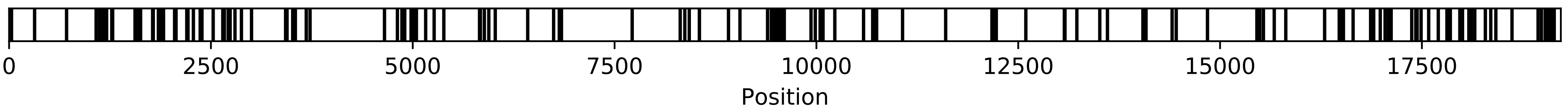

### Region: 66



CR1 vs CR2
Number of differences: 244

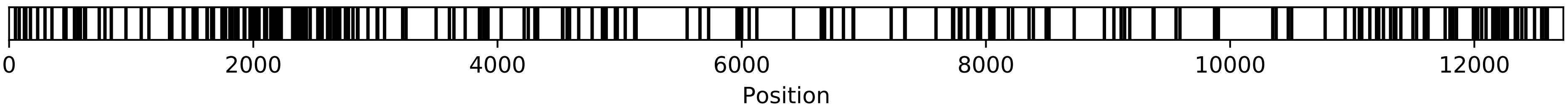


CR1 vs MA1
Number of differences: 88

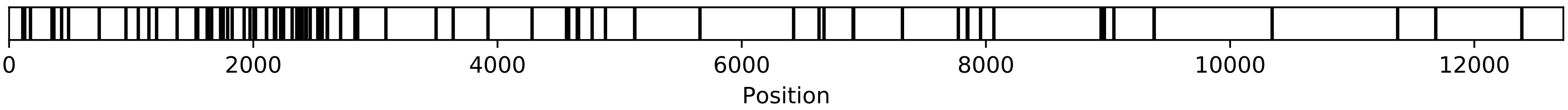


CR1 vs MA2
Number of differences: 248

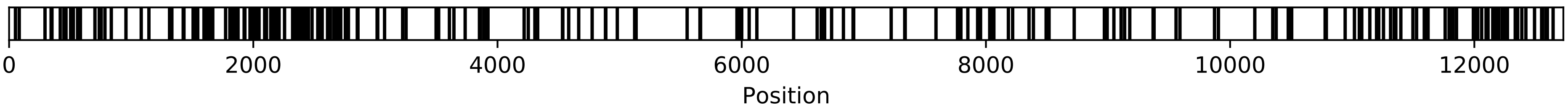


CR1 vs MM1
Number of differences: 88

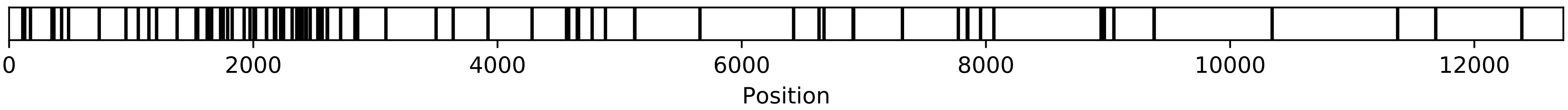


CR1 vs MM2
Number of differences: 81

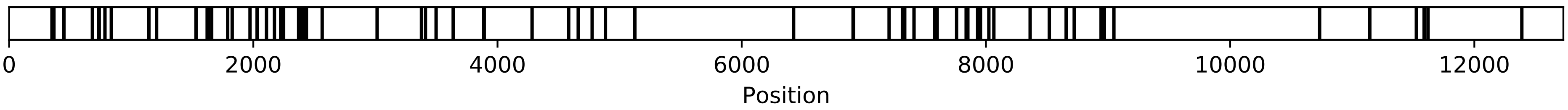


CR2 vs MA1
Number of differences: 237

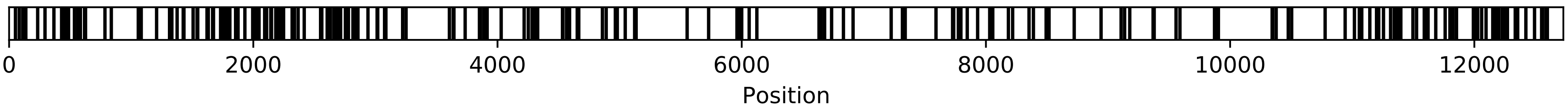


CR2 vs MA2
Number of differences: 61

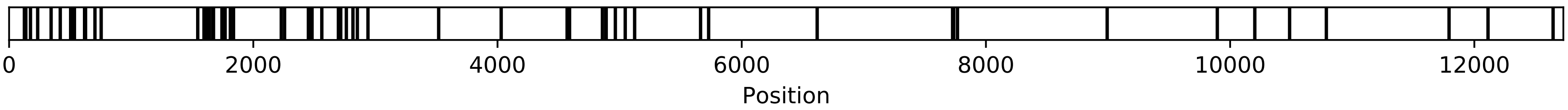


CR2 vs MM1
Number of differences: 237

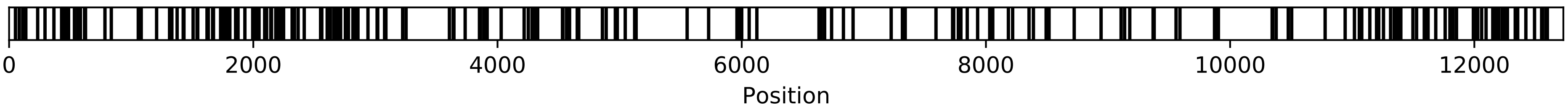


CR2 vs MM2
Number of differences: 244

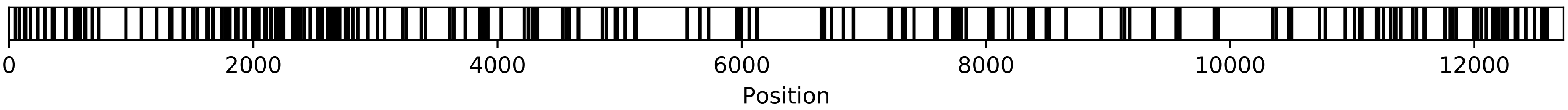


MA1 vs MA2
Number of differences: 242

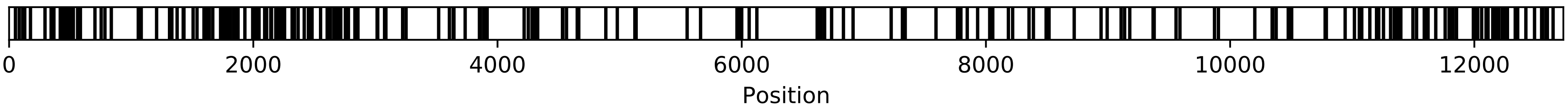


MA1 vs MM1
Number of differences: 0

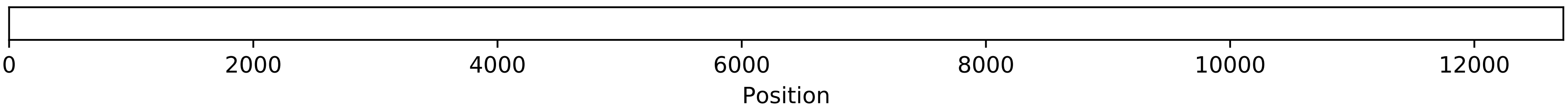


MA1 vs MM2
Number of differences: 88

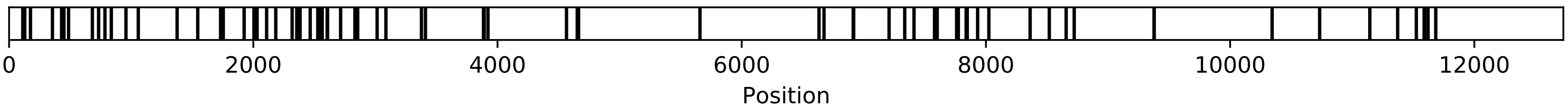


MA2 vs MM1
Number of differences: 242

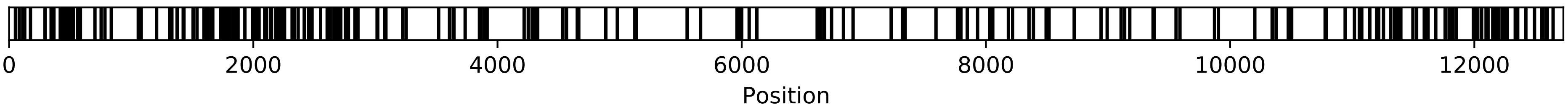


MA2 vs MM2
Number of differences: 247

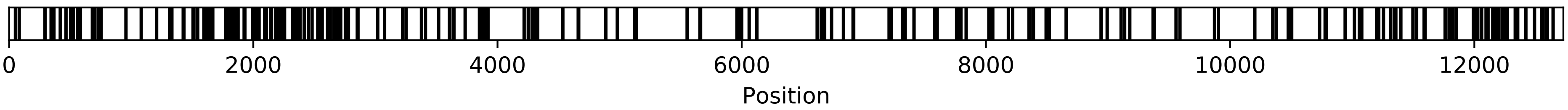


MM1 vs MM2
Number of differences: 88

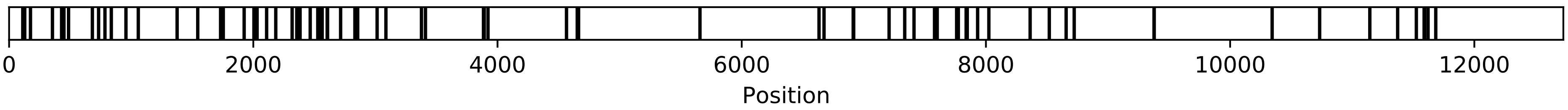

### Region: 68

CR1 vs CR2
Number of differences: 274

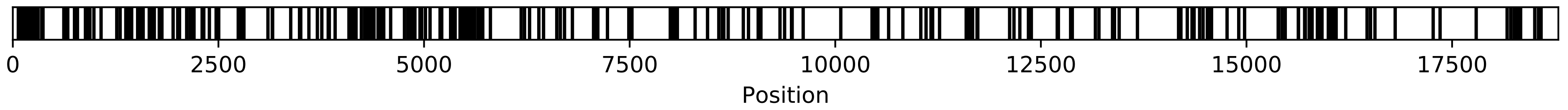


CR1 vs MA1
Number of differences: 265

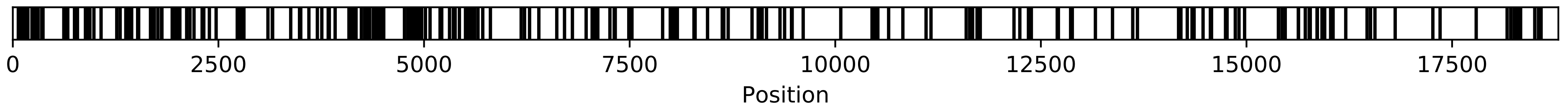


CR1 vs MA2
Number of differences: 74

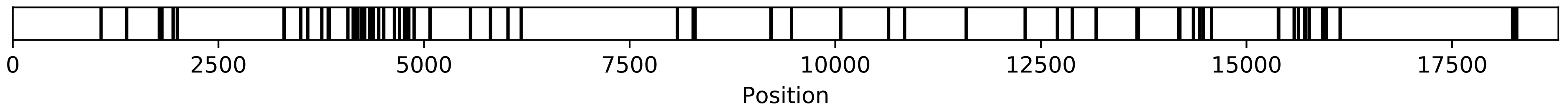


CR1 vs MM1
Number of differences: 259

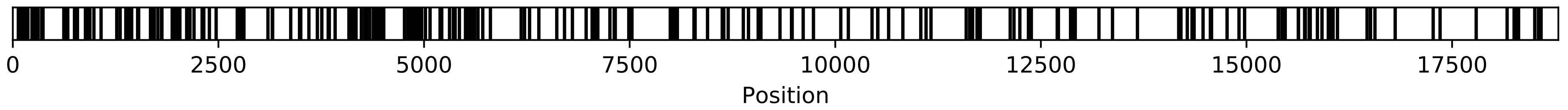


CR1 vs MM2
Number of differences: 265

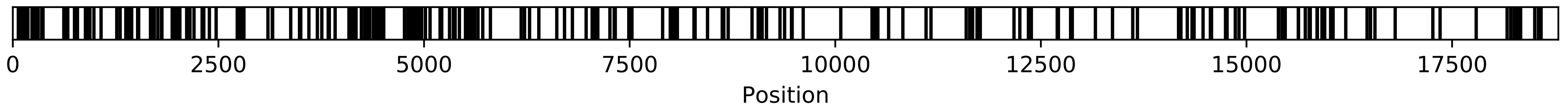


CR2 vs MA1
Number of differences: 88

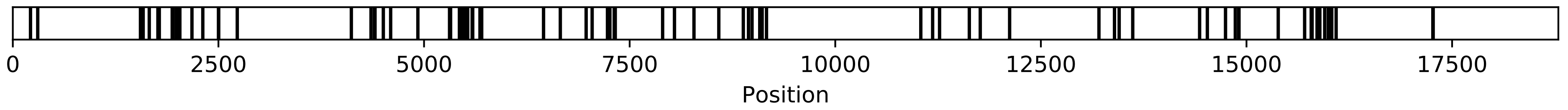


CR2 vs MA2
Number of differences: 265

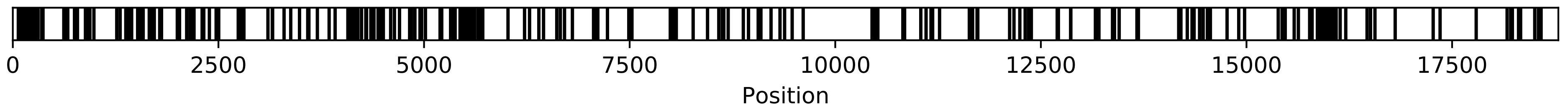


CR2 vs MM1
Number of differences: 82

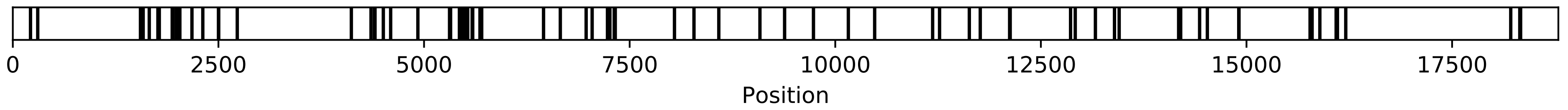


CR2 vs MM2
Number of differences: 88

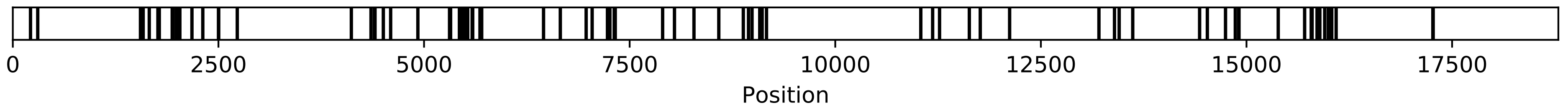


MA1 vs MA2
Number of differences: 258

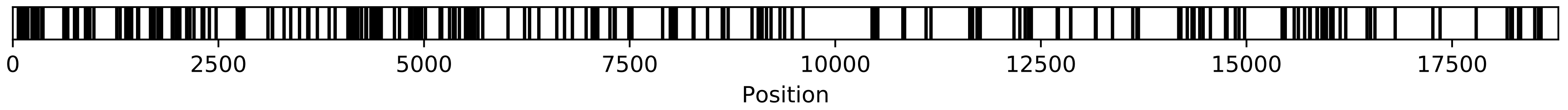


MA1 vs MM1
Number of differences: 39

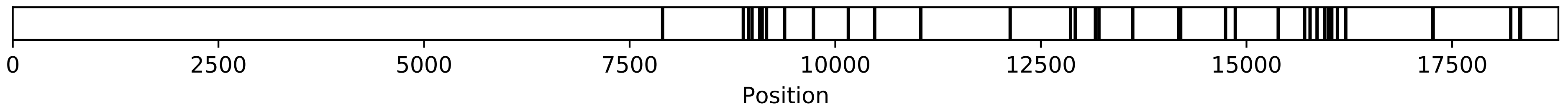


MA1 vs MM2
Number of differences: 0

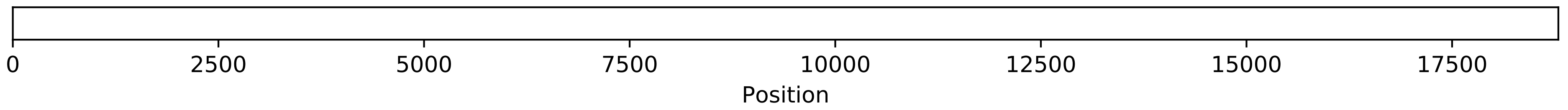


MA2 vs MM1
Number of differences: 254

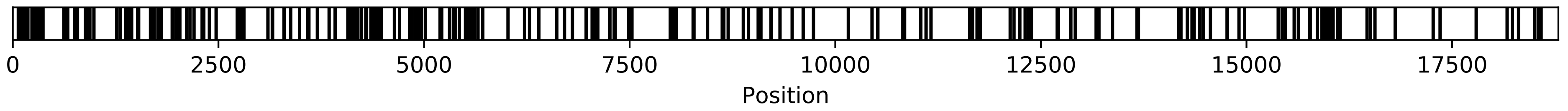


MA2 vs MM2
Number of differences: 258

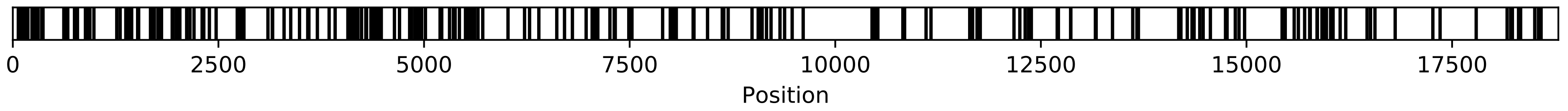


MM1 vs MM2
Number of differences: 39

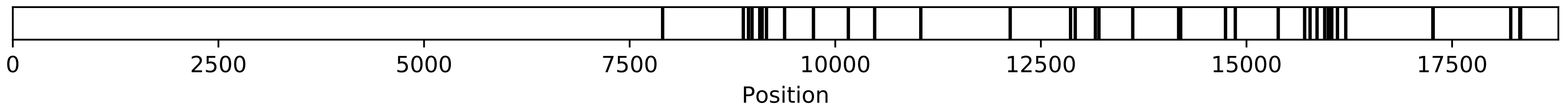

### Region: 70



CR1 vs CR2
Number of differences: 200

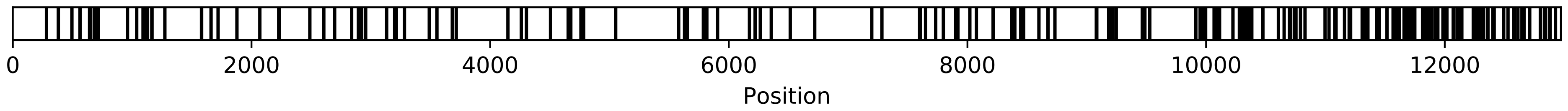


CR1 vs MA1
Number of differences: 204

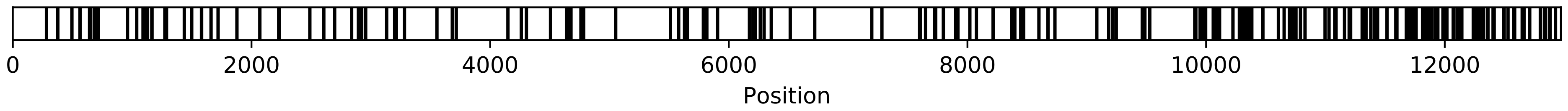


CR1 vs MA2
Number of differences: 129

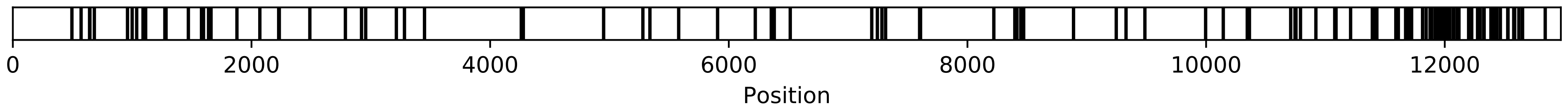


CR1 vs MM1
Number of differences: 129

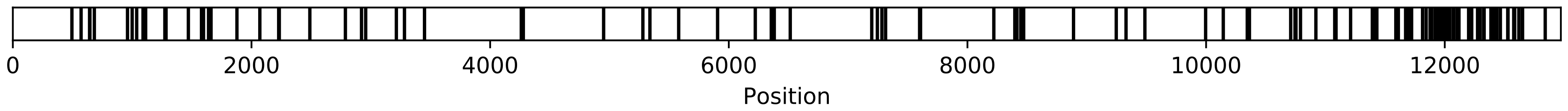


CR1 vs MM2
Number of differences: 119

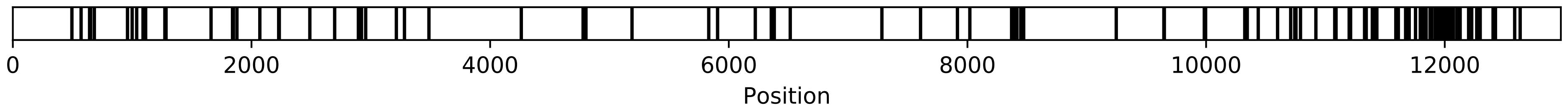


CR2 vs MA1
Number of differences: 39

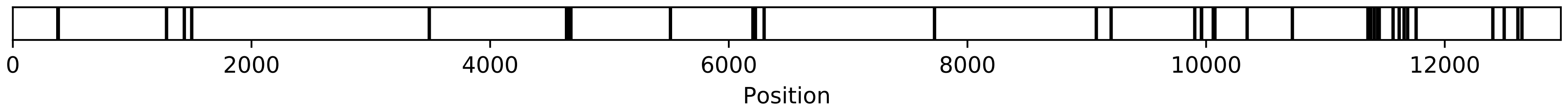


CR2 vs MA2
Number of differences: 181

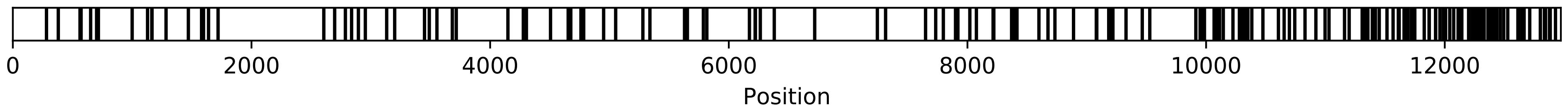


CR2 vs MM1
Number of differences: 181

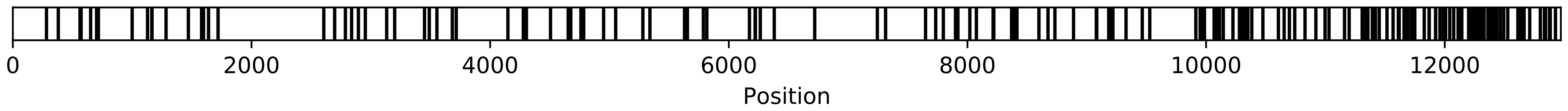


CR2 vs MM2
Number of differences: 171

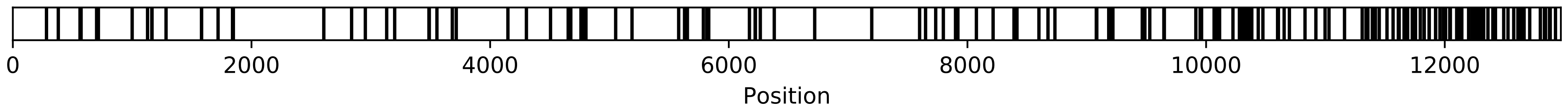


MA1 vs MA2
Number of differences: 187

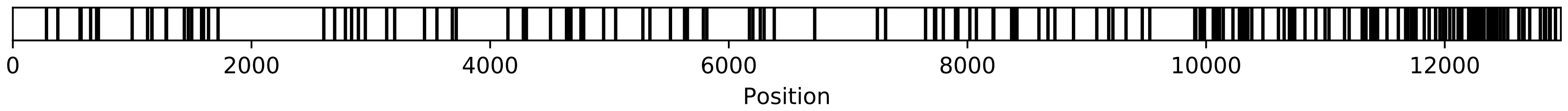


MA1 vs MM1
Number of differences: 187

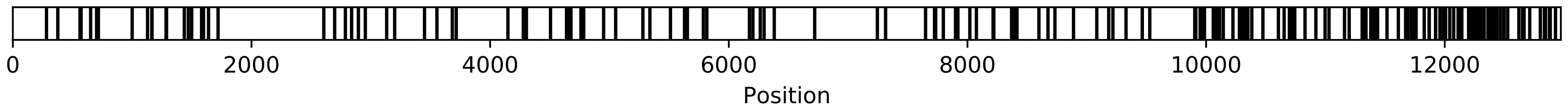


MA1 vs MM2
Number of differences: 175

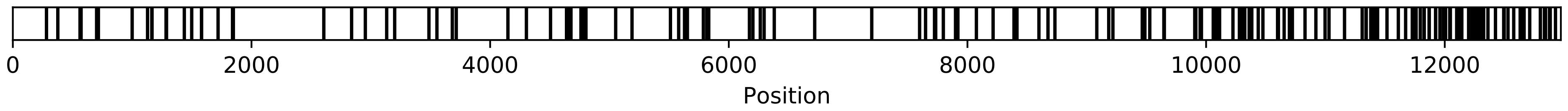


MA2 vs MM1
Number of differences: 0

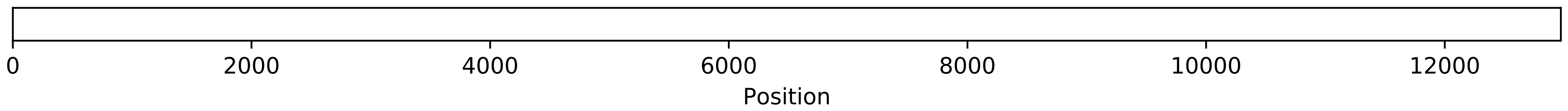


MA2 vs MM2
Number of differences: 78

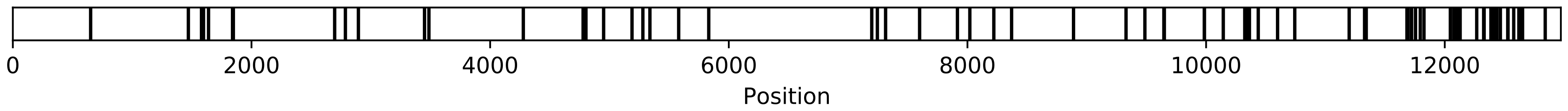


MM1 vs MM2
Number of differences: 78

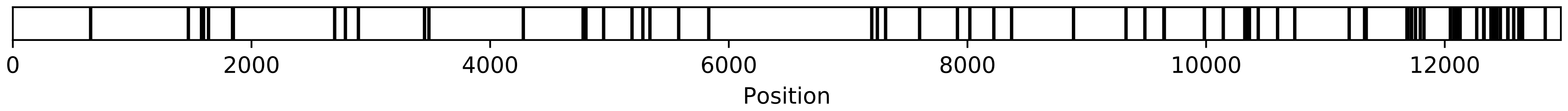

### Region: 71



CR1 vs CR2
Number of differences: 307

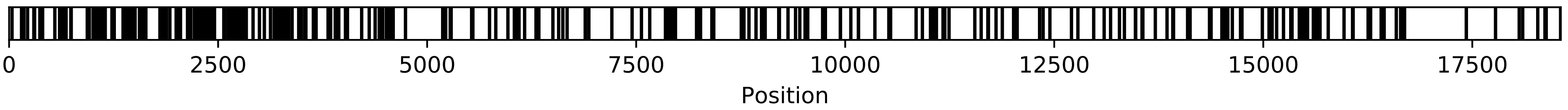


CR1 vs MA1
Number of differences: 145

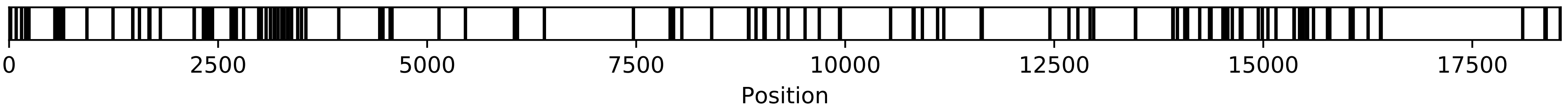


CR1 vs MA2
Number of differences: 299

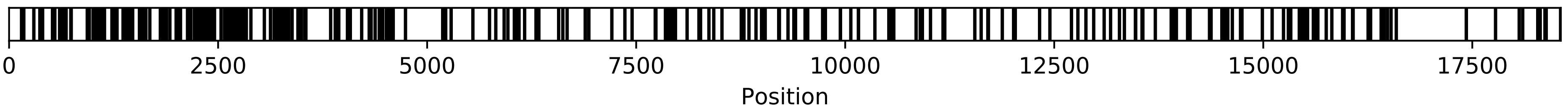


CR1 vs MM1
Number of differences: 134

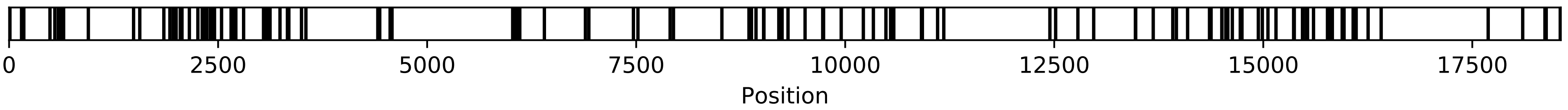


CR1 vs MM2
Number of differences: 126

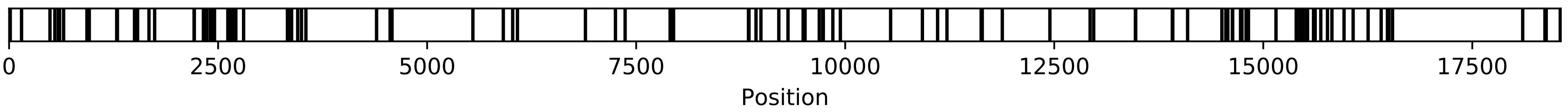


CR2 vs MA1
Number of differences: 305

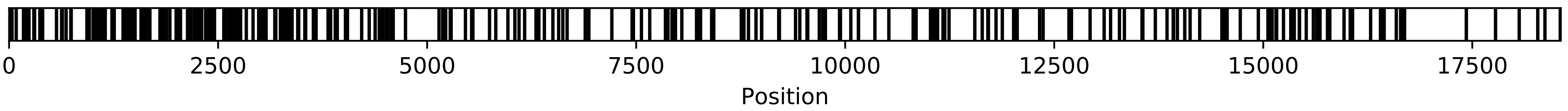


CR2 vs MA2
Number of differences: 101

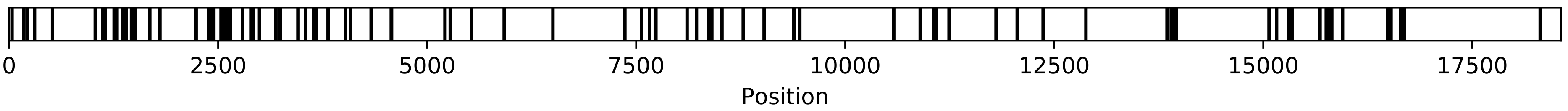


CR2 vs MM1
Number of differences: 309

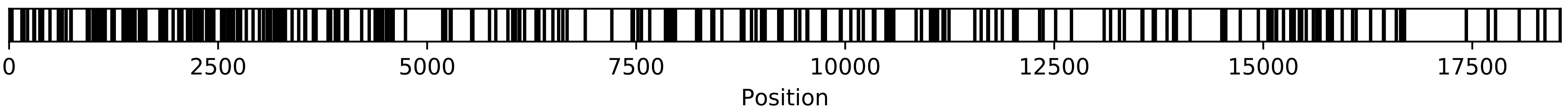


CR2 vs MM2
Number of differences: 299

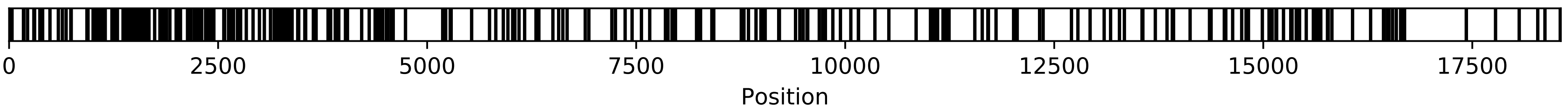


MA1 vs MA2
Number of differences: 297

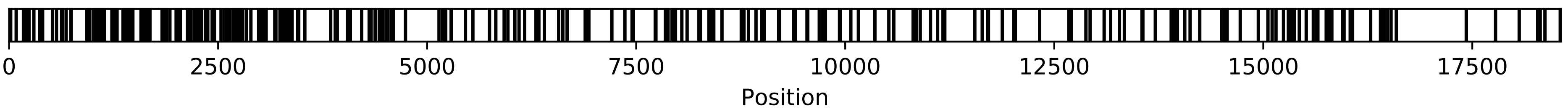


MA1 vs MM1
Number of differences: 146

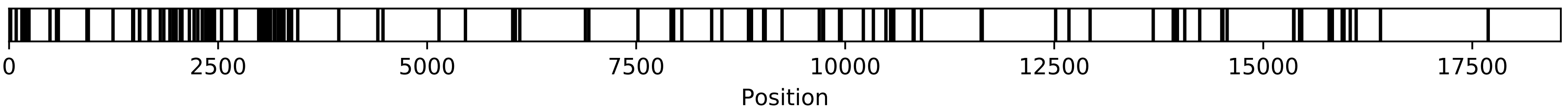


MA1 vs MM2
Number of differences: 123

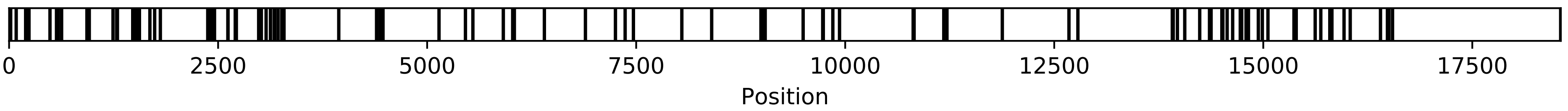


MA2 vs MM1
Number of differences: 295

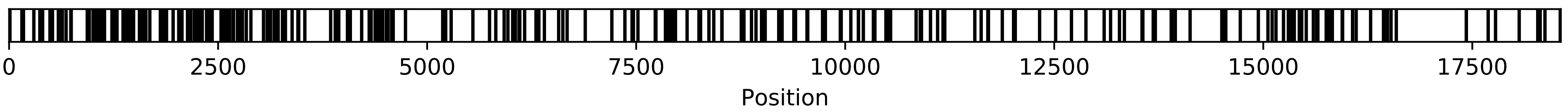


MA2 vs MM2
Number of differences: 288

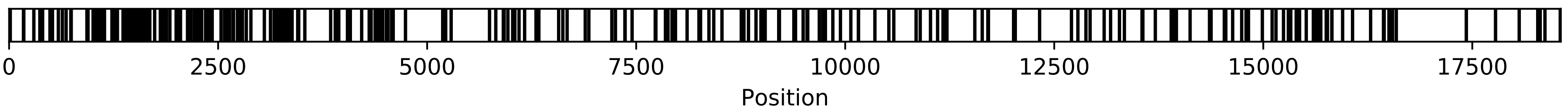


MM1 vs MM2
Number of differences: 127

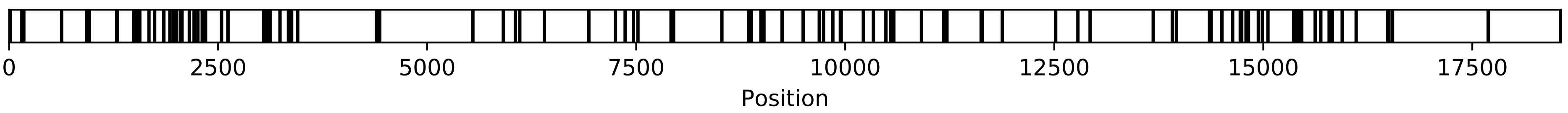

### Region: 72

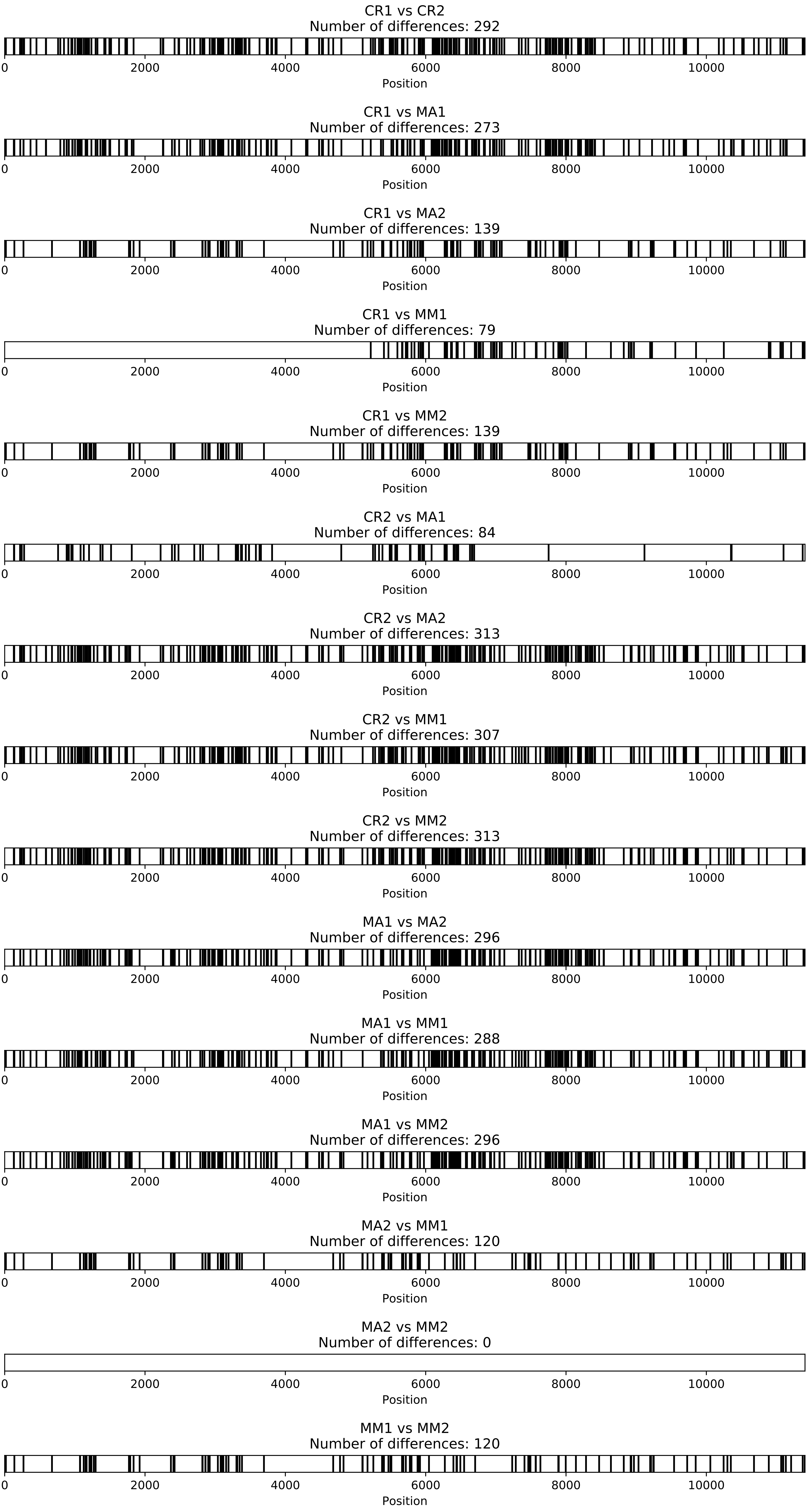
CR1 vs CR2
Number of differences: 292
CR1 vs MA1
Number of differences: 273
CR1 vs MA2
Number of differences: 139
CR1 vs MM1
Number of differences: 79
CR1 vs MM2
Number of differences: 139
CR2 vs MA1
Number of differences: 84
CR2 vs MA2
Number of differences: 313
CR2 vs MM1
Number of differences: 307
CR2 vs MM2
Number of differences: 313
MA1 vs MA2
Number of differences: 296
MA1 vs MM1
Number of differences: 288
MA1 vs MM2
Number of differences: 296
MA2 vs MM1
Number of differences: 120
MA2 vs MM2
Number of differences: 0
MM1 vs MM2
Number of differences: 120
0
2000
4000
6000
8000
10000
Position

### Region: 73



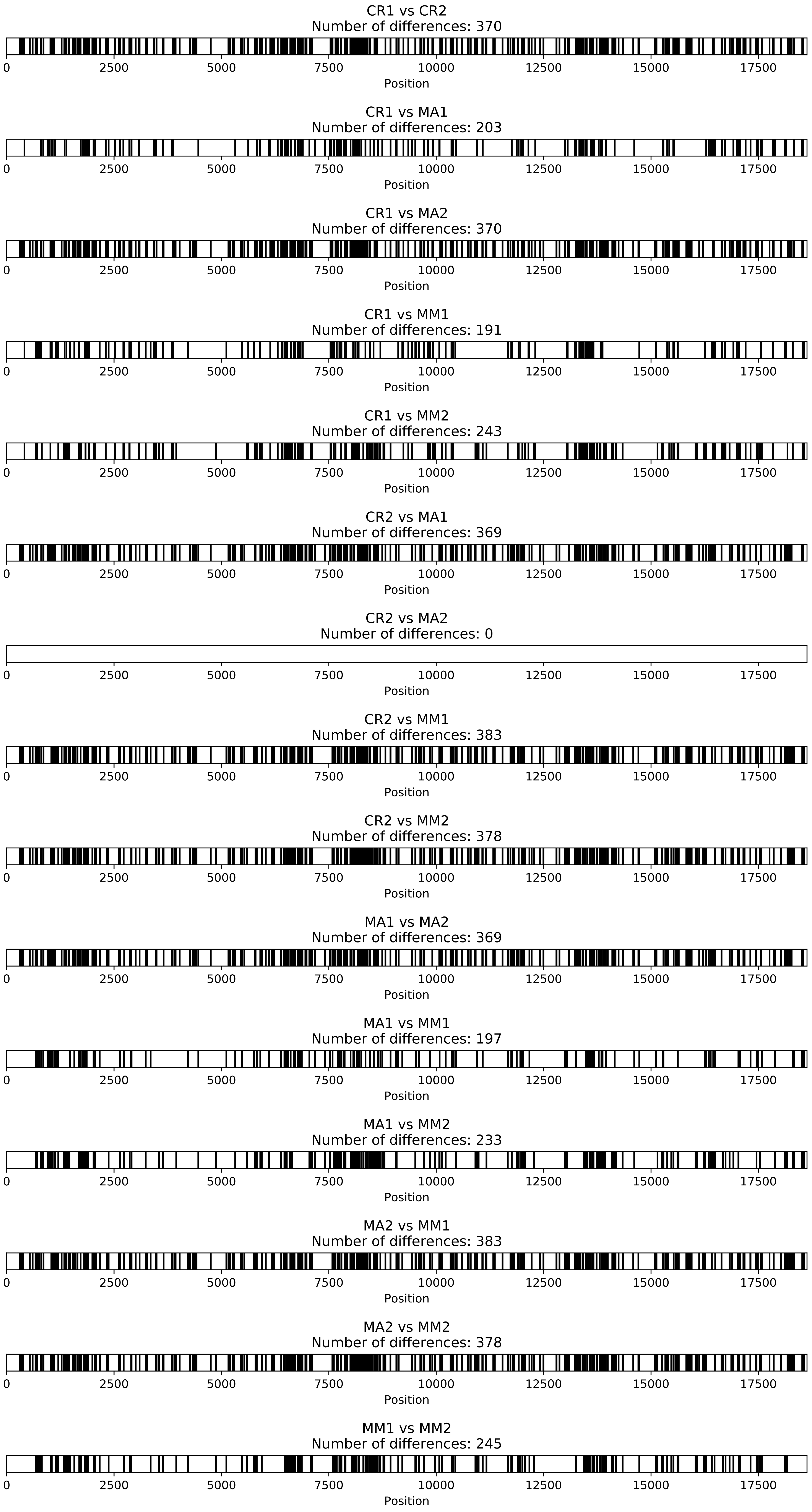
CR1 vs CR2
Number of differences: 370
CR1 vs MA1
Number of differences: 203
CR1 vs MA2
Number of differences: 370
CR1 vs MM1
Number of differences: 191
CR1 vs MM2
Number of differences: 243
CR2 vs MA1
Number of differences: 369
CR2 vs MA2
Number of differences: 0
CR2 vs MM1
Number of differences: 383
CR2 vs MM2
Number of differences: 378
MA1 vs MA2
Number of differences: 369
MA1 vs MM1
Number of differences: 197
MA1 vs MM2
Number of differences: 233
MA2 vs MM1
Number of differences: 383
MA2 vs MM2
Number of differences: 378
MM1 vs MM2
Number of differences: 245
0
2500
5000
7500
10000
12500
15000
17500
Position

### Region: 75



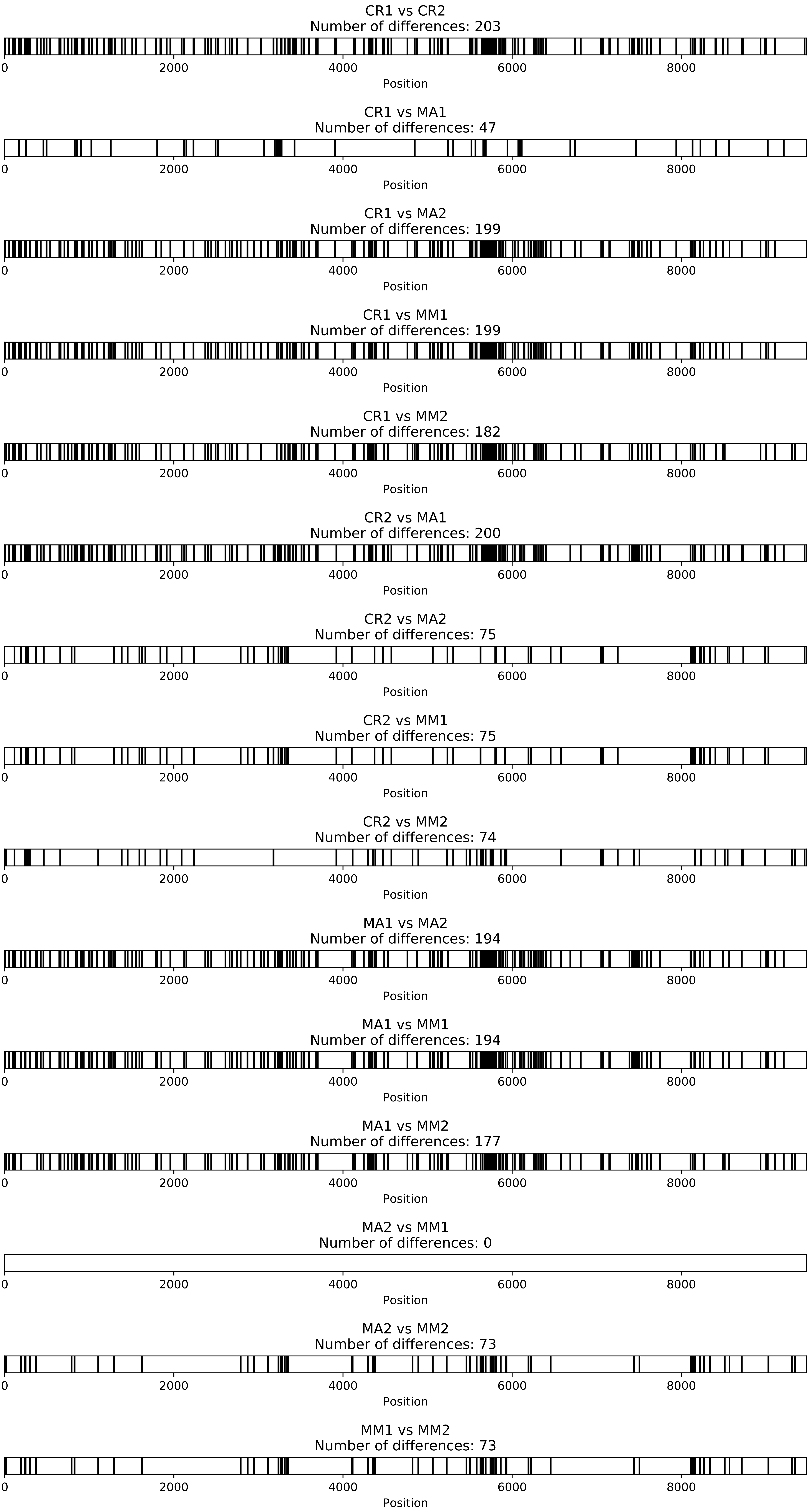
CR1 vs CR2
Number of differences: 203
0 2000 4000 6000 8000
Position
CR1 vs MA1
Number of differences: 47
0 2000 4000 6000 8000
Position
CR1 vs MA2
Number of differences: 199
0 2000 4000 6000 8000
Position
CR1 vs MM1
Number of differences: 199
0 2000 4000 6000 8000
Position
CR1 vs MM2
Number of differences: 182
0 2000 4000 6000 8000
Position
CR2 vs MA1
Number of differences: 200
0 2000 4000 6000 8000
Position
CR2 vs MA2
Number of differences: 75
0 2000 4000 6000 8000
Position
CR2 vs MM1
Number of differences: 75
0 2000 4000 6000 8000
Position
CR2 vs MM2
Number of differences: 74
0 2000 4000 6000 8000
Position
MA1 vs MA2
Number of differences: 194
0 2000 4000 6000 8000
Position
MA1 vs MM1
Number of differences: 194
0 2000 4000 6000 8000
Position
MA1 vs MM2
Number of differences: 177
0 2000 4000 6000 8000
Position
MA2 vs MM1
Number of differences: 0
0 2000 4000 6000 8000
Position
MA2 vs MM2
Number of differences: 73
0 2000 4000 6000 8000
Position
MM1 vs MM2
Number of differences: 73
0 2000 4000 6000 8000
Position

### Region: 76



CR1 vs CR2
Number of differences: 229

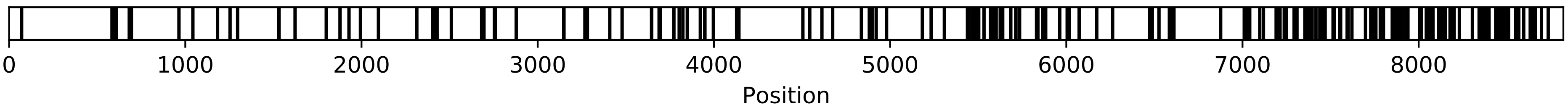

CR1 vs MA1
Number of differences: 143

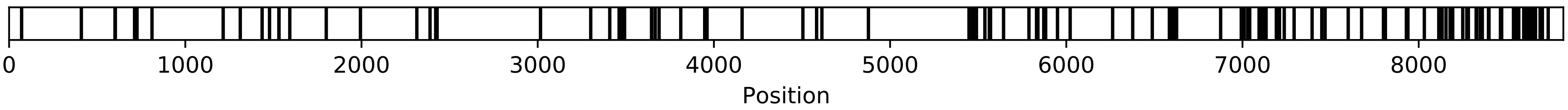

CR1 vs MA2
Number of differences: 225

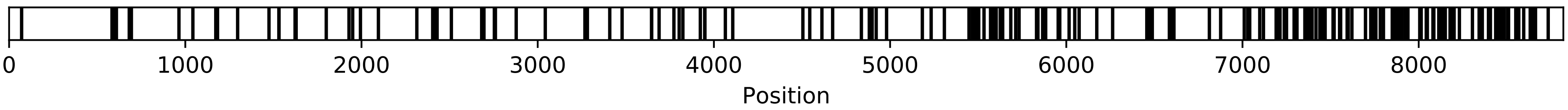

CR1 vs MM1
Number of differences: 110

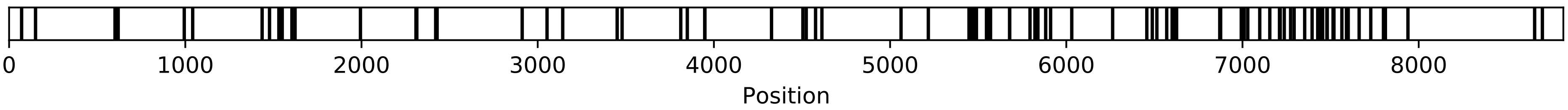

CR1 vs MM2
Number of differences: 143

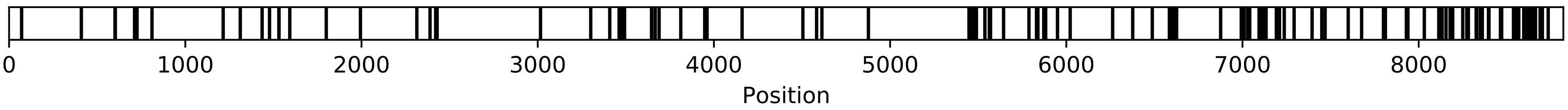

CR2 vs MA1
Number of differences: 212

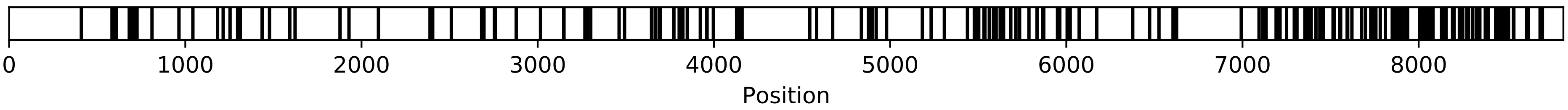

CR2 vs MA2
Number of differences: 31

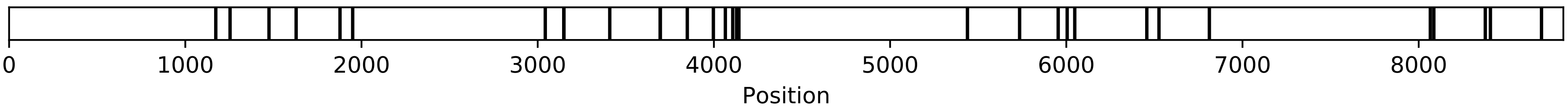

CR2 vs MM1
Number of differences: 230

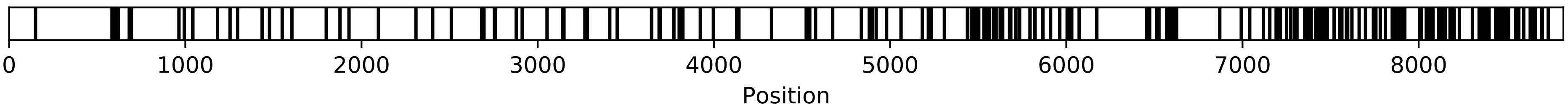

CR2 vs MM2
Number of differences: 212

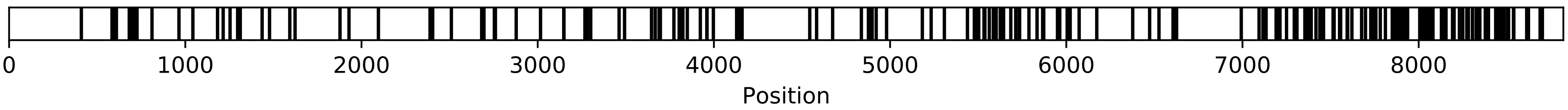

MA1 vs MA2
Number of differences: 209

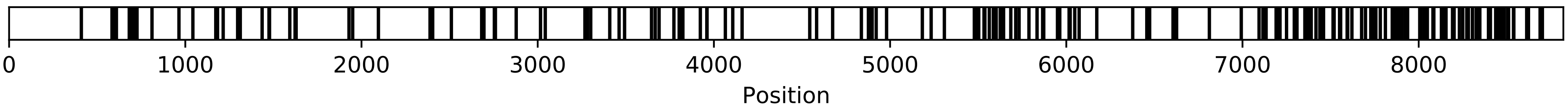

MA1 vs MM1
Number of differences: 140

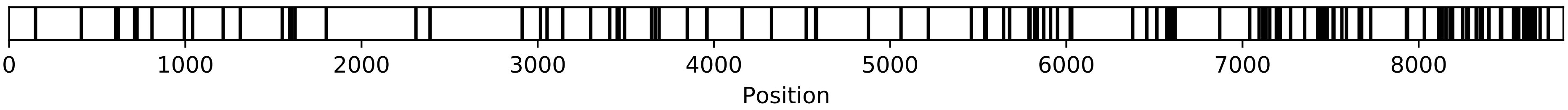

MA1 vs MM2
Number of differences: 0

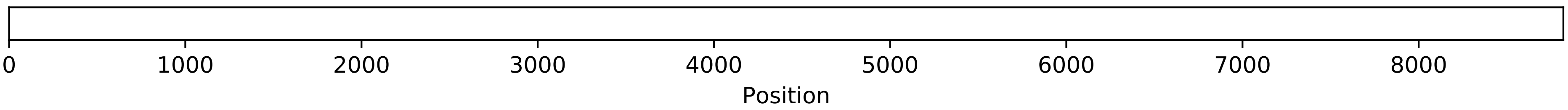

MA2 vs MM1
Number of differences: 226

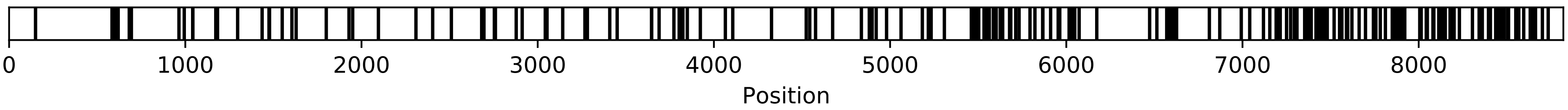

MA2 vs MM2
Number of differences: 209

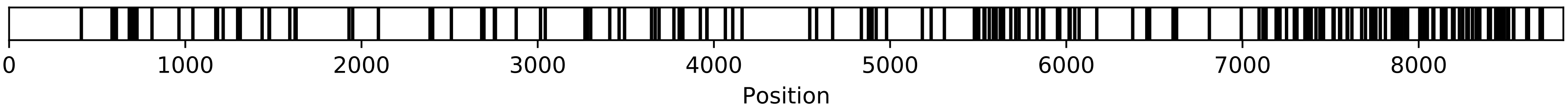

MM1 vs MM2
Number of differences: 140

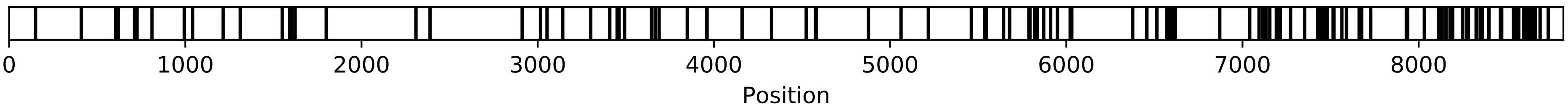

### Region: 77



CR1 vs CR2
Number of differences: 109

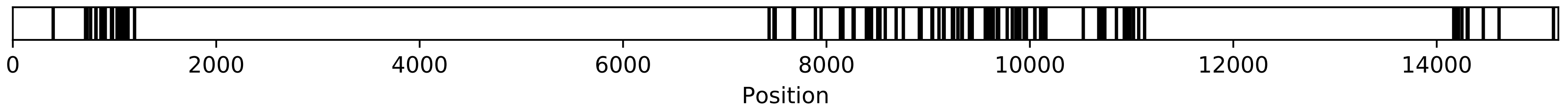


CR1 vs MA1
Number of differences: 310

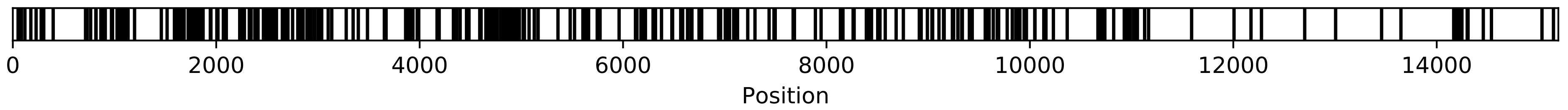


CR1 vs MA2
Number of differences: 191

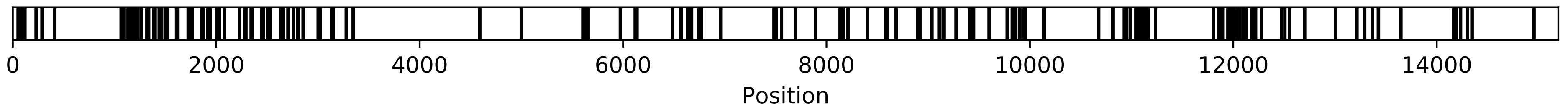


CR1 vs MM1
Number of differences: 226

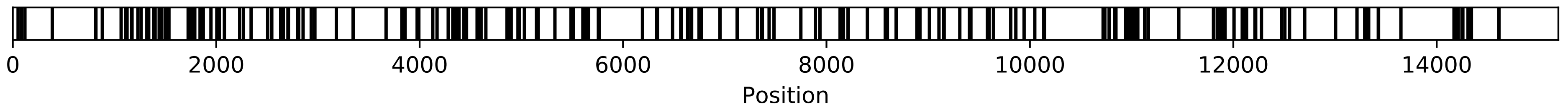


CR1 vs MM2
Number of differences: 191

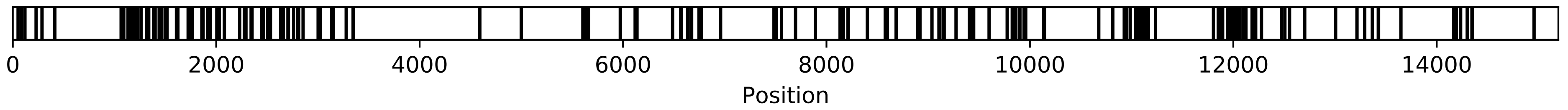


CR2 vs MA1
Number of differences: 213

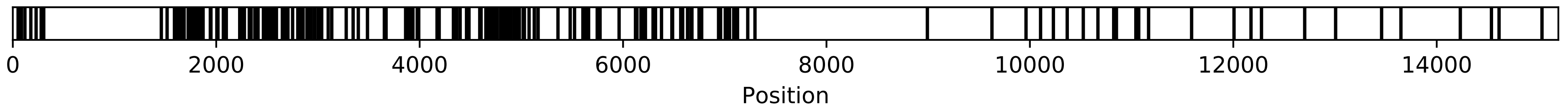


CR2 vs MA2
Number of differences: 229

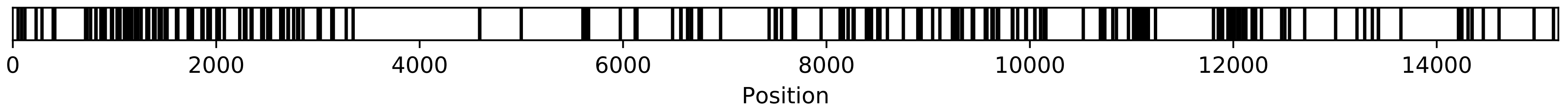


CR2 vs MM1
Number of differences: 274

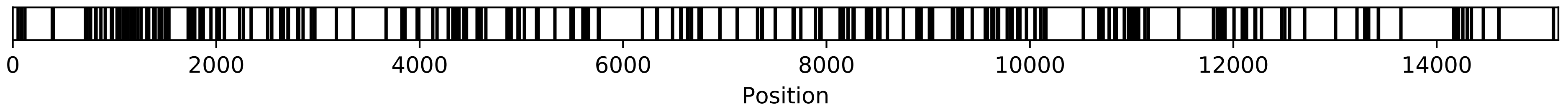


CR2 vs MM2
Number of differences: 229

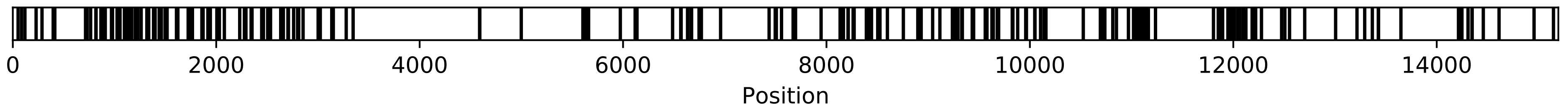


MA1 vs MA2
Number of differences: 327

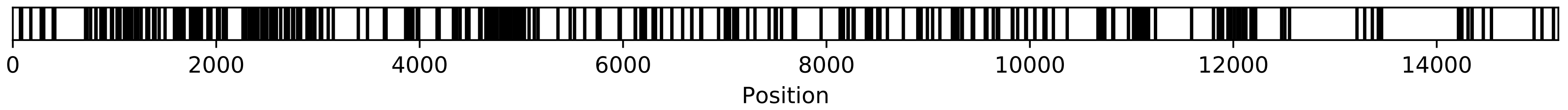


MA1 vs MM1
Number of differences: 342

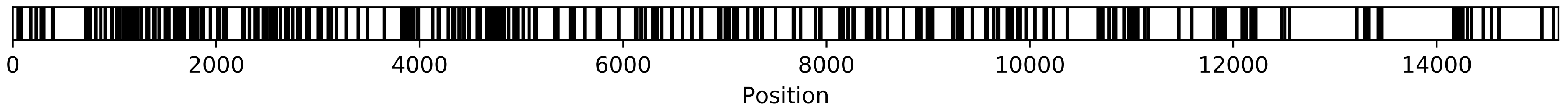


MA1 vs MM2
Number of differences: 327

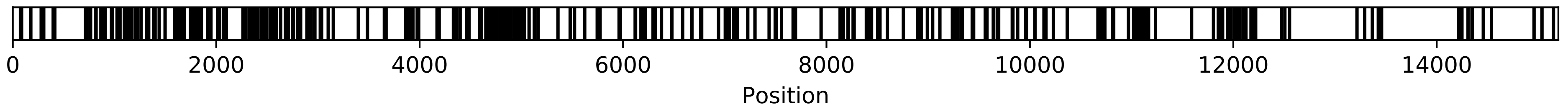


MA2 vs MM1
Number of differences: 196

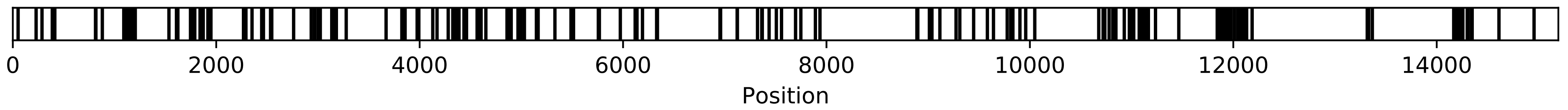


MA2 vs MM2
Number of differences: 0

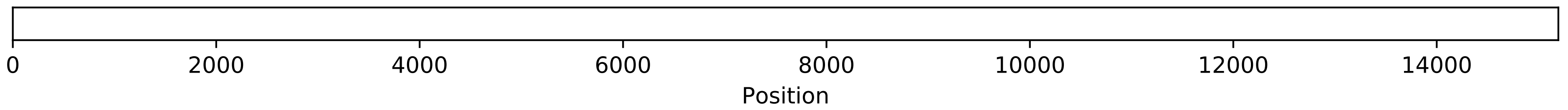


MM1 vs MM2
Number of differences: 196

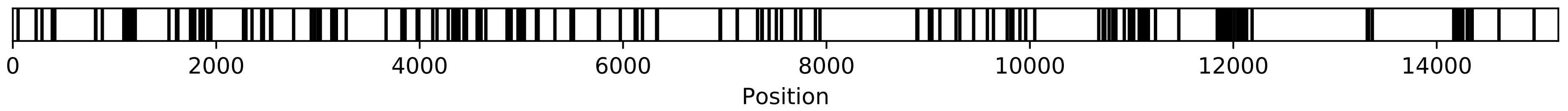

### Region: 78

CR1 vs CR2
Number of differences: 367

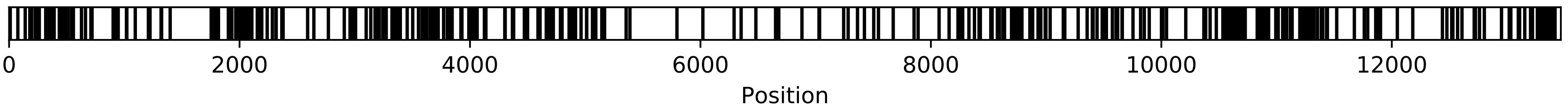


CR1 vs MA1
Number of differences: 191

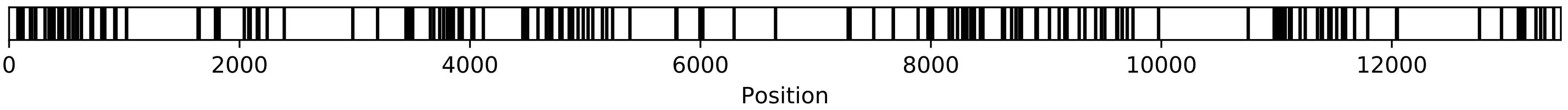


CR1 vs MA2
Number of differences: 326

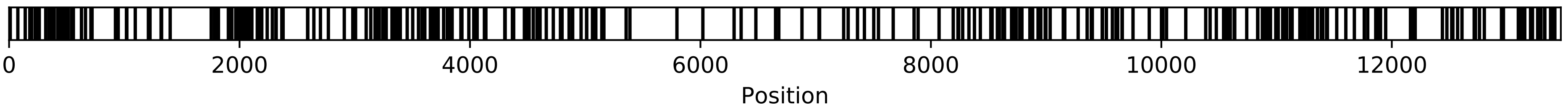


CR1 vs MM1
Number of differences: 191

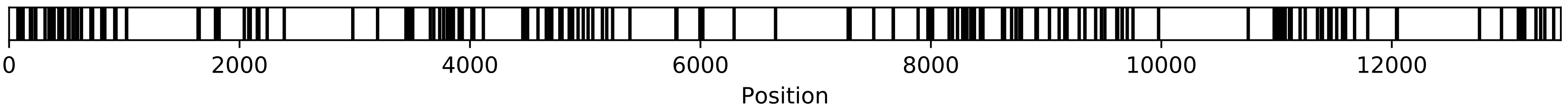


CR1 vs MM2
Number of differences: 220

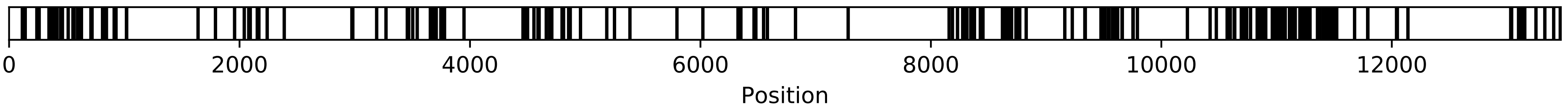


CR2 vs MA1
Number of differences: 373

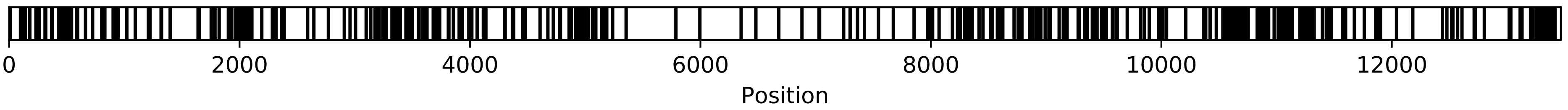


CR2 vs MA2
Number of differences: 102

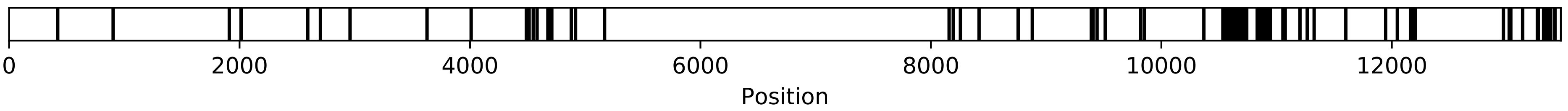


CR2 vs MM1
Number of differences: 373

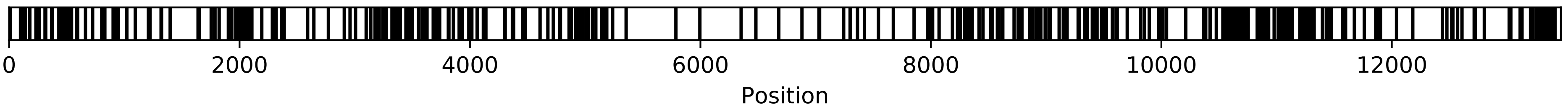


CR2 vs MM2
Number of differences: 400

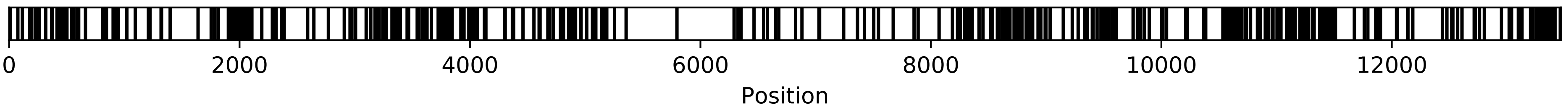


MA1 vs MA2
Number of differences: 339

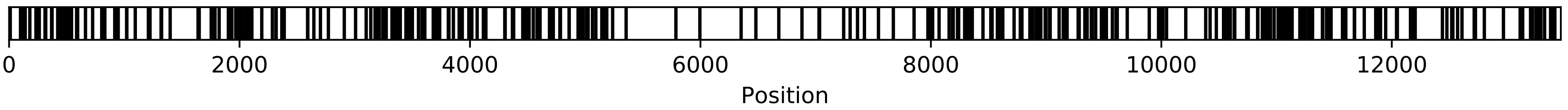


MA1 vs MM1
Number of differences: 0

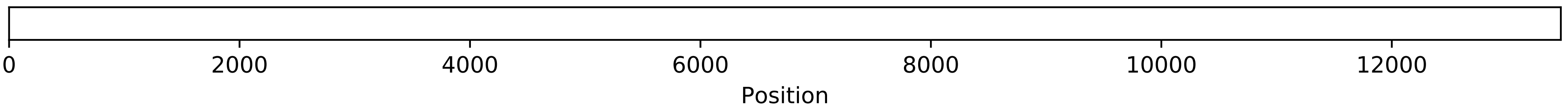


MA1 vs MM2
Number of differences: 233

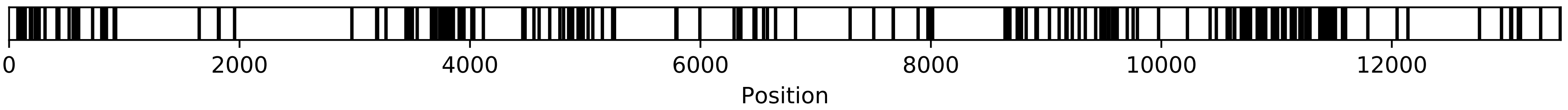


MA2 vs MM1
Number of differences: 339

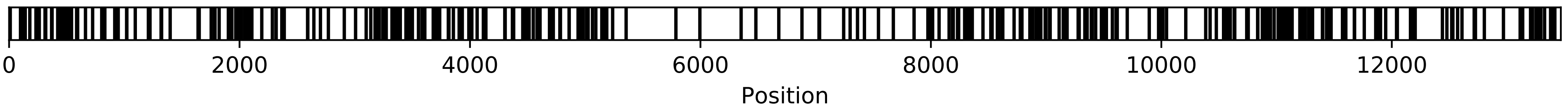


MA2 vs MM2
Number of differences: 383

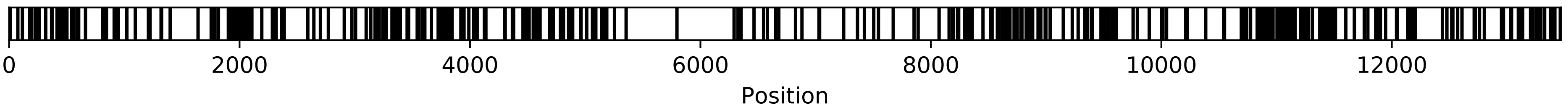


MM1 vs MM2
Number of differences: 233

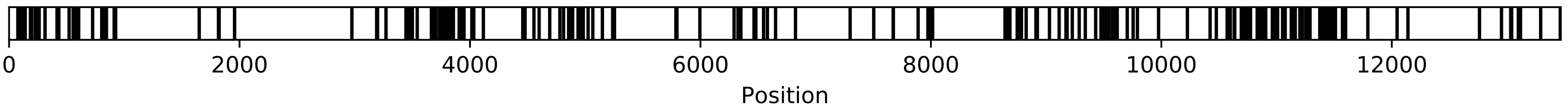

### Region: 79

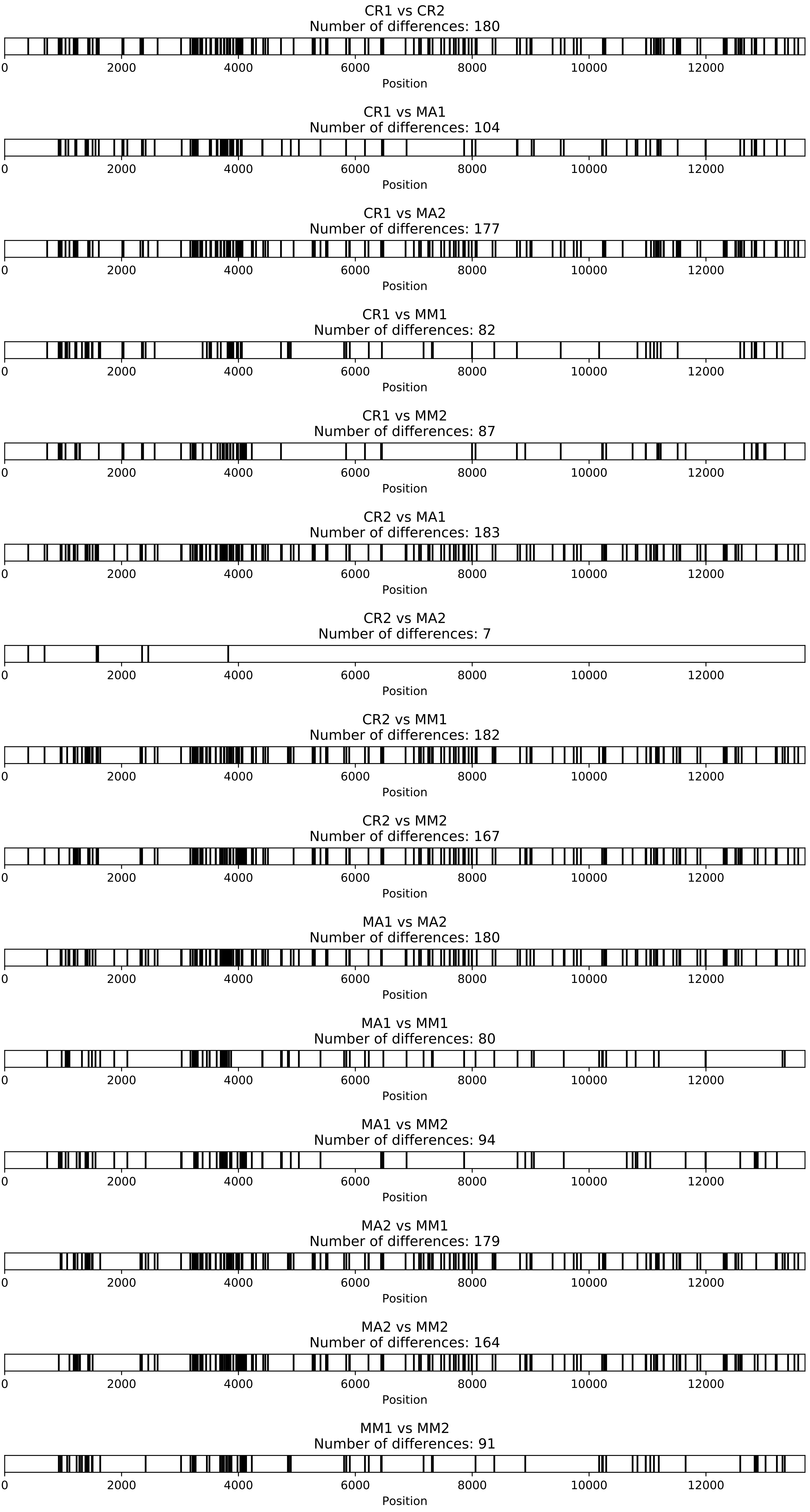
CR1 vs CR2
Number of differences: 180
CR1 vs MA1
Number of differences: 104
CR1 vs MA2
Number of differences: 177
CR1 vs MM1
Number of differences: 82
CR1 vs MM2
Number of differences: 87
CR2 vs MA1
Number of differences: 183
CR2 vs MA2
Number of differences: 7
CR2 vs MM1
Number of differences: 182
CR2 vs MM2
Number of differences: 167
MA1 vs MA2
Number of differences: 180
MA1 vs MM1
Number of differences: 80
MA1 vs MM2
Number of differences: 94
MA2 vs MM1
Number of differences: 179
MA2 vs MM2
Number of differences: 164
MM1 vs MM2
Number of differences: 91
0
2000
4000
6000
8000
10000
12000
Position

### Region: 81

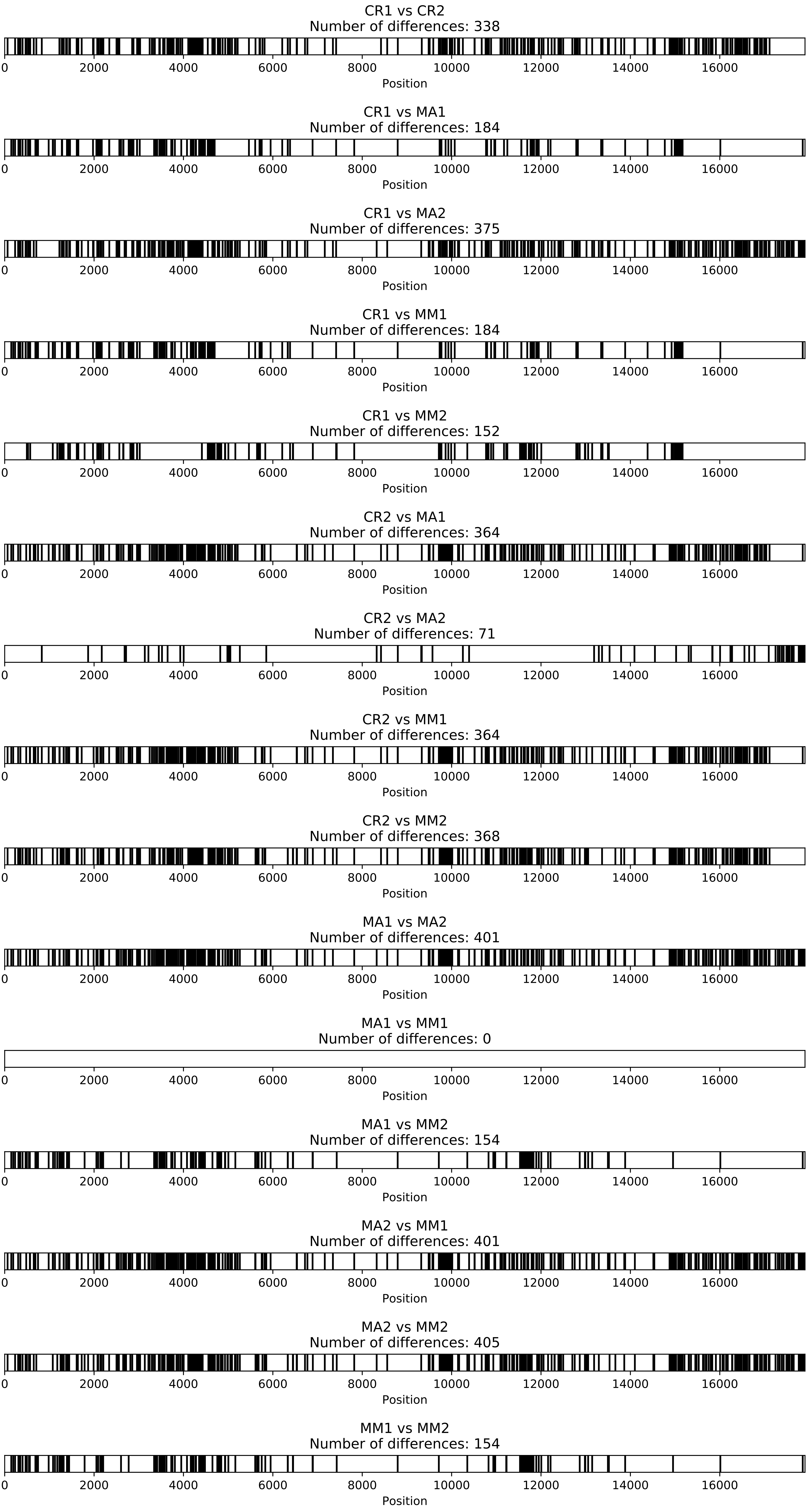
CR1 vs CR2
Number of differences: 338
CR1 vs MA1
Number of differences: 184
CR1 vs MA2
Number of differences: 375
CR1 vs MM1
Number of differences: 184
CR1 vs MM2
Number of differences: 152
CR2 vs MA1
Number of differences: 364
CR2 vs MA2
Number of differences: 71
CR2 vs MM1
Number of differences: 364
CR2 vs MM2
Number of differences: 368
MA1 vs MA2
Number of differences: 401
MA1 vs MM1
Number of differences: 0
MA1 vs MM2
Number of differences: 154
MA2 vs MM1
Number of differences: 401
MA2 vs MM2
Number of differences: 405
MM1 vs MM2
Number of differences: 154
0
2000
4000
6000
8000
10000
12000
14000
16000
Position

### Region: 82

CR1 vs CR2
Number of differences: 265

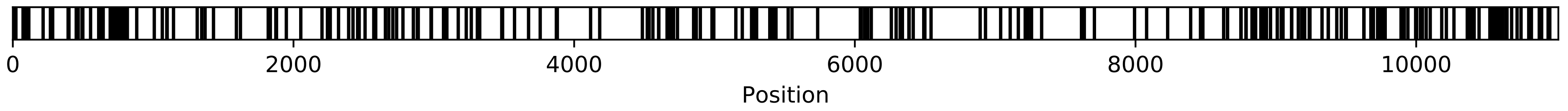


CR1 vs MA1
Number of differences: 272

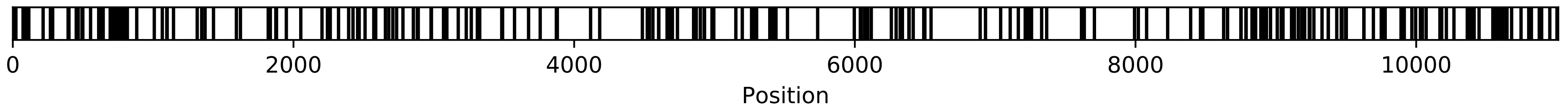


CR1 vs MA2
Number of differences: 159

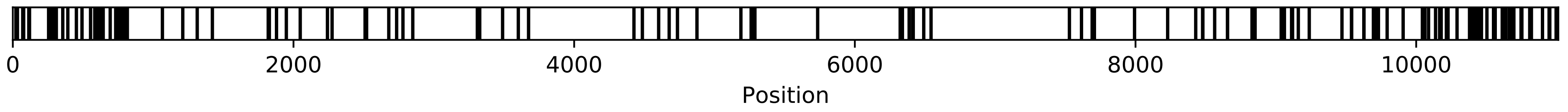


CR1 vs MM1
Number of differences: 159

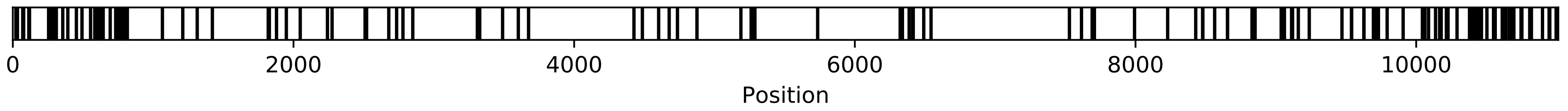


CR1 vs MM2
Number of differences: 134

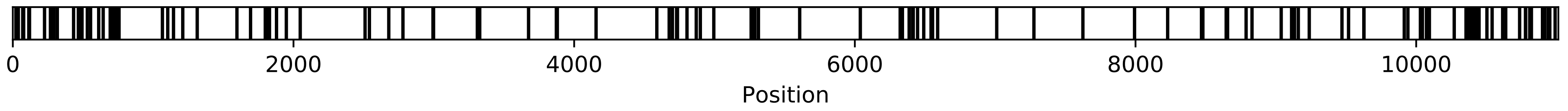


CR2 vs MA1
Number of differences: 38

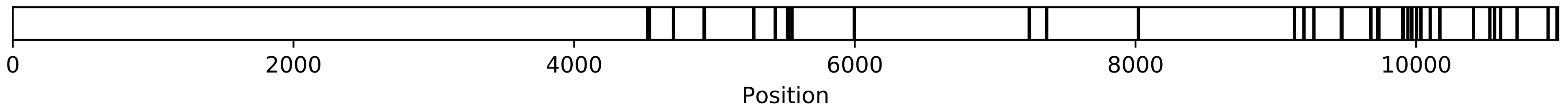


CR2 vs MA2
Number of differences: 277

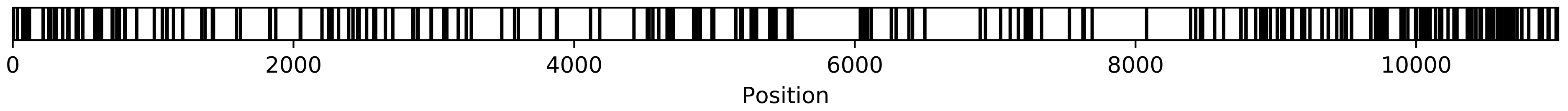


CR2 vs MM1
Number of differences: 277

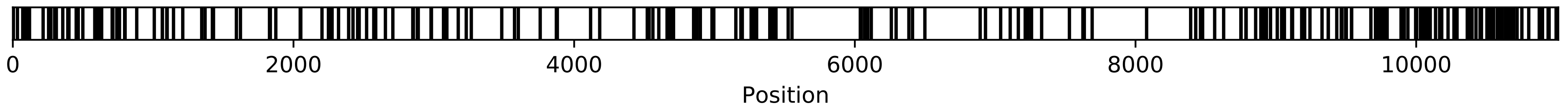


CR2 vs MM2
Number of differences: 258

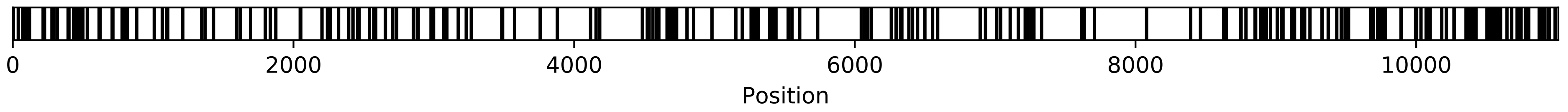


MA1 vs MA2
Number of differences: 282

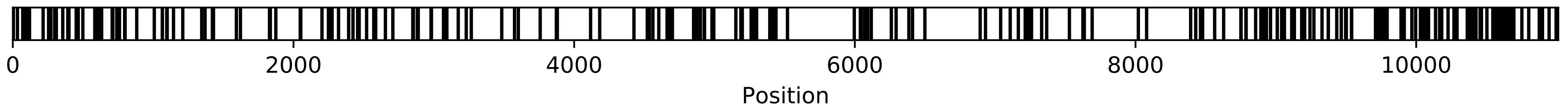


MA1 vs MM1
Number of differences: 282

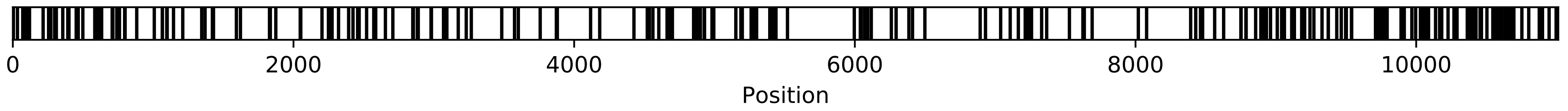


MA1 vs MM2
Number of differences: 266

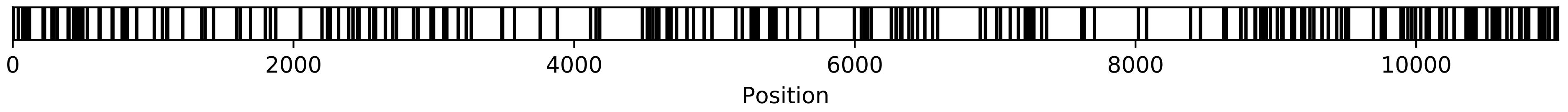


MA2 vs MM1
Number of differences: 0

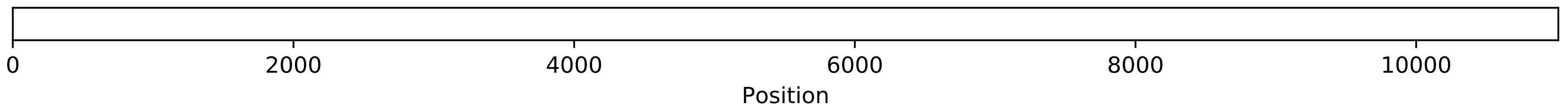


MA2 vs MM2
Number of differences: 151

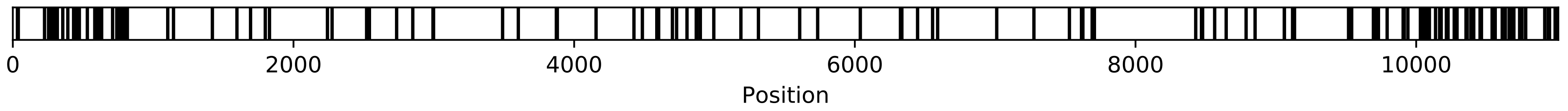


MM1 vs MM2
Number of differences: 151

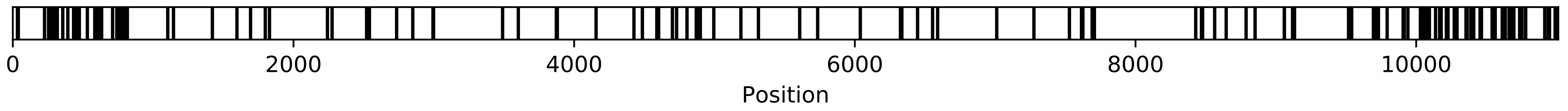

### Region: 86



CR1 vs CR2
Number of differences: 181

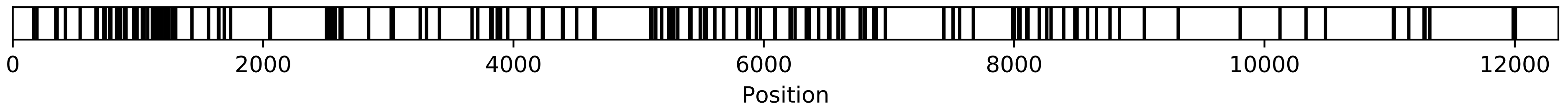


CR1 vs MA1
Number of differences: 90

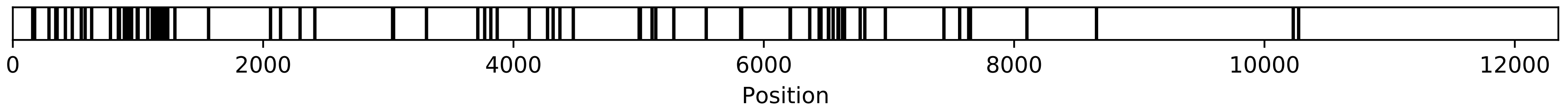


CR1 vs MA2
Number of differences: 188

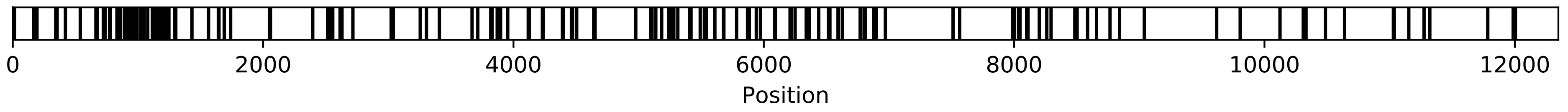


CR1 vs MM1
Number of differences: 90

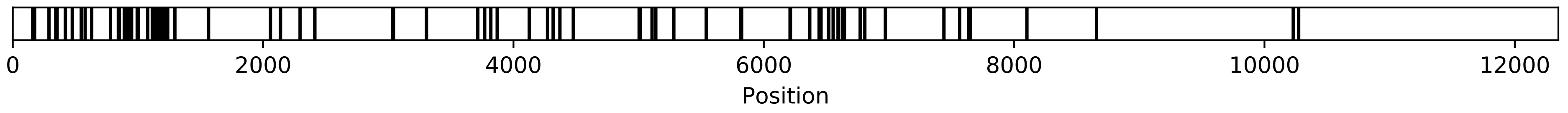


CR1 vs MM2
Number of differences: 87

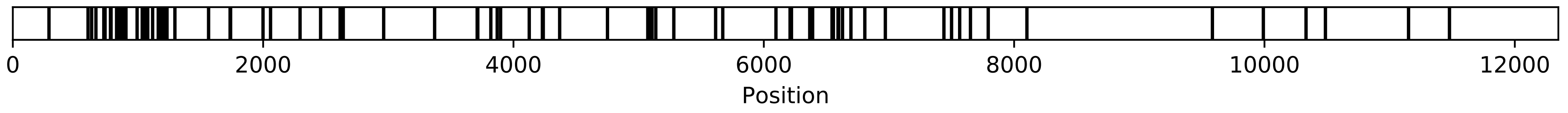


CR2 vs MA1
Number of differences: 181

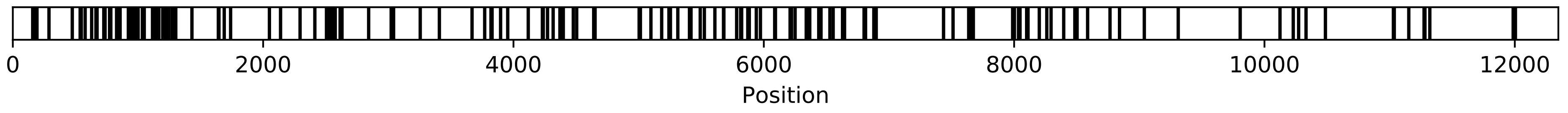


CR2 vs MA2
Number of differences: 41

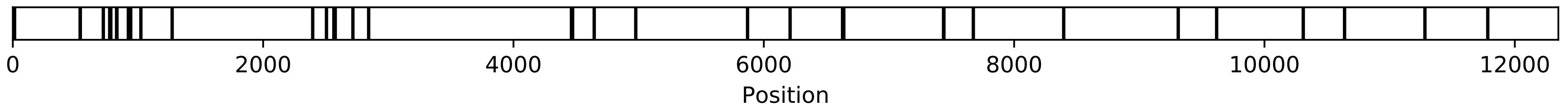


CR2 vs MM1
Number of differences: 181

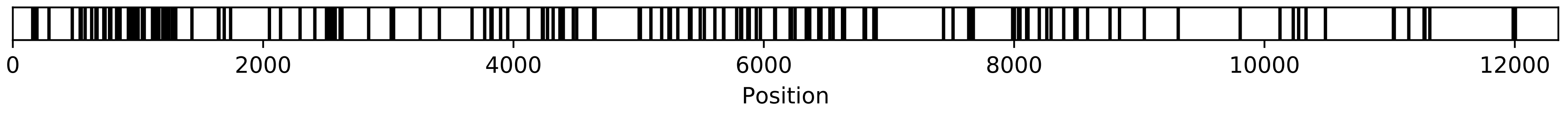


CR2 vs MM2
Number of differences: 169

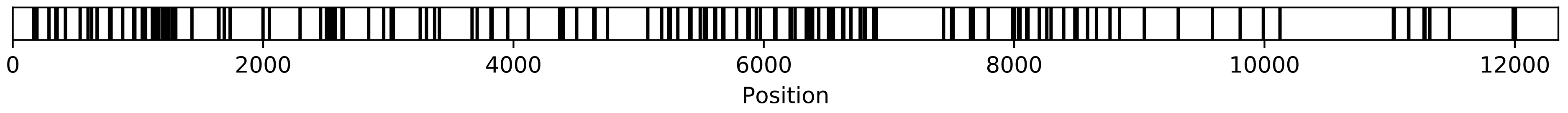


MA1 vs MA2
Number of differences: 186

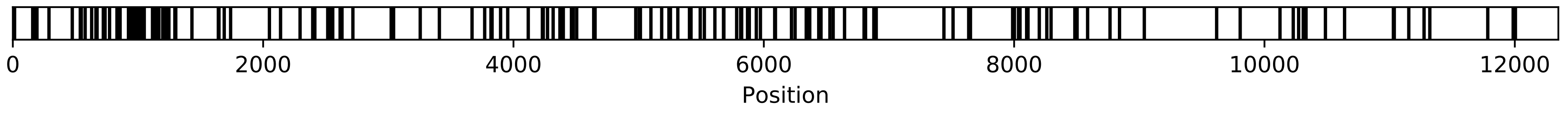


MA1 vs MM1
Number of differences: 0

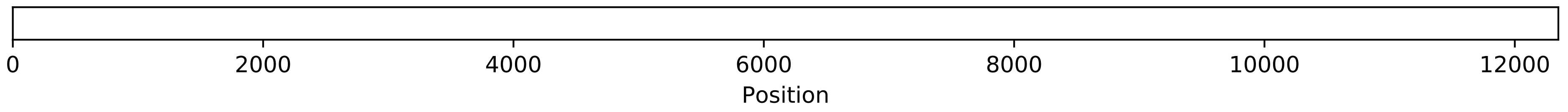


MA1 vs MM2
Number of differences: 99

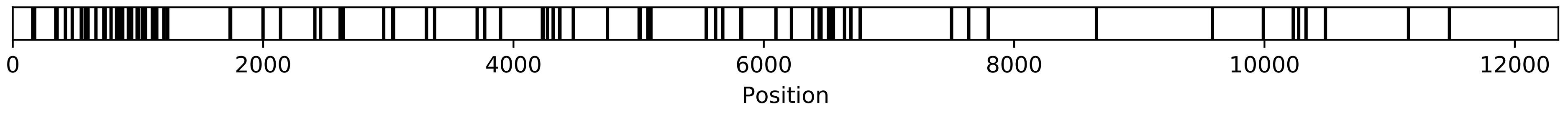


MA2 vs MM1
Number of differences: 186

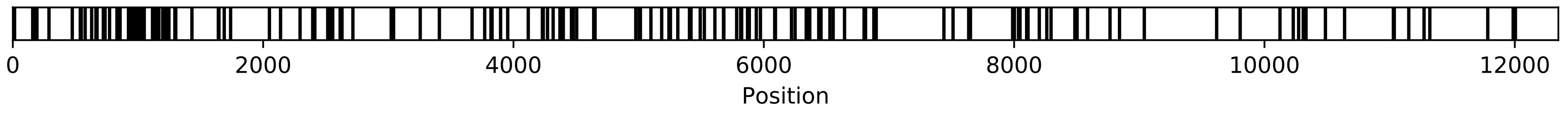


MA2 vs MM2
Number of differences: 178

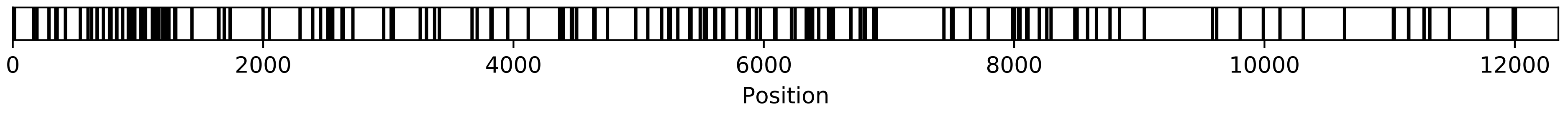


MM1 vs MM2
Number of differences: 99

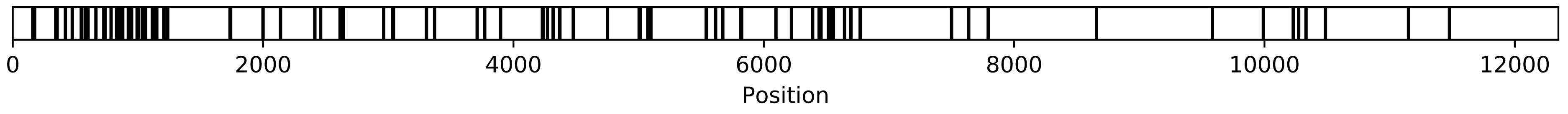

### Region: 87



CR1 vs CR2
Number of differences: 264

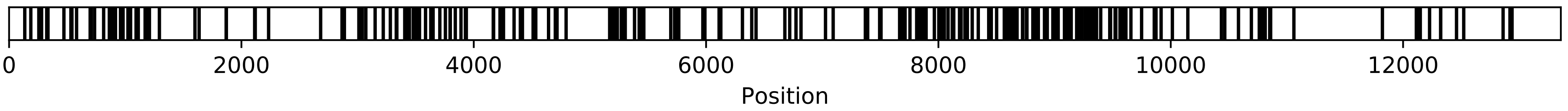


CR1 vs MA1
Number of differences: 137

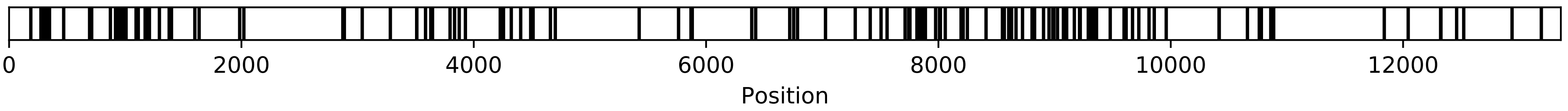


CR1 vs MA2
Number of differences: 255

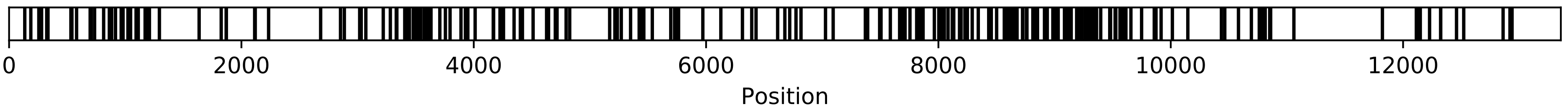


CR1 vs MM1
Number of differences: 132

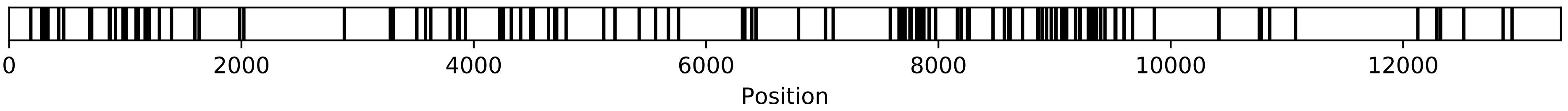


CR1 vs MM2
Number of differences: 137

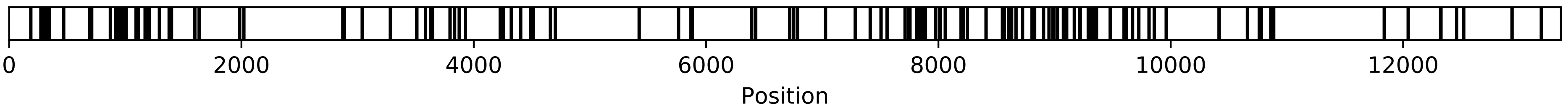


CR2 vs MA1
Number of differences: 287

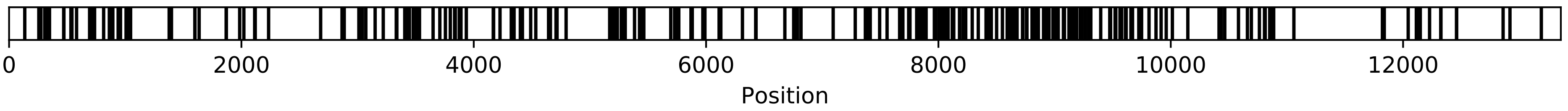


CR2 vs MA2
Number of differences: 41

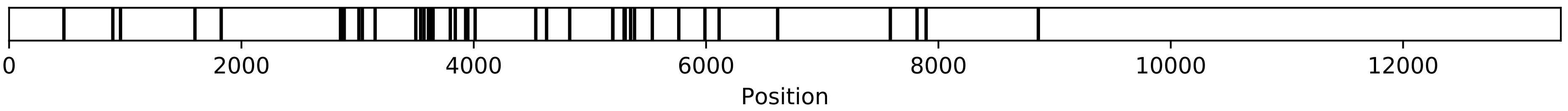


CR2 vs MM1
Number of differences: 257

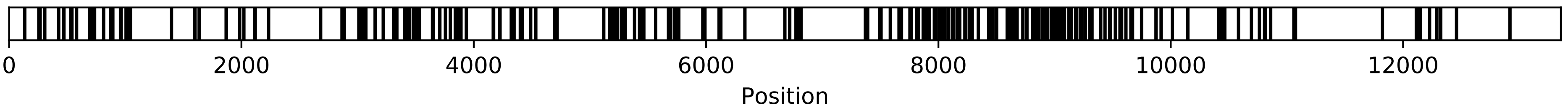


CR2 vs MM2
Number of differences: 287

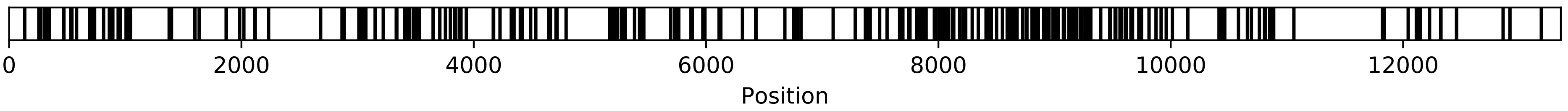


MA1 vs MA2
Number of differences: 280

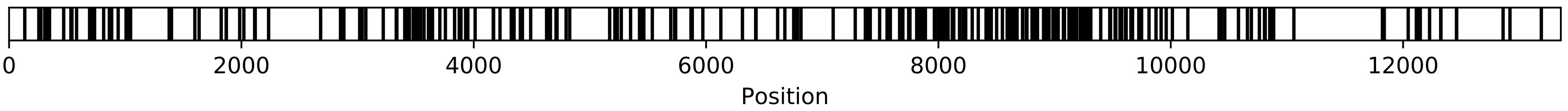


MA1 vs MM1
Number of differences: 130

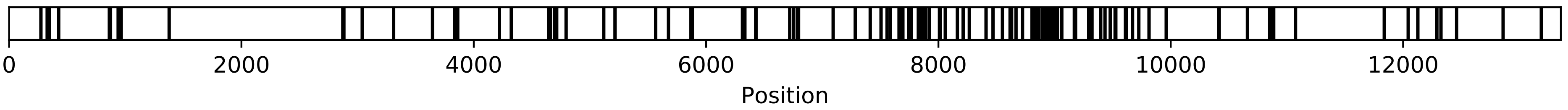


MA1 vs MM2
Number of differences: 0

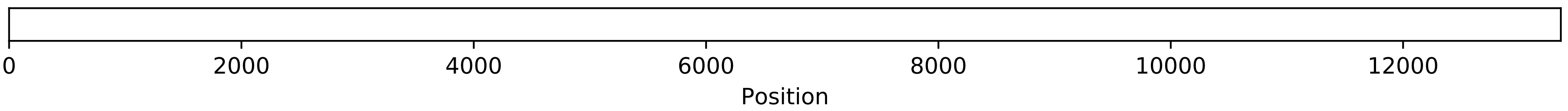


MA2 vs MM1
Number of differences: 246

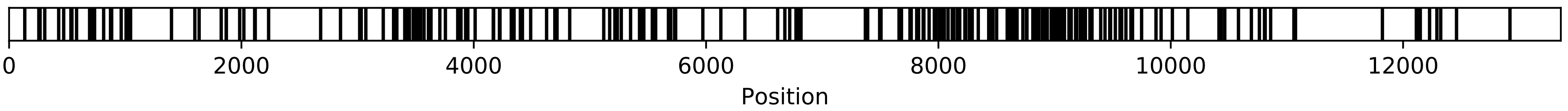


MA2 vs MM2
Number of differences: 280

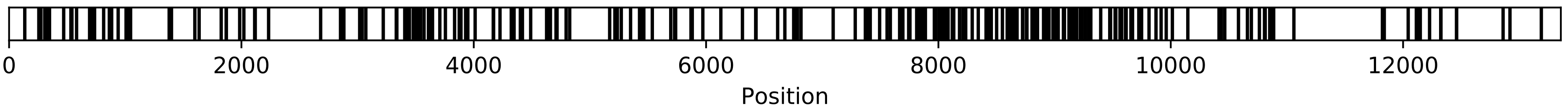


MM1 vs MM2
Number of differences: 130

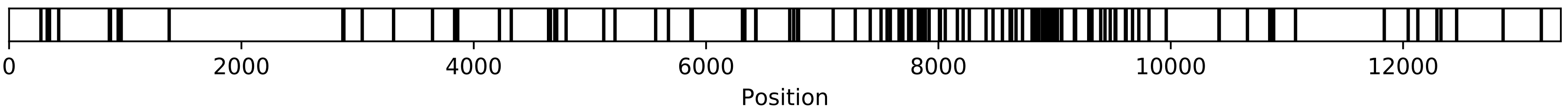

### Region: 88



CR1 vs CR2
Number of differences: 265

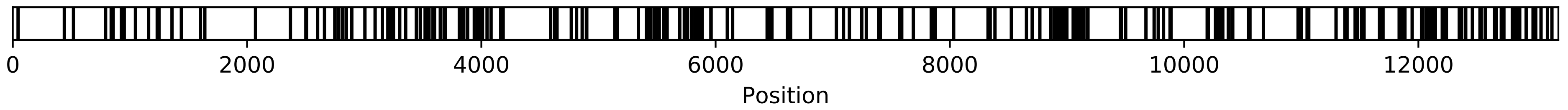


CR1 vs MA1
Number of differences: 93

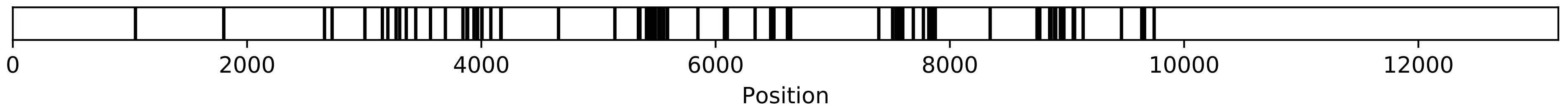


CR1 vs MA2
Number of differences: 260

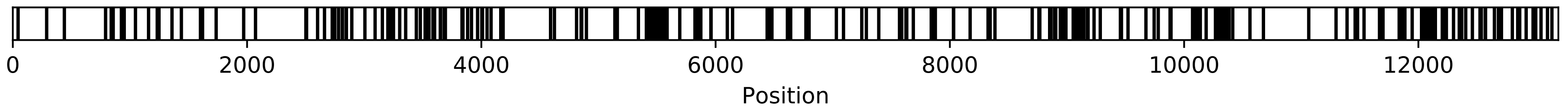


CR1 vs MM1
Number of differences: 144

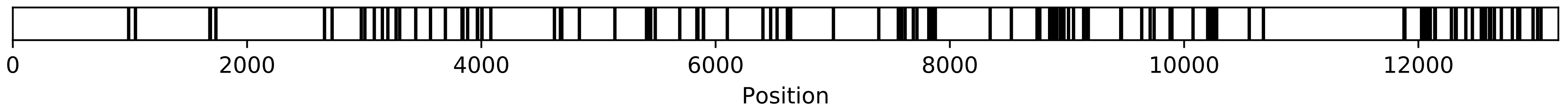


CR1 vs MM2
Number of differences: 93

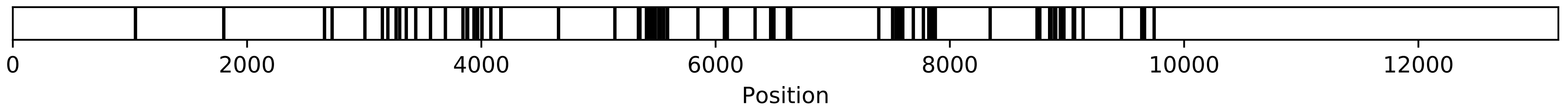


CR2 vs MA1
Number of differences: 264

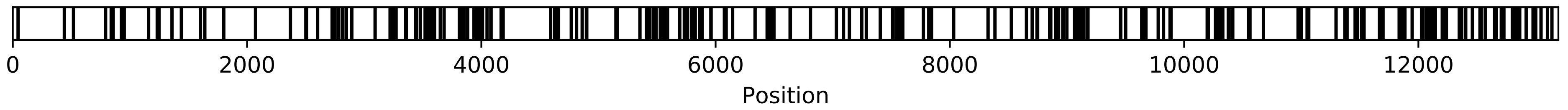


CR2 vs MA2
Number of differences: 93

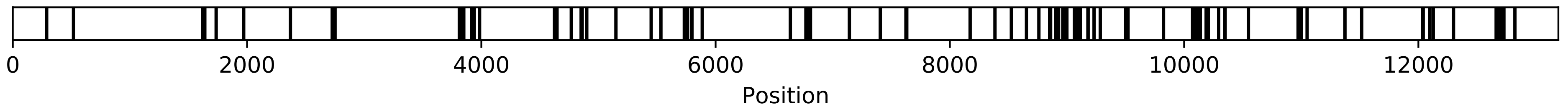


CR2 vs MM1
Number of differences: 262

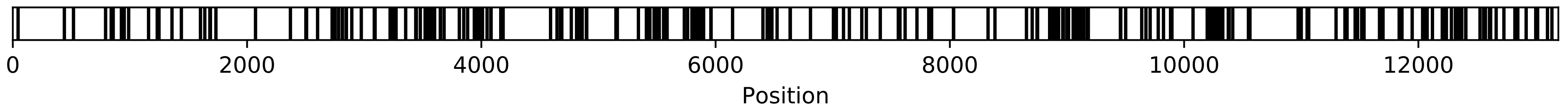


CR2 vs MM2
Number of differences: 264

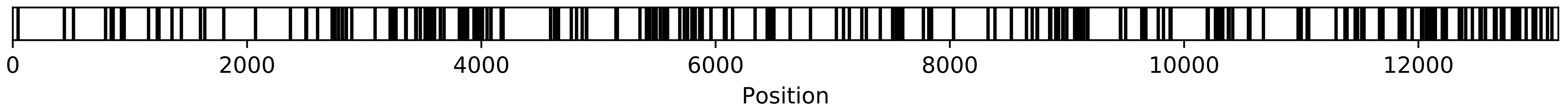


MA1 vs MA2
Number of differences: 255

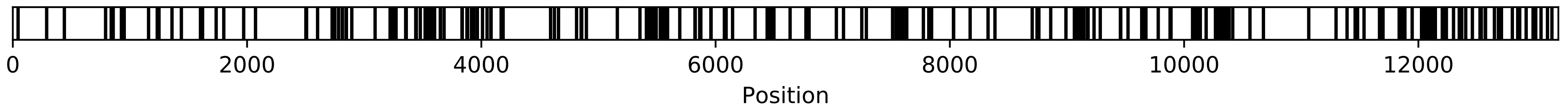


MA1 vs MM1
Number of differences: 123

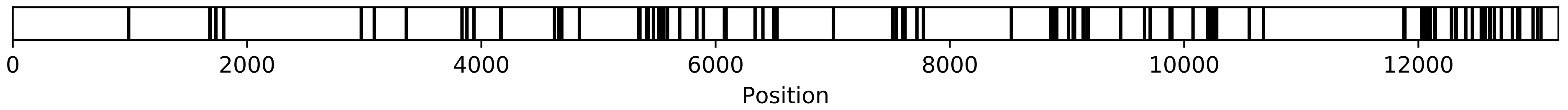


MA1 vs MM2
Number of differences: 0

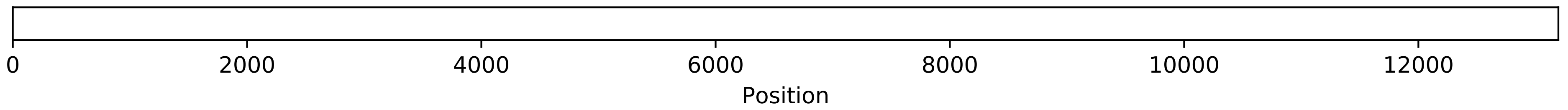


MA2 vs MM1
Number of differences: 258

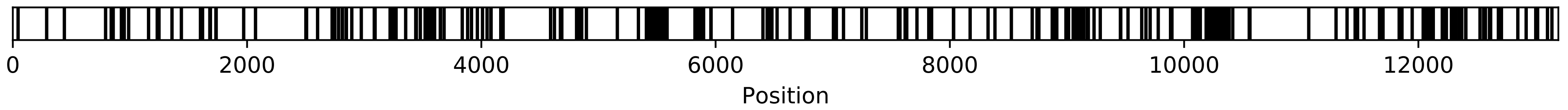


MA2 vs MM2
Number of differences: 255

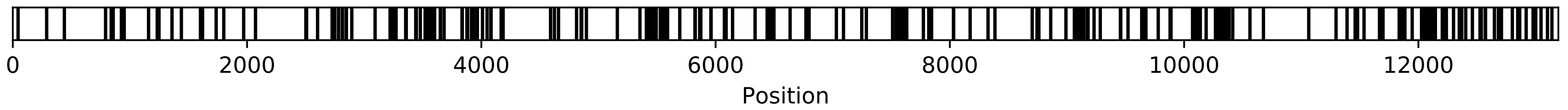


MM1 vs MM2
Number of differences: 123

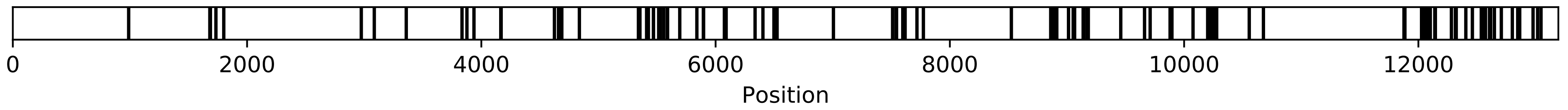

### Region: 90

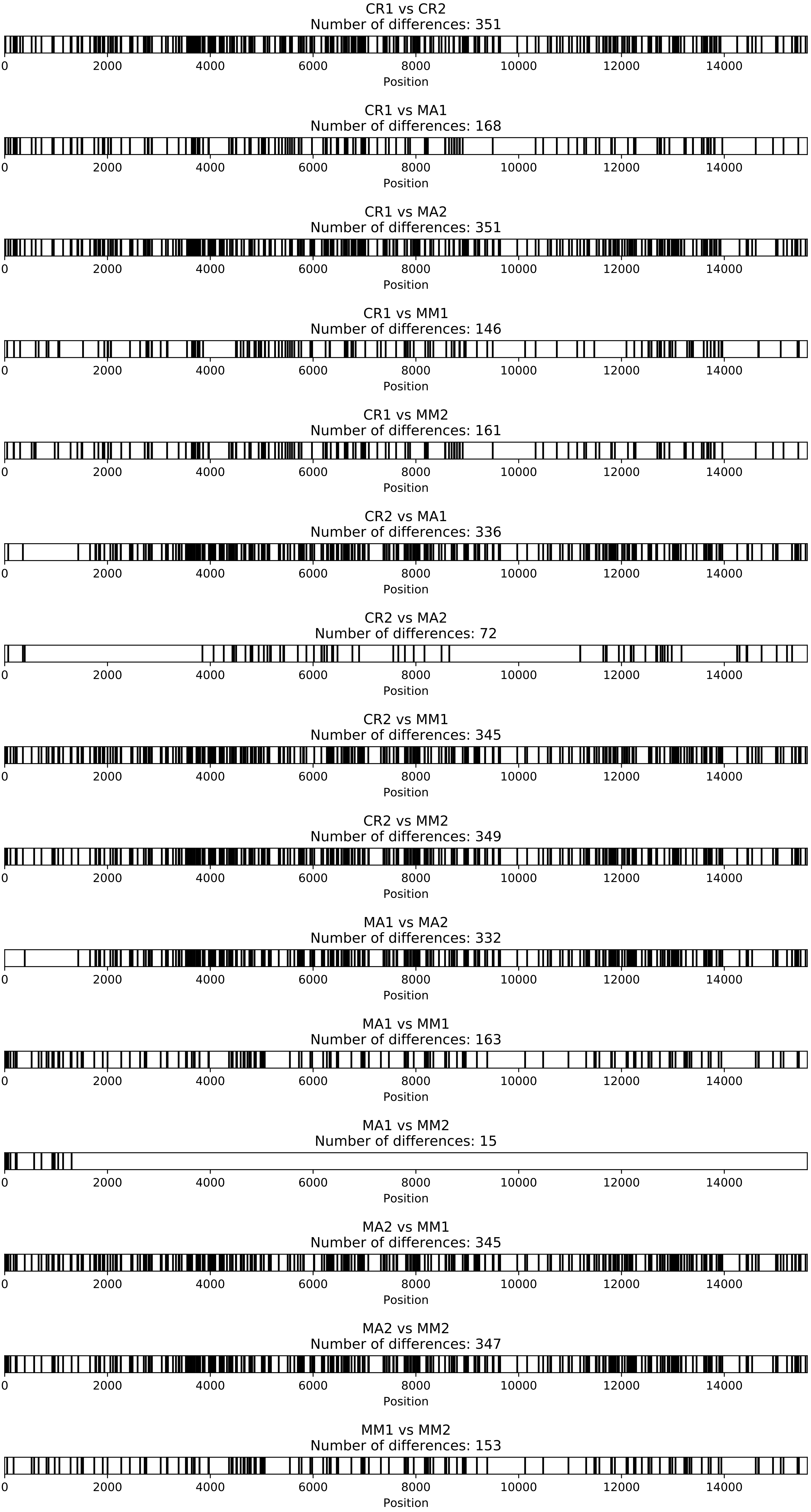
CR1 vs CR2
Number of differences: 351
CR1 vs MA1
Number of differences: 168
CR1 vs MA2
Number of differences: 351
CR1 vs MM1
Number of differences: 146
CR1 vs MM2
Number of differences: 161
CR2 vs MA1
Number of differences: 336
CR2 vs MA2
Number of differences: 72
CR2 vs MM1
Number of differences: 345
CR2 vs MM2
Number of differences: 349
MA1 vs MA2
Number of differences: 332
MA1 vs MM1
Number of differences: 163
MA1 vs MM2
Number of differences: 15
MA2 vs MM1
Number of differences: 345
MA2 vs MM2
Number of differences: 347
MM1 vs MM2
Number of differences: 153
0
2000
4000
6000
8000
10000
12000
14000
Position

### Region: 91



CR1 vs CR2
Number of differences: 121

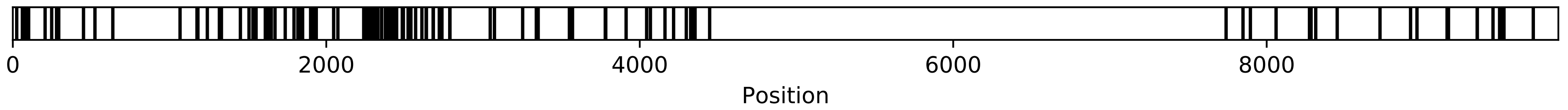


CR1 vs MA1
Number of differences: 152

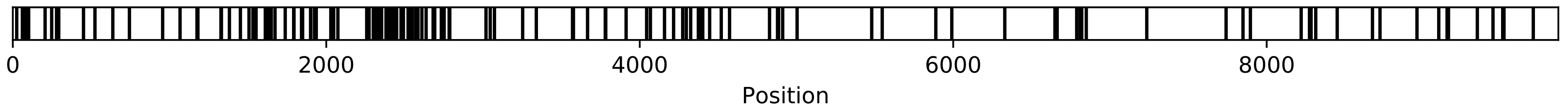


CR1 vs MA2
Number of differences: 79

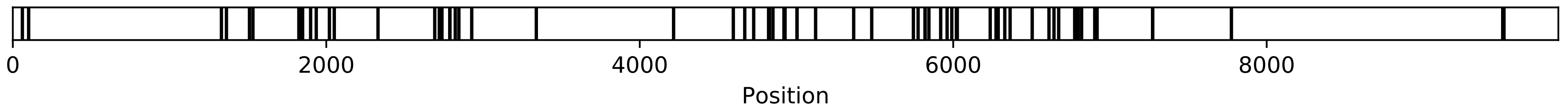


CR1 vs MM1
Number of differences: 139

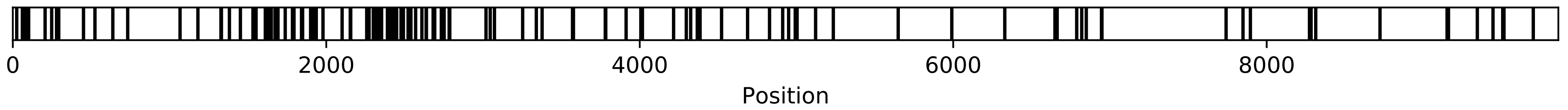


CR1 vs MM2
Number of differences: 152

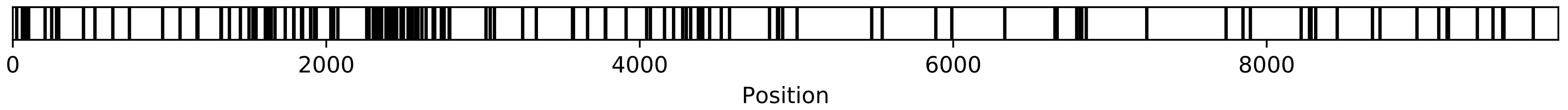


CR2 vs MA1
Number of differences: 92

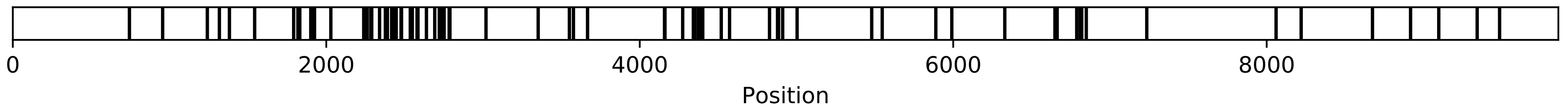


CR2 vs MA2
Number of differences: 166

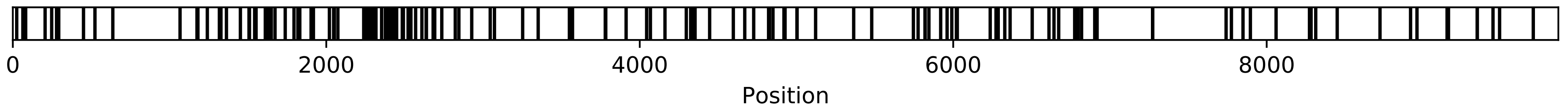


CR2 vs MM1
Number of differences: 88

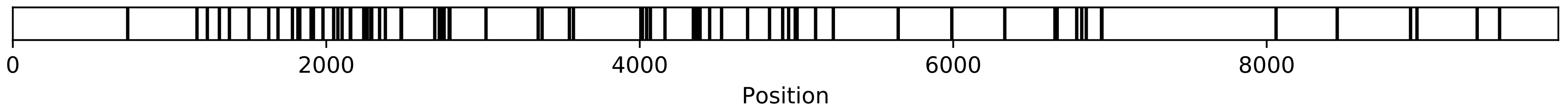


CR2 vs MM2
Number of differences: 92

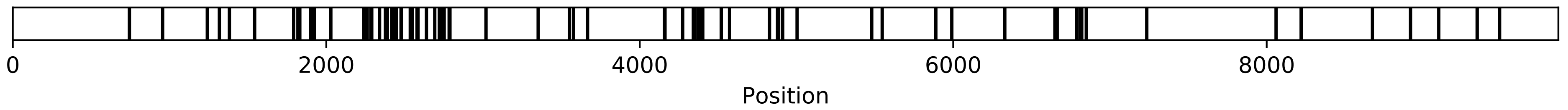


MA1 vs MA2
Number of differences: 189

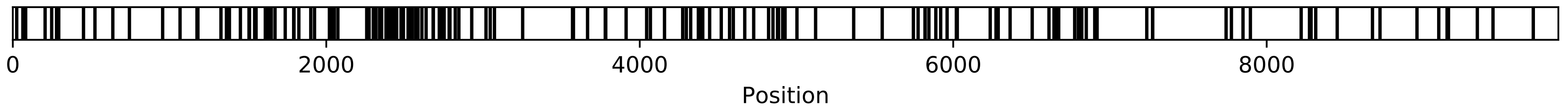


MA1 vs MM1
Number of differences: 70

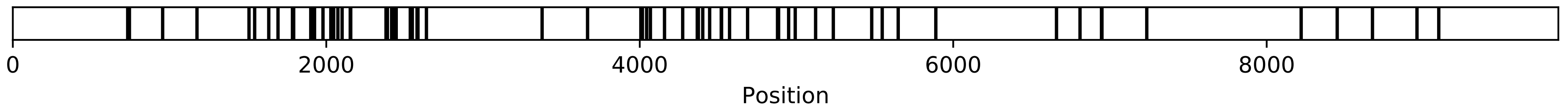


MA1 vs MM2
Number of differences: 0

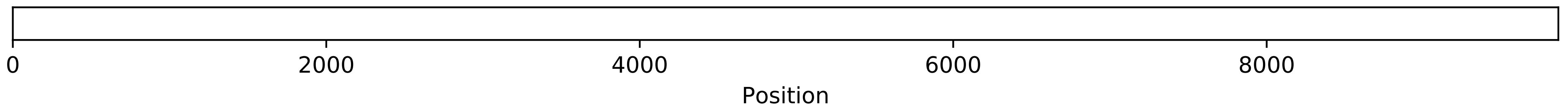


MA2 vs MM1
Number of differences: 176

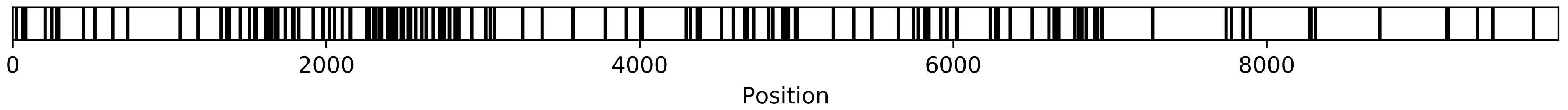


MA2 vs MM2
Number of differences: 189

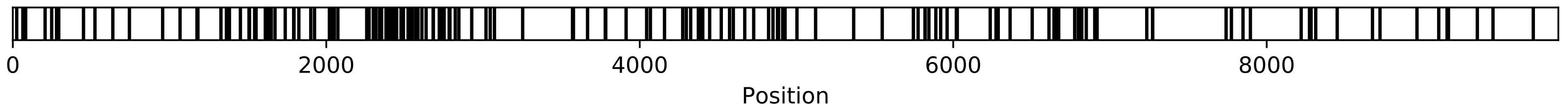


MM1 vs MM2
Number of differences: 70

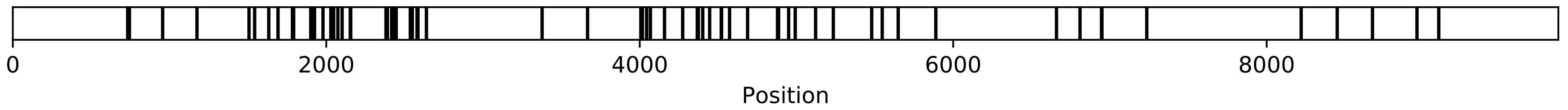

### Region: 92

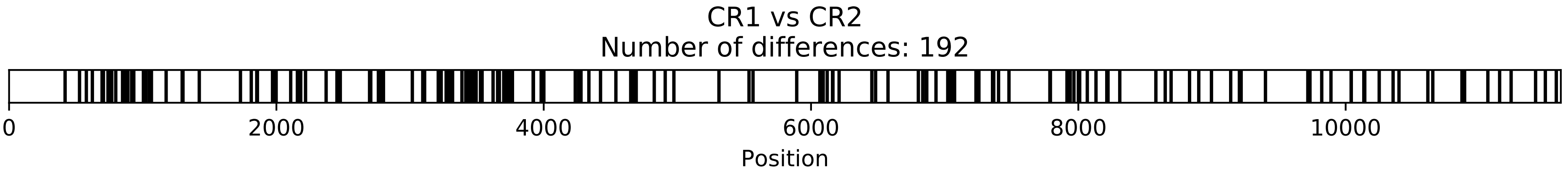
CR1 vs CR2
Number of differences: 192
0 2000 4000 6000 8000 10000
Position

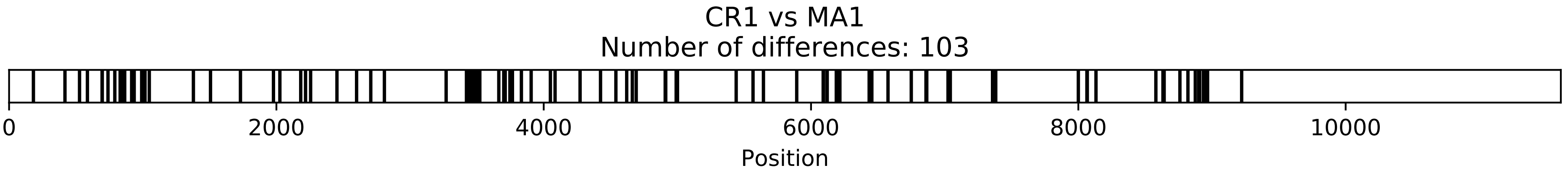
CR1 vs MA1
Number of differences: 103
0 2000 4000 6000 8000 10000
Position

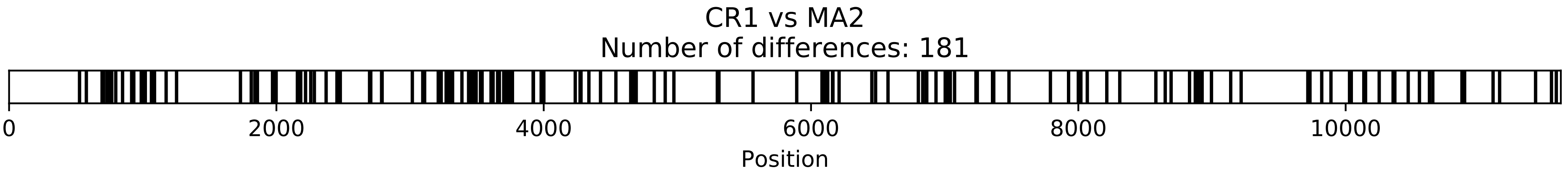
CR1 vs MA2
Number of differences: 181
0 2000 4000 6000 8000 10000
Position

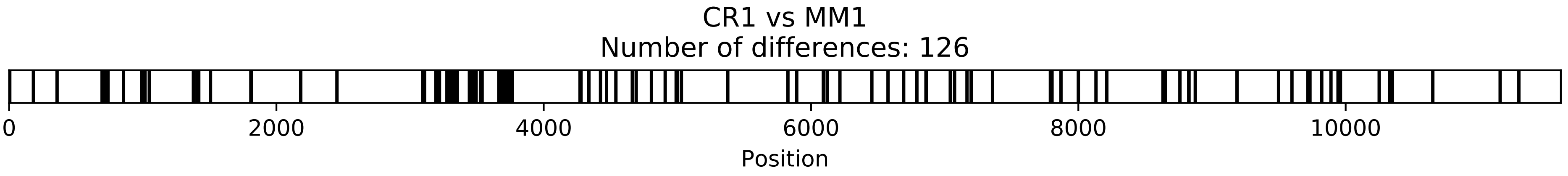
CR1 vs MM1
Number of differences: 126
0 2000 4000 6000 8000 10000
Position

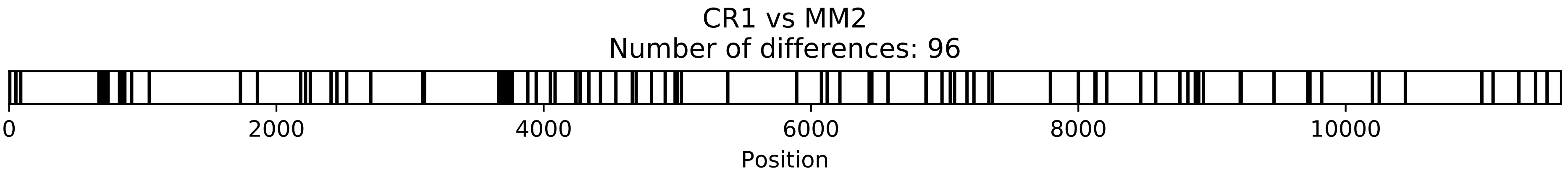
CR1 vs MM2
Number of differences: 96
0 2000 4000 6000 8000 10000
Position

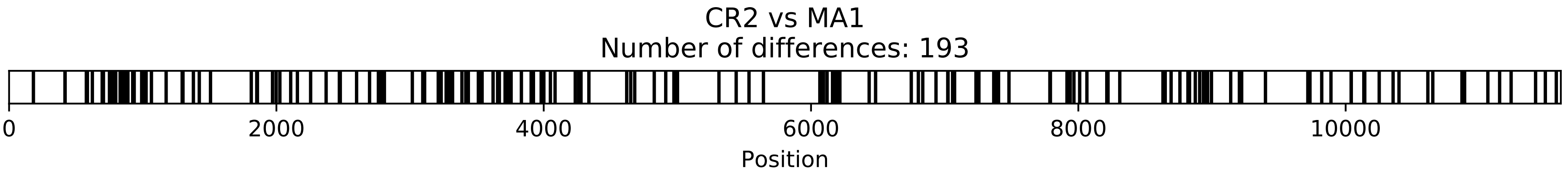
CR2 vs MA1
Number of differences: 193
0 2000 4000 6000 8000 10000
Position

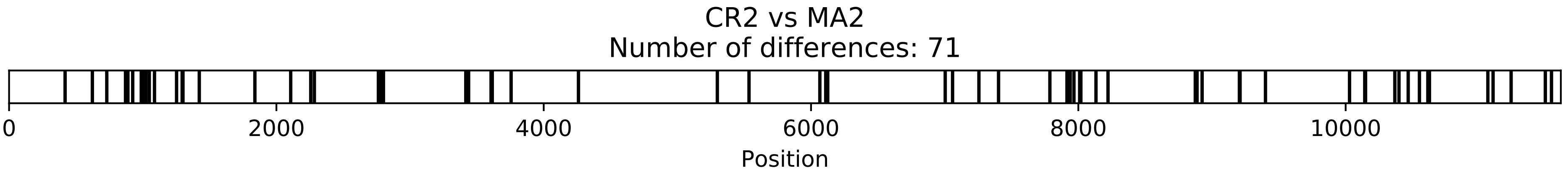
CR2 vs MA2
Number of differences: 71
0 2000 4000 6000 8000 10000
Position

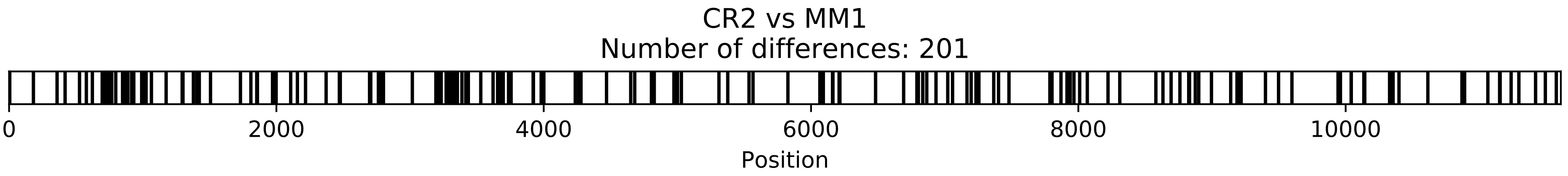
CR2 vs MM1
Number of differences: 201
0 2000 4000 6000 8000 10000
Position

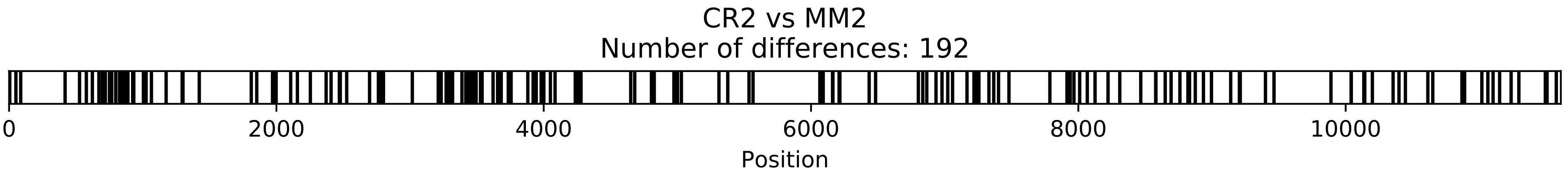
CR2 vs MM2
Number of differences: 192
0 2000 4000 6000 8000 10000
Position

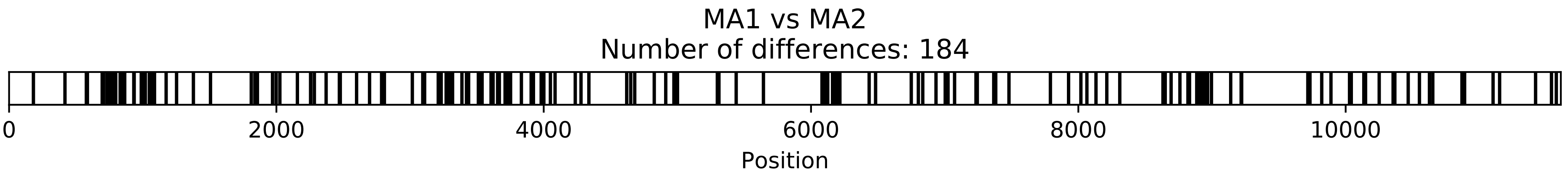
MA1 vs MA2
Number of differences: 184
0 2000 4000 6000 8000 10000
Position

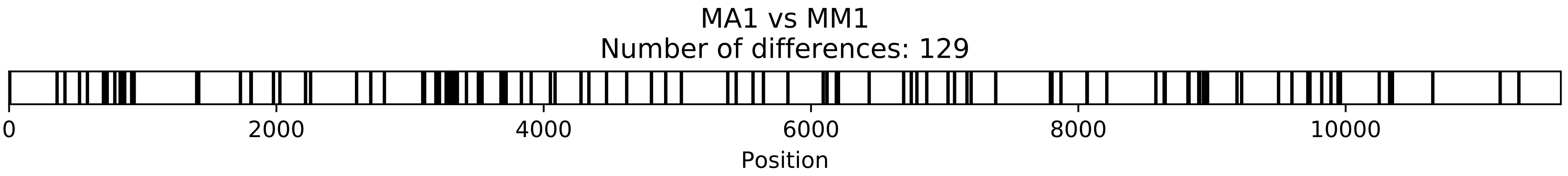
MA1 vs MM1
Number of differences: 129
0 2000 4000 6000 8000 10000
Position

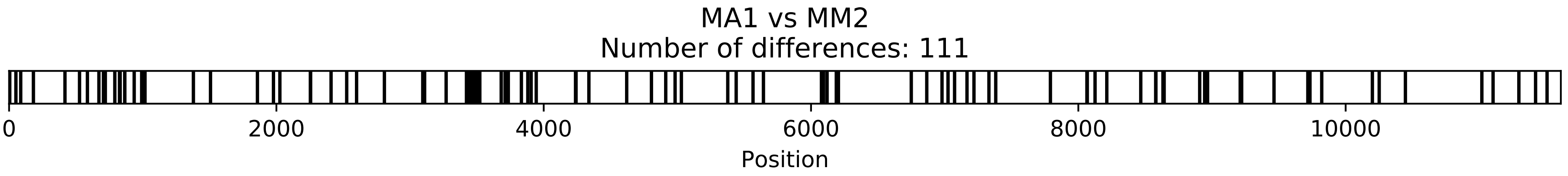
MA1 vs MM2
Number of differences: 111
0 2000 4000 6000 8000 10000
Position

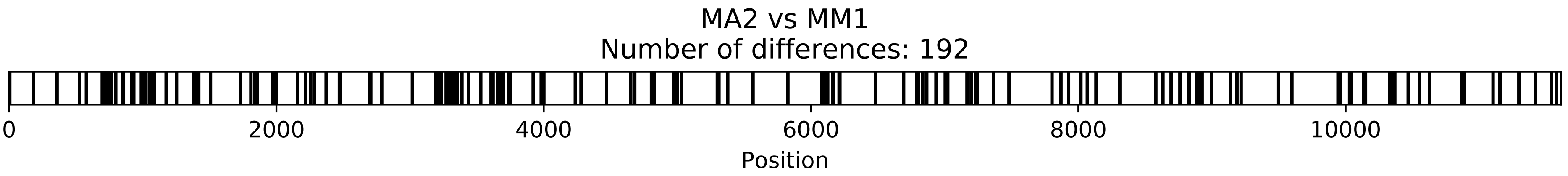
MA2 vs MM1
Number of differences: 192
0 2000 4000 6000 8000 10000
Position

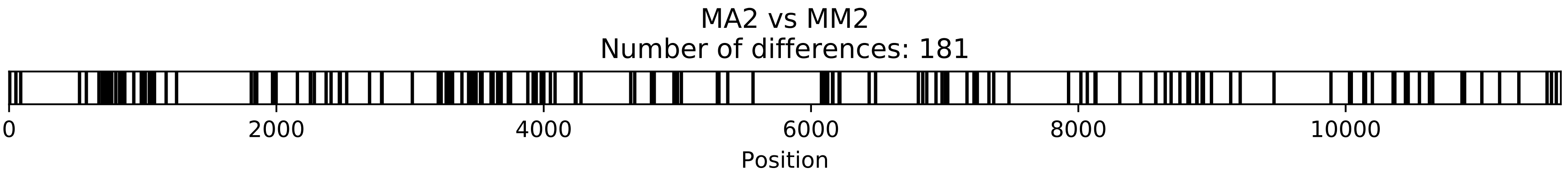
MA2 vs MM2
Number of differences: 181
0 2000 4000 6000 8000 10000
Position

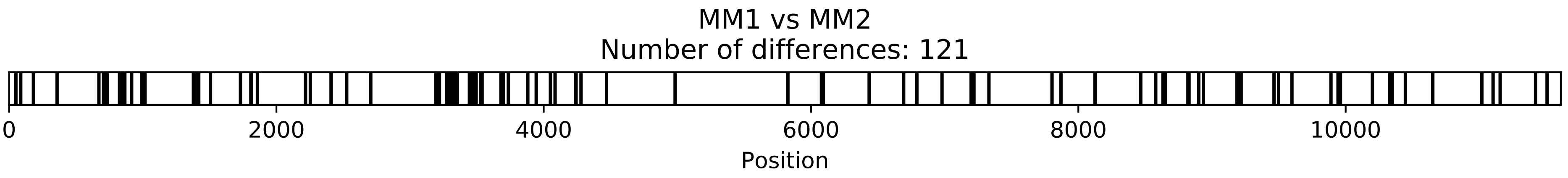
MM1 vs MM2
Number of differences: 121
0 2000 4000 6000 8000 10000
Position

### Region: 94



CR1 vs CR2
Number of differences: 119

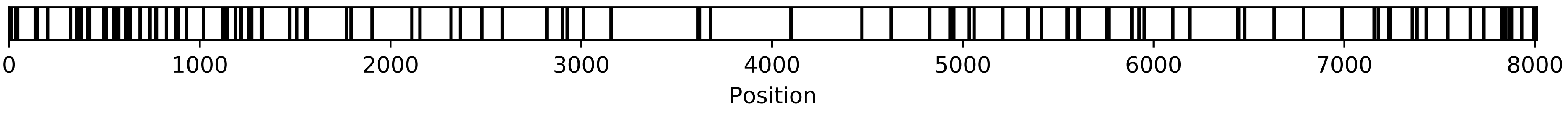


CR1 vs MA1
Number of differences: 78

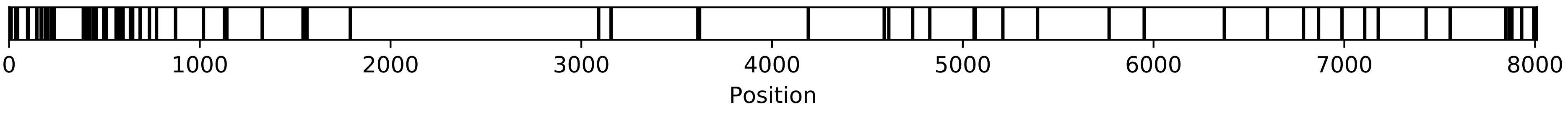


CR1 vs MA2
Number of differences: 118

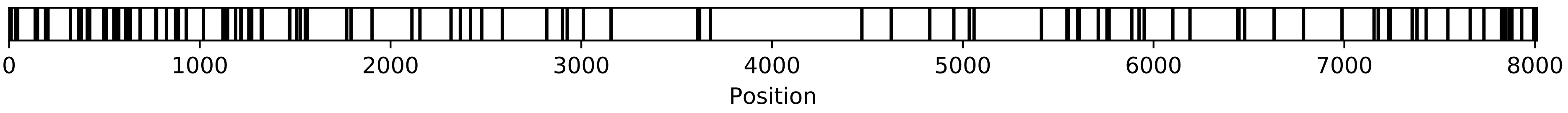


CR1 vs MM1
Number of differences: 78

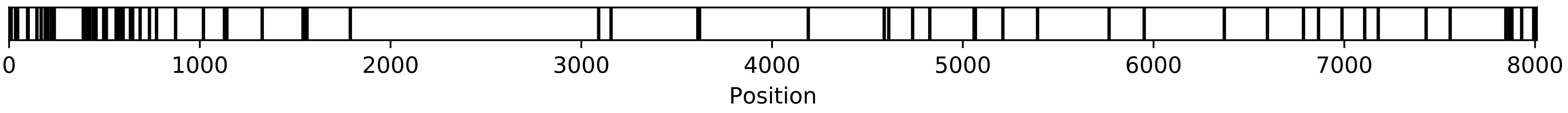


CR1 vs MM2
Number of differences: 83

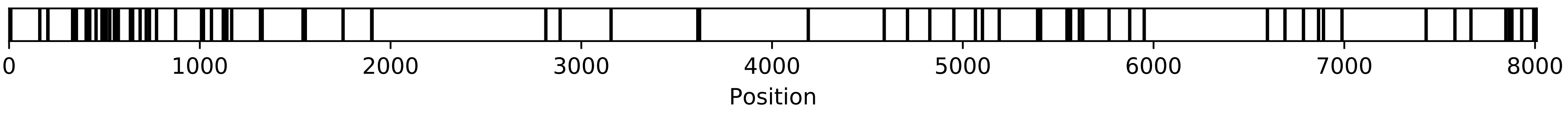


CR2 vs MA1
Number of differences: 112

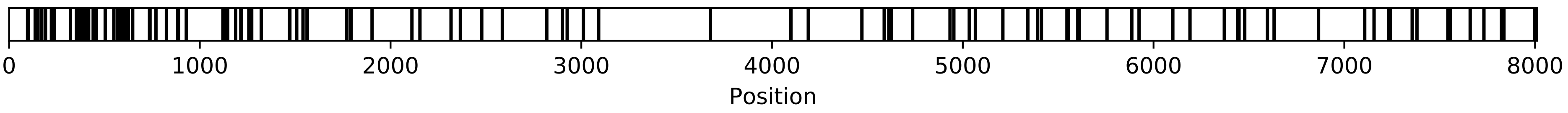


CR2 vs MA2
Number of differences: 11

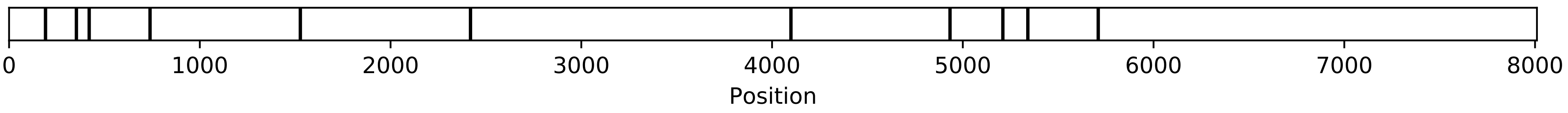


CR2 vs MM1
Number of differences: 112

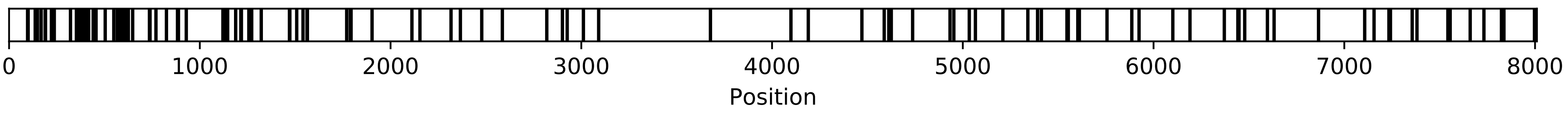


CR2 vs MM2
Number of differences: 116

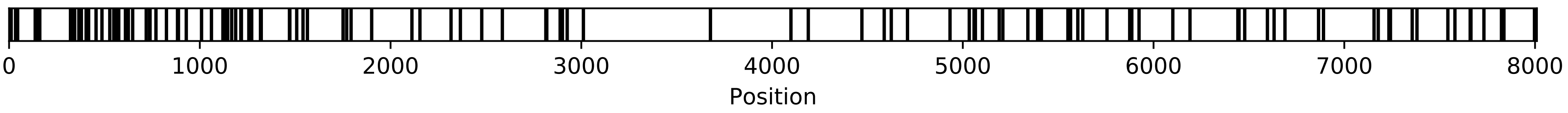


MA1 vs MA2
Number of differences: 110

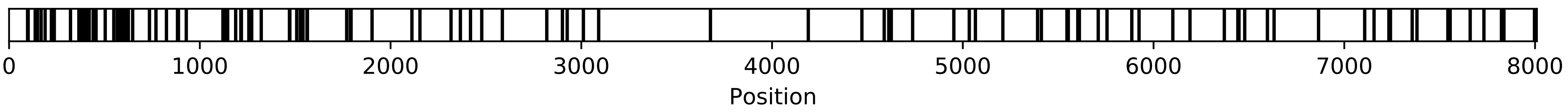


MA1 vs MM1
Number of differences: 0

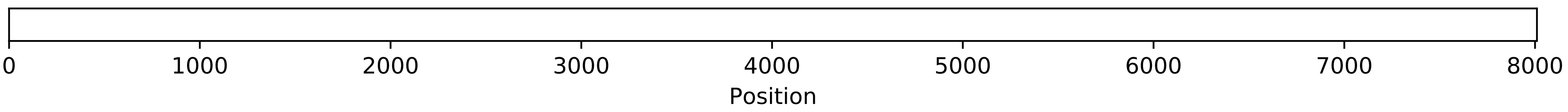


MA1 vs MM2
Number of differences: 61

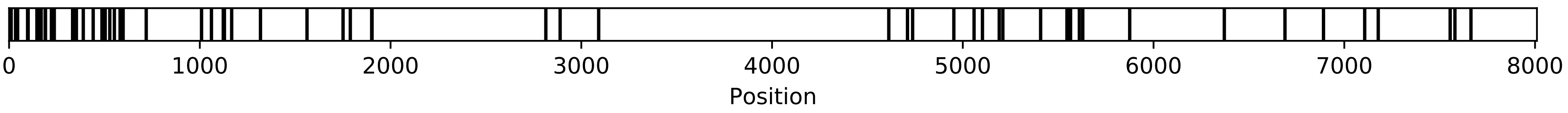


MA2 vs MM1
Number of differences: 110

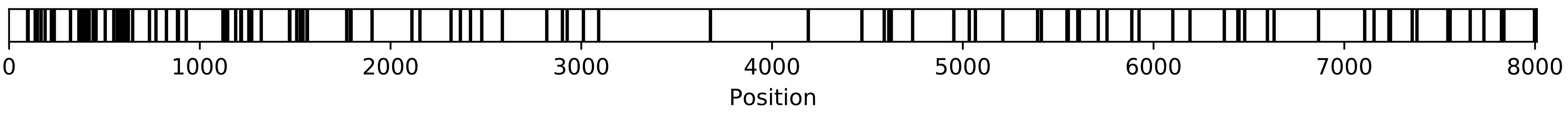


MA2 vs MM2
Number of differences: 117

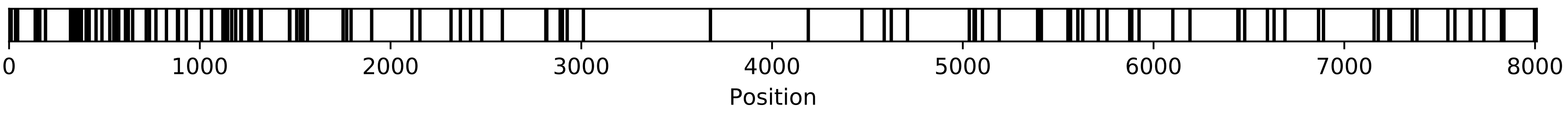


MM1 vs MM2
Number of differences: 61

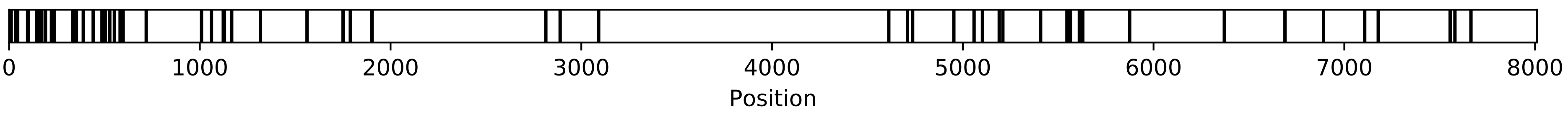

### Region: 96

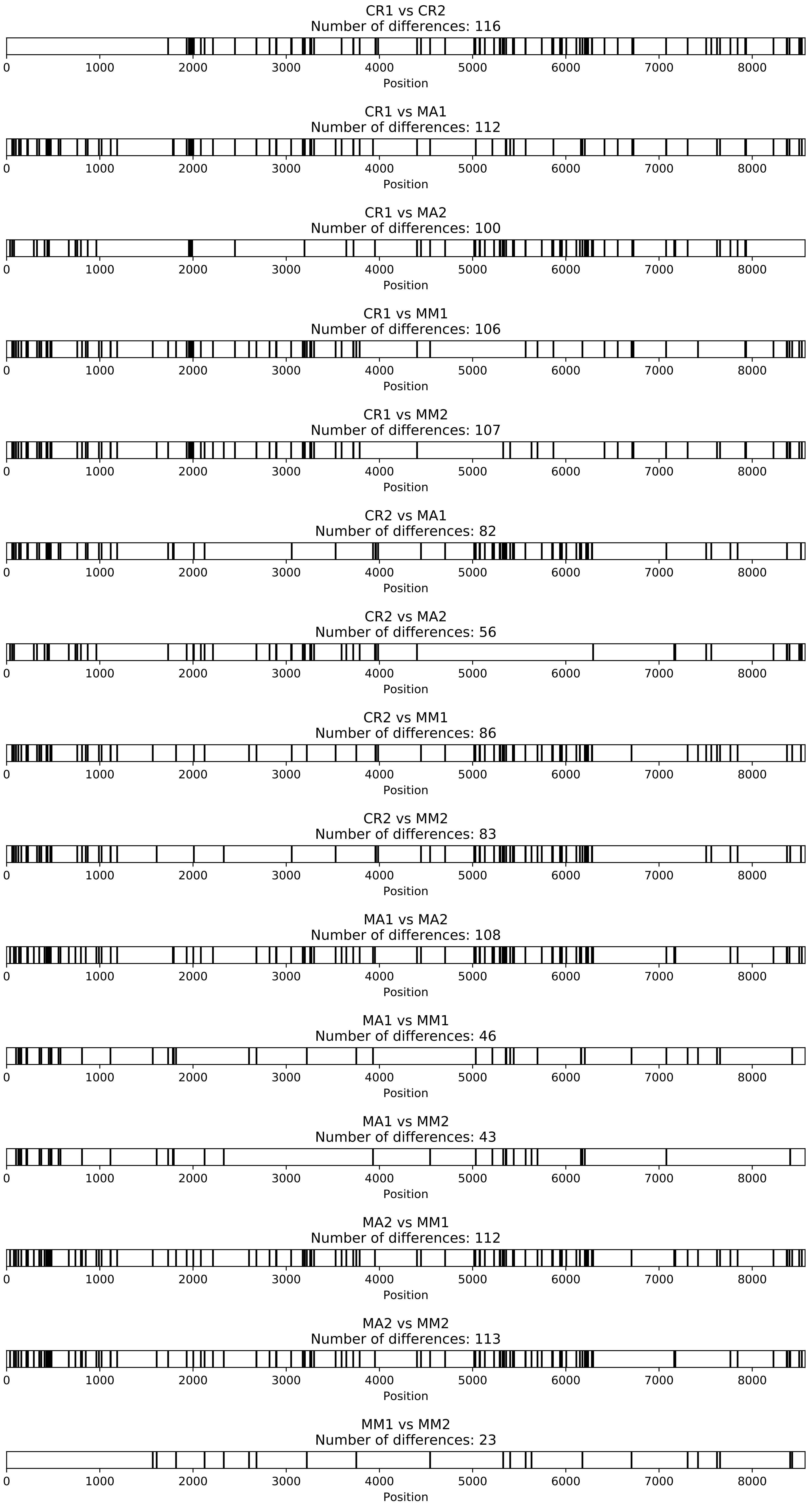
CR1 vs CR2
Number of differences: 116
CR1 vs MA1
Number of differences: 112
CR1 vs MA2
Number of differences: 100
CR1 vs MM1
Number of differences: 106
CR1 vs MM2
Number of differences: 107
CR2 vs MA1
Number of differences: 82
CR2 vs MA2
Number of differences: 56
CR2 vs MM1
Number of differences: 86
CR2 vs MM2
Number of differences: 83
MA1 vs MA2
Number of differences: 108
MA1 vs MM1
Number of differences: 46
MA1 vs MM2
Number of differences: 43
MA2 vs MM1
Number of differences: 112
MA2 vs MM2
Number of differences: 113
MM1 vs MM2
Number of differences: 23
0
1000
2000
3000
4000
5000
6000
7000
8000
Position

### Region: 97



CR1 vs CR2
Number of differences: 468

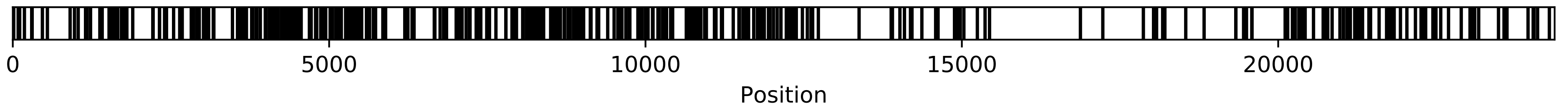


CR1 vs MA1
Number of differences: 238

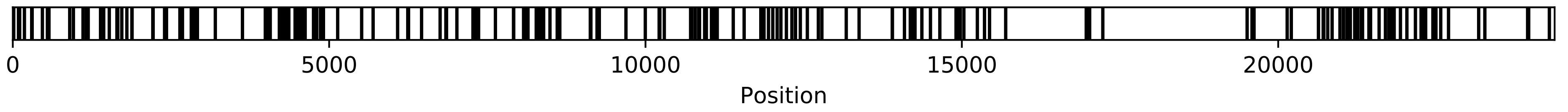


CR1 vs MA2
Number of differences: 237

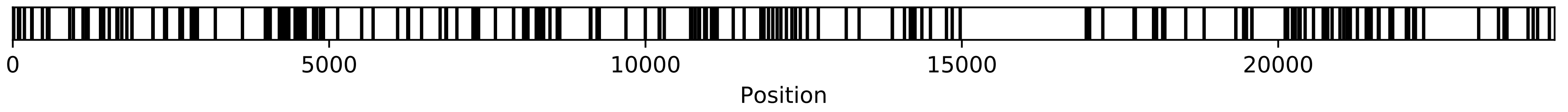


CR1 vs MM1
Number of differences: 209

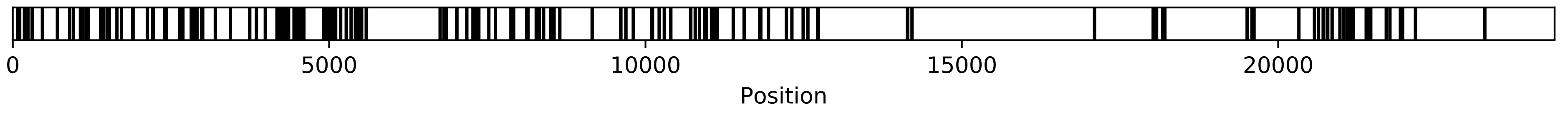


CR1 vs MM2
Number of differences: 212

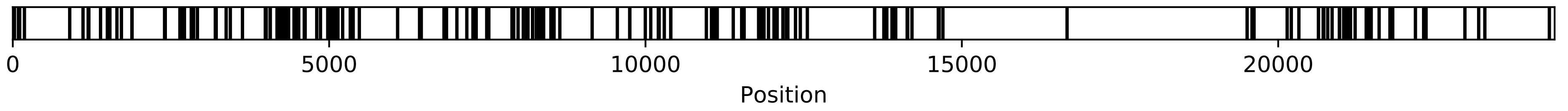


CR2 vs MA1
Number of differences: 420

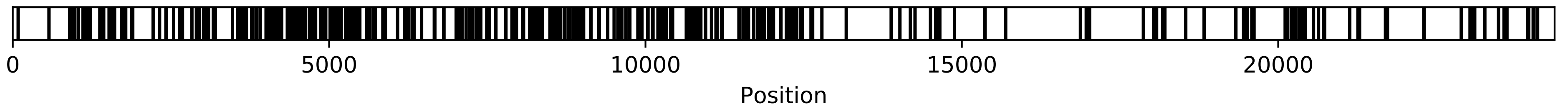


CR2 vs MA2
Number of differences: 419

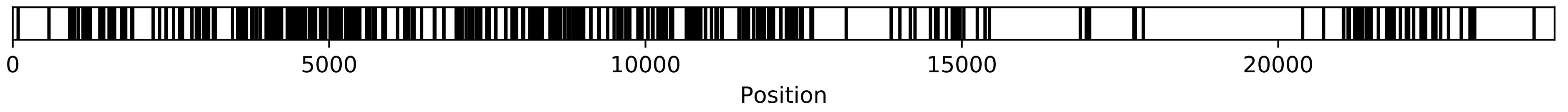


CR2 vs MM1
Number of differences: 456

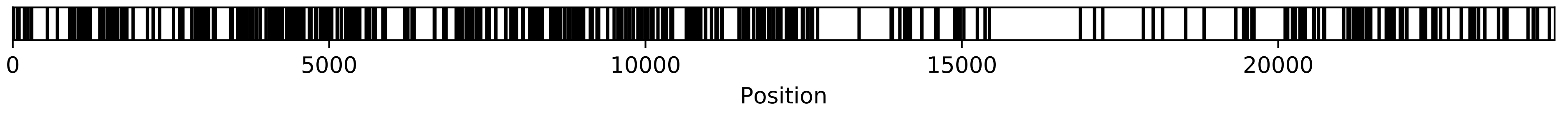


CR2 vs MM2
Number of differences: 478

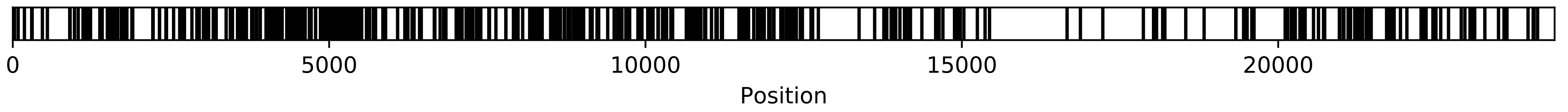


MA1 vs MA2
Number of differences: 89

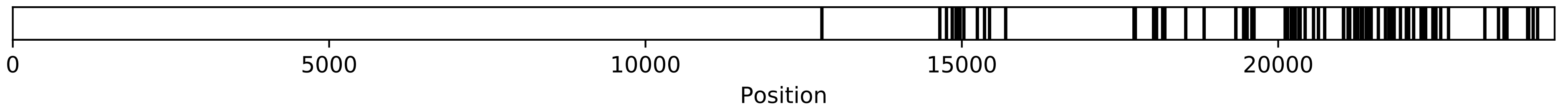


MA1 vs MM1
Number of differences: 245

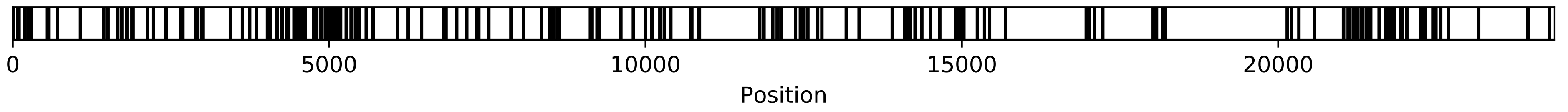


MA1 vs MM2
Number of differences: 244

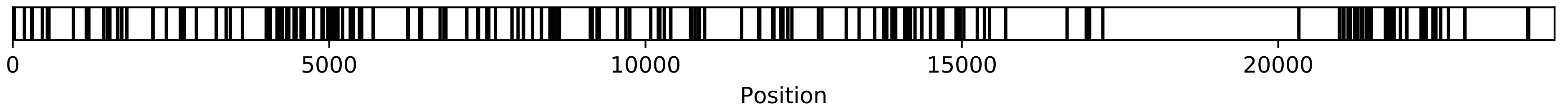


MA2 vs MM1
Number of differences: 240

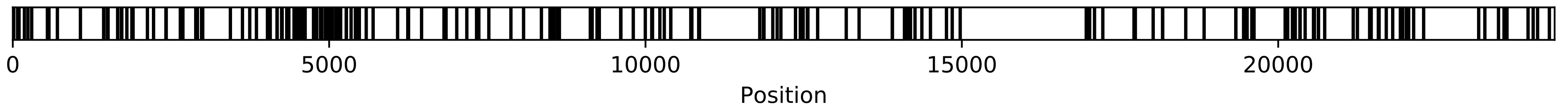


MA2 vs MM2
Number of differences: 251

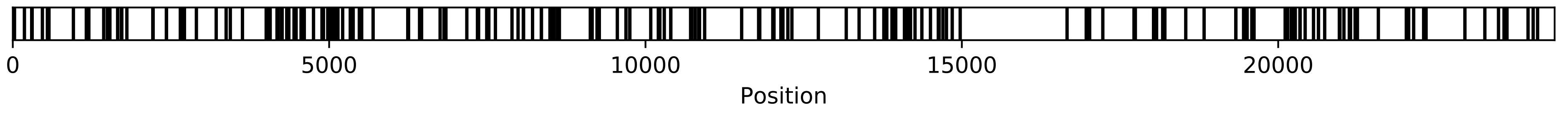


MM1 vs MM2
Number of differences: 201

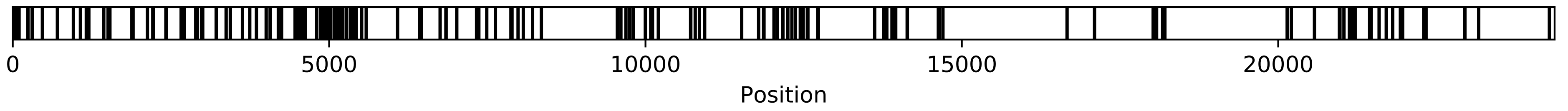

### Region: 98



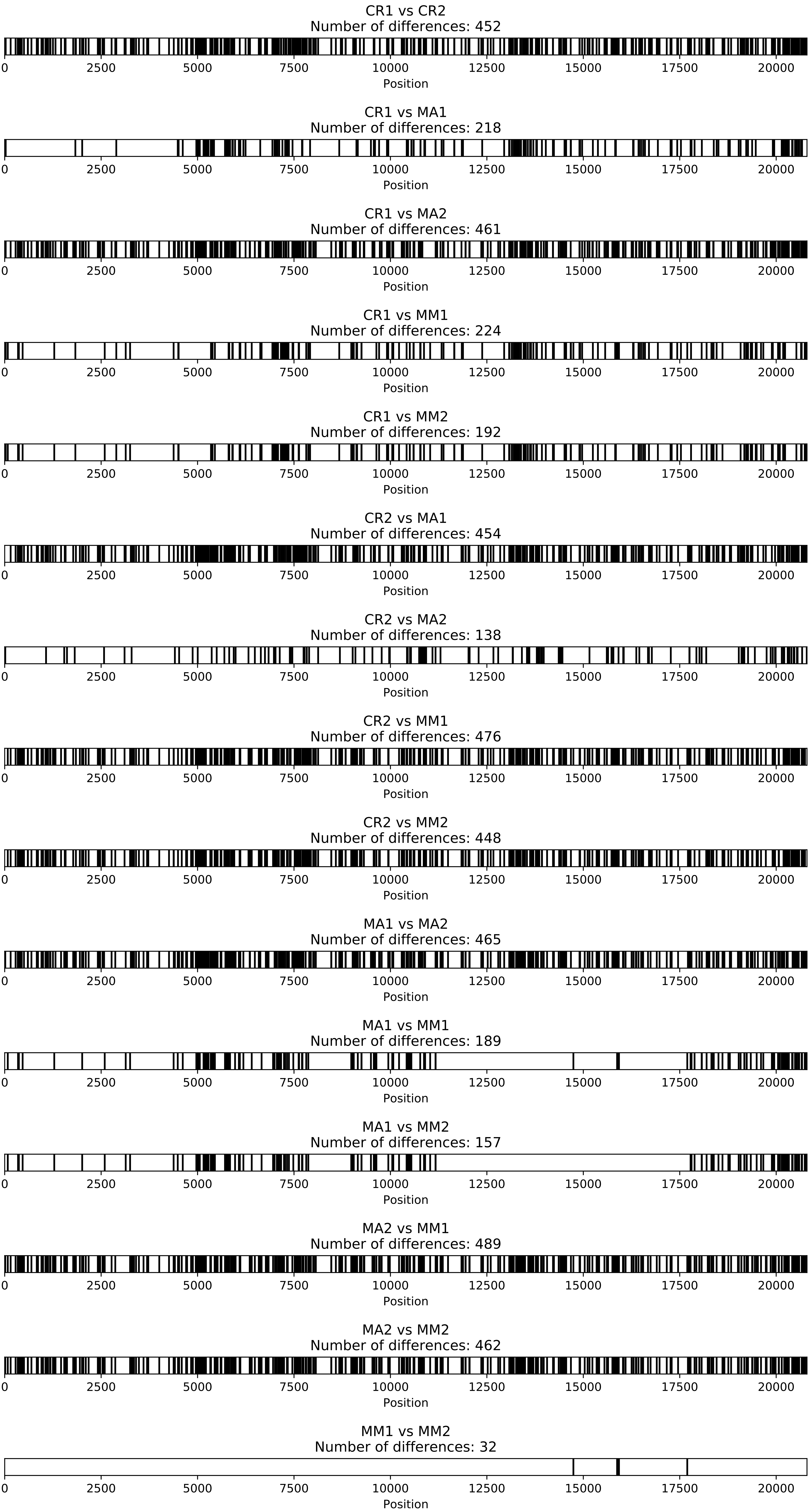
CR1 vs CR2
Number of differences: 452
CR1 vs MA1
Number of differences: 218
CR1 vs MA2
Number of differences: 461
CR1 vs MM1
Number of differences: 224
CR1 vs MM2
Number of differences: 192
CR2 vs MA1
Number of differences: 454
CR2 vs MA2
Number of differences: 138
CR2 vs MM1
Number of differences: 476
CR2 vs MM2
Number of differences: 448
MA1 vs MA2
Number of differences: 465
MA1 vs MM1
Number of differences: 189
MA1 vs MM2
Number of differences: 157
MA2 vs MM1
Number of differences: 489
MA2 vs MM2
Number of differences: 462
MM1 vs MM2
Number of differences: 32
0
2500
5000
7500
10000
12500
15000
17500
20000
Position

### Region: 99



CR1 vs CR2
Number of differences: 539

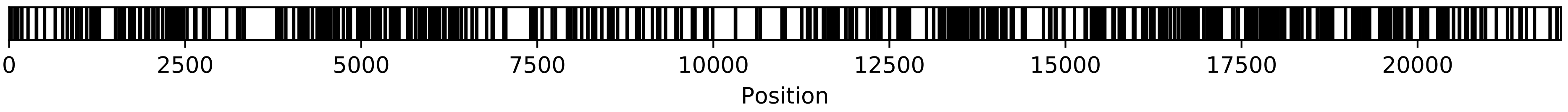


CR1 vs MA1
Number of differences: 131

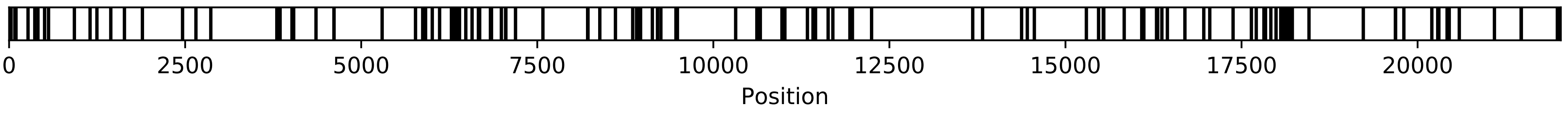


CR1 vs MA2
Number of differences: 512

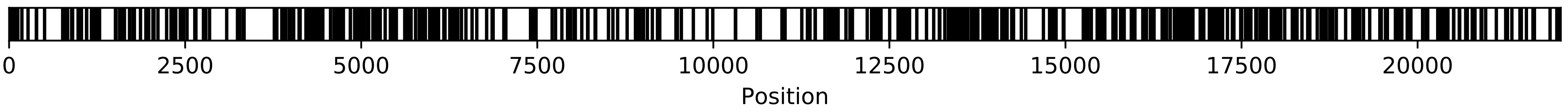


CR1 vs MM1
Number of differences: 548

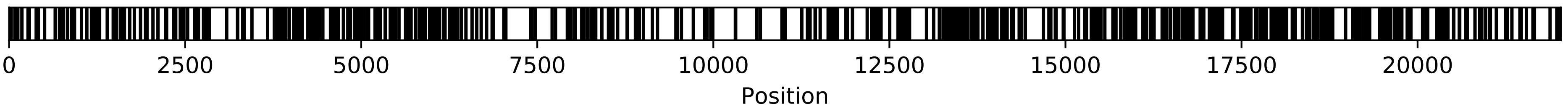


CR1 vs MM2
Number of differences: 512

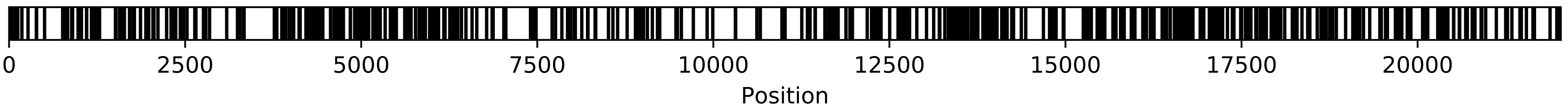


CR2 vs MA1
Number of differences: 550

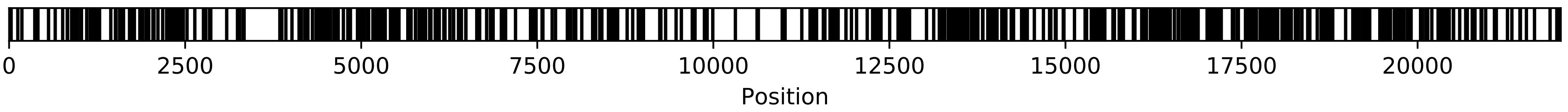


CR2 vs MA2
Number of differences: 215

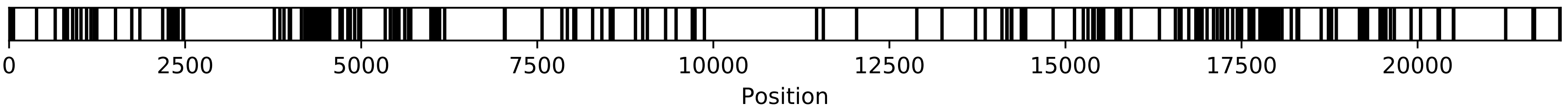


CR2 vs MM1
Number of differences: 231

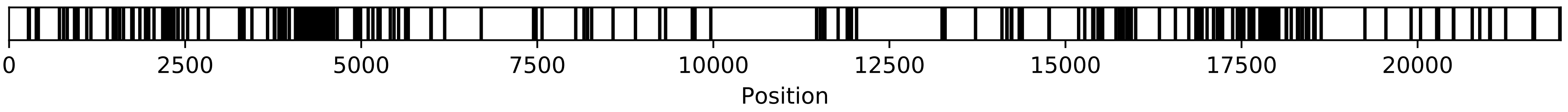


CR2 vs MM2
Number of differences: 215

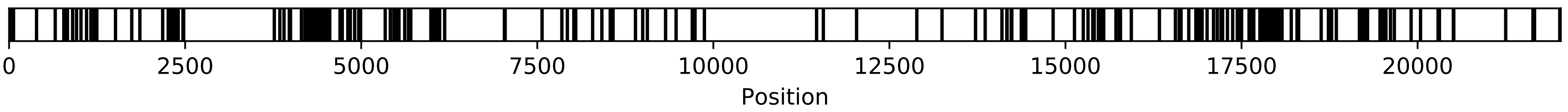


MA1 vs MA2
Number of differences: 522

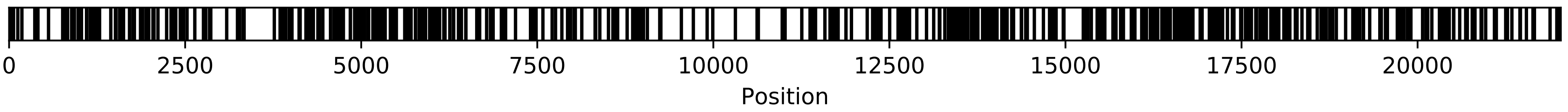


MA1 vs MM1
Number of differences: 561

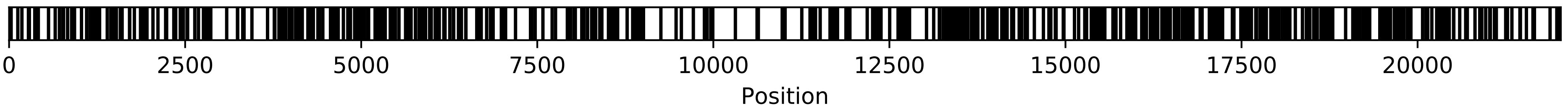


MA1 vs MM2
Number of differences: 522

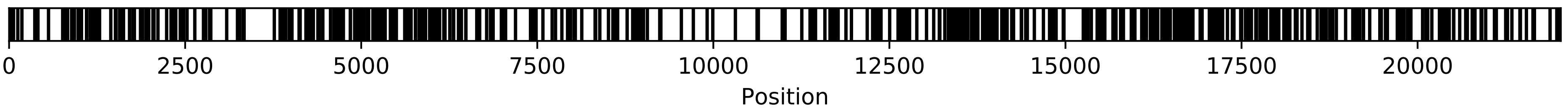


MA2 vs MM1
Number of differences: 198

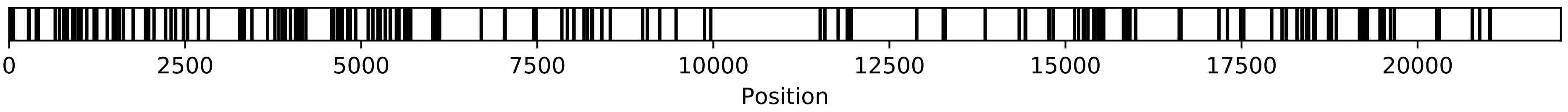


MA2 vs MM2
Number of differences: 0

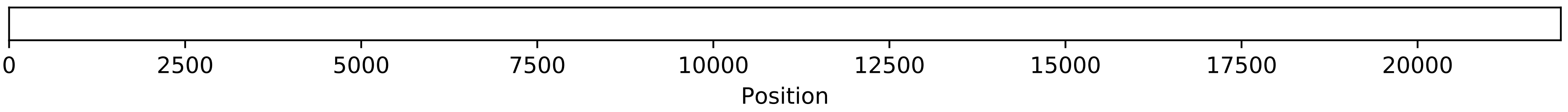


MM1 vs MM2
Number of differences: 198

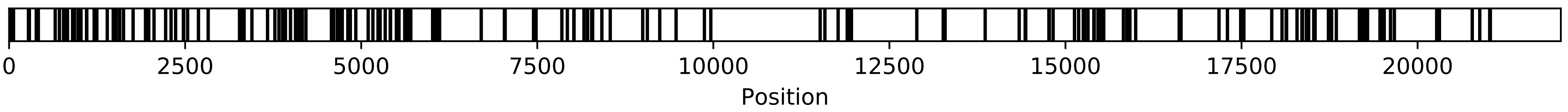

### Region: 101

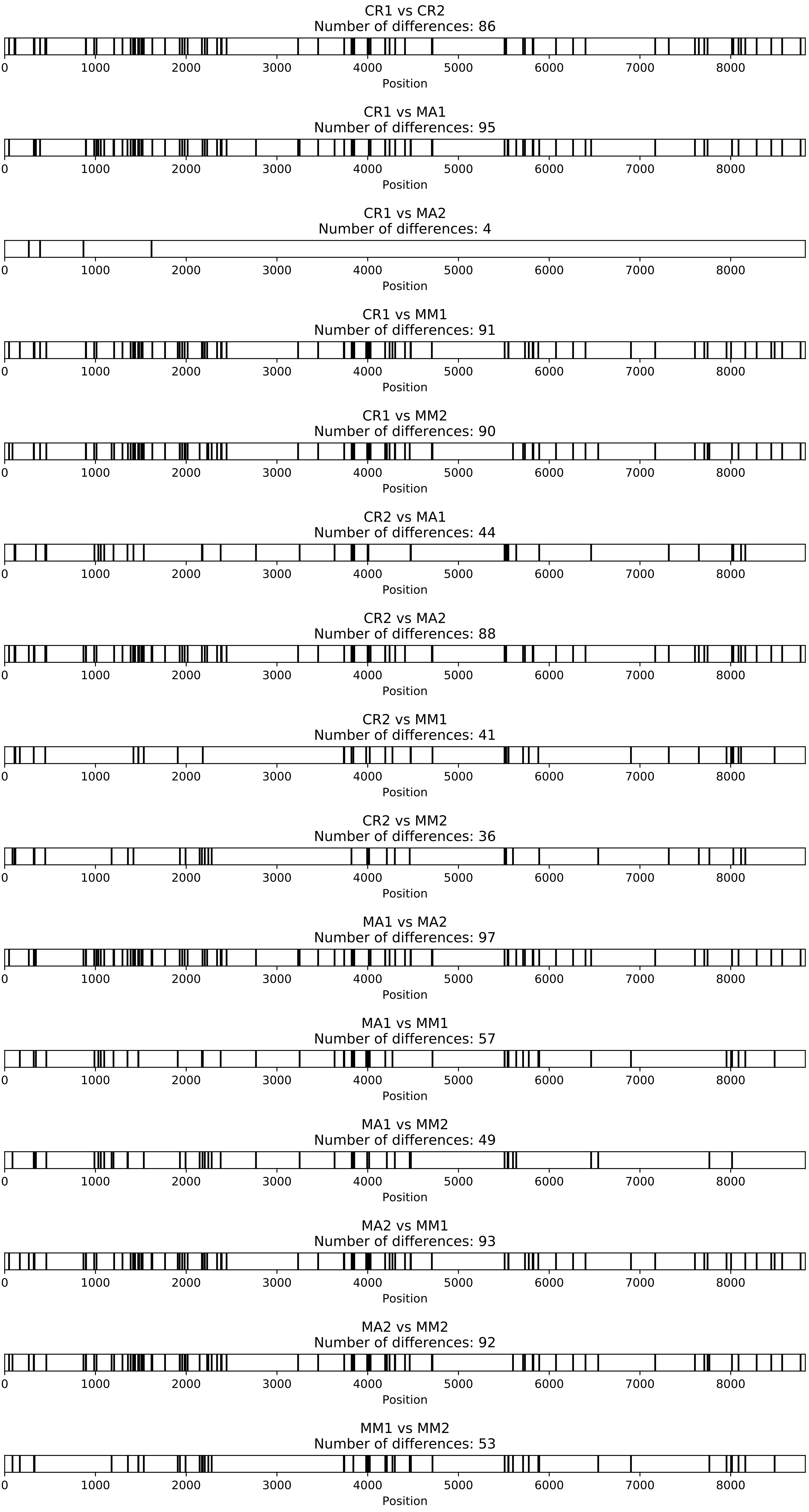
CR1 vs CR2
Number of differences: 86
CR1 vs MA1
Number of differences: 95
CR1 vs MA2
Number of differences: 4
CR1 vs MM1
Number of differences: 91
CR1 vs MM2
Number of differences: 90
CR2 vs MA1
Number of differences: 44
CR2 vs MA2
Number of differences: 88
CR2 vs MM1
Number of differences: 41
CR2 vs MM2
Number of differences: 36
MA1 vs MA2
Number of differences: 97
MA1 vs MM1
Number of differences: 57
MA1 vs MM2
Number of differences: 49
MA2 vs MM1
Number of differences: 93
MA2 vs MM2
Number of differences: 92
MM1 vs MM2
Number of differences: 53
0
1000
2000
3000
4000
5000
6000
7000
8000
Position

### Region: 102

CR1 vs CR2
Number of differences: 285

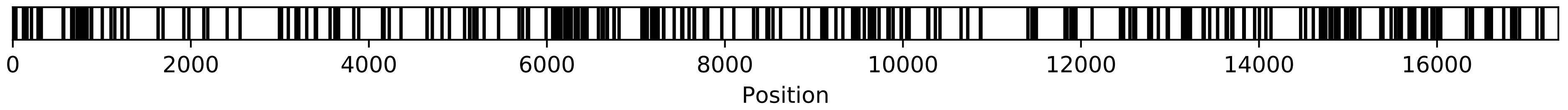


CR1 vs MA1
Number of differences: 269

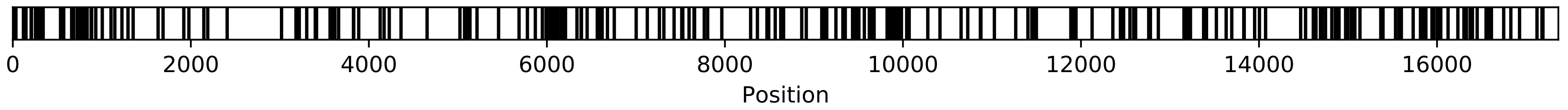


CR1 vs MA2
Number of differences: 170

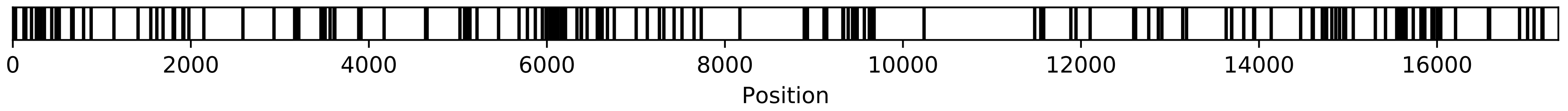


CR1 vs MM1
Number of differences: 101

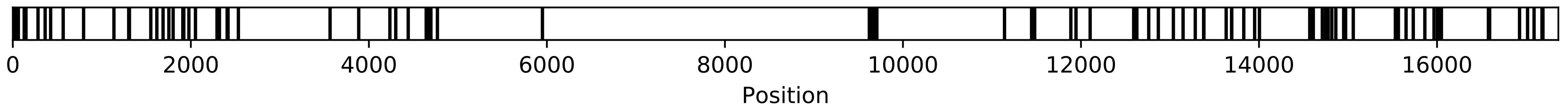


CR1 vs MM2
Number of differences: 163

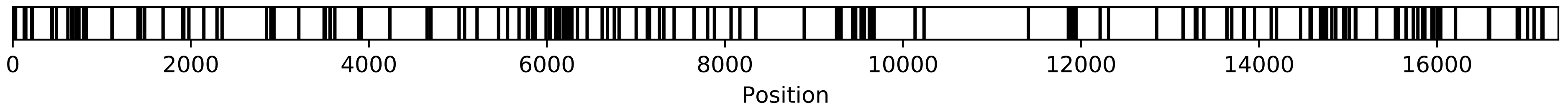


CR2 vs MA1
Number of differences: 158

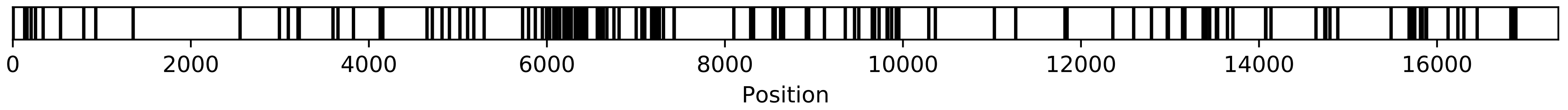


CR2 vs MA2
Number of differences: 279

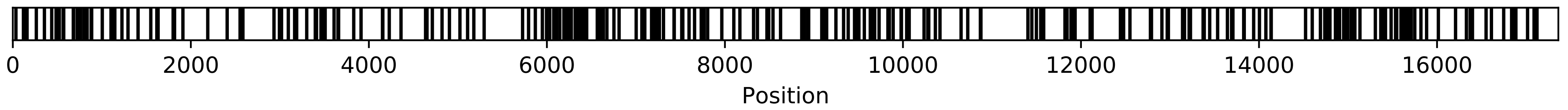


CR2 vs MM1
Number of differences: 280

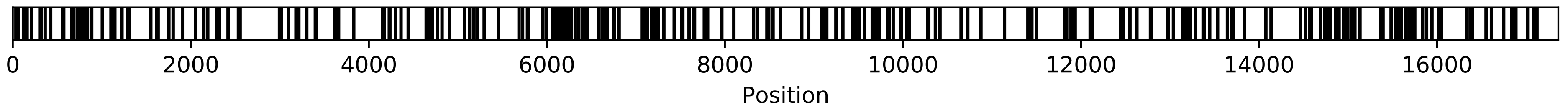


CR2 vs MM2
Number of differences: 298

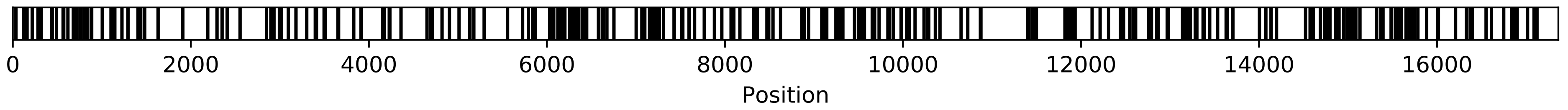


MA1 vs MA2
Number of differences: 224

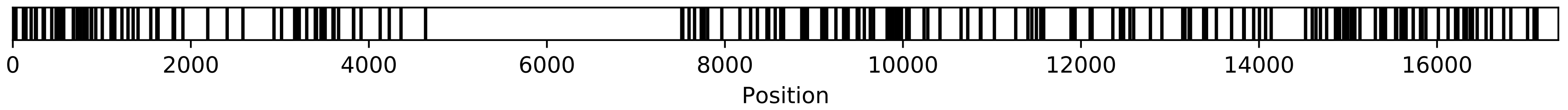


MA1 vs MM1
Number of differences: 268

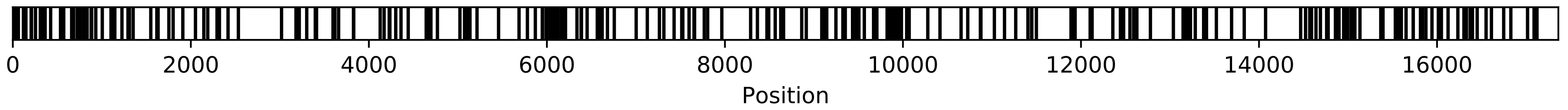


MA1 vs MM2
Number of differences: 285

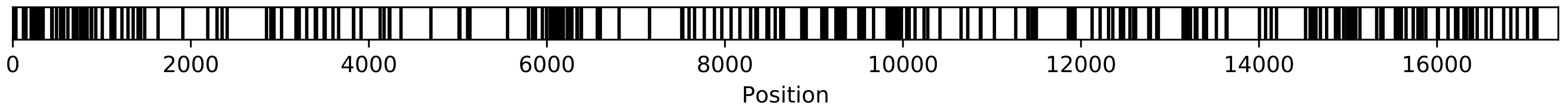


MA2 vs MM1
Number of differences: 155

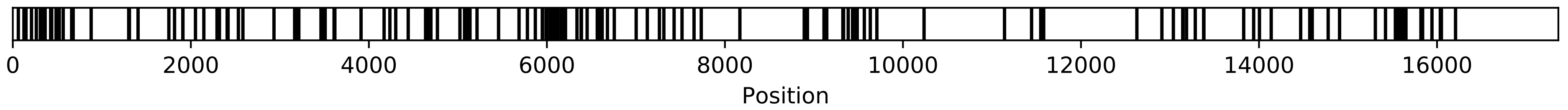


MA2 vs MM2
Number of differences: 162

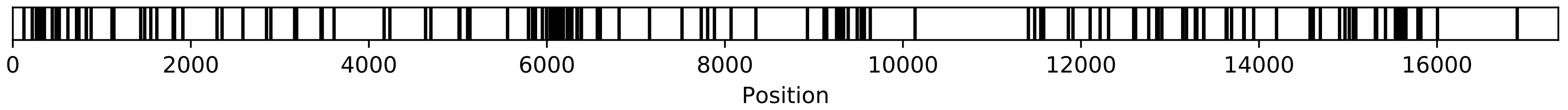


MM1 vs MM2
Number of differences: 158

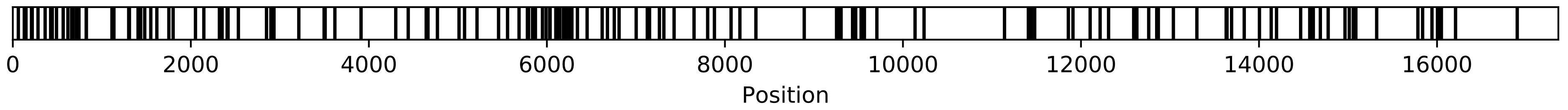

### Region: 103

CR1 vs CR2
Number of differences: 222

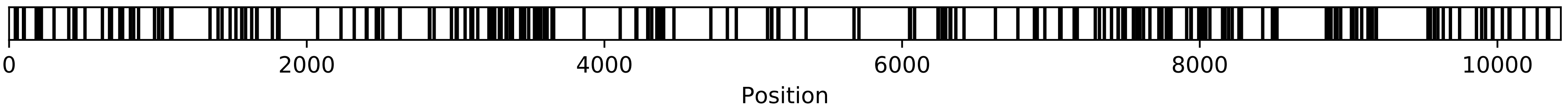


CR1 vs MA1
Number of differences: 223

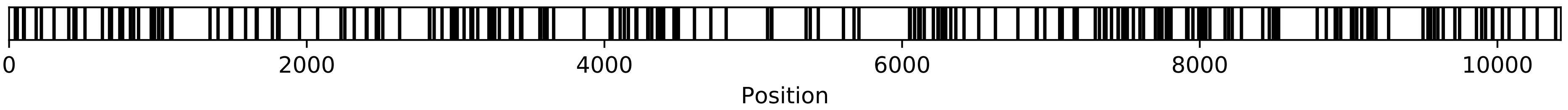


CR1 vs MA2
Number of differences: 175

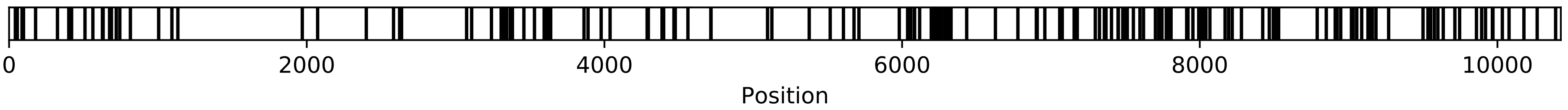


CR1 vs MM1
Number of differences: 128

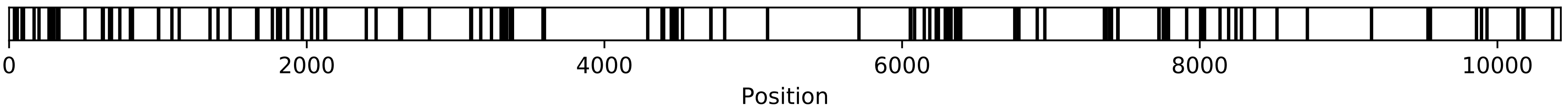


CR1 vs MM2
Number of differences: 120

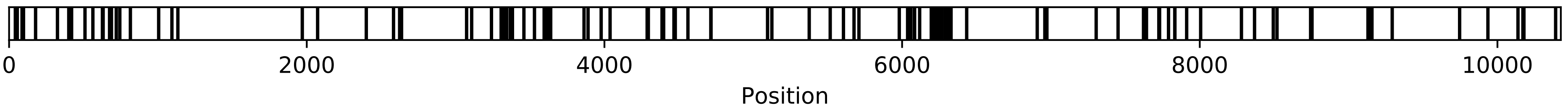


CR2 vs MA1
Number of differences: 101

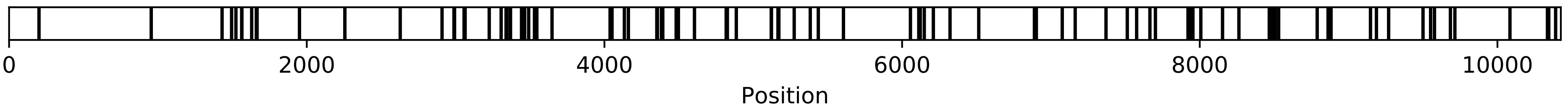


CR2 vs MA2
Number of differences: 170

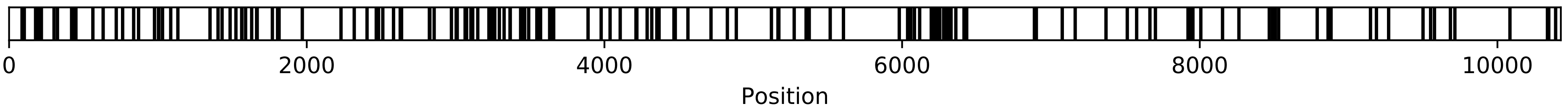


CR2 vs MM1
Number of differences: 220

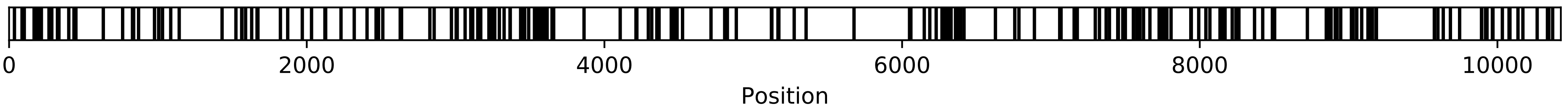


CR2 vs MM2
Number of differences: 233

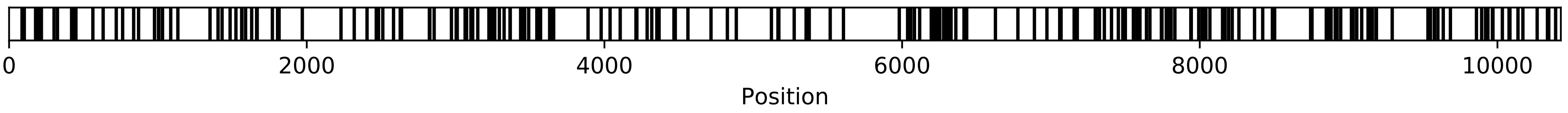


MA1 vs MA2
Number of differences: 139

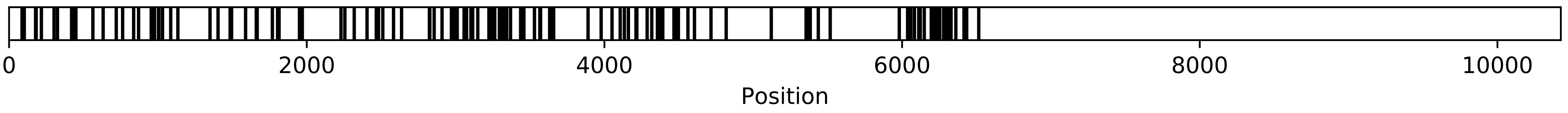


MA1 vs MM1
Number of differences: 224

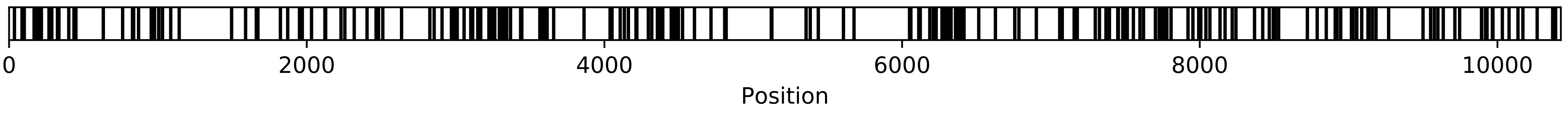


MA1 vs MM2
Number of differences: 234

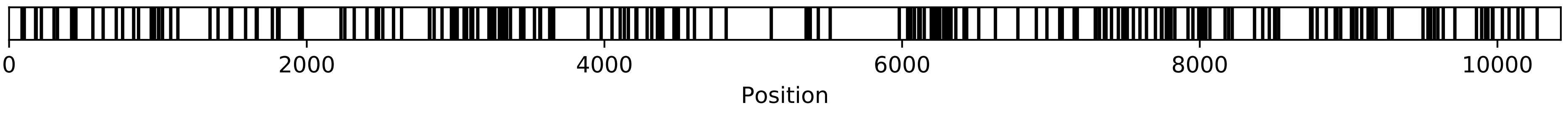


MA2 vs MM1
Number of differences: 183

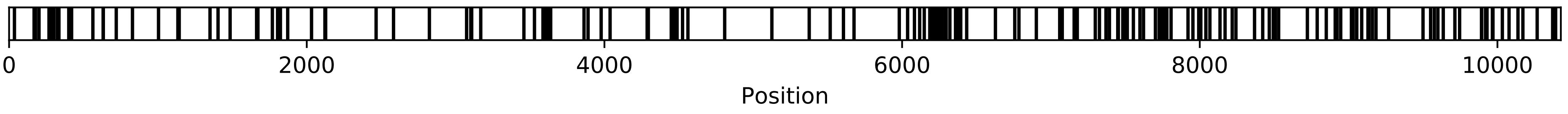


MA2 vs MM2
Number of differences: 95

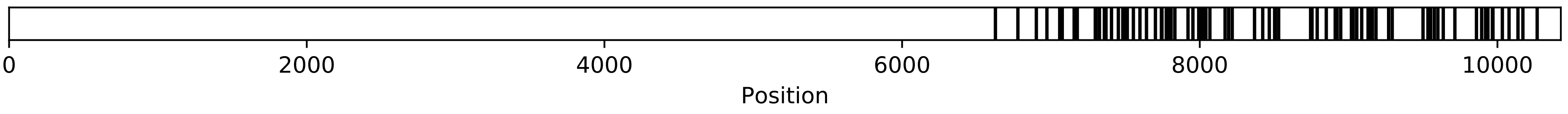


MM1 vs MM2
Number of differences: 143

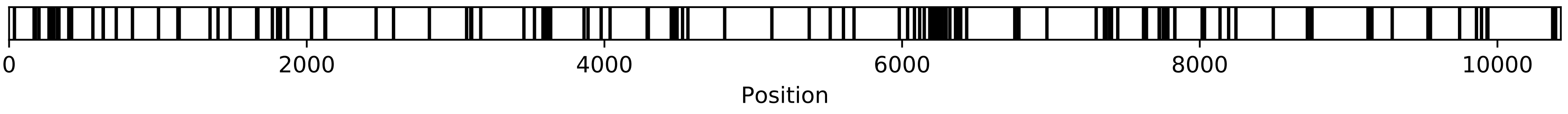

### Region: 104



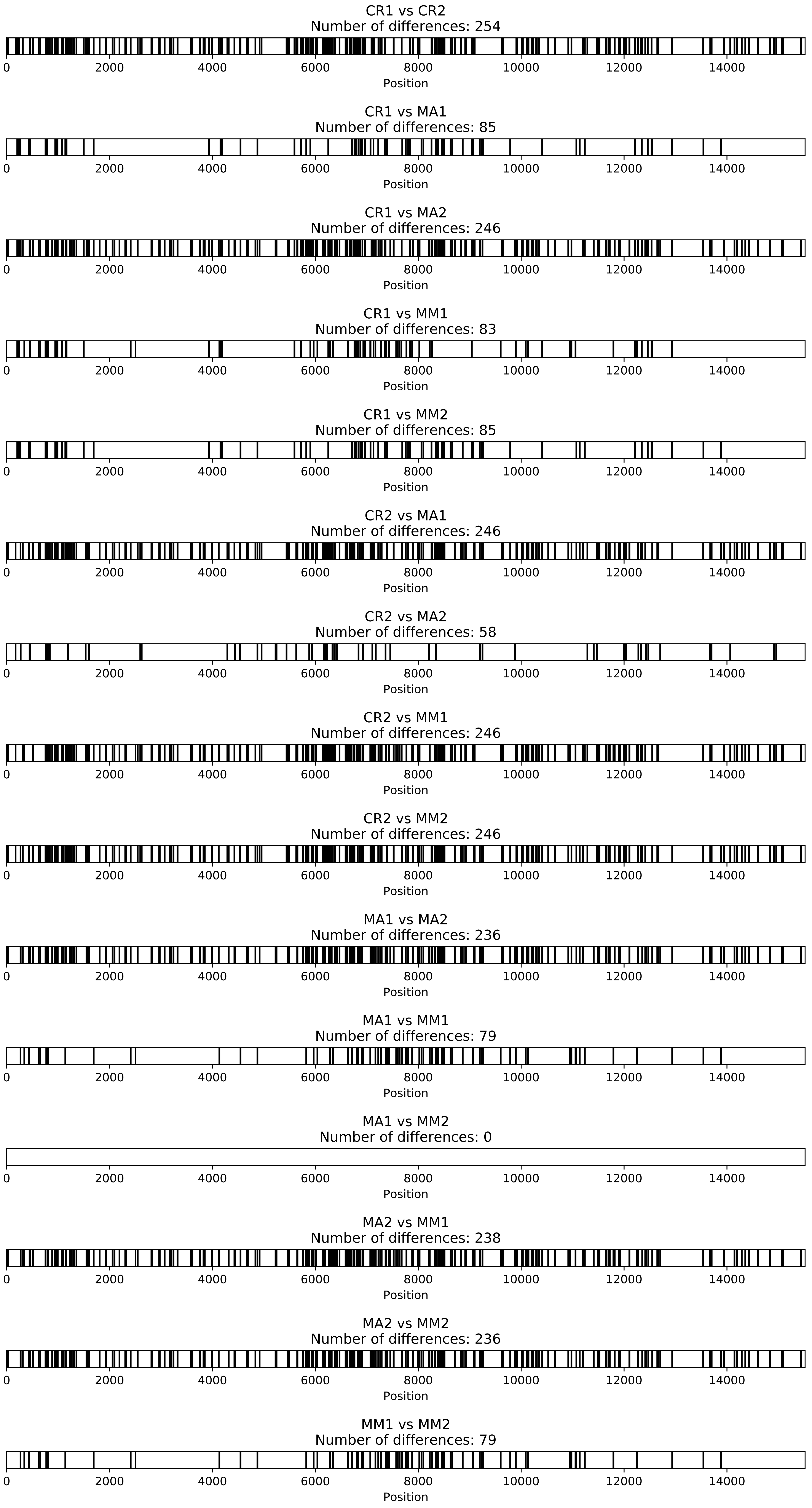
CR1 vs CR2
Number of differences: 254
CR1 vs MA1
Number of differences: 85
CR1 vs MA2
Number of differences: 246
CR1 vs MM1
Number of differences: 83
CR1 vs MM2
Number of differences: 85
CR2 vs MA1
Number of differences: 246
CR2 vs MA2
Number of differences: 58
CR2 vs MM1
Number of differences: 246
CR2 vs MM2
Number of differences: 246
MA1 vs MA2
Number of differences: 236
MA1 vs MM1
Number of differences: 79
MA1 vs MM2
Number of differences: 0
MA2 vs MM1
Number of differences: 238
MA2 vs MM2
Number of differences: 236
MM1 vs MM2
Number of differences: 79
0
2000
4000
6000
8000
10000
12000
14000
Position

### Region: 105



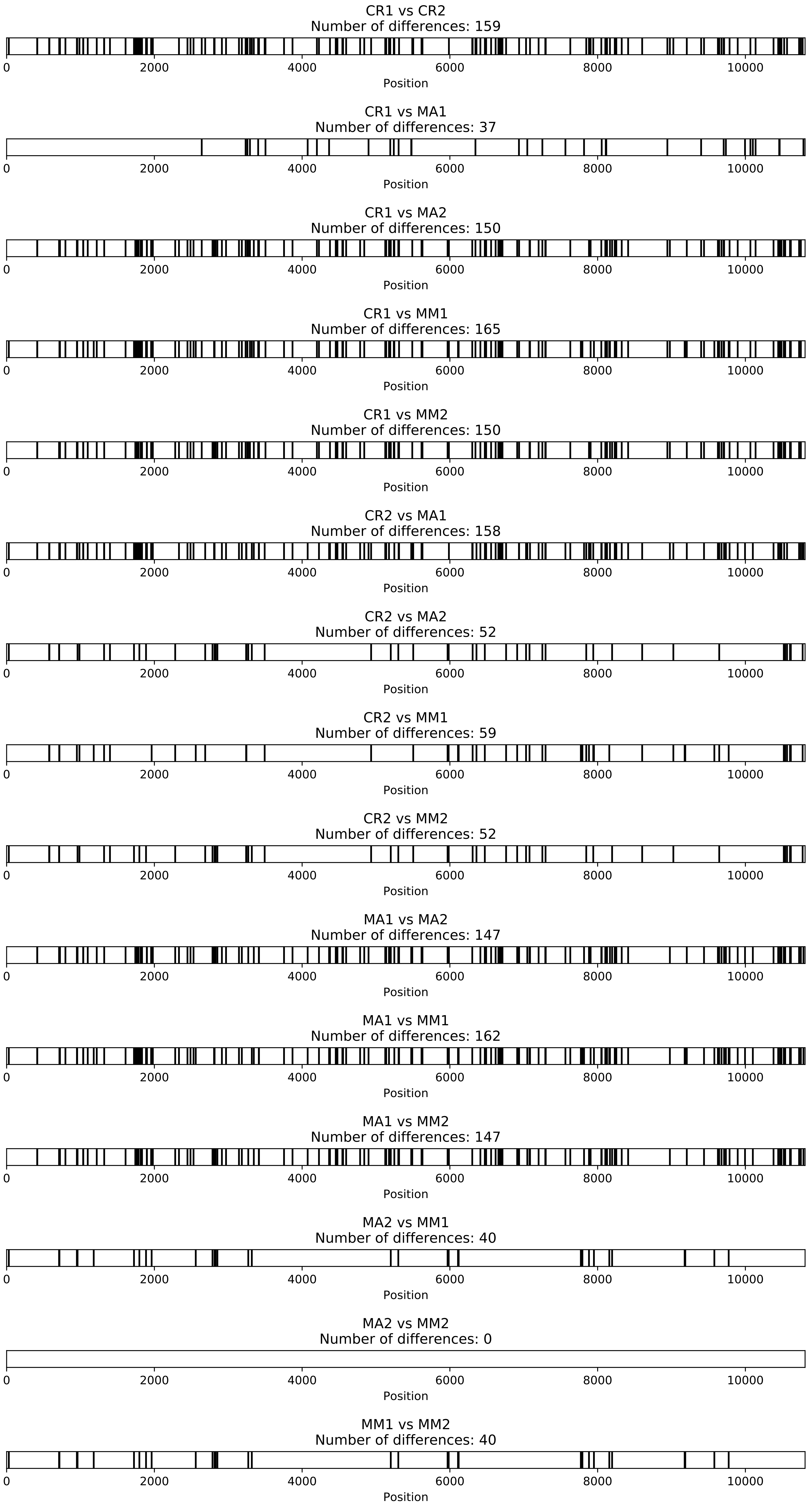
CR1 vs CR2
Number of differences: 159
CR1 vs MA1
Number of differences: 37
CR1 vs MA2
Number of differences: 150
CR1 vs MM1
Number of differences: 165
CR1 vs MM2
Number of differences: 150
CR2 vs MA1
Number of differences: 158
CR2 vs MA2
Number of differences: 52
CR2 vs MM1
Number of differences: 59
CR2 vs MM2
Number of differences: 52
MA1 vs MA2
Number of differences: 147
MA1 vs MM1
Number of differences: 162
MA1 vs MM2
Number of differences: 147
MA2 vs MM1
Number of differences: 40
MA2 vs MM2
Number of differences: 0
MM1 vs MM2
Number of differences: 40
0
2000
4000
6000
8000
10000
Position

### Region: 110



CR1 vs CR2
Number of differences: 293

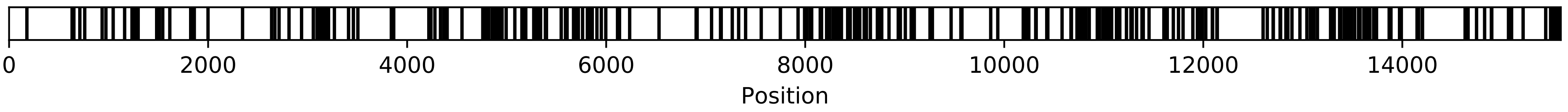


CR1 vs MA1
Number of differences: 186

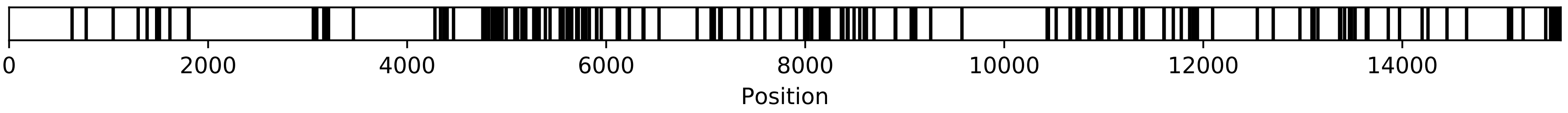


CR1 vs MA2
Number of differences: 277

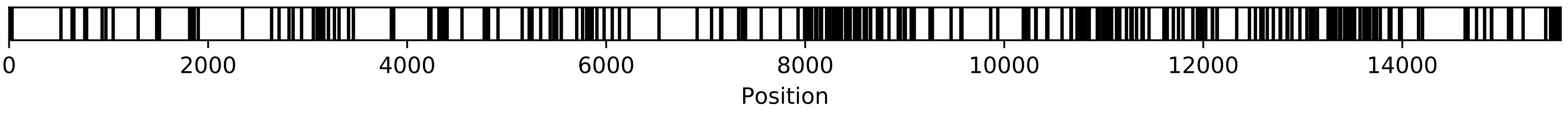


CR1 vs MM1
Number of differences: 164

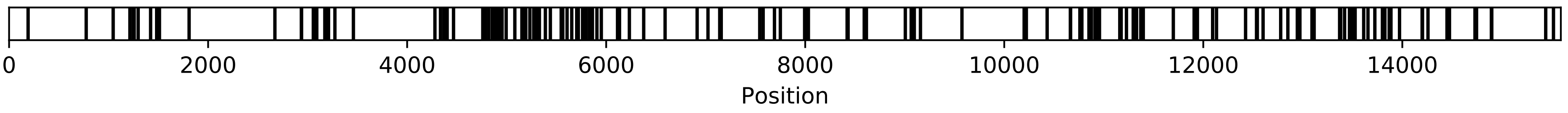


CR1 vs MM2
Number of differences: 191

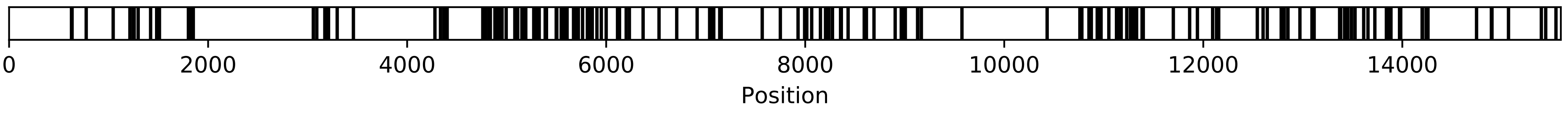


CR2 vs MA1
Number of differences: 261

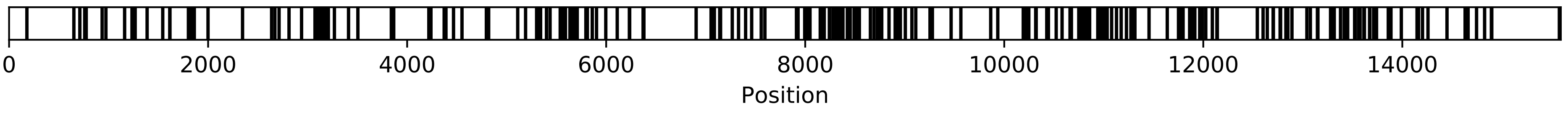


CR2 vs MA2
Number of differences: 111

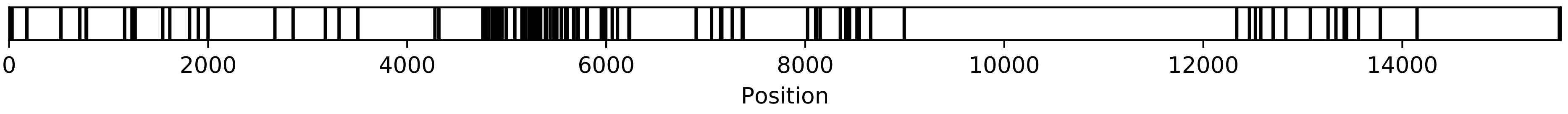


CR2 vs MM1
Number of differences: 271

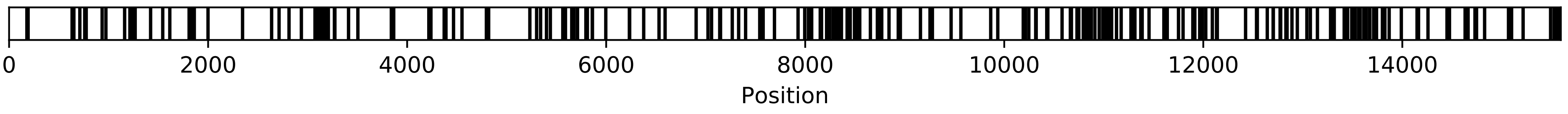


CR2 vs MM2
Number of differences: 279

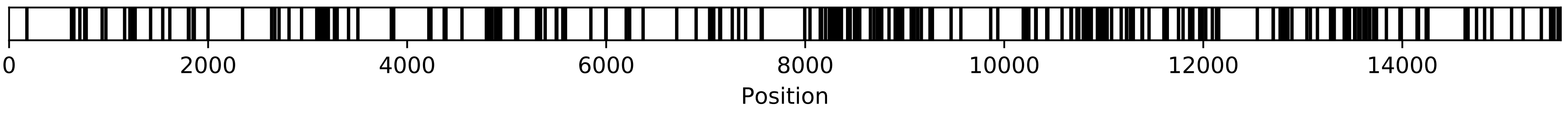


MA1 vs MA2
Number of differences: 285

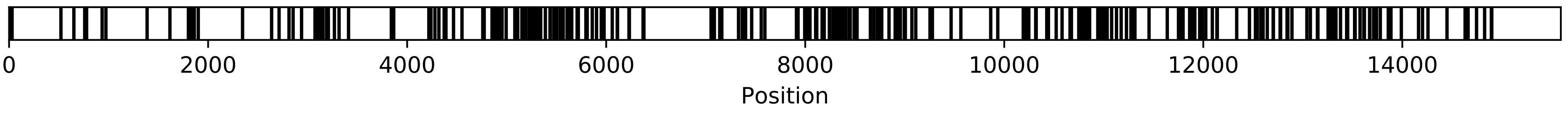


MA1 vs MM1
Number of differences: 143

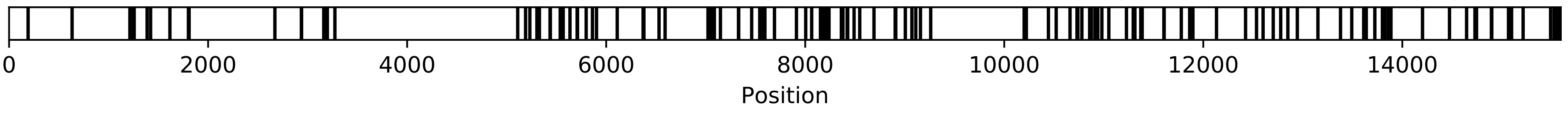


MA1 vs MM2
Number of differences: 160

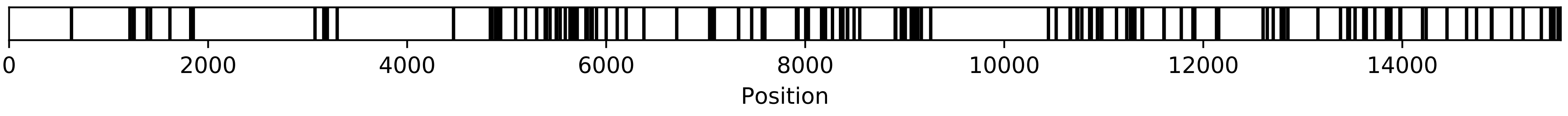


MA2 vs MM1
Number of differences: 303

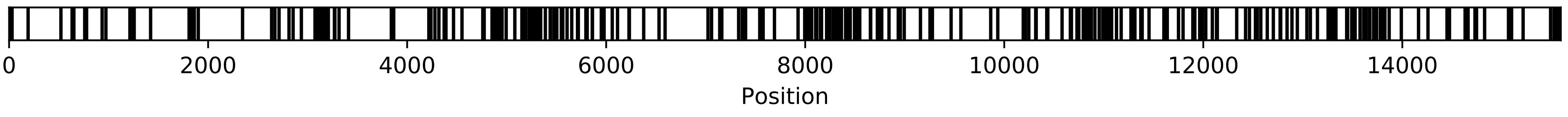


MA2 vs MM2
Number of differences: 314

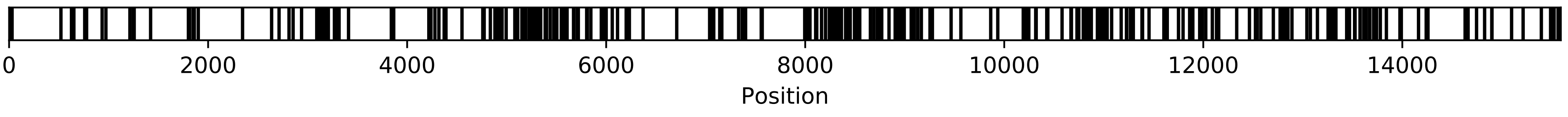


MM1 vs MM2
Number of differences: 146

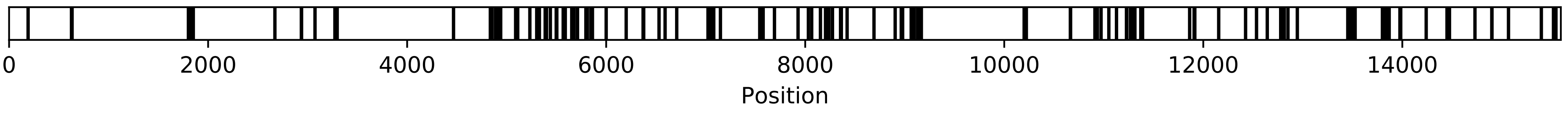

### Region: 111



CR1 vs CR2
Number of differences: 329

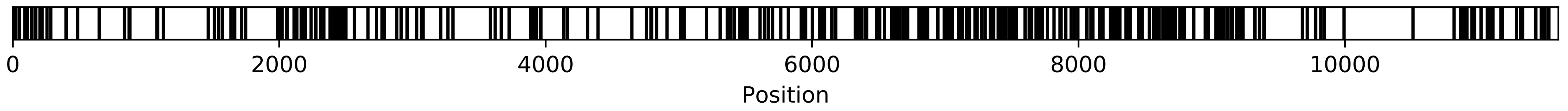


CR1 vs MA1
Number of differences: 116

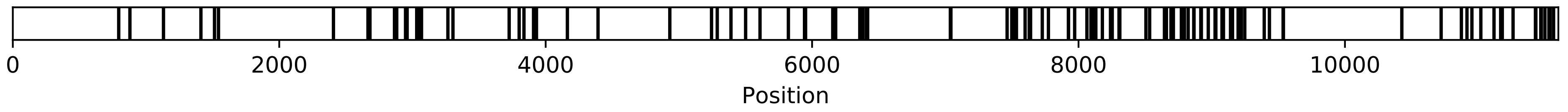


CR1 vs MA2
Number of differences: 308

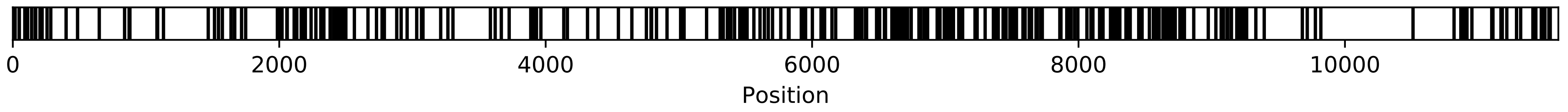


CR1 vs MM1
Number of differences: 116

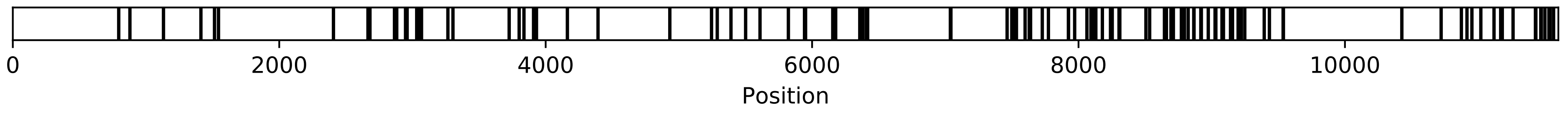


CR1 vs MM2
Number of differences: 155

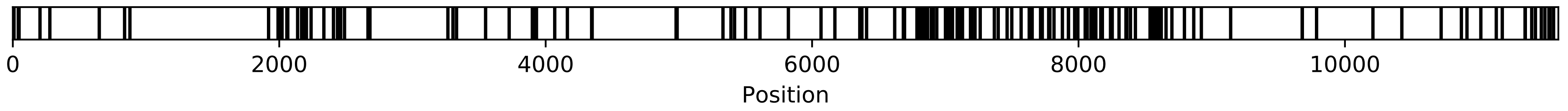


CR2 vs MA1
Number of differences: 323

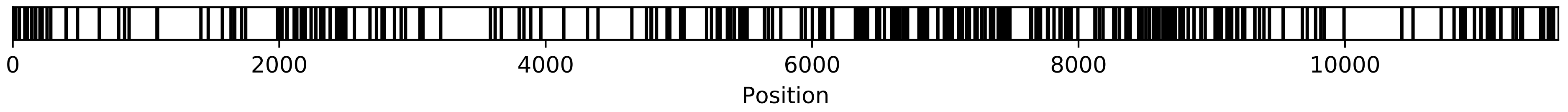


CR2 vs MA2
Number of differences: 74

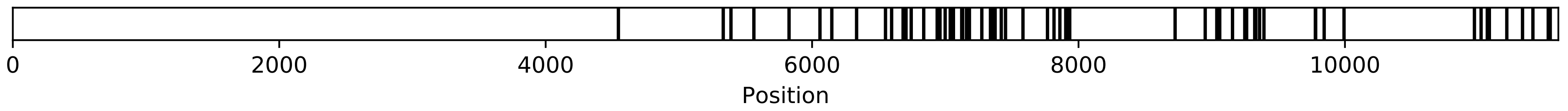


CR2 vs MM1
Number of differences: 323

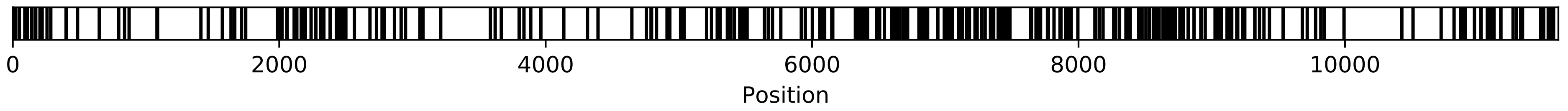


CR2 vs MM2
Number of differences: 330

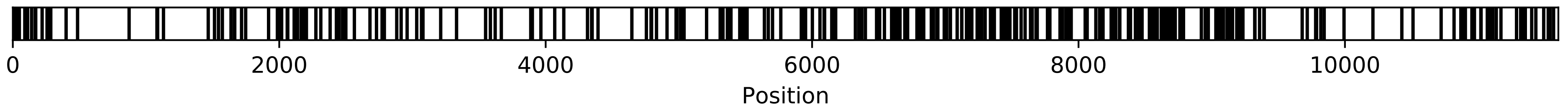


MA1 vs MA2
Number of differences: 299

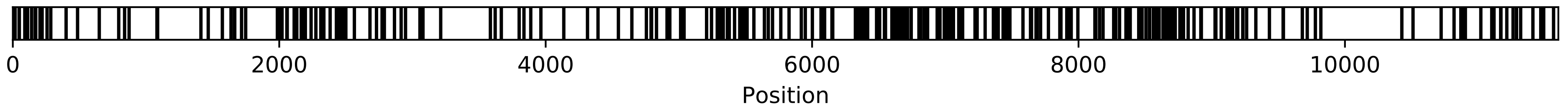


MA1 vs MM1
Number of differences: 0

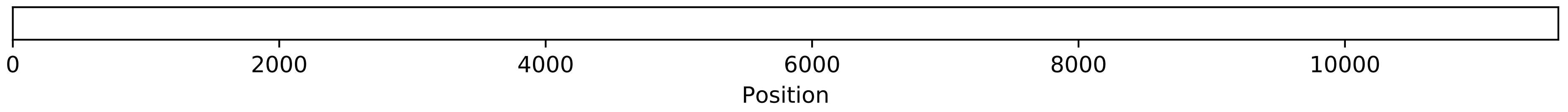


MA1 vs MM2
Number of differences: 175

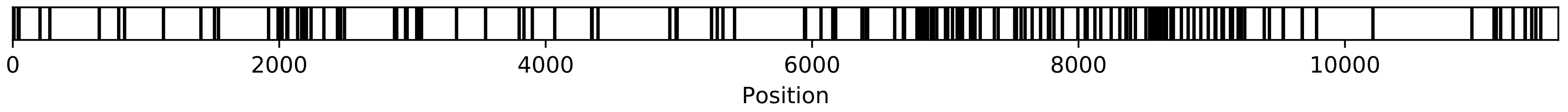


MA2 vs MM1
Number of differences: 299

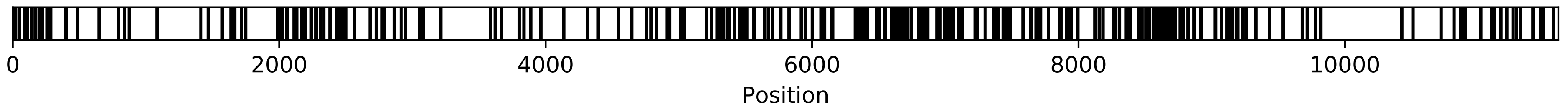


MA2 vs MM2
Number of differences: 308

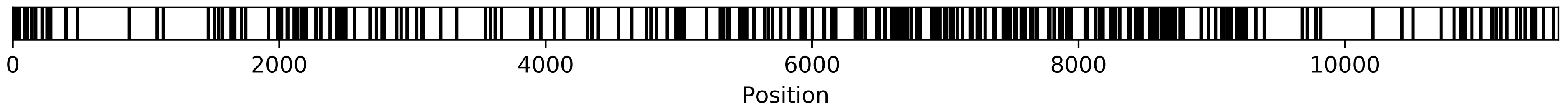


MM1 vs MM2
Number of differences: 175

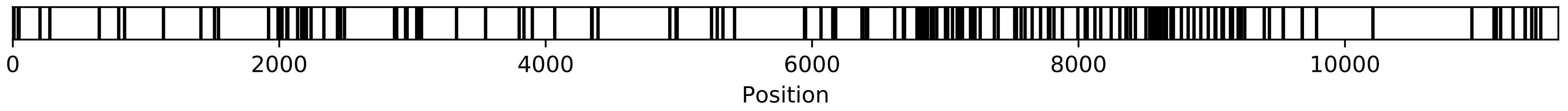

### Region: 112



CR1 vs CR2
Number of differences: 366

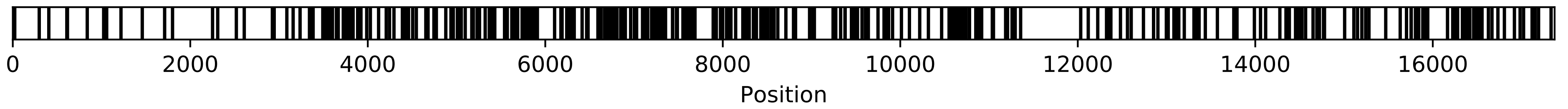


CR1 vs MA1
Number of differences: 369

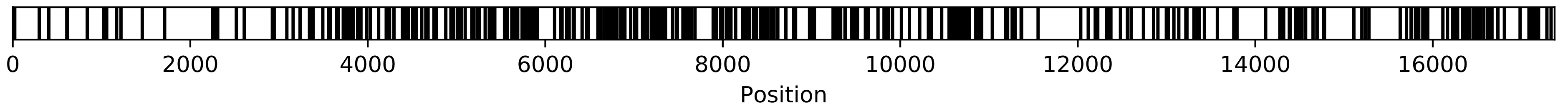


CR1 vs MA2
Number of differences: 250

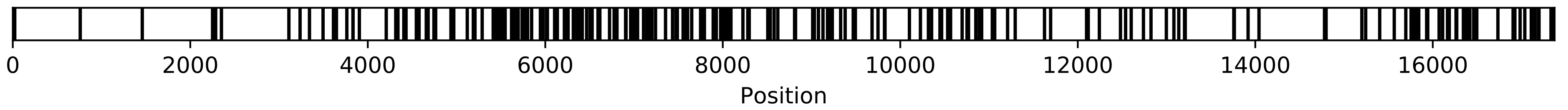


CR1 vs MM1
Number of differences: 242

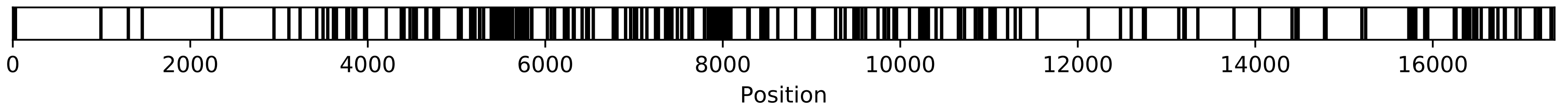


CR1 vs MM2
Number of differences: 232

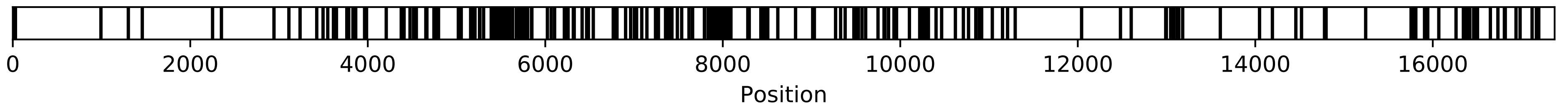


CR2 vs MA1
Number of differences: 65

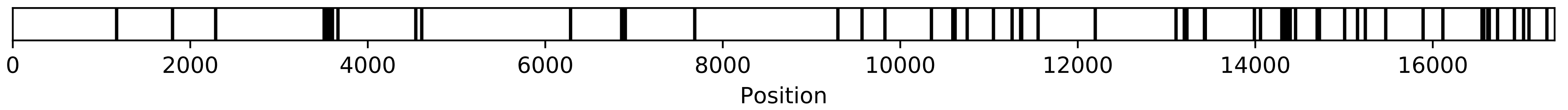


CR2 vs MA2
Number of differences: 386

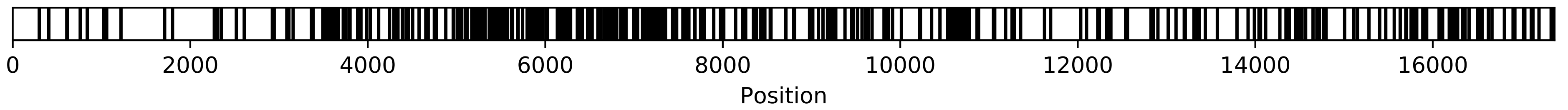


CR2 vs MM1
Number of differences: 387

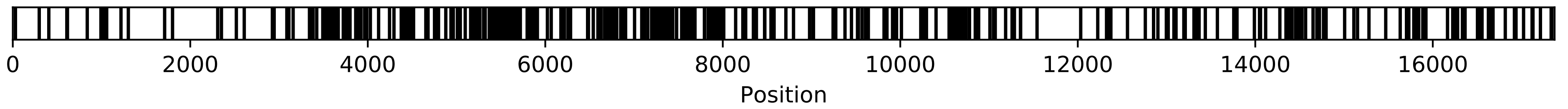


CR2 vs MM2
Number of differences: 383

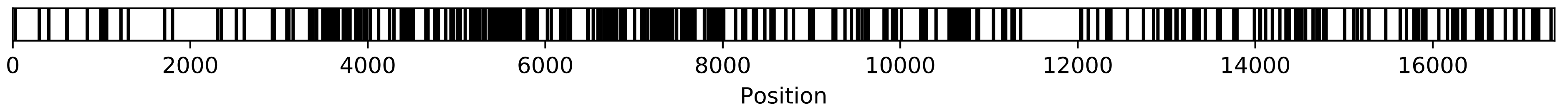


MA1 vs MA2
Number of differences: 387

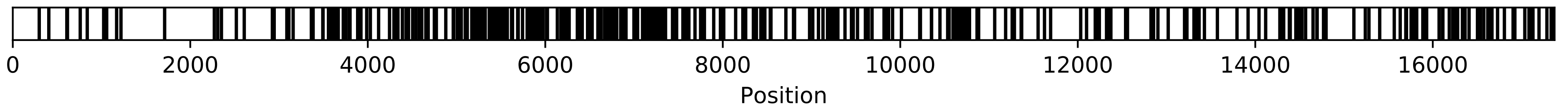


MA1 vs MM1
Number of differences: 391

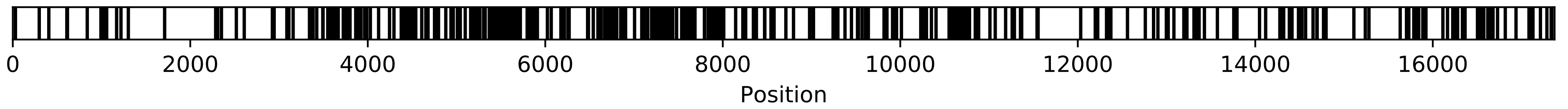


MA1 vs MM2
Number of differences: 387

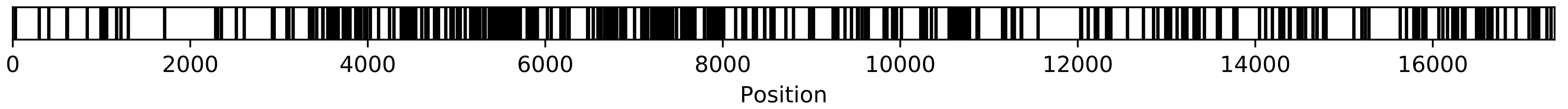


MA2 vs MM1
Number of differences: 245

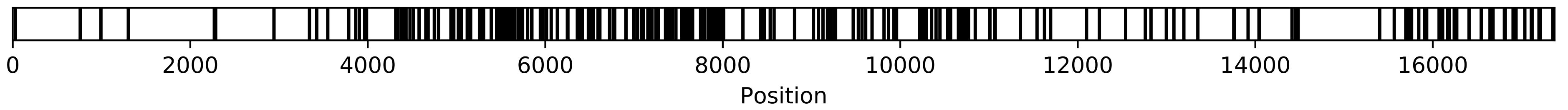


MA2 vs MM2
Number of differences: 247

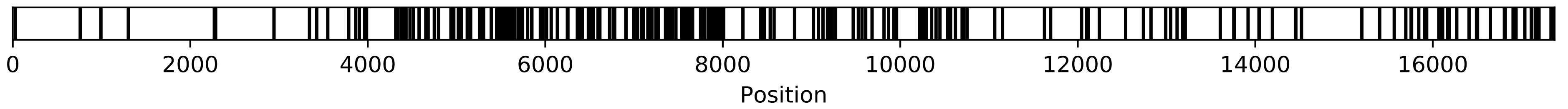


MM1 vs MM2
Number of differences: 56

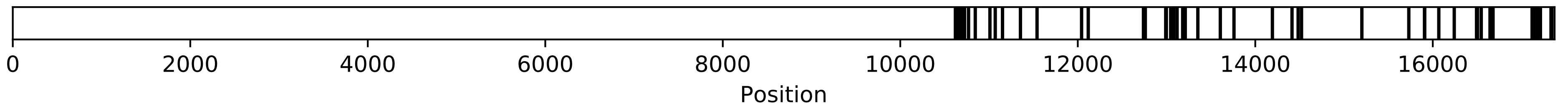

### Region: 114



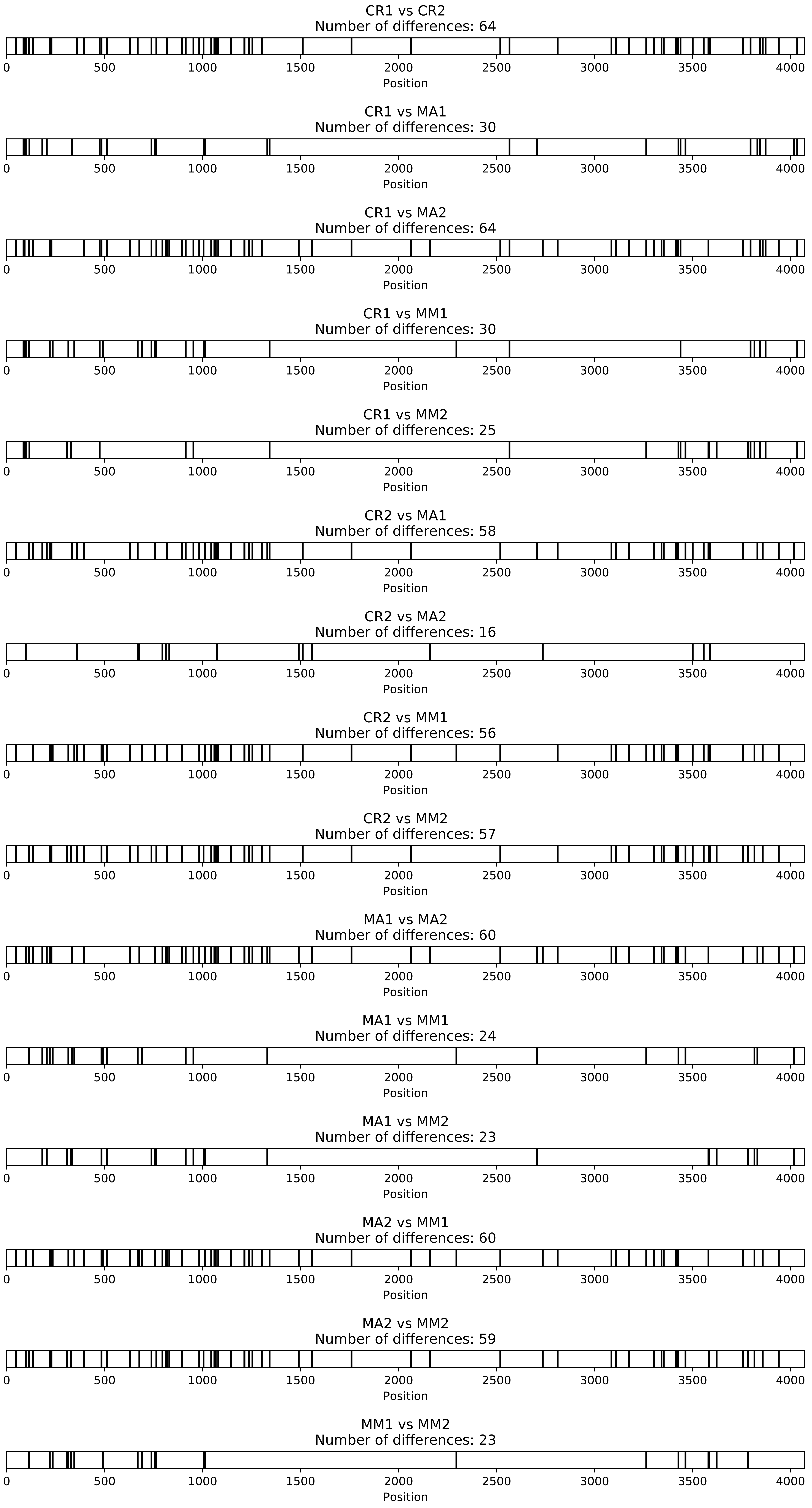
CR1 vs CR2
Number of differences: 64
CR1 vs MA1
Number of differences: 30
CR1 vs MA2
Number of differences: 64
CR1 vs MM1
Number of differences: 30
CR1 vs MM2
Number of differences: 25
CR2 vs MA1
Number of differences: 58
CR2 vs MA2
Number of differences: 16
CR2 vs MM1
Number of differences: 56
CR2 vs MM2
Number of differences: 57
MA1 vs MA2
Number of differences: 60
MA1 vs MM1
Number of differences: 24
MA1 vs MM2
Number of differences: 23
MA2 vs MM1
Number of differences: 60
MA2 vs MM2
Number of differences: 59
MM1 vs MM2
Number of differences: 23
0 500 1000 1500 2000 2500 3000 3500 4000
Position

### Region: 115

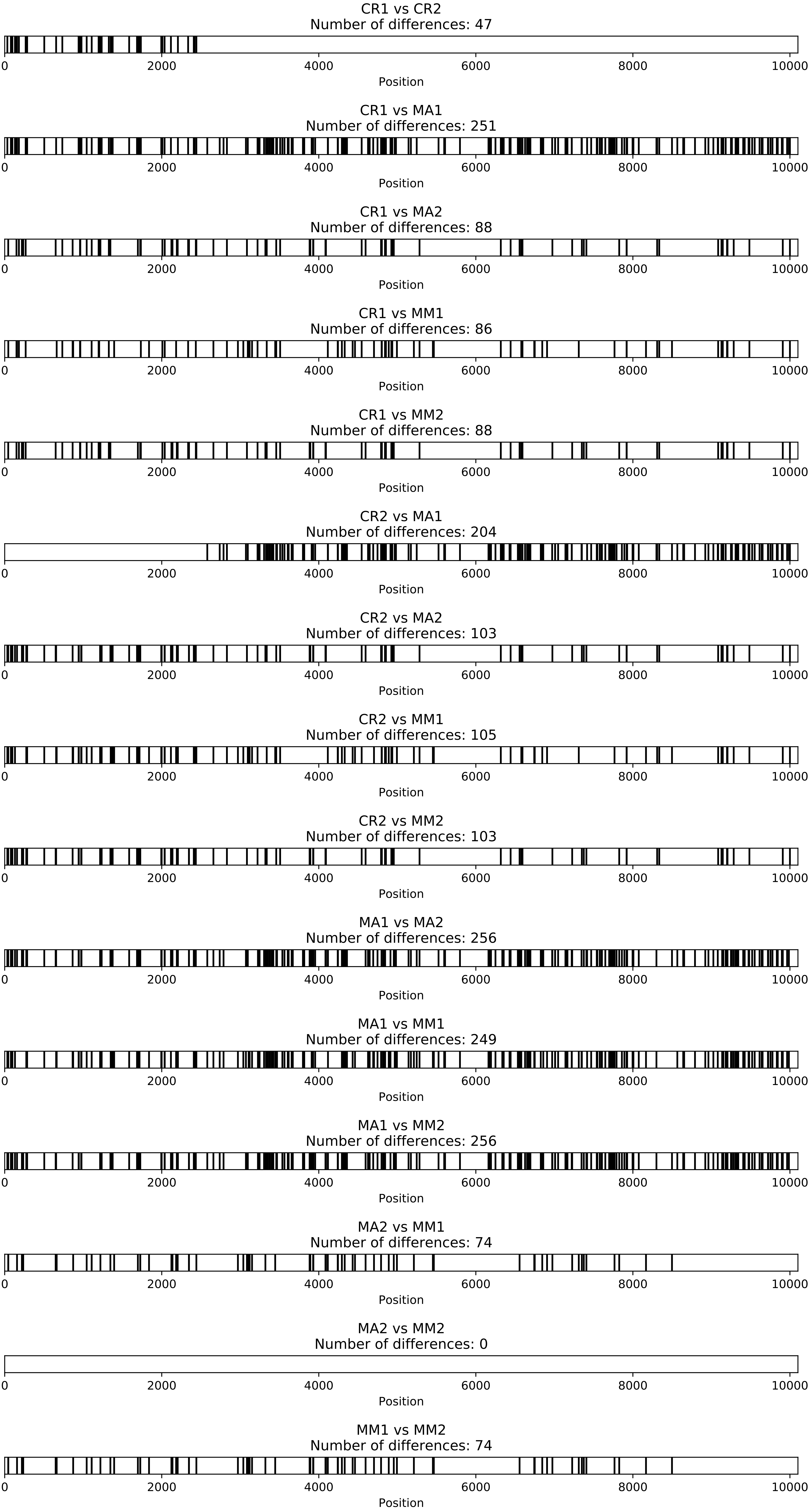
CR1 vs CR2
Number of differences: 47
CR1 vs MA1
Number of differences: 251
CR1 vs MA2
Number of differences: 88
CR1 vs MM1
Number of differences: 86
CR1 vs MM2
Number of differences: 88
CR2 vs MA1
Number of differences: 204
CR2 vs MA2
Number of differences: 103
CR2 vs MM1
Number of differences: 105
CR2 vs MM2
Number of differences: 103
MA1 vs MA2
Number of differences: 256
MA1 vs MM1
Number of differences: 249
MA1 vs MM2
Number of differences: 256
MA2 vs MM1
Number of differences: 74
MA2 vs MM2
Number of differences: 0
MM1 vs MM2
Number of differences: 74
0
2000
4000
6000
8000
10000
Position

### Region: 116



CR1 vs CR2
Number of differences: 212

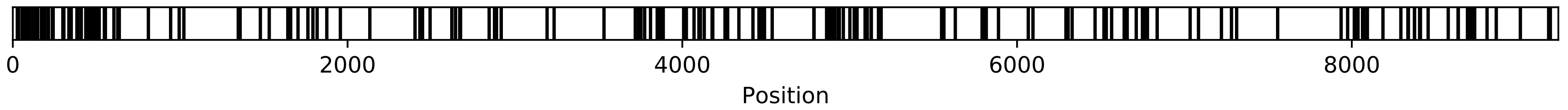


CR1 vs MA1
Number of differences: 81

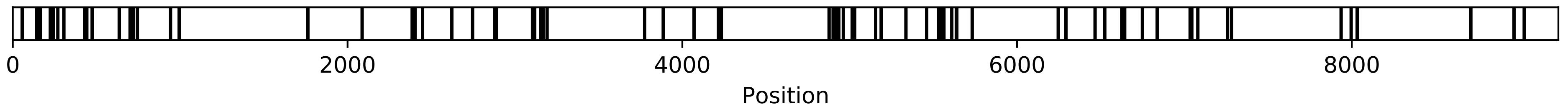


CR1 vs MA2
Number of differences: 194

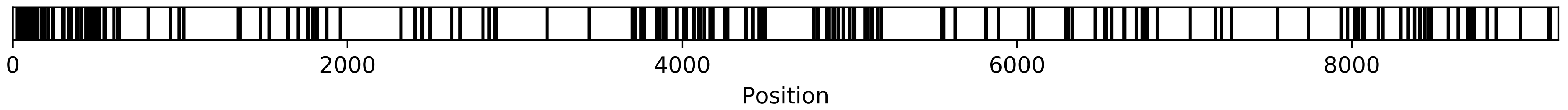


CR1 vs MM1
Number of differences: 81

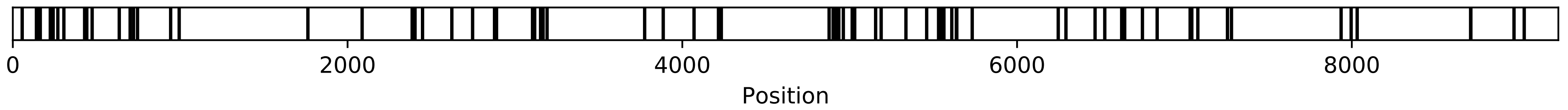


CR1 vs MM2
Number of differences: 78

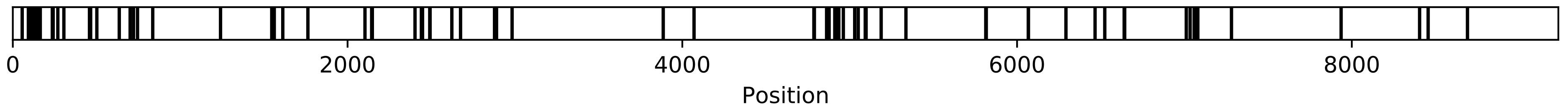


CR2 vs MA1
Number of differences: 218

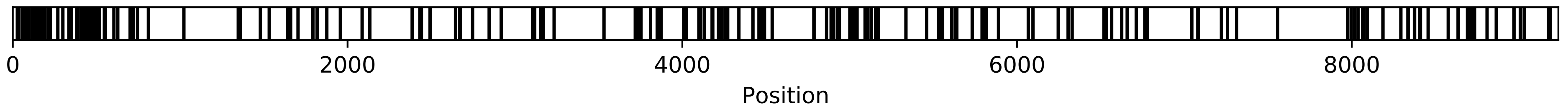


CR2 vs MA2
Number of differences: 67

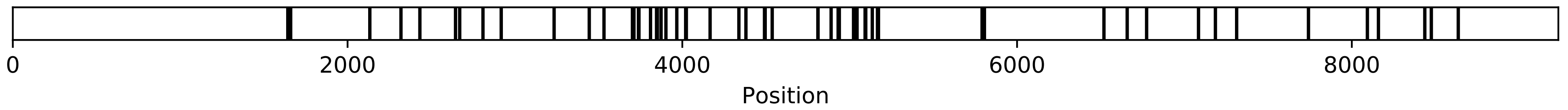


CR2 vs MM1
Number of differences: 218

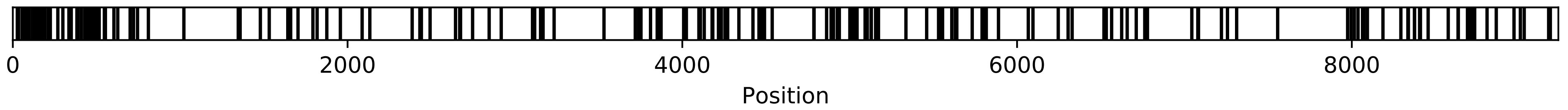


CR2 vs MM2
Number of differences: 201

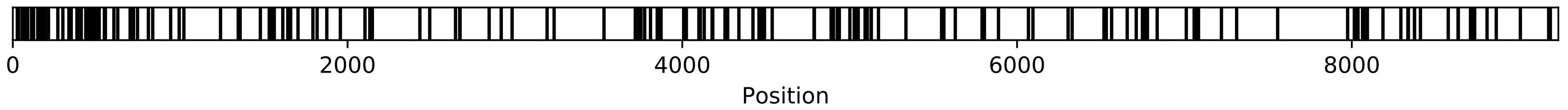


MA1 vs MA2
Number of differences: 200

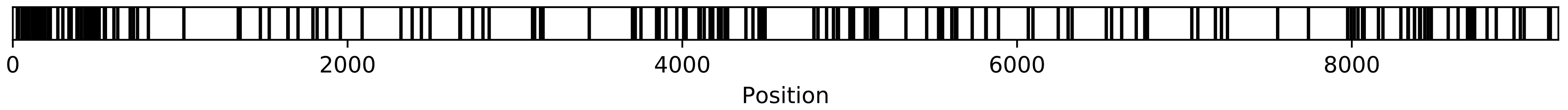


MA1 vs MM1
Number of differences: 0

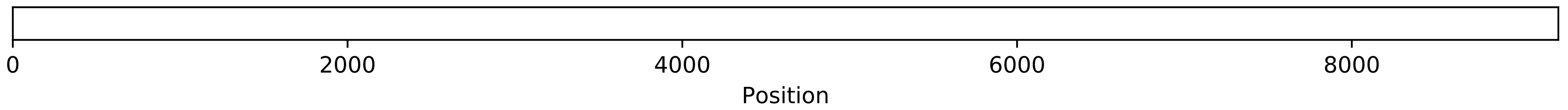


MA1 vs MM2
Number of differences: 85

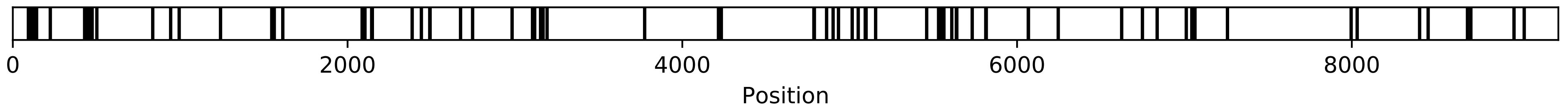


MA2 vs MM1
Number of differences: 200

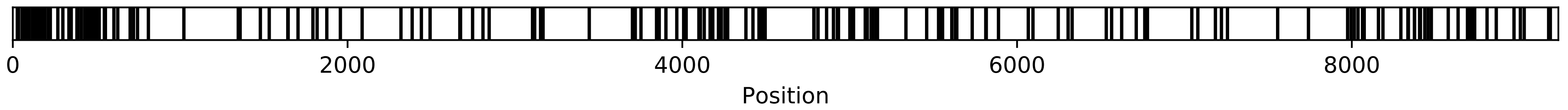


MA2 vs MM2
Number of differences: 182

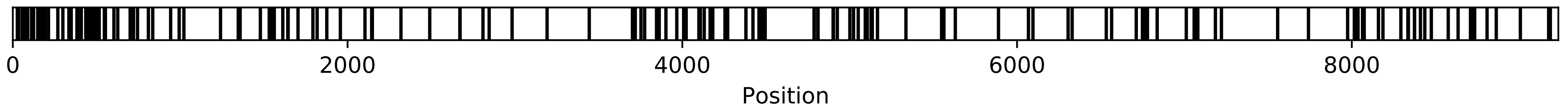


MM1 vs MM2
Number of differences: 85

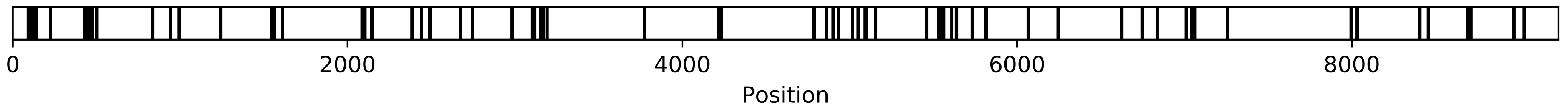

### Region: 119



CR1 vs CR2
Number of differences: 201

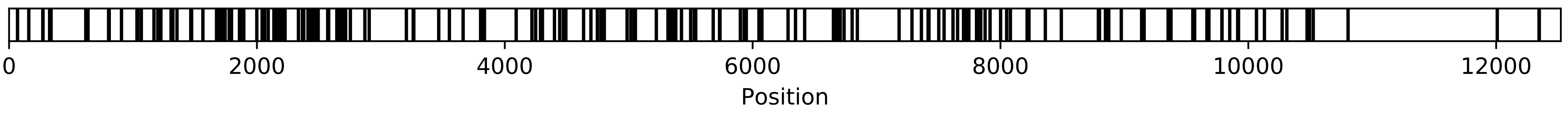


CR1 vs MA1
Number of differences: 107

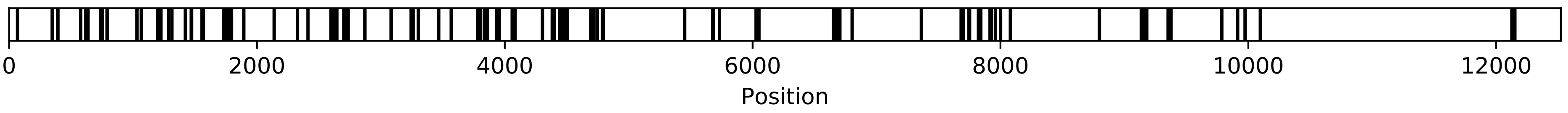


CR1 vs MA2
Number of differences: 198

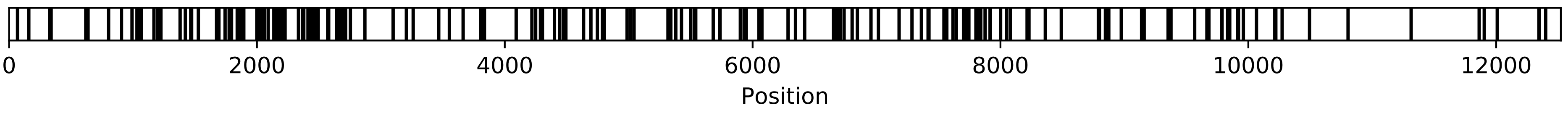


CR1 vs MM1
Number of differences: 88

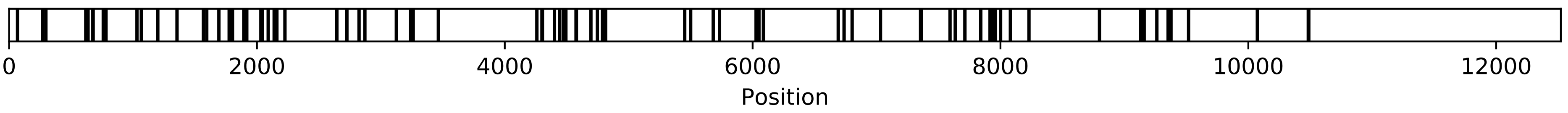


CR1 vs MM2
Number of differences: 107

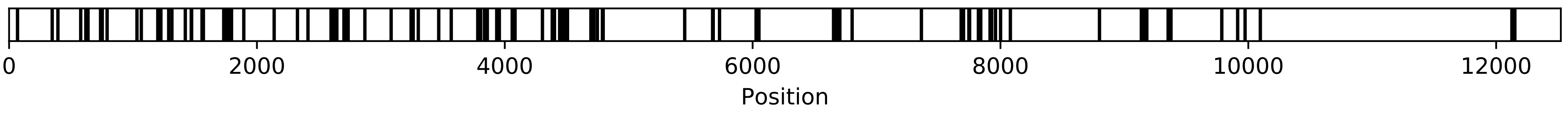


CR2 vs MA1
Number of differences: 205

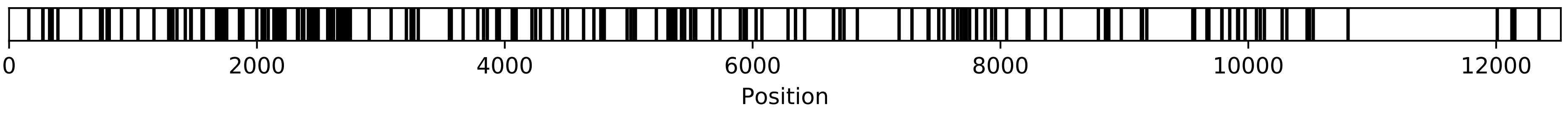


CR2 vs MA2
Number of differences: 52

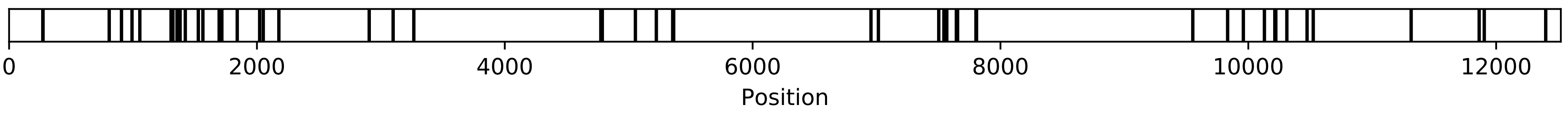


CR2 vs MM1
Number of differences: 196

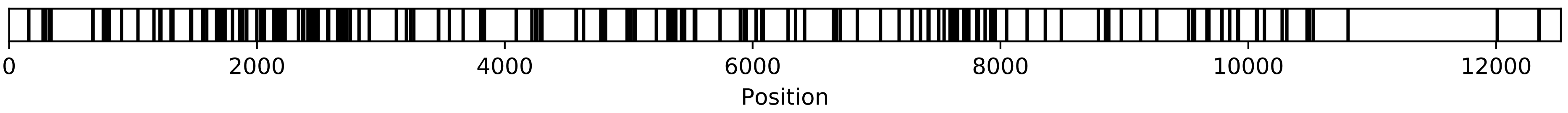


CR2 vs MM2
Number of differences: 205

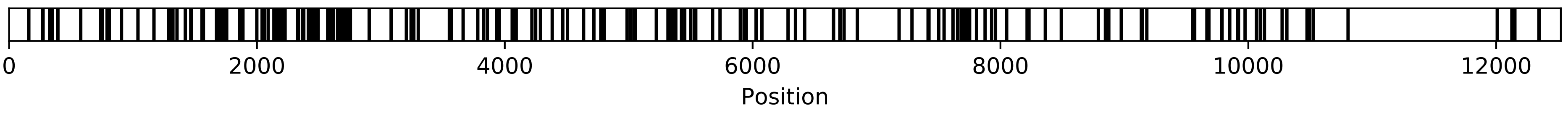


MA1 vs MA2
Number of differences: 201

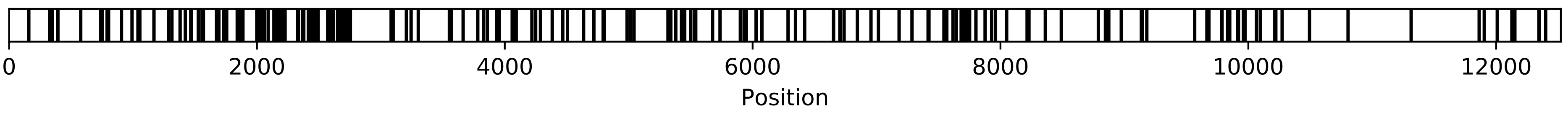


MA1 vs MM1
Number of differences: 109

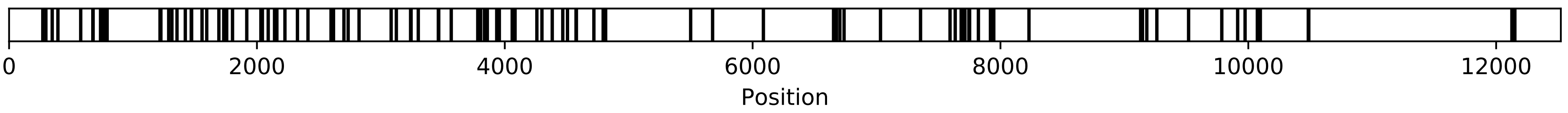


MA1 vs MM2
Number of differences: 0

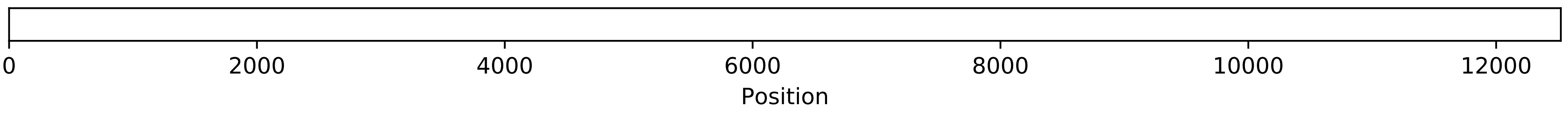


MA2 vs MM1
Number of differences: 197

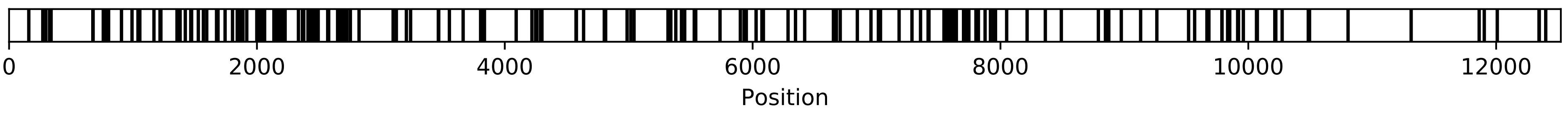


MA2 vs MM2
Number of differences: 201

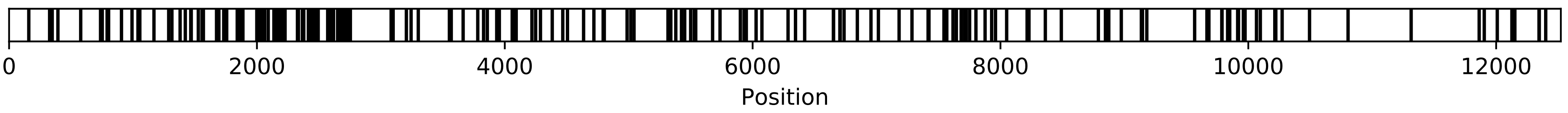


MM1 vs MM2
Number of differences: 109

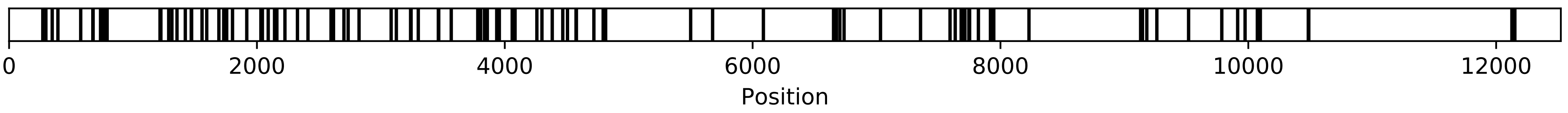

### Region: 120

CR1 vs CR2
Number of differences: 388

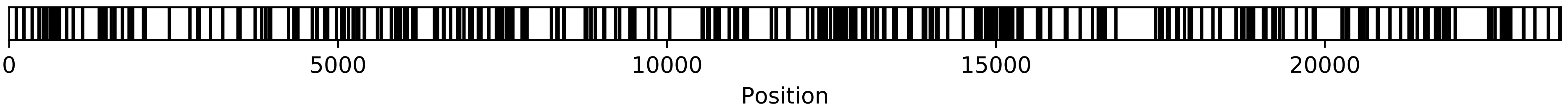


CR1 vs MA1
Number of differences: 227

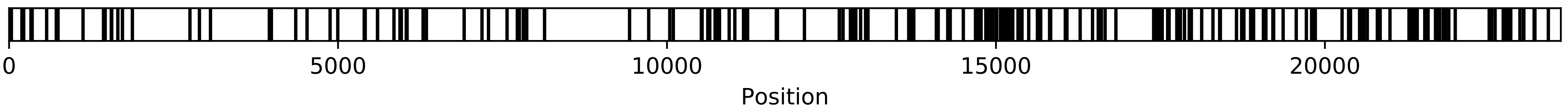


CR1 vs MA2
Number of differences: 364

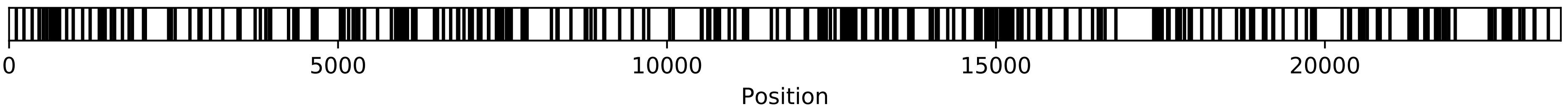


CR1 vs MM1
Number of differences: 183

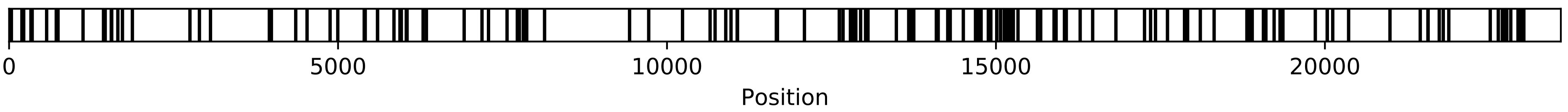


CR1 vs MM2
Number of differences: 183

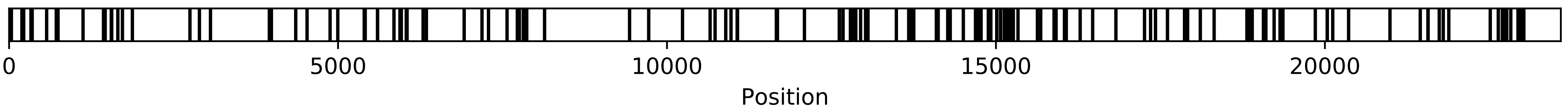


CR2 vs MA1
Number of differences: 288

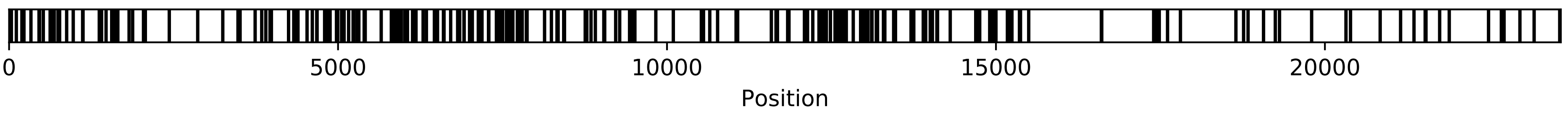


CR2 vs MA2
Number of differences: 144

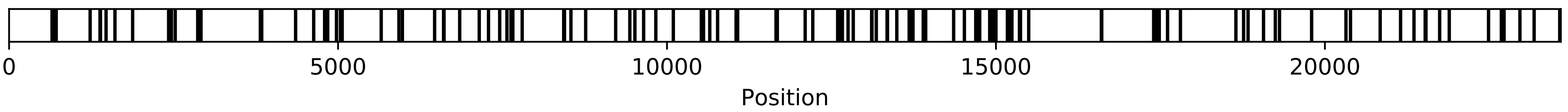


CR2 vs MM1
Number of differences: 397

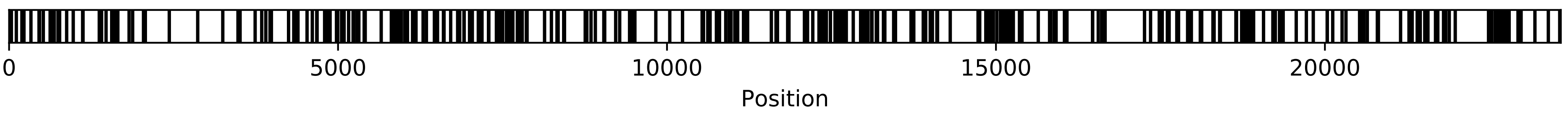


CR2 vs MM2
Number of differences: 397

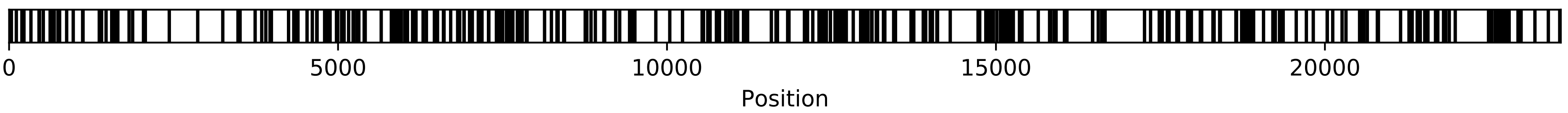


MA1 vs MA2
Number of differences: 218

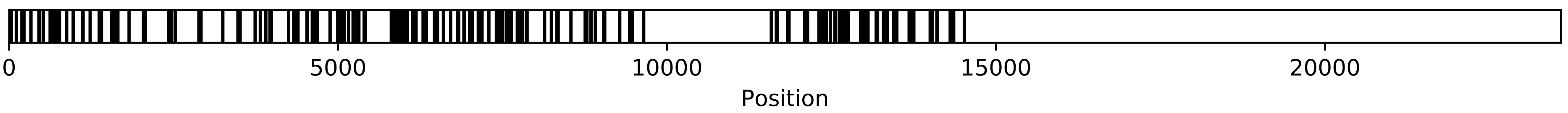


MA1 vs MM1
Number of differences: 165

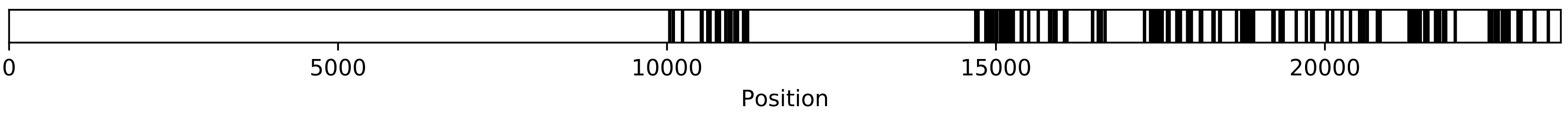


MA1 vs MM2
Number of differences: 165

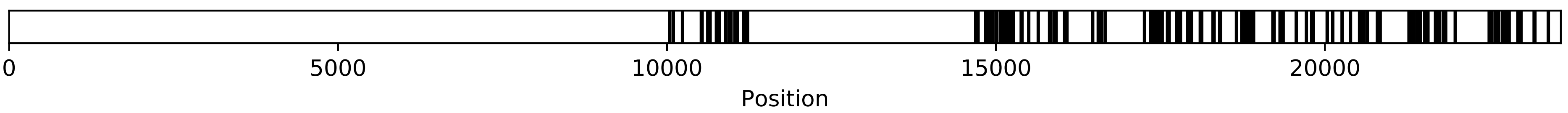


MA2 vs MM1
Number of differences: 383

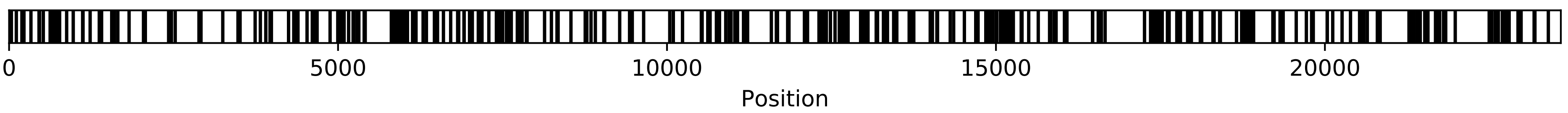


MA2 vs MM2
Number of differences: 383

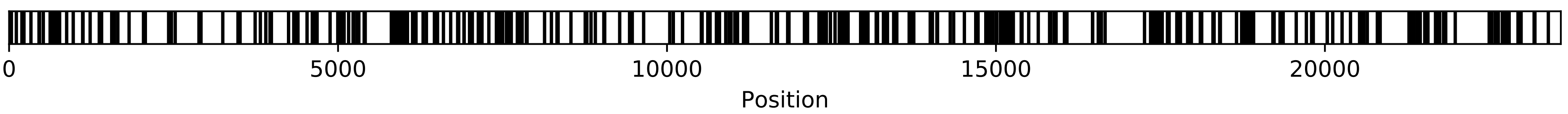


MM1 vs MM2
Number of differences: 0

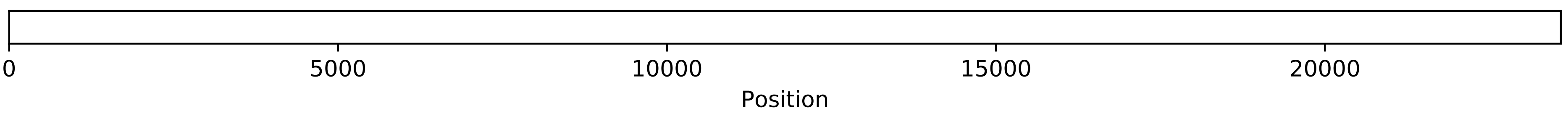

### Region: 122



CR1 vs CR2
Number of differences: 219

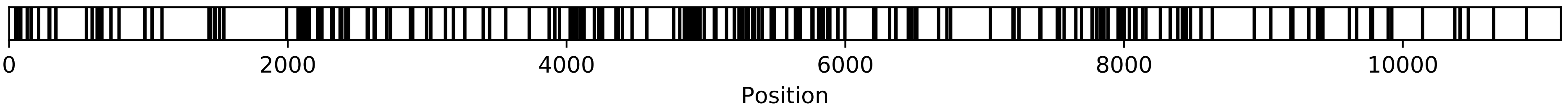


CR1 vs MA1
Number of differences: 89

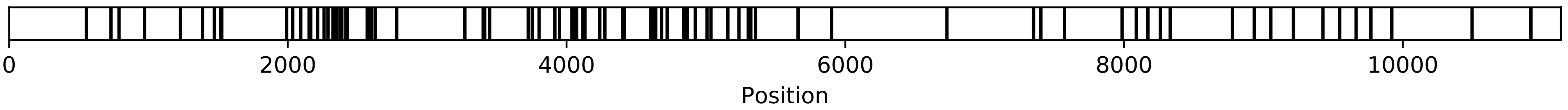


CR1 vs MA2
Number of differences: 218

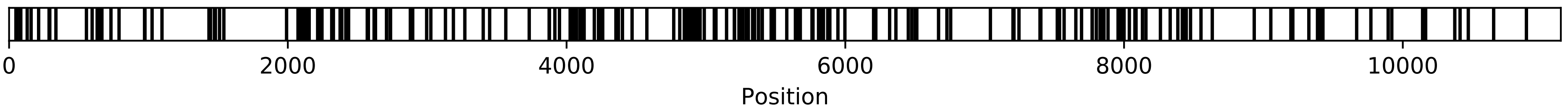


CR1 vs MM1
Number of differences: 218

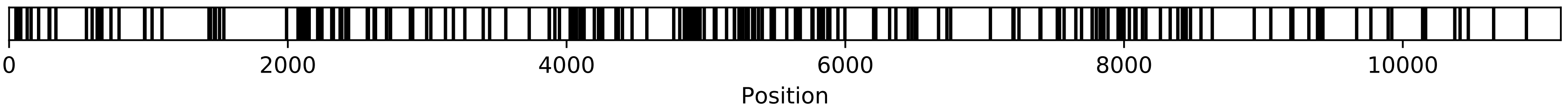


CR1 vs MM2
Number of differences: 243

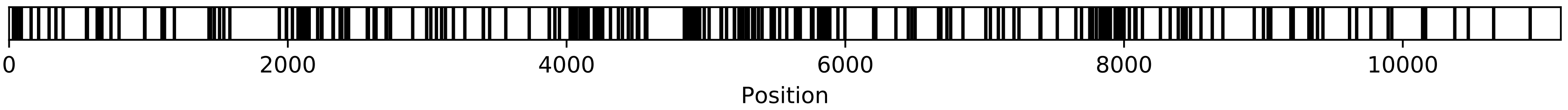


CR2 vs MA1
Number of differences: 221

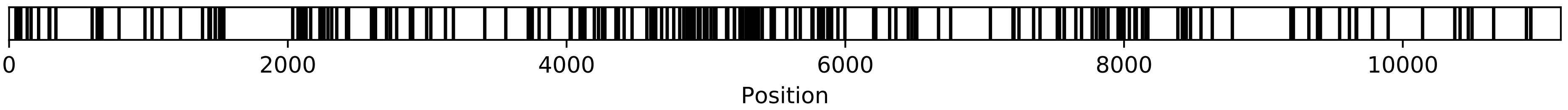


CR2 vs MA2
Number of differences: 3

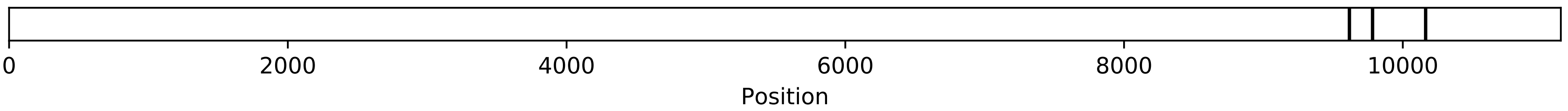


CR2 vs MM1
Number of differences: 3

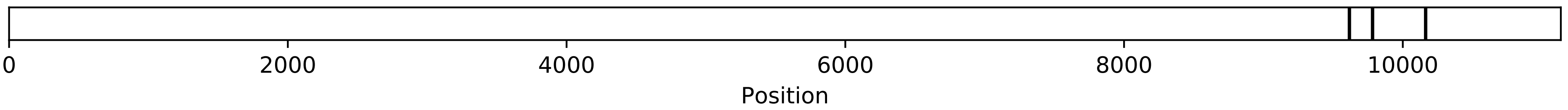


CR2 vs MM2
Number of differences: 96

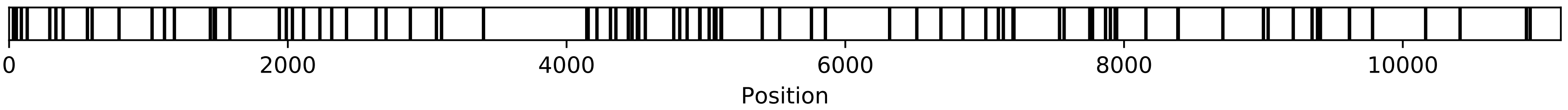


MA1 vs MA2
Number of differences: 220

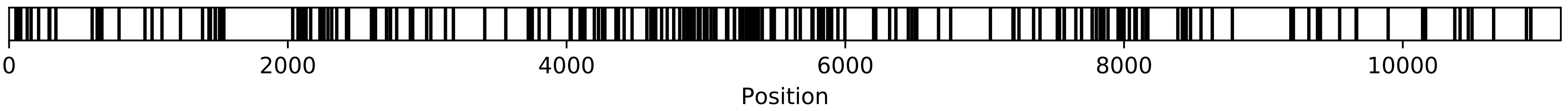


MA1 vs MM1
Number of differences: 220

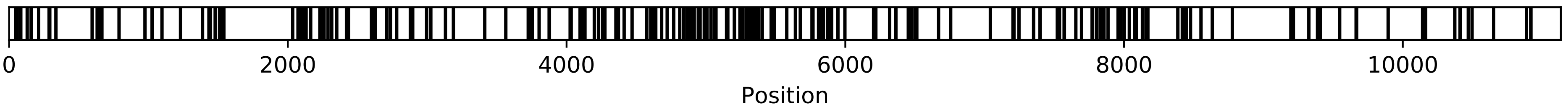


MA1 vs MM2
Number of differences: 244

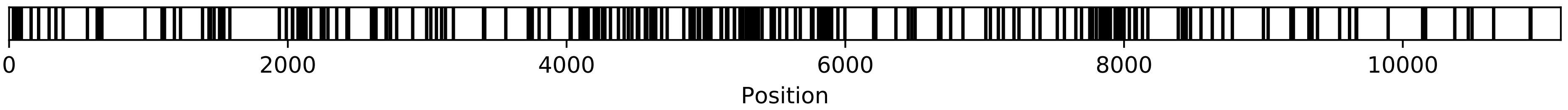


MA2 vs MM1
Number of differences: 0

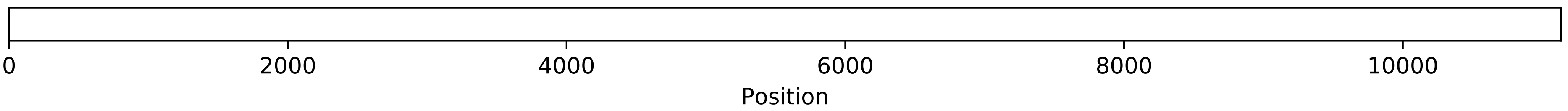


MA2 vs MM2
Number of differences: 93

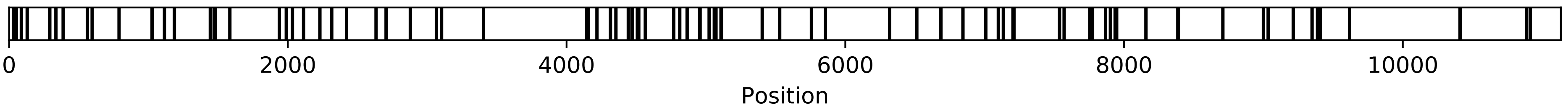


MM1 vs MM2
Number of differences: 93

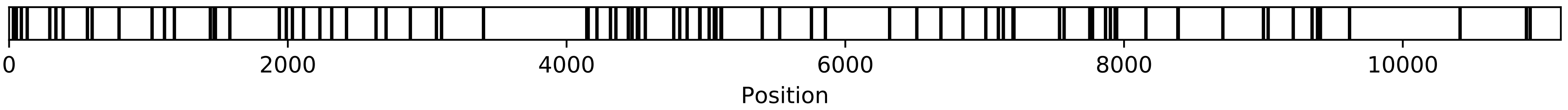

### Region: 123

CR1 vs CR2
Number of differences: 211

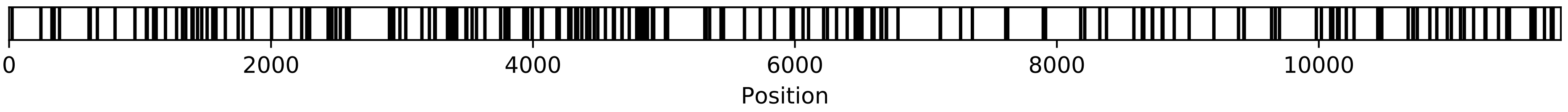


CR1 vs MA1
Number of differences: 152

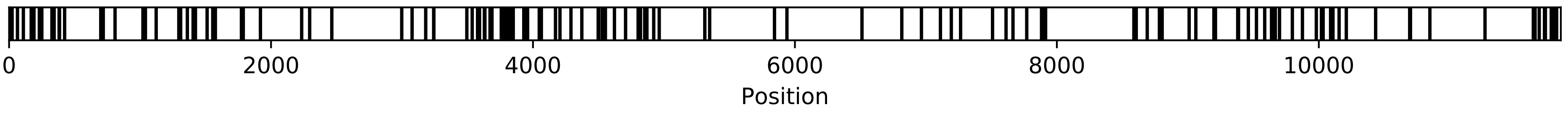


CR1 vs MA2
Number of differences: 207

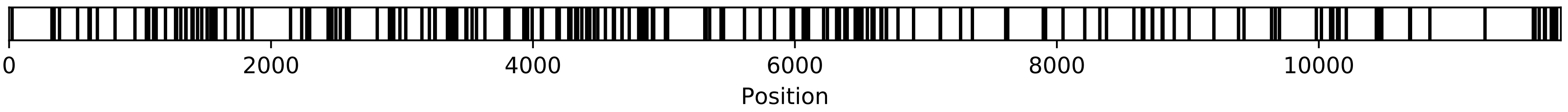


CR1 vs MM1
Number of differences: 109

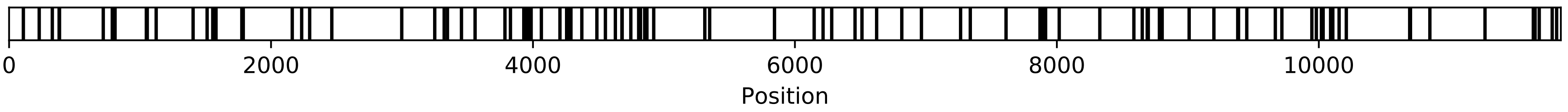


CR1 vs MM2
Number of differences: 156

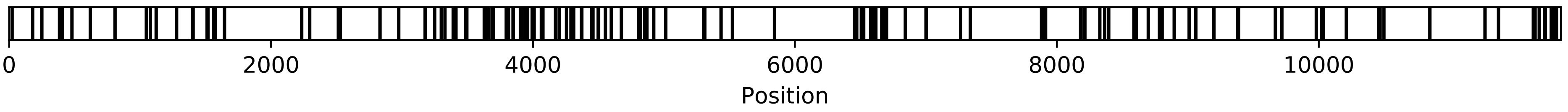


CR2 vs MA1
Number of differences: 231

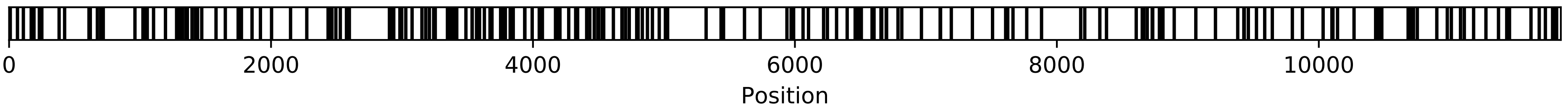


CR2 vs MA2
Number of differences: 52

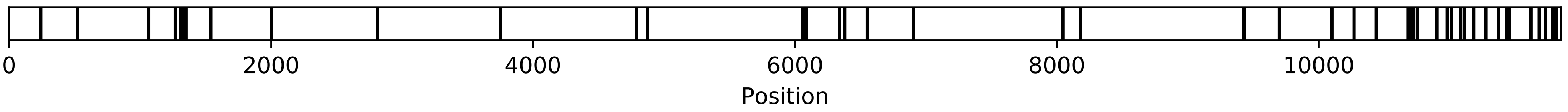


CR2 vs MM1
Number of differences: 199

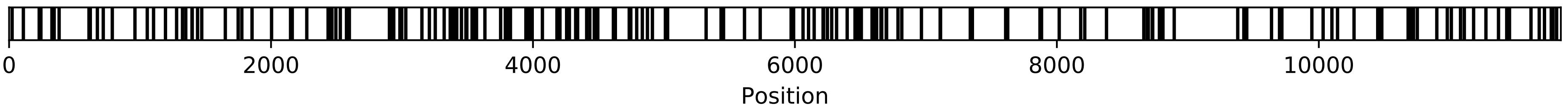


CR2 vs MM2
Number of differences: 223

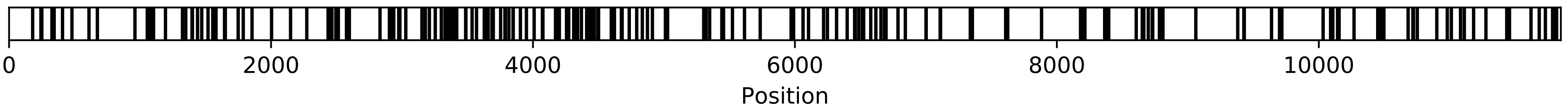


MA1 vs MA2
Number of differences: 211

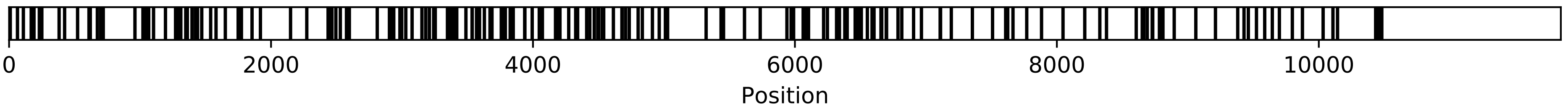


MA1 vs MM1
Number of differences: 121

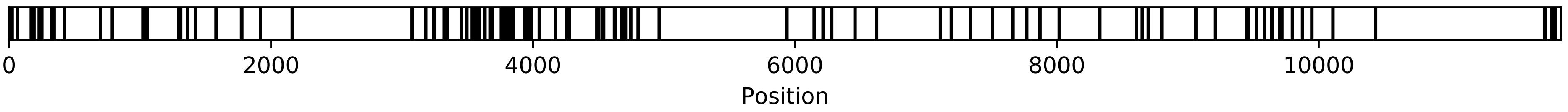


MA1 vs MM2
Number of differences: 166

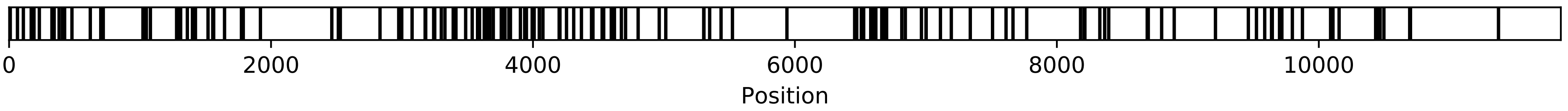


MA2 vs MM1
Number of differences: 187

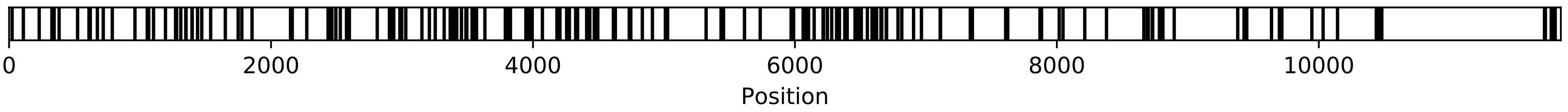


MA2 vs MM2
Number of differences: 209

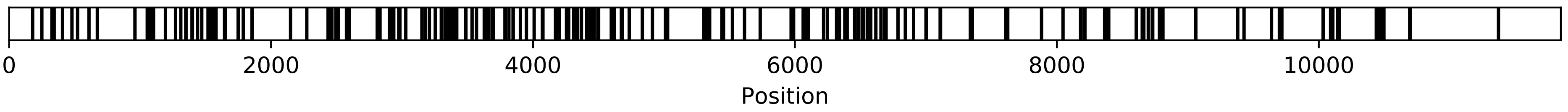


MM1 vs MM2
Number of differences: 153

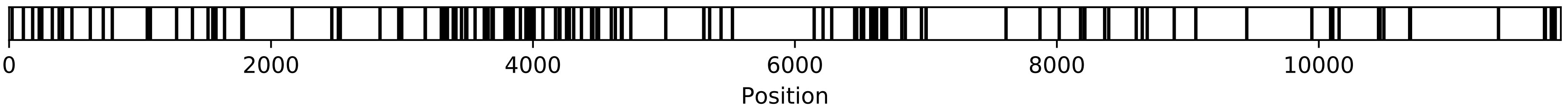

### Region: 124

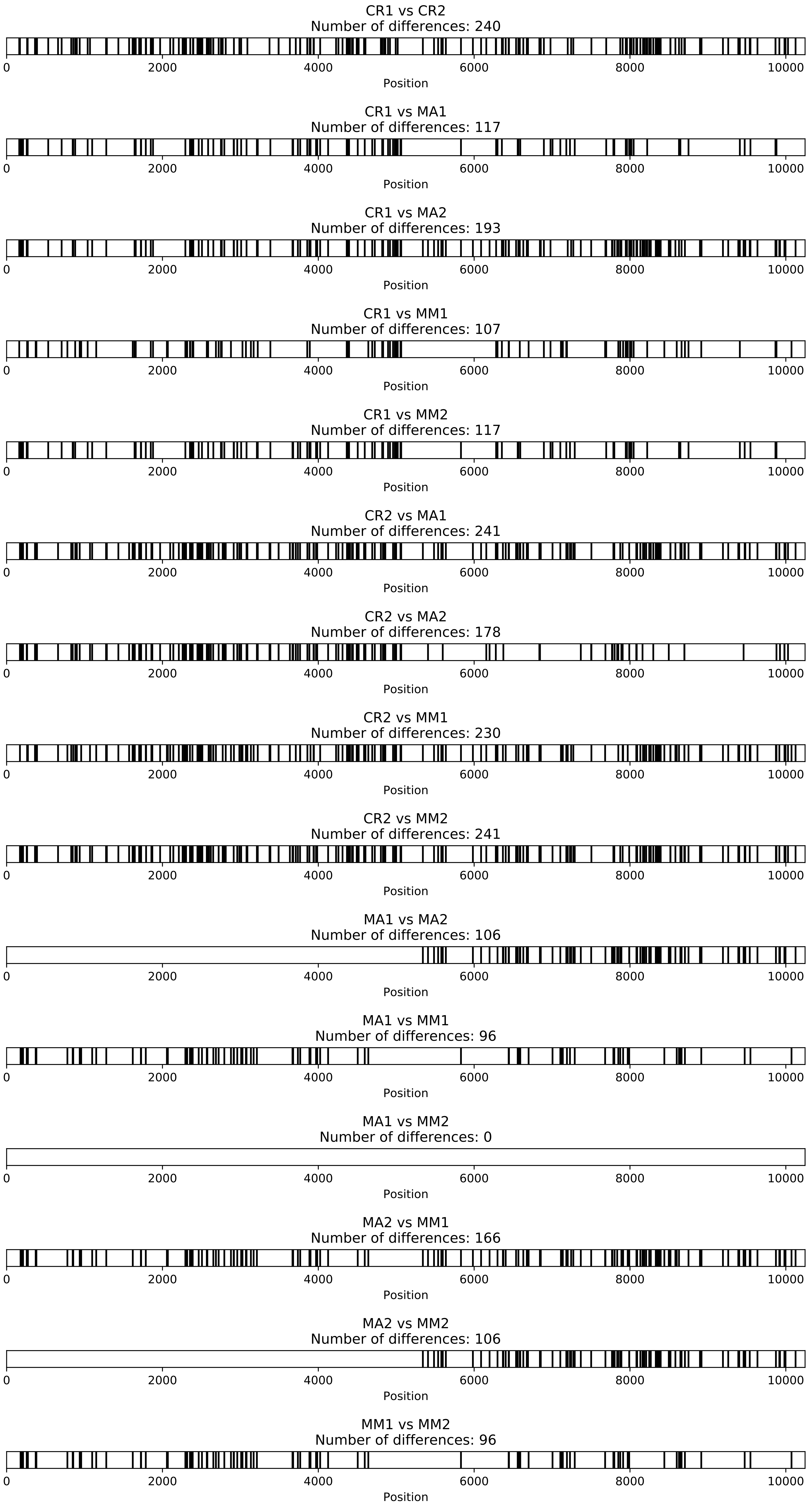
CR1 vs CR2
Number of differences: 240
0 2000 4000 6000 8000 10000
Position
CR1 vs MA1
Number of differences: 117
0 2000 4000 6000 8000 10000
Position
CR1 vs MA2
Number of differences: 193
0 2000 4000 6000 8000 10000
Position
CR1 vs MM1
Number of differences: 107
0 2000 4000 6000 8000 10000
Position
CR1 vs MM2
Number of differences: 117
0 2000 4000 6000 8000 10000
Position
CR2 vs MA1
Number of differences: 241
0 2000 4000 6000 8000 10000
Position
CR2 vs MA2
Number of differences: 178
0 2000 4000 6000 8000 10000
Position
CR2 vs MM1
Number of differences: 230
0 2000 4000 6000 8000 10000
Position
CR2 vs MM2
Number of differences: 241
0 2000 4000 6000 8000 10000
Position
MA1 vs MA2
Number of differences: 106
0 2000 4000 6000 8000 10000
Position
MA1 vs MM1
Number of differences: 96
0 2000 4000 6000 8000 10000
Position
MA1 vs MM2
Number of differences: 0
0 2000 4000 6000 8000 10000
Position
MA2 vs MM1
Number of differences: 166
0 2000 4000 6000 8000 10000
Position
MA2 vs MM2
Number of differences: 106
0 2000 4000 6000 8000 10000
Position
MM1 vs MM2
Number of differences: 96
0 2000 4000 6000 8000 10000
Position

### Region: 125



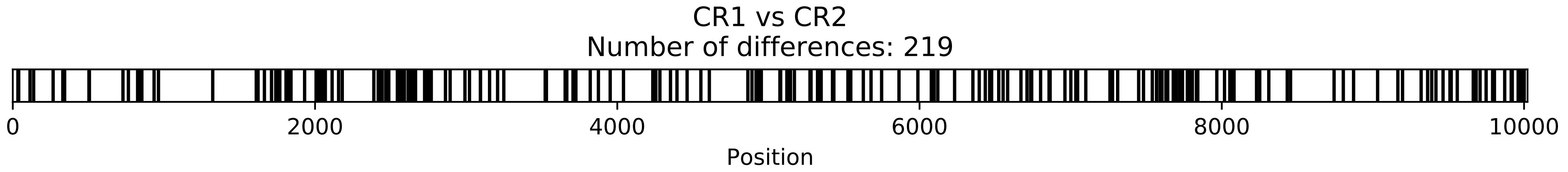
CR1 vs CR2
Number of differences: 219
0 2000 4000 6000 8000 10000
Position

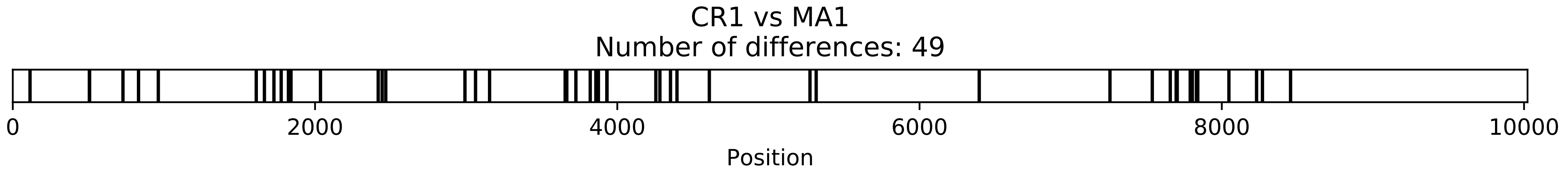
CR1 vs MA1
Number of differences: 49
0 2000 4000 6000 8000 10000
Position

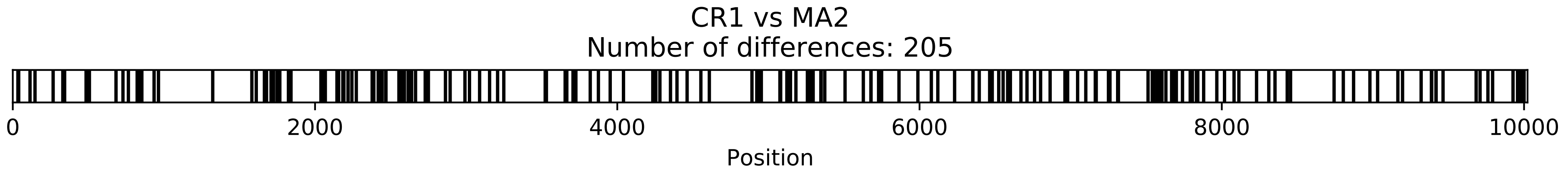
CR1 vs MA2
Number of differences: 205
0 2000 4000 6000 8000 10000
Position

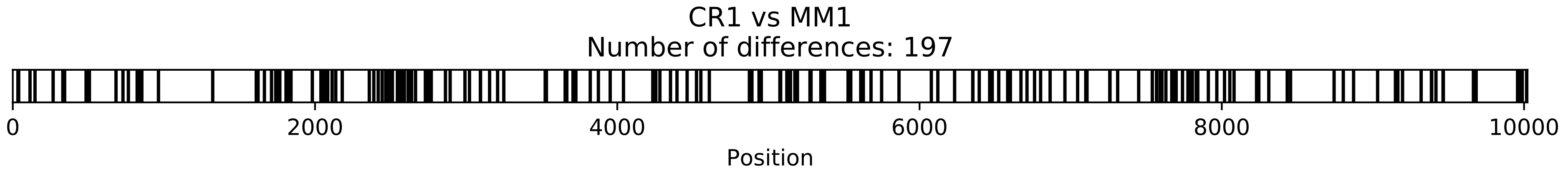
CR1 vs MM1
Number of differences: 197
0 2000 4000 6000 8000 10000
Position

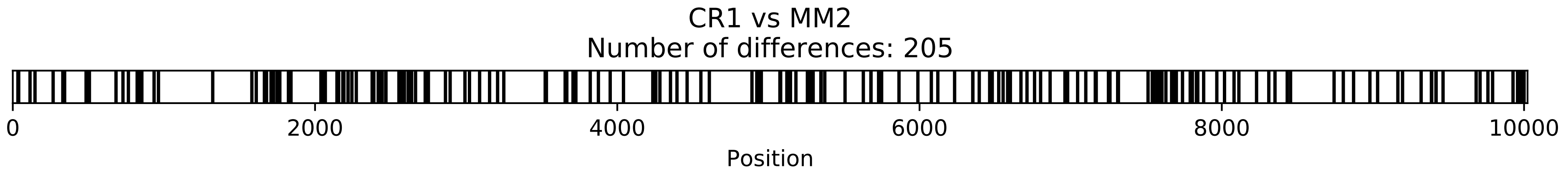
CR1 vs MM2
Number of differences: 205
0 2000 4000 6000 8000 10000
Position

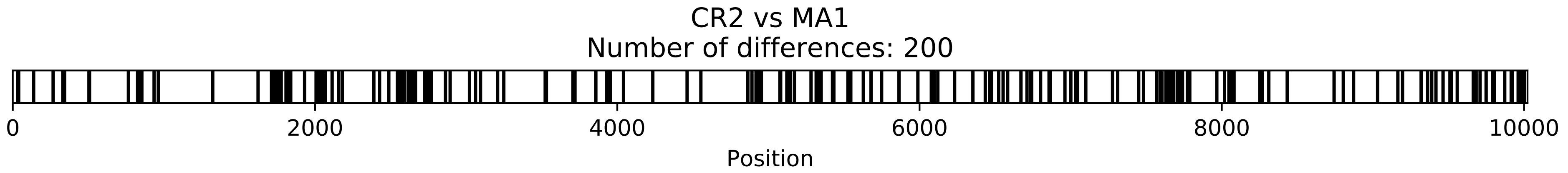
CR2 vs MA1
Number of differences: 200
0 2000 4000 6000 8000 10000
Position

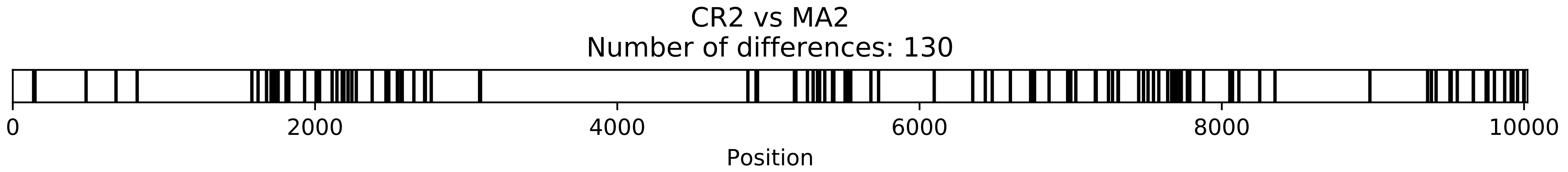
CR2 vs MA2
Number of differences: 130
0 2000 4000 6000 8000 10000
Position

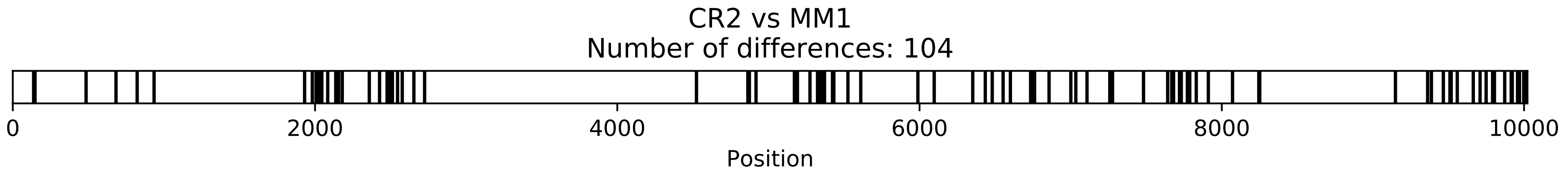
CR2 vs MM1
Number of differences: 104
0 2000 4000 6000 8000 10000
Position

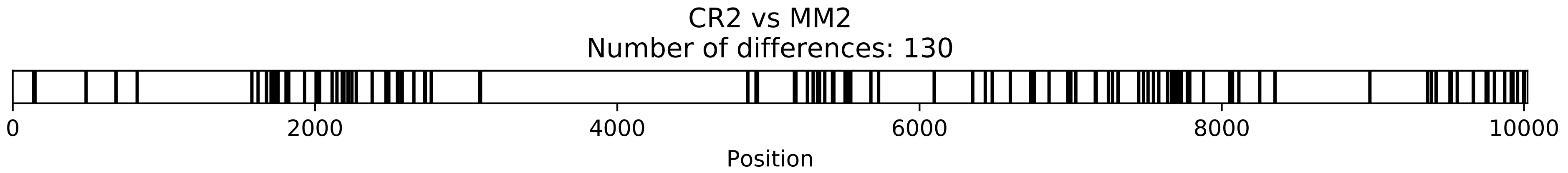
CR2 vs MM2
Number of differences: 130
0 2000 4000 6000 8000 10000
Position

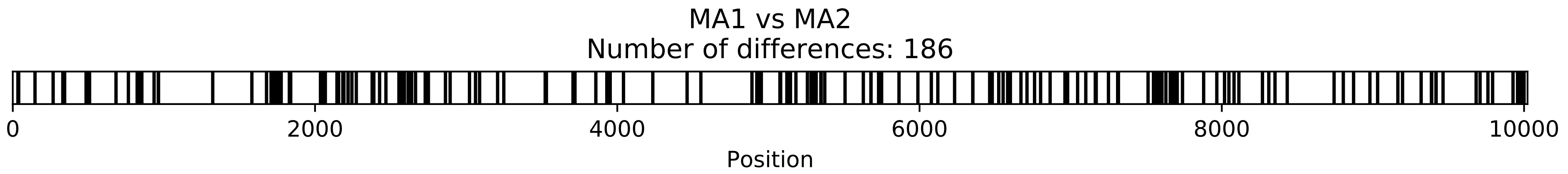
MA1 vs MA2
Number of differences: 186
0 2000 4000 6000 8000 10000
Position

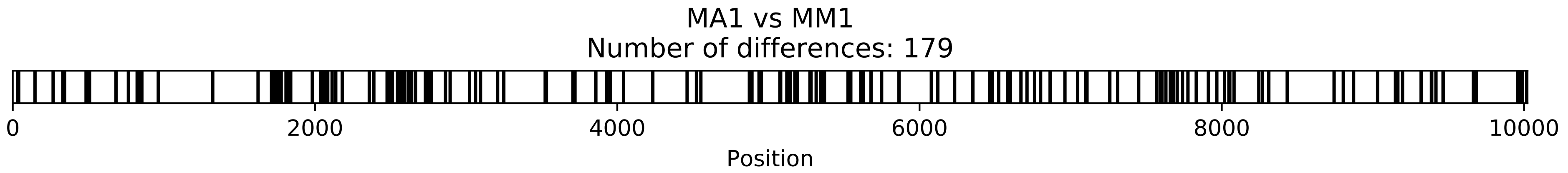
MA1 vs MM1
Number of differences: 179
0 2000 4000 6000 8000 10000
Position

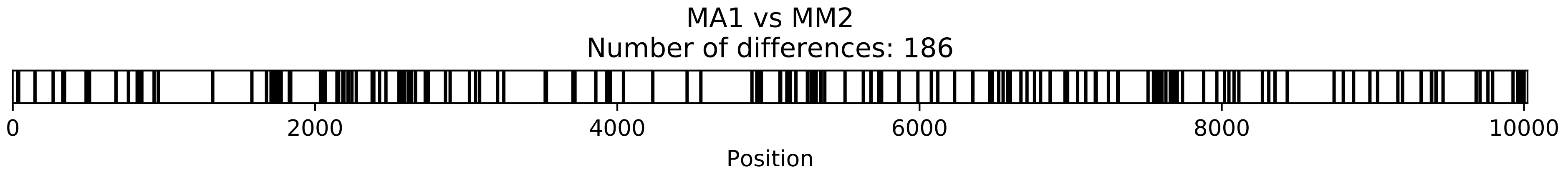
MA1 vs MM2
Number of differences: 186
0 2000 4000 6000 8000 10000
Position

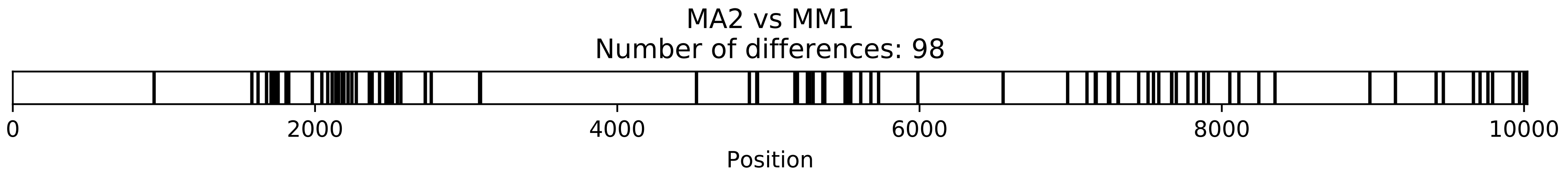
MA2 vs MM1
Number of differences: 98
0 2000 4000 6000 8000 10000
Position

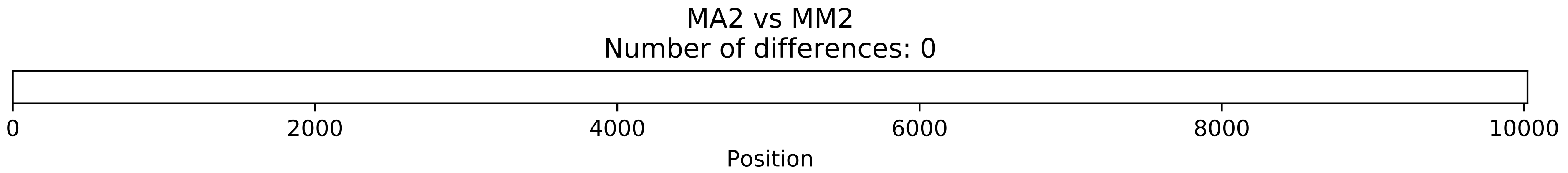
MA2 vs MM2
Number of differences: 0
0 2000 4000 6000 8000 10000
Position

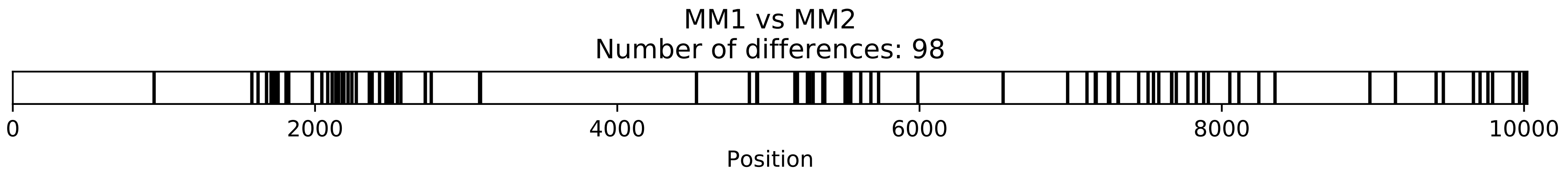
MM1 vs MM2
Number of differences: 98
0 2000 4000 6000 8000 10000
Position

### Region: 126

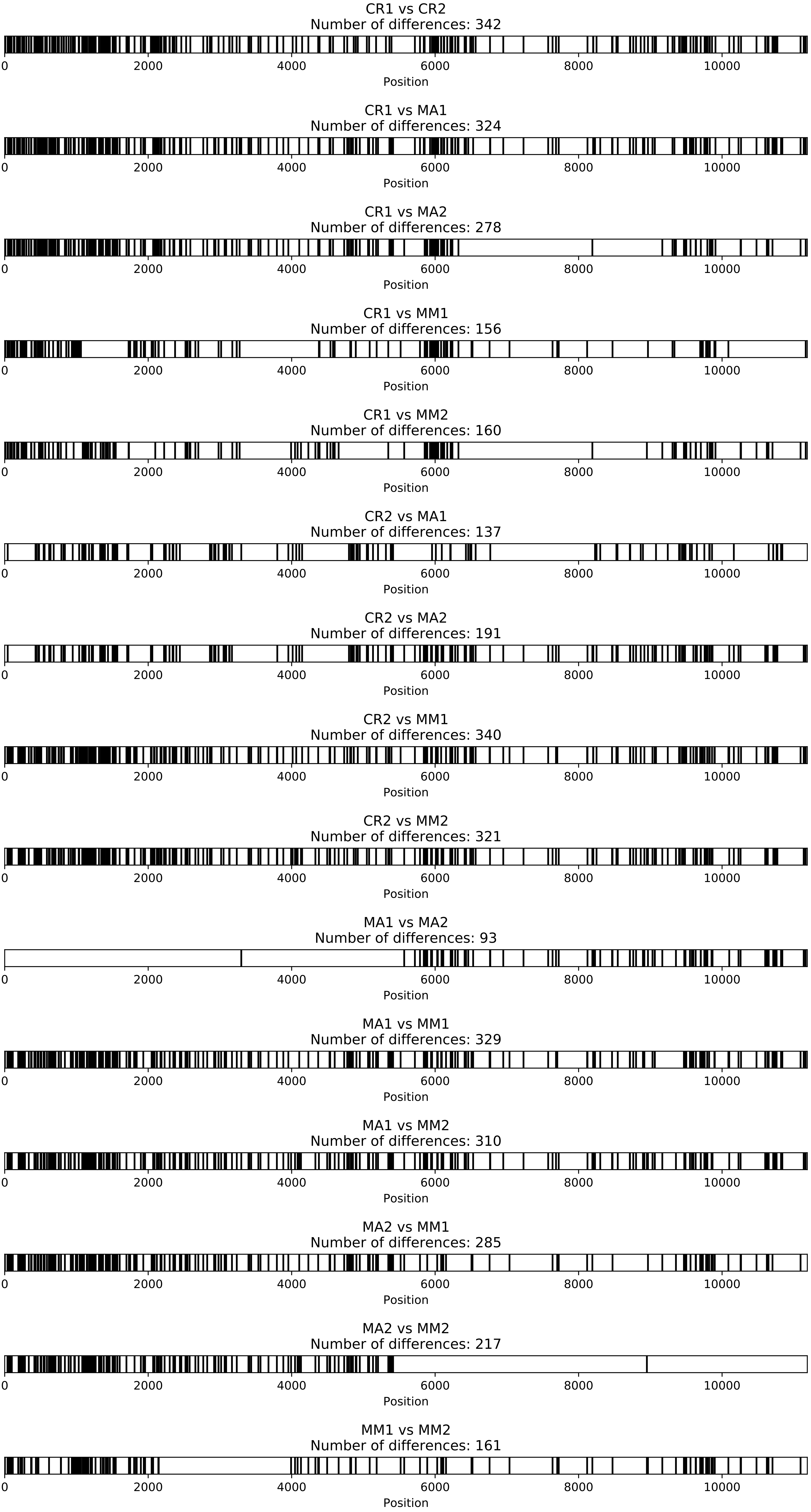
CR1 vs CR2
Number of differences: 342
CR1 vs MA1
Number of differences: 324
CR1 vs MA2
Number of differences: 278
CR1 vs MM1
Number of differences: 156
CR1 vs MM2
Number of differences: 160
CR2 vs MA1
Number of differences: 137
CR2 vs MA2
Number of differences: 191
CR2 vs MM1
Number of differences: 340
CR2 vs MM2
Number of differences: 321
MA1 vs MA2
Number of differences: 93
MA1 vs MM1
Number of differences: 329
MA1 vs MM2
Number of differences: 310
MA2 vs MM1
Number of differences: 285
MA2 vs MM2
Number of differences: 217
MM1 vs MM2
Number of differences: 161
0
2000
4000
6000
8000
10000
Position

### Region: 128



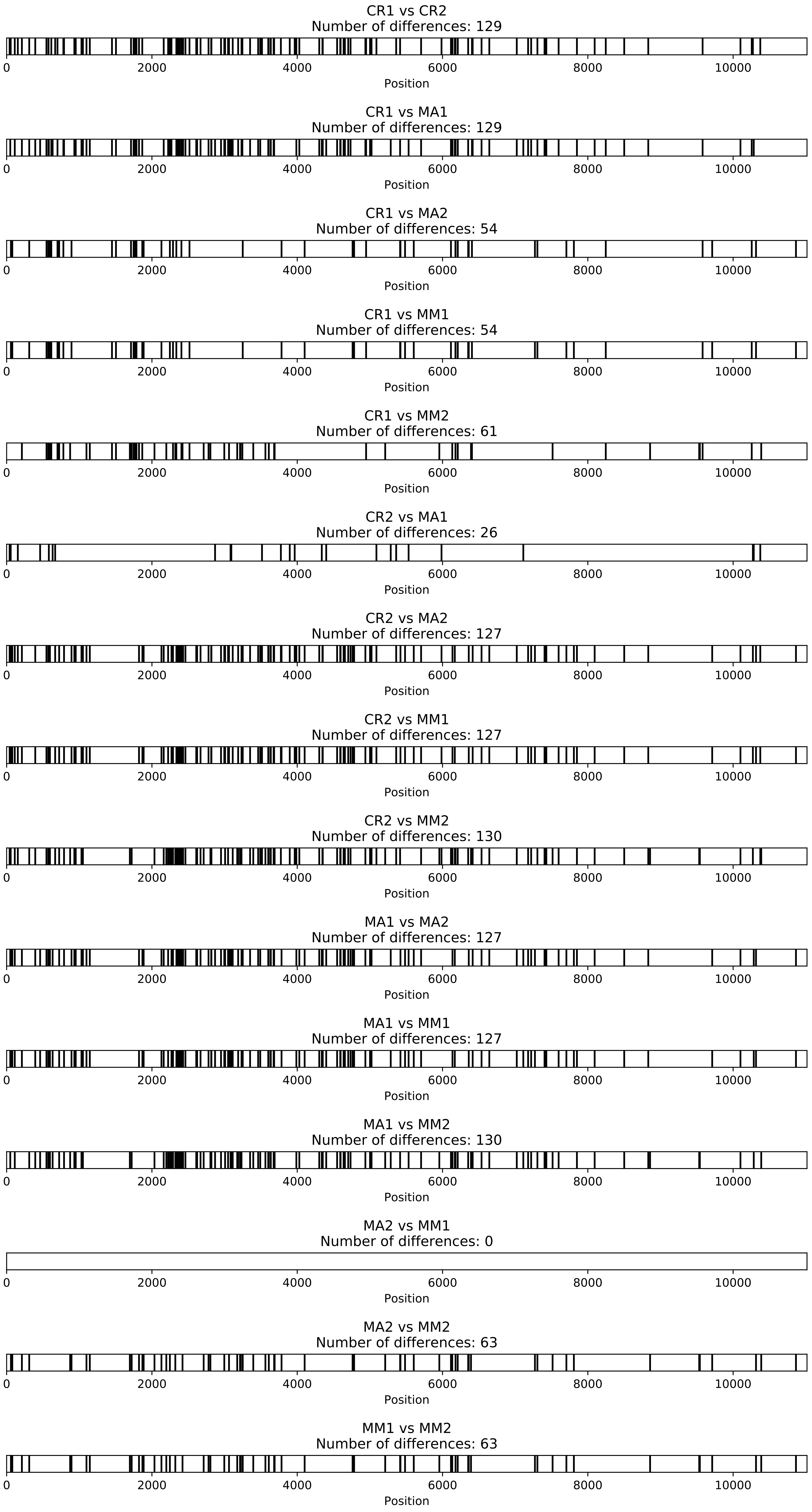
CR1 vs CR2
Number of differences: 129
0 2000 4000 6000 8000 10000
Position
CR1 vs MA1
Number of differences: 129
0 2000 4000 6000 8000 10000
Position
CR1 vs MA2
Number of differences: 54
0 2000 4000 6000 8000 10000
Position
CR1 vs MM1
Number of differences: 54
0 2000 4000 6000 8000 10000
Position
CR1 vs MM2
Number of differences: 61
0 2000 4000 6000 8000 10000
Position
CR2 vs MA1
Number of differences: 26
0 2000 4000 6000 8000 10000
Position
CR2 vs MA2
Number of differences: 127
0 2000 4000 6000 8000 10000
Position
CR2 vs MM1
Number of differences: 127
0 2000 4000 6000 8000 10000
Position
CR2 vs MM2
Number of differences: 130
0 2000 4000 6000 8000 10000
Position
MA1 vs MA2
Number of differences: 127
0 2000 4000 6000 8000 10000
Position
MA1 vs MM1
Number of differences: 127
0 2000 4000 6000 8000 10000
Position
MA1 vs MM2
Number of differences: 130
0 2000 4000 6000 8000 10000
Position
MA2 vs MM1
Number of differences: 0
0 2000 4000 6000 8000 10000
Position
MA2 vs MM2
Number of differences: 63
0 2000 4000 6000 8000 10000
Position
MM1 vs MM2
Number of differences: 63
0 2000 4000 6000 8000 10000
Position

### Region: 129



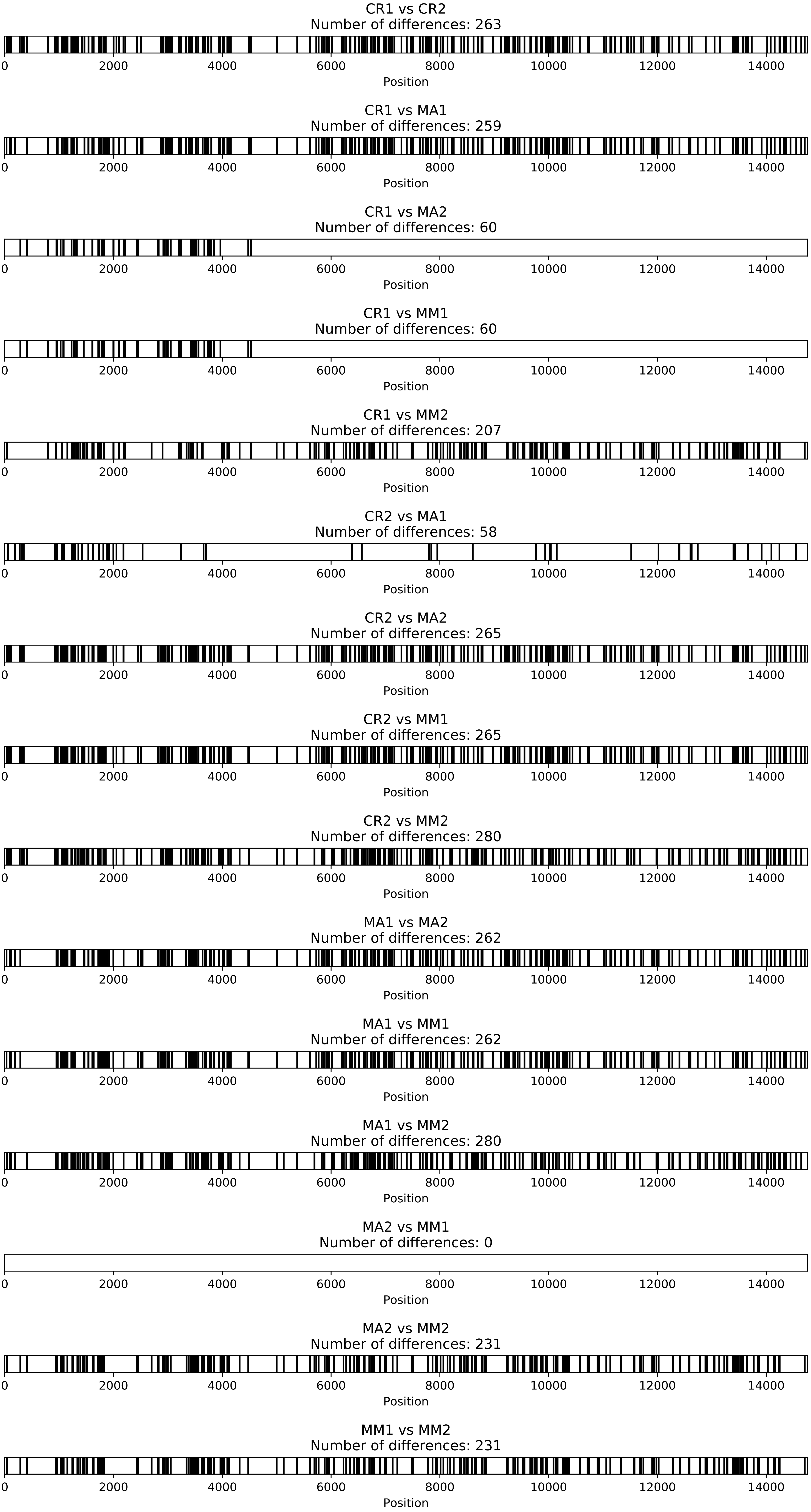
CR1 vs CR2
Number of differences: 263
CR1 vs MA1
Number of differences: 259
CR1 vs MA2
Number of differences: 60
CR1 vs MM1
Number of differences: 60
CR1 vs MM2
Number of differences: 207
CR2 vs MA1
Number of differences: 58
CR2 vs MA2
Number of differences: 265
CR2 vs MM1
Number of differences: 265
CR2 vs MM2
Number of differences: 280
MA1 vs MA2
Number of differences: 262
MA1 vs MM1
Number of differences: 262
MA1 vs MM2
Number of differences: 280
MA2 vs MM1
Number of differences: 0
MA2 vs MM2
Number of differences: 231
MM1 vs MM2
Number of differences: 231
0
2000
4000
6000
8000
10000
12000
14000
Position

### Region: 131

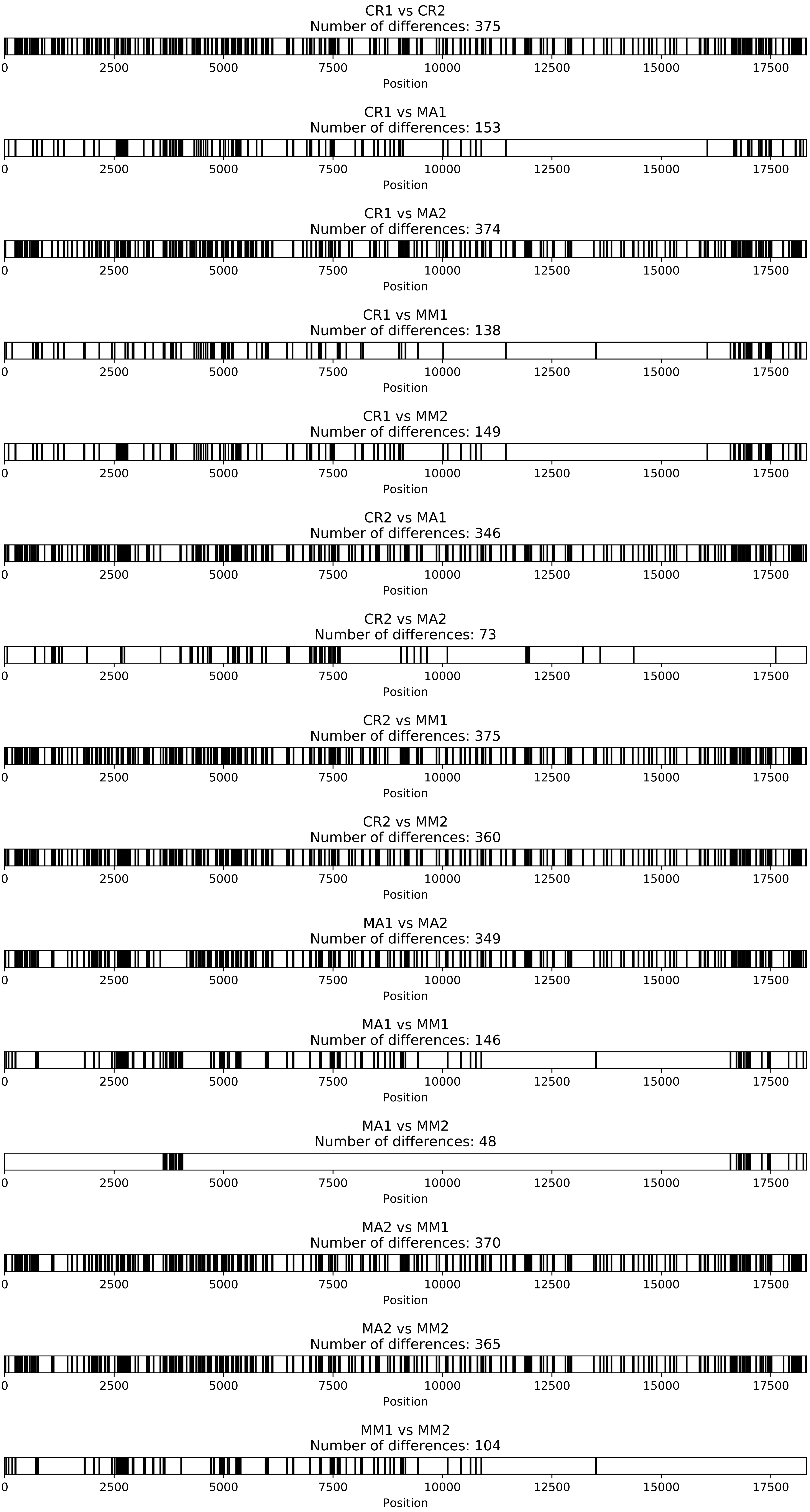
CR1 vs CR2
Number of differences: 375
CR1 vs MA1
Number of differences: 153
CR1 vs MA2
Number of differences: 374
CR1 vs MM1
Number of differences: 138
CR1 vs MM2
Number of differences: 149
CR2 vs MA1
Number of differences: 346
CR2 vs MA2
Number of differences: 73
CR2 vs MM1
Number of differences: 375
CR2 vs MM2
Number of differences: 360
MA1 vs MA2
Number of differences: 349
MA1 vs MM1
Number of differences: 146
MA1 vs MM2
Number of differences: 48
MA2 vs MM1
Number of differences: 370
MA2 vs MM2
Number of differences: 365
MM1 vs MM2
Number of differences: 104
0
2500
5000
7500
10000
12500
15000
17500
Position

### Region: 132



CR1 vs CR2
Number of differences: 353

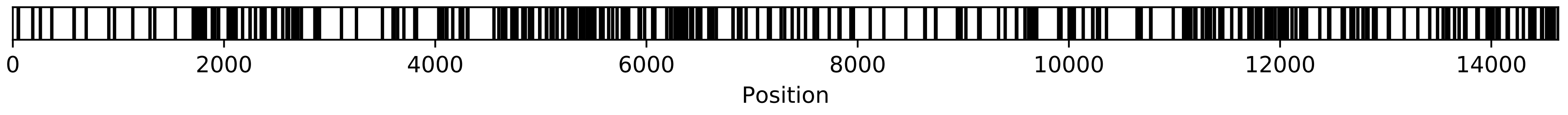


CR1 vs MA1
Number of differences: 186

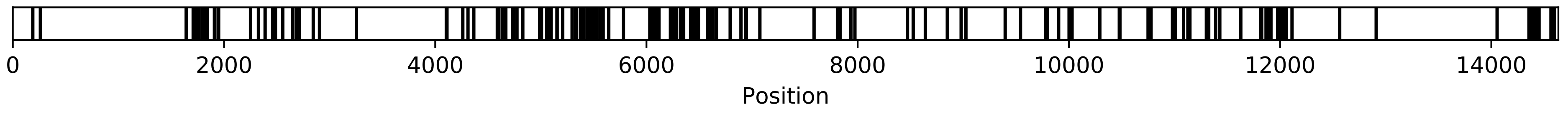


CR1 vs MA2
Number of differences: 353

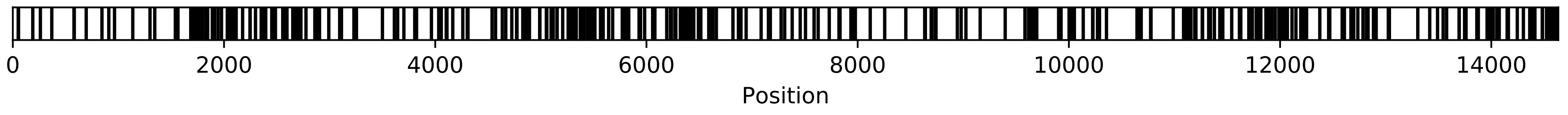


CR1 vs MM1
Number of differences: 186

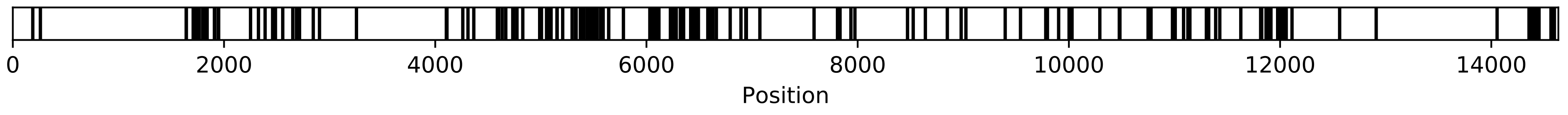


CR1 vs MM2
Number of differences: 169

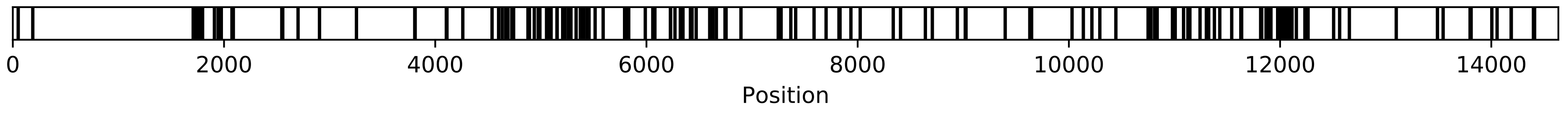


CR2 vs MA1
Number of differences: 324

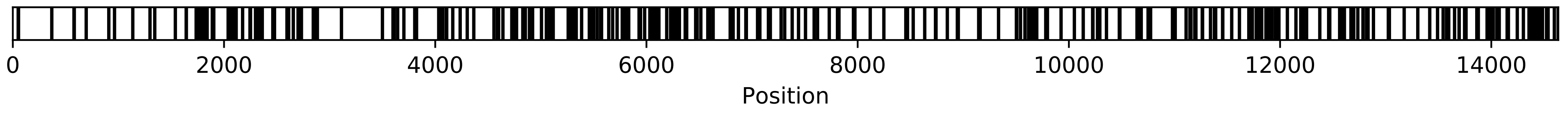


CR2 vs MA2
Number of differences: 83

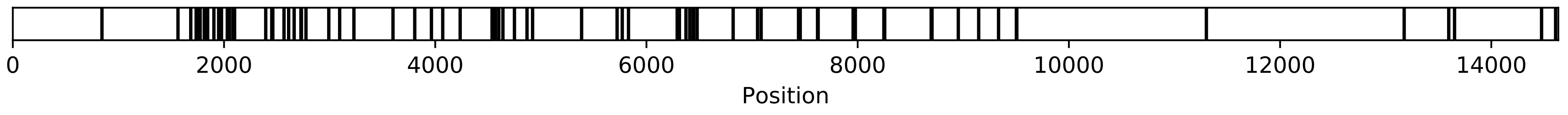


CR2 vs MM1
Number of differences: 324

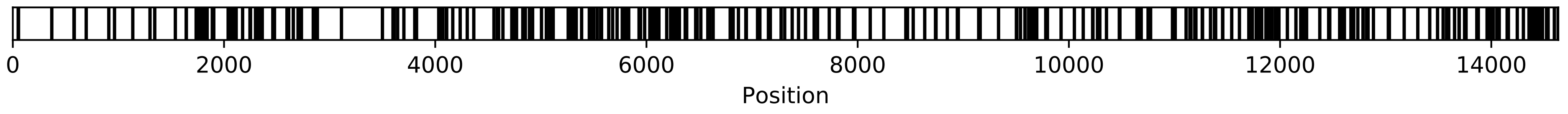


CR2 vs MM2
Number of differences: 339

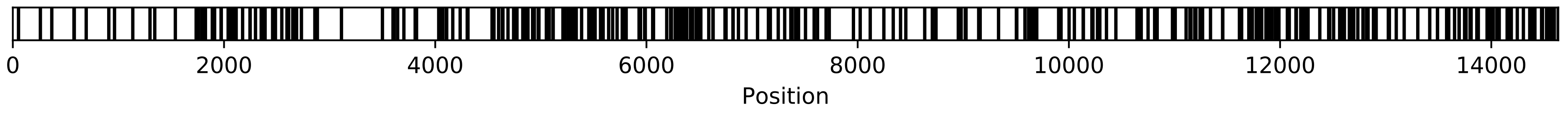


MA1 vs MA2
Number of differences: 333

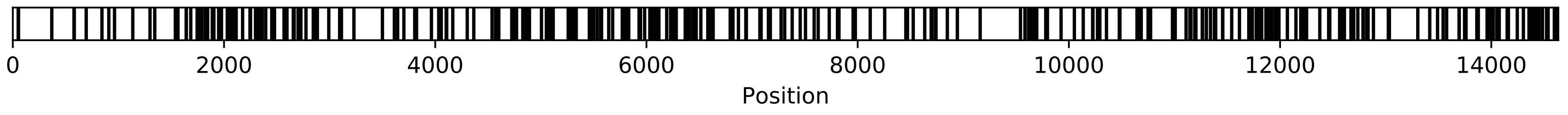


MA1 vs MM1
Number of differences: 0

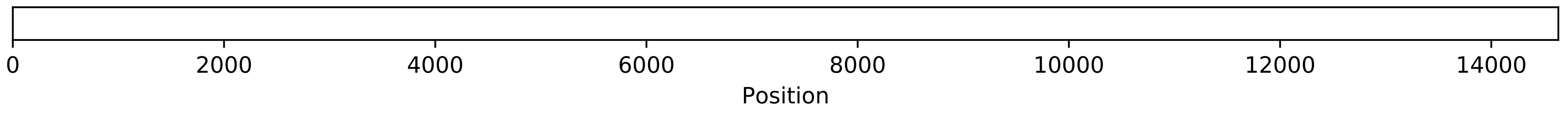


MA1 vs MM2
Number of differences: 184

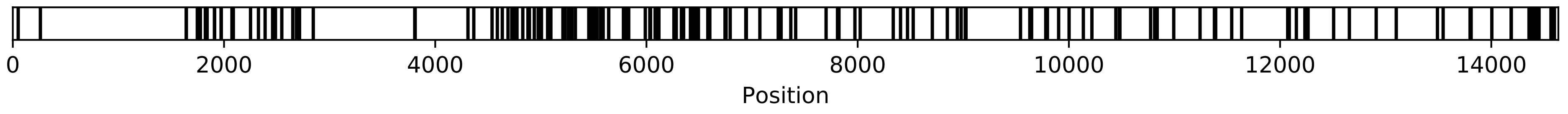


MA2 vs MM1
Number of differences: 333

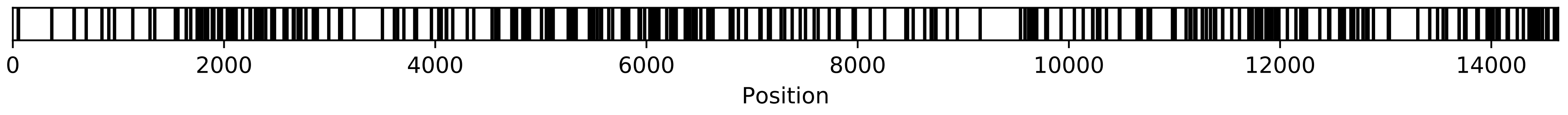


MA2 vs MM2
Number of differences: 345

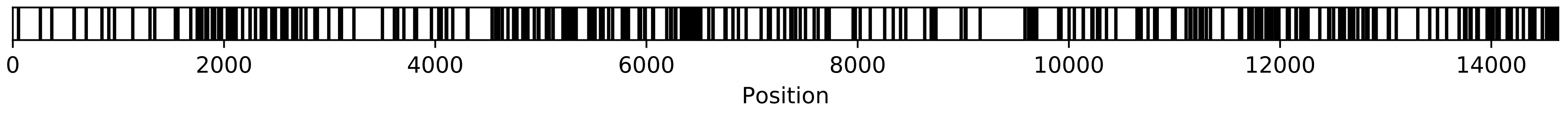


MM1 vs MM2
Number of differences: 184

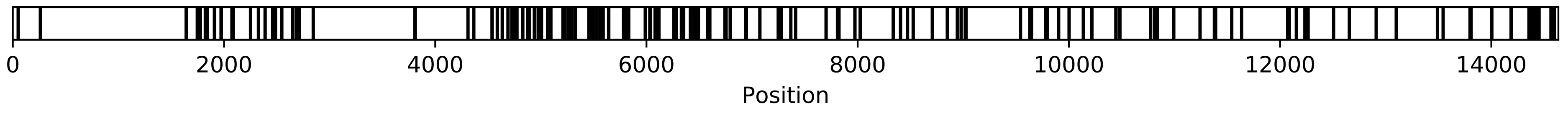

### Region: 133

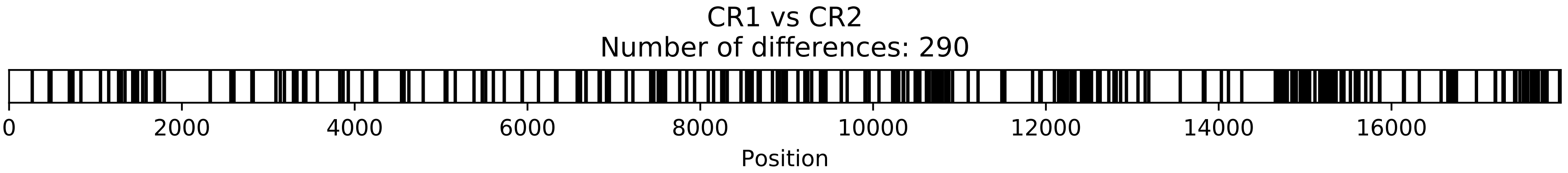
CR1 vs CR2
Number of differences: 290
0
2000
4000
6000
8000
10000
12000
14000
16000
Position

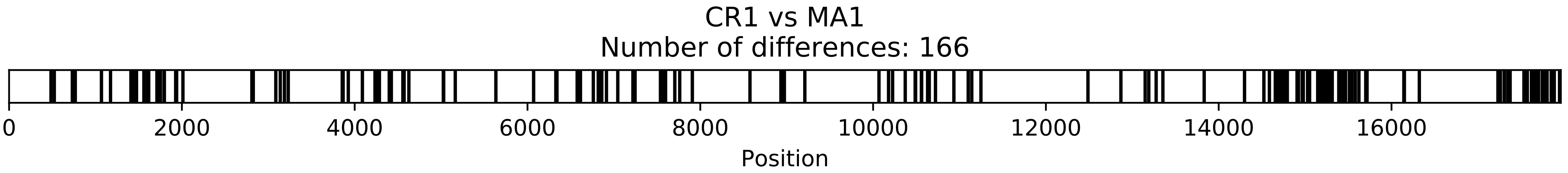
CR1 vs MA1
Number of differences: 166
0
2000
4000
6000
8000
10000
12000
14000
16000
Position

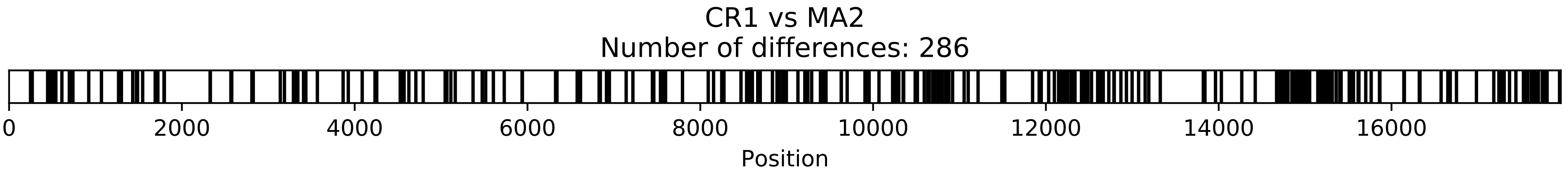
CR1 vs MA2
Number of differences: 286
0
2000
4000
6000
8000
10000
12000
14000
16000
Position

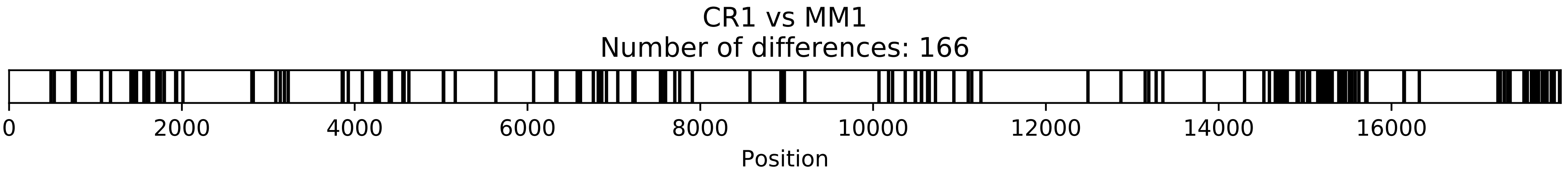
CR1 vs MM1
Number of differences: 166
0
2000
4000
6000
8000
10000
12000
14000
16000
Position

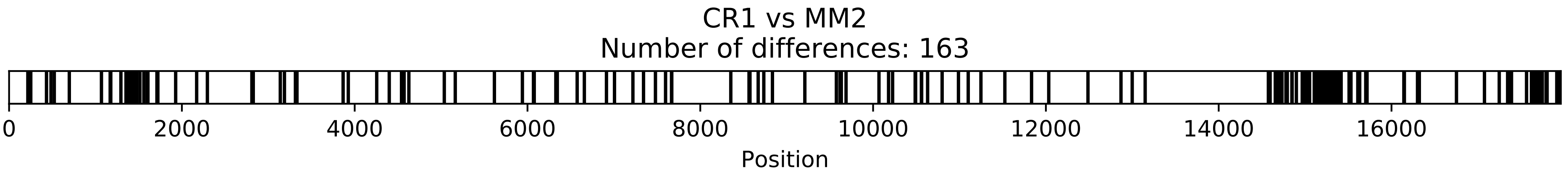
CR1 vs MM2
Number of differences: 163
0
2000
4000
6000
8000
10000
12000
14000
16000
Position

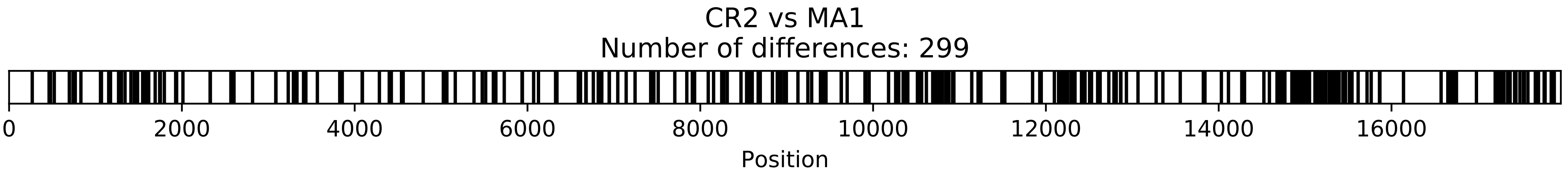
CR2 vs MA1
Number of differences: 299
0
2000
4000
6000
8000
10000
12000
14000
16000
Position

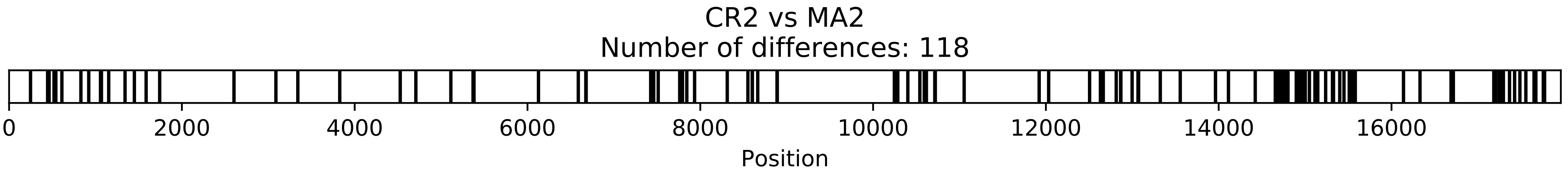
CR2 vs MA2
Number of differences: 118
0
2000
4000
6000
8000
10000
12000
14000
16000
Position

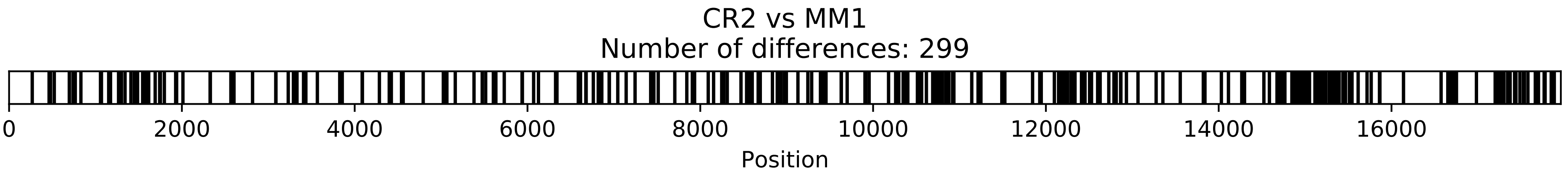
CR2 vs MM1
Number of differences: 299
0
2000
4000
6000
8000
10000
12000
14000
16000
Position

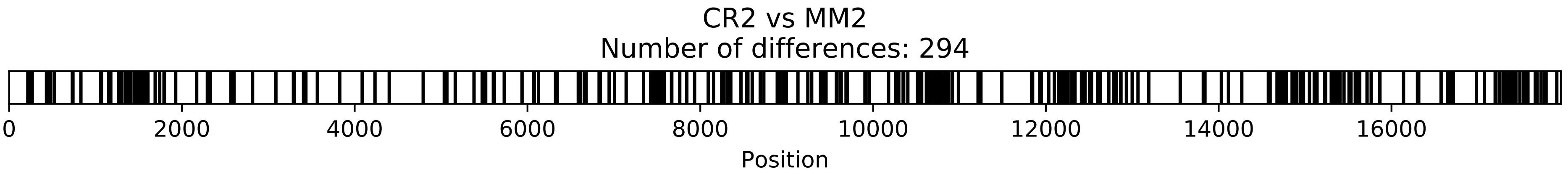
CR2 vs MM2
Number of differences: 294
0
2000
4000
6000
8000
10000
12000
14000
16000
Position

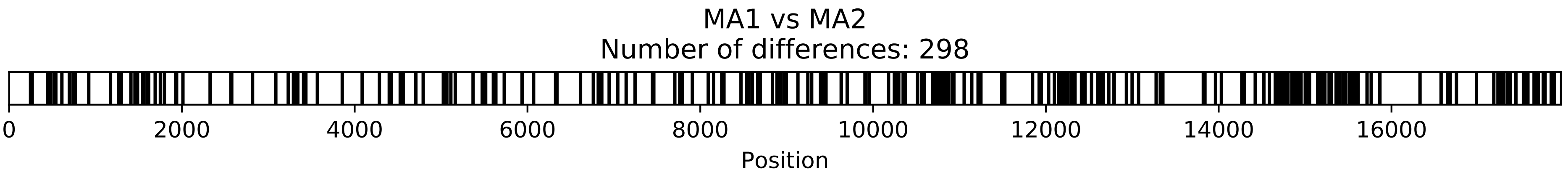
MA1 vs MA2
Number of differences: 298
0
2000
4000
6000
8000
10000
12000
14000
16000
Position

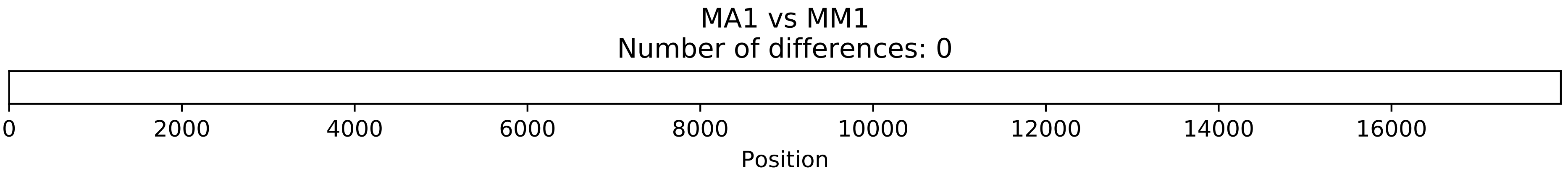
MA1 vs MM1
Number of differences: 0
0
2000
4000
6000
8000
10000
12000
14000
16000
Position

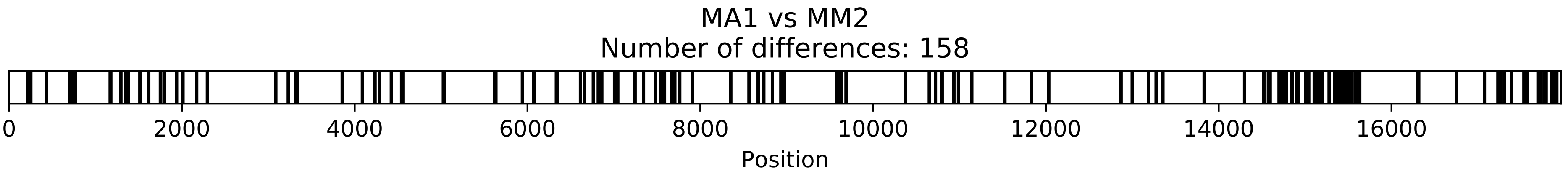
MA1 vs MM2
Number of differences: 158
0
2000
4000
6000
8000
10000
12000
14000
16000
Position

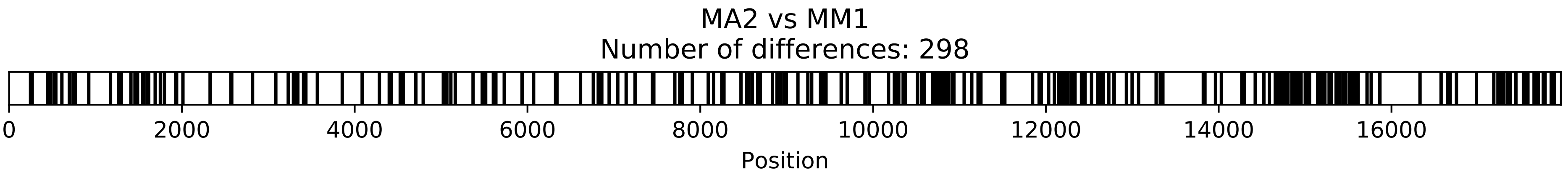
MA2 vs MM1
Number of differences: 298
0
2000
4000
6000
8000
10000
12000
14000
16000
Position

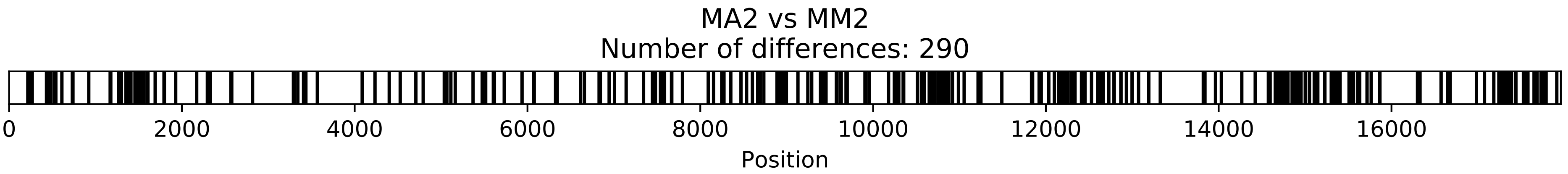
MA2 vs MM2
Number of differences: 290
0
2000
4000
6000
8000
10000
12000
14000
16000
Position

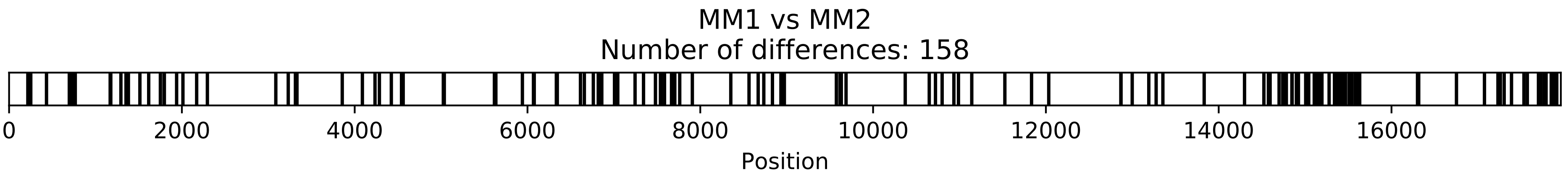
MM1 vs MM2
Number of differences: 158
0
2000
4000
6000
8000
10000
12000
14000
16000
Position

### Region: 134



CR1 vs CR2
Number of differences: 314

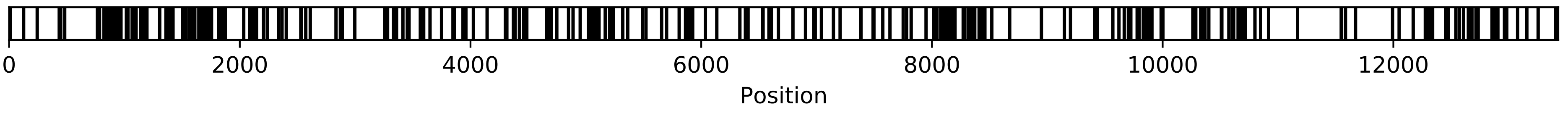


CR1 vs MA1
Number of differences: 11

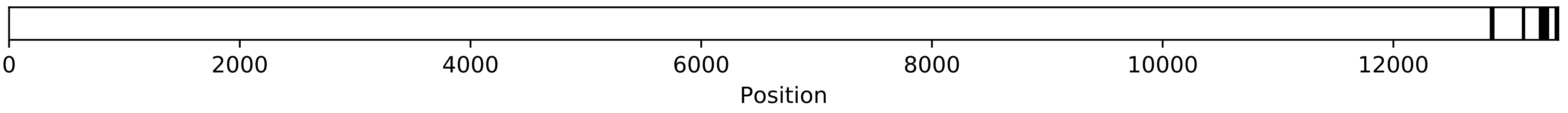


CR1 vs MA2
Number of differences: 321

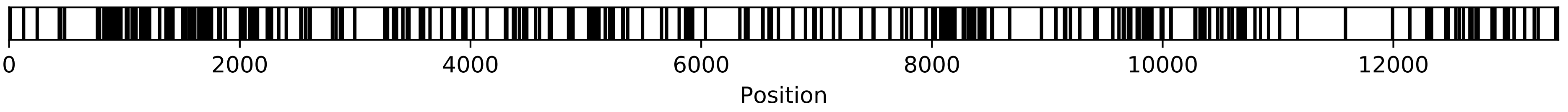


CR1 vs MM1
Number of differences: 152

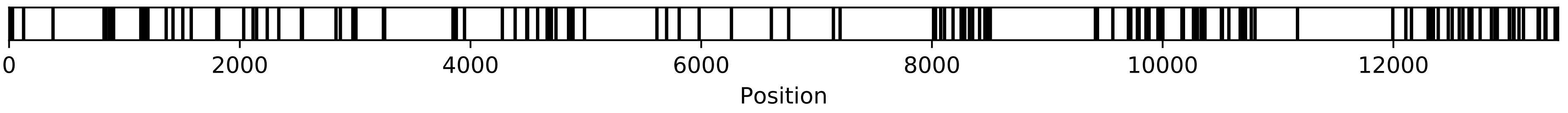


CR1 vs MM2
Number of differences: 36

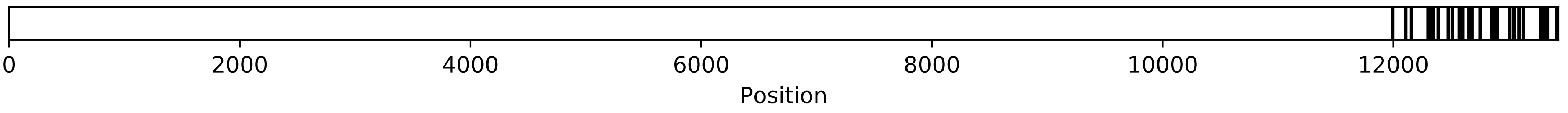


CR2 vs MA1
Number of differences: 321

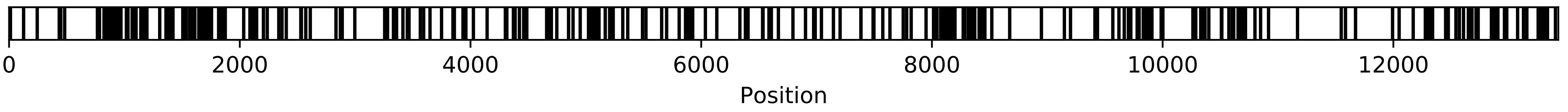


CR2 vs MA2
Number of differences: 80

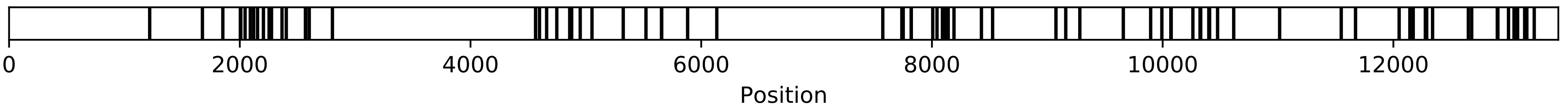


CR2 vs MM1
Number of differences: 298

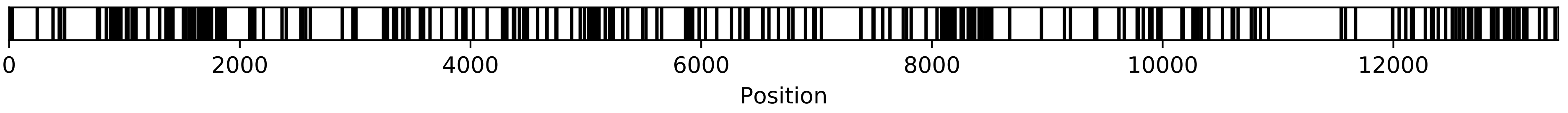


CR2 vs MM2
Number of differences: 326

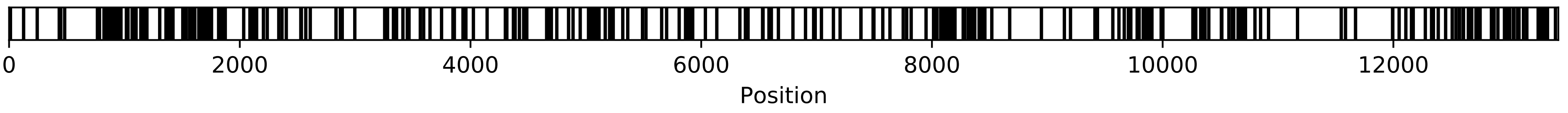


MA1 vs MA2
Number of differences: 328

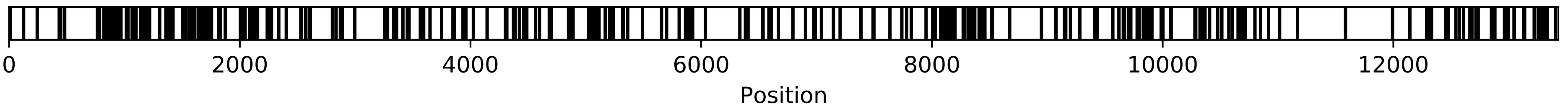


MA1 vs MM1
Number of differences: 149

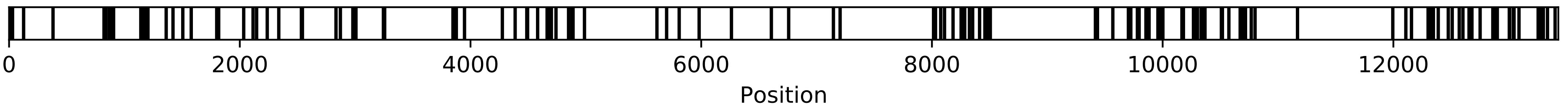


MA1 vs MM2
Number of differences: 27

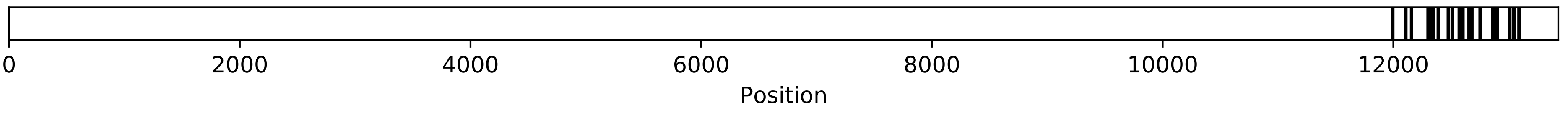


MA2 vs MM1
Number of differences: 303

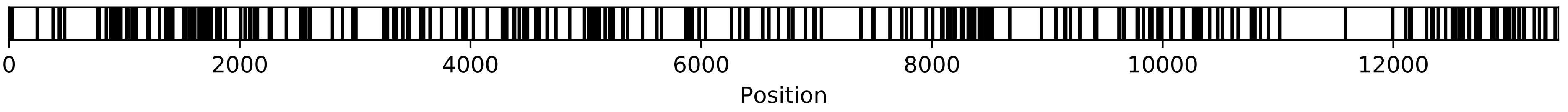


MA2 vs MM2
Number of differences: 333

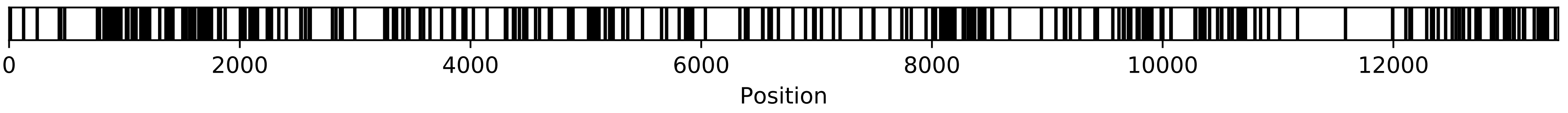


MM1 vs MM2
Number of differences: 122

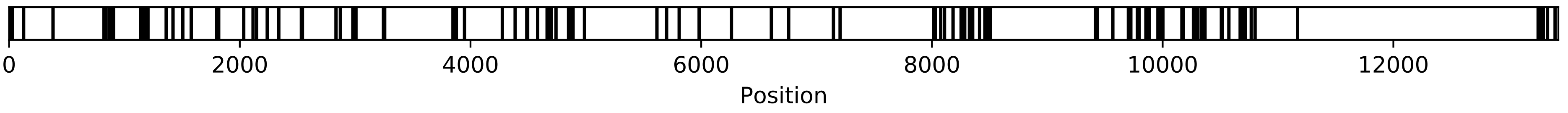

### Region: 137

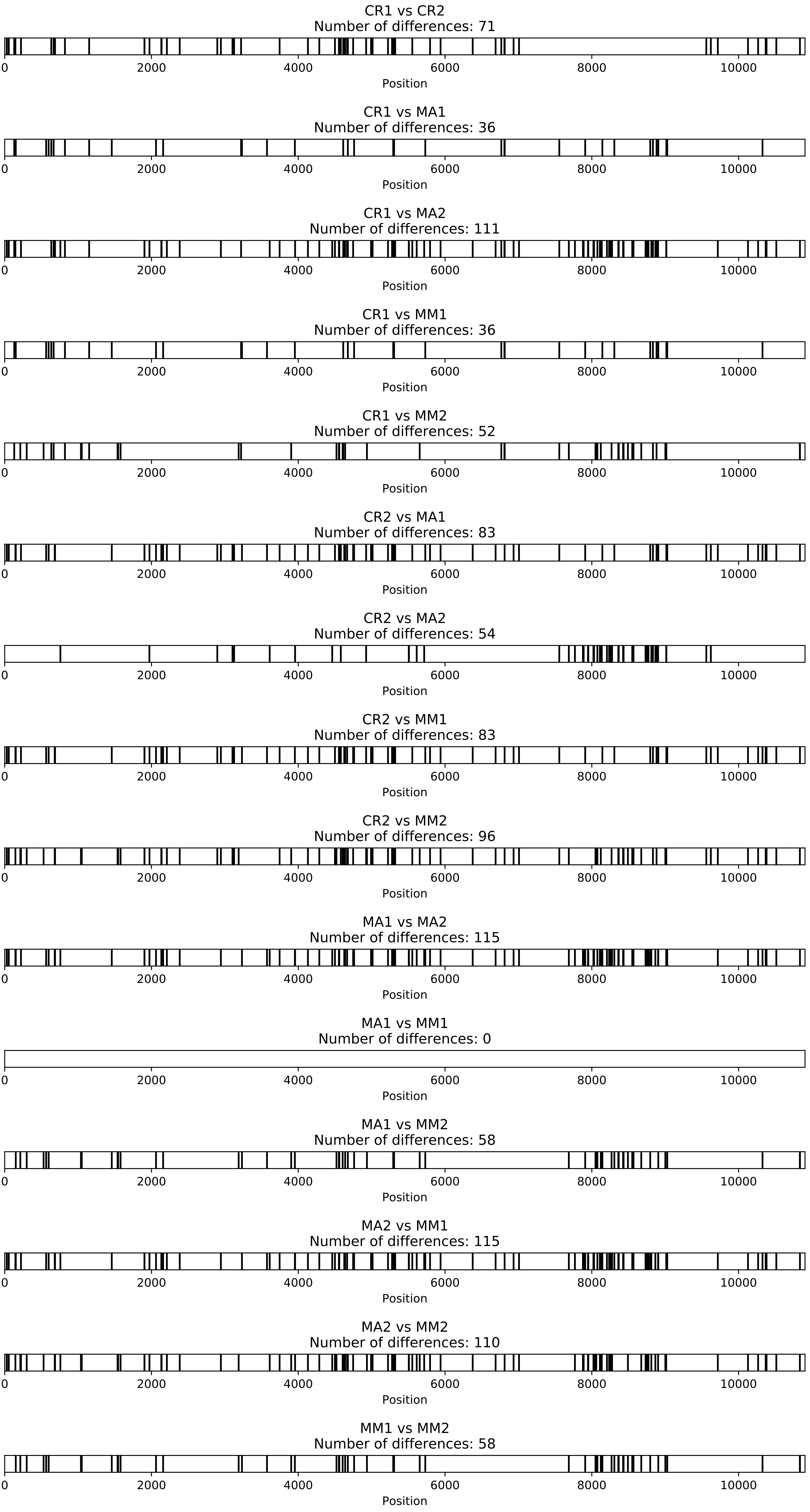
CR1 vs CR2
Number of differences: 71
CR1 vs MA1
Number of differences: 36
CR1 vs MA2
Number of differences: 111
CR1 vs MM1
Number of differences: 36
CR1 vs MM2
Number of differences: 52
CR2 vs MA1
Number of differences: 83
CR2 vs MA2
Number of differences: 54
CR2 vs MM1
Number of differences: 83
CR2 vs MM2
Number of differences: 96
MA1 vs MA2
Number of differences: 115
MA1 vs MM1
Number of differences: 0
MA1 vs MM2
Number of differences: 58
MA2 vs MM1
Number of differences: 115
MA2 vs MM2
Number of differences: 110
MM1 vs MM2
Number of differences: 58
0
2000
4000
6000
8000
10000
Position

### Region: 138



CR1 vs CR2
Number of differences: 292

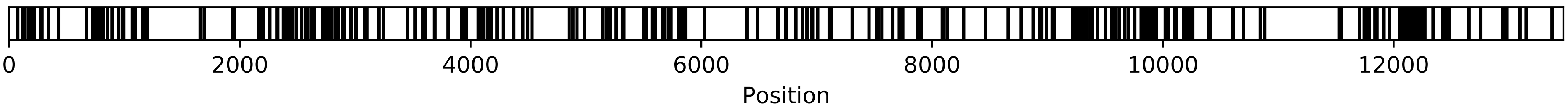


CR1 vs MA1
Number of differences: 150

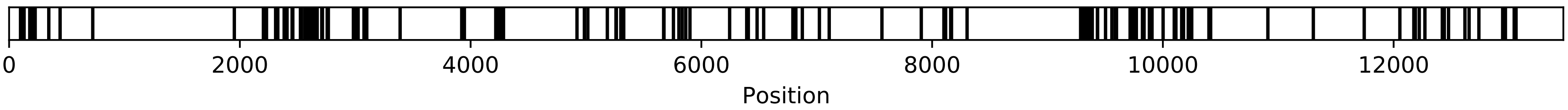


CR1 vs MA2
Number of differences: 298

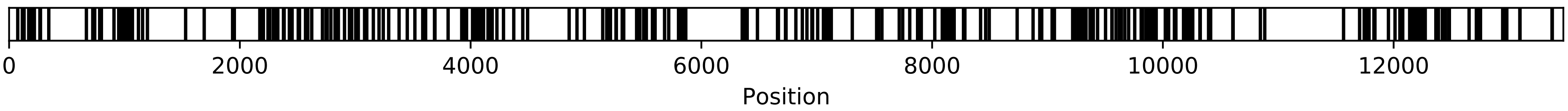


CR1 vs MM1
Number of differences: 298

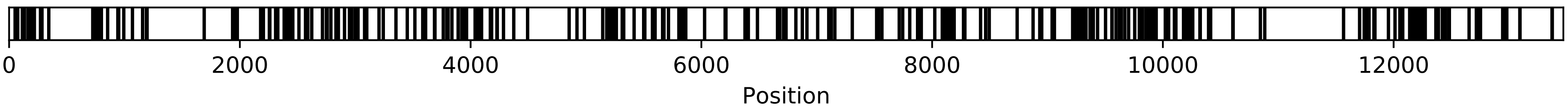


CR1 vs MM2
Number of differences: 298

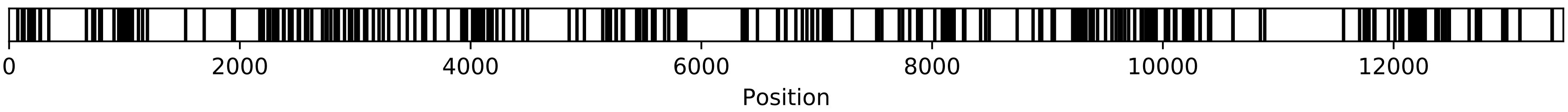


CR2 vs MA1
Number of differences: 295

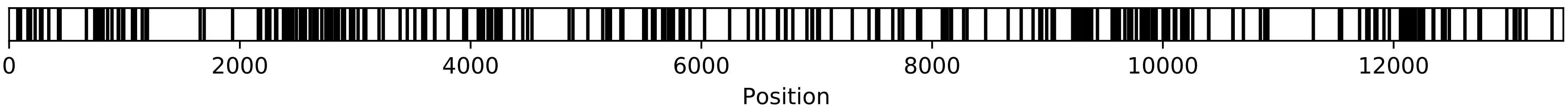


CR2 vs MA2
Number of differences: 166

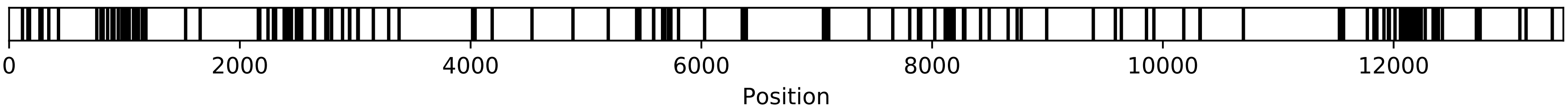


CR2 vs MM1
Number of differences: 174

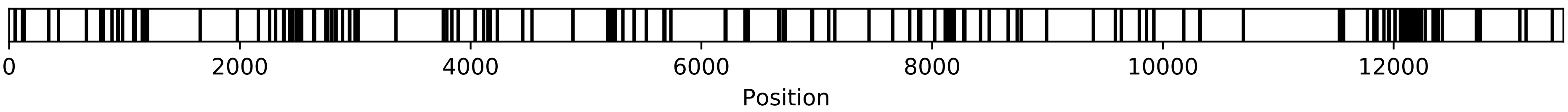


CR2 vs MM2
Number of differences: 166

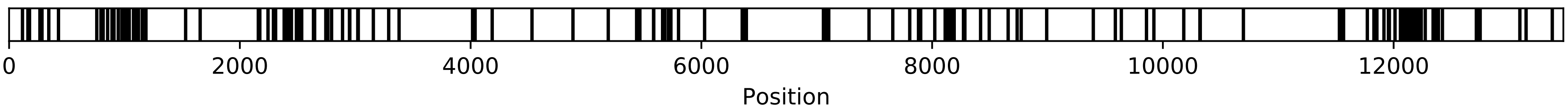


MA1 vs MA2
Number of differences: 297

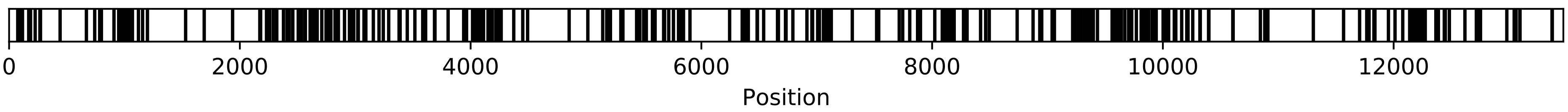


MA1 vs MM1
Number of differences: 291

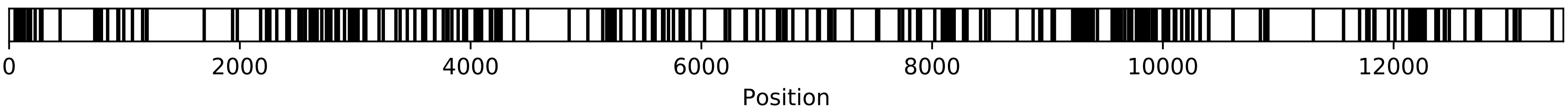


MA1 vs MM2
Number of differences: 297

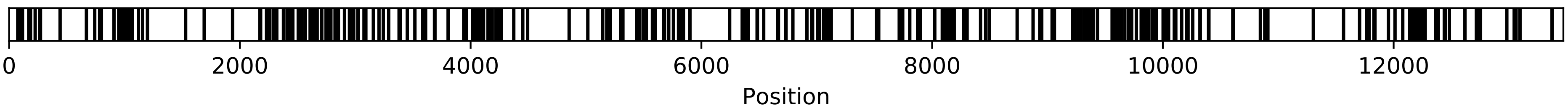


MA2 vs MM1
Number of differences: 96

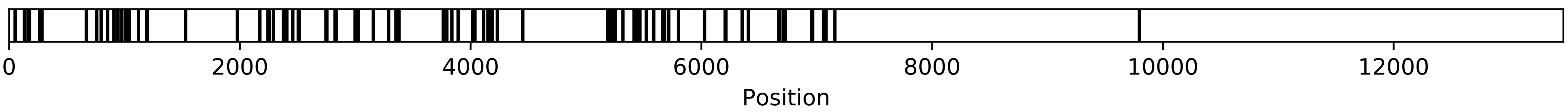


MA2 vs MM2
Number of differences: 0

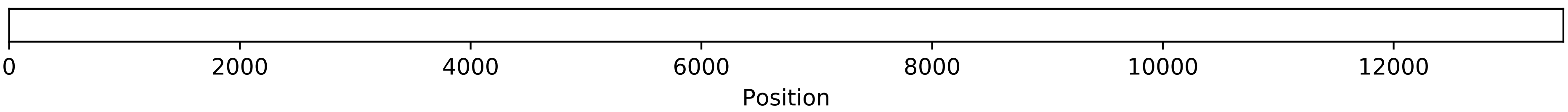


MM1 vs MM2
Number of differences: 96

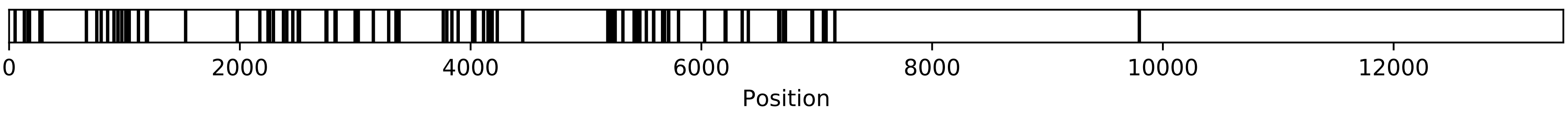

### Region: 139

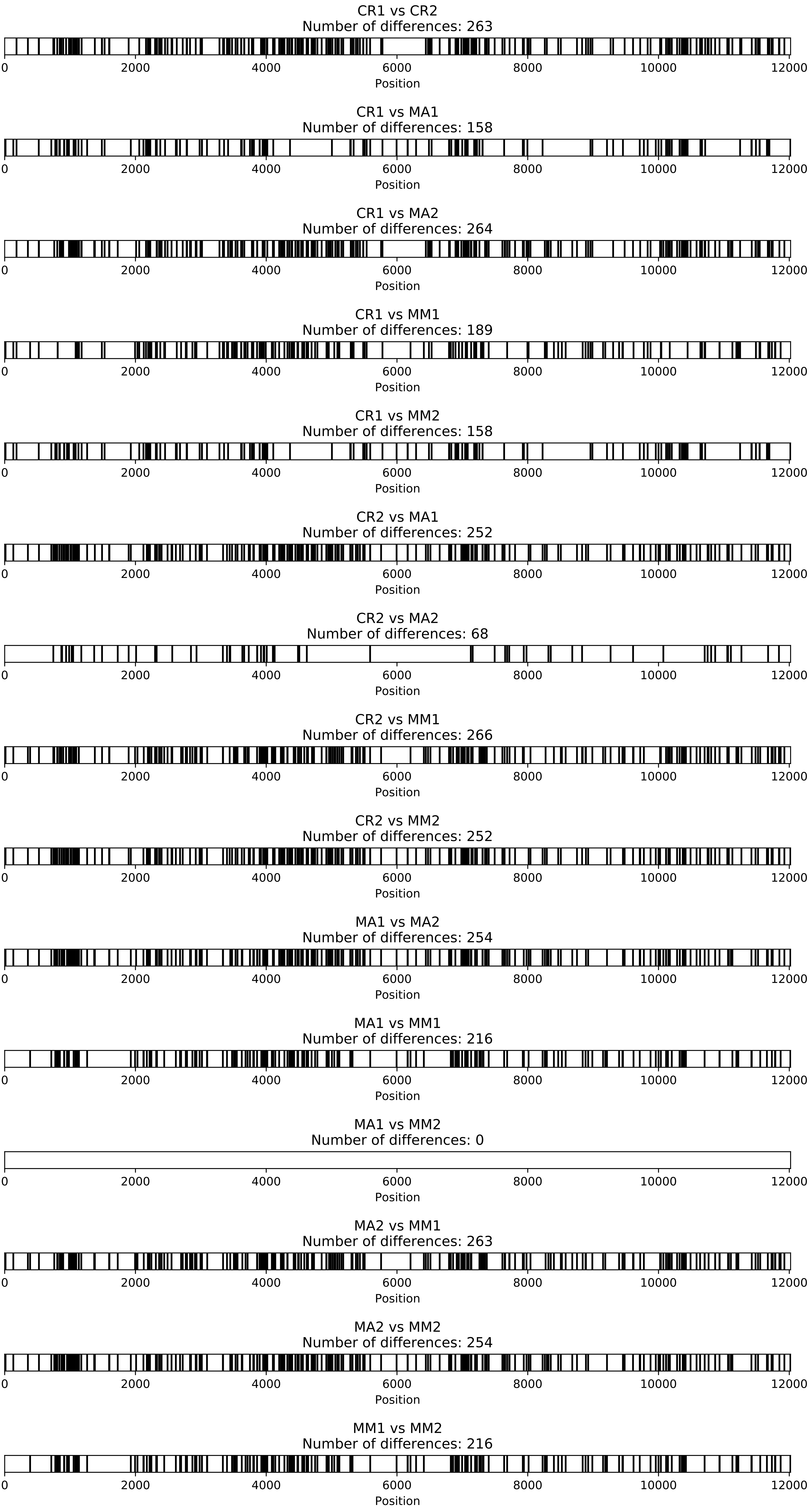
CR1 vs CR2
Number of differences: 263
CR1 vs MA1
Number of differences: 158
CR1 vs MA2
Number of differences: 264
CR1 vs MM1
Number of differences: 189
CR1 vs MM2
Number of differences: 158
CR2 vs MA1
Number of differences: 252
CR2 vs MA2
Number of differences: 68
CR2 vs MM1
Number of differences: 266
CR2 vs MM2
Number of differences: 252
MA1 vs MA2
Number of differences: 254
MA1 vs MM1
Number of differences: 216
MA1 vs MM2
Number of differences: 0
MA2 vs MM1
Number of differences: 263
MA2 vs MM2
Number of differences: 254
MM1 vs MM2
Number of differences: 216
0
2000
4000
6000
8000
10000
12000
Position

### Region: 140

CR1 vs CR2
Number of differences: 301

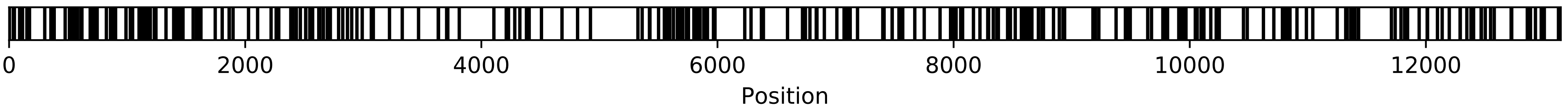


CR1 vs MA1
Number of differences: 177

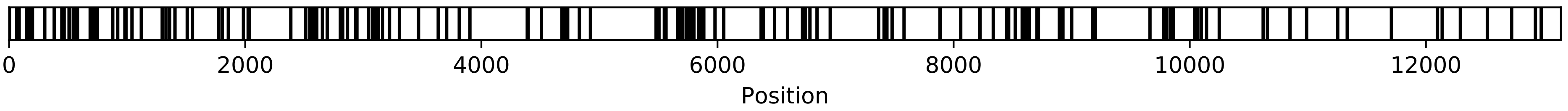


CR1 vs MA2
Number of differences: 306

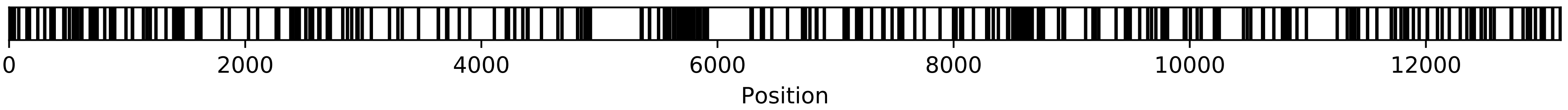


CR1 vs MM1
Number of differences: 178

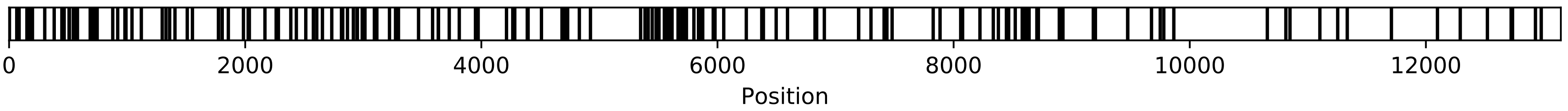


CR1 vs MM2
Number of differences: 155

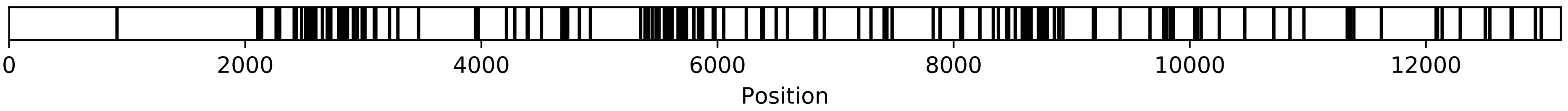


CR2 vs MA1
Number of differences: 300

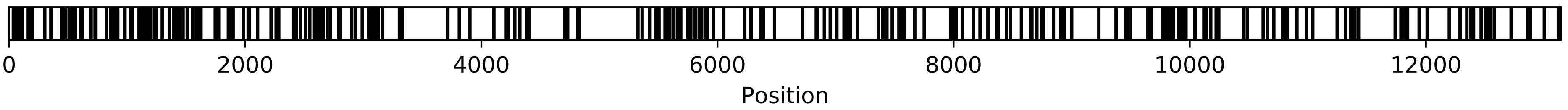


CR2 vs MA2
Number of differences: 146

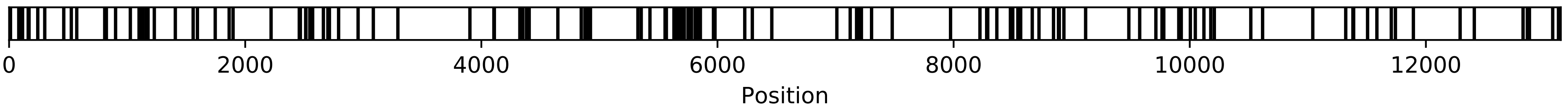


CR2 vs MM1
Number of differences: 302

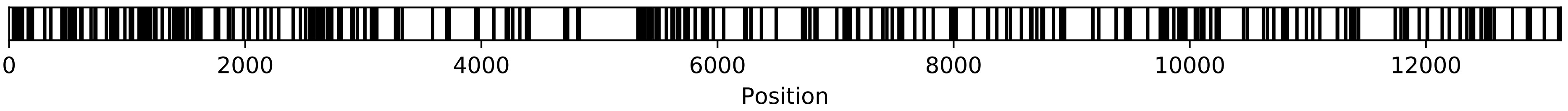


CR2 vs MM2
Number of differences: 304

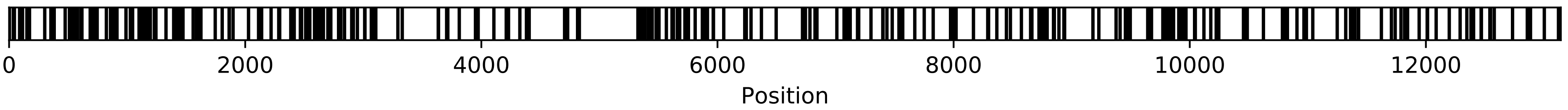


MA1 vs MA2
Number of differences: 302

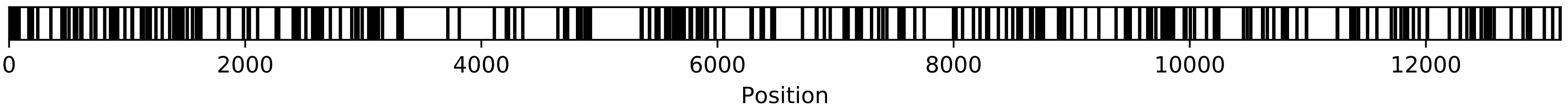


MA1 vs MM1
Number of differences: 117

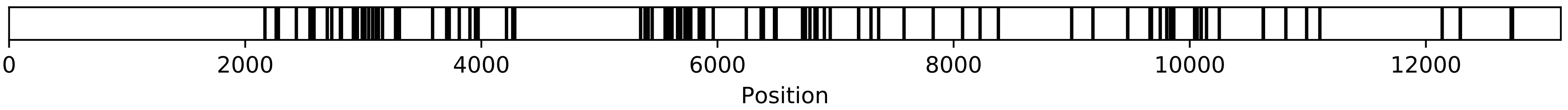


MA1 vs MM2
Number of differences: 179

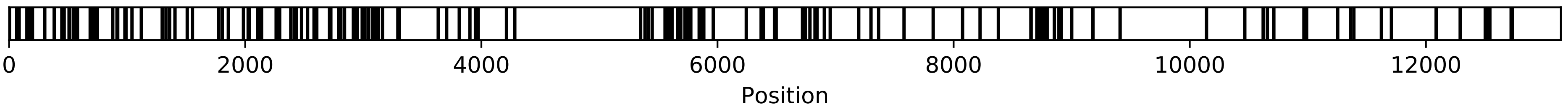


MA2 vs MM1
Number of differences: 309

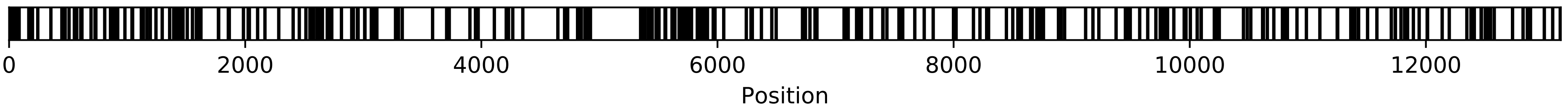


MA2 vs MM2
Number of differences: 311

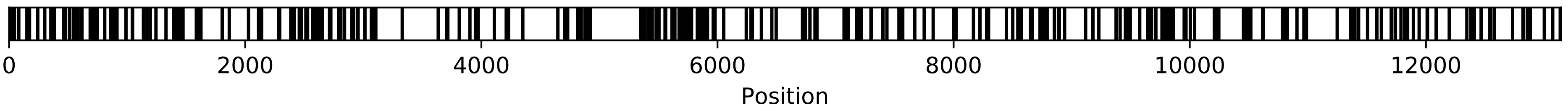


MM1 vs MM2
Number of differences: 127

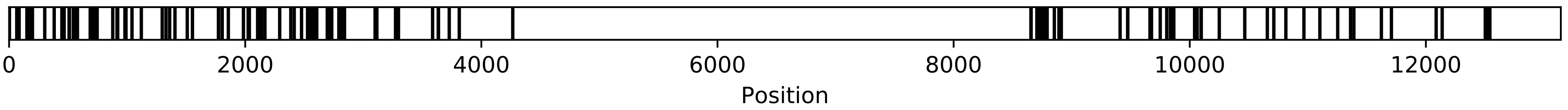

### Region: 141

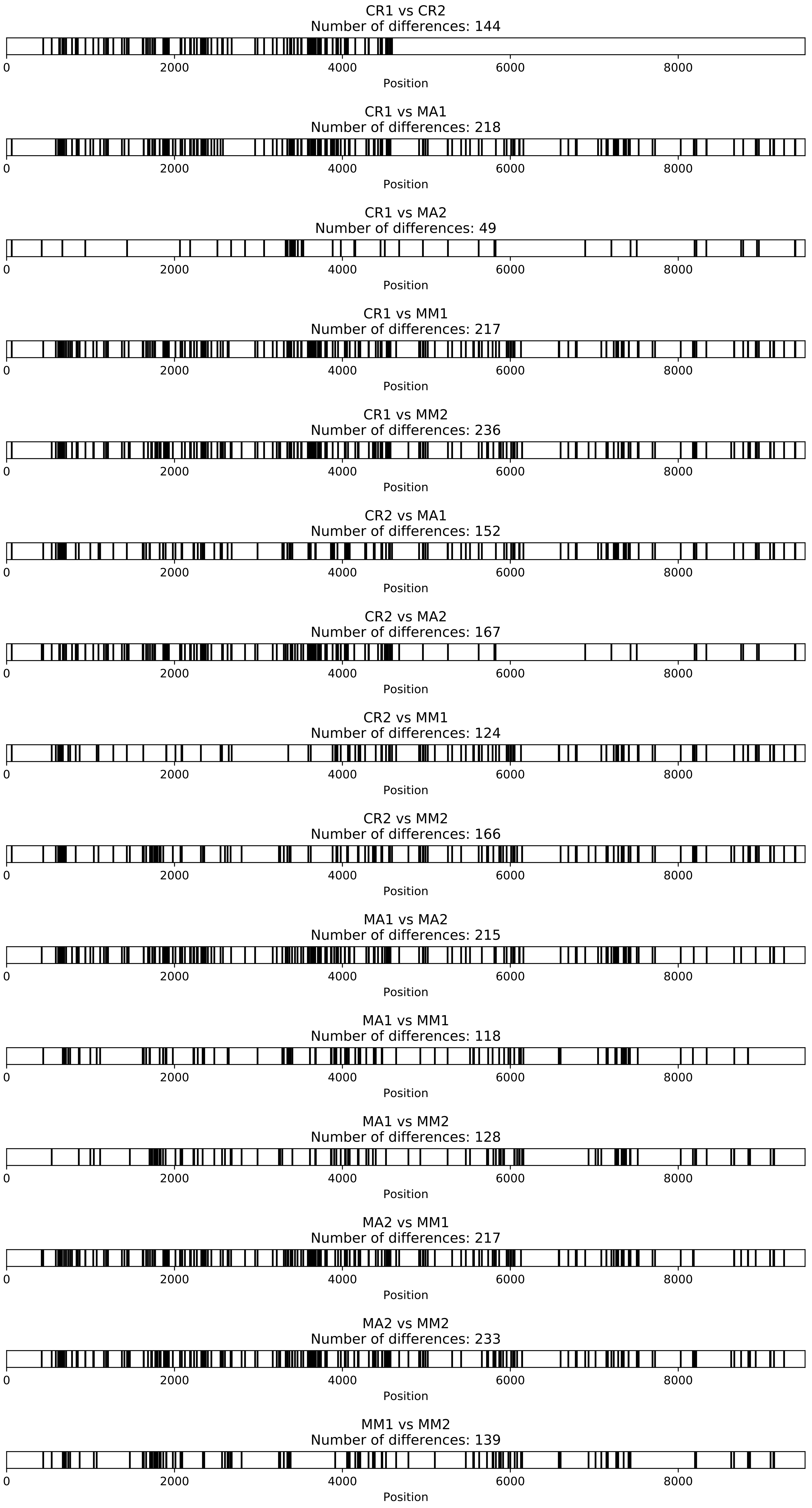
CR1 vs CR2
Number of differences: 144
CR1 vs MA1
Number of differences: 218
CR1 vs MA2
Number of differences: 49
CR1 vs MM1
Number of differences: 217
CR1 vs MM2
Number of differences: 236
CR2 vs MA1
Number of differences: 152
CR2 vs MA2
Number of differences: 167
CR2 vs MM1
Number of differences: 124
CR2 vs MM2
Number of differences: 166
MA1 vs MA2
Number of differences: 215
MA1 vs MM1
Number of differences: 118
MA1 vs MM2
Number of differences: 128
MA2 vs MM1
Number of differences: 217
MA2 vs MM2
Number of differences: 233
MM1 vs MM2
Number of differences: 139
0
2000
4000
6000
8000
Position

### Region: 142

CR1 vs CR2
Number of differences: 182

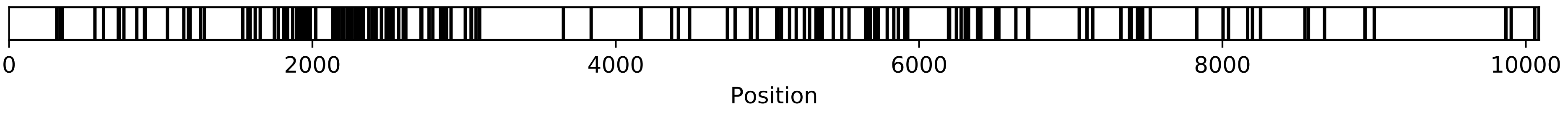


CR1 vs MA1
Number of differences: 82

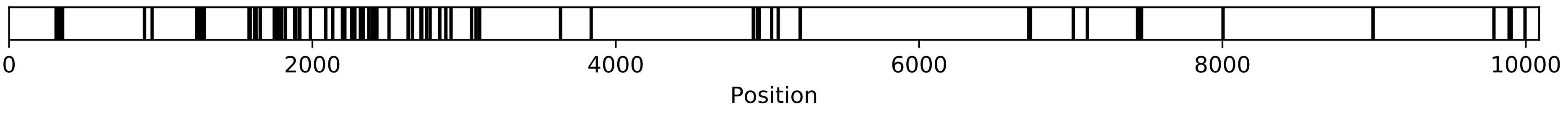


CR1 vs MA2
Number of differences: 169

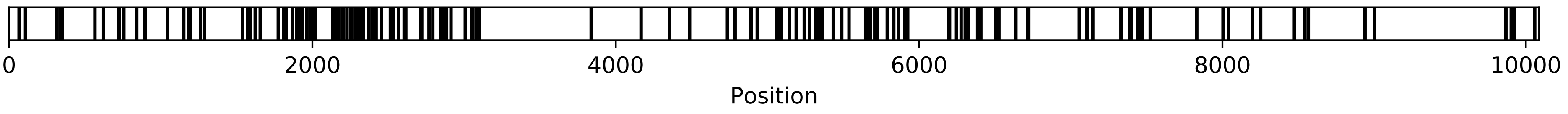


CR1 vs MM1
Number of differences: 82

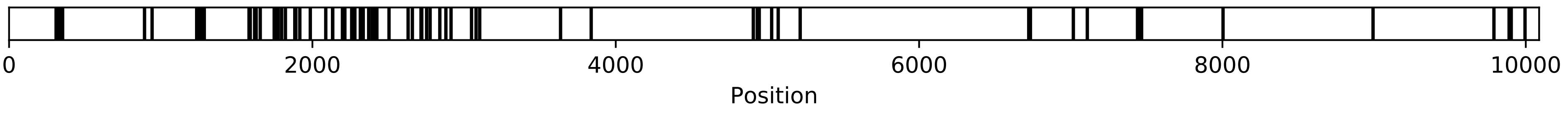


CR1 vs MM2
Number of differences: 84

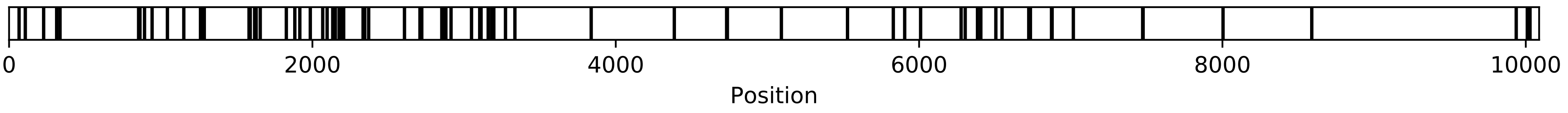


CR2 vs MA1
Number of differences: 190

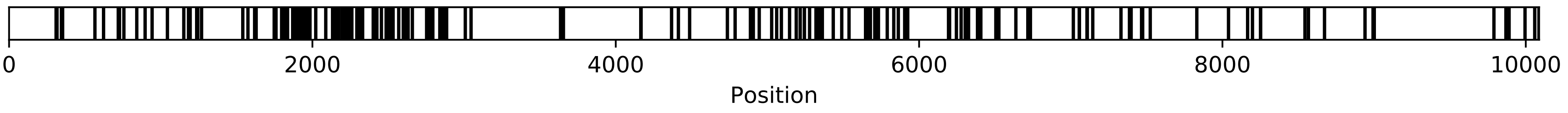


CR2 vs MA2
Number of differences: 41

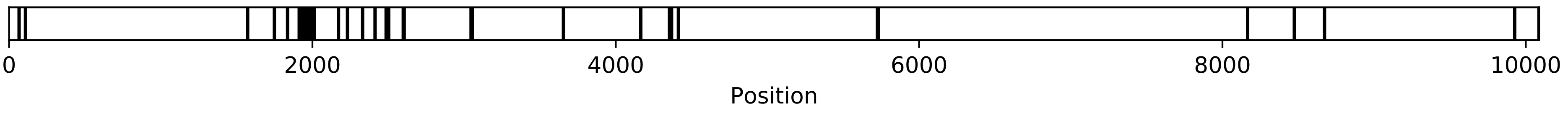


CR2 vs MM1
Number of differences: 190

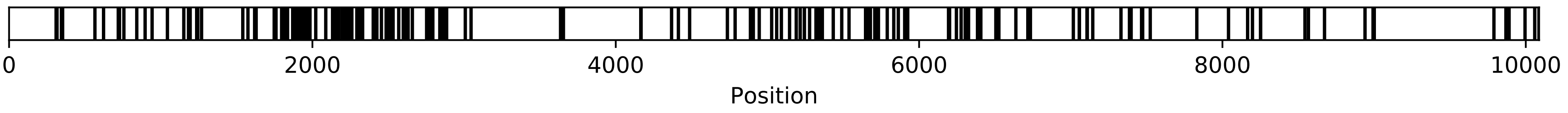


CR2 vs MM2
Number of differences: 188

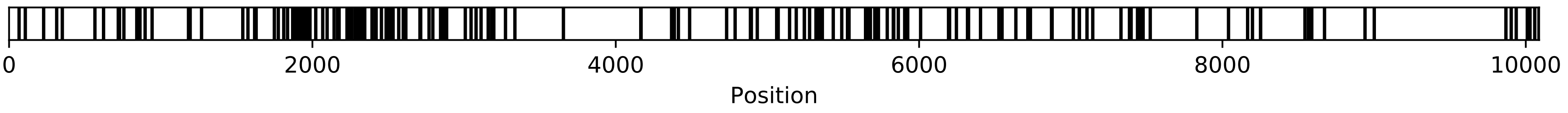


MA1 vs MA2
Number of differences: 177

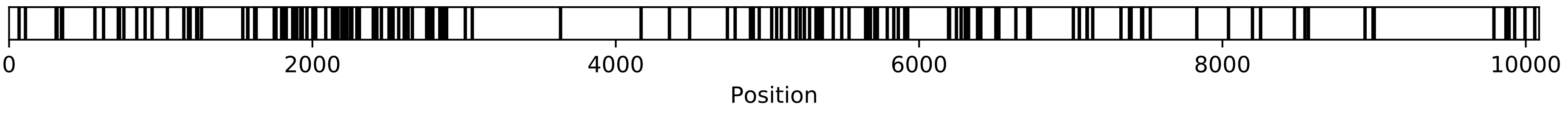


MA1 vs MM1
Number of differences: 0

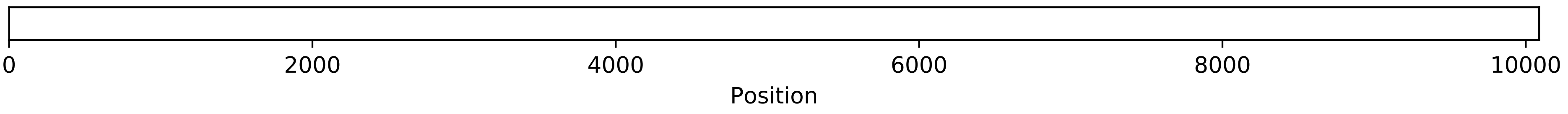


MA1 vs MM2
Number of differences: 102

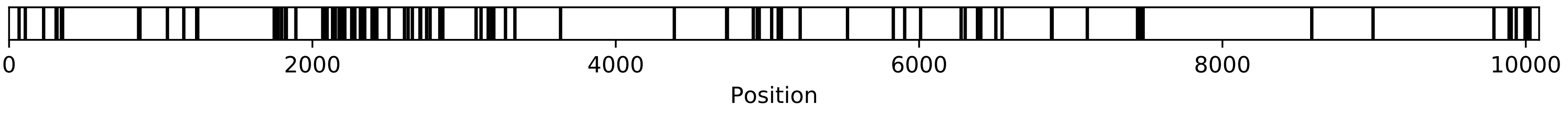


MA2 vs MM1
Number of differences: 177

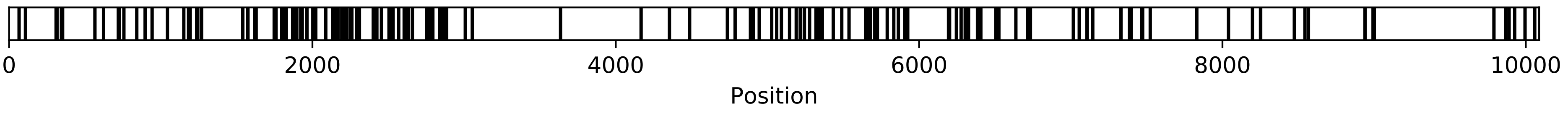


MA2 vs MM2
Number of differences: 173

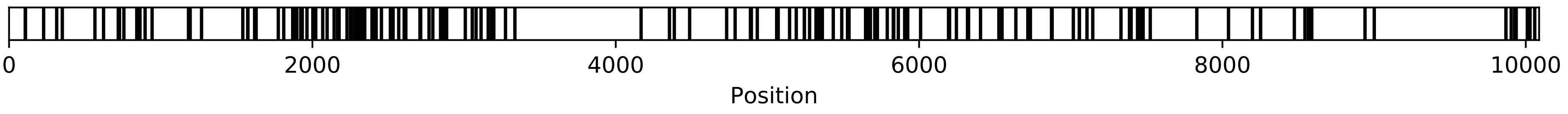


MM1 vs MM2
Number of differences: 102

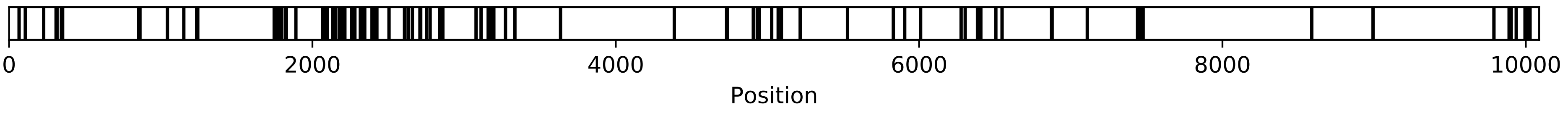

### Region: 143

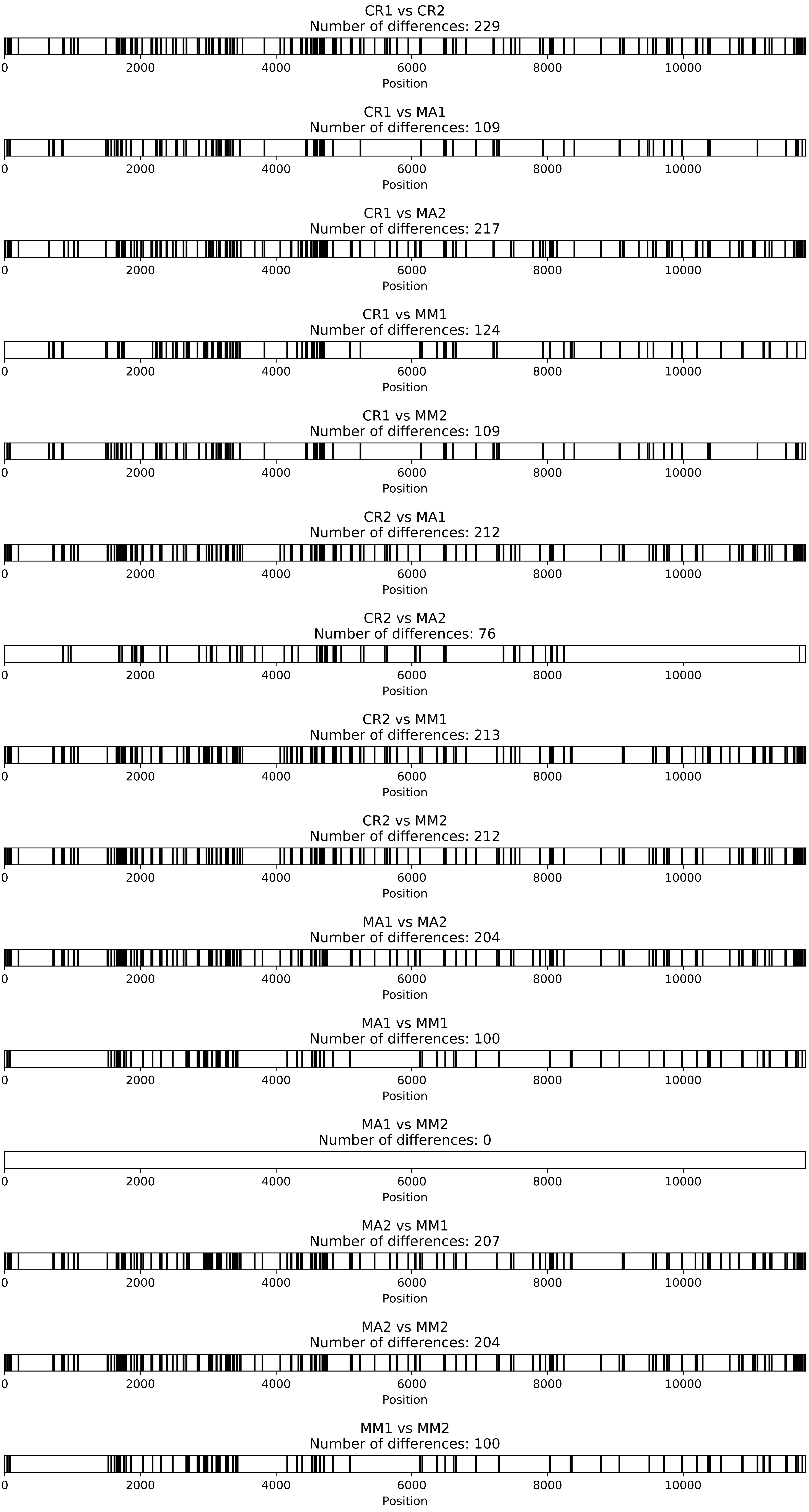
CR1 vs CR2
Number of differences: 229
CR1 vs MA1
Number of differences: 109
CR1 vs MA2
Number of differences: 217
CR1 vs MM1
Number of differences: 124
CR1 vs MM2
Number of differences: 109
CR2 vs MA1
Number of differences: 212
CR2 vs MA2
Number of differences: 76
CR2 vs MM1
Number of differences: 213
CR2 vs MM2
Number of differences: 212
MA1 vs MA2
Number of differences: 204
MA1 vs MM1
Number of differences: 100
MA1 vs MM2
Number of differences: 0
MA2 vs MM1
Number of differences: 207
MA2 vs MM2
Number of differences: 204
MM1 vs MM2
Number of differences: 100
0
2000
4000
6000
8000
10000
Position

### Region: 144

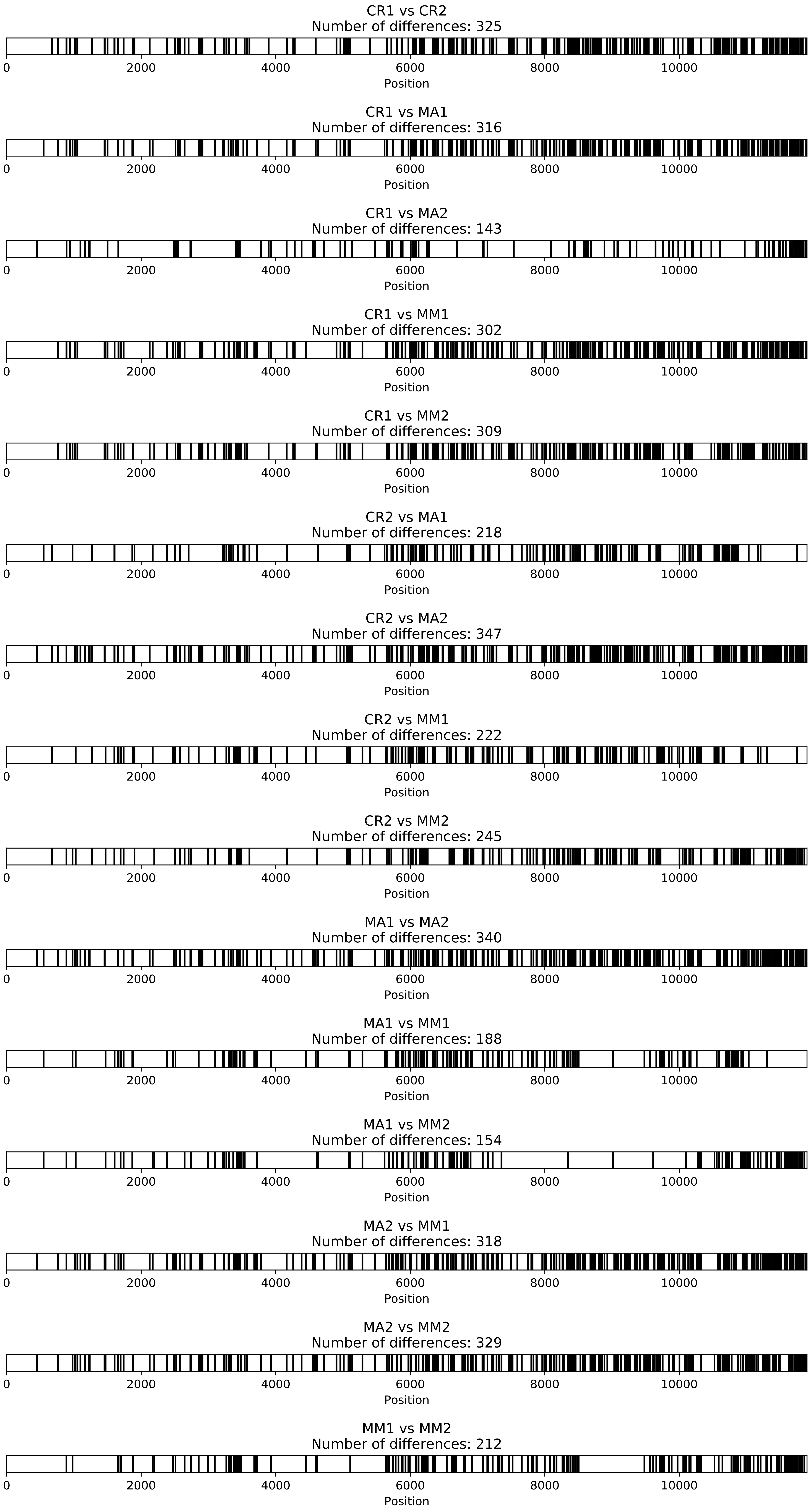
CR1 vs CR2
Number of differences: 325
0 2000 4000 6000 8000 10000
Position
CR1 vs MA1
Number of differences: 316
0 2000 4000 6000 8000 10000
Position
CR1 vs MA2
Number of differences: 143
0 2000 4000 6000 8000 10000
Position
CR1 vs MM1
Number of differences: 302
0 2000 4000 6000 8000 10000
Position
CR1 vs MM2
Number of differences: 309
0 2000 4000 6000 8000 10000
Position
CR2 vs MA1
Number of differences: 218
0 2000 4000 6000 8000 10000
Position
CR2 vs MA2
Number of differences: 347
0 2000 4000 6000 8000 10000
Position
CR2 vs MM1
Number of differences: 222
0 2000 4000 6000 8000 10000
Position
CR2 vs MM2
Number of differences: 245
0 2000 4000 6000 8000 10000
Position
MA1 vs MA2
Number of differences: 340
0 2000 4000 6000 8000 10000
Position
MA1 vs MM1
Number of differences: 188
0 2000 4000 6000 8000 10000
Position
MA1 vs MM2
Number of differences: 154
0 2000 4000 6000 8000 10000
Position
MA2 vs MM1
Number of differences: 318
0 2000 4000 6000 8000 10000
Position
MA2 vs MM2
Number of differences: 329
0 2000 4000 6000 8000 10000
Position
MM1 vs MM2
Number of differences: 212
0 2000 4000 6000 8000 10000
Position

### Region: 146



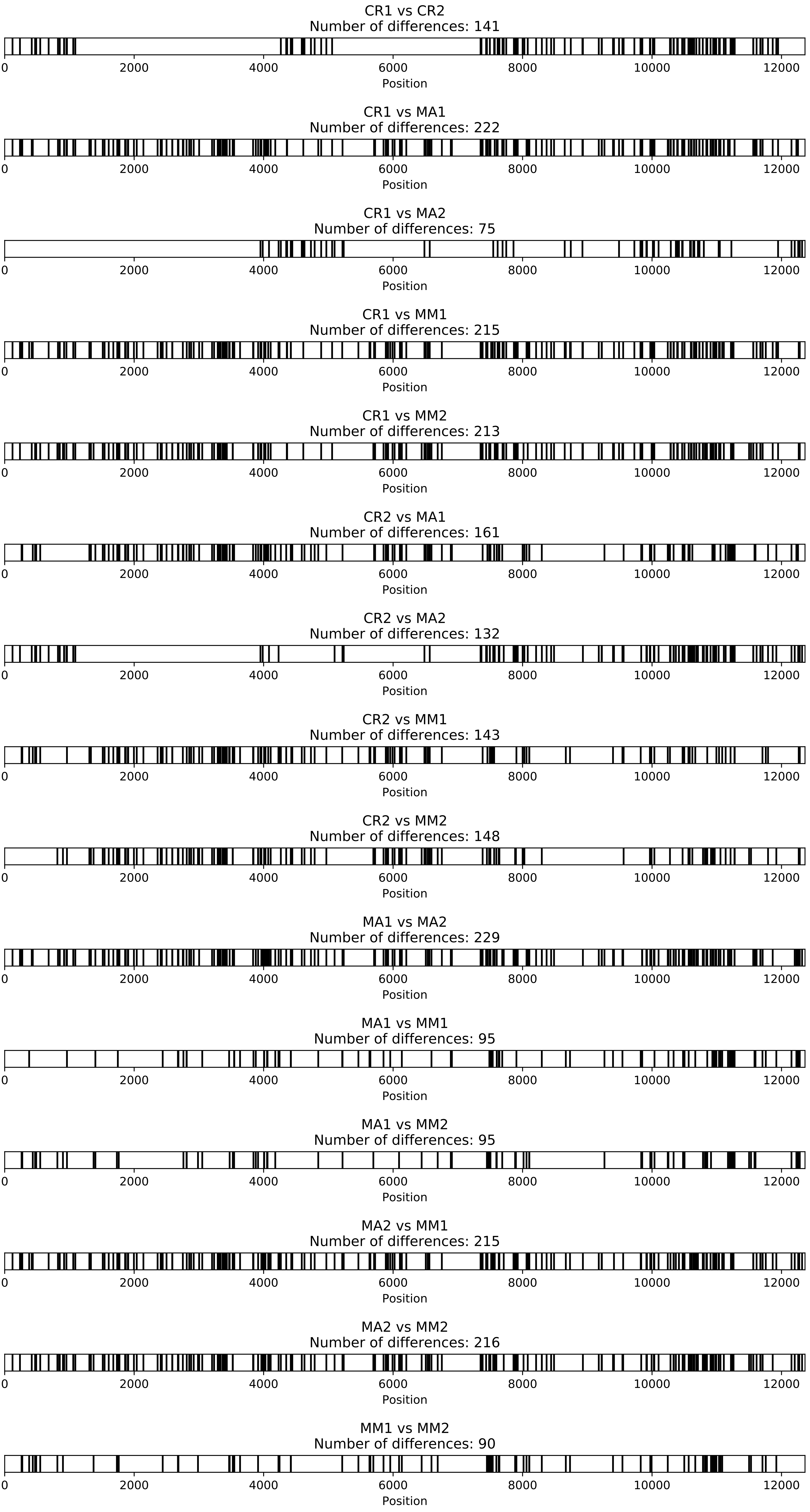
CR1 vs CR2
Number of differences: 141
CR1 vs MA1
Number of differences: 222
CR1 vs MA2
Number of differences: 75
CR1 vs MM1
Number of differences: 215
CR1 vs MM2
Number of differences: 213
CR2 vs MA1
Number of differences: 161
CR2 vs MA2
Number of differences: 132
CR2 vs MM1
Number of differences: 143
CR2 vs MM2
Number of differences: 148
MA1 vs MA2
Number of differences: 229
MA1 vs MM1
Number of differences: 95
MA1 vs MM2
Number of differences: 95
MA2 vs MM1
Number of differences: 215
MA2 vs MM2
Number of differences: 216
MM1 vs MM2
Number of differences: 90

### Region: 147



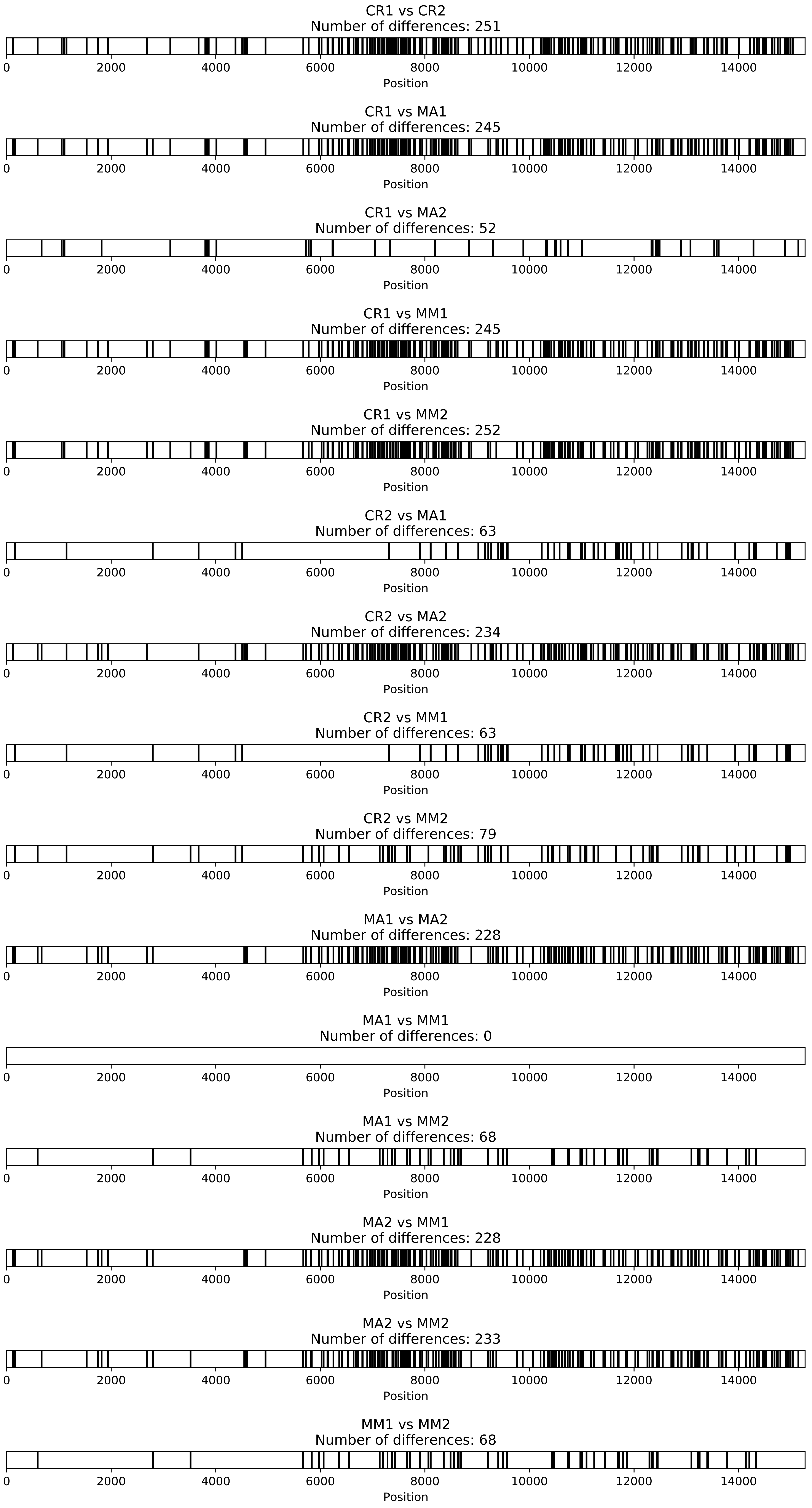
CR1 vs CR2
Number of differences: 251
0 2000 4000 6000 8000 10000 12000 14000
Position
CR1 vs MA1
Number of differences: 245
0 2000 4000 6000 8000 10000 12000 14000
Position
CR1 vs MA2
Number of differences: 52
0 2000 4000 6000 8000 10000 12000 14000
Position
CR1 vs MM1
Number of differences: 245
0 2000 4000 6000 8000 10000 12000 14000
Position
CR1 vs MM2
Number of differences: 252
0 2000 4000 6000 8000 10000 12000 14000
Position
CR2 vs MA1
Number of differences: 63
0 2000 4000 6000 8000 10000 12000 14000
Position
CR2 vs MA2
Number of differences: 234
0 2000 4000 6000 8000 10000 12000 14000
Position
CR2 vs MM1
Number of differences: 63
0 2000 4000 6000 8000 10000 12000 14000
Position
CR2 vs MM2
Number of differences: 79
0 2000 4000 6000 8000 10000 12000 14000
Position
MA1 vs MA2
Number of differences: 228
0 2000 4000 6000 8000 10000 12000 14000
Position
MA1 vs MM1
Number of differences: 0
0 2000 4000 6000 8000 10000 12000 14000
Position
MA1 vs MM2
Number of differences: 68
0 2000 4000 6000 8000 10000 12000 14000
Position
MA2 vs MM1
Number of differences: 228
0 2000 4000 6000 8000 10000 12000 14000
Position
MA2 vs MM2
Number of differences: 233
0 2000 4000 6000 8000 10000 12000 14000
Position
MM1 vs MM2
Number of differences: 68
0 2000 4000 6000 8000 10000 12000 14000
Position

### Region: 148



CR1 vs CR2
Number of differences: 331

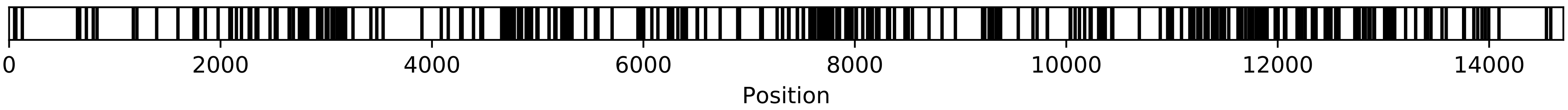


CR1 vs MA1
Number of differences: 168

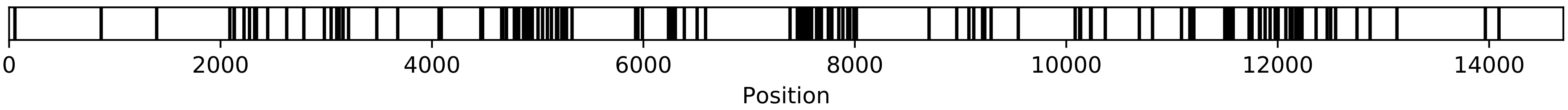


CR1 vs MA2
Number of differences: 338

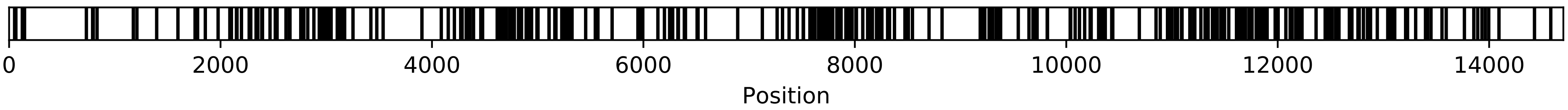


CR1 vs MM1
Number of differences: 162

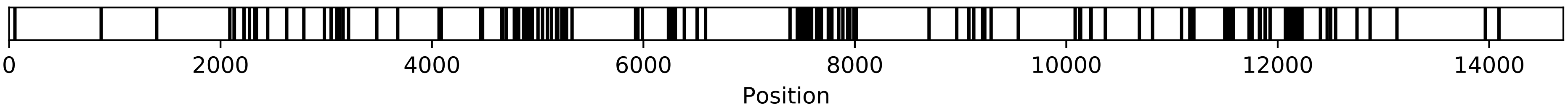


CR1 vs MM2
Number of differences: 189

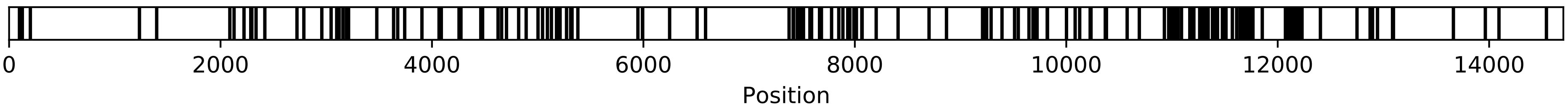


CR2 vs MA1
Number of differences: 324

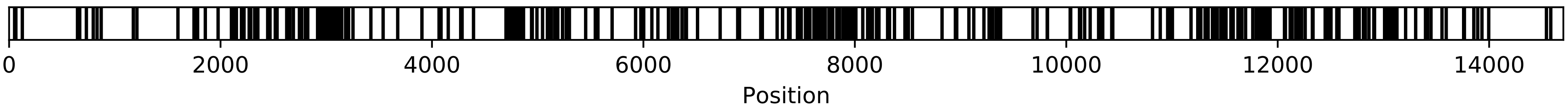


CR2 vs MA2
Number of differences: 116

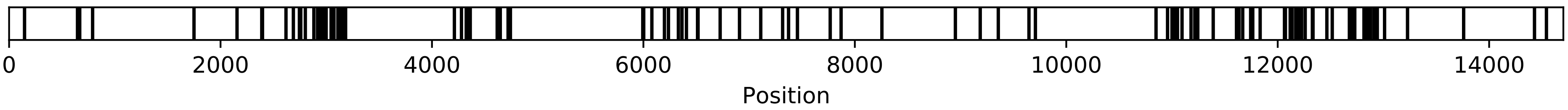


CR2 vs MM1
Number of differences: 326

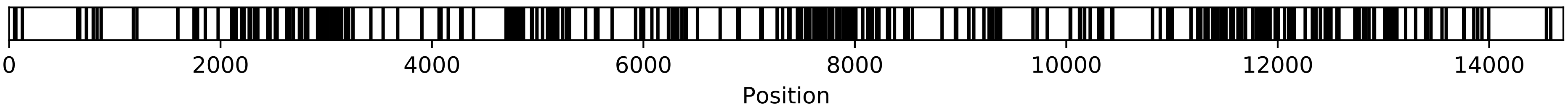


CR2 vs MM2
Number of differences: 341

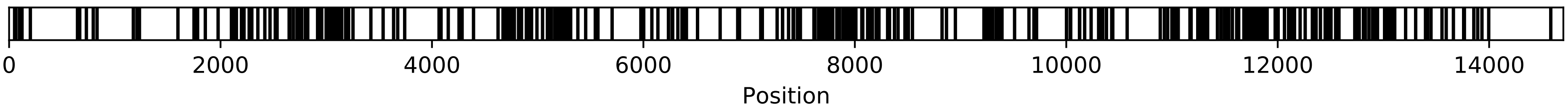


MA1 vs MA2
Number of differences: 318

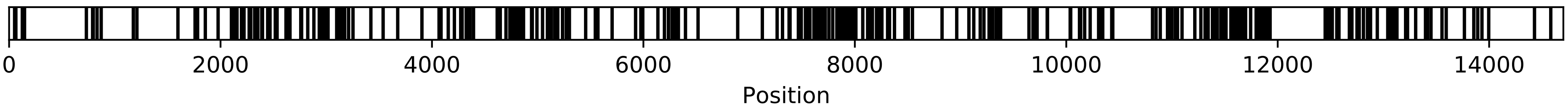


MA1 vs MM1
Number of differences: 16

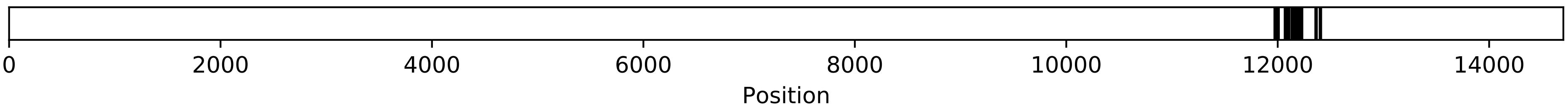


MA1 vs MM2
Number of differences: 181

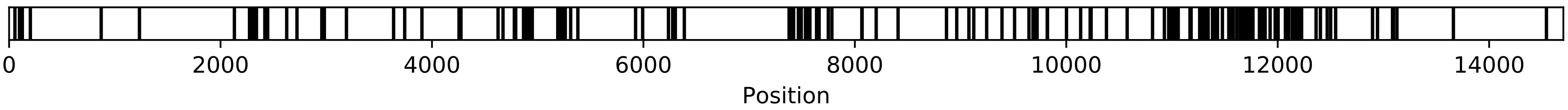


MA2 vs MM1
Number of differences: 334

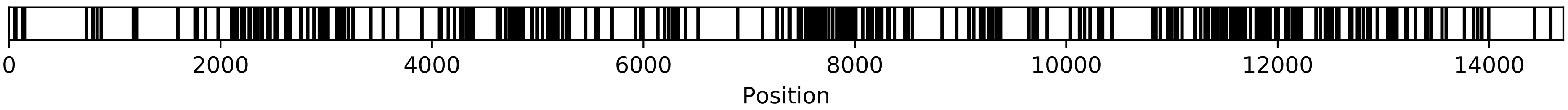


MA2 vs MM2
Number of differences: 347

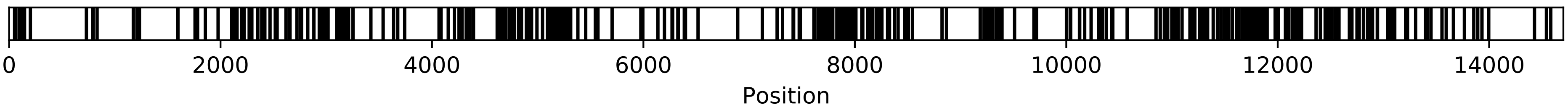


MM1 vs MM2
Number of differences: 165

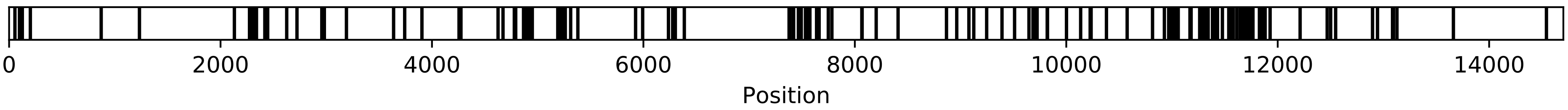

### Region: 149



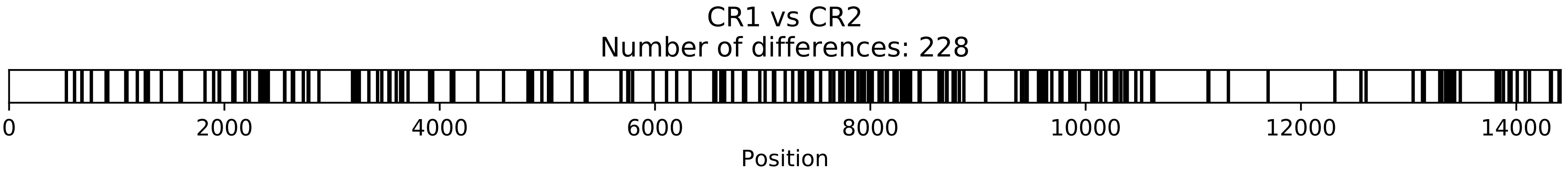
CR1 vs CR2
Number of differences: 228
0
2000
4000
6000
8000
10000
12000
14000
Position

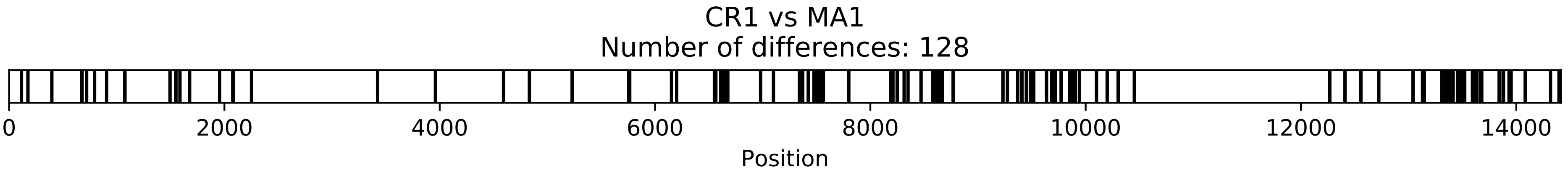
CR1 vs MA1
Number of differences: 128
0
2000
4000
6000
8000
10000
12000
14000
Position

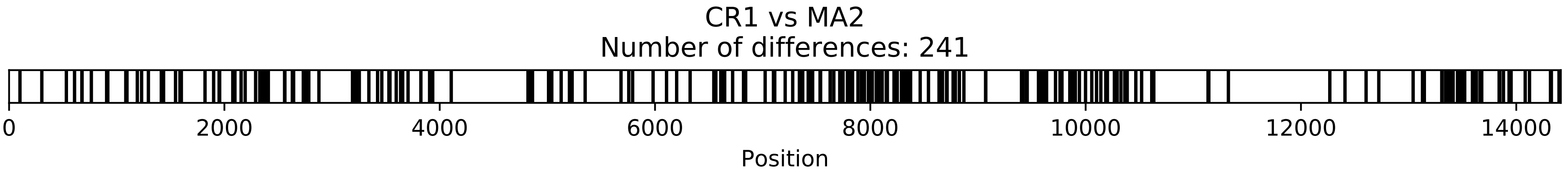
CR1 vs MA2
Number of differences: 241
0
2000
4000
6000
8000
10000
12000
14000
Position

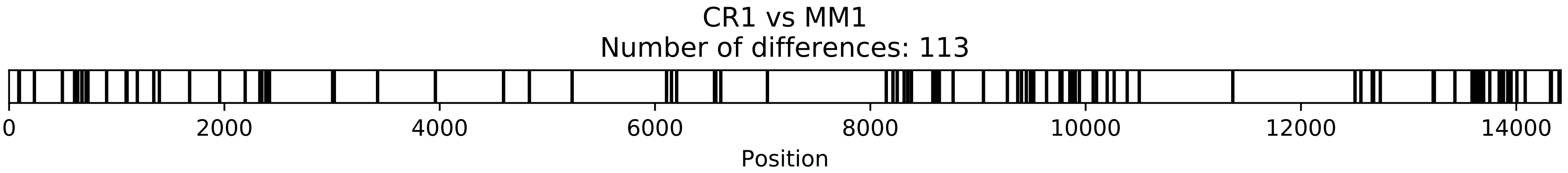
CR1 vs MM1
Number of differences: 113
0
2000
4000
6000
8000
10000
12000
14000
Position

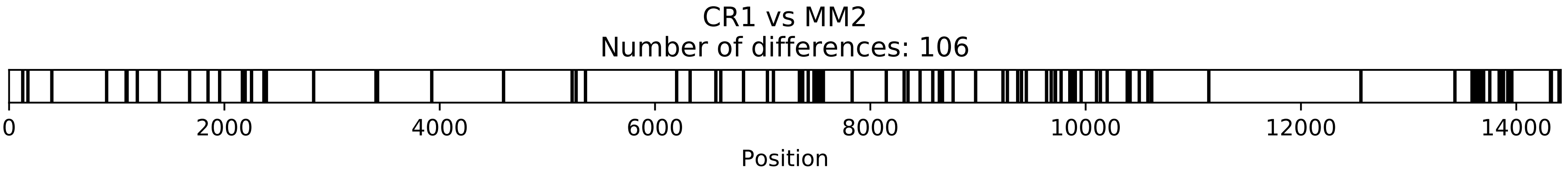
CR1 vs MM2
Number of differences: 106
0
2000
4000
6000
8000
10000
12000
14000
Position

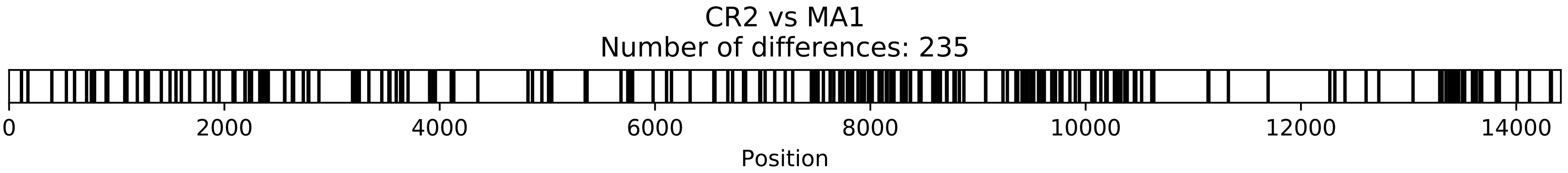
CR2 vs MA1
Number of differences: 235
0
2000
4000
6000
8000
10000
12000
14000
Position

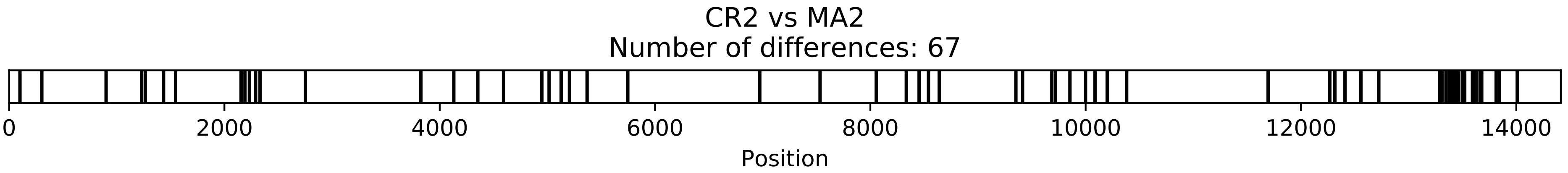
CR2 vs MA2
Number of differences: 67
0
2000
4000
6000
8000
10000
12000
14000
Position

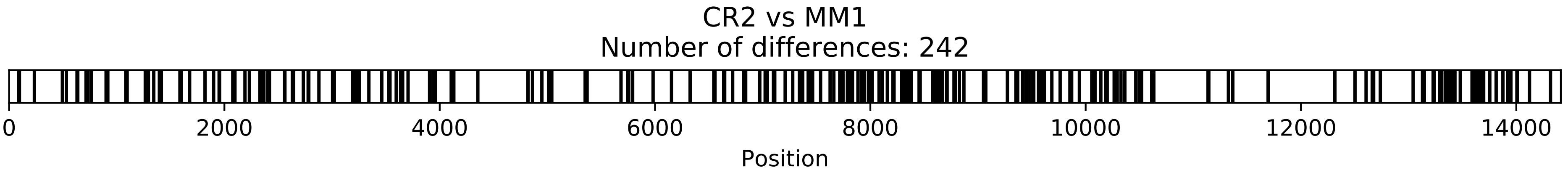
CR2 vs MM1
Number of differences: 242
0
2000
4000
6000
8000
10000
12000
14000
Position

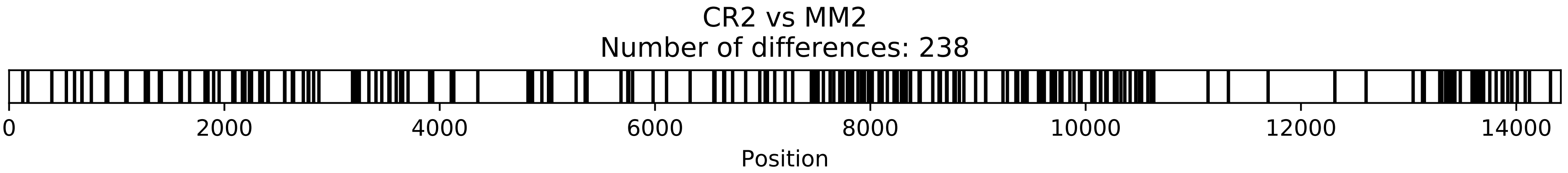
CR2 vs MM2
Number of differences: 238
0
2000
4000
6000
8000
10000
12000
14000
Position

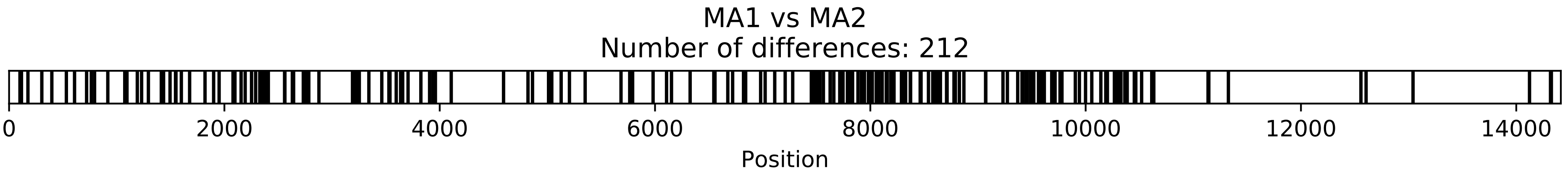
MA1 vs MA2
Number of differences: 212
0
2000
4000
6000
8000
10000
12000
14000
Position

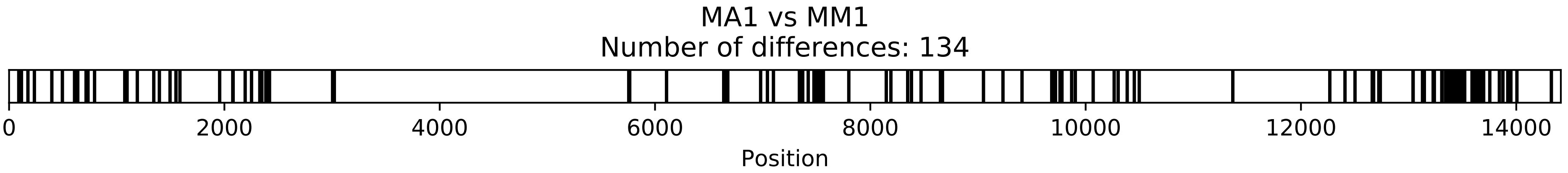
MA1 vs MM1
Number of differences: 134
0
2000
4000
6000
8000
10000
12000
14000
Position

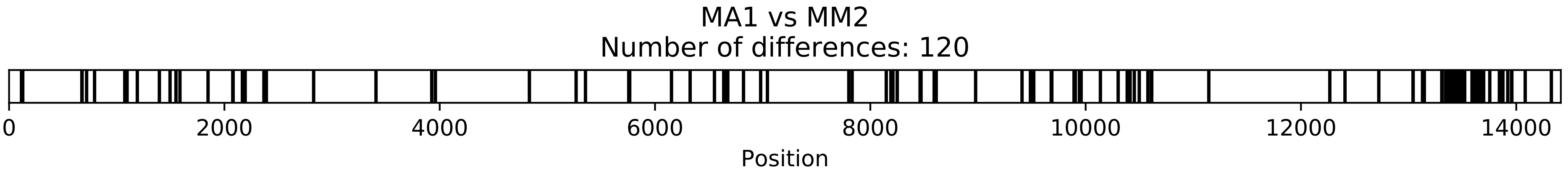
MA1 vs MM2
Number of differences: 120
0
2000
4000
6000
8000
10000
12000
14000
Position

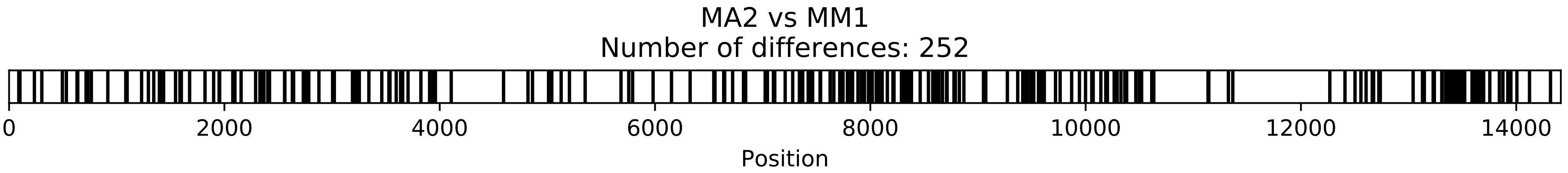
MA2 vs MM1
Number of differences: 252
0
2000
4000
6000
8000
10000
12000
14000
Position

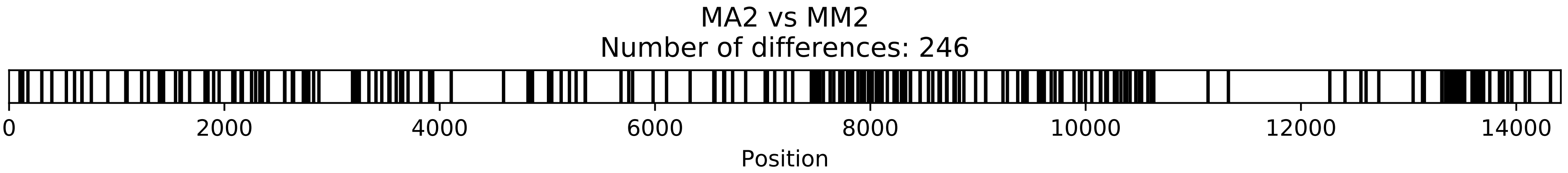
MA2 vs MM2
Number of differences: 246
0
2000
4000
6000
8000
10000
12000
14000
Position

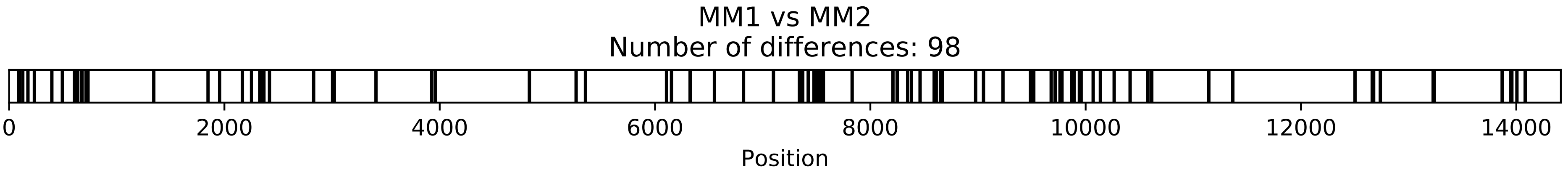
MM1 vs MM2
Number of differences: 98
0
2000
4000
6000
8000
10000
12000
14000
Position

### Region: 154

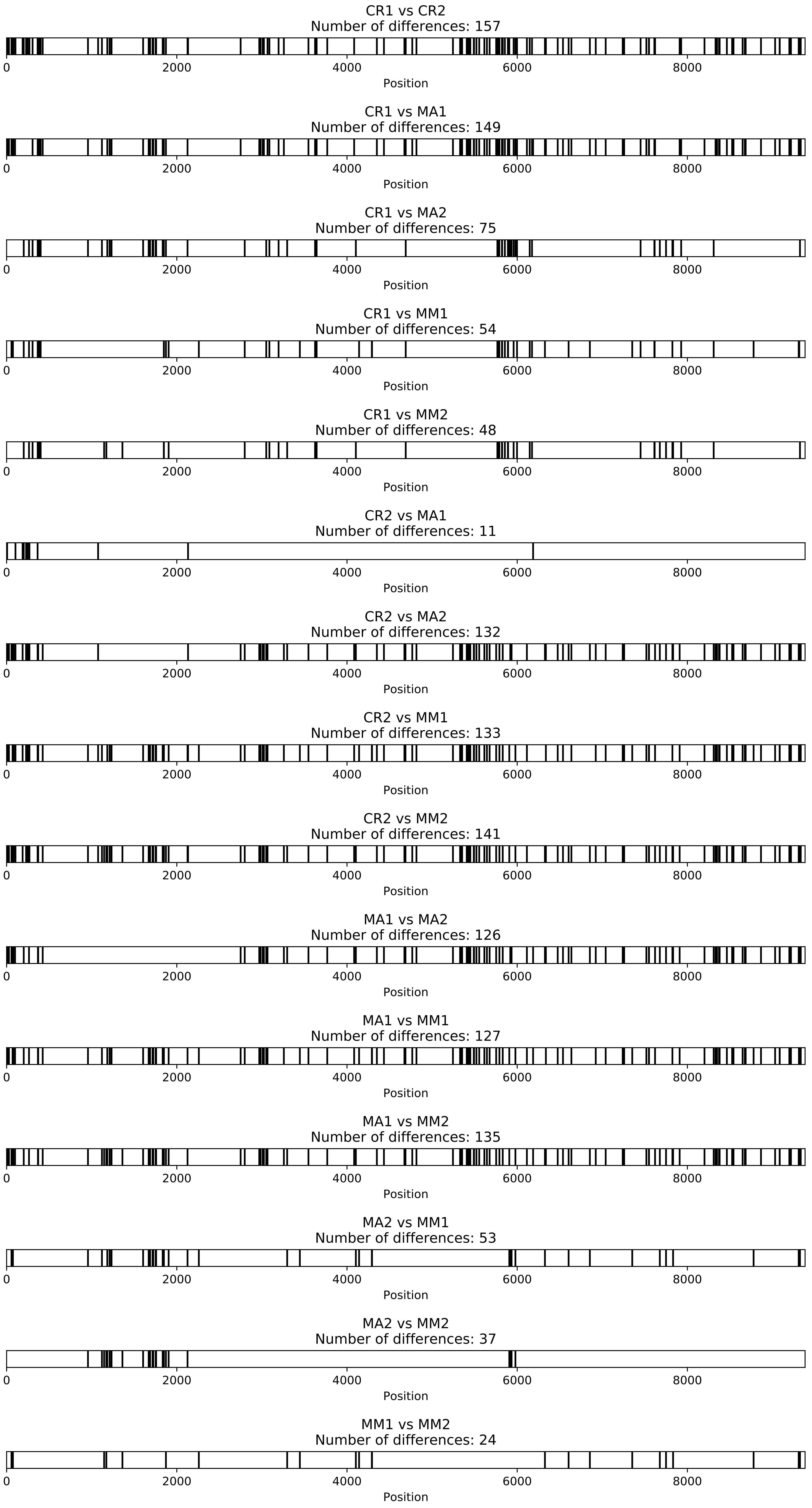
CR1 vs CR2
Number of differences: 157
CR1 vs MA1
Number of differences: 149
CR1 vs MA2
Number of differences: 75
CR1 vs MM1
Number of differences: 54
CR1 vs MM2
Number of differences: 48
CR2 vs MA1
Number of differences: 11
CR2 vs MA2
Number of differences: 132
CR2 vs MM1
Number of differences: 133
CR2 vs MM2
Number of differences: 141
MA1 vs MA2
Number of differences: 126
MA1 vs MM1
Number of differences: 127
MA1 vs MM2
Number of differences: 135
MA2 vs MM1
Number of differences: 53
MA2 vs MM2
Number of differences: 37
MM1 vs MM2
Number of differences: 24
0
2000
4000
6000
8000
Position

### Region: 156



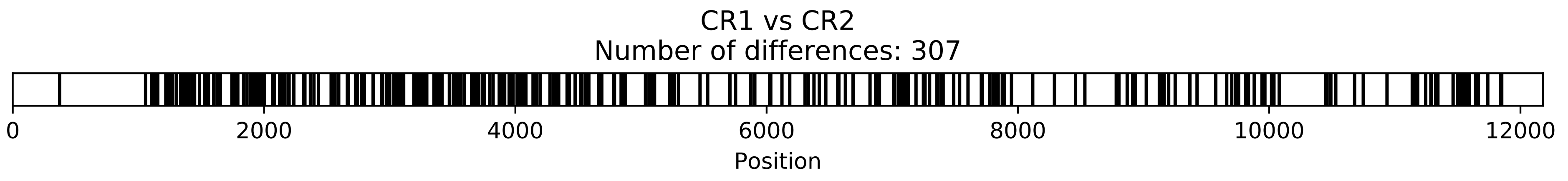
CR1 vs CR2
Number of differences: 307
0 2000 4000 6000 8000 10000 12000
Position

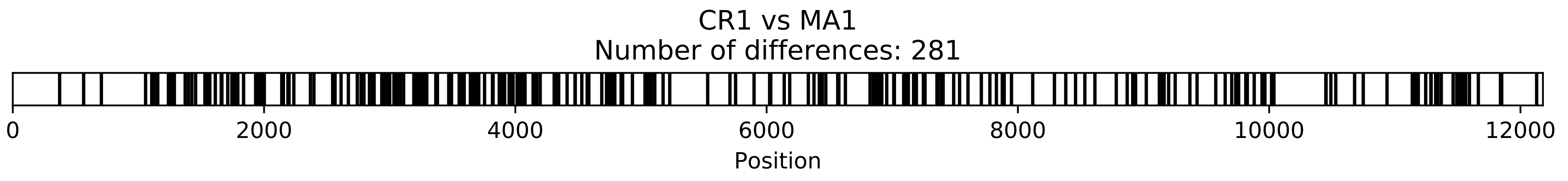
CR1 vs MA1
Number of differences: 281
0 2000 4000 6000 8000 10000 12000
Position

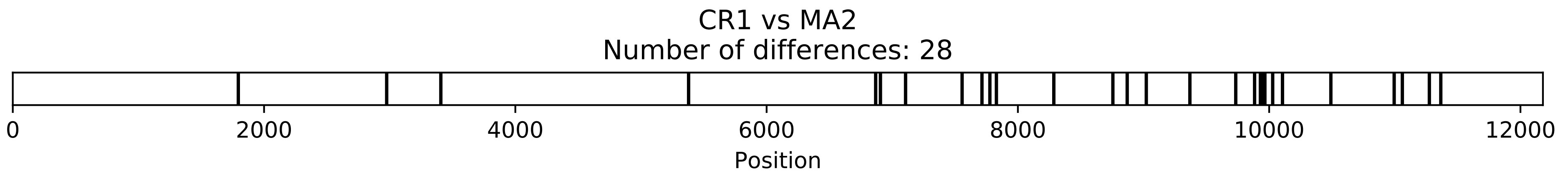
CR1 vs MA2
Number of differences: 28
0 2000 4000 6000 8000 10000 12000
Position

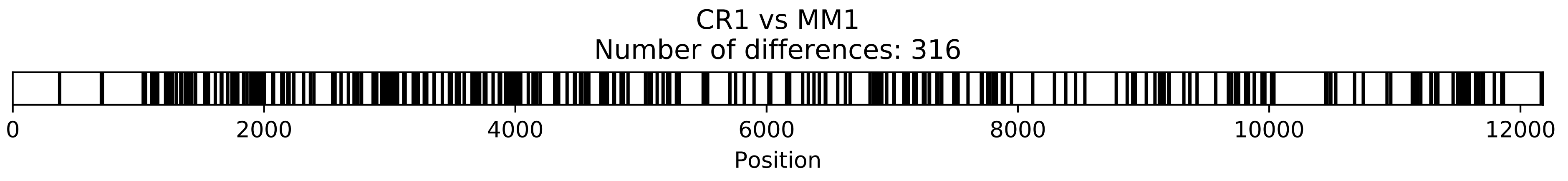
CR1 vs MM1
Number of differences: 316
0 2000 4000 6000 8000 10000 12000
Position

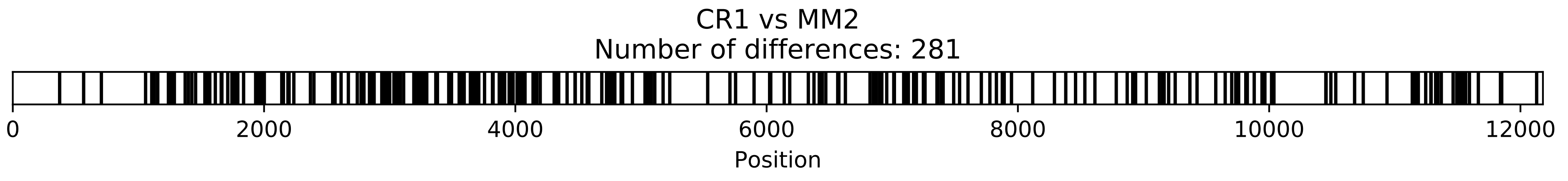
CR1 vs MM2
Number of differences: 281
0 2000 4000 6000 8000 10000 12000
Position

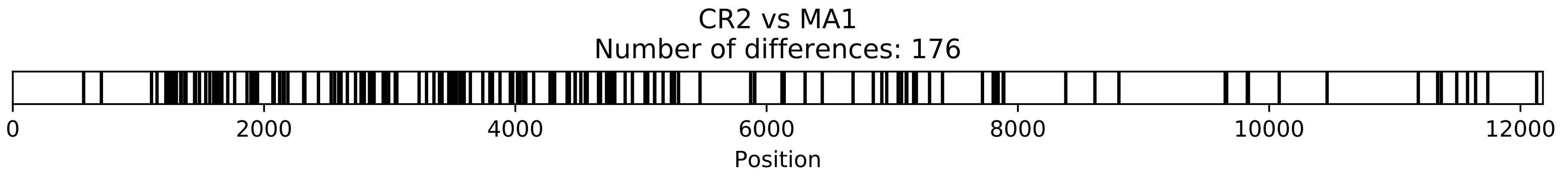
CR2 vs MA1
Number of differences: 176
0 2000 4000 6000 8000 10000 12000
Position

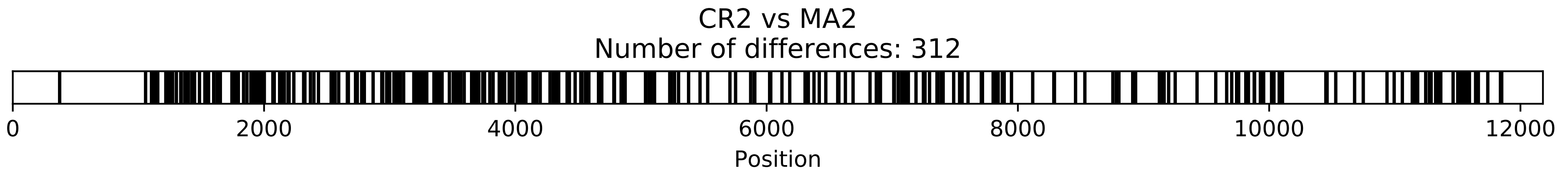
CR2 vs MA2
Number of differences: 312
0 2000 4000 6000 8000 10000 12000
Position

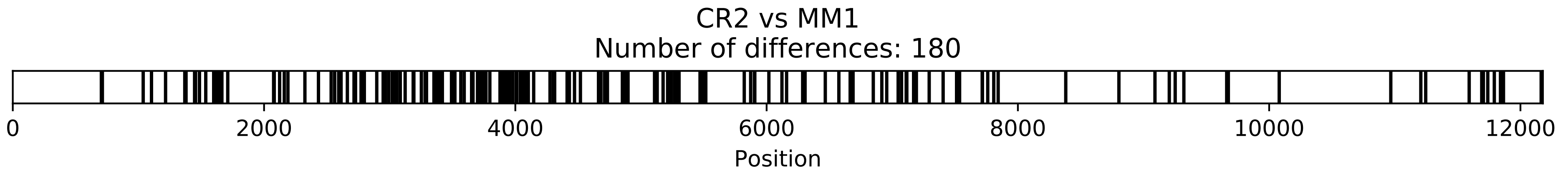
CR2 vs MM1
Number of differences: 180
0 2000 4000 6000 8000 10000 12000
Position

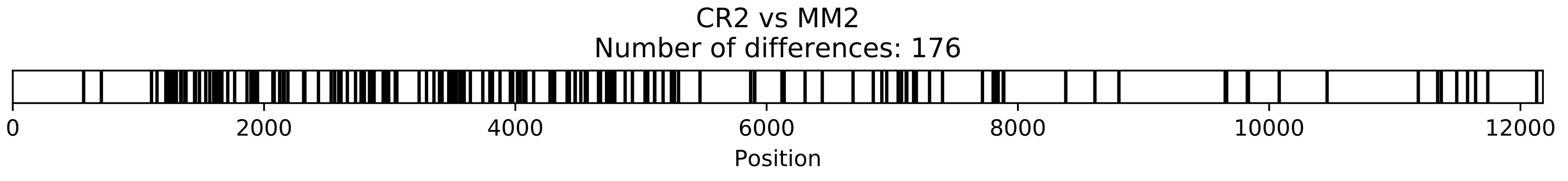
CR2 vs MM2
Number of differences: 176
0 2000 4000 6000 8000 10000 12000
Position

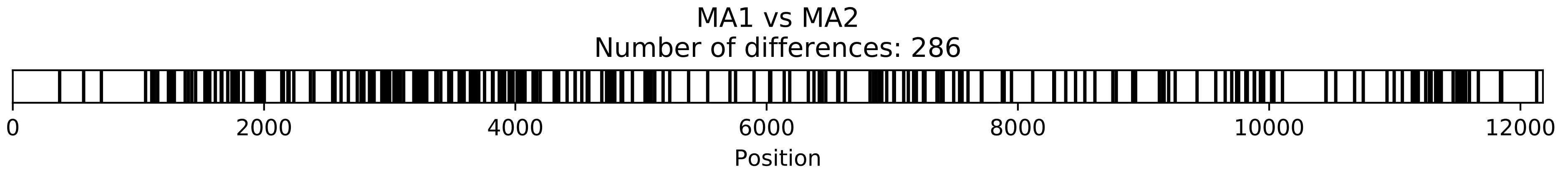
MA1 vs MA2
Number of differences: 286
0 2000 4000 6000 8000 10000 12000
Position

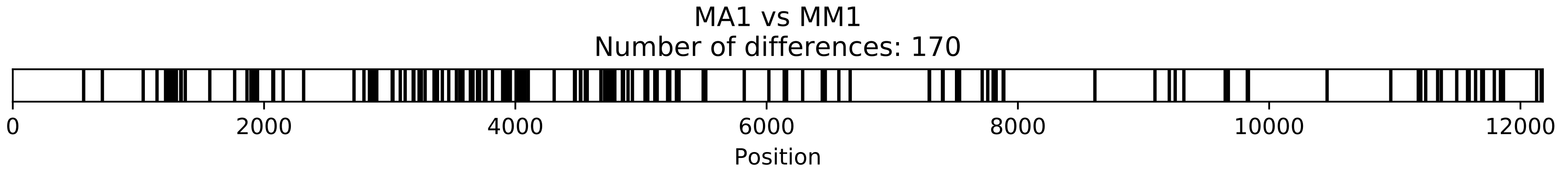
MA1 vs MM1
Number of differences: 170
0 2000 4000 6000 8000 10000 12000
Position

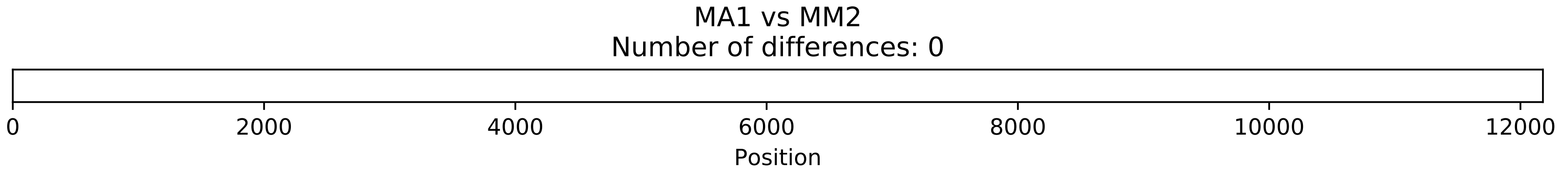
MA1 vs MM2
Number of differences: 0
0 2000 4000 6000 8000 10000 12000
Position

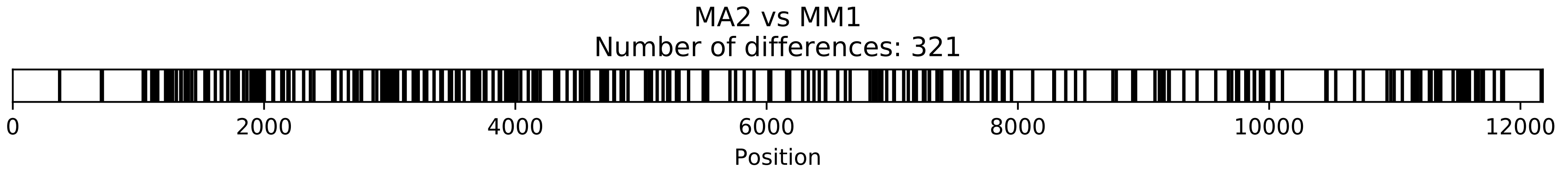
MA2 vs MM1
Number of differences: 321
0 2000 4000 6000 8000 10000 12000
Position

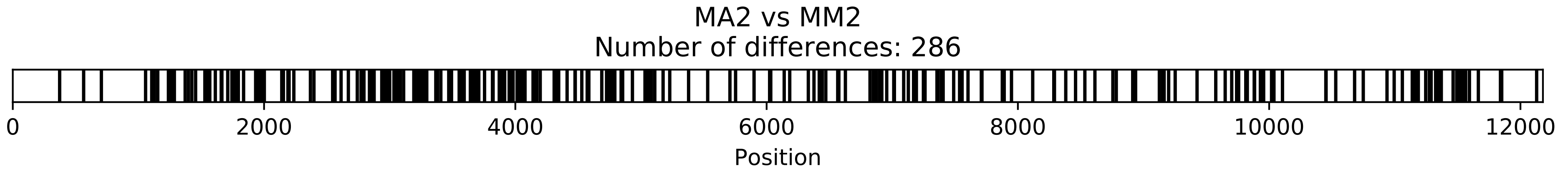
MA2 vs MM2
Number of differences: 286
0 2000 4000 6000 8000 10000 12000
Position

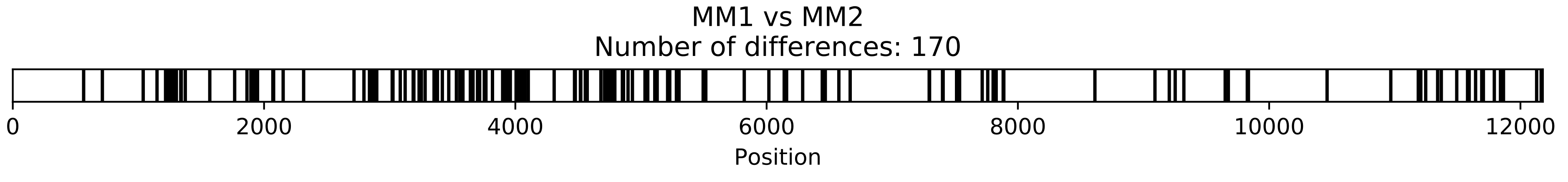
MM1 vs MM2
Number of differences: 170
0 2000 4000 6000 8000 10000 12000
Position

### Region: 159

CR1 vs CR2
Number of differences: 284

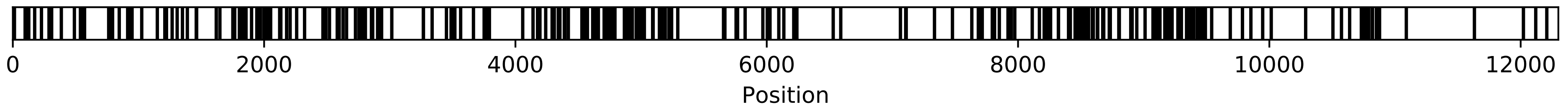


CR1 vs MA1
Number of differences: 59

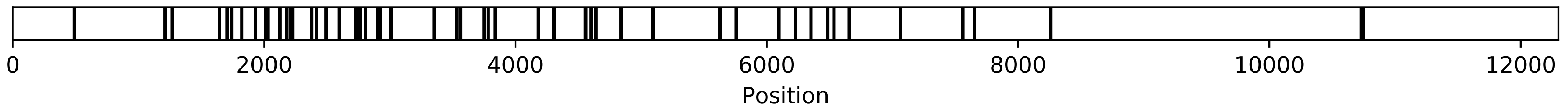


CR1 vs MA2
Number of differences: 266

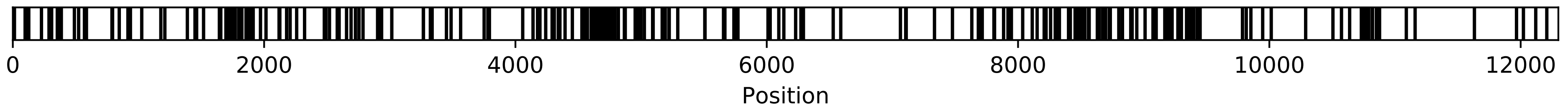


CR1 vs MM1
Number of differences: 260

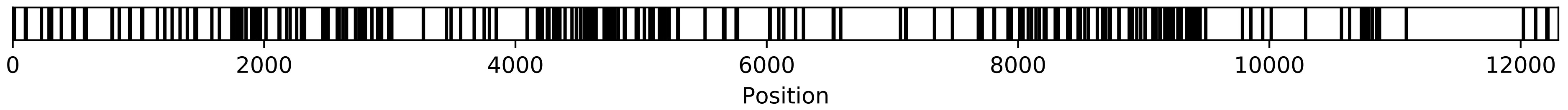


CR1 vs MM2
Number of differences: 261

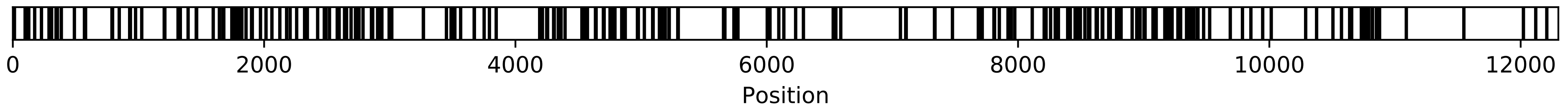


CR2 vs MA1
Number of differences: 285

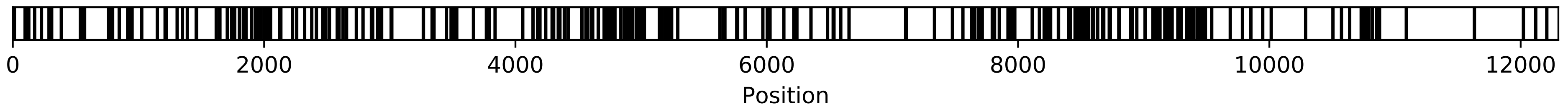


CR2 vs MA2
Number of differences: 154

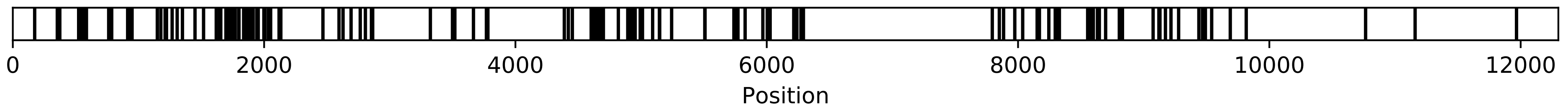


CR2 vs MM1
Number of differences: 182

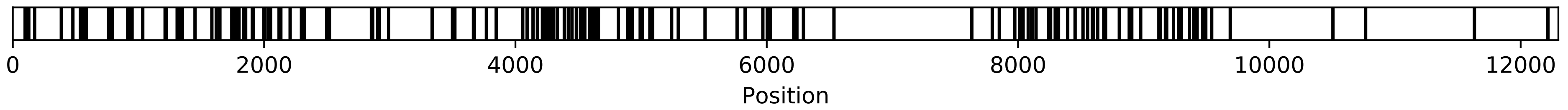


CR2 vs MM2
Number of differences: 169

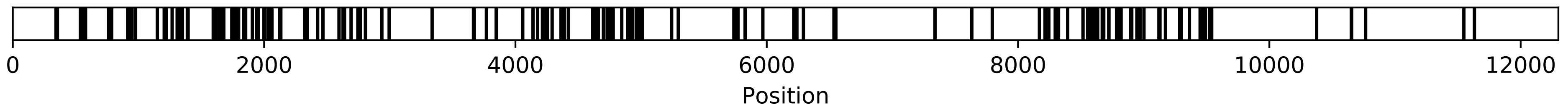


MA1 vs MA2
Number of differences: 273

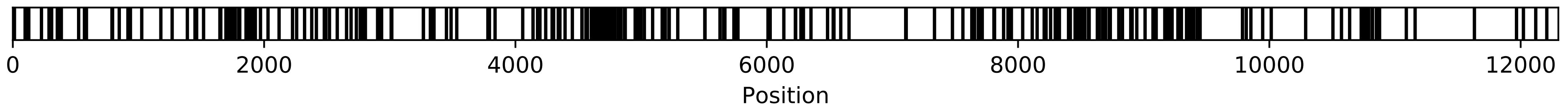


MA1 vs MM1
Number of differences: 257

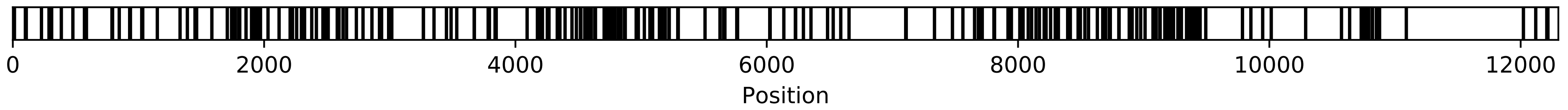


MA1 vs MM2
Number of differences: 264

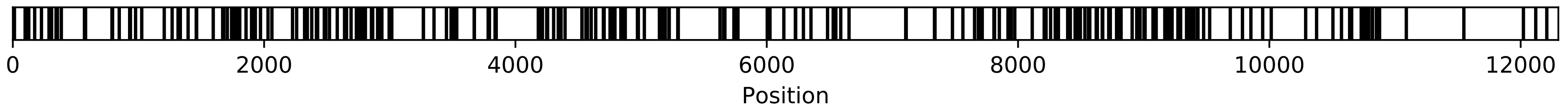


MA2 vs MM1
Number of differences: 146

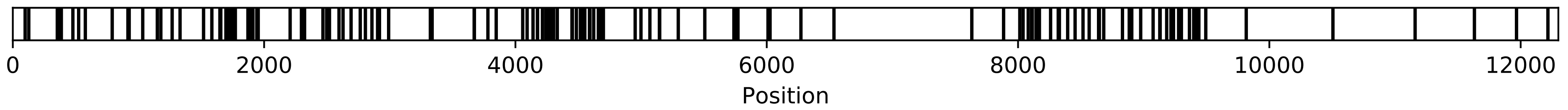


MA2 vs MM2
Number of differences: 148

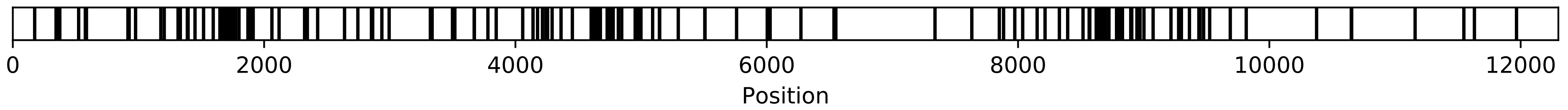


MM1 vs MM2
Number of differences: 157

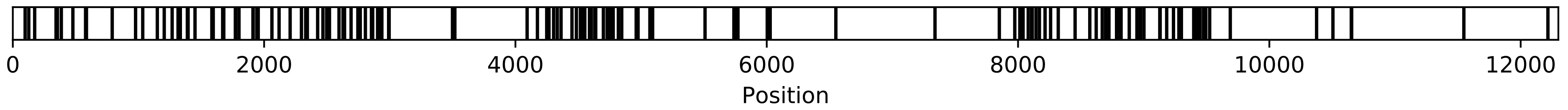

### Region: 160



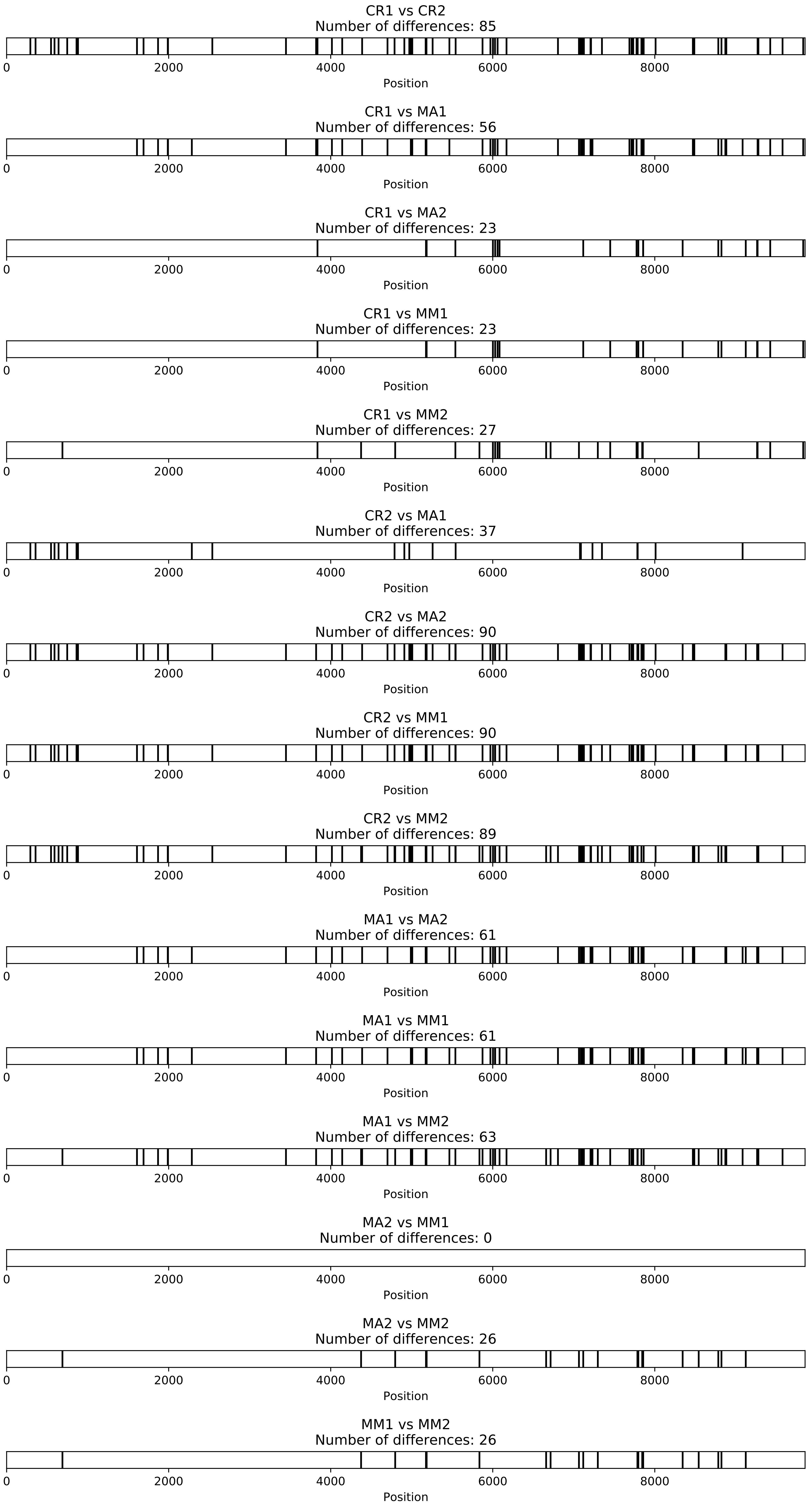
CR1 vs CR2
Number of differences: 85
CR1 vs MA1
Number of differences: 56
CR1 vs MA2
Number of differences: 23
CR1 vs MM1
Number of differences: 23
CR1 vs MM2
Number of differences: 27
CR2 vs MA1
Number of differences: 37
CR2 vs MA2
Number of differences: 90
CR2 vs MM1
Number of differences: 90
CR2 vs MM2
Number of differences: 89
MA1 vs MA2
Number of differences: 61
MA1 vs MM1
Number of differences: 61
MA1 vs MM2
Number of differences: 63
MA2 vs MM1
Number of differences: 0
MA2 vs MM2
Number of differences: 26
MM1 vs MM2
Number of differences: 26
0
2000
4000
6000
8000
Position

### Region: 161



CR1 vs CR2
Number of differences: 371

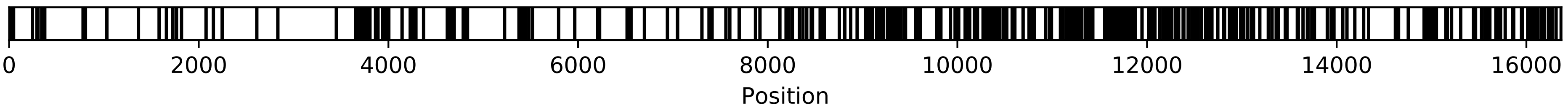


CR1 vs MA1
Number of differences: 384

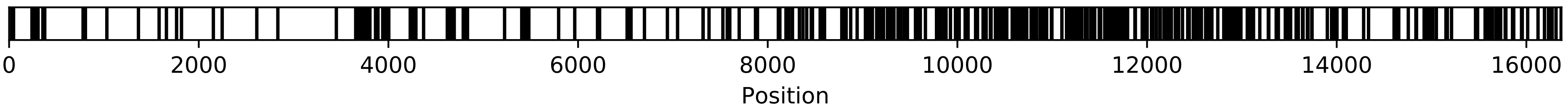


CR1 vs MA2
Number of differences: 160

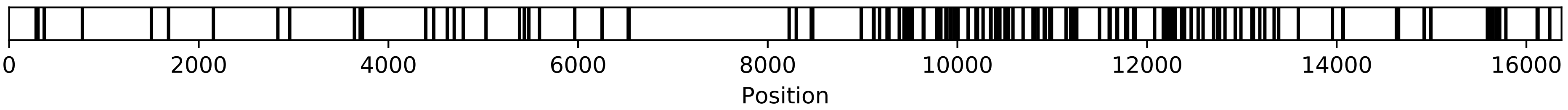


CR1 vs MM1
Number of differences: 386

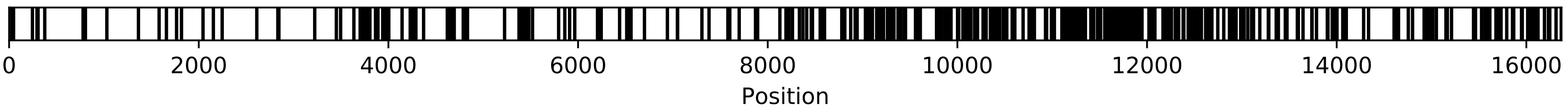


CR1 vs MM2
Number of differences: 391

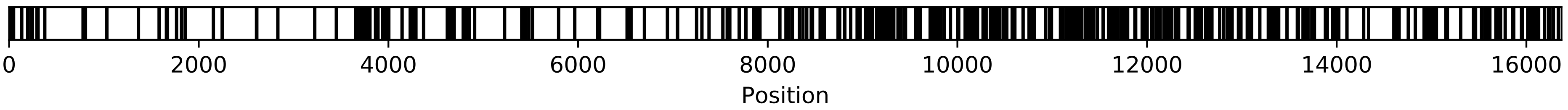


CR2 vs MA1
Number of differences: 191

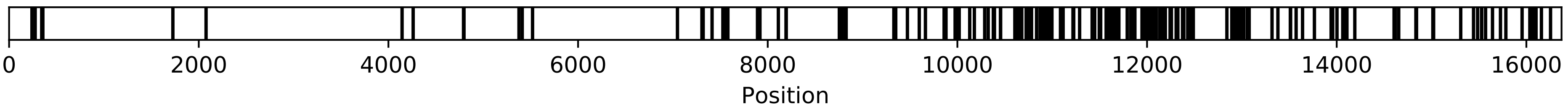


CR2 vs MA2
Number of differences: 393

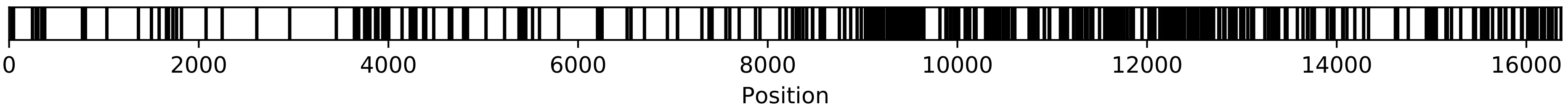


CR2 vs MM1
Number of differences: 123

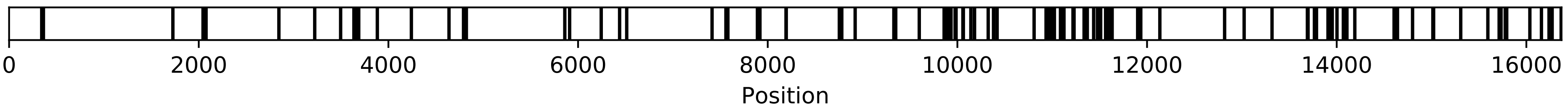


CR2 vs MM2
Number of differences: 187

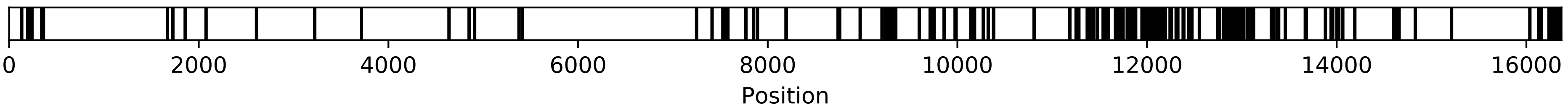


MA1 vs MA2
Number of differences: 401

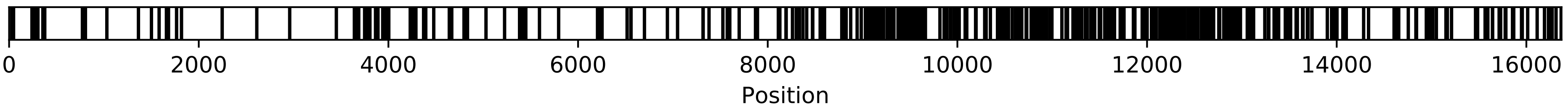


MA1 vs MM1
Number of differences: 204

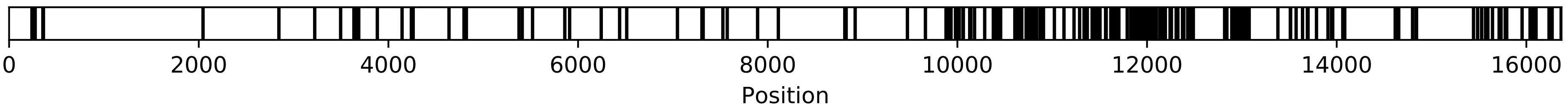


MA1 vs MM2
Number of differences: 206

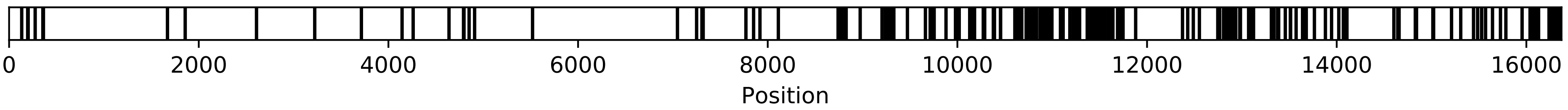


MA2 vs MM1
Number of differences: 409

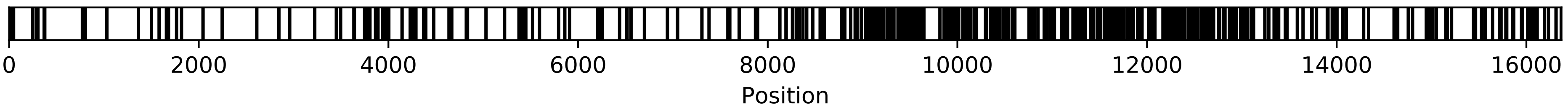


MA2 vs MM2
Number of differences: 415

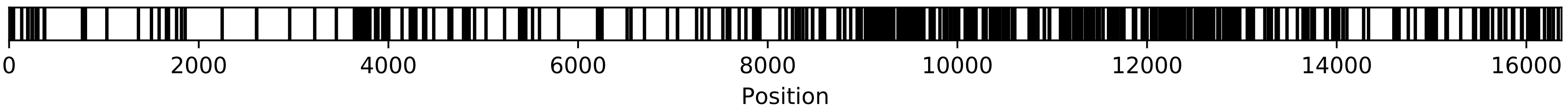


MM1 vs MM2
Number of differences: 228

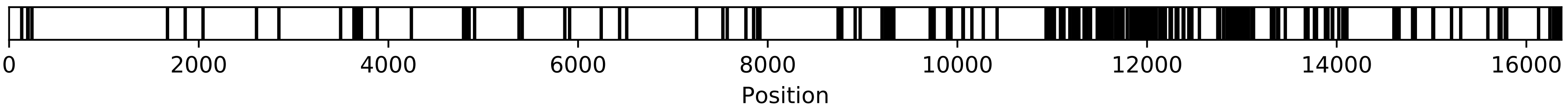

### Region: 165

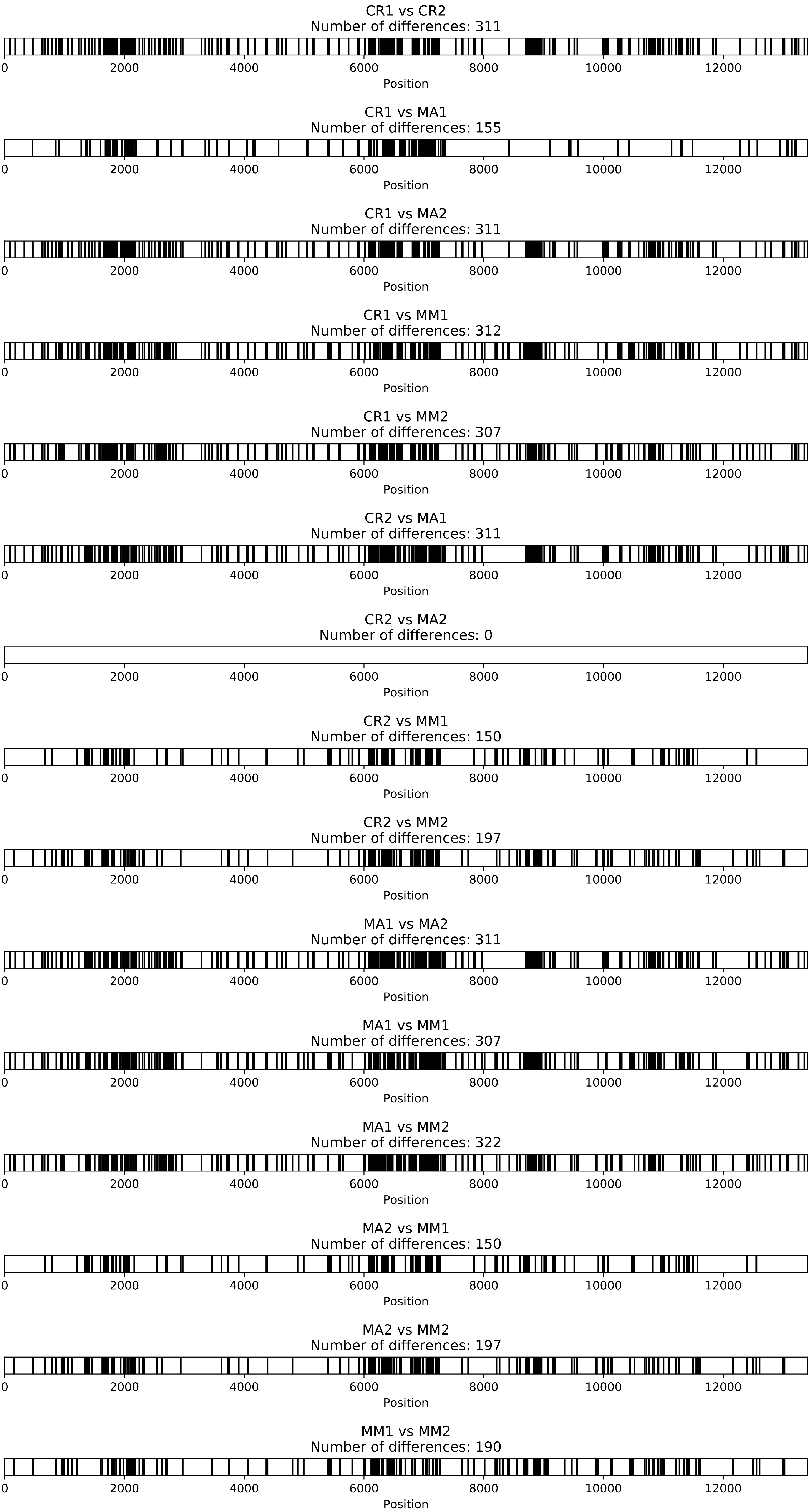
CR1 vs CR2
Number of differences: 311
CR1 vs MA1
Number of differences: 155
CR1 vs MA2
Number of differences: 311
CR1 vs MM1
Number of differences: 312
CR1 vs MM2
Number of differences: 307
CR2 vs MA1
Number of differences: 311
CR2 vs MA2
Number of differences: 0
CR2 vs MM1
Number of differences: 150
CR2 vs MM2
Number of differences: 197
MA1 vs MA2
Number of differences: 311
MA1 vs MM1
Number of differences: 307
MA1 vs MM2
Number of differences: 322
MA2 vs MM1
Number of differences: 150
MA2 vs MM2
Number of differences: 197
MM1 vs MM2
Number of differences: 190
0
2000
4000
6000
8000
10000
12000
Position

### Region: 167



CR1 vs CR2
Number of differences: 300

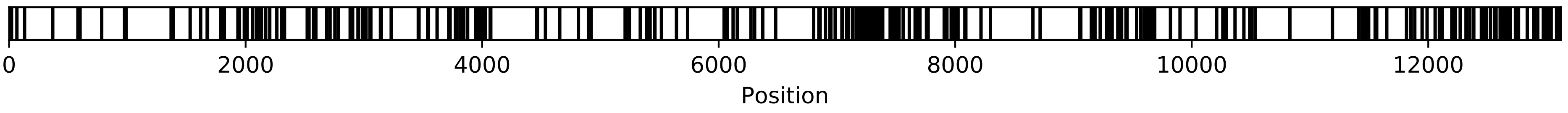


CR1 vs MA1
Number of differences: 0

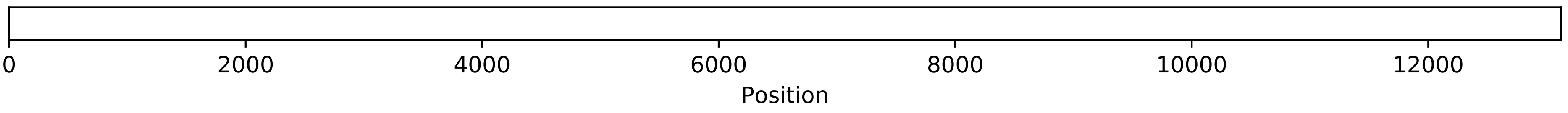


CR1 vs MA2
Number of differences: 297

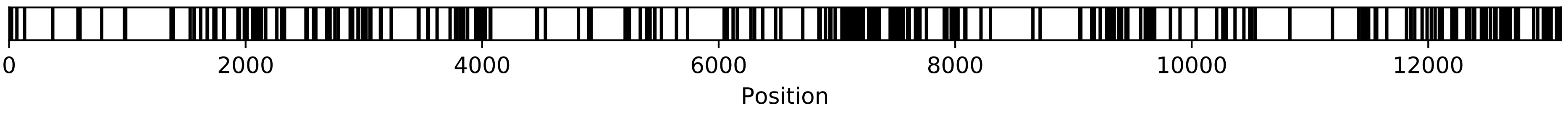


CR1 vs MM1
Number of differences: 106

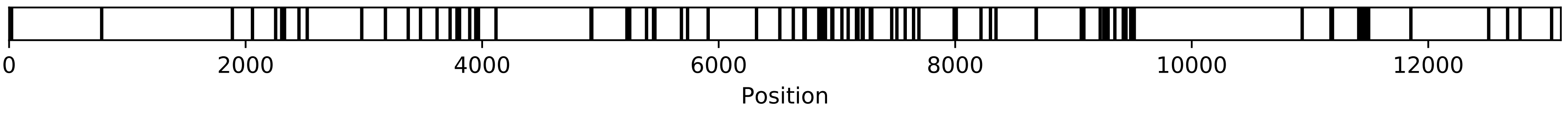


CR1 vs MM2
Number of differences: 0

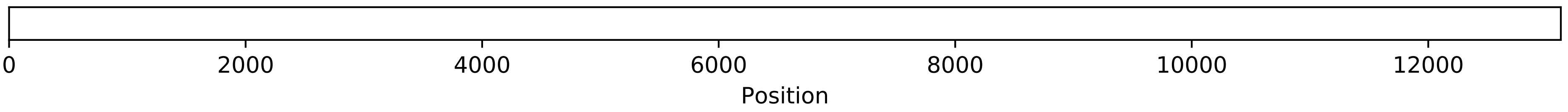


CR2 vs MA1
Number of differences: 300

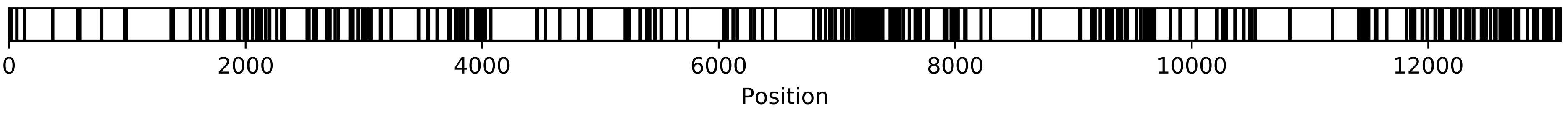


CR2 vs MA2
Number of differences: 55

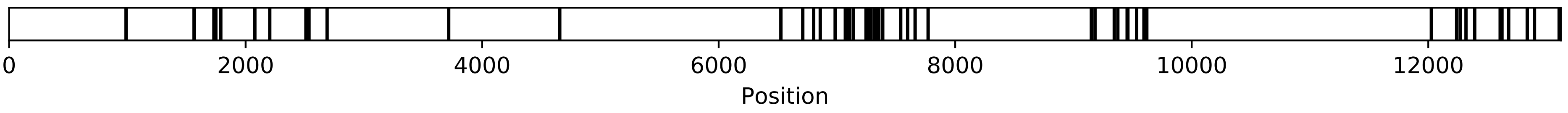


CR2 vs MM1
Number of differences: 303

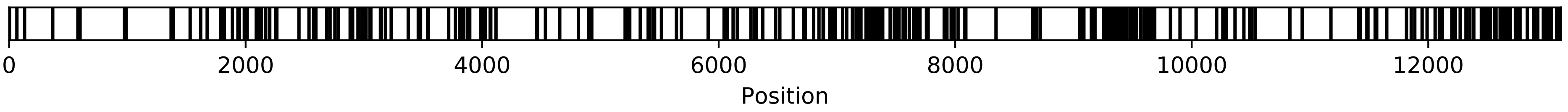


CR2 vs MM2
Number of differences: 300

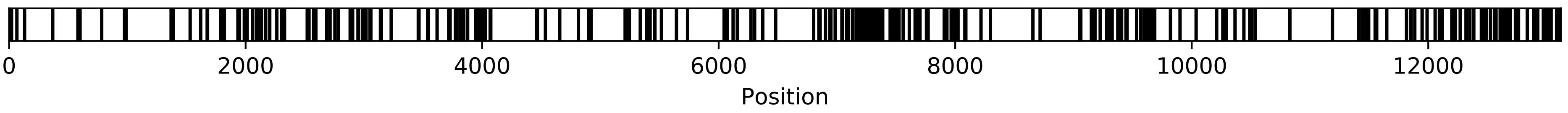


MA1 vs MA2
Number of differences: 297

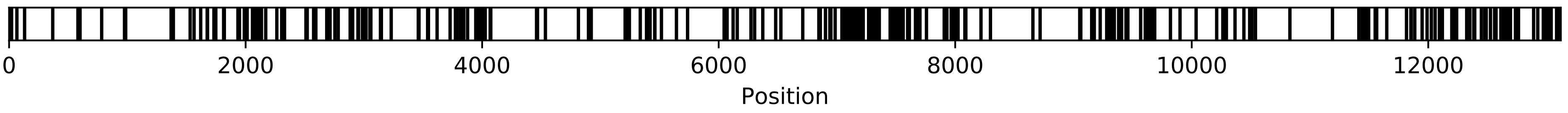


MA1 vs MM1
Number of differences: 106

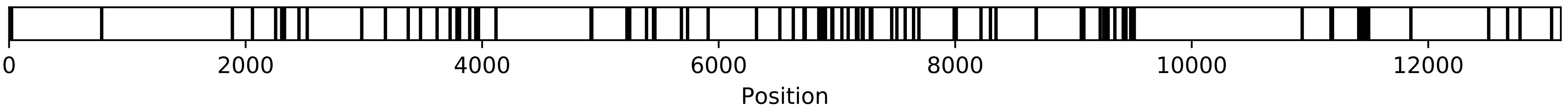


MA1 vs MM2
Number of differences: 0

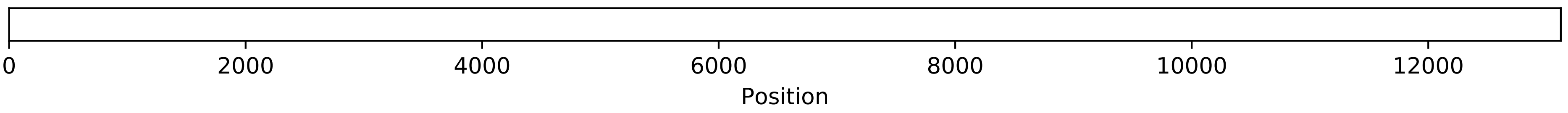


MA2 vs MM1
Number of differences: 300

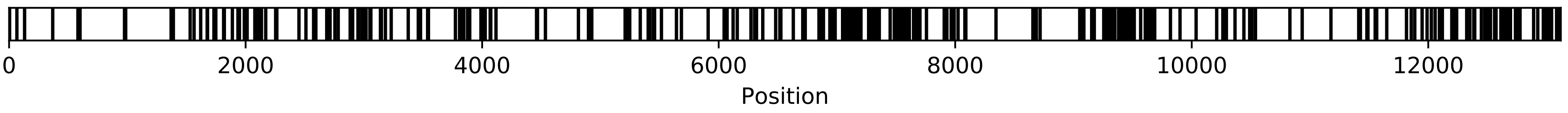


MA2 vs MM2
Number of differences: 297

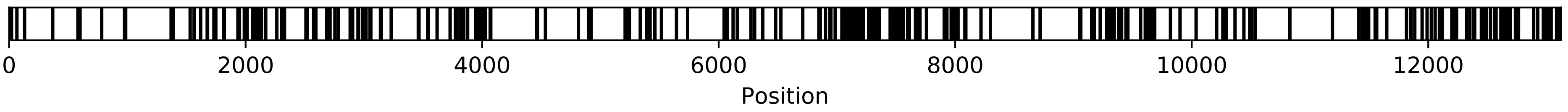


MM1 vs MM2
Number of differences: 106

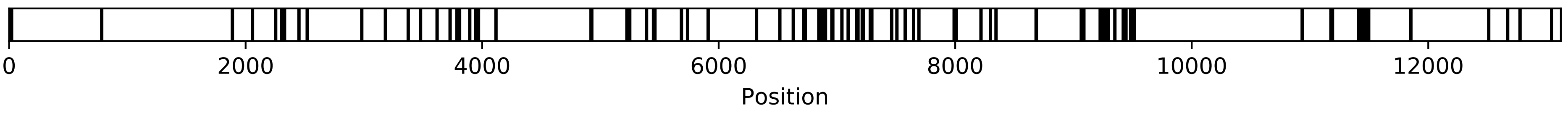

### Region: 168



CR1 vs CR2
Number of differences: 305

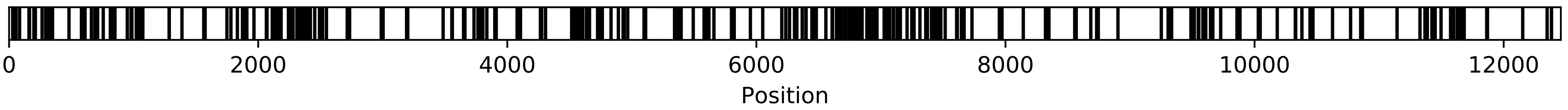


CR1 vs MA1
Number of differences: 207

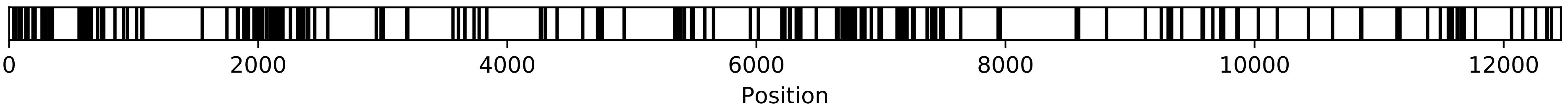


CR1 vs MA2
Number of differences: 307

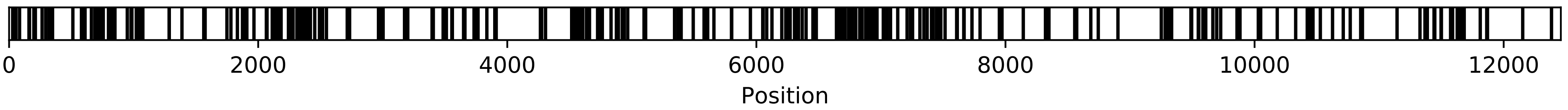


CR1 vs MM1
Number of differences: 189

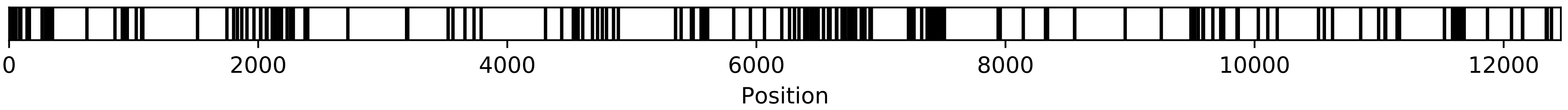


CR1 vs MM2
Number of differences: 228

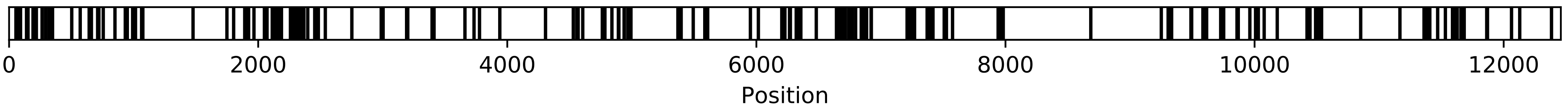


CR2 vs MA1
Number of differences: 294

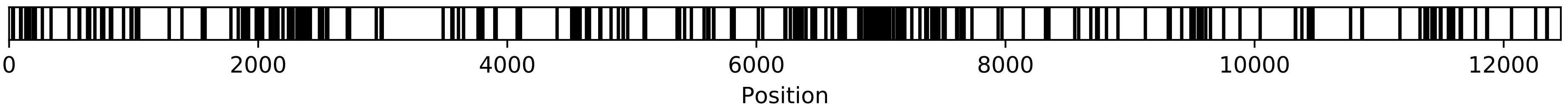


CR2 vs MA2
Number of differences: 99

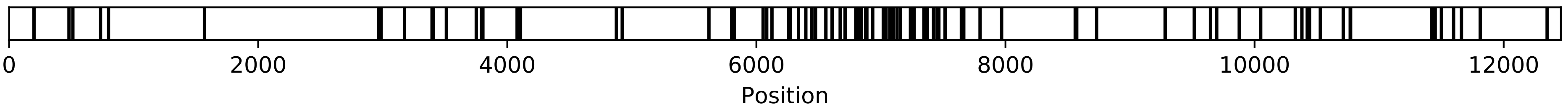


CR2 vs MM1
Number of differences: 296

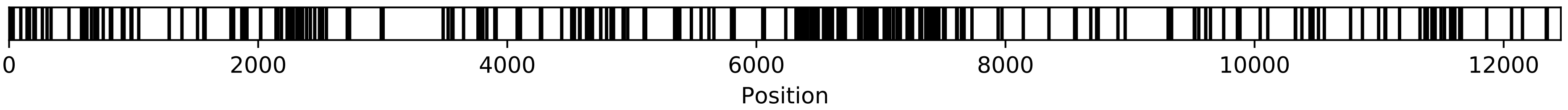


CR2 vs MM2
Number of differences: 319

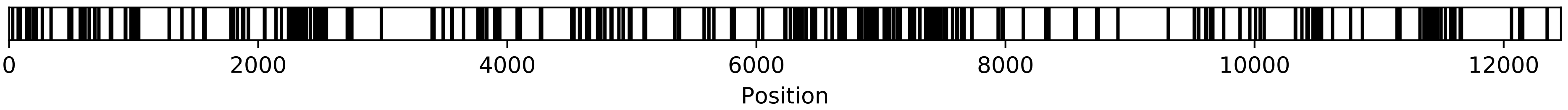


MA1 vs MA2
Number of differences: 297

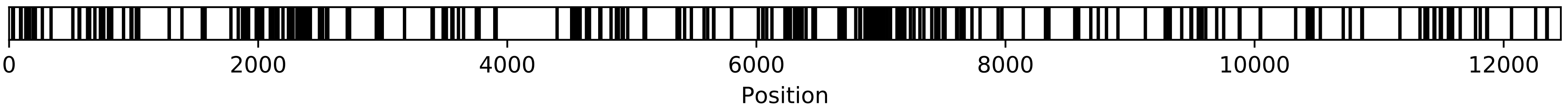


MA1 vs MM1
Number of differences: 207

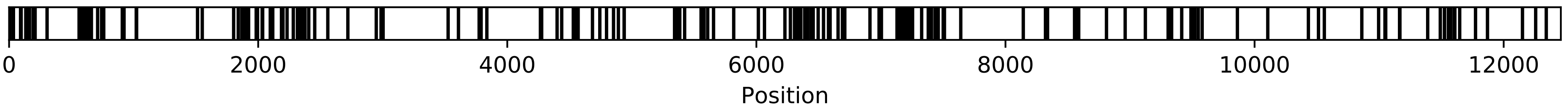


MA1 vs MM2
Number of differences: 203

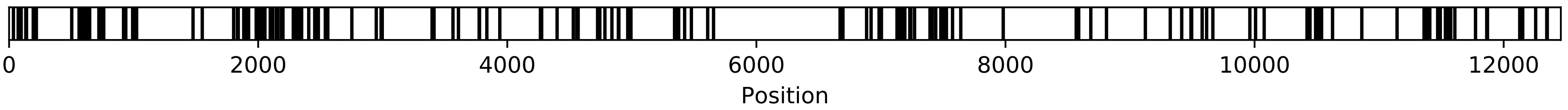


MA2 vs MM1
Number of differences: 300

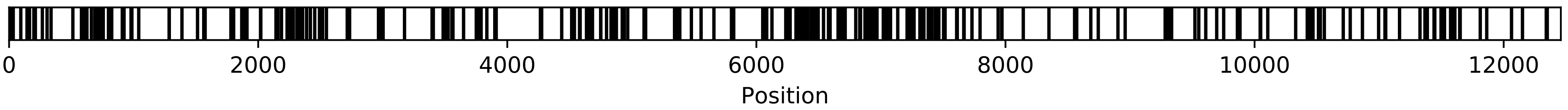


MA2 vs MM2
Number of differences: 317

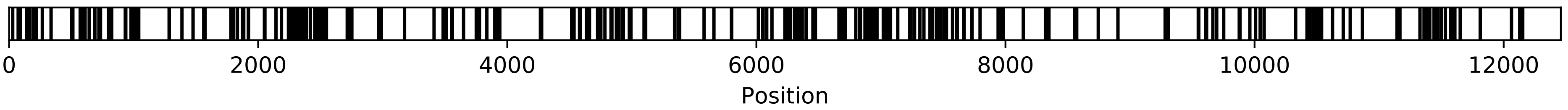


MM1 vs MM2
Number of differences: 230

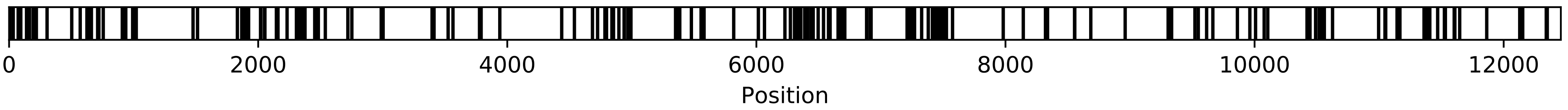

### Region: 170

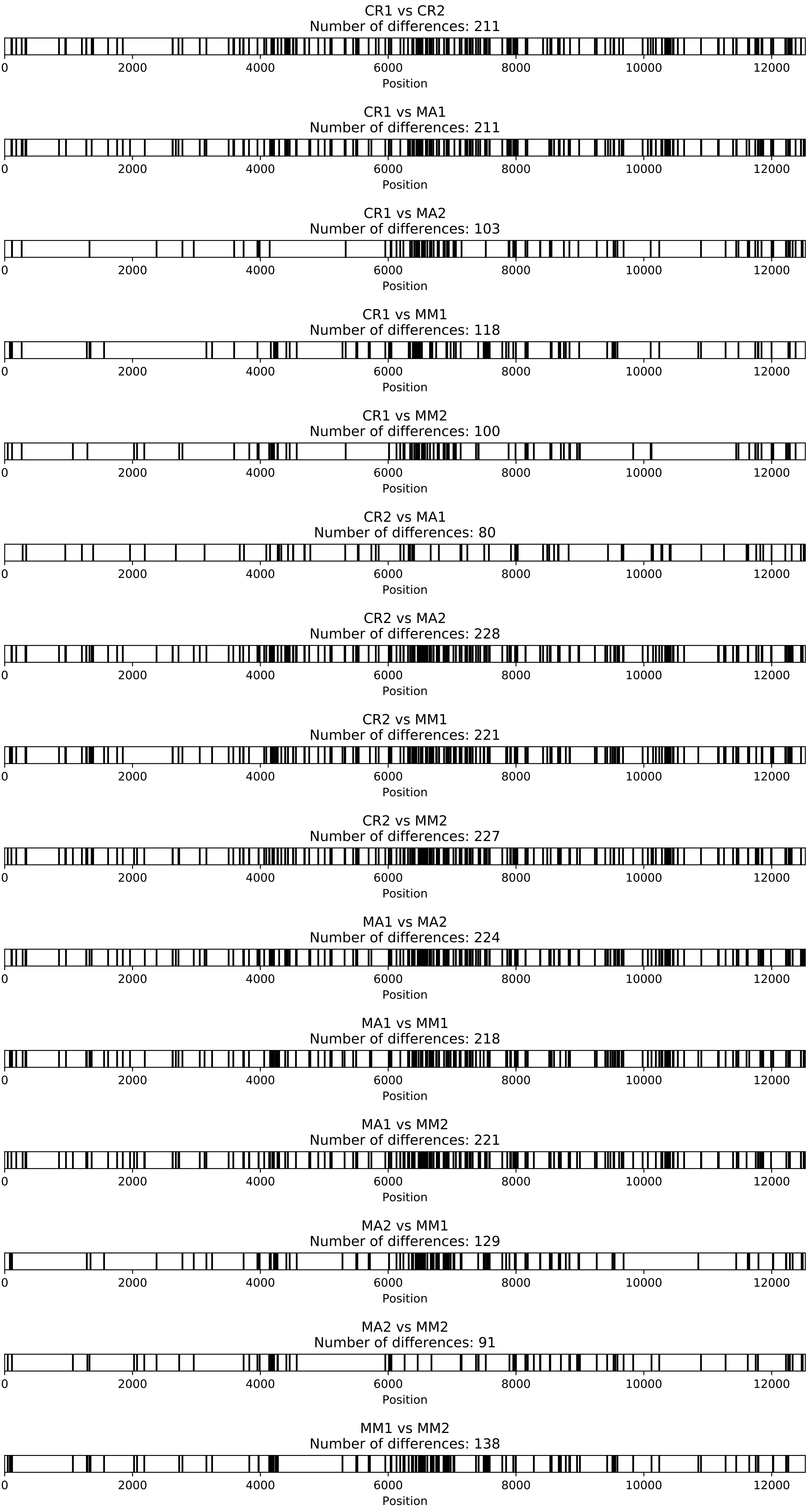
CR1 vs CR2
Number of differences: 211
CR1 vs MA1
Number of differences: 211
CR1 vs MA2
Number of differences: 103
CR1 vs MM1
Number of differences: 118
CR1 vs MM2
Number of differences: 100
CR2 vs MA1
Number of differences: 80
CR2 vs MA2
Number of differences: 228
CR2 vs MM1
Number of differences: 221
CR2 vs MM2
Number of differences: 227
MA1 vs MA2
Number of differences: 224
MA1 vs MM1
Number of differences: 218
MA1 vs MM2
Number of differences: 221
MA2 vs MM1
Number of differences: 129
MA2 vs MM2
Number of differences: 91
MM1 vs MM2
Number of differences: 138
0
2000
4000
6000
8000
10000
12000
Position

### Region: 172



CR1 vs CR2
Number of differences: 288

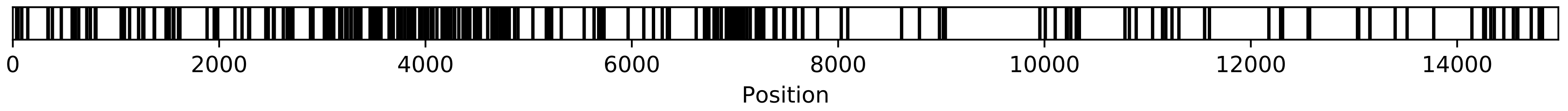


CR1 vs MA1
Number of differences: 162

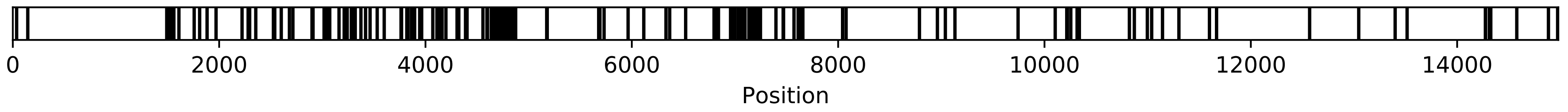


CR1 vs MA2
Number of differences: 279

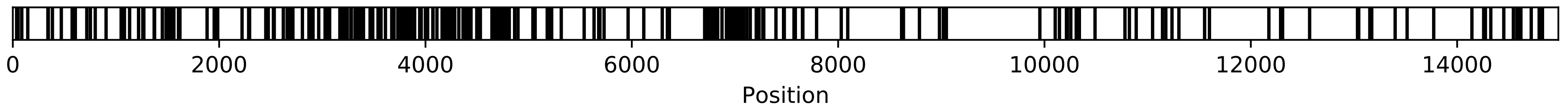


CR1 vs MM1
Number of differences: 162

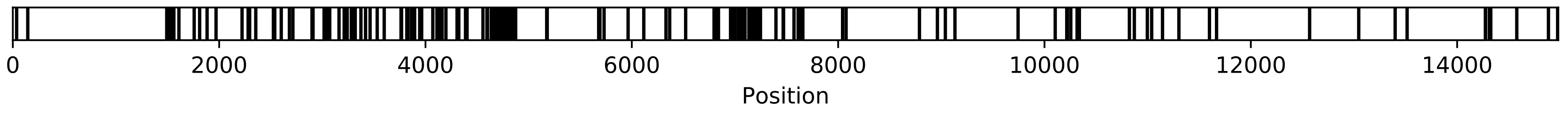


CR1 vs MM2
Number of differences: 141

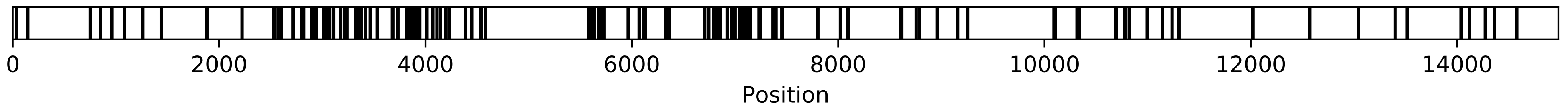


CR2 vs MA1
Number of differences: 282

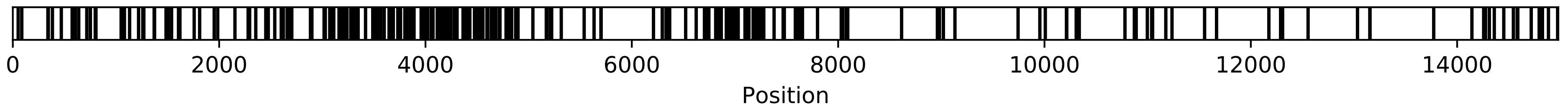


CR2 vs MA2
Number of differences: 107

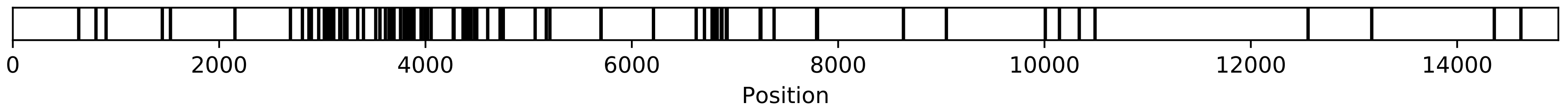


CR2 vs MM1
Number of differences: 282

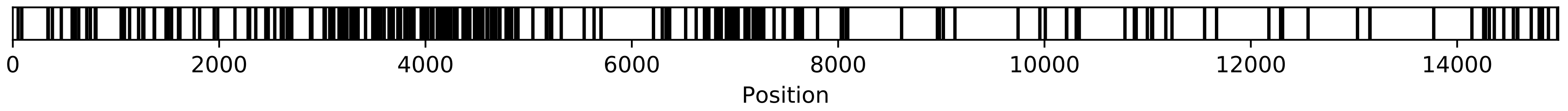


CR2 vs MM2
Number of differences: 281

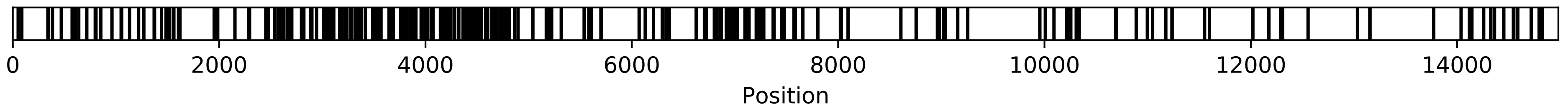


MA1 vs MA2
Number of differences: 281

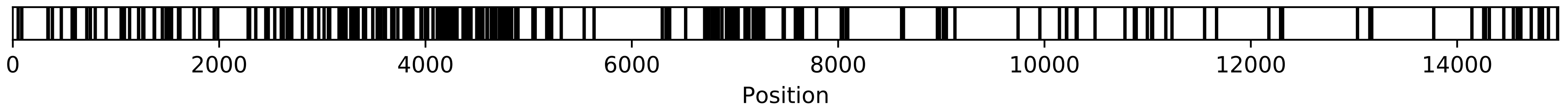


MA1 vs MM1
Number of differences: 0

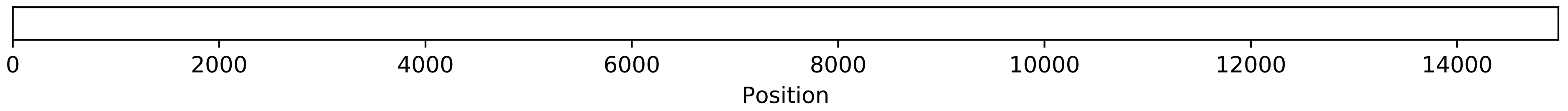


MA1 vs MM2
Number of differences: 178

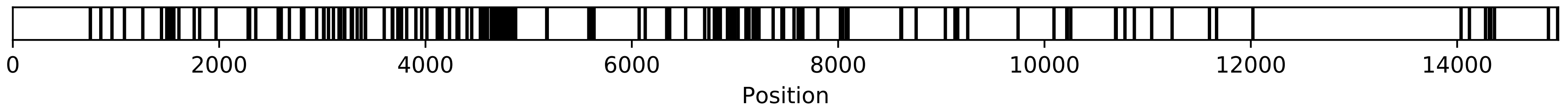


MA2 vs MM1
Number of differences: 281

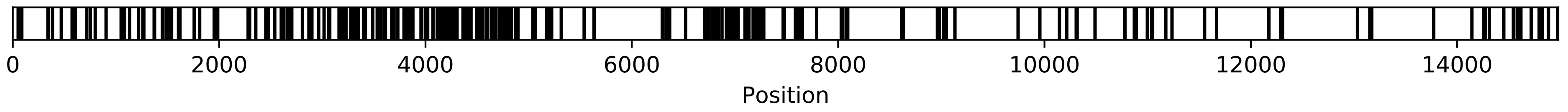


MA2 vs MM2
Number of differences: 276

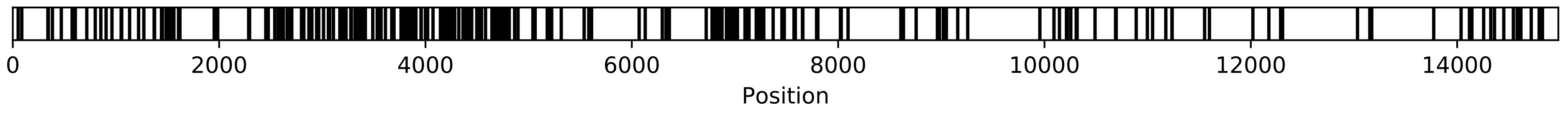


MM1 vs MM2
Number of differences: 178

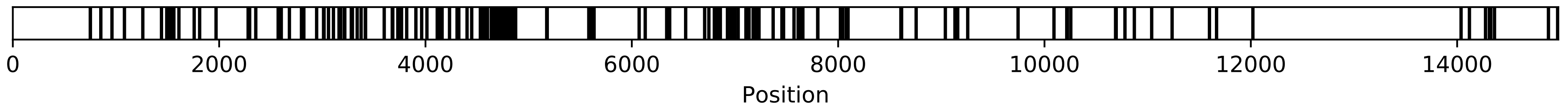

### Region: 173



CR1 vs CR2
Number of differences: 229

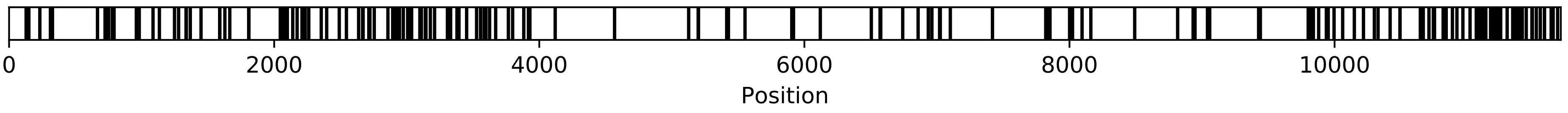


CR1 vs MA1
Number of differences: 113

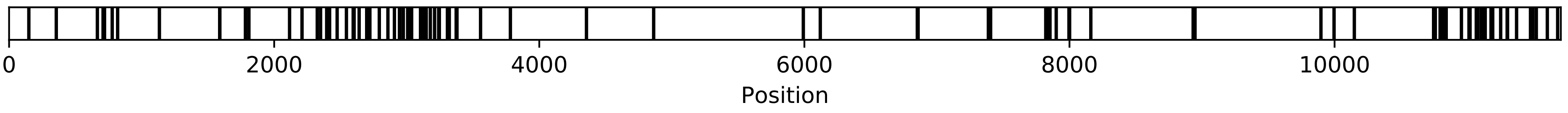


CR1 vs MA2
Number of differences: 234

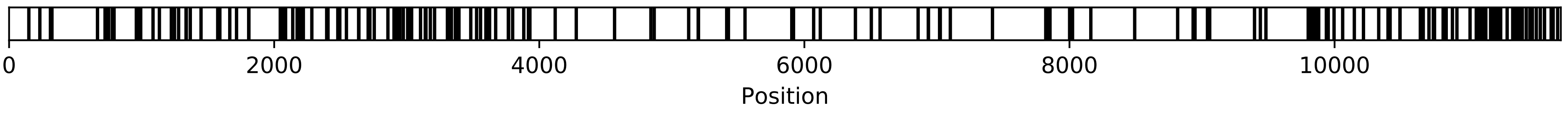


CR1 vs MM1
Number of differences: 113

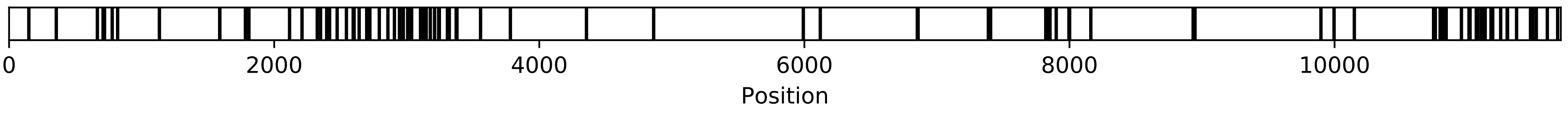


CR1 vs MM2
Number of differences: 108

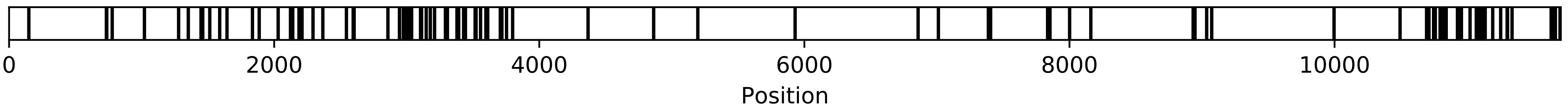


CR2 vs MA1
Number of differences: 220

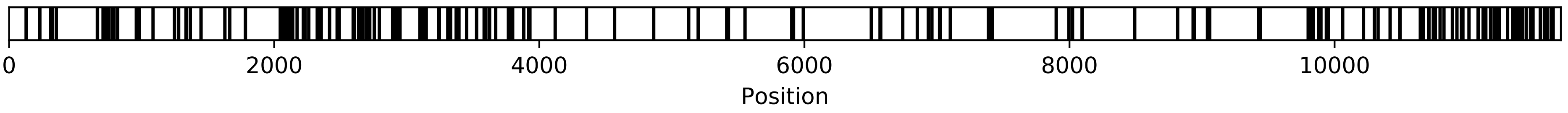


CR2 vs MA2
Number of differences: 65

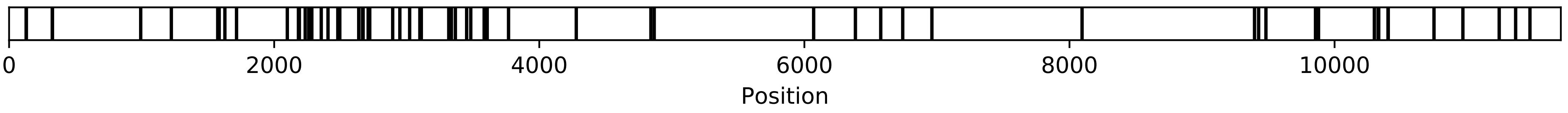


CR2 vs MM1
Number of differences: 220

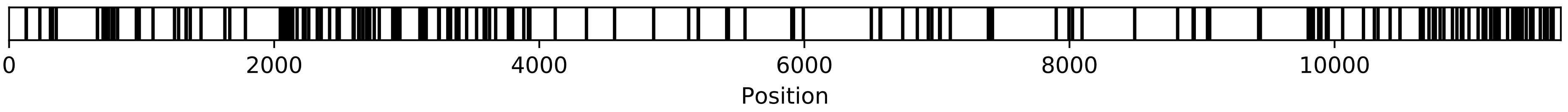


CR2 vs MM2
Number of differences: 231

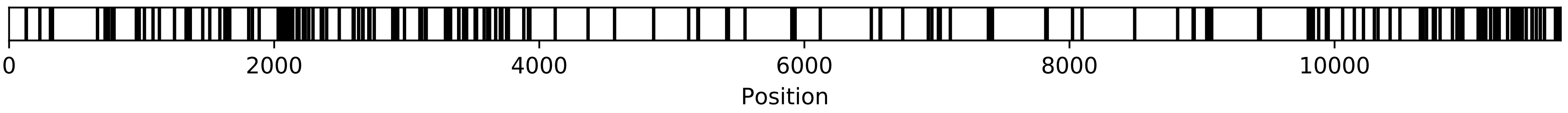


MA1 vs MA2
Number of differences: 223

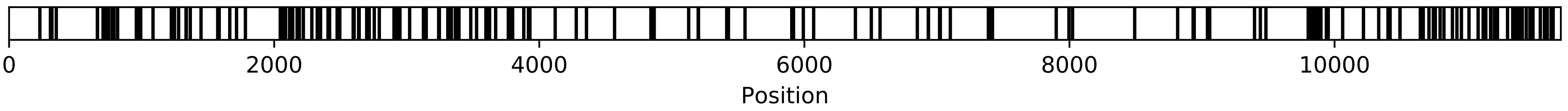


MA1 vs MM1
Number of differences: 0

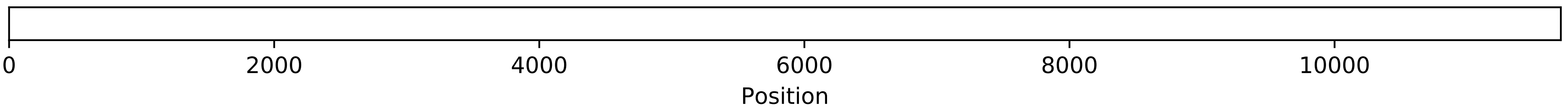


MA1 vs MM2
Number of differences: 103

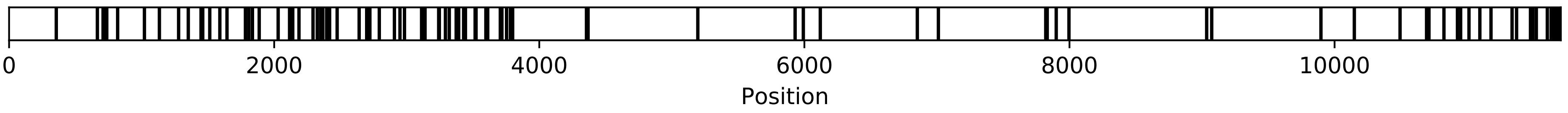


MA2 vs MM1
Number of differences: 223

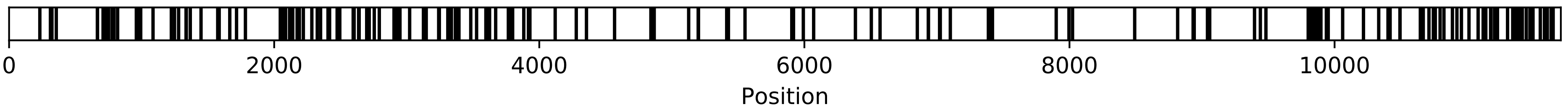


MA2 vs MM2
Number of differences: 234

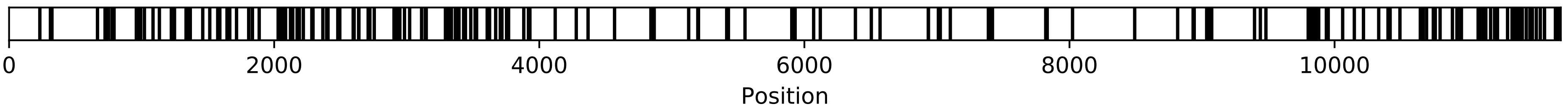


MM1 vs MM2
Number of differences: 103

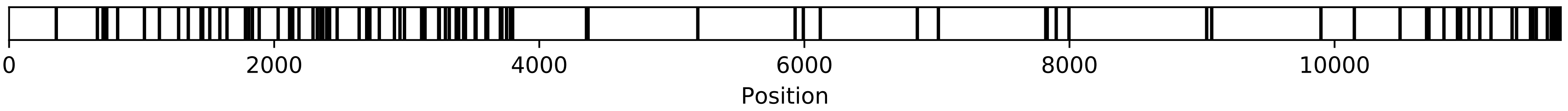

### Region: 174



CR1 vs CR2
Number of differences: 157

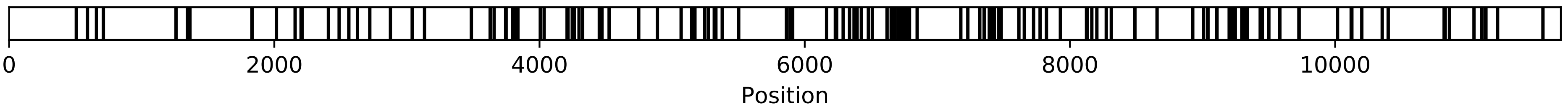


CR1 vs MA1
Number of differences: 163

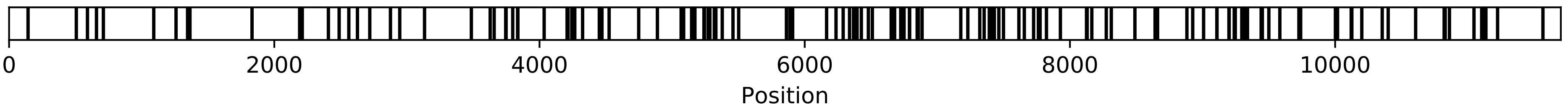


CR1 vs MA2
Number of differences: 47

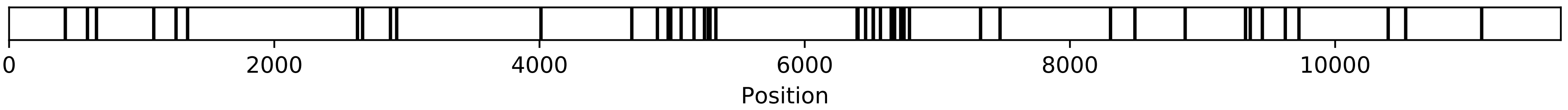


CR1 vs MM1
Number of differences: 157

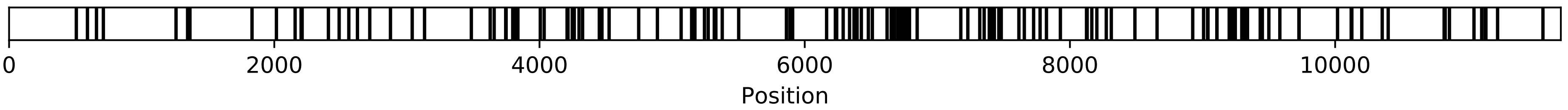


CR1 vs MM2
Number of differences: 163

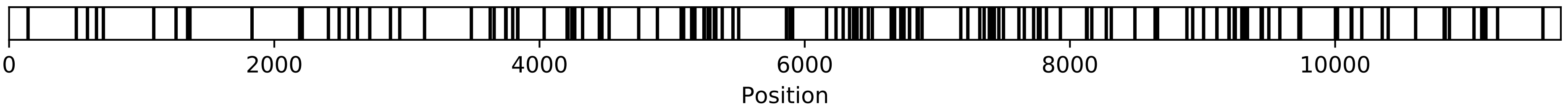


CR2 vs MA1
Number of differences: 71

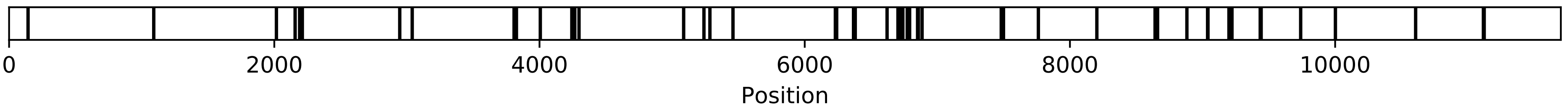


CR2 vs MA2
Number of differences: 156

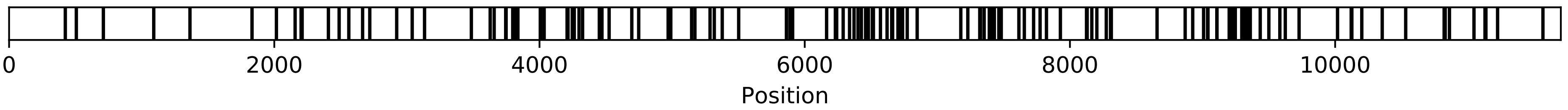


CR2 vs MM1
Number of differences: 0

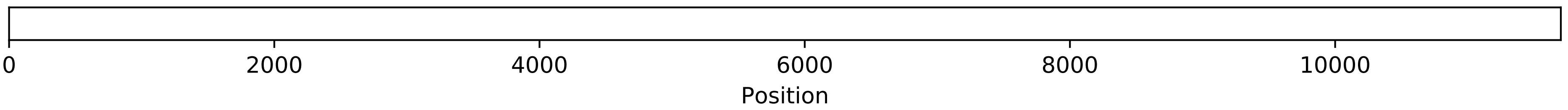


CR2 vs MM2
Number of differences: 71

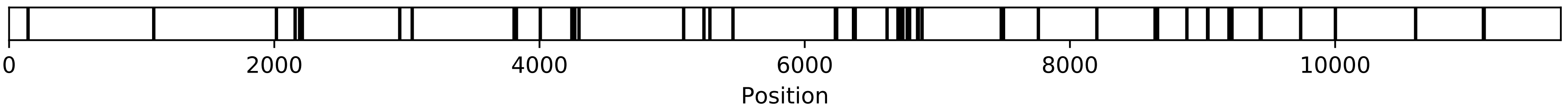


MA1 vs MA2
Number of differences: 160

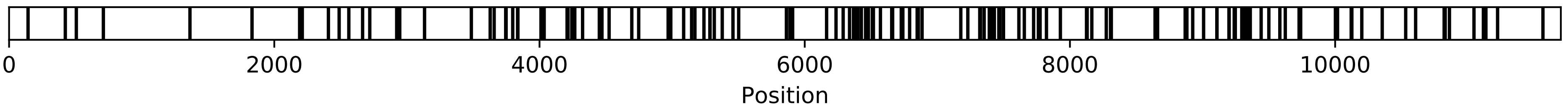


MA1 vs MM1
Number of differences: 71

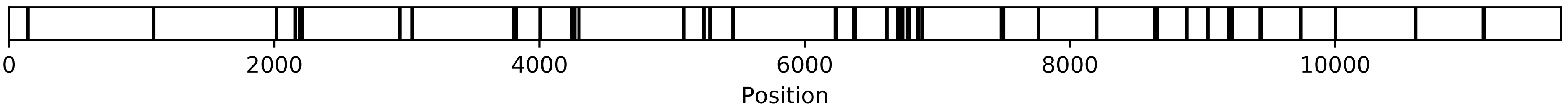


MA1 vs MM2
Number of differences: 0

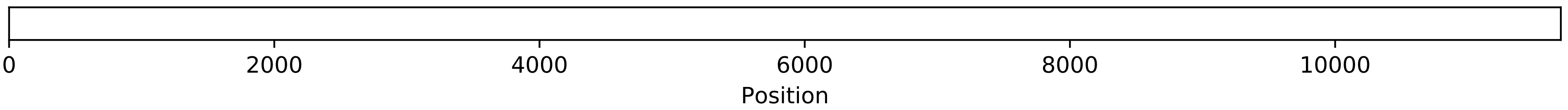


MA2 vs MM1
Number of differences: 156

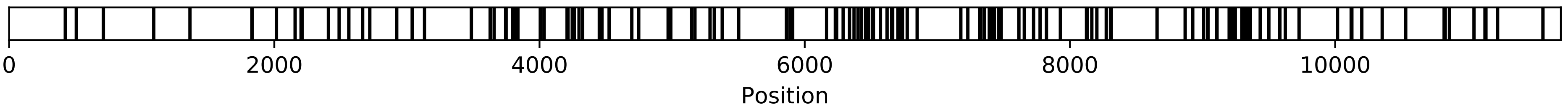


MA2 vs MM2
Number of differences: 160

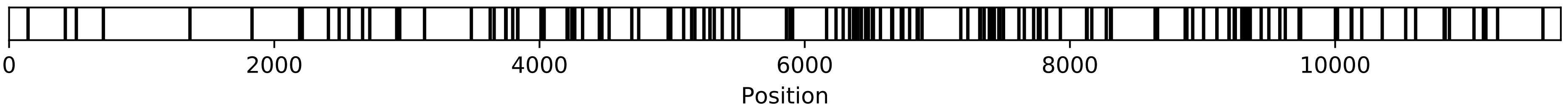


MM1 vs MM2
Number of differences: 71

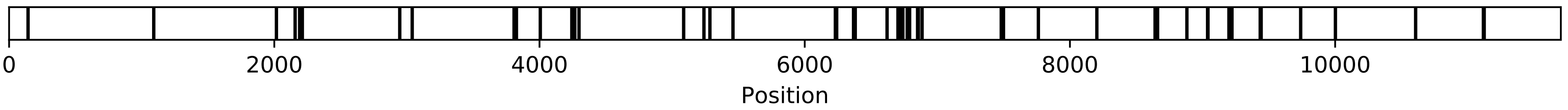

### Region: 175

CR1 vs CR2
Number of differences: 20

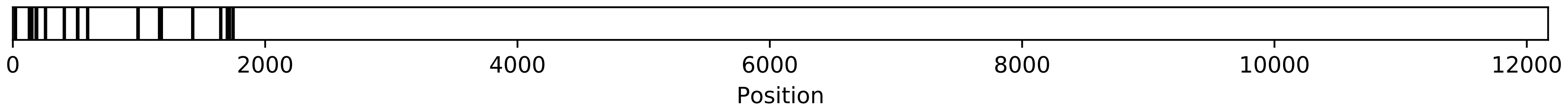


CR1 vs MA1
Number of differences: 270

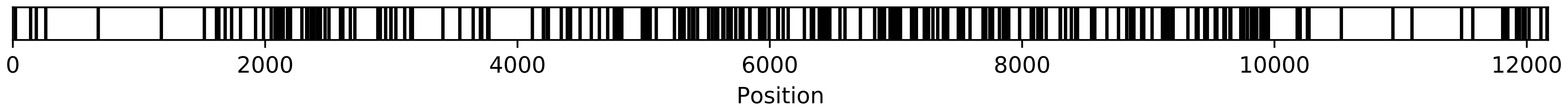


CR1 vs MA2
Number of differences: 126

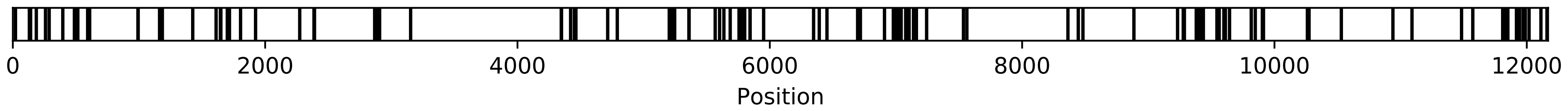


CR1 vs MM1
Number of differences: 111

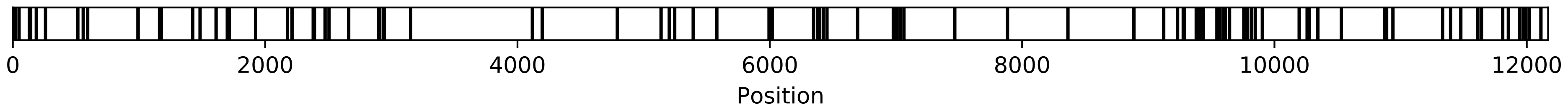


CR1 vs MM2
Number of differences: 126

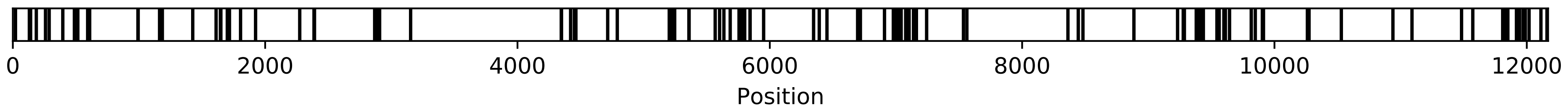


CR2 vs MA1
Number of differences: 284

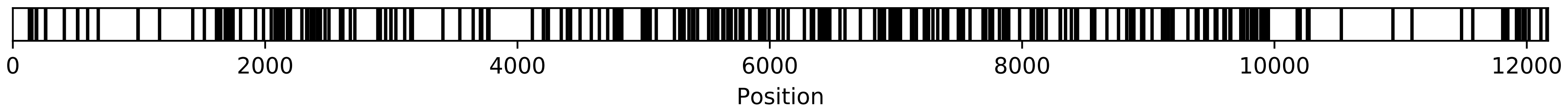


CR2 vs MA2
Number of differences: 120

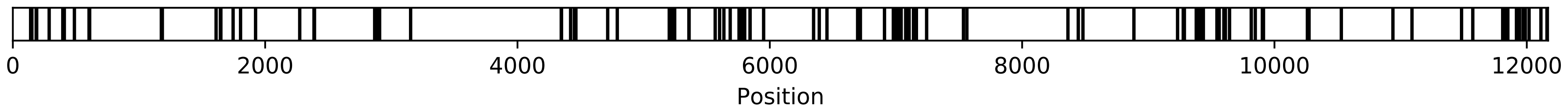


CR2 vs MM1
Number of differences: 103

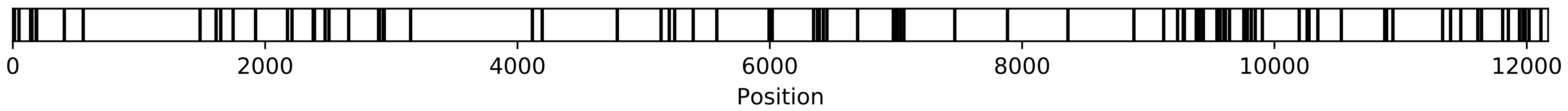


CR2 vs MM2
Number of differences: 120

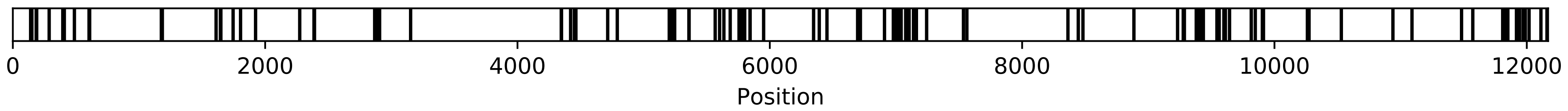


MA1 vs MA2
Number of differences: 271

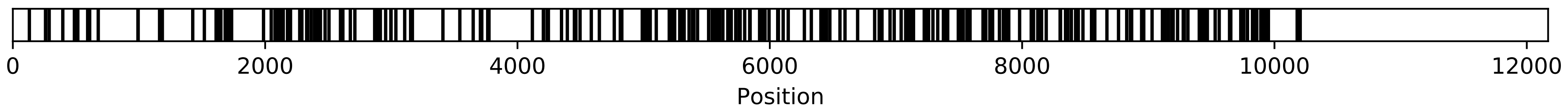


MA1 vs MM1
Number of differences: 287

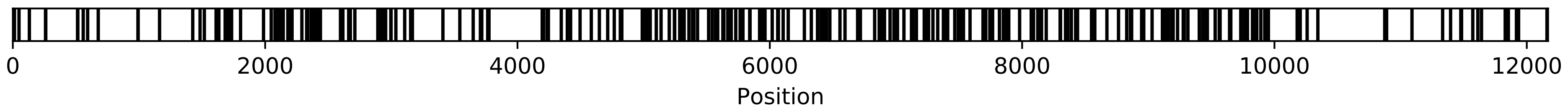


MA1 vs MM2
Number of differences: 271

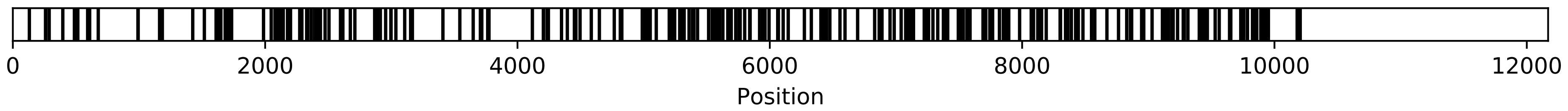


MA2 vs MM1
Number of differences: 104

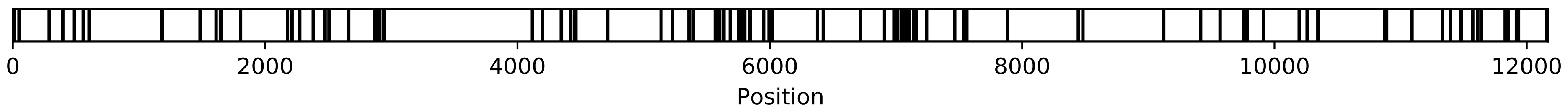


MA2 vs MM2
Number of differences: 0

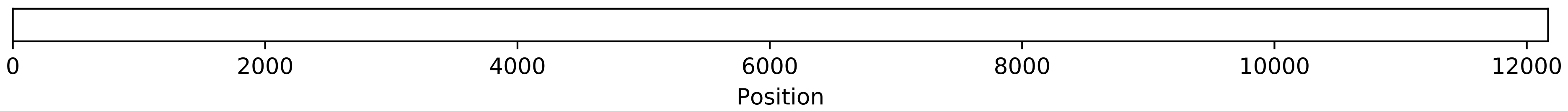


MM1 vs MM2
Number of differences: 104

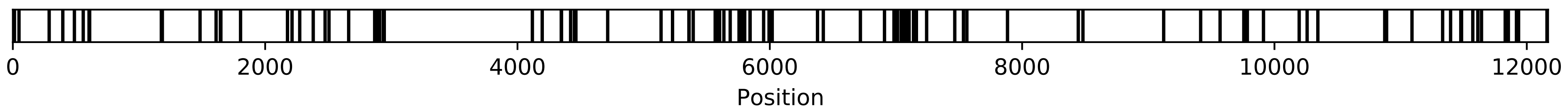

### Region: 176



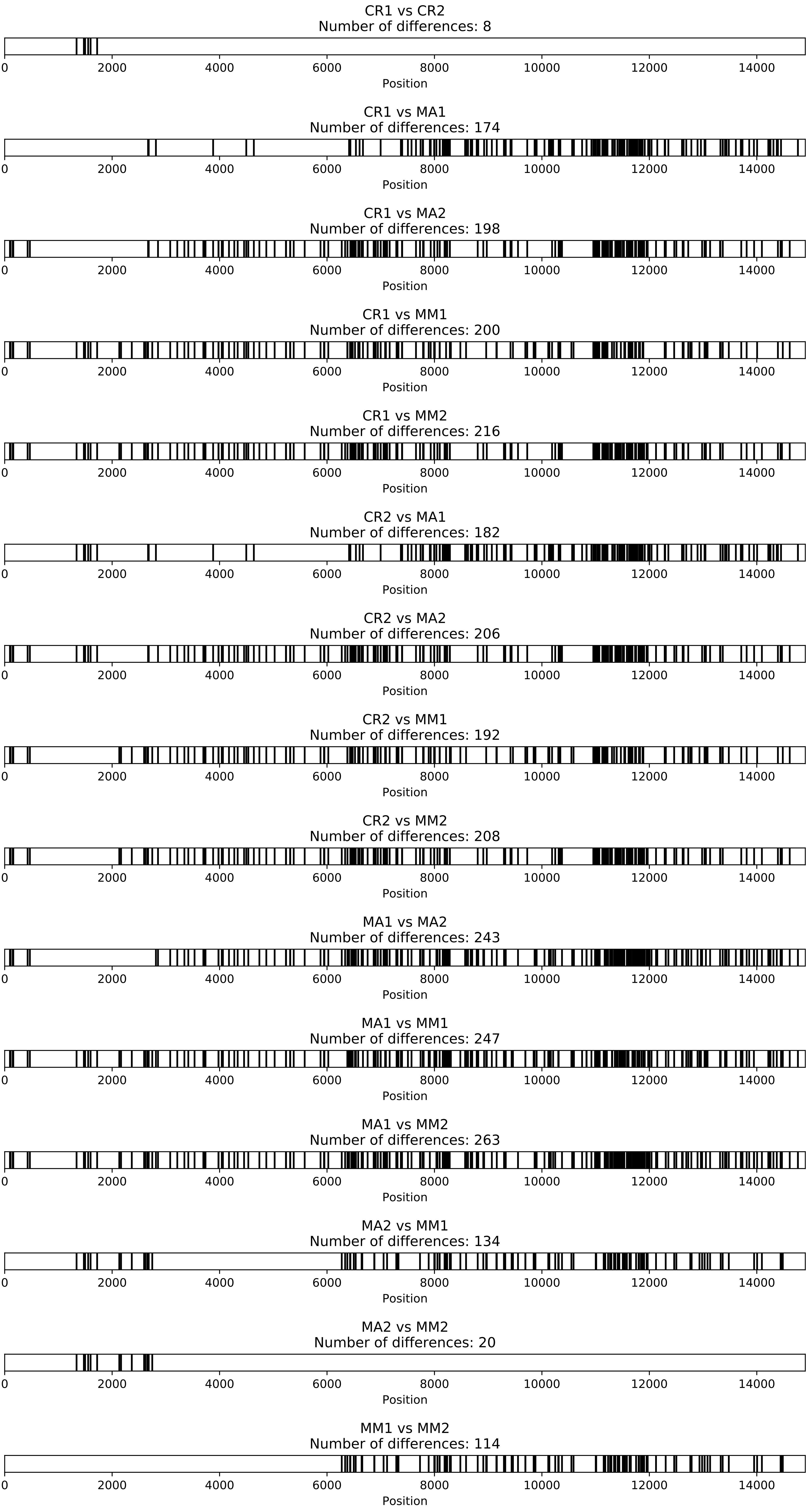
CR1 vs CR2
Number of differences: 8
CR1 vs MA1
Number of differences: 174
CR1 vs MA2
Number of differences: 198
CR1 vs MM1
Number of differences: 200
CR1 vs MM2
Number of differences: 216
CR2 vs MA1
Number of differences: 182
CR2 vs MA2
Number of differences: 206
CR2 vs MM1
Number of differences: 192
CR2 vs MM2
Number of differences: 208
MA1 vs MA2
Number of differences: 243
MA1 vs MM1
Number of differences: 247
MA1 vs MM2
Number of differences: 263
MA2 vs MM1
Number of differences: 134
MA2 vs MM2
Number of differences: 20
MM1 vs MM2
Number of differences: 114
0
2000
4000
6000
8000
10000
12000
14000
Position

### Region: 177



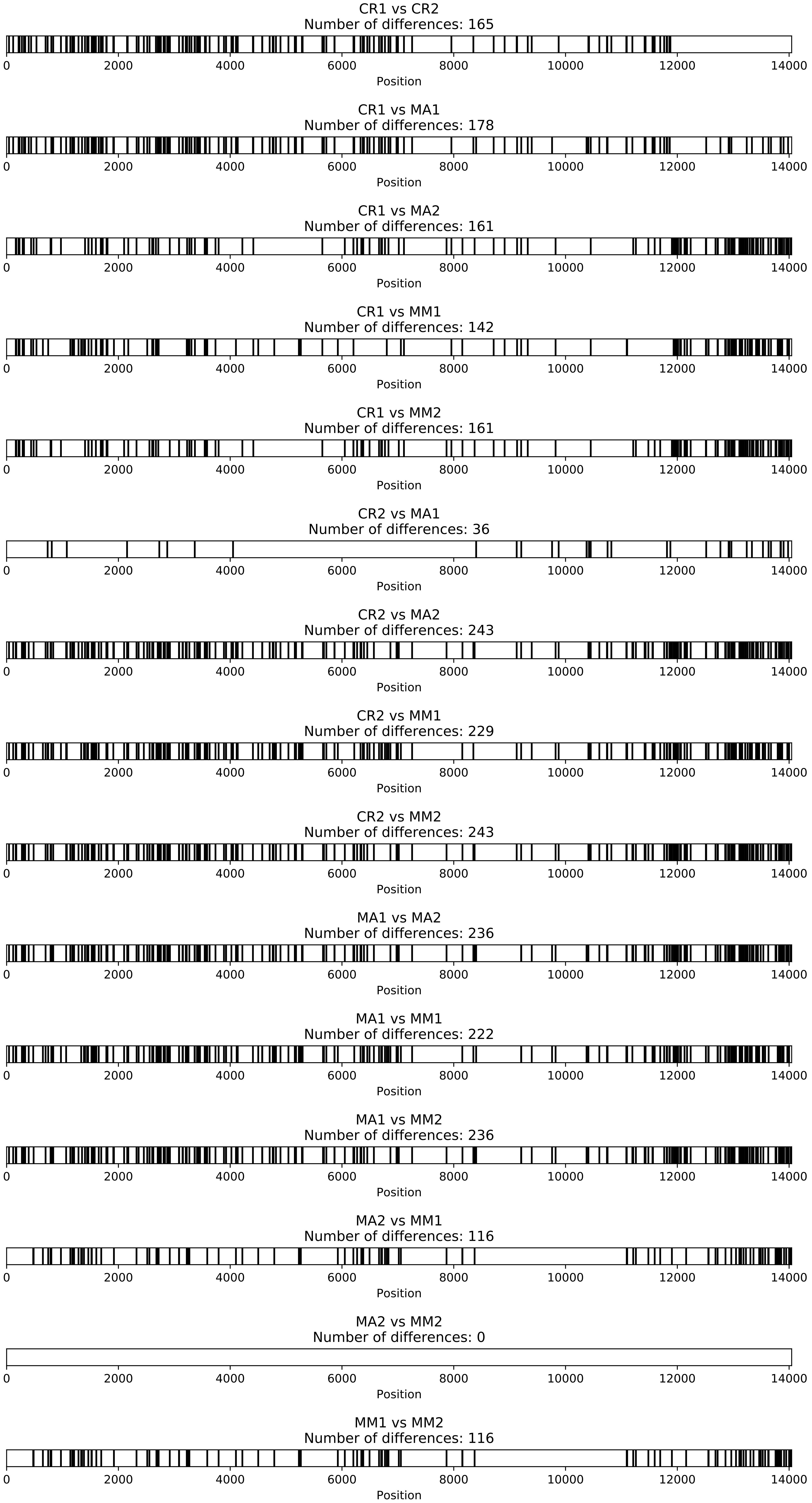
CR1 vs CR2
Number of differences: 165
CR1 vs MA1
Number of differences: 178
CR1 vs MA2
Number of differences: 161
CR1 vs MM1
Number of differences: 142
CR1 vs MM2
Number of differences: 161
CR2 vs MA1
Number of differences: 36
CR2 vs MA2
Number of differences: 243
CR2 vs MM1
Number of differences: 229
CR2 vs MM2
Number of differences: 243
MA1 vs MA2
Number of differences: 236
MA1 vs MM1
Number of differences: 222
MA1 vs MM2
Number of differences: 236
MA2 vs MM1
Number of differences: 116
MA2 vs MM2
Number of differences: 0
MM1 vs MM2
Number of differences: 116
Position

### Region: 178



CR1 vs CR2
Number of differences: 445

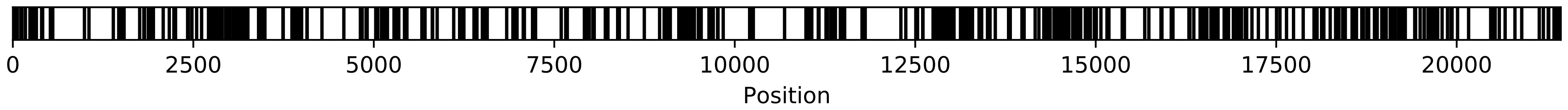


CR1 vs MA1
Number of differences: 192

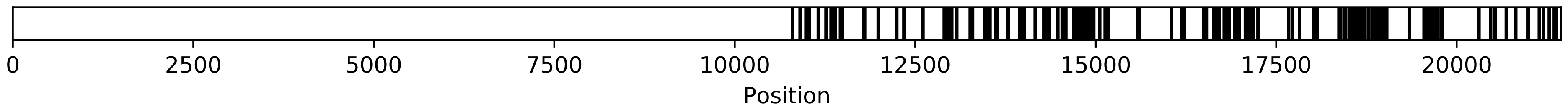


CR1 vs MA2
Number of differences: 465

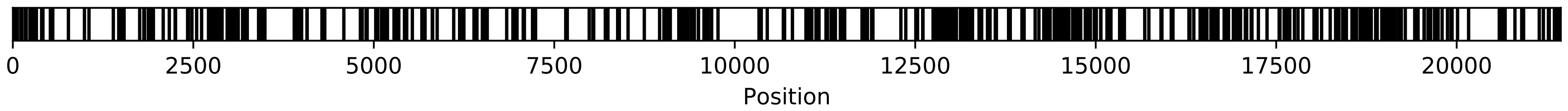


CR1 vs MM1
Number of differences: 260

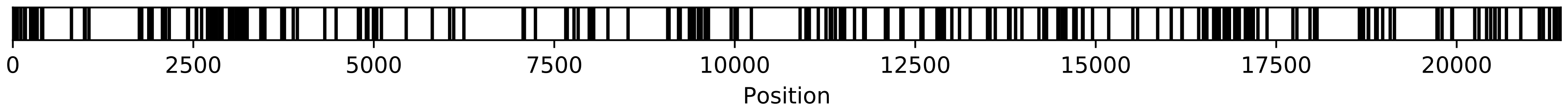


CR1 vs MM2
Number of differences: 245

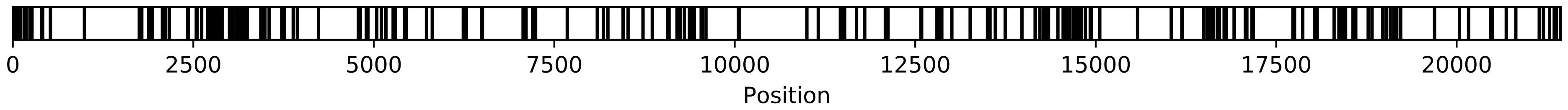


CR2 vs MA1
Number of differences: 437

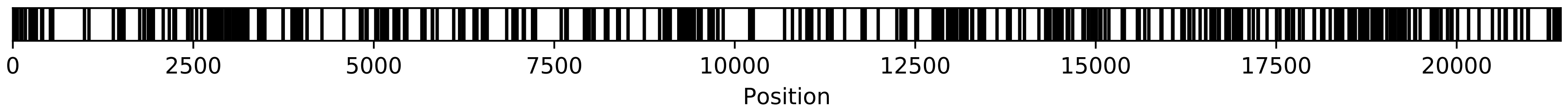


CR2 vs MA2
Number of differences: 149

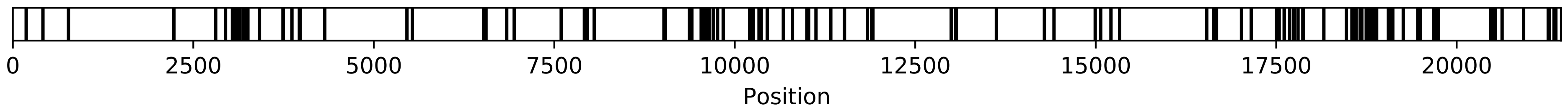


CR2 vs MM1
Number of differences: 447

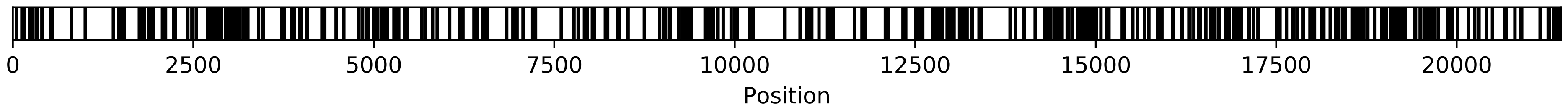


CR2 vs MM2
Number of differences: 446

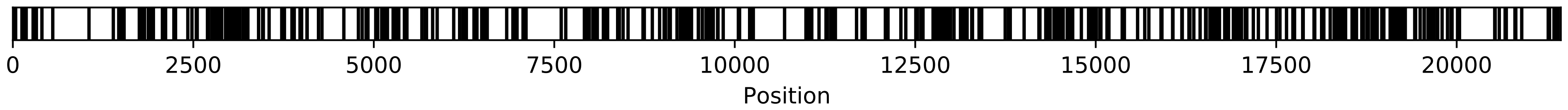


MA1 vs MA2
Number of differences: 463

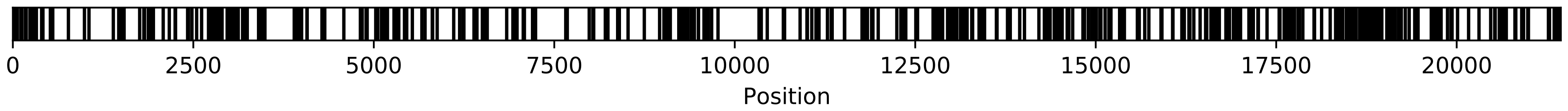


MA1 vs MM1
Number of differences: 282

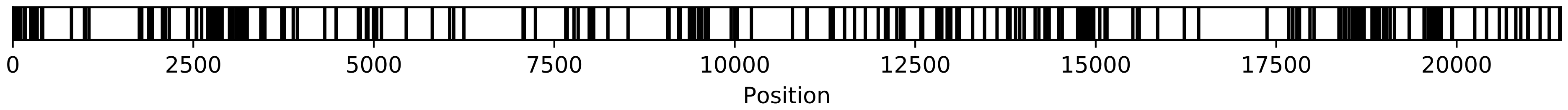


MA1 vs MM2
Number of differences: 311

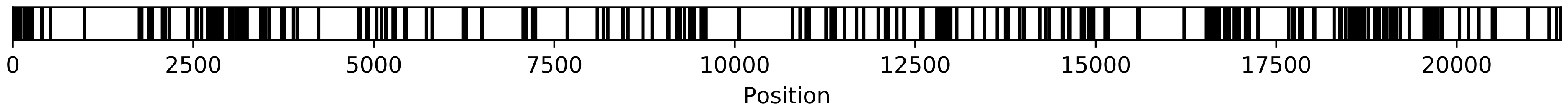


MA2 vs MM1
Number of differences: 475

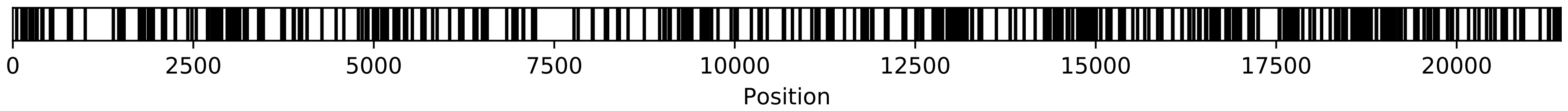


MA2 vs MM2
Number of differences: 469

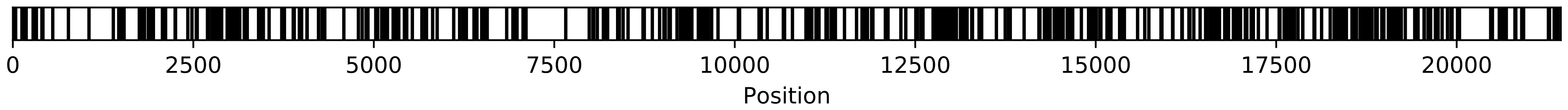


MM1 vs MM2
Number of differences: 223

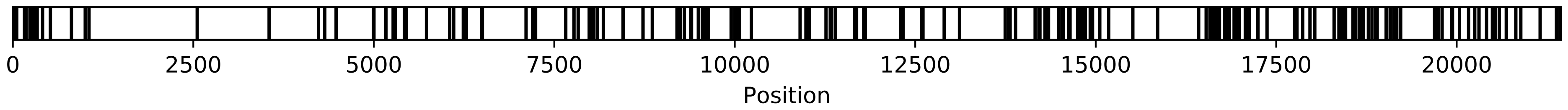

### Region: 179

CR1 vs CR2
Number of differences: 285

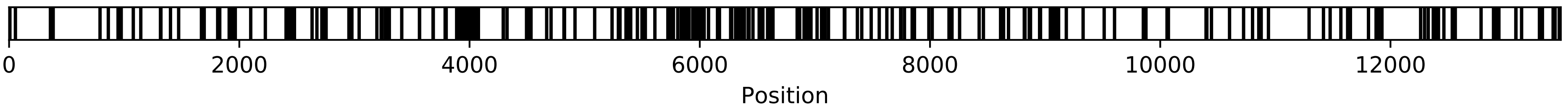


CR1 vs MA1
Number of differences: 151

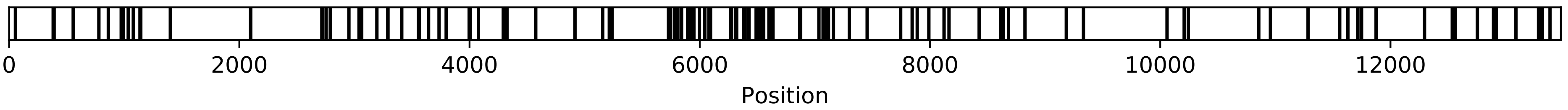


CR1 vs MA2
Number of differences: 276

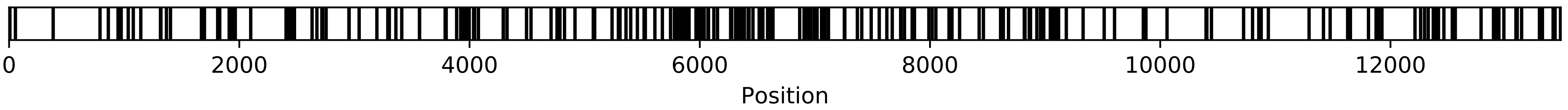


CR1 vs MM1
Number of differences: 151

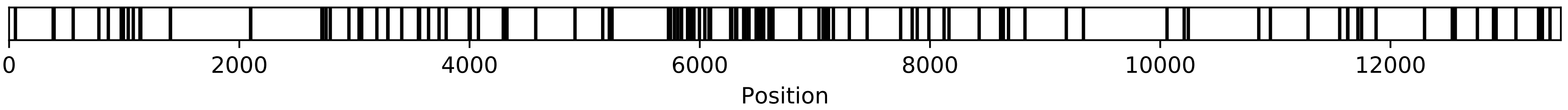


CR1 vs MM2
Number of differences: 137

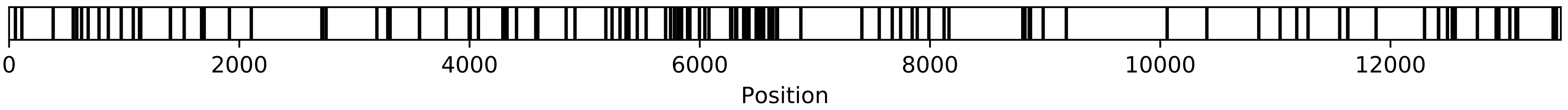


CR2 vs MA1
Number of differences: 292

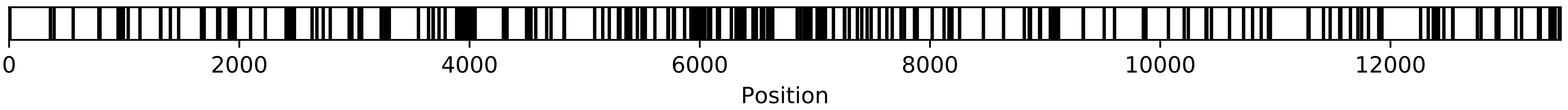


CR2 vs MA2
Number of differences: 65

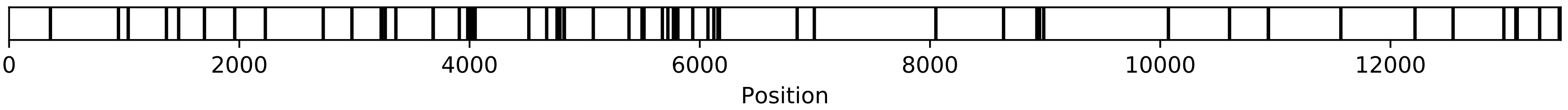


CR2 vs MM1
Number of differences: 292

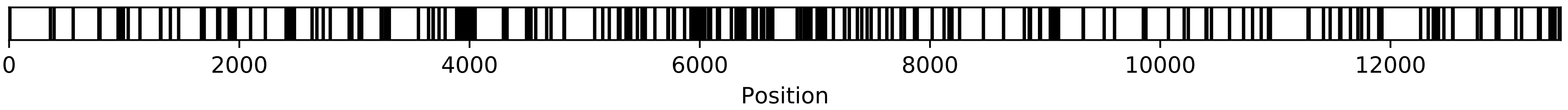


CR2 vs MM2
Number of differences: 275

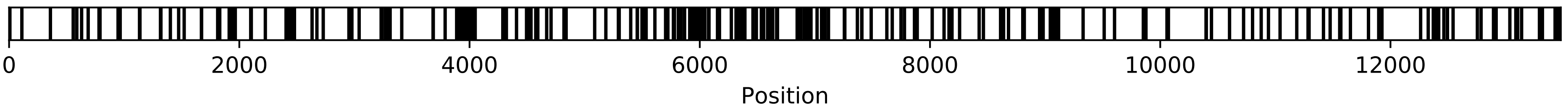


MA1 vs MA2
Number of differences: 279

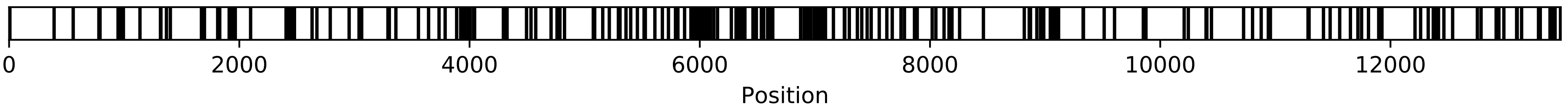


MA1 vs MM1
Number of differences: 0

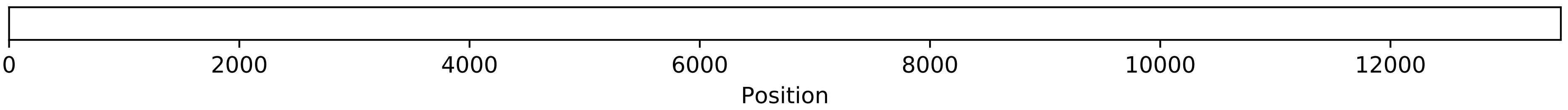


MA1 vs MM2
Number of differences: 127

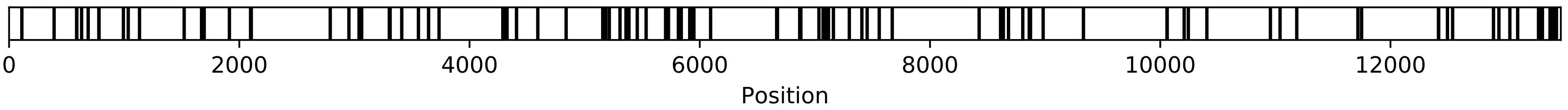


MA2 vs MM1
Number of differences: 279

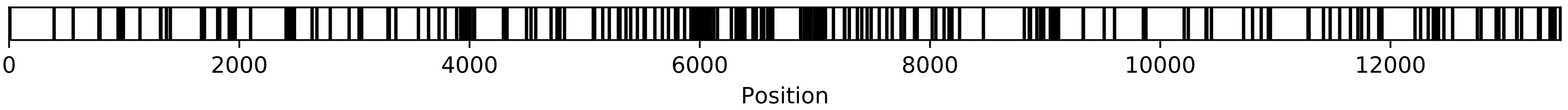


MA2 vs MM2
Number of differences: 266

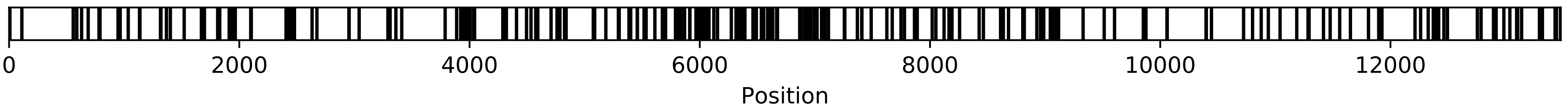


MM1 vs MM2
Number of differences: 127

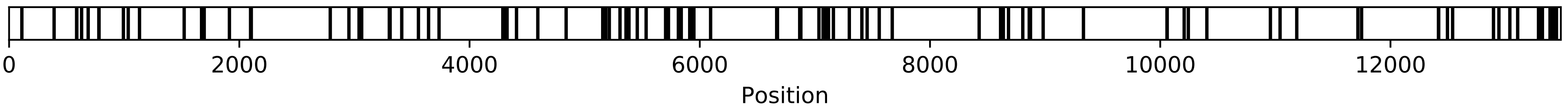

### Region: 180



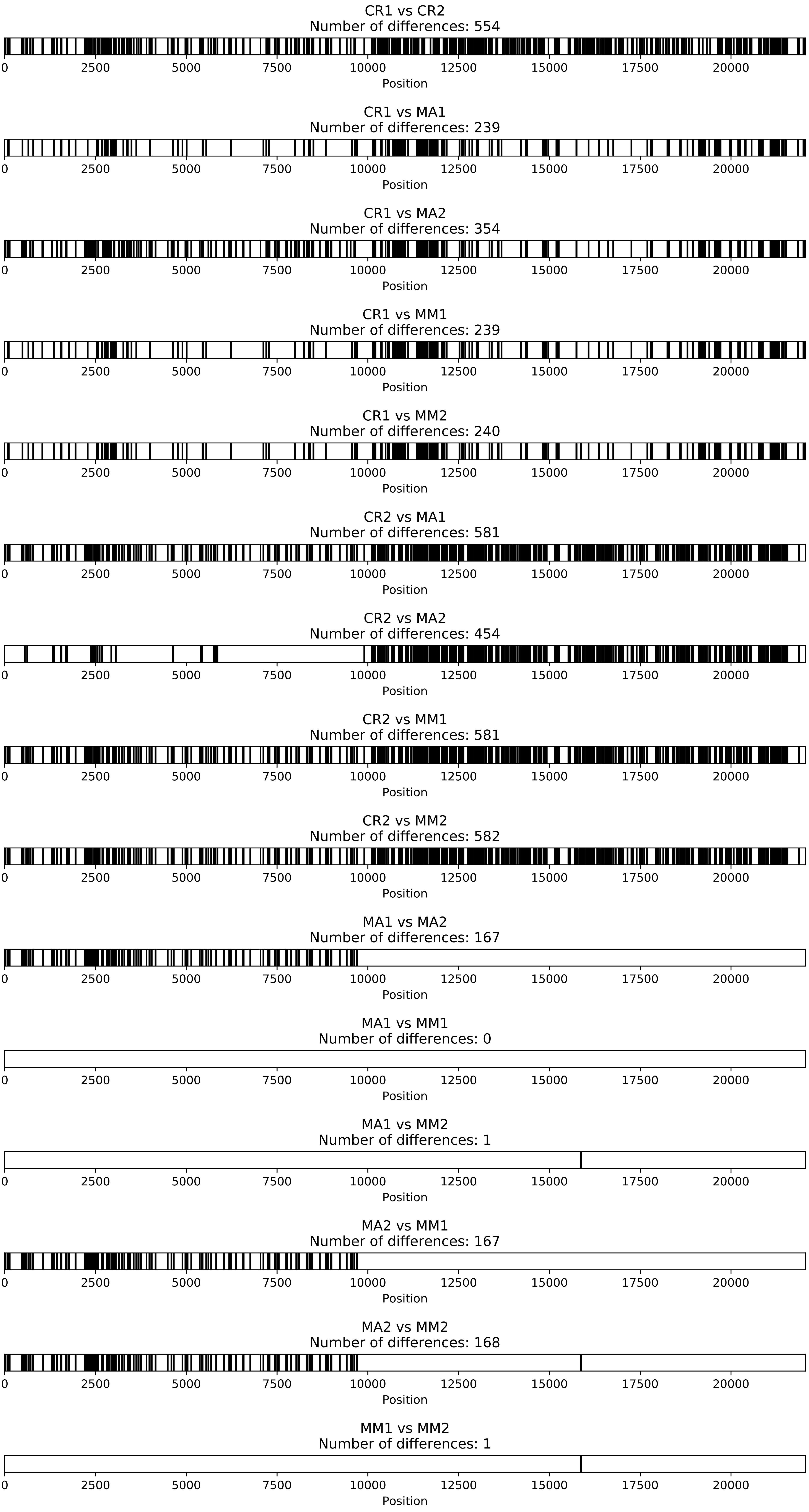
CR1 vs CR2
Number of differences: 554
CR1 vs MA1
Number of differences: 239
CR1 vs MA2
Number of differences: 354
CR1 vs MM1
Number of differences: 239
CR1 vs MM2
Number of differences: 240
CR2 vs MA1
Number of differences: 581
CR2 vs MA2
Number of differences: 454
CR2 vs MM1
Number of differences: 581
CR2 vs MM2
Number of differences: 582
MA1 vs MA2
Number of differences: 167
MA1 vs MM1
Number of differences: 0
MA1 vs MM2
Number of differences: 1
MA2 vs MM1
Number of differences: 167
MA2 vs MM2
Number of differences: 168
MM1 vs MM2
Number of differences: 1
0
2500
5000
7500
10000
12500
15000
17500
20000
Position

### Region: 181



CR1 vs CR2
Number of differences: 272

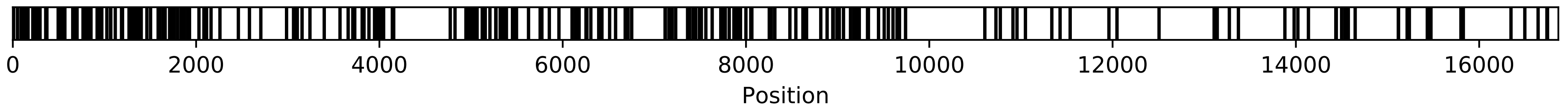


CR1 vs MA1
Number of differences: 275

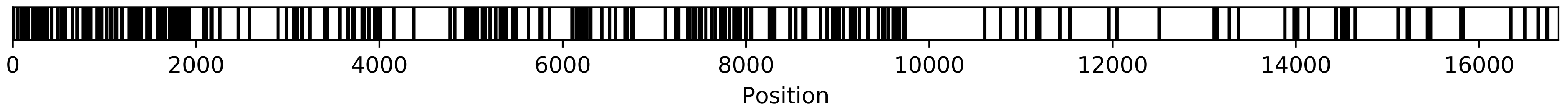


CR1 vs MA2
Number of differences: 120

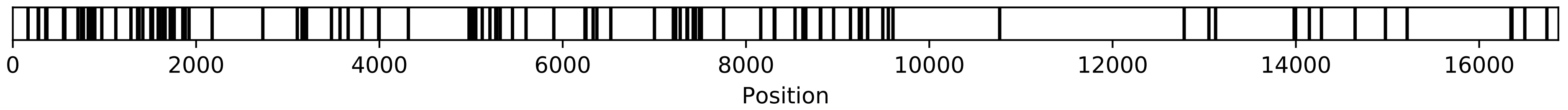


CR1 vs MM1
Number of differences: 0

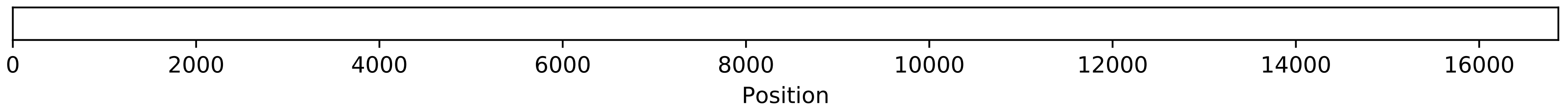


CR1 vs MM2
Number of differences: 129

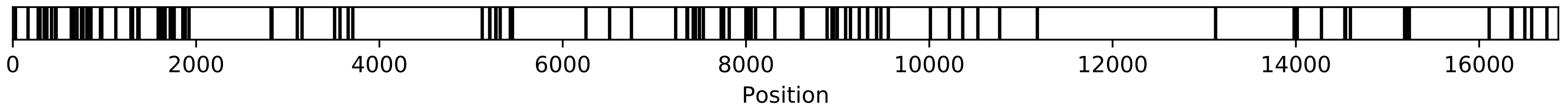


CR2 vs MA1
Number of differences: 55

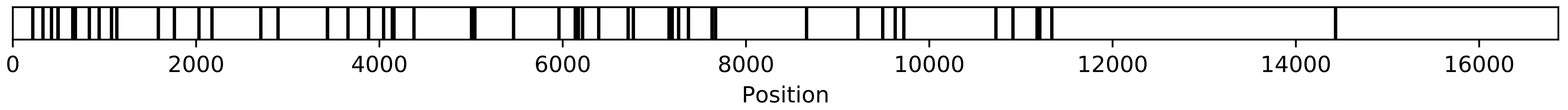


CR2 vs MA2
Number of differences: 259

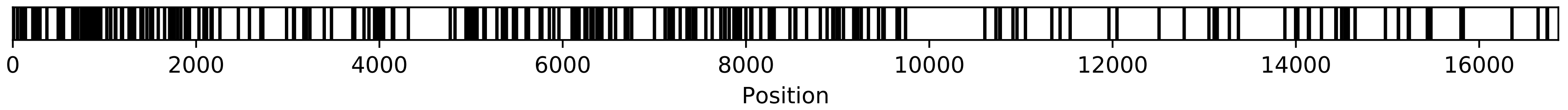


CR2 vs MM1
Number of differences: 272

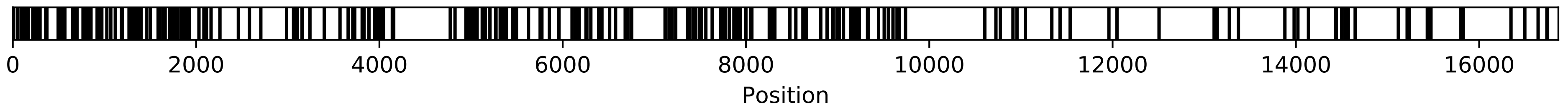


CR2 vs MM2
Number of differences: 267

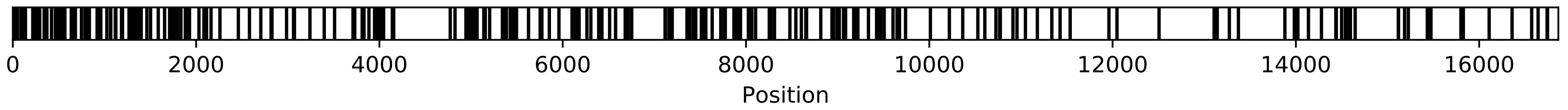


MA1 vs MA2
Number of differences: 261

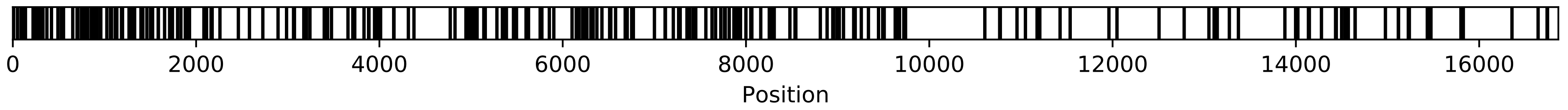


MA1 vs MM1
Number of differences: 275

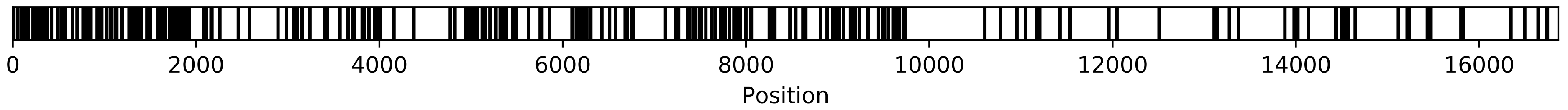


MA1 vs MM2
Number of differences: 267

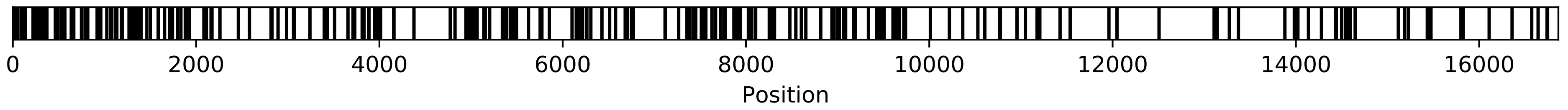


MA2 vs MM1
Number of differences: 120

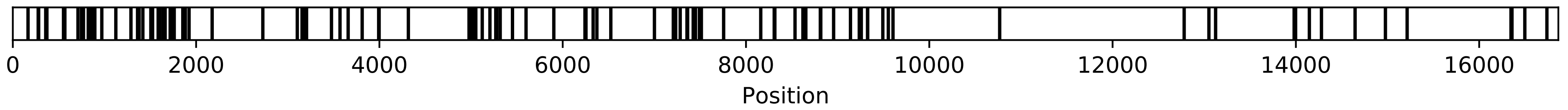


MA2 vs MM2
Number of differences: 120

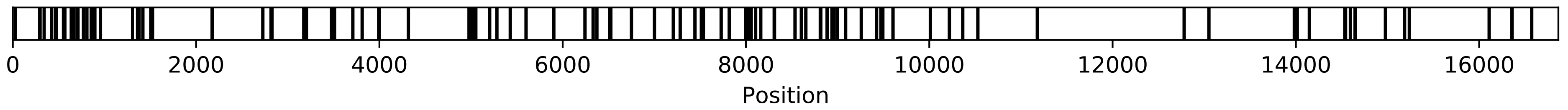


MM1 vs MM2
Number of differences: 129

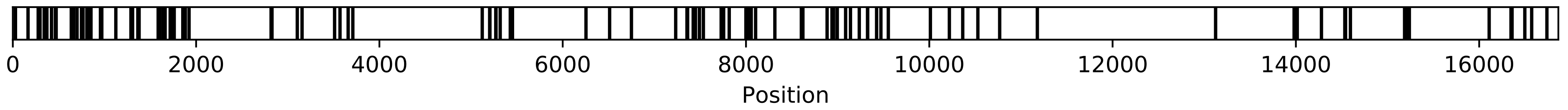

### Region: 183

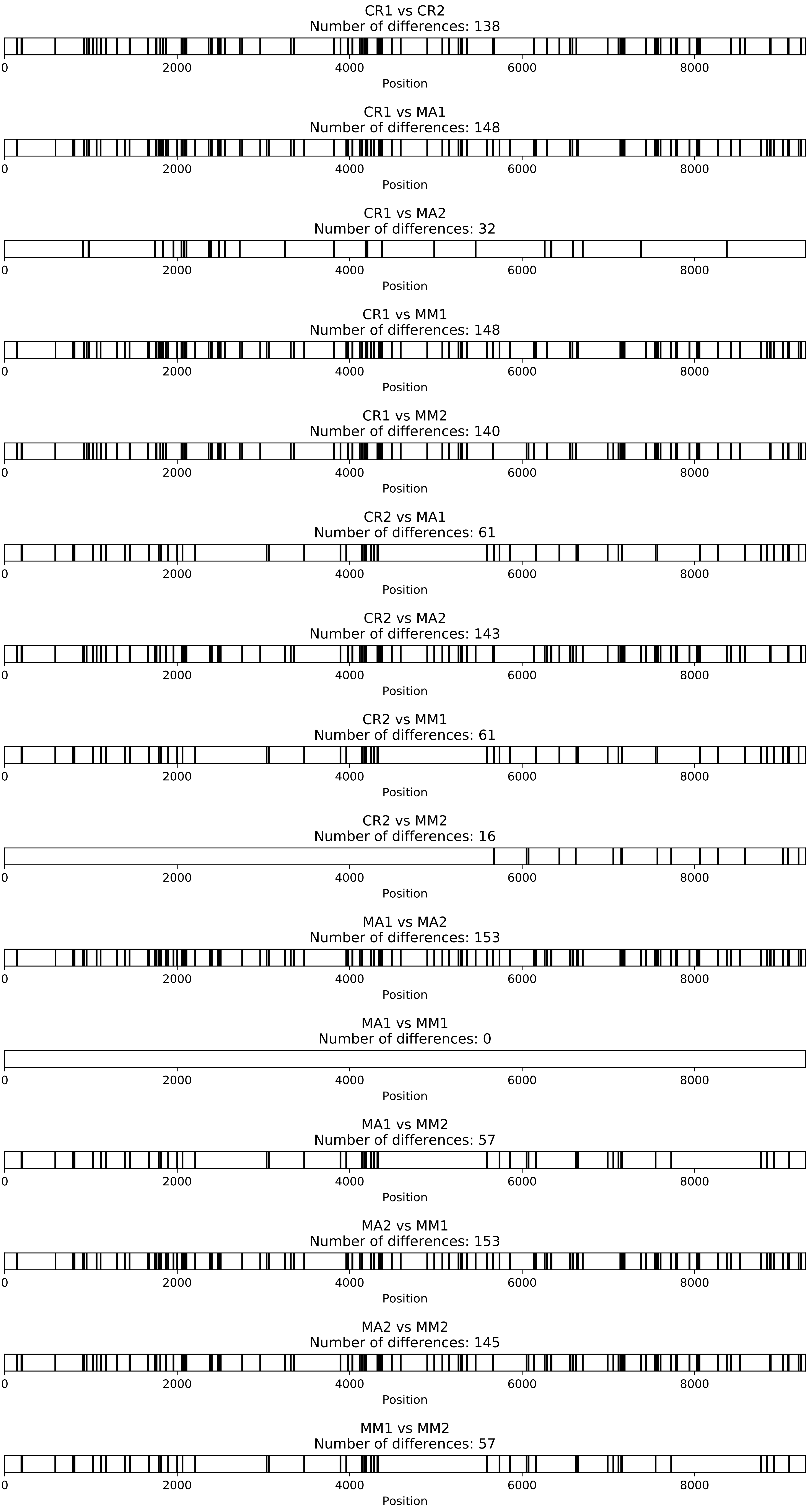
CR1 vs CR2
Number of differences: 138
CR1 vs MA1
Number of differences: 148
CR1 vs MA2
Number of differences: 32
CR1 vs MM1
Number of differences: 148
CR1 vs MM2
Number of differences: 140
CR2 vs MA1
Number of differences: 61
CR2 vs MA2
Number of differences: 143
CR2 vs MM1
Number of differences: 61
CR2 vs MM2
Number of differences: 16
MA1 vs MA2
Number of differences: 153
MA1 vs MM1
Number of differences: 0
MA1 vs MM2
Number of differences: 57
MA2 vs MM1
Number of differences: 153
MA2 vs MM2
Number of differences: 145
MM1 vs MM2
Number of differences: 57
0
2000
4000
6000
8000
Position

### Region: 186

CR1 vs CR2
Number of differences: 220

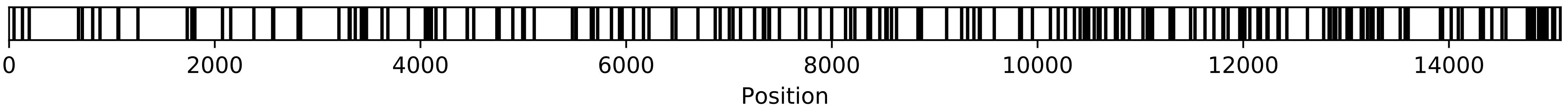


CR1 vs MA1
Number of differences: 73

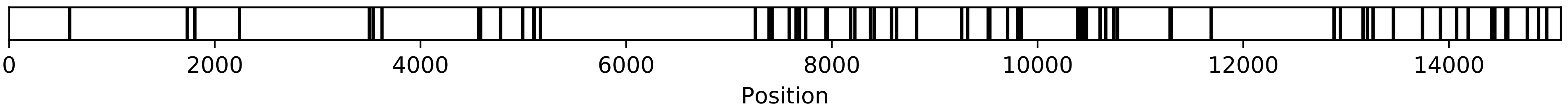


CR1 vs MA2
Number of differences: 226

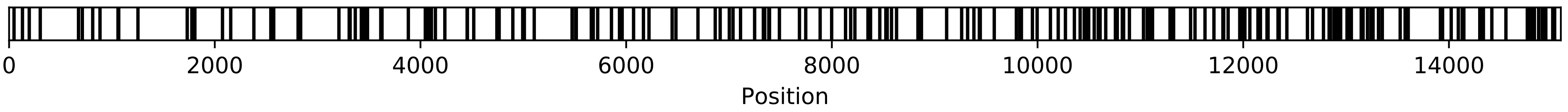


CR1 vs MM1
Number of differences: 73

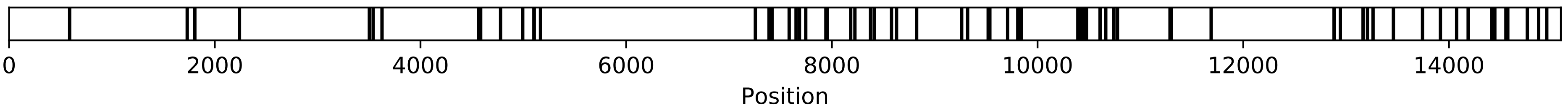


CR1 vs MM2
Number of differences: 61

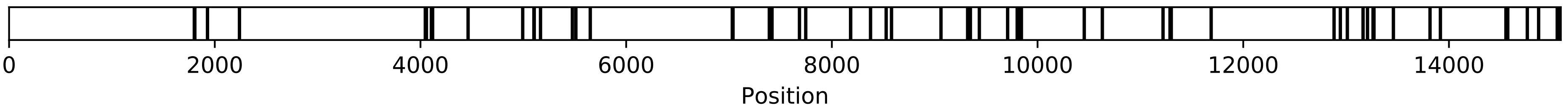


CR2 vs MA1
Number of differences: 223

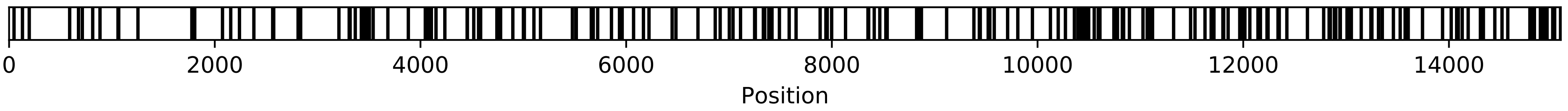


CR2 vs MA2
Number of differences: 32

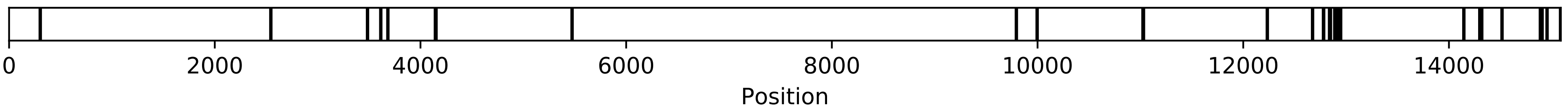


CR2 vs MM1
Number of differences: 223

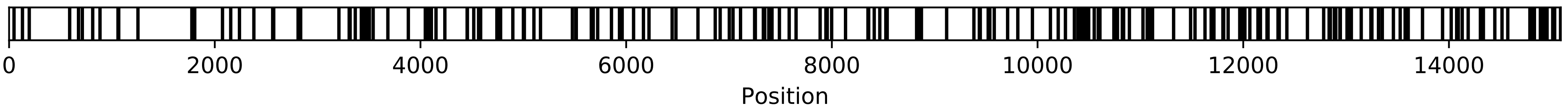


CR2 vs MM2
Number of differences: 214

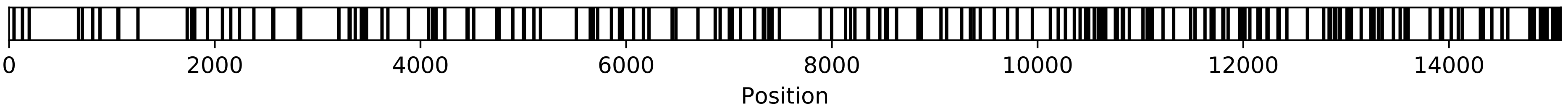


MA1 vs MA2
Number of differences: 229

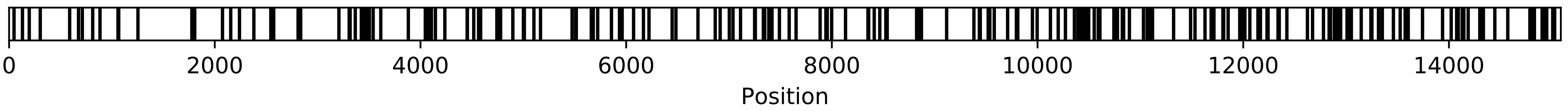


MA1 vs MM1
Number of differences: 0

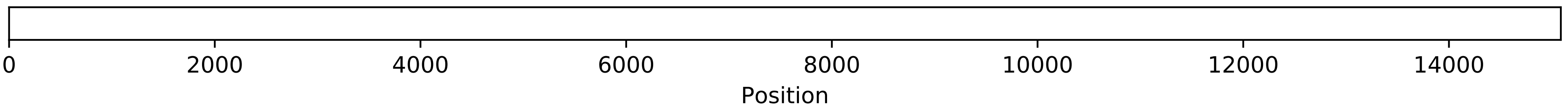


MA1 vs MM2
Number of differences: 66

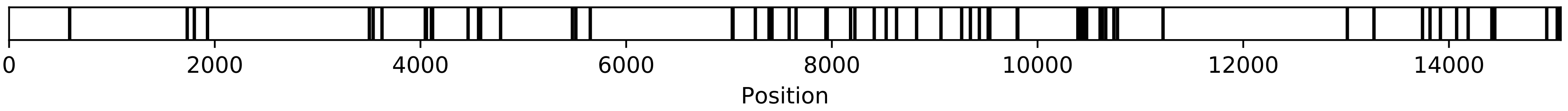


MA2 vs MM1
Number of differences: 229

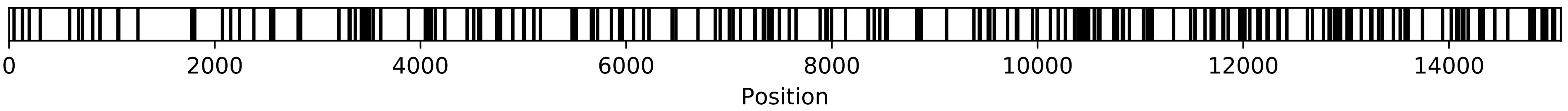


MA2 vs MM2
Number of differences: 220

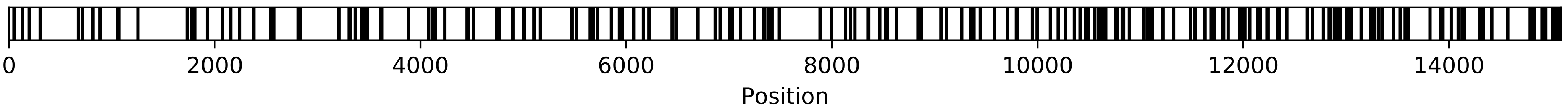


MM1 vs MM2
Number of differences: 66

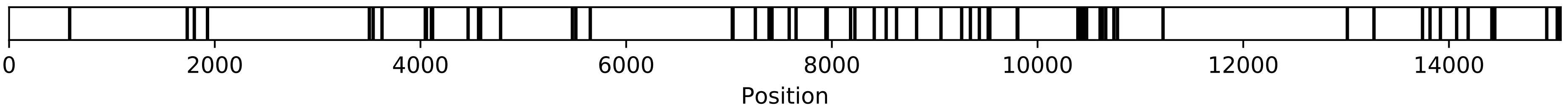

### Region: 187



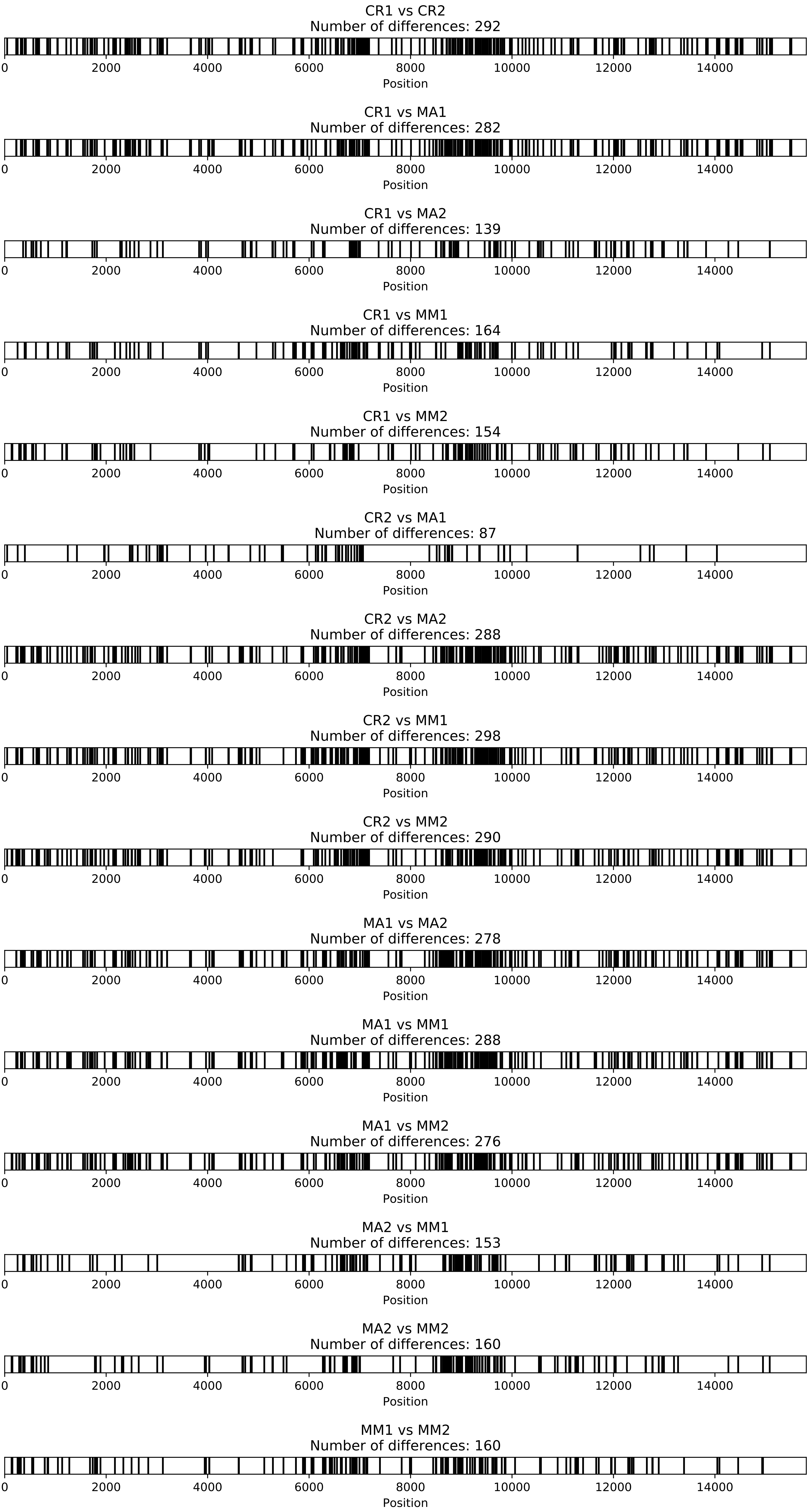
CR1 vs CR2
Number of differences: 292
CR1 vs MA1
Number of differences: 282
CR1 vs MA2
Number of differences: 139
CR1 vs MM1
Number of differences: 164
CR1 vs MM2
Number of differences: 154
CR2 vs MA1
Number of differences: 87
CR2 vs MA2
Number of differences: 288
CR2 vs MM1
Number of differences: 298
CR2 vs MM2
Number of differences: 290
MA1 vs MA2
Number of differences: 278
MA1 vs MM1
Number of differences: 288
MA1 vs MM2
Number of differences: 276
MA2 vs MM1
Number of differences: 153
MA2 vs MM2
Number of differences: 160
MM1 vs MM2
Number of differences: 160
0
2000
4000
6000
8000
10000
12000
14000
Position

### Region: 188



CR1 vs CR2
Number of differences: 110

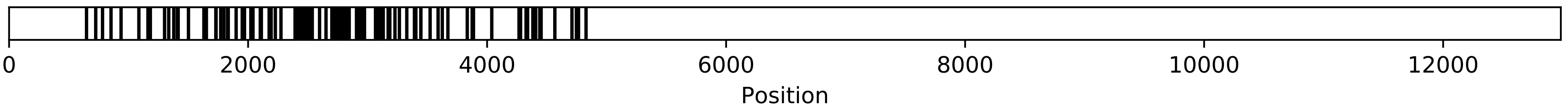

CR1 vs MA1
Number of differences: 169

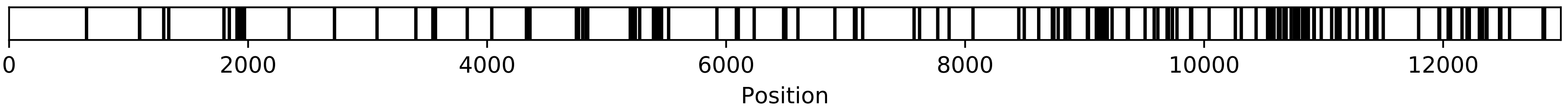

CR1 vs MA2
Number of differences: 169

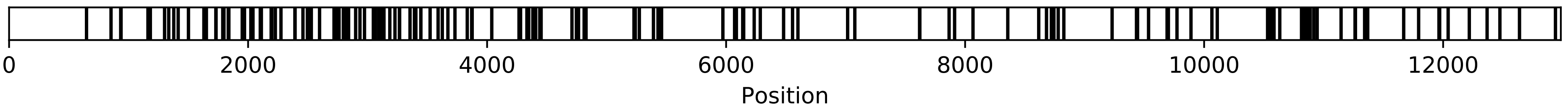

CR1 vs MM1
Number of differences: 169

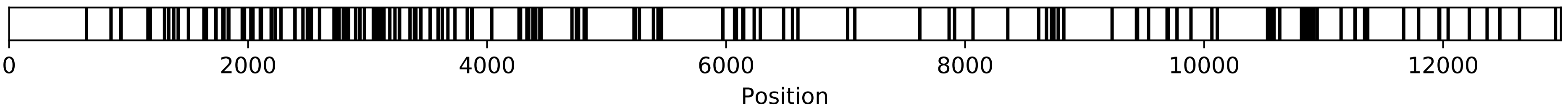

CR1 vs MM2
Number of differences: 177

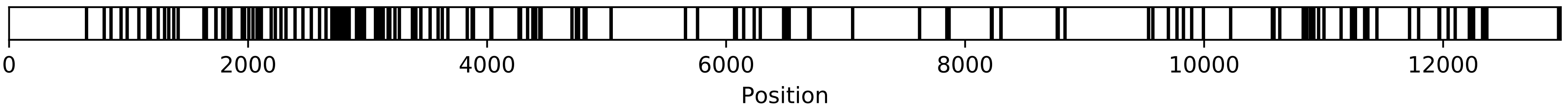

CR2 vs MA1
Number of differences: 253

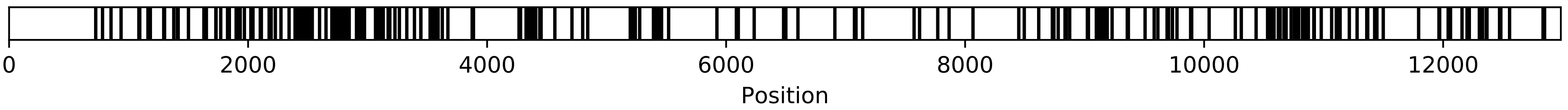

CR2 vs MA2
Number of differences: 129

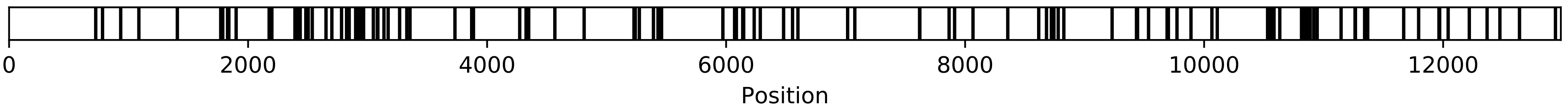

CR2 vs MM1
Number of differences: 129

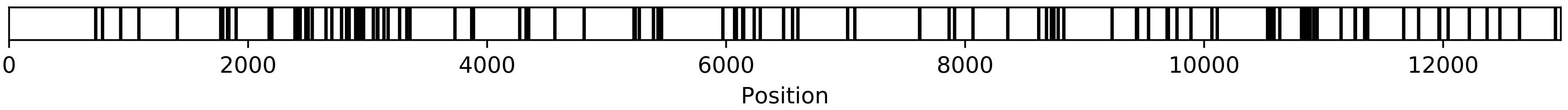

CR2 vs MM2
Number of differences: 104

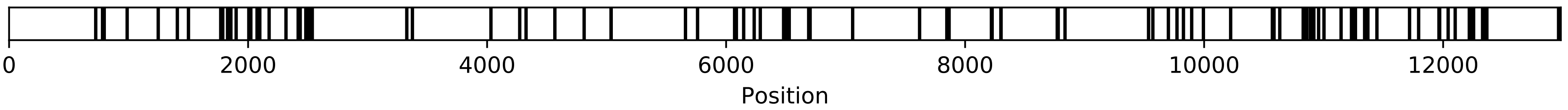

MA1 vs MA2
Number of differences: 241

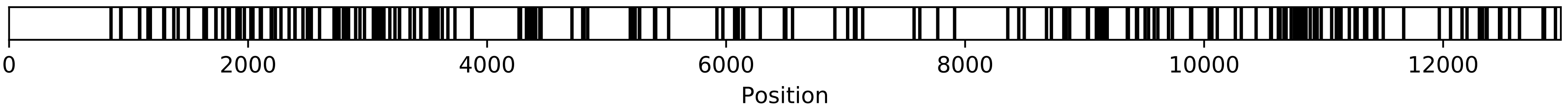

MA1 vs MM1
Number of differences: 241

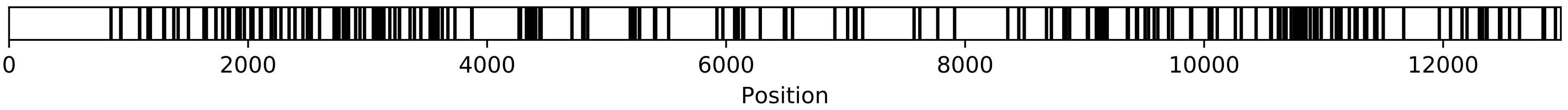

MA1 vs MM2
Number of differences: 270

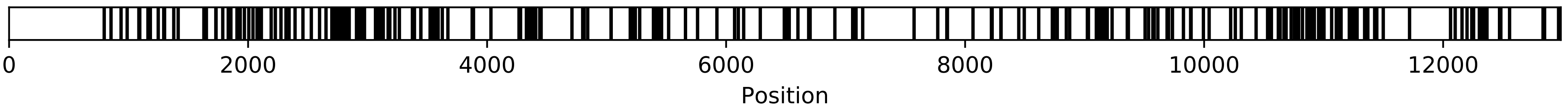

MA2 vs MM1
Number of differences: 0

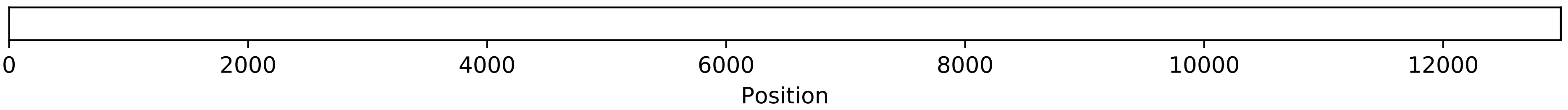

MA2 vs MM2
Number of differences: 139

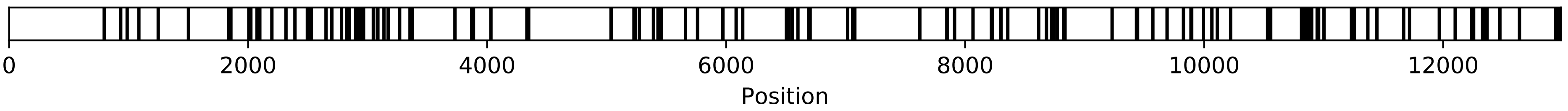

MM1 vs MM2
Number of differences: 139

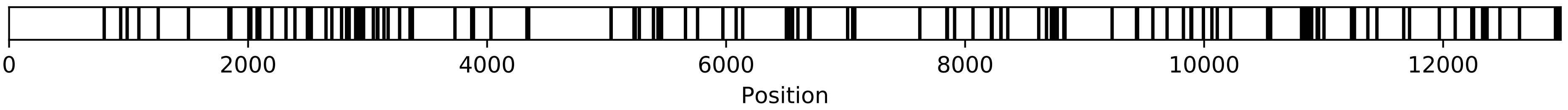

### Region: 189



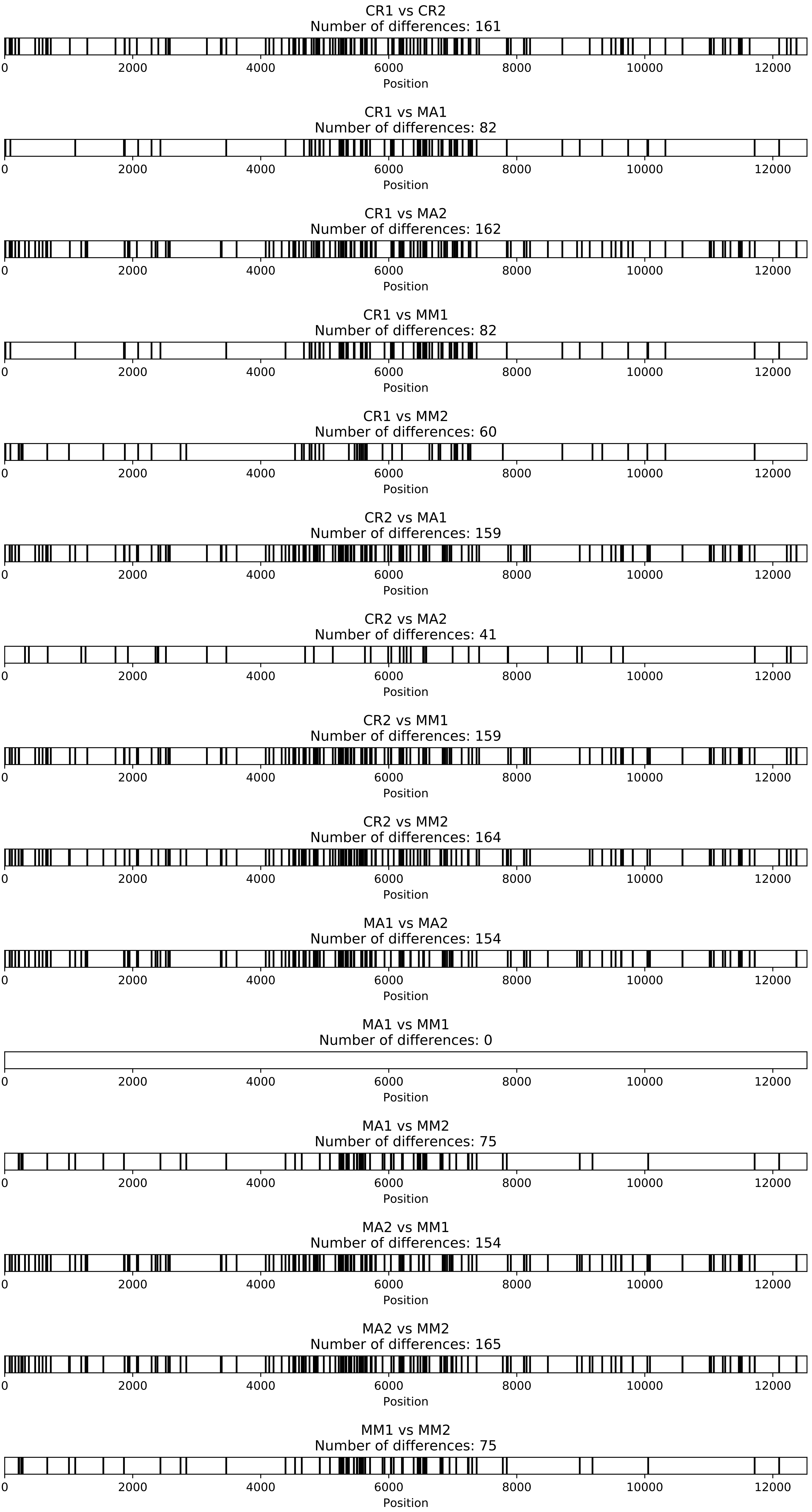
CR1 vs CR2
Number of differences: 161
CR1 vs MA1
Number of differences: 82
CR1 vs MA2
Number of differences: 162
CR1 vs MM1
Number of differences: 82
CR1 vs MM2
Number of differences: 60
CR2 vs MA1
Number of differences: 159
CR2 vs MA2
Number of differences: 41
CR2 vs MM1
Number of differences: 159
CR2 vs MM2
Number of differences: 164
MA1 vs MA2
Number of differences: 154
MA1 vs MM1
Number of differences: 0
MA1 vs MM2
Number of differences: 75
MA2 vs MM1
Number of differences: 154
MA2 vs MM2
Number of differences: 165
MM1 vs MM2
Number of differences: 75
0
2000
4000
6000
8000
10000
12000
Position

### Region: 197



CR1 vs CR2
Number of differences: 203

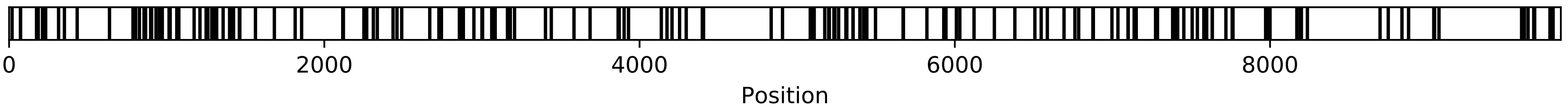


CR1 vs MA1
Number of differences: 206

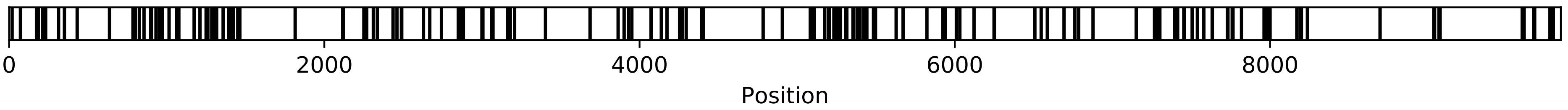


CR1 vs MA2
Number of differences: 84

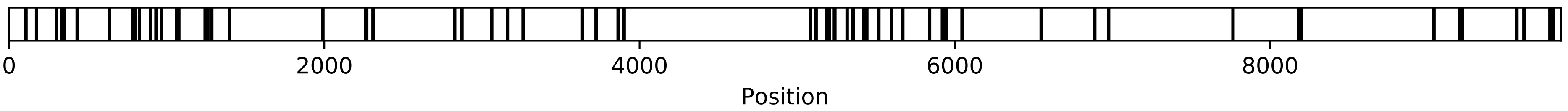


CR1 vs MM1
Number of differences: 103

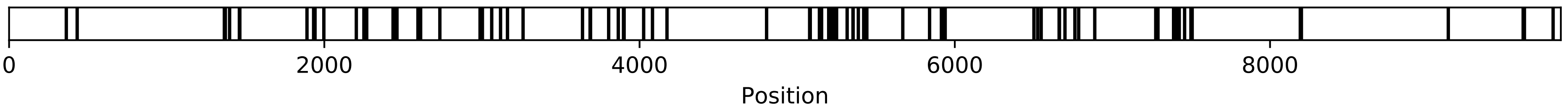


CR1 vs MM2
Number of differences: 84

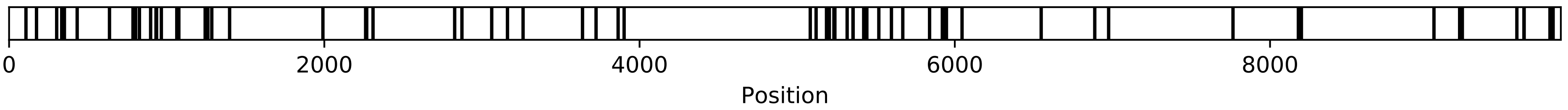


CR2 vs MA1
Number of differences: 100

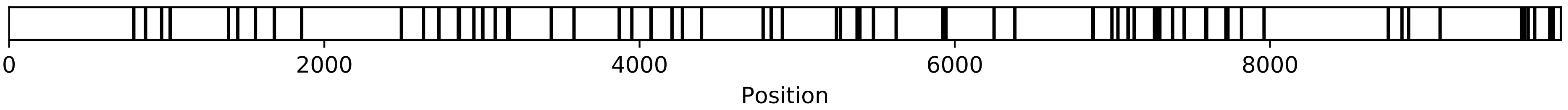


CR2 vs MA2
Number of differences: 199

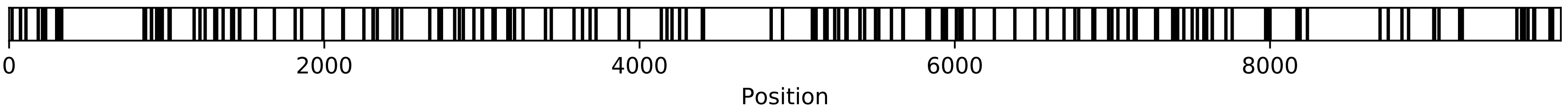


CR2 vs MM1
Number of differences: 217

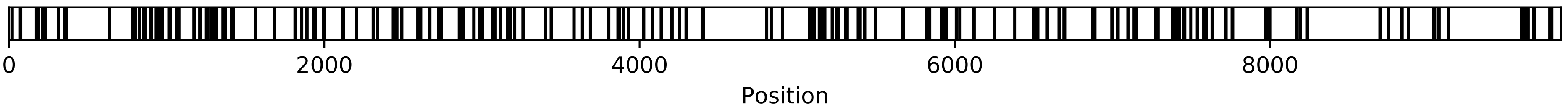


CR2 vs MM2
Number of differences: 199

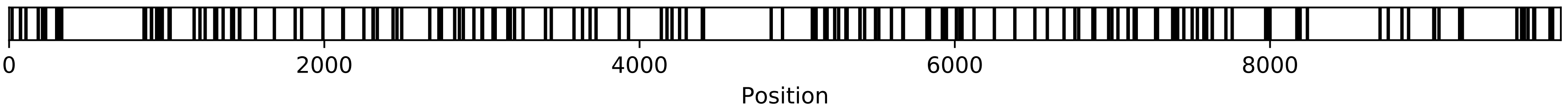


MA1 vs MA2
Number of differences: 198

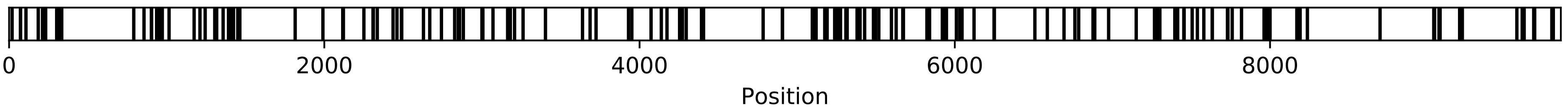


MA1 vs MM1
Number of differences: 215

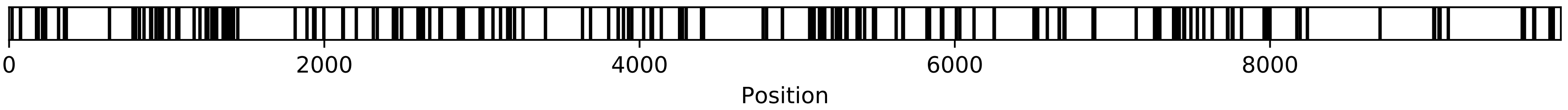


MA1 vs MM2
Number of differences: 198

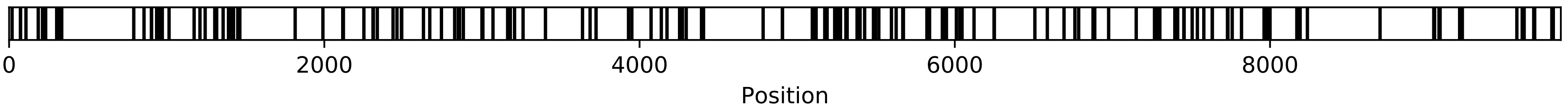


MA2 vs MM1
Number of differences: 123

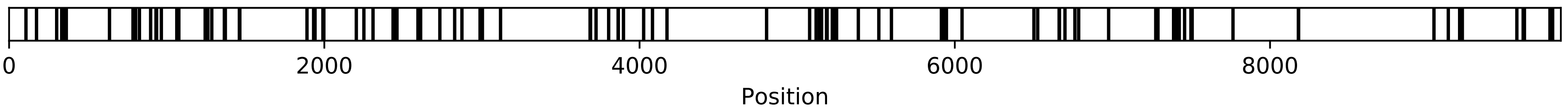


MA2 vs MM2
Number of differences: 0

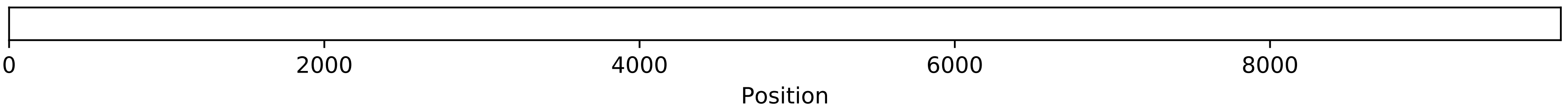


MM1 vs MM2
Number of differences: 123

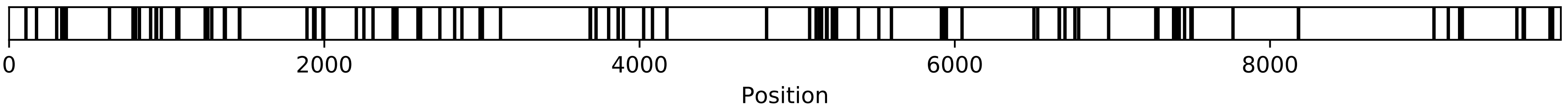

### Region: 198

CR1 vs CR2
Number of differences: 195

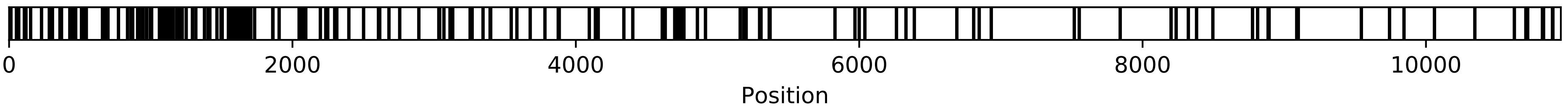


CR1 vs MA1
Number of differences: 83

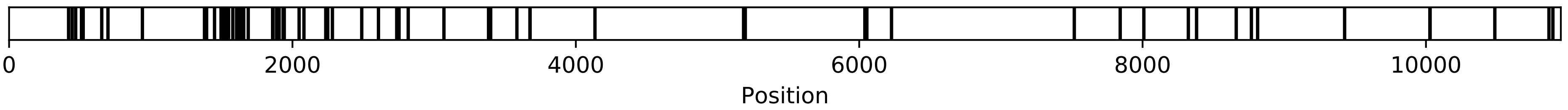


CR1 vs MA2
Number of differences: 194

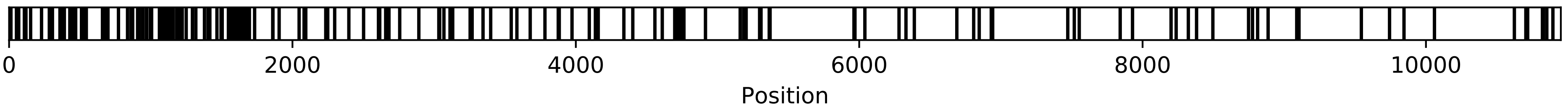


CR1 vs MM1
Number of differences: 52

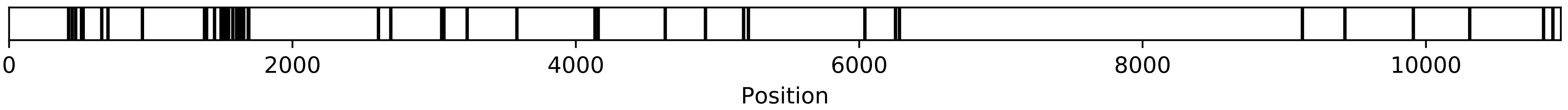


CR1 vs MM2
Number of differences: 83

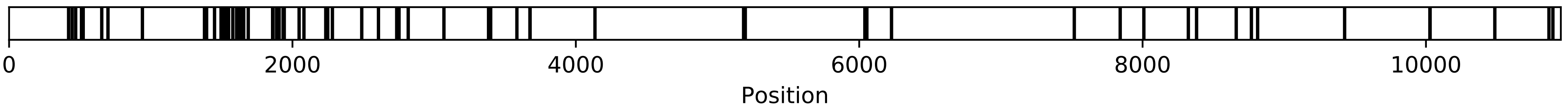


CR2 vs MA1
Number of differences: 207

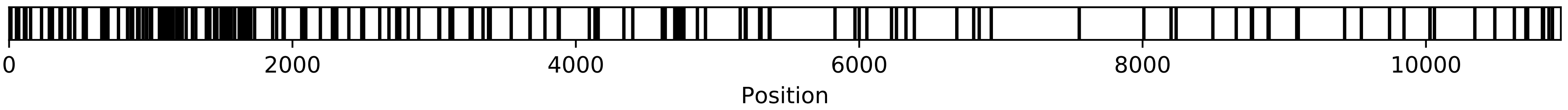


CR2 vs MA2
Number of differences: 25

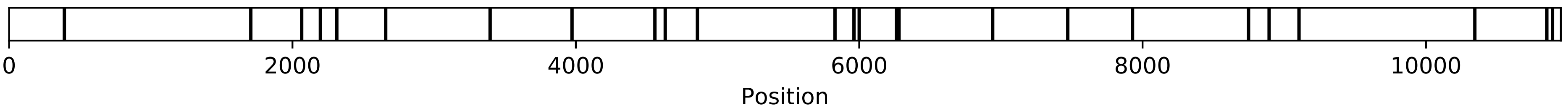


CR2 vs MM1
Number of differences: 193

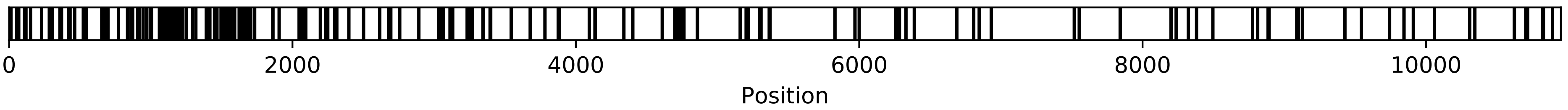


CR2 vs MM2
Number of differences: 207

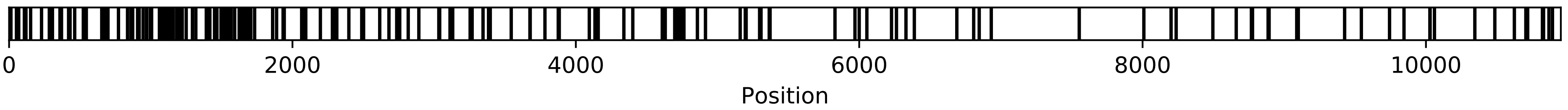


MA1 vs MA2
Number of differences: 206

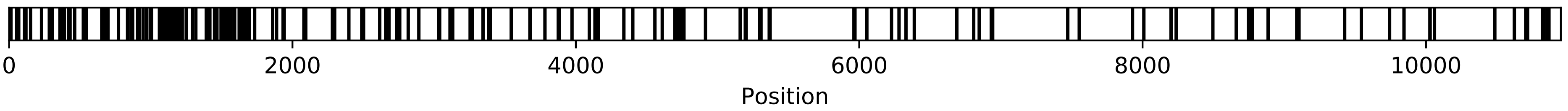


MA1 vs MM1
Number of differences: 63

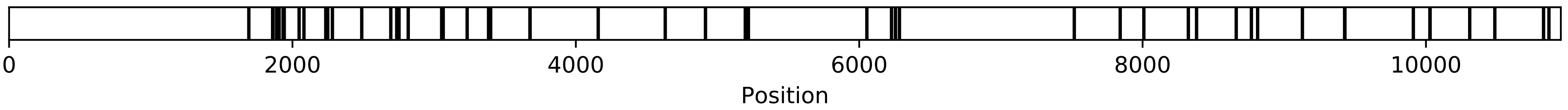


MA1 vs MM2
Number of differences: 0

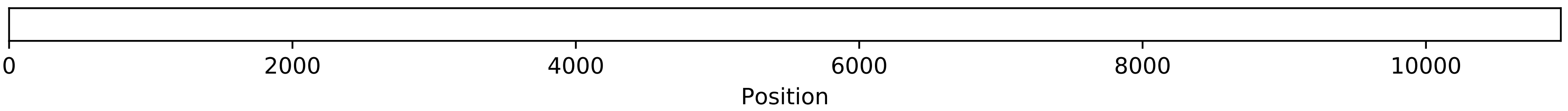


MA2 vs MM1
Number of differences: 194

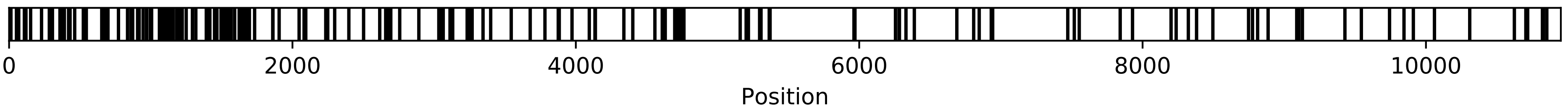


MA2 vs MM2
Number of differences: 206

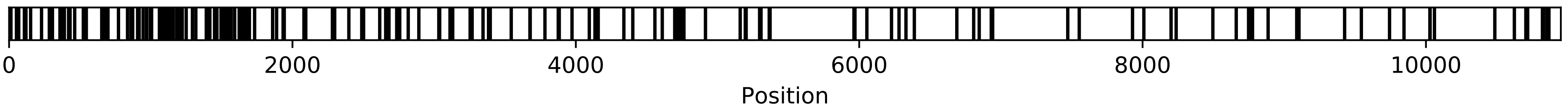


MM1 vs MM2
Number of differences: 63

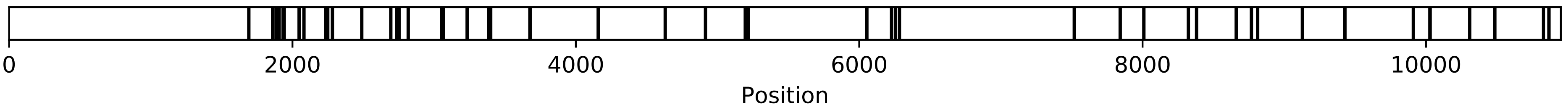

### Region: 199



CR1 vs CR2
Number of differences: 310

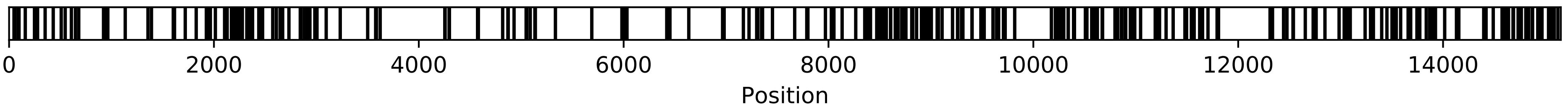


CR1 vs MA1
Number of differences: 309

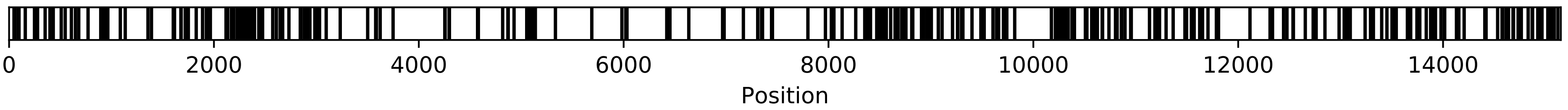


CR1 vs MA2
Number of differences: 139

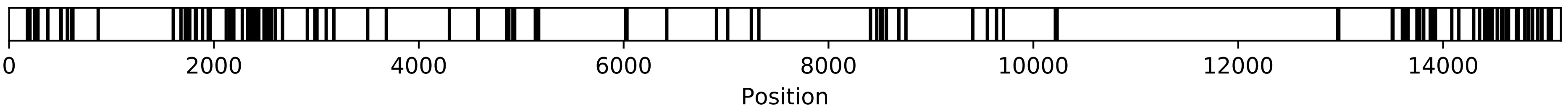


CR1 vs MM1
Number of differences: 169

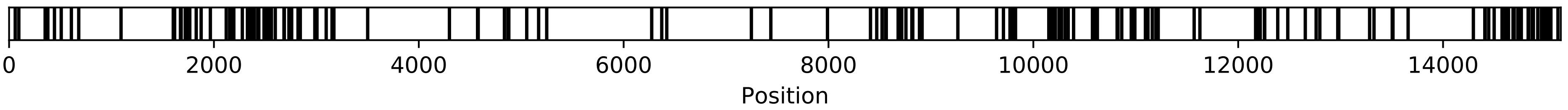


CR1 vs MM2
Number of differences: 141

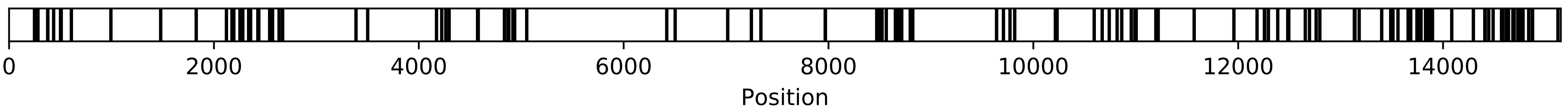


CR2 vs MA1
Number of differences: 81

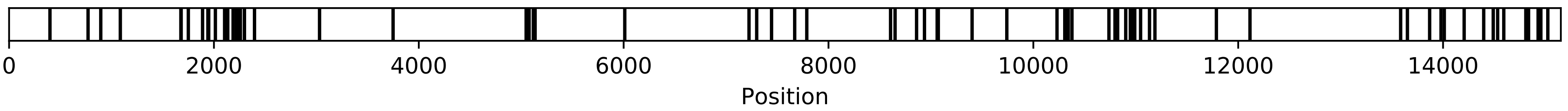


CR2 vs MA2
Number of differences: 322

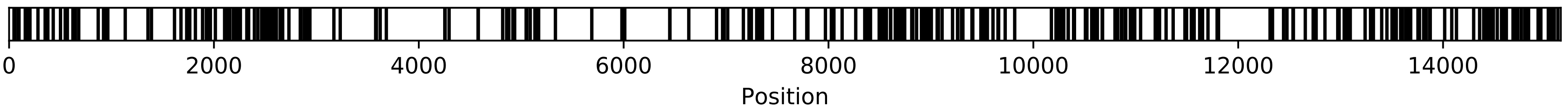


CR2 vs MM1
Number of differences: 323

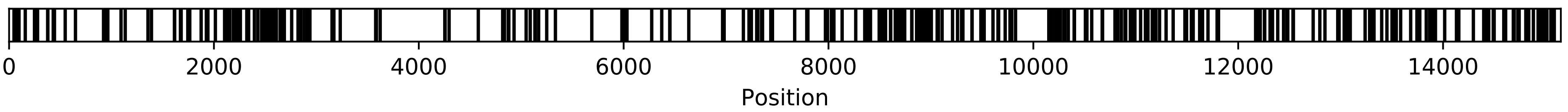


CR2 vs MM2
Number of differences: 325

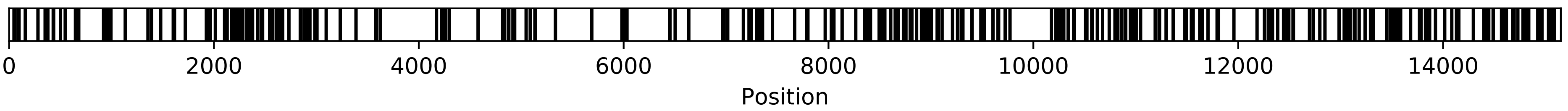


MA1 vs MA2
Number of differences: 319

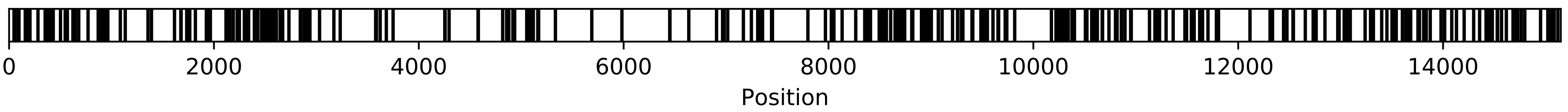


MA1 vs MM1
Number of differences: 320

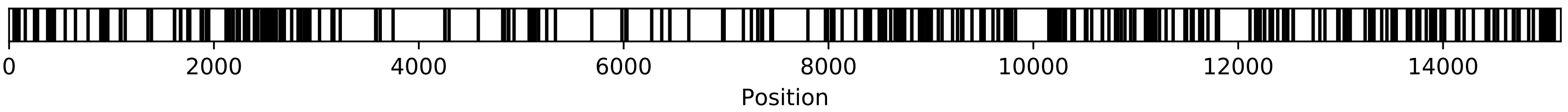


MA1 vs MM2
Number of differences: 324

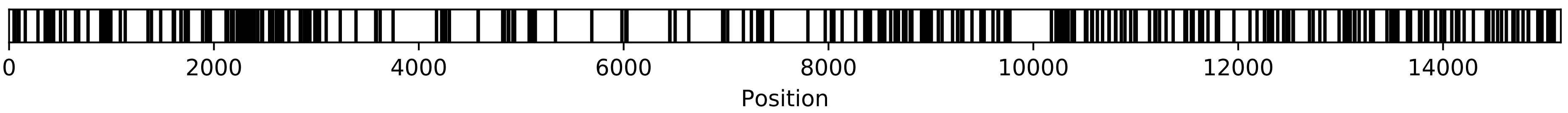


MA2 vs MM1
Number of differences: 166

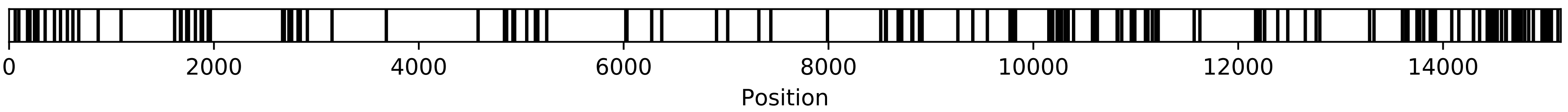


MA2 vs MM2
Number of differences: 161

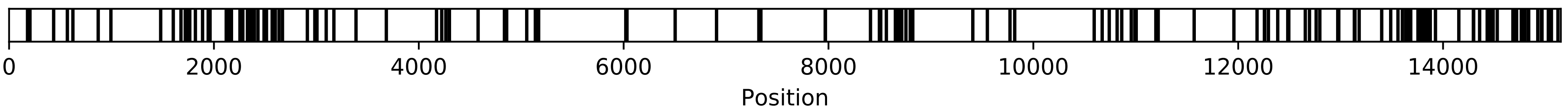


MM1 vs MM2
Number of differences: 161

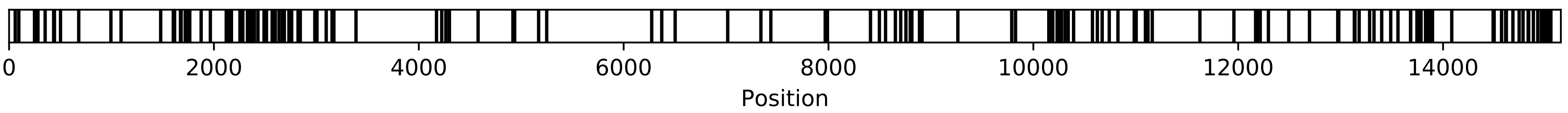

### Region: 201



CR1 vs CR2
Number of differences: 176

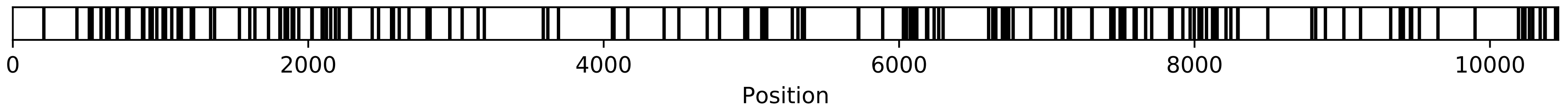


CR1 vs MA1
Number of differences: 63

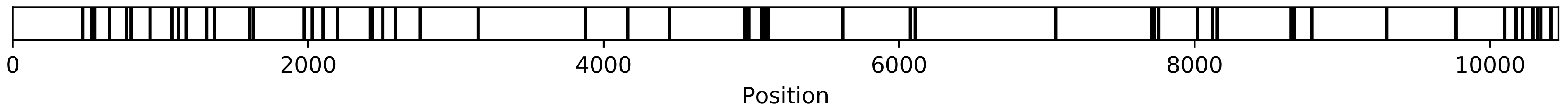


CR1 vs MA2
Number of differences: 180

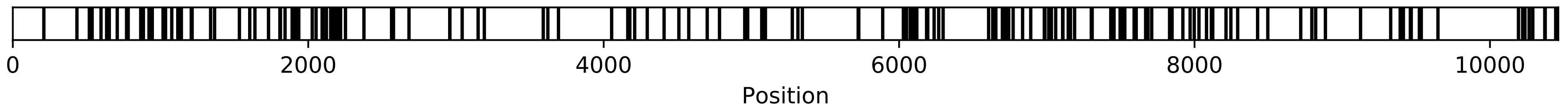


CR1 vs MM1
Number of differences: 63

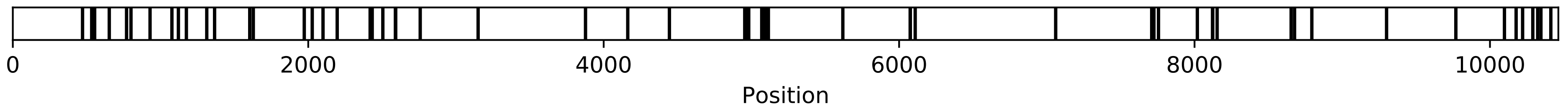


CR1 vs MM2
Number of differences: 84

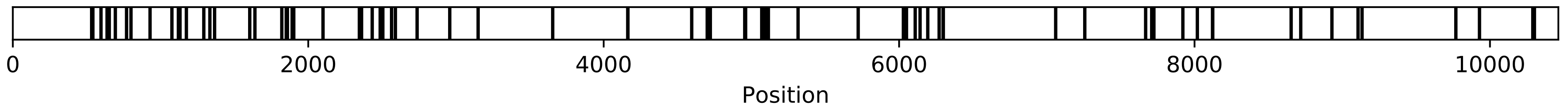


CR2 vs MA1
Number of differences: 185

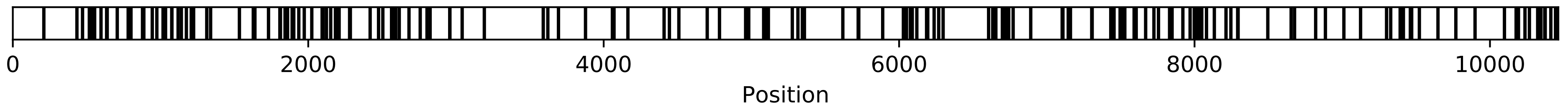


CR2 vs MA2
Number of differences: 62

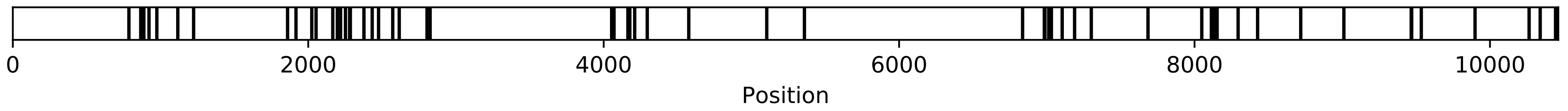


CR2 vs MM1
Number of differences: 185

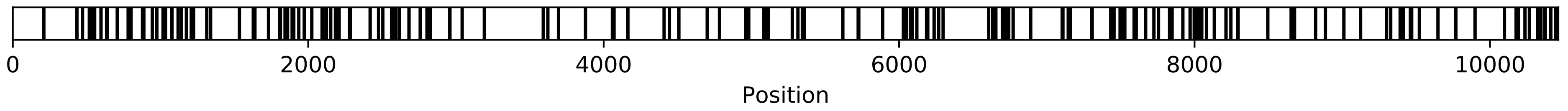


CR2 vs MM2
Number of differences: 180

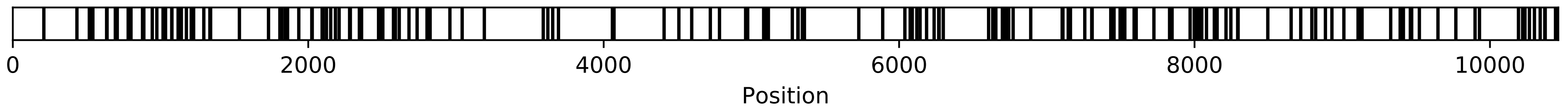


MA1 vs MA2
Number of differences: 193

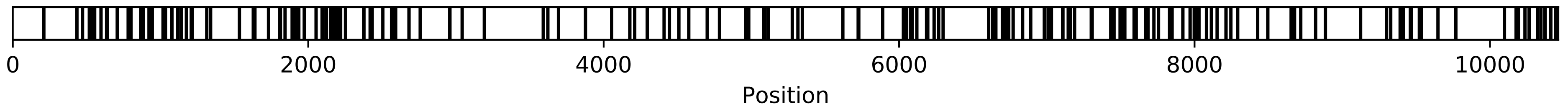


MA1 vs MM1
Number of differences: 0

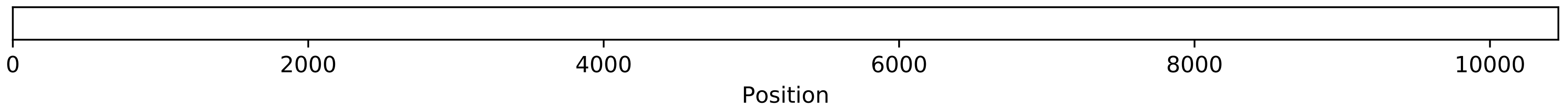


MA1 vs MM2
Number of differences: 77

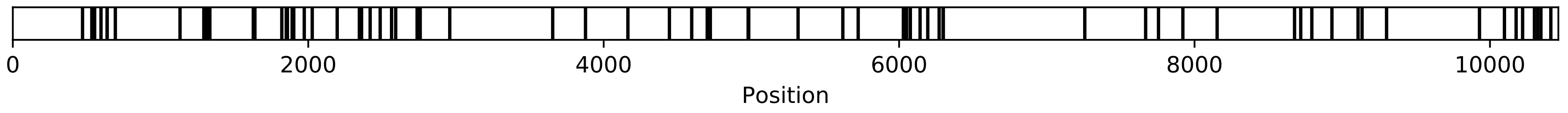


MA2 vs MM1
Number of differences: 193

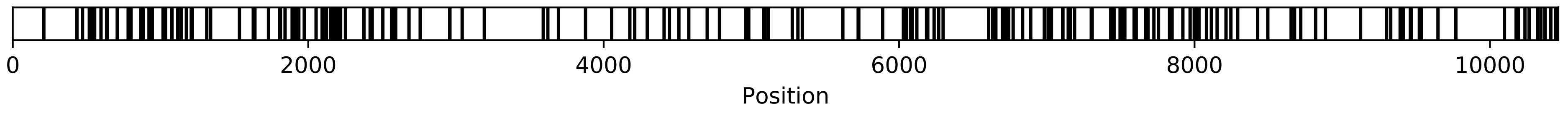


MA2 vs MM2
Number of differences: 186

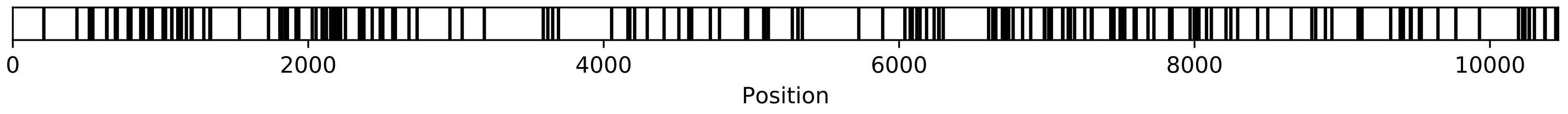


MM1 vs MM2
Number of differences: 77

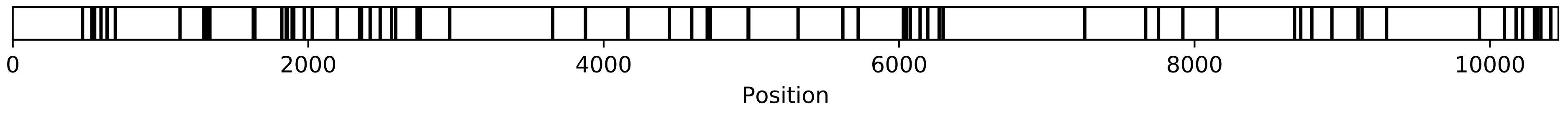

### Region: 203

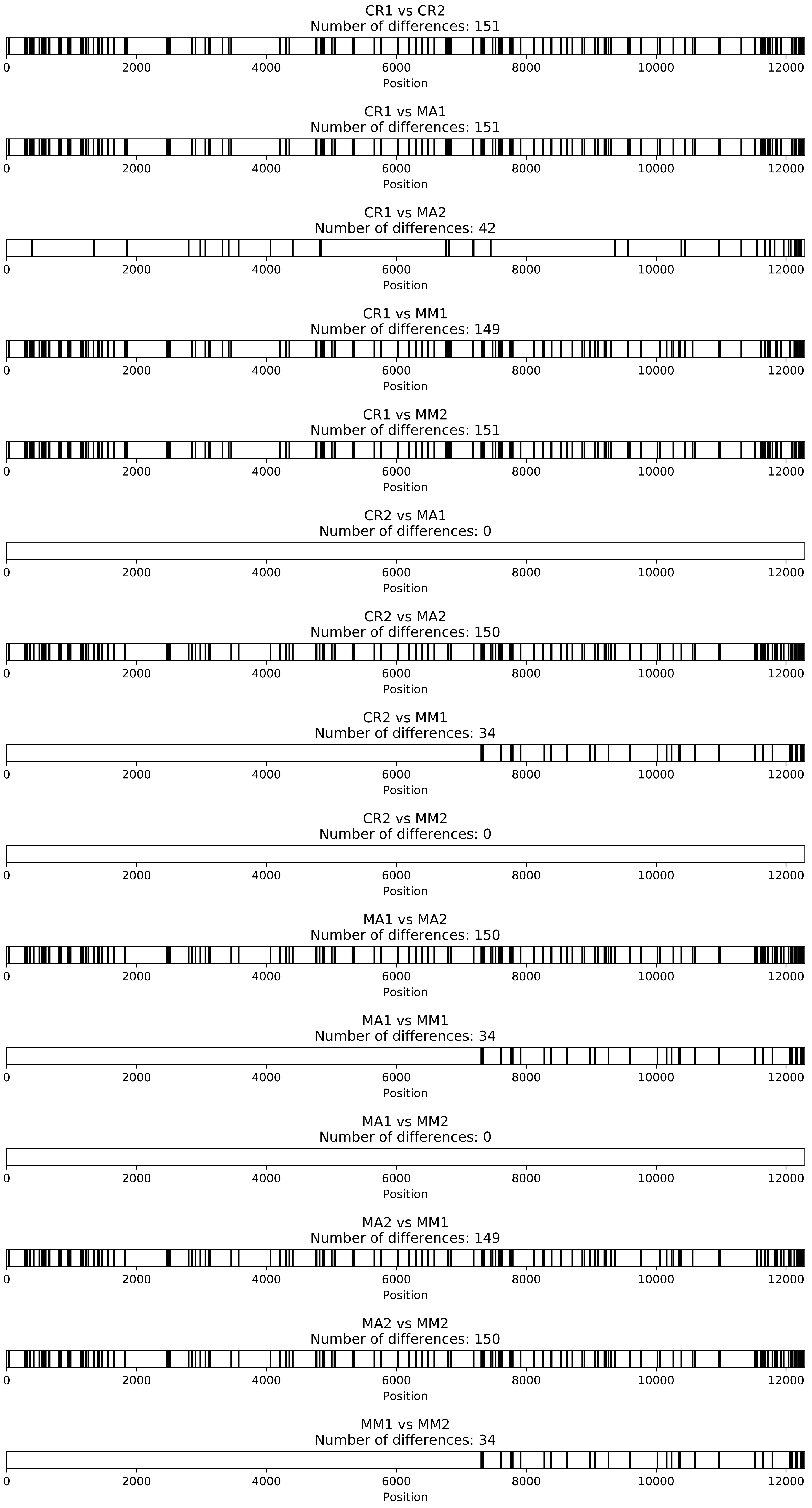
CR1 vs CR2
Number of differences: 151
CR1 vs MA1
Number of differences: 151
CR1 vs MA2
Number of differences: 42
CR1 vs MM1
Number of differences: 149
CR1 vs MM2
Number of differences: 151
CR2 vs MA1
Number of differences: 0
CR2 vs MA2
Number of differences: 150
CR2 vs MM1
Number of differences: 34
CR2 vs MM2
Number of differences: 0
MA1 vs MA2
Number of differences: 150
MA1 vs MM1
Number of differences: 34
MA1 vs MM2
Number of differences: 0
MA2 vs MM1
Number of differences: 149
MA2 vs MM2
Number of differences: 150
MM1 vs MM2
Number of differences: 34
0
2000
4000
6000
8000
10000
12000
Position

### Region: 204



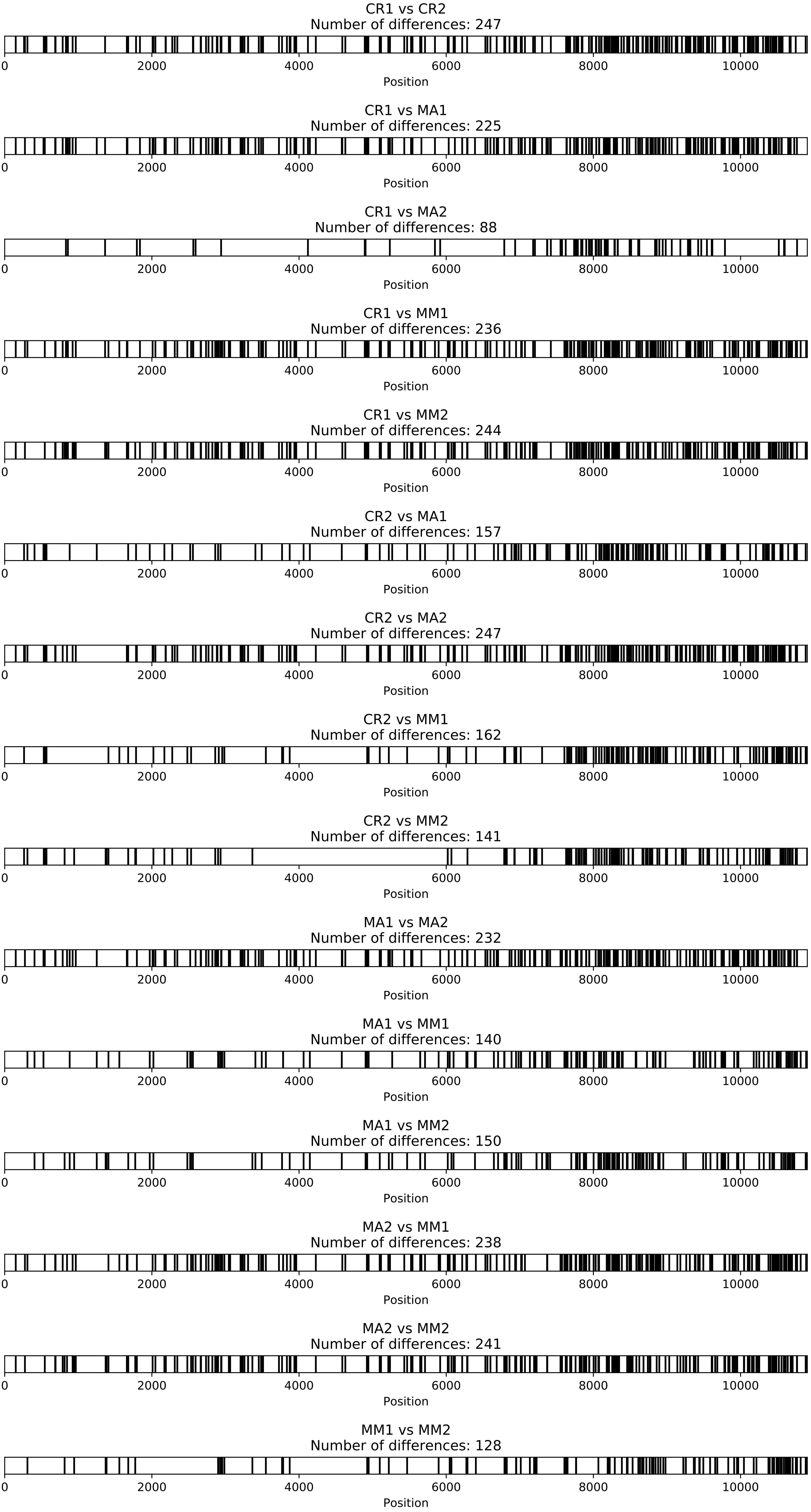
CR1 vs CR2
Number of differences: 247
CR1 vs MA1
Number of differences: 225
CR1 vs MA2
Number of differences: 88
CR1 vs MM1
Number of differences: 236
CR1 vs MM2
Number of differences: 244
CR2 vs MA1
Number of differences: 157
CR2 vs MA2
Number of differences: 247
CR2 vs MM1
Number of differences: 162
CR2 vs MM2
Number of differences: 141
MA1 vs MA2
Number of differences: 232
MA1 vs MM1
Number of differences: 140
MA1 vs MM2
Number of differences: 150
MA2 vs MM1
Number of differences: 238
MA2 vs MM2
Number of differences: 241
MM1 vs MM2
Number of differences: 128
0
2000
4000
6000
8000
10000
Position

### Region: 205

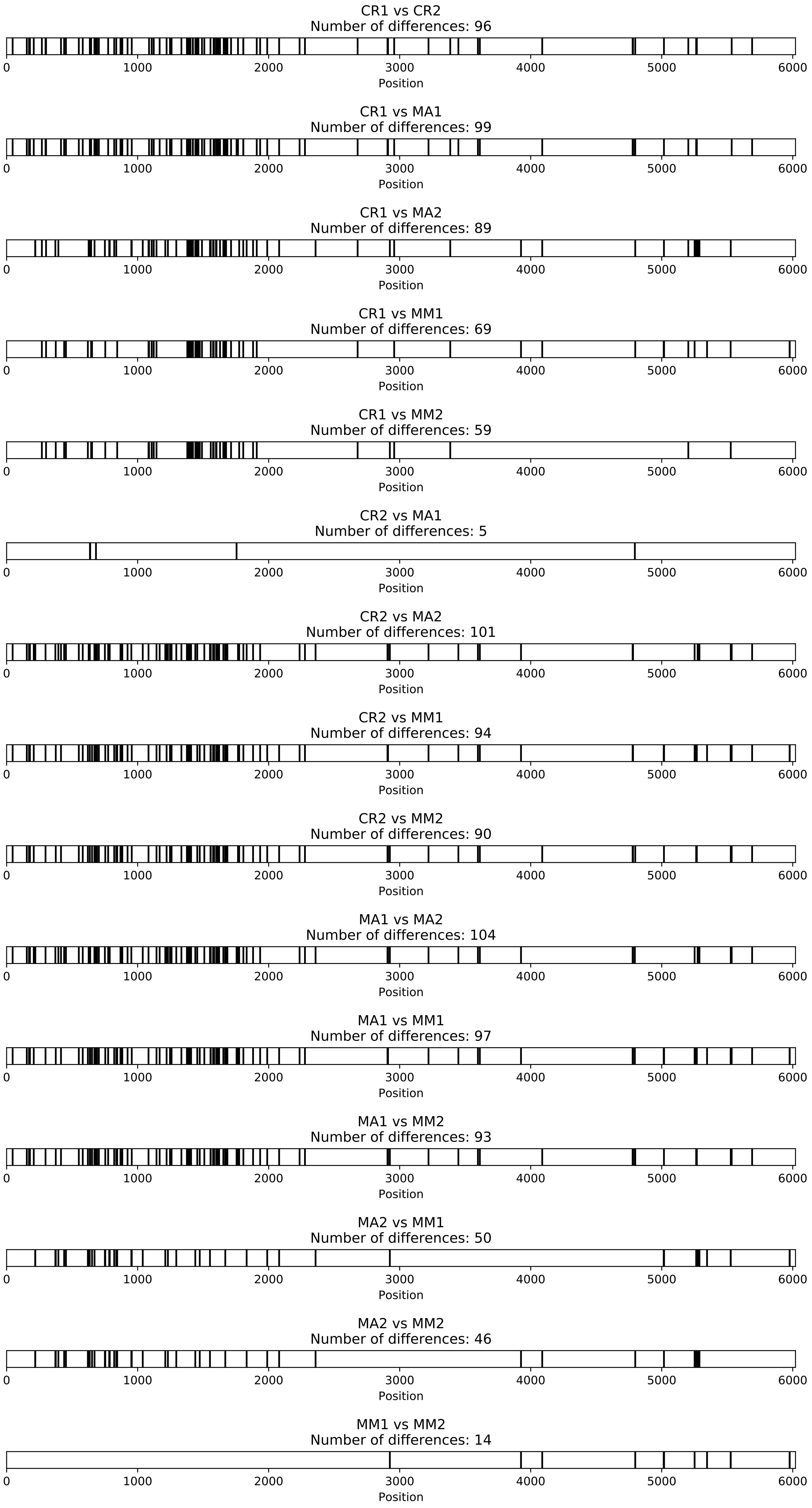
CR1 vs CR2
Number of differences: 96
CR1 vs MA1
Number of differences: 99
CR1 vs MA2
Number of differences: 89
CR1 vs MM1
Number of differences: 69
CR1 vs MM2
Number of differences: 59
CR2 vs MA1
Number of differences: 5
CR2 vs MA2
Number of differences: 101
CR2 vs MM1
Number of differences: 94
CR2 vs MM2
Number of differences: 90
MA1 vs MA2
Number of differences: 104
MA1 vs MM1
Number of differences: 97
MA1 vs MM2
Number of differences: 93
MA2 vs MM1
Number of differences: 50
MA2 vs MM2
Number of differences: 46
MM1 vs MM2
Number of differences: 14
0
1000
2000
3000
4000
5000
6000
Position

### Region: 206



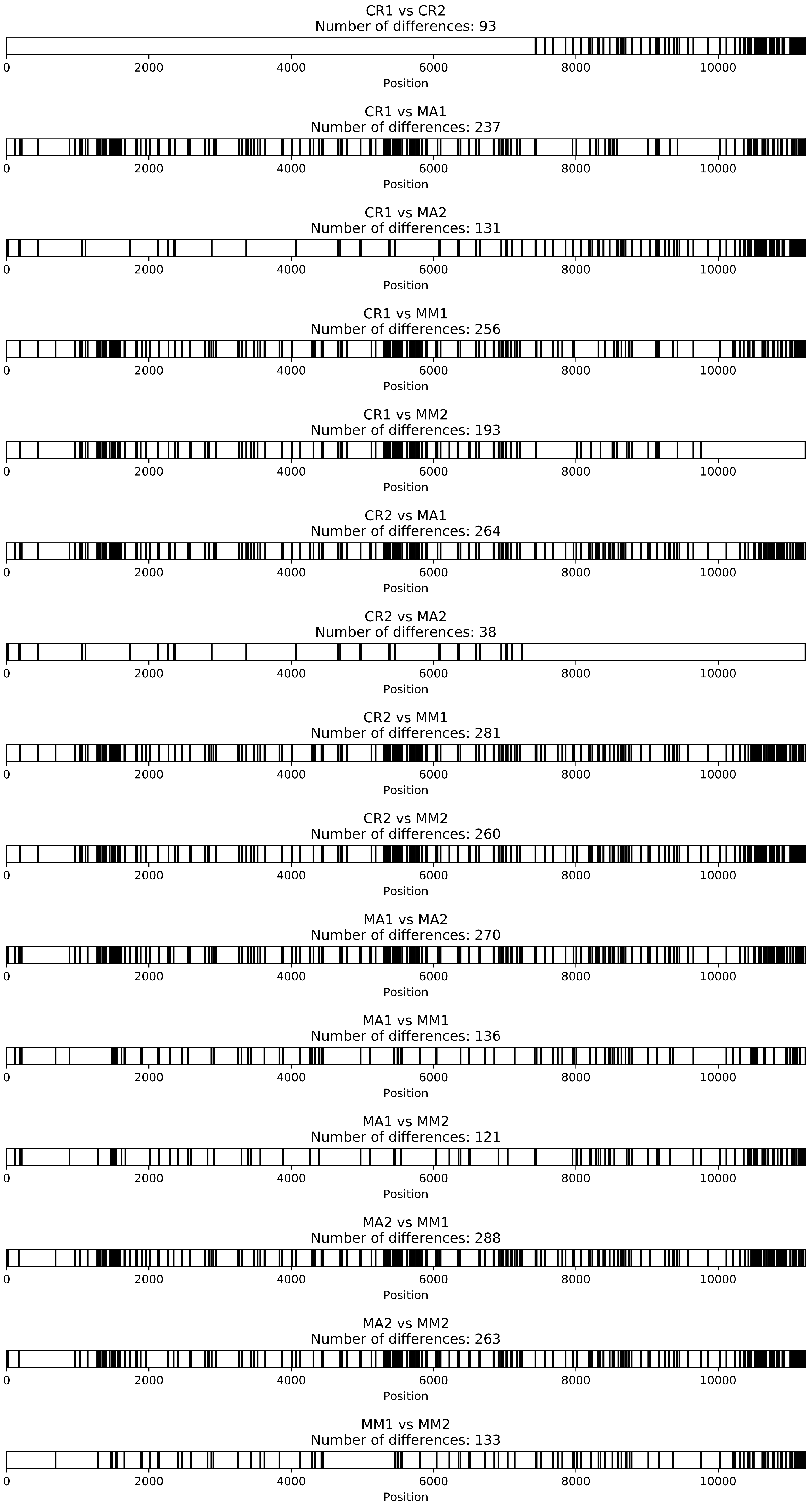
CR1 vs CR2
Number of differences: 93
CR1 vs MA1
Number of differences: 237
CR1 vs MA2
Number of differences: 131
CR1 vs MM1
Number of differences: 256
CR1 vs MM2
Number of differences: 193
CR2 vs MA1
Number of differences: 264
CR2 vs MA2
Number of differences: 38
CR2 vs MM1
Number of differences: 281
CR2 vs MM2
Number of differences: 260
MA1 vs MA2
Number of differences: 270
MA1 vs MM1
Number of differences: 136
MA1 vs MM2
Number of differences: 121
MA2 vs MM1
Number of differences: 288
MA2 vs MM2
Number of differences: 263
MM1 vs MM2
Number of differences: 133
0
2000
4000
6000
8000
10000
Position

### Region: 207



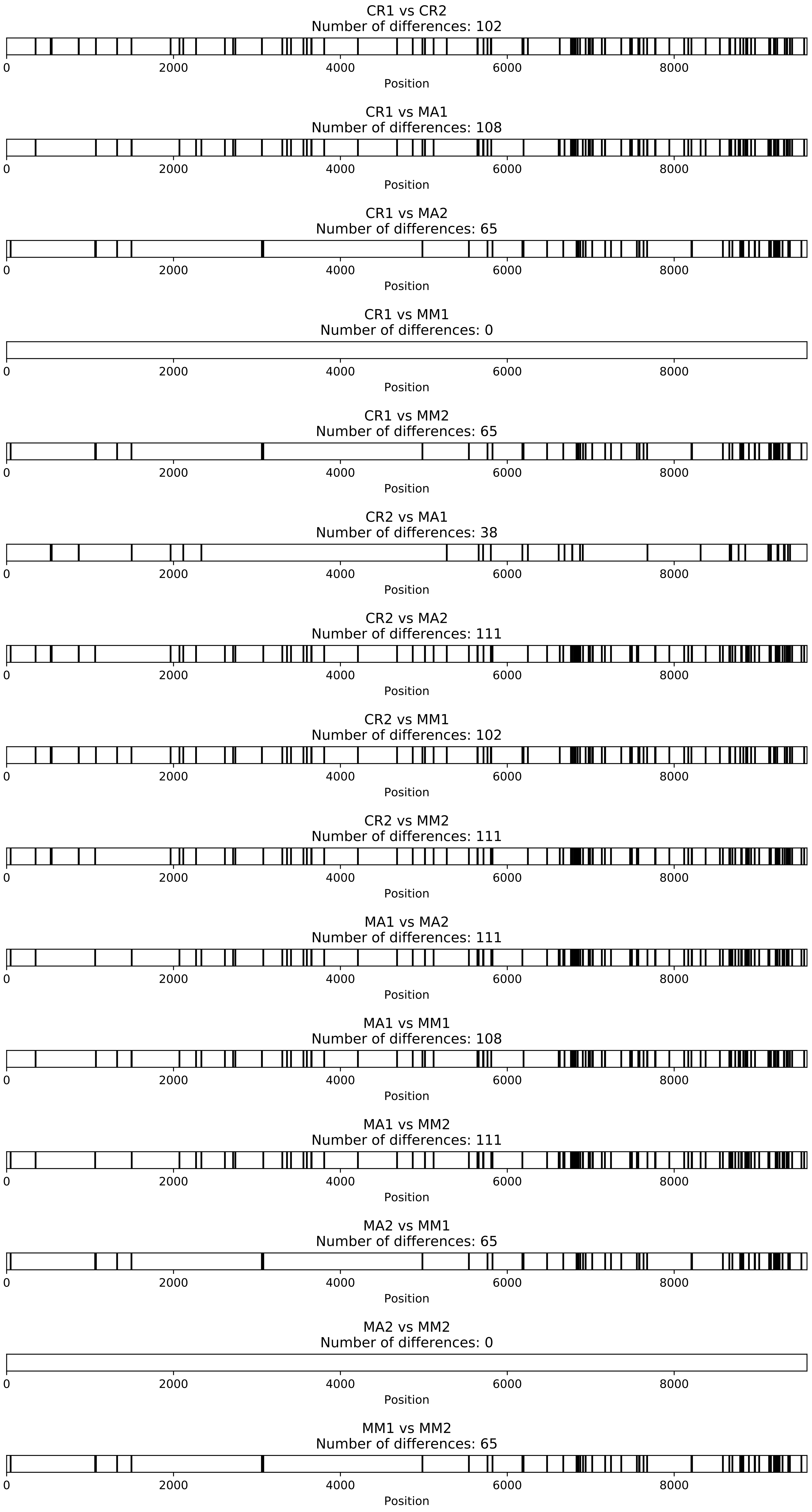
CR1 vs CR2
Number of differences: 102
CR1 vs MA1
Number of differences: 108
CR1 vs MA2
Number of differences: 65
CR1 vs MM1
Number of differences: 0
CR1 vs MM2
Number of differences: 65
CR2 vs MA1
Number of differences: 38
CR2 vs MA2
Number of differences: 111
CR2 vs MM1
Number of differences: 102
CR2 vs MM2
Number of differences: 111
MA1 vs MA2
Number of differences: 111
MA1 vs MM1
Number of differences: 108
MA1 vs MM2
Number of differences: 111
MA2 vs MM1
Number of differences: 65
MA2 vs MM2
Number of differences: 0
MM1 vs MM2
Number of differences: 65
0
2000
4000
6000
8000
Position

### Region: 208

CR1 vs CR2
Number of differences: 70

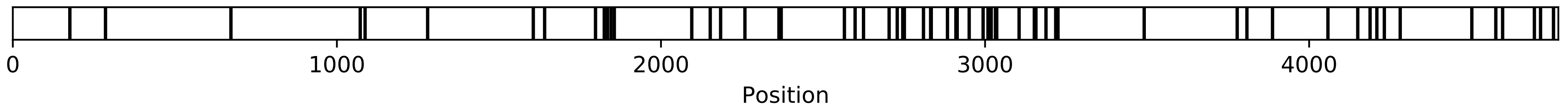


CR1 vs MA1
Number of differences: 33

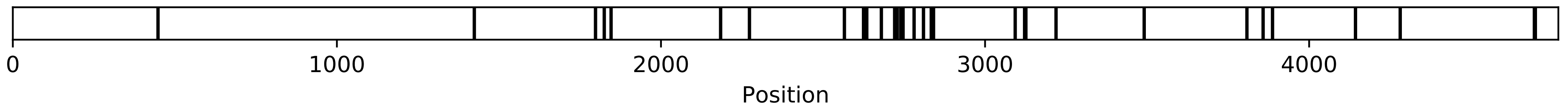


CR1 vs MA2
Number of differences: 74

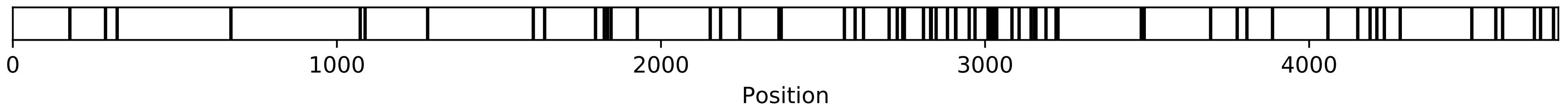


CR1 vs MM1
Number of differences: 36

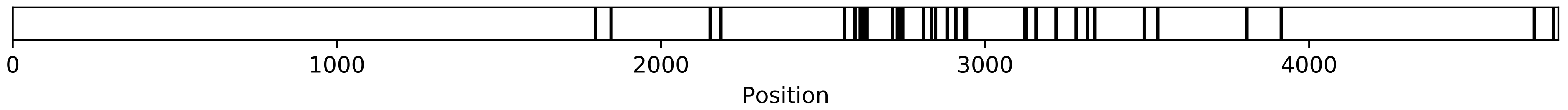


CR1 vs MM2
Number of differences: 31

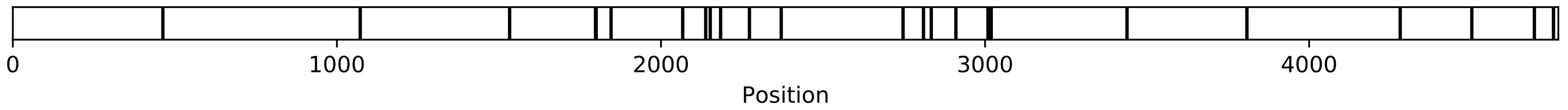


CR2 vs MA1
Number of differences: 70

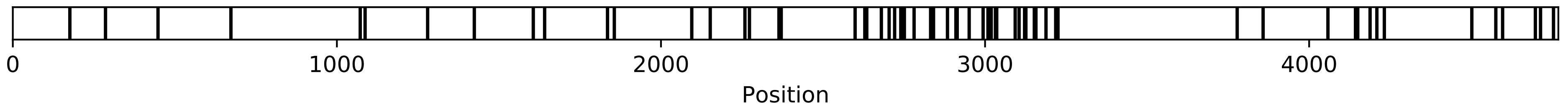


CR2 vs MA2
Number of differences: 18

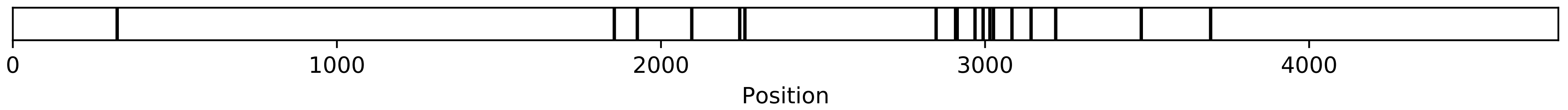


CR2 vs MM1
Number of differences: 66

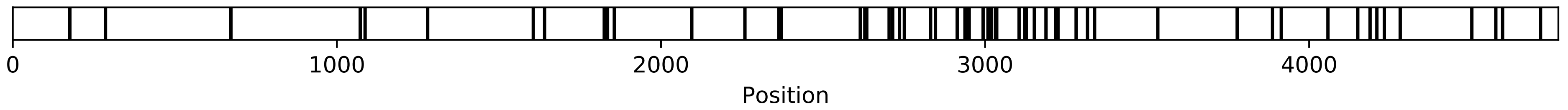


CR2 vs MM2
Number of differences: 55

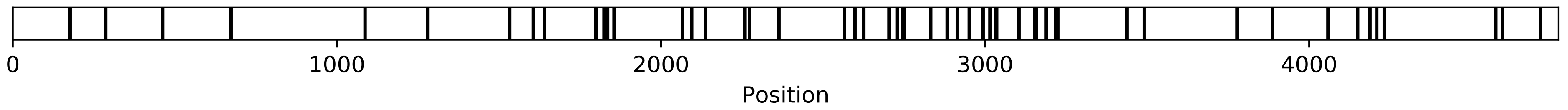


MA1 vs MA2
Number of differences: 74

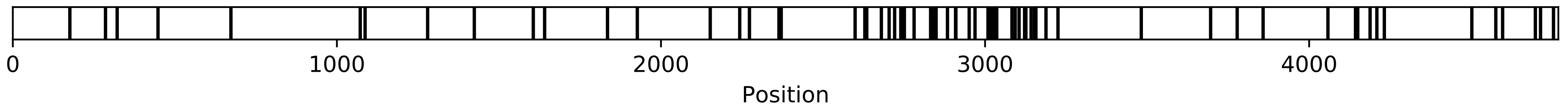


MA1 vs MM1
Number of differences: 34

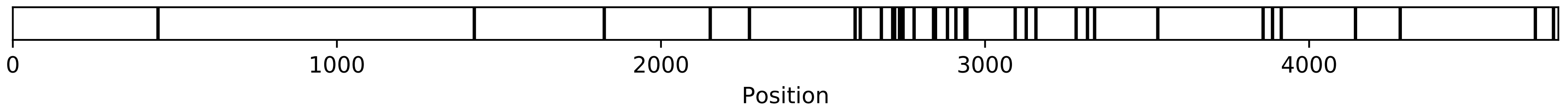


MA1 vs MM2
Number of differences: 45

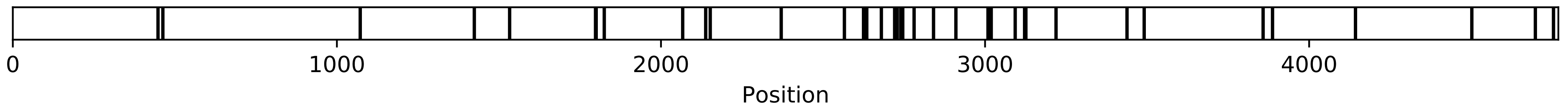


MA2 vs MM1
Number of differences: 70

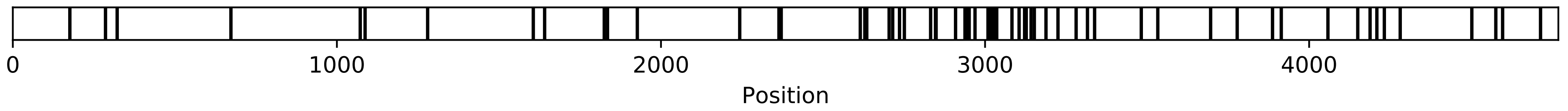


MA2 vs MM2
Number of differences: 59

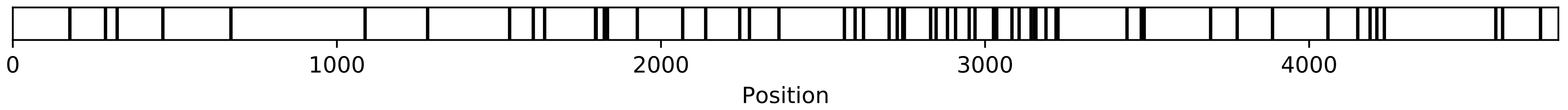


MM1 vs MM2
Number of differences: 47

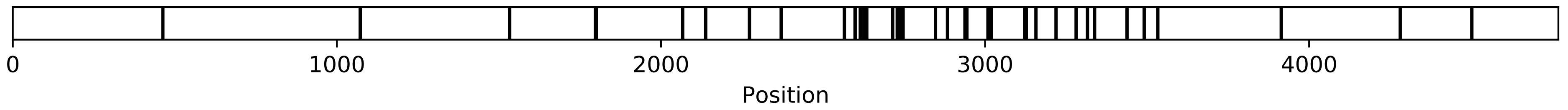

### Region: 209

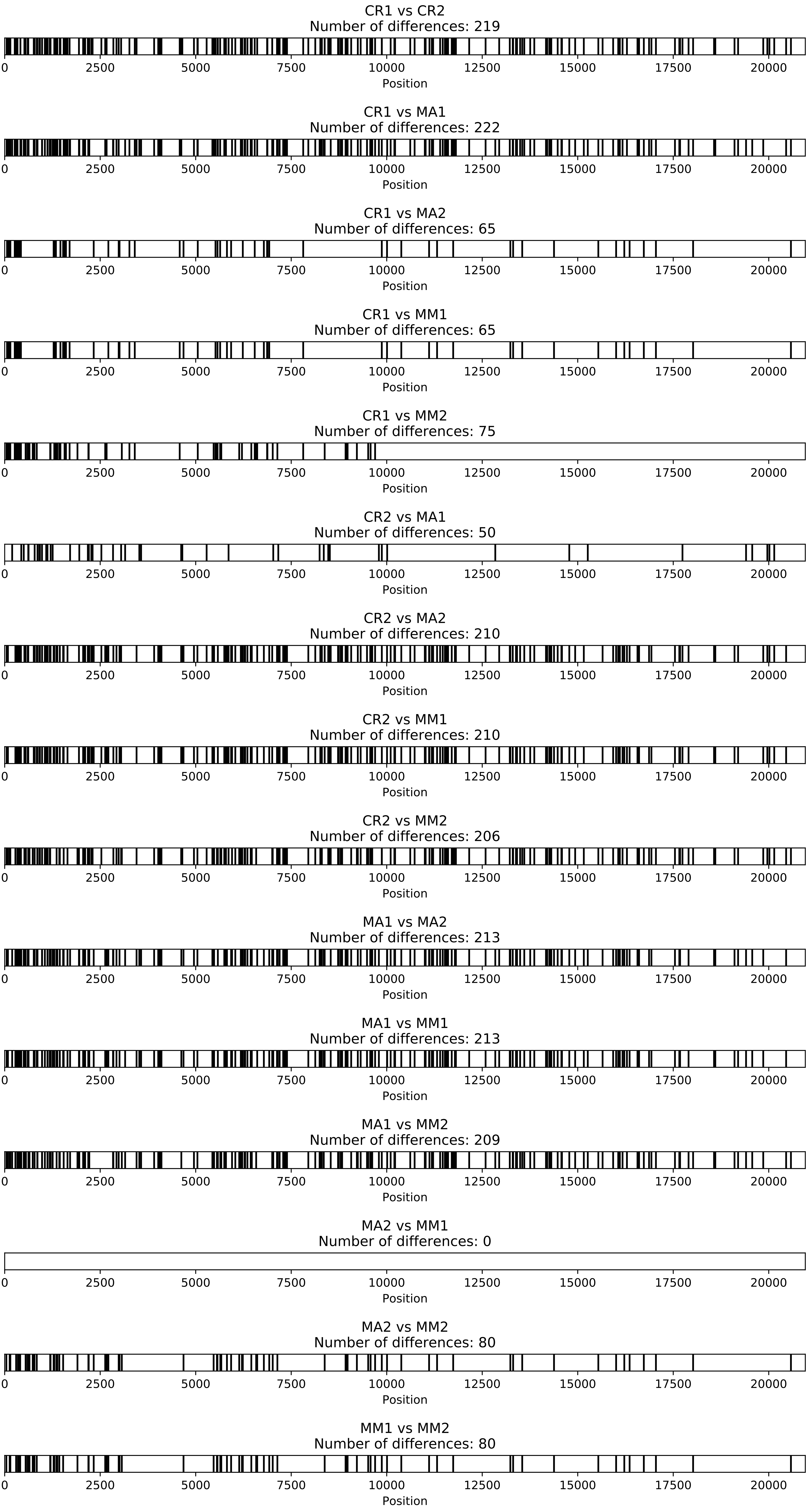
CR1 vs CR2
Number of differences: 219
CR1 vs MA1
Number of differences: 222
CR1 vs MA2
Number of differences: 65
CR1 vs MM1
Number of differences: 65
CR1 vs MM2
Number of differences: 75
CR2 vs MA1
Number of differences: 50
CR2 vs MA2
Number of differences: 210
CR2 vs MM1
Number of differences: 210
CR2 vs MM2
Number of differences: 206
MA1 vs MA2
Number of differences: 213
MA1 vs MM1
Number of differences: 213
MA1 vs MM2
Number of differences: 209
MA2 vs MM1
Number of differences: 0
MA2 vs MM2
Number of differences: 80
MM1 vs MM2
Number of differences: 80
0
2500
5000
7500
10000
12500
15000
17500
20000
Position

### Region: 210



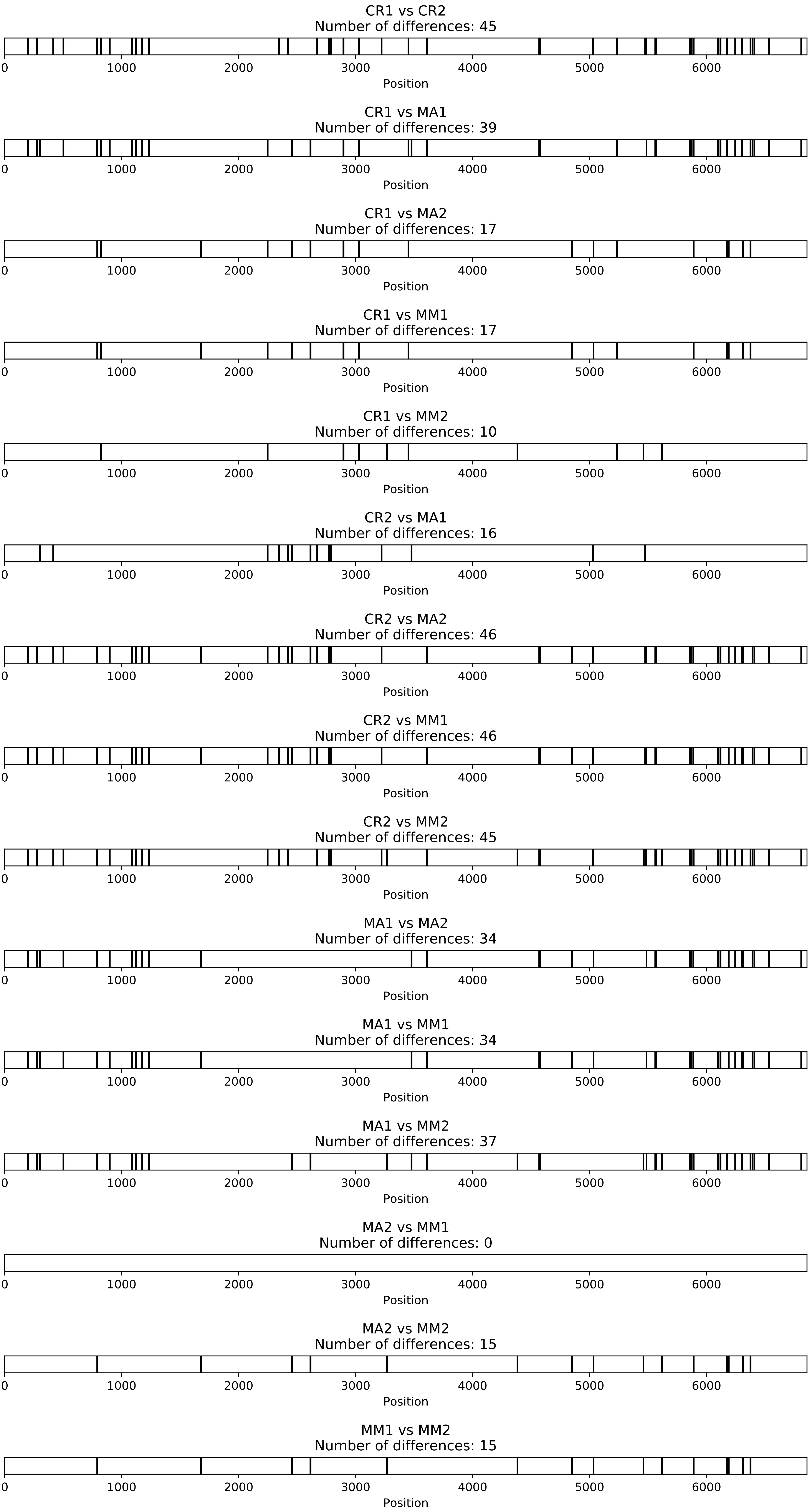
CR1 vs CR2
Number of differences: 45
CR1 vs MA1
Number of differences: 39
CR1 vs MA2
Number of differences: 17
CR1 vs MM1
Number of differences: 17
CR1 vs MM2
Number of differences: 10
CR2 vs MA1
Number of differences: 16
CR2 vs MA2
Number of differences: 46
CR2 vs MM1
Number of differences: 46
CR2 vs MM2
Number of differences: 45
MA1 vs MA2
Number of differences: 34
MA1 vs MM1
Number of differences: 34
MA1 vs MM2
Number of differences: 37
MA2 vs MM1
Number of differences: 0
MA2 vs MM2
Number of differences: 15
MM1 vs MM2
Number of differences: 15
0
1000
2000
3000
4000
5000
6000
Position

### Region: 211

CR1 vs CR2
Number of differences: 294

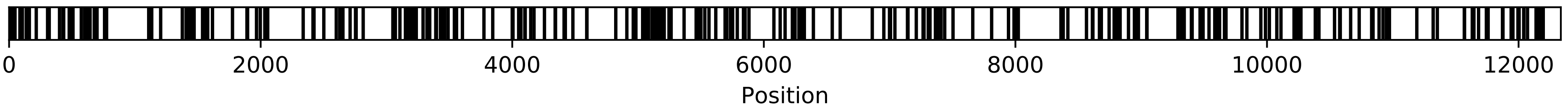


CR1 vs MA1
Number of differences: 132

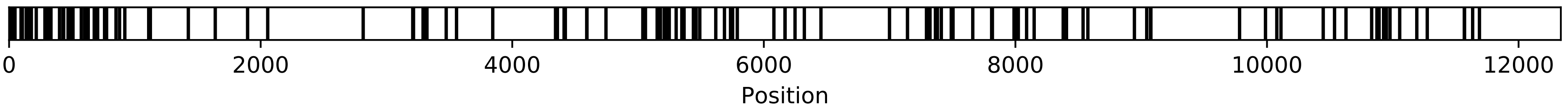


CR1 vs MA2
Number of differences: 291

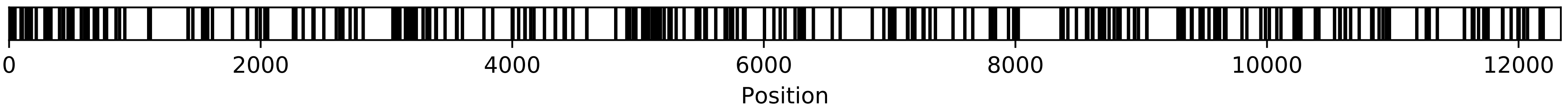


CR1 vs MM1
Number of differences: 119

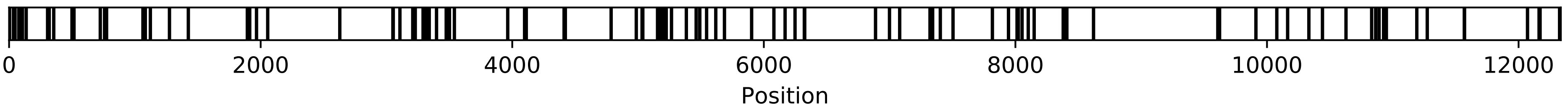


CR1 vs MM2
Number of differences: 123

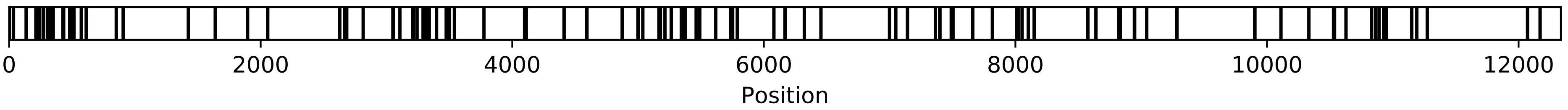


CR2 vs MA1
Number of differences: 256

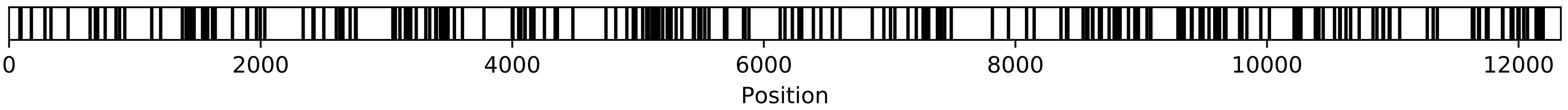


CR2 vs MA2
Number of differences: 89

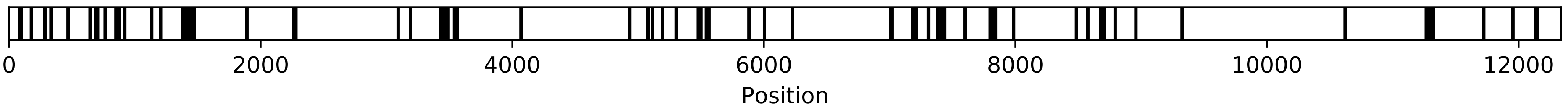


CR2 vs MM1
Number of differences: 289

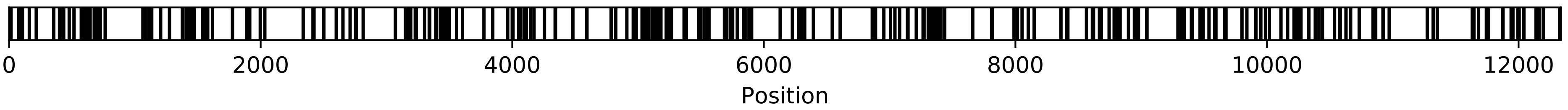


CR2 vs MM2
Number of differences: 287

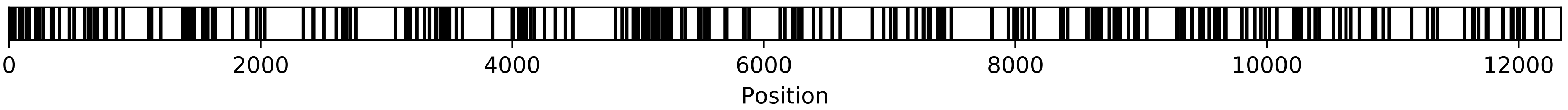


MA1 vs MA2
Number of differences: 233

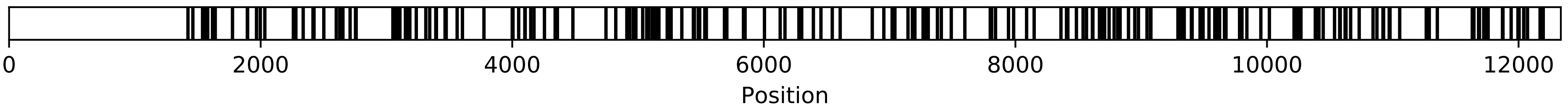


MA1 vs MM1
Number of differences: 142

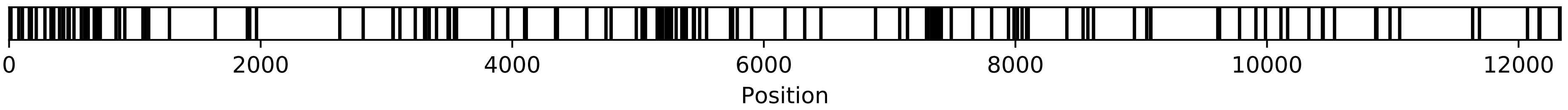


MA1 vs MM2
Number of differences: 130

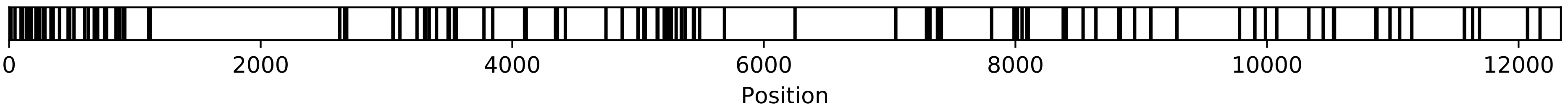


MA2 vs MM1
Number of differences: 288

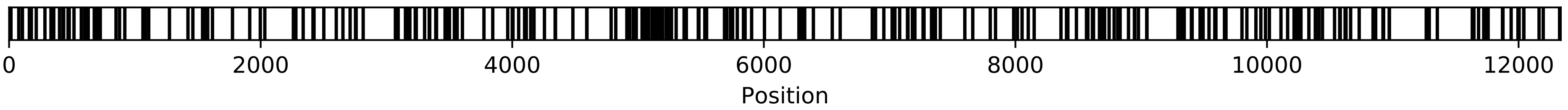


MA2 vs MM2
Number of differences: 284

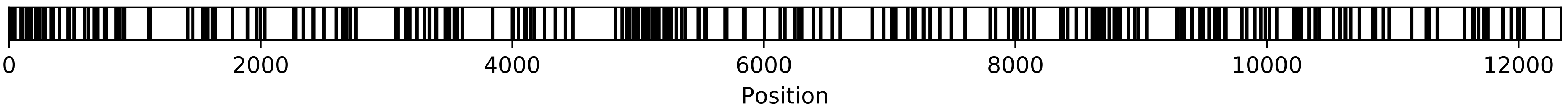


MM1 vs MM2
Number of differences: 113

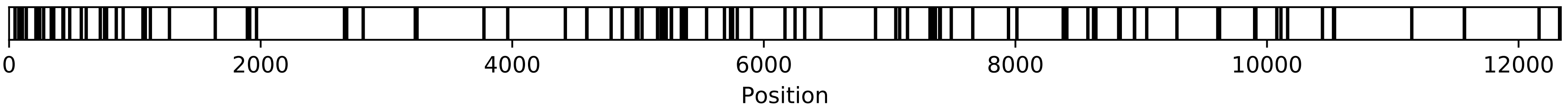

### Region: 212

CR1 vs CR2
Number of differences: 254

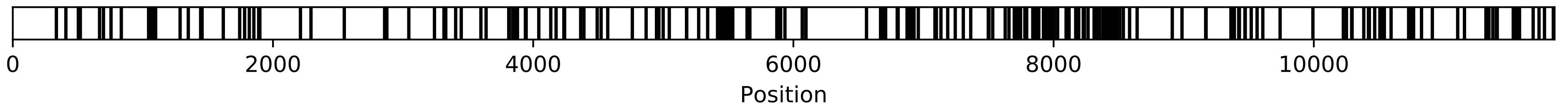


CR1 vs MA1
Number of differences: 195

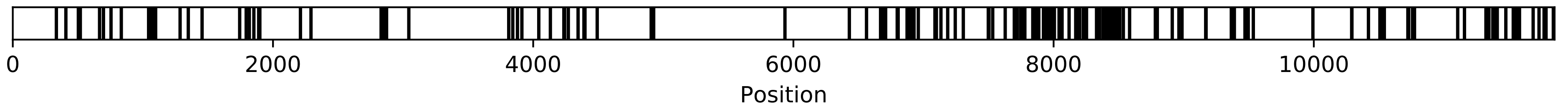


CR1 vs MA2
Number of differences: 188

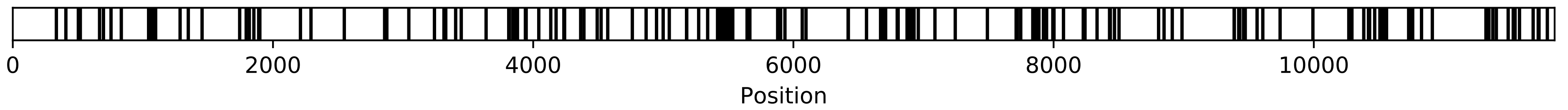


CR1 vs MM1
Number of differences: 158

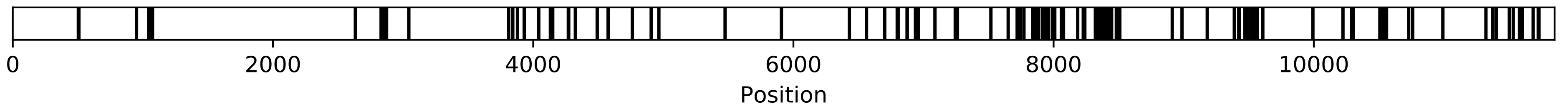


CR1 vs MM2
Number of differences: 130

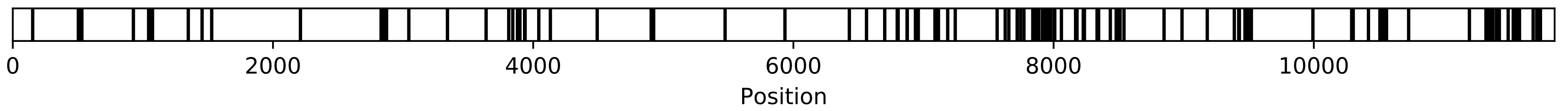


CR2 vs MA1
Number of differences: 127

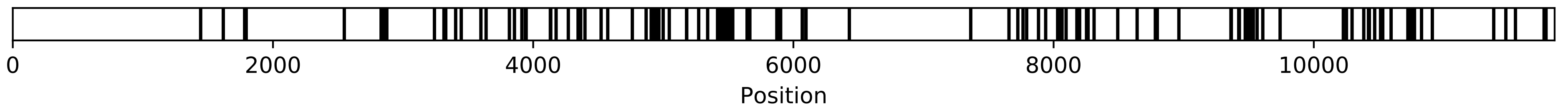


CR2 vs MA2
Number of differences: 121

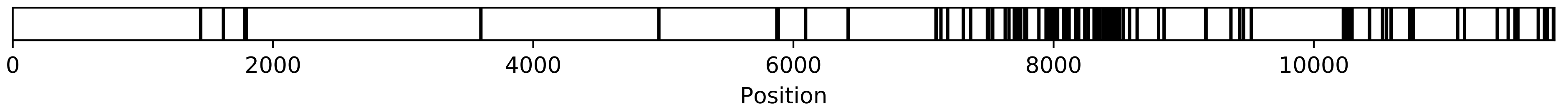


CR2 vs MM1
Number of differences: 237

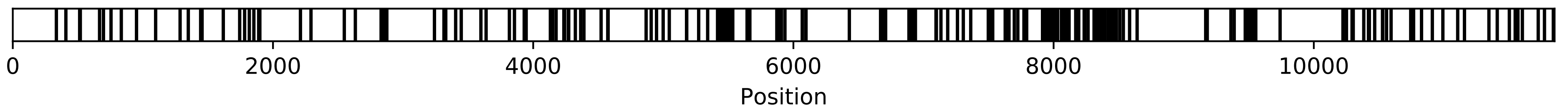


CR2 vs MM2
Number of differences: 228

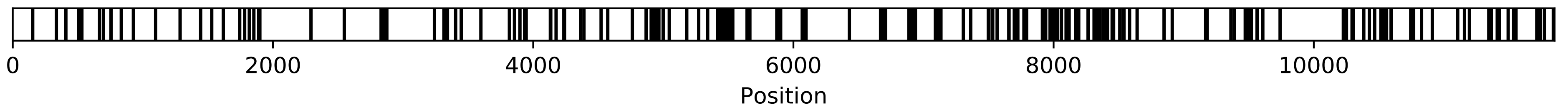


MA1 vs MA2
Number of differences: 180

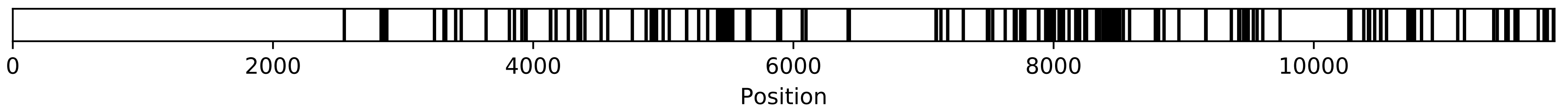


MA1 vs MM1
Number of differences: 179

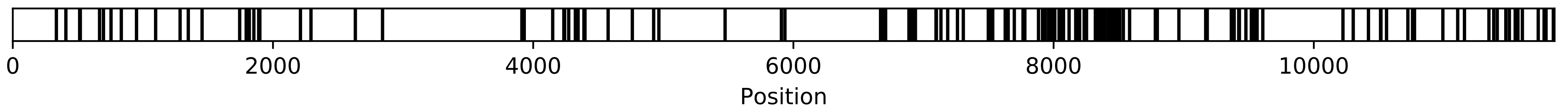


MA1 vs MM2
Number of differences: 162

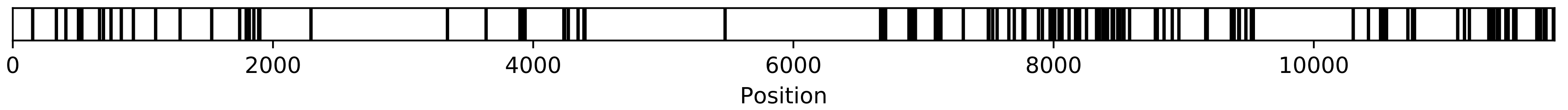


MA2 vs MM1
Number of differences: 202

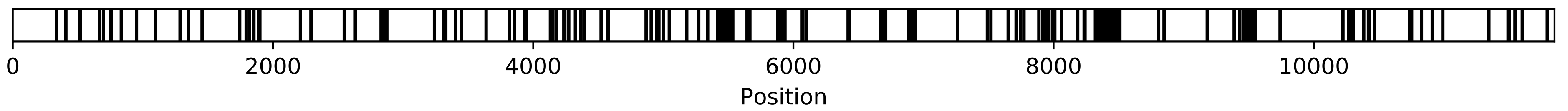


MA2 vs MM2
Number of differences: 175

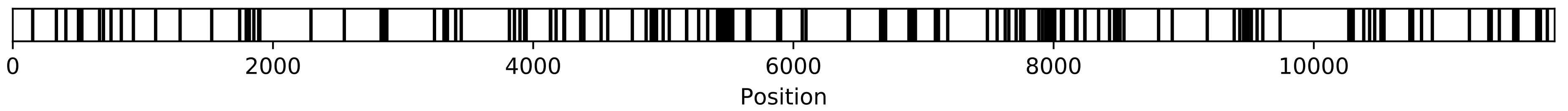


MM1 vs MM2
Number of differences: 132

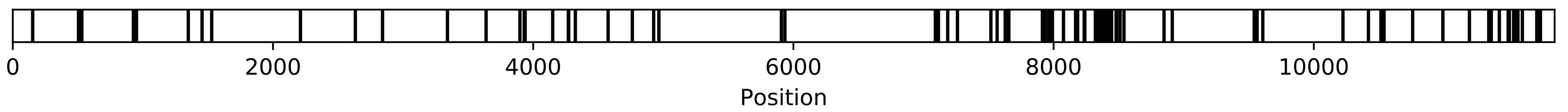

### Region: 213



CR1 vs CR2
Number of differences: 529

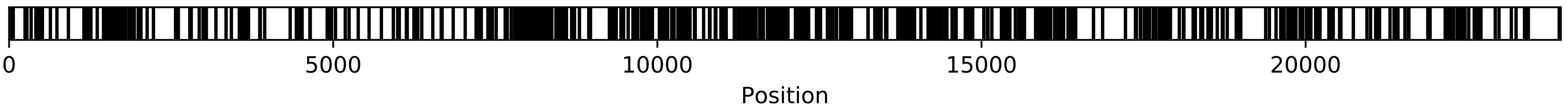


CR1 vs MA1
Number of differences: 298

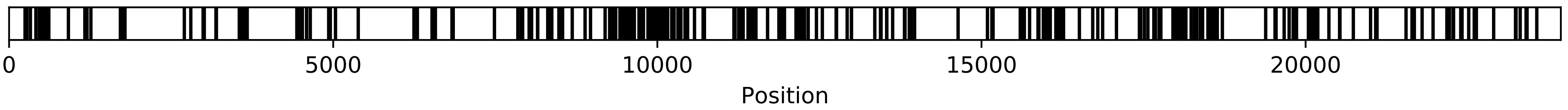


CR1 vs MA2
Number of differences: 549

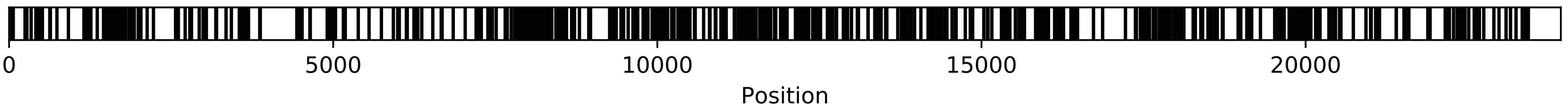


CR1 vs MM1
Number of differences: 299

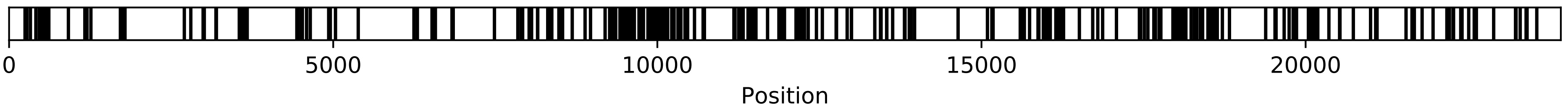


CR1 vs MM2
Number of differences: 335

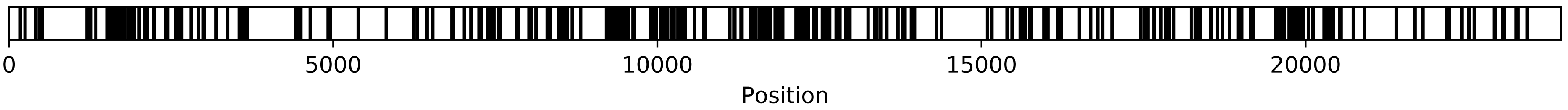


CR2 vs MA1
Number of differences: 510

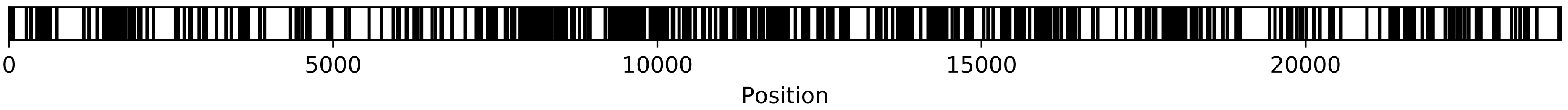


CR2 vs MA2
Number of differences: 138

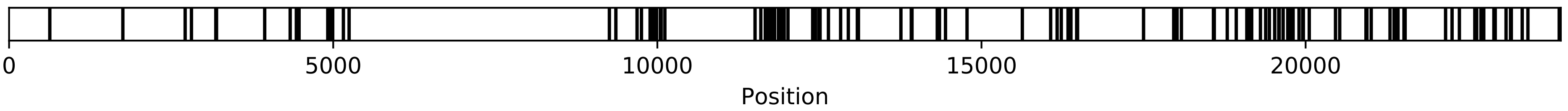


CR2 vs MM1
Number of differences: 511

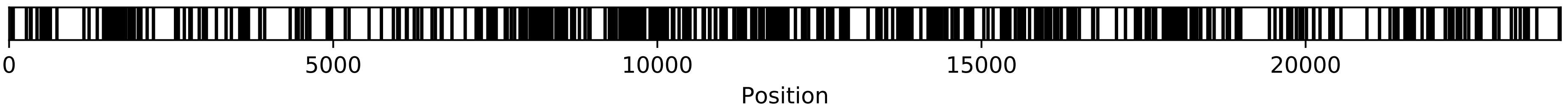


CR2 vs MM2
Number of differences: 516

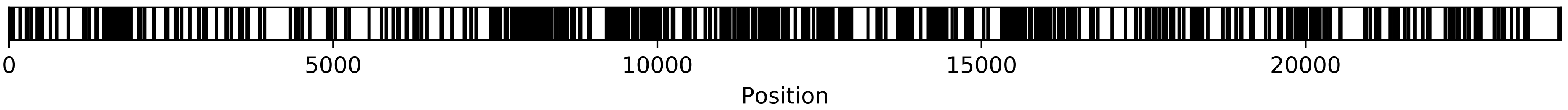


MA1 vs MA2
Number of differences: 520

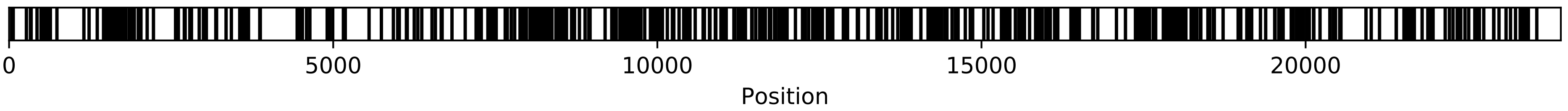


MA1 vs MM1
Number of differences: 1

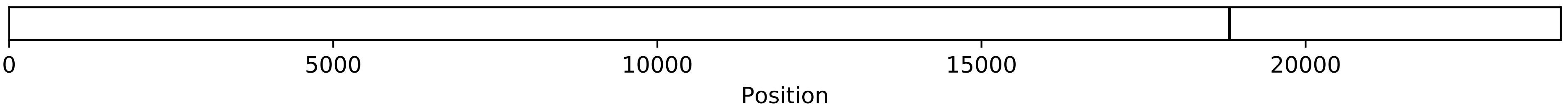


MA1 vs MM2
Number of differences: 320

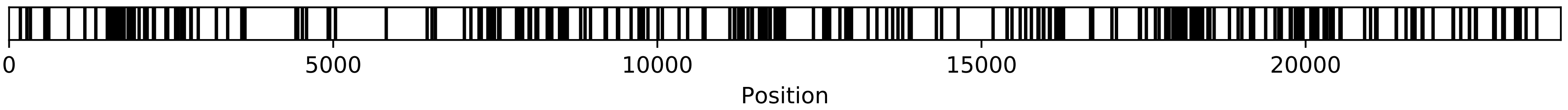


MA2 vs MM1
Number of differences: 521

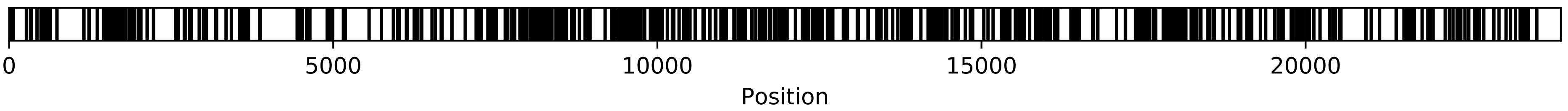


MA2 vs MM2
Number of differences: 527

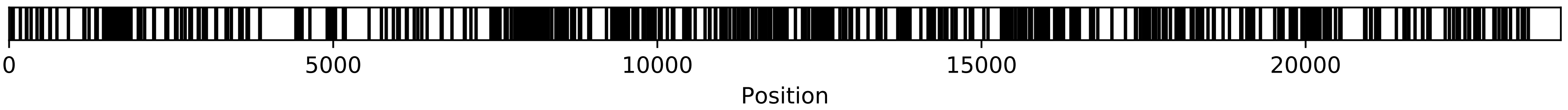


MM1 vs MM2
Number of differences: 319

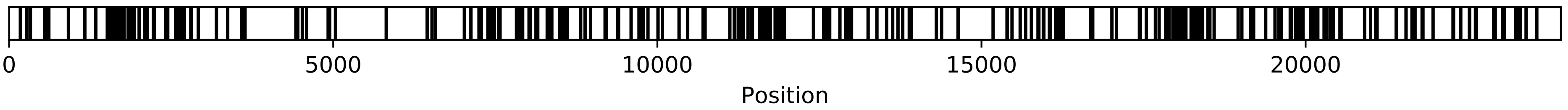

### Region: 216



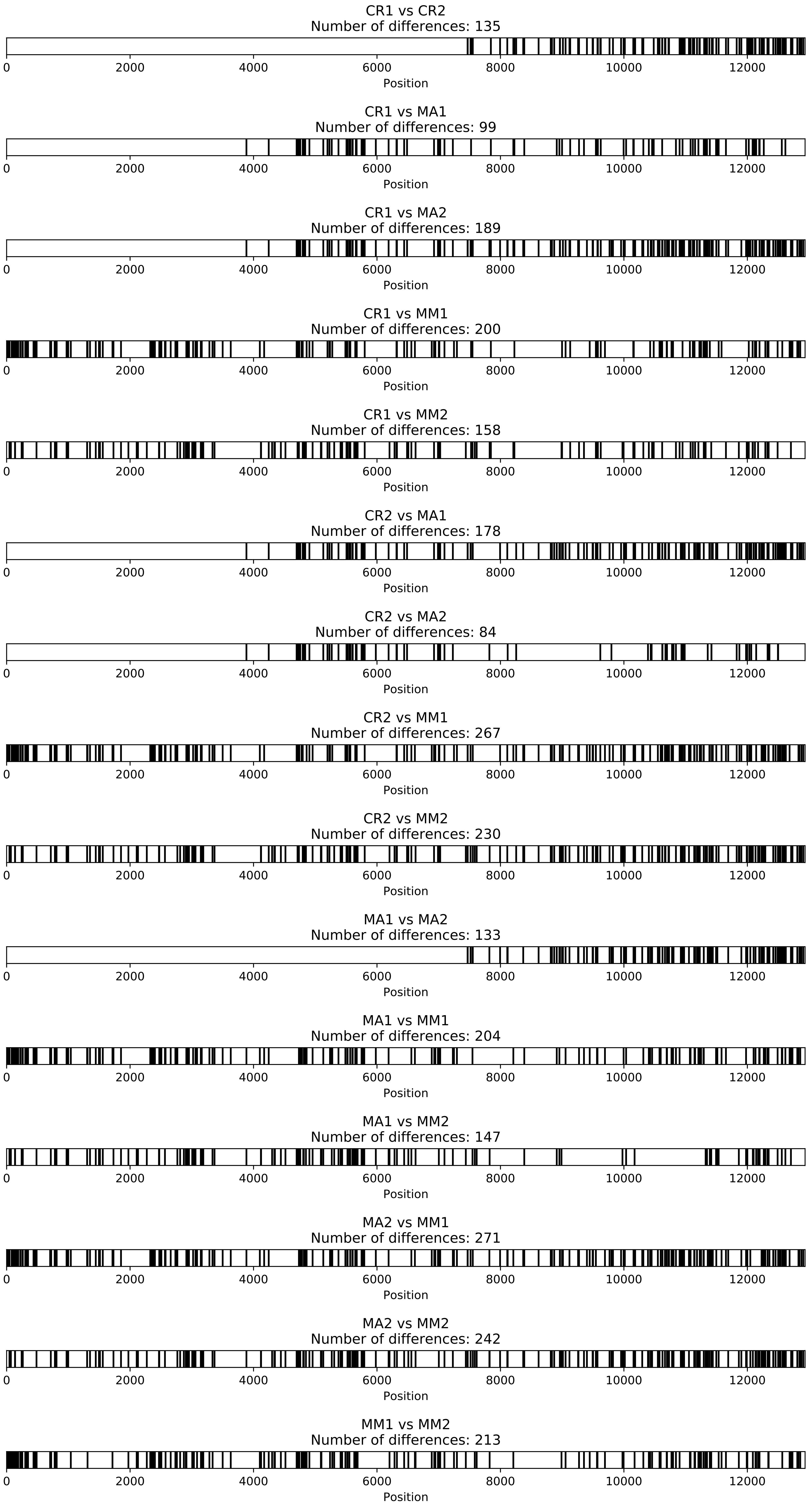
CR1 vs CR2
Number of differences: 135
CR1 vs MA1
Number of differences: 99
CR1 vs MA2
Number of differences: 189
CR1 vs MM1
Number of differences: 200
CR1 vs MM2
Number of differences: 158
CR2 vs MA1
Number of differences: 178
CR2 vs MA2
Number of differences: 84
CR2 vs MM1
Number of differences: 267
CR2 vs MM2
Number of differences: 230
MA1 vs MA2
Number of differences: 133
MA1 vs MM1
Number of differences: 204
MA1 vs MM2
Number of differences: 147
MA2 vs MM1
Number of differences: 271
MA2 vs MM2
Number of differences: 242
MM1 vs MM2
Number of differences: 213
0
2000
4000
6000
8000
10000
12000
Position

### Region: 217

CR1 vs CR2
Number of differences: 273

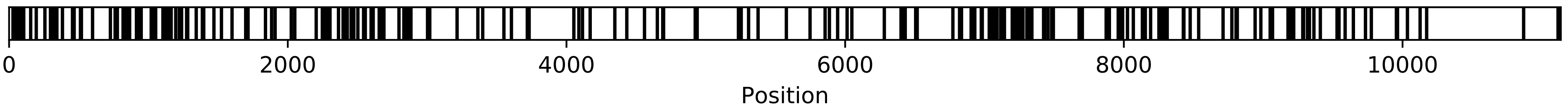


CR1 vs MA1
Number of differences: 48

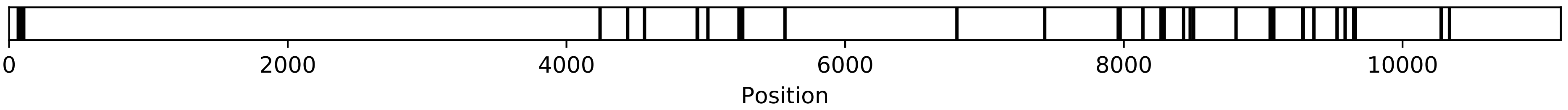


CR1 vs MA2
Number of differences: 266

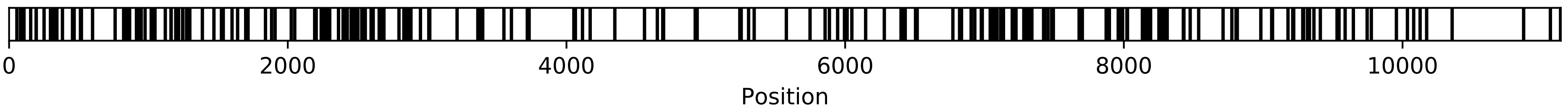


CR1 vs MM1
Number of differences: 141

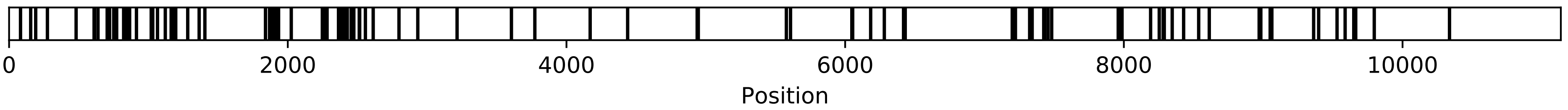


CR1 vs MM2
Number of differences: 48

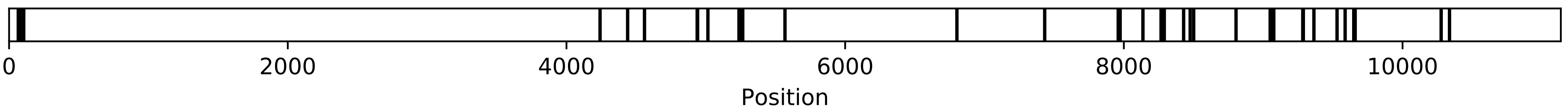


CR2 vs MA1
Number of differences: 262

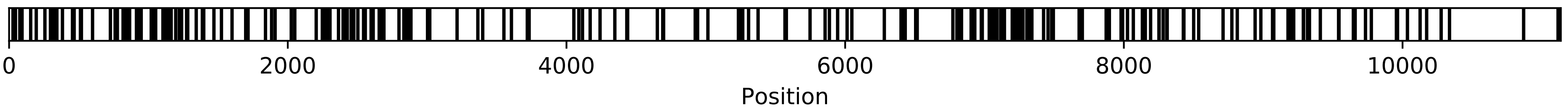


CR2 vs MA2
Number of differences: 86

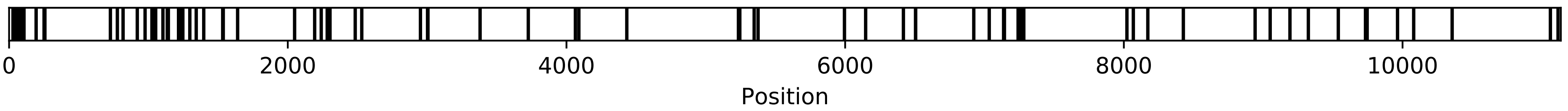


CR2 vs MM1
Number of differences: 255

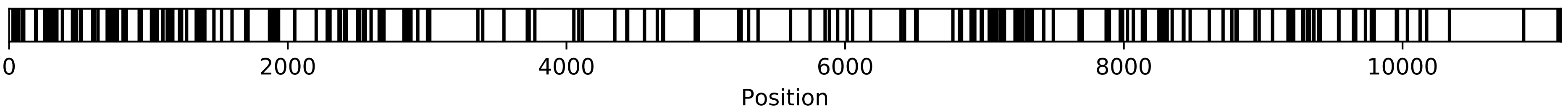


CR2 vs MM2
Number of differences: 262

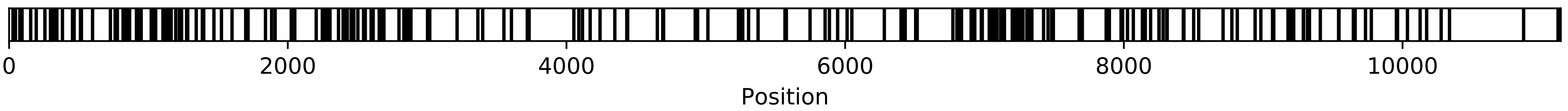


MA1 vs MA2
Number of differences: 259

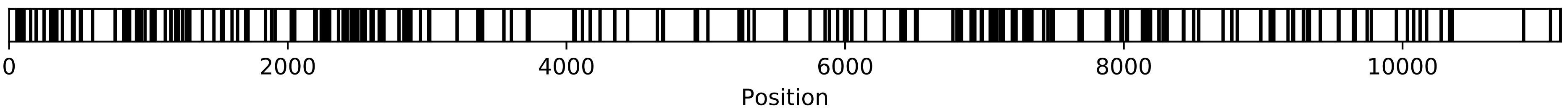


MA1 vs MM1
Number of differences: 138

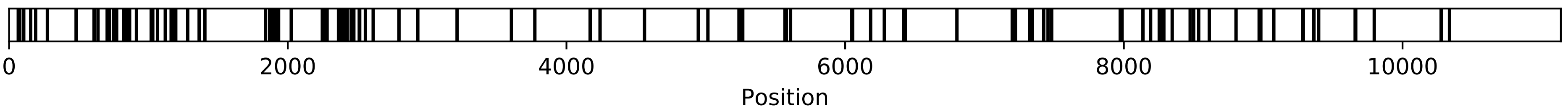


MA1 vs MM2
Number of differences: 0

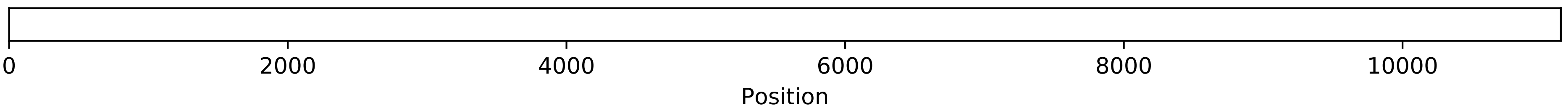


MA2 vs MM1
Number of differences: 250

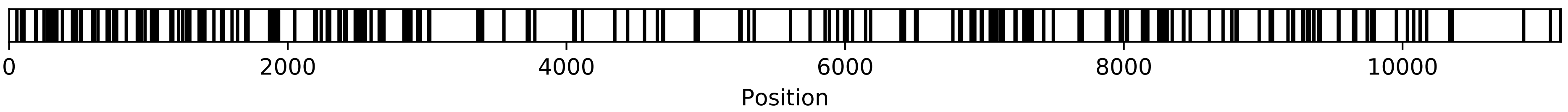


MA2 vs MM2
Number of differences: 259

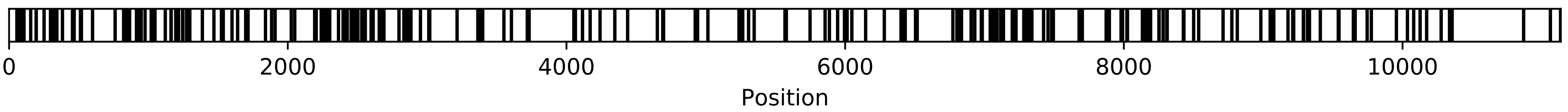


MM1 vs MM2
Number of differences: 138

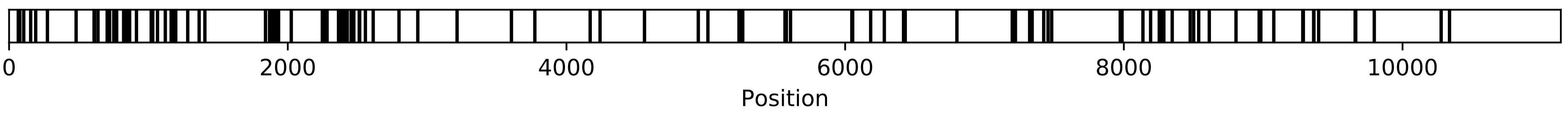

### Region: 218



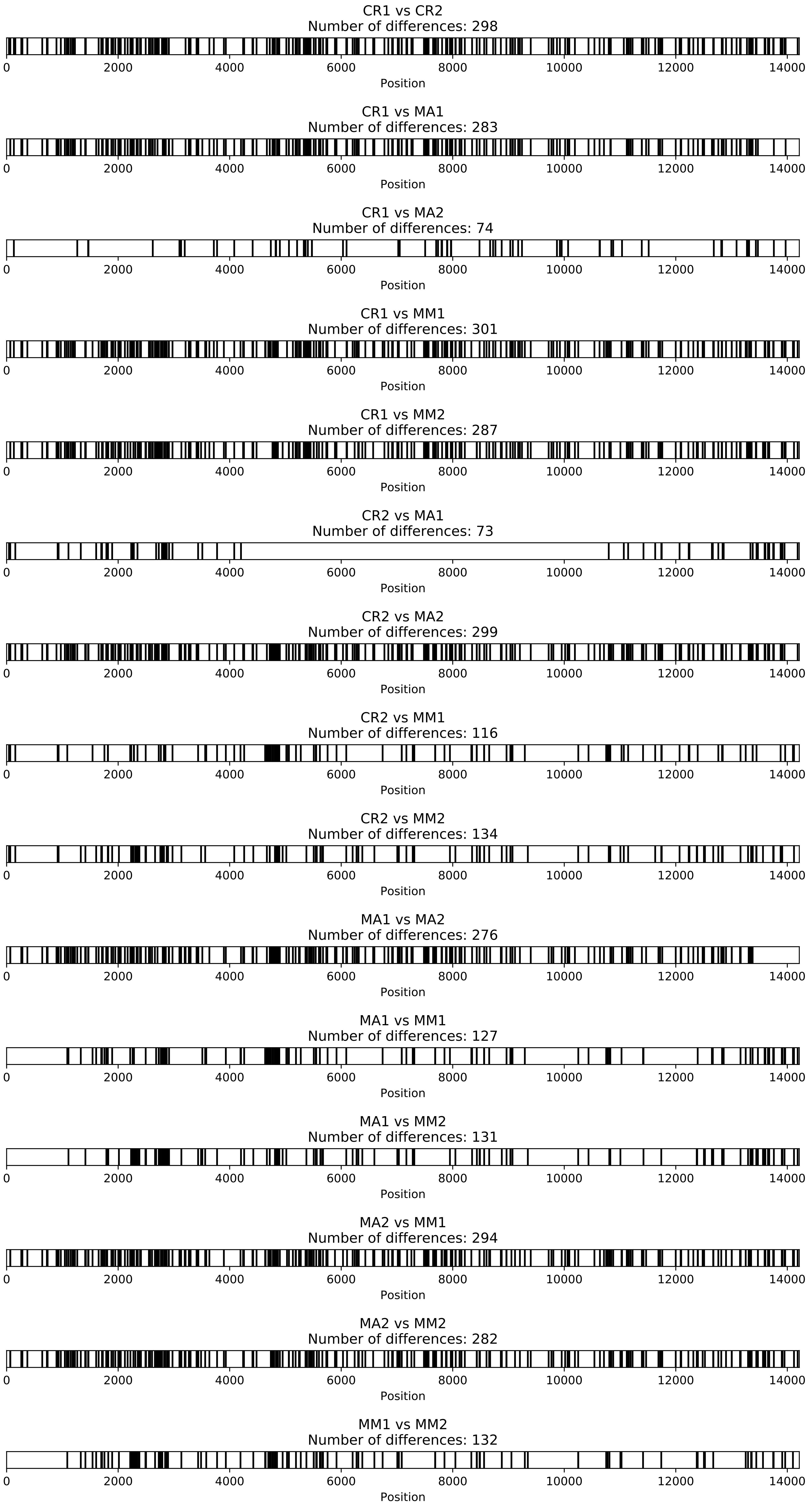
CR1 vs CR2
Number of differences: 298
CR1 vs MA1
Number of differences: 283
CR1 vs MA2
Number of differences: 74
CR1 vs MM1
Number of differences: 301
CR1 vs MM2
Number of differences: 287
CR2 vs MA1
Number of differences: 73
CR2 vs MA2
Number of differences: 299
CR2 vs MM1
Number of differences: 116
CR2 vs MM2
Number of differences: 134
MA1 vs MA2
Number of differences: 276
MA1 vs MM1
Number of differences: 127
MA1 vs MM2
Number of differences: 131
MA2 vs MM1
Number of differences: 294
MA2 vs MM2
Number of differences: 282
MM1 vs MM2
Number of differences: 132
0
2000
4000
6000
8000
10000
12000
14000
Position

### Region: 220

CR1 vs CR2
Number of differences: 307

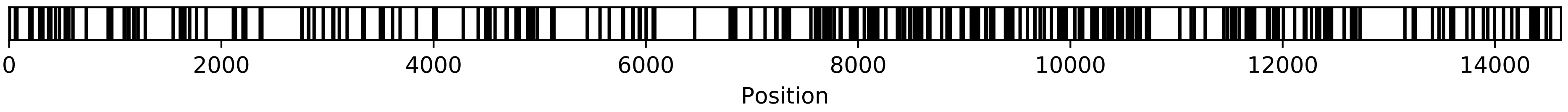


CR1 vs MA1
Number of differences: 0

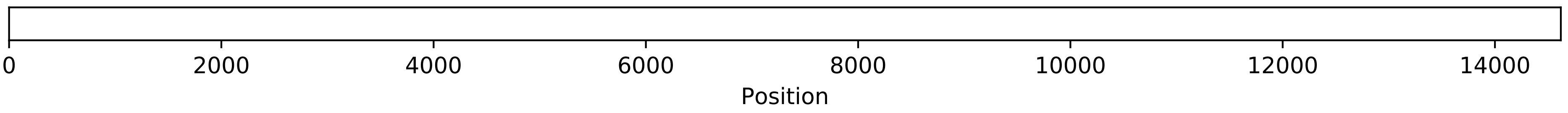


CR1 vs MA2
Number of differences: 305

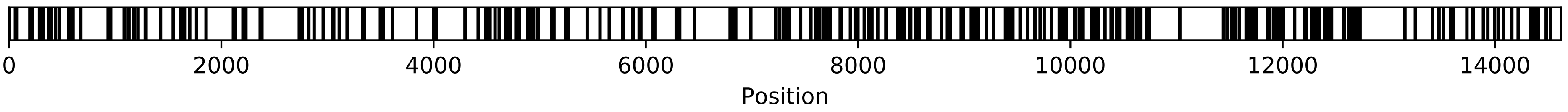


CR1 vs MM1
Number of differences: 0

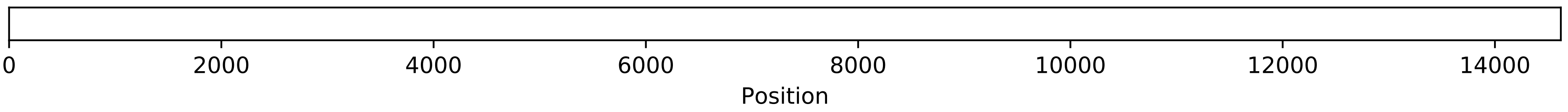


CR1 vs MM2
Number of differences: 123

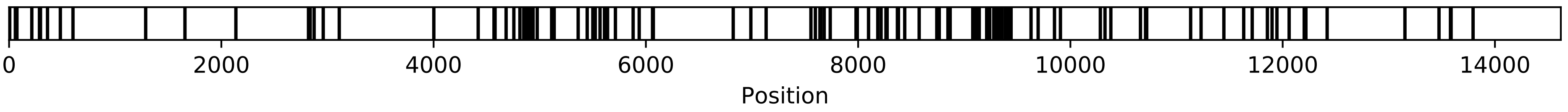


CR2 vs MA1
Number of differences: 307

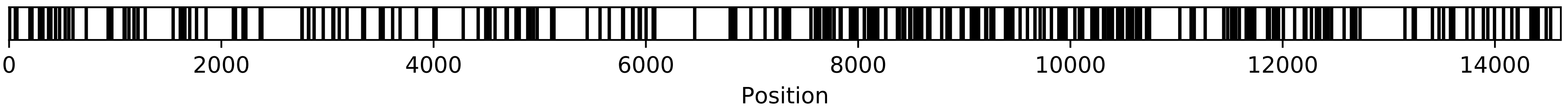


CR2 vs MA2
Number of differences: 73

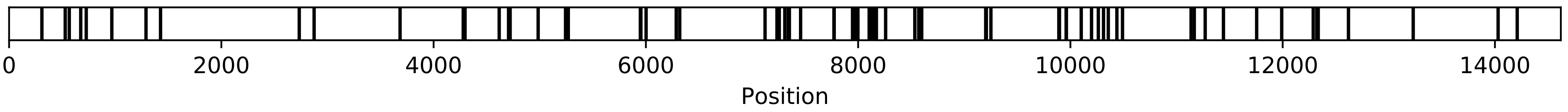


CR2 vs MM1
Number of differences: 307

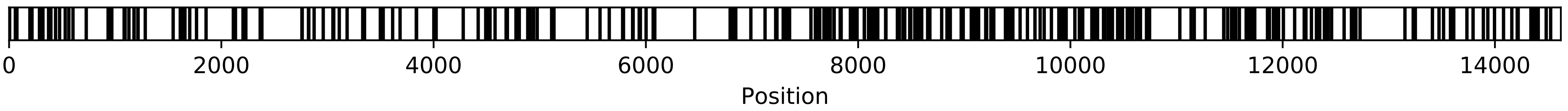


CR2 vs MM2
Number of differences: 302

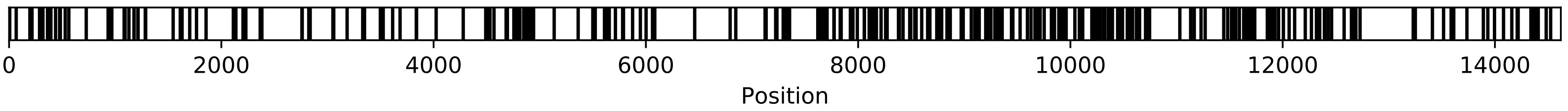


MA1 vs MA2
Number of differences: 305

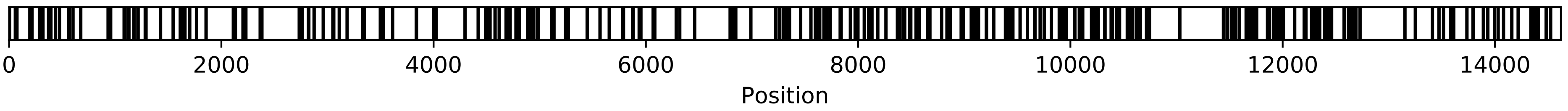


MA1 vs MM1
Number of differences: 0

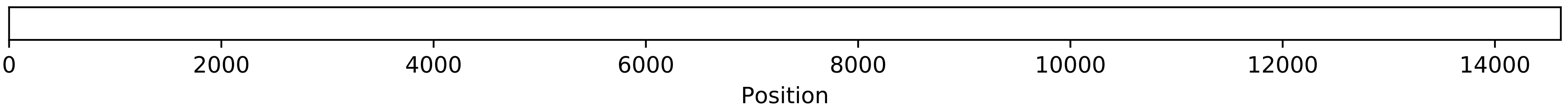


MA1 vs MM2
Number of differences: 123

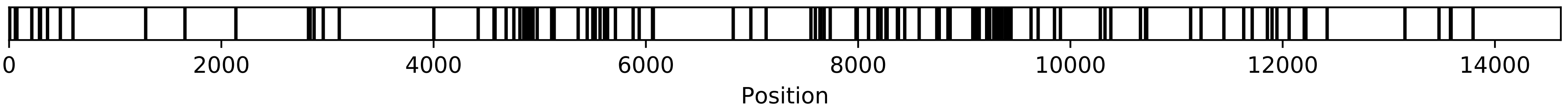


MA2 vs MM1
Number of differences: 305

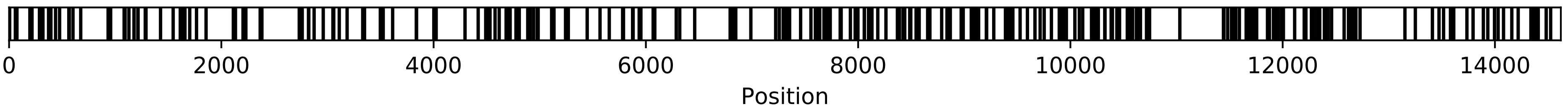


MA2 vs MM2
Number of differences: 301

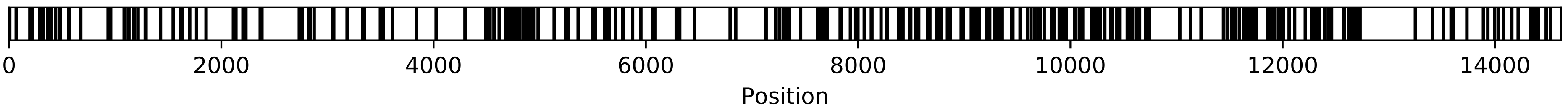


MM1 vs MM2
Number of differences: 123

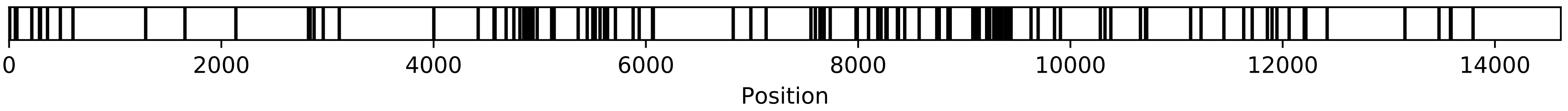

### Region: 222



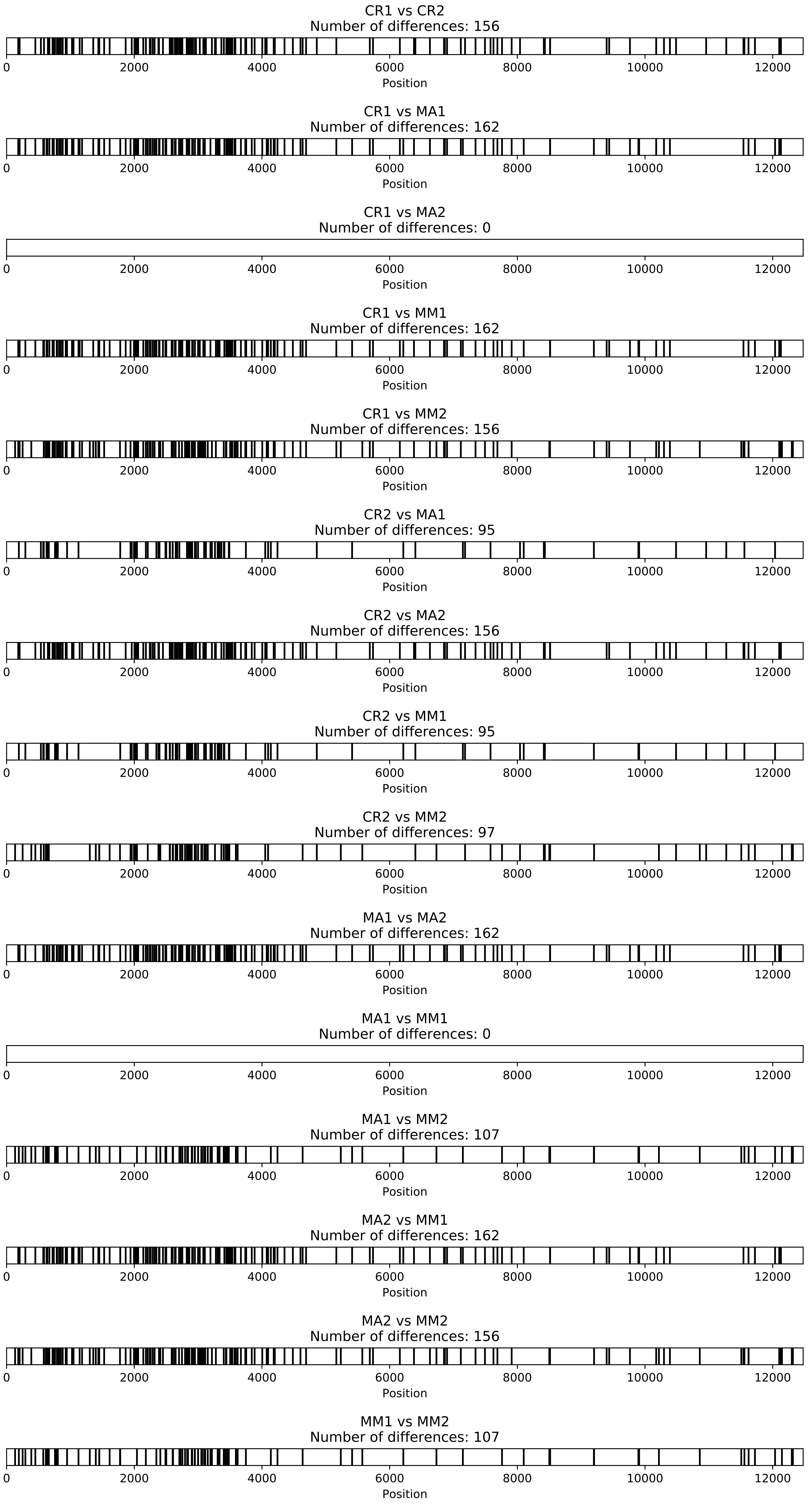
CR1 vs CR2
Number of differences: 156
CR1 vs MA1
Number of differences: 162
CR1 vs MA2
Number of differences: 0
CR1 vs MM1
Number of differences: 162
CR1 vs MM2
Number of differences: 156
CR2 vs MA1
Number of differences: 95
CR2 vs MA2
Number of differences: 156
CR2 vs MM1
Number of differences: 95
CR2 vs MM2
Number of differences: 97
MA1 vs MA2
Number of differences: 162
MA1 vs MM1
Number of differences: 0
MA1 vs MM2
Number of differences: 107
MA2 vs MM1
Number of differences: 162
MA2 vs MM2
Number of differences: 156
MM1 vs MM2
Number of differences: 107
0
2000
4000
6000
8000
10000
12000
Position

### Region: 223



CR1 vs CR2
Number of differences: 288

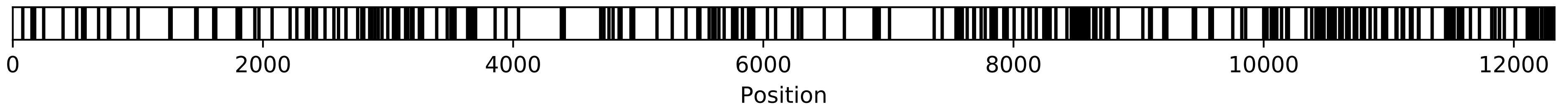


CR1 vs MA1
Number of differences: 141

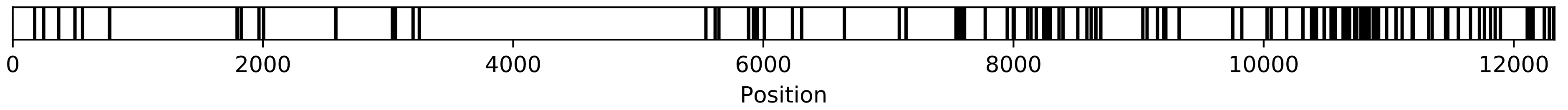


CR1 vs MA2
Number of differences: 280

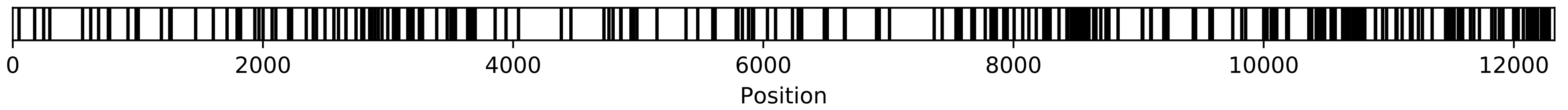


CR1 vs MM1
Number of differences: 163

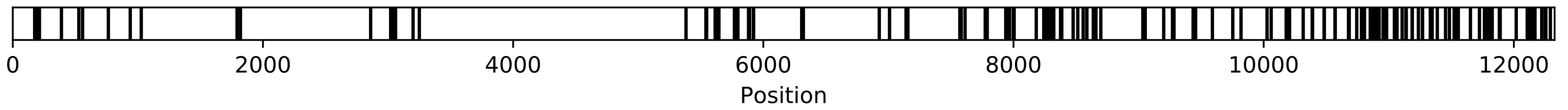


CR1 vs MM2
Number of differences: 139

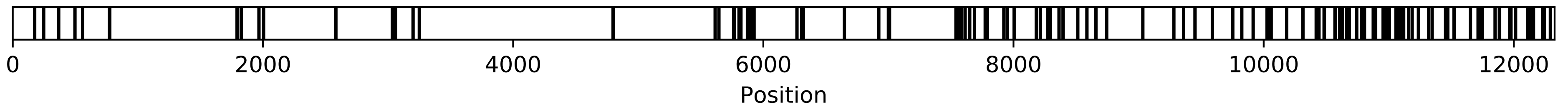


CR2 vs MA1
Number of differences: 272

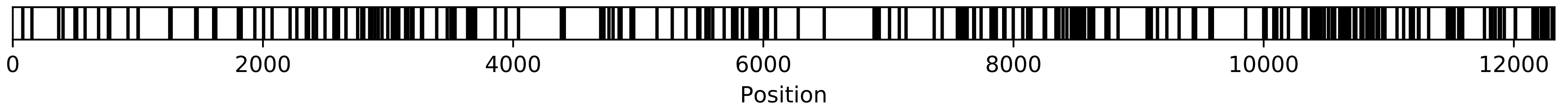


CR2 vs MA2
Number of differences: 111

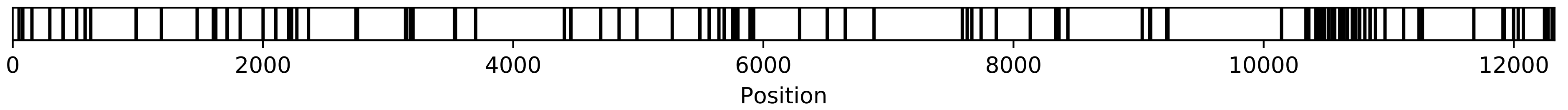


CR2 vs MM1
Number of differences: 290

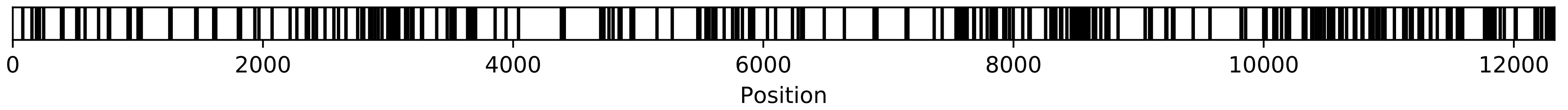


CR2 vs MM2
Number of differences: 264

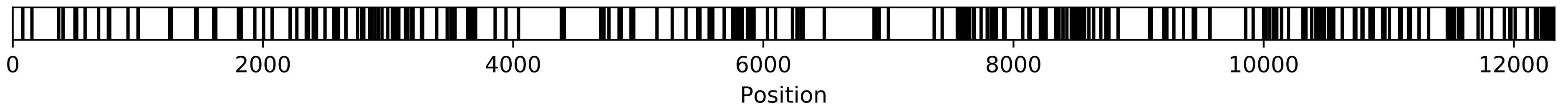


MA1 vs MA2
Number of differences: 266

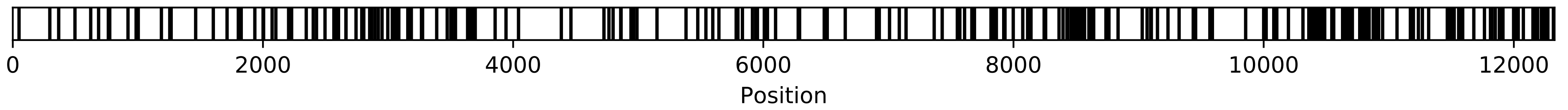


MA1 vs MM1
Number of differences: 156

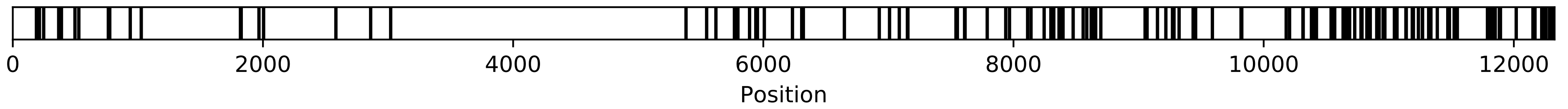


MA1 vs MM2
Number of differences: 113

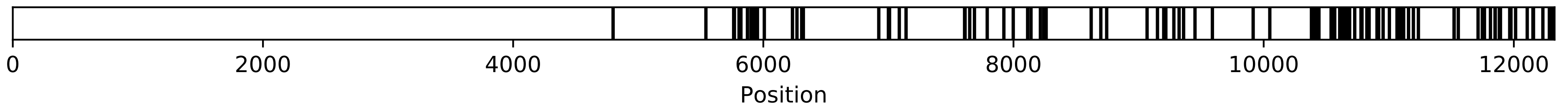


MA2 vs MM1
Number of differences: 279

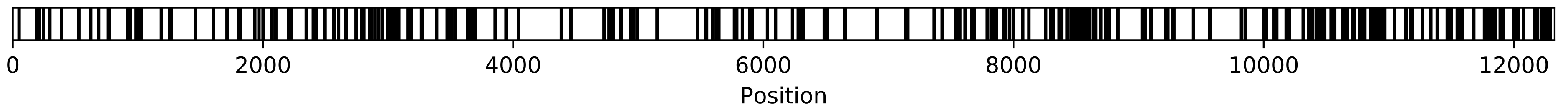


MA2 vs MM2
Number of differences: 262

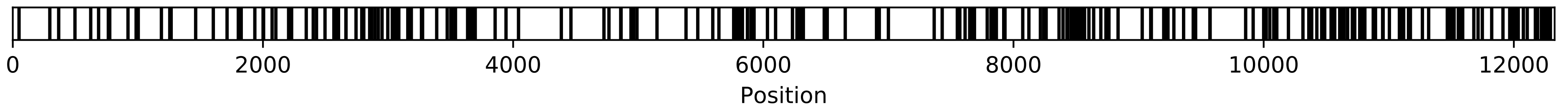


MM1 vs MM2
Number of differences: 148

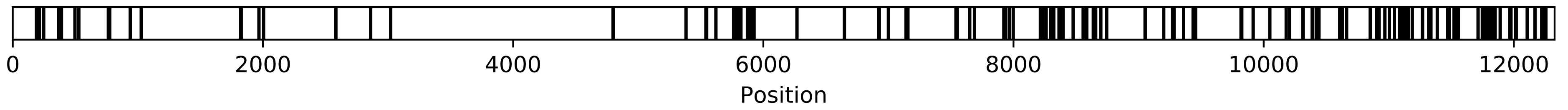

### Region: 224



CR1 vs CR2
Number of differences: 195

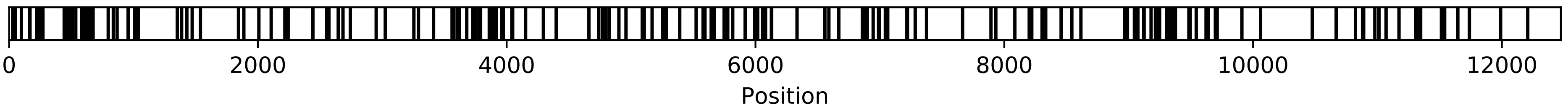


CR1 vs MA1
Number of differences: 209

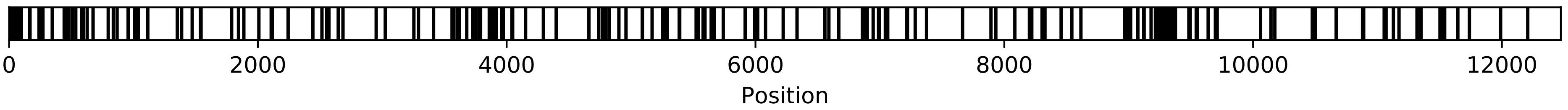


CR1 vs MA2
Number of differences: 125

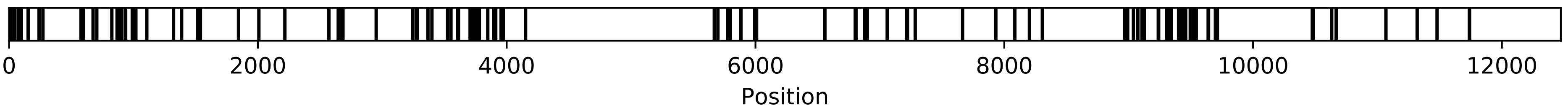


CR1 vs MM1
Number of differences: 148

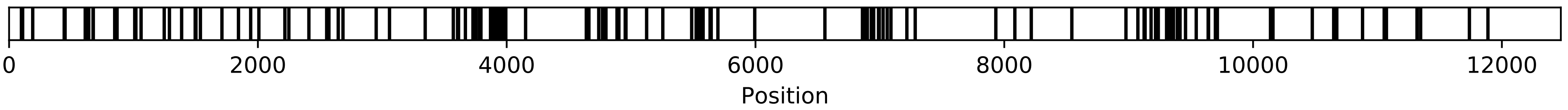


CR1 vs MM2
Number of differences: 125

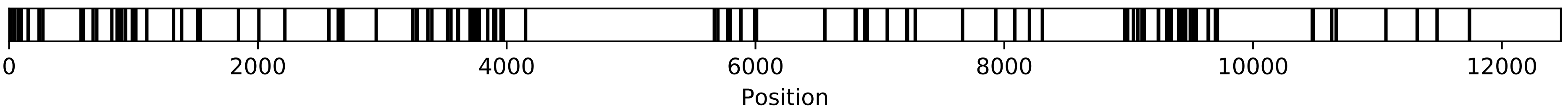


CR2 vs MA1
Number of differences: 72

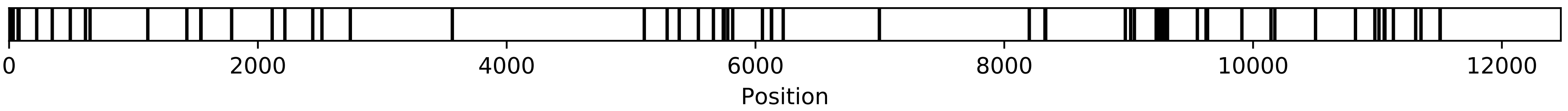


CR2 vs MA2
Number of differences: 198

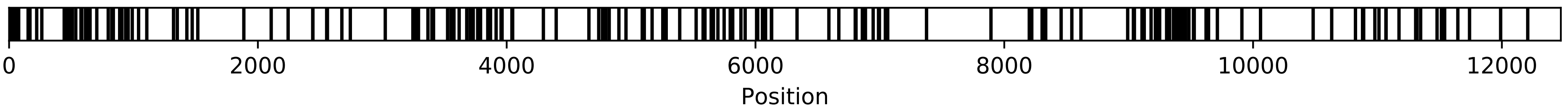


CR2 vs MM1
Number of differences: 192

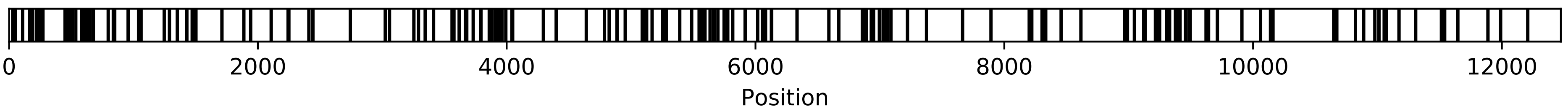


CR2 vs MM2
Number of differences: 198

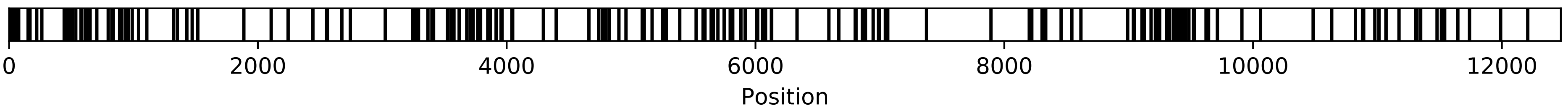


MA1 vs MA2
Number of differences: 210

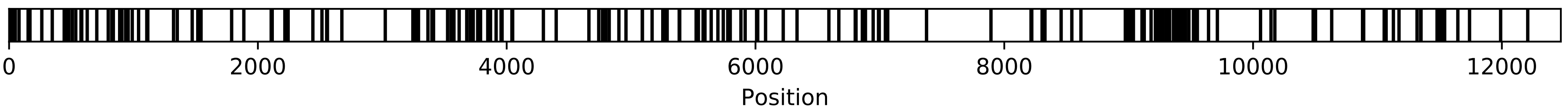


MA1 vs MM1
Number of differences: 198

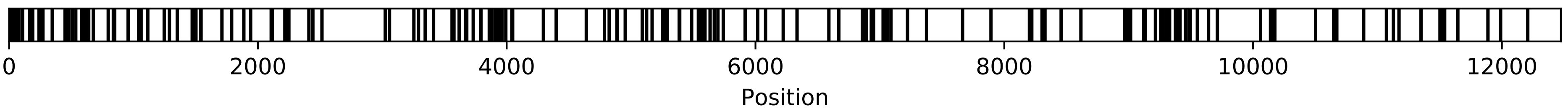


MA1 vs MM2
Number of differences: 210

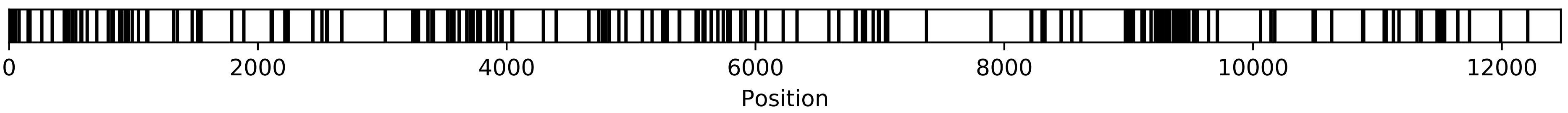


MA2 vs MM1
Number of differences: 159

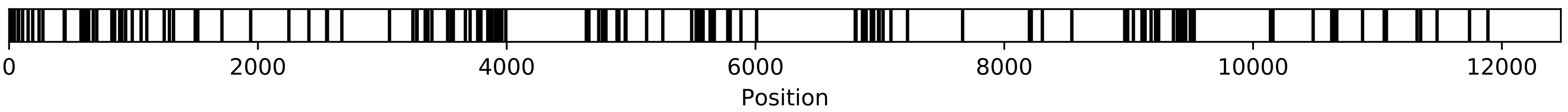


MA2 vs MM2
Number of differences: 0

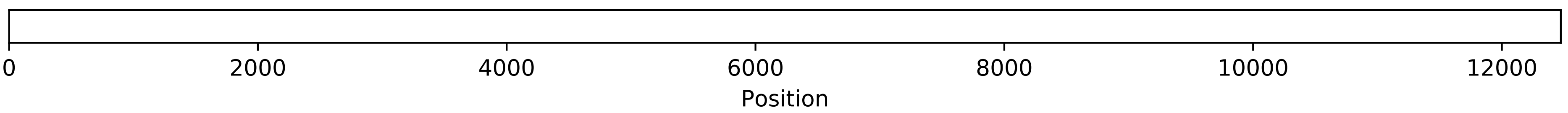


MM1 vs MM2
Number of differences: 159

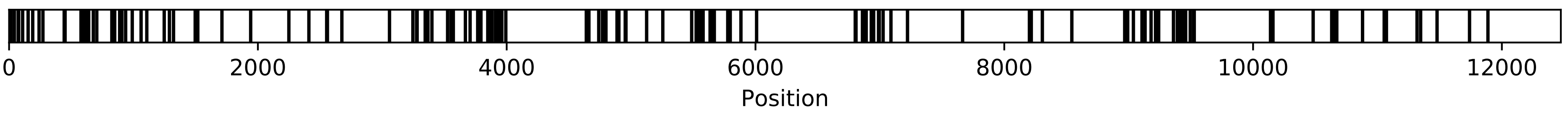

### Region: 227

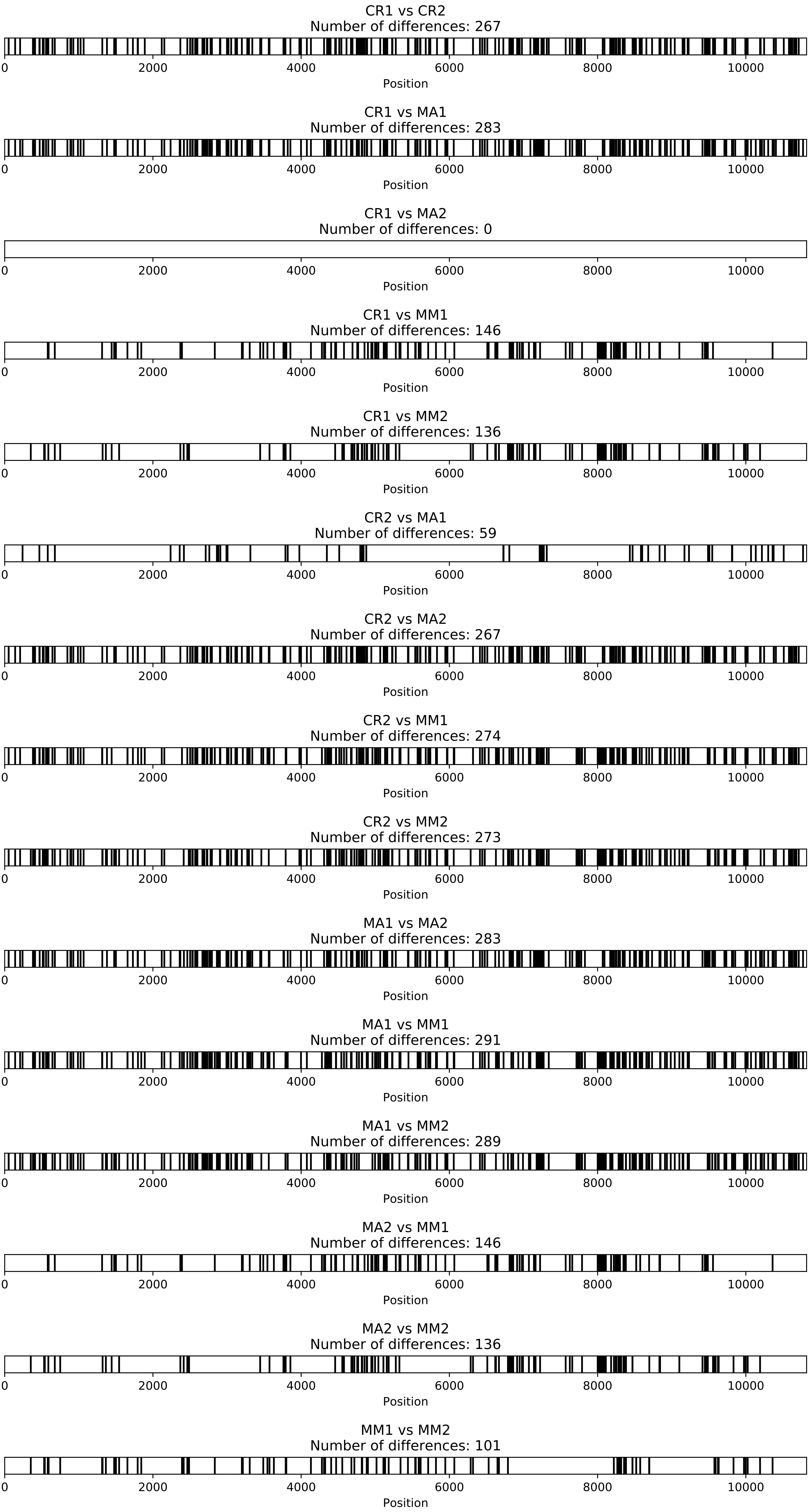
CR1 vs CR2
Number of differences: 267
CR1 vs MA1
Number of differences: 283
CR1 vs MA2
Number of differences: 0
CR1 vs MM1
Number of differences: 146
CR1 vs MM2
Number of differences: 136
CR2 vs MA1
Number of differences: 59
CR2 vs MA2
Number of differences: 267
CR2 vs MM1
Number of differences: 274
CR2 vs MM2
Number of differences: 273
MA1 vs MA2
Number of differences: 283
MA1 vs MM1
Number of differences: 291
MA1 vs MM2
Number of differences: 289
MA2 vs MM1
Number of differences: 146
MA2 vs MM2
Number of differences: 136
MM1 vs MM2
Number of differences: 101
0
2000
4000
6000
8000
10000
Position

### Region: 228



CR1 vs CR2
Number of differences: 244

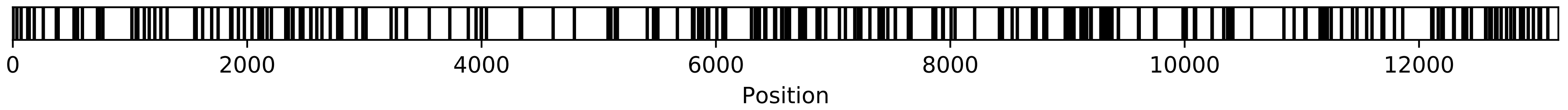


CR1 vs MA1
Number of differences: 101

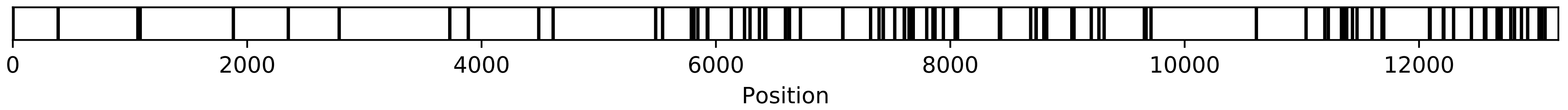


CR1 vs MA2
Number of differences: 242

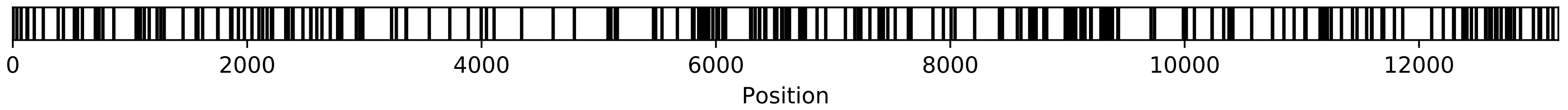


CR1 vs MM1
Number of differences: 116

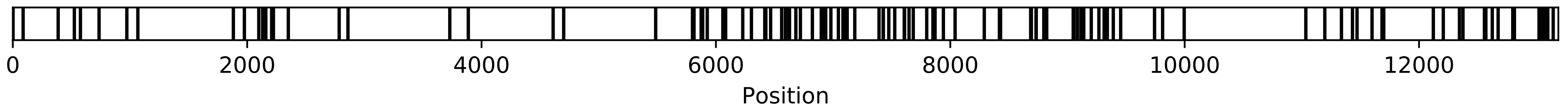


CR1 vs MM2
Number of differences: 101

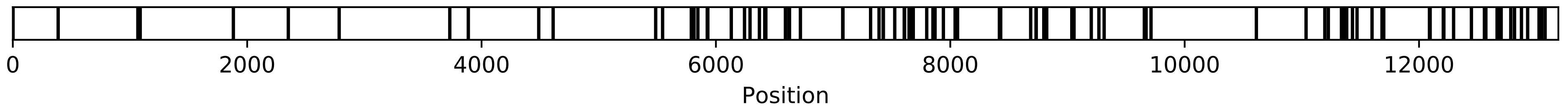


CR2 vs MA1
Number of differences: 245

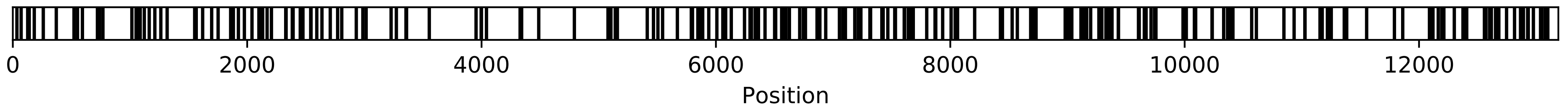


CR2 vs MA2
Number of differences: 83

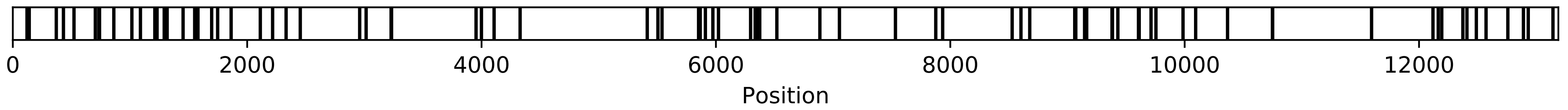


CR2 vs MM1
Number of differences: 238

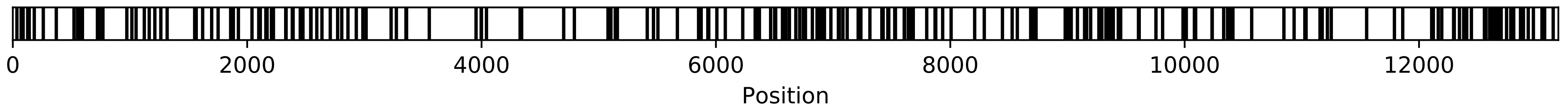


CR2 vs MM2
Number of differences: 245

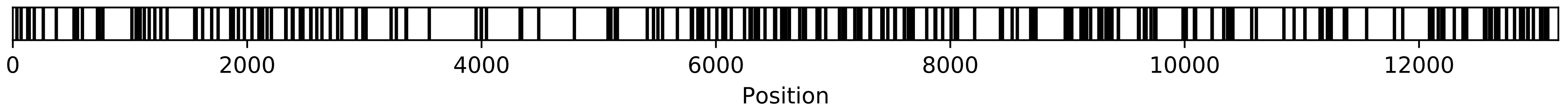


MA1 vs MA2
Number of differences: 241

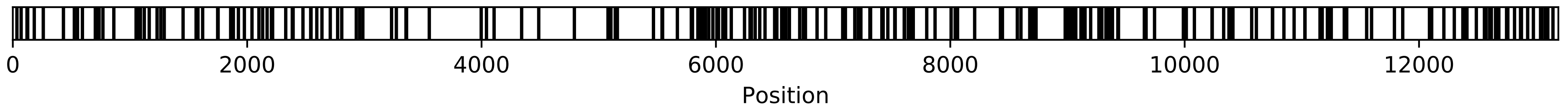


MA1 vs MM1
Number of differences: 101

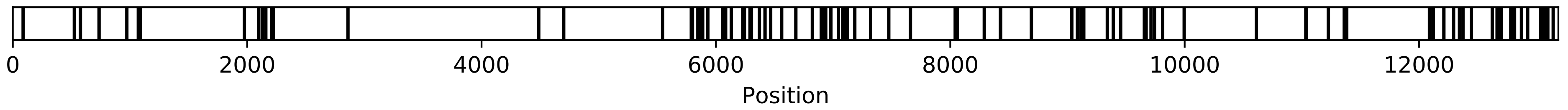


MA1 vs MM2
Number of differences: 0

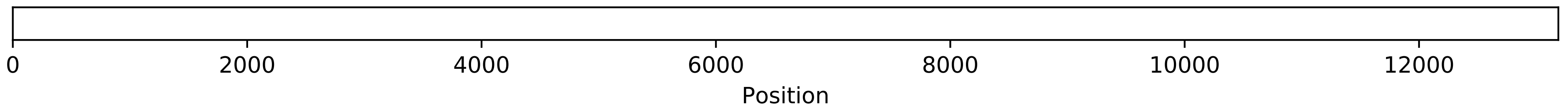


MA2 vs MM1
Number of differences: 236

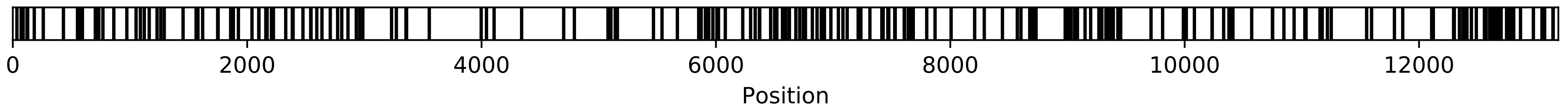


MA2 vs MM2
Number of differences: 241

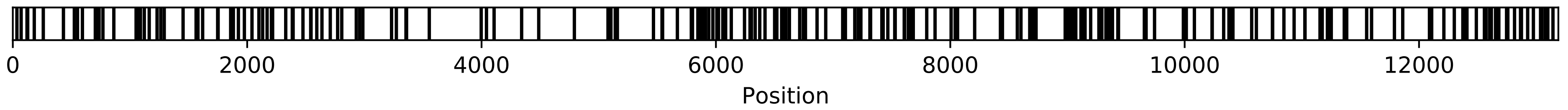


MM1 vs MM2
Number of differences: 101

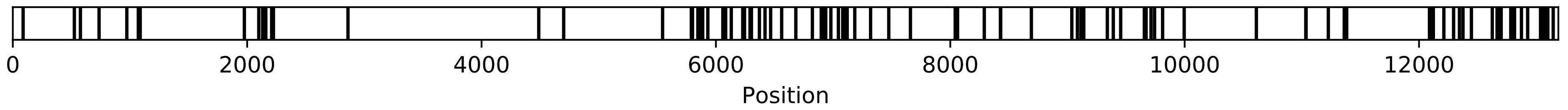

### Region: 231



CR1 vs CR2
Number of differences: 196

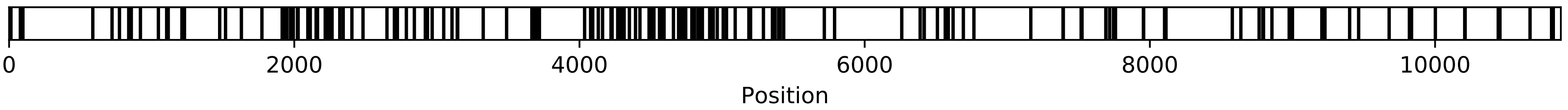


CR1 vs MA1
Number of differences: 203

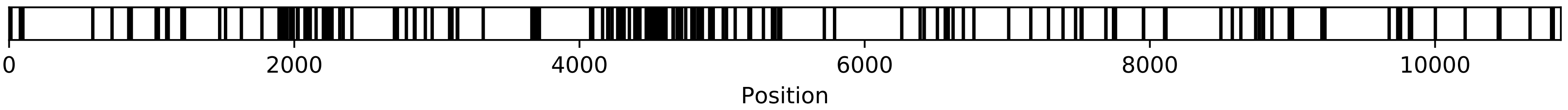


CR1 vs MA2
Number of differences: 204

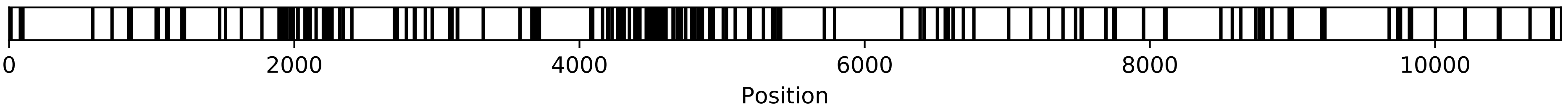


CR1 vs MM1
Number of differences: 38

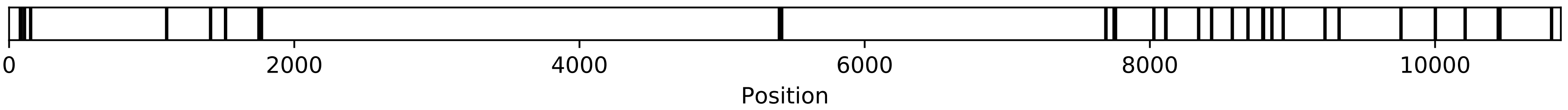


CR1 vs MM2
Number of differences: 118

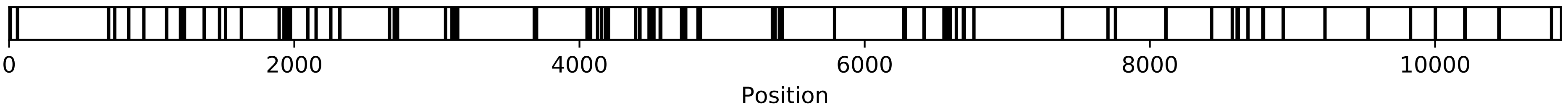


CR2 vs MA1
Number of differences: 55

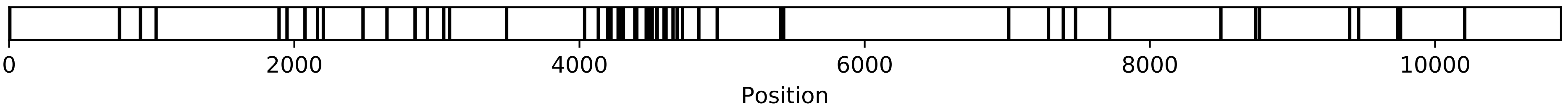


CR2 vs MA2
Number of differences: 53

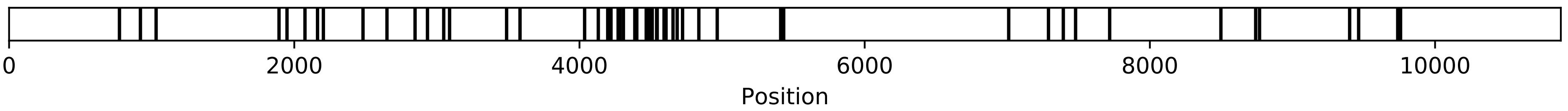


CR2 vs MM1
Number of differences: 189

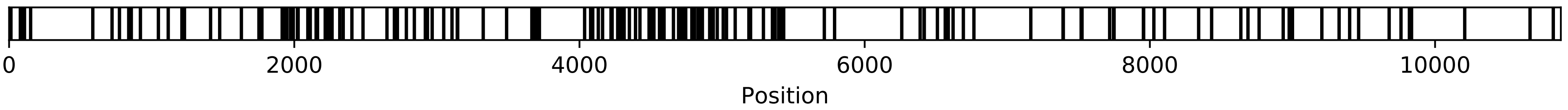


CR2 vs MM2
Number of differences: 185

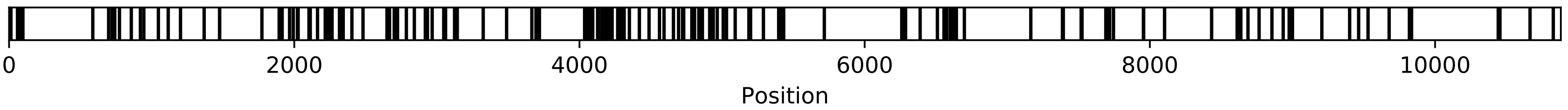


MA1 vs MA2
Number of differences: 4

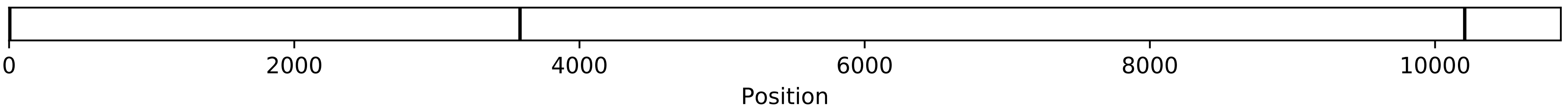


MA1 vs MM1
Number of differences: 194

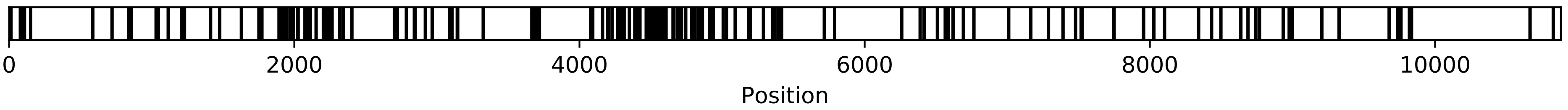


MA1 vs MM2
Number of differences: 191

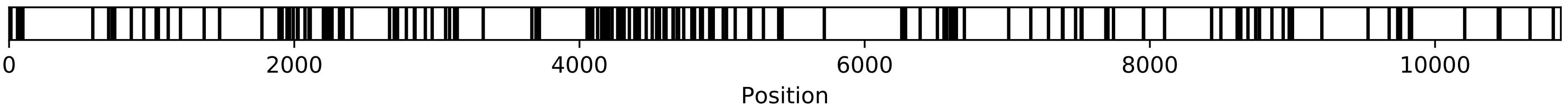


MA2 vs MM1
Number of differences: 195

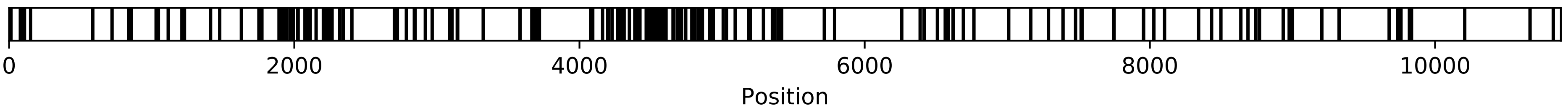


MA2 vs MM2
Number of differences: 190

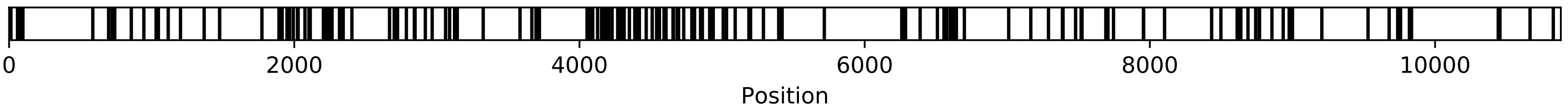


MM1 vs MM2
Number of differences: 116

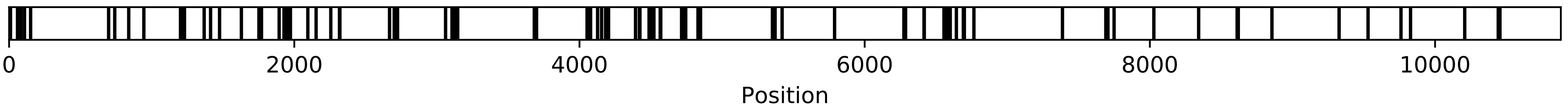

### Region: 232



CR1 vs CR2
Number of differences: 294

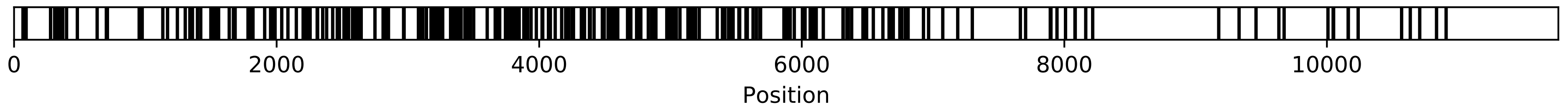


CR1 vs MA1
Number of differences: 188

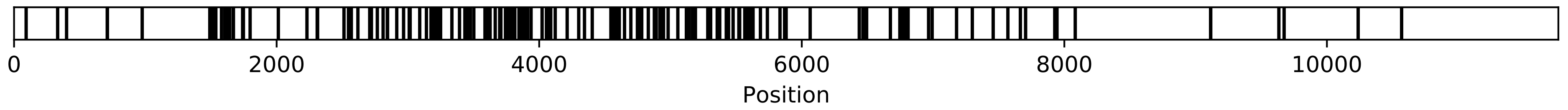


CR1 vs MA2
Number of differences: 312

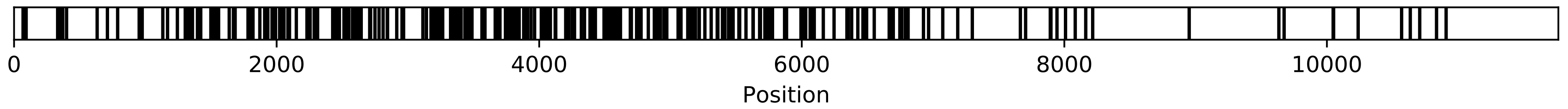


CR1 vs MM1
Number of differences: 188

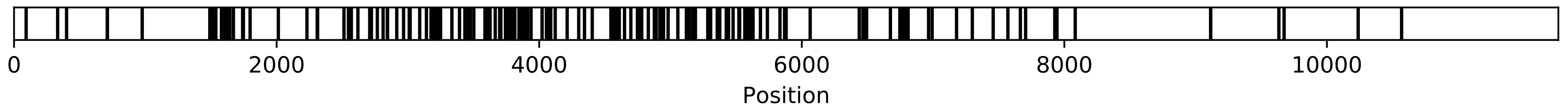


CR1 vs MM2
Number of differences: 121

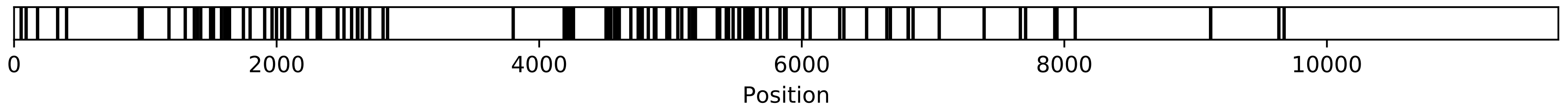


CR2 vs MA1
Number of differences: 293

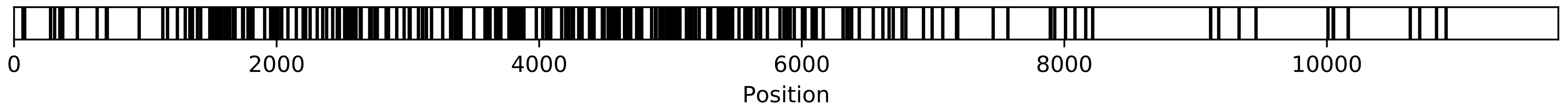


CR2 vs MA2
Number of differences: 161

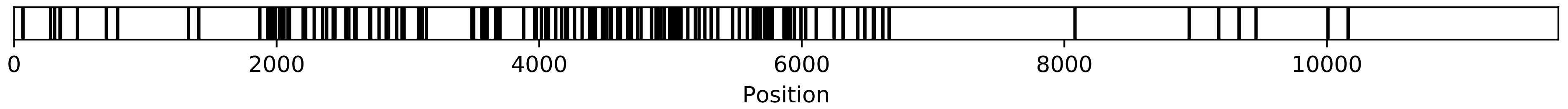


CR2 vs MM1
Number of differences: 293

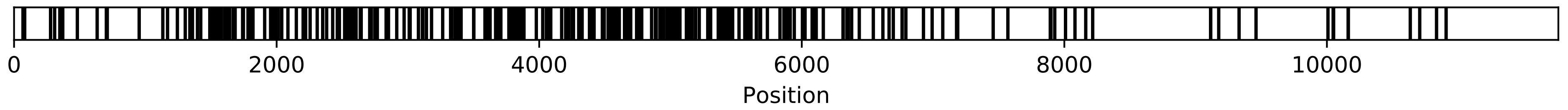


CR2 vs MM2
Number of differences: 304

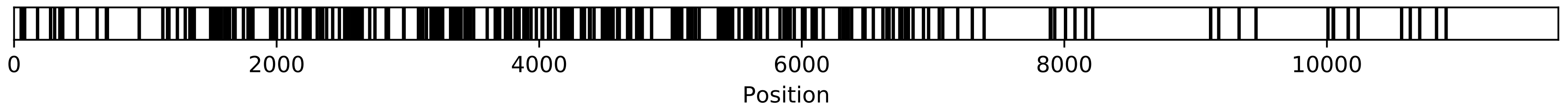


MA1 vs MA2
Number of differences: 303

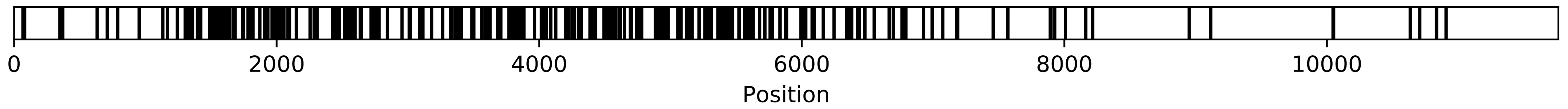


MA1 vs MM1
Number of differences: 0

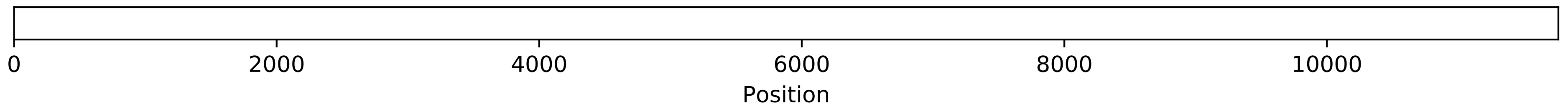


MA1 vs MM2
Number of differences: 169

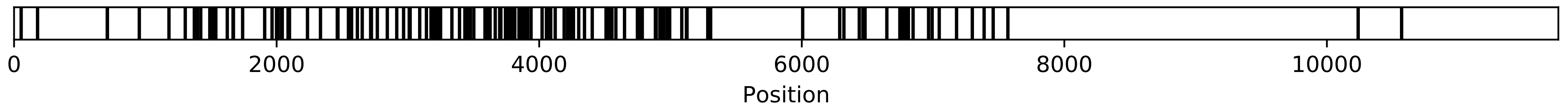


MA2 vs MM1
Number of differences: 303

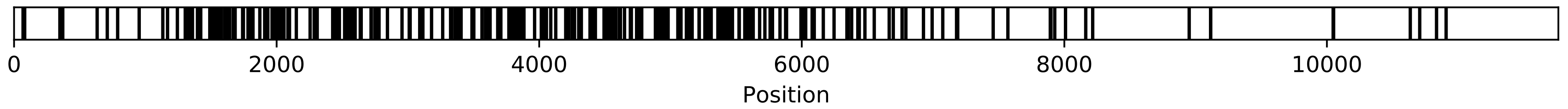


MA2 vs MM2
Number of differences: 312

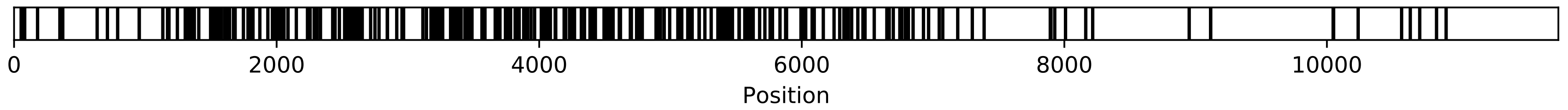


MM1 vs MM2
Number of differences: 169

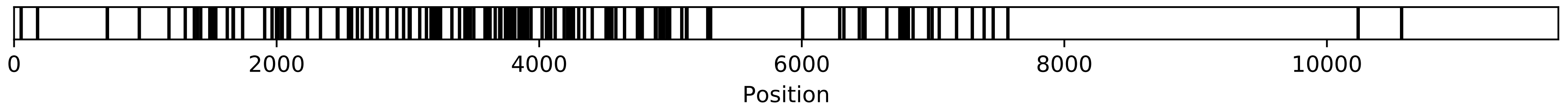

### Region: 233



CR1 vs CR2
Number of differences: 247

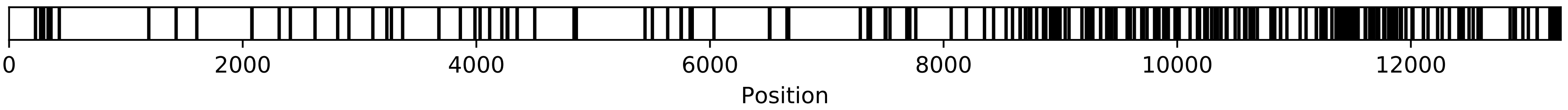


CR1 vs MA1
Number of differences: 96

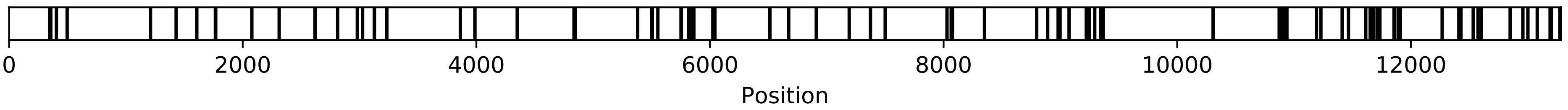


CR1 vs MA2
Number of differences: 241

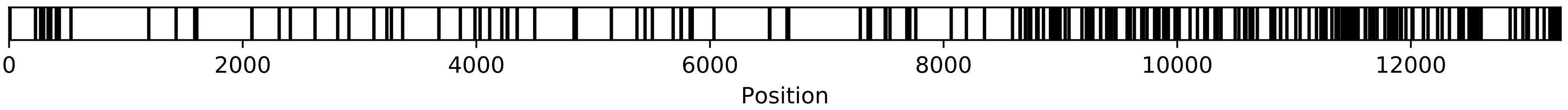


CR1 vs MM1
Number of differences: 37

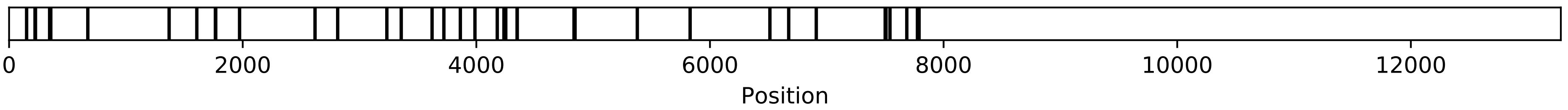


CR1 vs MM2
Number of differences: 96

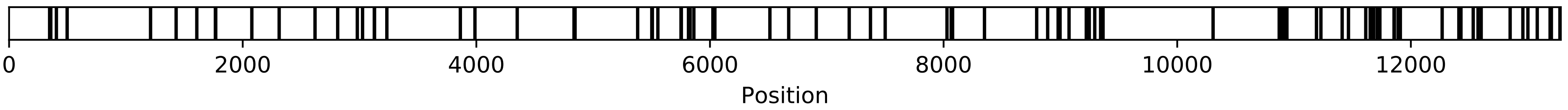


CR2 vs MA1
Number of differences: 236

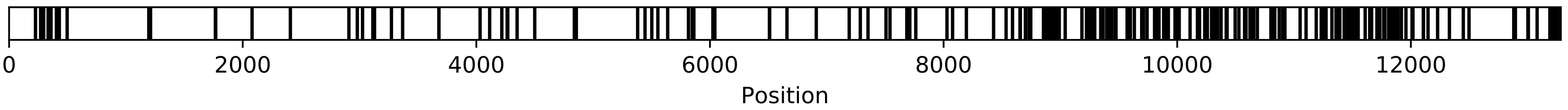


CR2 vs MA2
Number of differences: 46

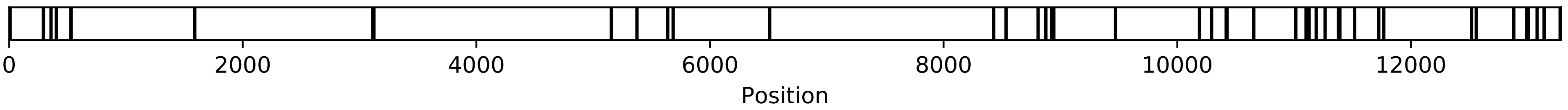


CR2 vs MM1
Number of differences: 245

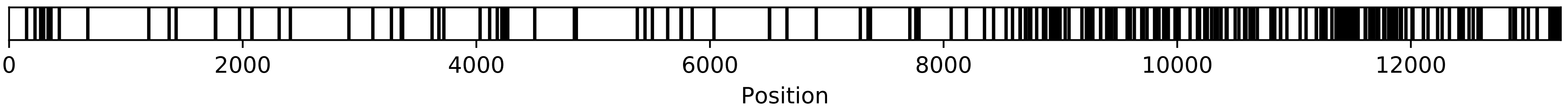


CR2 vs MM2
Number of differences: 236

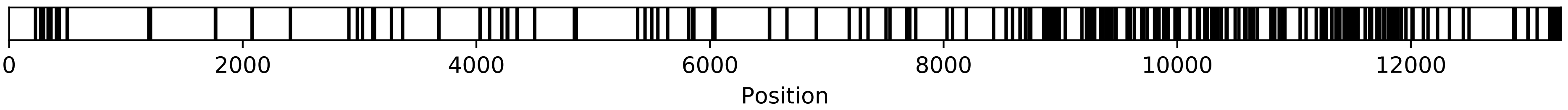


MA1 vs MA2
Number of differences: 230

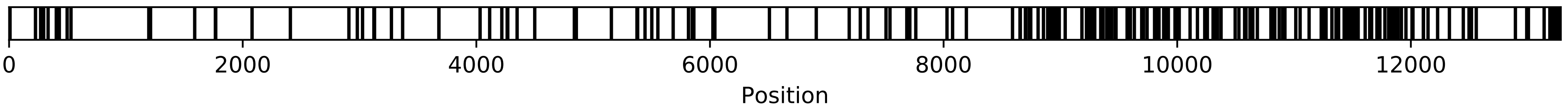


MA1 vs MM1
Number of differences: 97

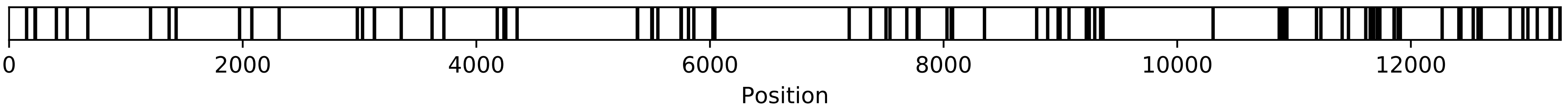


MA1 vs MM2
Number of differences: 0

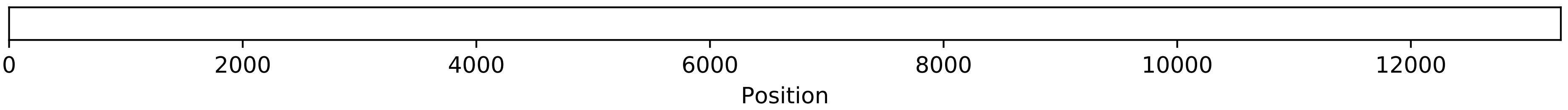


MA2 vs MM1
Number of differences: 239

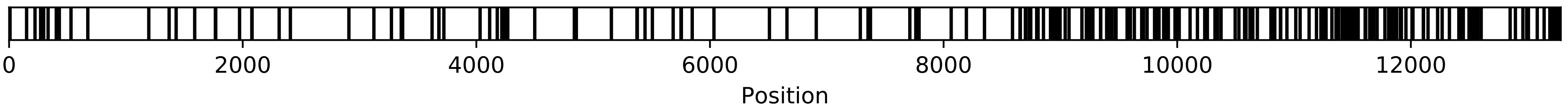


MA2 vs MM2
Number of differences: 230

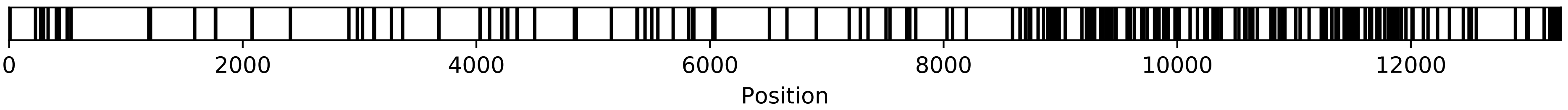


MM1 vs MM2
Number of differences: 97

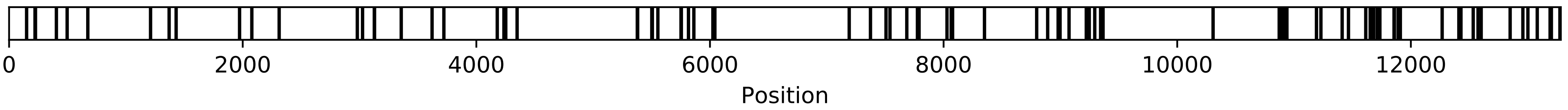

### Region: 235

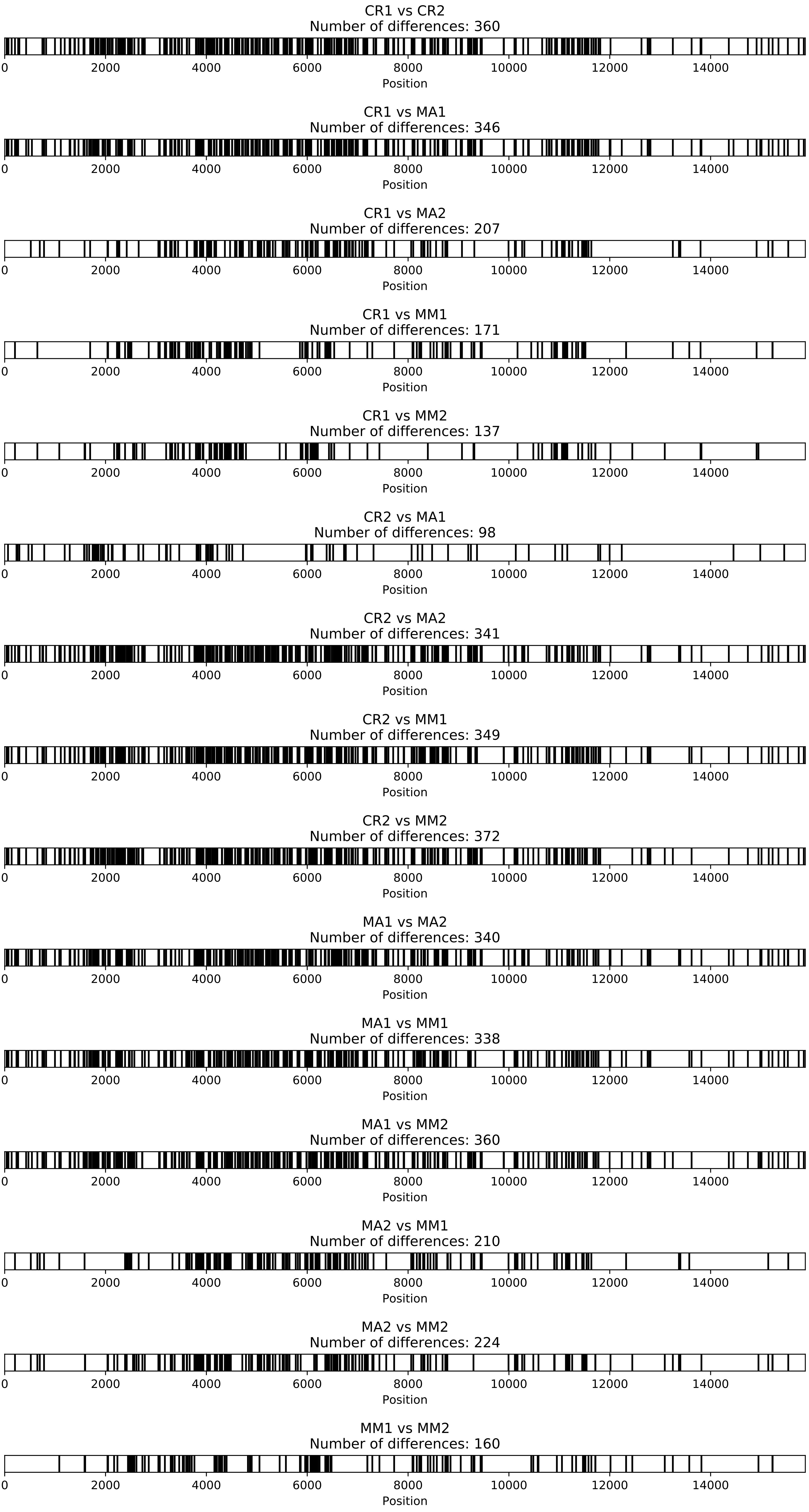
CR1 vs CR2
Number of differences: 360
CR1 vs MA1
Number of differences: 346
CR1 vs MA2
Number of differences: 207
CR1 vs MM1
Number of differences: 171
CR1 vs MM2
Number of differences: 137
CR2 vs MA1
Number of differences: 98
CR2 vs MA2
Number of differences: 341
CR2 vs MM1
Number of differences: 349
CR2 vs MM2
Number of differences: 372
MA1 vs MA2
Number of differences: 340
MA1 vs MM1
Number of differences: 338
MA1 vs MM2
Number of differences: 360
MA2 vs MM1
Number of differences: 210
MA2 vs MM2
Number of differences: 224
MM1 vs MM2
Number of differences: 160
0
2000
4000
6000
8000
10000
12000
14000
Position

### Region: 237



CR1 vs CR2
Number of differences: 234

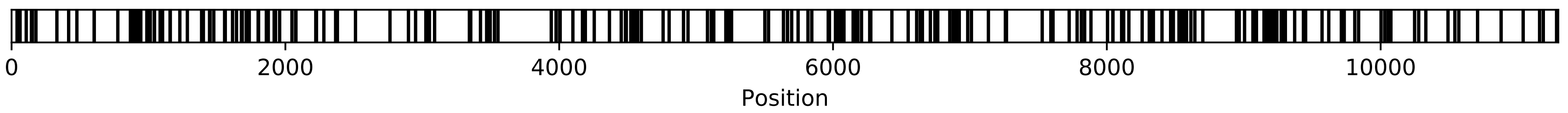


CR1 vs MA1
Number of differences: 77

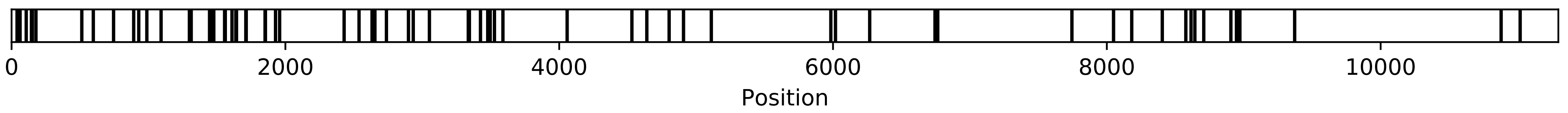


CR1 vs MA2
Number of differences: 238

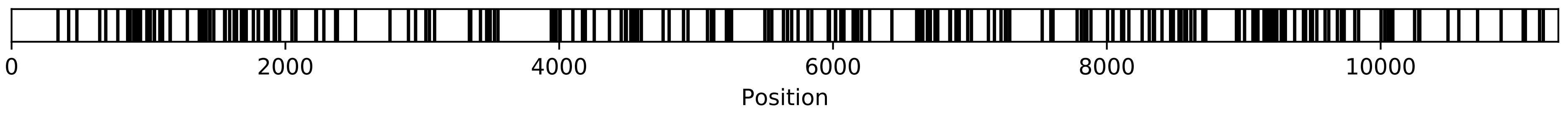


CR1 vs MM1
Number of differences: 233

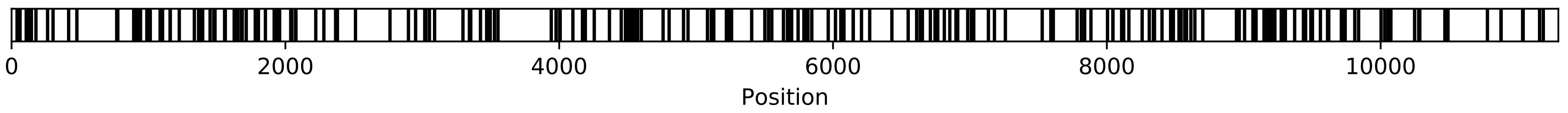


CR1 vs MM2
Number of differences: 230

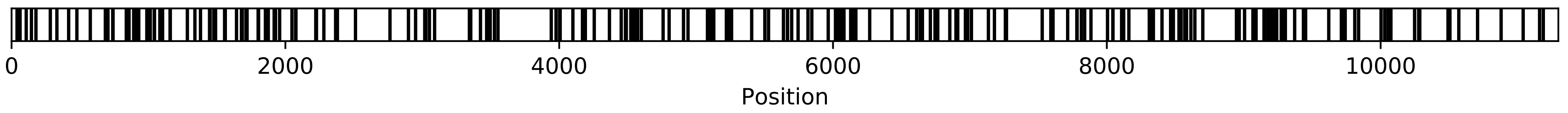


CR2 vs MA1
Number of differences: 228

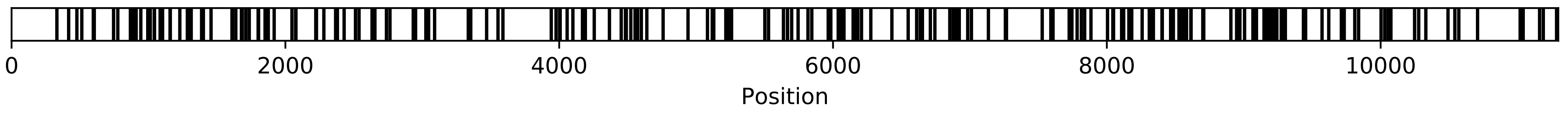


CR2 vs MA2
Number of differences: 74

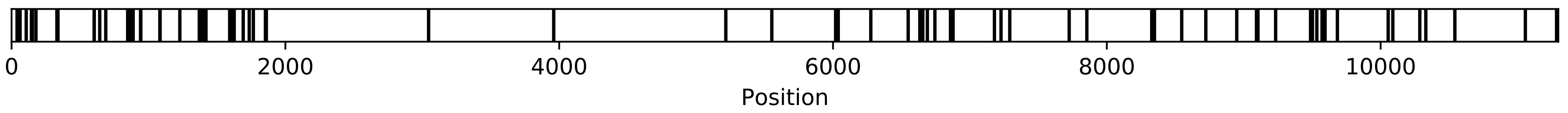


CR2 vs MM1
Number of differences: 91

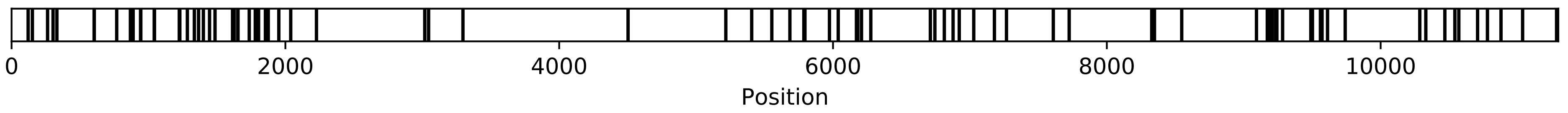


CR2 vs MM2
Number of differences: 74

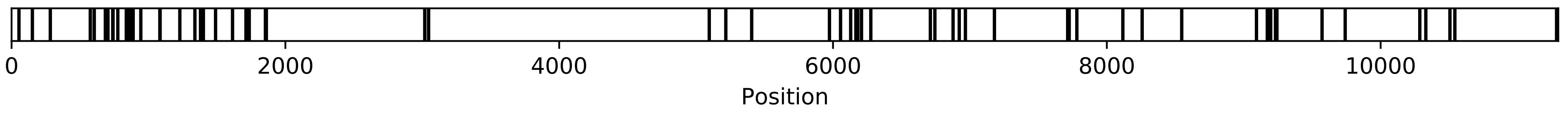


MA1 vs MA2
Number of differences: 244

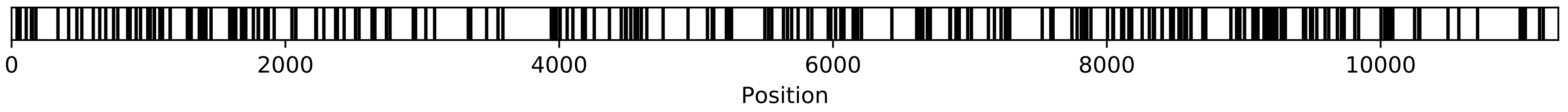


MA1 vs MM1
Number of differences: 229

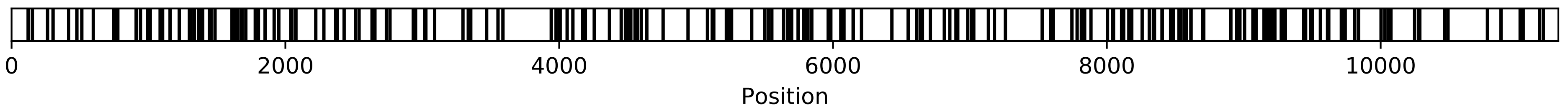


MA1 vs MM2
Number of differences: 225

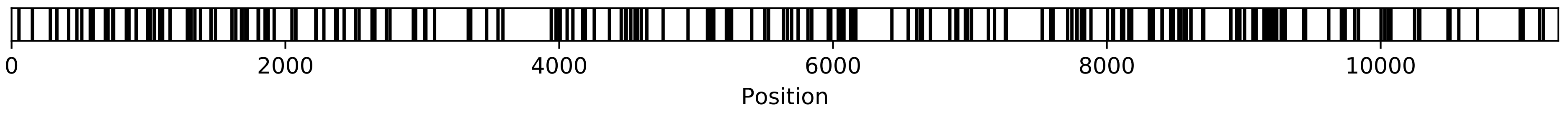


MA2 vs MM1
Number of differences: 95

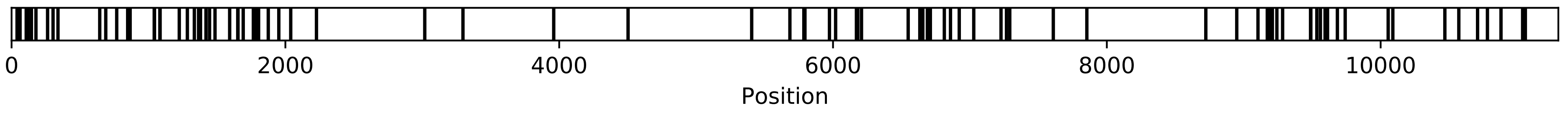


MA2 vs MM2
Number of differences: 87

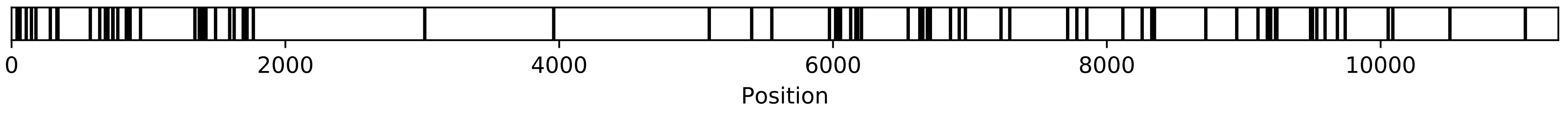


MM1 vs MM2
Number of differences: 87

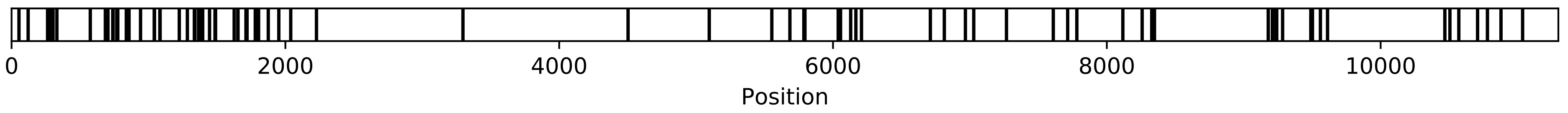

### Region: 239

CR1 vs CR2
Number of differences: 365

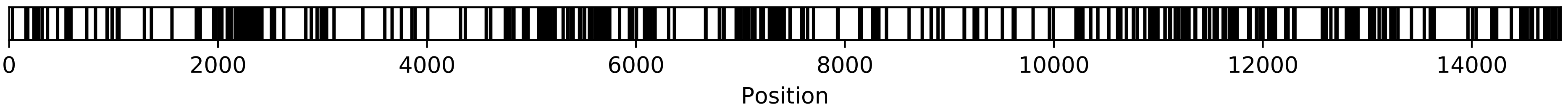


CR1 vs MA1
Number of differences: 180

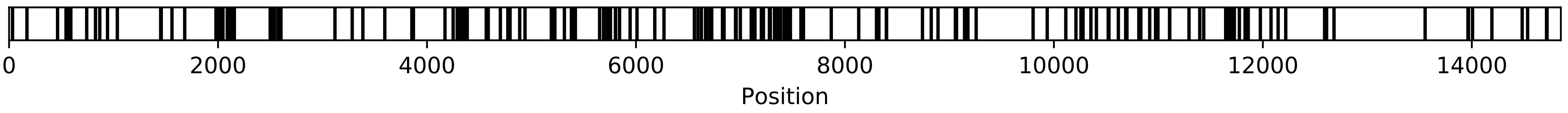


CR1 vs MA2
Number of differences: 352

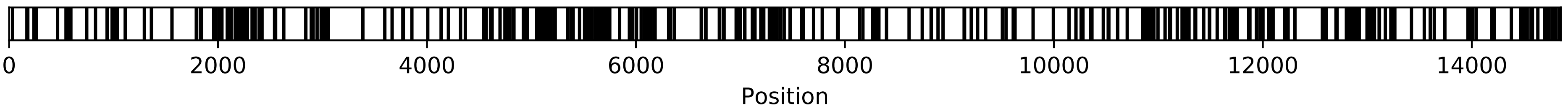


CR1 vs MM1
Number of differences: 190

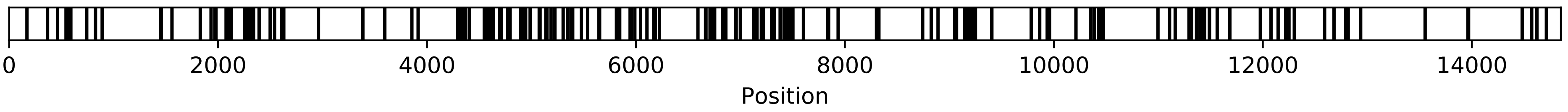


CR1 vs MM2
Number of differences: 155

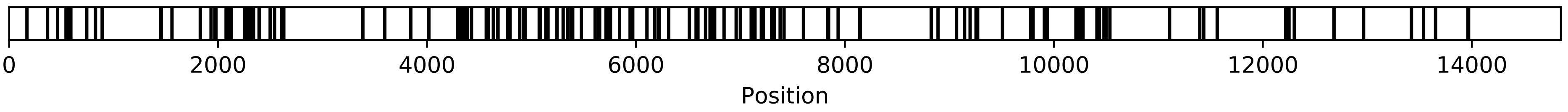


CR2 vs MA1
Number of differences: 379

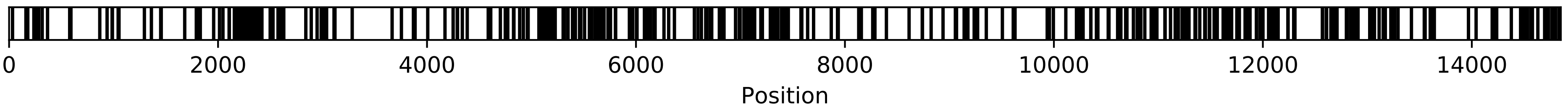


CR2 vs MA2
Number of differences: 133

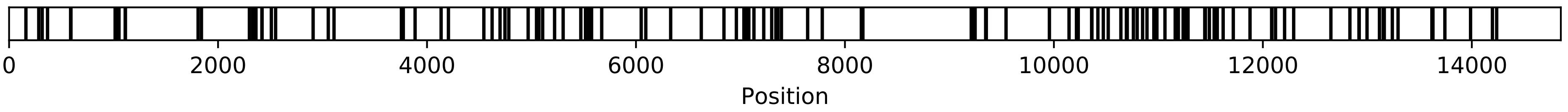


CR2 vs MM1
Number of differences: 387

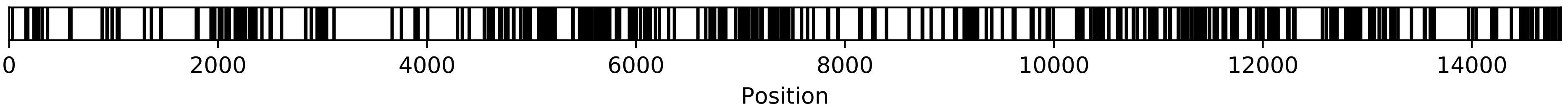


CR2 vs MM2
Number of differences: 366

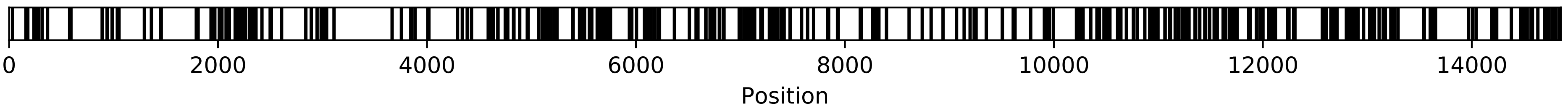


MA1 vs MA2
Number of differences: 365

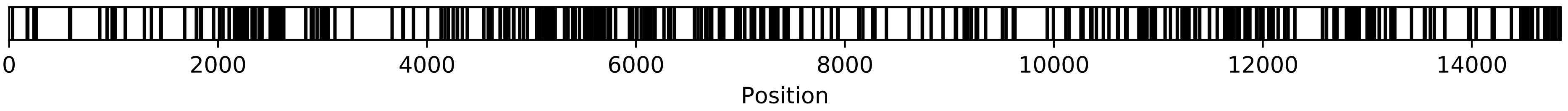


MA1 vs MM1
Number of differences: 191

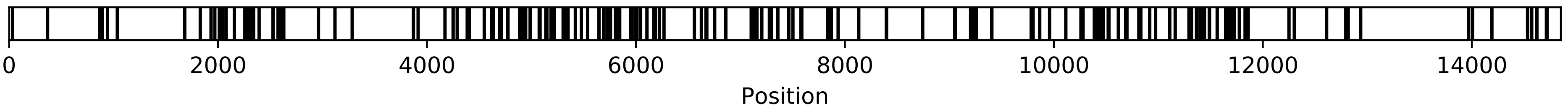


MA1 vs MM2
Number of differences: 169

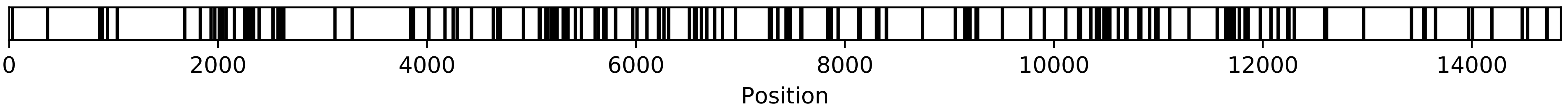


MA2 vs MM1
Number of differences: 374

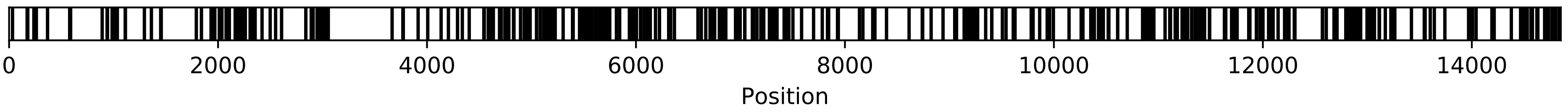


MA2 vs MM2
Number of differences: 353

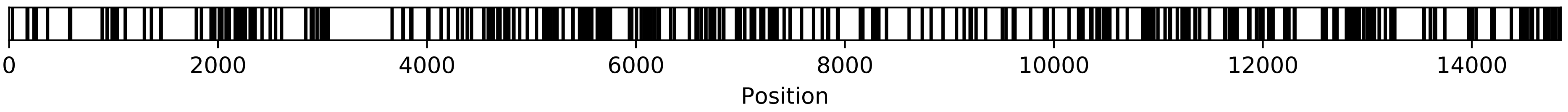


MM1 vs MM2
Number of differences: 147

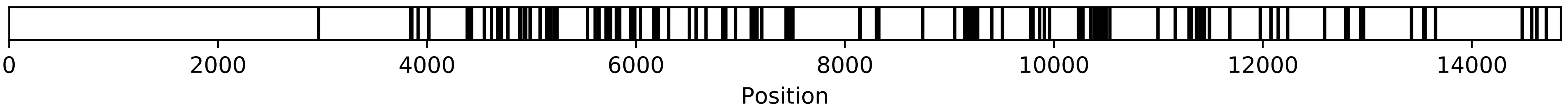

### Region: 240

CR1 vs CR2
Number of differences: 248

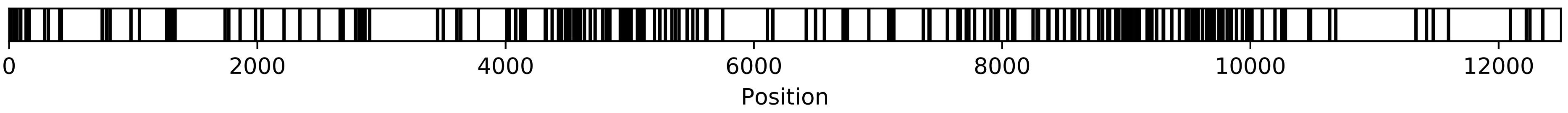

CR1 vs MA1
Number of differences: 124

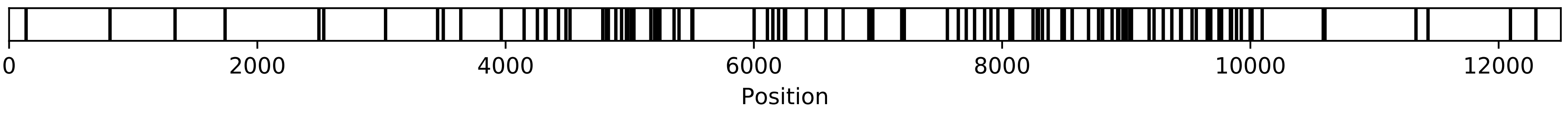

CR1 vs MA2
Number of differences: 241

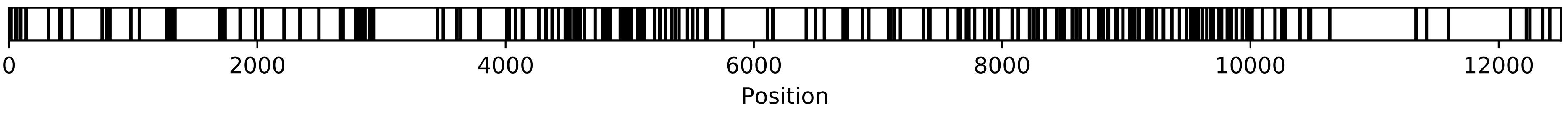

CR1 vs MM1
Number of differences: 148

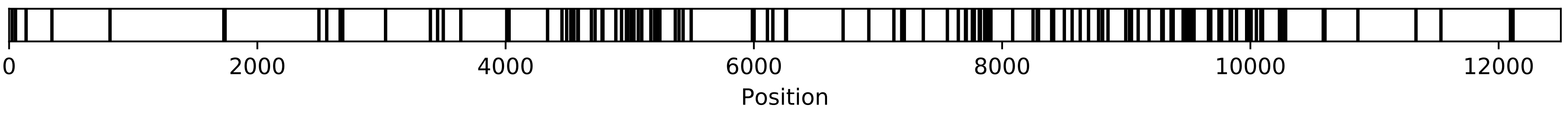

CR1 vs MM2
Number of differences: 151

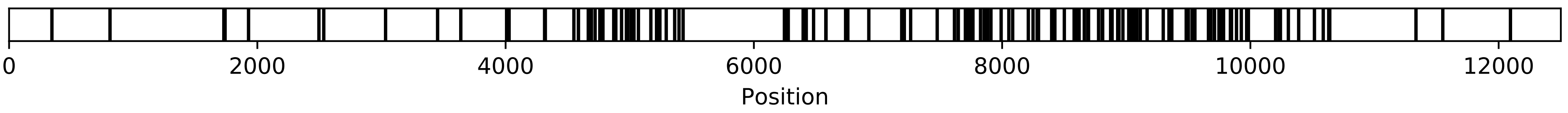

CR2 vs MA1
Number of differences: 227

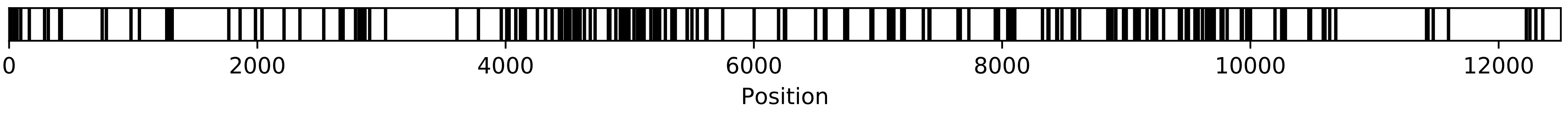

CR2 vs MA2
Number of differences: 68

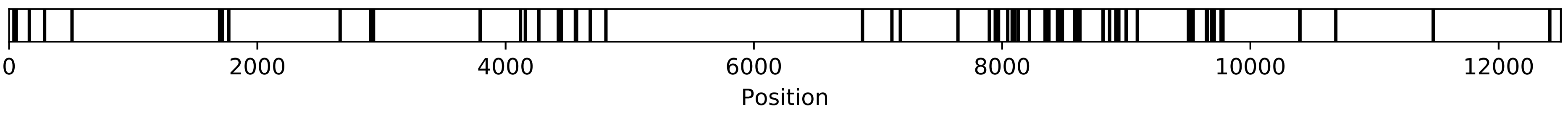

CR2 vs MM1
Number of differences: 240

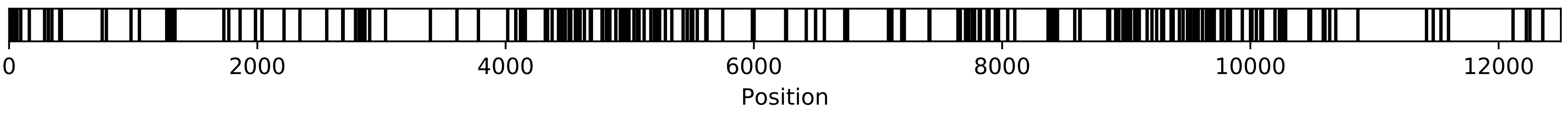

CR2 vs MM2
Number of differences: 262

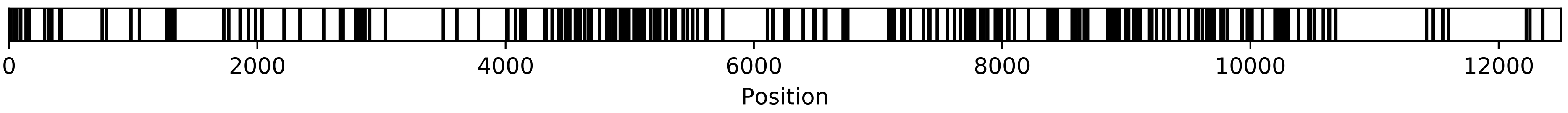

MA1 vs MA2
Number of differences: 222

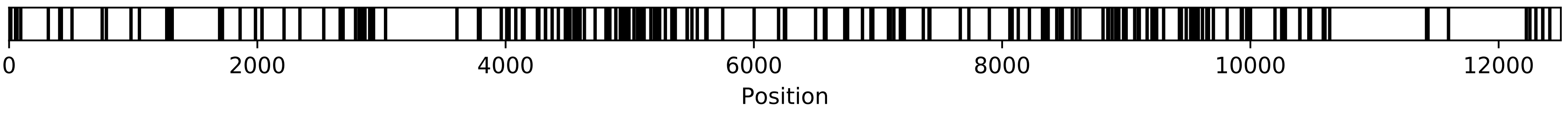

MA1 vs MM1
Number of differences: 130

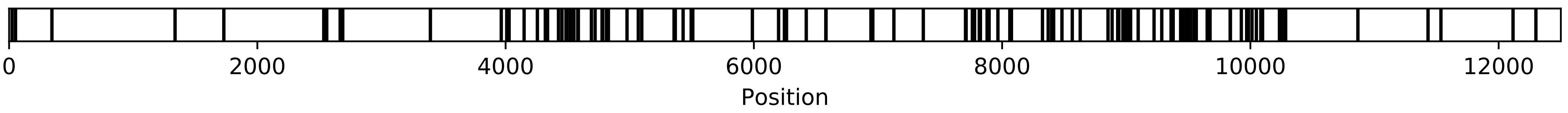

MA1 vs MM2
Number of differences: 139

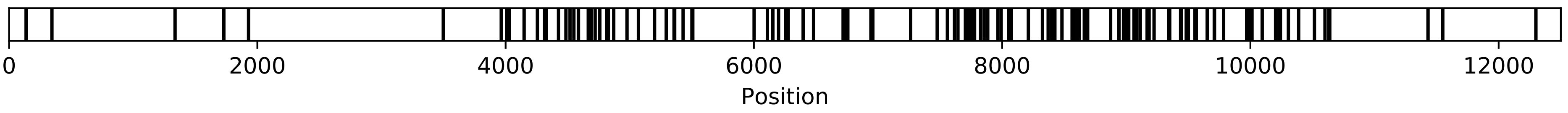

MA2 vs MM1
Number of differences: 234

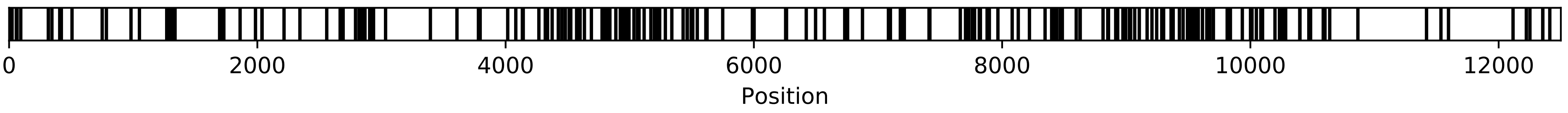

MA2 vs MM2
Number of differences: 259

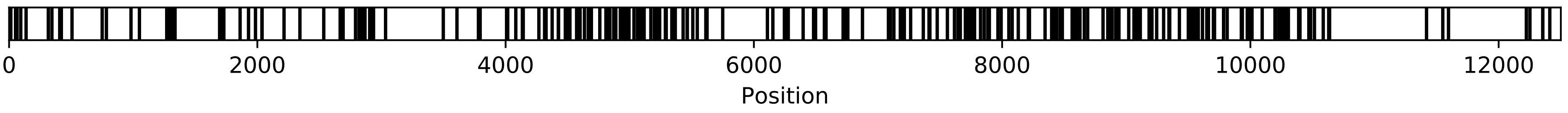

MM1 vs MM2
Number of differences: 148

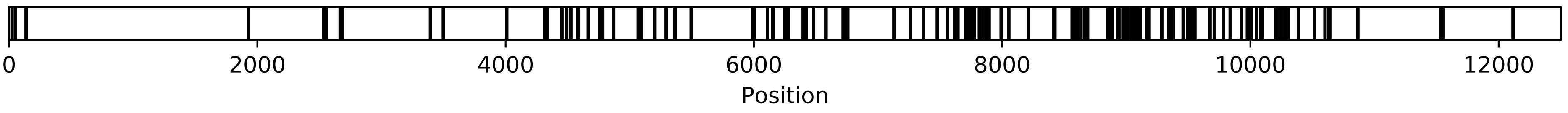

### Region: 242

CR1 vs CR2
Number of differences: 612

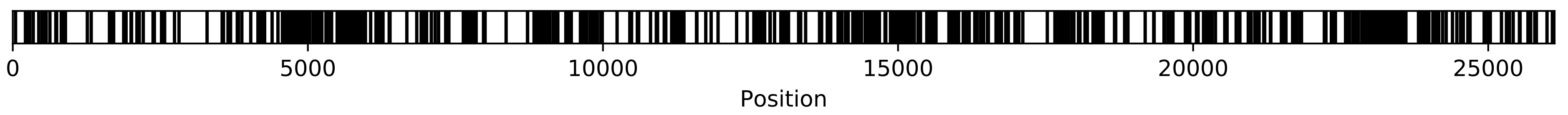


CR1 vs MA1
Number of differences: 602

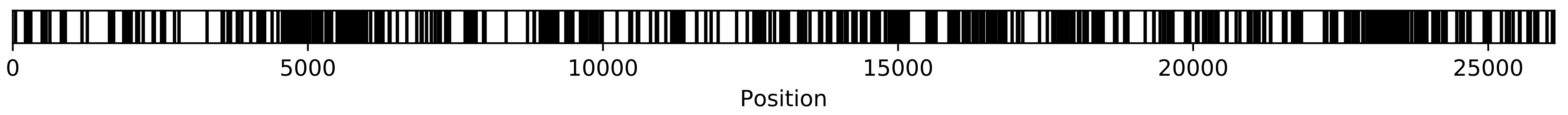


CR1 vs MA2
Number of differences: 397

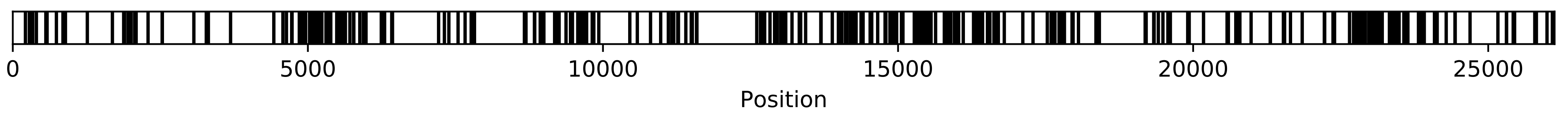


CR1 vs MM1
Number of differences: 346

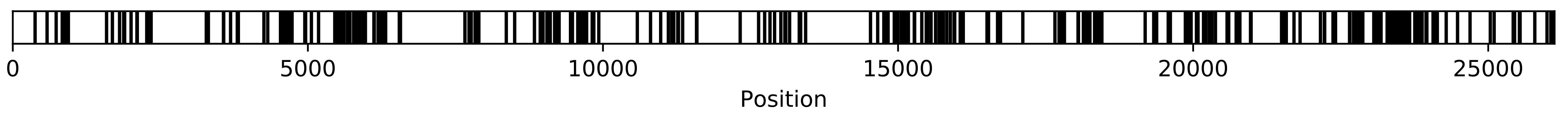


CR1 vs MM2
Number of differences: 379

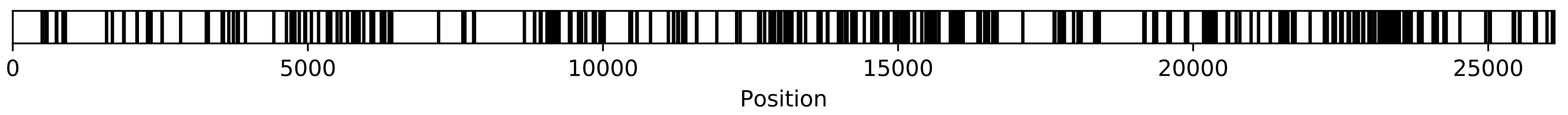


CR2 vs MA1
Number of differences: 201

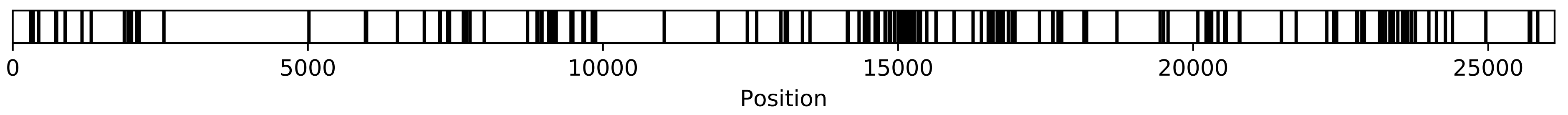


CR2 vs MA2
Number of differences: 607

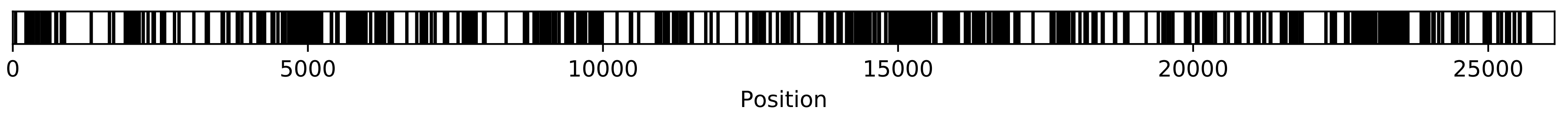


CR2 vs MM1
Number of differences: 593

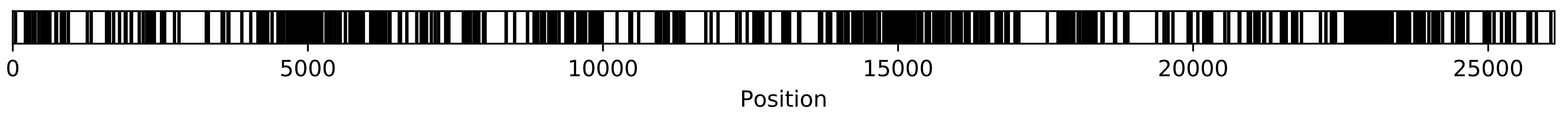


CR2 vs MM2
Number of differences: 608

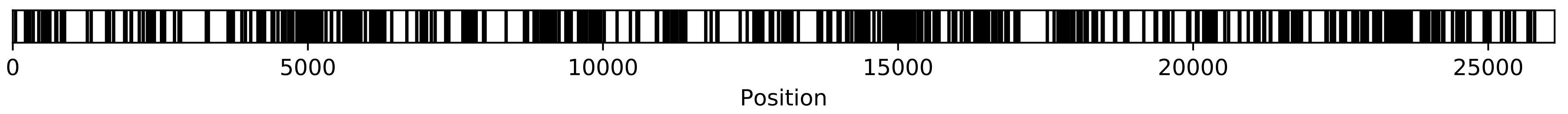


MA1 vs MA2
Number of differences: 617

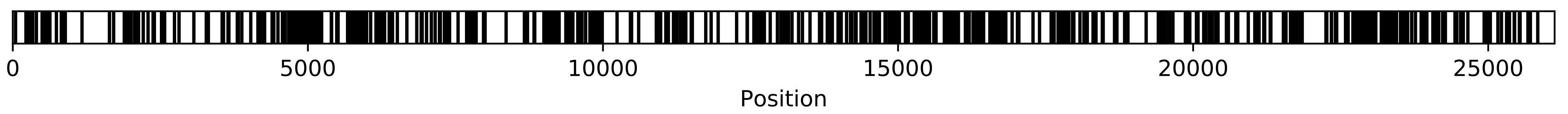


MA1 vs MM1
Number of differences: 600

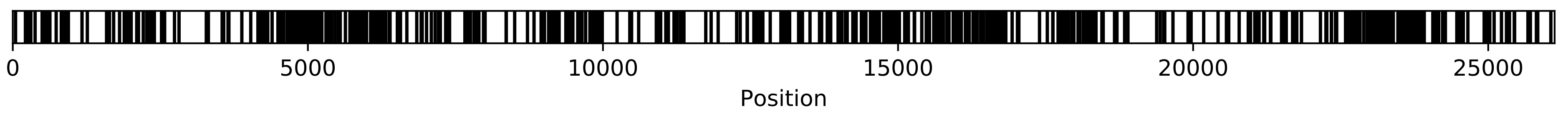


MA1 vs MM2
Number of differences: 613

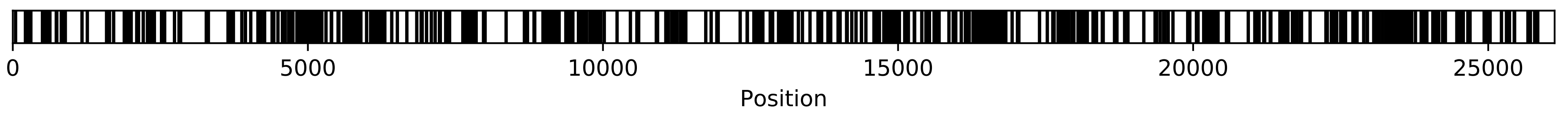


MA2 vs MM1
Number of differences: 418

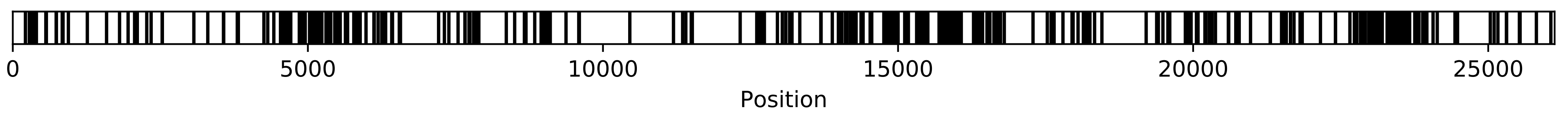


MA2 vs MM2
Number of differences: 409

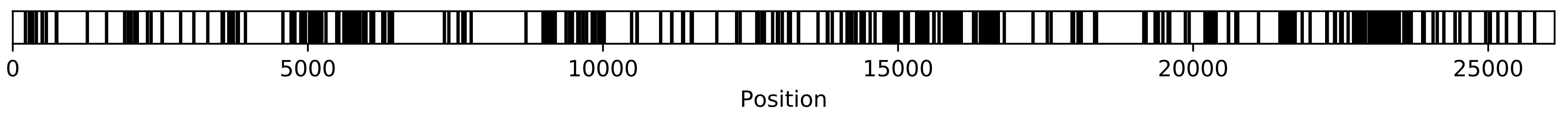


MM1 vs MM2
Number of differences: 327

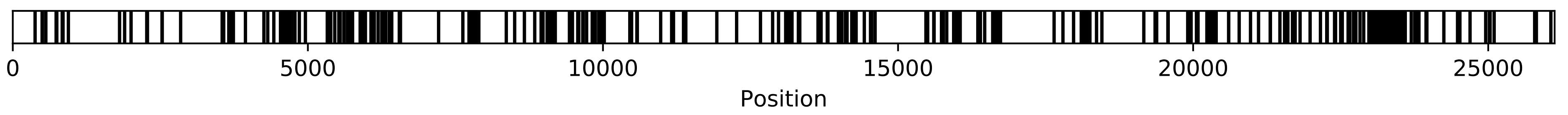

### Region: 243

CR1 vs CR2
Number of differences: 291

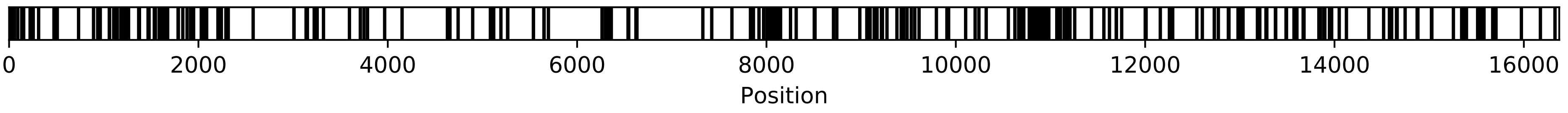


CR1 vs MA1
Number of differences: 172

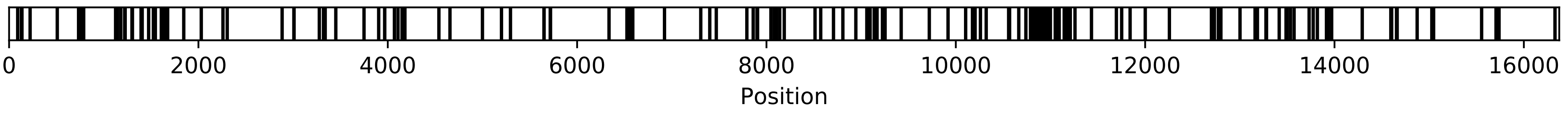


CR1 vs MA2
Number of differences: 288

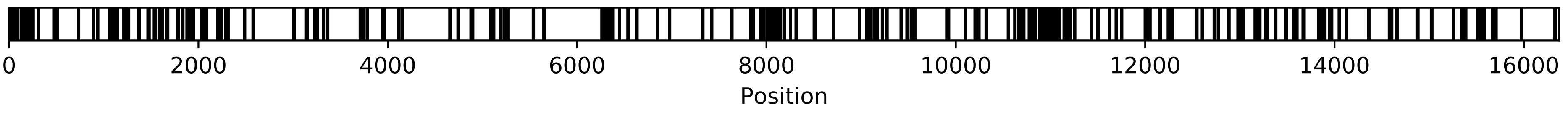


CR1 vs MM1
Number of differences: 172

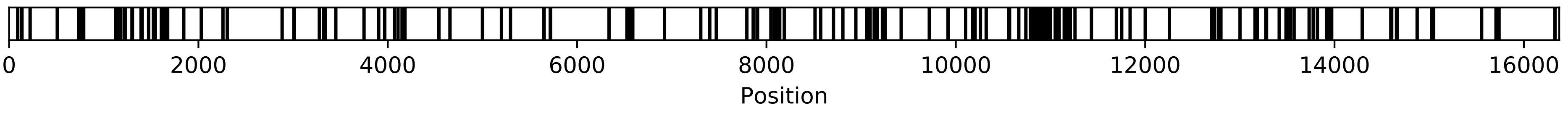


CR1 vs MM2
Number of differences: 175

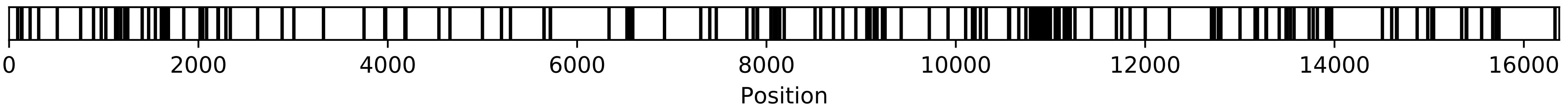


CR2 vs MA1
Number of differences: 286

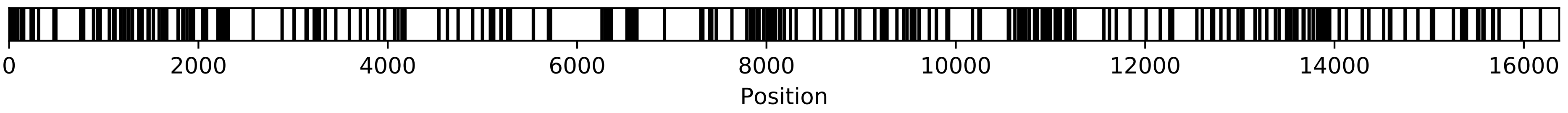


CR2 vs MA2
Number of differences: 86

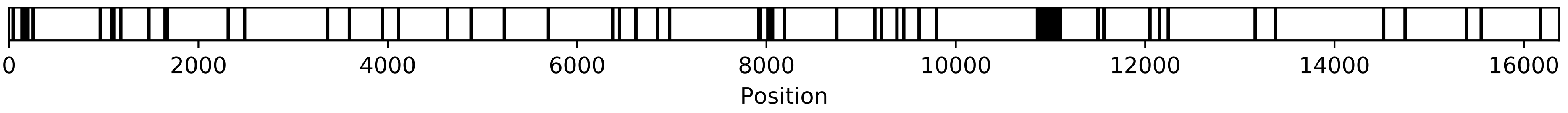


CR2 vs MM1
Number of differences: 286

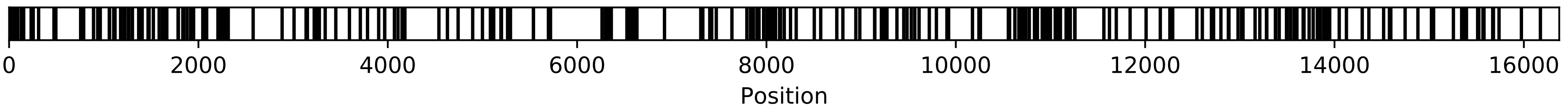


CR2 vs MM2
Number of differences: 278

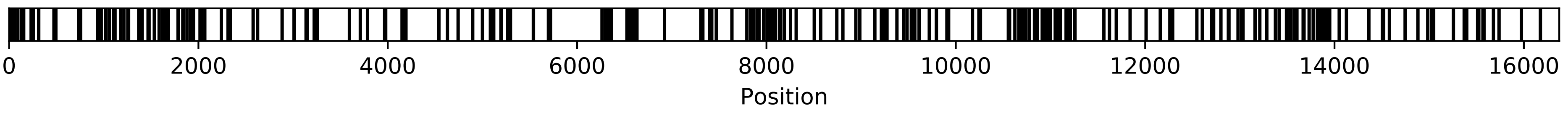


MA1 vs MA2
Number of differences: 292

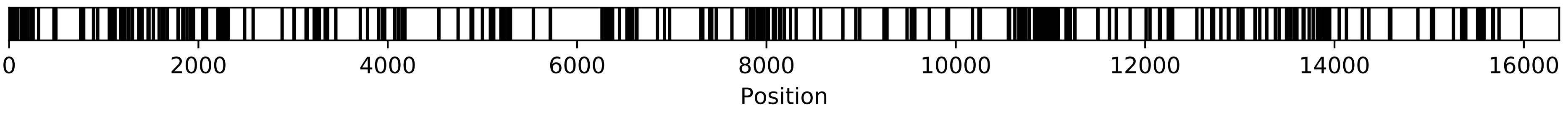


MA1 vs MM1
Number of differences: 0

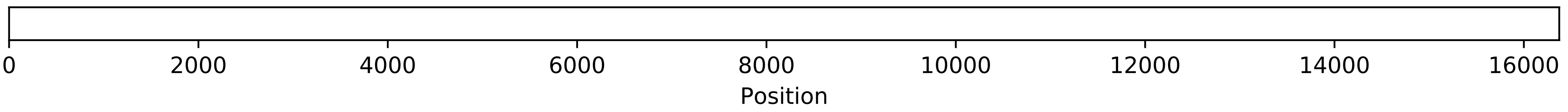


MA1 vs MM2
Number of differences: 45

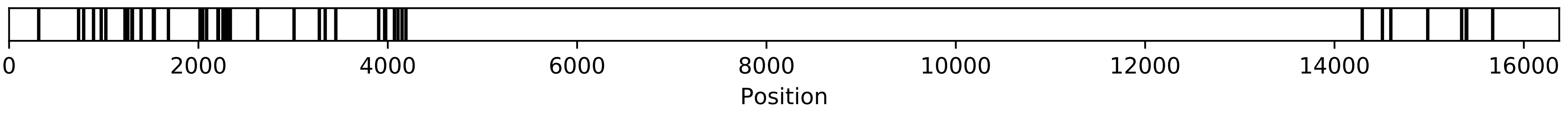


MA2 vs MM1
Number of differences: 292

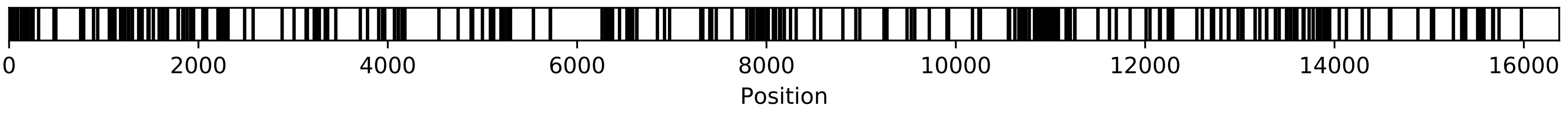


MA2 vs MM2
Number of differences: 284

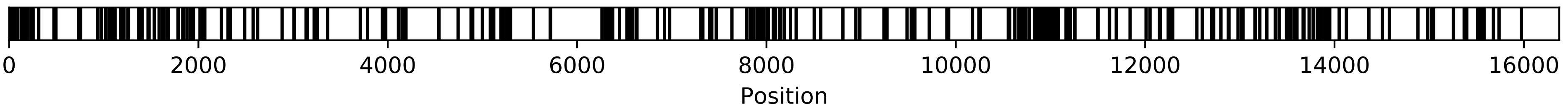


MM1 vs MM2
Number of differences: 45

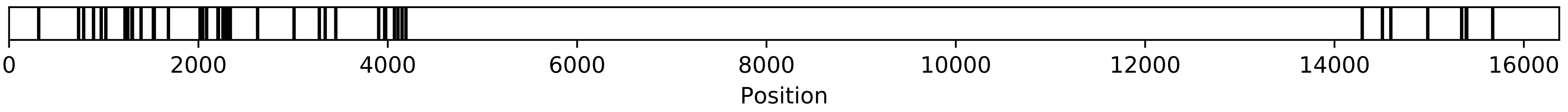

### Region: 245



CR1 vs CR2
Number of differences: 235

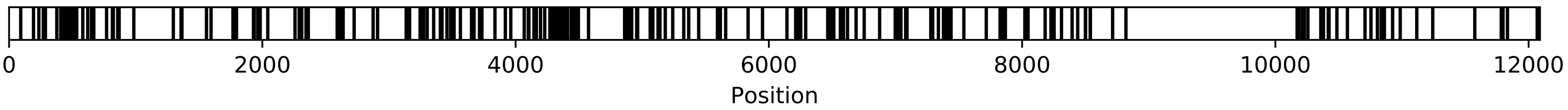


CR1 vs MA1
Number of differences: 254

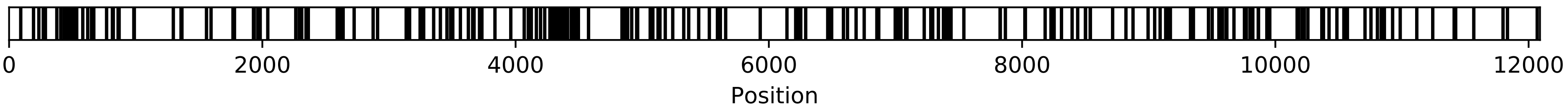


CR1 vs MA2
Number of differences: 149

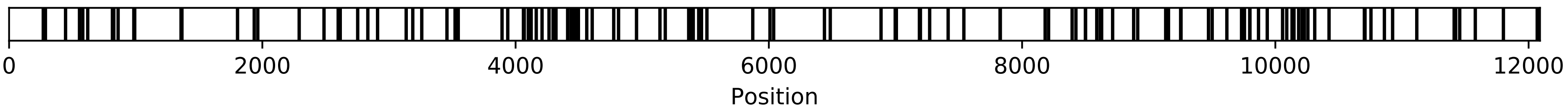


CR1 vs MM1
Number of differences: 142

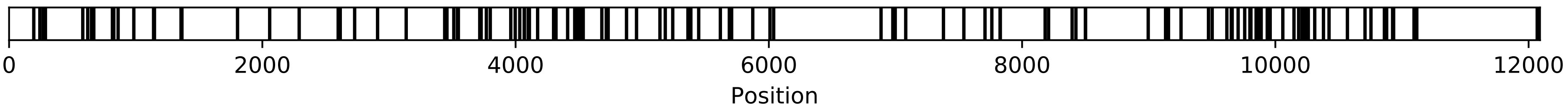


CR1 vs MM2
Number of differences: 149

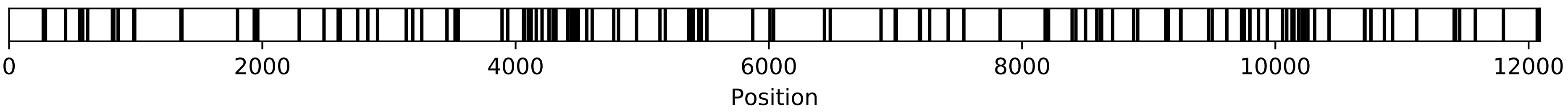


CR2 vs MA1
Number of differences: 87

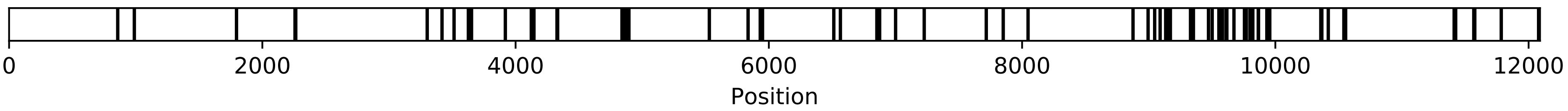


CR2 vs MA2
Number of differences: 252

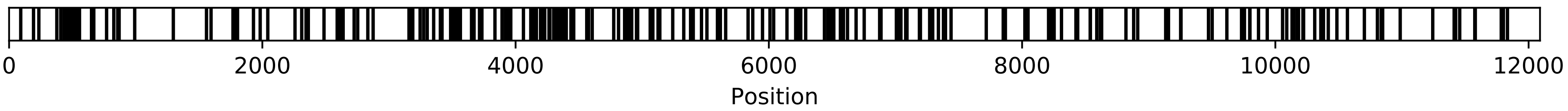


CR2 vs MM1
Number of differences: 248

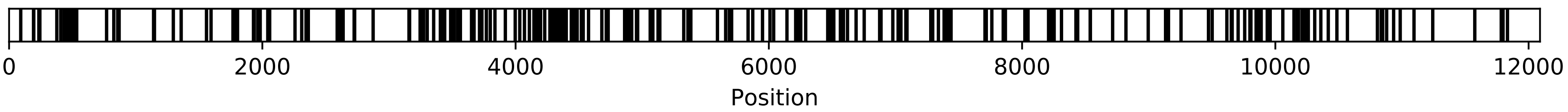


CR2 vs MM2
Number of differences: 252

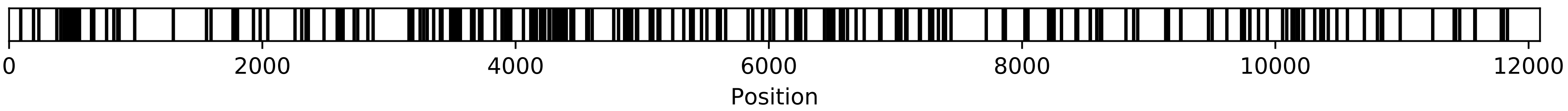


MA1 vs MA2
Number of differences: 257

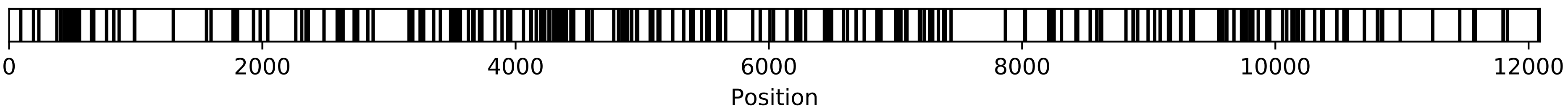


MA1 vs MM1
Number of differences: 263

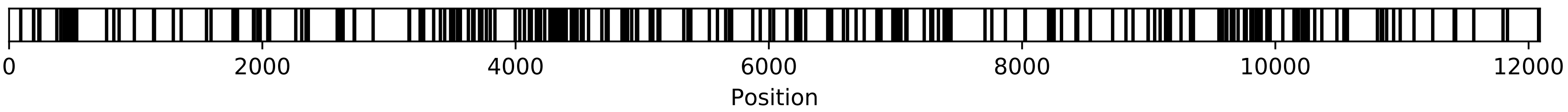


MA1 vs MM2
Number of differences: 257

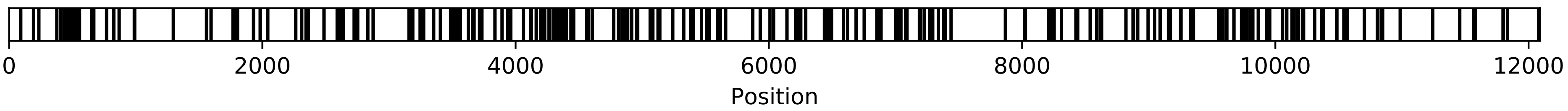


MA2 vs MM1
Number of differences: 133

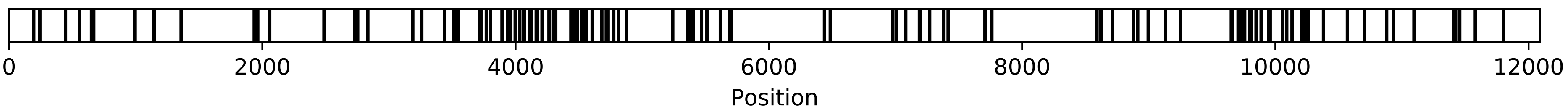


MA2 vs MM2
Number of differences: 0

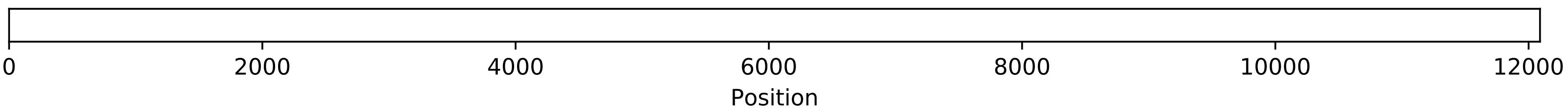


MM1 vs MM2
Number of differences: 133

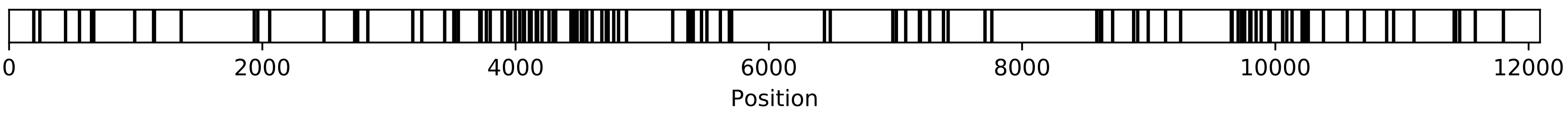

### Region: 246



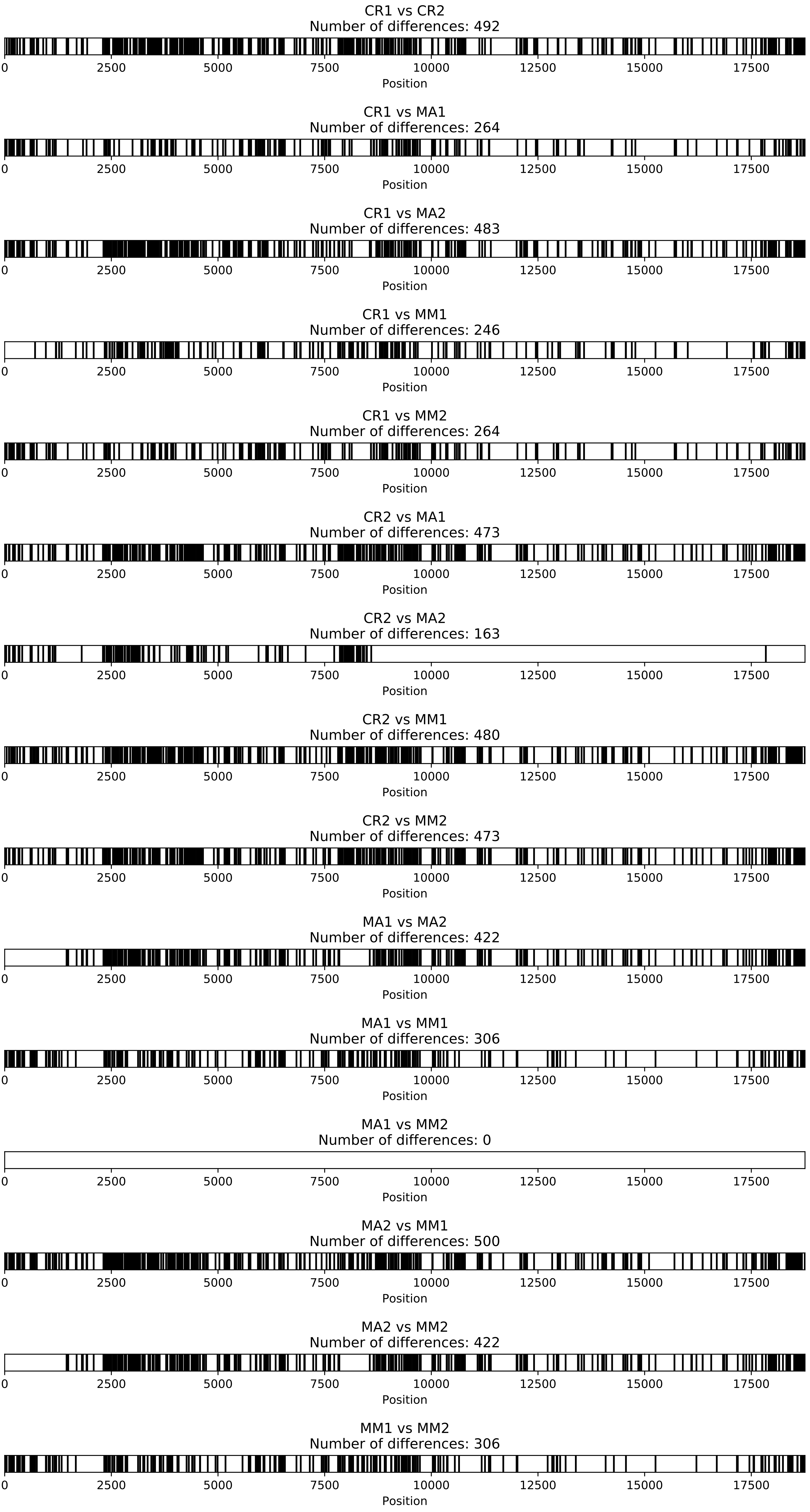
CR1 vs CR2
Number of differences: 492
CR1 vs MA1
Number of differences: 264
CR1 vs MA2
Number of differences: 483
CR1 vs MM1
Number of differences: 246
CR1 vs MM2
Number of differences: 264
CR2 vs MA1
Number of differences: 473
CR2 vs MA2
Number of differences: 163
CR2 vs MM1
Number of differences: 480
CR2 vs MM2
Number of differences: 473
MA1 vs MA2
Number of differences: 422
MA1 vs MM1
Number of differences: 306
MA1 vs MM2
Number of differences: 0
MA2 vs MM1
Number of differences: 500
MA2 vs MM2
Number of differences: 422
MM1 vs MM2
Number of differences: 306
0
2500
5000
7500
10000
12500
15000
17500
Position

### Region: 247

CR1 vs CR2
Number of differences: 267

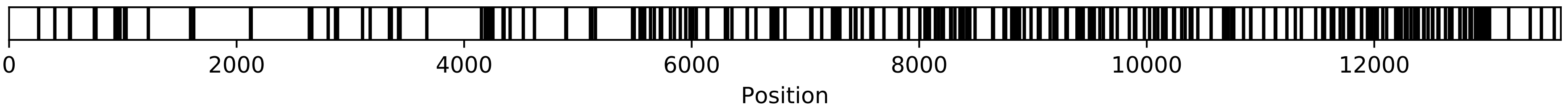


CR1 vs MA1
Number of differences: 245

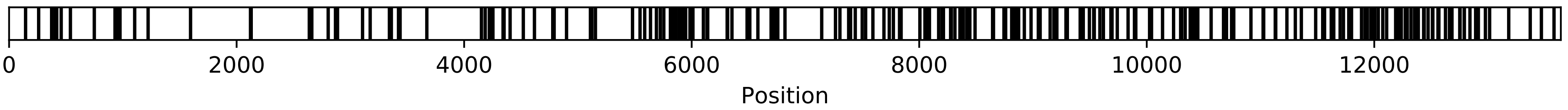


CR1 vs MA2
Number of differences: 47

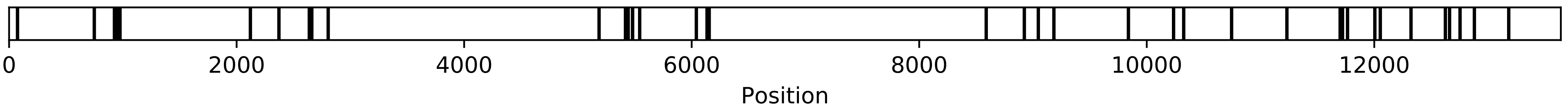


CR1 vs MM1
Number of differences: 274

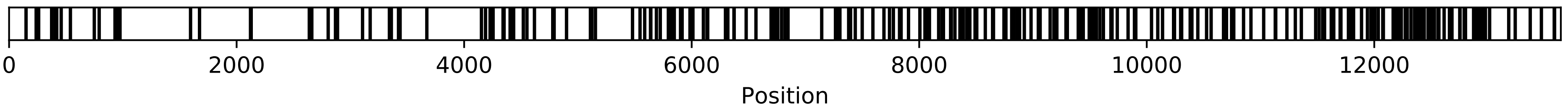


CR1 vs MM2
Number of differences: 255

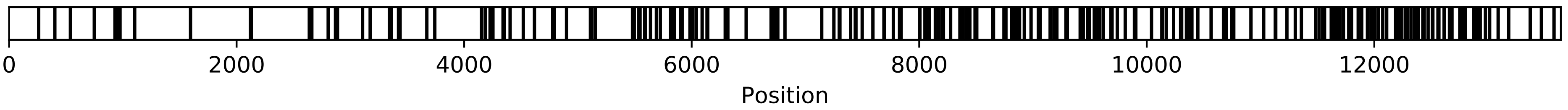


CR2 vs MA1
Number of differences: 95

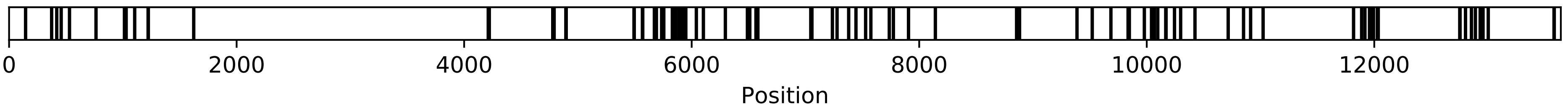


CR2 vs MA2
Number of differences: 258

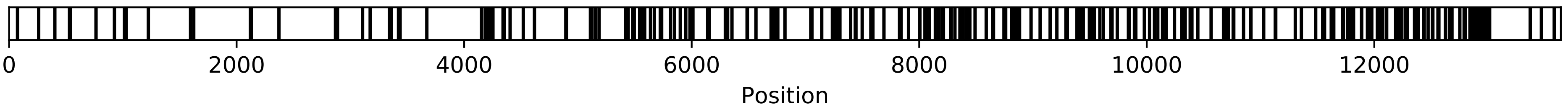


CR2 vs MM1
Number of differences: 122

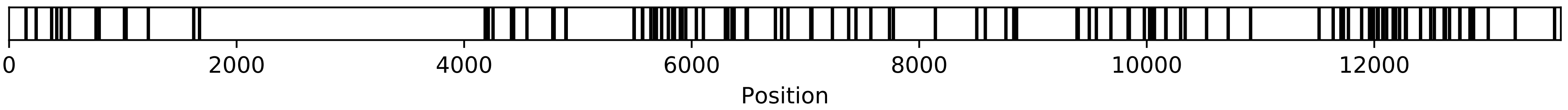


CR2 vs MM2
Number of differences: 107

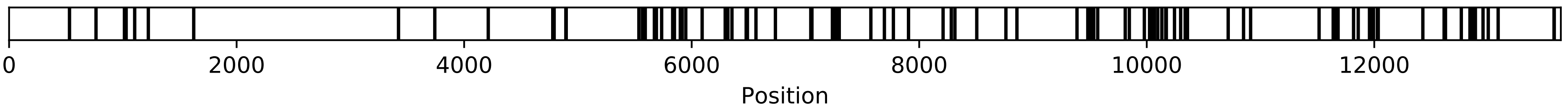


MA1 vs MA2
Number of differences: 239

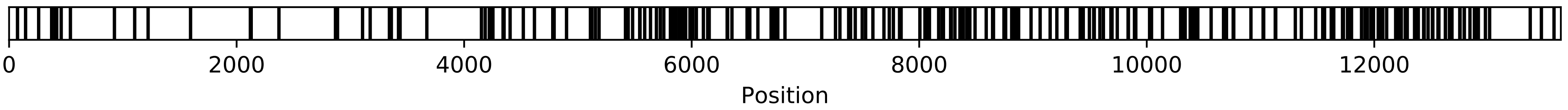


MA1 vs MM1
Number of differences: 95

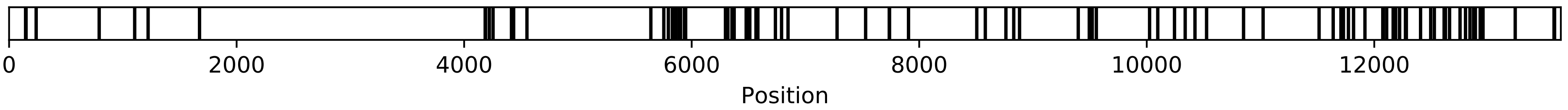


MA1 vs MM2
Number of differences: 87

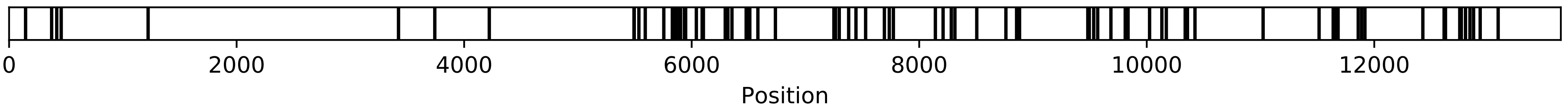


MA2 vs MM1
Number of differences: 266

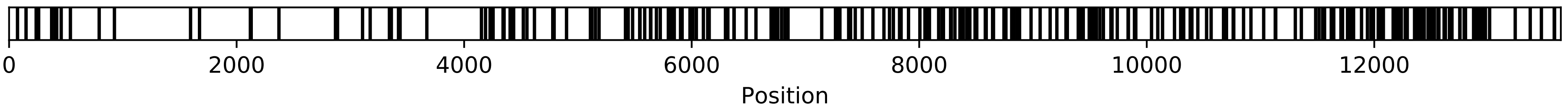


MA2 vs MM2
Number of differences: 246

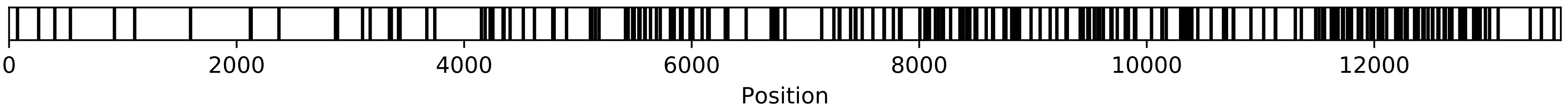


MM1 vs MM2
Number of differences: 100

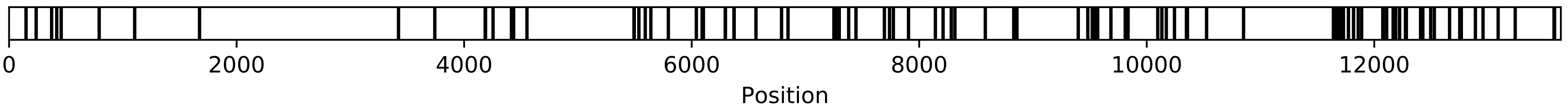

### Region: 248



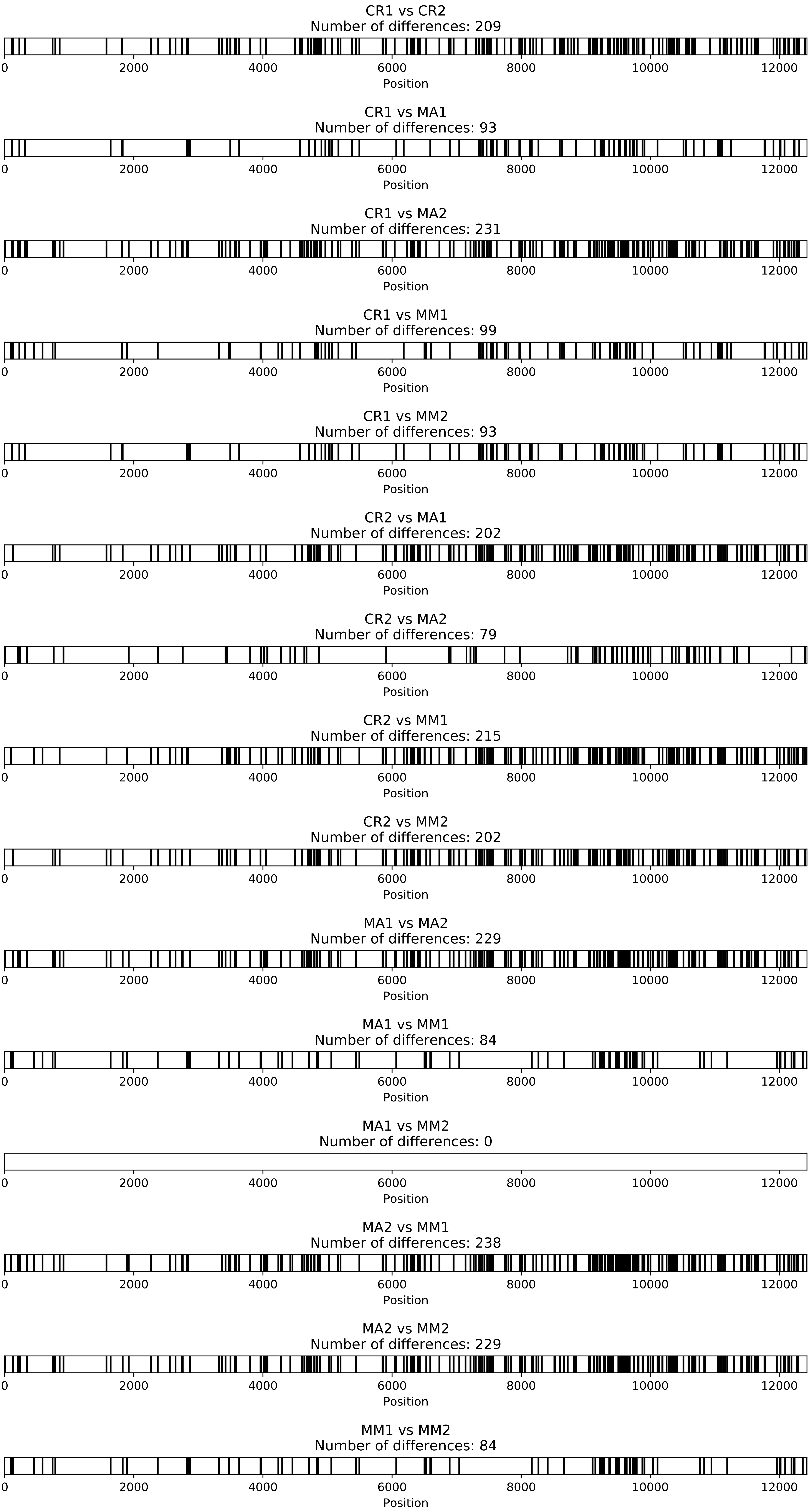
CR1 vs CR2
Number of differences: 209
CR1 vs MA1
Number of differences: 93
CR1 vs MA2
Number of differences: 231
CR1 vs MM1
Number of differences: 99
CR1 vs MM2
Number of differences: 93
CR2 vs MA1
Number of differences: 202
CR2 vs MA2
Number of differences: 79
CR2 vs MM1
Number of differences: 215
CR2 vs MM2
Number of differences: 202
MA1 vs MA2
Number of differences: 229
MA1 vs MM1
Number of differences: 84
MA1 vs MM2
Number of differences: 0
MA2 vs MM1
Number of differences: 238
MA2 vs MM2
Number of differences: 229
MM1 vs MM2
Number of differences: 84
0
2000
4000
6000
8000
10000
12000
Position

### Region: 249

CR1 vs CR2
Number of differences: 156

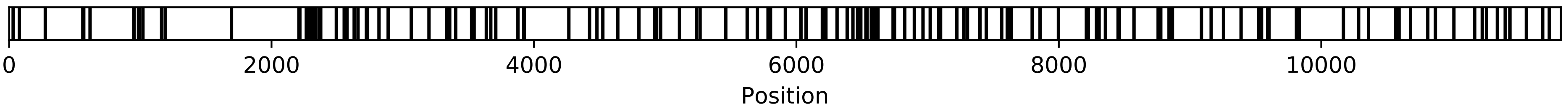


CR1 vs MA1
Number of differences: 166

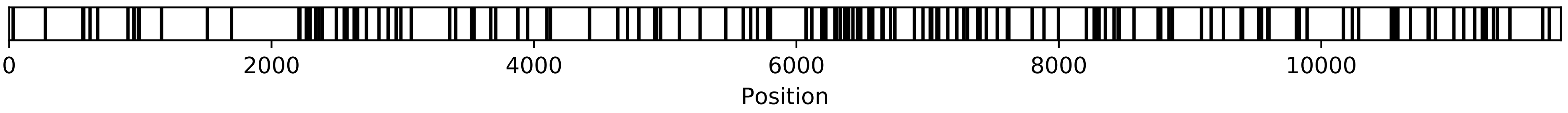


CR1 vs MA2
Number of differences: 65

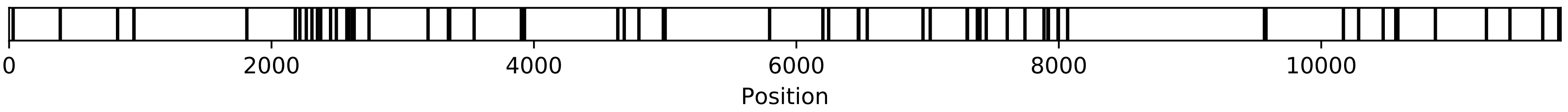


CR1 vs MM1
Number of differences: 166

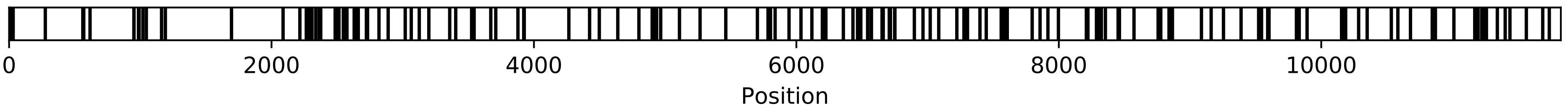


CR1 vs MM2
Number of differences: 166

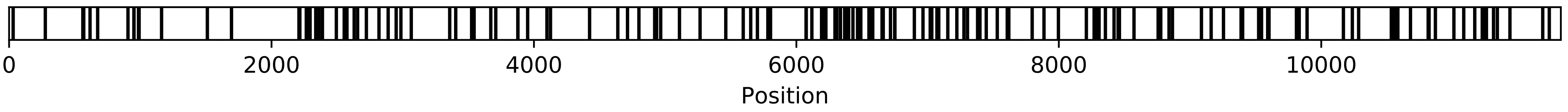


CR2 vs MA1
Number of differences: 102

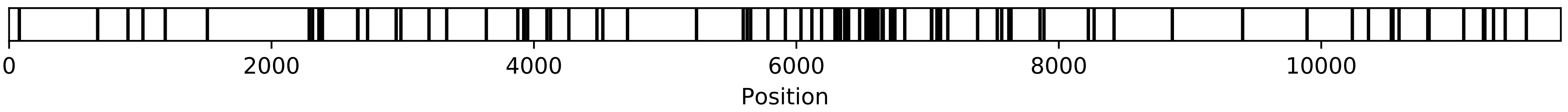


CR2 vs MA2
Number of differences: 161

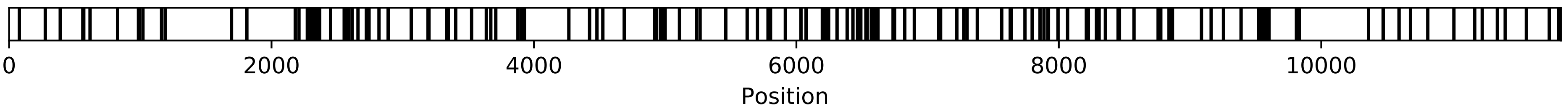


CR2 vs MM1
Number of differences: 88

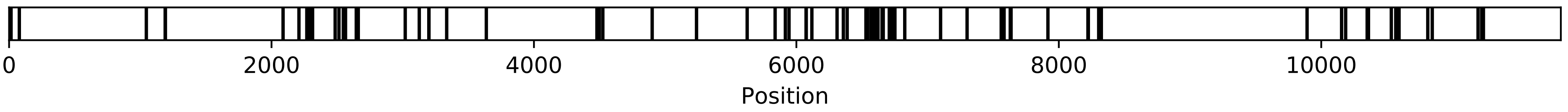


CR2 vs MM2
Number of differences: 102

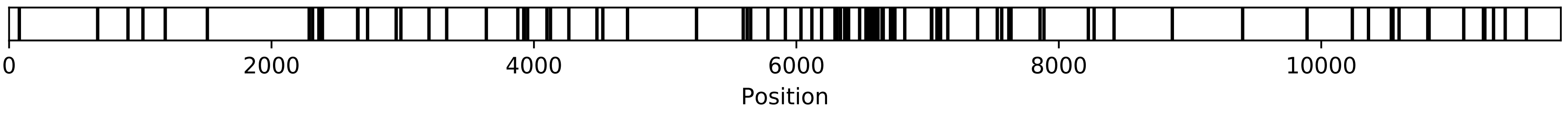


MA1 vs MA2
Number of differences: 171

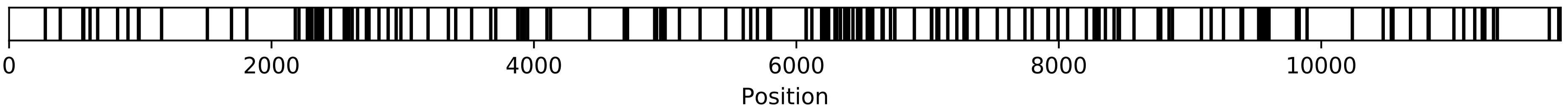


MA1 vs MM1
Number of differences: 102

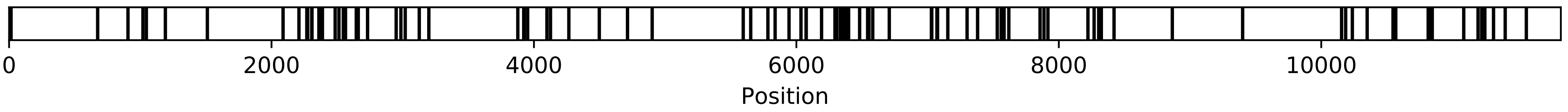


MA1 vs MM2
Number of differences: 0

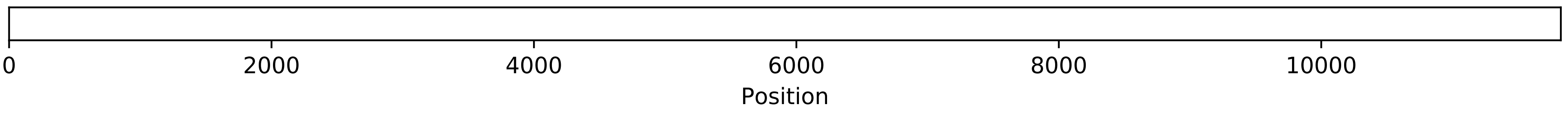


MA2 vs MM1
Number of differences: 168

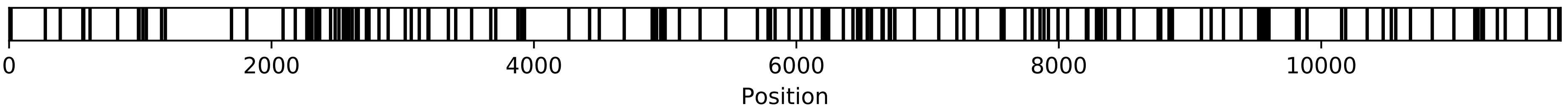


MA2 vs MM2
Number of differences: 171

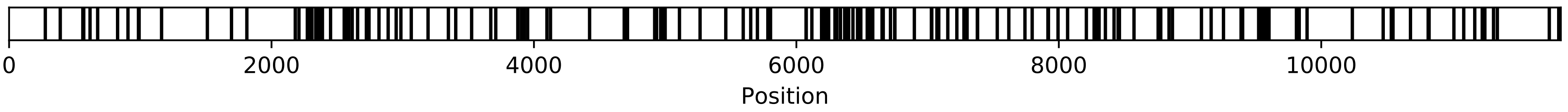


MM1 vs MM2
Number of differences: 102

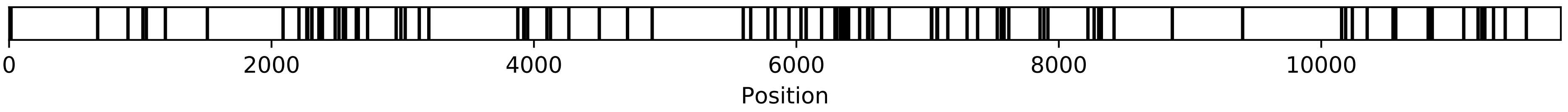

### Region: 250



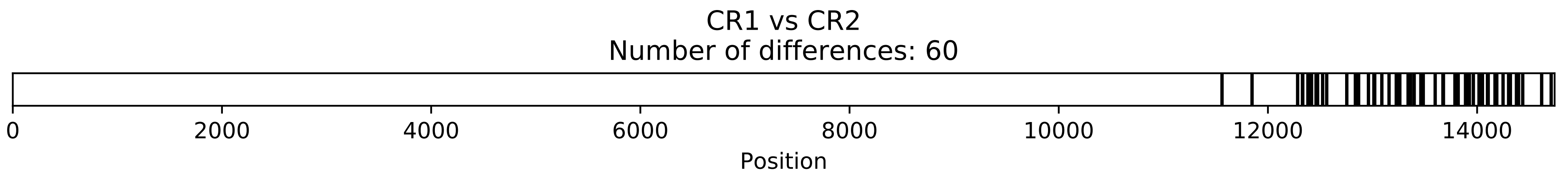
CR1 vs CR2
Number of differences: 60
0 2000 4000 6000 8000 10000 12000 14000
Position

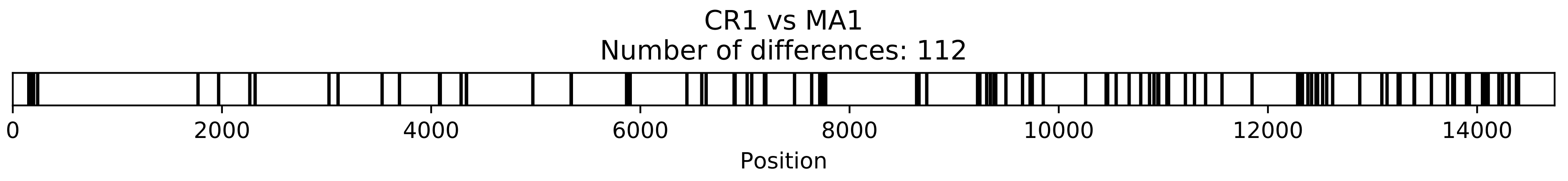
CR1 vs MA1
Number of differences: 112
0 2000 4000 6000 8000 10000 12000 14000
Position

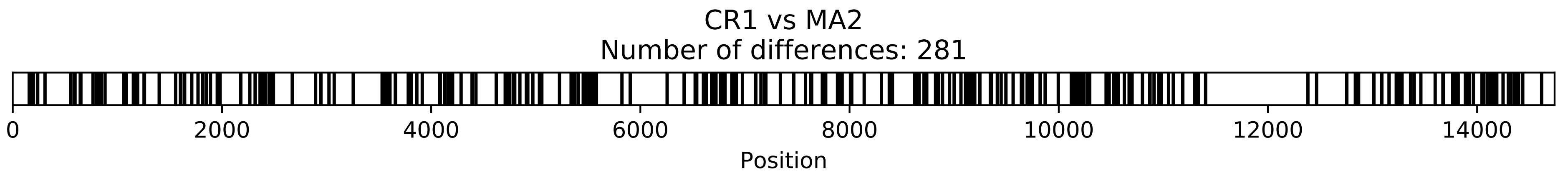
CR1 vs MA2
Number of differences: 281
0 2000 4000 6000 8000 10000 12000 14000
Position

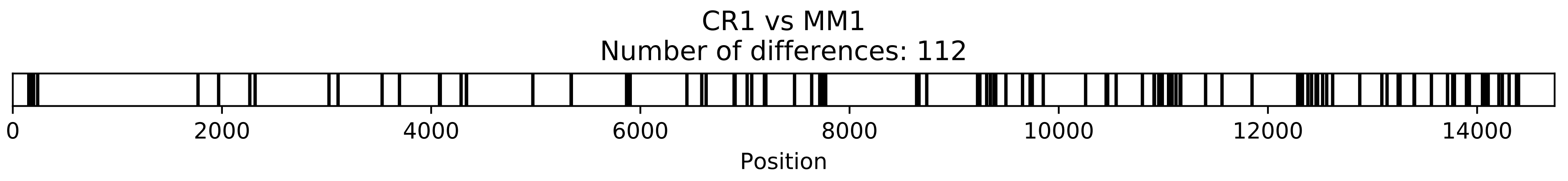
CR1 vs MM1
Number of differences: 112
0 2000 4000 6000 8000 10000 12000 14000
Position

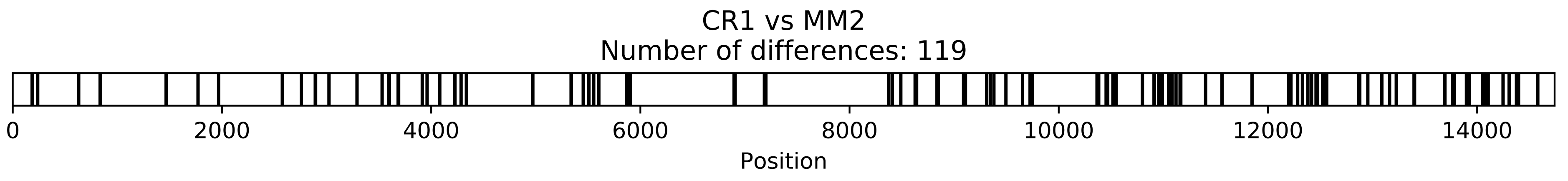
CR1 vs MM2
Number of differences: 119
0 2000 4000 6000 8000 10000 12000 14000
Position

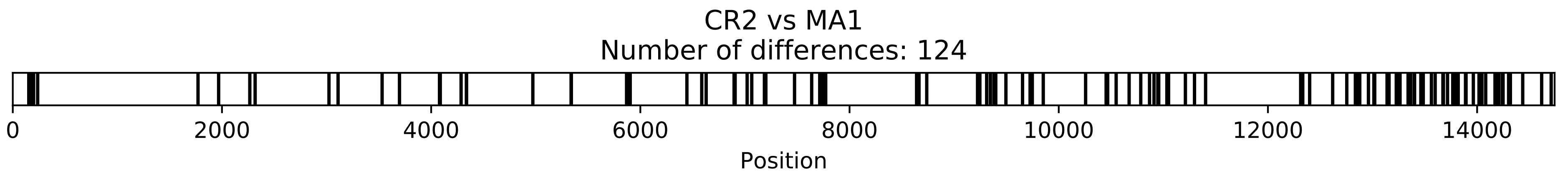
CR2 vs MA1
Number of differences: 124
0 2000 4000 6000 8000 10000 12000 14000
Position

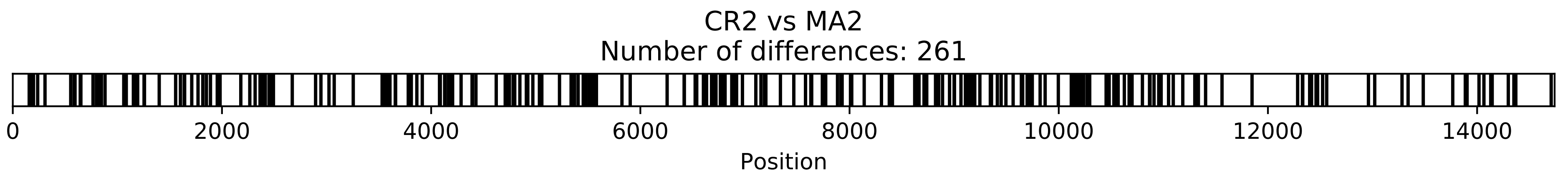
CR2 vs MA2
Number of differences: 261
0 2000 4000 6000 8000 10000 12000 14000
Position

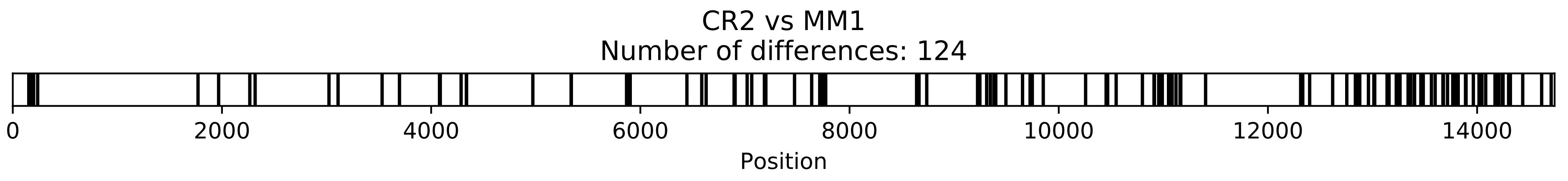
CR2 vs MM1
Number of differences: 124
0 2000 4000 6000 8000 10000 12000 14000
Position

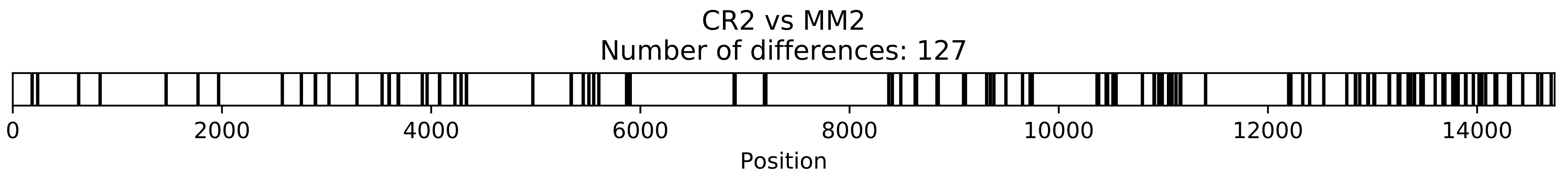
CR2 vs MM2
Number of differences: 127
0 2000 4000 6000 8000 10000 12000 14000
Position

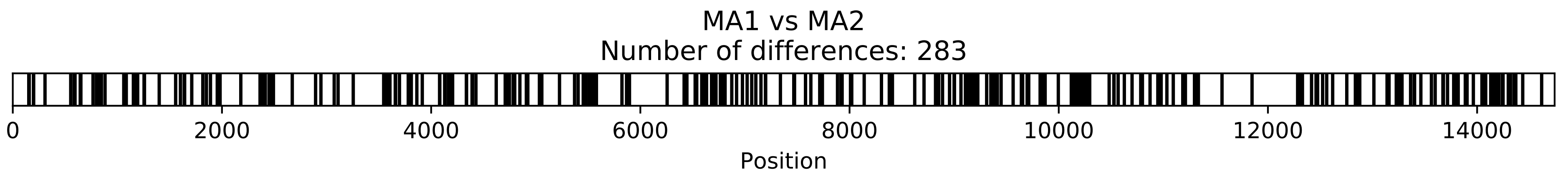
MA1 vs MA2
Number of differences: 283
0 2000 4000 6000 8000 10000 12000 14000
Position

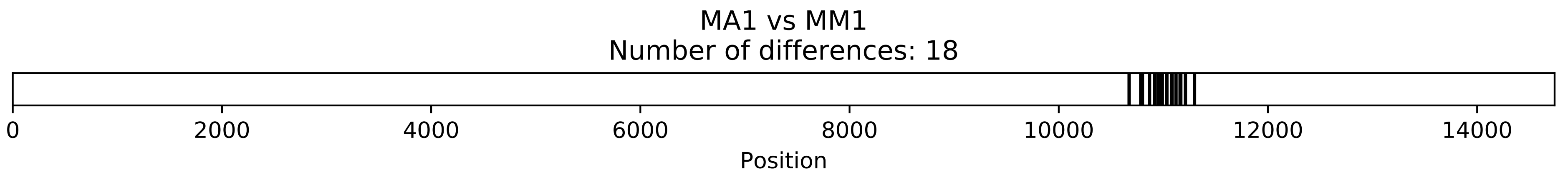
MA1 vs MM1
Number of differences: 18
0 2000 4000 6000 8000 10000 12000 14000
Position

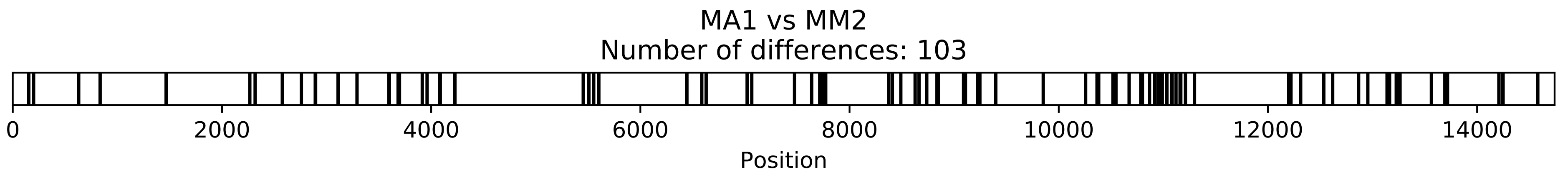
MA1 vs MM2
Number of differences: 103
0 2000 4000 6000 8000 10000 12000 14000
Position

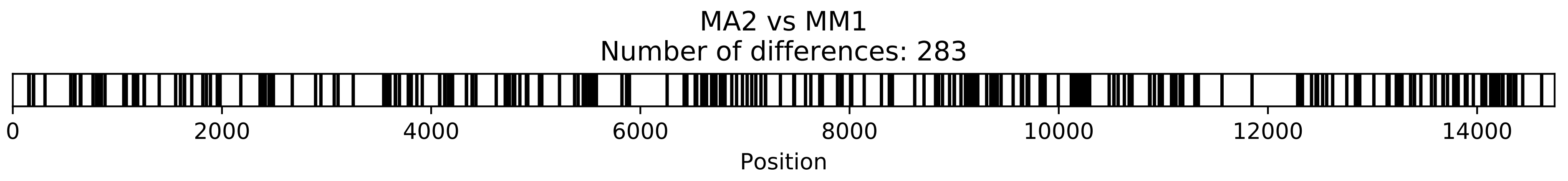
MA2 vs MM1
Number of differences: 283
0 2000 4000 6000 8000 10000 12000 14000
Position

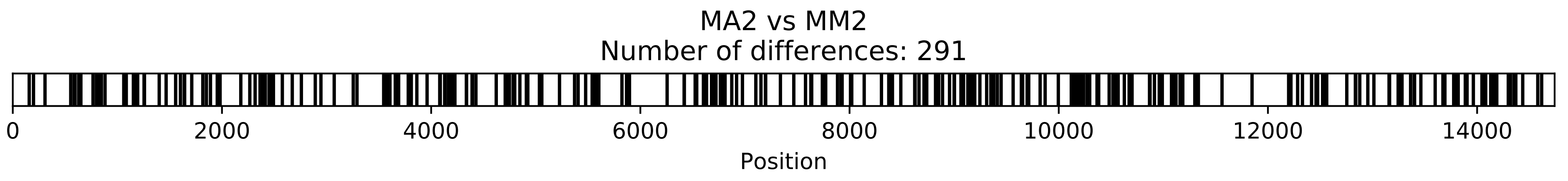
MA2 vs MM2
Number of differences: 291
0 2000 4000 6000 8000 10000 12000 14000
Position

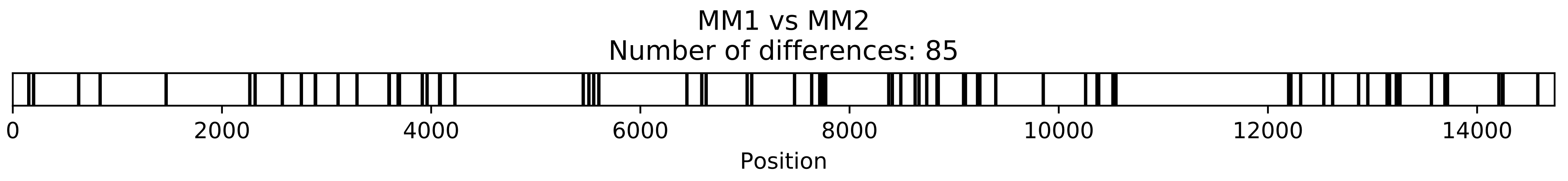
MM1 vs MM2
Number of differences: 85
0 2000 4000 6000 8000 10000 12000 14000
Position

### Region: 251

CR1 vs CR2
Number of differences: 262

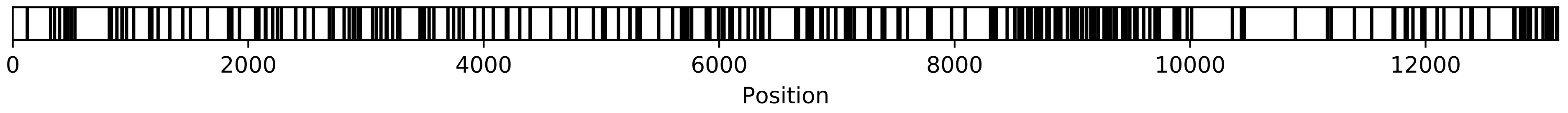


CR1 vs MA1
Number of differences: 145

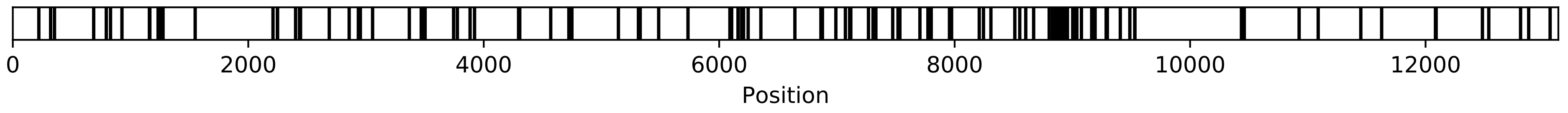


CR1 vs MA2
Number of differences: 272

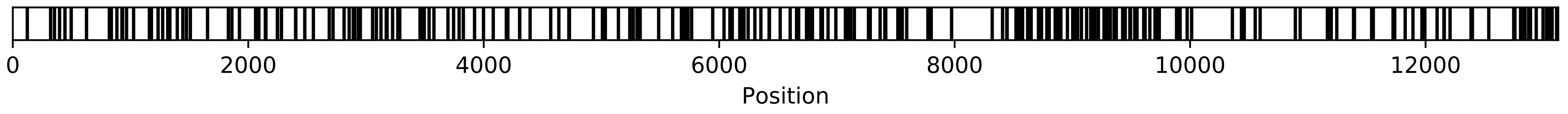


CR1 vs MM1
Number of differences: 155

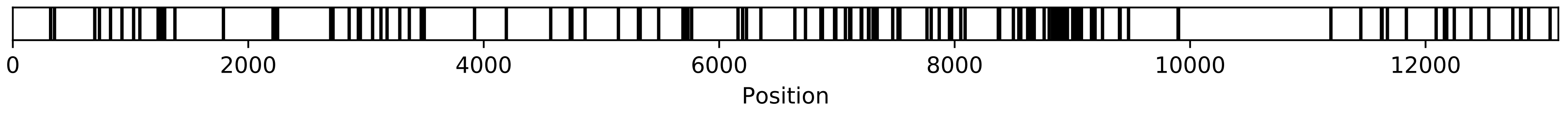


CR1 vs MM2
Number of differences: 150

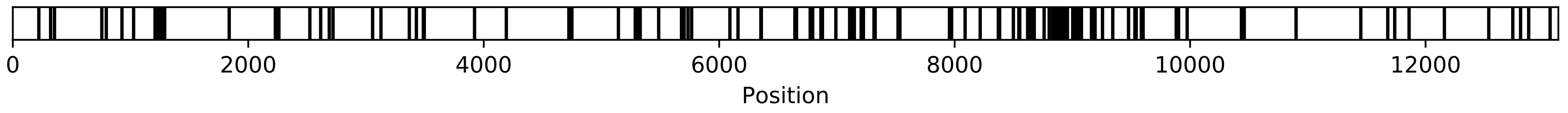


CR2 vs MA1
Number of differences: 284

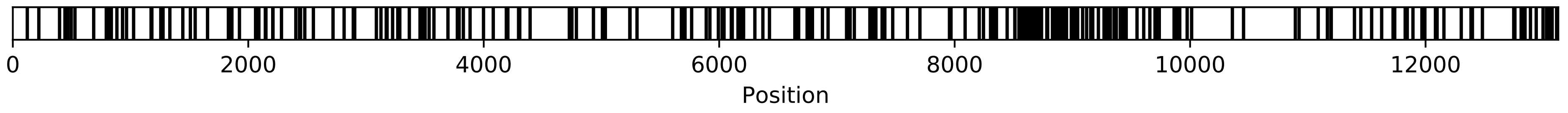


CR2 vs MA2
Number of differences: 82

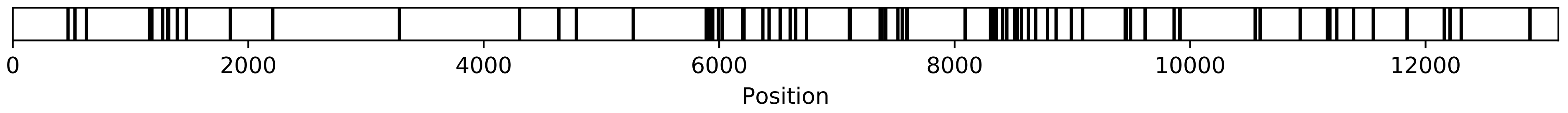


CR2 vs MM1
Number of differences: 291

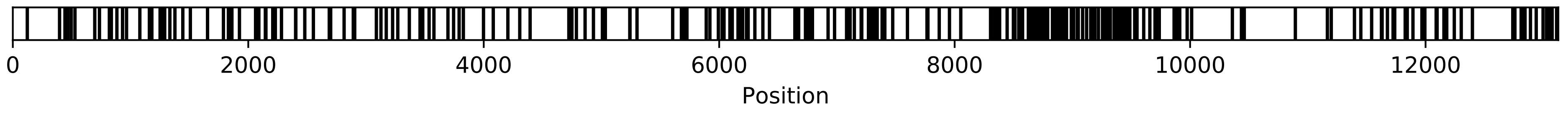


CR2 vs MM2
Number of differences: 290

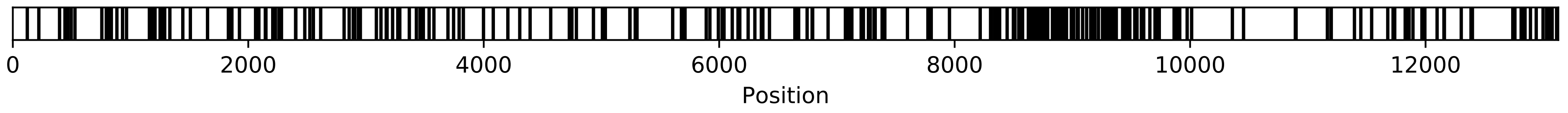


MA1 vs MA2
Number of differences: 297

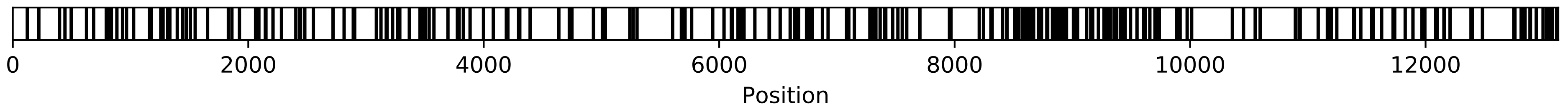


MA1 vs MM1
Number of differences: 127

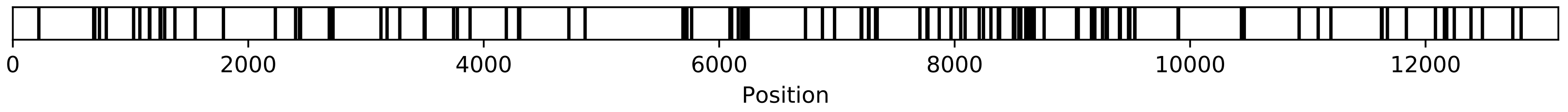


MA1 vs MM2
Number of differences: 133

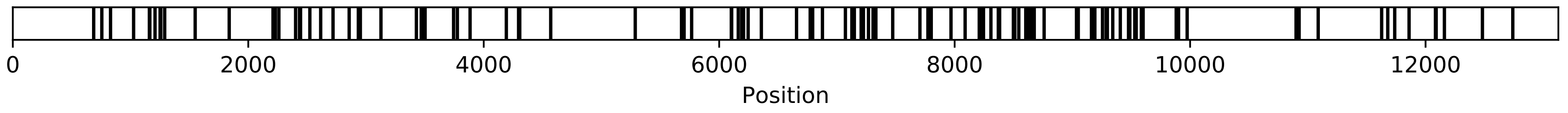


MA2 vs MM1
Number of differences: 304

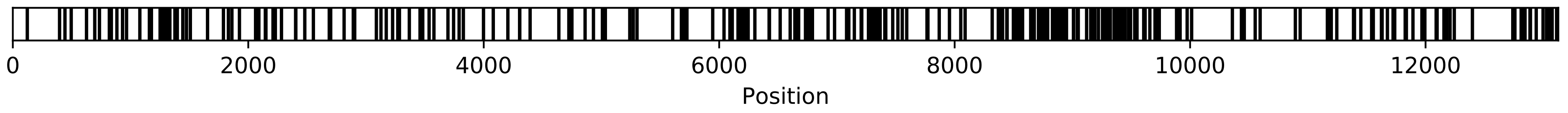


MA2 vs MM2
Number of differences: 304

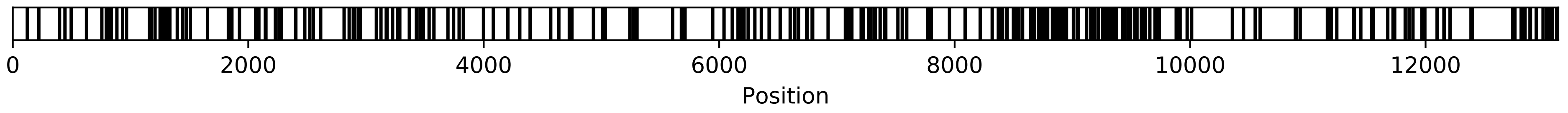


MM1 vs MM2
Number of differences: 85

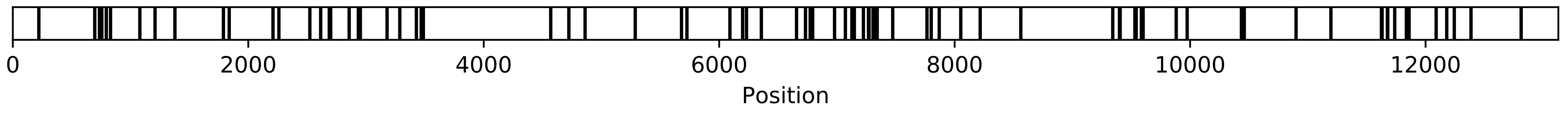

### Region: 252



CR1 vs CR2
Number of differences: 100

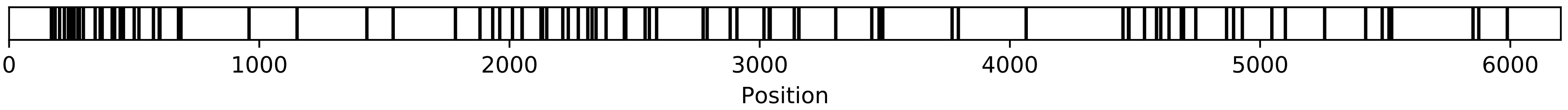


CR1 vs MA1
Number of differences: 29

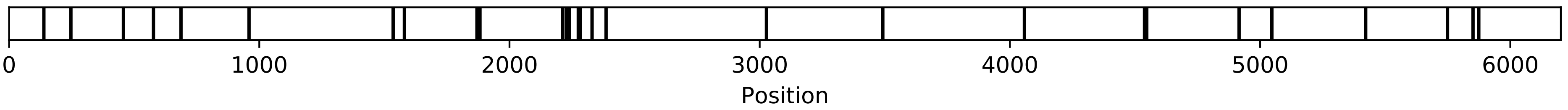


CR1 vs MA2
Number of differences: 98

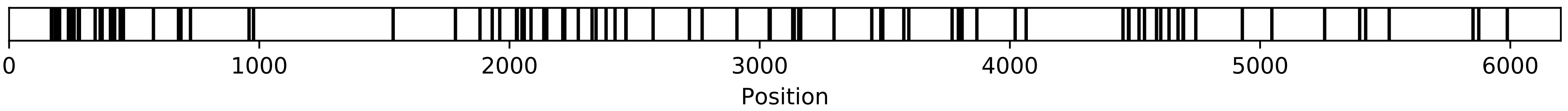


CR1 vs MM1
Number of differences: 109

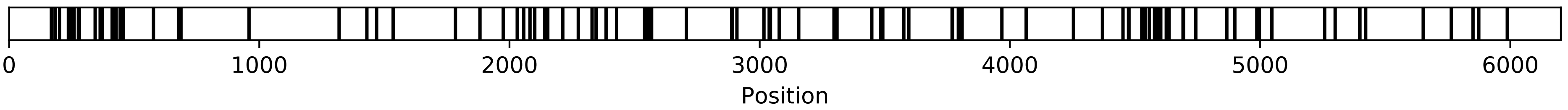


CR1 vs MM2
Number of differences: 108

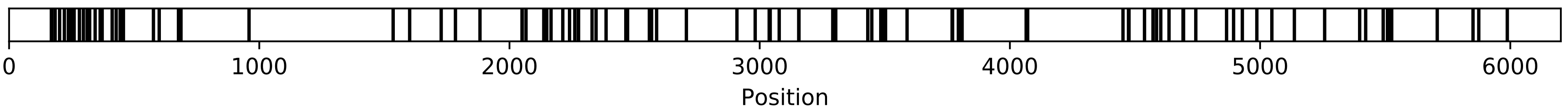


CR2 vs MA1
Number of differences: 95

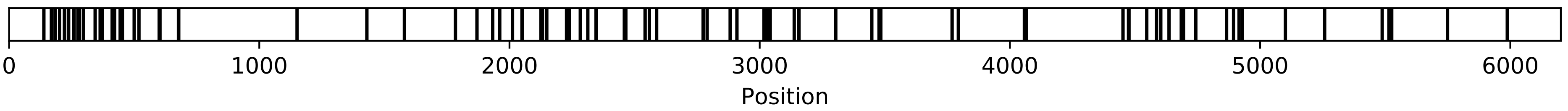


CR2 vs MA2
Number of differences: 66

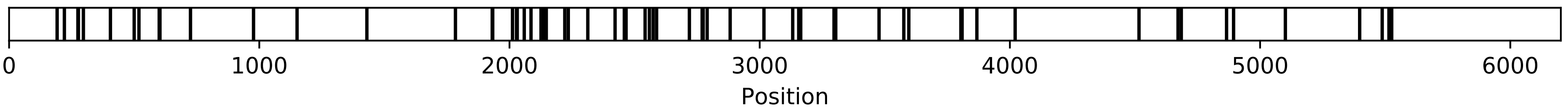


CR2 vs MM1
Number of differences: 85

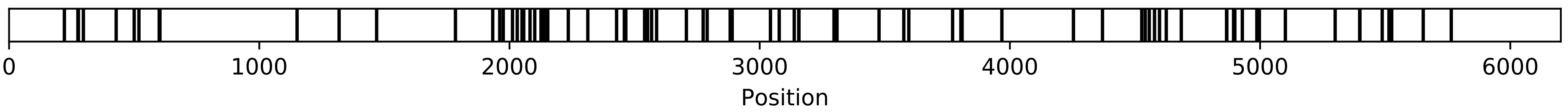


CR2 vs MM2
Number of differences: 66

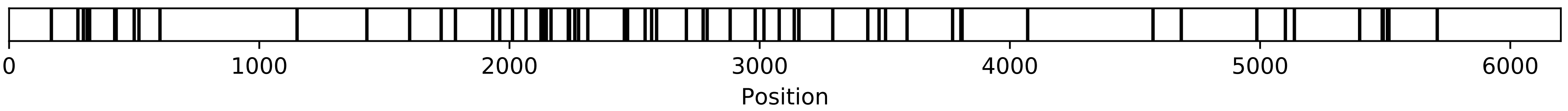


MA1 vs MA2
Number of differences: 93

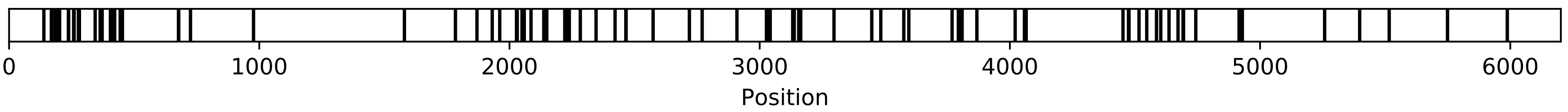


MA1 vs MM1
Number of differences: 104

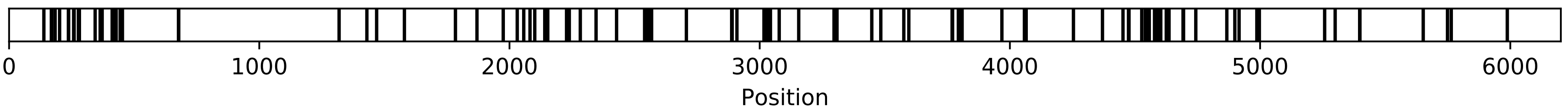


MA1 vs MM2
Number of differences: 103

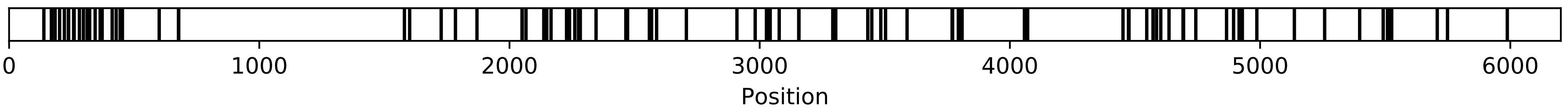


MA2 vs MM1
Number of differences: 69

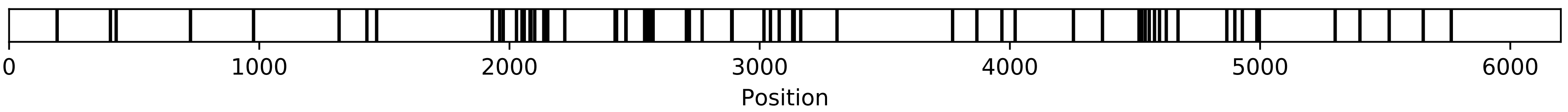


MA2 vs MM2
Number of differences: 65

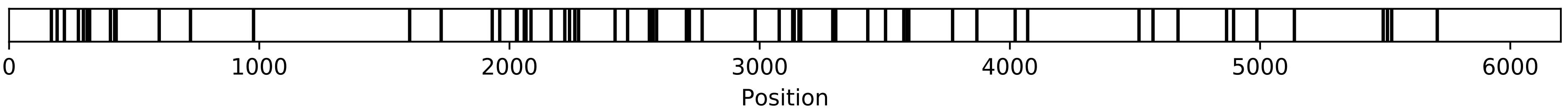


MM1 vs MM2
Number of differences: 78

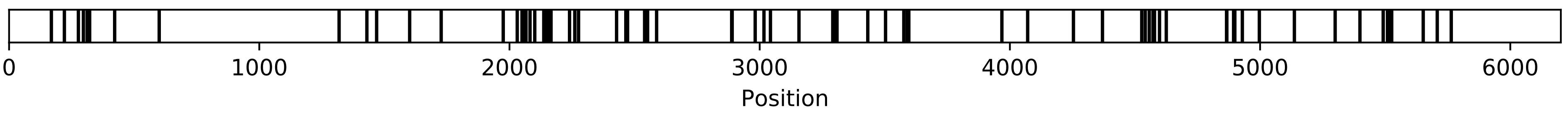

### Region: 253



CR1 vs CR2
Number of differences: 152

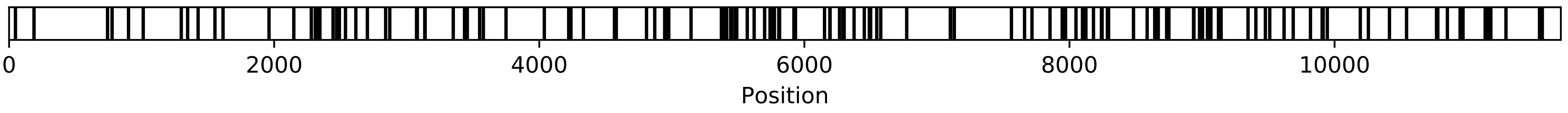


CR1 vs MA1
Number of differences: 141

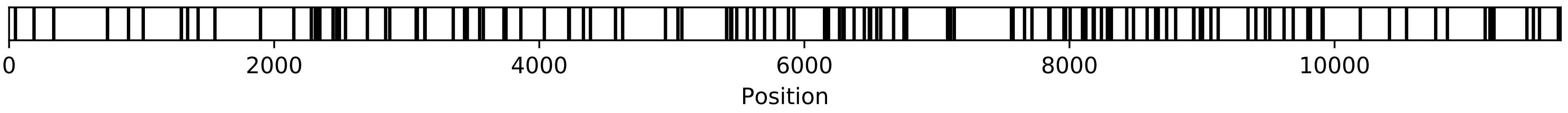


CR1 vs MA2
Number of differences: 55

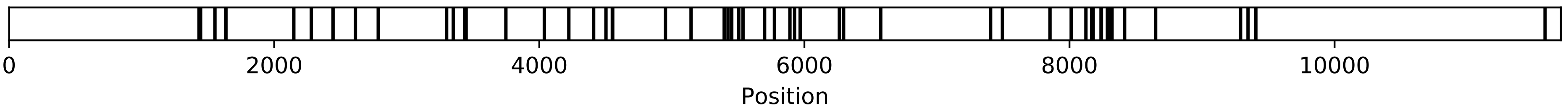


CR1 vs MM1
Number of differences: 143

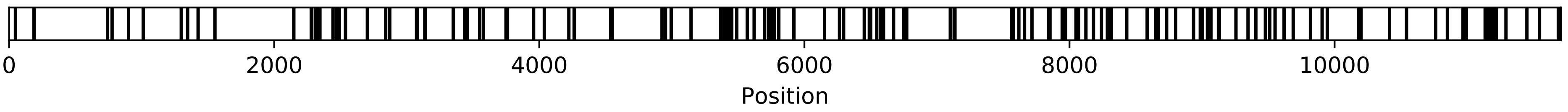


CR1 vs MM2
Number of differences: 142

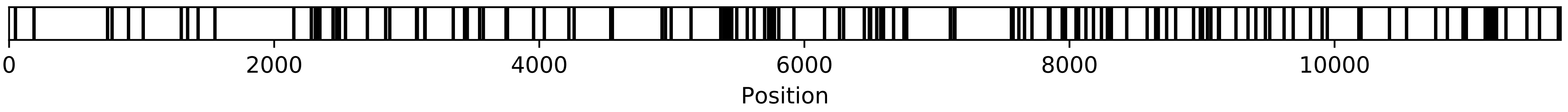


CR2 vs MA1
Number of differences: 88

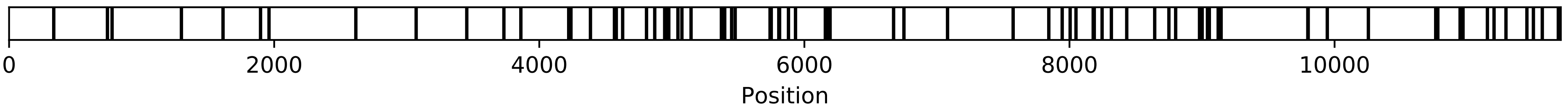


CR2 vs MA2
Number of differences: 159

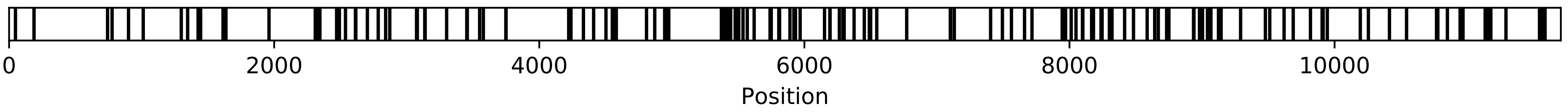


CR2 vs MM1
Number of differences: 82

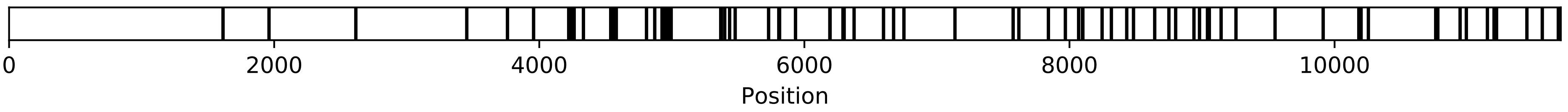


CR2 vs MM2
Number of differences: 81

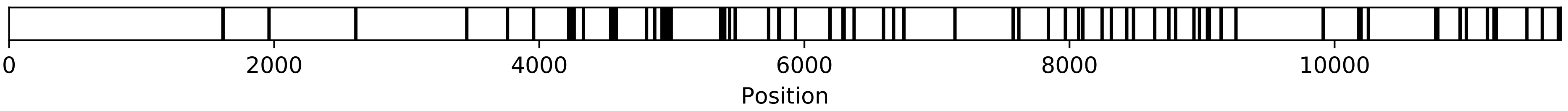


MA1 vs MA2
Number of differences: 146

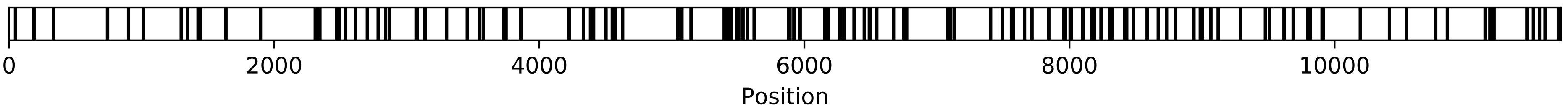


MA1 vs MM1
Number of differences: 80

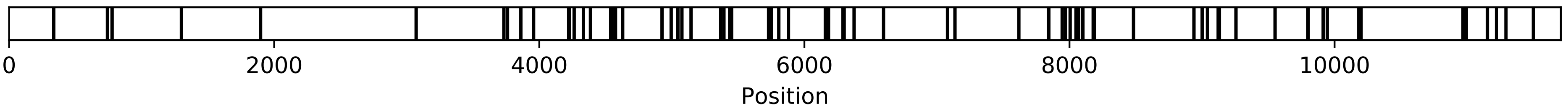


MA1 vs MM2
Number of differences: 79

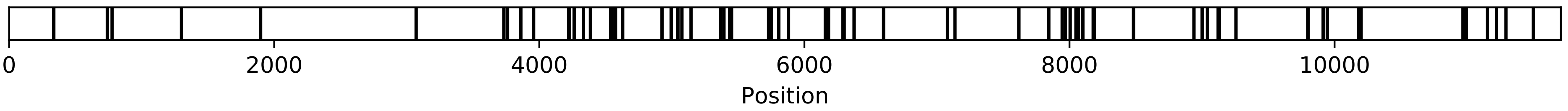


MA2 vs MM1
Number of differences: 146

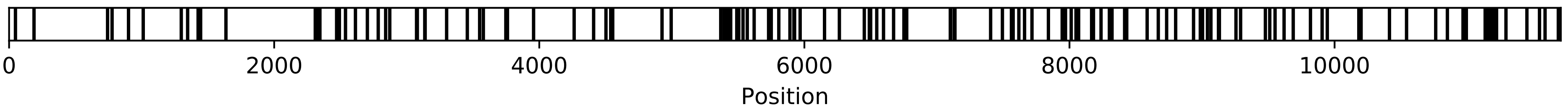


MA2 vs MM2
Number of differences: 145

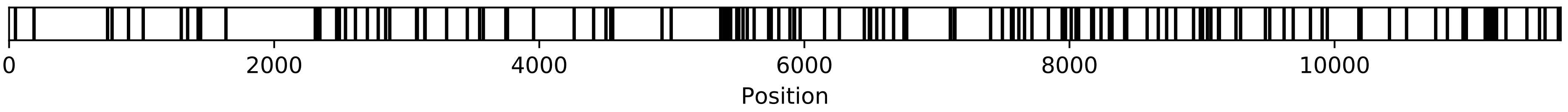


MM1 vs MM2
Number of differences: 1

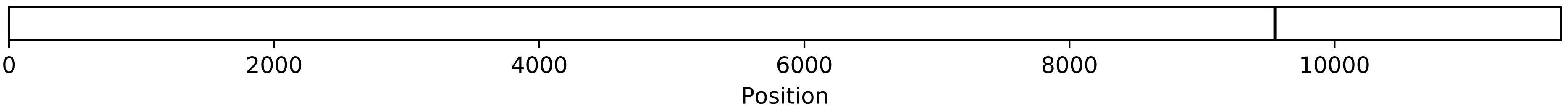

### Region: 256

CR1 vs CR2
Number of differences: 174

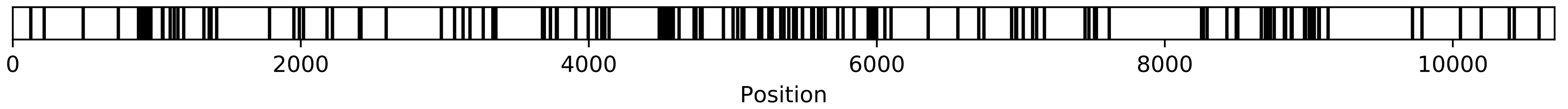

CR1 vs MA1
Number of differences: 71

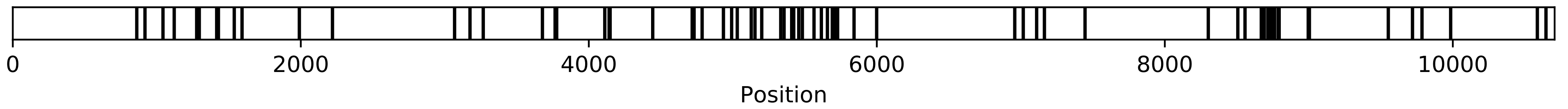

CR1 vs MA2
Number of differences: 177

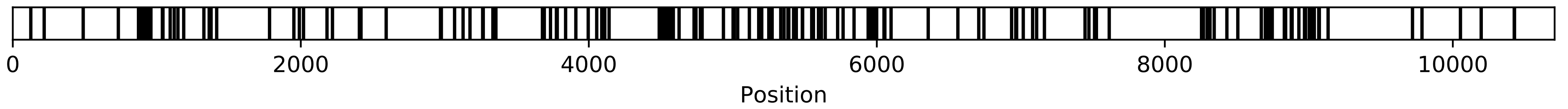

CR1 vs MM1
Number of differences: 61

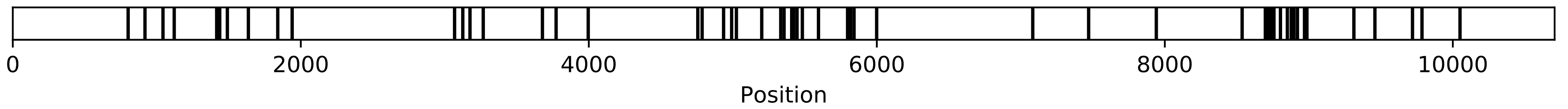

CR1 vs MM2
Number of differences: 71

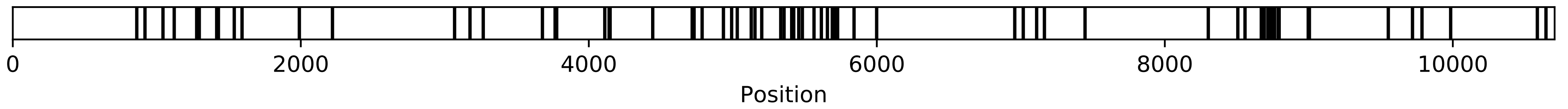

CR2 vs MA1
Number of differences: 179

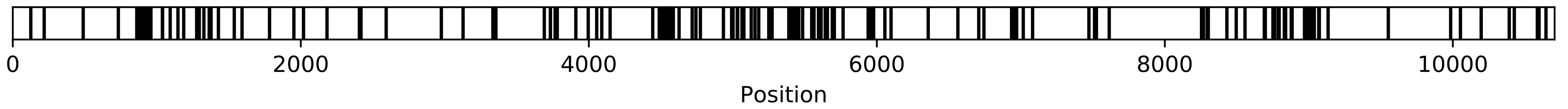

CR2 vs MA2
Number of differences: 21

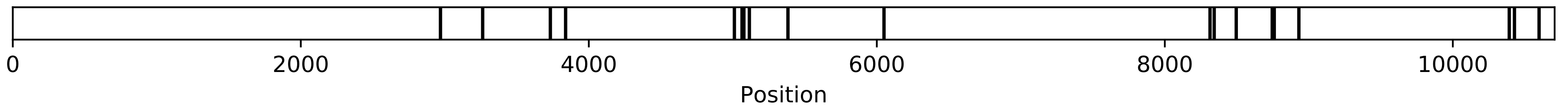

CR2 vs MM1
Number of differences: 183

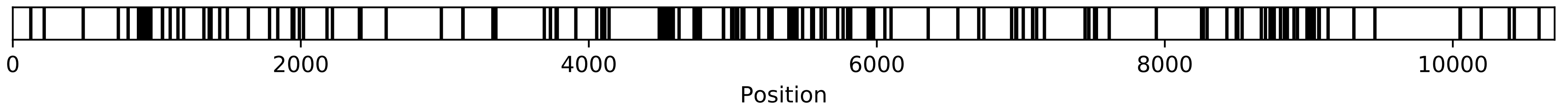

CR2 vs MM2
Number of differences: 179

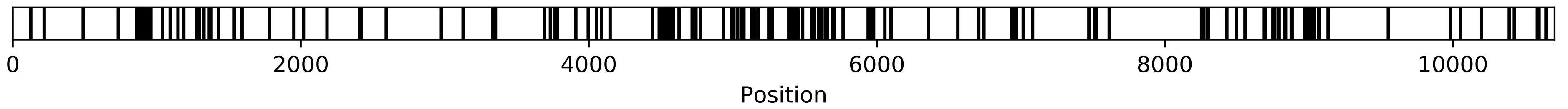

MA1 vs MA2
Number of differences: 182

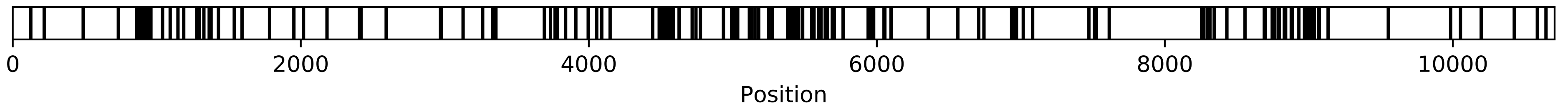

MA1 vs MM1
Number of differences: 84

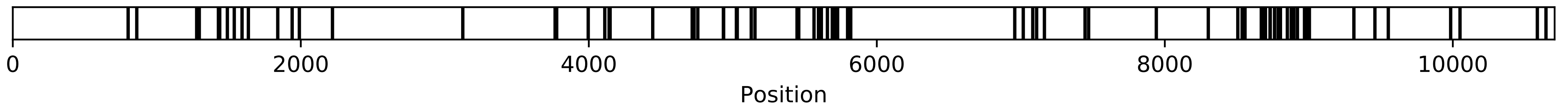

MA1 vs MM2
Number of differences: 0

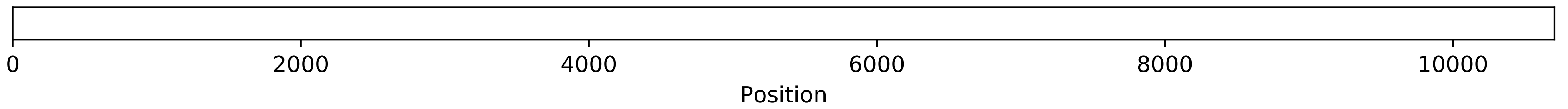

MA2 vs MM1
Number of differences: 186

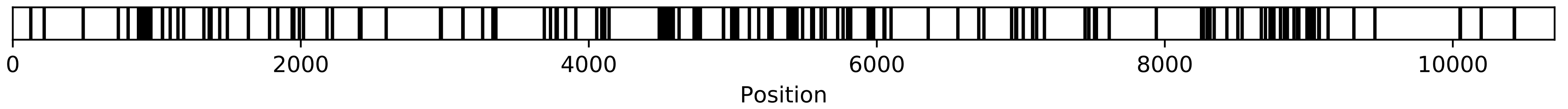

MA2 vs MM2
Number of differences: 182

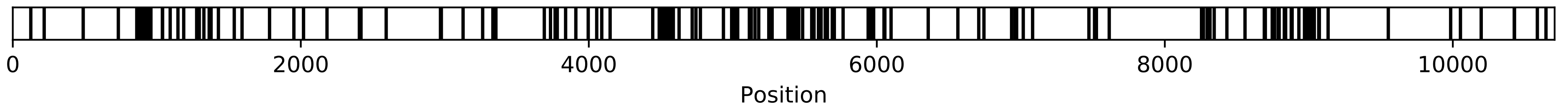

MM1 vs MM2
Number of differences: 84

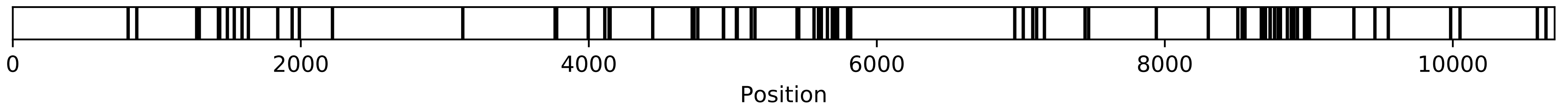

### Region: 257

CR1 vs CR2
Number of differences: 202

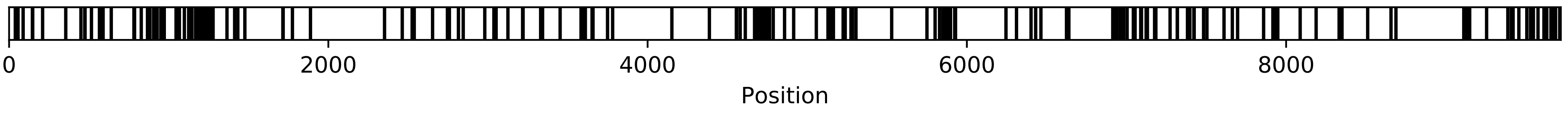


CR1 vs MA1
Number of differences: 152

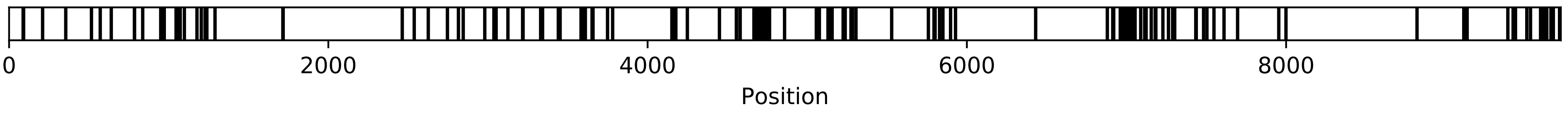


CR1 vs MA2
Number of differences: 180

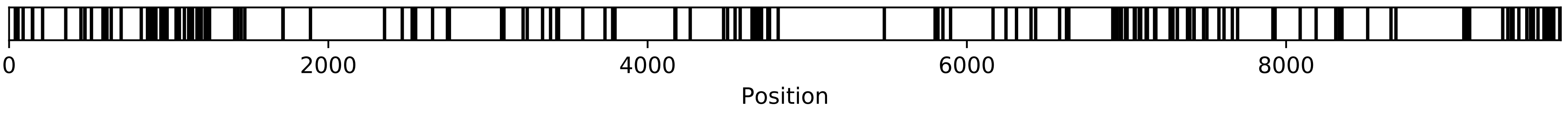


CR1 vs MM1
Number of differences: 148

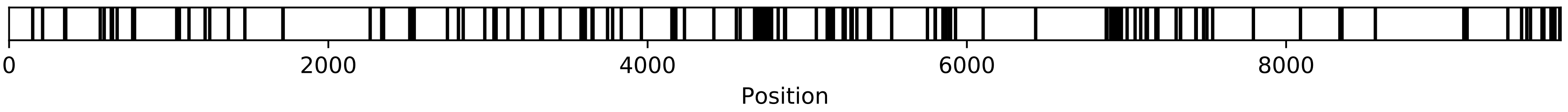


CR1 vs MM2
Number of differences: 152

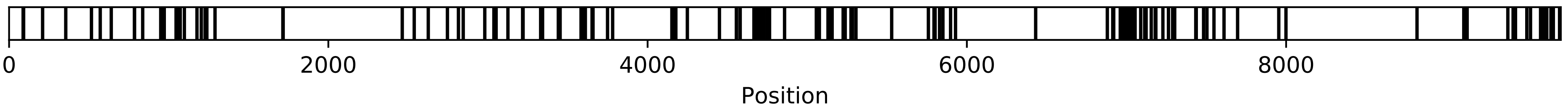


CR2 vs MA1
Number of differences: 150

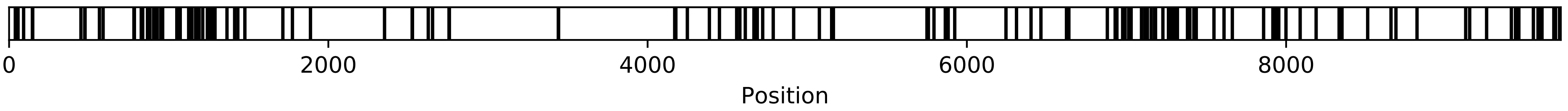


CR2 vs MA2
Number of differences: 118

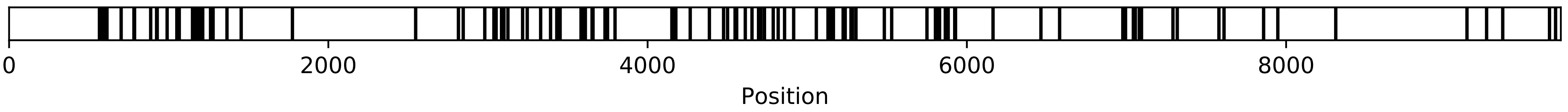


CR2 vs MM1
Number of differences: 166

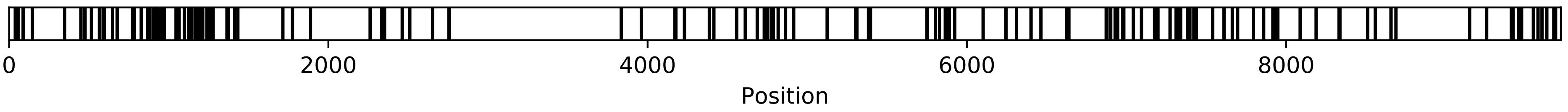


CR2 vs MM2
Number of differences: 150

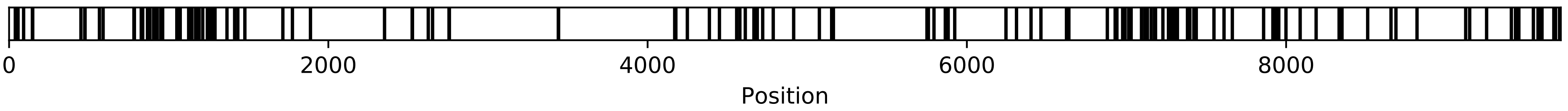


MA1 vs MA2
Number of differences: 195

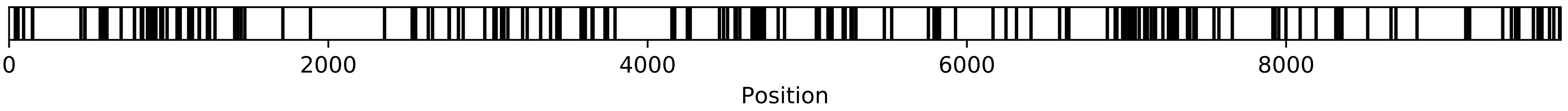


MA1 vs MM1
Number of differences: 123

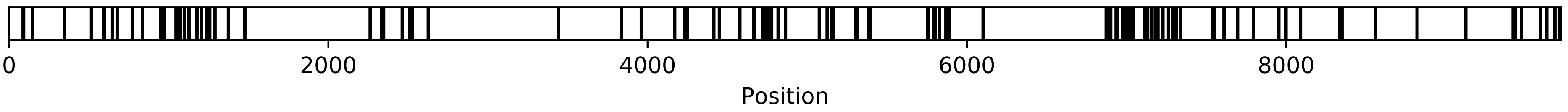


MA1 vs MM2
Number of differences: 0

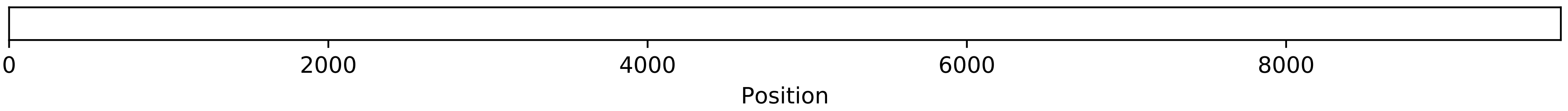


MA2 vs MM1
Number of differences: 202

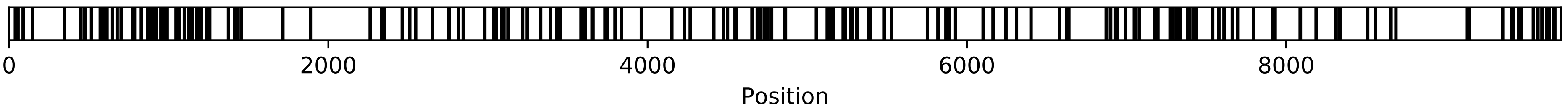


MA2 vs MM2
Number of differences: 195

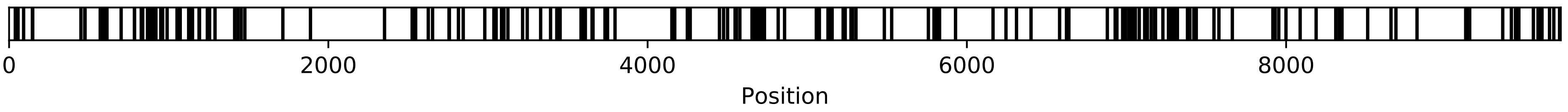


MM1 vs MM2
Number of differences: 123

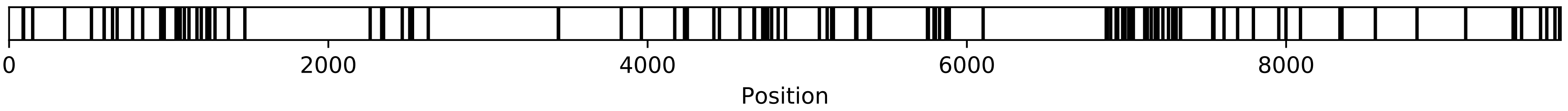

### Region: 258

CR1 vs CR2
Number of differences: 324

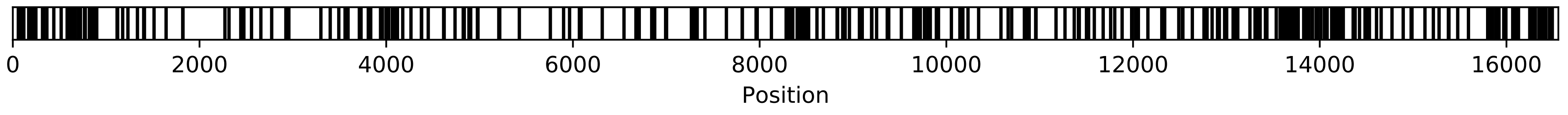


CR1 vs MA1
Number of differences: 325

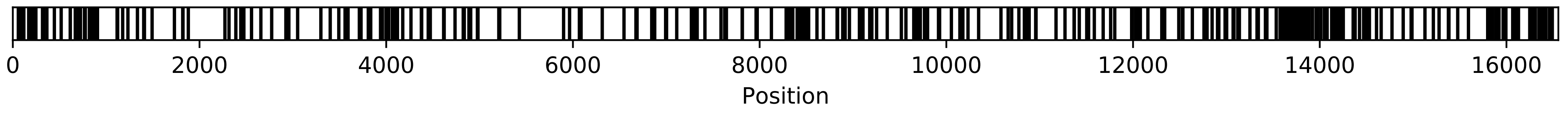


CR1 vs MA2
Number of differences: 156

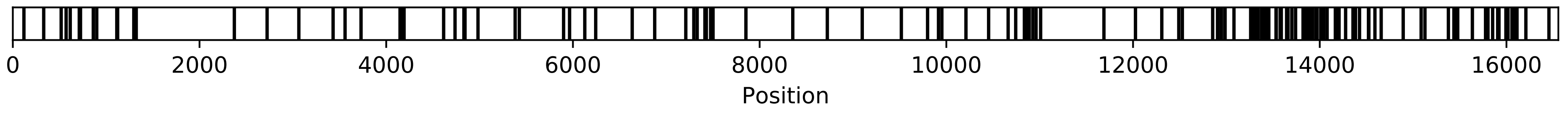


CR1 vs MM1
Number of differences: 139

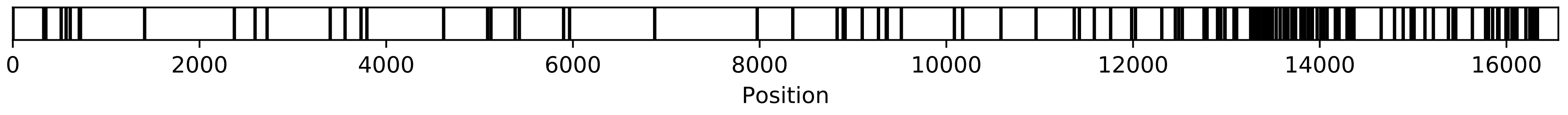


CR1 vs MM2
Number of differences: 0

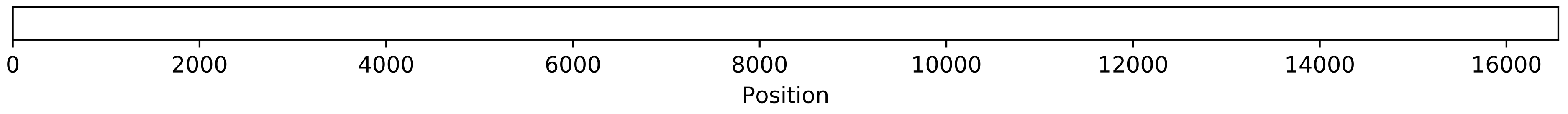


CR2 vs MA1
Number of differences: 56

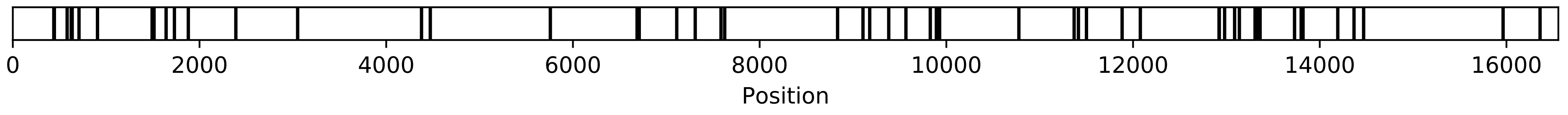


CR2 vs MA2
Number of differences: 315

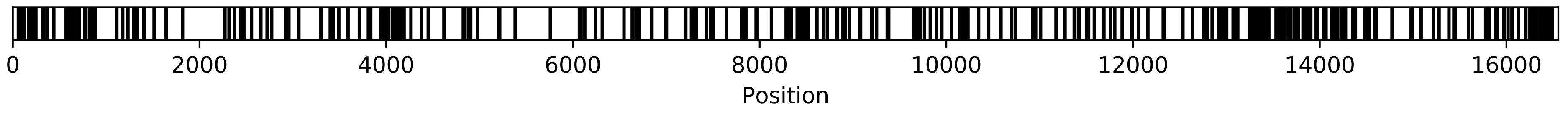


CR2 vs MM1
Number of differences: 307

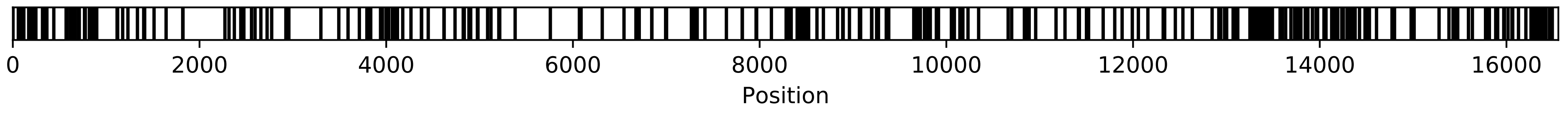


CR2 vs MM2
Number of differences: 324

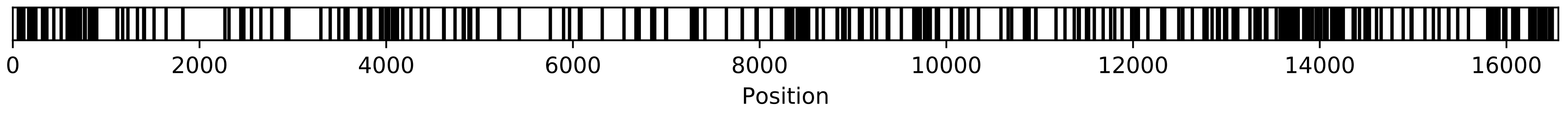


MA1 vs MA2
Number of differences: 317

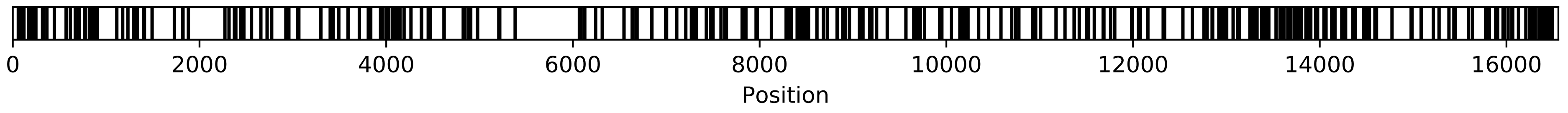


MA1 vs MM1
Number of differences: 306

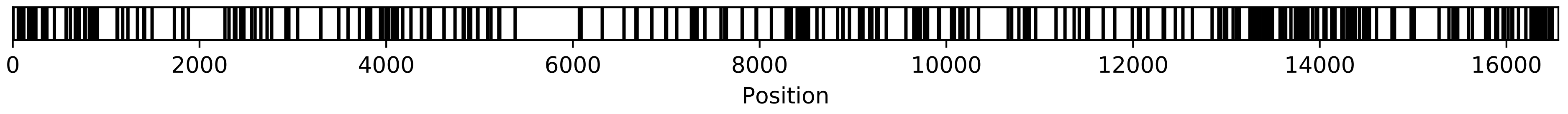


MA1 vs MM2
Number of differences: 325

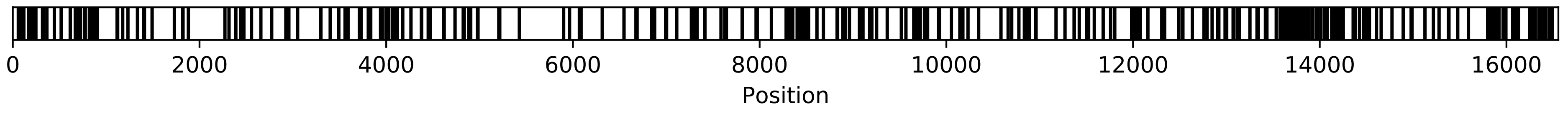


MA2 vs MM1
Number of differences: 147

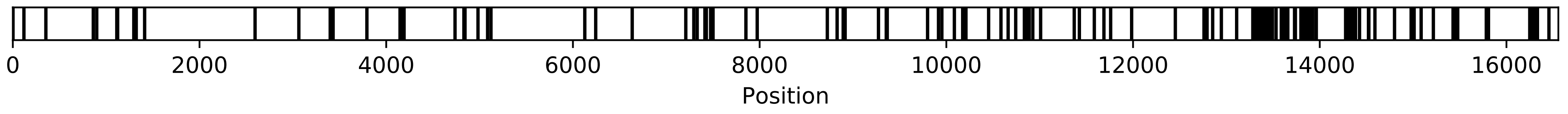


MA2 vs MM2
Number of differences: 156

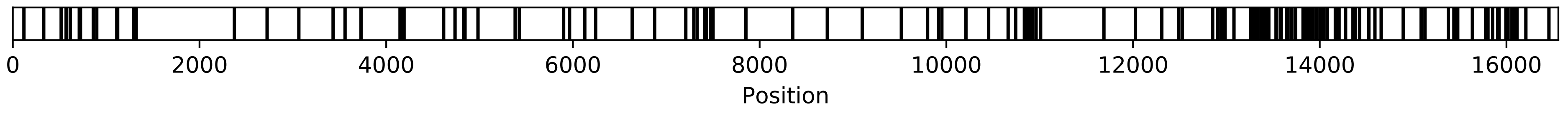


MM1 vs MM2
Number of differences: 139

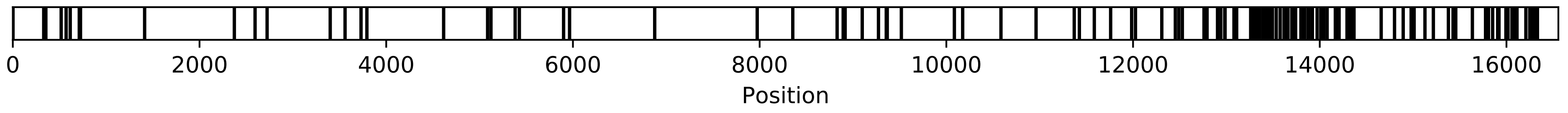

### Region: 259



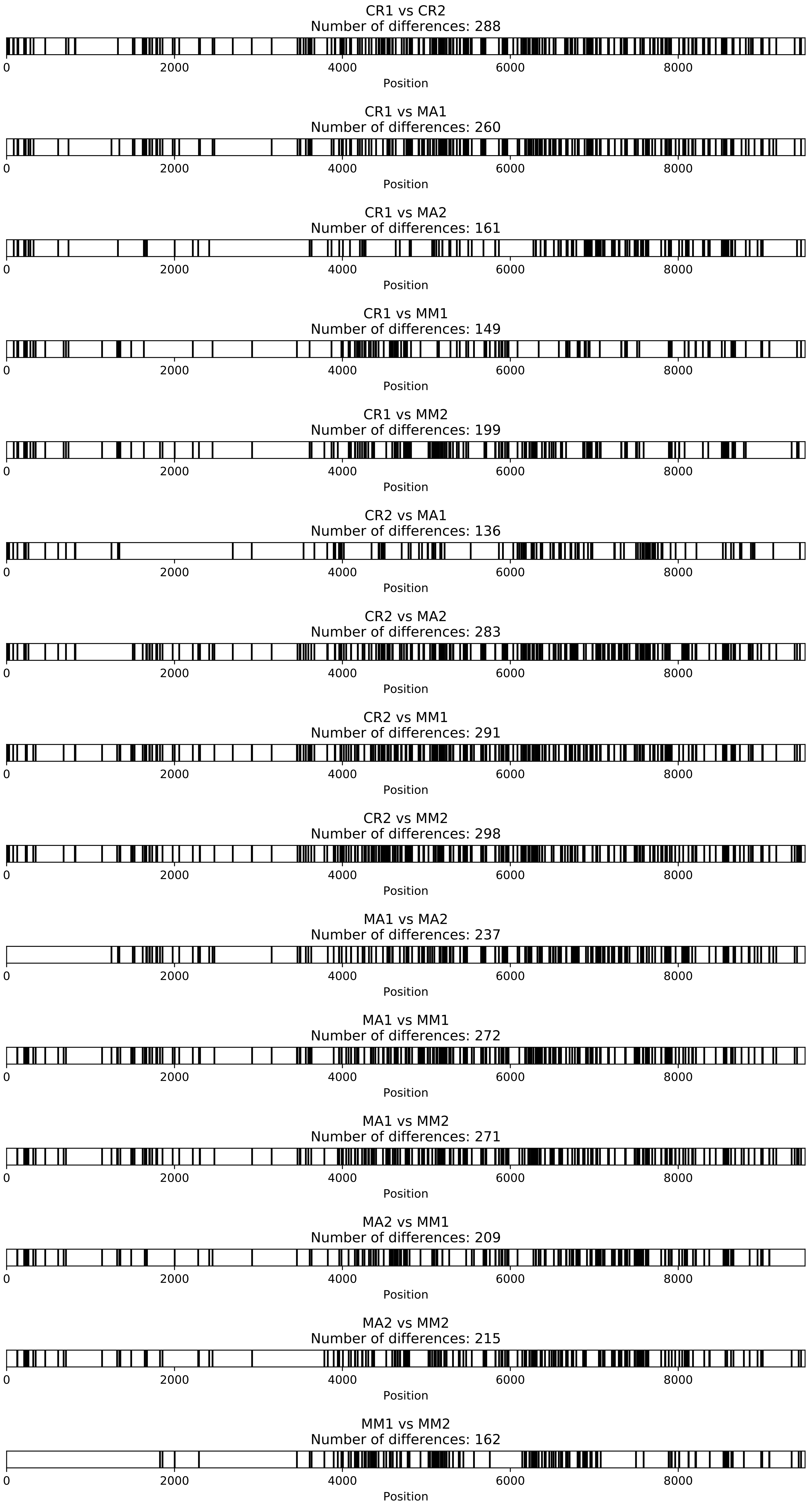
CR1 vs CR2
Number of differences: 288
0 2000 4000 6000 8000
Position
CR1 vs MA1
Number of differences: 260
0 2000 4000 6000 8000
Position
CR1 vs MA2
Number of differences: 161
0 2000 4000 6000 8000
Position
CR1 vs MM1
Number of differences: 149
0 2000 4000 6000 8000
Position
CR1 vs MM2
Number of differences: 199
0 2000 4000 6000 8000
Position
CR2 vs MA1
Number of differences: 136
0 2000 4000 6000 8000
Position
CR2 vs MA2
Number of differences: 283
0 2000 4000 6000 8000
Position
CR2 vs MM1
Number of differences: 291
0 2000 4000 6000 8000
Position
CR2 vs MM2
Number of differences: 298
0 2000 4000 6000 8000
Position
MA1 vs MA2
Number of differences: 237
0 2000 4000 6000 8000
Position
MA1 vs MM1
Number of differences: 272
0 2000 4000 6000 8000
Position
MA1 vs MM2
Number of differences: 271
0 2000 4000 6000 8000
Position
MA2 vs MM1
Number of differences: 209
0 2000 4000 6000 8000
Position
MA2 vs MM2
Number of differences: 215
0 2000 4000 6000 8000
Position
MM1 vs MM2
Number of differences: 162
0 2000 4000 6000 8000
Position

### Region: 261



CR1 vs CR2
Number of differences: 146

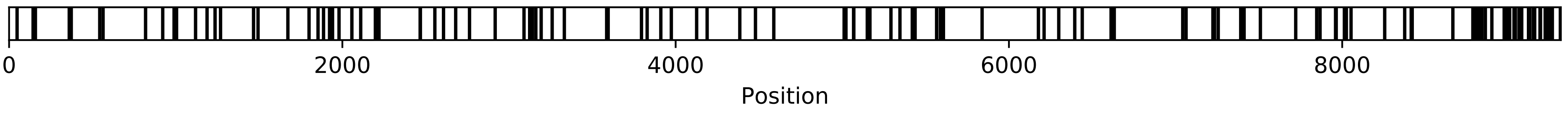


CR1 vs MA1
Number of differences: 77

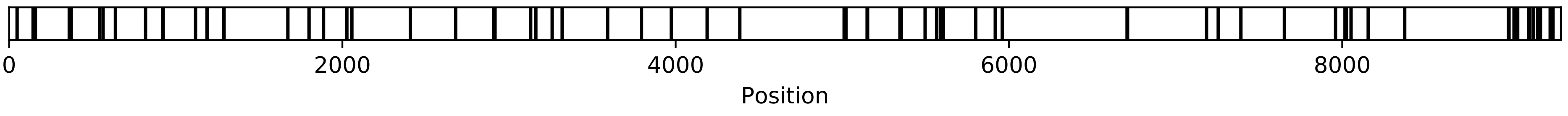


CR1 vs MA2
Number of differences: 138

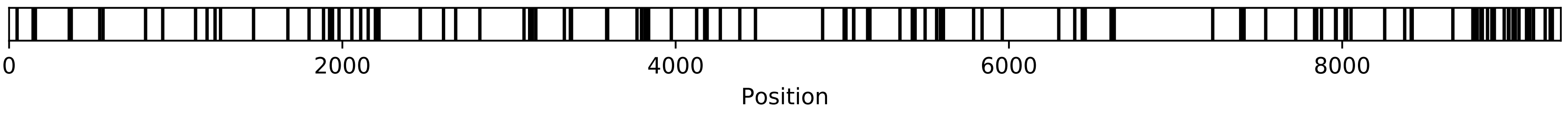


CR1 vs MM1
Number of differences: 78

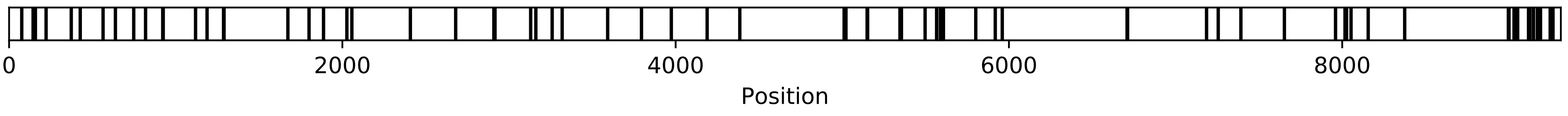


CR1 vs MM2
Number of differences: 78

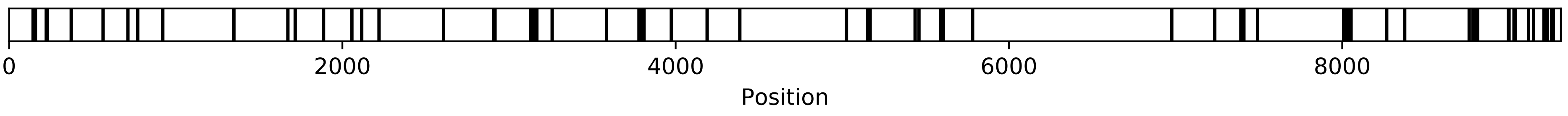


CR2 vs MA1
Number of differences: 122

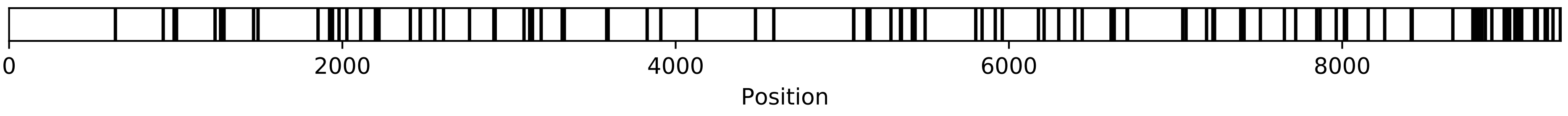


CR2 vs MA2
Number of differences: 75

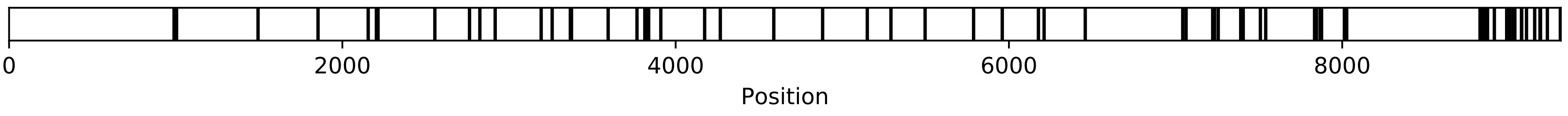


CR2 vs MM1
Number of differences: 129

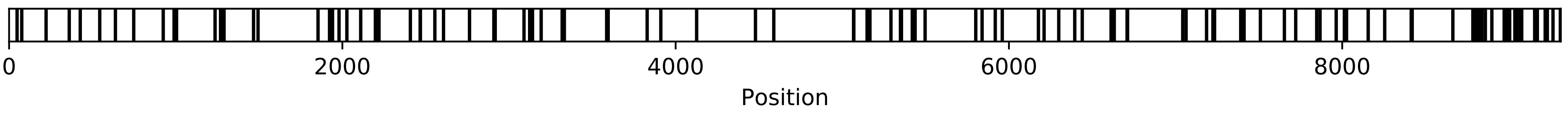


CR2 vs MM2
Number of differences: 136

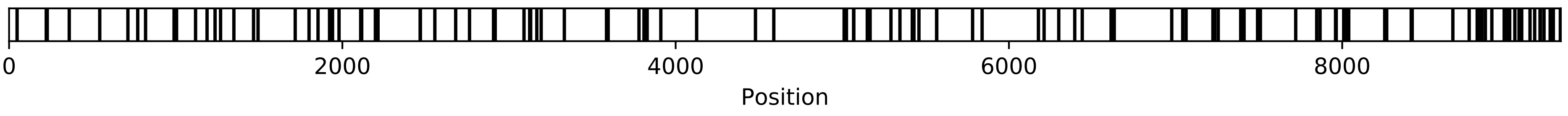


MA1 vs MA2
Number of differences: 116

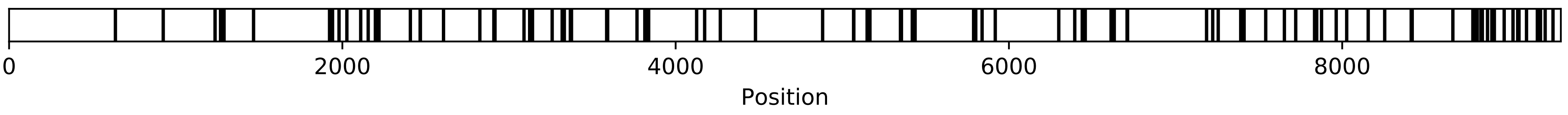


MA1 vs MM1
Number of differences: 7

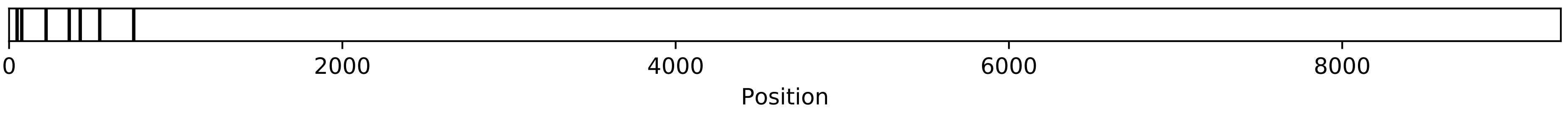


MA1 vs MM2
Number of differences: 89

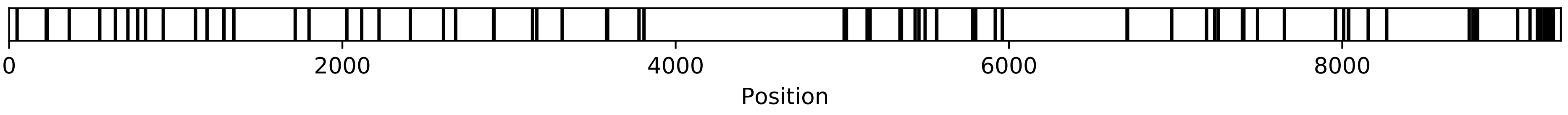


MA2 vs MM1
Number of differences: 123

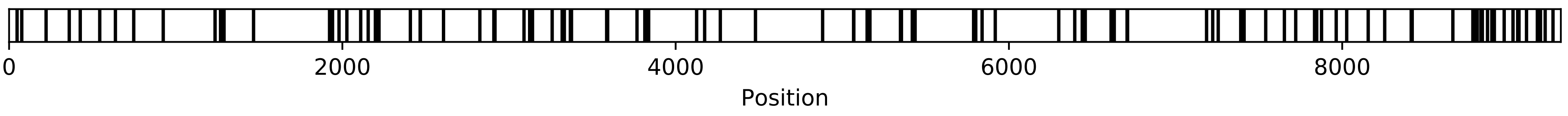


MA2 vs MM2
Number of differences: 132

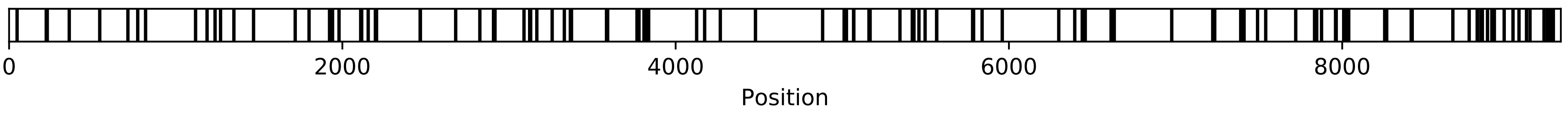


MM1 vs MM2
Number of differences: 90

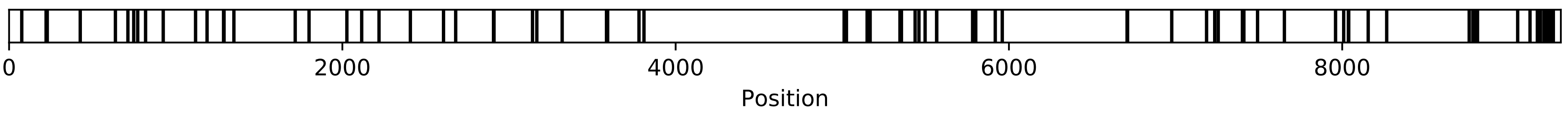

### Region: 262

CR1 vs CR2
Number of differences: 149

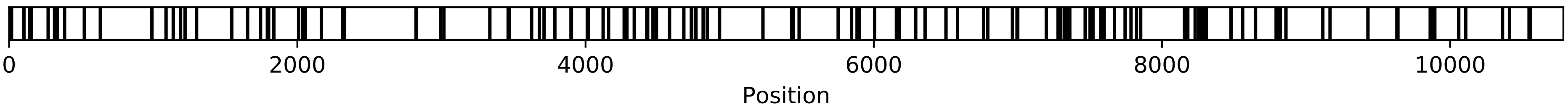


CR1 vs MA1
Number of differences: 62

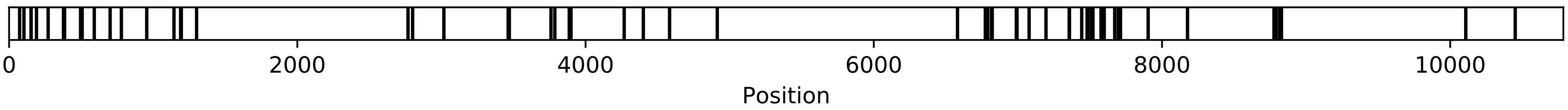


CR1 vs MA2
Number of differences: 153

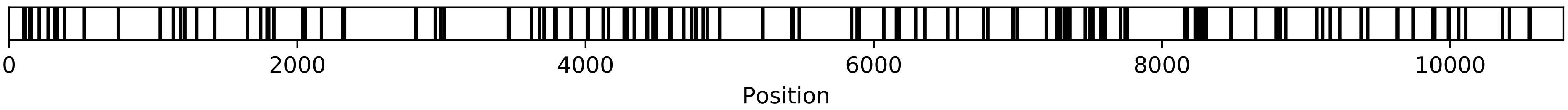


CR1 vs MM1
Number of differences: 55

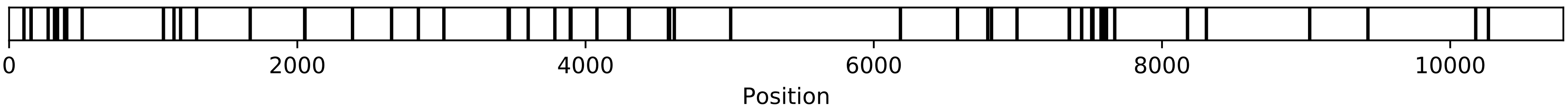


CR1 vs MM2
Number of differences: 62

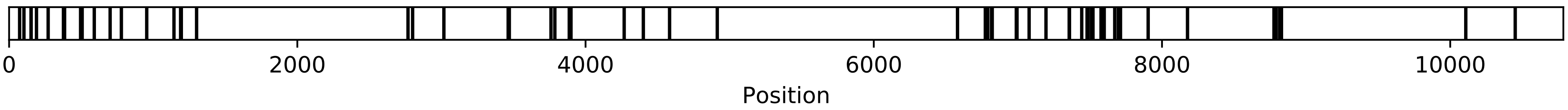


CR2 vs MA1
Number of differences: 159

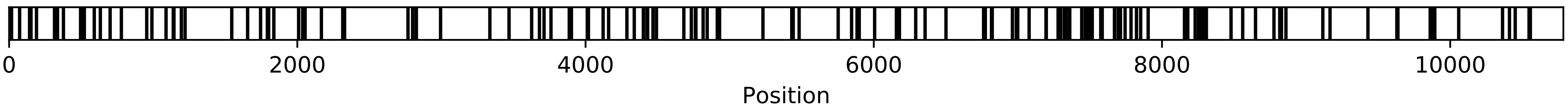


CR2 vs MA2
Number of differences: 54

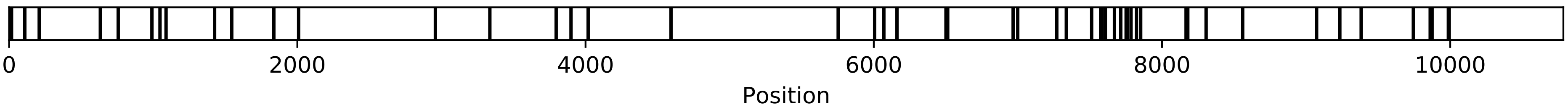


CR2 vs MM1
Number of differences: 153

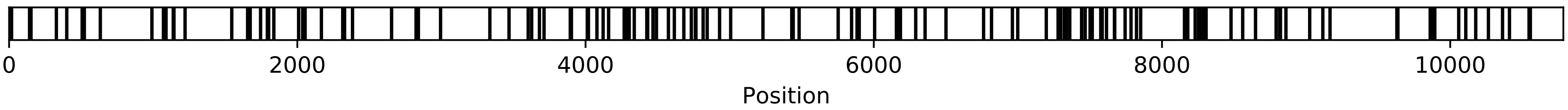


CR2 vs MM2
Number of differences: 159

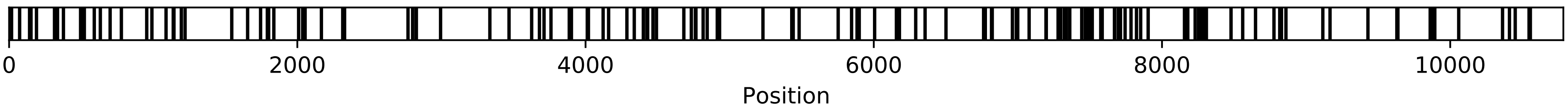


MA1 vs MA2
Number of differences: 162

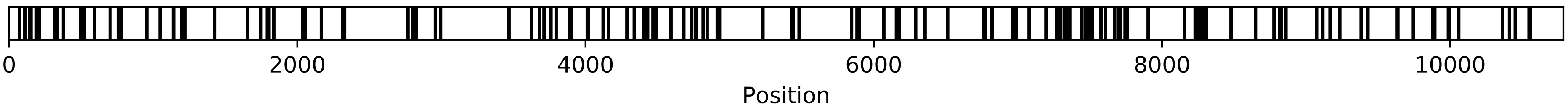


MA1 vs MM1
Number of differences: 63

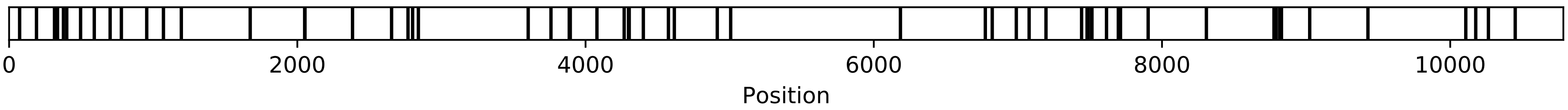


MA1 vs MM2
Number of differences: 0

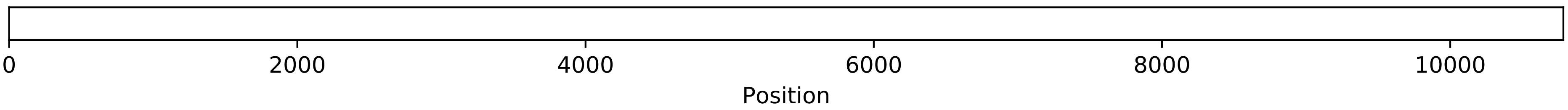


MA2 vs MM1
Number of differences: 154

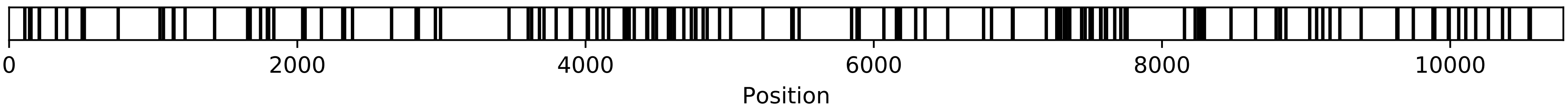


MA2 vs MM2
Number of differences: 162

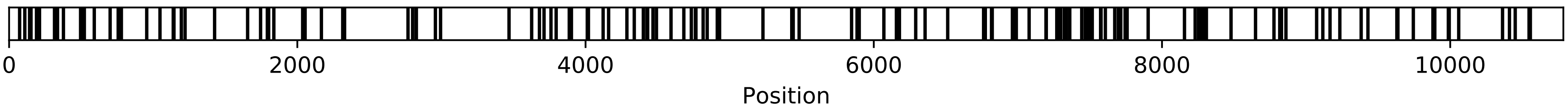


MM1 vs MM2
Number of differences: 63

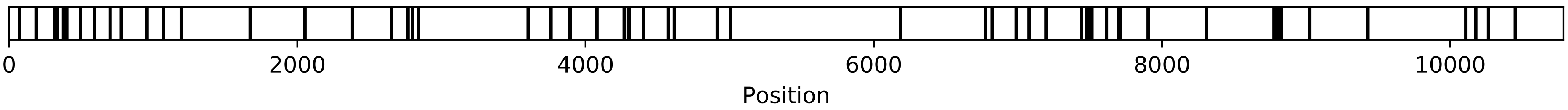

### Region: 264



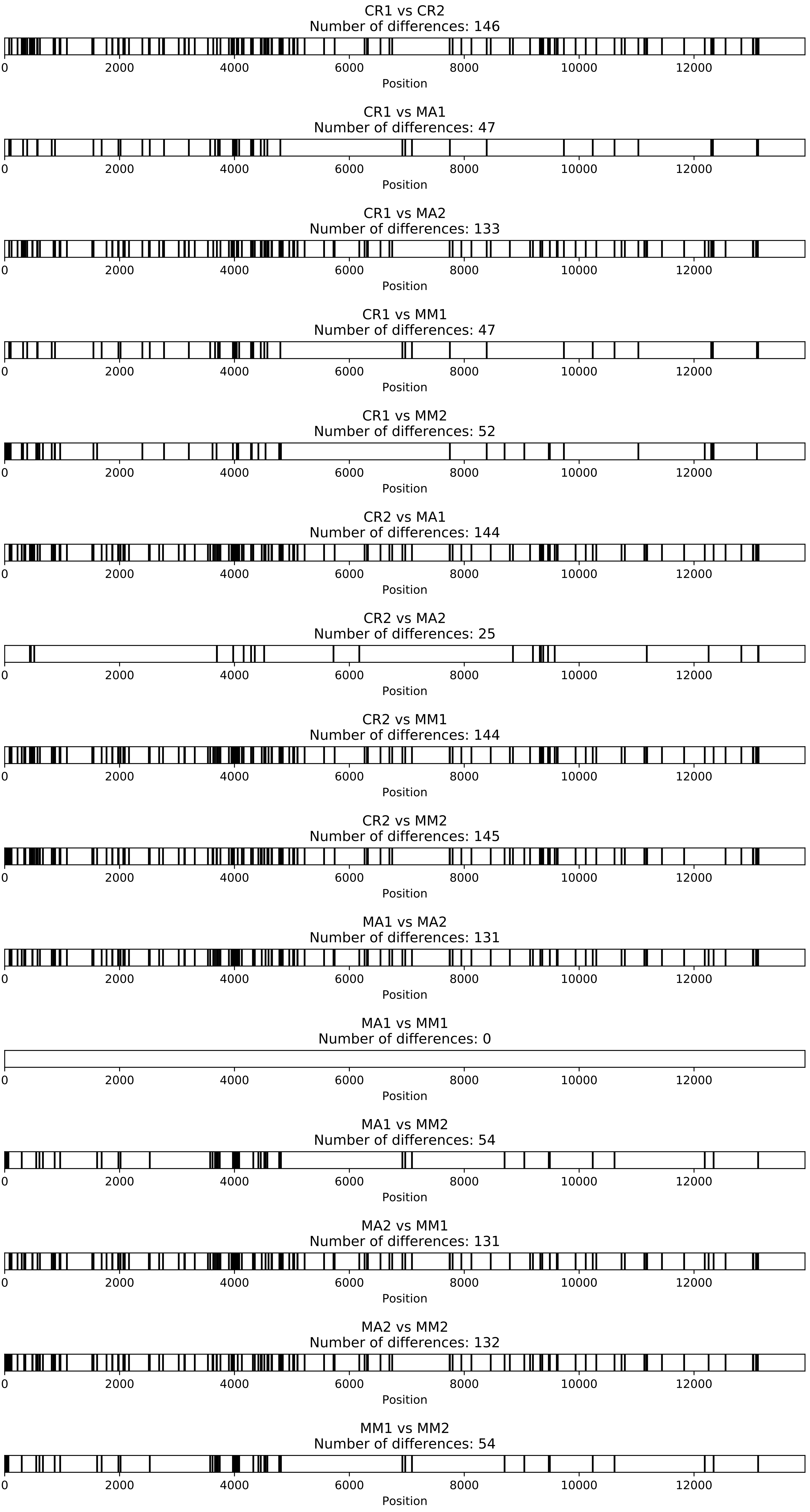
CR1 vs CR2
Number of differences: 146
CR1 vs MA1
Number of differences: 47
CR1 vs MA2
Number of differences: 133
CR1 vs MM1
Number of differences: 47
CR1 vs MM2
Number of differences: 52
CR2 vs MA1
Number of differences: 144
CR2 vs MA2
Number of differences: 25
CR2 vs MM1
Number of differences: 144
CR2 vs MM2
Number of differences: 145
MA1 vs MA2
Number of differences: 131
MA1 vs MM1
Number of differences: 0
MA1 vs MM2
Number of differences: 54
MA2 vs MM1
Number of differences: 131
MA2 vs MM2
Number of differences: 132
MM1 vs MM2
Number of differences: 54
0
2000
4000
6000
8000
10000
12000
Position

### Region: 265



CR1 vs CR2
Number of differences: 131

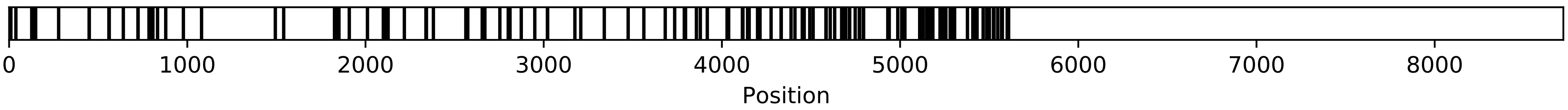


CR1 vs MA1
Number of differences: 158

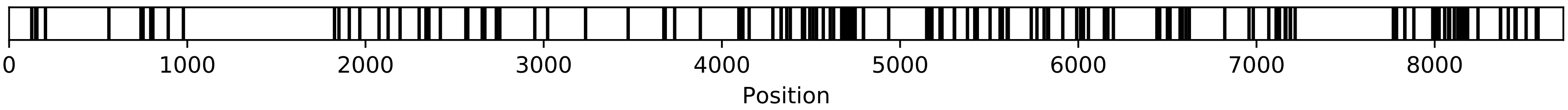


CR1 vs MA2
Number of differences: 249

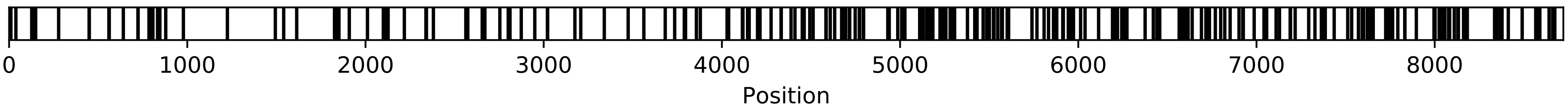


CR1 vs MM1
Number of differences: 125

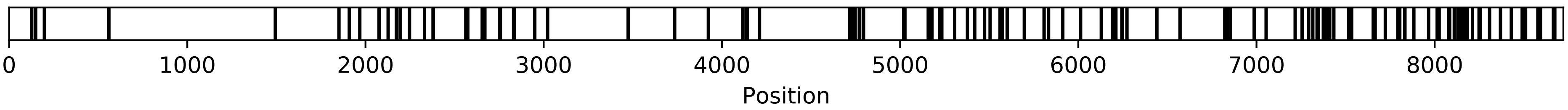


CR1 vs MM2
Number of differences: 122

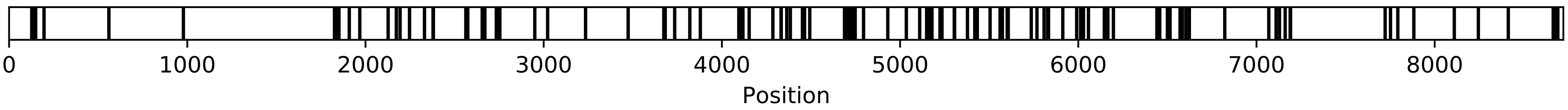


CR2 vs MA1
Number of differences: 186

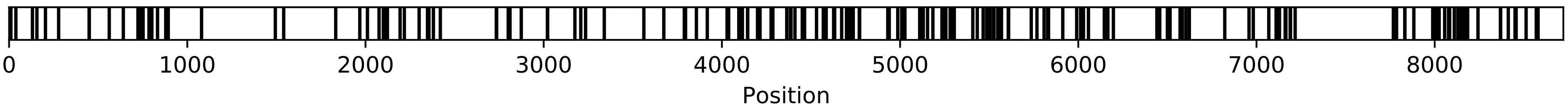


CR2 vs MA2
Number of differences: 124

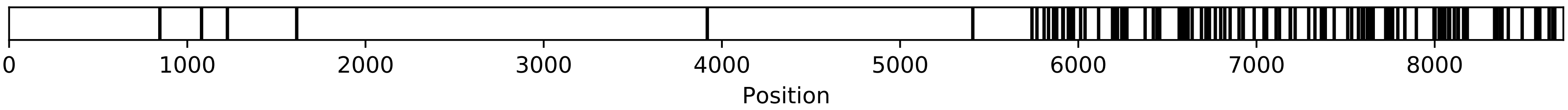


CR2 vs MM1
Number of differences: 191

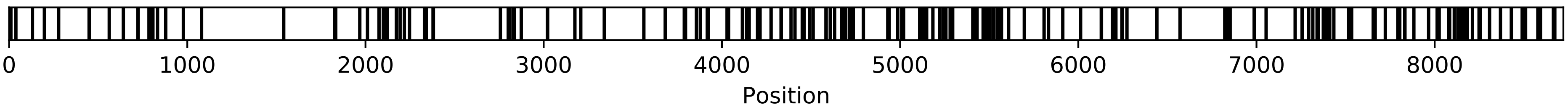


CR2 vs MM2
Number of differences: 162

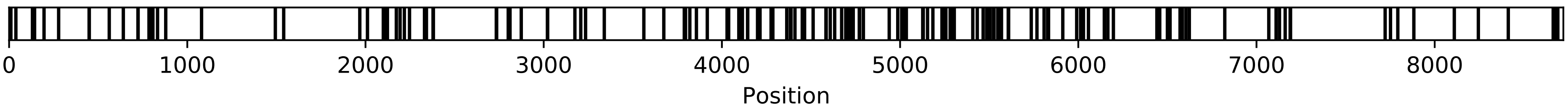


MA1 vs MA2
Number of differences: 246

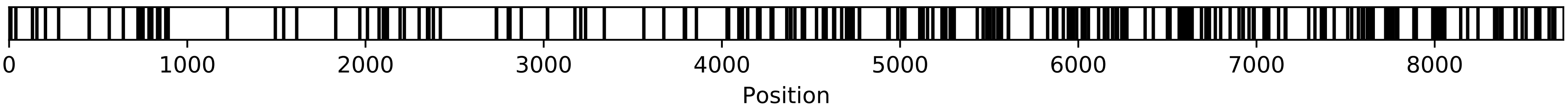


MA1 vs MM1
Number of differences: 161

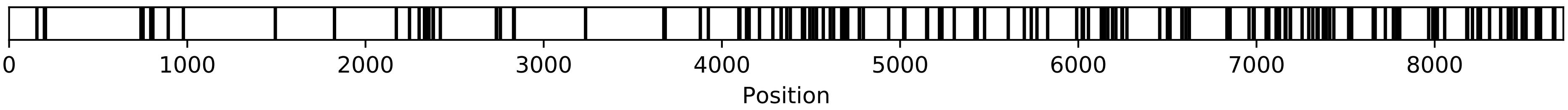


MA1 vs MM2
Number of differences: 78

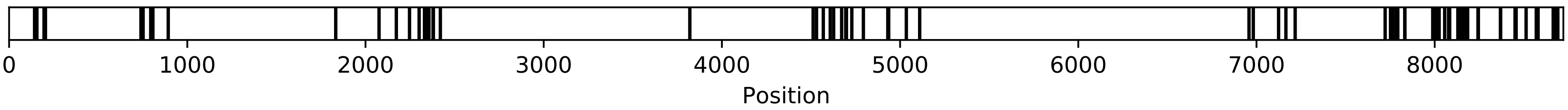


MA2 vs MM1
Number of differences: 234

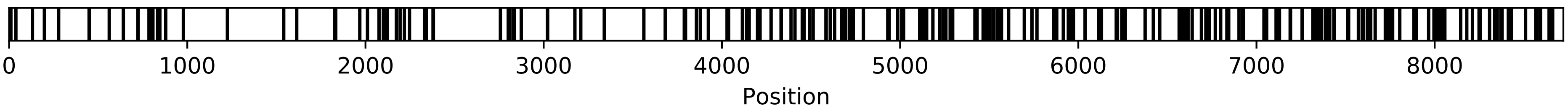


MA2 vs MM2
Number of differences: 242

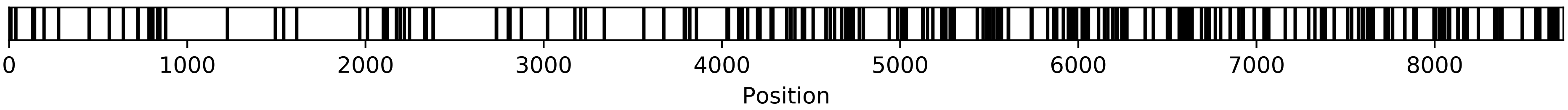


MM1 vs MM2
Number of differences: 143

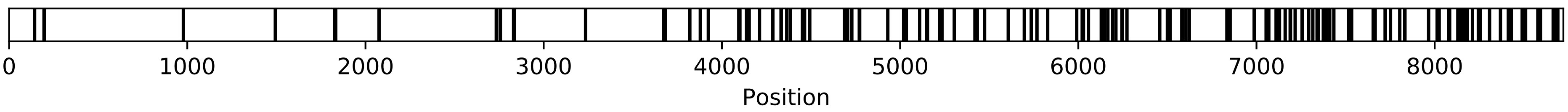

### Region: 266

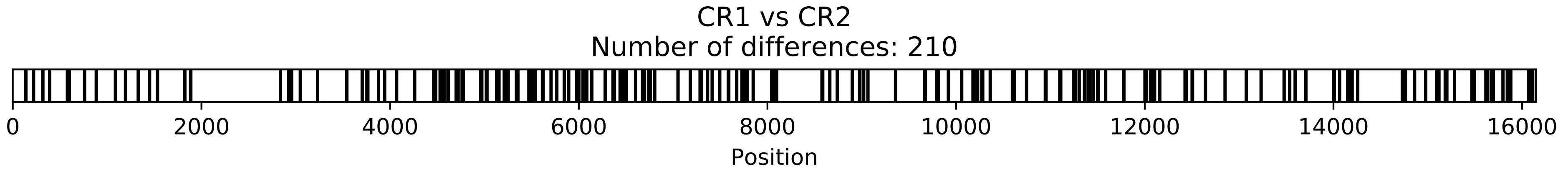
CR1 vs CR2
Number of differences: 210
0 2000 4000 6000 8000 10000 12000 14000 16000
Position

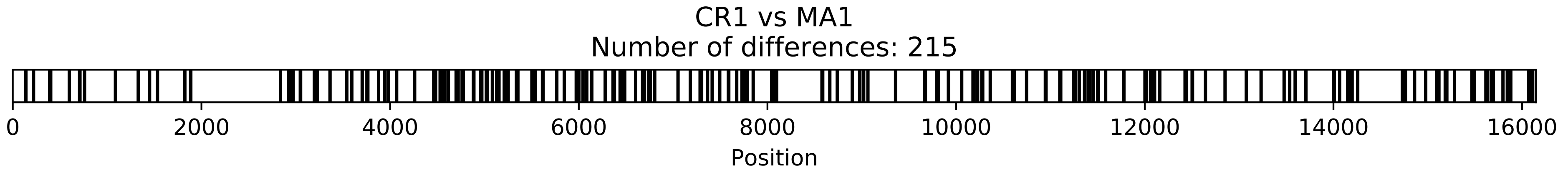
CR1 vs MA1
Number of differences: 215
0 2000 4000 6000 8000 10000 12000 14000 16000
Position

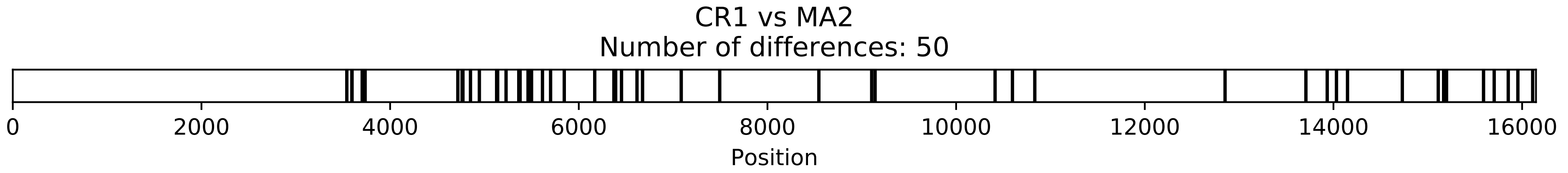
CR1 vs MA2
Number of differences: 50
0 2000 4000 6000 8000 10000 12000 14000 16000
Position

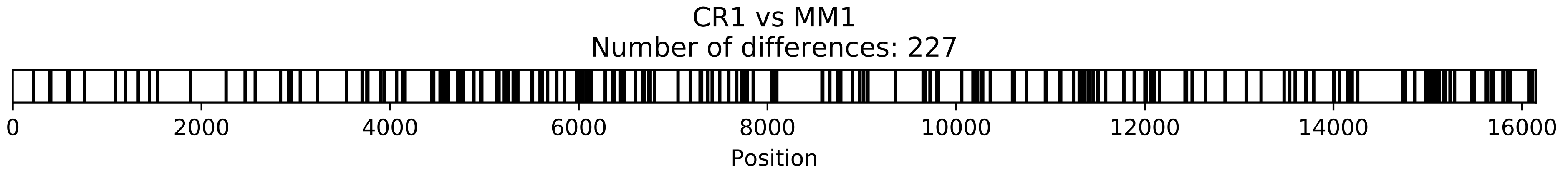
CR1 vs MM1
Number of differences: 227
0 2000 4000 6000 8000 10000 12000 14000 16000
Position

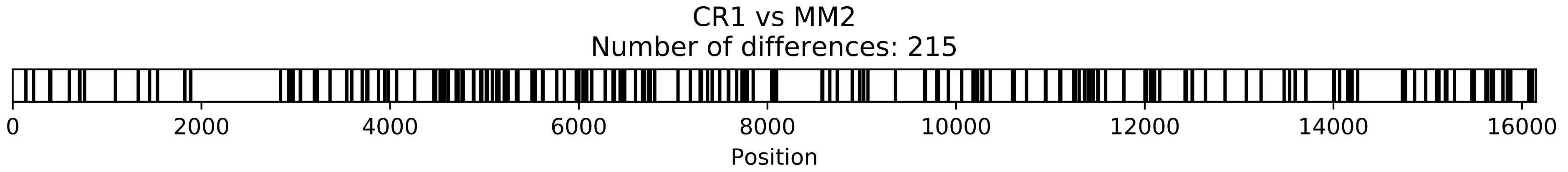
CR1 vs MM2
Number of differences: 215
0 2000 4000 6000 8000 10000 12000 14000 16000
Position

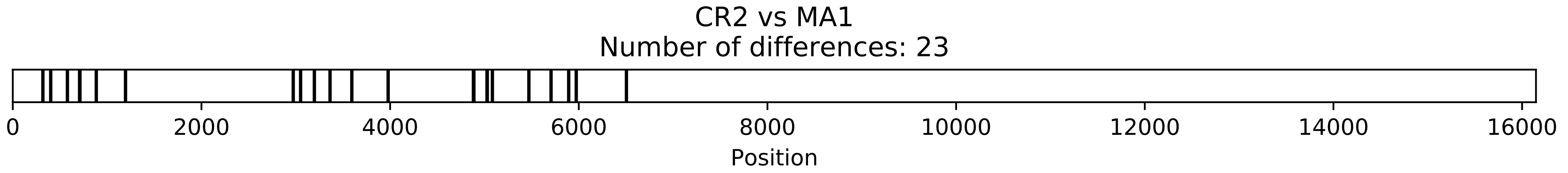
CR2 vs MA1
Number of differences: 23
0 2000 4000 6000 8000 10000 12000 14000 16000
Position

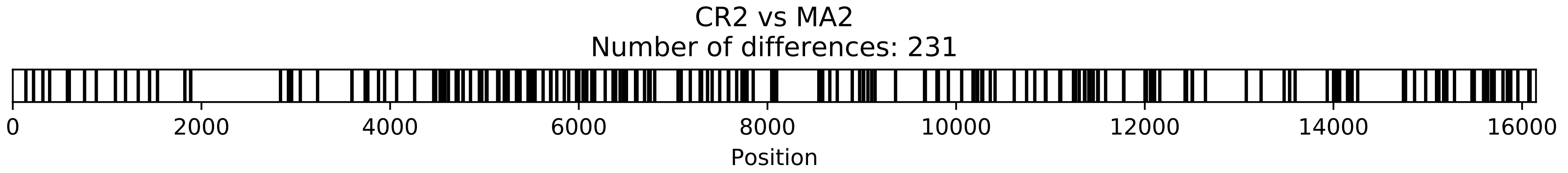
CR2 vs MA2
Number of differences: 231
0 2000 4000 6000 8000 10000 12000 14000 16000
Position

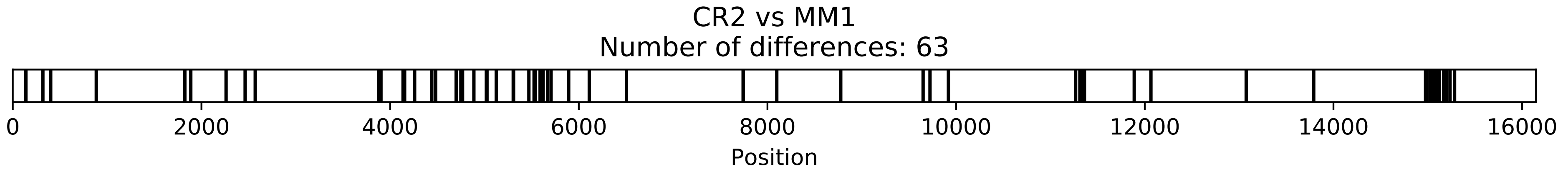
CR2 vs MM1
Number of differences: 63
0 2000 4000 6000 8000 10000 12000 14000 16000
Position

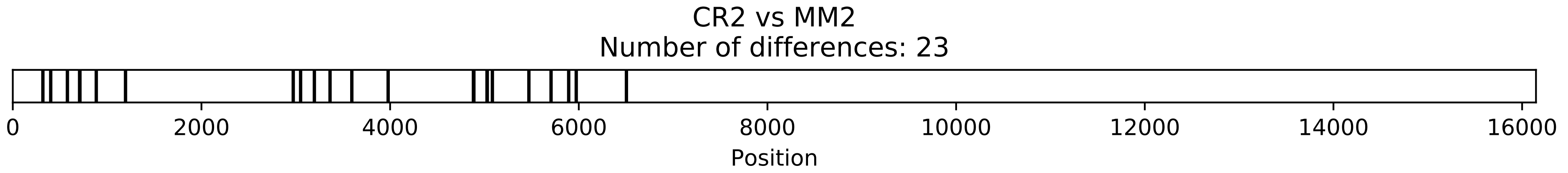
CR2 vs MM2
Number of differences: 23
0 2000 4000 6000 8000 10000 12000 14000 16000
Position

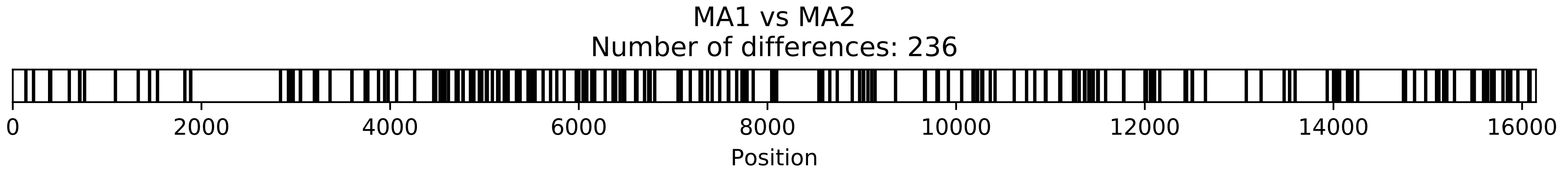
MA1 vs MA2
Number of differences: 236
0 2000 4000 6000 8000 10000 12000 14000 16000
Position

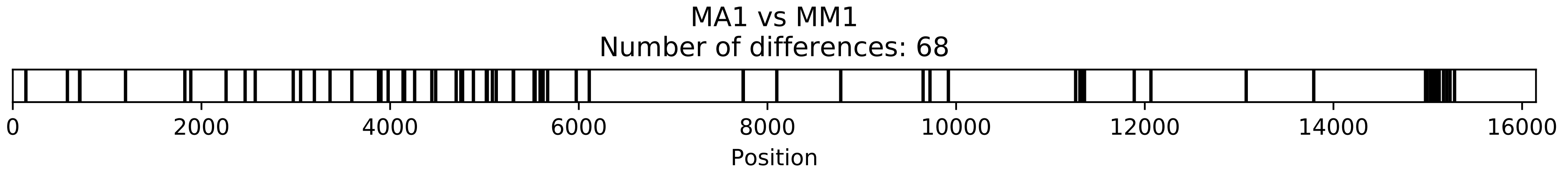
MA1 vs MM1
Number of differences: 68
0 2000 4000 6000 8000 10000 12000 14000 16000
Position

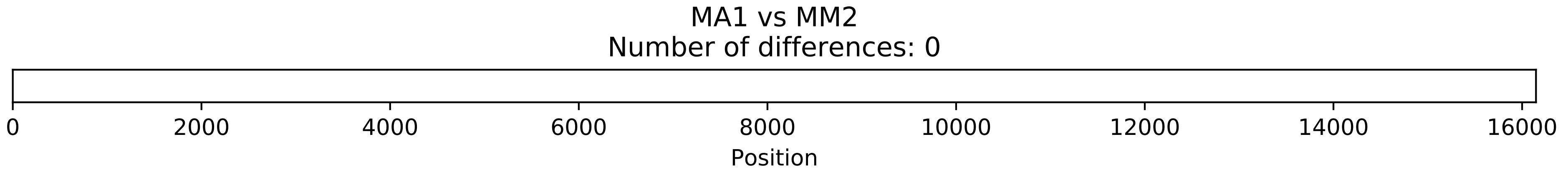
MA1 vs MM2
Number of differences: 0
0 2000 4000 6000 8000 10000 12000 14000 16000
Position

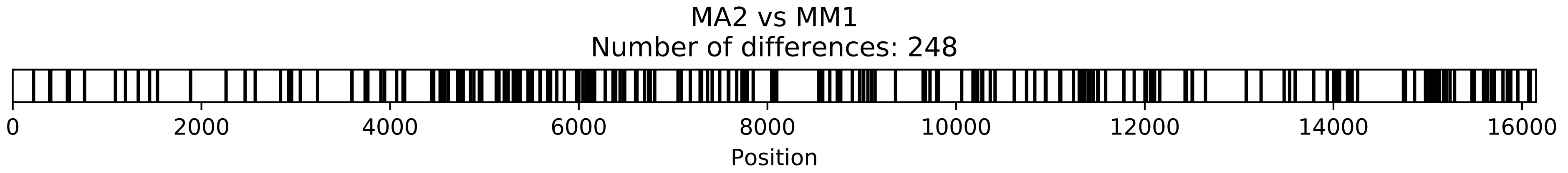
MA2 vs MM1
Number of differences: 248
0 2000 4000 6000 8000 10000 12000 14000 16000
Position

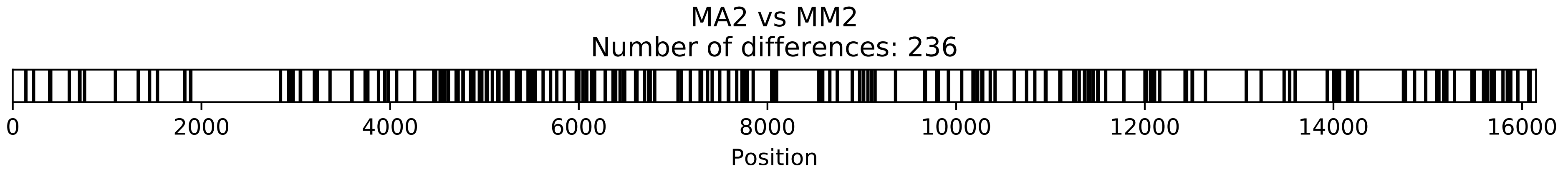
MA2 vs MM2
Number of differences: 236
0 2000 4000 6000 8000 10000 12000 14000 16000
Position

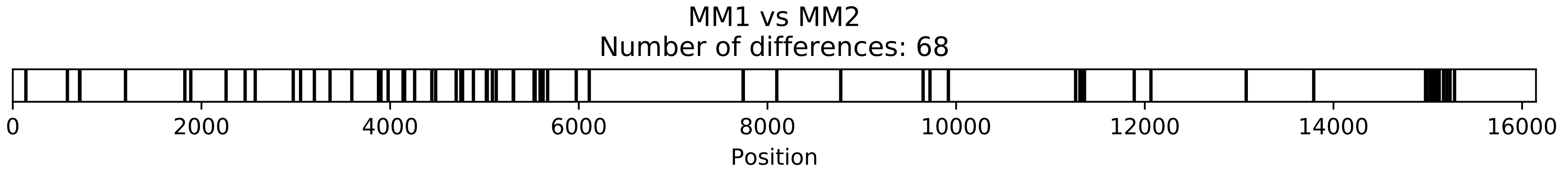
MM1 vs MM2
Number of differences: 68
0 2000 4000 6000 8000 10000 12000 14000 16000
Position

### Region: 267



CR1 vs CR2
Number of differences: 412

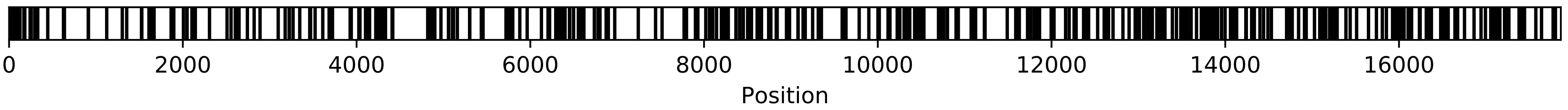


CR1 vs MA1
Number of differences: 439

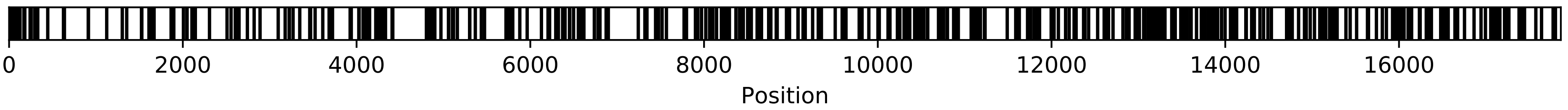


CR1 vs MA2
Number of differences: 202

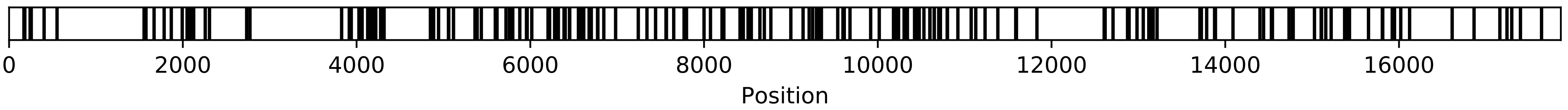


CR1 vs MM1
Number of differences: 193

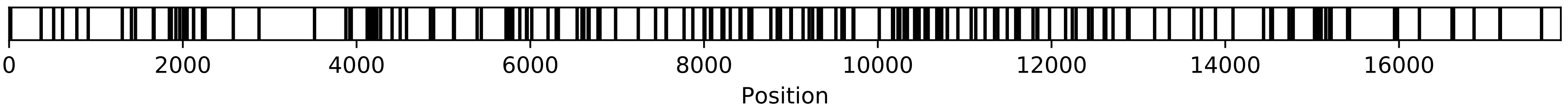


CR1 vs MM2
Number of differences: 184

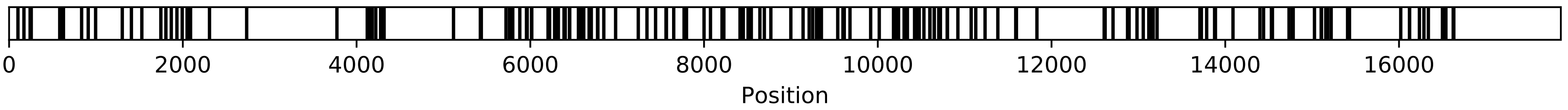


CR2 vs MA1
Number of differences: 54

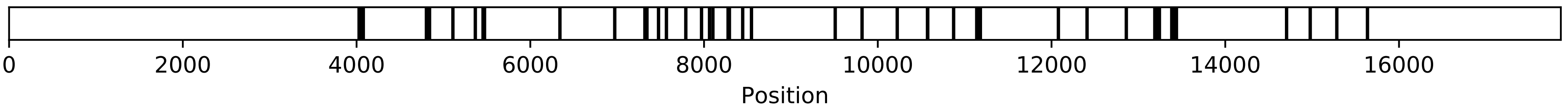


CR2 vs MA2
Number of differences: 427

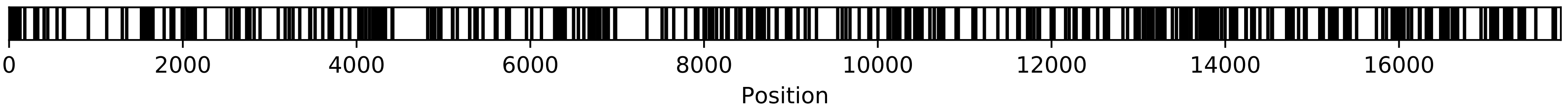


CR2 vs MM1
Number of differences: 427

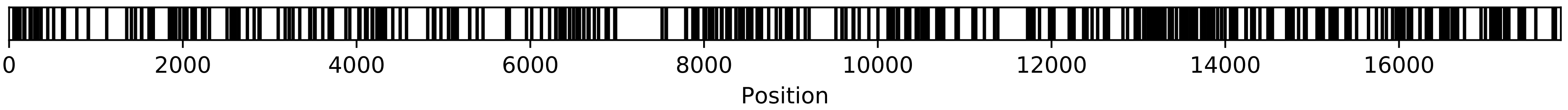


CR2 vs MM2
Number of differences: 426

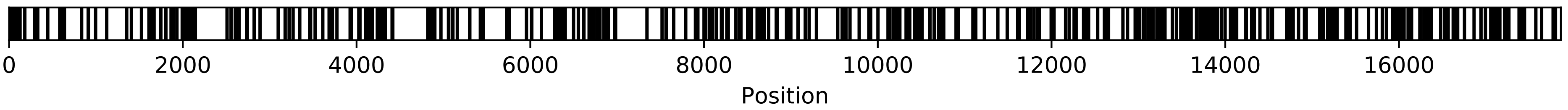


MA1 vs MA2
Number of differences: 450

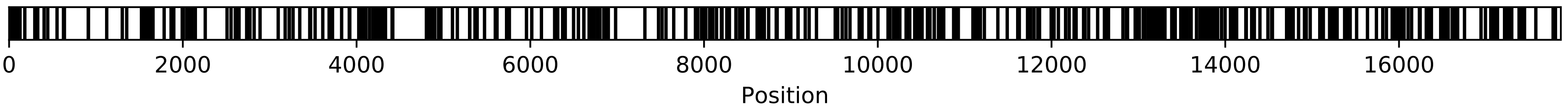


MA1 vs MM1
Number of differences: 452

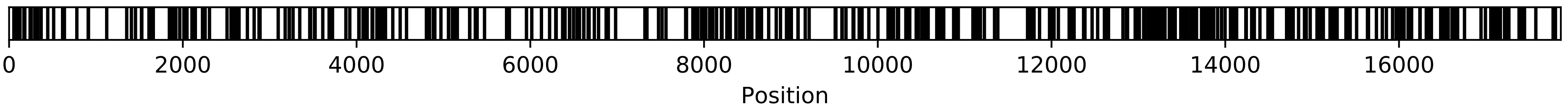


MA1 vs MM2
Number of differences: 449

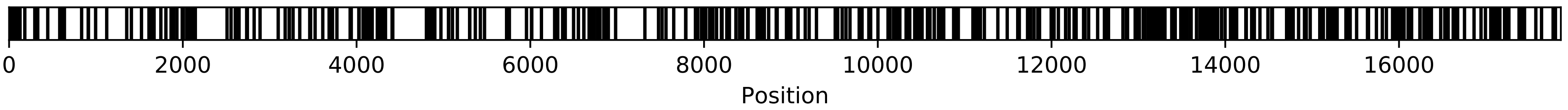


MA2 vs MM1
Number of differences: 190

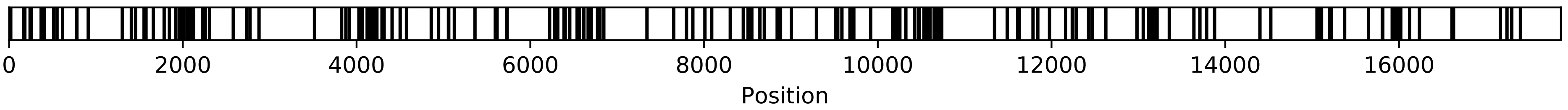


MA2 vs MM2
Number of differences: 78

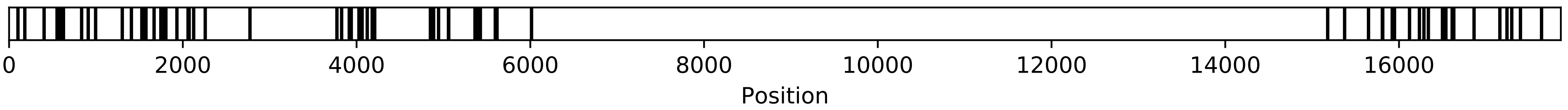


MM1 vs MM2
Number of differences: 190

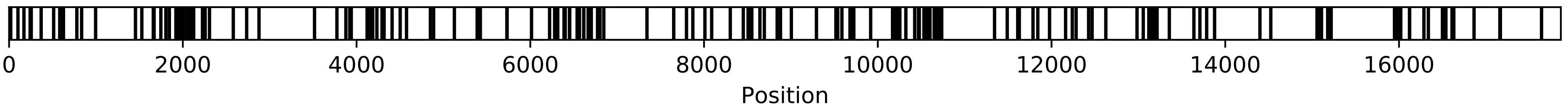

### Region: 268



CR1 vs CR2
Number of differences: 85

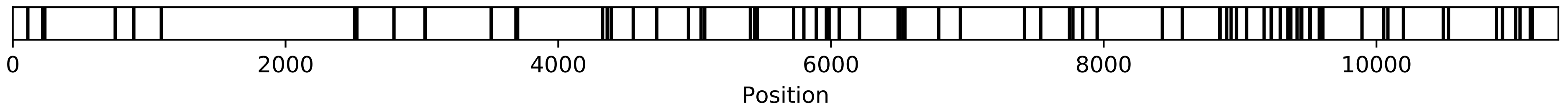

CR1 vs MA1
Number of differences: 42

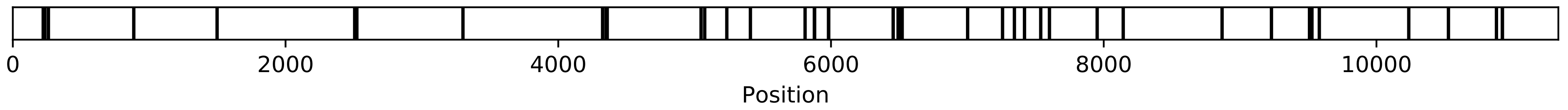

CR1 vs MA2
Number of differences: 90

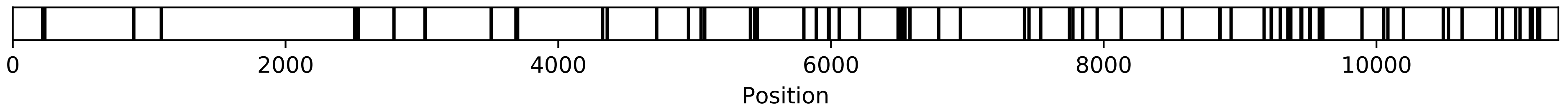

CR1 vs MM1
Number of differences: 42

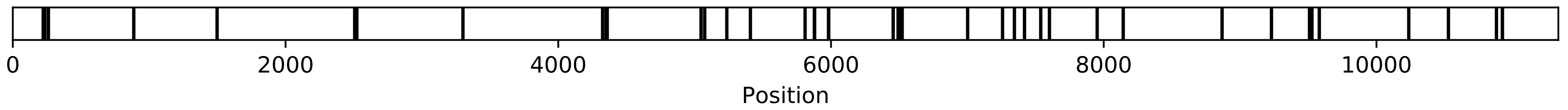

CR1 vs MM2
Number of differences: 33

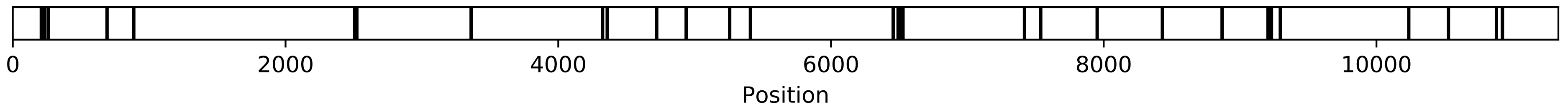

CR2 vs MA1
Number of differences: 88

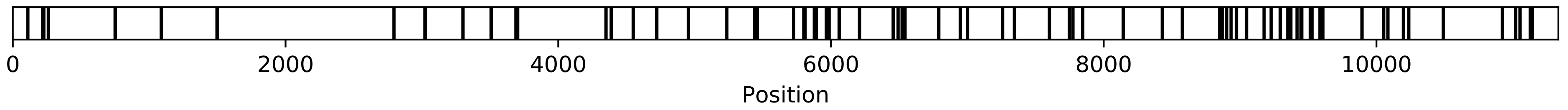

CR2 vs MA2
Number of differences: 27

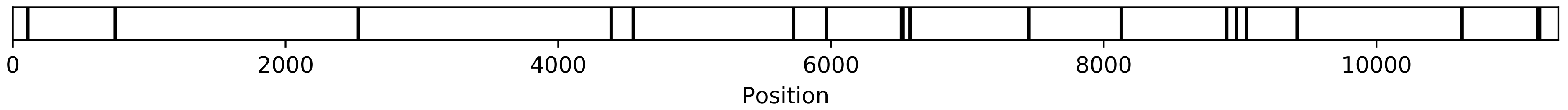

CR2 vs MM1
Number of differences: 88

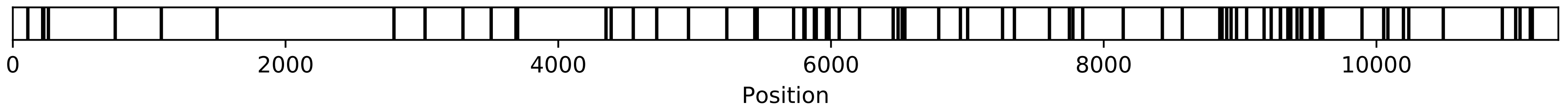

CR2 vs MM2
Number of differences: 81

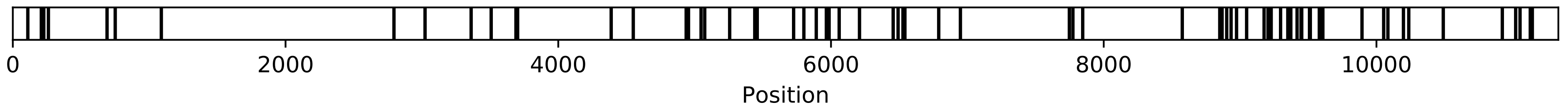

MA1 vs MA2
Number of differences: 93

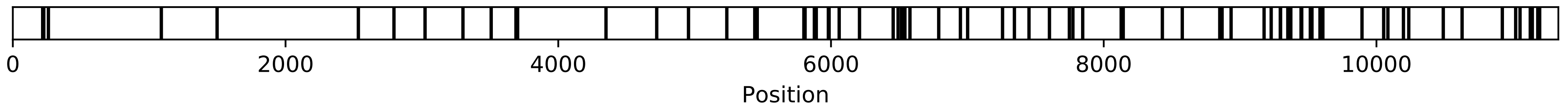

MA1 vs MM1
Number of differences: 0

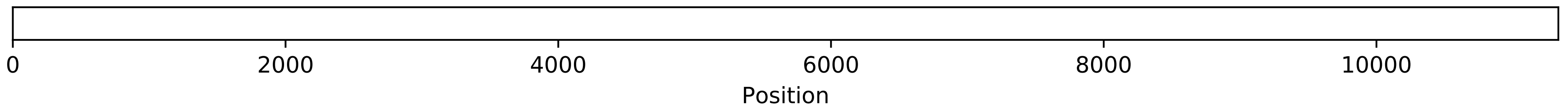

MA1 vs MM2
Number of differences: 33

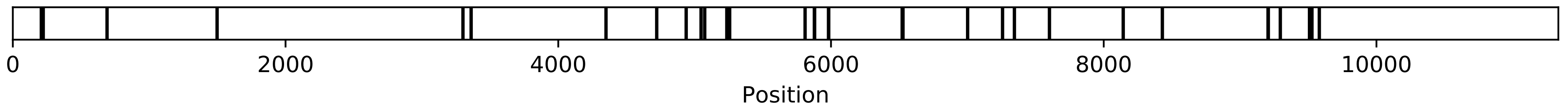

MA2 vs MM1
Number of differences: 93

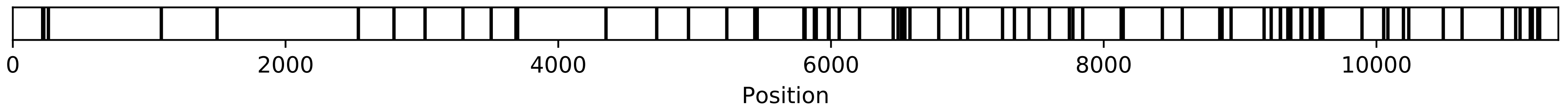

MA2 vs MM2
Number of differences: 86

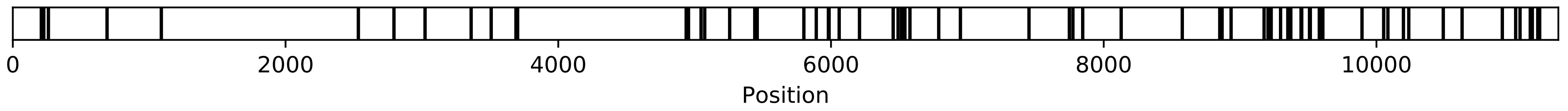

MM1 vs MM2
Number of differences: 33

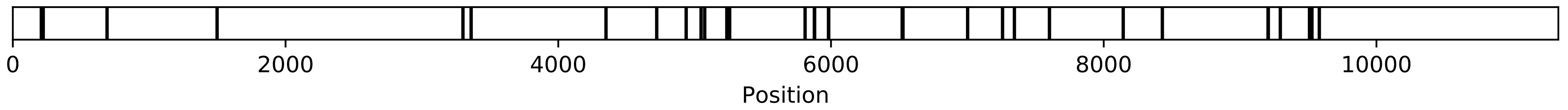

### Region: 270

CR1 vs CR2
Number of differences: 121

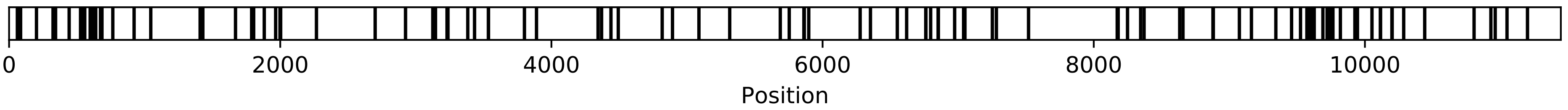


CR1 vs MA1
Number of differences: 39

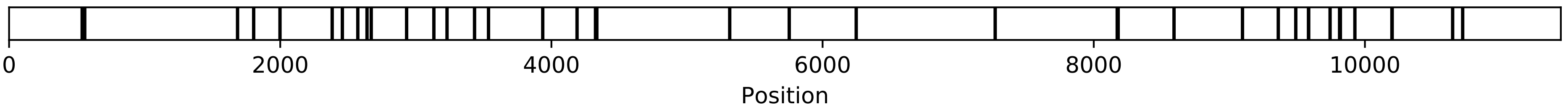


CR1 vs MA2
Number of differences: 124

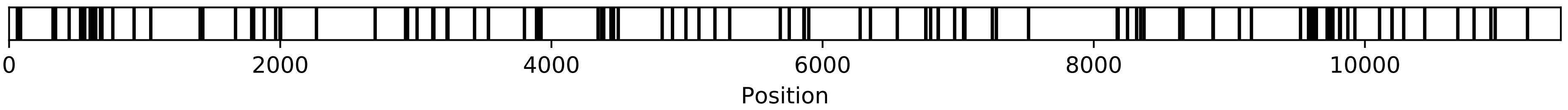


CR1 vs MM1
Number of differences: 59

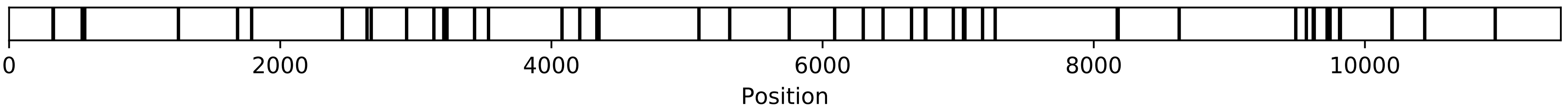


CR1 vs MM2
Number of differences: 39

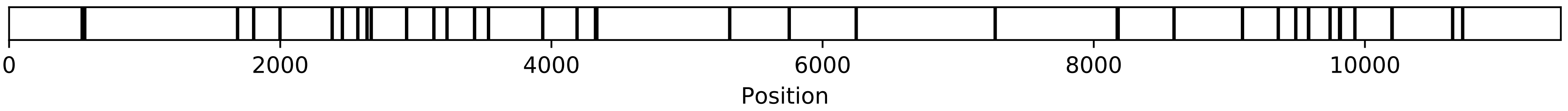


CR2 vs MA1
Number of differences: 127

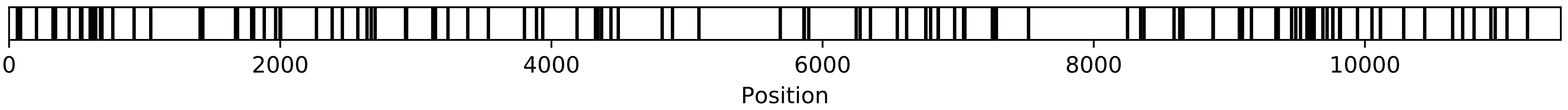


CR2 vs MA2
Number of differences: 32

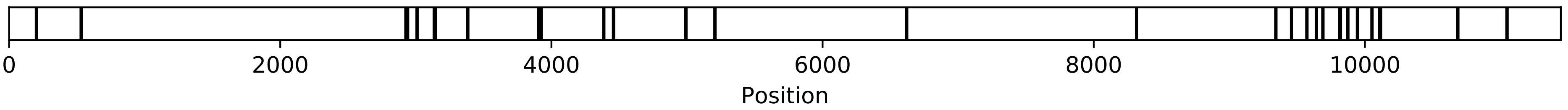


CR2 vs MM1
Number of differences: 126

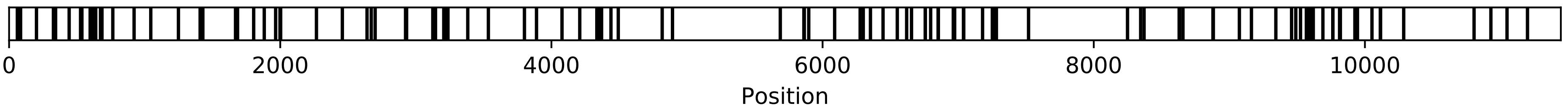


CR2 vs MM2
Number of differences: 127

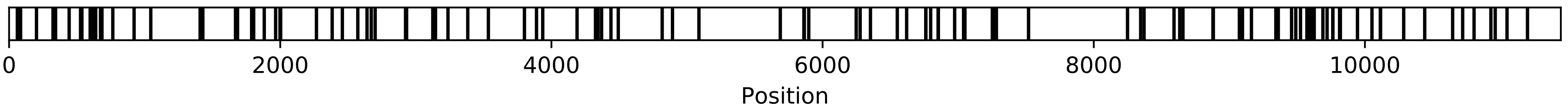


MA1 vs MA2
Number of differences: 128

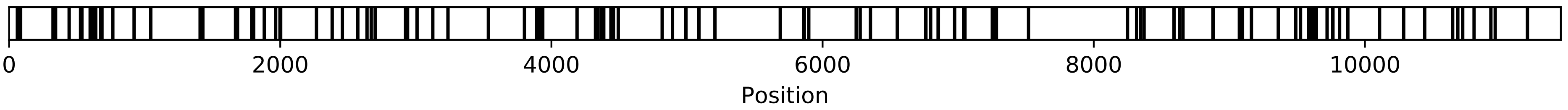


MA1 vs MM1
Number of differences: 52

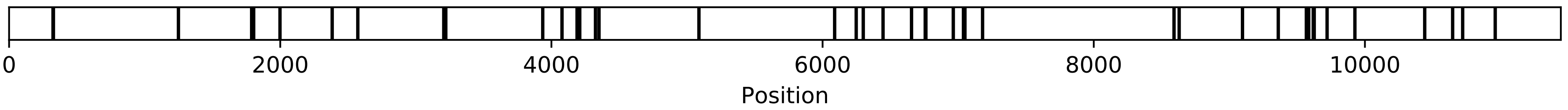


MA1 vs MM2
Number of differences: 0

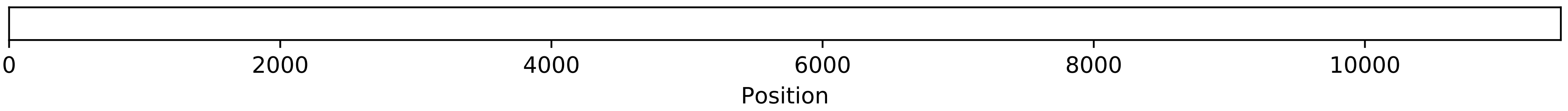


MA2 vs MM1
Number of differences: 127

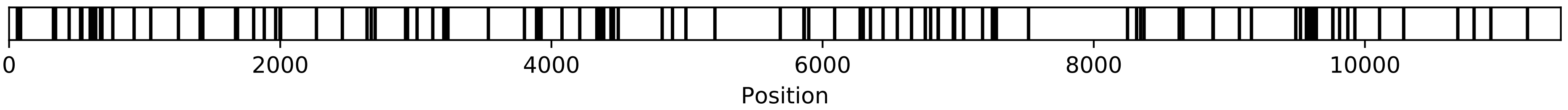


MA2 vs MM2
Number of differences: 128

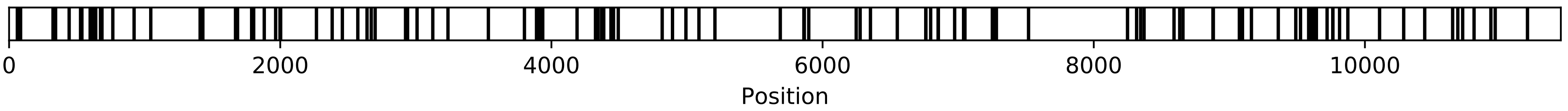


MM1 vs MM2
Number of differences: 52

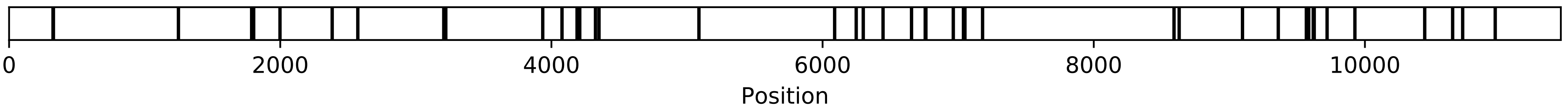

### Region: 273



CR1 vs CR2
Number of differences: 214

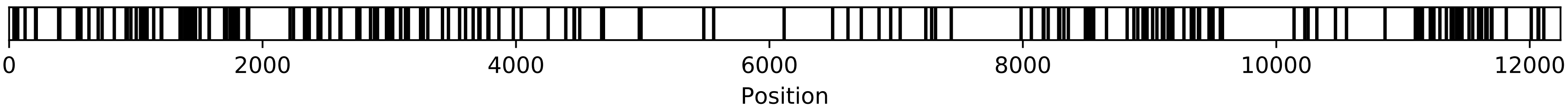


CR1 vs MA1
Number of differences: 95

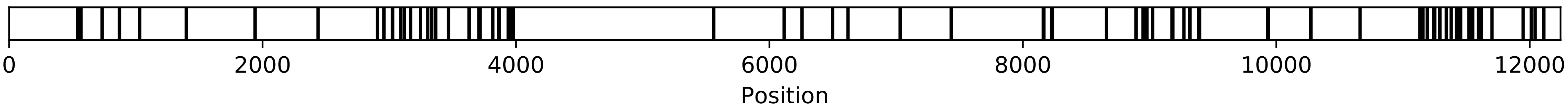


CR1 vs MA2
Number of differences: 212

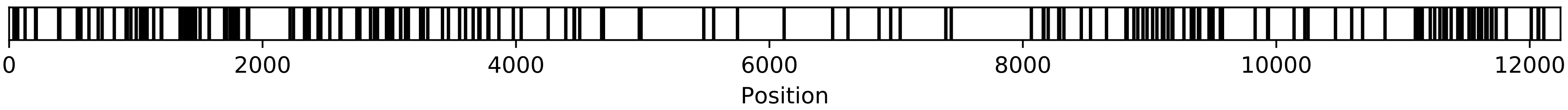


CR1 vs MM1
Number of differences: 95

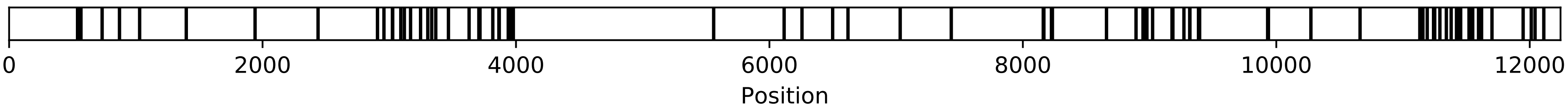


CR1 vs MM2
Number of differences: 85

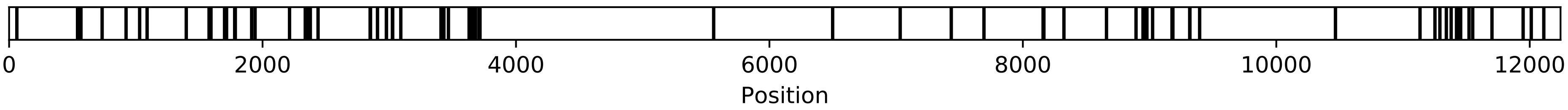


CR2 vs MA1
Number of differences: 214

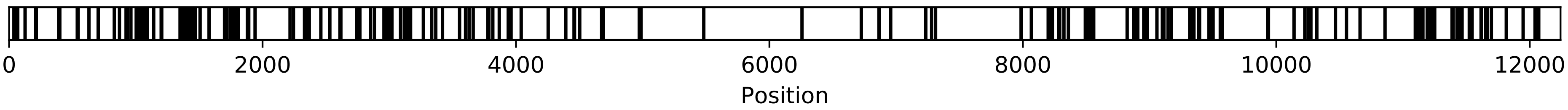


CR2 vs MA2
Number of differences: 40

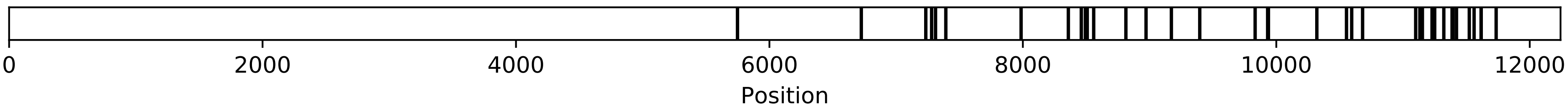


CR2 vs MM1
Number of differences: 214

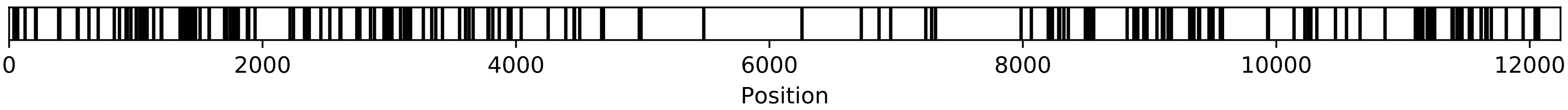


CR2 vs MM2
Number of differences: 184

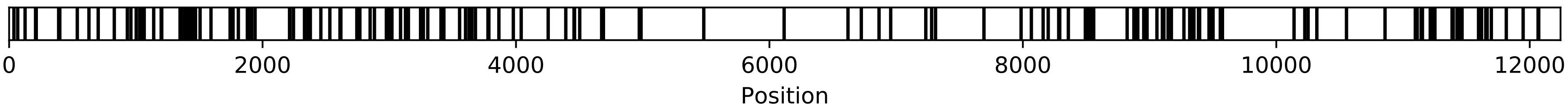


MA1 vs MA2
Number of differences: 201

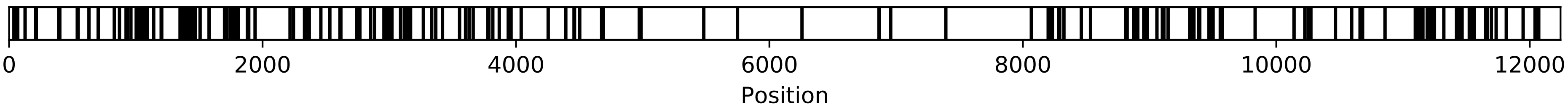


MA1 vs MM1
Number of differences: 0

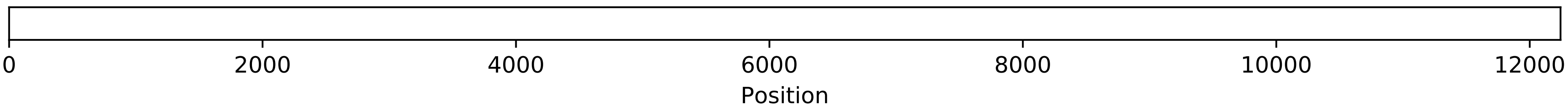


MA1 vs MM2
Number of differences: 80

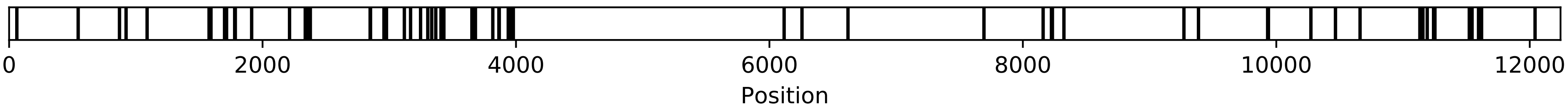


MA2 vs MM1
Number of differences: 201

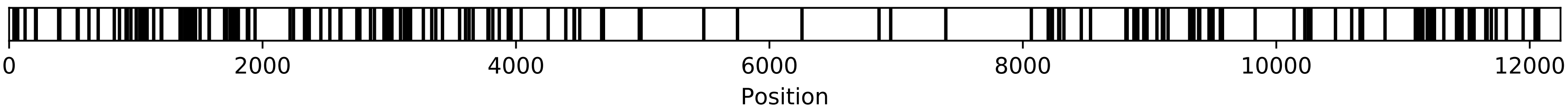


MA2 vs MM2
Number of differences: 184

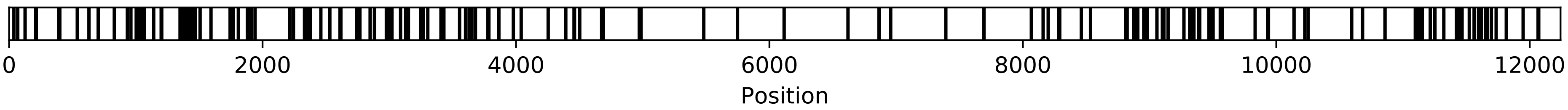


MM1 vs MM2
Number of differences: 80

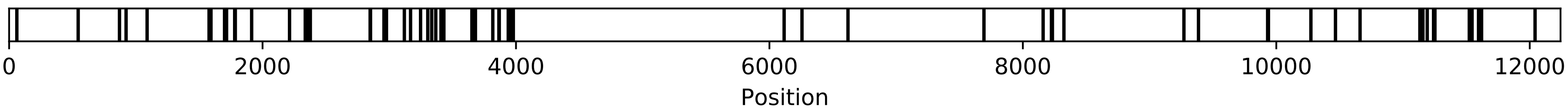

### Region: 275



CR1 vs CR2
Number of differences: 286

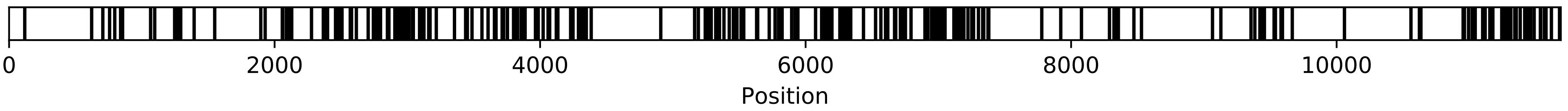


CR1 vs MA1
Number of differences: 156

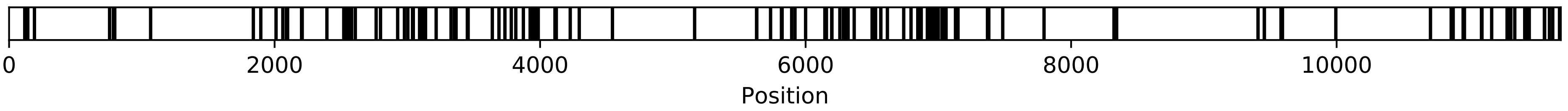


CR1 vs MA2
Number of differences: 286

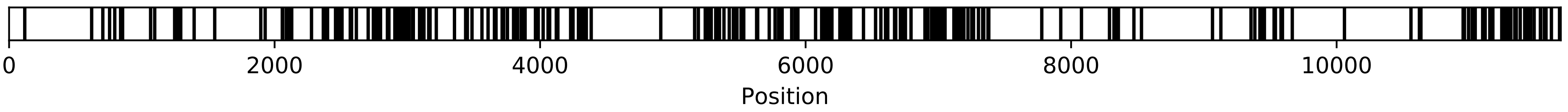


CR1 vs MM1
Number of differences: 149

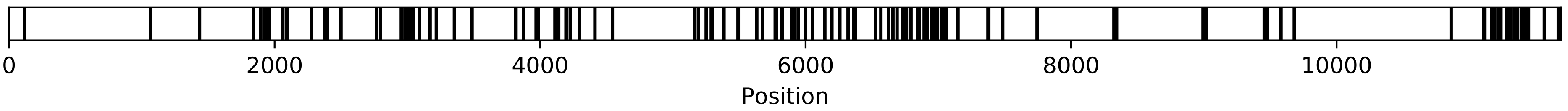


CR1 vs MM2
Number of differences: 156

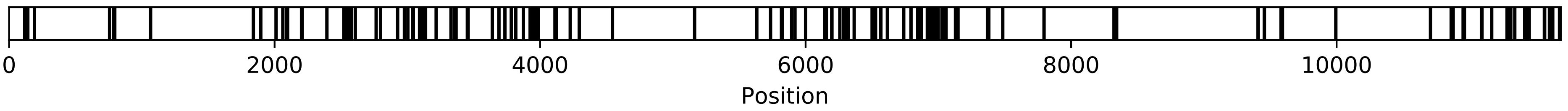


CR2 vs MA1
Number of differences: 290

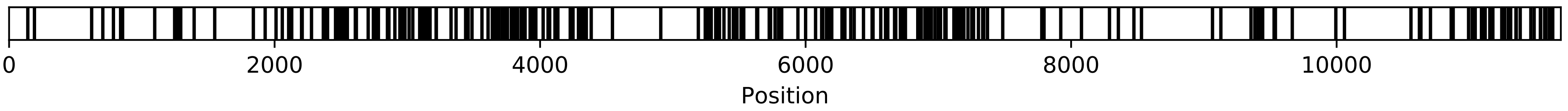


CR2 vs MA2
Number of differences: 0

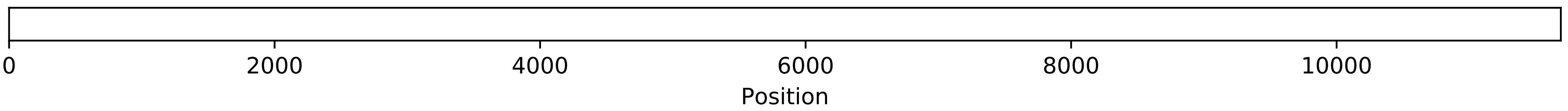


CR2 vs MM1
Number of differences: 284

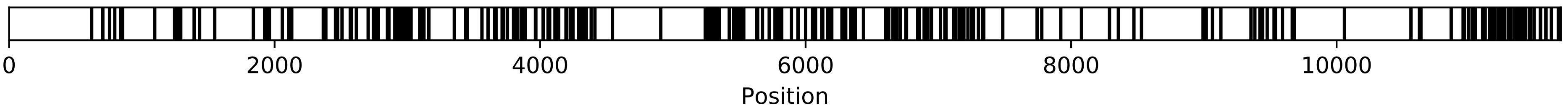


CR2 vs MM2
Number of differences: 290

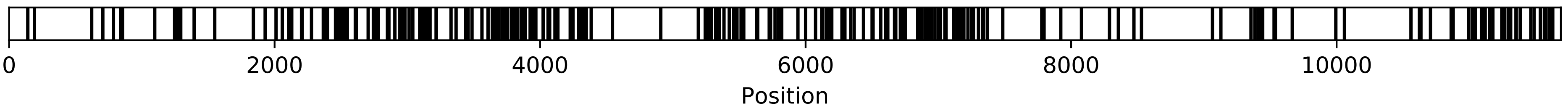


MA1 vs MA2
Number of differences: 290

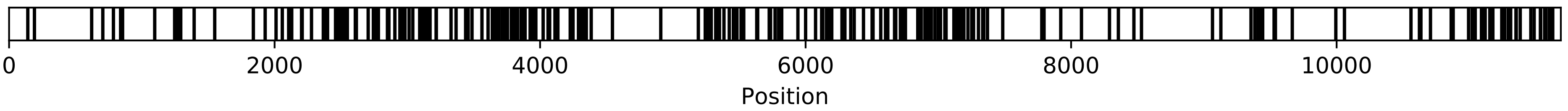


MA1 vs MM1
Number of differences: 165

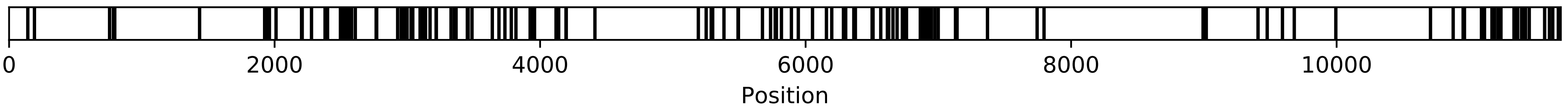


MA1 vs MM2
Number of differences: 0

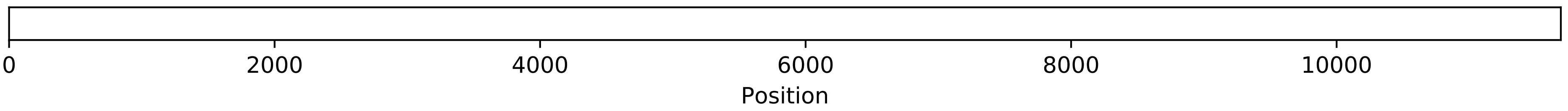


MA2 vs MM1
Number of differences: 284

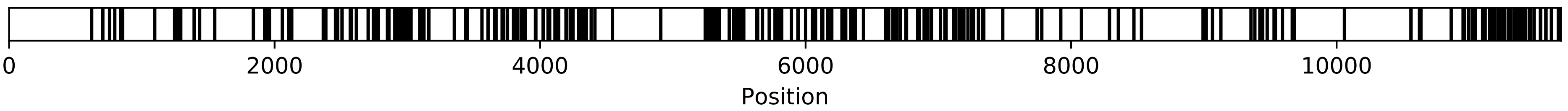


MA2 vs MM2
Number of differences: 290

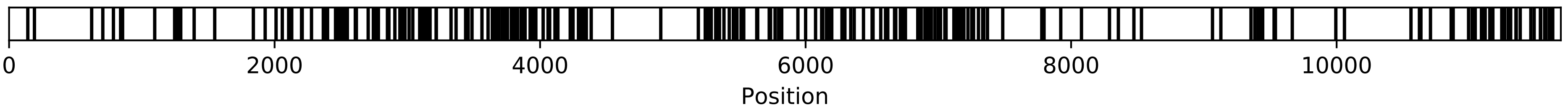


MM1 vs MM2
Number of differences: 165

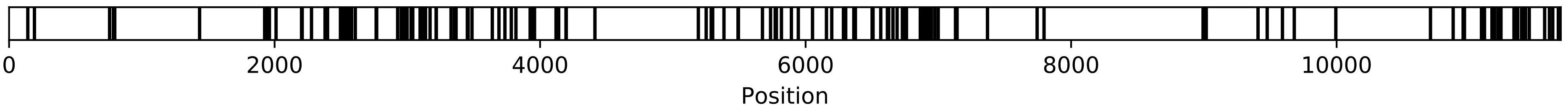

### Region: 276

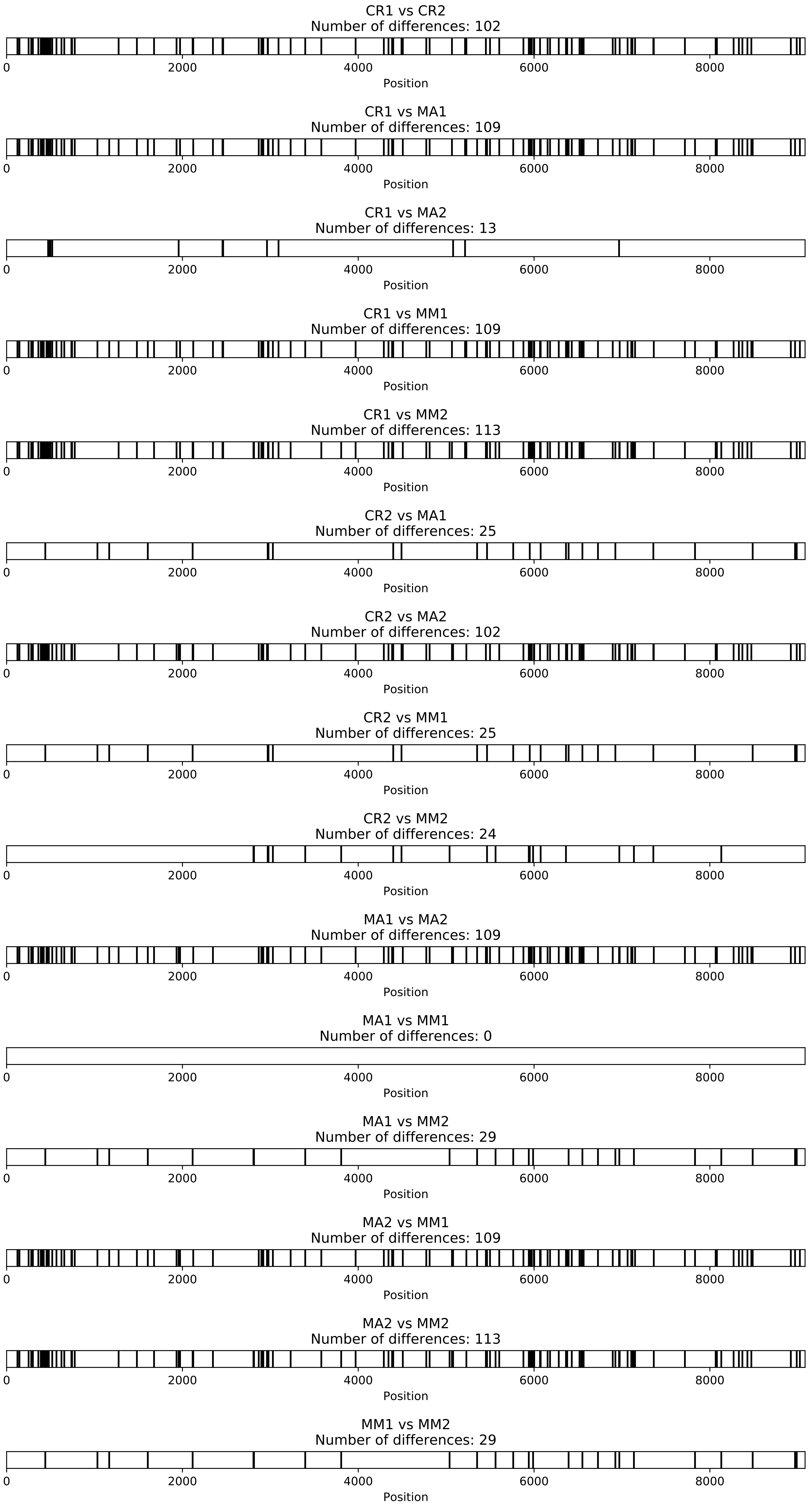
CR1 vs CR2
Number of differences: 102
CR1 vs MA1
Number of differences: 109
CR1 vs MA2
Number of differences: 13
CR1 vs MM1
Number of differences: 109
CR1 vs MM2
Number of differences: 113
CR2 vs MA1
Number of differences: 25
CR2 vs MA2
Number of differences: 102
CR2 vs MM1
Number of differences: 25
CR2 vs MM2
Number of differences: 24
MA1 vs MA2
Number of differences: 109
MA1 vs MM1
Number of differences: 0
MA1 vs MM2
Number of differences: 29
MA2 vs MM1
Number of differences: 109
MA2 vs MM2
Number of differences: 113
MM1 vs MM2
Number of differences: 29
0
2000
4000
6000
8000
Position

### Region: 277



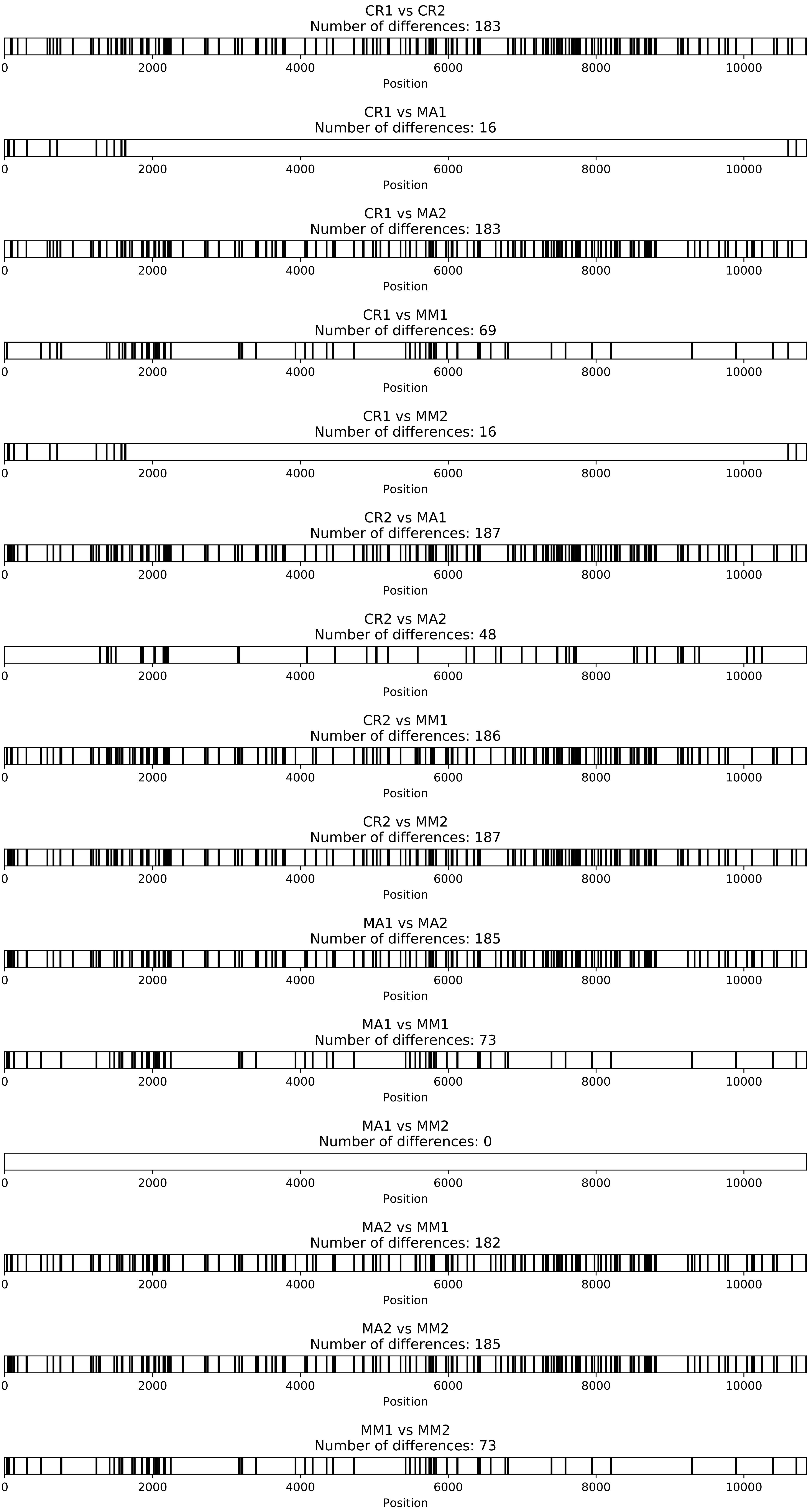
CR1 vs CR2
Number of differences: 183
CR1 vs MA1
Number of differences: 16
CR1 vs MA2
Number of differences: 183
CR1 vs MM1
Number of differences: 69
CR1 vs MM2
Number of differences: 16
CR2 vs MA1
Number of differences: 187
CR2 vs MA2
Number of differences: 48
CR2 vs MM1
Number of differences: 186
CR2 vs MM2
Number of differences: 187
MA1 vs MA2
Number of differences: 185
MA1 vs MM1
Number of differences: 73
MA1 vs MM2
Number of differences: 0
MA2 vs MM1
Number of differences: 182
MA2 vs MM2
Number of differences: 185
MM1 vs MM2
Number of differences: 73
0
2000
4000
6000
8000
10000
Position

### Region: 278

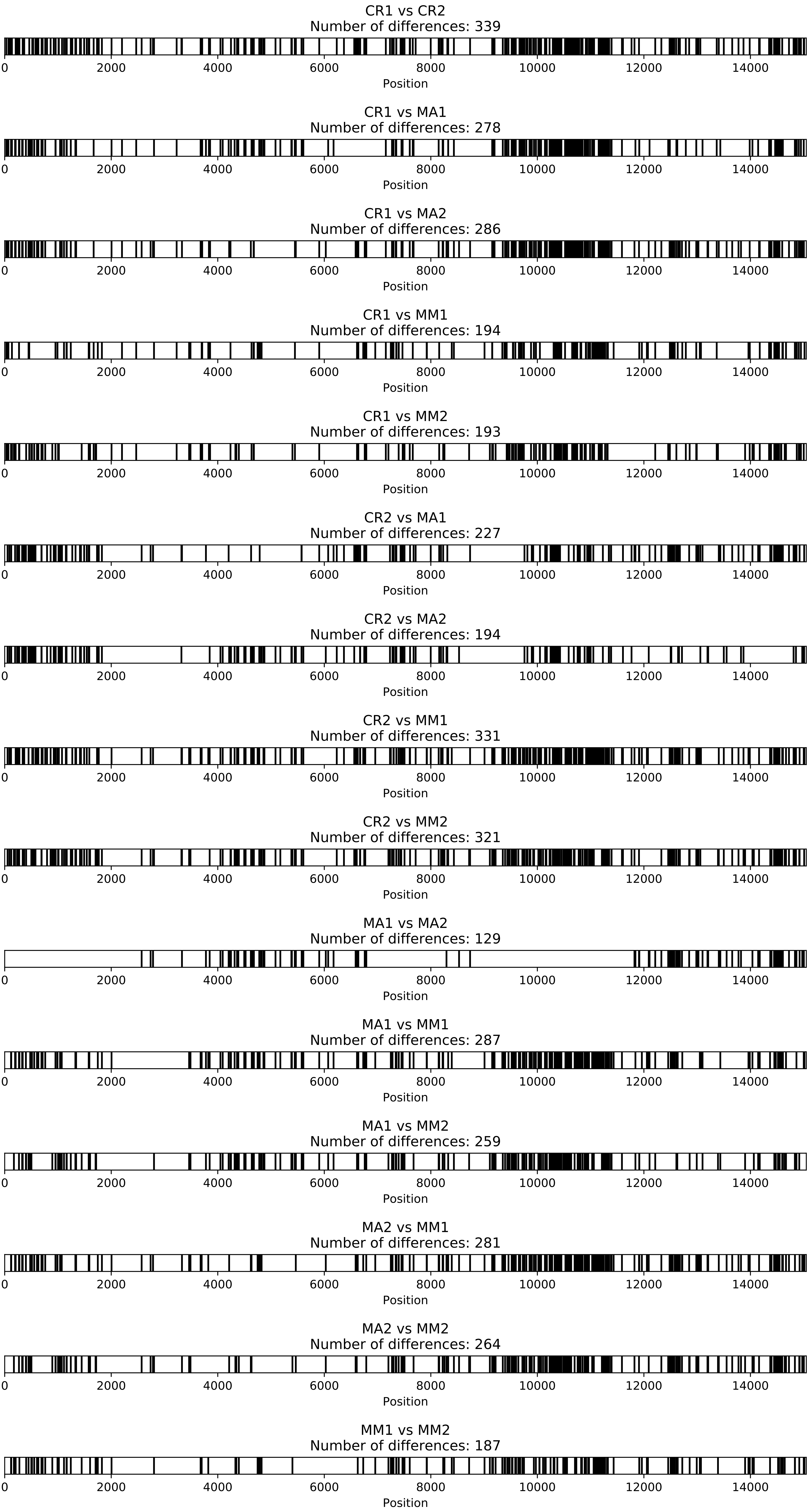
CR1 vs CR2
Number of differences: 339
CR1 vs MA1
Number of differences: 278
CR1 vs MA2
Number of differences: 286
CR1 vs MM1
Number of differences: 194
CR1 vs MM2
Number of differences: 193
CR2 vs MA1
Number of differences: 227
CR2 vs MA2
Number of differences: 194
CR2 vs MM1
Number of differences: 331
CR2 vs MM2
Number of differences: 321
MA1 vs MA2
Number of differences: 129
MA1 vs MM1
Number of differences: 287
MA1 vs MM2
Number of differences: 259
MA2 vs MM1
Number of differences: 281
MA2 vs MM2
Number of differences: 264
MM1 vs MM2
Number of differences: 187
0
2000
4000
6000
8000
10000
12000
14000
Position

### Region: 279



CR1 vs CR2
Number of differences: 264

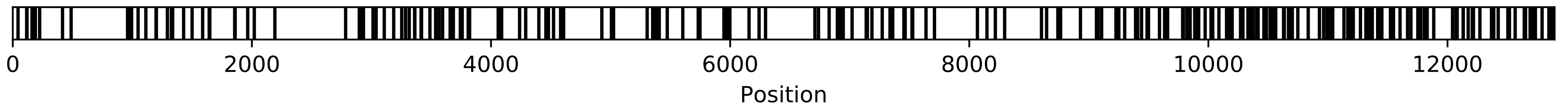


CR1 vs MA1
Number of differences: 264

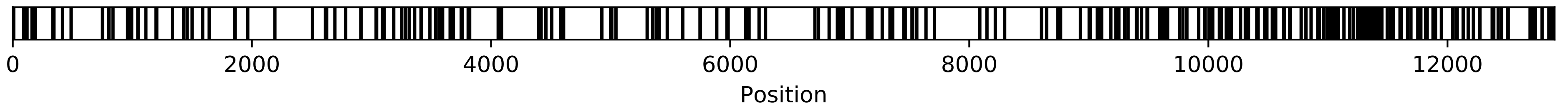


CR1 vs MA2
Number of differences: 93

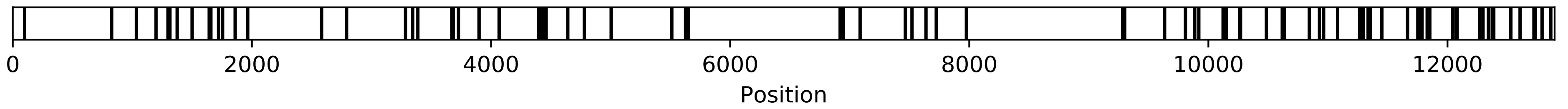


CR1 vs MM1
Number of differences: 254

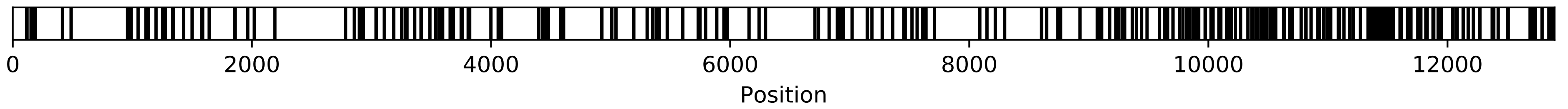


CR1 vs MM2
Number of differences: 248

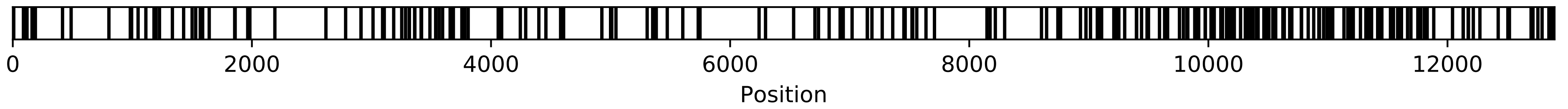


CR2 vs MA1
Number of differences: 155

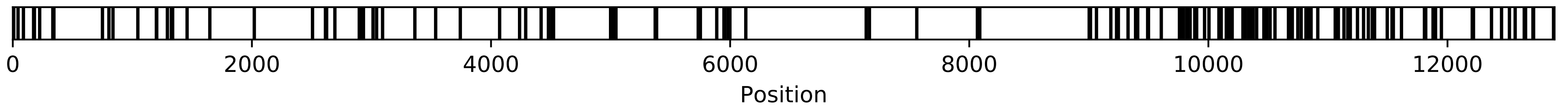


CR2 vs MA2
Number of differences: 277

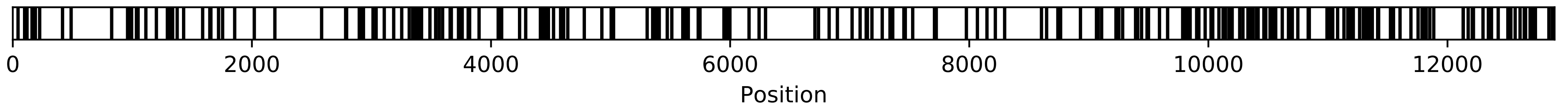


CR2 vs MM1
Number of differences: 125

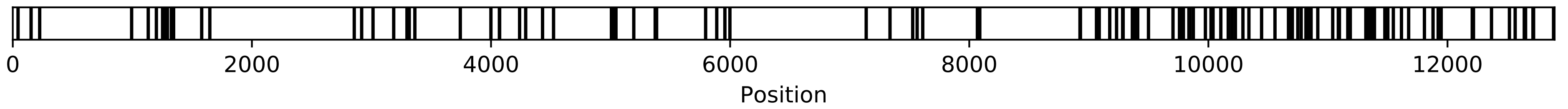


CR2 vs MM2
Number of differences: 138

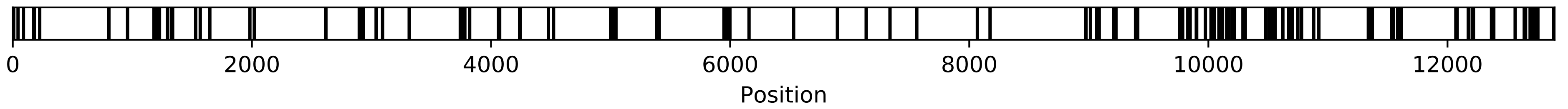


MA1 vs MA2
Number of differences: 279

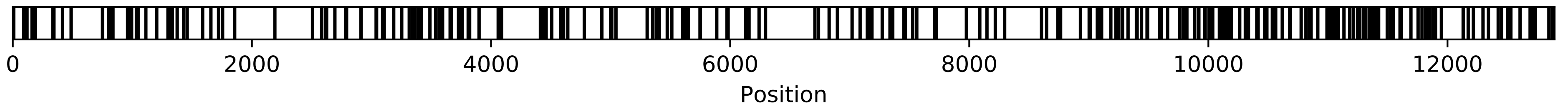


MA1 vs MM1
Number of differences: 130

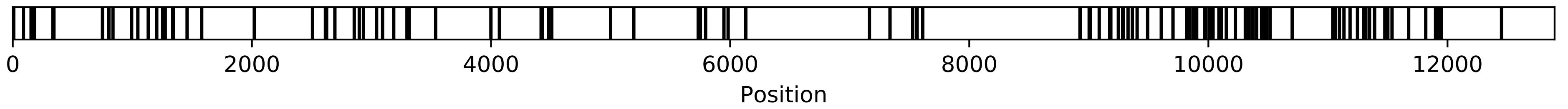


MA1 vs MM2
Number of differences: 153

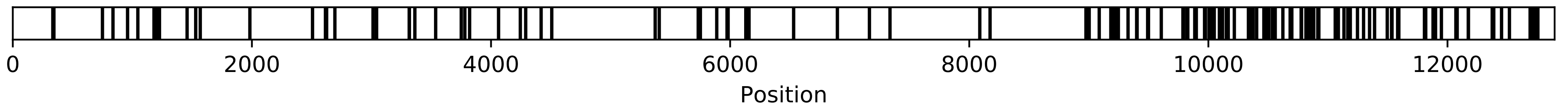


MA2 vs MM1
Number of differences: 265

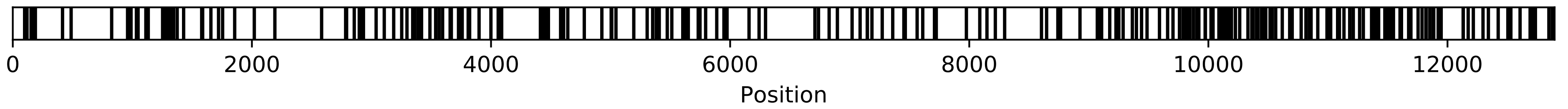


MA2 vs MM2
Number of differences: 269

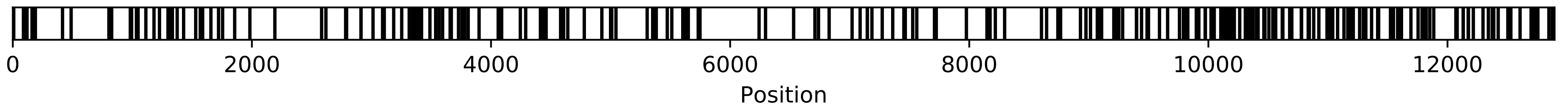


MM1 vs MM2
Number of differences: 150

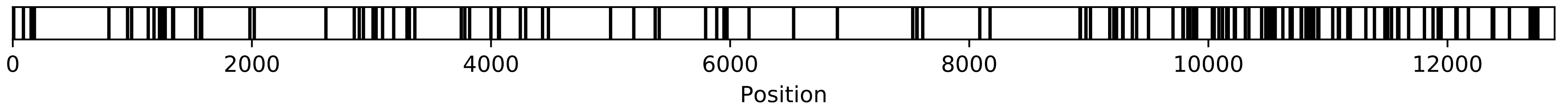

### Region: 280



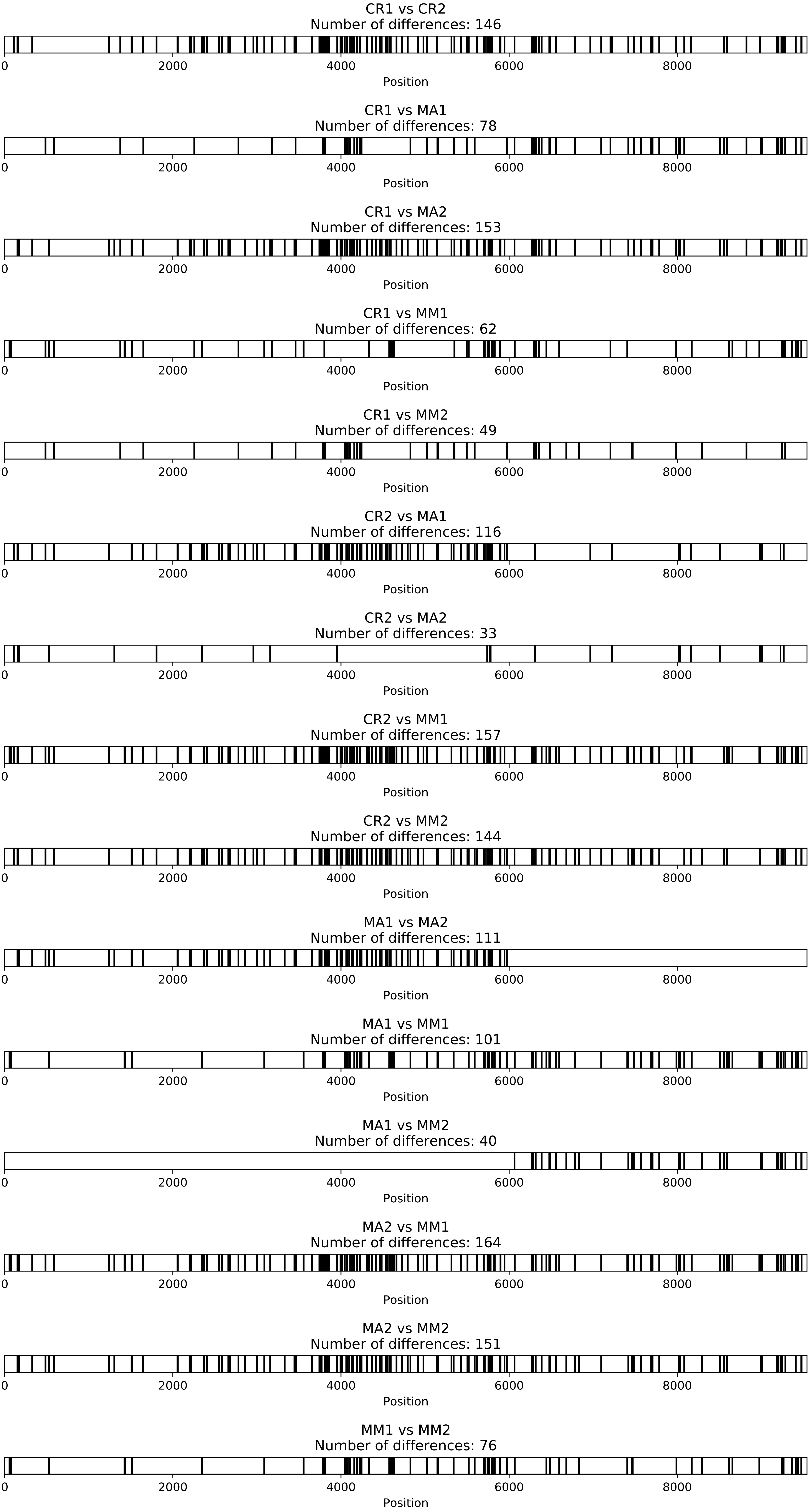
CR1 vs CR2
Number of differences: 146
CR1 vs MA1
Number of differences: 78
CR1 vs MA2
Number of differences: 153
CR1 vs MM1
Number of differences: 62
CR1 vs MM2
Number of differences: 49
CR2 vs MA1
Number of differences: 116
CR2 vs MA2
Number of differences: 33
CR2 vs MM1
Number of differences: 157
CR2 vs MM2
Number of differences: 144
MA1 vs MA2
Number of differences: 111
MA1 vs MM1
Number of differences: 101
MA1 vs MM2
Number of differences: 40
MA2 vs MM1
Number of differences: 164
MA2 vs MM2
Number of differences: 151
MM1 vs MM2
Number of differences: 76
0
2000
4000
6000
8000
Position

### Region: 281



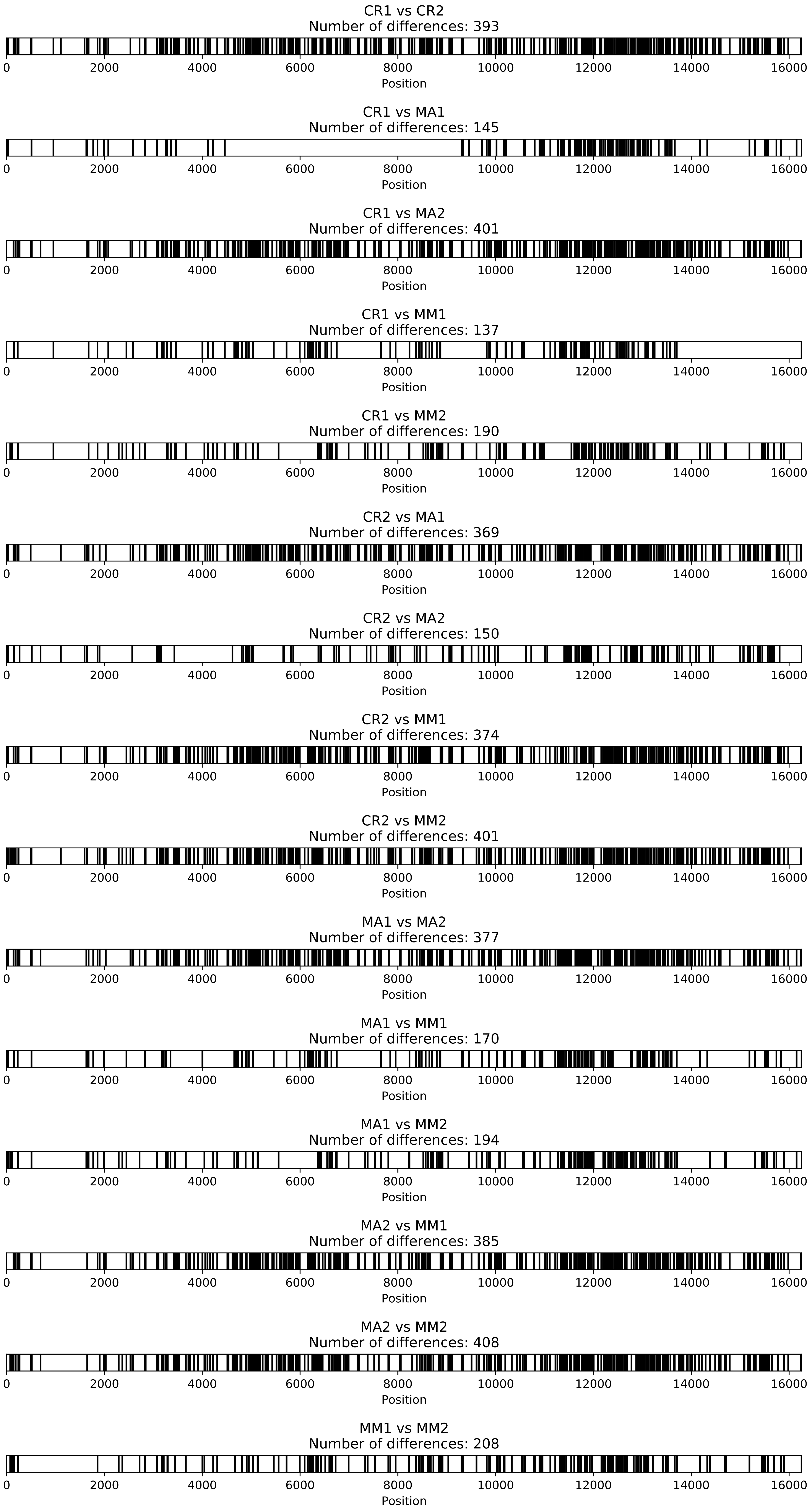
CR1 vs CR2
Number of differences: 393
CR1 vs MA1
Number of differences: 145
CR1 vs MA2
Number of differences: 401
CR1 vs MM1
Number of differences: 137
CR1 vs MM2
Number of differences: 190
CR2 vs MA1
Number of differences: 369
CR2 vs MA2
Number of differences: 150
CR2 vs MM1
Number of differences: 374
CR2 vs MM2
Number of differences: 401
MA1 vs MA2
Number of differences: 377
MA1 vs MM1
Number of differences: 170
MA1 vs MM2
Number of differences: 194
MA2 vs MM1
Number of differences: 385
MA2 vs MM2
Number of differences: 408
MM1 vs MM2
Number of differences: 208
0
2000
4000
6000
8000
10000
12000
14000
16000
Position

### Region: 282

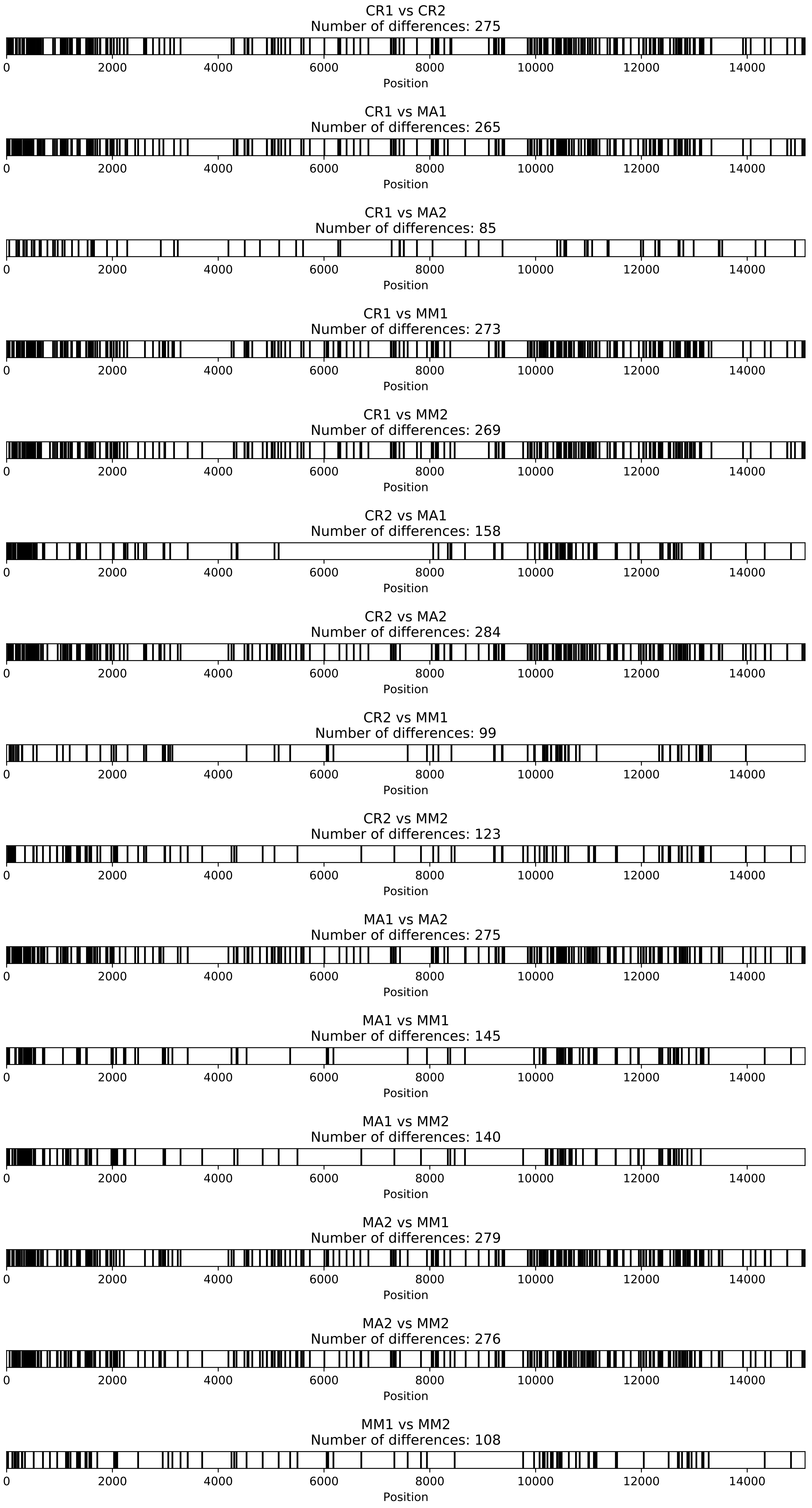
CR1 vs CR2
Number of differences: 275
CR1 vs MA1
Number of differences: 265
CR1 vs MA2
Number of differences: 85
CR1 vs MM1
Number of differences: 273
CR1 vs MM2
Number of differences: 269
CR2 vs MA1
Number of differences: 158
CR2 vs MA2
Number of differences: 284
CR2 vs MM1
Number of differences: 99
CR2 vs MM2
Number of differences: 123
MA1 vs MA2
Number of differences: 275
MA1 vs MM1
Number of differences: 145
MA1 vs MM2
Number of differences: 140
MA2 vs MM1
Number of differences: 279
MA2 vs MM2
Number of differences: 276
MM1 vs MM2
Number of differences: 108
0
2000
4000
6000
8000
10000
12000
14000
Position

### Region: 283



CR1 vs CR2
Number of differences: 471

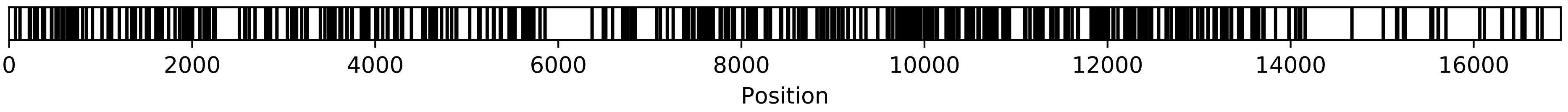


CR1 vs MA1
Number of differences: 471

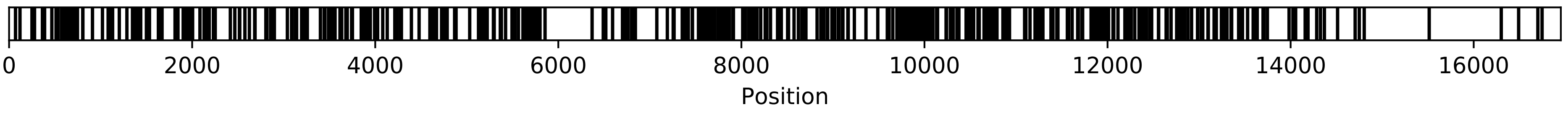


CR1 vs MA2
Number of differences: 413

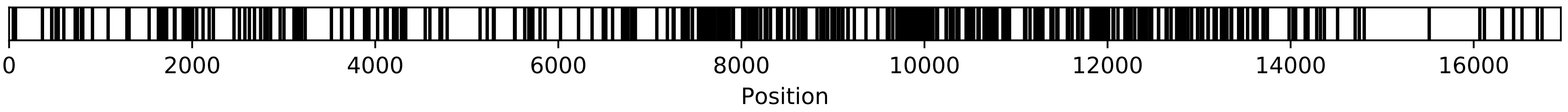


CR1 vs MM1
Number of differences: 325

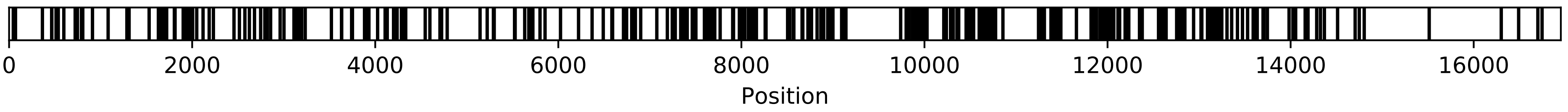


CR1 vs MM2
Number of differences: 340

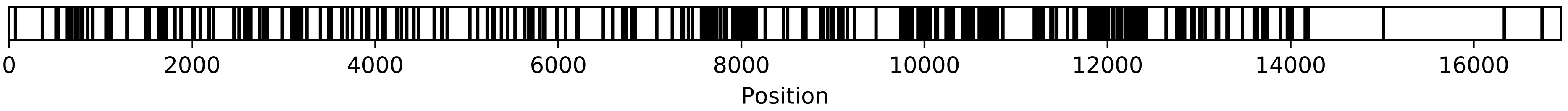


CR2 vs MA1
Number of differences: 180

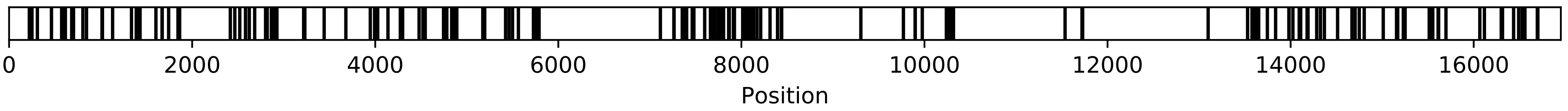


CR2 vs MA2
Number of differences: 232

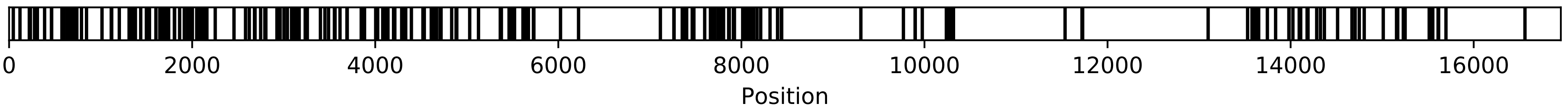


CR2 vs MM1
Number of differences: 444

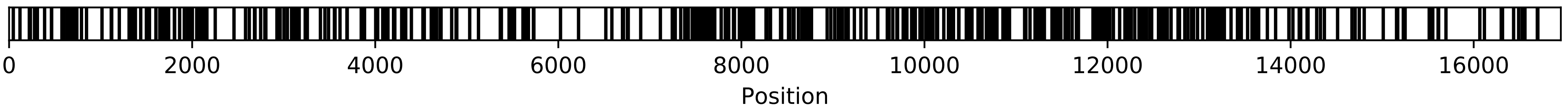


CR2 vs MM2
Number of differences: 430

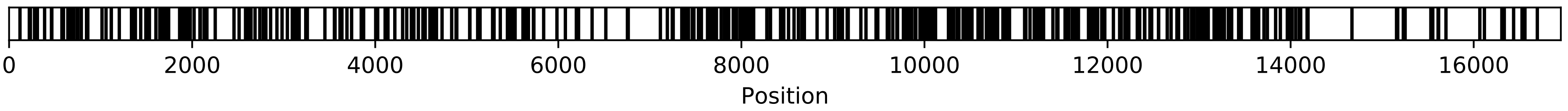


MA1 vs MA2
Number of differences: 170

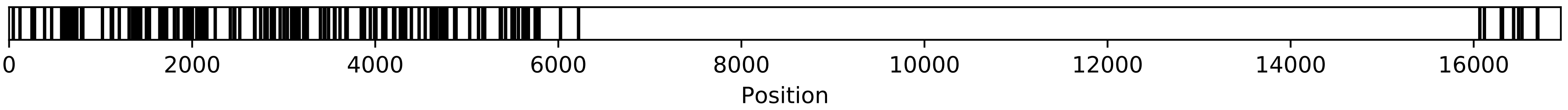


MA1 vs MM1
Number of differences: 404

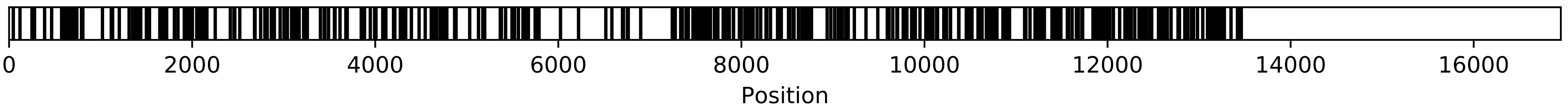


MA1 vs MM2
Number of differences: 431

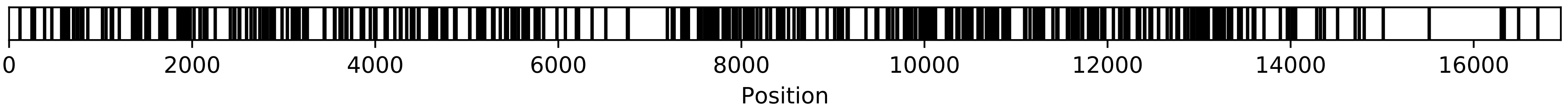


MA2 vs MM1
Number of differences: 262

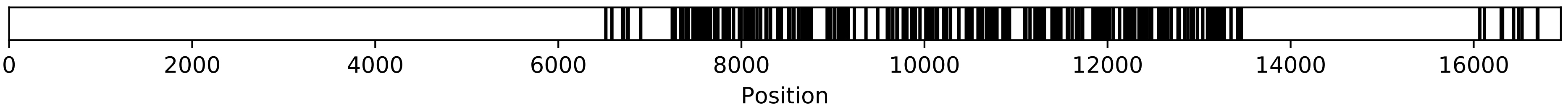


MA2 vs MM2
Number of differences: 363

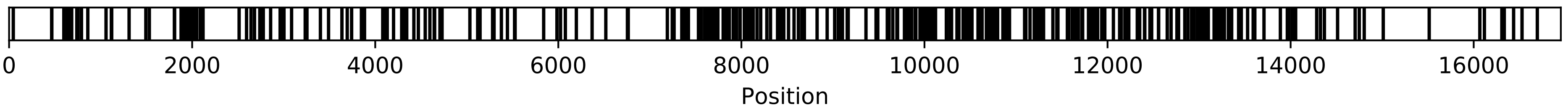


MM1 vs MM2
Number of differences: 300

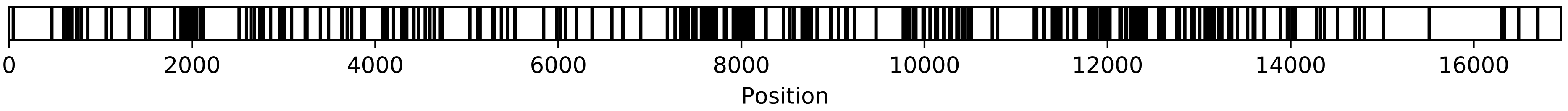

### Region: 284

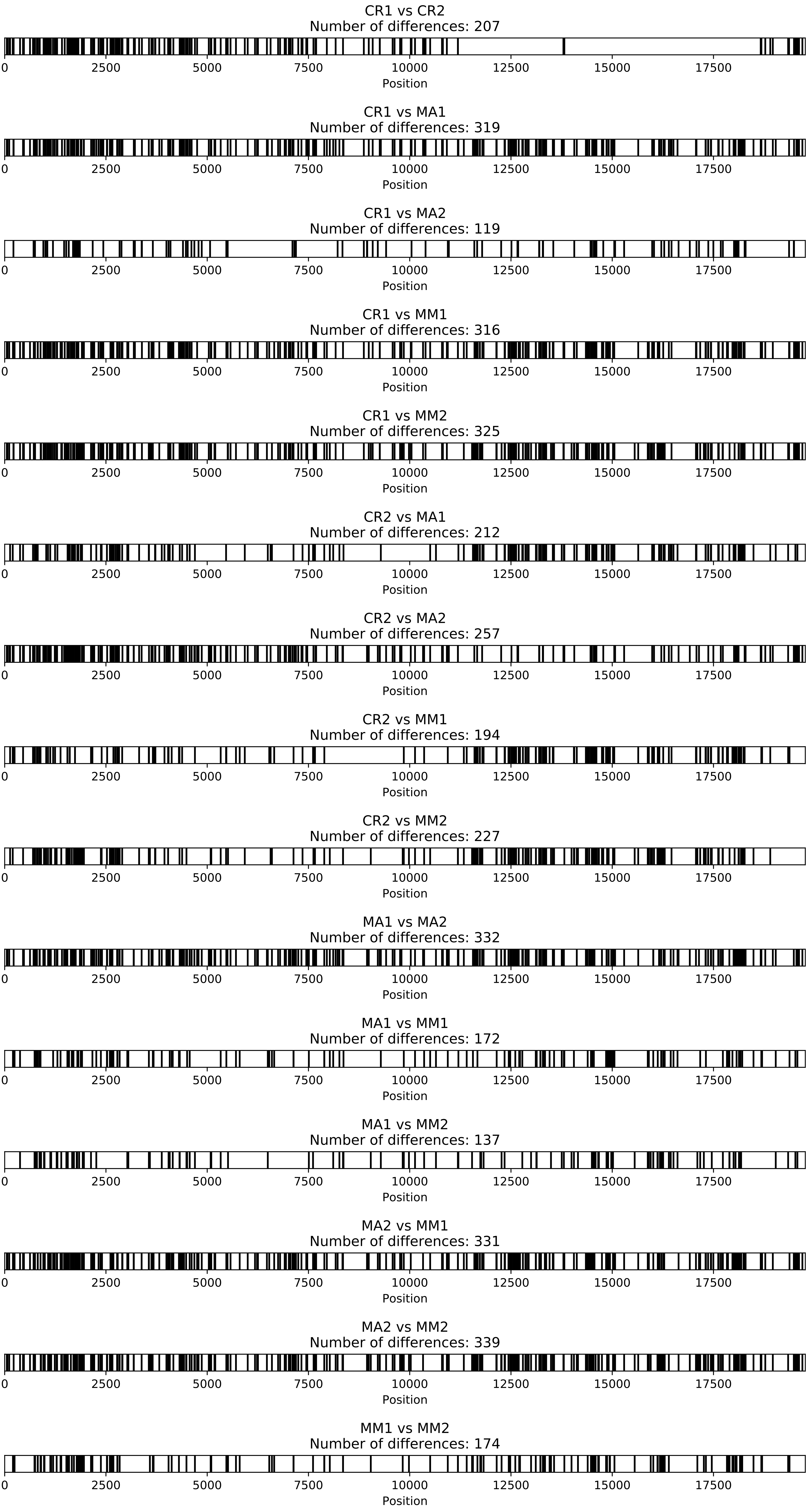
CR1 vs CR2
Number of differences: 207
CR1 vs MA1
Number of differences: 319
CR1 vs MA2
Number of differences: 119
CR1 vs MM1
Number of differences: 316
CR1 vs MM2
Number of differences: 325
CR2 vs MA1
Number of differences: 212
CR2 vs MA2
Number of differences: 257
CR2 vs MM1
Number of differences: 194
CR2 vs MM2
Number of differences: 227
MA1 vs MA2
Number of differences: 332
MA1 vs MM1
Number of differences: 172
MA1 vs MM2
Number of differences: 137
MA2 vs MM1
Number of differences: 331
MA2 vs MM2
Number of differences: 339
MM1 vs MM2
Number of differences: 174
0
2500
5000
7500
10000
12500
15000
17500
Position

### Region: 287



CR1 vs CR2
Number of differences: 539

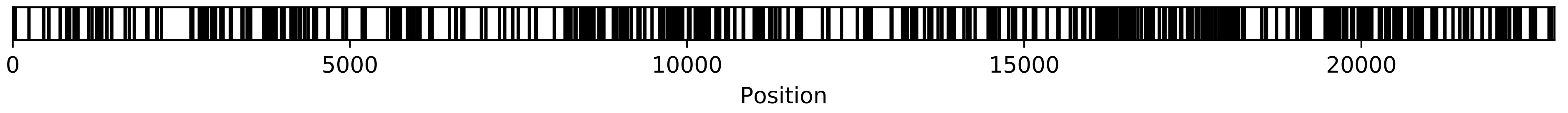


CR1 vs MA1
Number of differences: 250

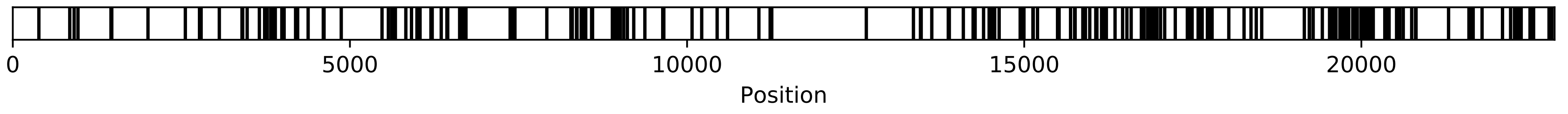


CR1 vs MA2
Number of differences: 537

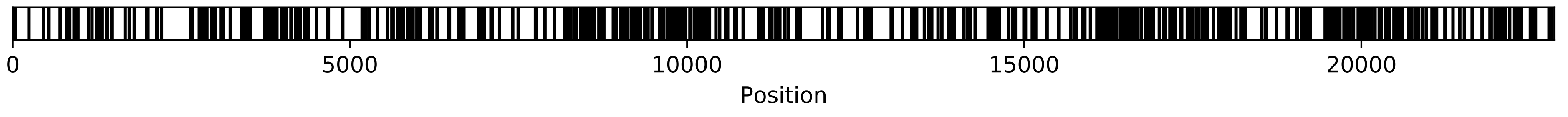


CR1 vs MM1
Number of differences: 251

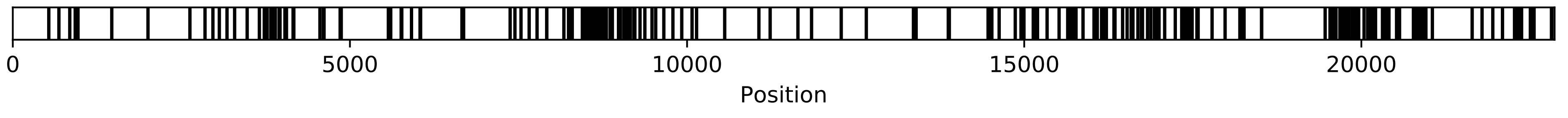


CR1 vs MM2
Number of differences: 240

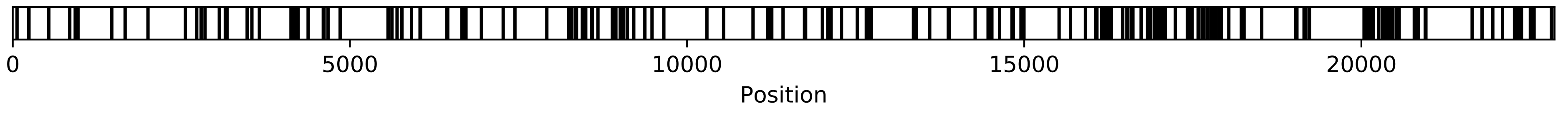


CR2 vs MA1
Number of differences: 541

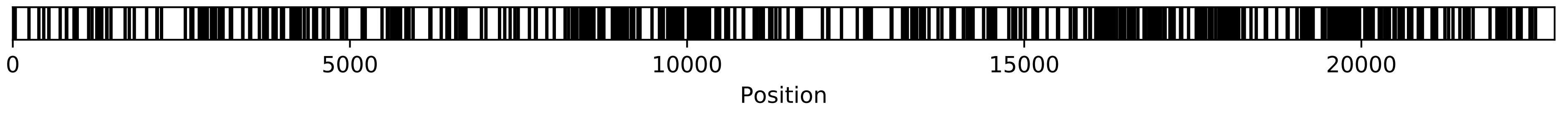


CR2 vs MA2
Number of differences: 162

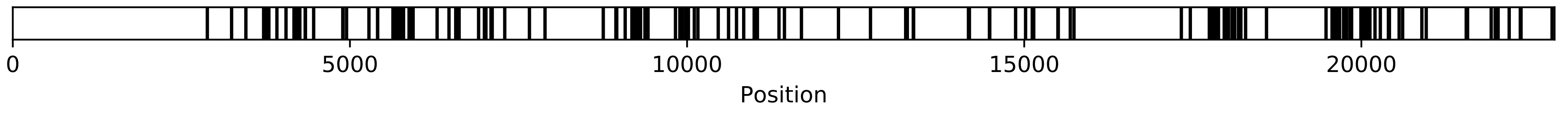


CR2 vs MM1
Number of differences: 569

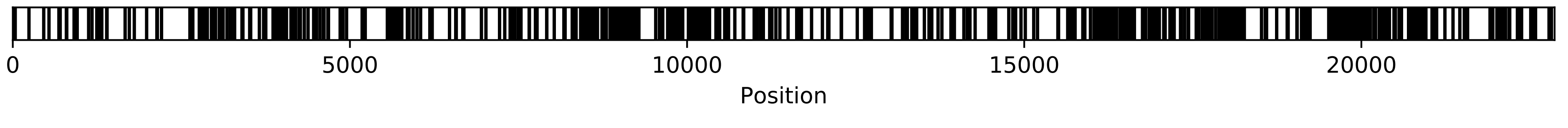


CR2 vs MM2
Number of differences: 554

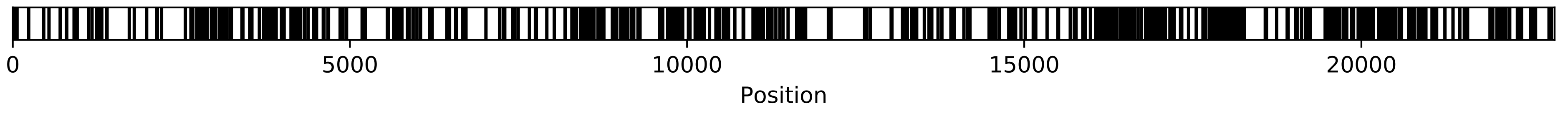


MA1 vs MA2
Number of differences: 535

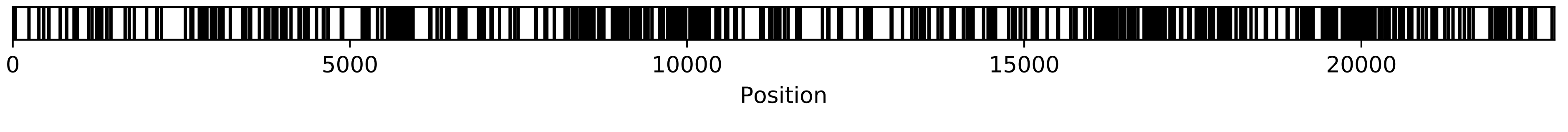


MA1 vs MM1
Number of differences: 261

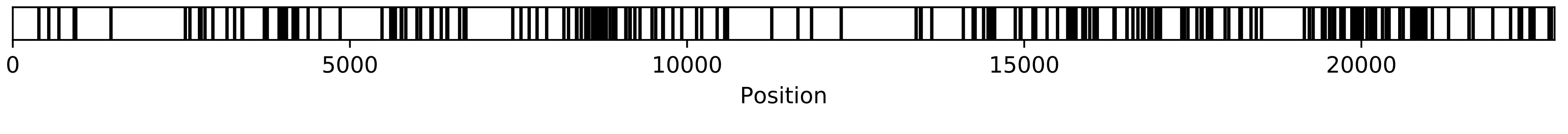


MA1 vs MM2
Number of differences: 242

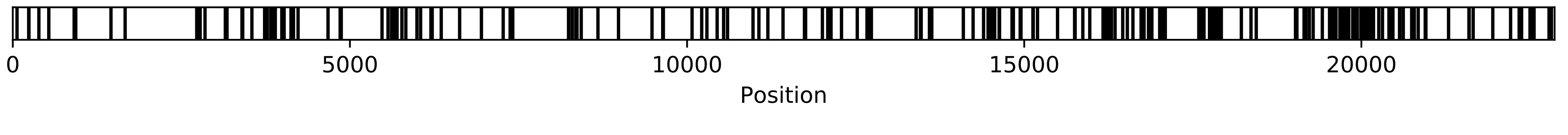


MA2 vs MM1
Number of differences: 562

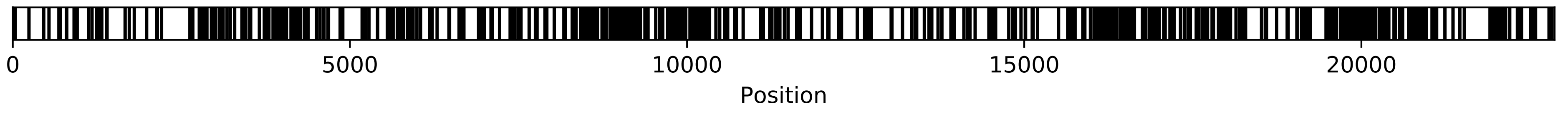


MA2 vs MM2
Number of differences: 550

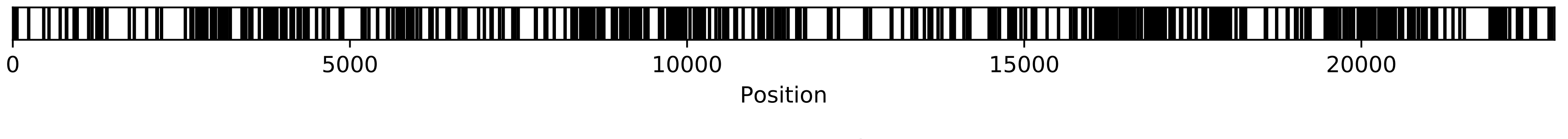


MM1 vs MM2
Number of differences: 267

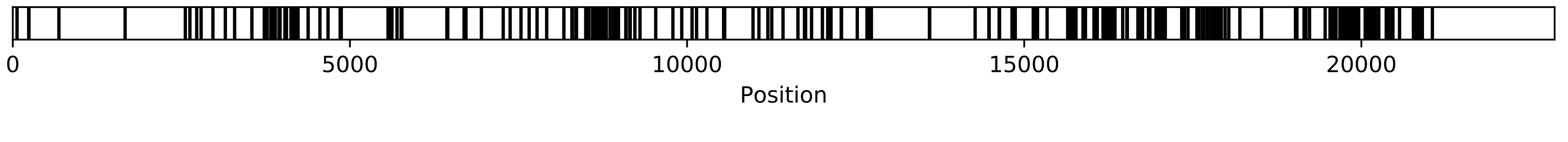

### Region: 290

CR1 vs CR2
Number of differences: 161

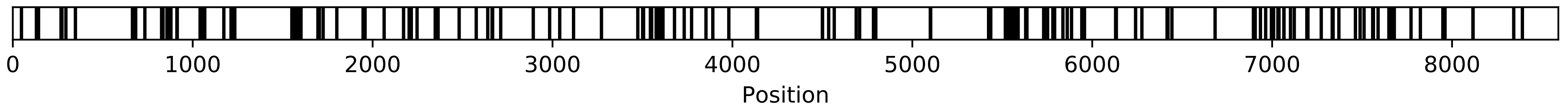


CR1 vs MA1
Number of differences: 38

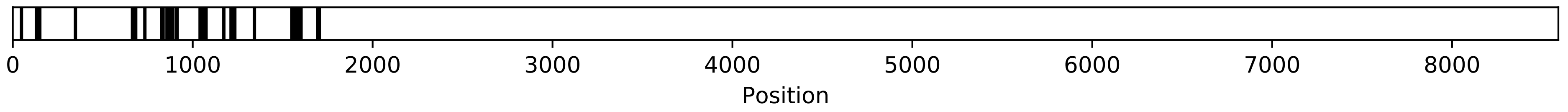


CR1 vs MA2
Number of differences: 35

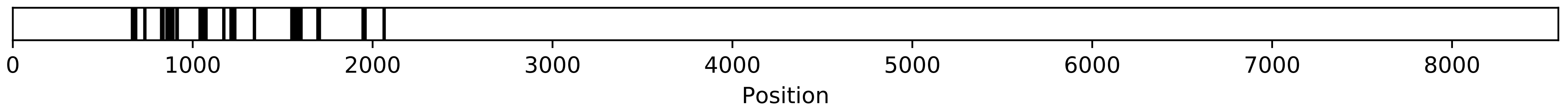


CR1 vs MM1
Number of differences: 74

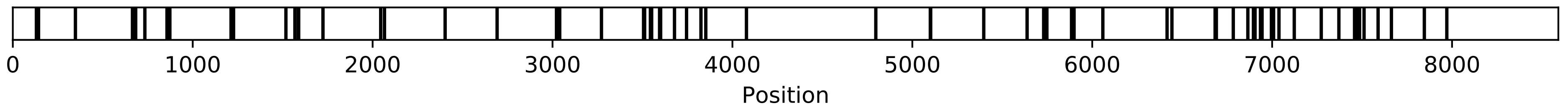


CR1 vs MM2
Number of differences: 81

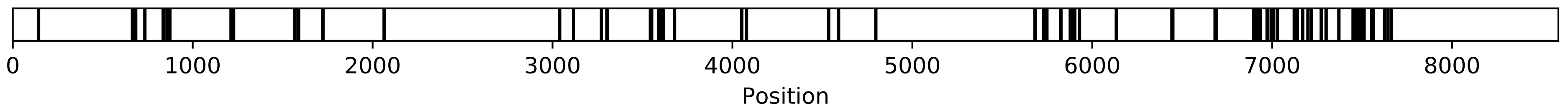


CR2 vs MA1
Number of differences: 132

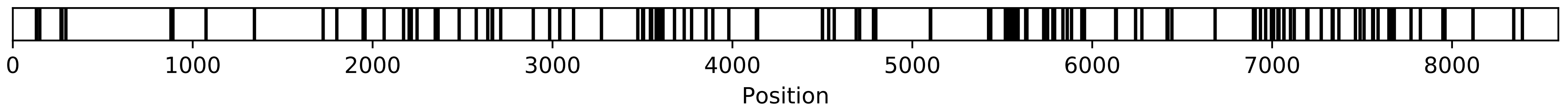


CR2 vs MA2
Number of differences: 132

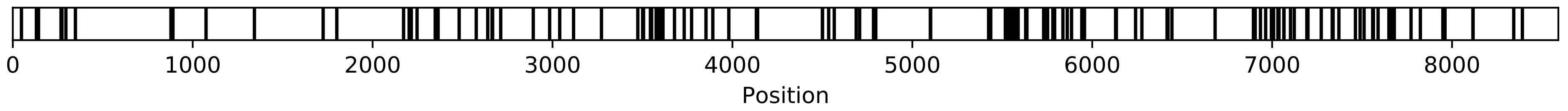


CR2 vs MM1
Number of differences: 146

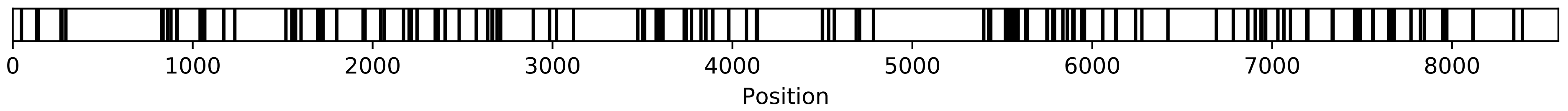


CR2 vs MM2
Number of differences: 156

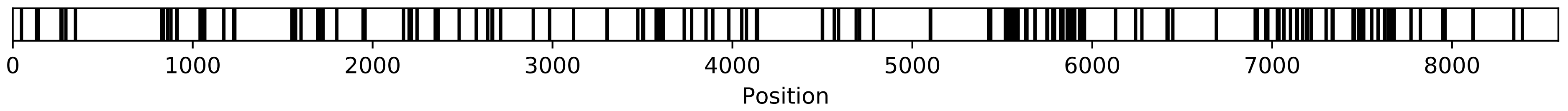


MA1 vs MA2
Number of differences: 9

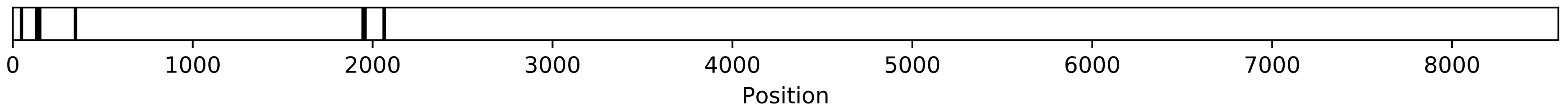


MA1 vs MM1
Number of differences: 82

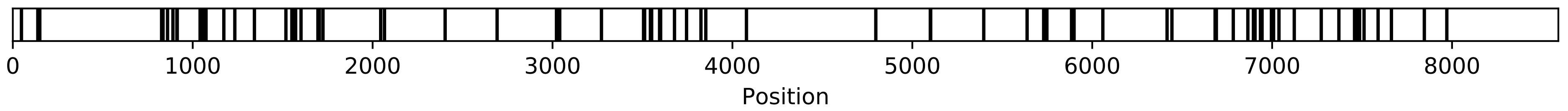


MA1 vs MM2
Number of differences: 94

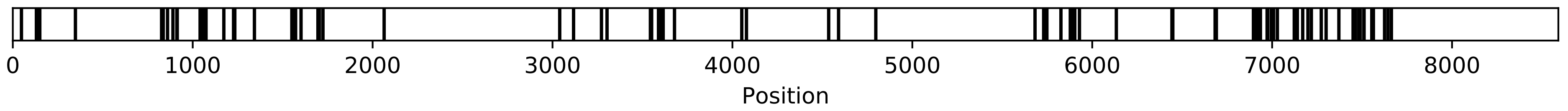


MA2 vs MM1
Number of differences: 85

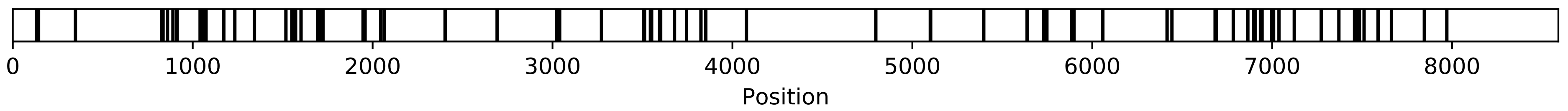


MA2 vs MM2
Number of differences: 91

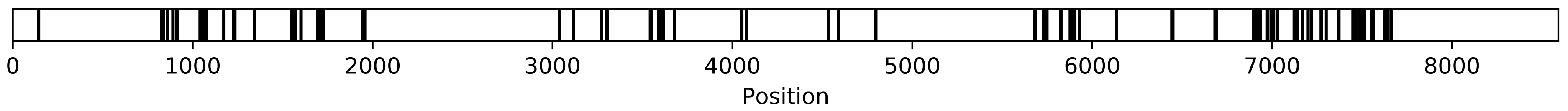


MM1 vs MM2
Number of differences: 71

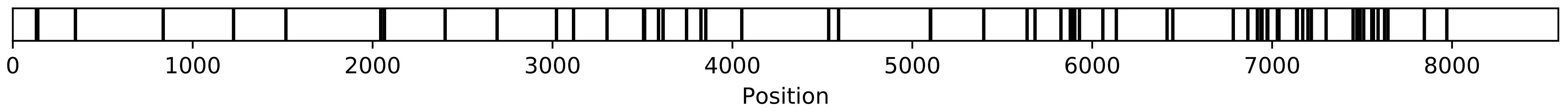

### Region: 291

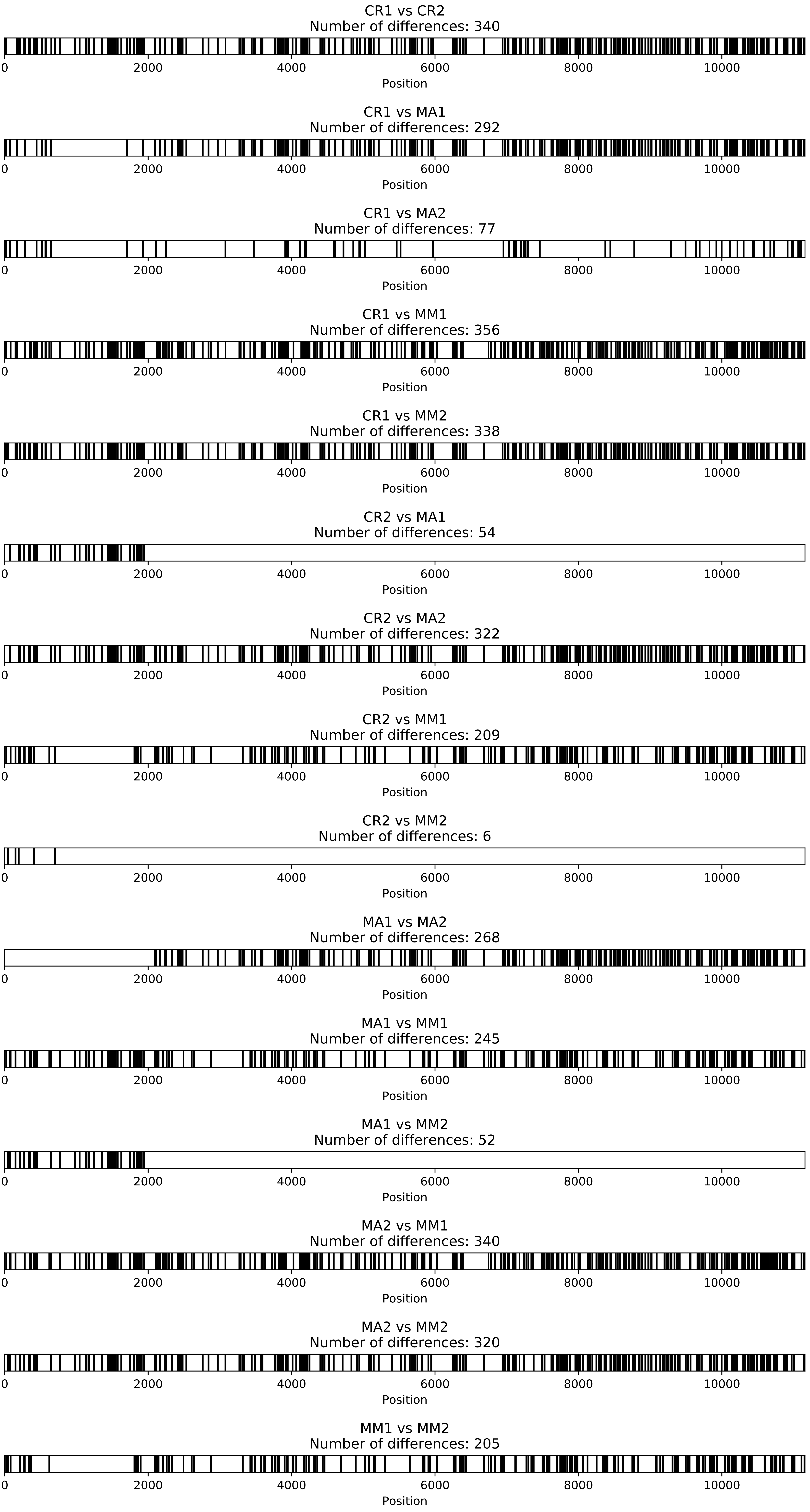
CR1 vs CR2
Number of differences: 340
0 2000 4000 6000 8000 10000
Position
CR1 vs MA1
Number of differences: 292
0 2000 4000 6000 8000 10000
Position
CR1 vs MA2
Number of differences: 77
0 2000 4000 6000 8000 10000
Position
CR1 vs MM1
Number of differences: 356
0 2000 4000 6000 8000 10000
Position
CR1 vs MM2
Number of differences: 338
0 2000 4000 6000 8000 10000
Position
CR2 vs MA1
Number of differences: 54
0 2000 4000 6000 8000 10000
Position
CR2 vs MA2
Number of differences: 322
0 2000 4000 6000 8000 10000
Position
CR2 vs MM1
Number of differences: 209
0 2000 4000 6000 8000 10000
Position
CR2 vs MM2
Number of differences: 6
0 2000 4000 6000 8000 10000
Position
MA1 vs MA2
Number of differences: 268
0 2000 4000 6000 8000 10000
Position
MA1 vs MM1
Number of differences: 245
0 2000 4000 6000 8000 10000
Position
MA1 vs MM2
Number of differences: 52
0 2000 4000 6000 8000 10000
Position
MA2 vs MM1
Number of differences: 340
0 2000 4000 6000 8000 10000
Position
MA2 vs MM2
Number of differences: 320
0 2000 4000 6000 8000 10000
Position
MM1 vs MM2
Number of differences: 205
0 2000 4000 6000 8000 10000
Position

### Region: 297



CR1 vs CR2
Number of differences: 292

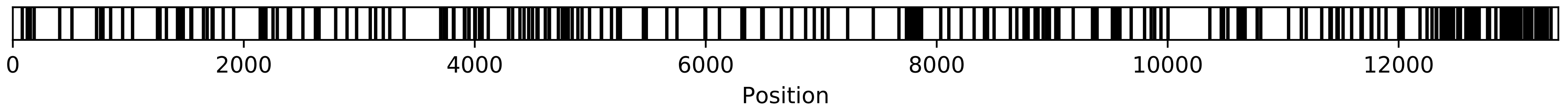


CR1 vs MA1
Number of differences: 100

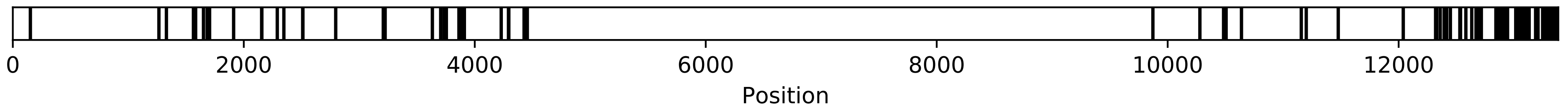


CR1 vs MA2
Number of differences: 294

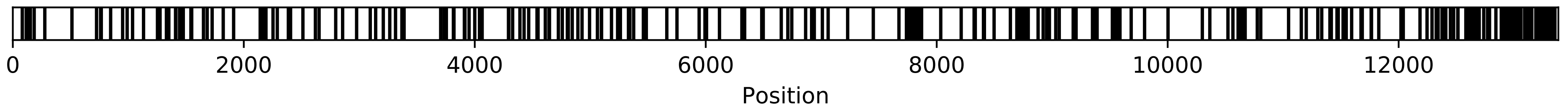


CR1 vs MM1
Number of differences: 100

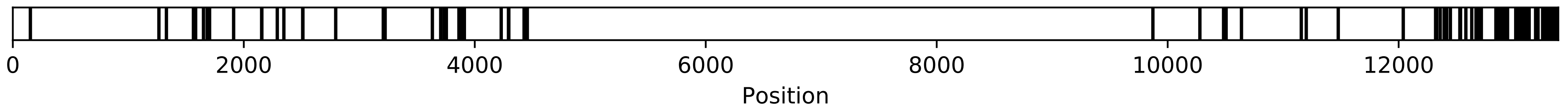


CR1 vs MM2
Number of differences: 83

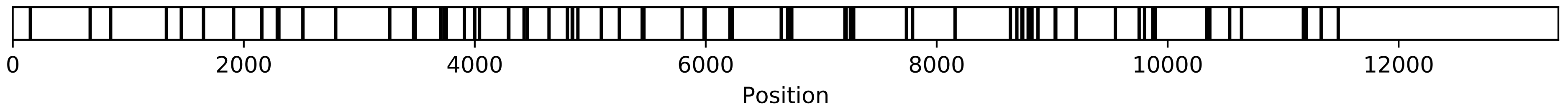


CR2 vs MA1
Number of differences: 290

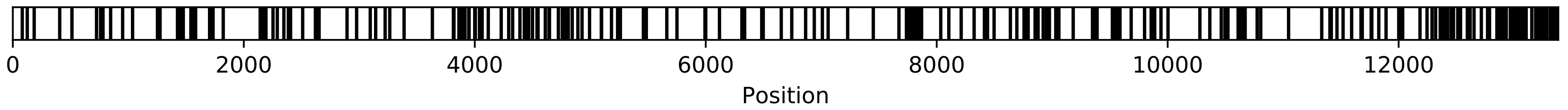


CR2 vs MA2
Number of differences: 89

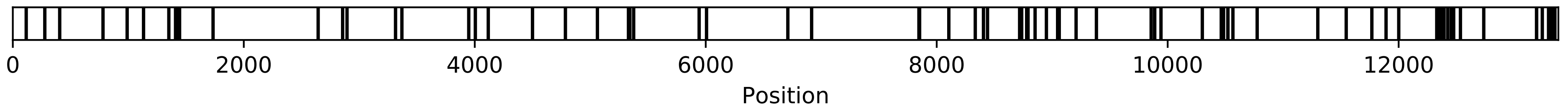


CR2 vs MM1
Number of differences: 290

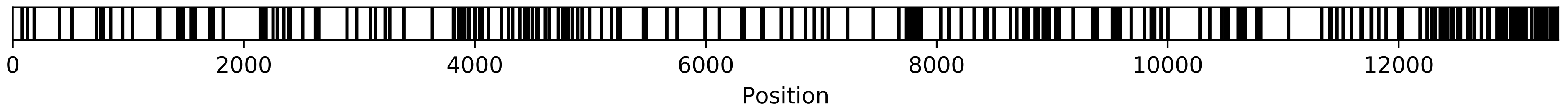


CR2 vs MM2
Number of differences: 290

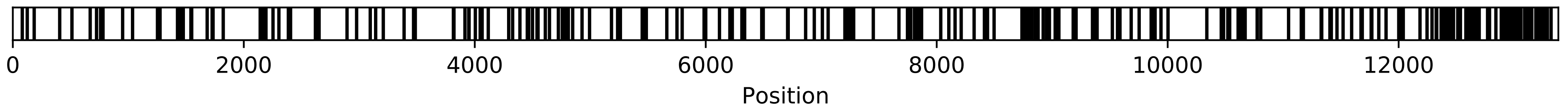


MA1 vs MA2
Number of differences: 293

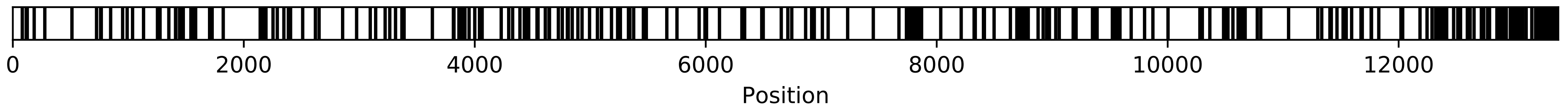


MA1 vs MM1
Number of differences: 0

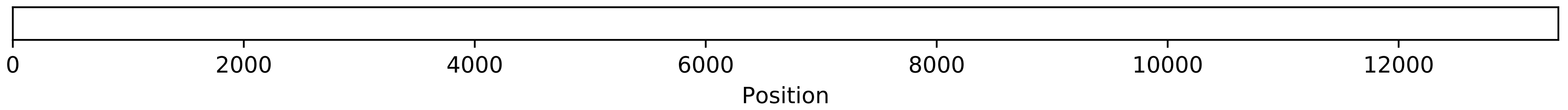


MA1 vs MM2
Number of differences: 142

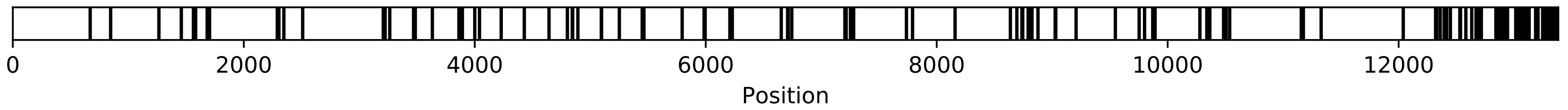


MA2 vs MM1
Number of differences: 293

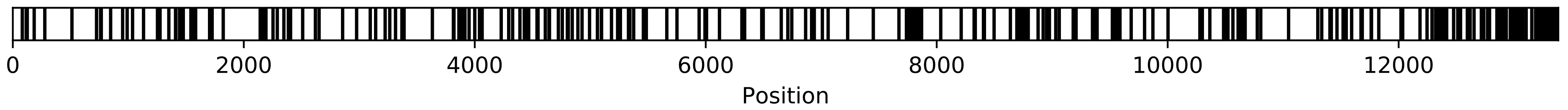


MA2 vs MM2
Number of differences: 293

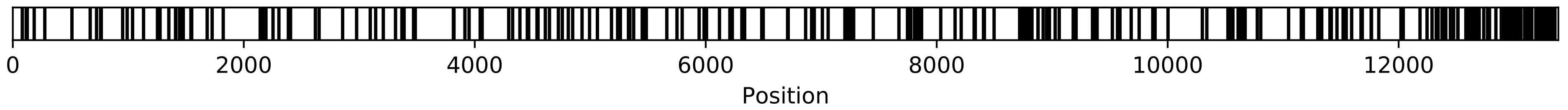


MM1 vs MM2
Number of differences: 142

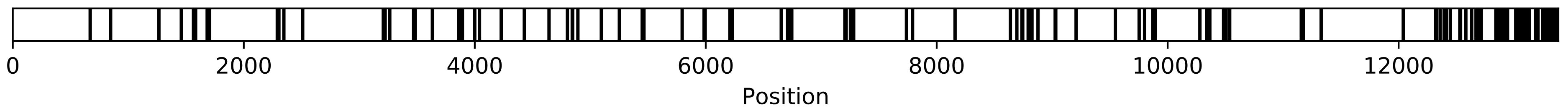

### Region: 300

CR1 vs CR2
Number of differences: 123

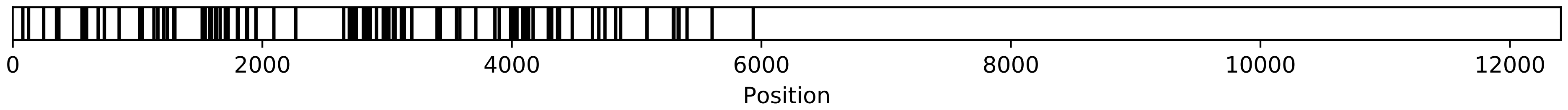


CR1 vs MA1
Number of differences: 159

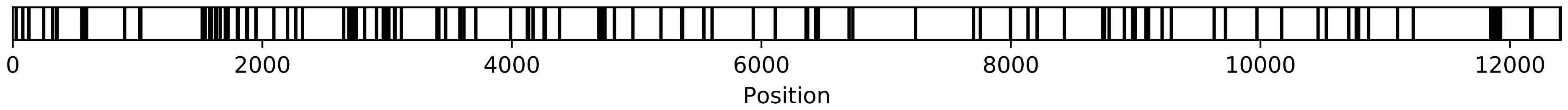


CR1 vs MA2
Number of differences: 167

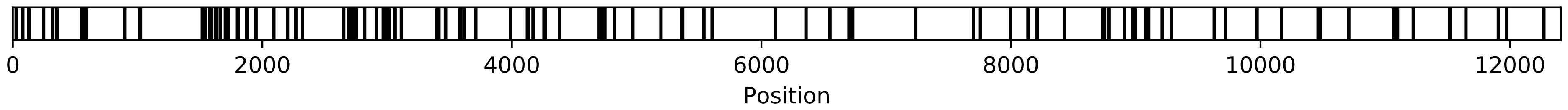


CR1 vs MM1
Number of differences: 0

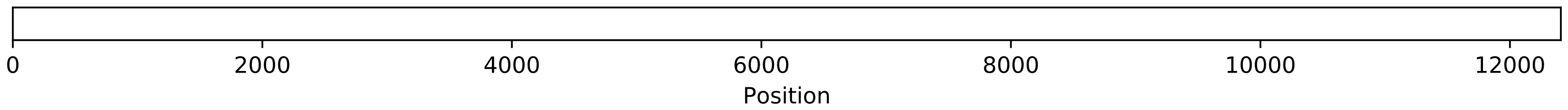


CR1 vs MM2
Number of differences: 99

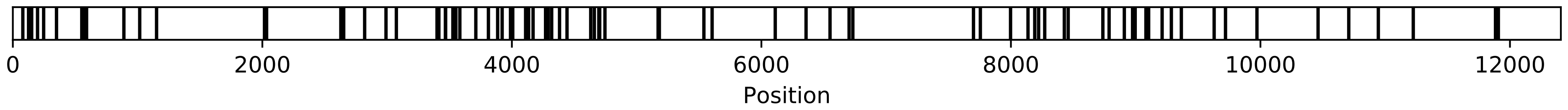


CR2 vs MA1
Number of differences: 134

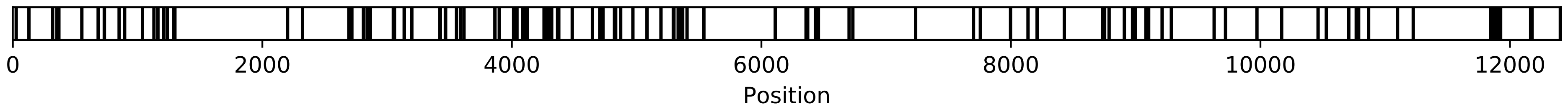


CR2 vs MA2
Number of differences: 144

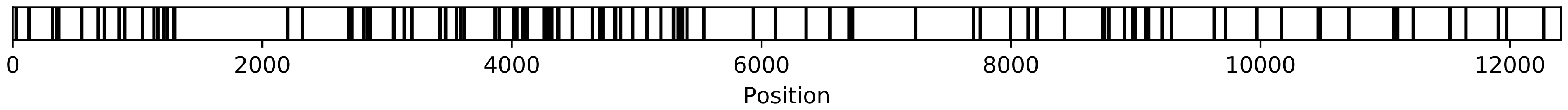


CR2 vs MM1
Number of differences: 123

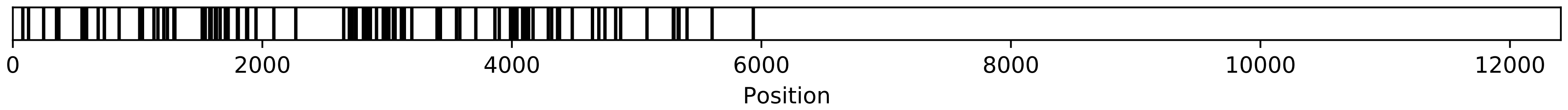


CR2 vs MM2
Number of differences: 154

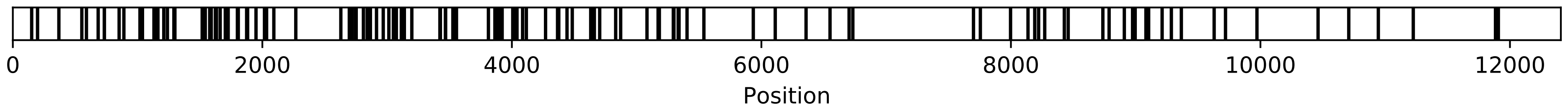


MA1 vs MA2
Number of differences: 45

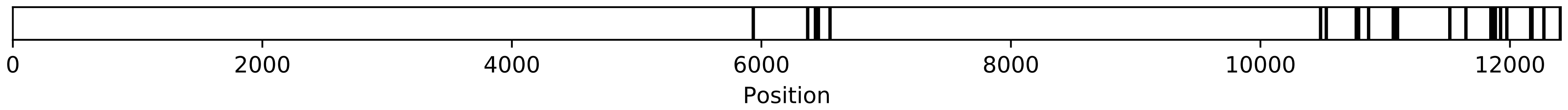


MA1 vs MM1
Number of differences: 159

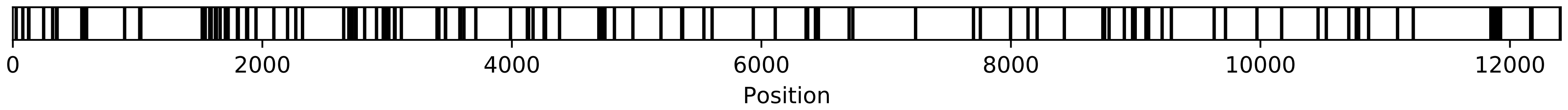


MA1 vs MM2
Number of differences: 121

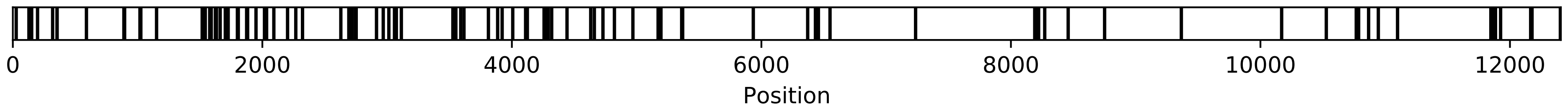


MA2 vs MM1
Number of differences: 167

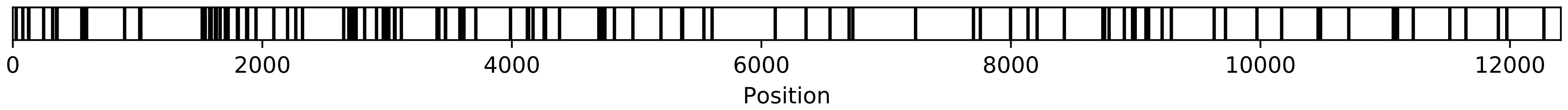


MA2 vs MM2
Number of differences: 127

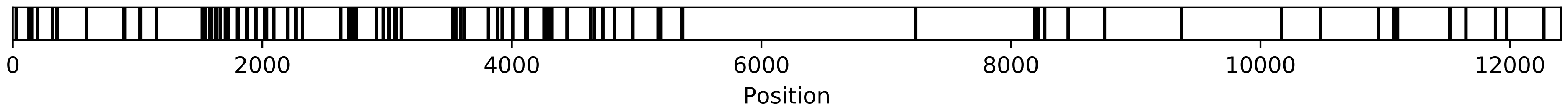


MM1 vs MM2
Number of differences: 99

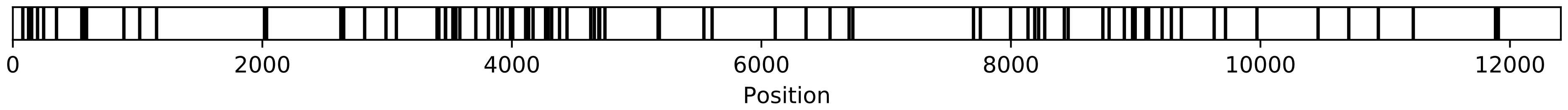

### Region: 302



CR1 vs CR2
Number of differences: 224

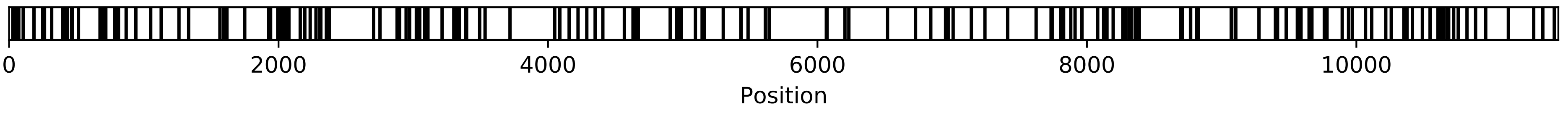


CR1 vs MA1
Number of differences: 142

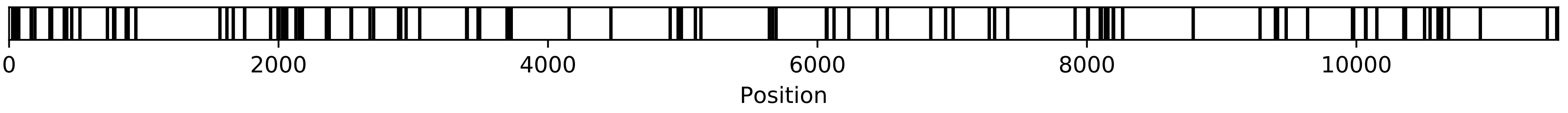


CR1 vs MA2
Number of differences: 223

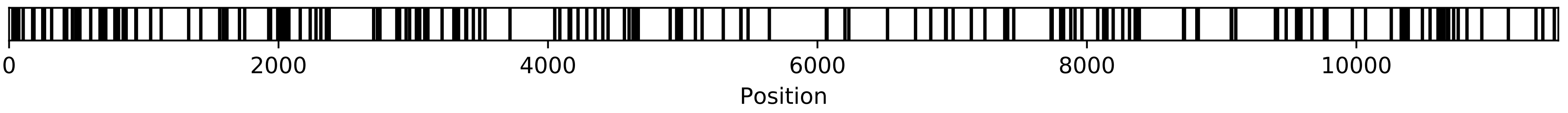


CR1 vs MM1
Number of differences: 142

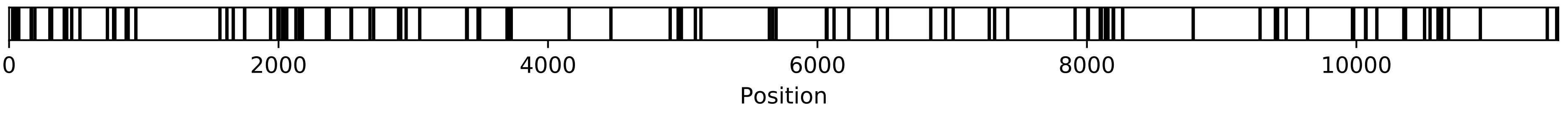


CR1 vs MM2
Number of differences: 137

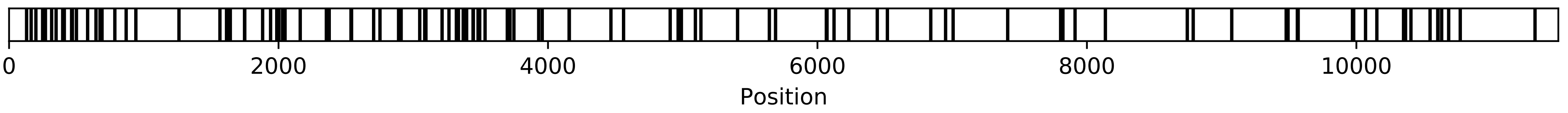


CR2 vs MA1
Number of differences: 225

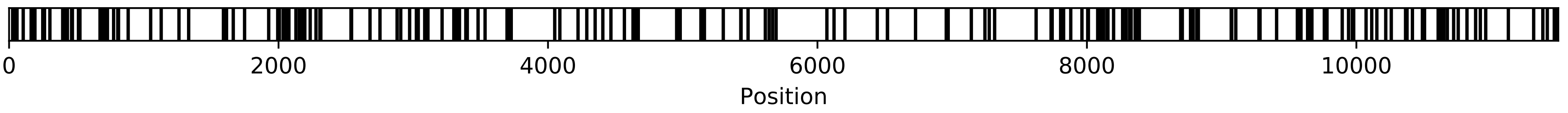


CR2 vs MA2
Number of differences: 63

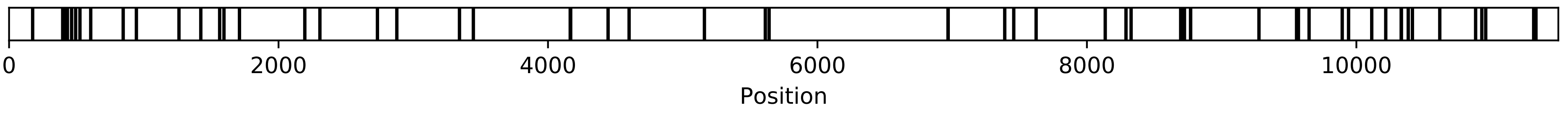


CR2 vs MM1
Number of differences: 225

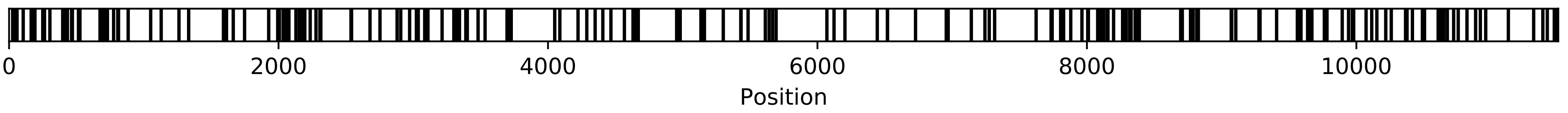


CR2 vs MM2
Number of differences: 208

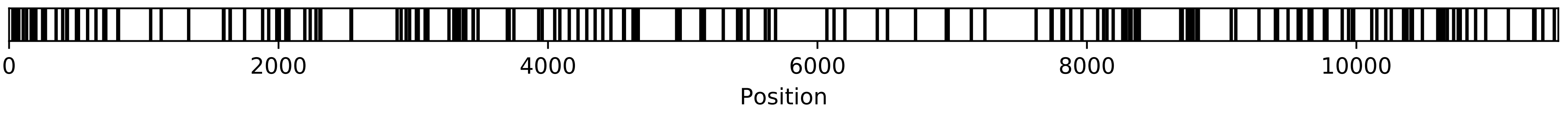


MA1 vs MA2
Number of differences: 220

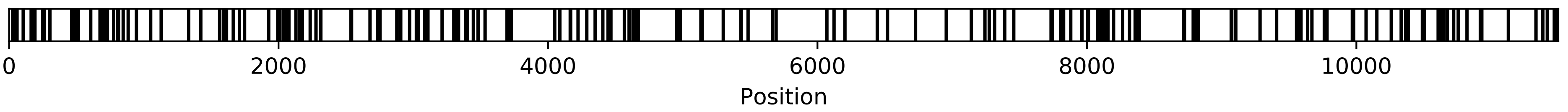


MA1 vs MM1
Number of differences: 0

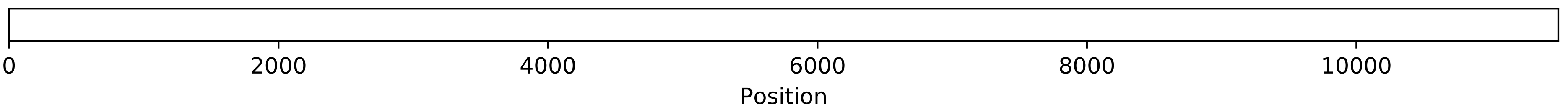


MA1 vs MM2
Number of differences: 115

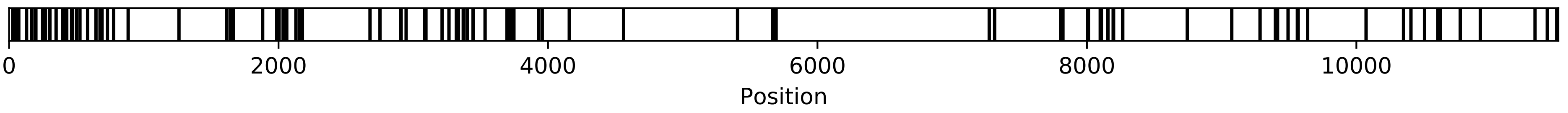


MA2 vs MM1
Number of differences: 220

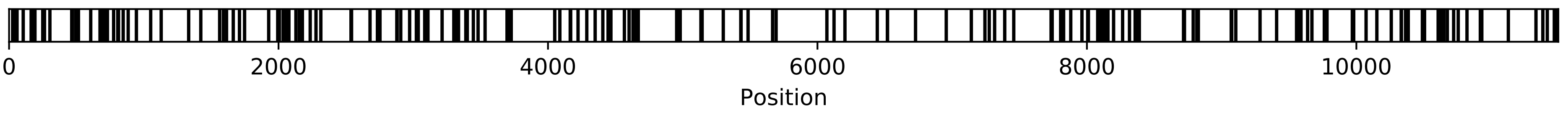


MA2 vs MM2
Number of differences: 208

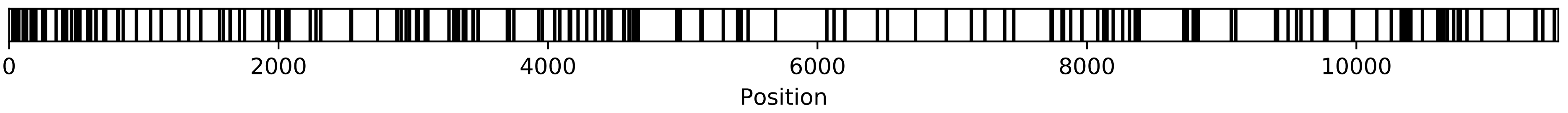


MM1 vs MM2
Number of differences: 115

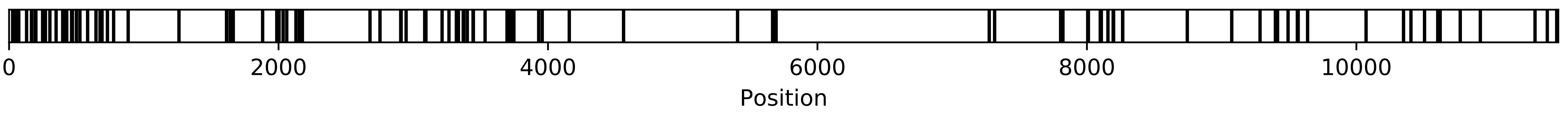

### Region: 303

CR1 vs CR2
Number of differences: 137

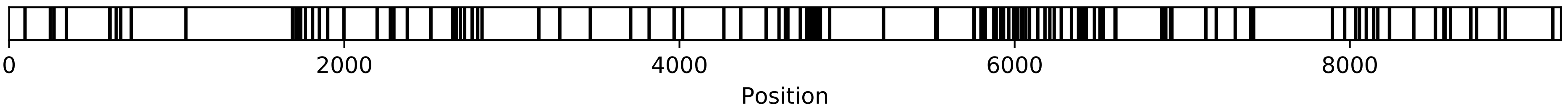


CR1 vs MA1
Number of differences: 27

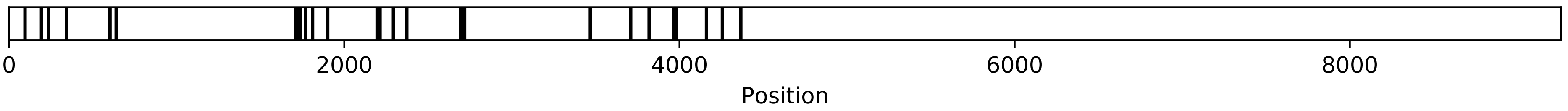


CR1 vs MA2
Number of differences: 132

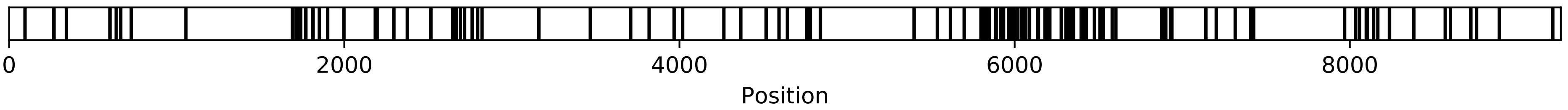


CR1 vs MM1
Number of differences: 27

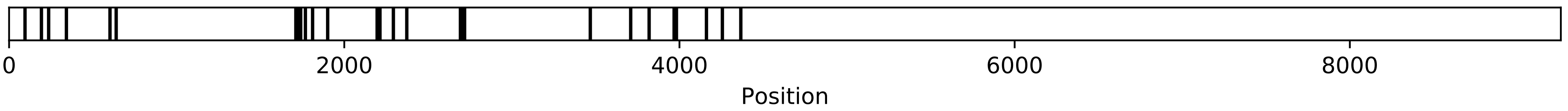


CR1 vs MM2
Number of differences: 53

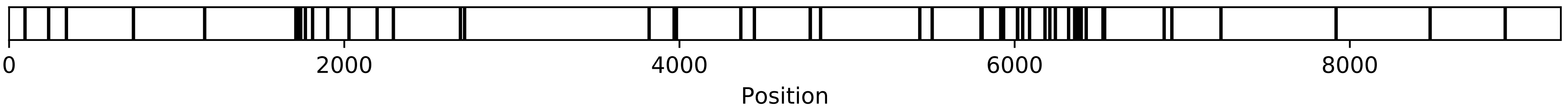


CR2 vs MA1
Number of differences: 128

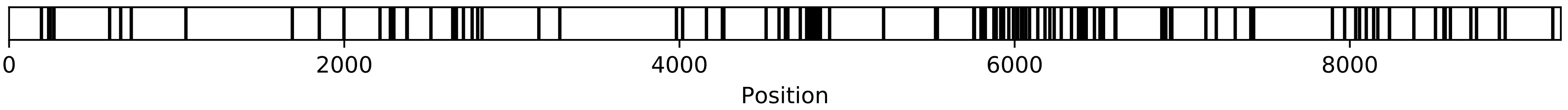


CR2 vs MA2
Number of differences: 48

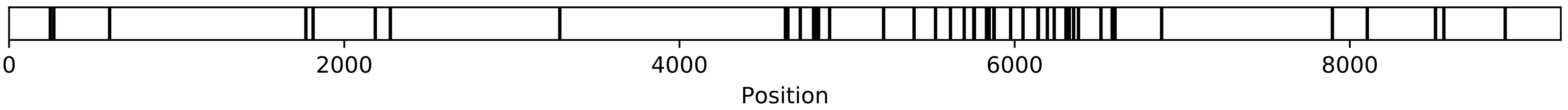


CR2 vs MM1
Number of differences: 128

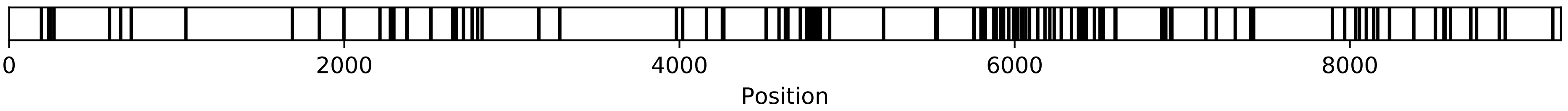


CR2 vs MM2
Number of differences: 128

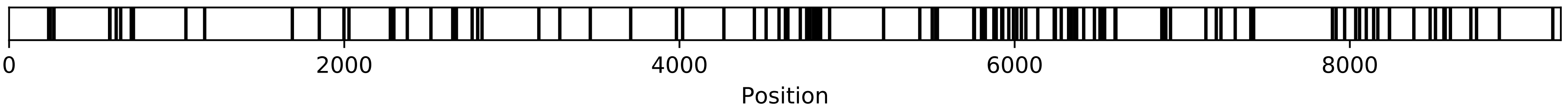


MA1 vs MA2
Number of differences: 123

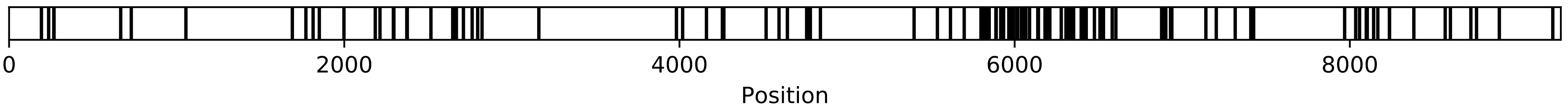


MA1 vs MM1
Number of differences: 0

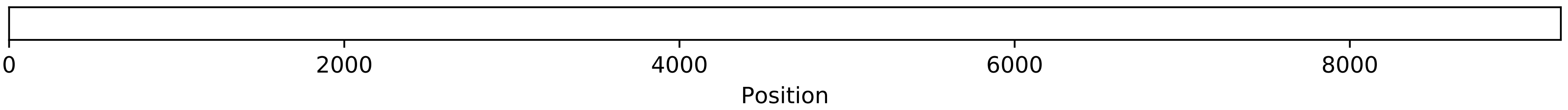


MA1 vs MM2
Number of differences: 46

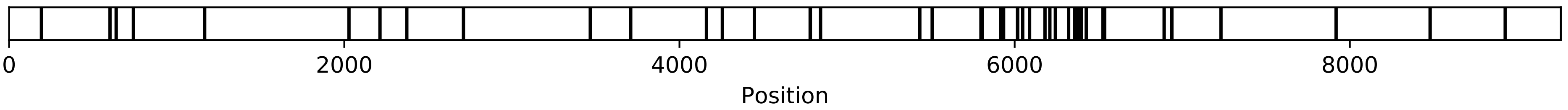


MA2 vs MM1
Number of differences: 123

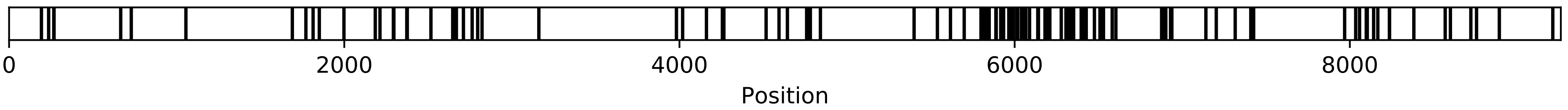


MA2 vs MM2
Number of differences: 127

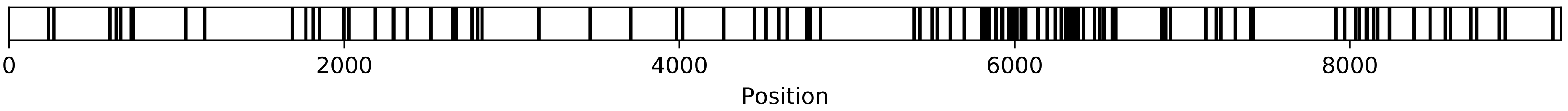


MM1 vs MM2
Number of differences: 46

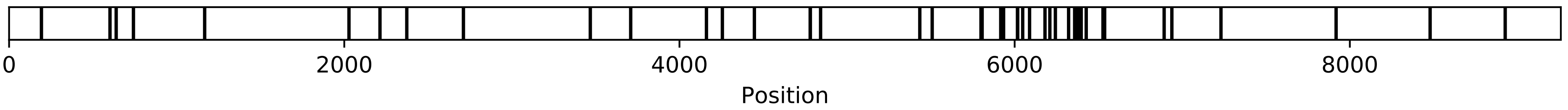

### Region: 304



CR1 vs CR2
Number of differences: 361

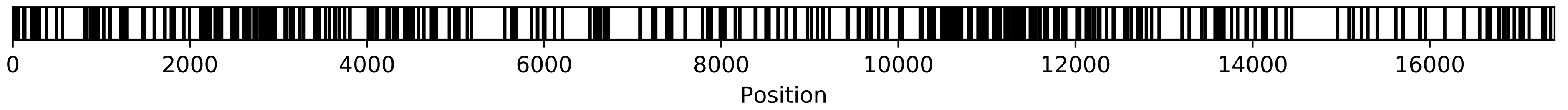


CR1 vs MA1
Number of differences: 198

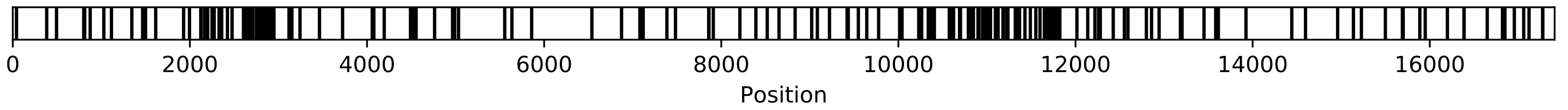


CR1 vs MA2
Number of differences: 355

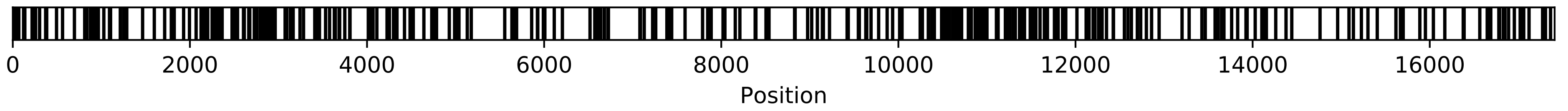


CR1 vs MM1
Number of differences: 200

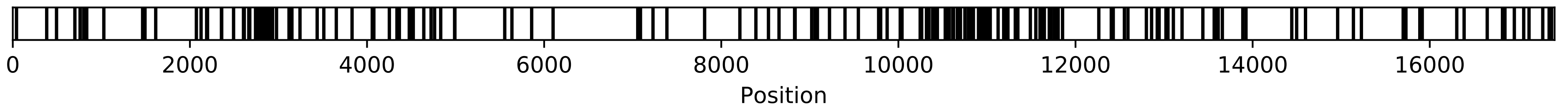


CR1 vs MM2
Number of differences: 198

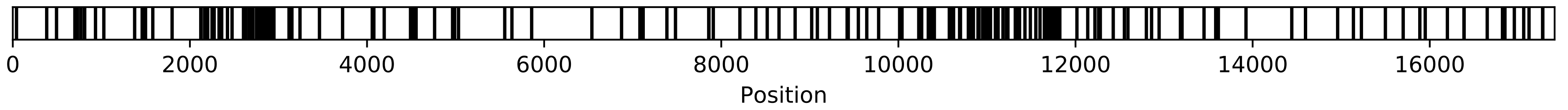


CR2 vs MA1
Number of differences: 325

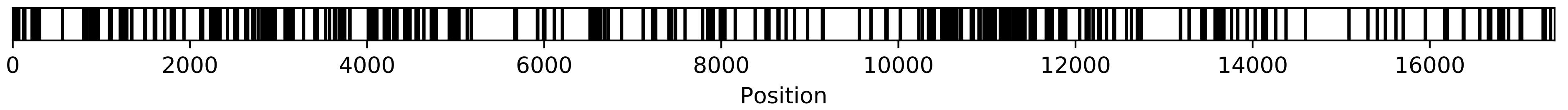


CR2 vs MA2
Number of differences: 44

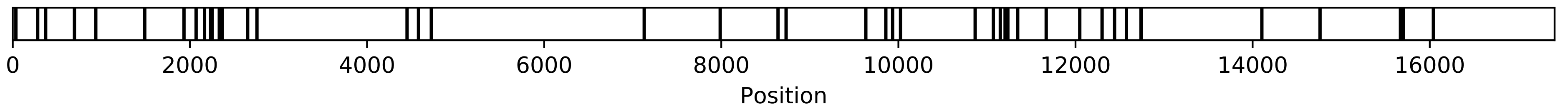


CR2 vs MM1
Number of differences: 346

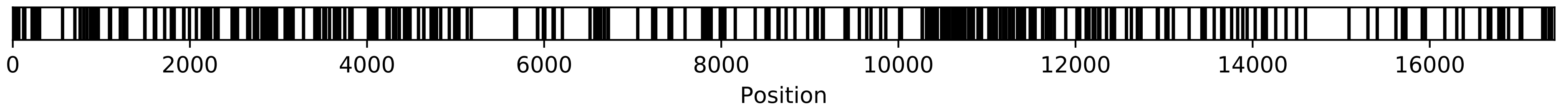


CR2 vs MM2
Number of differences: 329

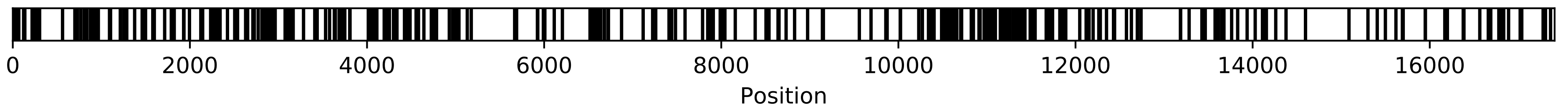


MA1 vs MA2
Number of differences: 319

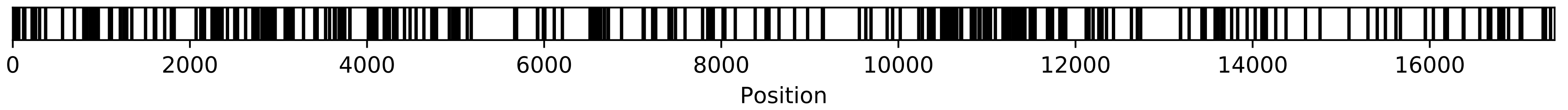


MA1 vs MM1
Number of differences: 178

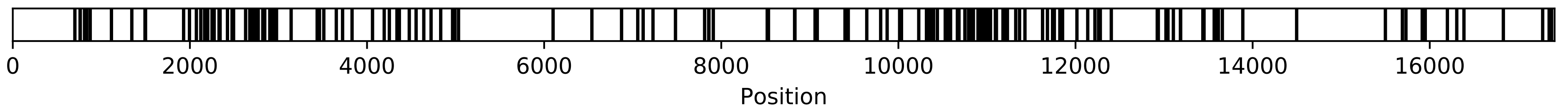


MA1 vs MM2
Number of differences: 18

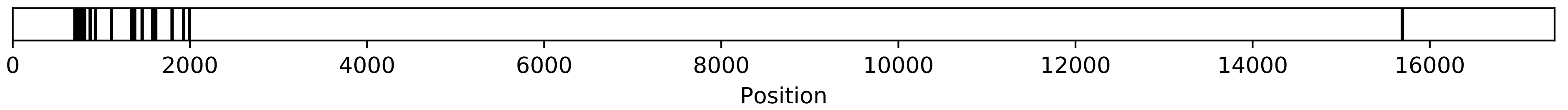


MA2 vs MM1
Number of differences: 341

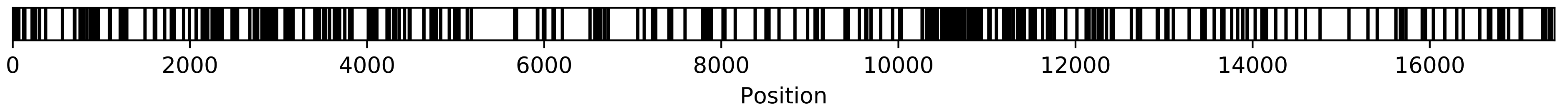


MA2 vs MM2
Number of differences: 323

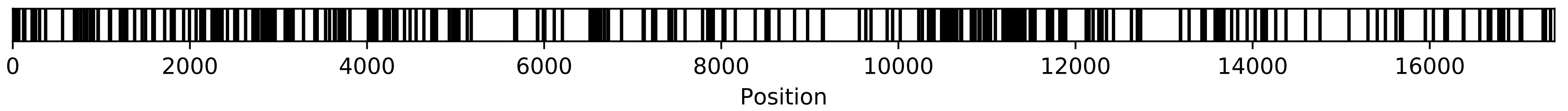


MM1 vs MM2
Number of differences: 180

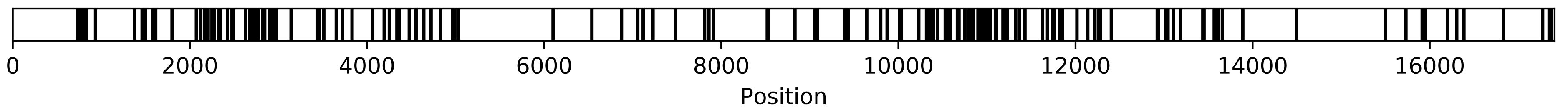

### Region: 308



CR1 vs CR2
Number of differences: 195

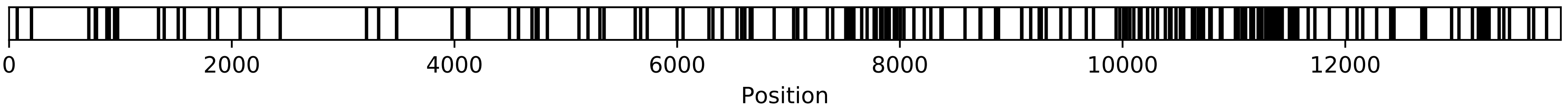


CR1 vs MA1
Number of differences: 104

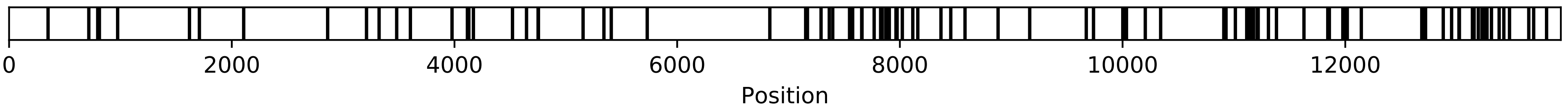


CR1 vs MA2
Number of differences: 141

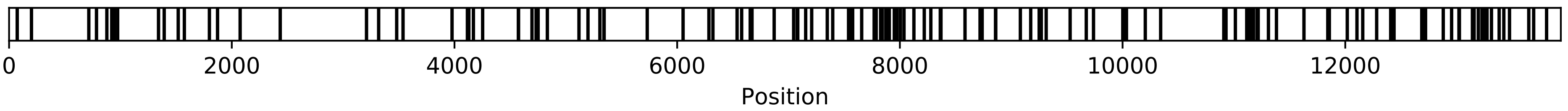


CR1 vs MM1
Number of differences: 31

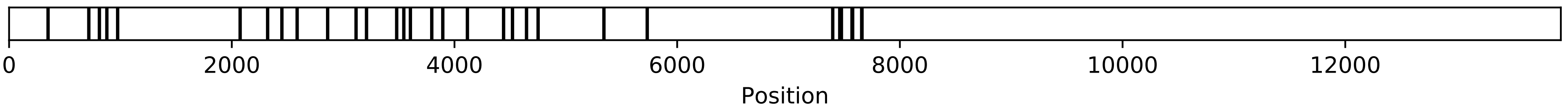


CR1 vs MM2
Number of differences: 86

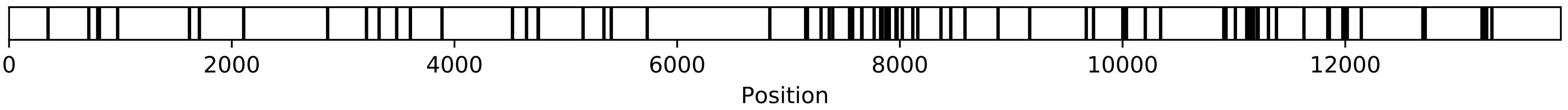


CR2 vs MA1
Number of differences: 193

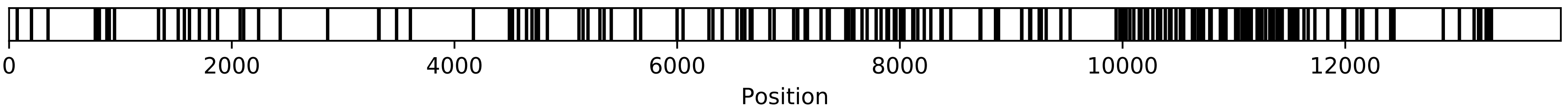


CR2 vs MA2
Number of differences: 107

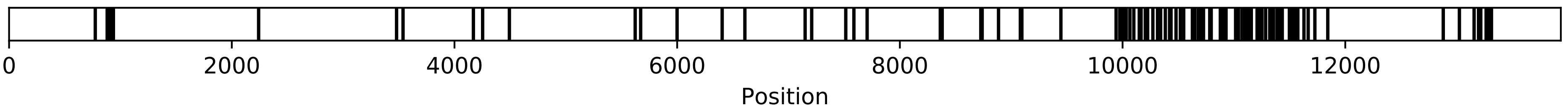


CR2 vs MM1
Number of differences: 201

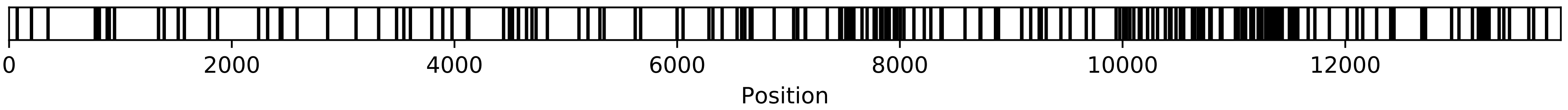


CR2 vs MM2
Number of differences: 205

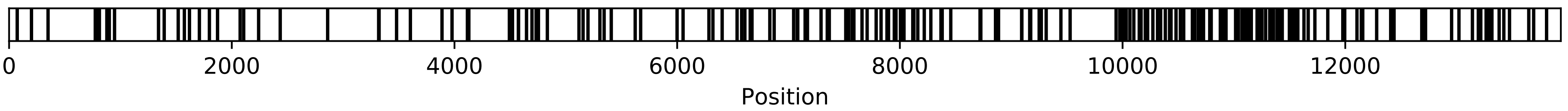


MA1 vs MA2
Number of differences: 102

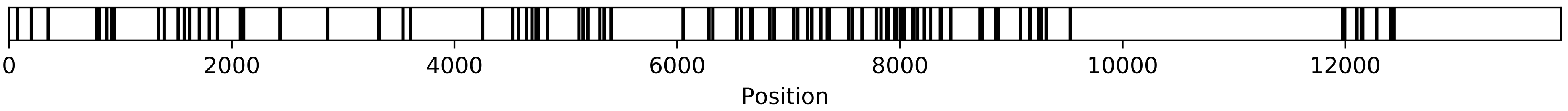


MA1 vs MM1
Number of differences: 98

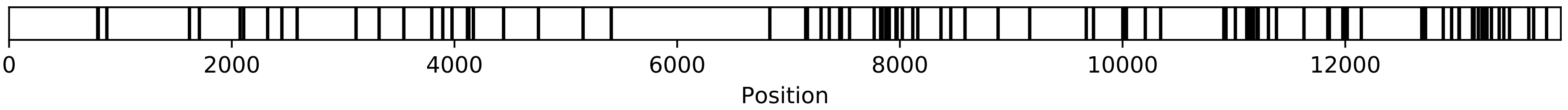


MA1 vs MM2
Number of differences: 24

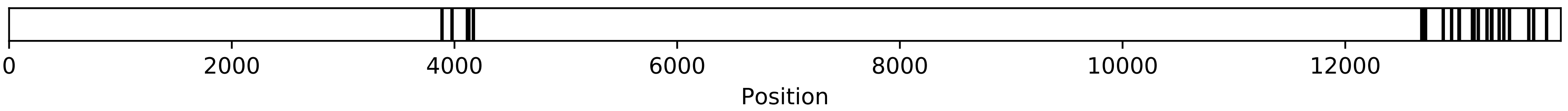


MA2 vs MM1
Number of differences: 147

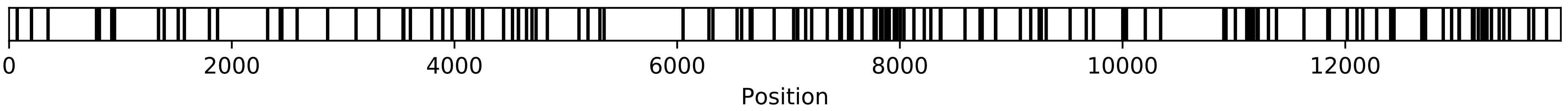


MA2 vs MM2
Number of differences: 126

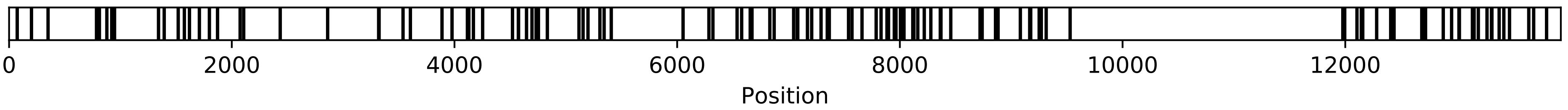


MM1 vs MM2
Number of differences: 81

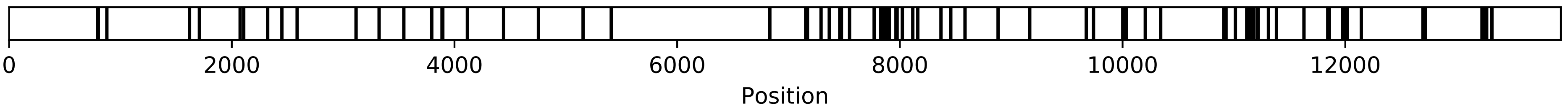

### Region: 311

CR1 vs CR2
Number of differences: 144

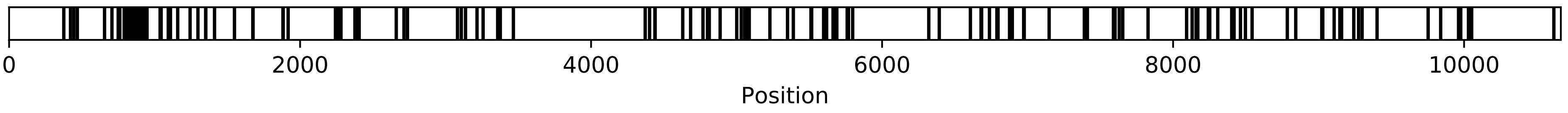


CR1 vs MA1
Number of differences: 48

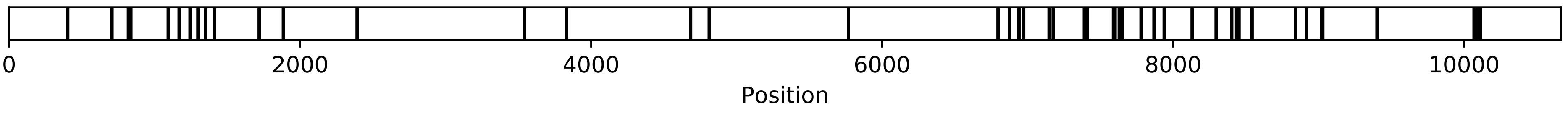


CR1 vs MA2
Number of differences: 148

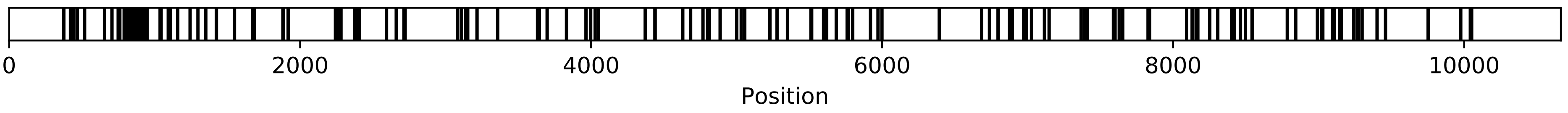


CR1 vs MM1
Number of differences: 49

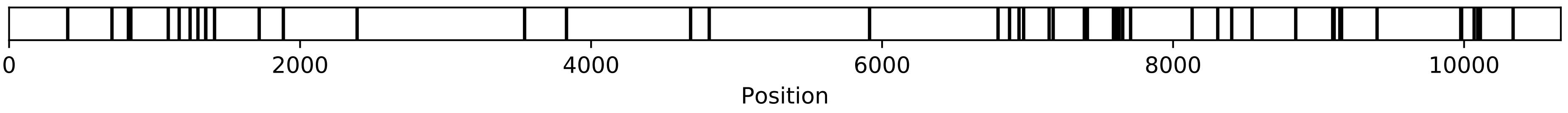


CR1 vs MM2
Number of differences: 48

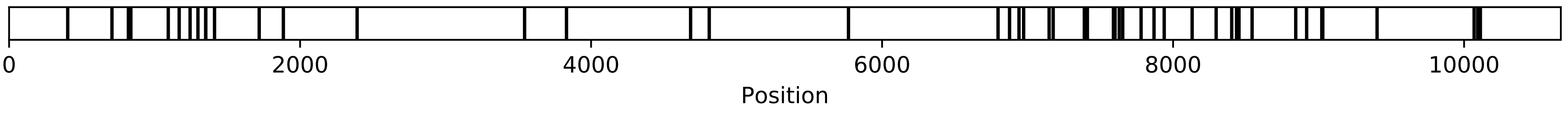


CR2 vs MA1
Number of differences: 135

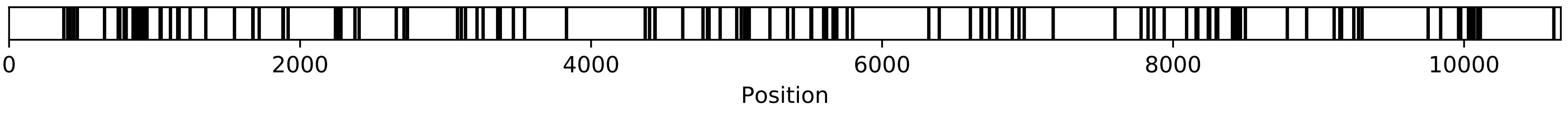


CR2 vs MA2
Number of differences: 53

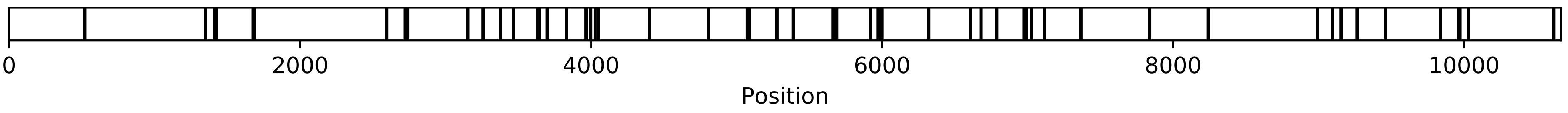


CR2 vs MM1
Number of differences: 132

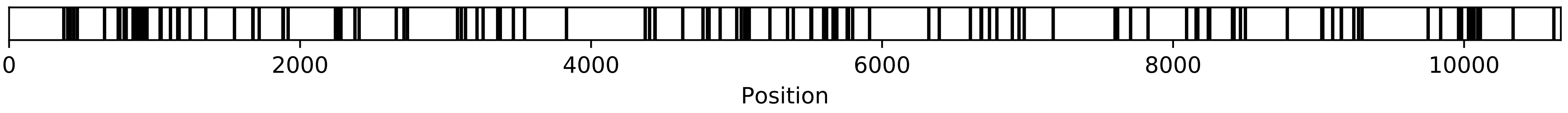


CR2 vs MM2
Number of differences: 135

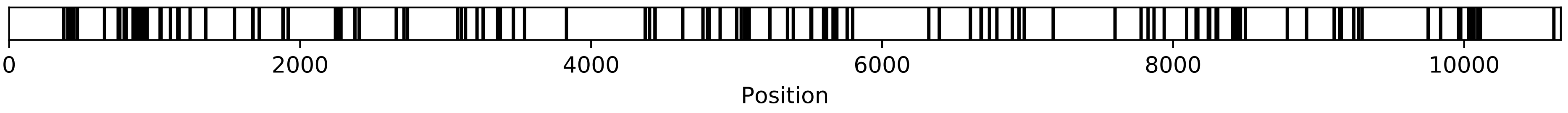


MA1 vs MA2
Number of differences: 138

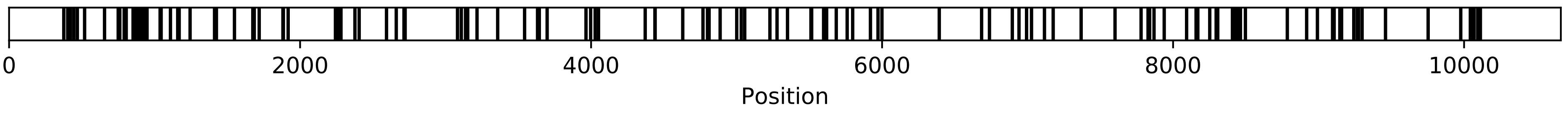


MA1 vs MM1
Number of differences: 21

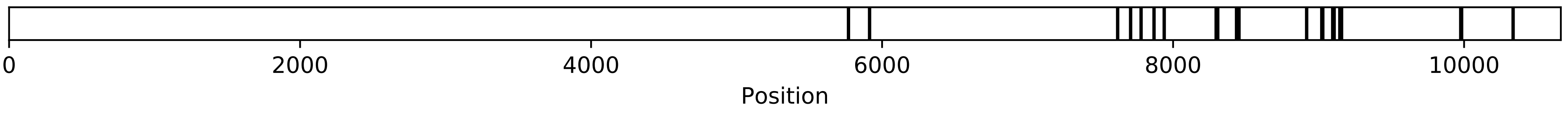


MA1 vs MM2
Number of differences: 0

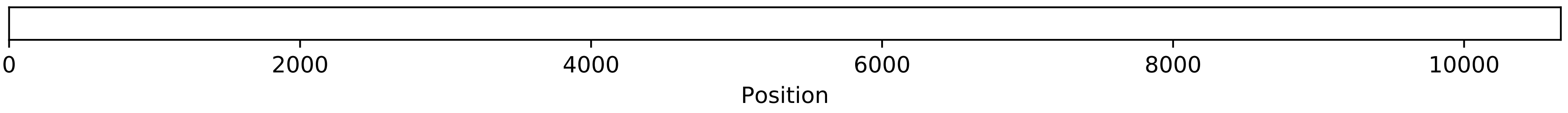


MA2 vs MM1
Number of differences: 135

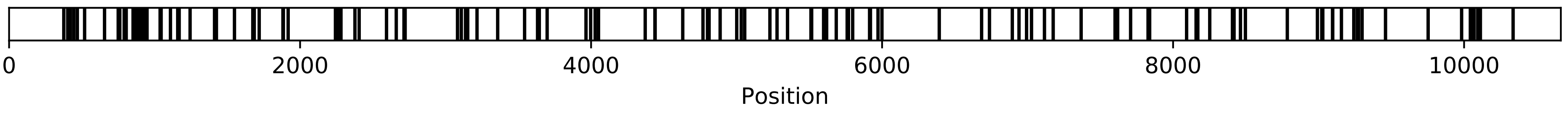


MA2 vs MM2
Number of differences: 138

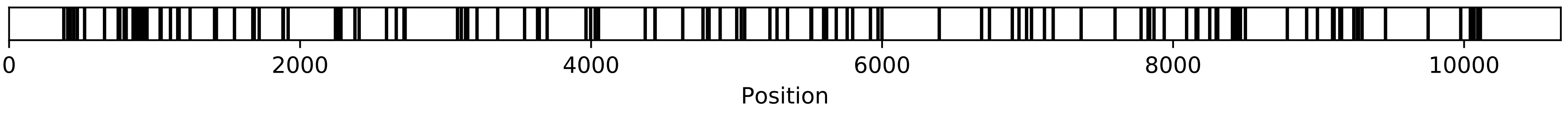


MM1 vs MM2
Number of differences: 21

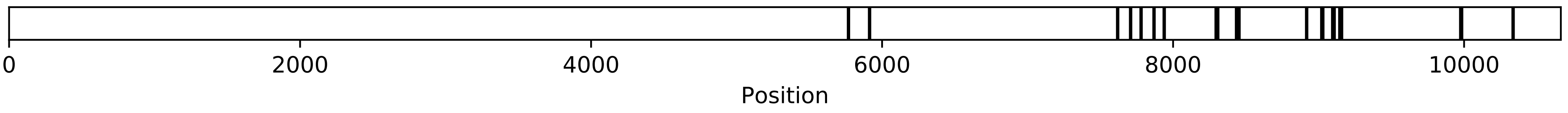

### Region: 312

CR1 vs CR2
Number of differences: 168

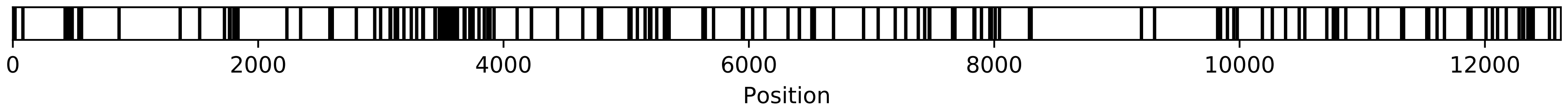


CR1 vs MA1
Number of differences: 70

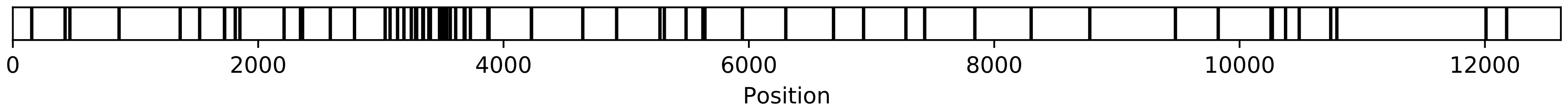


CR1 vs MA2
Number of differences: 171

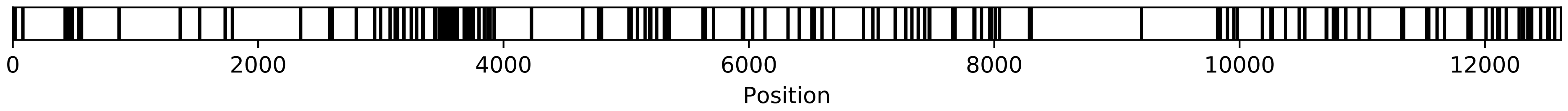


CR1 vs MM1
Number of differences: 70

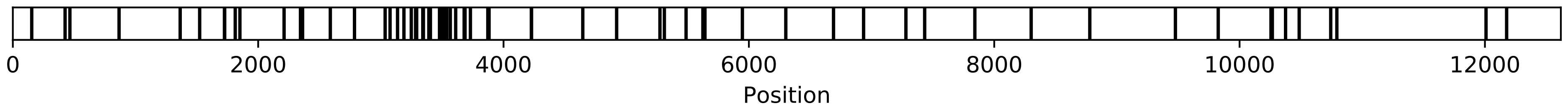


CR1 vs MM2
Number of differences: 75

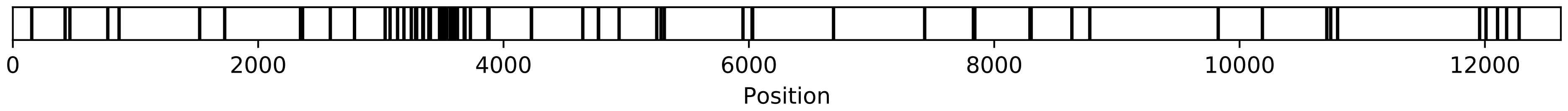


CR2 vs MA1
Number of differences: 155

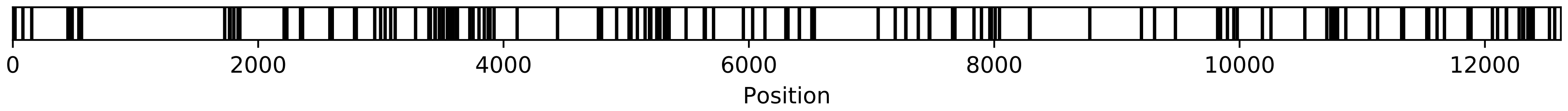


CR2 vs MA2
Number of differences: 35

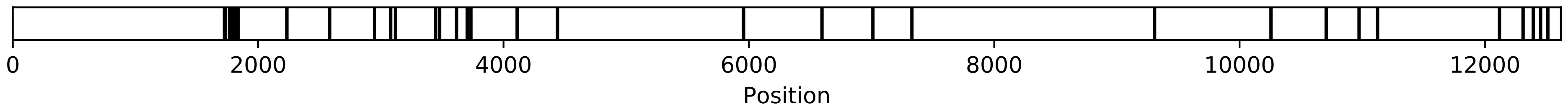


CR2 vs MM1
Number of differences: 155

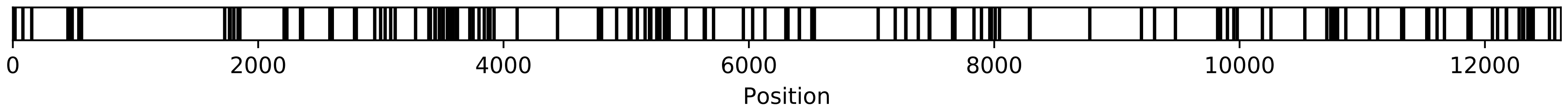


CR2 vs MM2
Number of differences: 153

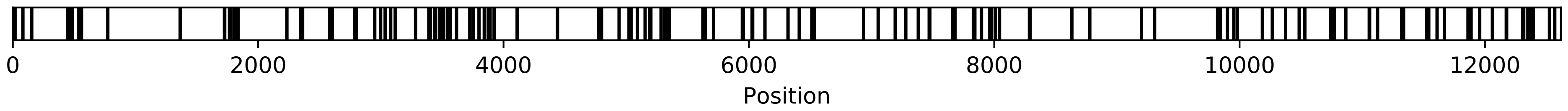


MA1 vs MA2
Number of differences: 160

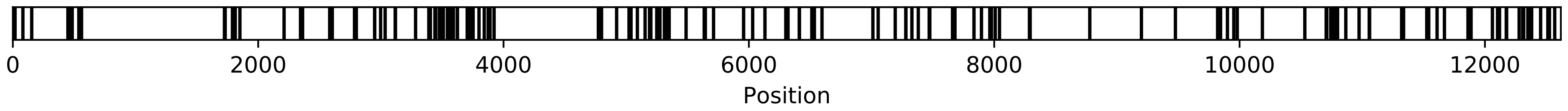


MA1 vs MM1
Number of differences: 0

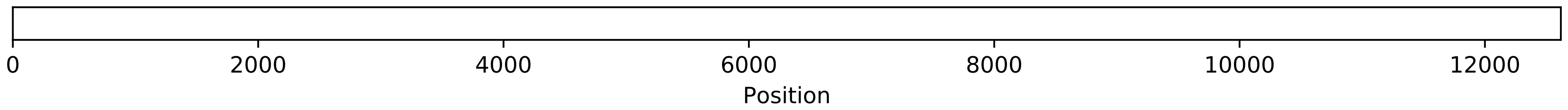


MA1 vs MM2
Number of differences: 49

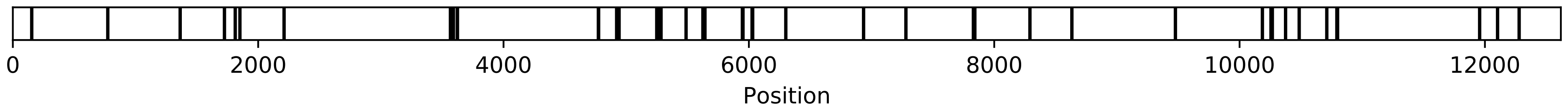


MA2 vs MM1
Number of differences: 160

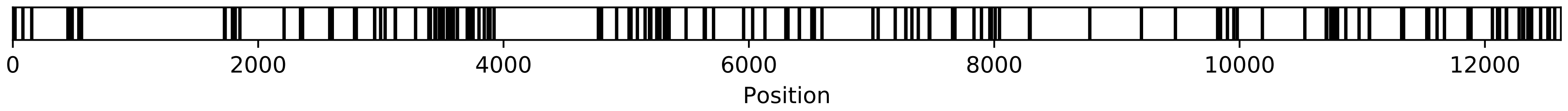


MA2 vs MM2
Number of differences: 156

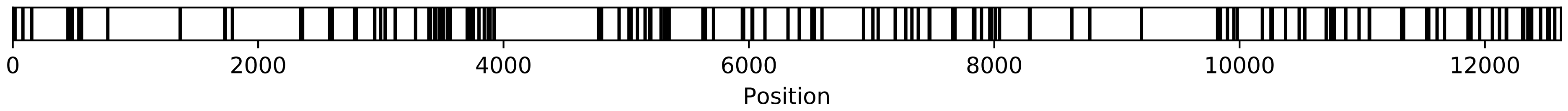


MM1 vs MM2
Number of differences: 49

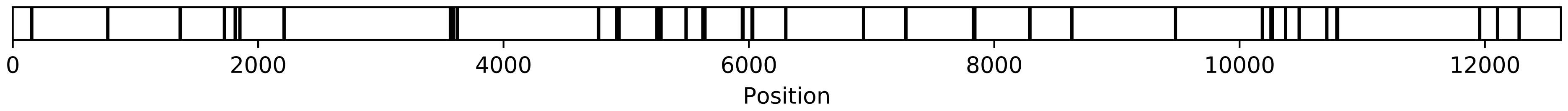

### Region: 313



CR1 vs CR2
Number of differences: 87

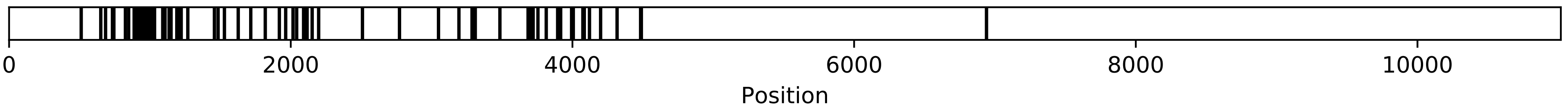


CR1 vs MA1
Number of differences: 145

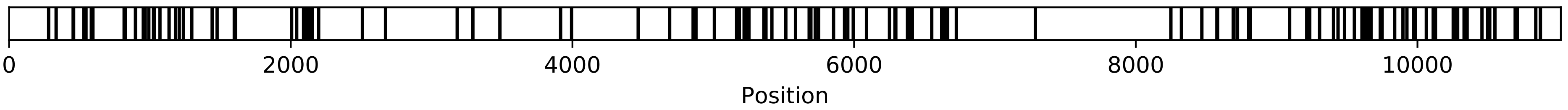


CR1 vs MA2
Number of differences: 162

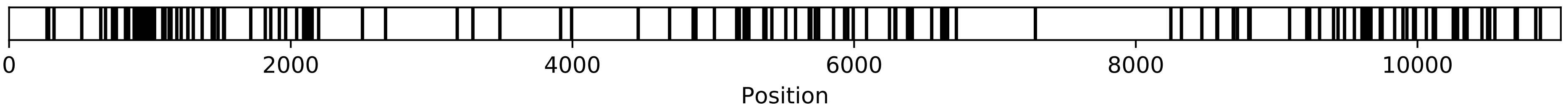


CR1 vs MM1
Number of differences: 221

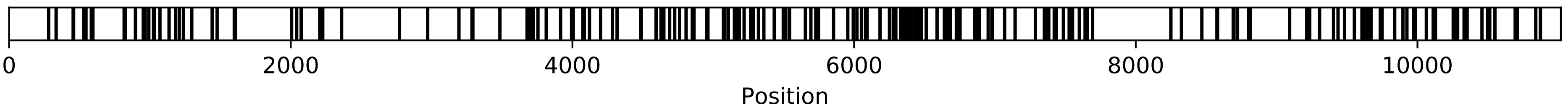


CR1 vs MM2
Number of differences: 238

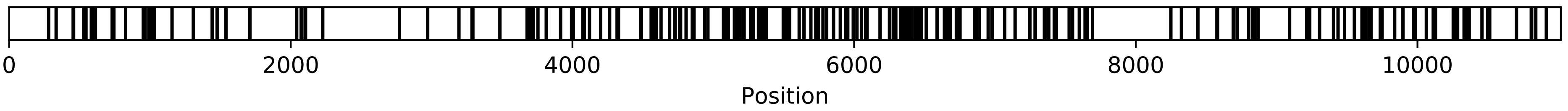


CR2 vs MA1
Number of differences: 195

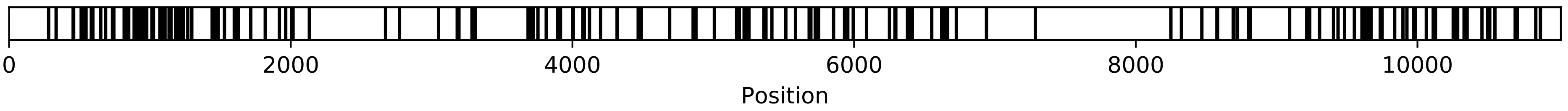


CR2 vs MA2
Number of differences: 144

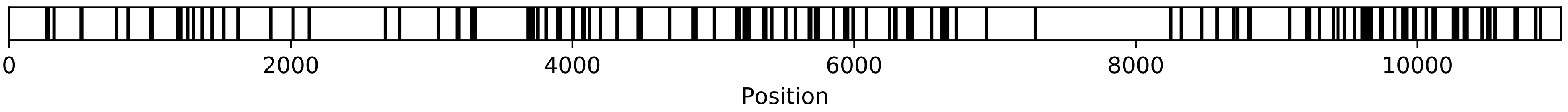


CR2 vs MM1
Number of differences: 239

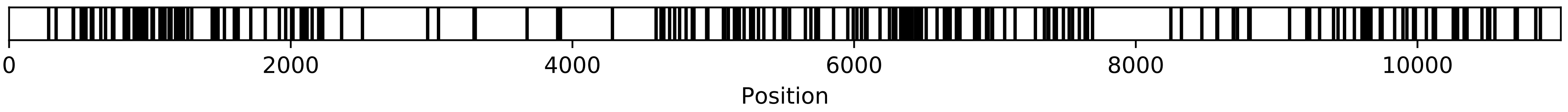


CR2 vs MM2
Number of differences: 251

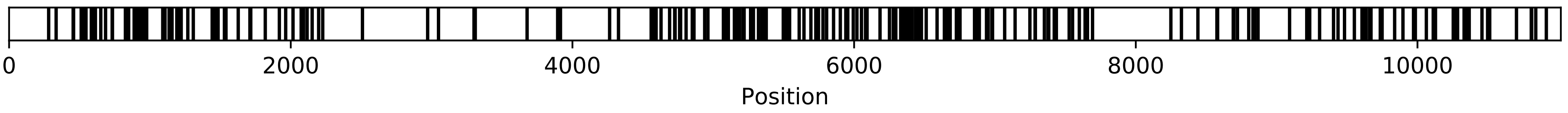


MA1 vs MA2
Number of differences: 74

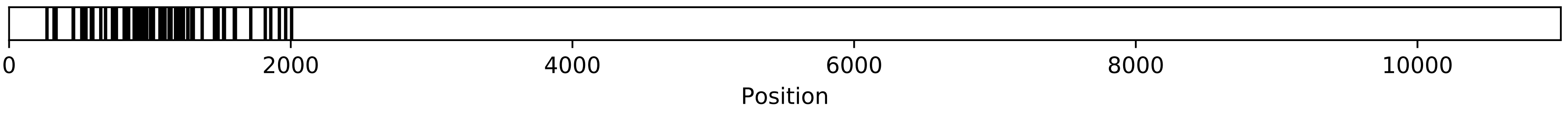


MA1 vs MM1
Number of differences: 141

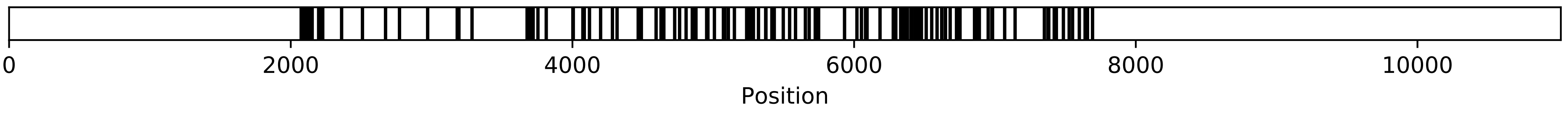


MA1 vs MM2
Number of differences: 199

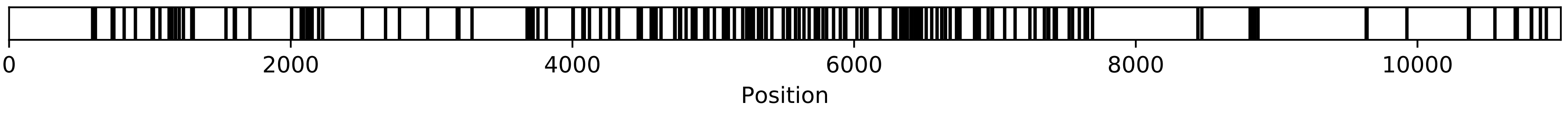


MA2 vs MM1
Number of differences: 215

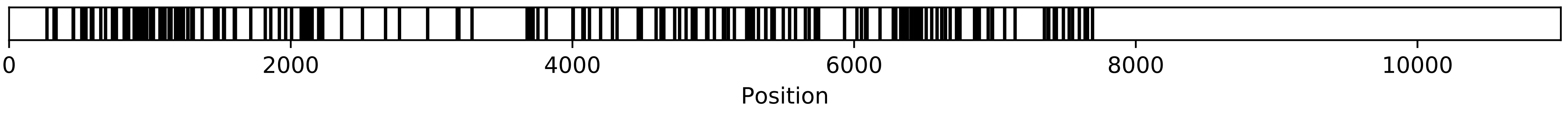


MA2 vs MM2
Number of differences: 243

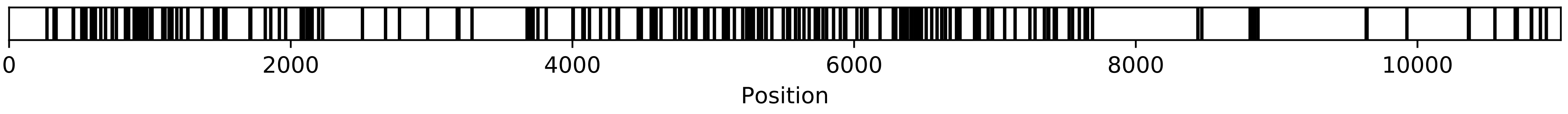


MM1 vs MM2
Number of differences: 78

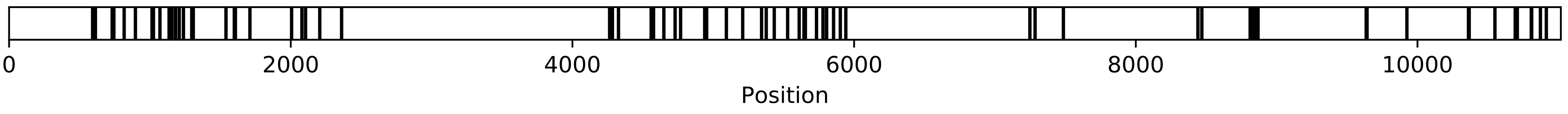

### Region: 315

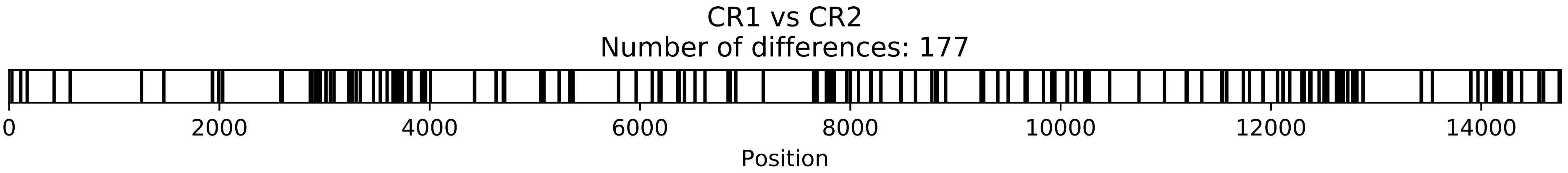
CR1 vs CR2
Number of differences: 177
0 2000 4000 6000 8000 10000 12000 14000
Position

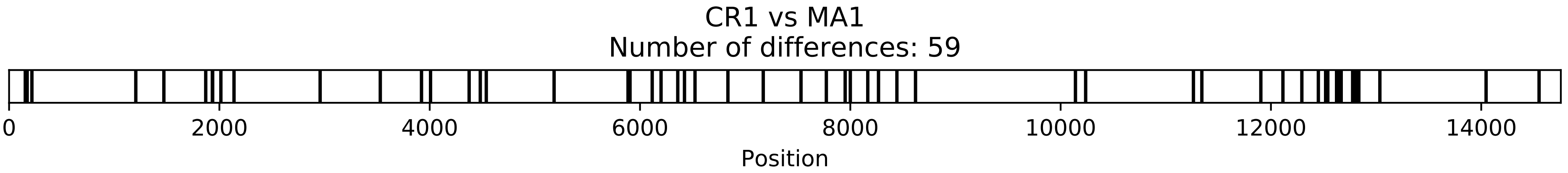
CR1 vs MA1
Number of differences: 59
0 2000 4000 6000 8000 10000 12000 14000
Position

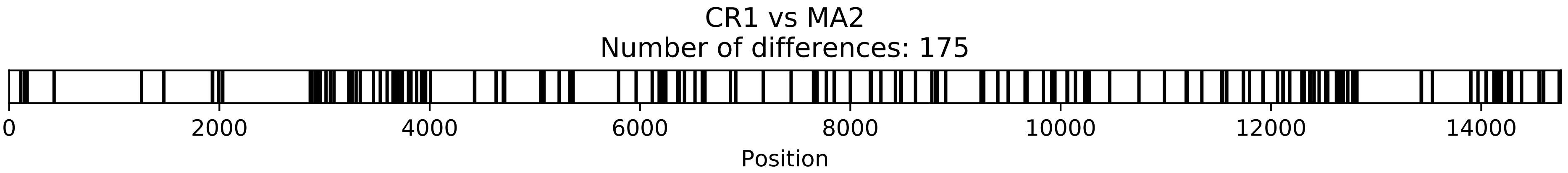
CR1 vs MA2
Number of differences: 175
0 2000 4000 6000 8000 10000 12000 14000
Position

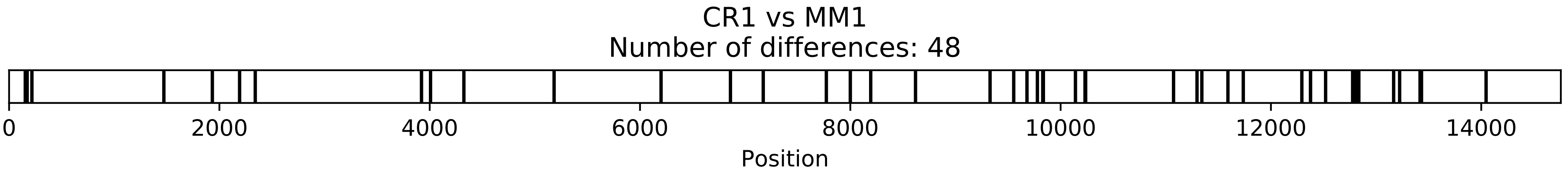
CR1 vs MM1
Number of differences: 48
0 2000 4000 6000 8000 10000 12000 14000
Position

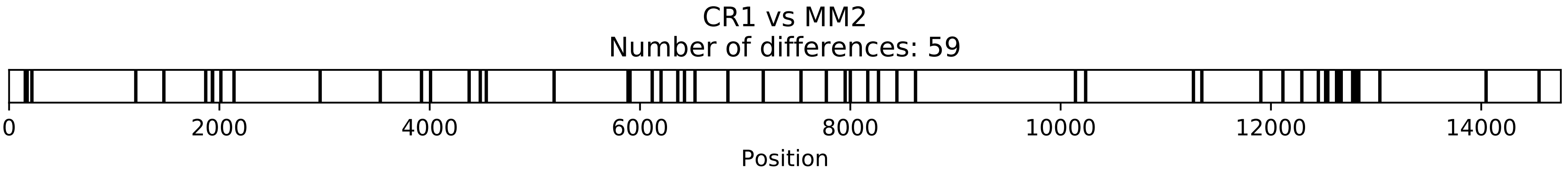
CR1 vs MM2
Number of differences: 59
0 2000 4000 6000 8000 10000 12000 14000
Position

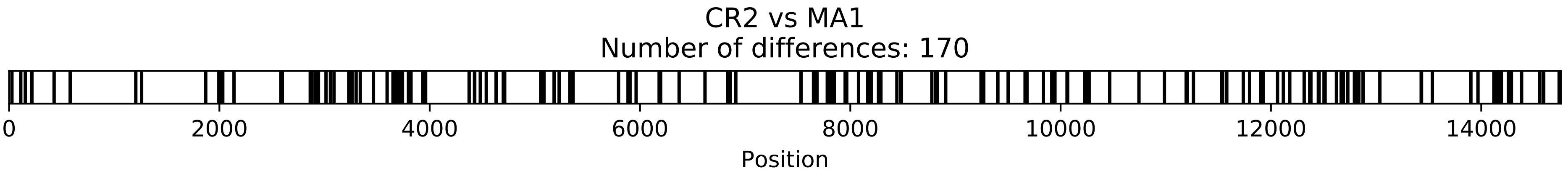
CR2 vs MA1
Number of differences: 170
0 2000 4000 6000 8000 10000 12000 14000
Position

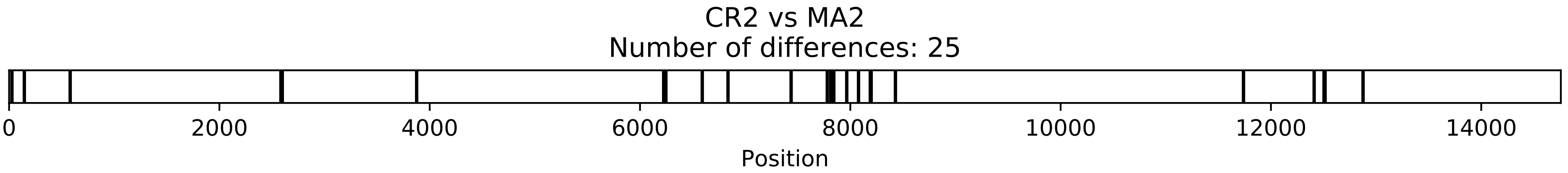
CR2 vs MA2
Number of differences: 25
0 2000 4000 6000 8000 10000 12000 14000
Position

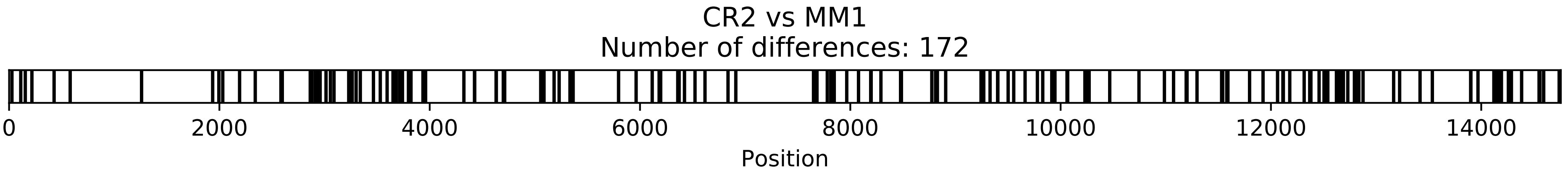
CR2 vs MM1
Number of differences: 172
0 2000 4000 6000 8000 10000 12000 14000
Position

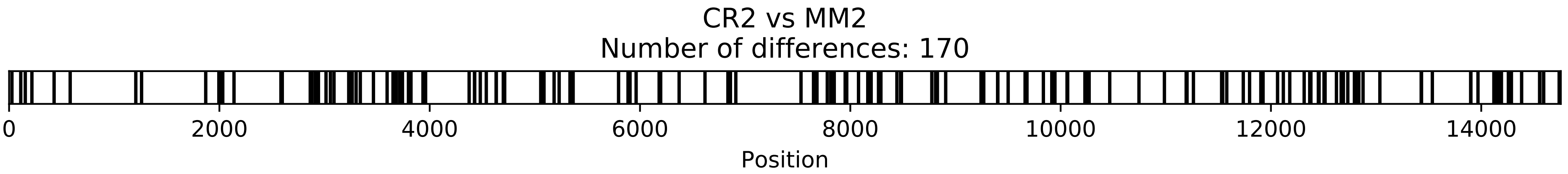
CR2 vs MM2
Number of differences: 170
0 2000 4000 6000 8000 10000 12000 14000
Position

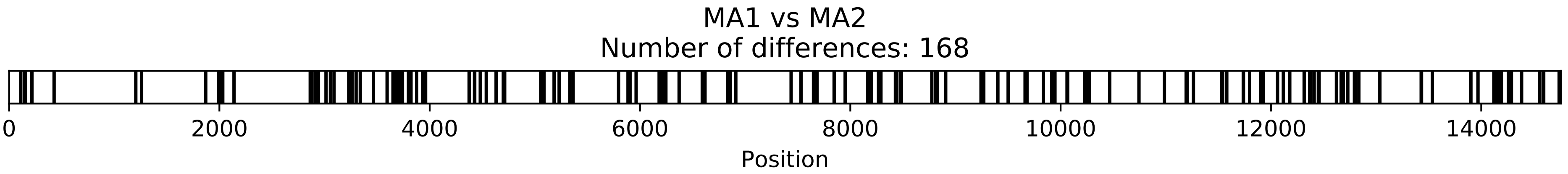
MA1 vs MA2
Number of differences: 168
0 2000 4000 6000 8000 10000 12000 14000
Position

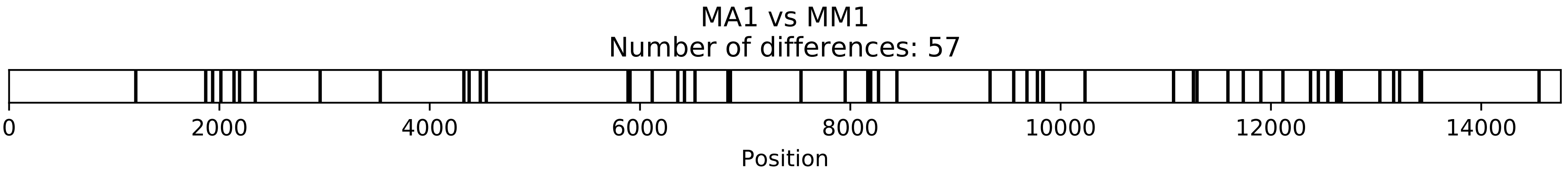
MA1 vs MM1
Number of differences: 57
0 2000 4000 6000 8000 10000 12000 14000
Position

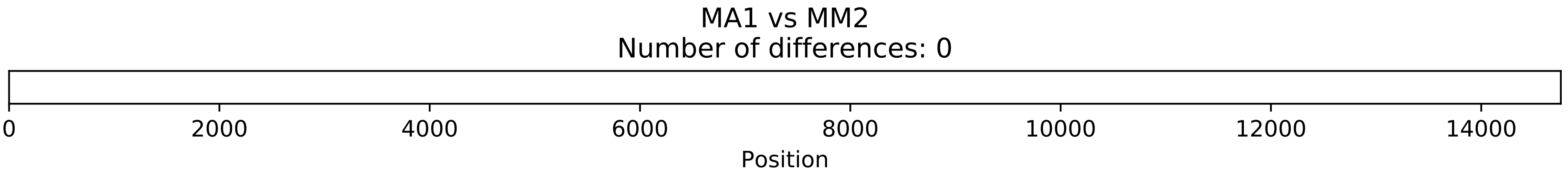
MA1 vs MM2
Number of differences: 0
0 2000 4000 6000 8000 10000 12000 14000
Position

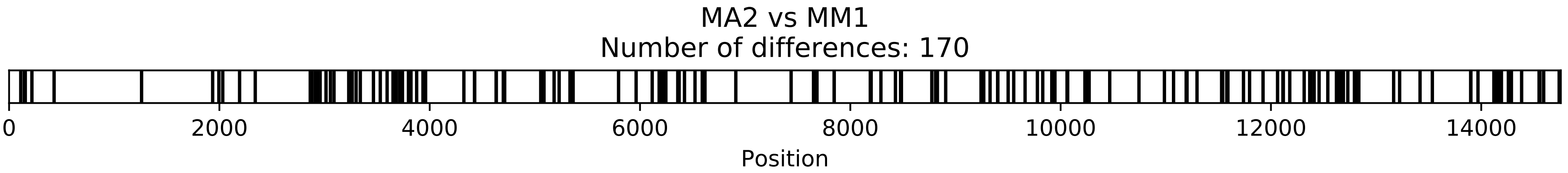
MA2 vs MM1
Number of differences: 170
0 2000 4000 6000 8000 10000 12000 14000
Position

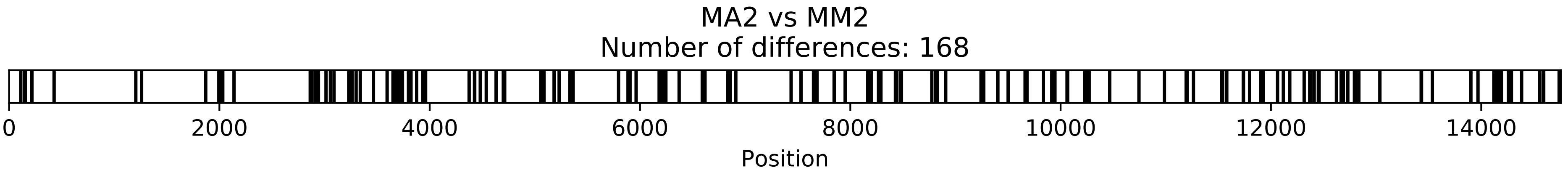
MA2 vs MM2
Number of differences: 168
0 2000 4000 6000 8000 10000 12000 14000
Position

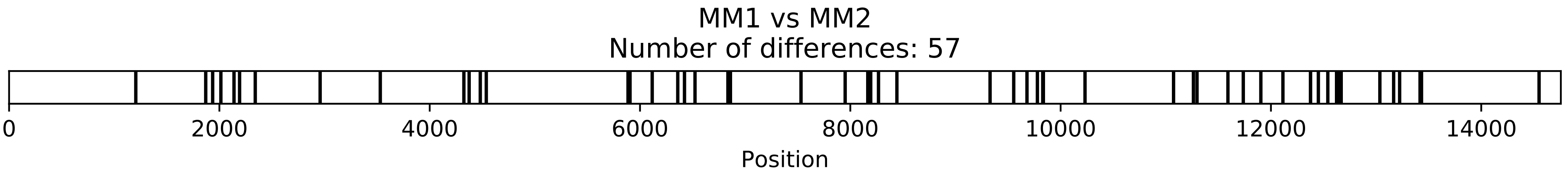
MM1 vs MM2
Number of differences: 57
0 2000 4000 6000 8000 10000 12000 14000
Position

### Region: 316

CR1 vs CR2
Number of differences: 184

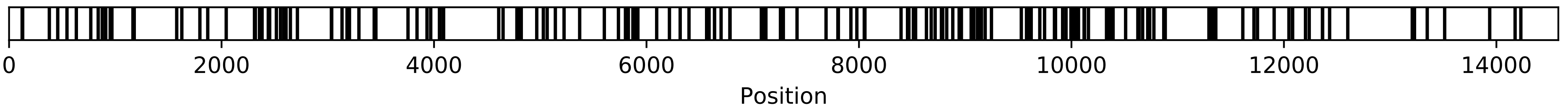


CR1 vs MA1
Number of differences: 54

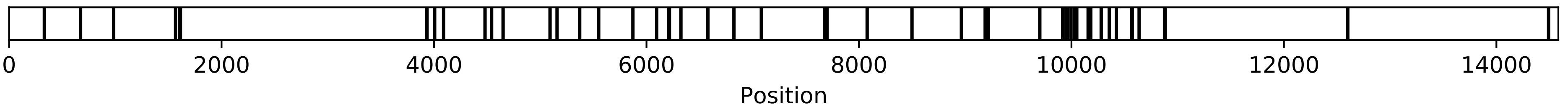


CR1 vs MA2
Number of differences: 183

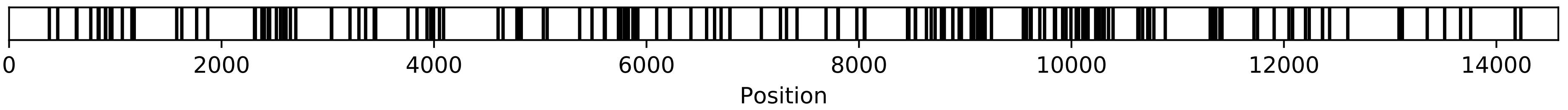


CR1 vs MM1
Number of differences: 194

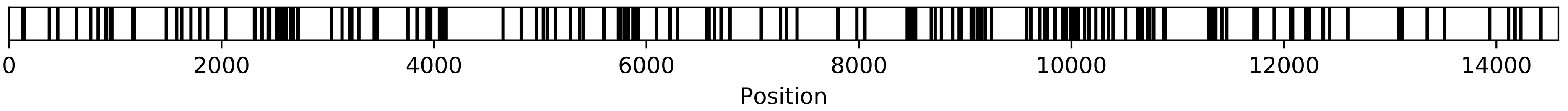


CR1 vs MM2
Number of differences: 183

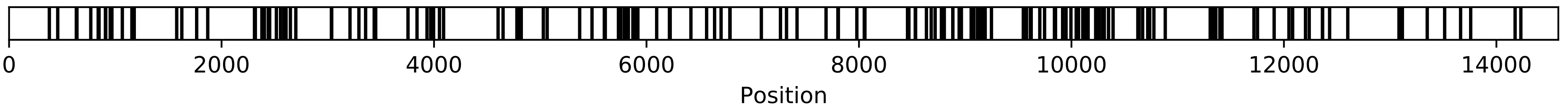


CR2 vs MA1
Number of differences: 198

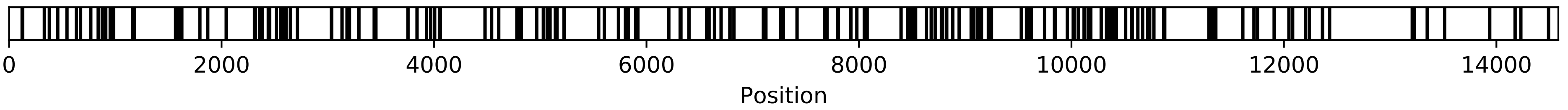


CR2 vs MA2
Number of differences: 94

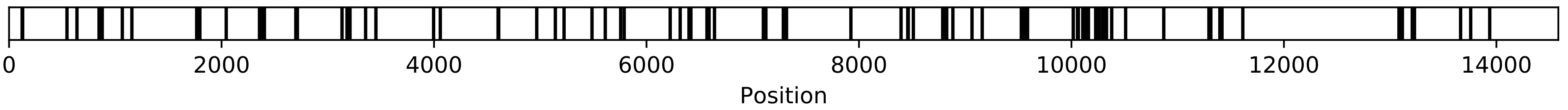


CR2 vs MM1
Number of differences: 82

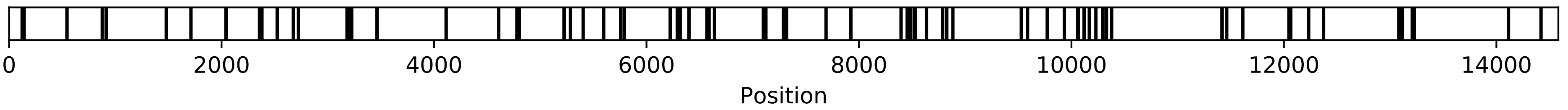


CR2 vs MM2
Number of differences: 94

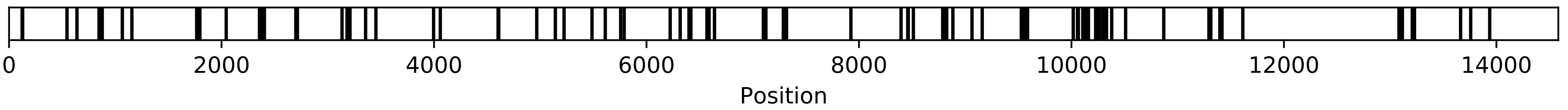


MA1 vs MA2
Number of differences: 195

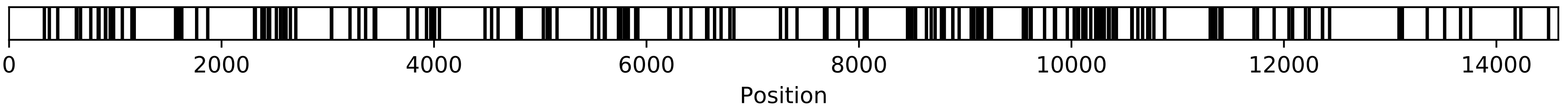


MA1 vs MM1
Number of differences: 208

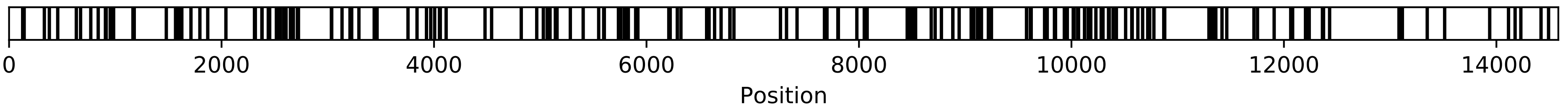


MA1 vs MM2
Number of differences: 195

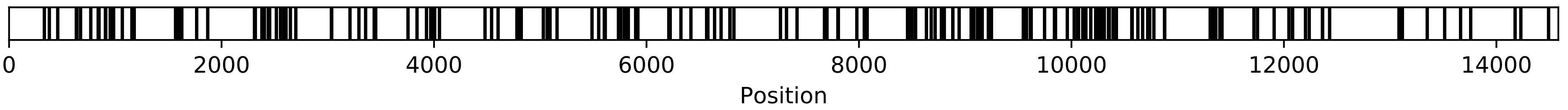


MA2 vs MM1
Number of differences: 89

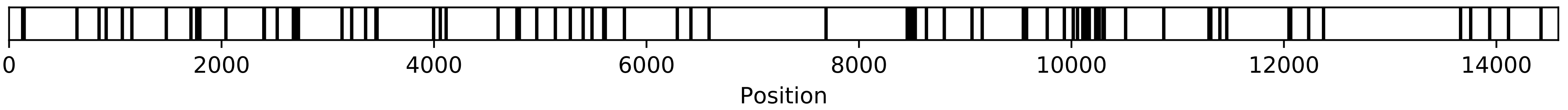


MA2 vs MM2
Number of differences: 0

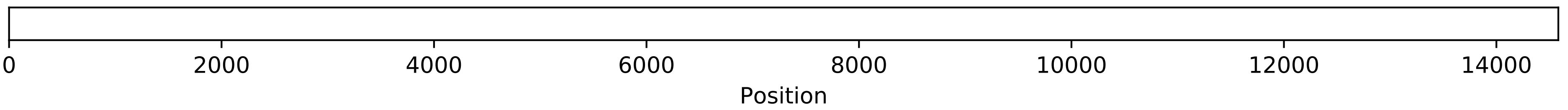


MM1 vs MM2
Number of differences: 89

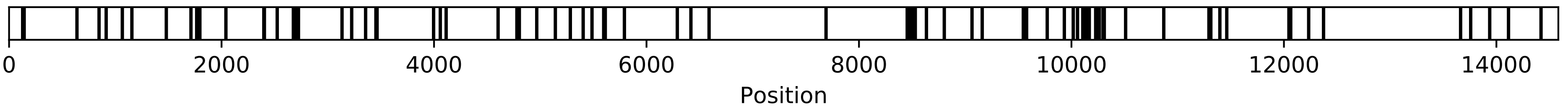

### Region: 317



CR1 vs CR2
Number of differences: 297

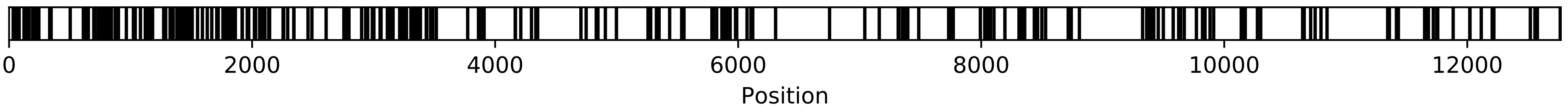


CR1 vs MA1
Number of differences: 152

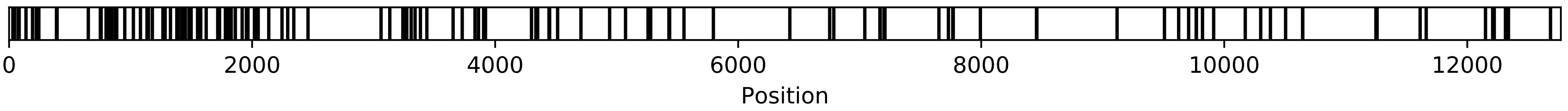


CR1 vs MA2
Number of differences: 303

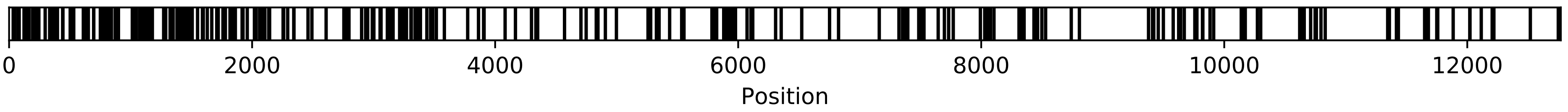


CR1 vs MM1
Number of differences: 165

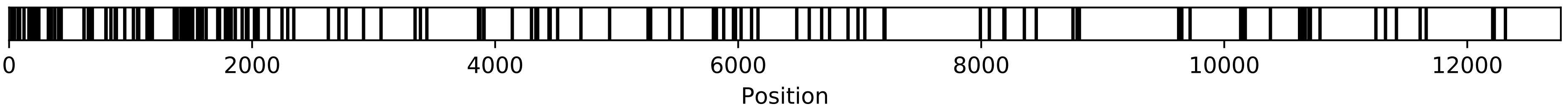


CR1 vs MM2
Number of differences: 152

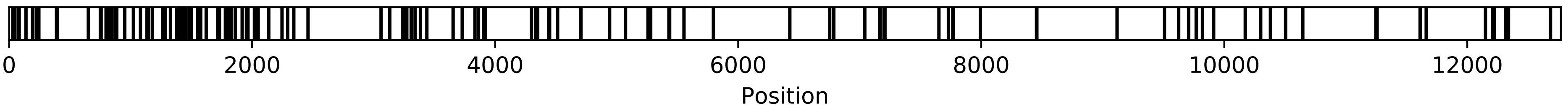


CR2 vs MA1
Number of differences: 295

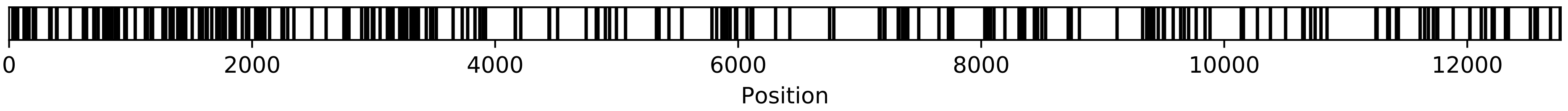


CR2 vs MA2
Number of differences: 88

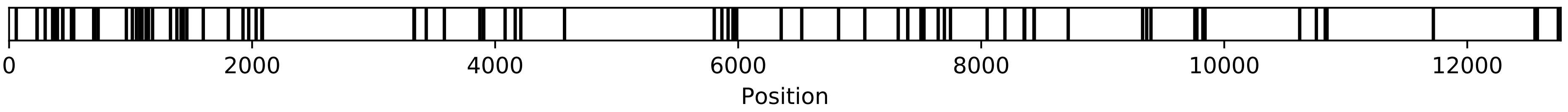


CR2 vs MM1
Number of differences: 315

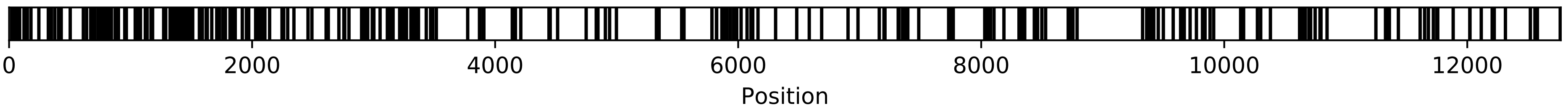


CR2 vs MM2
Number of differences: 295

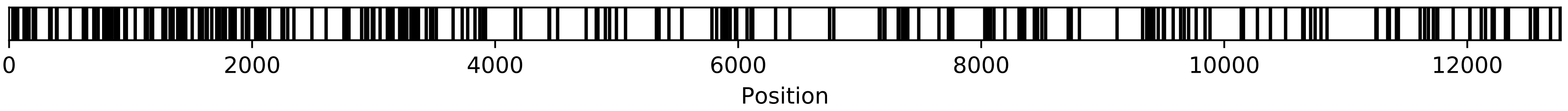


MA1 vs MA2
Number of differences: 310

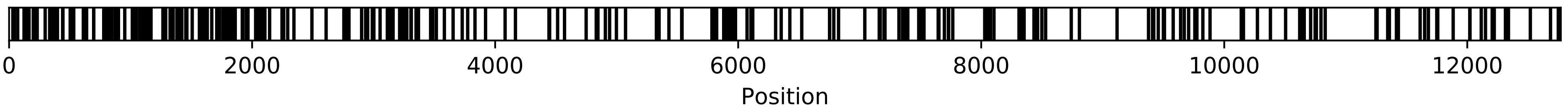


MA1 vs MM1
Number of differences: 152

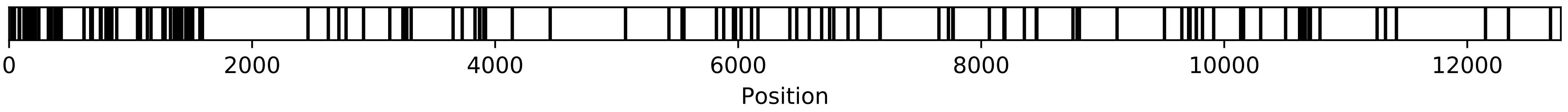


MA1 vs MM2
Number of differences: 0

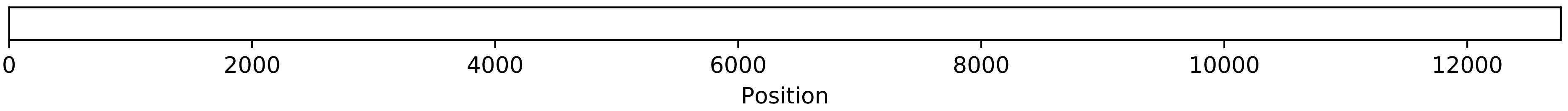


MA2 vs MM1
Number of differences: 324

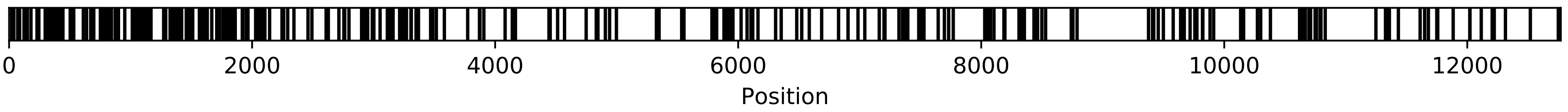


MA2 vs MM2
Number of differences: 310

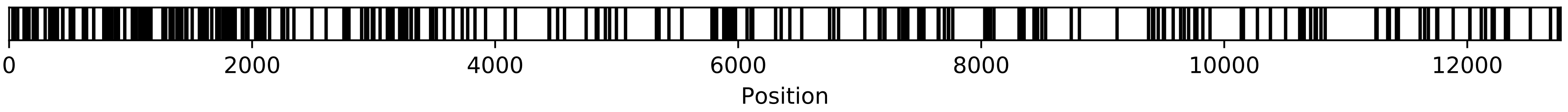


MM1 vs MM2
Number of differences: 152

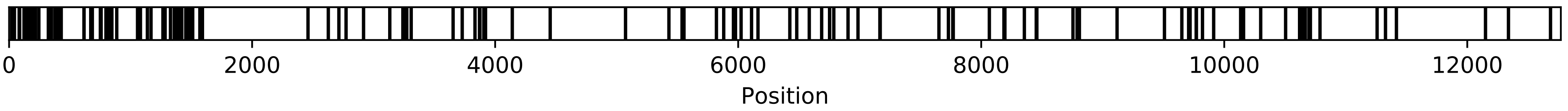

### Region: 318



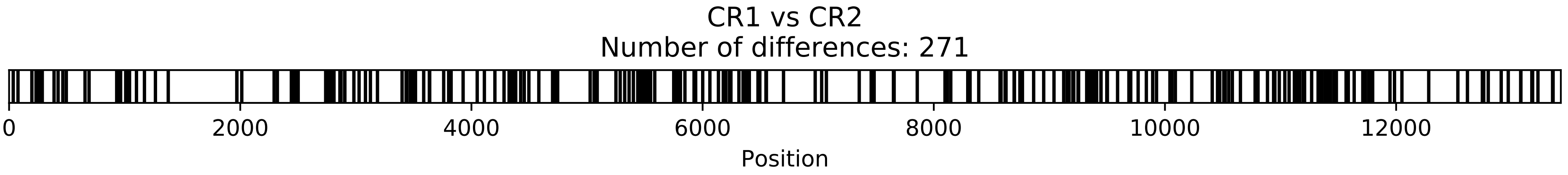
CR1 vs CR2
Number of differences: 271
0 2000 4000 6000 8000 10000 12000
Position

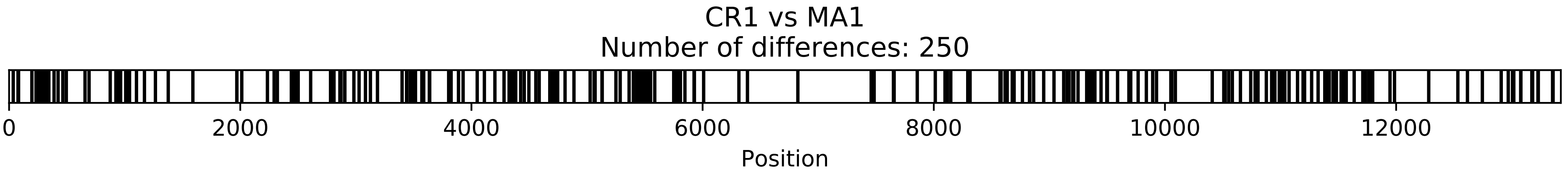
CR1 vs MA1
Number of differences: 250
0 2000 4000 6000 8000 10000 12000
Position

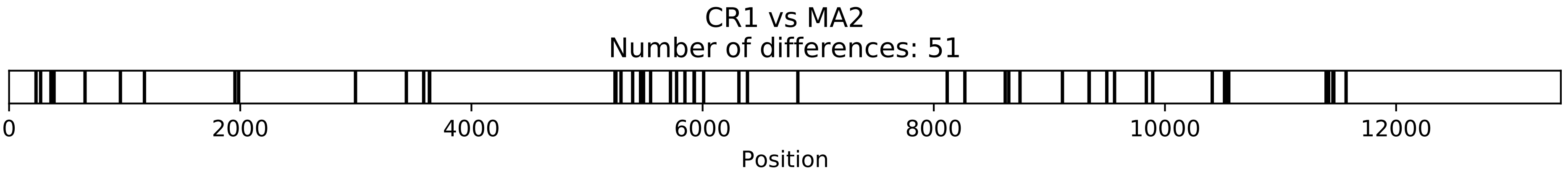
CR1 vs MA2
Number of differences: 51
0 2000 4000 6000 8000 10000 12000
Position

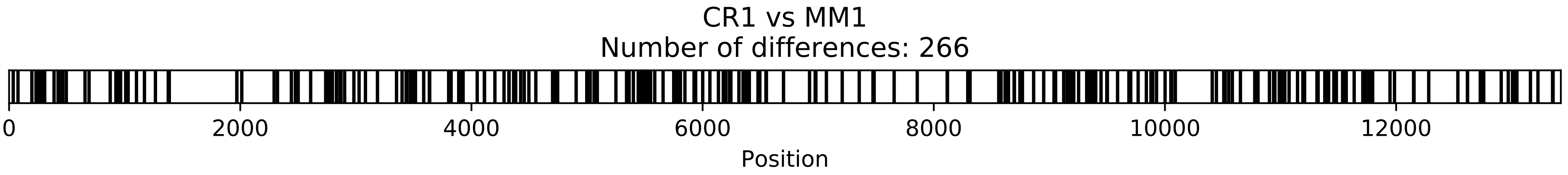
CR1 vs MM1
Number of differences: 266
0 2000 4000 6000 8000 10000 12000
Position

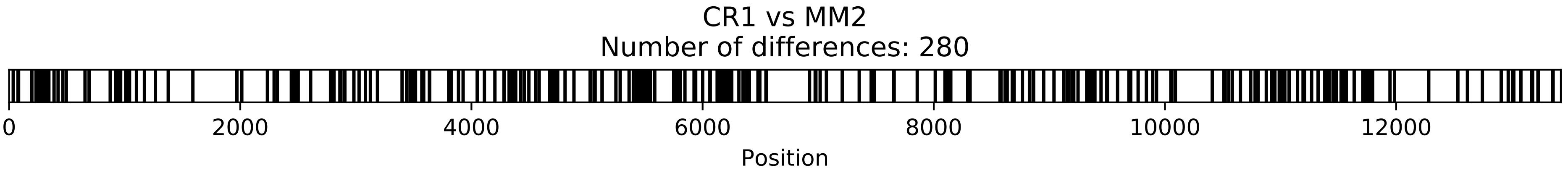
CR1 vs MM2
Number of differences: 280
0 2000 4000 6000 8000 10000 12000
Position

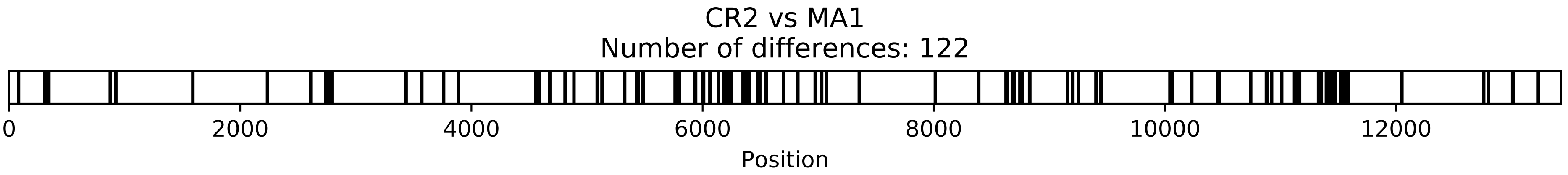
CR2 vs MA1
Number of differences: 122
0 2000 4000 6000 8000 10000 12000
Position

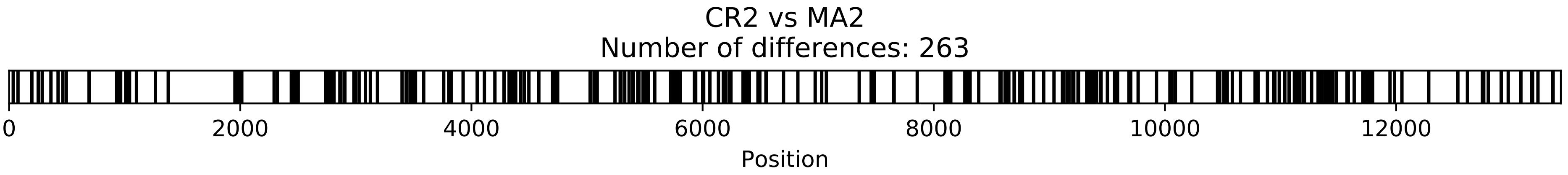
CR2 vs MA2
Number of differences: 263
0 2000 4000 6000 8000 10000 12000
Position

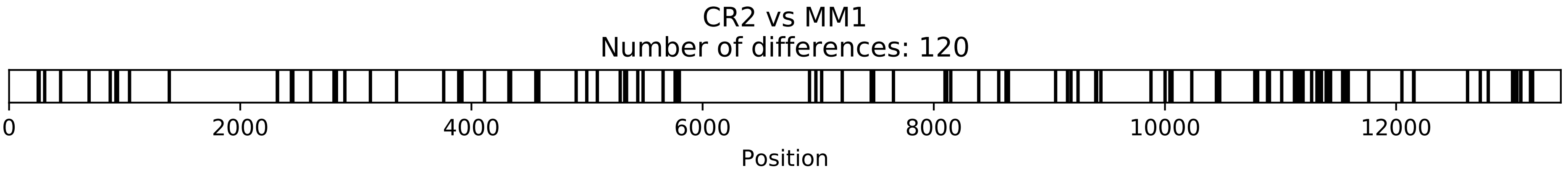
CR2 vs MM1
Number of differences: 120
0 2000 4000 6000 8000 10000 12000
Position

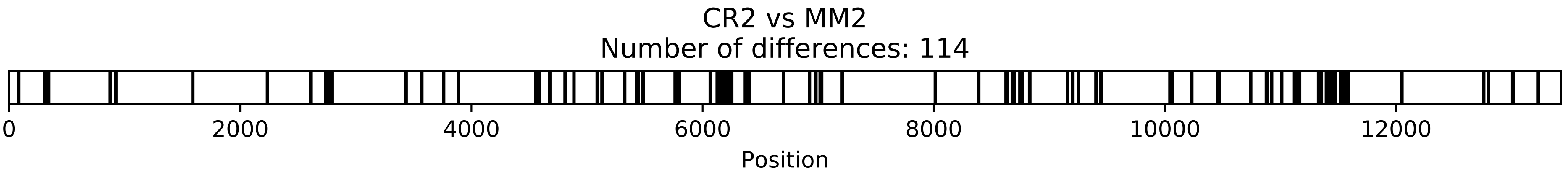
CR2 vs MM2
Number of differences: 114
0 2000 4000 6000 8000 10000 12000
Position

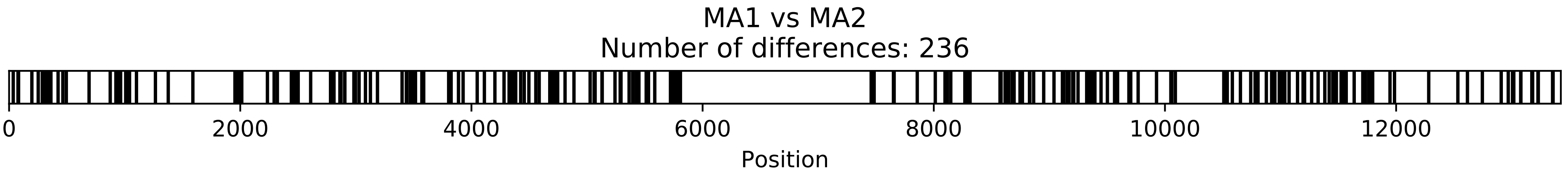
MA1 vs MA2
Number of differences: 236
0 2000 4000 6000 8000 10000 12000
Position

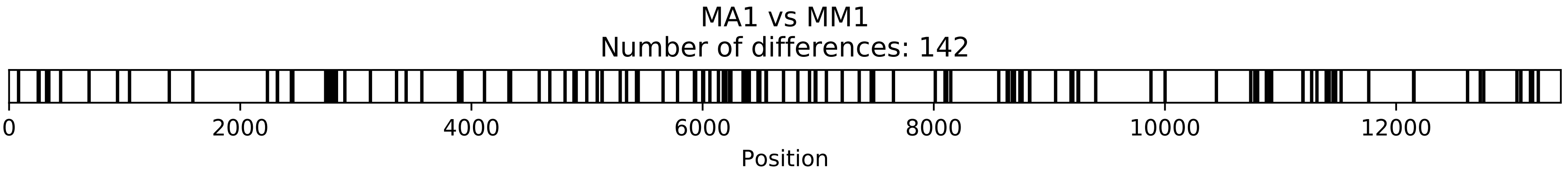
MA1 vs MM1
Number of differences: 142
0 2000 4000 6000 8000 10000 12000
Position

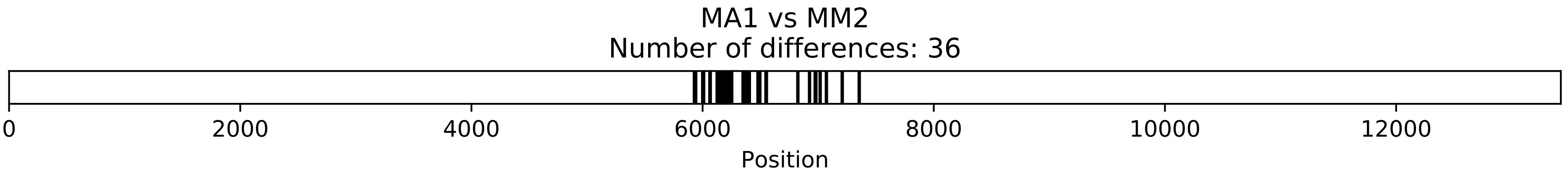
MA1 vs MM2
Number of differences: 36
0 2000 4000 6000 8000 10000 12000
Position

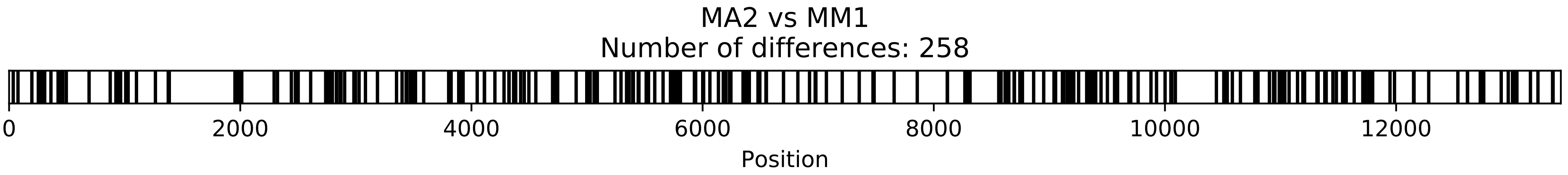
MA2 vs MM1
Number of differences: 258
0 2000 4000 6000 8000 10000 12000
Position

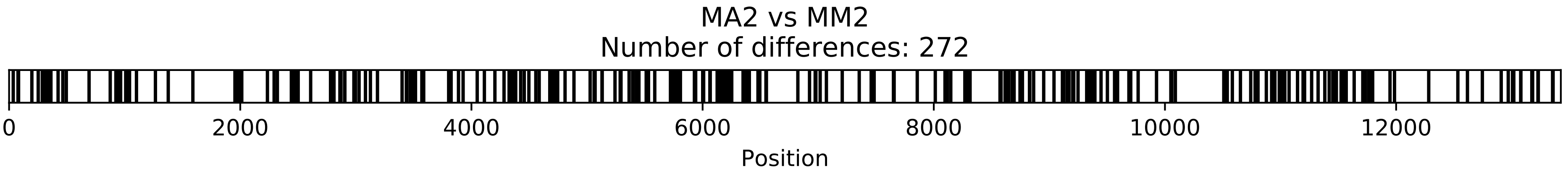
MA2 vs MM2
Number of differences: 272
0 2000 4000 6000 8000 10000 12000
Position

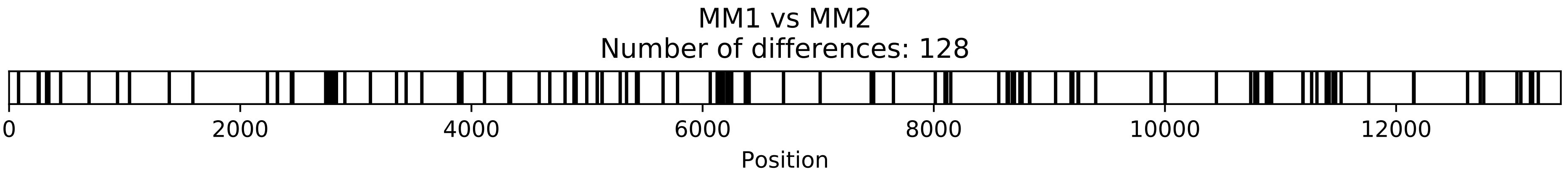
MM1 vs MM2
Number of differences: 128
0 2000 4000 6000 8000 10000 12000
Position

### Region: 319



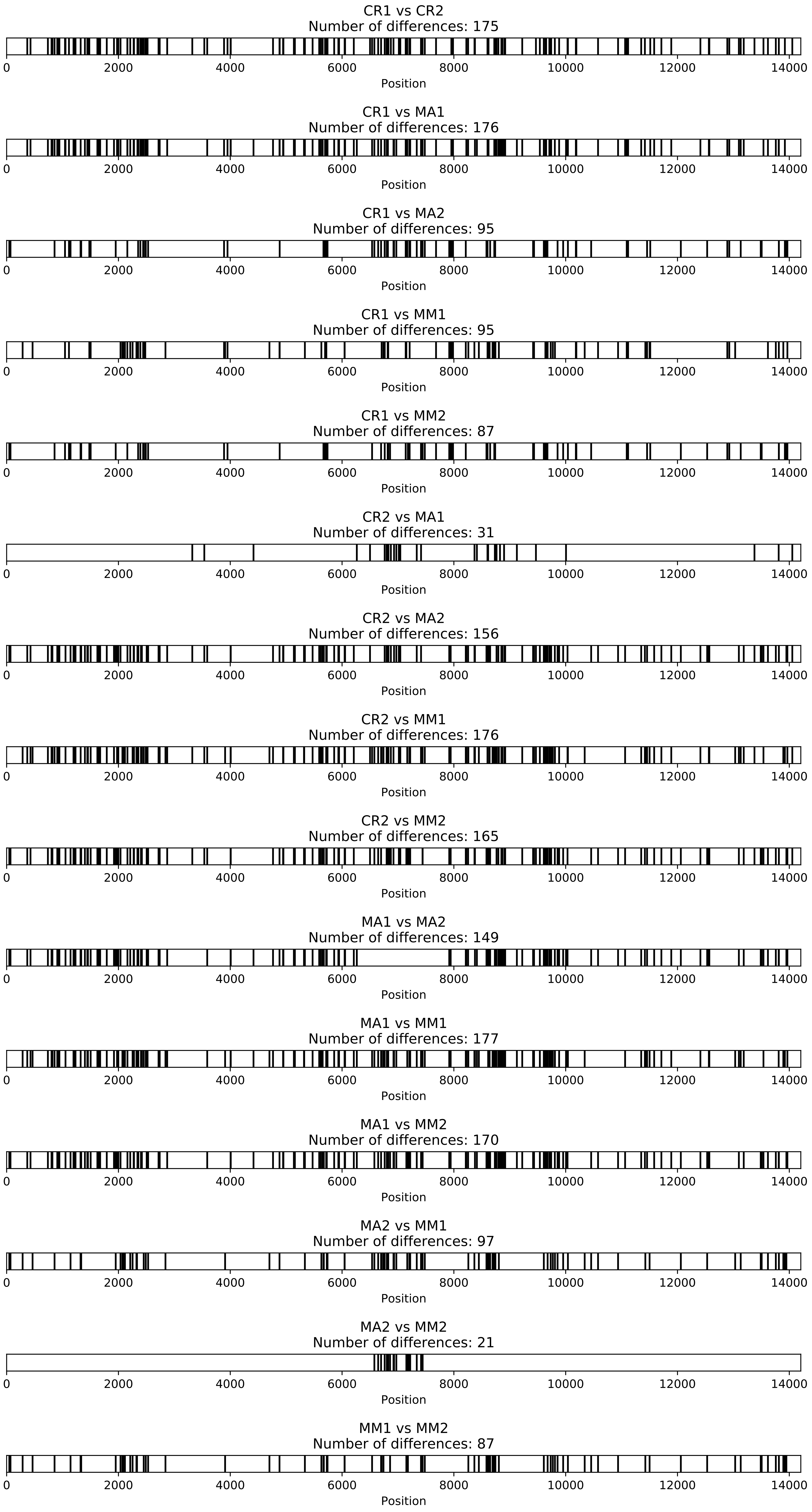
CR1 vs CR2
Number of differences: 175
CR1 vs MA1
Number of differences: 176
CR1 vs MA2
Number of differences: 95
CR1 vs MM1
Number of differences: 95
CR1 vs MM2
Number of differences: 87
CR2 vs MA1
Number of differences: 31
CR2 vs MA2
Number of differences: 156
CR2 vs MM1
Number of differences: 176
CR2 vs MM2
Number of differences: 165
MA1 vs MA2
Number of differences: 149
MA1 vs MM1
Number of differences: 177
MA1 vs MM2
Number of differences: 170
MA2 vs MM1
Number of differences: 97
MA2 vs MM2
Number of differences: 21
MM1 vs MM2
Number of differences: 87
0
2000
4000
6000
8000
10000
12000
14000
Position

### Region: 320

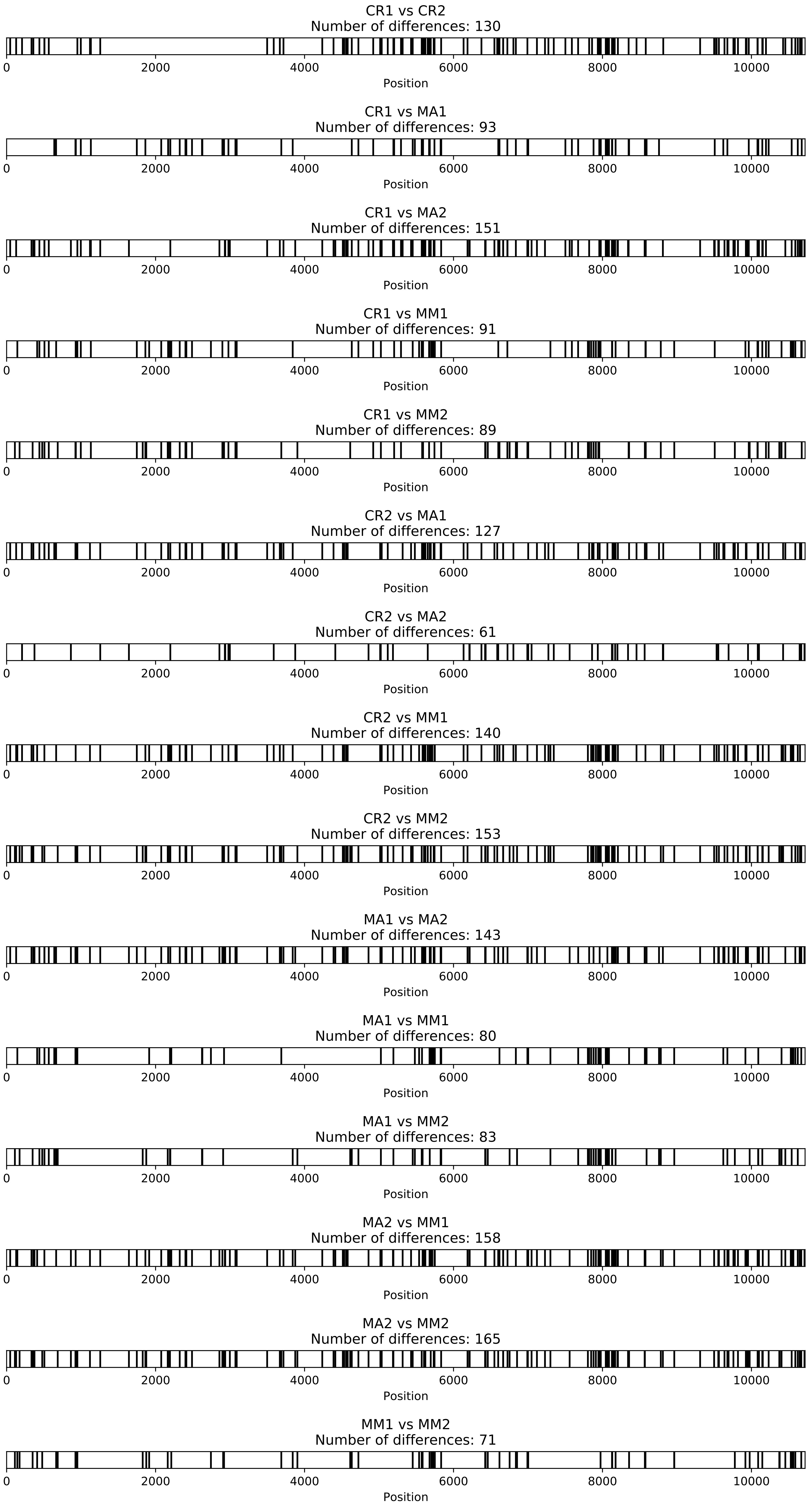
CR1 vs CR2
Number of differences: 130
CR1 vs MA1
Number of differences: 93
CR1 vs MA2
Number of differences: 151
CR1 vs MM1
Number of differences: 91
CR1 vs MM2
Number of differences: 89
CR2 vs MA1
Number of differences: 127
CR2 vs MA2
Number of differences: 61
CR2 vs MM1
Number of differences: 140
CR2 vs MM2
Number of differences: 153
MA1 vs MA2
Number of differences: 143
MA1 vs MM1
Number of differences: 80
MA1 vs MM2
Number of differences: 83
MA2 vs MM1
Number of differences: 158
MA2 vs MM2
Number of differences: 165
MM1 vs MM2
Number of differences: 71
0
2000
4000
6000
8000
10000
Position

### Region: 322



CR1 vs CR2
Number of differences: 164

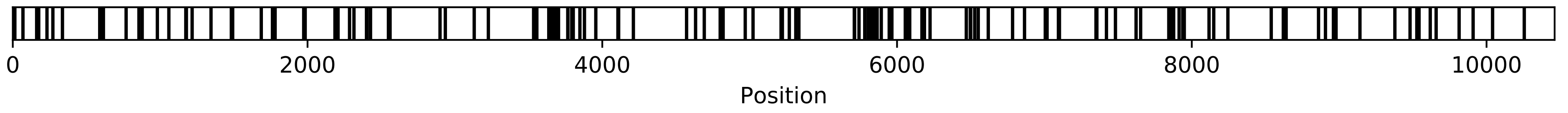


CR1 vs MA1
Number of differences: 173

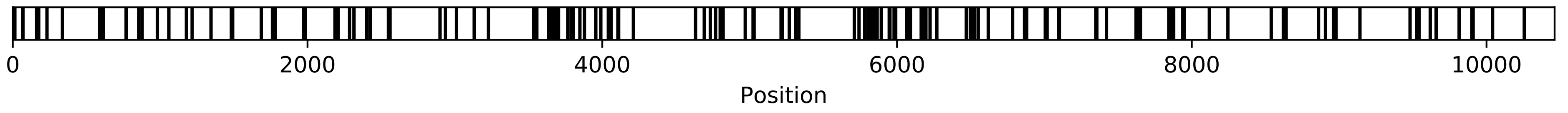


CR1 vs MA2
Number of differences: 101

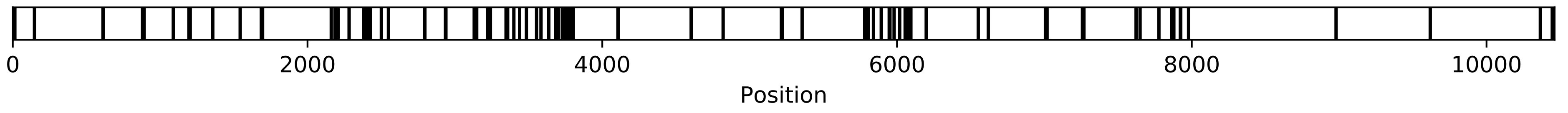


CR1 vs MM1
Number of differences: 101

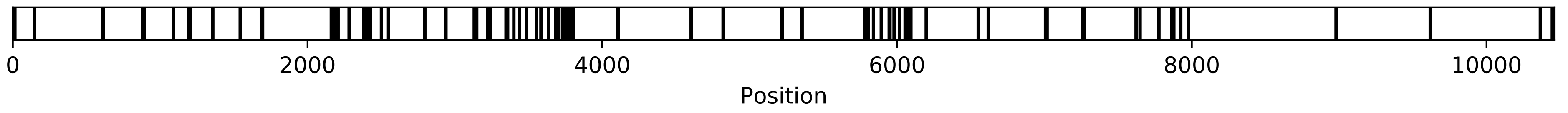


CR1 vs MM2
Number of differences: 116

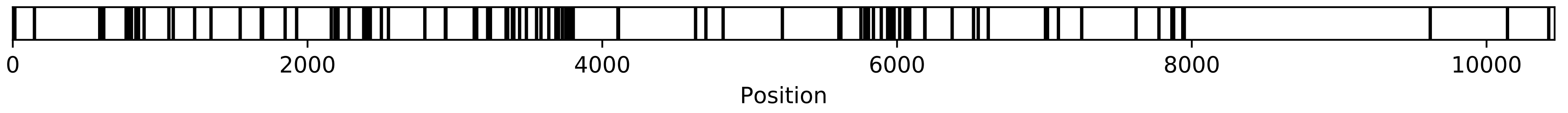


CR2 vs MA1
Number of differences: 39

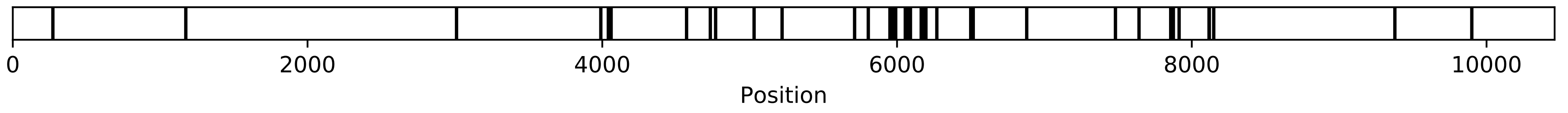


CR2 vs MA2
Number of differences: 172

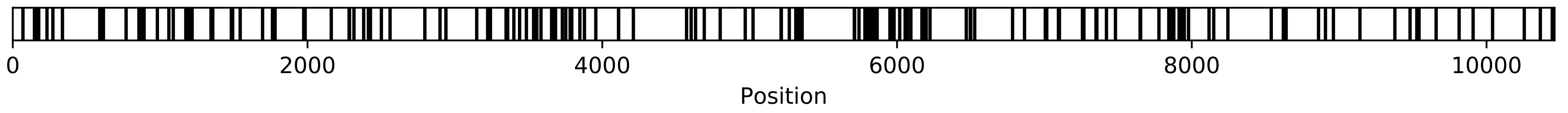


CR2 vs MM1
Number of differences: 172

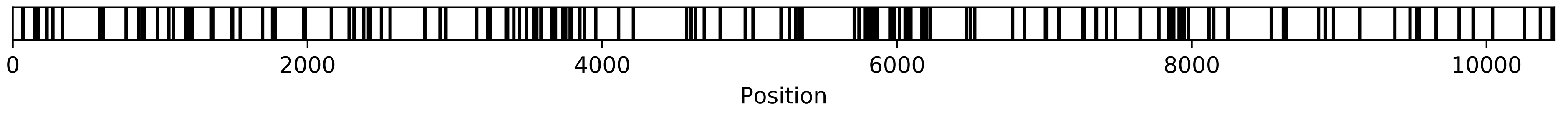


CR2 vs MM2
Number of differences: 170

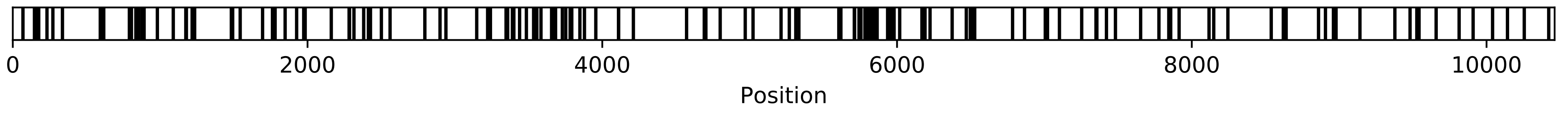


MA1 vs MA2
Number of differences: 175

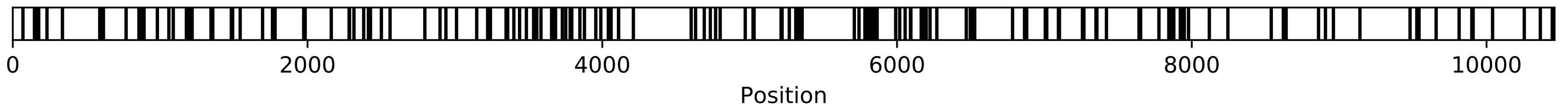


MA1 vs MM1
Number of differences: 175

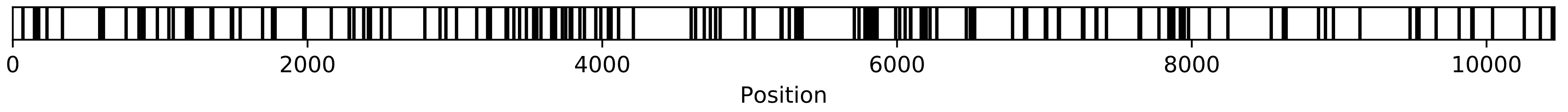


MA1 vs MM2
Number of differences: 177

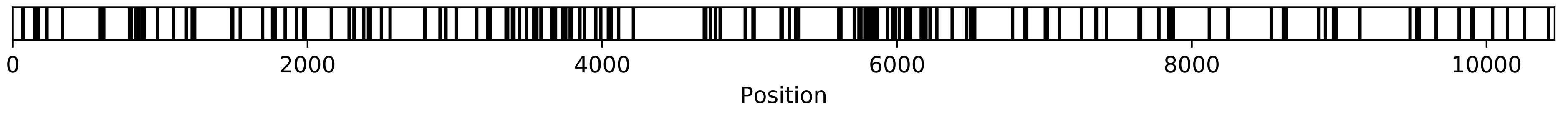


MA2 vs MM1
Number of differences: 0

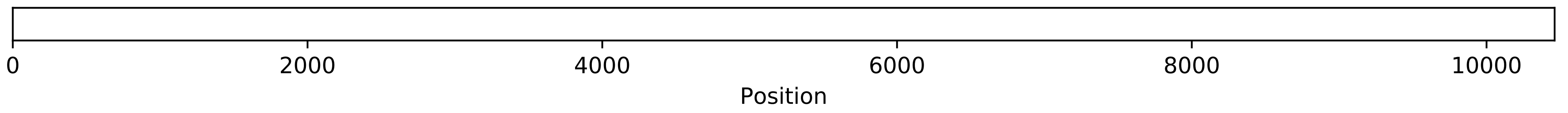


MA2 vs MM2
Number of differences: 58

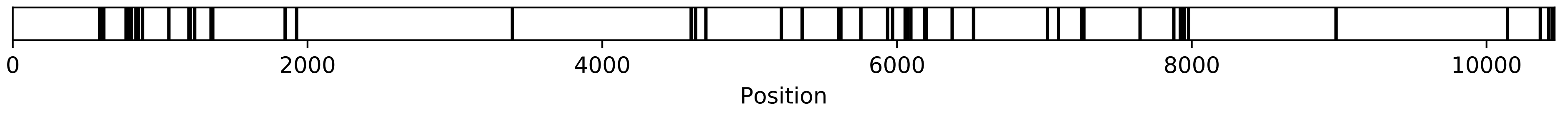


MM1 vs MM2
Number of differences: 58

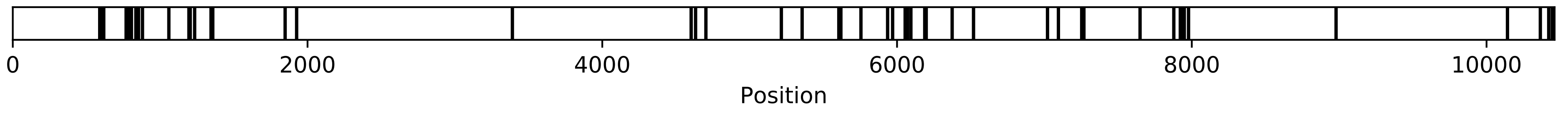

### Region: 324



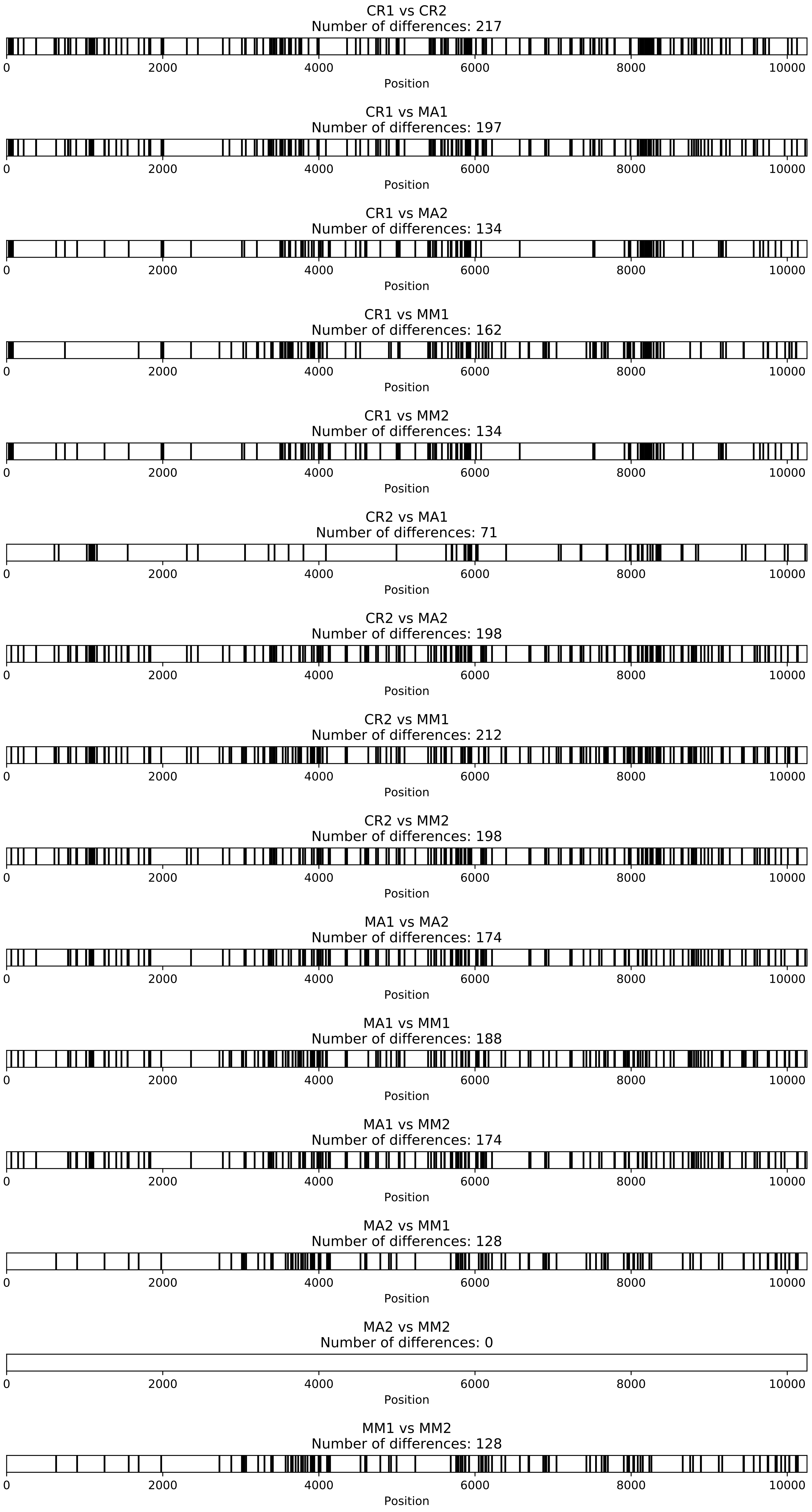
CR1 vs CR2
Number of differences: 217
CR1 vs MA1
Number of differences: 197
CR1 vs MA2
Number of differences: 134
CR1 vs MM1
Number of differences: 162
CR1 vs MM2
Number of differences: 134
CR2 vs MA1
Number of differences: 71
CR2 vs MA2
Number of differences: 198
CR2 vs MM1
Number of differences: 212
CR2 vs MM2
Number of differences: 198
MA1 vs MA2
Number of differences: 174
MA1 vs MM1
Number of differences: 188
MA1 vs MM2
Number of differences: 174
MA2 vs MM1
Number of differences: 128
MA2 vs MM2
Number of differences: 0
MM1 vs MM2
Number of differences: 128
0
2000
4000
6000
8000
10000
Position

### Region: 326



CR1 vs CR2
Number of differences: 201

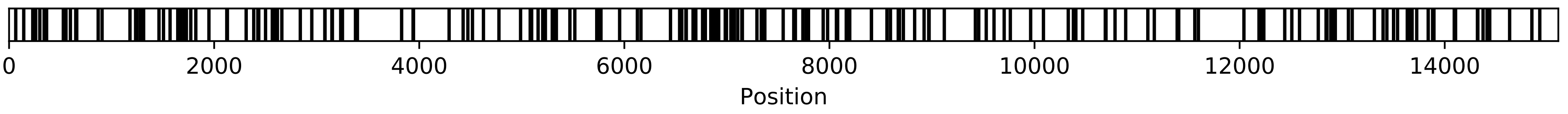


CR1 vs MA1
Number of differences: 67

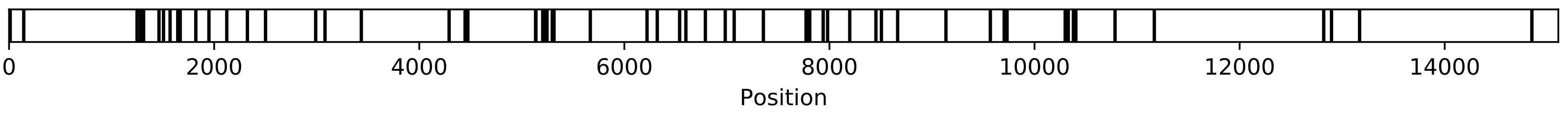


CR1 vs MA2
Number of differences: 200

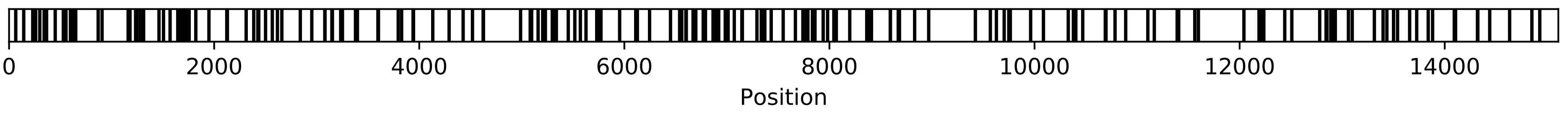


CR1 vs MM1
Number of differences: 199

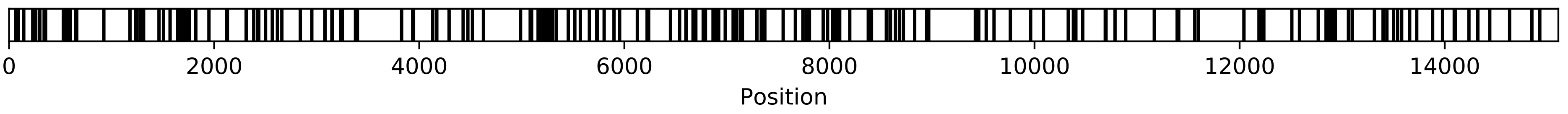


CR1 vs MM2
Number of differences: 200

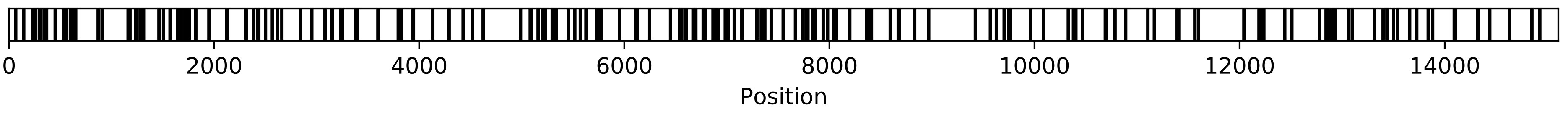


CR2 vs MA1
Number of differences: 186

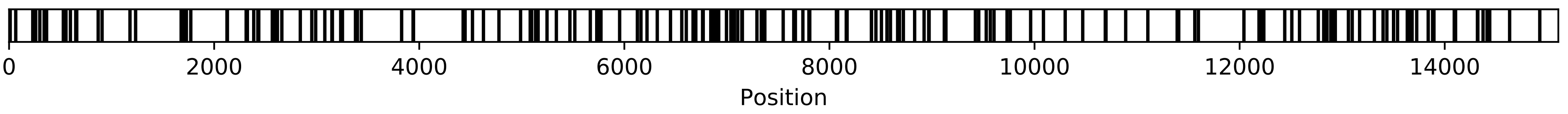


CR2 vs MA2
Number of differences: 72

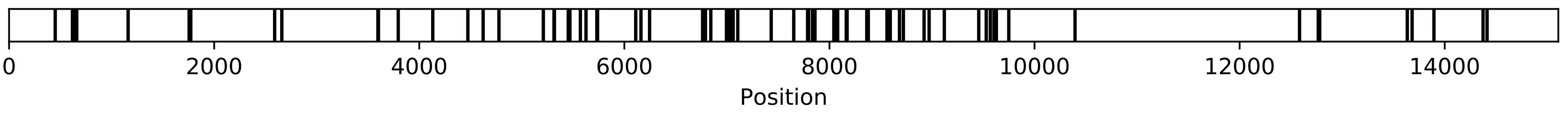


CR2 vs MM1
Number of differences: 73

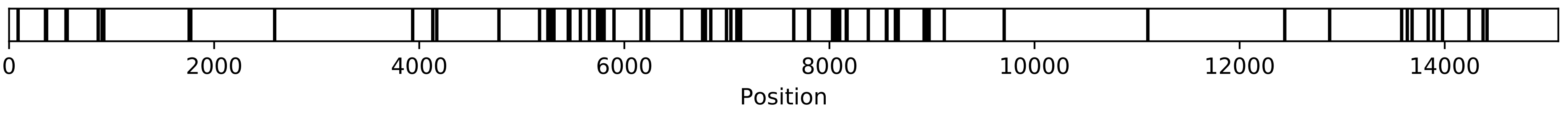


CR2 vs MM2
Number of differences: 72

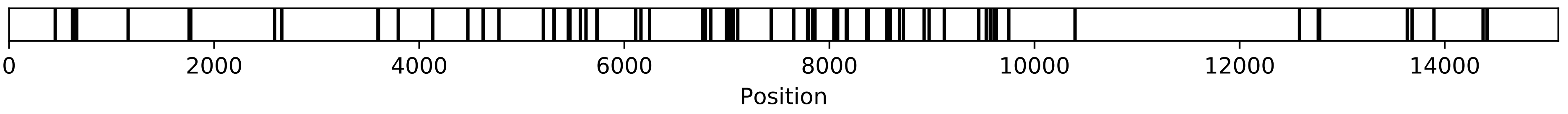


MA1 vs MA2
Number of differences: 189

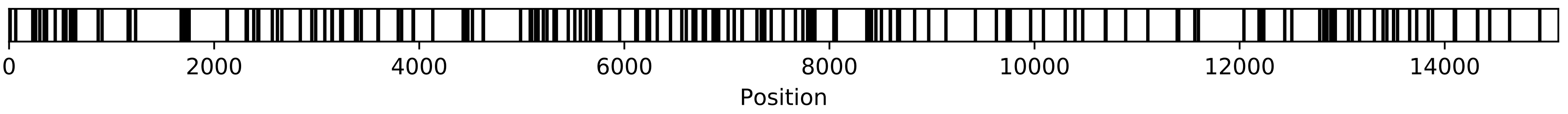


MA1 vs MM1
Number of differences: 188

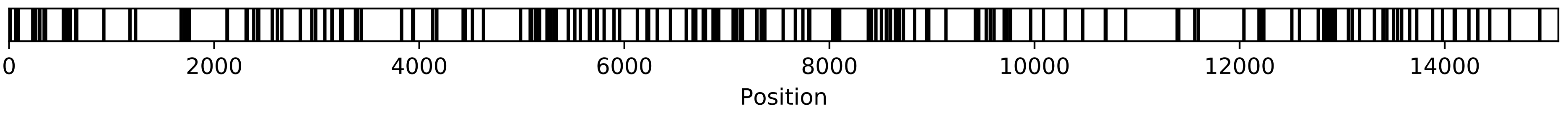


MA1 vs MM2
Number of differences: 189

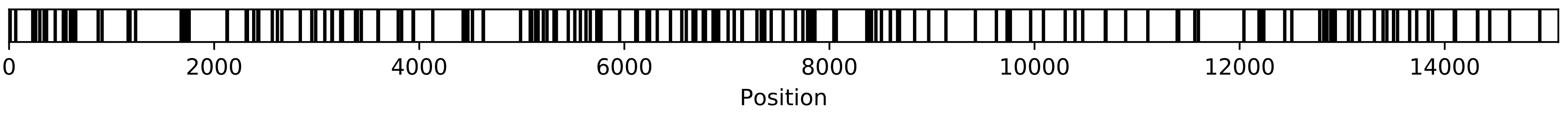


MA2 vs MM1
Number of differences: 81

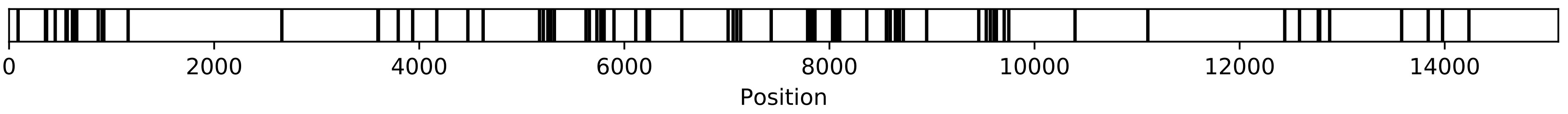


MA2 vs MM2
Number of differences: 0

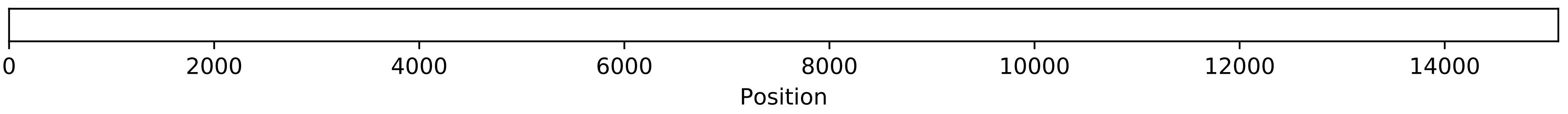


MM1 vs MM2
Number of differences: 81

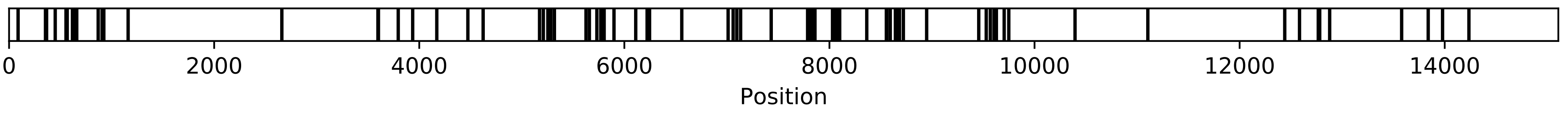

### Region: 329



CR1 vs CR2
Number of differences: 357

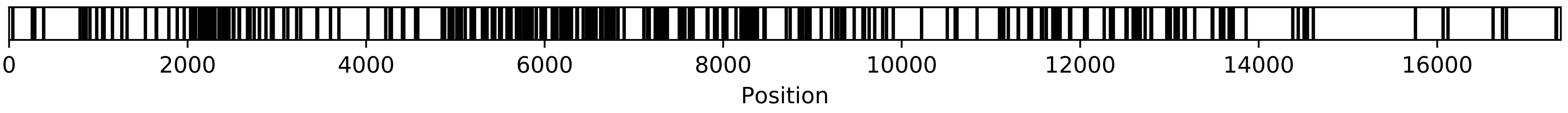


CR1 vs MA1
Number of differences: 213

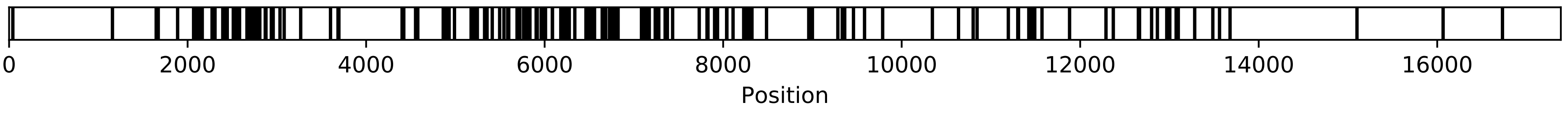


CR1 vs MA2
Number of differences: 353

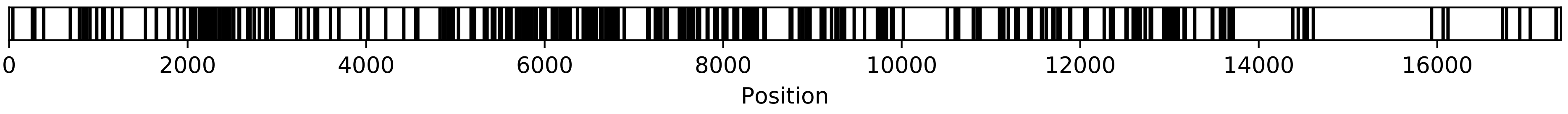


CR1 vs MM1
Number of differences: 231

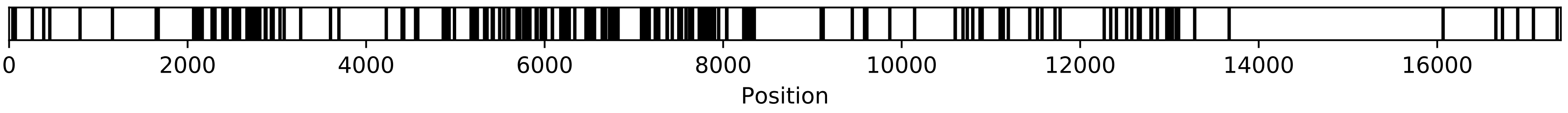


CR1 vs MM2
Number of differences: 226

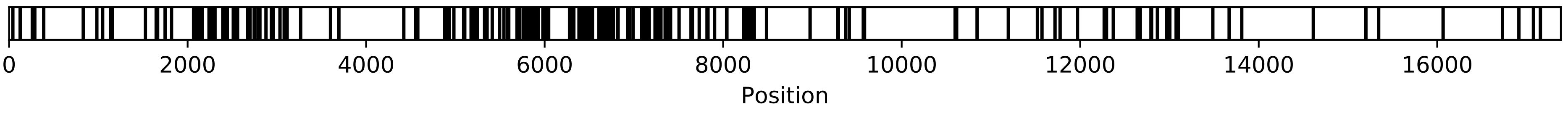


CR2 vs MA1
Number of differences: 337

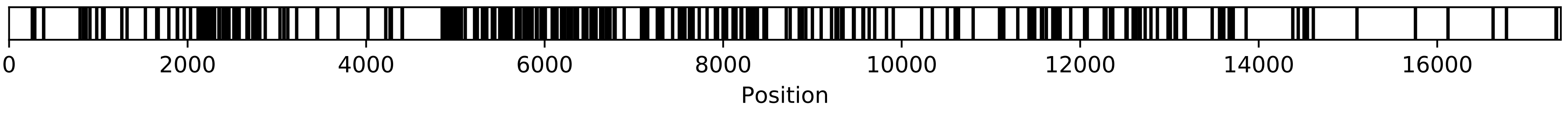


CR2 vs MA2
Number of differences: 100

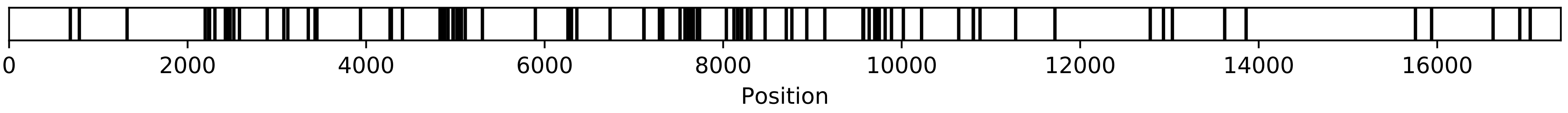


CR2 vs MM1
Number of differences: 346

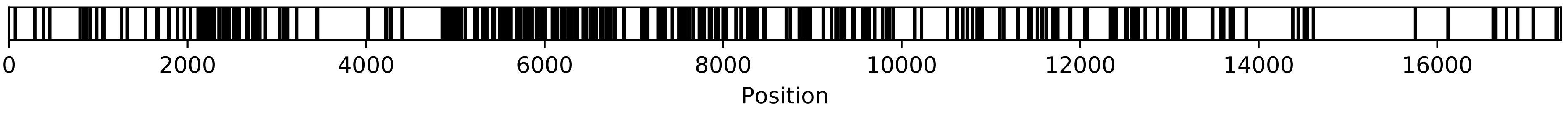


CR2 vs MM2
Number of differences: 344

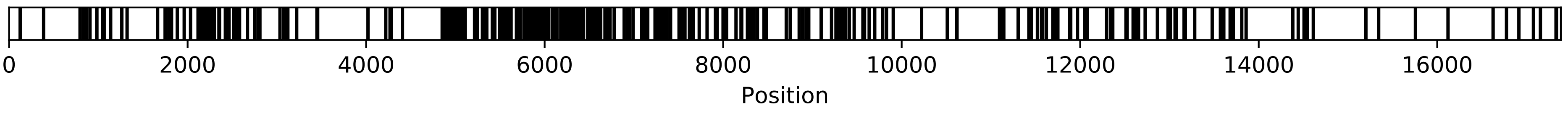


MA1 vs MA2
Number of differences: 331

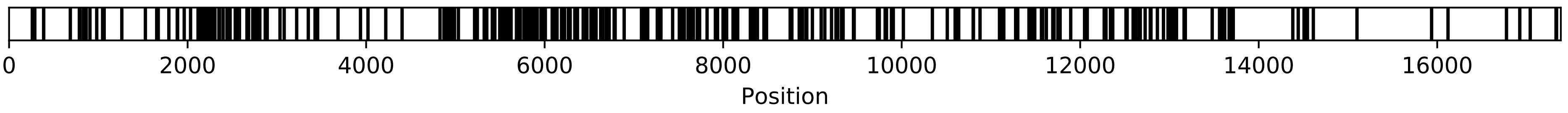


MA1 vs MM1
Number of differences: 102

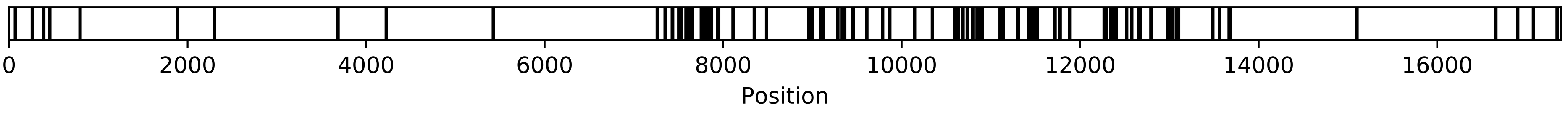


MA1 vs MM2
Number of differences: 175

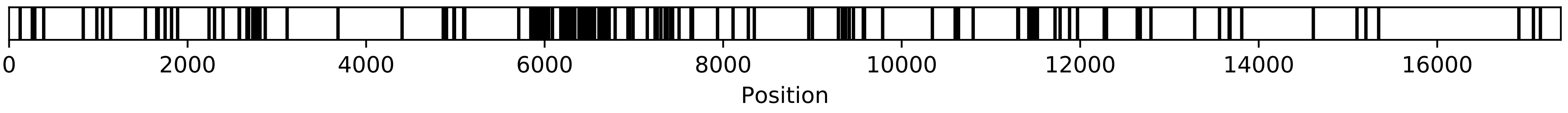


MA2 vs MM1
Number of differences: 335

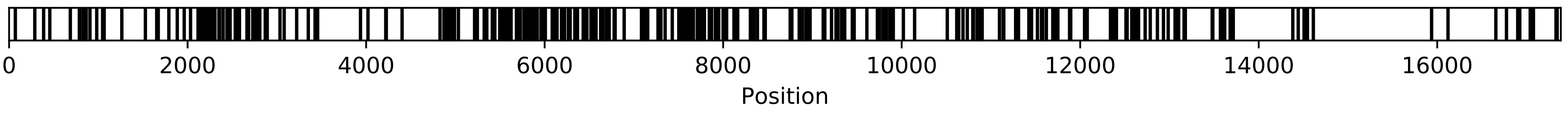


MA2 vs MM2
Number of differences: 344

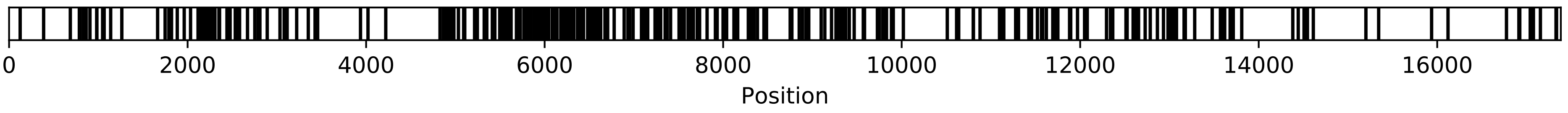


MM1 vs MM2
Number of differences: 176

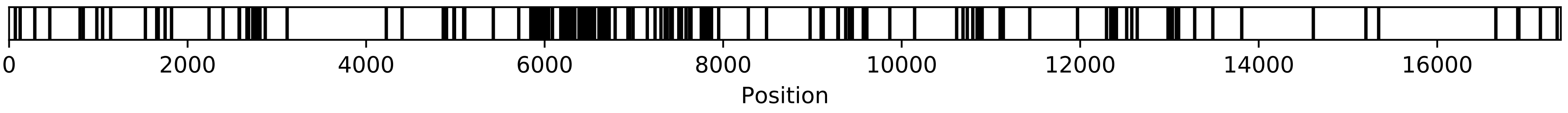

### Region: 334



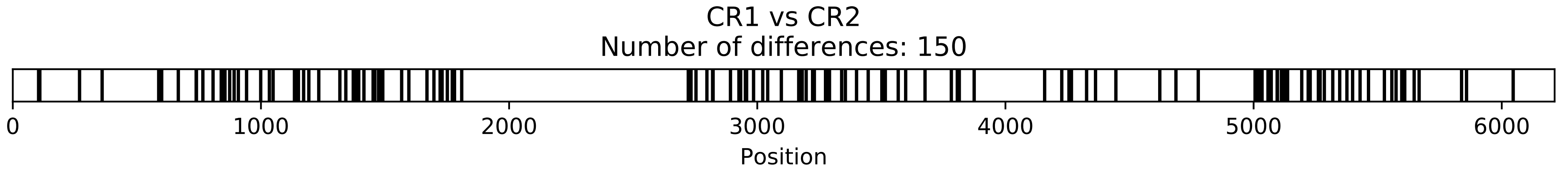
CR1 vs CR2
Number of differences: 150
0 1000 2000 3000 4000 5000 6000
Position

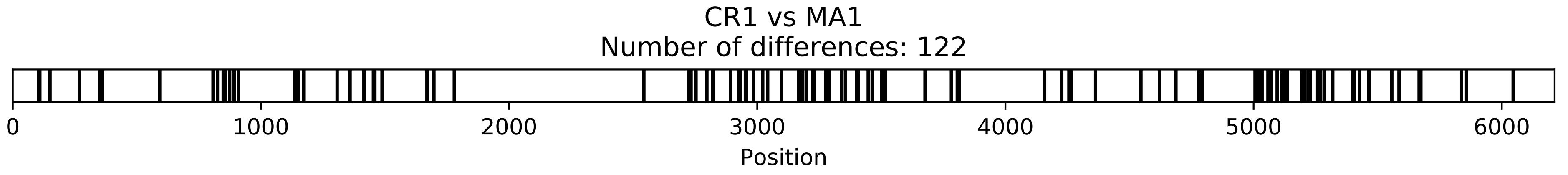
CR1 vs MA1
Number of differences: 122
0 1000 2000 3000 4000 5000 6000
Position

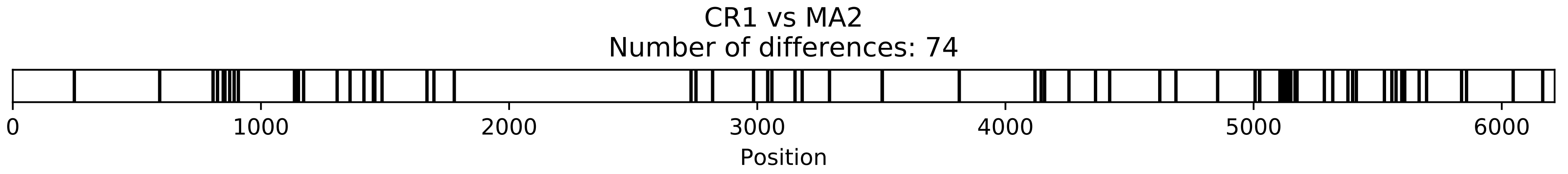
CR1 vs MA2
Number of differences: 74
0 1000 2000 3000 4000 5000 6000
Position

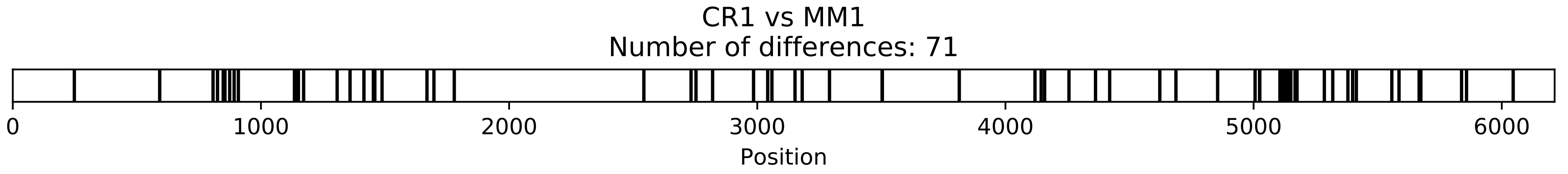
CR1 vs MM1
Number of differences: 71
0 1000 2000 3000 4000 5000 6000
Position

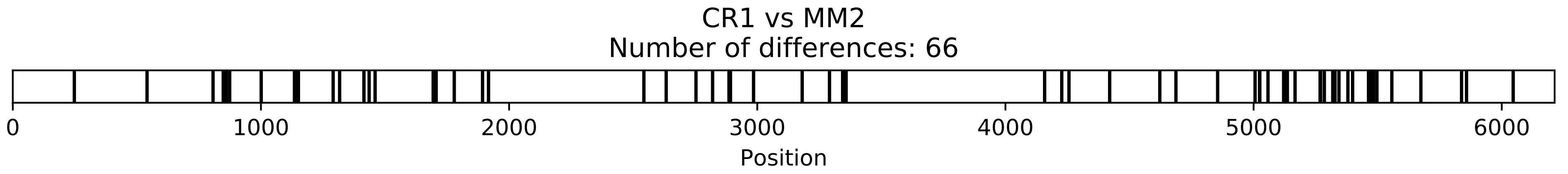
CR1 vs MM2
Number of differences: 66
0 1000 2000 3000 4000 5000 6000
Position

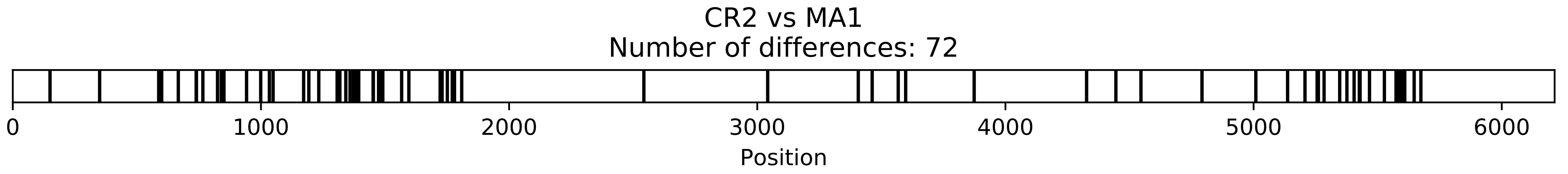
CR2 vs MA1
Number of differences: 72
0 1000 2000 3000 4000 5000 6000
Position

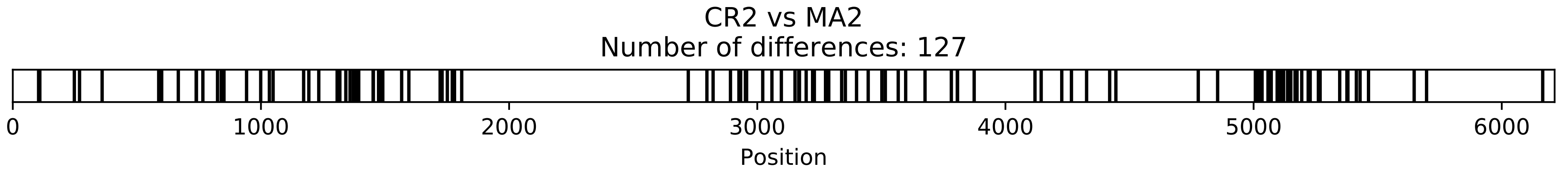
CR2 vs MA2
Number of differences: 127
0 1000 2000 3000 4000 5000 6000
Position

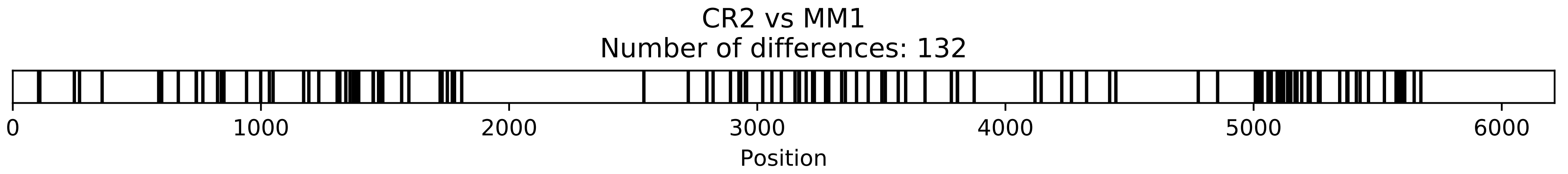
CR2 vs MM1
Number of differences: 132
0 1000 2000 3000 4000 5000 6000
Position

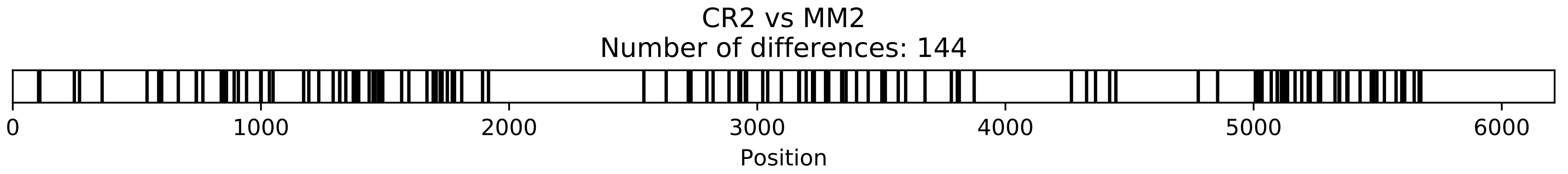
CR2 vs MM2
Number of differences: 144
0 1000 2000 3000 4000 5000 6000
Position

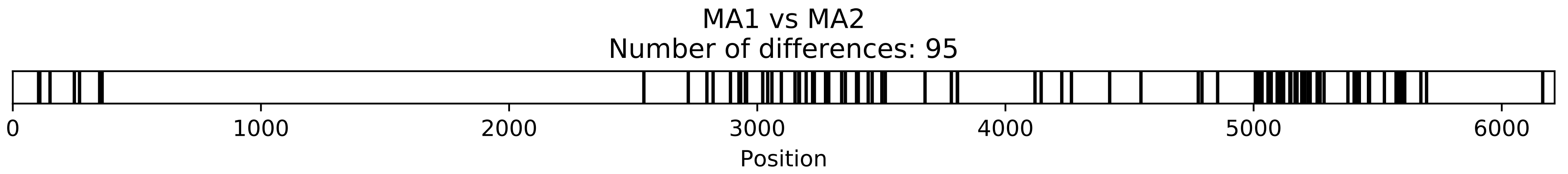
MA1 vs MA2
Number of differences: 95
0 1000 2000 3000 4000 5000 6000
Position

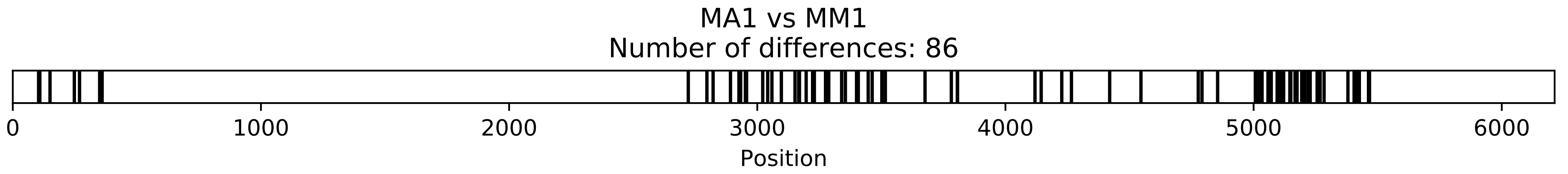
MA1 vs MM1
Number of differences: 86
0 1000 2000 3000 4000 5000 6000
Position

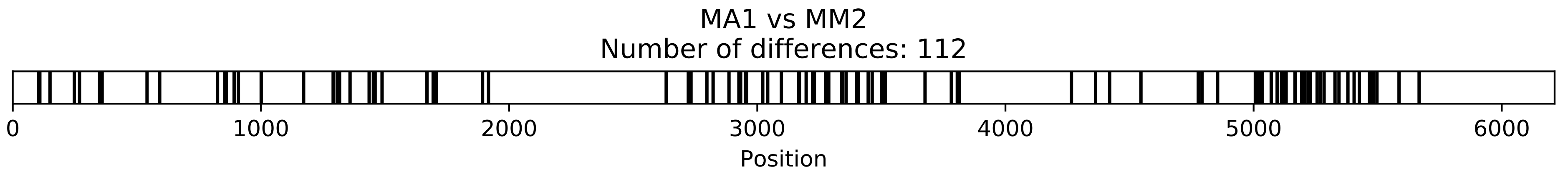
MA1 vs MM2
Number of differences: 112
0 1000 2000 3000 4000 5000 6000
Position

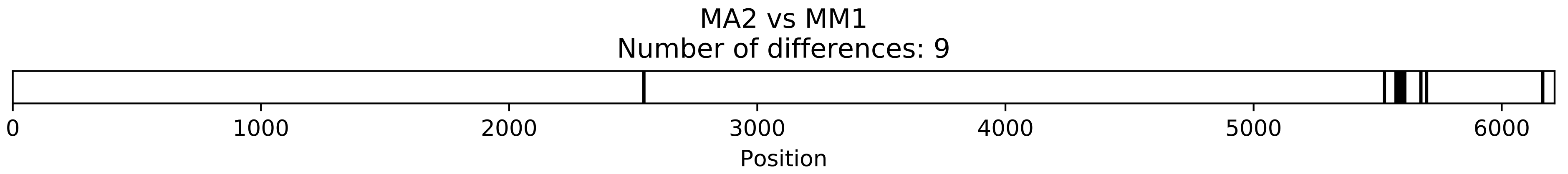
MA2 vs MM1
Number of differences: 9
0 1000 2000 3000 4000 5000 6000
Position

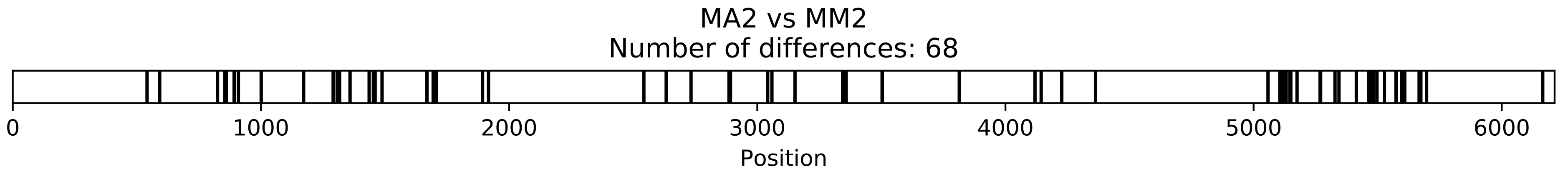
MA2 vs MM2
Number of differences: 68
0 1000 2000 3000 4000 5000 6000
Position

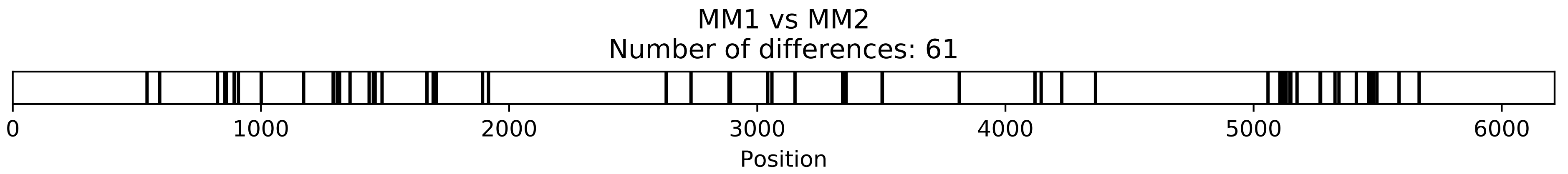
MM1 vs MM2
Number of differences: 61
0 1000 2000 3000 4000 5000 6000
Position

### Region: 335

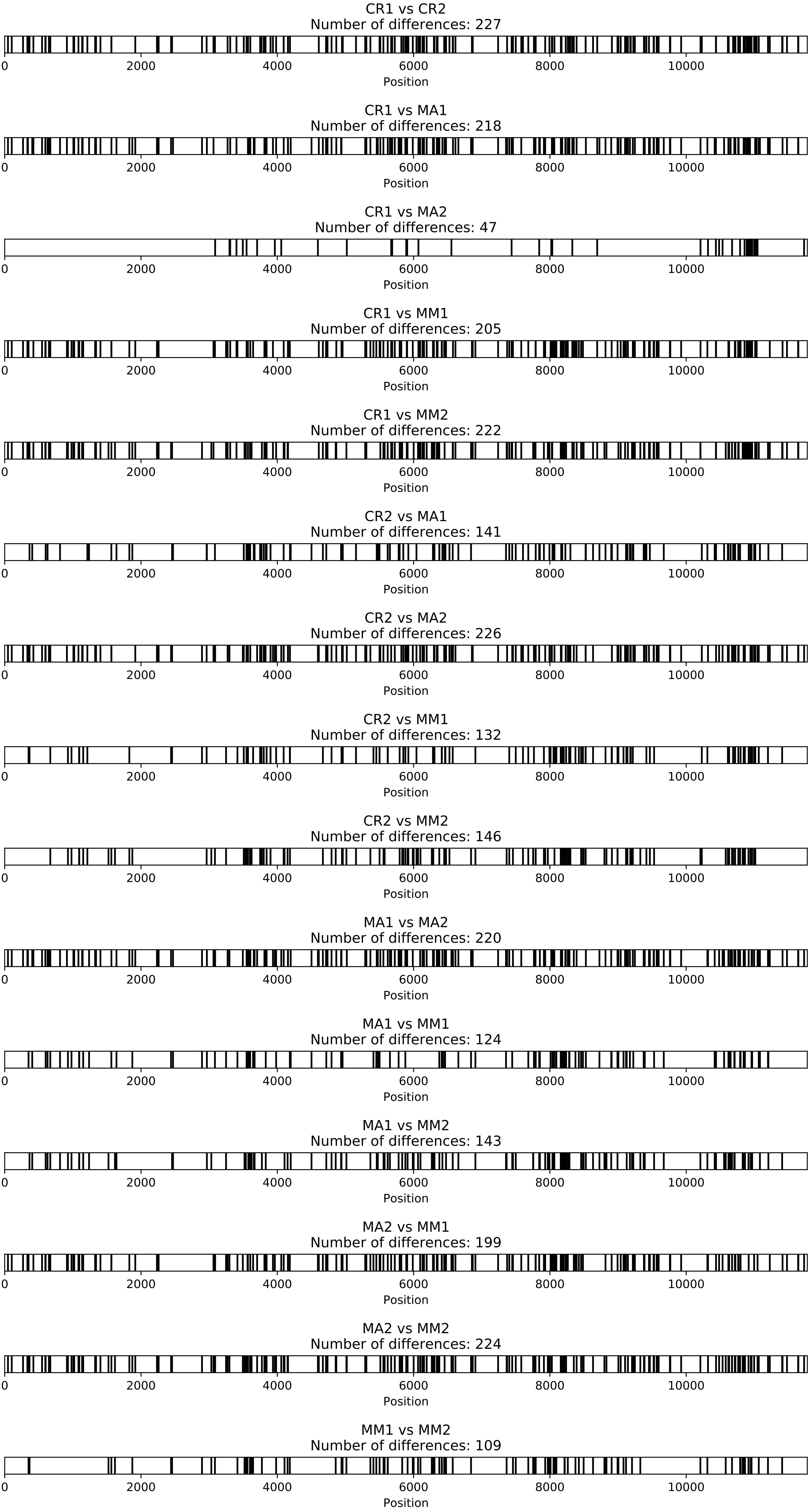
CR1 vs CR2
Number of differences: 227
CR1 vs MA1
Number of differences: 218
CR1 vs MA2
Number of differences: 47
CR1 vs MM1
Number of differences: 205
CR1 vs MM2
Number of differences: 222
CR2 vs MA1
Number of differences: 141
CR2 vs MA2
Number of differences: 226
CR2 vs MM1
Number of differences: 132
CR2 vs MM2
Number of differences: 146
MA1 vs MA2
Number of differences: 220
MA1 vs MM1
Number of differences: 124
MA1 vs MM2
Number of differences: 143
MA2 vs MM1
Number of differences: 199
MA2 vs MM2
Number of differences: 224
MM1 vs MM2
Number of differences: 109
0
2000
4000
6000
8000
10000
Position

### Region: 336



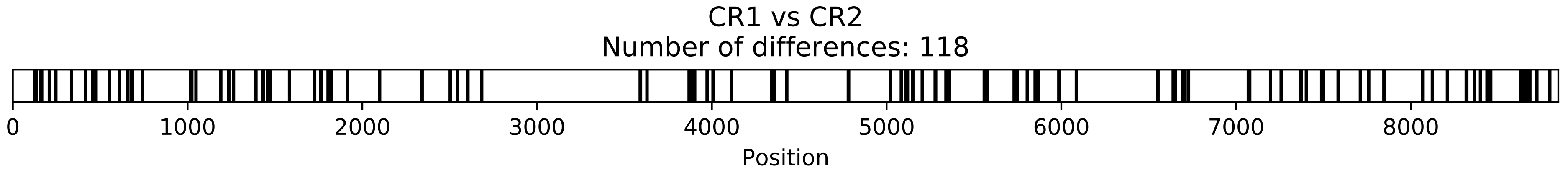
CR1 vs CR2
Number of differences: 118
0
1000
2000
3000
4000
5000
6000
7000
8000
Position

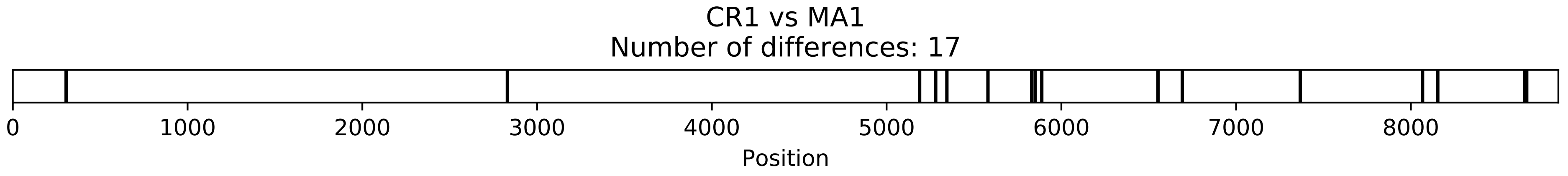
CR1 vs MA1
Number of differences: 17
0
1000
2000
3000
4000
5000
6000
7000
8000
Position

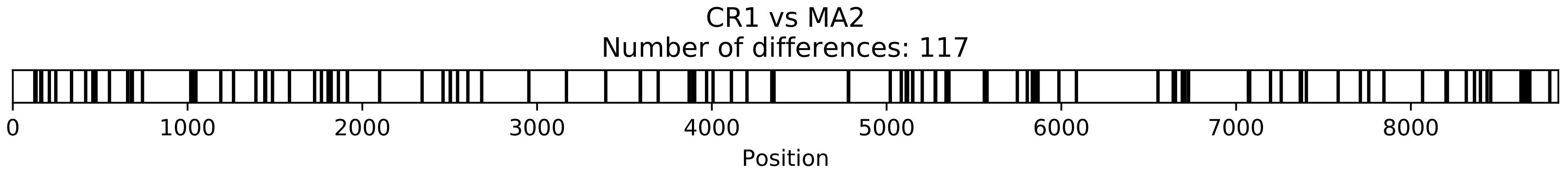
CR1 vs MA2
Number of differences: 117
0
1000
2000
3000
4000
5000
6000
7000
8000
Position

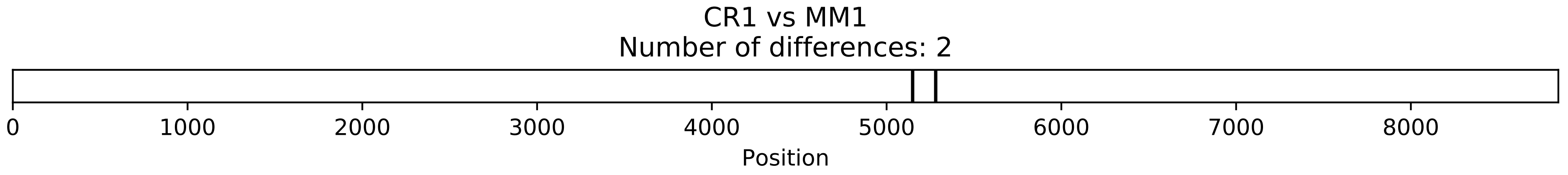
CR1 vs MM1
Number of differences: 2
0
1000
2000
3000
4000
5000
6000
7000
8000
Position

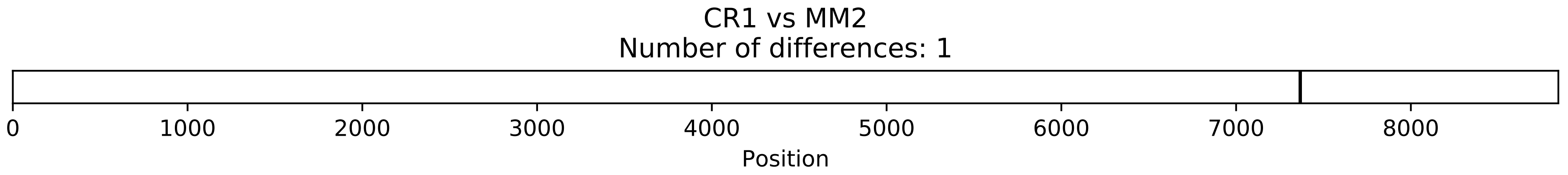
CR1 vs MM2
Number of differences: 1
0
1000
2000
3000
4000
5000
6000
7000
8000
Position

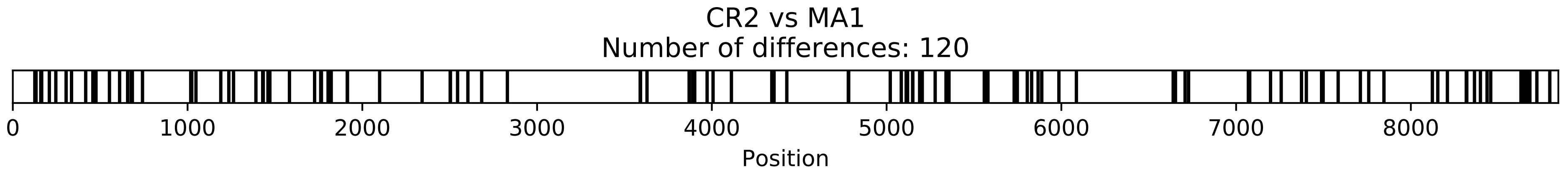
CR2 vs MA1
Number of differences: 120
0
1000
2000
3000
4000
5000
6000
7000
8000
Position

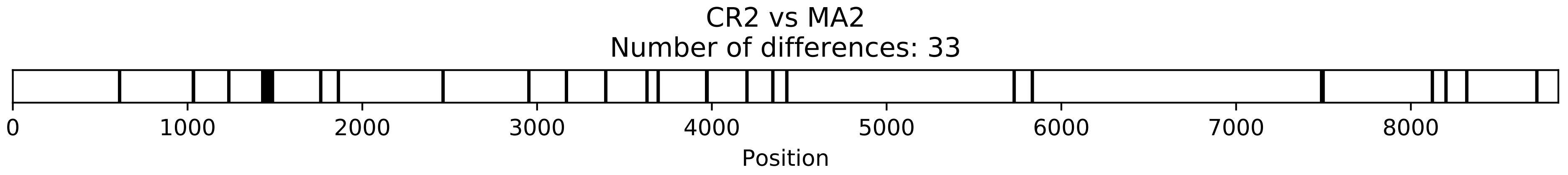
CR2 vs MA2
Number of differences: 33
0
1000
2000
3000
4000
5000
6000
7000
8000
Position

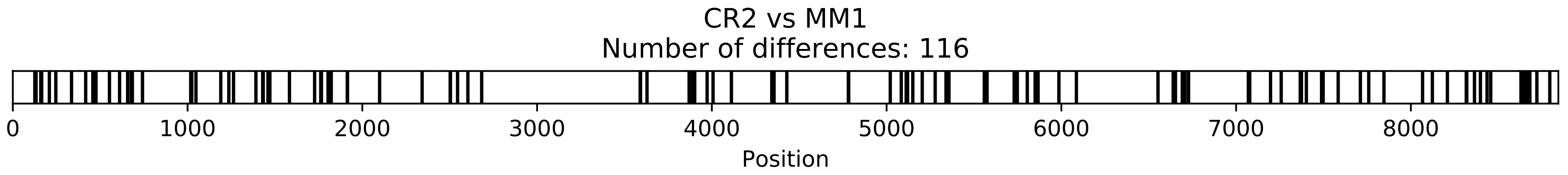
CR2 vs MM1
Number of differences: 116
0
1000
2000
3000
4000
5000
6000
7000
8000
Position

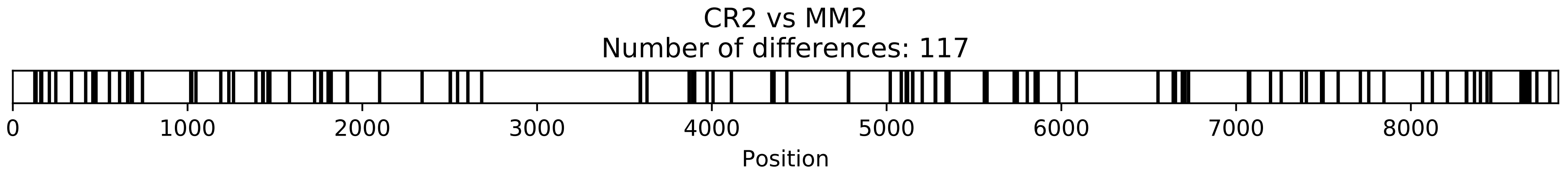
CR2 vs MM2
Number of differences: 117
0
1000
2000
3000
4000
5000
6000
7000
8000
Position

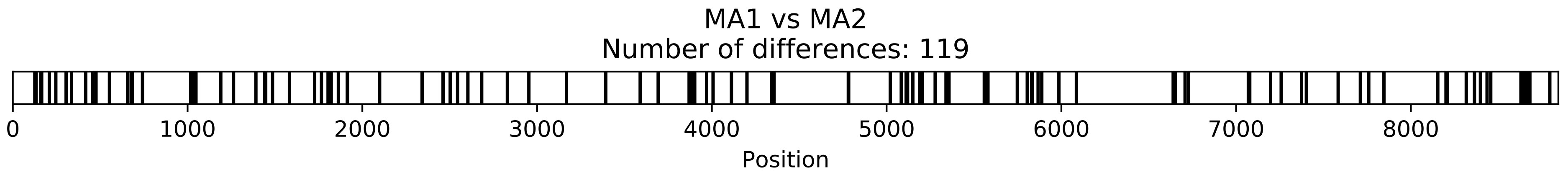
MA1 vs MA2
Number of differences: 119
0
1000
2000
3000
4000
5000
6000
7000
8000
Position

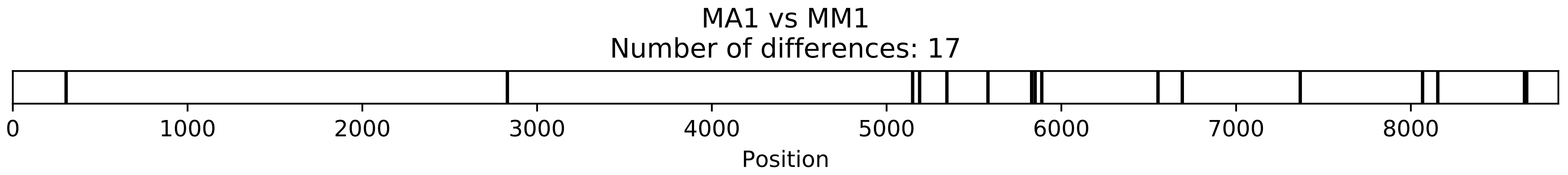
MA1 vs MM1
Number of differences: 17
0
1000
2000
3000
4000
5000
6000
7000
8000
Position

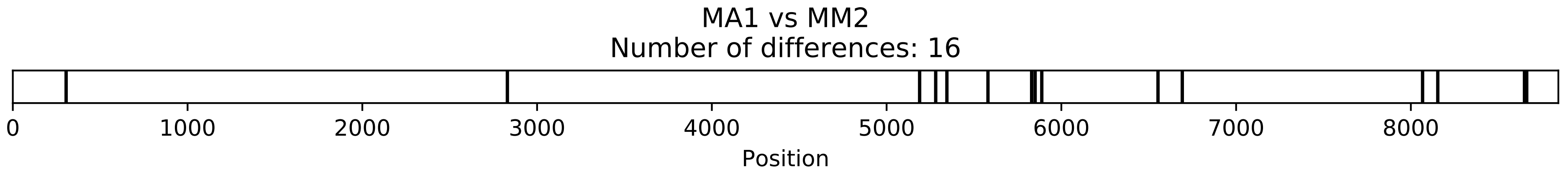
MA1 vs MM2
Number of differences: 16
0
1000
2000
3000
4000
5000
6000
7000
8000
Position

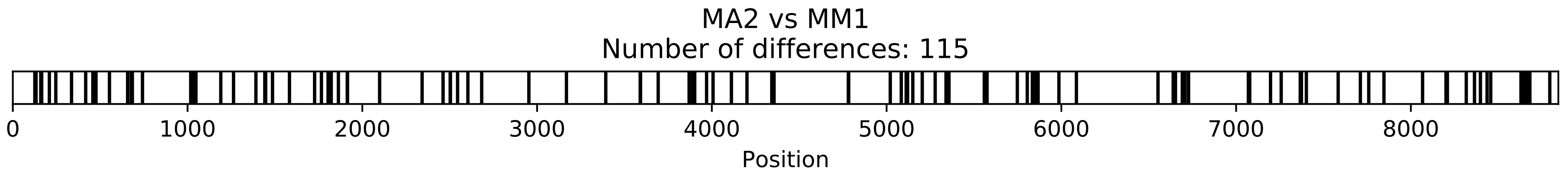
MA2 vs MM1
Number of differences: 115
0
1000
2000
3000
4000
5000
6000
7000
8000
Position

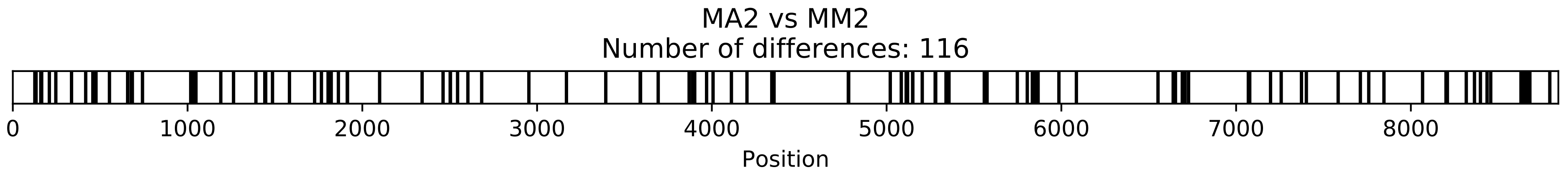
MA2 vs MM2
Number of differences: 116
0
1000
2000
3000
4000
5000
6000
7000
8000
Position

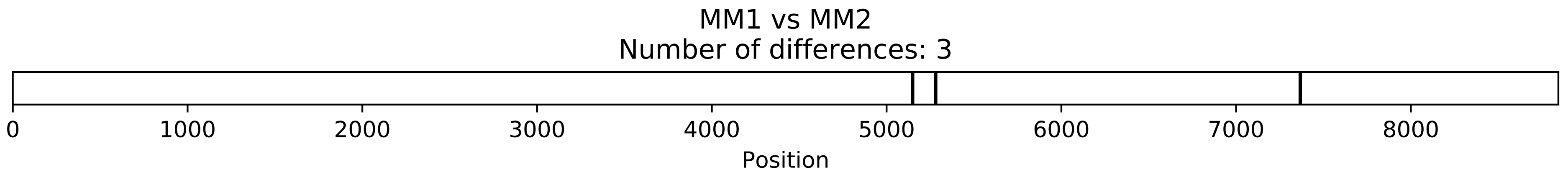
MM1 vs MM2
Number of differences: 3
0
1000
2000
3000
4000
5000
6000
7000
8000
Position

### Region: 337

CR1 vs CR2
Number of differences: 126

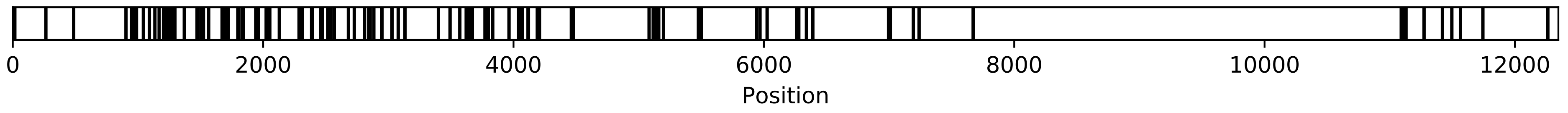


CR1 vs MA1
Number of differences: 173

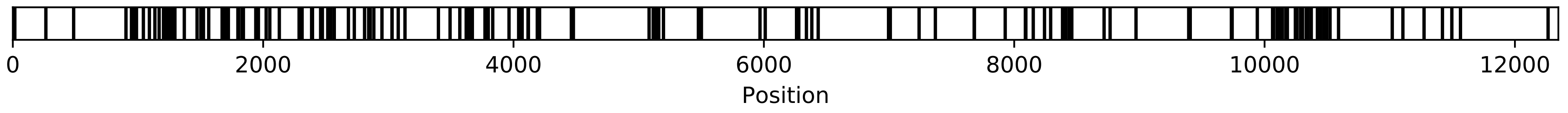


CR1 vs MA2
Number of differences: 97

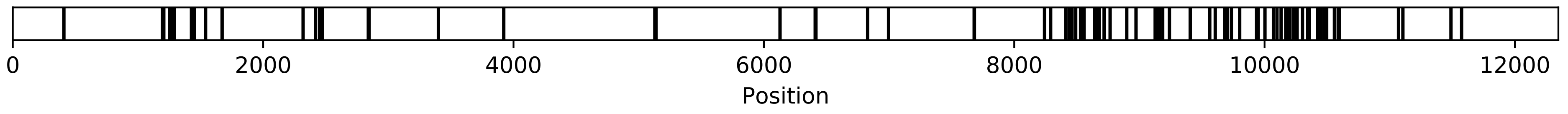


CR1 vs MM1
Number of differences: 165

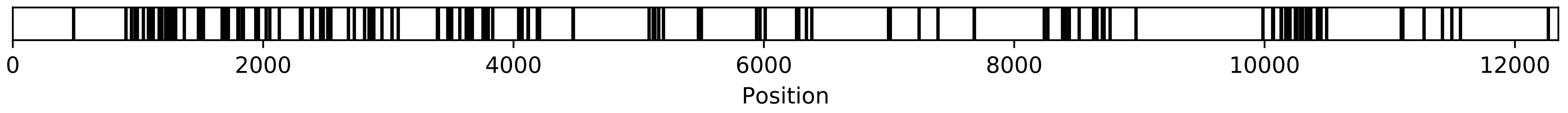


CR1 vs MM2
Number of differences: 173

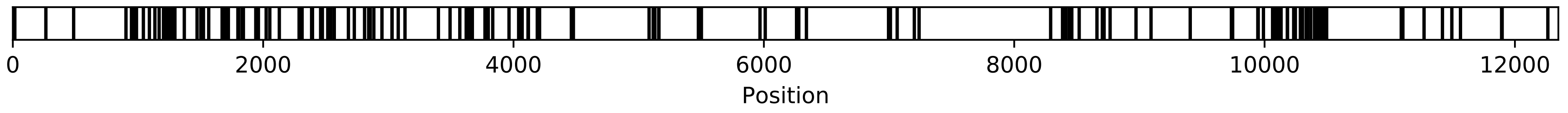


CR2 vs MA1
Number of differences: 67

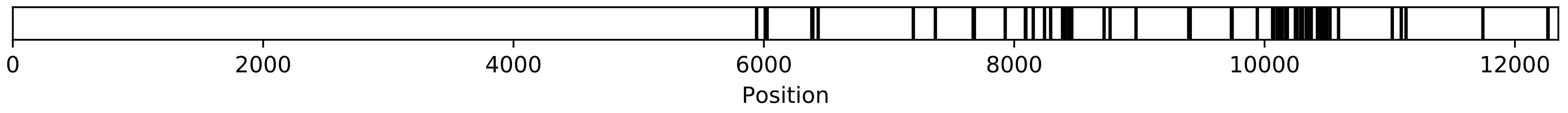


CR2 vs MA2
Number of differences: 201

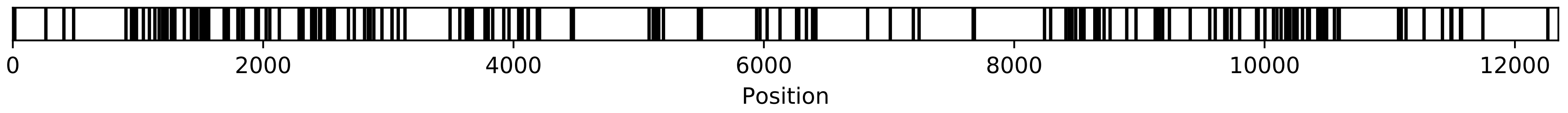


CR2 vs MM1
Number of differences: 106

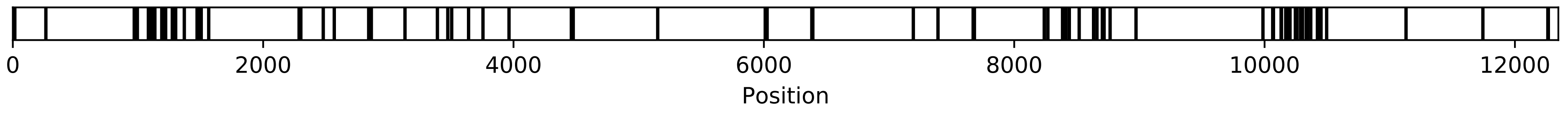


CR2 vs MM2
Number of differences: 69

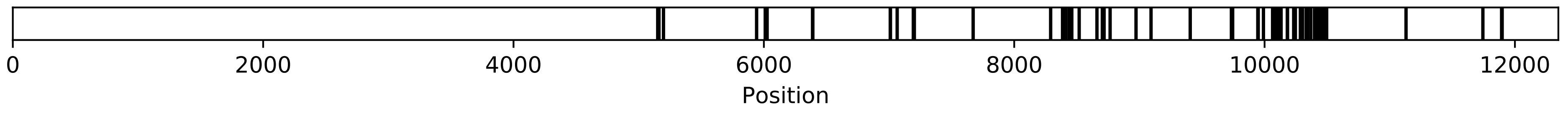


MA1 vs MA2
Number of differences: 190

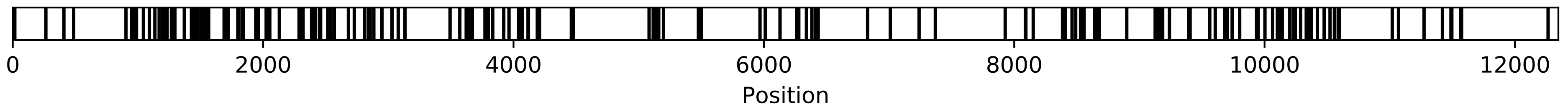


MA1 vs MM1
Number of differences: 97

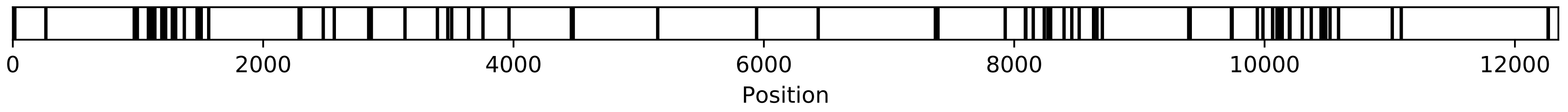


MA1 vs MM2
Number of differences: 64

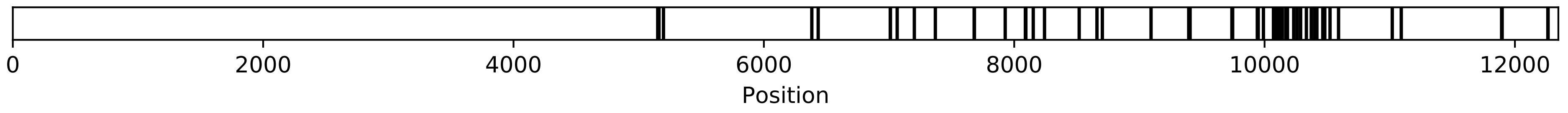


MA2 vs MM1
Number of differences: 192

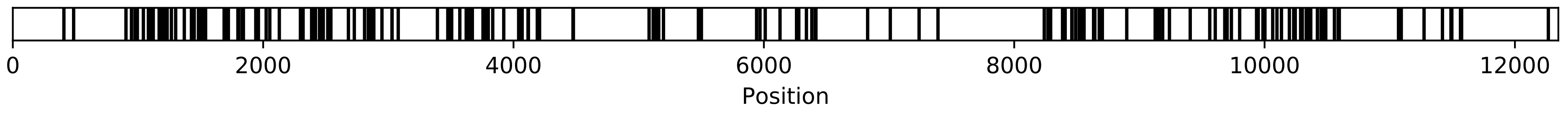


MA2 vs MM2
Number of differences: 199

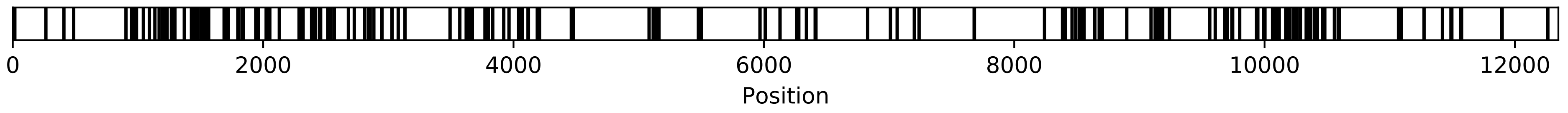


MM1 vs MM2
Number of differences: 113

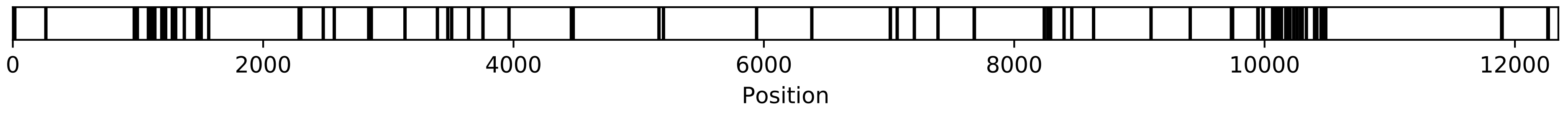

### Region: 342



CR1 vs CR2
Number of differences: 277

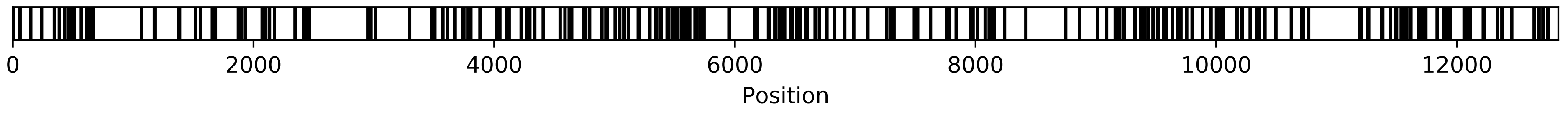


CR1 vs MA1
Number of differences: 172

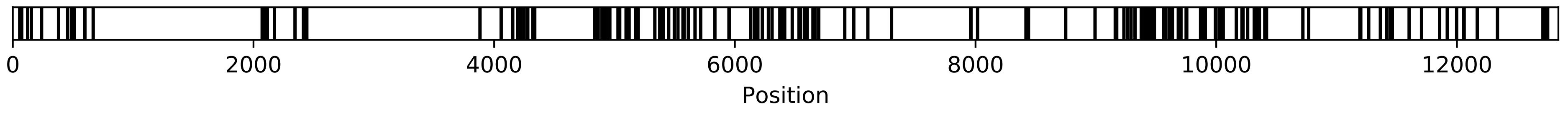


CR1 vs MA2
Number of differences: 271

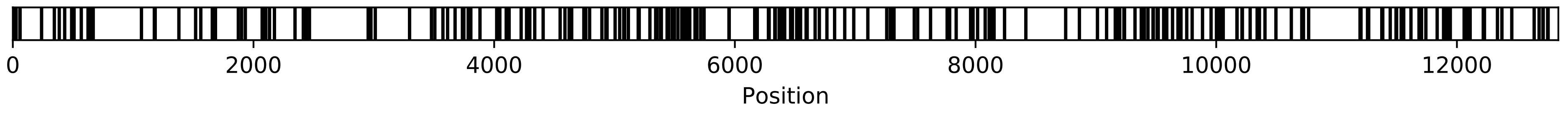


CR1 vs MM1
Number of differences: 164

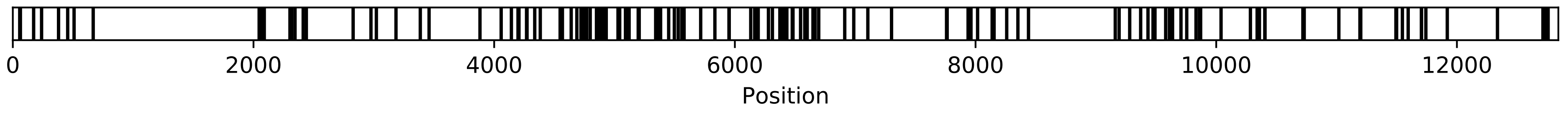


CR1 vs MM2
Number of differences: 185

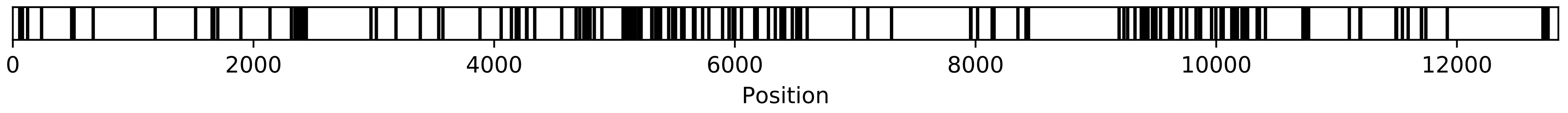


CR2 vs MA1
Number of differences: 283

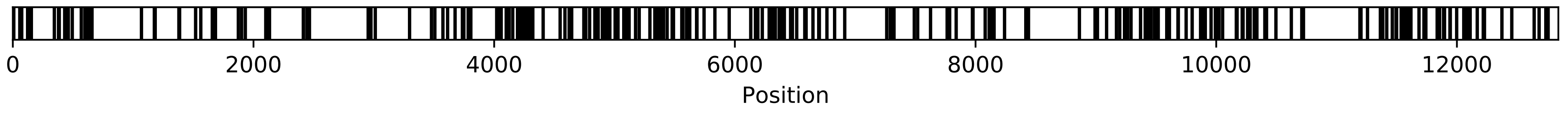


CR2 vs MA2
Number of differences: 10

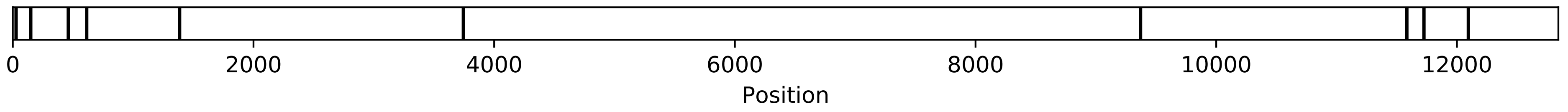


CR2 vs MM1
Number of differences: 290

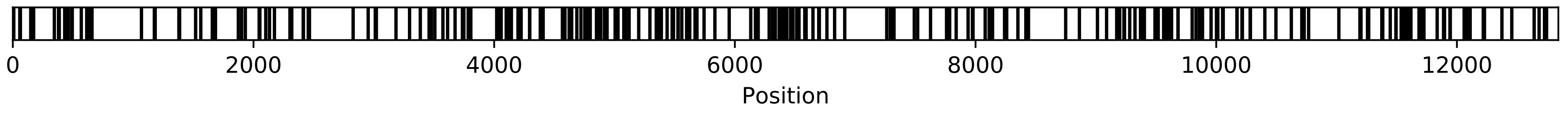


CR2 vs MM2
Number of differences: 302

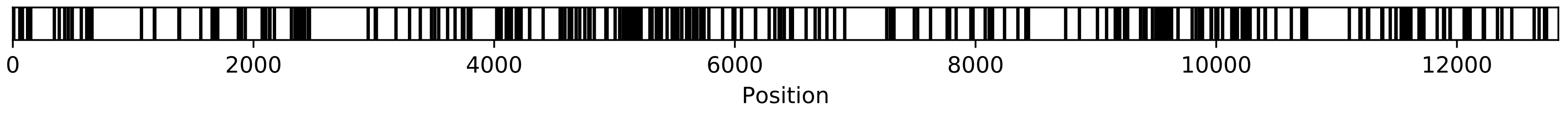


MA1 vs MA2
Number of differences: 277

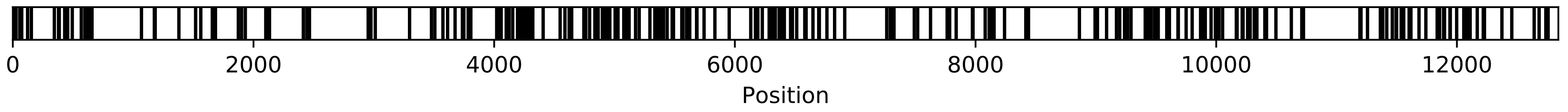


MA1 vs MM1
Number of differences: 163

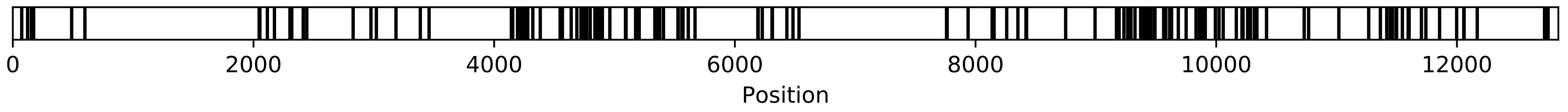


MA1 vs MM2
Number of differences: 207

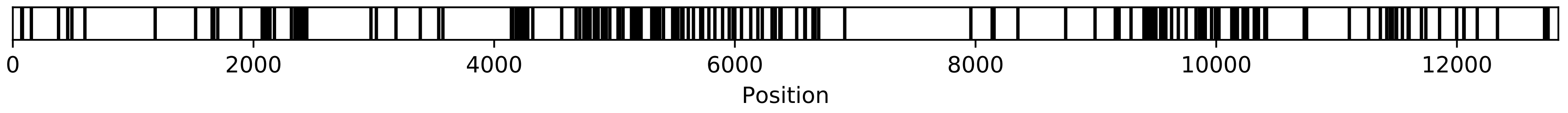


MA2 vs MM1
Number of differences: 284

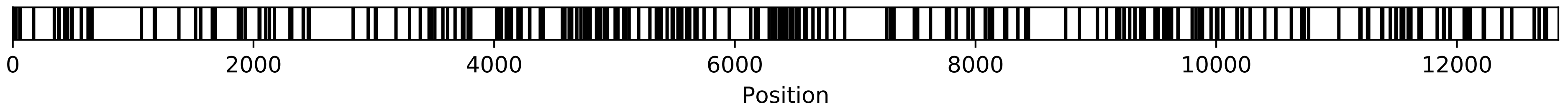


MA2 vs MM2
Number of differences: 296

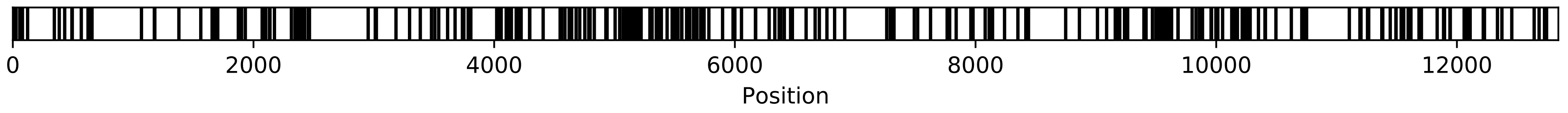


MM1 vs MM2
Number of differences: 152

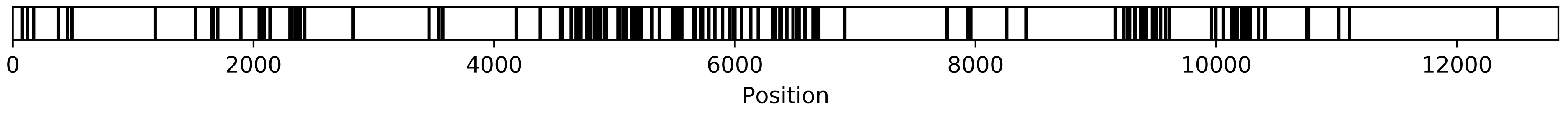

### Region: 343



CR1 vs CR2
Number of differences: 359

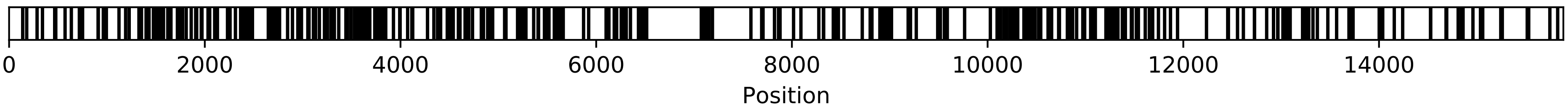


CR1 vs MA1
Number of differences: 179

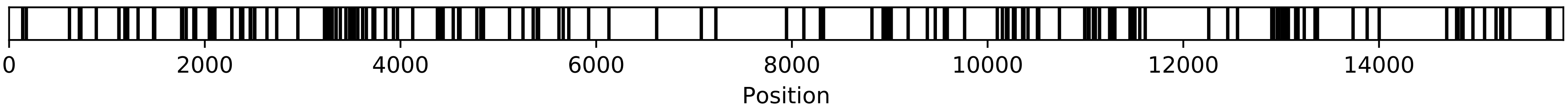


CR1 vs MA2
Number of differences: 358

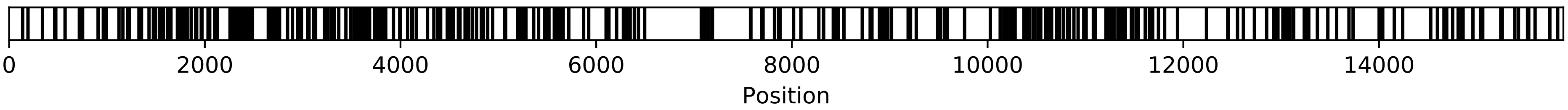


CR1 vs MM1
Number of differences: 167

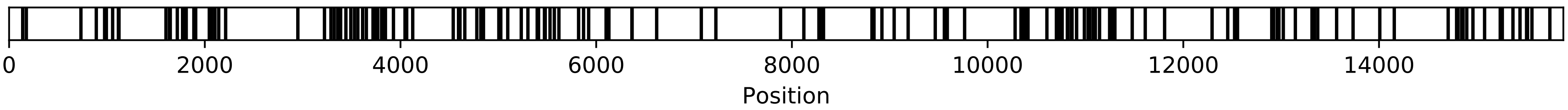


CR1 vs MM2
Number of differences: 179

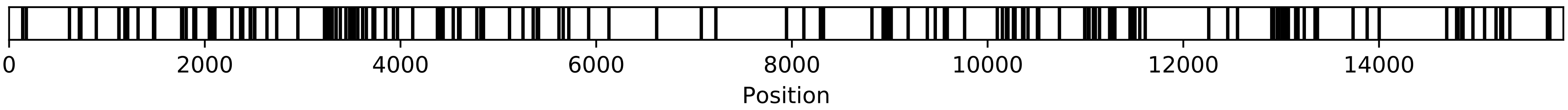


CR2 vs MA1
Number of differences: 370

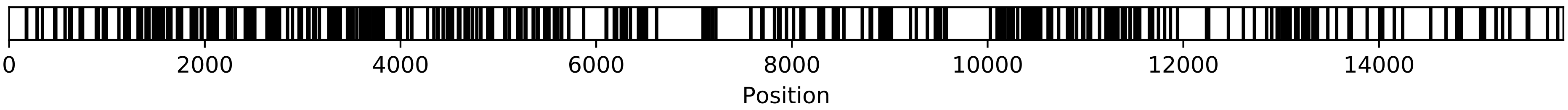


CR2 vs MA2
Number of differences: 96

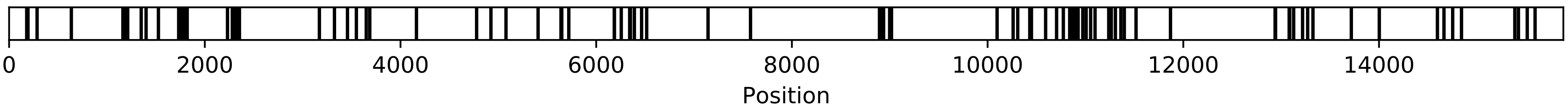


CR2 vs MM1
Number of differences: 372

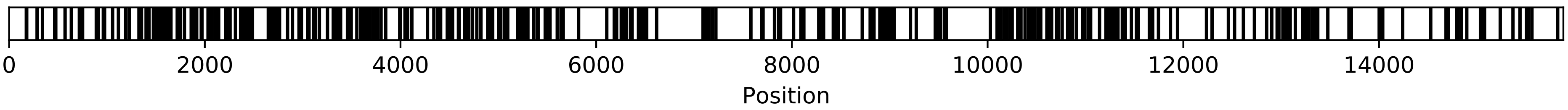


CR2 vs MM2
Number of differences: 370

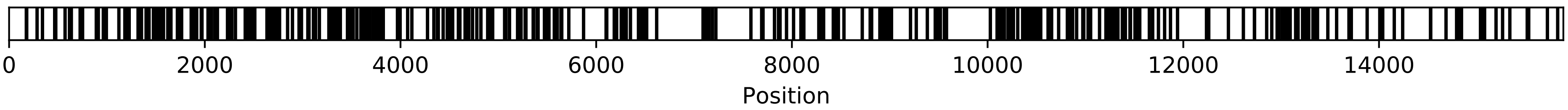


MA1 vs MA2
Number of differences: 369

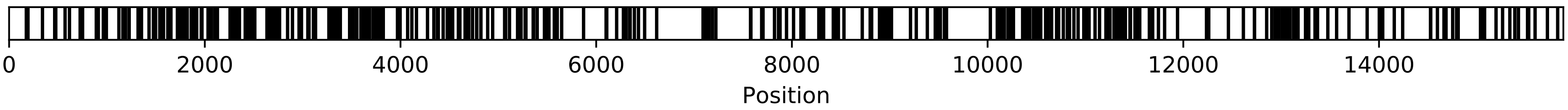


MA1 vs MM1
Number of differences: 172

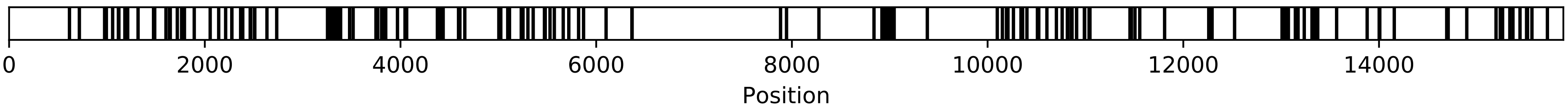


MA1 vs MM2
Number of differences: 0

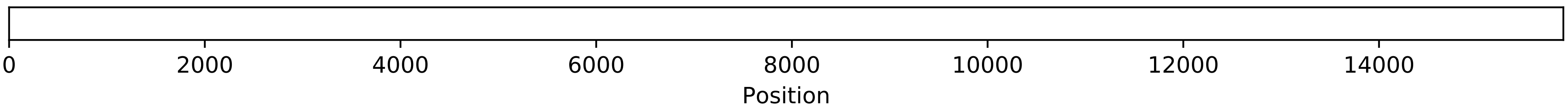


MA2 vs MM1
Number of differences: 373

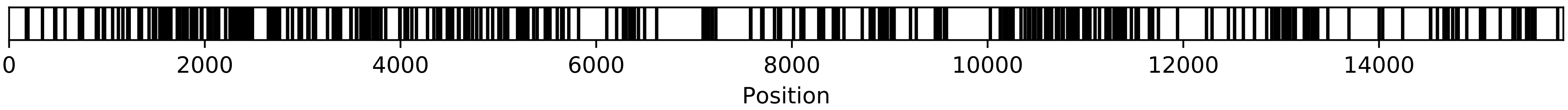


MA2 vs MM2
Number of differences: 369

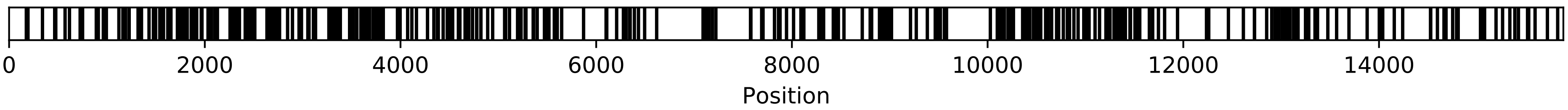


MM1 vs MM2
Number of differences: 172

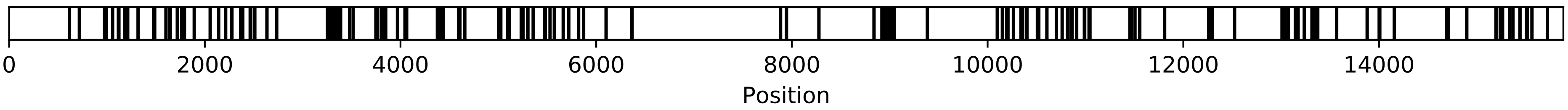

### Region: 344



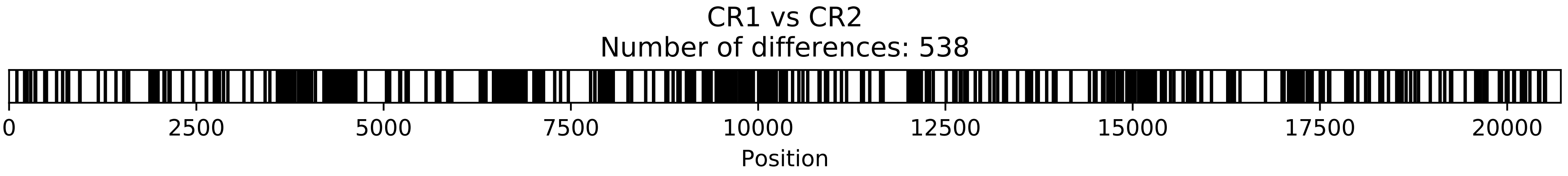
CR1 vs CR2
Number of differences: 538
0 2500 5000 7500 10000 12500 15000 17500 20000
Position

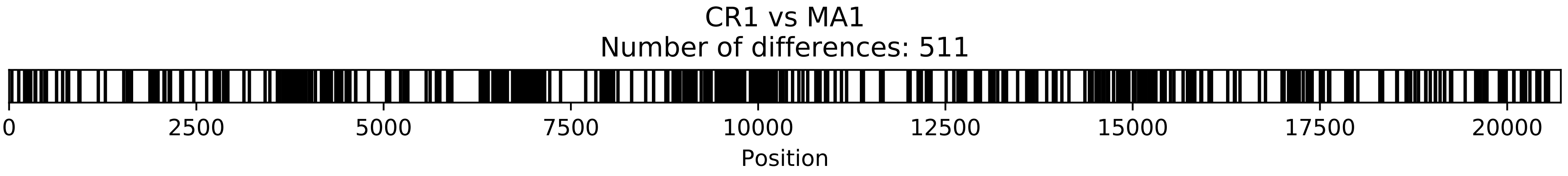
CR1 vs MA1
Number of differences: 511
0 2500 5000 7500 10000 12500 15000 17500 20000
Position

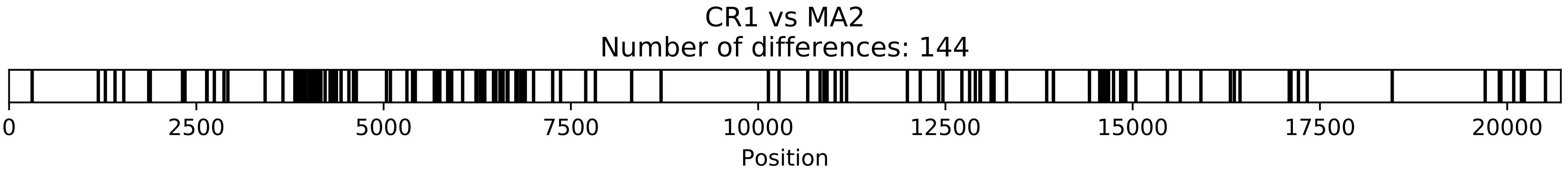
CR1 vs MA2
Number of differences: 144
0 2500 5000 7500 10000 12500 15000 17500 20000
Position

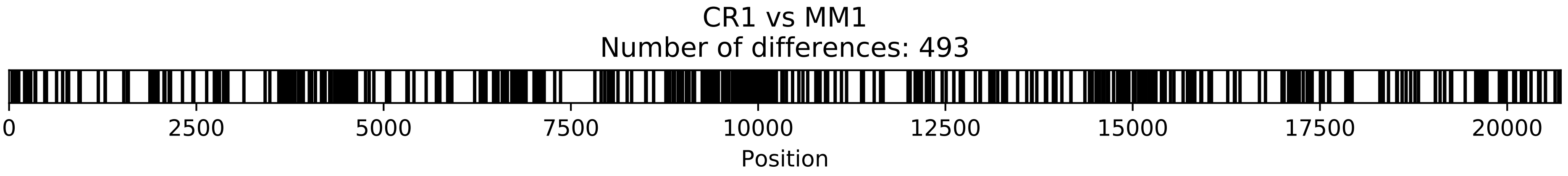
CR1 vs MM1
Number of differences: 493
0 2500 5000 7500 10000 12500 15000 17500 20000
Position

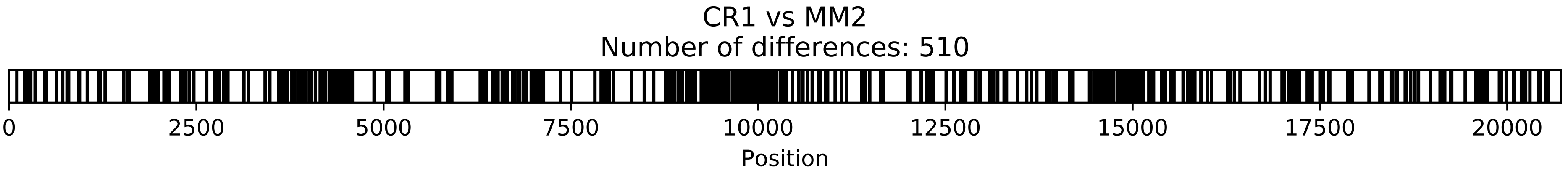
CR1 vs MM2
Number of differences: 510
0 2500 5000 7500 10000 12500 15000 17500 20000
Position

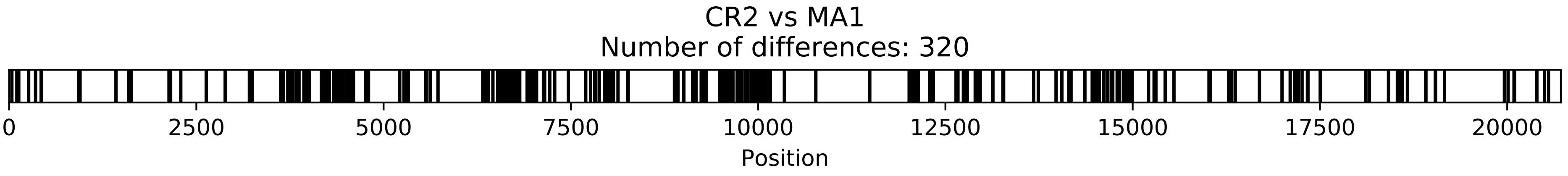
CR2 vs MA1
Number of differences: 320
0 2500 5000 7500 10000 12500 15000 17500 20000
Position

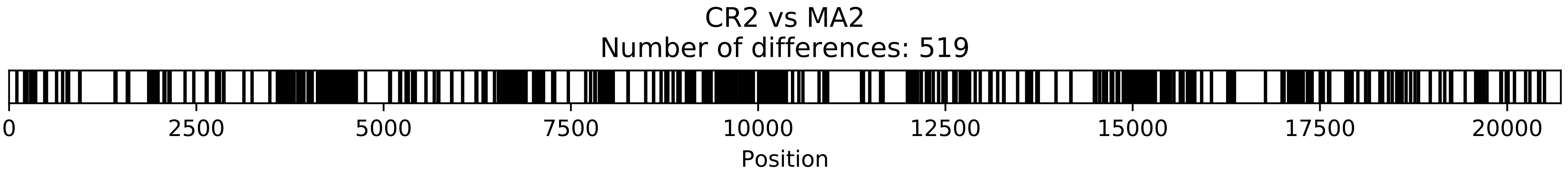
CR2 vs MA2
Number of differences: 519
0 2500 5000 7500 10000 12500 15000 17500 20000
Position

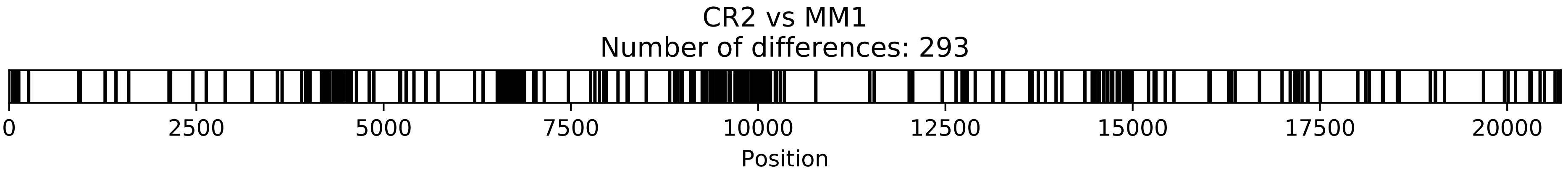
CR2 vs MM1
Number of differences: 293
0 2500 5000 7500 10000 12500 15000 17500 20000
Position

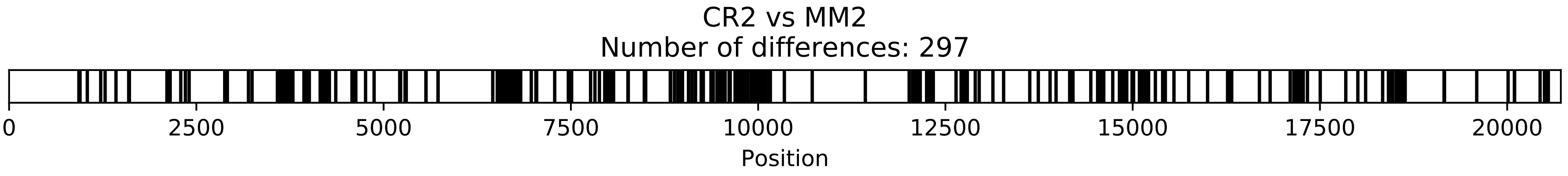
CR2 vs MM2
Number of differences: 297
0 2500 5000 7500 10000 12500 15000 17500 20000
Position

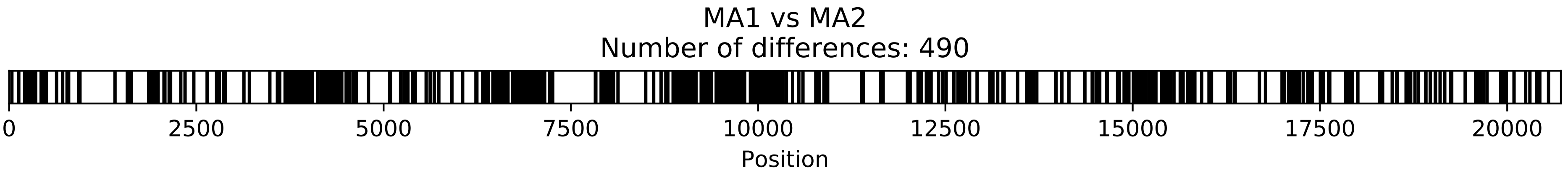
MA1 vs MA2
Number of differences: 490
0 2500 5000 7500 10000 12500 15000 17500 20000
Position

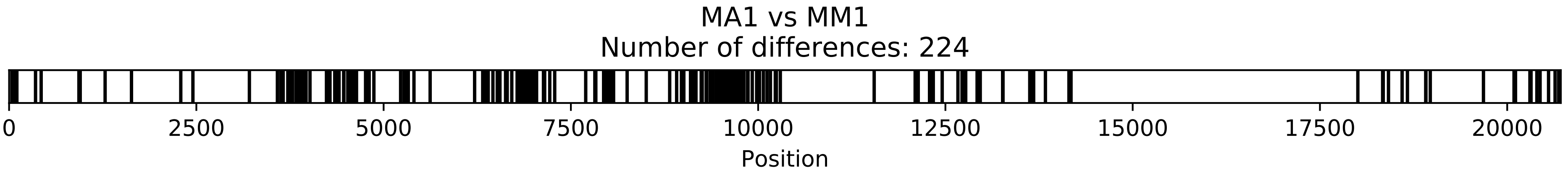
MA1 vs MM1
Number of differences: 224
0 2500 5000 7500 10000 12500 15000 17500 20000
Position

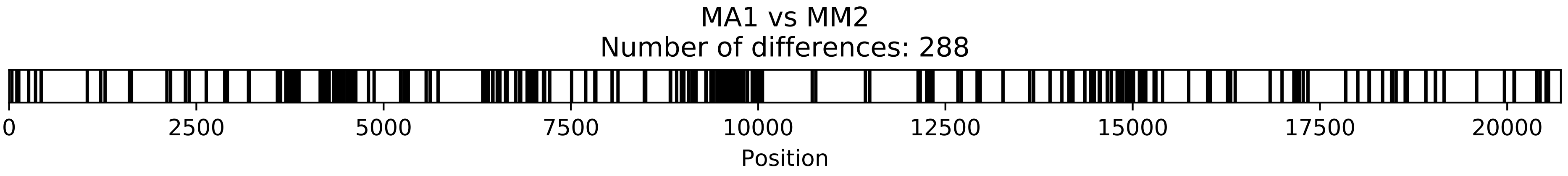
MA1 vs MM2
Number of differences: 288
0 2500 5000 7500 10000 12500 15000 17500 20000
Position

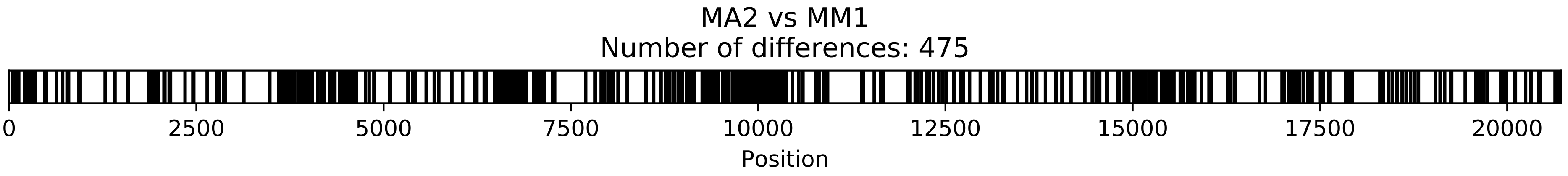
MA2 vs MM1
Number of differences: 475
0 2500 5000 7500 10000 12500 15000 17500 20000
Position

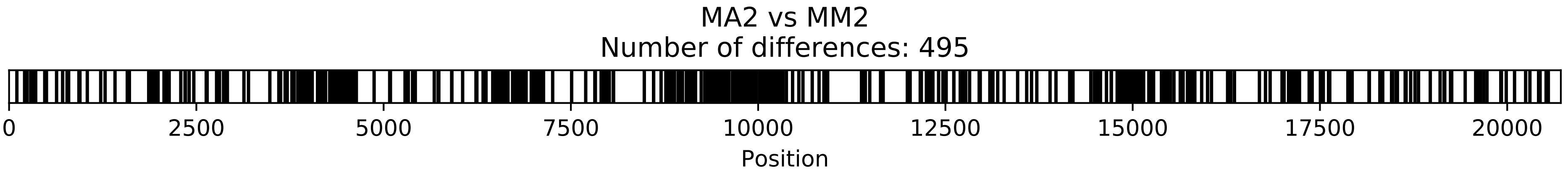
MA2 vs MM2
Number of differences: 495
0 2500 5000 7500 10000 12500 15000 17500 20000
Position

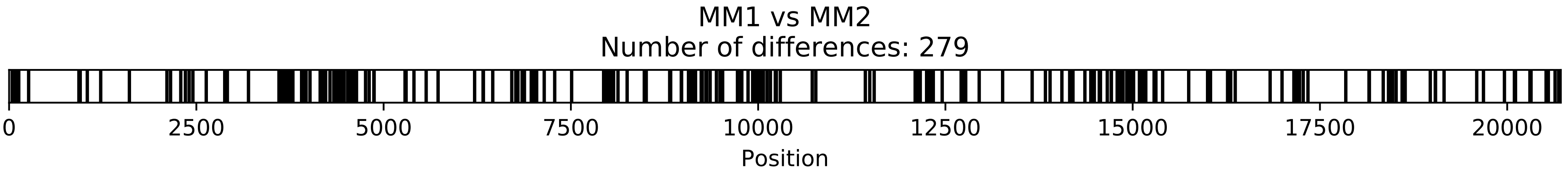
MM1 vs MM2
Number of differences: 279
0 2500 5000 7500 10000 12500 15000 17500 20000
Position

### Region: 346

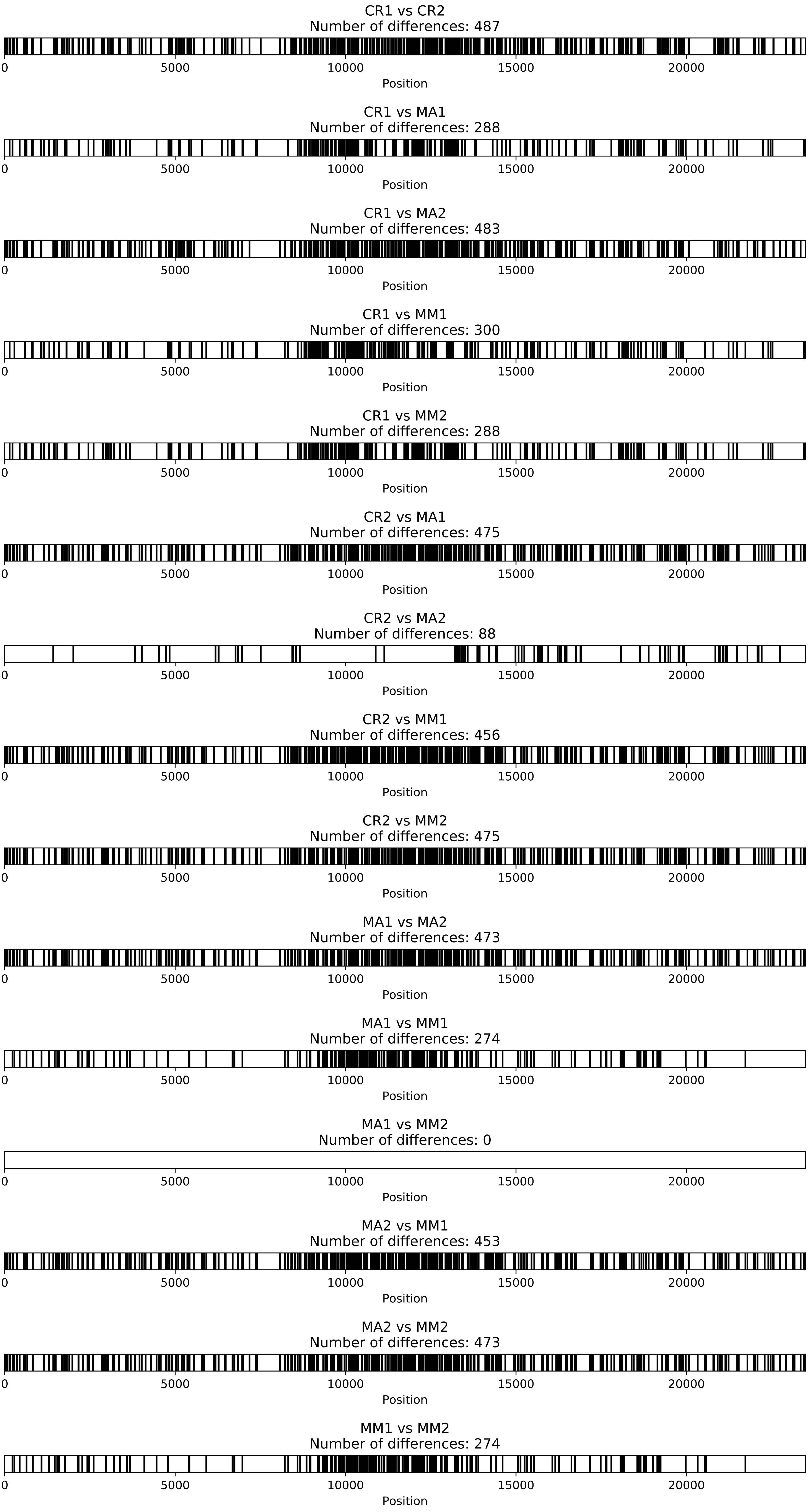
CR1 vs CR2
Number of differences: 487
CR1 vs MA1
Number of differences: 288
CR1 vs MA2
Number of differences: 483
CR1 vs MM1
Number of differences: 300
CR1 vs MM2
Number of differences: 288
CR2 vs MA1
Number of differences: 475
CR2 vs MA2
Number of differences: 88
CR2 vs MM1
Number of differences: 456
CR2 vs MM2
Number of differences: 475
MA1 vs MA2
Number of differences: 473
MA1 vs MM1
Number of differences: 274
MA1 vs MM2
Number of differences: 0
MA2 vs MM1
Number of differences: 453
MA2 vs MM2
Number of differences: 473
MM1 vs MM2
Number of differences: 274
0
5000
10000
15000
20000
Position

### Region: 348

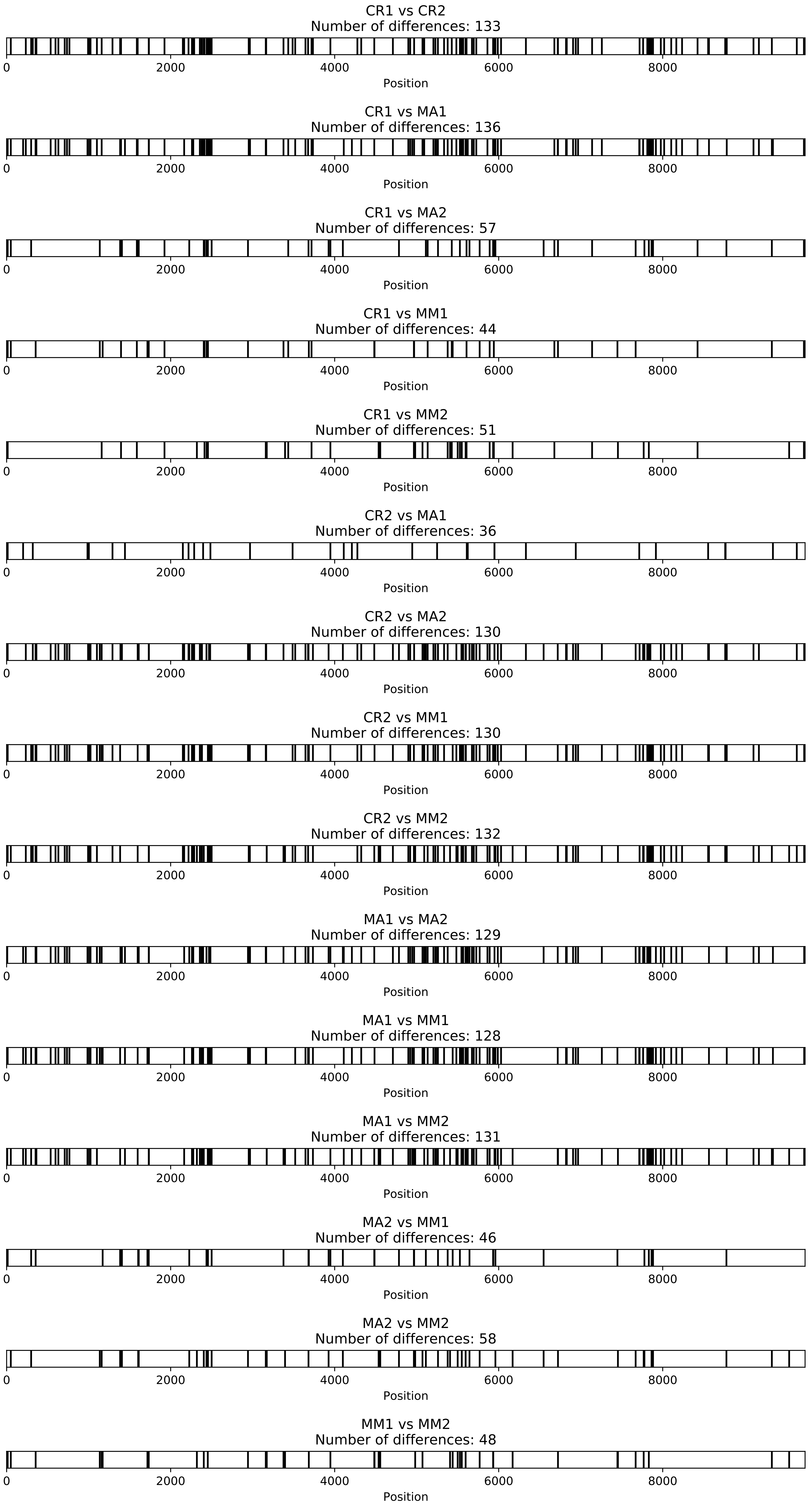
CR1 vs CR2
Number of differences: 133
CR1 vs MA1
Number of differences: 136
CR1 vs MA2
Number of differences: 57
CR1 vs MM1
Number of differences: 44
CR1 vs MM2
Number of differences: 51
CR2 vs MA1
Number of differences: 36
CR2 vs MA2
Number of differences: 130
CR2 vs MM1
Number of differences: 130
CR2 vs MM2
Number of differences: 132
MA1 vs MA2
Number of differences: 129
MA1 vs MM1
Number of differences: 128
MA1 vs MM2
Number of differences: 131
MA2 vs MM1
Number of differences: 46
MA2 vs MM2
Number of differences: 58
MM1 vs MM2
Number of differences: 48
0
2000
4000
6000
8000
Position

Laine et al.


### Region: 349

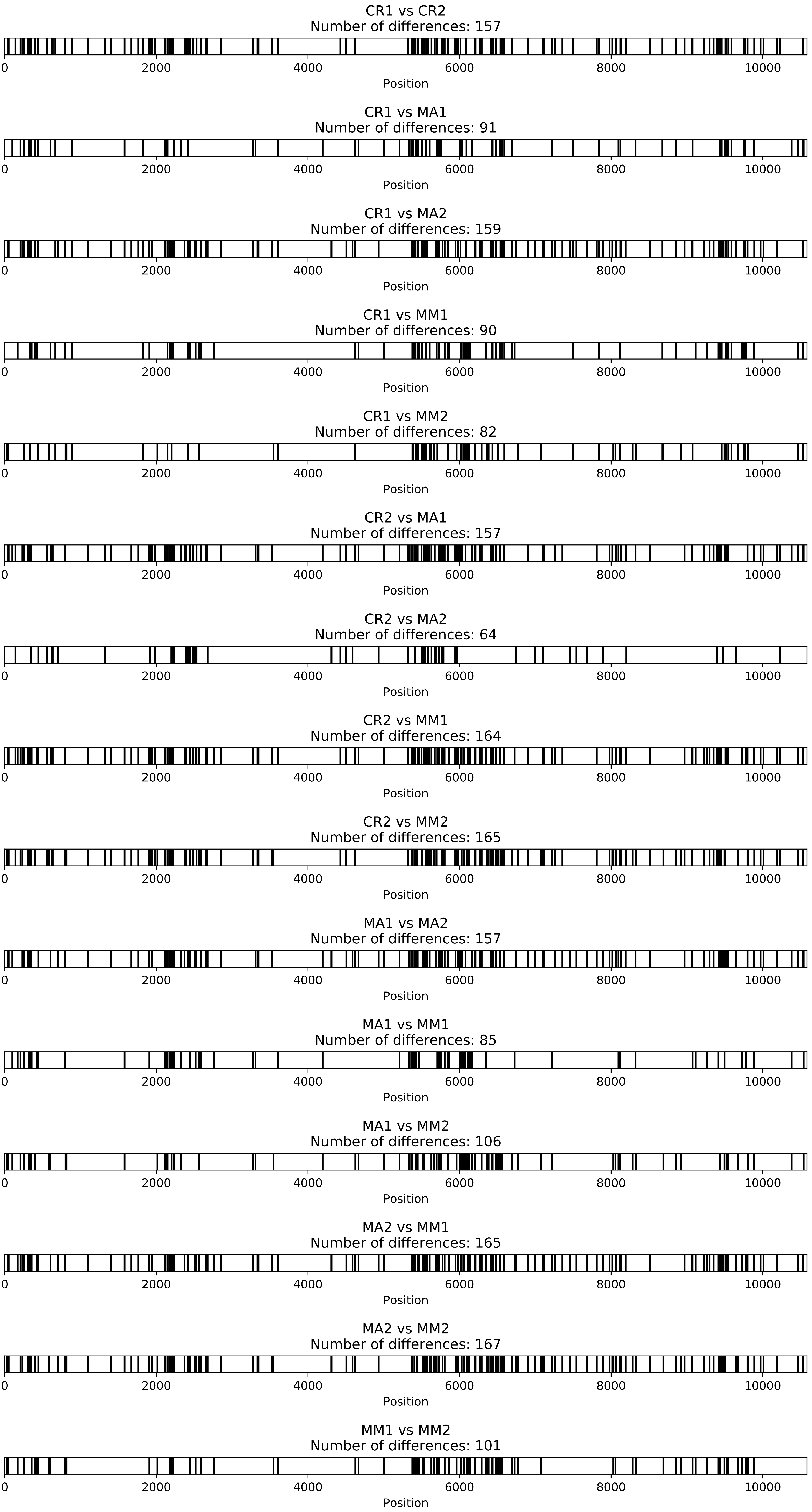

### Region: 352



CR1 vs CR2
Number of differences: 288

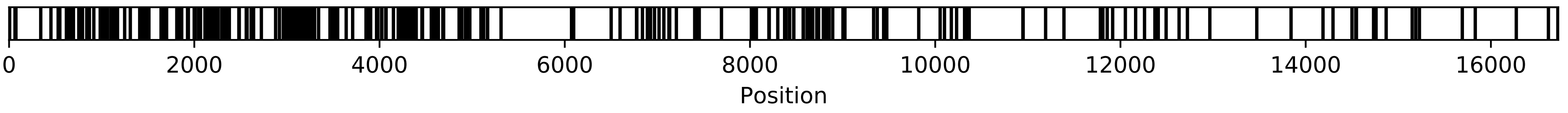


CR1 vs MA1
Number of differences: 139

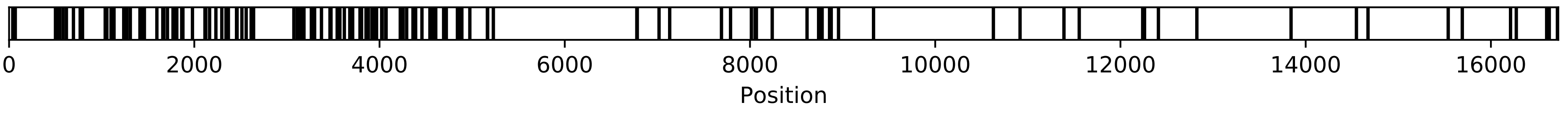


CR1 vs MA2
Number of differences: 285

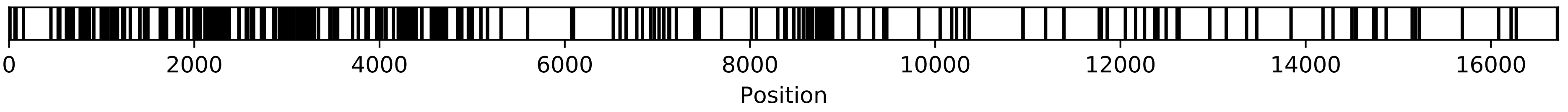


CR1 vs MM1
Number of differences: 141

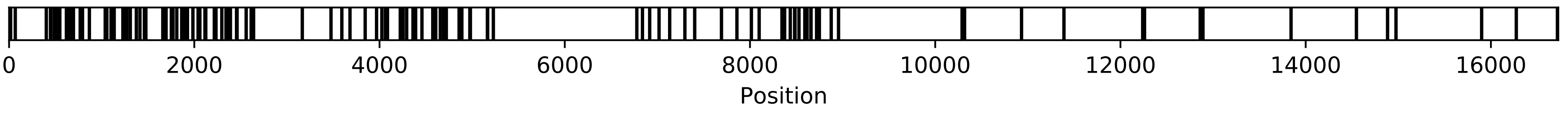


CR1 vs MM2
Number of differences: 139

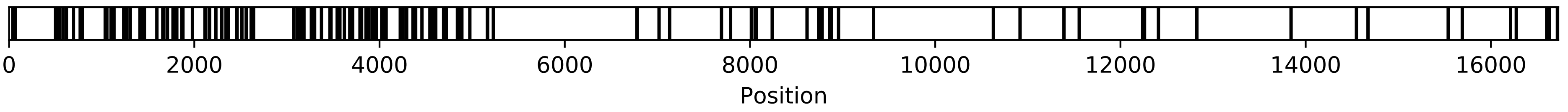


CR2 vs MA1
Number of differences: 289

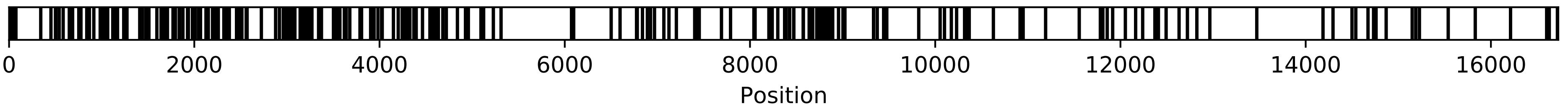


CR2 vs MA2
Number of differences: 111

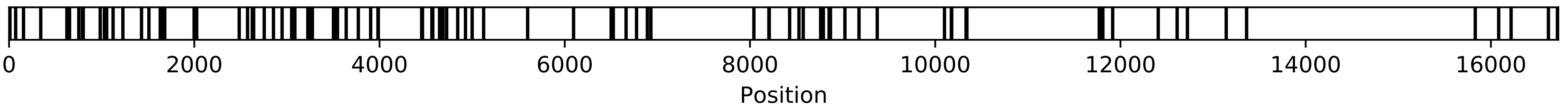


CR2 vs MM1
Number of differences: 295

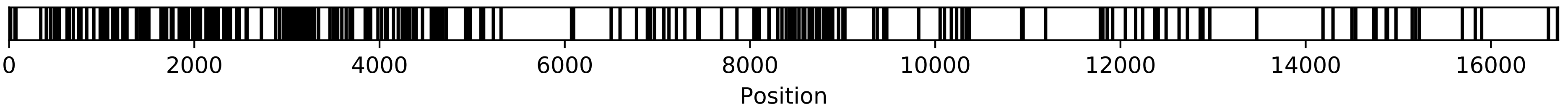


CR2 vs MM2
Number of differences: 289

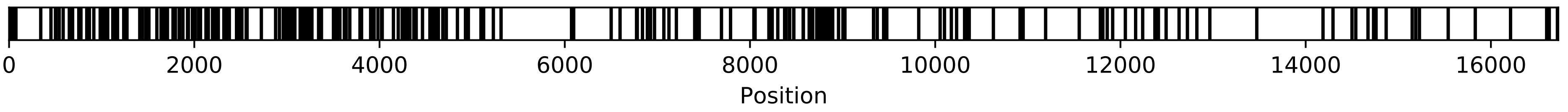


MA1 vs MA2
Number of differences: 278

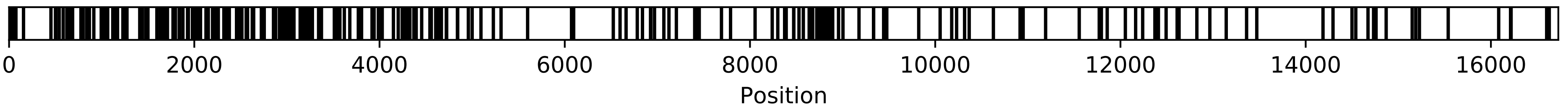


MA1 vs MM1
Number of differences: 136

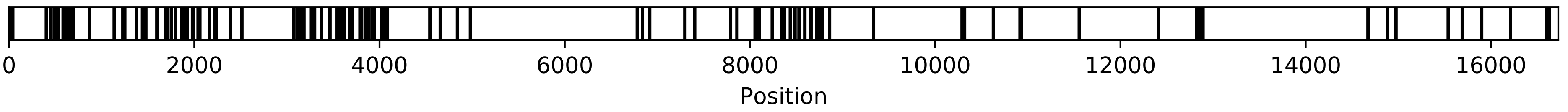


MA1 vs MM2
Number of differences: 0

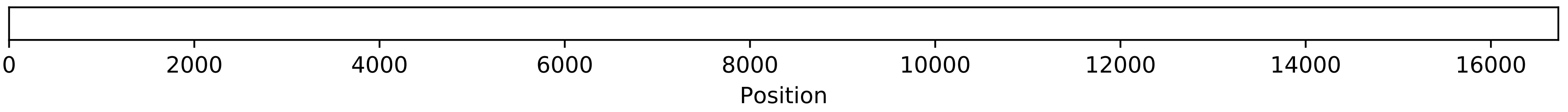


MA2 vs MM1
Number of differences: 278

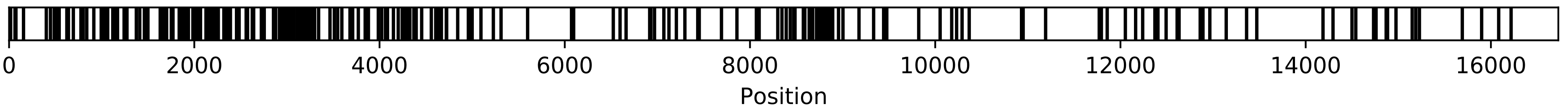


MA2 vs MM2
Number of differences: 278

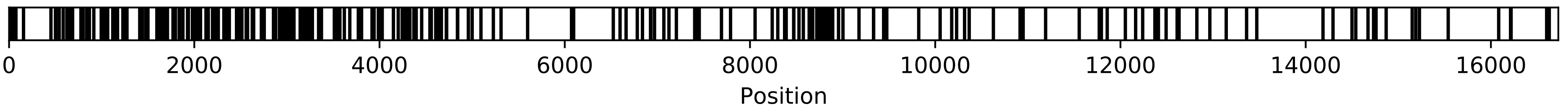


MM1 vs MM2
Number of differences: 136

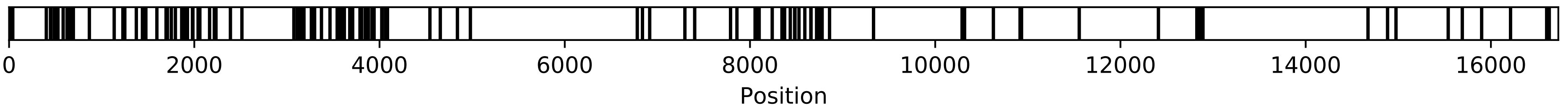

### Region: 354

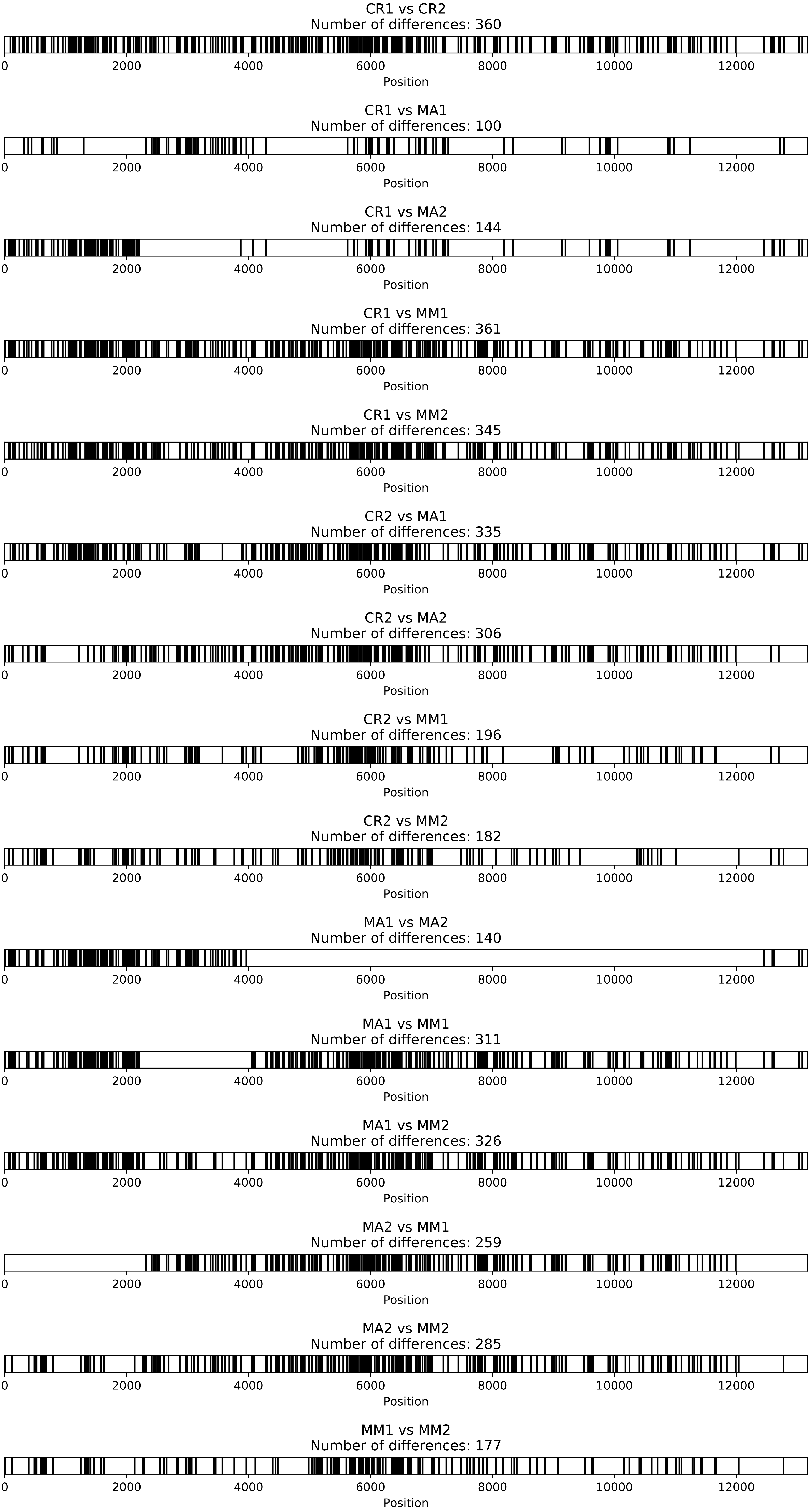
CR1 vs CR2
Number of differences: 360
0 2000 4000 6000 8000 10000 12000
Position
CR1 vs MA1
Number of differences: 100
0 2000 4000 6000 8000 10000 12000
Position
CR1 vs MA2
Number of differences: 144
0 2000 4000 6000 8000 10000 12000
Position
CR1 vs MM1
Number of differences: 361
0 2000 4000 6000 8000 10000 12000
Position
CR1 vs MM2
Number of differences: 345
0 2000 4000 6000 8000 10000 12000
Position
CR2 vs MA1
Number of differences: 335
0 2000 4000 6000 8000 10000 12000
Position
CR2 vs MA2
Number of differences: 306
0 2000 4000 6000 8000 10000 12000
Position
CR2 vs MM1
Number of differences: 196
0 2000 4000 6000 8000 10000 12000
Position
CR2 vs MM2
Number of differences: 182
0 2000 4000 6000 8000 10000 12000
Position
MA1 vs MA2
Number of differences: 140
0 2000 4000 6000 8000 10000 12000
Position
MA1 vs MM1
Number of differences: 311
0 2000 4000 6000 8000 10000 12000
Position
MA1 vs MM2
Number of differences: 326
0 2000 4000 6000 8000 10000 12000
Position
MA2 vs MM1
Number of differences: 259
0 2000 4000 6000 8000 10000 12000
Position
MA2 vs MM2
Number of differences: 285
0 2000 4000 6000 8000 10000 12000
Position
MM1 vs MM2
Number of differences: 177
0 2000 4000 6000 8000 10000 12000
Position

### Region: 356

CR1 vs CR2
Number of differences: 33

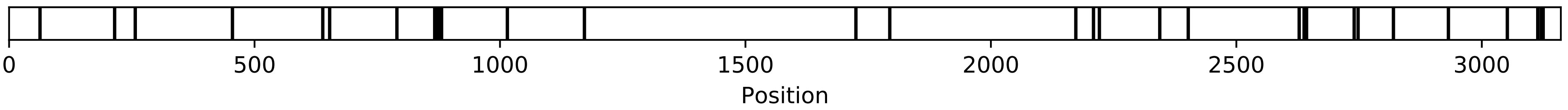


CR1 vs MA1
Number of differences: 16

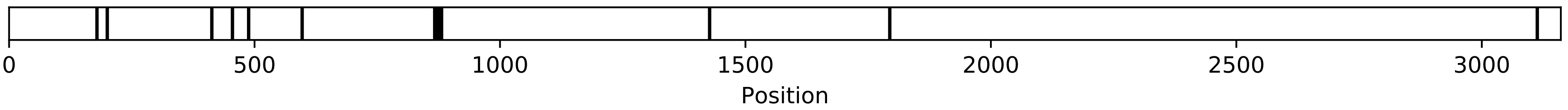


CR1 vs MA2
Number of differences: 35

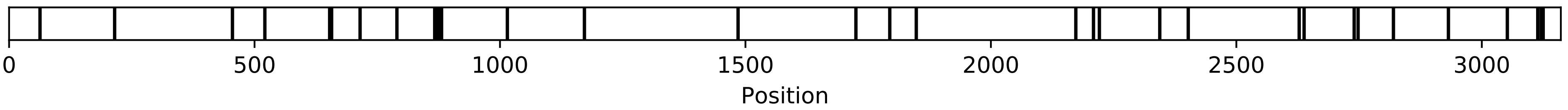


CR1 vs MM1
Number of differences: 18

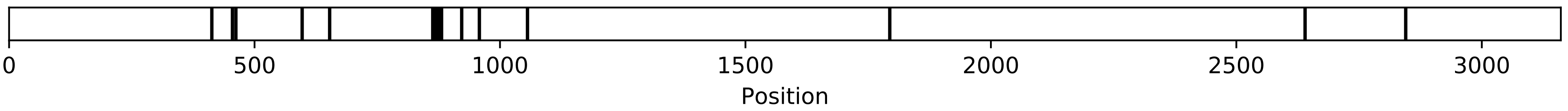


CR1 vs MM2
Number of differences: 19

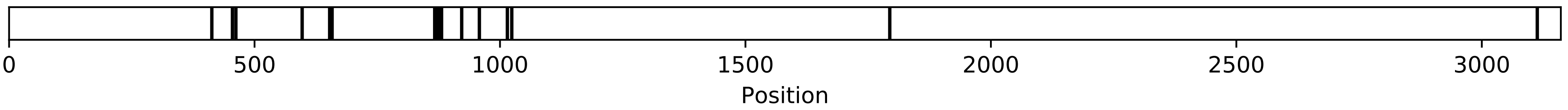


CR2 vs MA1
Number of differences: 33

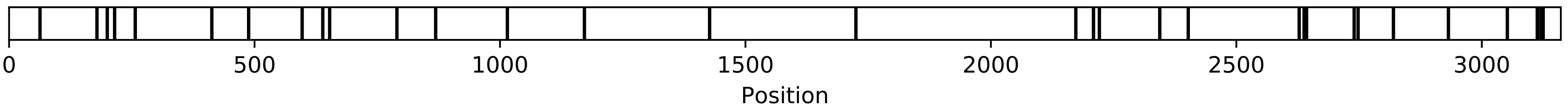


CR2 vs MA2
Number of differences: 8

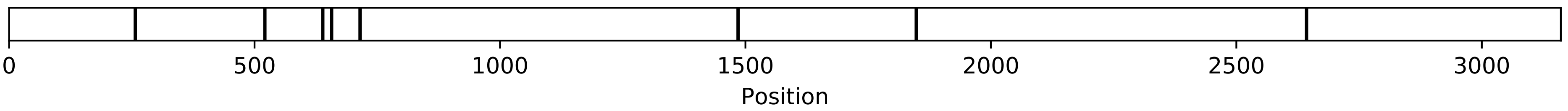


CR2 vs MM1
Number of differences: 33

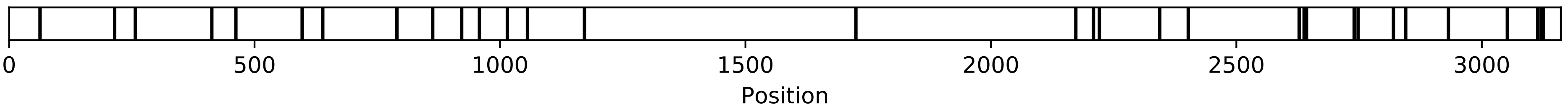


CR2 vs MM2
Number of differences: 32

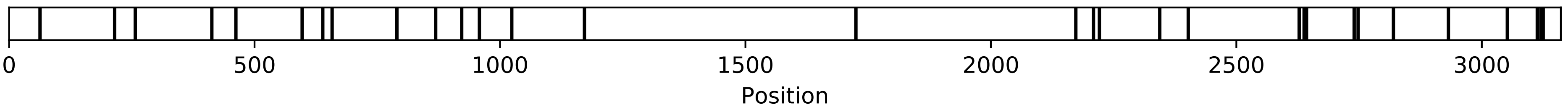


MA1 vs MA2
Number of differences: 35

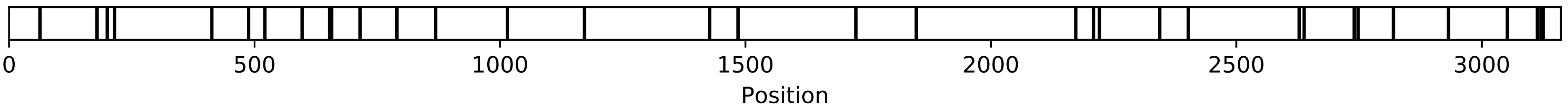


MA1 vs MM1
Number of differences: 14

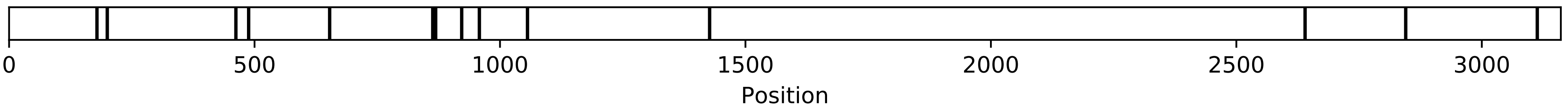


MA1 vs MM2
Number of differences: 11

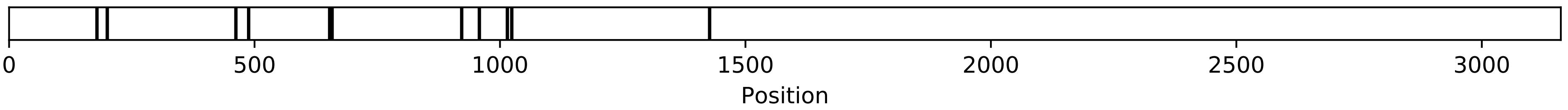


MA2 vs MM1
Number of differences: 35

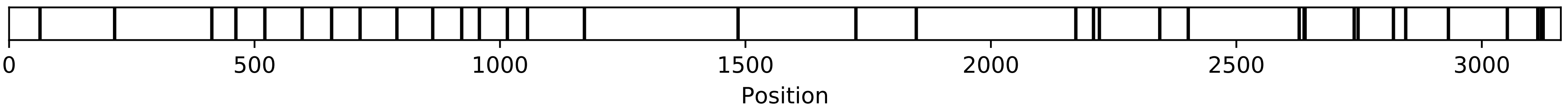


MA2 vs MM2
Number of differences: 34

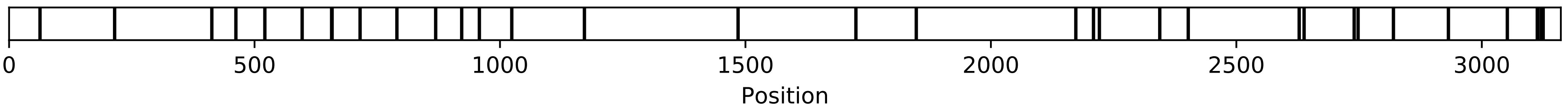


MM1 vs MM2
Number of differences: 9

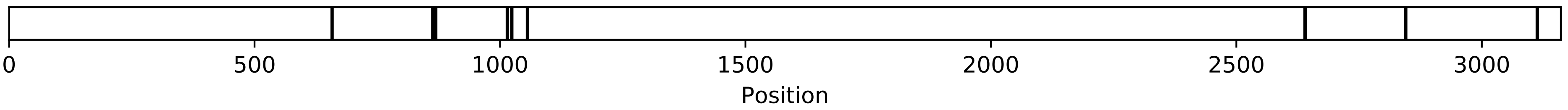

### Region: 358



CR1 vs CR2
Number of differences: 127

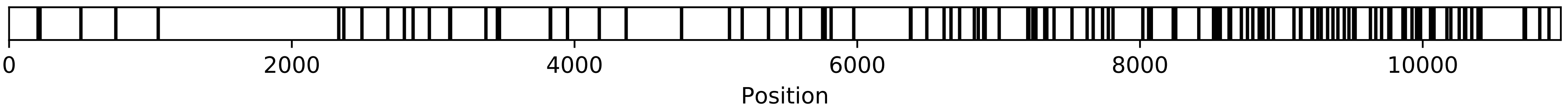

CR1 vs MA1
Number of differences: 79

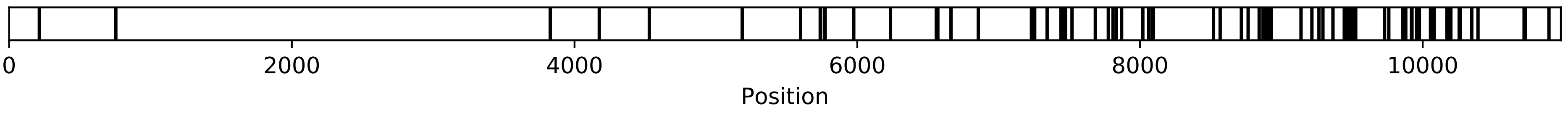

CR1 vs MA2
Number of differences: 87

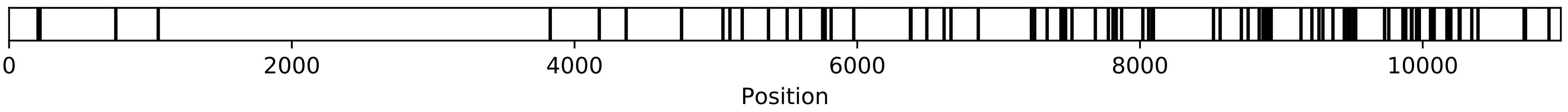

CR1 vs MM1
Number of differences: 56

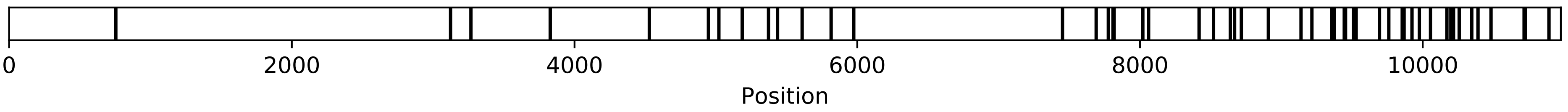

CR1 vs MM2
Number of differences: 70

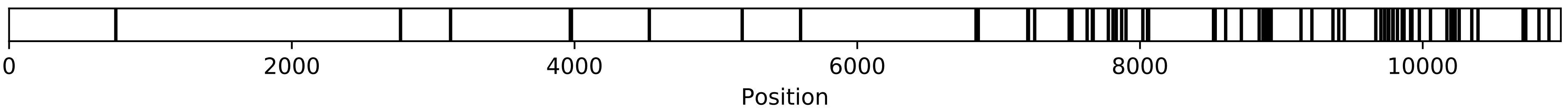

CR2 vs MA1
Number of differences: 121

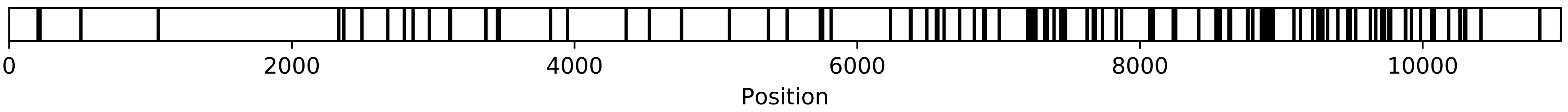

CR2 vs MA2
Number of differences: 103

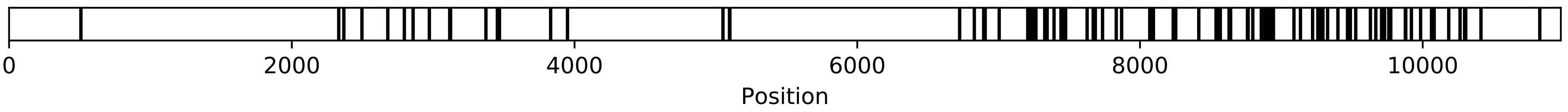

CR2 vs MM1
Number of differences: 114

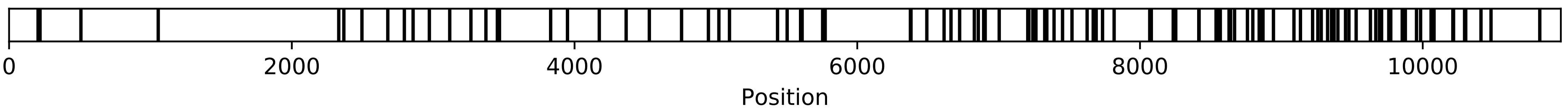

CR2 vs MM2
Number of differences: 124

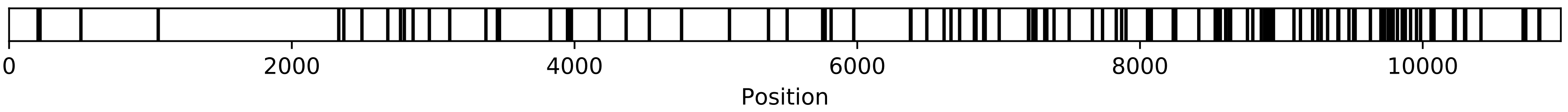

MA1 vs MA2
Number of differences: 22

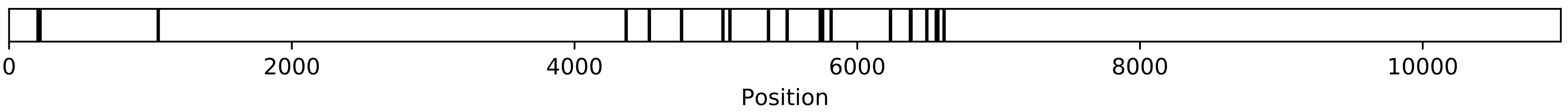

MA1 vs MM1
Number of differences: 70

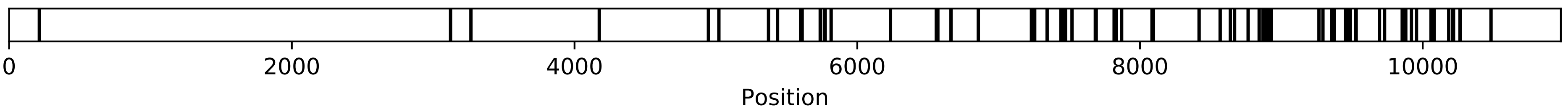

MA1 vs MM2
Number of differences: 67

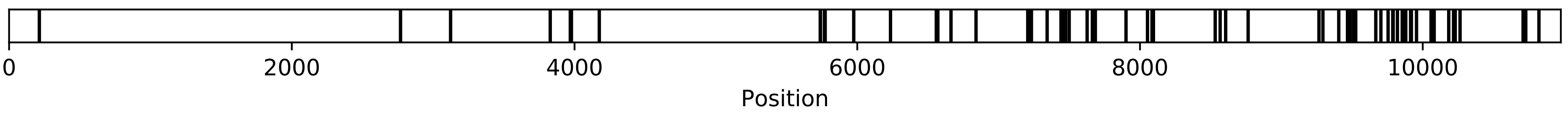

MA2 vs MM1
Number of differences: 76

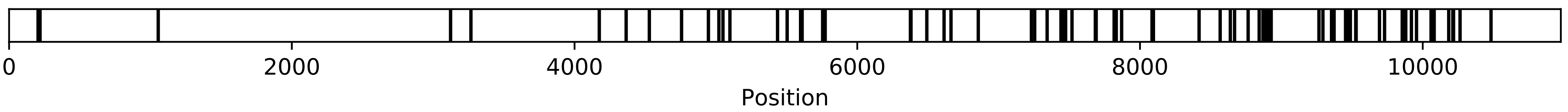

MA2 vs MM2
Number of differences: 77

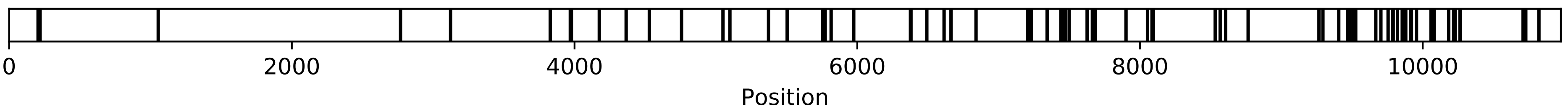

MM1 vs MM2
Number of differences: 64

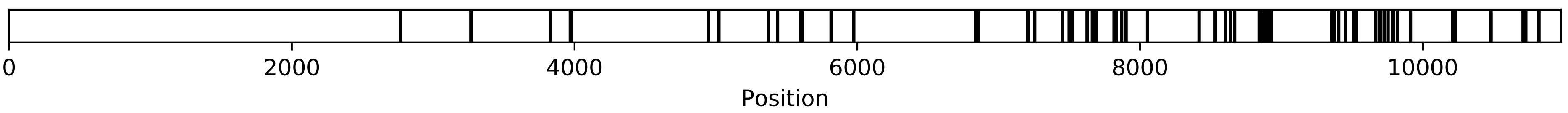

### Region: 361

CR1 vs CR2
Number of differences: 208

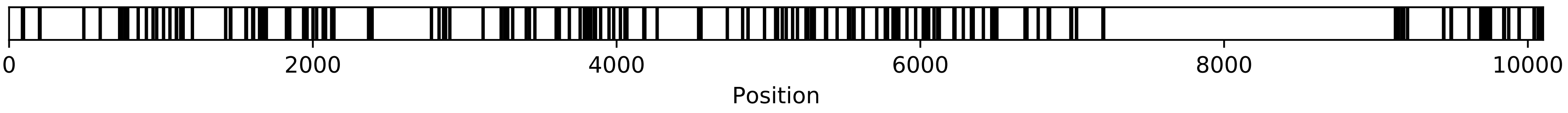


CR1 vs MA1
Number of differences: 197

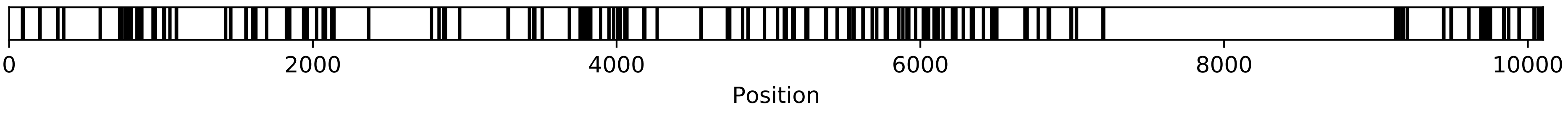


CR1 vs MA2
Number of differences: 158

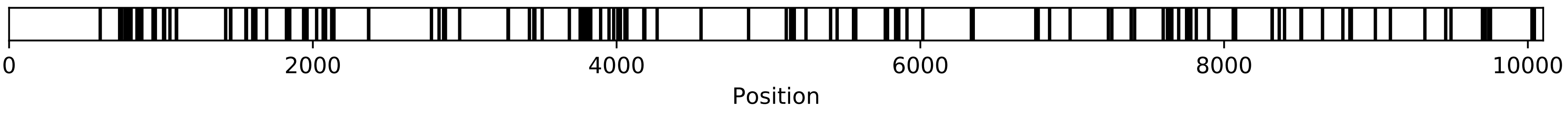


CR1 vs MM1
Number of differences: 197

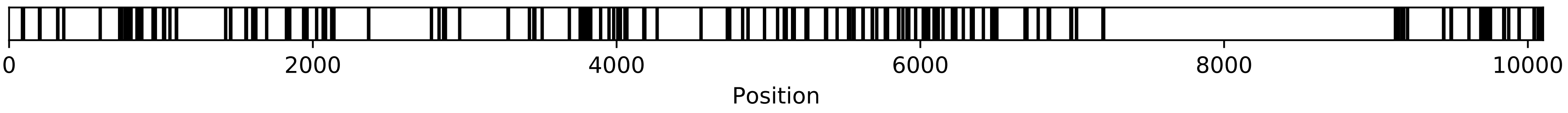


CR1 vs MM2
Number of differences: 228

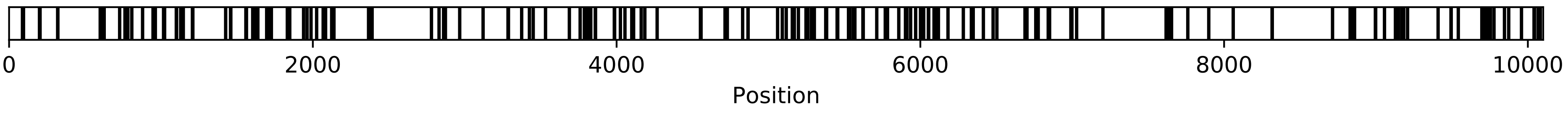


CR2 vs MA1
Number of differences: 86

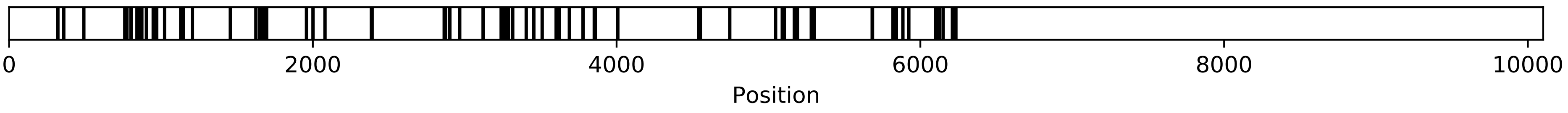


CR2 vs MA2
Number of differences: 203

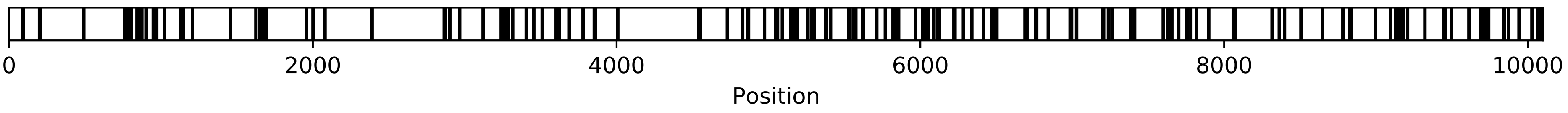


CR2 vs MM1
Number of differences: 86

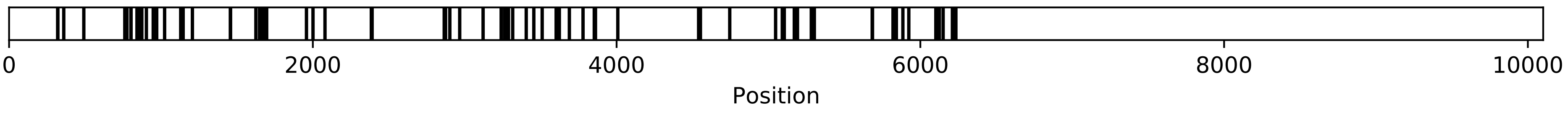


CR2 vs MM2
Number of differences: 157

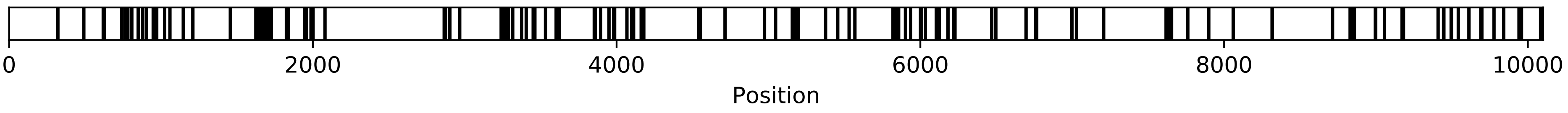


MA1 vs MA2
Number of differences: 145

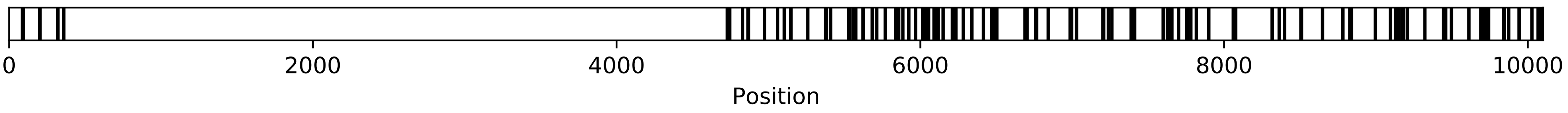


MA1 vs MM1
Number of differences: 0

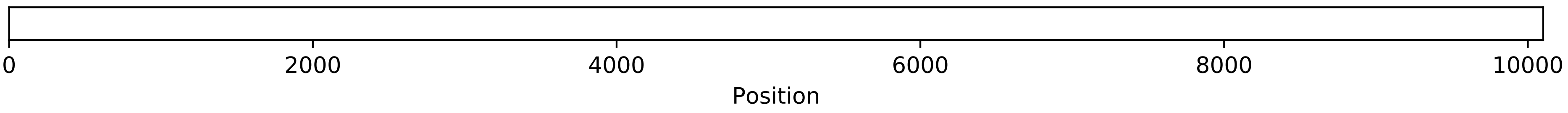


MA1 vs MM2
Number of differences: 144

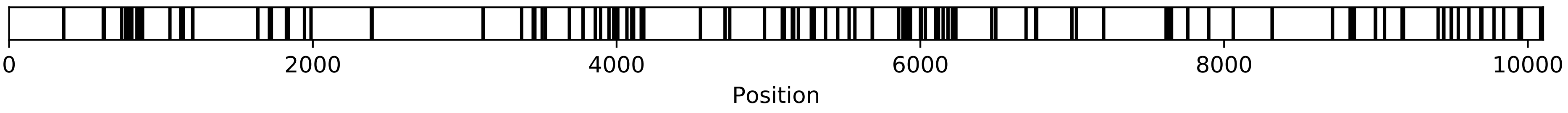


MA2 vs MM1
Number of differences: 145

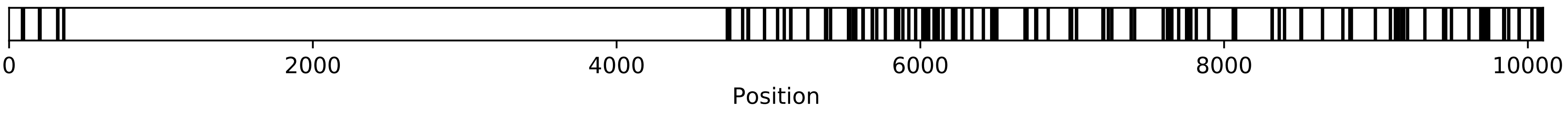


MA2 vs MM2
Number of differences: 192

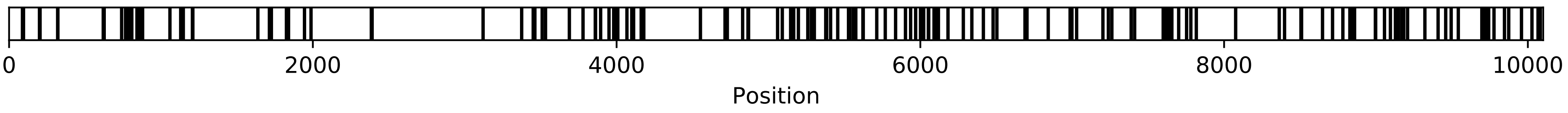


MM1 vs MM2
Number of differences: 144

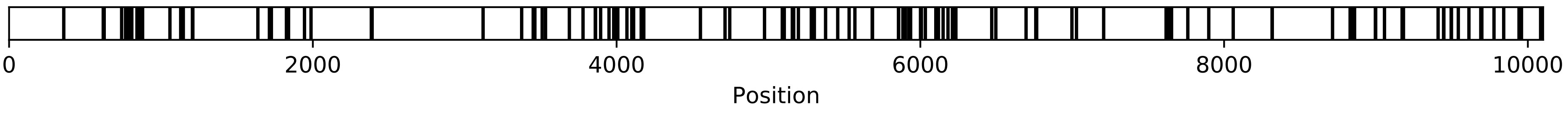

### Region: 362

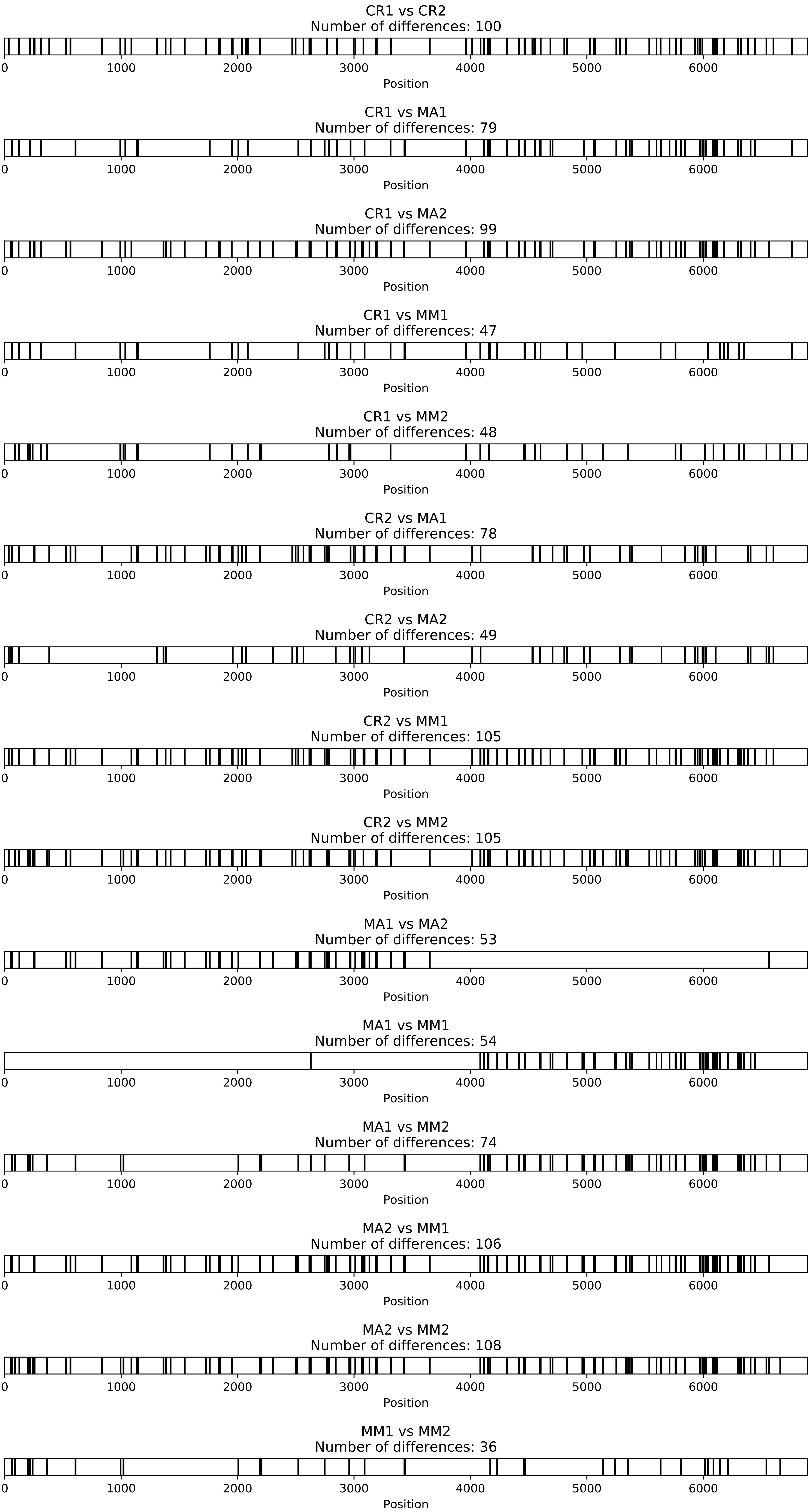
CR1 vs CR2
Number of differences: 100
CR1 vs MA1
Number of differences: 79
CR1 vs MA2
Number of differences: 99
CR1 vs MM1
Number of differences: 47
CR1 vs MM2
Number of differences: 48
CR2 vs MA1
Number of differences: 78
CR2 vs MA2
Number of differences: 49
CR2 vs MM1
Number of differences: 105
CR2 vs MM2
Number of differences: 105
MA1 vs MA2
Number of differences: 53
MA1 vs MM1
Number of differences: 54
MA1 vs MM2
Number of differences: 74
MA2 vs MM1
Number of differences: 106
MA2 vs MM2
Number of differences: 108
MM1 vs MM2
Number of differences: 36
0
1000
2000
3000
4000
5000
6000
Position

### Region: 364



CR1 vs CR2
Number of differences: 0

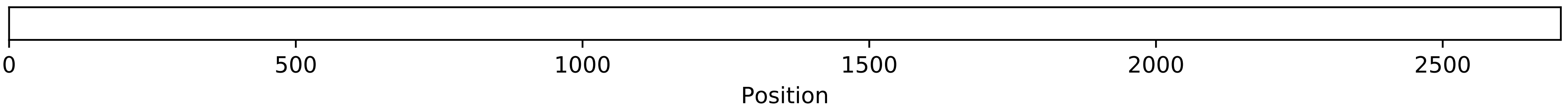


CR1 vs MA1
Number of differences: 58

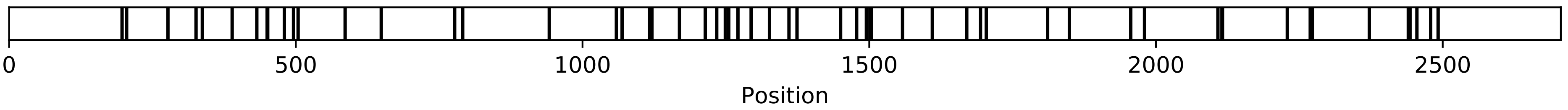


CR1 vs MA2
Number of differences: 23

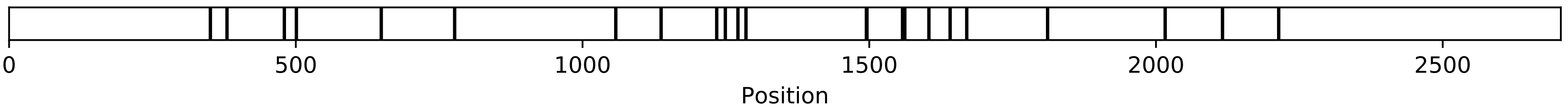


CR1 vs MM1
Number of differences: 24

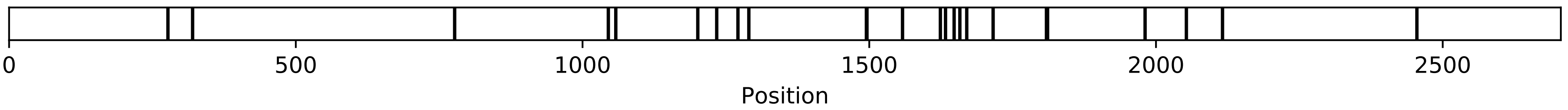


CR1 vs MM2
Number of differences: 14

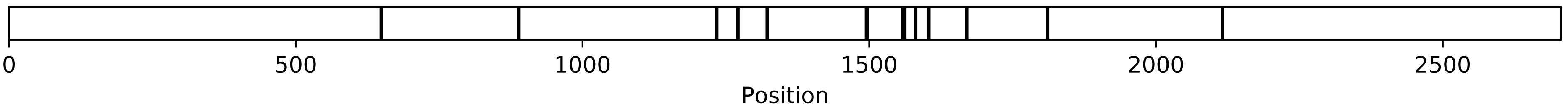


CR2 vs MA1
Number of differences: 58

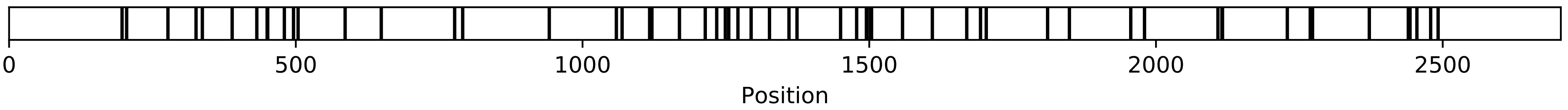


CR2 vs MA2
Number of differences: 23

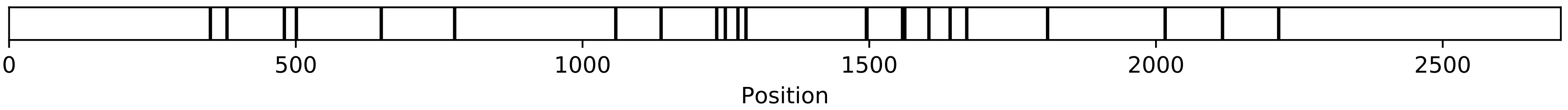


CR2 vs MM1
Number of differences: 24

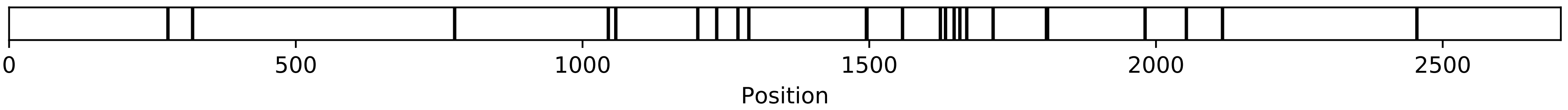


CR2 vs MM2
Number of differences: 14

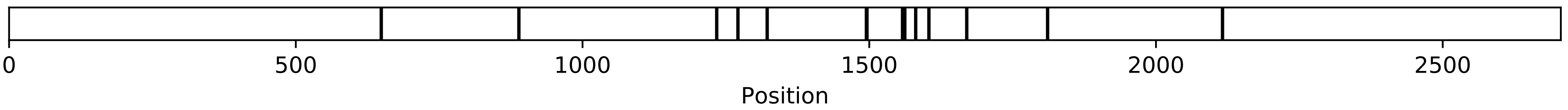


MA1 vs MA2
Number of differences: 57

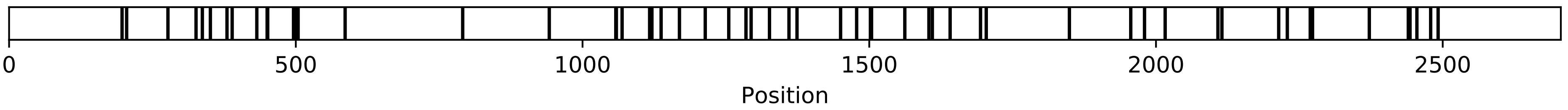


MA1 vs MM1
Number of differences: 60

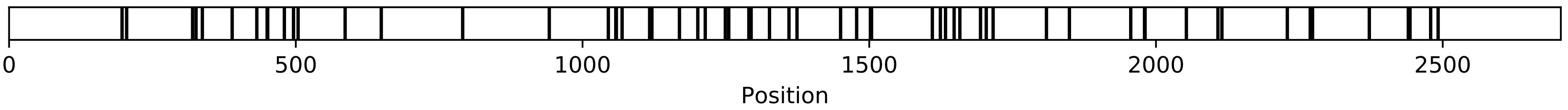


MA1 vs MM2
Number of differences: 54

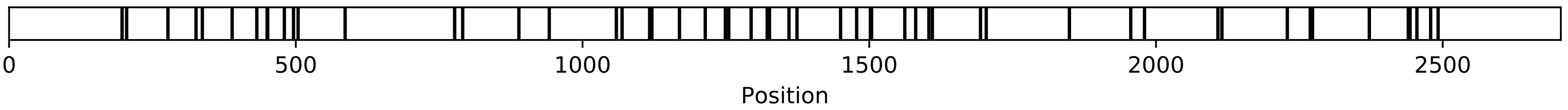


MA2 vs MM1
Number of differences: 27

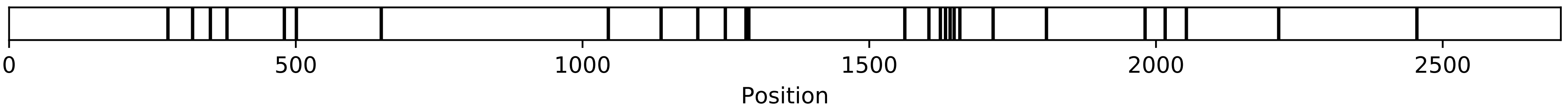


MA2 vs MM2
Number of differences: 15

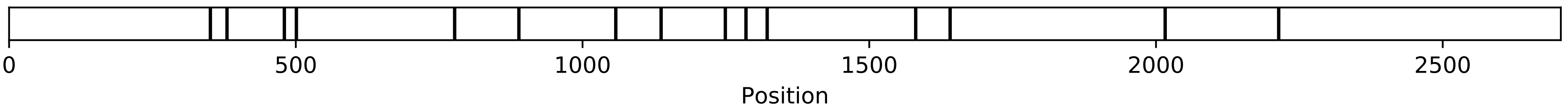


MM1 vs MM2
Number of differences: 22

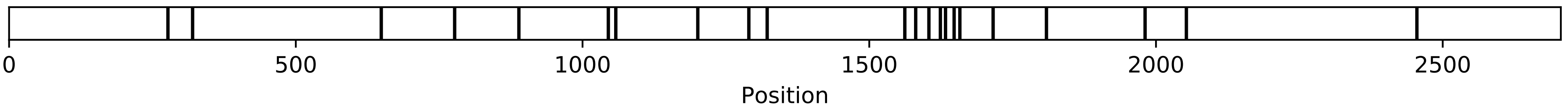

### Region: 366

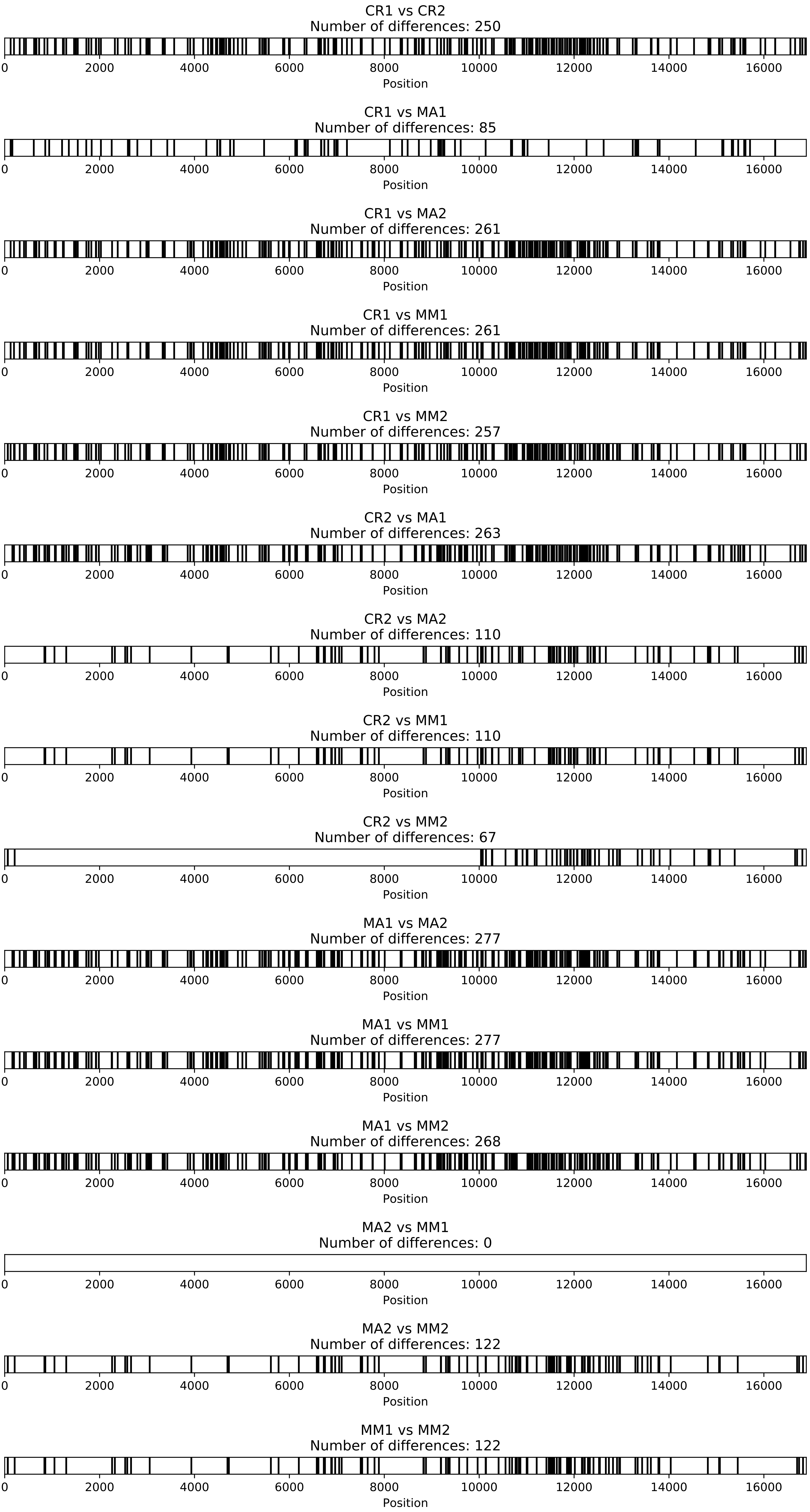
CR1 vs CR2
Number of differences: 250
CR1 vs MA1
Number of differences: 85
CR1 vs MA2
Number of differences: 261
CR1 vs MM1
Number of differences: 261
CR1 vs MM2
Number of differences: 257
CR2 vs MA1
Number of differences: 263
CR2 vs MA2
Number of differences: 110
CR2 vs MM1
Number of differences: 110
CR2 vs MM2
Number of differences: 67
MA1 vs MA2
Number of differences: 277
MA1 vs MM1
Number of differences: 277
MA1 vs MM2
Number of differences: 268
MA2 vs MM1
Number of differences: 0
MA2 vs MM2
Number of differences: 122
MM1 vs MM2
Number of differences: 122
0
2000
4000
6000
8000
10000
12000
14000
16000
Position

### Region: 367

CR1 vs CR2
Number of differences: 409

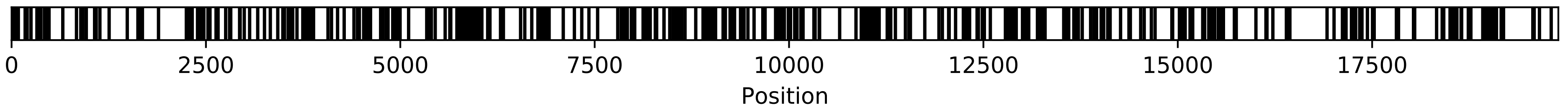


CR1 vs MA1
Number of differences: 173

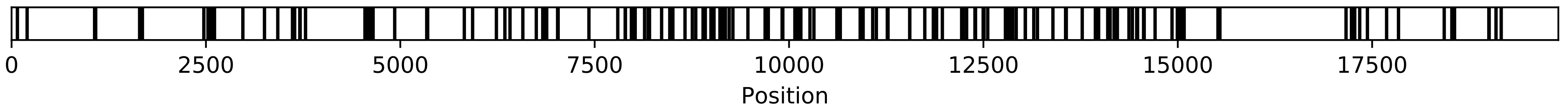


CR1 vs MA2
Number of differences: 401

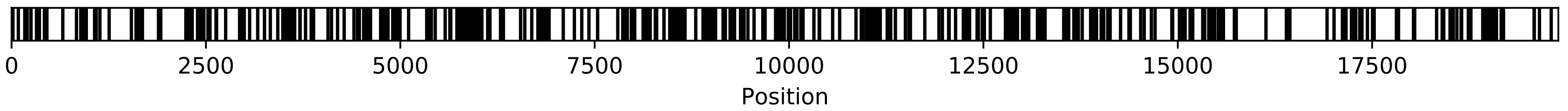


CR1 vs MM1
Number of differences: 184

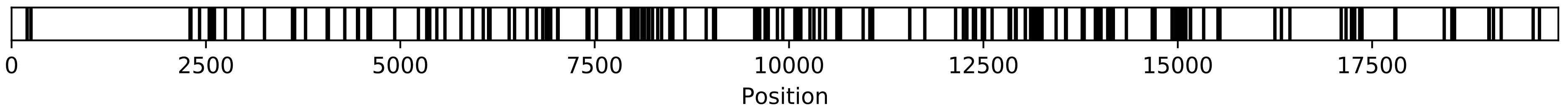


CR1 vs MM2
Number of differences: 197

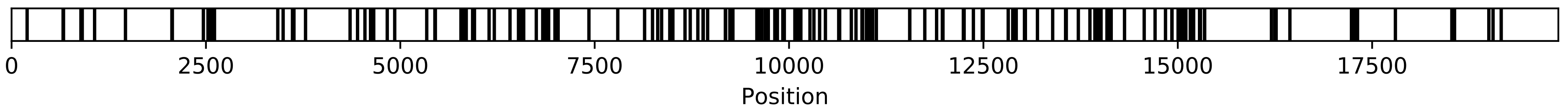


CR2 vs MA1
Number of differences: 392

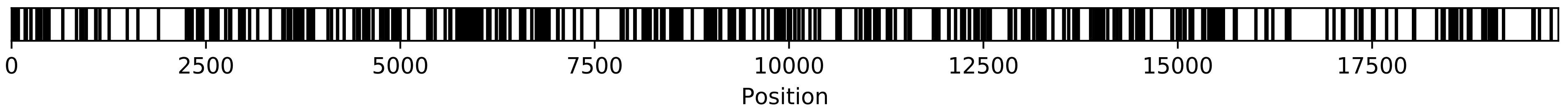


CR2 vs MA2
Number of differences: 56

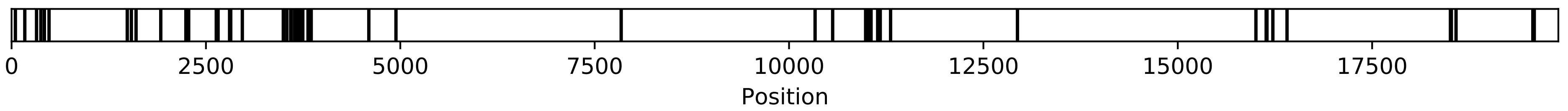


CR2 vs MM1
Number of differences: 444

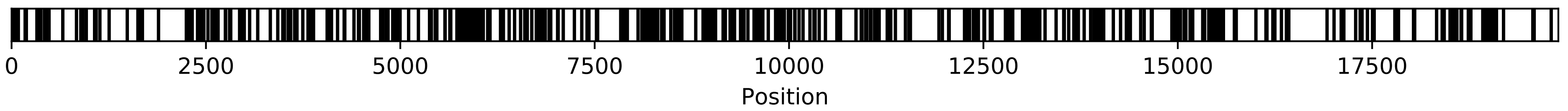


CR2 vs MM2
Number of differences: 443

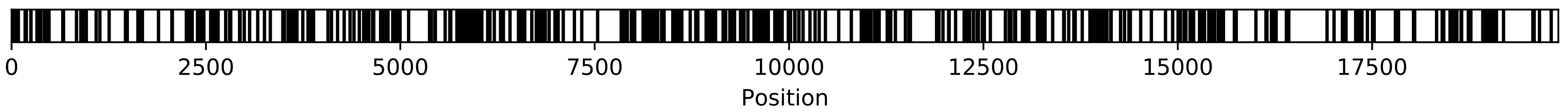


MA1 vs MA2
Number of differences: 378

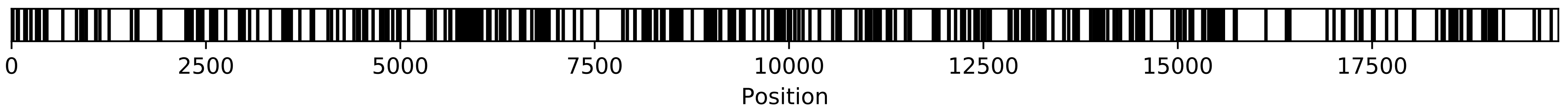


MA1 vs MM1
Number of differences: 184

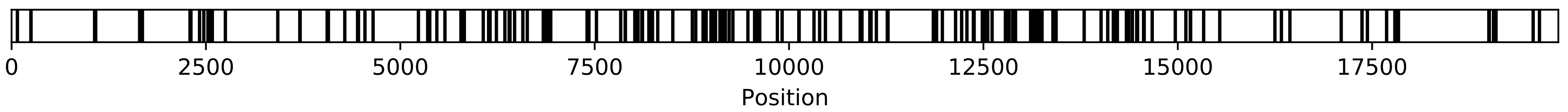


MA1 vs MM2
Number of differences: 202

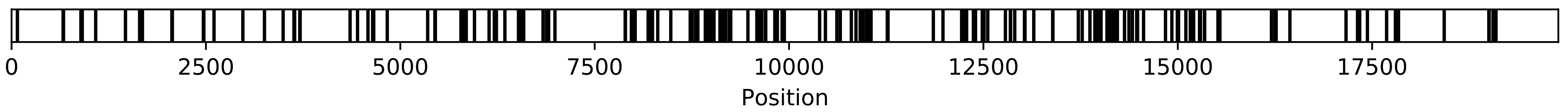


MA2 vs MM1
Number of differences: 430

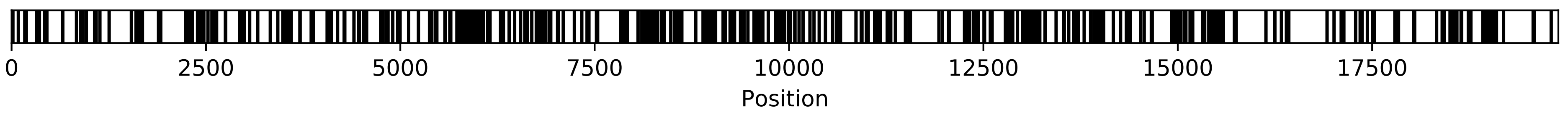


MA2 vs MM2
Number of differences: 431

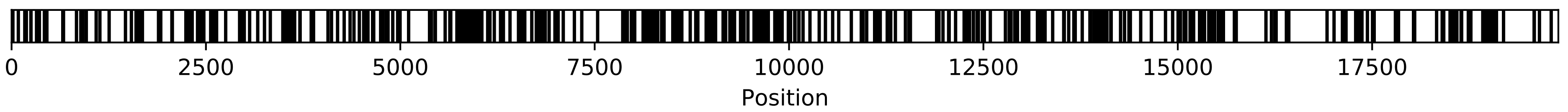


MM1 vs MM2
Number of differences: 215

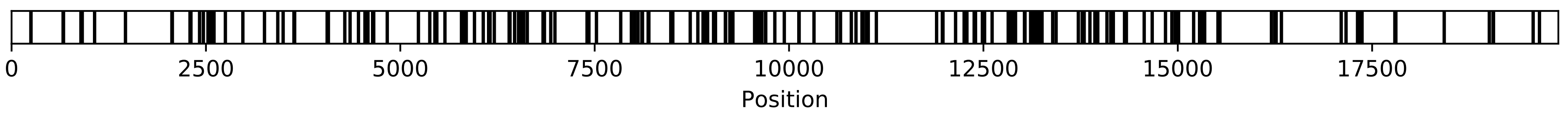

### Region: 368



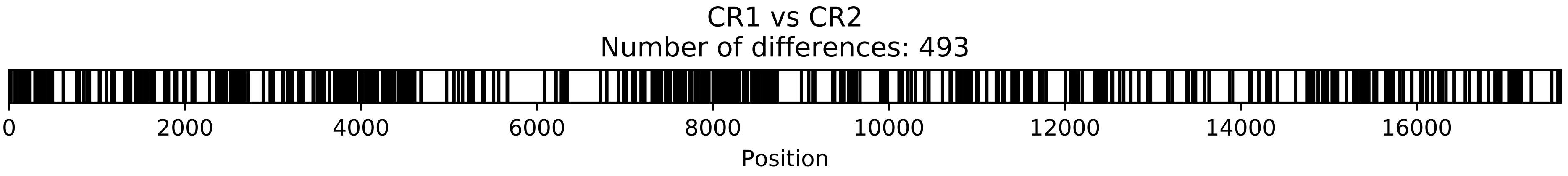
CR1 vs CR2
Number of differences: 493
0 2000 4000 6000 8000 10000 12000 14000 16000
Position

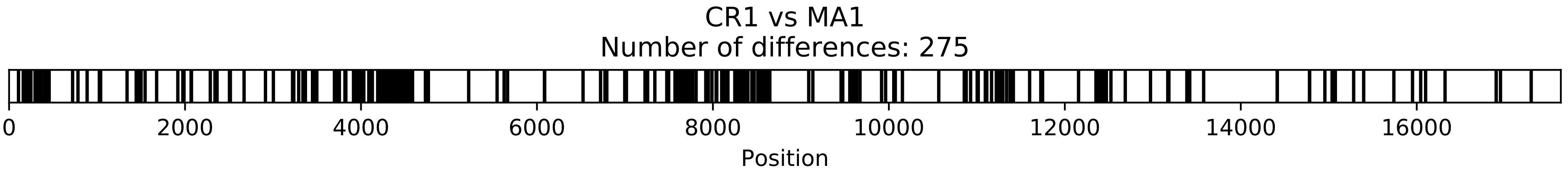
CR1 vs MA1
Number of differences: 275
0 2000 4000 6000 8000 10000 12000 14000 16000
Position

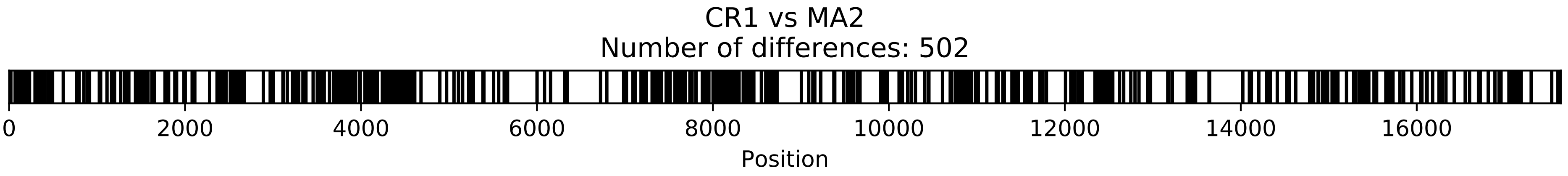
CR1 vs MA2
Number of differences: 502
0 2000 4000 6000 8000 10000 12000 14000 16000
Position

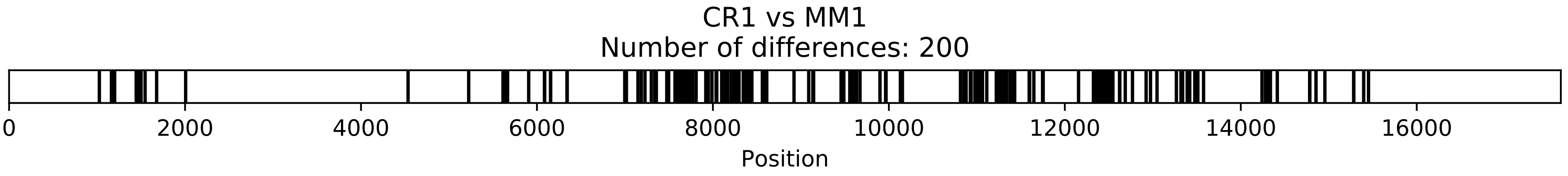
CR1 vs MM1
Number of differences: 200
0 2000 4000 6000 8000 10000 12000 14000 16000
Position

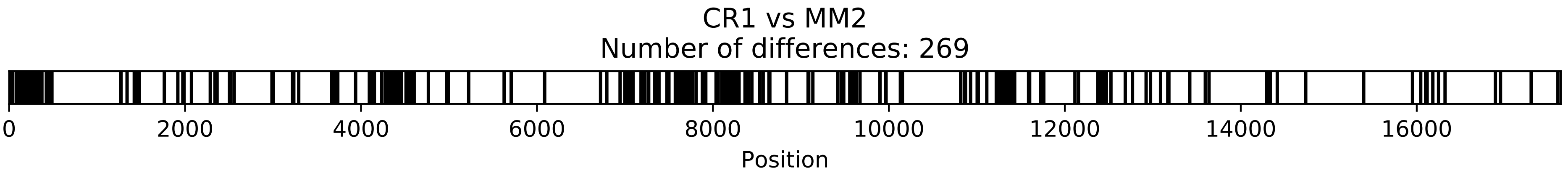
CR1 vs MM2
Number of differences: 269
0 2000 4000 6000 8000 10000 12000 14000 16000
Position

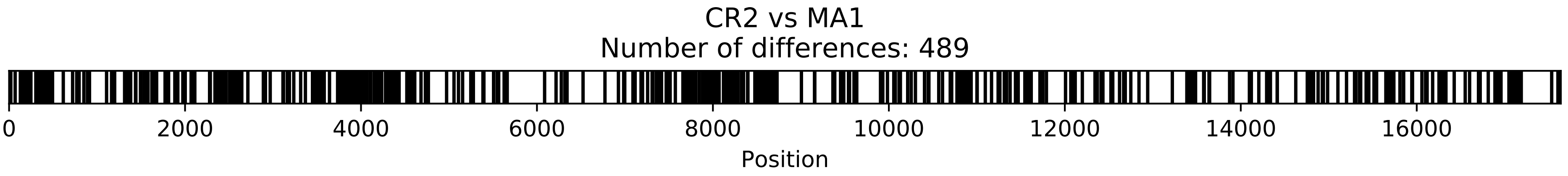
CR2 vs MA1
Number of differences: 489
0 2000 4000 6000 8000 10000 12000 14000 16000
Position

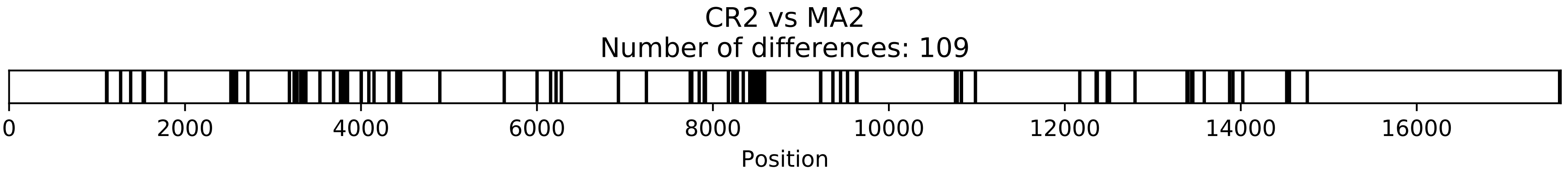
CR2 vs MA2
Number of differences: 109
0 2000 4000 6000 8000 10000 12000 14000 16000
Position

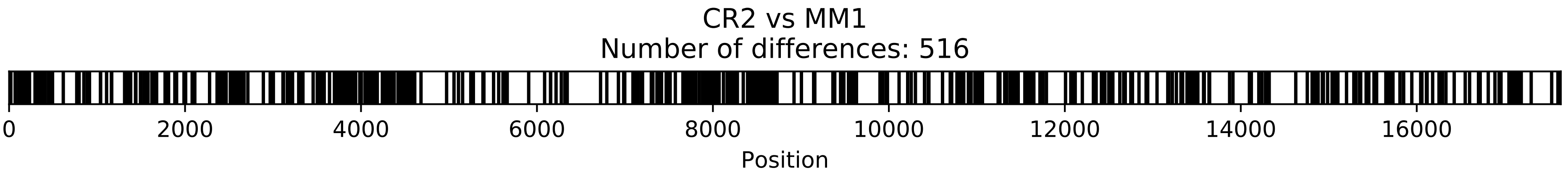
CR2 vs MM1
Number of differences: 516
0 2000 4000 6000 8000 10000 12000 14000 16000
Position

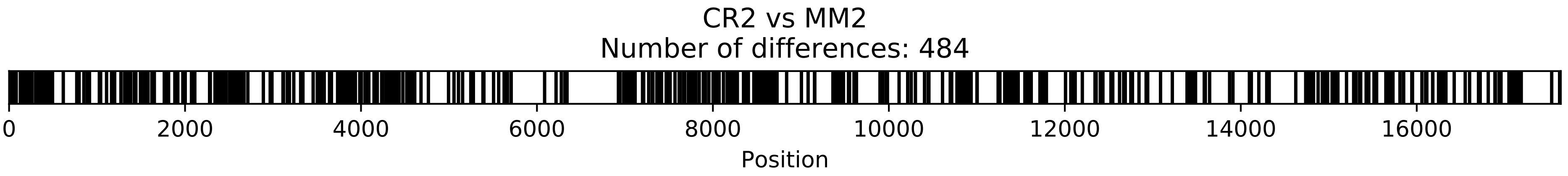
CR2 vs MM2
Number of differences: 484
0 2000 4000 6000 8000 10000 12000 14000 16000
Position

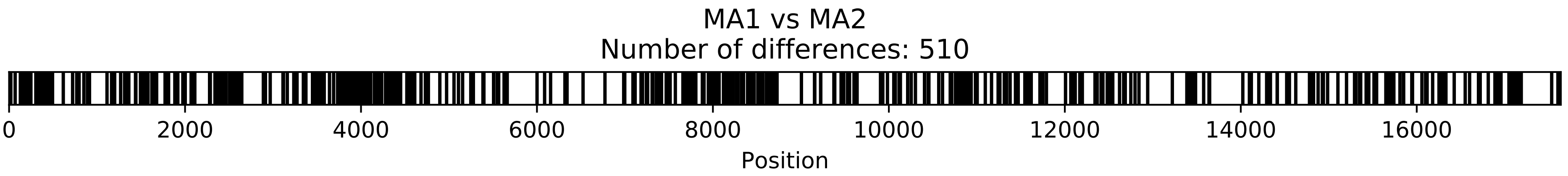
MA1 vs MA2
Number of differences: 510
0 2000 4000 6000 8000 10000 12000 14000 16000
Position

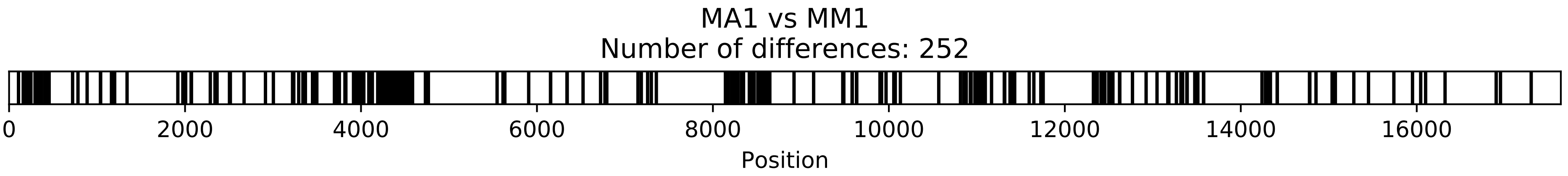
MA1 vs MM1
Number of differences: 252
0 2000 4000 6000 8000 10000 12000 14000 16000
Position

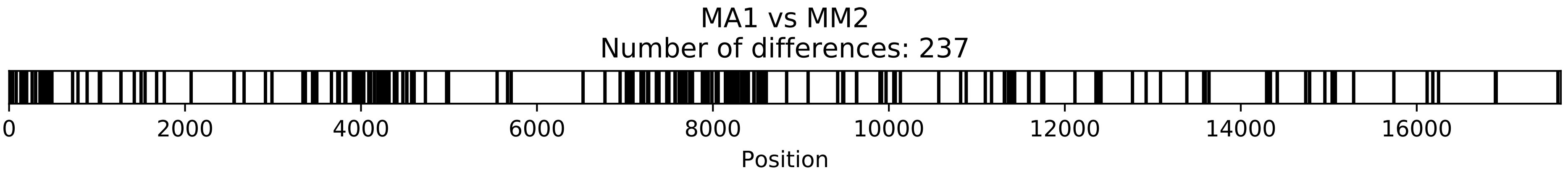
MA1 vs MM2
Number of differences: 237
0 2000 4000 6000 8000 10000 12000 14000 16000
Position

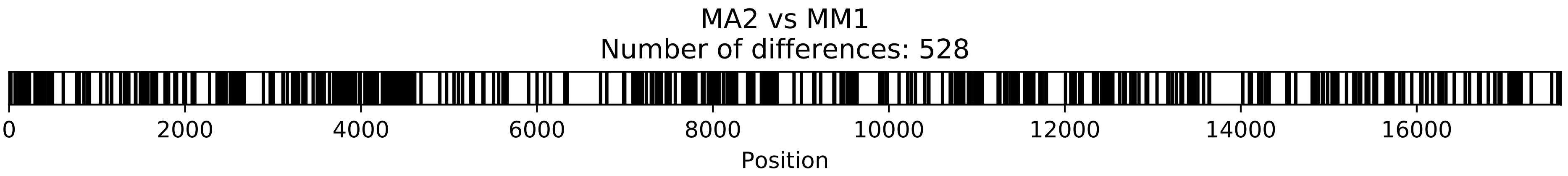
MA2 vs MM1
Number of differences: 528
0 2000 4000 6000 8000 10000 12000 14000 16000
Position

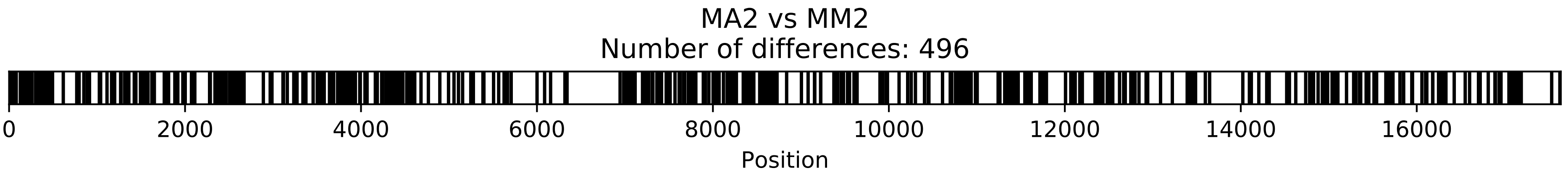
MA2 vs MM2
Number of differences: 496
0 2000 4000 6000 8000 10000 12000 14000 16000
Position

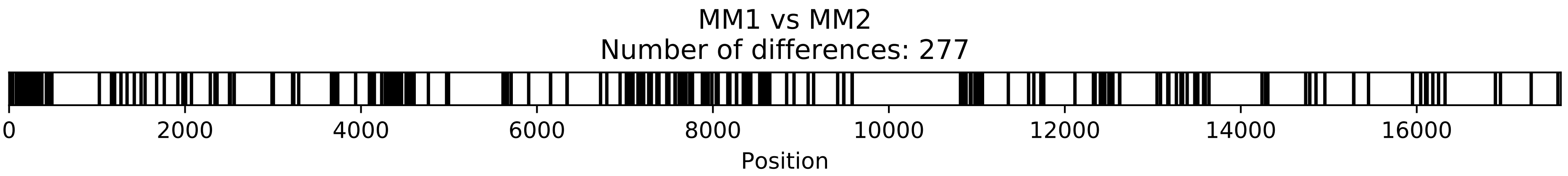
MM1 vs MM2
Number of differences: 277
0 2000 4000 6000 8000 10000 12000 14000 16000
Position

### Region: 370

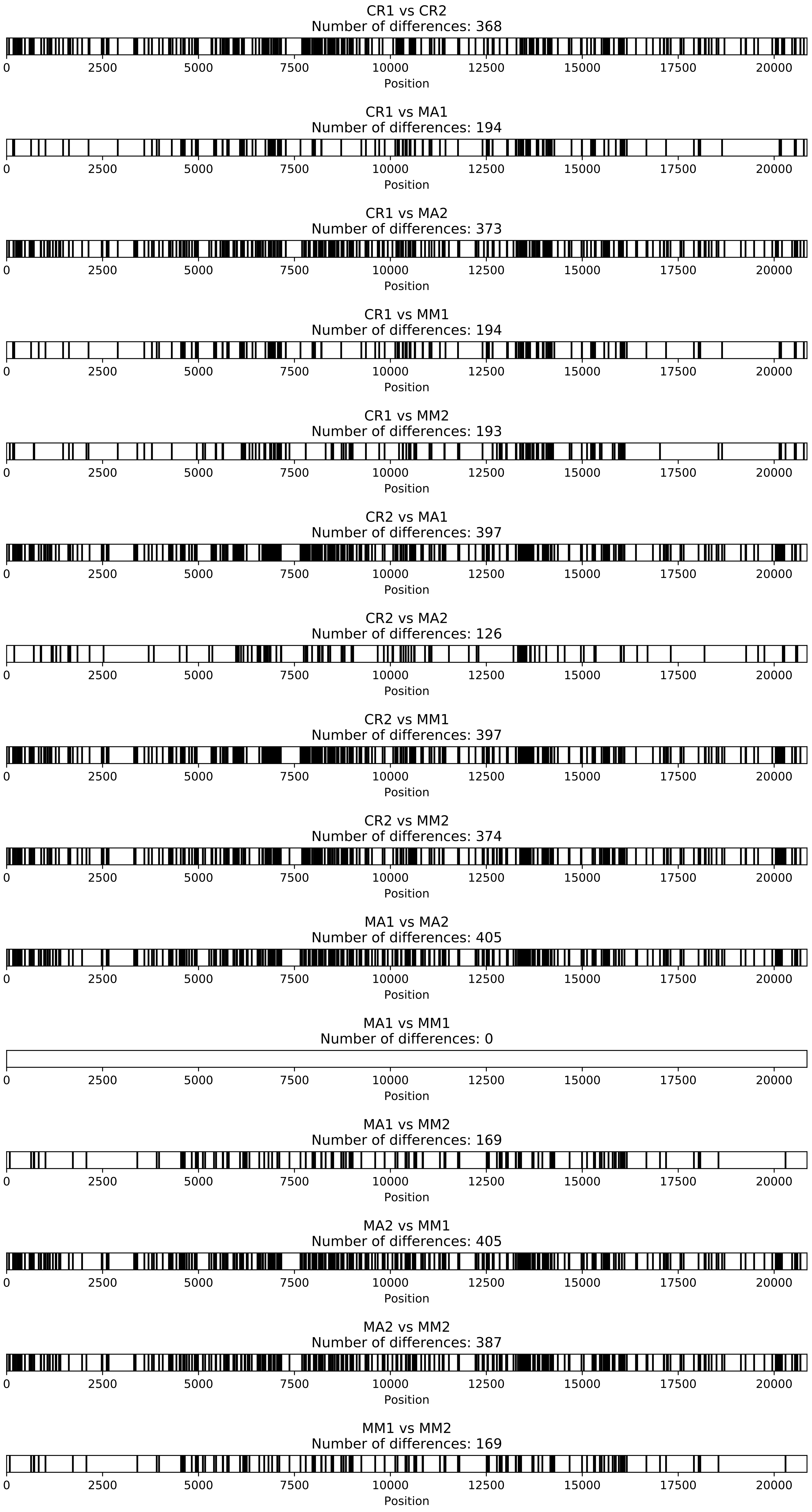
CR1 vs CR2
Number of differences: 368
CR1 vs MA1
Number of differences: 194
CR1 vs MA2
Number of differences: 373
CR1 vs MM1
Number of differences: 194
CR1 vs MM2
Number of differences: 193
CR2 vs MA1
Number of differences: 397
CR2 vs MA2
Number of differences: 126
CR2 vs MM1
Number of differences: 397
CR2 vs MM2
Number of differences: 374
MA1 vs MA2
Number of differences: 405
MA1 vs MM1
Number of differences: 0
MA1 vs MM2
Number of differences: 169
MA2 vs MM1
Number of differences: 405
MA2 vs MM2
Number of differences: 387
MM1 vs MM2
Number of differences: 169
0
2500
5000
7500
10000
12500
15000
17500
20000
Position

### Region: 371



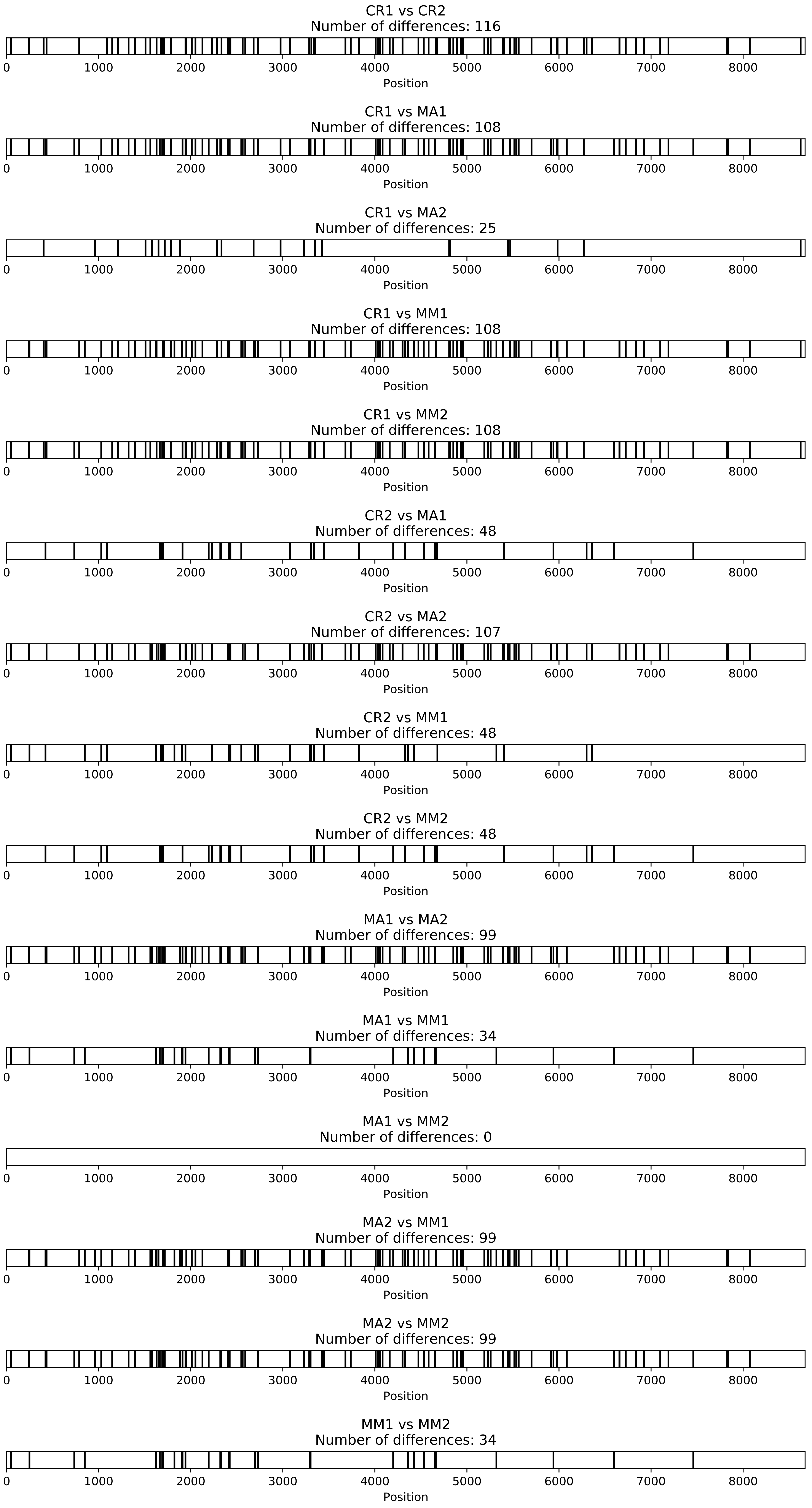
CR1 vs CR2
Number of differences: 116
CR1 vs MA1
Number of differences: 108
CR1 vs MA2
Number of differences: 25
CR1 vs MM1
Number of differences: 108
CR1 vs MM2
Number of differences: 108
CR2 vs MA1
Number of differences: 48
CR2 vs MA2
Number of differences: 107
CR2 vs MM1
Number of differences: 48
CR2 vs MM2
Number of differences: 48
MA1 vs MA2
Number of differences: 99
MA1 vs MM1
Number of differences: 34
MA1 vs MM2
Number of differences: 0
MA2 vs MM1
Number of differences: 99
MA2 vs MM2
Number of differences: 99
MM1 vs MM2
Number of differences: 34
0
1000
2000
3000
4000
5000
6000
7000
8000
Position

### Region: 372



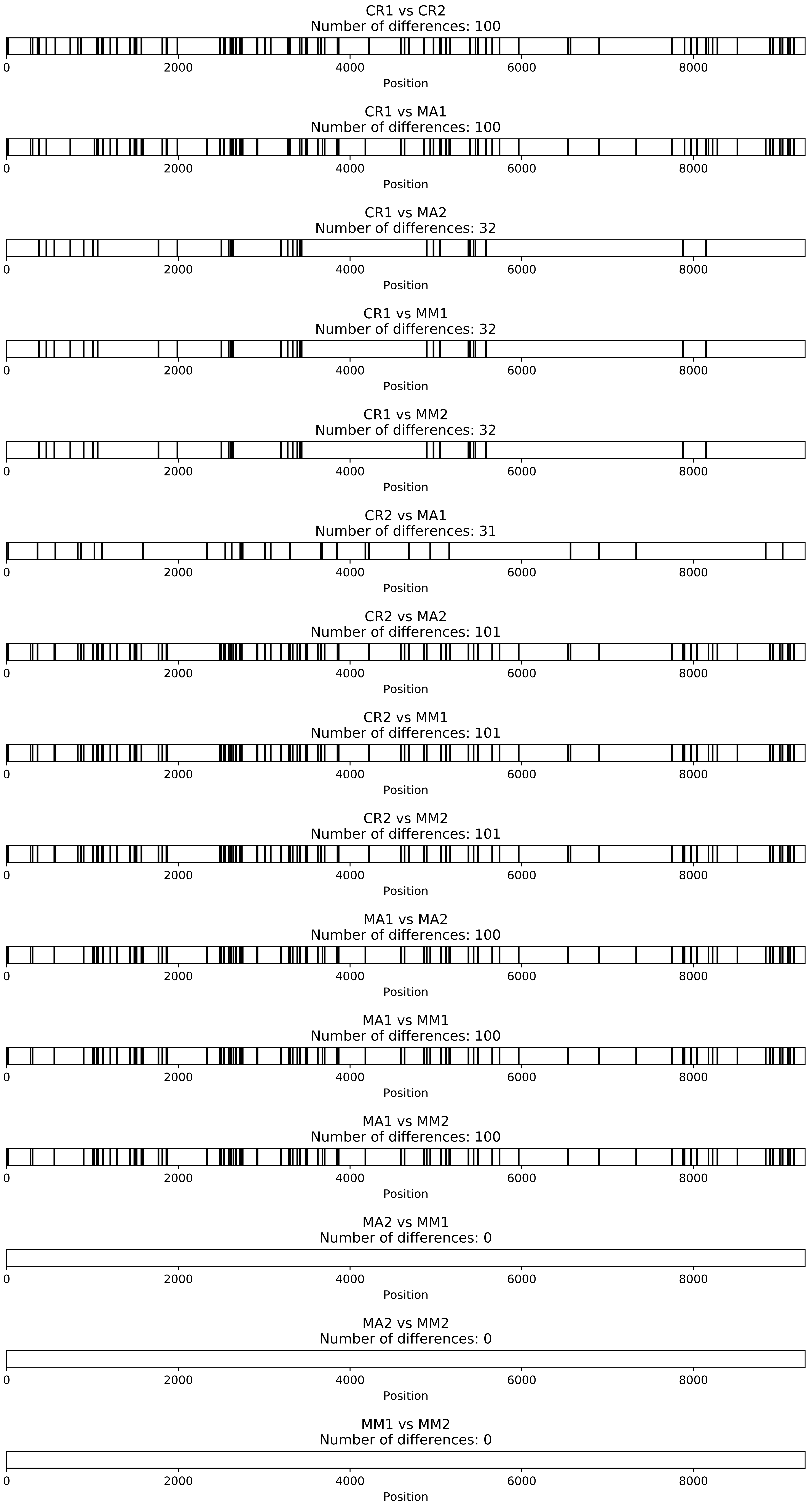
CR1 vs CR2
Number of differences: 100
0 2000 4000 6000 8000
Position
CR1 vs MA1
Number of differences: 100
0 2000 4000 6000 8000
Position
CR1 vs MA2
Number of differences: 32
0 2000 4000 6000 8000
Position
CR1 vs MM1
Number of differences: 32
0 2000 4000 6000 8000
Position
CR1 vs MM2
Number of differences: 32
0 2000 4000 6000 8000
Position
CR2 vs MA1
Number of differences: 31
0 2000 4000 6000 8000
Position
CR2 vs MA2
Number of differences: 101
0 2000 4000 6000 8000
Position
CR2 vs MM1
Number of differences: 101
0 2000 4000 6000 8000
Position
CR2 vs MM2
Number of differences: 101
0 2000 4000 6000 8000
Position
MA1 vs MA2
Number of differences: 100
0 2000 4000 6000 8000
Position
MA1 vs MM1
Number of differences: 100
0 2000 4000 6000 8000
Position
MA1 vs MM2
Number of differences: 100
0 2000 4000 6000 8000
Position
MA2 vs MM1
Number of differences: 0
0 2000 4000 6000 8000
Position
MA2 vs MM2
Number of differences: 0
0 2000 4000 6000 8000
Position
MM1 vs MM2
Number of differences: 0
0 2000 4000 6000 8000
Position

### Region: 373



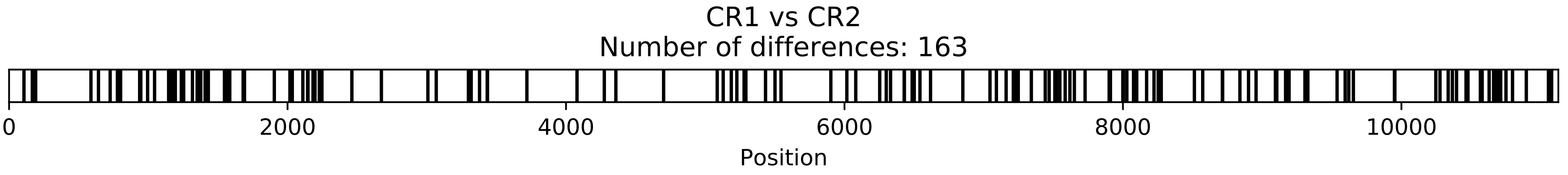
CR1 vs CR2
Number of differences: 163
0
2000
4000
6000
8000
10000
Position

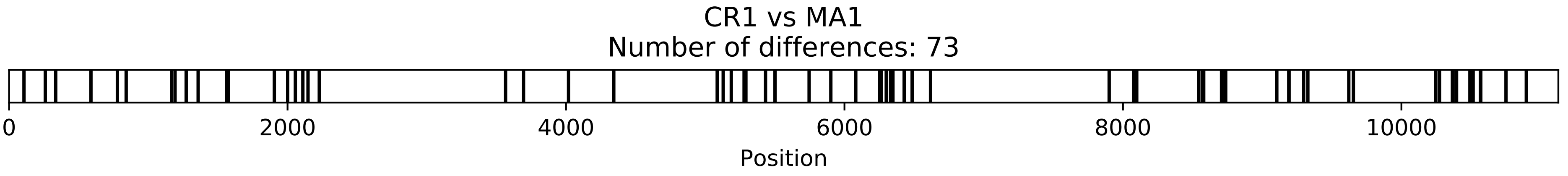
CR1 vs MA1
Number of differences: 73
0
2000
4000
6000
8000
10000
Position

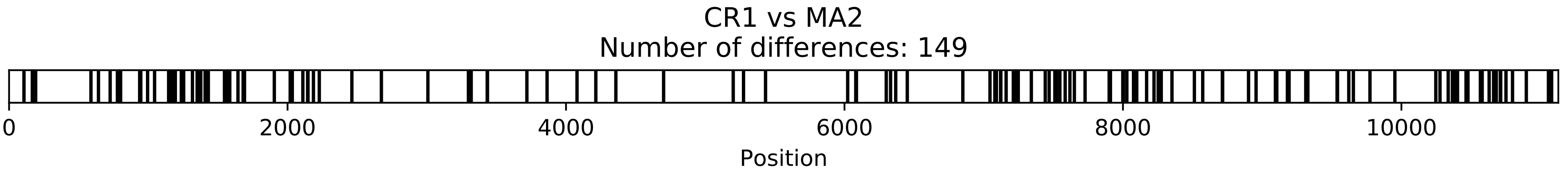
CR1 vs MA2
Number of differences: 149
0
2000
4000
6000
8000
10000
Position

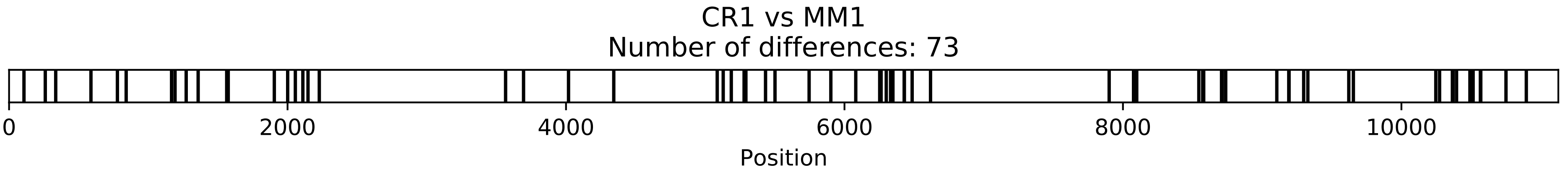
CR1 vs MM1
Number of differences: 73
0
2000
4000
6000
8000
10000
Position

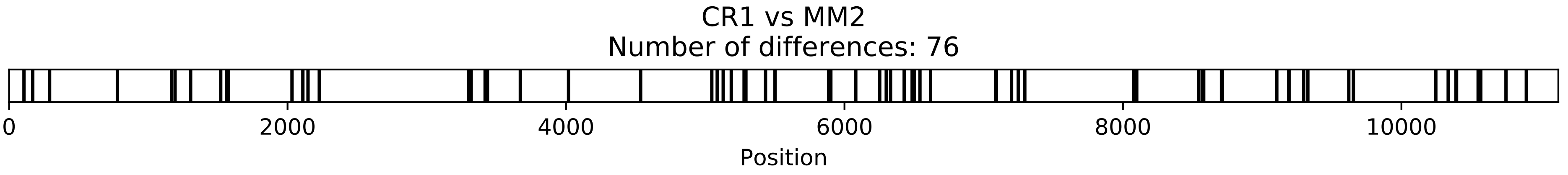
CR1 vs MM2
Number of differences: 76
0
2000
4000
6000
8000
10000
Position

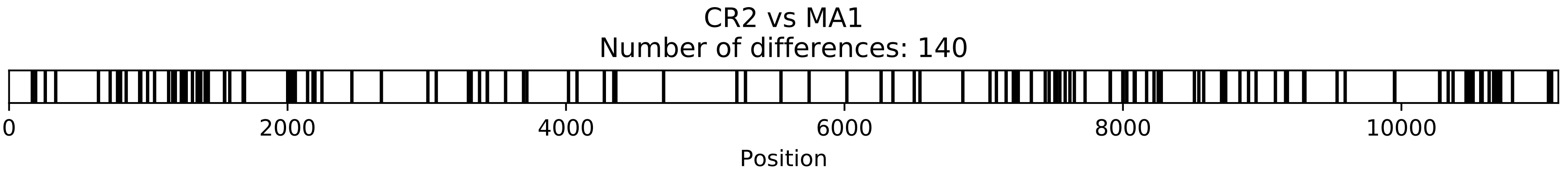
CR2 vs MA1
Number of differences: 140
0
2000
4000
6000
8000
10000
Position

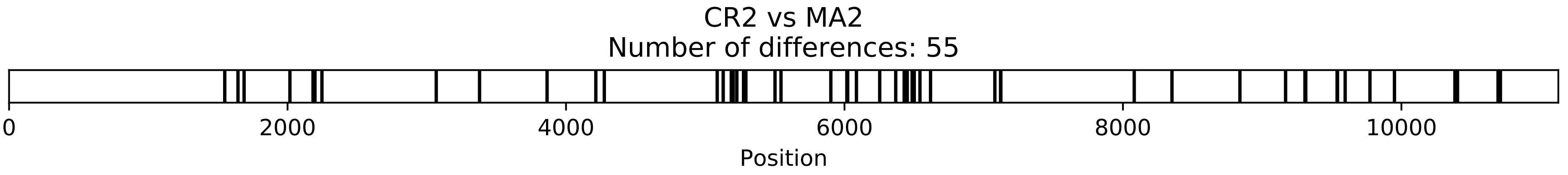
CR2 vs MA2
Number of differences: 55
0
2000
4000
6000
8000
10000
Position

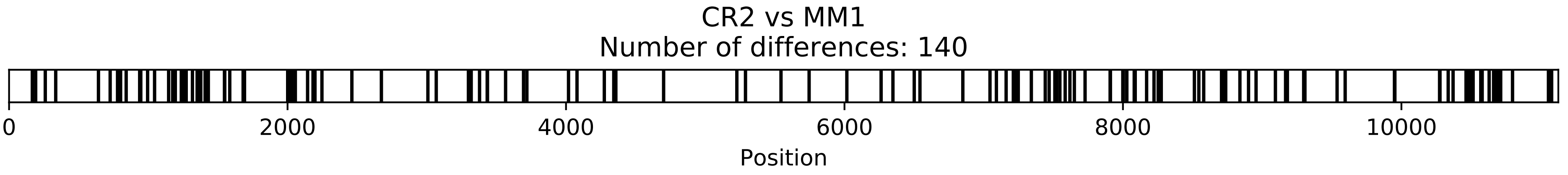
CR2 vs MM1
Number of differences: 140
0
2000
4000
6000
8000
10000
Position

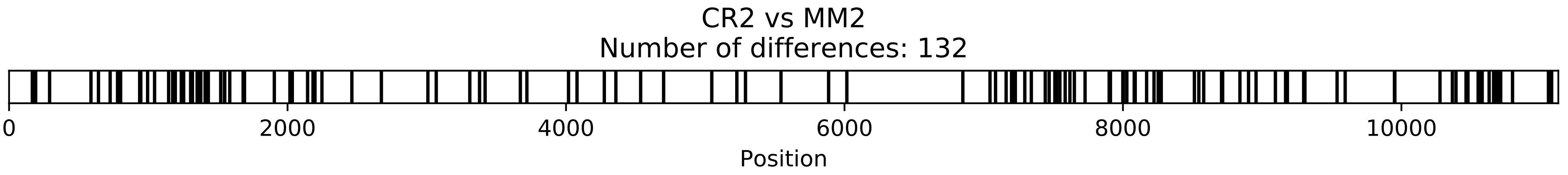
CR2 vs MM2
Number of differences: 132
0
2000
4000
6000
8000
10000
Position

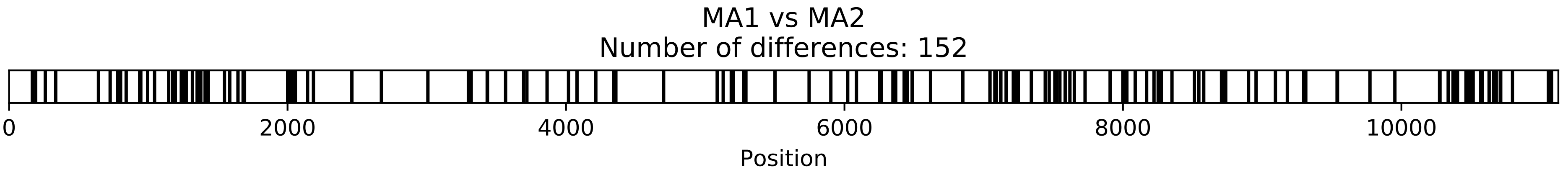
MA1 vs MA2
Number of differences: 152
0
2000
4000
6000
8000
10000
Position

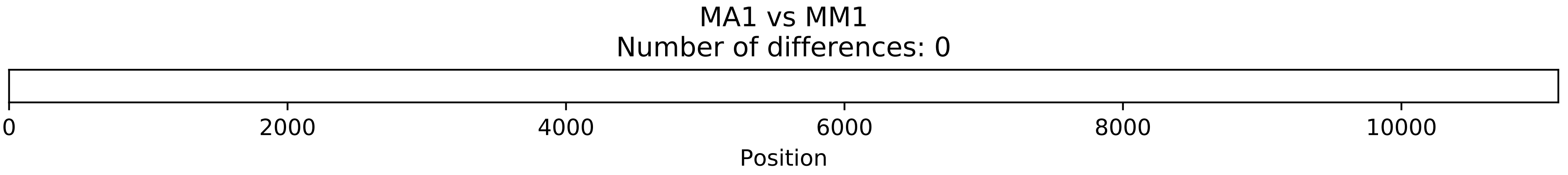
MA1 vs MM1
Number of differences: 0
0
2000
4000
6000
8000
10000
Position

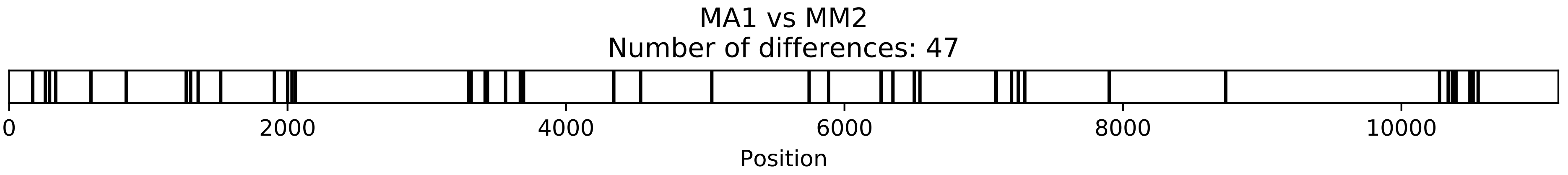
MA1 vs MM2
Number of differences: 47
0
2000
4000
6000
8000
10000
Position

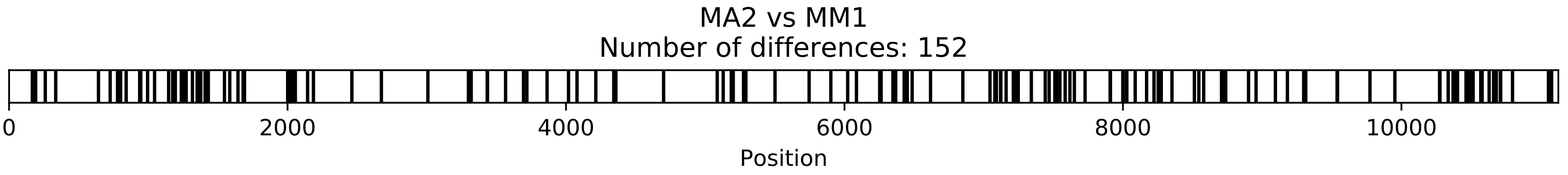
MA2 vs MM1
Number of differences: 152
0
2000
4000
6000
8000
10000
Position

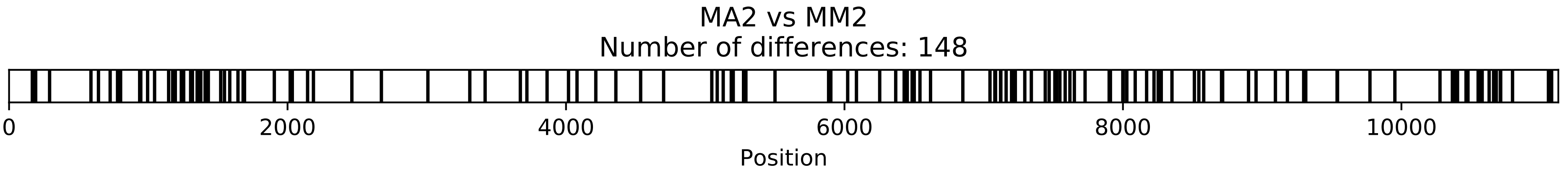
MA2 vs MM2
Number of differences: 148
0
2000
4000
6000
8000
10000
Position

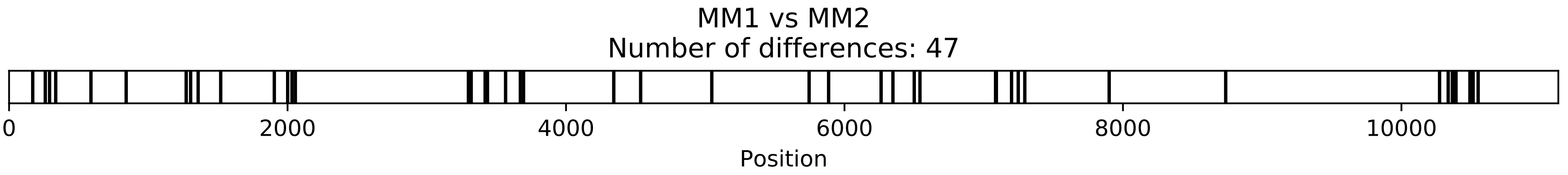
MM1 vs MM2
Number of differences: 47
0
2000
4000
6000
8000
10000
Position

### Region: 374

CR1 vs CR2
Number of differences: 185

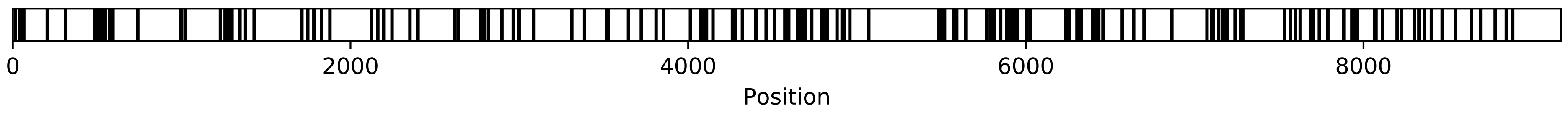


CR1 vs MA1
Number of differences: 9

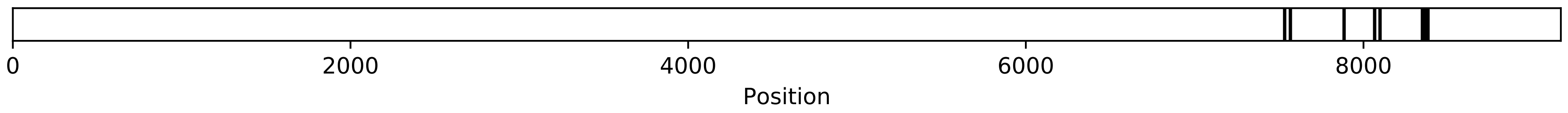


CR1 vs MA2
Number of differences: 180

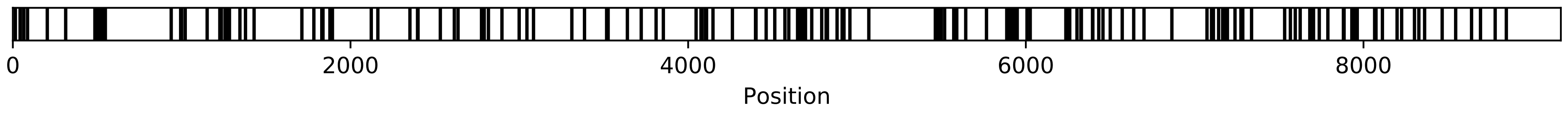


CR1 vs MM1
Number of differences: 9

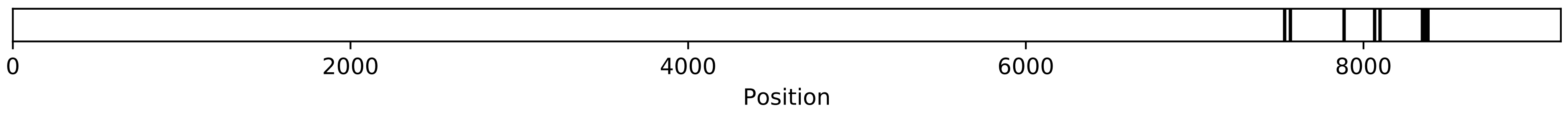


CR1 vs MM2
Number of differences: 110

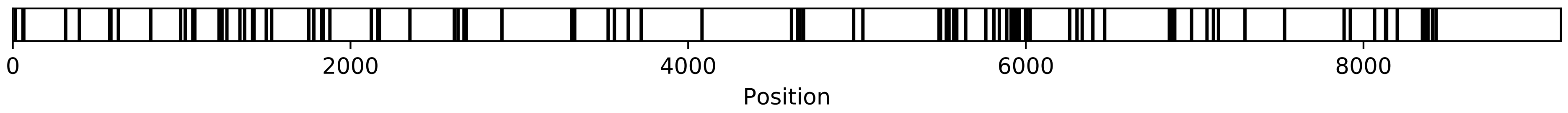


CR2 vs MA1
Number of differences: 184

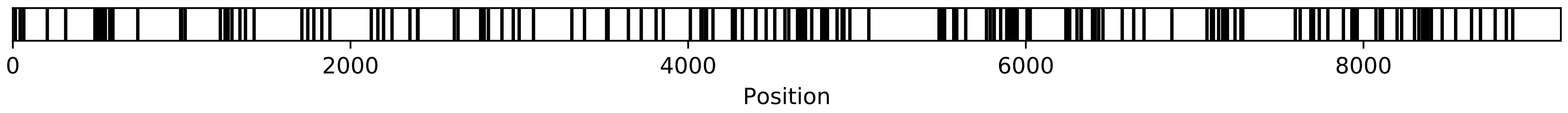


CR2 vs MA2
Number of differences: 55

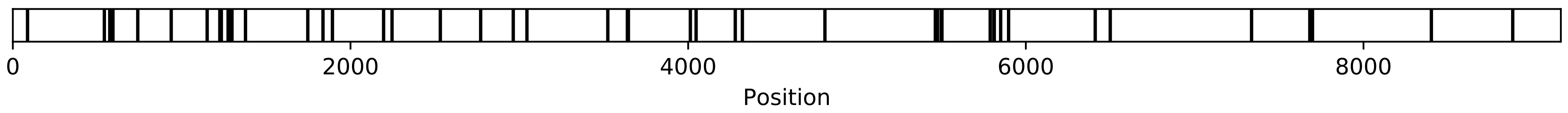


CR2 vs MM1
Number of differences: 184

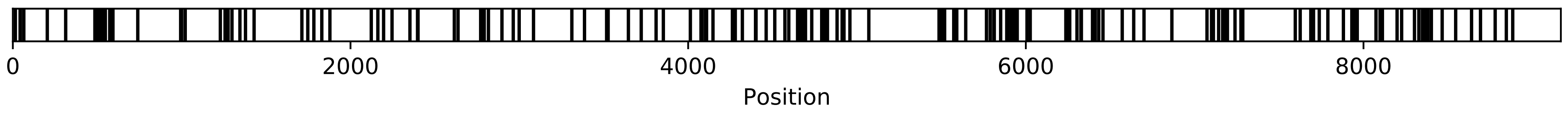


CR2 vs MM2
Number of differences: 189

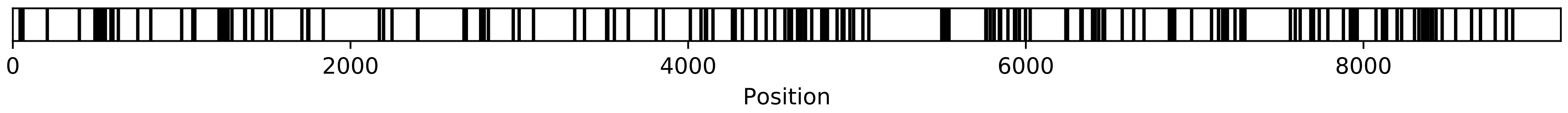


MA1 vs MA2
Number of differences: 179

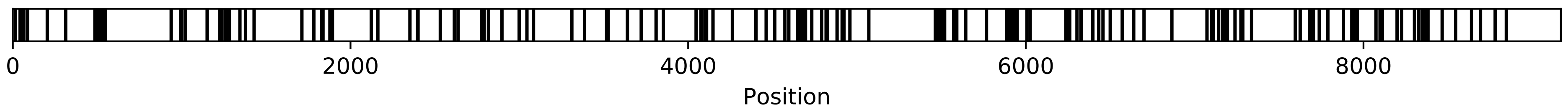


MA1 vs MM1
Number of differences: 0

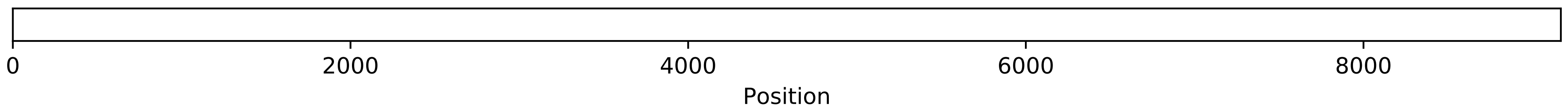


MA1 vs MM2
Number of differences: 105

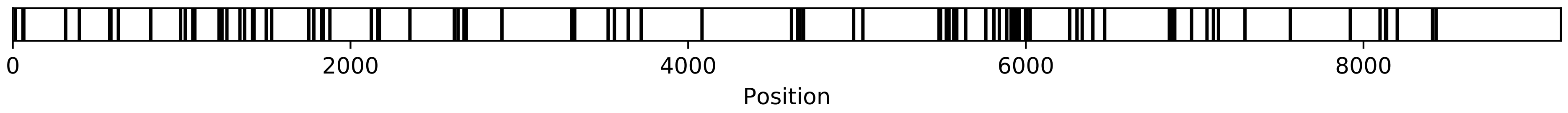


MA2 vs MM1
Number of differences: 179

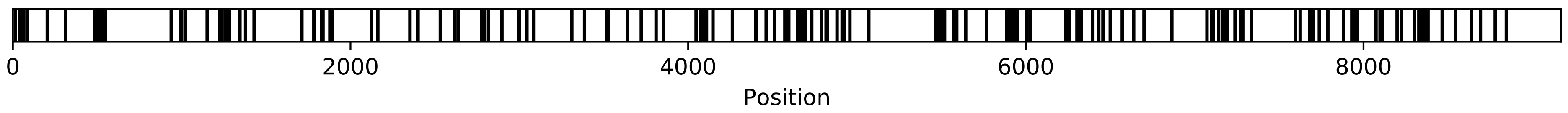


MA2 vs MM2
Number of differences: 188

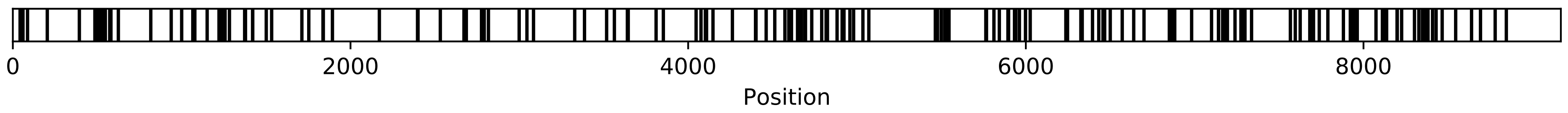


MM1 vs MM2
Number of differences: 105

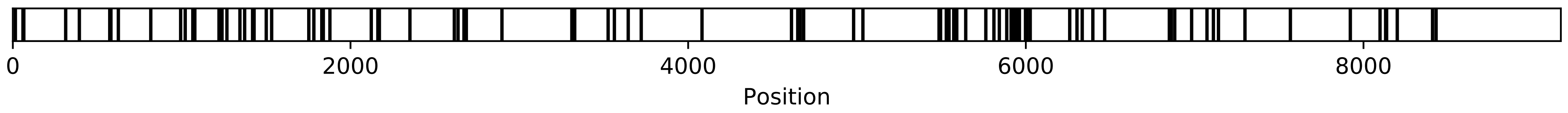

### Region: 376

CR1 vs CR2
Number of differences: 42

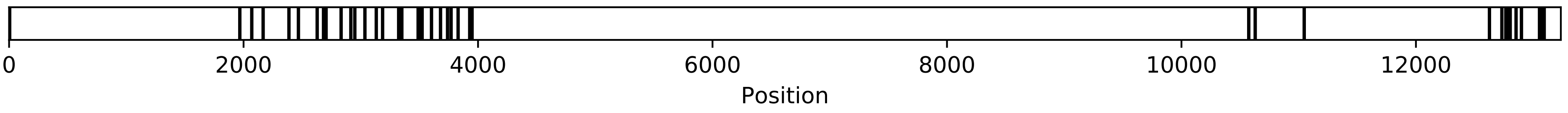

CR1 vs MA1
Number of differences: 263

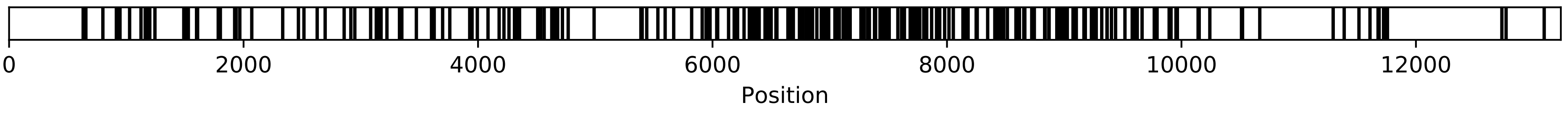

CR1 vs MA2
Number of differences: 154

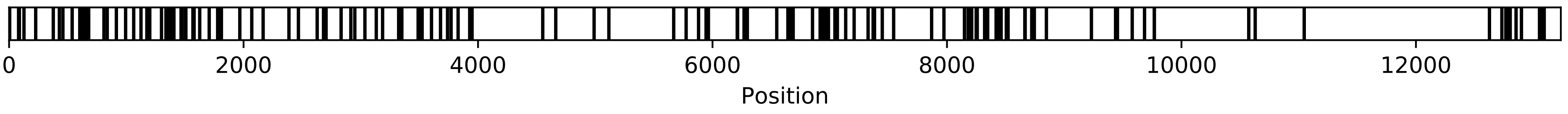

CR1 vs MM1
Number of differences: 263

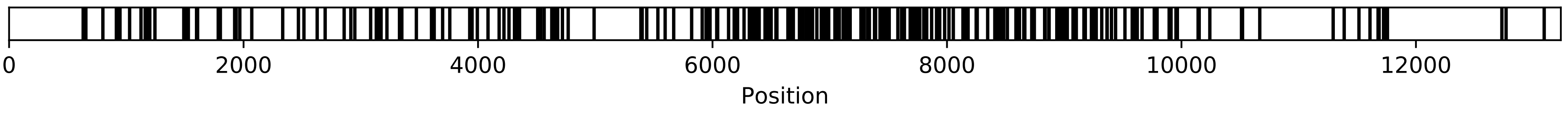

CR1 vs MM2
Number of differences: 251

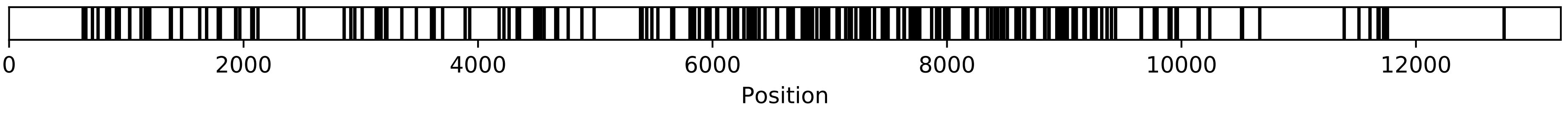

CR2 vs MA1
Number of differences: 277

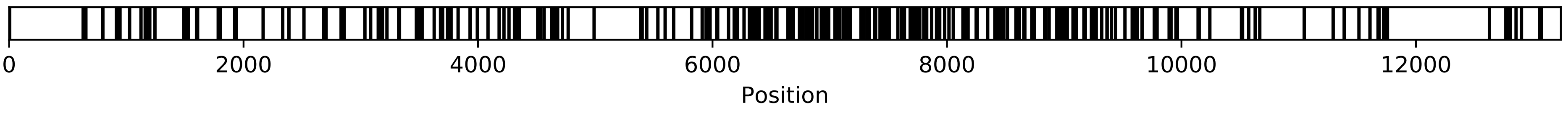

CR2 vs MA2
Number of differences: 112

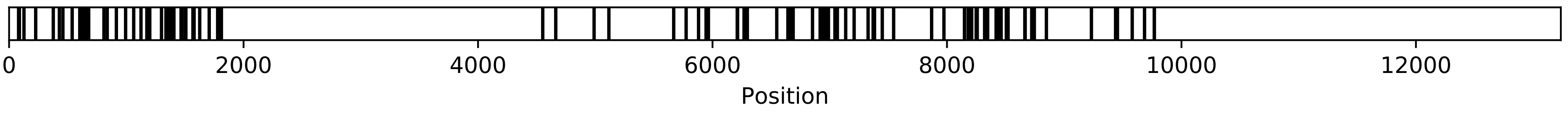

CR2 vs MM1
Number of differences: 277

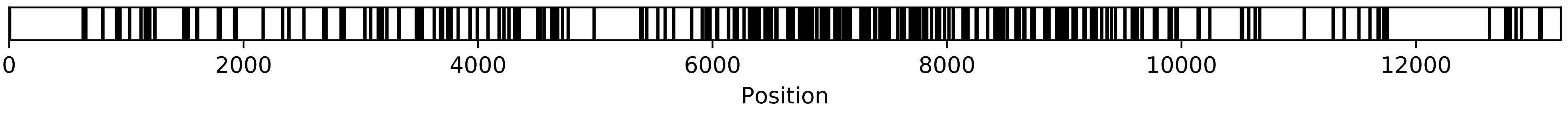

CR2 vs MM2
Number of differences: 275

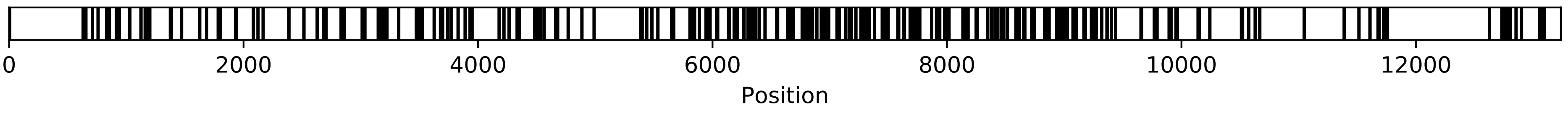

MA1 vs MA2
Number of differences: 296

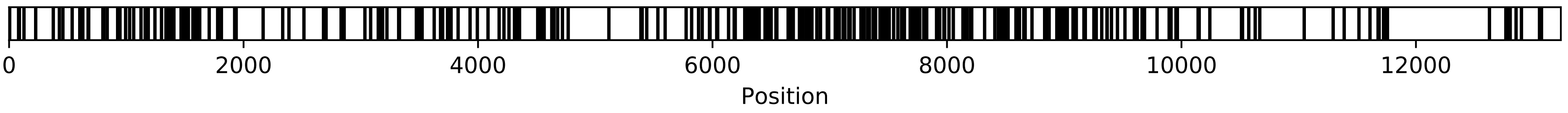

MA1 vs MM1
Number of differences: 0

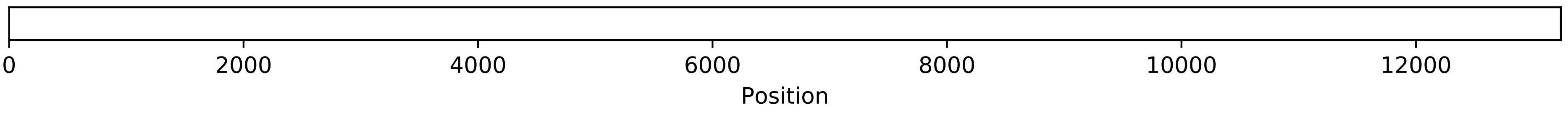

MA1 vs MM2
Number of differences: 120

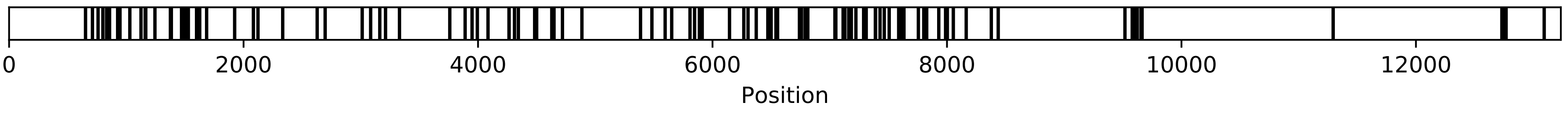

MA2 vs MM1
Number of differences: 296

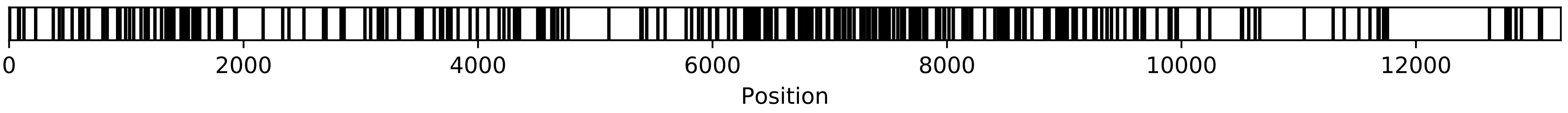

MA2 vs MM2
Number of differences: 293

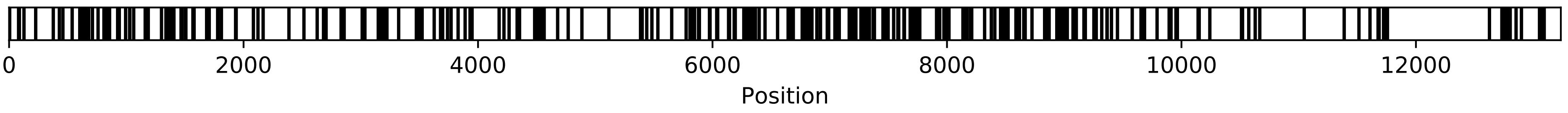

MM1 vs MM2
Number of differences: 120

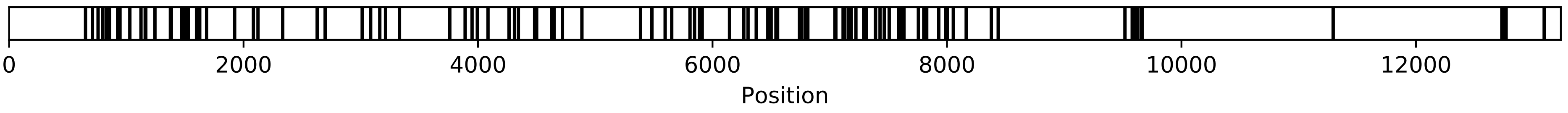

### Region: 378

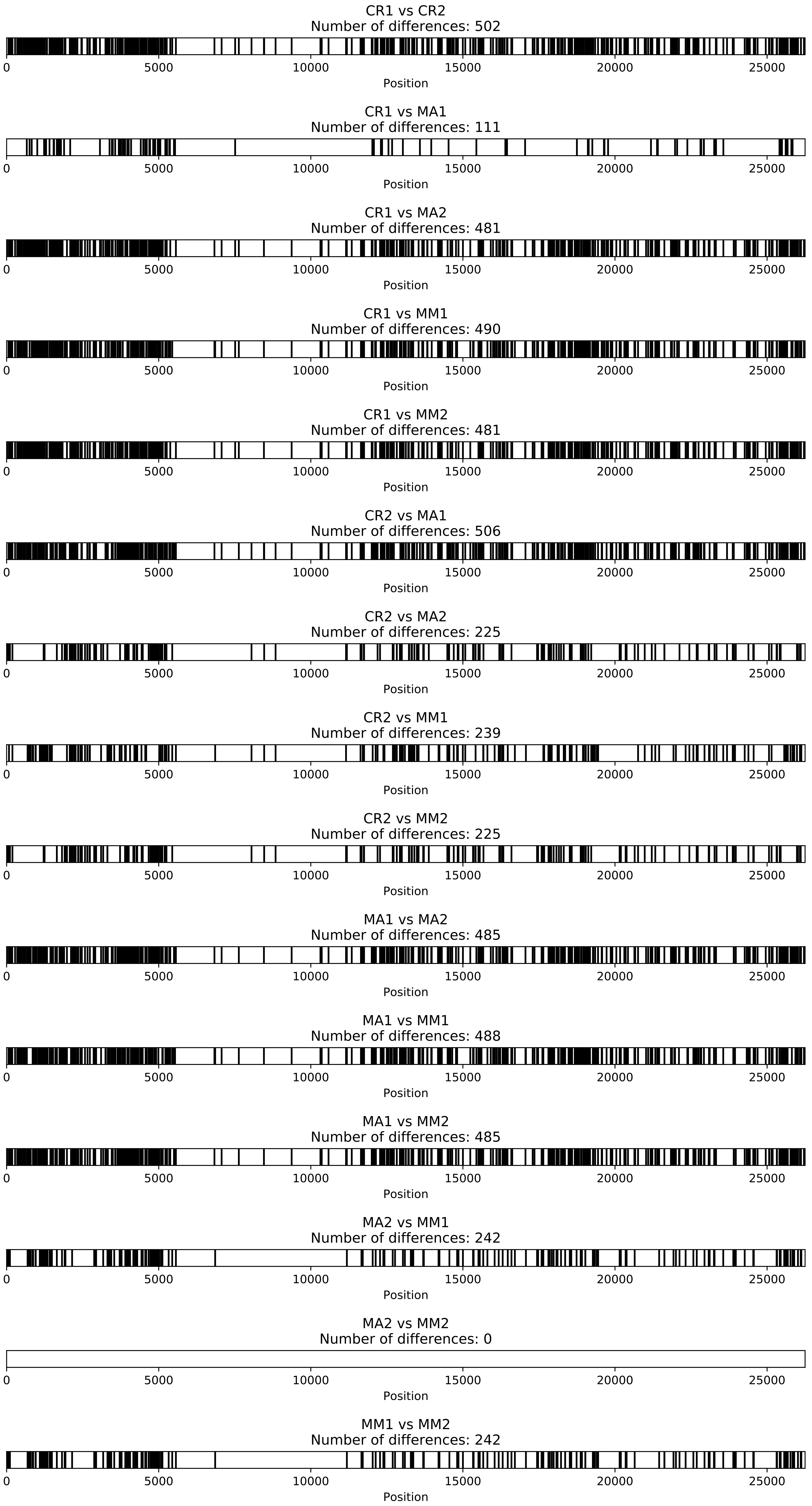
CR1 vs CR2
Number of differences: 502
CR1 vs MA1
Number of differences: 111
CR1 vs MA2
Number of differences: 481
CR1 vs MM1
Number of differences: 490
CR1 vs MM2
Number of differences: 481
CR2 vs MA1
Number of differences: 506
CR2 vs MA2
Number of differences: 225
CR2 vs MM1
Number of differences: 239
CR2 vs MM2
Number of differences: 225
MA1 vs MA2
Number of differences: 485
MA1 vs MM1
Number of differences: 488
MA1 vs MM2
Number of differences: 485
MA2 vs MM1
Number of differences: 242
MA2 vs MM2
Number of differences: 0
MM1 vs MM2
Number of differences: 242
0
5000
10000
15000
20000
25000
Position

### Region: 379



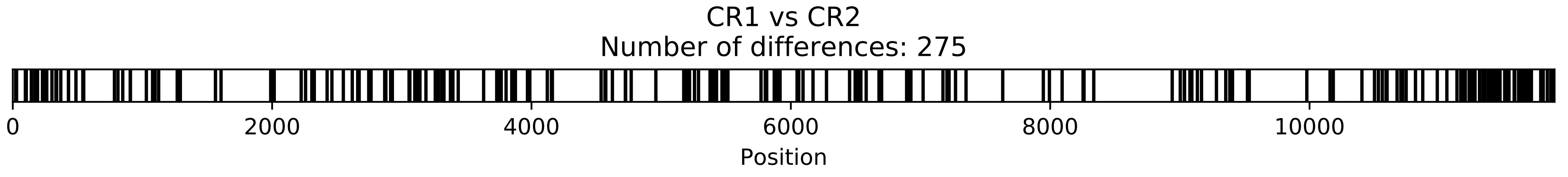
CR1 vs CR2
Number of differences: 275
0 2000 4000 6000 8000 10000
Position

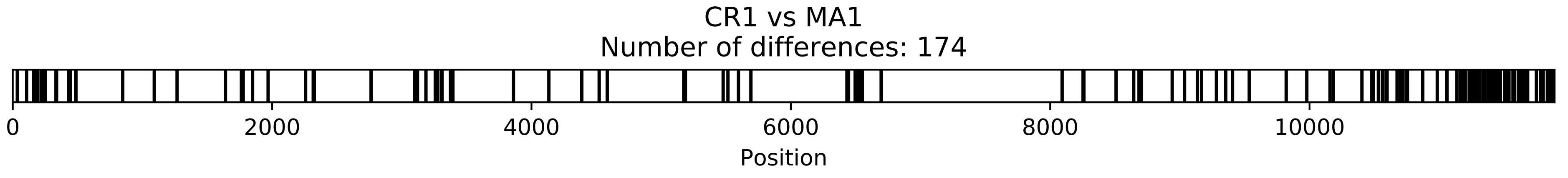
CR1 vs MA1
Number of differences: 174
0 2000 4000 6000 8000 10000
Position

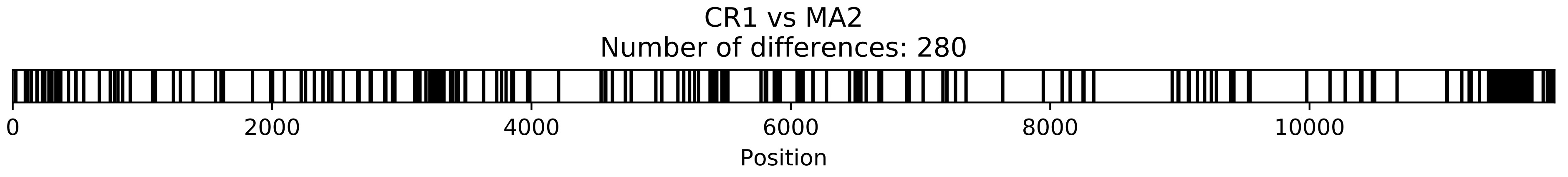
CR1 vs MA2
Number of differences: 280
0 2000 4000 6000 8000 10000
Position

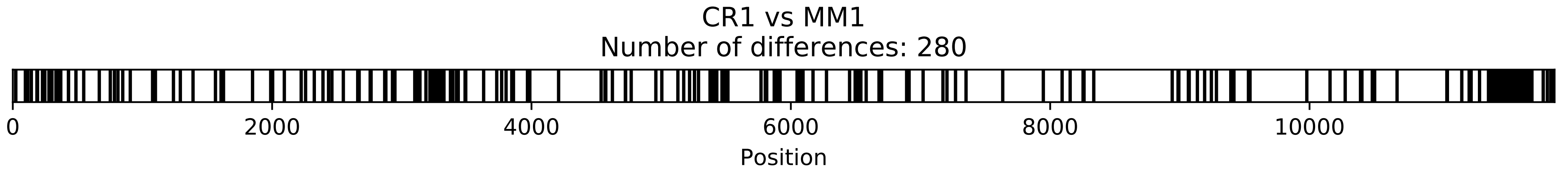
CR1 vs MM1
Number of differences: 280
0 2000 4000 6000 8000 10000
Position

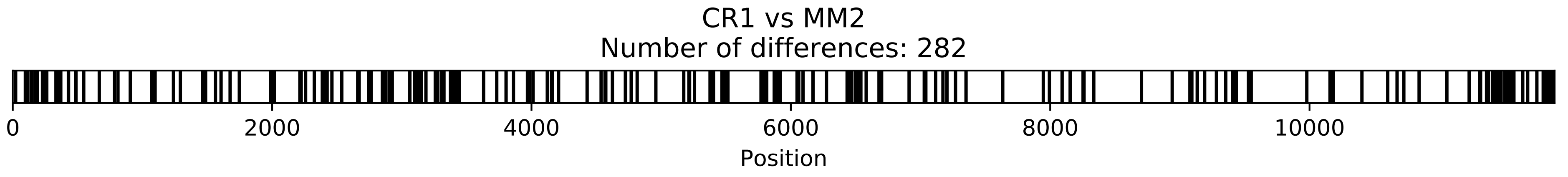
CR1 vs MM2
Number of differences: 282
0 2000 4000 6000 8000 10000
Position

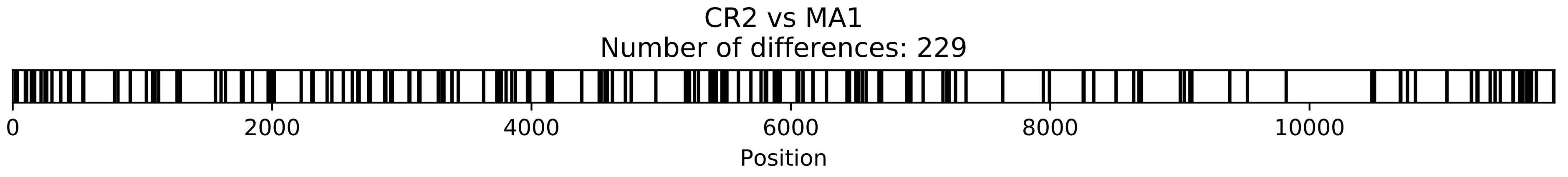
CR2 vs MA1
Number of differences: 229
0 2000 4000 6000 8000 10000
Position

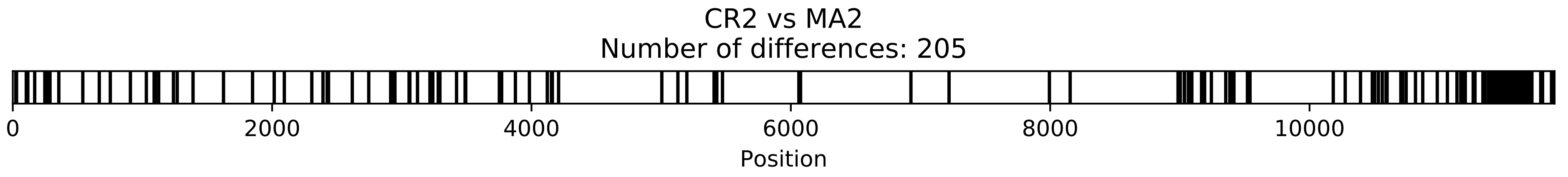
CR2 vs MA2
Number of differences: 205
0 2000 4000 6000 8000 10000
Position

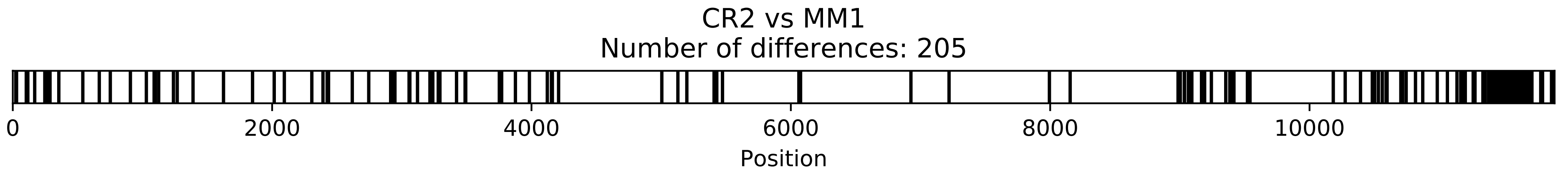
CR2 vs MM1
Number of differences: 205
0 2000 4000 6000 8000 10000
Position

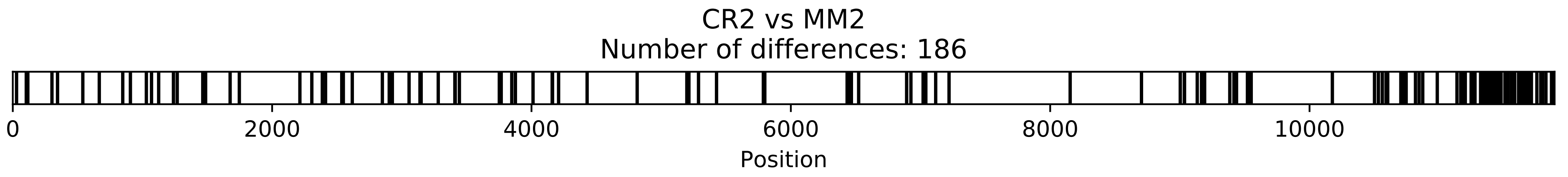
CR2 vs MM2
Number of differences: 186
0 2000 4000 6000 8000 10000
Position

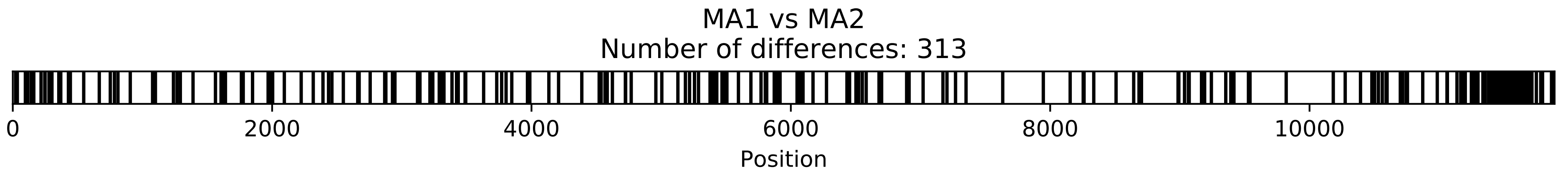
MA1 vs MA2
Number of differences: 313
0 2000 4000 6000 8000 10000
Position

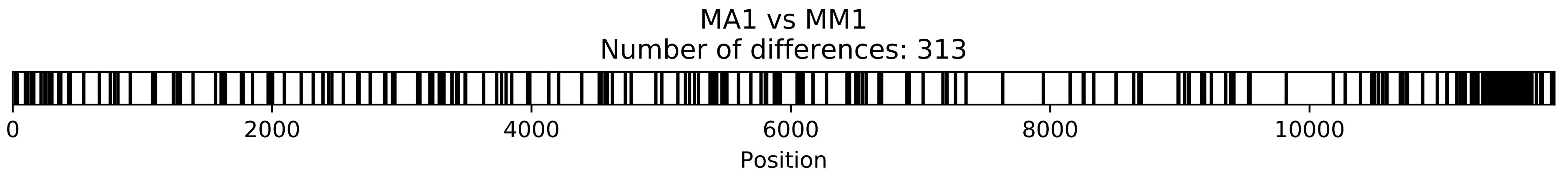
MA1 vs MM1
Number of differences: 313
0 2000 4000 6000 8000 10000
Position

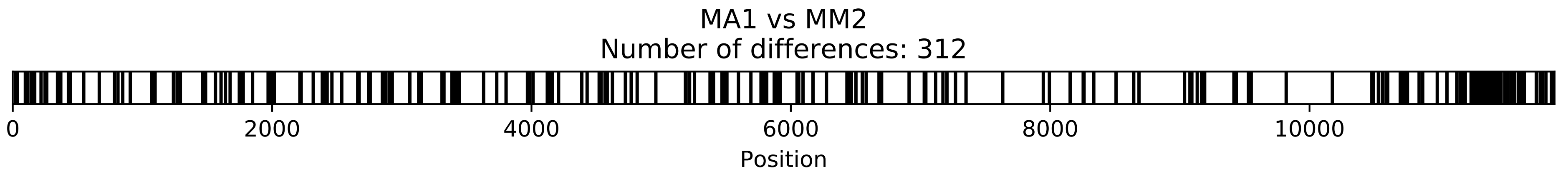
MA1 vs MM2
Number of differences: 312
0 2000 4000 6000 8000 10000
Position

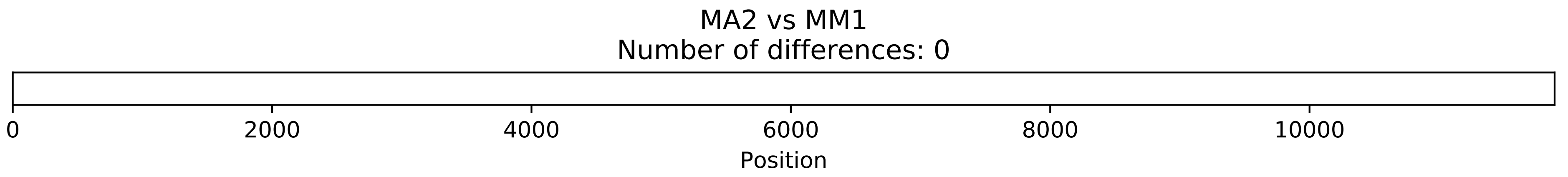
MA2 vs MM1
Number of differences: 0
0 2000 4000 6000 8000 10000
Position

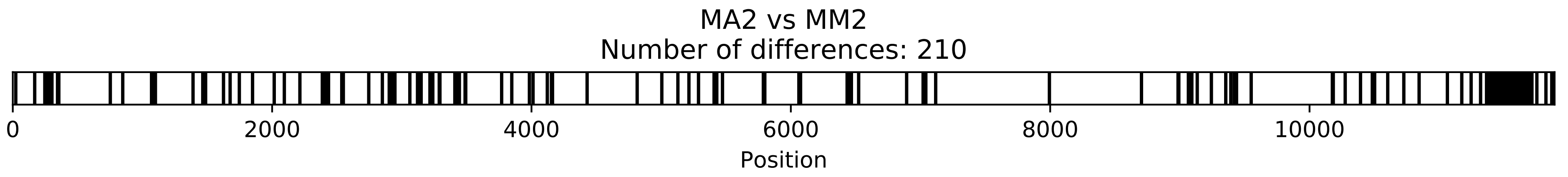
MA2 vs MM2
Number of differences: 210
0 2000 4000 6000 8000 10000
Position

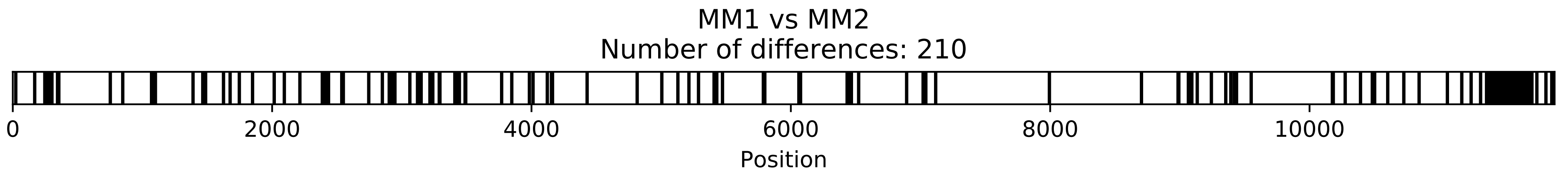
MM1 vs MM2
Number of differences: 210
0 2000 4000 6000 8000 10000
Position

### Region: 380



CR1 vs CR2
Number of differences: 230

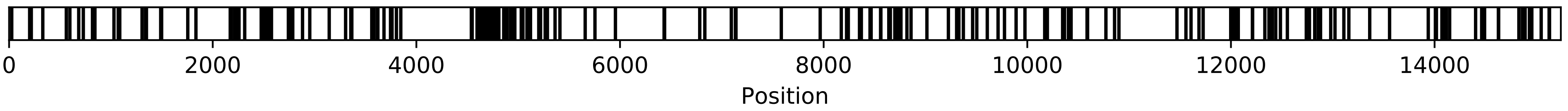


CR1 vs MA1
Number of differences: 115

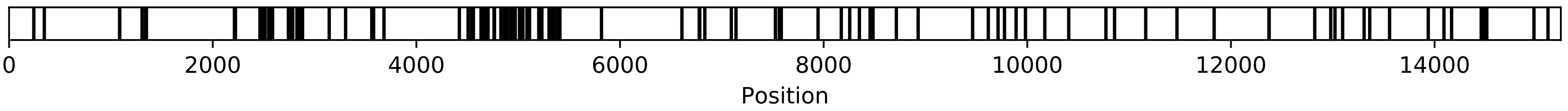


CR1 vs MA2
Number of differences: 227

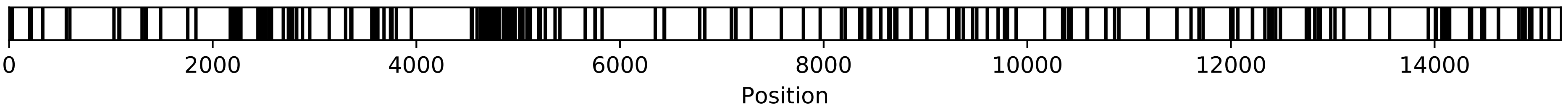


CR1 vs MM1
Number of differences: 79

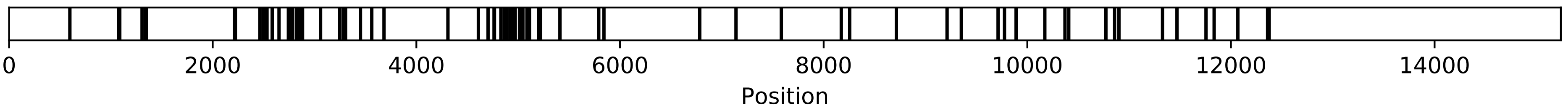


CR1 vs MM2
Number of differences: 115

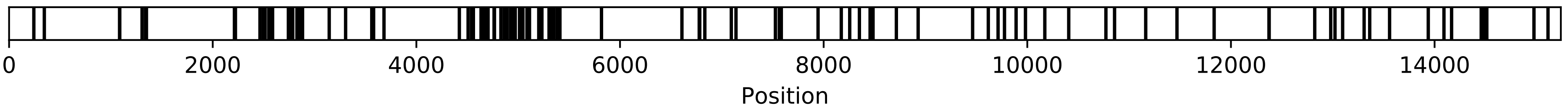


CR2 vs MA1
Number of differences: 226

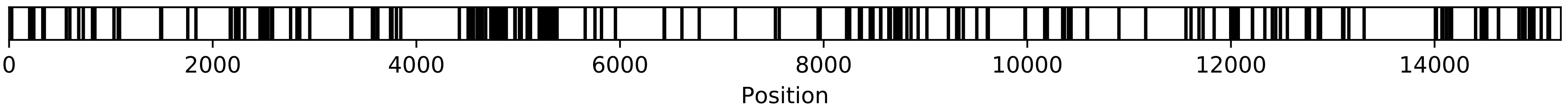


CR2 vs MA2
Number of differences: 67

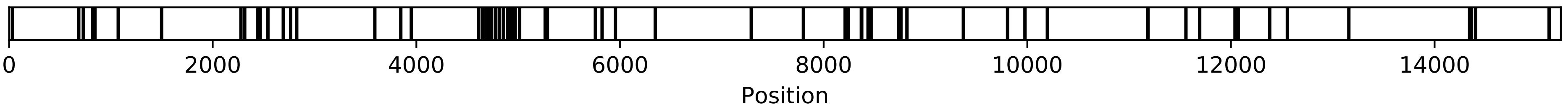


CR2 vs MM1
Number of differences: 219

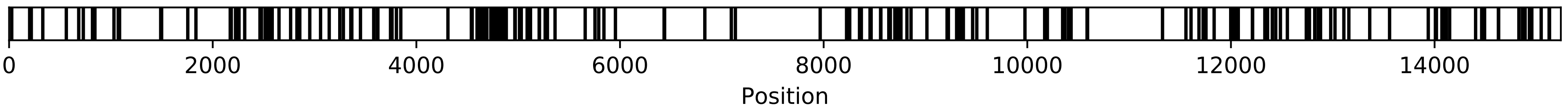


CR2 vs MM2
Number of differences: 226

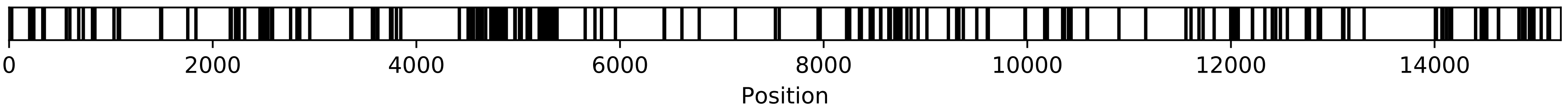


MA1 vs MA2
Number of differences: 221

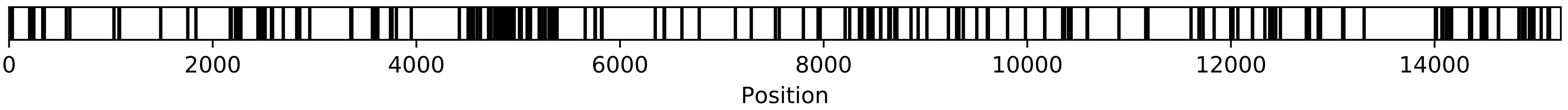


MA1 vs MM1
Number of differences: 78

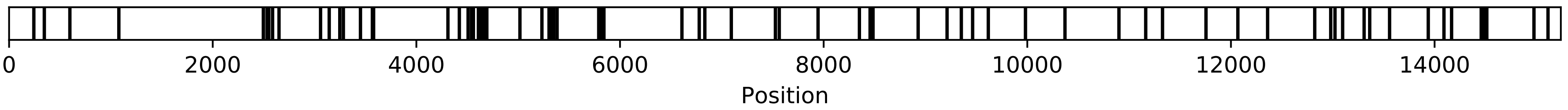


MA1 vs MM2
Number of differences: 0

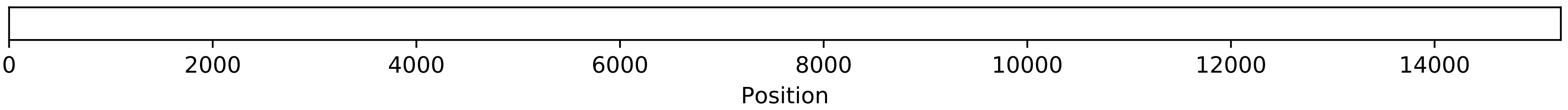


MA2 vs MM1
Number of differences: 214

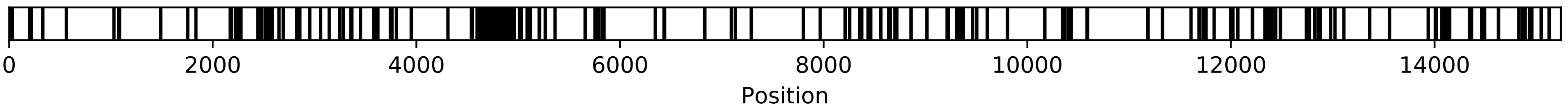


MA2 vs MM2
Number of differences: 221

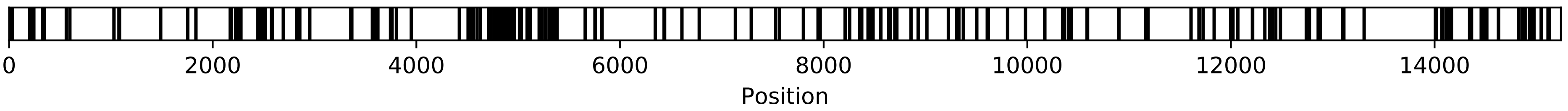


MM1 vs MM2
Number of differences: 78

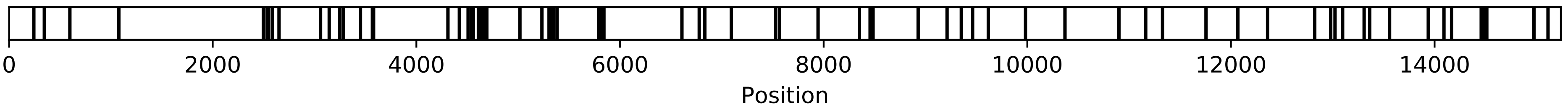

### Region: 382



CR1 vs CR2
Number of differences: 233

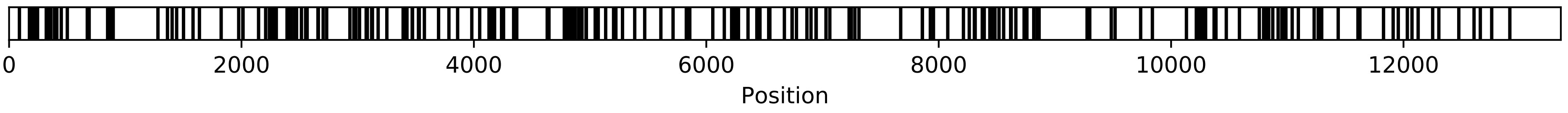


CR1 vs MA1
Number of differences: 109

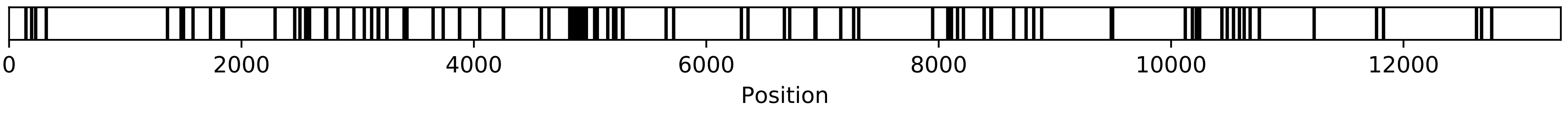


CR1 vs MA2
Number of differences: 222

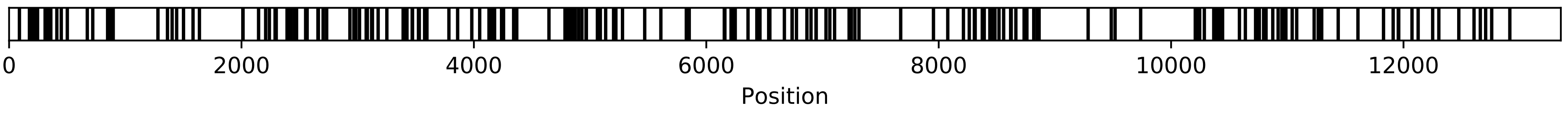


CR1 vs MM1
Number of differences: 0

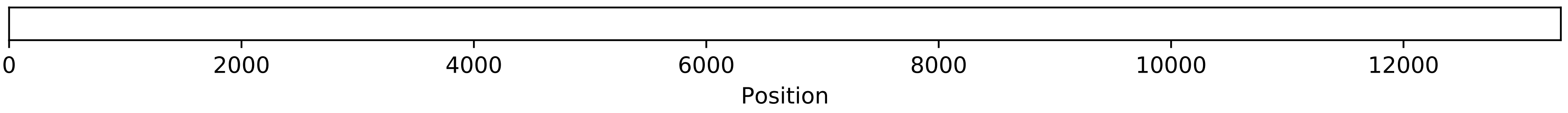


CR1 vs MM2
Number of differences: 119

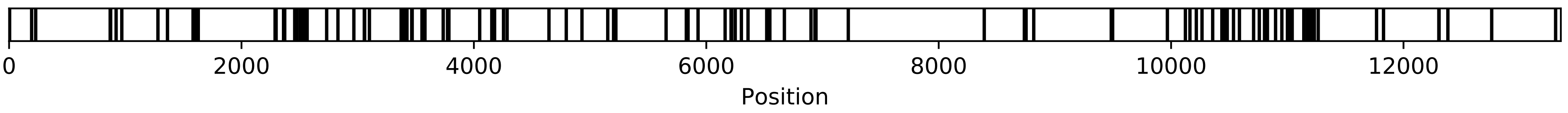


CR2 vs MA1
Number of differences: 248

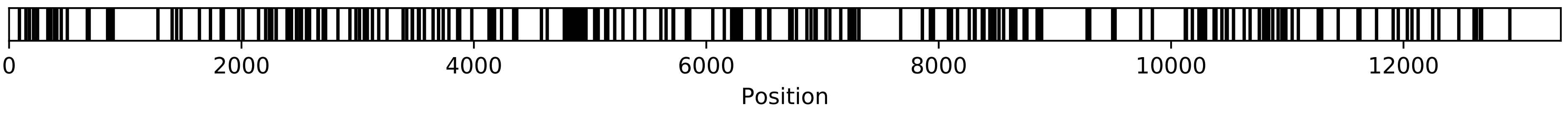


CR2 vs MA2
Number of differences: 65

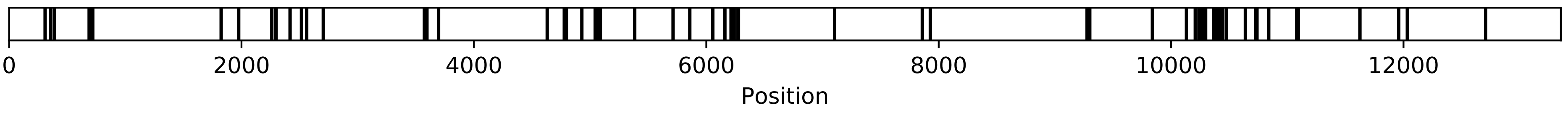


CR2 vs MM1
Number of differences: 233

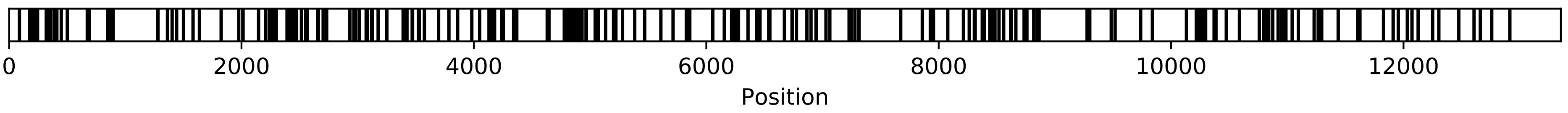


CR2 vs MM2
Number of differences: 251

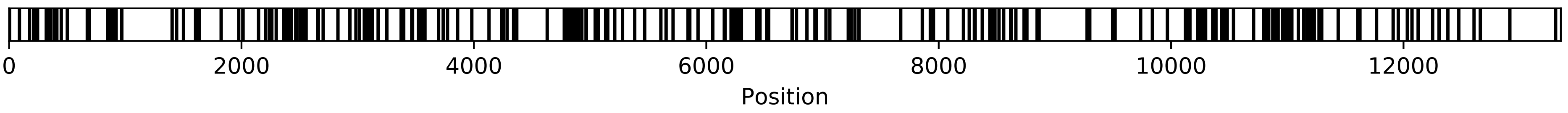


MA1 vs MA2
Number of differences: 236

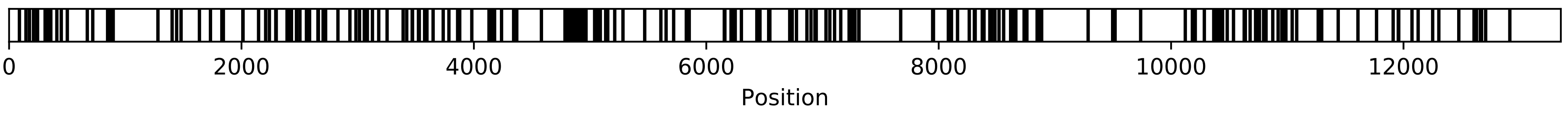


MA1 vs MM1
Number of differences: 109

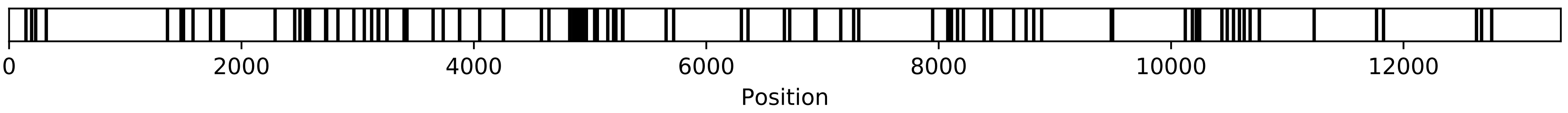


MA1 vs MM2
Number of differences: 143

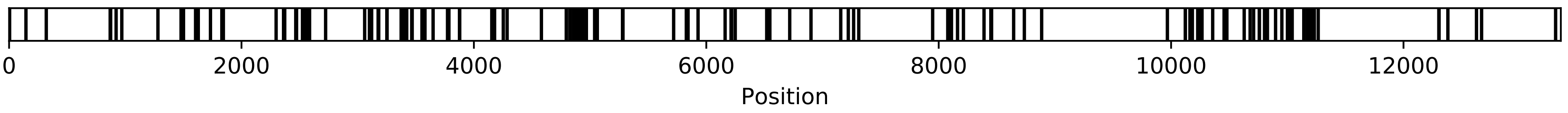


MA2 vs MM1
Number of differences: 222

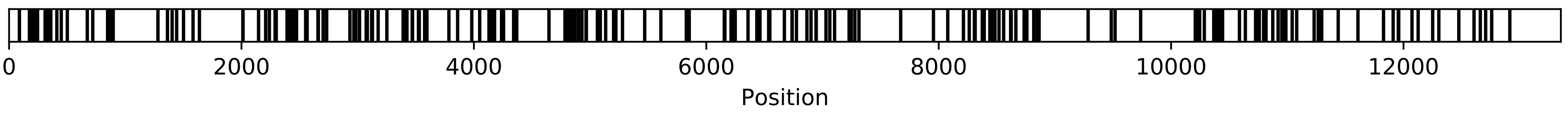


MA2 vs MM2
Number of differences: 239

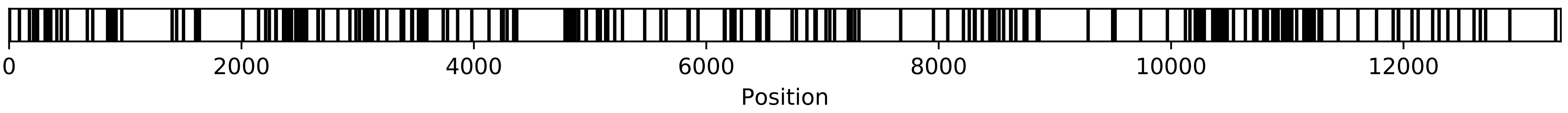


MM1 vs MM2
Number of differences: 119

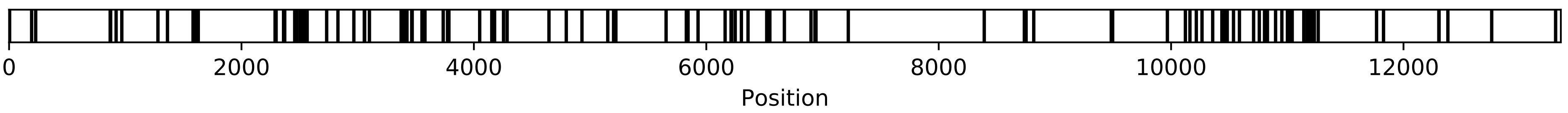

### Region: 384

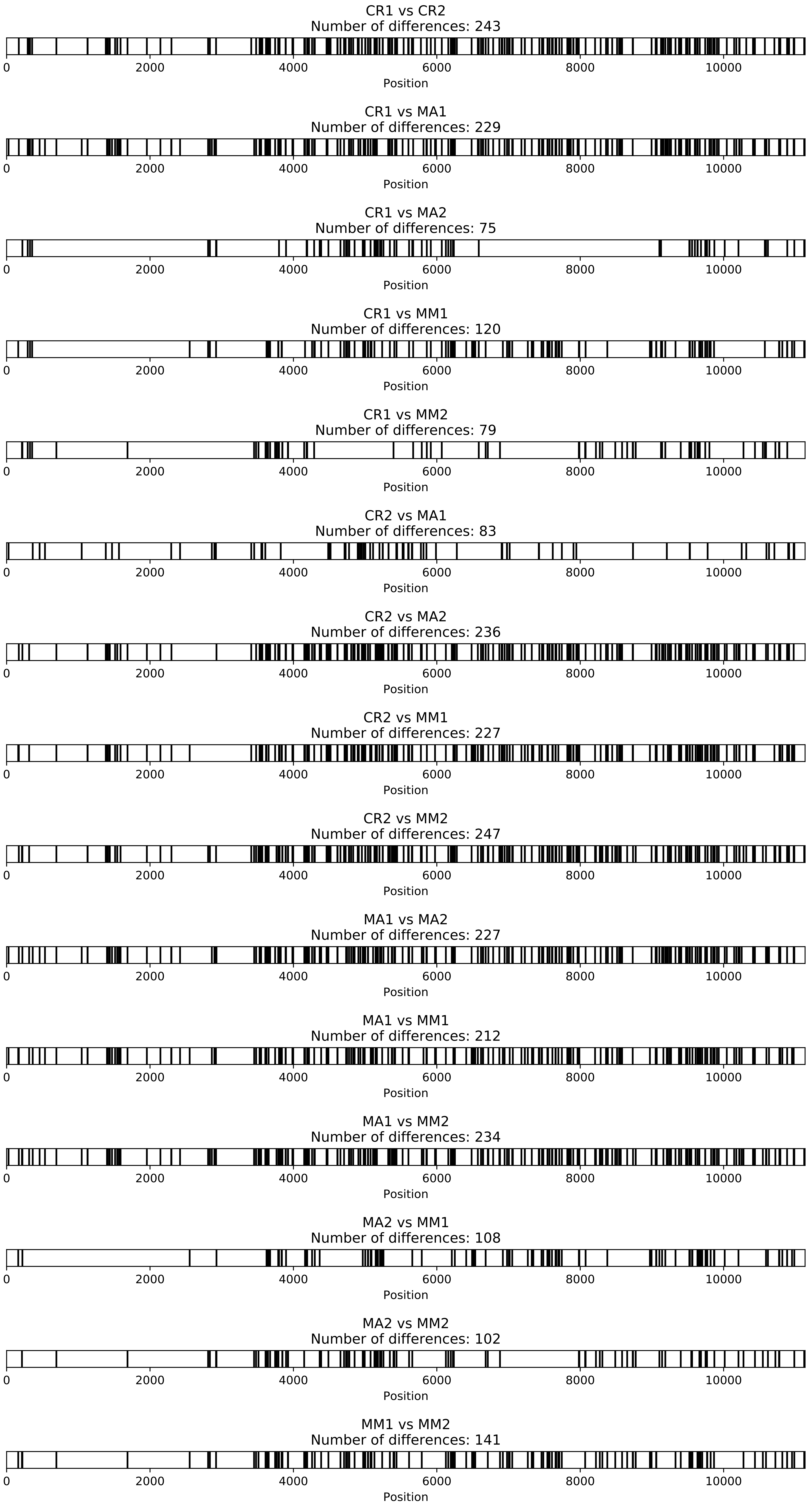
CR1 vs CR2
Number of differences: 243
CR1 vs MA1
Number of differences: 229
CR1 vs MA2
Number of differences: 75
CR1 vs MM1
Number of differences: 120
CR1 vs MM2
Number of differences: 79
CR2 vs MA1
Number of differences: 83
CR2 vs MA2
Number of differences: 236
CR2 vs MM1
Number of differences: 227
CR2 vs MM2
Number of differences: 247
MA1 vs MA2
Number of differences: 227
MA1 vs MM1
Number of differences: 212
MA1 vs MM2
Number of differences: 234
MA2 vs MM1
Number of differences: 108
MA2 vs MM2
Number of differences: 102
MM1 vs MM2
Number of differences: 141
0
2000
4000
6000
8000
10000
Position

### Region: 385

CR1 vs CR2
Number of differences: 151

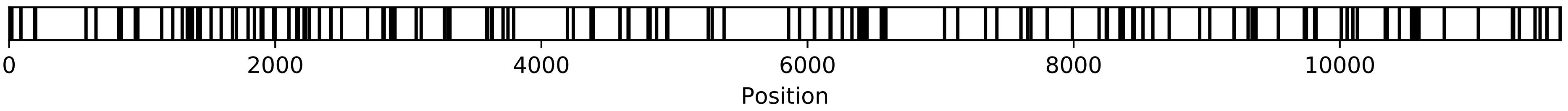


CR1 vs MA1
Number of differences: 62

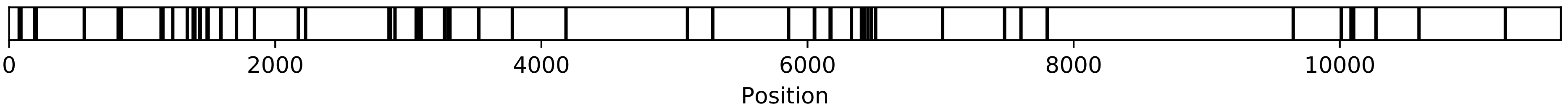


CR1 vs MA2
Number of differences: 157

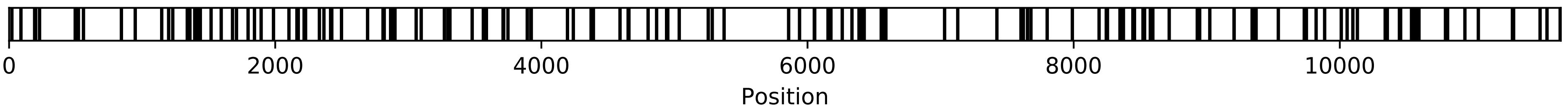


CR1 vs MM1
Number of differences: 62

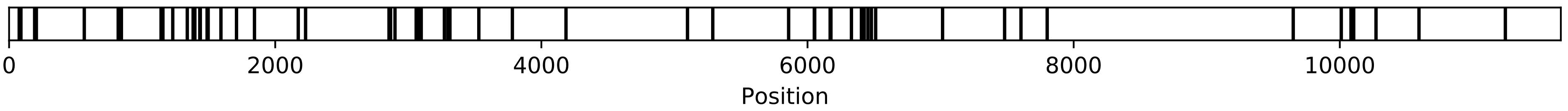


CR1 vs MM2
Number of differences: 75

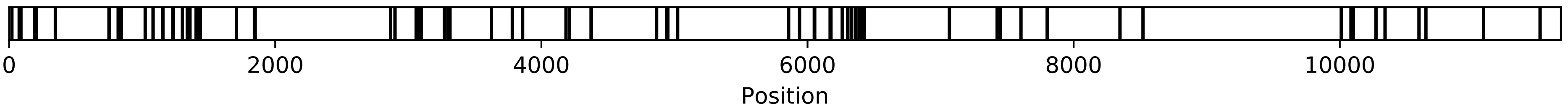


CR2 vs MA1
Number of differences: 154

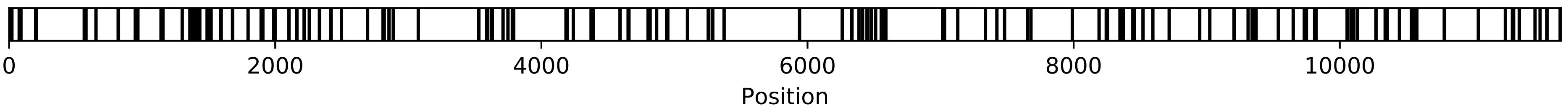


CR2 vs MA2
Number of differences: 61

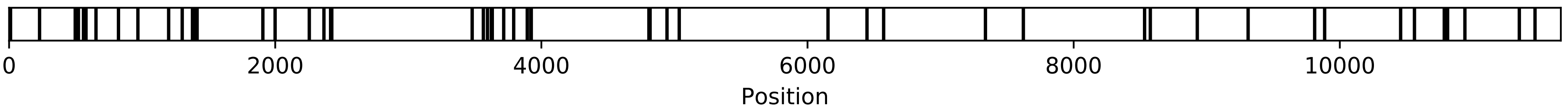


CR2 vs MM1
Number of differences: 154

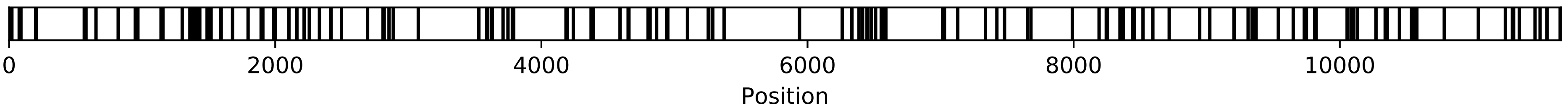


CR2 vs MM2
Number of differences: 147

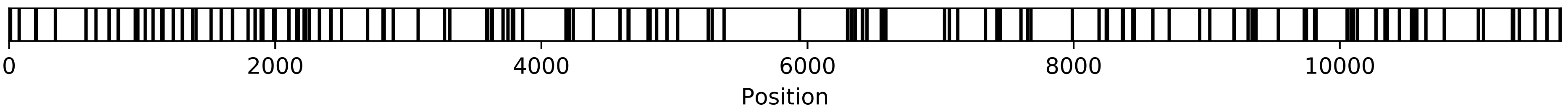


MA1 vs MA2
Number of differences: 160

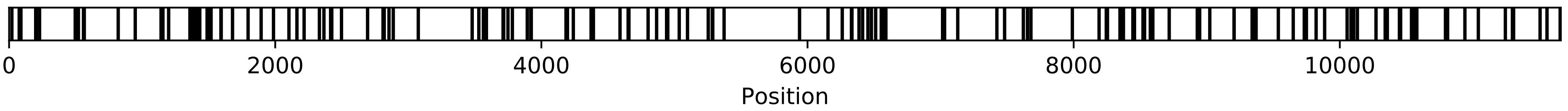


MA1 vs MM1
Number of differences: 0

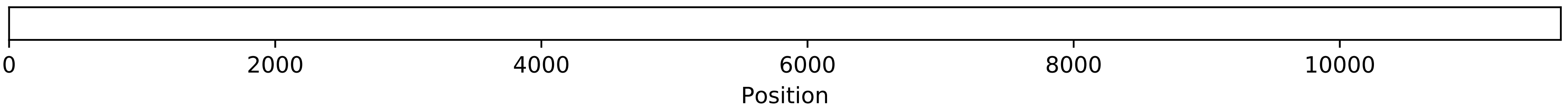


MA1 vs MM2
Number of differences: 69

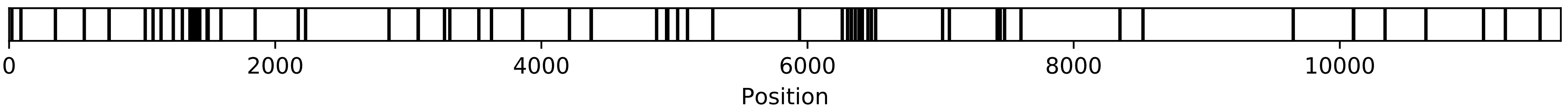


MA2 vs MM1
Number of differences: 160

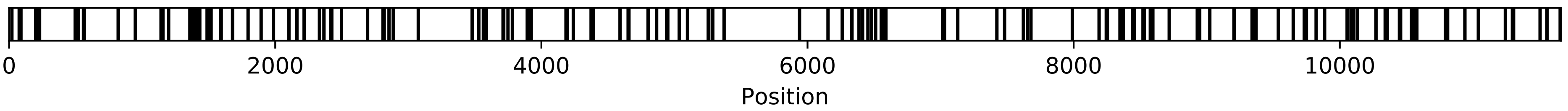


MA2 vs MM2
Number of differences: 153

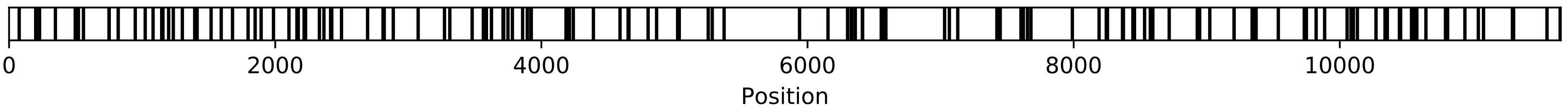


MM1 vs MM2
Number of differences: 69

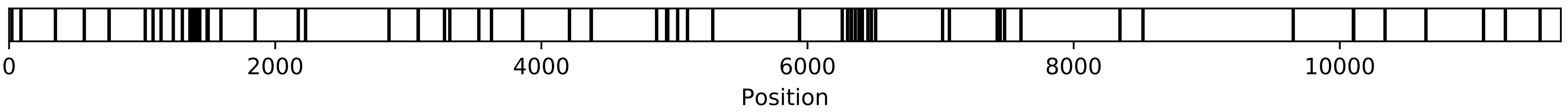

### Region: 386

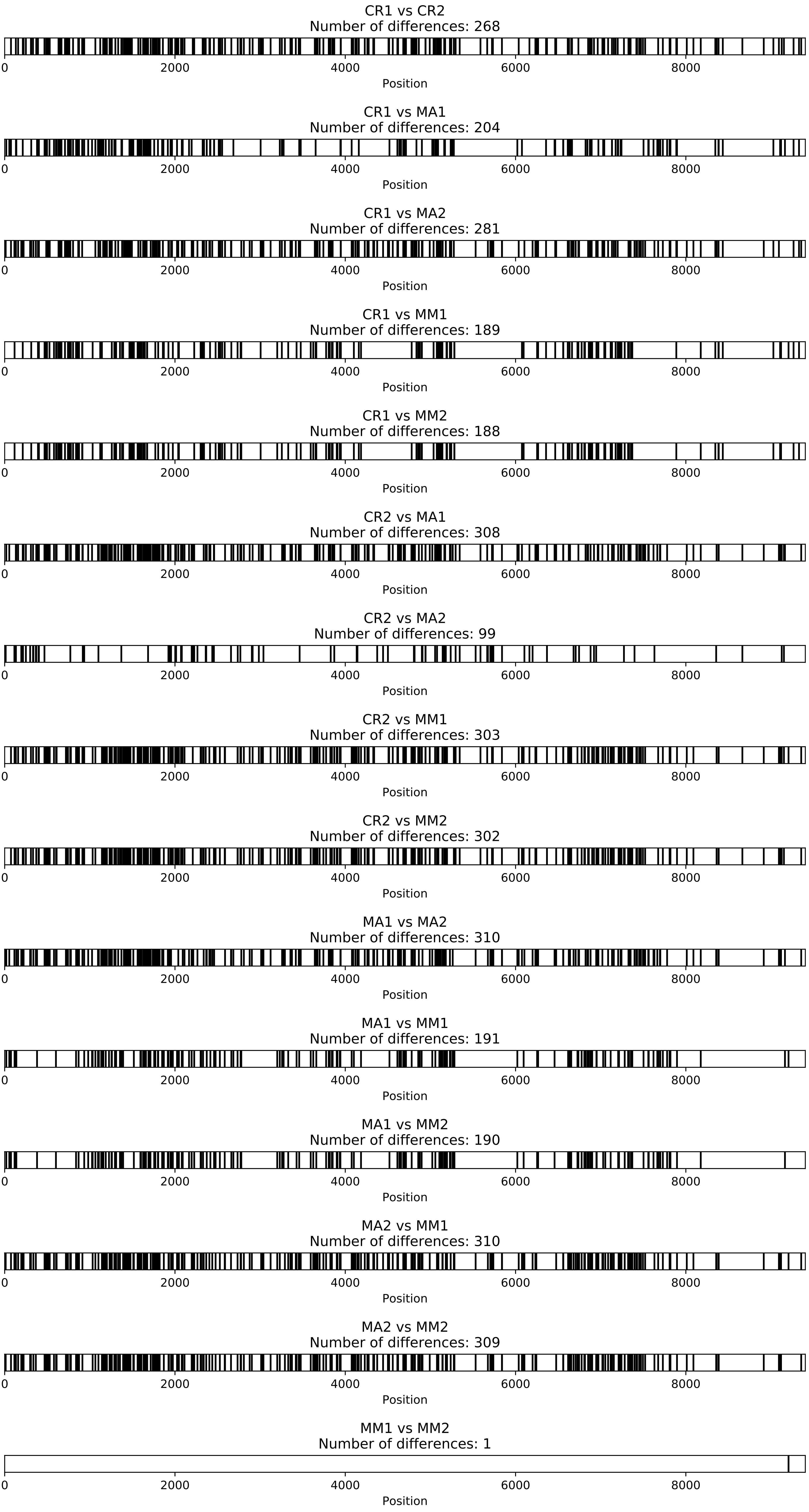
CR1 vs CR2
Number of differences: 268
CR1 vs MA1
Number of differences: 204
CR1 vs MA2
Number of differences: 281
CR1 vs MM1
Number of differences: 189
CR1 vs MM2
Number of differences: 188
CR2 vs MA1
Number of differences: 308
CR2 vs MA2
Number of differences: 99
CR2 vs MM1
Number of differences: 303
CR2 vs MM2
Number of differences: 302
MA1 vs MA2
Number of differences: 310
MA1 vs MM1
Number of differences: 191
MA1 vs MM2
Number of differences: 190
MA2 vs MM1
Number of differences: 310
MA2 vs MM2
Number of differences: 309
MM1 vs MM2
Number of differences: 1
0
2000
4000
6000
8000
Position

### Region: 387



CR1 vs CR2
Number of differences: 269

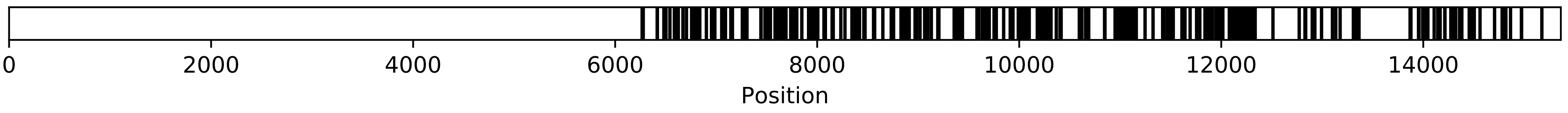


CR1 vs MA1
Number of differences: 270

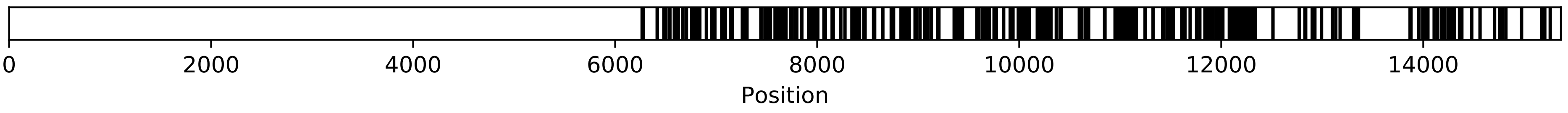


CR1 vs MA2
Number of differences: 297

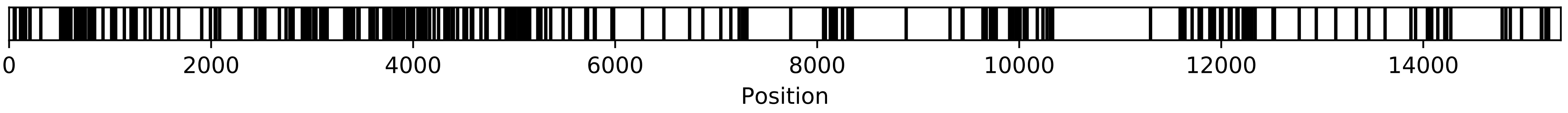


CR1 vs MM1
Number of differences: 387

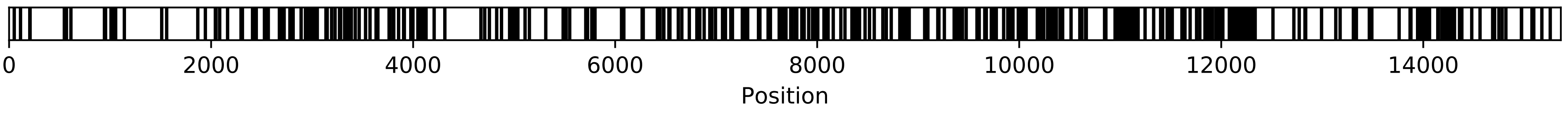


CR1 vs MM2
Number of differences: 387

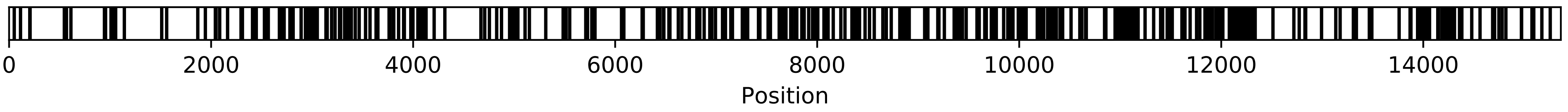


CR2 vs MA1
Number of differences: 15

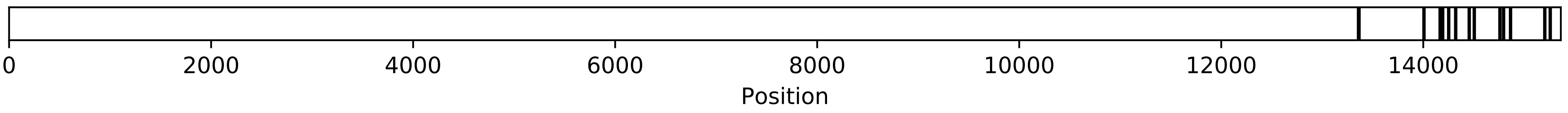


CR2 vs MA2
Number of differences: 456

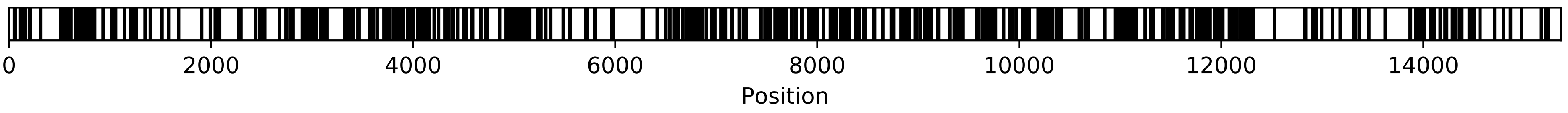


CR2 vs MM1
Number of differences: 252

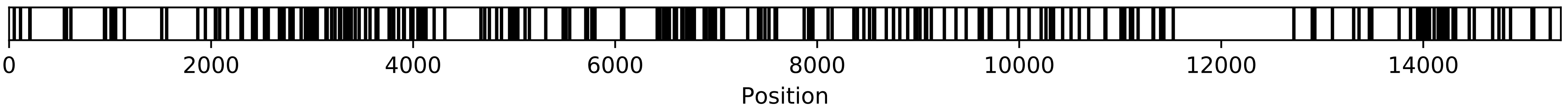


CR2 vs MM2
Number of differences: 252

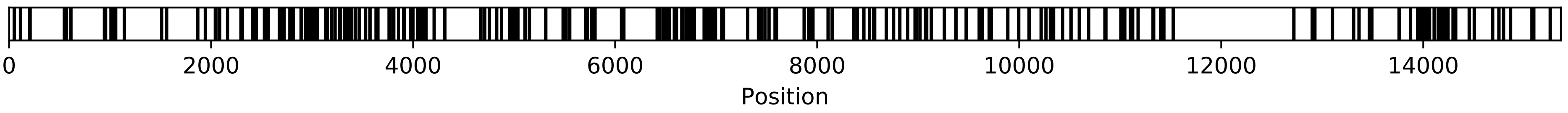


MA1 vs MA2
Number of differences: 457

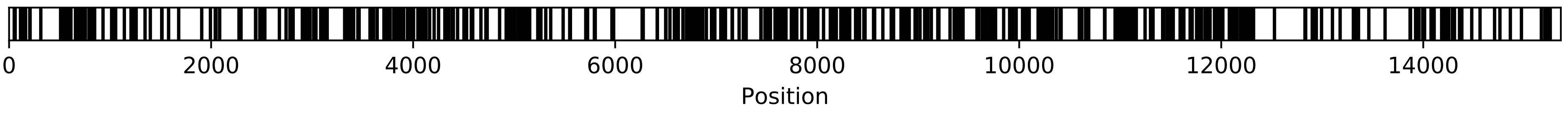


MA1 vs MM1
Number of differences: 245

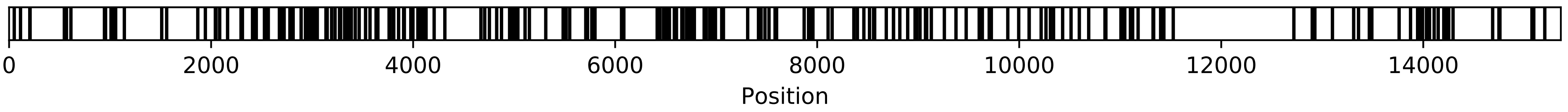


MA1 vs MM2
Number of differences: 245

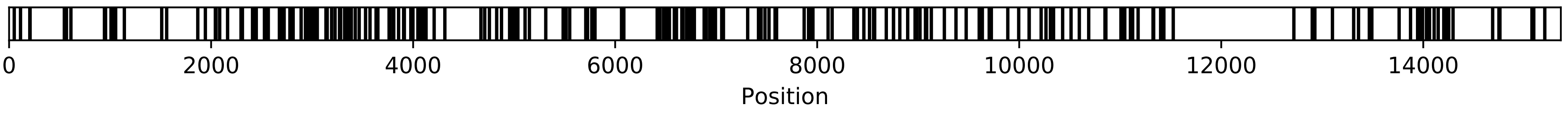


MA2 vs MM1
Number of differences: 452

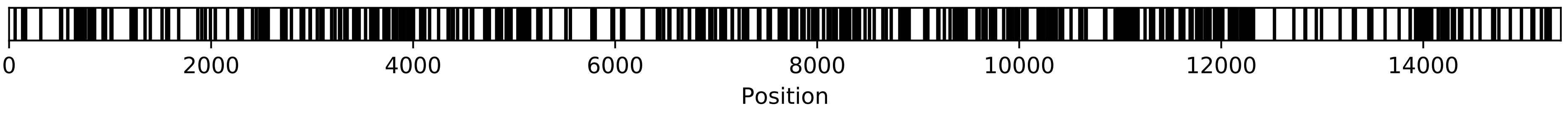


MA2 vs MM2
Number of differences: 452

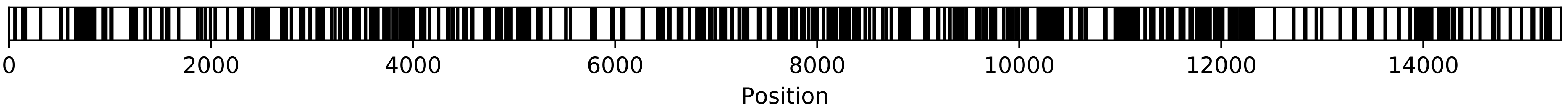


MM1 vs MM2
Number of differences: 0

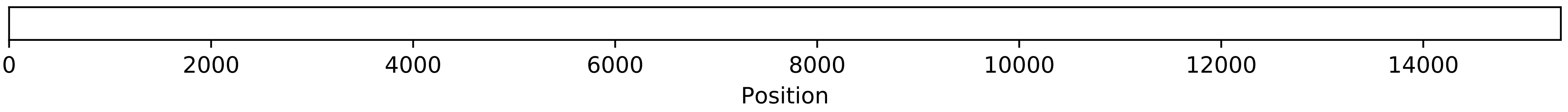

### Region: 388



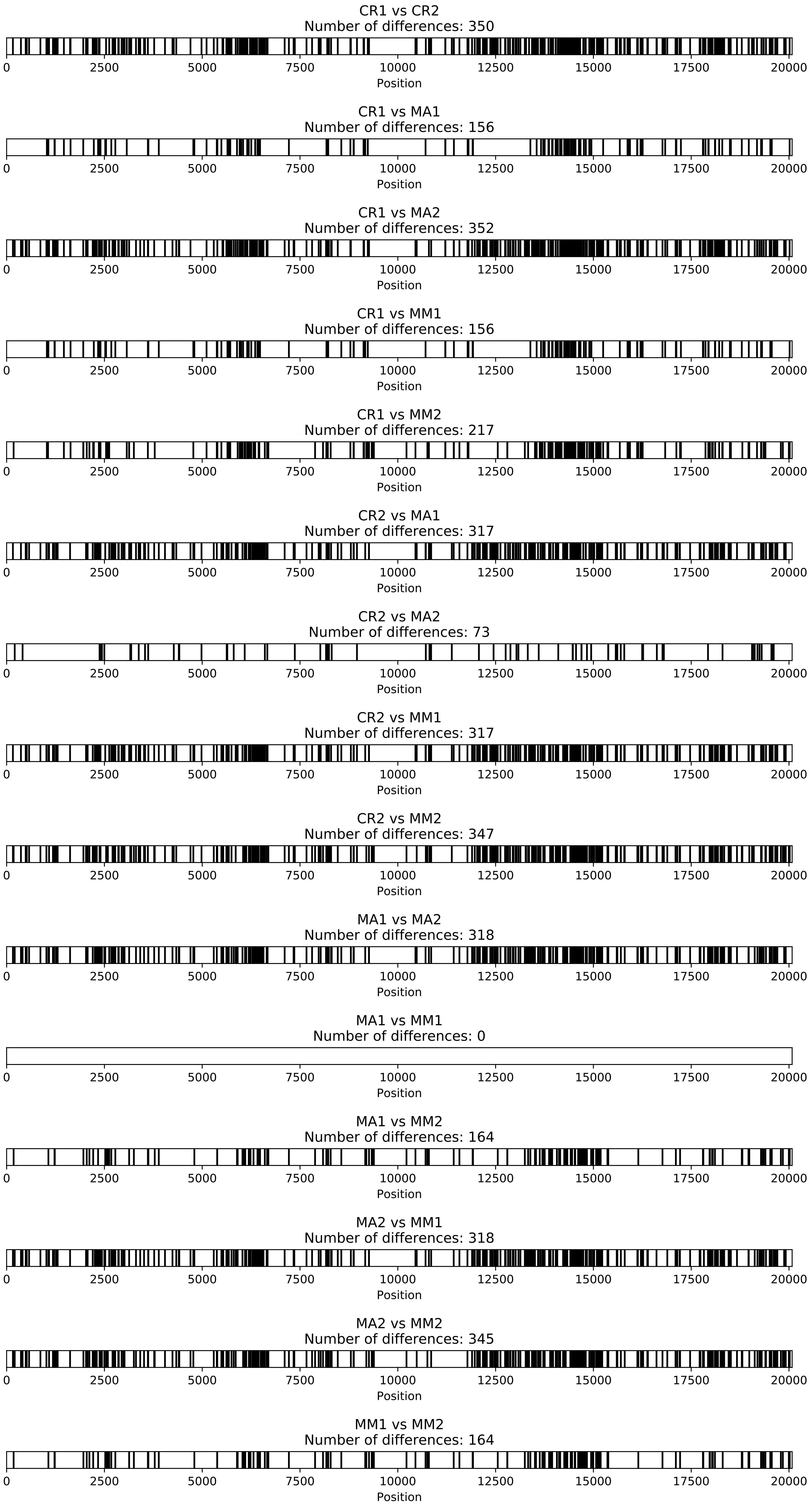
CR1 vs CR2
Number of differences: 350
CR1 vs MA1
Number of differences: 156
CR1 vs MA2
Number of differences: 352
CR1 vs MM1
Number of differences: 156
CR1 vs MM2
Number of differences: 217
CR2 vs MA1
Number of differences: 317
CR2 vs MA2
Number of differences: 73
CR2 vs MM1
Number of differences: 317
CR2 vs MM2
Number of differences: 347
MA1 vs MA2
Number of differences: 318
MA1 vs MM1
Number of differences: 0
MA1 vs MM2
Number of differences: 164
MA2 vs MM1
Number of differences: 318
MA2 vs MM2
Number of differences: 345
MM1 vs MM2
Number of differences: 164
0 2500 5000 7500 10000 12500 15000 17500 20000
Position

### Region: 391

CR1 vs CR2
Number of differences: 325

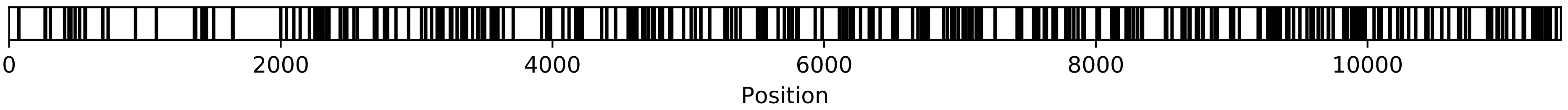


CR1 vs MA1
Number of differences: 232

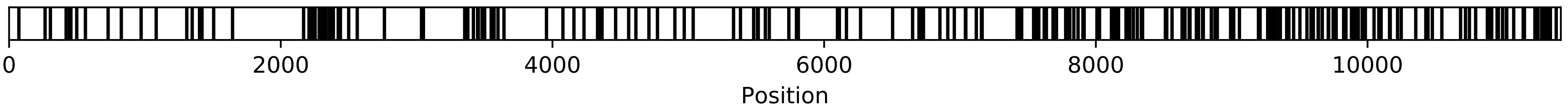


CR1 vs MA2
Number of differences: 242

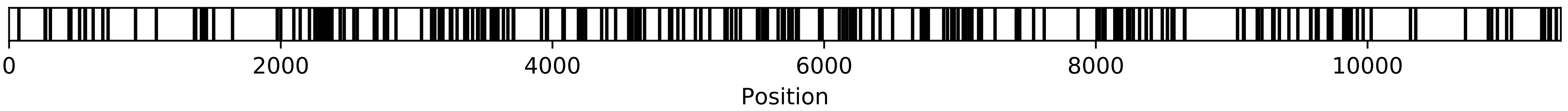


CR1 vs MM1
Number of differences: 166

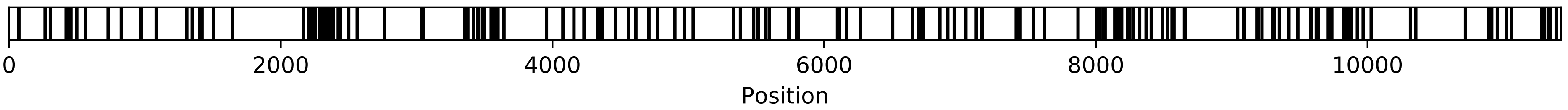


CR1 vs MM2
Number of differences: 169

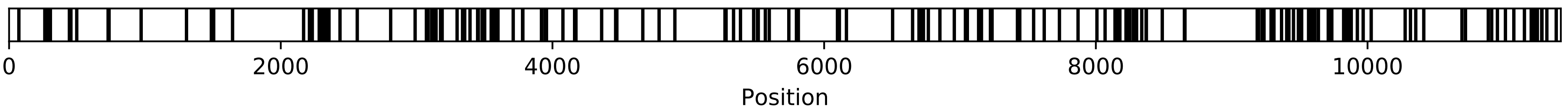


CR2 vs MA1
Number of differences: 194

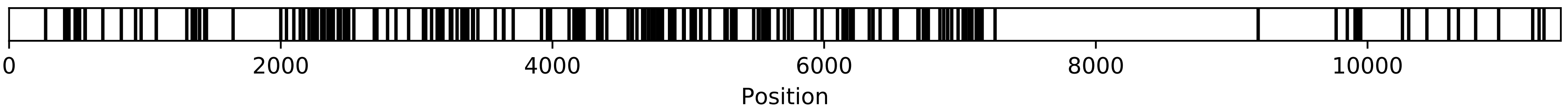


CR2 vs MA2
Number of differences: 206

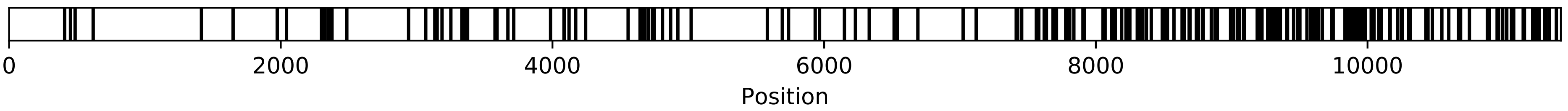


CR2 vs MM1
Number of differences: 319

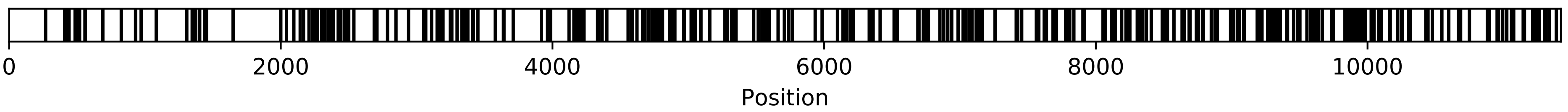


CR2 vs MM2
Number of differences: 321

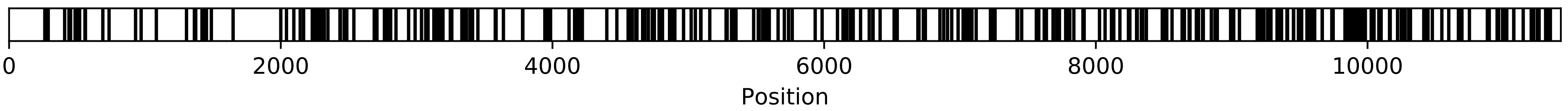


MA1 vs MA2
Number of differences: 301

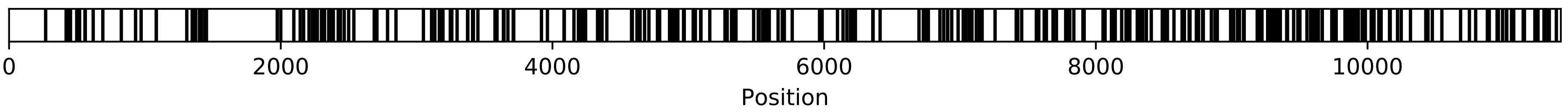


MA1 vs MM1
Number of differences: 137

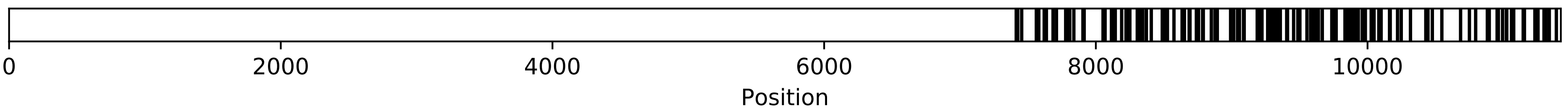


MA1 vs MM2
Number of differences: 227

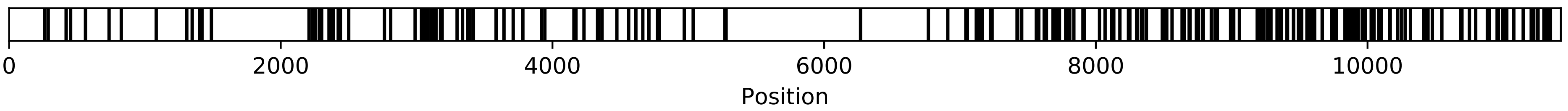


MA2 vs MM1
Number of differences: 164

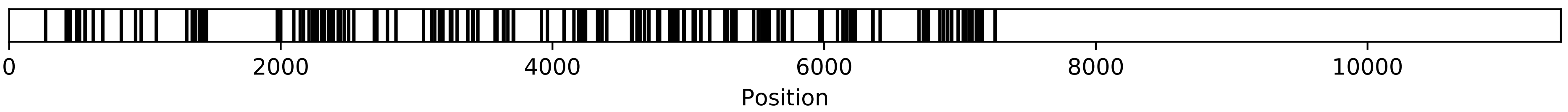


MA2 vs MM2
Number of differences: 223

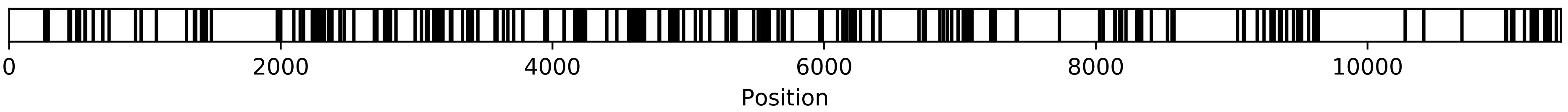


MM1 vs MM2
Number of differences: 143

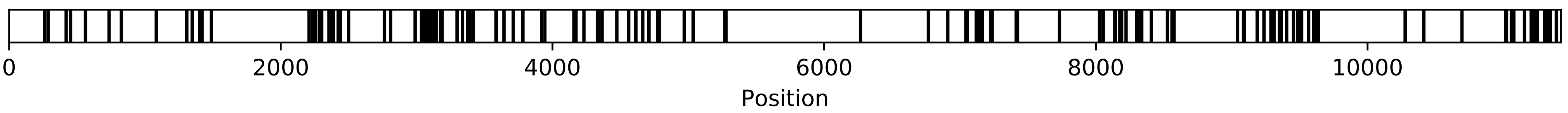

### Region: 392

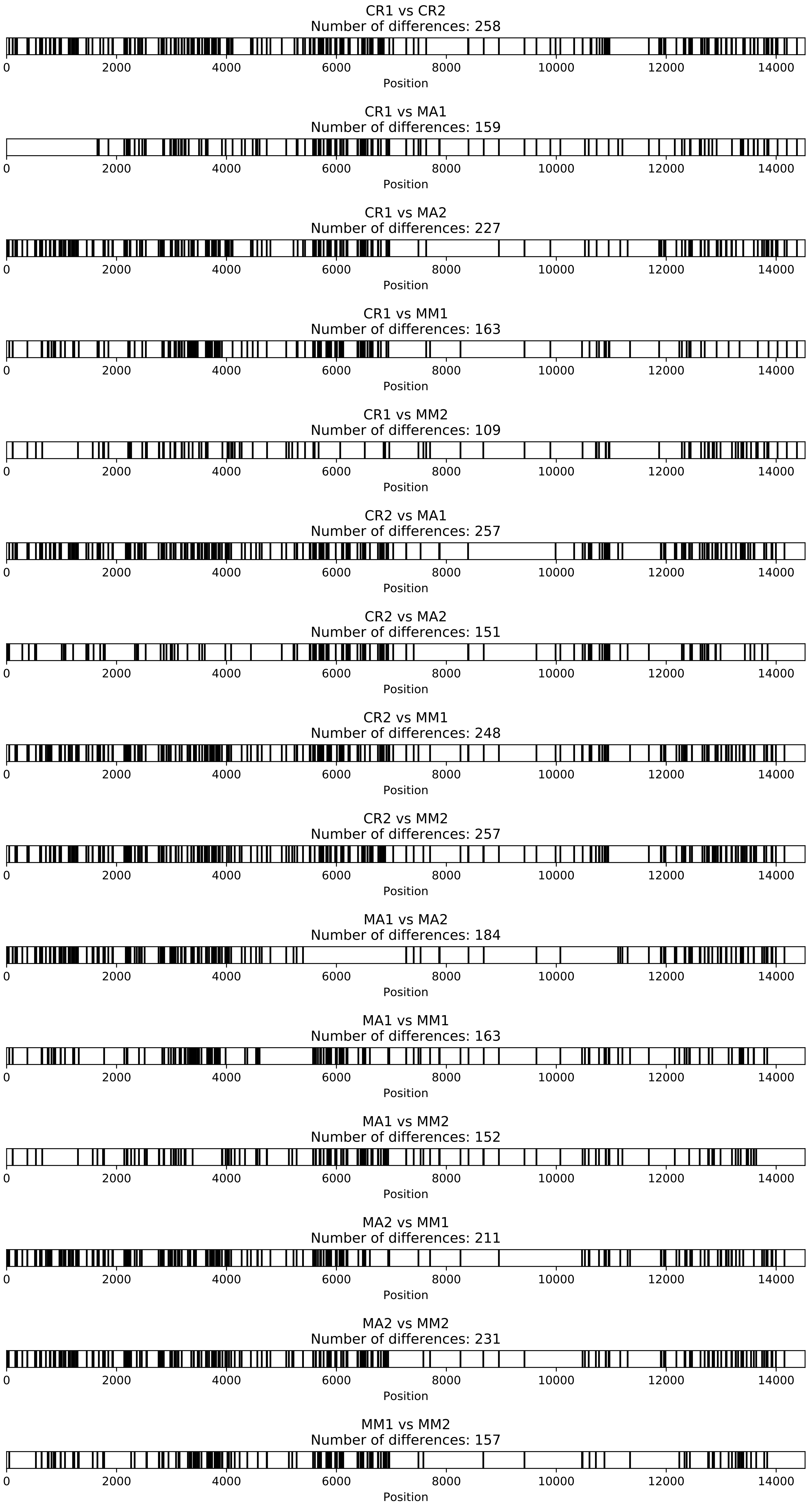
CR1 vs CR2
Number of differences: 258
0 2000 4000 6000 8000 10000 12000 14000
Position
CR1 vs MA1
Number of differences: 159
0 2000 4000 6000 8000 10000 12000 14000
Position
CR1 vs MA2
Number of differences: 227
0 2000 4000 6000 8000 10000 12000 14000
Position
CR1 vs MM1
Number of differences: 163
0 2000 4000 6000 8000 10000 12000 14000
Position
CR1 vs MM2
Number of differences: 109
0 2000 4000 6000 8000 10000 12000 14000
Position
CR2 vs MA1
Number of differences: 257
0 2000 4000 6000 8000 10000 12000 14000
Position
CR2 vs MA2
Number of differences: 151
0 2000 4000 6000 8000 10000 12000 14000
Position
CR2 vs MM1
Number of differences: 248
0 2000 4000 6000 8000 10000 12000 14000
Position
CR2 vs MM2
Number of differences: 257
0 2000 4000 6000 8000 10000 12000 14000
Position
MA1 vs MA2
Number of differences: 184
0 2000 4000 6000 8000 10000 12000 14000
Position
MA1 vs MM1
Number of differences: 163
0 2000 4000 6000 8000 10000 12000 14000
Position
MA1 vs MM2
Number of differences: 152
0 2000 4000 6000 8000 10000 12000 14000
Position
MA2 vs MM1
Number of differences: 211
0 2000 4000 6000 8000 10000 12000 14000
Position
MA2 vs MM2
Number of differences: 231
0 2000 4000 6000 8000 10000 12000 14000
Position
MM1 vs MM2
Number of differences: 157
0 2000 4000 6000 8000 10000 12000 14000
Position

### Region: 393



CR1 vs CR2
Number of differences: 349

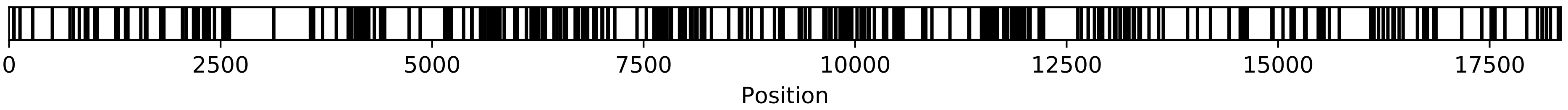


CR1 vs MA1
Number of differences: 329

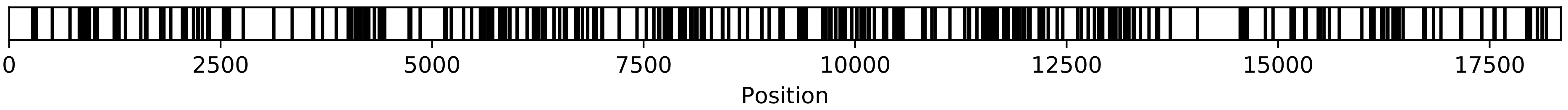


CR1 vs MA2
Number of differences: 122

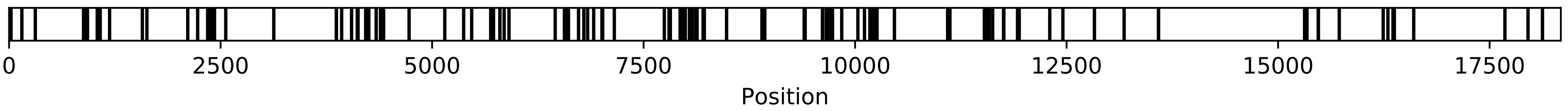


CR1 vs MM1
Number of differences: 343

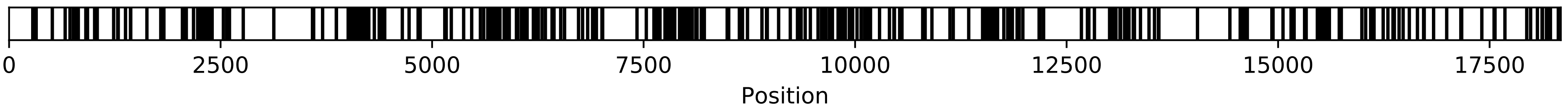


CR1 vs MM2
Number of differences: 350

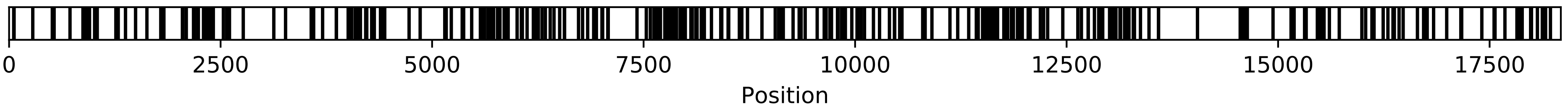


CR2 vs MA1
Number of differences: 174

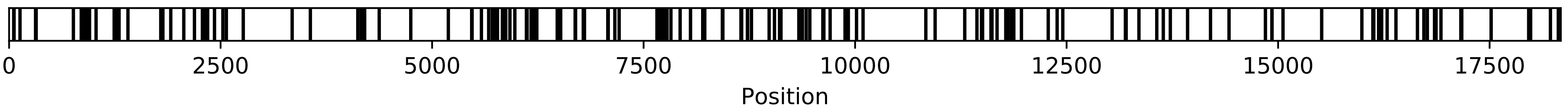


CR2 vs MA2
Number of differences: 336

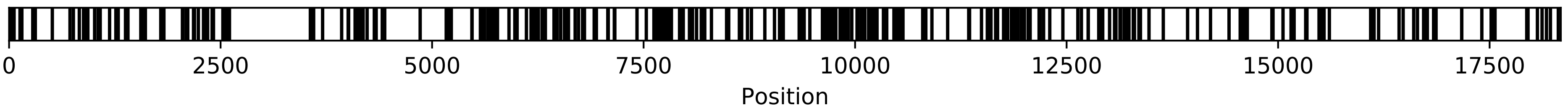


CR2 vs MM1
Number of differences: 219

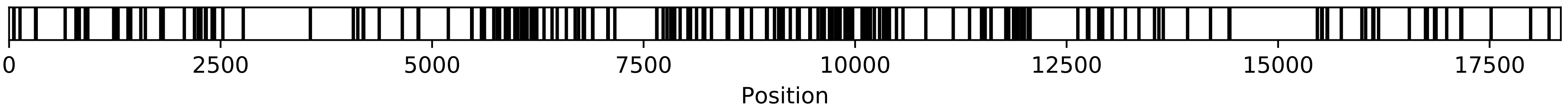


CR2 vs MM2
Number of differences: 203

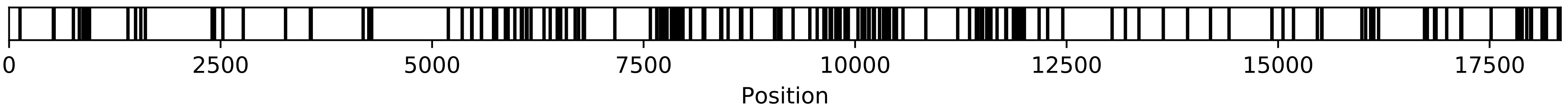


MA1 vs MA2
Number of differences: 317

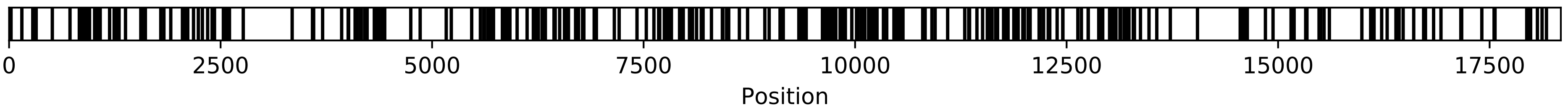


MA1 vs MM1
Number of differences: 216

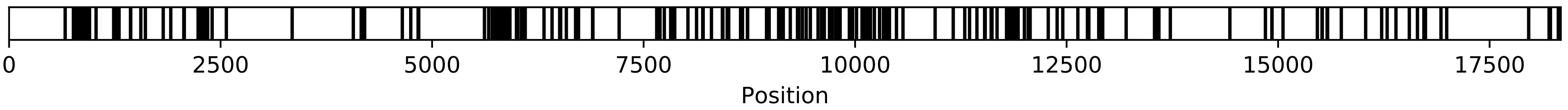


MA1 vs MM2
Number of differences: 202

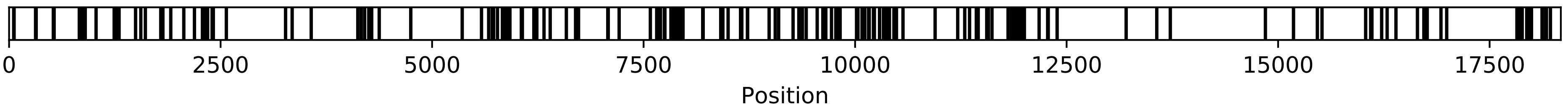


MA2 vs MM1
Number of differences: 331

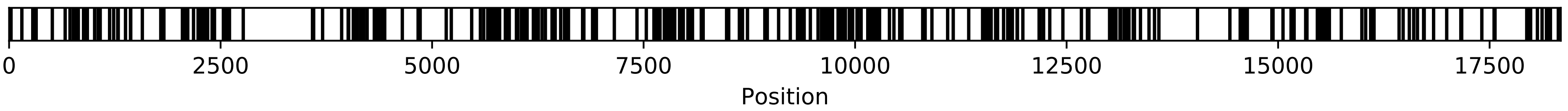


MA2 vs MM2
Number of differences: 338

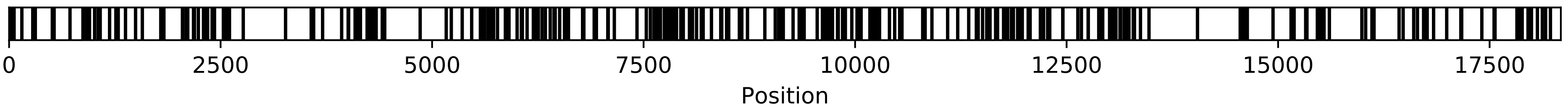


MM1 vs MM2
Number of differences: 203

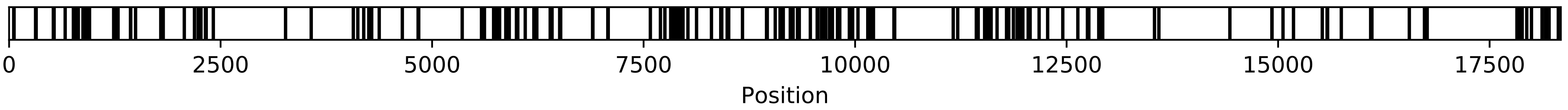

### Region: 394

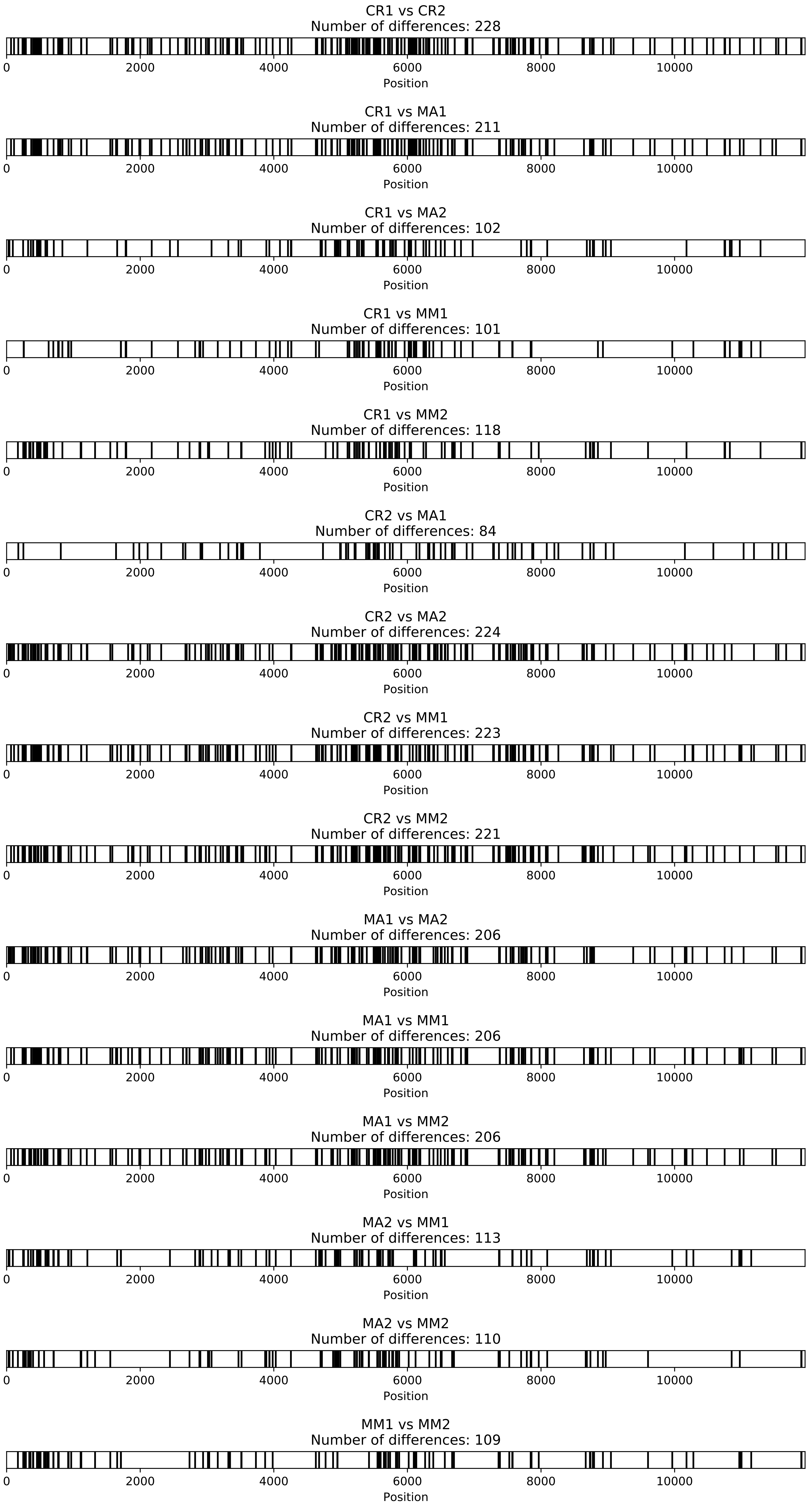
CR1 vs CR2
Number of differences: 228
CR1 vs MA1
Number of differences: 211
CR1 vs MA2
Number of differences: 102
CR1 vs MM1
Number of differences: 101
CR1 vs MM2
Number of differences: 118
CR2 vs MA1
Number of differences: 84
CR2 vs MA2
Number of differences: 224
CR2 vs MM1
Number of differences: 223
CR2 vs MM2
Number of differences: 221
MA1 vs MA2
Number of differences: 206
MA1 vs MM1
Number of differences: 206
MA1 vs MM2
Number of differences: 206
MA2 vs MM1
Number of differences: 113
MA2 vs MM2
Number of differences: 110
MM1 vs MM2
Number of differences: 109
0
2000
4000
6000
8000
10000
Position

### Region: 395



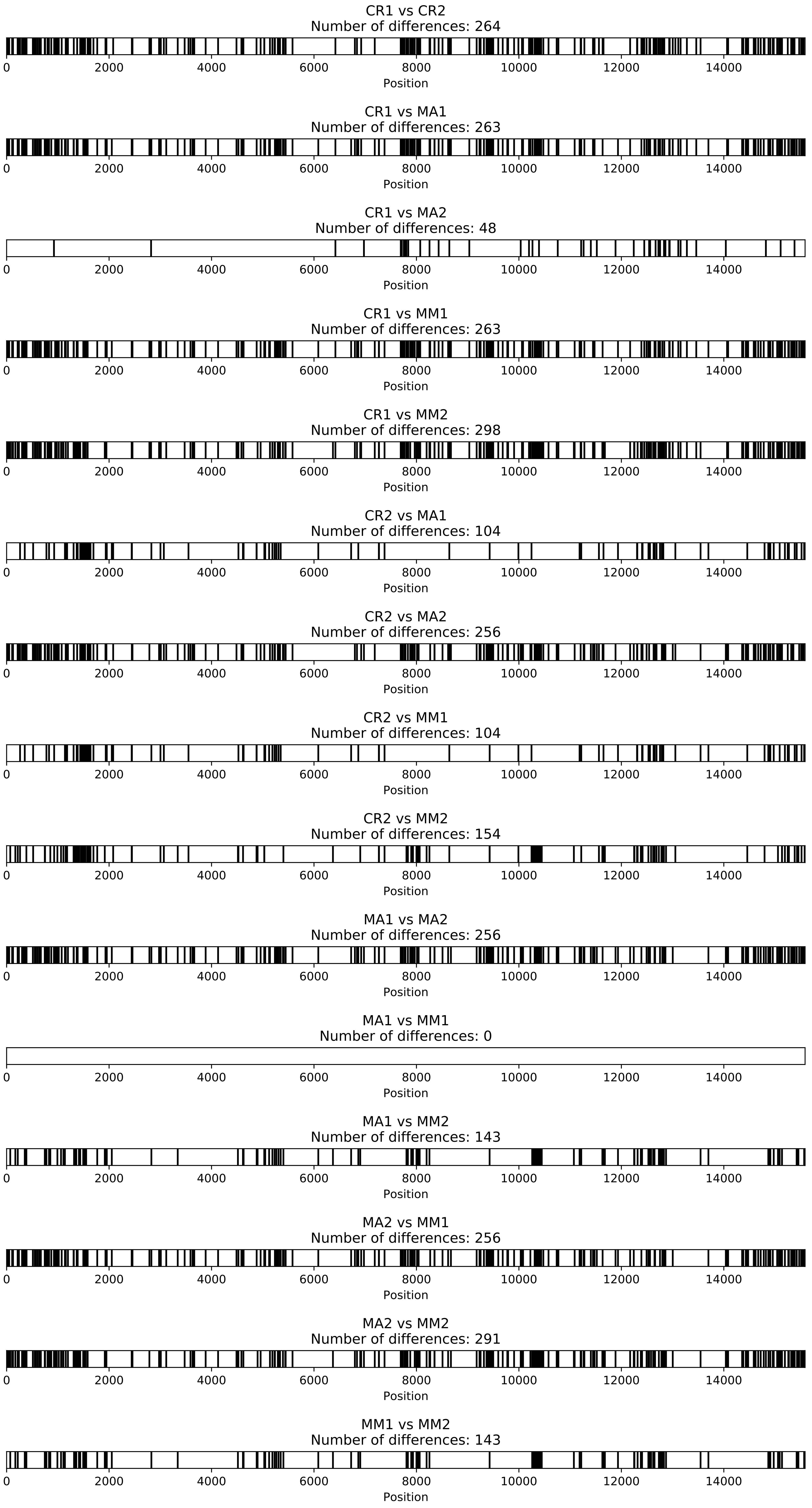
CR1 vs CR2
Number of differences: 264
CR1 vs MA1
Number of differences: 263
CR1 vs MA2
Number of differences: 48
CR1 vs MM1
Number of differences: 263
CR1 vs MM2
Number of differences: 298
CR2 vs MA1
Number of differences: 104
CR2 vs MA2
Number of differences: 256
CR2 vs MM1
Number of differences: 104
CR2 vs MM2
Number of differences: 154
MA1 vs MA2
Number of differences: 256
MA1 vs MM1
Number of differences: 0
MA1 vs MM2
Number of differences: 143
MA2 vs MM1
Number of differences: 256
MA2 vs MM2
Number of differences: 291
MM1 vs MM2
Number of differences: 143
0
2000
4000
6000
8000
10000
12000
14000
Position

### Region: 396

CR1 vs CR2
Number of differences: 276

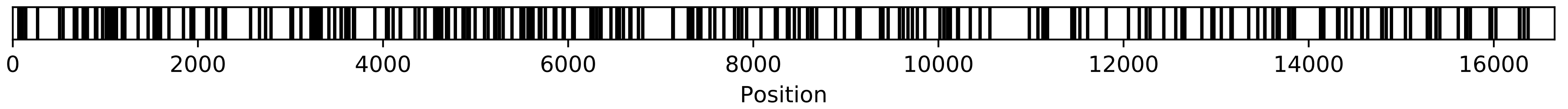


CR1 vs MA1
Number of differences: 288

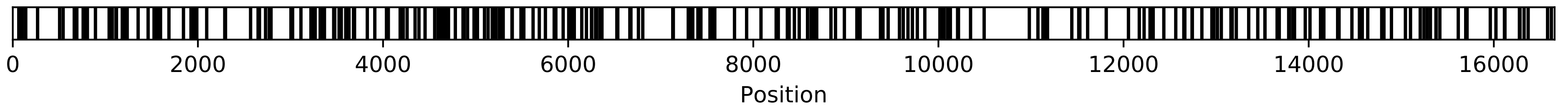


CR1 vs MA2
Number of differences: 47

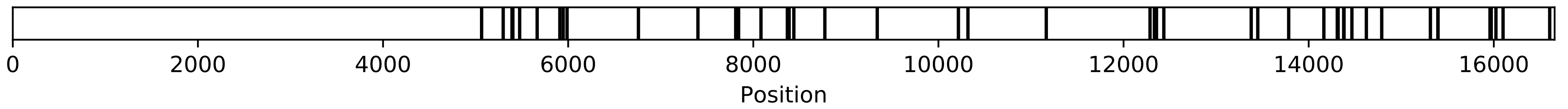


CR1 vs MM1
Number of differences: 288

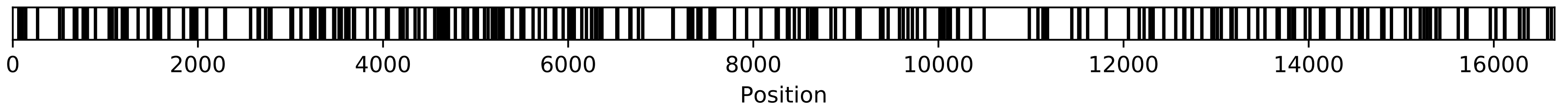


CR1 vs MM2
Number of differences: 287

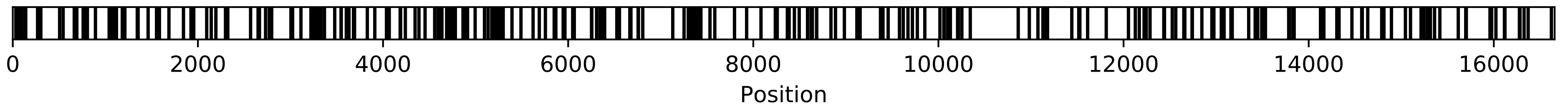


CR2 vs MA1
Number of differences: 121

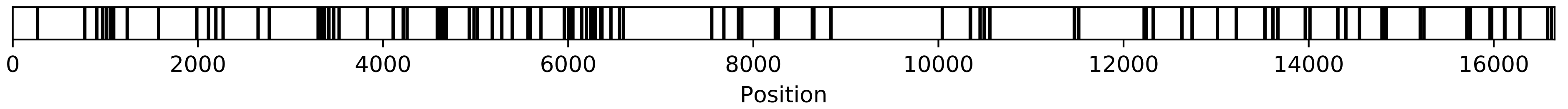


CR2 vs MA2
Number of differences: 279

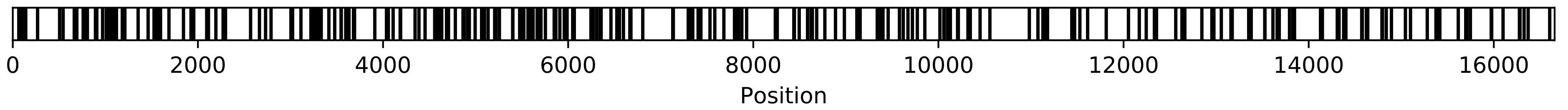


CR2 vs MM1
Number of differences: 121

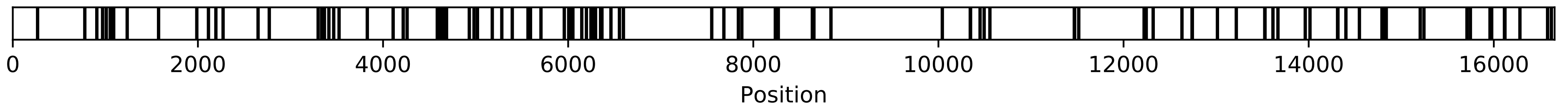


CR2 vs MM2
Number of differences: 132

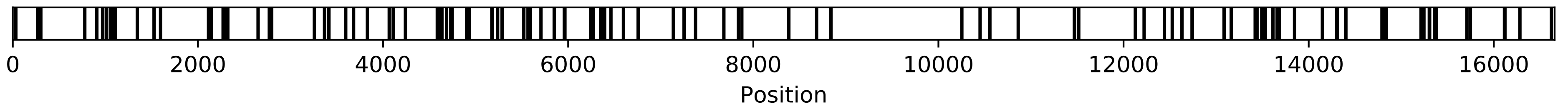


MA1 vs MA2
Number of differences: 293

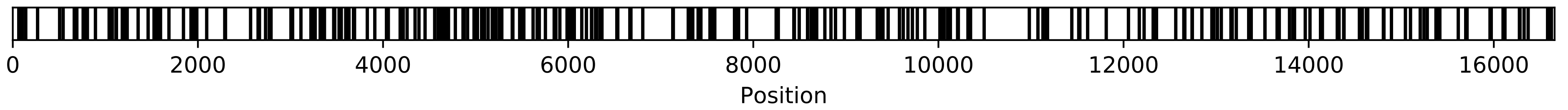


MA1 vs MM1
Number of differences: 0

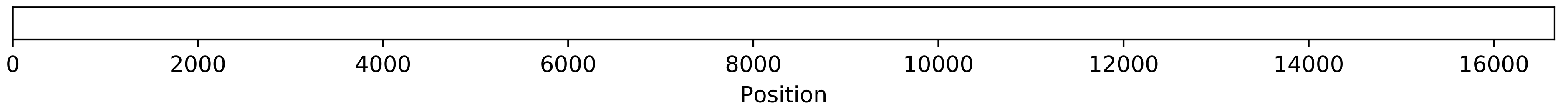


MA1 vs MM2
Number of differences: 105

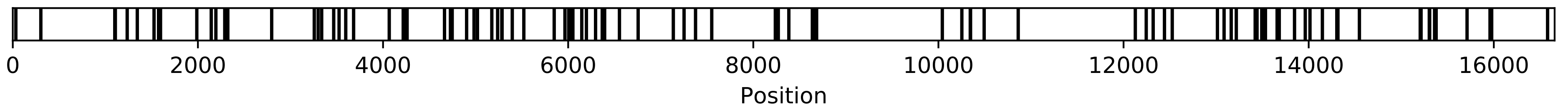


MA2 vs MM1
Number of differences: 293

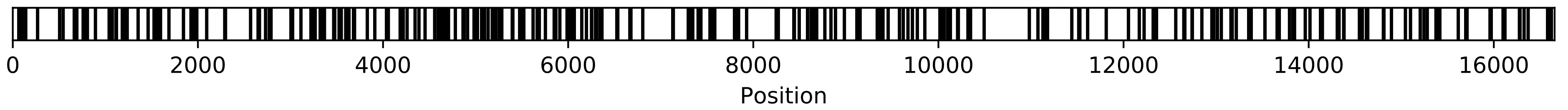


MA2 vs MM2
Number of differences: 290

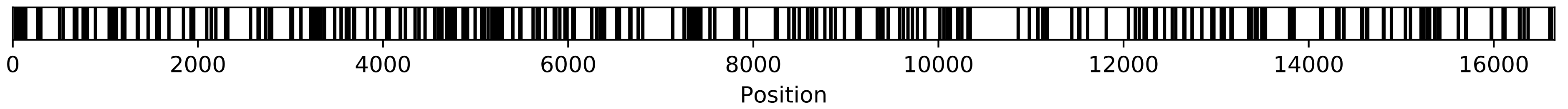


MM1 vs MM2
Number of differences: 105

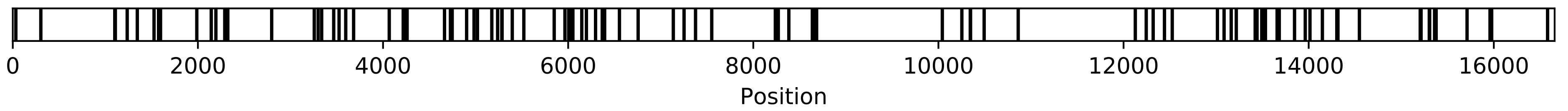

### Region: 397

CR1 vs CR2
Number of differences: 160

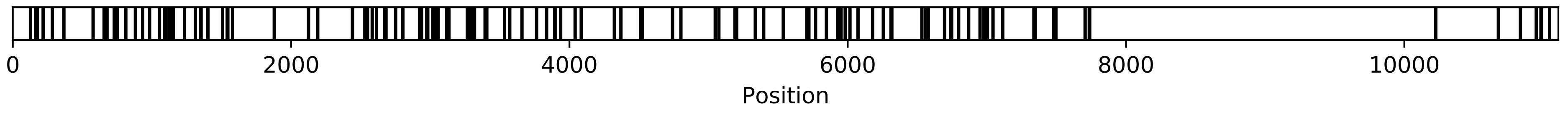


CR1 vs MA1
Number of differences: 106

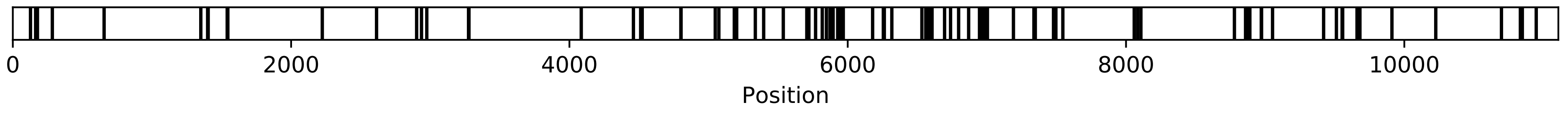


CR1 vs MA2
Number of differences: 187

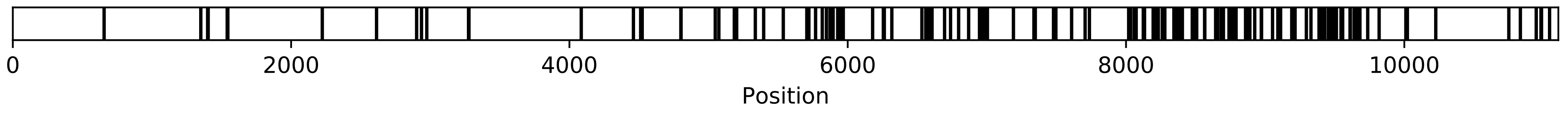


CR1 vs MM1
Number of differences: 269

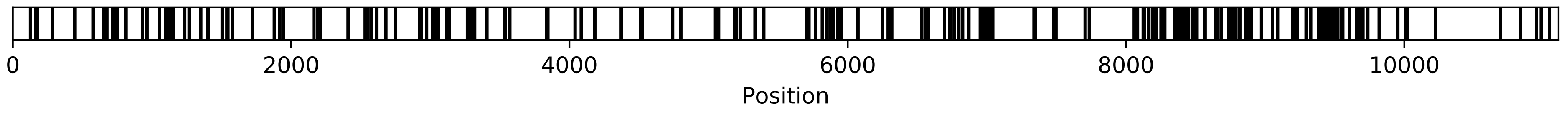


CR1 vs MM2
Number of differences: 254

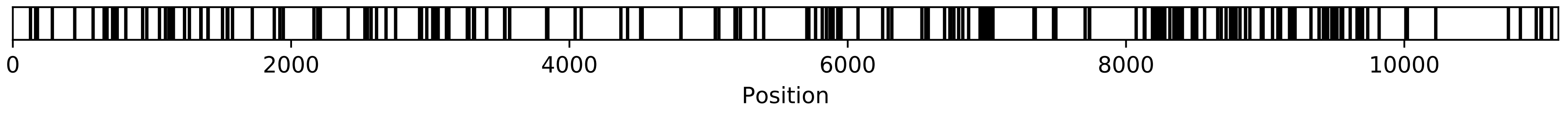


CR2 vs MA1
Number of differences: 136

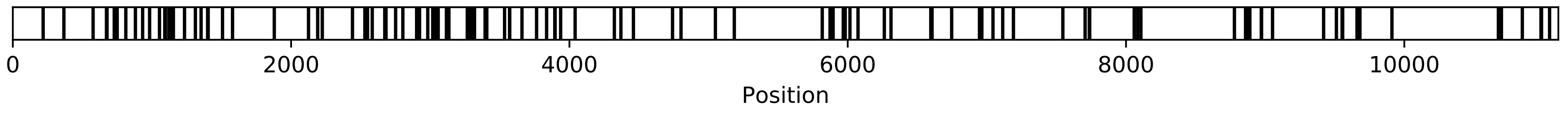


CR2 vs MA2
Number of differences: 223

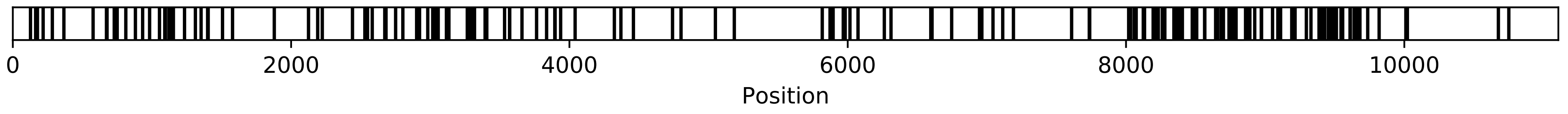


CR2 vs MM1
Number of differences: 178

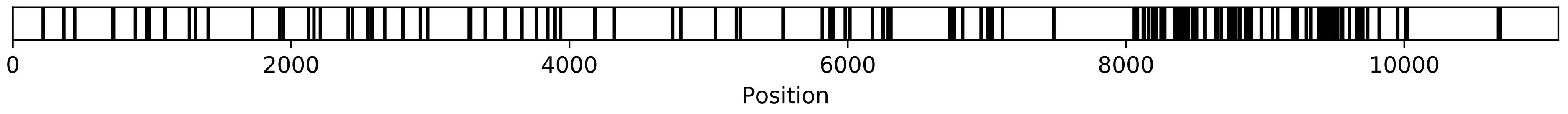


CR2 vs MM2
Number of differences: 165

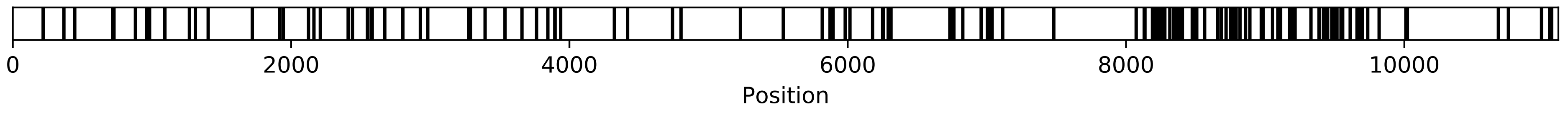


MA1 vs MA2
Number of differences: 109

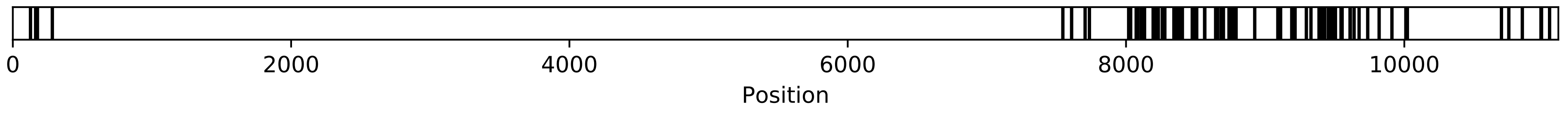


MA1 vs MM1
Number of differences: 210

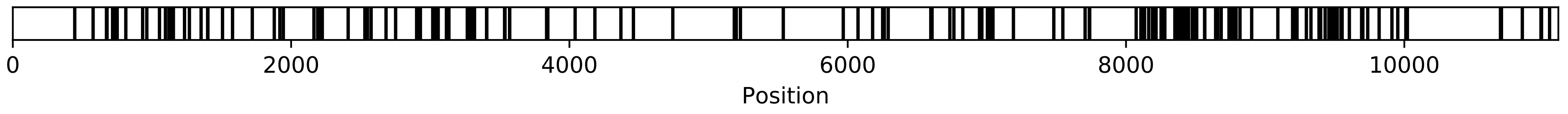


MA1 vs MM2
Number of differences: 200

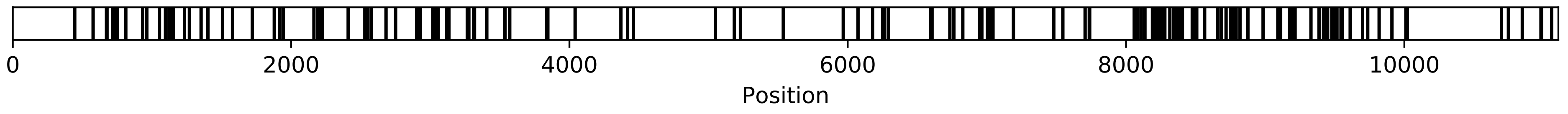


MA2 vs MM1
Number of differences: 183

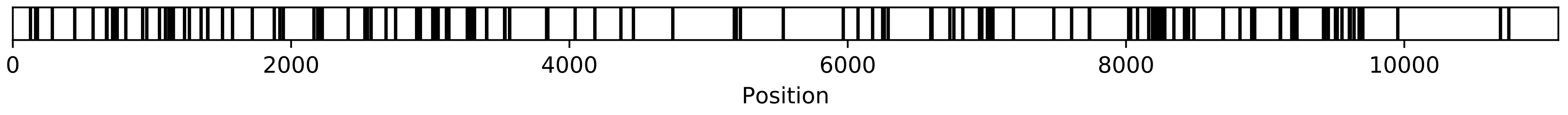


MA2 vs MM2
Number of differences: 181

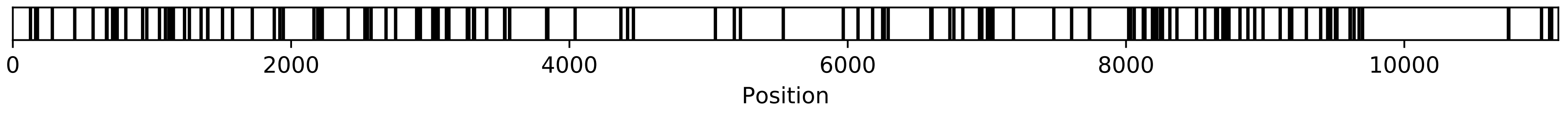


MM1 vs MM2
Number of differences: 68

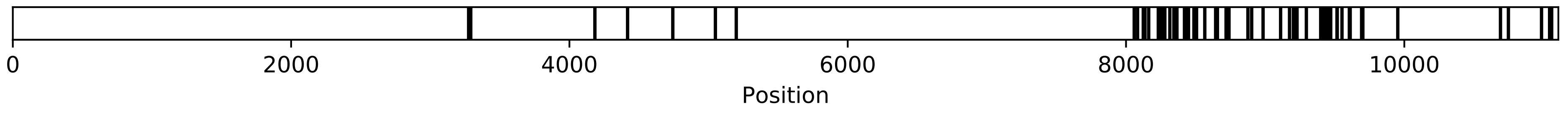

### Region: 398



CR1 vs CR2
Number of differences: 170

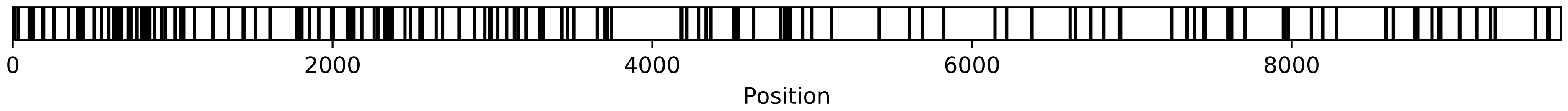


CR1 vs MA1
Number of differences: 102

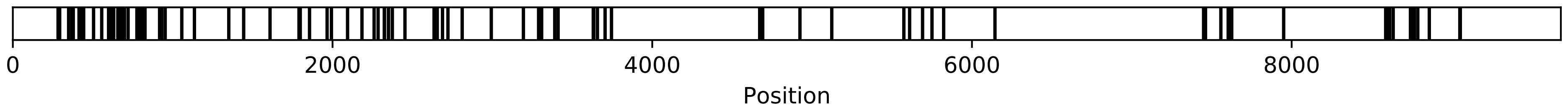


CR1 vs MA2
Number of differences: 171

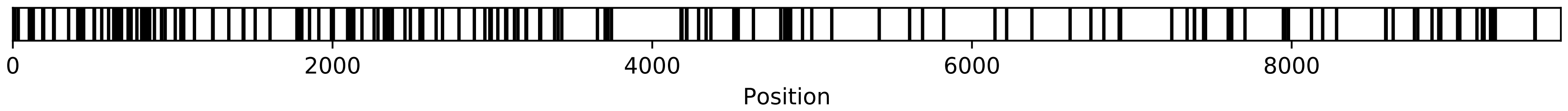


CR1 vs MM1
Number of differences: 90

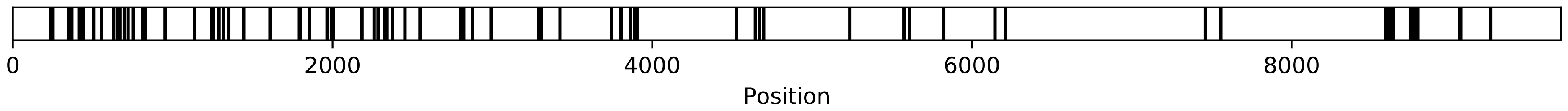


CR1 vs MM2
Number of differences: 102

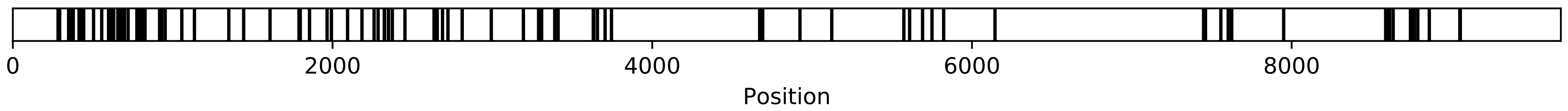


CR2 vs MA1
Number of differences: 163

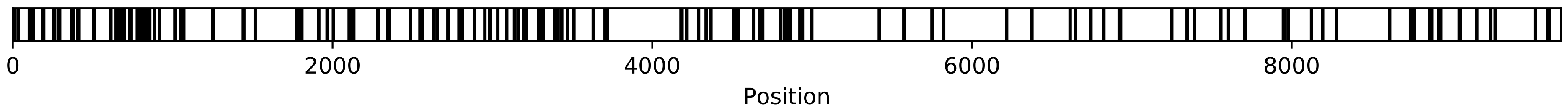


CR2 vs MA2
Number of differences: 17

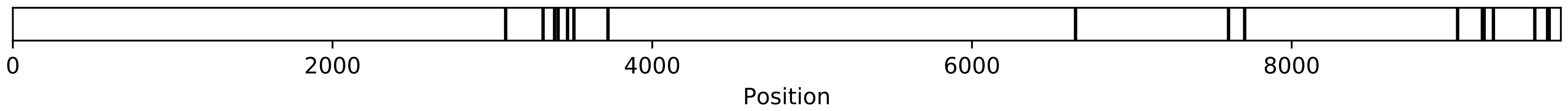


CR2 vs MM1
Number of differences: 171

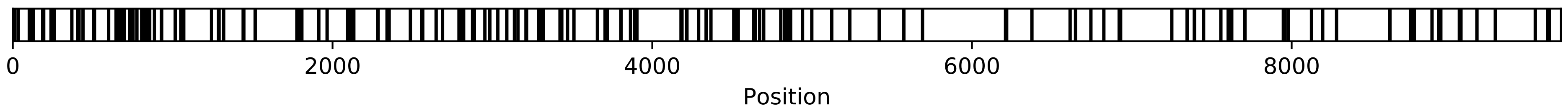


CR2 vs MM2
Number of differences: 163

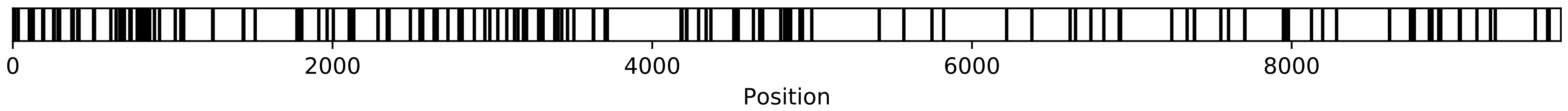


MA1 vs MA2
Number of differences: 162

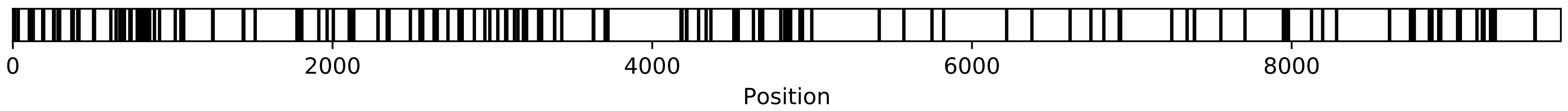


MA1 vs MM1
Number of differences: 79

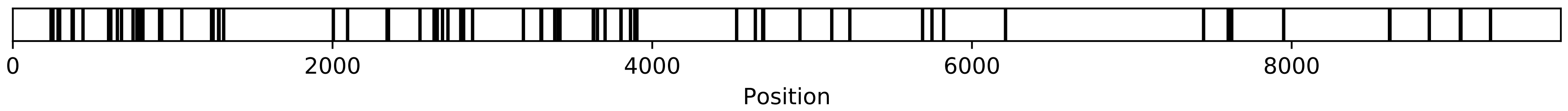


MA1 vs MM2
Number of differences: 0

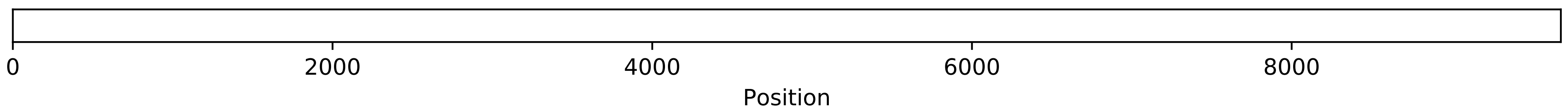


MA2 vs MM1
Number of differences: 172

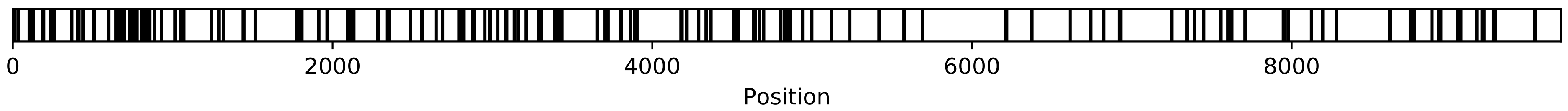


MA2 vs MM2
Number of differences: 162

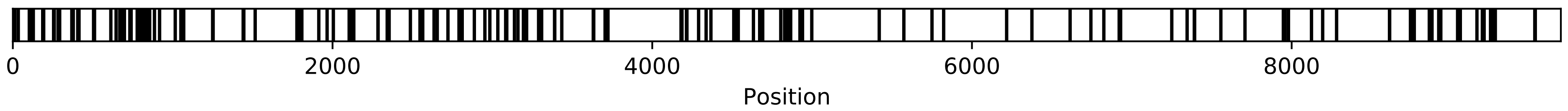


MM1 vs MM2
Number of differences: 79

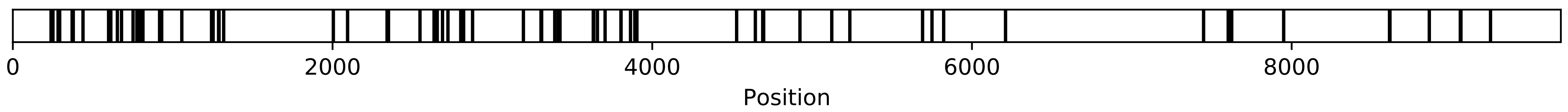

### Region: 400



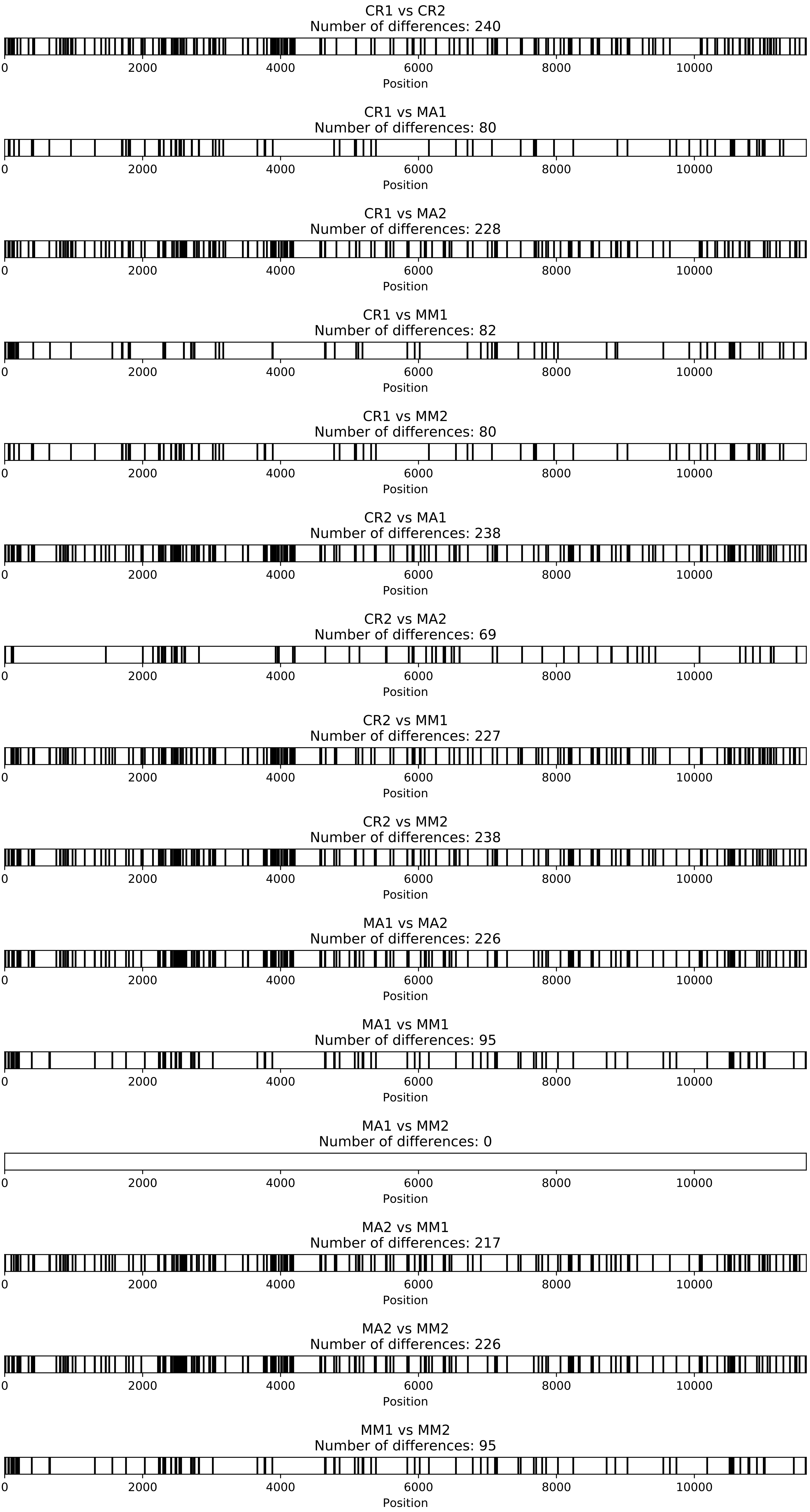
CR1 vs CR2
Number of differences: 240
CR1 vs MA1
Number of differences: 80
CR1 vs MA2
Number of differences: 228
CR1 vs MM1
Number of differences: 82
CR1 vs MM2
Number of differences: 80
CR2 vs MA1
Number of differences: 238
CR2 vs MA2
Number of differences: 69
CR2 vs MM1
Number of differences: 227
CR2 vs MM2
Number of differences: 238
MA1 vs MA2
Number of differences: 226
MA1 vs MM1
Number of differences: 95
MA1 vs MM2
Number of differences: 0
MA2 vs MM1
Number of differences: 217
MA2 vs MM2
Number of differences: 226
MM1 vs MM2
Number of differences: 95
0
2000
4000
6000
8000
10000
Position

### Region: 402



CR1 vs CR2
Number of differences: 279

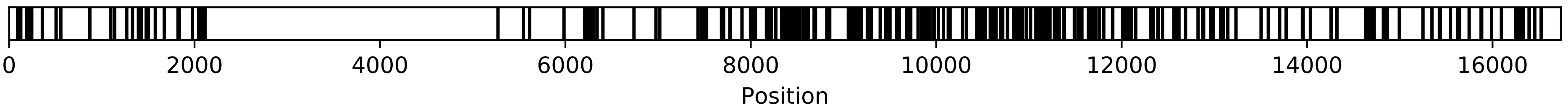


CR1 vs MA1
Number of differences: 277

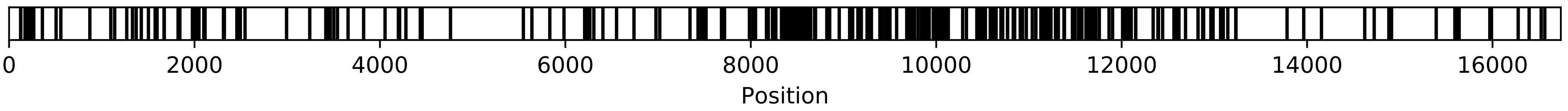


CR1 vs MA2
Number of differences: 274

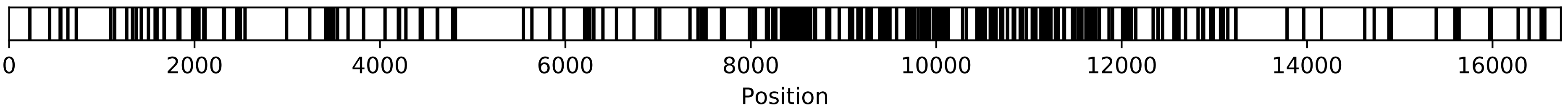


CR1 vs MM1
Number of differences: 324

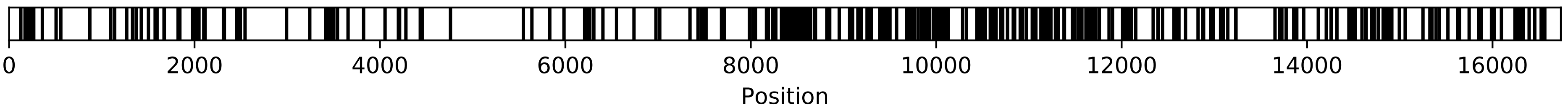


CR1 vs MM2
Number of differences: 344

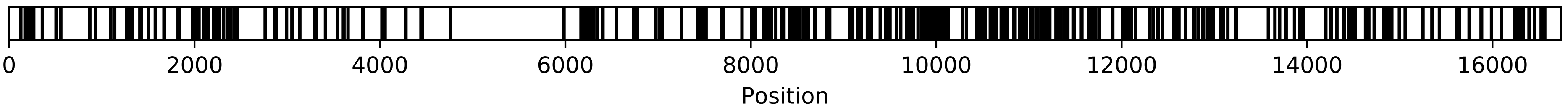


CR2 vs MA1
Number of differences: 201

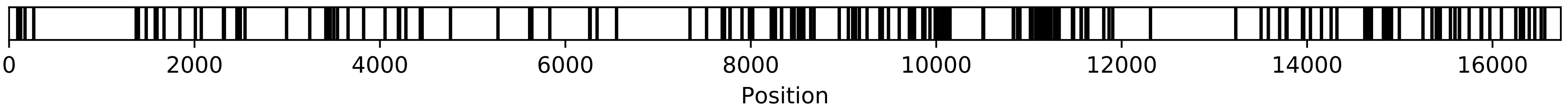


CR2 vs MA2
Number of differences: 212

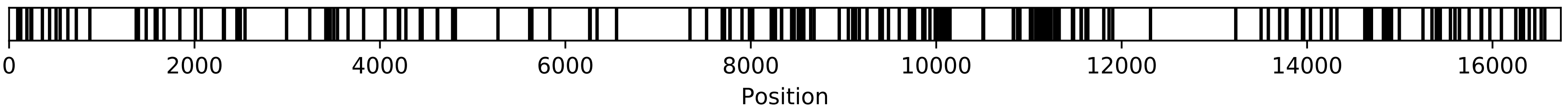


CR2 vs MM1
Number of differences: 194

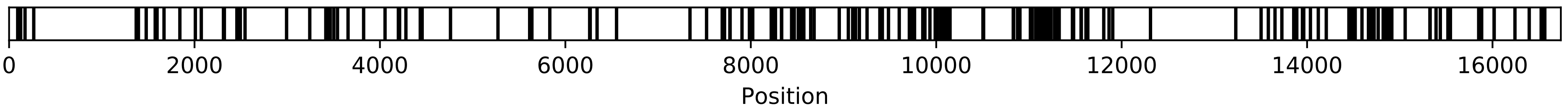


CR2 vs MM2
Number of differences: 186

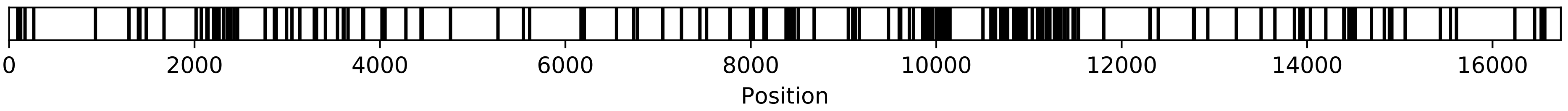


MA1 vs MA2
Number of differences: 19

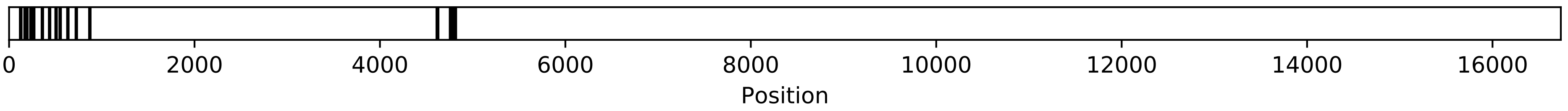


MA1 vs MM1
Number of differences: 66

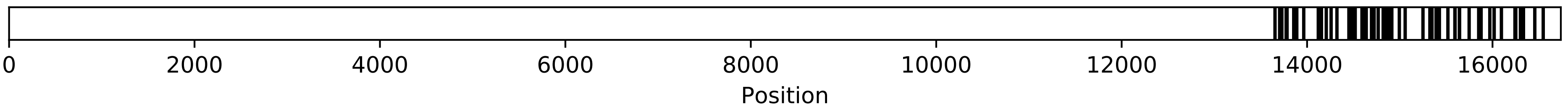


MA1 vs MM2
Number of differences: 229

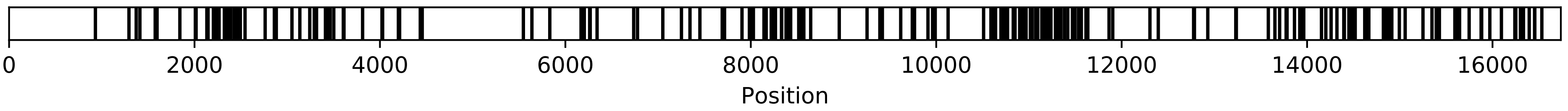


MA2 vs MM1
Number of differences: 85

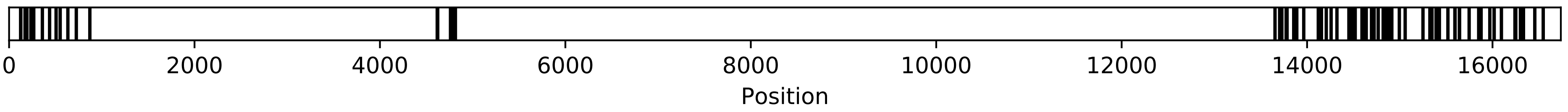


MA2 vs MM2
Number of differences: 248

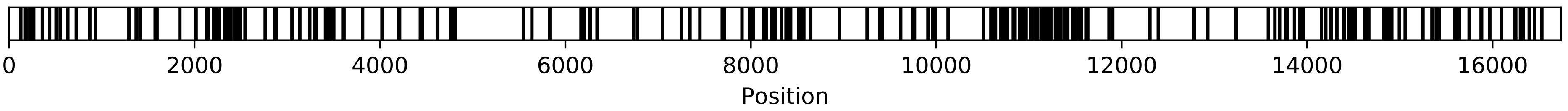


MM1 vs MM2
Number of differences: 206

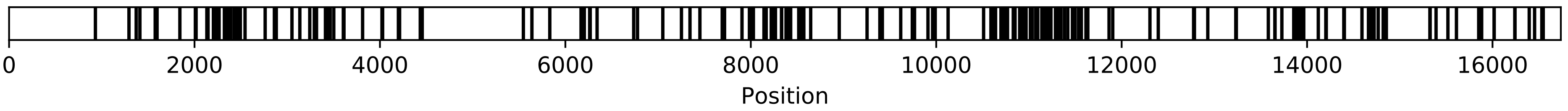

### Region: 404

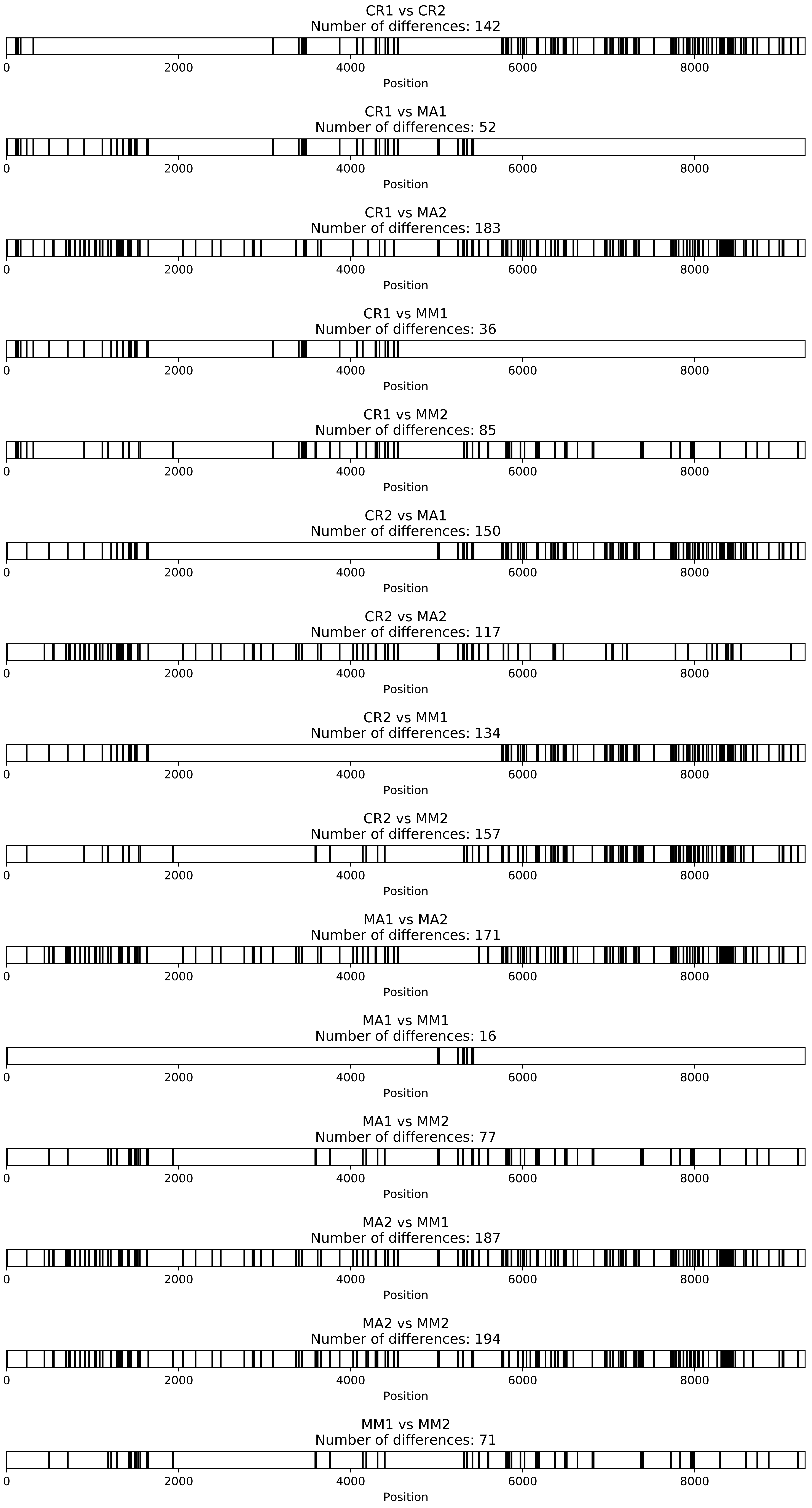
CR1 vs CR2
Number of differences: 142
CR1 vs MA1
Number of differences: 52
CR1 vs MA2
Number of differences: 183
CR1 vs MM1
Number of differences: 36
CR1 vs MM2
Number of differences: 85
CR2 vs MA1
Number of differences: 150
CR2 vs MA2
Number of differences: 117
CR2 vs MM1
Number of differences: 134
CR2 vs MM2
Number of differences: 157
MA1 vs MA2
Number of differences: 171
MA1 vs MM1
Number of differences: 16
MA1 vs MM2
Number of differences: 77
MA2 vs MM1
Number of differences: 187
MA2 vs MM2
Number of differences: 194
MM1 vs MM2
Number of differences: 71
0
2000
4000
6000
8000
Position

### Region: 407



CR1 vs CR2
Number of differences: 151

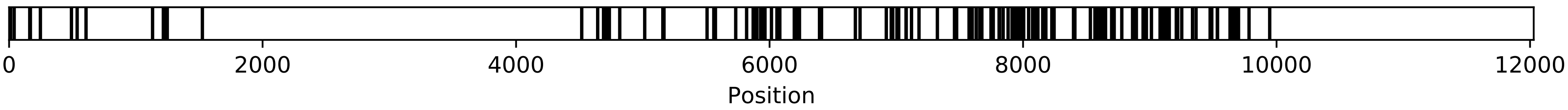


CR1 vs MA1
Number of differences: 284

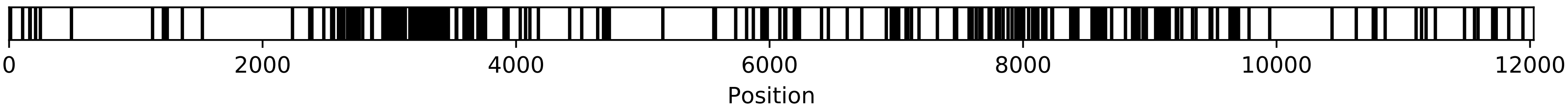


CR1 vs MA2
Number of differences: 126

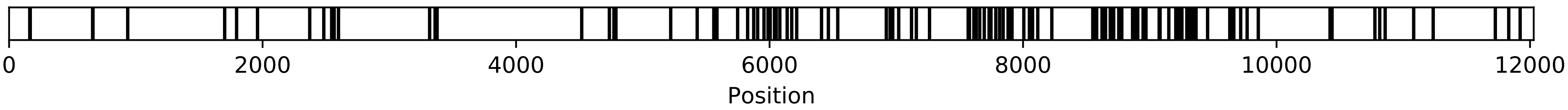


CR1 vs MM1
Number of differences: 121

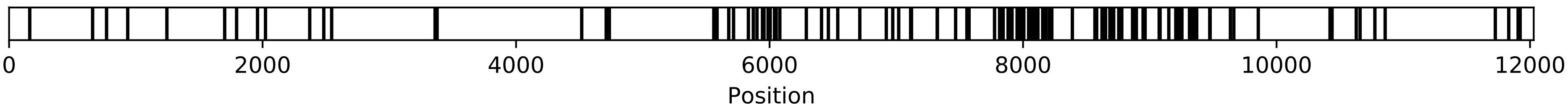


CR1 vs MM2
Number of differences: 126

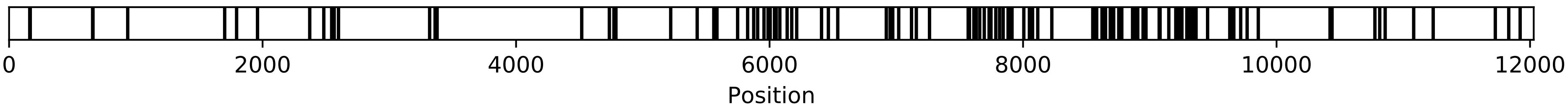


CR2 vs MA1
Number of differences: 199

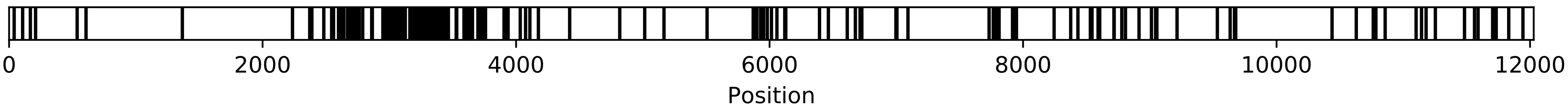


CR2 vs MA2
Number of differences: 187

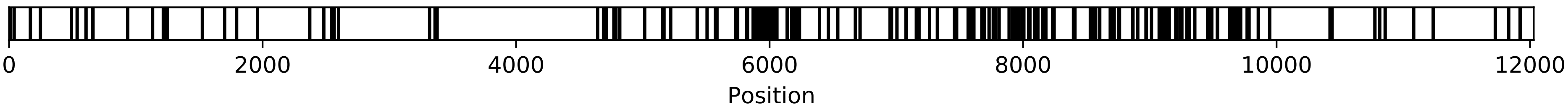


CR2 vs MM1
Number of differences: 170

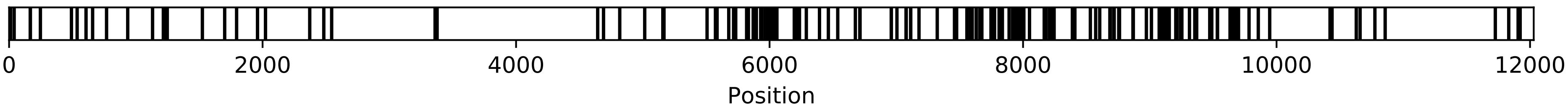


CR2 vs MM2
Number of differences: 187

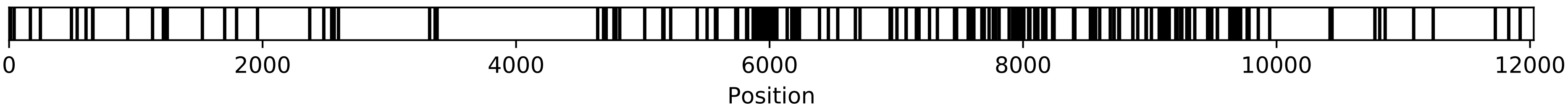


MA1 vs MA2
Number of differences: 296

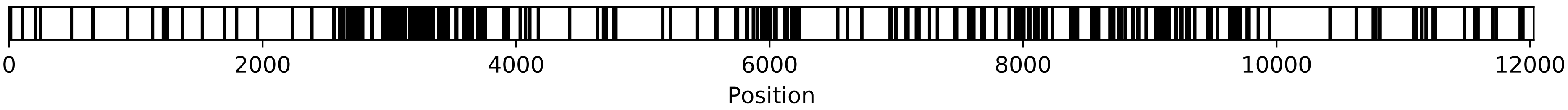


MA1 vs MM1
Number of differences: 284

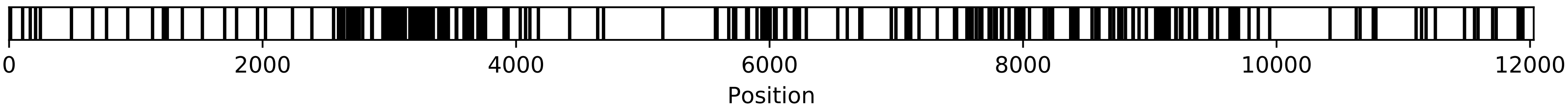


MA1 vs MM2
Number of differences: 296

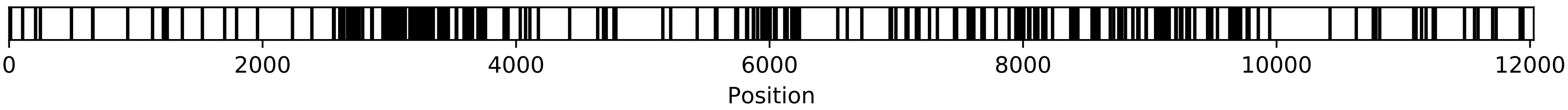


MA2 vs MM1
Number of differences: 92

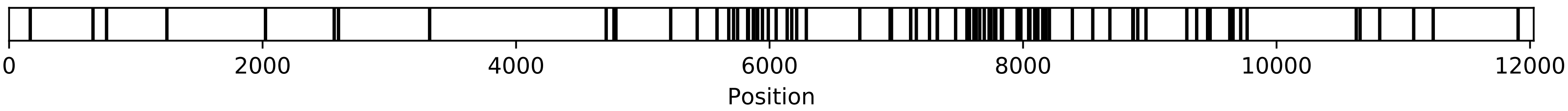


MA2 vs MM2
Number of differences: 0

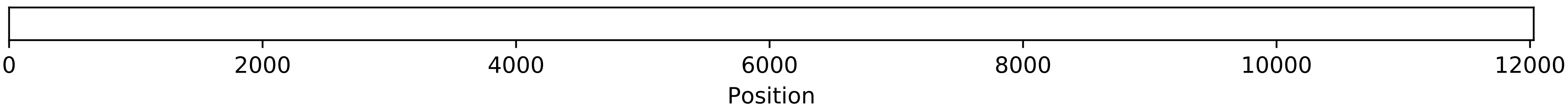


MM1 vs MM2
Number of differences: 92

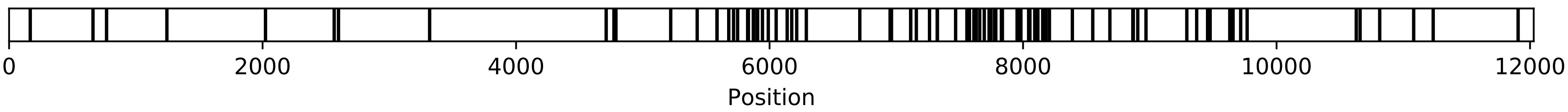

### Region: 408

CR1 vs CR2
Number of differences: 277

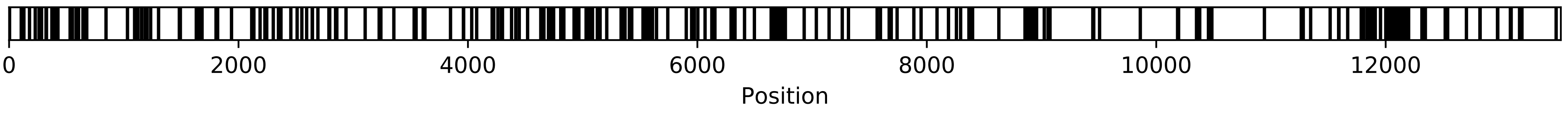


CR1 vs MA1
Number of differences: 56

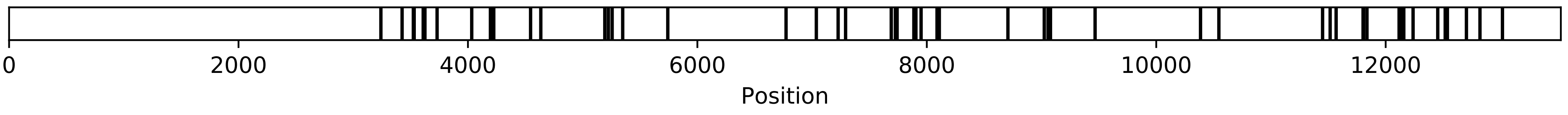


CR1 vs MA2
Number of differences: 251

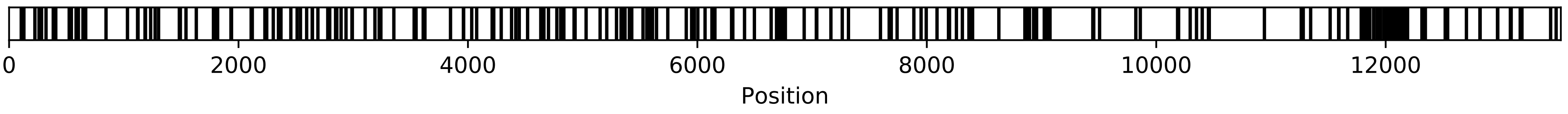


CR1 vs MM1
Number of differences: 251

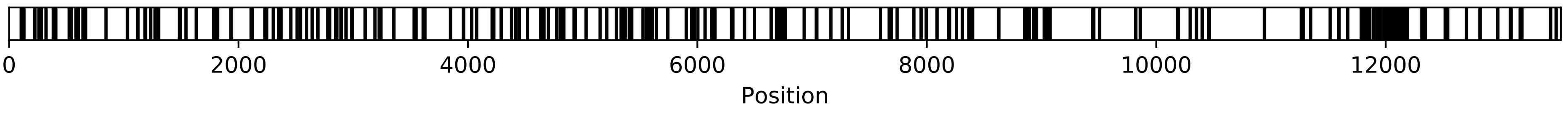


CR1 vs MM2
Number of differences: 238

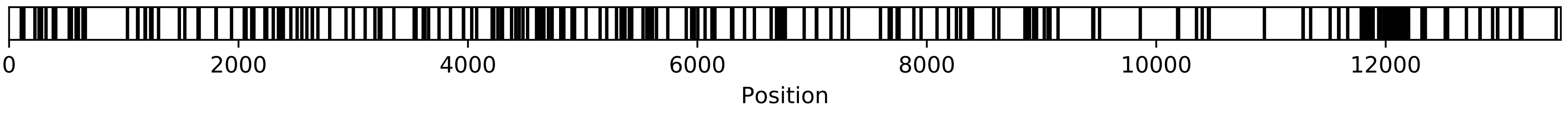


CR2 vs MA1
Number of differences: 278

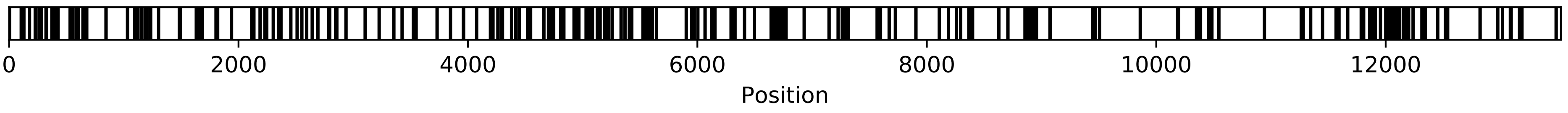


CR2 vs MA2
Number of differences: 127

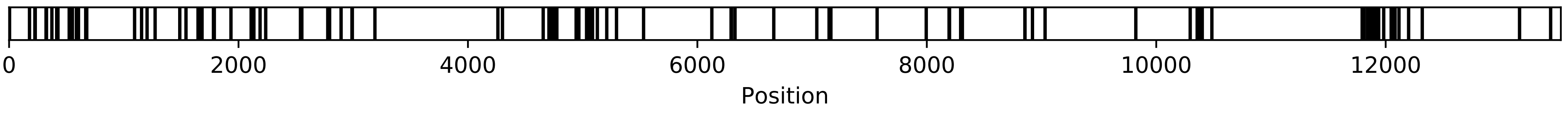


CR2 vs MM1
Number of differences: 127

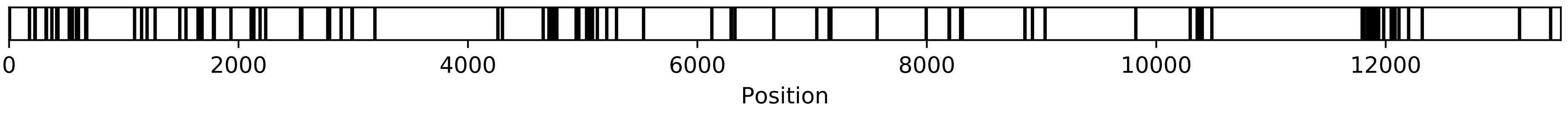


CR2 vs MM2
Number of differences: 127

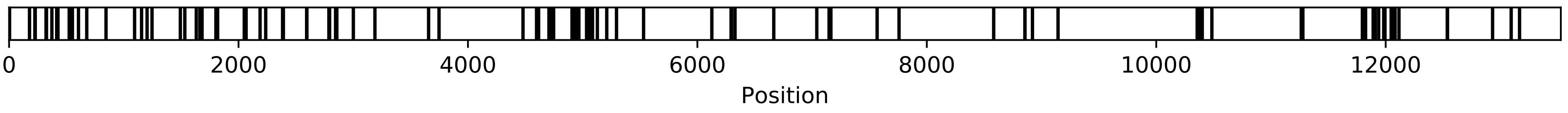


MA1 vs MA2
Number of differences: 251

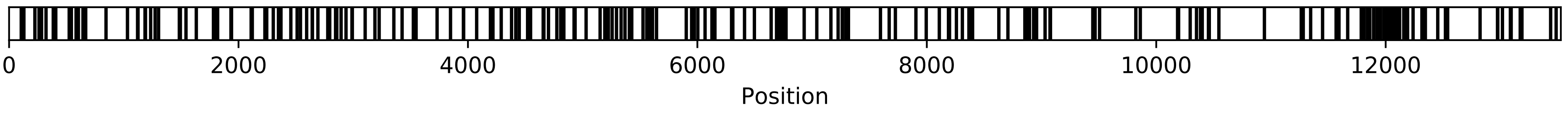


MA1 vs MM1
Number of differences: 251

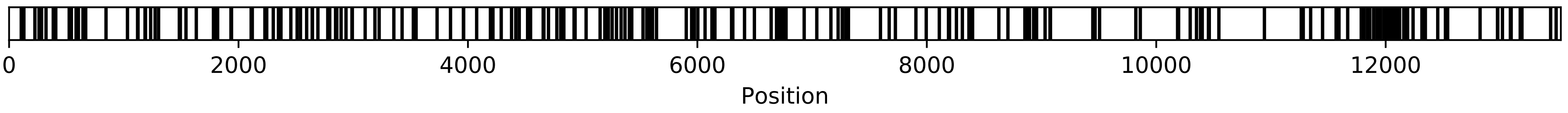


MA1 vs MM2
Number of differences: 239

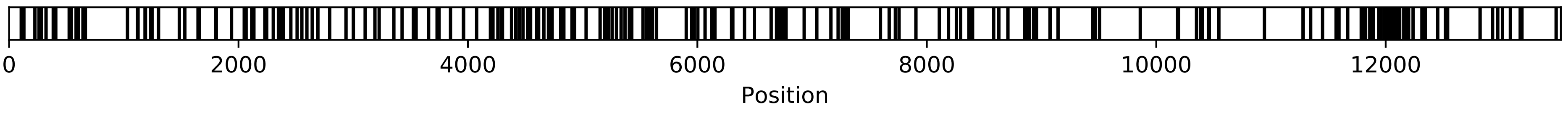


MA2 vs MM1
Number of differences: 0

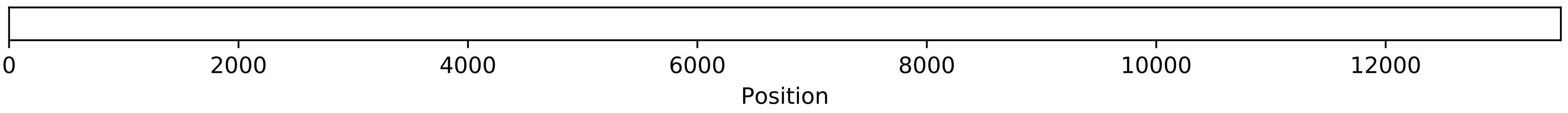


MA2 vs MM2
Number of differences: 95

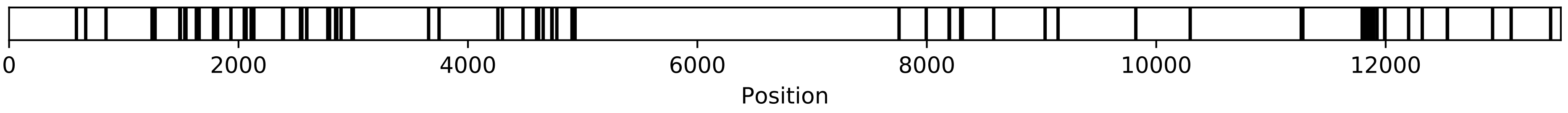


MM1 vs MM2
Number of differences: 95

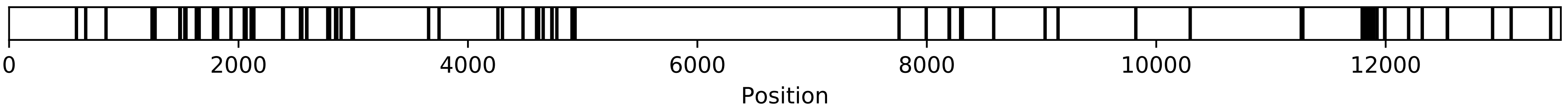

### Region: 409



CR1 vs CR2
Number of differences: 288

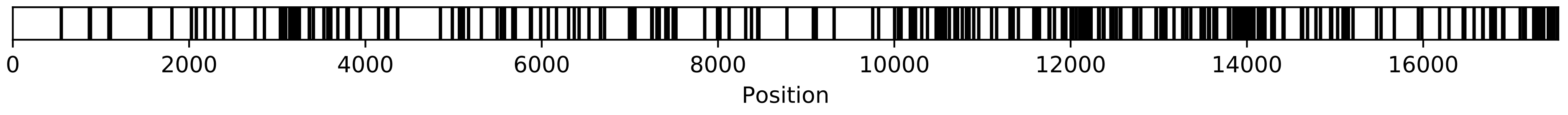


CR1 vs MA1
Number of differences: 278

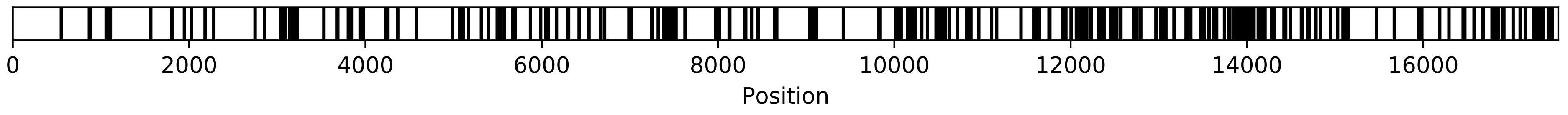


CR1 vs MA2
Number of differences: 94

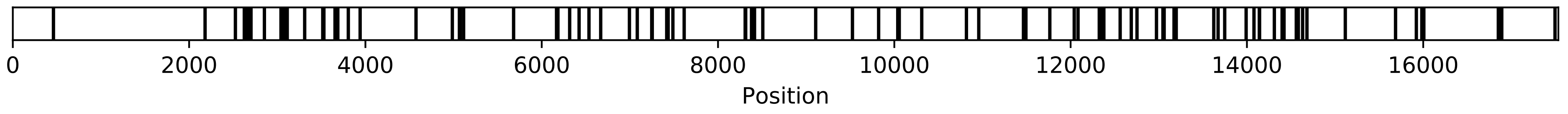


CR1 vs MM1
Number of differences: 294

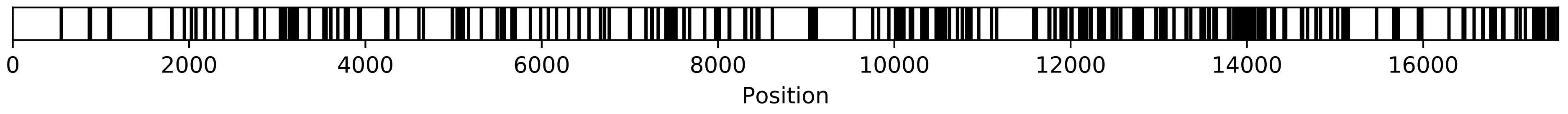


CR1 vs MM2
Number of differences: 293

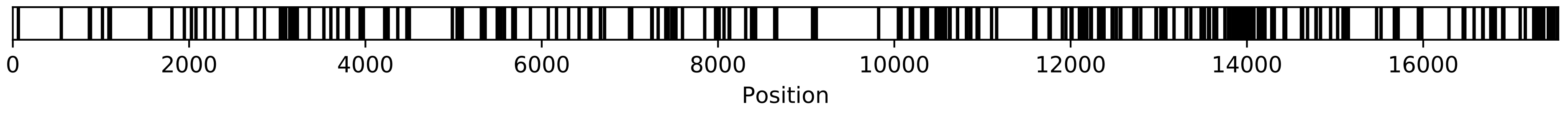


CR2 vs MA1
Number of differences: 128

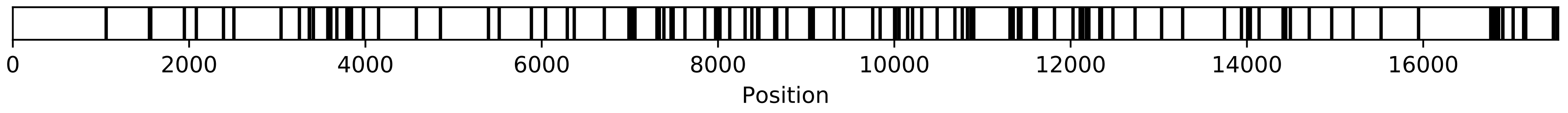


CR2 vs MA2
Number of differences: 294

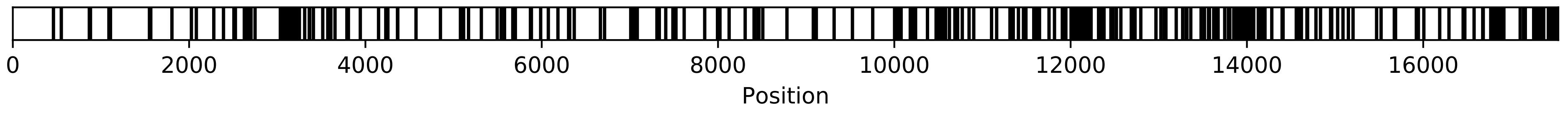


CR2 vs MM1
Number of differences: 124

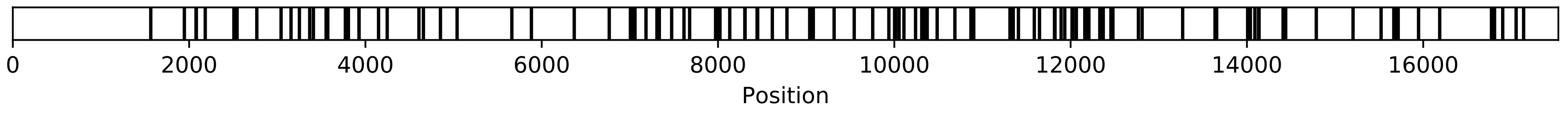


CR2 vs MM2
Number of differences: 130

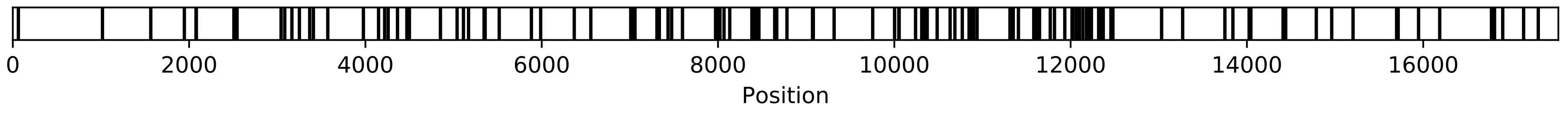


MA1 vs MA2
Number of differences: 282

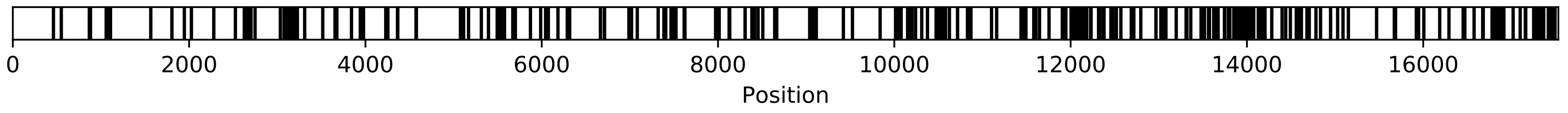


MA1 vs MM1
Number of differences: 116

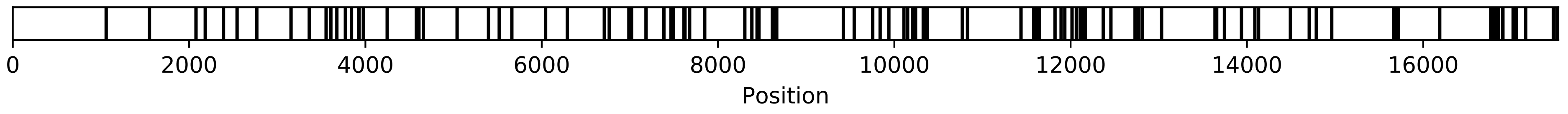


MA1 vs MM2
Number of differences: 131

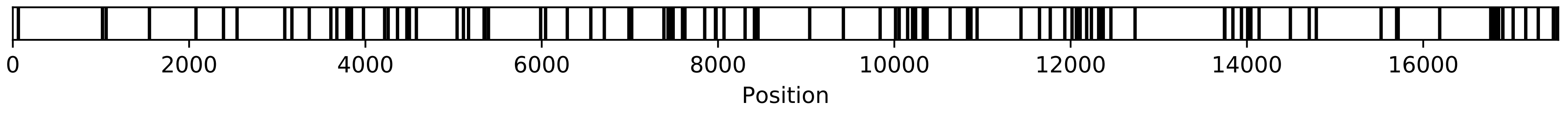


MA2 vs MM1
Number of differences: 299

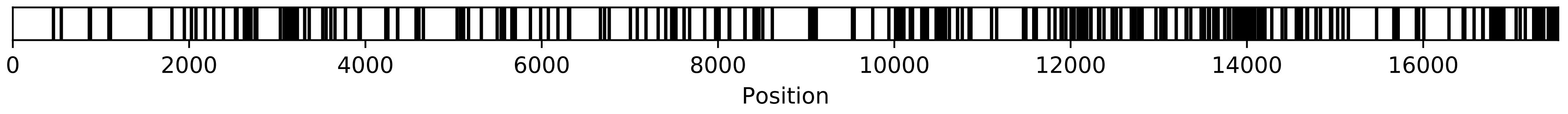


MA2 vs MM2
Number of differences: 298

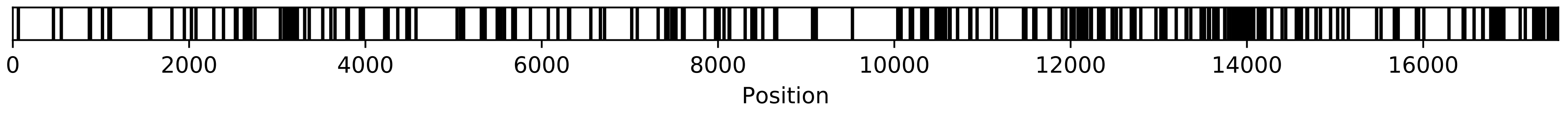


MM1 vs MM2
Number of differences: 116

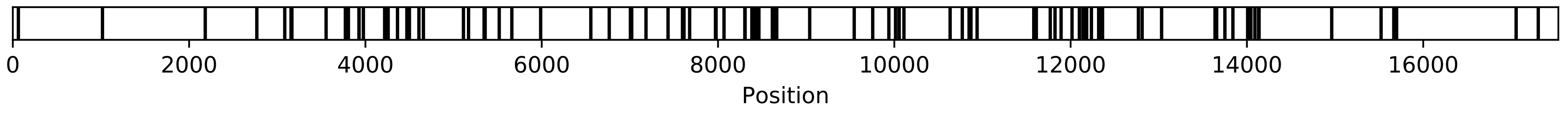

### Region: 410



CR1 vs CR2
Number of differences: 228

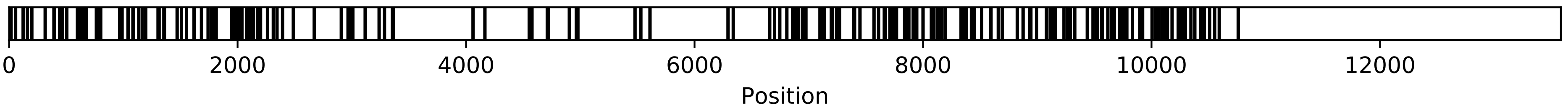


CR1 vs MA1
Number of differences: 377

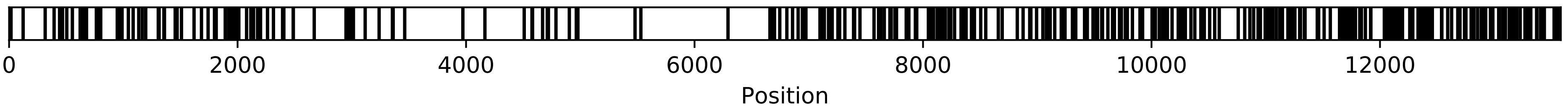


CR1 vs MA2
Number of differences: 77

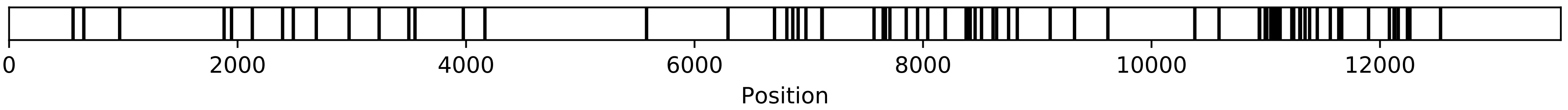


CR1 vs MM1
Number of differences: 350

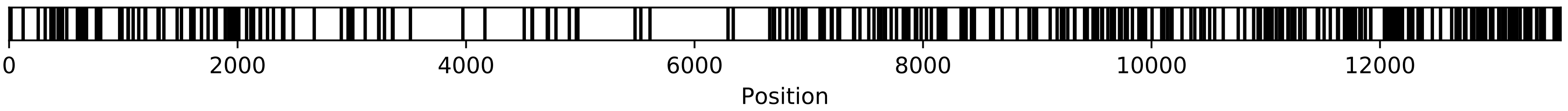


CR1 vs MM2
Number of differences: 368

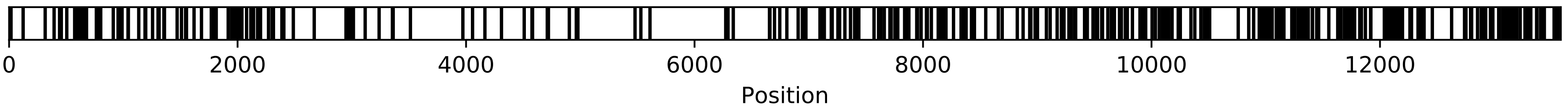


CR2 vs MA1
Number of differences: 257

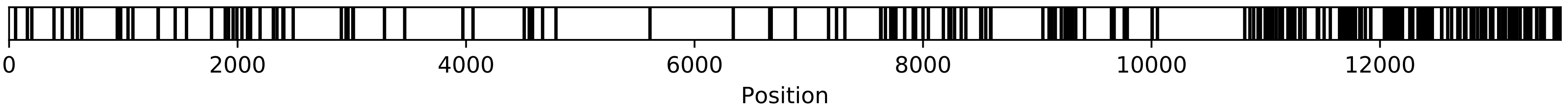


CR2 vs MA2
Number of differences: 251

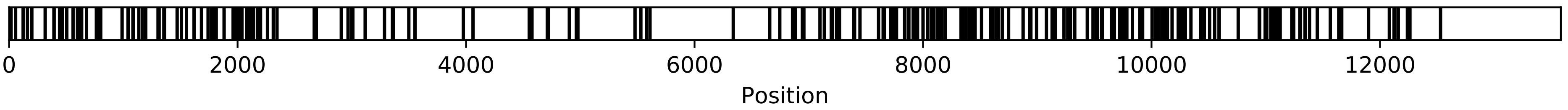


CR2 vs MM1
Number of differences: 256

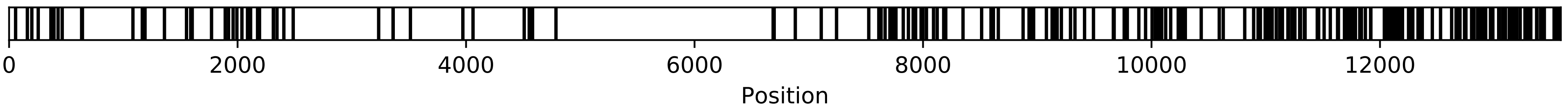


CR2 vs MM2
Number of differences: 259

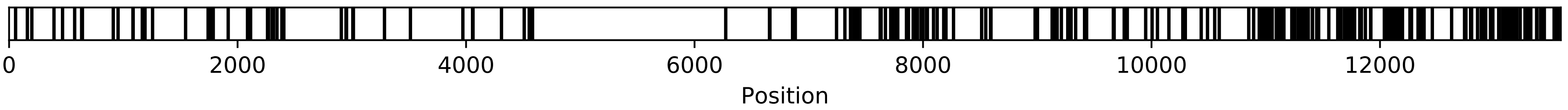


MA1 vs MA2
Number of differences: 368

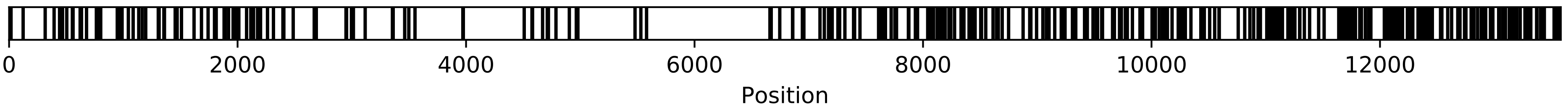


MA1 vs MM1
Number of differences: 165

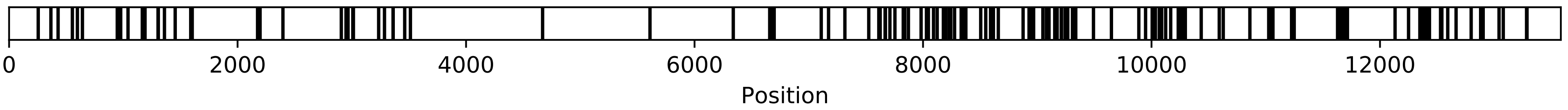


MA1 vs MM2
Number of differences: 196

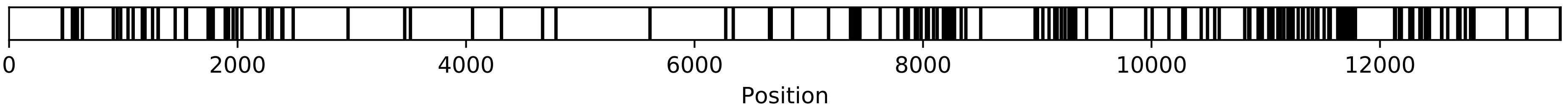


MA2 vs MM1
Number of differences: 339

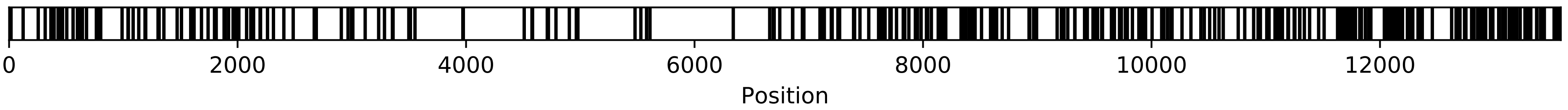


MA2 vs MM2
Number of differences: 361

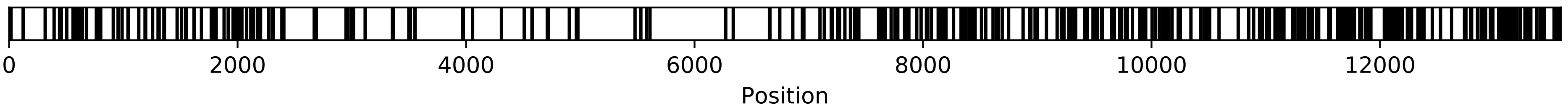


MM1 vs MM2
Number of differences: 185

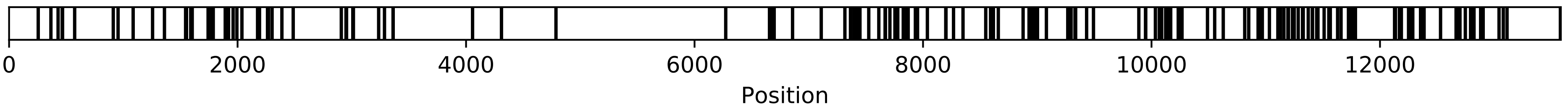

### Region: 413



CR1 vs CR2
Number of differences: 173

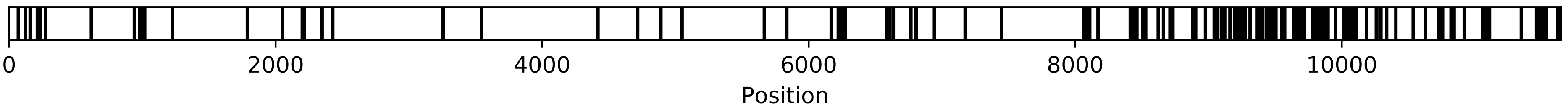


CR1 vs MA1
Number of differences: 167

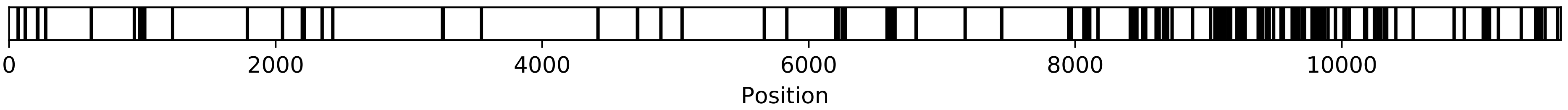


CR1 vs MA2
Number of differences: 75

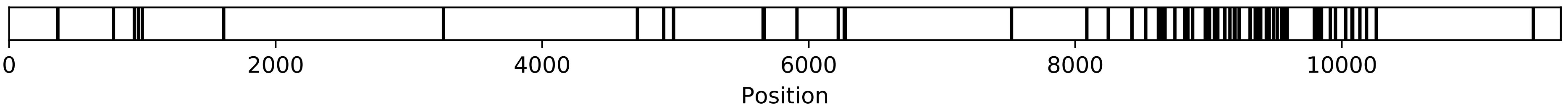


CR1 vs MM1
Number of differences: 176

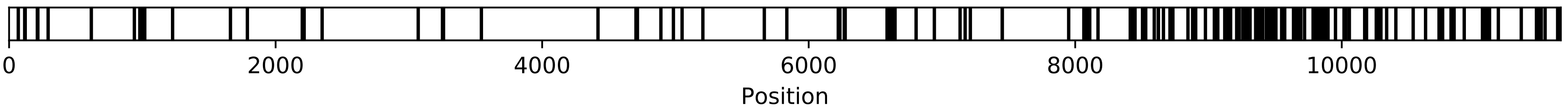


CR1 vs MM2
Number of differences: 182

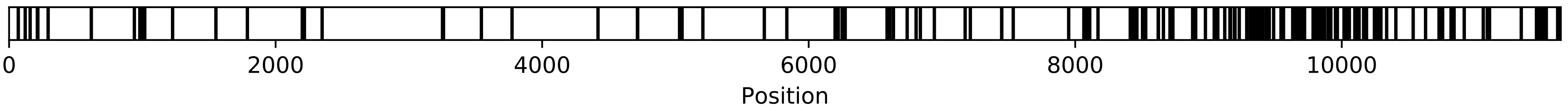


CR2 vs MA1
Number of differences: 90

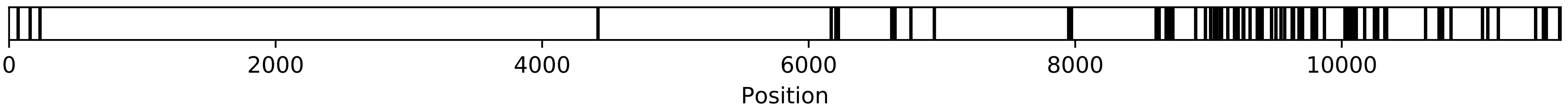


CR2 vs MA2
Number of differences: 178

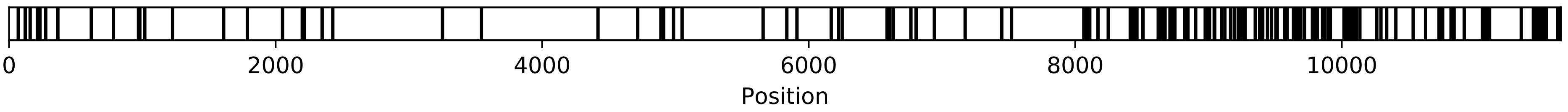


CR2 vs MM1
Number of differences: 64

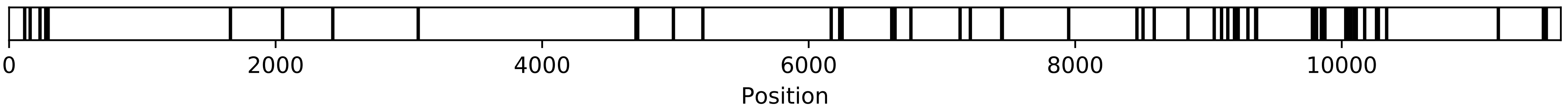


CR2 vs MM2
Number of differences: 84

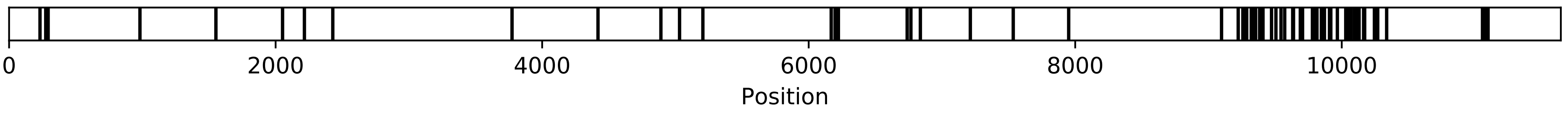


MA1 vs MA2
Number of differences: 179

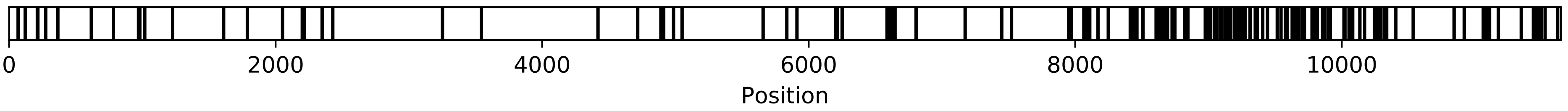


MA1 vs MM1
Number of differences: 92

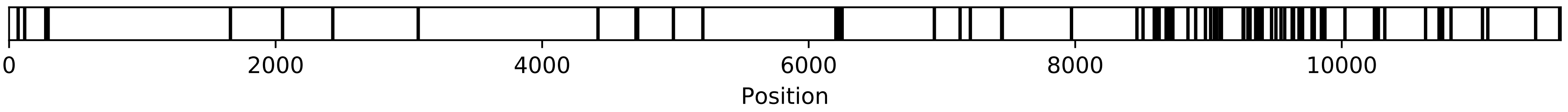


MA1 vs MM2
Number of differences: 111

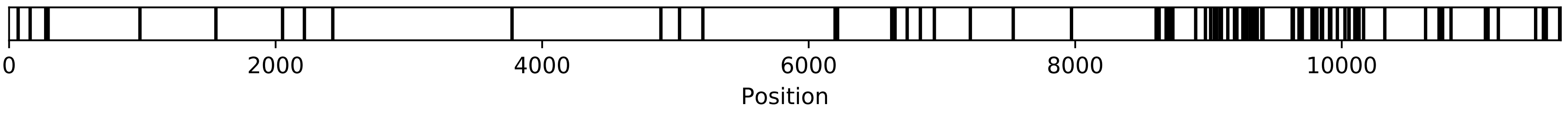


MA2 vs MM1
Number of differences: 181

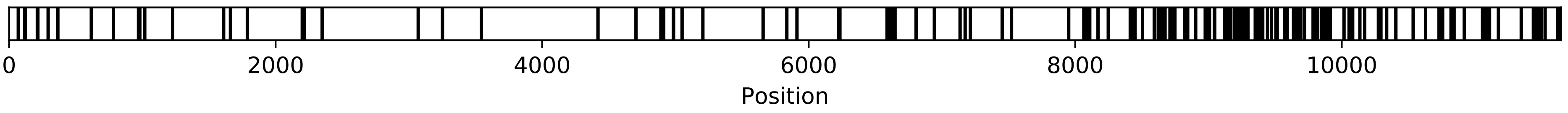


MA2 vs MM2
Number of differences: 187

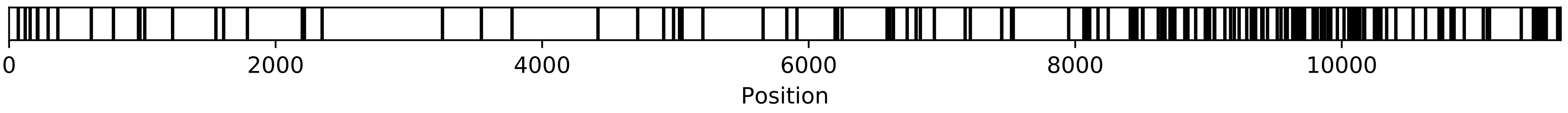


MM1 vs MM2
Number of differences: 93

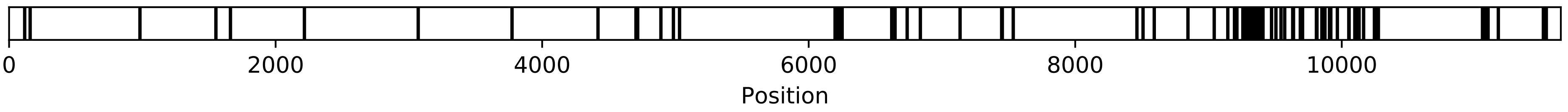

### Region: 414

CR1 vs CR2
Number of differences: 125

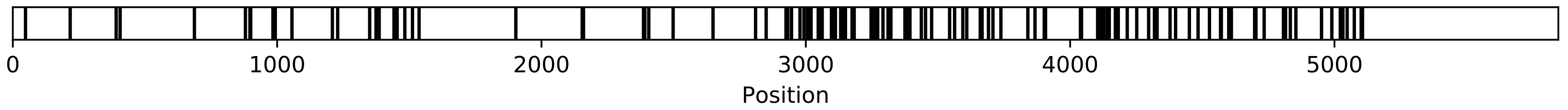


CR1 vs MA1
Number of differences: 54

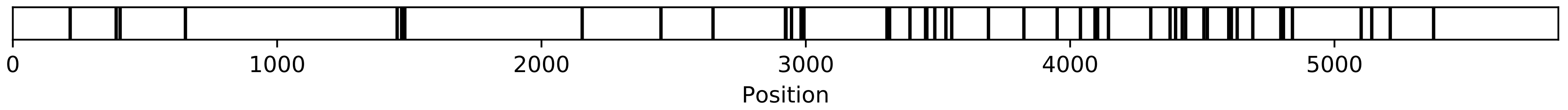


CR1 vs MA2
Number of differences: 128

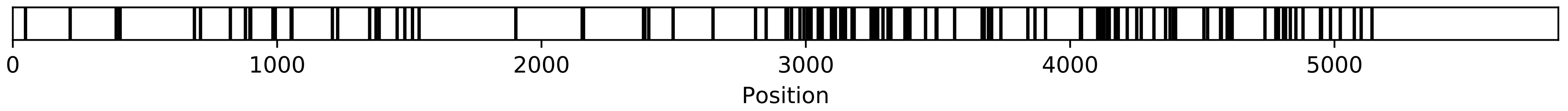


CR1 vs MM1
Number of differences: 141

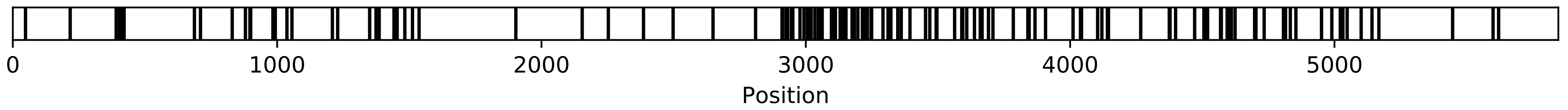


CR1 vs MM2
Number of differences: 128

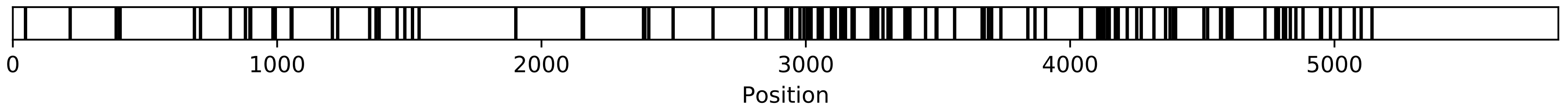


CR2 vs MA1
Number of differences: 135

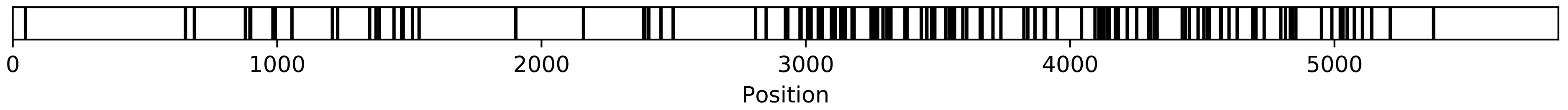


CR2 vs MA2
Number of differences: 52

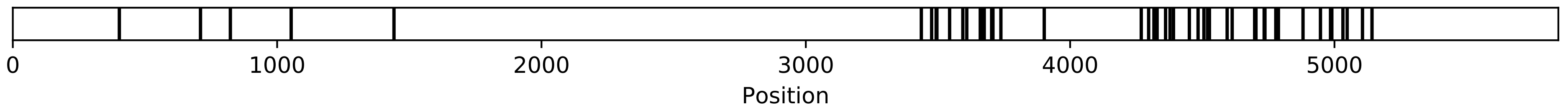


CR2 vs MM1
Number of differences: 79

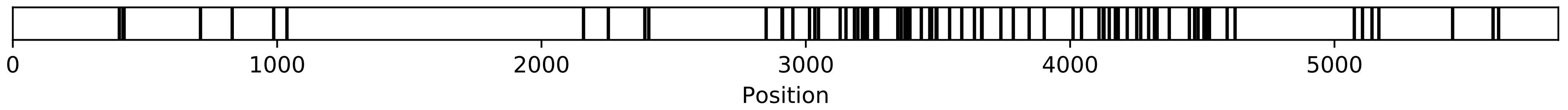


CR2 vs MM2
Number of differences: 52

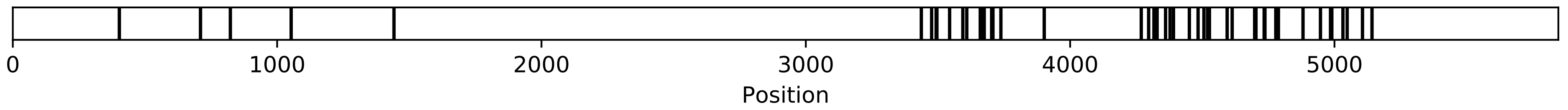


MA1 vs MA2
Number of differences: 137

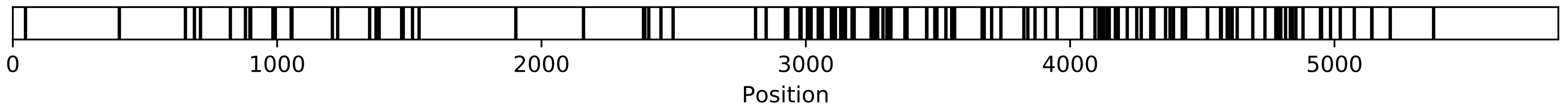


MA1 vs MM1
Number of differences: 150

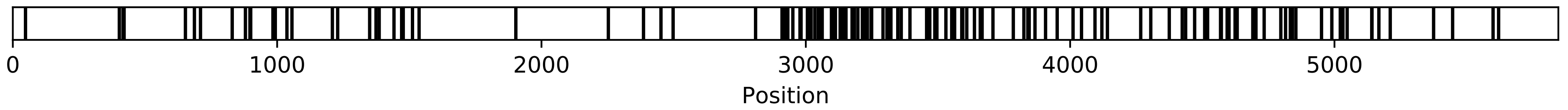


MA1 vs MM2
Number of differences: 137

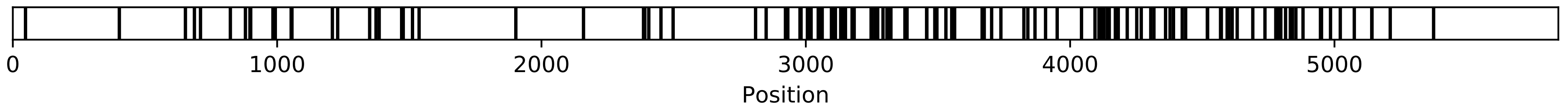


MA2 vs MM1
Number of differences: 86

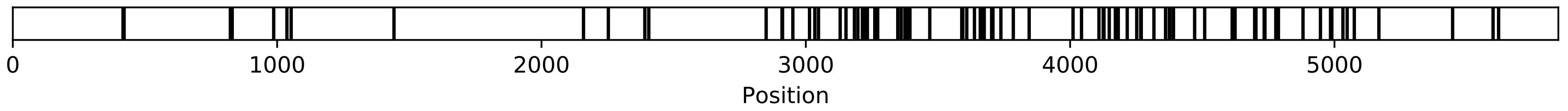


MA2 vs MM2
Number of differences: 0

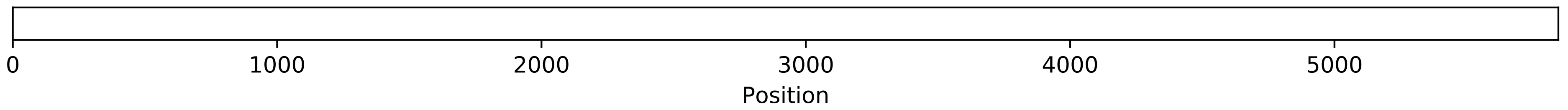


MM1 vs MM2
Number of differences: 86

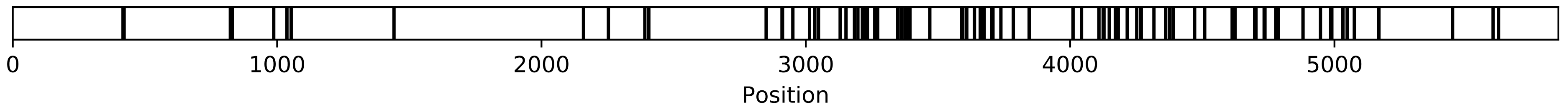

### Region: 417



CR1 vs CR2
Number of differences: 184

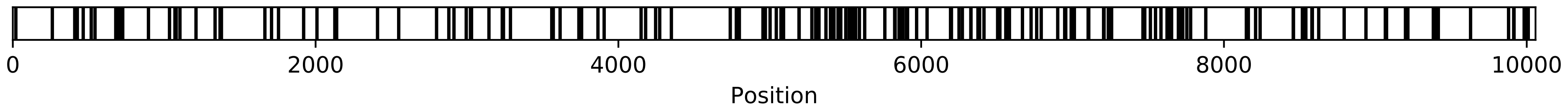


CR1 vs MA1
Number of differences: 173

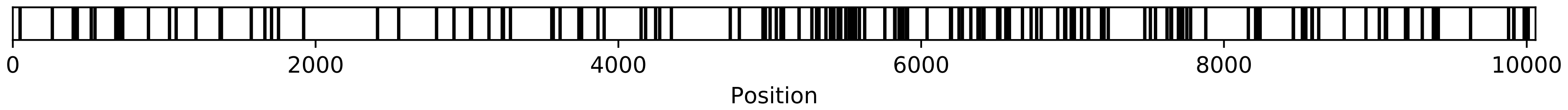


CR1 vs MA2
Number of differences: 51

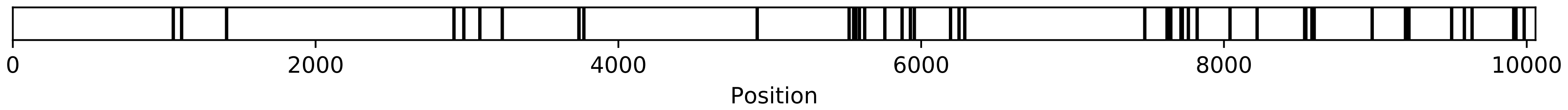


CR1 vs MM1
Number of differences: 66

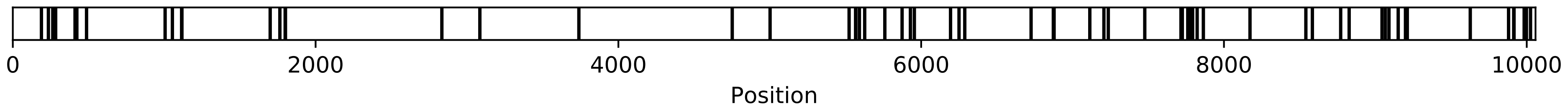


CR1 vs MM2
Number of differences: 51

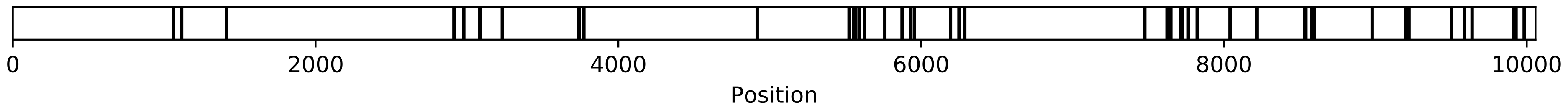


CR2 vs MA1
Number of differences: 31

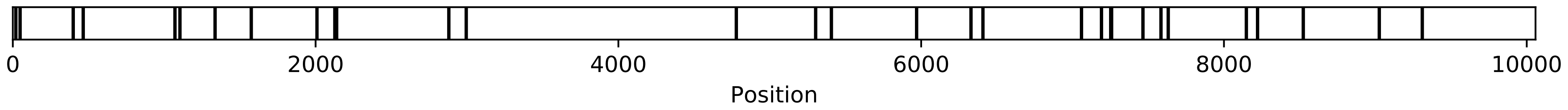


CR2 vs MA2
Number of differences: 190

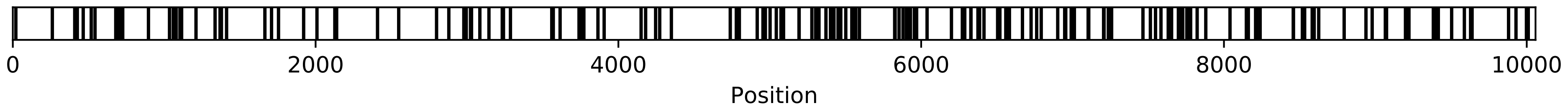


CR2 vs MM1
Number of differences: 197

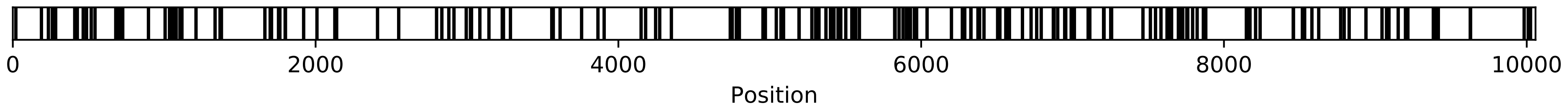


CR2 vs MM2
Number of differences: 190

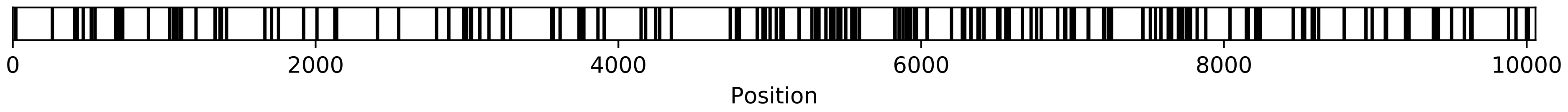


MA1 vs MA2
Number of differences: 179

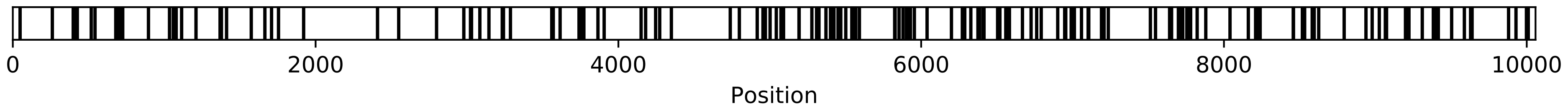


MA1 vs MM1
Number of differences: 186

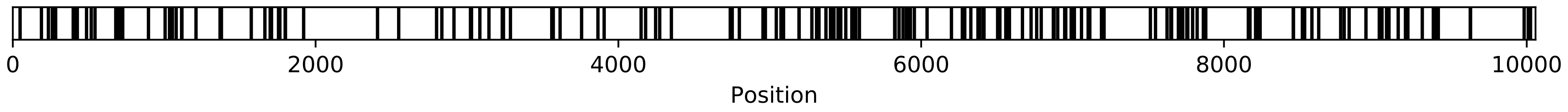


MA1 vs MM2
Number of differences: 179

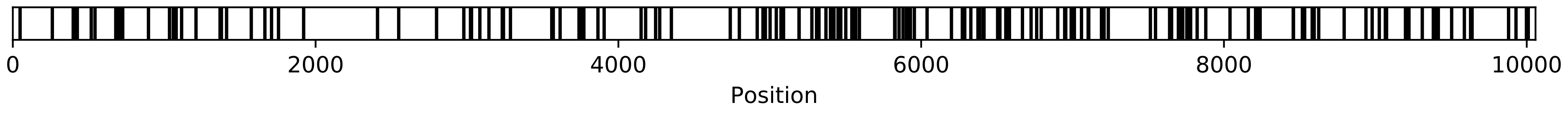


MA2 vs MM1
Number of differences: 67

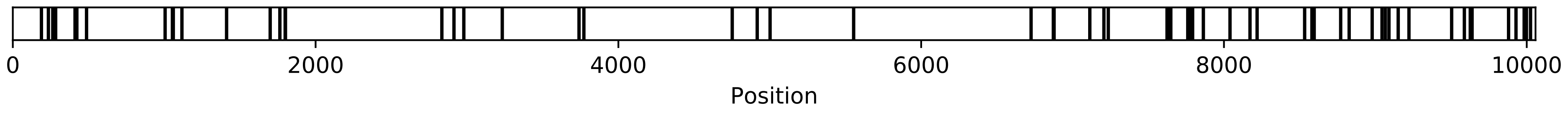


MA2 vs MM2
Number of differences: 0

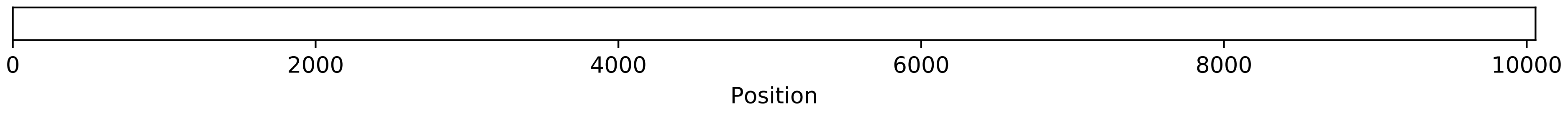


MM1 vs MM2
Number of differences: 67

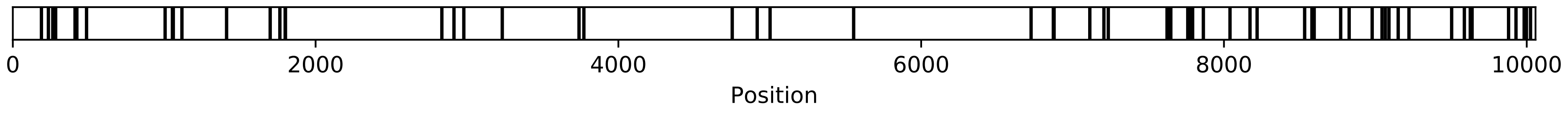

### Region: 418

CR1 vs CR2
Number of differences: 220

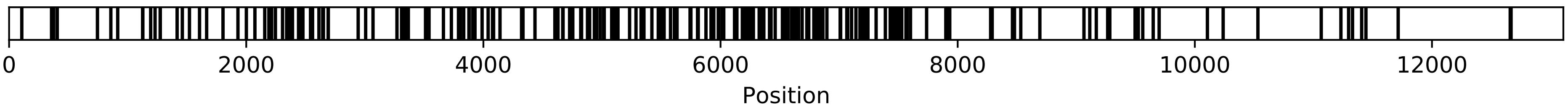


CR1 vs MA1
Number of differences: 105

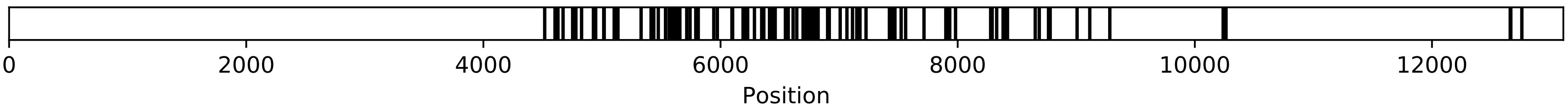


CR1 vs MA2
Number of differences: 211

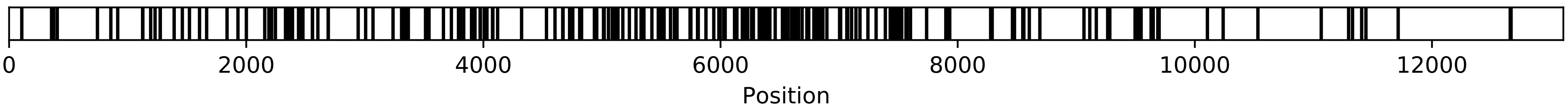


CR1 vs MM1
Number of differences: 135

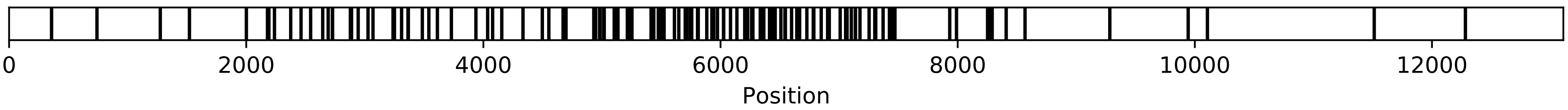


CR1 vs MM2
Number of differences: 127

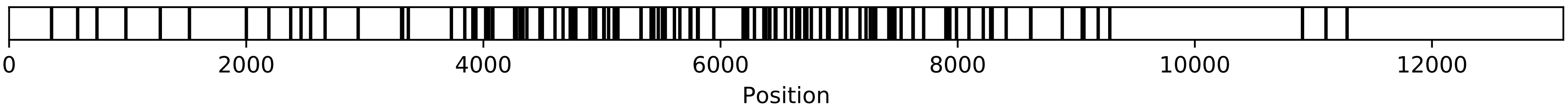


CR2 vs MA1
Number of differences: 231

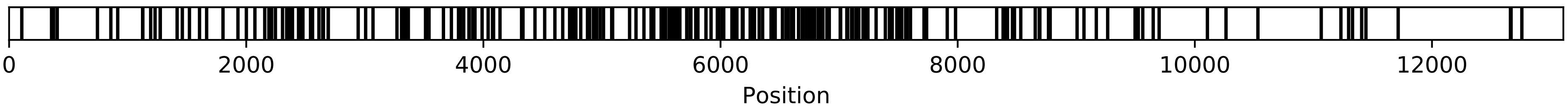


CR2 vs MA2
Number of differences: 70

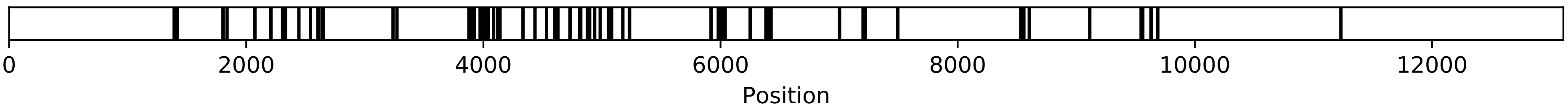


CR2 vs MM1
Number of differences: 233

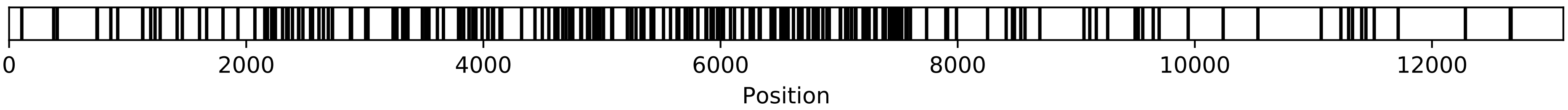


CR2 vs MM2
Number of differences: 237

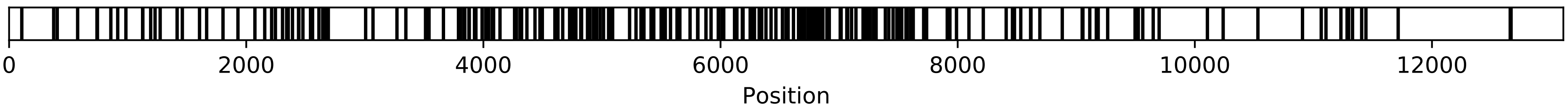


MA1 vs MA2
Number of differences: 225

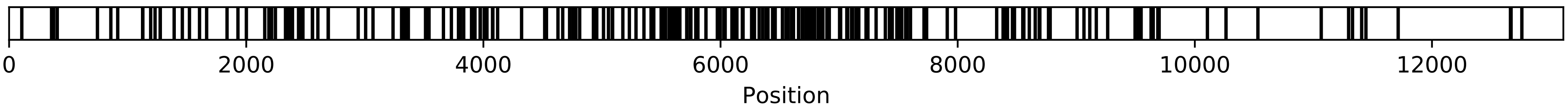


MA1 vs MM1
Number of differences: 153

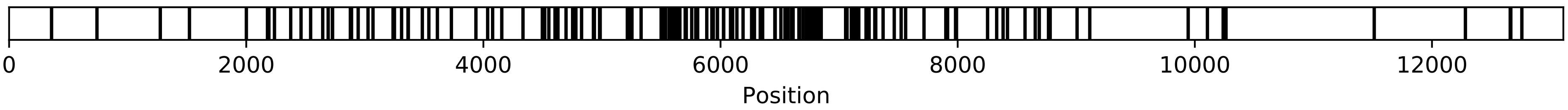


MA1 vs MM2
Number of differences: 128

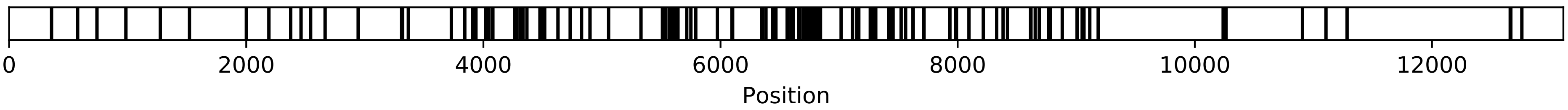


MA2 vs MM1
Number of differences: 229

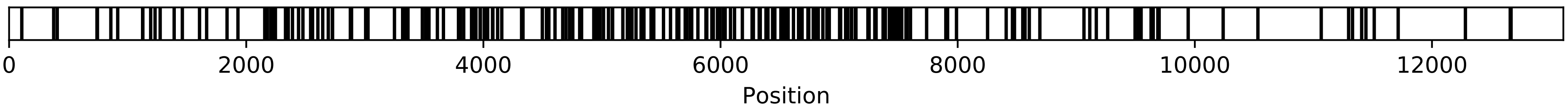


MA2 vs MM2
Number of differences: 226

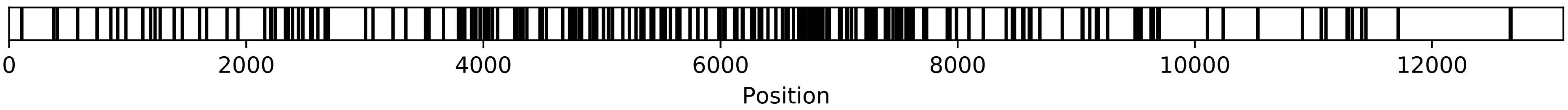


MM1 vs MM2
Number of differences: 145

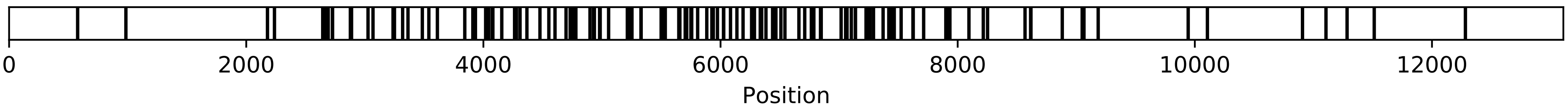

### Region: 422

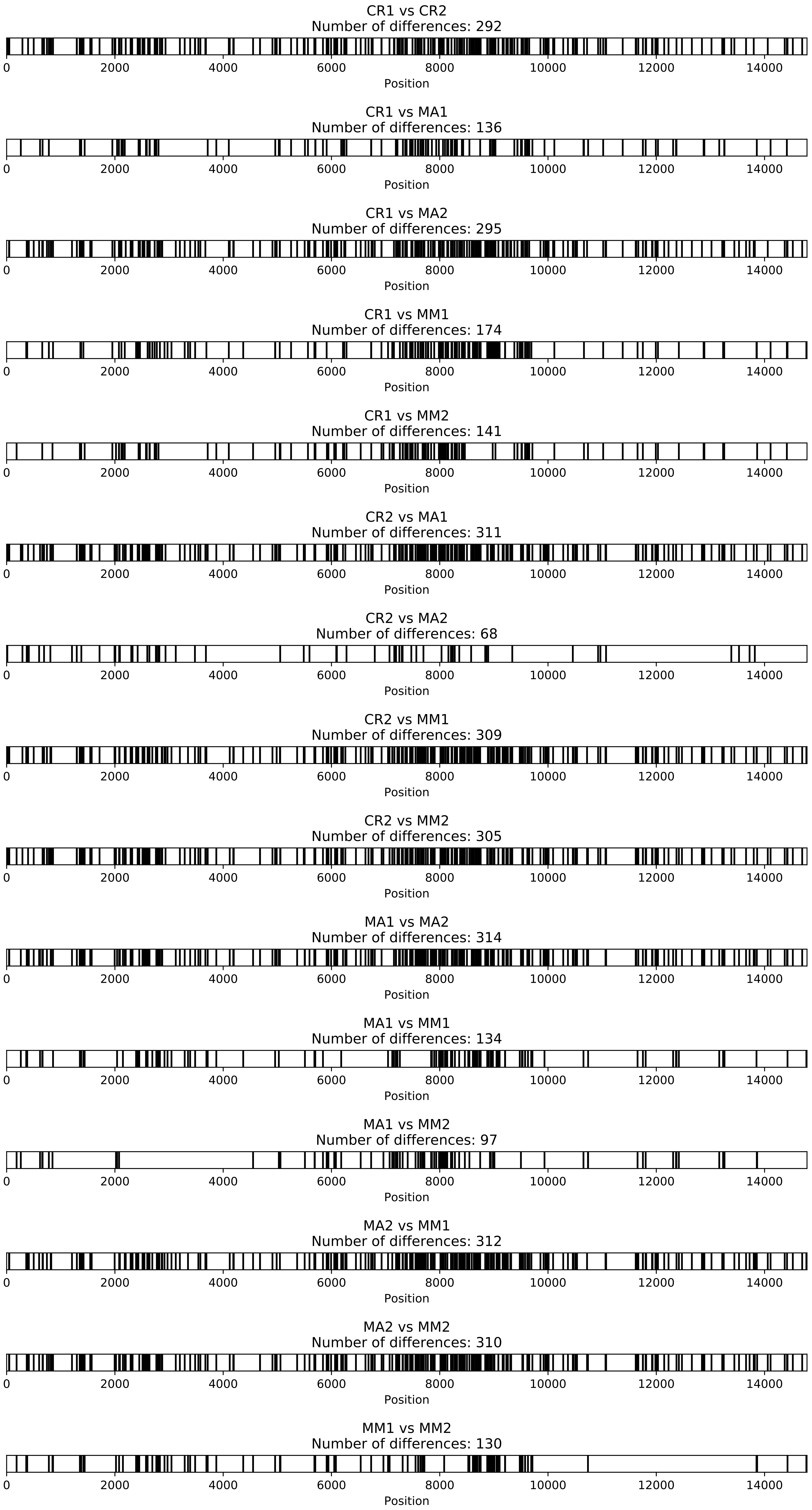
CR1 vs CR2
Number of differences: 292
CR1 vs MA1
Number of differences: 136
CR1 vs MA2
Number of differences: 295
CR1 vs MM1
Number of differences: 174
CR1 vs MM2
Number of differences: 141
CR2 vs MA1
Number of differences: 311
CR2 vs MA2
Number of differences: 68
CR2 vs MM1
Number of differences: 309
CR2 vs MM2
Number of differences: 305
MA1 vs MA2
Number of differences: 314
MA1 vs MM1
Number of differences: 134
MA1 vs MM2
Number of differences: 97
MA2 vs MM1
Number of differences: 312
MA2 vs MM2
Number of differences: 310
MM1 vs MM2
Number of differences: 130
0
2000
4000
6000
8000
10000
12000
14000
Position

### Region: 424

CR1 vs CR2
Number of differences: 295

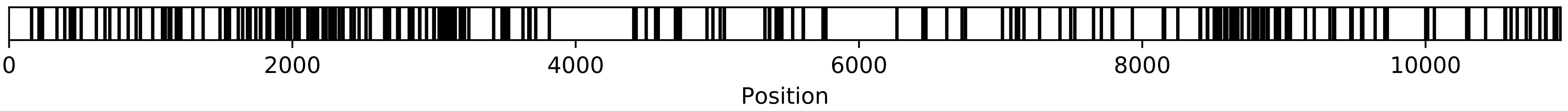


CR1 vs MA1
Number of differences: 189

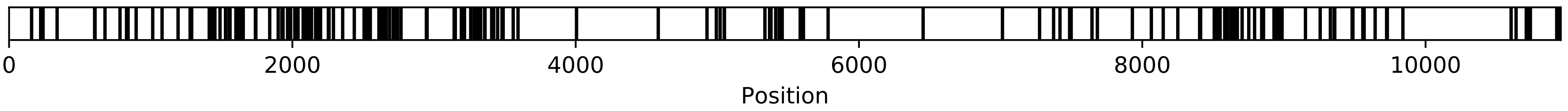


CR1 vs MA2
Number of differences: 253

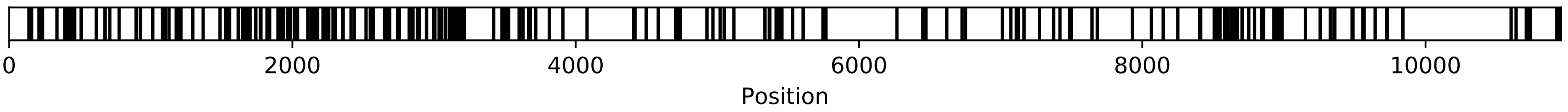


CR1 vs MM1
Number of differences: 145

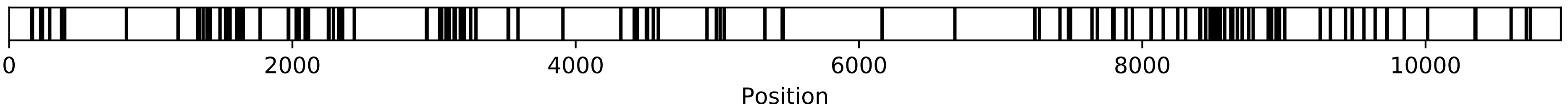


CR1 vs MM2
Number of differences: 139

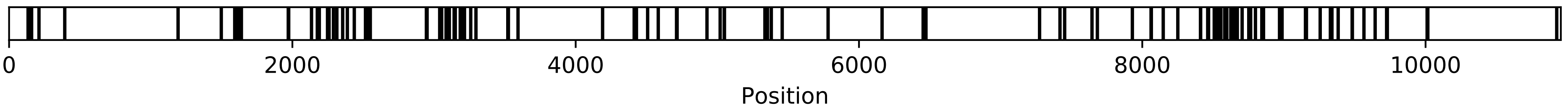


CR2 vs MA1
Number of differences: 316

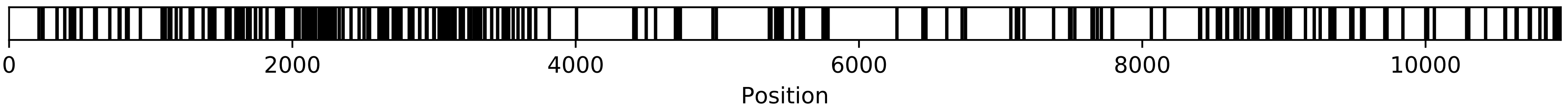


CR2 vs MA2
Number of differences: 159

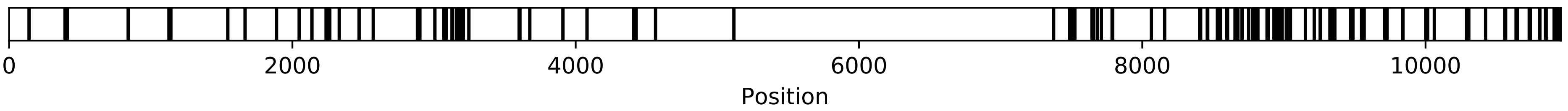


CR2 vs MM1
Number of differences: 313

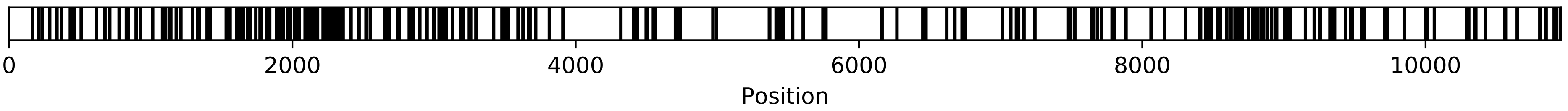


CR2 vs MM2
Number of differences: 298

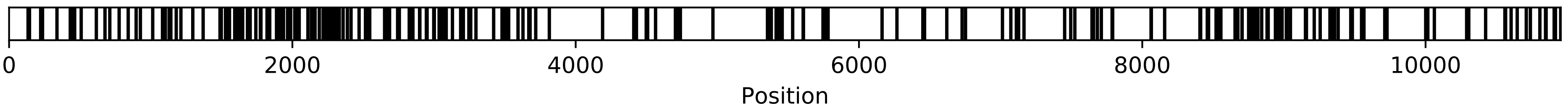


MA1 vs MA2
Number of differences: 211

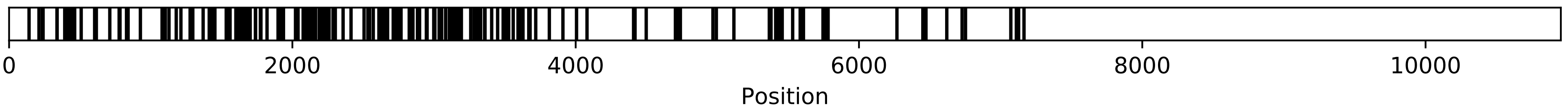


MA1 vs MM1
Number of differences: 174

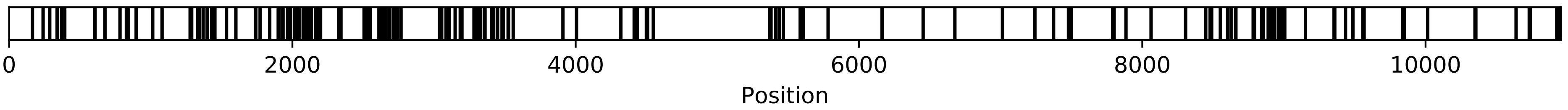


MA1 vs MM2
Number of differences: 176

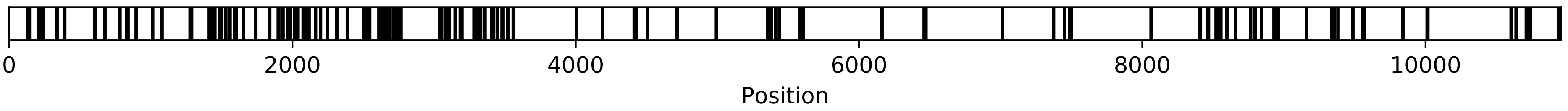


MA2 vs MM1
Number of differences: 257

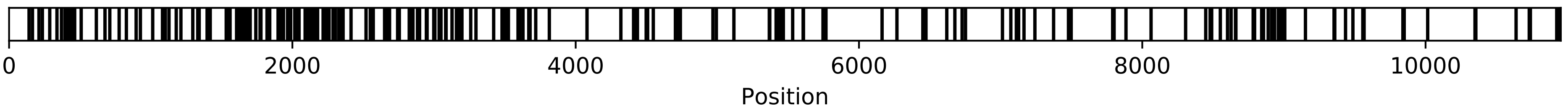


MA2 vs MM2
Number of differences: 245

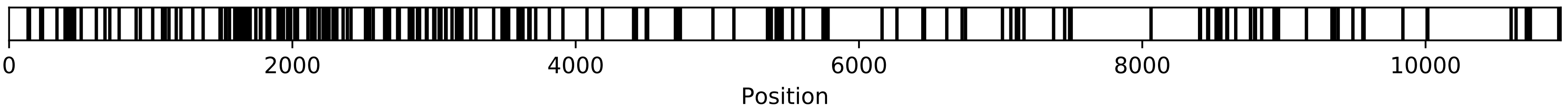


MM1 vs MM2
Number of differences: 131

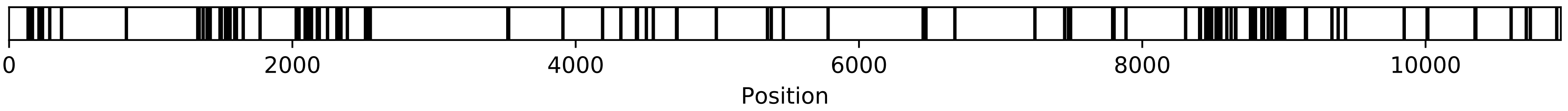

### Region: 427



CR1 vs CR2
Number of differences: 296

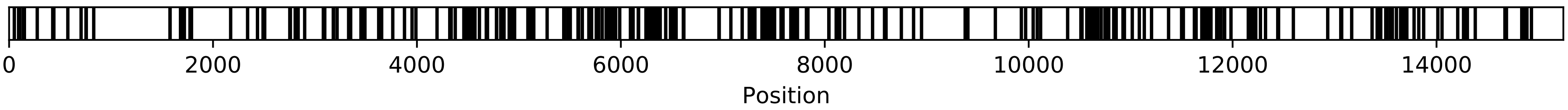


CR1 vs MA1
Number of differences: 152

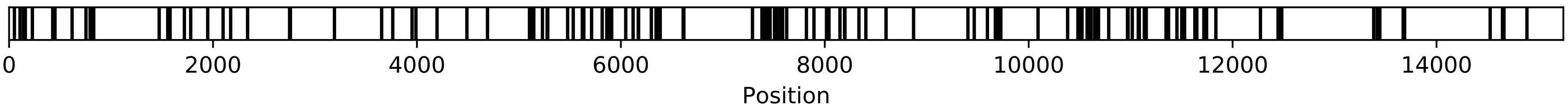


CR1 vs MA2
Number of differences: 286

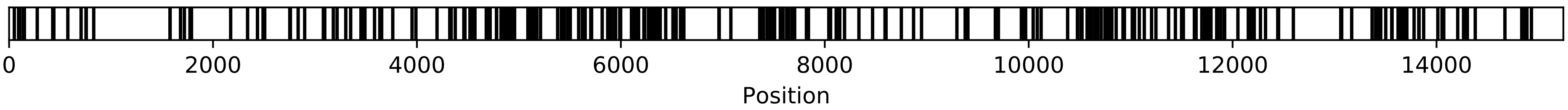


CR1 vs MM1
Number of differences: 150

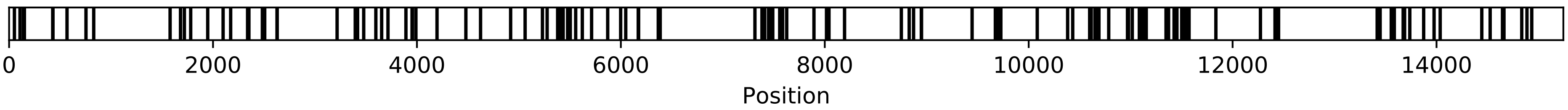


CR1 vs MM2
Number of differences: 150

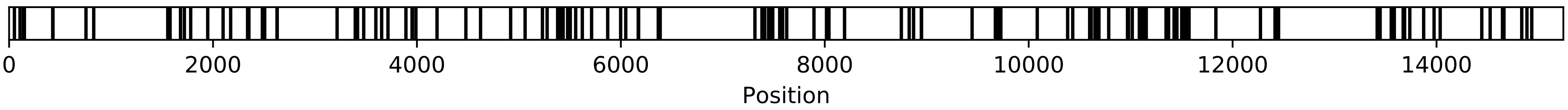


CR2 vs MA1
Number of differences: 281

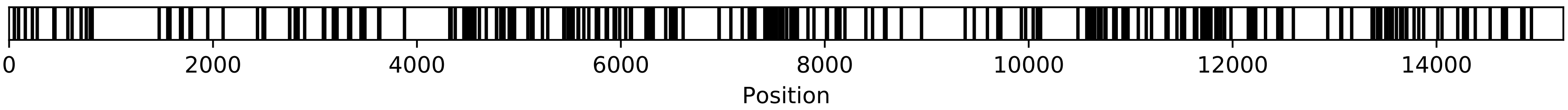


CR2 vs MA2
Number of differences: 114

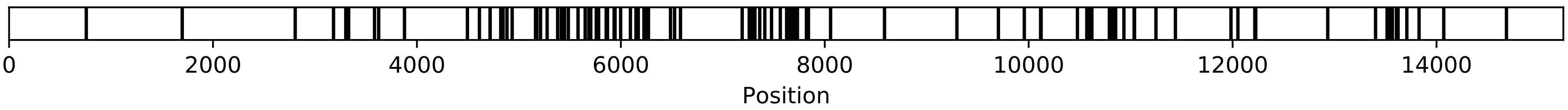


CR2 vs MM1
Number of differences: 293

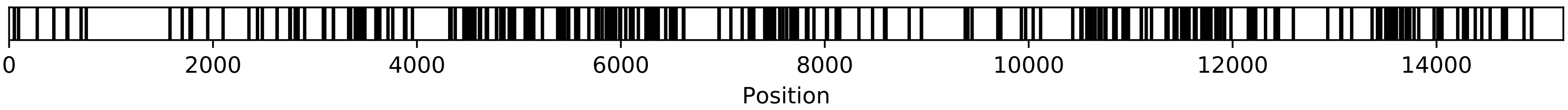


CR2 vs MM2
Number of differences: 293

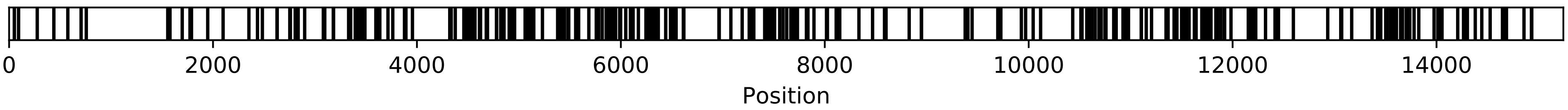


MA1 vs MA2
Number of differences: 278

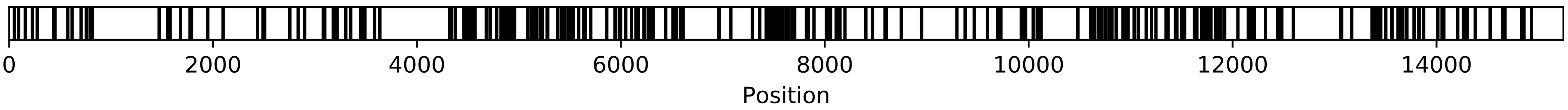


MA1 vs MM1
Number of differences: 132

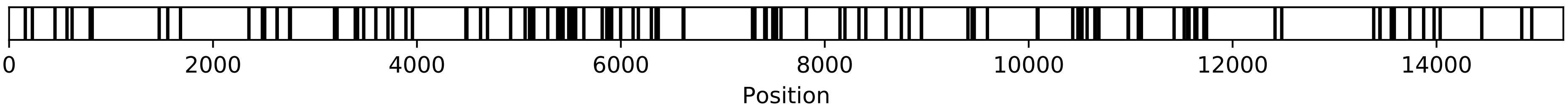


MA1 vs MM2
Number of differences: 130

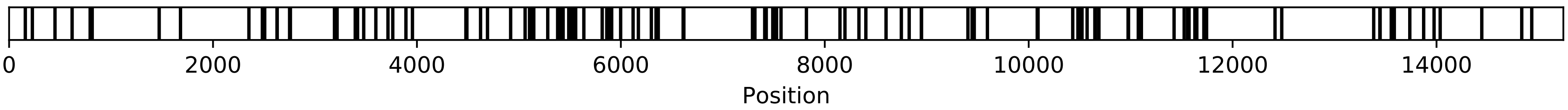


MA2 vs MM1
Number of differences: 287

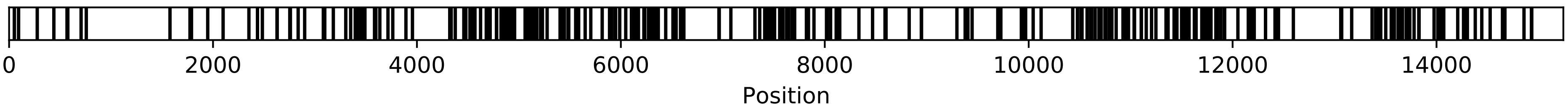


MA2 vs MM2
Number of differences: 287

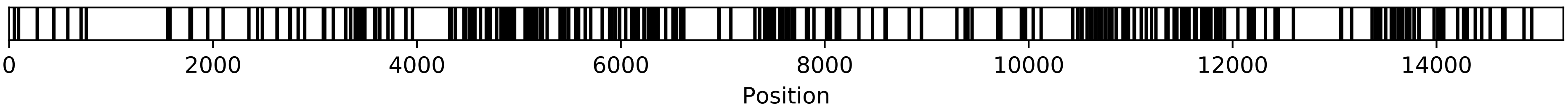


MM1 vs MM2
Number of differences: 2

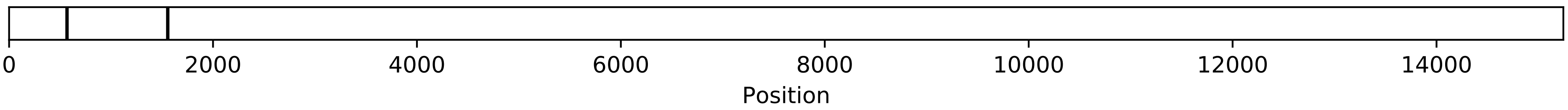

### Region: 428

CR1 vs CR2
Number of differences: 304

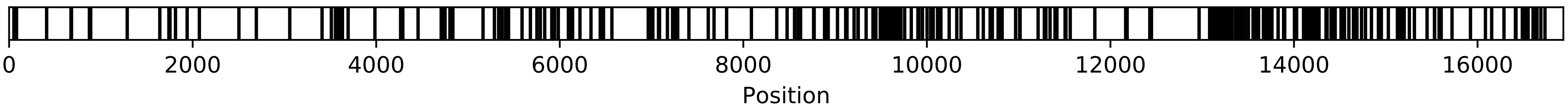


CR1 vs MA1
Number of differences: 303

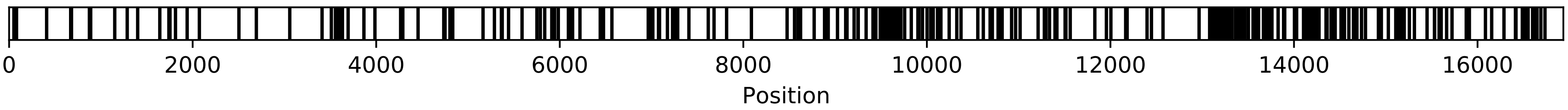


CR1 vs MA2
Number of differences: 33

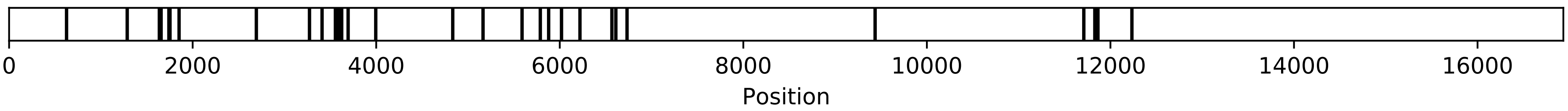


CR1 vs MM1
Number of differences: 163

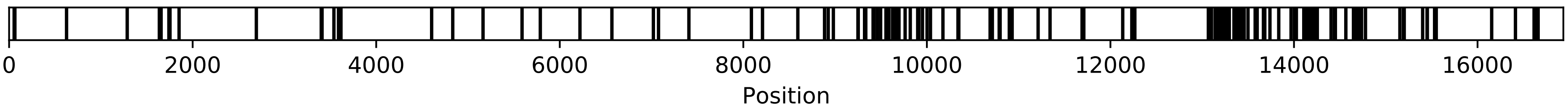


CR1 vs MM2
Number of differences: 27

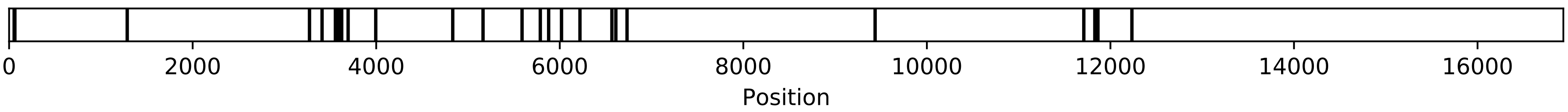


CR2 vs MA1
Number of differences: 33

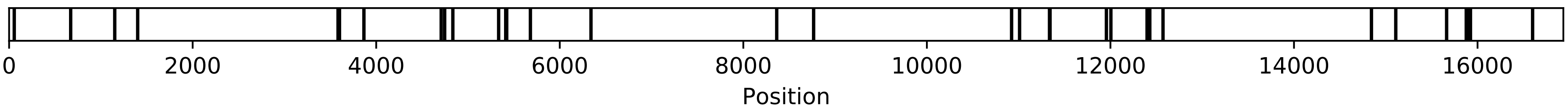


CR2 vs MA2
Number of differences: 304

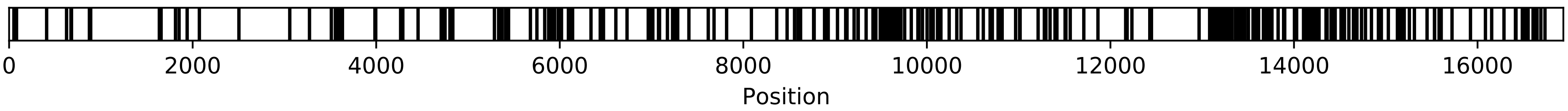


CR2 vs MM1
Number of differences: 292

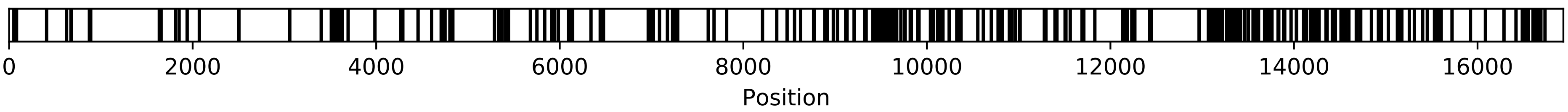


CR2 vs MM2
Number of differences: 306

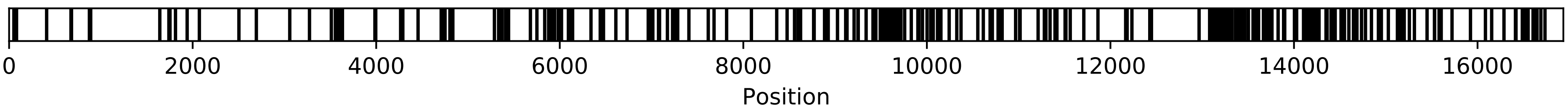


MA1 vs MA2
Number of differences: 303

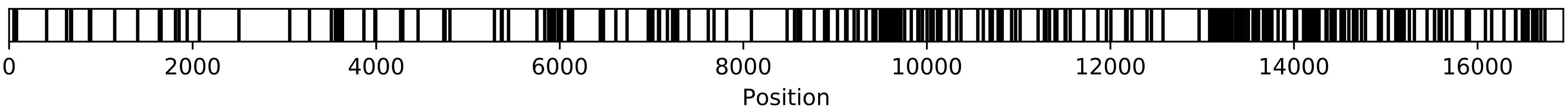


MA1 vs MM1
Number of differences: 293

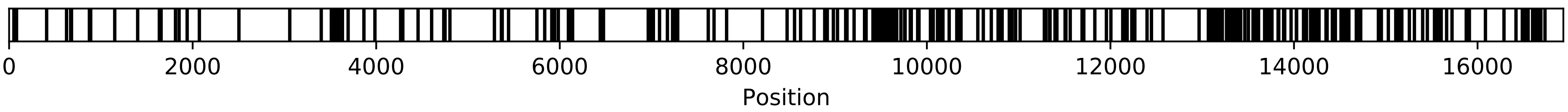


MA1 vs MM2
Number of differences: 305

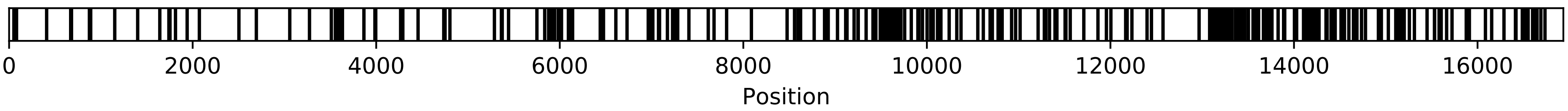


MA2 vs MM1
Number of differences: 158

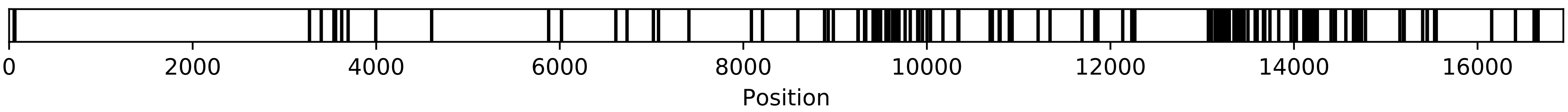


MA2 vs MM2
Number of differences: 10

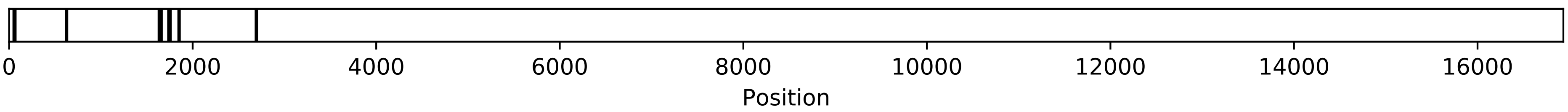


MM1 vs MM2
Number of differences: 164

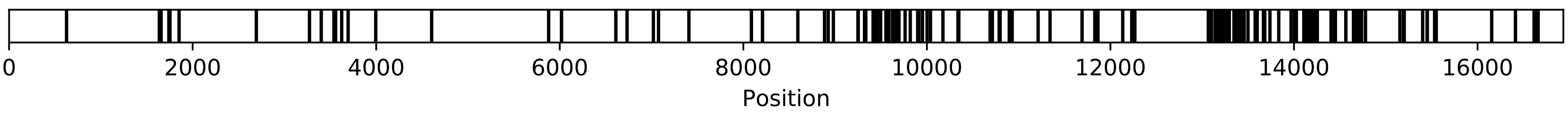

### Region: 429



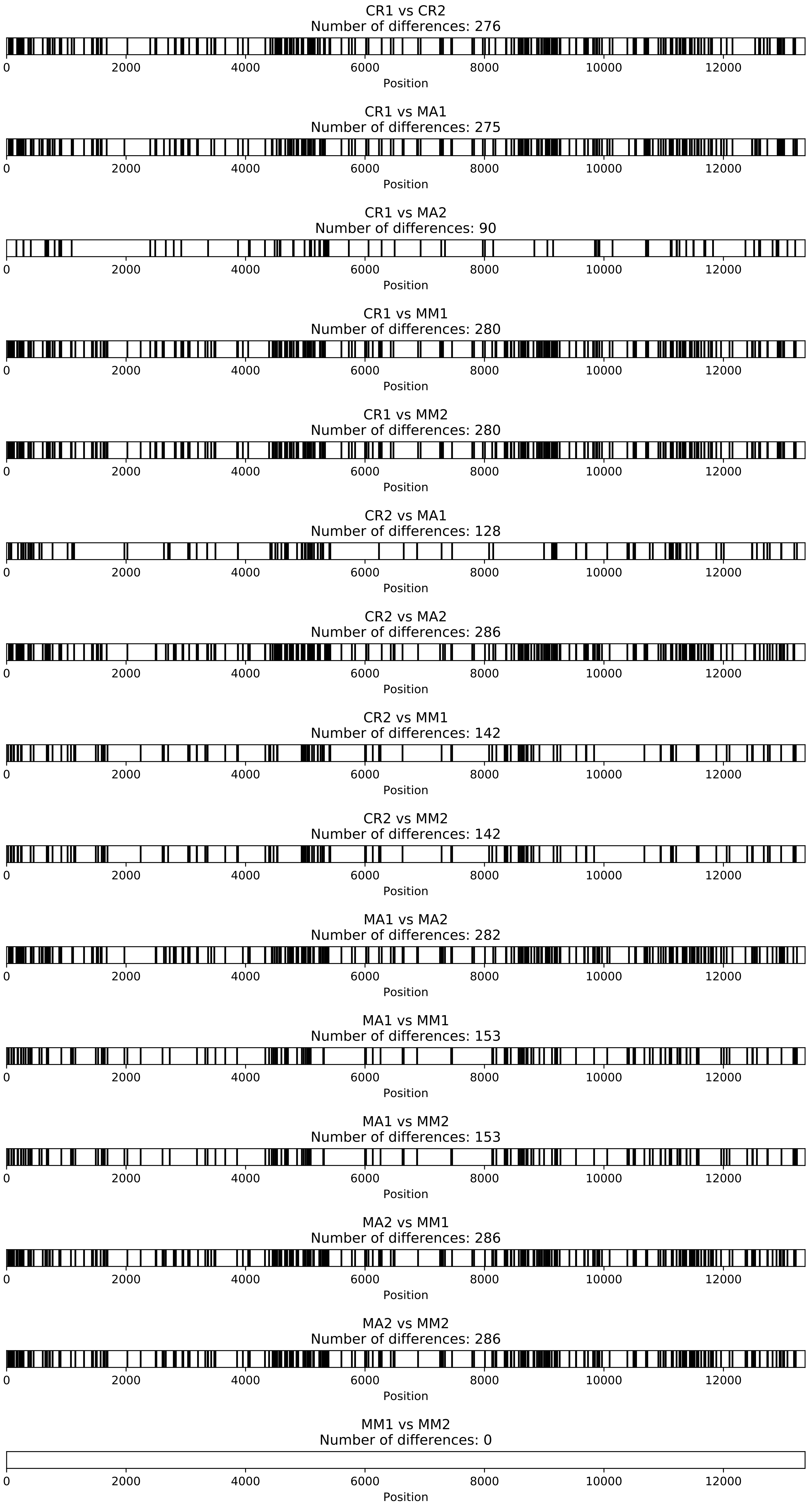
CR1 vs CR2
Number of differences: 276
CR1 vs MA1
Number of differences: 275
CR1 vs MA2
Number of differences: 90
CR1 vs MM1
Number of differences: 280
CR1 vs MM2
Number of differences: 280
CR2 vs MA1
Number of differences: 128
CR2 vs MA2
Number of differences: 286
CR2 vs MM1
Number of differences: 142
CR2 vs MM2
Number of differences: 142
MA1 vs MA2
Number of differences: 282
MA1 vs MM1
Number of differences: 153
MA1 vs MM2
Number of differences: 153
MA2 vs MM1
Number of differences: 286
MA2 vs MM2
Number of differences: 286
MM1 vs MM2
Number of differences: 0
0
2000
4000
6000
8000
10000
12000
Position

### Region: 432



CR1 vs CR2
Number of differences: 112

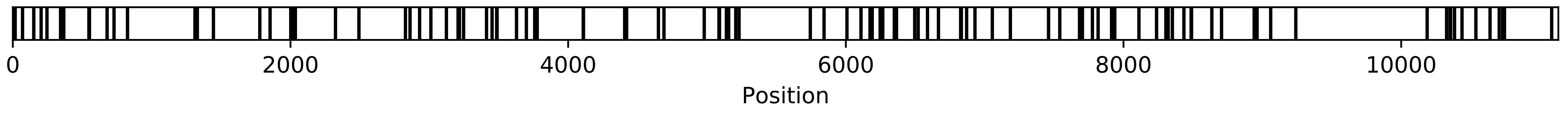


CR1 vs MA1
Number of differences: 102

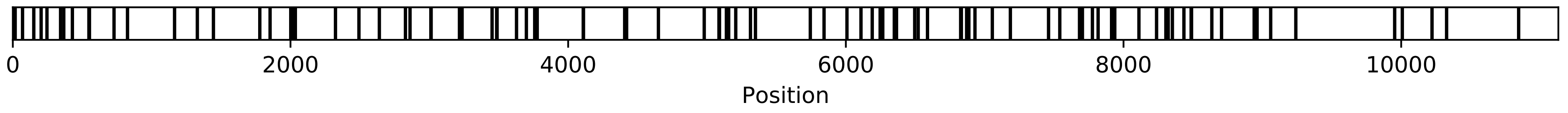


CR1 vs MA2
Number of differences: 58

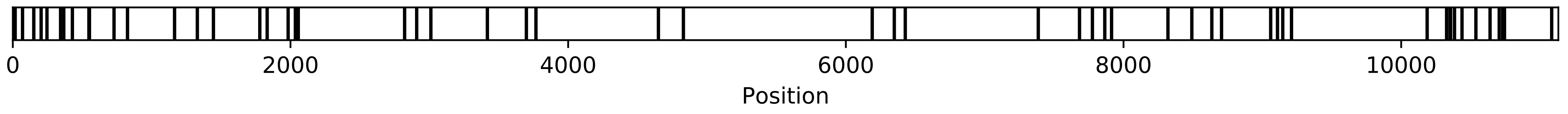


CR1 vs MM1
Number of differences: 32

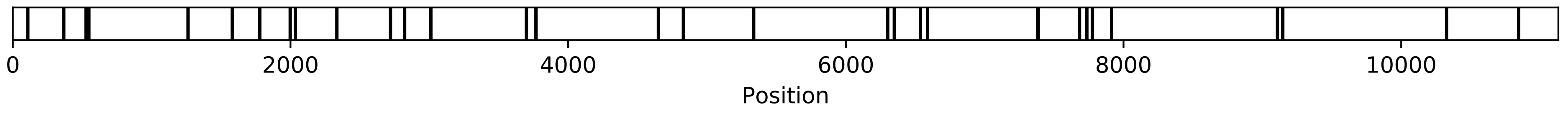


CR1 vs MM2
Number of differences: 41

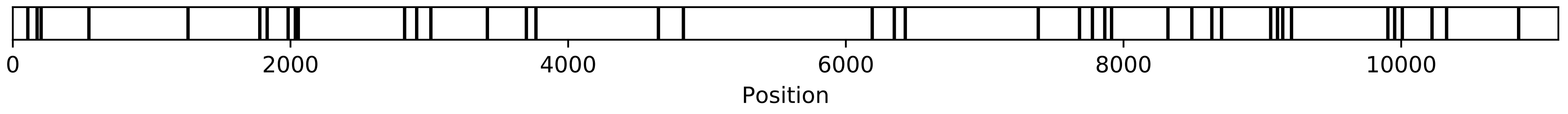


CR2 vs MA1
Number of differences: 34

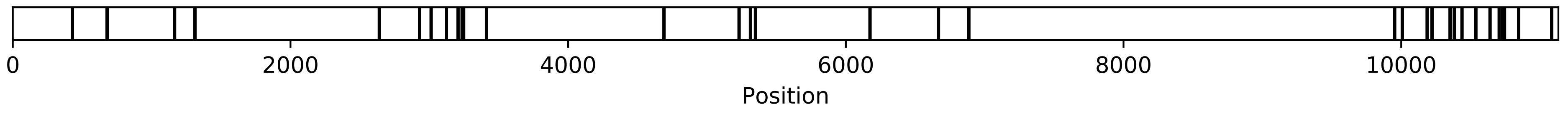


CR2 vs MA2
Number of differences: 88

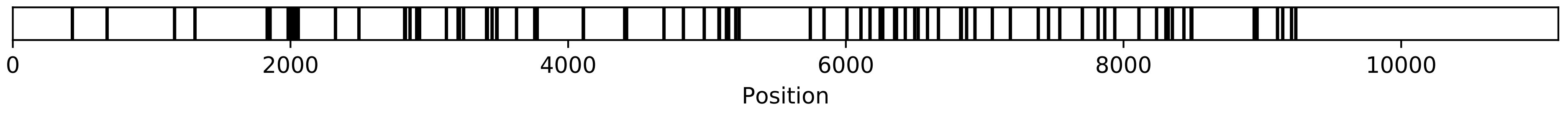


CR2 vs MM1
Number of differences: 116

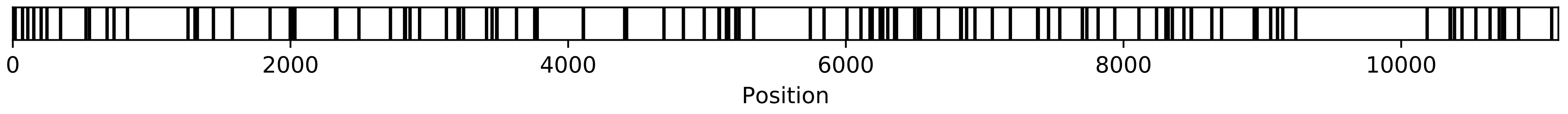


CR2 vs MM2
Number of differences: 117

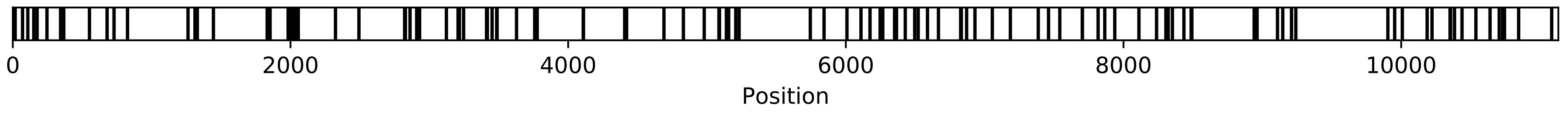


MA1 vs MA2
Number of differences: 96

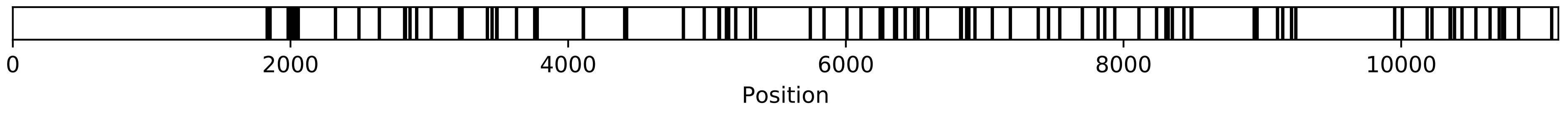


MA1 vs MM1
Number of differences: 104

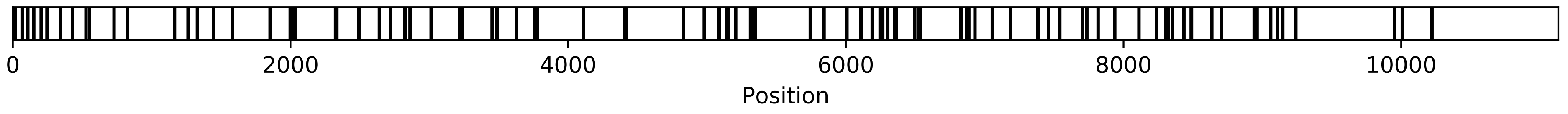


MA1 vs MM2
Number of differences: 99

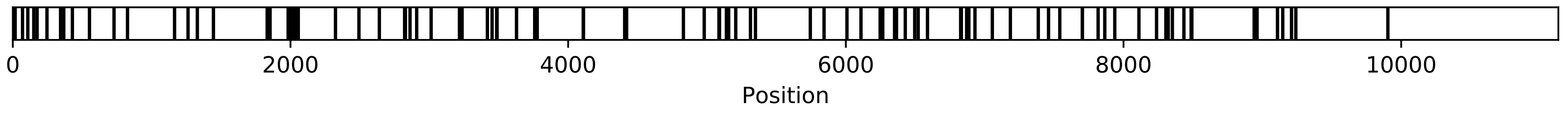


MA2 vs MM1
Number of differences: 54

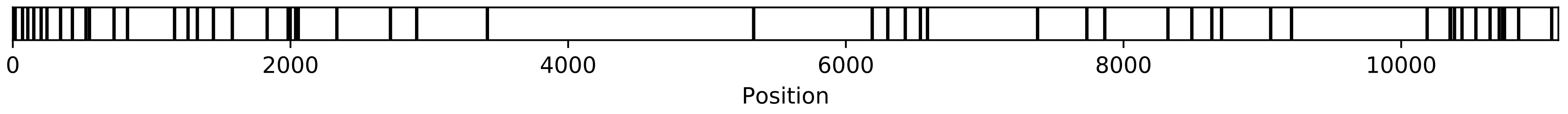


MA2 vs MM2
Number of differences: 33

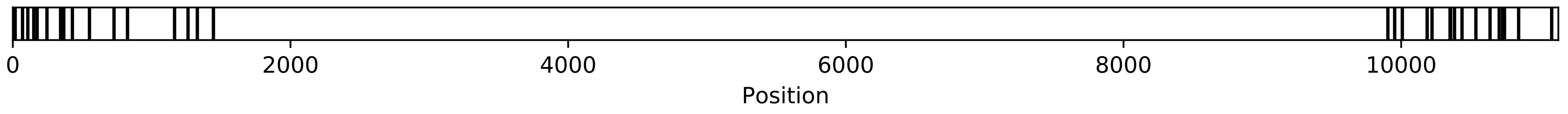


MM1 vs MM2
Number of differences: 33

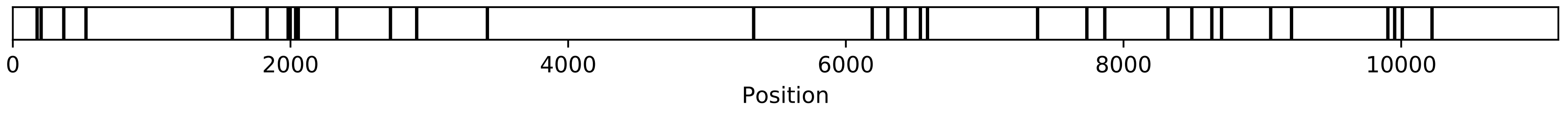

### Region: 433



CR1 vs CR2
Number of differences: 172

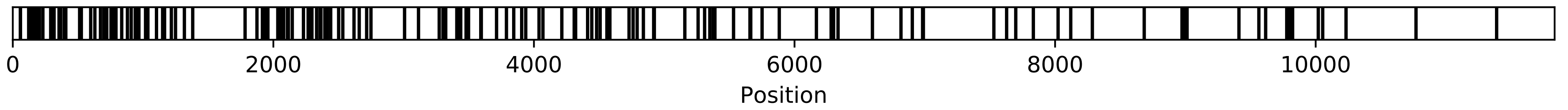


CR1 vs MA1
Number of differences: 80

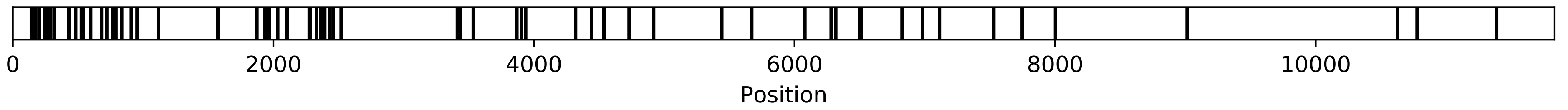


CR1 vs MA2
Number of differences: 172

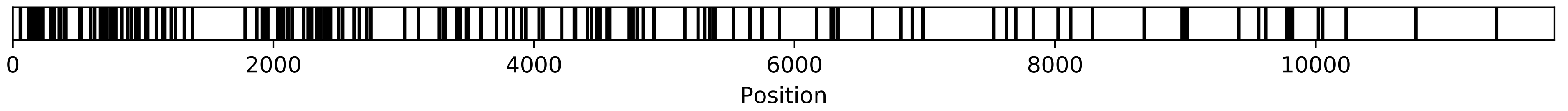


CR1 vs MM1
Number of differences: 98

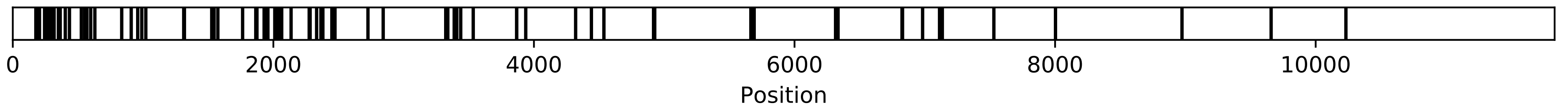


CR1 vs MM2
Number of differences: 80

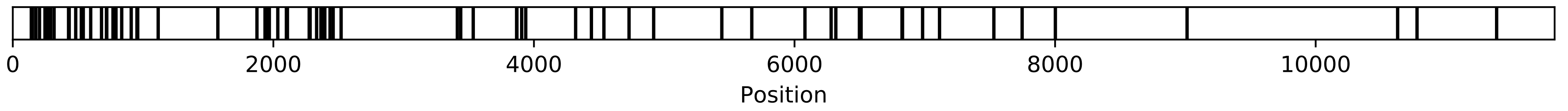


CR2 vs MA1
Number of differences: 172

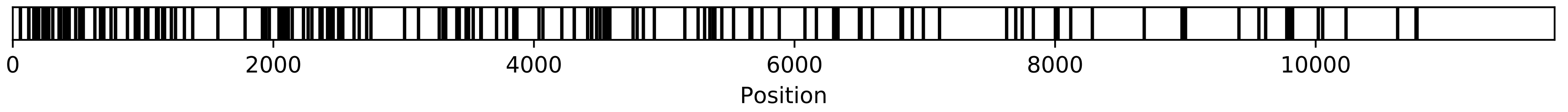


CR2 vs MA2
Number of differences: 0

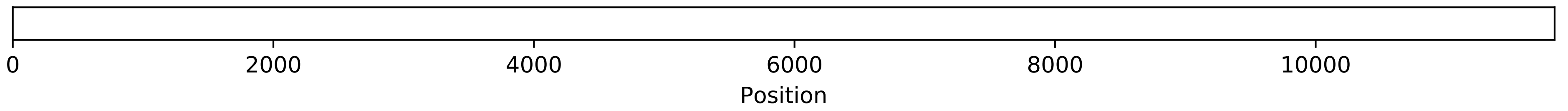


CR2 vs MM1
Number of differences: 203

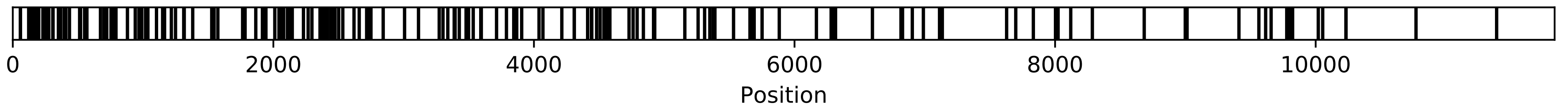


CR2 vs MM2
Number of differences: 172

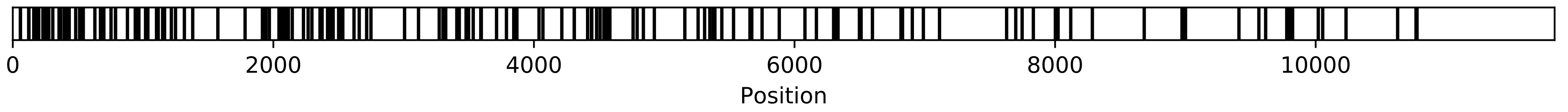


MA1 vs MA2
Number of differences: 172

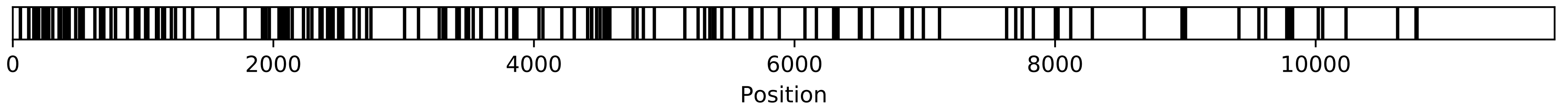


MA1 vs MM1
Number of differences: 100

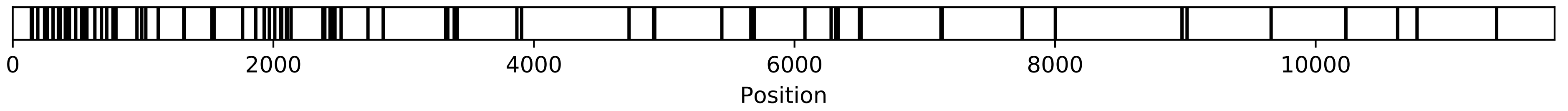


MA1 vs MM2
Number of differences: 0

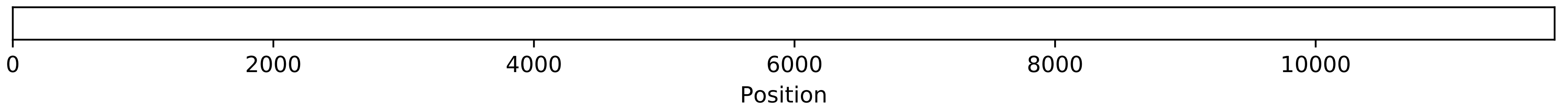


MA2 vs MM1
Number of differences: 203

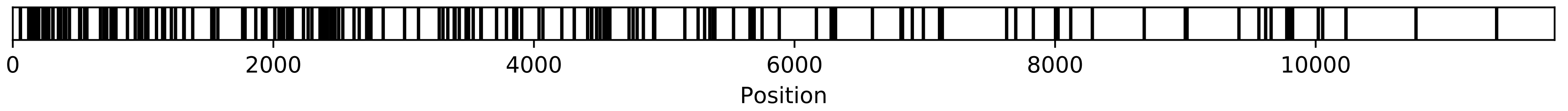


MA2 vs MM2
Number of differences: 172

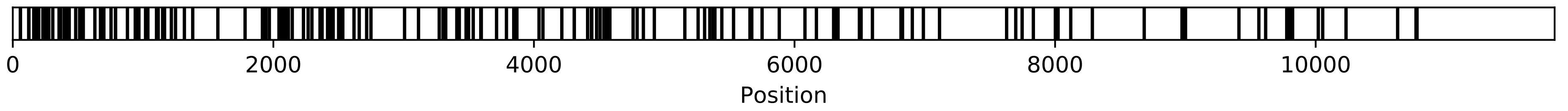


MM1 vs MM2
Number of differences: 100

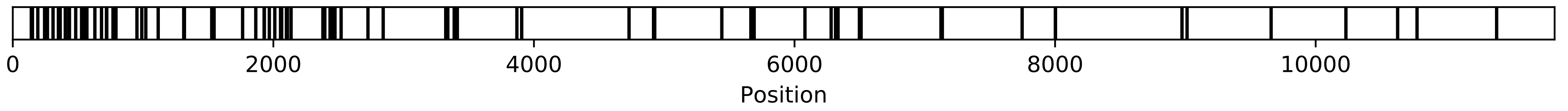

### Region: 435



CR1 vs CR2
Number of differences: 633

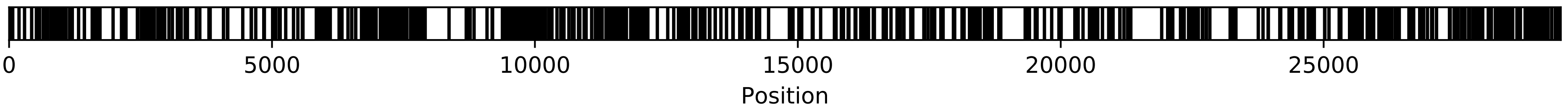


CR1 vs MA1
Number of differences: 625

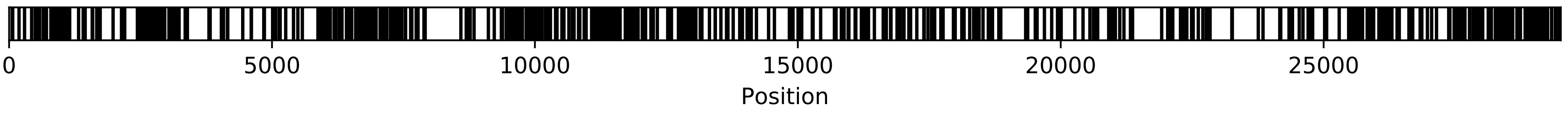


CR1 vs MA2
Number of differences: 176

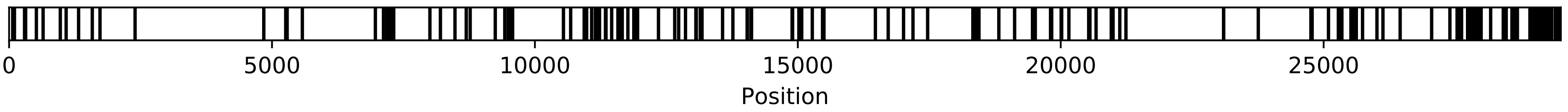


CR1 vs MM1
Number of differences: 620

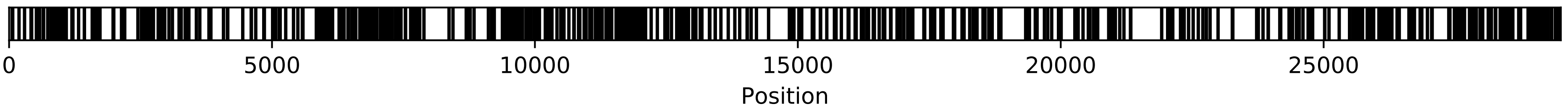


CR1 vs MM2
Number of differences: 625

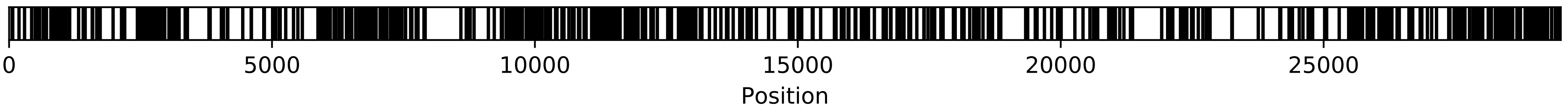


CR2 vs MA1
Number of differences: 299

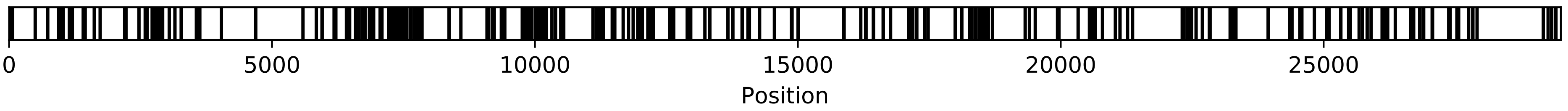


CR2 vs MA2
Number of differences: 639

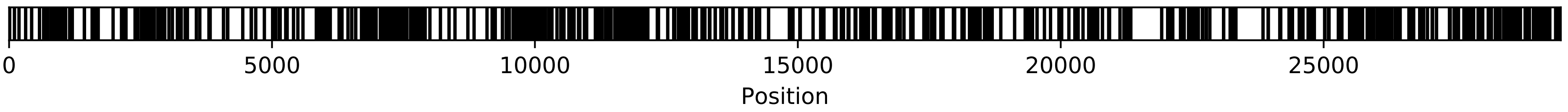


CR2 vs MM1
Number of differences: 320

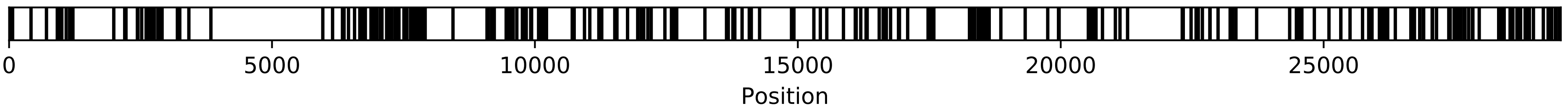


CR2 vs MM2
Number of differences: 299

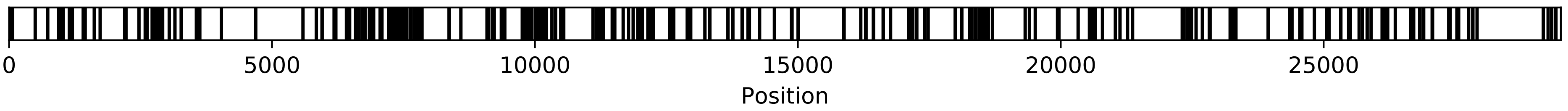


MA1 vs MA2
Number of differences: 634

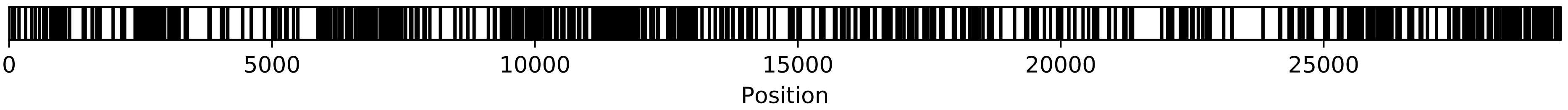


MA1 vs MM1
Number of differences: 314

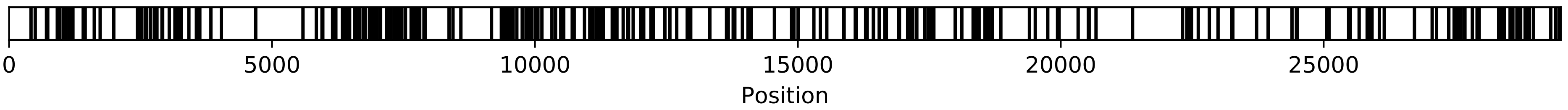


MA1 vs MM2
Number of differences: 0

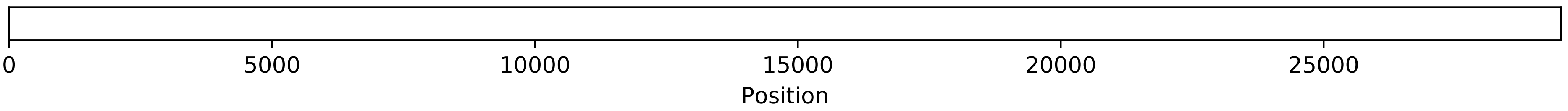


MA2 vs MM1
Number of differences: 630

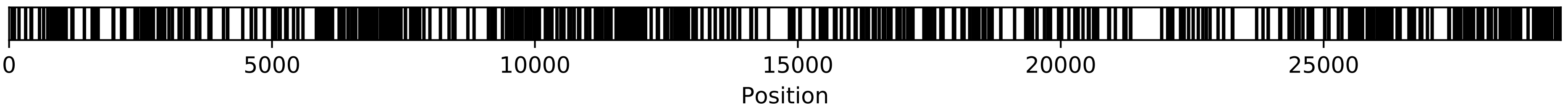


MA2 vs MM2
Number of differences: 634

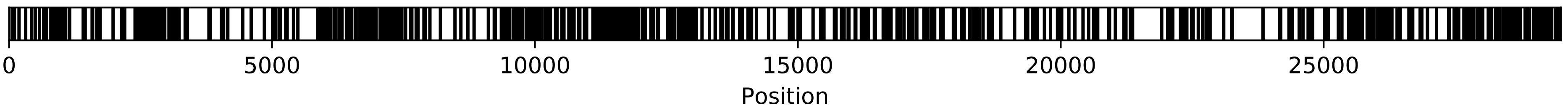


MM1 vs MM2
Number of differences: 314

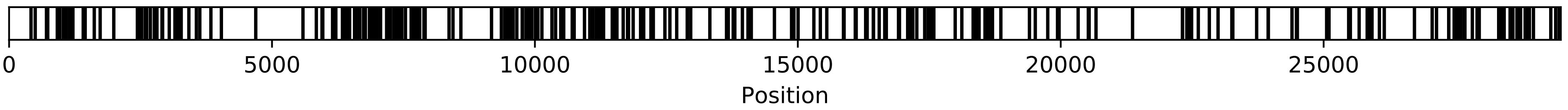

### Region: 436



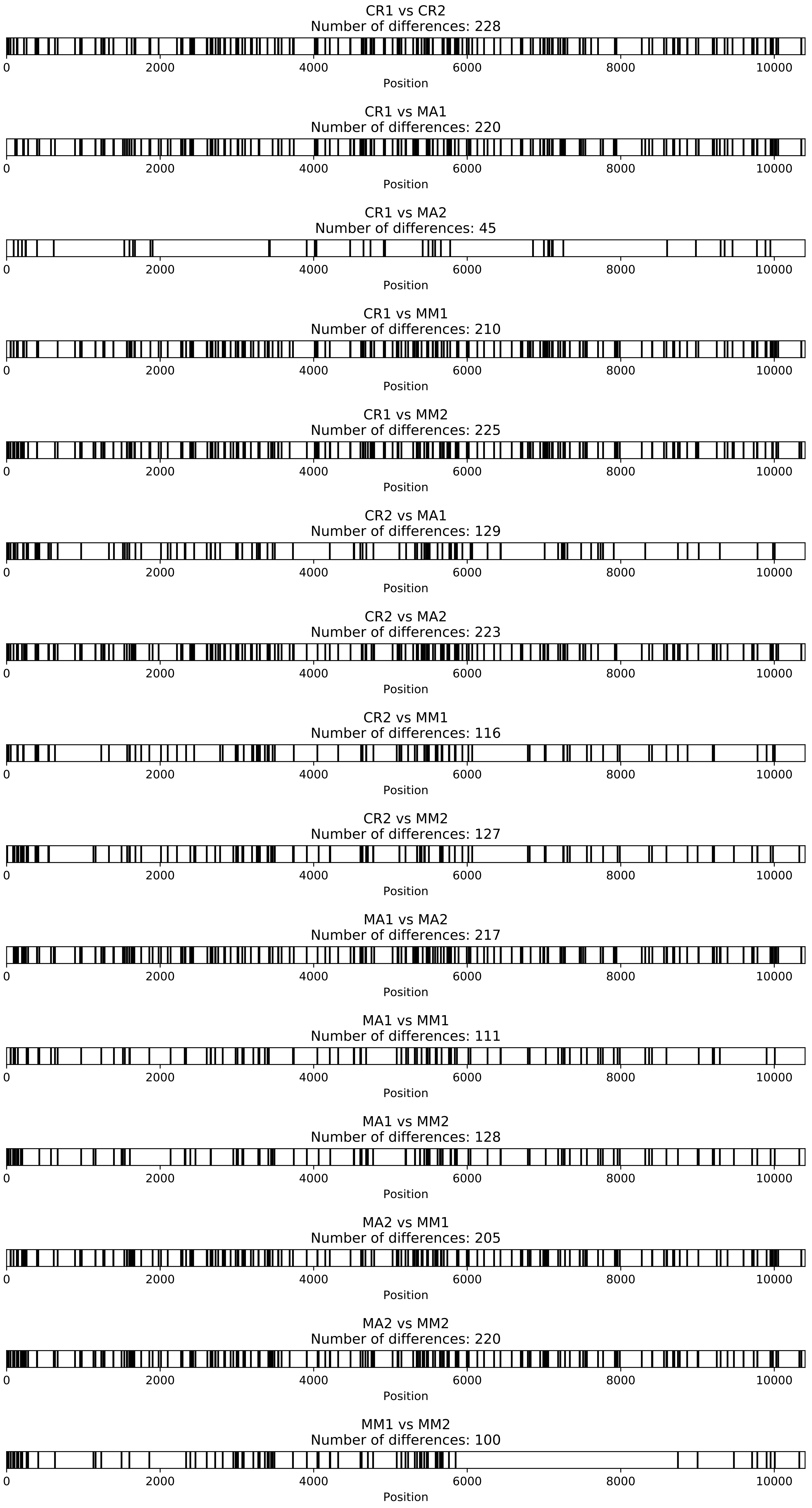
CR1 vs CR2
Number of differences: 228
CR1 vs MA1
Number of differences: 220
CR1 vs MA2
Number of differences: 45
CR1 vs MM1
Number of differences: 210
CR1 vs MM2
Number of differences: 225
CR2 vs MA1
Number of differences: 129
CR2 vs MA2
Number of differences: 223
CR2 vs MM1
Number of differences: 116
CR2 vs MM2
Number of differences: 127
MA1 vs MA2
Number of differences: 217
MA1 vs MM1
Number of differences: 111
MA1 vs MM2
Number of differences: 128
MA2 vs MM1
Number of differences: 205
MA2 vs MM2
Number of differences: 220
MM1 vs MM2
Number of differences: 100
0
2000
4000
6000
8000
10000
Position

### Region: 437



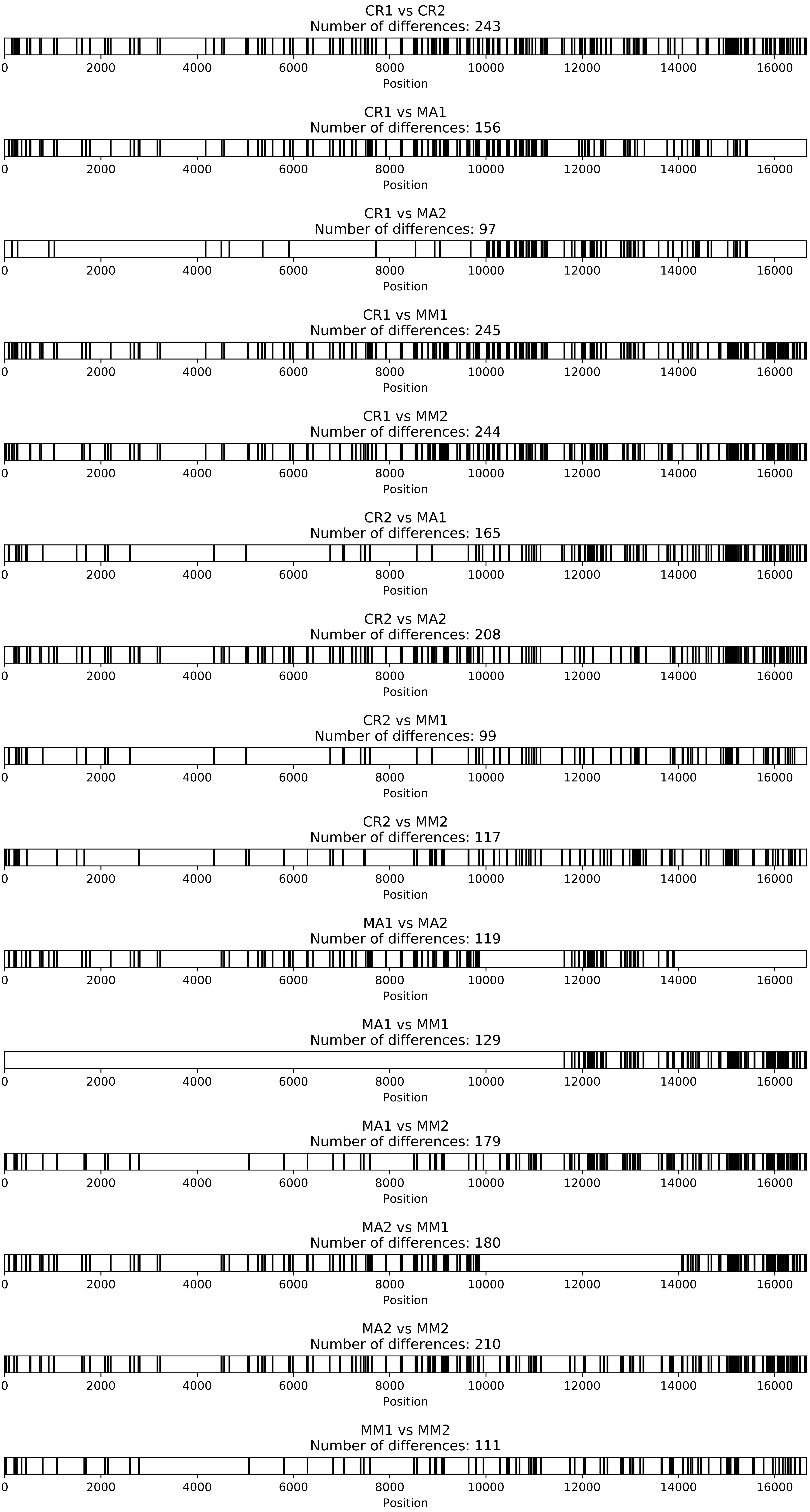
CR1 vs CR2
Number of differences: 243
CR1 vs MA1
Number of differences: 156
CR1 vs MA2
Number of differences: 97
CR1 vs MM1
Number of differences: 245
CR1 vs MM2
Number of differences: 244
CR2 vs MA1
Number of differences: 165
CR2 vs MA2
Number of differences: 208
CR2 vs MM1
Number of differences: 99
CR2 vs MM2
Number of differences: 117
MA1 vs MA2
Number of differences: 119
MA1 vs MM1
Number of differences: 129
MA1 vs MM2
Number of differences: 179
MA2 vs MM1
Number of differences: 180
MA2 vs MM2
Number of differences: 210
MM1 vs MM2
Number of differences: 111
0
2000
4000
6000
8000
10000
12000
14000
16000
Position

### Region: 441



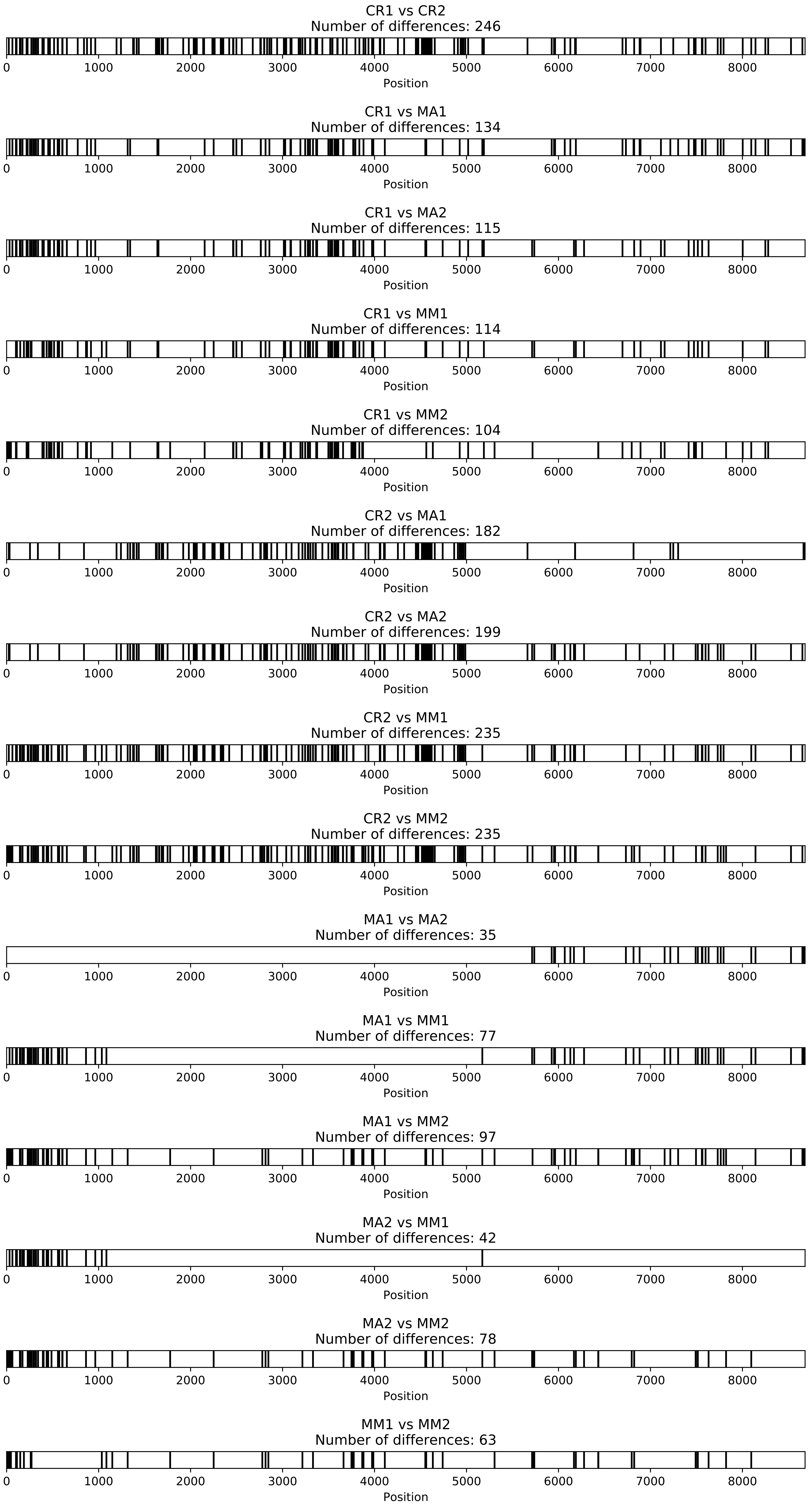
CR1 vs CR2
Number of differences: 246
CR1 vs MA1
Number of differences: 134
CR1 vs MA2
Number of differences: 115
CR1 vs MM1
Number of differences: 114
CR1 vs MM2
Number of differences: 104
CR2 vs MA1
Number of differences: 182
CR2 vs MA2
Number of differences: 199
CR2 vs MM1
Number of differences: 235
CR2 vs MM2
Number of differences: 235
MA1 vs MA2
Number of differences: 35
MA1 vs MM1
Number of differences: 77
MA1 vs MM2
Number of differences: 97
MA2 vs MM1
Number of differences: 42
MA2 vs MM2
Number of differences: 78
MM1 vs MM2
Number of differences: 63
0
1000
2000
3000
4000
5000
6000
7000
8000
Position

### Region: 442

CR1 vs CR2
Number of differences: 293

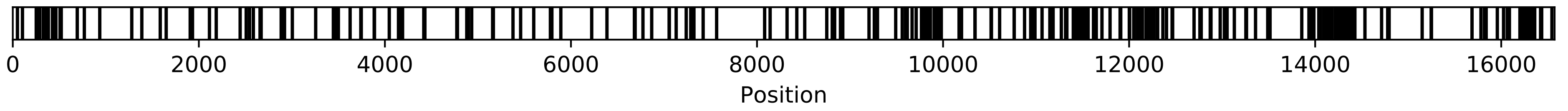

CR1 vs MA1
Number of differences: 289

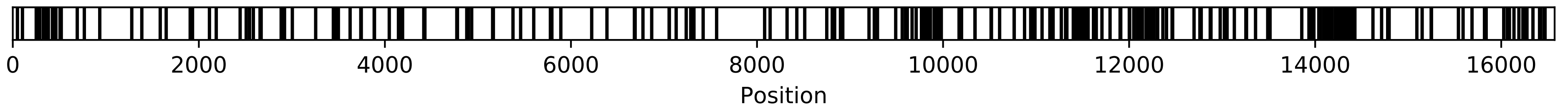

CR1 vs MA2
Number of differences: 239

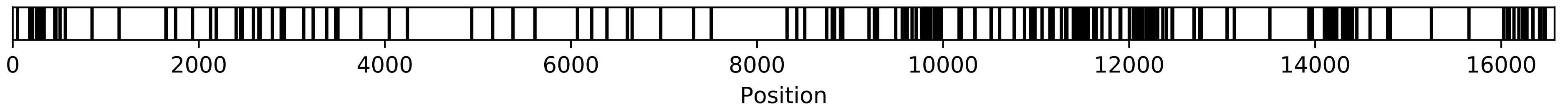

CR1 vs MM1
Number of differences: 186

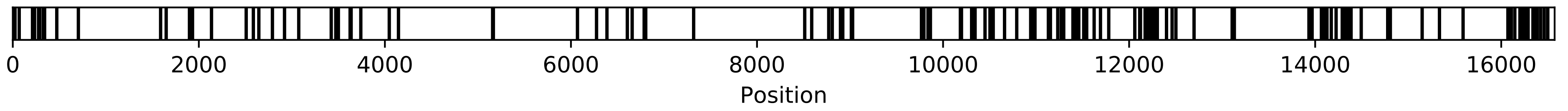

CR1 vs MM2
Number of differences: 175

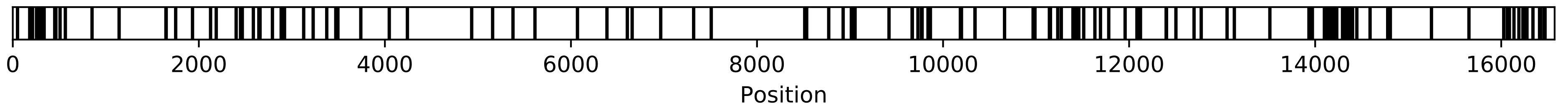

CR2 vs MA1
Number of differences: 42

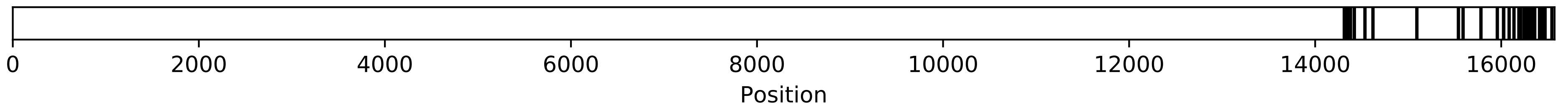

CR2 vs MA2
Number of differences: 194

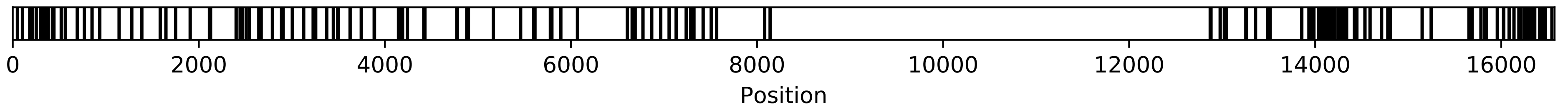

CR2 vs MM1
Number of differences: 304

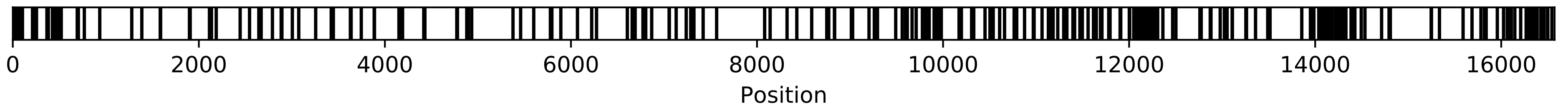

CR2 vs MM2
Number of differences: 304

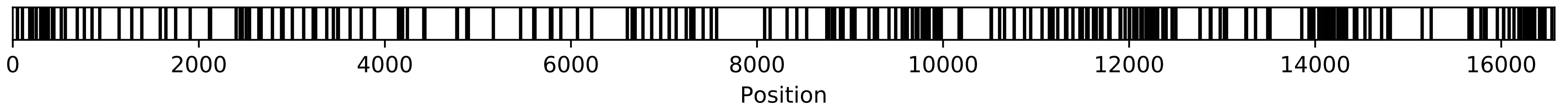

MA1 vs MA2
Number of differences: 172

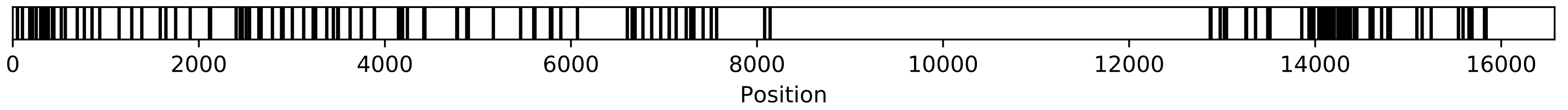

MA1 vs MM1
Number of differences: 302

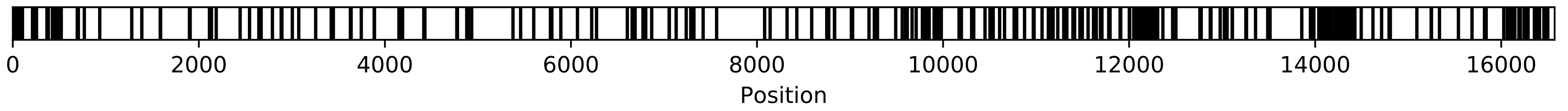

MA1 vs MM2
Number of differences: 282

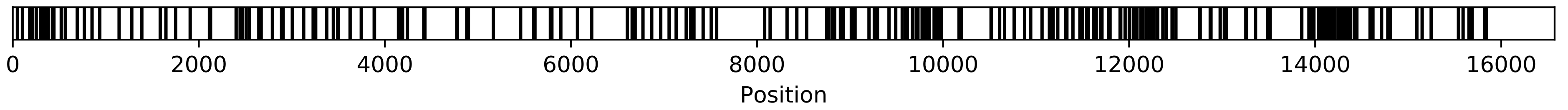

MA2 vs MM1
Number of differences: 264

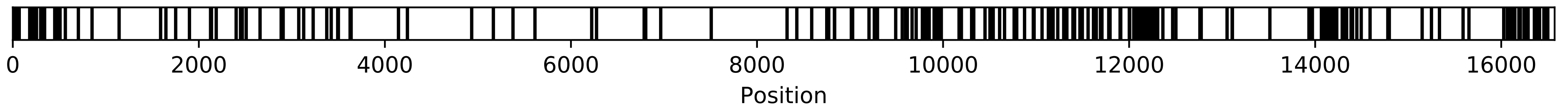

MA2 vs MM2
Number of differences: 110

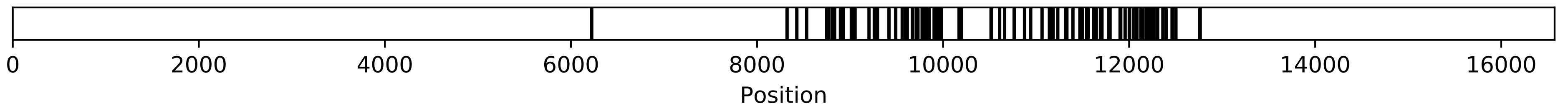

MM1 vs MM2
Number of differences: 219

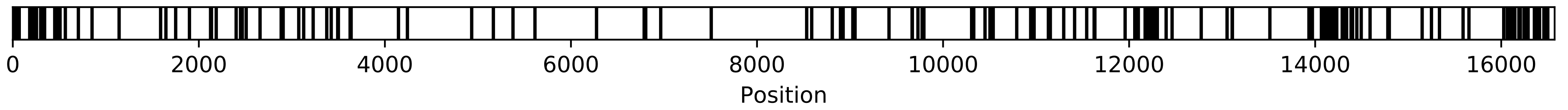

### Region: 445

CR1 vs CR2
Number of differences: 338

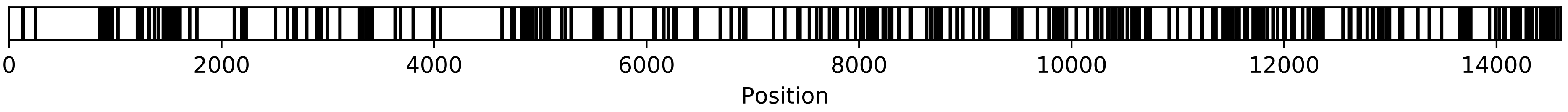


CR1 vs MA1
Number of differences: 198

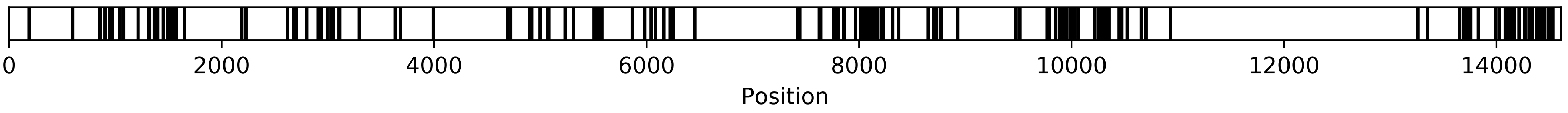


CR1 vs MA2
Number of differences: 344

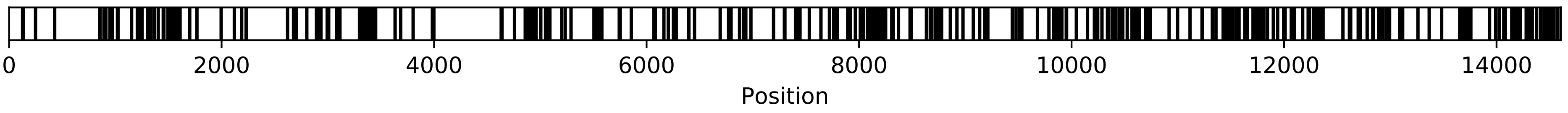


CR1 vs MM1
Number of differences: 197

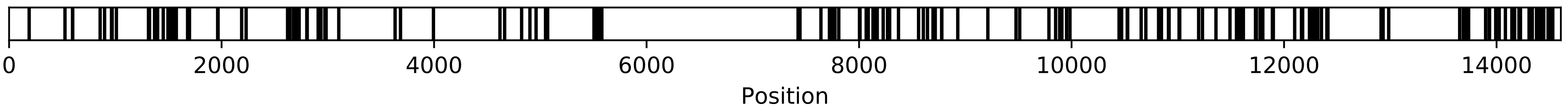


CR1 vs MM2
Number of differences: 221

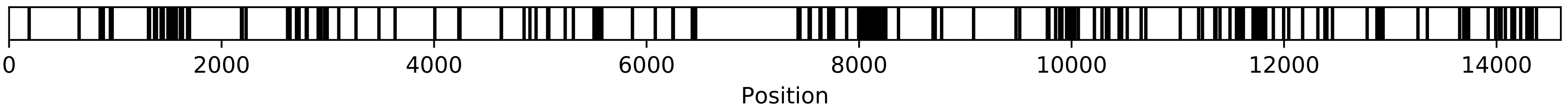


CR2 vs MA1
Number of differences: 292

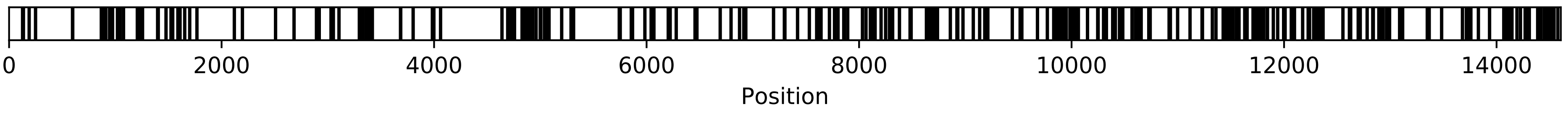


CR2 vs MA2
Number of differences: 52

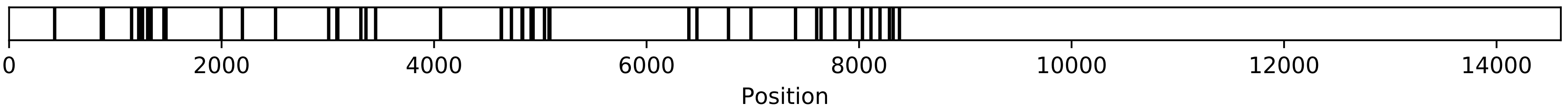


CR2 vs MM1
Number of differences: 301

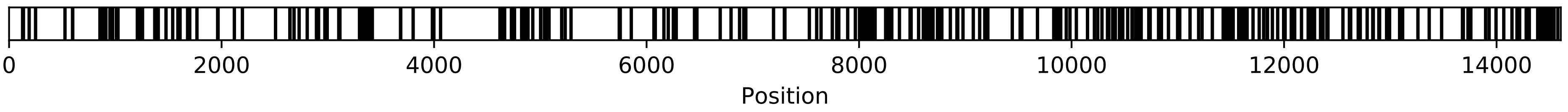


CR2 vs MM2
Number of differences: 297

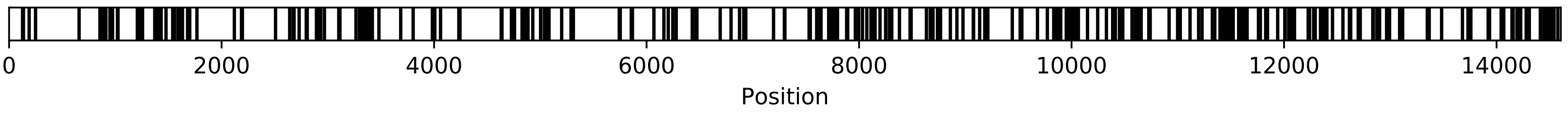


MA1 vs MA2
Number of differences: 298

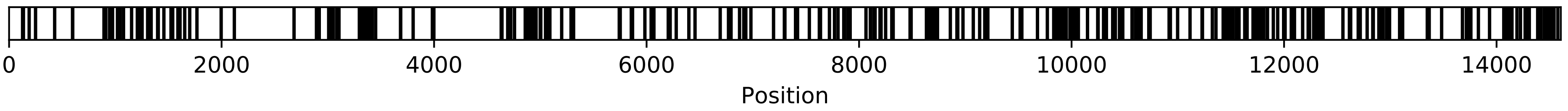


MA1 vs MM1
Number of differences: 184

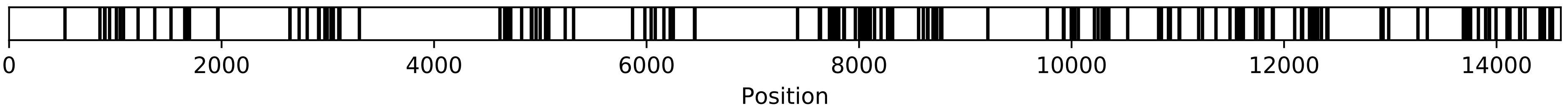


MA1 vs MM2
Number of differences: 176

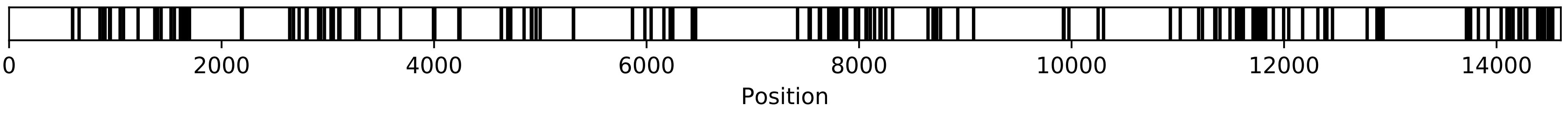


MA2 vs MM1
Number of differences: 309

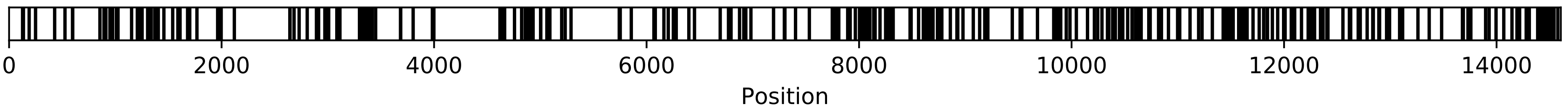


MA2 vs MM2
Number of differences: 299

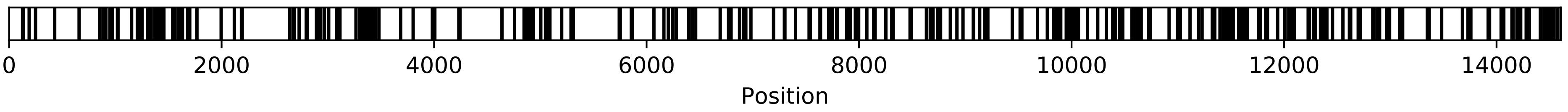


MM1 vs MM2
Number of differences: 164

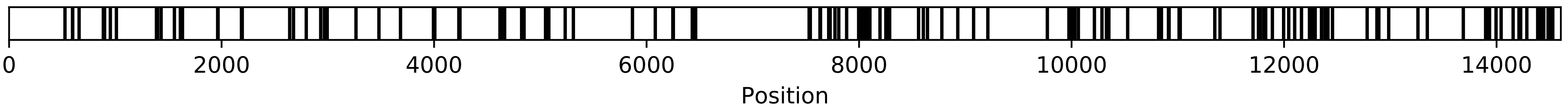

### Region: 447



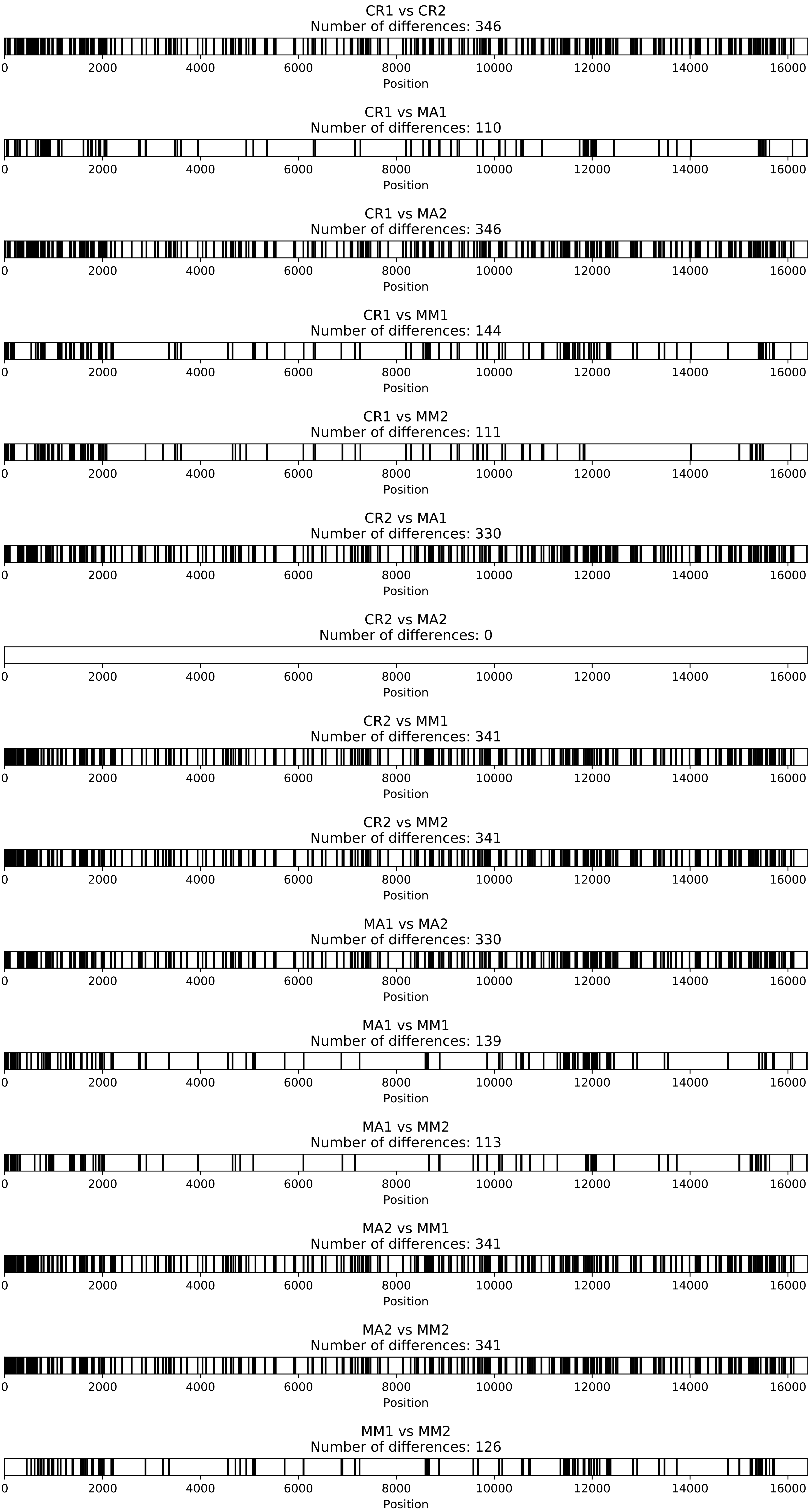
CR1 vs CR2
Number of differences: 346
CR1 vs MA1
Number of differences: 110
CR1 vs MA2
Number of differences: 346
CR1 vs MM1
Number of differences: 144
CR1 vs MM2
Number of differences: 111
CR2 vs MA1
Number of differences: 330
CR2 vs MA2
Number of differences: 0
CR2 vs MM1
Number of differences: 341
CR2 vs MM2
Number of differences: 341
MA1 vs MA2
Number of differences: 330
MA1 vs MM1
Number of differences: 139
MA1 vs MM2
Number of differences: 113
MA2 vs MM1
Number of differences: 341
MA2 vs MM2
Number of differences: 341
MM1 vs MM2
Number of differences: 126
0
2000
4000
6000
8000
10000
12000
14000
16000
Position

### Region: 448

CR1 vs CR2
Number of differences: 259

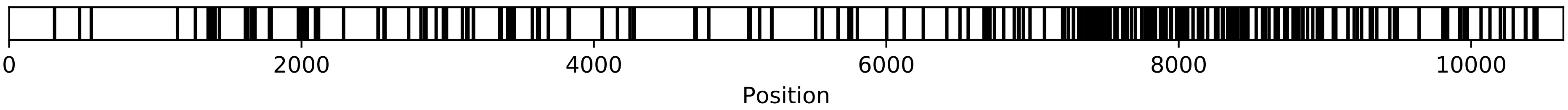


CR1 vs MA1
Number of differences: 240

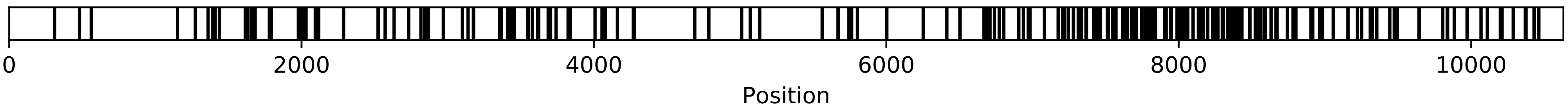


CR1 vs MA2
Number of differences: 123

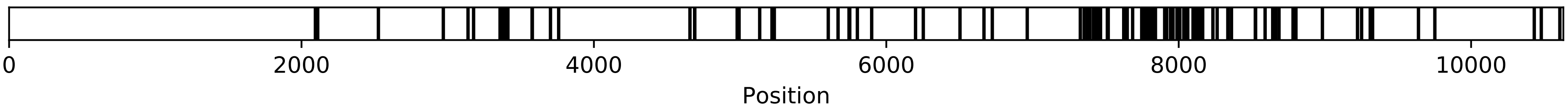


CR1 vs MM1
Number of differences: 242

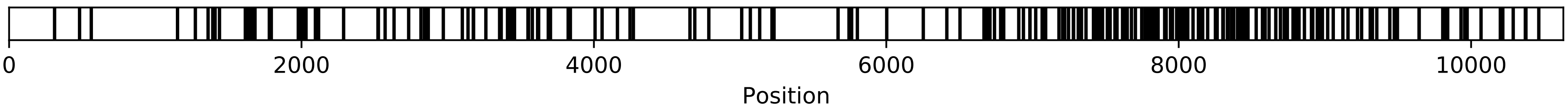


CR1 vs MM2
Number of differences: 240

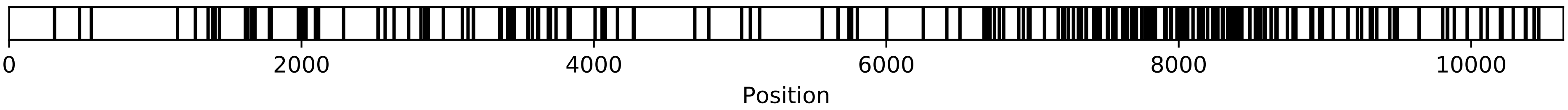


CR2 vs MA1
Number of differences: 120

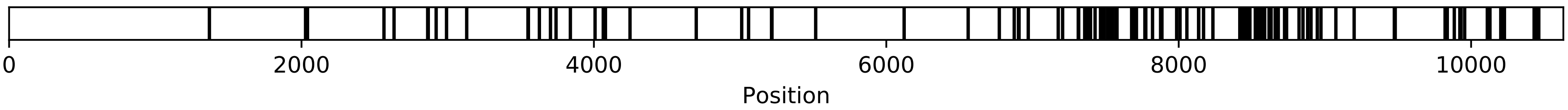


CR2 vs MA2
Number of differences: 238

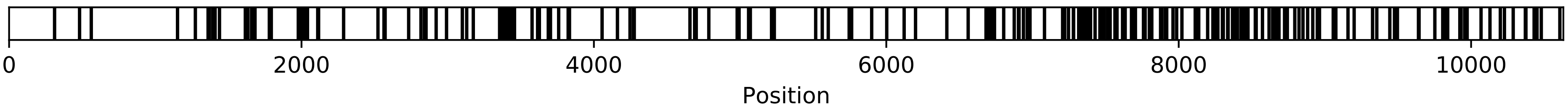


CR2 vs MM1
Number of differences: 106

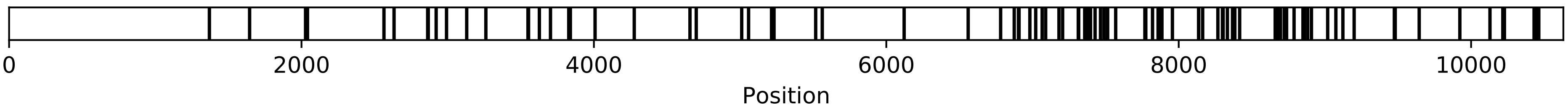


CR2 vs MM2
Number of differences: 120

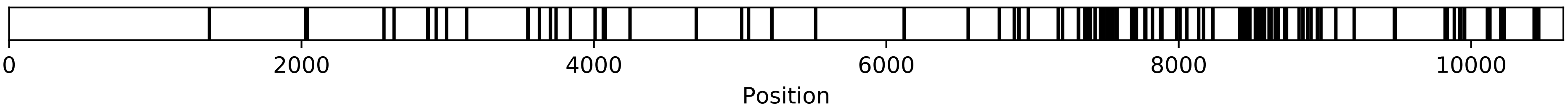


MA1 vs MA2
Number of differences: 215

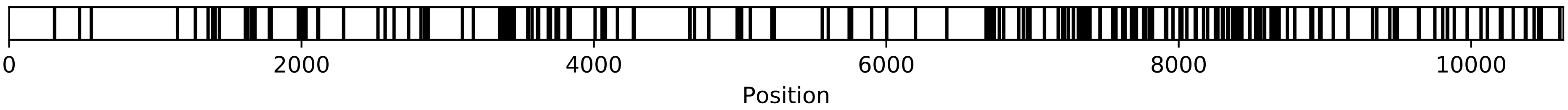


MA1 vs MM1
Number of differences: 95

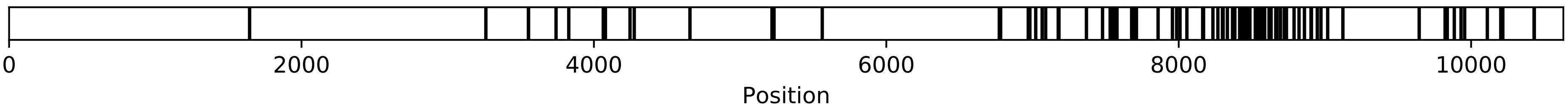


MA1 vs MM2
Number of differences: 0

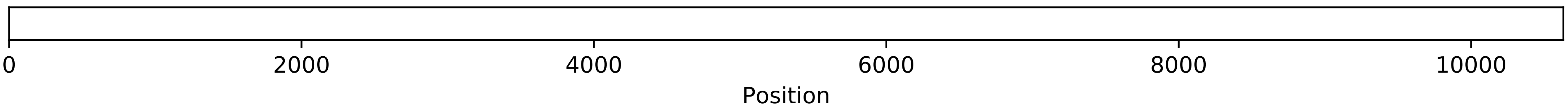


MA2 vs MM1
Number of differences: 220

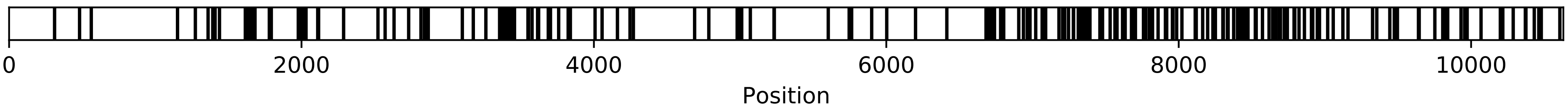


MA2 vs MM2
Number of differences: 215

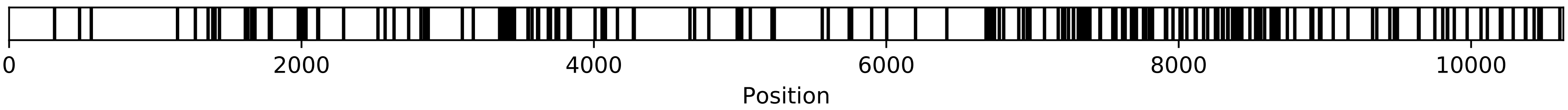


MM1 vs MM2
Number of differences: 95

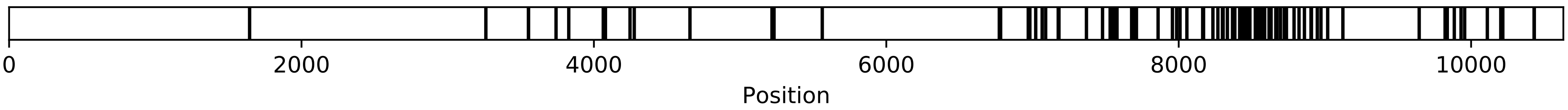

### Region: 452



CR1 vs CR2
Number of differences: 164

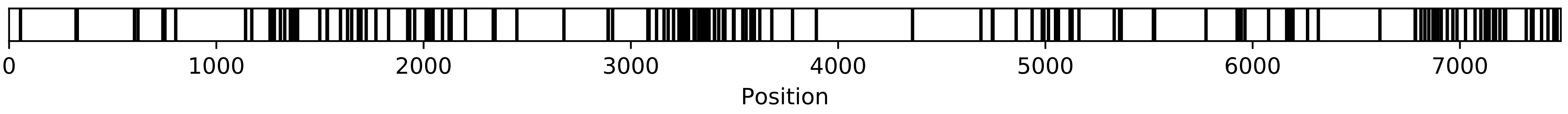


CR1 vs MA1
Number of differences: 166

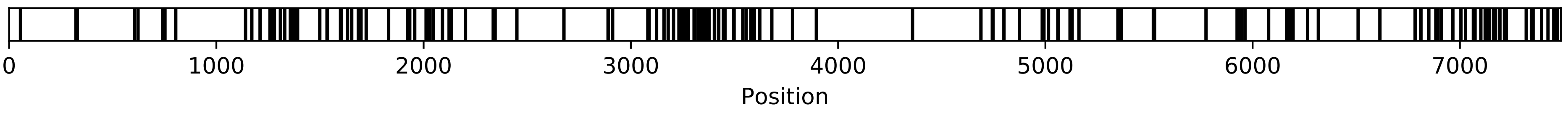


CR1 vs MA2
Number of differences: 105

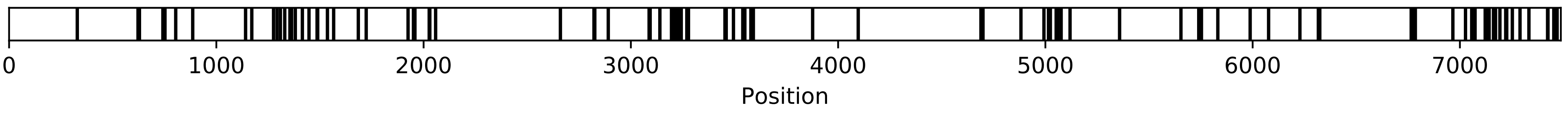


CR1 vs MM1
Number of differences: 105

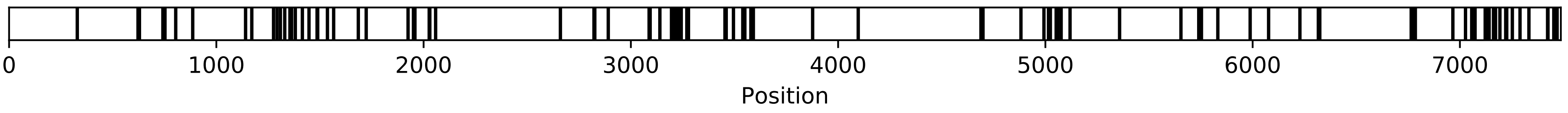


CR1 vs MM2
Number of differences: 88

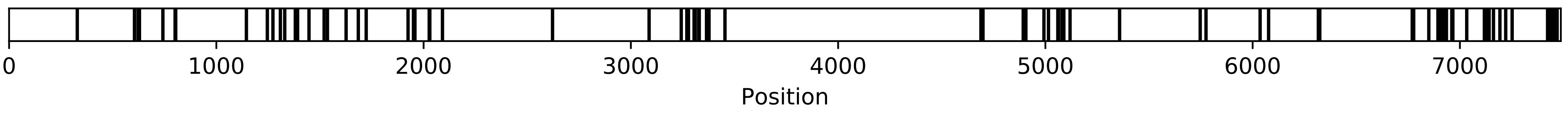


CR2 vs MA1
Number of differences: 21

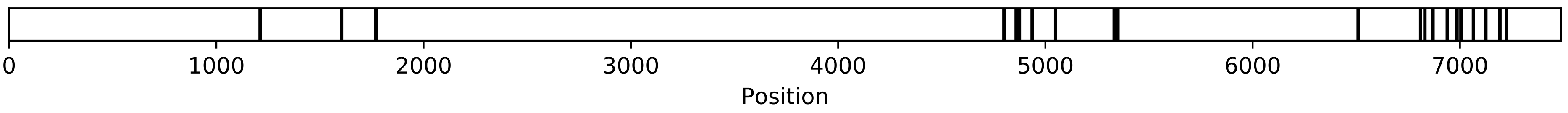


CR2 vs MA2
Number of differences: 169

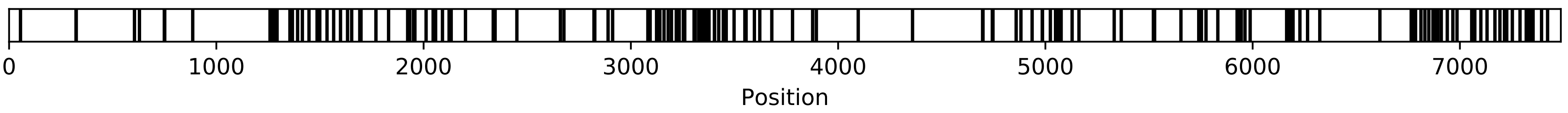


CR2 vs MM1
Number of differences: 169

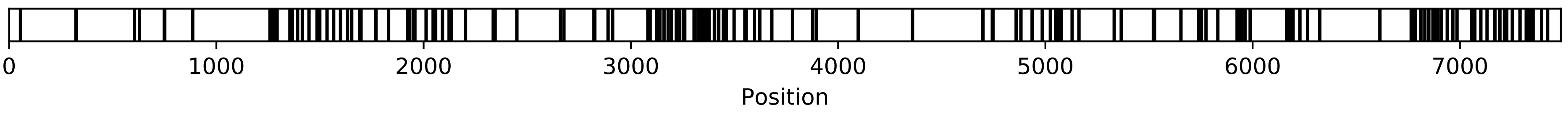


CR2 vs MM2
Number of differences: 143

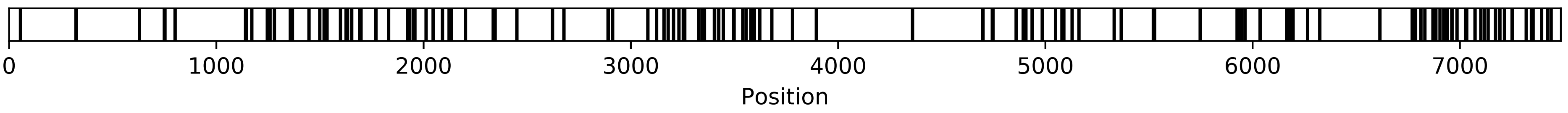


MA1 vs MA2
Number of differences: 170

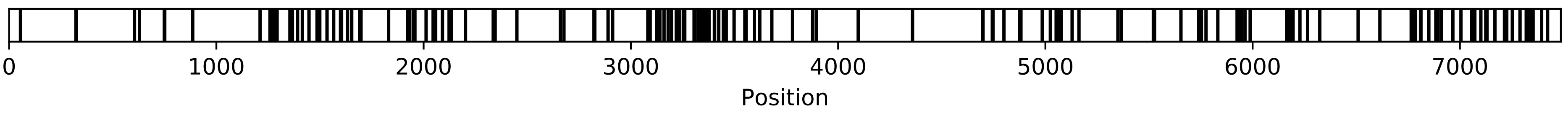


MA1 vs MM1
Number of differences: 170

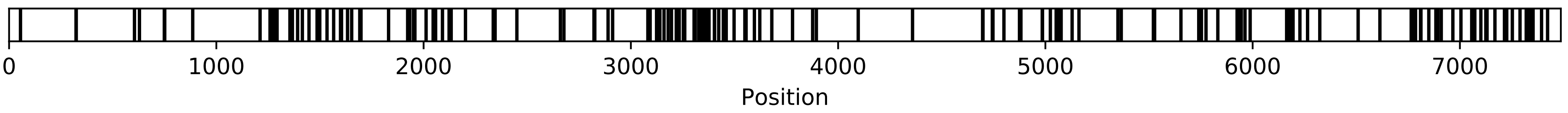


MA1 vs MM2
Number of differences: 144

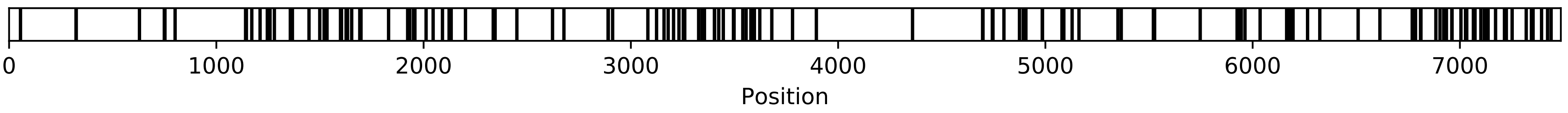


MA2 vs MM1
Number of differences: 0

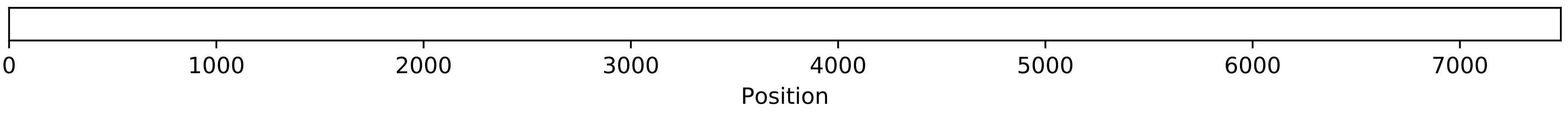


MA2 vs MM2
Number of differences: 99

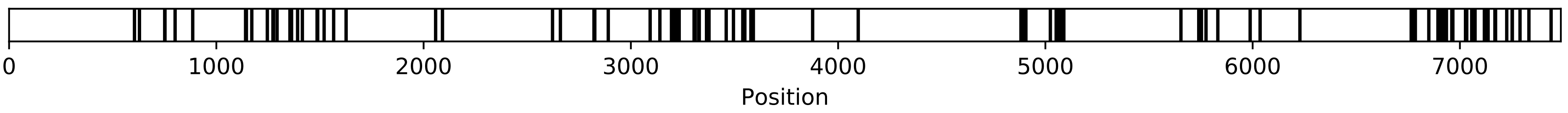


MM1 vs MM2
Number of differences: 99

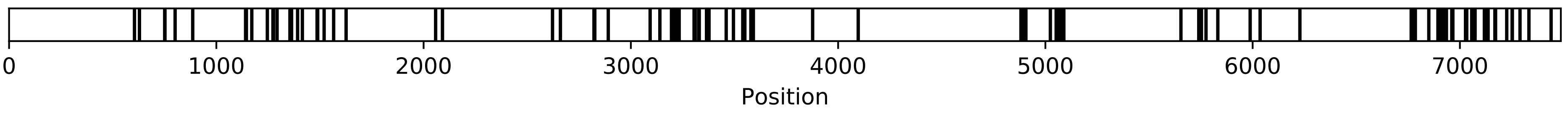

### Region: 453



CR1 vs CR2
Number of differences: 342

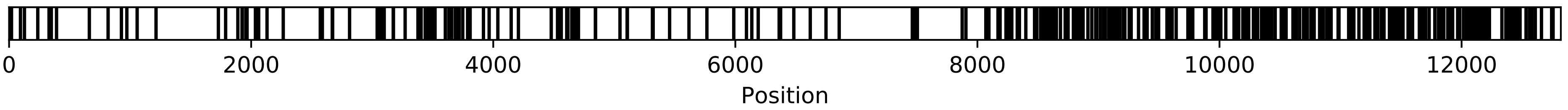


CR1 vs MA1
Number of differences: 237

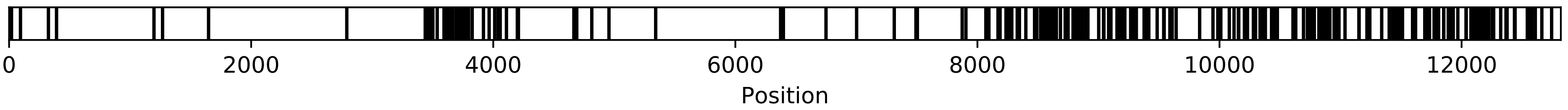


CR1 vs MA2
Number of differences: 349

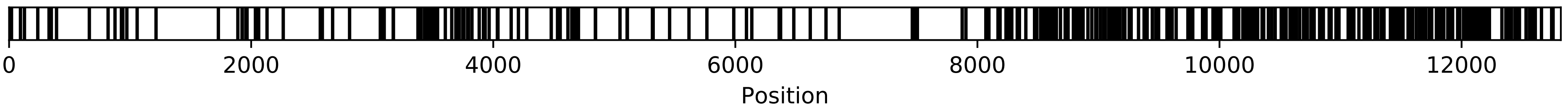


CR1 vs MM1
Number of differences: 225

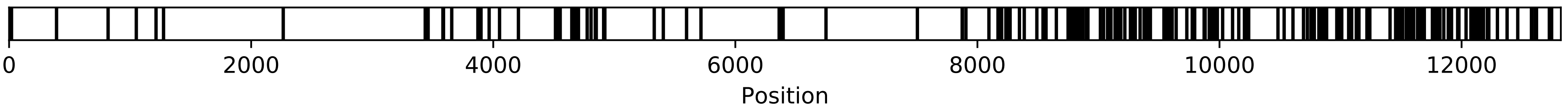


CR1 vs MM2
Number of differences: 219

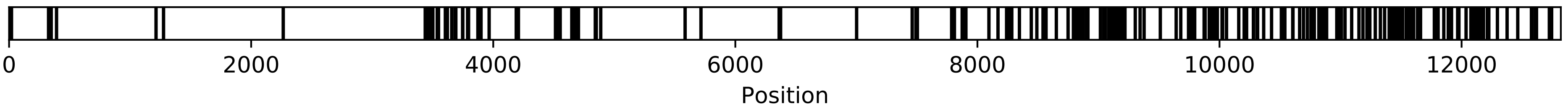


CR2 vs MA1
Number of differences: 309

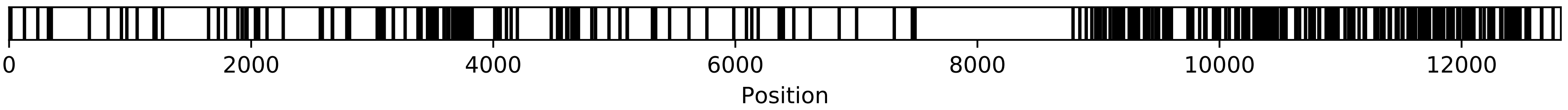


CR2 vs MA2
Number of differences: 54

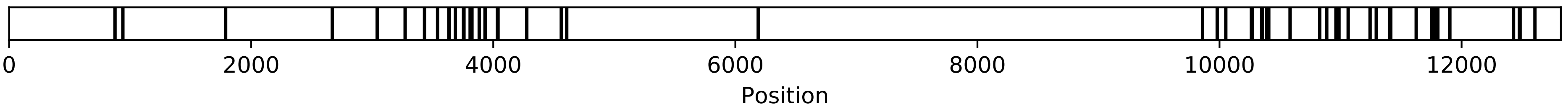


CR2 vs MM1
Number of differences: 333

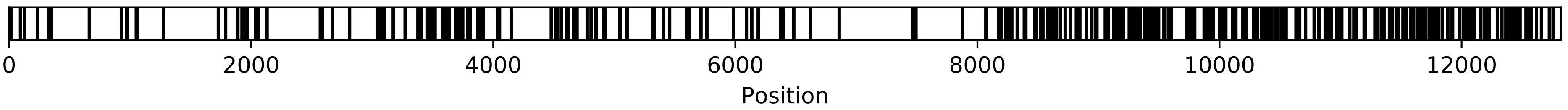


CR2 vs MM2
Number of differences: 314

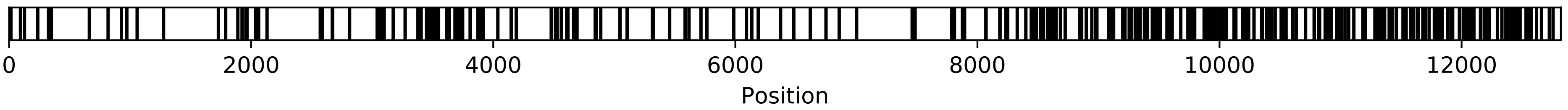


MA1 vs MA2
Number of differences: 316

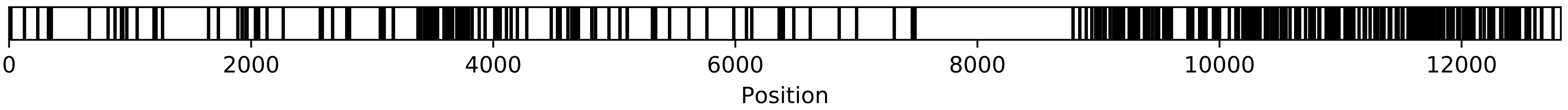


MA1 vs MM1
Number of differences: 263

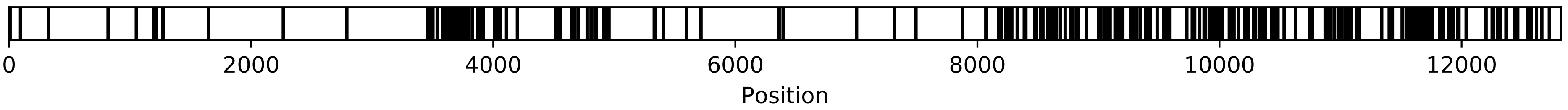


MA1 vs MM2
Number of differences: 248

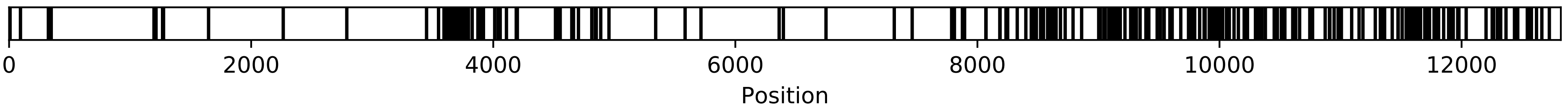


MA2 vs MM1
Number of differences: 339

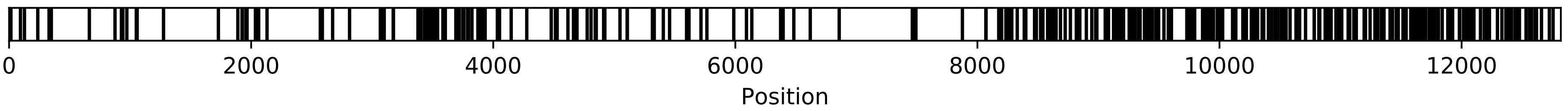


MA2 vs MM2
Number of differences: 322

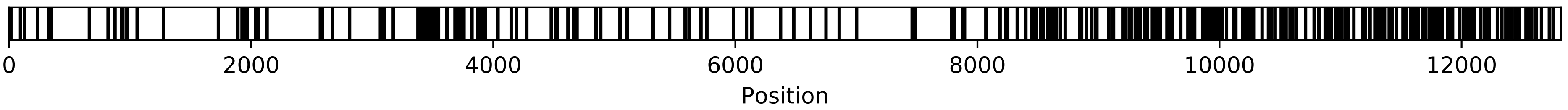


MM1 vs MM2
Number of differences: 159

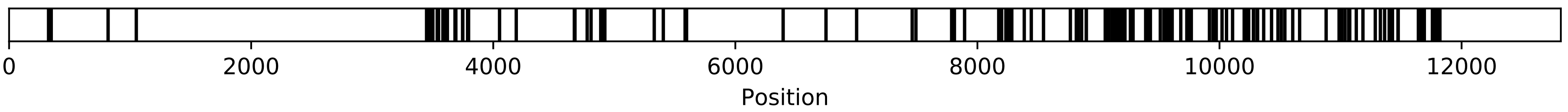

### Region: 454

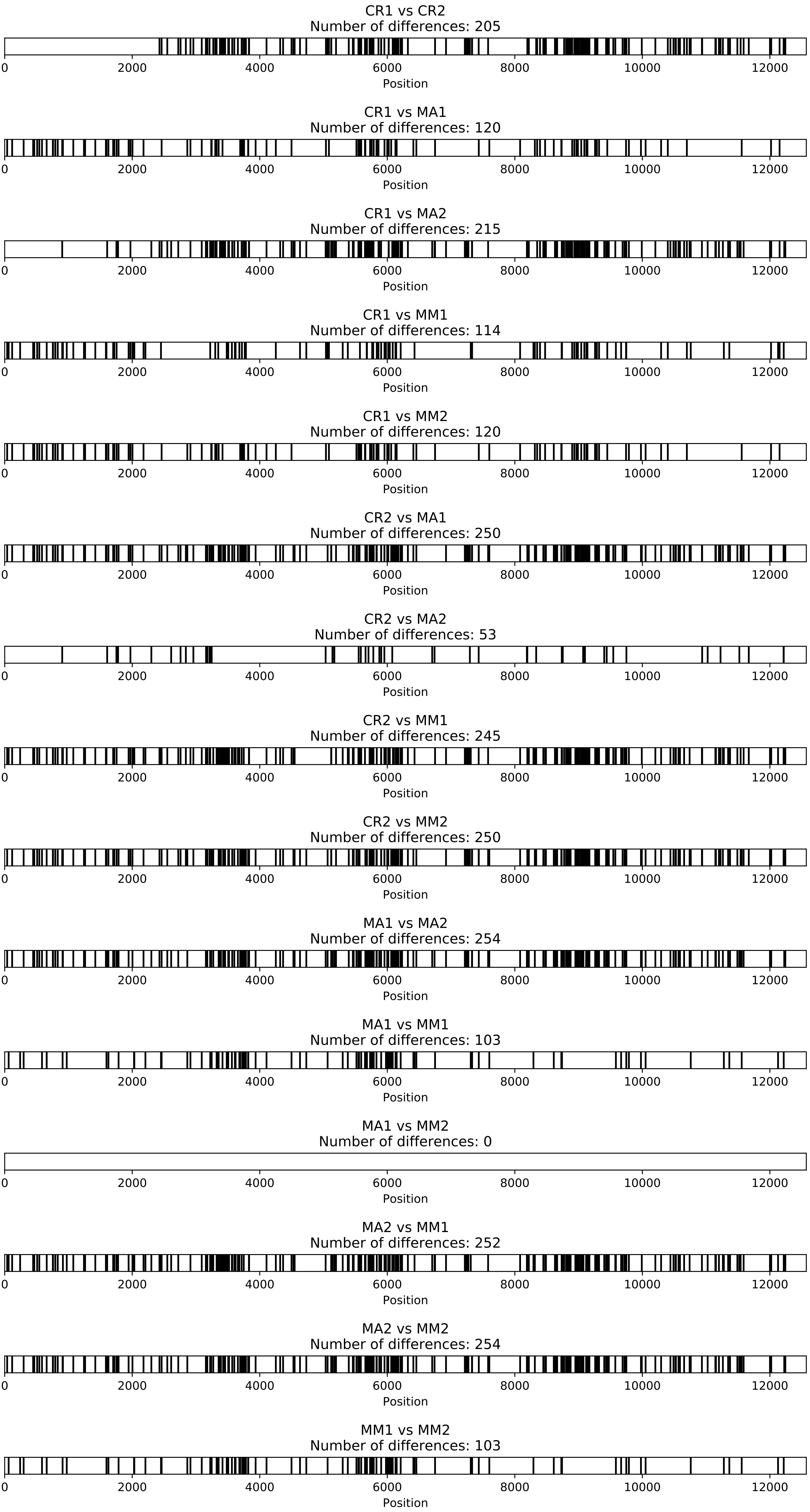
CR1 vs CR2
Number of differences: 205
CR1 vs MA1
Number of differences: 120
CR1 vs MA2
Number of differences: 215
CR1 vs MM1
Number of differences: 114
CR1 vs MM2
Number of differences: 120
CR2 vs MA1
Number of differences: 250
CR2 vs MA2
Number of differences: 53
CR2 vs MM1
Number of differences: 245
CR2 vs MM2
Number of differences: 250
MA1 vs MA2
Number of differences: 254
MA1 vs MM1
Number of differences: 103
MA1 vs MM2
Number of differences: 0
MA2 vs MM1
Number of differences: 252
MA2 vs MM2
Number of differences: 254
MM1 vs MM2
Number of differences: 103
0
2000
4000
6000
8000
10000
12000
Position

### Region: 456



CR1 vs CR2
Number of differences: 230

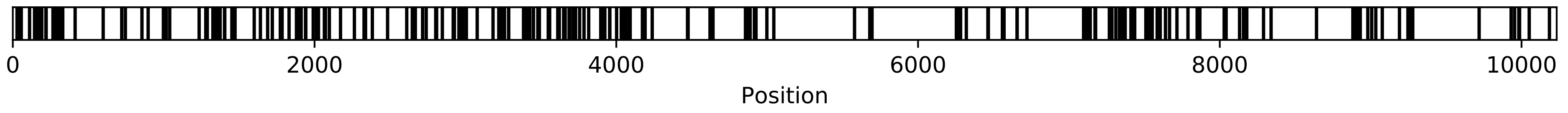


CR1 vs MA1
Number of differences: 143

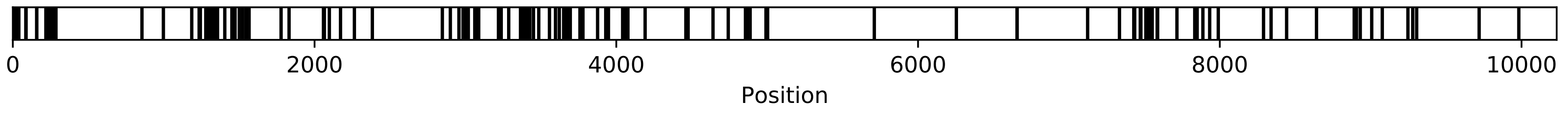


CR1 vs MA2
Number of differences: 252

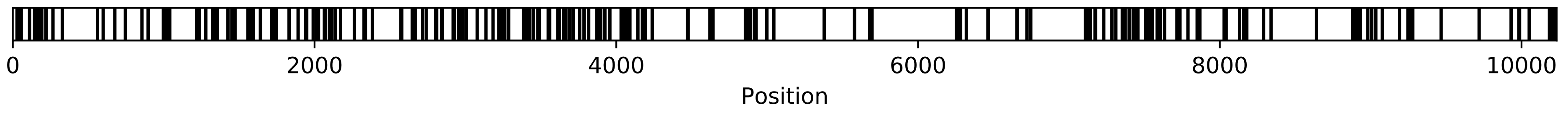


CR1 vs MM1
Number of differences: 143

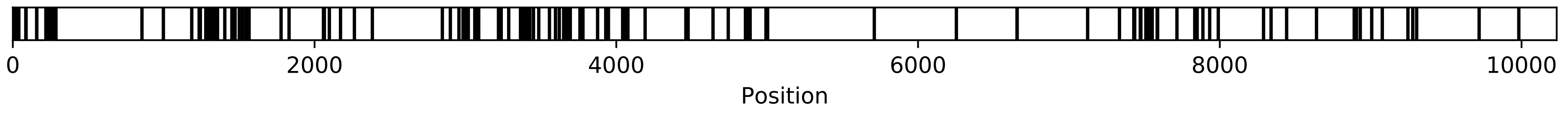


CR1 vs MM2
Number of differences: 127

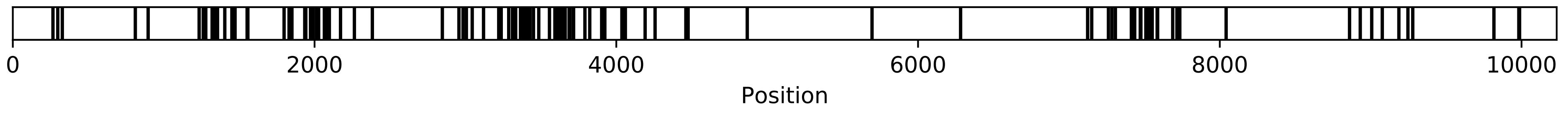


CR2 vs MA1
Number of differences: 205

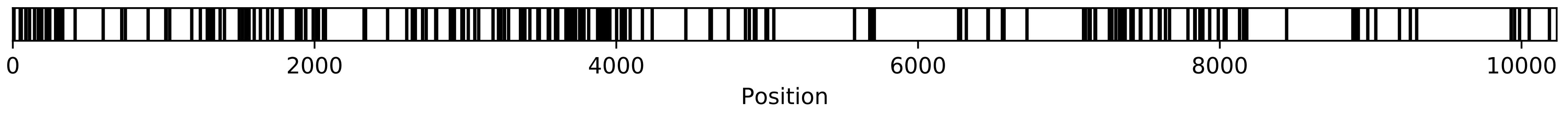


CR2 vs MA2
Number of differences: 97

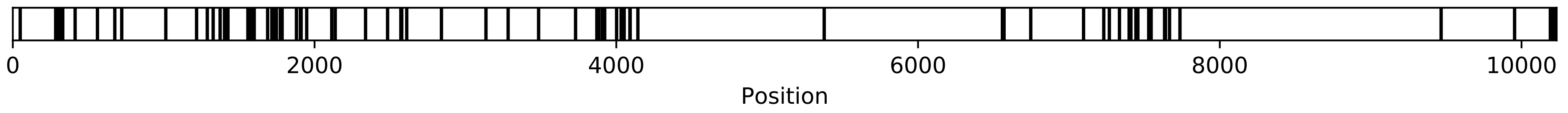


CR2 vs MM1
Number of differences: 205

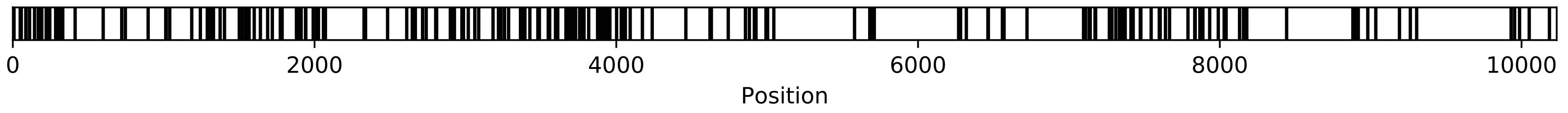


CR2 vs MM2
Number of differences: 201

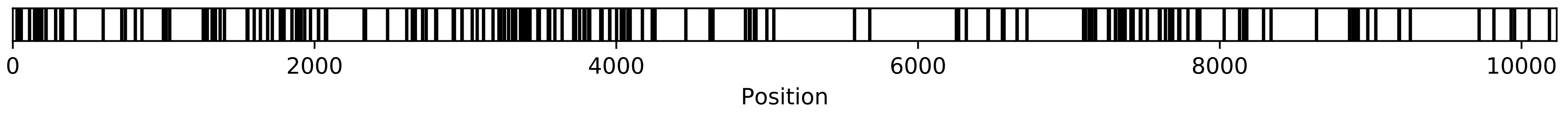


MA1 vs MA2
Number of differences: 226

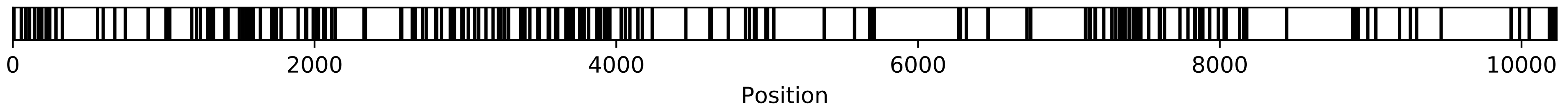


MA1 vs MM1
Number of differences: 0

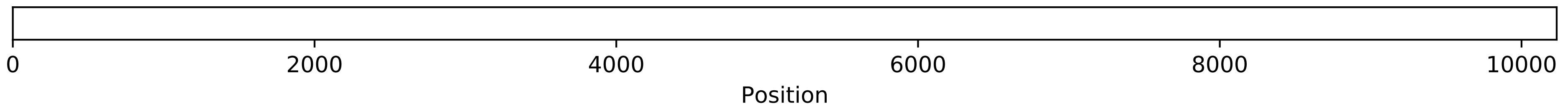


MA1 vs MM2
Number of differences: 134

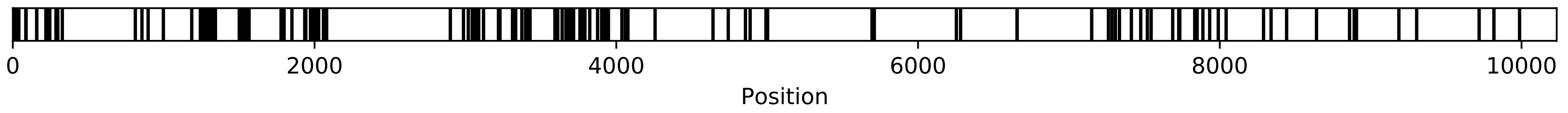


MA2 vs MM1
Number of differences: 226

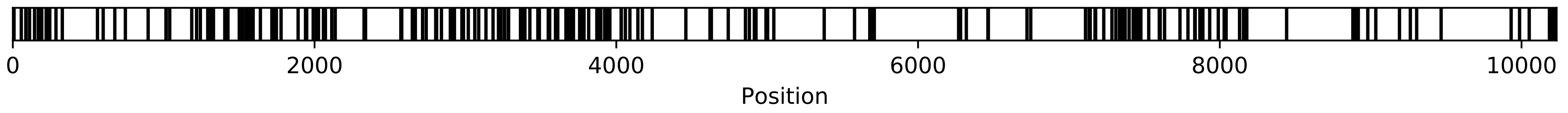


MA2 vs MM2
Number of differences: 223

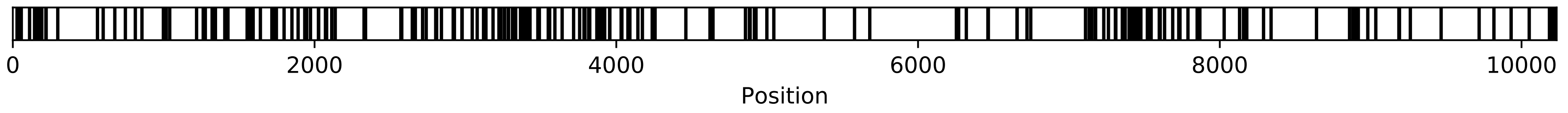


MM1 vs MM2
Number of differences: 134

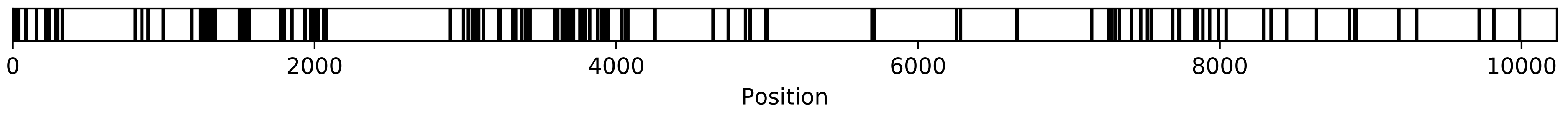

### Region: 457

CR1 vs CR2
Number of differences: 124

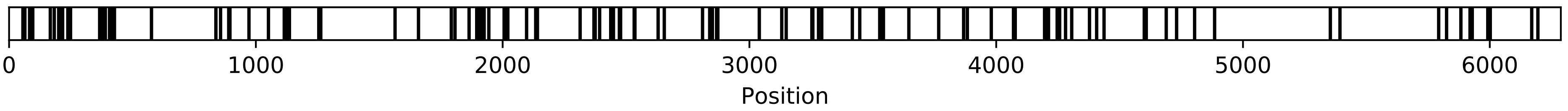


CR1 vs MA1
Number of differences: 59

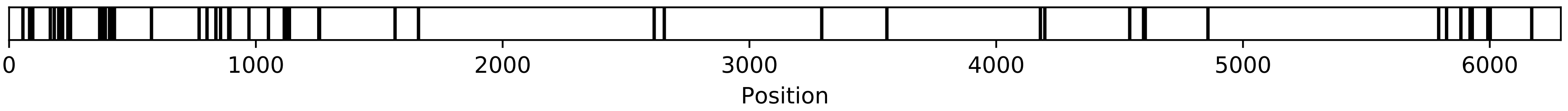


CR1 vs MA2
Number of differences: 90

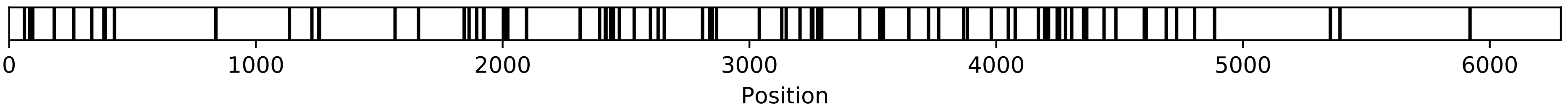


CR1 vs MM1
Number of differences: 82

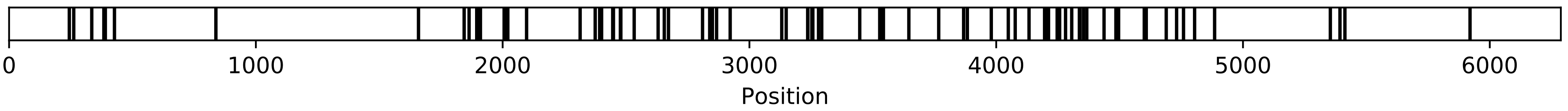


CR1 vs MM2
Number of differences: 90

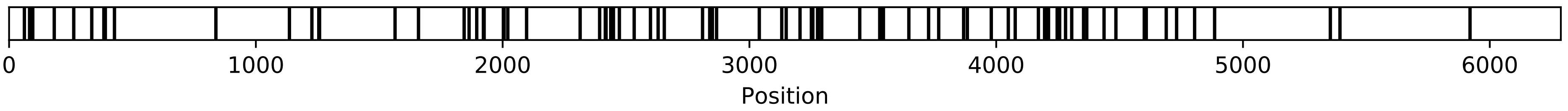


CR2 vs MA1
Number of differences: 83

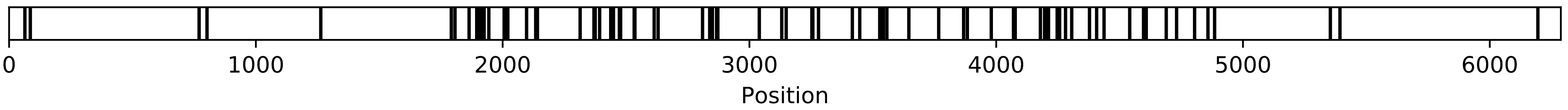


CR2 vs MA2
Number of differences: 74

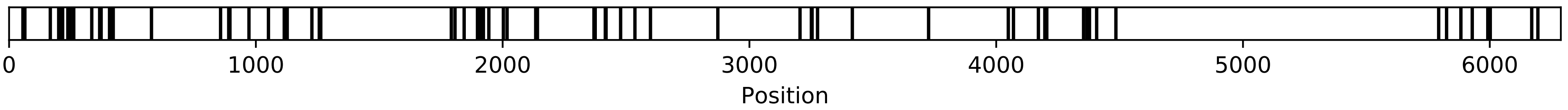


CR2 vs MM1
Number of differences: 88

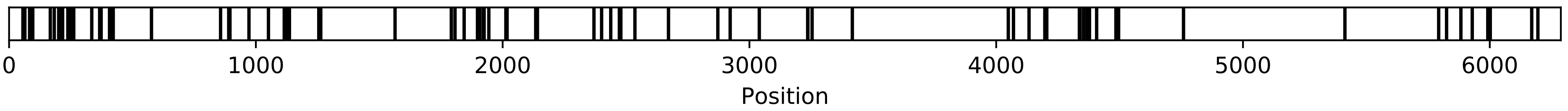


CR2 vs MM2
Number of differences: 74

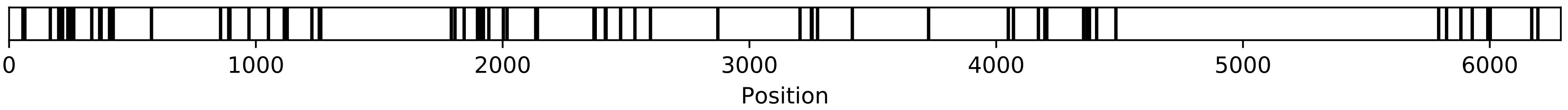


MA1 vs MA2
Number of differences: 112

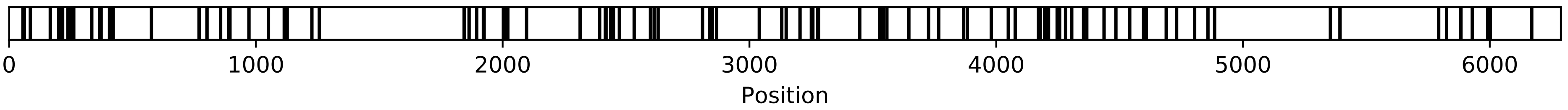


MA1 vs MM1
Number of differences: 121

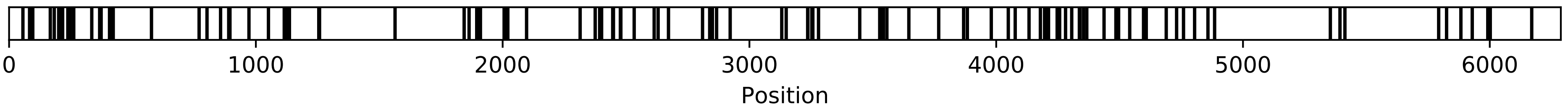


MA1 vs MM2
Number of differences: 112

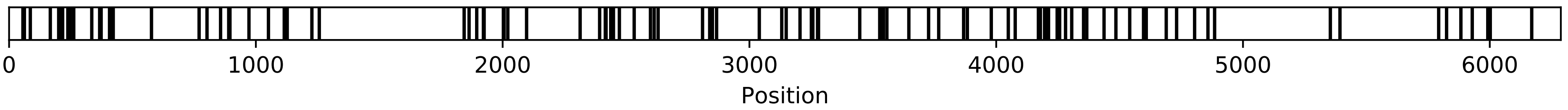


MA2 vs MM1
Number of differences: 44

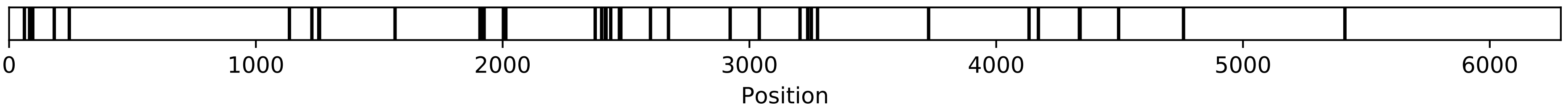


MA2 vs MM2
Number of differences: 0

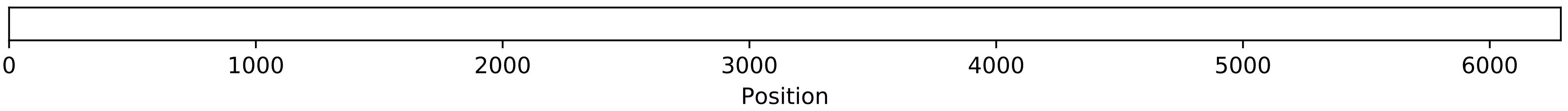


MM1 vs MM2
Number of differences: 44

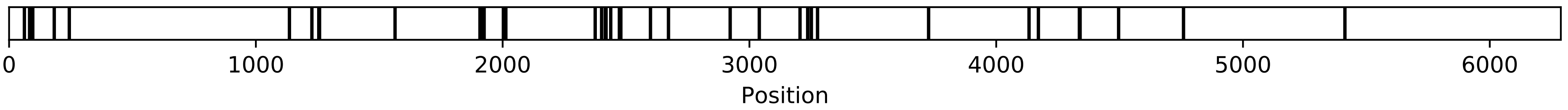

### Region: 458

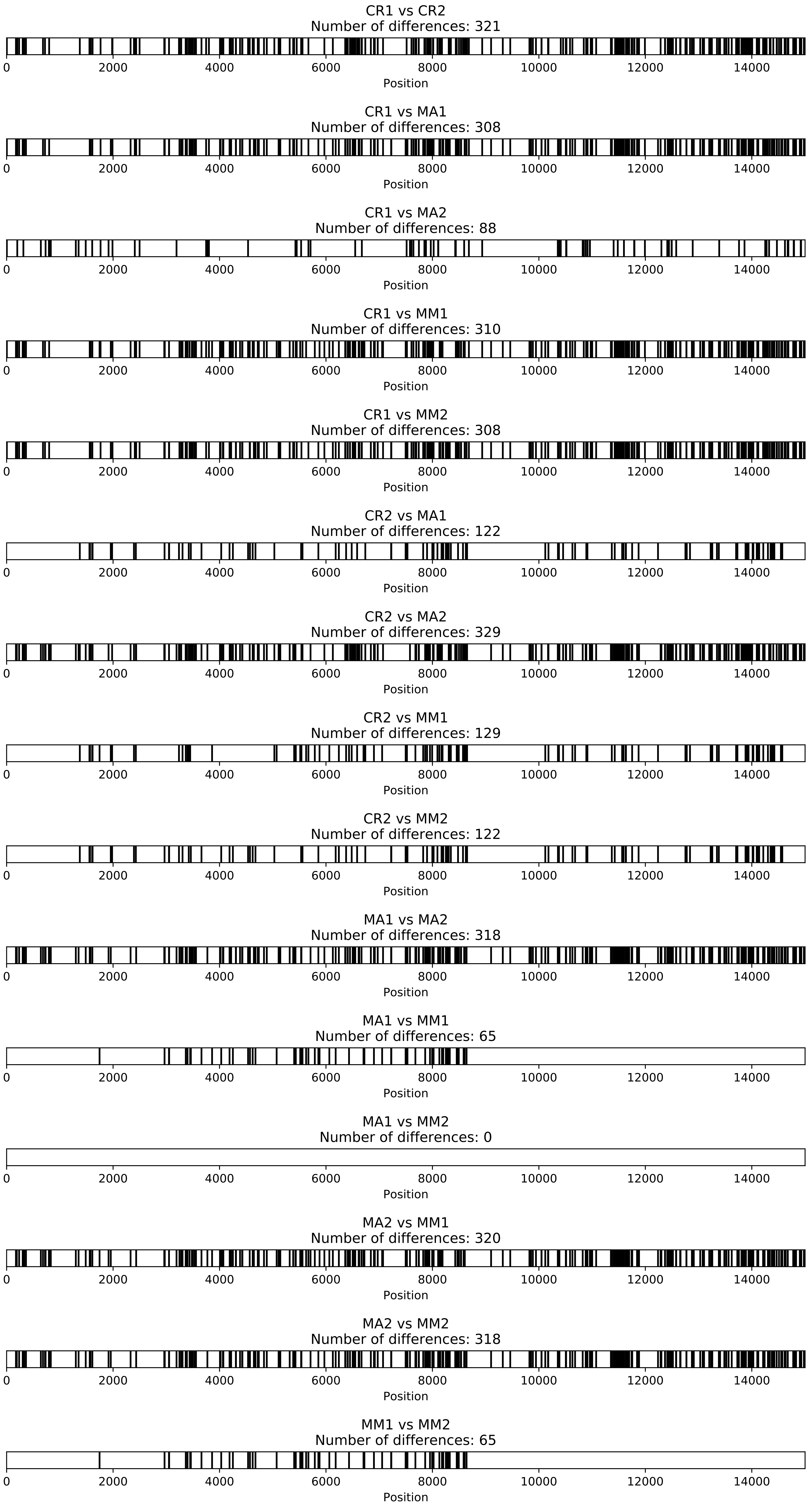
CR1 vs CR2
Number of differences: 321
CR1 vs MA1
Number of differences: 308
CR1 vs MA2
Number of differences: 88
CR1 vs MM1
Number of differences: 310
CR1 vs MM2
Number of differences: 308
CR2 vs MA1
Number of differences: 122
CR2 vs MA2
Number of differences: 329
CR2 vs MM1
Number of differences: 129
CR2 vs MM2
Number of differences: 122
MA1 vs MA2
Number of differences: 318
MA1 vs MM1
Number of differences: 65
MA1 vs MM2
Number of differences: 0
MA2 vs MM1
Number of differences: 320
MA2 vs MM2
Number of differences: 318
MM1 vs MM2
Number of differences: 65
0
2000
4000
6000
8000
10000
12000
14000
Position

### Region: 459



CR1 vs CR2
Number of differences: 283

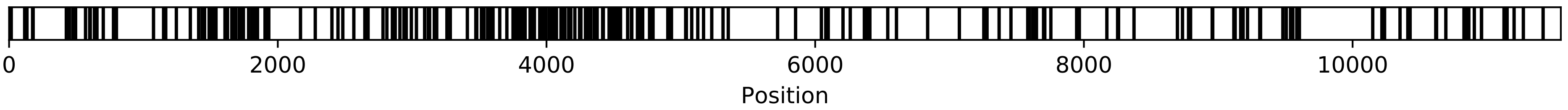


CR1 vs MA1
Number of differences: 151

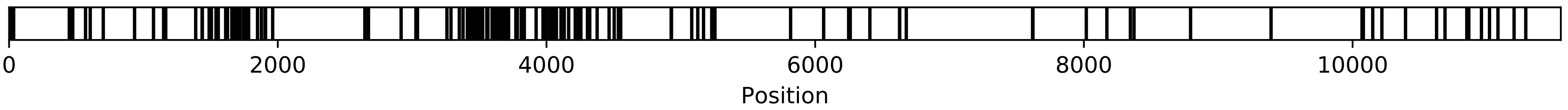


CR1 vs MA2
Number of differences: 274

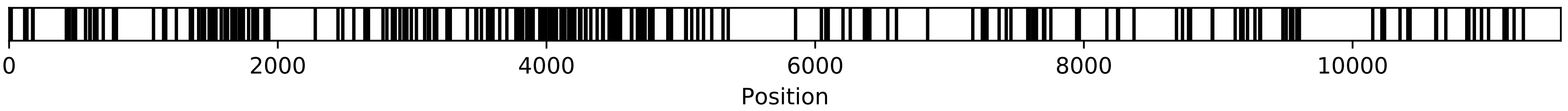


CR1 vs MM1
Number of differences: 161

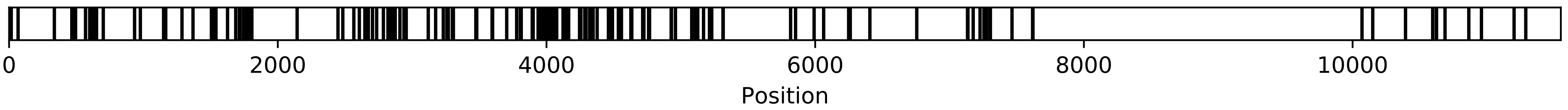


CR1 vs MM2
Number of differences: 141

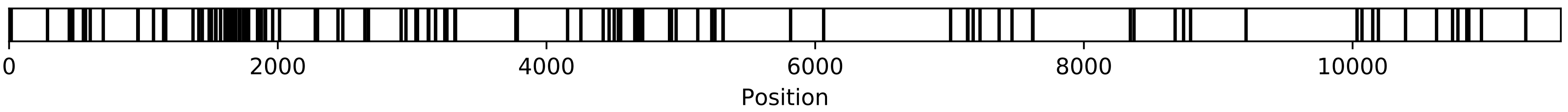


CR2 vs MA1
Number of differences: 270

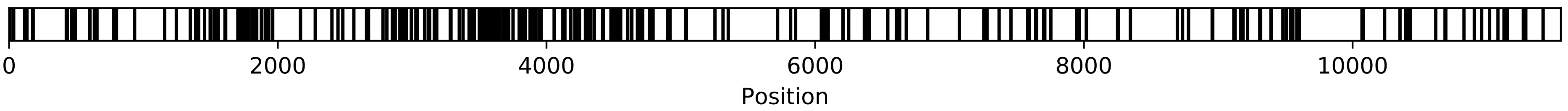


CR2 vs MA2
Number of differences: 53

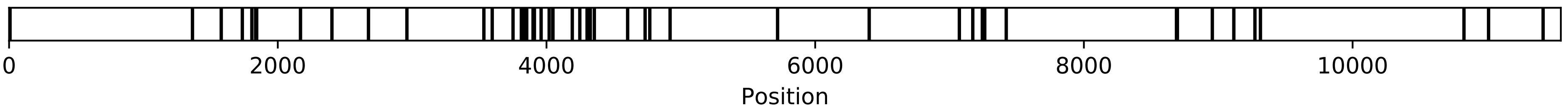


CR2 vs MM1
Number of differences: 287

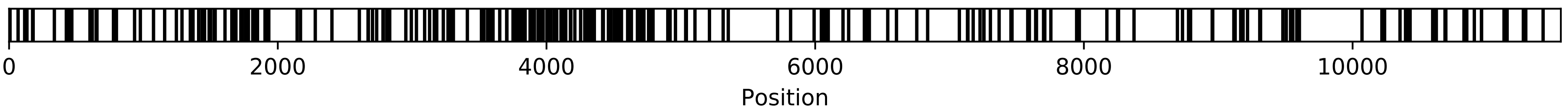


CR2 vs MM2
Number of differences: 287

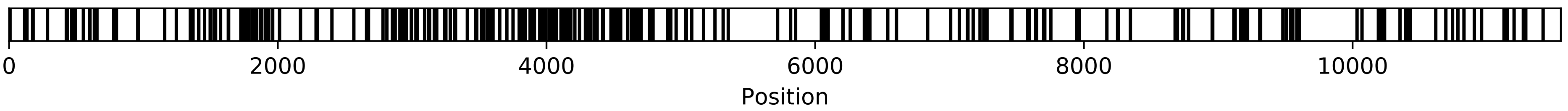


MA1 vs MA2
Number of differences: 263

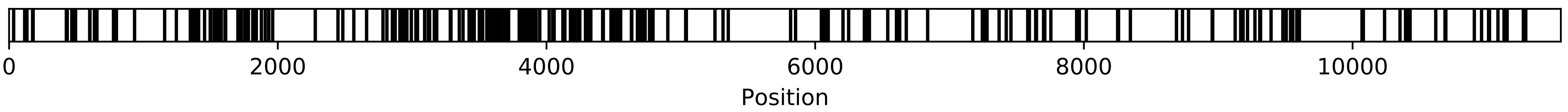


MA1 vs MM1
Number of differences: 189

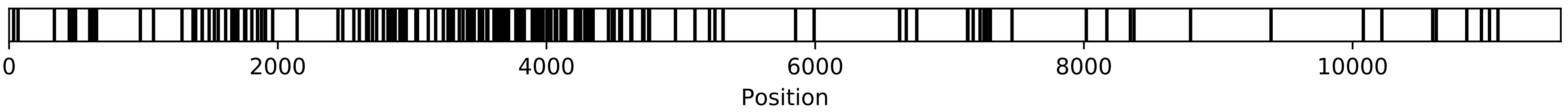


MA1 vs MM2
Number of differences: 138

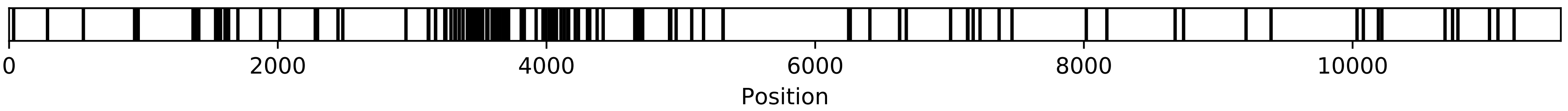


MA2 vs MM1
Number of differences: 281

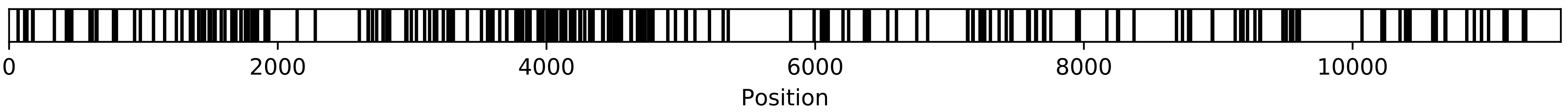


MA2 vs MM2
Number of differences: 277

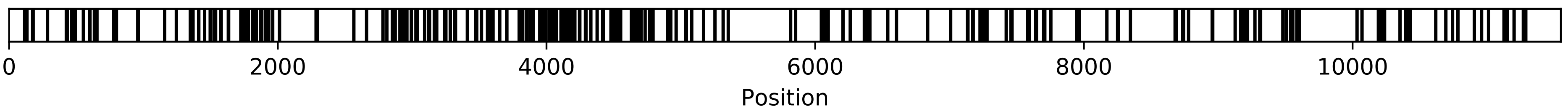


MM1 vs MM2
Number of differences: 191

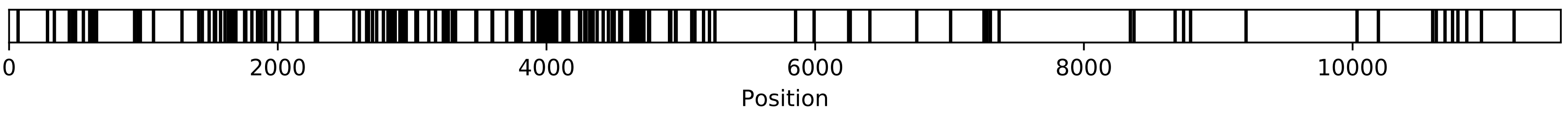

### Region: 461

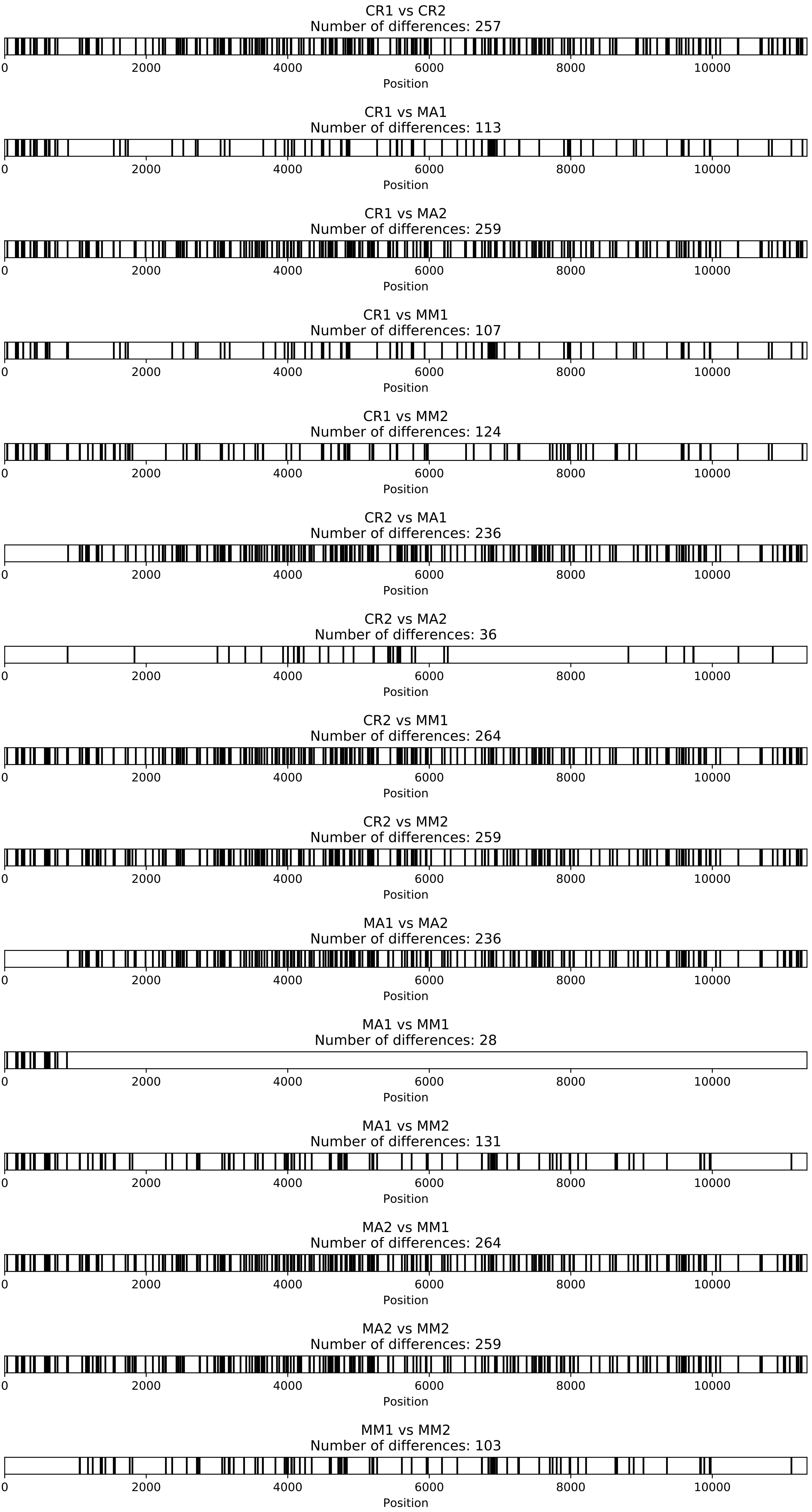
CR1 vs CR2
Number of differences: 257
CR1 vs MA1
Number of differences: 113
CR1 vs MA2
Number of differences: 259
CR1 vs MM1
Number of differences: 107
CR1 vs MM2
Number of differences: 124
CR2 vs MA1
Number of differences: 236
CR2 vs MA2
Number of differences: 36
CR2 vs MM1
Number of differences: 264
CR2 vs MM2
Number of differences: 259
MA1 vs MA2
Number of differences: 236
MA1 vs MM1
Number of differences: 28
MA1 vs MM2
Number of differences: 131
MA2 vs MM1
Number of differences: 264
MA2 vs MM2
Number of differences: 259
MM1 vs MM2
Number of differences: 103
0
2000
4000
6000
8000
10000
Position

### Region: 462



CR1 vs CR2
Number of differences: 322

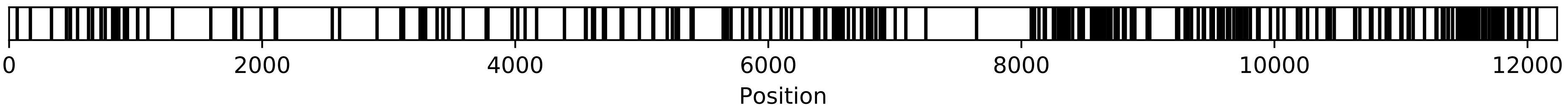


CR1 vs MA1
Number of differences: 282

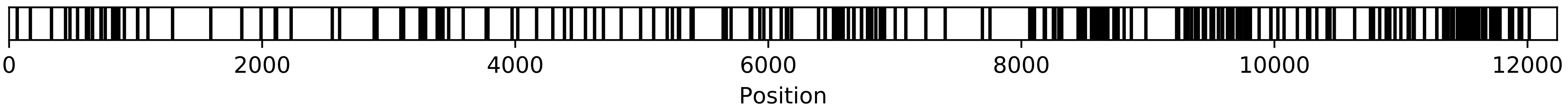


CR1 vs MA2
Number of differences: 132

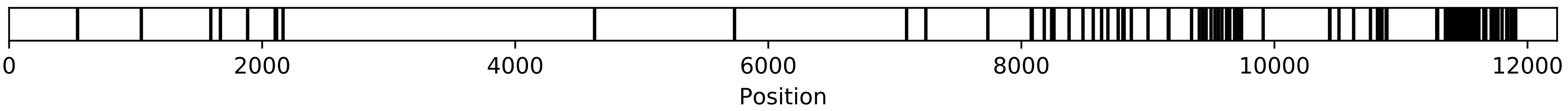


CR1 vs MM1
Number of differences: 292

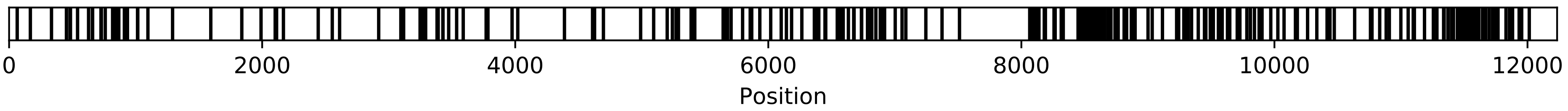


CR1 vs MM2
Number of differences: 293

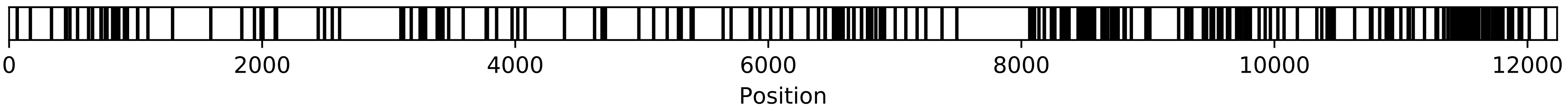


CR2 vs MA1
Number of differences: 150

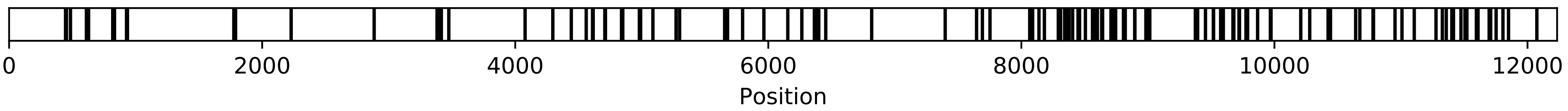


CR2 vs MA2
Number of differences: 318

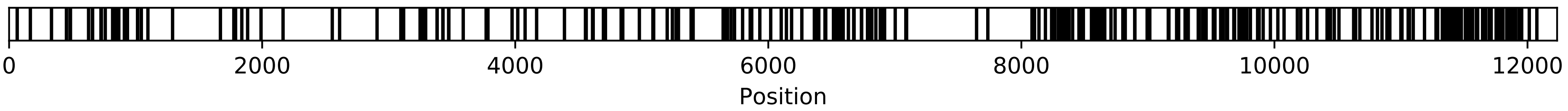


CR2 vs MM1
Number of differences: 133

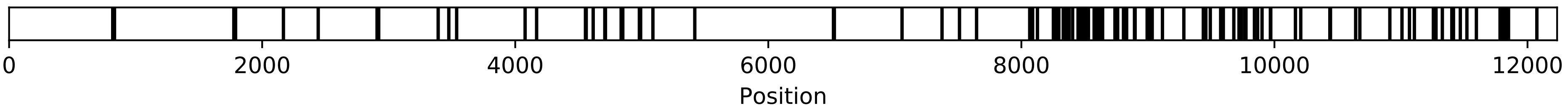


CR2 vs MM2
Number of differences: 141

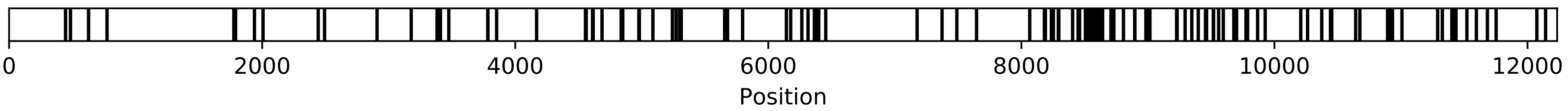


MA1 vs MA2
Number of differences: 281

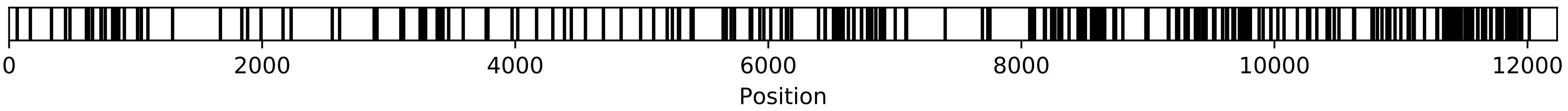


MA1 vs MM1
Number of differences: 132

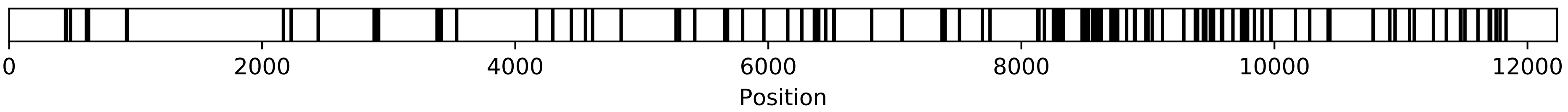


MA1 vs MM2
Number of differences: 124

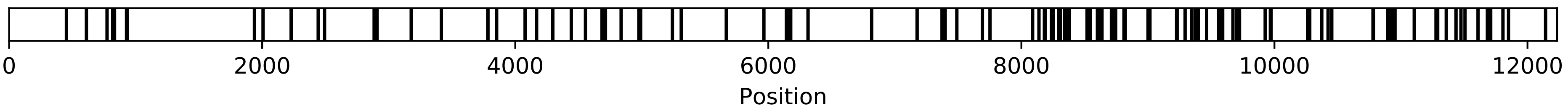


MA2 vs MM1
Number of differences: 286

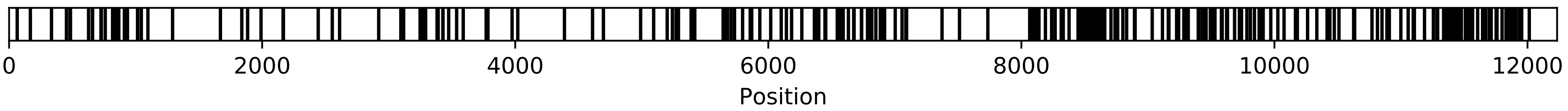


MA2 vs MM2
Number of differences: 284

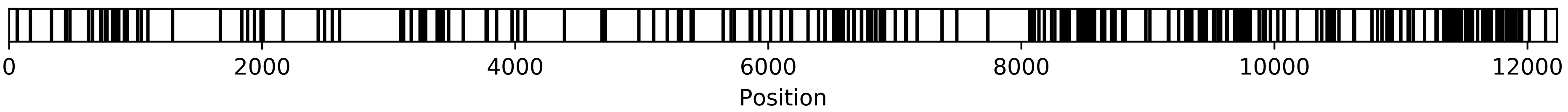


MM1 vs MM2
Number of differences: 148

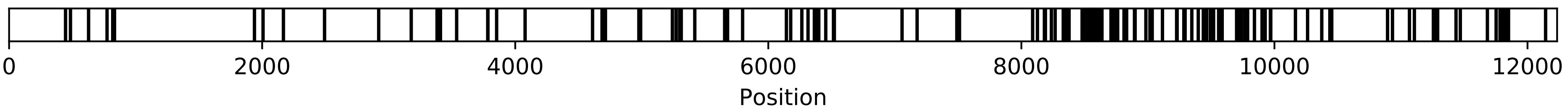

### Region: 464

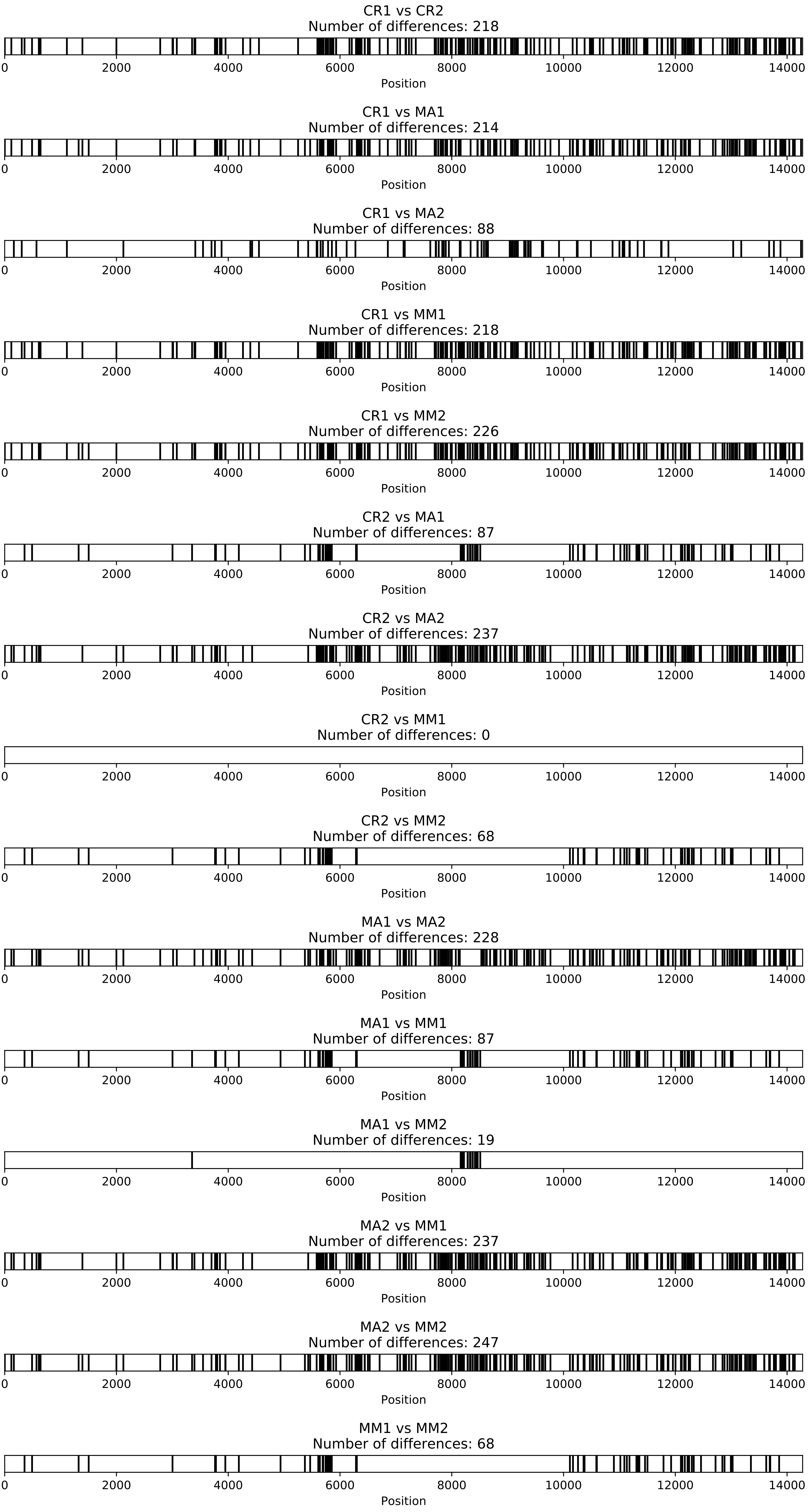
CR1 vs CR2
Number of differences: 218
CR1 vs MA1
Number of differences: 214
CR1 vs MA2
Number of differences: 88
CR1 vs MM1
Number of differences: 218
CR1 vs MM2
Number of differences: 226
CR2 vs MA1
Number of differences: 87
CR2 vs MA2
Number of differences: 237
CR2 vs MM1
Number of differences: 0
CR2 vs MM2
Number of differences: 68
MA1 vs MA2
Number of differences: 228
MA1 vs MM1
Number of differences: 87
MA1 vs MM2
Number of differences: 19
MA2 vs MM1
Number of differences: 237
MA2 vs MM2
Number of differences: 247
MM1 vs MM2
Number of differences: 68
0
2000
4000
6000
8000
10000
12000
14000
Position

### Region: 465



CR1 vs CR2
Number of differences: 221

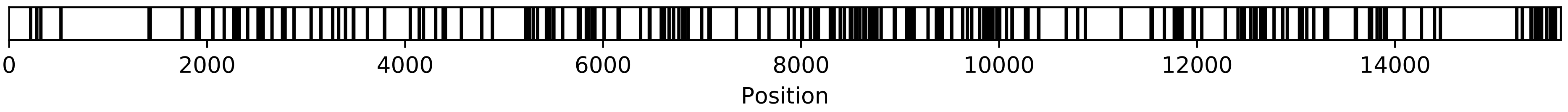


CR1 vs MA1
Number of differences: 216

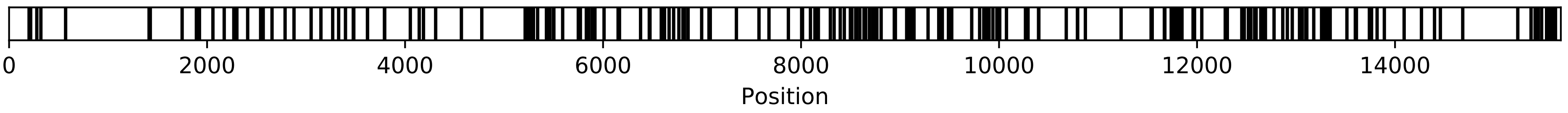


CR1 vs MA2
Number of differences: 76

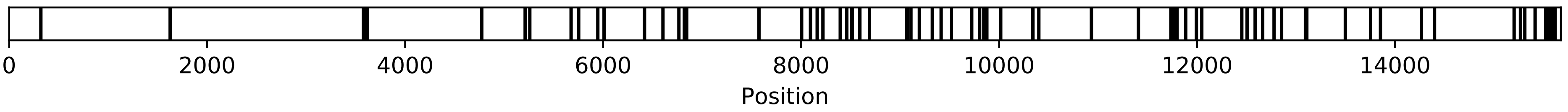


CR1 vs MM1
Number of differences: 76

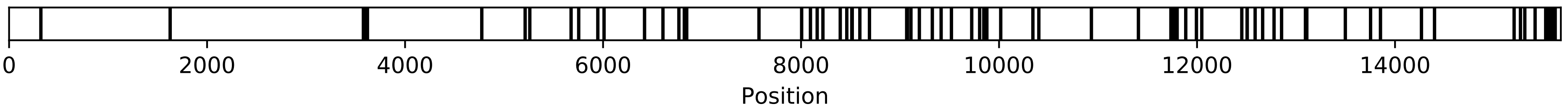


CR1 vs MM2
Number of differences: 85

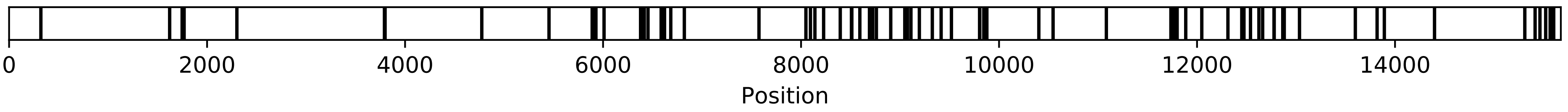


CR2 vs MA1
Number of differences: 49

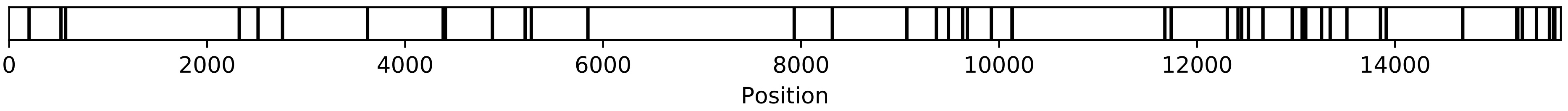


CR2 vs MA2
Number of differences: 217

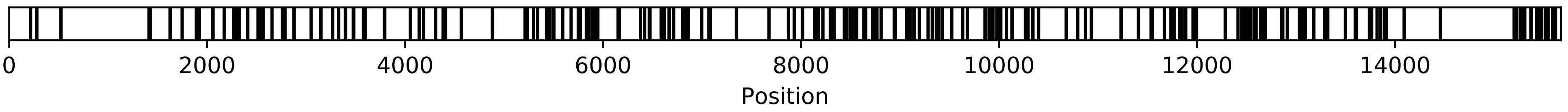


CR2 vs MM1
Number of differences: 217

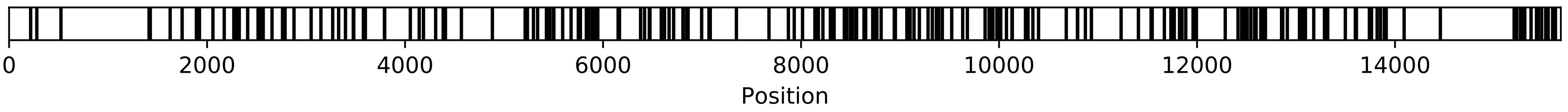


CR2 vs MM2
Number of differences: 209

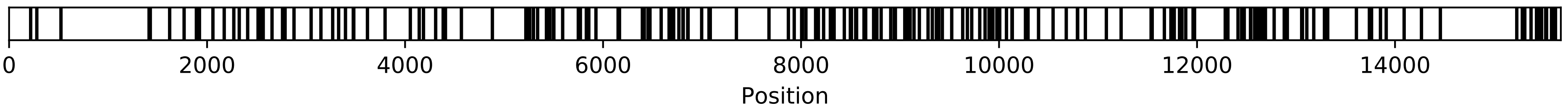


MA1 vs MA2
Number of differences: 211

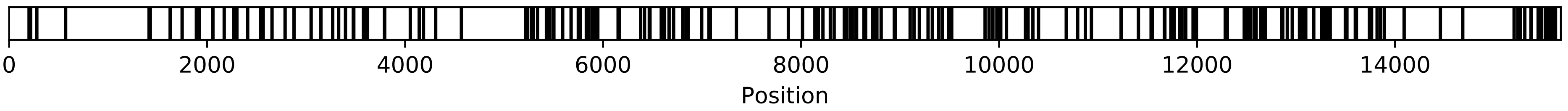


MA1 vs MM1
Number of differences: 211

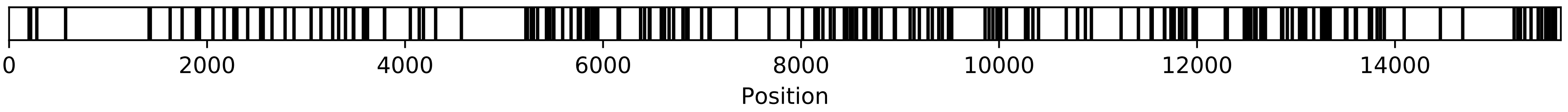


MA1 vs MM2
Number of differences: 204

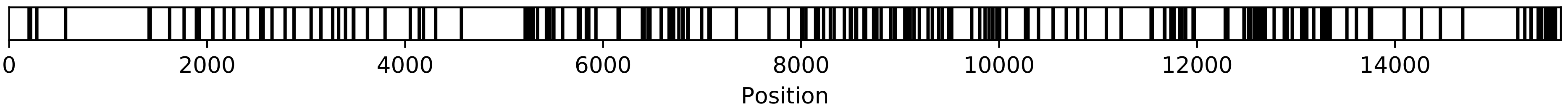


MA2 vs MM1
Number of differences: 0

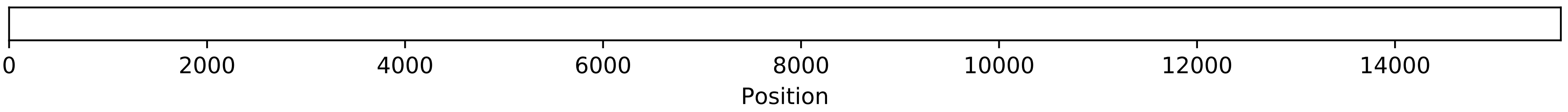


MA2 vs MM2
Number of differences: 83

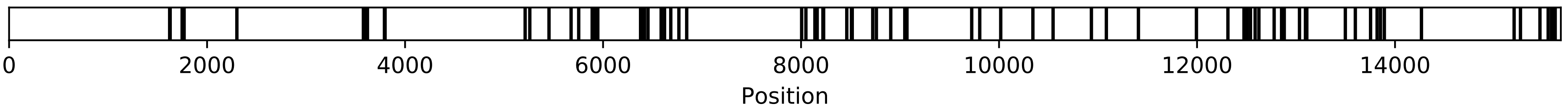


MM1 vs MM2
Number of differences: 83

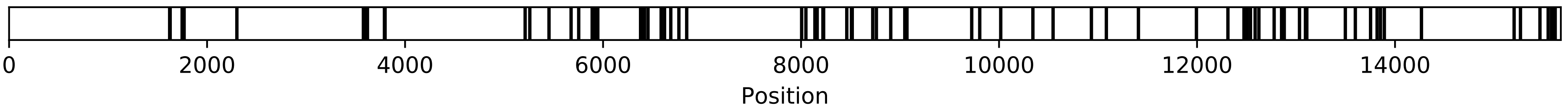

### Region: 467



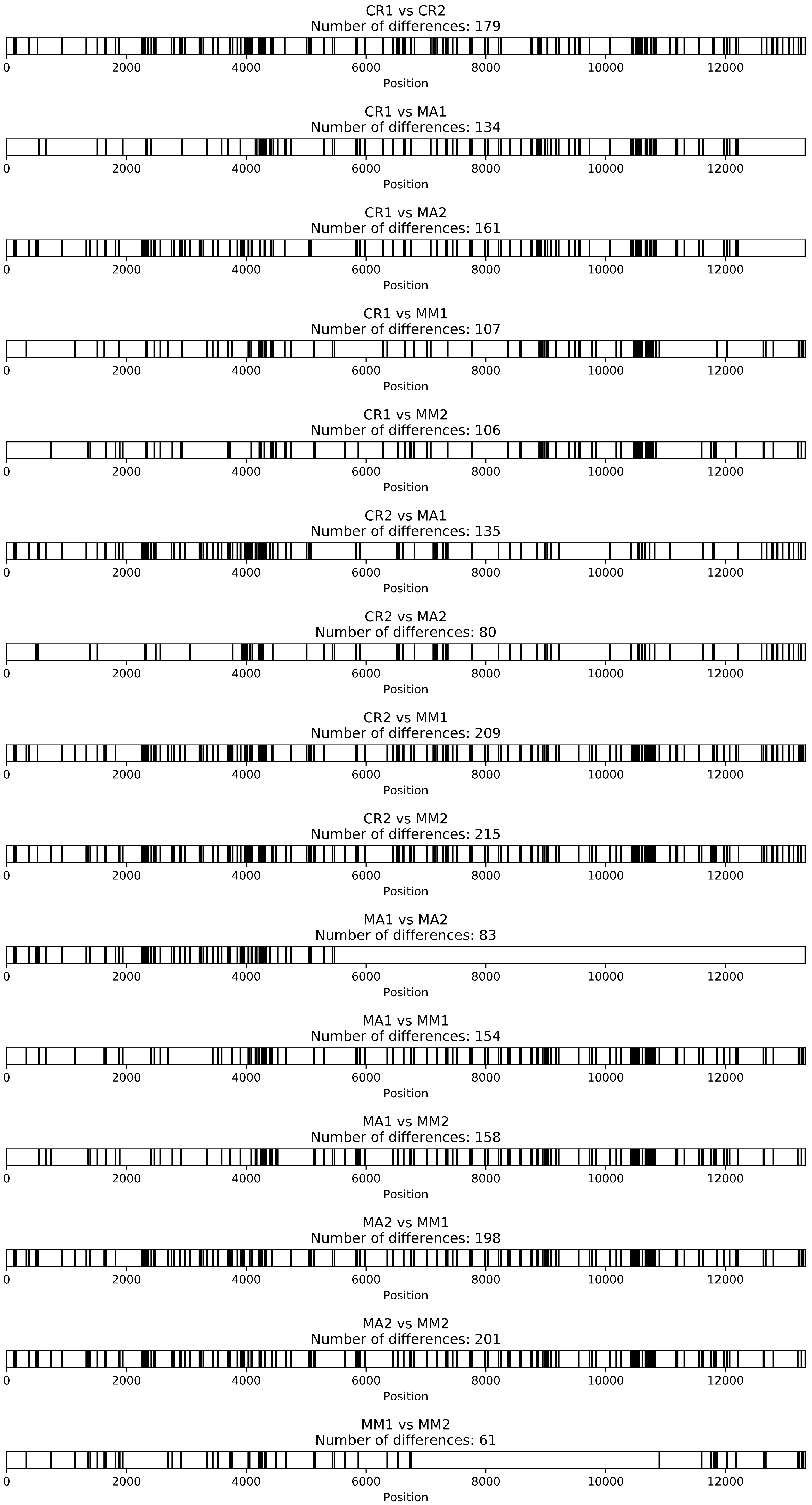
CR1 vs CR2
Number of differences: 179
CR1 vs MA1
Number of differences: 134
CR1 vs MA2
Number of differences: 161
CR1 vs MM1
Number of differences: 107
CR1 vs MM2
Number of differences: 106
CR2 vs MA1
Number of differences: 135
CR2 vs MA2
Number of differences: 80
CR2 vs MM1
Number of differences: 209
CR2 vs MM2
Number of differences: 215
MA1 vs MA2
Number of differences: 83
MA1 vs MM1
Number of differences: 154
MA1 vs MM2
Number of differences: 158
MA2 vs MM1
Number of differences: 198
MA2 vs MM2
Number of differences: 201
MM1 vs MM2
Number of differences: 61
Position
0
2000
4000
6000
8000
10000
12000

### Region: 472

CR1 vs CR2
Number of differences: 262

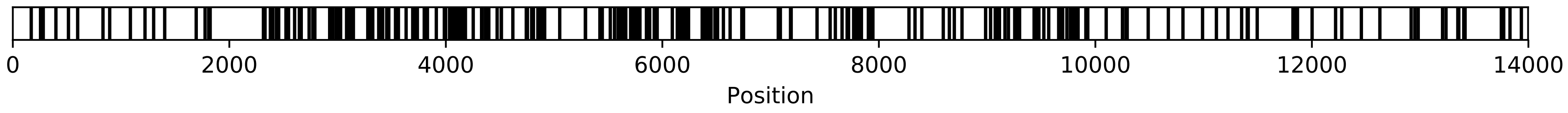


CR1 vs MA1
Number of differences: 250

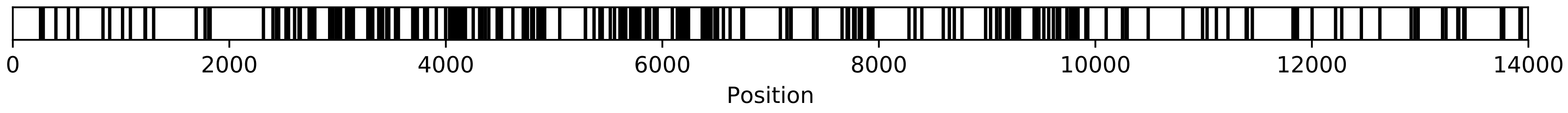


CR1 vs MA2
Number of differences: 124

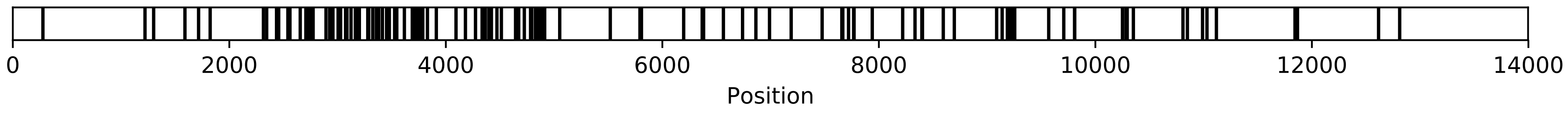


CR1 vs MM1
Number of differences: 124

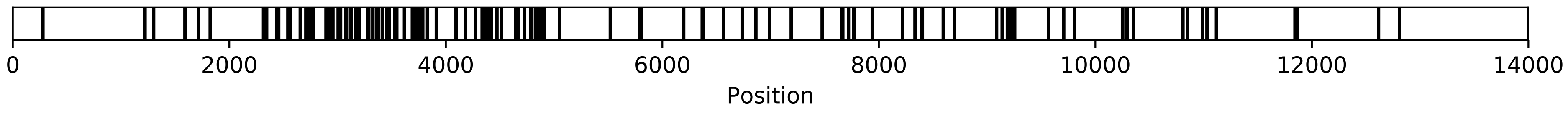


CR1 vs MM2
Number of differences: 129

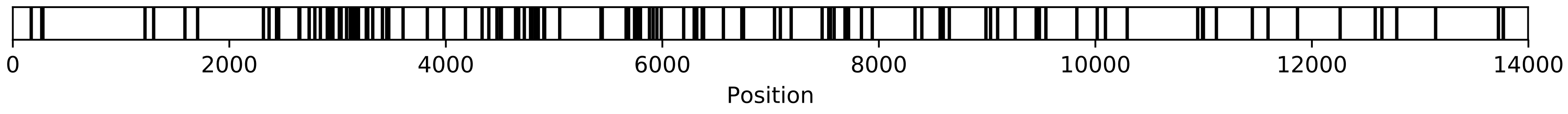


CR2 vs MA1
Number of differences: 68

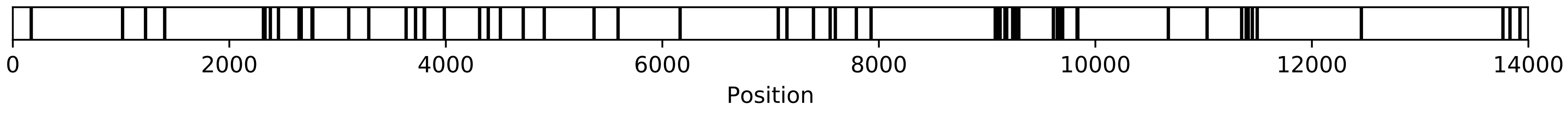


CR2 vs MA2
Number of differences: 260

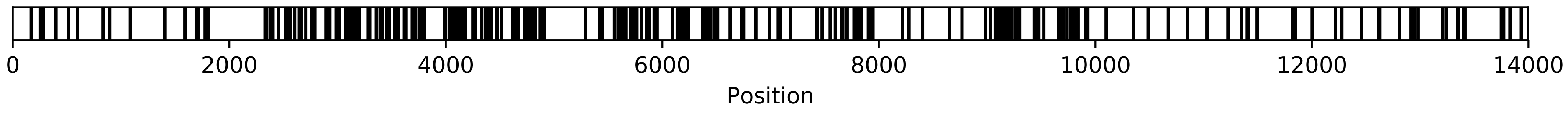


CR2 vs MM1
Number of differences: 260

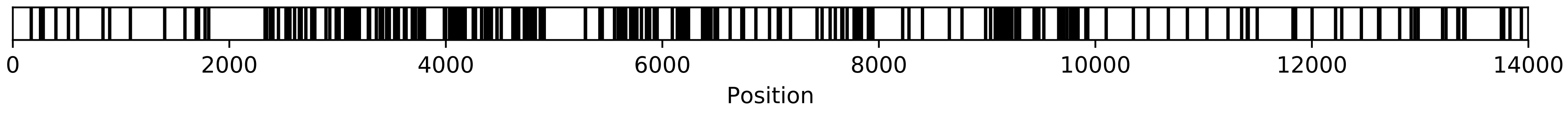


CR2 vs MM2
Number of differences: 262

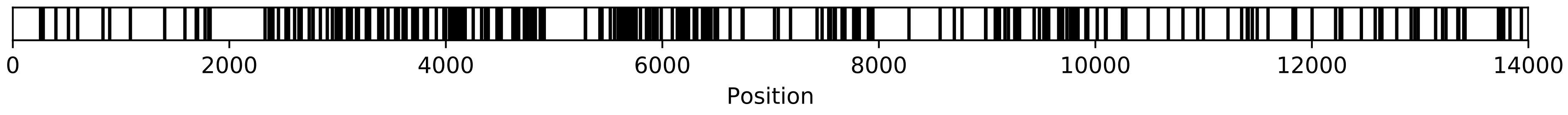


MA1 vs MA2
Number of differences: 247

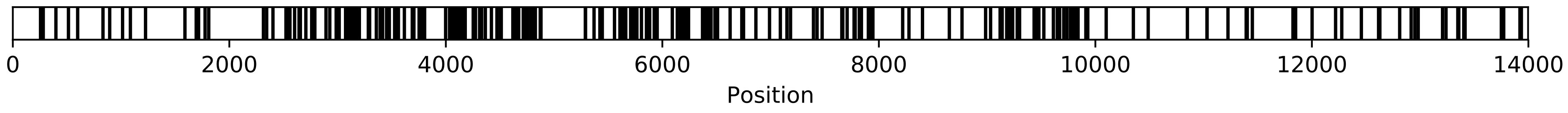


MA1 vs MM1
Number of differences: 247

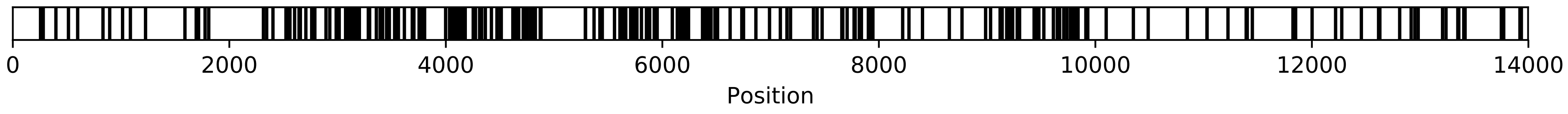


MA1 vs MM2
Number of differences: 251

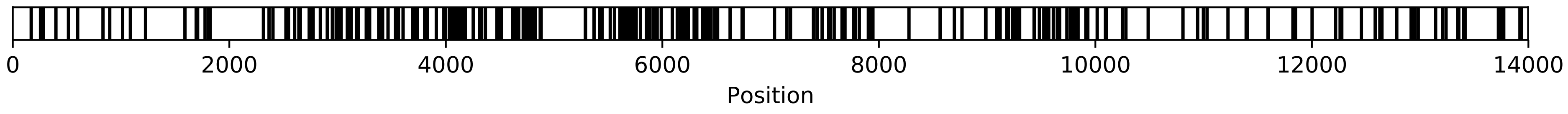


MA2 vs MM1
Number of differences: 0

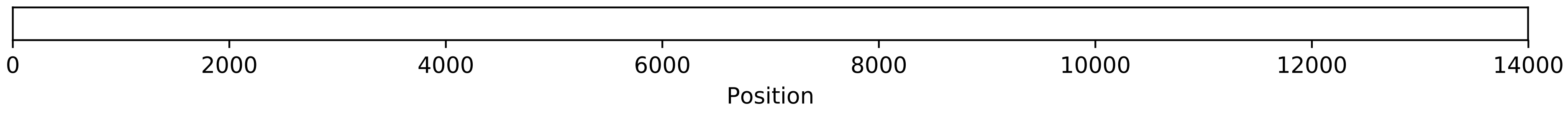


MA2 vs MM2
Number of differences: 149

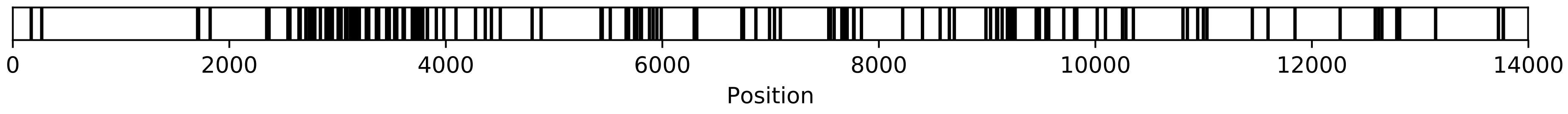


MM1 vs MM2
Number of differences: 149

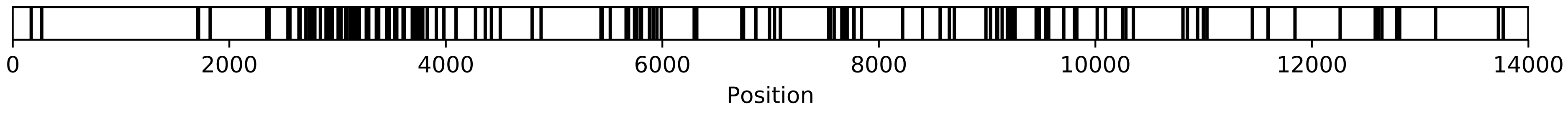

### Region: 473



CR1 vs CR2
Number of differences: 102

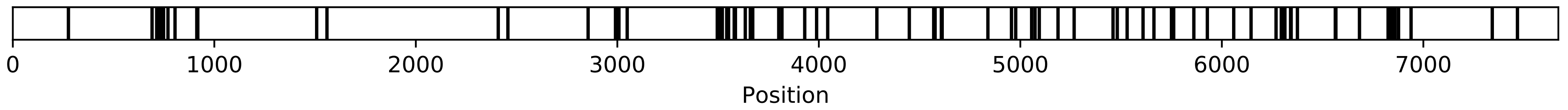


CR1 vs MA1
Number of differences: 101

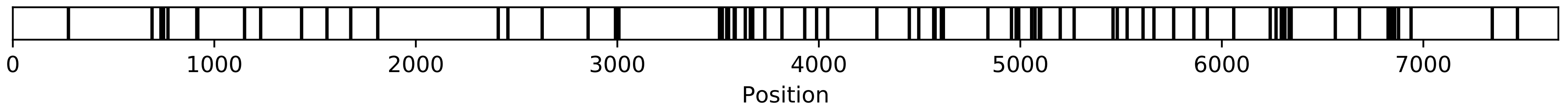


CR1 vs MA2
Number of differences: 6

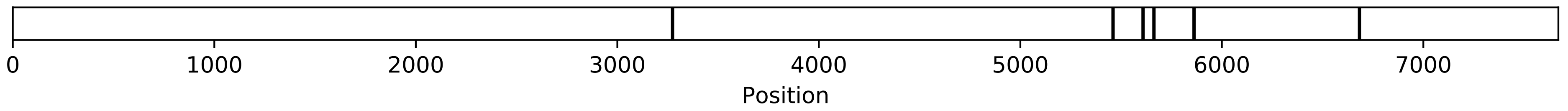


CR1 vs MM1
Number of differences: 99

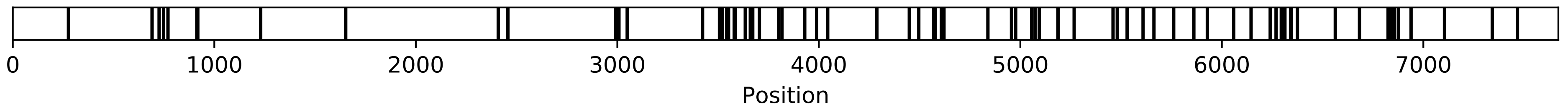


CR1 vs MM2
Number of differences: 96

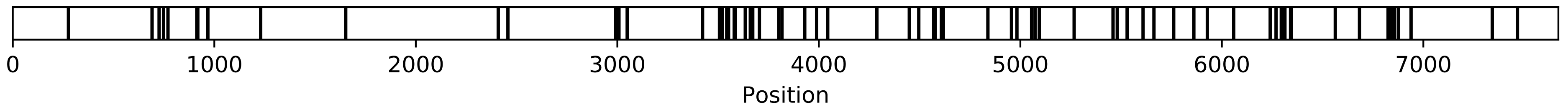


CR2 vs MA1
Number of differences: 33

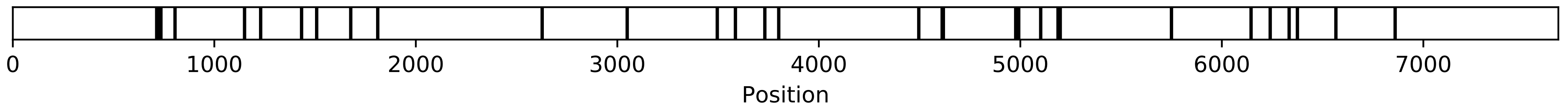


CR2 vs MA2
Number of differences: 100

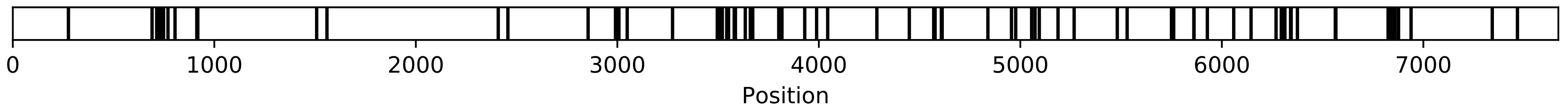


CR2 vs MM1
Number of differences: 19

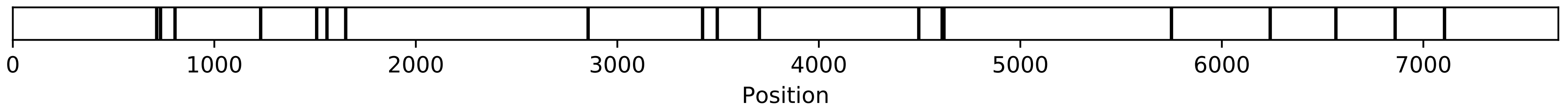


CR2 vs MM2
Number of differences: 24

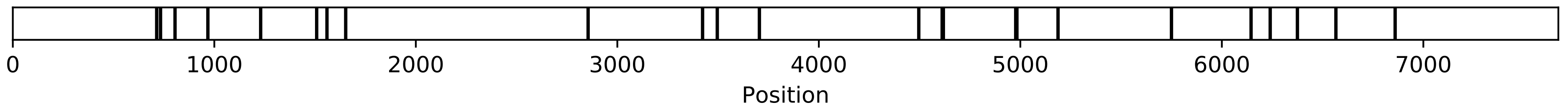


MA1 vs MA2
Number of differences: 99

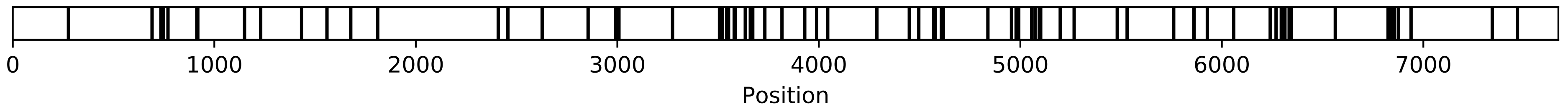


MA1 vs MM1
Number of differences: 28

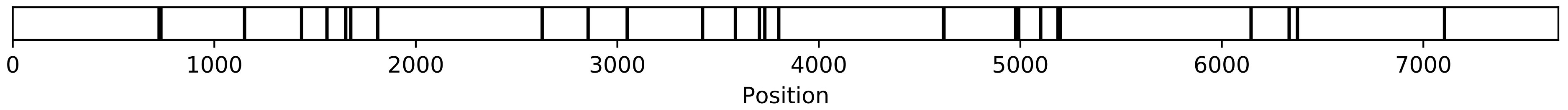


MA1 vs MM2
Number of differences: 23

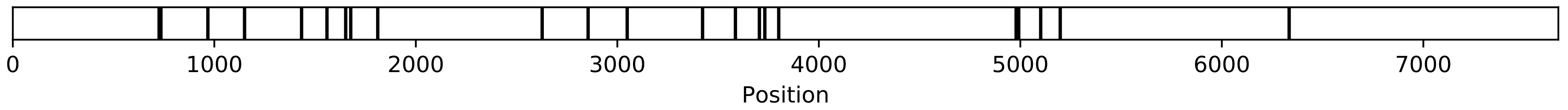


MA2 vs MM1
Number of differences: 97

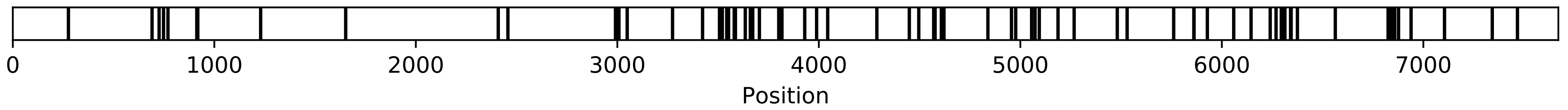


MA2 vs MM2
Number of differences: 94

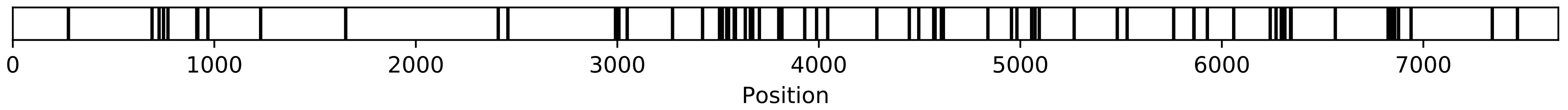


MM1 vs MM2
Number of differences: 9

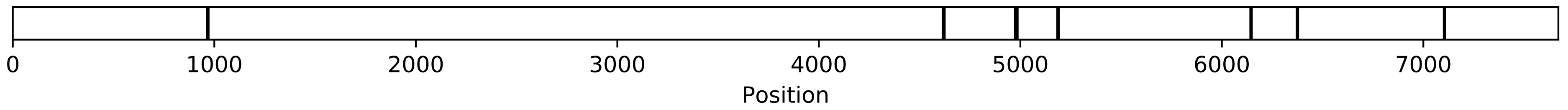

### Region: 475



CR1 vs CR2
Number of differences: 100

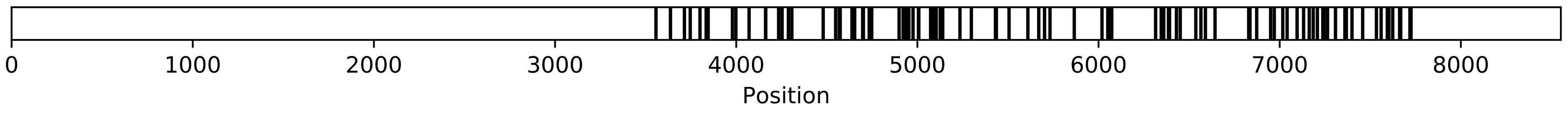


CR1 vs MA1
Number of differences: 34

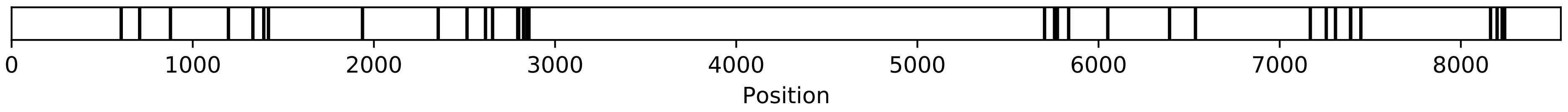


CR1 vs MA2
Number of differences: 178

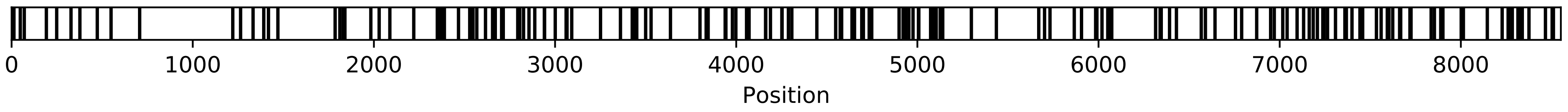


CR1 vs MM1
Number of differences: 180

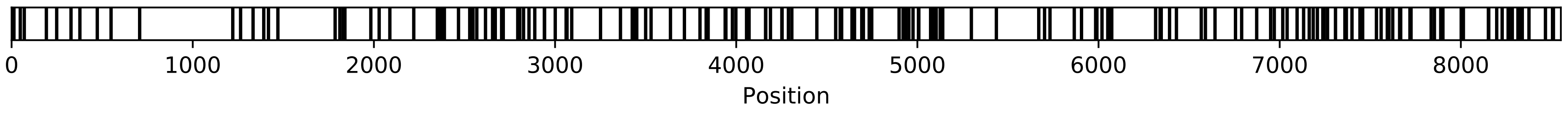


CR1 vs MM2
Number of differences: 189

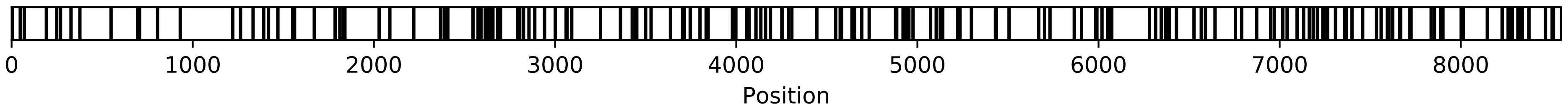


CR2 vs MA1
Number of differences: 126

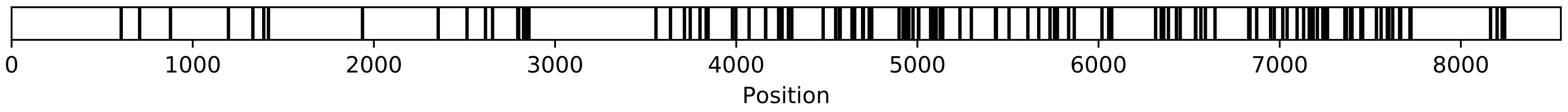


CR2 vs MA2
Number of differences: 118

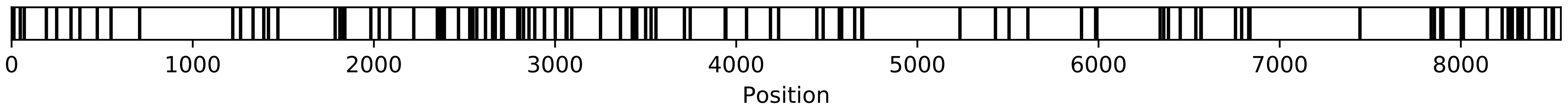


CR2 vs MM1
Number of differences: 118

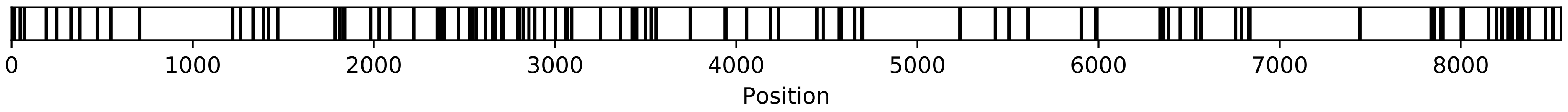


CR2 vs MM2
Number of differences: 133

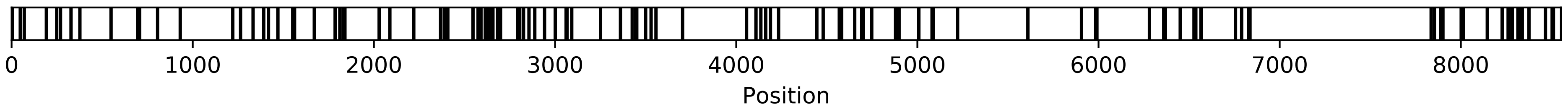


MA1 vs MA2
Number of differences: 185

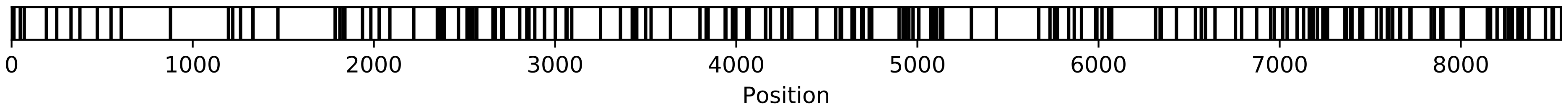


MA1 vs MM1
Number of differences: 187

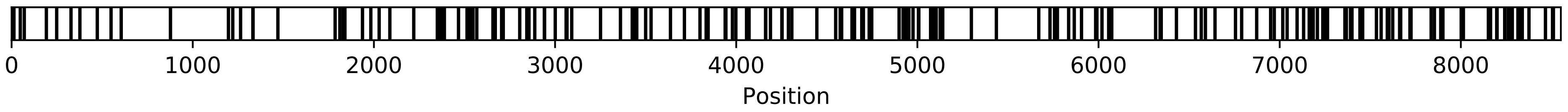


MA1 vs MM2
Number of differences: 196

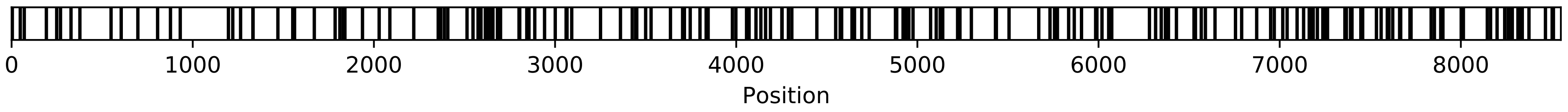


MA2 vs MM1
Number of differences: 6

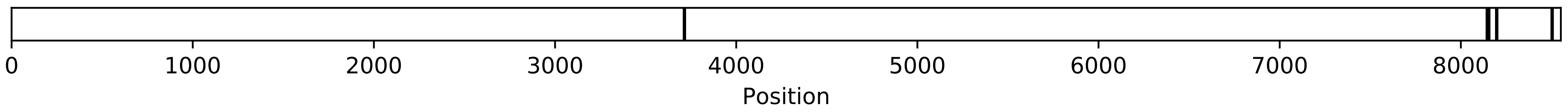


MA2 vs MM2
Number of differences: 63

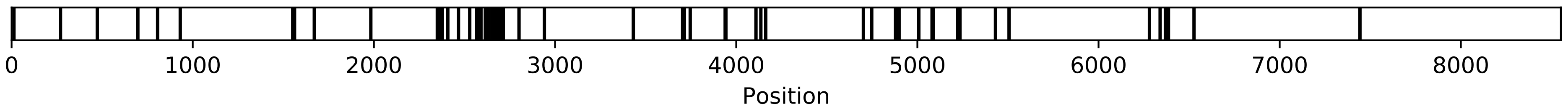


MM1 vs MM2
Number of differences: 67

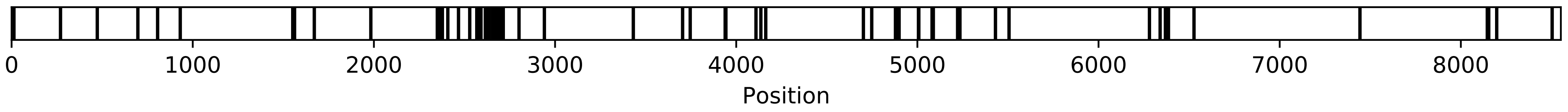

### Region: 476

CR1 vs CR2
Number of differences: 239

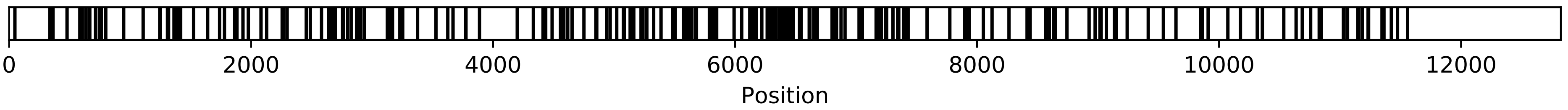


CR1 vs MA1
Number of differences: 154

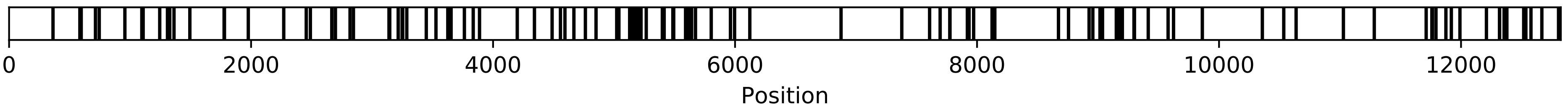


CR1 vs MA2
Number of differences: 241

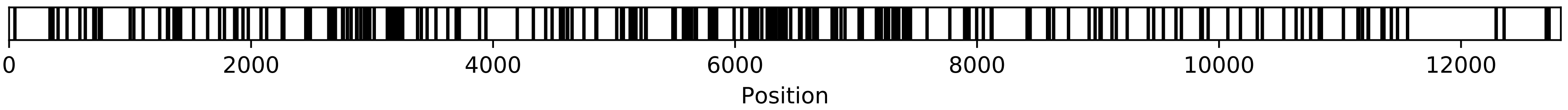


CR1 vs MM1
Number of differences: 174

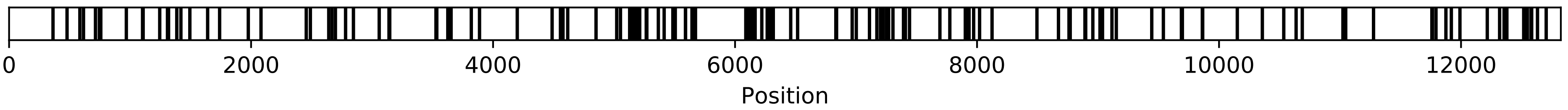


CR1 vs MM2
Number of differences: 167

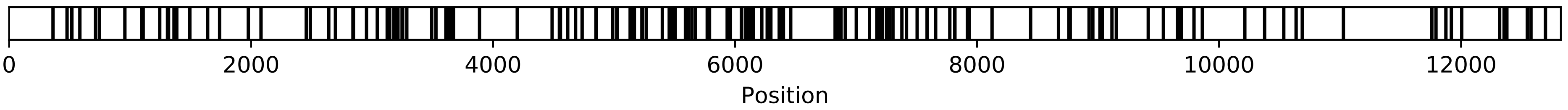


CR2 vs MA1
Number of differences: 290

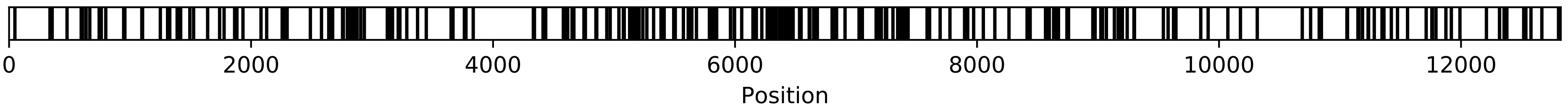


CR2 vs MA2
Number of differences: 95

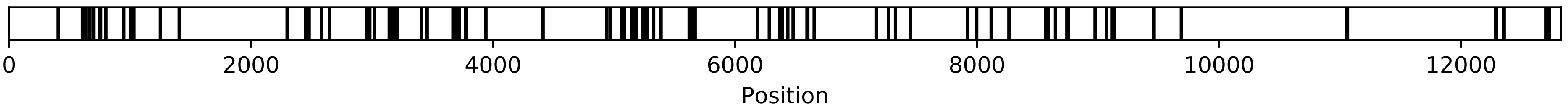


CR2 vs MM1
Number of differences: 287

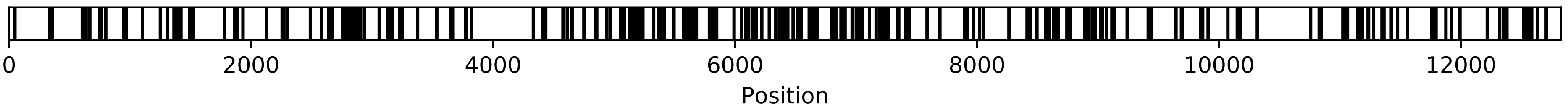


CR2 vs MM2
Number of differences: 257

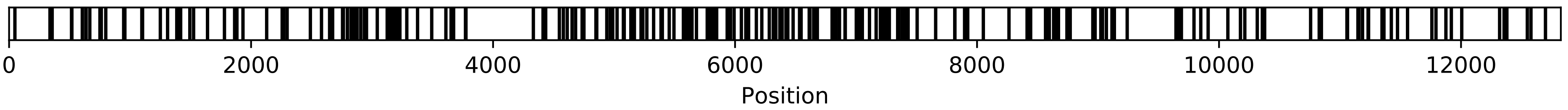


MA1 vs MA2
Number of differences: 285

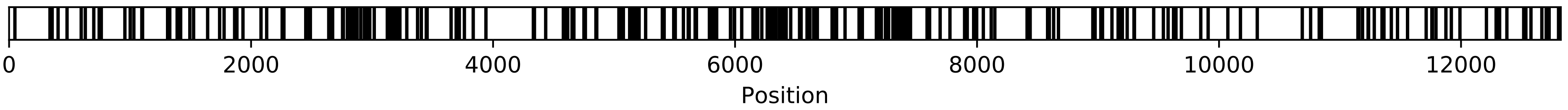


MA1 vs MM1
Number of differences: 174

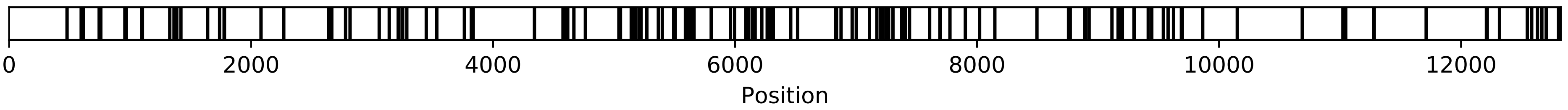


MA1 vs MM2
Number of differences: 181

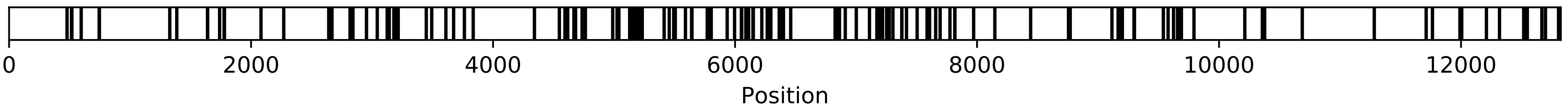


MA2 vs MM1
Number of differences: 280

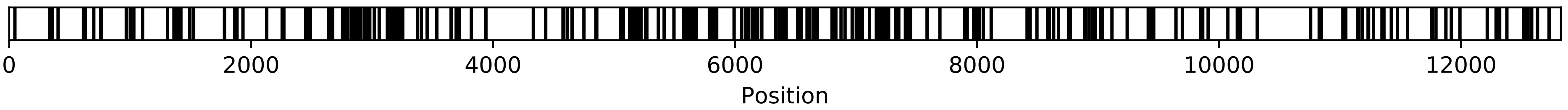


MA2 vs MM2
Number of differences: 250

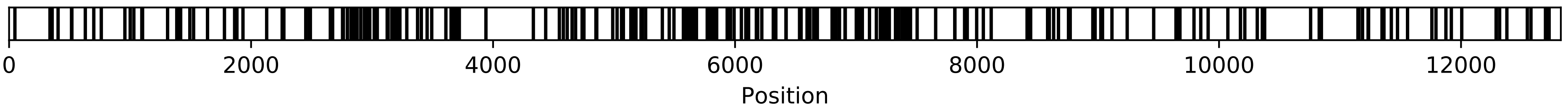


MM1 vs MM2
Number of differences: 156

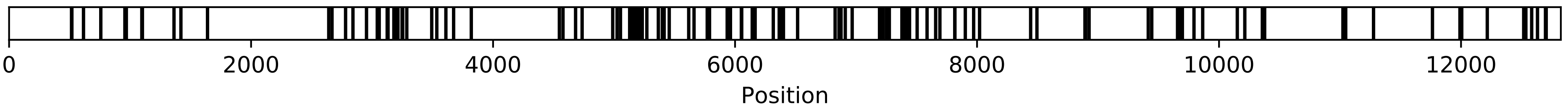

### Region: 477

CR1 vs CR2
Number of differences: 284

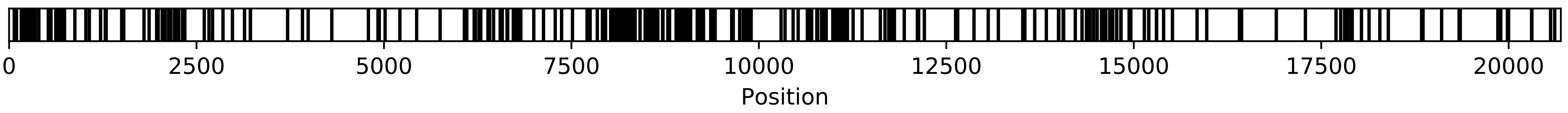


CR1 vs MA1
Number of differences: 132

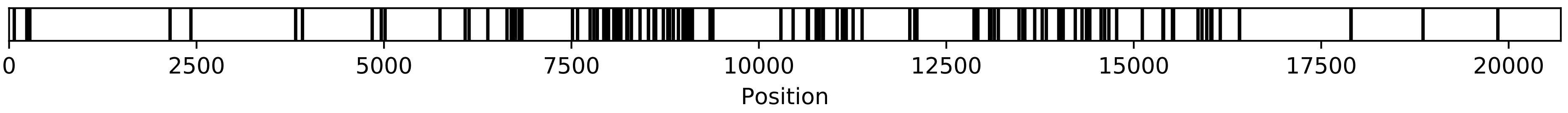


CR1 vs MA2
Number of differences: 261

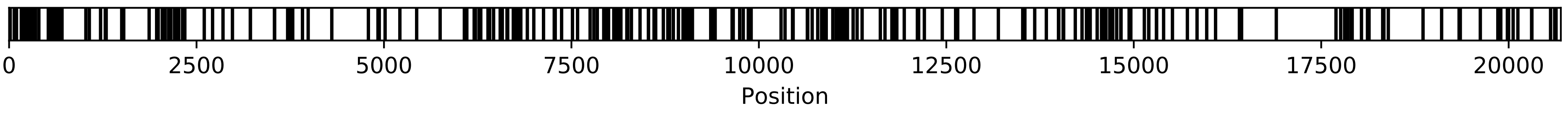


CR1 vs MM1
Number of differences: 139

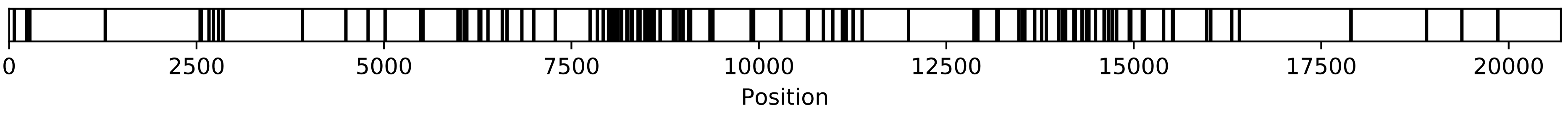


CR1 vs MM2
Number of differences: 128

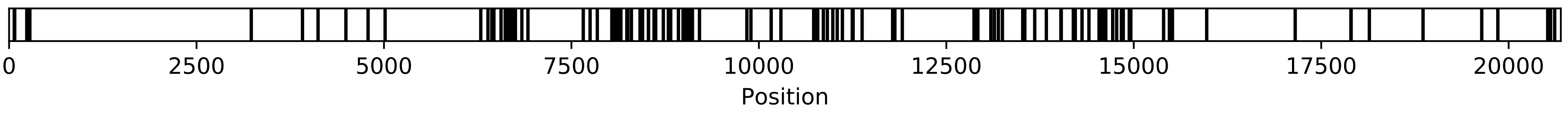


CR2 vs MA1
Number of differences: 280

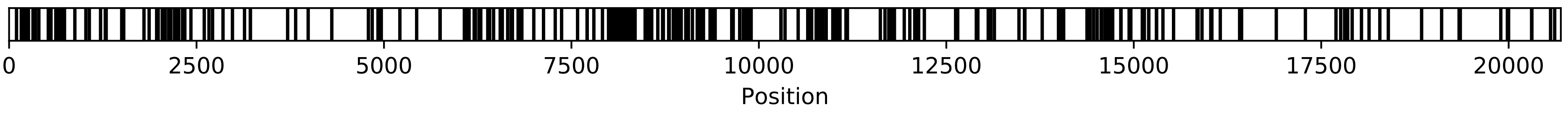


CR2 vs MA2
Number of differences: 109

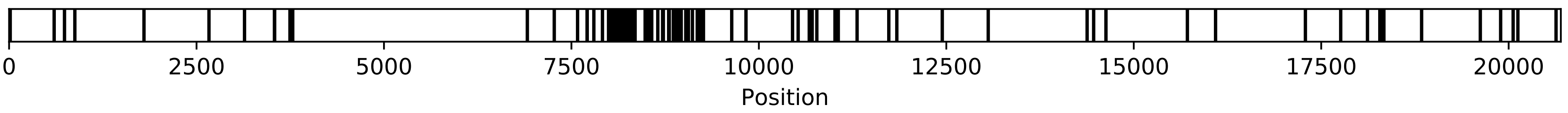


CR2 vs MM1
Number of differences: 266

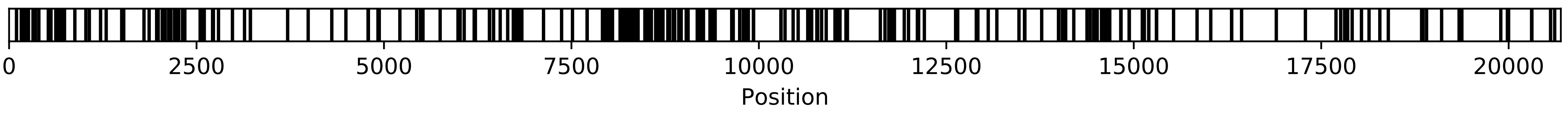


CR2 vs MM2
Number of differences: 292

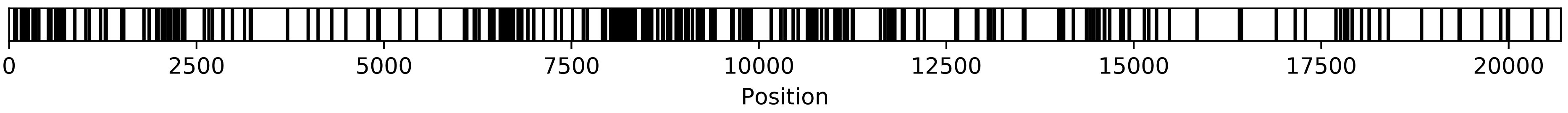


MA1 vs MA2
Number of differences: 224

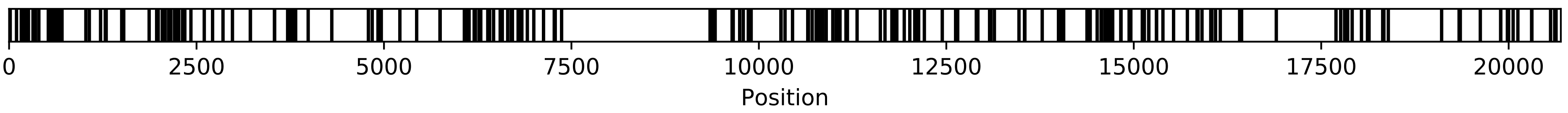


MA1 vs MM1
Number of differences: 138

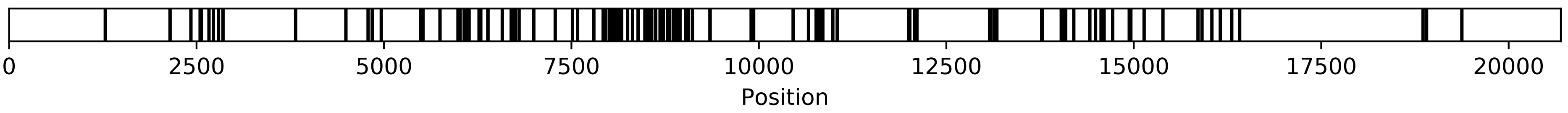


MA1 vs MM2
Number of differences: 119

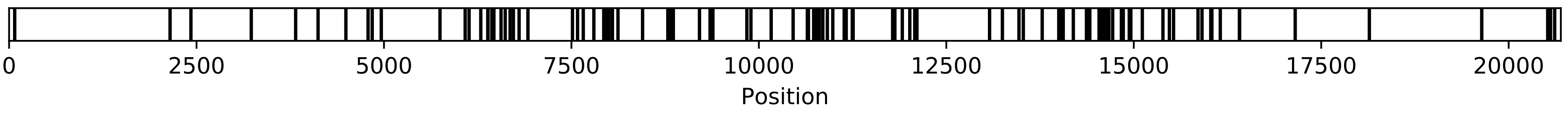


MA2 vs MM1
Number of differences: 259

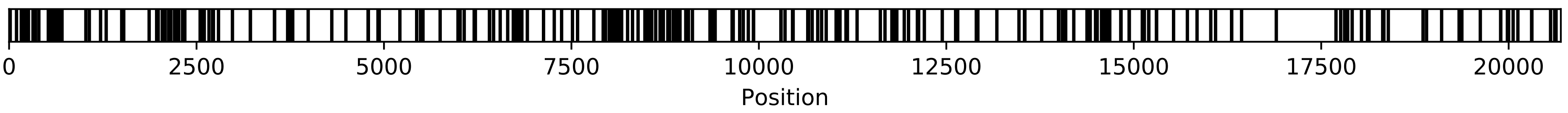


MA2 vs MM2
Number of differences: 247

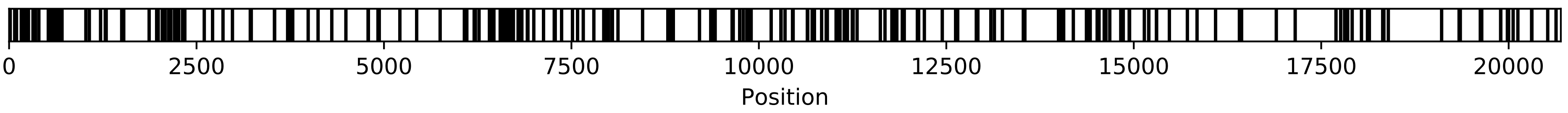


MM1 vs MM2
Number of differences: 155

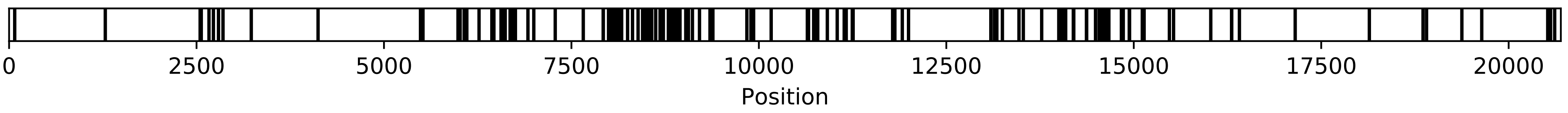

### Region: 478

CR1 vs CR2
Number of differences: 401

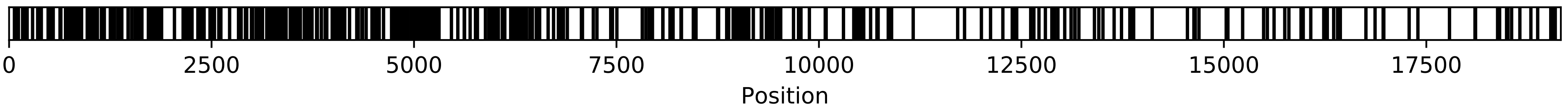


CR1 vs MA1
Number of differences: 394

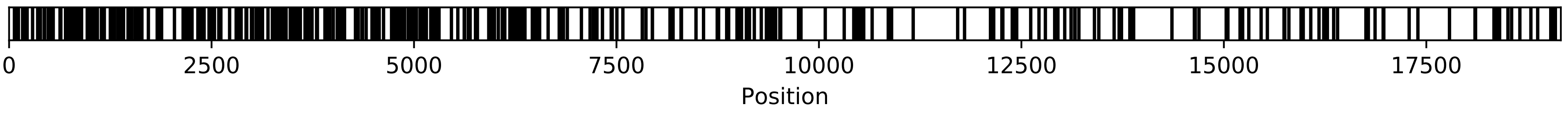


CR1 vs MA2
Number of differences: 158

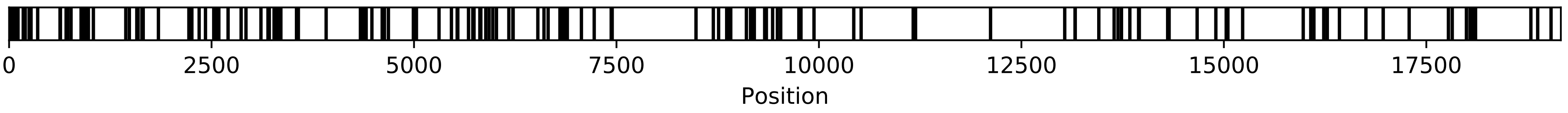


CR1 vs MM1
Number of differences: 394

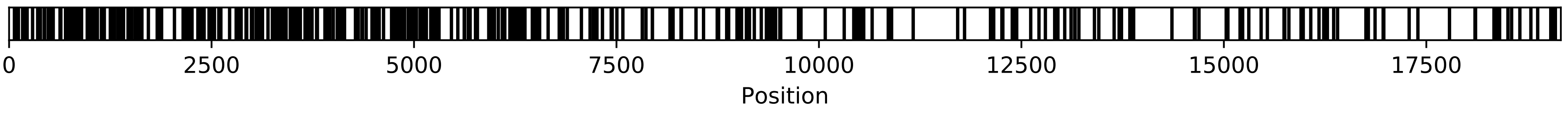


CR1 vs MM2
Number of differences: 393

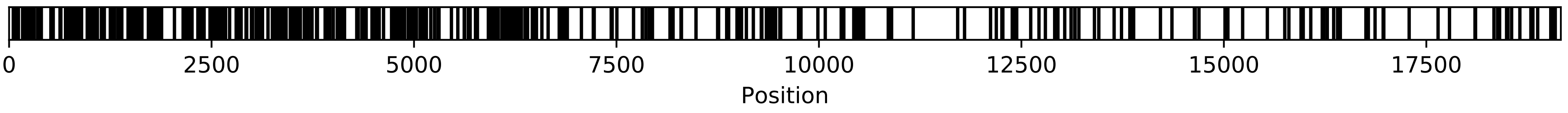


CR2 vs MA1
Number of differences: 166

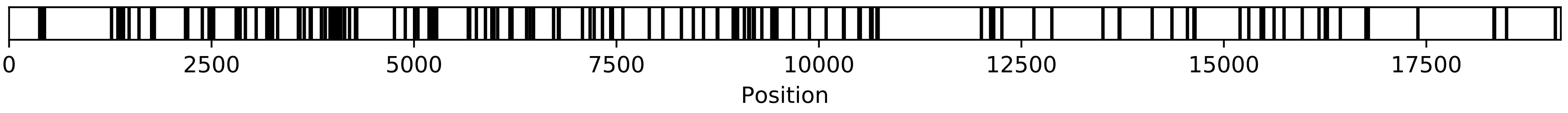


CR2 vs MA2
Number of differences: 410

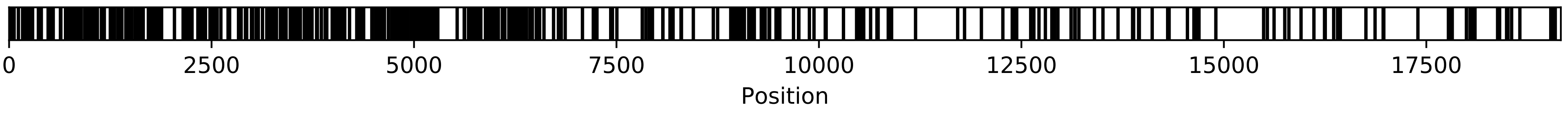


CR2 vs MM1
Number of differences: 166

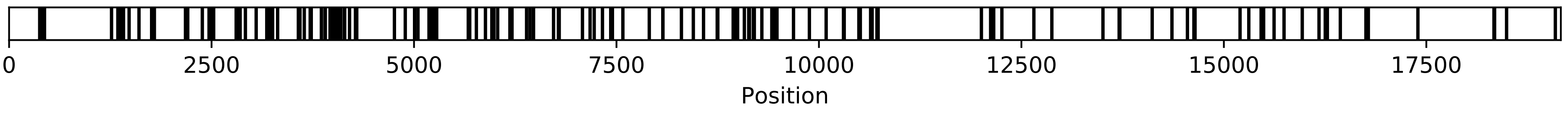


CR2 vs MM2
Number of differences: 144

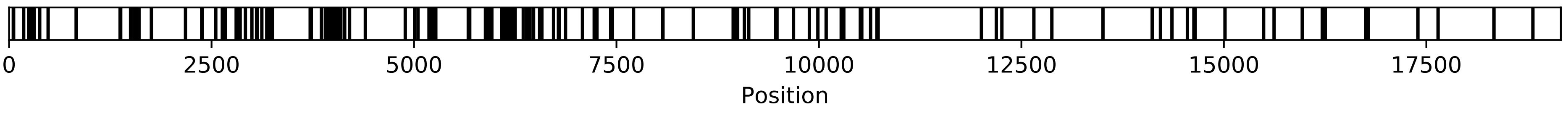


MA1 vs MA2
Number of differences: 399

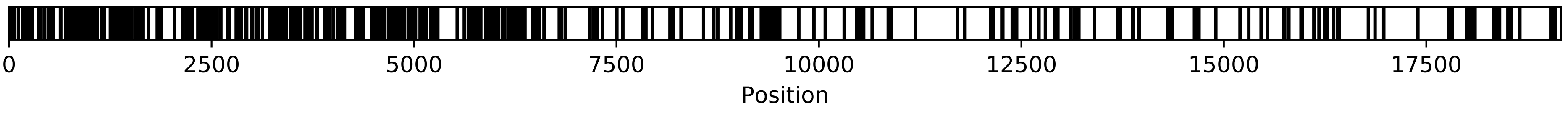


MA1 vs MM1
Number of differences: 0

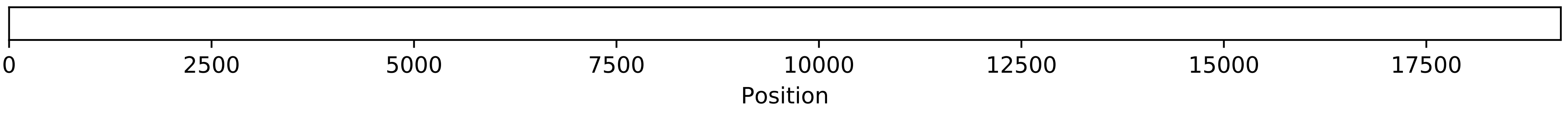


MA1 vs MM2
Number of differences: 125

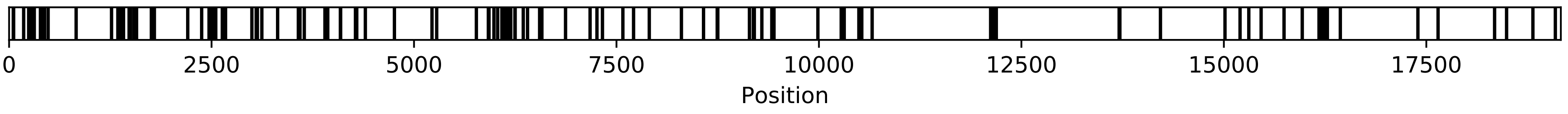


MA2 vs MM1
Number of differences: 399

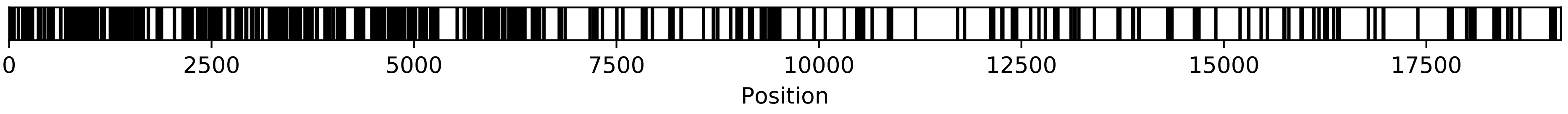


MA2 vs MM2
Number of differences: 400

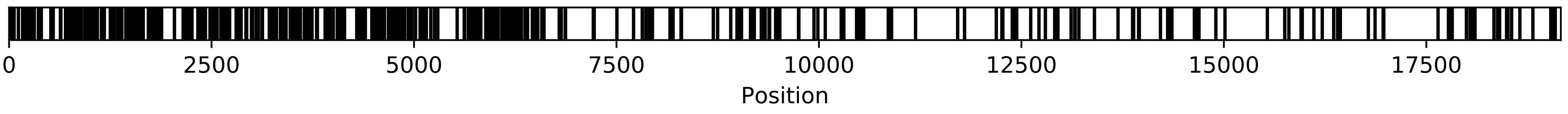


MM1 vs MM2
Number of differences: 125

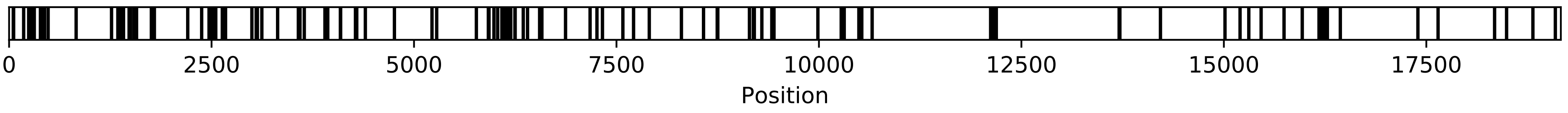

### Region: 480



## CR1 vs CR2
Number of differences: 75

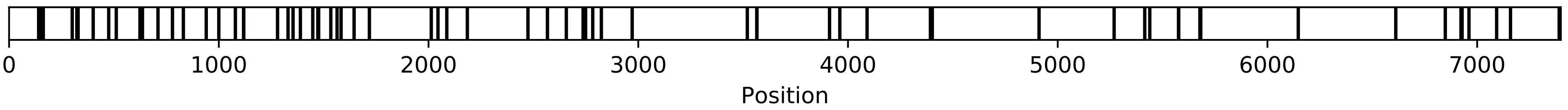


## CR1 vs MA1
Number of differences: 20

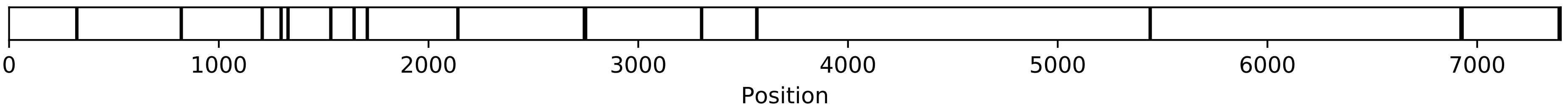


## CR1 vs MA2
Number of differences: 76

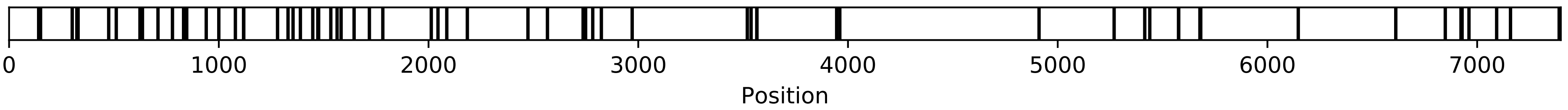


## CR1 vs MM1
Number of differences: 18

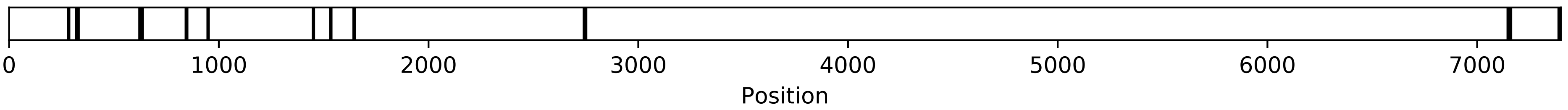


## CR1 vs MM2
Number of differences: 20

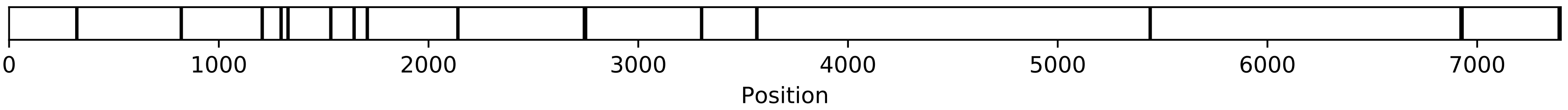


## CR2 vs MA1
Number of differences: 72

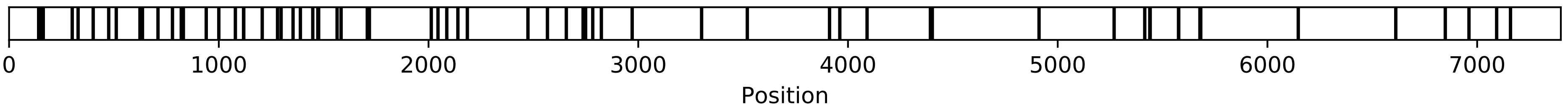


## CR2 vs MA2
Number of differences: 19

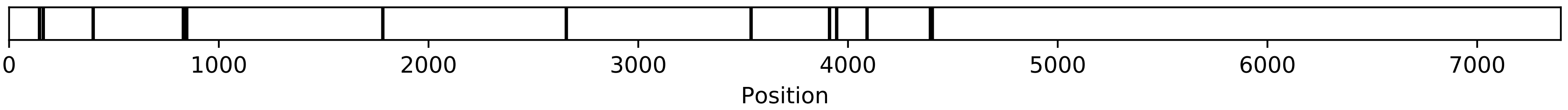


## CR2 vs MM1
Number of differences: 74

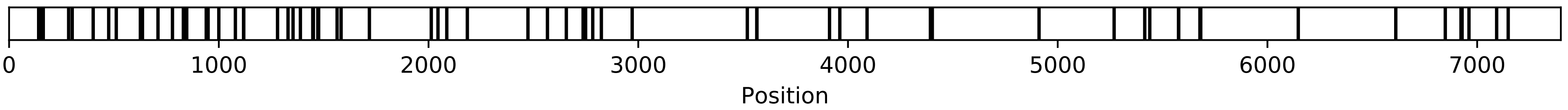


## CR2 vs MM2
Number of differences: 72

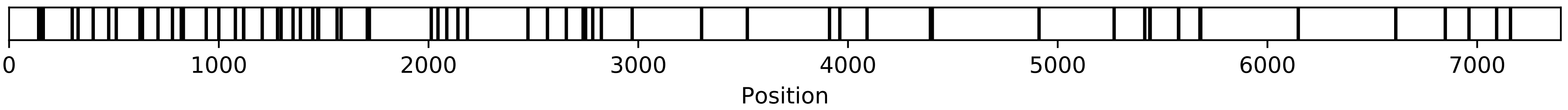


## MA1 vs MA2
Number of differences: 73

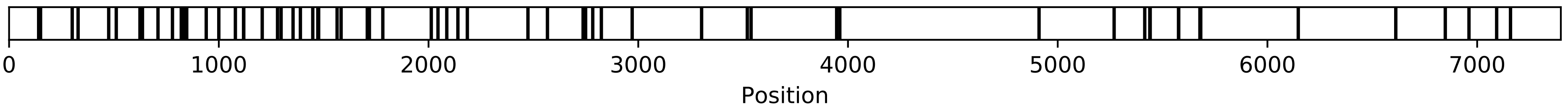


## MA1 vs MM1
Number of differences: 26

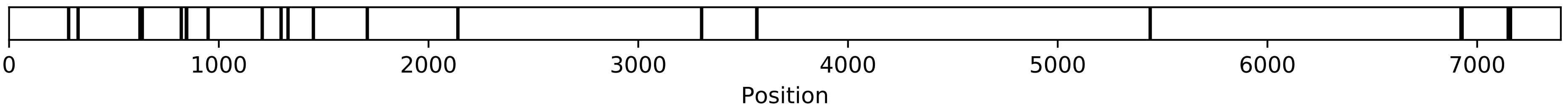


## MA1 vs MM2
Number of differences: 0

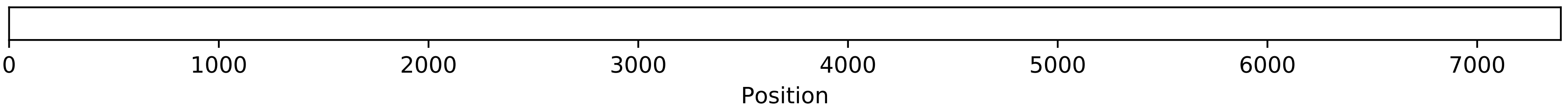


## MA2 vs MM1
Number of differences: 74

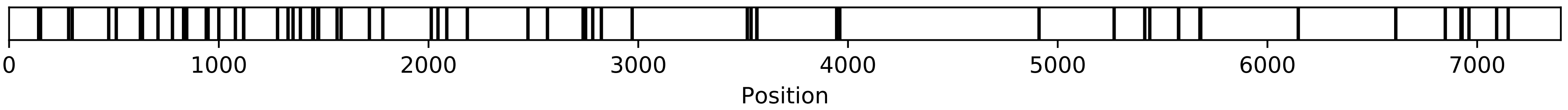


## MA2 vs MM2
Number of differences: 73

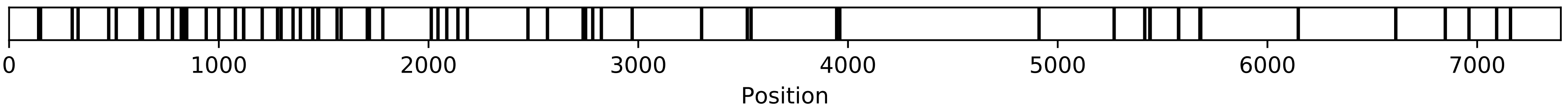


## MM1 vs MM2
Number of differences: 26

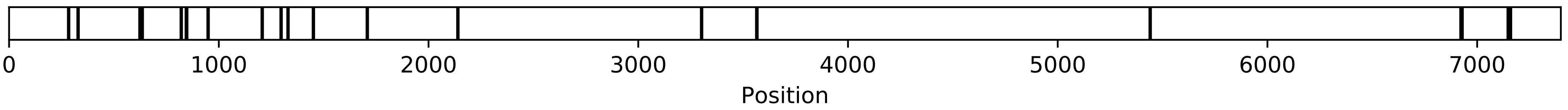

### Region: 481



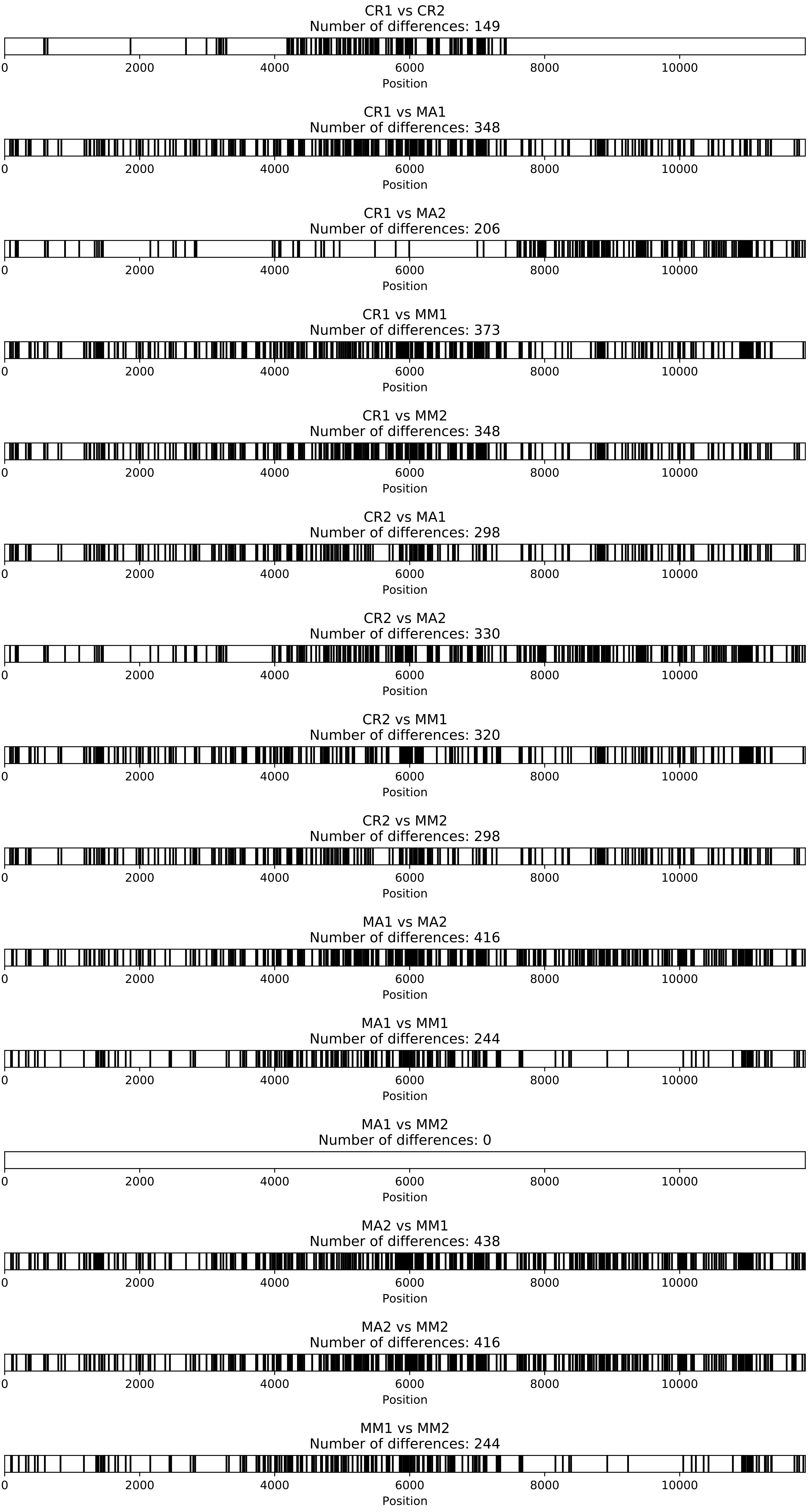
CR1 vs CR2
Number of differences: 149
CR1 vs MA1
Number of differences: 348
CR1 vs MA2
Number of differences: 206
CR1 vs MM1
Number of differences: 373
CR1 vs MM2
Number of differences: 348
CR2 vs MA1
Number of differences: 298
CR2 vs MA2
Number of differences: 330
CR2 vs MM1
Number of differences: 320
CR2 vs MM2
Number of differences: 298
MA1 vs MA2
Number of differences: 416
MA1 vs MM1
Number of differences: 244
MA1 vs MM2
Number of differences: 0
MA2 vs MM1
Number of differences: 438
MA2 vs MM2
Number of differences: 416
MM1 vs MM2
Number of differences: 244
0
2000
4000
6000
8000
10000
Position

### Region: 482

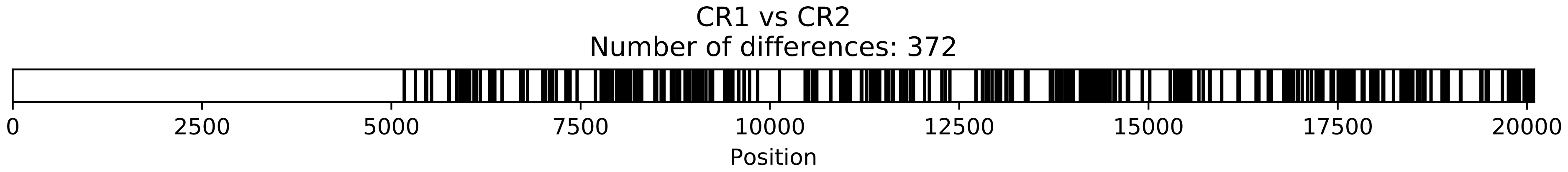
CR1 vs CR2
Number of differences: 372
0
2500
5000
7500
10000
12500
15000
17500
20000
Position

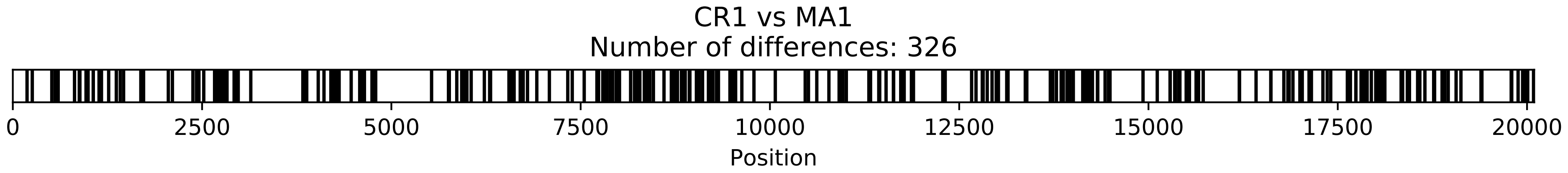
CR1 vs MA1
Number of differences: 326
Position

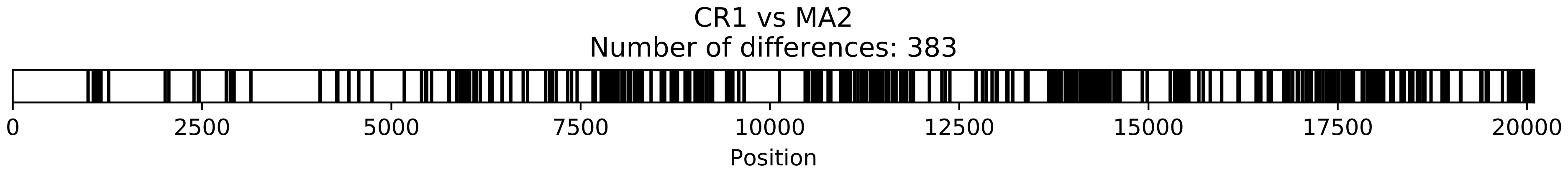
CR1 vs MA2
Number of differences: 383
Position

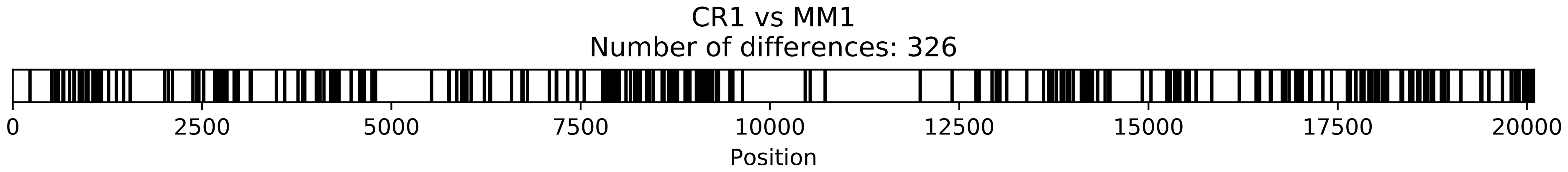
CR1 vs MM1
Number of differences: 326
Position

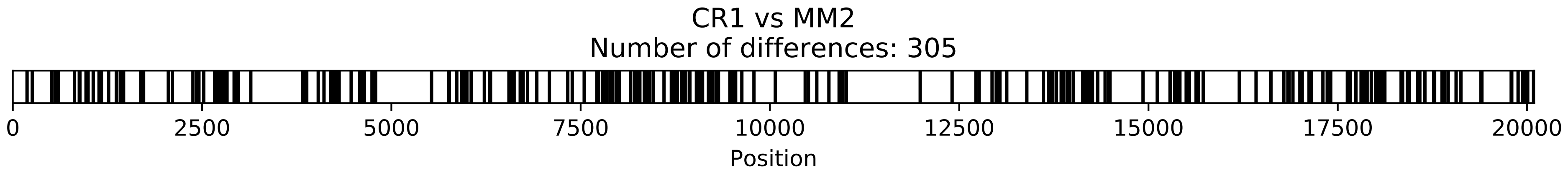
CR1 vs MM2
Number of differences: 305
Position

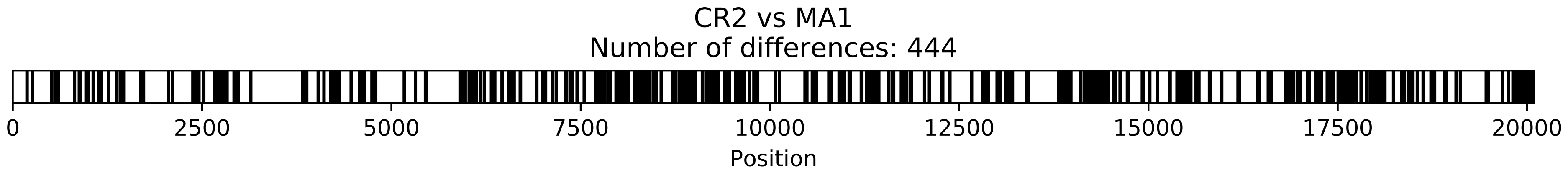
CR2 vs MA1
Number of differences: 444
Position

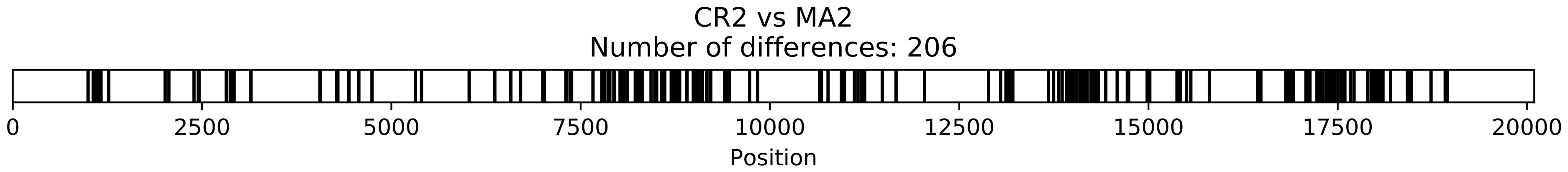
CR2 vs MA2
Number of differences: 206
Position

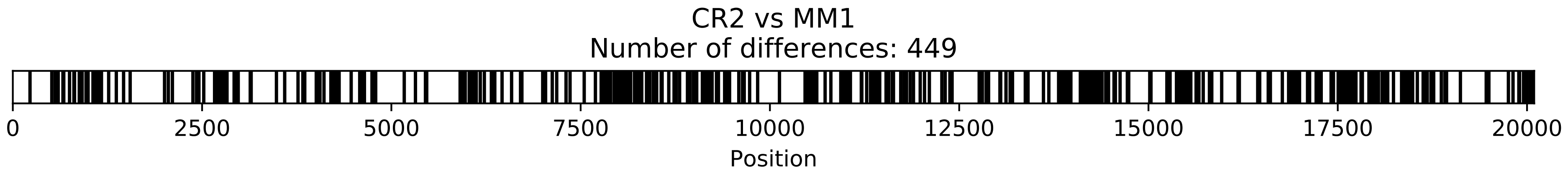
CR2 vs MM1
Number of differences: 449
Position

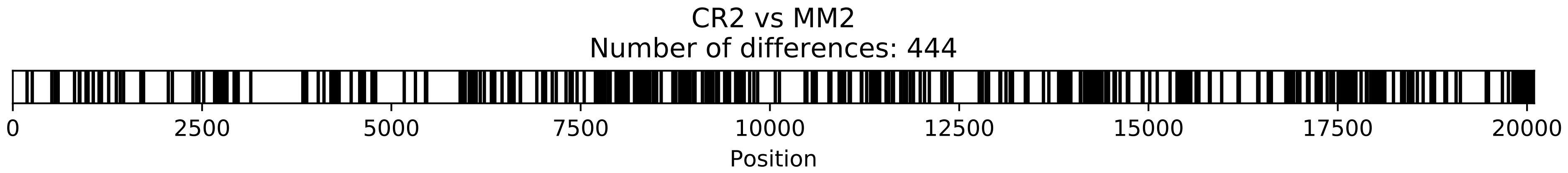
CR2 vs MM2
Number of differences: 444
Position

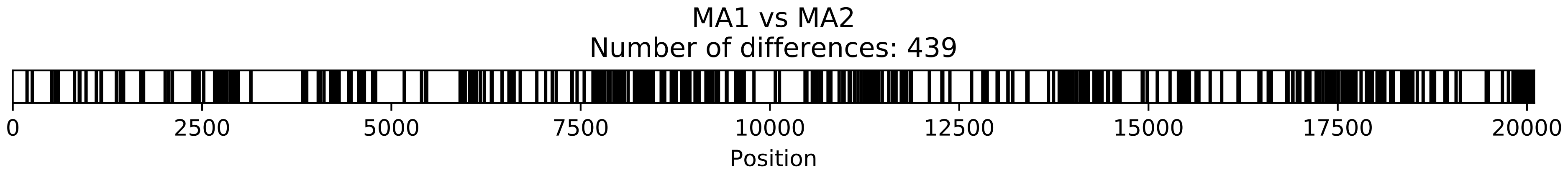
MA1 vs MA2
Number of differences: 439
Position

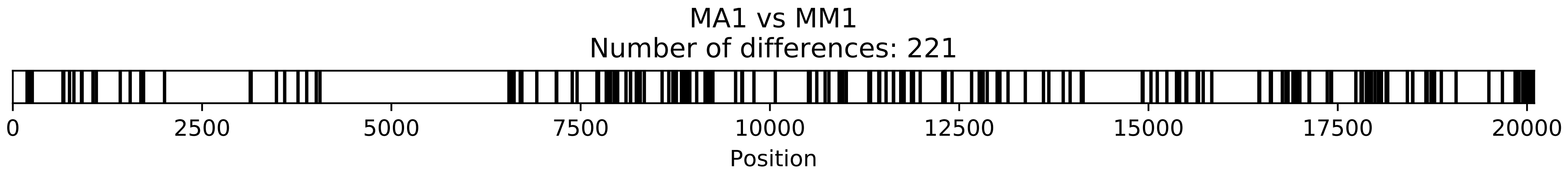
MA1 vs MM1
Number of differences: 221
Position

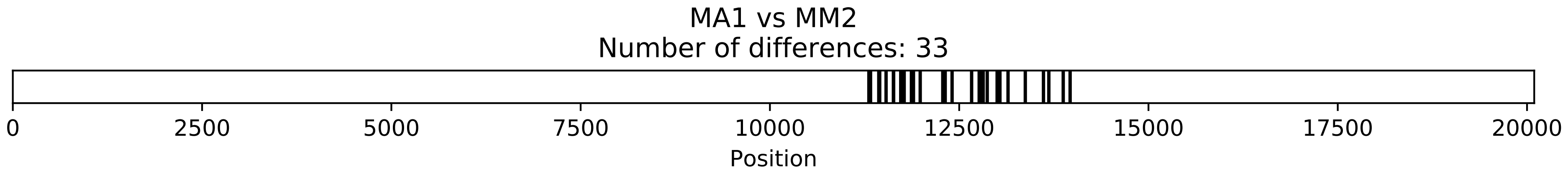
MA1 vs MM2
Number of differences: 33
Position

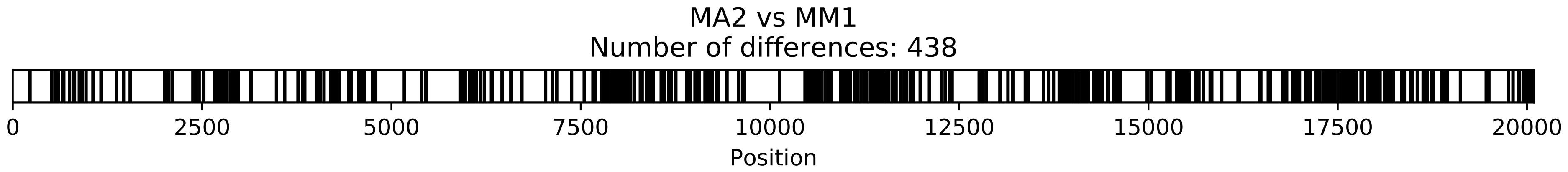
MA2 vs MM1
Number of differences: 438
Position

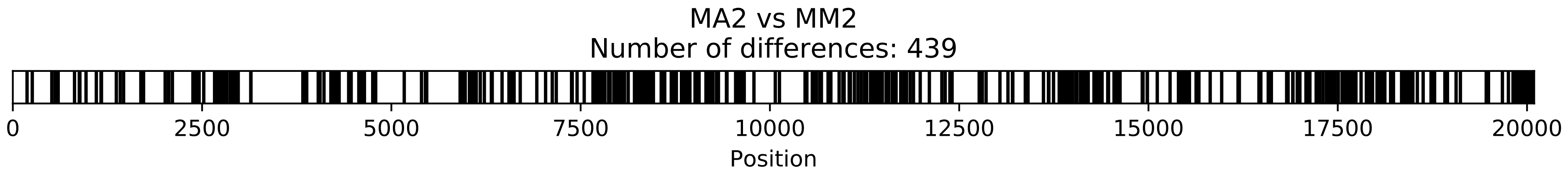
MA2 vs MM2
Number of differences: 439
Position

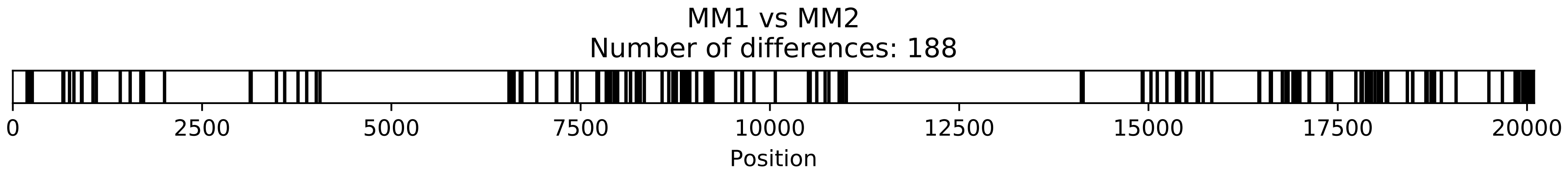
MM1 vs MM2
Number of differences: 188
Position

### Region: 483



CR1 vs CR2
Number of differences: 140

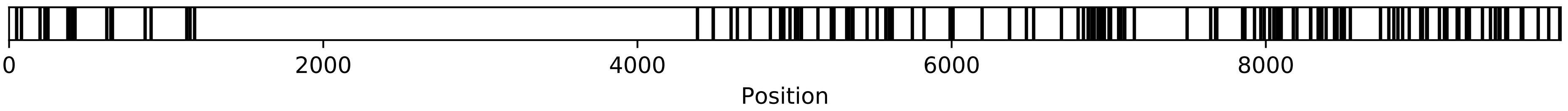


CR1 vs MA1
Number of differences: 159

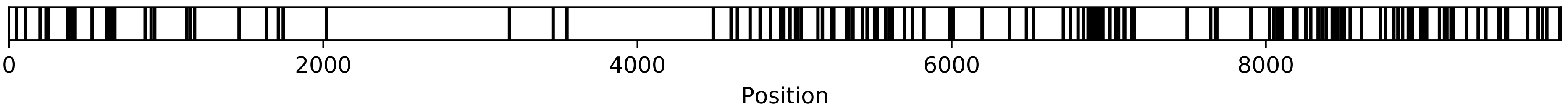


CR1 vs MA2
Number of differences: 92

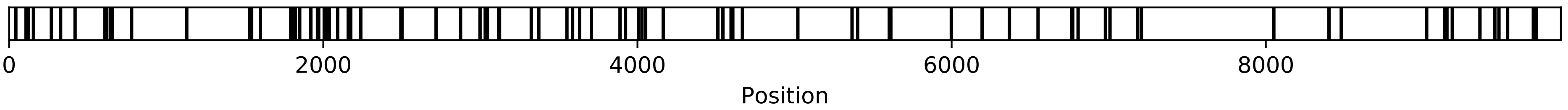


CR1 vs MM1
Number of differences: 166

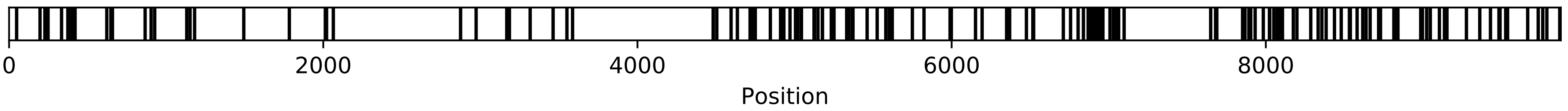


CR1 vs MM2
Number of differences: 159

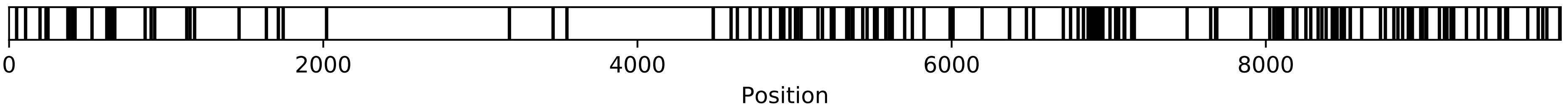


CR2 vs MA1
Number of differences: 81

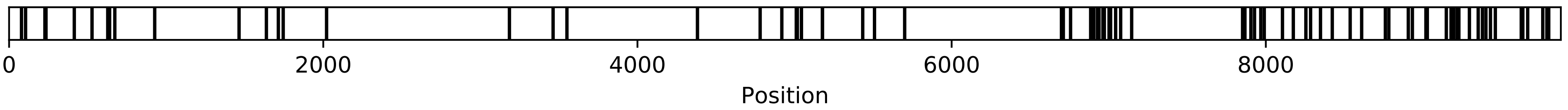


CR2 vs MA2
Number of differences: 188

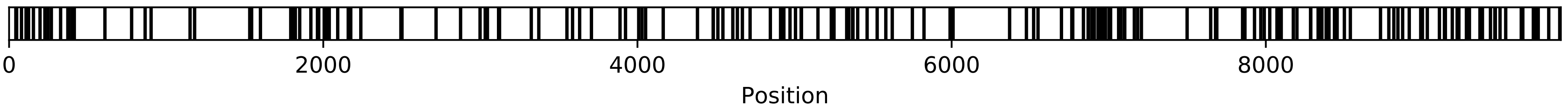


CR2 vs MM1
Number of differences: 92

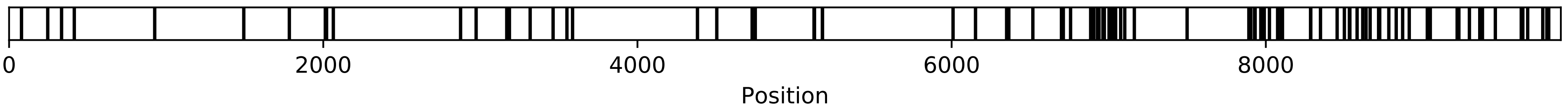


CR2 vs MM2
Number of differences: 81

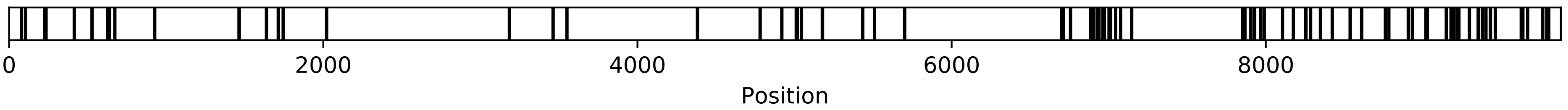


MA1 vs MA2
Number of differences: 203

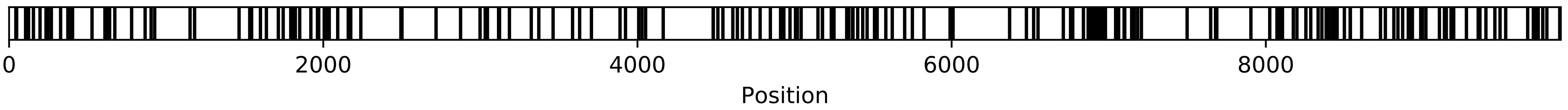


MA1 vs MM1
Number of differences: 87

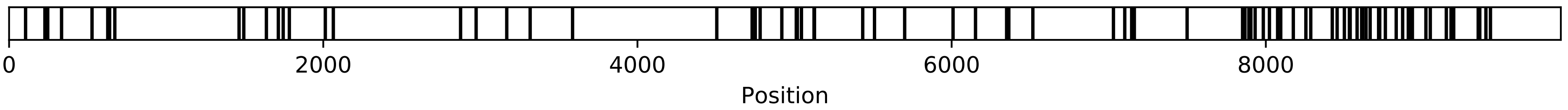


MA1 vs MM2
Number of differences: 0

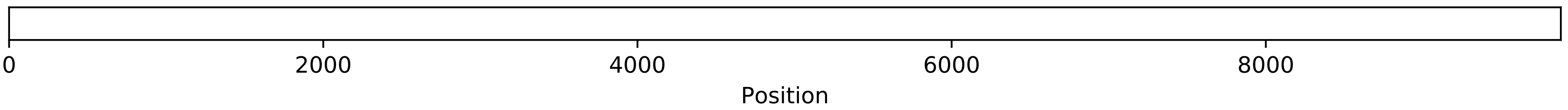


MA2 vs MM1
Number of differences: 206

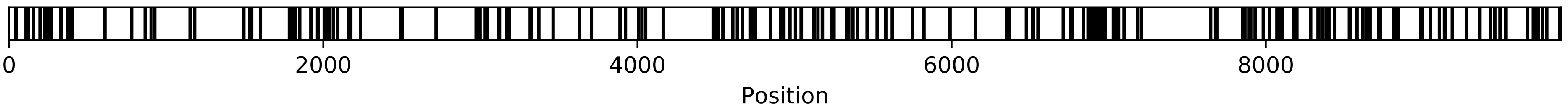


MA2 vs MM2
Number of differences: 203

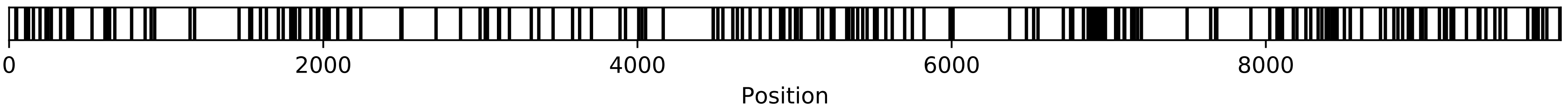


MM1 vs MM2
Number of differences: 87

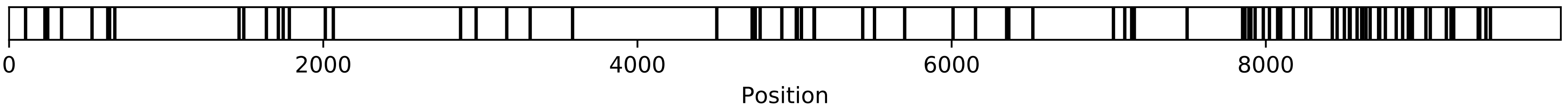

### Region: 484



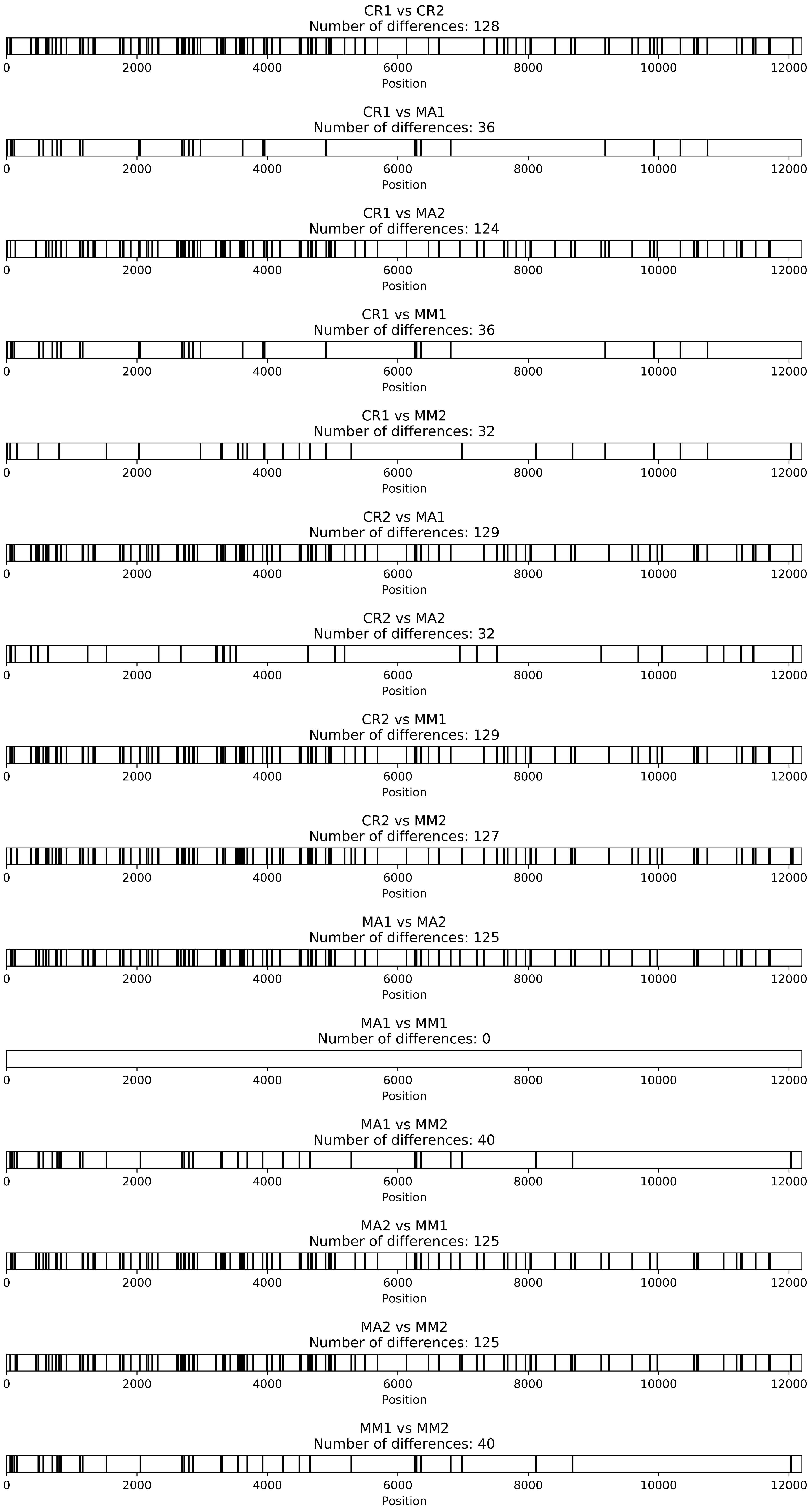
CR1 vs CR2
Number of differences: 128
CR1 vs MA1
Number of differences: 36
CR1 vs MA2
Number of differences: 124
CR1 vs MM1
Number of differences: 36
CR1 vs MM2
Number of differences: 32
CR2 vs MA1
Number of differences: 129
CR2 vs MA2
Number of differences: 32
CR2 vs MM1
Number of differences: 129
CR2 vs MM2
Number of differences: 127
MA1 vs MA2
Number of differences: 125
MA1 vs MM1
Number of differences: 0
MA1 vs MM2
Number of differences: 40
MA2 vs MM1
Number of differences: 125
MA2 vs MM2
Number of differences: 125
MM1 vs MM2
Number of differences: 40
0
2000
4000
6000
8000
10000
12000
Position

### Region: 485

CR1 vs CR2
Number of differences: 184

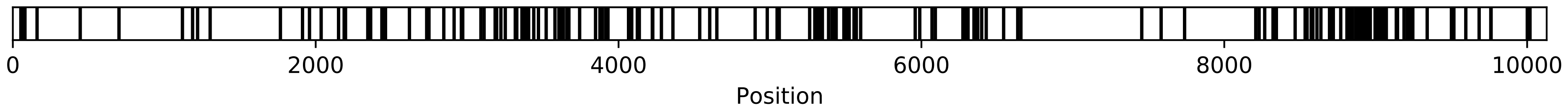


CR1 vs MA1
Number of differences: 95

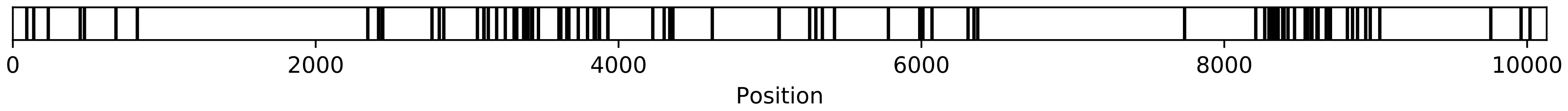


CR1 vs MA2
Number of differences: 187

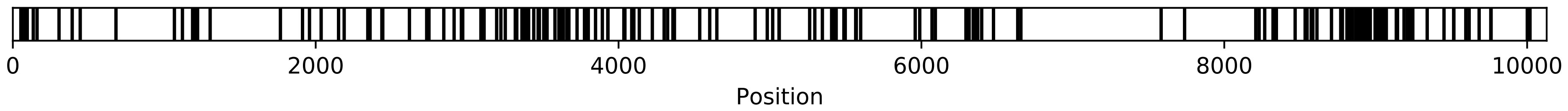


CR1 vs MM1
Number of differences: 199

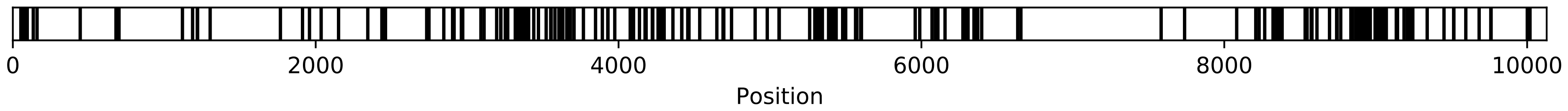


CR1 vs MM2
Number of differences: 187

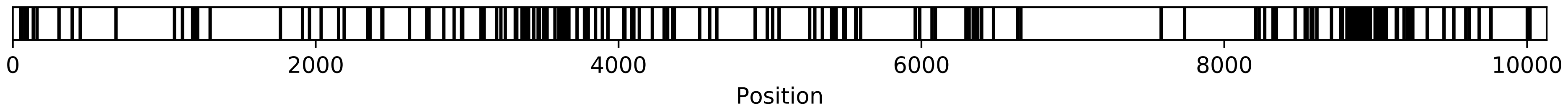


CR2 vs MA1
Number of differences: 193

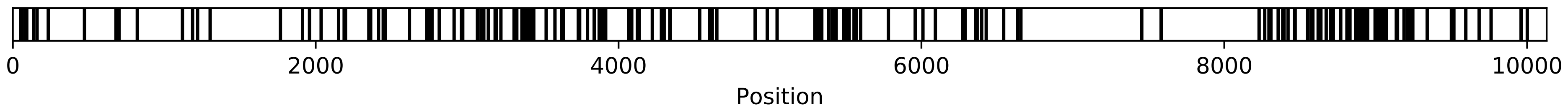


CR2 vs MA2
Number of differences: 78

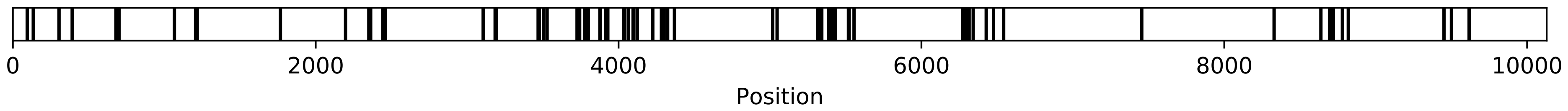


CR2 vs MM1
Number of differences: 90

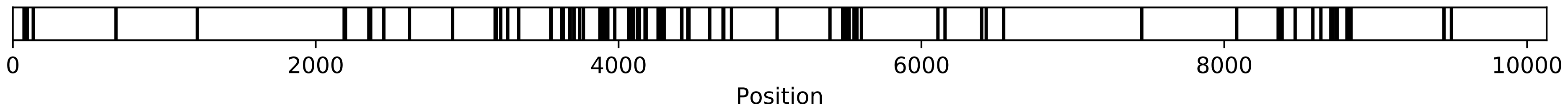


CR2 vs MM2
Number of differences: 78

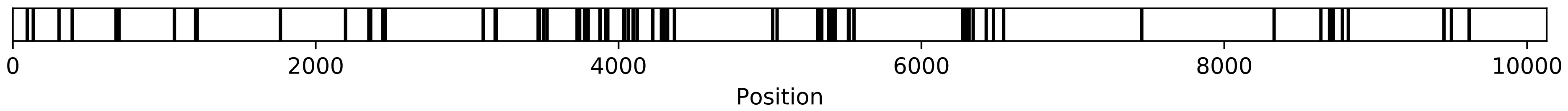


MA1 vs MA2
Number of differences: 198

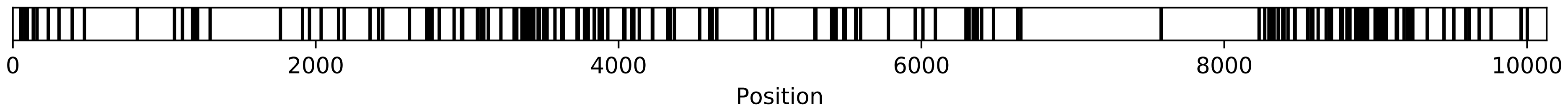


MA1 vs MM1
Number of differences: 203

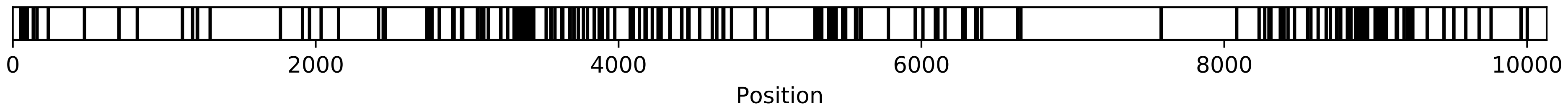


MA1 vs MM2
Number of differences: 198

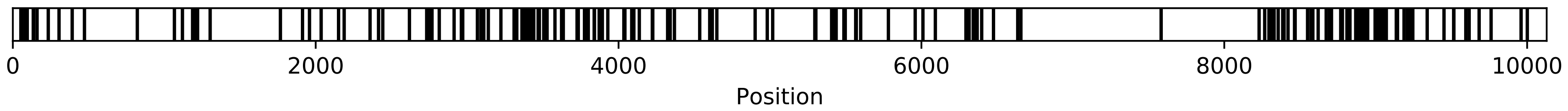


MA2 vs MM1
Number of differences: 106

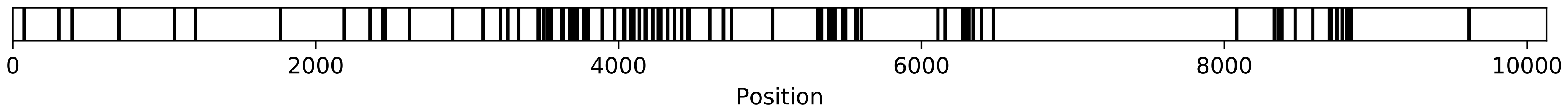


MA2 vs MM2
Number of differences: 0

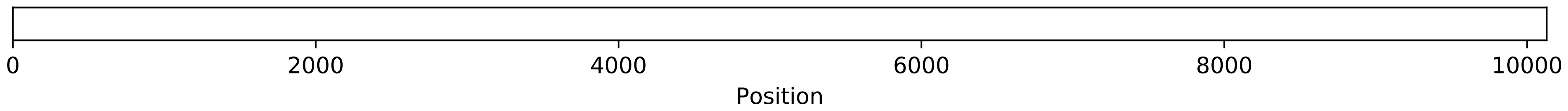


MM1 vs MM2
Number of differences: 106

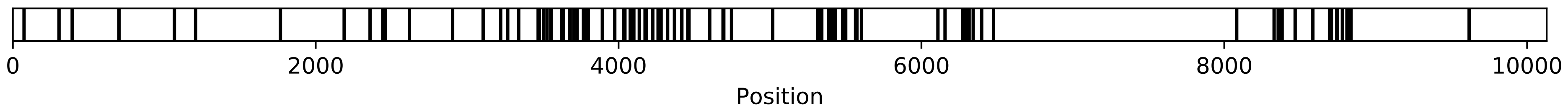

### Region: 486

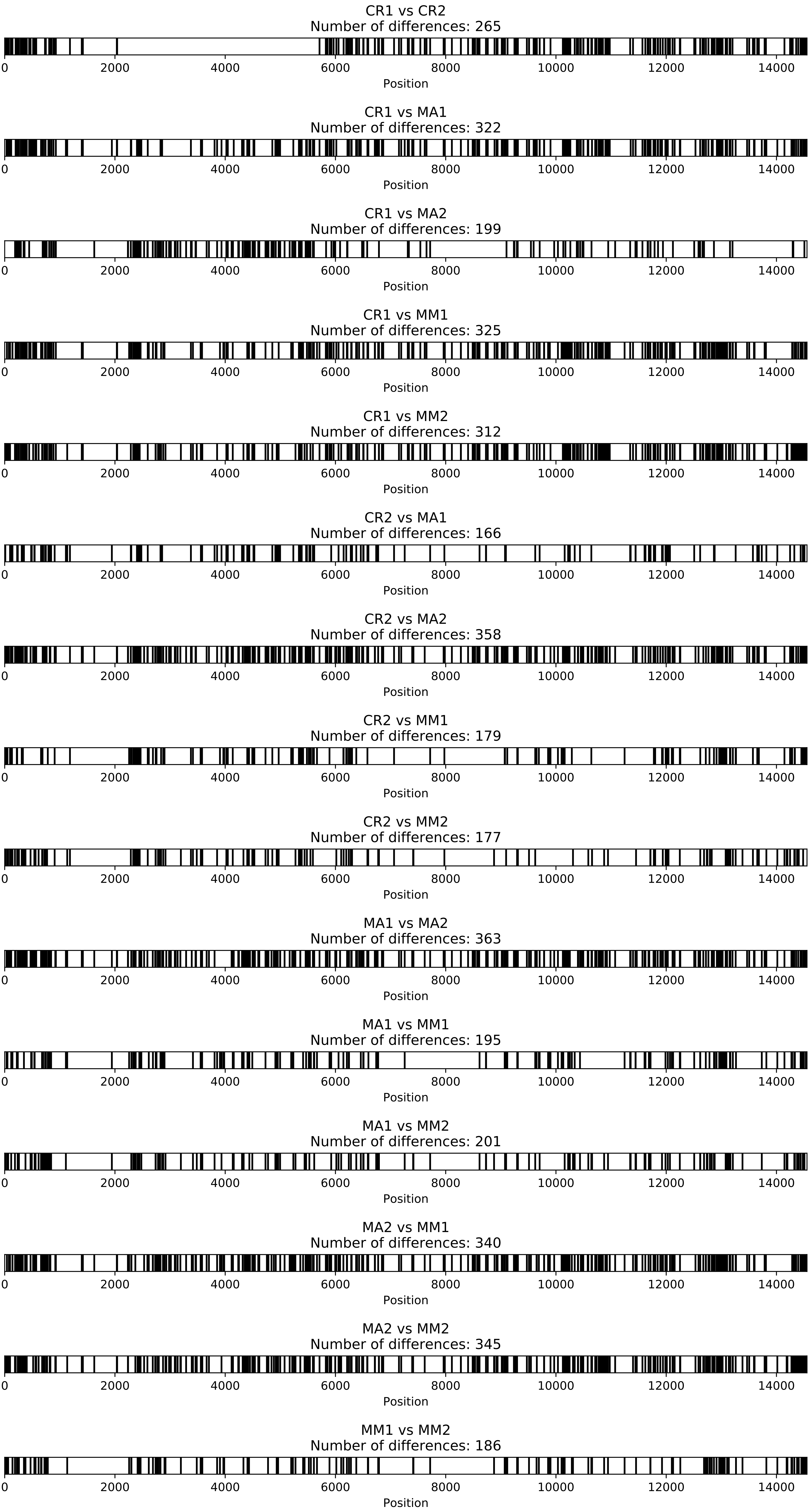
CR1 vs CR2
Number of differences: 265
CR1 vs MA1
Number of differences: 322
CR1 vs MA2
Number of differences: 199
CR1 vs MM1
Number of differences: 325
CR1 vs MM2
Number of differences: 312
CR2 vs MA1
Number of differences: 166
CR2 vs MA2
Number of differences: 358
CR2 vs MM1
Number of differences: 179
CR2 vs MM2
Number of differences: 177
MA1 vs MA2
Number of differences: 363
MA1 vs MM1
Number of differences: 195
MA1 vs MM2
Number of differences: 201
MA2 vs MM1
Number of differences: 340
MA2 vs MM2
Number of differences: 345
MM1 vs MM2
Number of differences: 186
0
2000
4000
6000
8000
10000
12000
14000
Position

### Region: 487



CR1 vs CR2
Number of differences: 310

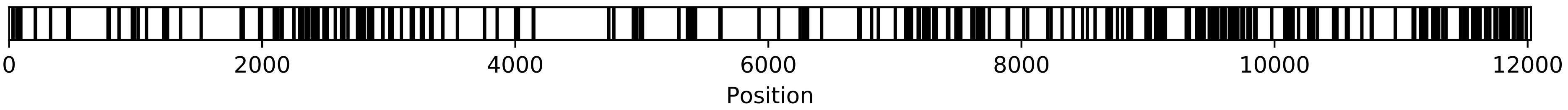


CR1 vs MA1
Number of differences: 227

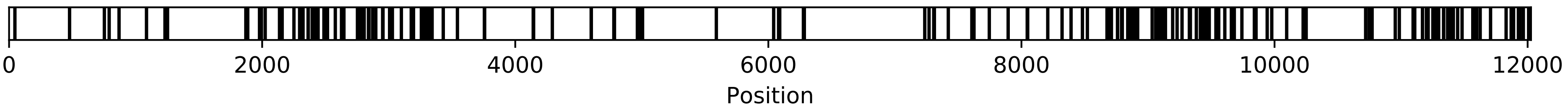


CR1 vs MA2
Number of differences: 256

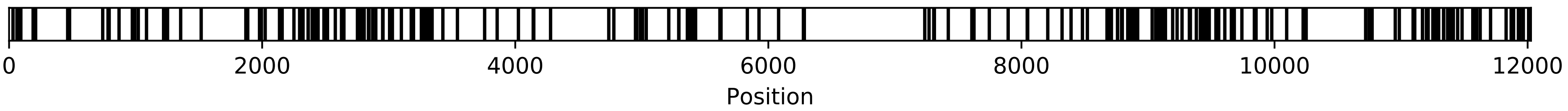


CR1 vs MM1
Number of differences: 148

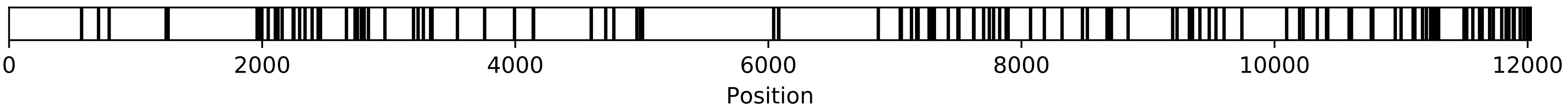


CR1 vs MM2
Number of differences: 190

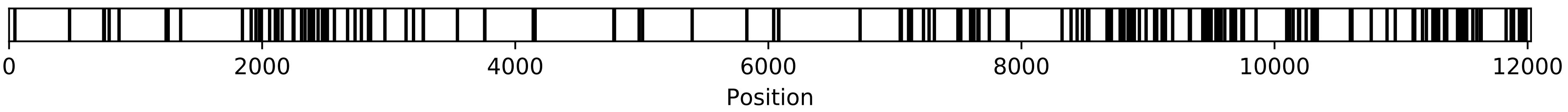


CR2 vs MA1
Number of differences: 256

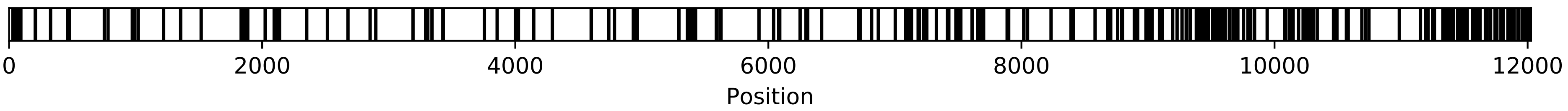


CR2 vs MA2
Number of differences: 212

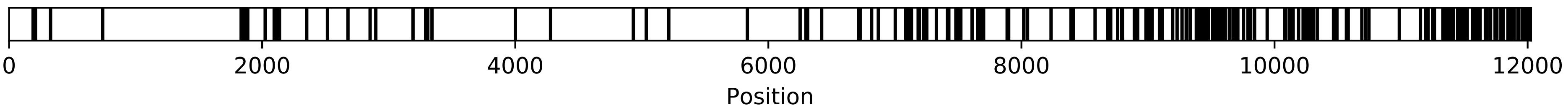


CR2 vs MM1
Number of differences: 303

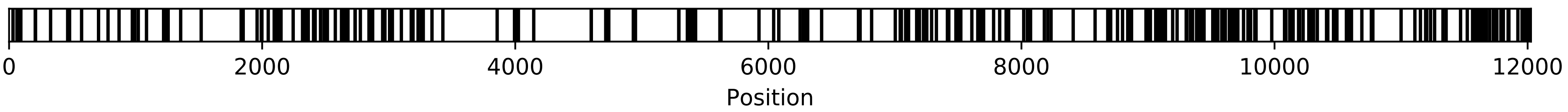


CR2 vs MM2
Number of differences: 308

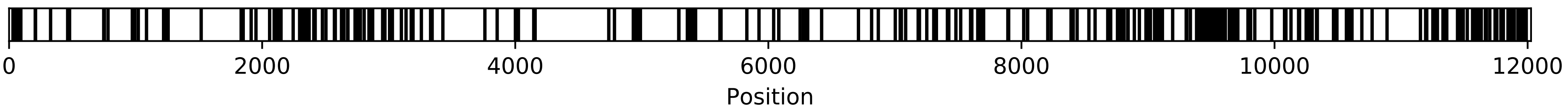


MA1 vs MA2
Number of differences: 58

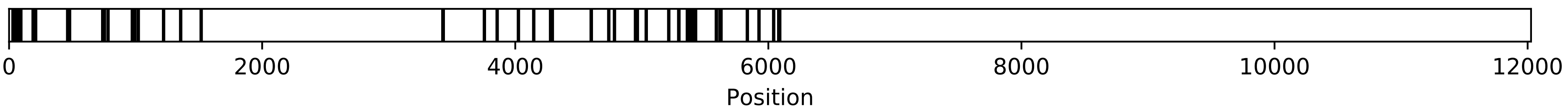


MA1 vs MM1
Number of differences: 221

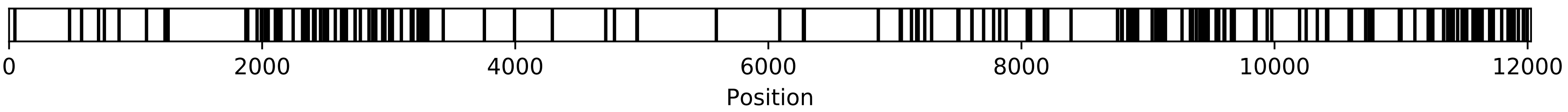


MA1 vs MM2
Number of differences: 206

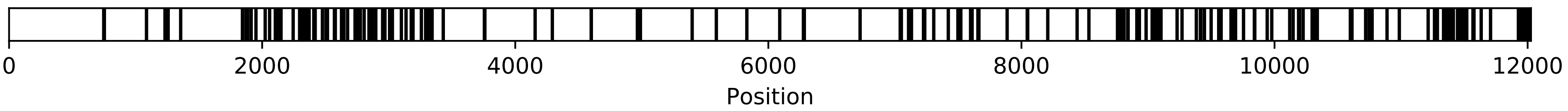


MA2 vs MM1
Number of differences: 258

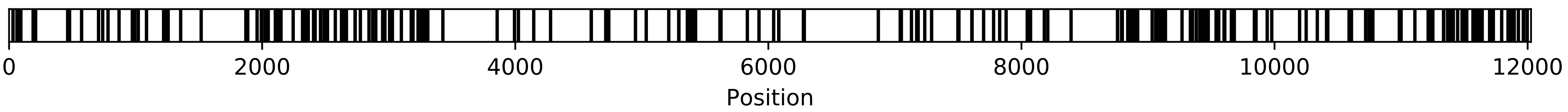


MA2 vs MM2
Number of differences: 243

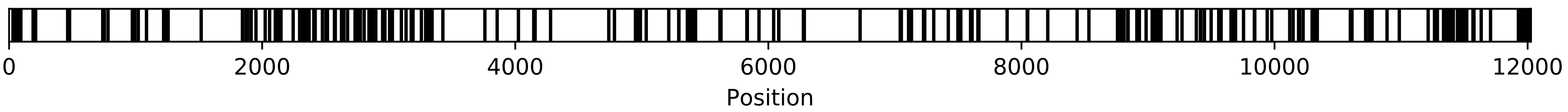


MM1 vs MM2
Number of differences: 184

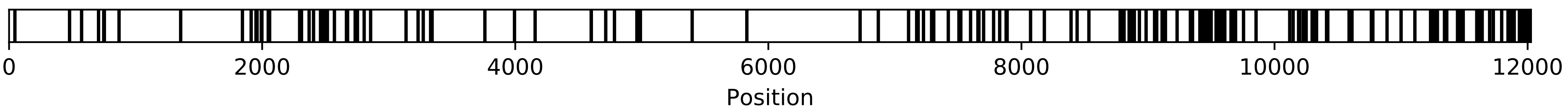

### Region: 488

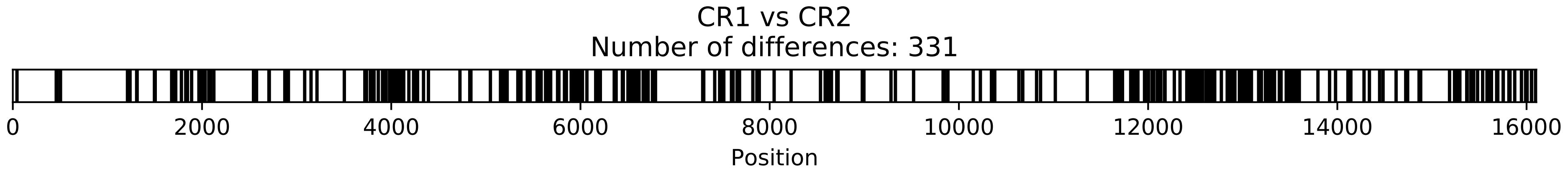
CR1 vs CR2
Number of differences: 331
0 2000 4000 6000 8000 10000 12000 14000 16000
Position

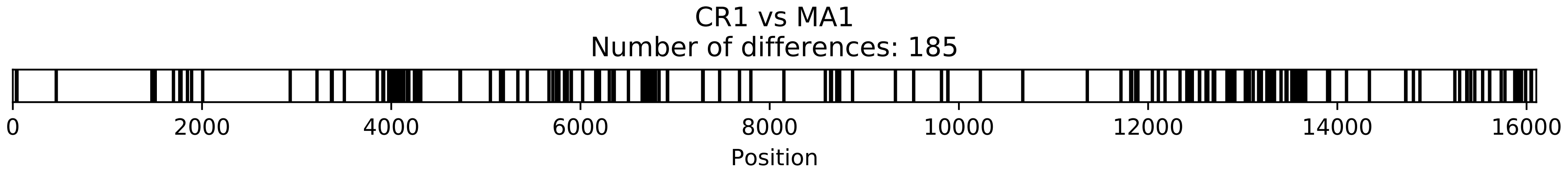
CR1 vs MA1
Number of differences: 185
0 2000 4000 6000 8000 10000 12000 14000 16000
Position

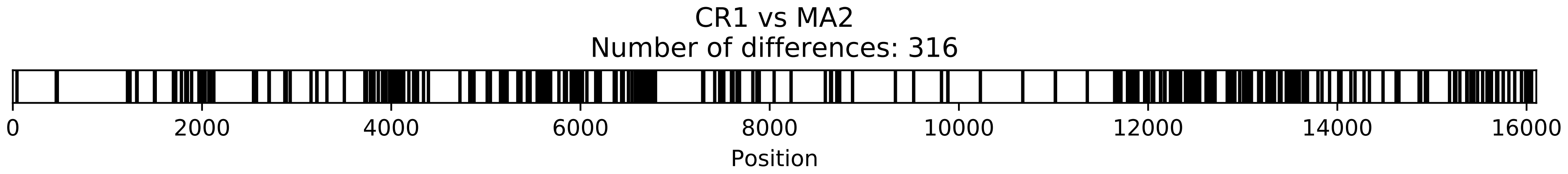
CR1 vs MA2
Number of differences: 316
0 2000 4000 6000 8000 10000 12000 14000 16000
Position

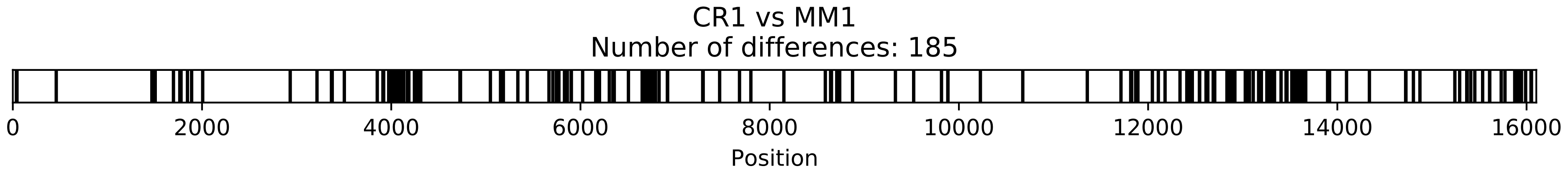
CR1 vs MM1
Number of differences: 185
0 2000 4000 6000 8000 10000 12000 14000 16000
Position

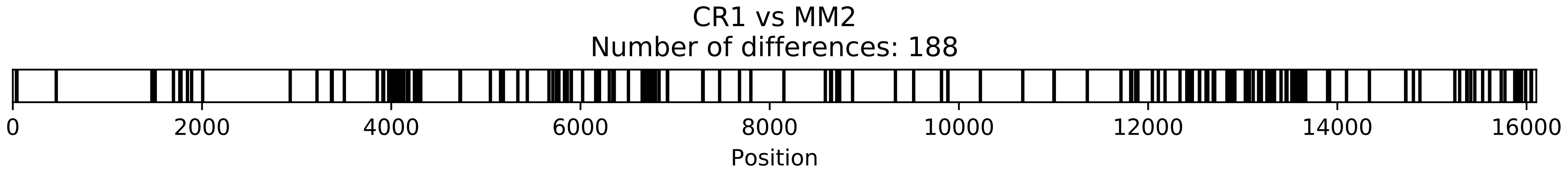
CR1 vs MM2
Number of differences: 188
0 2000 4000 6000 8000 10000 12000 14000 16000
Position

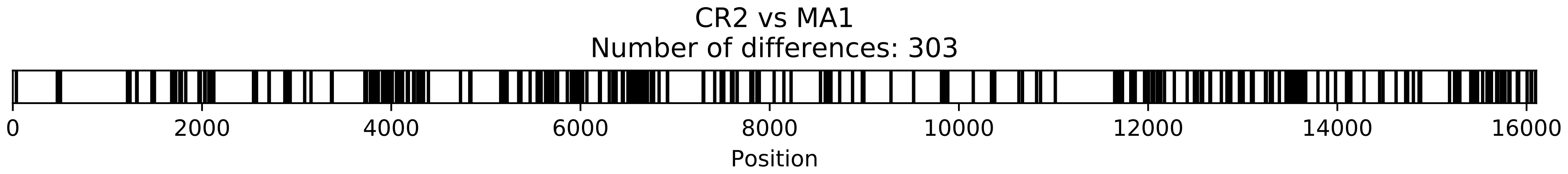
CR2 vs MA1
Number of differences: 303
0 2000 4000 6000 8000 10000 12000 14000 16000
Position

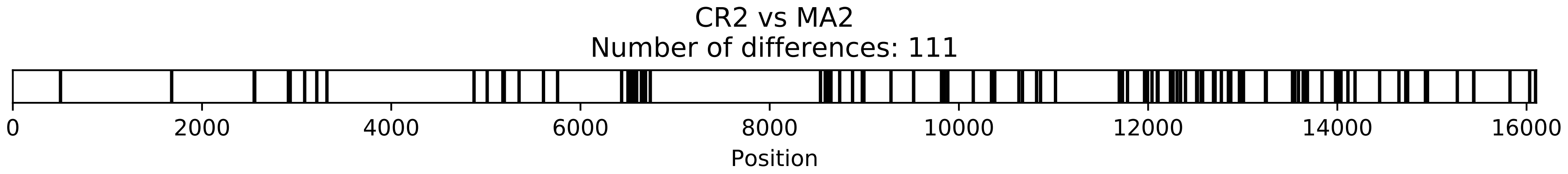
CR2 vs MA2
Number of differences: 111
0 2000 4000 6000 8000 10000 12000 14000 16000
Position

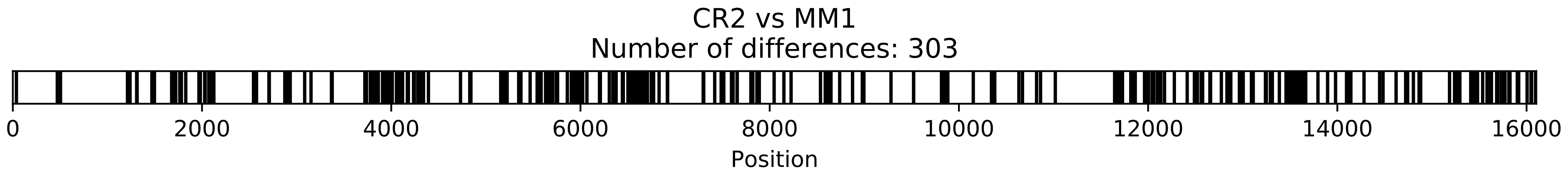
CR2 vs MM1
Number of differences: 303
0 2000 4000 6000 8000 10000 12000 14000 16000
Position

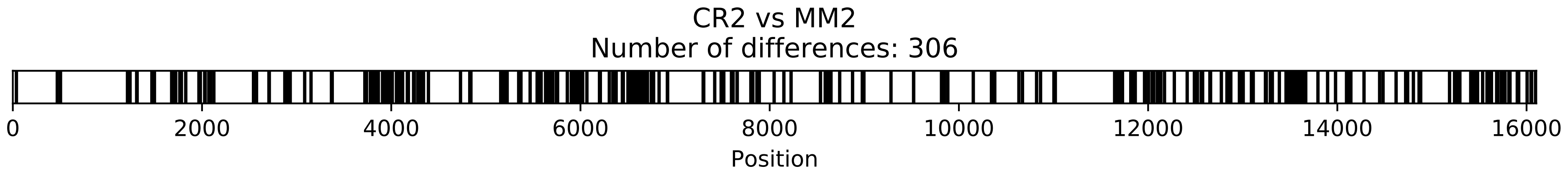
CR2 vs MM2
Number of differences: 306
0 2000 4000 6000 8000 10000 12000 14000 16000
Position

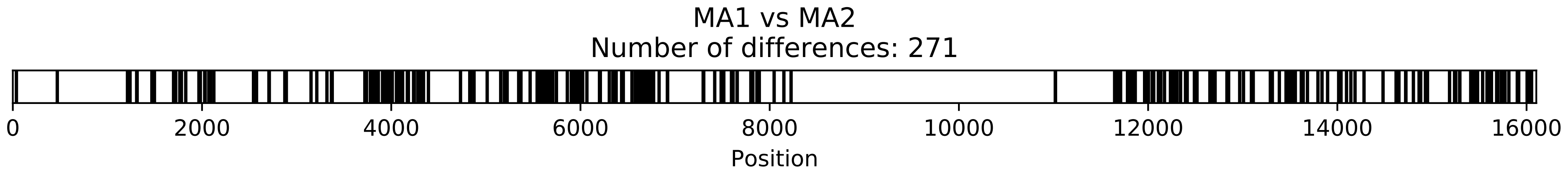
MA1 vs MA2
Number of differences: 271
0 2000 4000 6000 8000 10000 12000 14000 16000
Position

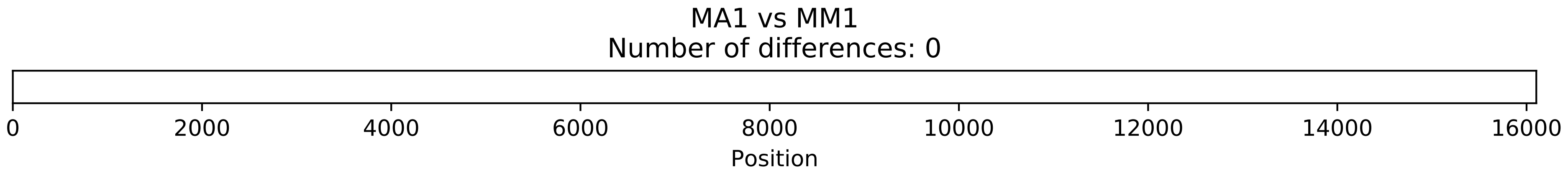
MA1 vs MM1
Number of differences: 0
0 2000 4000 6000 8000 10000 12000 14000 16000
Position

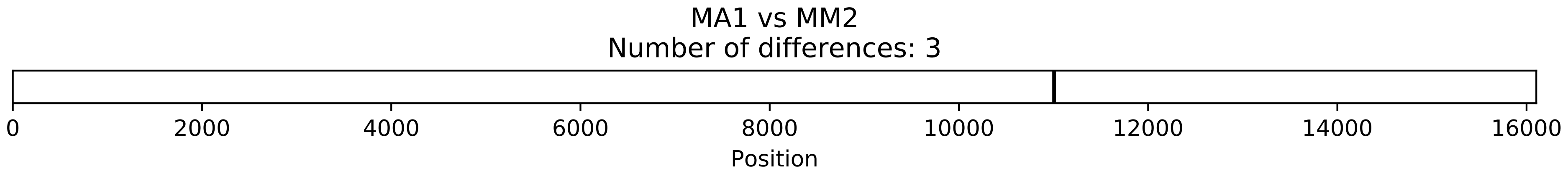
MA1 vs MM2
Number of differences: 3
0 2000 4000 6000 8000 10000 12000 14000 16000
Position

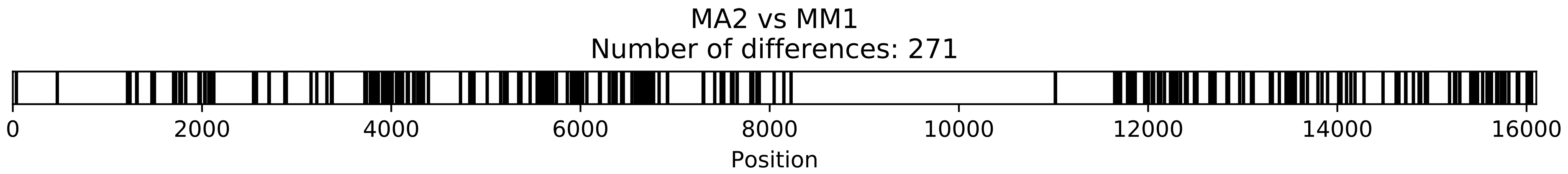
MA2 vs MM1
Number of differences: 271
0 2000 4000 6000 8000 10000 12000 14000 16000
Position

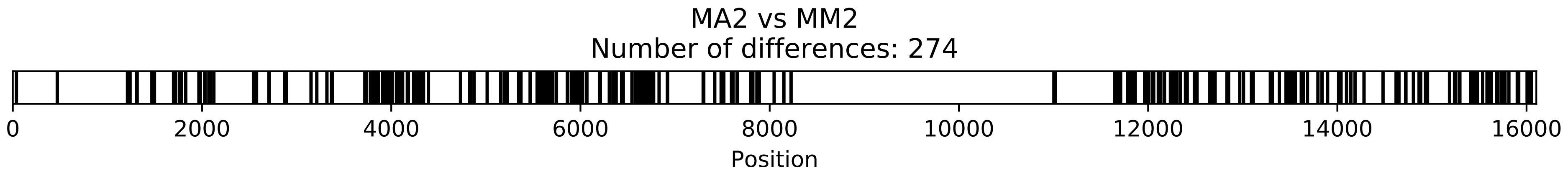
MA2 vs MM2
Number of differences: 274
0 2000 4000 6000 8000 10000 12000 14000 16000
Position

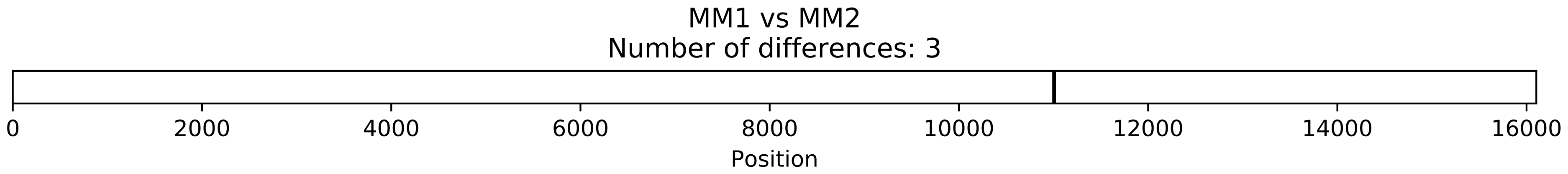
MM1 vs MM2
Number of differences: 3
0 2000 4000 6000 8000 10000 12000 14000 16000
Position

### Region: 491



CR1 vs CR2
Number of differences: 246

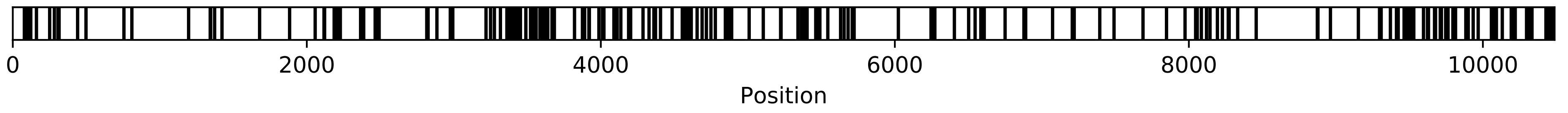


CR1 vs MA1
Number of differences: 220

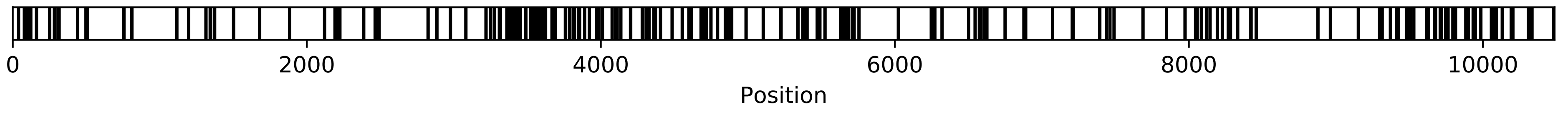


CR1 vs MA2
Number of differences: 107

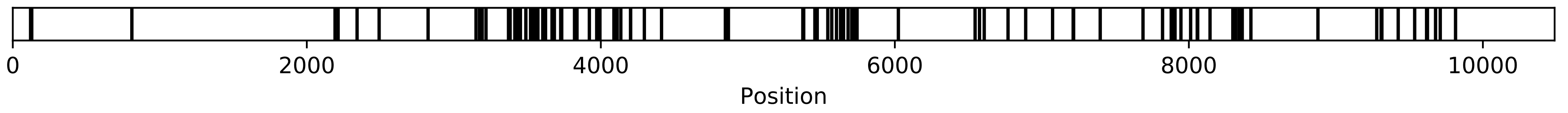


CR1 vs MM1
Number of differences: 248

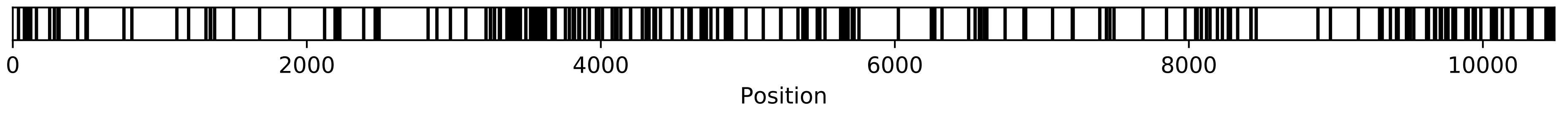


CR1 vs MM2
Number of differences: 253

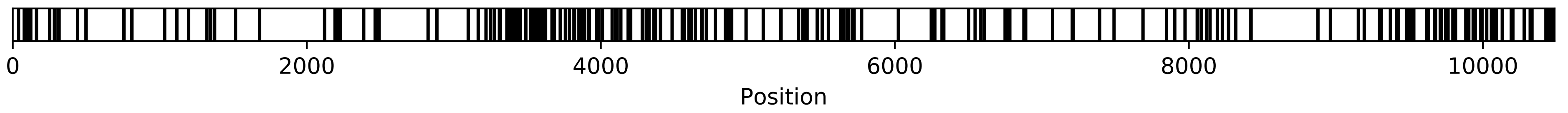


CR2 vs MA1
Number of differences: 127

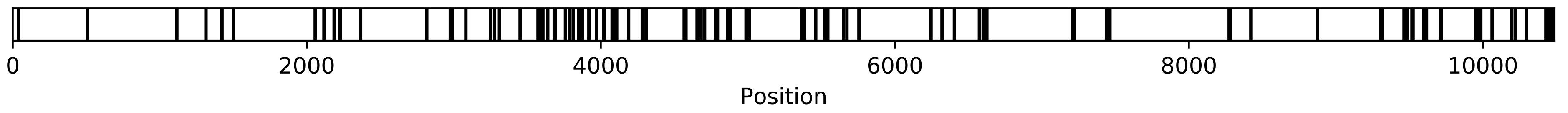


CR2 vs MA2
Number of differences: 235

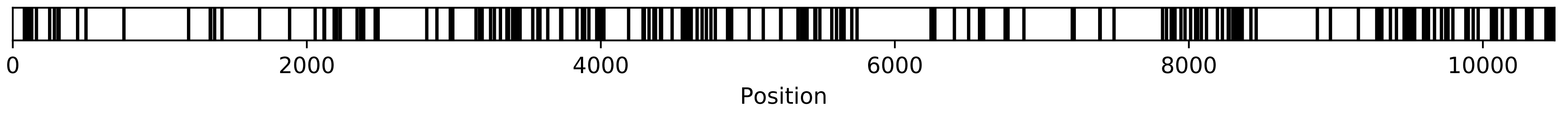


CR2 vs MM1
Number of differences: 98

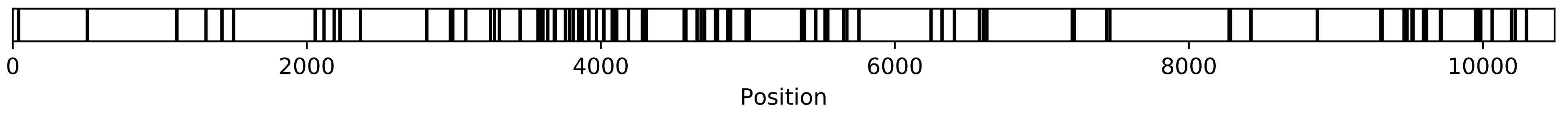


CR2 vs MM2
Number of differences: 112

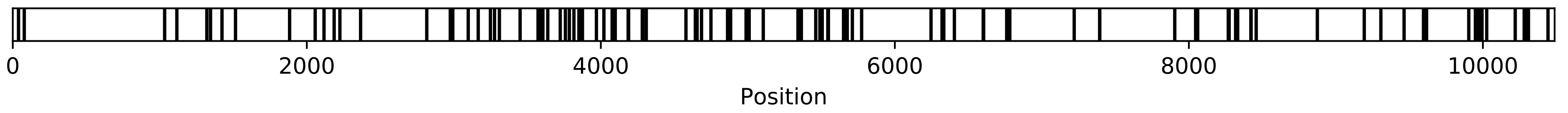


MA1 vs MA2
Number of differences: 209

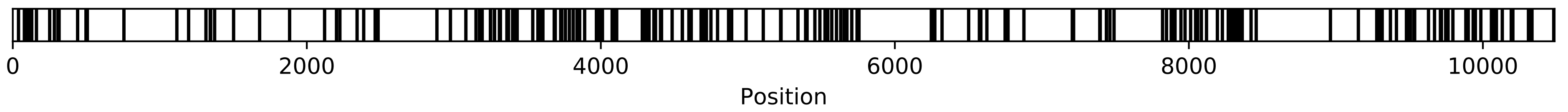


MA1 vs MM1
Number of differences: 29

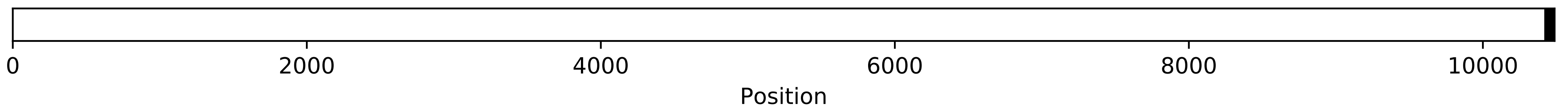


MA1 vs MM2
Number of differences: 117

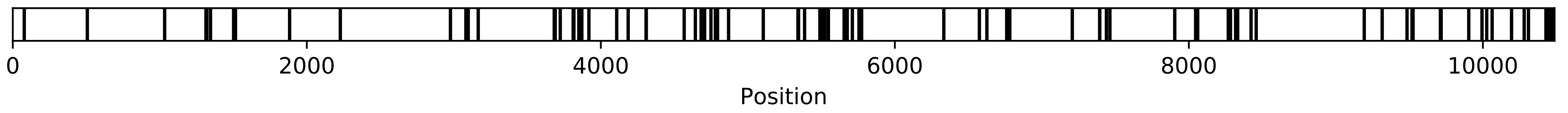


MA2 vs MM1
Number of differences: 237

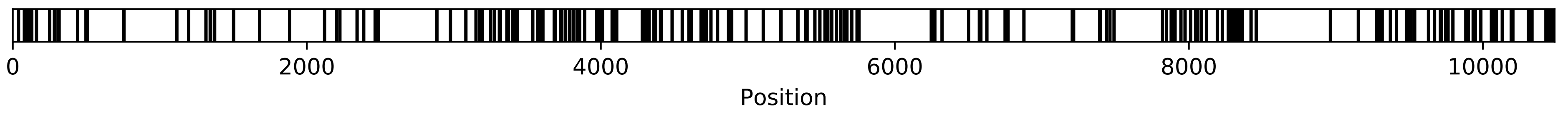


MA2 vs MM2
Number of differences: 238

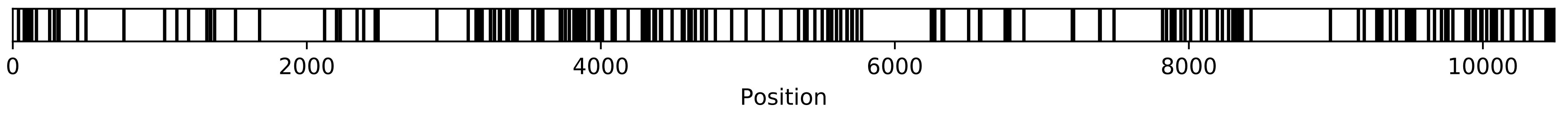


MM1 vs MM2
Number of differences: 89

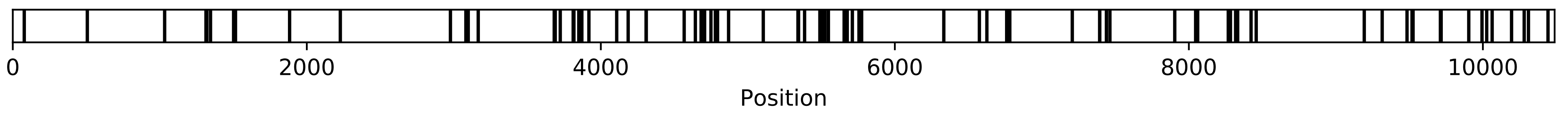

### Region: 492



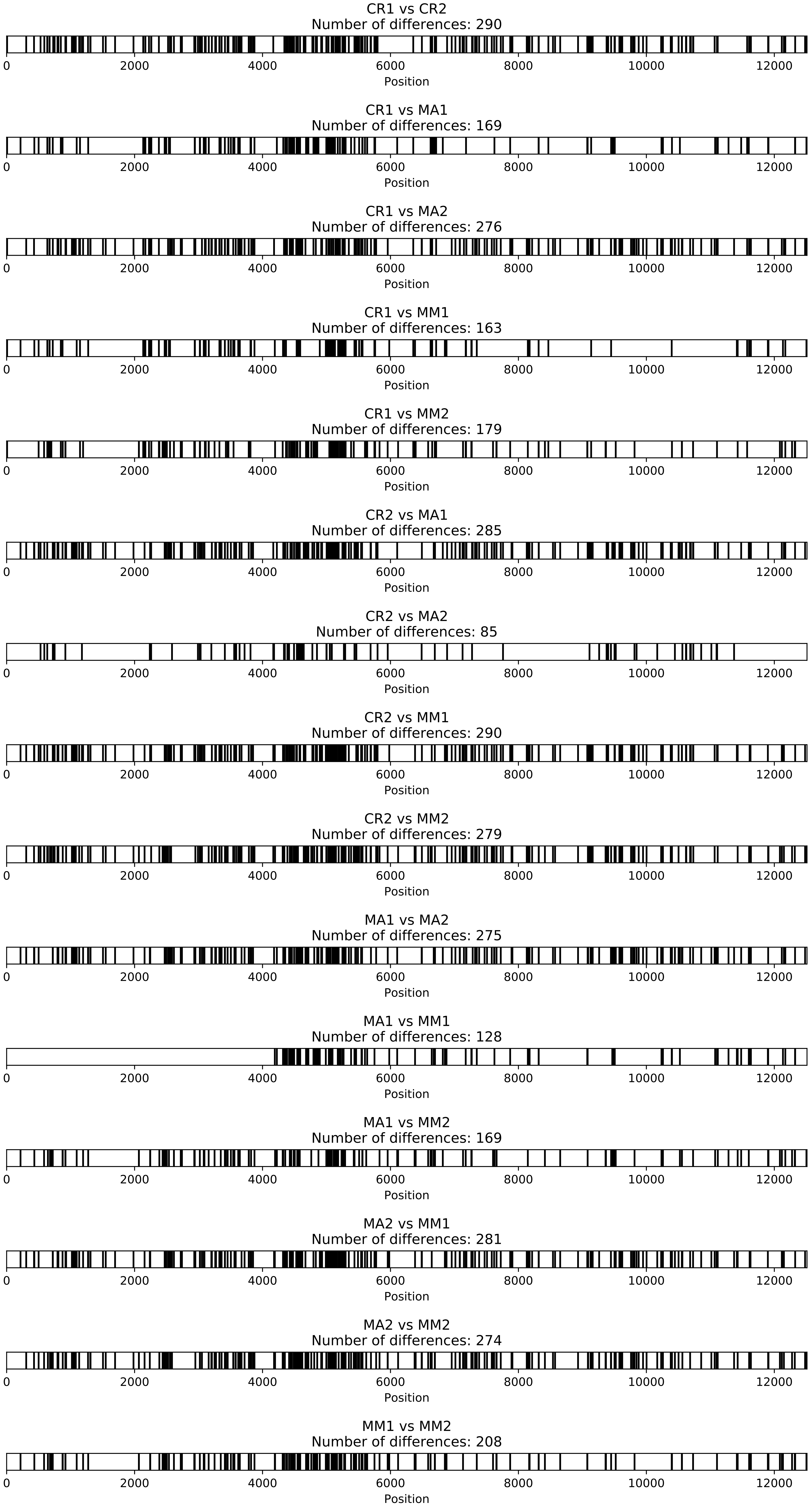
CR1 vs CR2
Number of differences: 290
CR1 vs MA1
Number of differences: 169
CR1 vs MA2
Number of differences: 276
CR1 vs MM1
Number of differences: 163
CR1 vs MM2
Number of differences: 179
CR2 vs MA1
Number of differences: 285
CR2 vs MA2
Number of differences: 85
CR2 vs MM1
Number of differences: 290
CR2 vs MM2
Number of differences: 279
MA1 vs MA2
Number of differences: 275
MA1 vs MM1
Number of differences: 128
MA1 vs MM2
Number of differences: 169
MA2 vs MM1
Number of differences: 281
MA2 vs MM2
Number of differences: 274
MM1 vs MM2
Number of differences: 208
0
2000
4000
6000
8000
10000
12000
Position

### Region: 493

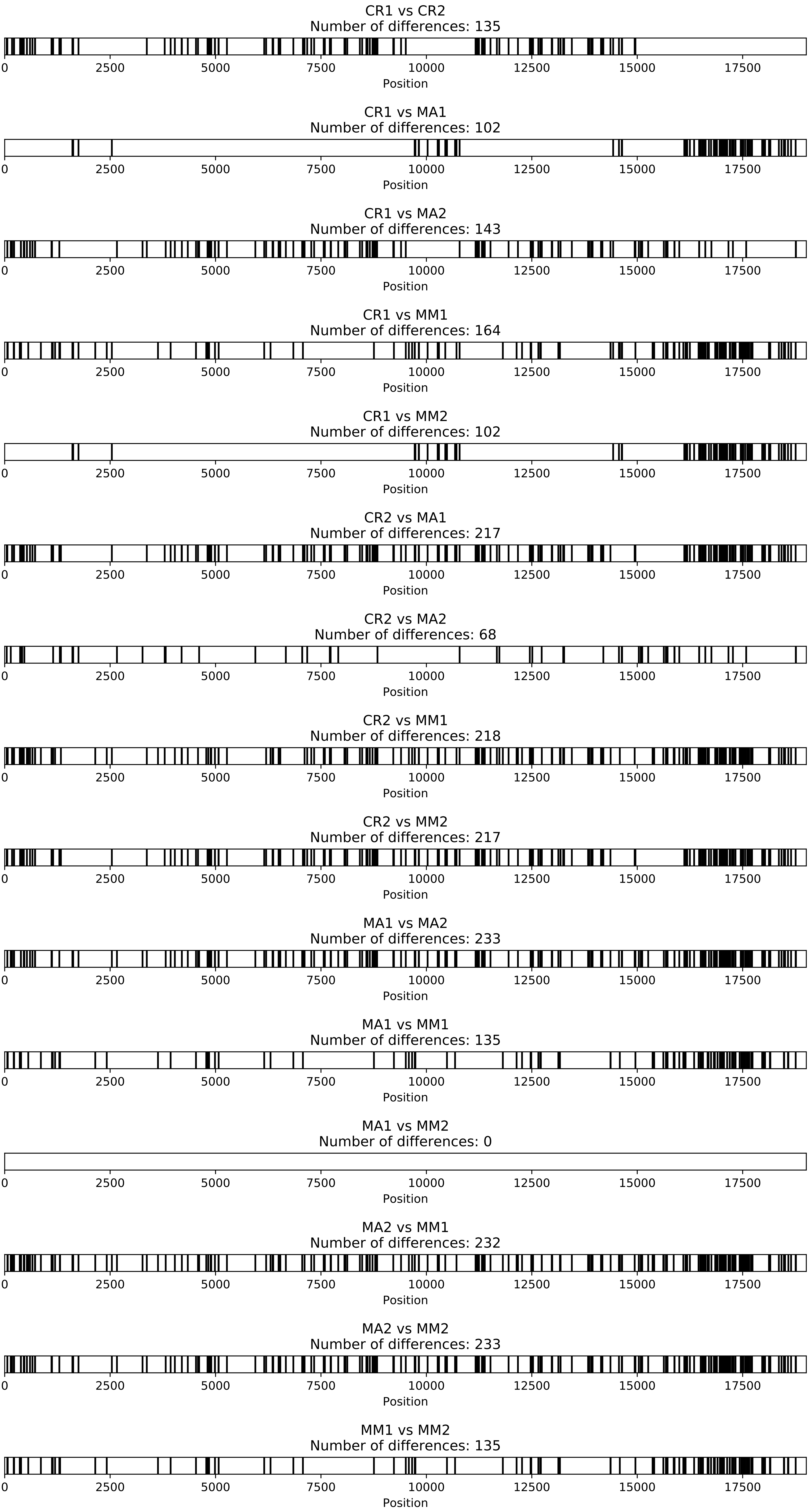
CR1 vs CR2
Number of differences: 135
CR1 vs MA1
Number of differences: 102
CR1 vs MA2
Number of differences: 143
CR1 vs MM1
Number of differences: 164
CR1 vs MM2
Number of differences: 102
CR2 vs MA1
Number of differences: 217
CR2 vs MA2
Number of differences: 68
CR2 vs MM1
Number of differences: 218
CR2 vs MM2
Number of differences: 217
MA1 vs MA2
Number of differences: 233
MA1 vs MM1
Number of differences: 135
MA1 vs MM2
Number of differences: 0
MA2 vs MM1
Number of differences: 232
MA2 vs MM2
Number of differences: 233
MM1 vs MM2
Number of differences: 135
0
2500
5000
7500
10000
12500
15000
17500
Position

### Region: 494

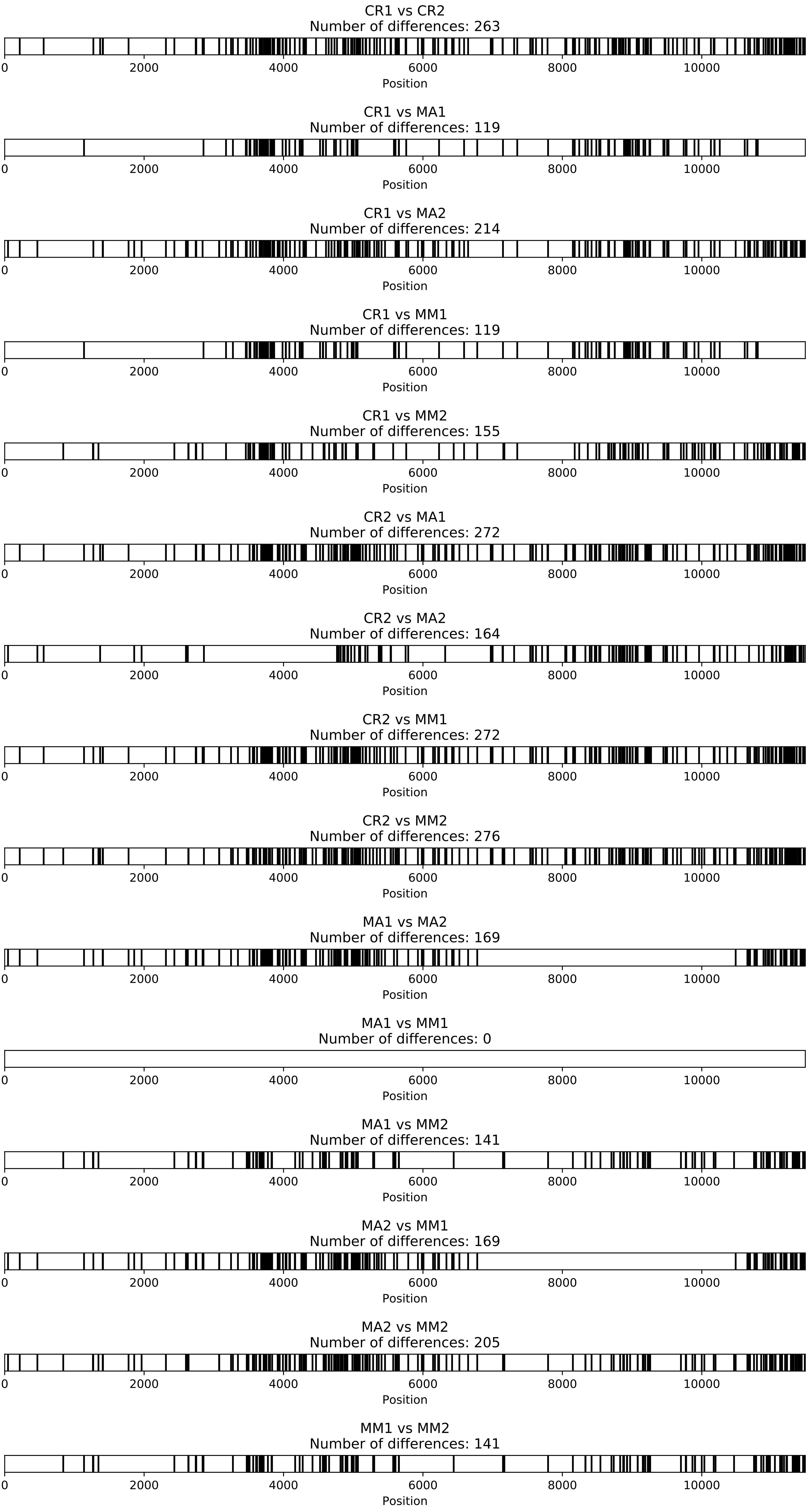
CR1 vs CR2
Number of differences: 263
CR1 vs MA1
Number of differences: 119
CR1 vs MA2
Number of differences: 214
CR1 vs MM1
Number of differences: 119
CR1 vs MM2
Number of differences: 155
CR2 vs MA1
Number of differences: 272
CR2 vs MA2
Number of differences: 164
CR2 vs MM1
Number of differences: 272
CR2 vs MM2
Number of differences: 276
MA1 vs MA2
Number of differences: 169
MA1 vs MM1
Number of differences: 0
MA1 vs MM2
Number of differences: 141
MA2 vs MM1
Number of differences: 169
MA2 vs MM2
Number of differences: 205
MM1 vs MM2
Number of differences: 141
0
2000
4000
6000
8000
10000
Position

### Region: 496



CR1 vs CR2
Number of differences: 77

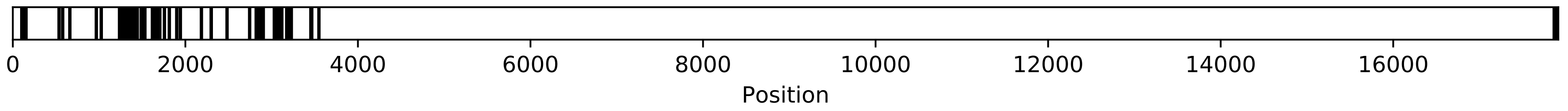


CR1 vs MA1
Number of differences: 238

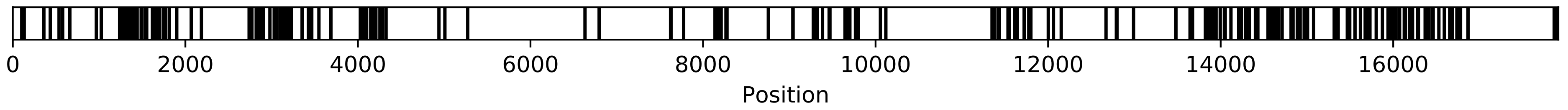


CR1 vs MA2
Number of differences: 427

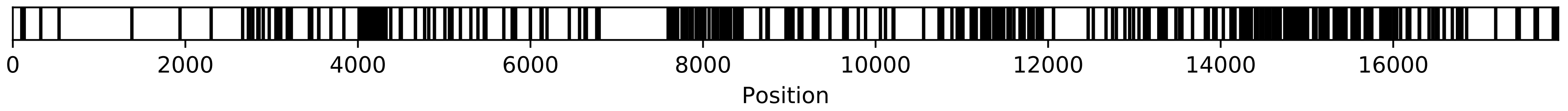


CR1 vs MM1
Number of differences: 434

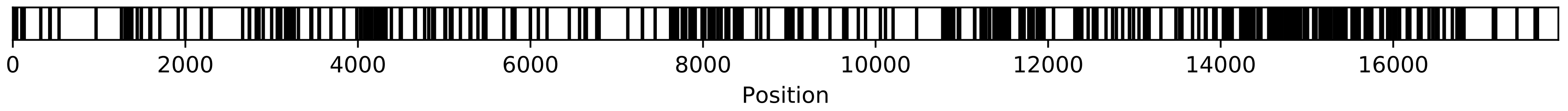


CR1 vs MM2
Number of differences: 457

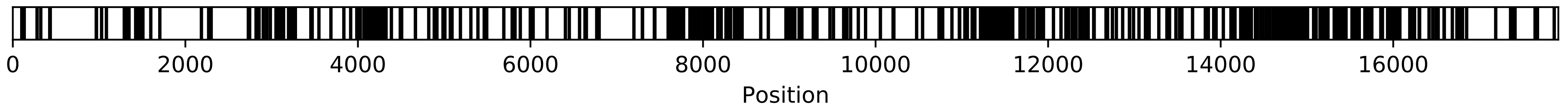


CR2 vs MA1
Number of differences: 207

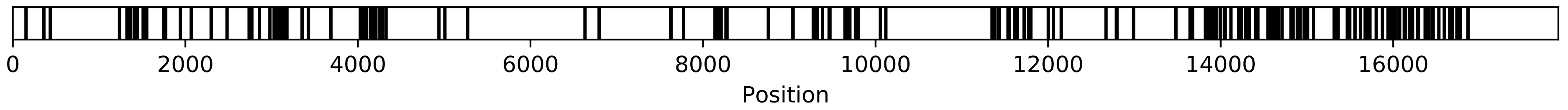


CR2 vs MA2
Number of differences: 470

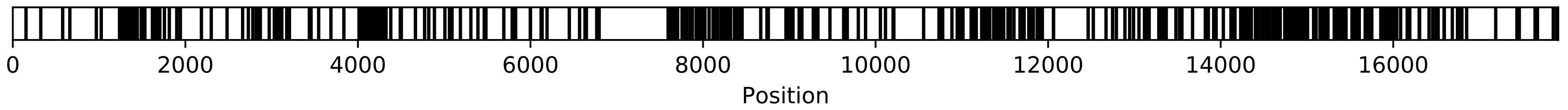


CR2 vs MM1
Number of differences: 458

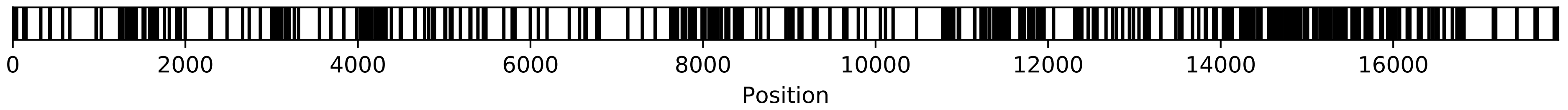


CR2 vs MM2
Number of differences: 485

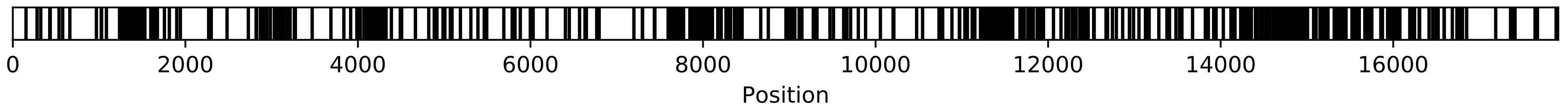


MA1 vs MA2
Number of differences: 478

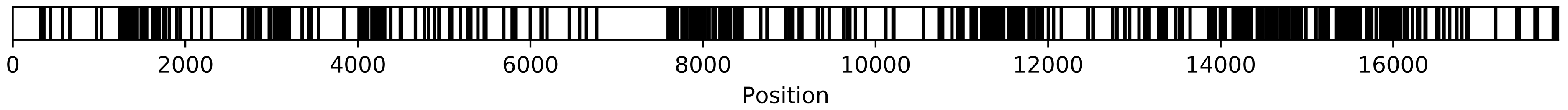


MA1 vs MM1
Number of differences: 455

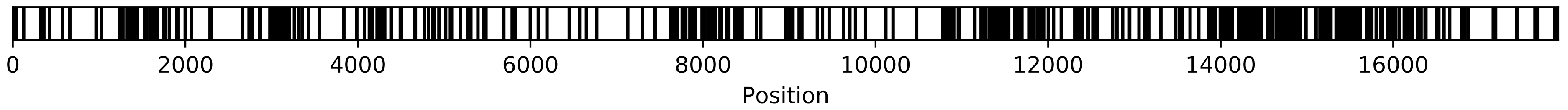


MA1 vs MM2
Number of differences: 488

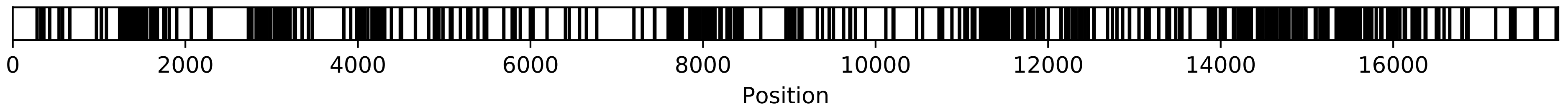


MA2 vs MM1
Number of differences: 282

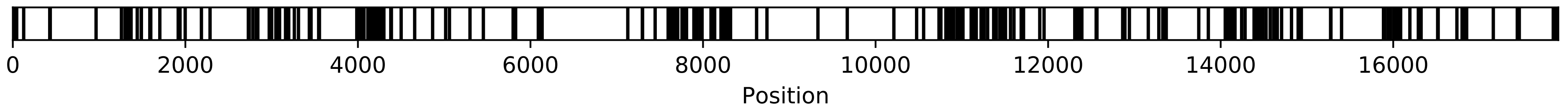


MA2 vs MM2
Number of differences: 248

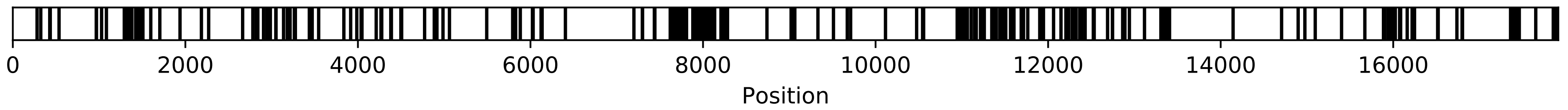


MM1 vs MM2
Number of differences: 272

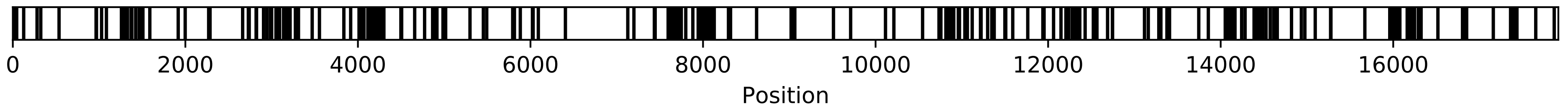

### Region: 500

CR1 vs CR2
Number of differences: 0

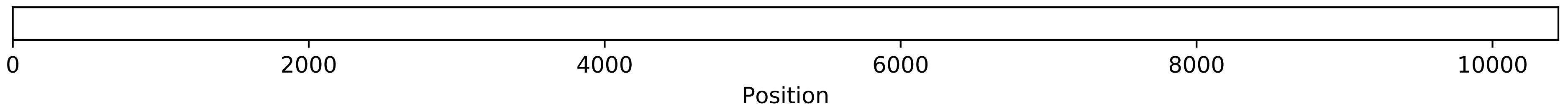


CR1 vs MA1
Number of differences: 158

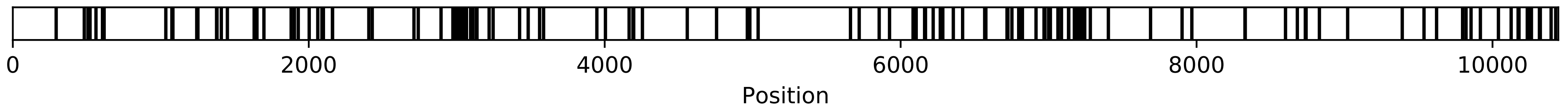


CR1 vs MA2
Number of differences: 211

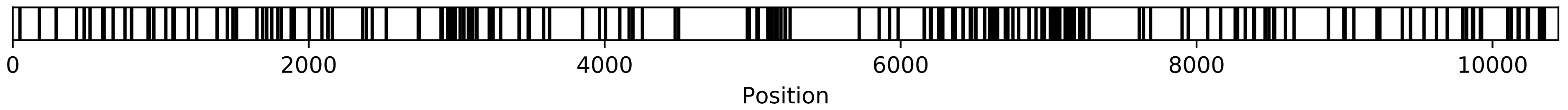


CR1 vs MM1
Number of differences: 158

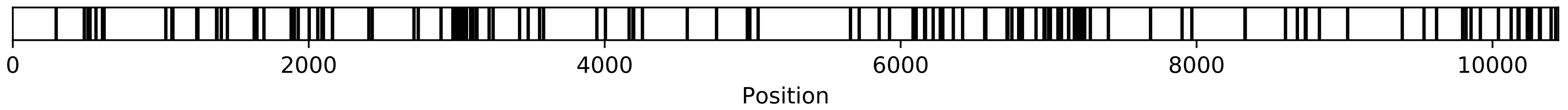


CR1 vs MM2
Number of differences: 102

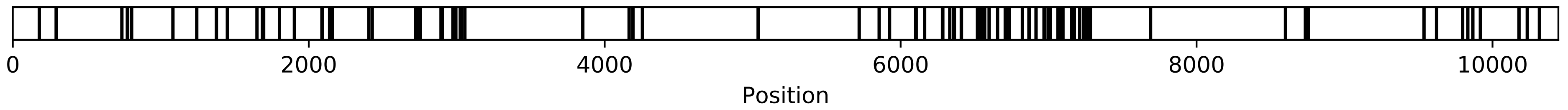


CR2 vs MA1
Number of differences: 158

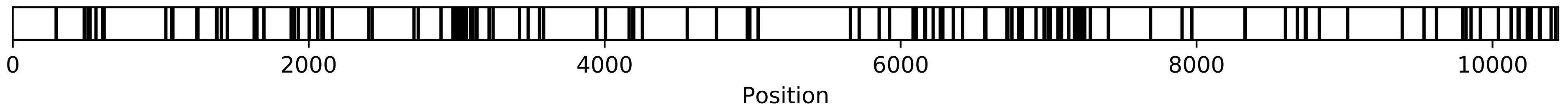


CR2 vs MA2
Number of differences: 211

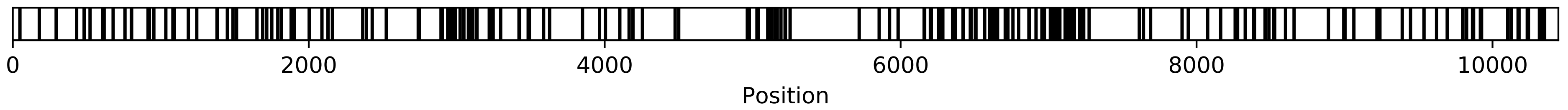


CR2 vs MM1
Number of differences: 158

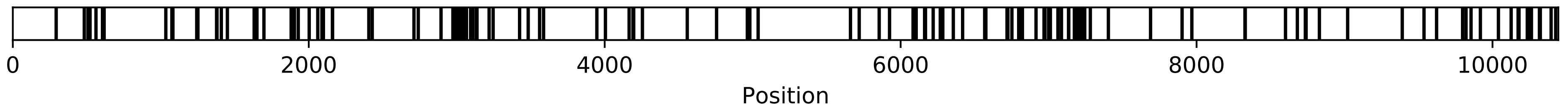


CR2 vs MM2
Number of differences: 102

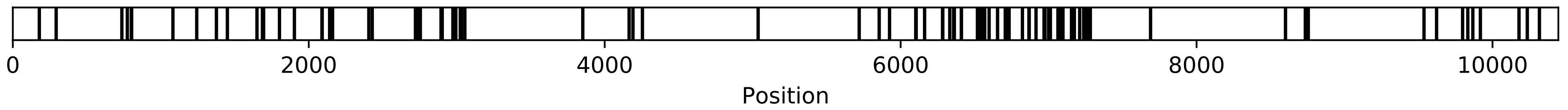


MA1 vs MA2
Number of differences: 222

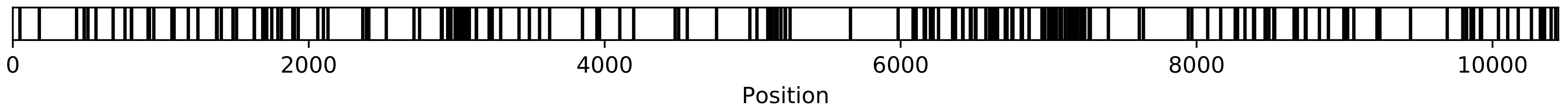


MA1 vs MM1
Number of differences: 0

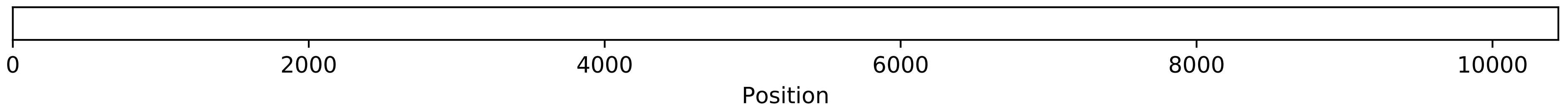


MA1 vs MM2
Number of differences: 147

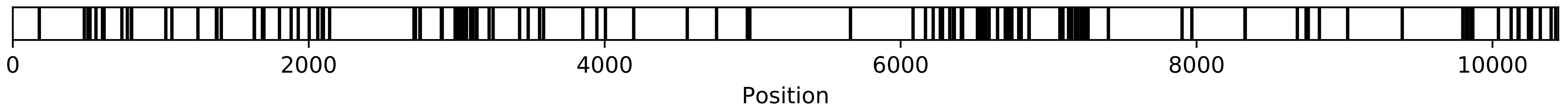


MA2 vs MM1
Number of differences: 222

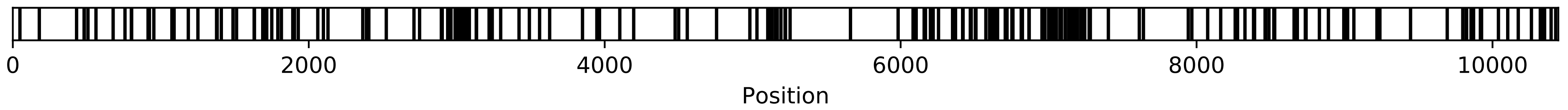


MA2 vs MM2
Number of differences: 209

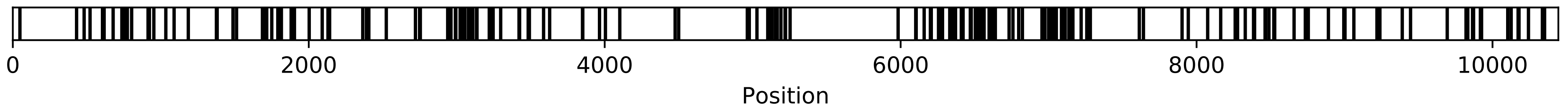


MM1 vs MM2
Number of differences: 147

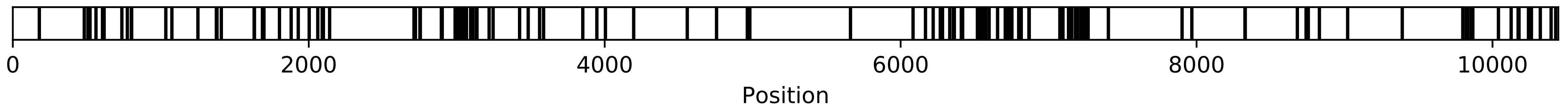

### Region: 501

CR1 vs CR2
Number of differences: 324

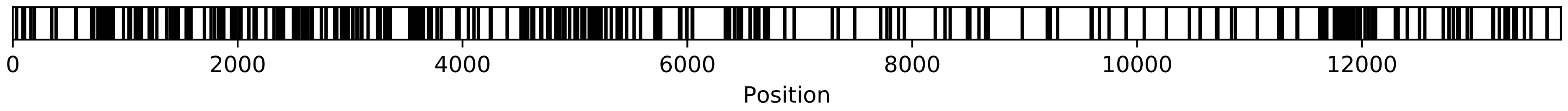


CR1 vs MA1
Number of differences: 132

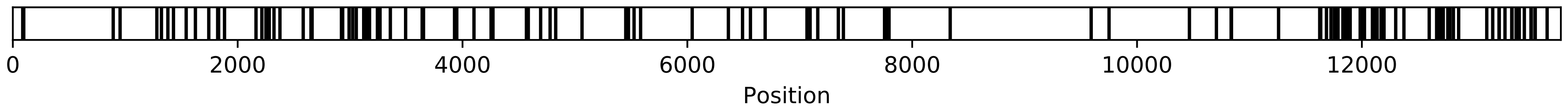


CR1 vs MA2
Number of differences: 329

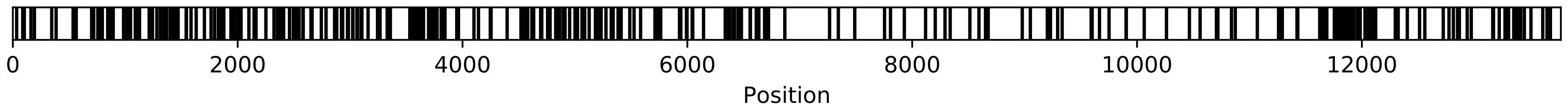


CR1 vs MM1
Number of differences: 102

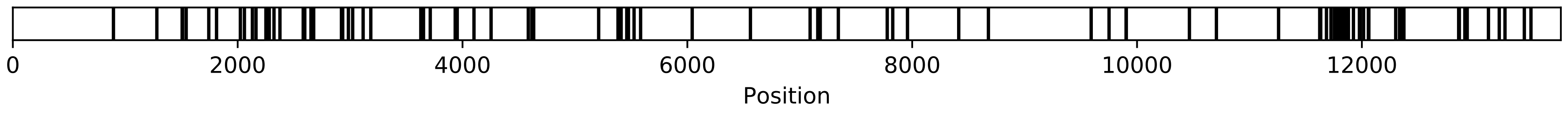


CR1 vs MM2
Number of differences: 134

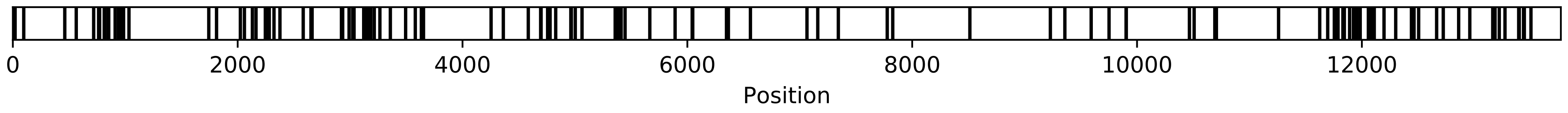


CR2 vs MA1
Number of differences: 332

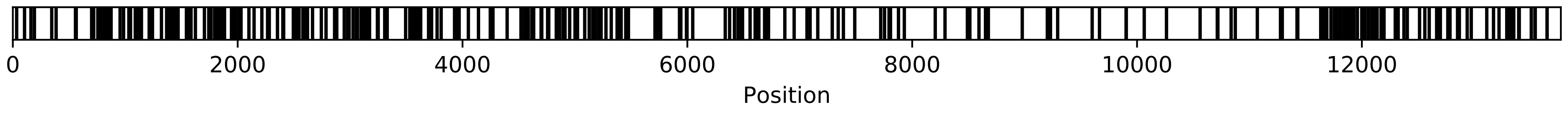


CR2 vs MA2
Number of differences: 73

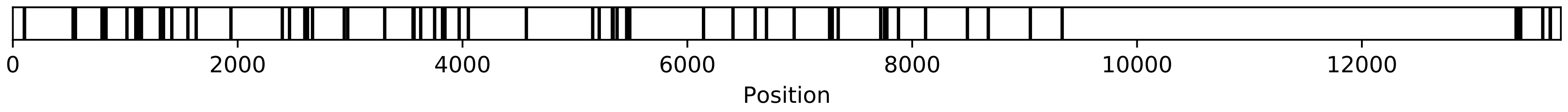


CR2 vs MM1
Number of differences: 314

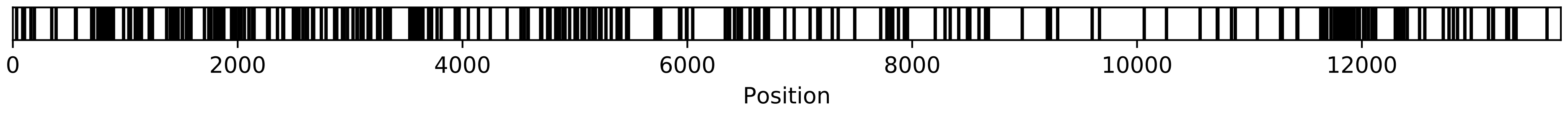


CR2 vs MM2
Number of differences: 327

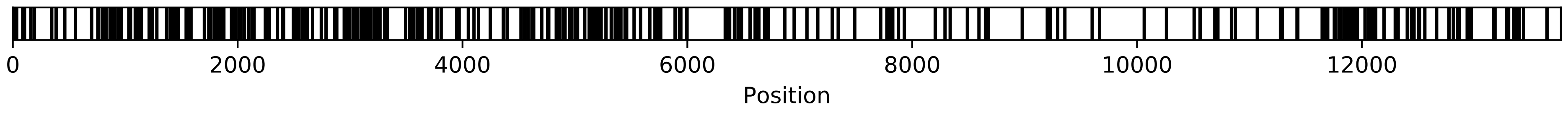


MA1 vs MA2
Number of differences: 337

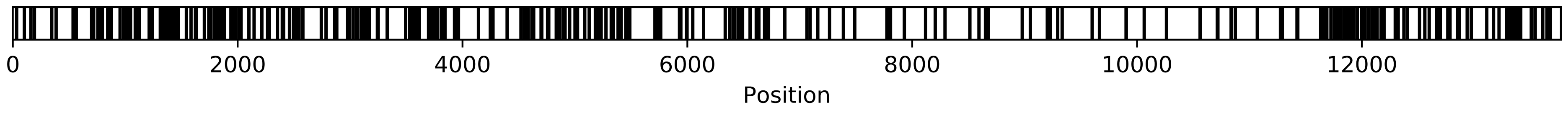


MA1 vs MM1
Number of differences: 126

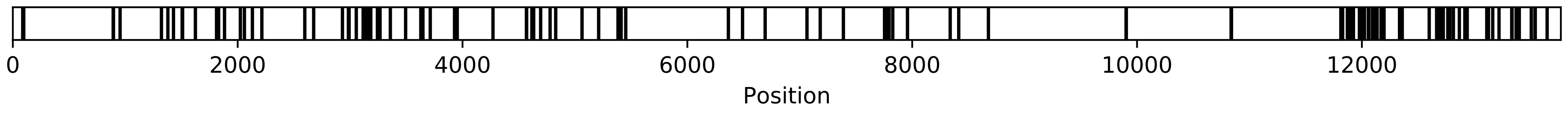


MA1 vs MM2
Number of differences: 142

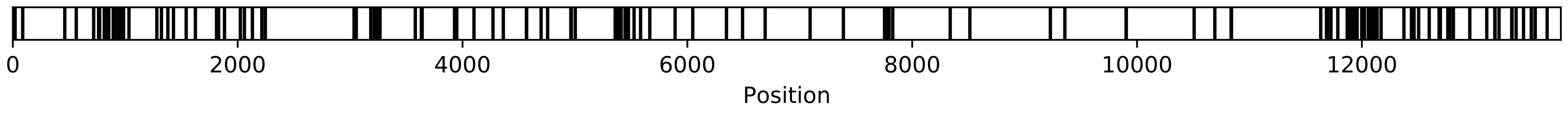


MA2 vs MM1
Number of differences: 321

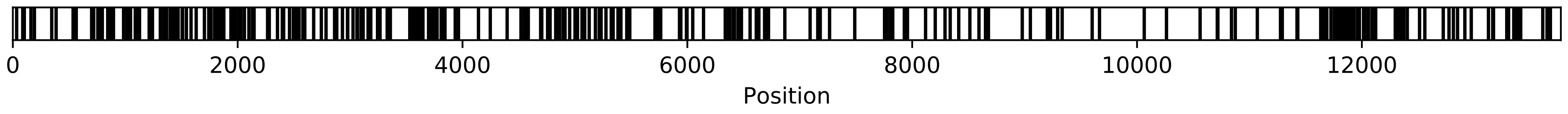


MA2 vs MM2
Number of differences: 332

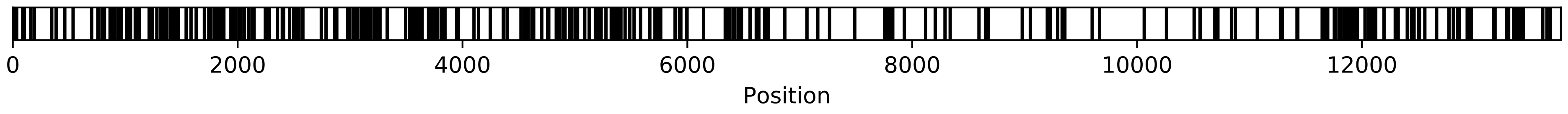


MM1 vs MM2
Number of differences: 126

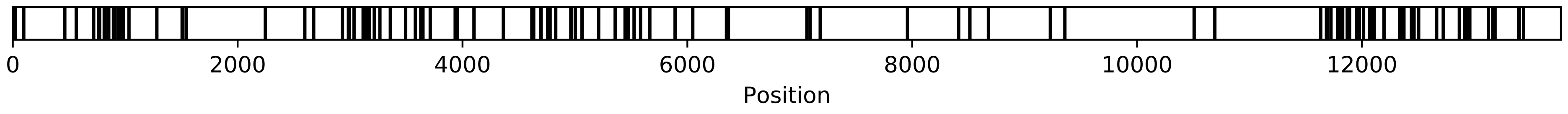

### Region: 502



CR1 vs CR2
Number of differences: 211

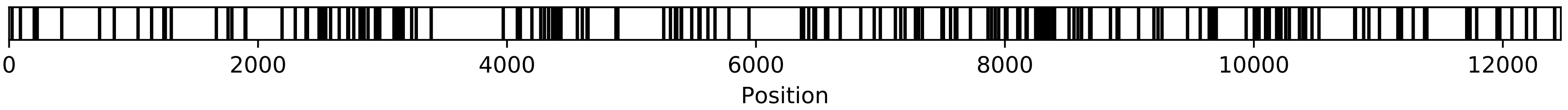


CR1 vs MA1
Number of differences: 214

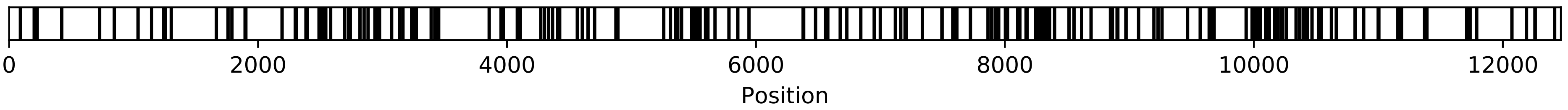


CR1 vs MA2
Number of differences: 49

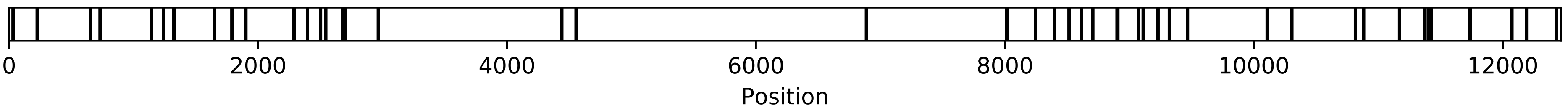


CR1 vs MM1
Number of differences: 211

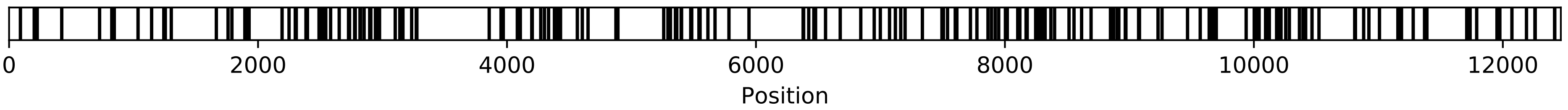


CR1 vs MM2
Number of differences: 214

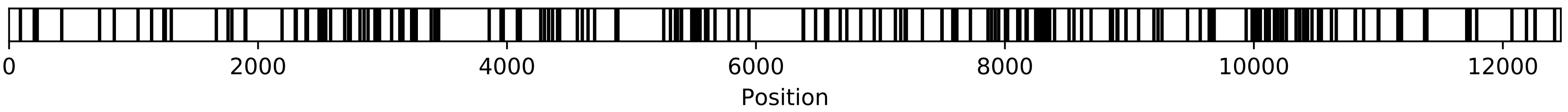


CR2 vs MA1
Number of differences: 95

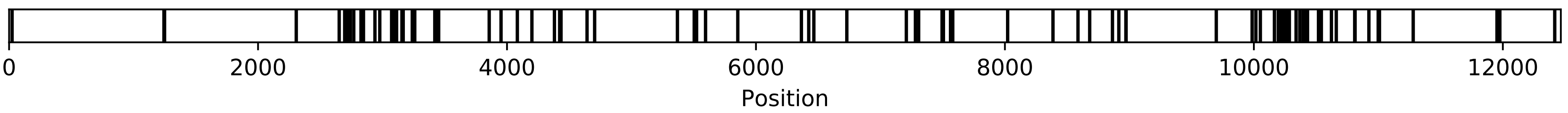


CR2 vs MA2
Number of differences: 205

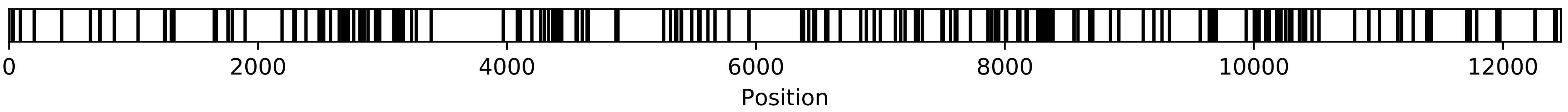


CR2 vs MM1
Number of differences: 66

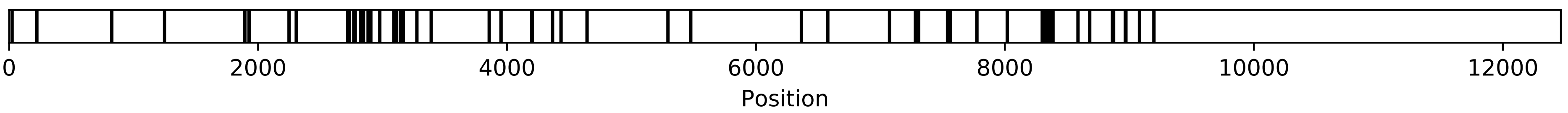


CR2 vs MM2
Number of differences: 95

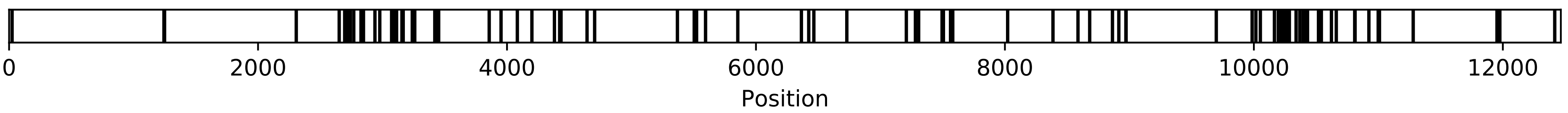


MA1 vs MA2
Number of differences: 208

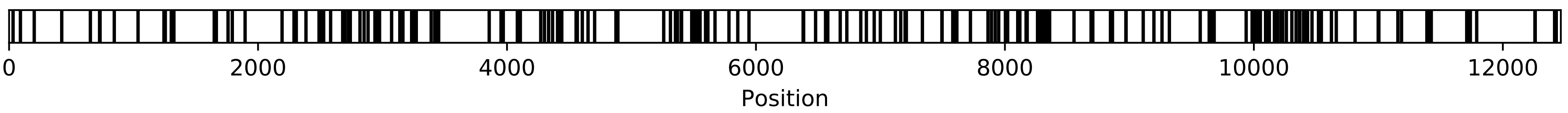


MA1 vs MM1
Number of differences: 105

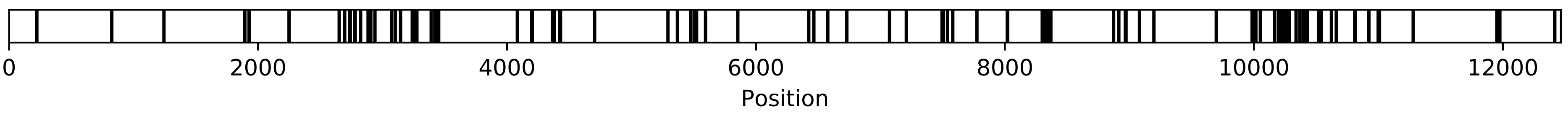


MA1 vs MM2
Number of differences: 0

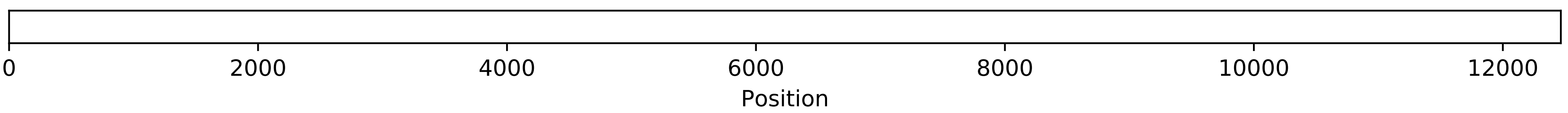


MA2 vs MM1
Number of differences: 205

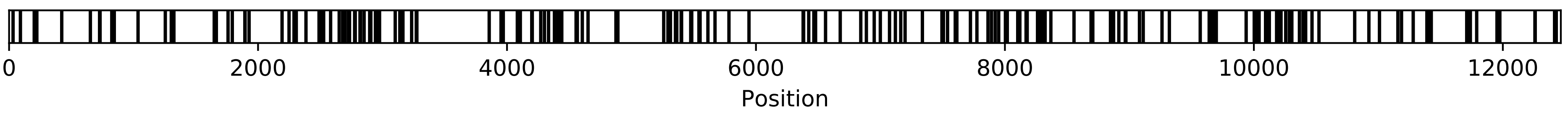


MA2 vs MM2
Number of differences: 208

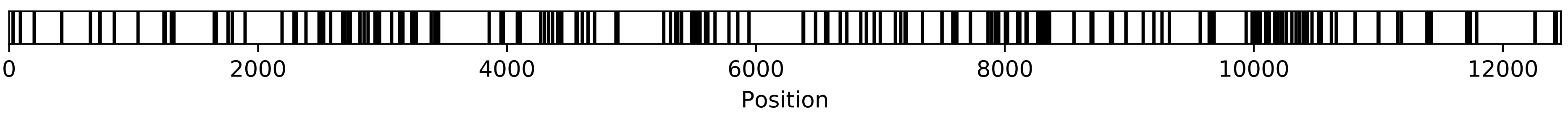


MM1 vs MM2
Number of differences: 105

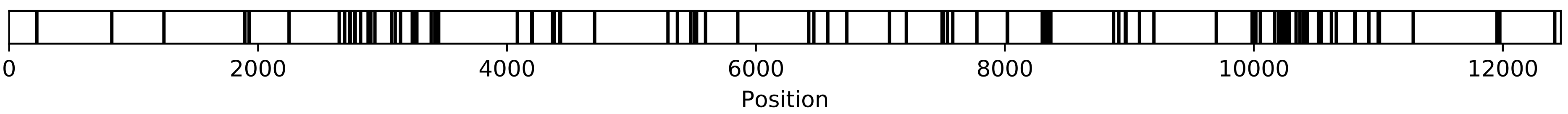

### Region: 503



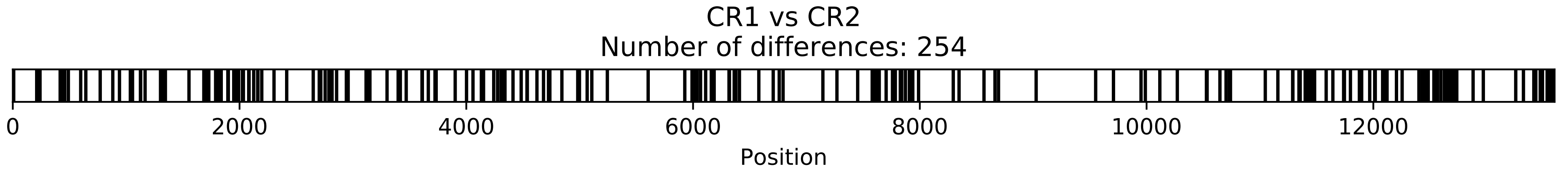
CR1 vs CR2
Number of differences: 254
0
2000
4000
6000
8000
10000
12000
Position

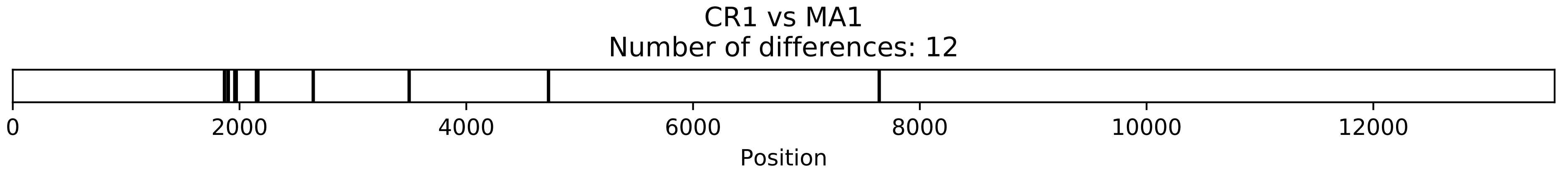
CR1 vs MA1
Number of differences: 12
0
2000
4000
6000
8000
10000
12000
Position

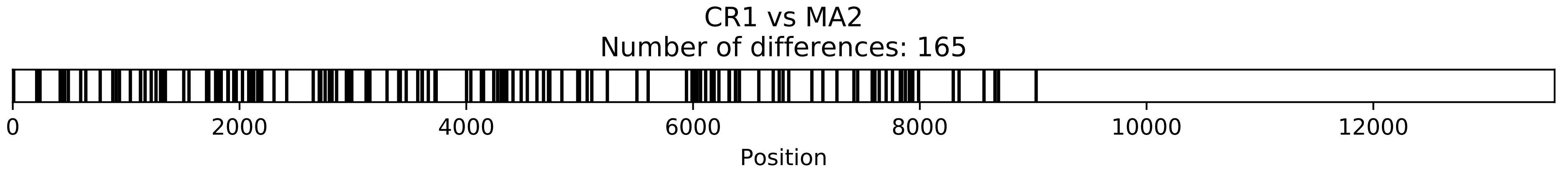
CR1 vs MA2
Number of differences: 165
0
2000
4000
6000
8000
10000
12000
Position

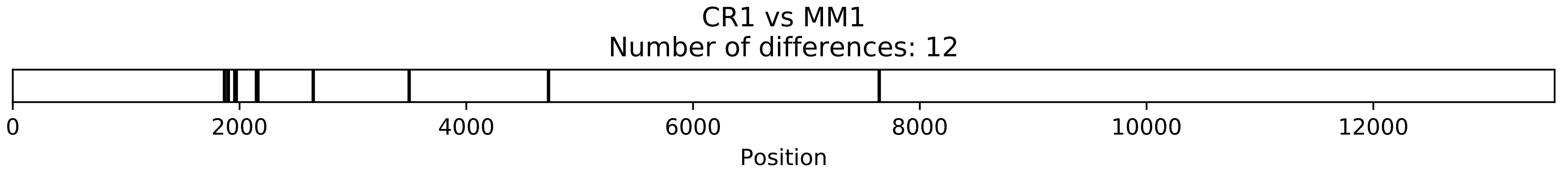
CR1 vs MM1
Number of differences: 12
0
2000
4000
6000
8000
10000
12000
Position

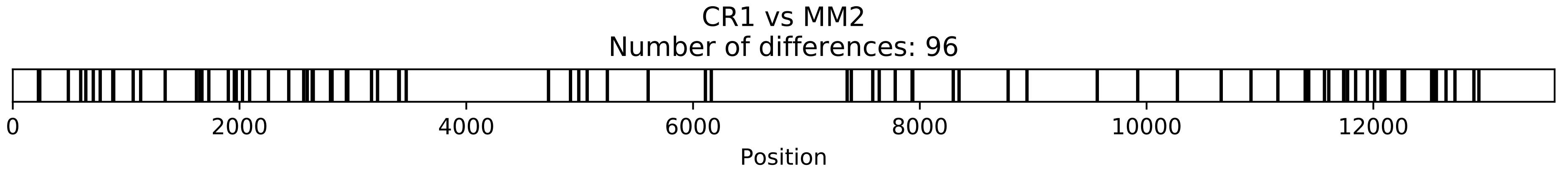
CR1 vs MM2
Number of differences: 96
0
2000
4000
6000
8000
10000
12000
Position

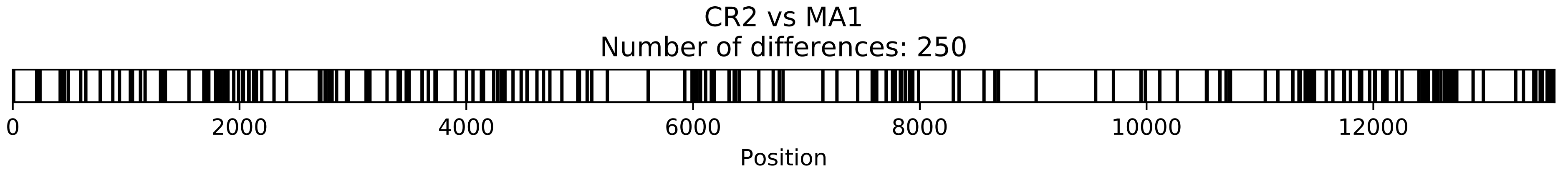
CR2 vs MA1
Number of differences: 250
0
2000
4000
6000
8000
10000
12000
Position

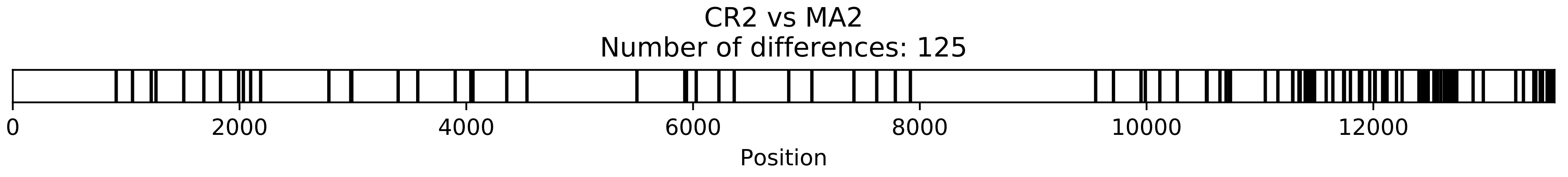
CR2 vs MA2
Number of differences: 125
0
2000
4000
6000
8000
10000
12000
Position

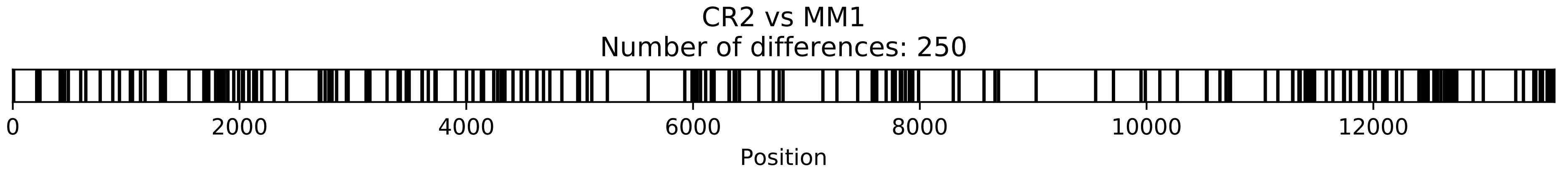
CR2 vs MM1
Number of differences: 250
0
2000
4000
6000
8000
10000
12000
Position

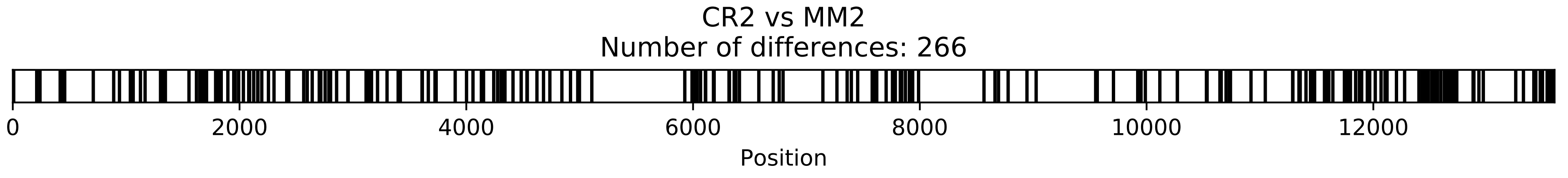
CR2 vs MM2
Number of differences: 266
0
2000
4000
6000
8000
10000
12000
Position

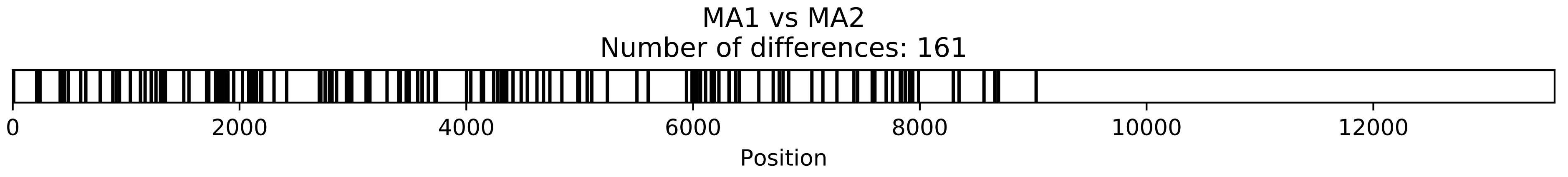
MA1 vs MA2
Number of differences: 161
0
2000
4000
6000
8000
10000
12000
Position

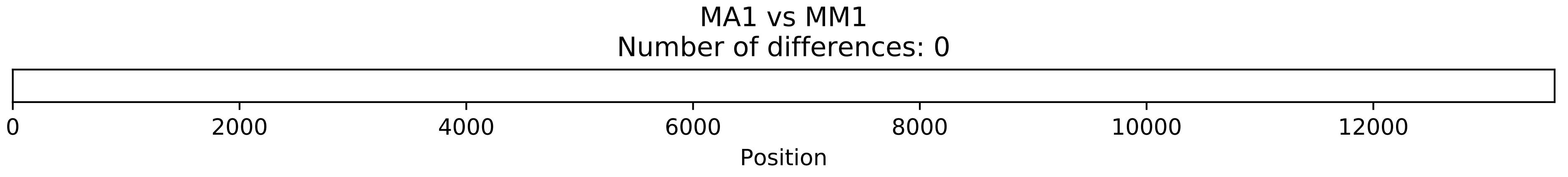
MA1 vs MM1
Number of differences: 0
0
2000
4000
6000
8000
10000
12000
Position

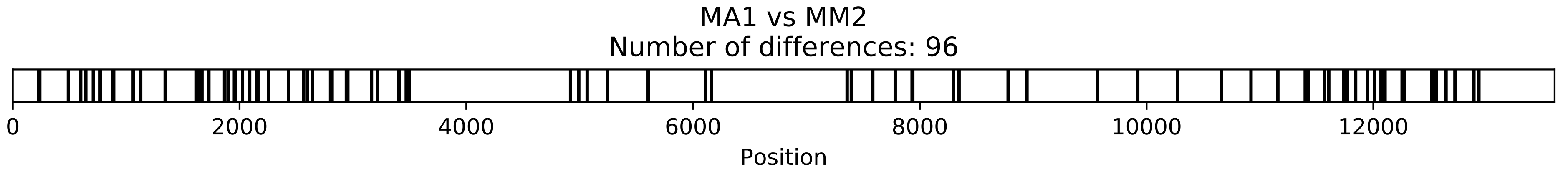
MA1 vs MM2
Number of differences: 96
0
2000
4000
6000
8000
10000
12000
Position

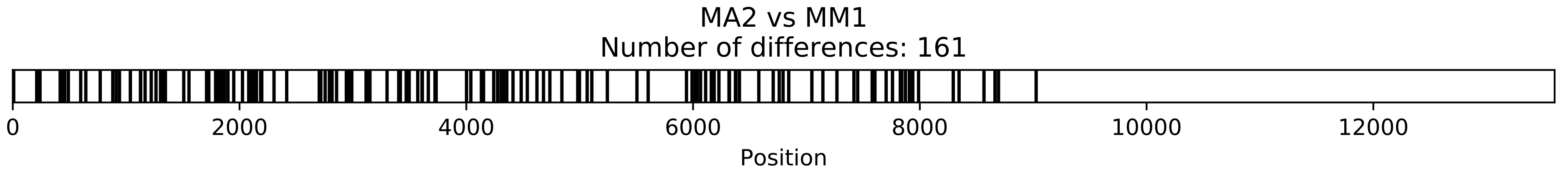
MA2 vs MM1
Number of differences: 161
0
2000
4000
6000
8000
10000
12000
Position

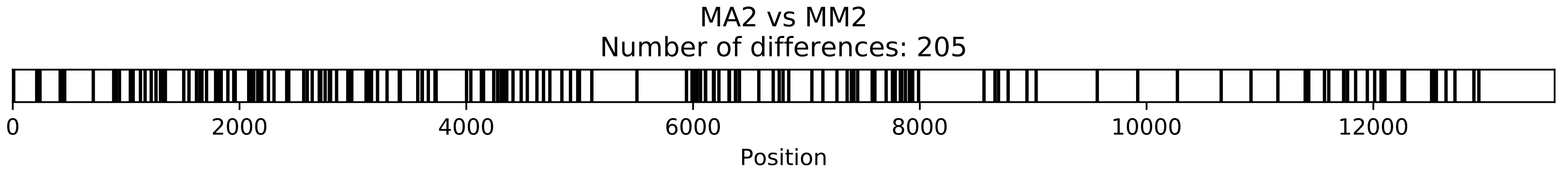
MA2 vs MM2
Number of differences: 205
0
2000
4000
6000
8000
10000
12000
Position

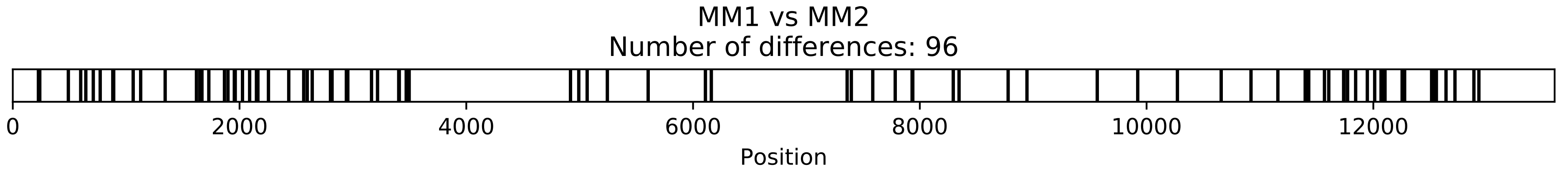
MM1 vs MM2
Number of differences: 96
0
2000
4000
6000
8000
10000
12000
Position

### Region: 505

CR1 vs CR2
Number of differences: 283

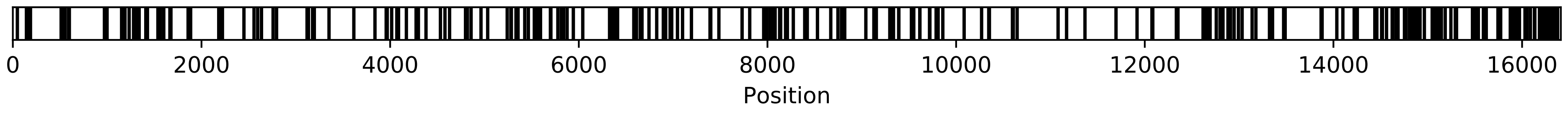


CR1 vs MA1
Number of differences: 261

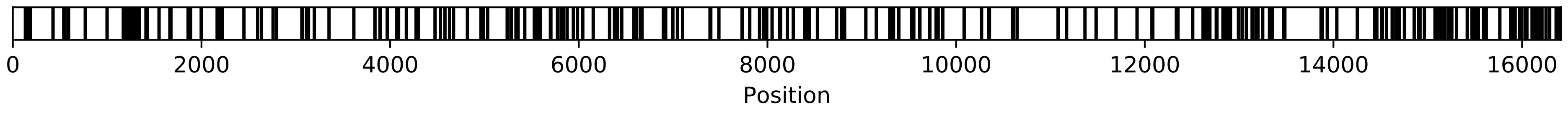


CR1 vs MA2
Number of differences: 82

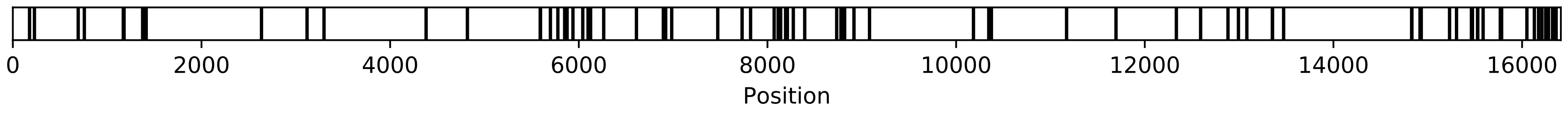


CR1 vs MM1
Number of differences: 261

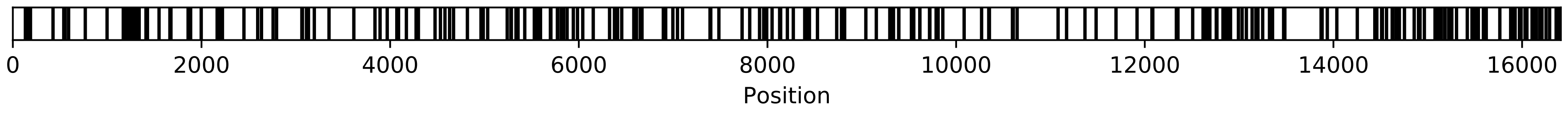


CR1 vs MM2
Number of differences: 292

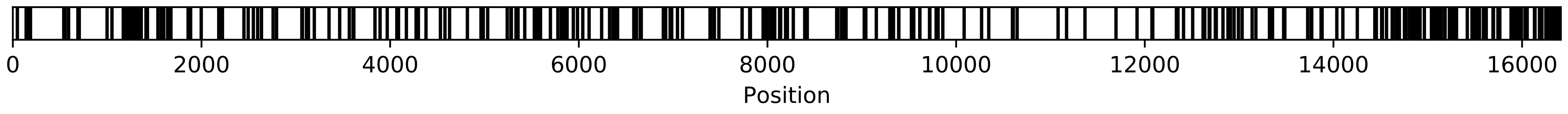


CR2 vs MA1
Number of differences: 144

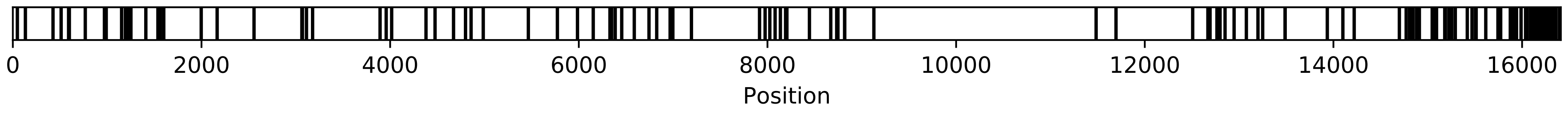


CR2 vs MA2
Number of differences: 288

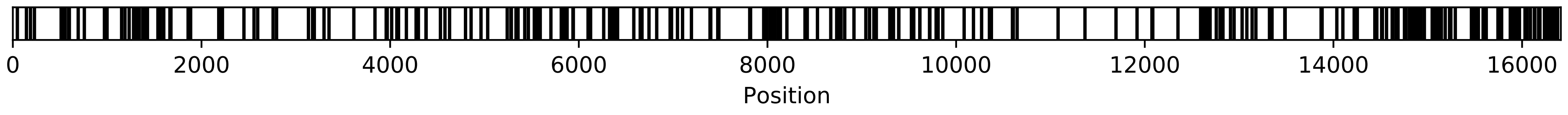


CR2 vs MM1
Number of differences: 144

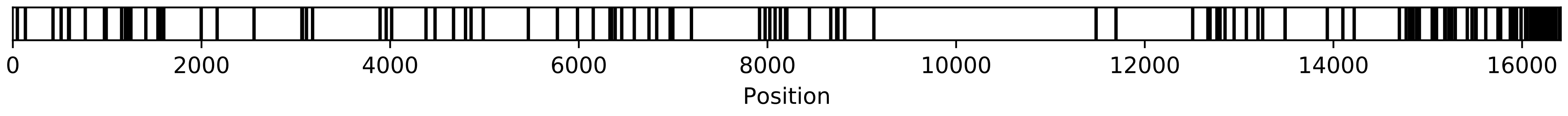


CR2 vs MM2
Number of differences: 129

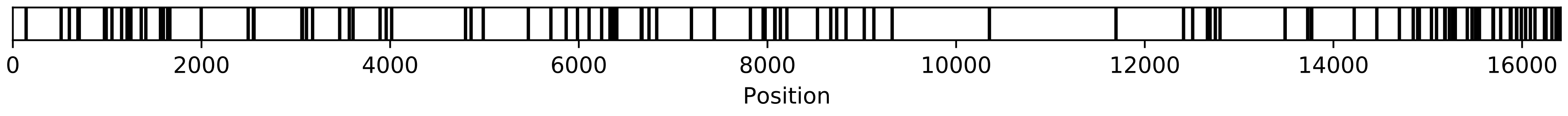


MA1 vs MA2
Number of differences: 266

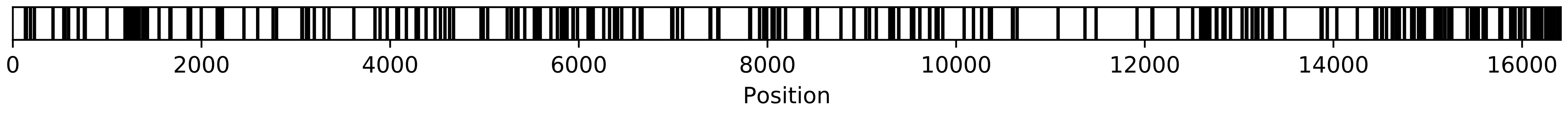


MA1 vs MM1
Number of differences: 0

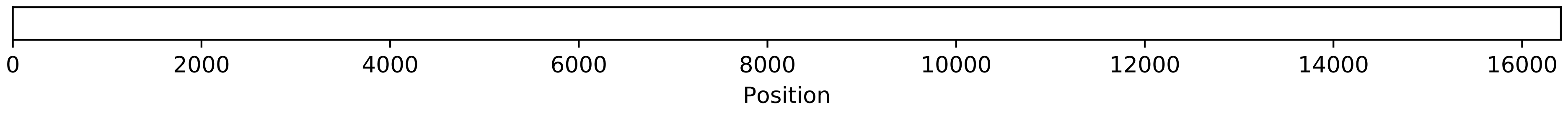


MA1 vs MM2
Number of differences: 129

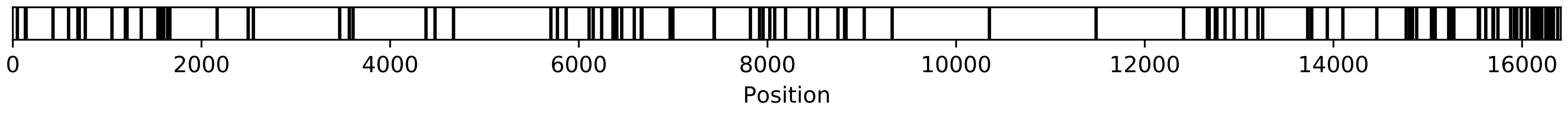


MA2 vs MM1
Number of differences: 266

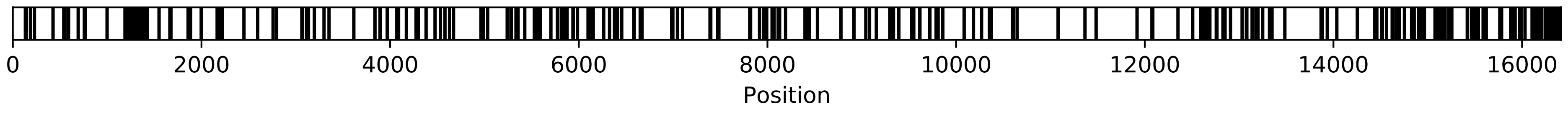


MA2 vs MM2
Number of differences: 294

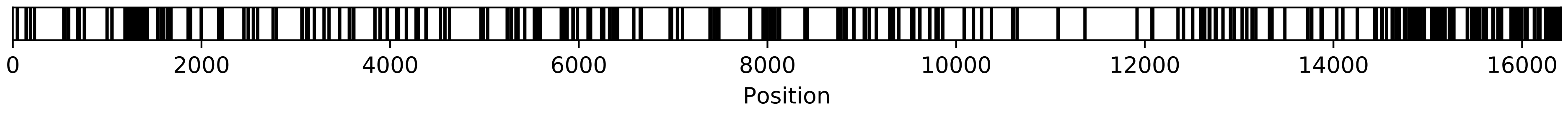


MM1 vs MM2
Number of differences: 129

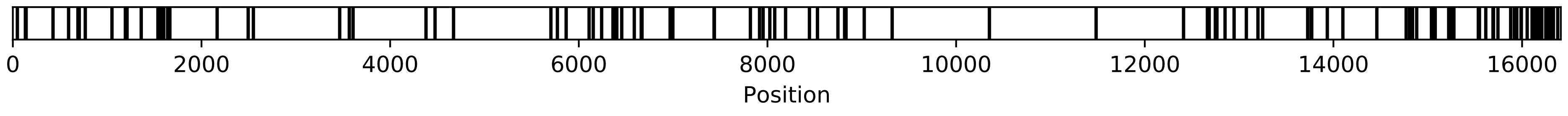

### Region: 507

CR1 vs CR2
Number of differences: 1

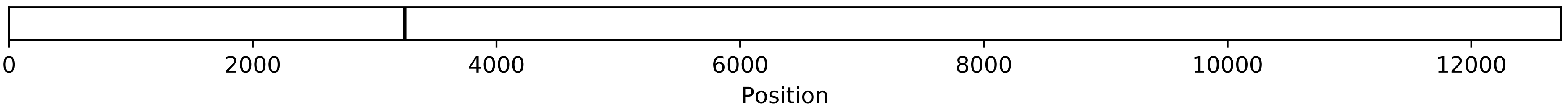


CR1 vs MA1
Number of differences: 237

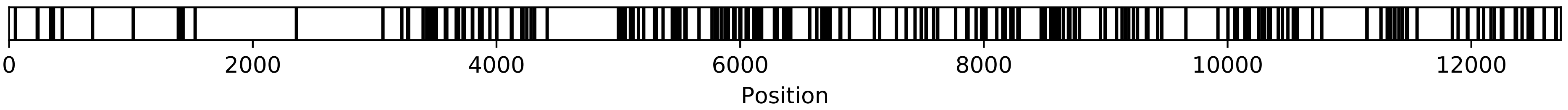


CR1 vs MA2
Number of differences: 237

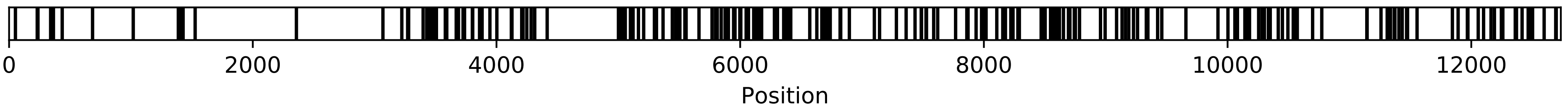


CR1 vs MM1
Number of differences: 252

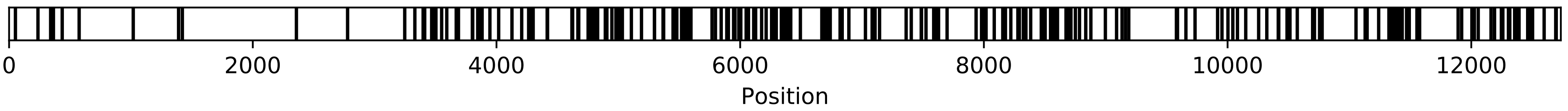


CR1 vs MM2
Number of differences: 218

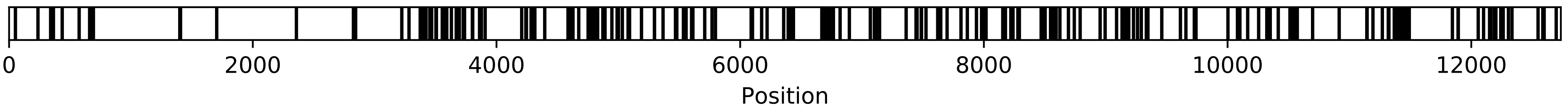


CR2 vs MA1
Number of differences: 238

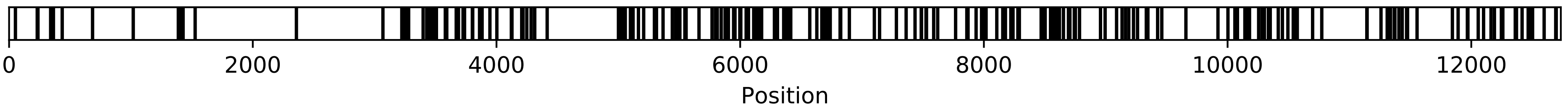


CR2 vs MA2
Number of differences: 238

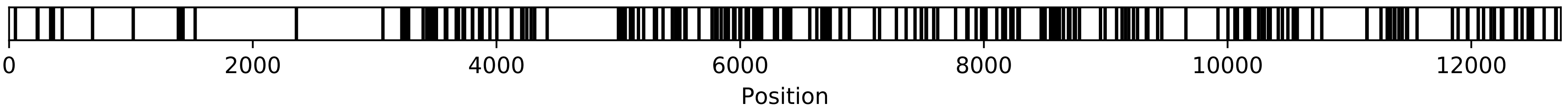


CR2 vs MM1
Number of differences: 251

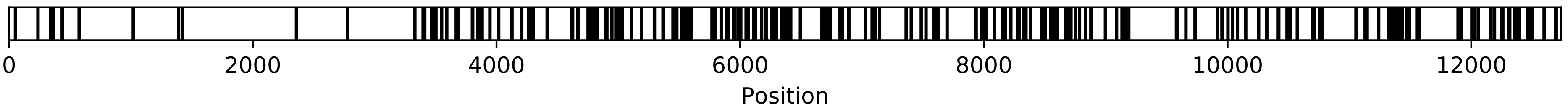


CR2 vs MM2
Number of differences: 219

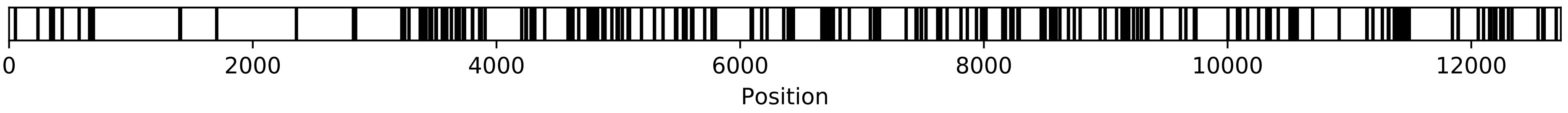


MA1 vs MA2
Number of differences: 0

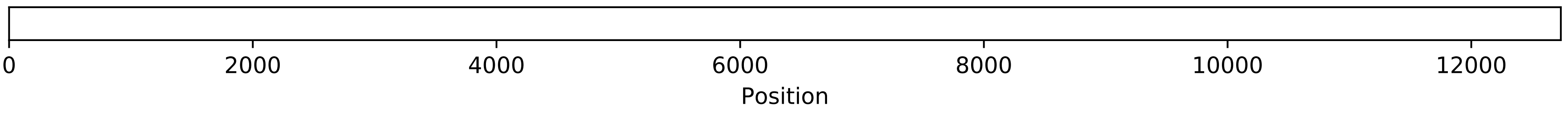


MA1 vs MM1
Number of differences: 227

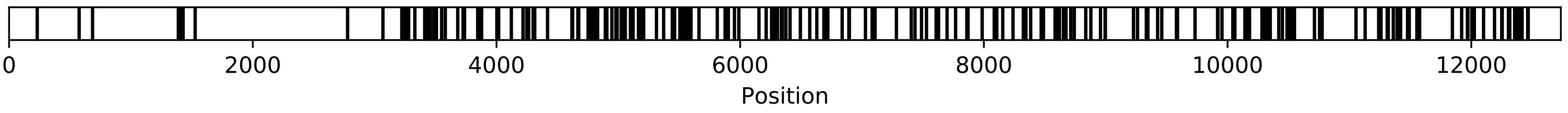


MA1 vs MM2
Number of differences: 216

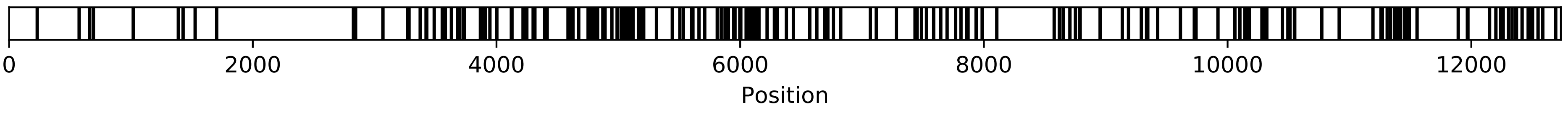


MA2 vs MM1
Number of differences: 227

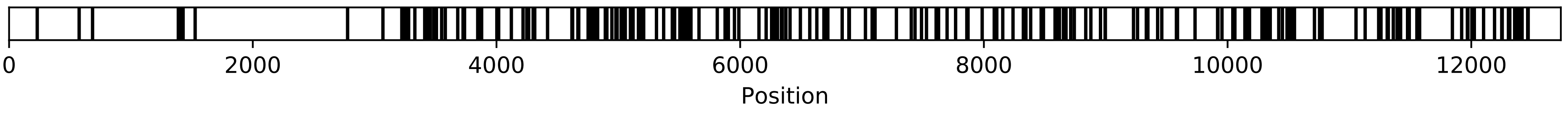


MA2 vs MM2
Number of differences: 216

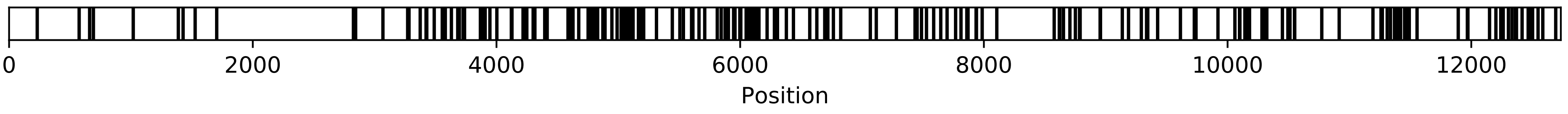


MM1 vs MM2
Number of differences: 235

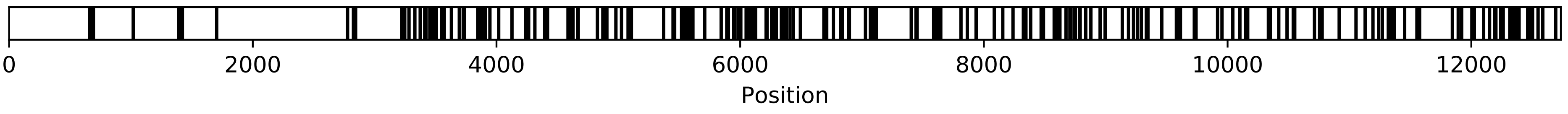

### Region: 510



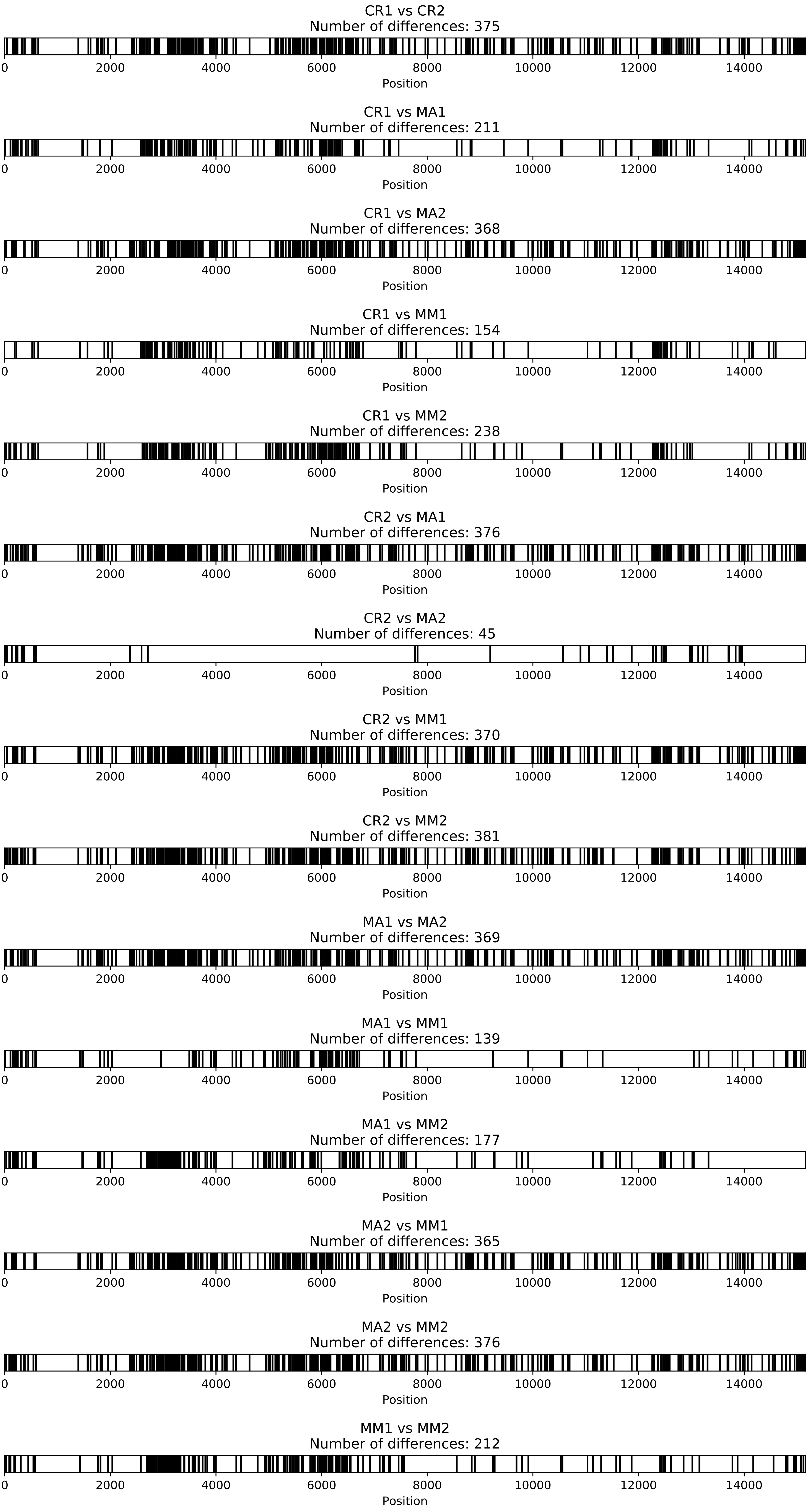
CR1 vs CR2
Number of differences: 375
CR1 vs MA1
Number of differences: 211
CR1 vs MA2
Number of differences: 368
CR1 vs MM1
Number of differences: 154
CR1 vs MM2
Number of differences: 238
CR2 vs MA1
Number of differences: 376
CR2 vs MA2
Number of differences: 45
CR2 vs MM1
Number of differences: 370
CR2 vs MM2
Number of differences: 381
MA1 vs MA2
Number of differences: 369
MA1 vs MM1
Number of differences: 139
MA1 vs MM2
Number of differences: 177
MA2 vs MM1
Number of differences: 365
MA2 vs MM2
Number of differences: 376
MM1 vs MM2
Number of differences: 212
0
2000
4000
6000
8000
10000
12000
14000
Position

### Region: 511

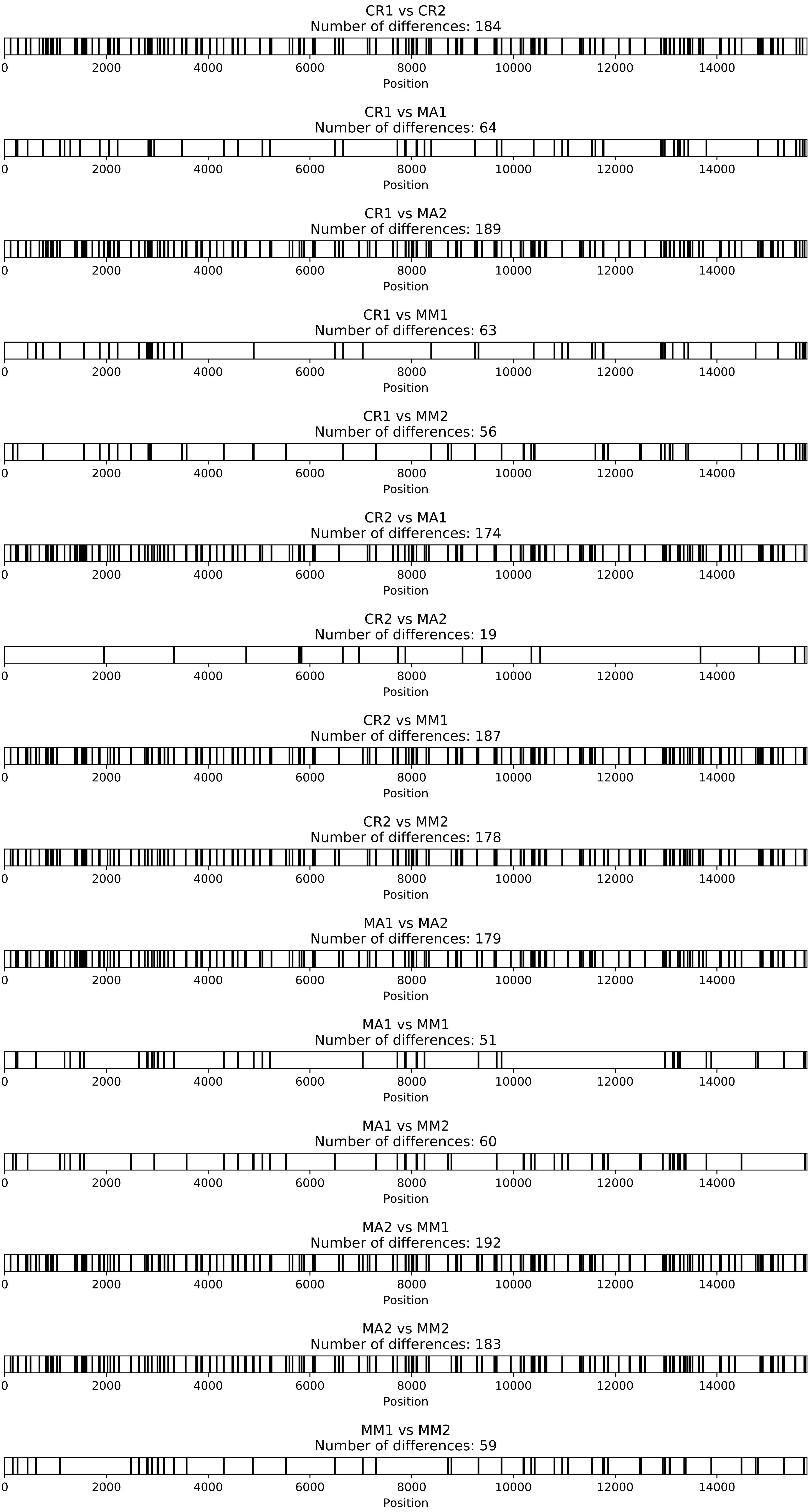
CR1 vs CR2
Number of differences: 184
CR1 vs MA1
Number of differences: 64
CR1 vs MA2
Number of differences: 189
CR1 vs MM1
Number of differences: 63
CR1 vs MM2
Number of differences: 56
CR2 vs MA1
Number of differences: 174
CR2 vs MA2
Number of differences: 19
CR2 vs MM1
Number of differences: 187
CR2 vs MM2
Number of differences: 178
MA1 vs MA2
Number of differences: 179
MA1 vs MM1
Number of differences: 51
MA1 vs MM2
Number of differences: 60
MA2 vs MM1
Number of differences: 192
MA2 vs MM2
Number of differences: 183
MM1 vs MM2
Number of differences: 59
0
2000
4000
6000
8000
10000
12000
14000
Position

### Region: 512



CR1 vs CR2
Number of differences: 356

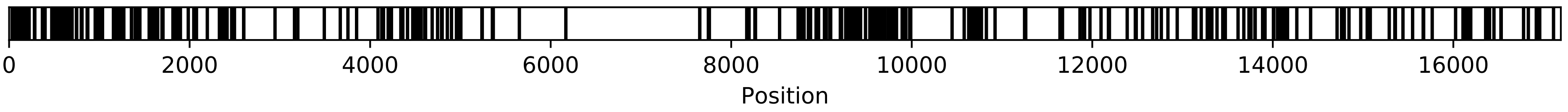


CR1 vs MA1
Number of differences: 280

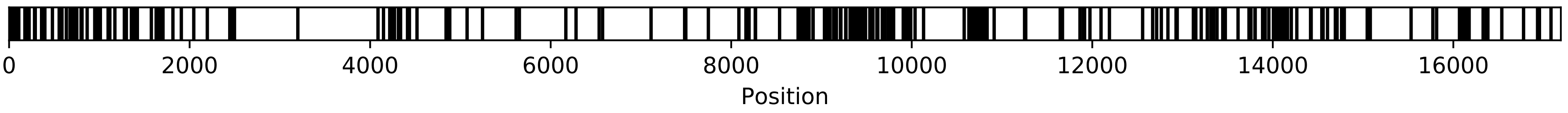


CR1 vs MA2
Number of differences: 295

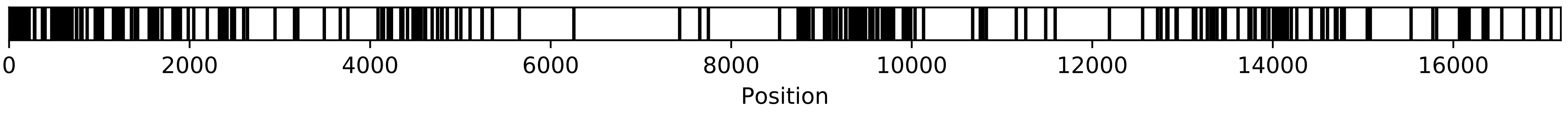


CR1 vs MM1
Number of differences: 201

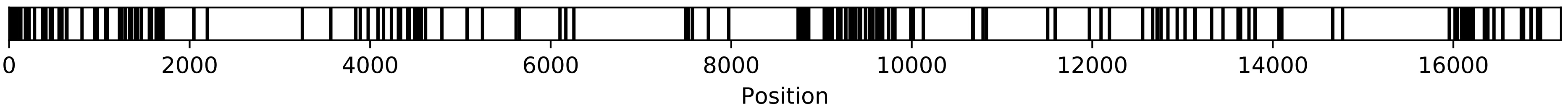


CR1 vs MM2
Number of differences: 210

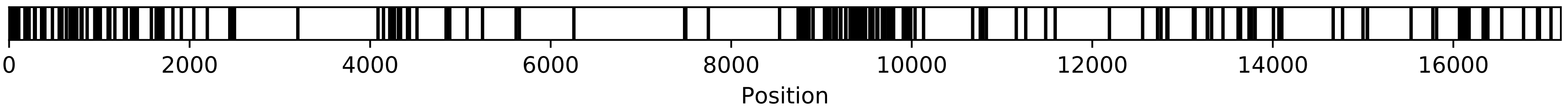


CR2 vs MA1
Number of differences: 329

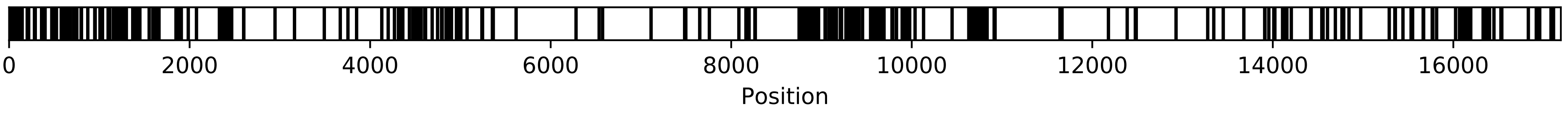


CR2 vs MA2
Number of differences: 219

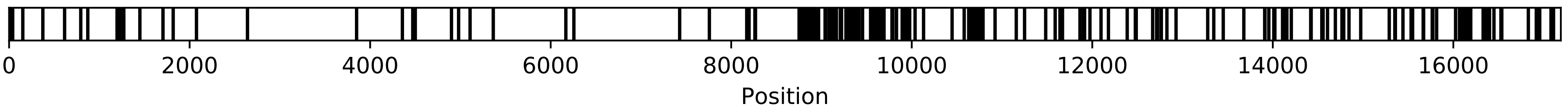


CR2 vs MM1
Number of differences: 375

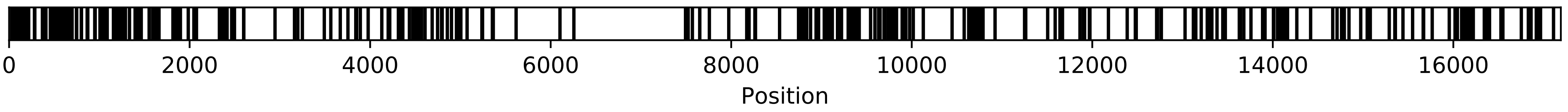


CR2 vs MM2
Number of differences: 362

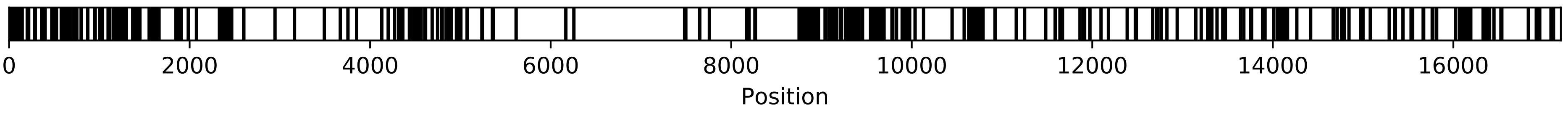


MA1 vs MA2
Number of differences: 203

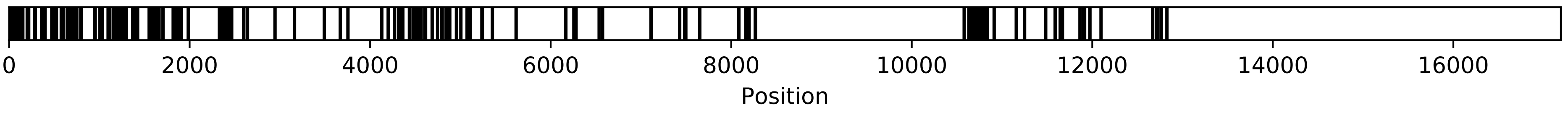


MA1 vs MM1
Number of differences: 272

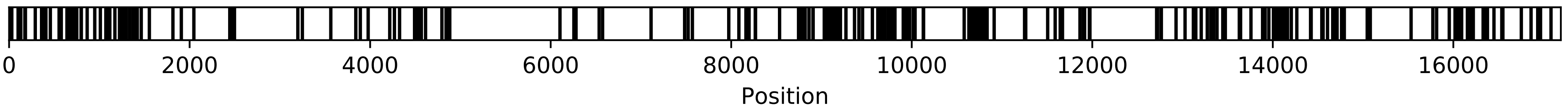


MA1 vs MM2
Number of differences: 109

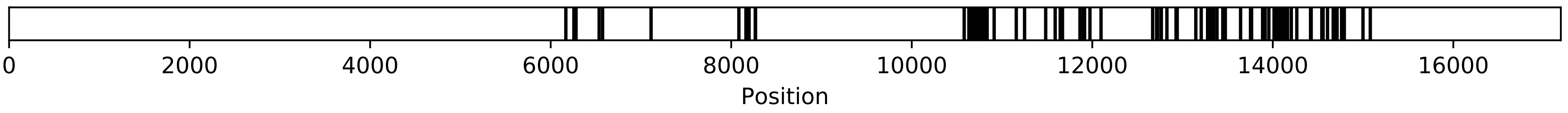


MA2 vs MM1
Number of differences: 306

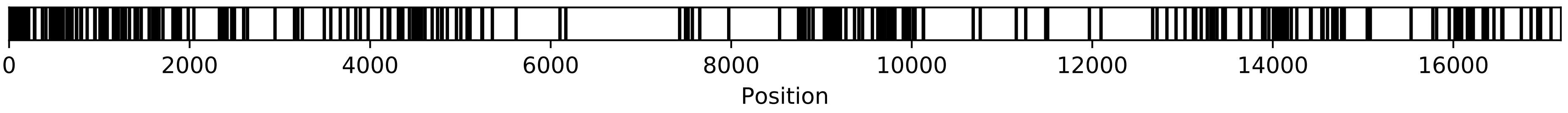


MA2 vs MM2
Number of differences: 202

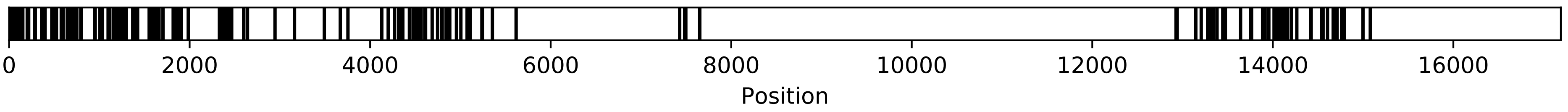


MM1 vs MM2
Number of differences: 195

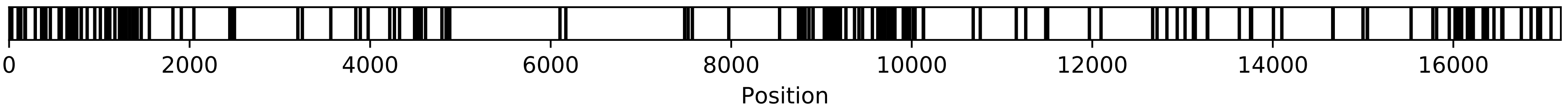

### Region: 515



CR1 vs CR2
Number of differences: 258

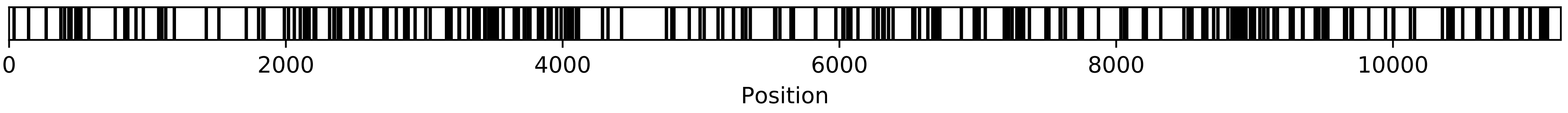


CR1 vs MA1
Number of differences: 76

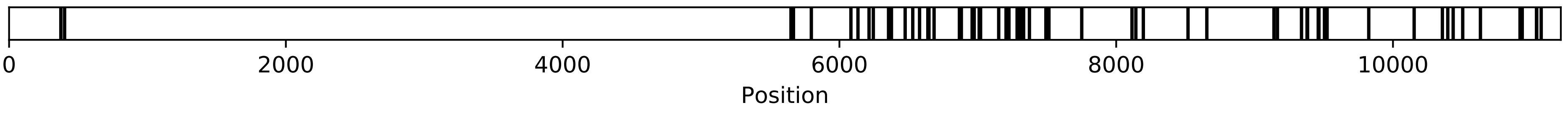


CR1 vs MA2
Number of differences: 255

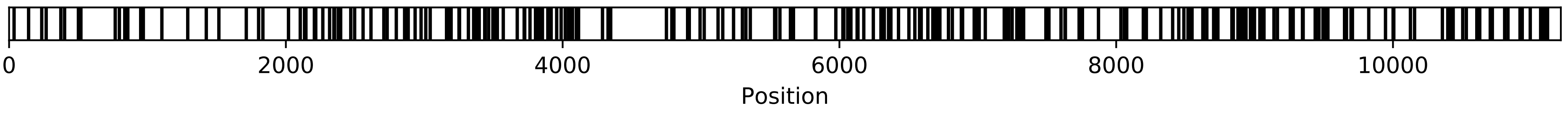


CR1 vs MM1
Number of differences: 153

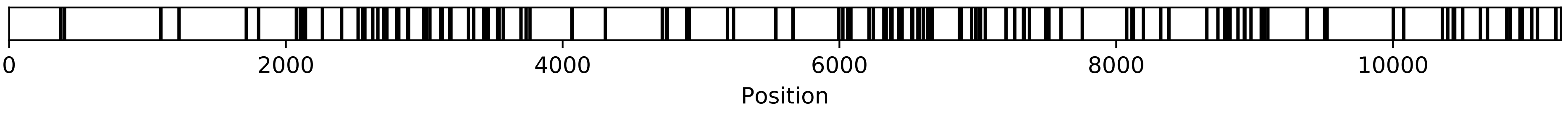


CR1 vs MM2
Number of differences: 149

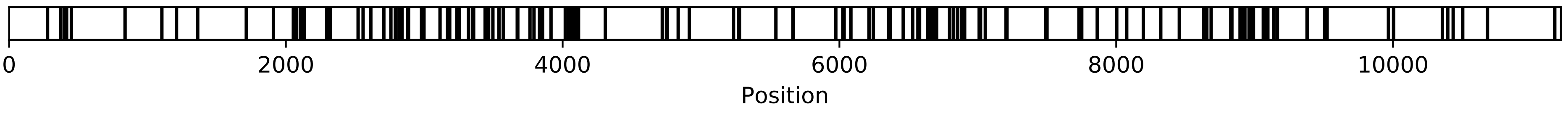


CR2 vs MA1
Number of differences: 261

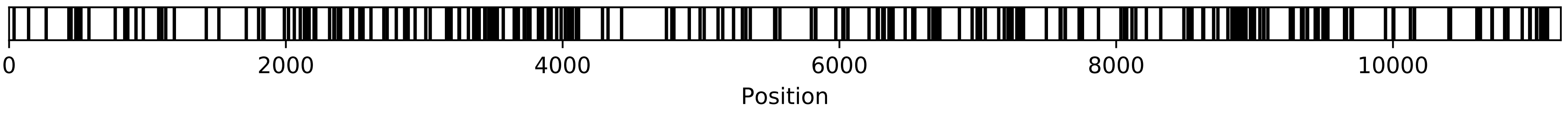


CR2 vs MA2
Number of differences: 74

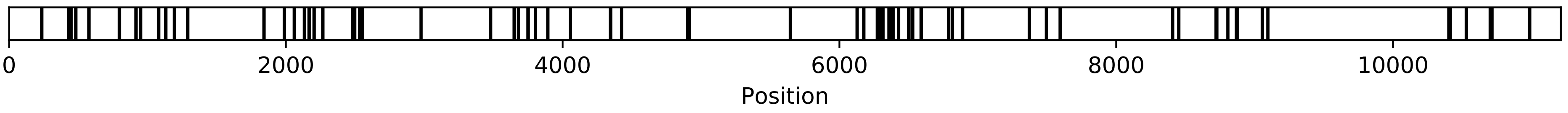


CR2 vs MM1
Number of differences: 290

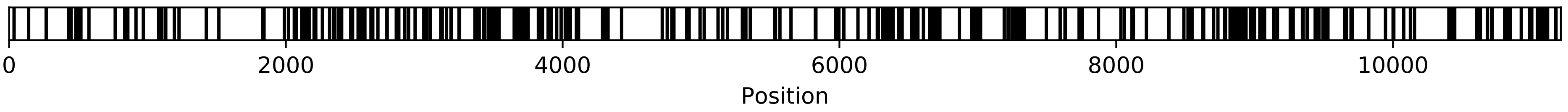


CR2 vs MM2
Number of differences: 269

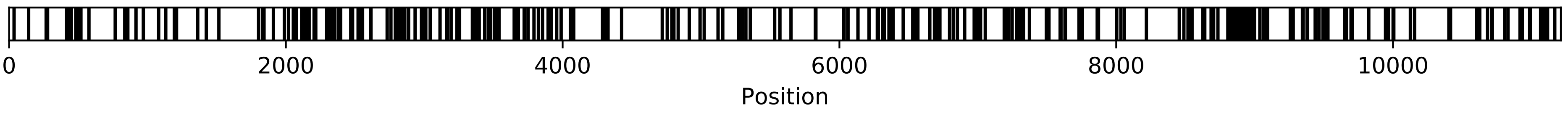


MA1 vs MA2
Number of differences: 259

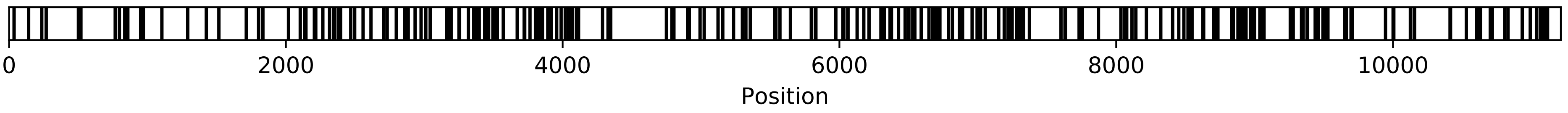


MA1 vs MM1
Number of differences: 161

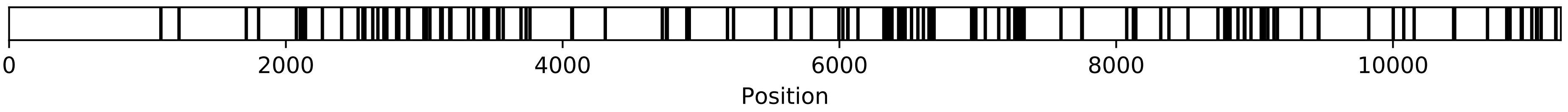


MA1 vs MM2
Number of differences: 163

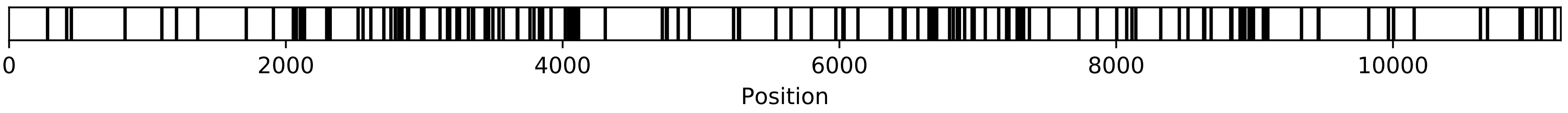


MA2 vs MM1
Number of differences: 295

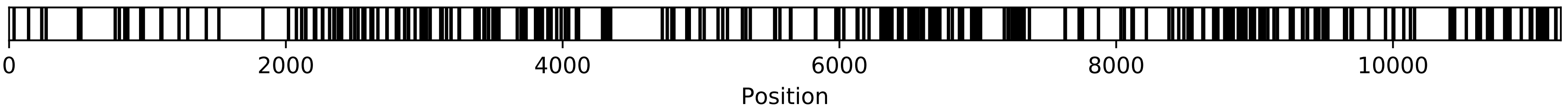


MA2 vs MM2
Number of differences: 264

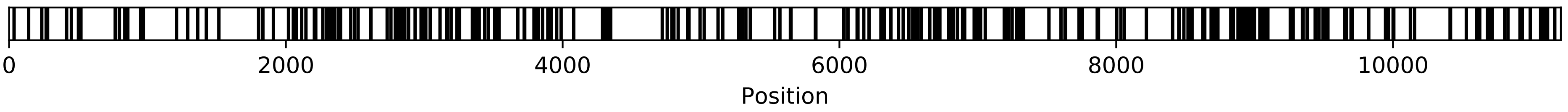


MM1 vs MM2
Number of differences: 172

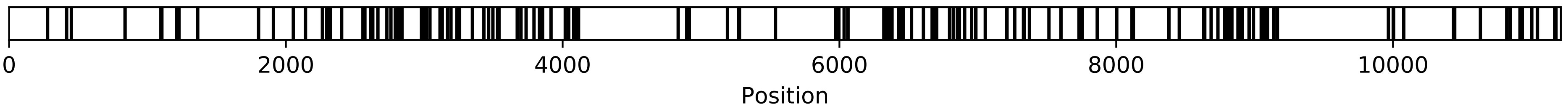

### Region: 516



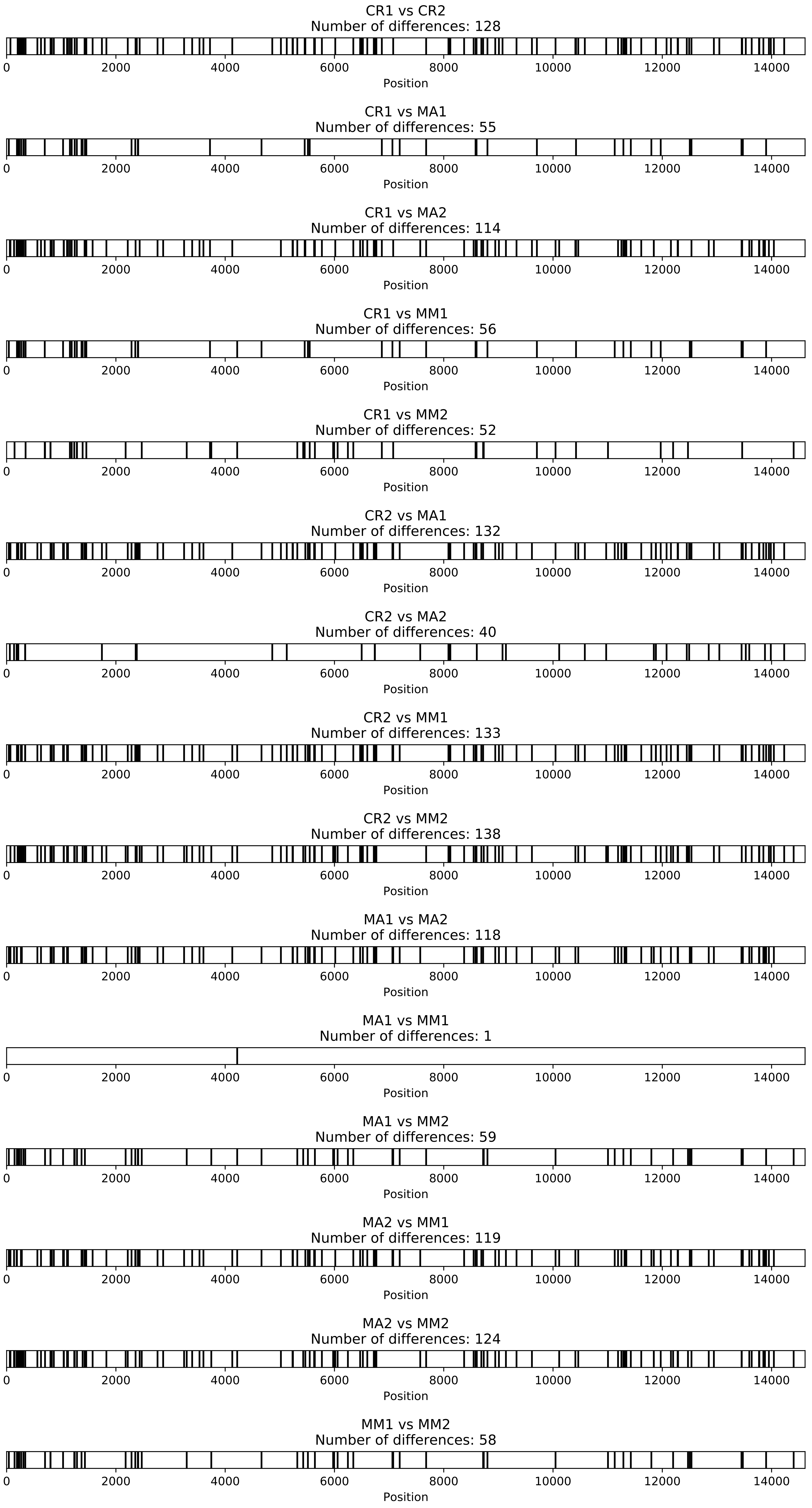
CR1 vs CR2
Number of differences: 128
0 2000 4000 6000 8000 10000 12000 14000
Position
CR1 vs MA1
Number of differences: 55
0 2000 4000 6000 8000 10000 12000 14000
Position
CR1 vs MA2
Number of differences: 114
0 2000 4000 6000 8000 10000 12000 14000
Position
CR1 vs MM1
Number of differences: 56
0 2000 4000 6000 8000 10000 12000 14000
Position
CR1 vs MM2
Number of differences: 52
0 2000 4000 6000 8000 10000 12000 14000
Position
CR2 vs MA1
Number of differences: 132
0 2000 4000 6000 8000 10000 12000 14000
Position
CR2 vs MA2
Number of differences: 40
0 2000 4000 6000 8000 10000 12000 14000
Position
CR2 vs MM1
Number of differences: 133
0 2000 4000 6000 8000 10000 12000 14000
Position
CR2 vs MM2
Number of differences: 138
0 2000 4000 6000 8000 10000 12000 14000
Position
MA1 vs MA2
Number of differences: 118
0 2000 4000 6000 8000 10000 12000 14000
Position
MA1 vs MM1
Number of differences: 1
0 2000 4000 6000 8000 10000 12000 14000
Position
MA1 vs MM2
Number of differences: 59
0 2000 4000 6000 8000 10000 12000 14000
Position
MA2 vs MM1
Number of differences: 119
0 2000 4000 6000 8000 10000 12000 14000
Position
MA2 vs MM2
Number of differences: 124
0 2000 4000 6000 8000 10000 12000 14000
Position
MM1 vs MM2
Number of differences: 58
0 2000 4000 6000 8000 10000 12000 14000
Position

### Region: 517



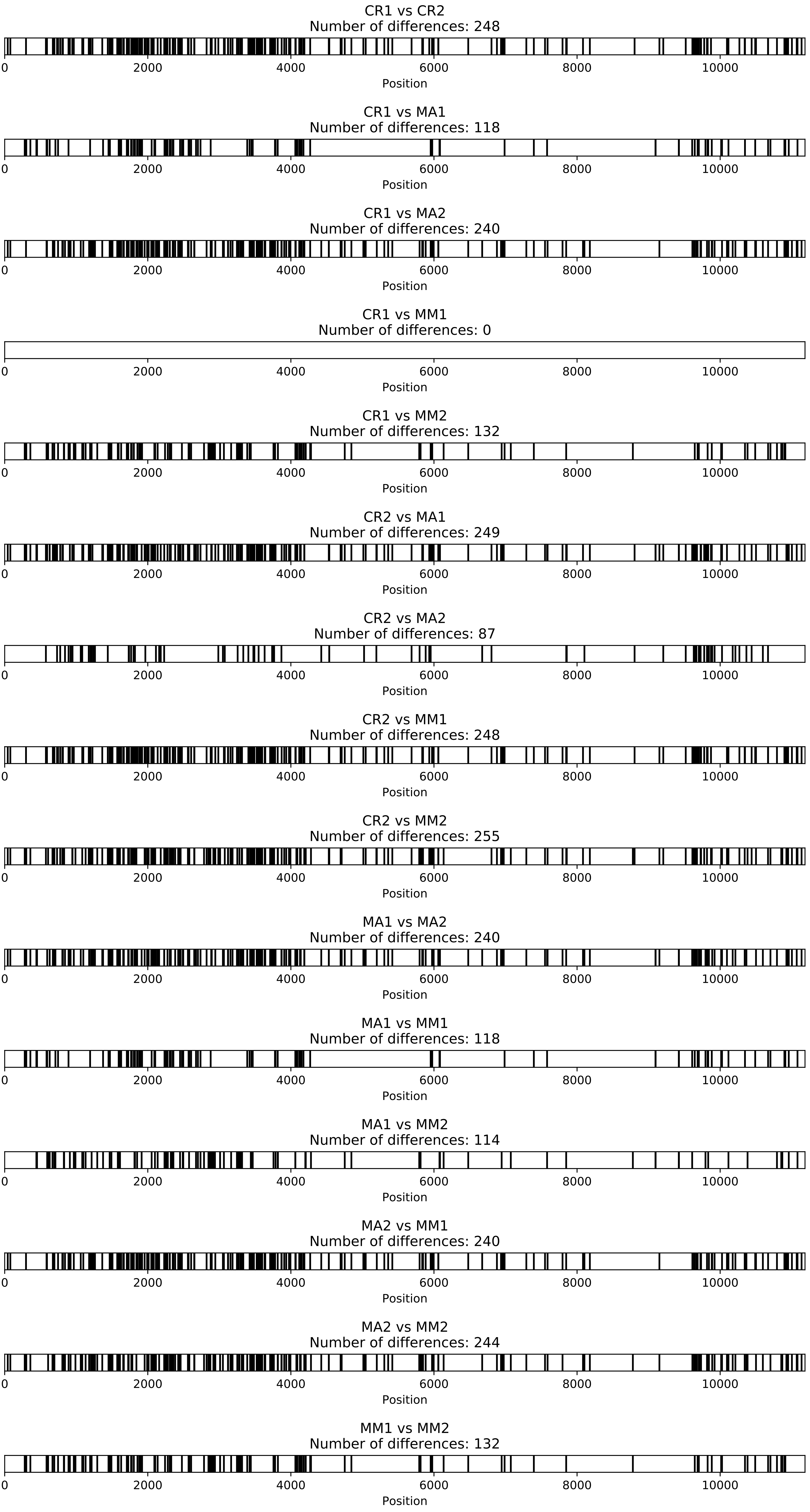
CR1 vs CR2
Number of differences: 248
CR1 vs MA1
Number of differences: 118
CR1 vs MA2
Number of differences: 240
CR1 vs MM1
Number of differences: 0
CR1 vs MM2
Number of differences: 132
CR2 vs MA1
Number of differences: 249
CR2 vs MA2
Number of differences: 87
CR2 vs MM1
Number of differences: 248
CR2 vs MM2
Number of differences: 255
MA1 vs MA2
Number of differences: 240
MA1 vs MM1
Number of differences: 118
MA1 vs MM2
Number of differences: 114
MA2 vs MM1
Number of differences: 240
MA2 vs MM2
Number of differences: 244
MM1 vs MM2
Number of differences: 132
0
2000
4000
6000
8000
10000
Position

### Region: 519

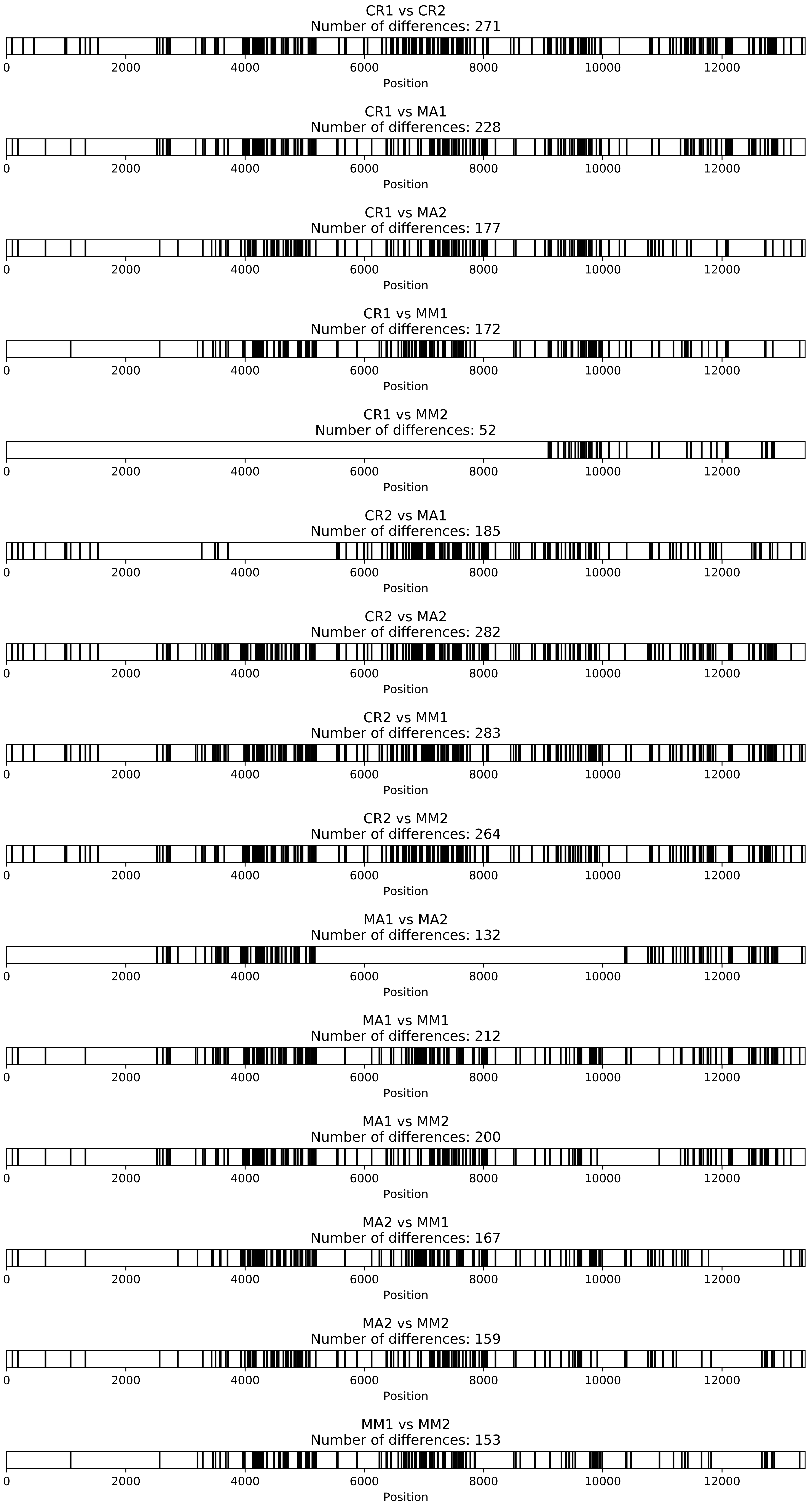
CR1 vs CR2
Number of differences: 271
CR1 vs MA1
Number of differences: 228
CR1 vs MA2
Number of differences: 177
CR1 vs MM1
Number of differences: 172
CR1 vs MM2
Number of differences: 52
CR2 vs MA1
Number of differences: 185
CR2 vs MA2
Number of differences: 282
CR2 vs MM1
Number of differences: 283
CR2 vs MM2
Number of differences: 264
MA1 vs MA2
Number of differences: 132
MA1 vs MM1
Number of differences: 212
MA1 vs MM2
Number of differences: 200
MA2 vs MM1
Number of differences: 167
MA2 vs MM2
Number of differences: 159
MM1 vs MM2
Number of differences: 153

### Region: 520



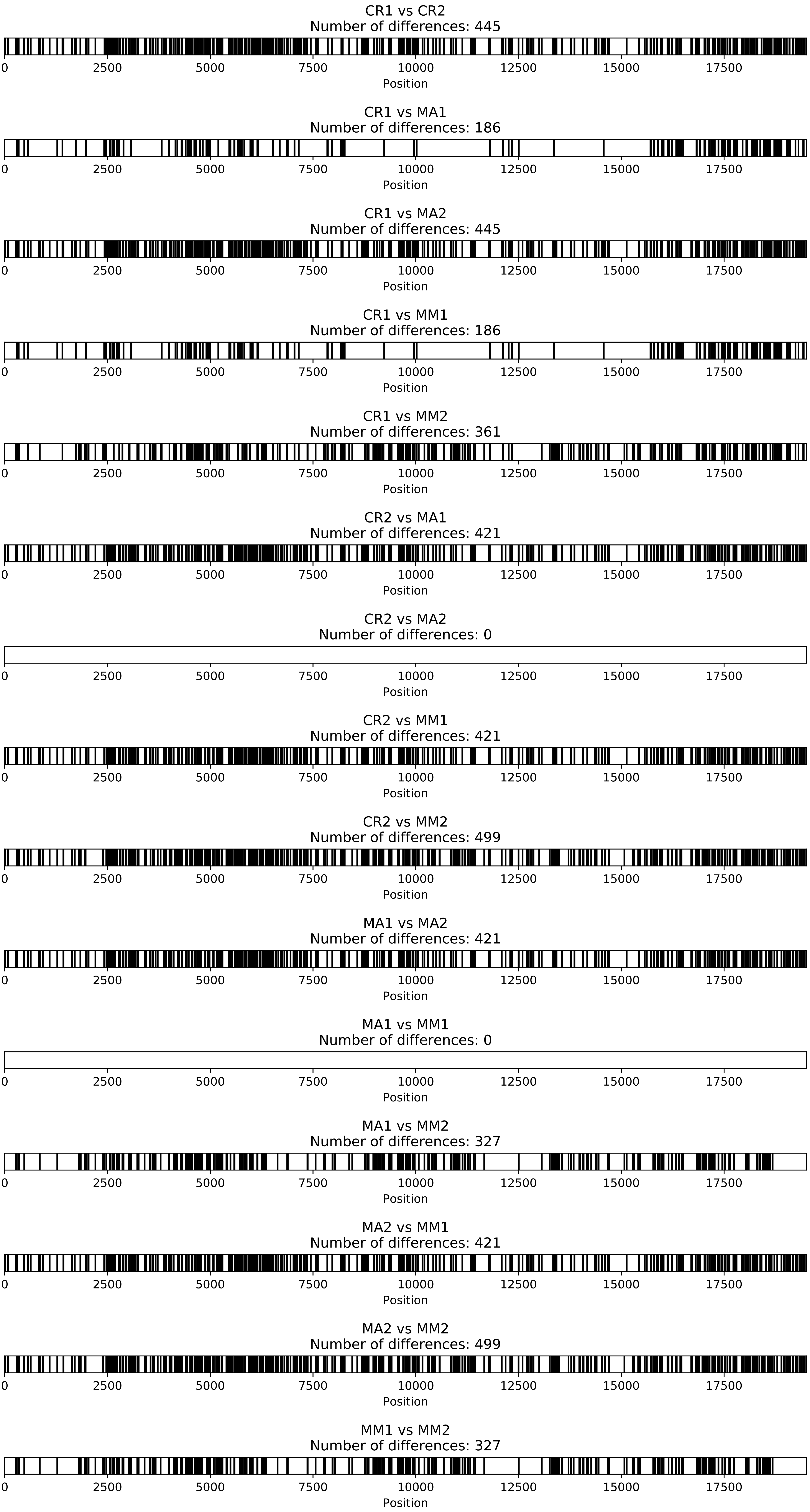
CR1 vs CR2
Number of differences: 445
CR1 vs MA1
Number of differences: 186
CR1 vs MA2
Number of differences: 445
CR1 vs MM1
Number of differences: 186
CR1 vs MM2
Number of differences: 361
CR2 vs MA1
Number of differences: 421
CR2 vs MA2
Number of differences: 0
CR2 vs MM1
Number of differences: 421
CR2 vs MM2
Number of differences: 499
MA1 vs MA2
Number of differences: 421
MA1 vs MM1
Number of differences: 0
MA1 vs MM2
Number of differences: 327
MA2 vs MM1
Number of differences: 421
MA2 vs MM2
Number of differences: 499
MM1 vs MM2
Number of differences: 327
0
2500
5000
7500
10000
12500
15000
17500
Position

### Region: 521



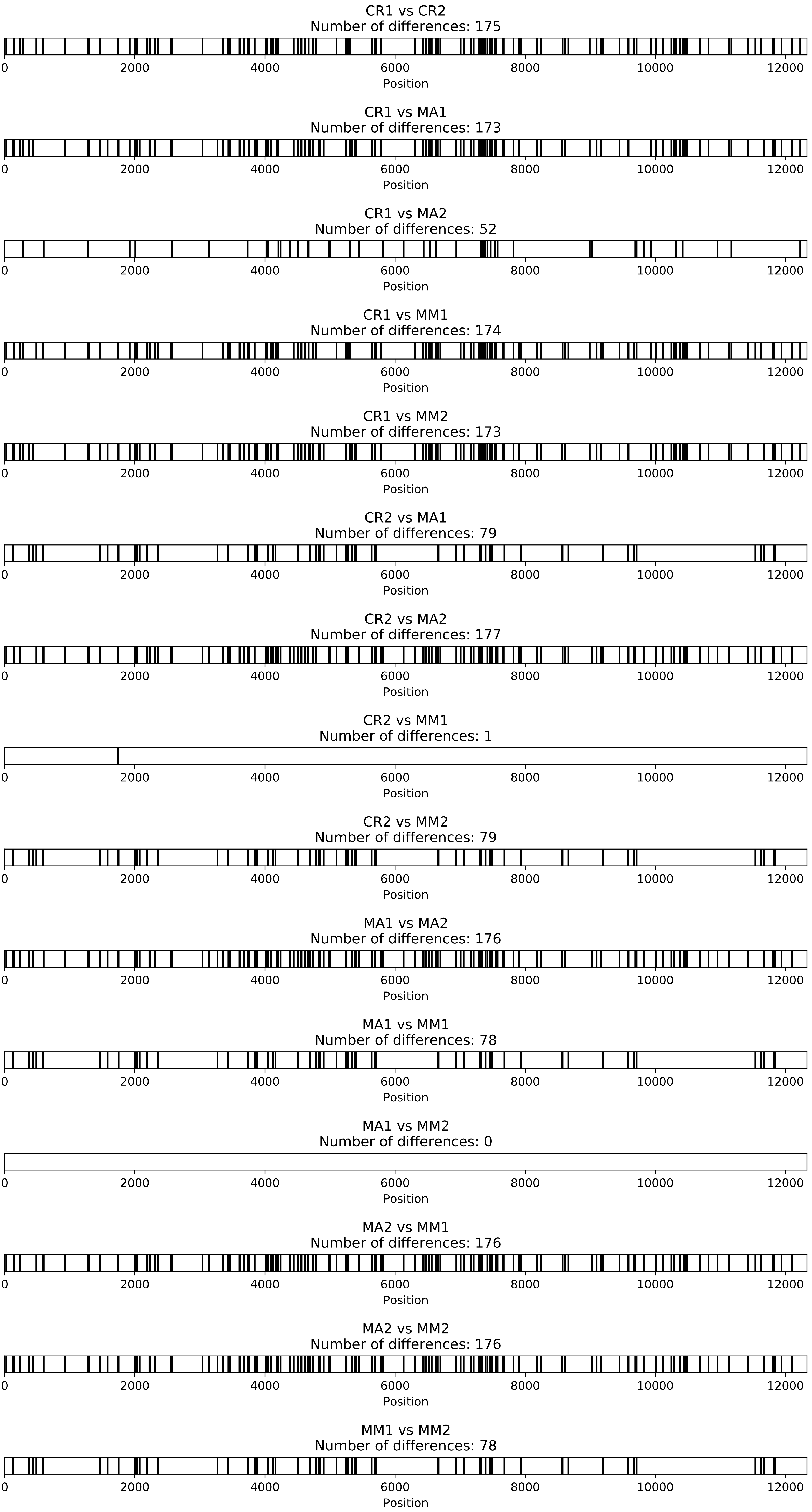
CR1 vs CR2
Number of differences: 175
CR1 vs MA1
Number of differences: 173
CR1 vs MA2
Number of differences: 52
CR1 vs MM1
Number of differences: 174
CR1 vs MM2
Number of differences: 173
CR2 vs MA1
Number of differences: 79
CR2 vs MA2
Number of differences: 177
CR2 vs MM1
Number of differences: 1
CR2 vs MM2
Number of differences: 79
MA1 vs MA2
Number of differences: 176
MA1 vs MM1
Number of differences: 78
MA1 vs MM2
Number of differences: 0
MA2 vs MM1
Number of differences: 176
MA2 vs MM2
Number of differences: 176
MM1 vs MM2
Number of differences: 78
0
2000
4000
6000
8000
10000
12000
Position

### Region: 522



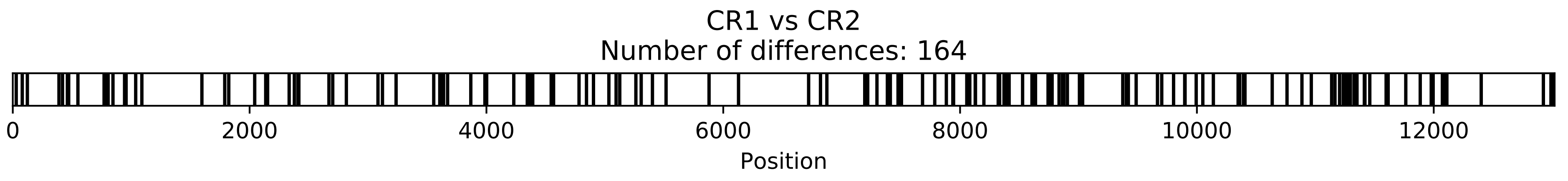
CR1 vs CR2
Number of differences: 164
0 2000 4000 6000 8000 10000 12000
Position

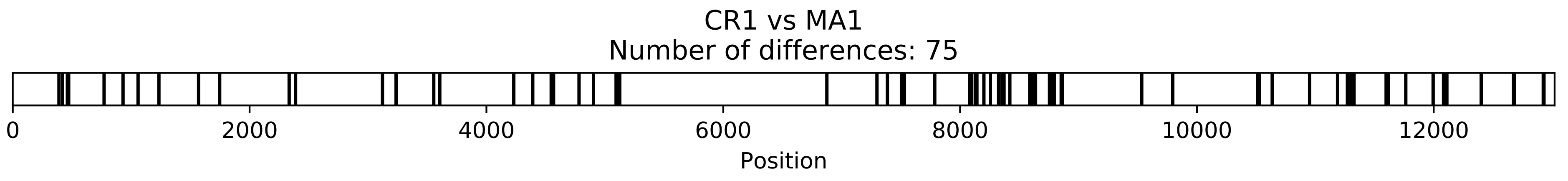
CR1 vs MA1
Number of differences: 75
0 2000 4000 6000 8000 10000 12000
Position

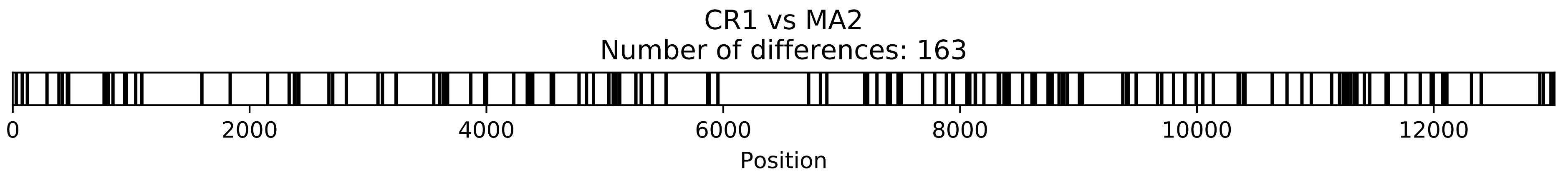
CR1 vs MA2
Number of differences: 163
0 2000 4000 6000 8000 10000 12000
Position

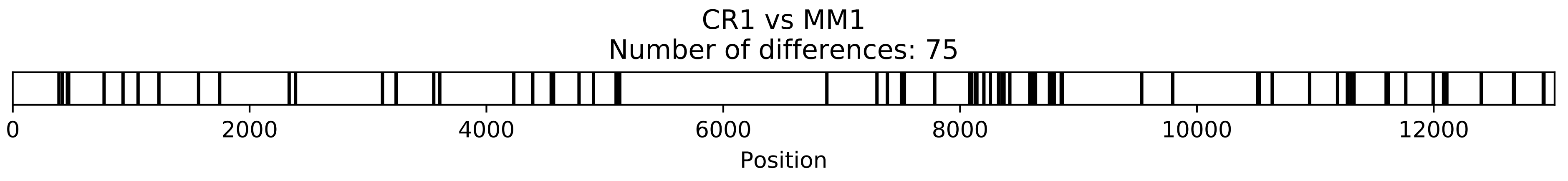
CR1 vs MM1
Number of differences: 75
0 2000 4000 6000 8000 10000 12000
Position

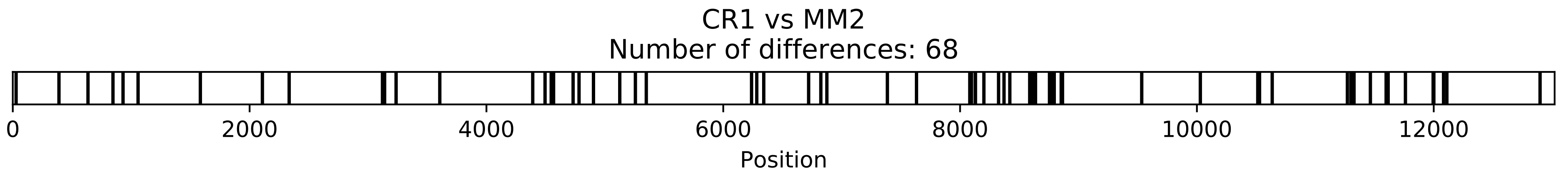
CR1 vs MM2
Number of differences: 68
0 2000 4000 6000 8000 10000 12000
Position

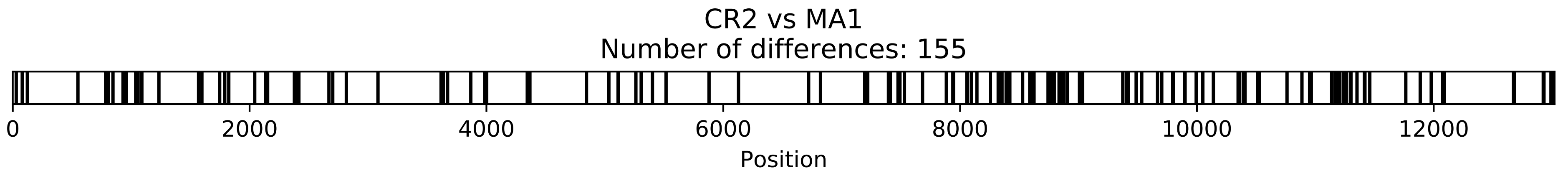
CR2 vs MA1
Number of differences: 155
0 2000 4000 6000 8000 10000 12000
Position

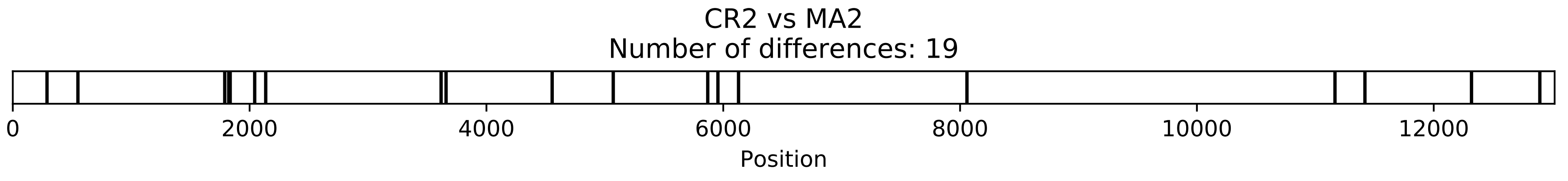
CR2 vs MA2
Number of differences: 19
0 2000 4000 6000 8000 10000 12000
Position

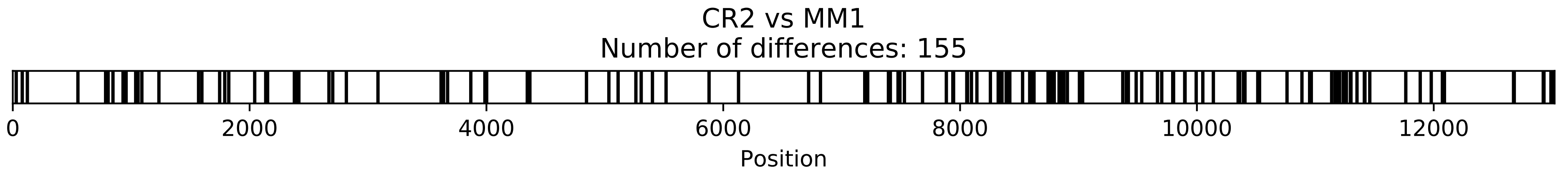
CR2 vs MM1
Number of differences: 155
0 2000 4000 6000 8000 10000 12000
Position

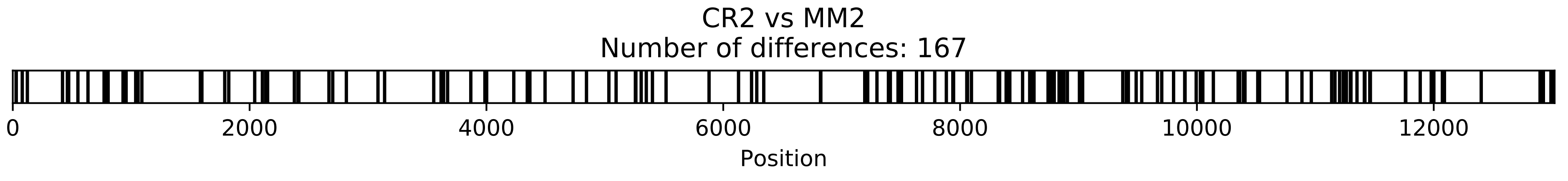
CR2 vs MM2
Number of differences: 167
0 2000 4000 6000 8000 10000 12000
Position

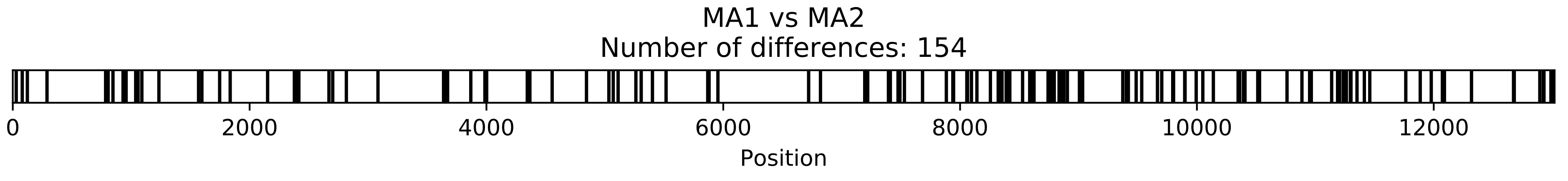
MA1 vs MA2
Number of differences: 154
0 2000 4000 6000 8000 10000 12000
Position

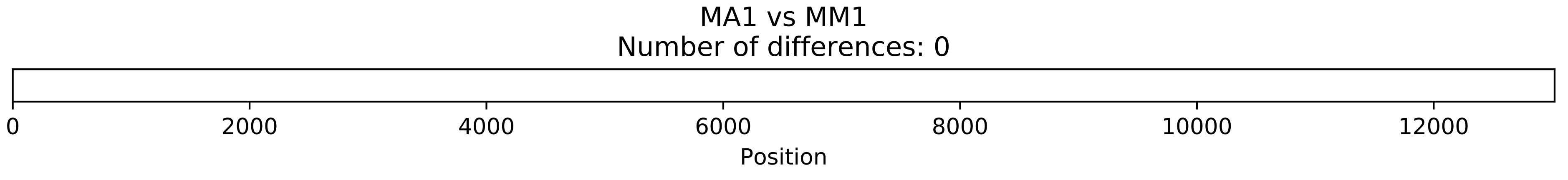
MA1 vs MM1
Number of differences: 0
0 2000 4000 6000 8000 10000 12000
Position

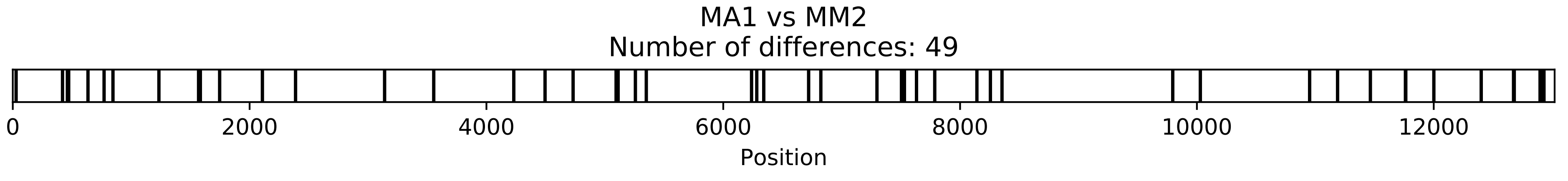
MA1 vs MM2
Number of differences: 49
0 2000 4000 6000 8000 10000 12000
Position

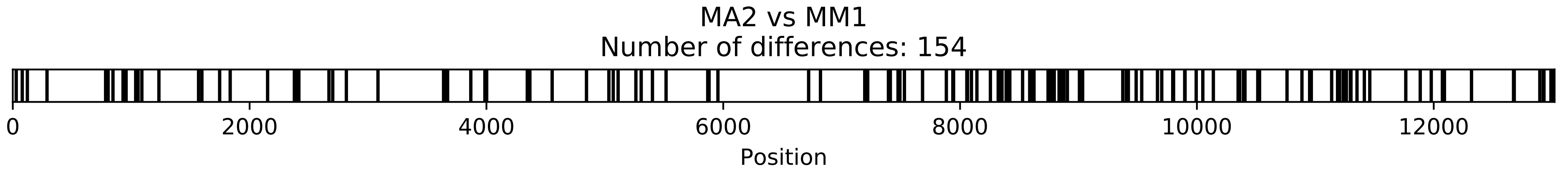
MA2 vs MM1
Number of differences: 154
0 2000 4000 6000 8000 10000 12000
Position

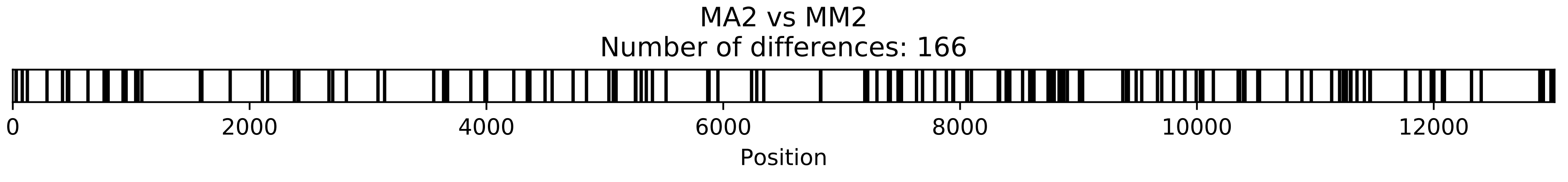
MA2 vs MM2
Number of differences: 166
0 2000 4000 6000 8000 10000 12000
Position

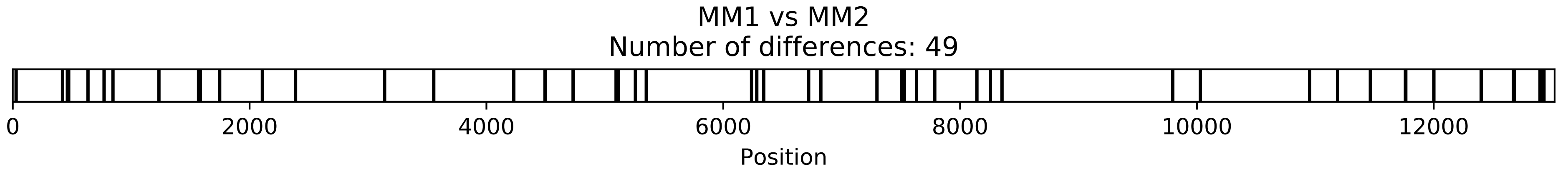
MM1 vs MM2
Number of differences: 49
0 2000 4000 6000 8000 10000 12000
Position

### Region: 524

CR1 vs CR2
Number of differences: 213

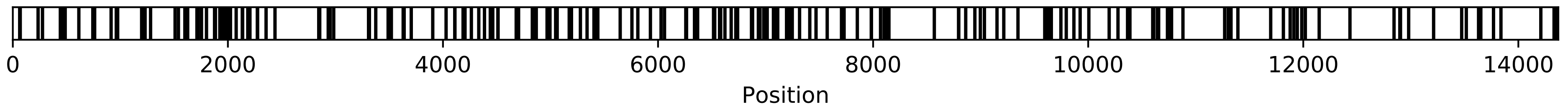


CR1 vs MA1
Number of differences: 124

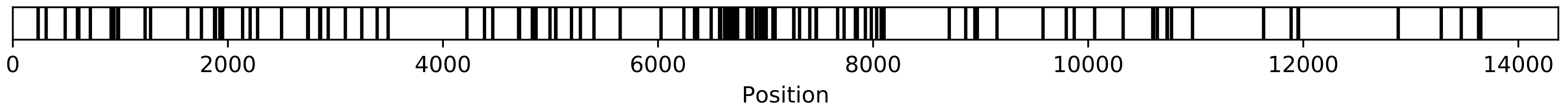


CR1 vs MA2
Number of differences: 205

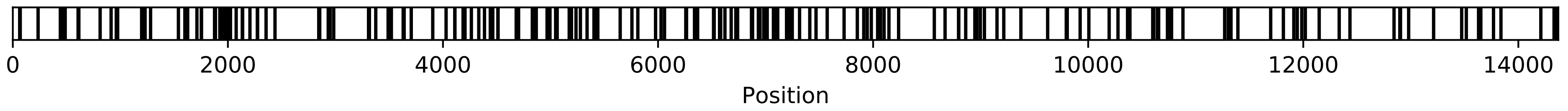


CR1 vs MM1
Number of differences: 59

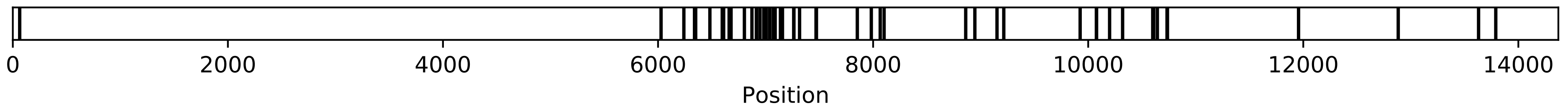


CR1 vs MM2
Number of differences: 124

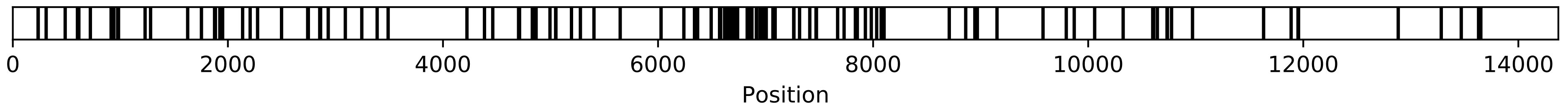


CR2 vs MA1
Number of differences: 212

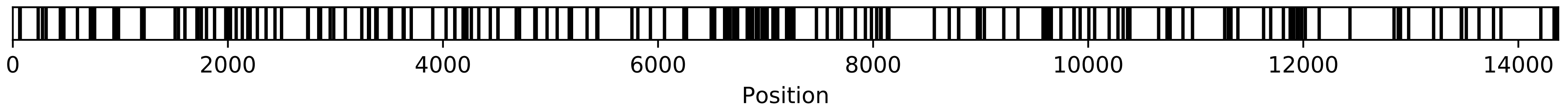


CR2 vs MA2
Number of differences: 34

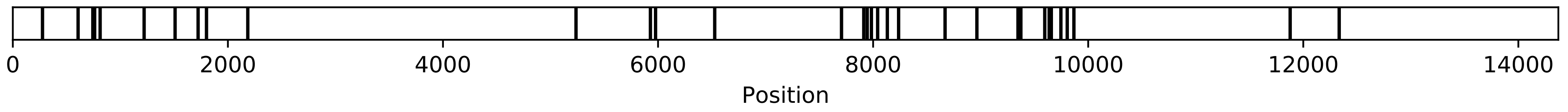


CR2 vs MM1
Number of differences: 225

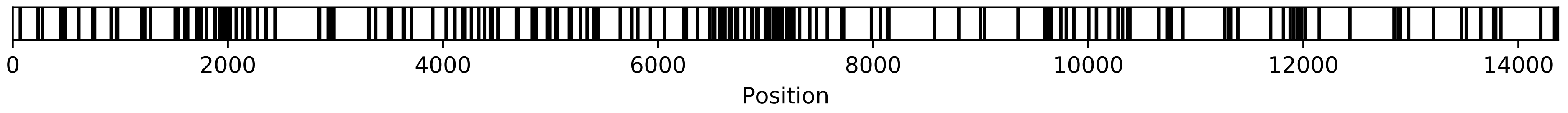


CR2 vs MM2
Number of differences: 212

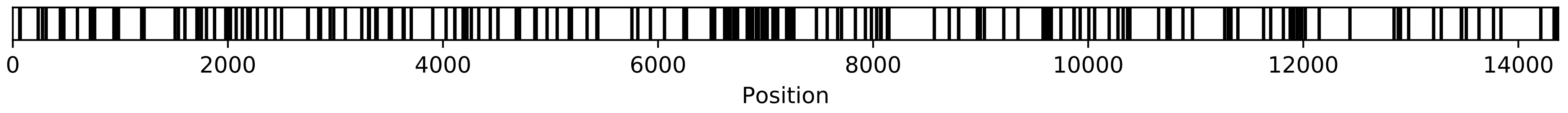


MA1 vs MA2
Number of differences: 204

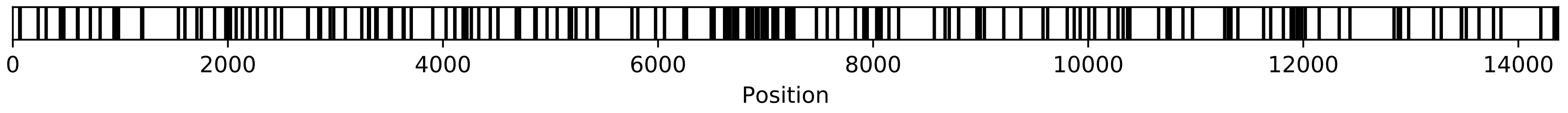


MA1 vs MM1
Number of differences: 118

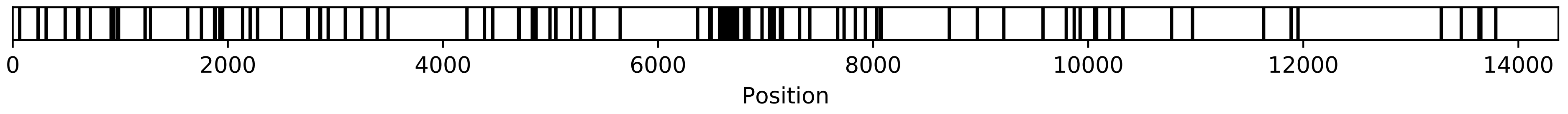


MA1 vs MM2
Number of differences: 0

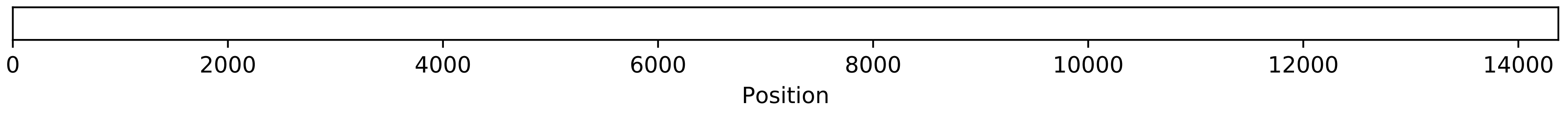


MA2 vs MM1
Number of differences: 217

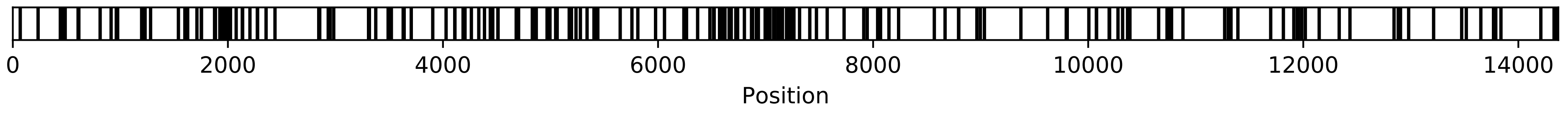


MA2 vs MM2
Number of differences: 204

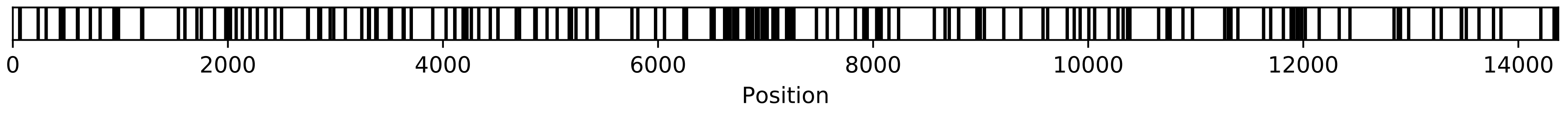


MM1 vs MM2
Number of differences: 118

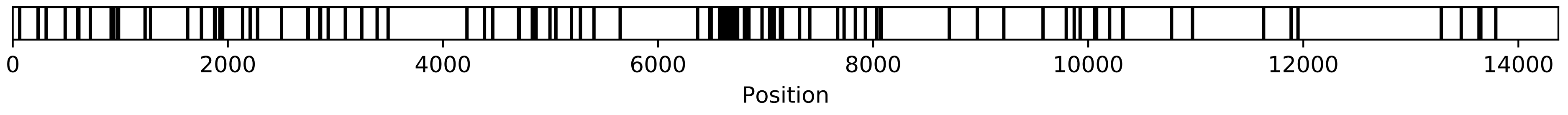

### Region: 525

CR1 vs CR2
Number of differences: 135

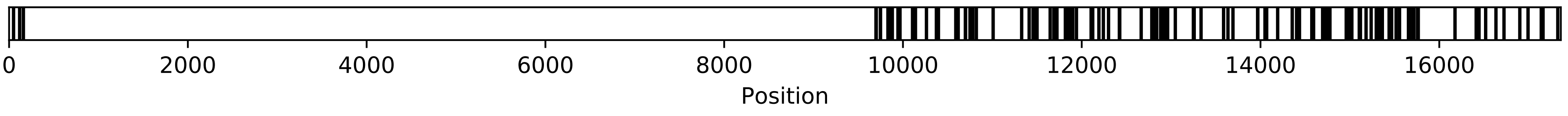


CR1 vs MA1
Number of differences: 421

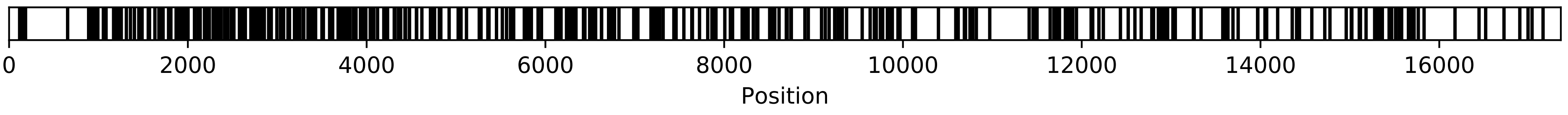


CR1 vs MA2
Number of differences: 193

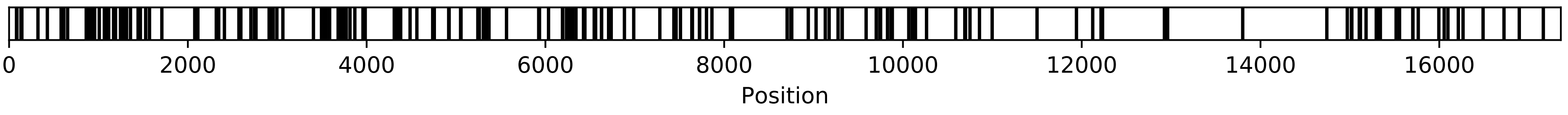


CR1 vs MM1
Number of differences: 202

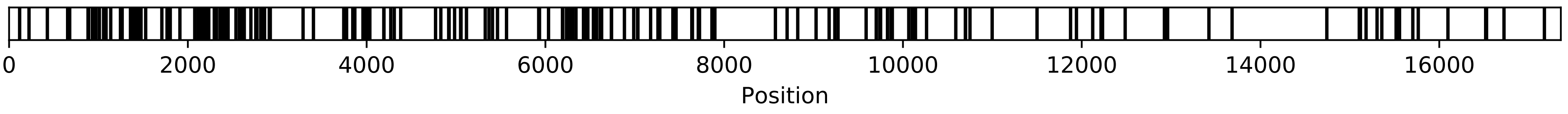


CR1 vs MM2
Number of differences: 158

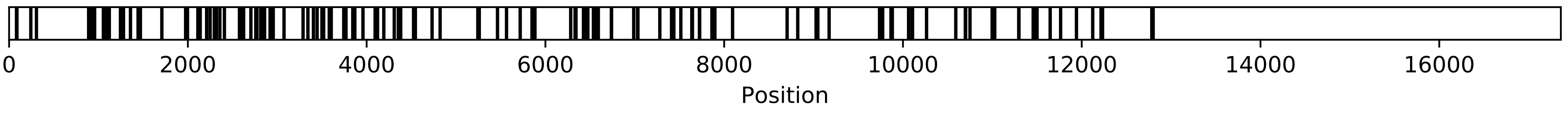


CR2 vs MA1
Number of differences: 337

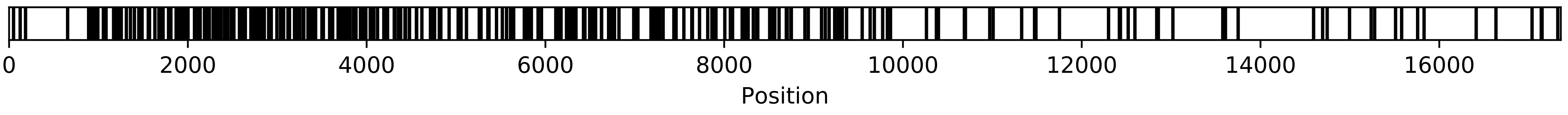


CR2 vs MA2
Number of differences: 276

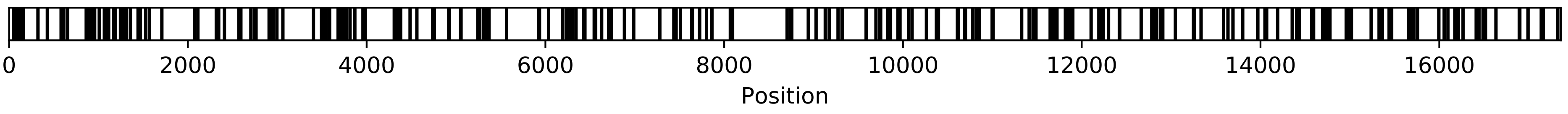


CR2 vs MM1
Number of differences: 285

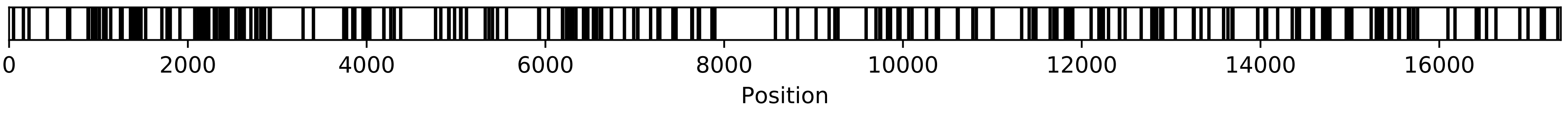


CR2 vs MM2
Number of differences: 264

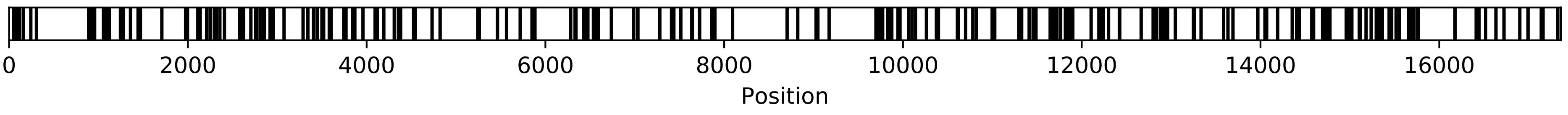


MA1 vs MA2
Number of differences: 403

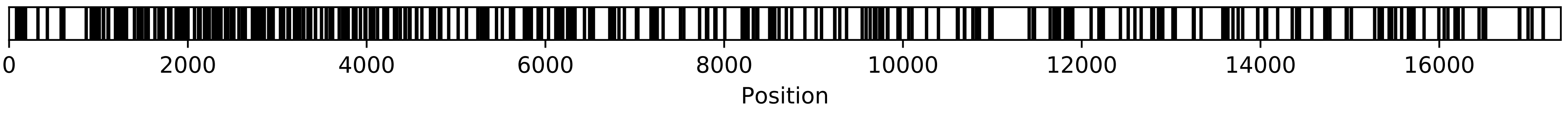


MA1 vs MM1
Number of differences: 410

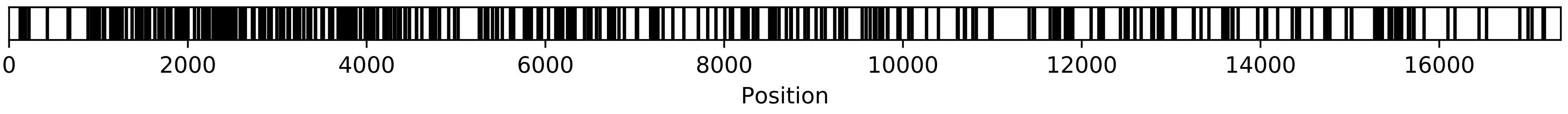


MA1 vs MM2
Number of differences: 413

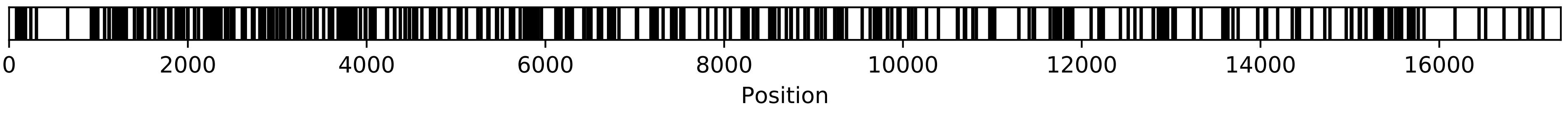


MA2 vs MM1
Number of differences: 189

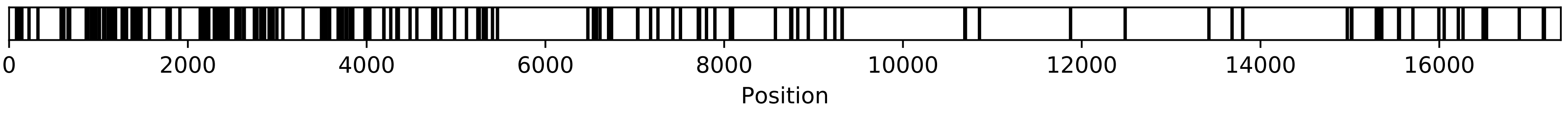


MA2 vs MM2
Number of differences: 208

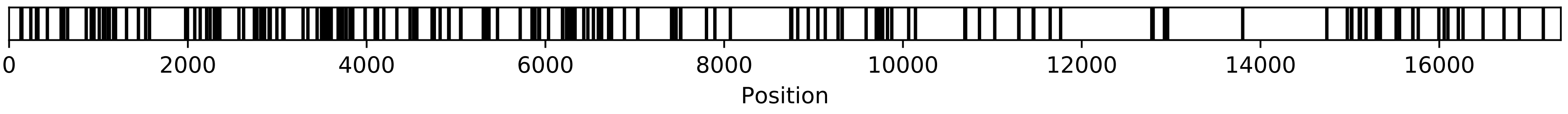


MM1 vs MM2
Number of differences: 176

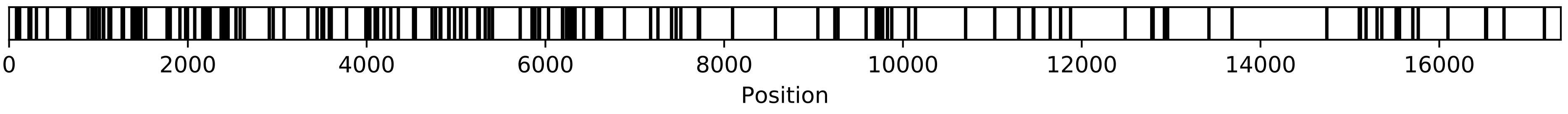

### Region: 526



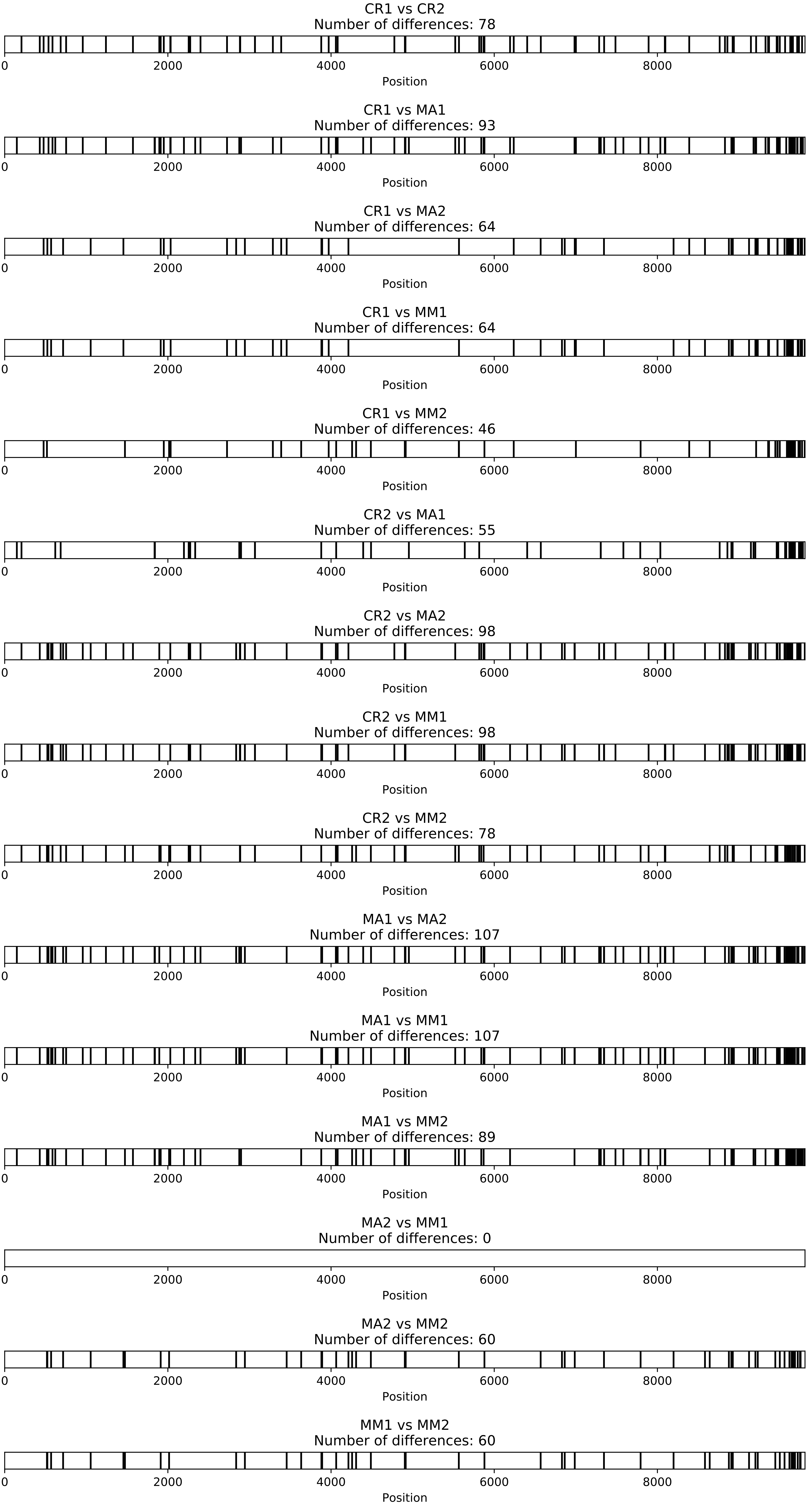
CR1 vs CR2
Number of differences: 78
CR1 vs MA1
Number of differences: 93
CR1 vs MA2
Number of differences: 64
CR1 vs MM1
Number of differences: 64
CR1 vs MM2
Number of differences: 46
CR2 vs MA1
Number of differences: 55
CR2 vs MA2
Number of differences: 98
CR2 vs MM1
Number of differences: 98
CR2 vs MM2
Number of differences: 78
MA1 vs MA2
Number of differences: 107
MA1 vs MM1
Number of differences: 107
MA1 vs MM2
Number of differences: 89
MA2 vs MM1
Number of differences: 0
MA2 vs MM2
Number of differences: 60
MM1 vs MM2
Number of differences: 60

### Region: 527

CR1 vs CR2
Number of differences: 234

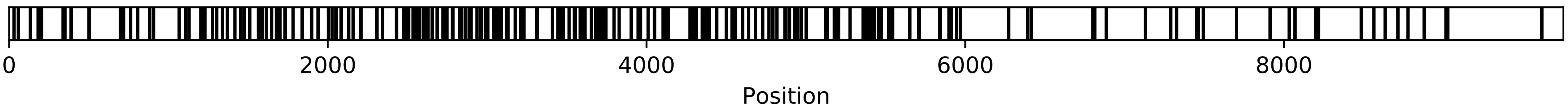


CR1 vs MA1
Number of differences: 219

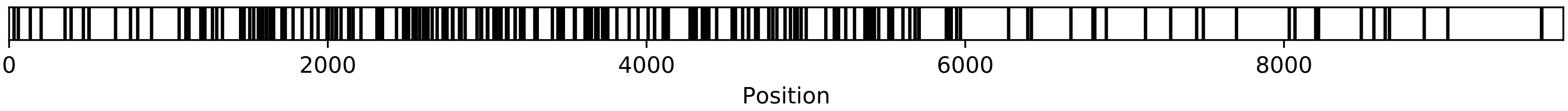


CR1 vs MA2
Number of differences: 141

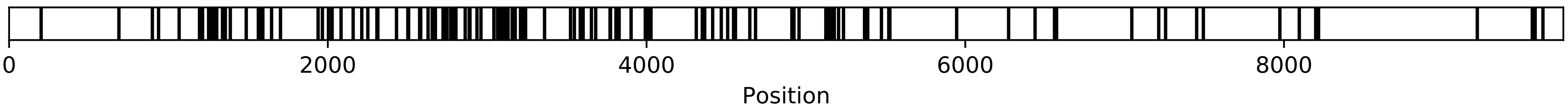


CR1 vs MM1
Number of differences: 144

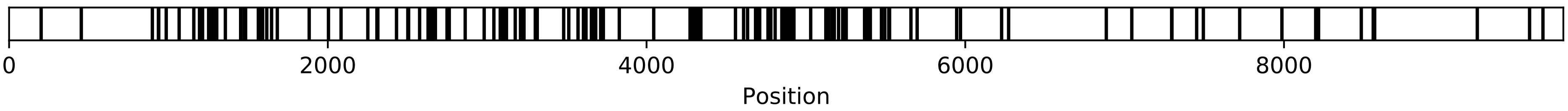


CR1 vs MM2
Number of differences: 123

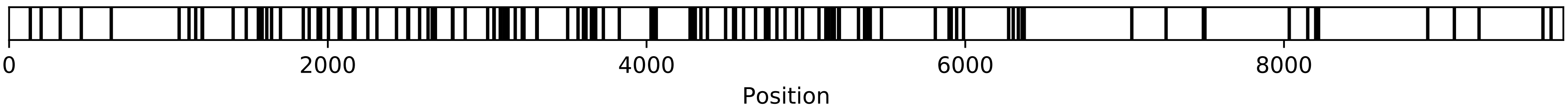


CR2 vs MA1
Number of differences: 88

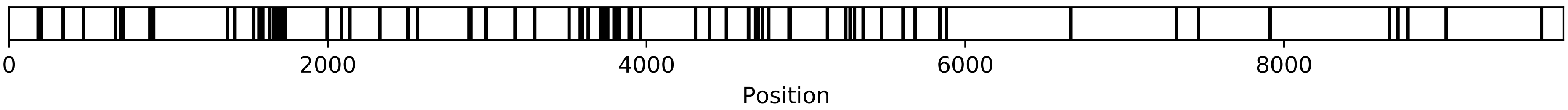


CR2 vs MA2
Number of differences: 246

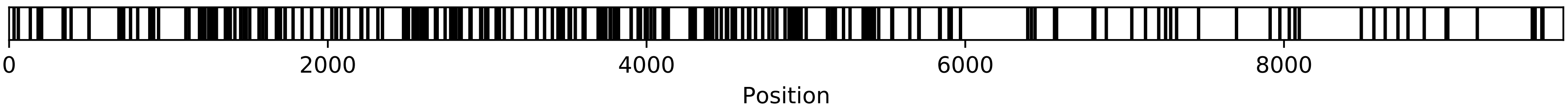


CR2 vs MM1
Number of differences: 253

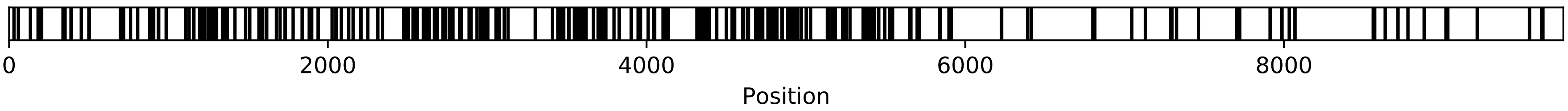


CR2 vs MM2
Number of differences: 241

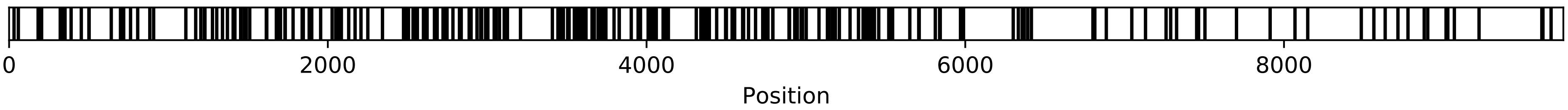


MA1 vs MA2
Number of differences: 236

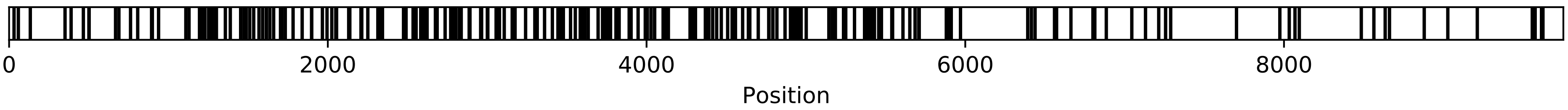


MA1 vs MM1
Number of differences: 242

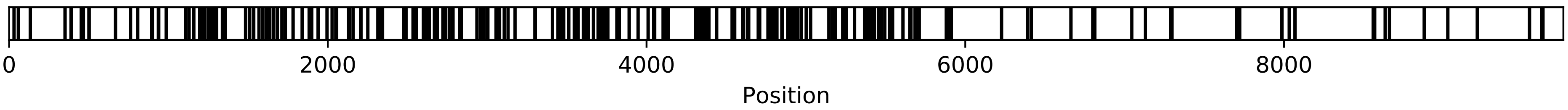


MA1 vs MM2
Number of differences: 227

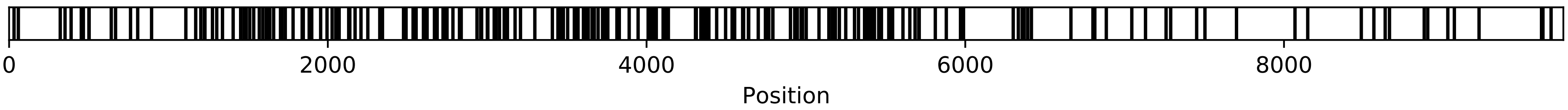


MA2 vs MM1
Number of differences: 142

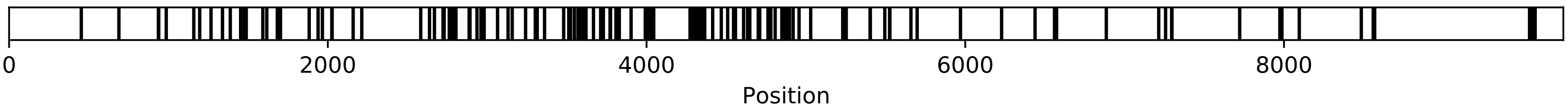


MA2 vs MM2
Number of differences: 151

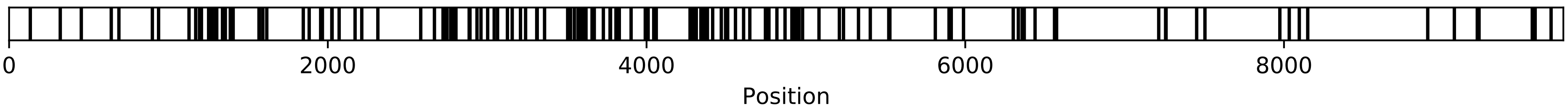


MM1 vs MM2
Number of differences: 129

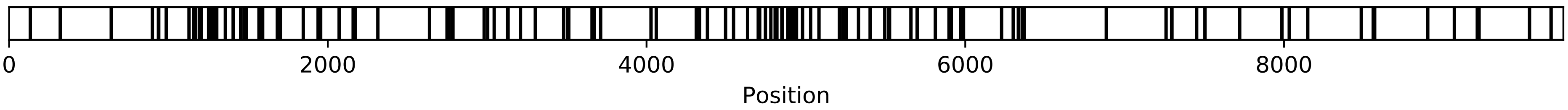

### Region: 529

CR1 vs CR2
Number of differences: 258

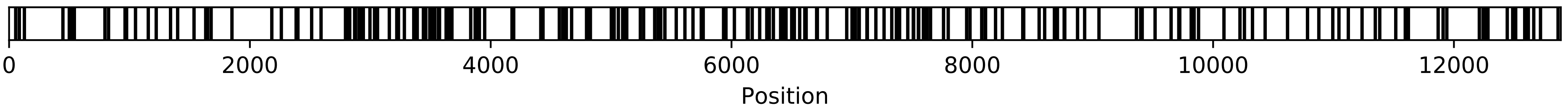


CR1 vs MA1
Number of differences: 278

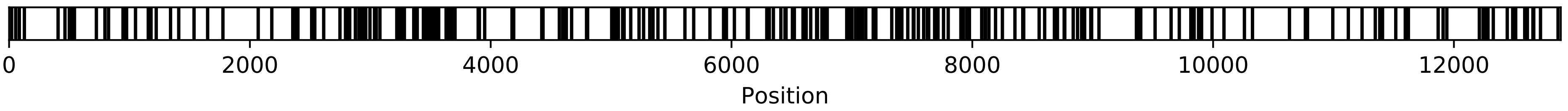


CR1 vs MA2
Number of differences: 81

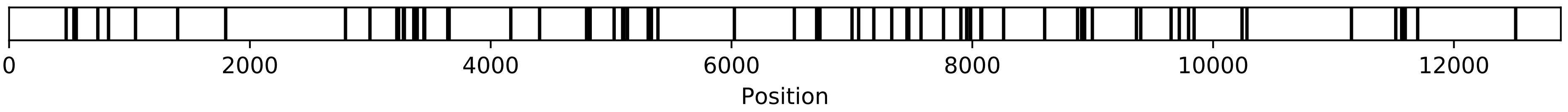


CR1 vs MM1
Number of differences: 274

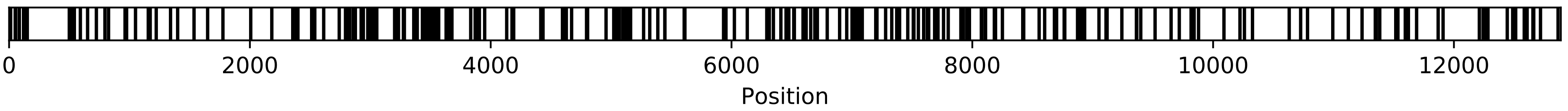


CR1 vs MM2
Number of differences: 277

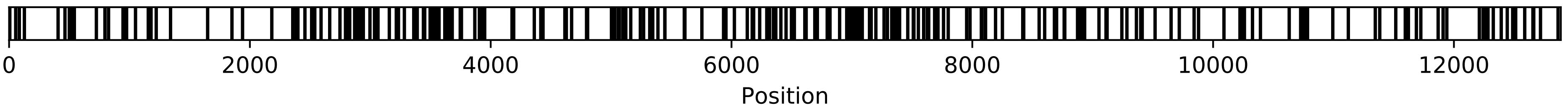


CR2 vs MA1
Number of differences: 157

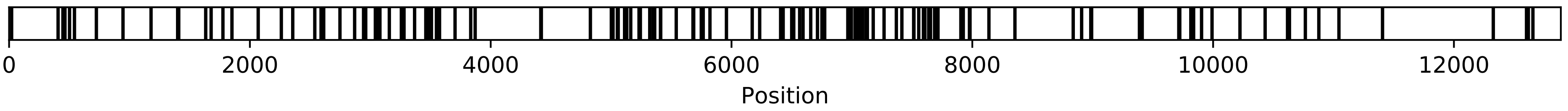


CR2 vs MA2
Number of differences: 252

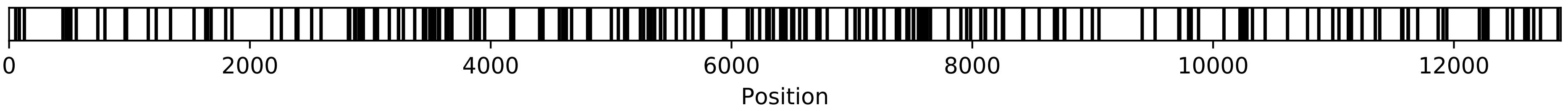


CR2 vs MM1
Number of differences: 174

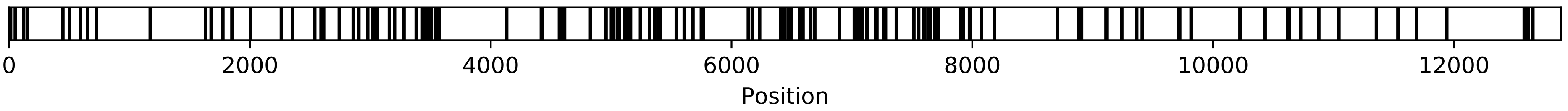


CR2 vs MM2
Number of differences: 167

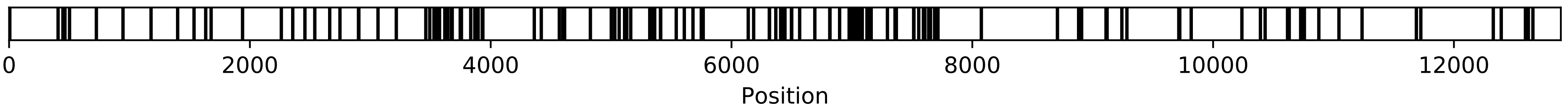


MA1 vs MA2
Number of differences: 271

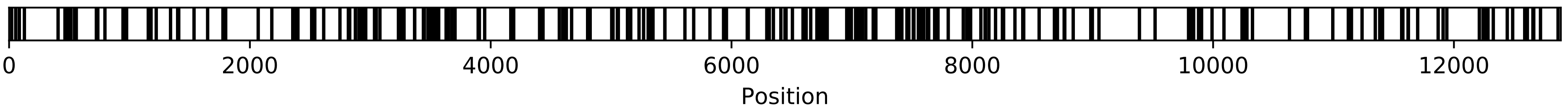


MA1 vs MM1
Number of differences: 151

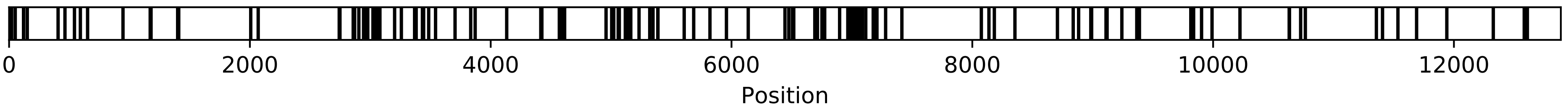


MA1 vs MM2
Number of differences: 183

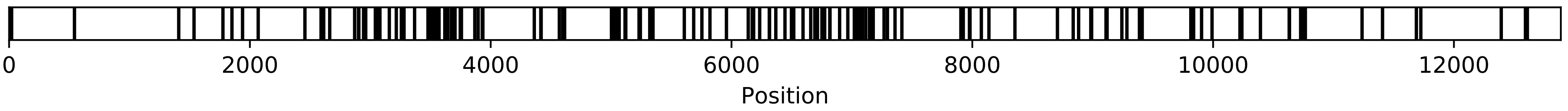


MA2 vs MM1
Number of differences: 268

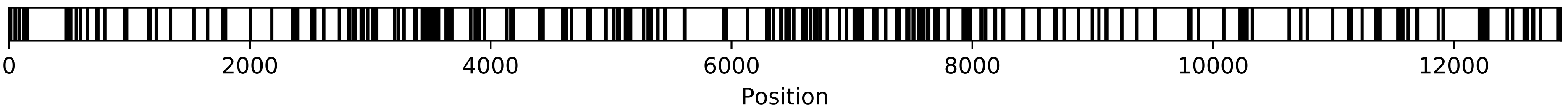


MA2 vs MM2
Number of differences: 271

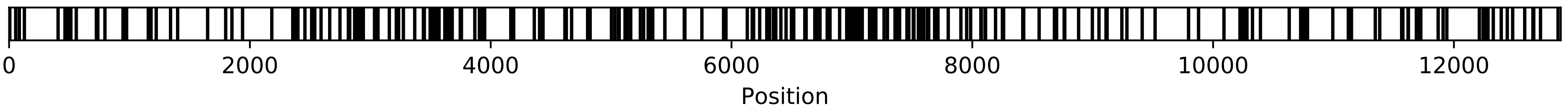


MM1 vs MM2
Number of differences: 151

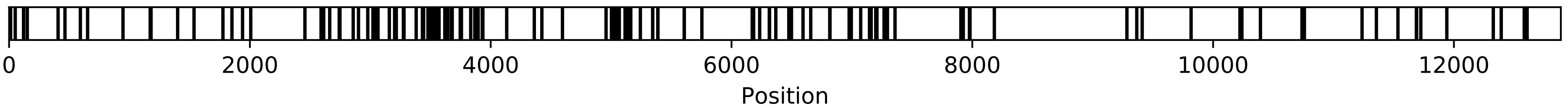

### Region: 530

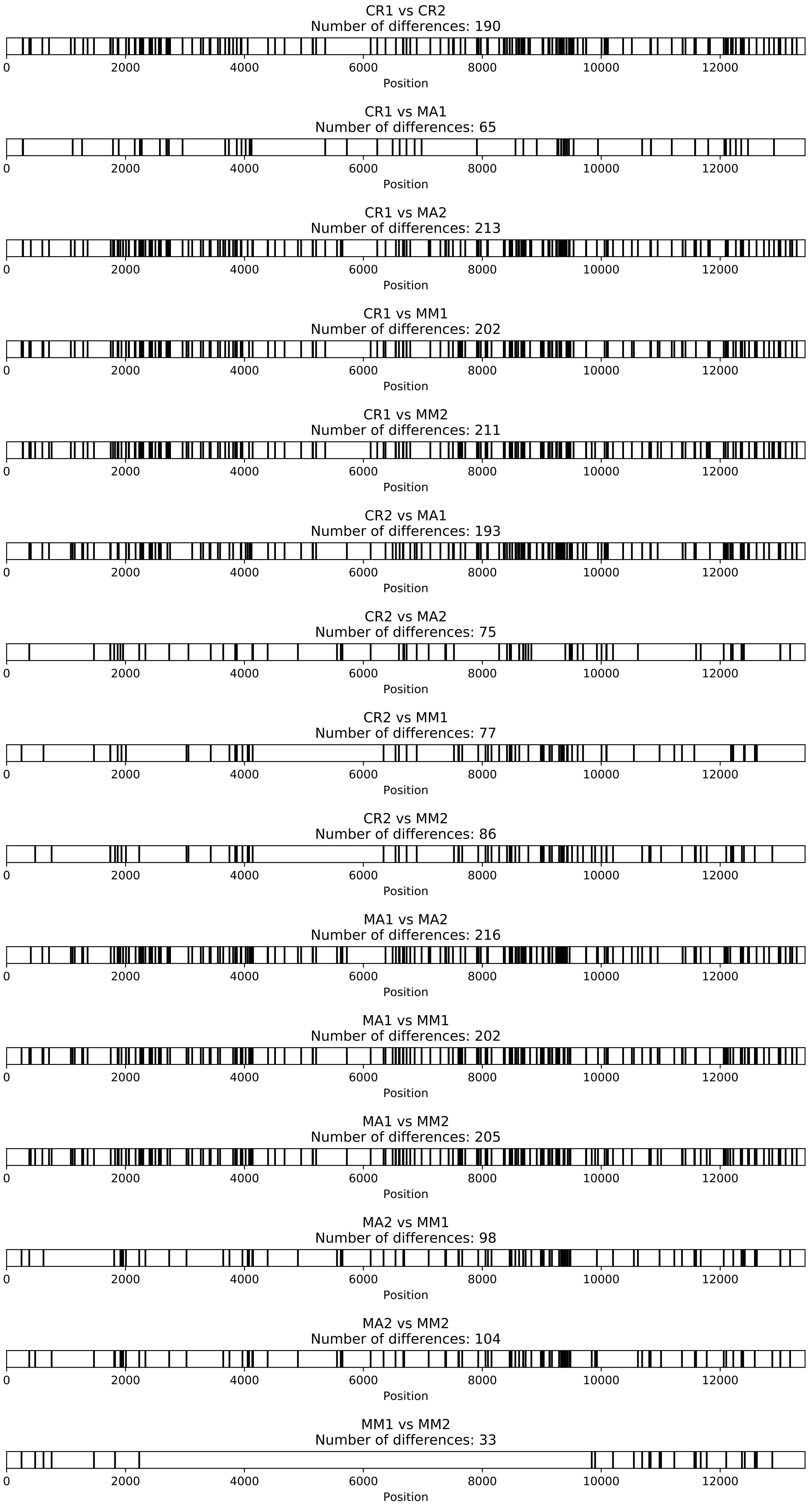
CR1 vs CR2
Number of differences: 190
CR1 vs MA1
Number of differences: 65
CR1 vs MA2
Number of differences: 213
CR1 vs MM1
Number of differences: 202
CR1 vs MM2
Number of differences: 211
CR2 vs MA1
Number of differences: 193
CR2 vs MA2
Number of differences: 75
CR2 vs MM1
Number of differences: 77
CR2 vs MM2
Number of differences: 86
MA1 vs MA2
Number of differences: 216
MA1 vs MM1
Number of differences: 202
MA1 vs MM2
Number of differences: 205
MA2 vs MM1
Number of differences: 98
MA2 vs MM2
Number of differences: 104
MM1 vs MM2
Number of differences: 33
0
2000
4000
6000
8000
10000
12000
Position
