## Supplementary material for "Genomic Signature of Sexual Reproduction in the Bdelloid Rotifer *Macrotrachella quadricornifera*": Table S5

### Region 6



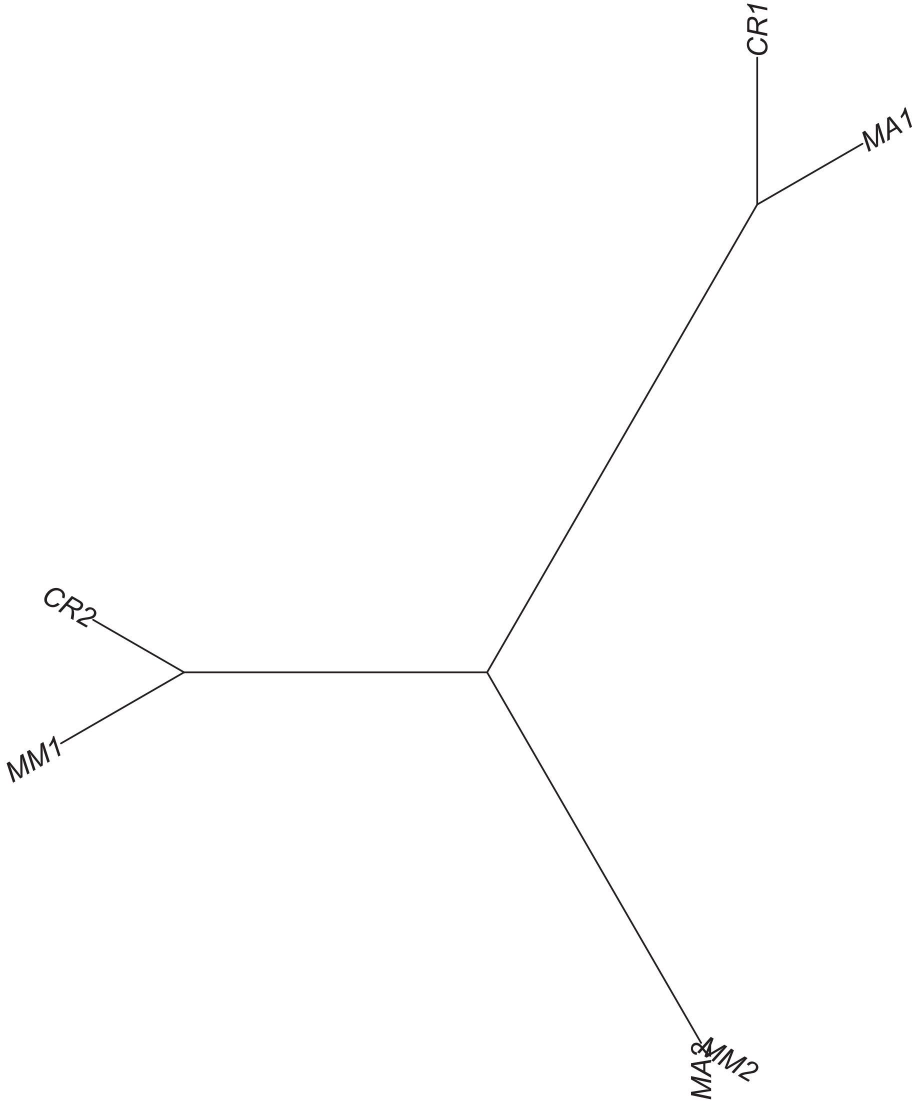

### Region 7



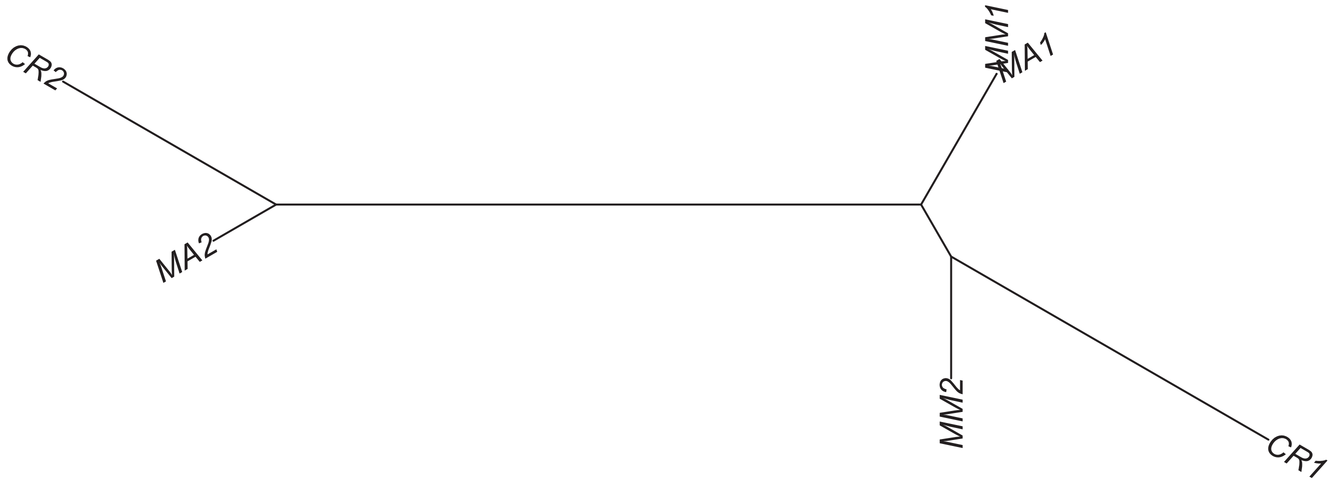

Region 8

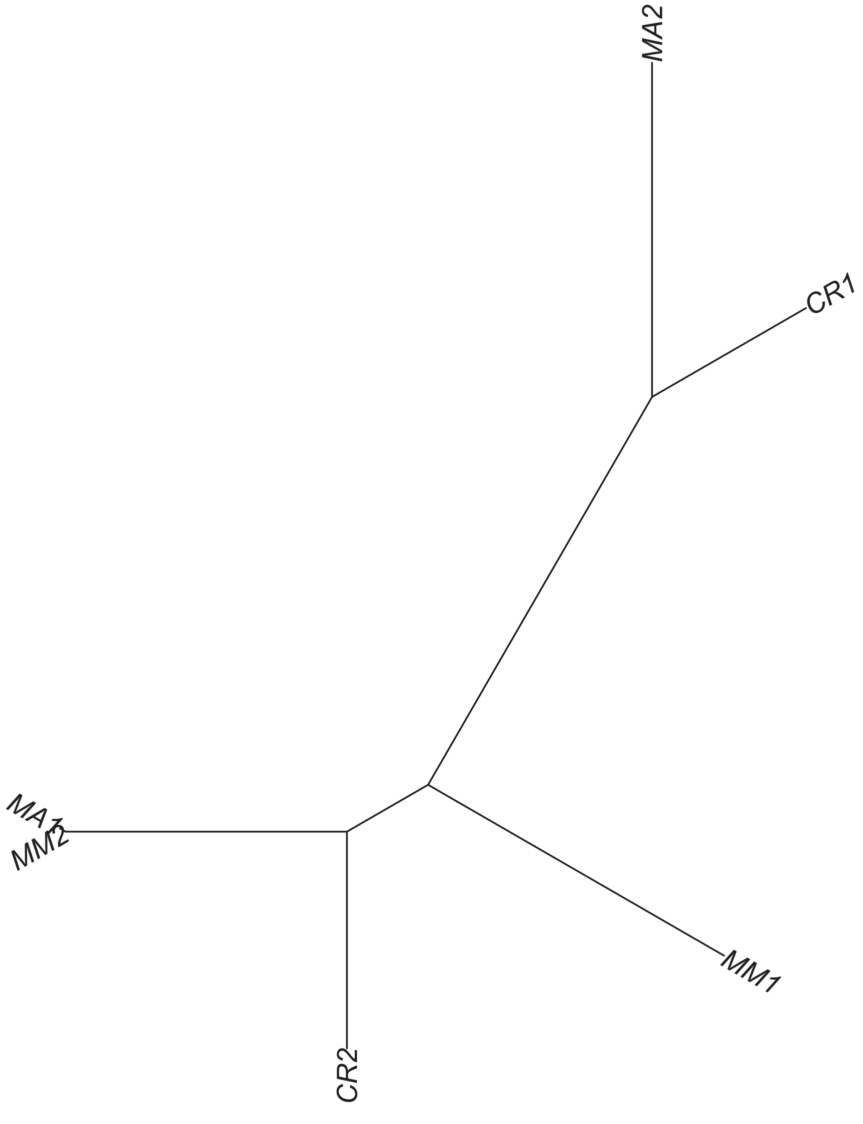

Region 10

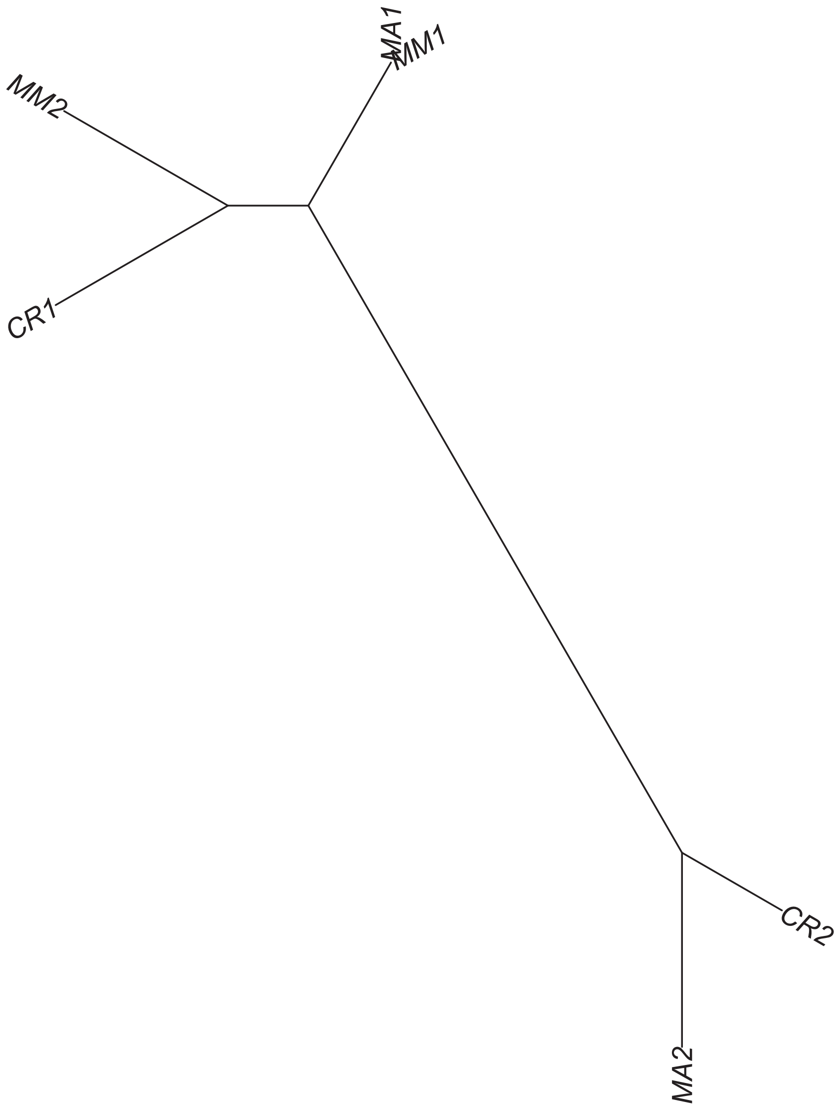

### Region 12



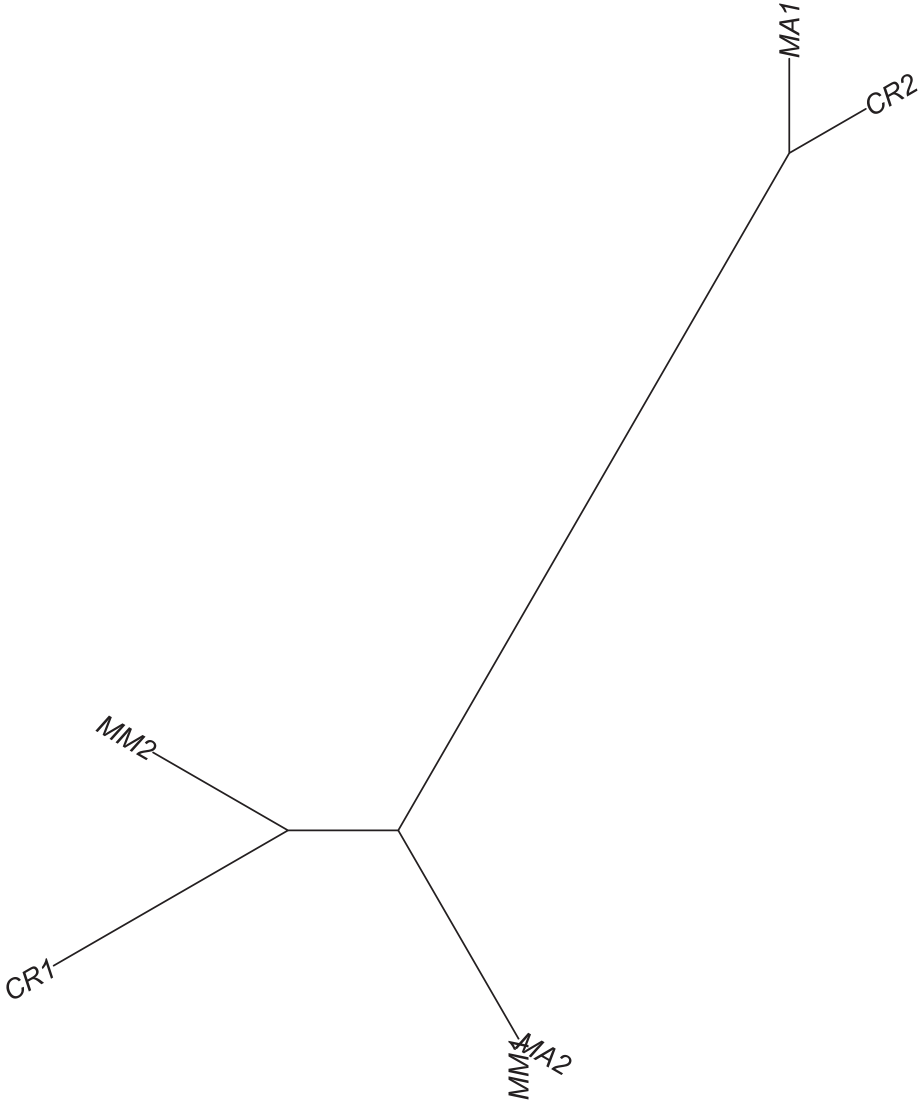

#### Region 13

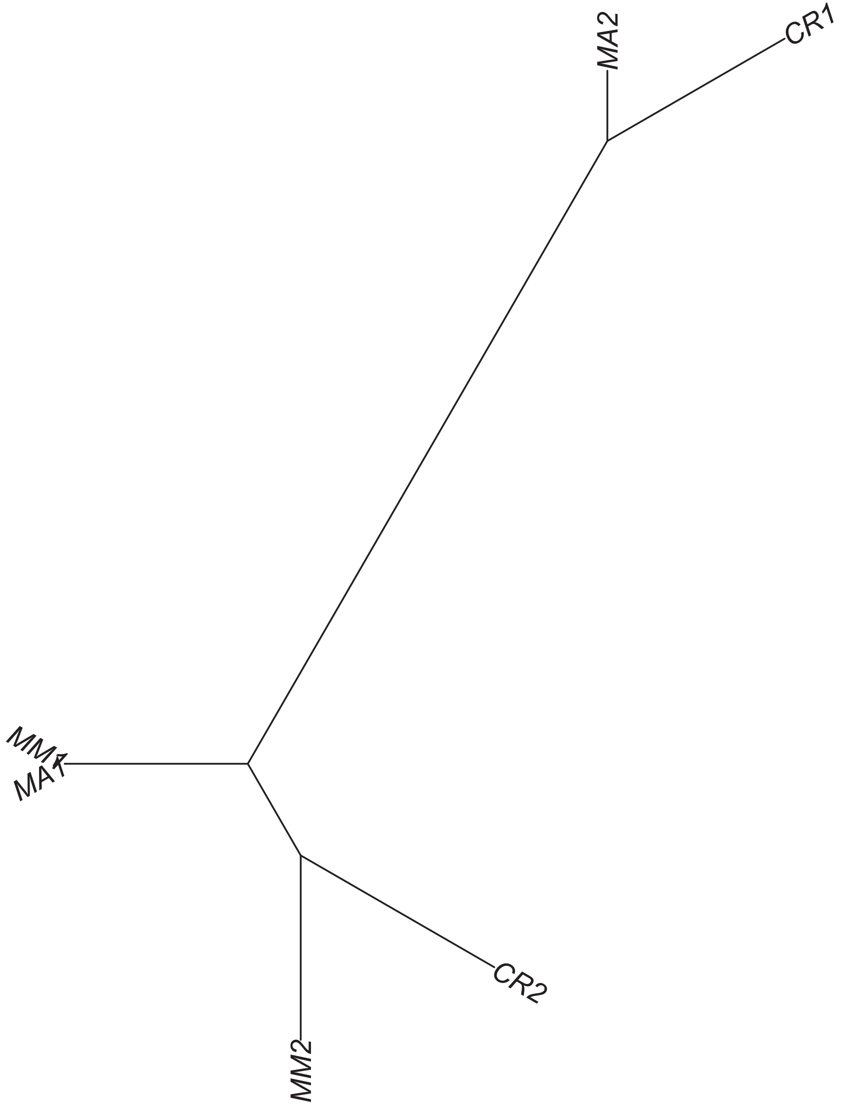

Region 14



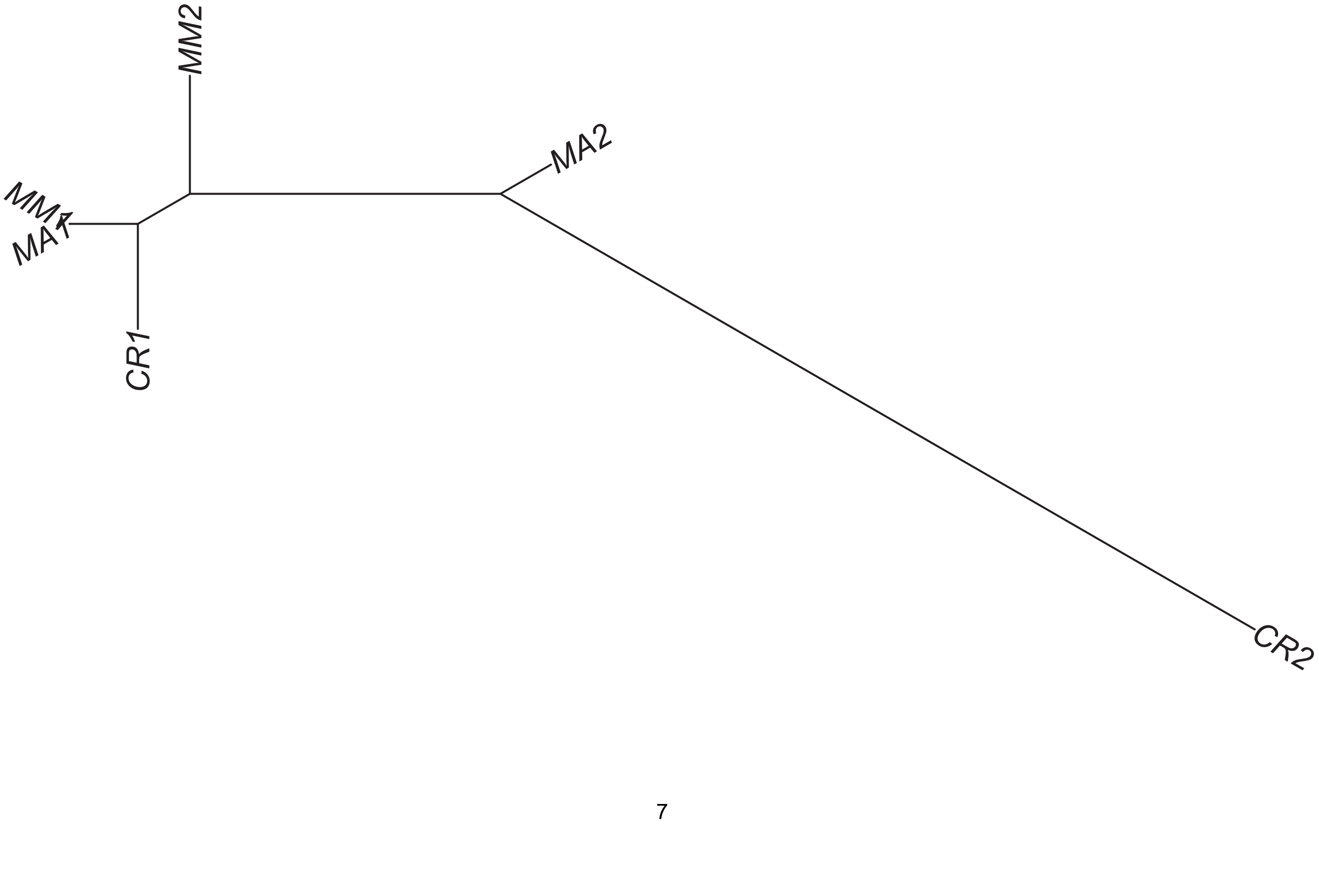

### Region 15

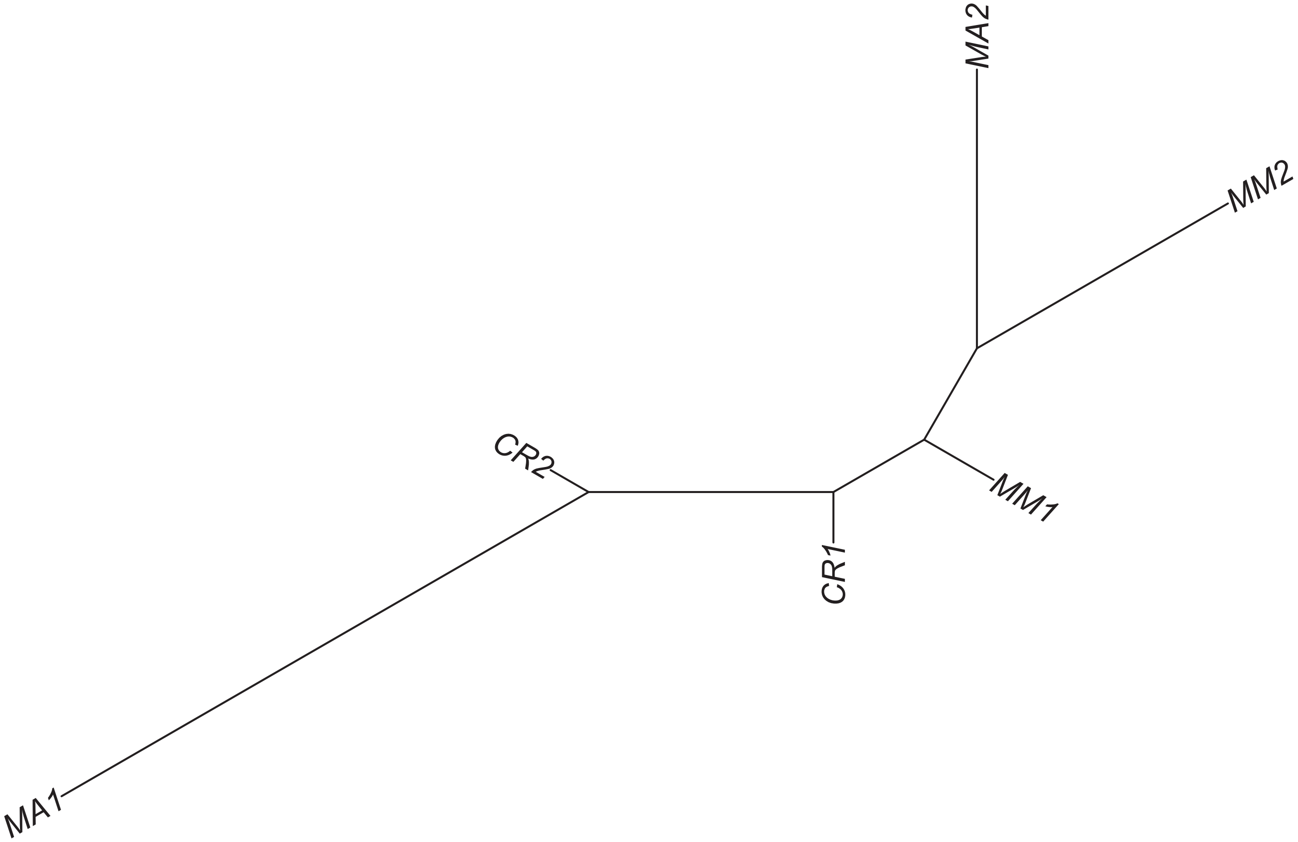

Region 18

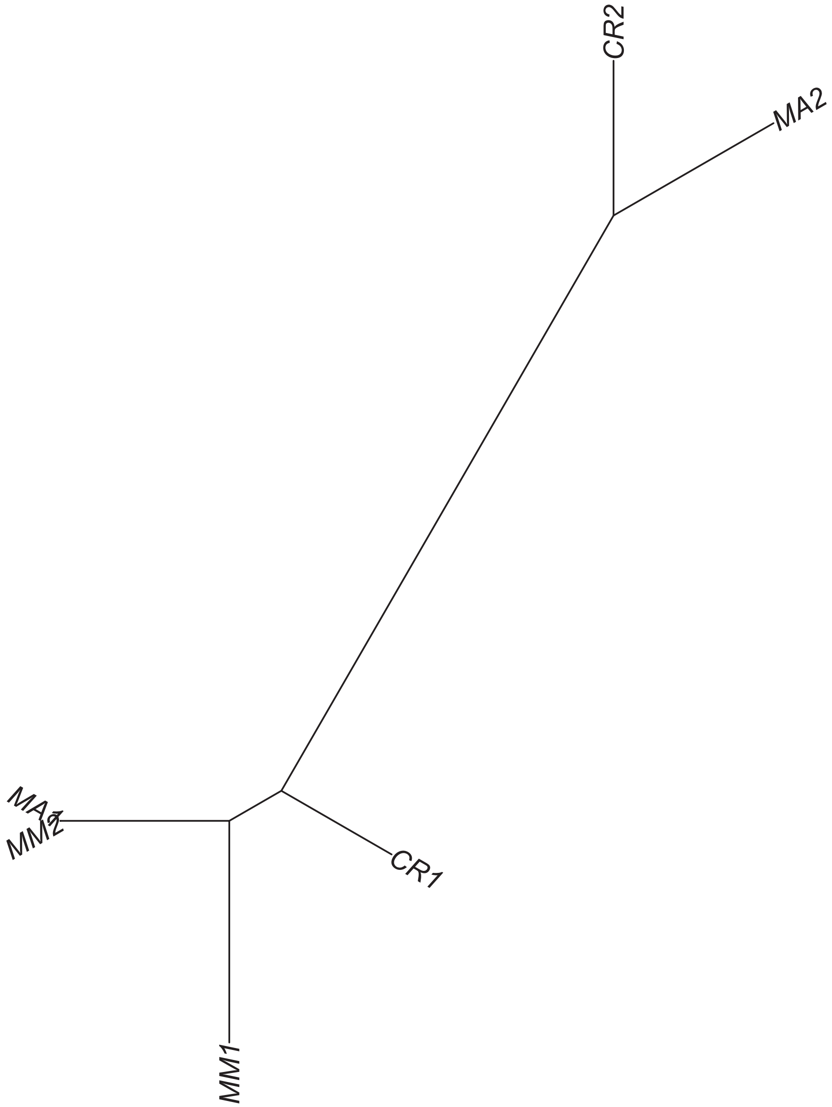

### Region 20

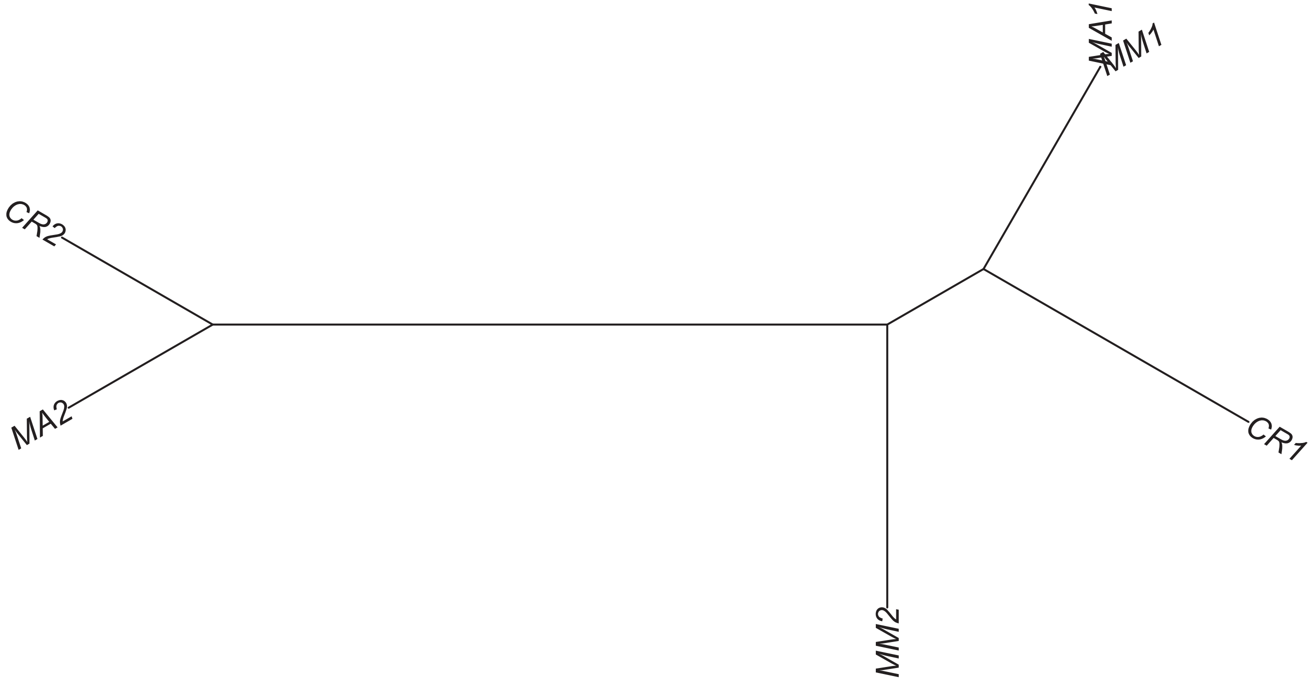

### Region 23

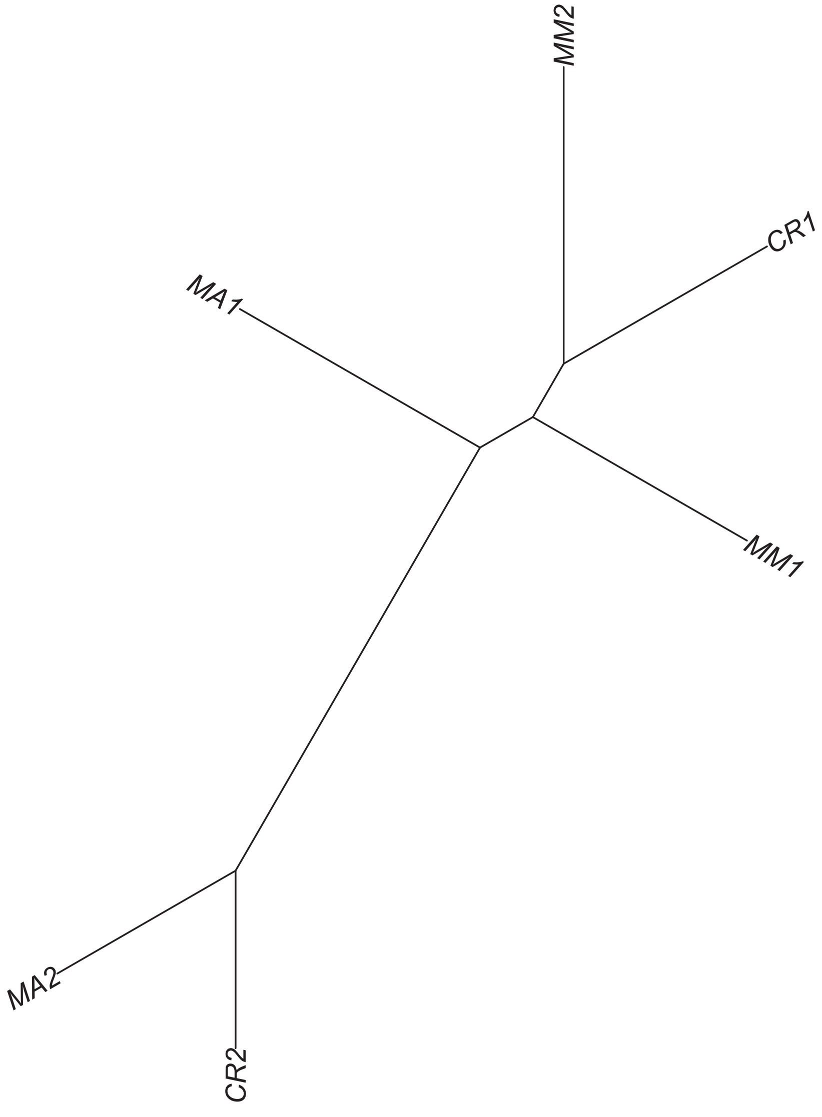

### Region 25



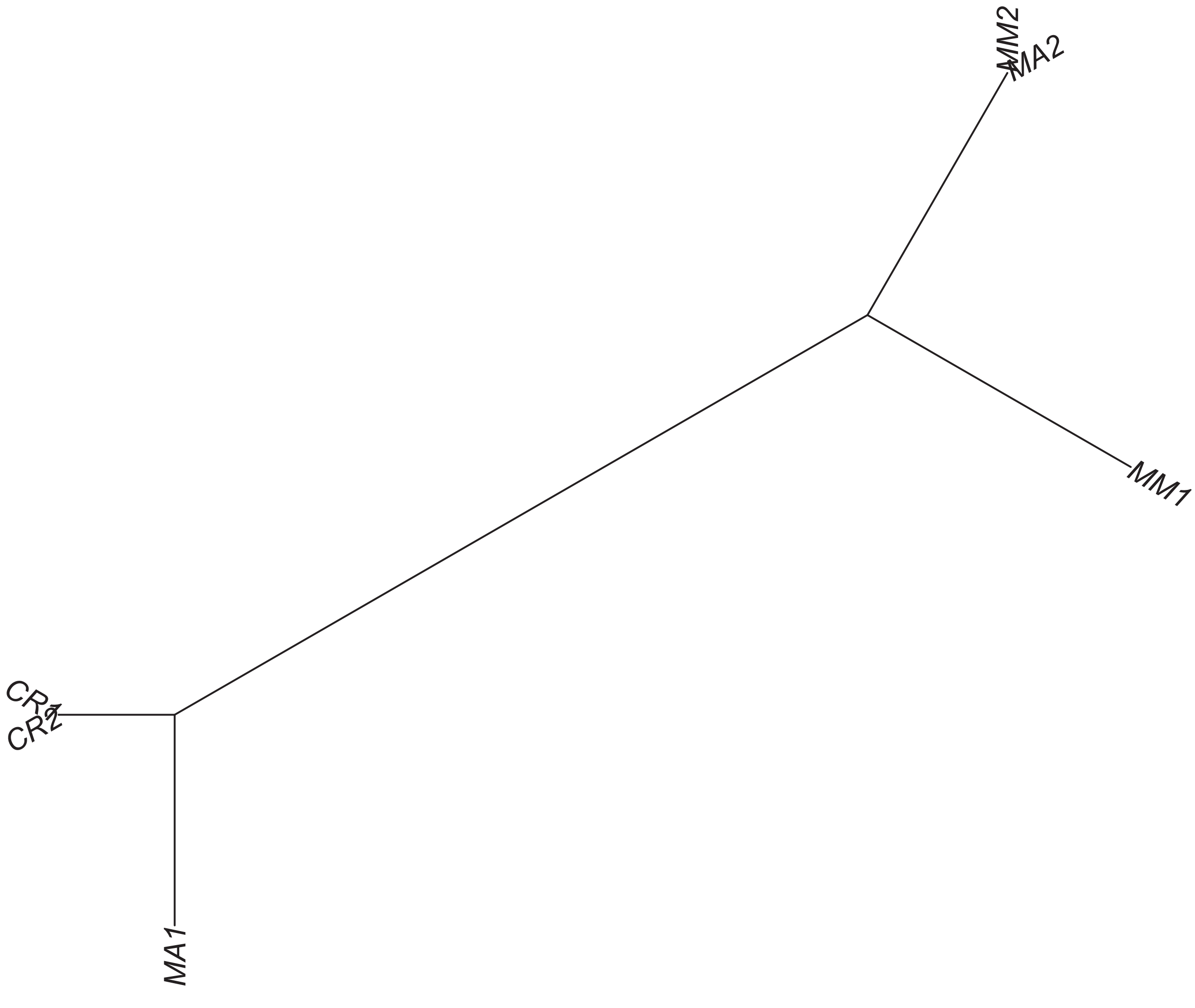

### Region 27



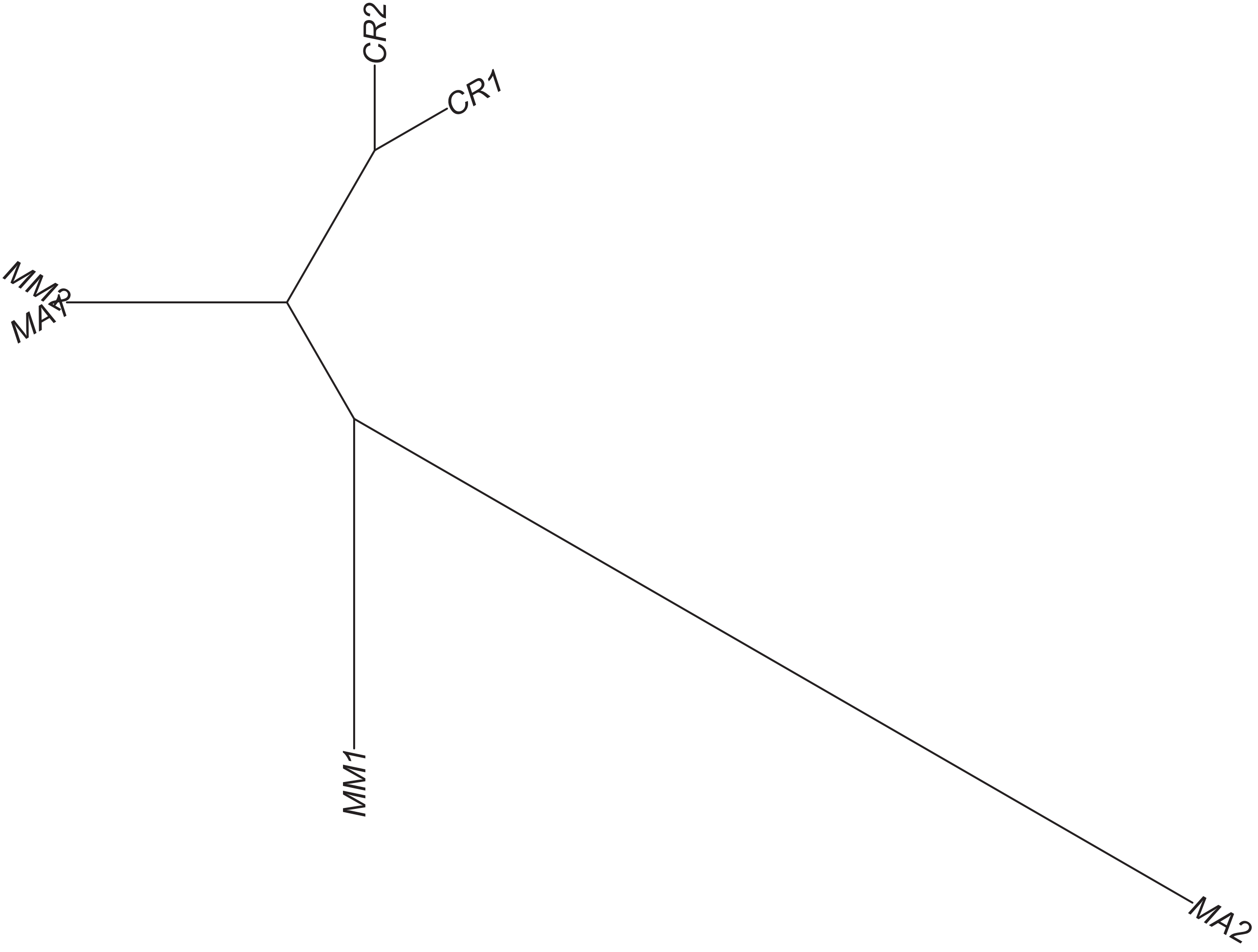

Region 28

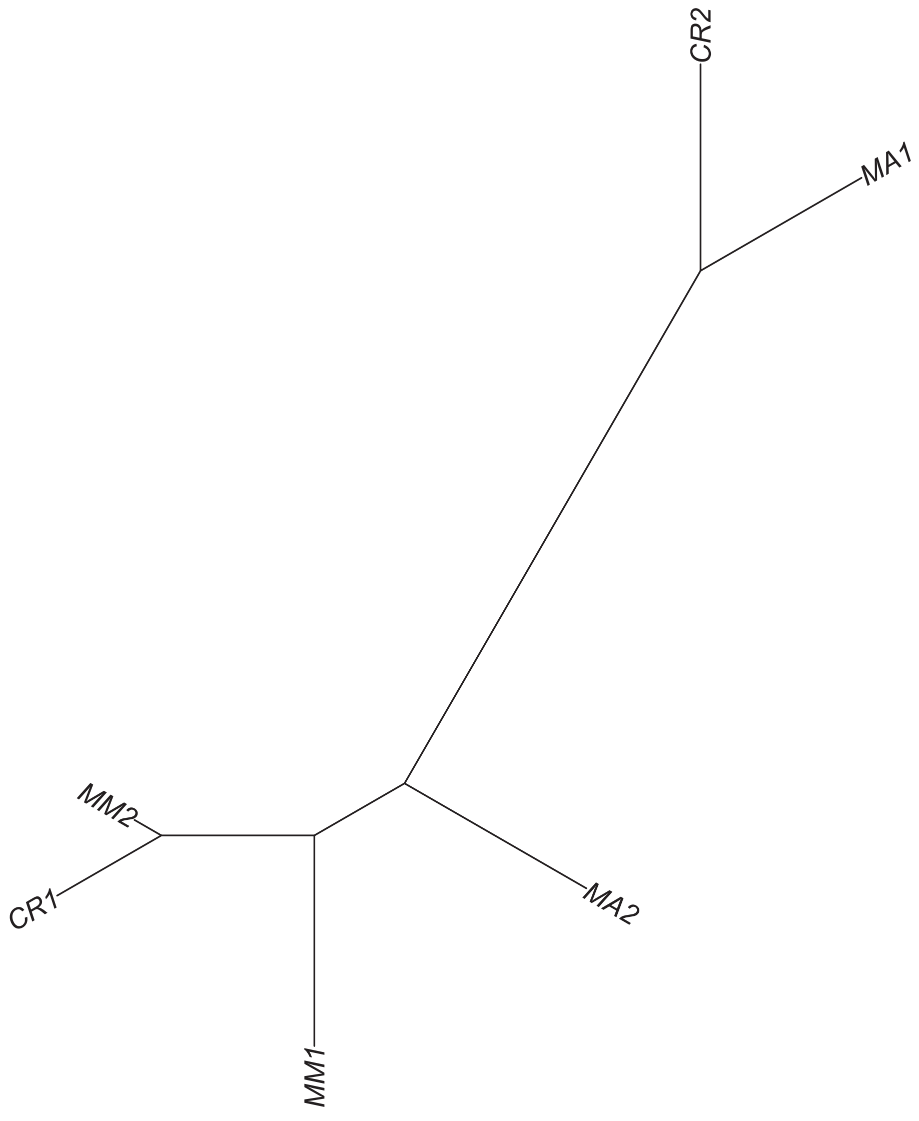

### Region 30

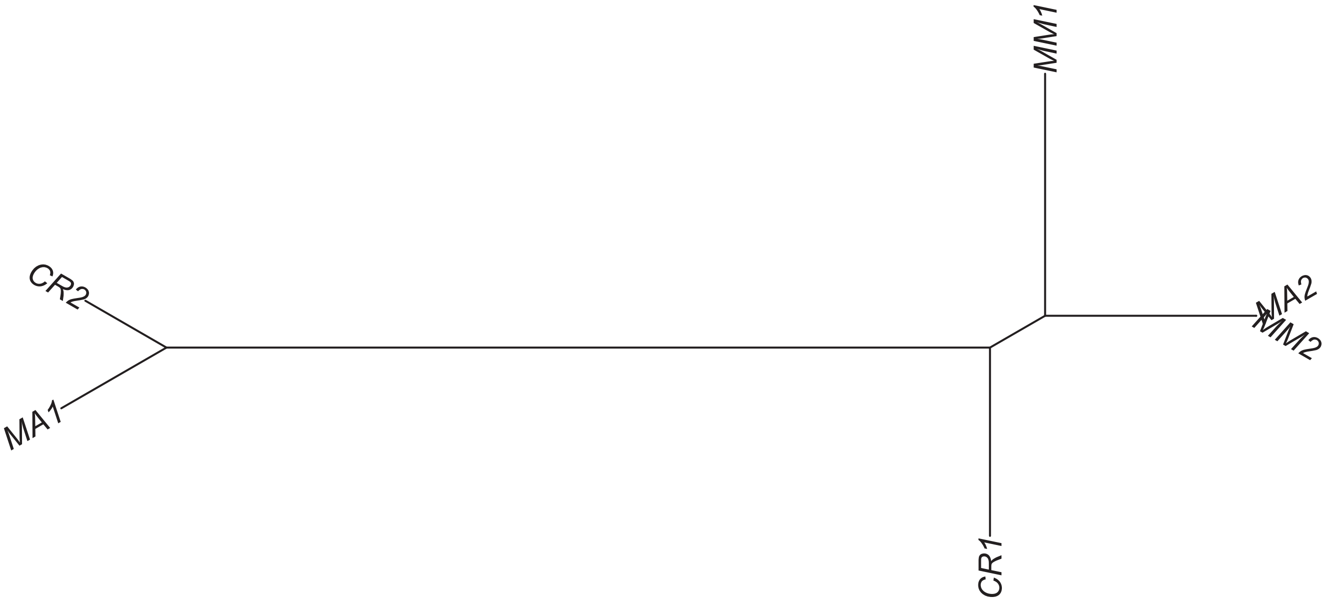

### Region 31

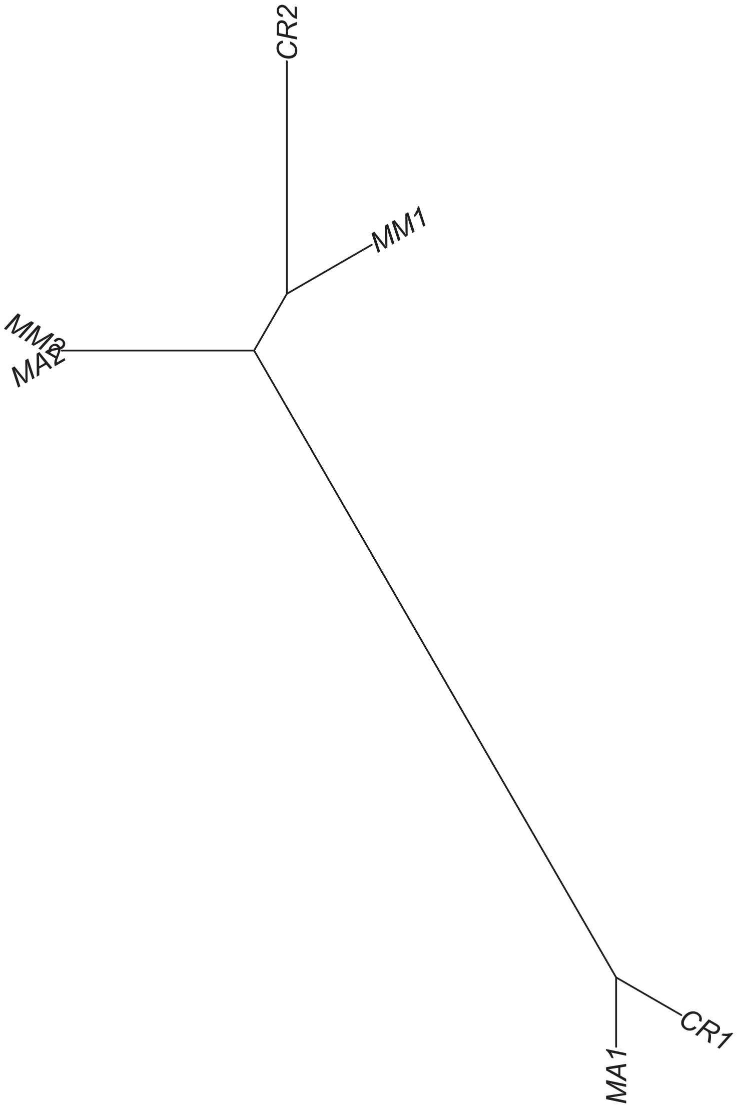

### Region 33



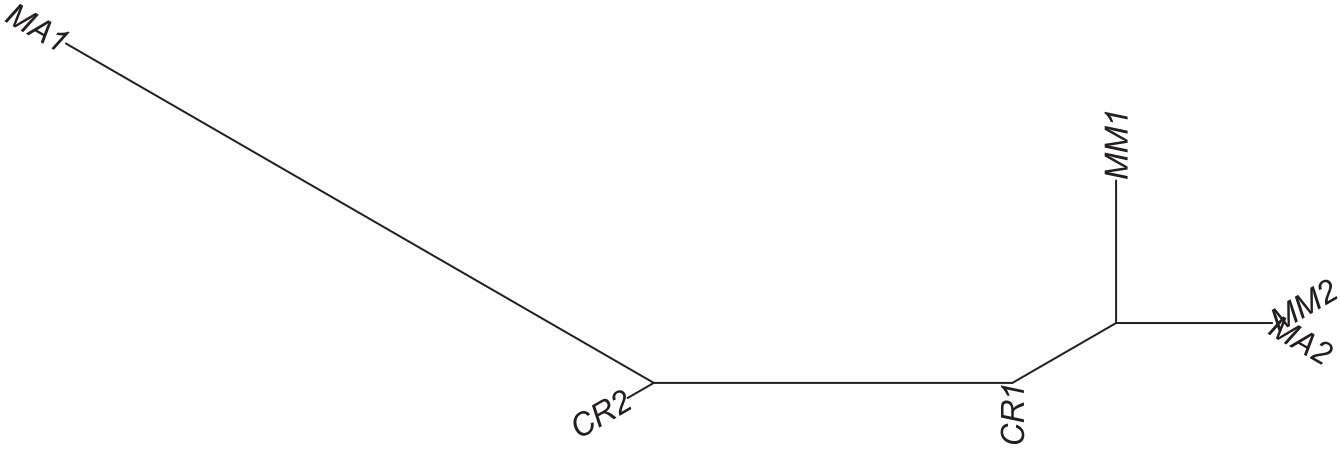

Region 34

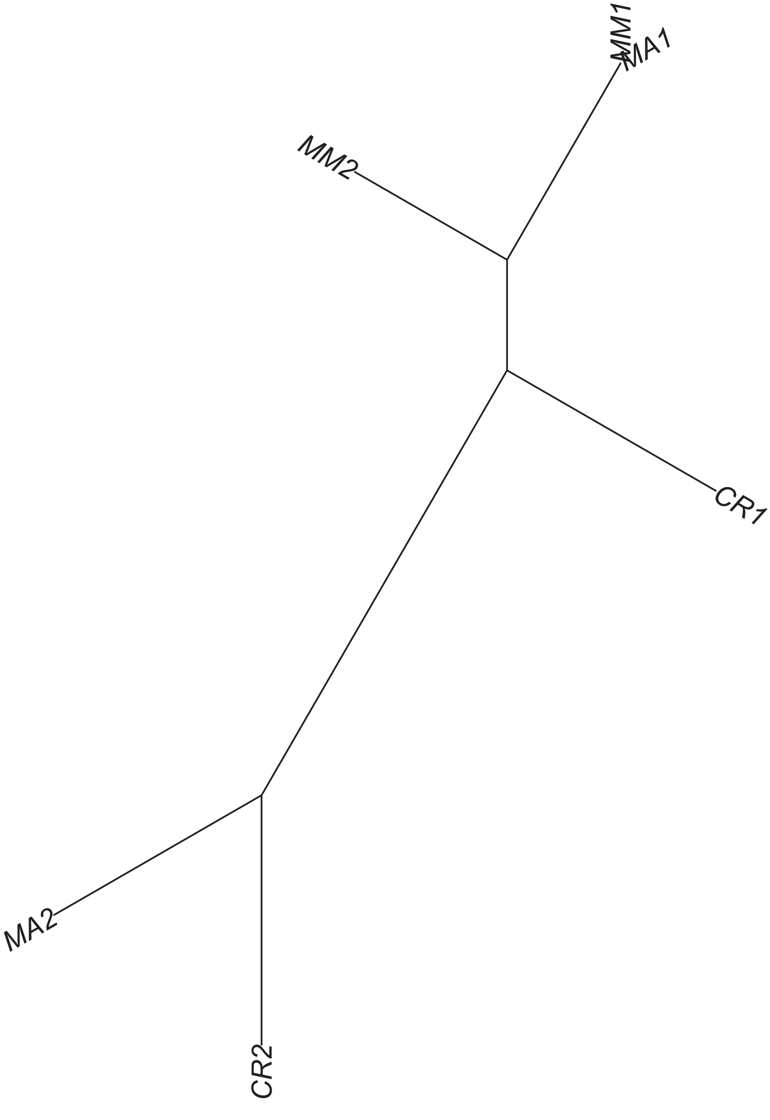

Region 36

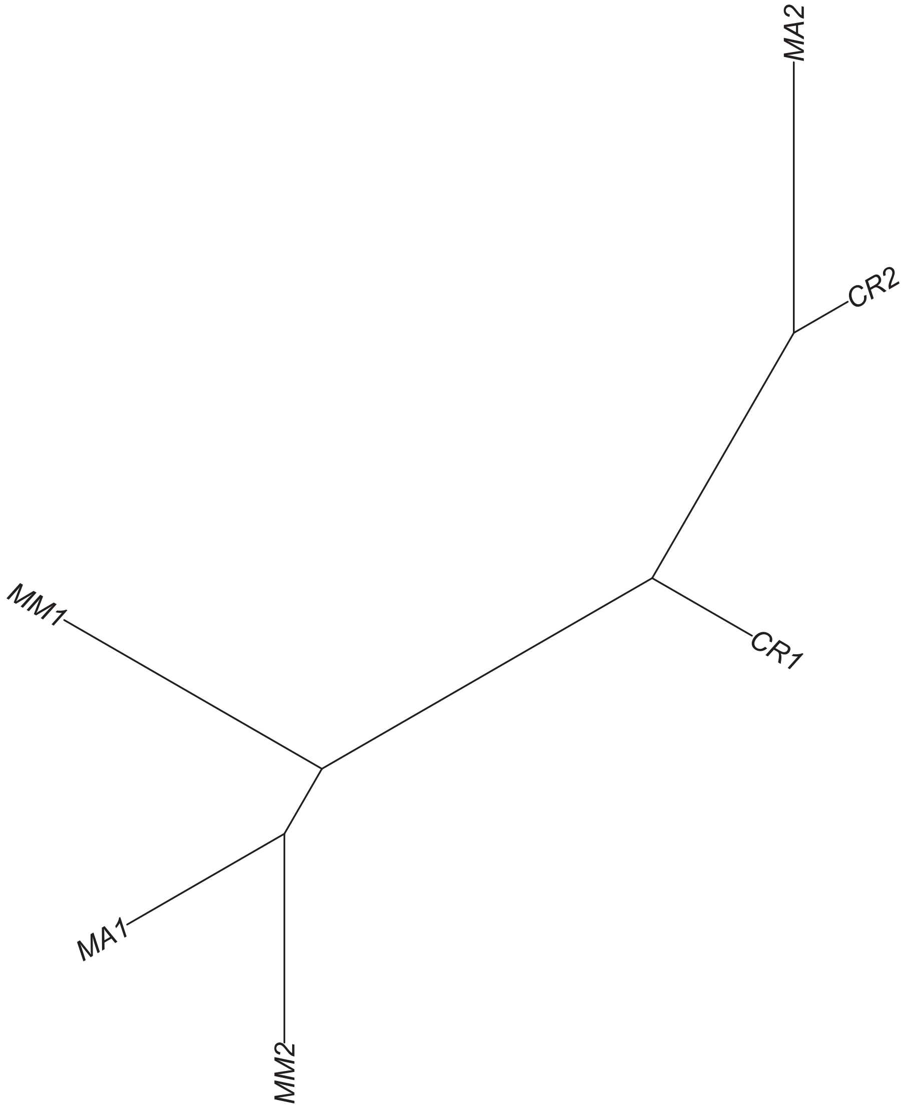

Region 37

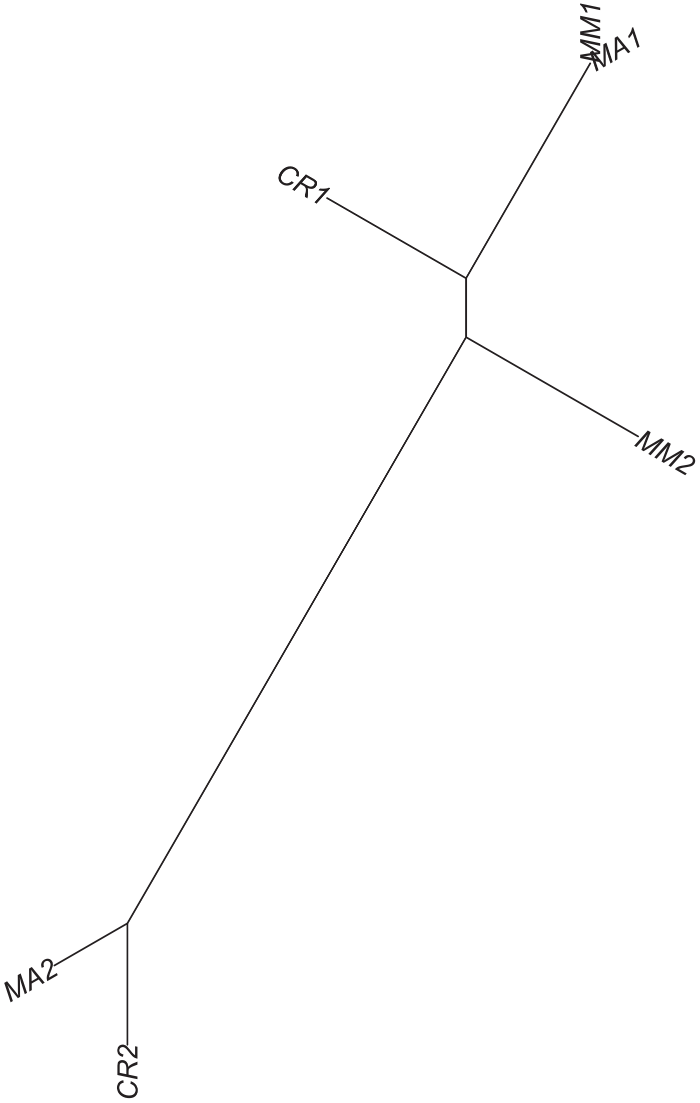

Region 38

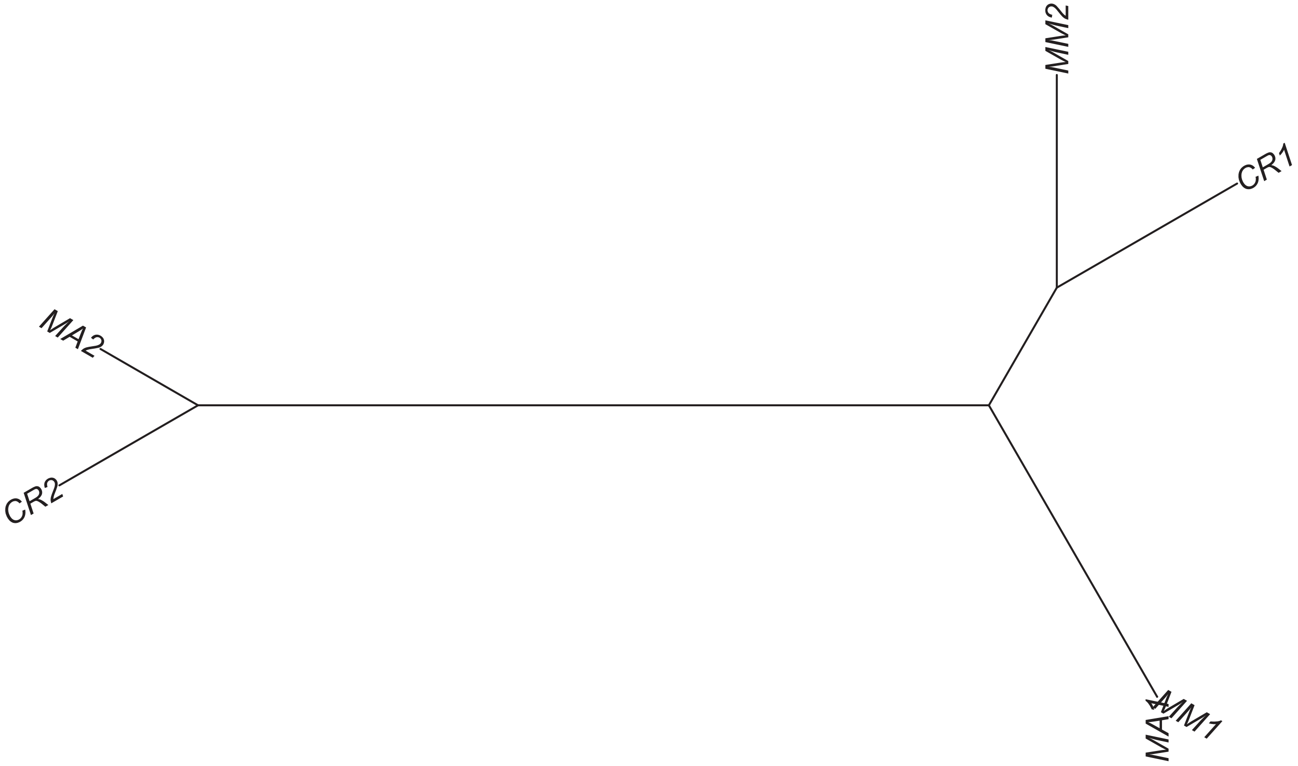

### Region 39



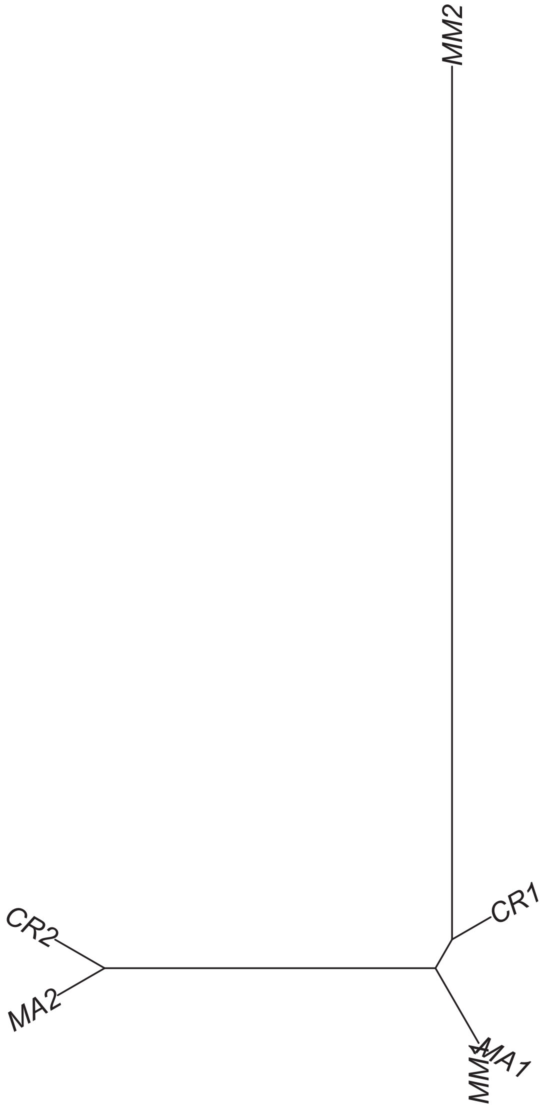

#### Region 40



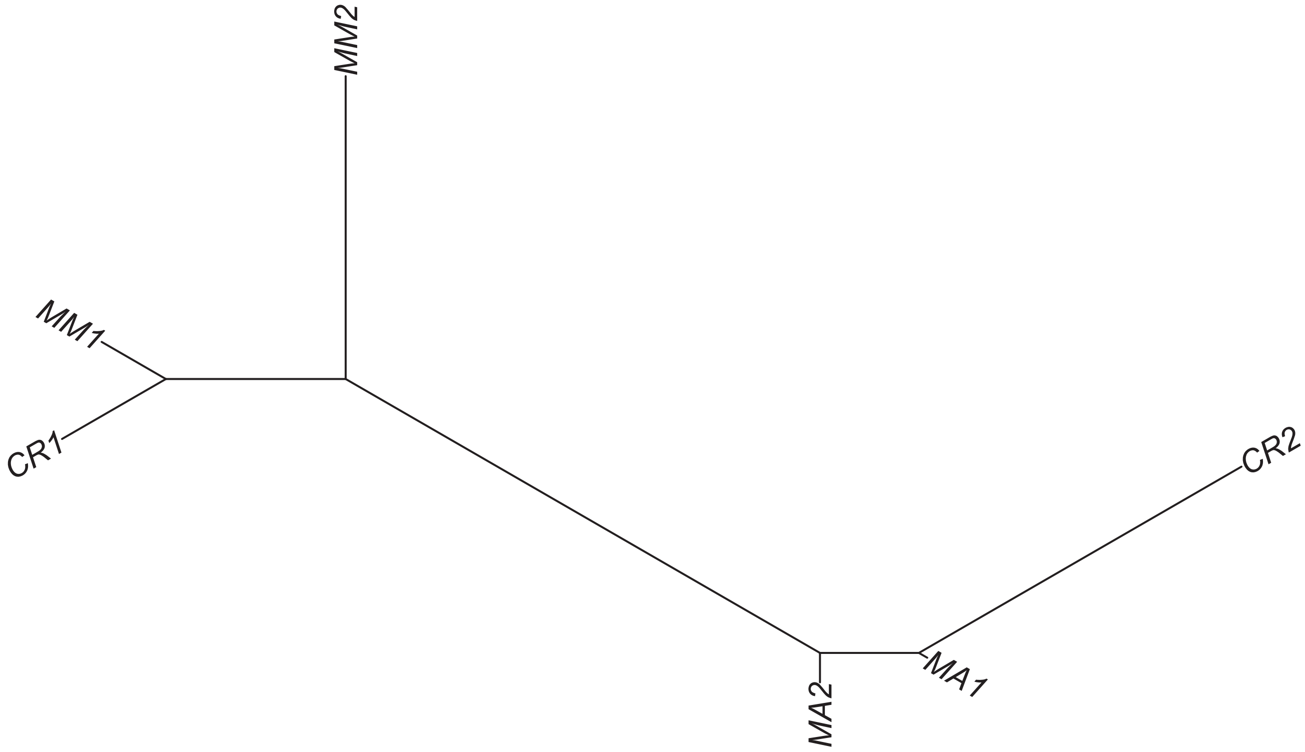

Region 42

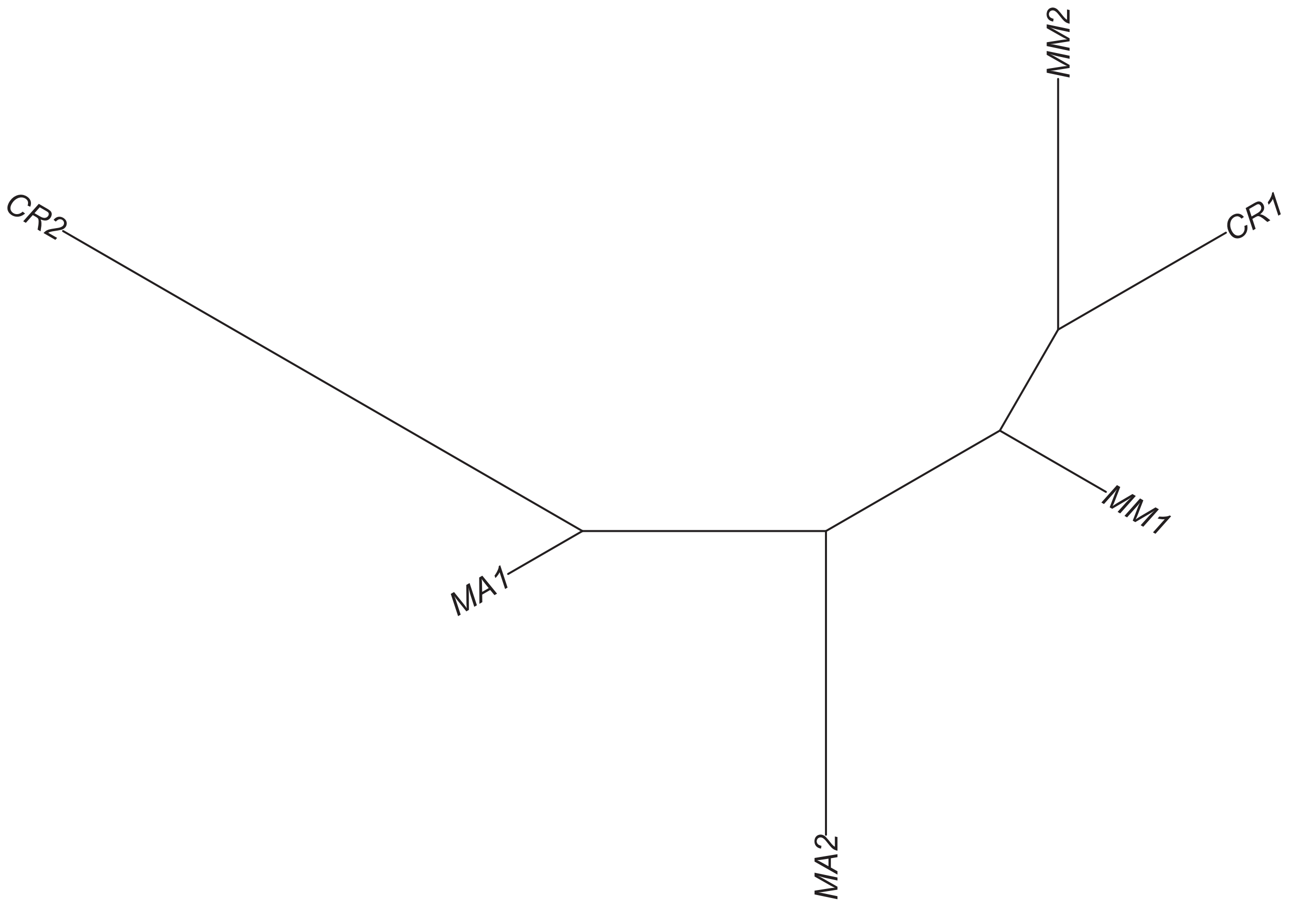

### Region 44



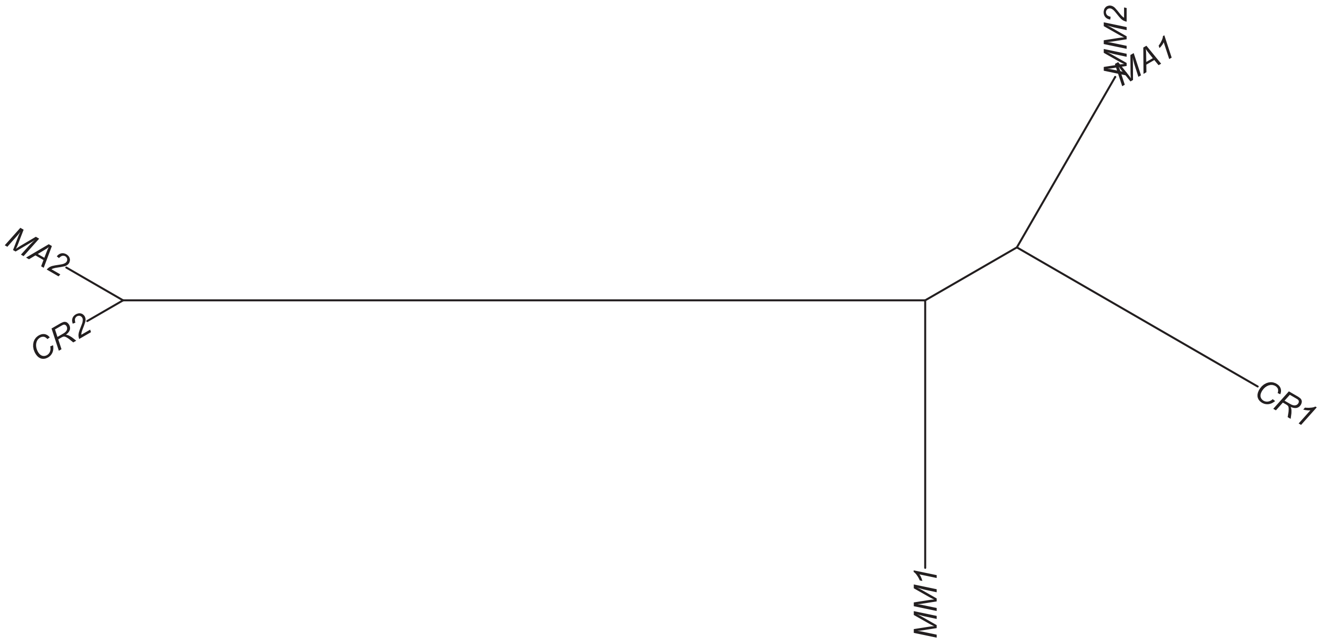

### Region 46



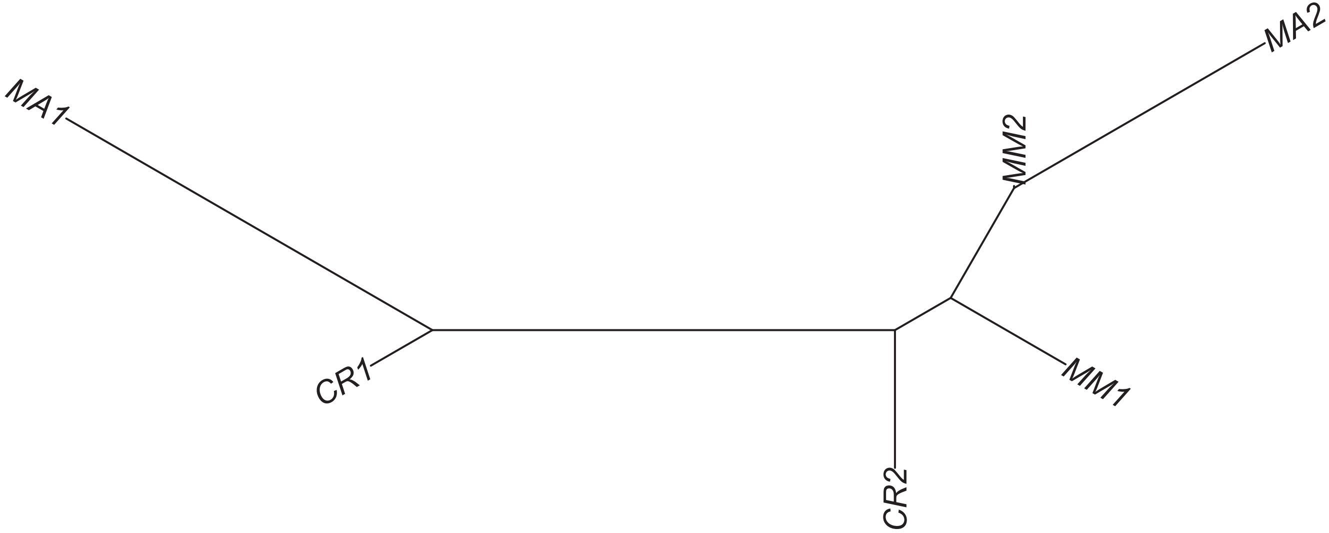

Region 47

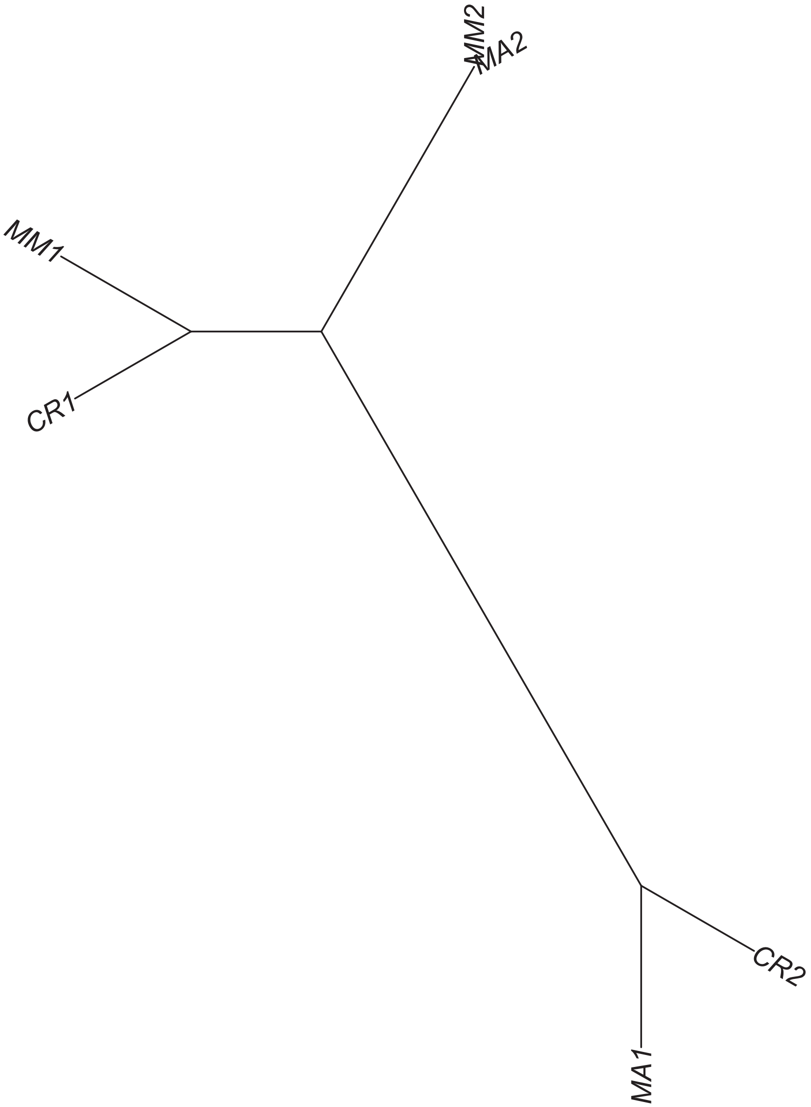

### Region 49

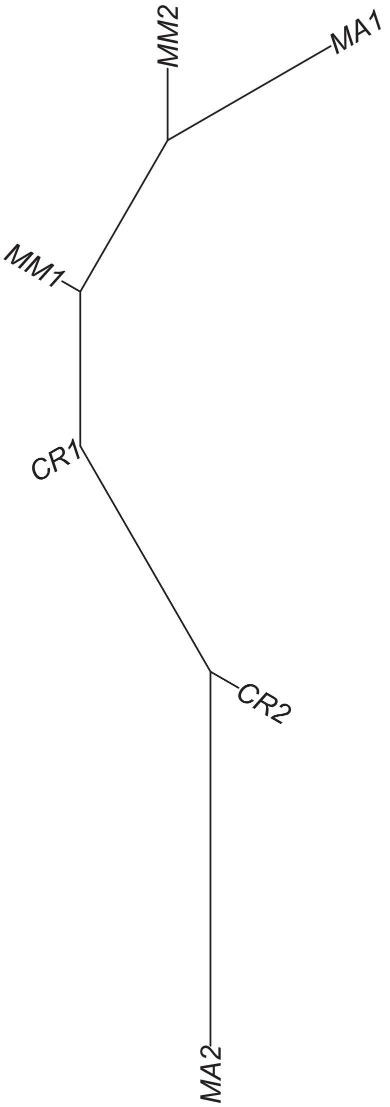

Region 50



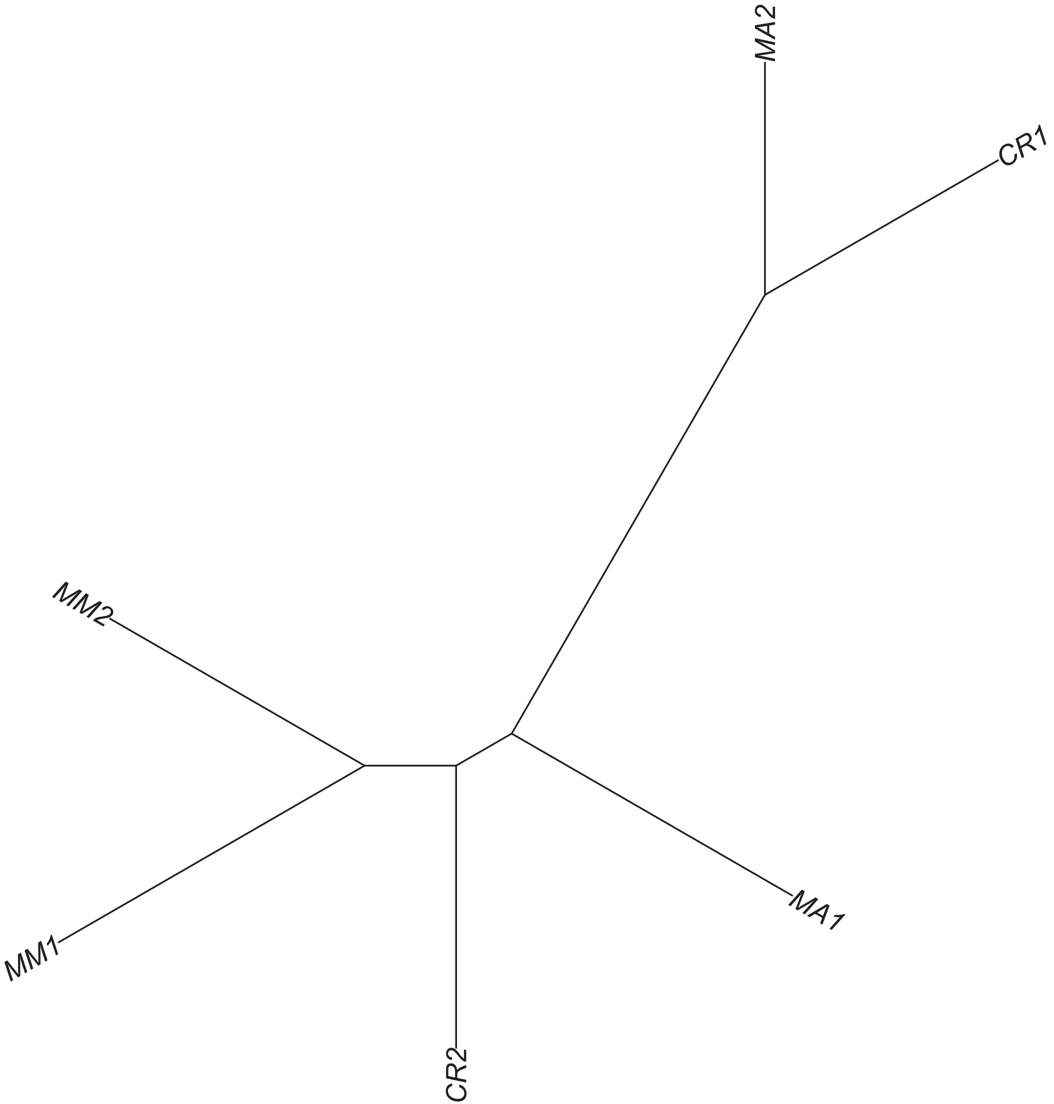

Region 51

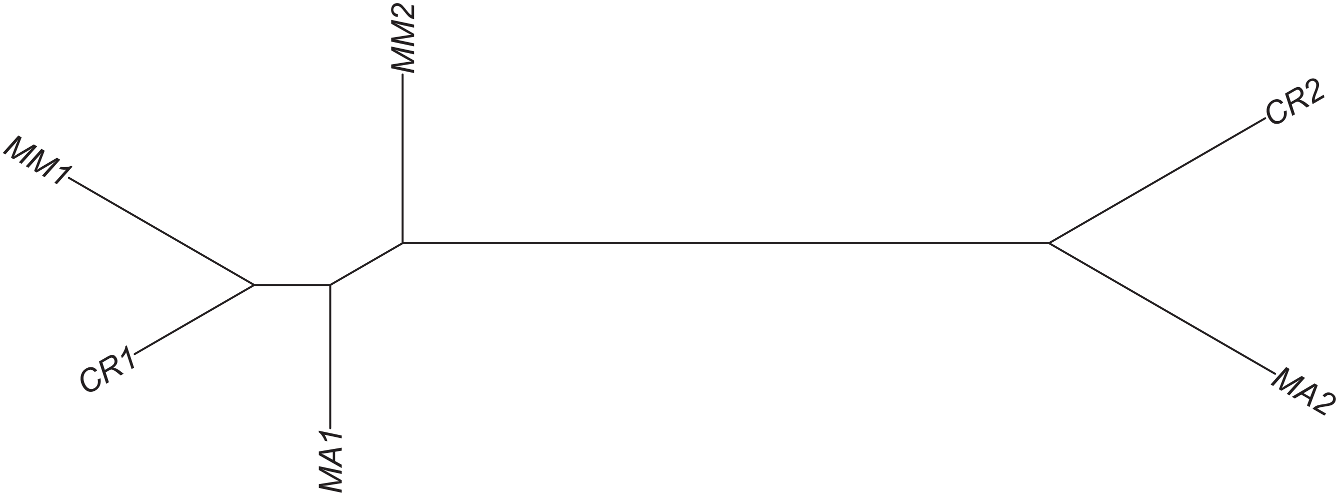

### Region 52

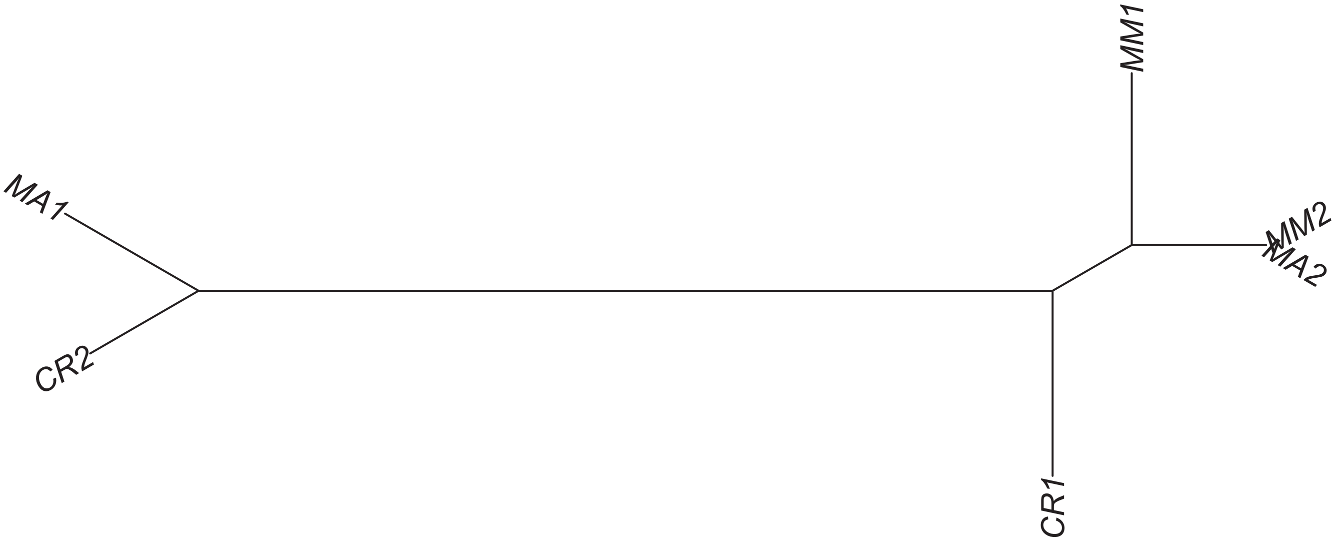

### Region 53

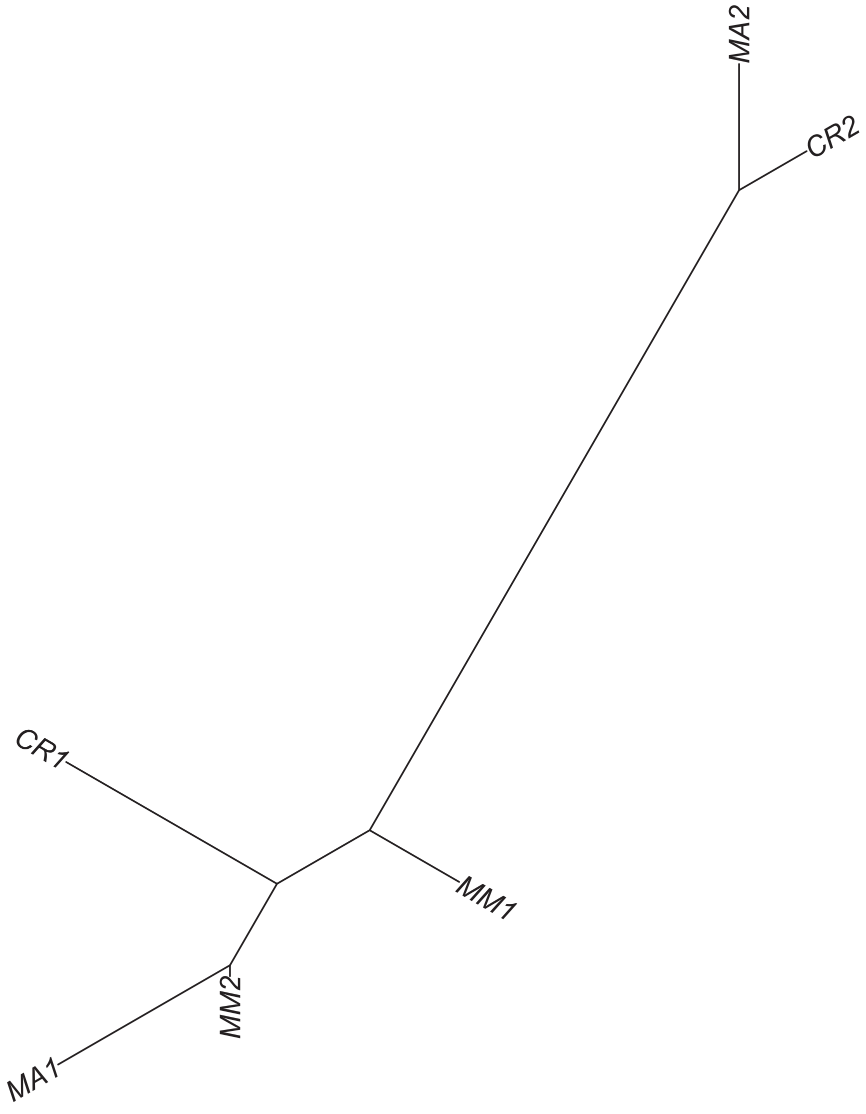

### Region 54

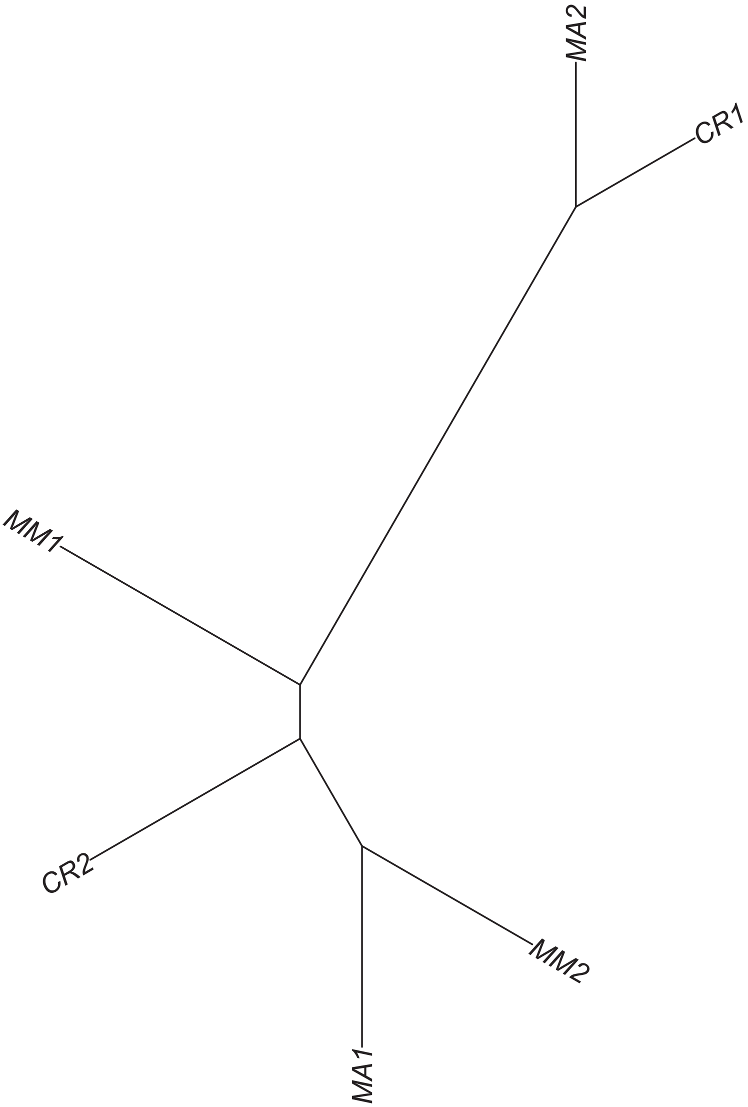

### Region 55



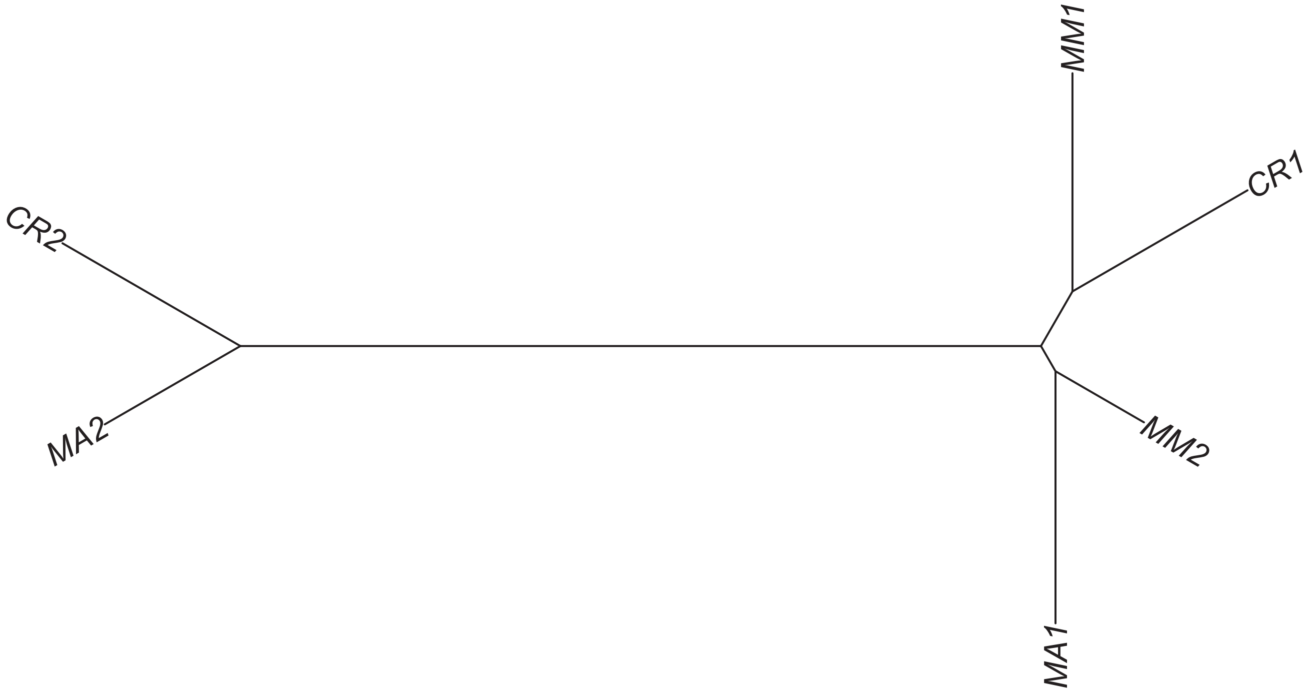

### Region 57

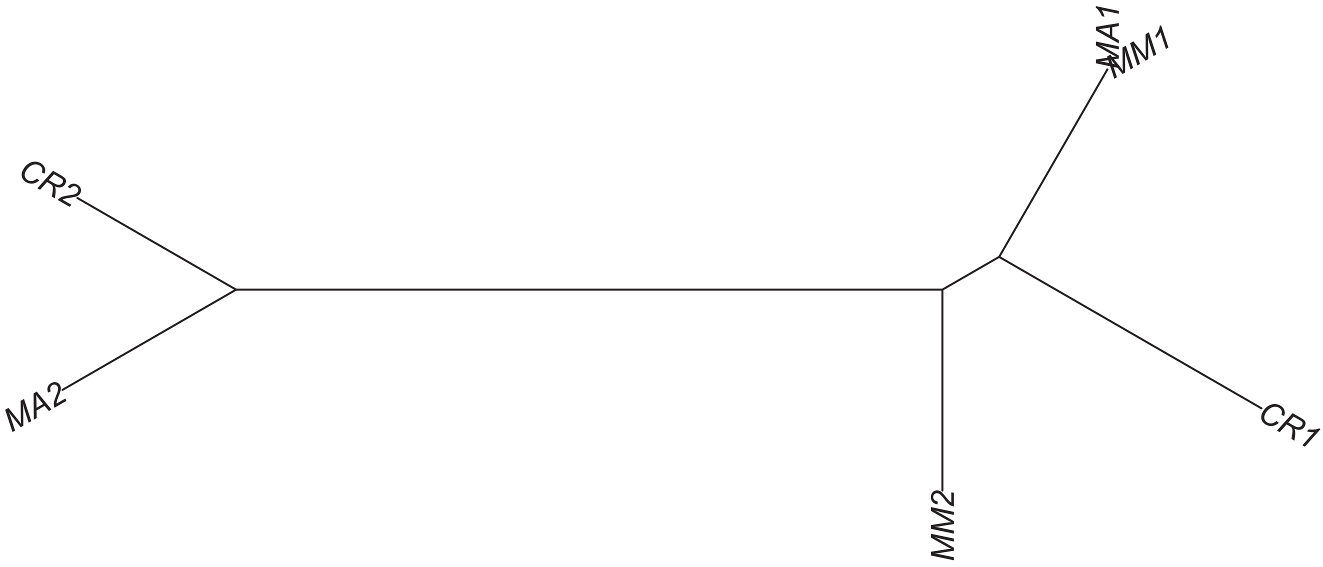

Region 64

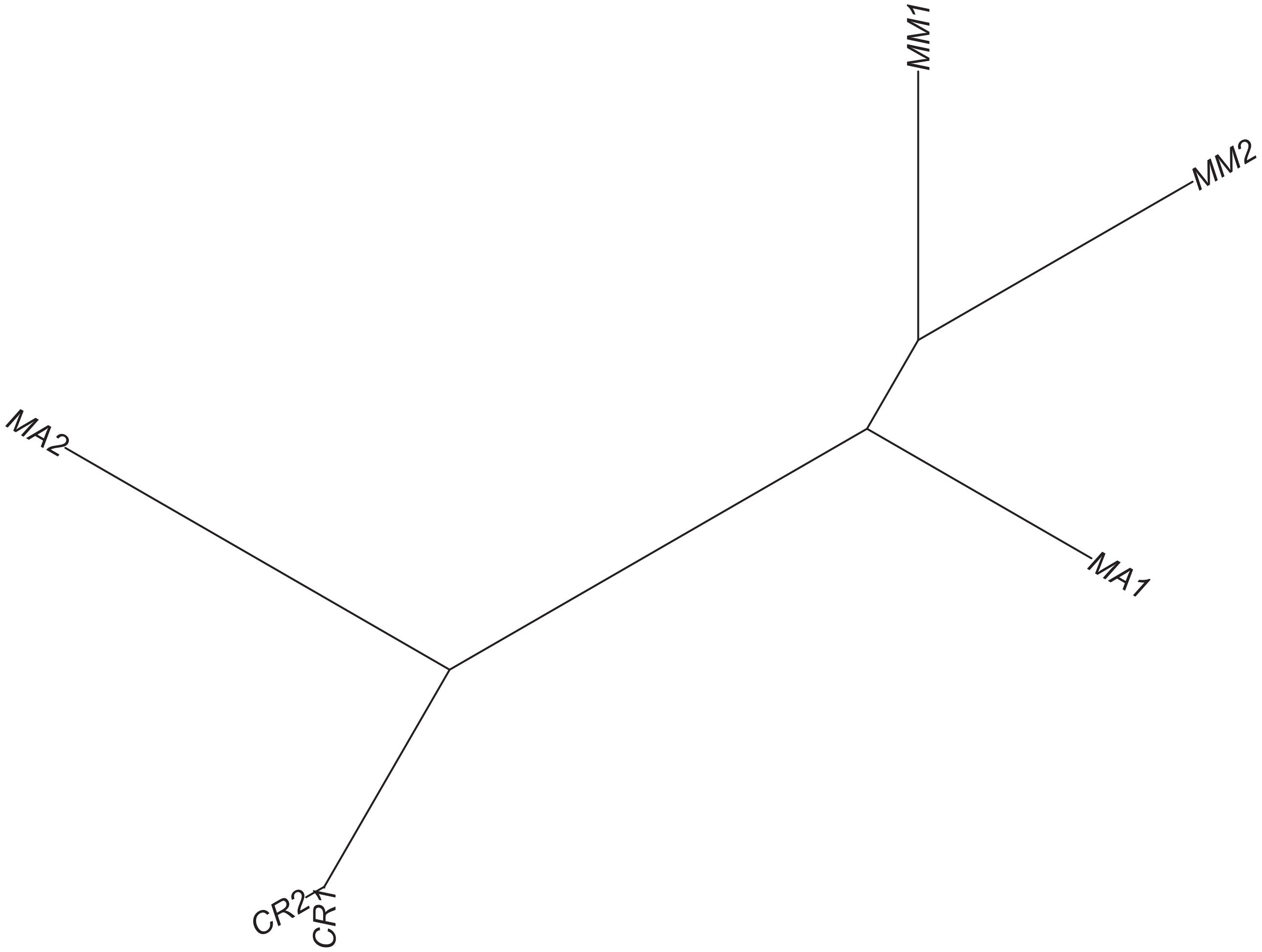

Region 65

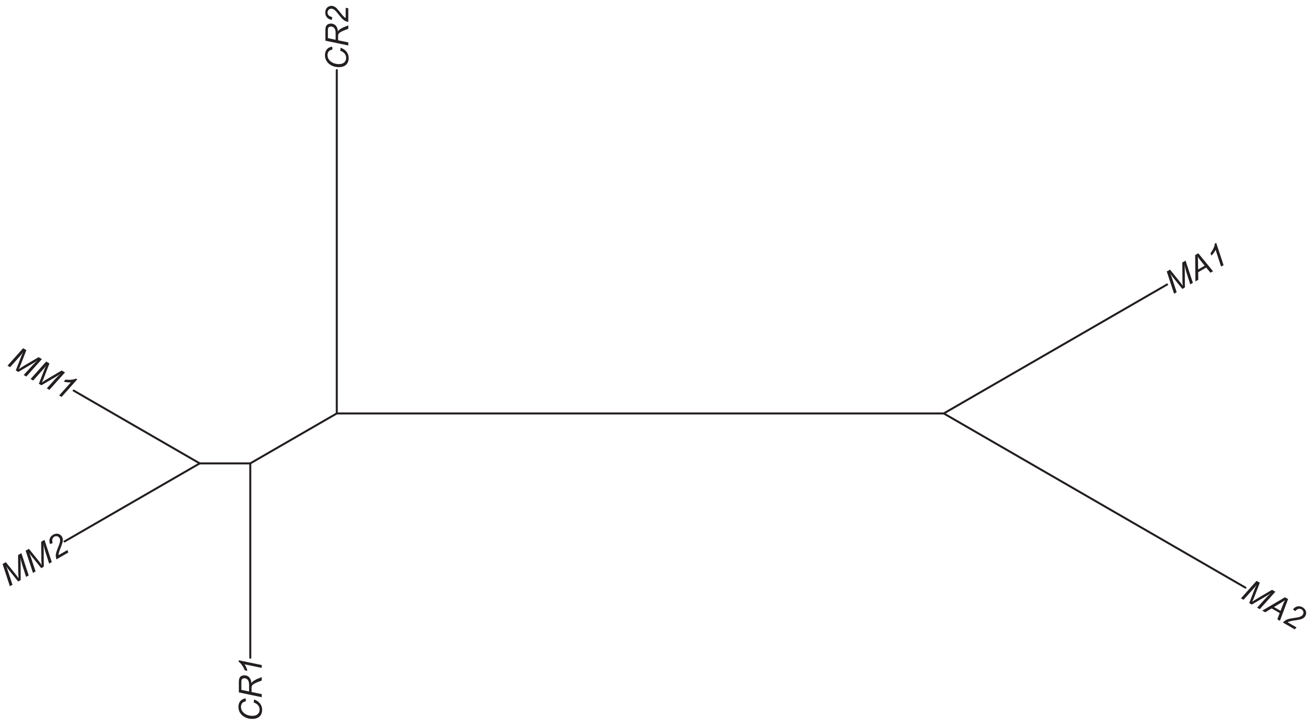

Region 66

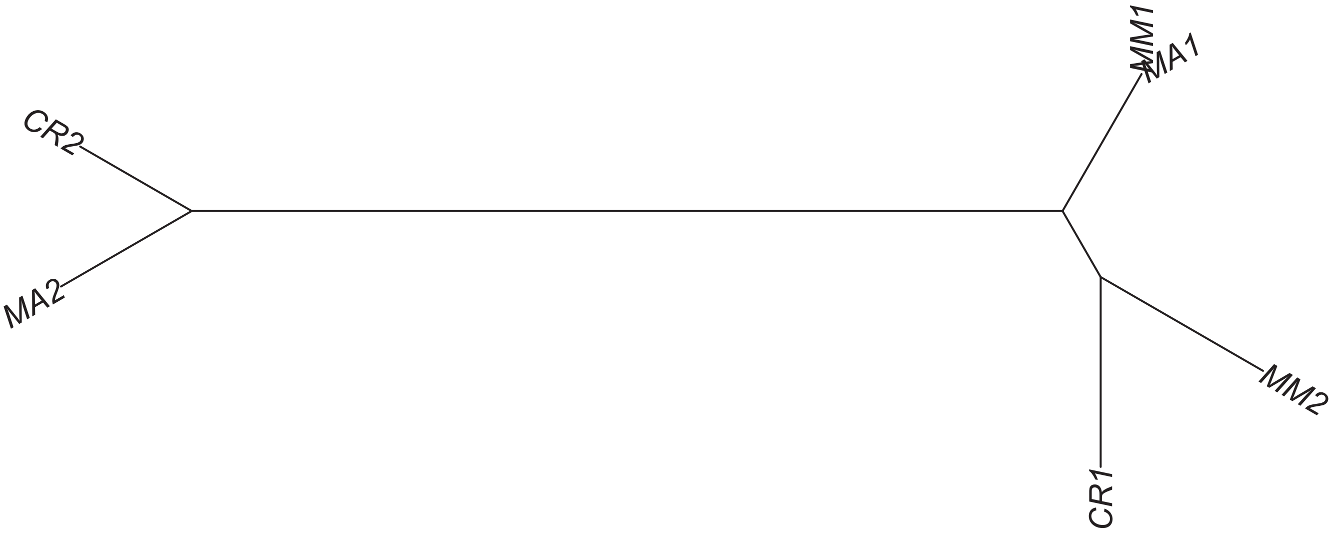

Region 68

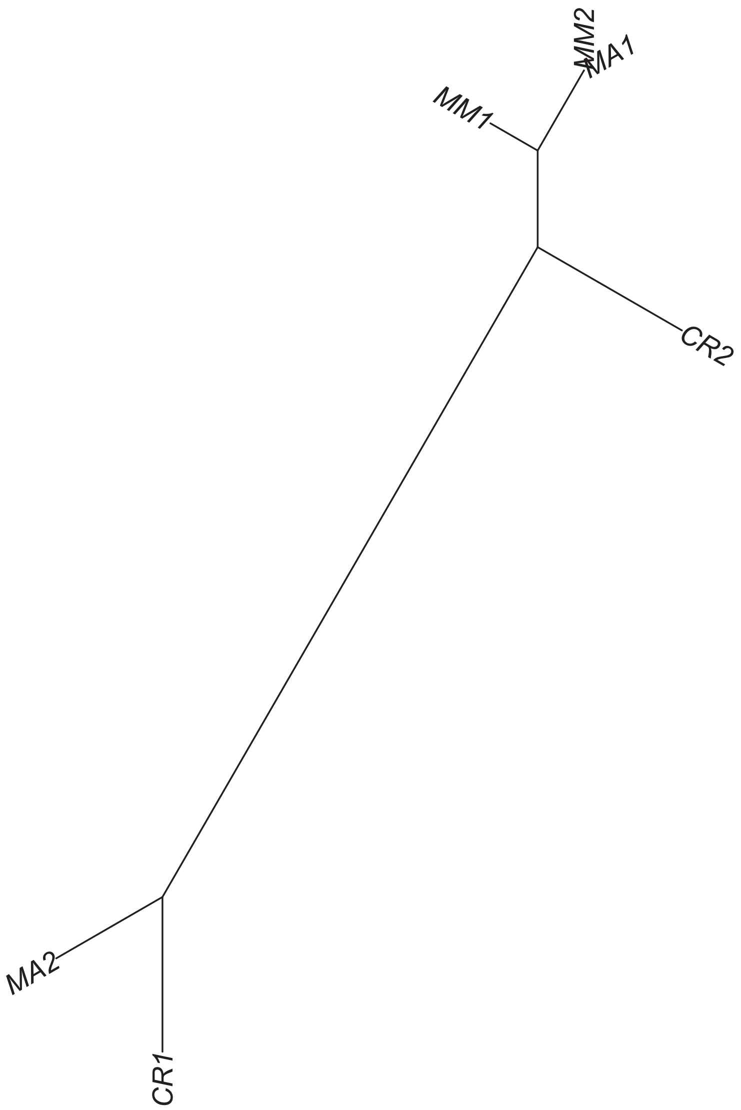

Region 70

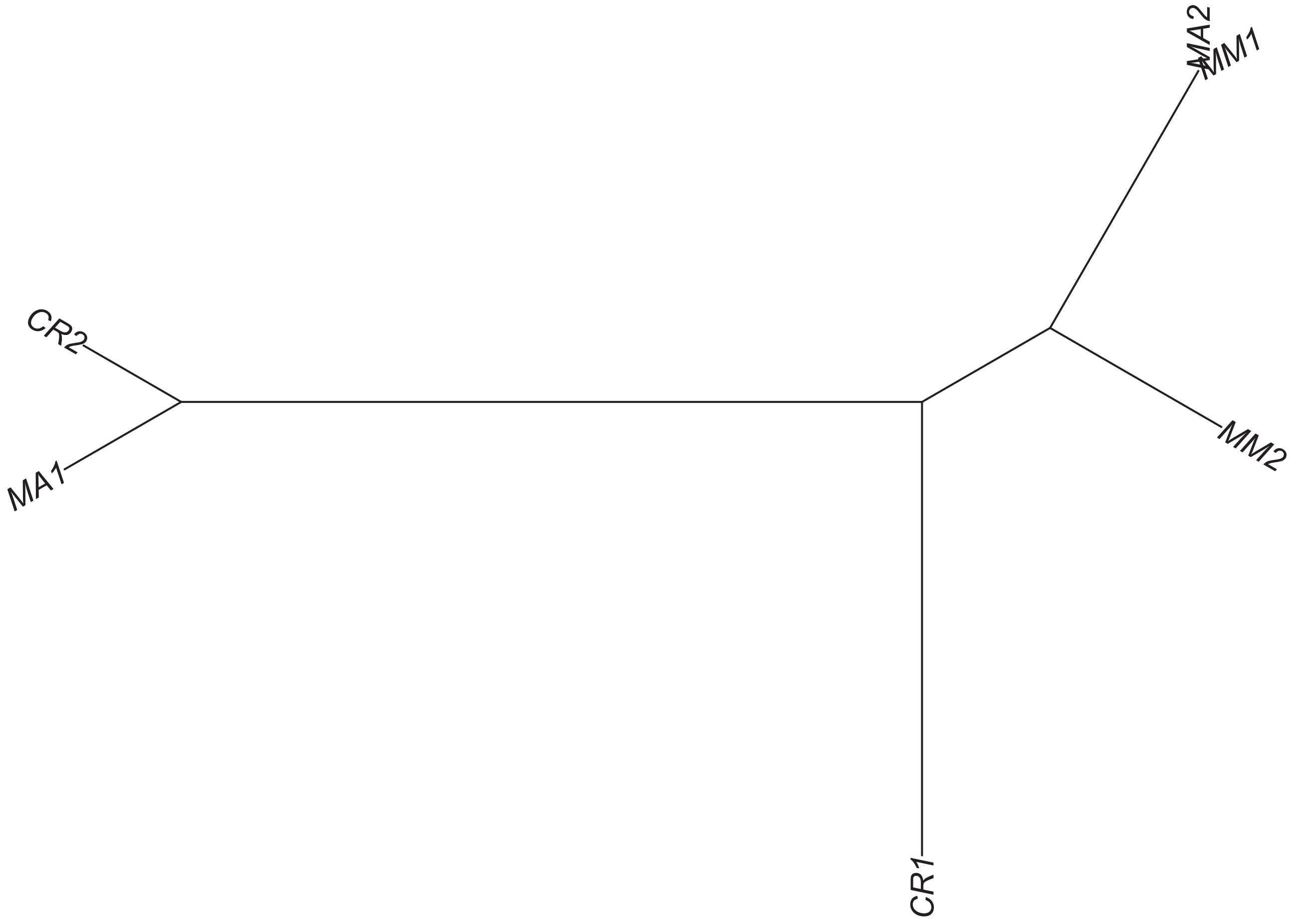

Region 71

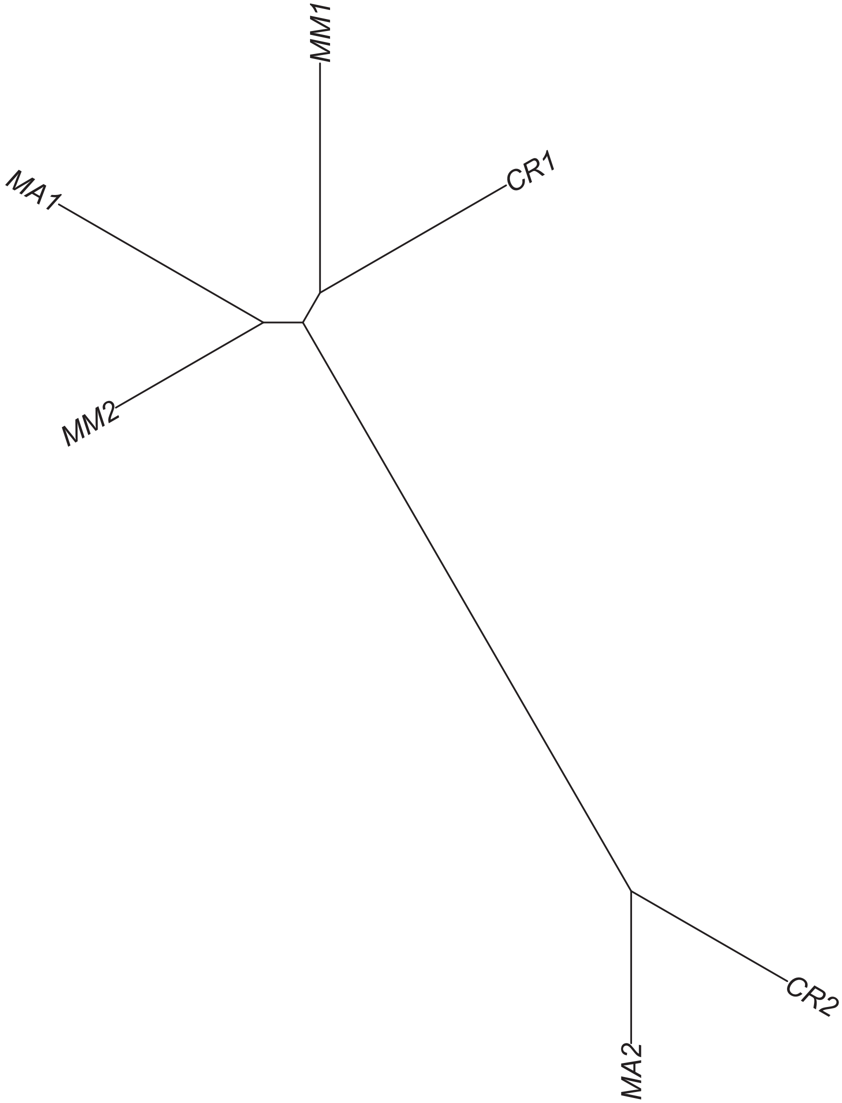

### Region 72

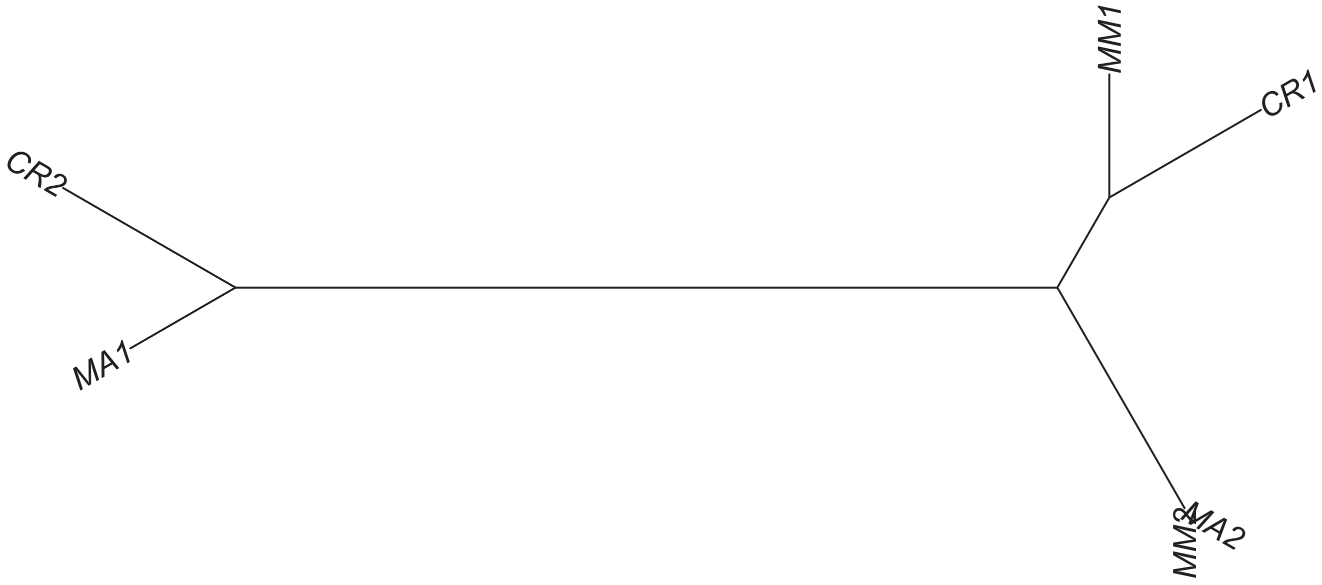

Region 73

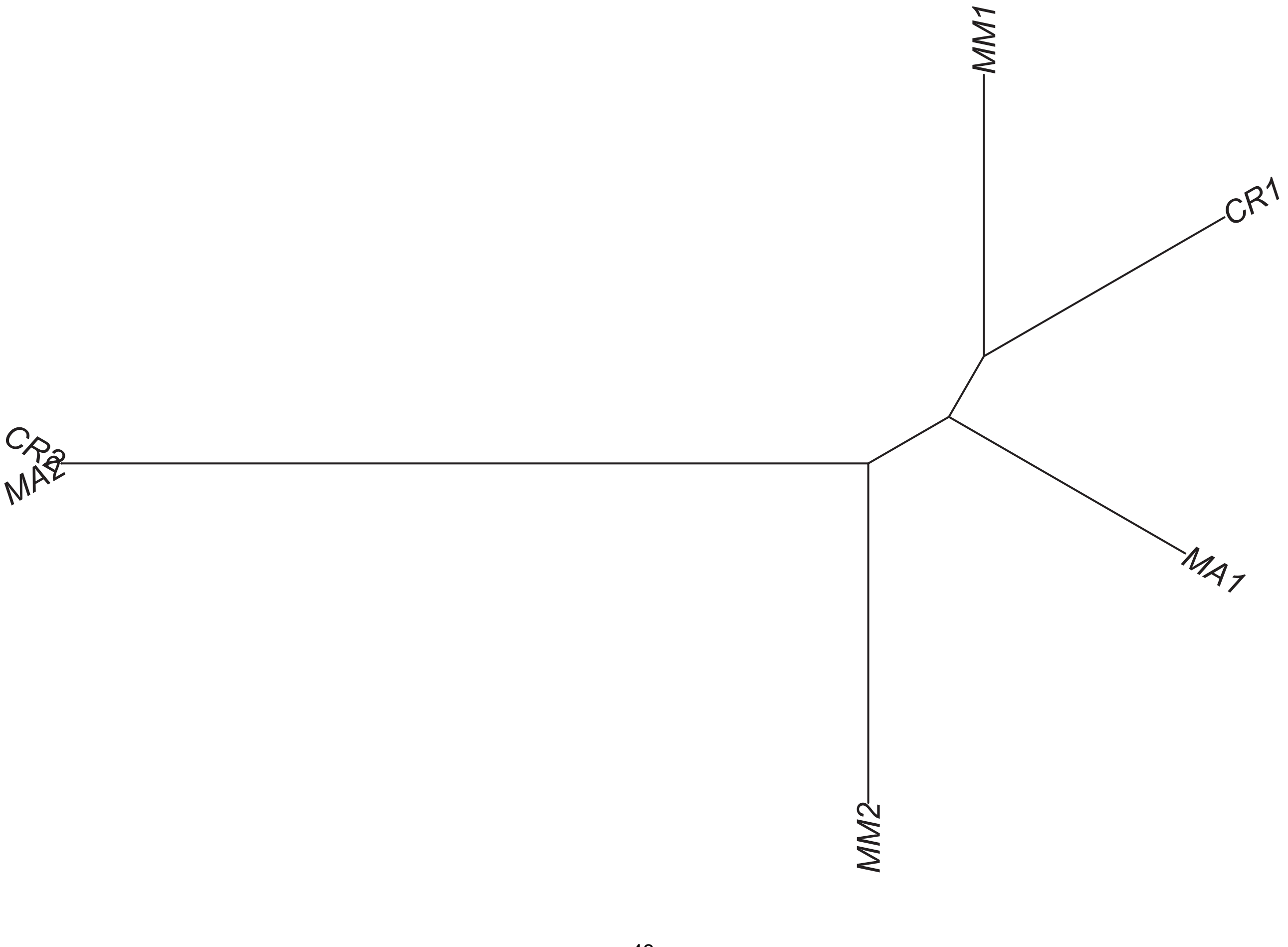

Region 75

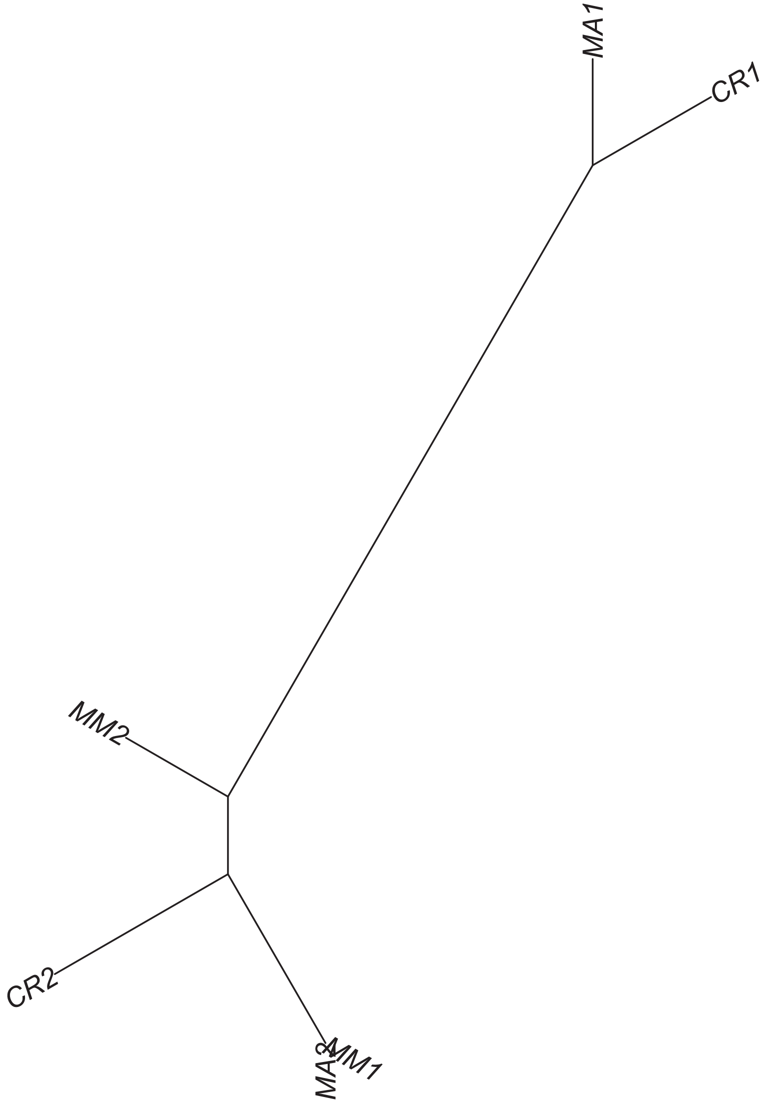

Region 76

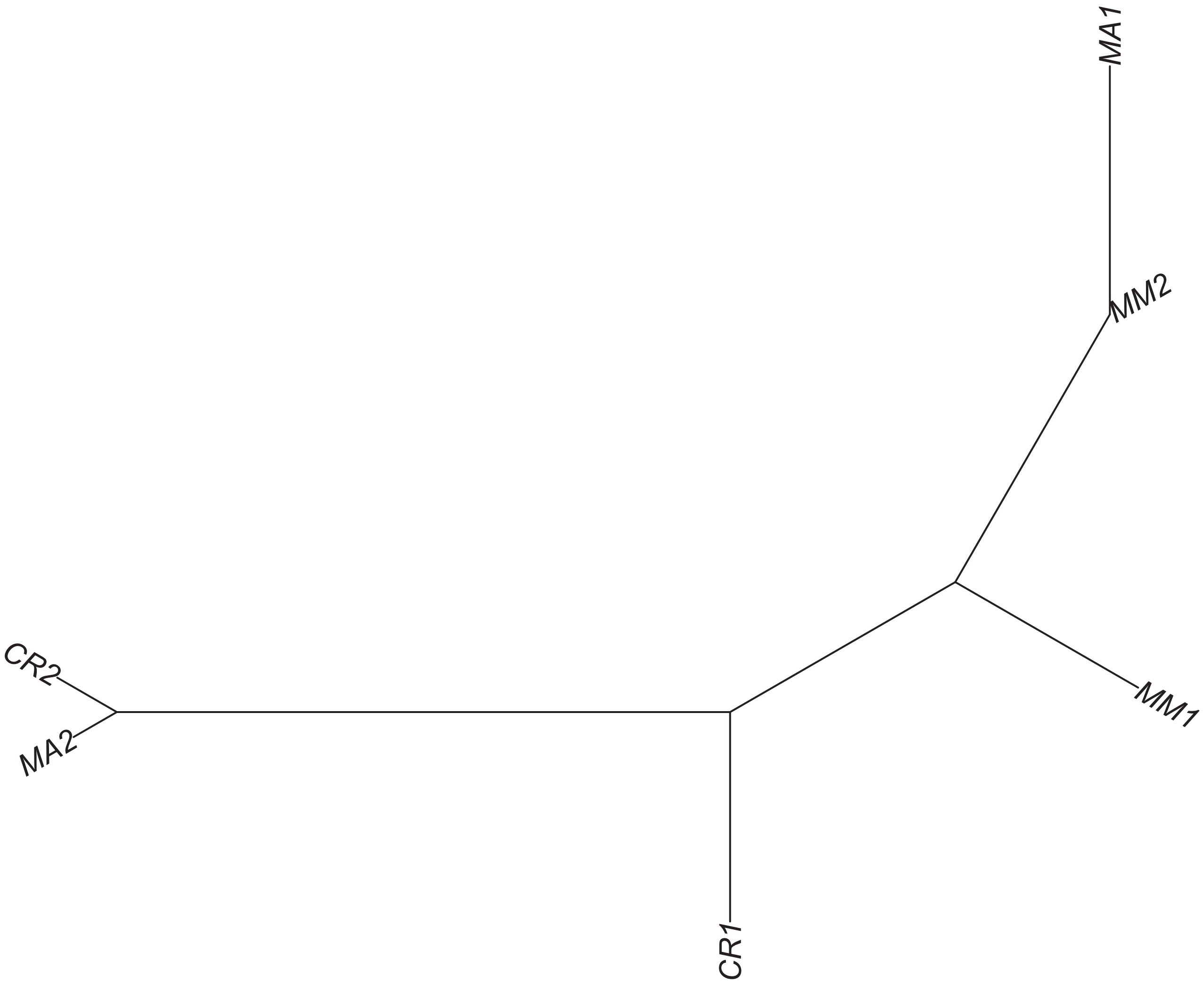

### Region 77



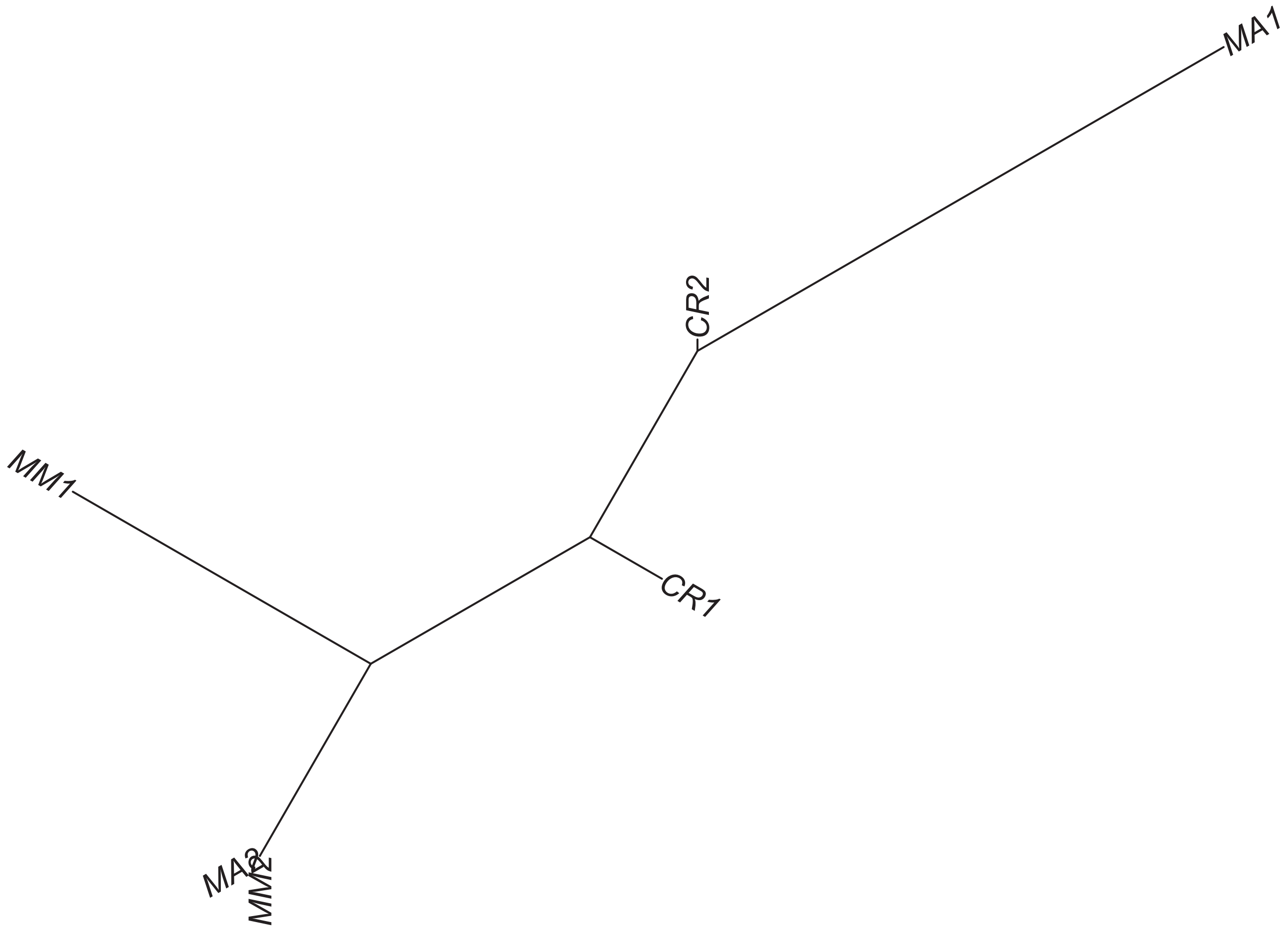

### Region 78

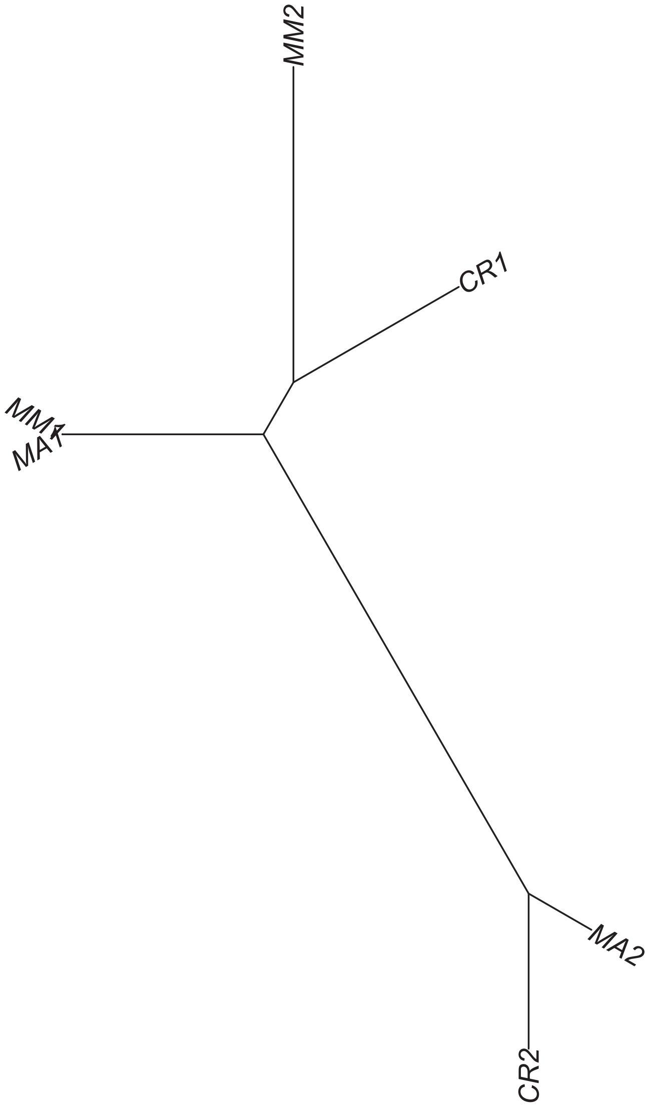

Region 79

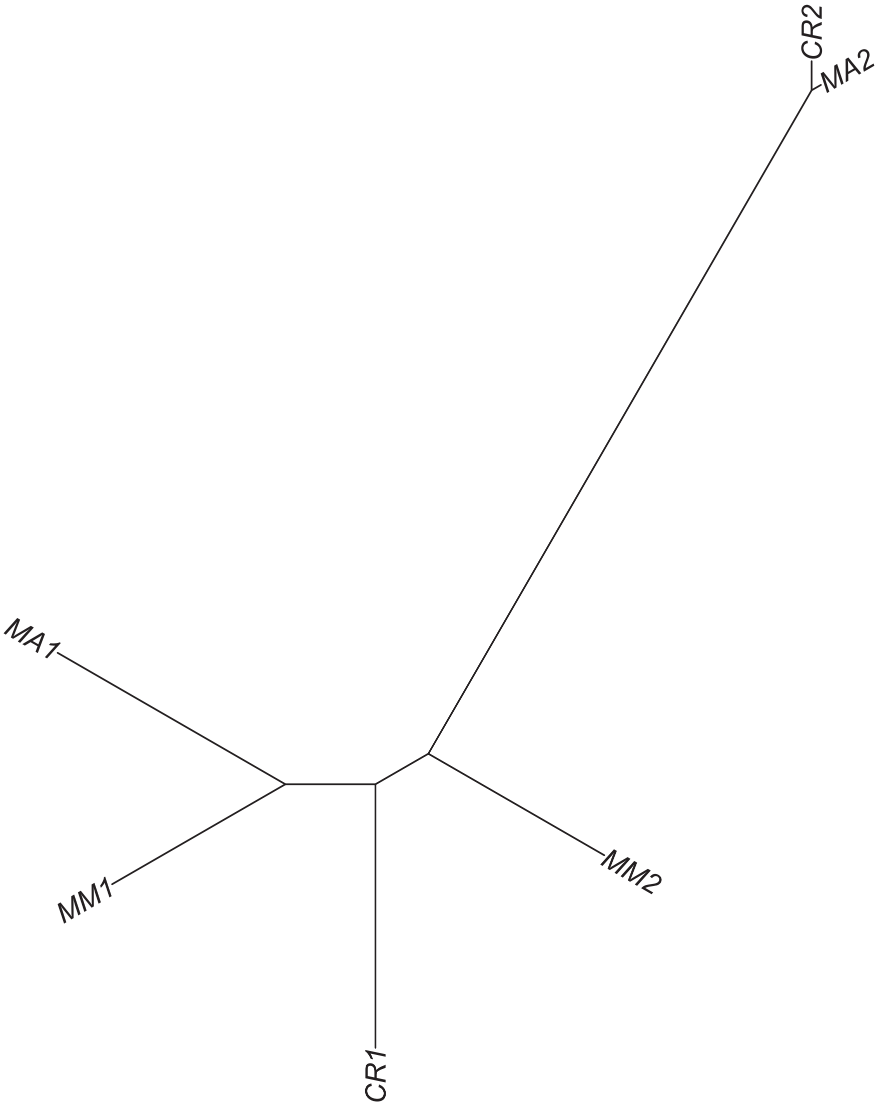

Region 81

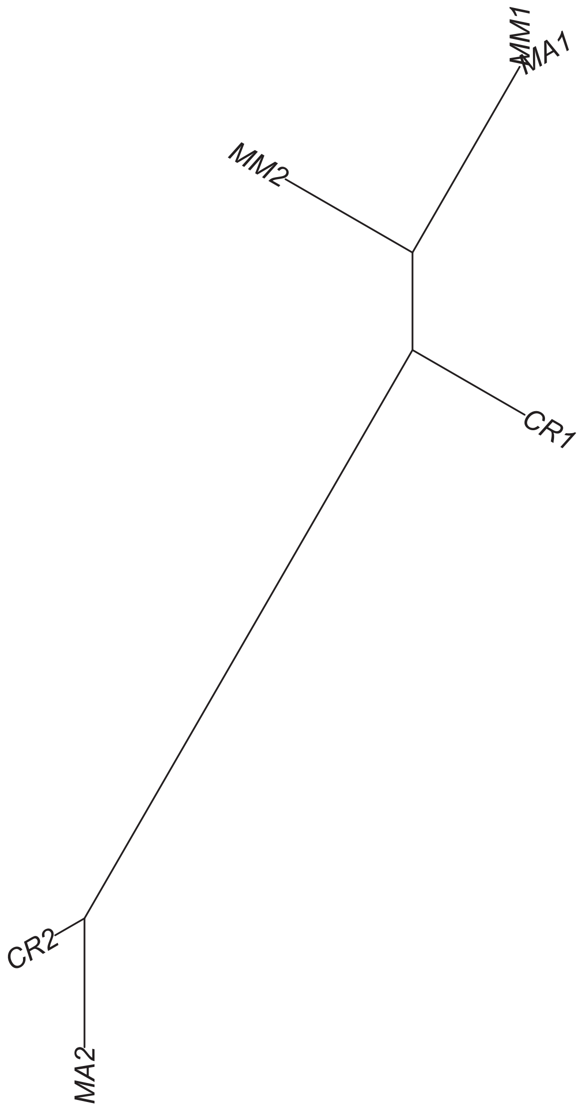

### Region 82

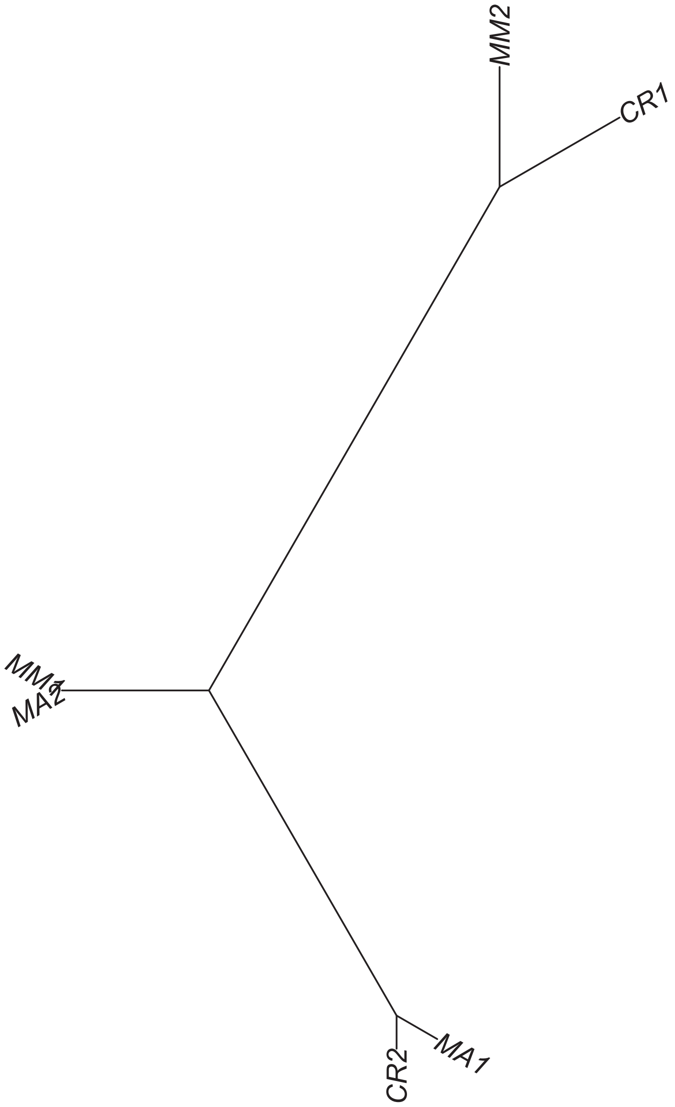

### Region 86



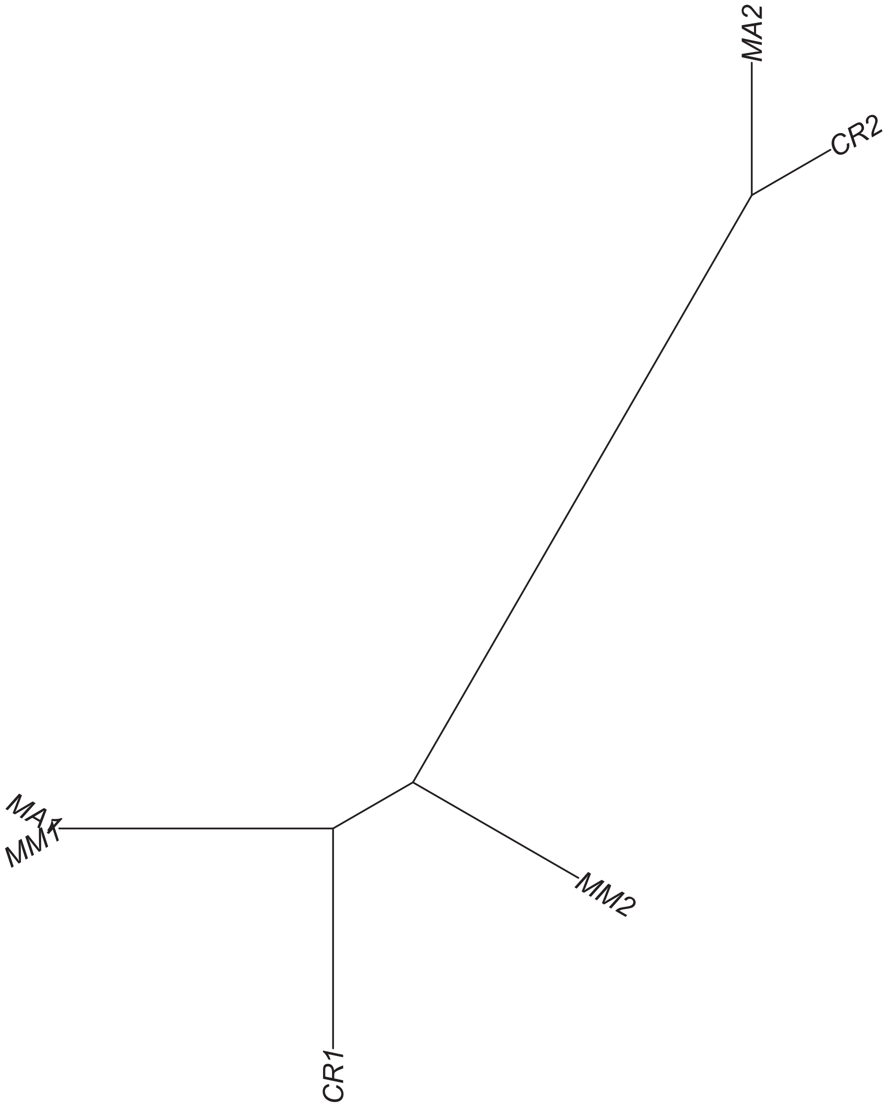

Region 87

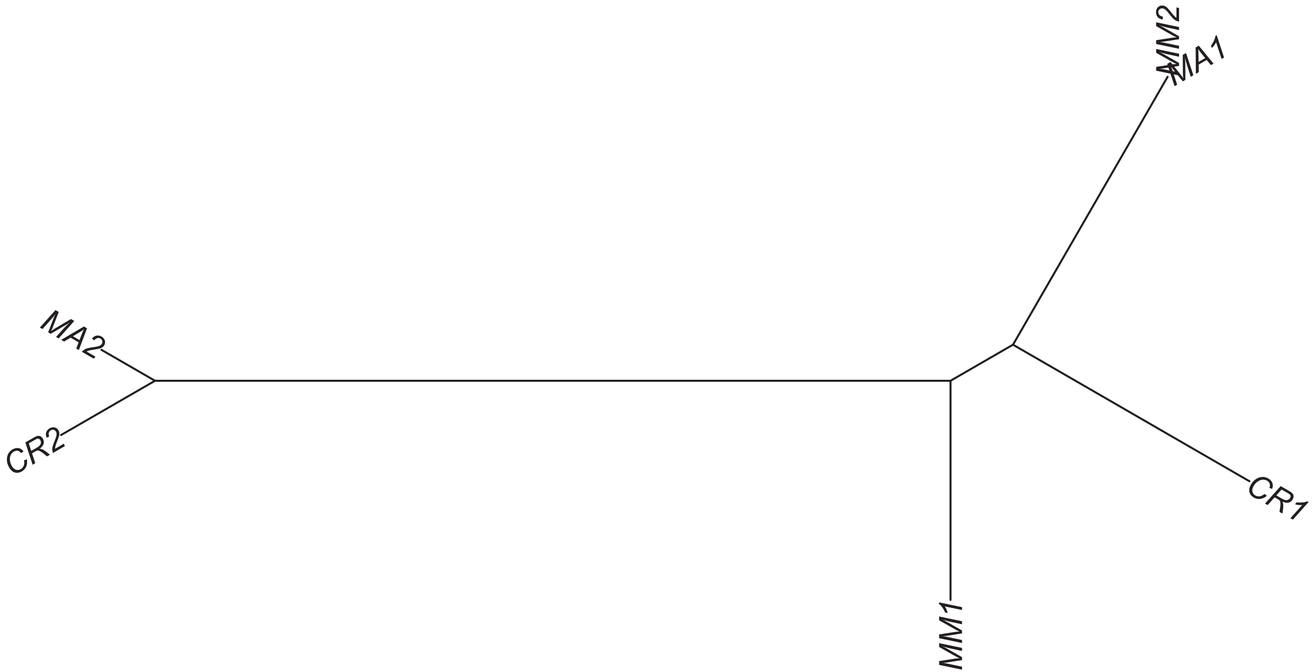

### Region 88

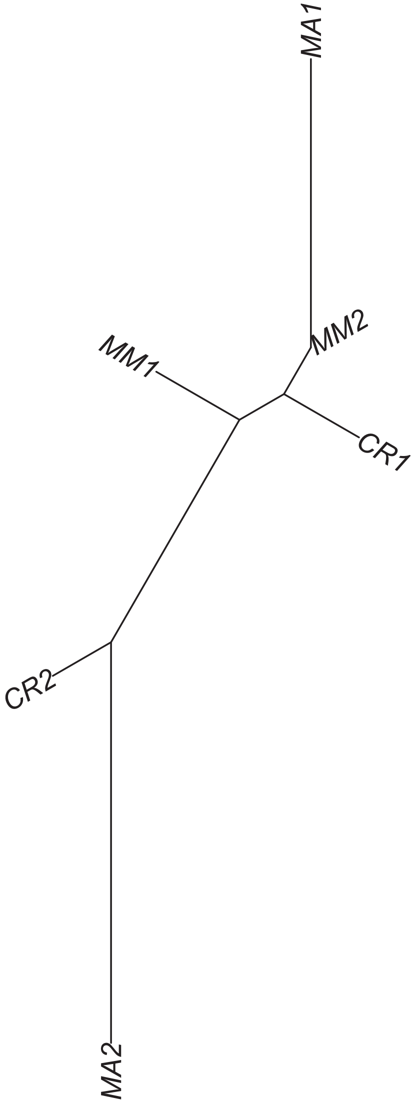

### Region 90

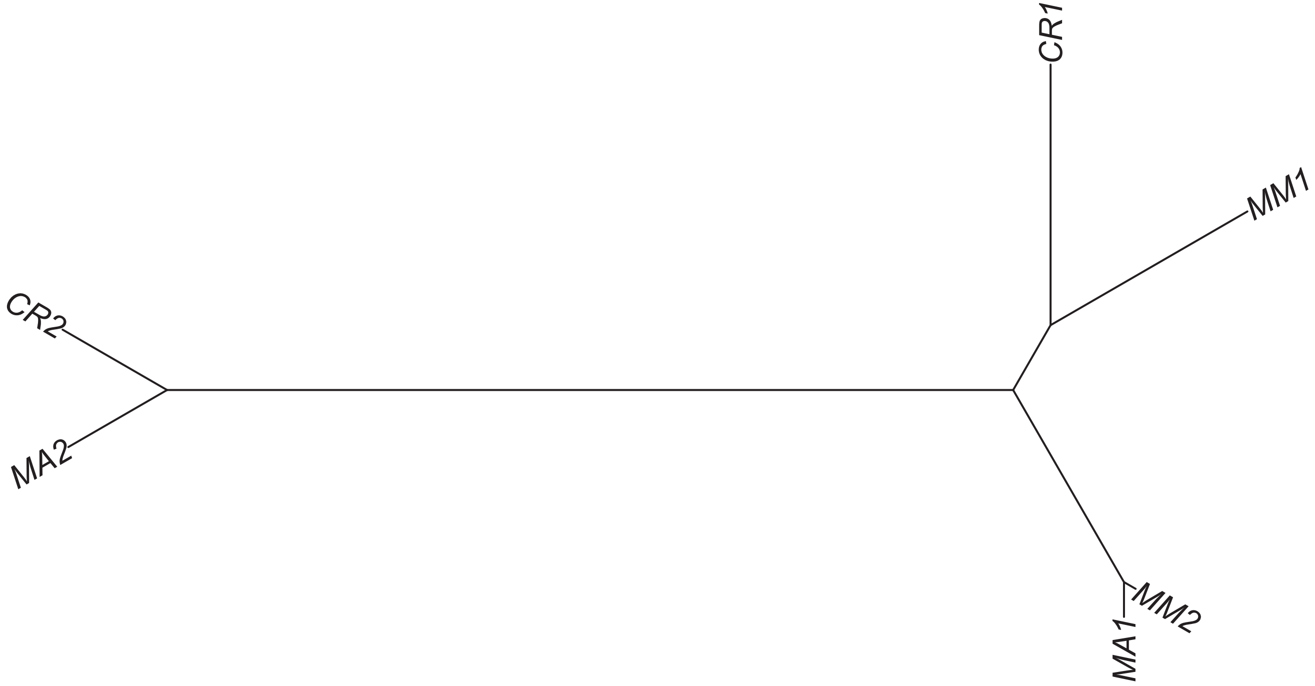

### Region 91

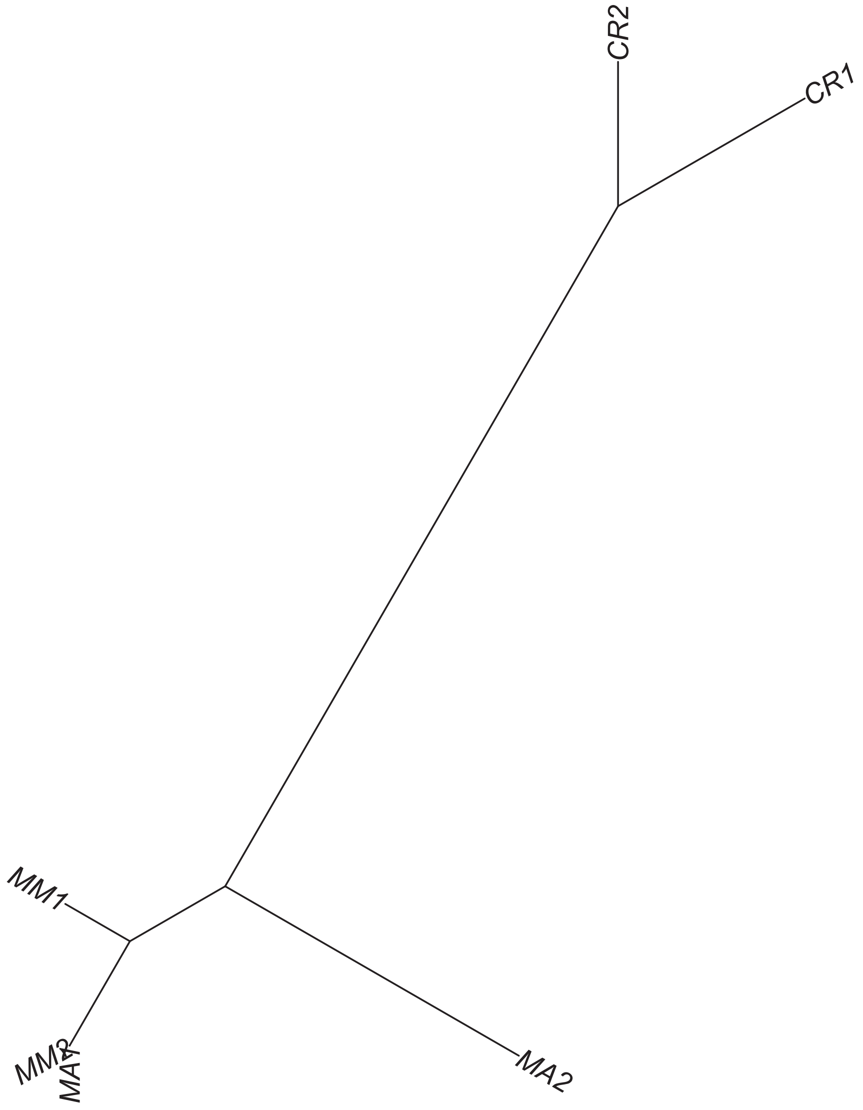

### Region 92

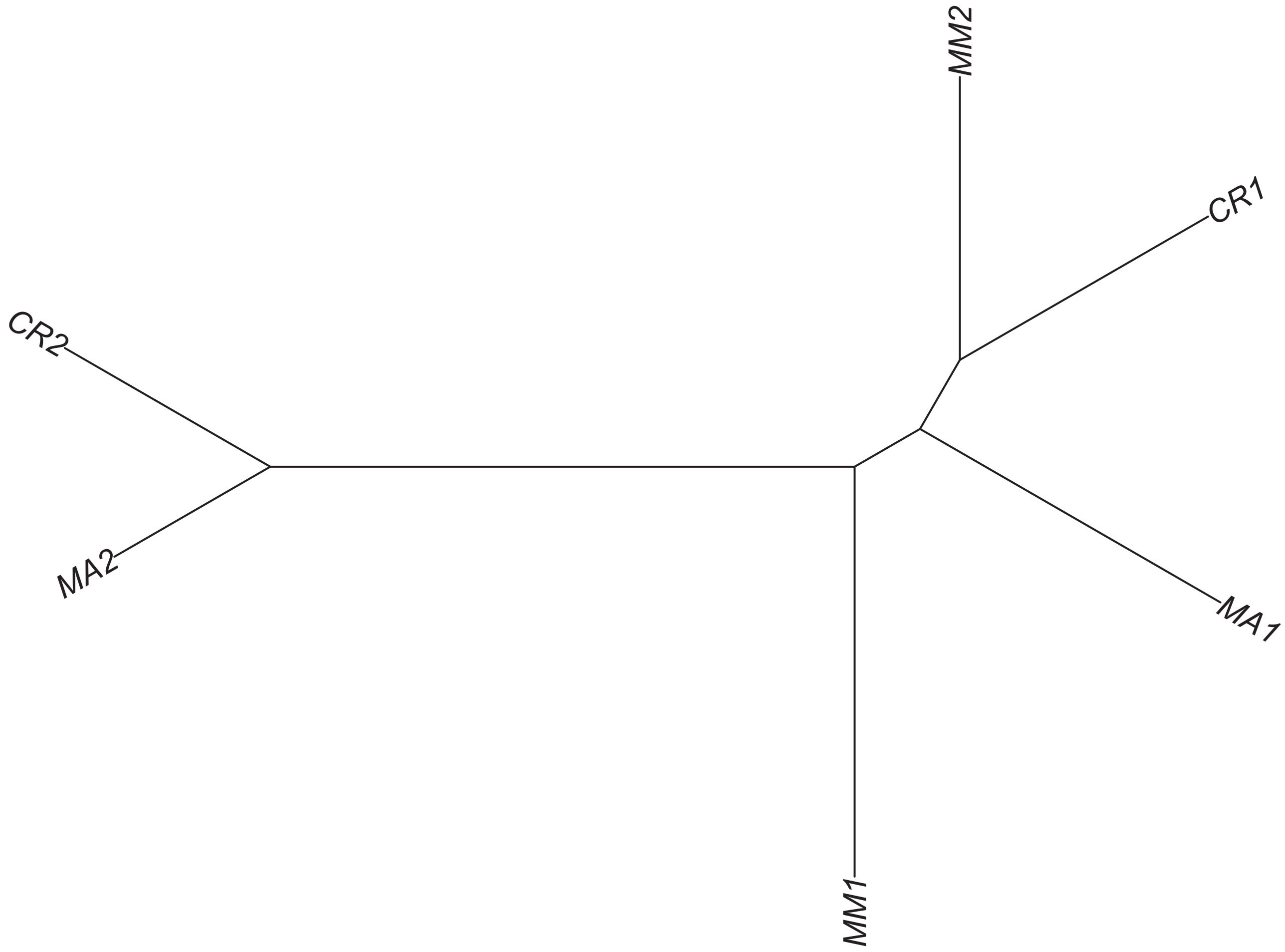

#### Region 94

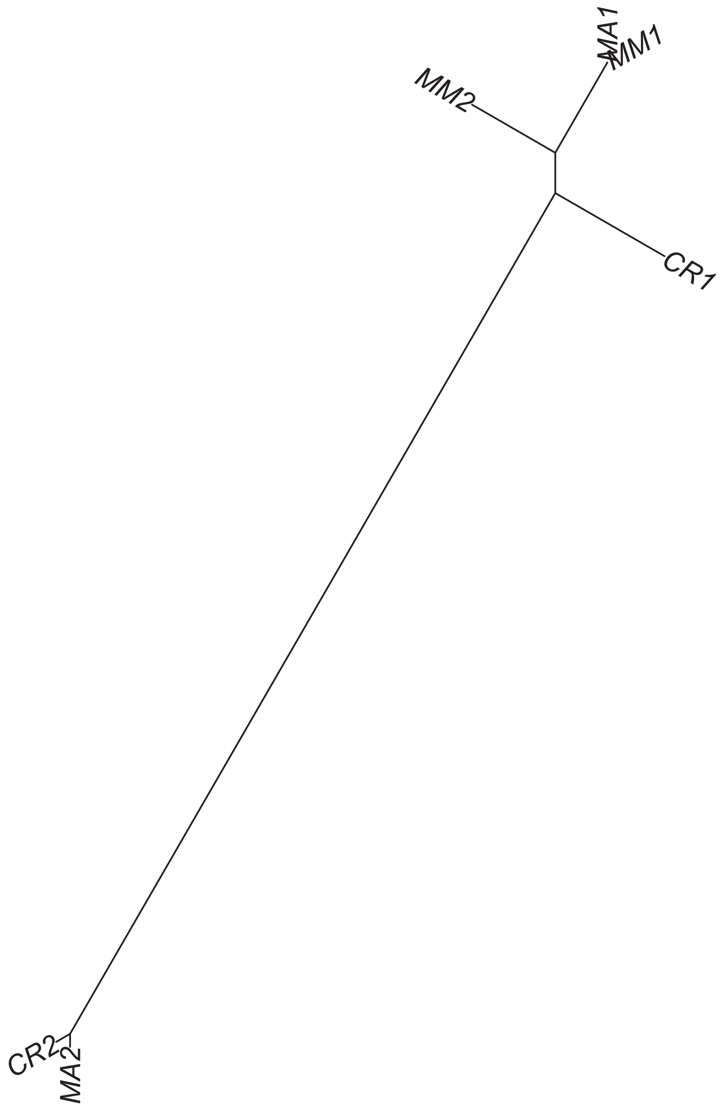

Region 96

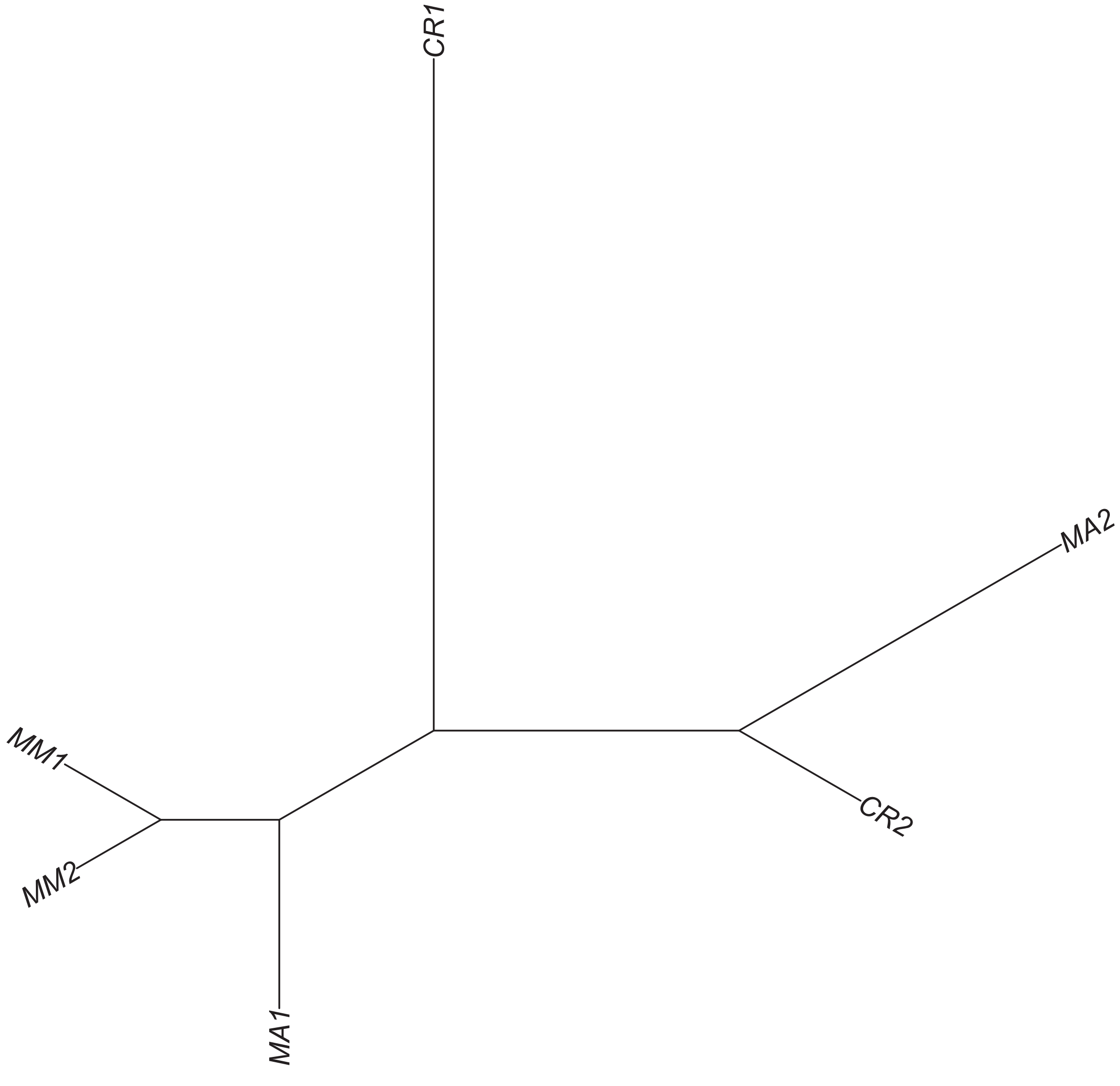

Region 97

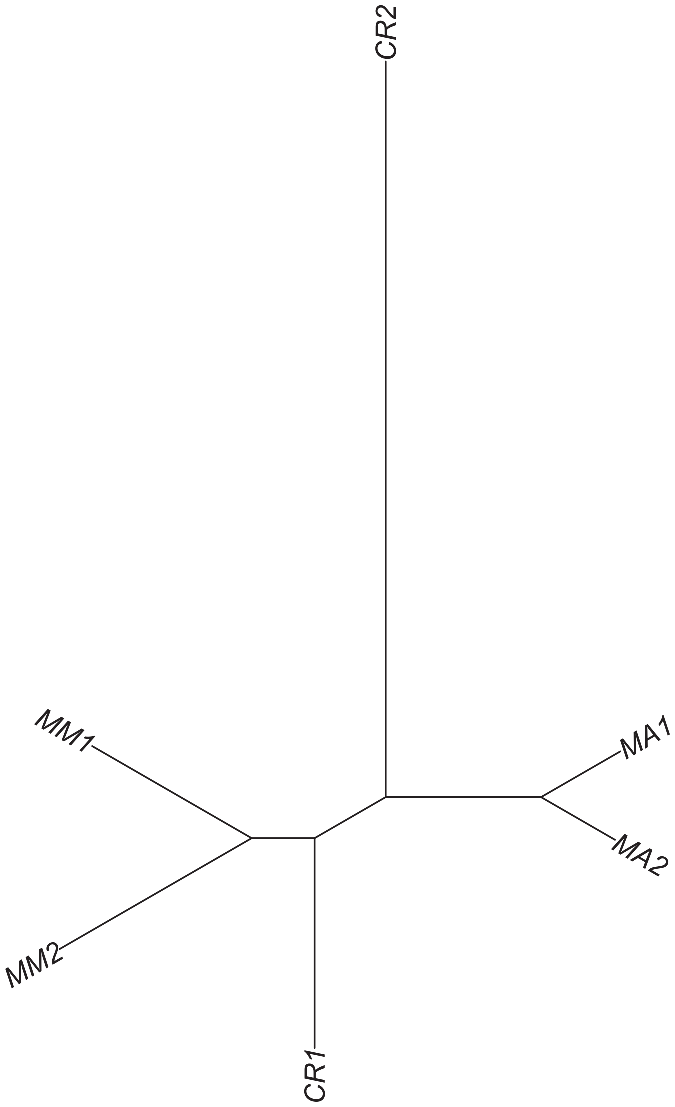

### Region 98

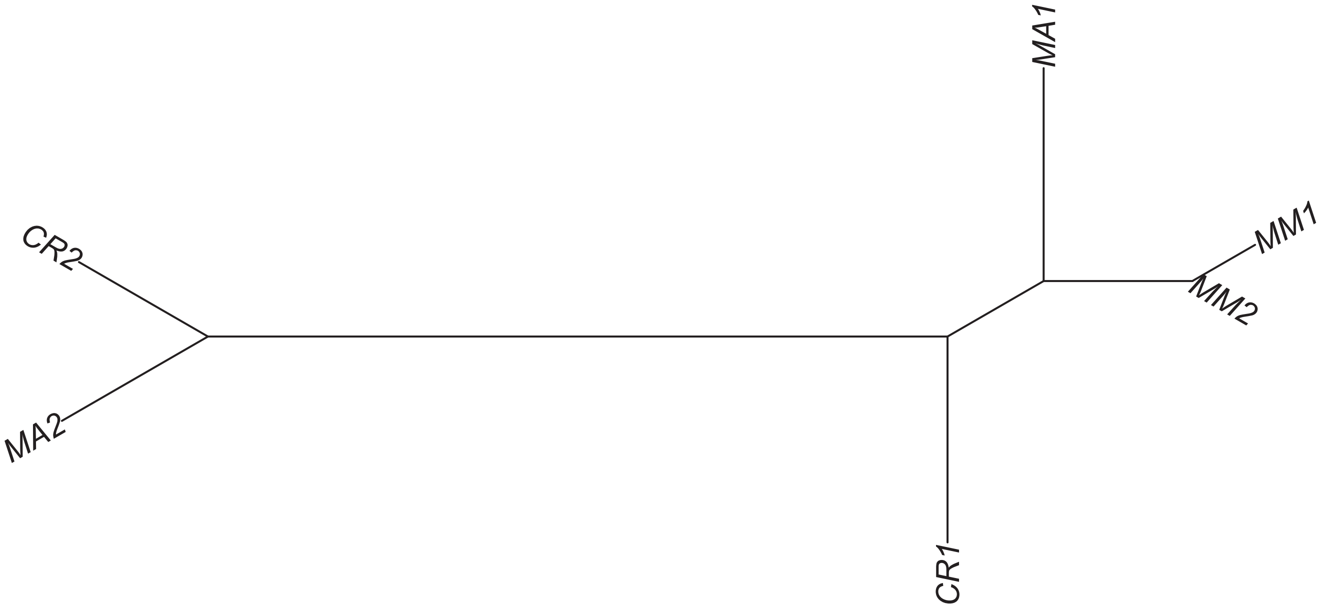

### Region 99



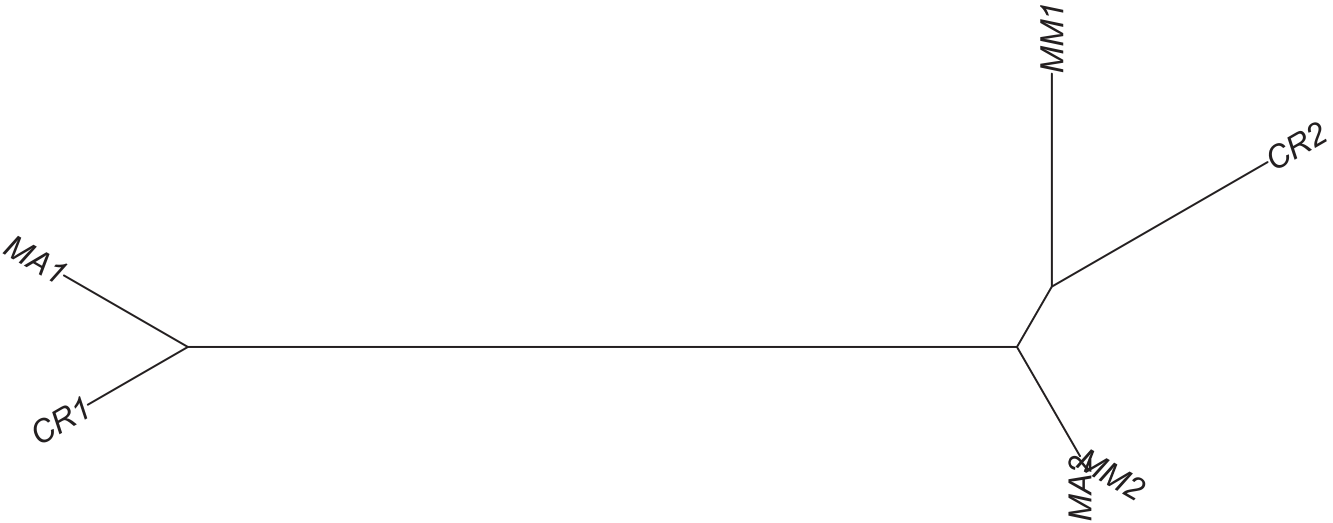

### Region 101

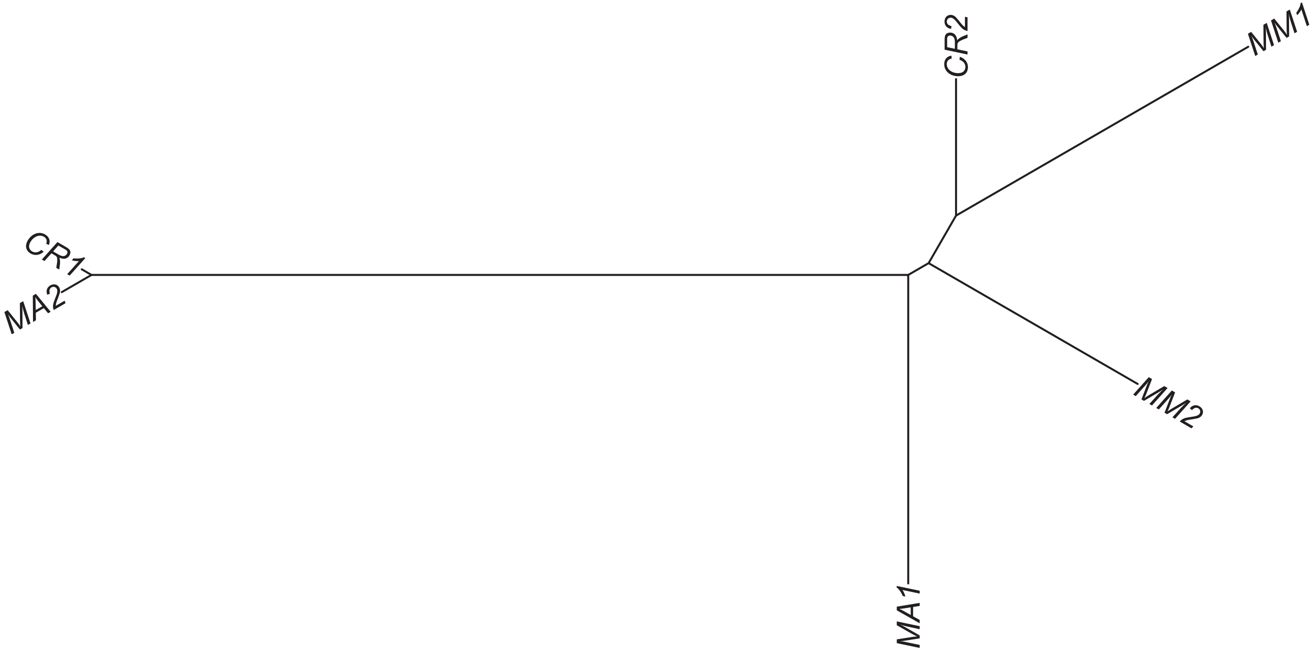

Region 102



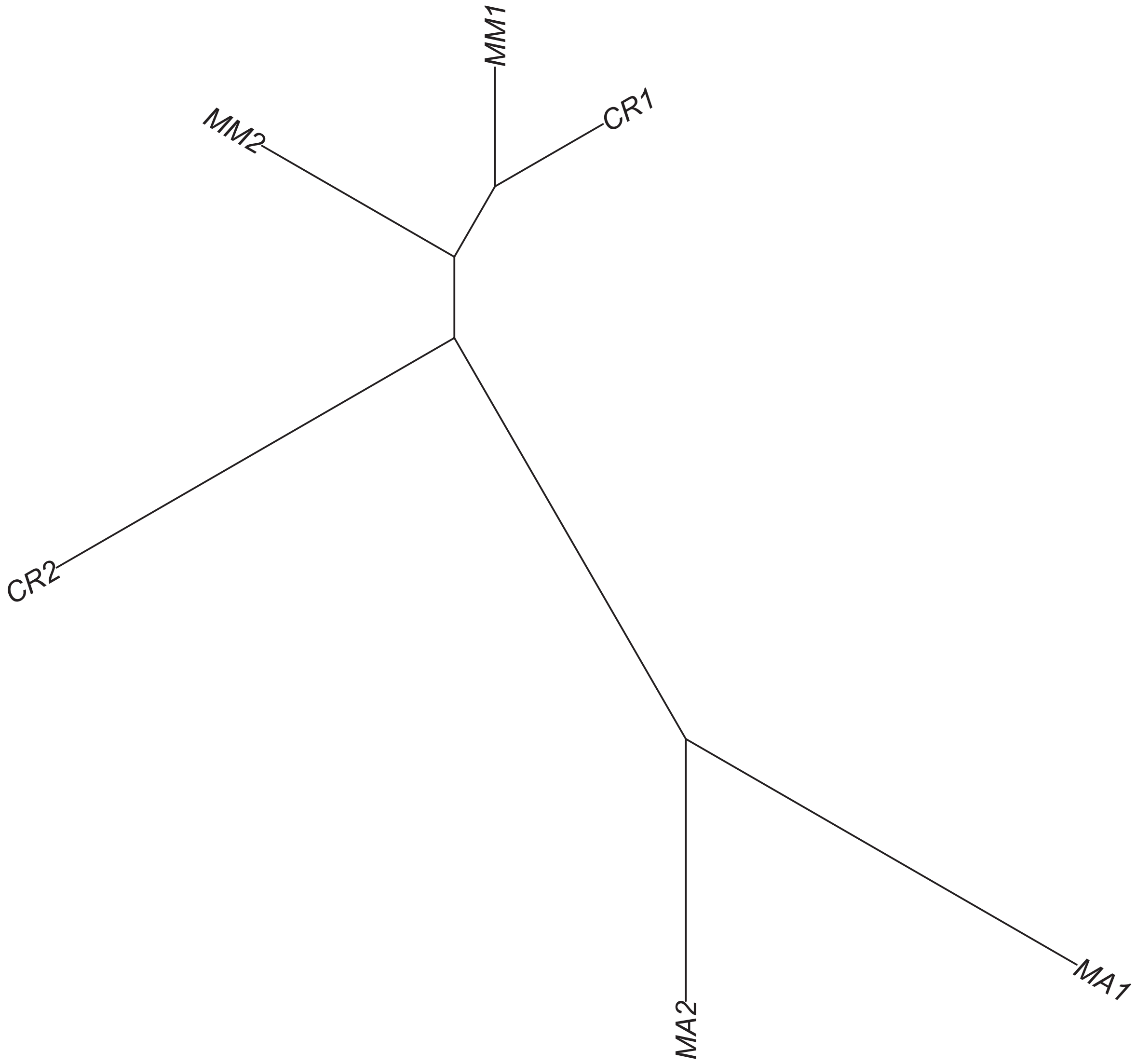

Region 103

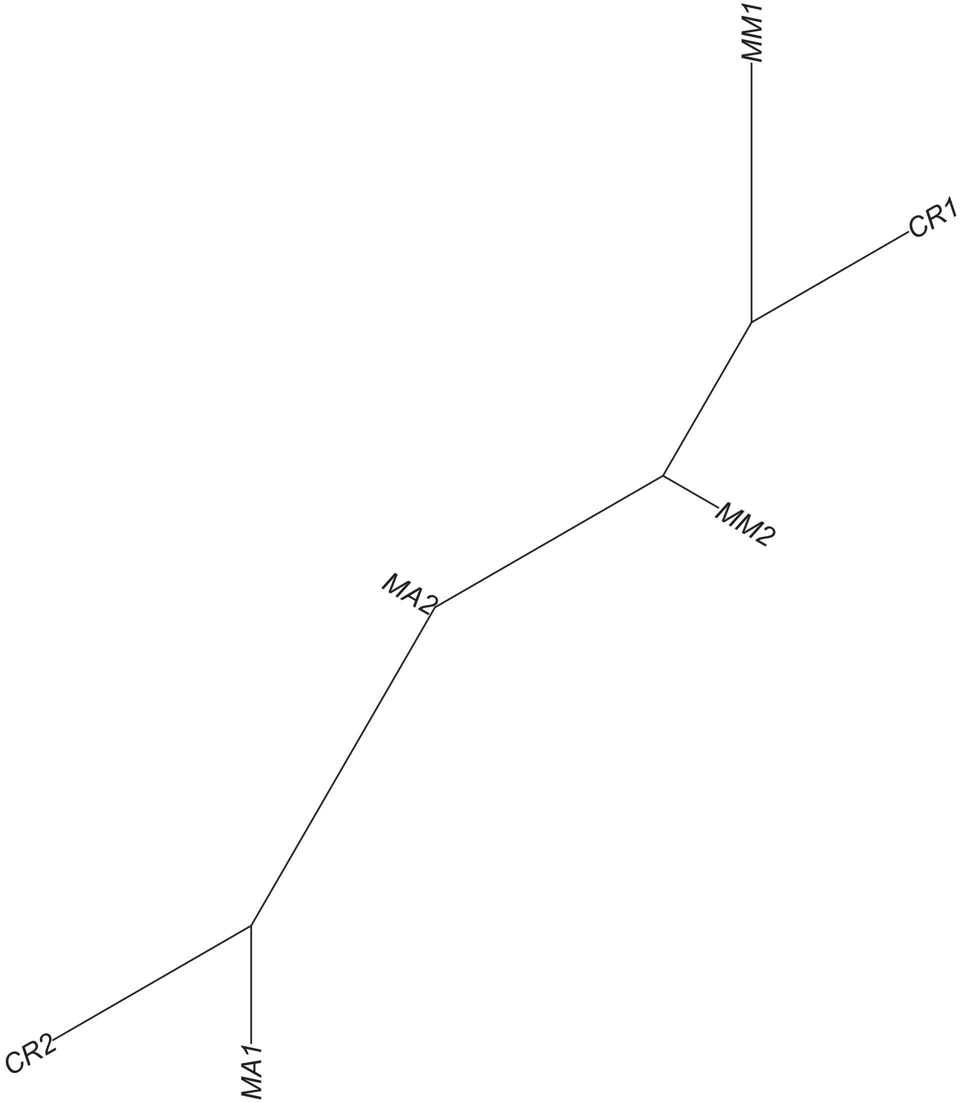

### Region 104



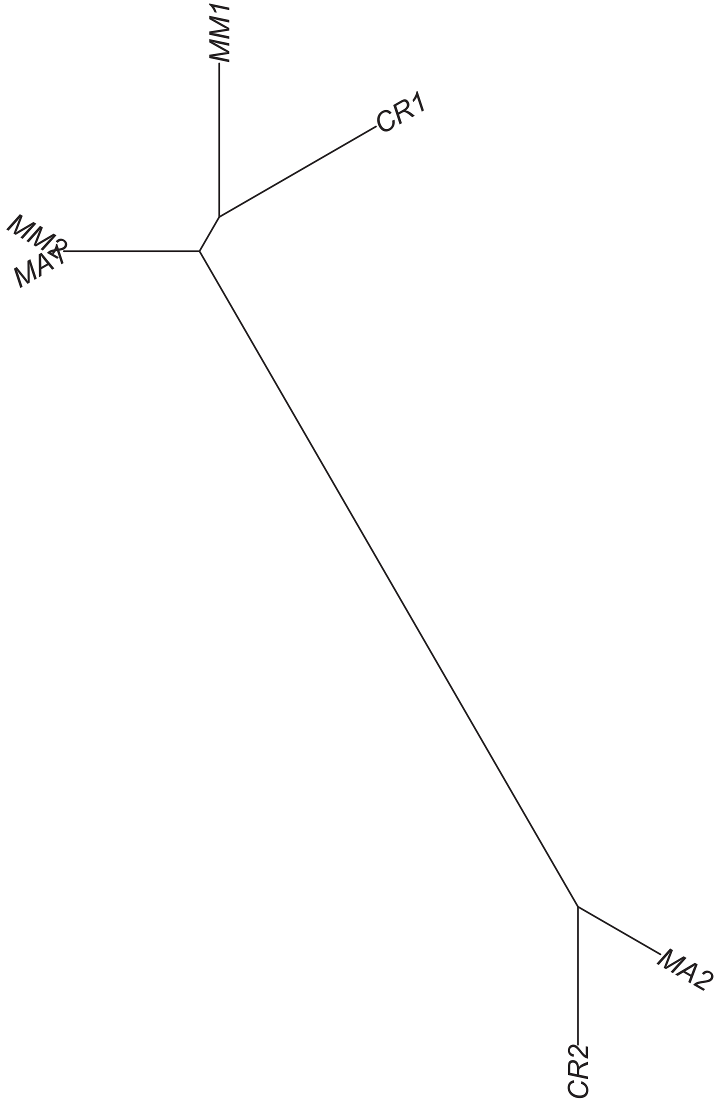

Region 105

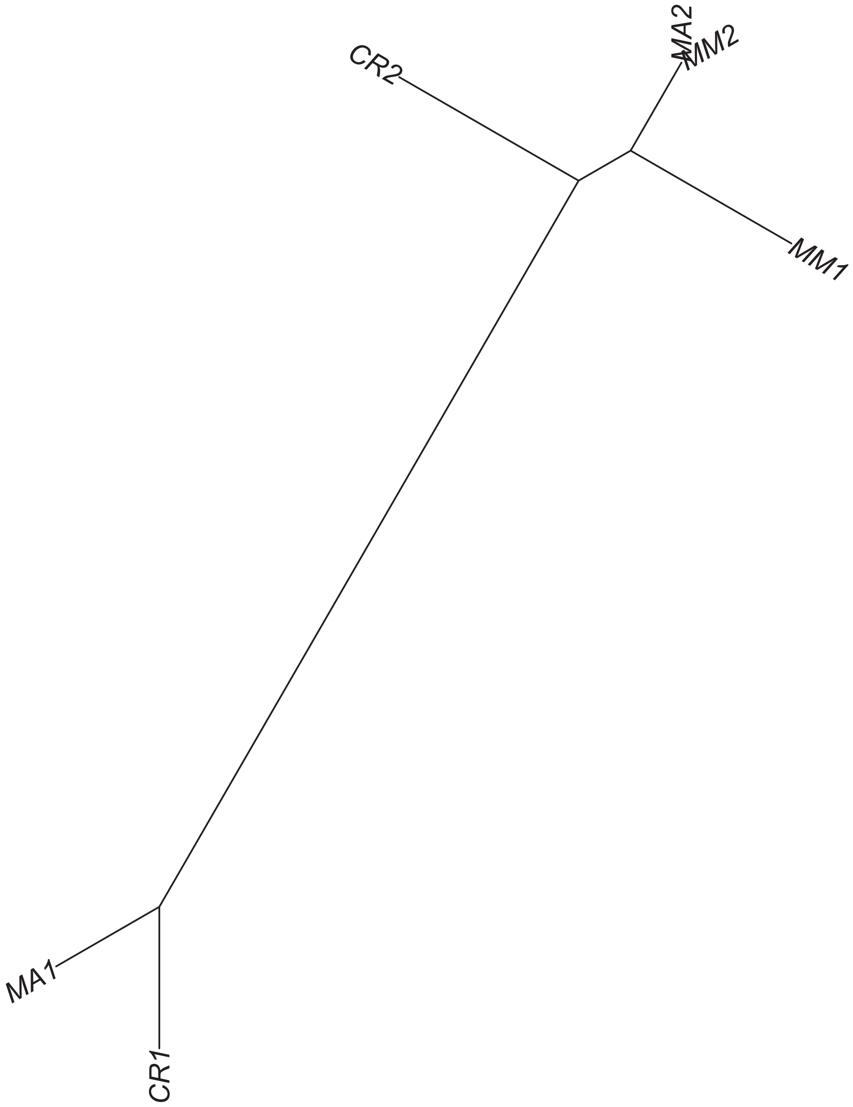

### Region 110



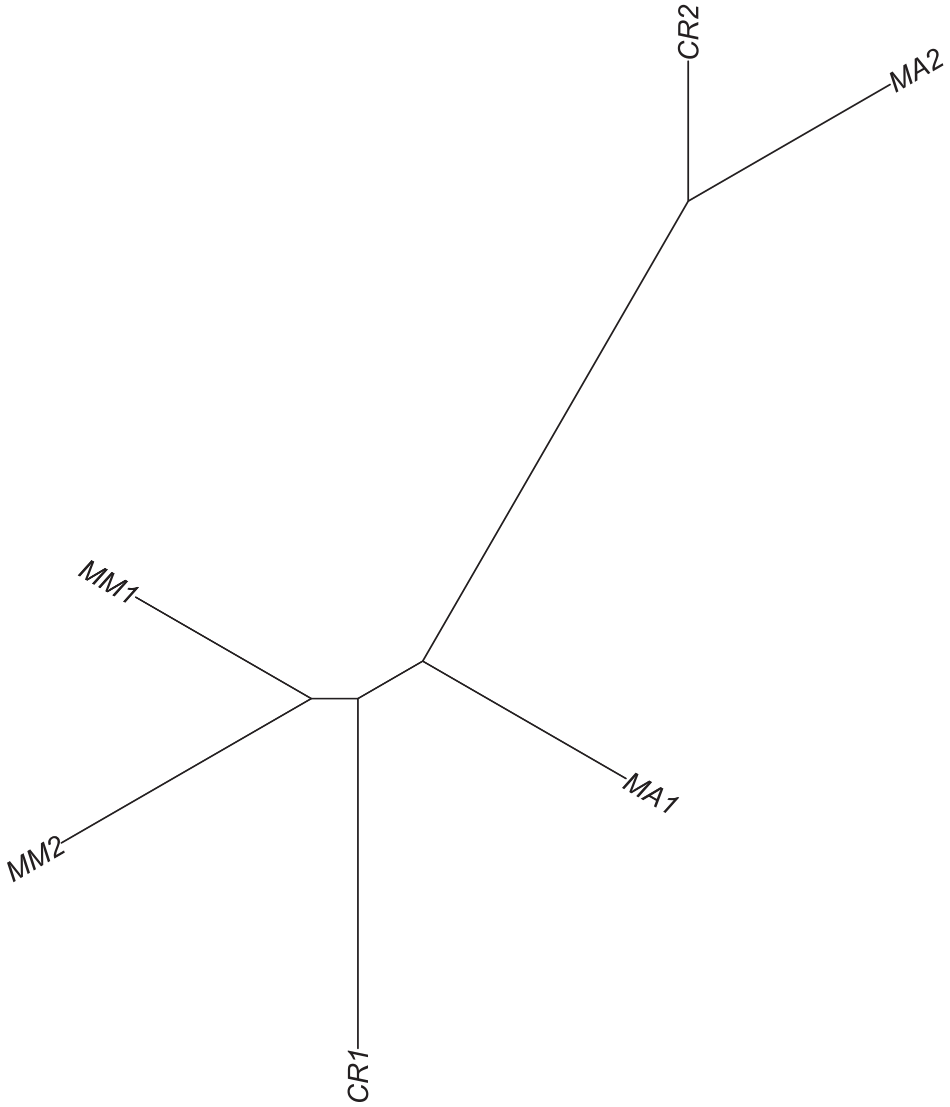

Region 111

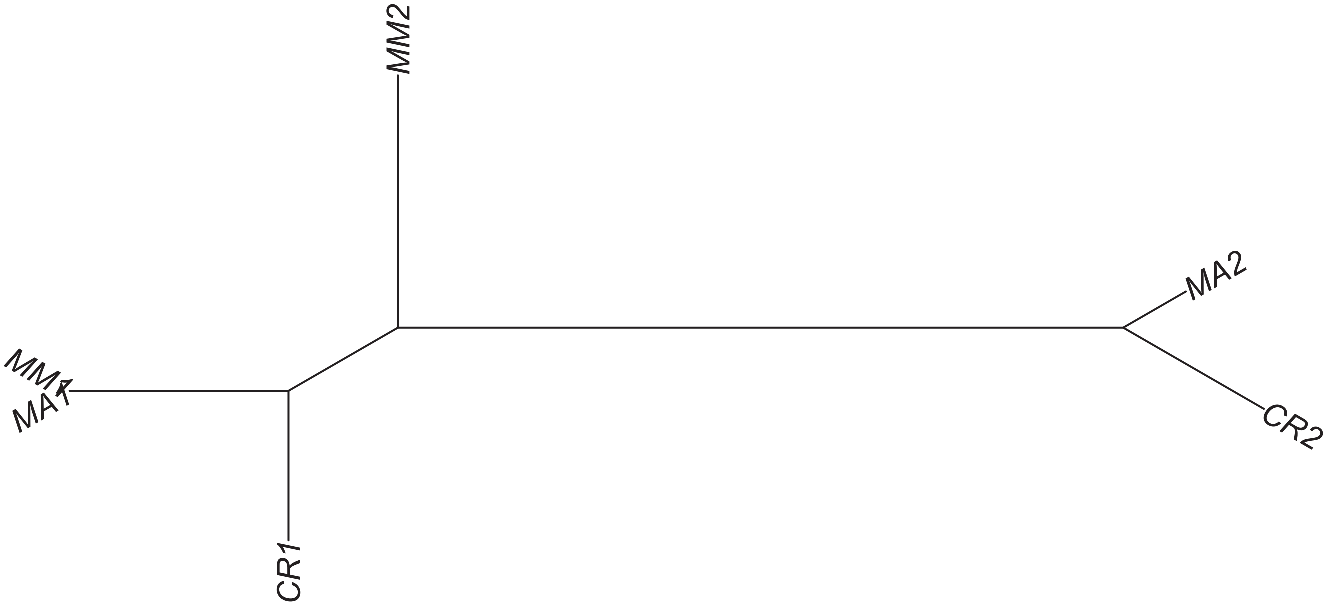

### Region 112



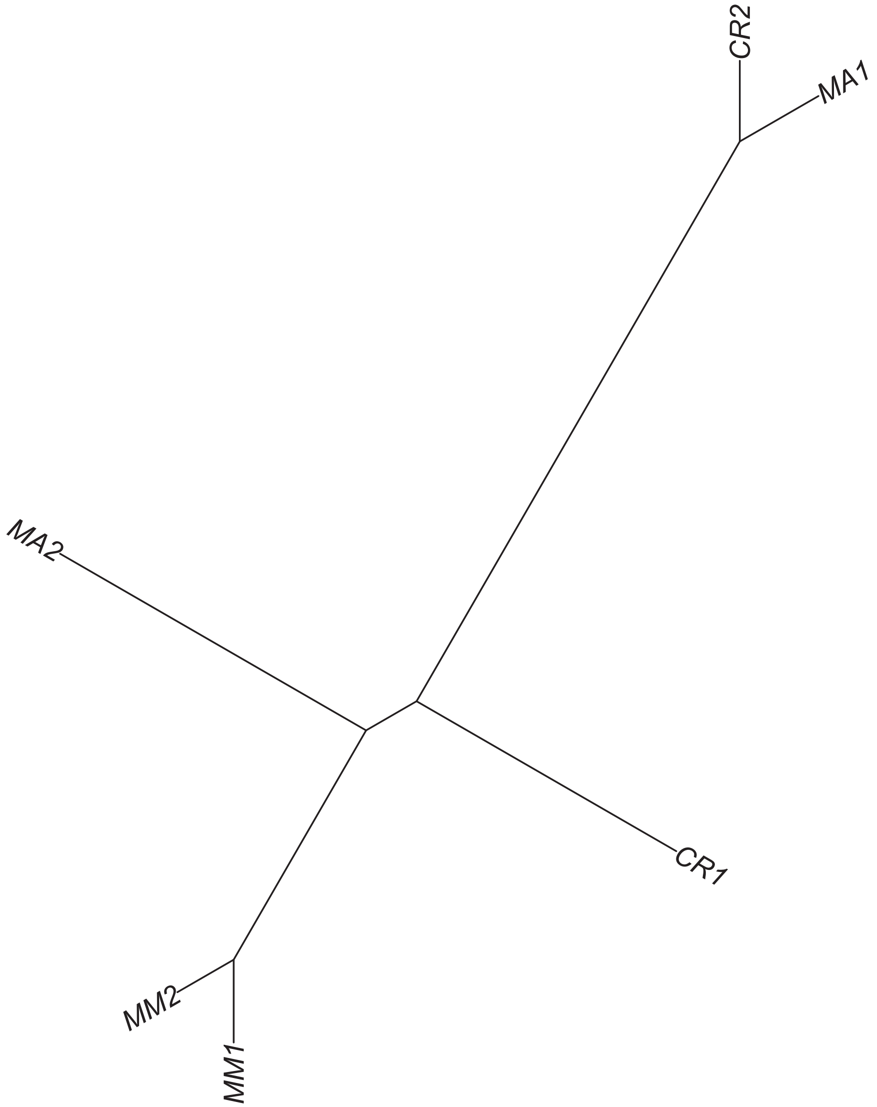

### Region 114



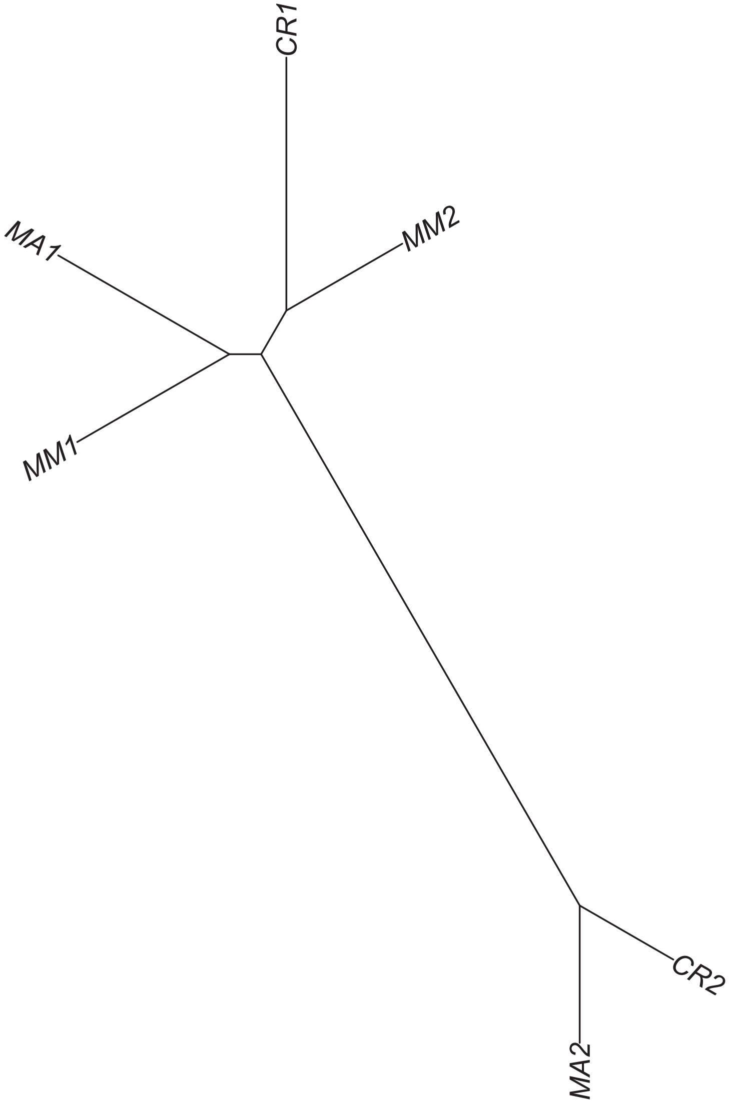

### Region 115

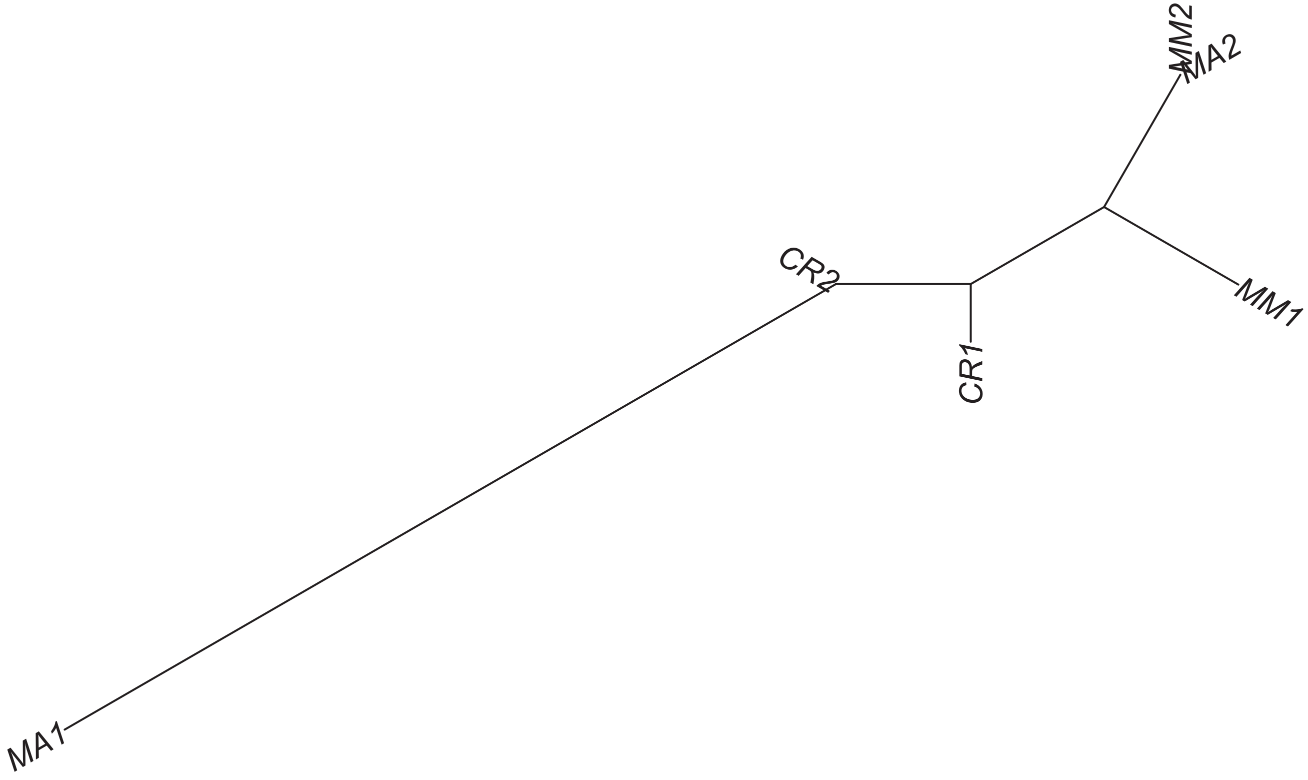

### Region 116

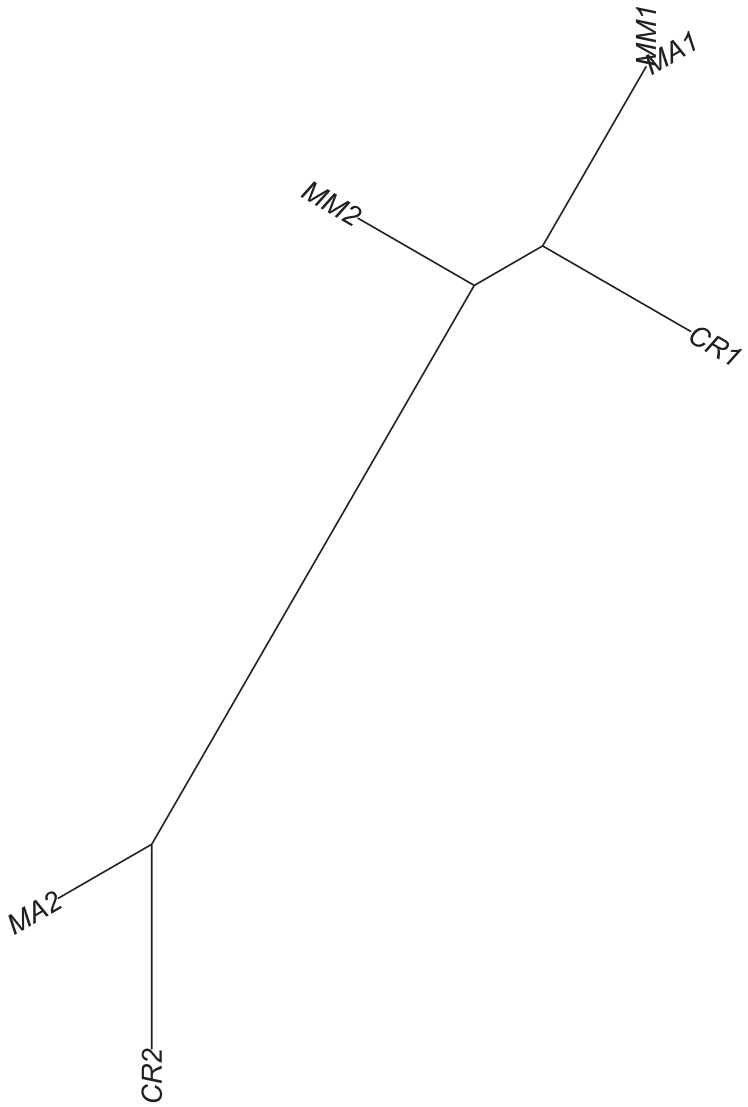

Region 119



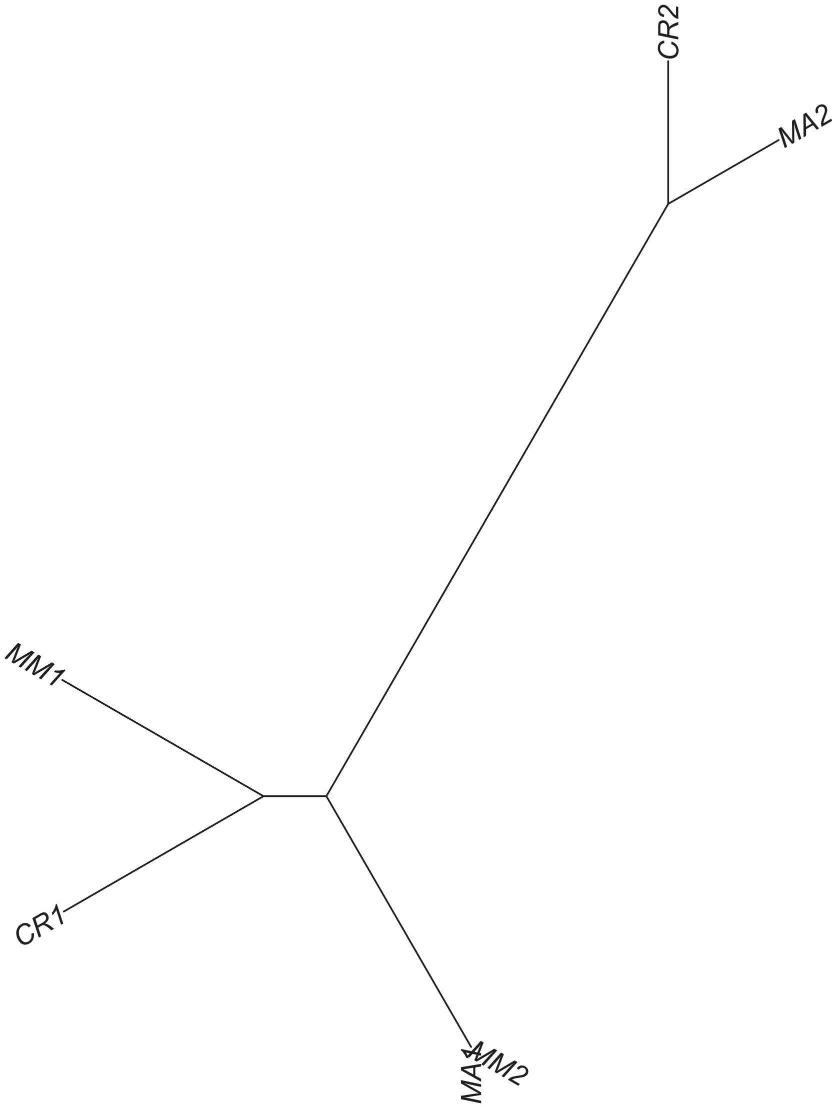

Region 120

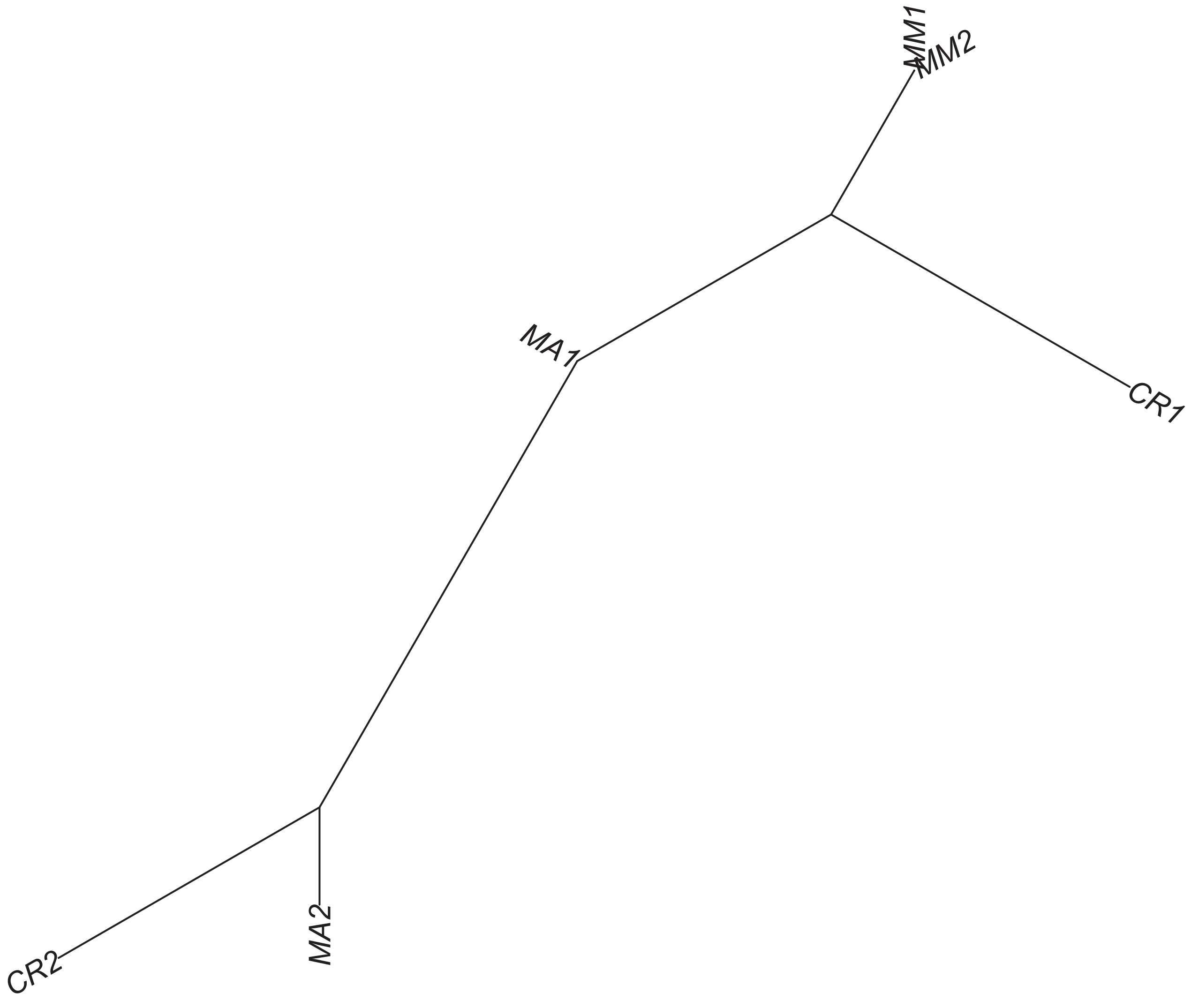

Region 122

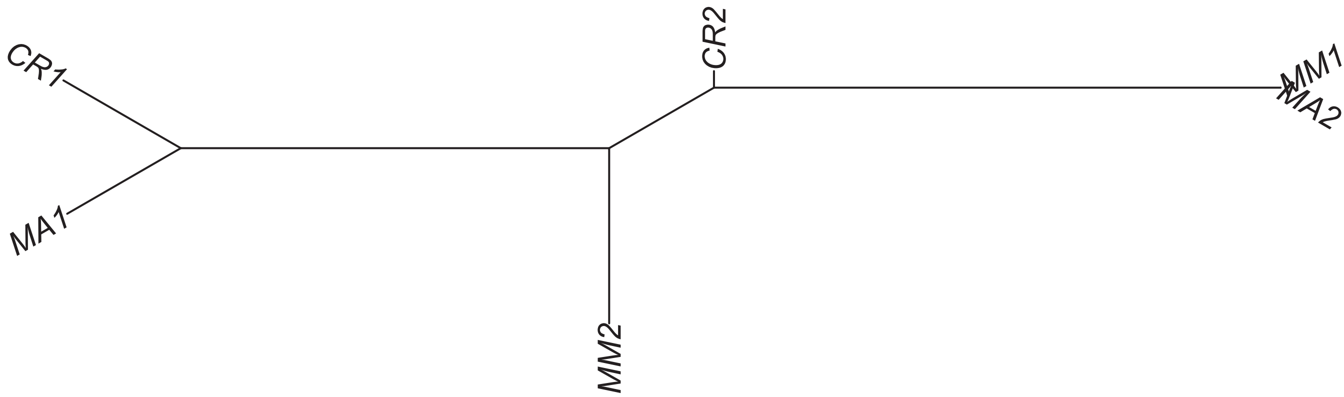

### Region 123

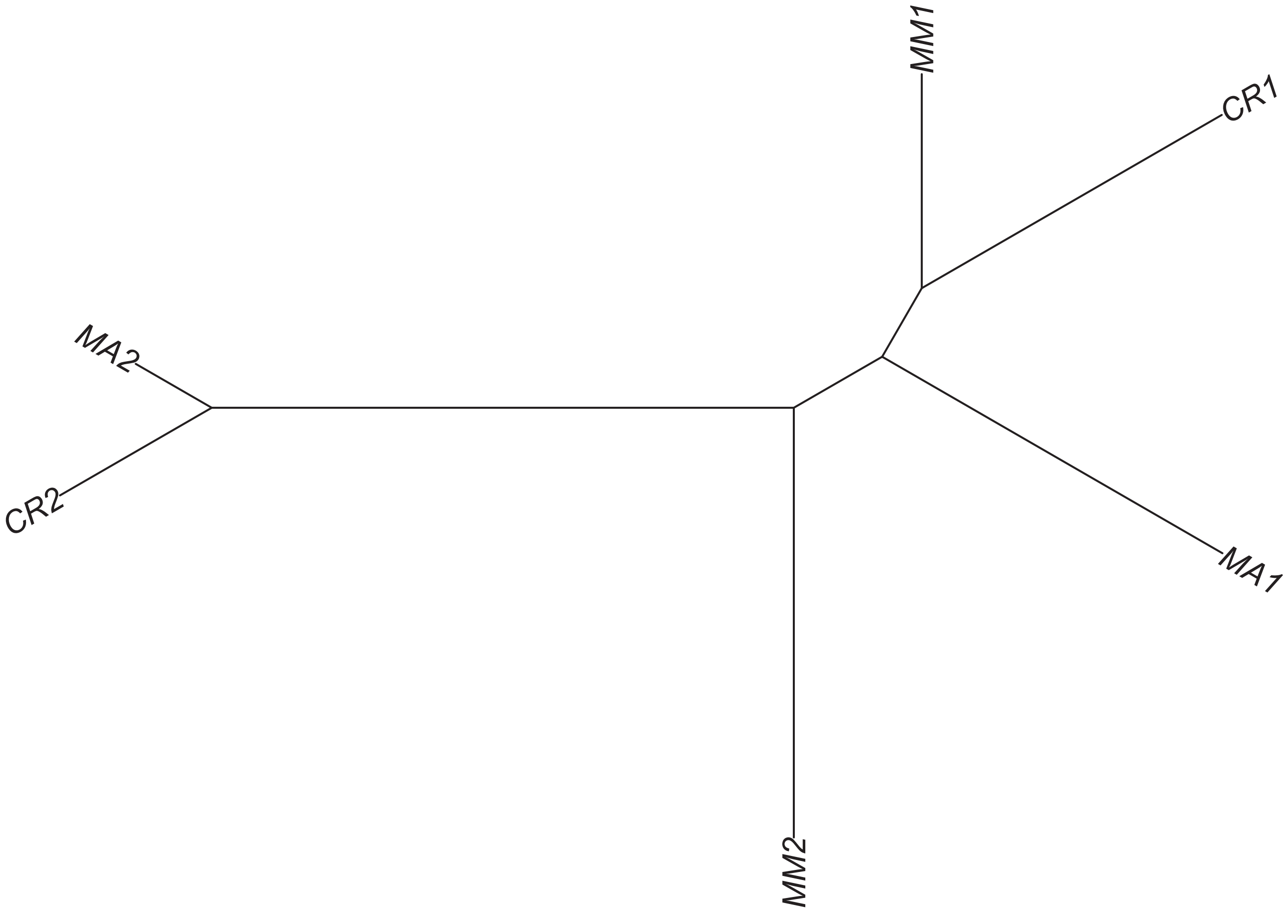

#### Region 124



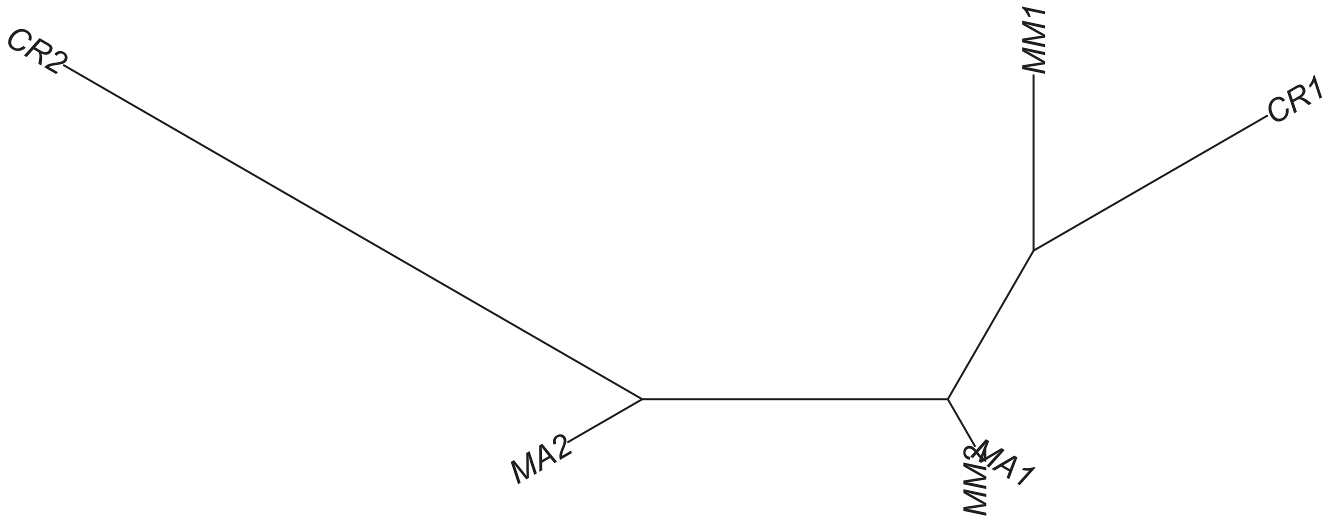

### Region 125

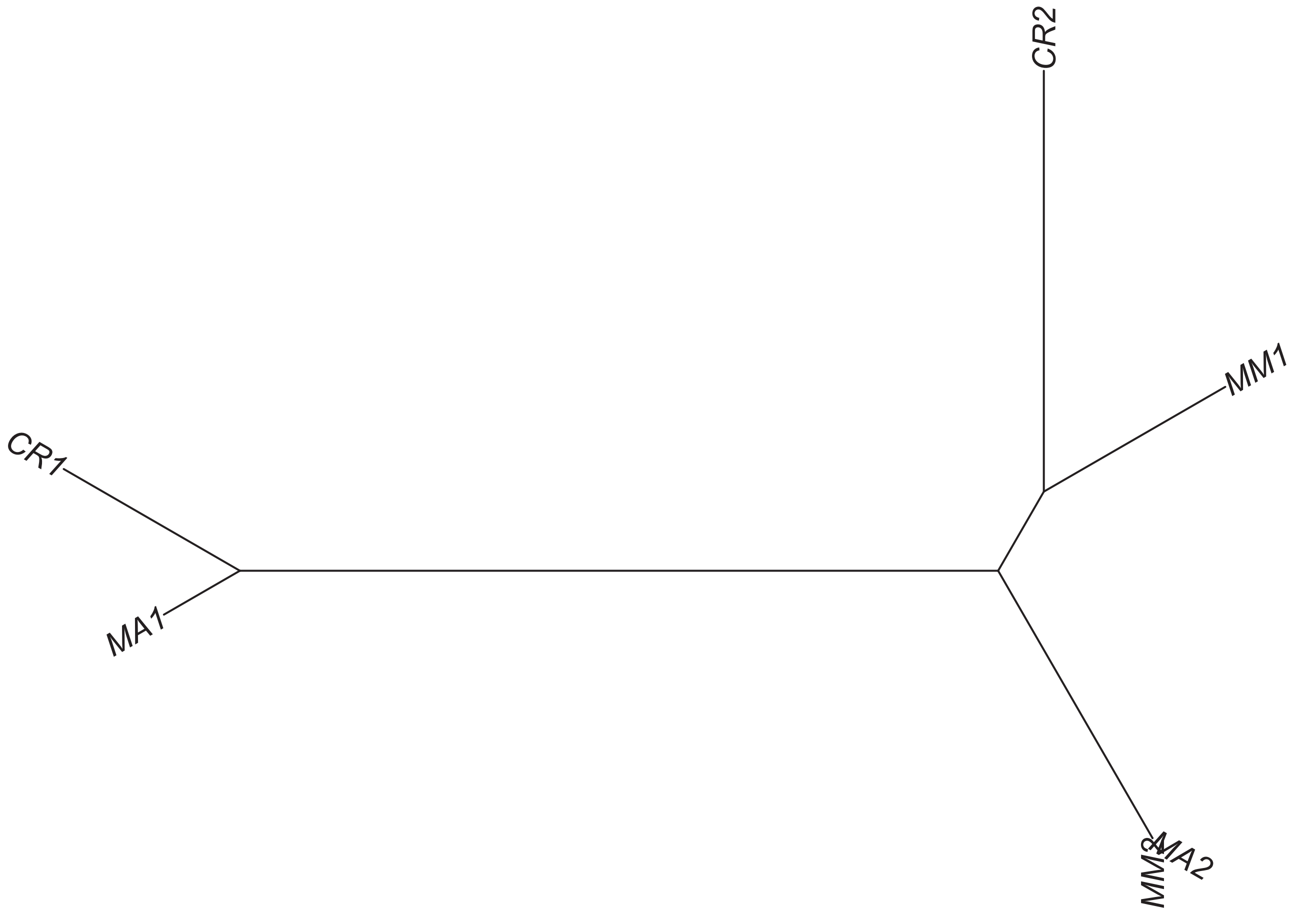

### Region 126

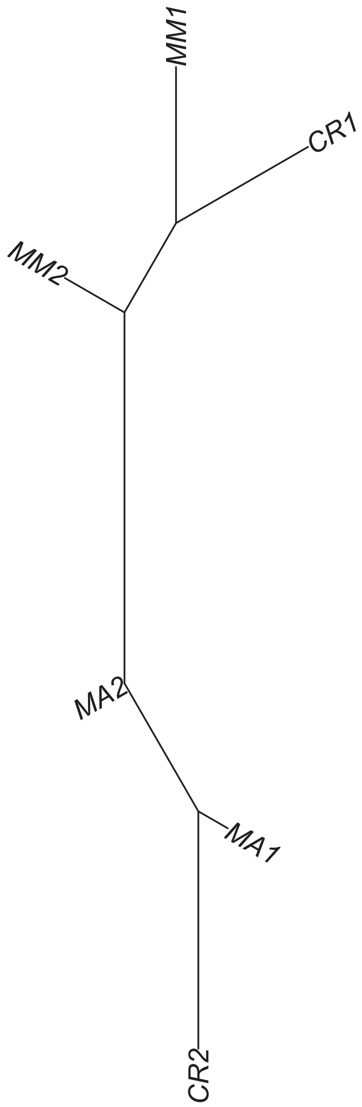

### Region 128

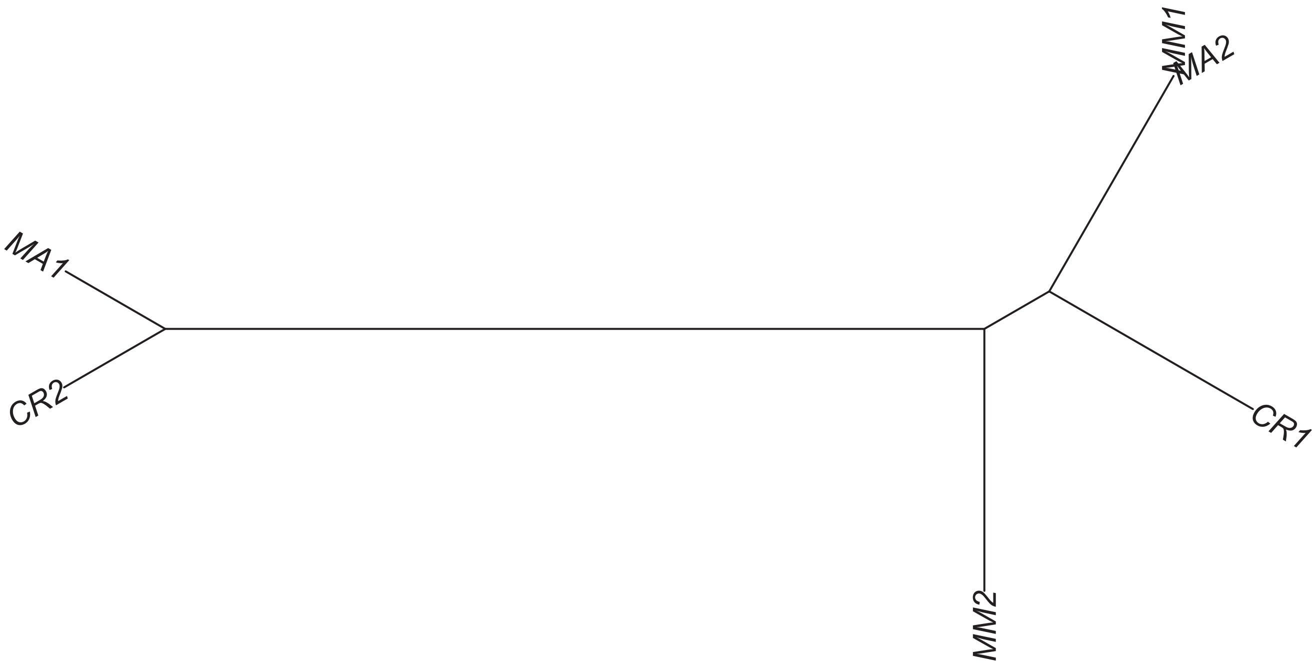

Region 129

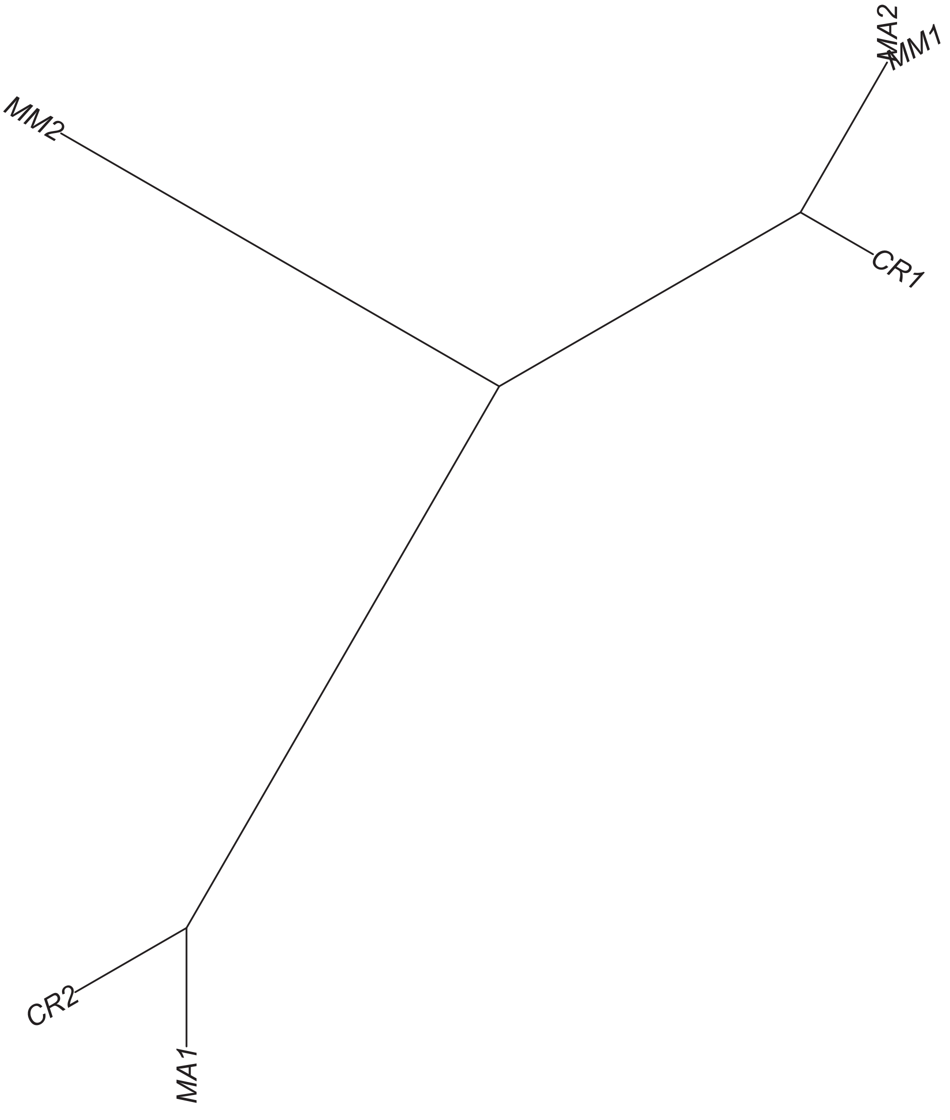

### Region 131

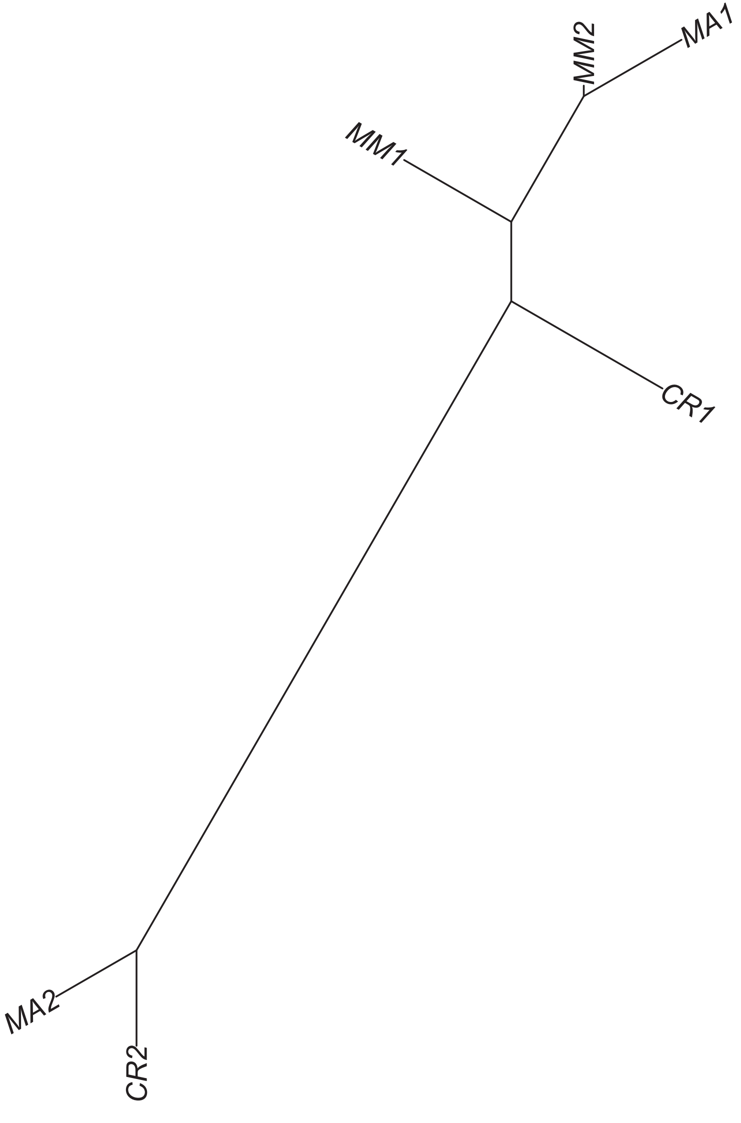

Region 132



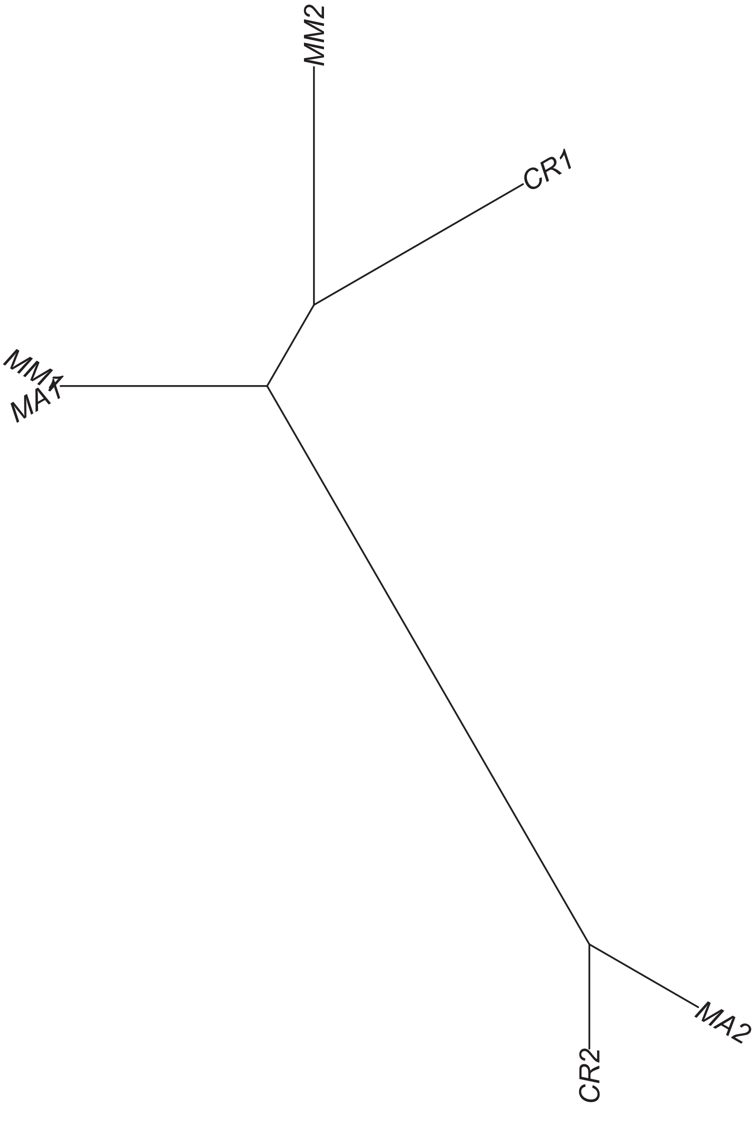

### Region 133

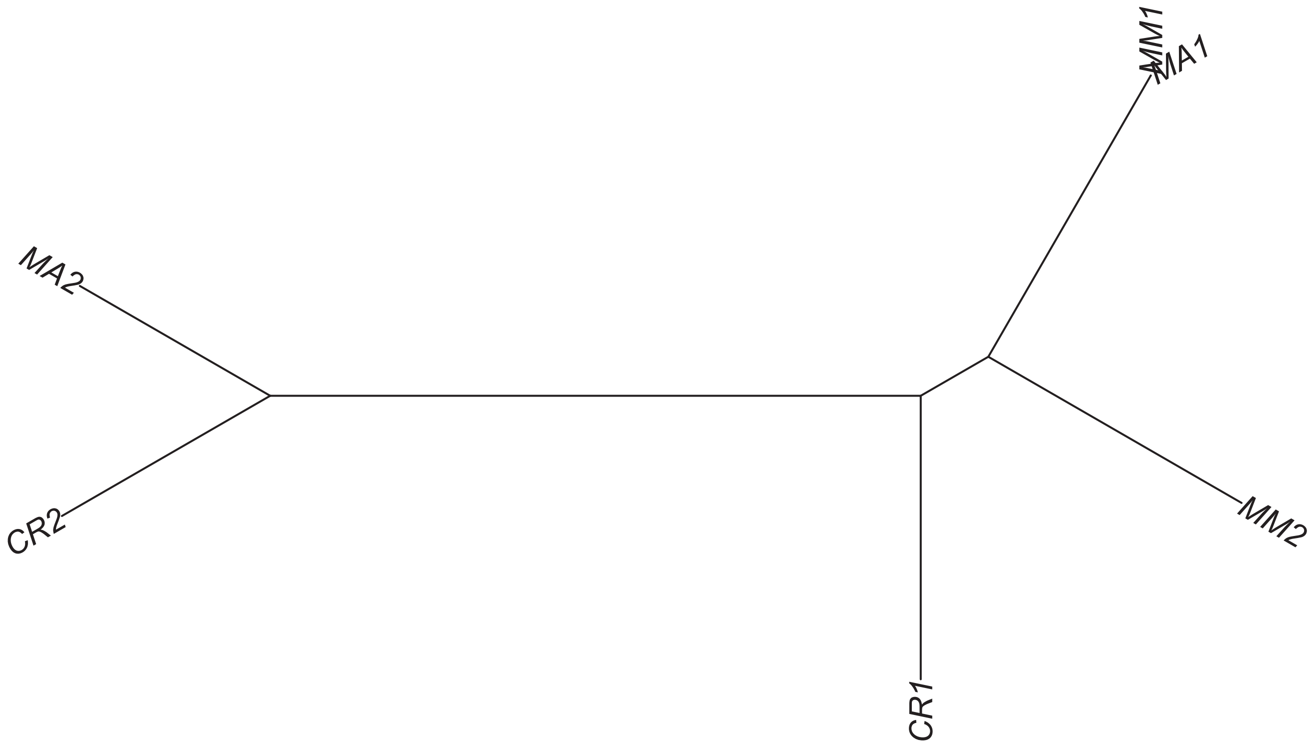

Region 134

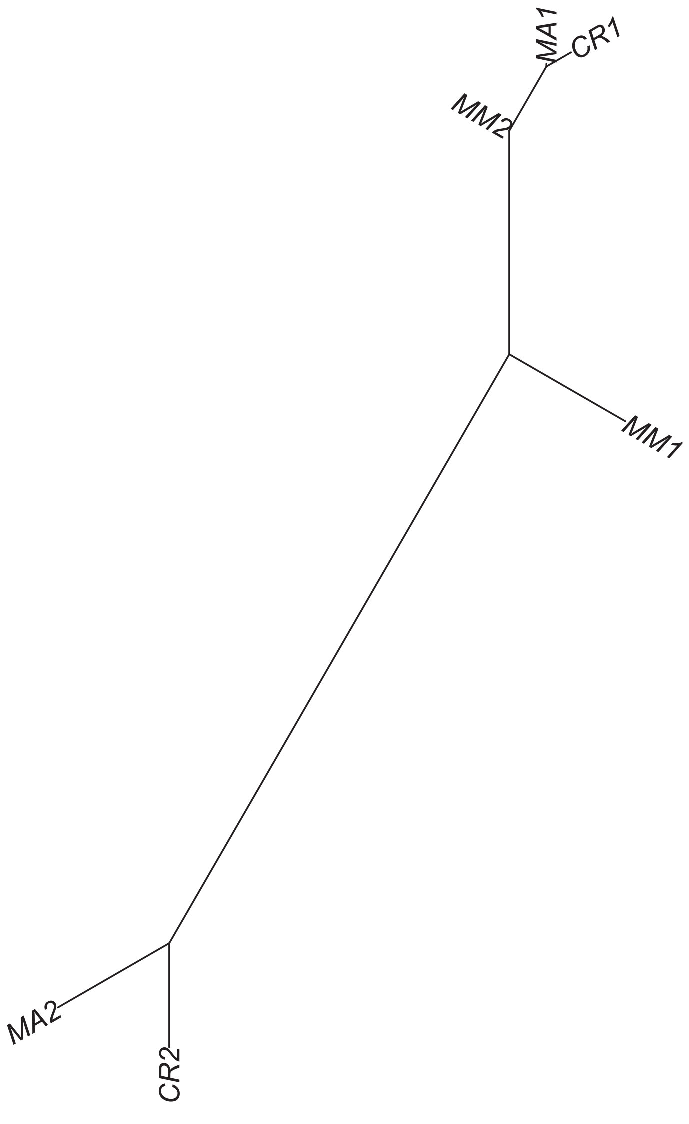

### Region 137

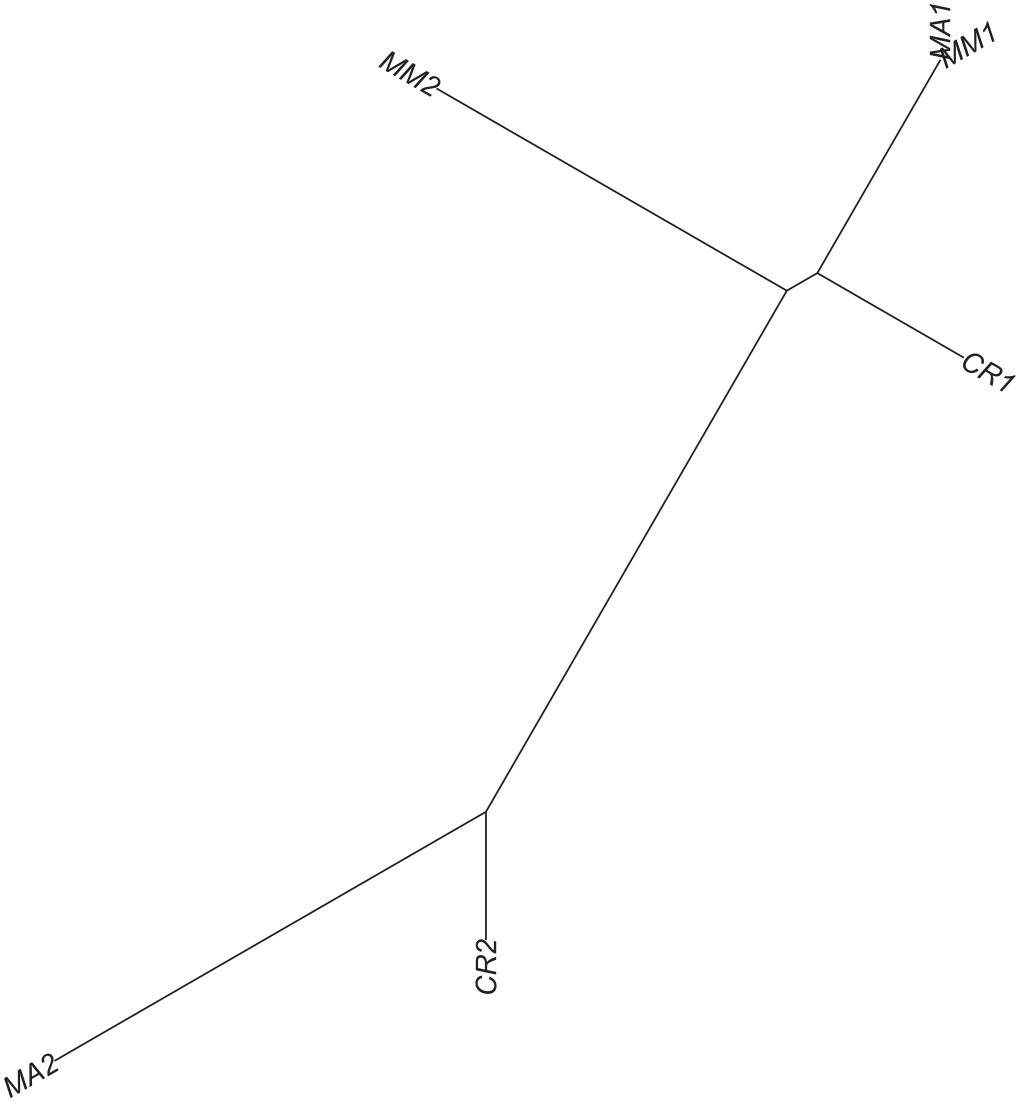

### Region 138

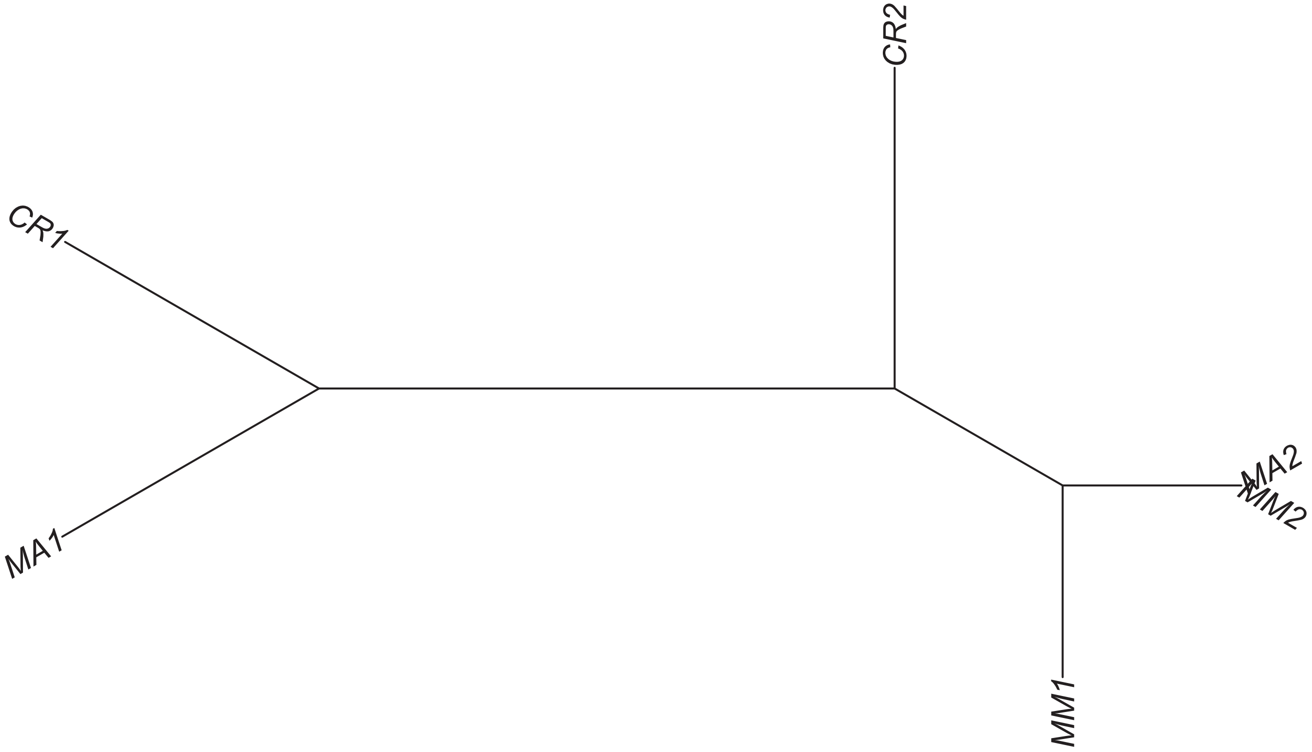

### Region 139

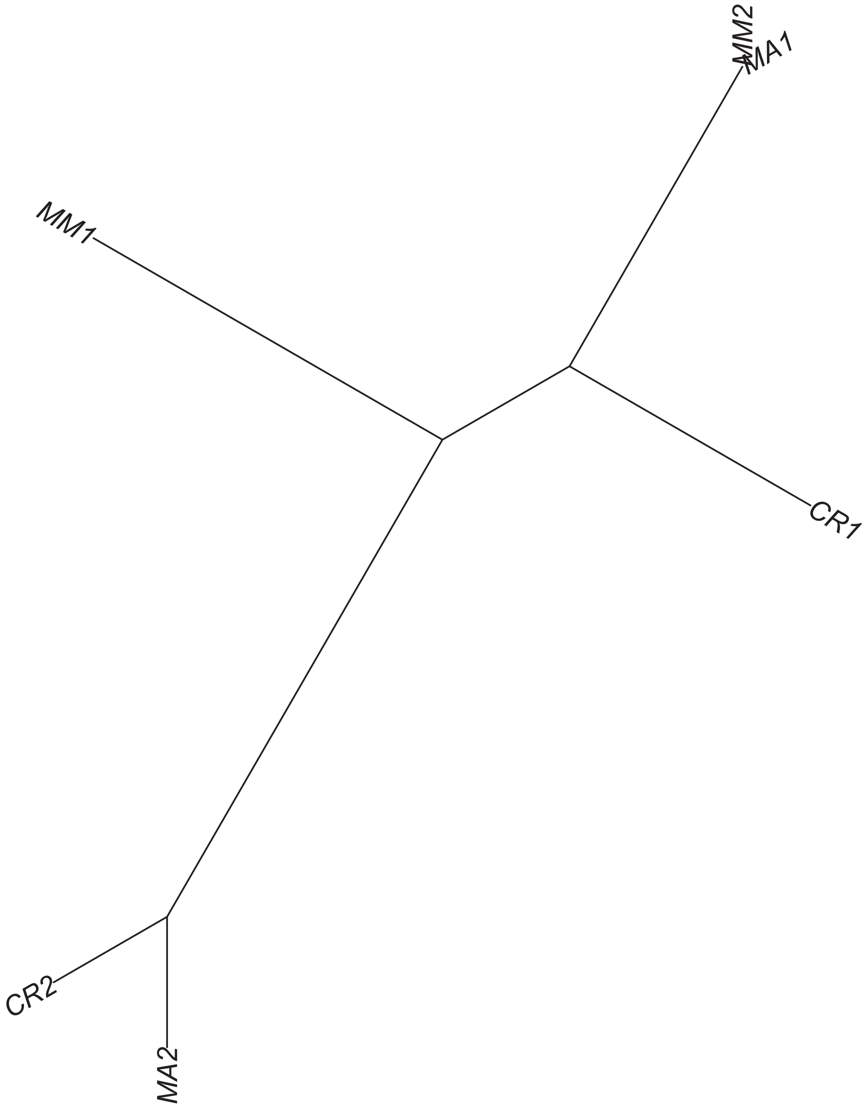

Region 140

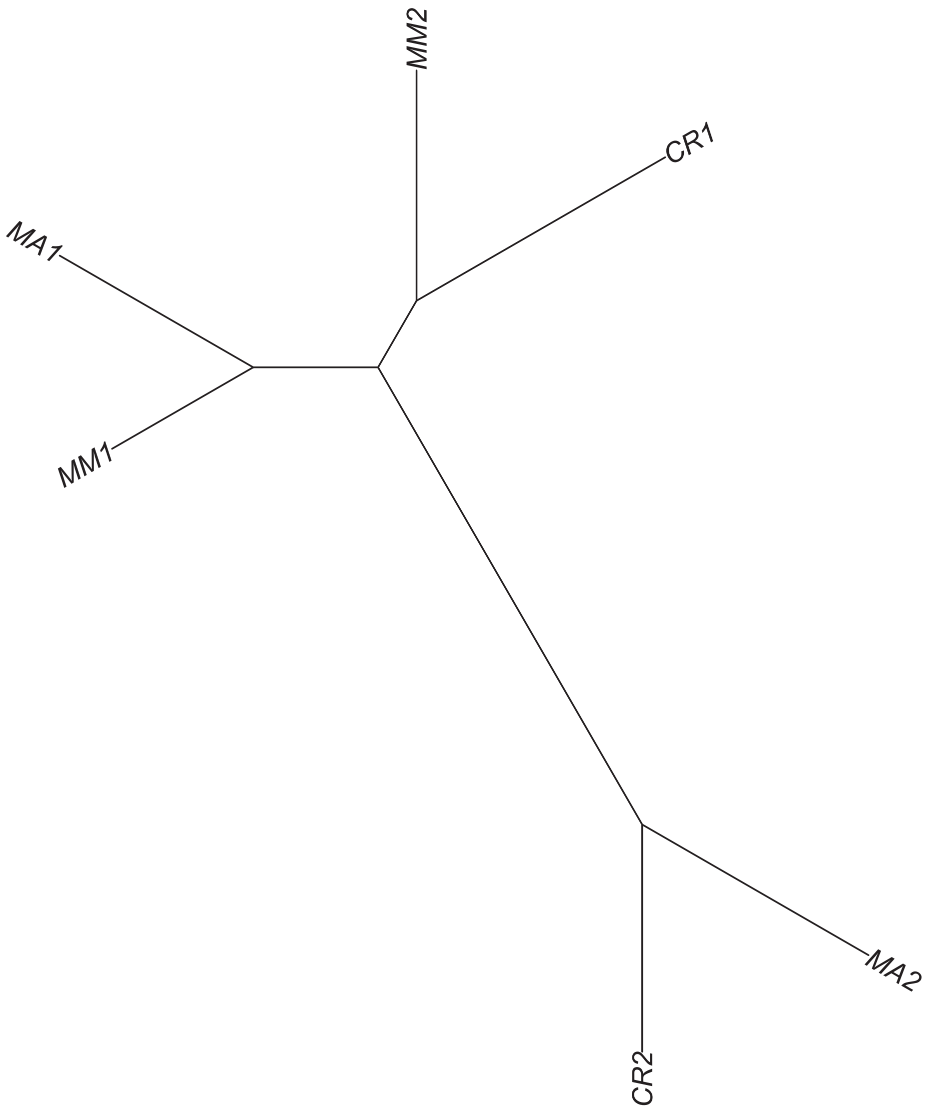

### Region 141



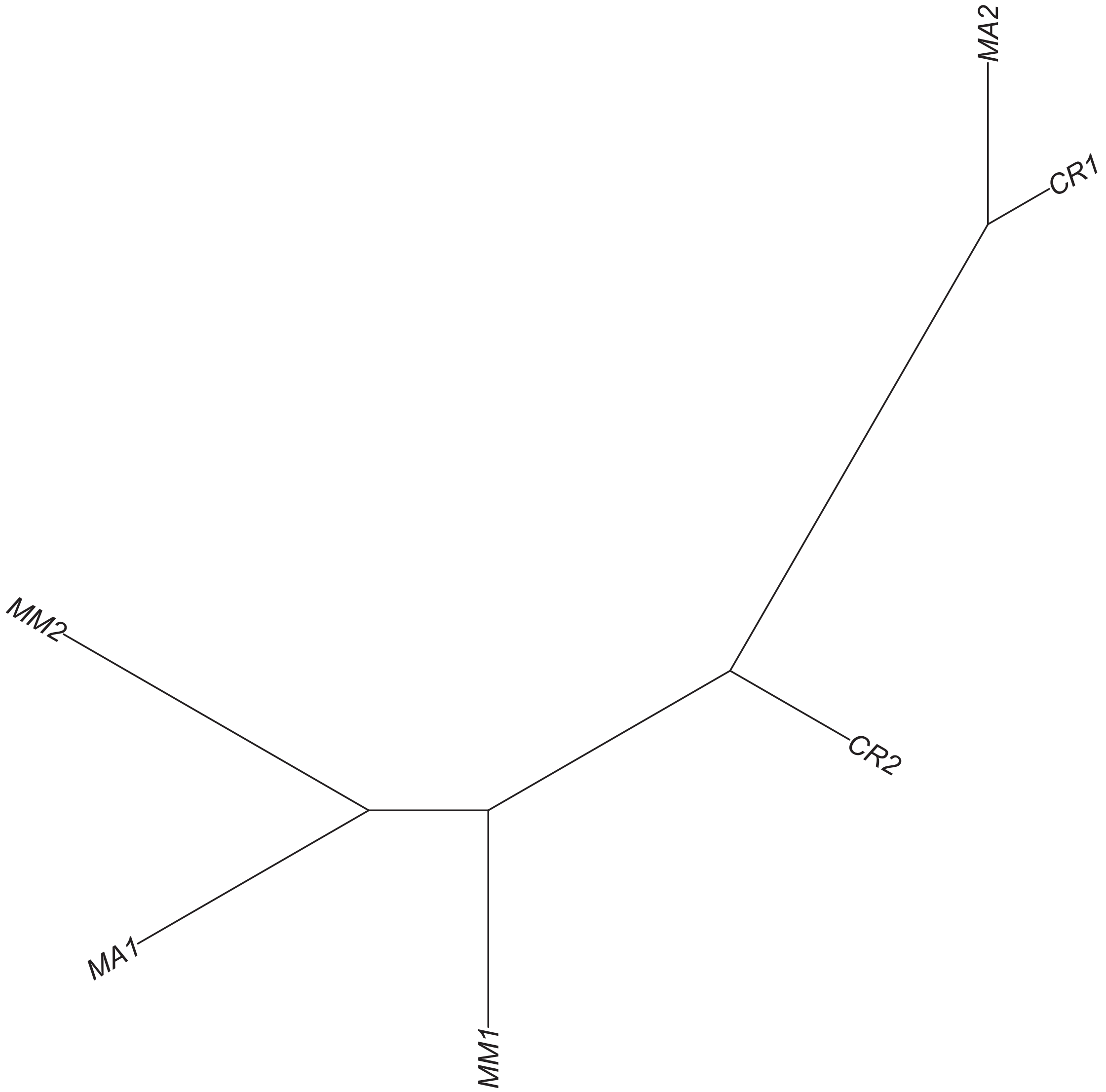

### Region 142

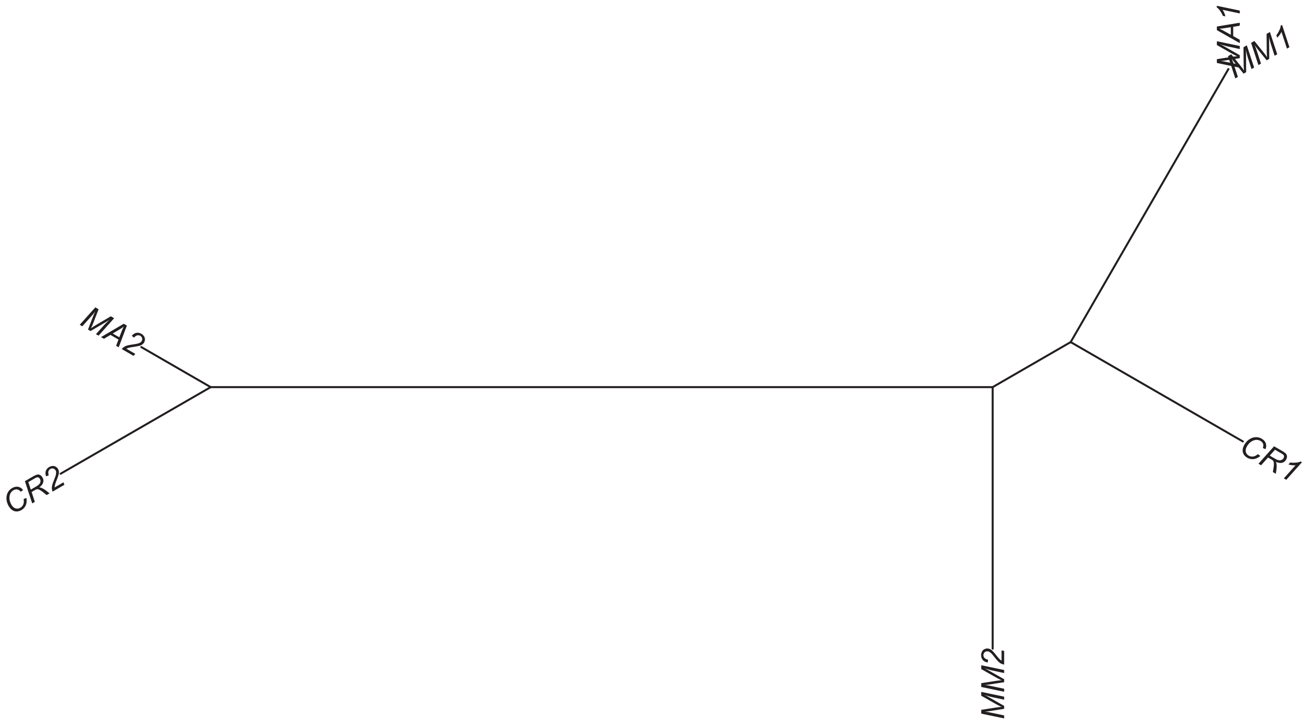

### Region 143

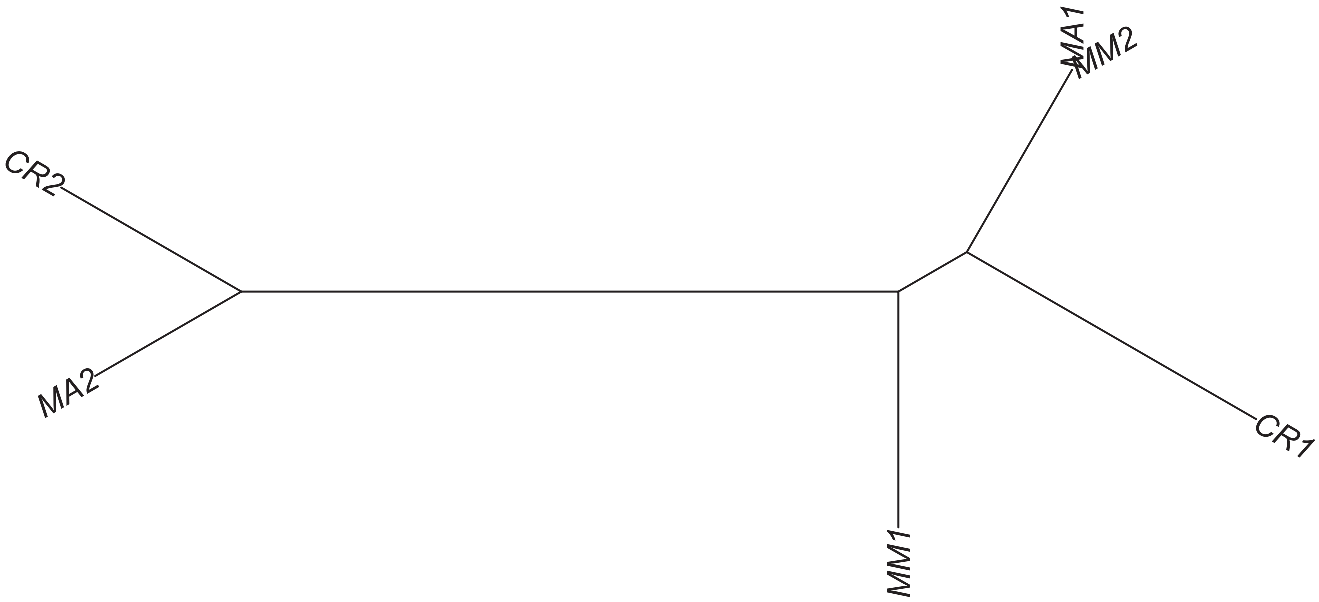

Region 144

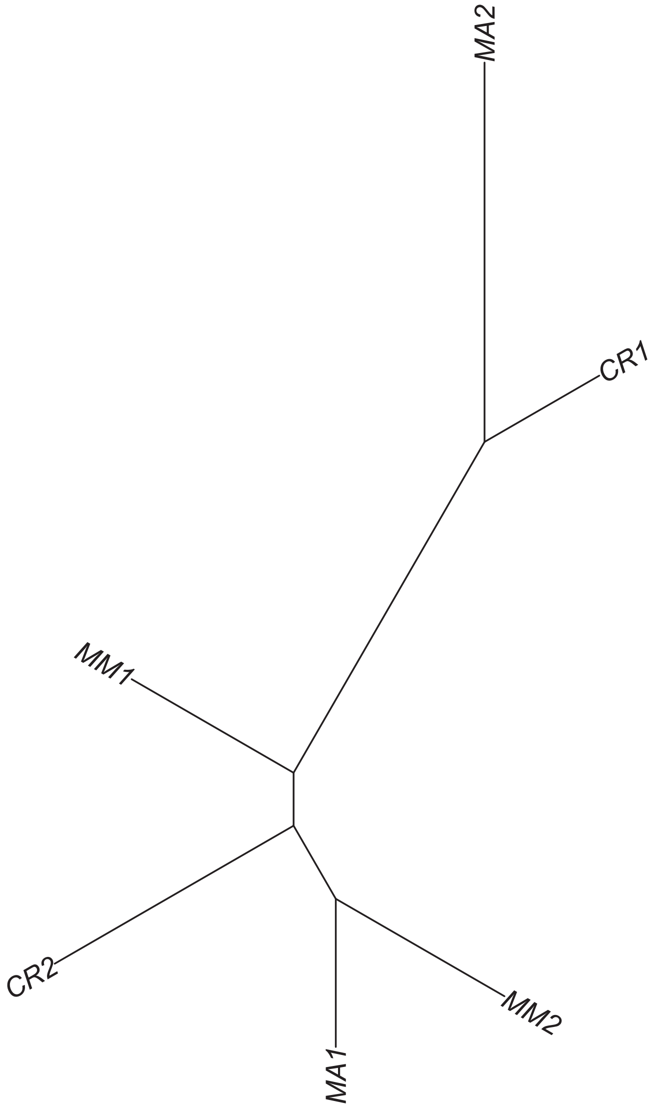

### Region 146



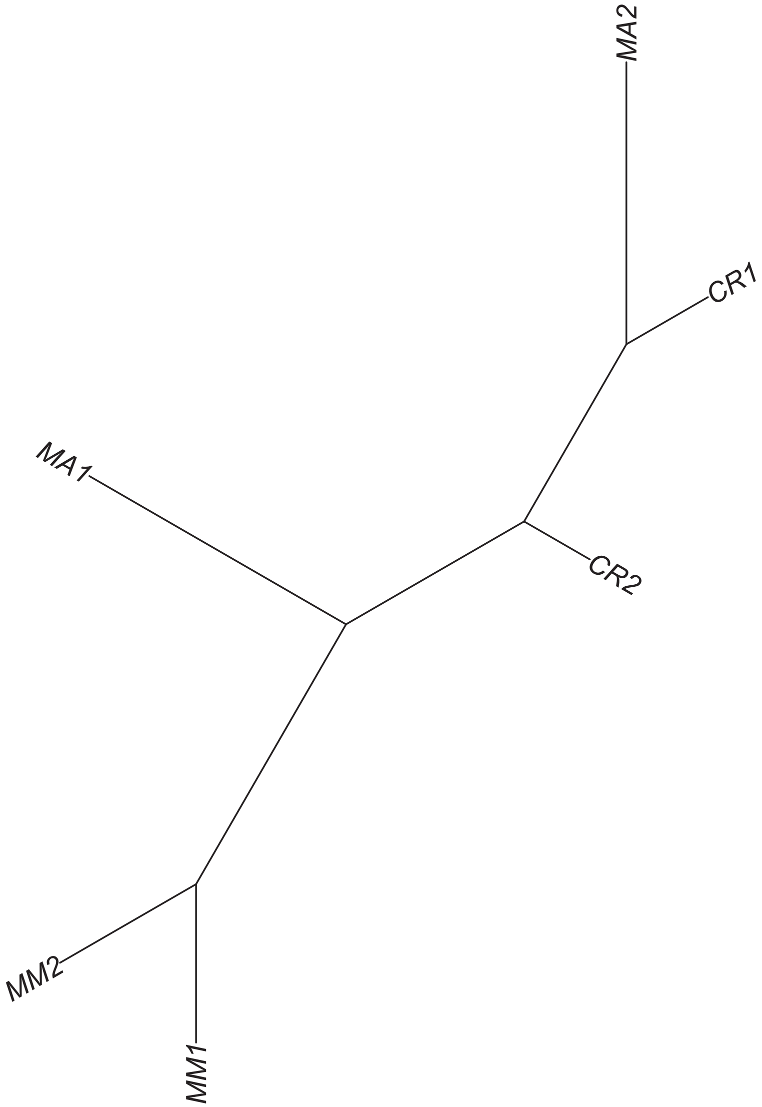

### Region 147

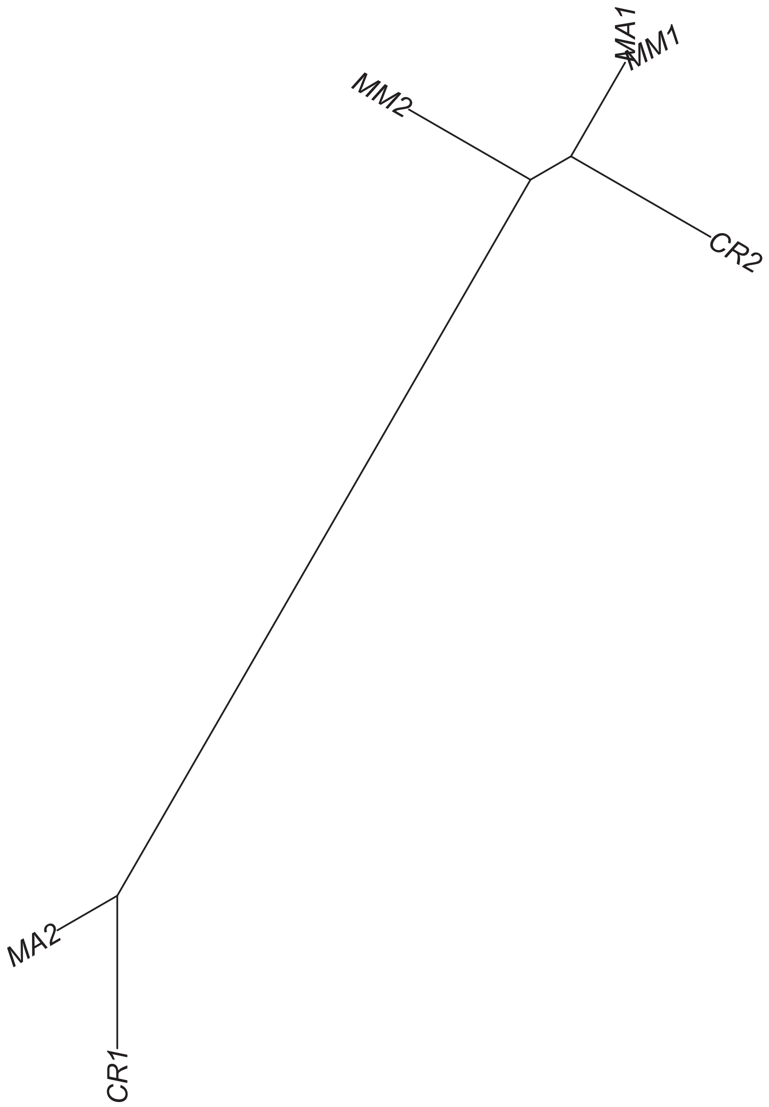

### Region 148

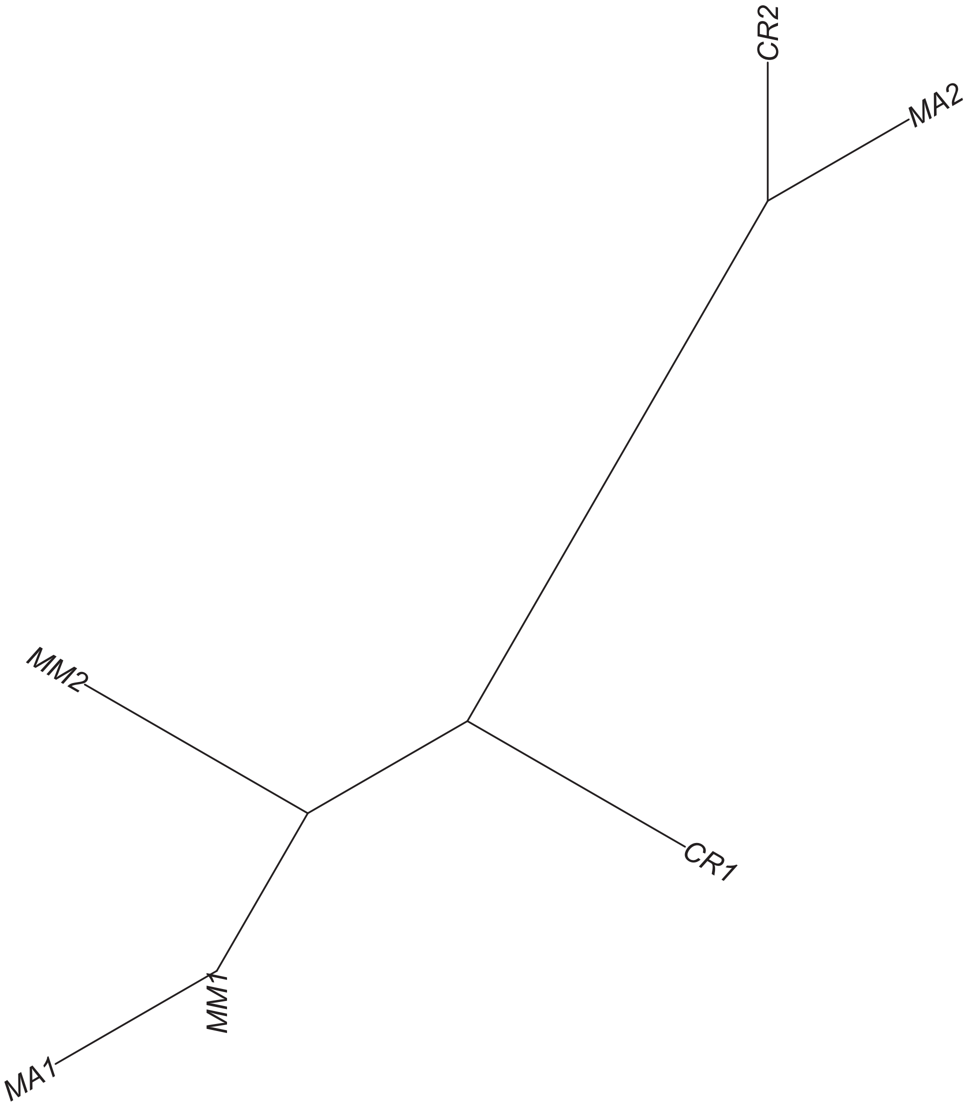

#### Region 149



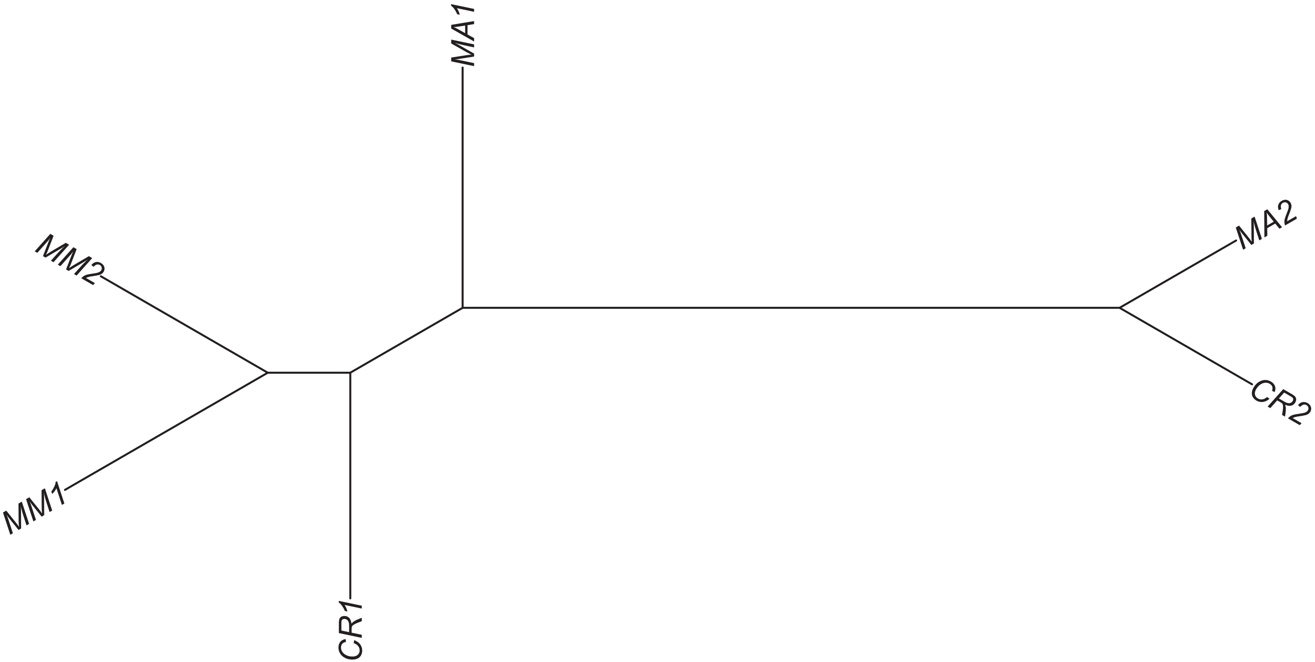

Region 154

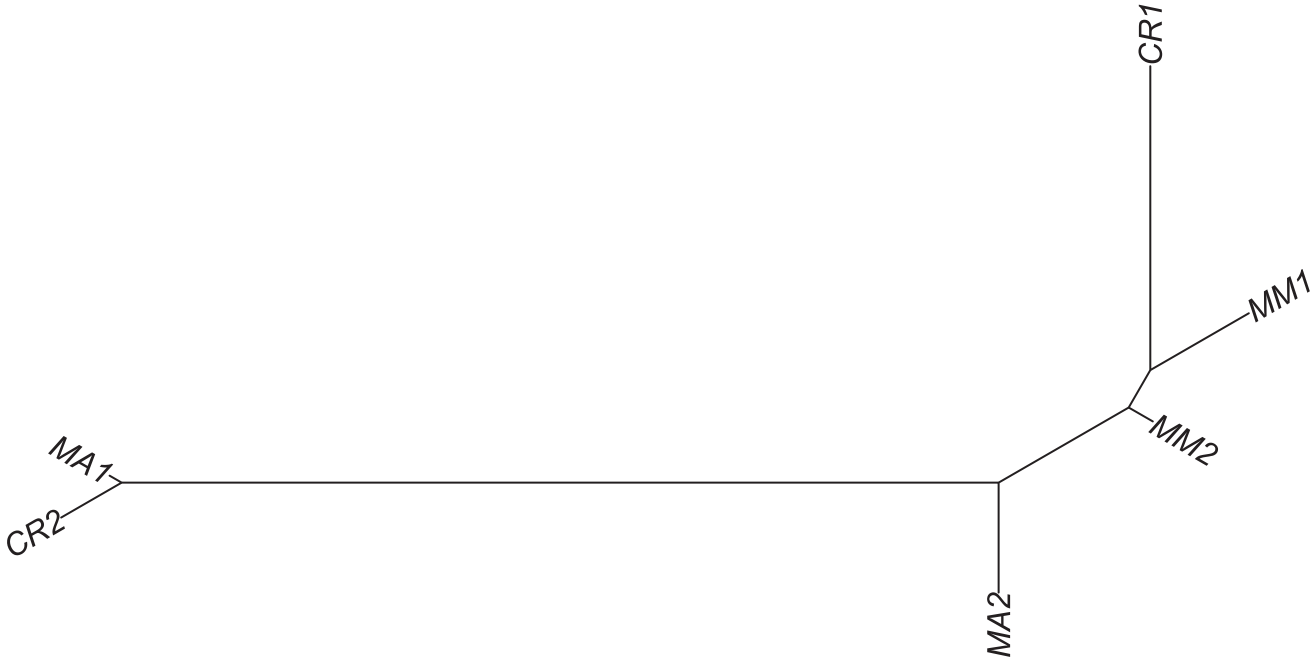

Region 156

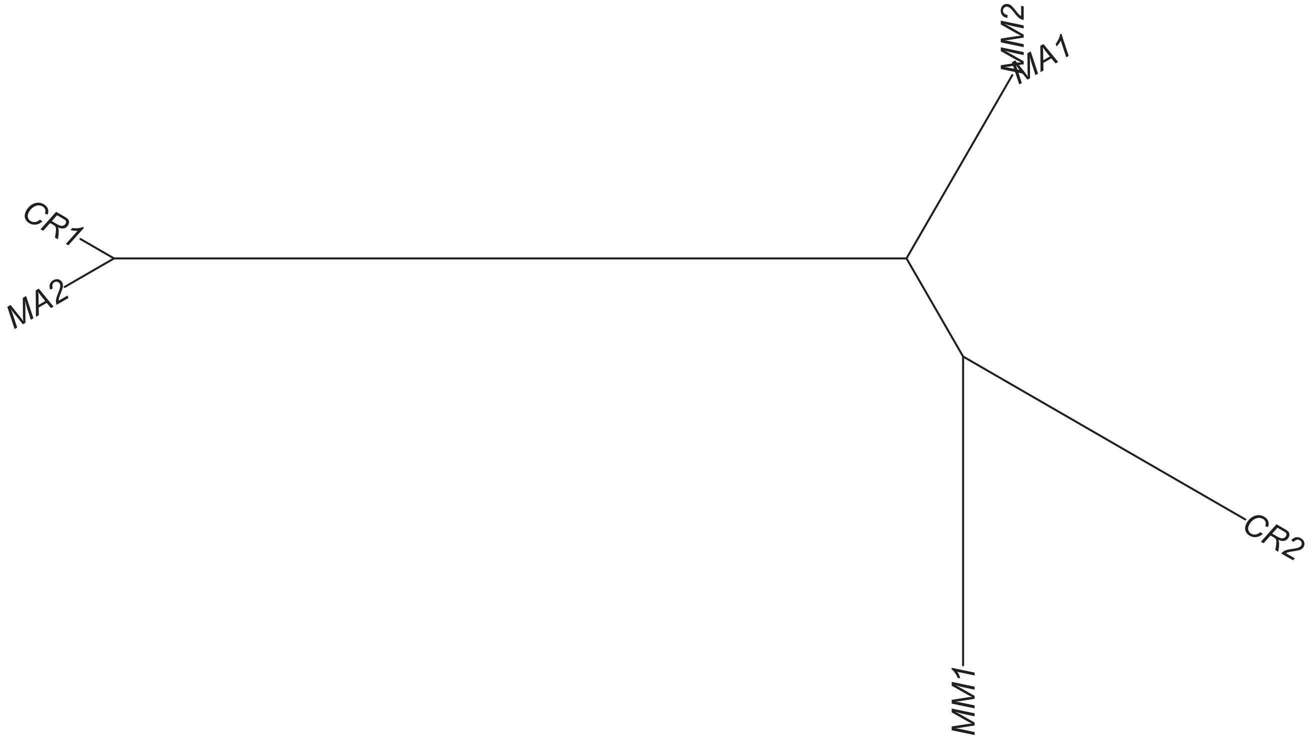

#### Region 159

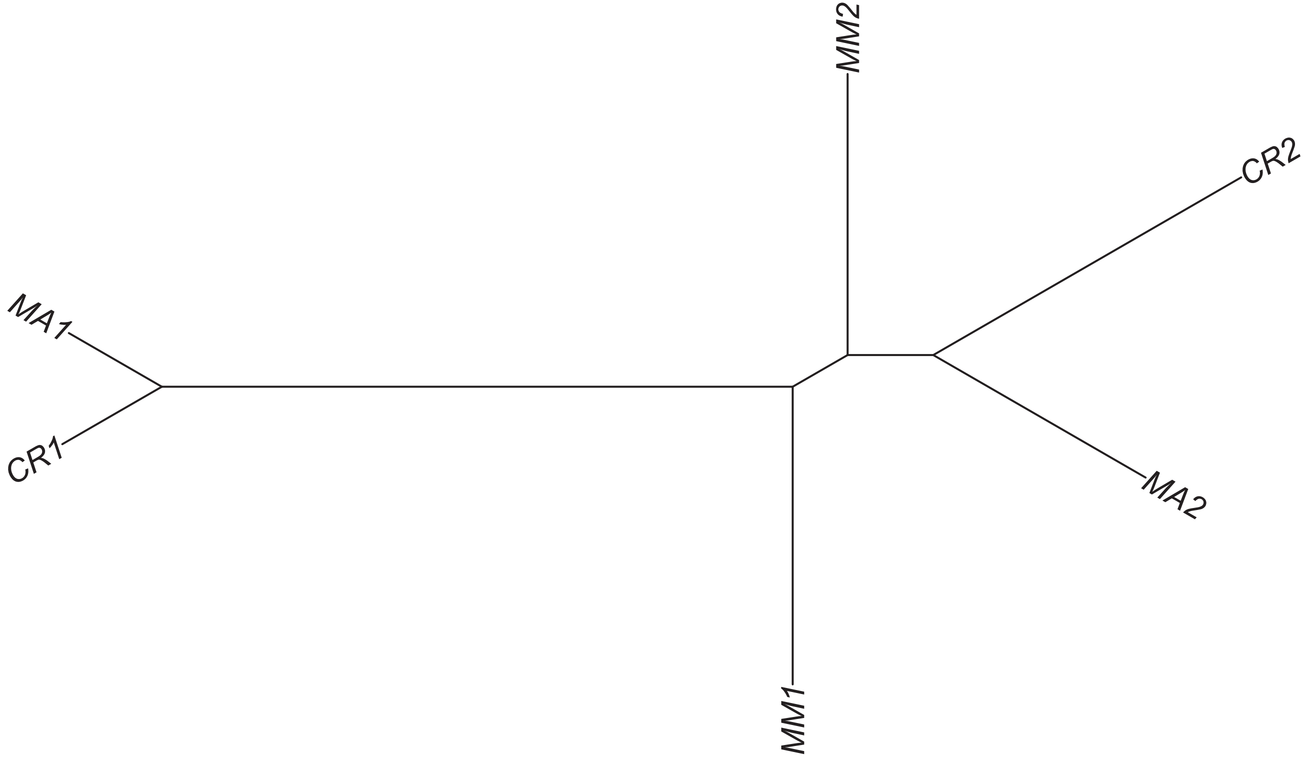

Region 160

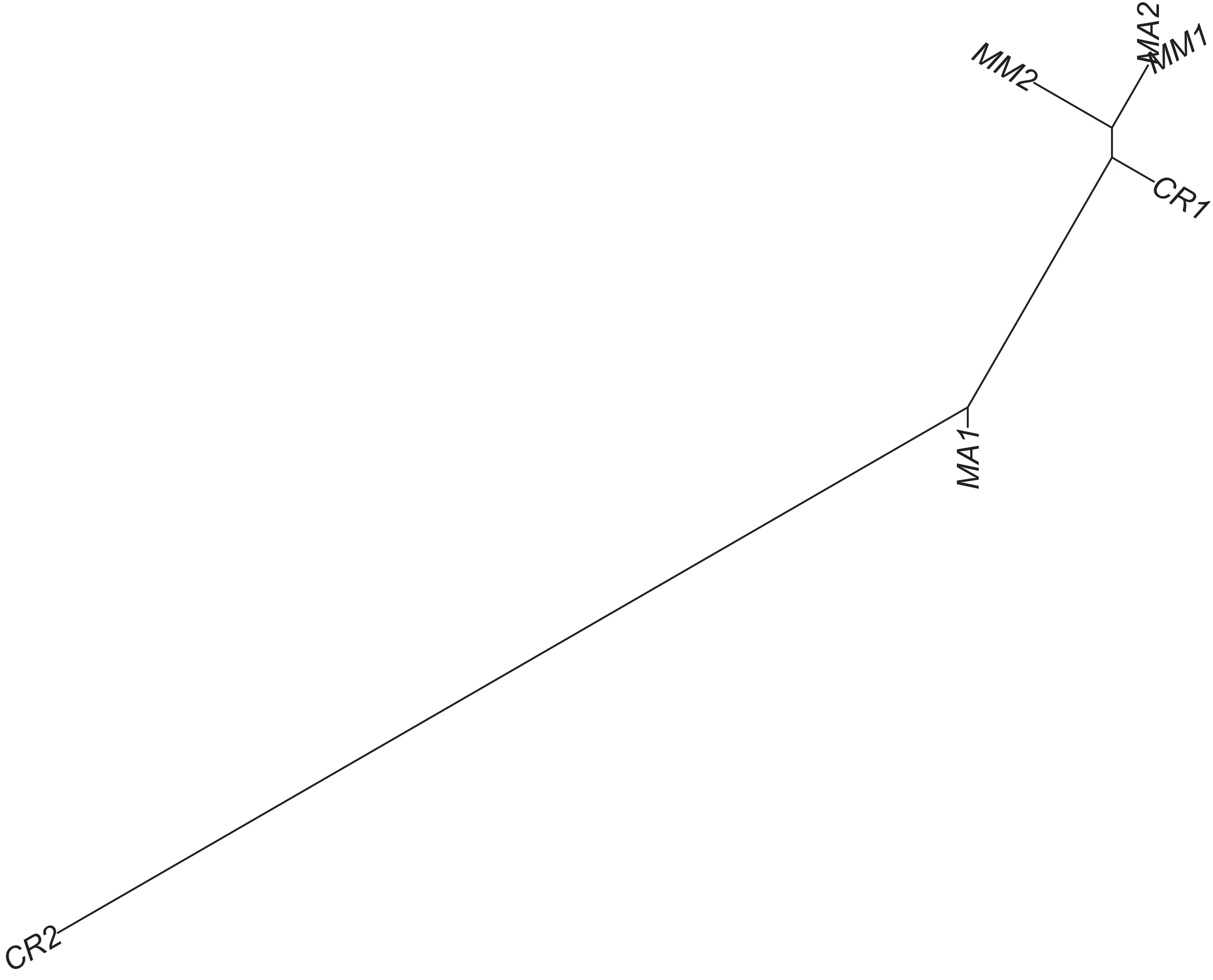

### Region 161



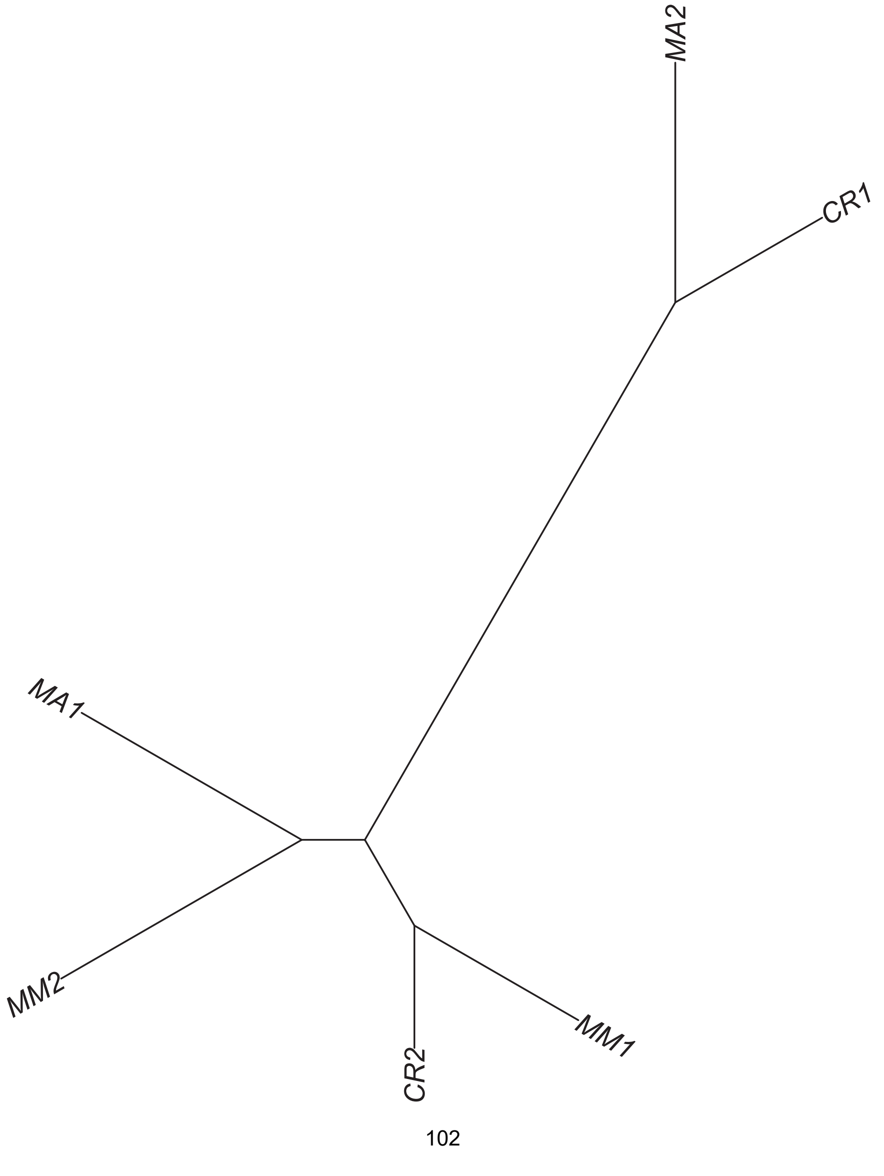

#### Region 165



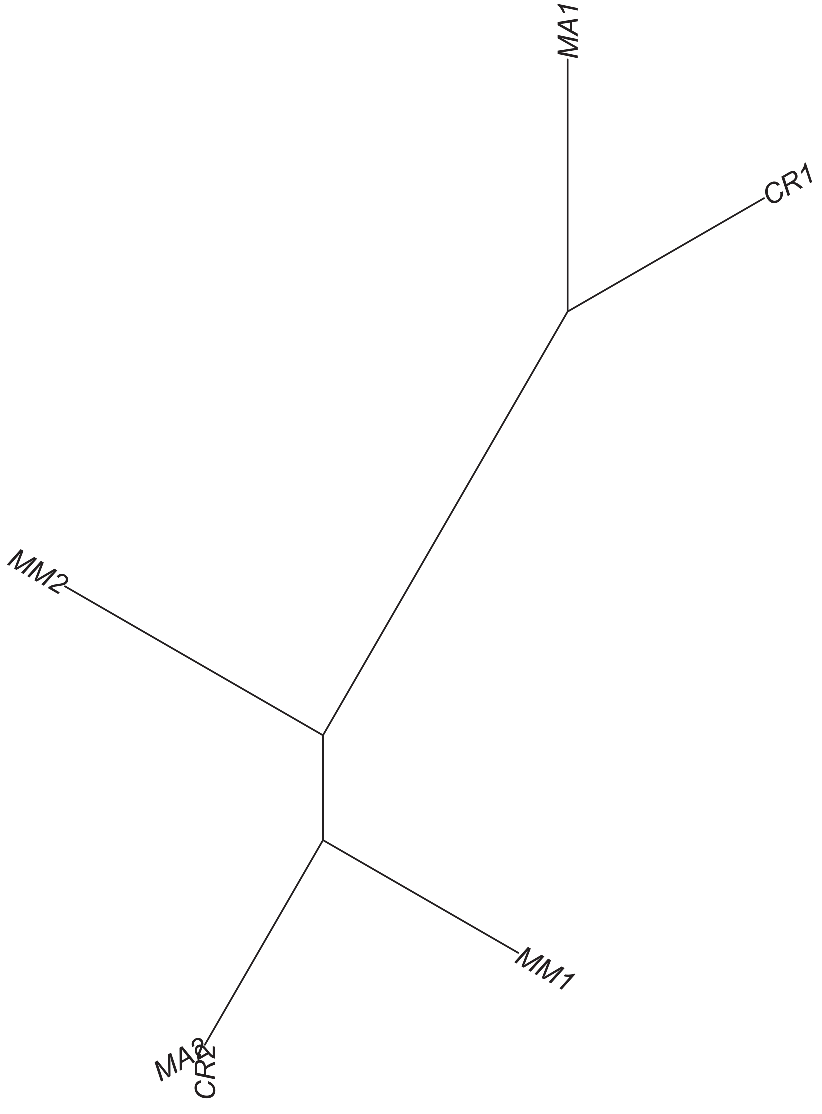

### Region 167

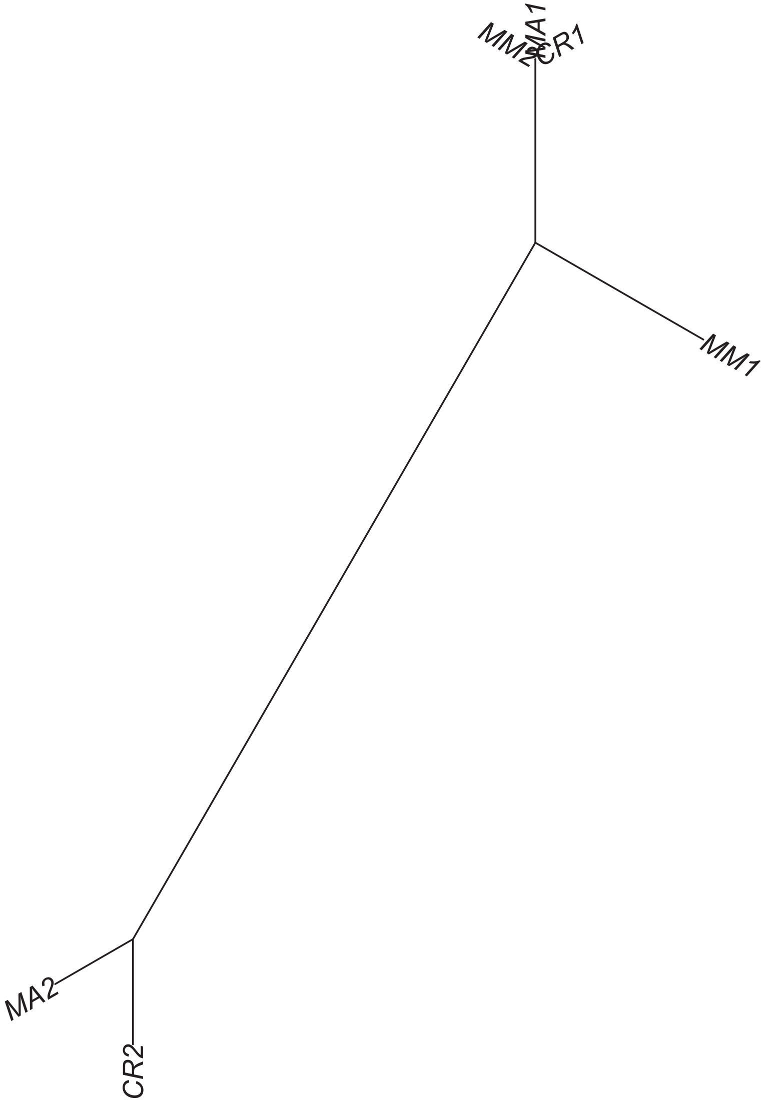

Region 168

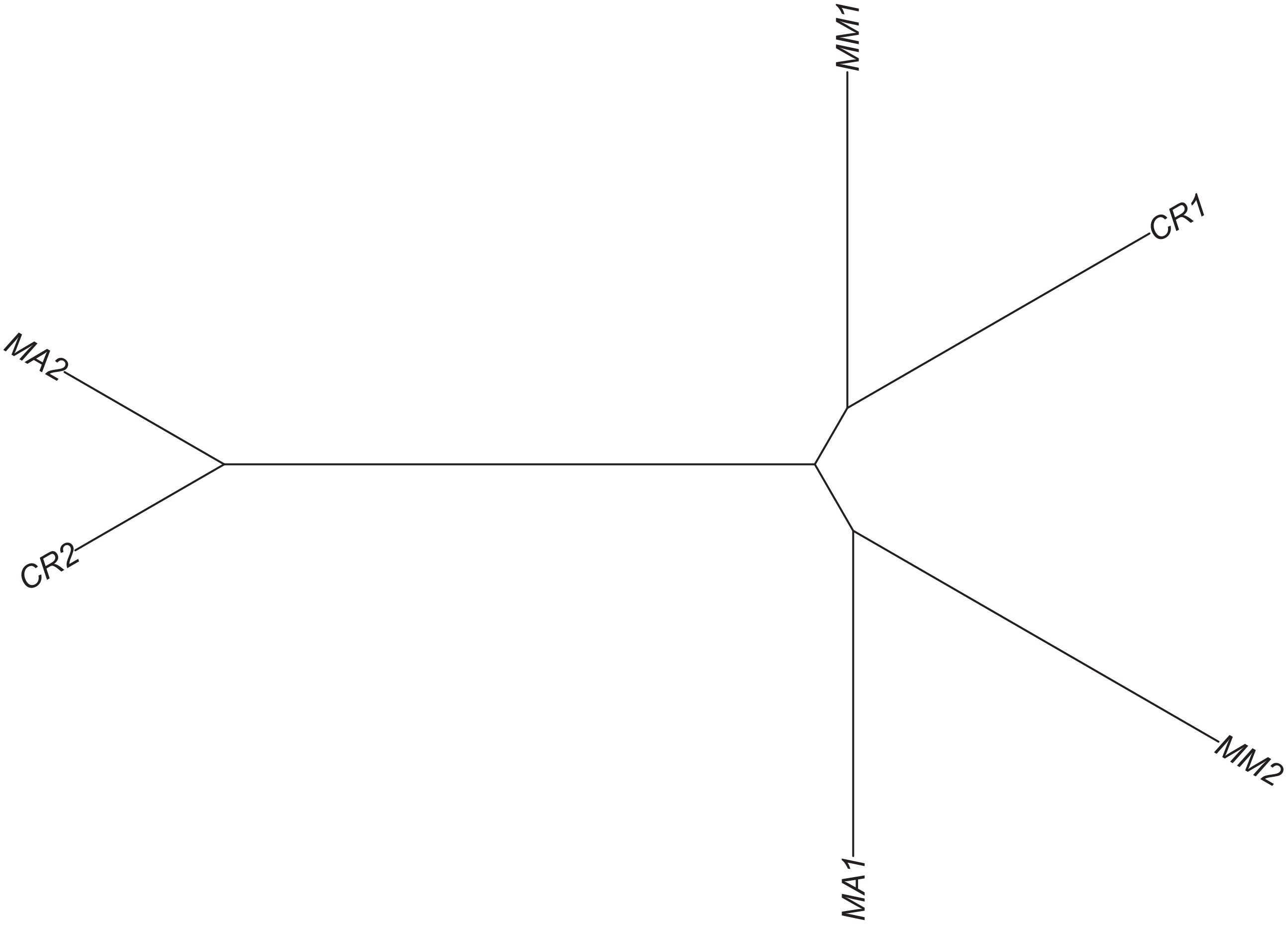

Region 170

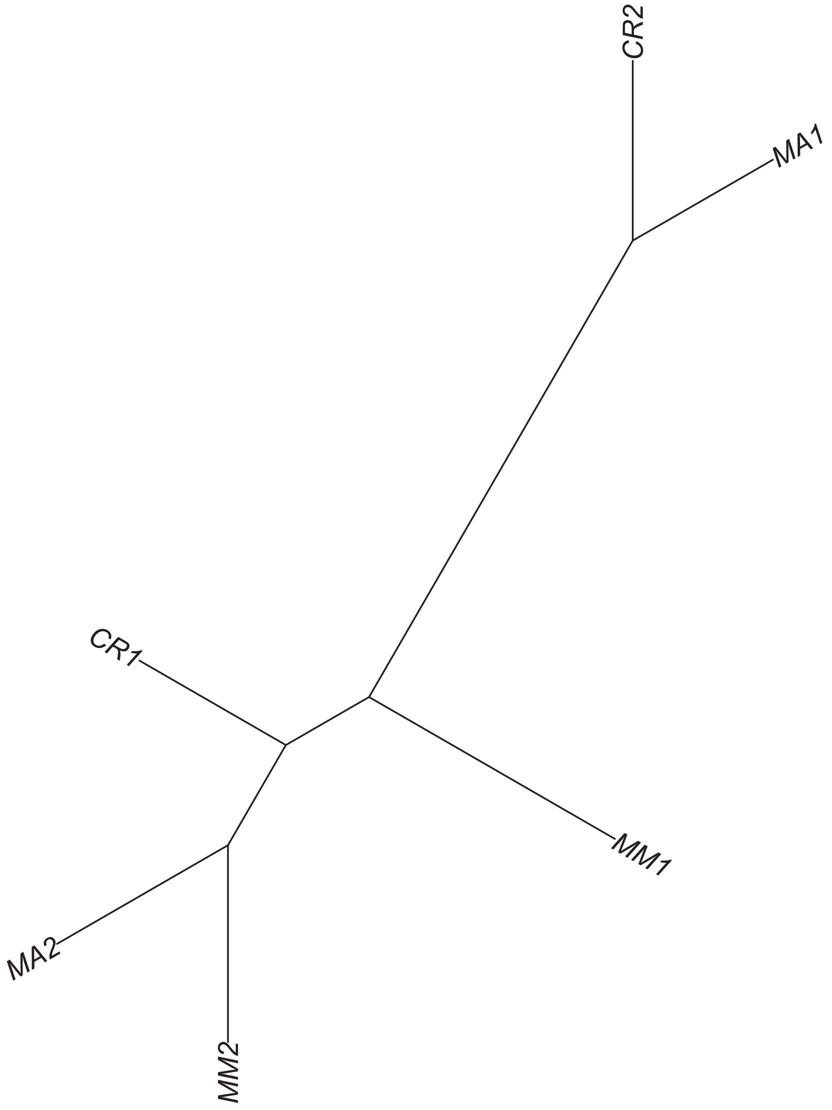

### Region 172



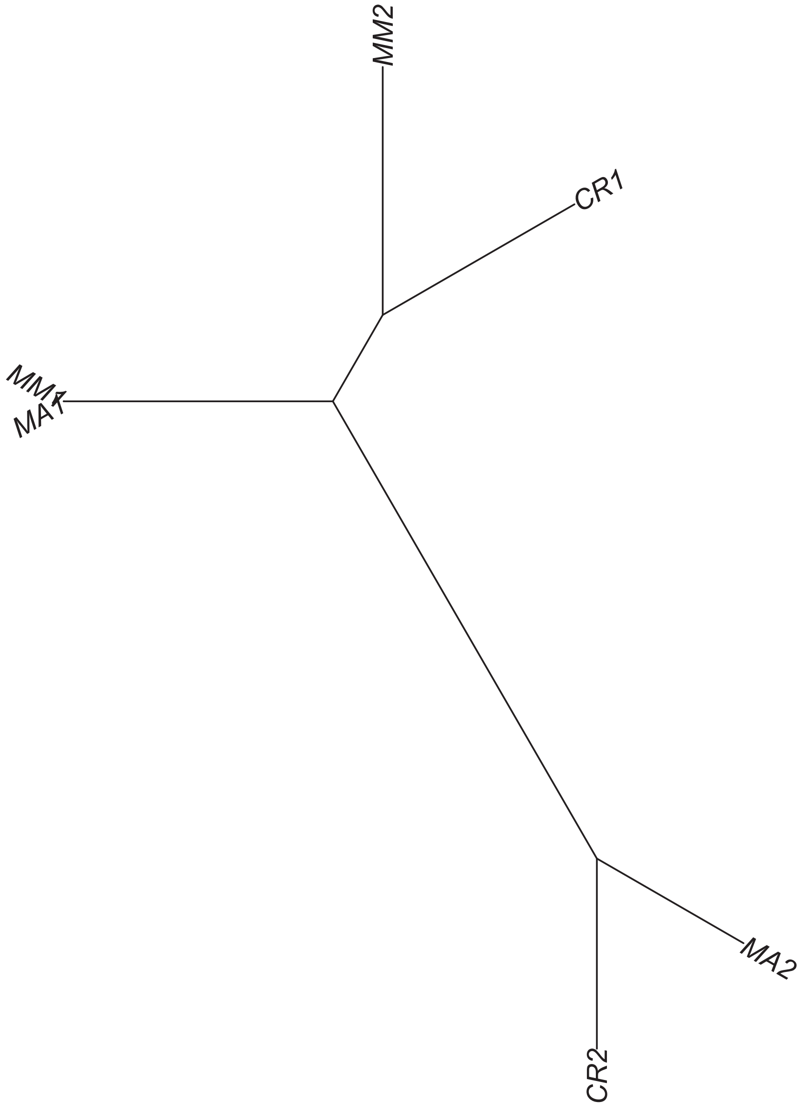

Region 173

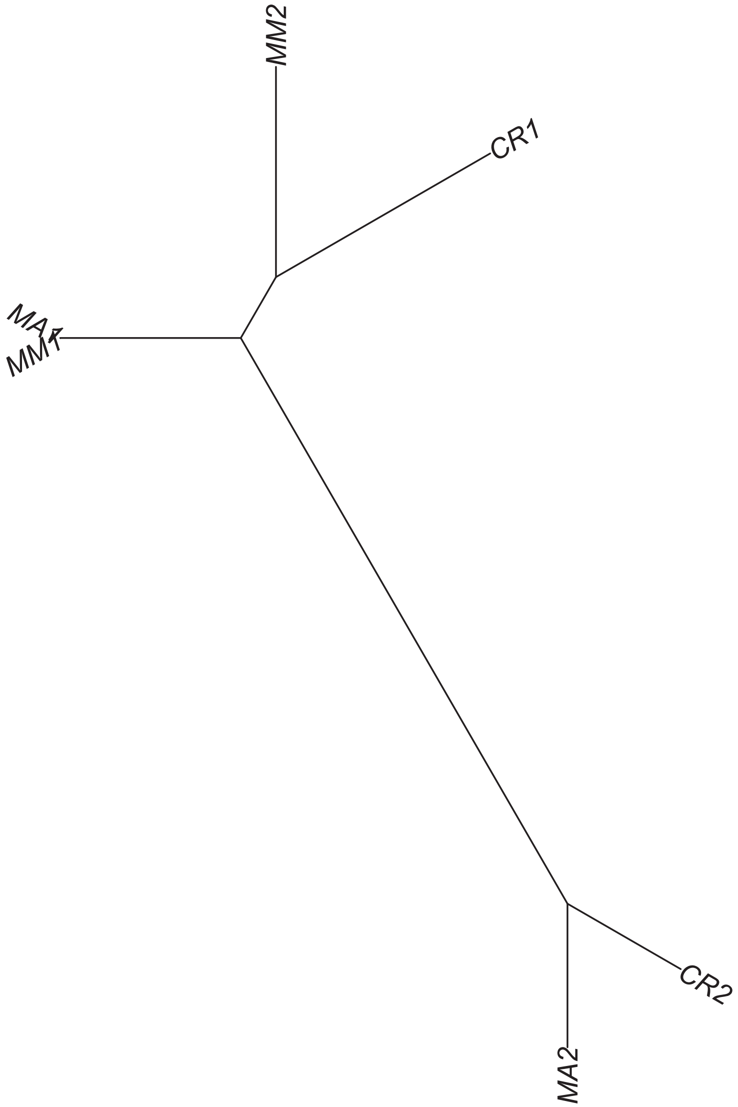

### Region 174

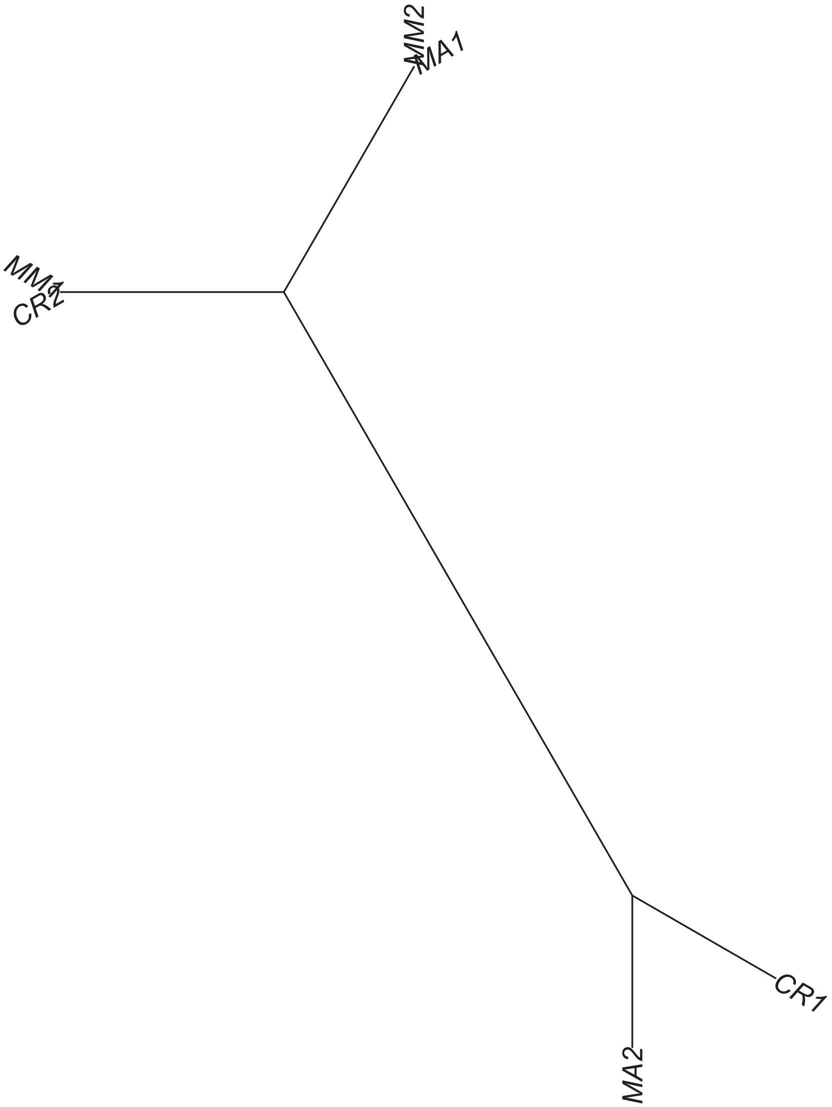

### Region 175

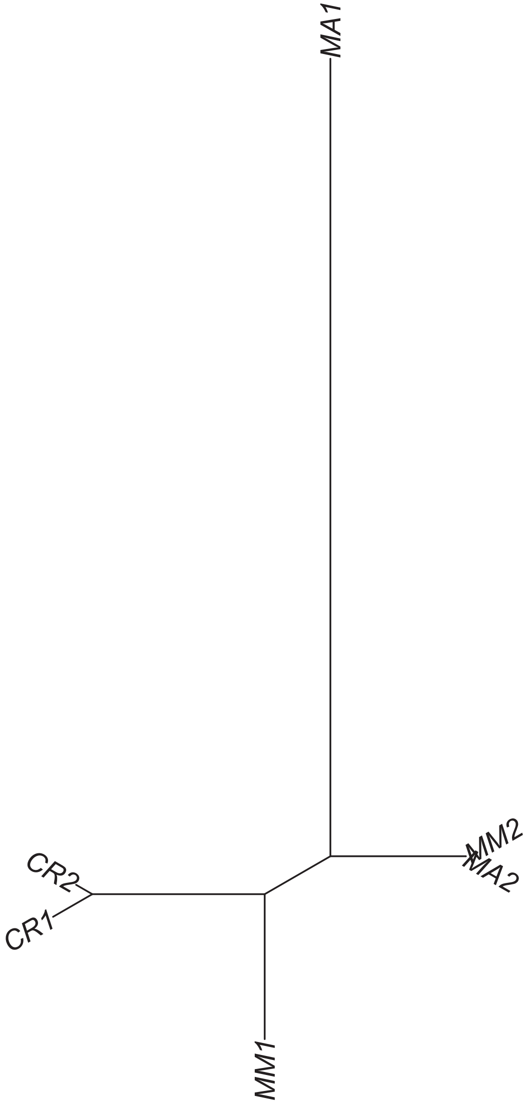

### Region 176

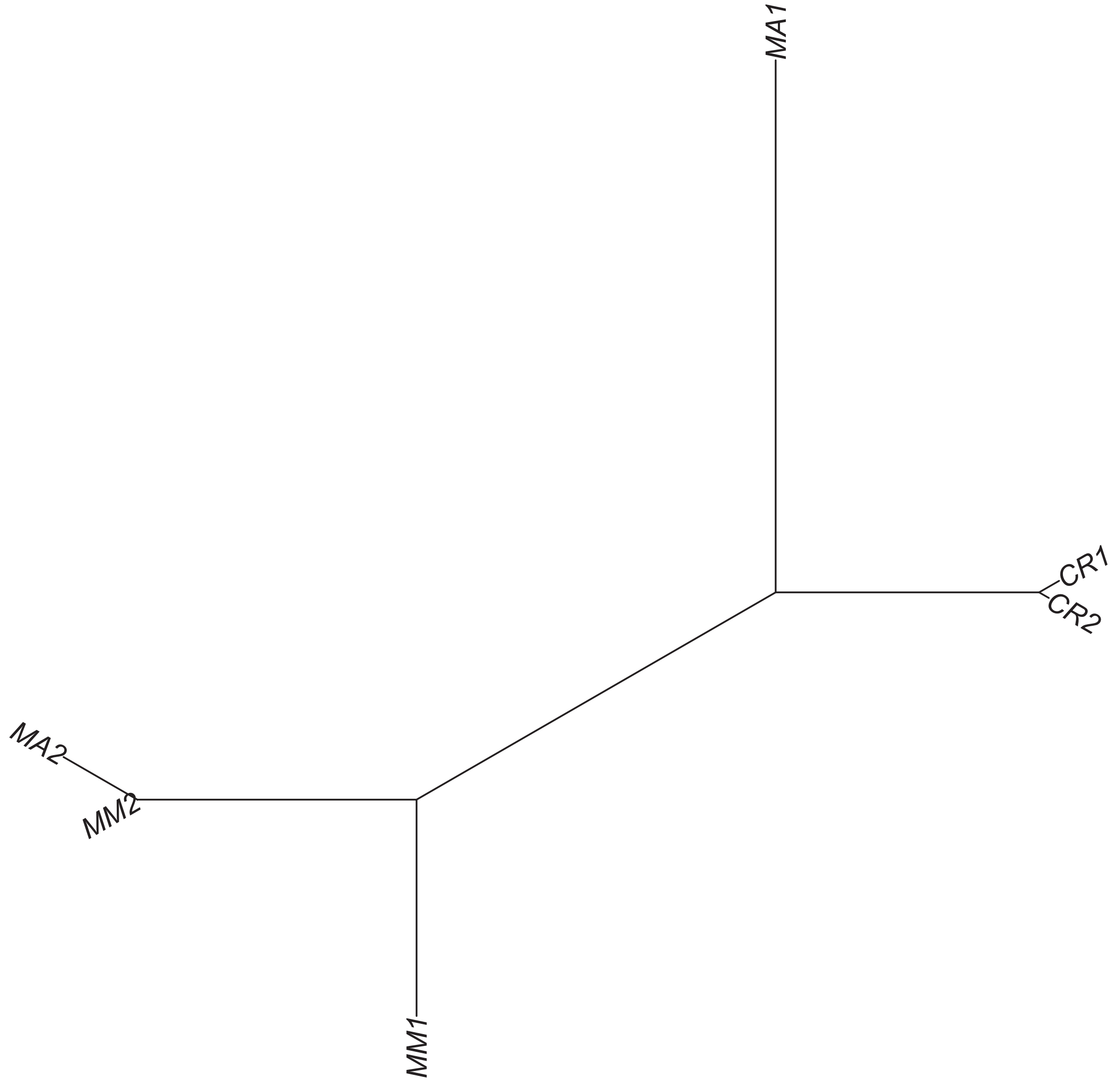

Region 177

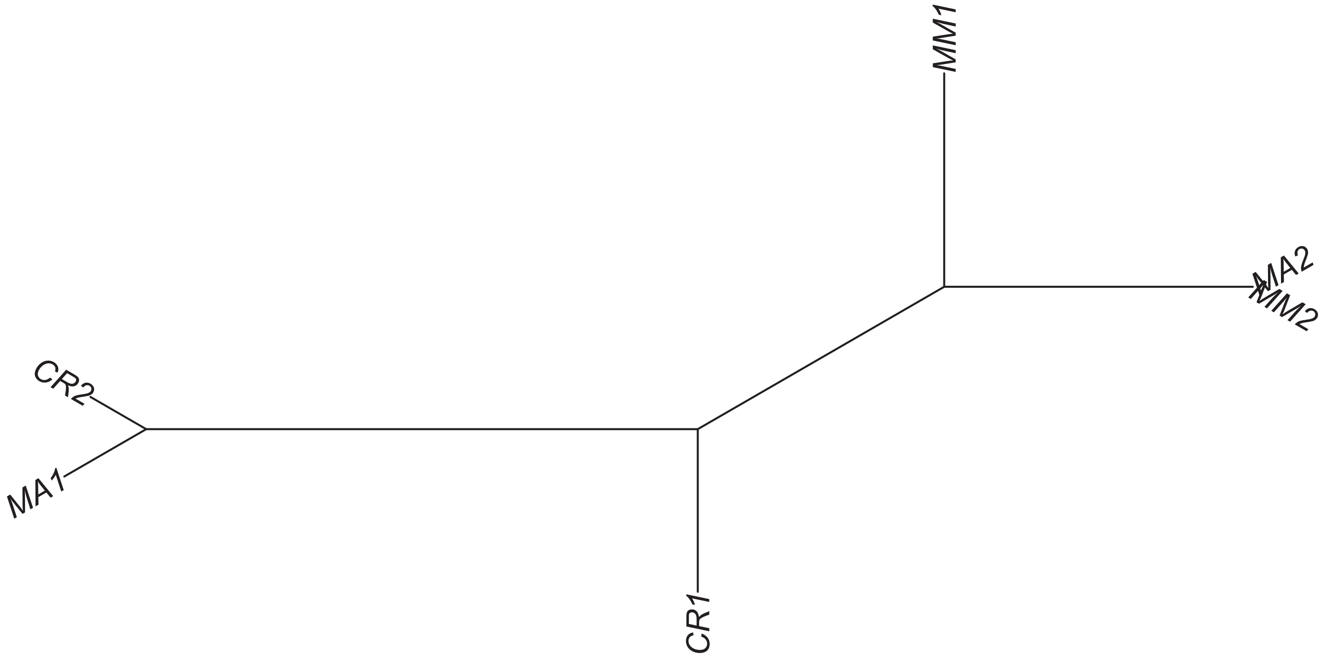

### Region 178

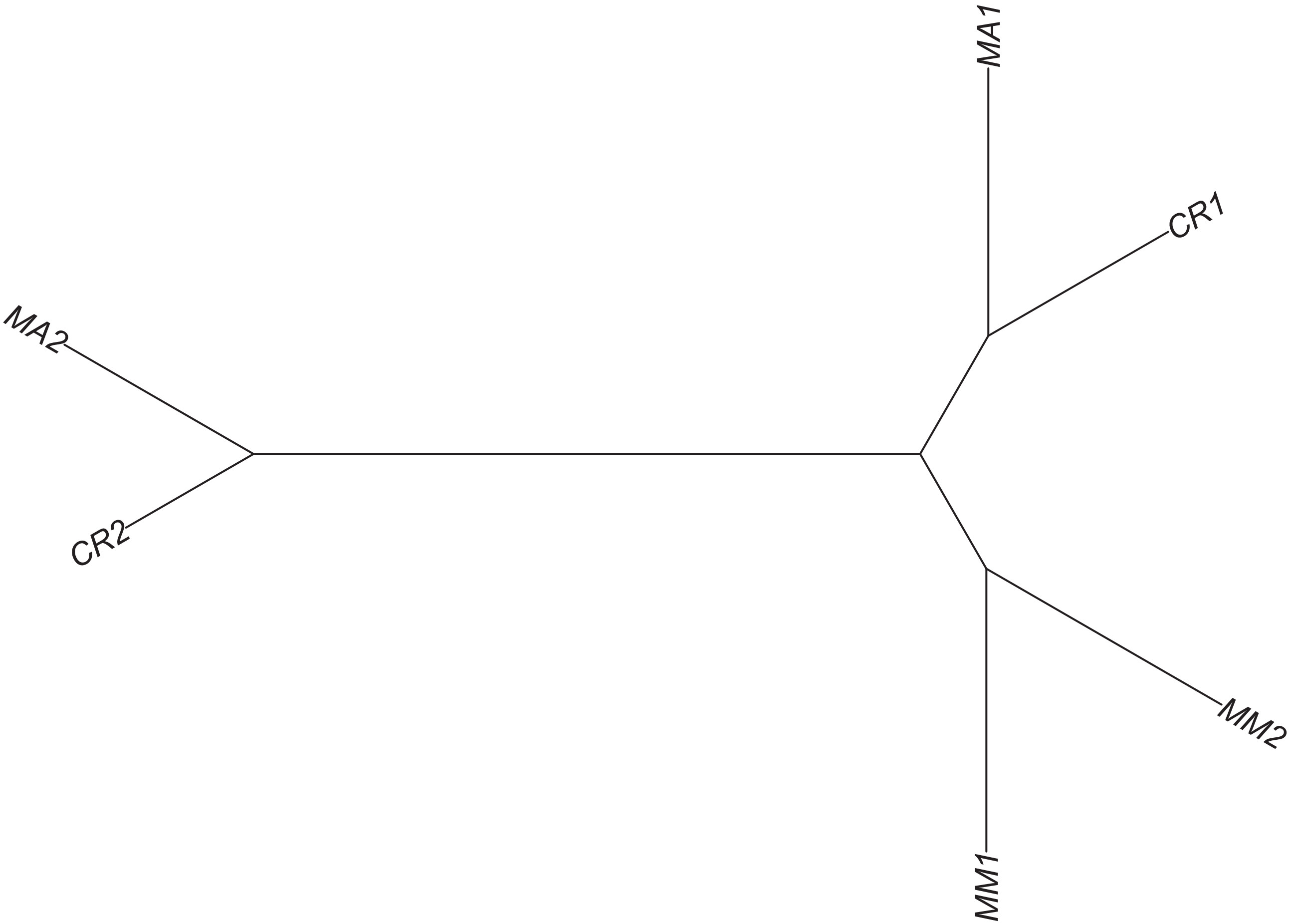

### Region 179



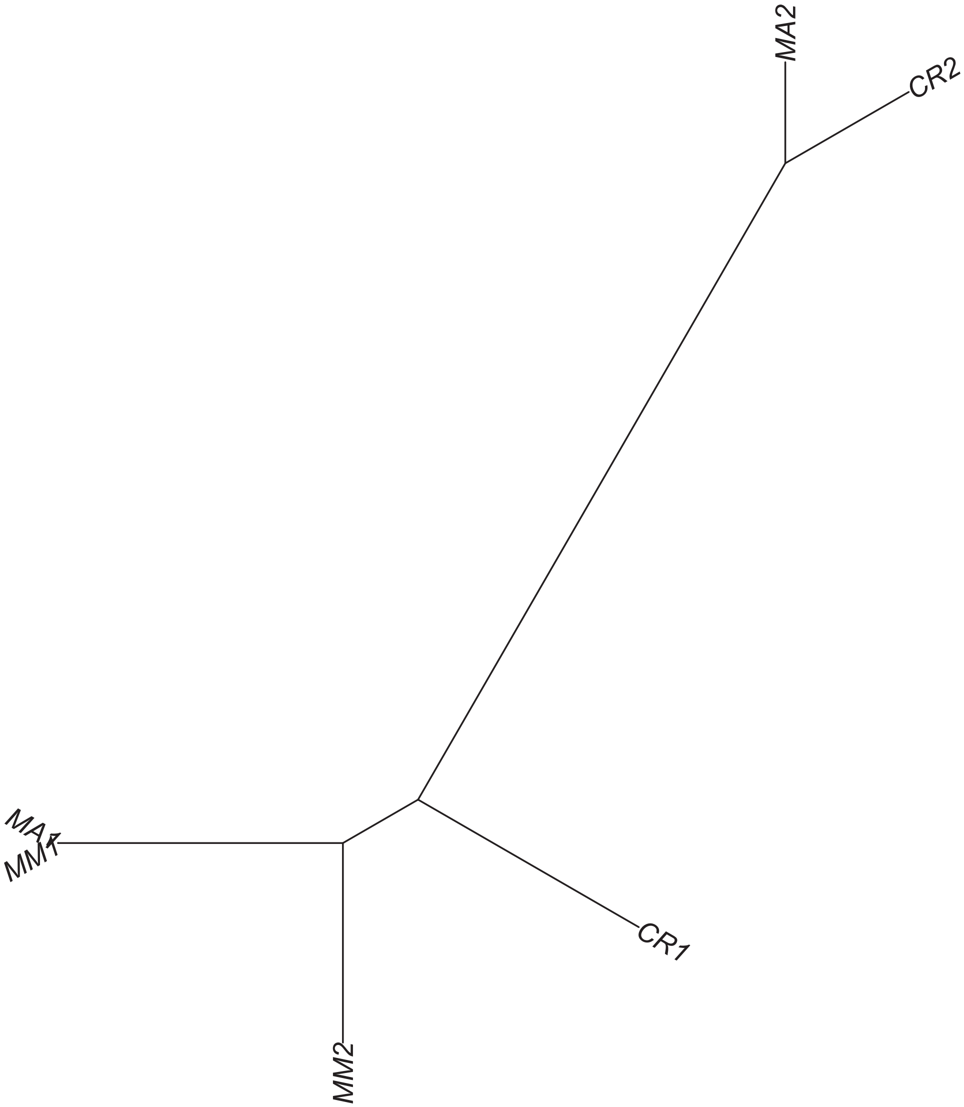

### Region 180

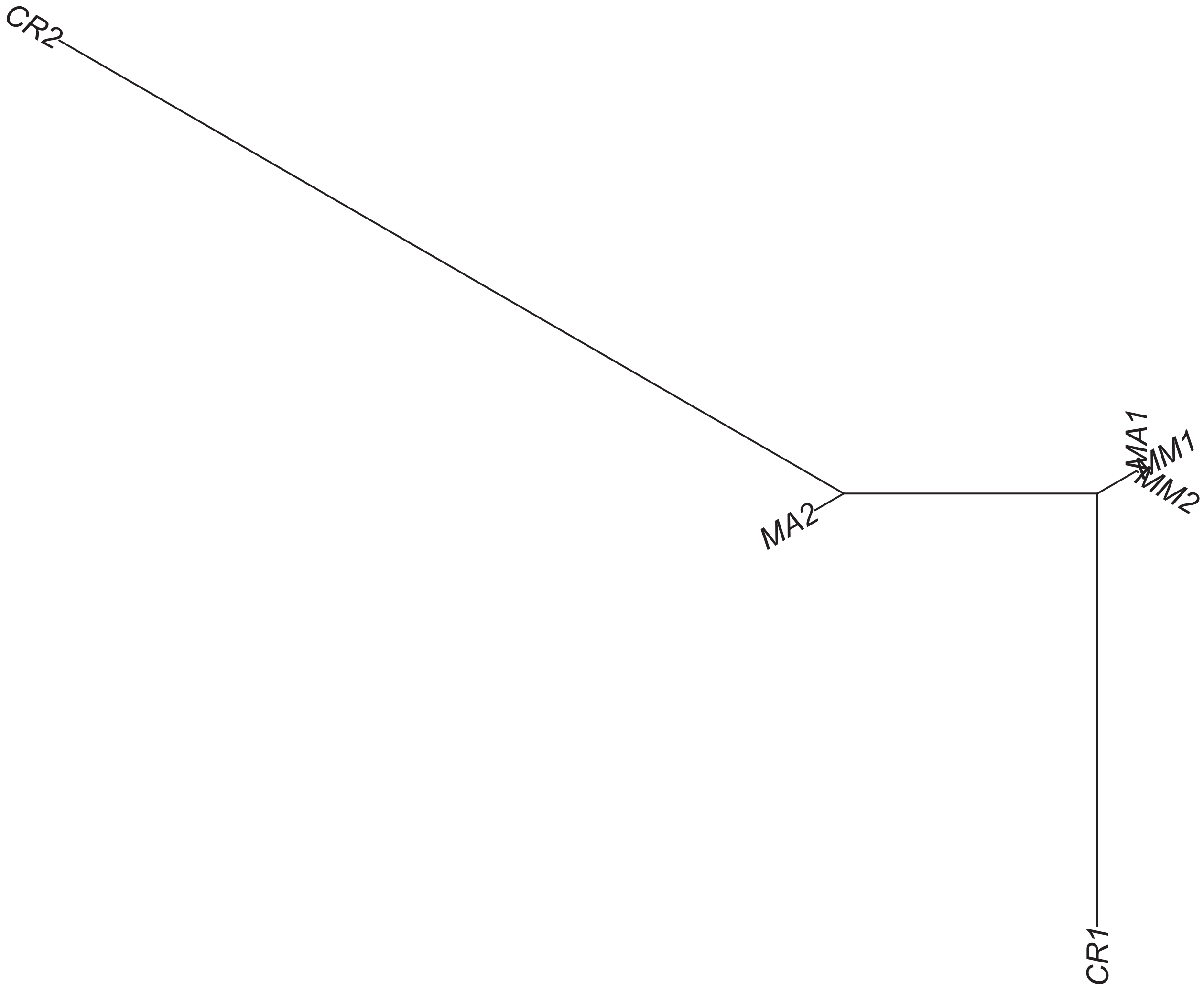

### Region 181



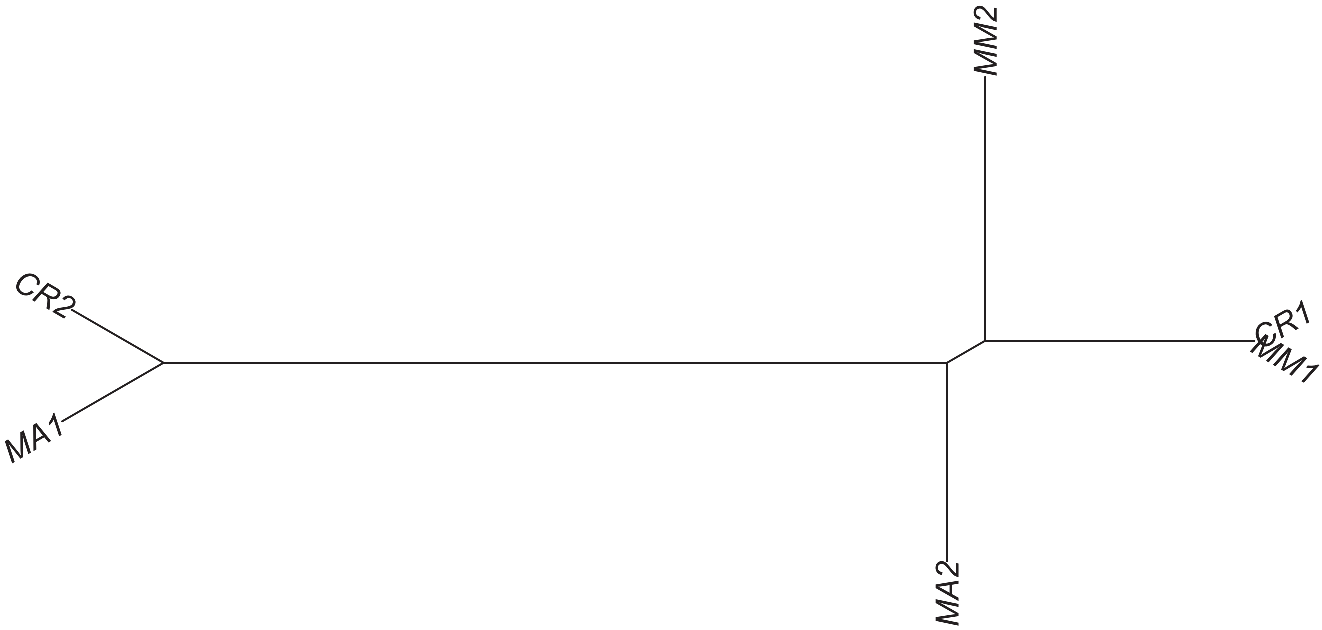

### Region 183

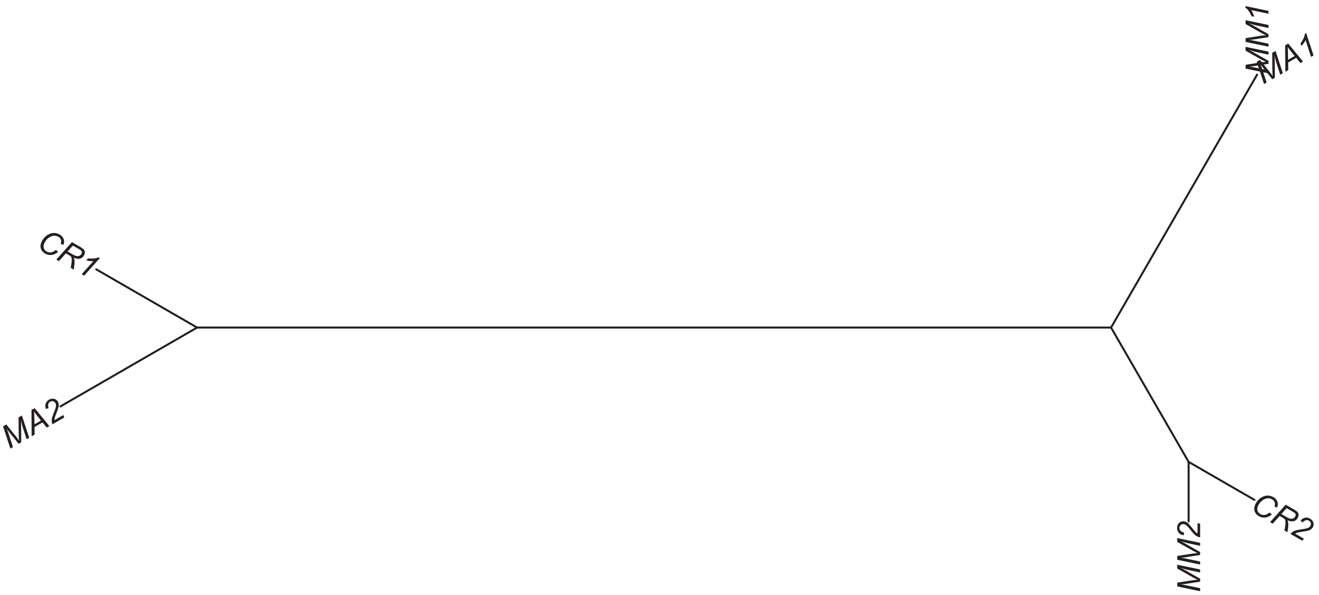

### Region 186



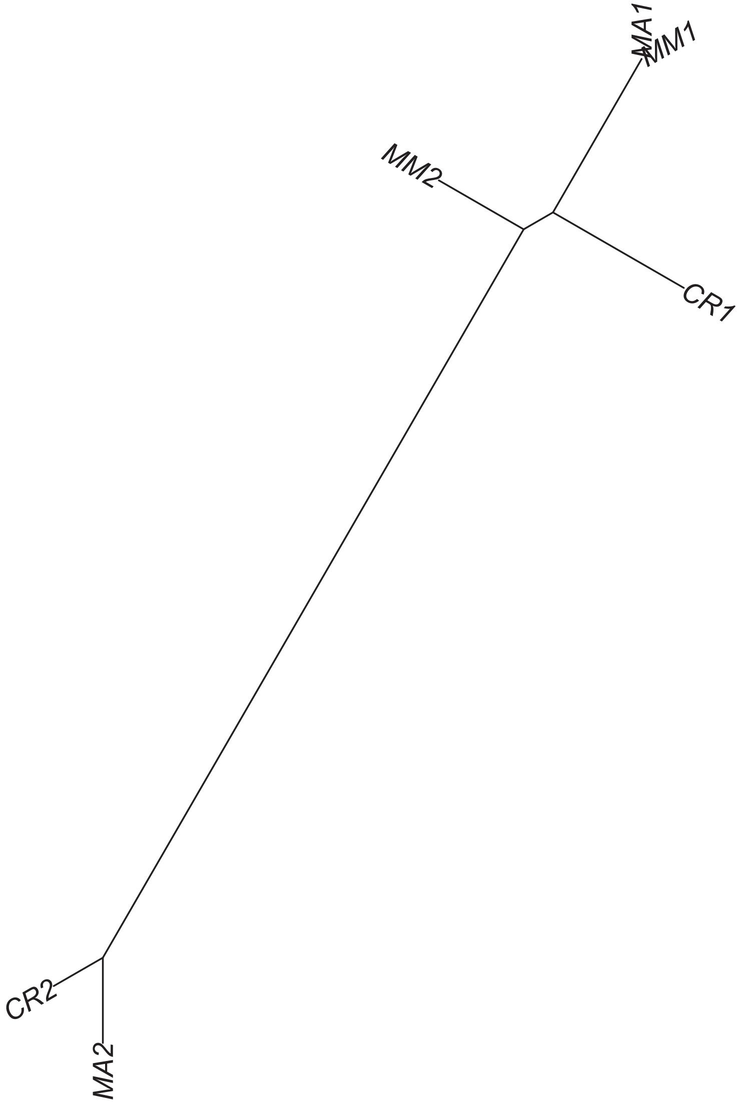

Region 187

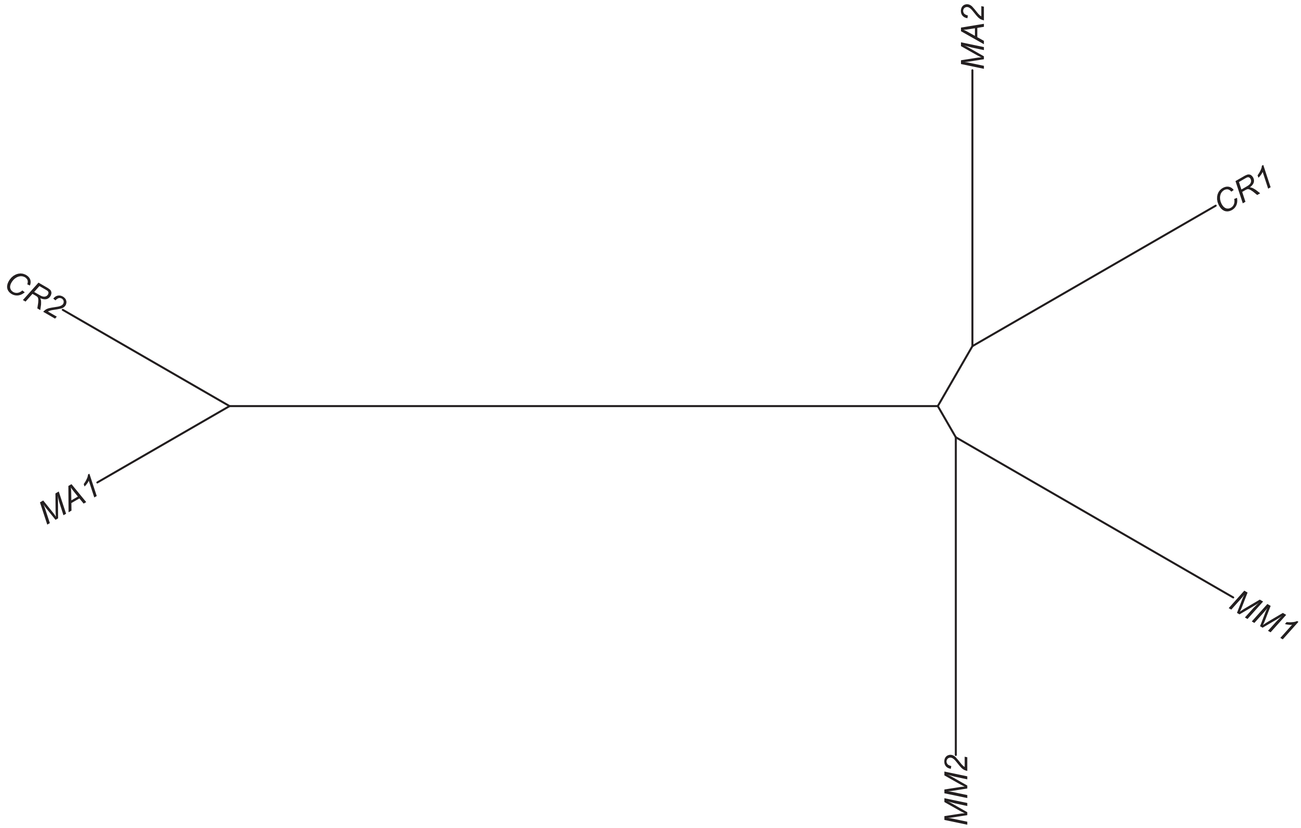

### Region 188

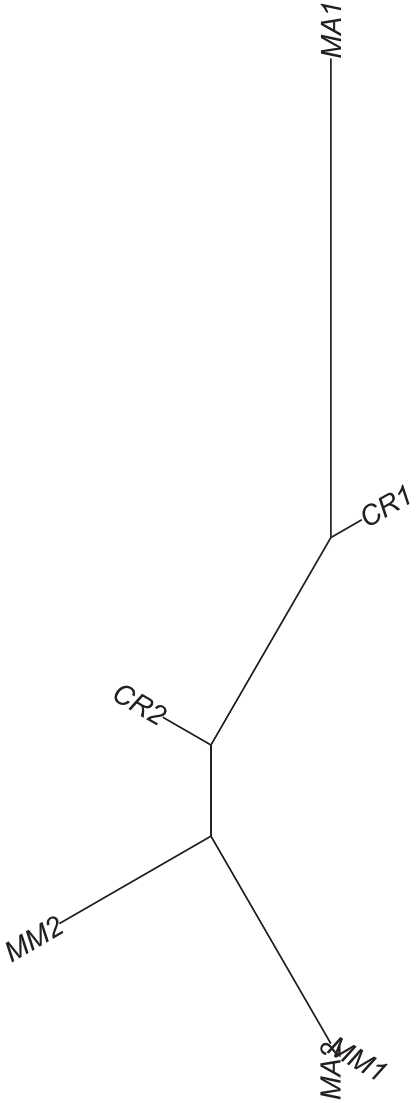

### Region 189

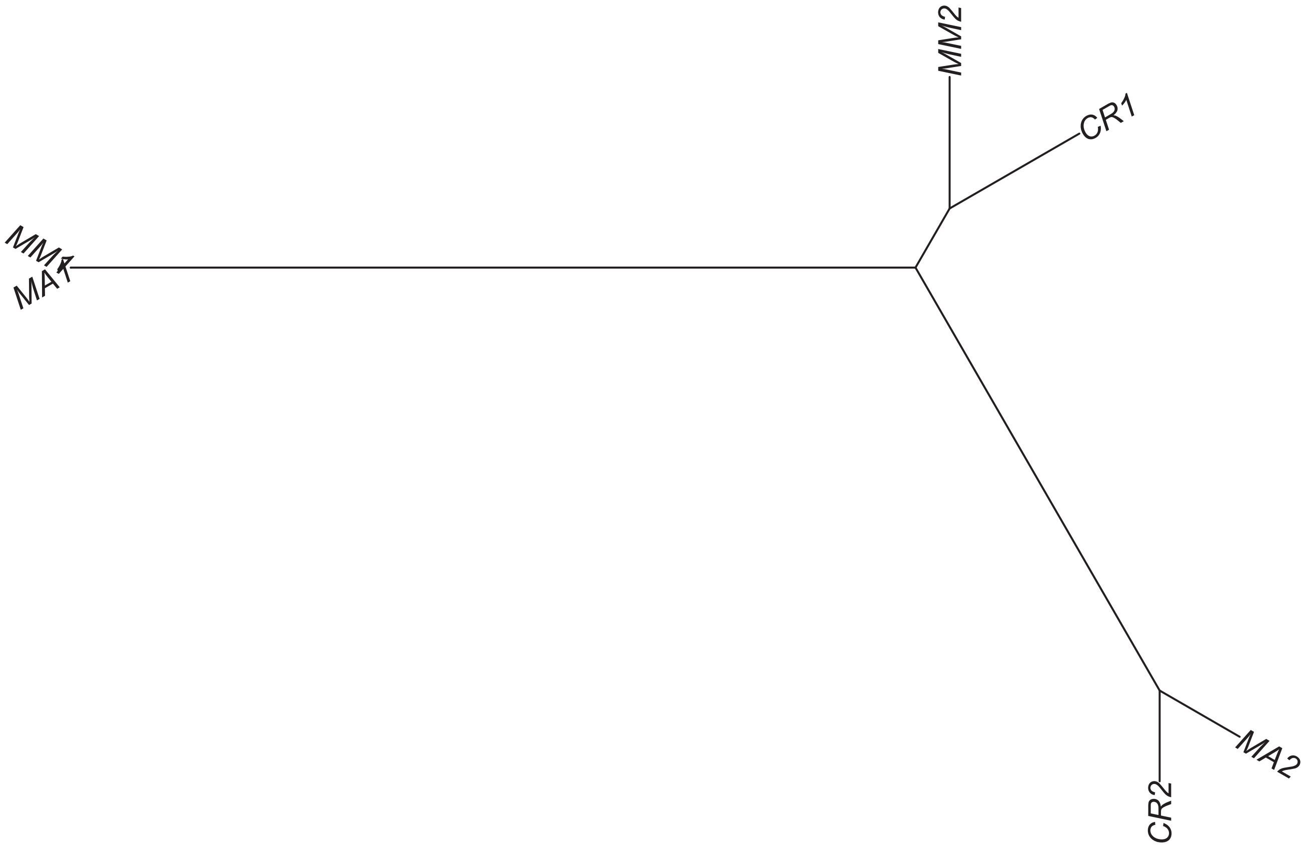

Region 197

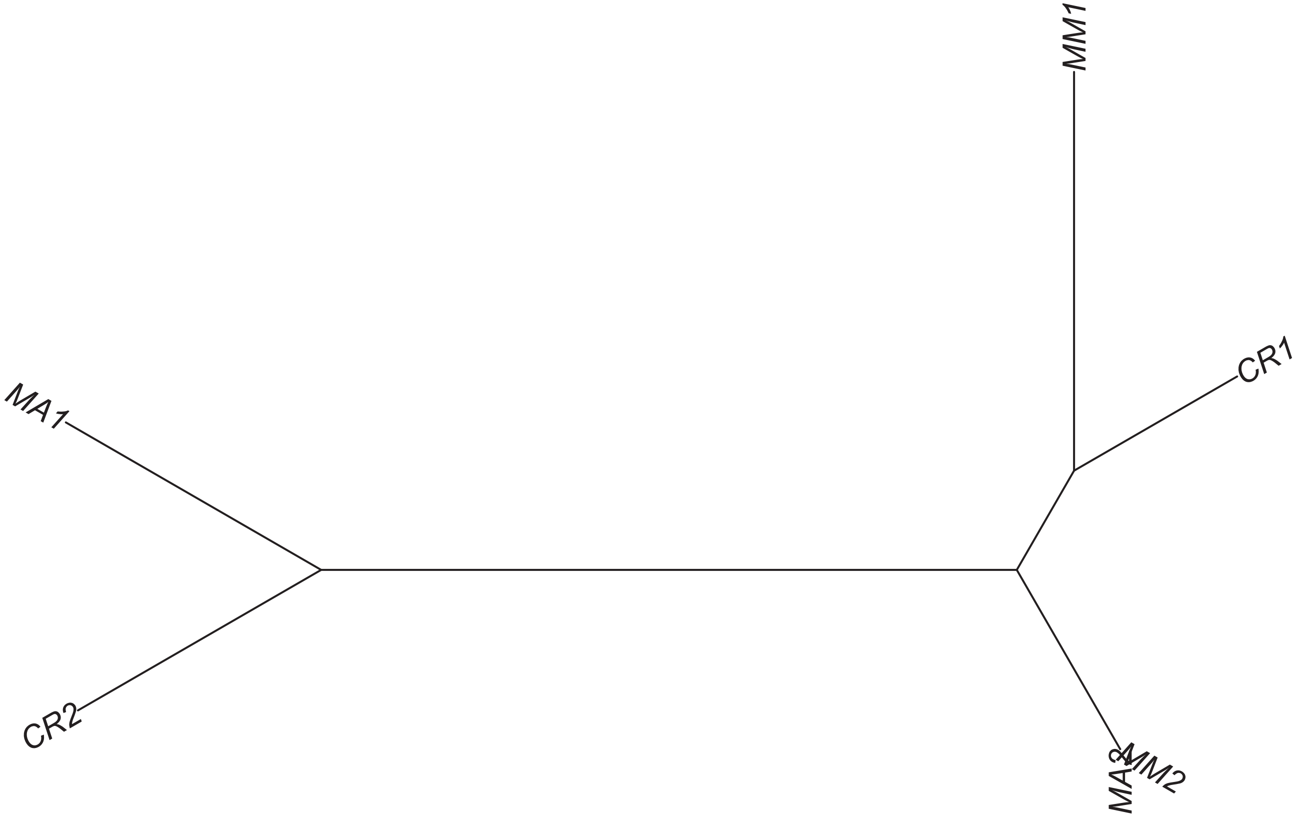

### Region 198

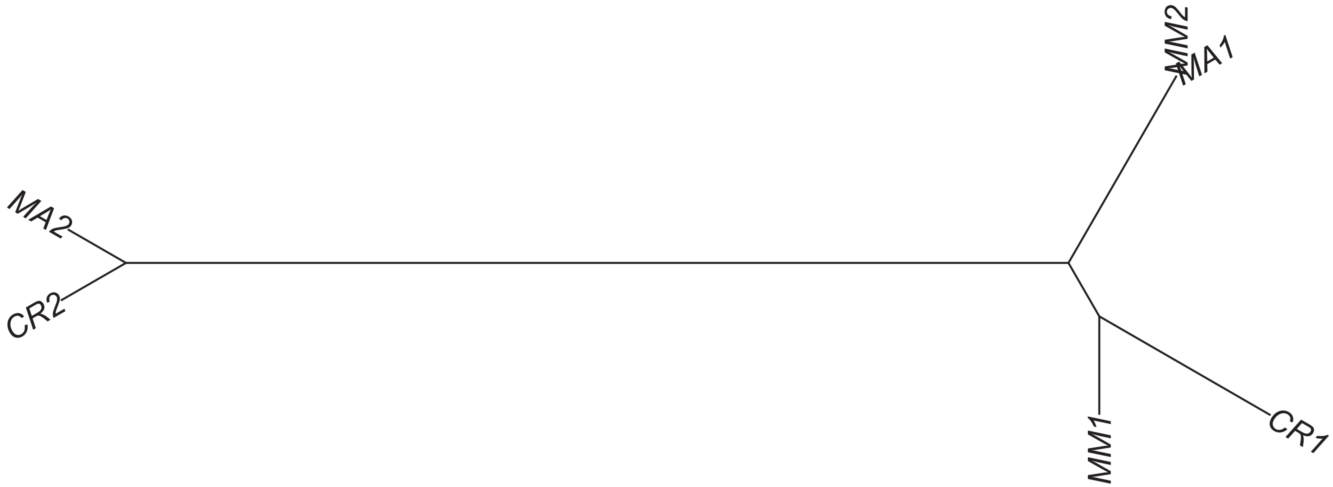

### Region 199

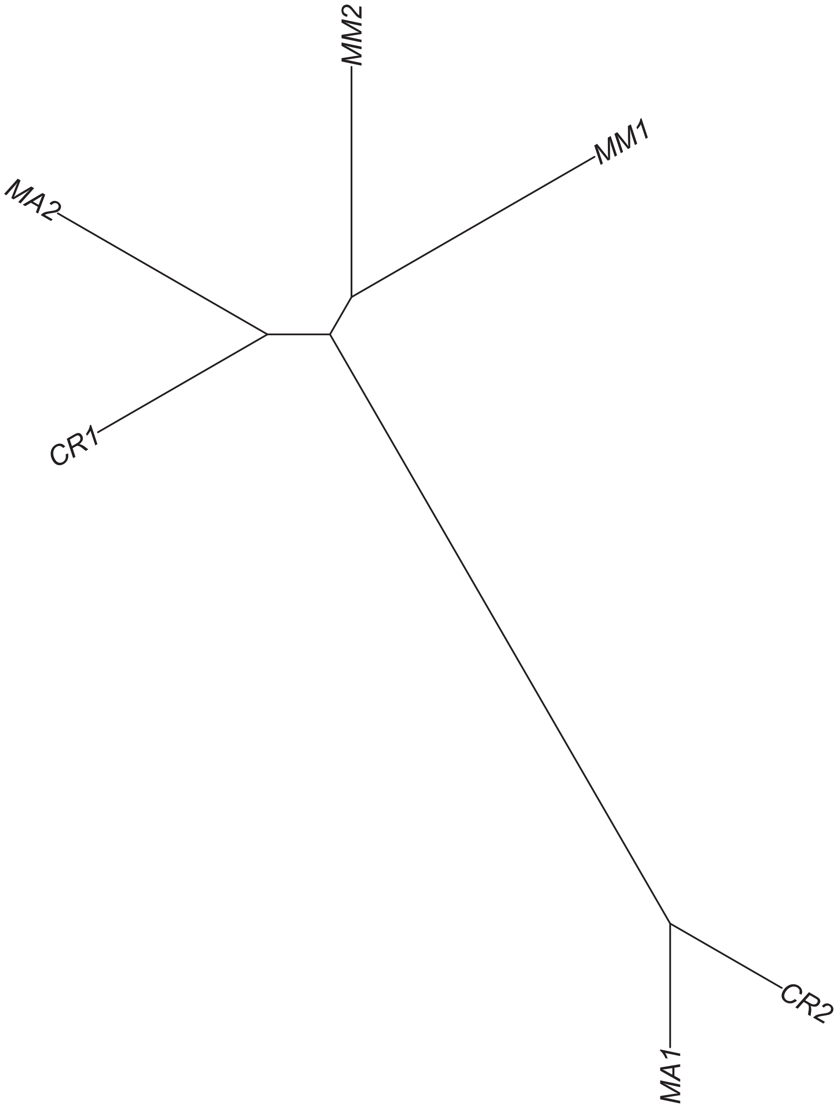

Region 201

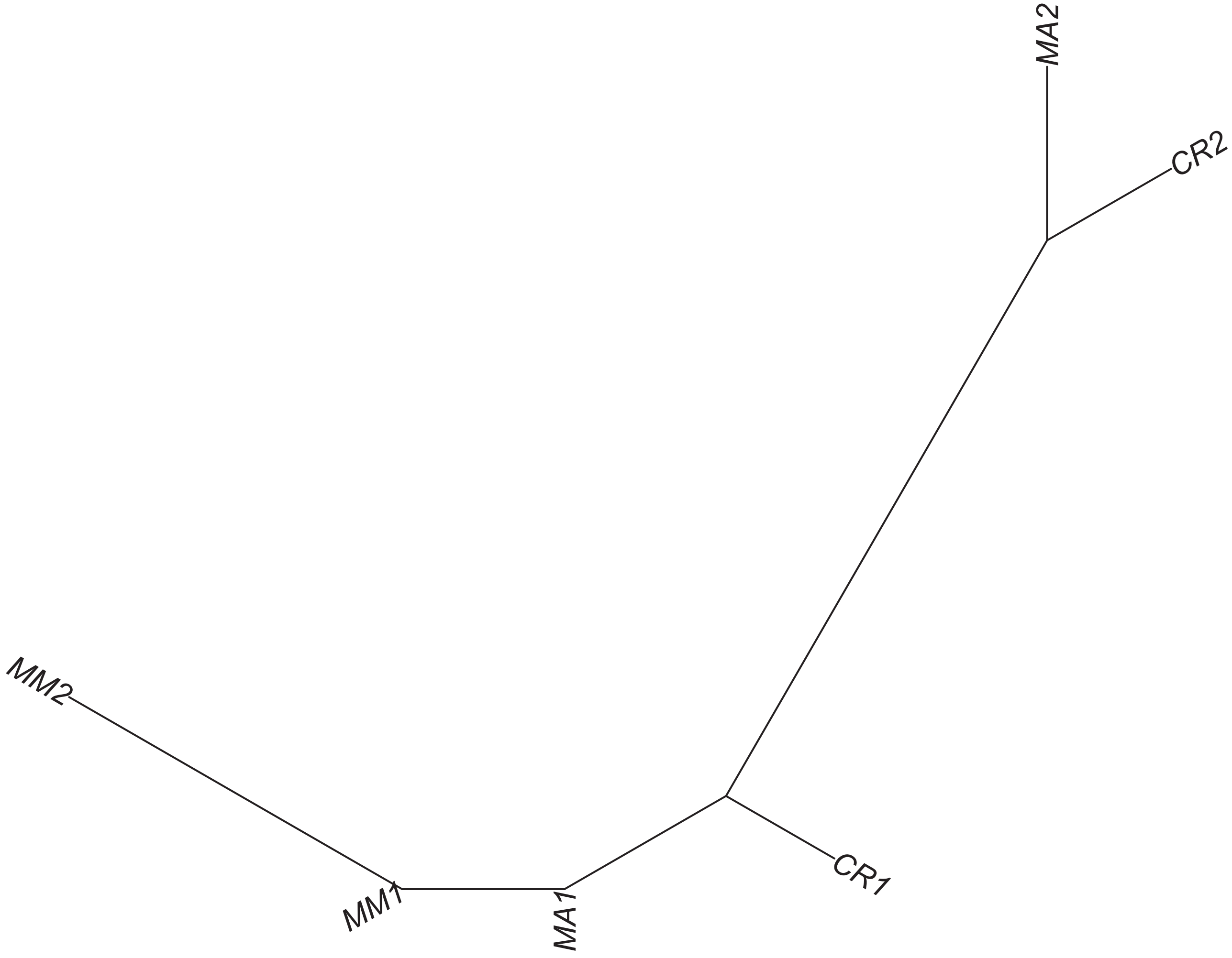

Region 203

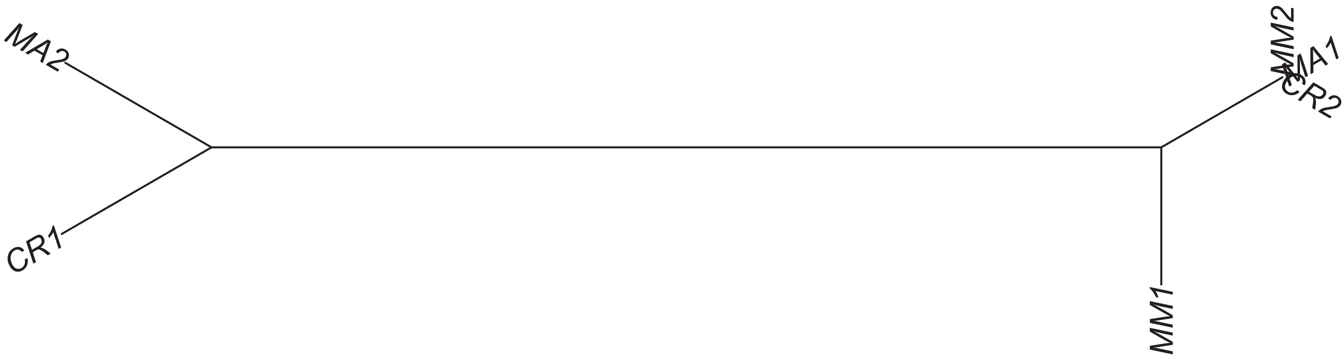

### Region 204

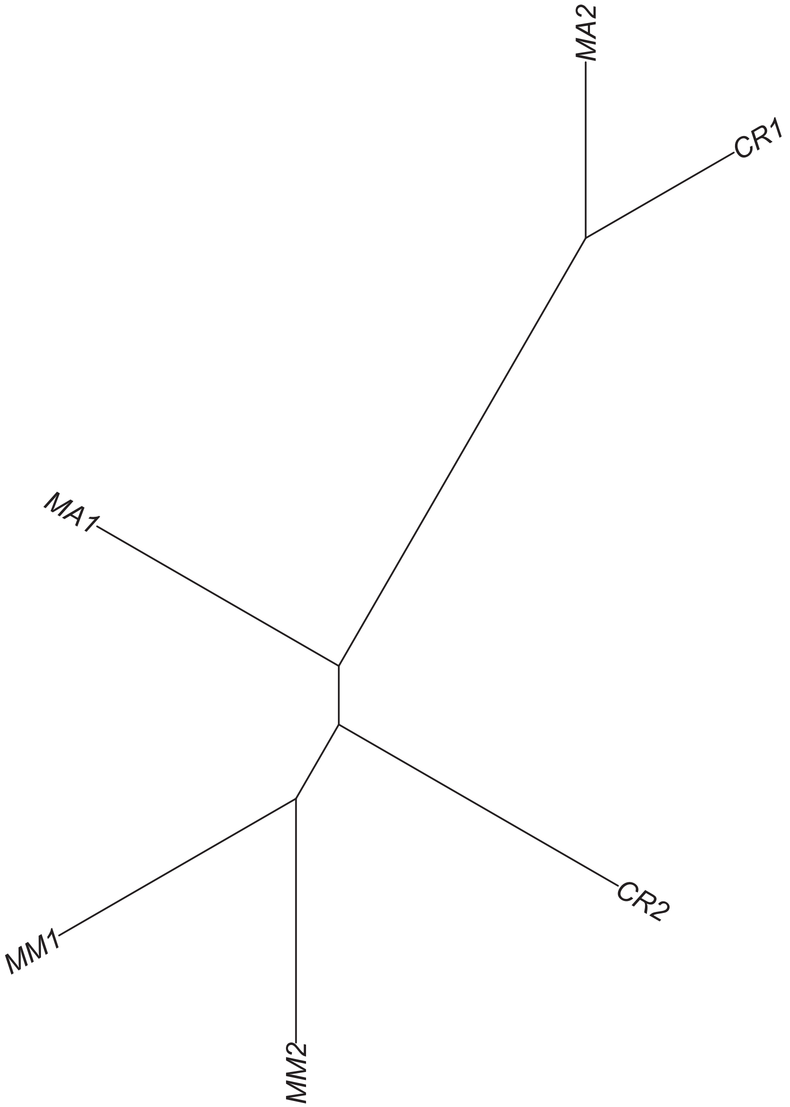

### Region 205

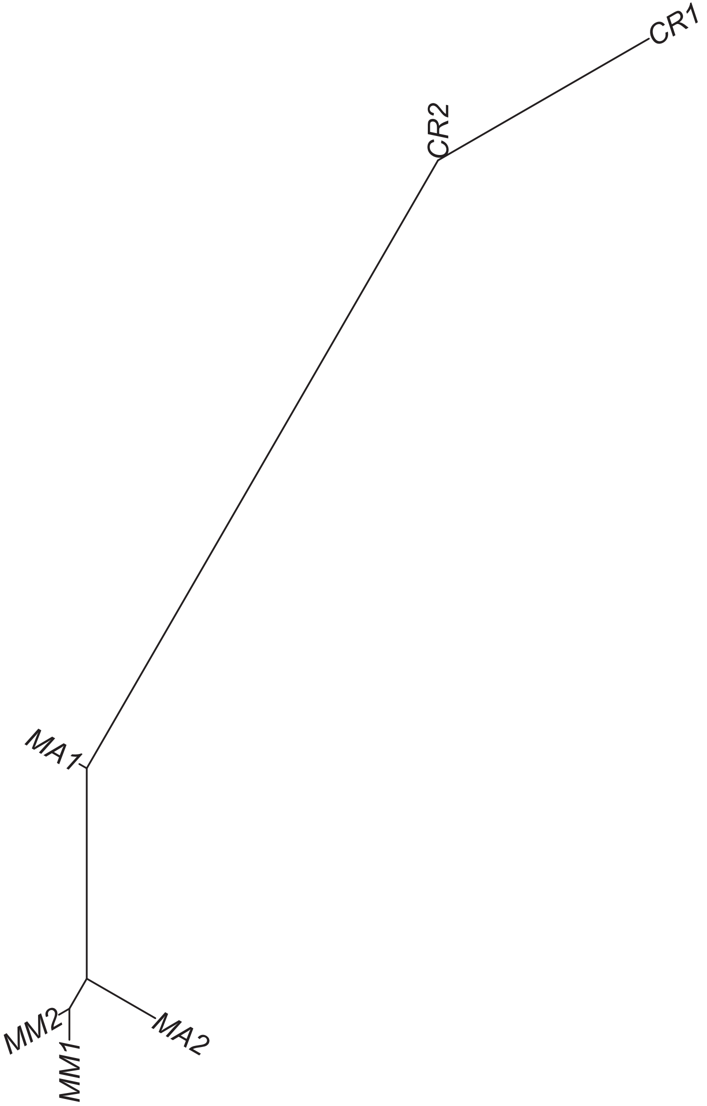

### Region 206

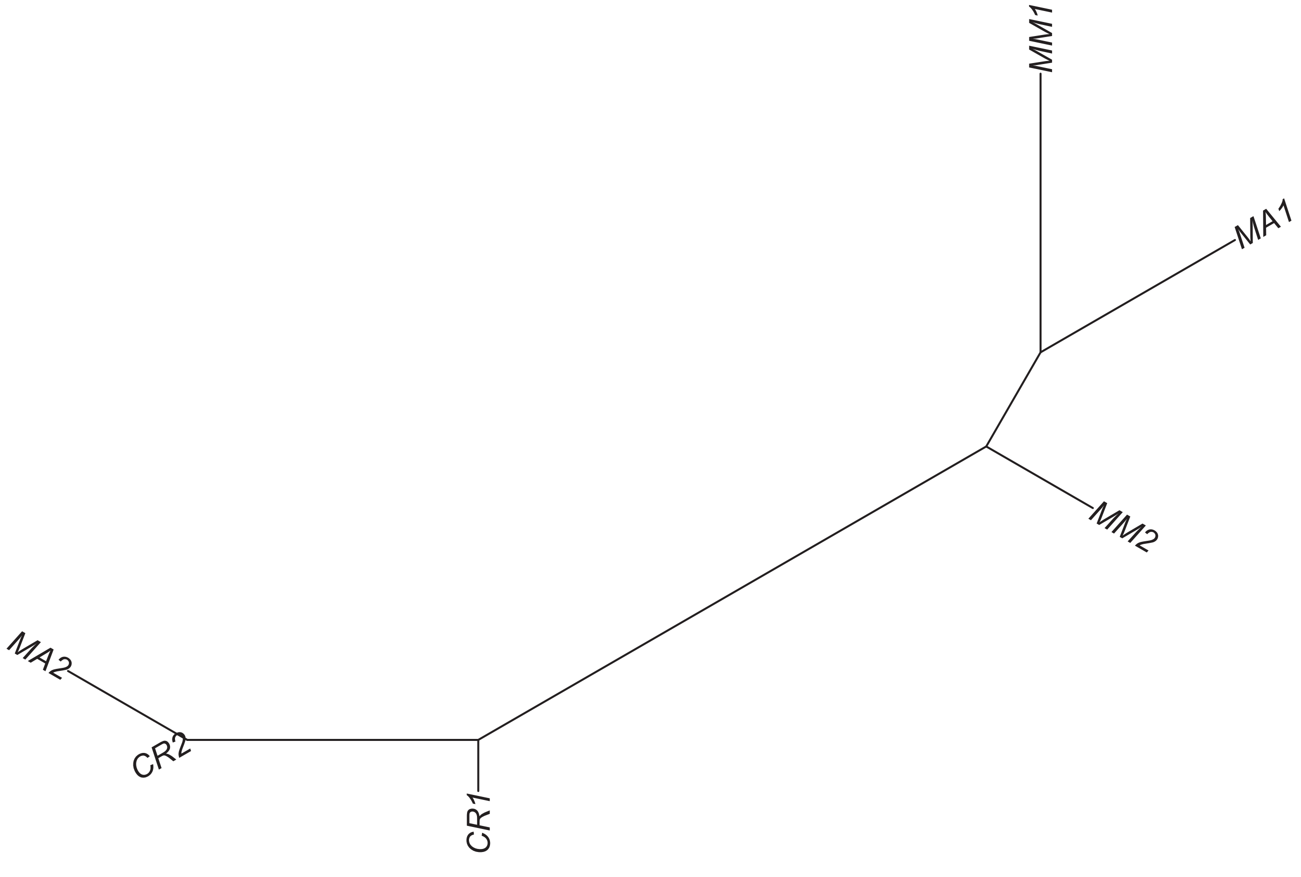

### Region 207

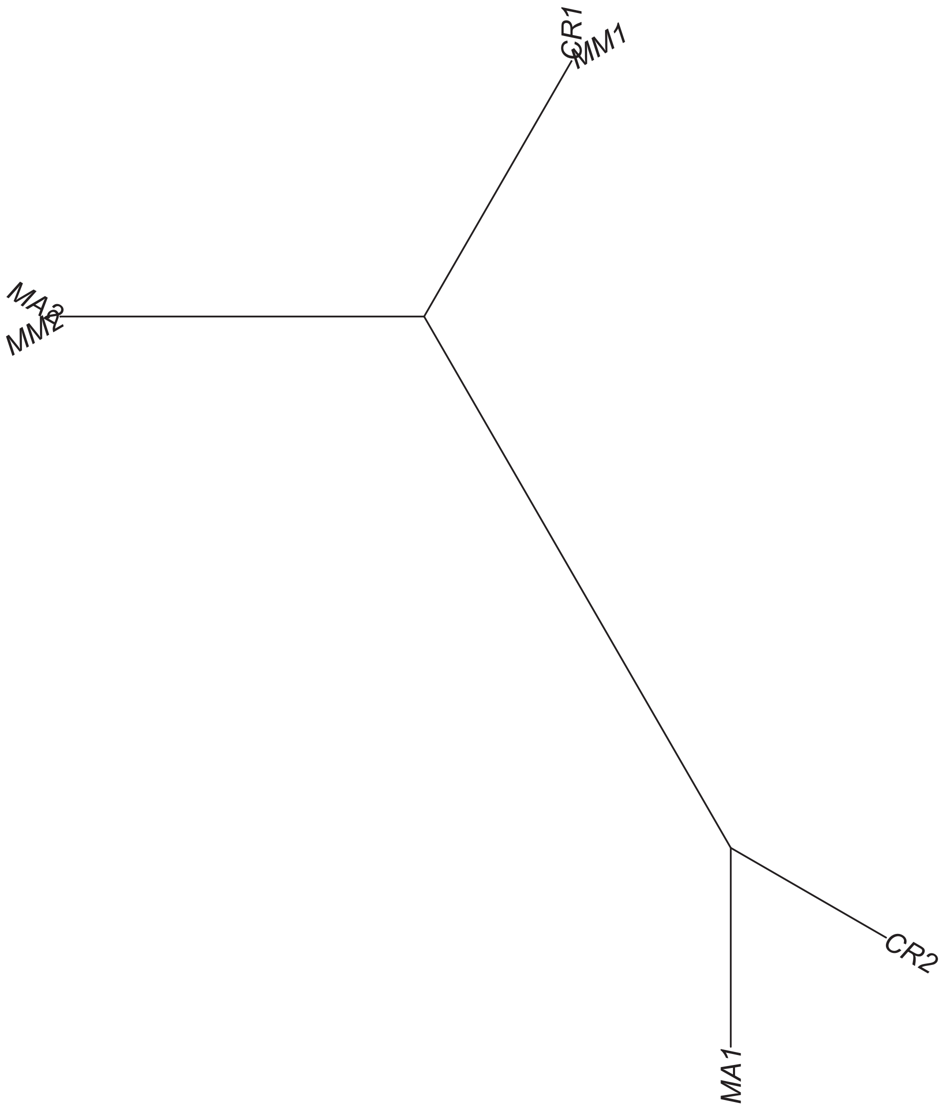

### Region 208

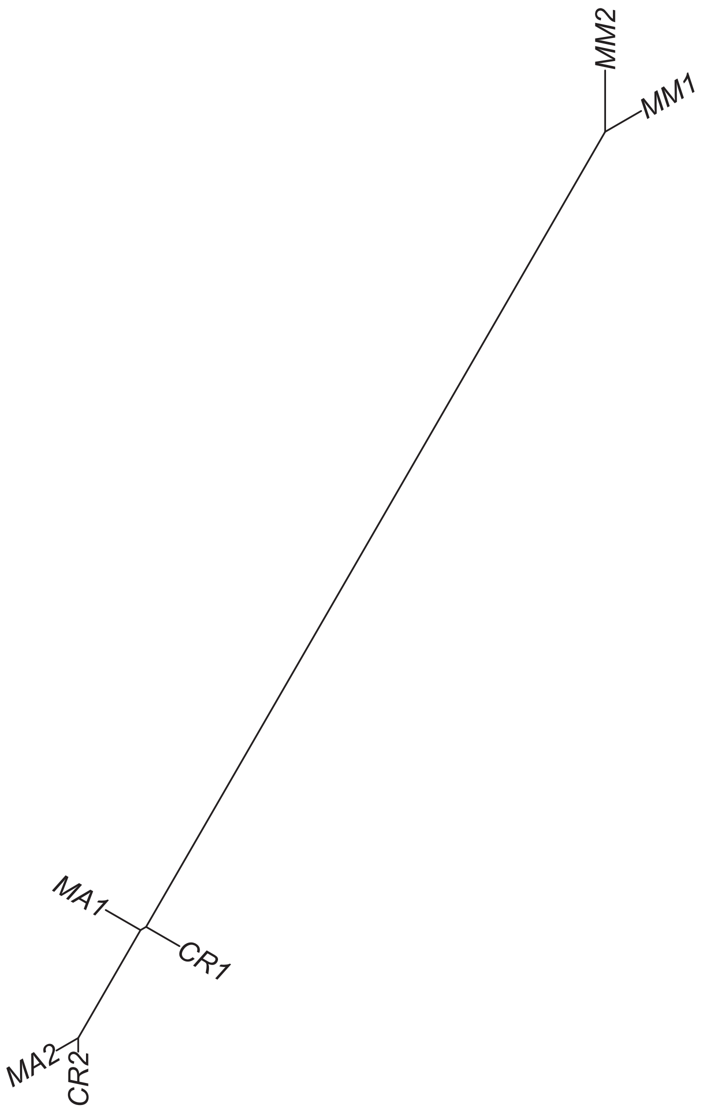

### Region 209

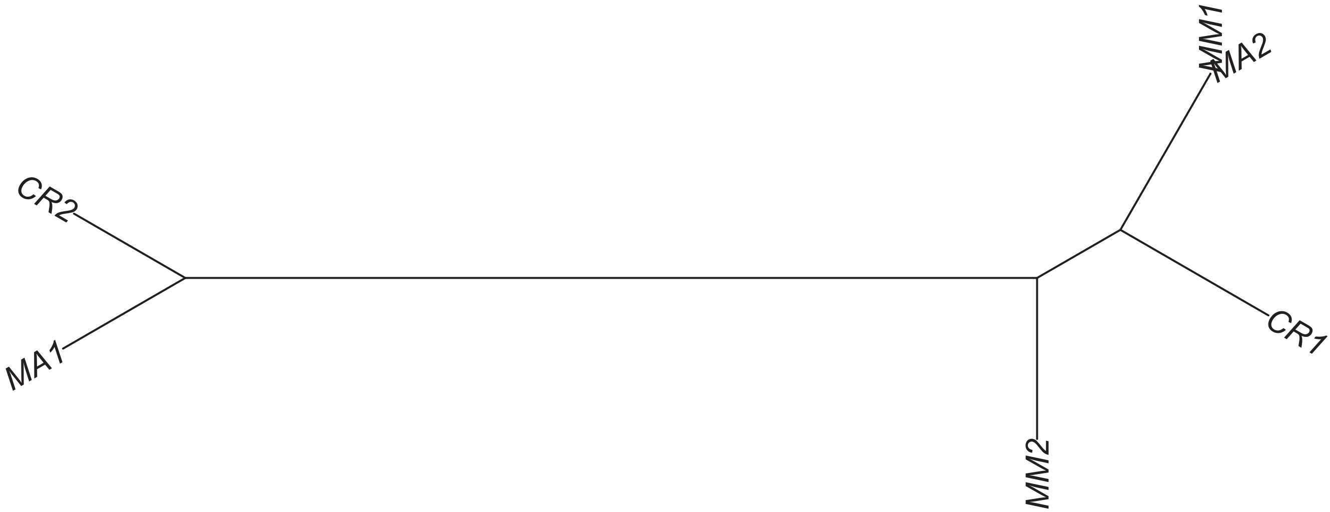

### Region 210



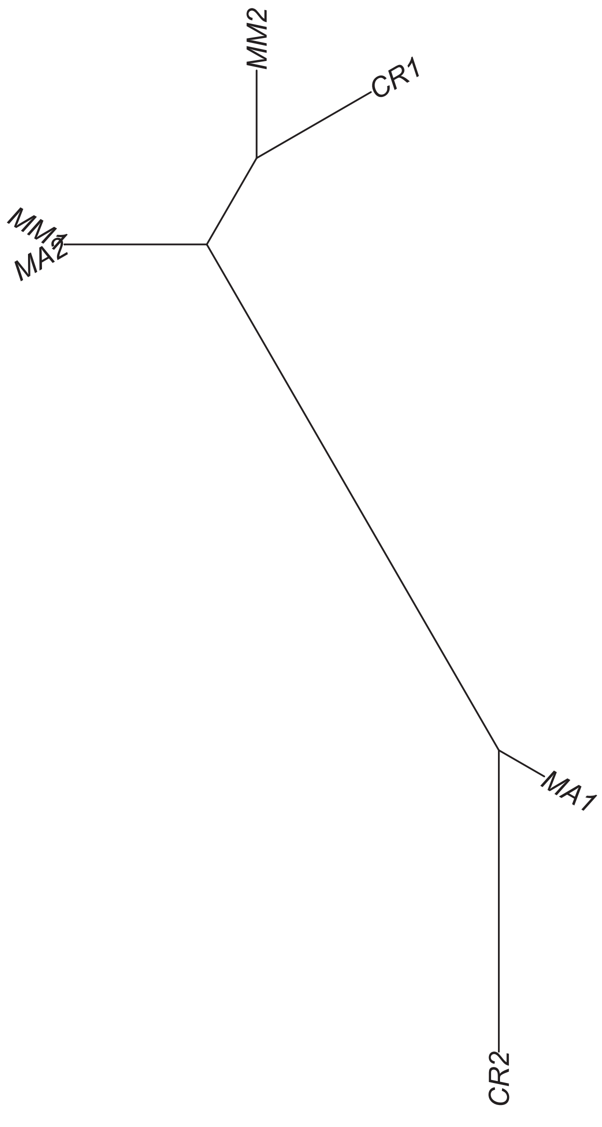

### Region 211

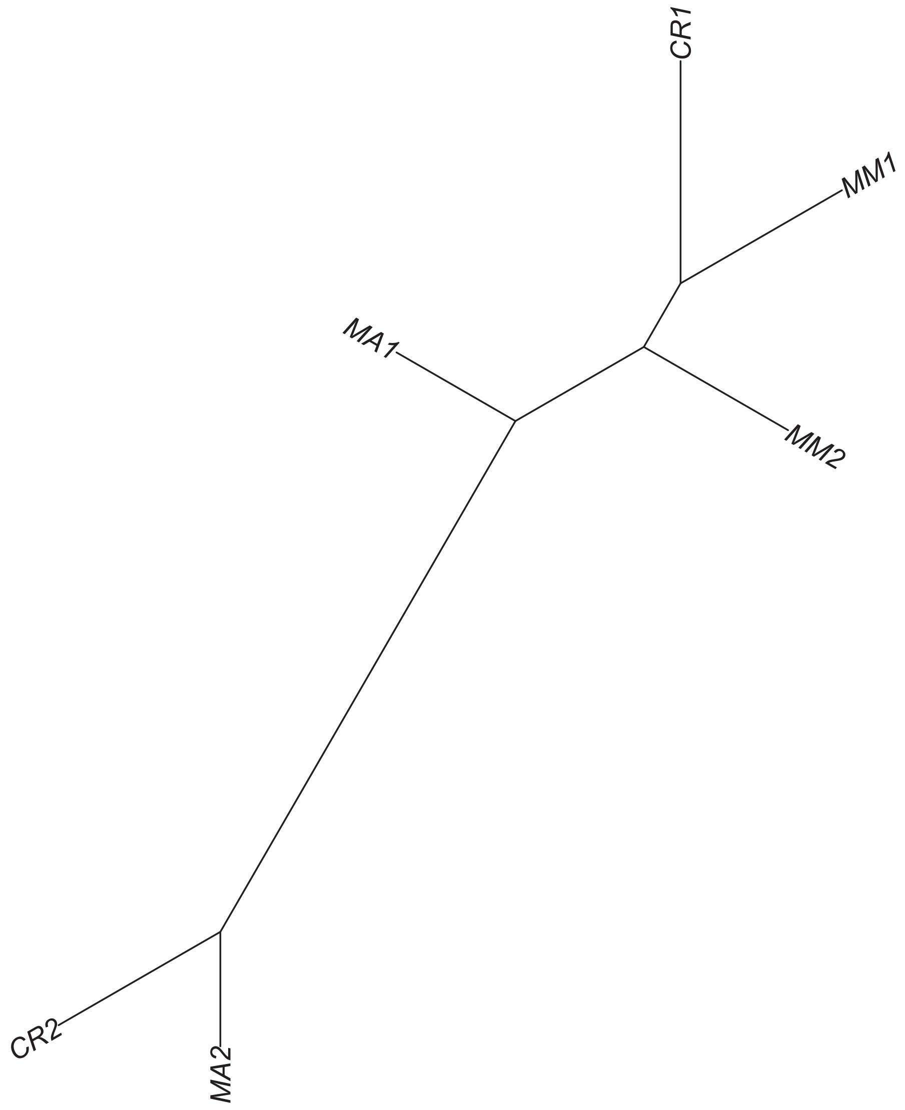

### Region 212

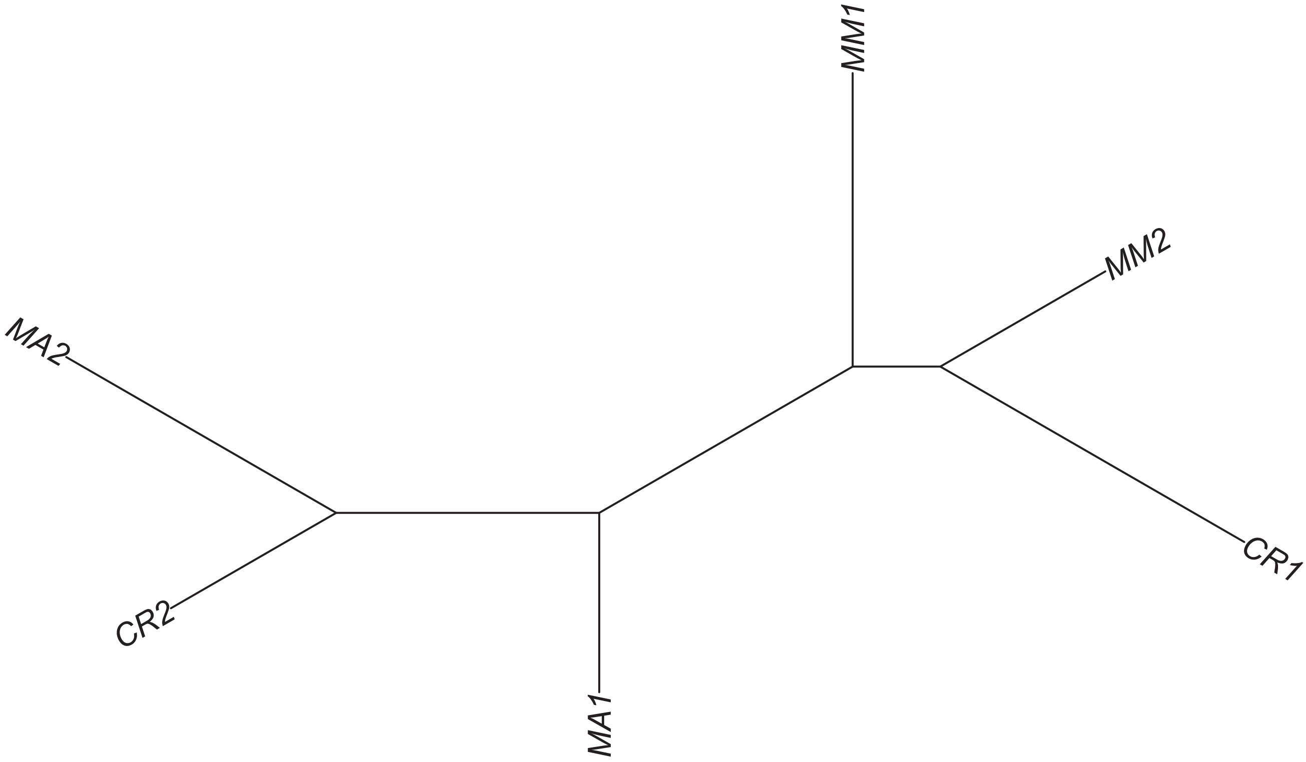

Region 213

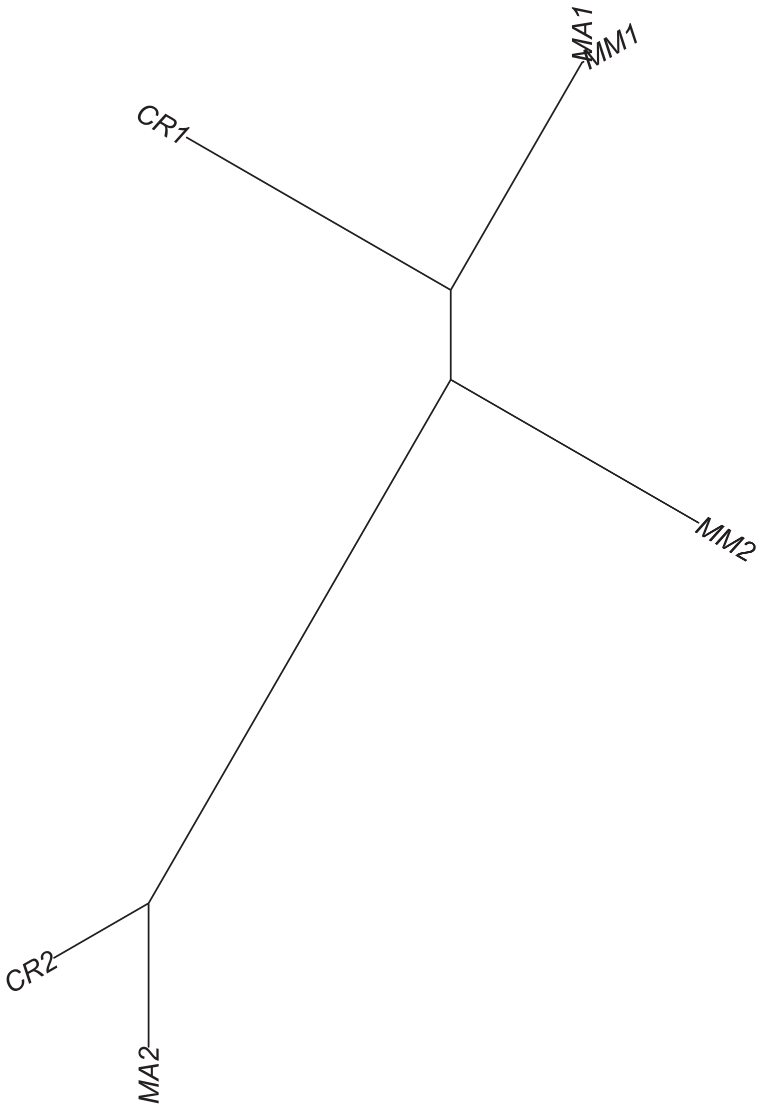

### Region 216

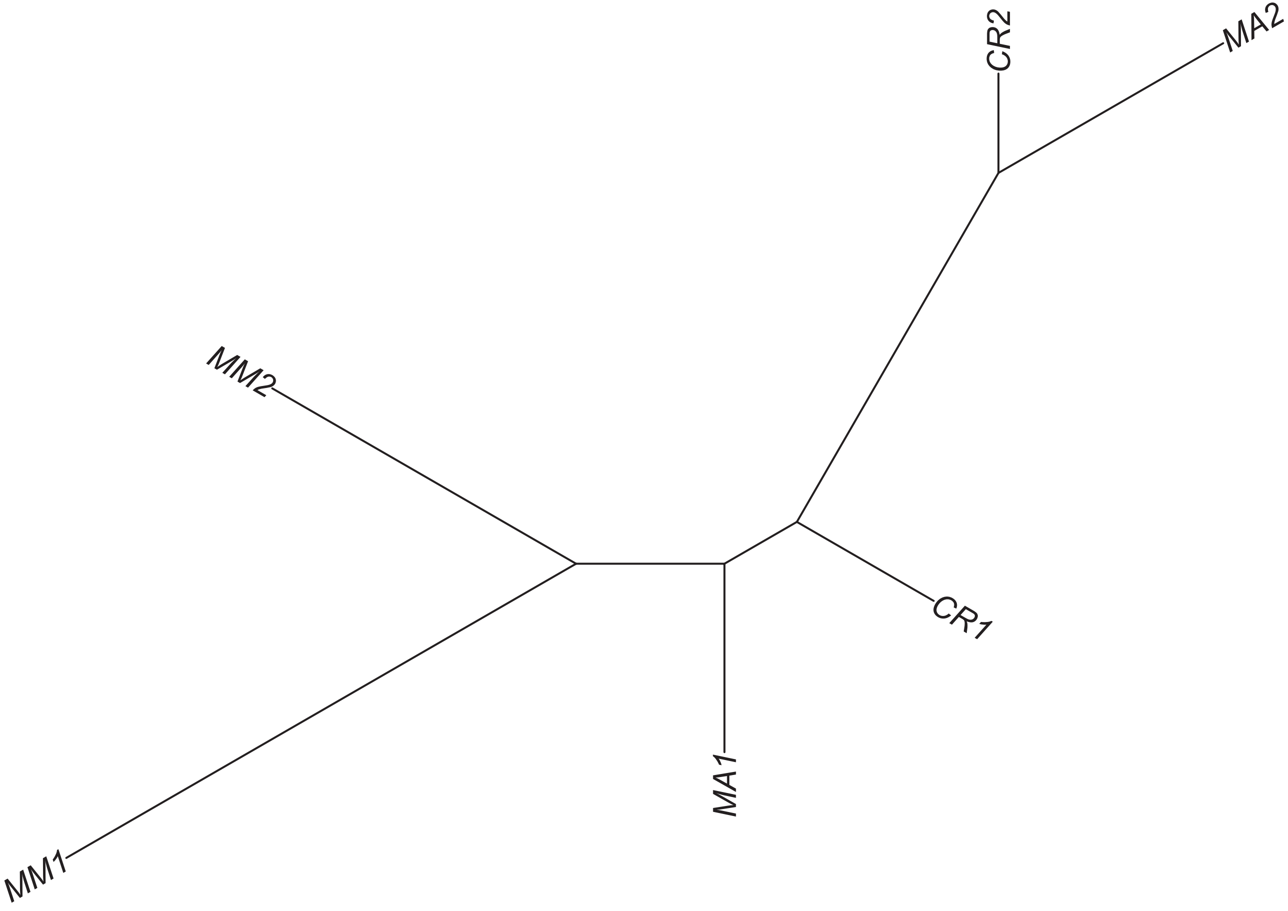

### Region 217



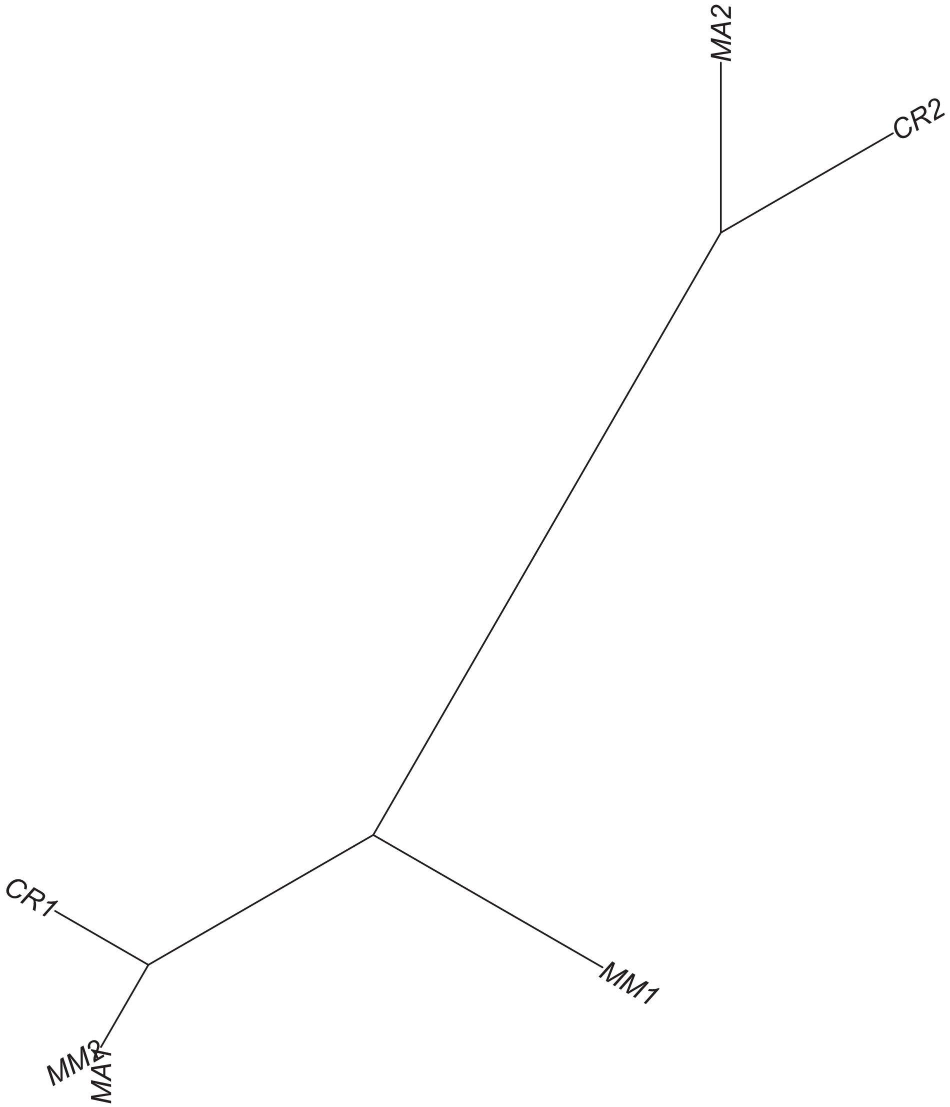

### Region 218

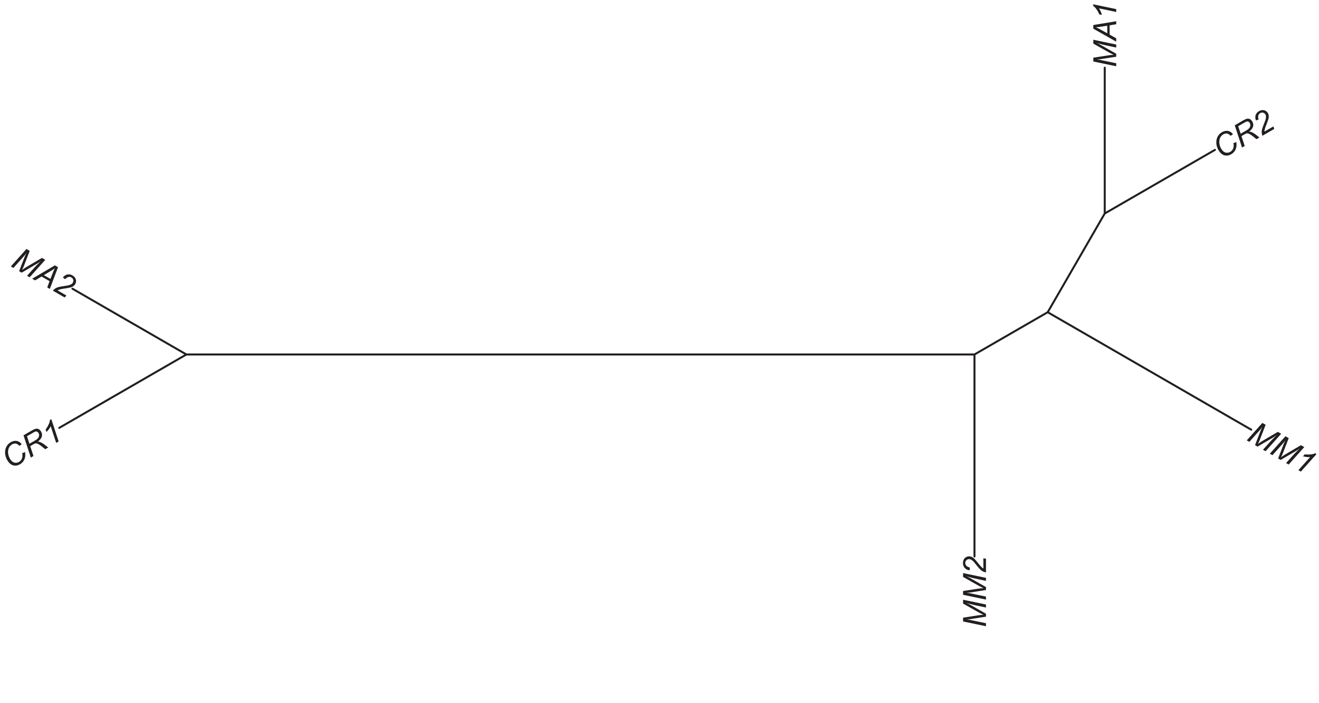

Region 220

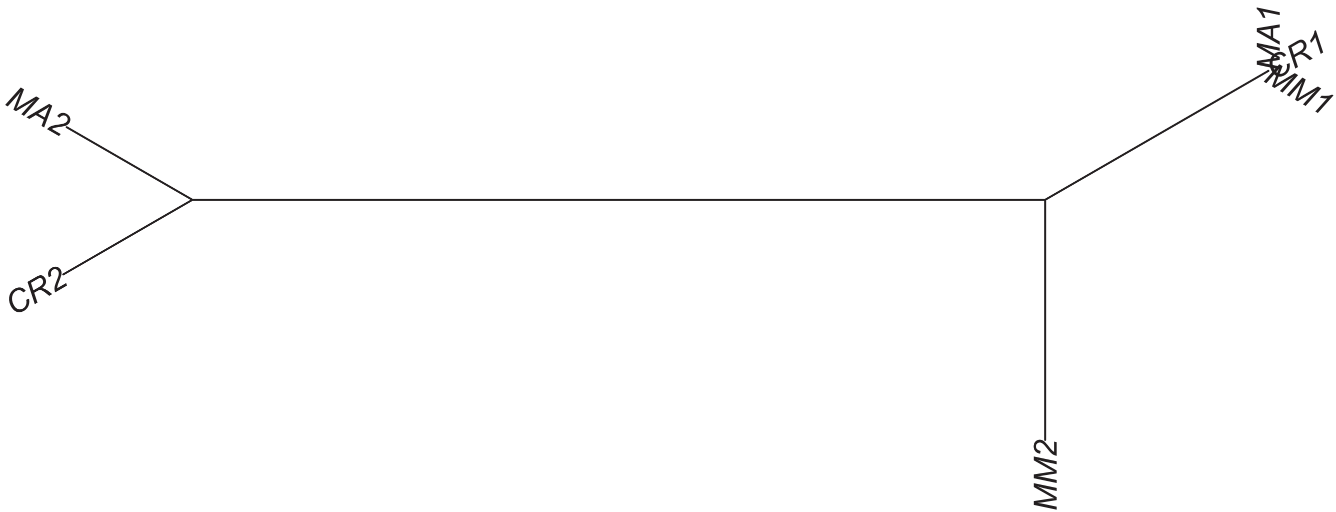

### Region 222



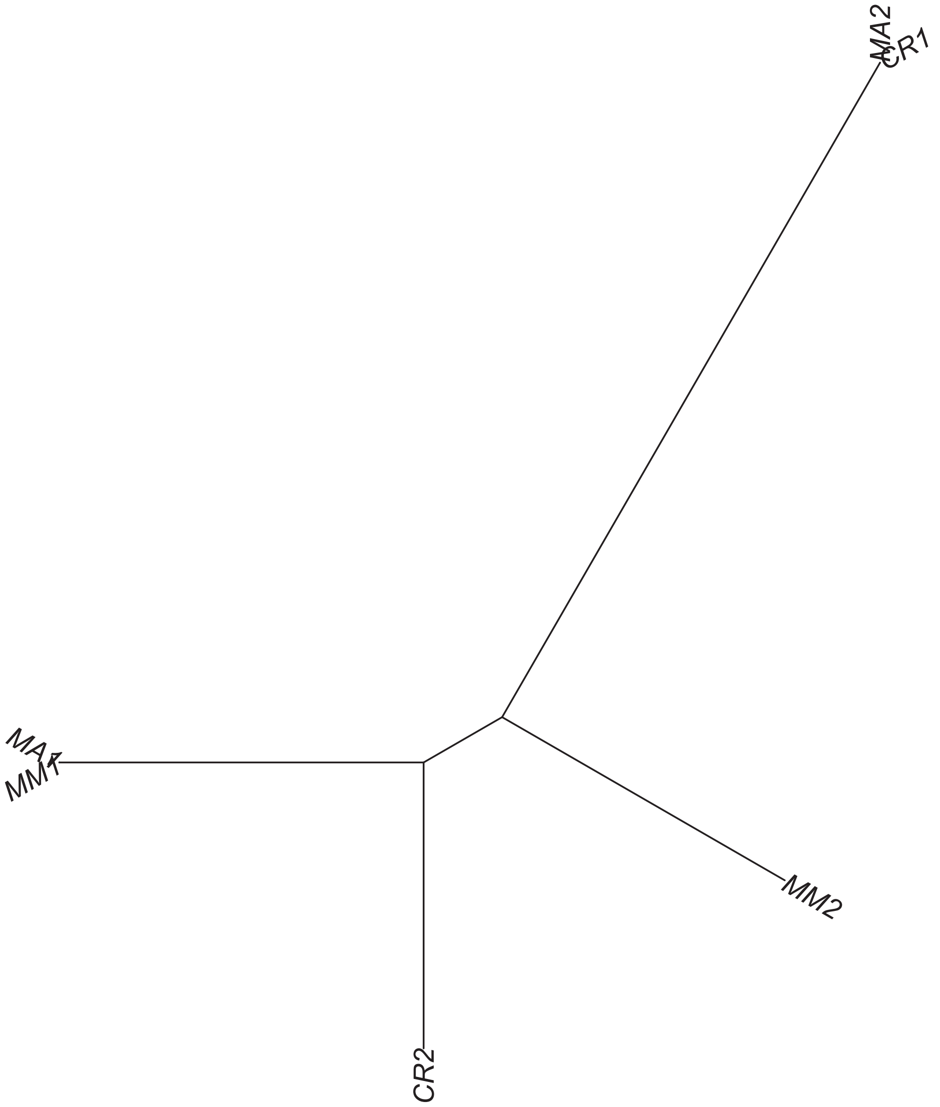

Region 223

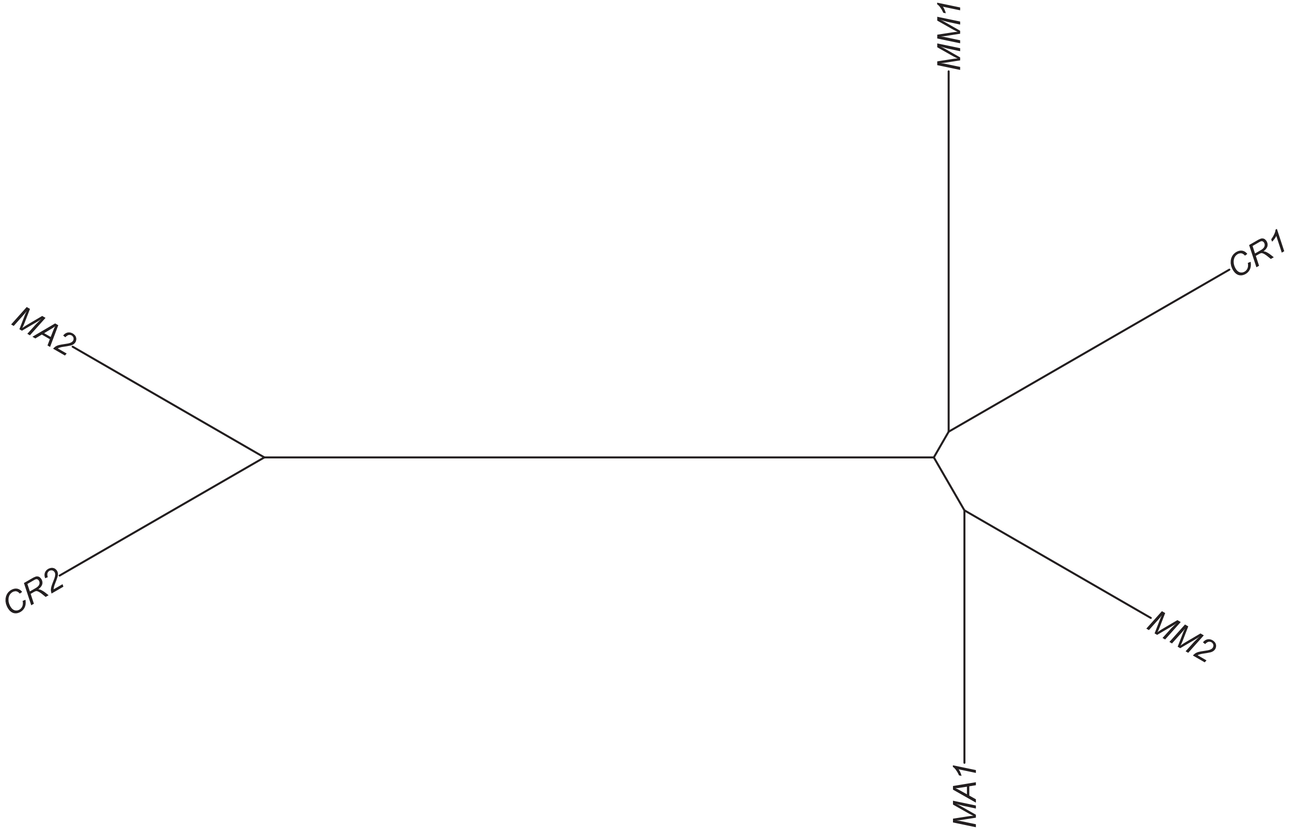

### Region 224

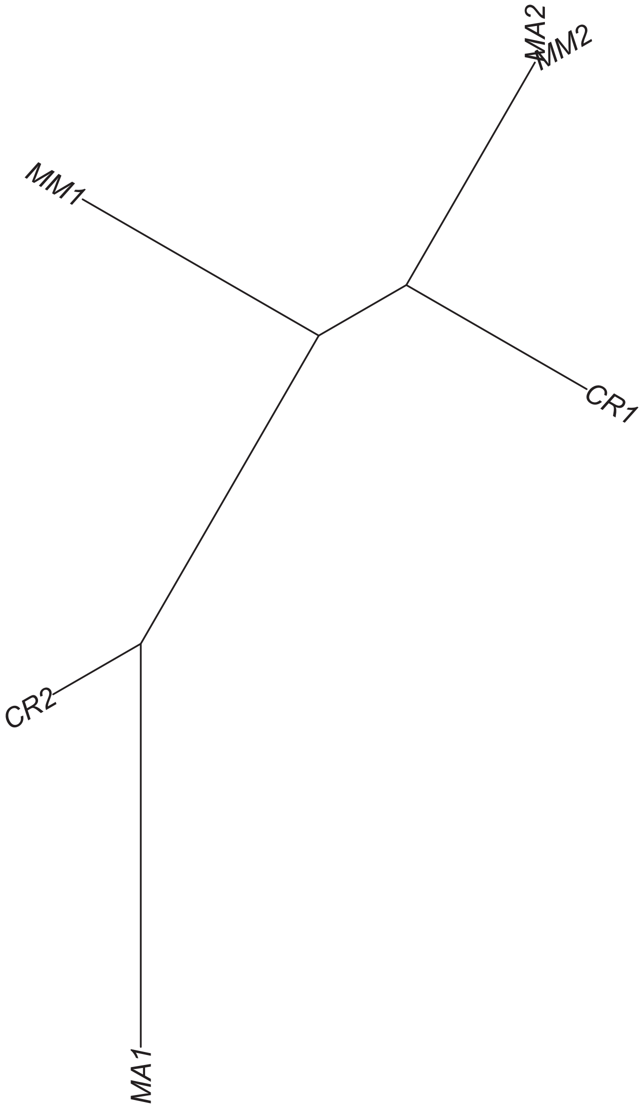

#### Region 227



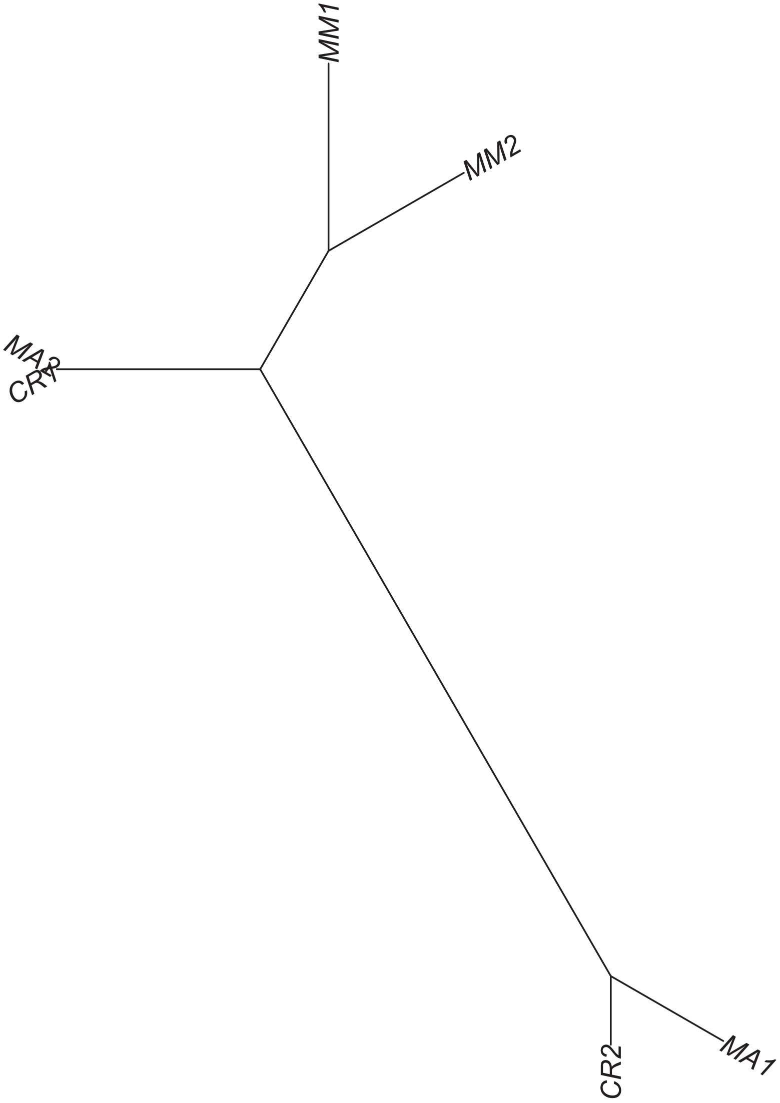

### Region 228

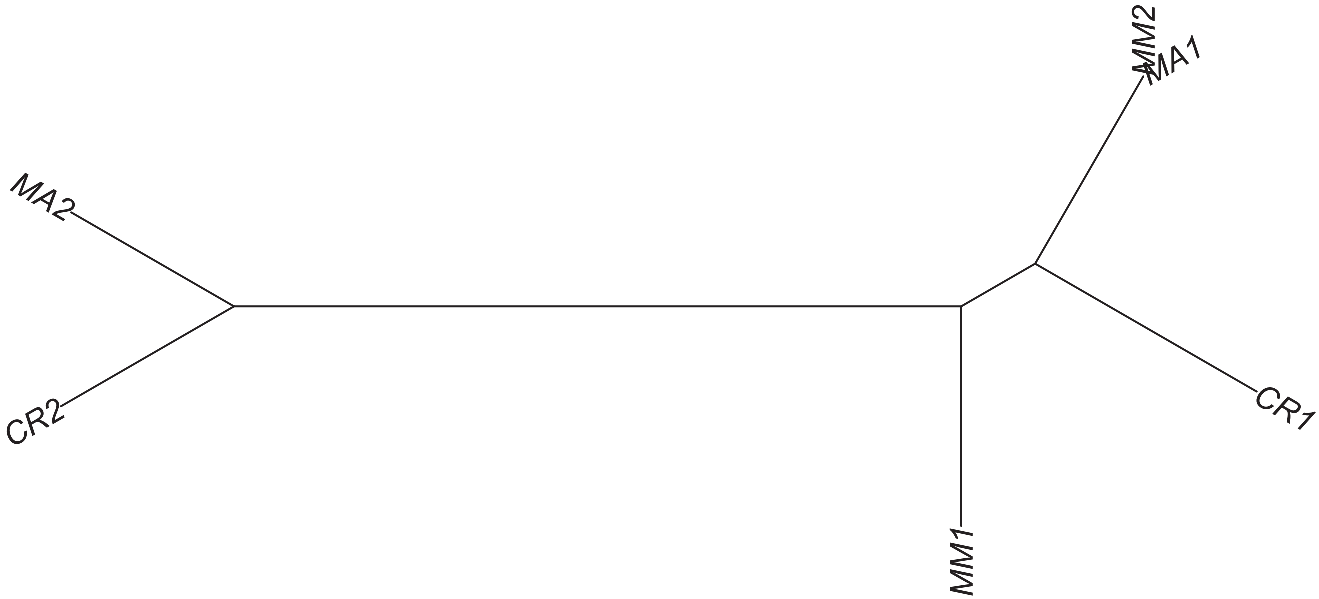

Region 231

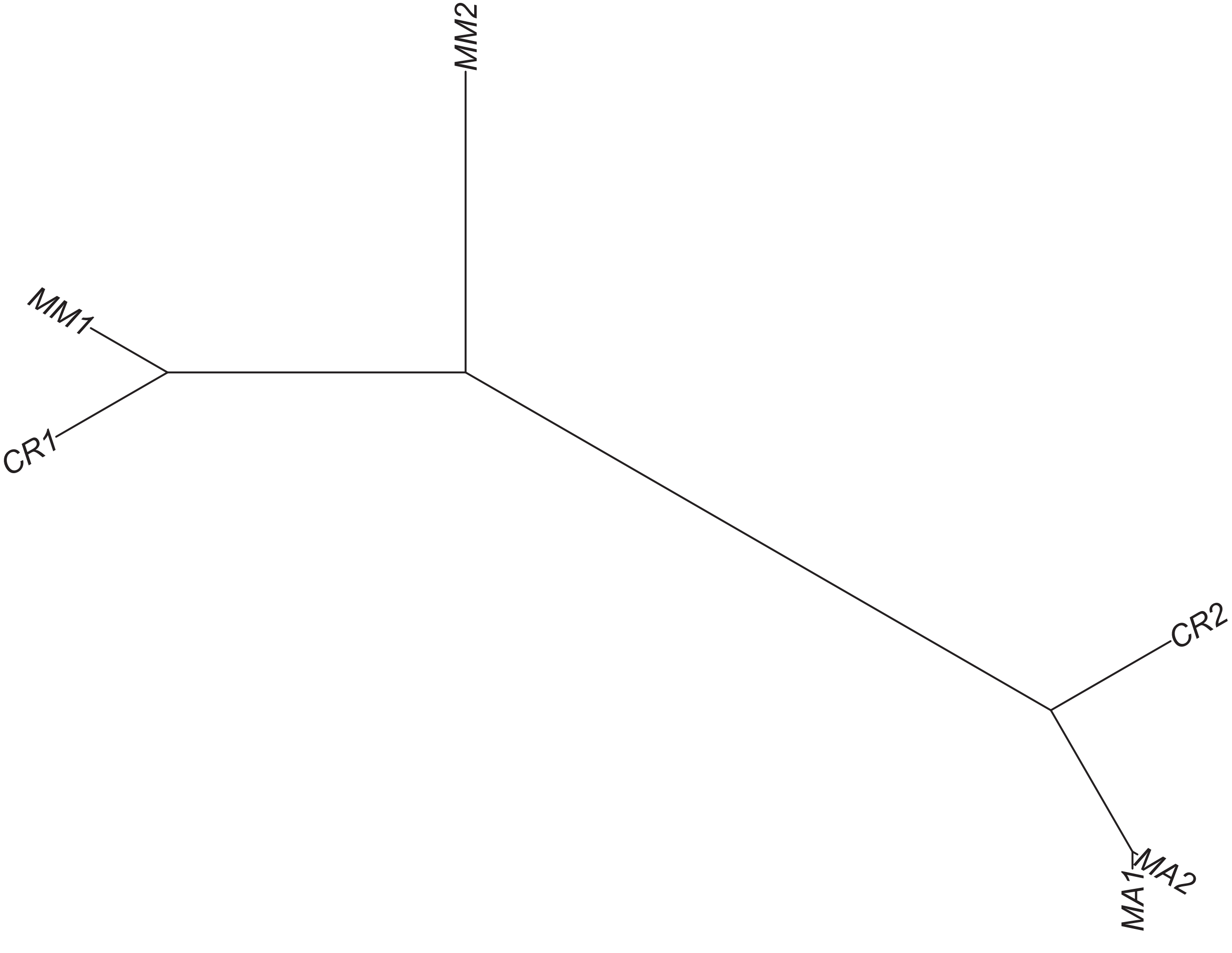

### Region 232

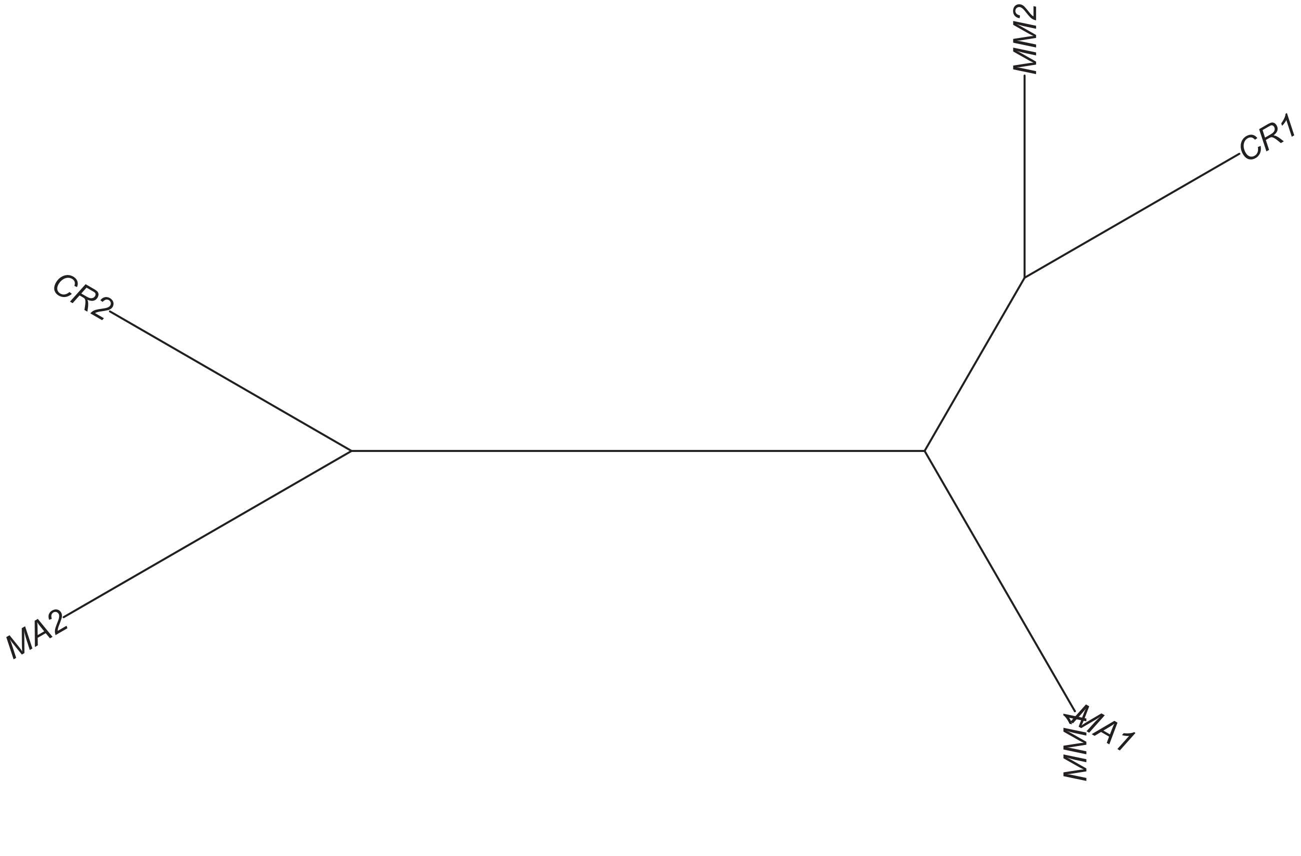

### Region 233



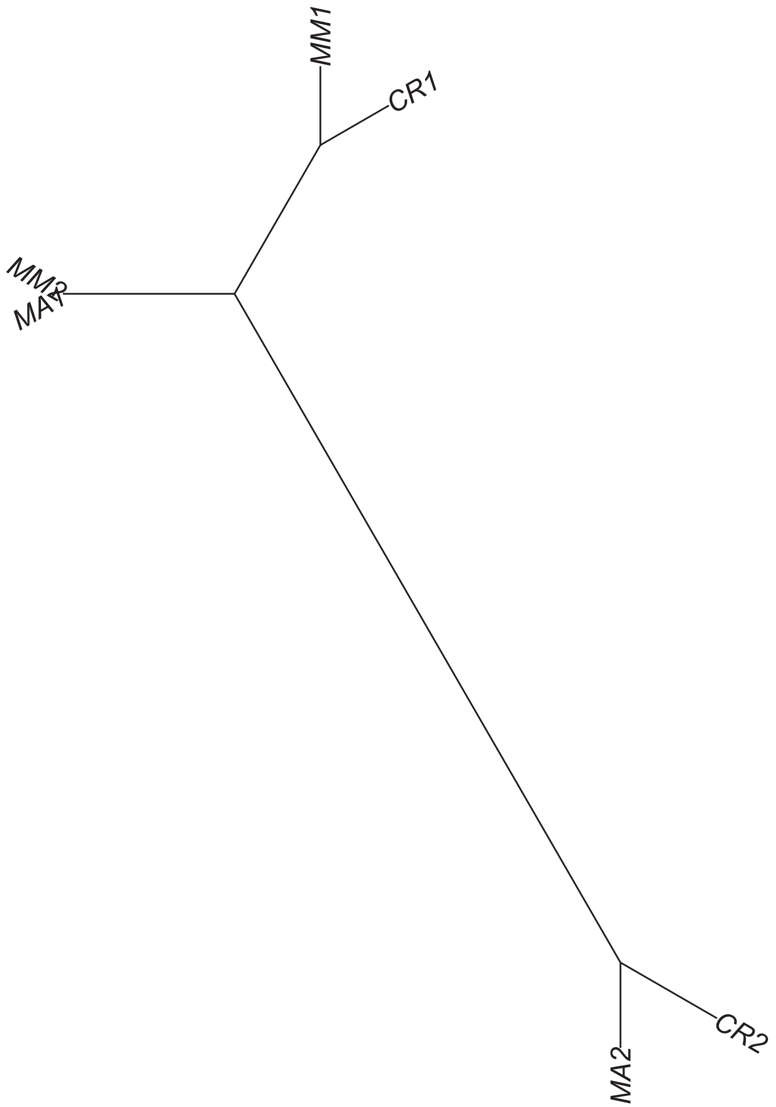

### Region 235



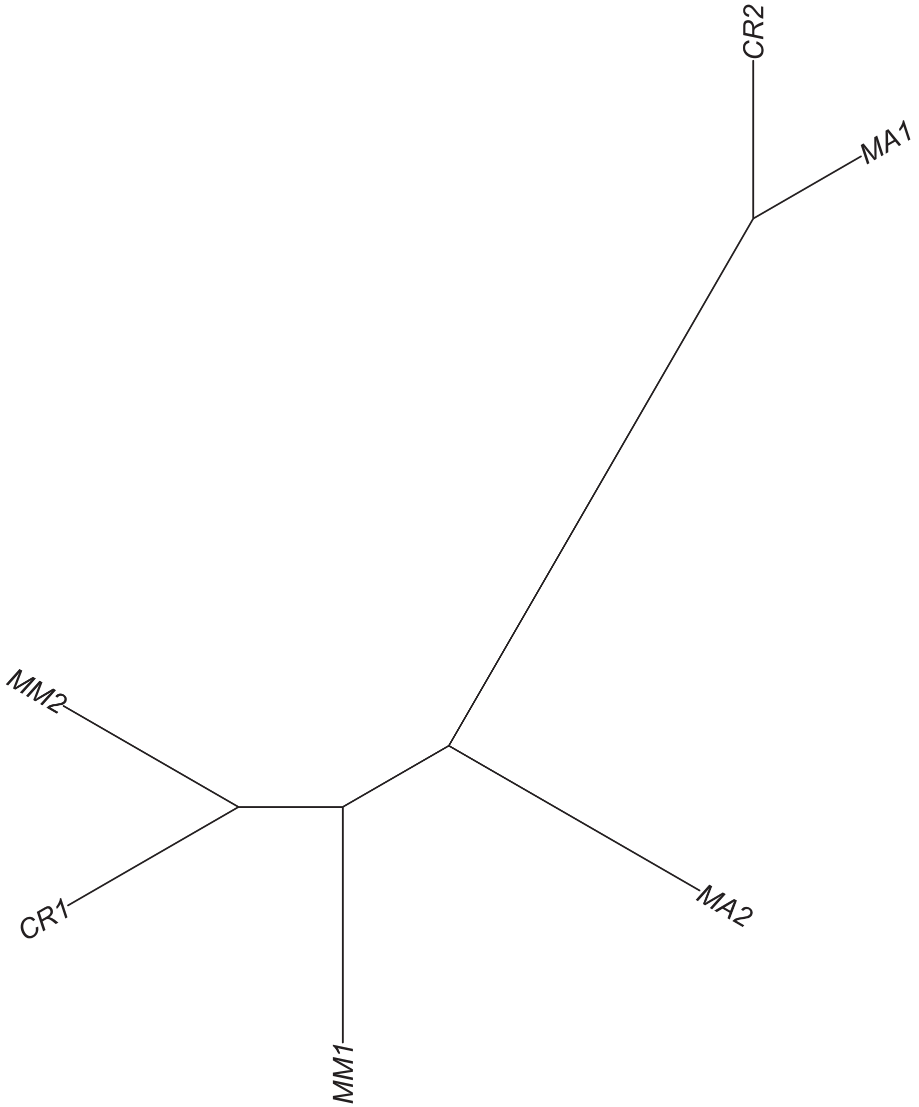

### Region 237

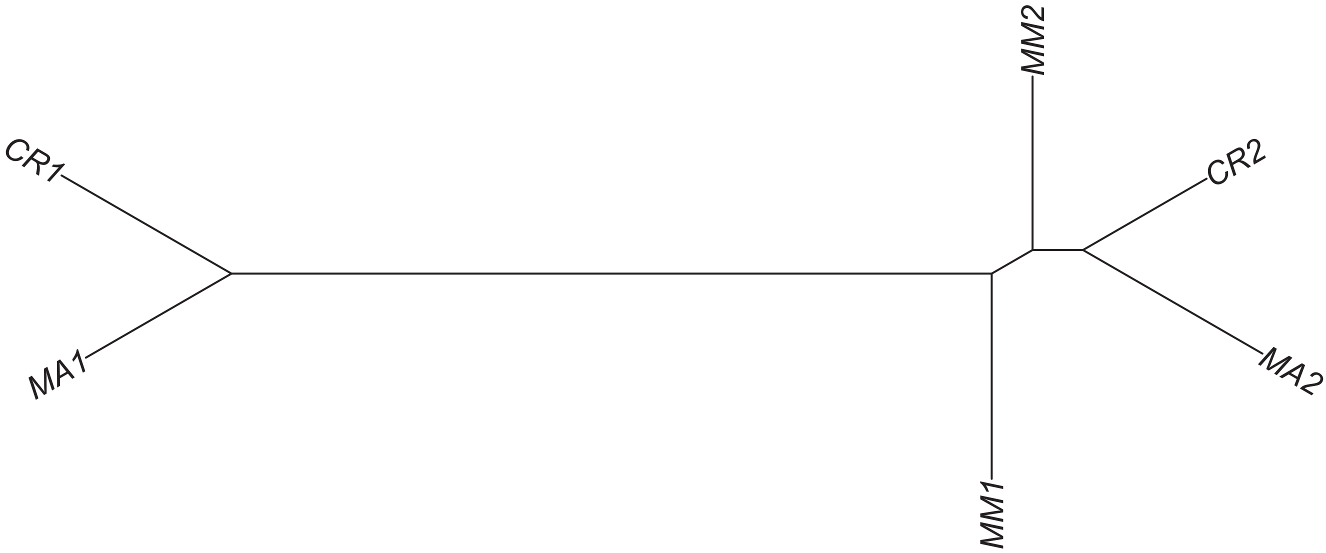

### Region 239

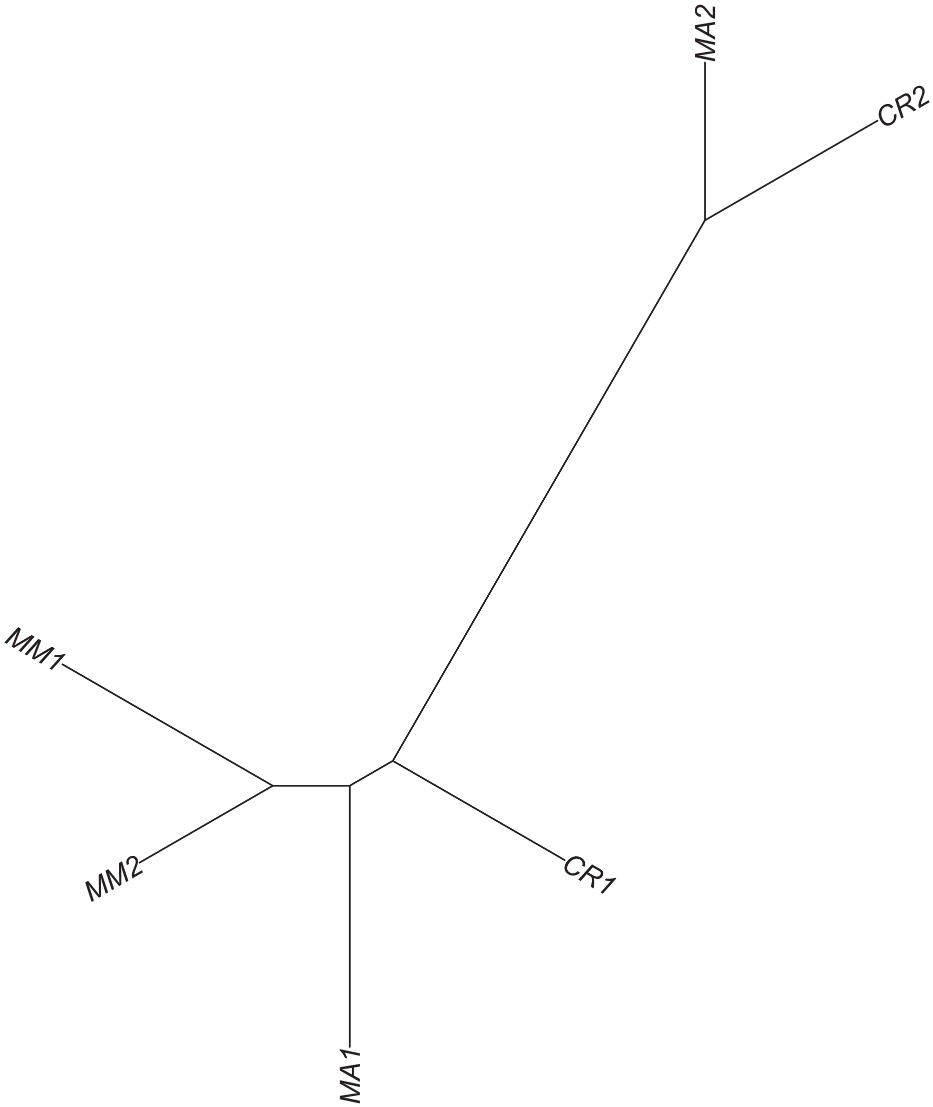

### Region 240

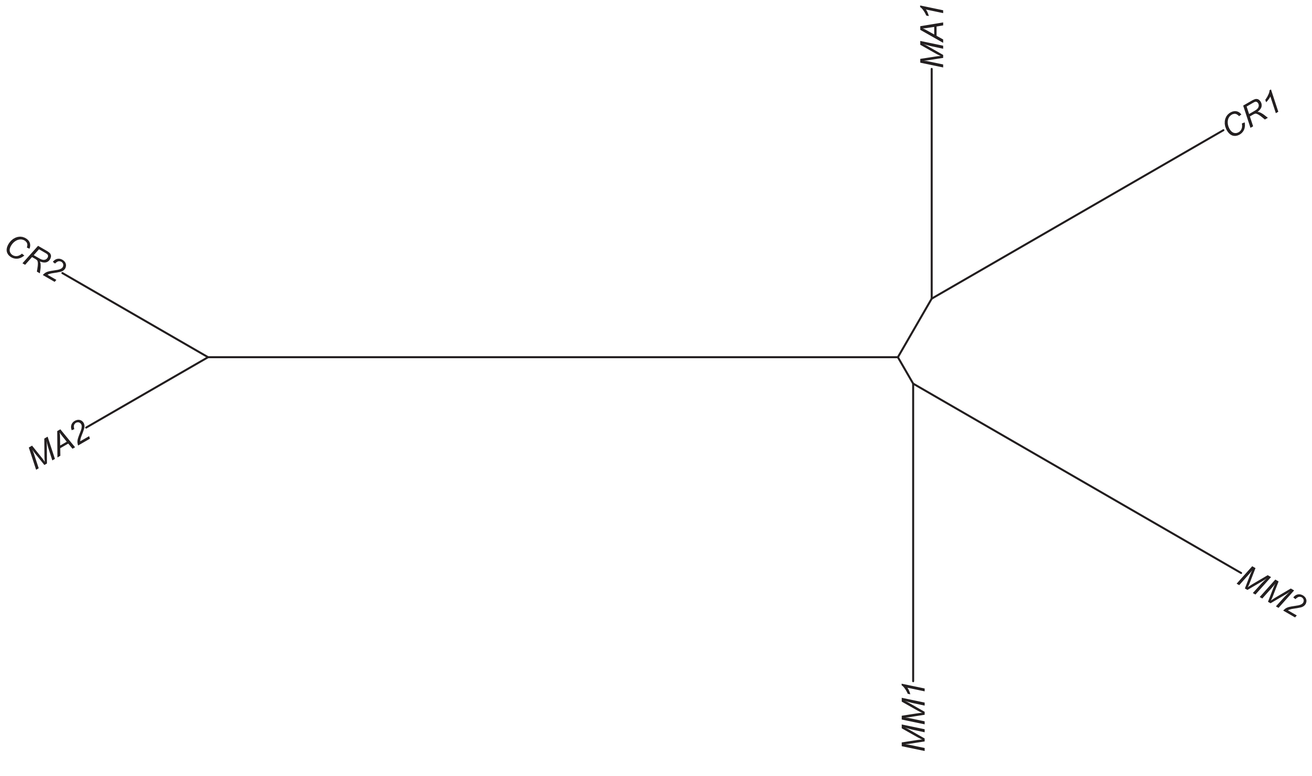

### Region 242



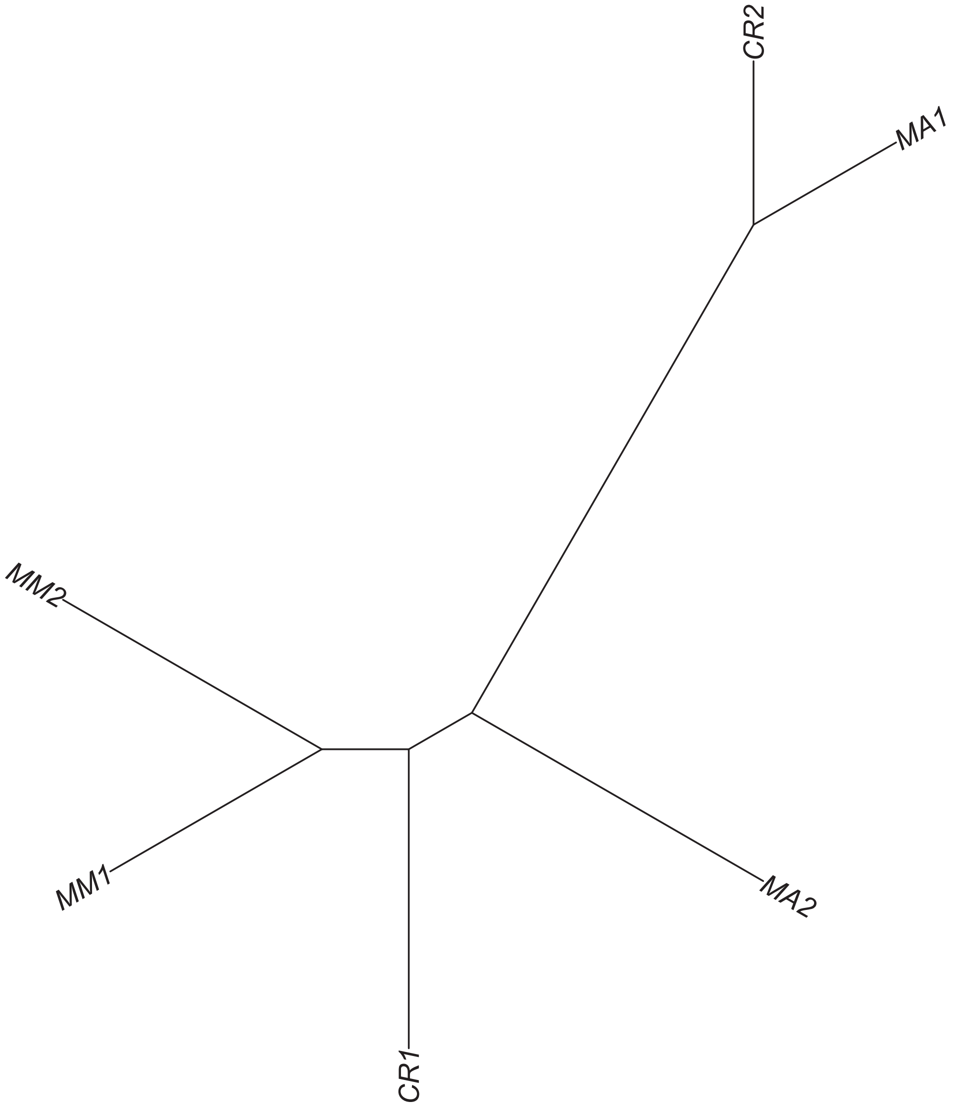

Region 243

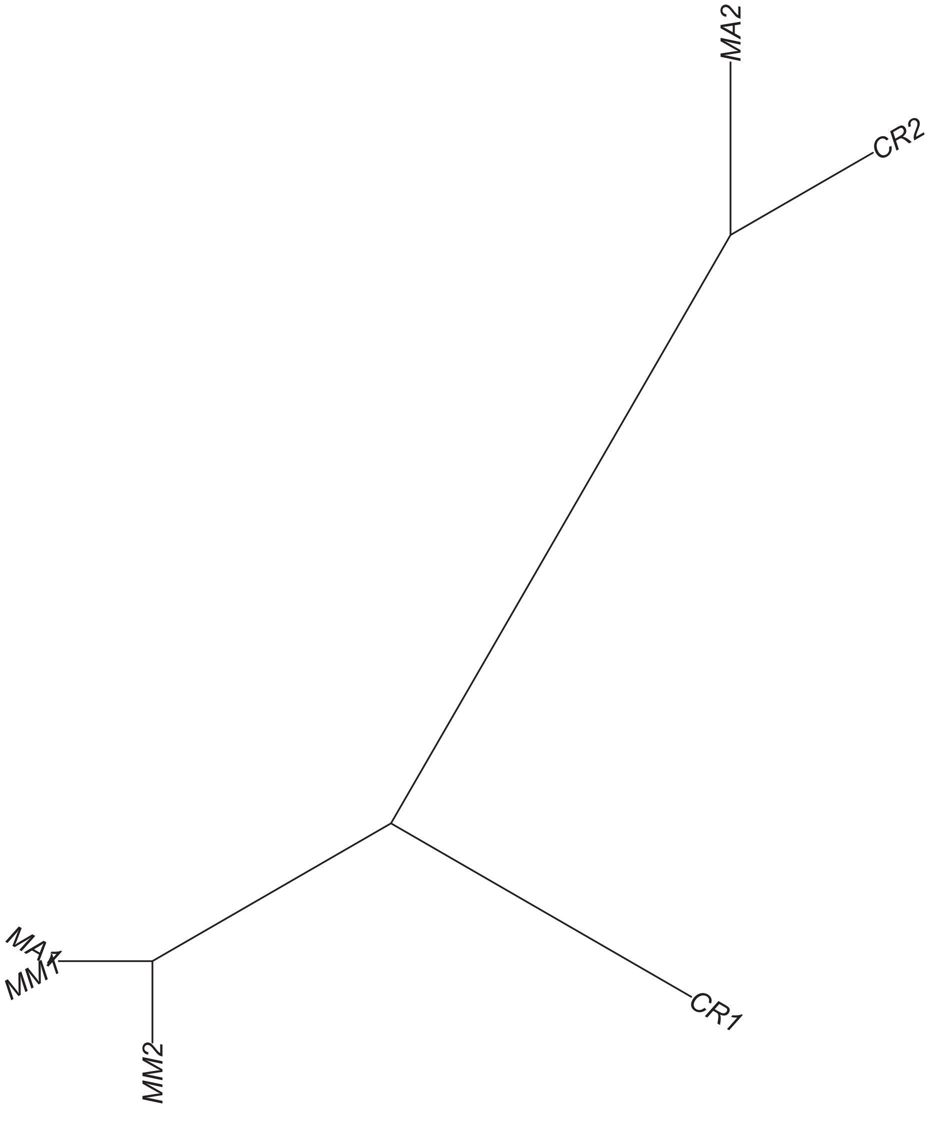

### Region 245

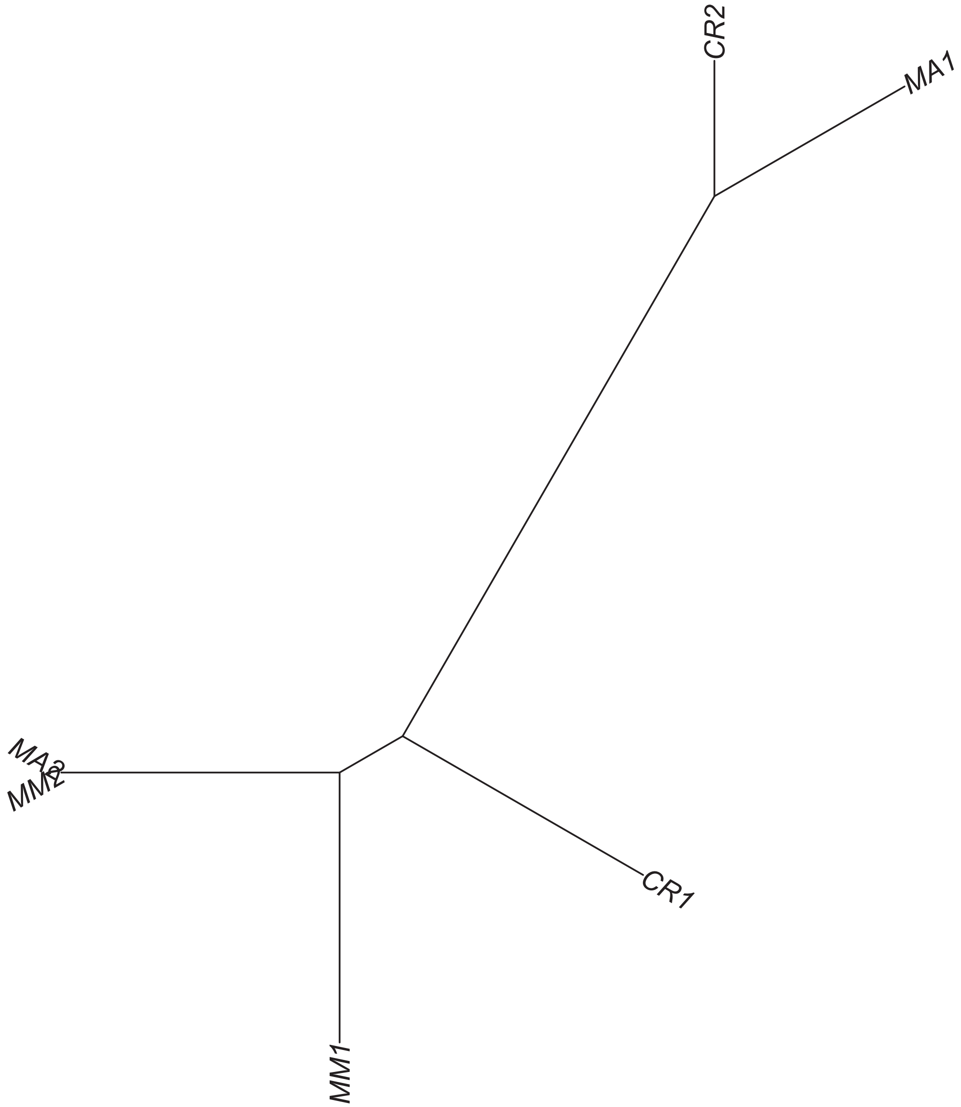

### Region 246

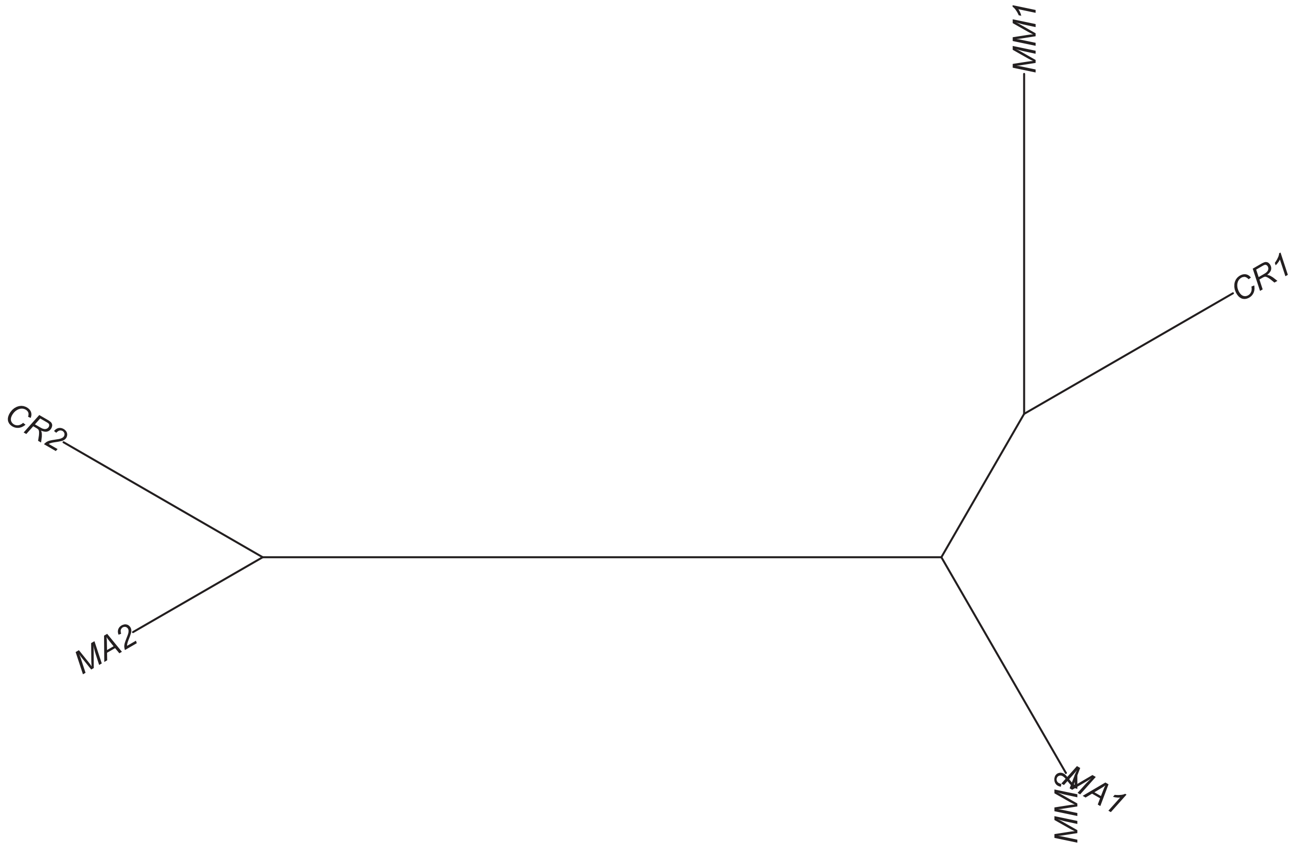

Region 247

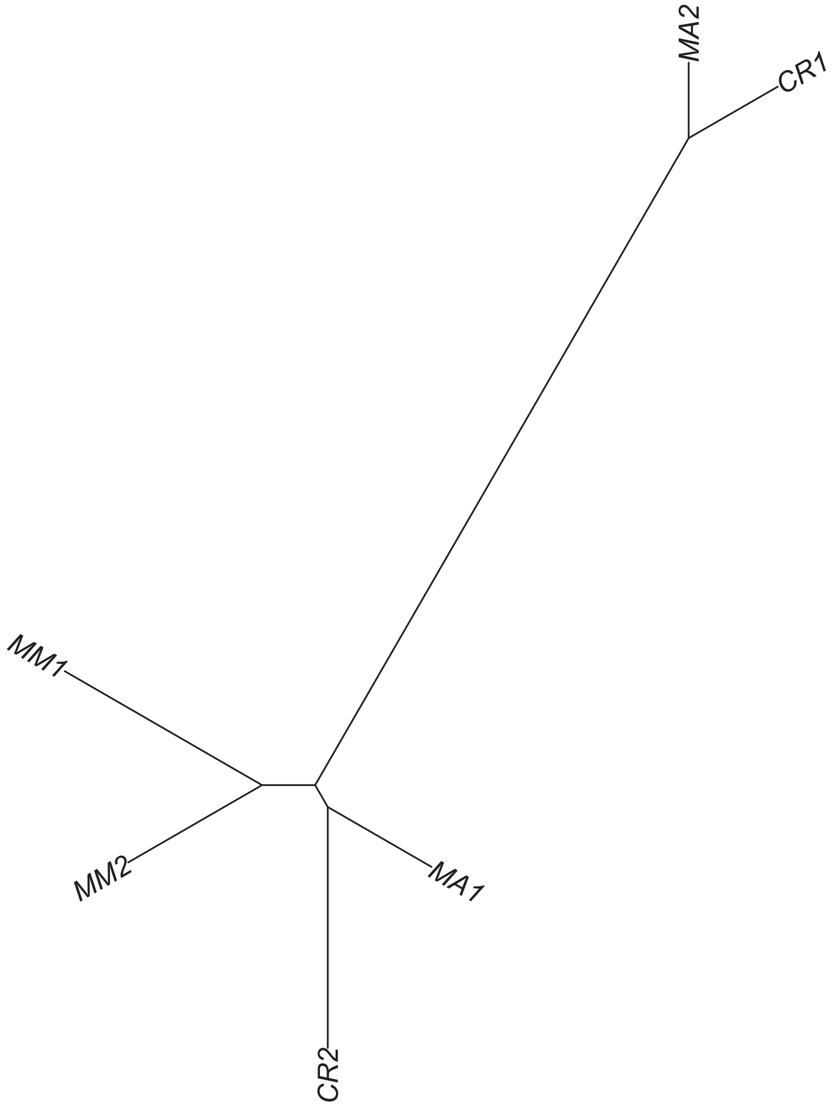

### Region 248

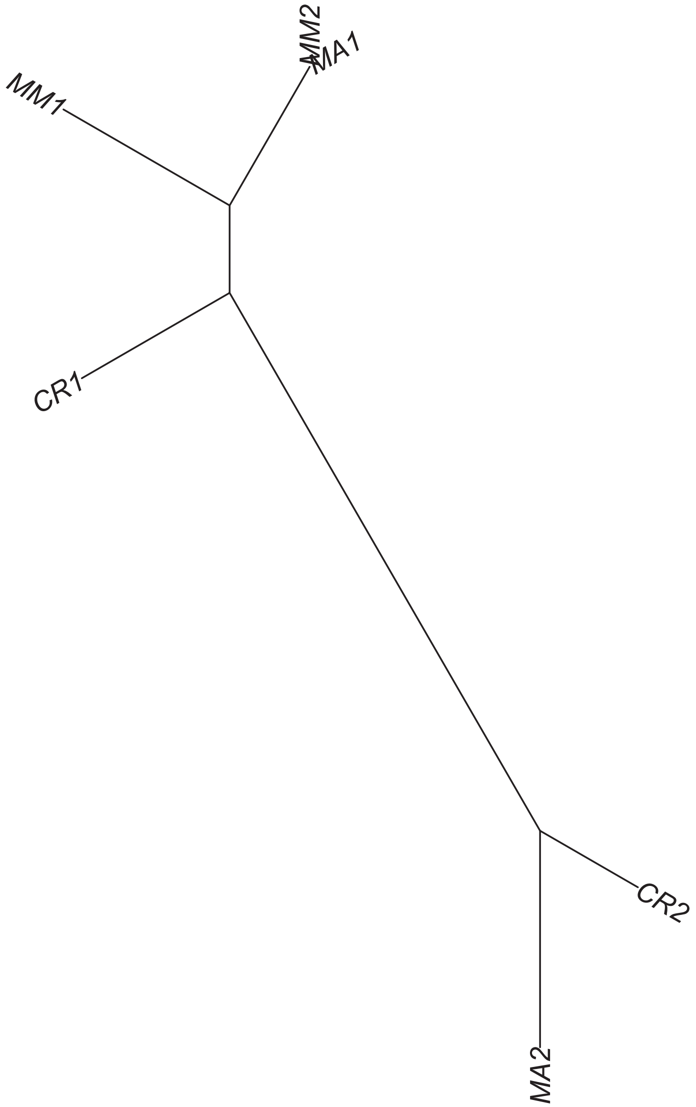

### Region 249

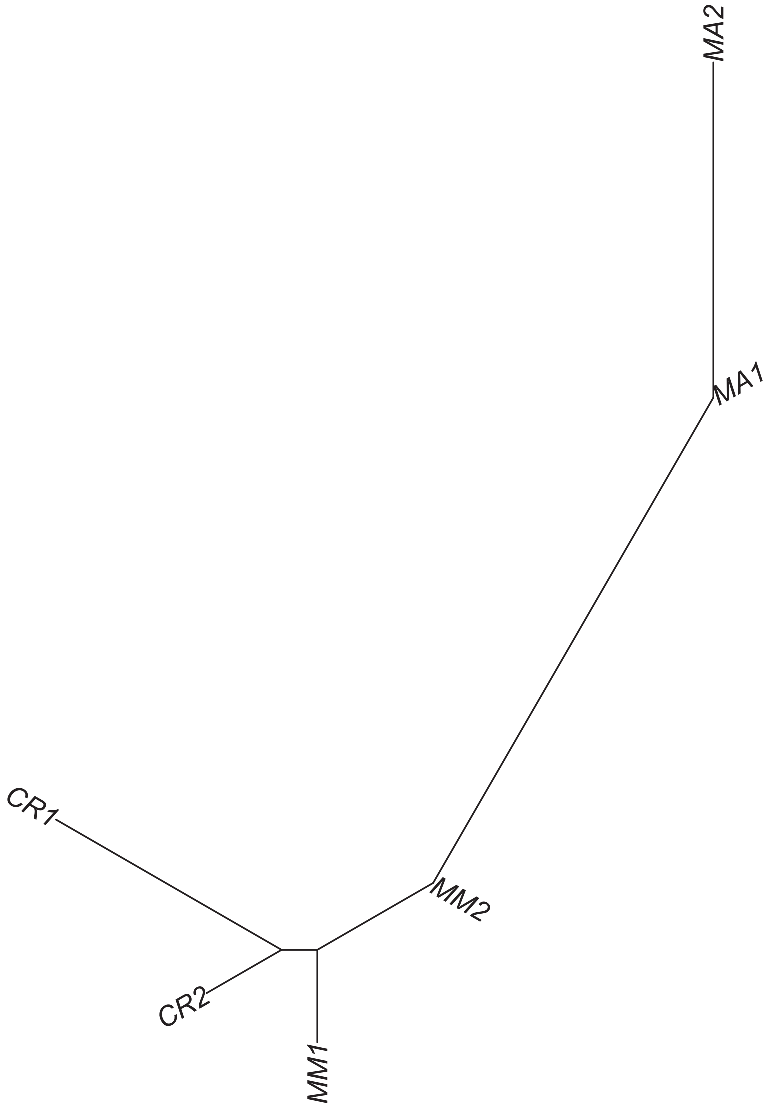

Region 250

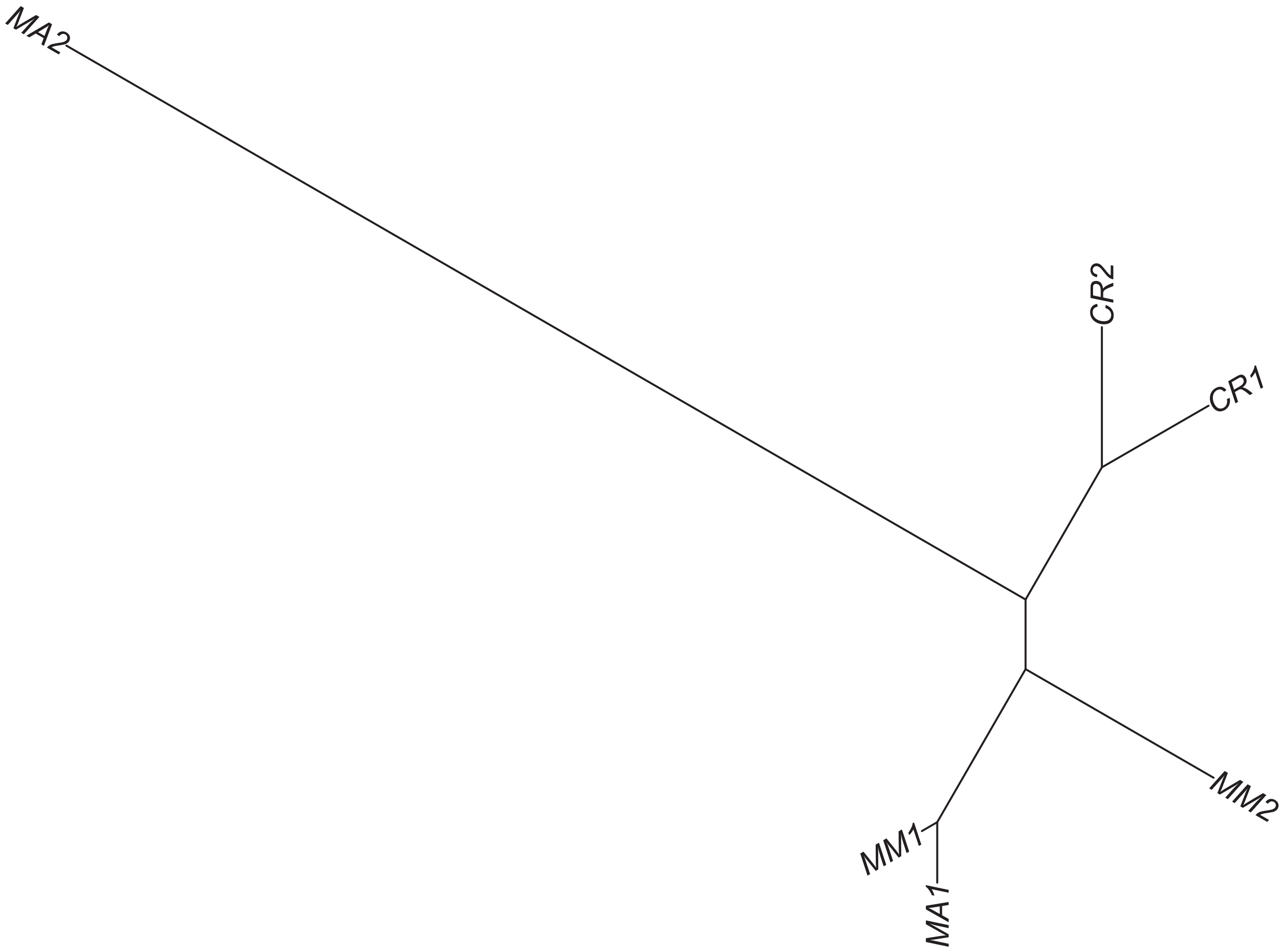

### Region 251

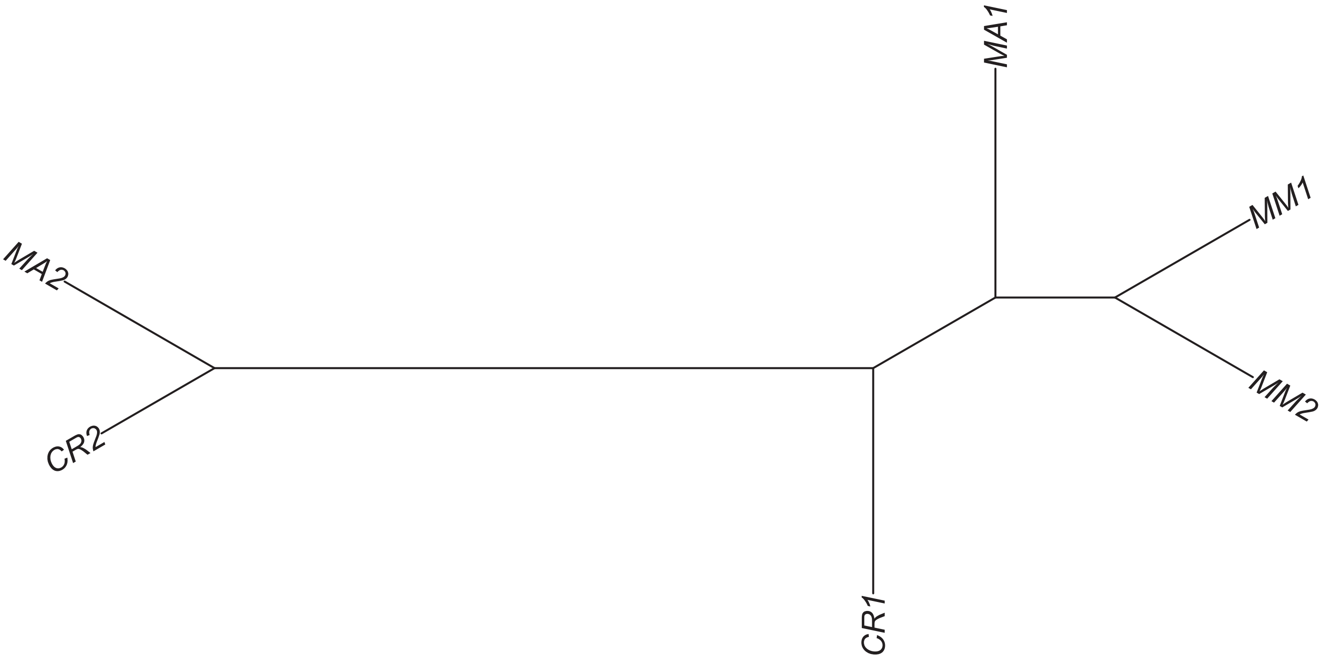

Region 252

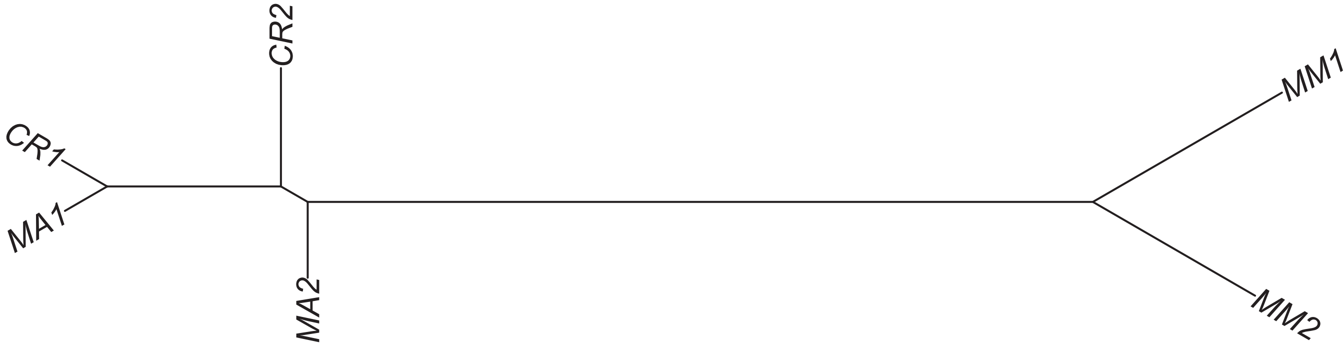

### Region 253

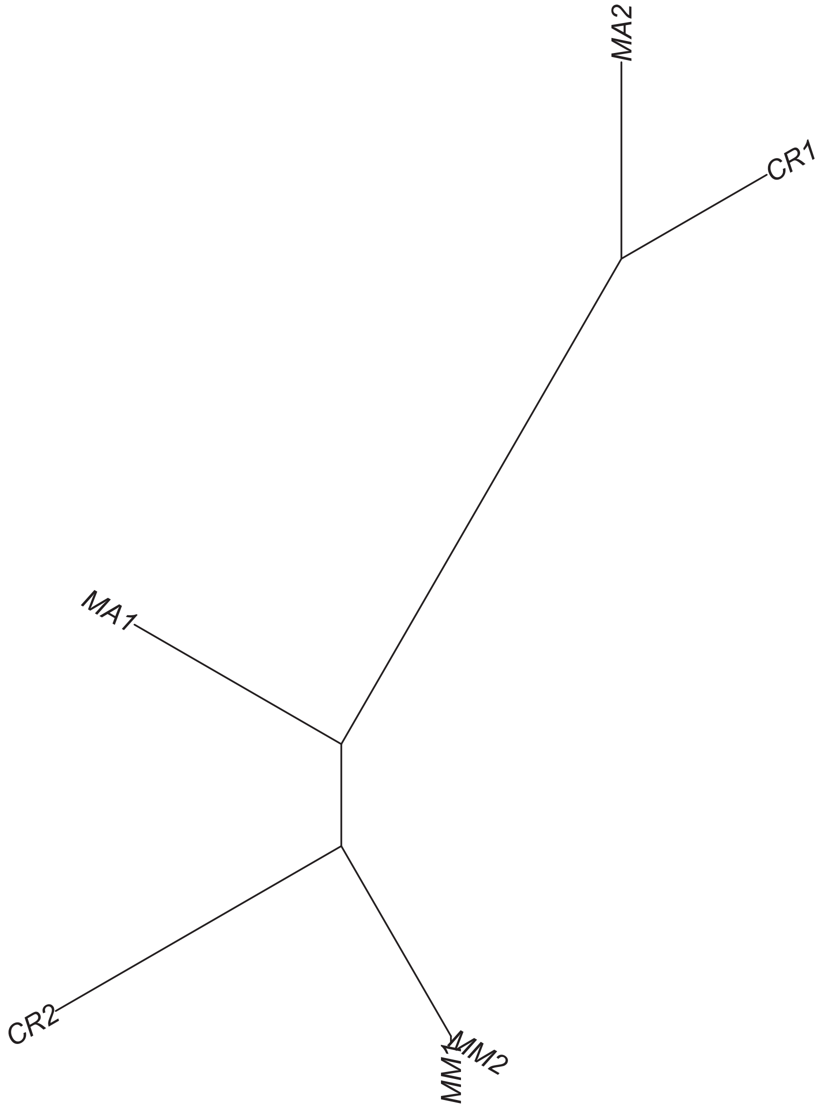

### Region 256



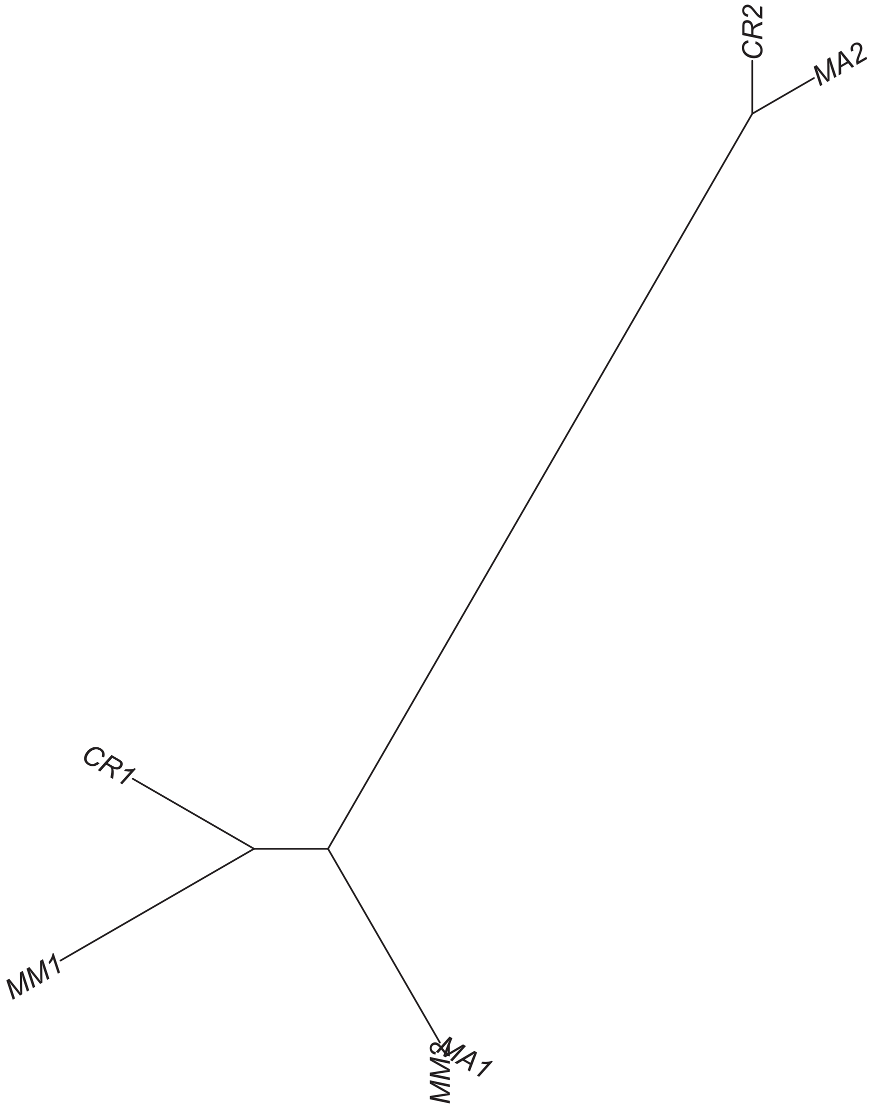

Region 257

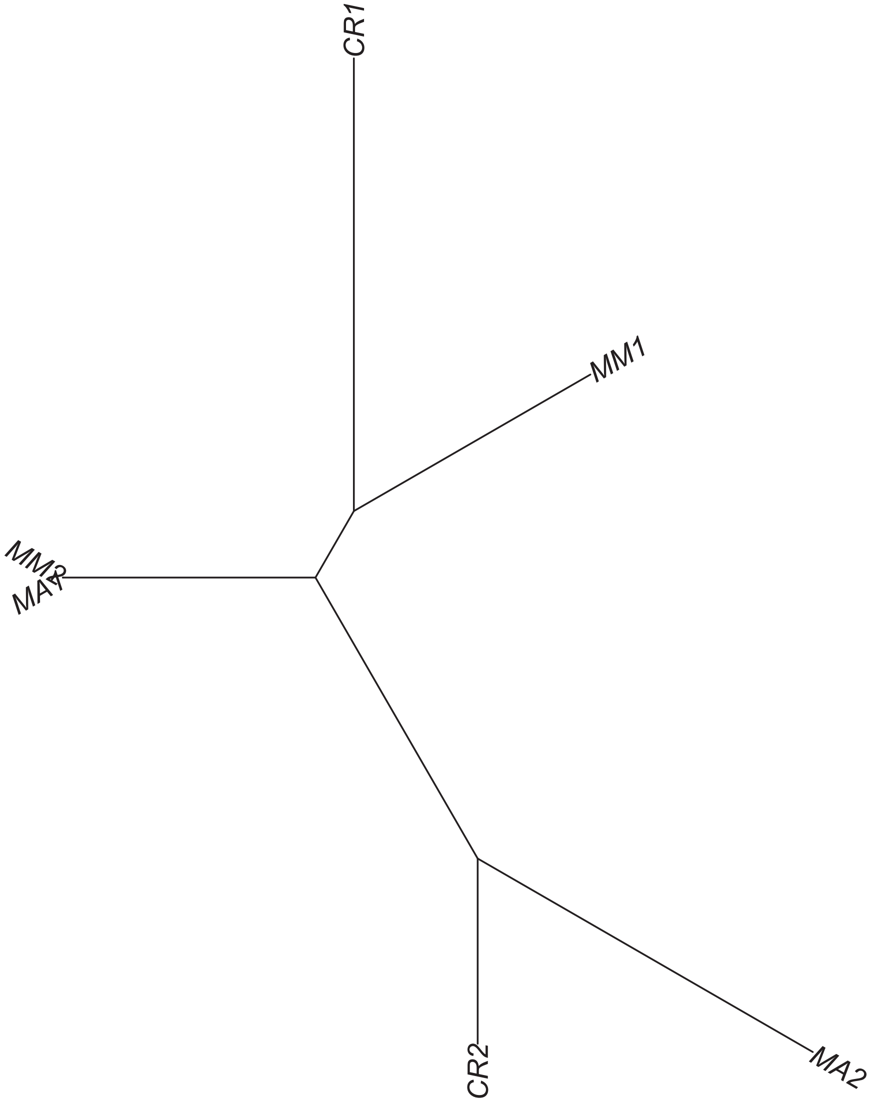

### Region 258

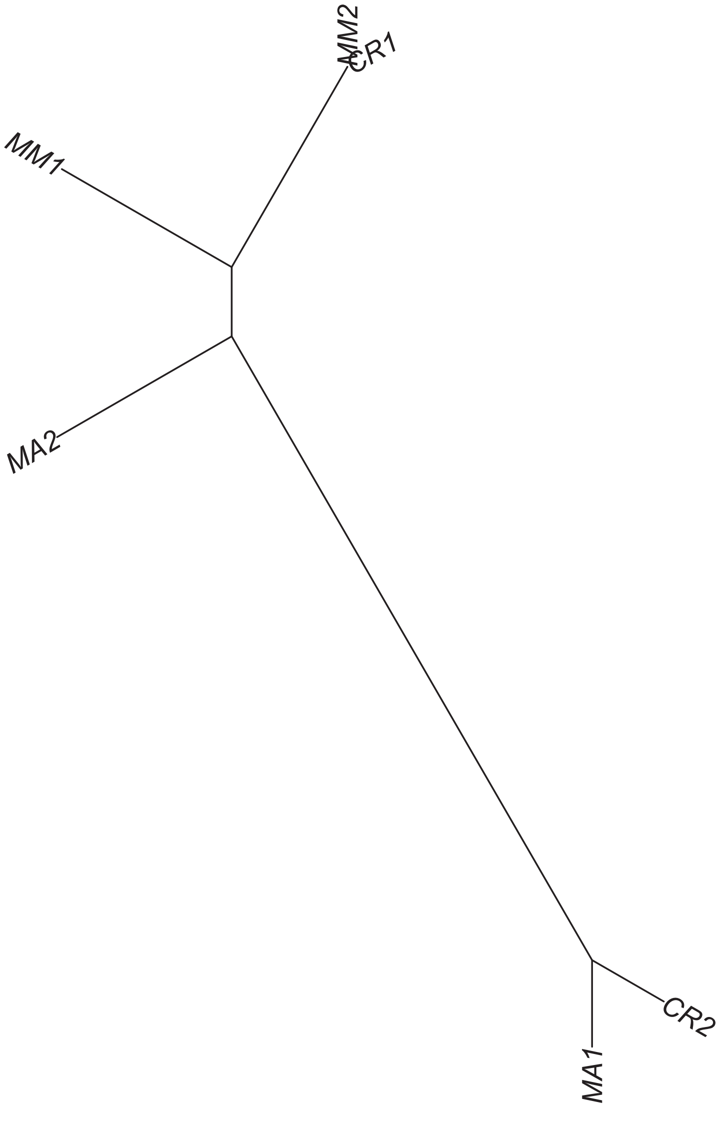

### Region 259



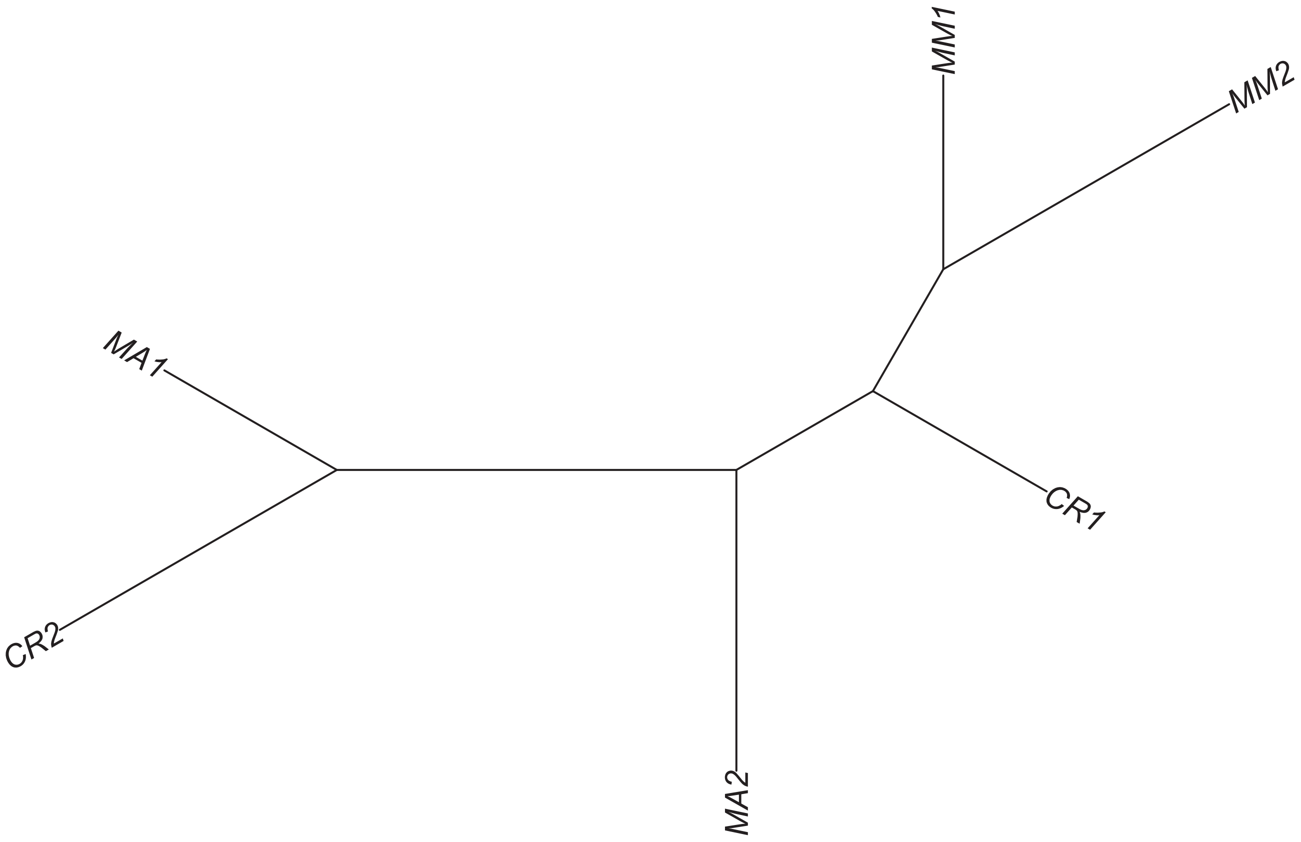

### Region 261

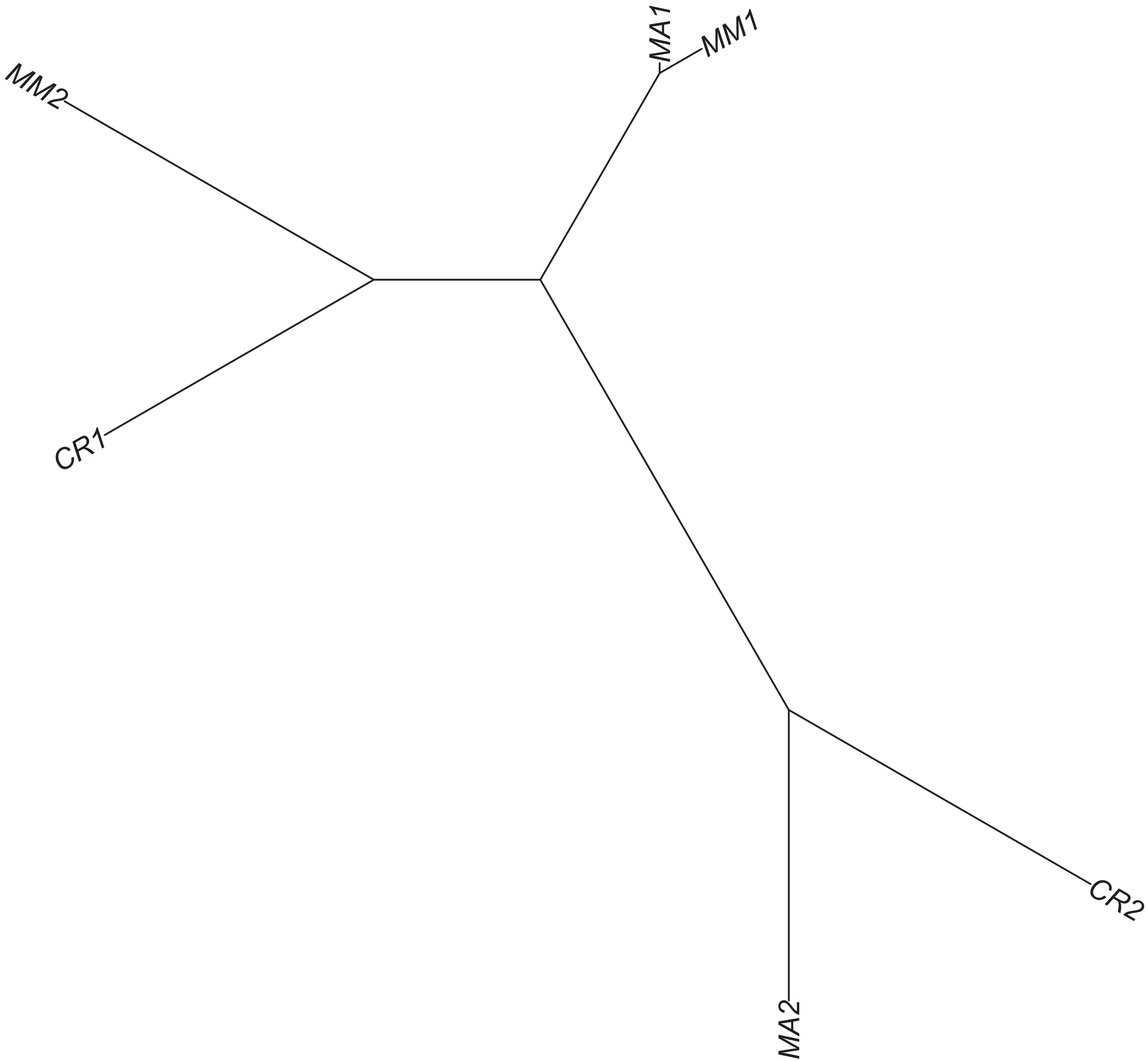

### Region 262



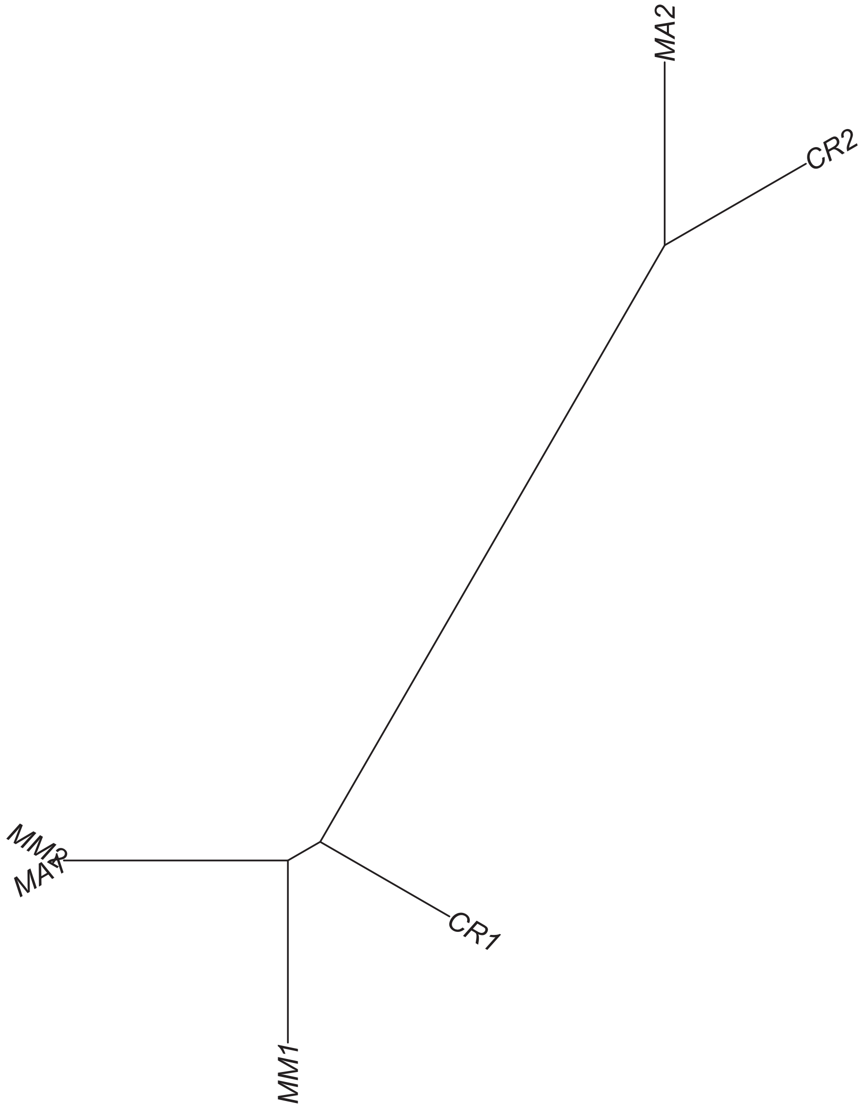

### Region 264



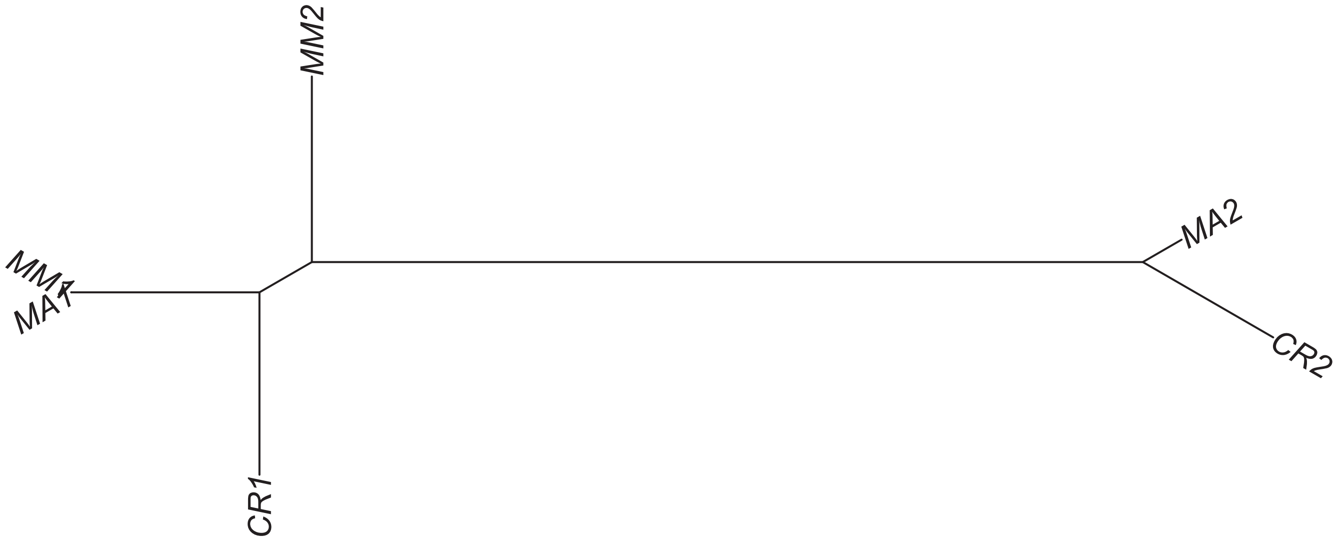

Region 265

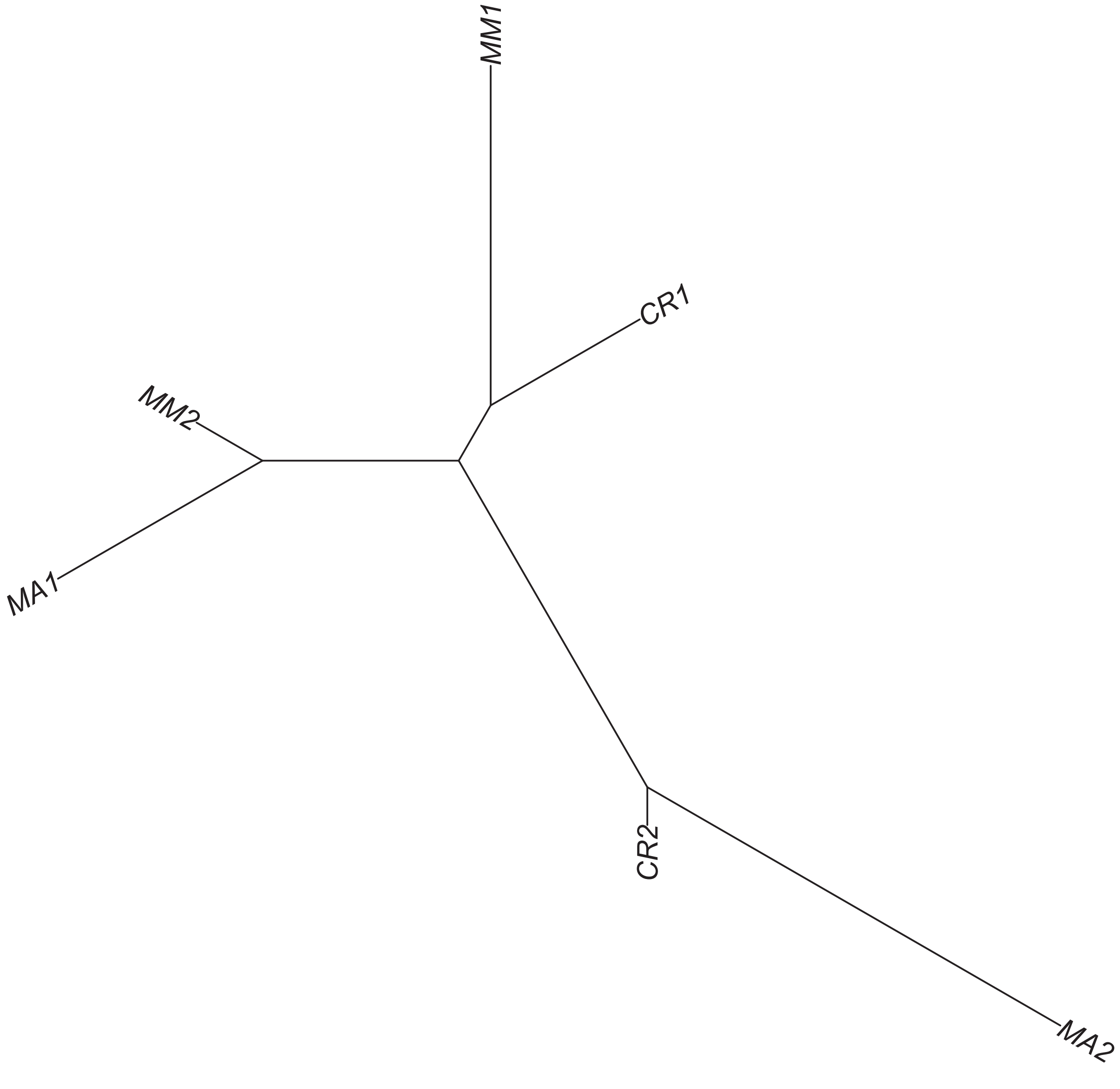

### Region 266

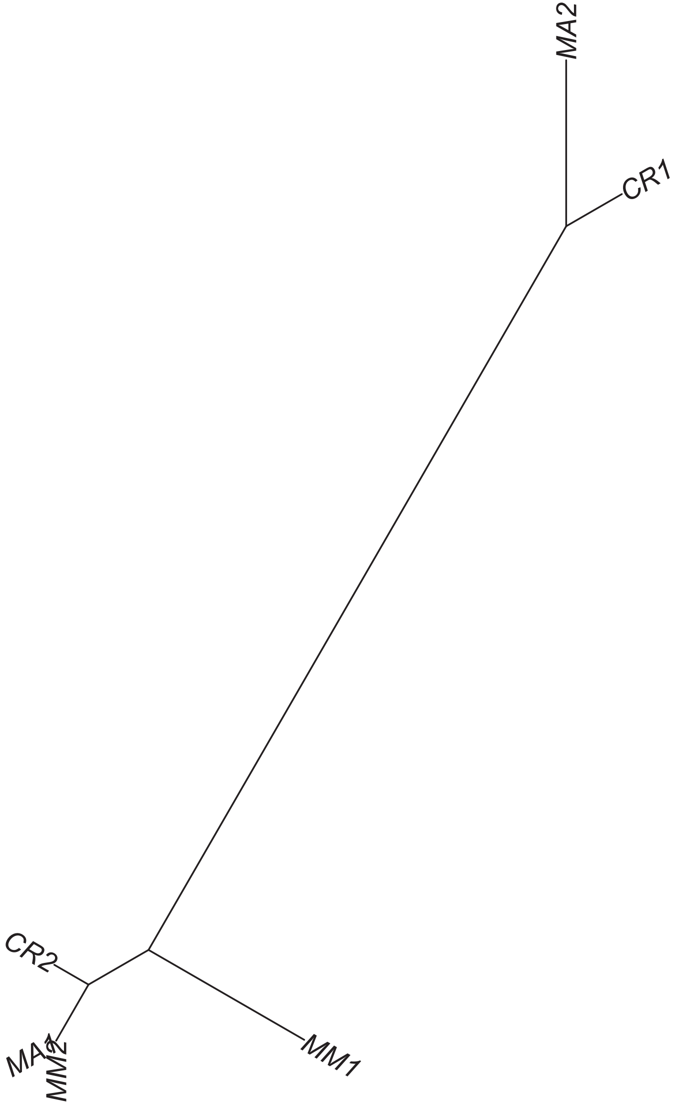

### Region 267

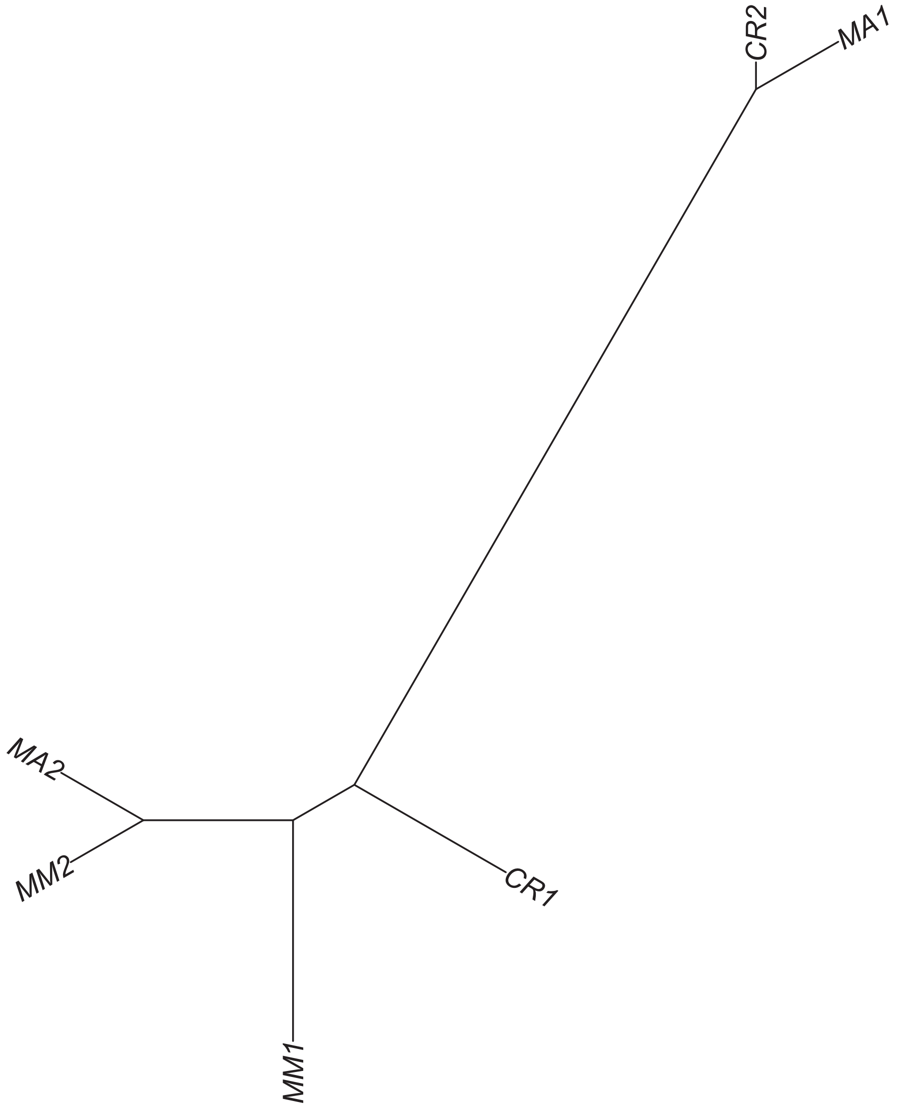

#### Region 268



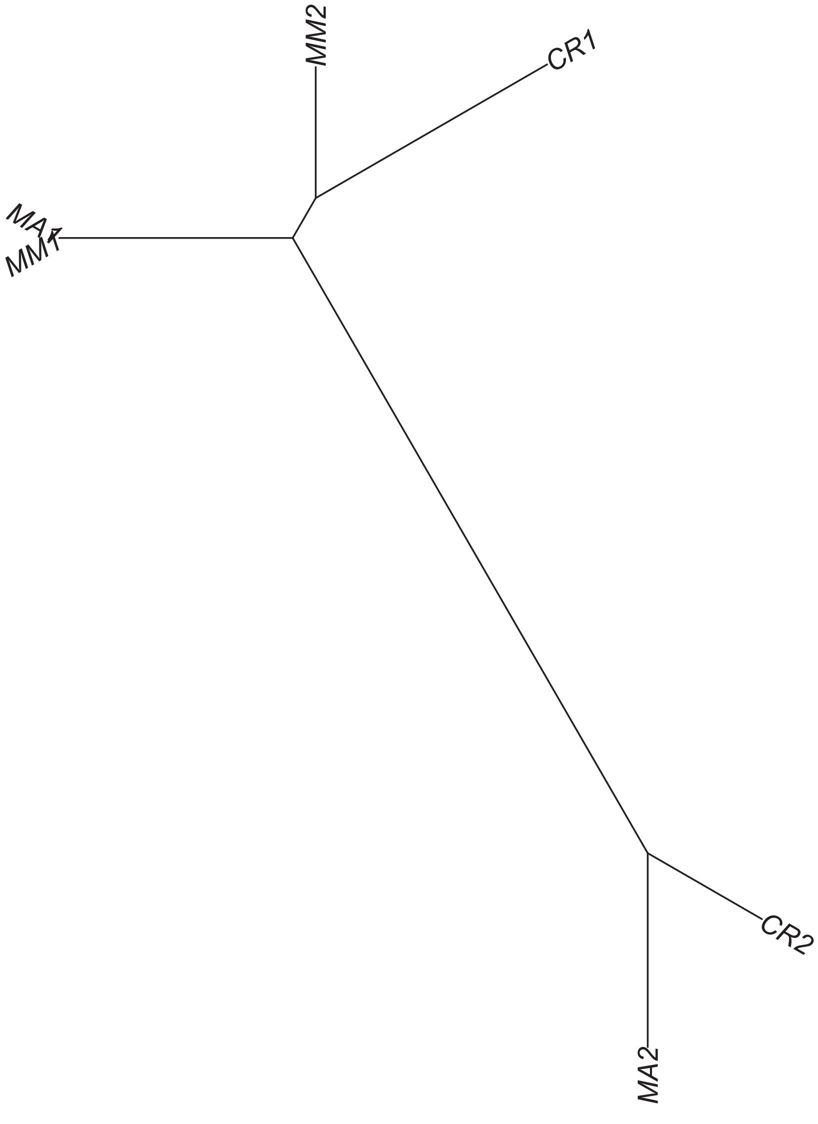

### Region 270



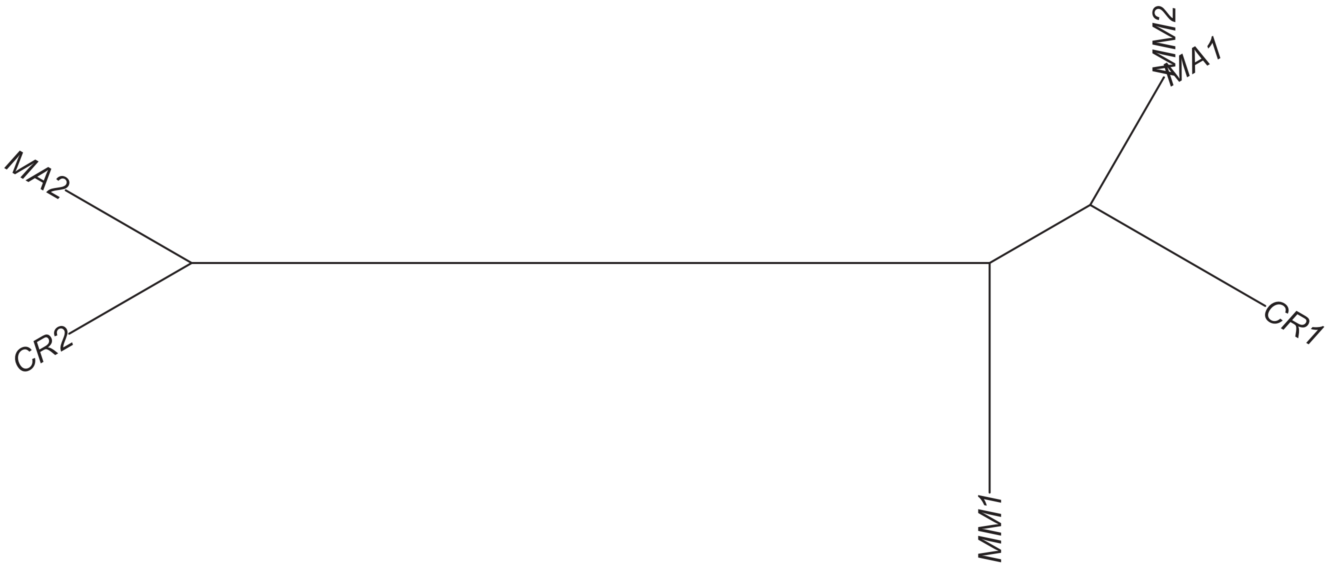

### Region 273

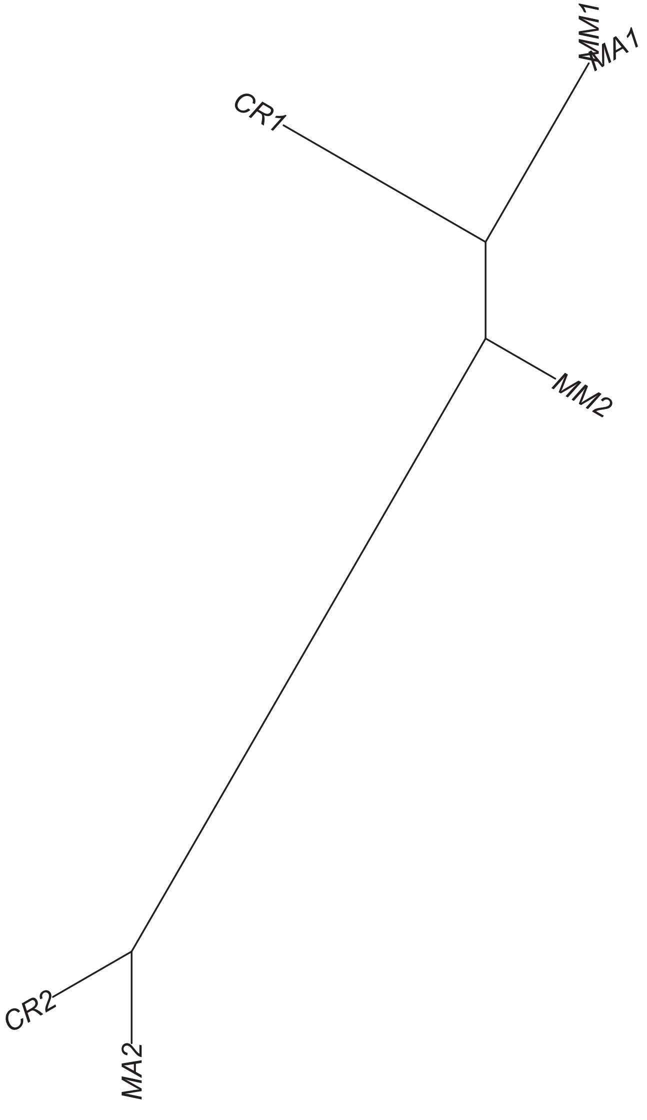

### Region 275

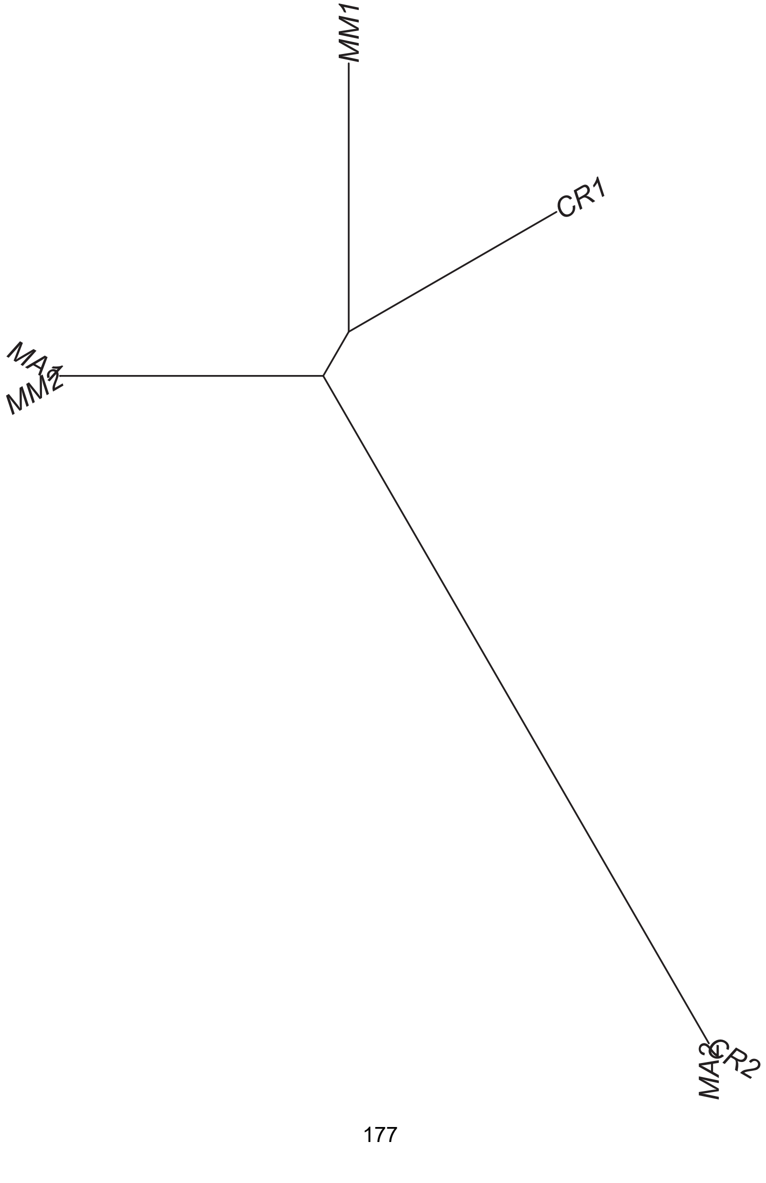

### Region 276

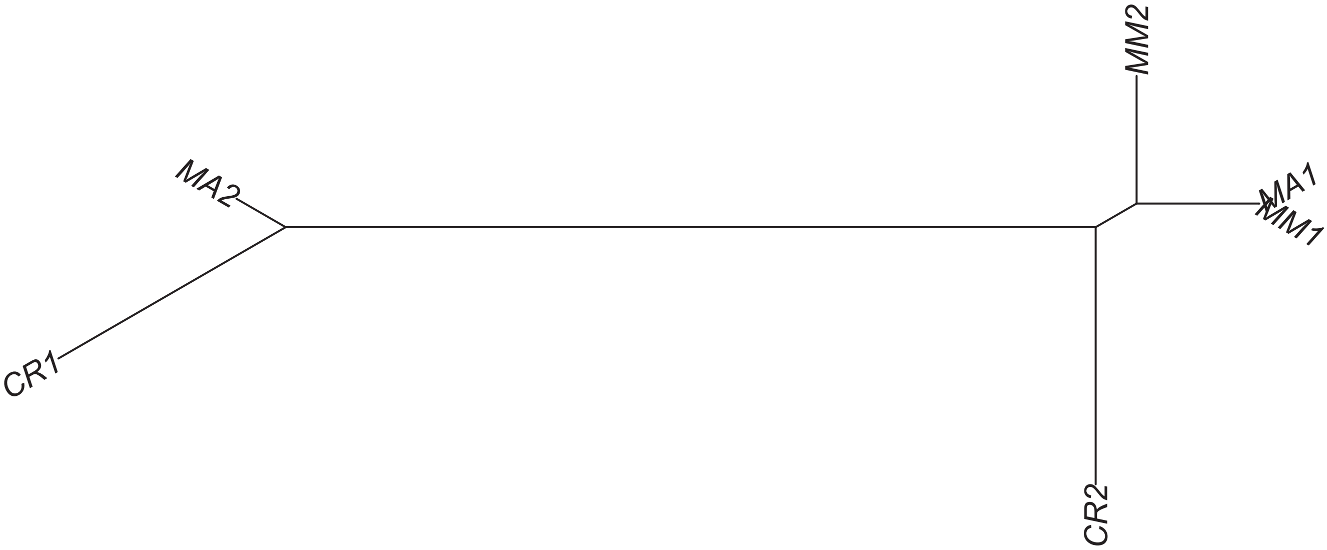

### Region 277



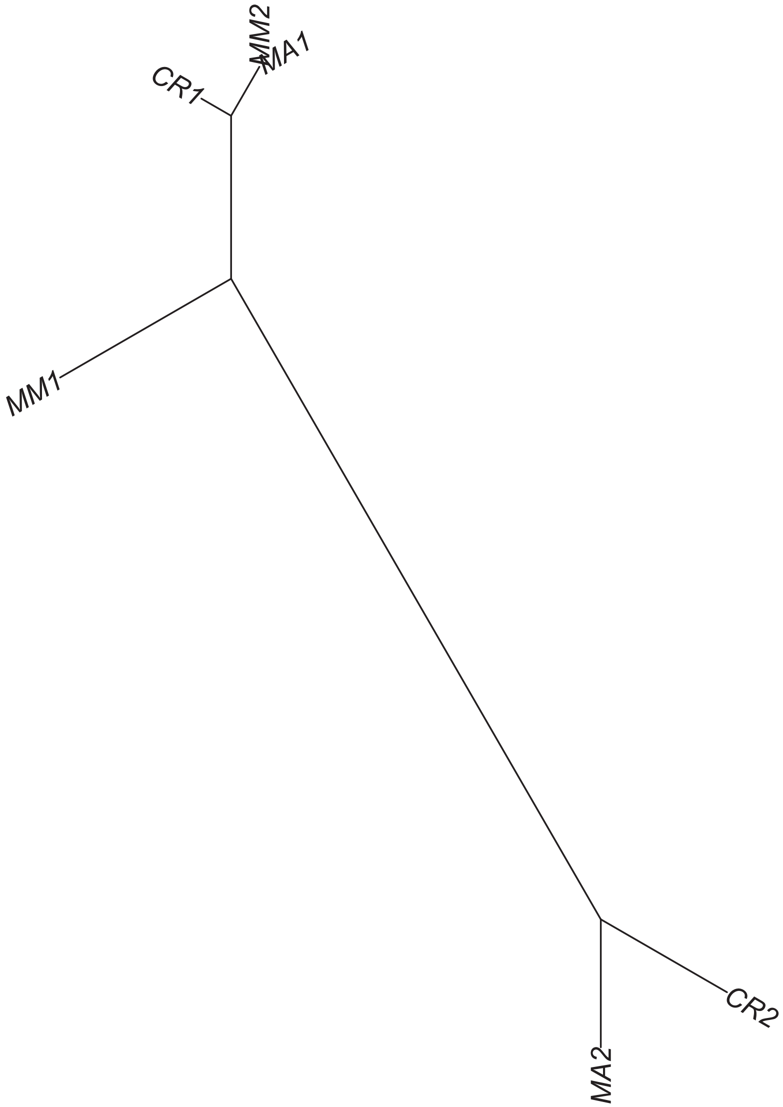

### Region 278

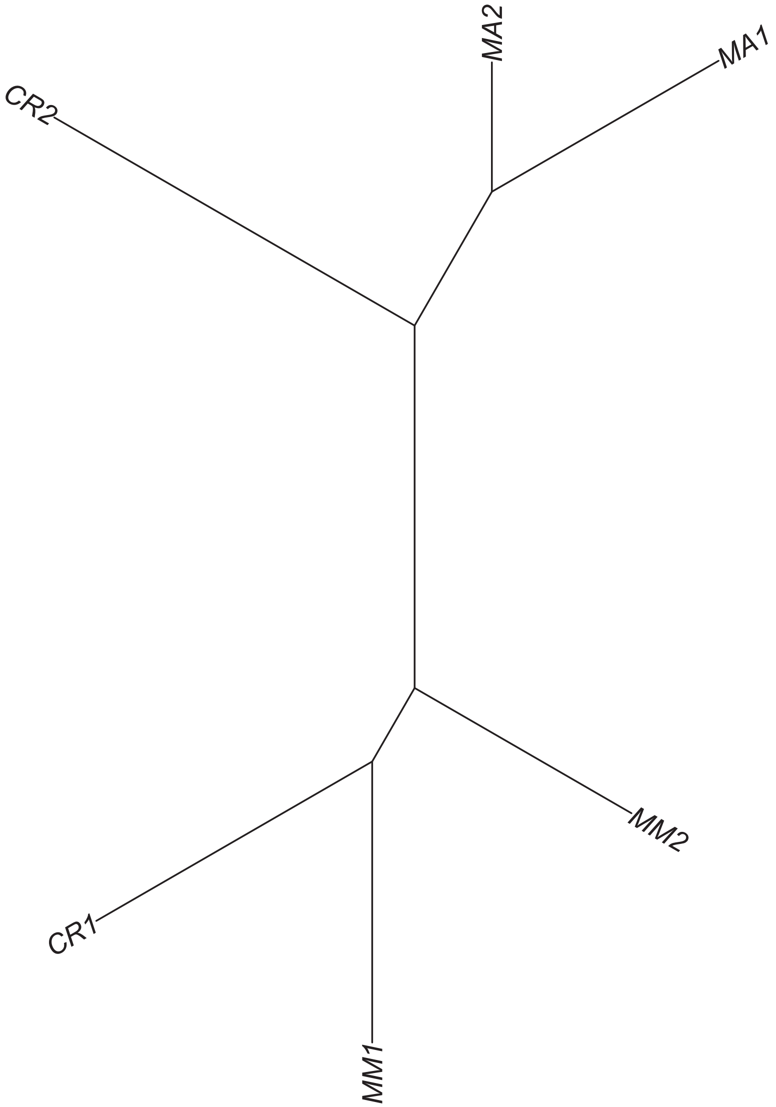

### Region 279

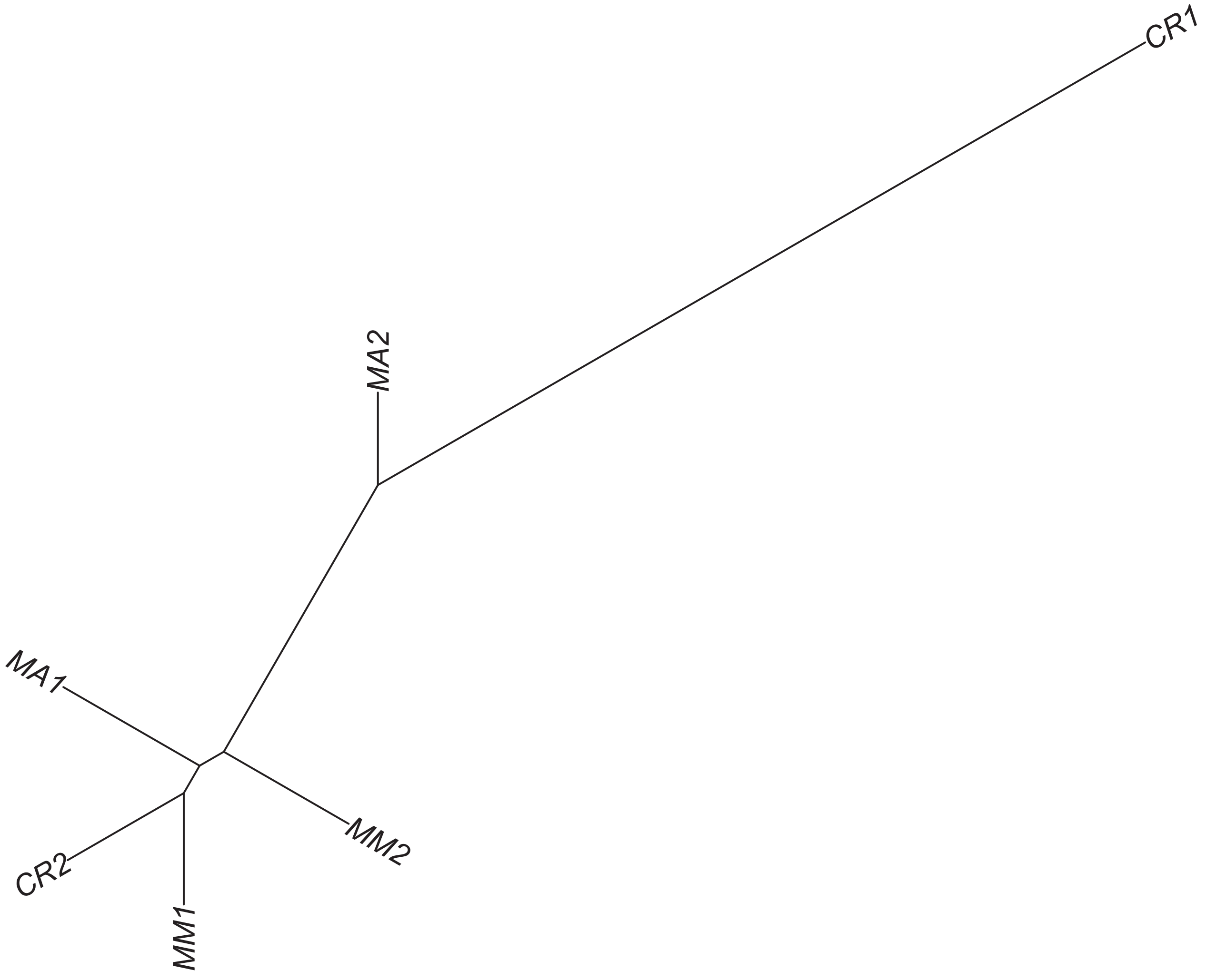

### Region 280

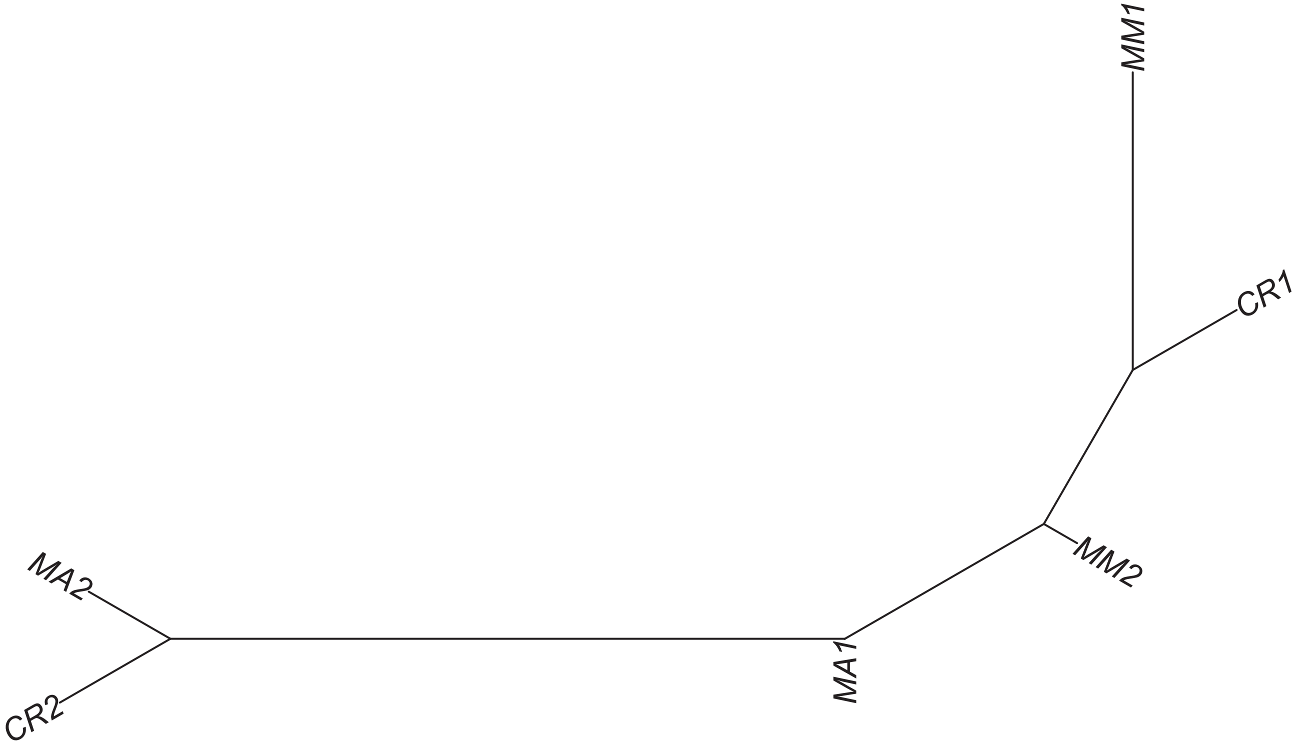

### Region 281



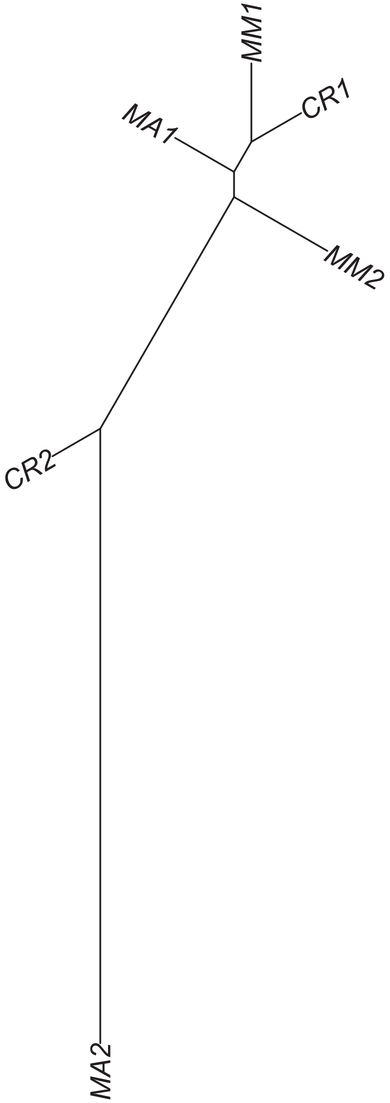

Region 282



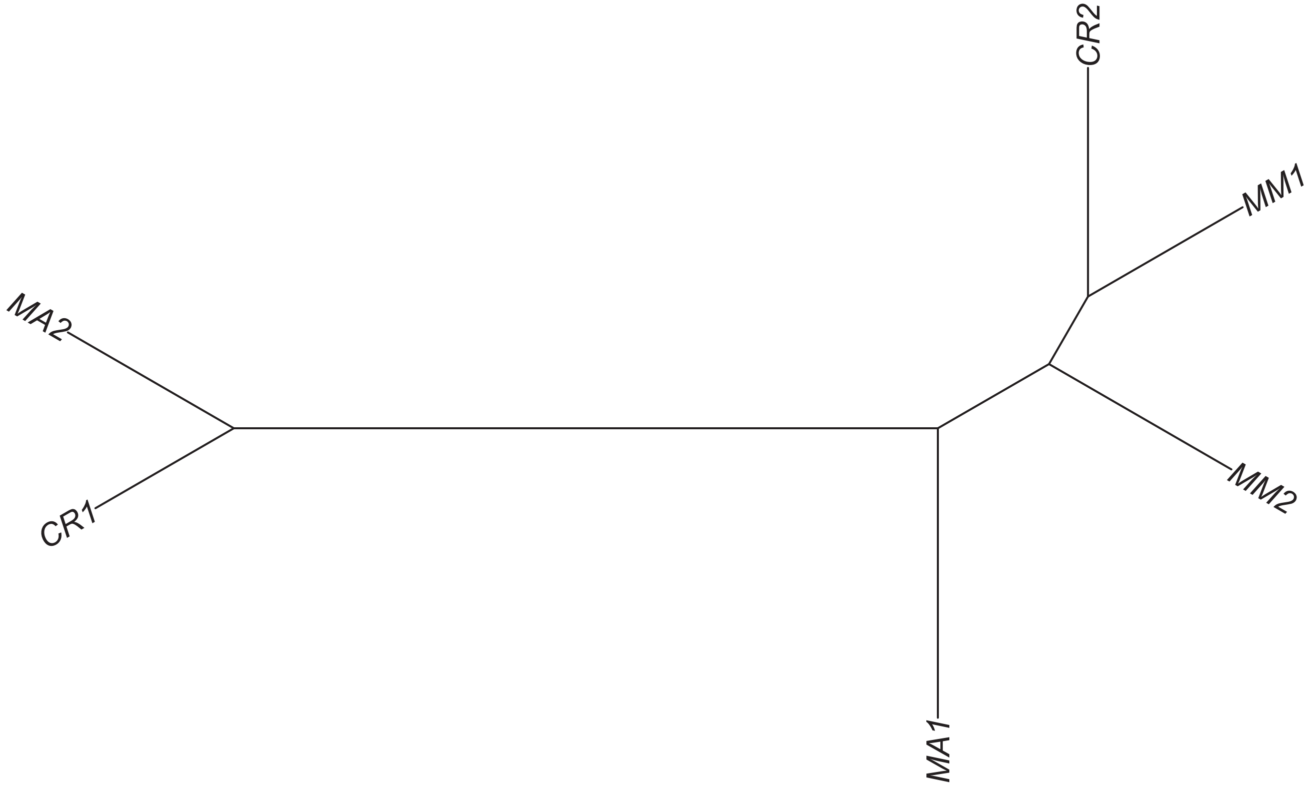

### Region 283

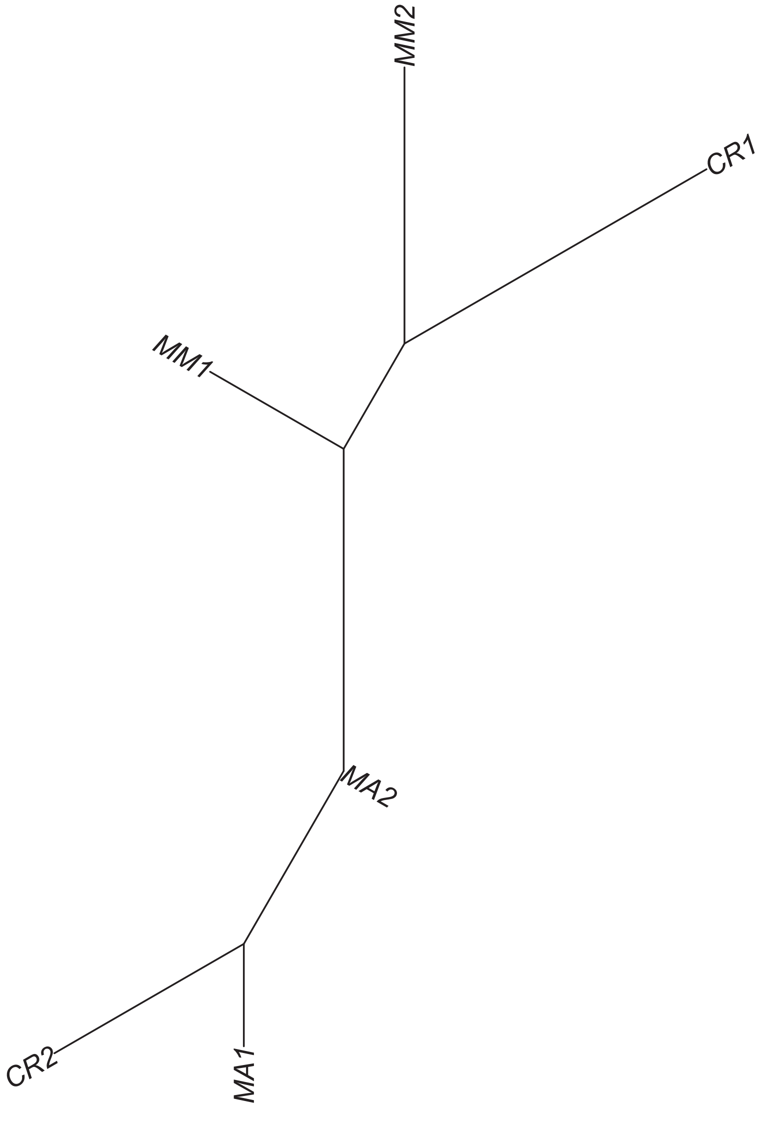

### Region 284

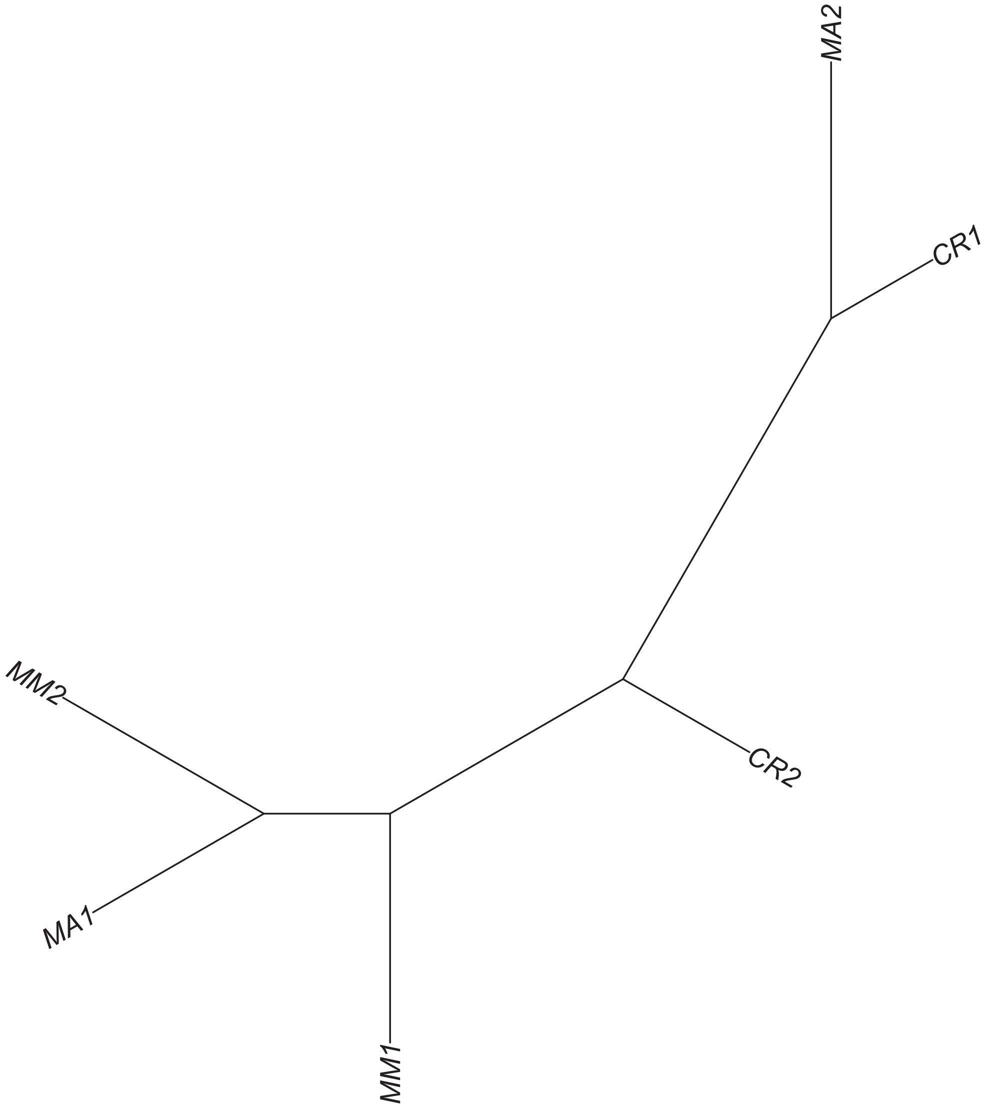

### Region 287



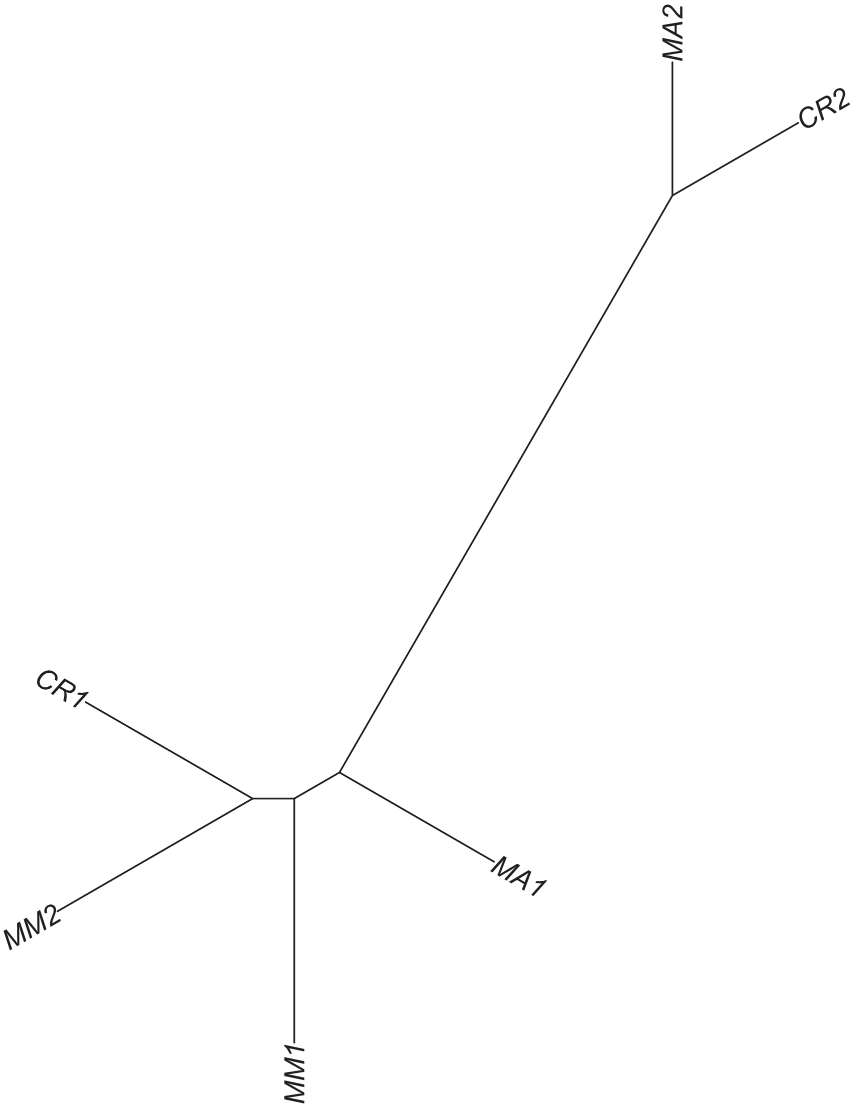

### Region 290



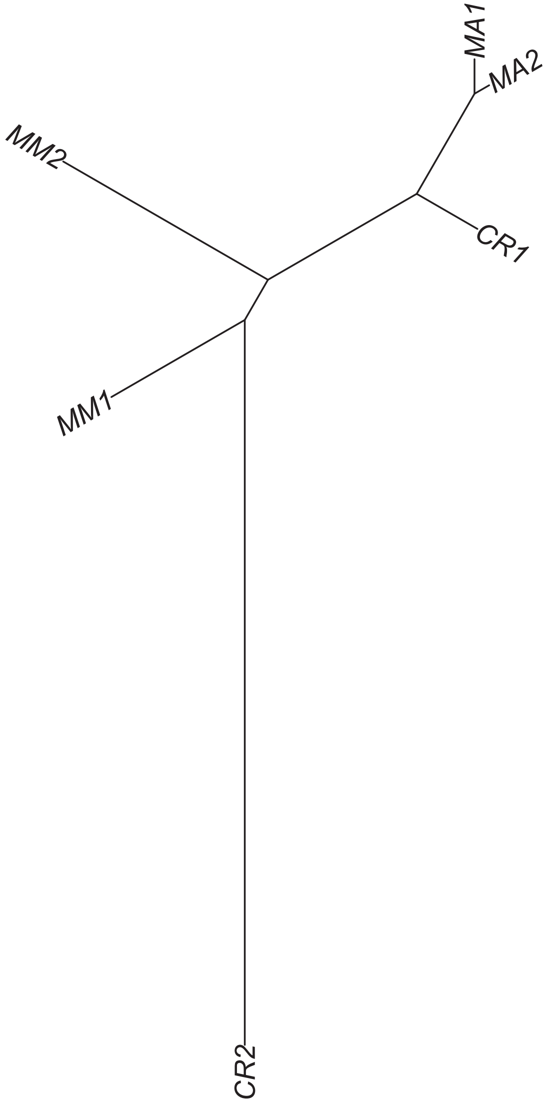

### Region 291

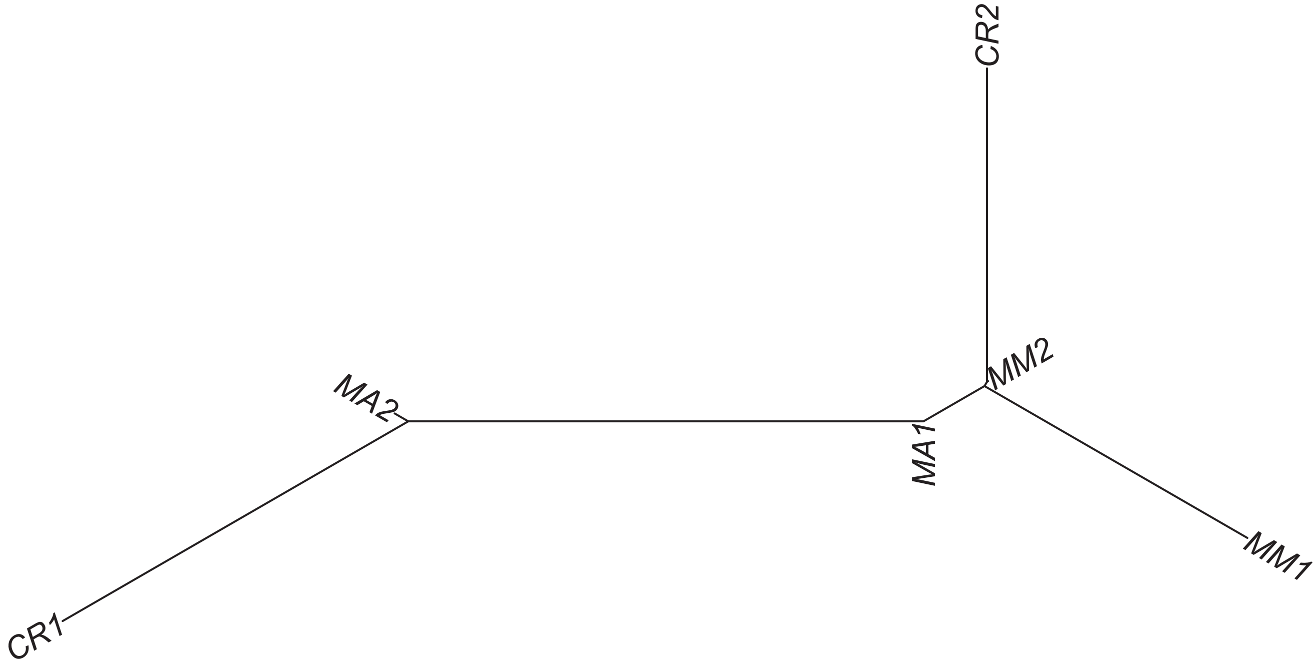

### Region 297

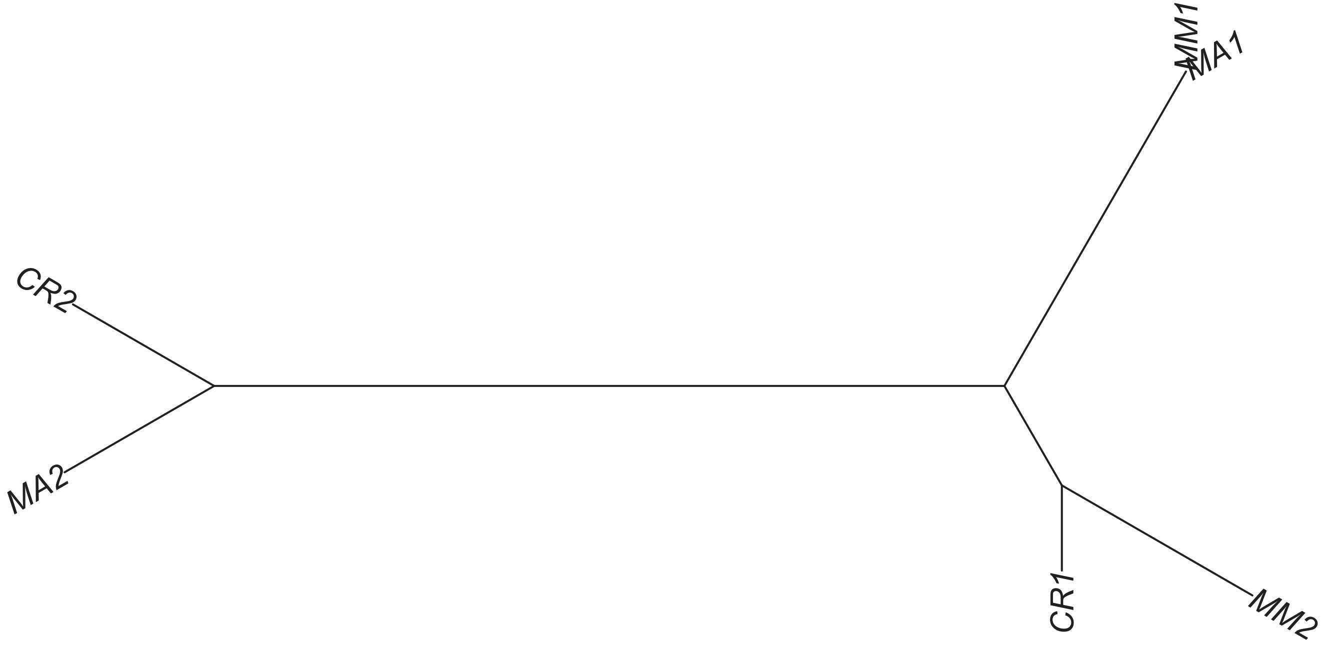

### Region 300



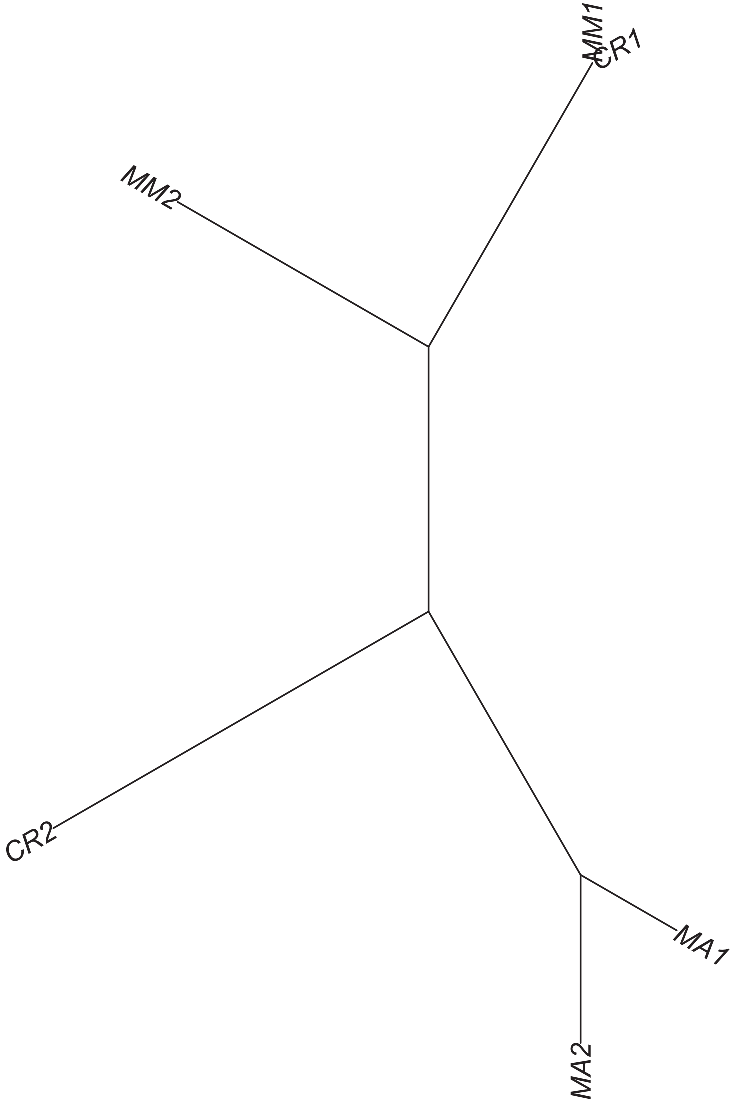

### Region 302

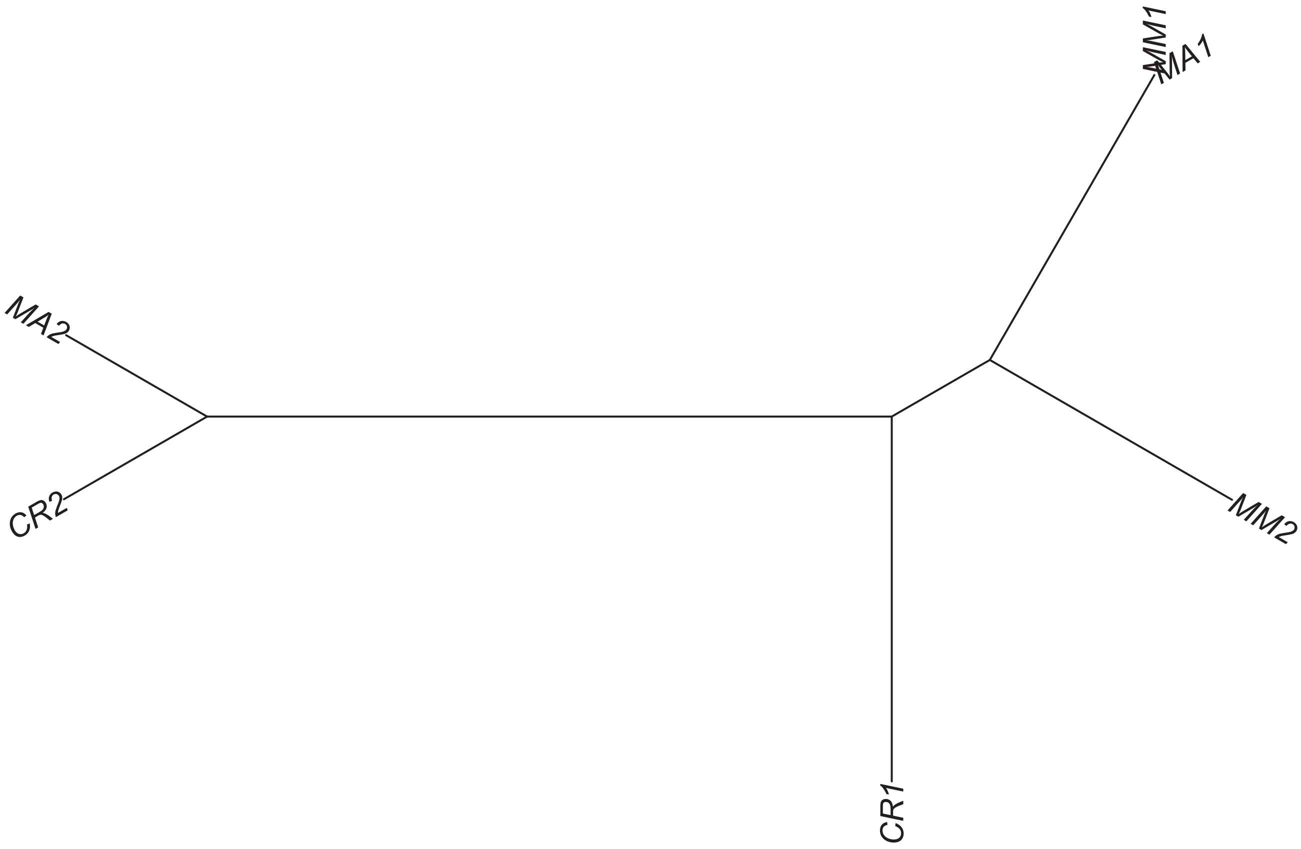

### Region 303

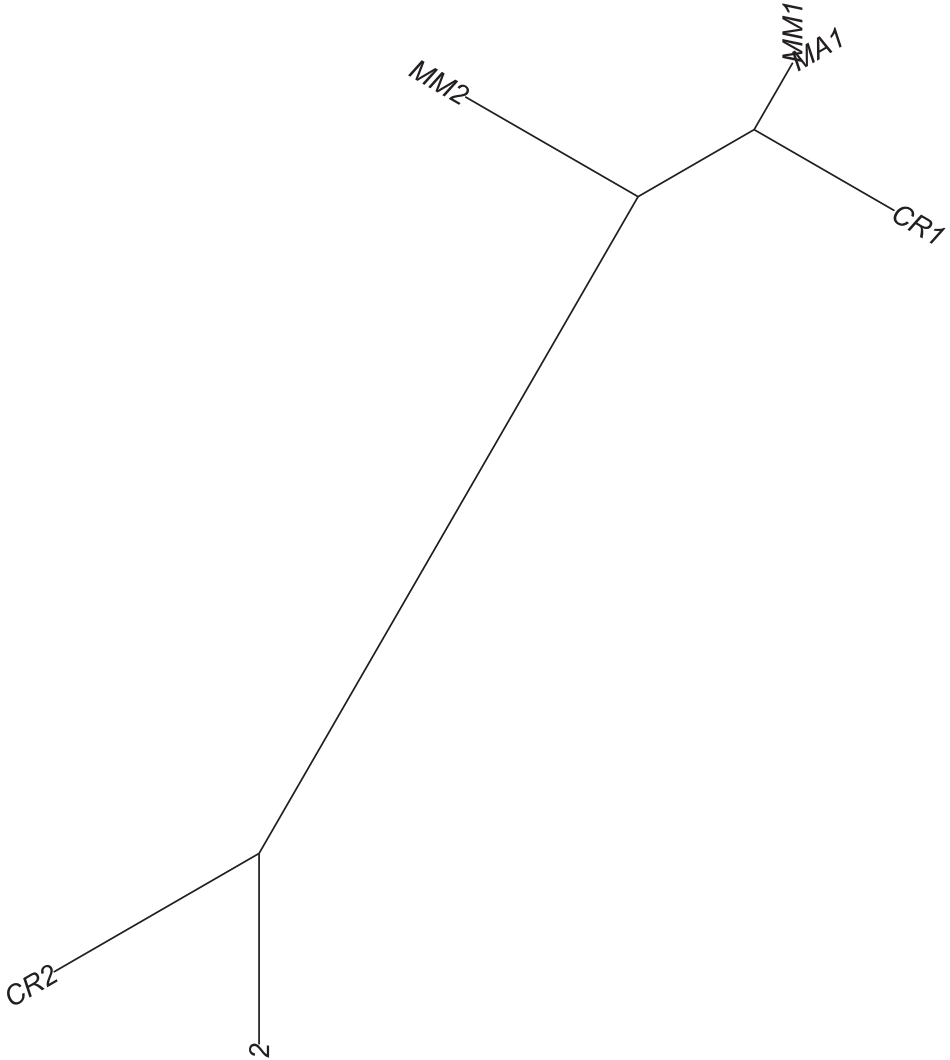

Region 304

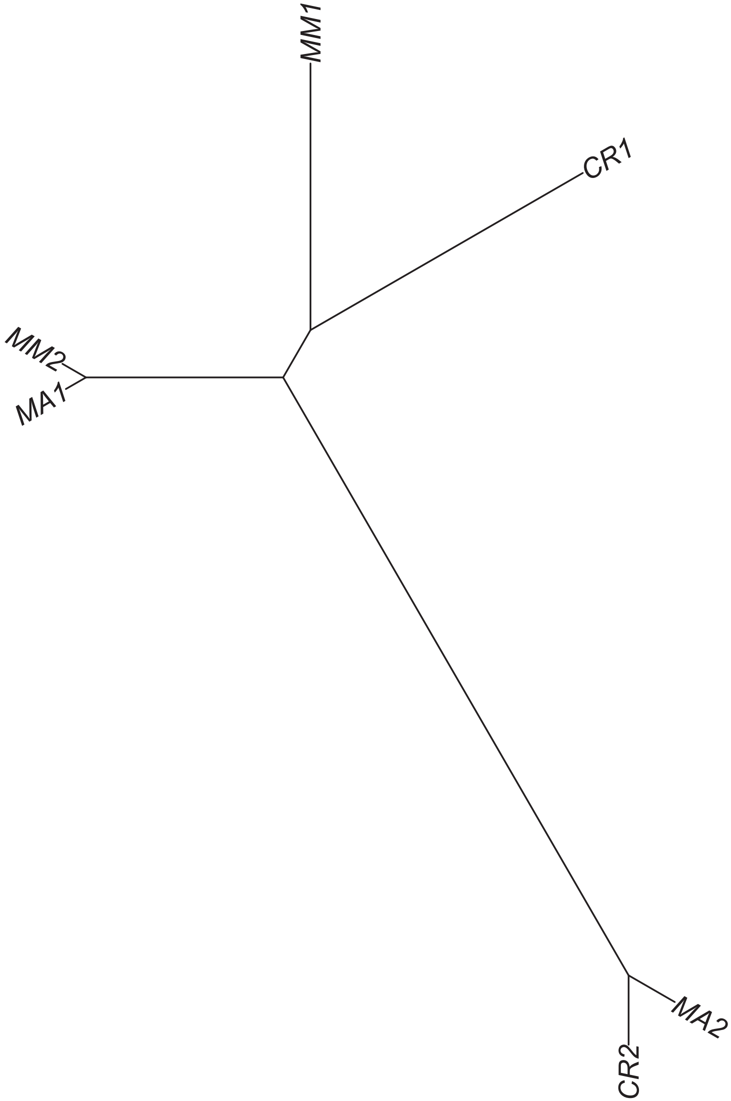

### Region 308

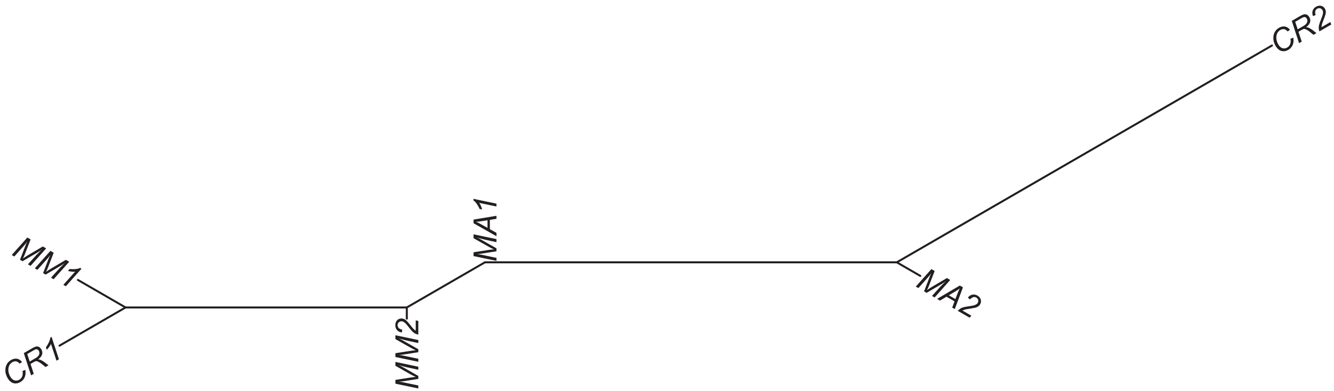

### Region 311

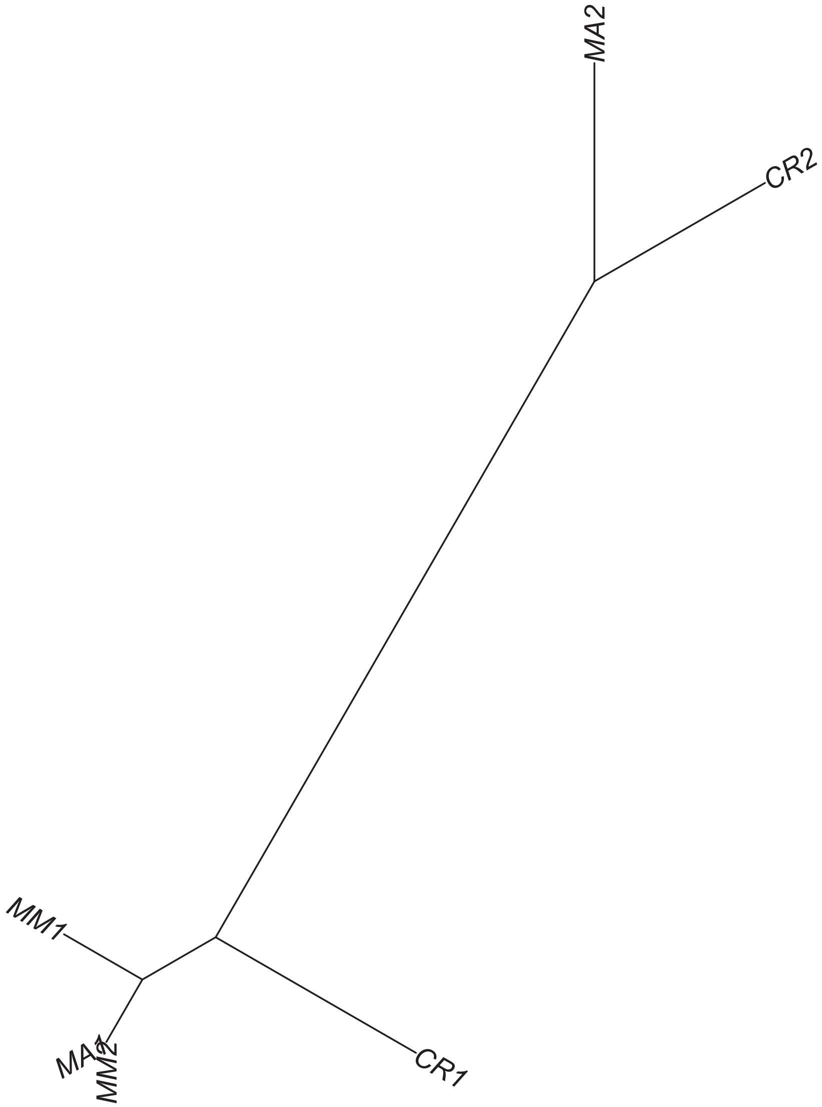

### Region 312



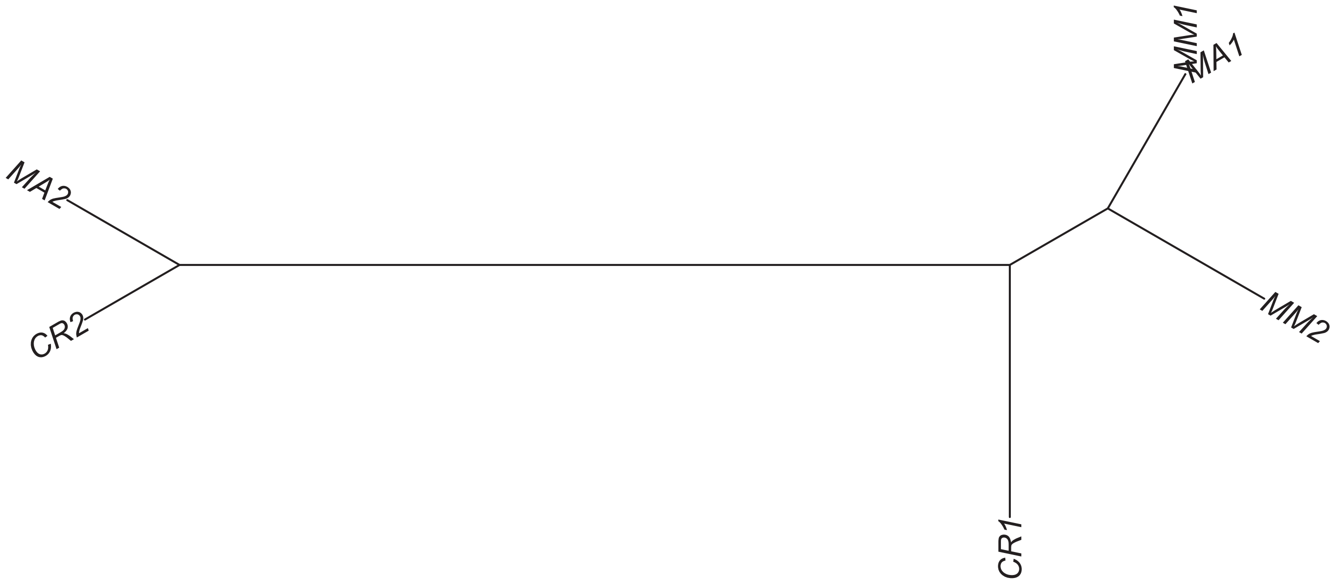

### Region 313

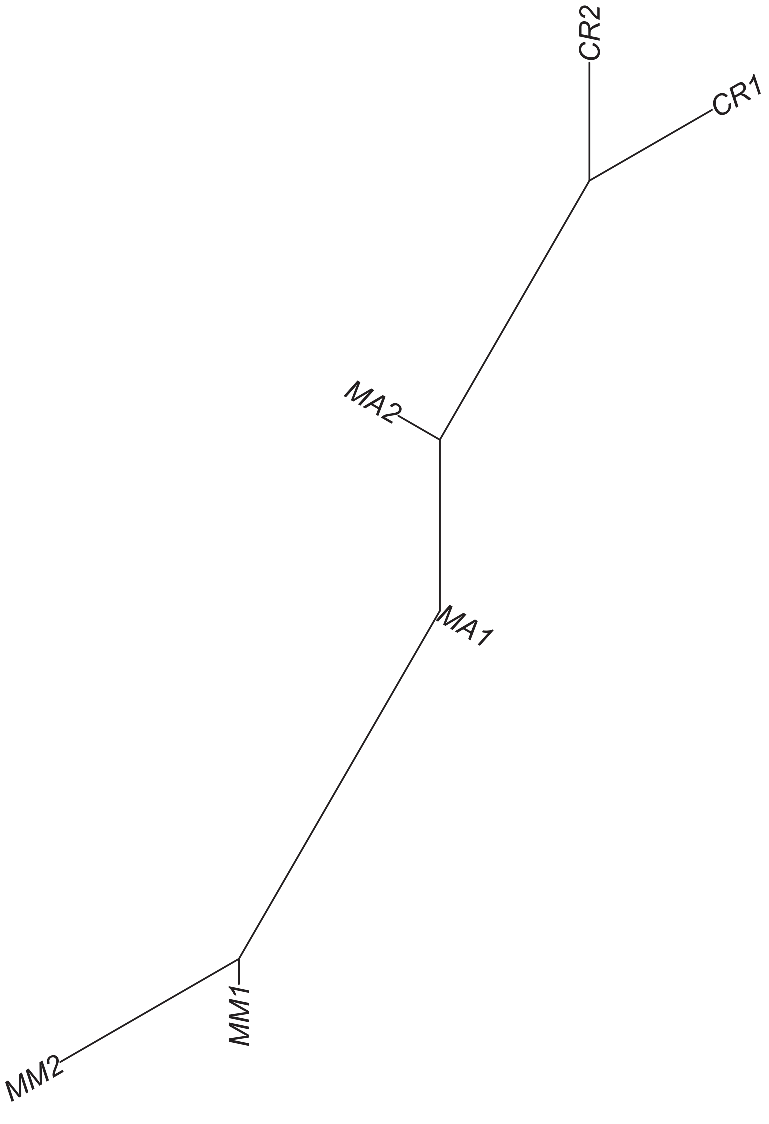

### Region 315

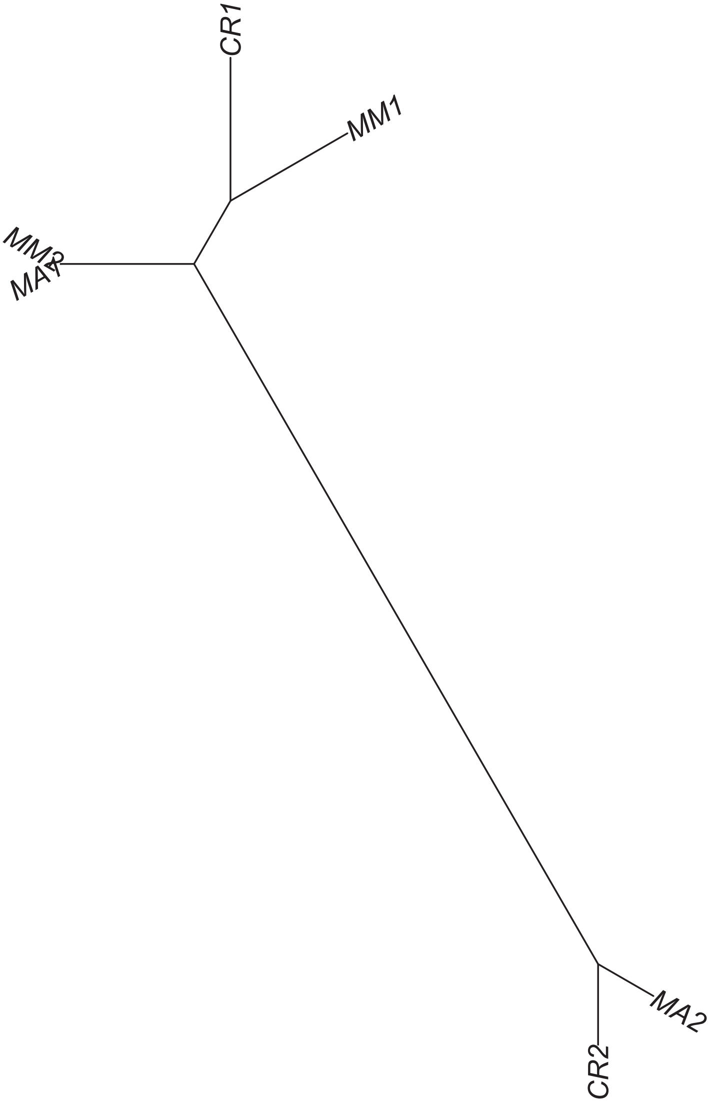

### Region 316



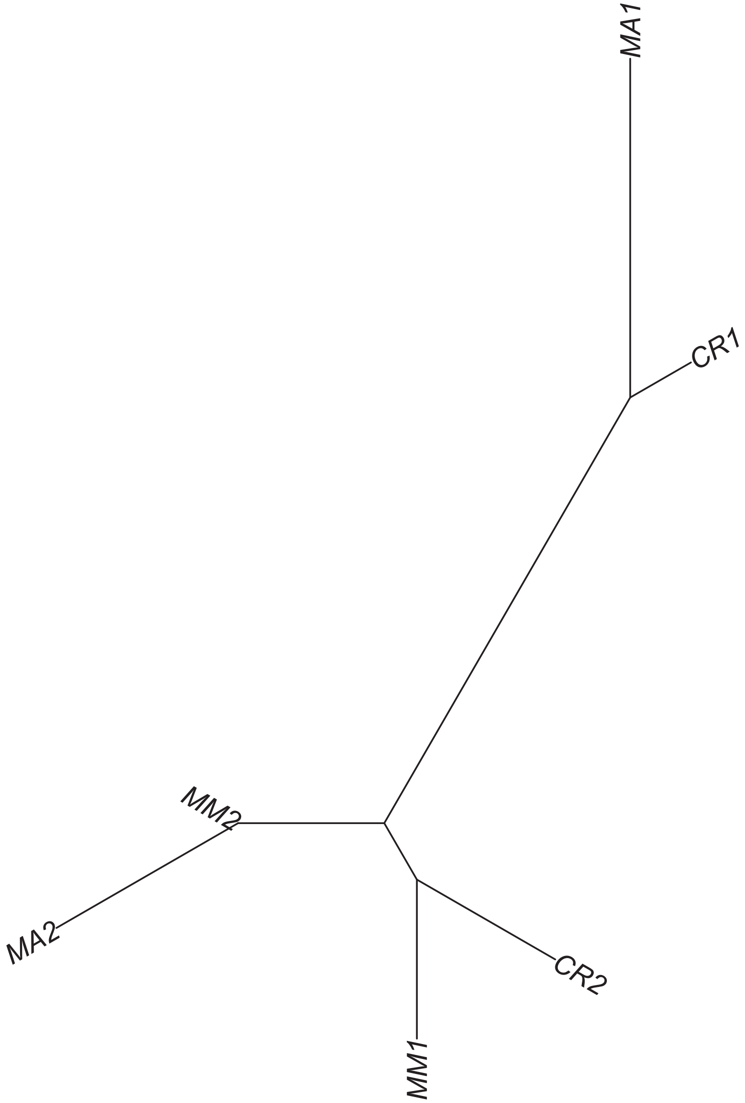

### Region 317

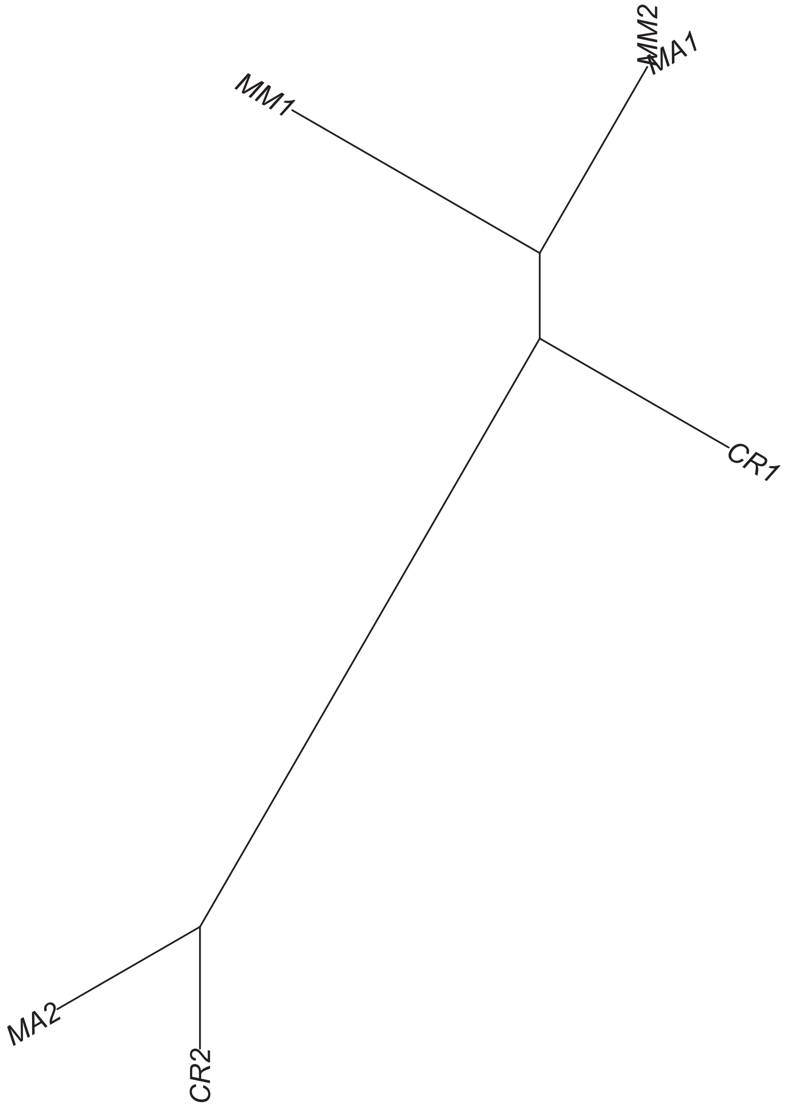

### Region 318



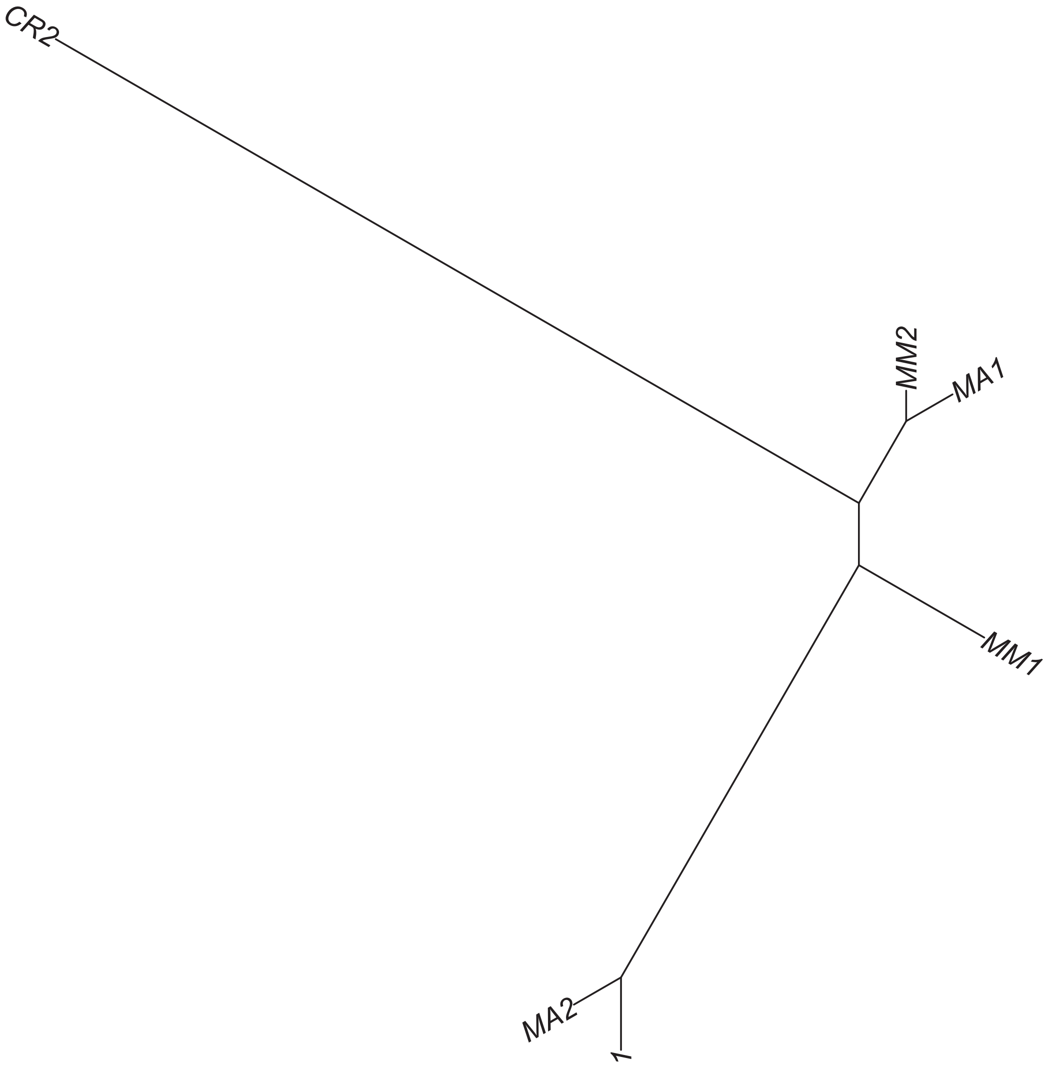

### Region 319



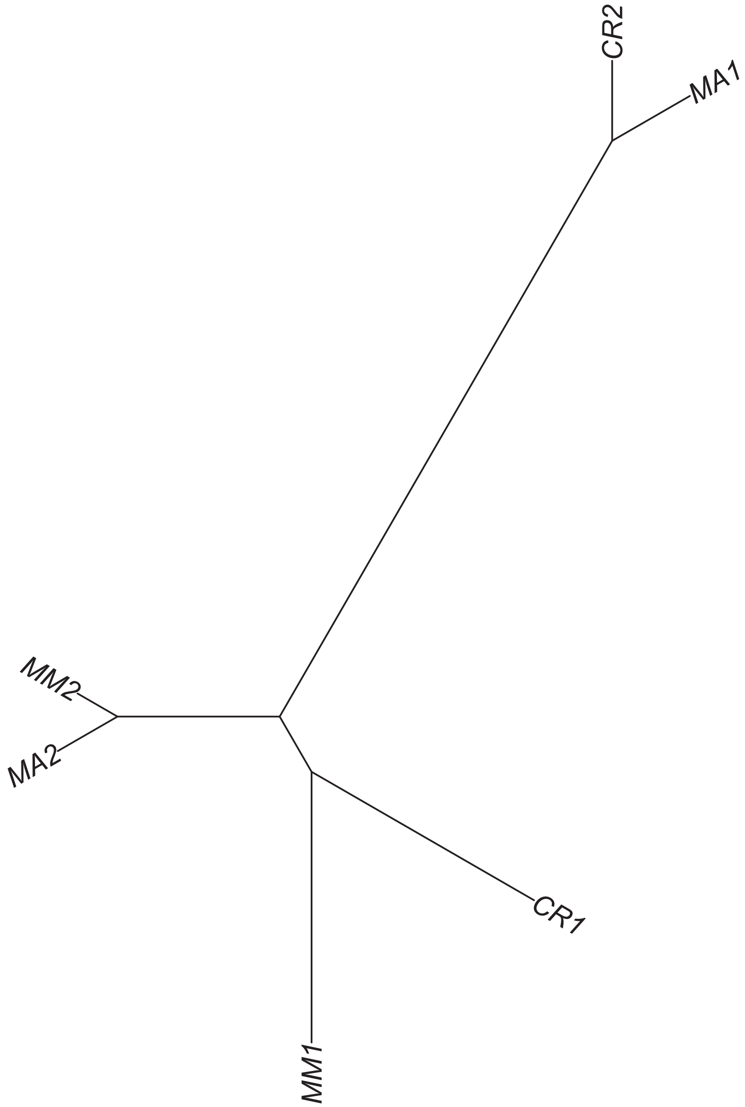

### Region 320

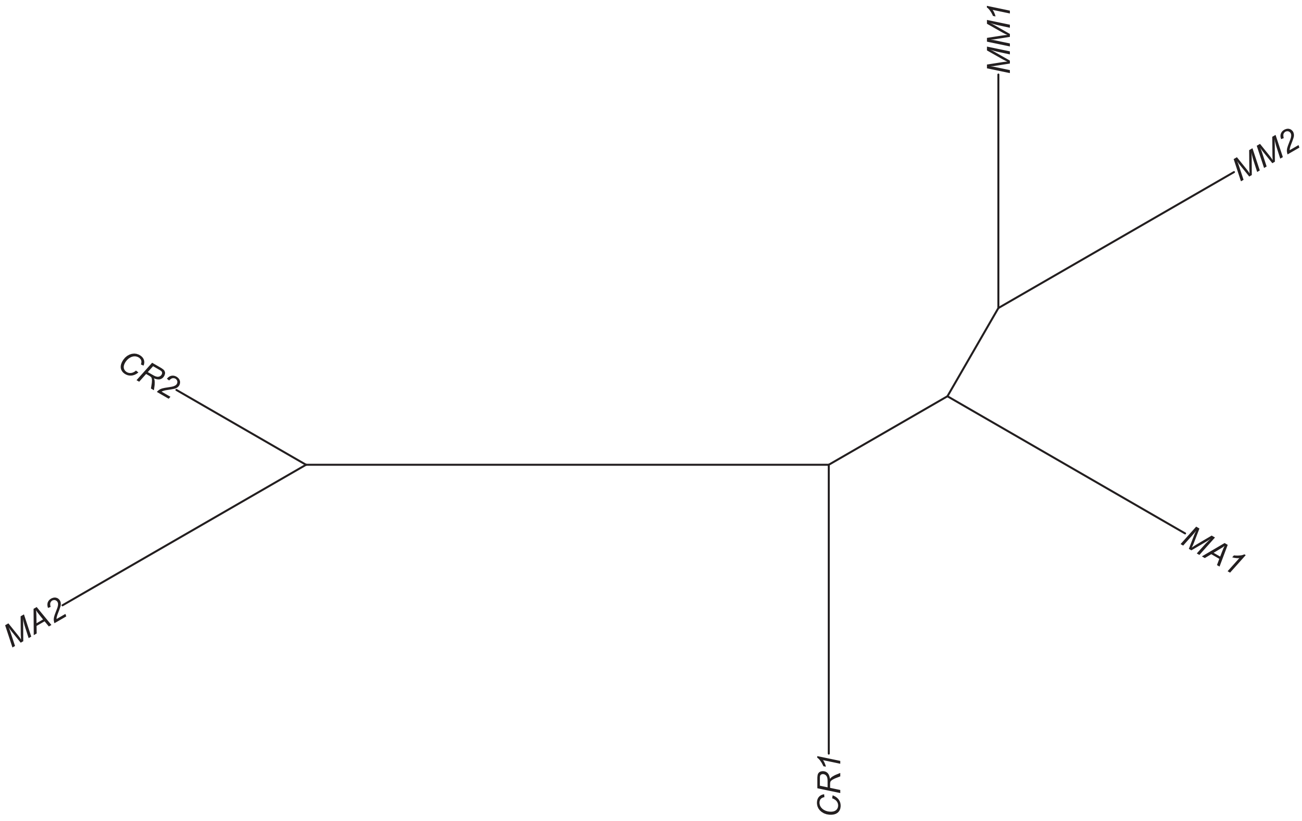

### Region 322

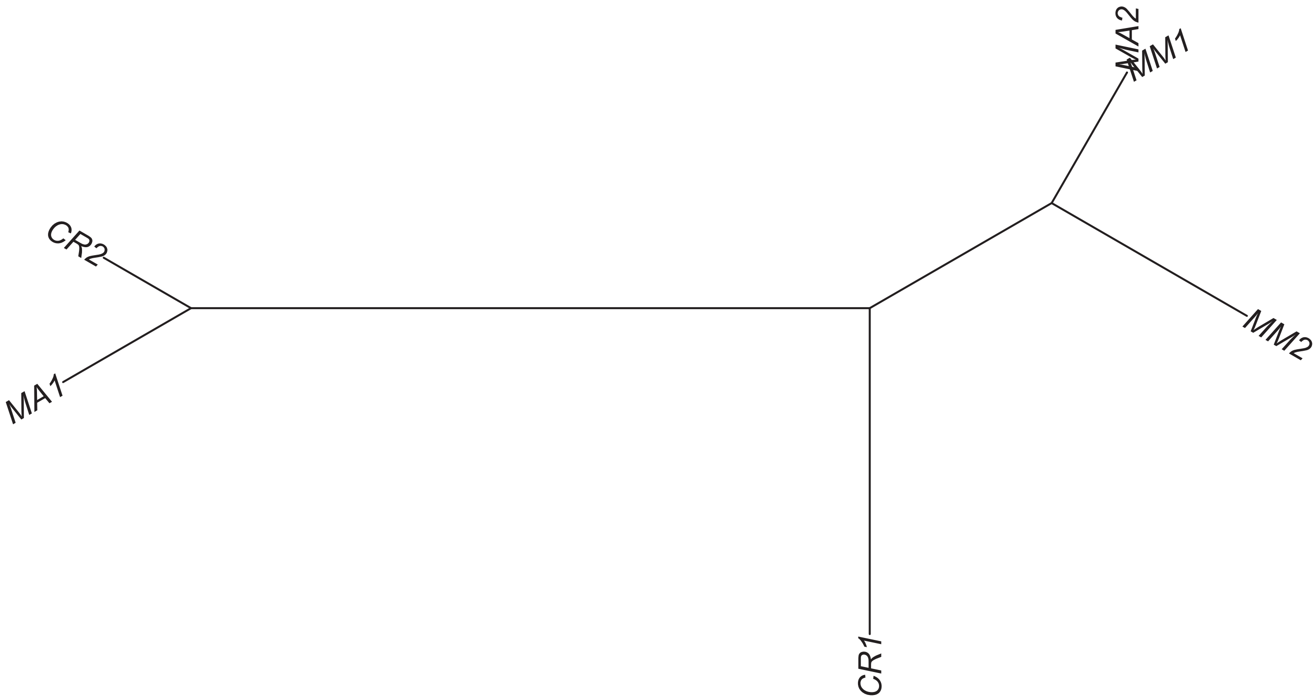

### Region 324



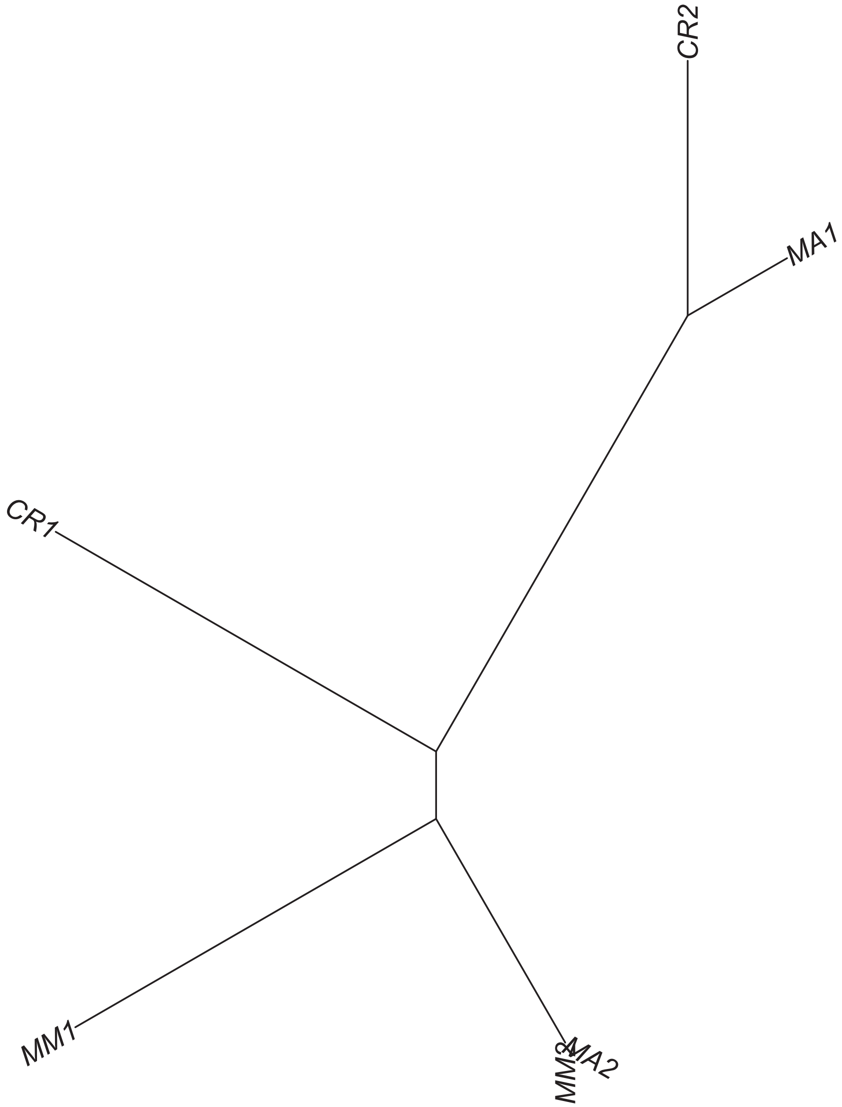

### Region 326



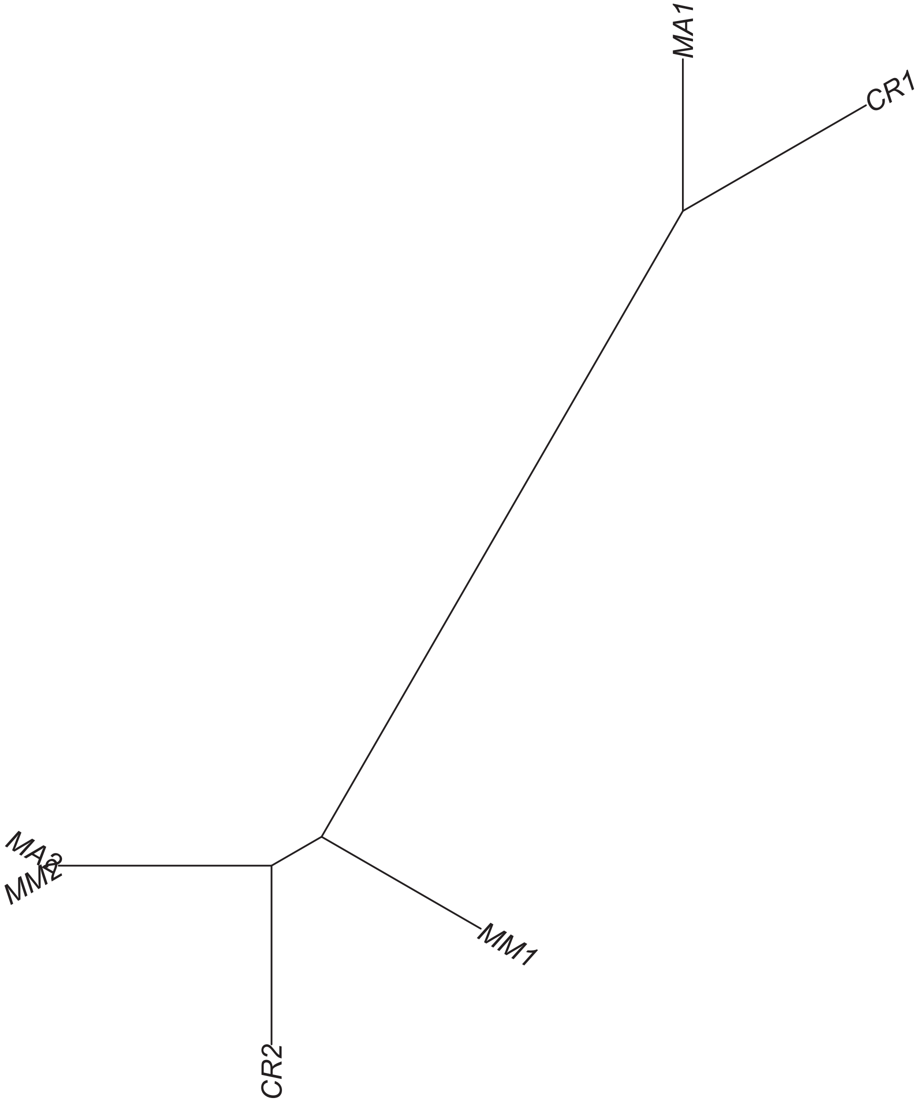

### Region 329

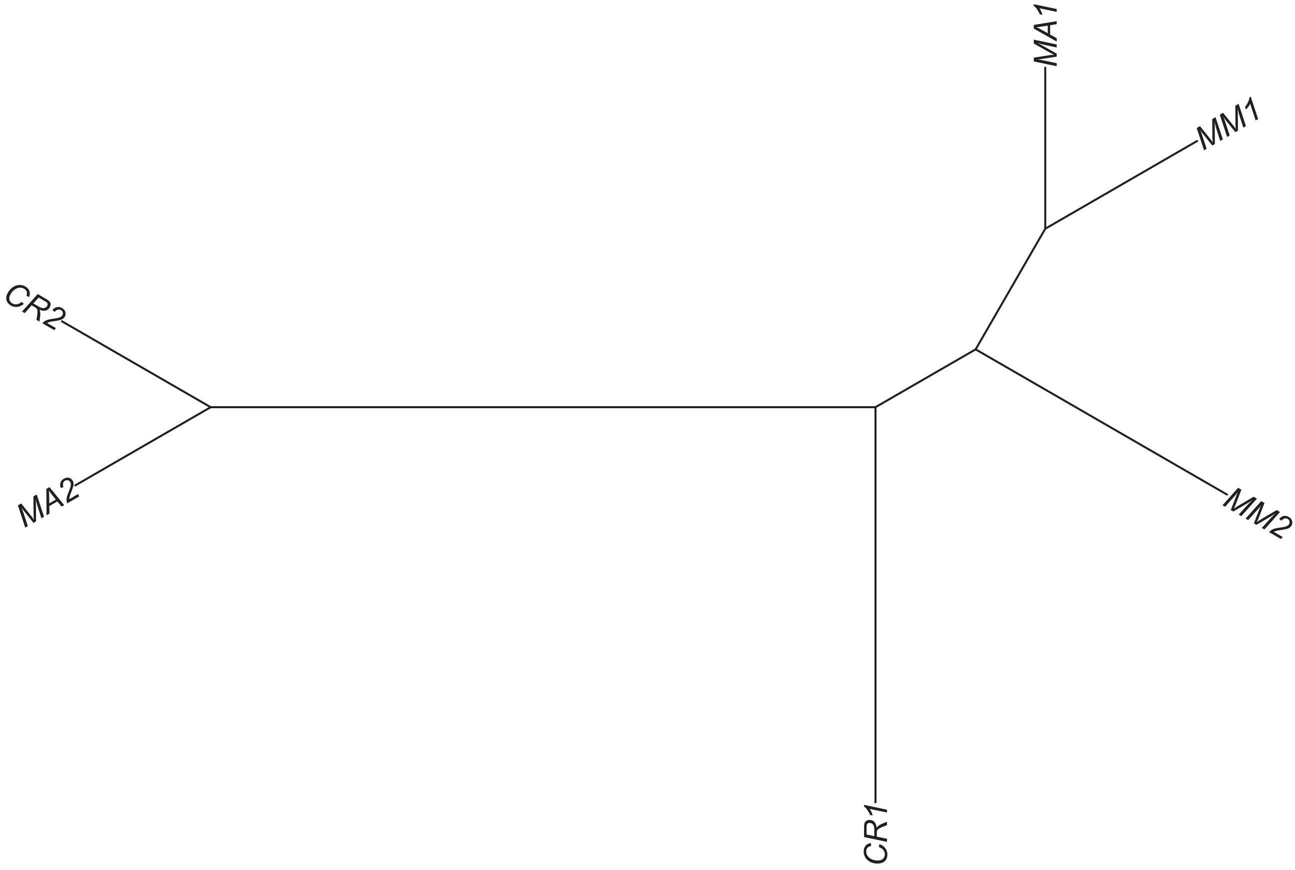

Region 334

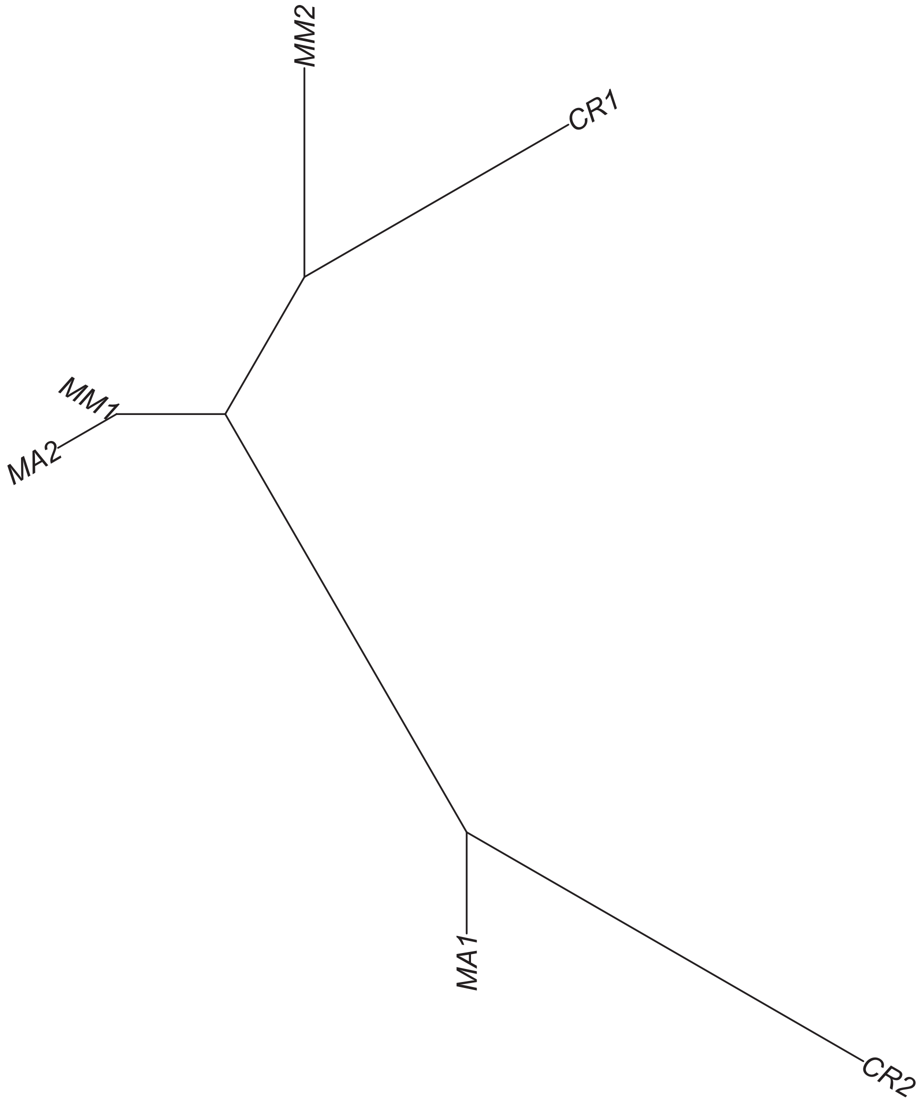

Region 335

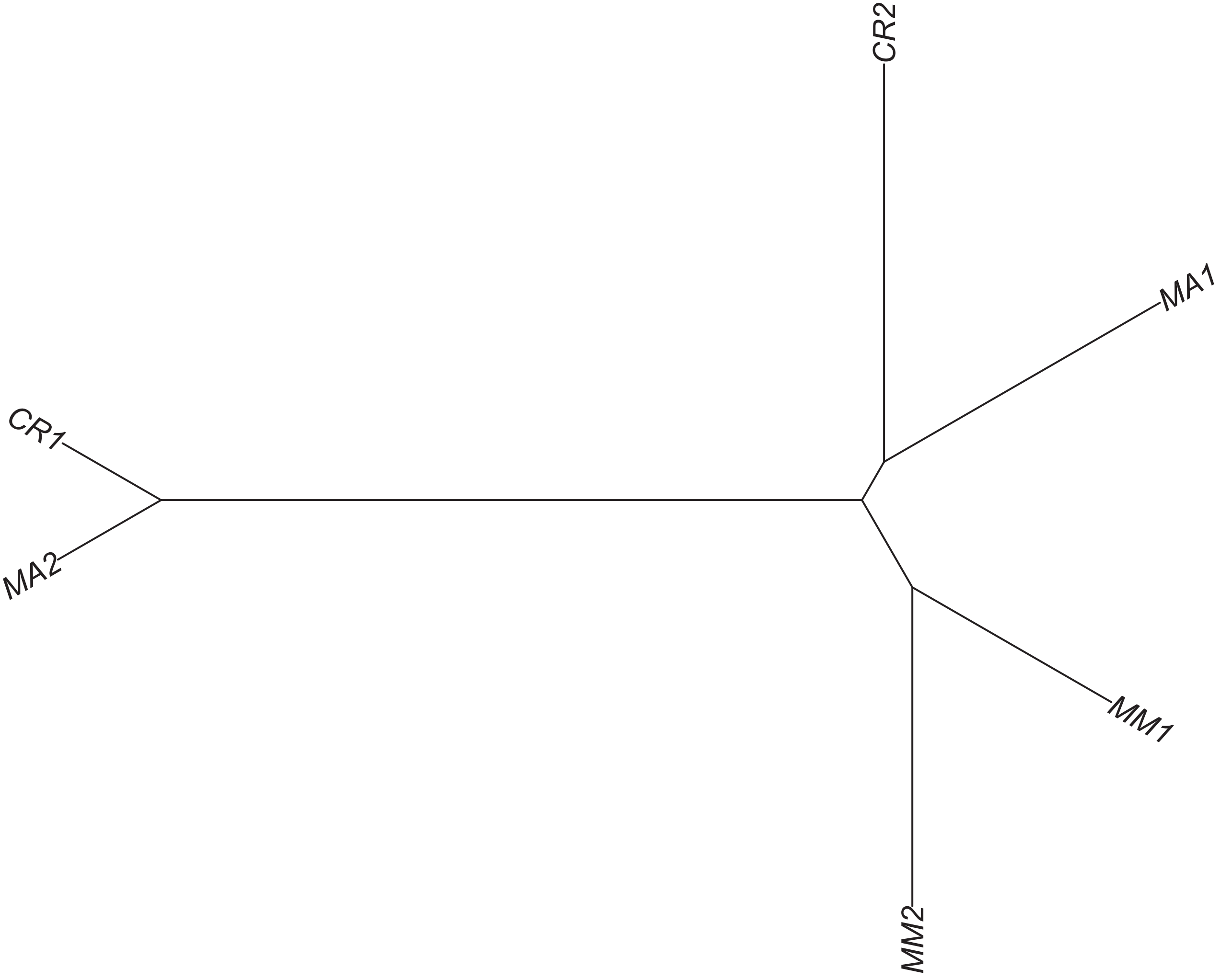

Region 336

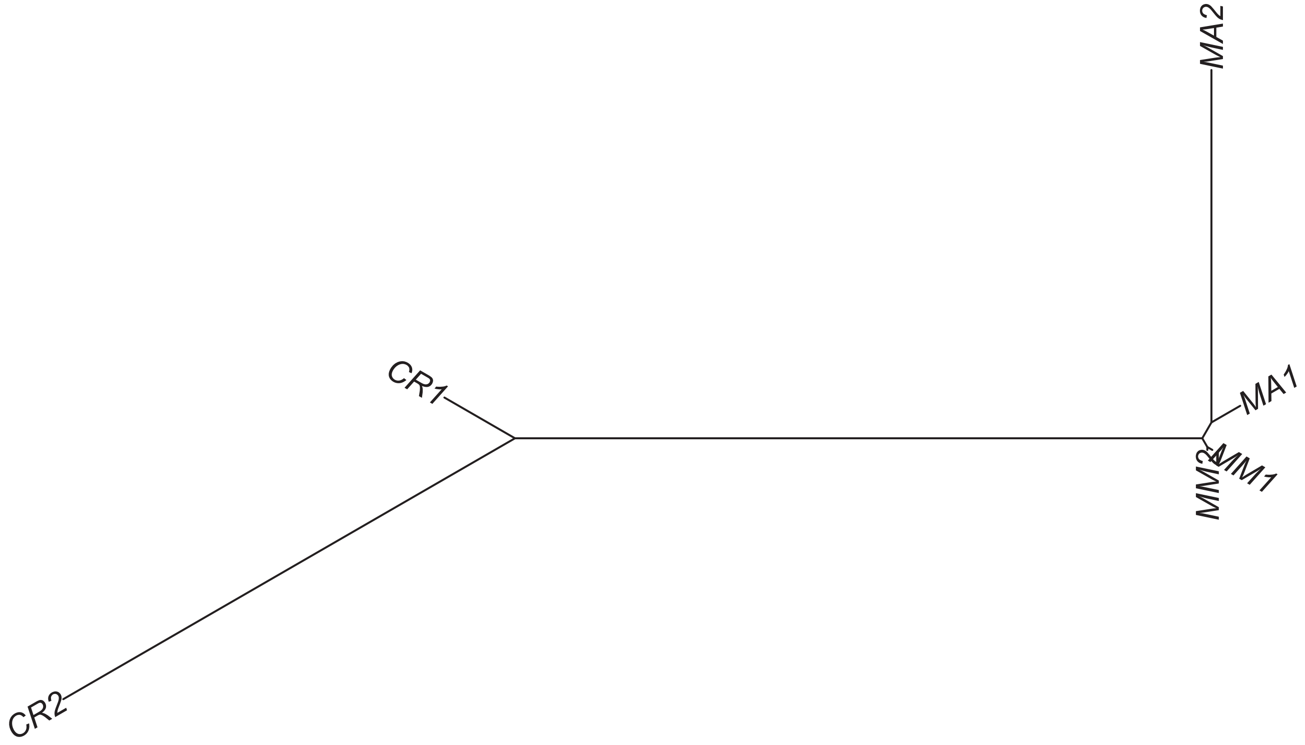

### Region 337



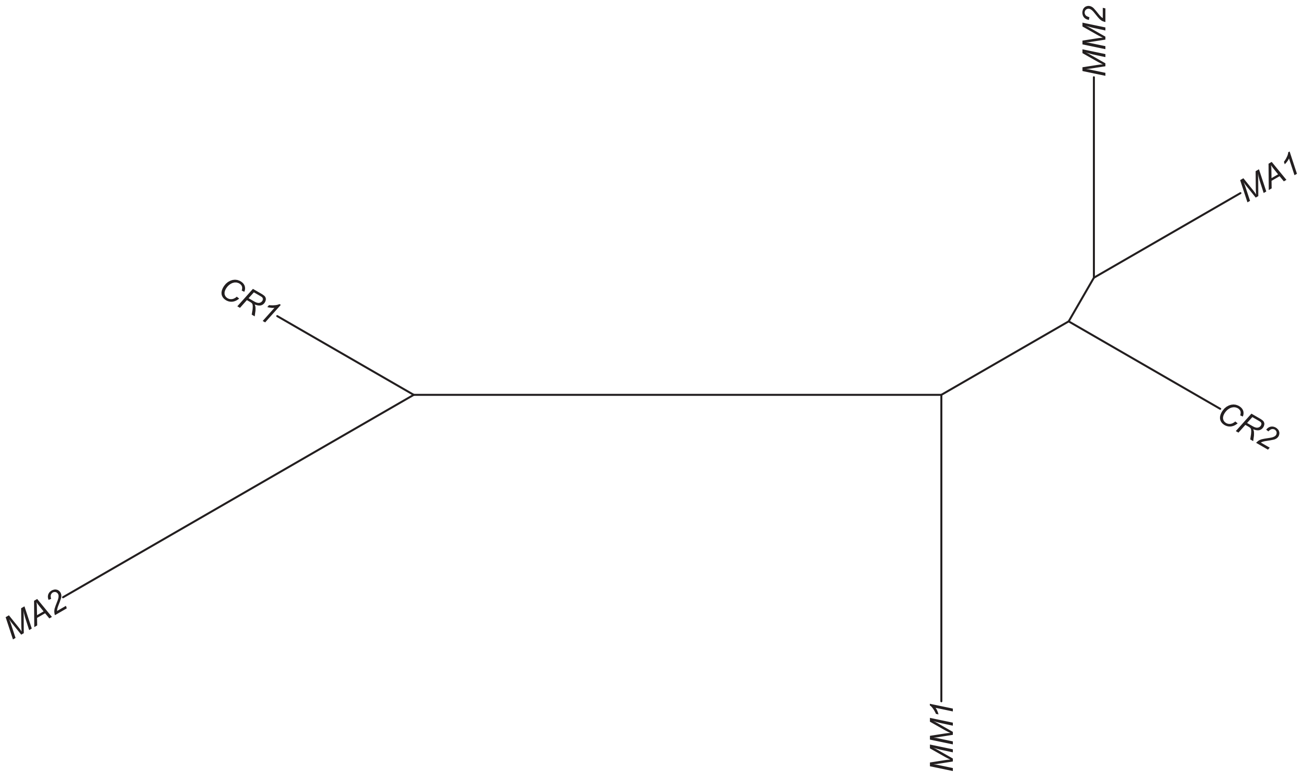

### Region 342

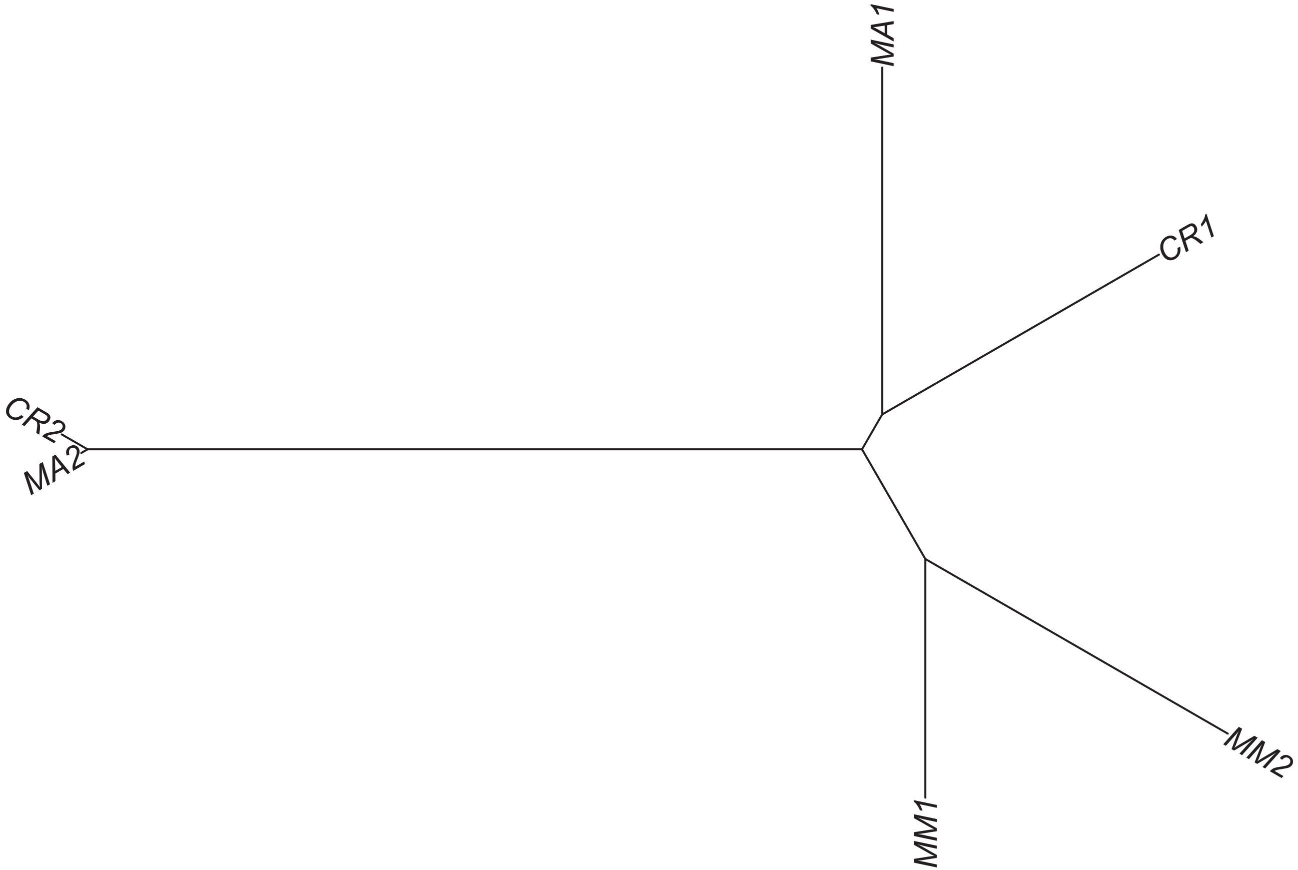

### Region 343



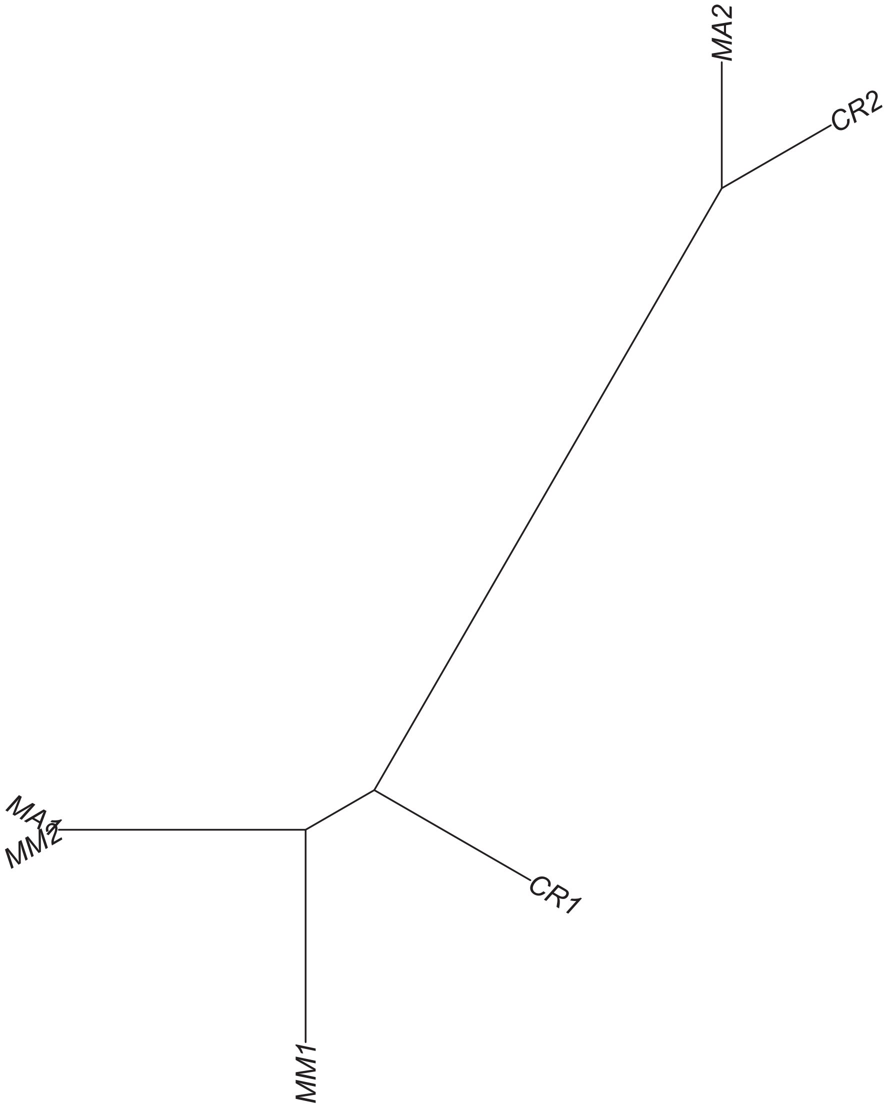

### Region 344

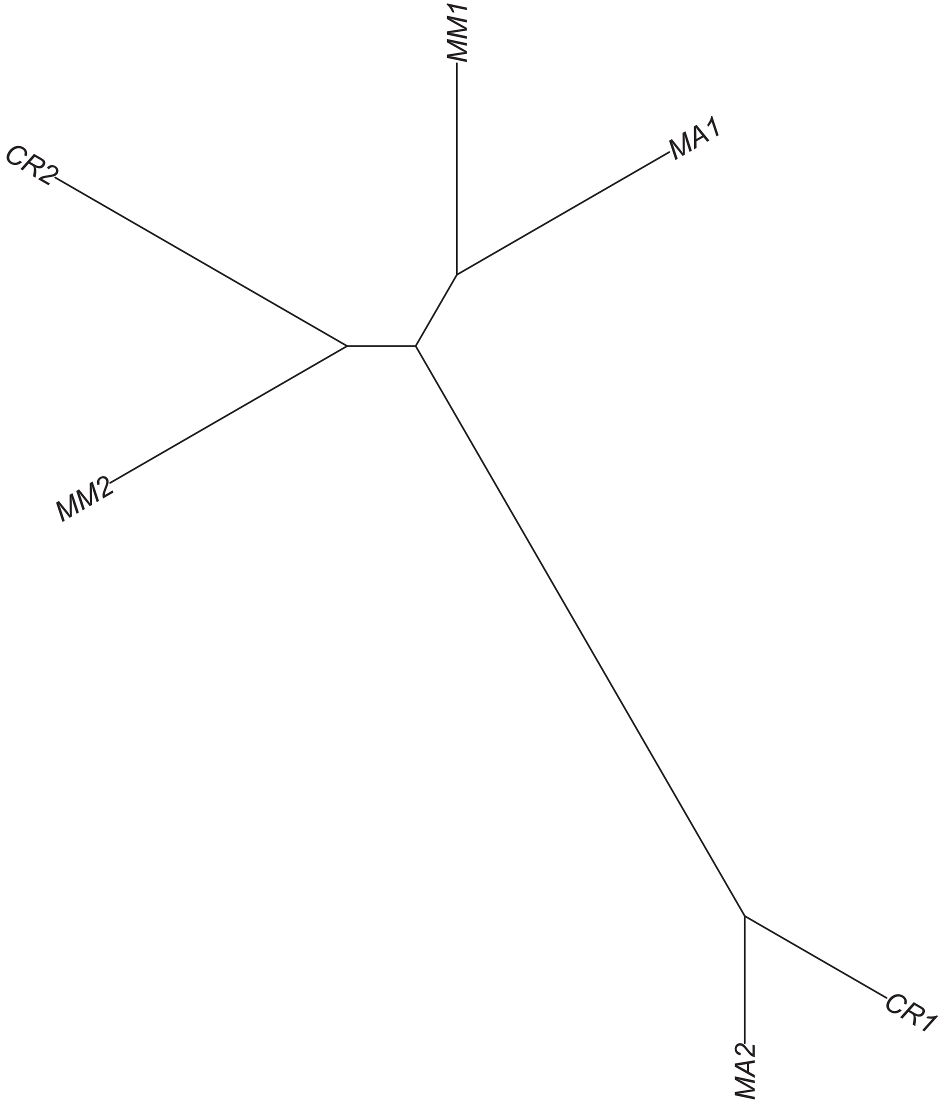

### Region 346

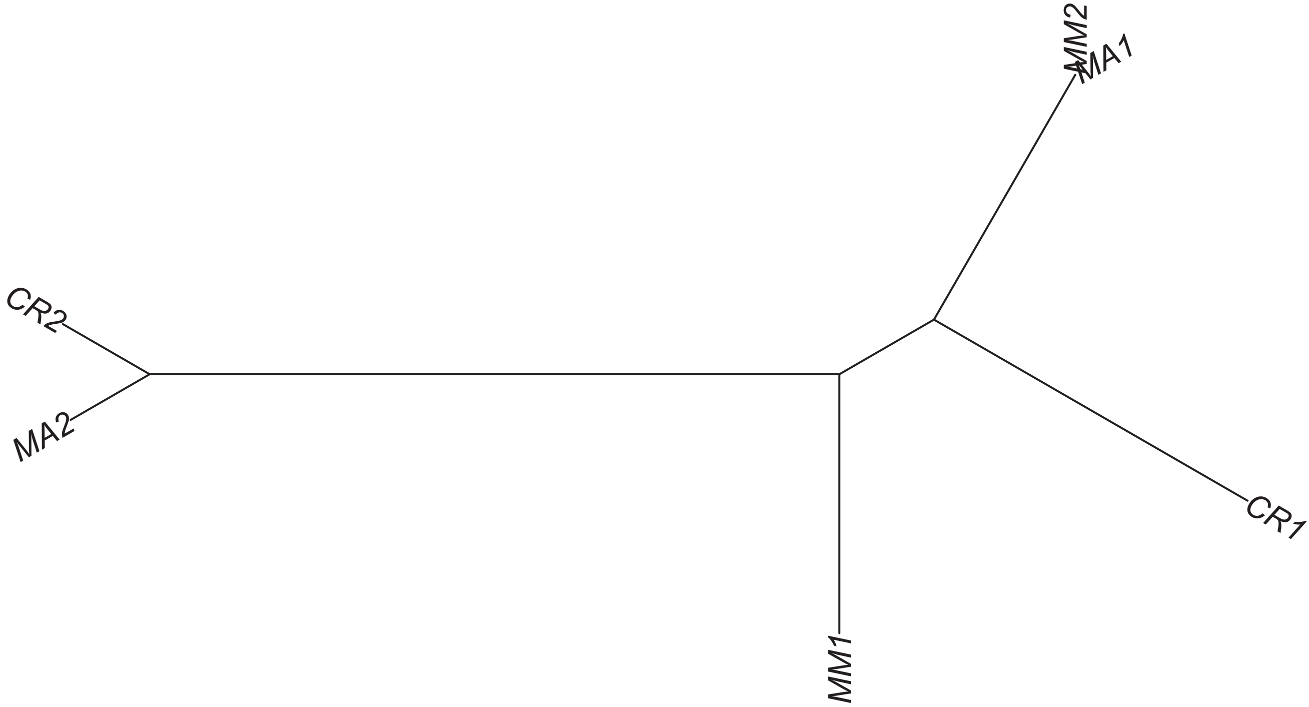

### Region 348



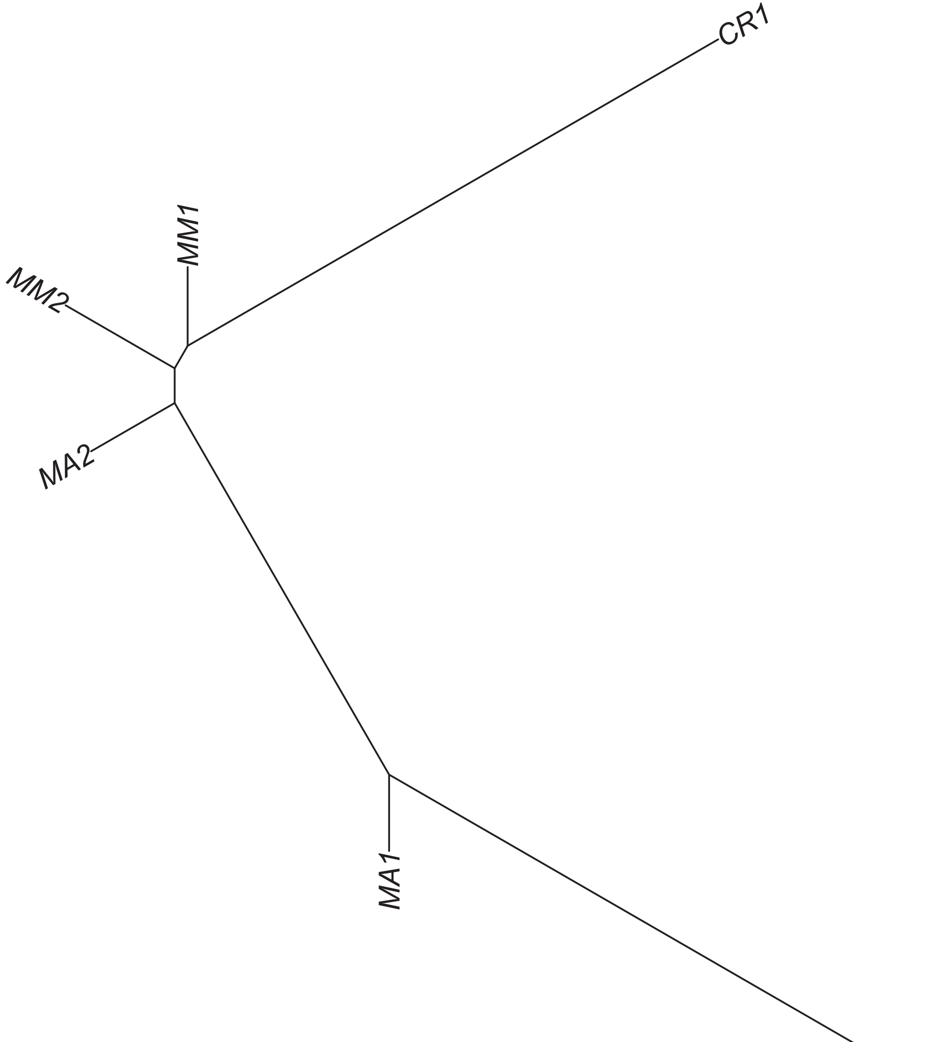

### Region 349

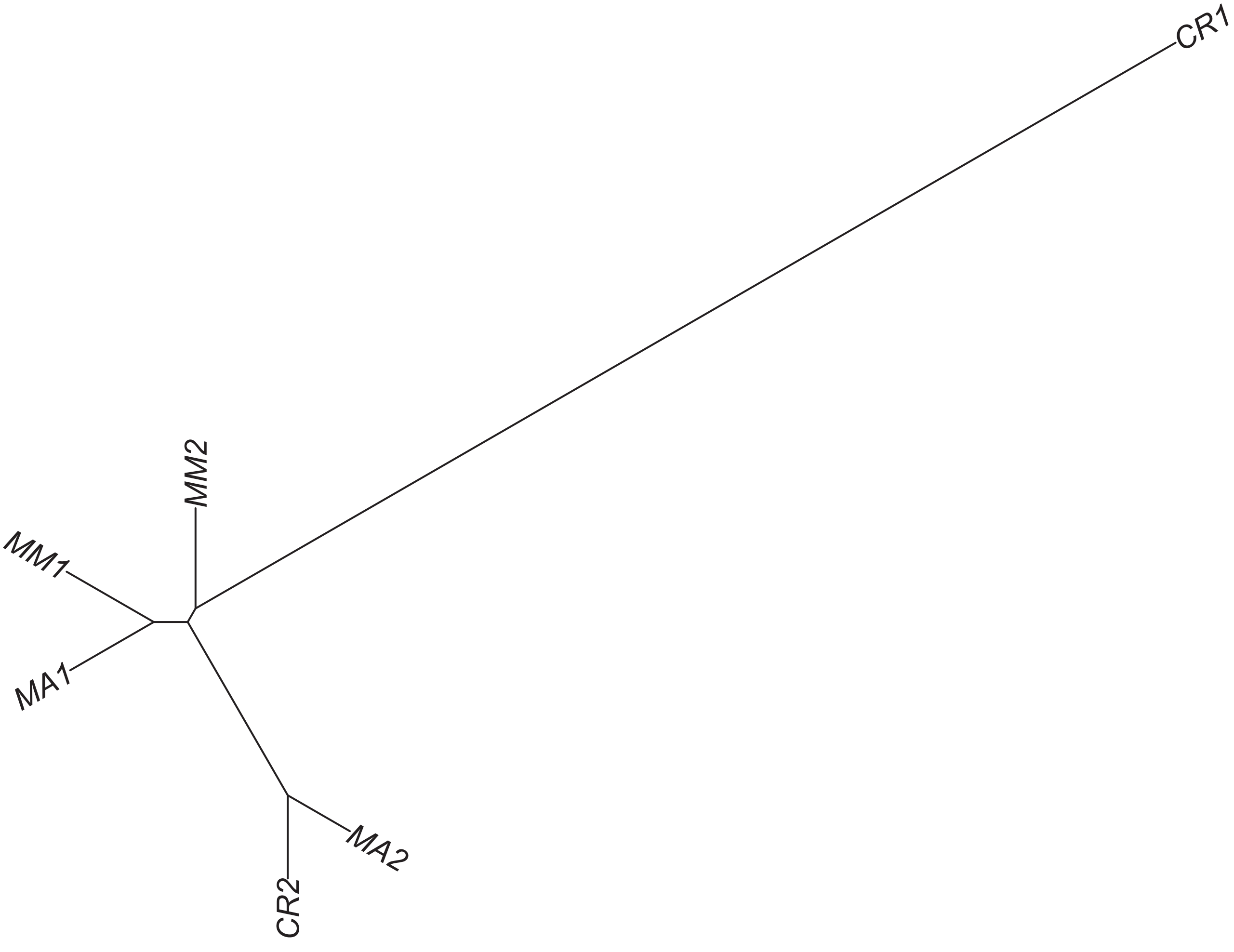

### Region 352



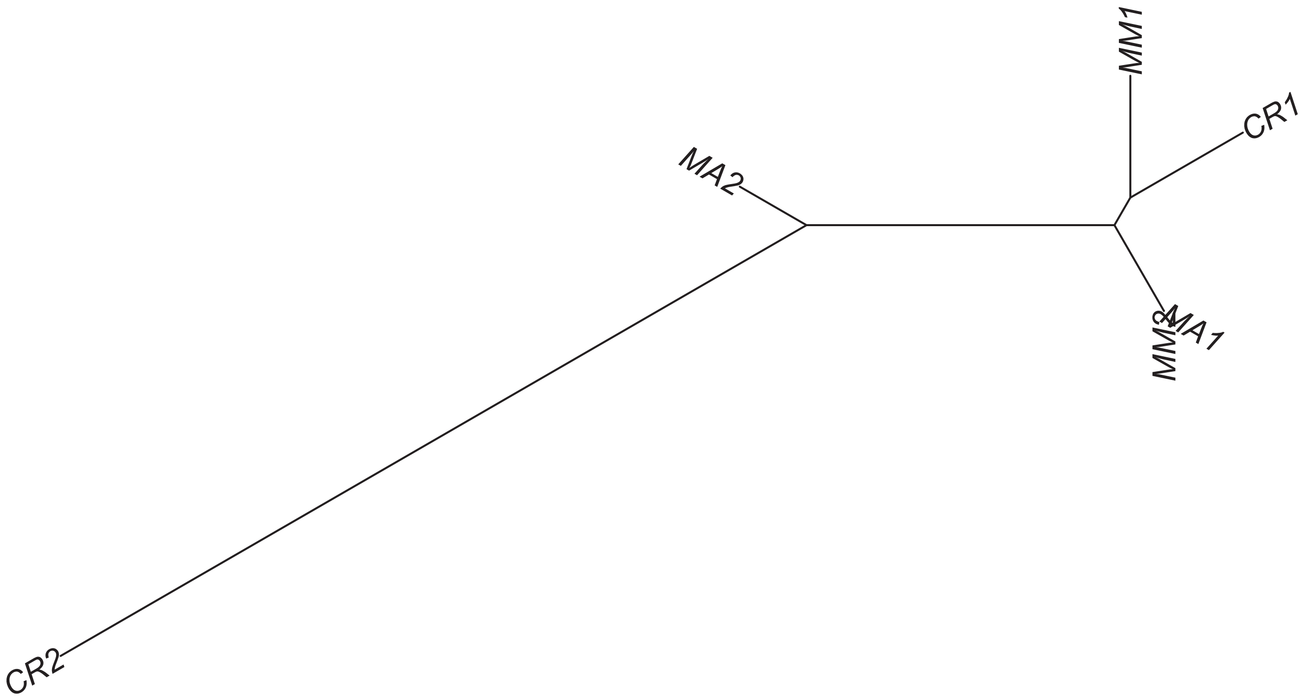

### Region 354

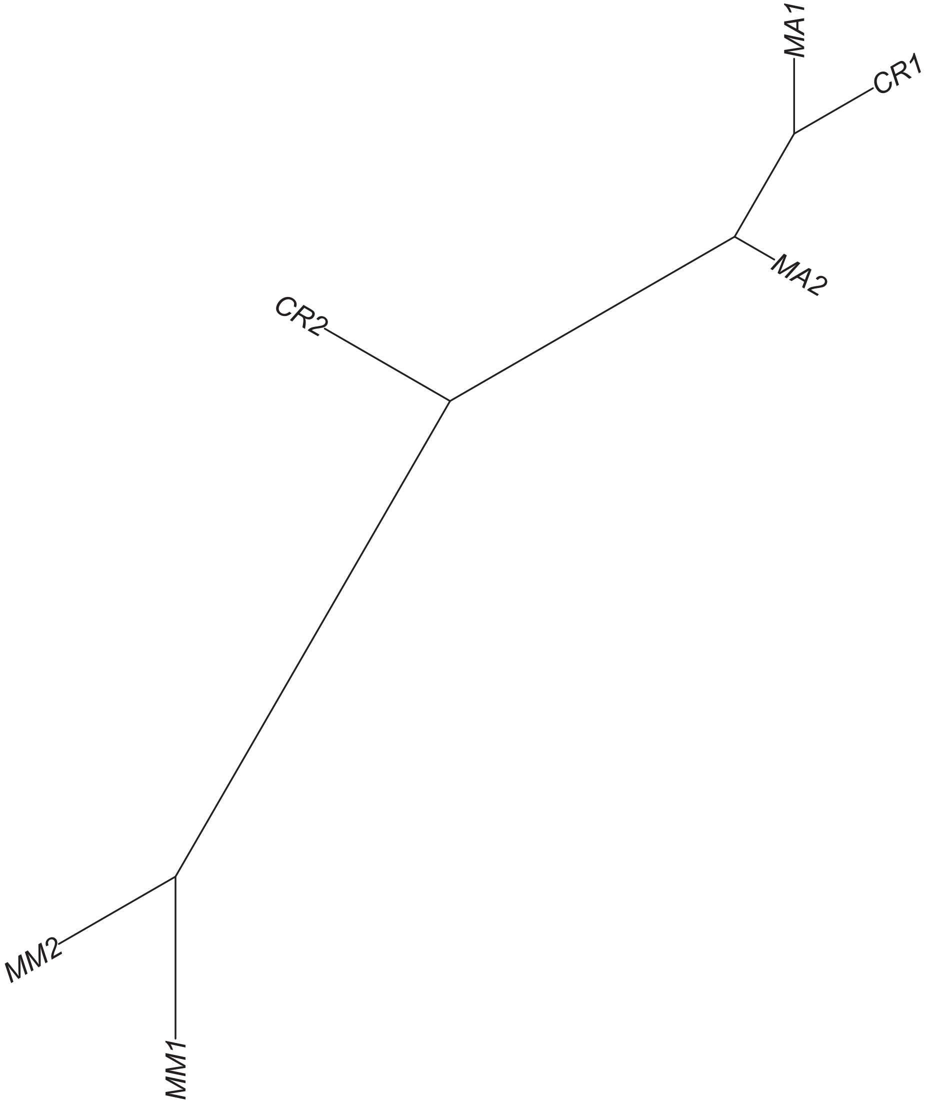

### Region 356

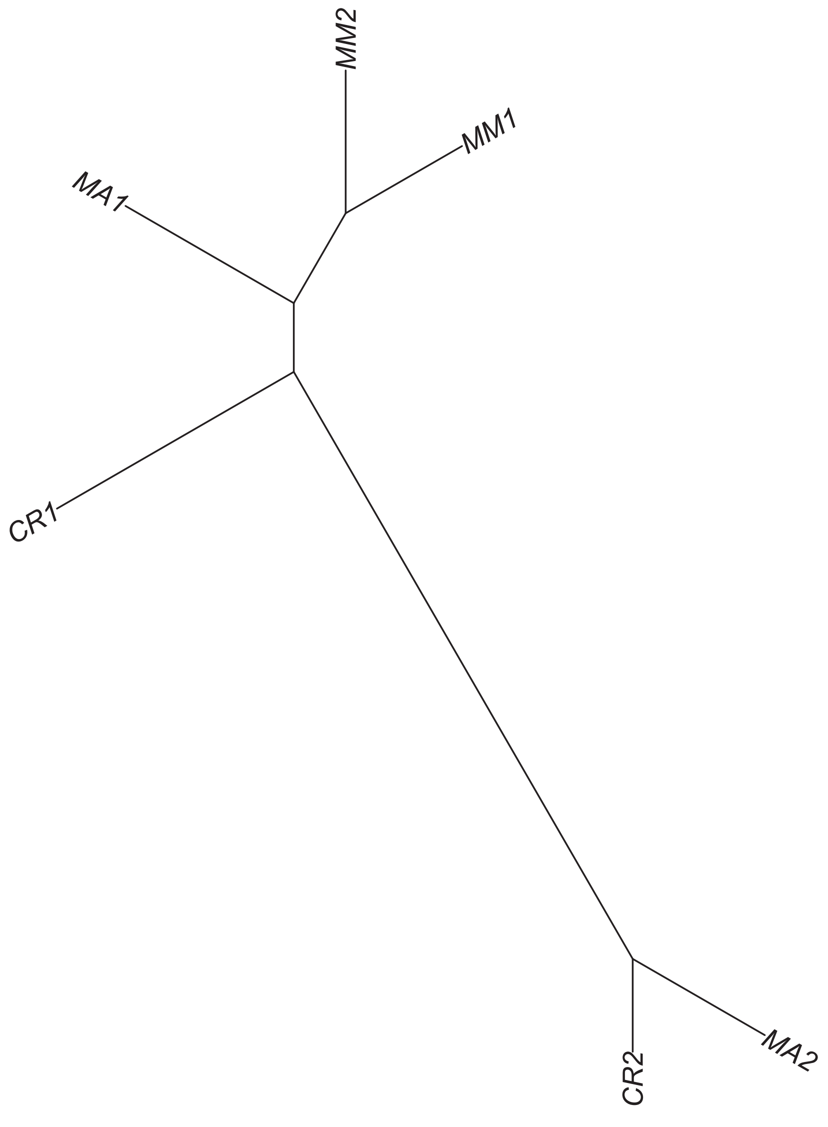

### Region 358



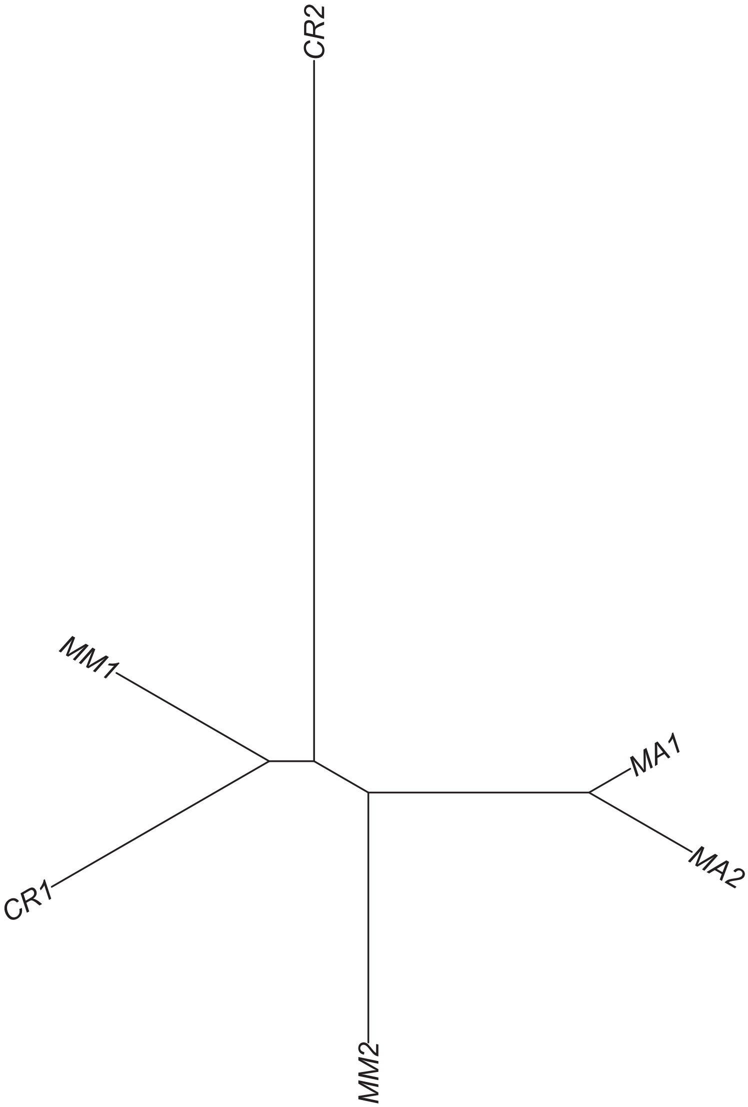

### Region 361

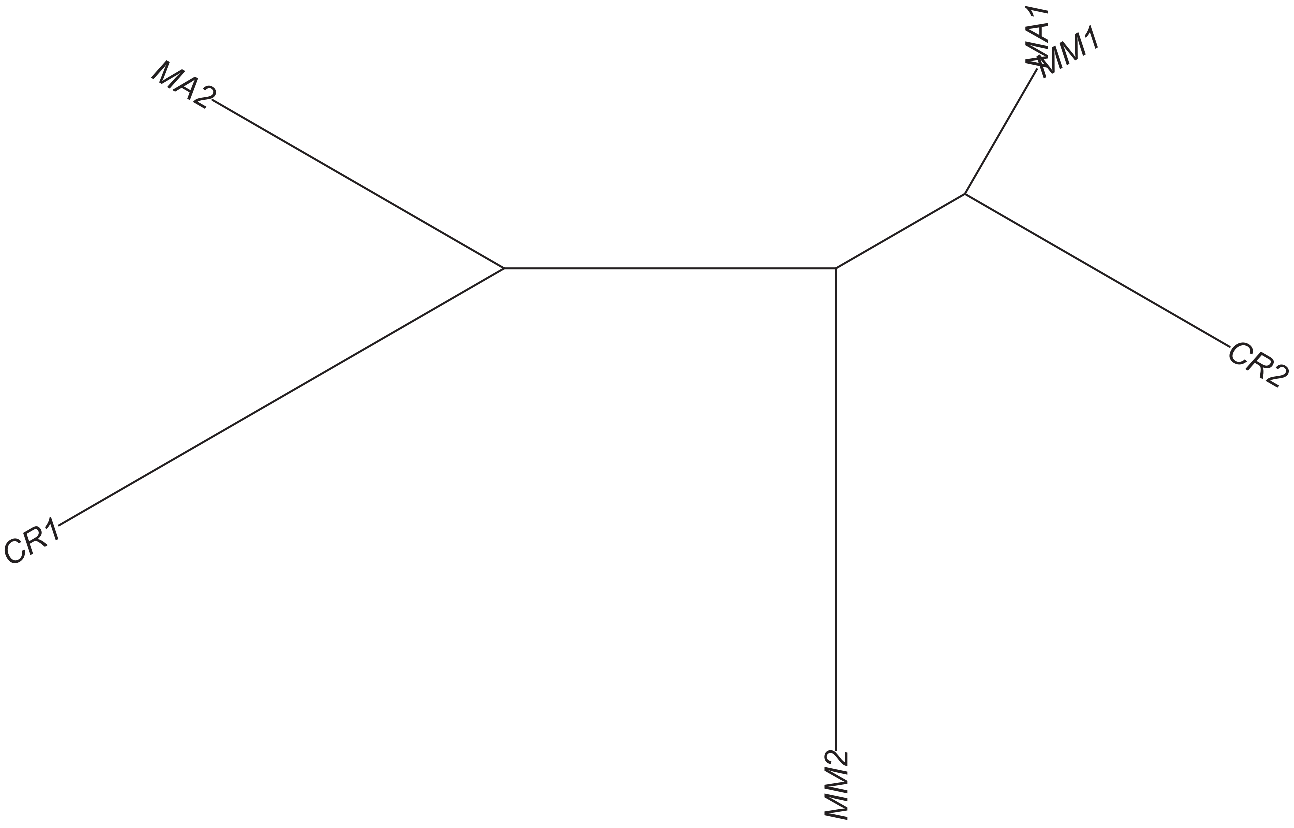

### Region 362

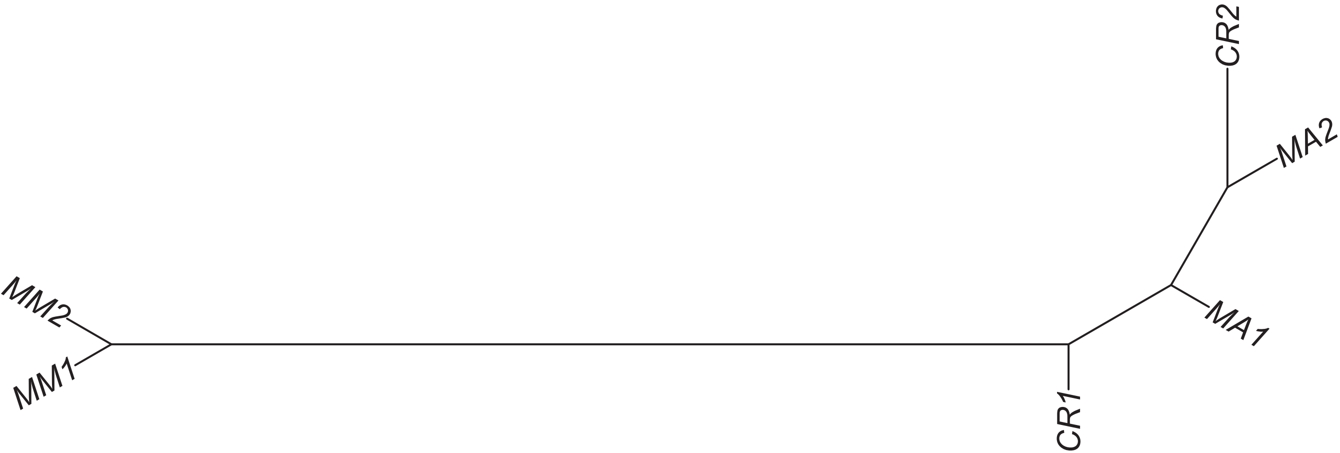

### Region 364



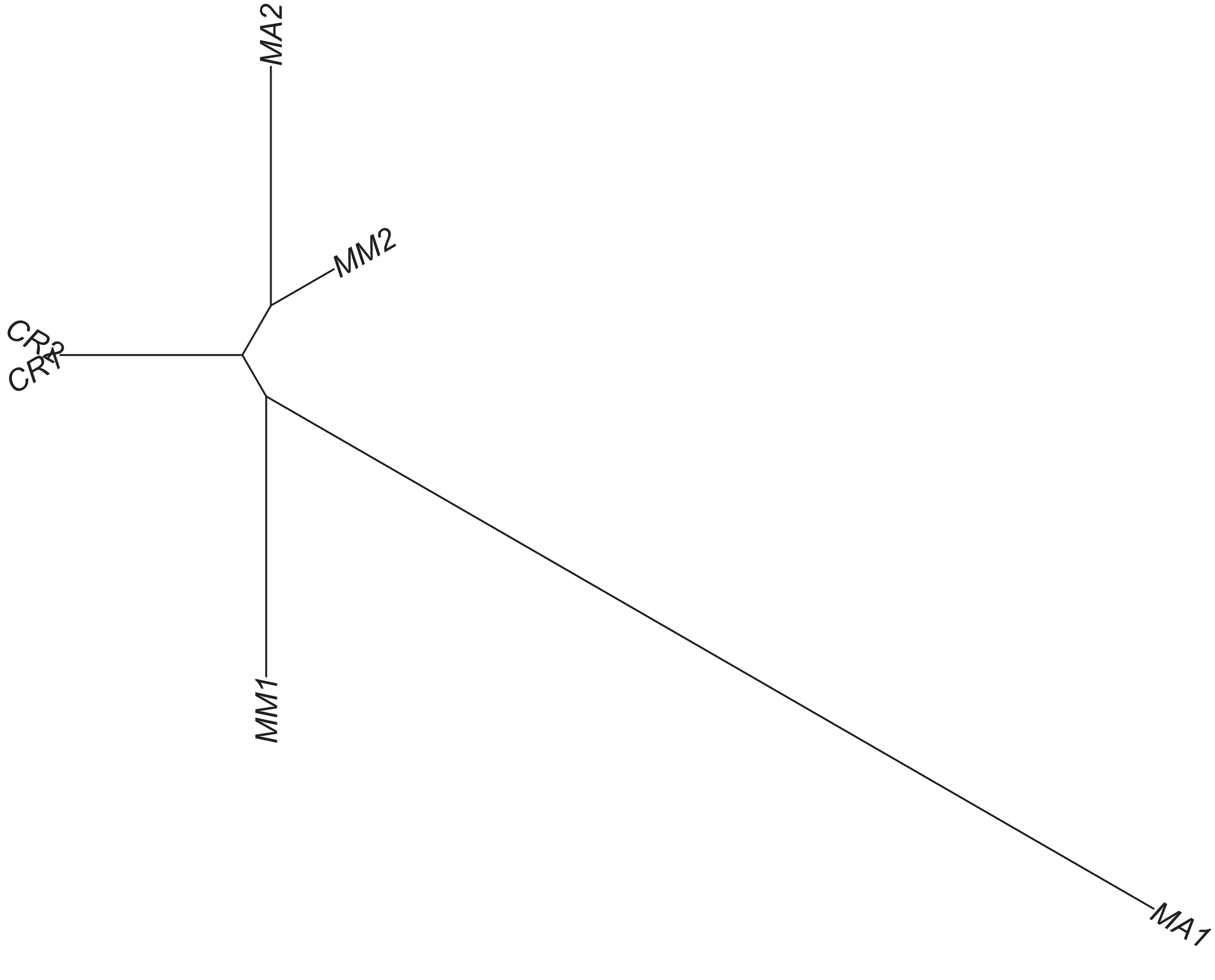

### Region 366

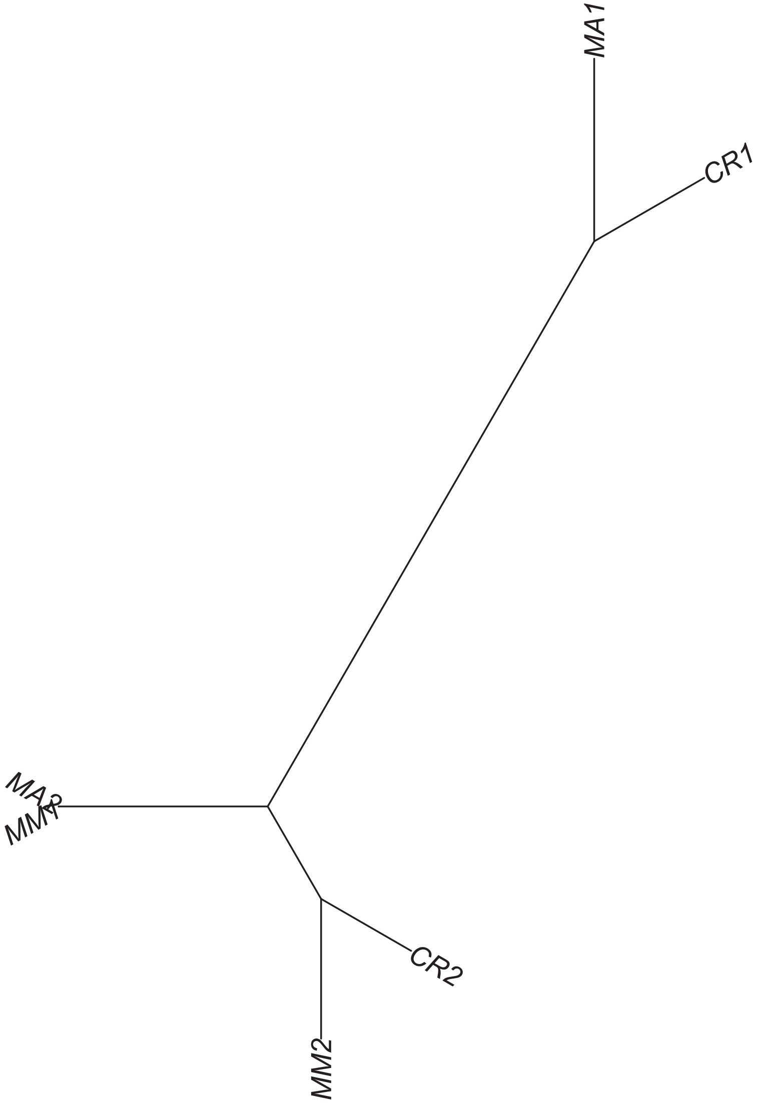

### Region 367

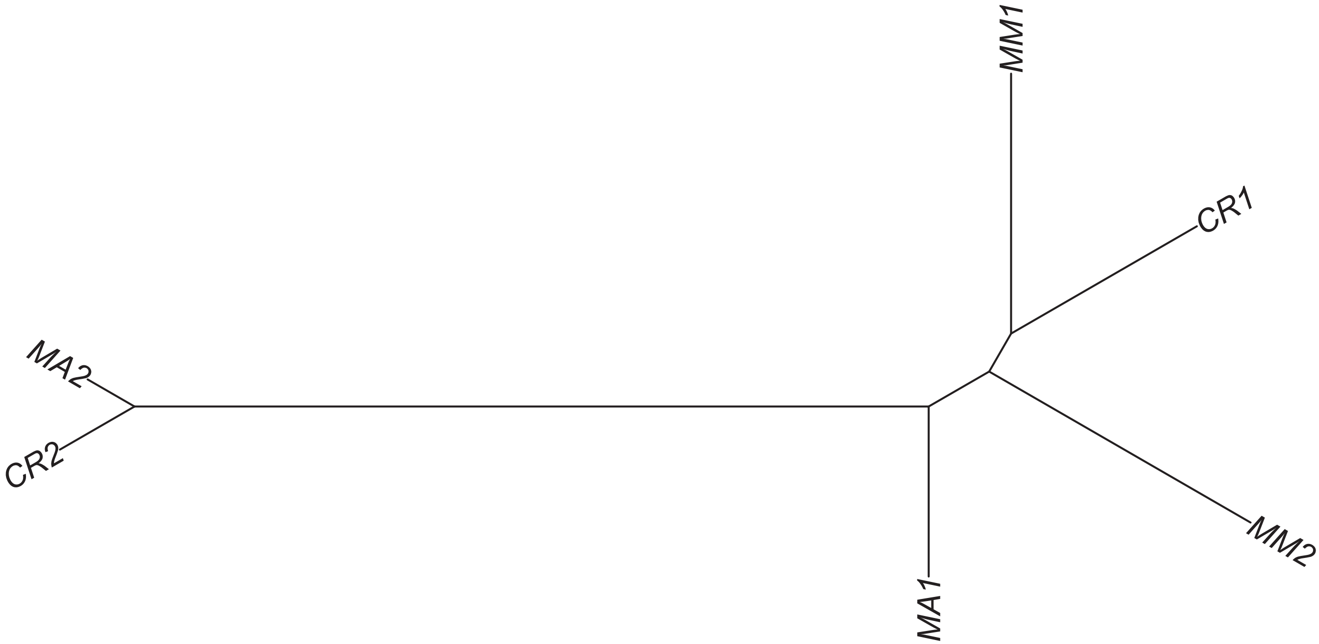

### Region 368



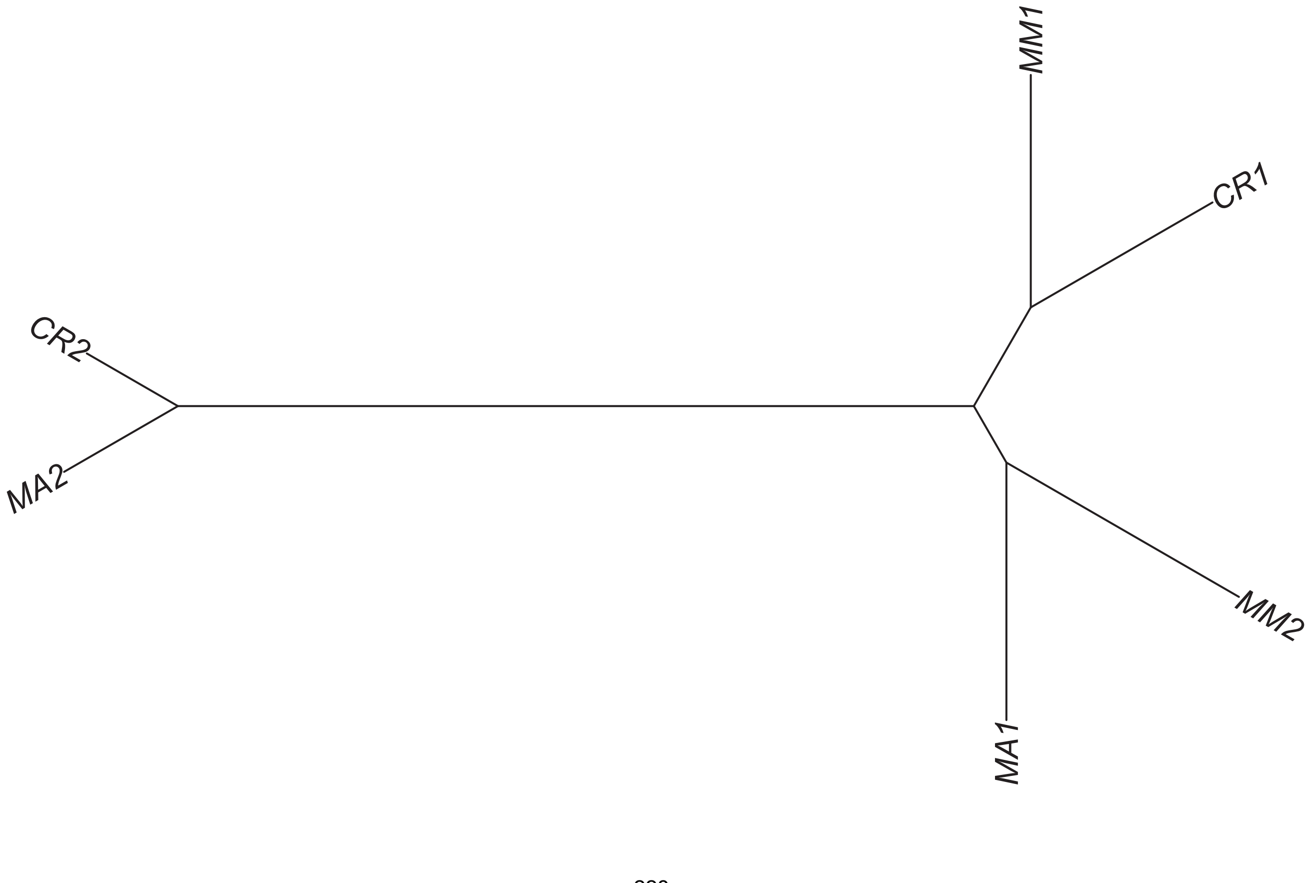

### Region 370



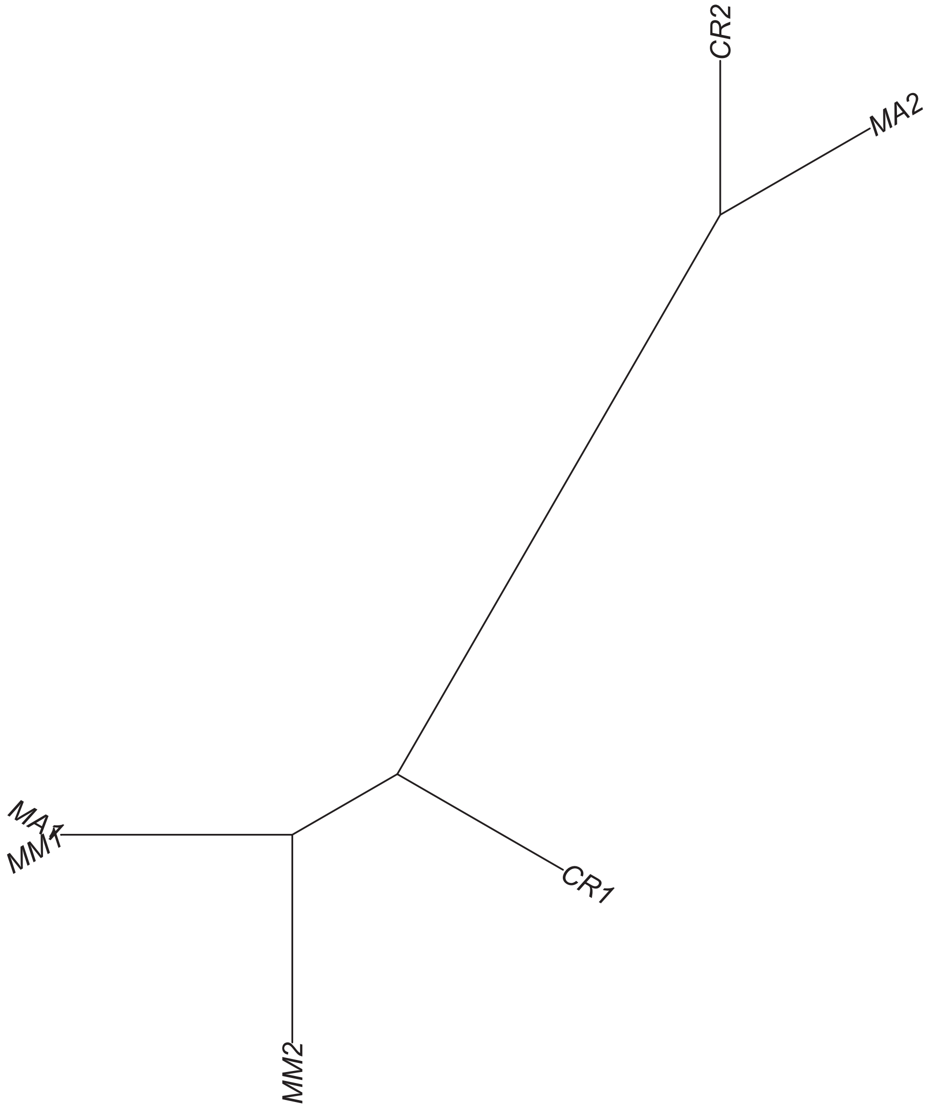

### Region 371

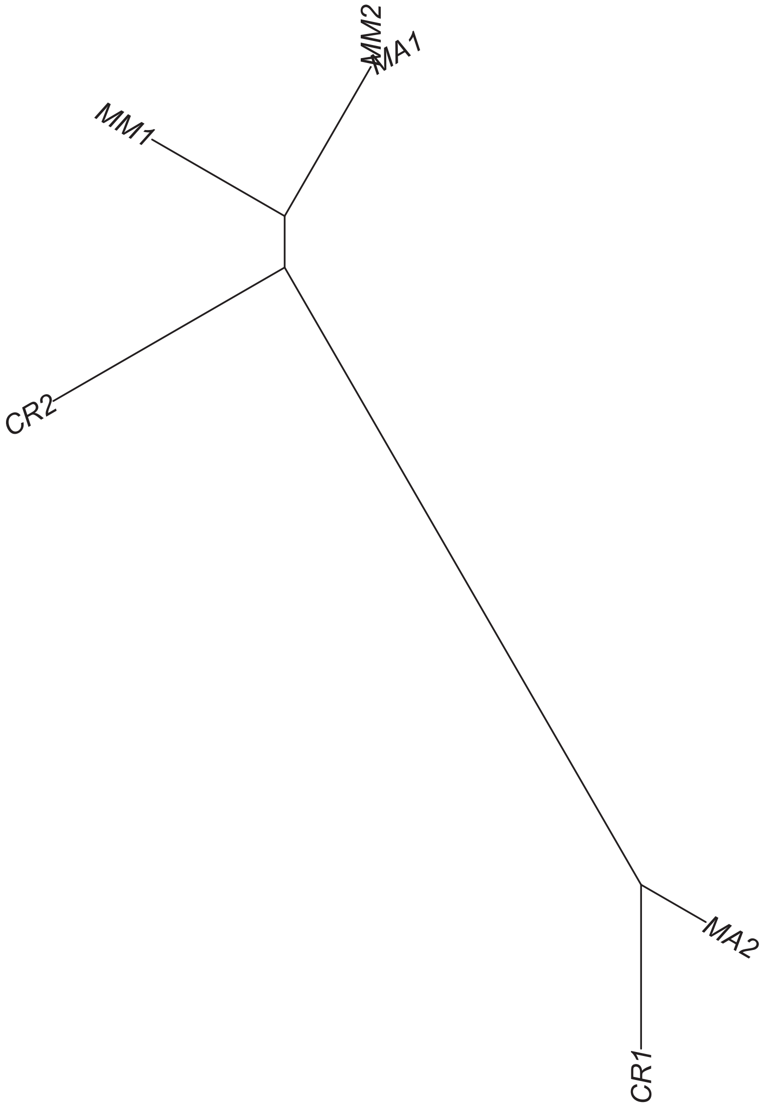

### Region 372



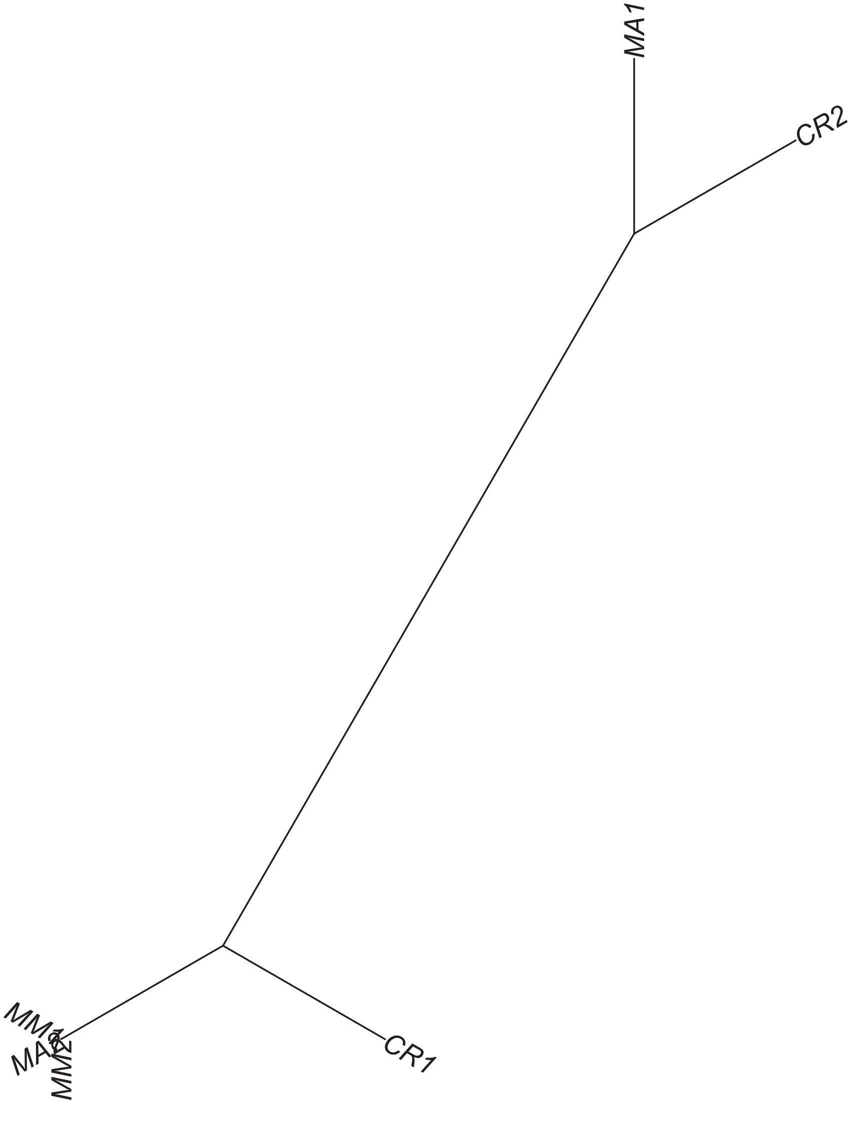

### Region 373

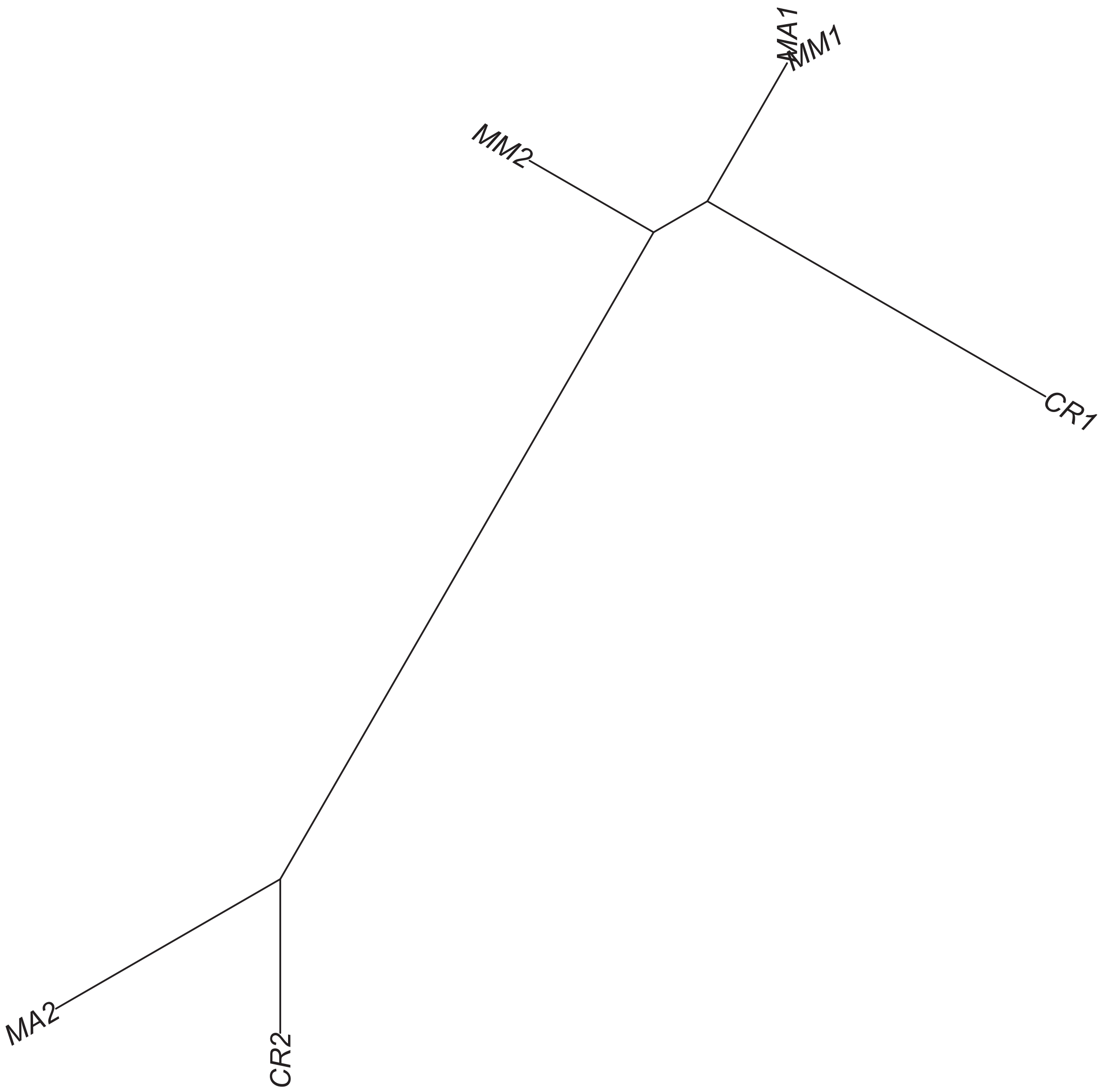

### Region 374



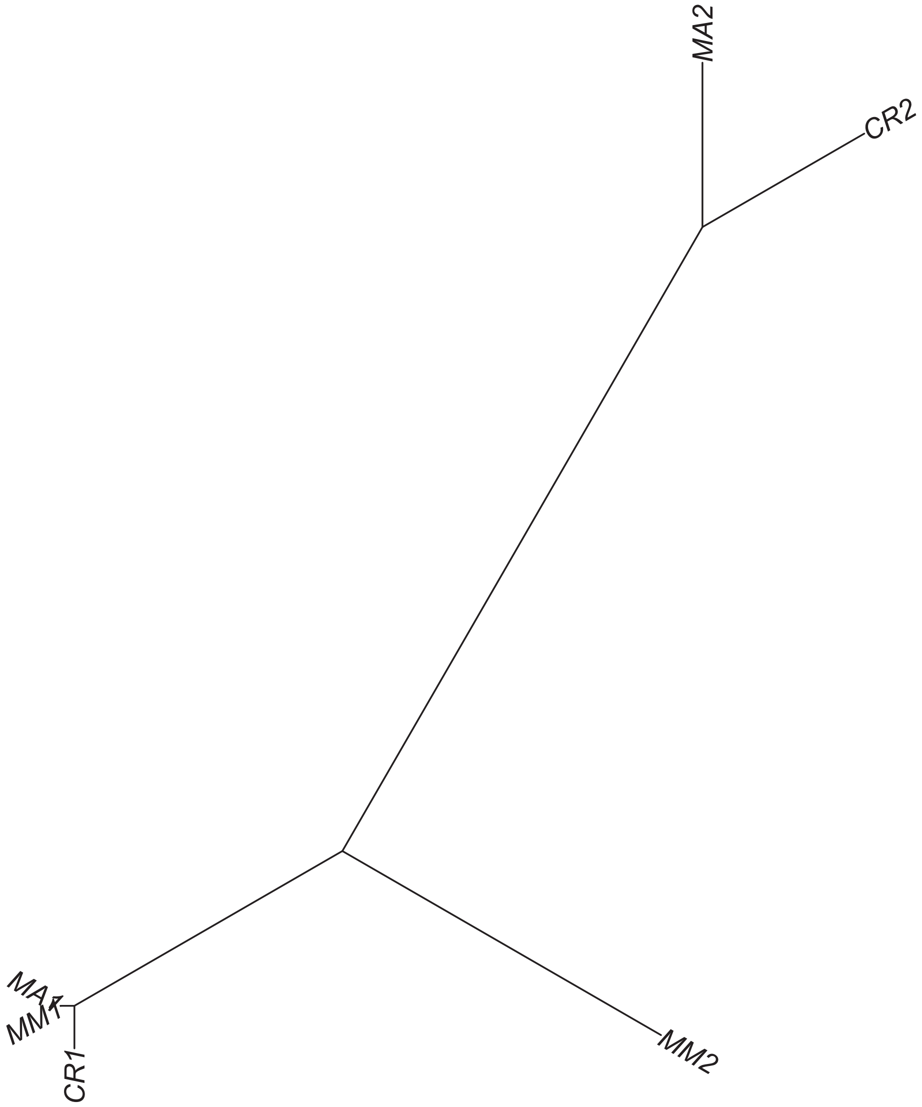

Region 376



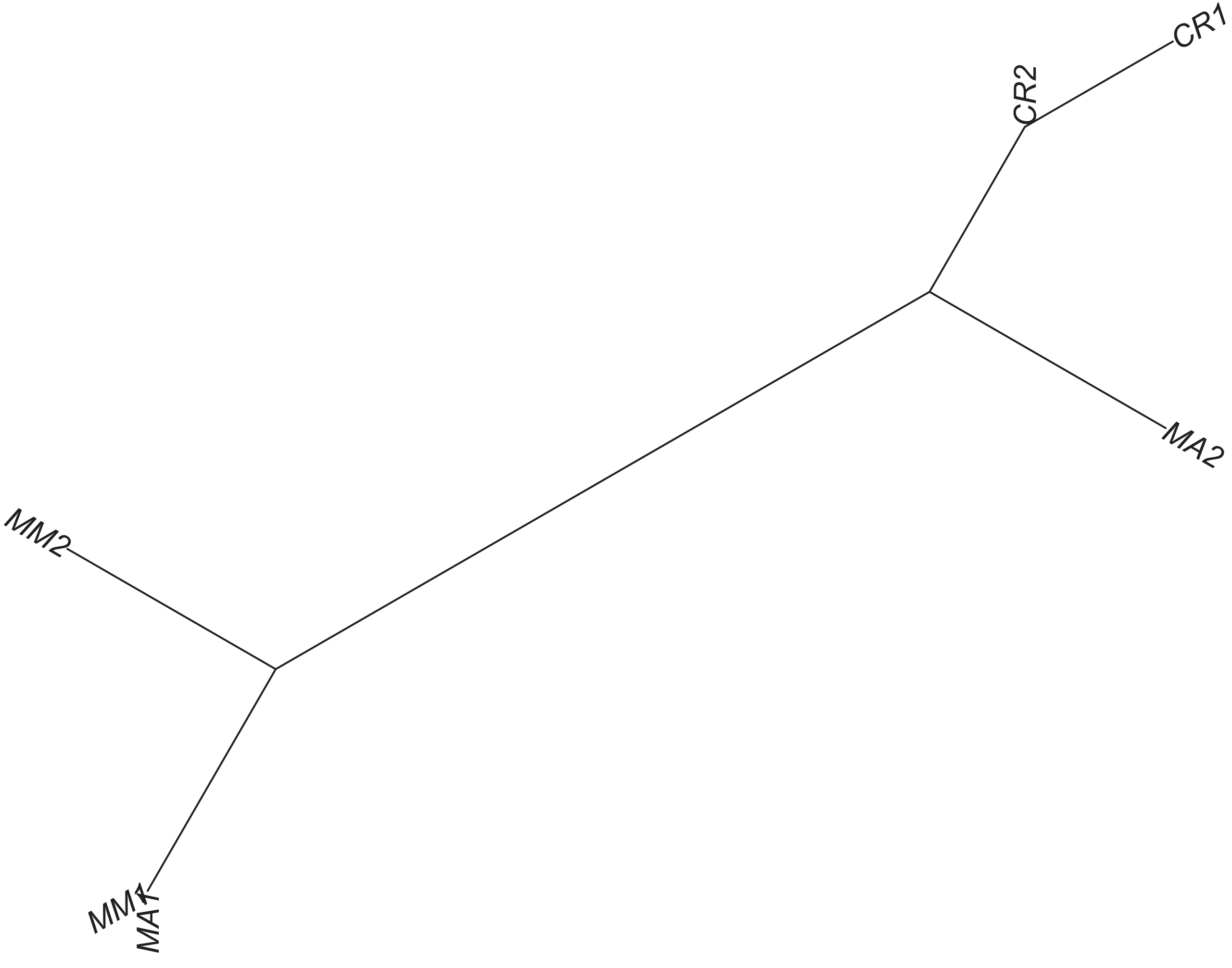

### Region 378

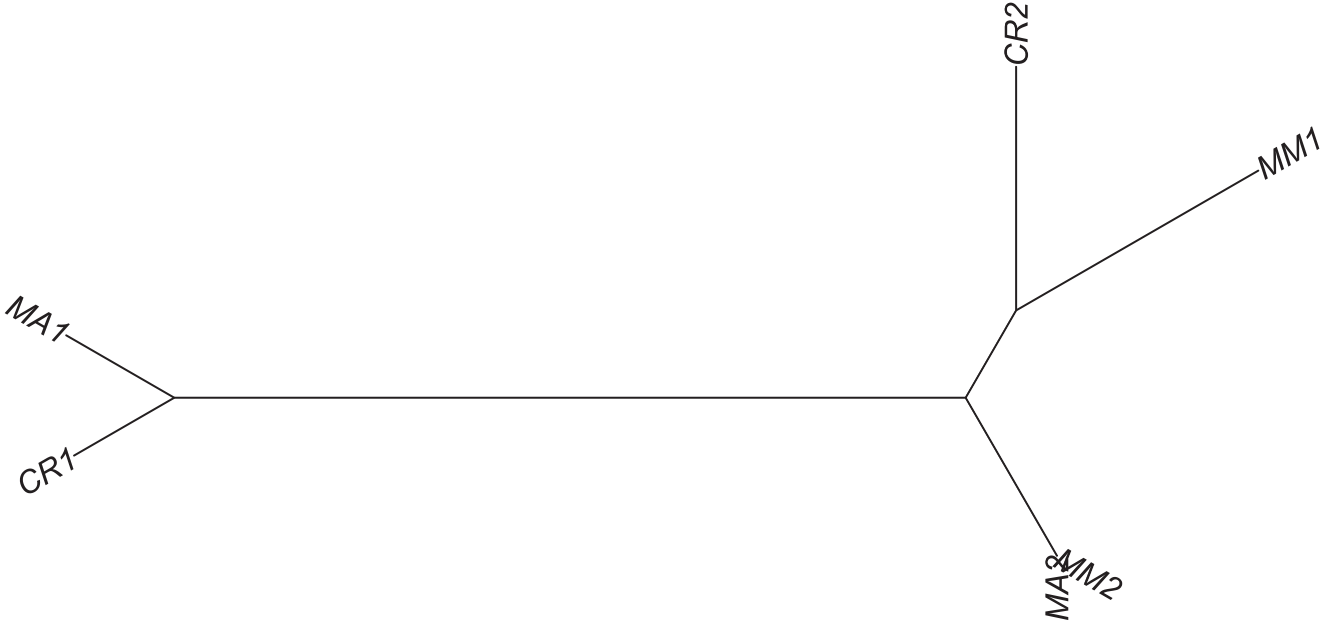

### Region 379



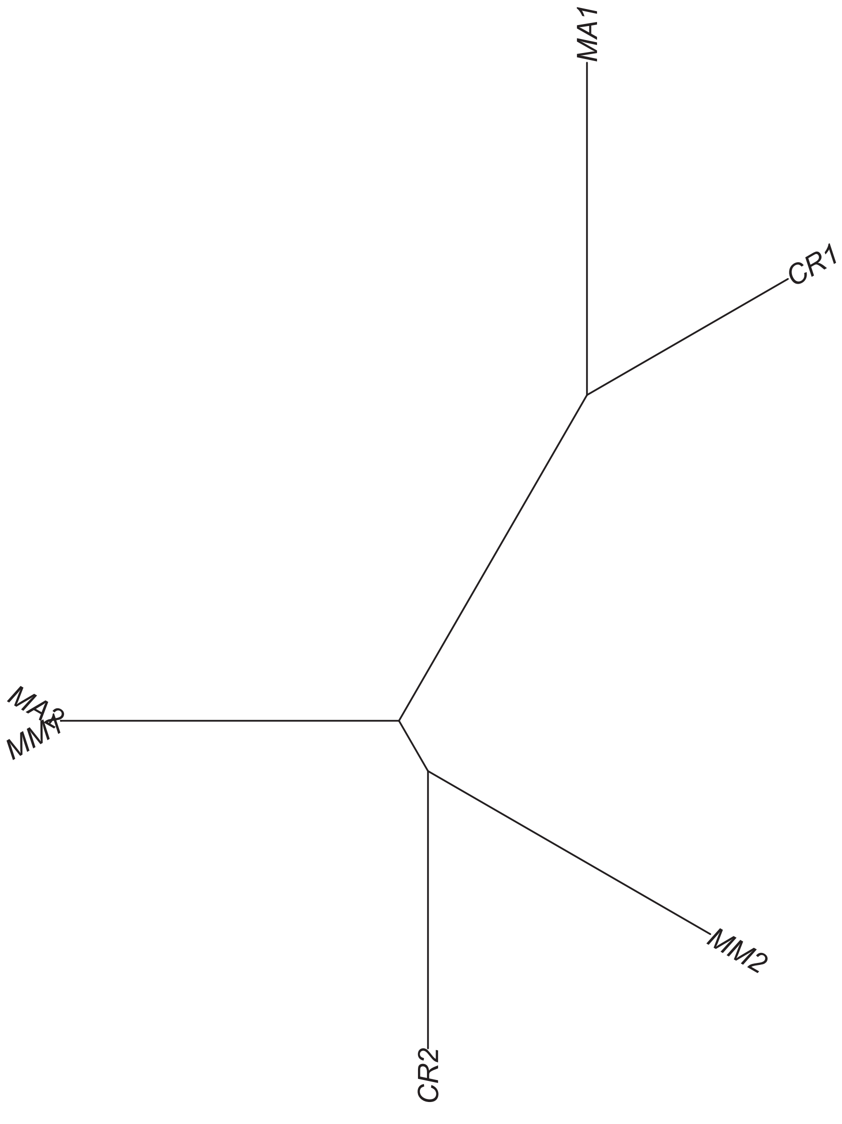

### Region 380

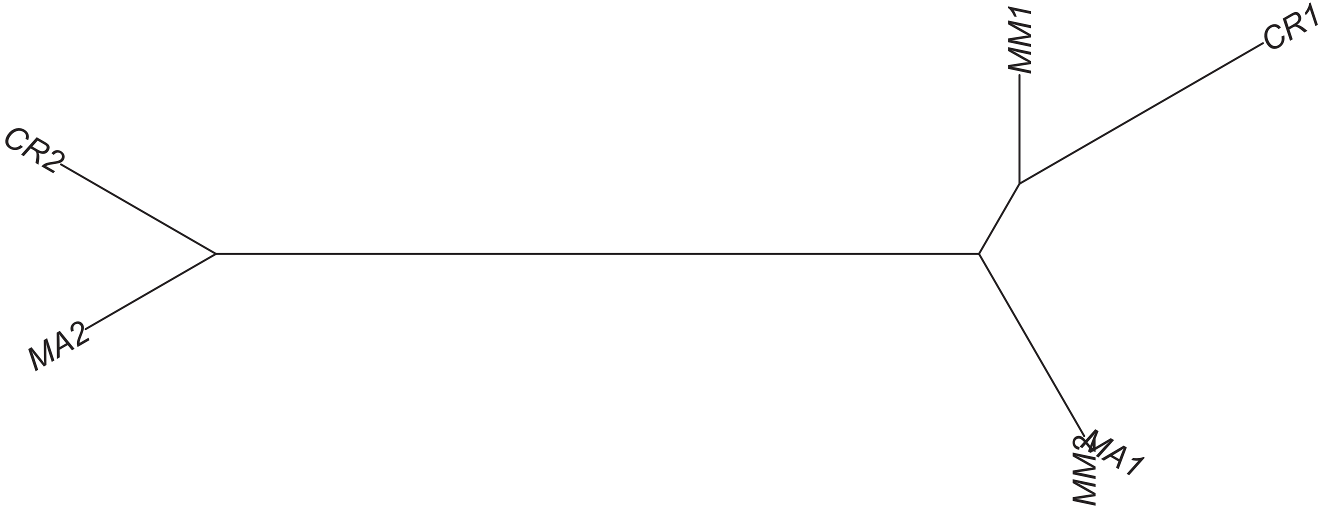

### Region 382

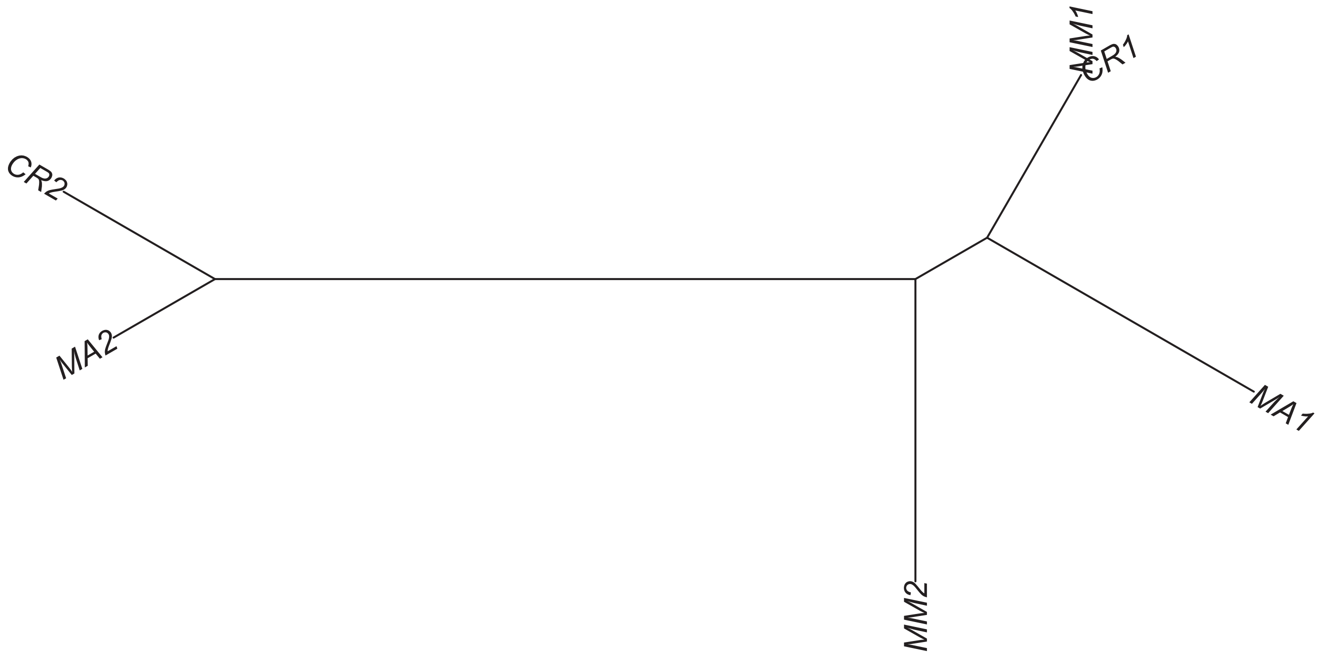

### Region 384

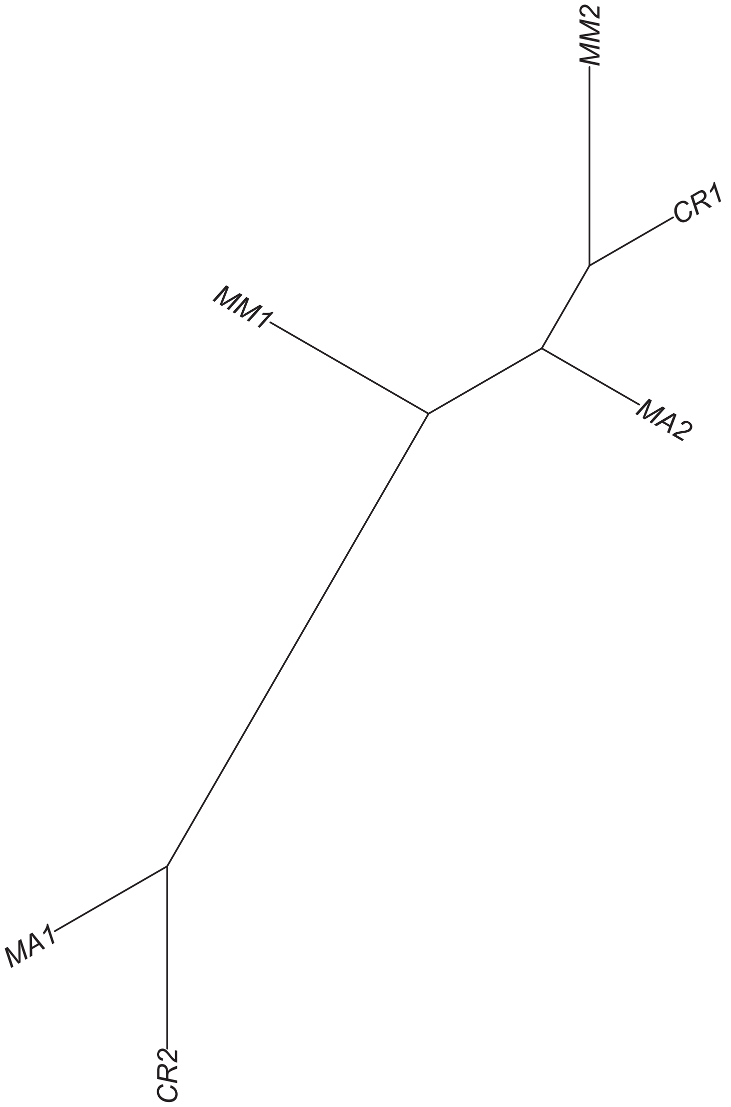

### Region 385

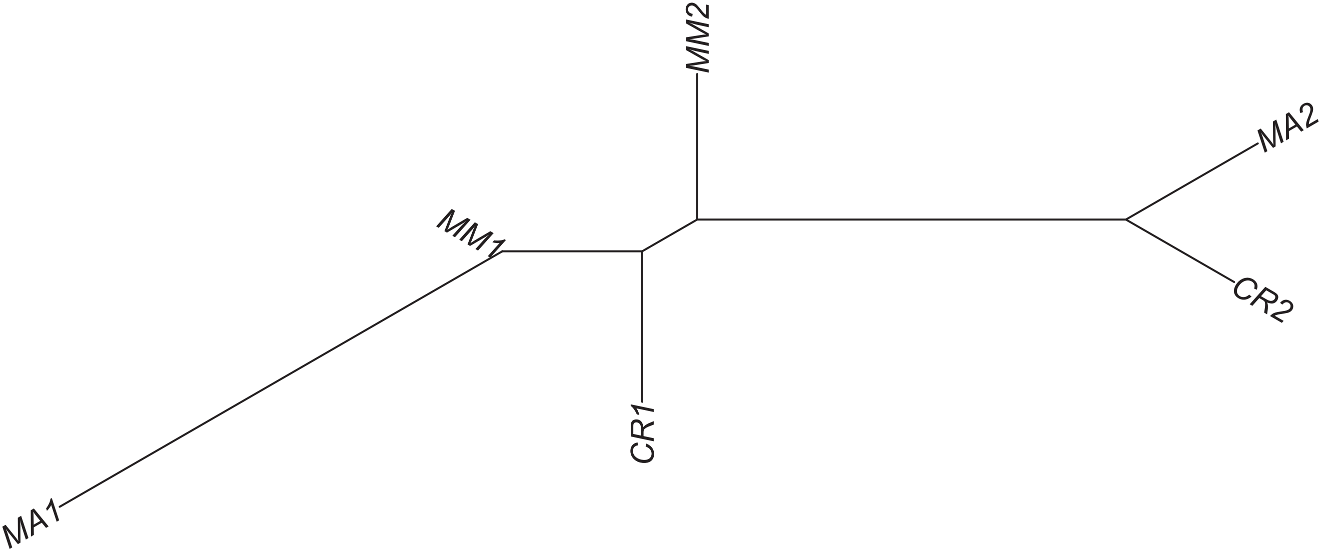

Region 386



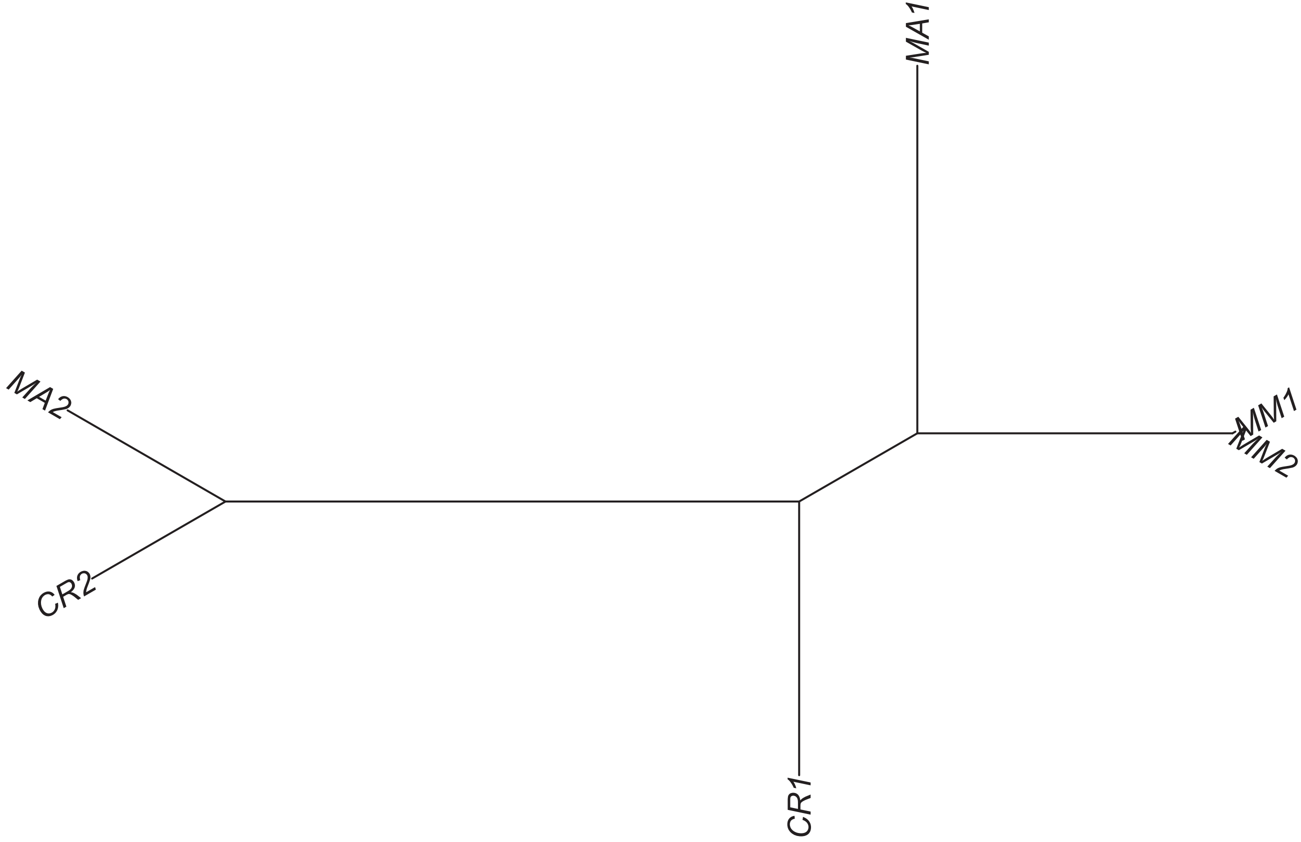

### Region 387

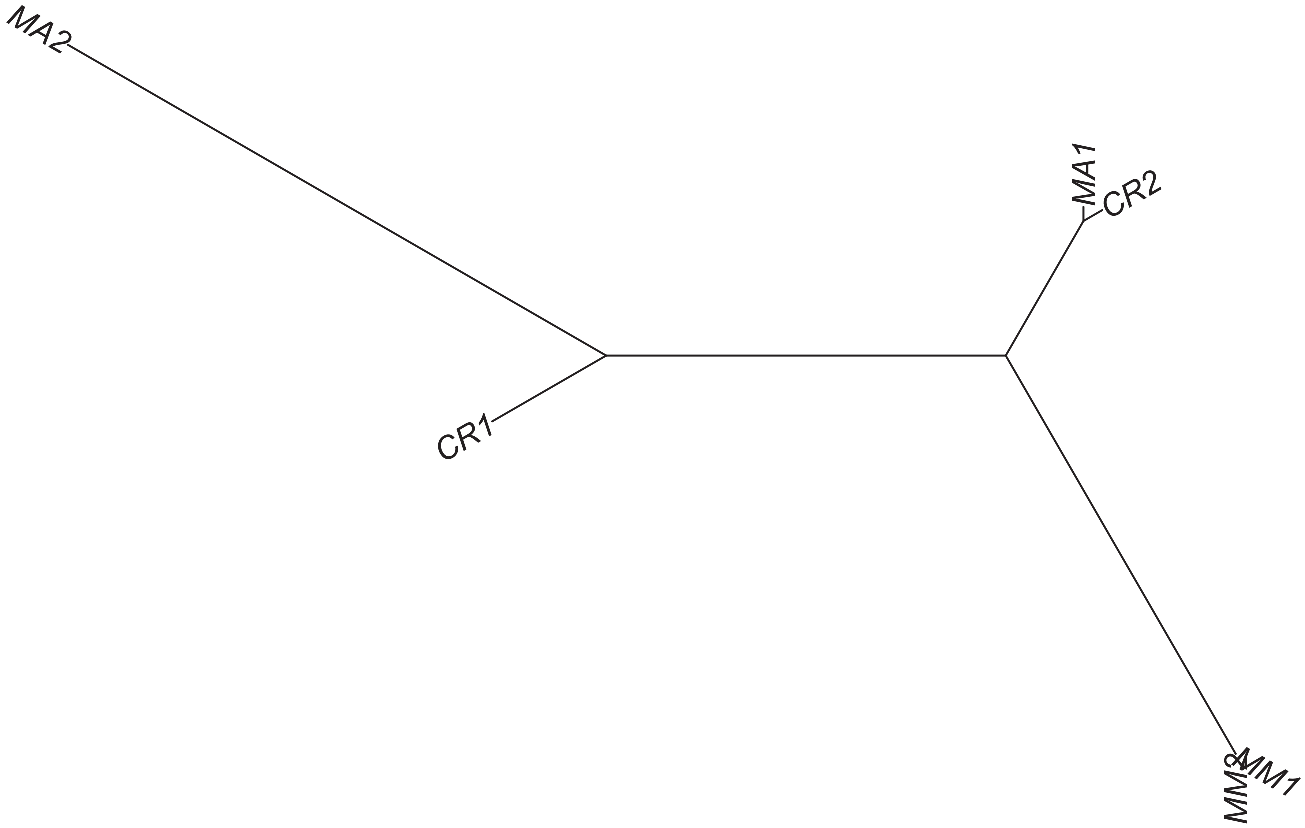

### Region 388



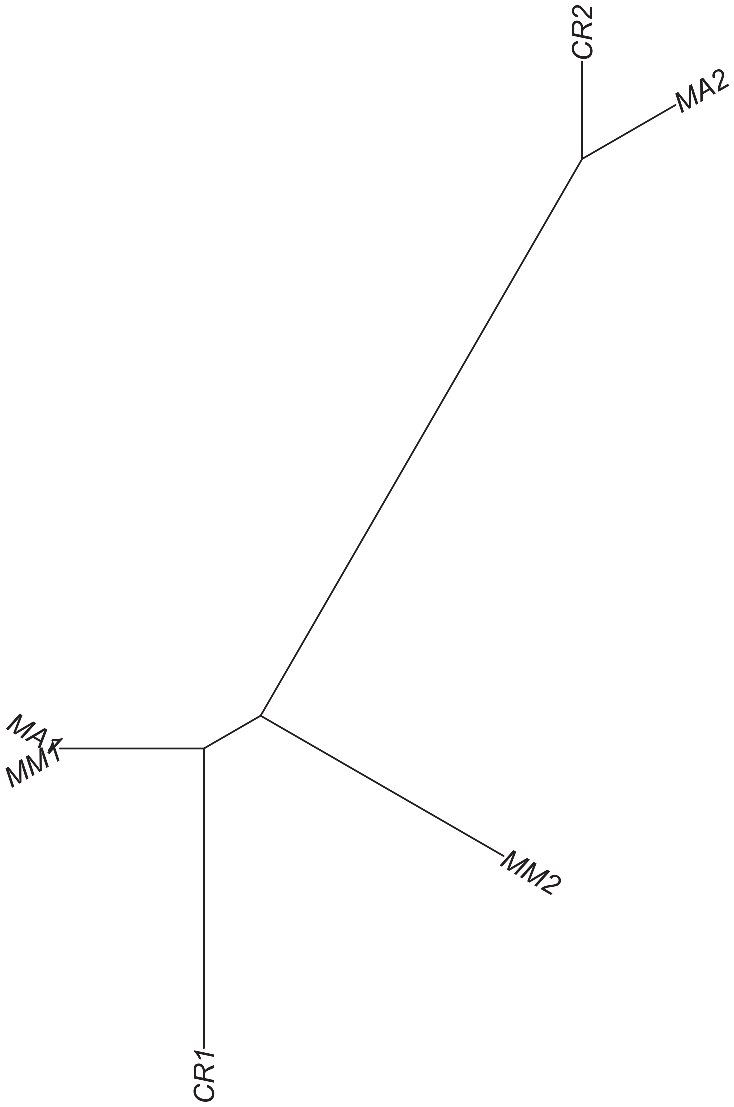

### Region 391

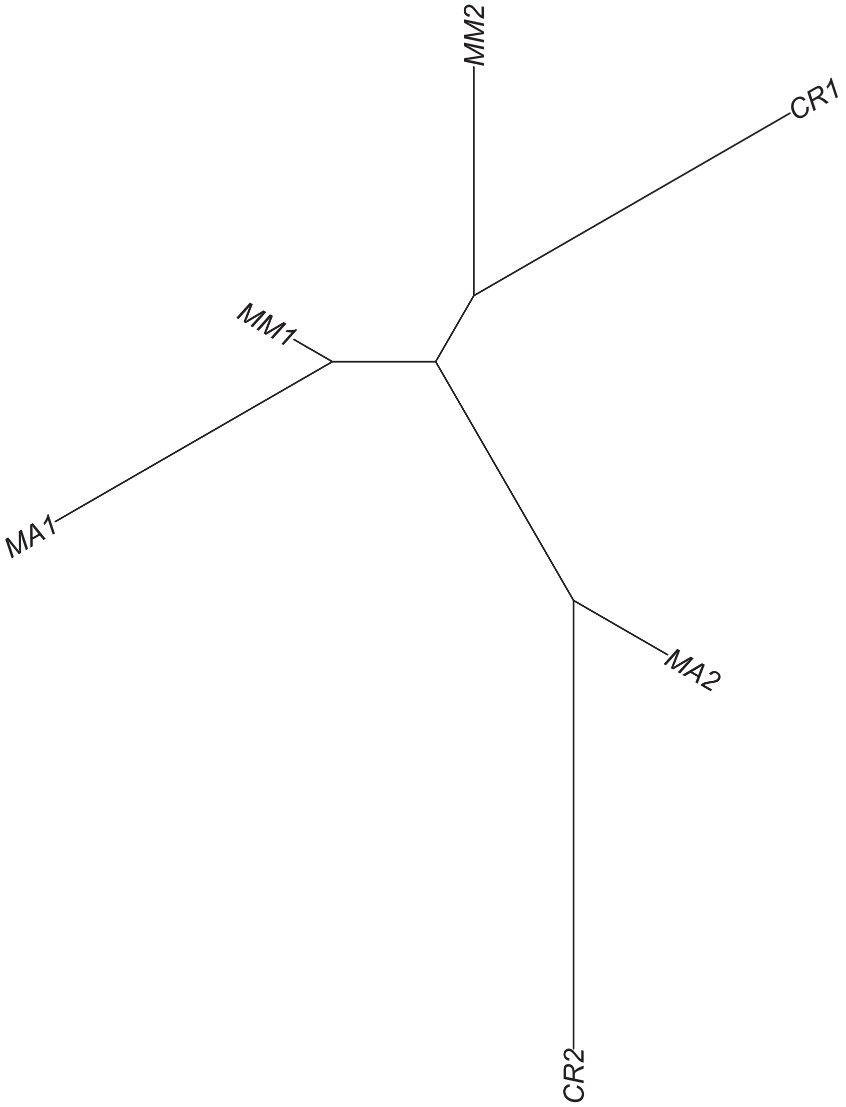

### Region 392



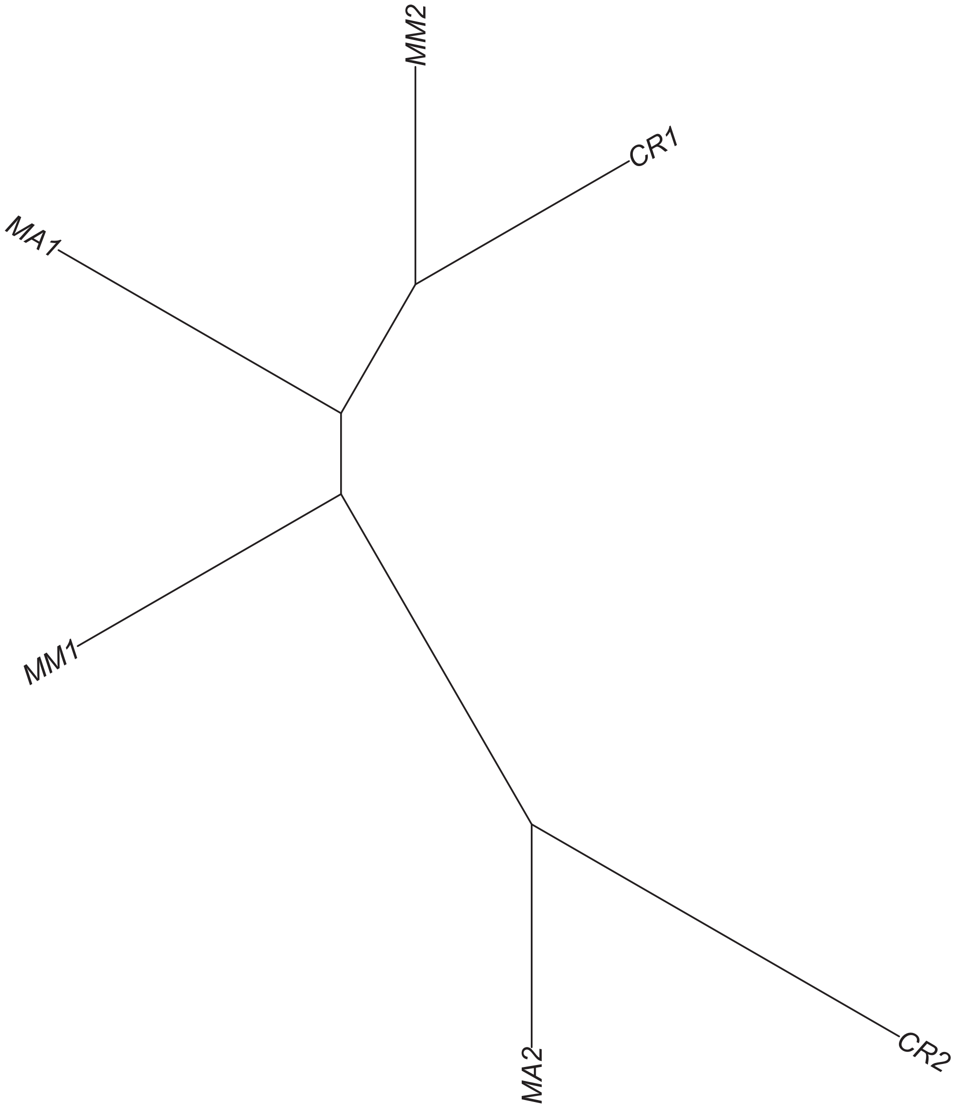

### Region 393

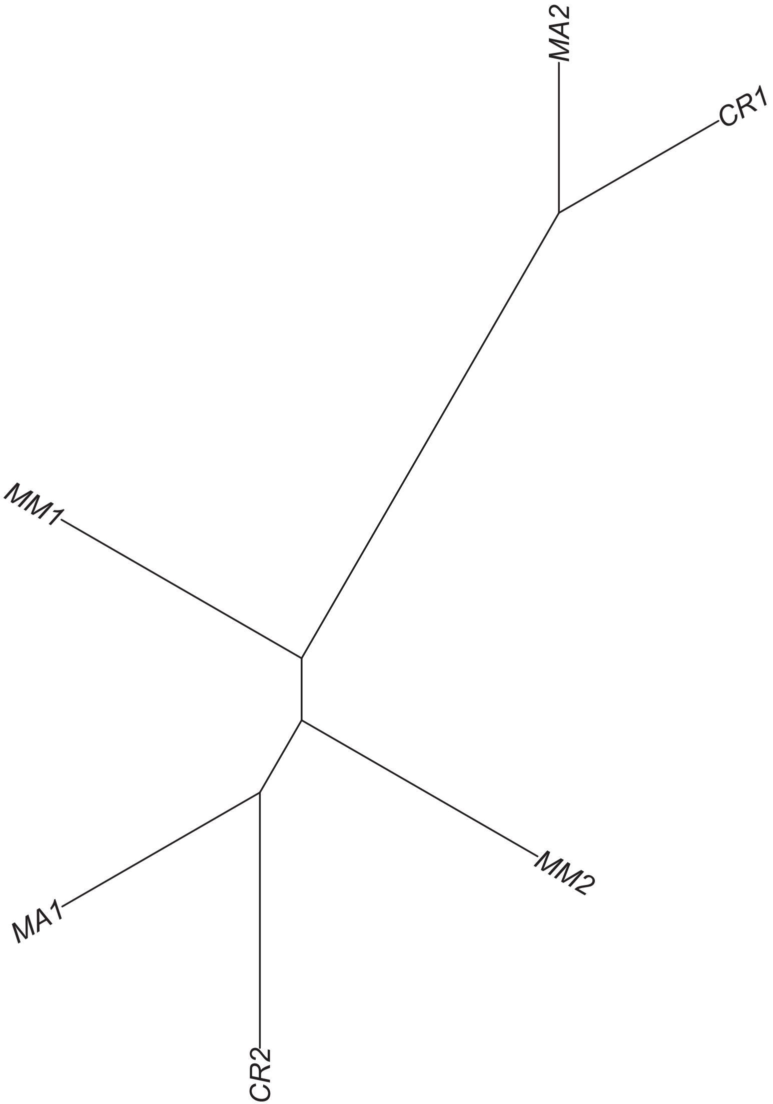

### Region 394

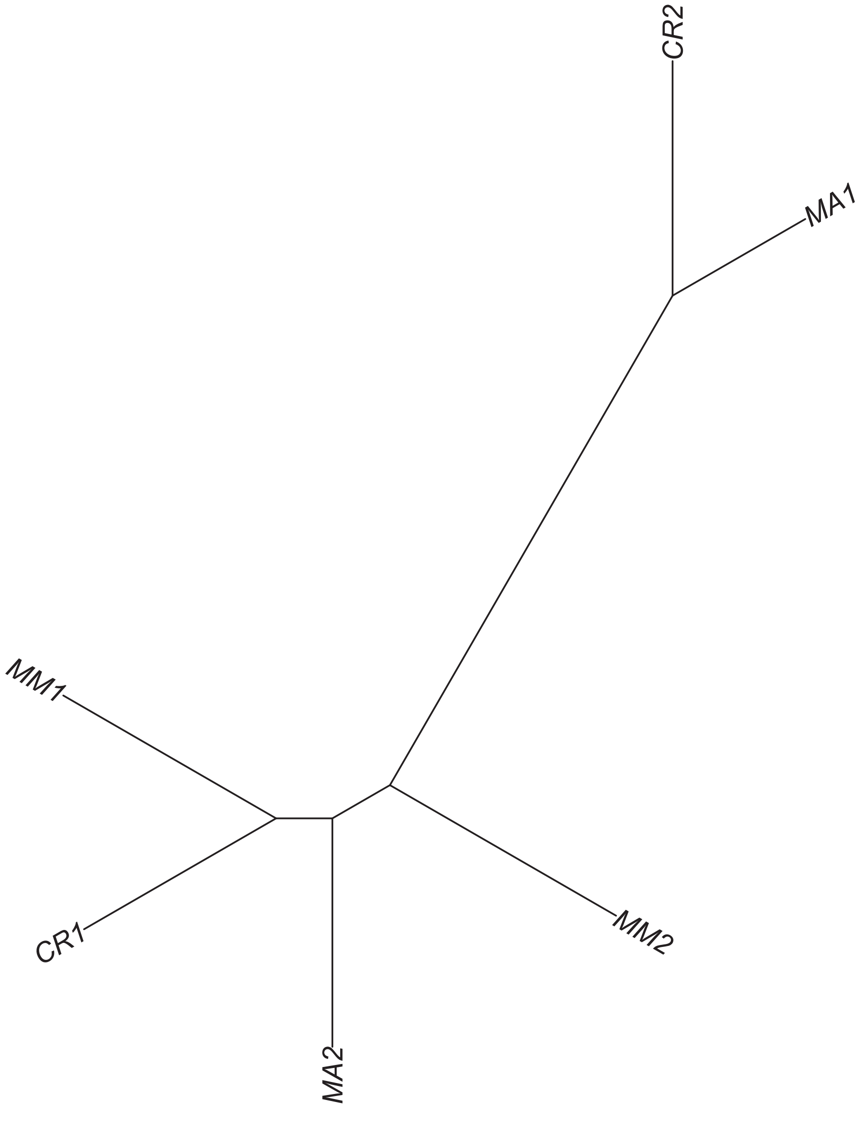

### Region 395

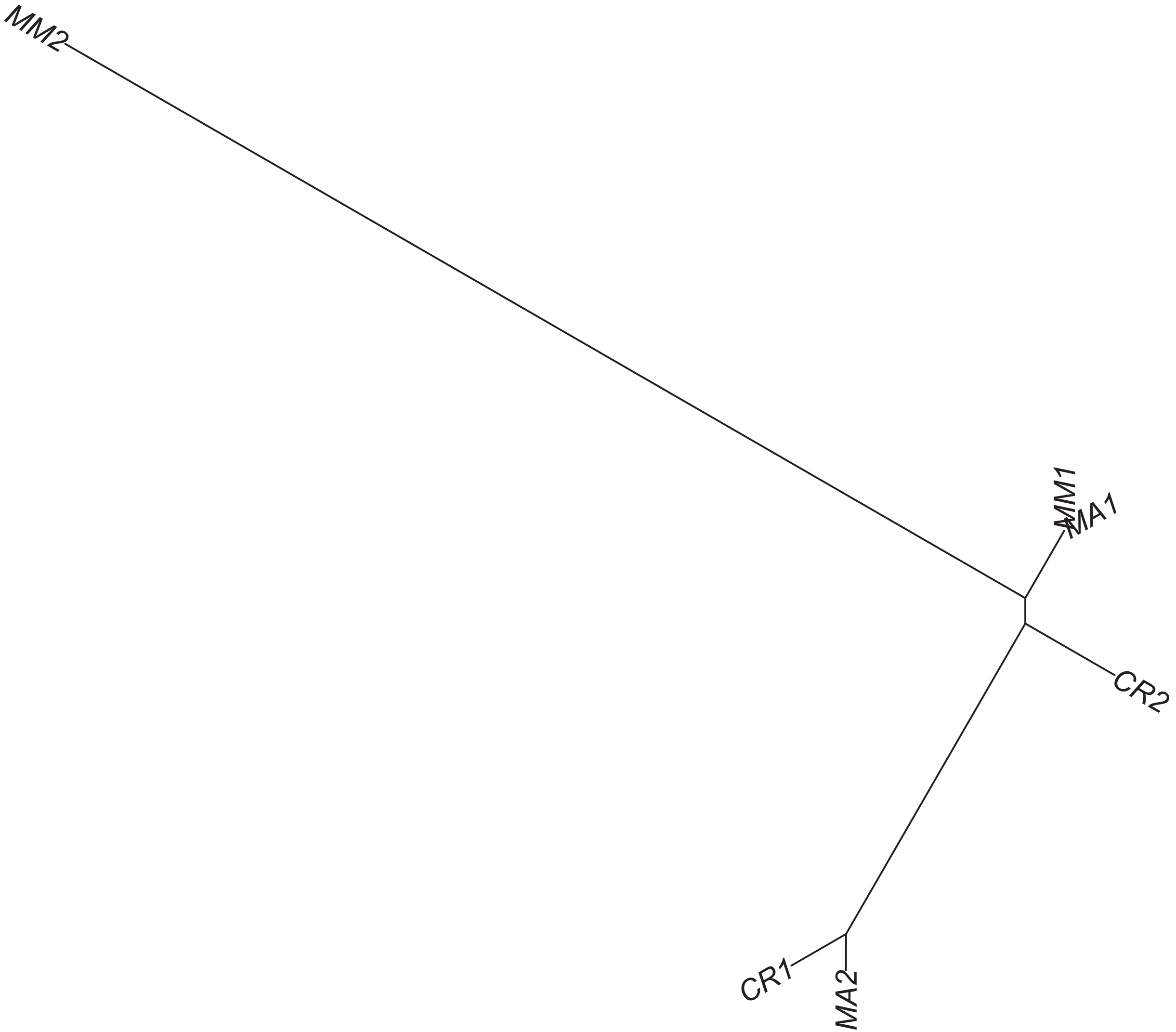

### Region 396

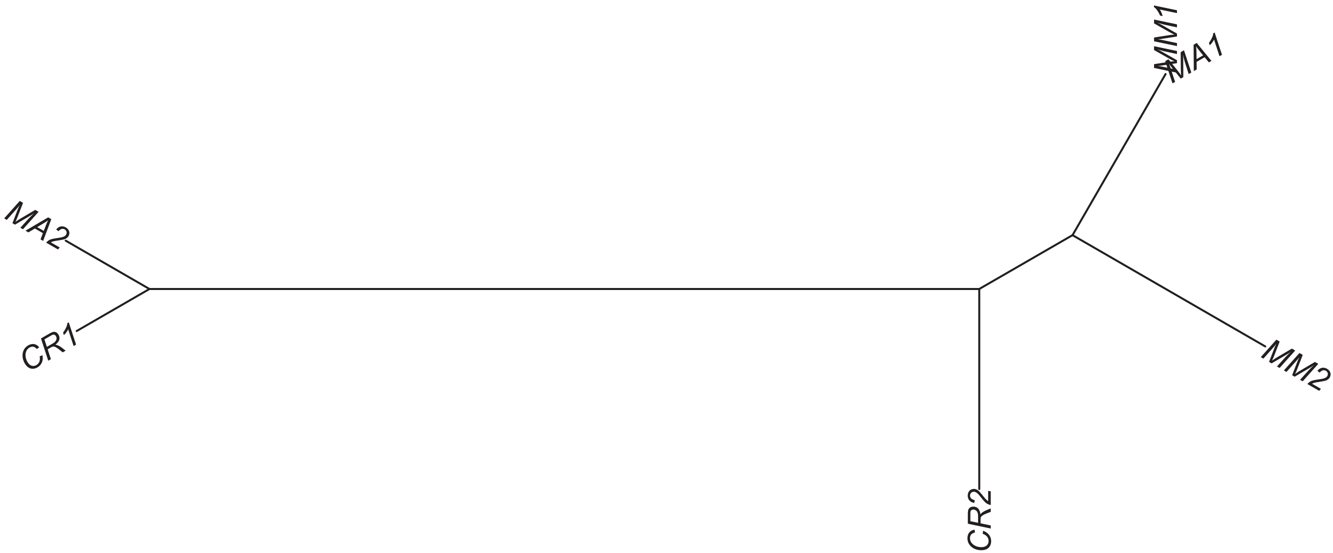

### Region 397

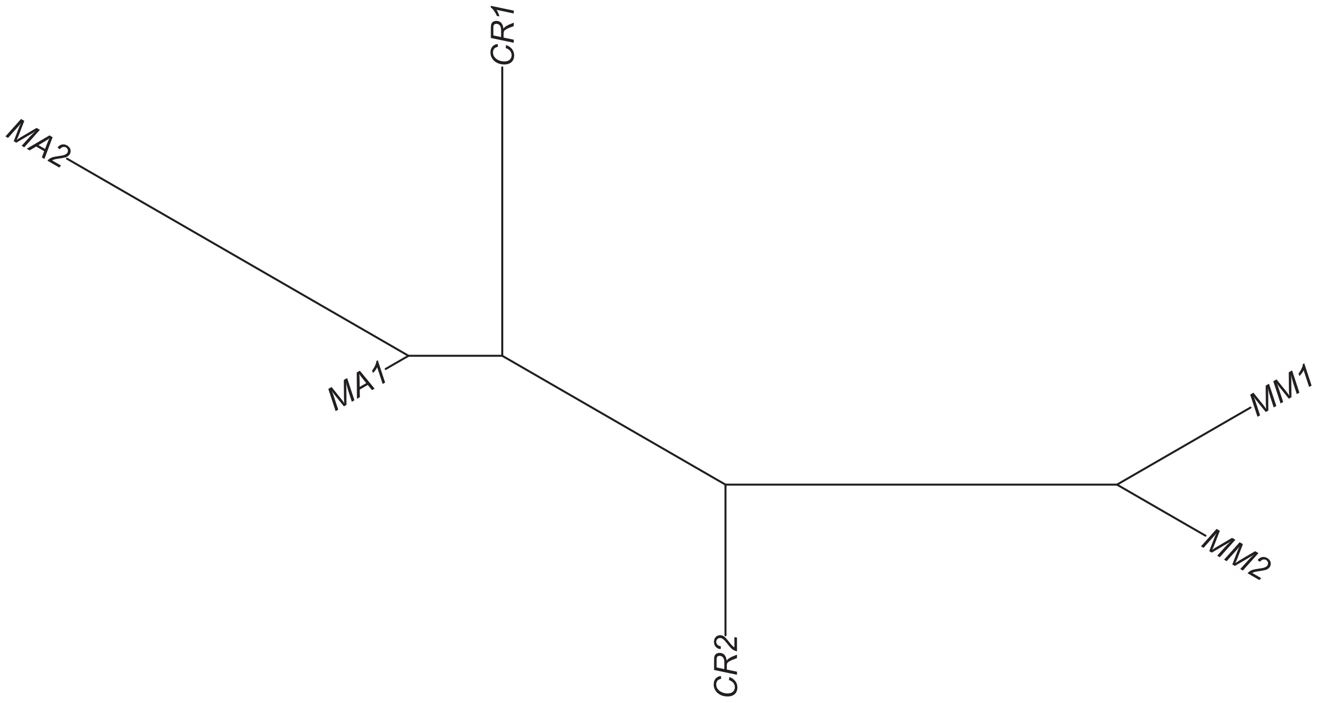

### Region 398

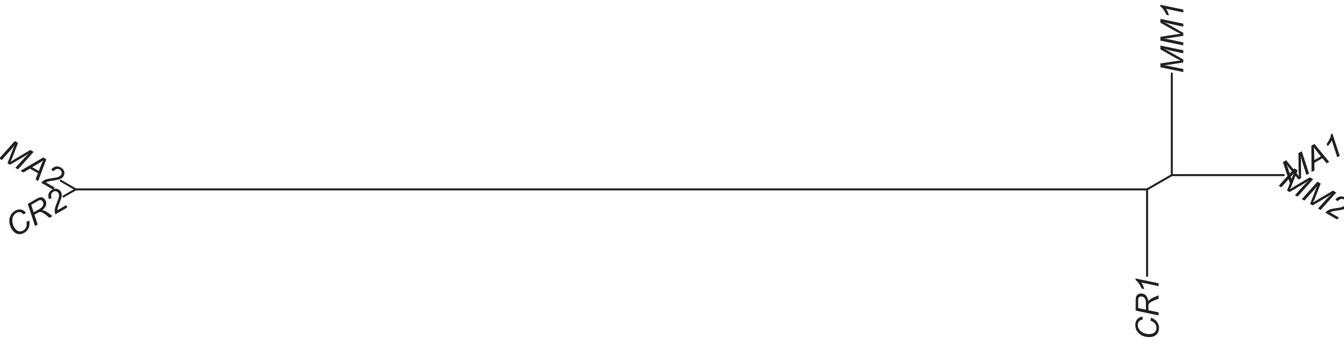

Region 400

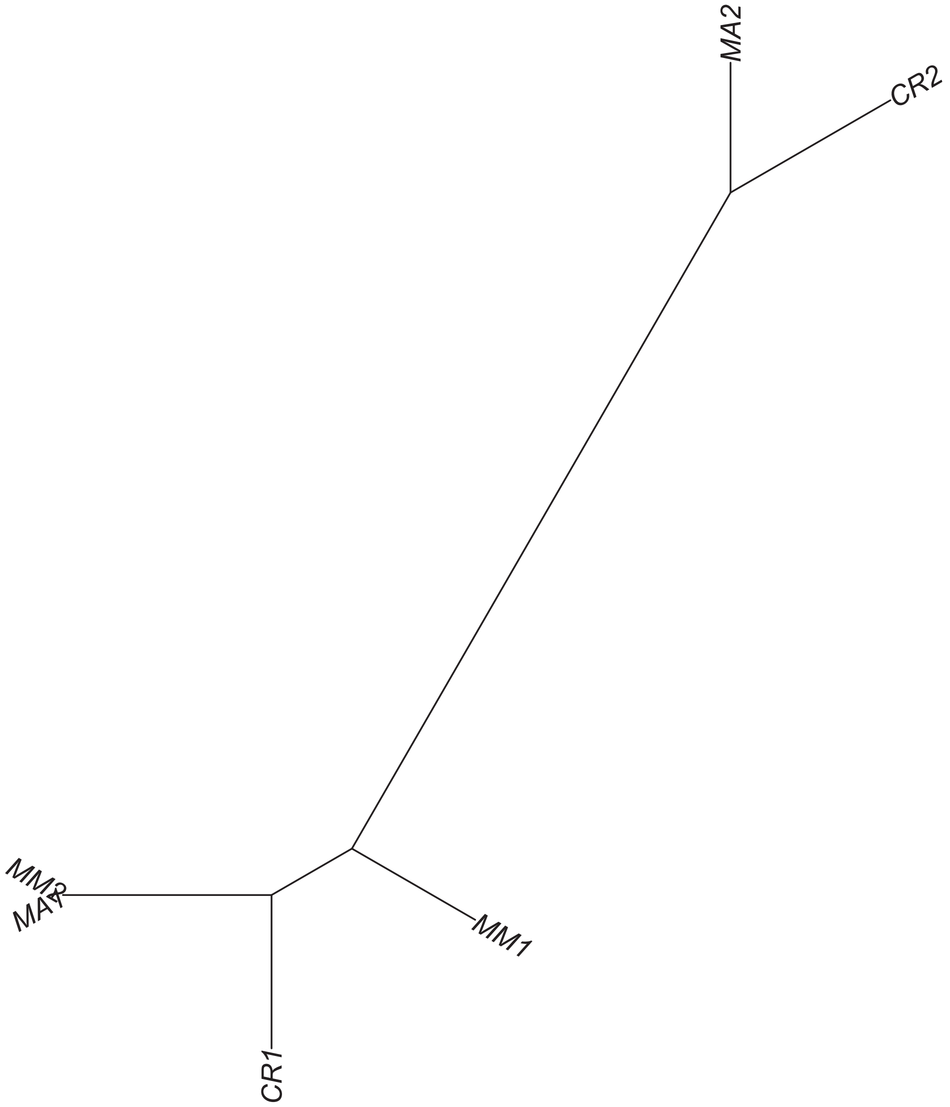

### Region 402

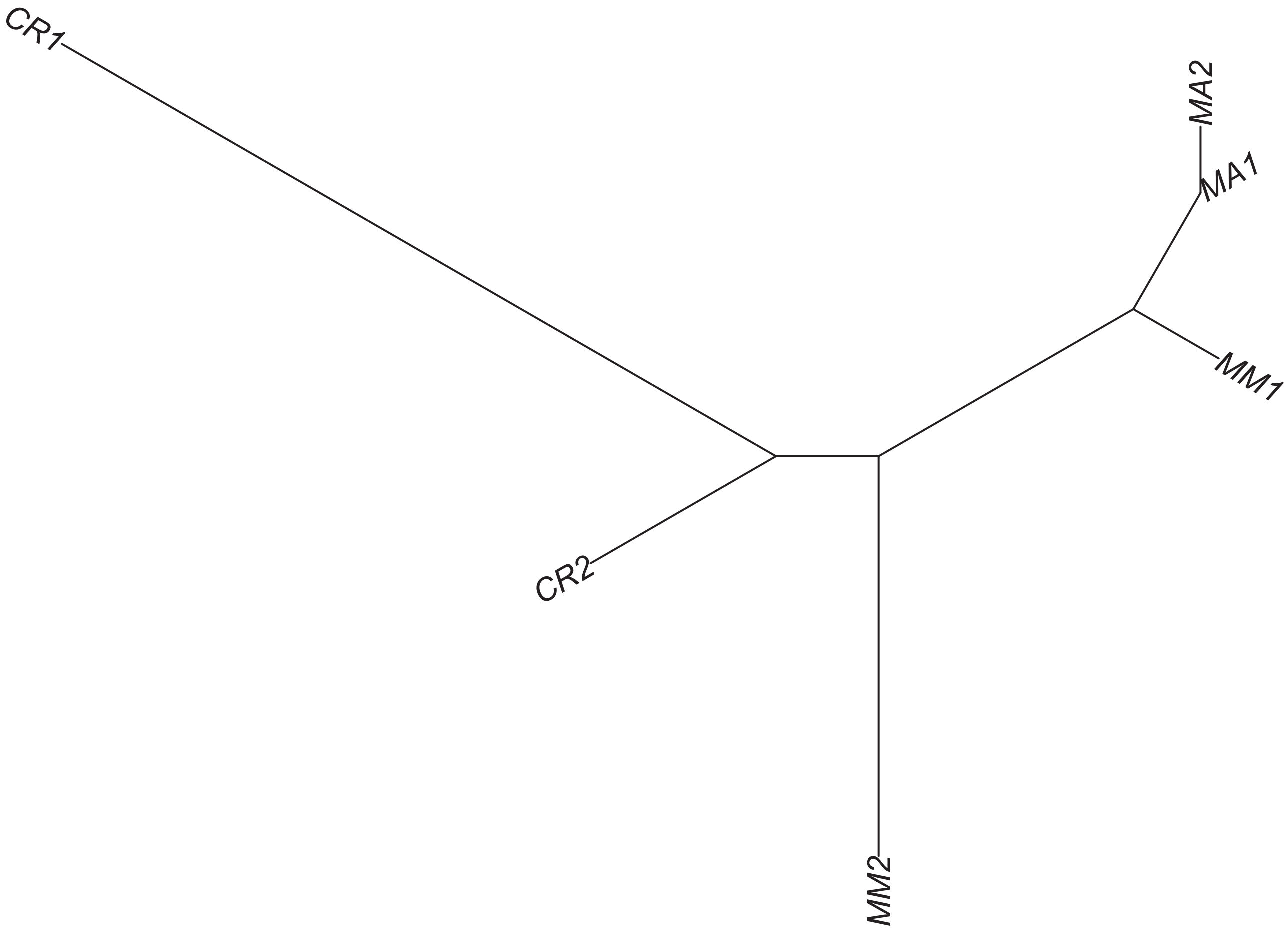

### Region 404



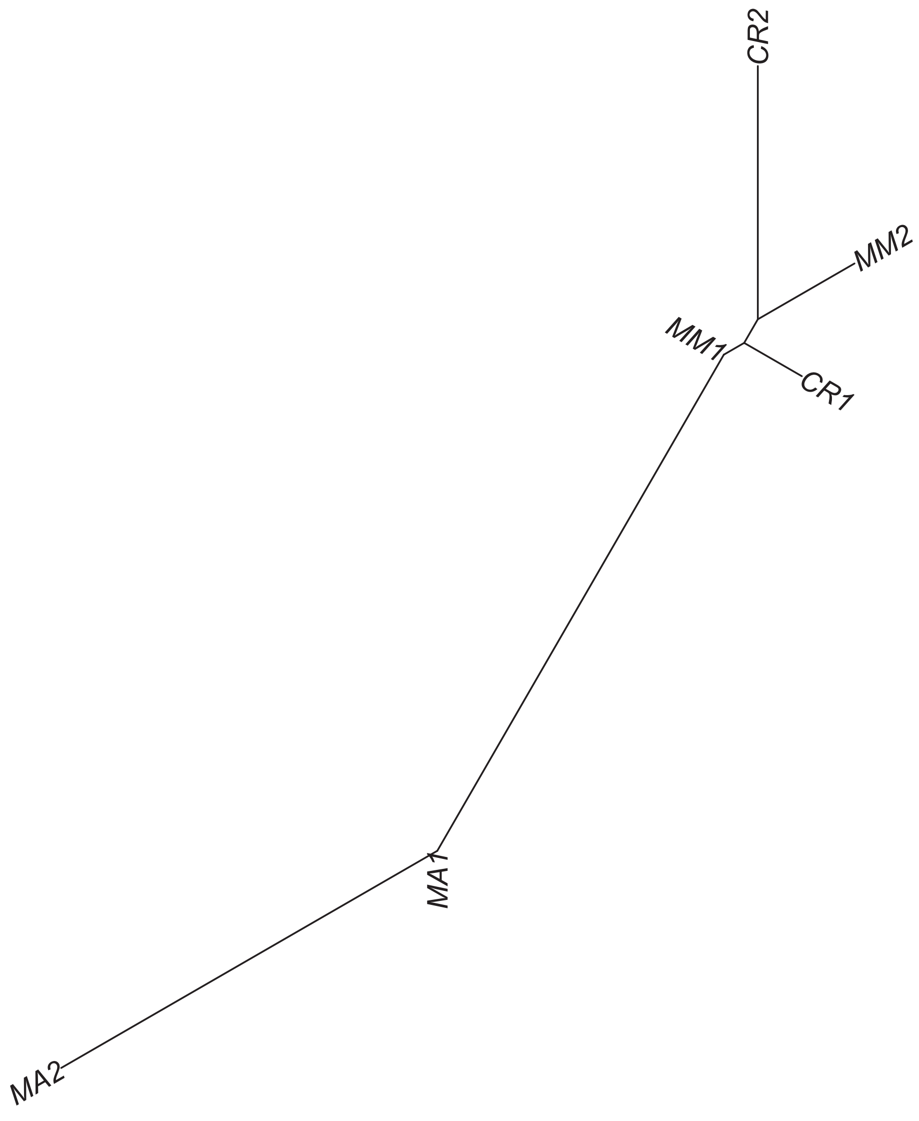

Region 407

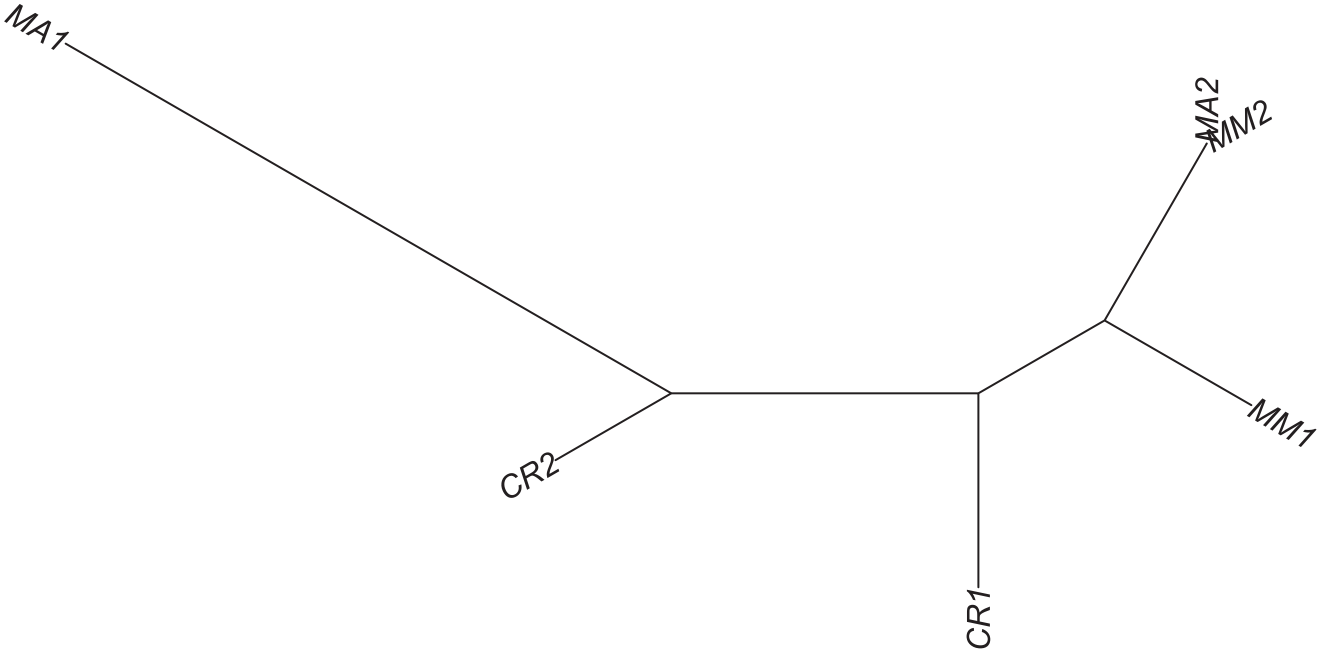

### Region 408

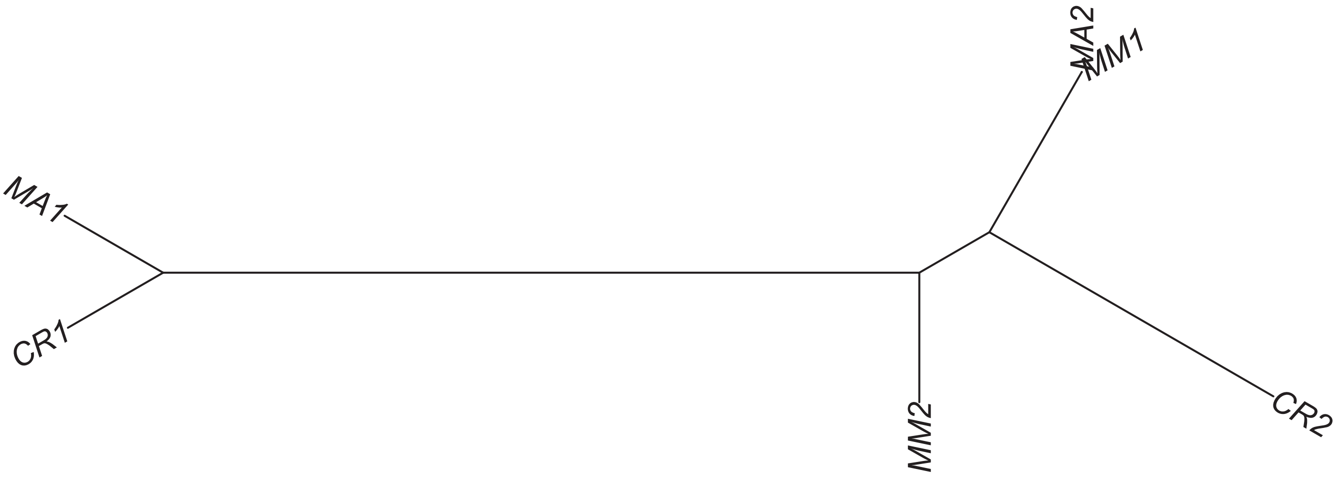

### Region 409

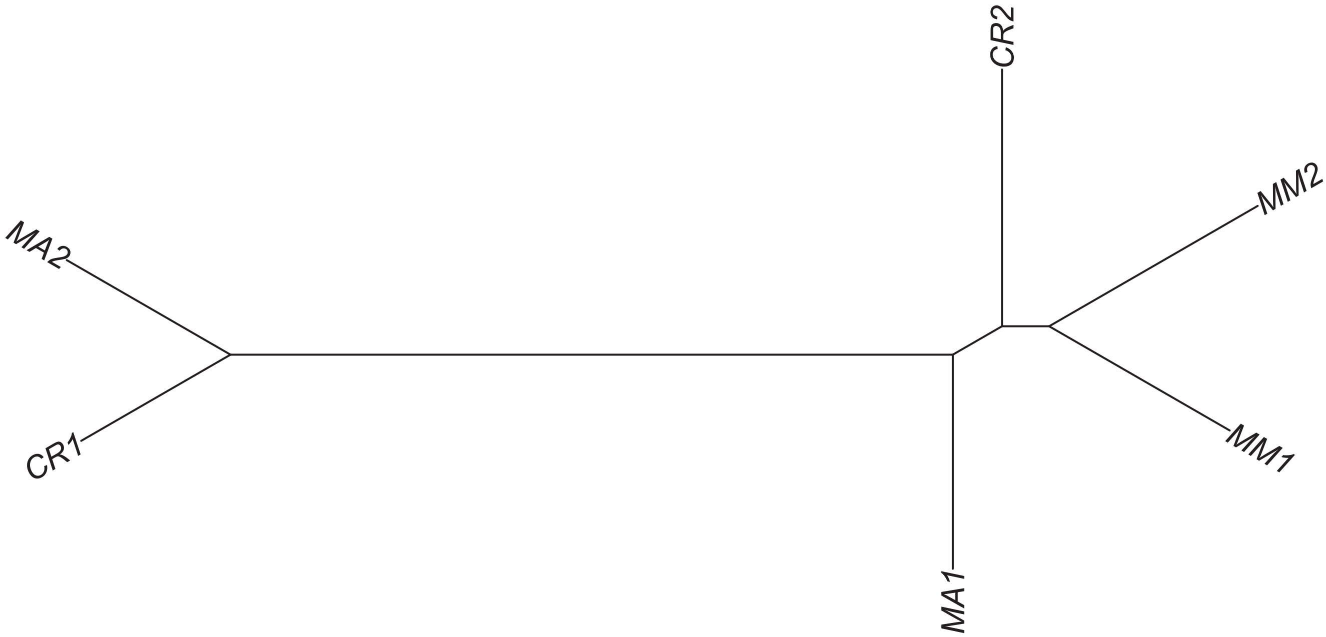

#### Region 410

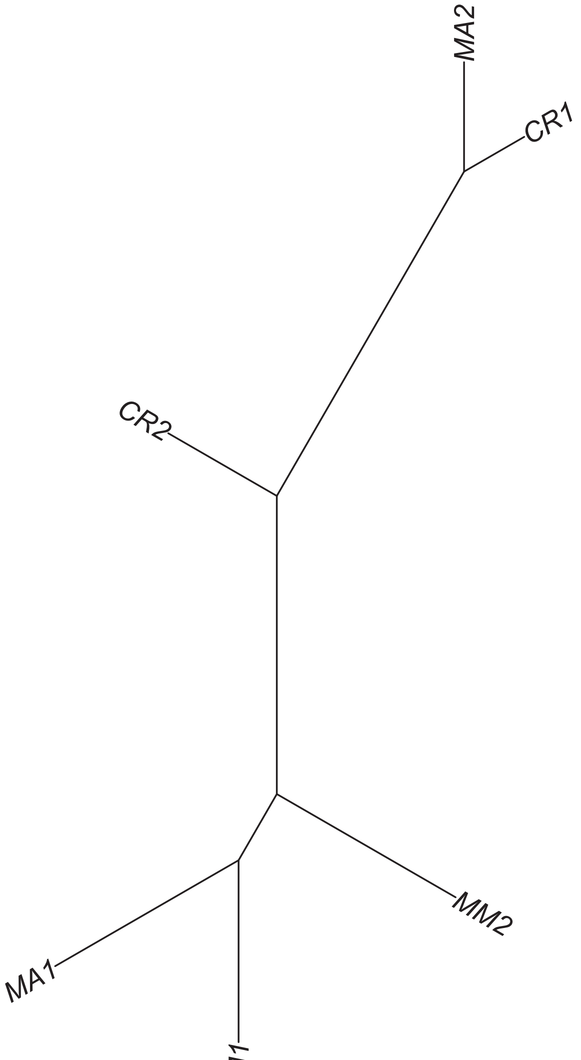

### Region 413



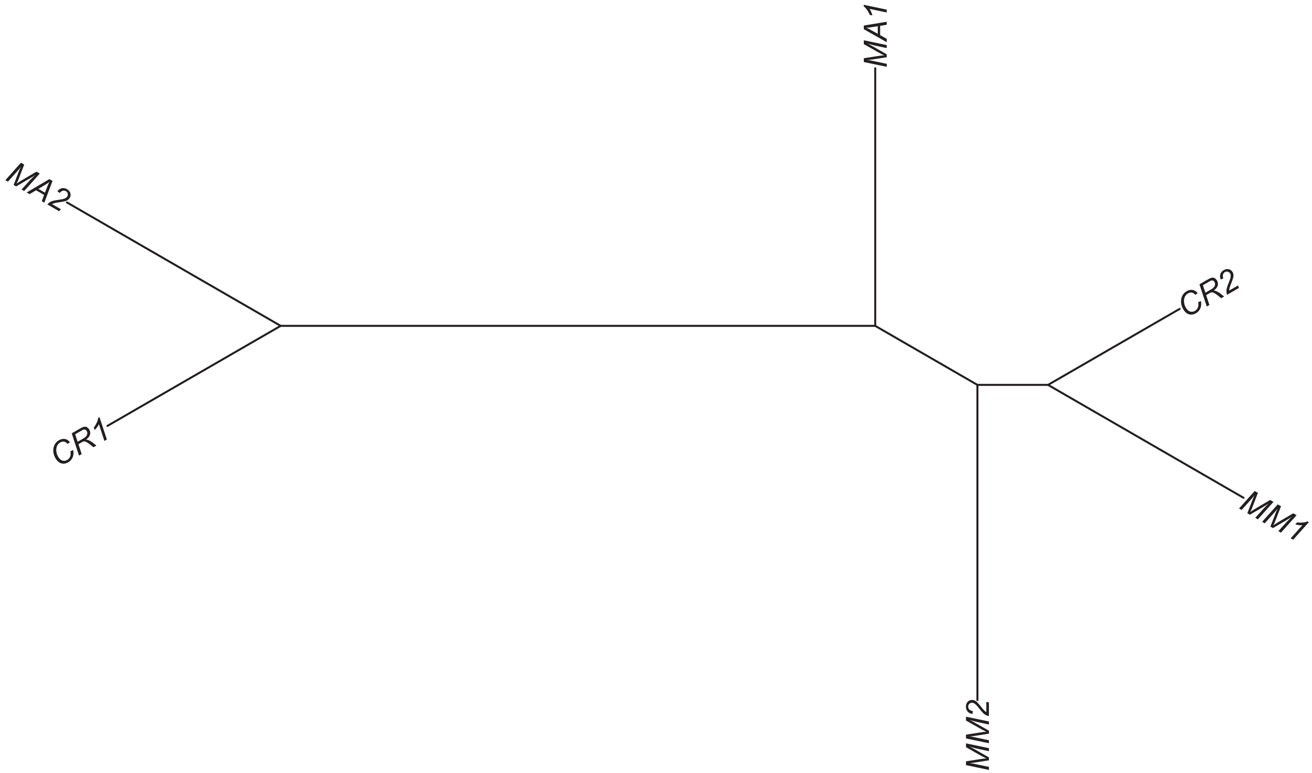

### Region 414

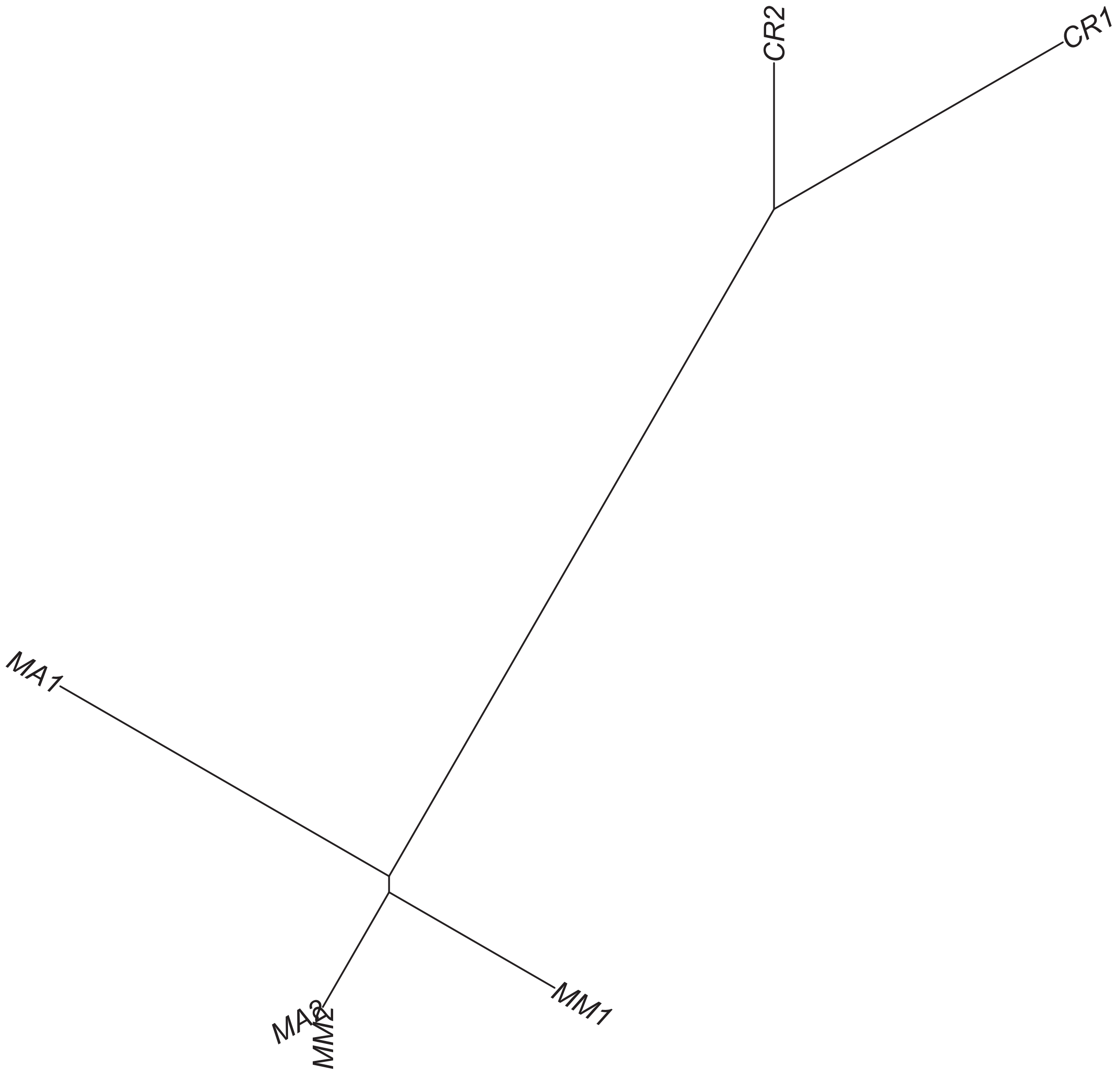

### Region 417

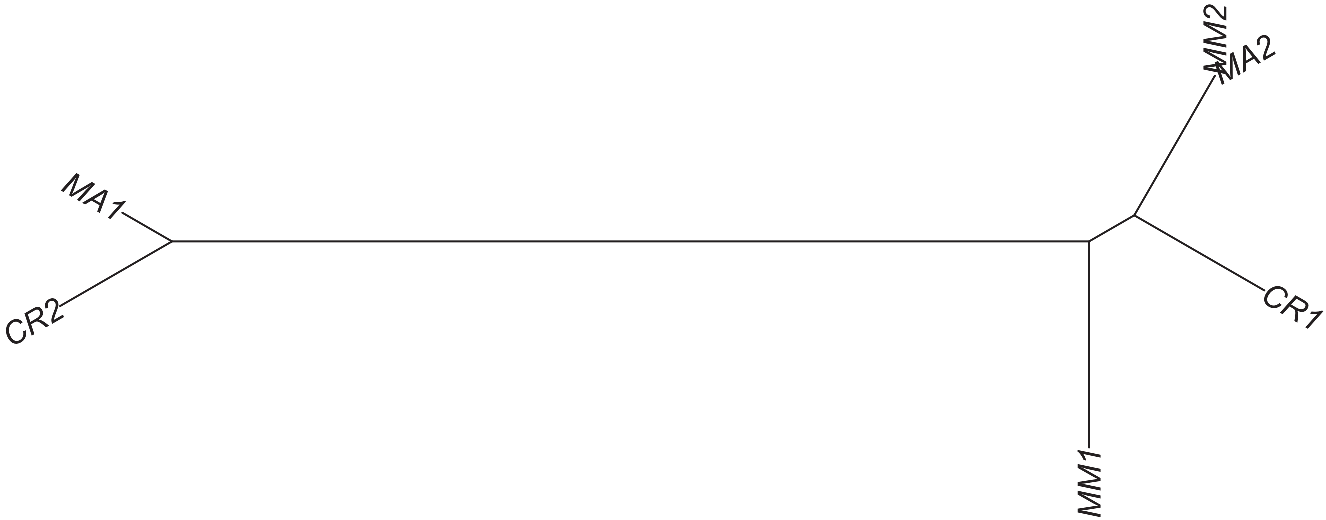

Region 418

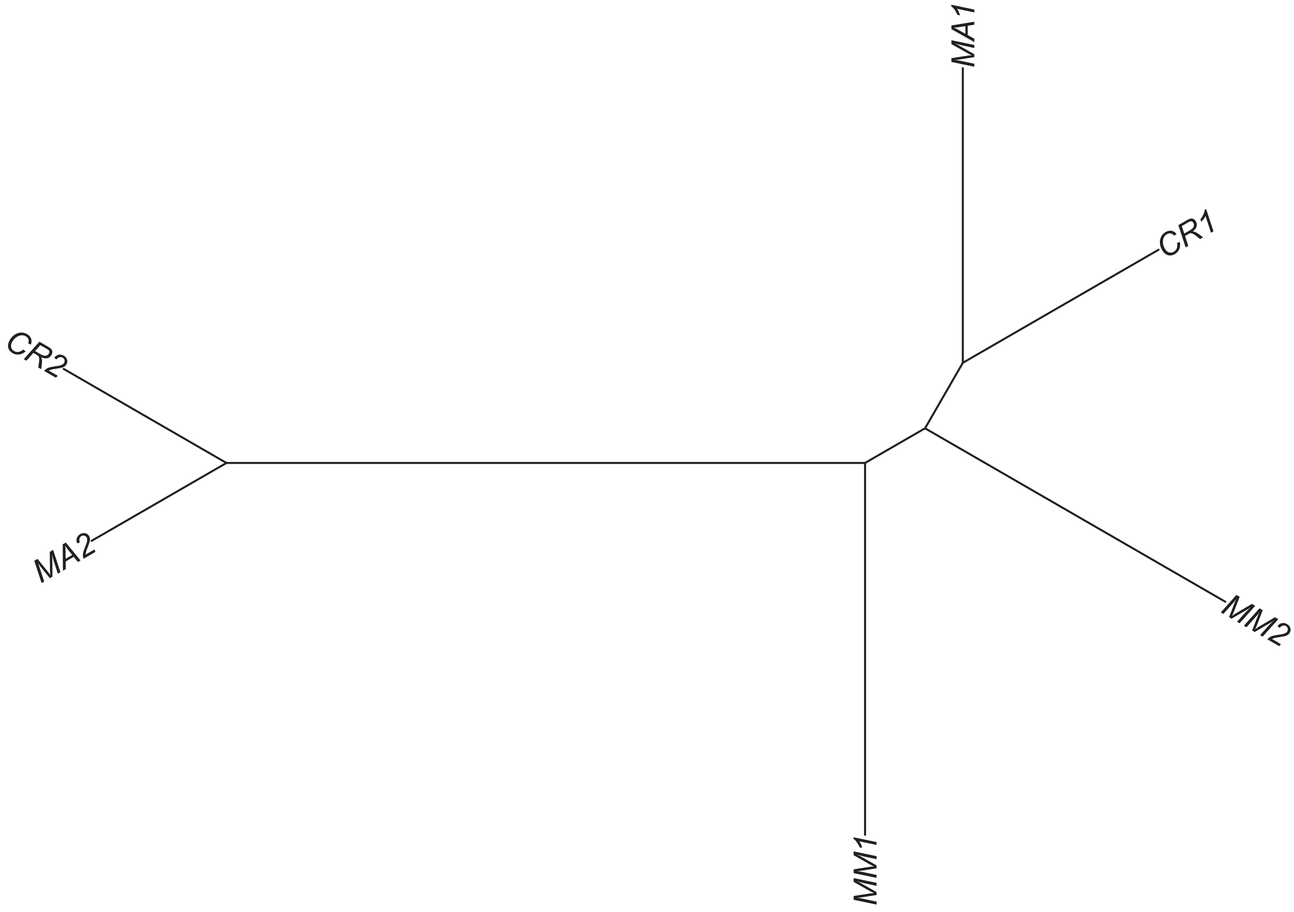

Region 422



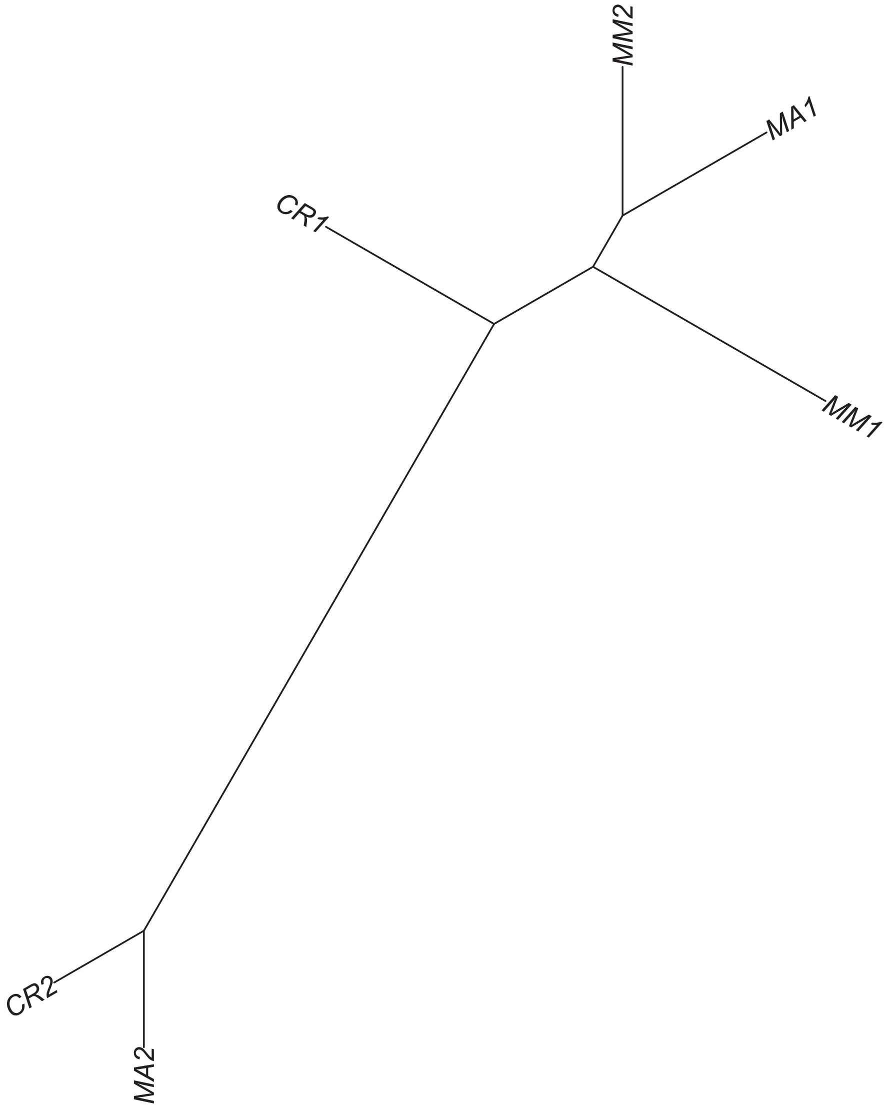

### Region 424

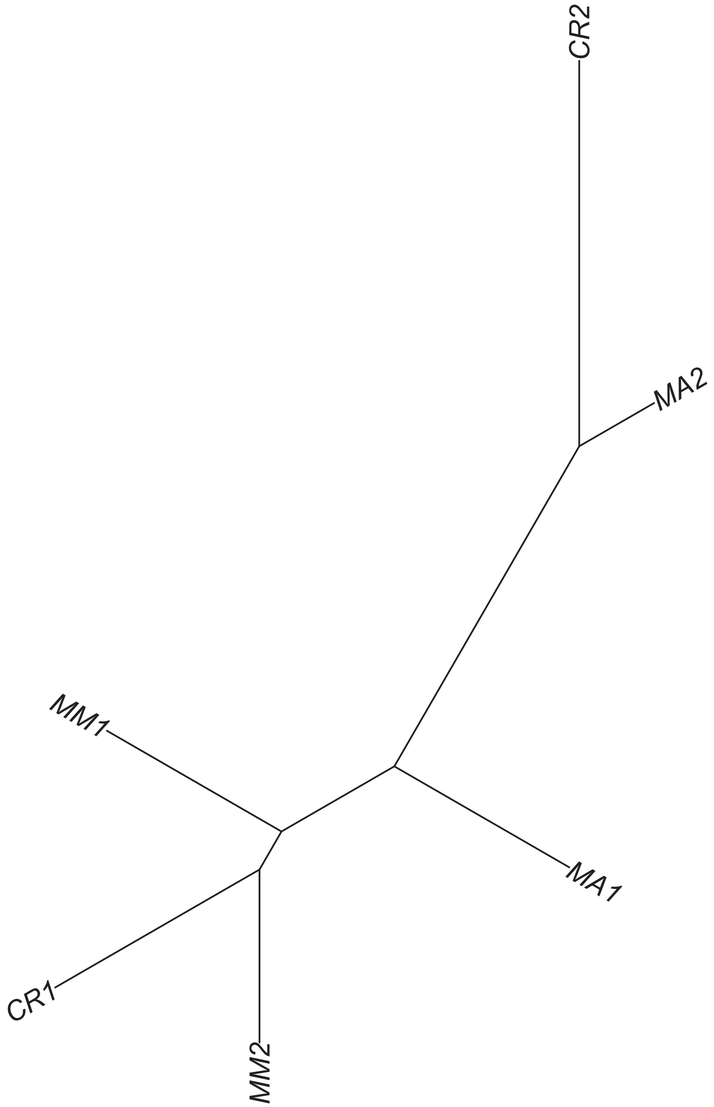

### Region 427

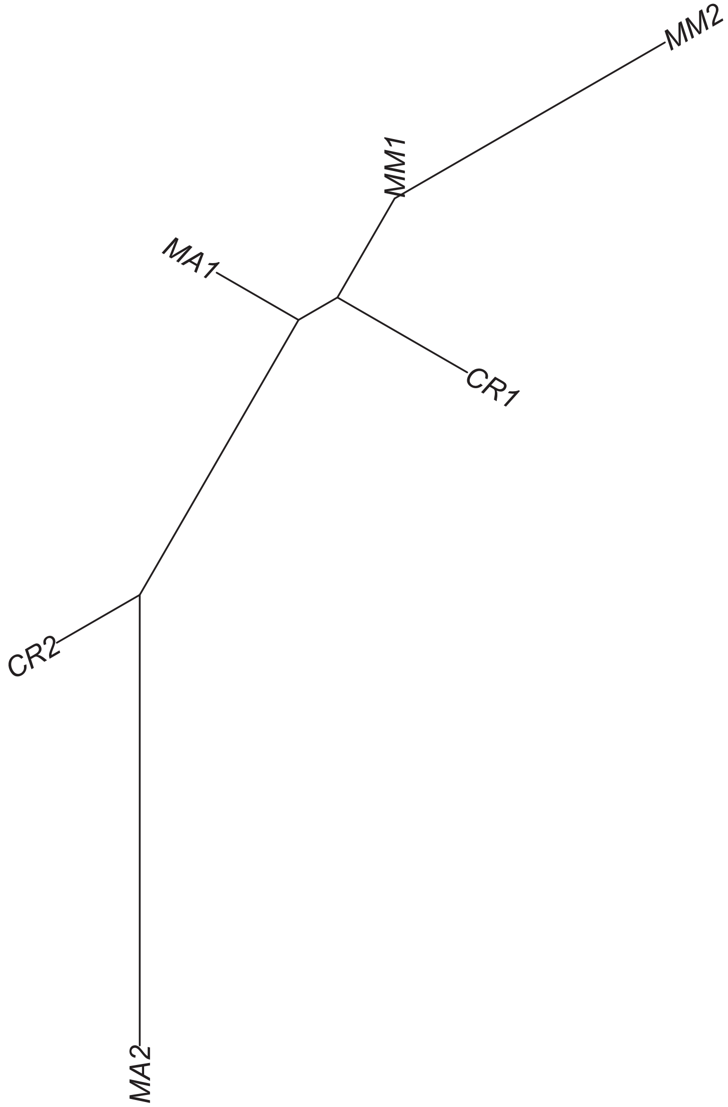

### Region 428



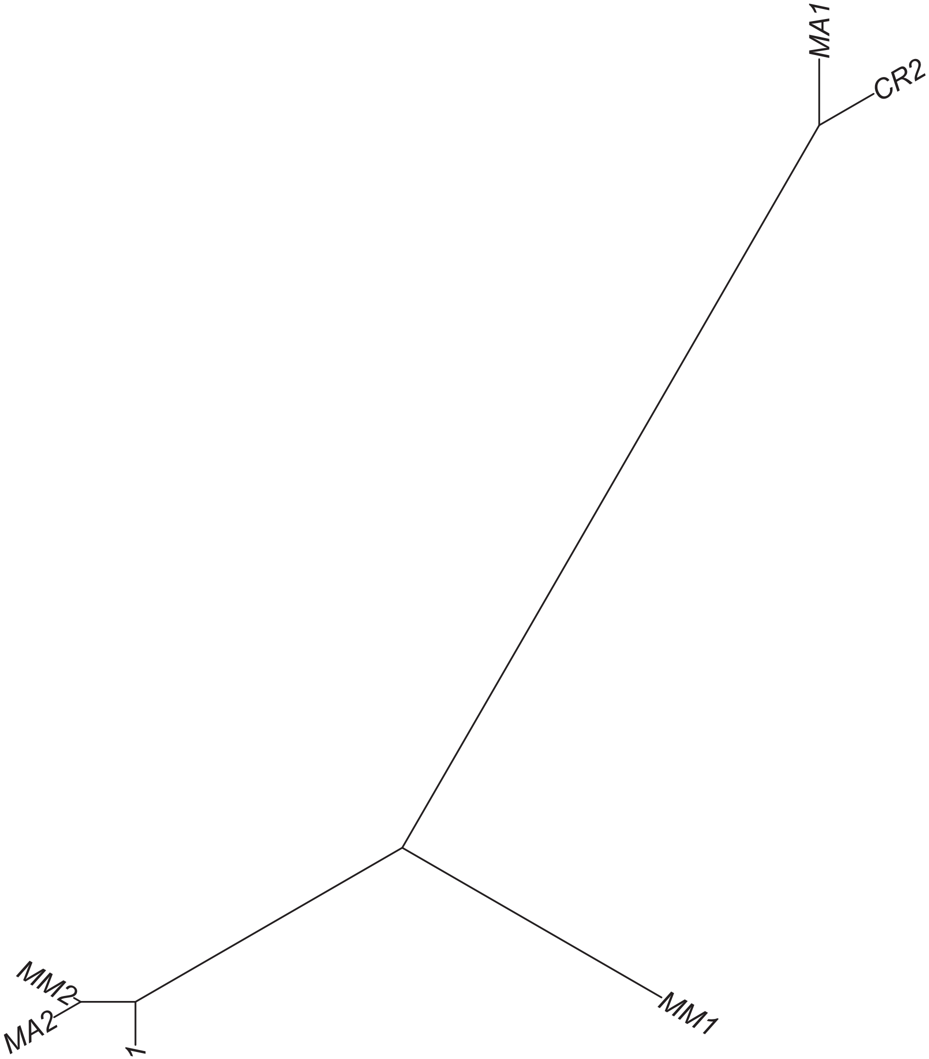

### Region 429

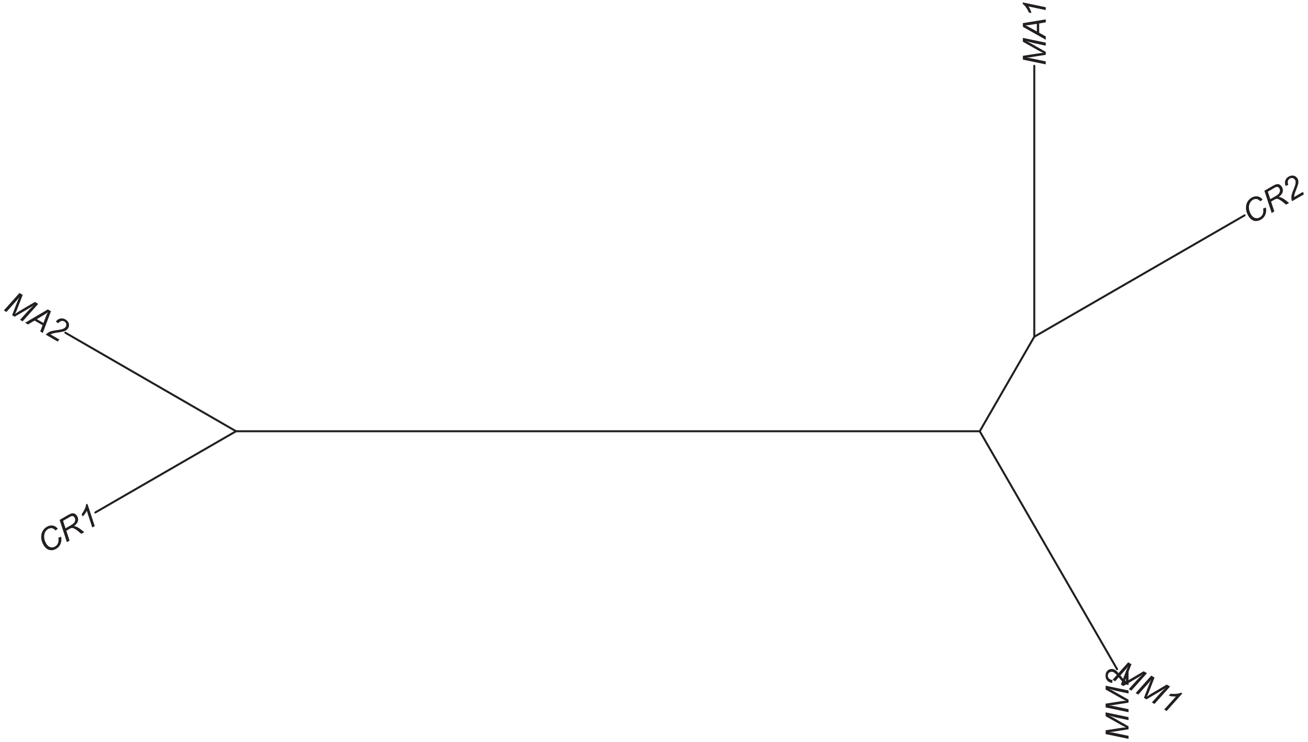

### Region 432

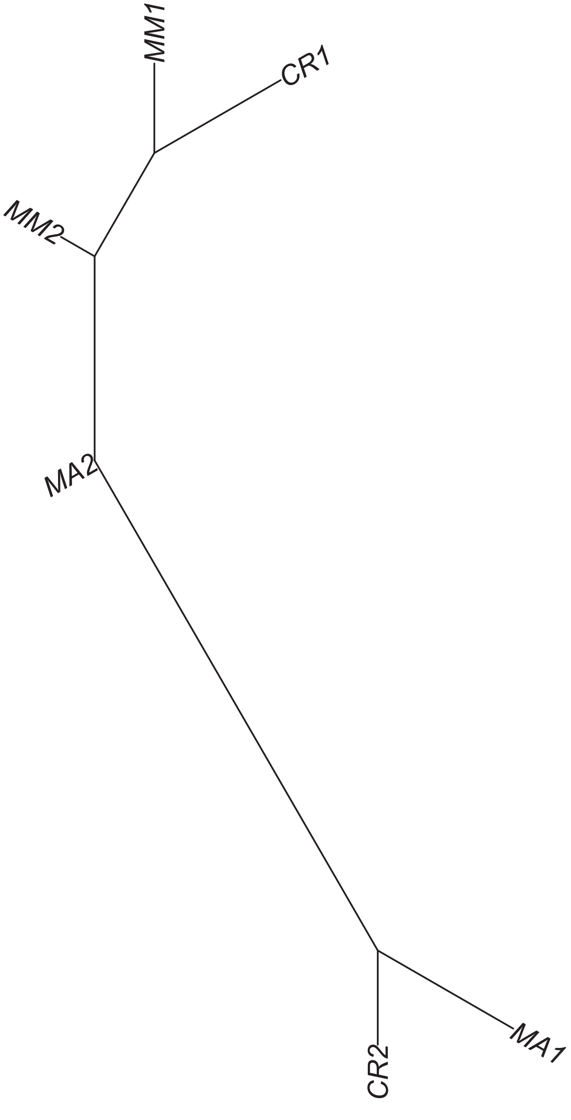

### Region 433

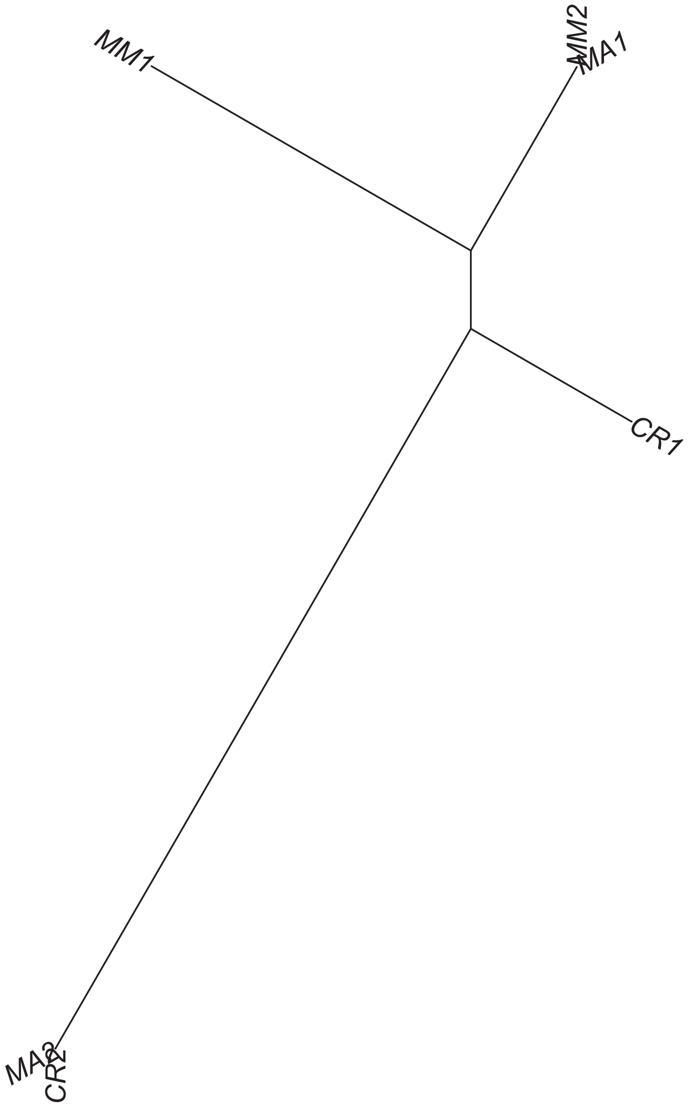

### Region 435



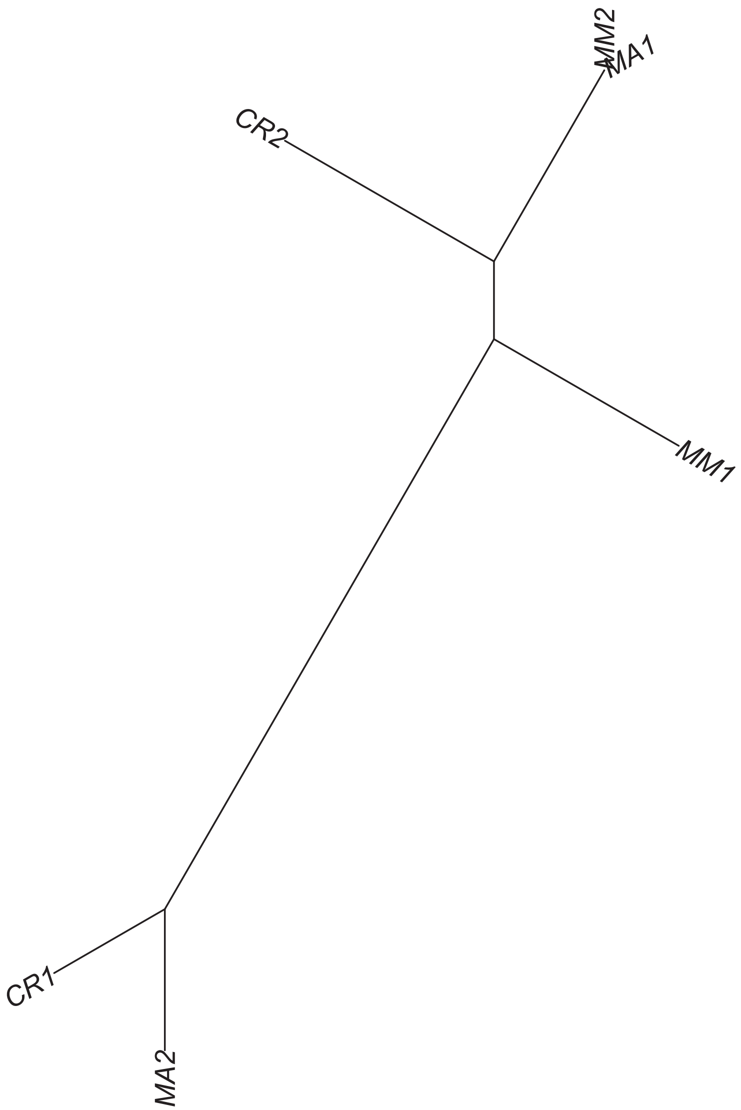

### Region 436



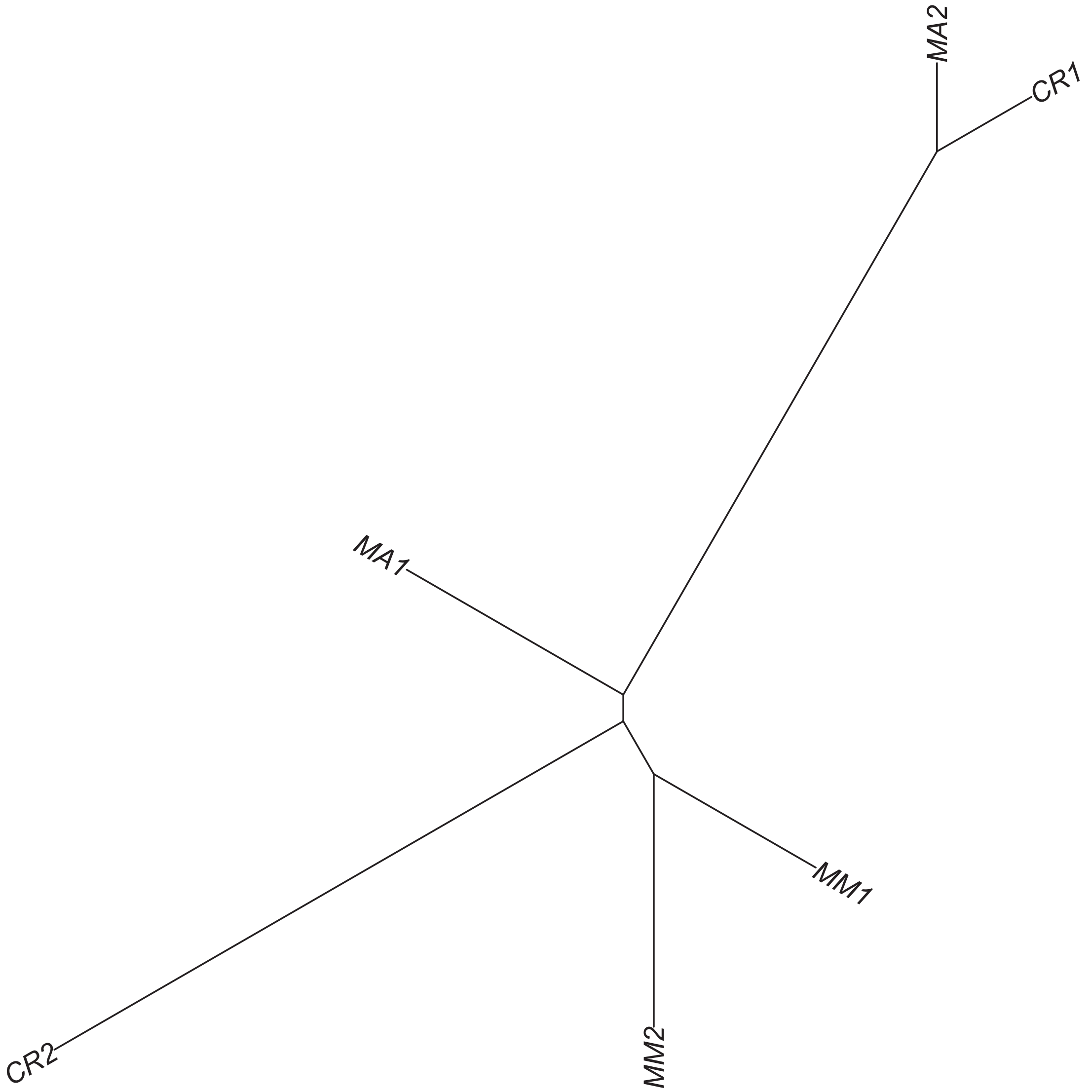

### Region 437



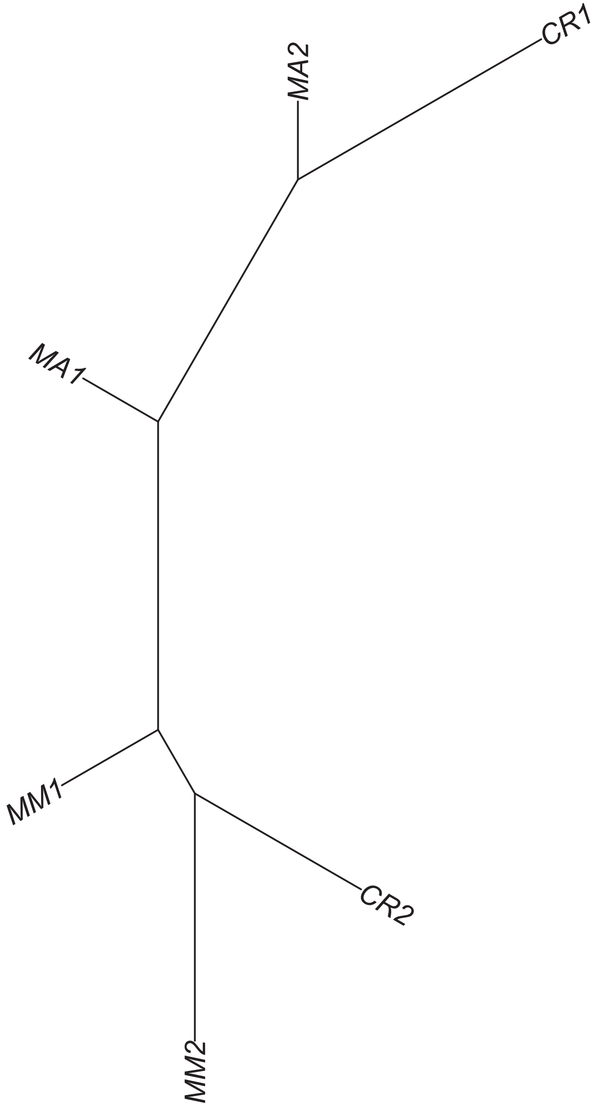

Region 441

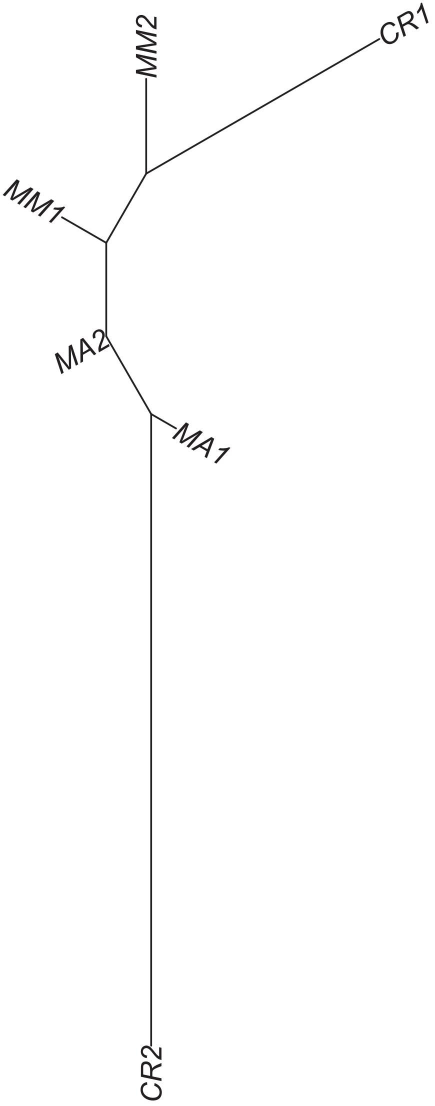

### Region 442

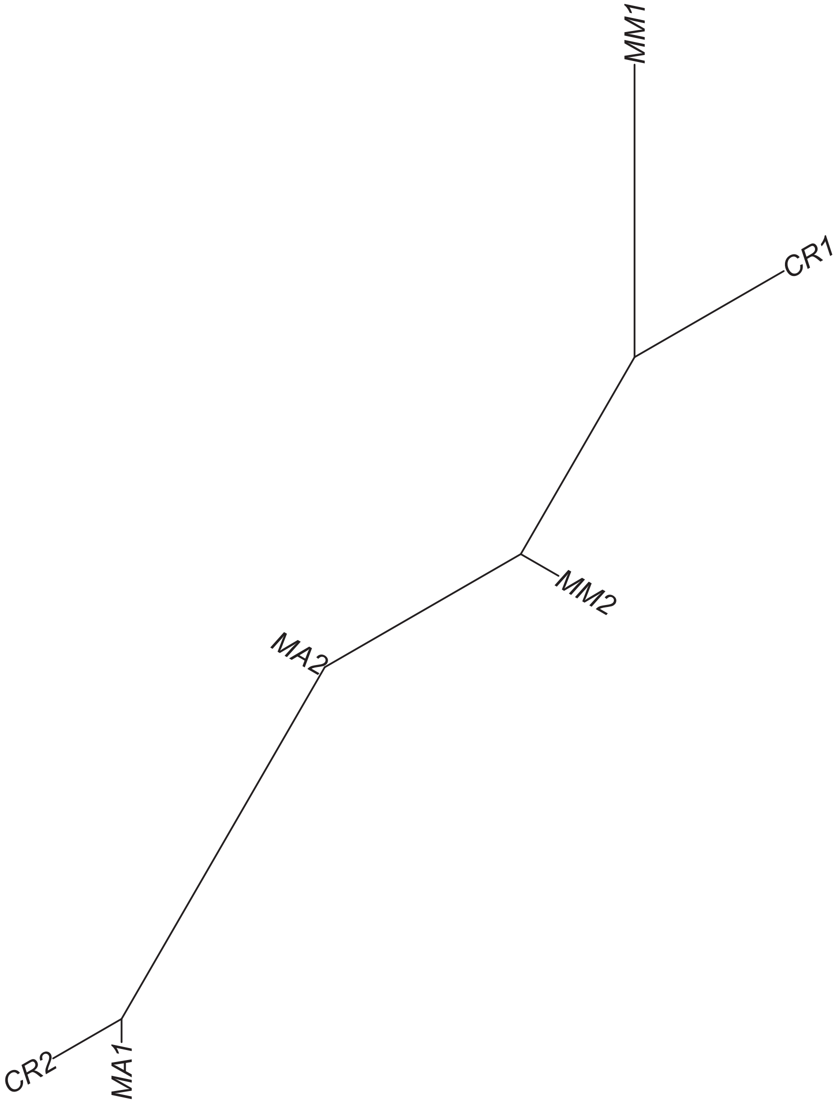

### Region 445

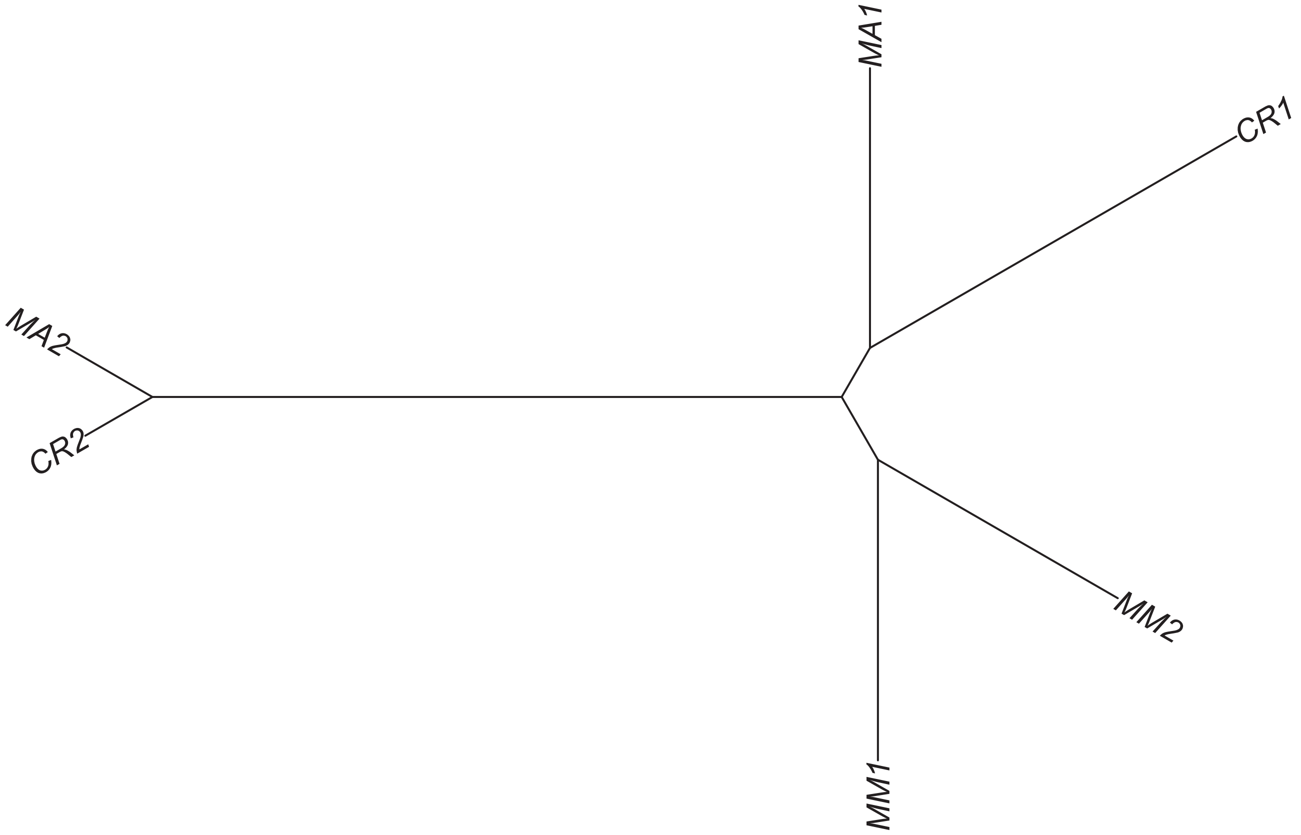

### Region 447



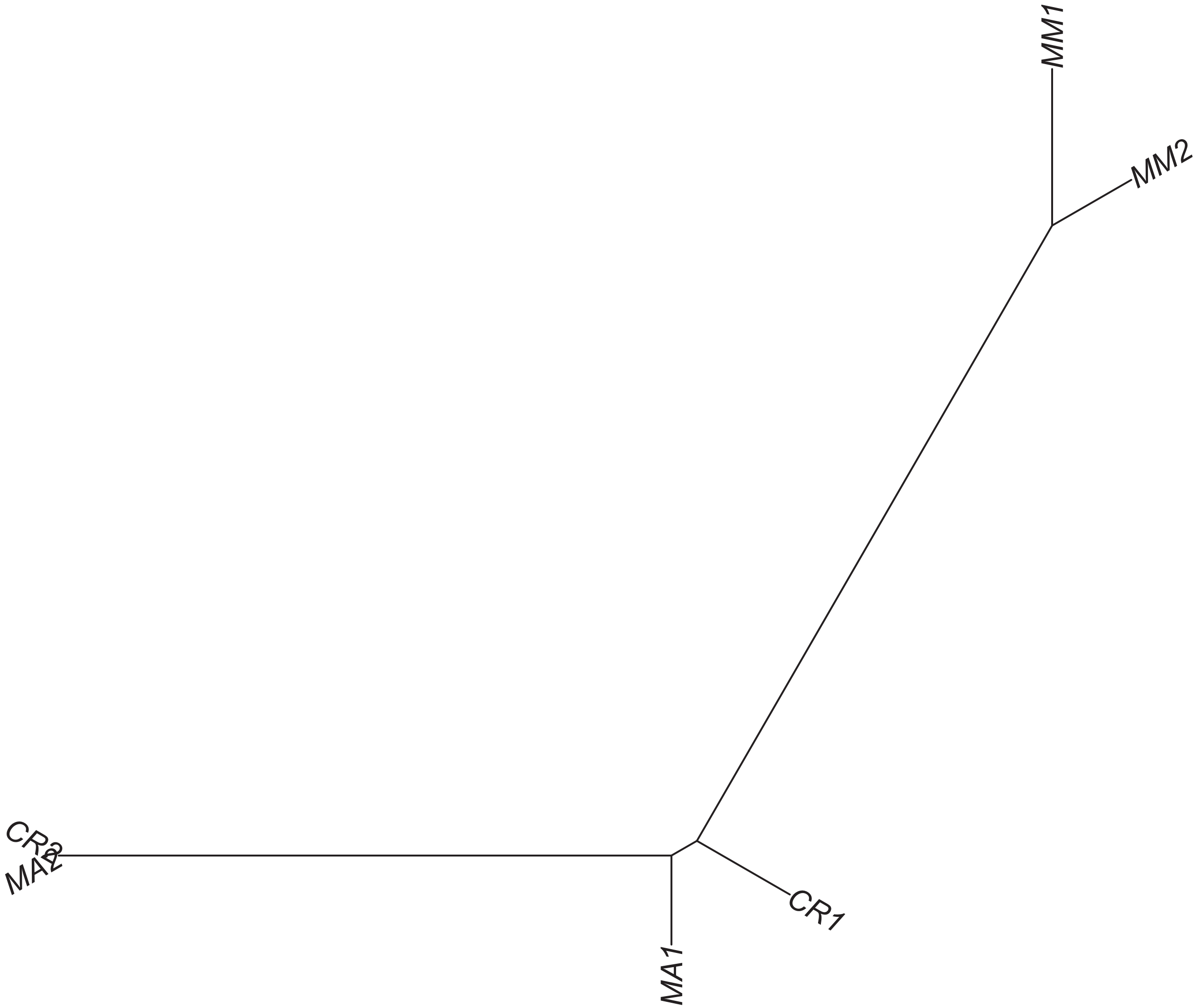

### Region 448

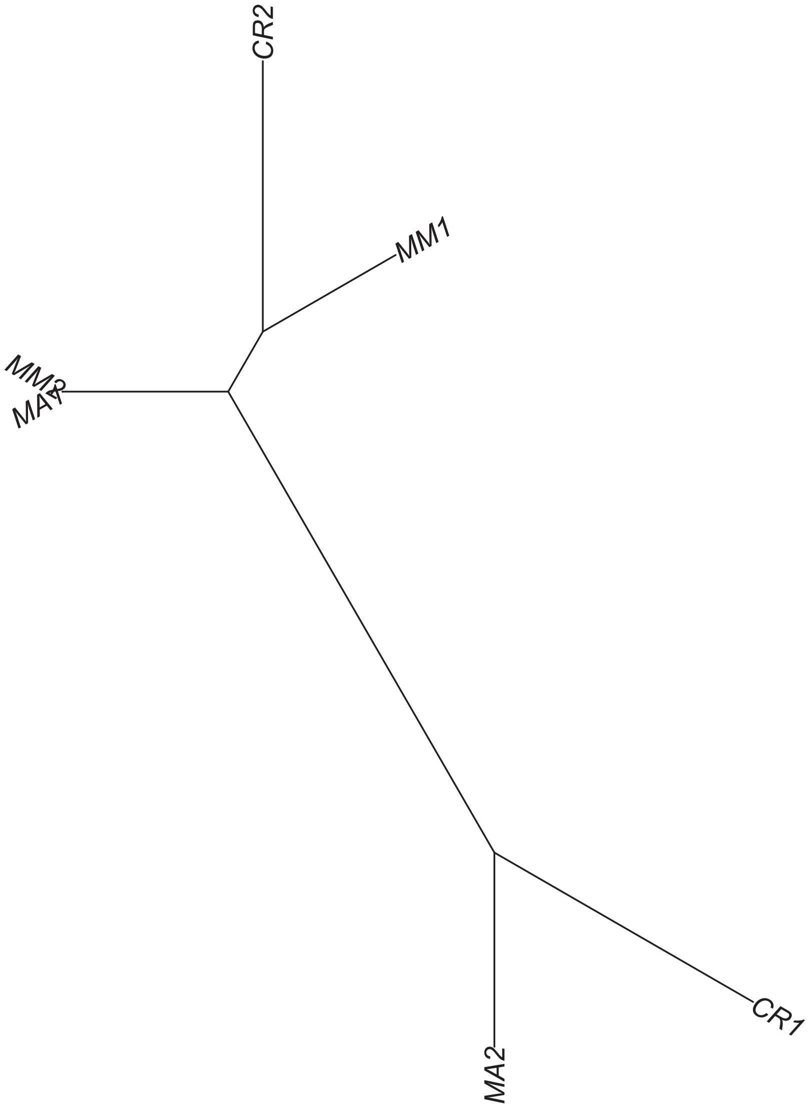

### Region 452

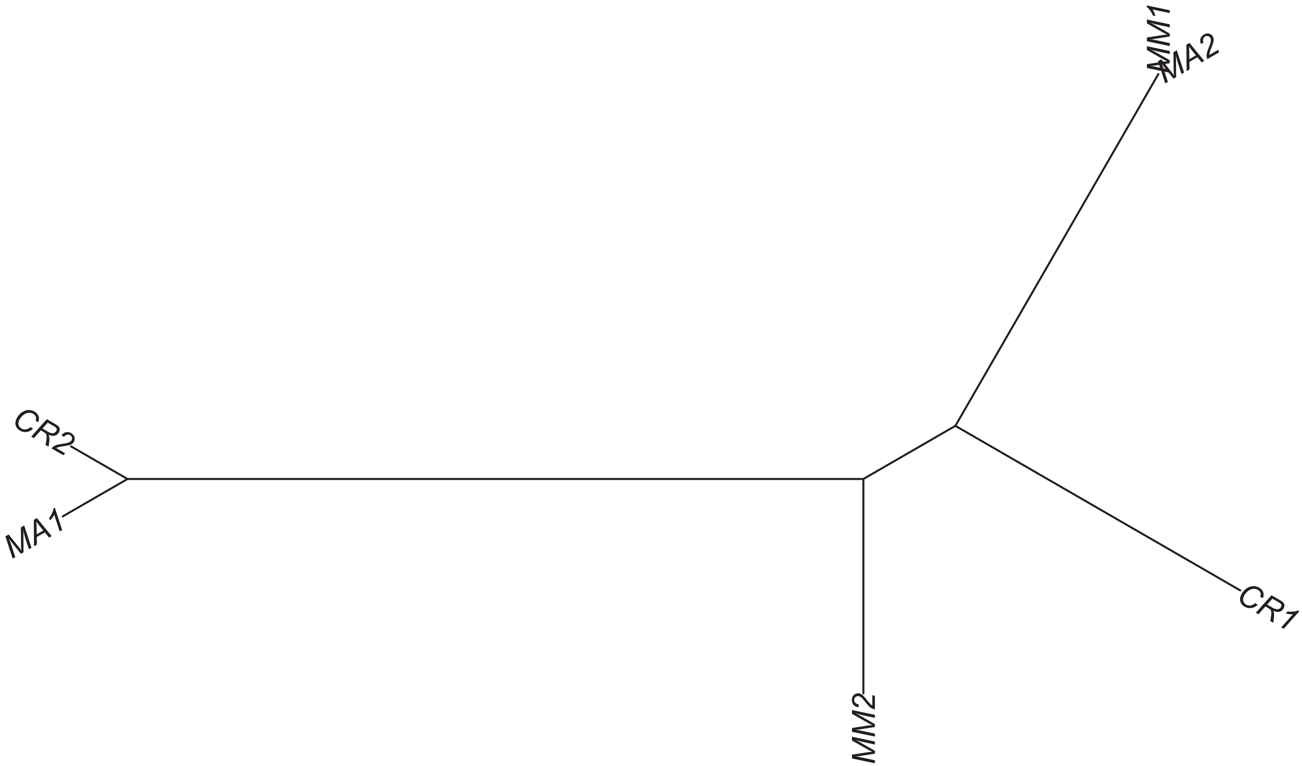

### Region 453



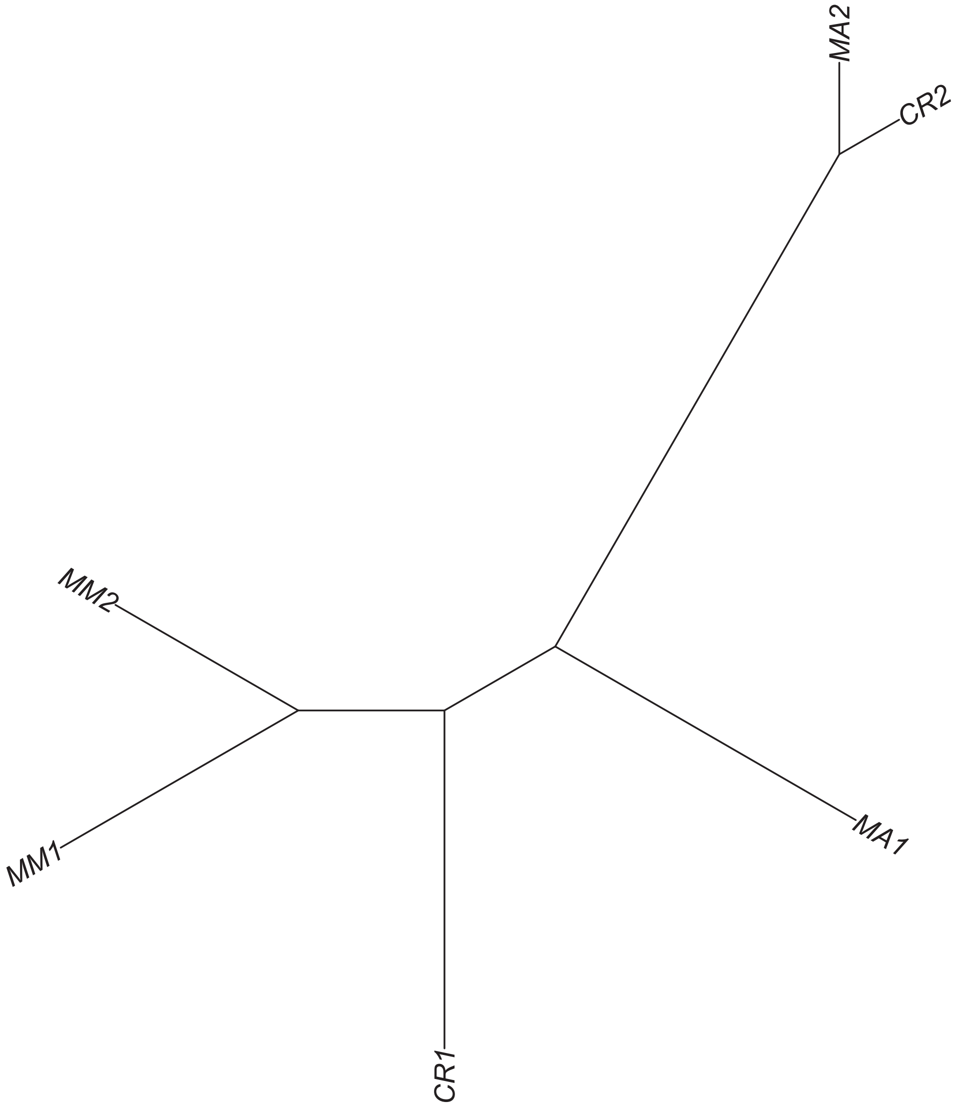

### Region 454



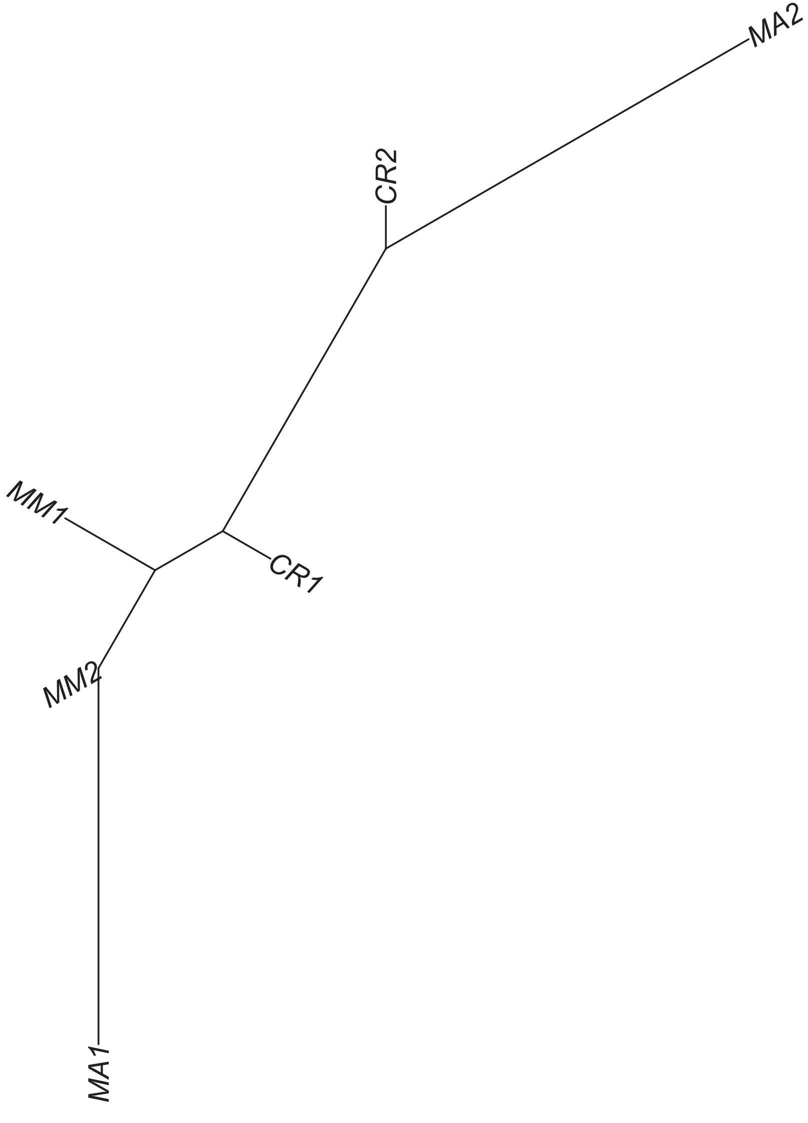

Region 456

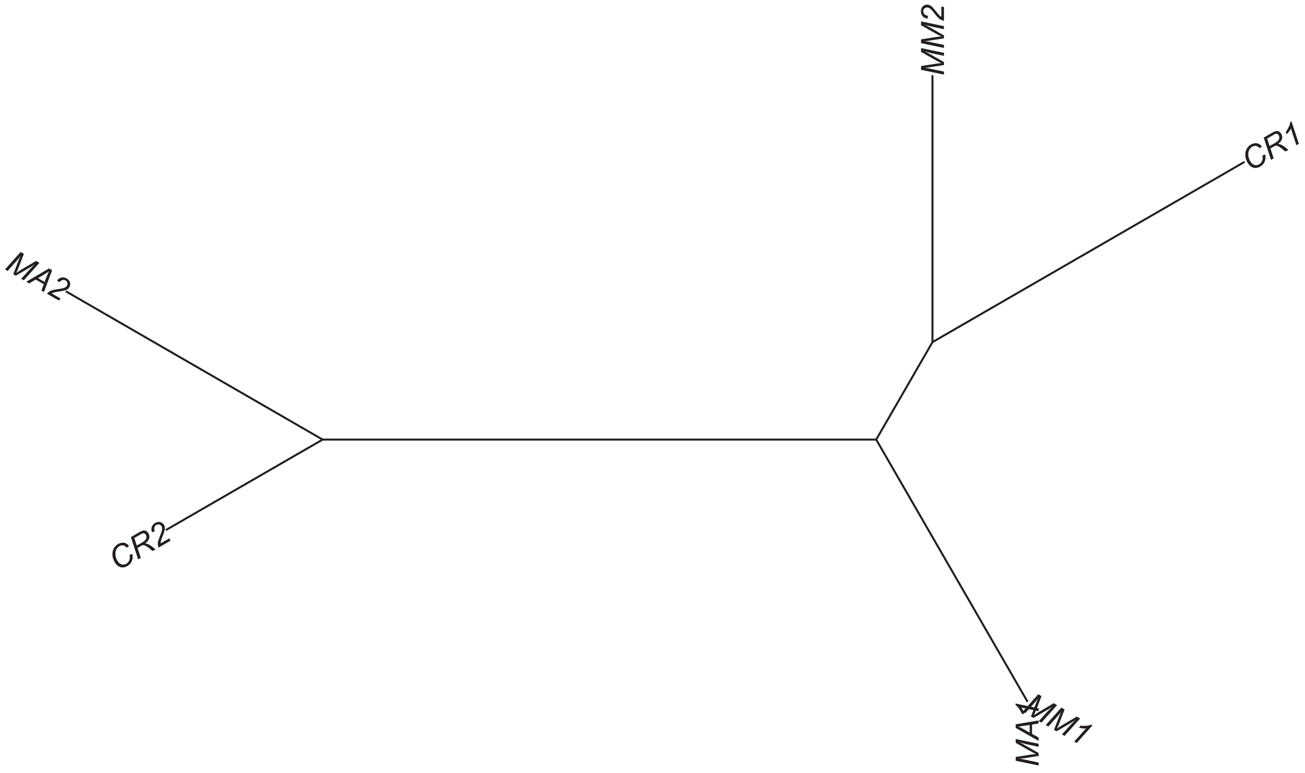

Region 457

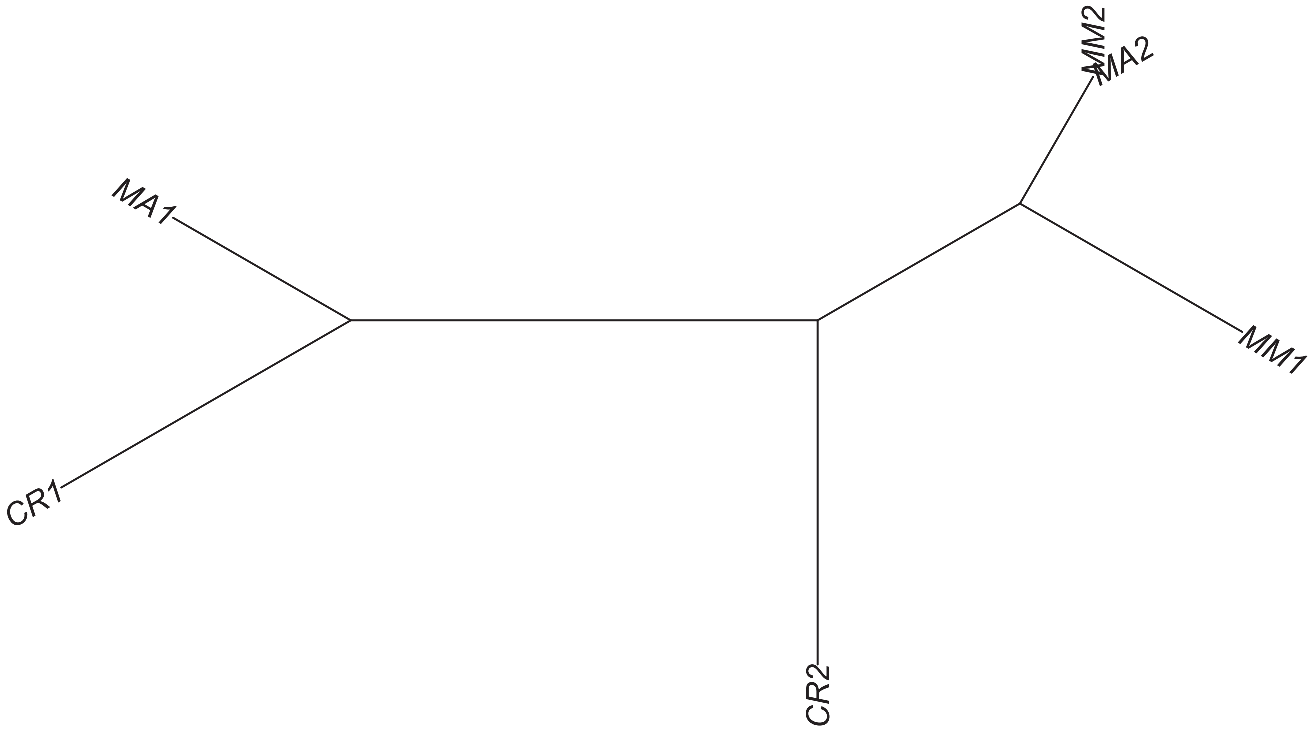

### Region 458



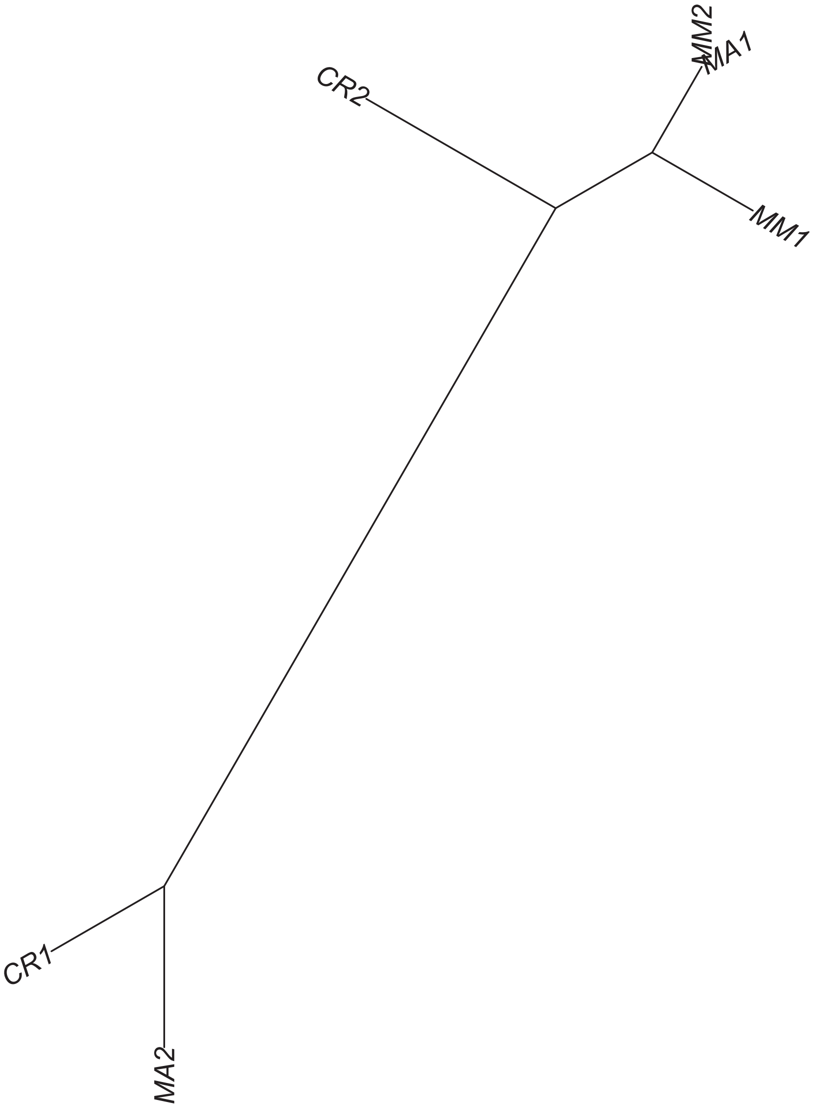

### Region 459



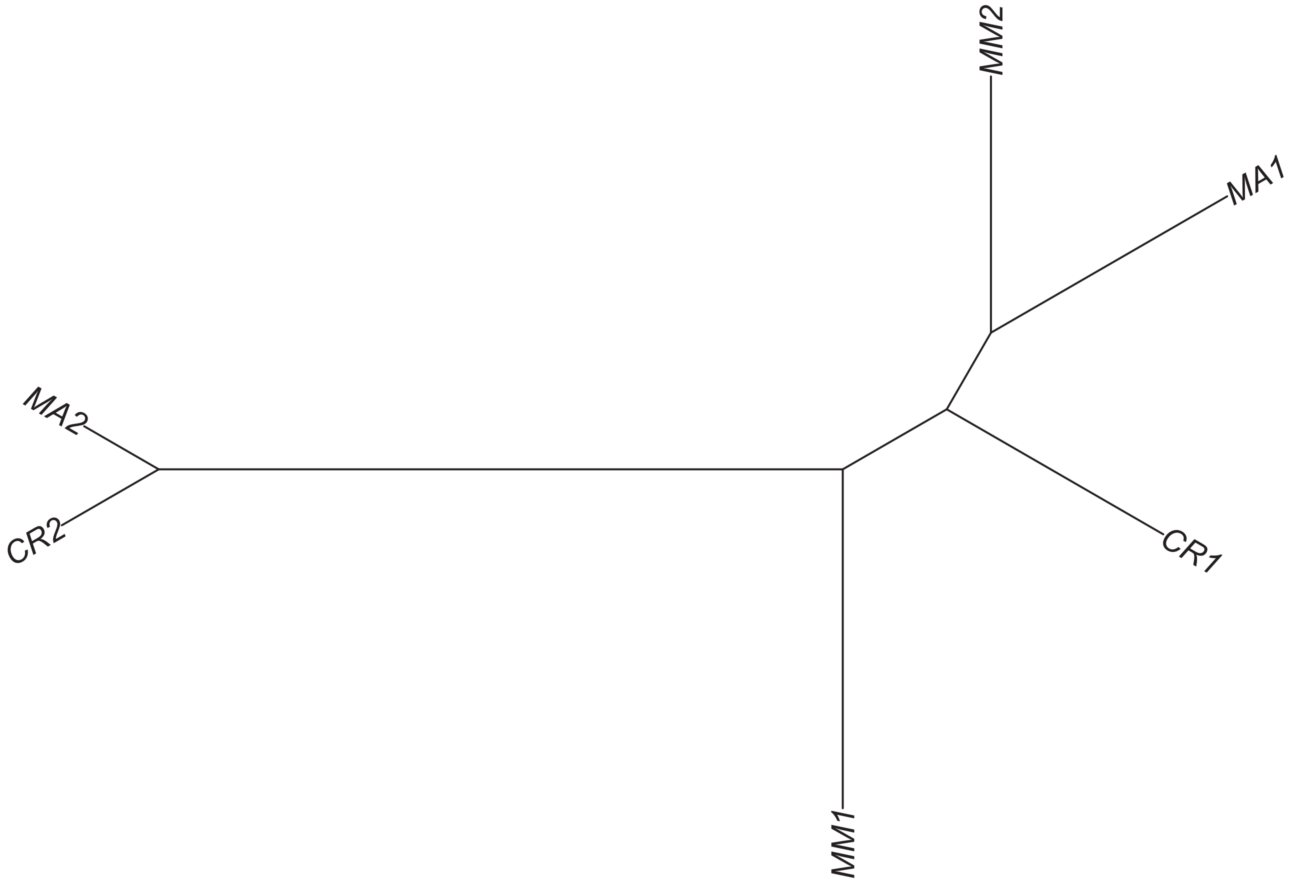

### Region 461



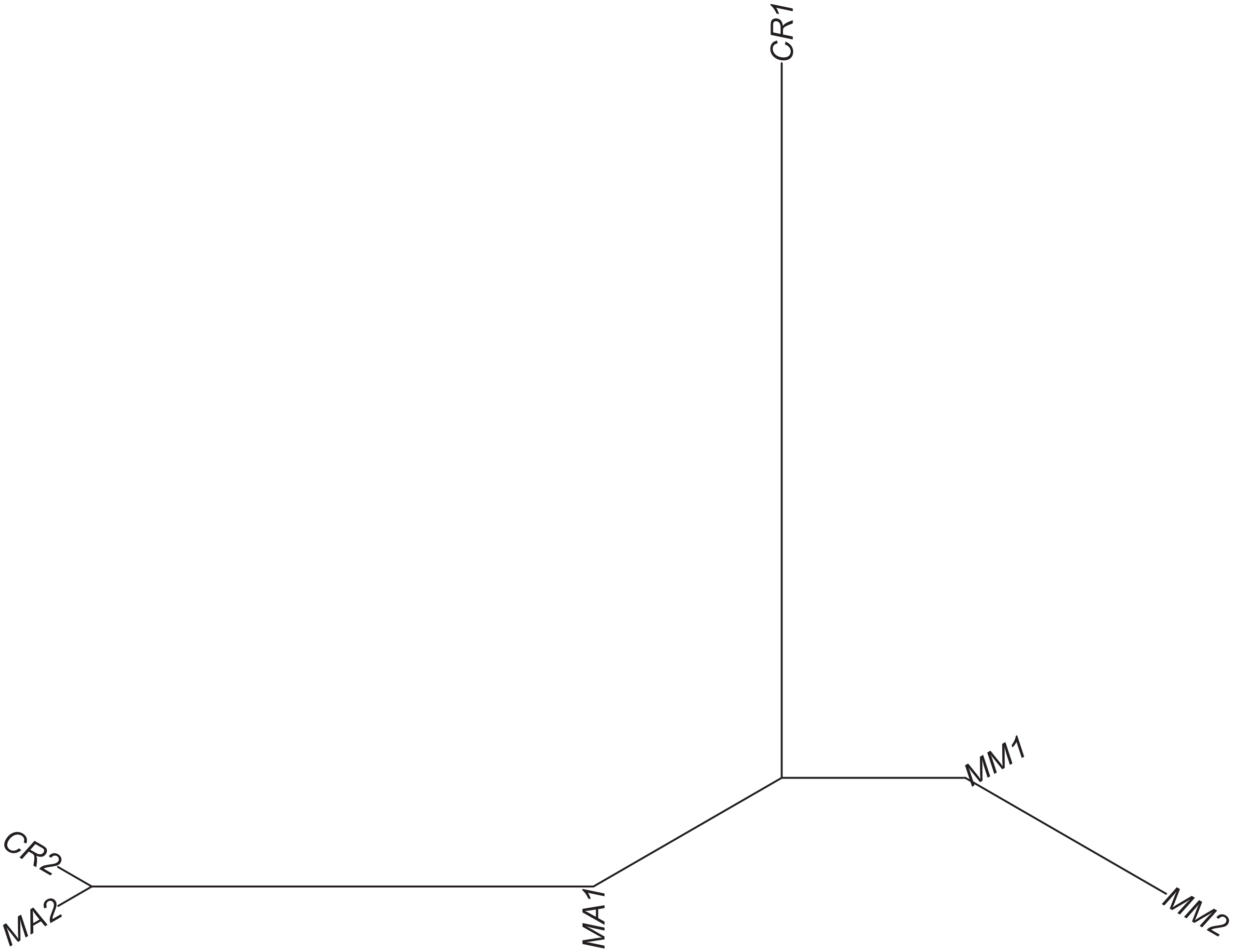

### Region 462

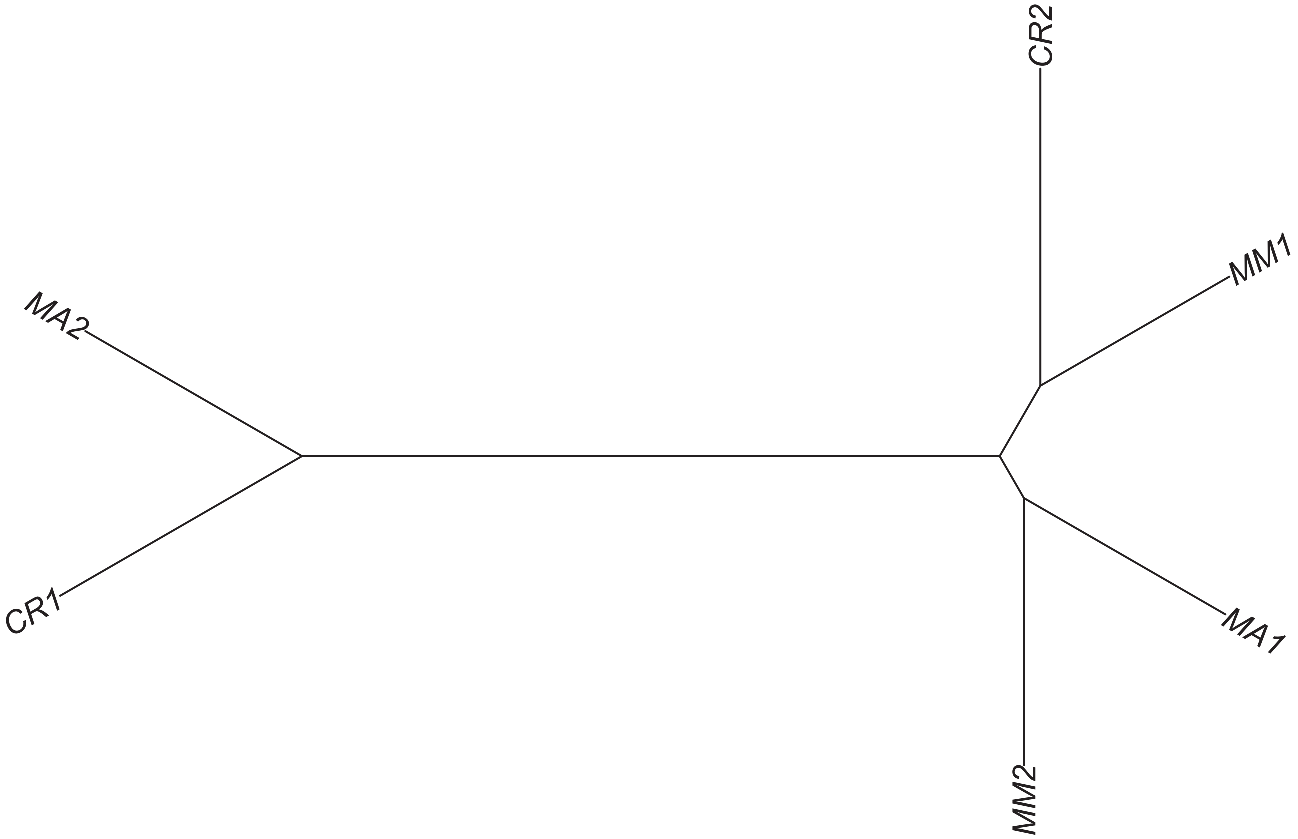

### Region 464



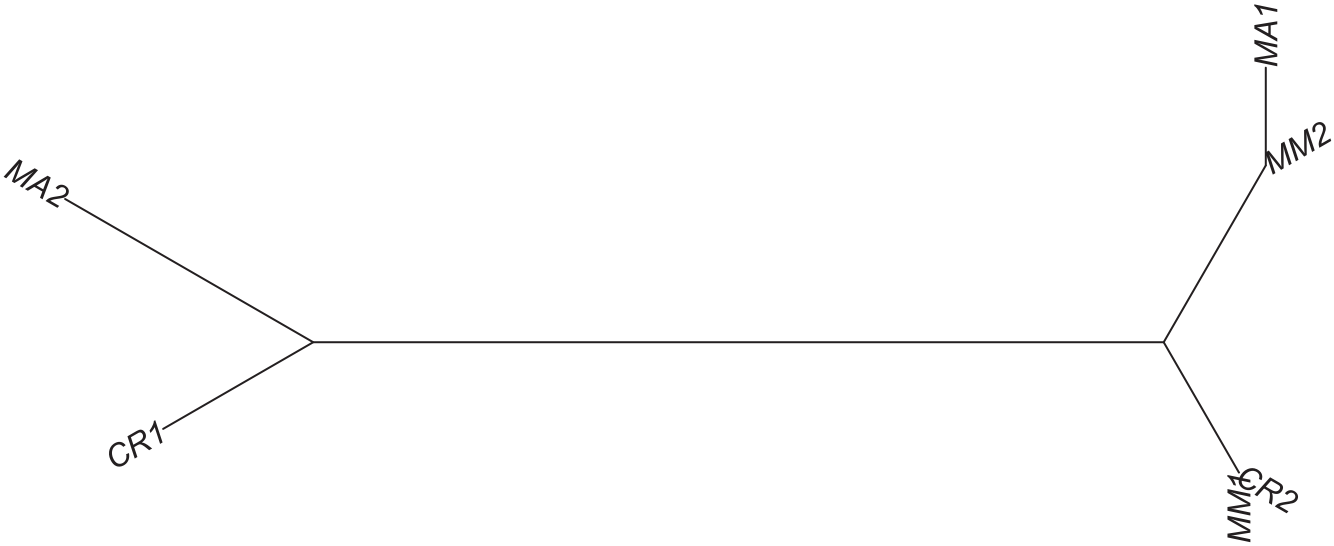

### Region 465

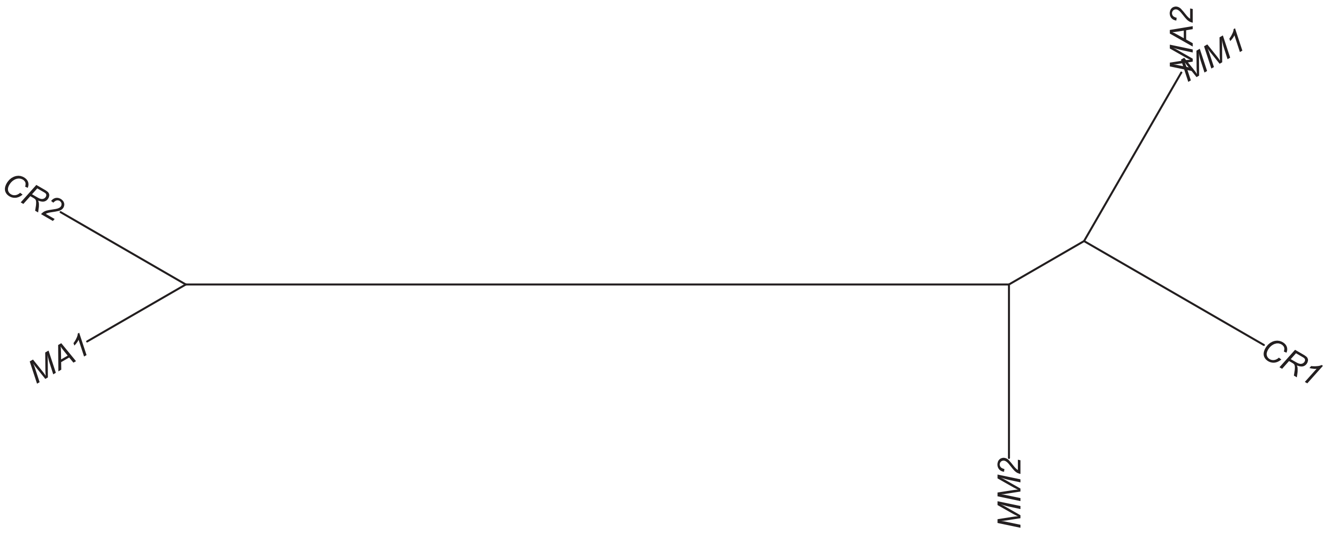

### Region 467



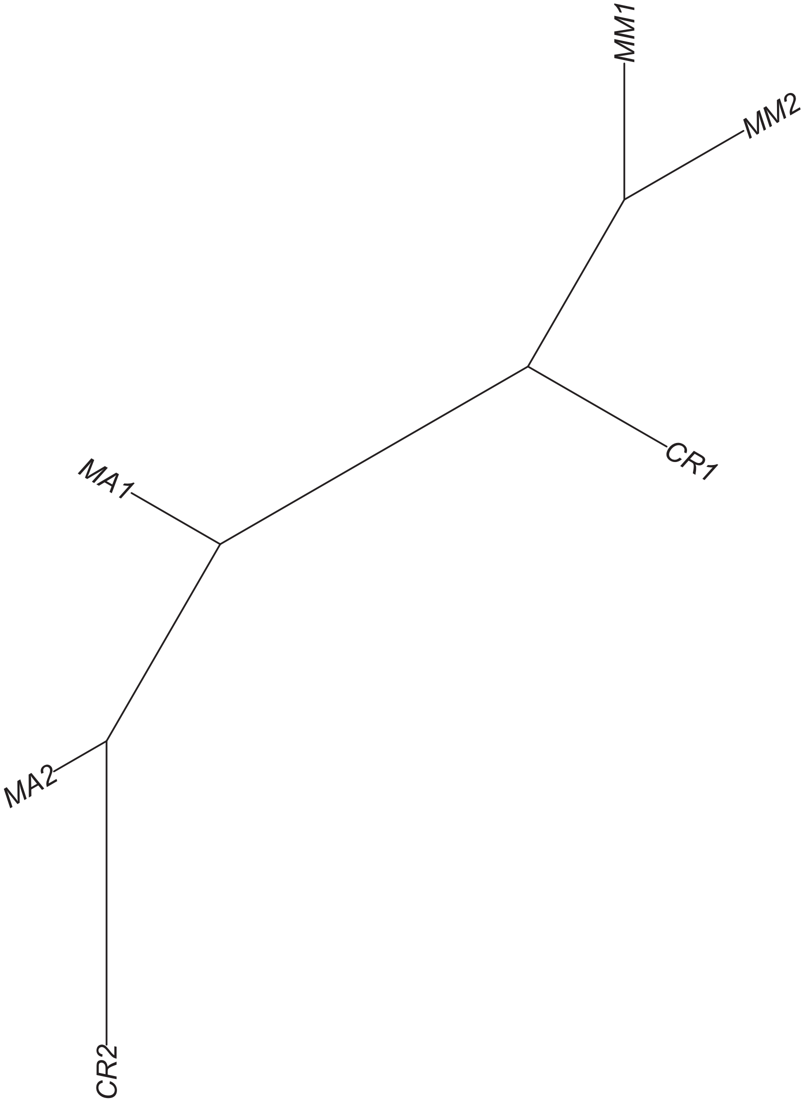

### Region 472

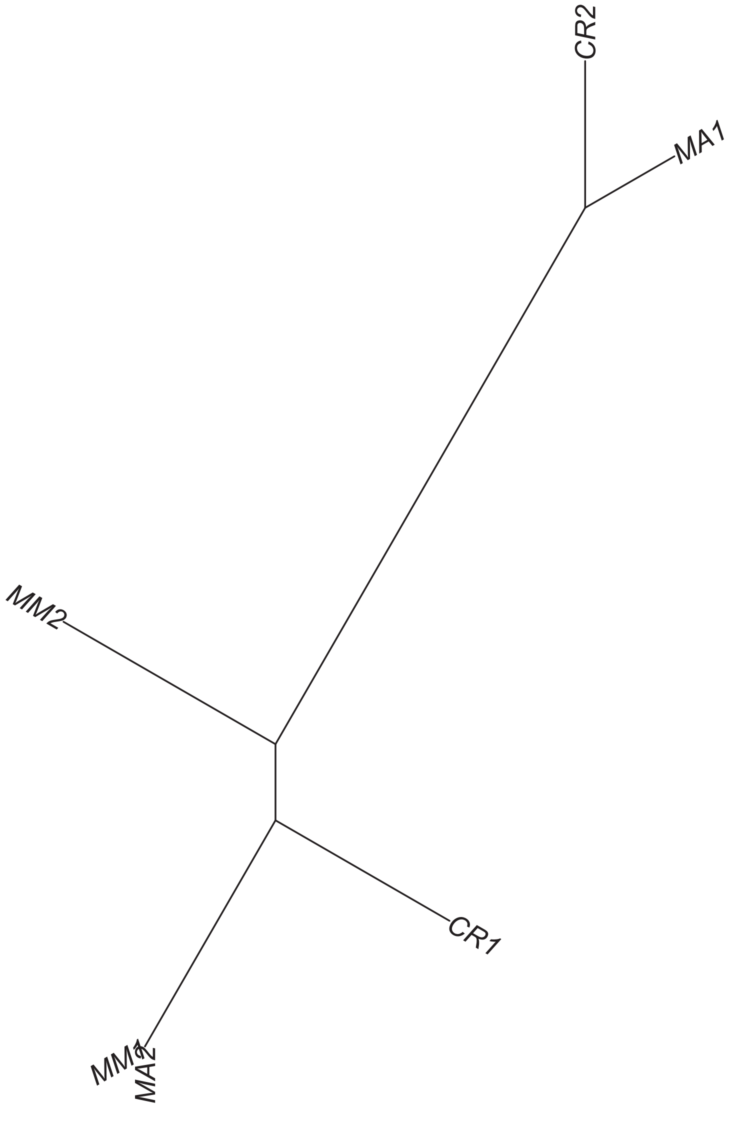

### Region 473

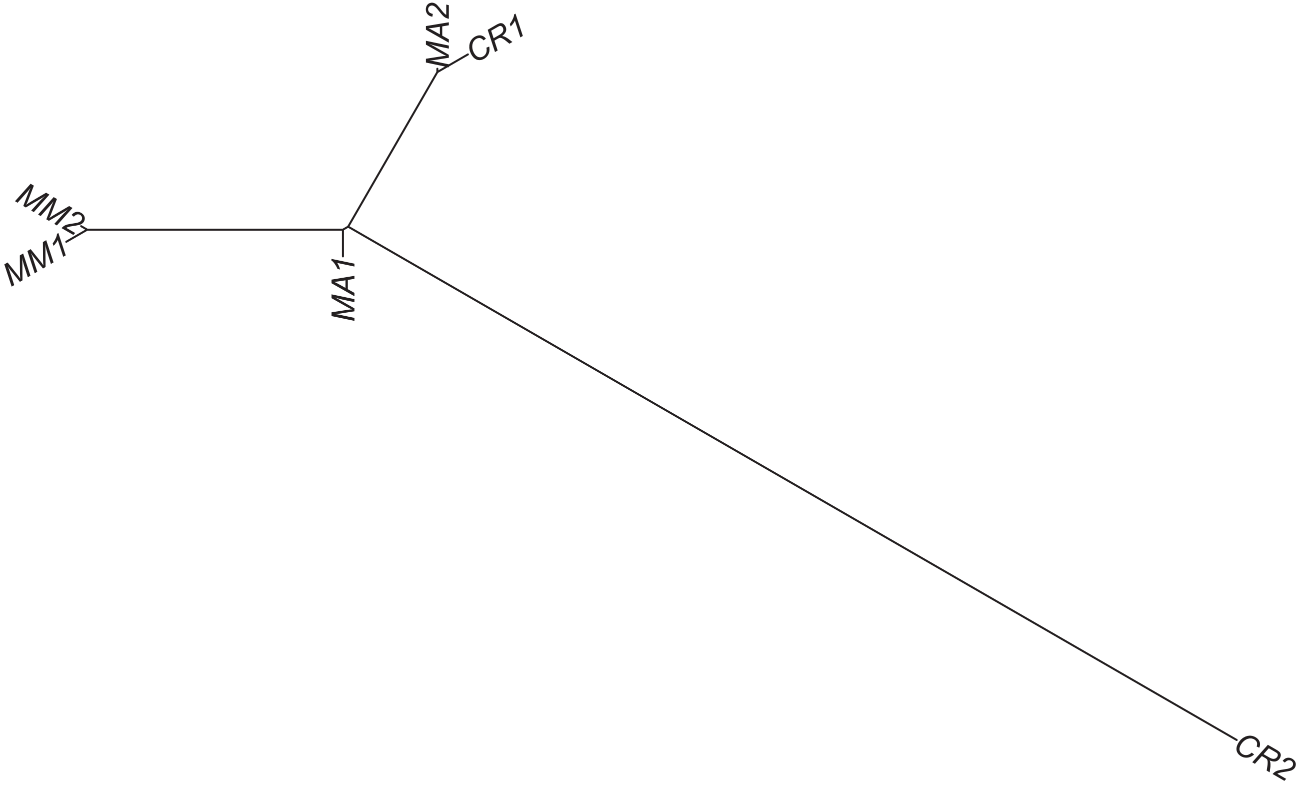

Region 475



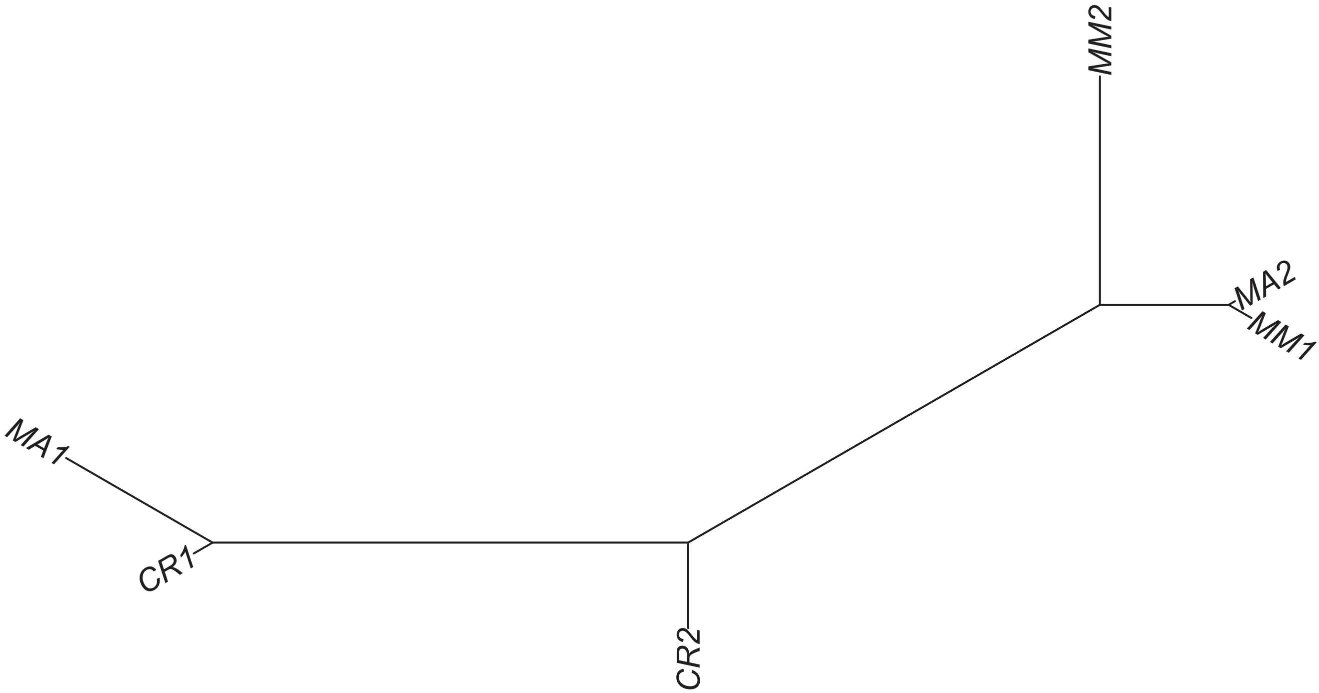

Region 476

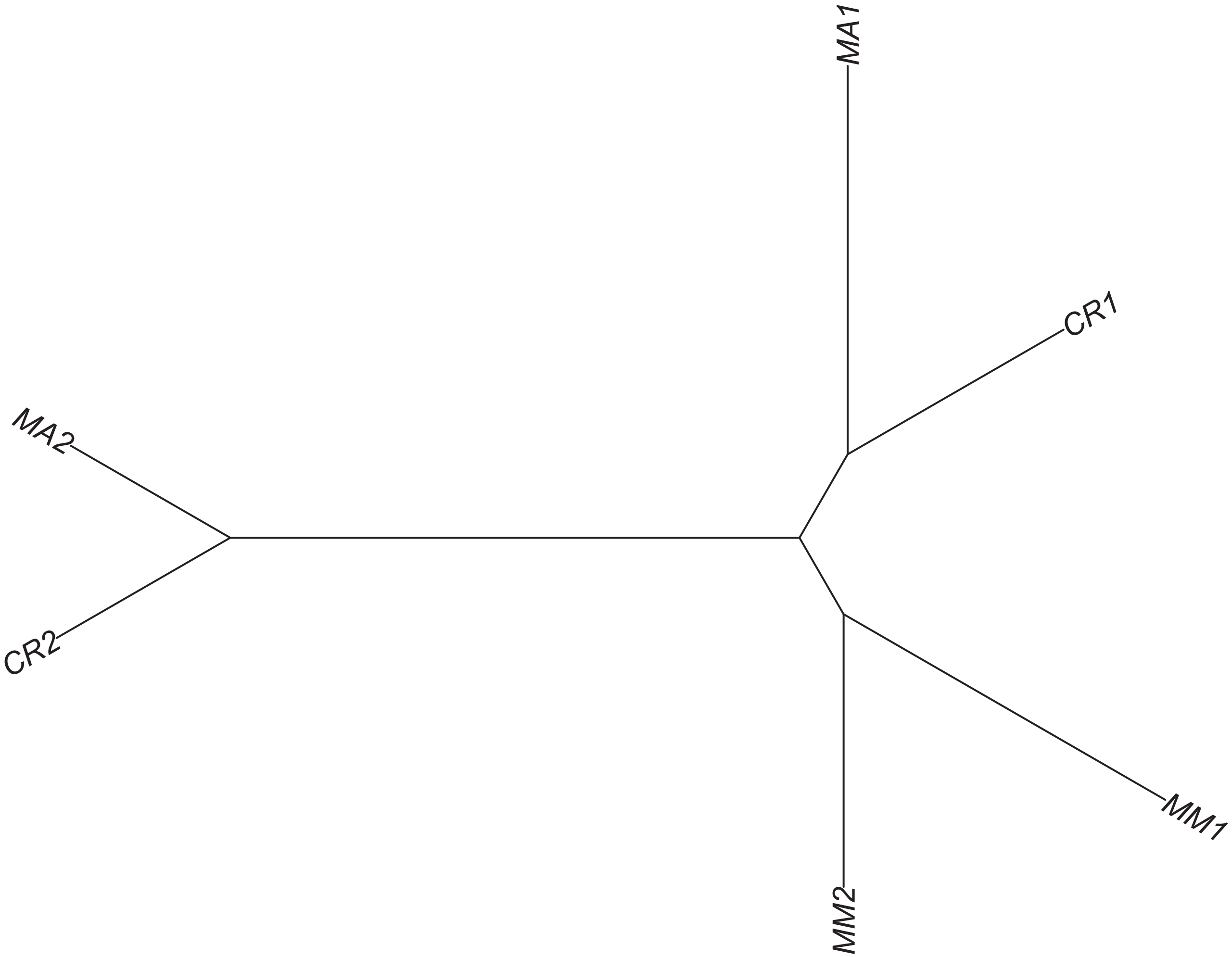

### Region 477

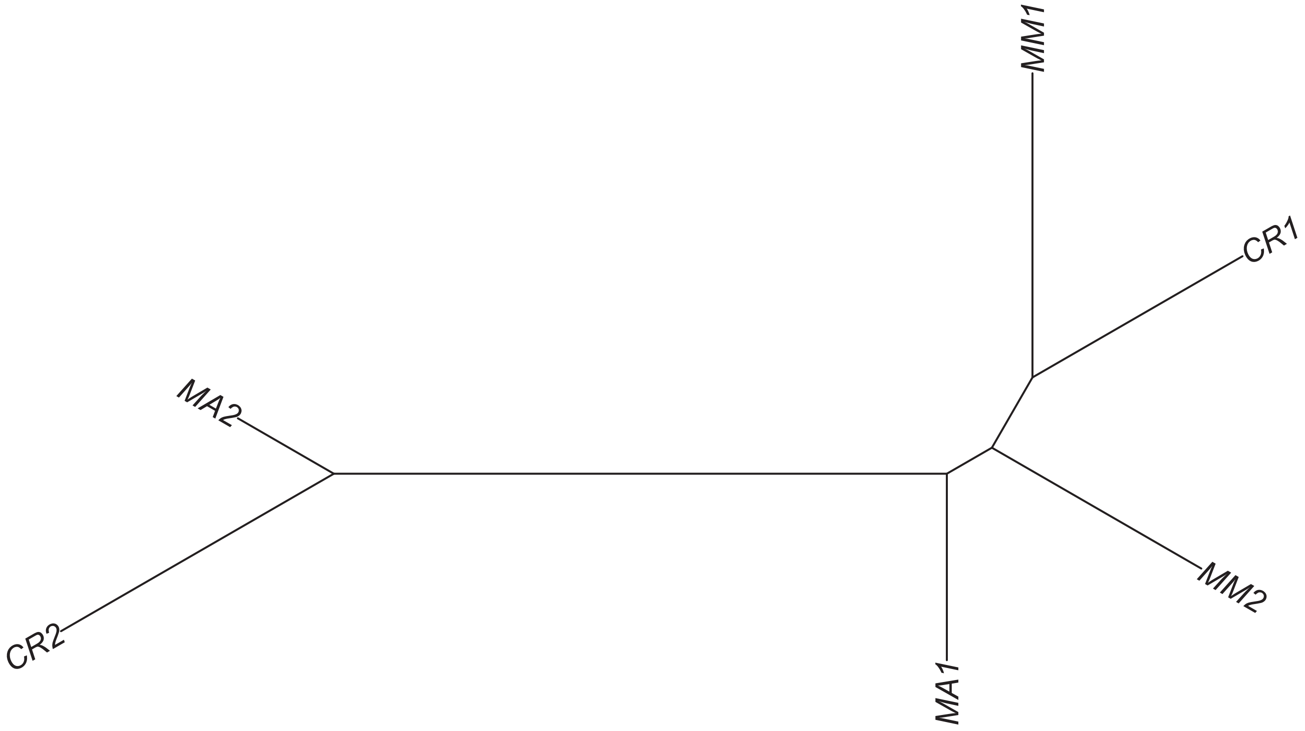

#### Region 478

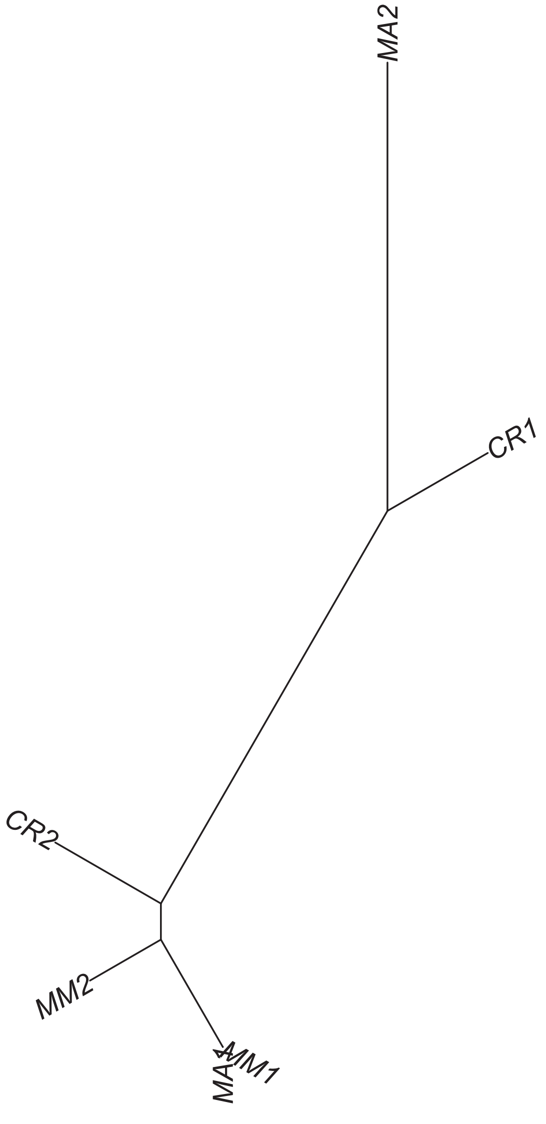

### Region 480



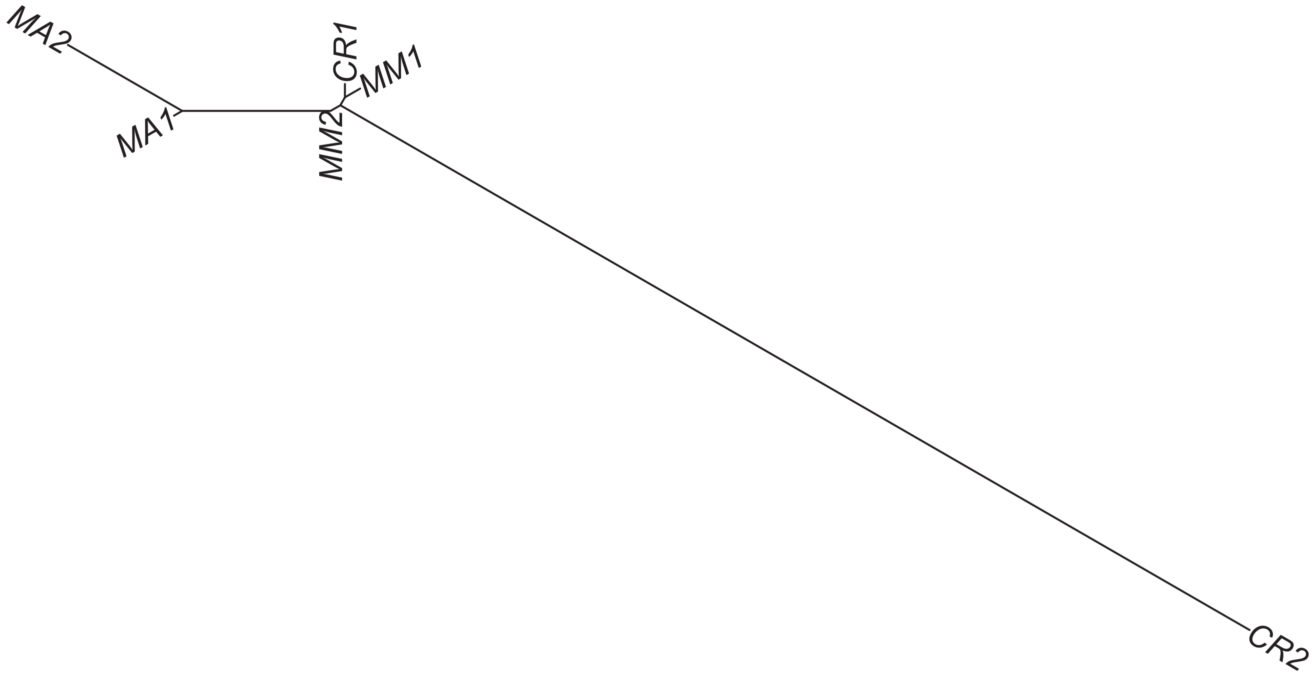

Region 481

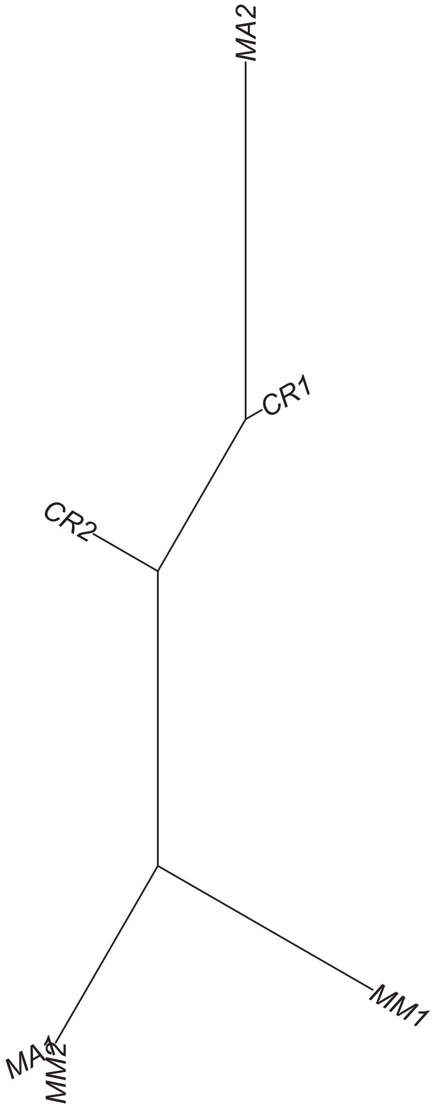

### Region 482



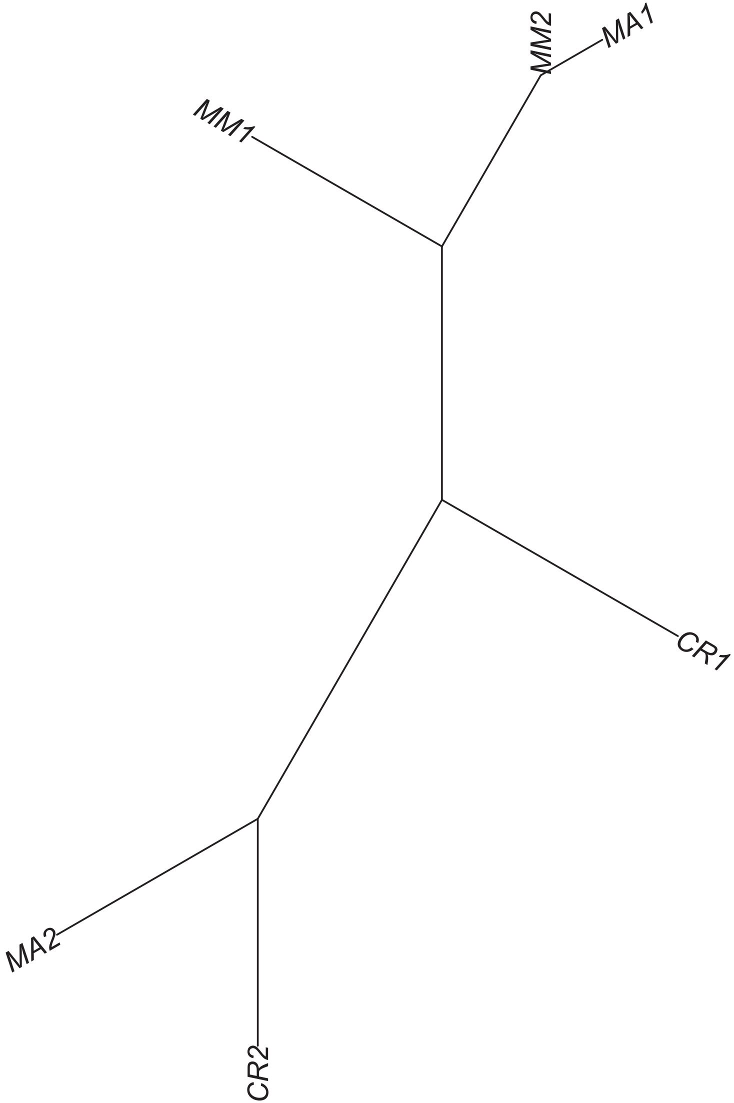

### Region 483



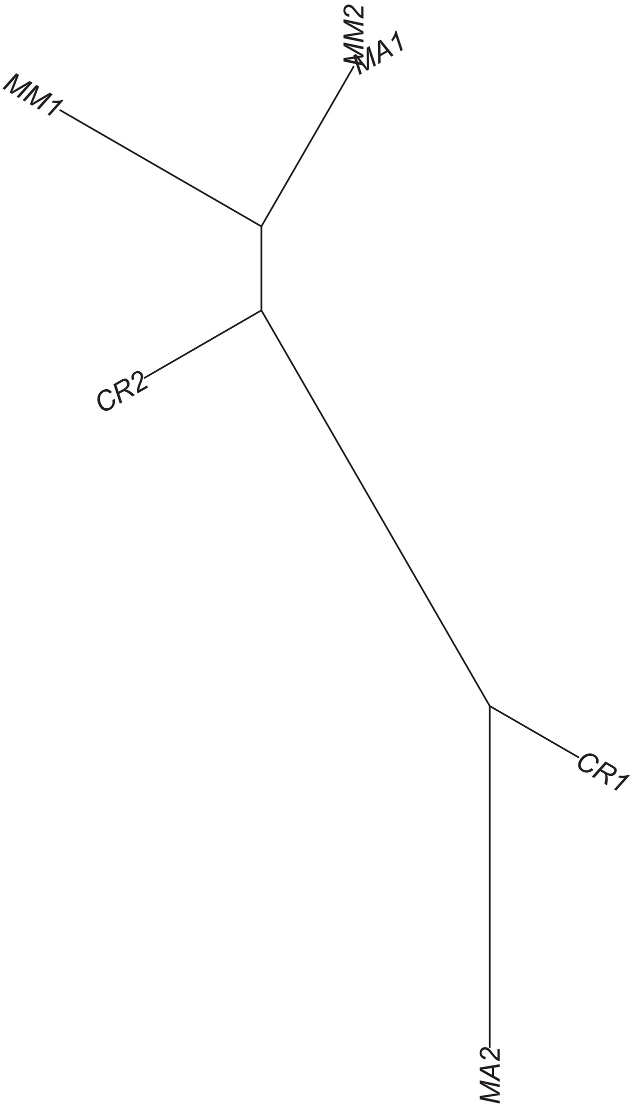

### Region 484



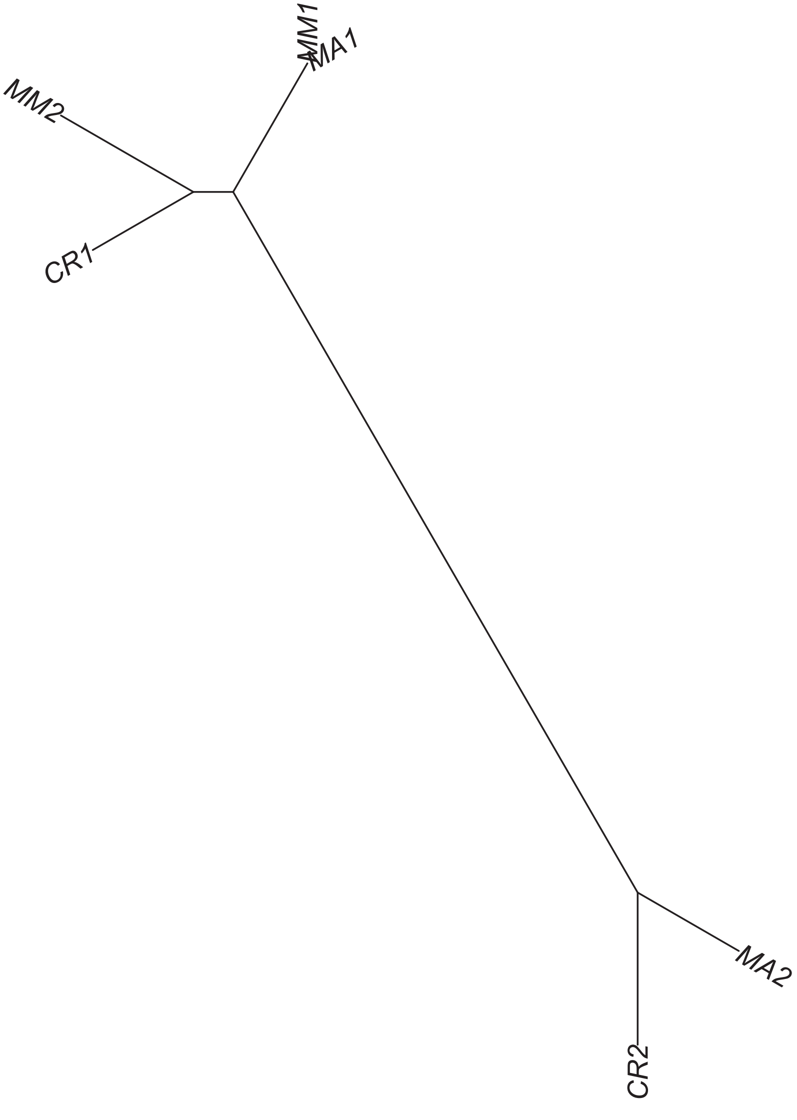

### Region 485



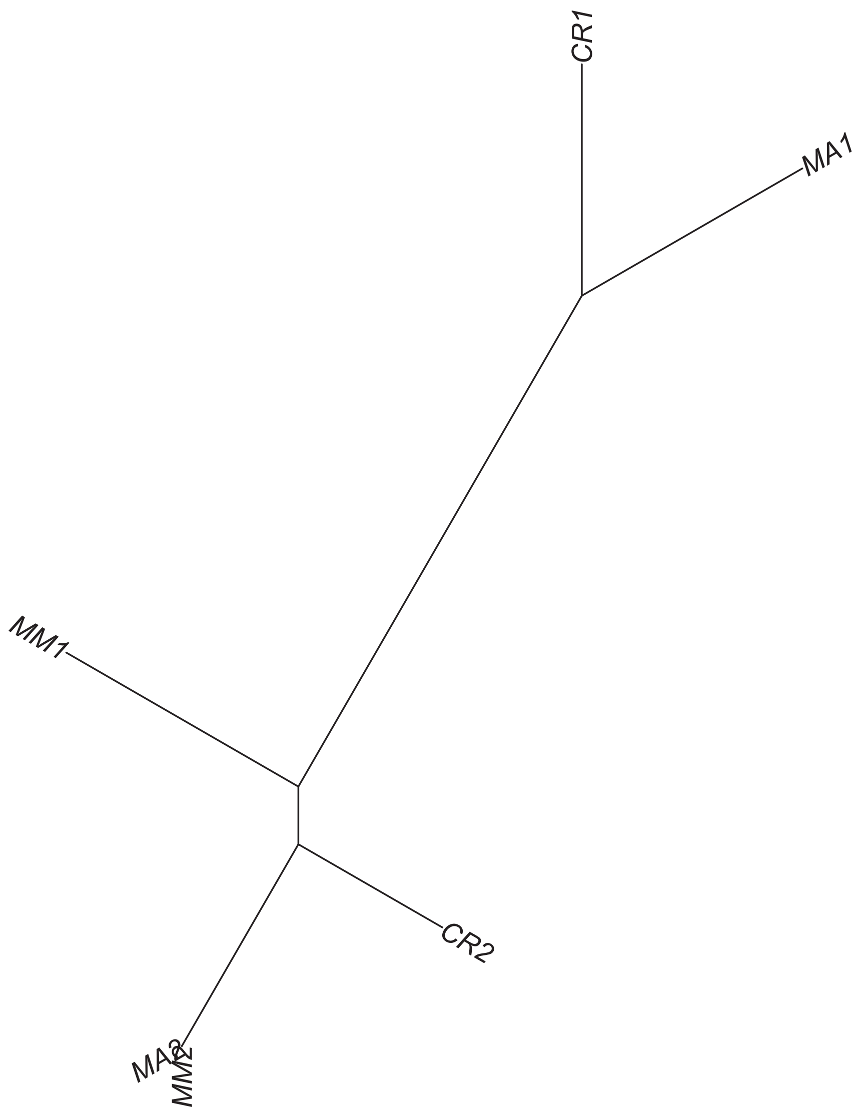

Region 486



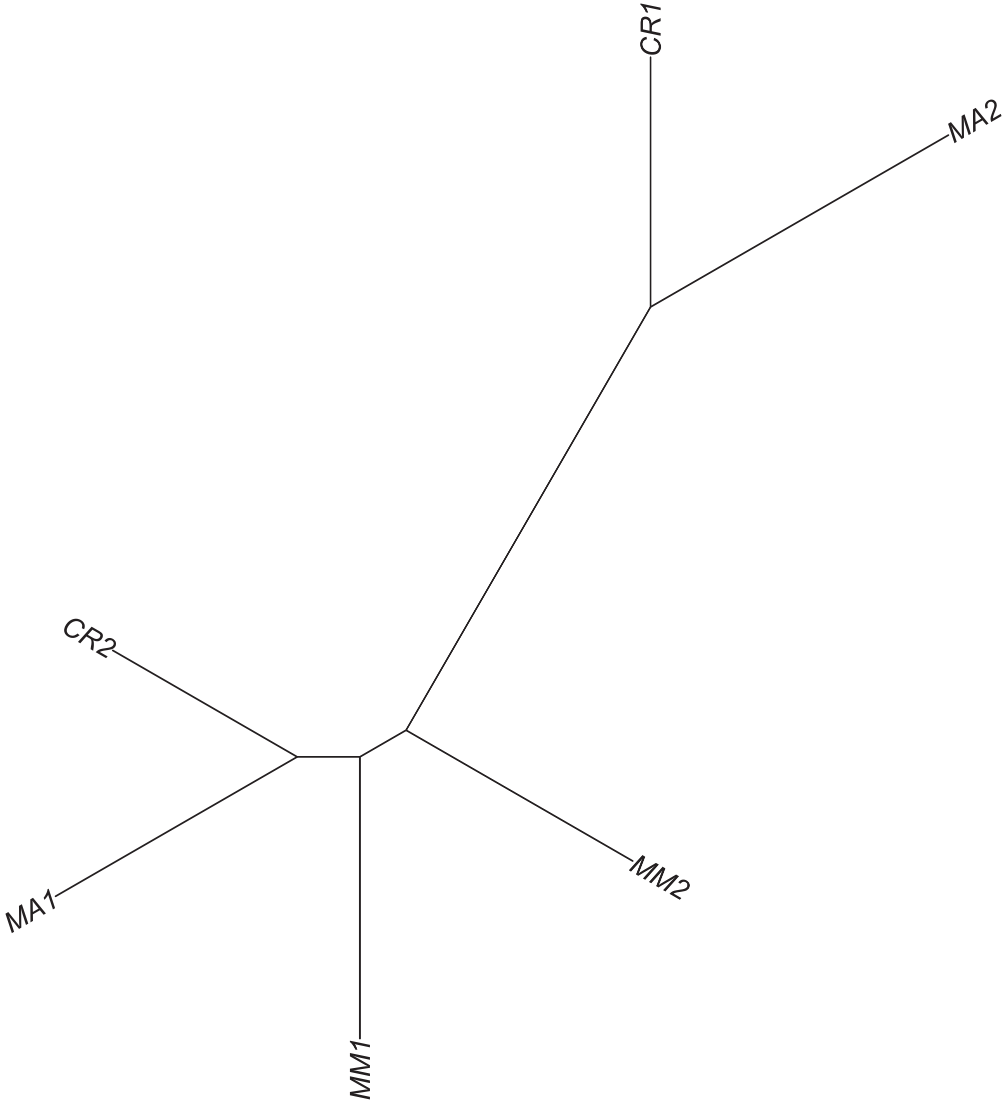

### Region 487

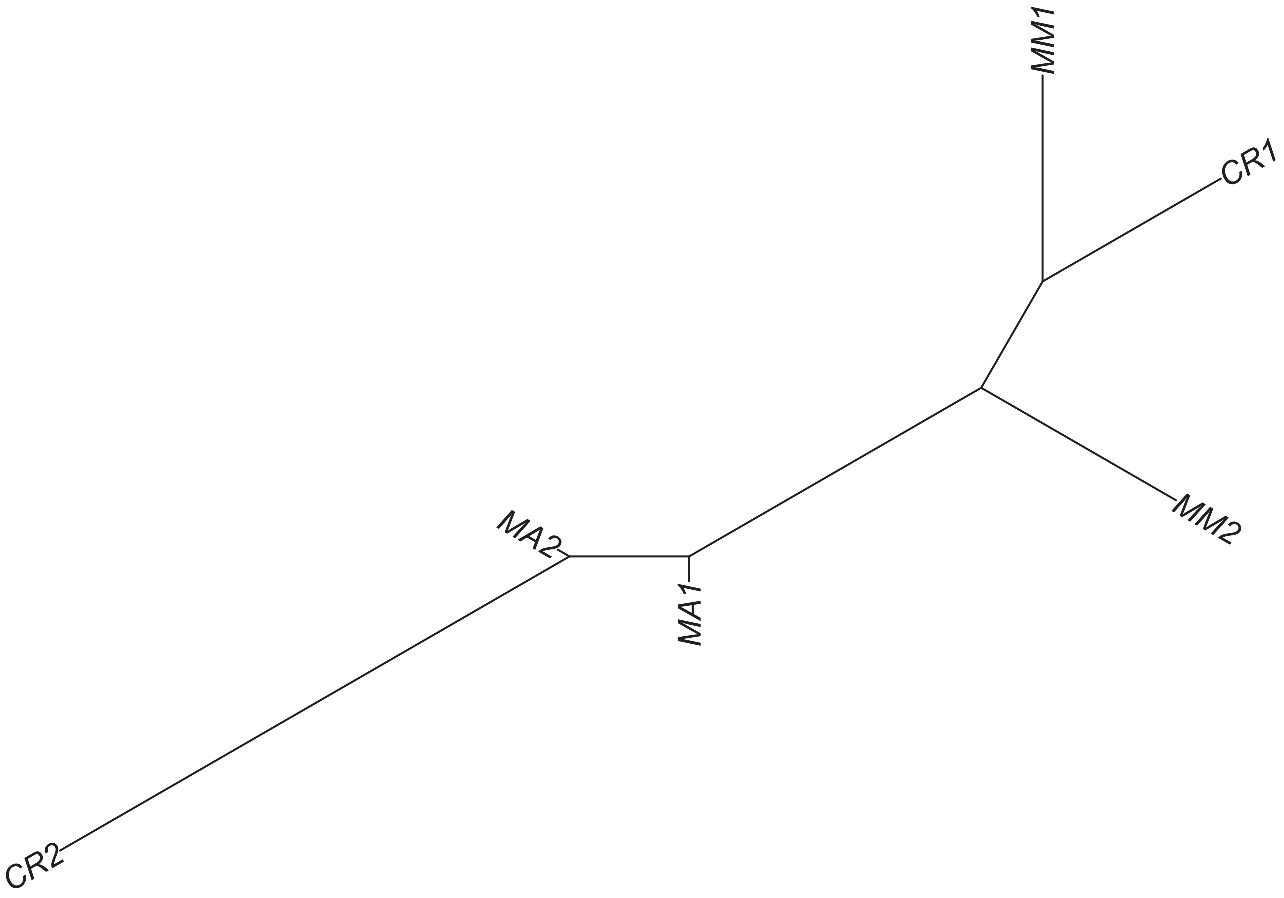

### Region 488



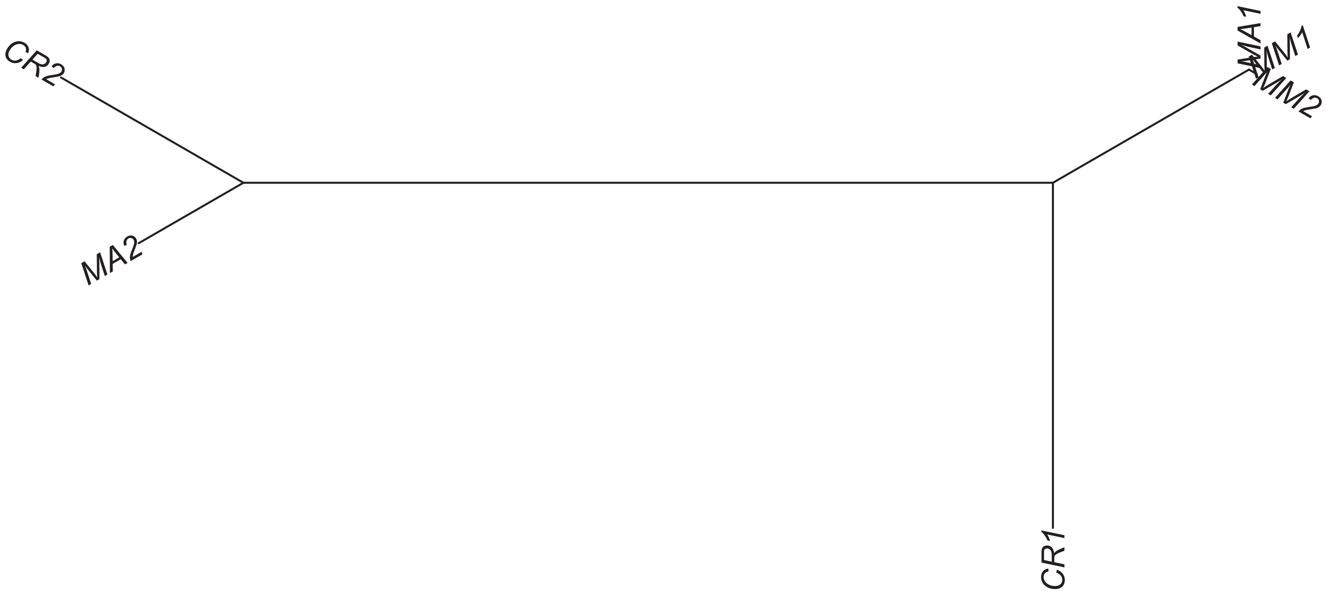

### Region 491

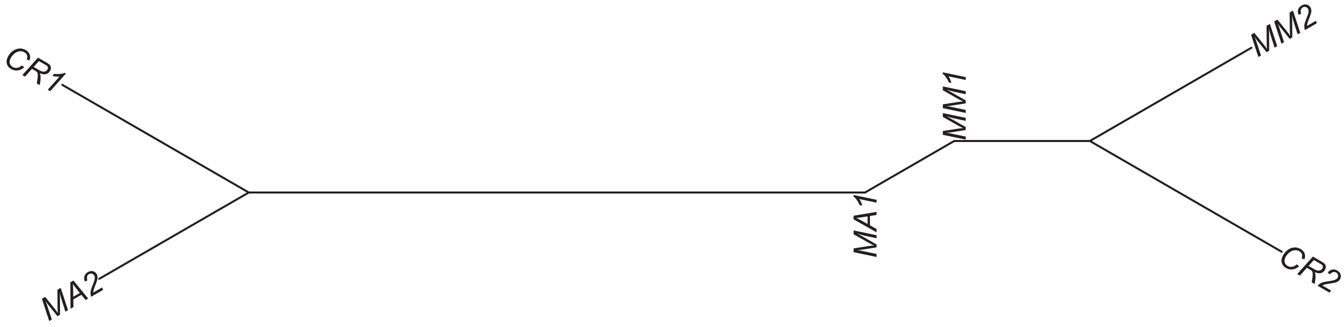

### Region 492

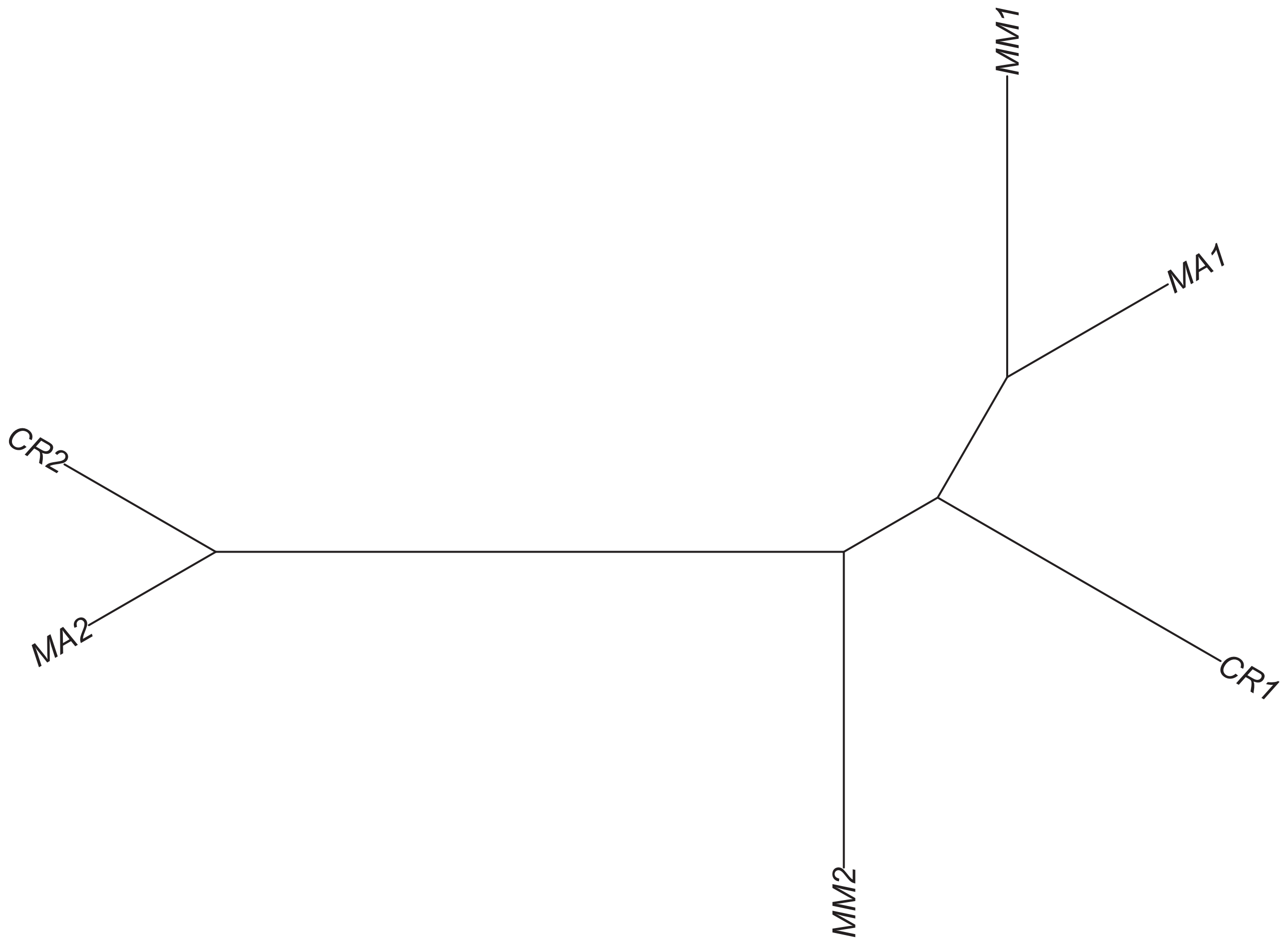

### Region 493



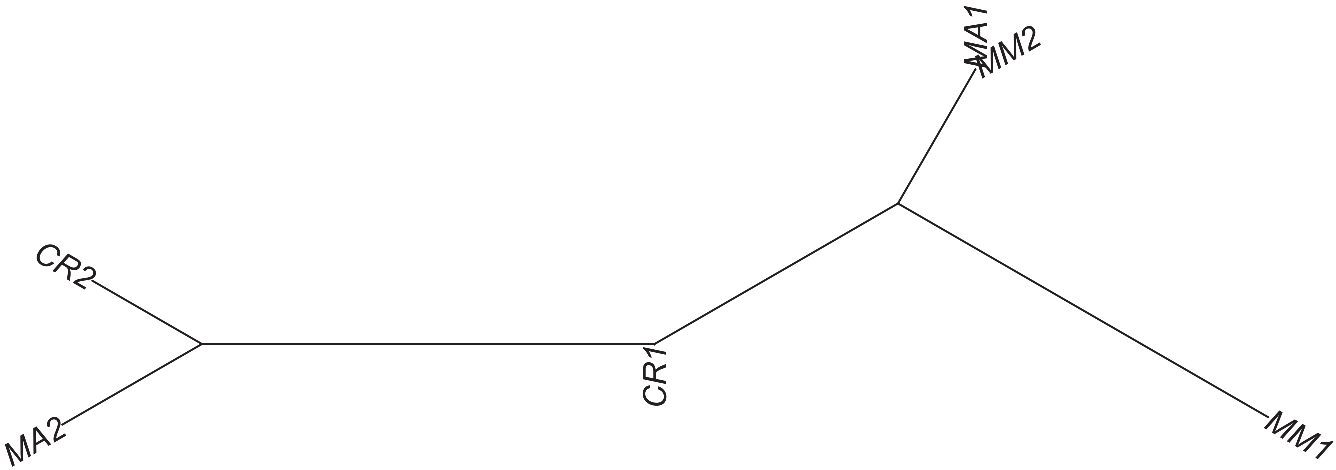

### Region 494



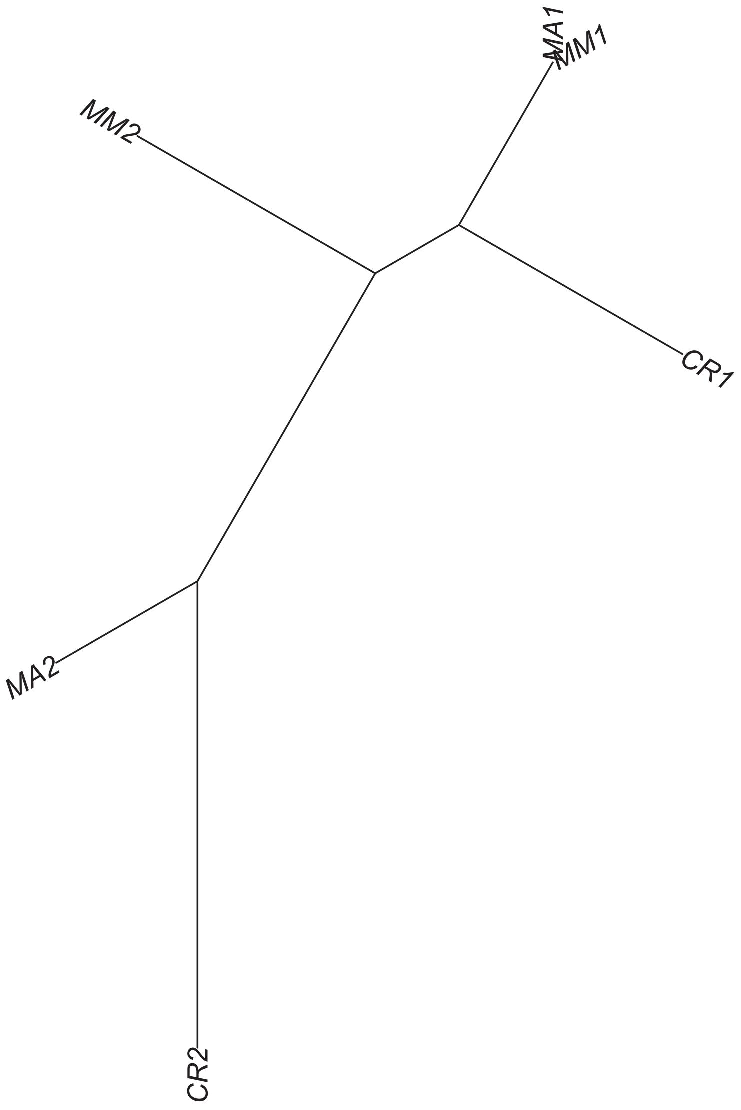

### Region 496



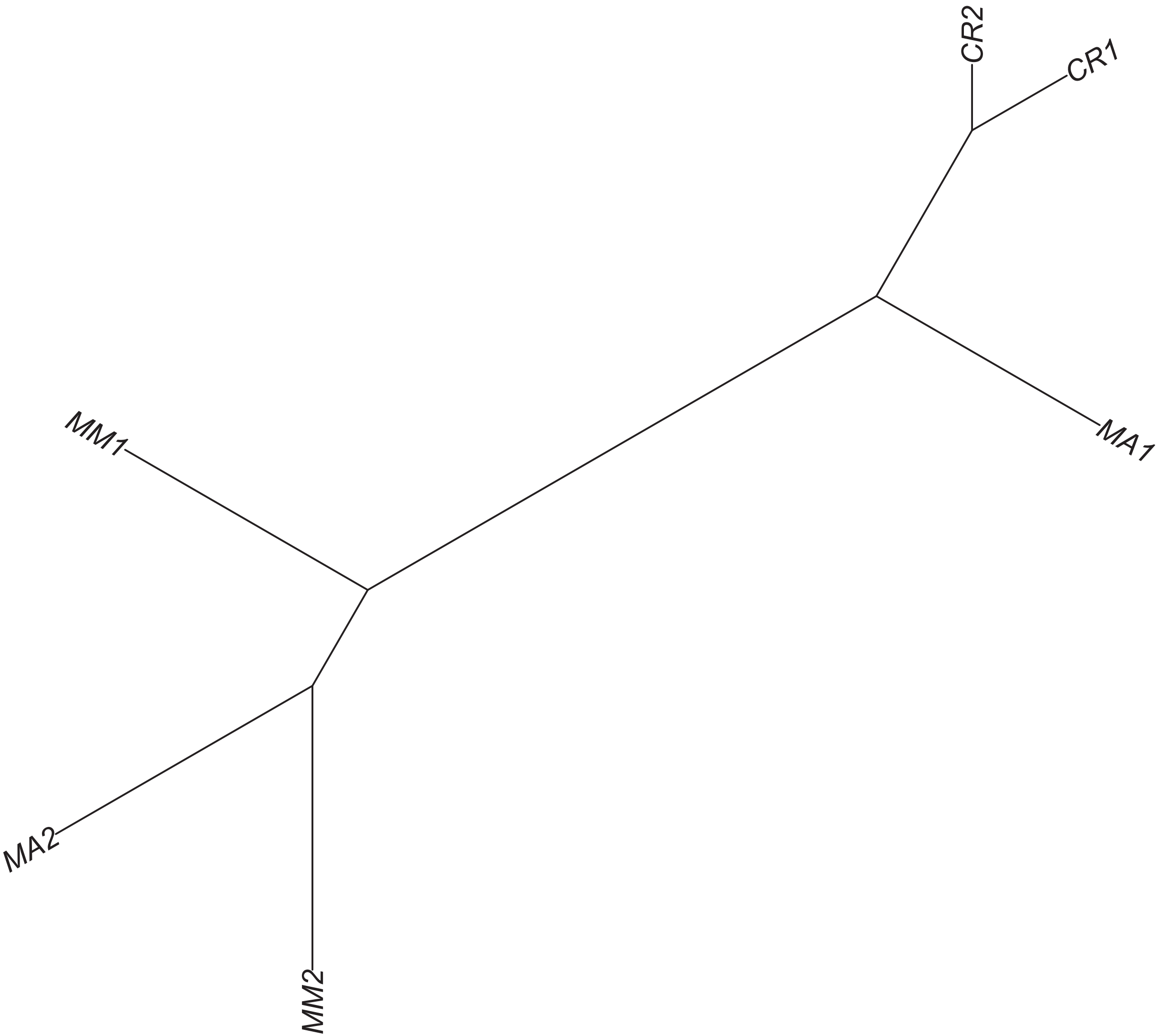

### Region 500

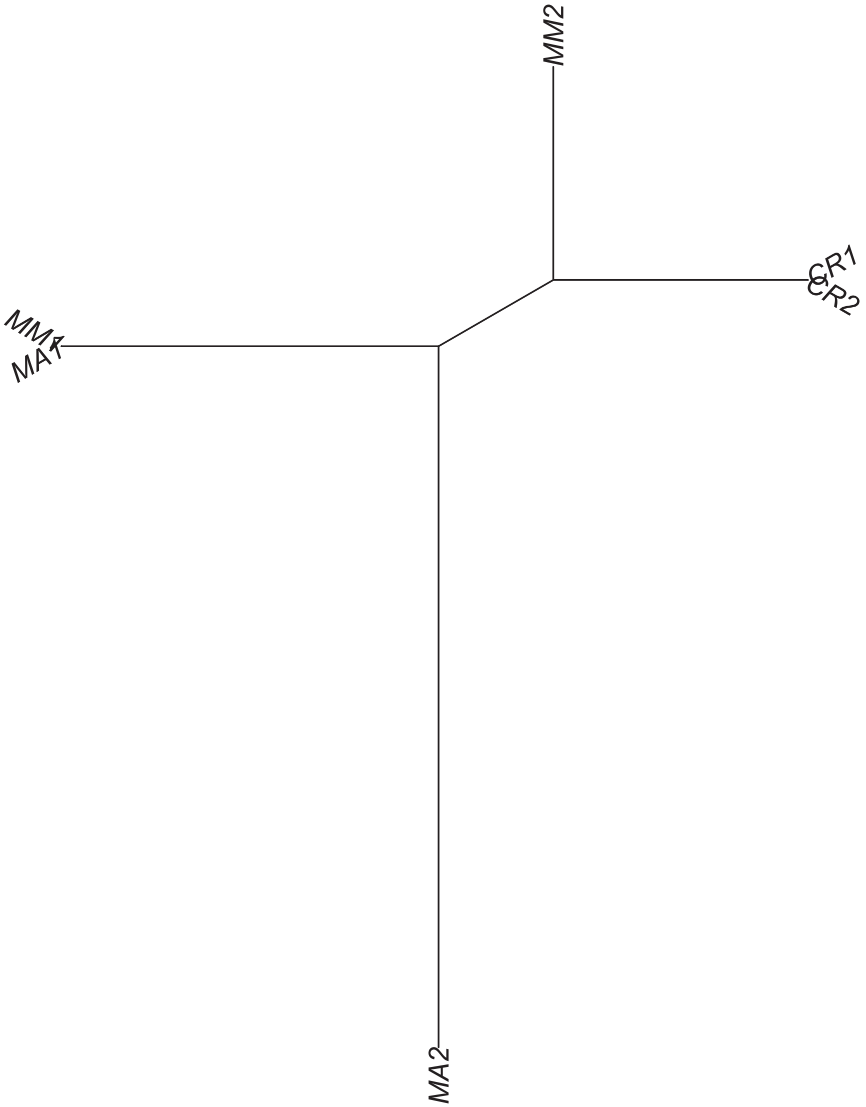

### Region 501



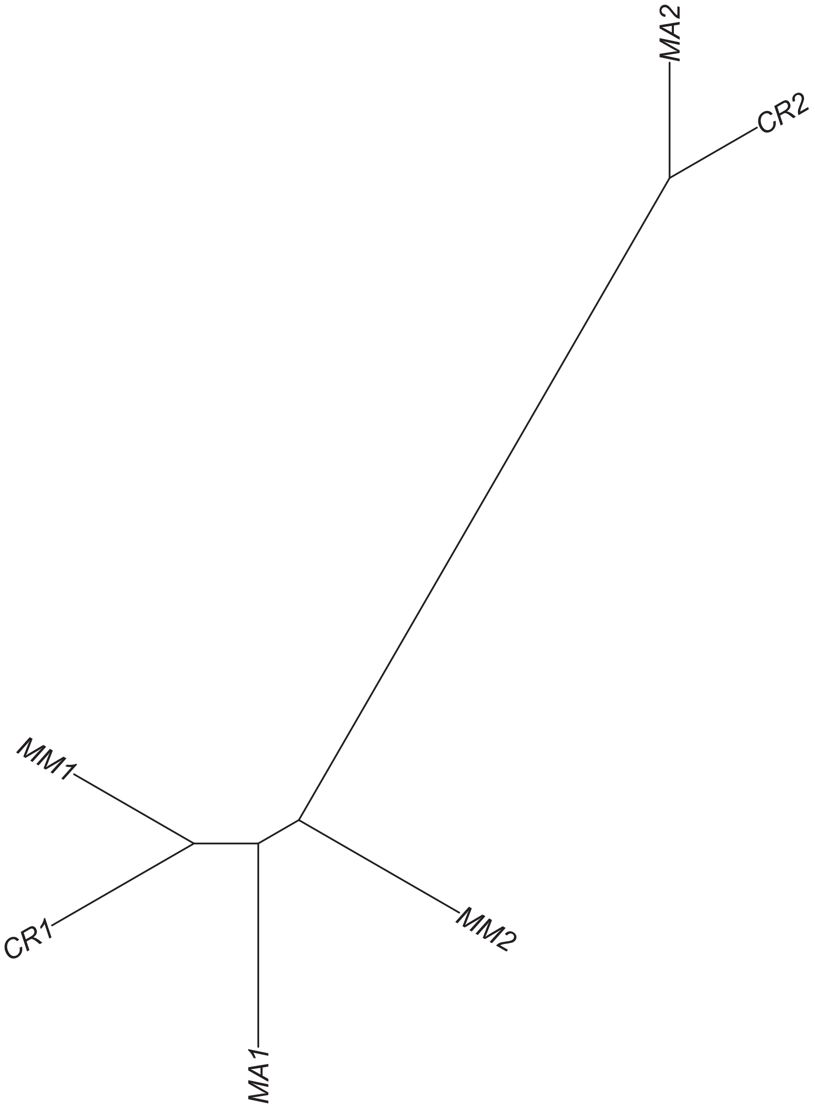

Region 502

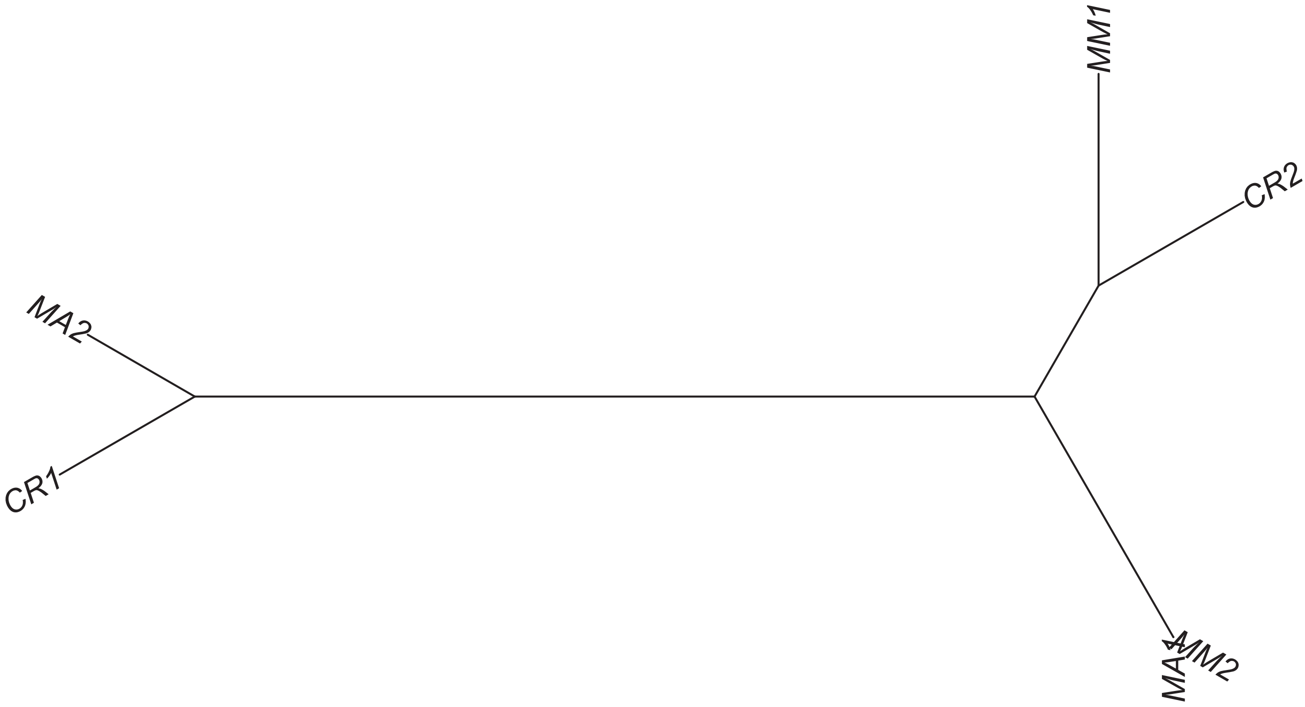

### Region 503



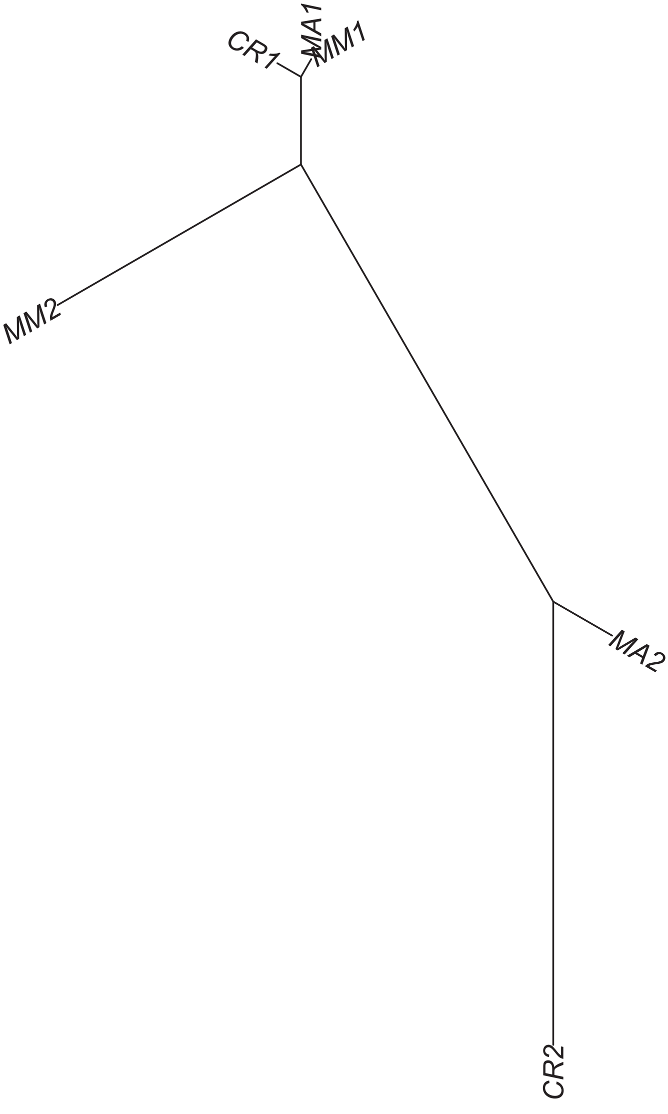

### Region 505

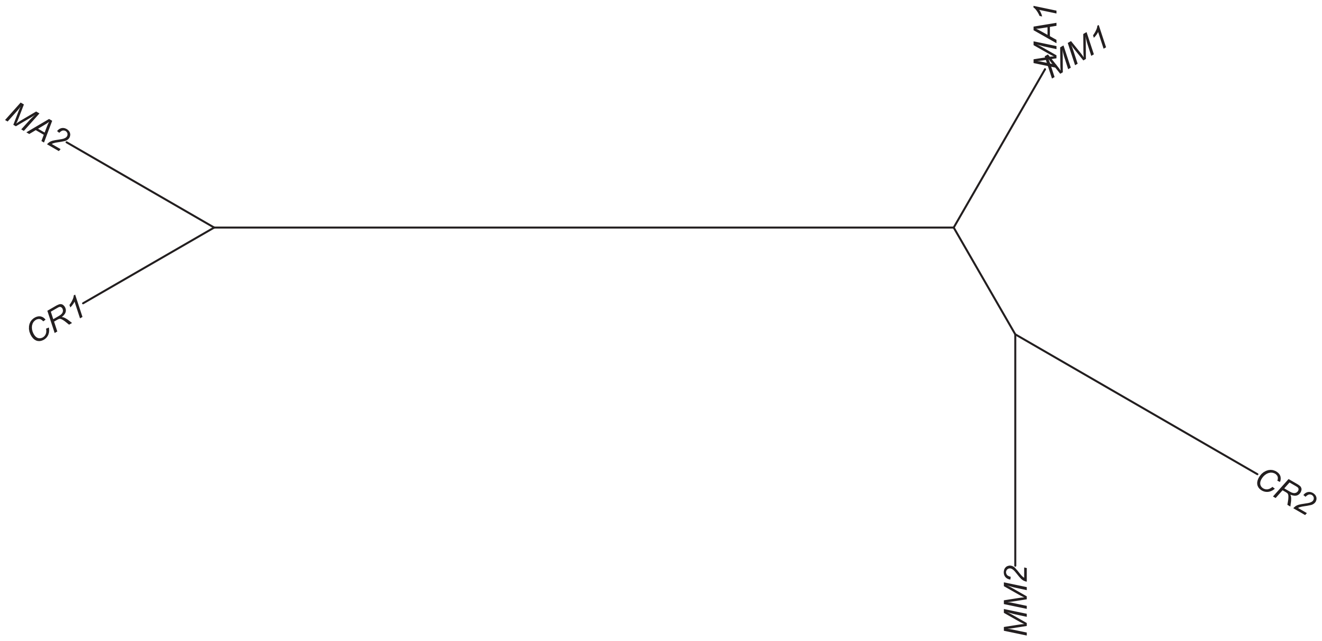

### Region 507



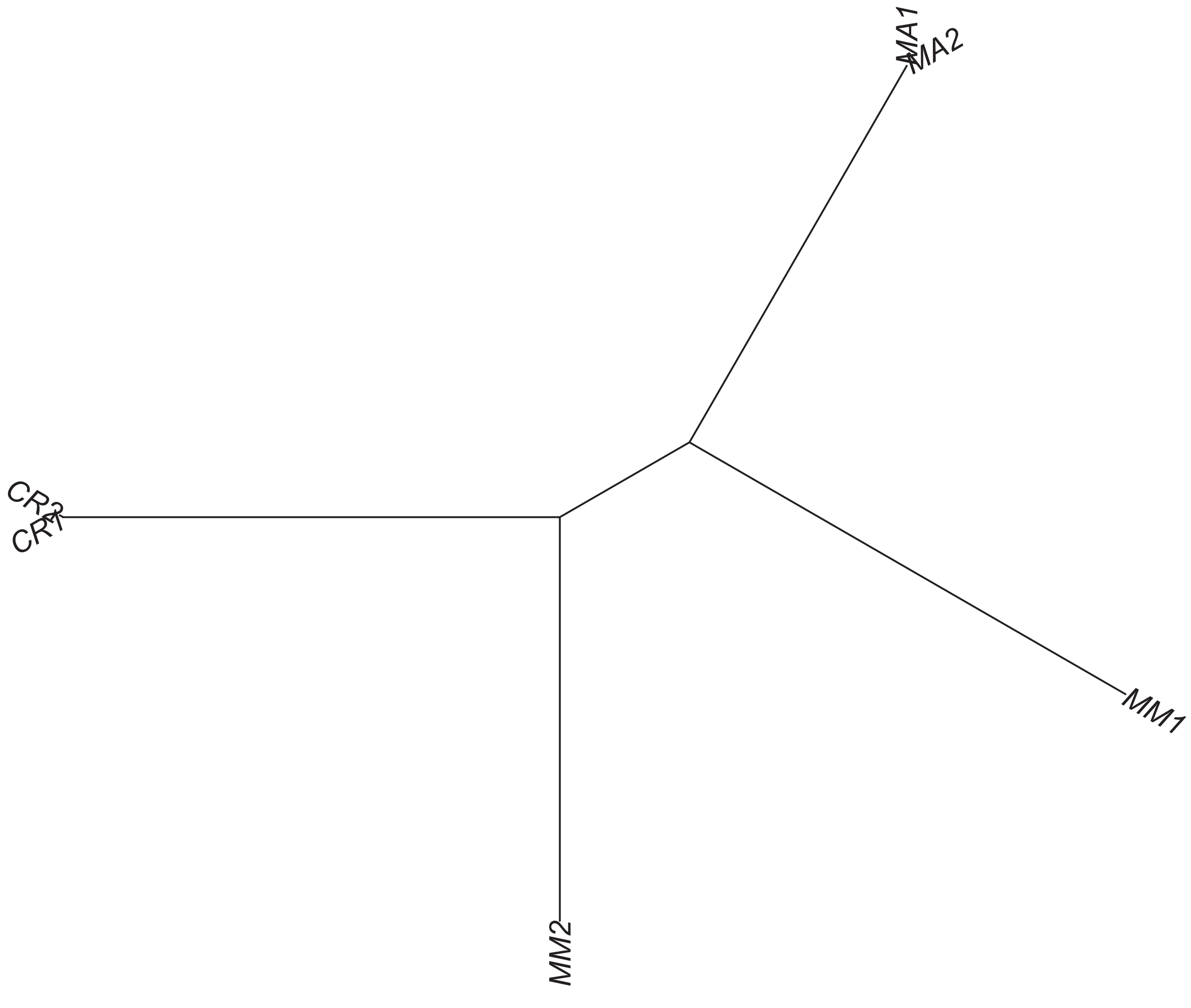

### Region 510

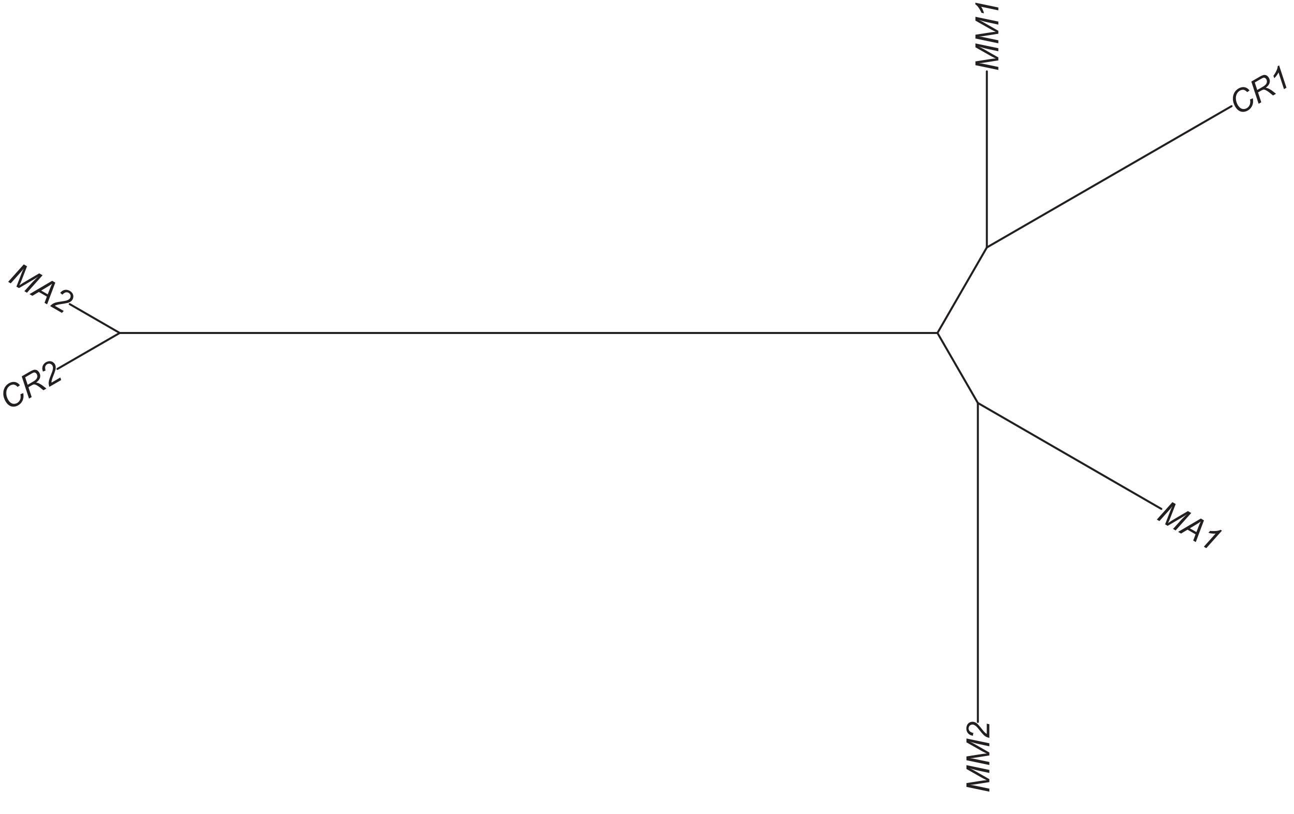

### Region 511



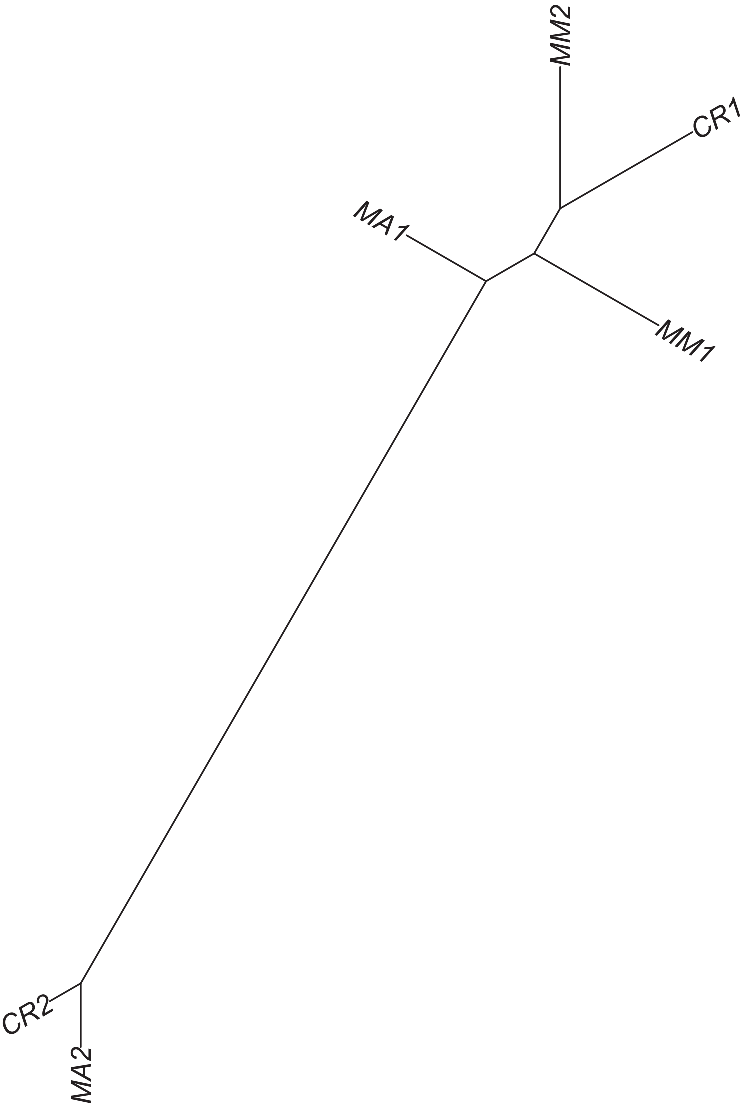

### Region 512



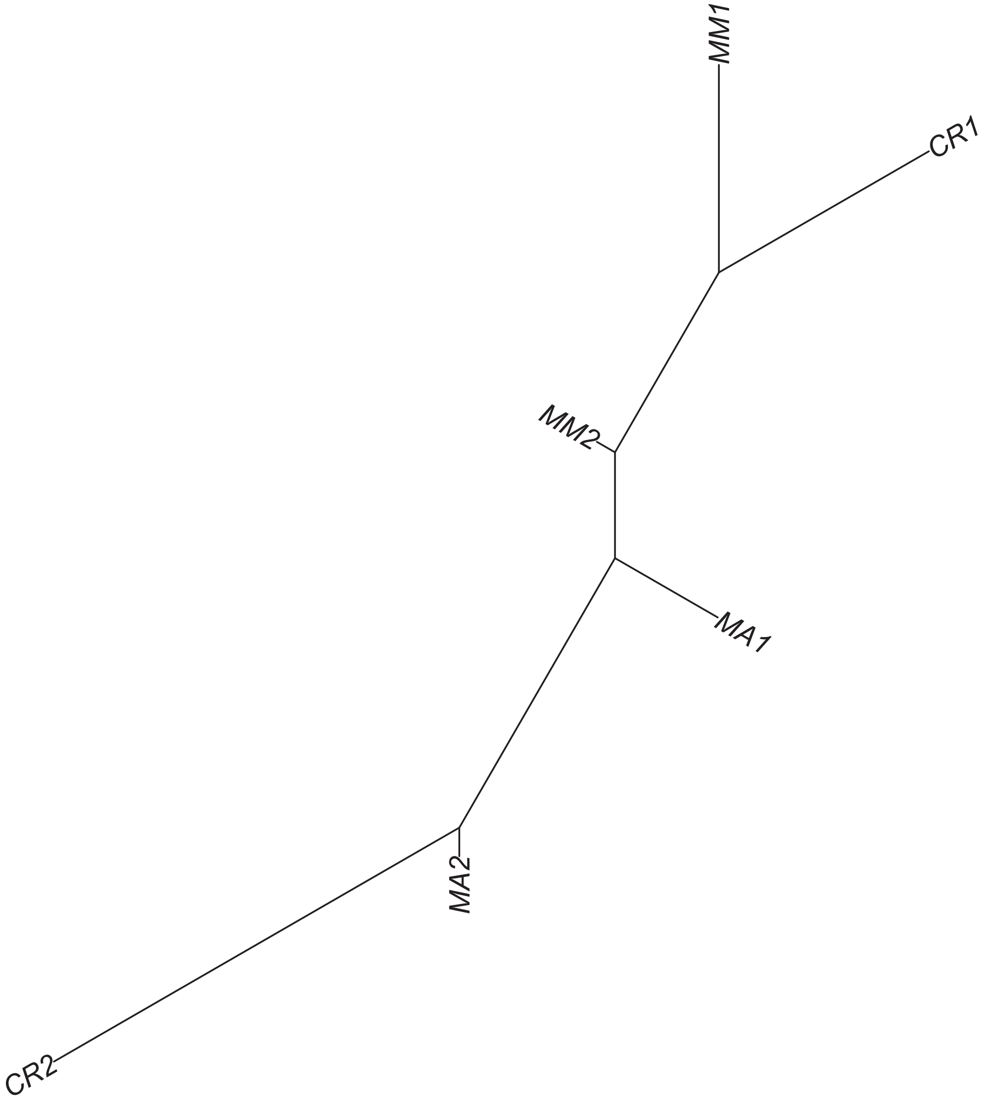

### Region 515



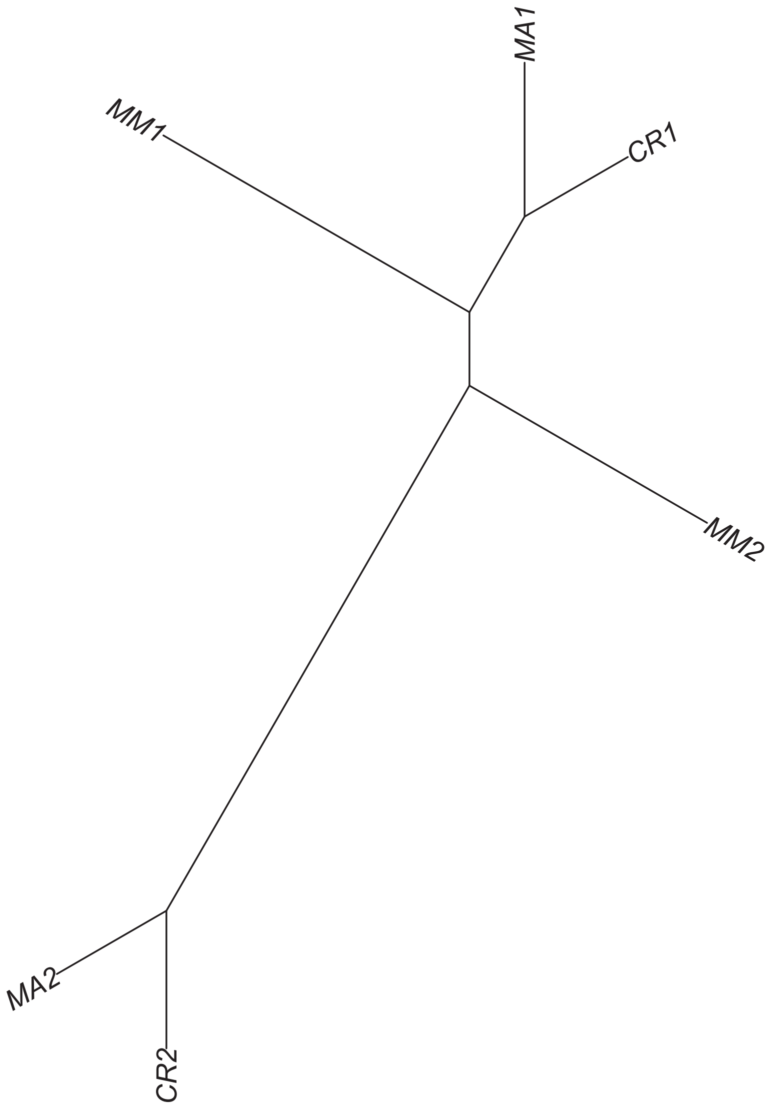

Region 516

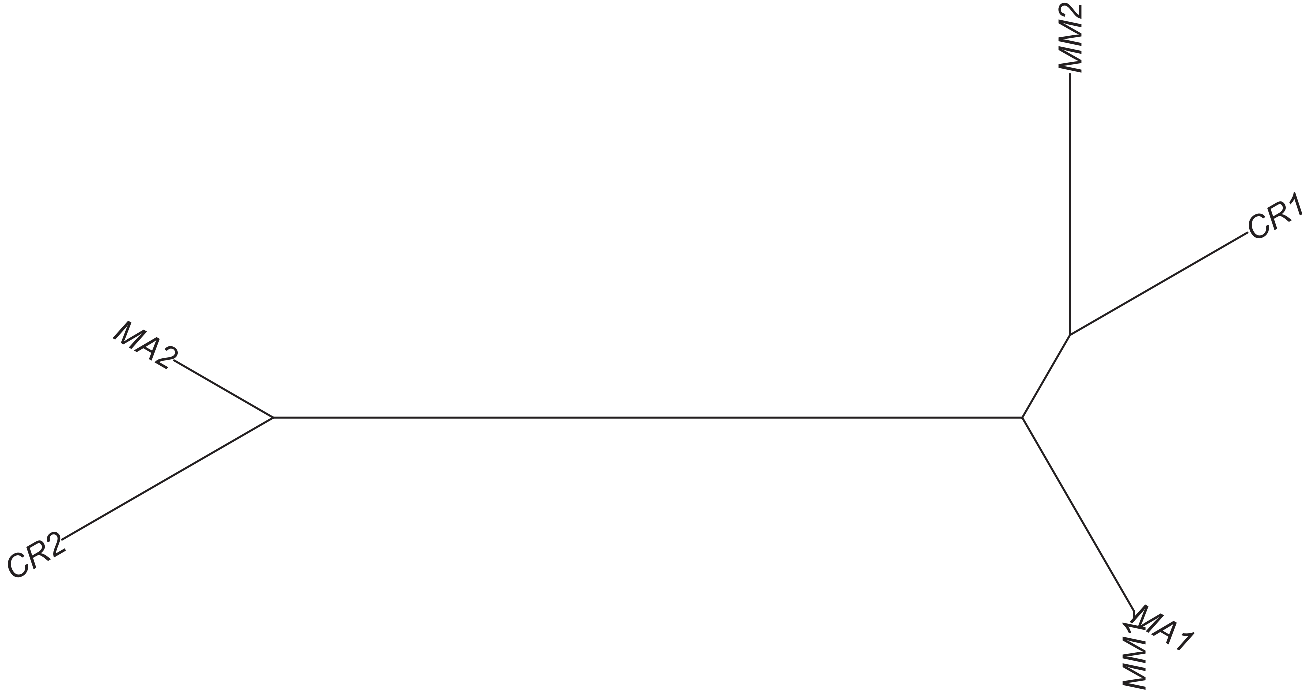

### Region 517

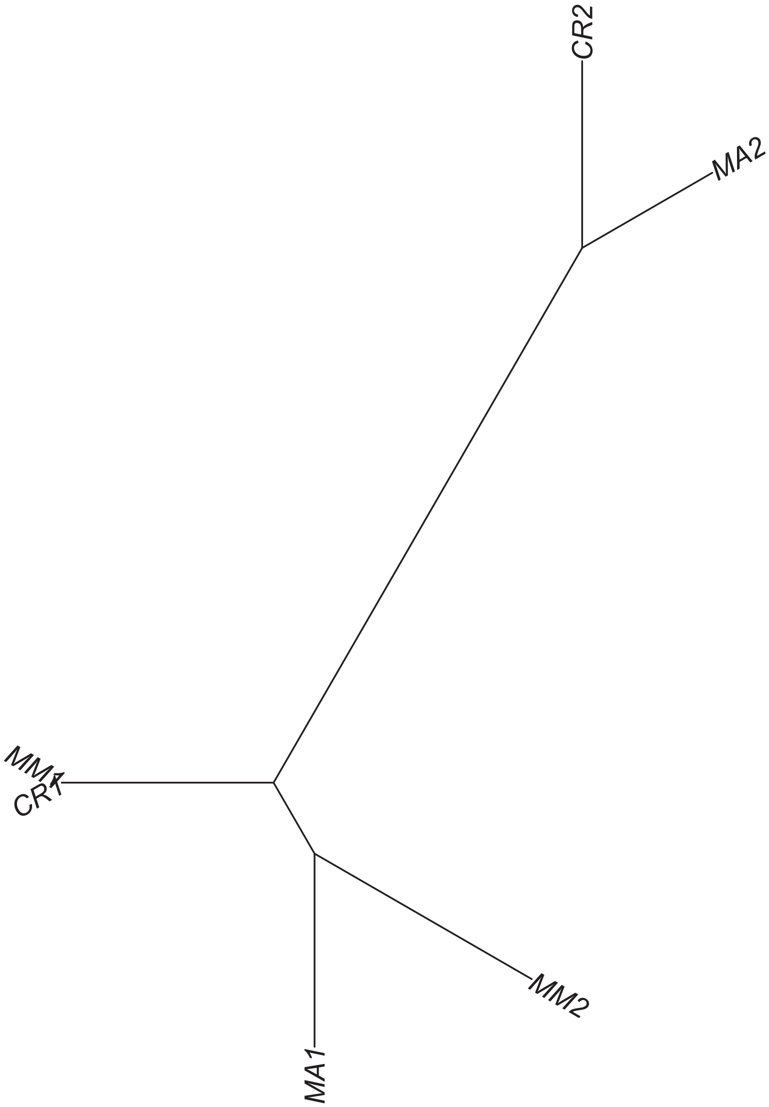

### Region 519

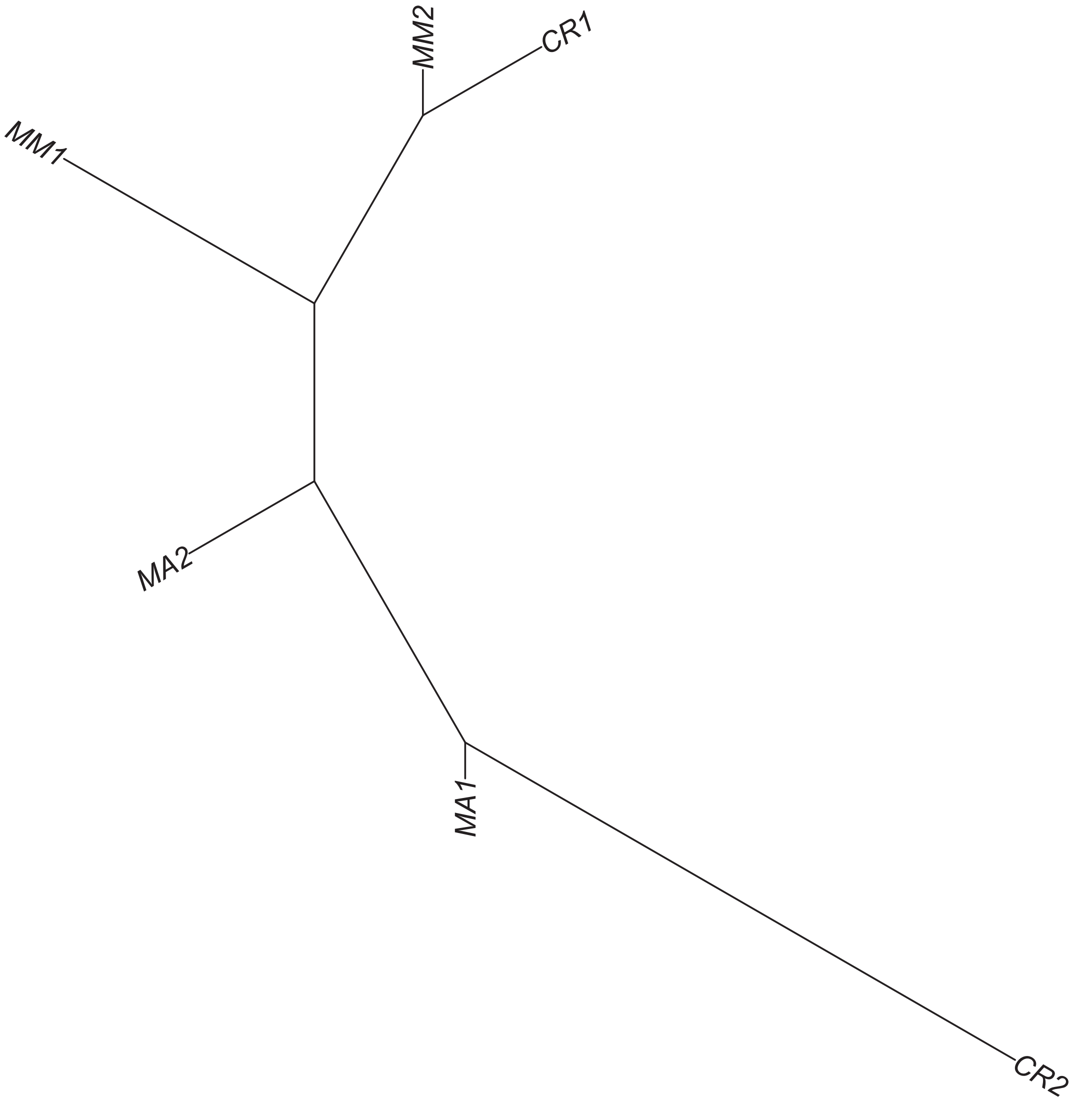

### Region 520



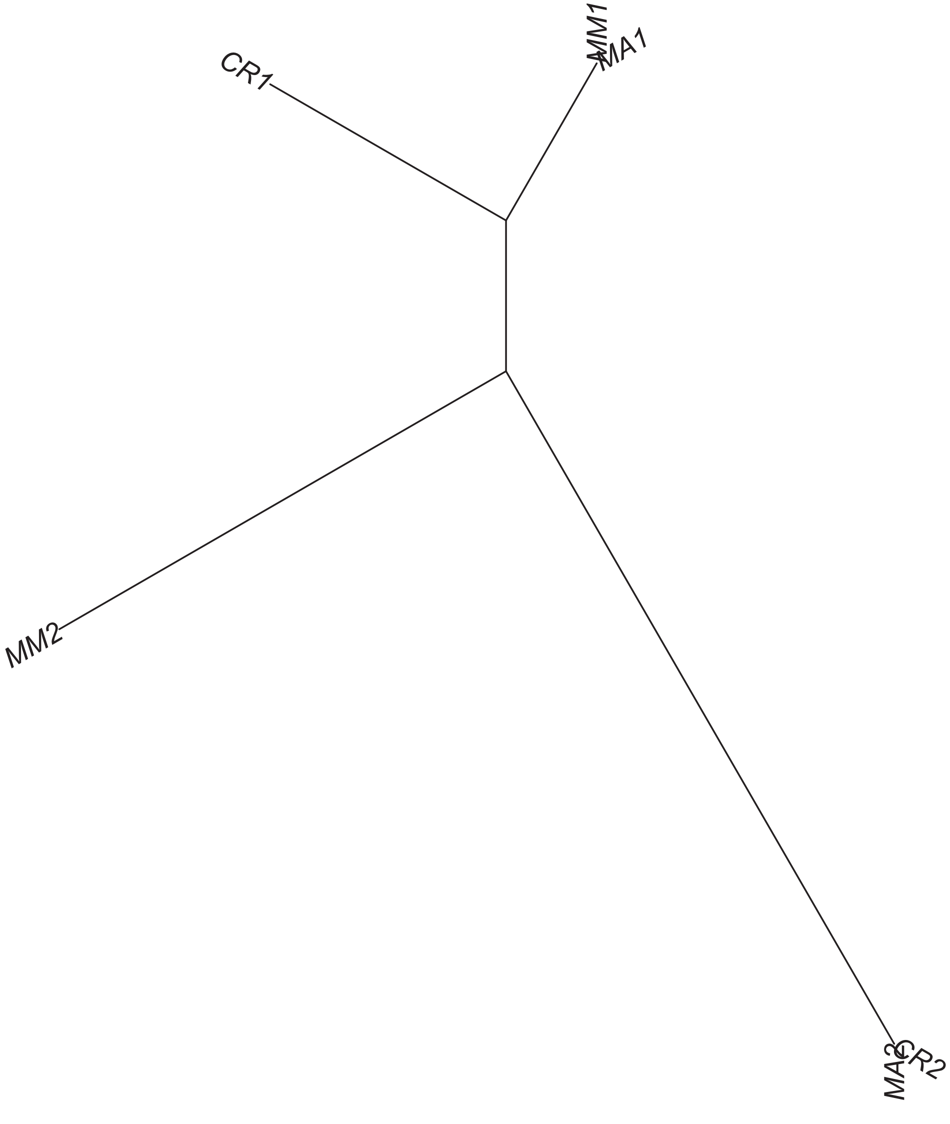

### Region 521



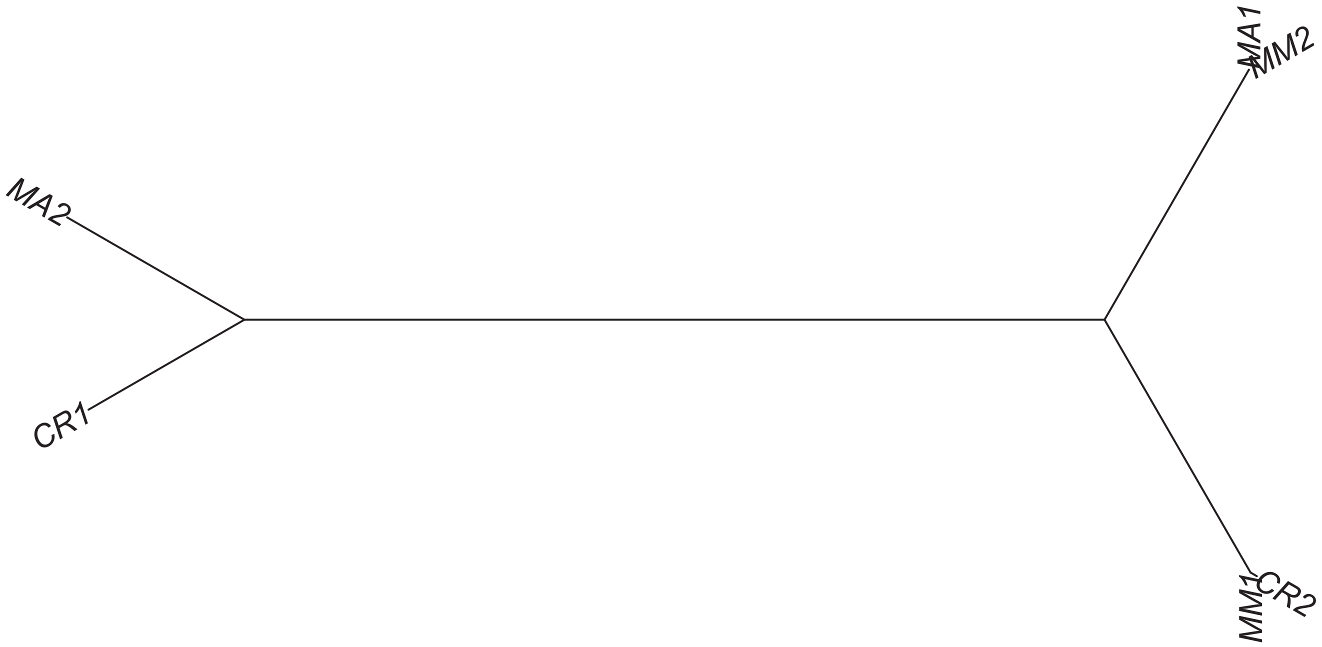

### Region 522

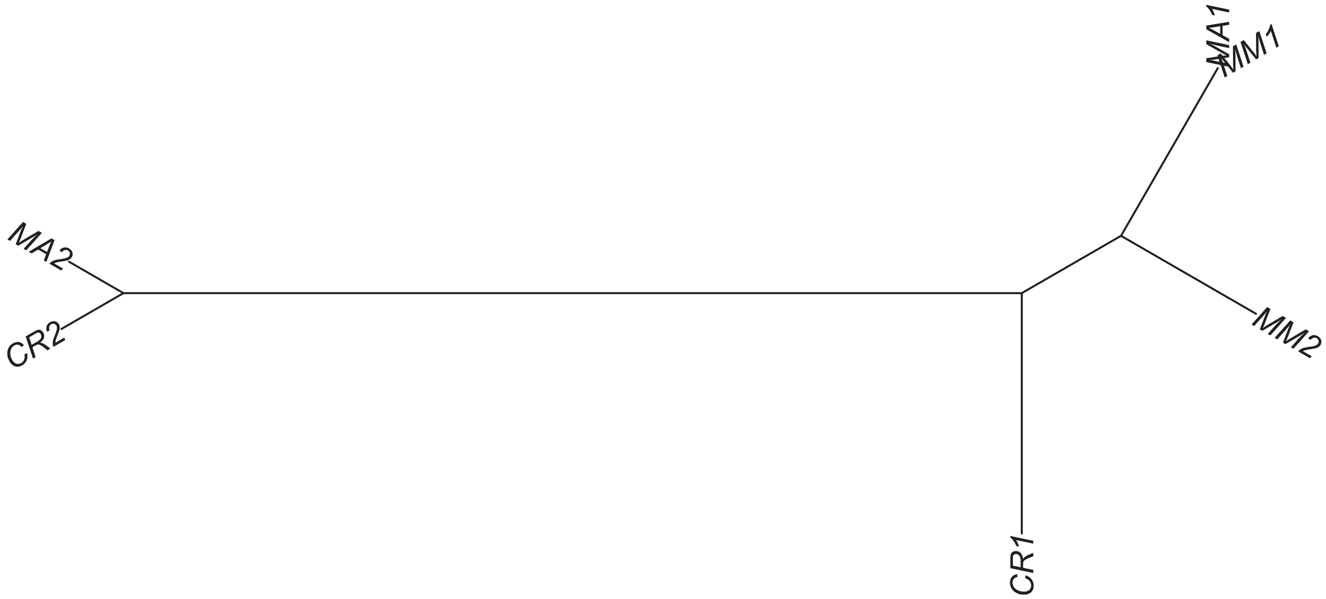

### Region 524



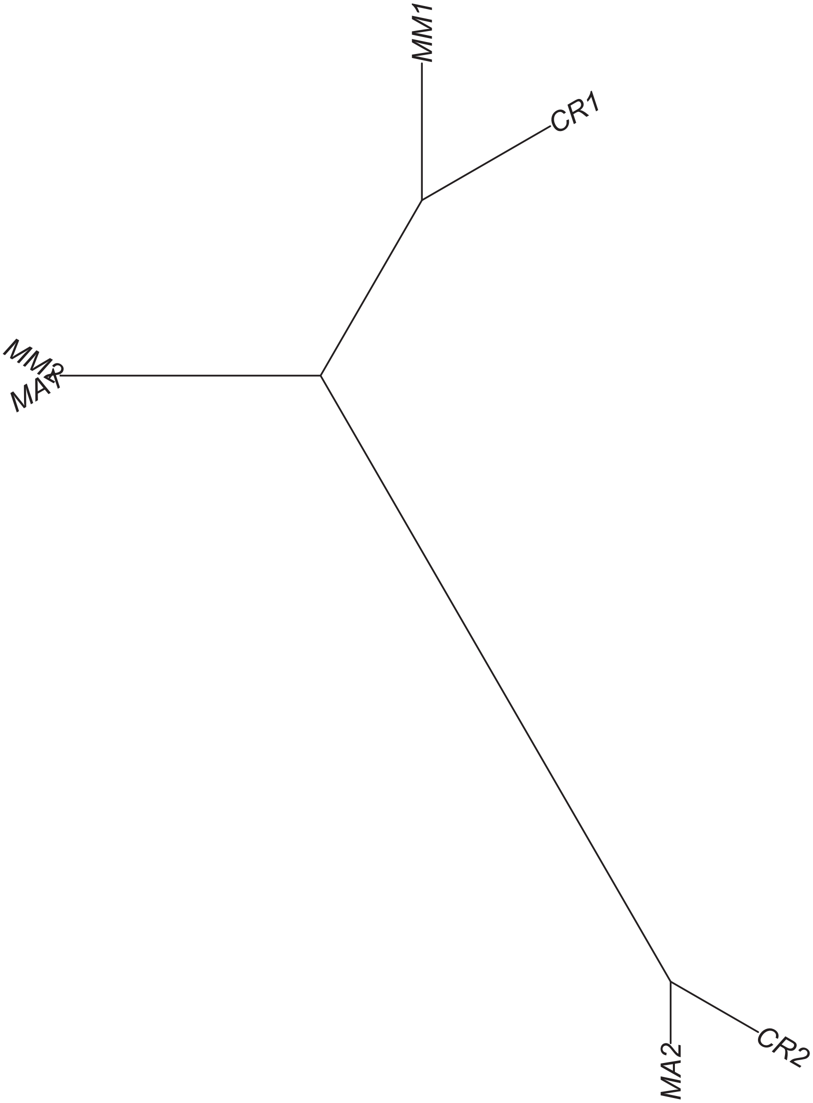

### Region 525



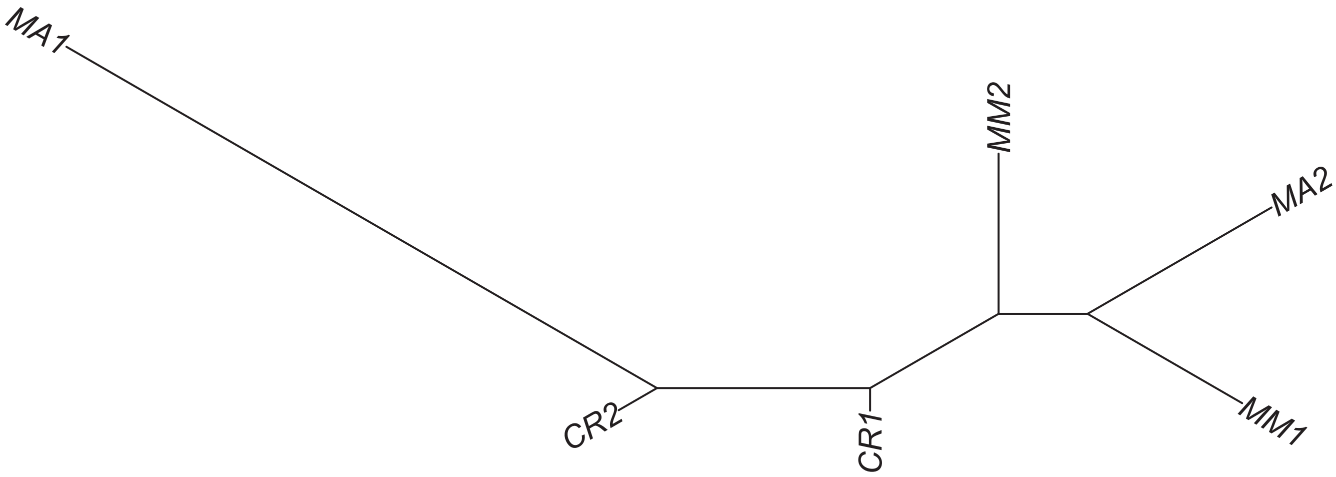

### Region 526

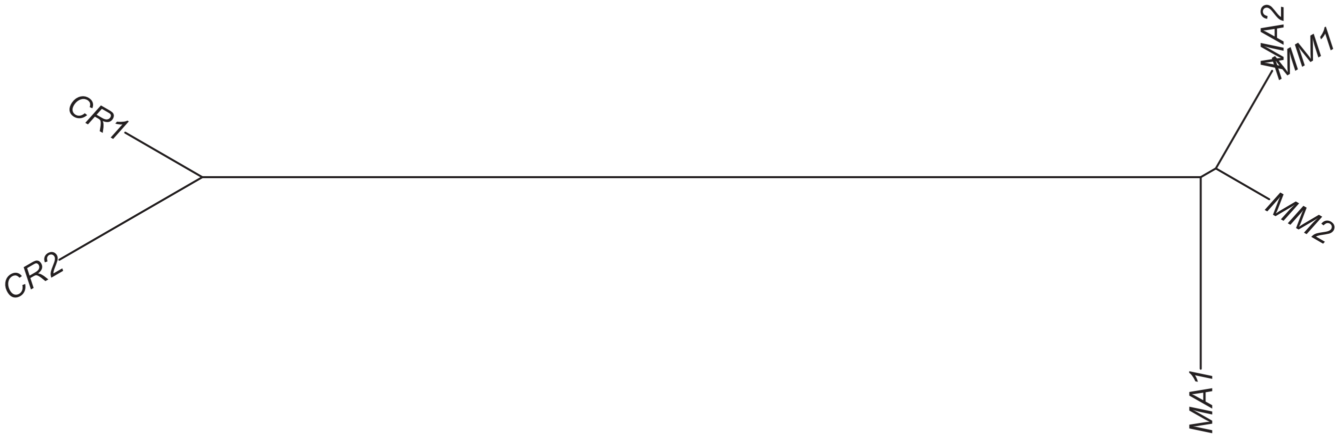

### Region 527



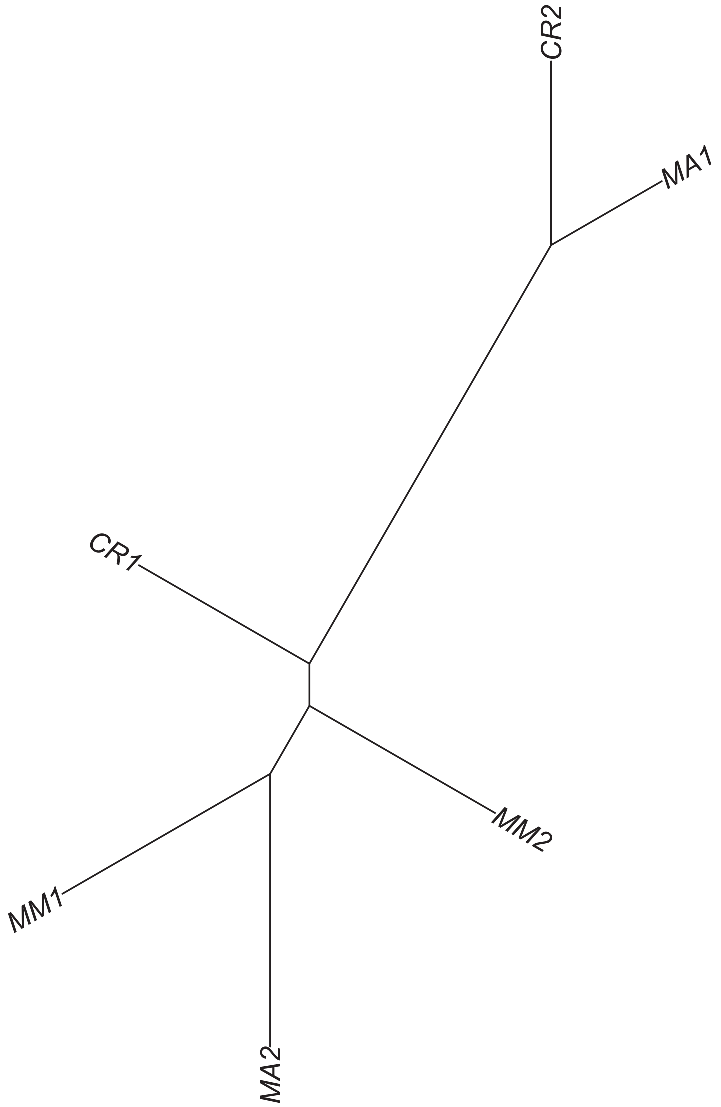

### Region 529

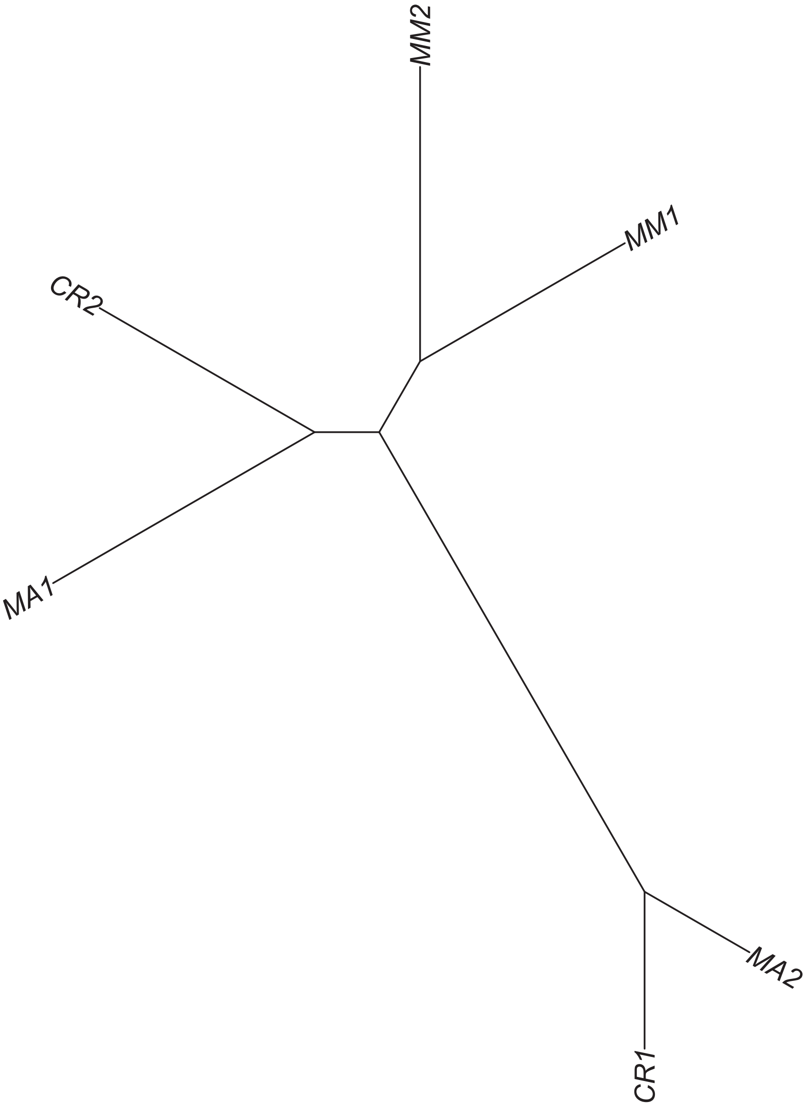

### Region 530



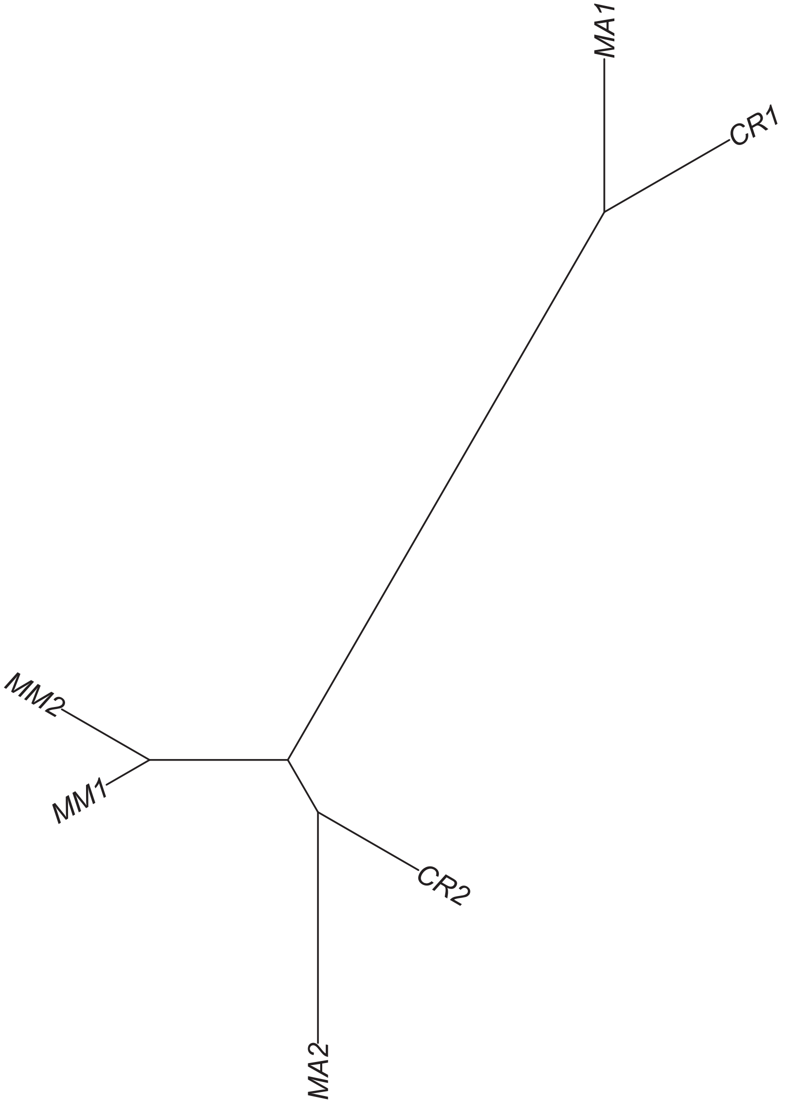
