## Supplementary material for "Genomic Signature of Sexual Reproduction in the Bdelloid Rotifer *Macrotrachella quadricornifera*": Table S6

| 15 | | | | | 11778 |
|---|---|---|---|---|---|
| | CR1 | CR2 | MA1 | MA2 | |
| CR1 | 0 | 214 | 208 | 77 | |
| CR2 | 0.0182 | 0 | 49 | 217 | |
| MA1 | 0.0177 | 0.0042 | 0 | 214 | |
| MA2 | 0.0065 | 0.0184 | 0.0182 | 0 | |
| 18 | | | | | 19954 |
| | CR1 | CR2 | MA1 | MA2 | |
| CR1 | 0 | 263 | 275 | 96 | |
| CR2 | 0.0132 | 0 | 168 | 302 | |
| MA1 | 0.0138 | 0.0084 | 0 | 295 | |
| MA2 | 0.0048 | 0.0151 | 0.0148 | 0 | |
| 20 | | | | | 8051 |
| | CR1 | CR2 | MA1 | MA2 | |
| CR1 | 0 | 143 | 143 | 96 | |
| CR2 | 0.0178 | 0 | 0 | 146 | |
| MA1 | 0.0178 | 0.0000 | 0 | 146 | |
| MA2 | 0.0119 | 0.0181 | 0.0181 | 0 | |
| 22 | | | | | 7638 |
| | CR1 | CR2 | MA1 | MA2 | |
| CR1 | 0 | 146 | 149 | 84 | |
| CR2 | 0.0191 | 0 | 41 | 149 | |
| MA1 | 0.0195 | 0.0054 | 0 | 150 | |
| MA2 | 0.0110 | 0.0195 | 0.0196 | 0 | |
| 29 | | | | | 3190 |
| | CR1 | CR2 | MA1 | MA2 | |
| CR1 | 0 | 52 | 27 | 63 | |
| CR2 | 0.0163 | 0 | 59 | 45 | |
| MA1 | 0.0085 | 0.0185 | 0 | 68 | |
| MA2 | 0.0197 | 0.0141 | 0.0213 | 0 | |
| 31 | | | | | 5807 |
| | CR1 | CR2 | MA1 | MA2 | |
| CR1 | 0 | 129 | 85 | 69 | |
| CR2 | 0.0222 | 0 | 116 | 126 | |
| MA1 | 0.0146 | 0.0200 | 0 | 24 | |
| MA2 | 0.0119 | 0.0217 | 0.0041 | 0 | |
| 42 | | | | | 11535 |
| | CR1 | CR2 | MA1 | MA2 | |
| CR1 | 0 | 238 | 69 | 237 | |
| CR2 | 0.0206 | 0 | 223 | 25 | |
| MA1 | 0.0060 | 0.0193 | 0 | 222 | |
| MA2 | 0.0205 | 0.0022 | 0.0192 | 0 | |

Table S7: Difference Matrices.



| 44 | | | | | 13086 |
|---|---|---|---|---|---|
| | CR1 | CR2 | MA1 | MA2 | |
| CR1 | 0 | 162 | 221 | 102 | |
| CR2 | 0.0124 | 0 | 105 | 192 | |
| MA1 | 0.0169 | 0.0080 | 0 | 245 | |
| MA2 | 0.0078 | 0.0147 | 0.0187 | 0 | |

| 48 | | | | | 20424 |
|---|---|---|---|---|---|
| | CR1 | CR2 | MA1 | MA2 | |
| CR1 | 0 | 549 | 192 | 537 | |
| CR2 | 0.0269 | 0 | 553 | 205 | |
| MA1 | 0.0094 | 0.0271 | 0 | 548 | |
| MA2 | 0.0263 | 0.0100 | 0.0268 | 0 | |

| 49 | | | | | 12319 |
|---|---|---|---|---|---|
| | CR1 | CR2 | MA1 | MA2 | |
| CR1 | 0 | 281 | 274 | 117 | |
| CR2 | 0.0228 | 0 | 65 | 292 | |
| MA1 | 0.0222 | 0.0053 | 0 | 285 | |
| MA2 | 0.0095 | 0.0237 | 0.0231 | 0 | |

| 55 | | | | | 13907 |
|---|---|---|---|---|---|
| | CR1 | CR2 | MA1 | MA2 | |
| CR1 | 0 | 61 | 212 | 223 | |
| CR2 | 0.0044 | 0 | 224 | 185 | |
| MA1 | 0.0152 | 0.0161 | 0 | 266 | |
| MA2 | 0.0160 | 0.0133 | 0.0191 | 0 | |

| 59 | | | | | 12720 |
|---|---|---|---|---|---|
| | CR1 | CR2 | MA1 | MA2 | |
| CR1 | 0 | 144 | 48 | 152 | |
| CR2 | 0.0113 | 0 | 145 | 62 | |
| MA1 | 0.0038 | 0.0114 | 0 | 154 | |
| MA2 | 0.0119 | 0.0049 | 0.0121 | 0 | |

| 73 | | | | | 9604 |
|---|---|---|---|---|---|
| | CR1 | CR2 | MA1 | MA2 | |
| CR1 | 0 | 141 | 147 | 40 | |
| CR2 | 0.0147 | 0 | 55 | 147 | |
| MA1 | 0.0153 | 0.0057 | 0 | 154 | |
| MA2 | 0.0042 | 0.0153 | 0.0160 | 0 | |

| 84 | | | | | 11206 |
|---|---|---|---|---|---|
| | CR1 | CR2 | MA1 | MA2 | |
| CR1 | 0 | 56 | 130 | 283 | |
| CR2 | 0.0050 | 0 | 97 | 299 | |
| MA1 | 0.0116 | 0.0087 | 0 | 296 | |
| MA2 | 0.0253 | 0.0267 | 0.0264 | 0 | |

| 109 | | | | | 14720 |
|---|---|---|---|---|---|
| | CR1 | CR2 | MA1 | MA2 | |
| CR1 | 0 | 227 | 288 | 297 | |
| CR2 | 0.0154 | 0 | 155 | 390 | |
| MA1 | 0.0196 | 0.0105 | 0 | 388 | |
| MA2 | 0.0202 | 0.0265 | 0.0264 | 0 | |

| 111 | | | | | 12175 |
|---|---|---|---|---|---|
| | CR1 | CR2 | MA1 | MA2 | |
| CR1 | 0 | 212 | 165 | 133 | |
| CR2 | 0.0174 | 0 | 134 | 203 | |
| MA1 | 0.0136 | 0.0110 | 0 | 101 | |
| MA2 | 0.0109 | 0.0167 | 0.0083 | 0 | |

| 122 | | | | | 9796 |
|---|---|---|---|---|---|
| | CR1 | CR2 | MA1 | MA2 | |
| CR1 | 0 | 133 | 91 | 134 | |
| CR2 | 0.0136 | 0 | 133 | 13 | |
| MA1 | 0.0093 | 0.0136 | 0 | 134 | |
| MA2 | 0.0137 | 0.0013 | 0.0137 | 0 | |

| 127 | | | | | 7683 |
|---|---|---|---|---|---|
| | CR1 | CR2 | MA1 | MA2 | |
| CR1 | 0 | 65 | 75 | 24 | |
| CR2 | 0.0085 | 0 | 26 | 69 | |
| MA1 | 0.0098 | 0.0034 | 0 | 77 | |
| MA2 | 0.0031 | 0.0090 | 0.0100 | 0 | |

| 129 | | | | | 19591 |
|---|---|---|---|---|---|
| | CR1 | CR2 | MA1 | MA2 | |
| CR1 | 0 | 307 | 311 | 64 | |
| CR2 | 0.0157 | 0 | 119 | 318 | |
| MA1 | 0.0159 | 0.0061 | 0 | 320 | |
| MA2 | 0.0033 | 0.0162 | 0.0163 | 0 | |

| 132 | | | | | 14102 |
|---|---|---|---|---|---|
| | CR1 | CR2 | MA1 | MA2 | |
| CR1 | 0 | 81 | 246 | 128 | |
| CR2 | 0.0057 | 0 | 302 | 126 | |
| MA1 | 0.0174 | 0.0214 | 0 | 248 | |
| MA2 | 0.0091 | 0.0089 | 0.0176 | 0 | |

| 142 | | | | | 7781 |
|---|---|---|---|---|---|
| | CR1 | CR2 | MA1 | MA2 | |
| CR1 | 0 | 104 | 36 | 60 | |
| CR2 | 0.0134 | 0 | 96 | 84 | |
| MA1 | 0.0046 | 0.0123 | 0 | 44 | |
| MA2 | 0.0077 | 0.0108 | 0.0057 | 0 | |

| 144 | | | | | 7654 |
|---|---|---|---|---|---|
| | CR1 | CR2 | MA1 | MA2 | |
| CR1 | 0 | 104 | 65 | 105 | |
| CR2 | 0.0136 | 0 | 53 | 6 | |
| MA1 | 0.0085 | 0.0069 | 0 | 54 | |
| MA2 | 0.0137 | 0.0008 | 0.0071 | 0 | |
| 153 | | | | | 29791 |
| | CR1 | CR2 | MA1 | MA2 | |
| CR1 | 0 | 570 | 568 | 90 | |
| CR2 | 0.0191 | 0 | 161 | 567 | |
| MA1 | 0.0191 | 0.0054 | 0 | 565 | |
| MA2 | 0.0030 | 0.0190 | 0.0190 | 0 | |
| 161 | | | | | 7944 |
| | CR1 | CR2 | MA1 | MA2 | |
| CR1 | 0 | 216 | 109 | 195 | |
| CR2 | 0.0272 | 0 | 204 | 55 | |
| MA1 | 0.0137 | 0.0257 | 0 | 183 | |
| MA2 | 0.0245 | 0.0069 | 0.0230 | 0 | |
| 165 | | | | | 13822 |
| | CR1 | CR2 | MA1 | MA2 | |
| CR1 | 0 | 207 | 61 | 233 | |
| CR2 | 0.0150 | 0 | 198 | 102 | |
| MA1 | 0.0044 | 0.0143 | 0 | 226 | |
| MA2 | 0.0169 | 0.0074 | 0.0164 | 0 | |
| 181 | | | | | 7304 |
| | CR1 | CR2 | MA1 | MA2 | |
| CR1 | 0 | 162 | 133 | 112 | |
| CR2 | 0.0222 | 0 | 135 | 172 | |
| MA1 | 0.0182 | 0.0185 | 0 | 65 | |
| MA2 | 0.0153 | 0.0235 | 0.0089 | 0 | |
| 184 | | | | | 9639 |
| | CR1 | CR2 | MA1 | MA2 | |
| CR1 | 0 | 140 | 142 | 60 | |
| CR2 | 0.0145 | 0 | 20 | 154 | |
| MA1 | 0.0147 | 0.0021 | 0 | 156 | |
| MA2 | 0.0062 | 0.0160 | 0.0162 | 0 | |
| 185 | | | | | 16433 |
| | CR1 | CR2 | MA1 | MA2 | |
| CR1 | 0 | 557 | 552 | 442 | |
| CR2 | 0.0339 | 0 | 270 | 521 | |
| MA1 | 0.0336 | 0.0164 | 0 | 520 | |
| MA2 | 0.0269 | 0.0317 | 0.0316 | 0 | |

| 195 | | | | | 14586 |
|---|---|---|---|---|---|
| | CR1 | CR2 | MA1 | MA2 | |
| CR1 | 0 | 360 | 333 | 283 | |
| CR2 | 0.0247 | 0 | 109 | 172 | |
| MA1 | 0.0228 | 0.0075 | 0 | 79 | |
| MA2 | 0.0194 | 0.0118 | 0.0054 | 0 | |
| 205 | | | | | 8176 |
| | CR1 | CR2 | MA1 | MA2 | |
| CR1 | 0 | 21 | 156 | 111 | |
| CR2 | 0.0026 | 0 | 135 | 114 | |
| MA1 | 0.0191 | 0.0165 | 0 | 167 | |
| MA2 | 0.0136 | 0.0139 | 0.0204 | 0 | |
| 207 | | | | | 13994 |
| | CR1 | CR2 | MA1 | MA2 | |
| CR1 | 0 | 451 | 271 | 440 | |
| CR2 | 0.0322 | 0 | 456 | 91 | |
| MA1 | 0.0194 | 0.0326 | 0 | 450 | |
| MA2 | 0.0314 | 0.0065 | 0.0322 | 0 | |
| 218 | | | | | 11867 |
| | CR1 | CR2 | MA1 | MA2 | |
| CR1 | 0 | 103 | 196 | 190 | |
| CR2 | 0.0087 | 0 | 196 | 225 | |
| MA1 | 0.0165 | 0.0165 | 0 | 304 | |
| MA2 | 0.0160 | 0.0190 | 0.0256 | 0 | |
| 221 | | | | | 15921 |
| | CR1 | CR2 | MA1 | MA2 | |
| CR1 | 0 | 299 | 301 | 129 | |
| CR2 | 0.0188 | 0 | 64 | 280 | |
| MA1 | 0.0189 | 0.0040 | 0 | 279 | |
| MA2 | 0.0081 | 0.0176 | 0.0175 | 0 | |
| 225 | | | | | 7650 |
| | CR1 | CR2 | MA1 | MA2 | |
| CR1 | 0 | 143 | 139 | 59 | |
| CR2 | 0.0187 | 0 | 16 | 152 | |
| MA1 | 0.0182 | 0.0021 | 0 | 148 | |
| MA2 | 0.0077 | 0.0199 | 0.0193 | 0 | |
| 234 | | | | | 14603 |
| | CR1 | CR2 | MA1 | MA2 | |
| CR1 | 0 | 178 | 132 | 88 | |
| CR2 | 0.0122 | 0 | 120 | 180 | |
| MA1 | 0.0090 | 0.0082 | 0 | 118 | |
| MA2 | 0.0060 | 0.0123 | 0.0081 | 0 | |

| 240 | | | | | 18071 |
|---|---|---|---|---|---|
| | CR1 | CR2 | MA1 | MA2 | |
| CR1 | 0 | 429 | 423 | 204 | |
| CR2 | 0.0237 | 0 | 137 | 461 | |
| MA1 | 0.0234 | 0.0076 | 0 | 455 | |
| MA2 | 0.0113 | 0.0255 | 0.0252 | 0 | |
| 241 | | | | | 18876 |
| | CR1 | CR2 | MA1 | MA2 | |
| CR1 | 0 | 510 | 503 | 317 | |
| CR2 | 0.0270 | 0 | 164 | 538 | |
| MA1 | 0.0266 | 0.0087 | 0 | 526 | |
| MA2 | 0.0168 | 0.0285 | 0.0279 | 0 | |
| 249 | | | | | 22839 |
| | CR1 | CR2 | MA1 | MA2 | |
| CR1 | 0 | 319 | 328 | 98 | |
| CR2 | 0.0140 | 0 | 89 | 317 | |
| MA1 | 0.0144 | 0.0039 | 0 | 327 | |
| MA2 | 0.0043 | 0.0139 | 0.0143 | 0 | |
| 251 | | | | | 9582 |
| | CR1 | CR2 | MA1 | MA2 | |
| CR1 | 0 | 183 | 84 | 184 | |
| CR2 | 0.0191 | 0 | 174 | 17 | |
| MA1 | 0.0088 | 0.0182 | 0 | 175 | |
| MA2 | 0.0192 | 0.0018 | 0.0183 | 0 | |
| 263 | | | | | 13288 |
| | CR1 | CR2 | MA1 | MA2 | |
| CR1 | 0 | 226 | 234 | 176 | |
| CR2 | 0.0170 | 0 | 182 | 249 | |
| MA1 | 0.0176 | 0.0137 | 0 | 112 | |
| MA2 | 0.0132 | 0.0187 | 0.0084 | 0 | |
| 271 | | | | | 7680 |
| | CR1 | CR2 | MA1 | MA2 | |
| CR1 | 0 | 69 | 88 | 51 | |
| CR2 | 0.0090 | 0 | 93 | 59 | |
| MA1 | 0.0115 | 0.0121 | 0 | 105 | |
| MA2 | 0.0066 | 0.0077 | 0.0137 | 0 | |
| 274 | | | | | 13200 |
| | CR1 | CR2 | MA1 | MA2 | |
| CR1 | 0 | 331 | 351 | 122 | |
| CR2 | 0.0251 | 0 | 144 | 311 | |
| MA1 | 0.0266 | 0.0109 | 0 | 334 | |
| MA2 | 0.0092 | 0.0236 | 0.0253 | 0 | |

| 278 | | | | | 10142 |
|---|---|---|---|---|---|
| | CR1 | CR2 | MA1 | MA2 | |
| CR1 | 0 | 240 | 230 | 110 | |
| CR2 | 0.0237 | 0 | 59 | 235 | |
| MA1 | 0.0227 | 0.0058 | 0 | 227 | |
| MA2 | 0.0108 | 0.0232 | 0.0224 | 0 | |
| 284 | | | | | 6665 |
| | CR1 | CR2 | MA1 | MA2 | |
| CR1 | 0 | 150 | 166 | 226 | |
| CR2 | 0.0225 | 0 | 198 | 132 | |
| MA1 | 0.0249 | 0.0297 | 0 | 236 | |
| MA2 | 0.0339 | 0.0198 | 0.0354 | 0 | |
| 287 | | | | | 8507 |
| | CR1 | CR2 | MA1 | MA2 | |
| CR1 | 0 | 352 | 153 | 328 | |
| CR2 | 0.0414 | 0 | 334 | 320 | |
| MA1 | 0.0180 | 0.0393 | 0 | 349 | |
| MA2 | 0.0386 | 0.0376 | 0.0410 | 0 | |
| 290 | | | | | 9884 |
| | CR1 | CR2 | MA1 | MA2 | |
| CR1 | 0 | 203 | 217 | 102 | |
| CR2 | 0.0205 | 0 | 44 | 193 | |
| MA1 | 0.0220 | 0.0045 | 0 | 207 | |
| MA2 | 0.0103 | 0.0195 | 0.0209 | 0 | |
| 296 | | | | | 7069 |
| | CR1 | CR2 | MA1 | MA2 | |
| CR1 | 0 | 176 | 65 | 177 | |
| CR2 | 0.0249 | 0 | 178 | 11 | |
| MA1 | 0.0092 | 0.0252 | 0 | 181 | |
| MA2 | 0.0250 | 0.0016 | 0.0256 | 0 | |
| 298 | | | | | 14124 |
| | CR1 | CR2 | MA1 | MA2 | |
| CR1 | 0 | 322 | 248 | 232 | |
| CR2 | 0.0228 | 0 | 323 | 336 | |
| MA1 | 0.0176 | 0.0229 | 0 | 18 | |
| MA2 | 0.0164 | 0.0238 | 0.0013 | 0 | |
| 299 | | | | | 10195 |
| | CR1 | CR2 | MA1 | MA2 | |
| CR1 | 0 | 125 | 119 | 65 | |
| CR2 | 0.0123 | 0 | 24 | 125 | |
| MA1 | 0.0117 | 0.0024 | 0 | 121 | |
| MA2 | 0.0064 | 0.0123 | 0.0119 | 0 | |

| 303 | | | | | 9223 |
|---|---|---|---|---|---|
| | CR1 | CR2 | MA1 | MA2 | |
| CR1 | 0 | 185 | 178 | 105 | |
| CR2 | 0.0201 | 0 | 72 | 187 | |
| MA1 | 0.0193 | 0.0078 | 0 | 185 | |
| MA2 | 0.0114 | 0.0203 | 0.0201 | 0 | |
| 309 | | | | | 7145 |
| | CR1 | CR2 | MA1 | MA2 | |
| CR1 | 0 | 145 | 78 | 147 | |
| CR2 | 0.0203 | 0 | 135 | 50 | |
| MA1 | 0.0109 | 0.0189 | 0 | 139 | |
| MA2 | 0.0206 | 0.0070 | 0.0195 | 0 | |
| 312 | | | | | 8014 |
| | CR1 | CR2 | MA1 | MA2 | |
| CR1 | 0 | 271 | 157 | 180 | |
| CR2 | 0.0338 | 0 | 282 | 256 | |
| MA1 | 0.0196 | 0.0352 | 0 | 48 | |
| MA2 | 0.0225 | 0.0319 | 0.0060 | 0 | |
| 313 | | | | | 9547 |
| | CR1 | CR2 | MA1 | MA2 | |
| CR1 | 0 | 255 | 263 | 61 | |
| CR2 | 0.0267 | 0 | 131 | 258 | |
| MA1 | 0.0275 | 0.0137 | 0 | 266 | |
| MA2 | 0.0064 | 0.0270 | 0.0279 | 0 | |
| 329 | | | | | 9864 |
| | CR1 | CR2 | MA1 | MA2 | |
| CR1 | 0 | 184 | 28 | 197 | |
| CR2 | 0.0187 | 0 | 186 | 129 | |
| MA1 | 0.0028 | 0.0189 | 0 | 199 | |
| MA2 | 0.0200 | 0.0131 | 0.0202 | 0 | |
| 332 | | | | | 8092 |
| | CR1 | CR2 | MA1 | MA2 | |
| CR1 | 0 | 172 | 179 | 67 | |
| CR2 | 0.0213 | 0 | 74 | 157 | |
| MA1 | 0.0221 | 0.0091 | 0 | 164 | |
| MA2 | 0.0083 | 0.0194 | 0.0203 | 0 | |
| 334 | | | | | 13665 |
| | CR1 | CR2 | MA1 | MA2 | |
| CR1 | 0 | 214 | 133 | 231 | |
| CR2 | 0.0157 | 0 | 228 | 55 | |
| MA1 | 0.0097 | 0.0167 | 0 | 241 | |
| MA2 | 0.0169 | 0.0040 | 0.0176 | 0 | |

| 340 | | | | | 9883 |
|---|---|---|---|---|---|
| | CR1 | CR2 | MA1 | MA2 | |
| CR1 | 0 | 182 | 183 | 13 | |
| CR2 | 0.0184 | 0 | 54 | 177 | |
| MA1 | 0.0185 | 0.0055 | 0 | 178 | |
| MA2 | 0.0013 | 0.0179 | 0.0180 | 0 | |
| 341 | | | | | 10840 |
| | CR1 | CR2 | MA1 | MA2 | |
| CR1 | 0 | 135 | 159 | 79 | |
| CR2 | 0.0125 | 0 | 113 | 166 | |
| MA1 | 0.0147 | 0.0104 | 0 | 110 | |
| MA2 | 0.0073 | 0.0153 | 0.0101 | 0 | |
| 344 | | | | | 9263 |
| | CR1 | CR2 | MA1 | MA2 | |
| CR1 | 0 | 193 | 110 | 194 | |
| CR2 | 0.0208 | 0 | 193 | 71 | |
| MA1 | 0.0119 | 0.0208 | 0 | 195 | |
| MA2 | 0.0209 | 0.0077 | 0.0211 | 0 | |
| 346 | | | | | 13000 |
| | CR1 | CR2 | MA1 | MA2 | |
| CR1 | 0 | 247 | 100 | 243 | |
| CR2 | 0.0190 | 0 | 243 | 90 | |
| MA1 | 0.0077 | 0.0187 | 0 | 237 | |
| MA2 | 0.0187 | 0.0069 | 0.0182 | 0 | |
| 349 | | | | | 13085 |
| | CR1 | CR2 | MA1 | MA2 | |
| CR1 | 0 | 271 | 158 | 270 | |
| CR2 | 0.0207 | 0 | 241 | 127 | |
| MA1 | 0.0121 | 0.0184 | 0 | 286 | |
| MA2 | 0.0206 | 0.0097 | 0.0219 | 0 | |
| 353 | | | | | 11272 |
| | CR1 | CR2 | MA1 | MA2 | |
| CR1 | 0 | 152 | 259 | 279 | |
| CR2 | 0.0135 | 0 | 375 | 203 | |
| MA1 | 0.0230 | 0.0333 | 0 | 375 | |
| MA2 | 0.0248 | 0.0180 | 0.0333 | 0 | |
| 360 | | | | | 7794 |
| | CR1 | CR2 | MA1 | MA2 | |
| CR1 | 0 | 156 | 62 | 161 | |
| CR2 | 0.0200 | 0 | 167 | 70 | |
| MA1 | 0.0080 | 0.0214 | 0 | 172 | |
| MA2 | 0.0207 | 0.0090 | 0.0221 | 0 | |

| 361 | | | | | 12630 |
|---|---|---|---|---|---|
| | CR1 | CR2 | MA1 | MA2 | |
| CR1 | 0 | 130 | 194 | 156 | |
| CR2 | 0.0103 | 0 | 240 | 102 | |
| MA1 | 0.0154 | 0.0190 | 0 | 241 | |
| MA2 | 0.0124 | 0.0081 | 0.0191 | 0 | |
| 365 | | | | | 14018 |
| | CR1 | CR2 | MA1 | MA2 | |
| CR1 | 0 | 271 | 237 | 144 | |
| CR2 | 0.0193 | 0 | 94 | 231 | |
| MA1 | 0.0169 | 0.0067 | 0 | 182 | |
| MA2 | 0.0103 | 0.0165 | 0.0130 | 0 | |
| 366 | | | | | 15282 |
| | CR1 | CR2 | MA1 | MA2 | |
| CR1 | 0 | 476 | 221 | 476 | |
| CR2 | 0.0311 | 0 | 471 | 44 | |
| MA1 | 0.0145 | 0.0308 | 0 | 471 | |
| MA2 | 0.0311 | 0.0029 | 0.0308 | 0 | |
| 372 | | | | | 9496 |
| | CR1 | CR2 | MA1 | MA2 | |
| CR1 | 0 | 57 | 163 | 116 | |
| CR2 | 0.0060 | 0 | 194 | 63 | |
| MA1 | 0.0172 | 0.0204 | 0 | 189 | |
| MA2 | 0.0122 | 0.0066 | 0.0199 | 0 | |
| 374 | | | | | 10100 |
| | CR1 | CR2 | MA1 | MA2 | |
| CR1 | 0 | 142 | 107 | 62 | |
| CR2 | 0.0141 | 0 | 84 | 114 | |
| MA1 | 0.0106 | 0.0083 | 0 | 104 | |
| MA2 | 0.0061 | 0.0113 | 0.0103 | 0 | |
| 377 | | | | | 17247 |
| | CR1 | CR2 | MA1 | MA2 | |
| CR1 | 0 | 1 | 353 | 331 | |
| CR2 | 0.0001 | 0 | 354 | 332 | |
| MA1 | 0.0205 | 0.0205 | 0 | 26 | |
| MA2 | 0.0192 | 0.0192 | 0.0015 | 0 | |
| 389 | | | | | 10405 |
| | CR1 | CR2 | MA1 | MA2 | |
| CR1 | 0 | 216 | 228 | 73 | |
| CR2 | 0.0208 | 0 | 84 | 227 | |
| MA1 | 0.0219 | 0.0081 | 0 | 237 | |
| MA2 | 0.0070 | 0.0218 | 0.0228 | 0 | |

| 410 | | | | | 2051 |
|---|---|---|---|---|---|
| | CR1 | CR2 | MA1 | MA2 | |
| CR1 | 0 | 16 | 6 | 14 | |
| CR2 | 0.0078 | 0 | 18 | 2 | |
| MA1 | 0.0029 | 0.0088 | 0 | 16 | |
| MA2 | 0.0068 | 0.0010 | 0.0078 | 0 | |
| 417 | | | | | 13923 |
| | CR1 | CR2 | MA1 | MA2 | |
| CR1 | 0 | 238 | 67 | 239 | |
| CR2 | 0.0171 | 0 | 227 | 56 | |
| MA1 | 0.0048 | 0.0163 | 0 | 227 | |
| MA2 | 0.0172 | 0.0040 | 0.0163 | 0 | |
| 421 | | | | | 7773 |
| | CR1 | CR2 | MA1 | MA2 | |
| CR1 | 0 | 148 | 91 | 153 | |
| CR2 | 0.0190 | 0 | 144 | 15 | |
| MA1 | 0.0117 | 0.0185 | 0 | 149 | |
| MA2 | 0.0197 | 0.0019 | 0.0192 | 0 | |
| 422 | | | | | 11551 |
| | CR1 | CR2 | MA1 | MA2 | |
| CR1 | 0 | 84 | 145 | 73 | |
| CR2 | 0.0073 | 0 | 125 | 141 | |
| MA1 | 0.0126 | 0.0108 | 0 | 172 | |
| MA2 | 0.0063 | 0.0122 | 0.0149 | 0 | |
| 442 | | | | | 16212 |
| | CR1 | CR2 | MA1 | MA2 | |
| CR1 | 0 | 226 | 114 | 241 | |
| CR2 | 0.0139 | 0 | 218 | 70 | |
| MA1 | 0.0070 | 0.0134 | 0 | 234 | |
| MA2 | 0.0149 | 0.0043 | 0.0144 | 0 | |
| 450 | | | | | 24883 |
| | CR1 | CR2 | MA1 | MA2 | |
| CR1 | 0 | 556 | 295 | 556 | |
| CR2 | 0.0223 | 0 | 587 | 141 | |
| MA1 | 0.0119 | 0.0236 | 0 | 587 | |
| MA2 | 0.0223 | 0.0057 | 0.0236 | 0 | |
| 451 | | | | | 12893 |
| | CR1 | CR2 | MA1 | MA2 | |
| CR1 | 0 | 159 | 154 | 67 | |
| CR2 | 0.0123 | 0 | 40 | 161 | |
| MA1 | 0.0119 | 0.0031 | 0 | 156 | |
| MA2 | 0.0052 | 0.0125 | 0.0121 | 0 | |

| 471 | | | | | 20260 |
|---|---|---|---|---|---|
| | CR1 | CR2 | MA1 | MA2 | |
| CR1 | 0 | 96 | 474 | 426 | |
| CR2 | 0.0047 | 0 | 437 | 486 | |
| MA1 | 0.0234 | 0.0216 | 0 | 99 | |
| MA2 | 0.0210 | 0.0240 | 0.0049 | 0 | |
| 492 | | | | | 11301 |
| | CR1 | CR2 | MA1 | MA2 | |
| CR1 | 0 | 301 | 138 | 300 | |
| CR2 | 0.0266 | 0 | 279 | 86 | |
| MA1 | 0.0122 | 0.0247 | 0 | 283 | |
| MA2 | 0.0265 | 0.0076 | 0.0250 | 0 | |
| 494 | | | | | 8728 |
| | CR1 | CR2 | MA1 | MA2 | |
| CR1 | 0 | 136 | 45 | 83 | |
| CR2 | 0.0156 | 0 | 145 | 97 | |
| MA1 | 0.0052 | 0.0166 | 0 | 70 | |
| MA2 | 0.0095 | 0.0111 | 0.0080 | 0 | |
| 500 | | | | | 10350 |
| | CR1 | CR2 | MA1 | MA2 | |
| CR1 | 0 | 68 | 142 | 158 | |
| CR2 | 0.0066 | 0 | 166 | 116 | |
| MA1 | 0.0137 | 0.0160 | 0 | 74 | |
| MA2 | 0.0153 | 0.0112 | 0.0071 | 0 | |
| 502 | | | | | 10569 |
| | CR1 | CR2 | MA1 | MA2 | |
| CR1 | 0 | 166 | 77 | 166 | |
| CR2 | 0.0157 | 0 | 180 | 48 | |
| MA1 | 0.0073 | 0.0170 | 0 | 182 | |
| MA2 | 0.0157 | 0.0045 | 0.0172 | 0 | |
| 506 | | | | | 8597 |
| | CR1 | CR2 | MA1 | MA2 | |
| CR1 | 0 | 167 | 148 | 48 | |
| CR2 | 0.0194 | 0 | 103 | 168 | |
| MA1 | 0.0172 | 0.0120 | 0 | 155 | |
| MA2 | 0.0056 | 0.0195 | 0.0180 | 0 | |
| 511 | | | | | 14480 |
| | CR1 | CR2 | MA1 | MA2 | |
| CR1 | 0 | 317 | 114 | 306 | |
| CR2 | 0.0219 | 0 | 314 | 180 | |
| MA1 | 0.0079 | 0.0217 | 0 | 306 | |
| MA2 | 0.0211 | 0.0124 | 0.0211 | 0 | |

| 515 | | | | | 17506 |
|---|---|---|---|---|---|
| | CR1 | CR2 | MA1 | MA2 | |
| CR1 | 0 | 82 | 251 | 441 | |
| CR2 | 0.0047 | 0 | 181 | 485 | |
| MA1 | 0.0143 | 0.0103 | 0 | 499 | |
| MA2 | 0.0252 | 0.0277 | 0.0285 | 0 | |

| 521 | | | | | 10935 |
|---|---|---|---|---|---|
| | CR1 | CR2 | MA1 | MA2 | |
| CR1 | 0 | 133 | 118 | 41 | |
| CR2 | 0.0122 | 0 | 50 | 140 | |
| MA1 | 0.0108 | 0.0046 | 0 | 125 | |
| MA2 | 0.0037 | 0.0128 | 0.0114 | 0 | |

| 527 | | | | | 17079 |
|---|---|---|---|---|---|
| | CR1 | CR2 | MA1 | MA2 | |
| CR1 | 0 | 369 | 338 | 187 | |
| CR2 | 0.0216 | 0 | 119 | 370 | |
| MA1 | 0.0198 | 0.0070 | 0 | 339 | |
| MA2 | 0.0109 | 0.0217 | 0.0198 | 0 | |

| 539 | | | | | 14831 |
|---|---|---|---|---|---|
| | CR1 | CR2 | MA1 | MA2 | |
| CR1 | 0 | 59 | 140 | 137 | |
| CR2 | 0.0040 | 0 | 164 | 106 | |
| MA1 | 0.0094 | 0.0111 | 0 | 74 | |
| MA2 | 0.0092 | 0.0071 | 0.0050 | 0 | |

| 562 | | | | | 12038 |
|---|---|---|---|---|---|
| | CR1 | CR2 | MA1 | MA2 | |
| CR1 | 0 | 213 | 64 | 216 | |
| CR2 | 0.0177 | 0 | 211 | 68 | |
| MA1 | 0.0053 | 0.0175 | 0 | 214 | |
| MA2 | 0.0179 | 0.0056 | 0.0178 | 0 | |

| 563 | | | | | 5430 |
|---|---|---|---|---|---|
| | CR1 | CR2 | MA1 | MA2 | |
| CR1 | 0 | 144 | 219 | 219 | |
| CR2 | 0.0265 | 0 | 179 | 179 | |
| MA1 | 0.0403 | 0.0330 | 0 | 0 | |
| MA2 | 0.0403 | 0.0330 | 0.0000 | 0 | |

| 566 | | | | | 7948 |
|---|---|---|---|---|---|
| | CR1 | CR2 | MA1 | MA2 | |
| CR1 | 0 | 204 | 199 | 117 | |
| CR2 | 0.0257 | 0 | 63 | 196 | |
| MA1 | 0.0250 | 0.0079 | 0 | 191 | |
| MA2 | 0.0147 | 0.0247 | 0.0240 | 0 | |

| 573 | | | | | 17006 |
|---|---|---|---|---|---|
| | CR1 | CR2 | MA1 | MA2 | |
| CR1 | 0 | 216 | 119 | 218 | |
| CR2 | 0.0127 | 0 | 233 | 81 | |
| MA1 | 0.0070 | 0.0137 | 0 | 238 | |
| MA2 | 0.0128 | 0.0048 | 0.0140 | 0 | |

| 590 | | | | | 8906 |
|---|---|---|---|---|---|
| | CR1 | CR2 | MA1 | MA2 | |
| CR1 | 0 | 145 | 94 | 158 | |
| CR2 | 0.0163 | 0 | 150 | 48 | |
| MA1 | 0.0106 | 0.0168 | 0 | 155 | |
| MA2 | 0.0177 | 0.0054 | 0.0174 | 0 | |

| 594 | | | | | 10321 |
|---|---|---|---|---|---|
| | CR1 | CR2 | MA1 | MA2 | |
| CR1 | 0 | 303 | 312 | 99 | |
| CR2 | 0.0294 | 0 | 174 | 318 | |
| MA1 | 0.0302 | 0.0169 | 0 | 329 | |
| MA2 | 0.0096 | 0.0308 | 0.0319 | 0 | |

| 604 | | | | | 13188 |
|---|---|---|---|---|---|
| | CR1 | CR2 | MA1 | MA2 | |
| CR1 | 0 | 392 | 408 | 173 | |
| CR2 | 0.0297 | 0 | 254 | 392 | |
| MA1 | 0.0309 | 0.0193 | 0 | 409 | |
| MA2 | 0.0131 | 0.0297 | 0.0310 | 0 | |

| 617 | | | | | 7367 |
|---|---|---|---|---|---|
| | CR1 | CR2 | MA1 | MA2 | |
| CR1 | 0 | 204 | 104 | 209 | |
| CR2 | 0.0277 | 0 | 217 | 36 | |
| MA1 | 0.0141 | 0.0295 | 0 | 224 | |
| MA2 | 0.0284 | 0.0049 | 0.0304 | 0 | |

| 620 | | | | | 9255 |
|---|---|---|---|---|---|
| | CR1 | CR2 | MA1 | MA2 | |
| CR1 | 0 | 127 | 119 | 21 | |
| CR2 | 0.0137 | 0 | 83 | 125 | |
| MA1 | 0.0129 | 0.0090 | 0 | 116 | |
| MA2 | 0.0023 | 0.0135 | 0.0125 | 0 | |

| 622 | | | | | 11142 |
|---|---|---|---|---|---|
| | CR1 | CR2 | MA1 | MA2 | |
| CR1 | 0 | 133 | 107 | 114 | |
| CR2 | 0.0119 | 0 | 94 | 115 | |
| MA1 | 0.0096 | 0.0084 | 0 | 80 | |
| MA2 | 0.0102 | 0.0103 | 0.0072 | 0 | |

| 636 | | | | | 13351 |
|---|---|---|---|---|---|
| | CR1 | CR2 | MA1 | MA2 | |
| CR1 | 0 | 360 | 224 | 294 | |
| CR2 | 0.0270 | 0 | 326 | 207 | |
| MA1 | 0.0168 | 0.0244 | 0 | 180 | |
| MA2 | 0.0220 | 0.0155 | 0.0135 | 0 | |
| 637 | | | | | 10975 |
| | CR1 | CR2 | MA1 | MA2 | |
| CR1 | 0 | 199 | 217 | 102 | |
| CR2 | 0.0181 | 0 | 103 | 190 | |
| MA1 | 0.0198 | 0.0094 | 0 | 196 | |
| MA2 | 0.0093 | 0.0173 | 0.0179 | 0 | |
| 643 | | | | | 14941 |
| | CR1 | CR2 | MA1 | MA2 | |
| CR1 | 0 | 289 | 72 | 279 | |
| CR2 | 0.0193 | 0 | 282 | 162 | |
| MA1 | 0.0048 | 0.0189 | 0 | 270 | |
| MA2 | 0.0187 | 0.0108 | 0.0181 | 0 | |
| 647 | | | | | 11881 |
| | CR1 | CR2 | MA1 | MA2 | |
| CR1 | 0 | 308 | 155 | 291 | |
| CR2 | 0.0259 | 0 | 304 | 205 | |
| MA1 | 0.0130 | 0.0256 | 0 | 268 | |
| MA2 | 0.0245 | 0.0173 | 0.0226 | 0 | |
| 653 | | | | | 8670 |
| | CR1 | CR2 | MA1 | MA2 | |
| CR1 | 0 | 179 | 113 | 120 | |
| CR2 | 0.0206 | 0 | 190 | 144 | |
| MA1 | 0.0130 | 0.0219 | 0 | 63 | |
| MA2 | 0.0138 | 0.0166 | 0.0073 | 0 | |
| 656 | | | | | 13356 |
| | CR1 | CR2 | MA1 | MA2 | |
| CR1 | 0 | 236 | 271 | 393 | |
| CR2 | 0.0177 | 0 | 340 | 293 | |
| MA1 | 0.0203 | 0.0255 | 0 | 435 | |
| MA2 | 0.0294 | 0.0219 | 0.0326 | 0 | |
| 660 | | | | | 7420 |
| | CR1 | CR2 | MA1 | MA2 | |
| CR1 | 0 | 118 | 96 | 122 | |
| CR2 | 0.0159 | 0 | 137 | 44 | |
| MA1 | 0.0129 | 0.0185 | 0 | 140 | |
| MA2 | 0.0164 | 0.0059 | 0.0189 | 0 | |

| 664 | | | | | 11284 |
|---|---|---|---|---|---|
| | CR1 | CR2 | MA1 | MA2 | |
| CR1 | 0 | 211 | 217 | 124 | |
| CR2 | 0.0187 | 0 | 76 | 202 | |
| MA1 | 0.0192 | 0.0067 | 0 | 210 | |
| MA2 | 0.0110 | 0.0179 | 0.0186 | 0 | |
| 665 | | | | | 20031 |
| | CR1 | CR2 | MA1 | MA2 | |
| CR1 | 0 | 289 | 43 | 294 | |
| CR2 | 0.0144 | 0 | 276 | 128 | |
| MA1 | 0.0021 | 0.0138 | 0 | 284 | |
| MA2 | 0.0147 | 0.0064 | 0.0142 | 0 | |
| 666 | | | | | 8051 |
| | CR1 | CR2 | MA1 | MA2 | |
| CR1 | 0 | 184 | 174 | 114 | |
| CR2 | 0.0229 | 0 | 63 | 171 | |
| MA1 | 0.0216 | 0.0078 | 0 | 150 | |
| MA2 | 0.0142 | 0.0212 | 0.0186 | 0 | |
| 675 | | | | | 11012 |
| | CR1 | CR2 | MA1 | MA2 | |
| CR1 | 0 | 191 | 180 | 48 | |
| CR2 | 0.0173 | 0 | 54 | 202 | |
| MA1 | 0.0163 | 0.0049 | 0 | 191 | |
| MA2 | 0.0044 | 0.0183 | 0.0173 | 0 | |
| 680 | | | | | 10411 |
| | CR1 | CR2 | MA1 | MA2 | |
| CR1 | 0 | 275 | 291 | 131 | |
| CR2 | 0.0264 | 0 | 101 | 281 | |
| MA1 | 0.0280 | 0.0097 | 0 | 293 | |
| MA2 | 0.0126 | 0.0270 | 0.0281 | 0 | |
| 683 | | | | | 7837 |
| | CR1 | CR2 | MA1 | MA2 | |
| CR1 | 0 | 116 | 91 | 29 | |
| CR2 | 0.0148 | 0 | 53 | 111 | |
| MA1 | 0.0116 | 0.0068 | 0 | 80 | |
| MA2 | 0.0037 | 0.0142 | 0.0102 | 0 | |

| 5 | | | | | 18450 |
|---|---|---|---|---|---|
| | MA1 | MA2 | MM1 | MM2 | |
| MA1 | 0 | 526 | 525 | 538 | |
| MA2 | 0.0285 | 0 | 201 | 297 | |
| MM1 | 0.0285 | 0.0109 | 0 | 286 | |
| MM2 | 0.0292 | 0.0161 | 0.0155 | 0 | |
| 12 | | | | | 16923 |
| | MA1 | MA2 | MM1 | MM2 | |
| MA1 | 0 | 87 | 88 | 87 | |
| MA2 | 0.0051 | 0 | 1 | 0 | |
| MM1 | 0.0052 | 0.0001 | 0 | 1 | |
| MM2 | 0.0051 | 0.0000 | 0.0001 | 0 | |
| 15 | | | | | 12230 |
| | MA1 | MA2 | MM1 | MM2 | |
| MA1 | 0 | 192 | 0 | 80 | |
| MA2 | 0.0157 | 0 | 192 | 191 | |
| MM1 | 0.0000 | 0.0157 | 0 | 80 | |
| MM2 | 0.0065 | 0.0156 | 0.0065 | 0 | |
| 20 | | | | | 6921 |
| | MA1 | MA2 | MM1 | MM2 | |
| MA1 | 0 | 59 | 62 | 59 | |
| MA2 | 0.0085 | 0 | 24 | 0 | |
| MM1 | 0.0090 | 0.0035 | 0 | 24 | |
| MM2 | 0.0085 | 0.0000 | 0.0035 | 0 | |
| 22 | | | | | 10243 |
| | MA1 | MA2 | MM1 | MM2 | |
| MA1 | 0 | 193 | 193 | 235 | |
| MA2 | 0.0188 | 0 | 0 | 117 | |
| MM1 | 0.0188 | 0.0000 | 0 | 117 | |
| MM2 | 0.0229 | 0.0114 | 0.0114 | 0 | |
| 24 | | | | | 11530 |
| | MA1 | MA2 | MM1 | MM2 | |
| MA1 | 0 | 188 | 7 | 7 | |
| MA2 | 0.0163 | 0 | 187 | 187 | |
| MM1 | 0.0006 | 0.0162 | 0 | 0 | |
| MM2 | 0.0006 | 0.0162 | 0.0000 | 0 | |
| 26 | | | | | 9747 |
| | MA1 | MA2 | MM1 | MM2 | |
| MA1 | 0 | 96 | 232 | 153 | |
| MA2 | 0.0098 | 0 | 286 | 249 | |
| MM1 | 0.0238 | 0.0293 | 0 | 177 | |
| MM2 | 0.0157 | 0.0255 | 0.0182 | 0 | |

| 27 | | | | | 16186 |
|---|---|---|---|---|---|
| | MA1 | MA2 | MM1 | MM2 | |
| MA1 | 0 | 0 | 384 | 384 | |
| MA2 | 0.0000 | 0 | 384 | 384 | |
| MM1 | 0.0237 | 0.0237 | 0 | 0 | |
| MM2 | 0.0237 | 0.0237 | 0.0000 | 0 | |
| 31 | | | | | 15749 |
| | MA1 | MA2 | MM1 | MM2 | |
| MA1 | 0 | 295 | 295 | 311 | |
| MA2 | 0.0187 | 0 | 0 | 118 | |
| MM1 | 0.0187 | 0.0000 | 0 | 118 | |
| MM2 | 0.0197 | 0.0075 | 0.0075 | 0 | |
| 32 | | | | | 8159 |
| | MA1 | MA2 | MM1 | MM2 | |
| MA1 | 0 | 115 | 57 | 0 | |
| MA2 | 0.0141 | 0 | 103 | 115 | |
| MM1 | 0.0070 | 0.0126 | 0 | 57 | |
| MM2 | 0.0000 | 0.0141 | 0.0070 | 0 | |
| 33 | | | | | 12165 |
| | MA1 | MA2 | MM1 | MM2 | |
| MA1 | 0 | 222 | 233 | 245 | |
| MA2 | 0.0182 | 0 | 112 | 102 | |
| MM1 | 0.0192 | 0.0092 | 0 | 128 | |
| MM2 | 0.0201 | 0.0084 | 0.0105 | 0 | |
| 49 | | | | | 10570 |
| | MA1 | MA2 | MM1 | MM2 | |
| MA1 | 0 | 198 | 79 | 7 | |
| MA2 | 0.0187 | 0 | 208 | 199 | |
| MM1 | 0.0075 | 0.0197 | 0 | 72 | |
| MM2 | 0.0007 | 0.0188 | 0.0068 | 0 | |
| 50 | | | | | 10128 |
| | MA1 | MA2 | MM1 | MM2 | |
| MA1 | 0 | 228 | 234 | 228 | |
| MA2 | 0.0225 | 0 | 113 | 0 | |
| MM1 | 0.0231 | 0.0112 | 0 | 113 | |
| MM2 | 0.0225 | 0.0000 | 0.0112 | 0 | |
| 59 | | | | | 9328 |
| | MA1 | MA2 | MM1 | MM2 | |
| MA1 | 0 | 90 | 211 | 9 | |
| MA2 | 0.0096 | 0 | 250 | 85 | |
| MM1 | 0.0226 | 0.0268 | 0 | 210 | |
| MM2 | 0.0010 | 0.0091 | 0.0225 | 0 | |

| 61 | | | | | 17874 |
|---|---|---|---|---|---|
| | MA1 | MA2 | MM1 | MM2 | |
| MA1 | 0 | 371 | 371 | 373 | |
| MA2 | 0.0208 | 0 | 120 | 153 | |
| MM1 | 0.0208 | 0.0067 | 0 | 150 | |
| MM2 | 0.0209 | 0.0086 | 0.0084 | 0 | |
| 65 | | | | | 20700 |
| | MA1 | MA2 | MM1 | MM2 | |
| MA1 | 0 | 440 | 236 | 229 | |
| MA2 | 0.0213 | 0 | 442 | 437 | |
| MM1 | 0.0114 | 0.0214 | 0 | 27 | |
| MM2 | 0.0111 | 0.0211 | 0.0013 | 0 | |
| 67 | | | | | 13724 |
| | MA1 | MA2 | MM1 | MM2 | |
| MA1 | 0 | 162 | 154 | 163 | |
| MA2 | 0.0118 | 0 | 60 | 62 | |
| MM1 | 0.0112 | 0.0044 | 0 | 62 | |
| MM2 | 0.0119 | 0.0045 | 0.0045 | 0 | |
| 71 | | | | | 4589 |
| | MA1 | MA2 | MM1 | MM2 | |
| MA1 | 0 | 6 | 6 | 6 | |
| MA2 | 0.0013 | 0 | 0 | 0 | |
| MM1 | 0.0013 | 0.0000 | 0 | 0 | |
| MM2 | 0.0013 | 0.0000 | 0.0000 | 0 | |
| 72 | | | | | 10511 |
| | MA1 | MA2 | MM1 | MM2 | |
| MA1 | 0 | 209 | 209 | 198 | |
| MA2 | 0.0199 | 0 | 0 | 129 | |
| MM1 | 0.0199 | 0.0000 | 0 | 129 | |
| MM2 | 0.0188 | 0.0123 | 0.0123 | 0 | |
| 73 | | | | | 14785 |
| | MA1 | MA2 | MM1 | MM2 | |
| MA1 | 0 | 175 | 97 | 0 | |
| MA2 | 0.0118 | 0 | 177 | 175 | |
| MM1 | 0.0066 | 0.0120 | 0 | 97 | |
| MM2 | 0.0000 | 0.0118 | 0.0066 | 0 | |
| 83 | | | | | 11604 |
| | MA1 | MA2 | MM1 | MM2 | |
| MA1 | 0 | 279 | 271 | 279 | |
| MA2 | 0.0240 | 0 | 93 | 0 | |
| MM1 | 0.0234 | 0.0080 | 0 | 93 | |
| MM2 | 0.0240 | 0.0000 | 0.0080 | 0 | |

| 87 | | | | | 17781 |
|---|---|---|---|---|---|
| | MA1 | MA2 | MM1 | MM2 | |
| MA1 | 0 | 431 | 418 | 431 | |
| MA2 | 0.0242 | 0 | 111 | 0 | |
| MM1 | 0.0235 | 0.0062 | 0 | 111 | |
| MM2 | 0.0242 | 0.0000 | 0.0062 | 0 | |
| 92 | | | | | 11201 |
| | MA1 | MA2 | MM1 | MM2 | |
| MA1 | 0 | 191 | 193 | 191 | |
| MA2 | 0.0171 | 0 | 68 | 0 | |
| MM1 | 0.0172 | 0.0061 | 0 | 68 | |
| MM2 | 0.0171 | 0.0000 | 0.0061 | 0 | |
| 95 | | | | | 8930 |
| | MA1 | MA2 | MM1 | MM2 | |
| MA1 | 0 | 122 | 165 | 122 | |
| MA2 | 0.0137 | 0 | 87 | 0 | |
| MM1 | 0.0185 | 0.0097 | 0 | 87 | |
| MM2 | 0.0137 | 0.0000 | 0.0097 | 0 | |
| 101 | | | | | 12248 |
| | MA1 | MA2 | MM1 | MM2 | |
| MA1 | 0 | 281 | 0 | 209 | |
| MA2 | 0.0229 | 0 | 281 | 290 | |
| MM1 | 0.0000 | 0.0229 | 0 | 209 | |
| MM2 | 0.0171 | 0.0237 | 0.0171 | 0 | |
| 102 | | | | | 9338 |
| | MA1 | MA2 | MM1 | MM2 | |
| MA1 | 0 | 127 | 139 | 127 | |
| MA2 | 0.0136 | 0 | 51 | 0 | |
| MM1 | 0.0149 | 0.0055 | 0 | 51 | |
| MM2 | 0.0136 | 0.0000 | 0.0055 | 0 | |
| 107 | | | | | 9663 |
| | MA1 | MA2 | MM1 | MM2 | |
| MA1 | 0 | 236 | 118 | 122 | |
| MA2 | 0.0244 | 0 | 236 | 253 | |
| MM1 | 0.0122 | 0.0244 | 0 | 116 | |
| MM2 | 0.0126 | 0.0262 | 0.0120 | 0 | |
| 108 | | | | | 12313 |
| | MA1 | MA2 | MM1 | MM2 | |
| MA1 | 0 | 352 | 362 | 349 | |
| MA2 | 0.0286 | 0 | 232 | 216 | |
| MM1 | 0.0294 | 0.0188 | 0 | 227 | |
| MM2 | 0.0283 | 0.0175 | 0.0184 | 0 | |

| 110 | | | | | 7340 |
|---|---|---|---|---|---|
| | MA1 | MA2 | MM1 | MM2 | |
| MA1 | 0 | 111 | 0 | 52 | |
| MA2 | 0.0151 | 0 | 111 | 120 | |
| MM1 | 0.0000 | 0.0151 | 0 | 52 | |
| MM2 | 0.0071 | 0.0163 | 0.0071 | 0 | |
| 112 | | | | | 8278 |
| | MA1 | MA2 | MM1 | MM2 | |
| MA1 | 0 | 177 | 0 | 50 | |
| MA2 | 0.0214 | 0 | 177 | 181 | |
| MM1 | 0.0000 | 0.0214 | 0 | 50 | |
| MM2 | 0.0060 | 0.0219 | 0.0060 | 0 | |
| 121 | | | | | 2149 |
| | MA1 | MA2 | MM1 | MM2 | |
| MA1 | 0 | 23 | 13 | 12 | |
| MA2 | 0.0107 | 0 | 16 | 17 | |
| MM1 | 0.0060 | 0.0074 | 0 | 3 | |
| MM2 | 0.0056 | 0.0079 | 0.0014 | 0 | |
| 125 | | | | | 9061 |
| | MA1 | MA2 | MM1 | MM2 | |
| MA1 | 0 | 219 | 219 | 217 | |
| MA2 | 0.0242 | 0 | 0 | 93 | |
| MM1 | 0.0242 | 0.0000 | 0 | 93 | |
| MM2 | 0.0239 | 0.0103 | 0.0103 | 0 | |
| 127 | | | | | 11815 |
| | MA1 | MA2 | MM1 | MM2 | |
| MA1 | 0 | 250 | 127 | 95 | |
| MA2 | 0.0212 | 0 | 238 | 228 | |
| MM1 | 0.0107 | 0.0201 | 0 | 114 | |
| MM2 | 0.0080 | 0.0193 | 0.0096 | 0 | |
| 131 | | | | | 12171 |
| | MA1 | MA2 | MM1 | MM2 | |
| MA1 | 0 | 0 | 35 | 99 | |
| MA2 | 0.0000 | 0 | 35 | 99 | |
| MM1 | 0.0029 | 0.0029 | 0 | 132 | |
| MM2 | 0.0081 | 0.0081 | 0.0108 | 0 | |
| 133 | | | | | 8383 |
| | MA1 | MA2 | MM1 | MM2 | |
| MA1 | 0 | 66 | 60 | 103 | |
| MA2 | 0.0079 | 0 | 6 | 76 | |
| MM1 | 0.0072 | 0.0007 | 0 | 70 | |
| MM2 | 0.0123 | 0.0091 | 0.0084 | 0 | |

| 141 | | | | | 6949 |
|---|---|---|---|---|---|
| | MA1 | MA2 | MM1 | MM2 | |
| MA1 | 0 | 145 | 145 | 145 | |
| MA2 | 0.0209 | 0 | 0 | 73 | |
| MM1 | 0.0209 | 0.0000 | 0 | 73 | |
| MM2 | 0.0209 | 0.0105 | 0.0105 | 0 | |
| 146 | | | | | 21096 |
| | MA1 | MA2 | MM1 | MM2 | |
| MA1 | 0 | 514 | 508 | 514 | |
| MA2 | 0.0244 | 0 | 358 | 0 | |
| MM1 | 0.0241 | 0.0170 | 0 | 358 | |
| MM2 | 0.0244 | 0.0000 | 0.0170 | 0 | |
| 147 | | | | | 8173 |
| | MA1 | MA2 | MM1 | MM2 | |
| MA1 | 0 | 0 | 109 | 126 | |
| MA2 | 0.0000 | 0 | 109 | 126 | |
| MM1 | 0.0133 | 0.0133 | 0 | 60 | |
| MM2 | 0.0154 | 0.0154 | 0.0073 | 0 | |
| 149 | | | | | 15239 |
| | MA1 | MA2 | MM1 | MM2 | |
| MA1 | 0 | 346 | 0 | 174 | |
| MA2 | 0.0227 | 0 | 346 | 352 | |
| MM1 | 0.0000 | 0.0227 | 0 | 174 | |
| MM2 | 0.0114 | 0.0231 | 0.0114 | 0 | |
| 150 | | | | | 3171 |
| | MA1 | MA2 | MM1 | MM2 | |
| MA1 | 0 | 77 | 70 | 77 | |
| MA2 | 0.0243 | 0 | 36 | 0 | |
| MM1 | 0.0221 | 0.0114 | 0 | 36 | |
| MM2 | 0.0243 | 0.0000 | 0.0114 | 0 | |
| 153 | | | | | 7908 |
| | MA1 | MA2 | MM1 | MM2 | |
| MA1 | 0 | 149 | 151 | 146 | |
| MA2 | 0.0188 | 0 | 2 | 63 | |
| MM1 | 0.0191 | 0.0003 | 0 | 61 | |
| MM2 | 0.0185 | 0.0080 | 0.0077 | 0 | |
| 158 | | | | | 17511 |
| | MA1 | MA2 | MM1 | MM2 | |
| MA1 | 0 | 529 | 529 | 510 | |
| MA2 | 0.0302 | 0 | 0 | 260 | |
| MM1 | 0.0302 | 0.0000 | 0 | 260 | |
| MM2 | 0.0291 | 0.0148 | 0.0148 | 0 | |

| 165 | | | | | 6918 |
|---|---|---|---|---|---|
| | MA1 | MA2 | MM1 | MM2 | |
| MA1 | 0 | 147 | 147 | 153 | |
| MA2 | 0.0212 | 0 | 0 | 62 | |
| MM1 | 0.0212 | 0.0000 | 0 | 62 | |
| MM2 | 0.0221 | 0.0090 | 0.0090 | 0 | |
| 167 | | | | | 11542 |
| | MA1 | MA2 | MM1 | MM2 | |
| MA1 | 0 | 220 | 233 | 220 | |
| MA2 | 0.0191 | 0 | 130 | 0 | |
| MM1 | 0.0202 | 0.0113 | 0 | 130 | |
| MM2 | 0.0191 | 0.0000 | 0.0113 | 0 | |
| 173 | | | | | 9180 |
| | MA1 | MA2 | MM1 | MM2 | |
| MA1 | 0 | 191 | 191 | 191 | |
| MA2 | 0.0208 | 0 | 68 | 0 | |
| MM1 | 0.0208 | 0.0074 | 0 | 68 | |
| MM2 | 0.0208 | 0.0000 | 0.0074 | 0 | |
| 176 | | | | | 10626 |
| | MA1 | MA2 | MM1 | MM2 | |
| MA1 | 0 | 252 | 231 | 274 | |
| MA2 | 0.0237 | 0 | 168 | 102 | |
| MM1 | 0.0217 | 0.0158 | 0 | 141 | |
| MM2 | 0.0258 | 0.0096 | 0.0133 | 0 | |
| 178 | | | | | 7766 |
| | MA1 | MA2 | MM1 | MM2 | |
| MA1 | 0 | 74 | 128 | 164 | |
| MA2 | 0.0095 | 0 | 202 | 202 | |
| MM1 | 0.0165 | 0.0260 | 0 | 78 | |
| MM2 | 0.0211 | 0.0260 | 0.0100 | 0 | |
| 179 | | | | | 13073 |
| | MA1 | MA2 | MM1 | MM2 | |
| MA1 | 0 | 261 | 155 | 0 | |
| MA2 | 0.0200 | 0 | 275 | 261 | |
| MM1 | 0.0119 | 0.0210 | 0 | 155 | |
| MM2 | 0.0000 | 0.0200 | 0.0119 | 0 | |
| 181 | | | | | 13000 |
| | MA1 | MA2 | MM1 | MM2 | |
| MA1 | 0 | 225 | 140 | 0 | |
| MA2 | 0.0173 | 0 | 223 | 225 | |
| MM1 | 0.0108 | 0.0172 | 0 | 140 | |
| MM2 | 0.0000 | 0.0173 | 0.0108 | 0 | |

| 183 | | | | | 13698 |
|---|---|---|---|---|---|
| | MA1 | MA2 | MM1 | MM2 | |
| MA1 | 0 | 15 | 168 | 15 | |
| MA2 | 0.0011 | 0 | 155 | 0 | |
| MM1 | 0.0123 | 0.0113 | 0 | 155 | |
| MM2 | 0.0011 | 0.0000 | 0.0113 | 0 | |
| 184 | | | | | 14196 |
| | MA1 | MA2 | MM1 | MM2 | |
| MA1 | 0 | 296 | 296 | 293 | |
| MA2 | 0.0209 | 0 | 0 | 168 | |
| MM1 | 0.0209 | 0.0000 | 0 | 168 | |
| MM2 | 0.0206 | 0.0118 | 0.0118 | 0 | |
| 189 | | | | | 12182 |
| | MA1 | MA2 | MM1 | MM2 | |
| MA1 | 0 | 229 | 220 | 229 | |
| MA2 | 0.0188 | 0 | 142 | 0 | |
| MM1 | 0.0181 | 0.0117 | 0 | 142 | |
| MM2 | 0.0188 | 0.0000 | 0.0117 | 0 | |
| 191 | | | | | 10068 |
| | MA1 | MA2 | MM1 | MM2 | |
| MA1 | 0 | 243 | 108 | 115 | |
| MA2 | 0.0241 | 0 | 235 | 246 | |
| MM1 | 0.0107 | 0.0233 | 0 | 130 | |
| MM2 | 0.0114 | 0.0244 | 0.0129 | 0 | |
| 196 | | | | | 7081 |
| | MA1 | MA2 | MM1 | MM2 | |
| MA1 | 0 | 167 | 195 | 179 | |
| MA2 | 0.0236 | 0 | 92 | 97 | |
| MM1 | 0.0275 | 0.0130 | 0 | 106 | |
| MM2 | 0.0253 | 0.0137 | 0.0150 | 0 | |
| 198 | | | | | 15581 |
| | MA1 | MA2 | MM1 | MM2 | |
| MA1 | 0 | 403 | 411 | 403 | |
| MA2 | 0.0259 | 0 | 270 | 0 | |
| MM1 | 0.0264 | 0.0173 | 0 | 270 | |
| MM2 | 0.0259 | 0.0000 | 0.0173 | 0 | |
| 199 | | | | | 7382 |
| | MA1 | MA2 | MM1 | MM2 | |
| MA1 | 0 | 117 | 0 | 51 | |
| MA2 | 0.0158 | 0 | 117 | 121 | |
| MM1 | 0.0000 | 0.0158 | 0 | 51 | |
| MM2 | 0.0069 | 0.0164 | 0.0069 | 0 | |

| 203 | | | | | 21330 |
|---|---|---|---|---|---|
| | MA1 | MA2 | MM1 | MM2 | |
| MA1 | 0 | 445 | 244 | 241 | |
| MA2 | 0.0209 | 0 | 471 | 461 | |
| MM1 | 0.0114 | 0.0221 | 0 | 235 | |
| MM2 | 0.0113 | 0.0216 | 0.0110 | 0 | |
| 206 | | | | | 11994 |
| | MA1 | MA2 | MM1 | MM2 | |
| MA1 | 0 | 265 | 265 | 265 | |
| MA2 | 0.0221 | 0 | 0 | 144 | |
| MM1 | 0.0221 | 0.0000 | 0 | 144 | |
| MM2 | 0.0221 | 0.0120 | 0.0120 | 0 | |
| 212 | | | | | 13556 |
| | MA1 | MA2 | MM1 | MM2 | |
| MA1 | 0 | 196 | 200 | 196 | |
| MA2 | 0.0145 | 0 | 78 | 0 | |
| MM1 | 0.0148 | 0.0058 | 0 | 78 | |
| MM2 | 0.0145 | 0.0000 | 0.0058 | 0 | |
| 213 | | | | | 10560 |
| | MA1 | MA2 | MM1 | MM2 | |
| MA1 | 0 | 184 | 95 | 0 | |
| MA2 | 0.0174 | 0 | 183 | 184 | |
| MM1 | 0.0090 | 0.0173 | 0 | 95 | |
| 215 | | | | | 26343 |
| | MA1 | MA2 | MM1 | MM2 | |
| MA1 | 0 | 598 | 364 | 351 | |
| MA2 | 0.0227 | 0 | 588 | 625 | |
| MM1 | 0.0138 | 0.0223 | 0 | 340 | |
| MM2 | 0.0133 | 0.0237 | 0.0129 | 0 | |
| 218 | | | | | 9450 |
| | MA1 | MA2 | MM1 | MM2 | |
| MA1 | 0 | 291 | 198 | 194 | |
| MA2 | 0.0308 | 0 | 323 | 327 | |
| MM1 | 0.0210 | 0.0342 | 0 | 56 | |
| MM2 | 0.0205 | 0.0346 | 0.0059 | 0 | |
| 219 | | | | | 14751 |
| | MA1 | MA2 | MM1 | MM2 | |
| MA1 | 0 | 0 | 734 | 734 | |
| MA2 | 0.0000 | 0 | 734 | 734 | |
| MM1 | 0.0498 | 0.0498 | 0 | 0 | |
| MM2 | 0.0498 | 0.0498 | 0.0000 | 0 | |
| 223 | | | | | 13504 |

| | MA1 | MA2 | MM1 | MM2 | |
|---|---|---|---|---|---|
| MA1 | 0 | 277 | 109 | 128 | |
| MA2 | 0.0205 | 0 | 283 | 283 | |
| MM1 | 0.0081 | 0.0210 | 0 | 121 | |
| MM2 | 0.0095 | 0.0210 | 0.0090 | 0 | |
| 225 | | | | | 10566 |
| | MA1 | MA2 | MM1 | MM2 | |
| MA1 | 0 | 231 | 225 | 225 | |
| MA2 | 0.0219 | 0 | 42 | 83 | |
| MM1 | 0.0213 | 0.0040 | 0 | 81 | |
| MM2 | 0.0213 | 0.0079 | 0.0077 | 0 | |
| 226 | | | | | 12080 |
| | MA1 | MA2 | MM1 | MM2 | |
| MA1 | 0 | 0 | 117 | 259 | |
| MA2 | 0.0000 | 0 | 117 | 259 | |
| MM1 | 0.0097 | 0.0097 | 0 | 142 | |
| MM2 | 0.0214 | 0.0214 | 0.0118 | 0 | |
| 227 | | | | | 8565 |
| | MA1 | MA2 | MM1 | MM2 | |
| MA1 | 0 | 43 | 148 | 176 | |
| MA2 | 0.0050 | 0 | 171 | 201 | |
| MM1 | 0.0173 | 0.0200 | 0 | 148 | |
| MM2 | 0.0205 | 0.0235 | 0.0173 | 0 | |
| 228 | | | | | 12250 |
| | MA1 | MA2 | MM1 | MM2 | |
| MA1 | 0 | 340 | 346 | 331 | |
| MA2 | 0.0278 | 0 | 98 | 143 | |
| MM1 | 0.0282 | 0.0080 | 0 | 157 | |
| MM2 | 0.0270 | 0.0117 | 0.0128 | 0 | |
| 237 | | | | | 15897 |
| | MA1 | MA2 | MM1 | MM2 | |
| MA1 | 0 | 369 | 271 | 0 | |
| MA2 | 0.0232 | 0 | 337 | 369 | |
| MM1 | 0.0170 | 0.0212 | 0 | 271 | |
| MM2 | 0.0000 | 0.0232 | 0.0170 | 0 | |
| 238 | | | | | 8873 |
| | MA1 | MA2 | MM1 | MM2 | |
| MA1 | 0 | 210 | 0 | 112 | |
| MA2 | 0.0237 | 0 | 210 | 215 | |
| MM1 | 0.0000 | 0.0237 | 0 | 112 | |
| MM2 | 0.0126 | 0.0242 | 0.0126 | 0 | |
| 240 | | | | | 12228 |

| | MA1 | MA2 | MM1 | MM2 | |
|---|---|---|---|---|---|
| MA1 | 0 | 271 | 130 | 0 | |
| MA2 | 0.0222 | 0 | 275 | 271 | |
| MM1 | 0.0106 | 0.0225 | 0 | 130 | |
| MM2 | 0.0000 | 0.0222 | 0.0106 | 0 | |
| 248 | | | | | 8518 |
| | MA1 | MA2 | MM1 | MM2 | |
| MA1 | 0 | 189 | 200 | 212 | |
| MA2 | 0.0222 | 0 | 99 | 110 | |
| MM1 | 0.0235 | 0.0116 | 0 | 76 | |
| MM2 | 0.0249 | 0.0129 | 0.0089 | 0 | |
| 251 | | | | | 15270 |
| | MA1 | MA2 | MM1 | MM2 | |
| MA1 | 0 | 453 | 453 | 451 | |
| MA2 | 0.0297 | 0 | 0 | 268 | |
| MM1 | 0.0297 | 0.0000 | 0 | 268 | |
| MM2 | 0.0295 | 0.0176 | 0.0176 | 0 | |
| 256 | | | | | 10867 |
| | MA1 | MA2 | MM1 | MM2 | |
| MA1 | 0 | 188 | 183 | 188 | |
| MA2 | 0.0173 | 0 | 64 | 0 | |
| MM1 | 0.0168 | 0.0059 | 0 | 64 | |
| MM2 | 0.0173 | 0.0000 | 0.0059 | 0 | |
| 261 | | | | | 9635 |
| | MA1 | MA2 | MM1 | MM2 | |
| MA1 | 0 | 226 | 218 | 226 | |
| MA2 | 0.0235 | 0 | 92 | 0 | |
| MM1 | 0.0226 | 0.0095 | 0 | 92 | |
| MM2 | 0.0235 | 0.0000 | 0.0095 | 0 | |
| 264 | | | | | 5077 |
| | MA1 | MA2 | MM1 | MM2 | |
| MA1 | 0 | 220 | 237 | 233 | |
| MA2 | 0.0433 | 0 | 167 | 140 | |
| MM1 | 0.0467 | 0.0329 | 0 | 27 | |
| MM2 | 0.0459 | 0.0276 | 0.0053 | 0 | |
| 267 | | | | | 12084 |
| | MA1 | MA2 | MM1 | MM2 | |
| MA1 | 0 | 114 | 109 | 95 | |
| MA2 | 0.0094 | 0 | 121 | 123 | |
| MM1 | 0.0090 | 0.0100 | 0 | 67 | |
| MM2 | 0.0079 | 0.0102 | 0.0055 | 0 | |
| 272 | | | | | 8749 |

| | MA1 | MA2 | MM1 | MM2 |
|---|---|---|---|---|
| MA1 | 0 | 164 | 169 | 164 |
| MA2 | 0.0187 | 0 | 75 | 0 |
| MM1 | 0.0193 | 0.0086 | 0 | 75 |
| MM2 | 0.0187 | 0.0000 | 0.0086 | 0 |

277 14114

| | MA1 | MA2 | MM1 | MM2 |
|---|---|---|---|---|
| MA1 | 0 | 222 | 228 | 221 |
| MA2 | 0.0157 | 0 | 63 | 109 |
| MM1 | 0.0162 | 0.0045 | 0 | 127 |
| MM2 | 0.0157 | 0.0077 | 0.0090 | 0 |

280 6919

| | MA1 | MA2 | MM1 | MM2 |
|---|---|---|---|---|
| MA1 | 0 | 91 | 104 | 97 |
| MA2 | 0.0132 | 0 | 56 | 33 |
| MM1 | 0.0150 | 0.0081 | 0 | 58 |
| MM2 | 0.0140 | 0.0048 | 0.0084 | 0 |

282 7450

| | MA1 | MA2 | MM1 | MM2 |
|---|---|---|---|---|
| MA1 | 0 | 176 | 122 | 104 |
| MA2 | 0.0236 | 0 | 196 | 161 |
| MM1 | 0.0164 | 0.0263 | 0 | 107 |
| MM2 | 0.0140 | 0.0216 | 0.0144 | 0 |

291 18560

| | MA1 | MA2 | MM1 | MM2 |
|---|---|---|---|---|
| MA1 | 0 | 367 | 386 | 371 |
| MA2 | 0.0198 | 0 | 171 | 193 |
| MM1 | 0.0208 | 0.0092 | 0 | 149 |
| MM2 | 0.0200 | 0.0104 | 0.0080 | 0 |

293 11376

| | MA1 | MA2 | MM1 | MM2 |
|---|---|---|---|---|
| MA1 | 0 | 223 | 223 | 217 |
| MA2 | 0.0196 | 0 | 0 | 147 |
| MM1 | 0.0196 | 0.0000 | 0 | 147 |
| MM2 | 0.0191 | 0.0129 | 0.0129 | 0 |

295 27334

| | MA1 | MA2 | MM1 | MM2 |
|---|---|---|---|---|
| MA1 | 0 | 303 | 0 | 282 |
| MA2 | 0.0111 | 0 | 303 | 442 |
| MM1 | 0.0000 | 0.0111 | 0 | 282 |
| MM2 | 0.0103 | 0.0162 | 0.0103 | 0 |

297 8864

| | MA1 | MA2 | MM1 | MM2 |
|---|---|---|---|---|
| MA1 | 0 | 254 | 180 | 0 |
| MA2 | 0.0287 | 0 | 256 | 254 |
| MM1 | 0.0203 | 0.0289 | 0 | 180 |
| MM2 | 0.0000 | 0.0287 | 0.0203 | 0 |

299 16716

| | MA1 | MA2 | MM1 | MM2 |
|---|---|---|---|---|
| MA1 | 0 | 348 | 333 | 344 |
| MA2 | 0.0208 | 0 | 202 | 205 |
| MM1 | 0.0199 | 0.0121 | 0 | 188 |
| MM2 | 0.0206 | 0.0123 | 0.0112 | 0 |

301 9426

| | MA1 | MA2 | MM1 | MM2 |
|---|---|---|---|---|
| MA1 | 0 | 183 | 190 | 192 |
| MA2 | 0.0194 | 0 | 93 | 104 |
| MM1 | 0.0202 | 0.0099 | 0 | 79 |
| MM2 | 0.0204 | 0.0110 | 0.0084 | 0 |

304 27288

| | MA1 | MA2 | MM1 | MM2 |
|---|---|---|---|---|
| MA1 | 0 | 603 | 617 | 603 |
| MA2 | 0.0221 | 0 | 302 | 0 |
| MM1 | 0.0226 | 0.0111 | 0 | 302 |
| MM2 | 0.0221 | 0.0000 | 0.0111 | 0 |

305 18319

| | MA1 | MA2 | MM1 | MM2 |
|---|---|---|---|---|
| MA1 | 0 | 502 | 514 | 500 |
| MA2 | 0.0274 | 0 | 240 | 218 |
| MM1 | 0.0281 | 0.0131 | 0 | 171 |
| MM2 | 0.0273 | 0.0119 | 0.0093 | 0 |

313 10254

| | MA1 | MA2 | MM1 | MM2 |
|---|---|---|---|---|
| MA1 | 0 | 253 | 0 | 99 |
| MA2 | 0.0247 | 0 | 253 | 265 |
| MM1 | 0.0000 | 0.0247 | 0 | 99 |
| MM2 | 0.0097 | 0.0258 | 0.0097 | 0 |

314 9821

| | MA1 | MA2 | MM1 | MM2 |
|---|---|---|---|---|
| MA1 | 0 | 168 | 0 | 115 |
| MA2 | 0.0171 | 0 | 168 | 153 |
| MM1 | 0.0000 | 0.0171 | 0 | 115 |
| MM2 | 0.0117 | 0.0156 | 0.0117 | 0 |

315 10166

| | MA1 | MA2 | MM1 | MM2 | |
|---|---|---|---|---|---|
| MA1 | 0 | 108 | 45 | 65 | |
| MA2 | 0.0106 | 0 | 113 | 101 | |
| MM1 | 0.0044 | 0.0111 | 0 | 68 | |
| MM2 | 0.0064 | 0.0099 | 0.0067 | 0 | |
| 318 | | | | | 11241 |
| | MA1 | MA2 | MM1 | MM2 | |
| MA1 | 0 | 204 | 207 | 204 | |
| MA2 | 0.0181 | 0 | 88 | 0 | |
| MM1 | 0.0184 | 0.0078 | 0 | 88 | |
| MM2 | 0.0181 | 0.0000 | 0.0078 | 0 | |
| 322 | | | | | 8176 |
| | MA1 | MA2 | MM1 | MM2 | |
| MA1 | 0 | 112 | 108 | 118 | |
| MA2 | 0.0137 | 0 | 37 | 40 | |
| MM1 | 0.0132 | 0.0045 | 0 | 43 | |
| MM2 | 0.0144 | 0.0049 | 0.0053 | 0 | |
| 324 | | | | | 18535 |
| | MA1 | MA2 | MM1 | MM2 | |
| MA1 | 0 | 555 | 355 | 285 | |
| MA2 | 0.0299 | 0 | 578 | 555 | |
| MM1 | 0.0192 | 0.0312 | 0 | 333 | |
| MM2 | 0.0154 | 0.0299 | 0.0180 | 0 | |
| 325 | | | | | 12373 |
| | MA1 | MA2 | MM1 | MM2 | |
| MA1 | 0 | 305 | 0 | 92 | |
| MA2 | 0.0247 | 0 | 305 | 310 | |
| MM1 | 0.0000 | 0.0247 | 0 | 92 | |
| MM2 | 0.0074 | 0.0251 | 0.0074 | 0 | |
| 331 | | | | | 10557 |
| | MA1 | MA2 | MM1 | MM2 | |
| MA1 | 0 | 217 | 69 | 0 | |
| MA2 | 0.0206 | 0 | 222 | 217 | |
| MM1 | 0.0065 | 0.0210 | 0 | 69 | |
| MM2 | 0.0000 | 0.0206 | 0.0065 | 0 | |
| 333 | | | | | 14395 |
| | MA1 | MA2 | MM1 | MM2 | |
| MA1 | 0 | 339 | 340 | 337 | |
| MA2 | 0.0235 | 0 | 3 | 124 | |
| MM1 | 0.0236 | 0.0002 | 0 | 121 | |
| MM2 | 0.0234 | 0.0086 | 0.0084 | 0 | |
| 334 | | | | | 15613 |

| | MA1 | MA2 | MM1 | MM2 | |
|---|---|---|---|---|---|
| MA1 | 0 | 479 | 479 | 466 | |
| MA2 | 0.0307 | 0 | 0 | 155 | |
| MM1 | 0.0307 | 0.0000 | 0 | 155 | |
| MM2 | 0.0298 | 0.0099 | 0.0099 | 0 | |
| 341 | | | | | 11160 |
| | MA1 | MA2 | MM1 | MM2 | |
| MA1 | 0 | 261 | 252 | 264 | |
| MA2 | 0.0234 | 0 | 97 | 142 | |
| MM1 | 0.0226 | 0.0087 | 0 | 149 | |
| MM2 | 0.0237 | 0.0127 | 0.0134 | 0 | |
| 343 | | | | | 15745 |
| | MA1 | MA2 | MM1 | MM2 | |
| MA1 | 0 | 280 | 0 | 100 | |
| MA2 | 0.0178 | 0 | 280 | 305 | |
| MM1 | 0.0000 | 0.0178 | 0 | 100 | |
| MM2 | 0.0064 | 0.0194 | 0.0064 | 0 | |
| 352 | | | | | 9213 |
| | MA1 | MA2 | MM1 | MM2 | |
| MA1 | 0 | 0 | 288 | 220 | |
| MA2 | 0.0000 | 0 | 288 | 220 | |
| MM1 | 0.0313 | 0.0313 | 0 | 189 | |
| MM2 | 0.0239 | 0.0239 | 0.0205 | 0 | |
| 353 | | | | | 10113 |
| | MA1 | MA2 | MM1 | MM2 | |
| MA1 | 0 | 196 | 196 | 203 | |
| MA2 | 0.0194 | 0 | 97 | 113 | |
| MM1 | 0.0194 | 0.0096 | 0 | 102 | |
| MM2 | 0.0201 | 0.0112 | 0.0101 | 0 | |
| 356 | | | | | 11976 |
| | MA1 | MA2 | MM1 | MM2 | |
| MA1 | 0 | 229 | 230 | 245 | |
| MA2 | 0.0191 | 0 | 79 | 123 | |
| MM1 | 0.0192 | 0.0066 | 0 | 90 | |
| MM2 | 0.0205 | 0.0103 | 0.0075 | 0 | |
| 357 | | | | | 9415 |
| | MA1 | MA2 | MM1 | MM2 | |
| MA1 | 0 | 214 | 120 | 120 | |
| MA2 | 0.0227 | 0 | 216 | 217 | |
| MM1 | 0.0127 | 0.0229 | 0 | 99 | |
| MM2 | 0.0127 | 0.0230 | 0.0105 | 0 | |
| 363 | | | | | 9876 |

| | MA1 | MA2 | MM1 | MM2 | |
|---|---|---|---|---|---|
| MA1 | 0 | 0 | 208 | 180 | |
| MA2 | 0.0000 | 0 | 208 | 180 | |
| MM1 | 0.0211 | 0.0211 | 0 | 126 | |
| MM2 | 0.0182 | 0.0182 | 0.0128 | 0 | |
| 365 | | | | | 10627 |
| | MA1 | MA2 | MM1 | MM2 | |
| MA1 | 0 | 281 | 113 | 0 | |
| MA2 | 0.0264 | 0 | 269 | 281 | |
| MM1 | 0.0106 | 0.0253 | 0 | 113 | |
| MM2 | 0.0000 | 0.0264 | 0.0106 | 0 | |
| 367 | | | | | 11533 |
| | MA1 | MA2 | MM1 | MM2 | |
| MA1 | 0 | 171 | 85 | 62 | |
| MA2 | 0.0148 | 0 | 197 | 172 | |
| MM1 | 0.0074 | 0.0171 | 0 | 92 | |
| MM2 | 0.0054 | 0.0149 | 0.0080 | 0 | |
| 368 | | | | | 14466 |
| | MA1 | MA2 | MM1 | MM2 | |
| MA1 | 0 | 318 | 130 | 0 | |
| MA2 | 0.0220 | 0 | 323 | 318 | |
| MM1 | 0.0090 | 0.0223 | 0 | 130 | |
| MM2 | 0.0000 | 0.0220 | 0.0090 | 0 | |
| 372 | | | | | 11051 |
| | MA1 | MA2 | MM1 | MM2 | |
| MA1 | 0 | 0 | 129 | 0 | |
| MA2 | 0.0000 | 0 | 129 | 0 | |
| MM1 | 0.0117 | 0.0117 | 0 | 129 | |
| MM2 | 0.0000 | 0.0000 | 0.0117 | 0 | |
| 373 | | | | | 7207 |
| | MA1 | MA2 | MM1 | MM2 | |
| MA1 | 0 | 192 | 67 | 115 | |
| MA2 | 0.0266 | 0 | 186 | 201 | |
| MM1 | 0.0093 | 0.0258 | 0 | 118 | |
| MM2 | 0.0160 | 0.0279 | 0.0164 | 0 | |
| 374 | | | | | 14227 |
| | MA1 | MA2 | MM1 | MM2 | |
| MA1 | 0 | 343 | 167 | 146 | |
| MA2 | 0.0241 | 0 | 344 | 322 | |
| MM1 | 0.0117 | 0.0242 | 0 | 182 | |
| MM2 | 0.0103 | 0.0226 | 0.0128 | 0 | |
| 379 | | | | | 8037 |

| | MA1 | MA2 | MM1 | MM2 | |
|---|---|---|---|---|---|
| MA1 | 0 | 393 | 387 | 393 | |
| MA2 | 0.0489 | 0 | 110 | 0 | |
| MM1 | 0.0482 | 0.0137 | 0 | 110 | |
| MM2 | 0.0489 | 0.0000 | 0.0137 | 0 | |
| 382 | | | | | 14446 |
| | MA1 | MA2 | MM1 | MM2 | |
| MA1 | 0 | 334 | 280 | 334 | |
| MA2 | 0.0231 | 0 | 196 | 0 | |
| MM1 | 0.0194 | 0.0136 | 0 | 196 | |
| MM2 | 0.0231 | 0.0000 | 0.0136 | 0 | |
| 383 | | | | | 9559 |
| | MA1 | MA2 | MM1 | MM2 | |
| MA1 | 0 | 249 | 95 | 0 | |
| MA2 | 0.0260 | 0 | 249 | 249 | |
| MM1 | 0.0099 | 0.0260 | 0 | 95 | |
| MM2 | 0.0000 | 0.0260 | 0.0099 | 0 | |
| 386 | | | | | 14656 |
| | MA1 | MA2 | MM1 | MM2 | |
| MA1 | 0 | 143 | 166 | 143 | |
| MA2 | 0.0098 | 0 | 67 | 0 | |
| MM1 | 0.0113 | 0.0046 | 0 | 67 | |
| MM2 | 0.0098 | 0.0000 | 0.0046 | 0 | |
| 388 | | | | | 9821 |
| | MA1 | MA2 | MM1 | MM2 | |
| MA1 | 0 | 108 | 8 | 8 | |
| MA2 | 0.0110 | 0 | 116 | 116 | |
| MM1 | 0.0008 | 0.0118 | 0 | 0 | |
| MM2 | 0.0008 | 0.0118 | 0.0000 | 0 | |
| 389 | | | | | 13805 |
| | MA1 | MA2 | MM1 | MM2 | |
| MA1 | 0 | 288 | 288 | 269 | |
| MA2 | 0.0209 | 0 | 0 | 110 | |
| MM1 | 0.0209 | 0.0000 | 0 | 110 | |
| MM2 | 0.0195 | 0.0080 | 0.0080 | 0 | |
| 392 | | | | | 7787 |
| | MA1 | MA2 | MM1 | MM2 | |
| MA1 | 0 | 197 | 218 | 195 | |
| MA2 | 0.0253 | 0 | 137 | 91 | |
| MM1 | 0.0280 | 0.0176 | 0 | 123 | |
| MM2 | 0.0250 | 0.0117 | 0.0158 | 0 | |
| 393 | | | | | 19247 |

| | MA1 | MA2 | MM1 | MM2 | |
|---|---|---|---|---|---|
| MA1 | 0 | 371 | 4 | 186 | |
| MA2 | 0.0193 | 0 | 373 | 404 | |
| MM1 | 0.0002 | 0.0194 | 0 | 190 | |
| MM2 | 0.0097 | 0.0210 | 0.0099 | 0 | |
| 396 | | | | | 11067 |
| | MA1 | MA2 | MM1 | MM2 | |
| MA1 | 0 | 3 | 259 | 241 | |
| MA2 | 0.0003 | 0 | 260 | 242 | |
| MM1 | 0.0234 | 0.0235 | 0 | 116 | |
| MM2 | 0.0218 | 0.0219 | 0.0105 | 0 | |
| 398 | | | | | 10345 |
| | MA1 | MA2 | MM1 | MM2 | |
| MA1 | 0 | 197 | 127 | 123 | |
| MA2 | 0.0190 | 0 | 218 | 205 | |
| MM1 | 0.0123 | 0.0211 | 0 | 143 | |
| MM2 | 0.0119 | 0.0198 | 0.0138 | 0 | |
| 404 | | | | | 18959 |
| | MA1 | MA2 | MM1 | MM2 | |
| MA1 | 0 | 327 | 150 | 0 | |
| MA2 | 0.0172 | 0 | 335 | 327 | |
| MM1 | 0.0079 | 0.0177 | 0 | 150 | |
| MM2 | 0.0000 | 0.0172 | 0.0079 | 0 | |
| 408 | | | | | 7635 |
| | MA1 | MA2 | MM1 | MM2 | |
| MA1 | 0 | 349 | 303 | 303 | |
| MA2 | 0.0457 | 0 | 229 | 200 | |
| MM1 | 0.0397 | 0.0300 | 0 | 29 | |
| MM2 | 0.0397 | 0.0262 | 0.0038 | 0 | |
| 413 | | | | | 10286 |
| | MA1 | MA2 | MM1 | MM2 | |
| MA1 | 0 | 105 | 113 | 105 | |
| MA2 | 0.0102 | 0 | 76 | 0 | |
| MM1 | 0.0110 | 0.0074 | 0 | 76 | |
| MM2 | 0.0102 | 0.0000 | 0.0074 | 0 | |
| 416 | | | | | 6241 |
| | MA1 | MA2 | MM1 | MM2 | |
| MA1 | 0 | 155 | 158 | 138 | |
| MA2 | 0.0248 | 0 | 77 | 71 | |
| MM1 | 0.0253 | 0.0123 | 0 | 95 | |
| MM2 | 0.0221 | 0.0114 | 0.0152 | 0 | |
| 420 | | | | | 7937 |

| | MA1 | MA2 | MM1 | MM2 | |
|---|---|---|---|---|---|
| MA1 | 0 | 137 | 94 | 0 | |
| MA2 | 0.0173 | 0 | 147 | 137 | |
| MM1 | 0.0118 | 0.0185 | 0 | 94 | |
| MM2 | 0.0000 | 0.0173 | 0.0118 | 0 | |
| 421 | | | | | 7167 |

| | MA1 | MA2 | MM1 | MM2 | |
|---|---|---|---|---|---|
| MA1 | 0 | 150 | 58 | 52 | |
| MA2 | 0.0209 | 0 | 138 | 135 | |
| MM1 | 0.0081 | 0.0193 | 0 | 52 | |
| MM2 | 0.0073 | 0.0188 | 0.0073 | 0 | |
| 424 | | | | | 10708 |

| | MA1 | MA2 | MM1 | MM2 | |
|---|---|---|---|---|---|
| MA1 | 0 | 237 | 255 | 237 | |
| MA2 | 0.0221 | 0 | 131 | 0 | |
| MM1 | 0.0238 | 0.0122 | 0 | 131 | |
| MM2 | 0.0221 | 0.0000 | 0.0122 | 0 | |
| 429 | | | | | 25264 |

| | MA1 | MA2 | MM1 | MM2 | |
|---|---|---|---|---|---|
| MA1 | 0 | 725 | 386 | 362 | |
| MA2 | 0.0287 | 0 | 718 | 701 | |
| MM1 | 0.0153 | 0.0284 | 0 | 360 | |
| MM2 | 0.0143 | 0.0277 | 0.0142 | 0 | |
| 430 | | | | | 6031 |

| | MA1 | MA2 | MM1 | MM2 | |
|---|---|---|---|---|---|
| MA1 | 0 | 110 | 175 | 121 | |
| MA2 | 0.0182 | 0 | 107 | 51 | |
| MM1 | 0.0290 | 0.0177 | 0 | 70 | |
| MM2 | 0.0201 | 0.0085 | 0.0116 | 0 | |
| 433 | | | | | 8284 |

| | MA1 | MA2 | MM1 | MM2 | |
|---|---|---|---|---|---|
| MA1 | 0 | 187 | 187 | 189 | |
| MA2 | 0.0226 | 0 | 0 | 58 | |
| MM1 | 0.0226 | 0.0000 | 0 | 58 | |
| MM2 | 0.0228 | 0.0070 | 0.0070 | 0 | |
| 435 | | | | | 11589 |

| | MA1 | MA2 | MM1 | MM2 | |
|---|---|---|---|---|---|
| MA1 | 0 | 184 | 184 | 177 | |
| MA2 | 0.0159 | 0 | 0 | 48 | |
| MM1 | 0.0159 | 0.0000 | 0 | 48 | |
| MM2 | 0.0153 | 0.0041 | 0.0041 | 0 | |
| 436 | | | | | 16692 |

Table S7: Difference Matrices.



| | MA1 | MA2 | MM1 | MM2 | |
|---|---|---|---|---|---|
| MA1 | 0 | 353 | 355 | 349 | |
| MA2 | 0.0211 | 0 | 196 | 208 | |
| MM1 | 0.0213 | 0.0117 | 0 | 20 | |
| MM2 | 0.0209 | 0.0125 | 0.0012 | 0 | |
| 438 | | | | | 7986 |
| | MA1 | MA2 | MM1 | MM2 | |
| MA1 | 0 | 178 | 195 | 178 | |
| MA2 | 0.0223 | 0 | 82 | 0 | |
| MM1 | 0.0244 | 0.0103 | 0 | 82 | |
| MM2 | 0.0223 | 0.0000 | 0.0103 | 0 | |
| 439 | | | | | 25973 |
| | MA1 | MA2 | MM1 | MM2 | |
| MA1 | 0 | 631 | 279 | 285 | |
| MA2 | 0.0243 | 0 | 613 | 624 | |
| MM1 | 0.0107 | 0.0236 | 0 | 278 | |
| MM2 | 0.0110 | 0.0240 | 0.0107 | 0 | |
| 441 | | | | | 9595 |
| | MA1 | MA2 | MM1 | MM2 | |
| MA1 | 0 | 175 | 175 | 182 | |
| MA2 | 0.0182 | 0 | 0 | 83 | |
| MM1 | 0.0182 | 0.0000 | 0 | 83 | |
| MM2 | 0.0190 | 0.0087 | 0.0087 | 0 | |
| 442 | | | | | 14357 |
| | MA1 | MA2 | MM1 | MM2 | |
| MA1 | 0 | 269 | 260 | 269 | |
| MA2 | 0.0187 | 0 | 126 | 0 | |
| MM1 | 0.0181 | 0.0088 | 0 | 126 | |
| MM2 | 0.0187 | 0.0000 | 0.0088 | 0 | |
| 453 | | | | | 8596 |
| | MA1 | MA2 | MM1 | MM2 | |
| MA1 | 0 | 0 | 354 | 351 | |
| MA2 | 0.0000 | 0 | 354 | 351 | |
| MM1 | 0.0412 | 0.0412 | 0 | 13 | |
| MM2 | 0.0408 | 0.0408 | 0.0015 | 0 | |
| 456 | | | | | 7080 |
| | MA1 | MA2 | MM1 | MM2 | |
| MA1 | 0 | 88 | 0 | 0 | |
| MA2 | 0.0124 | 0 | 88 | 88 | |
| MM1 | 0.0000 | 0.0124 | 0 | 0 | |
| MM2 | 0.0000 | 0.0124 | 0.0000 | 0 | |
| 464 | | | | | 12034 |

| | MA1 | MA2 | MM1 | MM2 | |
|---|---|---|---|---|---|
| MA1 | 0 | 258 | 243 | 251 | |
| MA2 | 0.0214 | 0 | 98 | 117 | |
| MM1 | 0.0202 | 0.0081 | 0 | 104 | |
| MM2 | 0.0209 | 0.0097 | 0.0086 | 0 | |
| 469 | | | | | 10254 |

| | MA1 | MA2 | MM1 | MM2 | |
|---|---|---|---|---|---|
| MA1 | 0 | 255 | 255 | 237 | |
| MA2 | 0.0249 | 0 | 0 | 105 | |
| MM1 | 0.0249 | 0.0000 | 0 | 105 | |
| MM2 | 0.0231 | 0.0102 | 0.0102 | 0 | |
| 470 | | | | | 21569 |

| | MA1 | MA2 | MM1 | MM2 | |
|---|---|---|---|---|---|
| MA1 | 0 | 32 | 320 | 32 | |
| MA2 | 0.0015 | 0 | 288 | 0 | |
| MM1 | 0.0148 | 0.0134 | 0 | 288 | |
| MM2 | 0.0015 | 0.0000 | 0.0134 | 0 | |
| 471 | | | | | 14756 |

| | MA1 | MA2 | MM1 | MM2 | |
|---|---|---|---|---|---|
| MA1 | 0 | 148 | 72 | 0 | |
| MA2 | 0.0100 | 0 | 149 | 148 | |
| MM1 | 0.0049 | 0.0101 | 0 | 72 | |
| MM2 | 0.0000 | 0.0100 | 0.0049 | 0 | |
| 473 | | | | | 13723 |

| | MA1 | MA2 | MM1 | MM2 | |
|---|---|---|---|---|---|
| MA1 | 0 | 227 | 238 | 242 | |
| MA2 | 0.0165 | 0 | 98 | 100 | |
| MM1 | 0.0173 | 0.0071 | 0 | 86 | |
| MM2 | 0.0176 | 0.0073 | 0.0063 | 0 | |
| 480 | | | | | 10830 |

| | MA1 | MA2 | MM1 | MM2 | |
|---|---|---|---|---|---|
| MA1 | 0 | 359 | 165 | 164 | |
| MA2 | 0.0331 | 0 | 354 | 350 | |
| MM1 | 0.0152 | 0.0327 | 0 | 64 | |
| MM2 | 0.0151 | 0.0323 | 0.0059 | 0 | |
| 483 | | | | | 14603 |

| | MA1 | MA2 | MM1 | MM2 | |
|---|---|---|---|---|---|
| MA1 | 0 | 405 | 0 | 193 | |
| MA2 | 0.0277 | 0 | 405 | 410 | |
| MM1 | 0.0000 | 0.0277 | 0 | 193 | |
| MM2 | 0.0132 | 0.0281 | 0.0132 | 0 | |
| 485 | | | | | 9818 |

| | MA1 | MA2 | MM1 | MM2 | |
|---|---|---|---|---|---|
| MA1 | 0 | 99 | 35 | 38 | |
| MA2 | 0.0101 | 0 | 100 | 100 | |
| MM1 | 0.0036 | 0.0102 | 0 | 15 | |
| MM2 | 0.0039 | 0.0102 | 0.0015 | 0 | |
| 488 | | | | | 19336 |

| | MA1 | MA2 | MM1 | MM2 | |
|---|---|---|---|---|---|
| MA1 | 0 | 65 | 315 | 65 | |
| MA2 | 0.0034 | 0 | 282 | 0 | |
| MM1 | 0.0163 | 0.0146 | 0 | 282 | |
| MM2 | 0.0034 | 0.0000 | 0.0146 | 0 | |
| 489 | | | | | 11697 |

| | MA1 | MA2 | MM1 | MM2 | |
|---|---|---|---|---|---|
| MA1 | 0 | 127 | 145 | 127 | |
| MA2 | 0.0109 | 0 | 81 | 0 | |
| MM1 | 0.0124 | 0.0069 | 0 | 81 | |
| MM2 | 0.0109 | 0.0000 | 0.0069 | 0 | |
| 491 | | | | | 12310 |

| | MA1 | MA2 | MM1 | MM2 | |
|---|---|---|---|---|---|
| MA1 | 0 | 273 | 109 | 102 | |
| MA2 | 0.0222 | 0 | 260 | 258 | |
| MM1 | 0.0089 | 0.0211 | 0 | 96 | |
| MM2 | 0.0083 | 0.0210 | 0.0078 | 0 | |
| 493 | | | | | 14885 |

| | MA1 | MA2 | MM1 | MM2 | |
|---|---|---|---|---|---|
| MA1 | 0 | 200 | 103 | 89 | |
| MA2 | 0.0134 | 0 | 217 | 212 | |
| MM1 | 0.0069 | 0.0146 | 0 | 91 | |
| MM2 | 0.0060 | 0.0142 | 0.0061 | 0 | |
| 495 | | | | | 13969 |

| | MA1 | MA2 | MM1 | MM2 | |
|---|---|---|---|---|---|
| MA1 | 0 | 308 | 296 | 308 | |
| MA2 | 0.0220 | 0 | 141 | 0 | |
| MM1 | 0.0212 | 0.0101 | 0 | 141 | |
| MM2 | 0.0220 | 0.0000 | 0.0101 | 0 | |
| 497 | | | | | 15397 |

| | MA1 | MA2 | MM1 | MM2 | |
|---|---|---|---|---|---|
| MA1 | 0 | 407 | 248 | 0 | |
| MA2 | 0.0264 | 0 | 425 | 407 | |
| MM1 | 0.0161 | 0.0276 | 0 | 248 | |
| MM2 | 0.0000 | 0.0264 | 0.0161 | 0 | |
| 499 | | | | | 7182 |

| | MA1 | MA2 | MM1 | MM2 | |
|---|---|---|---|---|---|
| MA1 | 0 | 119 | 119 | 128 | |
| MA2 | 0.0166 | 0 | 0 | 66 | |
| MM1 | 0.0166 | 0.0000 | 0 | 66 | |
| MM2 | 0.0178 | 0.0092 | 0.0092 | 0 | |
| 500 | | | | | 24012 |

| | MA1 | MA2 | MM1 | MM2 | |
|---|---|---|---|---|---|
| MA1 | 0 | 384 | 280 | 0 | |
| MA2 | 0.0160 | 0 | 505 | 384 | |
| MM1 | 0.0117 | 0.0210 | 0 | 280 | |
| MM2 | 0.0000 | 0.0160 | 0.0117 | 0 | |
| 501 | | | | | 14169 |

| | MA1 | MA2 | MM1 | MM2 | |
|---|---|---|---|---|---|
| MA1 | 0 | 257 | 265 | 237 | |
| MA2 | 0.0181 | 0 | 59 | 80 | |
| MM1 | 0.0187 | 0.0042 | 0 | 139 | |
| MM2 | 0.0167 | 0.0056 | 0.0098 | 0 | |
| 503 | | | | | 10712 |

| | MA1 | MA2 | MM1 | MM2 | |
|---|---|---|---|---|---|
| MA1 | 0 | 277 | 0 | 147 | |
| MA2 | 0.0259 | 0 | 277 | 272 | |
| MM1 | 0.0000 | 0.0259 | 0 | 147 | |
| MM2 | 0.0137 | 0.0254 | 0.0137 | 0 | |
| 508 | | | | | 12746 |

| | MA1 | MA2 | MM1 | MM2 | |
|---|---|---|---|---|---|
| MA1 | 0 | 328 | 324 | 337 | |
| MA2 | 0.0257 | 0 | 152 | 153 | |
| MM1 | 0.0254 | 0.0119 | 0 | 192 | |
| MM2 | 0.0264 | 0.0120 | 0.0151 | 0 | |
| 509 | | | | | 12335 |

| | MA1 | MA2 | MM1 | MM2 | |
|---|---|---|---|---|---|
| MA1 | 0 | 210 | 89 | 102 | |
| MA2 | 0.0170 | 0 | 200 | 195 | |
| MM1 | 0.0072 | 0.0162 | 0 | 79 | |
| MM2 | 0.0083 | 0.0158 | 0.0064 | 0 | |
| 519 | | | | | 13495 |

| | MA1 | MA2 | MM1 | MM2 | |
|---|---|---|---|---|---|
| MA1 | 0 | 240 | 0 | 44 | |
| MA2 | 0.0178 | 0 | 240 | 229 | |
| MM1 | 0.0000 | 0.0178 | 0 | 44 | |
| MM2 | 0.0033 | 0.0170 | 0.0033 | 0 | |
| 520 | | | | | 7272 |

| | MA1 | MA2 | MM1 | MM2 | |
|---|---|---|---|---|---|
| MA1 | 0 | 74 | 74 | 82 | |
| MA2 | 0.0102 | 0 | 0 | 26 | |
| MM1 | 0.0102 | 0.0000 | 0 | 26 | |
| MM2 | 0.0113 | 0.0036 | 0.0036 | 0 | |
| 521 | | | | | 9317 |
| | MA1 | MA2 | MM1 | MM2 | |
| MA1 | 0 | 187 | 51 | 0 | |
| MA2 | 0.0201 | 0 | 185 | 187 | |
| MM1 | 0.0055 | 0.0199 | 0 | 51 | |
| MM2 | 0.0000 | 0.0201 | 0.0055 | 0 | |
| 524 | | | | | 12858 |
| | MA1 | MA2 | MM1 | MM2 | |
| MA1 | 0 | 232 | 232 | 229 | |
| MA2 | 0.0180 | 0 | 0 | 123 | |
| MM1 | 0.0180 | 0.0000 | 0 | 123 | |
| MM2 | 0.0178 | 0.0096 | 0.0096 | 0 | |
| 530 | | | | | 14424 |
| | MA1 | MA2 | MM1 | MM2 | |
| MA1 | 0 | 153 | 153 | 151 | |
| MA2 | 0.0106 | 0 | 0 | 50 | |
| MM1 | 0.0106 | 0.0000 | 0 | 50 | |
| MM2 | 0.0105 | 0.0035 | 0.0035 | 0 | |
| 533 | | | | | 8587 |
| | MA1 | MA2 | MM1 | MM2 | |
| MA1 | 0 | 0 | 0 | 146 | |
| MA2 | 0.0000 | 0 | 0 | 146 | |
| MM1 | 0.0000 | 0.0000 | 0 | 146 | |
| MM2 | 0.0170 | 0.0170 | 0.0170 | 0 | |
| 535 | | | | | 12535 |
| | MA1 | MA2 | MM1 | MM2 | |
| MA1 | 0 | 66 | 376 | 279 | |
| MA2 | 0.0053 | 0 | 396 | 345 | |
| MM1 | 0.0300 | 0.0316 | 0 | 192 | |
| MM2 | 0.0223 | 0.0275 | 0.0153 | 0 | |
| 537 | | | | | 21065 |
| | MA1 | MA2 | MM1 | MM2 | |
| MA1 | 0 | 18 | 425 | 149 | |
| MA2 | 0.0009 | 0 | 413 | 131 | |
| MM1 | 0.0202 | 0.0196 | 0 | 337 | |
| MM2 | 0.0071 | 0.0062 | 0.0160 | 0 | |
| 541 | | | | | 16528 |

| | MA1 | MA2 | MM1 | MM2 | |
|---|---|---|---|---|---|
| MA1 | 0 | 334 | 315 | 334 | |
| MA2 | 0.0202 | 0 | 168 | 0 | |
| MM1 | 0.0191 | 0.0102 | 0 | 168 | |
| MM2 | 0.0202 | 0.0000 | 0.0102 | 0 | |
| 542 | | | | | 26787 |

| | MA1 | MA2 | MM1 | MM2 | |
|---|---|---|---|---|---|
| MA1 | 0 | 569 | 569 | 593 | |
| MA2 | 0.0212 | 0 | 0 | 289 | |
| MM1 | 0.0212 | 0.0000 | 0 | 289 | |
| MM2 | 0.0221 | 0.0108 | 0.0108 | 0 | |
| 543 | | | | | 13021 |

| | MA1 | MA2 | MM1 | MM2 | |
|---|---|---|---|---|---|
| MA1 | 0 | 302 | 137 | 0 | |
| MA2 | 0.0232 | 0 | 319 | 302 | |
| MM1 | 0.0105 | 0.0245 | 0 | 137 | |
| MM2 | 0.0000 | 0.0232 | 0.0105 | 0 | |
| 547 | | | | | 17591 |

| | MA1 | MA2 | MM1 | MM2 | |
|---|---|---|---|---|---|
| MA1 | 0 | 409 | 378 | 390 | |
| MA2 | 0.0233 | 0 | 211 | 192 | |
| MM1 | 0.0215 | 0.0120 | 0 | 187 | |
| MM2 | 0.0222 | 0.0109 | 0.0106 | 0 | |
| 548 | | | | | 9119 |

| | MA1 | MA2 | MM1 | MM2 | |
|---|---|---|---|---|---|
| MA1 | 0 | 227 | 118 | 110 | |
| MA2 | 0.0249 | 0 | 201 | 211 | |
| MM1 | 0.0129 | 0.0220 | 0 | 92 | |
| MM2 | 0.0121 | 0.0231 | 0.0101 | 0 | |
| 552 | | | | | 14204 |

| | MA1 | MA2 | MM1 | MM2 | |
|---|---|---|---|---|---|
| MA1 | 0 | 239 | 251 | 251 | |
| MA2 | 0.0168 | 0 | 131 | 143 | |
| MM1 | 0.0177 | 0.0092 | 0 | 118 | |
| MM2 | 0.0177 | 0.0101 | 0.0083 | 0 | |
| 556 | | | | | 10624 |

| | MA1 | MA2 | MM1 | MM2 | |
|---|---|---|---|---|---|
| MA1 | 0 | 22 | 138 | 0 | |
| MA2 | 0.0021 | 0 | 154 | 22 | |
| MM1 | 0.0130 | 0.0145 | 0 | 138 | |
| MM2 | 0.0000 | 0.0021 | 0.0130 | 0 | |
| 557 | | | | | 17506 |

Table S7: Difference Matrices.

Matrices MA-MM



| | MA1 | MA2 | MM1 | MM2 | |
|---|---|---|---|---|---|
| MA1 | 0 | 296 | 256 | 246 | |
| MA2 | 0.0169 | 0 | 413 | 394 | |
| MM1 | 0.0146 | 0.0236 | 0 | 206 | |
| MM2 | 0.0141 | 0.0225 | 0.0118 | 0 | |
| 562 | | | | | 14765 |
| | MA1 | MA2 | MM1 | MM2 | |
| MA1 | 0 | 176 | 0 | 97 | |
| MA2 | 0.0119 | 0 | 176 | 174 | |
| MM1 | 0.0000 | 0.0119 | 0 | 97 | |
| MM2 | 0.0066 | 0.0118 | 0.0066 | 0 | |
| 566 | | | | | 13422 |
| | MA1 | MA2 | MM1 | MM2 | |
| MA1 | 0 | 21 | 179 | 255 | |
| MA2 | 0.0016 | 0 | 194 | 271 | |
| MM1 | 0.0133 | 0.0145 | 0 | 206 | |
| MM2 | 0.0190 | 0.0202 | 0.0153 | 0 | |
| 569 | | | | | 10205 |
| | MA1 | MA2 | MM1 | MM2 | |
| MA1 | 0 | 212 | 212 | 217 | |
| MA2 | 0.0208 | 0 | 0 | 164 | |
| MM1 | 0.0208 | 0.0000 | 0 | 164 | |
| MM2 | 0.0213 | 0.0161 | 0.0161 | 0 | |
| 571 | | | | | 9467 |
| | MA1 | MA2 | MM1 | MM2 | |
| MA1 | 0 | 201 | 205 | 201 | |
| MA2 | 0.0212 | 0 | 94 | 0 | |
| MM1 | 0.0217 | 0.0099 | 0 | 94 | |
| MM2 | 0.0212 | 0.0000 | 0.0099 | 0 | |
| 577 | | | | | 10902 |
| | MA1 | MA2 | MM1 | MM2 | |
| MA1 | 0 | 246 | 236 | 241 | |
| MA2 | 0.0226 | 0 | 132 | 131 | |
| MM1 | 0.0216 | 0.0121 | 0 | 133 | |
| MM2 | 0.0221 | 0.0120 | 0.0122 | 0 | |
| 578 | | | | | 17850 |
| | MA1 | MA2 | MM1 | MM2 | |
| MA1 | 0 | 352 | 362 | 342 | |
| MA2 | 0.0197 | 0 | 186 | 189 | |
| MM1 | 0.0203 | 0.0104 | 0 | 181 | |
| MM2 | 0.0192 | 0.0106 | 0.0101 | 0 | |
| 584 | | | | | 11545 |

| | MA1 | MA2 | MM1 | MM2 | |
|---|---|---|---|---|---|
| MA1 | 0 | 375 | 0 | 178 | |
| MA2 | 0.0325 | 0 | 375 | 350 | |
| MM1 | 0.0000 | 0.0325 | 0 | 178 | |
| MM2 | 0.0154 | 0.0303 | 0.0154 | 0 | |
| 585 | | | | | 16227 |
| | MA1 | MA2 | MM1 | MM2 | |
| MA1 | 0 | 246 | 106 | 103 | |
| MA2 | 0.0152 | 0 | 249 | 273 | |
| MM1 | 0.0065 | 0.0153 | 0 | 114 | |
| MM2 | 0.0063 | 0.0168 | 0.0070 | 0 | |
| 586 | | | | | 13337 |
| | MA1 | MA2 | MM1 | MM2 | |
| MA1 | 0 | 235 | 235 | 232 | |
| MA2 | 0.0176 | 0 | 0 | 112 | |
| MM1 | 0.0176 | 0.0000 | 0 | 112 | |
| MM2 | 0.0174 | 0.0084 | 0.0084 | 0 | |
| 588 | | | | | 12021 |
| | MA1 | MA2 | MM1 | MM2 | |
| MA1 | 0 | 337 | 0 | 194 | |
| MA2 | 0.0280 | 0 | 337 | 358 | |
| MM1 | 0.0000 | 0.0280 | 0 | 194 | |
| MM2 | 0.0161 | 0.0298 | 0.0161 | 0 | |
| 589 | | | | | 12404 |
| | MA1 | MA2 | MM1 | MM2 | |
| MA1 | 0 | 223 | 60 | 73 | |
| MA2 | 0.0180 | 0 | 214 | 219 | |
| MM1 | 0.0048 | 0.0173 | 0 | 66 | |
| MM2 | 0.0059 | 0.0177 | 0.0053 | 0 | |
| 592 | | | | | 8725 |
| | MA1 | MA2 | MM1 | MM2 | |
| MA1 | 0 | 148 | 151 | 158 | |
| MA2 | 0.0170 | 0 | 19 | 99 | |
| MM1 | 0.0173 | 0.0022 | 0 | 80 | |
| MM2 | 0.0181 | 0.0113 | 0.0092 | 0 | |
| 596 | | | | | 8342 |
| | MA1 | MA2 | MM1 | MM2 | |
| MA1 | 0 | 135 | 135 | 135 | |
| MA2 | 0.0162 | 0 | 0 | 0 | |
| MM1 | 0.0162 | 0.0000 | 0 | 0 | |
| MM2 | 0.0162 | 0.0000 | 0.0000 | 0 | |
| 599 | | | | | 10767 |

Table S7: Difference Matrices.



| | MA1 | MA2 | MM1 | MM2 | |
|---|---|---|---|---|---|
| MA1 | 0 | 178 | 172 | 162 | |
| MA2 | 0.0165 | 0 | 86 | 88 | |
| MM1 | 0.0160 | 0.0080 | 0 | 86 | |
| MM2 | 0.0150 | 0.0082 | 0.0080 | 0 | |
| 605 | | | | | 7245 |
| | MA1 | MA2 | MM1 | MM2 | |
| MA1 | 0 | 214 | 135 | 127 | |
| MA2 | 0.0295 | 0 | 263 | 266 | |
| MM1 | 0.0186 | 0.0363 | 0 | 114 | |
| MM2 | 0.0175 | 0.0367 | 0.0157 | 0 | |
| 608 | | | | | 10383 |
| | MA1 | MA2 | MM1 | MM2 | |
| MA1 | 0 | 178 | 173 | 168 | |
| MA2 | 0.0171 | 0 | 98 | 93 | |
| MM1 | 0.0167 | 0.0094 | 0 | 61 | |
| MM2 | 0.0162 | 0.0090 | 0.0059 | 0 | |
| 610 | | | | | 12622 |
| | MA1 | MA2 | MM1 | MM2 | |
| MA1 | 0 | 202 | 202 | 201 | |
| MA2 | 0.0160 | 0 | 0 | 98 | |
| MM1 | 0.0160 | 0.0000 | 0 | 98 | |
| MM2 | 0.0159 | 0.0078 | 0.0078 | 0 | |
| 613 | | | | | 9528 |
| | MA1 | MA2 | MM1 | MM2 | |
| MA1 | 0 | 120 | 177 | 120 | |
| MA2 | 0.0126 | 0 | 107 | 0 | |
| MM1 | 0.0186 | 0.0112 | 0 | 107 | |
| MM2 | 0.0126 | 0.0000 | 0.0112 | 0 | |
| 622 | | | | | 7369 |
| | MA1 | MA2 | MM1 | MM2 | |
| MA1 | 0 | 162 | 162 | 166 | |
| MA2 | 0.0220 | 0 | 0 | 40 | |
| MM1 | 0.0220 | 0.0000 | 0 | 40 | |
| MM2 | 0.0225 | 0.0054 | 0.0054 | 0 | |
| 626 | | | | | 7386 |
| | MA1 | MA2 | MM1 | MM2 | |
| MA1 | 0 | 195 | 206 | 207 | |
| MA2 | 0.0264 | 0 | 142 | 143 | |
| MM1 | 0.0279 | 0.0192 | 0 | 106 | |
| MM2 | 0.0280 | 0.0194 | 0.0144 | 0 | |
| 635 | | | | | 13178 |

| | MA1 | MA2 | MM1 | MM2 | |
|---|---|---|---|---|---|
| MA1 | 0 | 246 | 214 | 246 | |
| MA2 | 0.0187 | 0 | 126 | 0 | |
| MM1 | 0.0162 | 0.0096 | 0 | 126 | |
| MM2 | 0.0187 | 0.0000 | 0.0096 | 0 | |
| 638 | | | | | 11079 |
| | MA1 | MA2 | MM1 | MM2 | |
| MA1 | 0 | 267 | 268 | 272 | |
| MA2 | 0.0241 | 0 | 144 | 152 | |
| MM1 | 0.0242 | 0.0130 | 0 | 148 | |
| MM2 | 0.0246 | 0.0137 | 0.0134 | 0 | |
| 639 | | | | | 12570 |
| | MA1 | MA2 | MM1 | MM2 | |
| MA1 | 0 | 456 | 290 | 249 | |
| MA2 | 0.0363 | 0 | 495 | 455 | |
| MM1 | 0.0231 | 0.0394 | 0 | 221 | |
| MM2 | 0.0198 | 0.0362 | 0.0176 | 0 | |
| 649 | | | | | 14893 |
| | MA1 | MA2 | MM1 | MM2 | |
| MA1 | 0 | 300 | 301 | 300 | |
| MA2 | 0.0201 | 0 | 140 | 0 | |
| MM1 | 0.0202 | 0.0094 | 0 | 140 | |
| MM2 | 0.0201 | 0.0000 | 0.0094 | 0 | |
| 650 | | | | | 15188 |
| | MA1 | MA2 | MM1 | MM2 | |
| MA1 | 0 | 241 | 0 | 134 | |
| MA2 | 0.0159 | 0 | 241 | 271 | |
| MM1 | 0.0000 | 0.0159 | 0 | 134 | |
| MM2 | 0.0088 | 0.0178 | 0.0088 | 0 | |
| 652 | | | | | 7452 |
| | MA1 | MA2 | MM1 | MM2 | |
| MA1 | 0 | 162 | 98 | 19 | |
| MA2 | 0.0217 | 0 | 177 | 165 | |
| MM1 | 0.0132 | 0.0238 | 0 | 92 | |
| MM2 | 0.0025 | 0.0221 | 0.0123 | 0 | |
| 661 | | | | | 13212 |
| | MA1 | MA2 | MM1 | MM2 | |
| MA1 | 0 | 337 | 337 | 351 | |
| MA2 | 0.0255 | 0 | 0 | 129 | |
| MM1 | 0.0255 | 0.0000 | 0 | 129 | |
| MM2 | 0.0266 | 0.0098 | 0.0098 | 0 | |
| 662 | | | | | 11457 |

| | MA1 | MA2 | MM1 | MM2 | |
|---|---|---|---|---|---|
| MA1 | 0 | 204 | 200 | 195 | |
| MA2 | 0.0178 | 0 | 92 | 76 | |
| MM1 | 0.0175 | 0.0080 | 0 | 81 | |
| MM2 | 0.0170 | 0.0066 | 0.0071 | 0 | |
| 665 | | | | | 12105 |

| | MA1 | MA2 | MM1 | MM2 | |
|---|---|---|---|---|---|
| MA1 | 0 | 85 | 86 | 173 | |
| MA2 | 0.0070 | 0 | 1 | 108 | |
| MM1 | 0.0071 | 0.0001 | 0 | 107 | |
| MM2 | 0.0143 | 0.0089 | 0.0088 | 0 | |
| 668 | | | | | 14222 |

| | MA1 | MA2 | MM1 | MM2 | |
|---|---|---|---|---|---|
| MA1 | 0 | 176 | 168 | 176 | |
| MA2 | 0.0124 | 0 | 48 | 57 | |
| MM1 | 0.0118 | 0.0034 | 0 | 39 | |
| MM2 | 0.0124 | 0.0040 | 0.0027 | 0 | |
| 669 | | | | | 32937 |

| | MA1 | MA2 | MM1 | MM2 | |
|---|---|---|---|---|---|
| MA1 | 0 | 775 | 249 | 404 | |
| MA2 | 0.0235 | 0 | 783 | 782 | |
| MM1 | 0.0076 | 0.0238 | 0 | 431 | |
| MM2 | 0.0123 | 0.0237 | 0.0131 | 0 | |
| 671 | | | | | 11380 |

| | MA1 | MA2 | MM1 | MM2 | |
|---|---|---|---|---|---|
| MA1 | 0 | 262 | 0 | 10 | |
| MA2 | 0.0230 | 0 | 262 | 255 | |
| MM1 | 0.0000 | 0.0230 | 0 | 10 | |
| MM2 | 0.0009 | 0.0224 | 0.0009 | 0 | |
| 672 | | | | | 9285 |

| | MA1 | MA2 | MM1 | MM2 | |
|---|---|---|---|---|---|
| MA1 | 0 | 227 | 133 | 49 | |
| MA2 | 0.0244 | 0 | 241 | 213 | |
| MM1 | 0.0143 | 0.0260 | 0 | 152 | |
| MM2 | 0.0053 | 0.0229 | 0.0164 | 0 | |
| 677 | | | | | 14185 |

| | MA1 | MA2 | MM1 | MM2 | |
|---|---|---|---|---|---|
| MA1 | 0 | 378 | 375 | 365 | |
| MA2 | 0.0266 | 0 | 213 | 222 | |
| MM1 | 0.0264 | 0.0150 | 0 | 213 | |
| MM2 | 0.0257 | 0.0157 | 0.0150 | 0 | |
| 685 | | | | | 15438 |

| | MA1 | MA2 | MM1 | MM2 | |
|---|---|---|---|---|---|
| MA1 | 0 | 471 | 471 | 472 | |
| MA2 | 0.0305 | 0 | 0 | 221 | |
| MM1 | 0.0305 | 0.0000 | 0 | 221 | |
| MM2 | 0.0306 | 0.0143 | 0.0143 | 0 | |
| 691 | | | | | 9840 |
| | MA1 | MA2 | MM1 | MM2 | |
| MA1 | 0 | 63 | 256 | 265 | |
| MA2 | 0.0064 | 0 | 222 | 202 | |
| MM1 | 0.0260 | 0.0226 | 0 | 141 | |
| MM2 | 0.0269 | 0.0205 | 0.0143 | 0 | |
| 692 | | | | | 12149 |
| | MA1 | MA2 | MM1 | MM2 | |
| MA1 | 0 | 258 | 116 | 0 | |
| MA2 | 0.0212 | 0 | 246 | 258 | |
| MM1 | 0.0095 | 0.0202 | 0 | 116 | |
| MM2 | 0.0000 | 0.0212 | 0.0095 | 0 | |
| 693 | | | | | 12220 |
| | MA1 | MA2 | MM1 | MM2 | |
| MA1 | 0 | 312 | 182 | 178 | |
| MA2 | 0.0255 | 0 | 337 | 333 | |
| MM1 | 0.0149 | 0.0276 | 0 | 4 | |
| MM2 | 0.0146 | 0.0273 | 0.0003 | 0 | |
| 694 | | | | | 18952 |
| | MA1 | MA2 | MM1 | MM2 | |
| MA1 | 0 | 483 | 487 | 502 | |
| MA2 | 0.0255 | 0 | 213 | 188 | |
| MM1 | 0.0257 | 0.0112 | 0 | 202 | |
| MM2 | 0.0265 | 0.0099 | 0.0107 | 0 | |
| 697 | | | | | 9490 |
| | MA1 | MA2 | MM1 | MM2 | |
| MA1 | 0 | 210 | 210 | 210 | |
| MA2 | 0.0221 | 0 | 96 | 0 | |
| MM1 | 0.0221 | 0.0101 | 0 | 96 | |
| MM2 | 0.0221 | 0.0000 | 0.0101 | 0 | |
| 699 | | | | | 19690 |
| | MA1 | MA2 | MM1 | MM2 | |
| MA1 | 0 | 407 | 444 | 457 | |
| MA2 | 0.0207 | 0 | 315 | 332 | |
| MM1 | 0.0225 | 0.0160 | 0 | 281 | |
| MM2 | 0.0232 | 0.0169 | 0.0143 | 0 | |
| 700 | | | | | 11327 |

| | MA1 | MA2 | MM1 | MM2 | |
|---|---|---|---|---|---|
| MA1 | 0 | 281 | 281 | 263 | |
| MA2 | 0.0248 | 0 | 0 | 126 | |
| MM1 | 0.0248 | 0.0000 | 0 | 126 | |
| MM2 | 0.0232 | 0.0111 | 0.0111 | 0 | |
| 701 | | | | | 7632 |
| | MA1 | MA2 | MM1 | MM2 | |
| MA1 | 0 | 184 | 47 | 160 | |
| MA2 | 0.0241 | 0 | 231 | 221 | |
| MM1 | 0.0062 | 0.0303 | 0 | 113 | |
| MM2 | 0.0210 | 0.0290 | 0.0148 | 0 | |
| 704 | | | | | 10009 |
| | MA1 | MA2 | MM1 | MM2 | |
| MA1 | 0 | 71 | 284 | 261 | |
| MA2 | 0.0071 | 0 | 289 | 275 | |
| MM1 | 0.0284 | 0.0289 | 0 | 162 | |
| MM2 | 0.0261 | 0.0275 | 0.0162 | 0 | |
| 718 | | | | | 15765 |
| | MA1 | MA2 | MM1 | MM2 | |
| MA1 | 0 | 334 | 330 | 334 | |
| MA2 | 0.0212 | 0 | 80 | 0 | |
| MM1 | 0.0209 | 0.0051 | 0 | 80 | |
| MM2 | 0.0212 | 0.0000 | 0.0051 | 0 | |
| 719 | | | | | 10962 |
| | MA1 | MA2 | MM1 | MM2 | |
| MA1 | 0 | 158 | 162 | 166 | |
| MA2 | 0.0144 | 0 | 51 | 46 | |
| MM1 | 0.0148 | 0.0047 | 0 | 65 | |
| MM2 | 0.0151 | 0.0042 | 0.0059 | 0 | |
| 722 | | | | | 6616 |
| | MA1 | MA2 | MM1 | MM2 | |
| MA1 | 0 | 31 | 22 | 8 | |
| MA2 | 0.0047 | 0 | 34 | 25 | |
| MM1 | 0.0033 | 0.0051 | 0 | 18 | |
| MM2 | 0.0012 | 0.0038 | 0.0027 | 0 | |
| 723 | | | | | 10336 |
| | MA1 | MA2 | MM1 | MM2 | |
| MA1 | 0 | 214 | 214 | 214 | |
| MA2 | 0.0207 | 0 | 64 | 0 | |
| MM1 | 0.0207 | 0.0062 | 0 | 64 | |
| MM2 | 0.0207 | 0.0000 | 0.0062 | 0 | |
| 725 | | | | | 10454 |

Table S7: Difference Matrices.



| | MA1 | MA2 | MM1 | MM2 | |
|---|---|---|---|---|---|
| MA1 | 0 | 306 | 264 | 215 | |
| MA2 | 0.0293 | 0 | 233 | 93 | |
| MM1 | 0.0253 | 0.0223 | 0 | 166 | |
| MM2 | 0.0206 | 0.0089 | 0.0159 | 0 | |
| 726 | | | | | 10441 |
| | MA1 | MA2 | MM1 | MM2 | |
| MA1 | 0 | 249 | 3 | 104 | |
| MA2 | 0.0238 | 0 | 246 | 256 | |
| MM1 | 0.0003 | 0.0236 | 0 | 101 | |
| MM2 | 0.0100 | 0.0245 | 0.0097 | 0 | |
| 732 | | | | | 9962 |
| | MA1 | MA2 | MM1 | MM2 | |
| MA1 | 0 | 181 | 189 | 181 | |
| MA2 | 0.0182 | 0 | 46 | 0 | |
| MM1 | 0.0190 | 0.0046 | 0 | 46 | |
| MM2 | 0.0182 | 0.0000 | 0.0046 | 0 | |
| 738 | | | | | 9735 |
| | MA1 | MA2 | MM1 | MM2 | |
| MA1 | 0 | 110 | 107 | 102 | |
| MA2 | 0.0113 | 0 | 164 | 181 | |
| MM1 | 0.0110 | 0.0168 | 0 | 77 | |
| MM2 | 0.0105 | 0.0186 | 0.0079 | 0 | |
| 740 | | | | | 12001 |
| | MA1 | MA2 | MM1 | MM2 | |
| MA1 | 0 | 193 | 0 | 112 | |
| MA2 | 0.0161 | 0 | 193 | 179 | |
| MM1 | 0.0000 | 0.0161 | 0 | 112 | |
| MM2 | 0.0093 | 0.0149 | 0.0093 | 0 | |
| 743 | | | | | 22003 |
| | MA1 | MA2 | MM1 | MM2 | |
| MA1 | 0 | 553 | 553 | 548 | |
| MA2 | 0.0251 | 0 | 0 | 261 | |
| MM1 | 0.0251 | 0.0000 | 0 | 261 | |
| MM2 | 0.0249 | 0.0119 | 0.0119 | 0 | |
| 746 | | | | | 10403 |
| | MA1 | MA2 | MM1 | MM2 | |
| MA1 | 0 | 227 | 227 | 217 | |
| MA2 | 0.0218 | 0 | 0 | 125 | |
| MM1 | 0.0218 | 0.0000 | 0 | 125 | |
| MM2 | 0.0209 | 0.0120 | 0.0120 | 0 | |
| 747 | | | | | 11830 |

| | MA1 | MA2 | MM1 | MM2 | |
|---|---|---|---|---|---|
| MA1 | 0 | 216 | 115 | 115 | |
| MA2 | 0.0183 | 0 | 217 | 217 | |
| MM1 | 0.0097 | 0.0183 | 0 | 0 | |
| MM2 | 0.0097 | 0.0183 | 0.0000 | 0 | |
| 761 | | | | | 13562 |
| | MA1 | MA2 | MM1 | MM2 | |
| MA1 | 0 | 245 | 245 | 227 | |
| MA2 | 0.0181 | 0 | 0 | 136 | |
| MM1 | 0.0181 | 0.0000 | 0 | 136 | |
| MM2 | 0.0167 | 0.0100 | 0.0100 | 0 | |
| 762 | | | | | 15898 |
| | MA1 | MA2 | MM1 | MM2 | |
| MA1 | 0 | 259 | 259 | 238 | |
| MA2 | 0.0163 | 0 | 0 | 103 | |
| MM1 | 0.0163 | 0.0000 | 0 | 103 | |
| MM2 | 0.0150 | 0.0065 | 0.0065 | 0 | |
| 769 | | | | | 9372 |
| | MA1 | MA2 | MM1 | MM2 | |
| MA1 | 0 | 182 | 12 | 0 | |
| MA2 | 0.0194 | 0 | 186 | 182 | |
| MM1 | 0.0013 | 0.0198 | 0 | 12 | |
| MM2 | 0.0000 | 0.0194 | 0.0013 | 0 | |
| 773 | | | | | 11866 |
| | MA1 | MA2 | MM1 | MM2 | |
| MA1 | 0 | 190 | 107 | 0 | |
| MA2 | 0.0160 | 0 | 179 | 190 | |
| MM1 | 0.0090 | 0.0151 | 0 | 107 | |
| MM2 | 0.0000 | 0.0160 | 0.0090 | 0 | |
| 776 | | | | | 12698 |
| | MA1 | MA2 | MM1 | MM2 | |
| MA1 | 0 | 110 | 165 | 129 | |
| MA2 | 0.0087 | 0 | 129 | 103 | |
| MM1 | 0.0130 | 0.0102 | 0 | 118 | |
| MM2 | 0.0102 | 0.0081 | 0.0093 | 0 | |
| 779 | | | | | 9973 |
| | MA1 | MA2 | MM1 | MM2 | |
| MA1 | 0 | 176 | 0 | 76 | |
| MA2 | 0.0176 | 0 | 176 | 175 | |
| MM1 | 0.0000 | 0.0176 | 0 | 76 | |
| MM2 | 0.0076 | 0.0175 | 0.0076 | 0 | |
| 780 | | | | | 11554 |

| | MA1 | MA2 | MM1 | MM2 | |
|---|---|---|---|---|---|
| MA1 | 0 | 221 | 221 | 214 | |
| MA2 | 0.0191 | 0 | 0 | 106 | |
| MM1 | 0.0191 | 0.0000 | 0 | 106 | |
| MM2 | 0.0185 | 0.0092 | 0.0092 | 0 | |
| 794 | | | | | 9830 |

| | MA1 | MA2 | MM1 | MM2 | |
|---|---|---|---|---|---|
| MA1 | 0 | 238 | 127 | 125 | |
| MA2 | 0.0242 | 0 | 235 | 233 | |
| MM1 | 0.0129 | 0.0239 | 0 | 2 | |
| MM2 | 0.0127 | 0.0237 | 0.0002 | 0 | |
| 796 | | | | | 11370 |

| | MA1 | MA2 | MM1 | MM2 | |
|---|---|---|---|---|---|
| MA1 | 0 | 293 | 300 | 293 | |
| MA2 | 0.0258 | 0 | 126 | 0 | |
| MM1 | 0.0264 | 0.0111 | 0 | 126 | |
| MM2 | 0.0258 | 0.0000 | 0.0111 | 0 | |
| 797 | | | | | 11724 |

| | MA1 | MA2 | MM1 | MM2 | |
|---|---|---|---|---|---|
| MA1 | 0 | 325 | 159 | 0 | |
| MA2 | 0.0277 | 0 | 305 | 325 | |
| MM1 | 0.0136 | 0.0260 | 0 | 159 | |
| MM2 | 0.0000 | 0.0277 | 0.0136 | 0 | |
| 802 | | | | | 9375 |

| | MA1 | MA2 | MM1 | MM2 | |
|---|---|---|---|---|---|
| MA1 | 0 | 282 | 287 | 286 | |
| MA2 | 0.0301 | 0 | 158 | 167 | |
| MM1 | 0.0306 | 0.0169 | 0 | 176 | |
| MM2 | 0.0305 | 0.0178 | 0.0188 | 0 | |
| 808 | | | | | 13526 |

| | MA1 | MA2 | MM1 | MM2 | |
|---|---|---|---|---|---|
| MA1 | 0 | 142 | 67 | 79 | |
| MA2 | 0.0105 | 0 | 146 | 145 | |
| MM1 | 0.0050 | 0.0108 | 0 | 70 | |
| MM2 | 0.0058 | 0.0107 | 0.0052 | 0 | |
| 810 | | | | | 21178 |

| | MA1 | MA2 | MM1 | MM2 | |
|---|---|---|---|---|---|
| MA1 | 0 | 468 | 460 | 476 | |
| MA2 | 0.0221 | 0 | 222 | 200 | |
| MM1 | 0.0217 | 0.0105 | 0 | 254 | |
| MM2 | 0.0225 | 0.0094 | 0.0120 | 0 | |
| 814 | | | | | 6658 |

| | MA1 | MA2 | MM1 | MM2 |
|---|---|---|---|---|
| MA1 | 0 | 267 | 247 | 237 |
| MA2 | 0.0401 | 0 | 162 | 154 |
| MM1 | 0.0371 | 0.0243 | 0 | 170 |
| MM2 | 0.0356 | 0.0231 | 0.0255 | 0 |

817 18542

| | MA1 | MA2 | MM1 | MM2 |
|---|---|---|---|---|
| MA1 | 0 | 264 | 262 | 264 |
| MA2 | 0.0142 | 0 | 110 | 0 |
| MM1 | 0.0141 | 0.0059 | 0 | 110 |
| MM2 | 0.0142 | 0.0000 | 0.0059 | 0 |

819 8582

| | MA1 | MA2 | MM1 | MM2 |
|---|---|---|---|---|
| MA1 | 0 | 172 | 172 | 164 |
| MA2 | 0.0200 | 0 | 0 | 34 |
| MM1 | 0.0200 | 0.0000 | 0 | 34 |
| MM2 | 0.0191 | 0.0040 | 0.0040 | 0 |

820 12475

| | MA1 | MA2 | MM1 | MM2 |
|---|---|---|---|---|
| MA1 | 0 | 267 | 241 | 270 |
| MA2 | 0.0214 | 0 | 118 | 149 |
| MM1 | 0.0193 | 0.0095 | 0 | 137 |
| MM2 | 0.0216 | 0.0119 | 0.0110 | 0 |

826 11646

| | MA1 | MA2 | MM1 | MM2 |
|---|---|---|---|---|
| MA1 | 0 | 238 | 238 | 239 |
| MA2 | 0.0204 | 0 | 0 | 94 |
| MM1 | 0.0204 | 0.0000 | 0 | 94 |
| MM2 | 0.0205 | 0.0081 | 0.0081 | 0 |

827 21803

| | MA1 | MA2 | MM1 | MM2 |
|---|---|---|---|---|
| MA1 | 0 | 470 | 470 | 468 |
| MA2 | 0.0216 | 0 | 0 | 225 |
| MM1 | 0.0216 | 0.0000 | 0 | 225 |
| MM2 | 0.0215 | 0.0103 | 0.0103 | 0 |

832 12725

| | MA1 | MA2 | MM1 | MM2 |
|---|---|---|---|---|
| MA1 | 0 | 47 | 177 | 158 |
| MA2 | 0.0037 | 0 | 195 | 170 |
| MM1 | 0.0139 | 0.0153 | 0 | 149 |
| MM2 | 0.0124 | 0.0134 | 0.0117 | 0 |

842 12159

| | MA1 | MA2 | MM1 | MM2 | |
|---|---|---|---|---|---|
| MA1 | 0 | 0 | 195 | 215 | |
| MA2 | 0.0000 | 0 | 195 | 215 | |
| MM1 | 0.0160 | 0.0160 | 0 | 127 | |
| MM2 | 0.0177 | 0.0177 | 0.0104 | 0 | |
| 845 | | | | | 12896 |
| | MA1 | MA2 | MM1 | MM2 | |
| MA1 | 0 | 215 | 214 | 212 | |
| MA2 | 0.0167 | 0 | 88 | 93 | |
| MM1 | 0.0166 | 0.0068 | 0 | 94 | |
| MM2 | 0.0164 | 0.0072 | 0.0073 | 0 | |
| 852 | | | | | 16323 |
| | MA1 | MA2 | MM1 | MM2 | |
| MA1 | 0 | 390 | 201 | 235 | |
| MA2 | 0.0239 | 0 | 392 | 408 | |
| MM1 | 0.0123 | 0.0240 | 0 | 250 | |
| MM2 | 0.0144 | 0.0250 | 0.0153 | 0 | |
| 855 | | | | | 19179 |
| | MA1 | MA2 | MM1 | MM2 | |
| MA1 | 0 | 350 | 372 | 363 | |
| MA2 | 0.0182 | 0 | 445 | 470 | |
| MM1 | 0.0194 | 0.0232 | 0 | 278 | |
| MM2 | 0.0189 | 0.0245 | 0.0145 | 0 | |
| 864 | | | | | 14312 |
| | MA1 | MA2 | MM1 | MM2 | |
| MA1 | 0 | 268 | 0 | 139 | |
| MA2 | 0.0187 | 0 | 268 | 264 | |
| MM1 | 0.0000 | 0.0187 | 0 | 139 | |
| MM2 | 0.0097 | 0.0184 | 0.0097 | 0 | |
| 869 | | | | | 9900 |
| | MA1 | MA2 | MM1 | MM2 | |
| MA1 | 0 | 0 | 206 | 0 | |
| MA2 | 0.0000 | 0 | 206 | 0 | |
| MM1 | 0.0208 | 0.0208 | 0 | 206 | |
| MM2 | 0.0000 | 0.0000 | 0.0208 | 0 | |
| 880 | | | | | 10735 |
| | MA1 | MA2 | MM1 | MM2 | |
| MA1 | 0 | 180 | 0 | 96 | |
| MA2 | 0.0168 | 0 | 180 | 183 | |
| MM1 | 0.0000 | 0.0168 | 0 | 96 | |
| MM2 | 0.0089 | 0.0170 | 0.0089 | 0 | |
| 894 | | | | | 7418 |

| | MA1 | MA2 | MM1 | MM2 |
|---|---|---|---|---|
| MA1 | 0 | 102 | 75 | 81 |
| MA2 | 0.0138 | 0 | 144 | 167 |
| MM1 | 0.0101 | 0.0194 | 0 | 105 |
| MM2 | 0.0109 | 0.0225 | 0.0142 | 0 |

902 19505

| | MA1 | MA2 | MM1 | MM2 |
|---|---|---|---|---|
| MA1 | 0 | 614 | 561 | 614 |
| MA2 | 0.0315 | 0 | 377 | 0 |
| MM1 | 0.0288 | 0.0193 | 0 | 377 |
| MM2 | 0.0315 | 0.0000 | 0.0193 | 0 |

905 19023

| | MA1 | MA2 | MM1 | MM2 |
|---|---|---|---|---|
| MA1 | 0 | 324 | 0 | 166 |
| MA2 | 0.0170 | 0 | 324 | 324 |
| MM1 | 0.0000 | 0.0170 | 0 | 166 |
| MM2 | 0.0087 | 0.0170 | 0.0087 | 0 |

906 17557

| | MA1 | MA2 | MM1 | MM2 |
|---|---|---|---|---|
| MA1 | 0 | 305 | 67 | 157 |
| MA2 | 0.0174 | 0 | 290 | 276 |
| MM1 | 0.0038 | 0.0165 | 0 | 136 |
| MM2 | 0.0089 | 0.0157 | 0.0077 | 0 |

907 10075

| | MA1 | MA2 | MM1 | MM2 |
|---|---|---|---|---|
| MA1 | 0 | 159 | 104 | 0 |
| MA2 | 0.0158 | 0 | 181 | 159 |
| MM1 | 0.0103 | 0.0180 | 0 | 104 |
| MM2 | 0.0000 | 0.0158 | 0.0103 | 0 |

908 15802

| | MA1 | MA2 | MM1 | MM2 |
|---|---|---|---|---|
| MA1 | 0 | 366 | 0 | 208 |
| MA2 | 0.0232 | 0 | 366 | 373 |
| MM1 | 0.0000 | 0.0232 | 0 | 208 |
| MM2 | 0.0132 | 0.0236 | 0.0132 | 0 |

917 6524

| | MA1 | MA2 | MM1 | MM2 |
|---|---|---|---|---|
| MA1 | 0 | 78 | 0 | 16 |
| MA2 | 0.0120 | 0 | 78 | 75 |
| MM1 | 0.0000 | 0.0120 | 0 | 16 |
| MM2 | 0.0025 | 0.0115 | 0.0025 | 0 |

919 19987

| | MA1 | MA2 | MM1 | MM2 | |
|---|---|---|---|---|---|
| MA1 | 0 | 65 | 267 | 0 | |
| MA2 | 0.0033 | 0 | 304 | 65 | |
| MM1 | 0.0134 | 0.0152 | 0 | 267 | |
| MM2 | 0.0000 | 0.0033 | 0.0134 | 0 | |
| 920 | | | | | 8976 |
| | MA1 | MA2 | MM1 | MM2 | |
| MA1 | 0 | 88 | 111 | 147 | |
| MA2 | 0.0098 | 0 | 23 | 86 | |
| MM1 | 0.0124 | 0.0026 | 0 | 76 | |
| MM2 | 0.0164 | 0.0096 | 0.0085 | 0 | |
| 922 | | | | | 14140 |
| | MA1 | MA2 | MM1 | MM2 | |
| MA1 | 0 | 364 | 248 | 196 | |
| MA2 | 0.0257 | 0 | 392 | 386 | |
| MM1 | 0.0175 | 0.0277 | 0 | 249 | |
| MM2 | 0.0139 | 0.0273 | 0.0176 | 0 | |
| 924 | | | | | 11711 |
| | MA1 | MA2 | MM1 | MM2 | |
| MA1 | 0 | 193 | 18 | 94 | |
| MA2 | 0.0165 | 0 | 211 | 207 | |
| MM1 | 0.0015 | 0.0180 | 0 | 76 | |
| MM2 | 0.0080 | 0.0177 | 0.0065 | 0 | |
| 926 | | | | | 12517 |
| | MA1 | MA2 | MM1 | MM2 | |
| MA1 | 0 | 219 | 201 | 219 | |
| MA2 | 0.0175 | 0 | 98 | 0 | |
| MM1 | 0.0161 | 0.0078 | 0 | 98 | |
| MM2 | 0.0175 | 0.0000 | 0.0078 | 0 | |
| 927 | | | | | 12300 |
| | MA1 | MA2 | MM1 | MM2 | |
| MA1 | 0 | 208 | 91 | 86 | |
| MA2 | 0.0169 | 0 | 209 | 222 | |
| MM1 | 0.0074 | 0.0170 | 0 | 61 | |
| MM2 | 0.0070 | 0.0180 | 0.0050 | 0 | |
| 928 | | | | | 9292 |
| | MA1 | MA2 | MM1 | MM2 | |
| MA1 | 0 | 137 | 137 | 124 | |
| MA2 | 0.0147 | 0 | 70 | 68 | |
| MM1 | 0.0147 | 0.0075 | 0 | 76 | |
| MM2 | 0.0133 | 0.0073 | 0.0082 | 0 | |
| 932 | | | | | 7794 |

Table S7: Difference Matrices.



| | MA1 | MA2 | MM1 | MM2 | |
|---|---|---|---|---|---|
| MA1 | 0 | 174 | 104 | 0 | |
| MA2 | 0.0223 | 0 | 178 | 174 | |
| MM1 | 0.0133 | 0.0228 | 0 | 104 | |
| MM2 | 0.0000 | 0.0223 | 0.0133 | 0 | |
| 933 | | | | | 18375 |

| | MA1 | MA2 | MM1 | MM2 | |
|---|---|---|---|---|---|
| MA1 | 0 | 339 | 179 | 0 | |
| MA2 | 0.0184 | 0 | 350 | 339 | |
| MM1 | 0.0097 | 0.0190 | 0 | 179 | |
| MM2 | 0.0000 | 0.0184 | 0.0097 | 0 | |
| 937 | | | | | 12005 |

| | MA1 | MA2 | MM1 | MM2 | |
|---|---|---|---|---|---|
| MA1 | 0 | 120 | 0 | 49 | |
| MA2 | 0.0100 | 0 | 120 | 123 | |
| MM1 | 0.0000 | 0.0100 | 0 | 49 | |
| MM2 | 0.0041 | 0.0102 | 0.0041 | 0 | |
| 939 | | | | | 13370 |

| | MA1 | MA2 | MM1 | MM2 | |
|---|---|---|---|---|---|
| MA1 | 0 | 247 | 238 | 248 | |
| MA2 | 0.0185 | 0 | 56 | 3 | |
| MM1 | 0.0178 | 0.0042 | 0 | 53 | |
| MM2 | 0.0185 | 0.0002 | 0.0040 | 0 | |
| 941 | | | | | 13748 |

| | MA1 | MA2 | MM1 | MM2 | |
|---|---|---|---|---|---|
| MA1 | 0 | 81 | 269 | 294 | |
| MA2 | 0.0059 | 0 | 188 | 247 | |
| MM1 | 0.0196 | 0.0137 | 0 | 178 | |
| MM2 | 0.0214 | 0.0180 | 0.0129 | 0 | |
| 943 | | | | | 8548 |

| | MA1 | MA2 | MM1 | MM2 | |
|---|---|---|---|---|---|
| MA1 | 0 | 189 | 188 | 188 | |
| MA2 | 0.0221 | 0 | 52 | 51 | |
| MM1 | 0.0220 | 0.0061 | 0 | 51 | |
| MM2 | 0.0220 | 0.0060 | 0.0060 | 0 | |
| 945 | | | | | 10924 |

| | MA1 | MA2 | MM1 | MM2 | |
|---|---|---|---|---|---|
| MA1 | 0 | 351 | 189 | 190 | |
| MA2 | 0.0321 | 0 | 364 | 350 | |
| MM1 | 0.0173 | 0.0333 | 0 | 217 | |
| MM2 | 0.0174 | 0.0320 | 0.0199 | 0 | |
| 953 | | | | | 10594 |

| | MA1 | MA2 | MM1 | MM2 | |
|---|---|---|---|---|---|
| MA1 | 0 | 225 | 234 | 218 | |
| MA2 | 0.0212 | 0 | 95 | 42 | |
| MM1 | 0.0221 | 0.0090 | 0 | 94 | |
| MM2 | 0.0206 | 0.0040 | 0.0089 | 0 | |
| 954 | | | | | 8375 |

| | MA1 | MA2 | MM1 | MM2 | |
|---|---|---|---|---|---|
| MA1 | 0 | 152 | 94 | 0 | |
| MA2 | 0.0181 | 0 | 150 | 152 | |
| MM1 | 0.0112 | 0.0179 | 0 | 94 | |
| MM2 | 0.0000 | 0.0181 | 0.0112 | 0 | |
| 957 | | | | | 11988 |

| | MA1 | MA2 | MM1 | MM2 | |
|---|---|---|---|---|---|
| MA1 | 0 | 241 | 209 | 114 | |
| MA2 | 0.0201 | 0 | 261 | 257 | |
| MM1 | 0.0174 | 0.0218 | 0 | 176 | |
| MM2 | 0.0095 | 0.0214 | 0.0147 | 0 | |
| 958 | | | | | 7665 |

| | MA1 | MA2 | MM1 | MM2 | |
|---|---|---|---|---|---|
| MA1 | 0 | 126 | 126 | 168 | |
| MA2 | 0.0164 | 0 | 0 | 116 | |
| MM1 | 0.0164 | 0.0000 | 0 | 116 | |
| MM2 | 0.0219 | 0.0151 | 0.0151 | 0 | |
| 964 | | | | | 15721 |

| | MA1 | MA2 | MM1 | MM2 | |
|---|---|---|---|---|---|
| MA1 | 0 | 325 | 325 | 314 | |
| MA2 | 0.0207 | 0 | 0 | 171 | |
| MM1 | 0.0207 | 0.0000 | 0 | 171 | |
| MM2 | 0.0200 | 0.0109 | 0.0109 | 0 | |
| 967 | | | | | 12529 |

| | MA1 | MA2 | MM1 | MM2 | |
|---|---|---|---|---|---|
| MA1 | 0 | 206 | 197 | 206 | |
| MA2 | 0.0164 | 0 | 74 | 0 | |
| MM1 | 0.0157 | 0.0059 | 0 | 74 | |
| MM2 | 0.0164 | 0.0000 | 0.0059 | 0 | |
| 974 | | | | | 15197 |

| | MA1 | MA2 | MM1 | MM2 | |
|---|---|---|---|---|---|
| MA1 | 0 | 391 | 407 | 414 | |
| MA2 | 0.0257 | 0 | 217 | 209 | |
| MM1 | 0.0268 | 0.0143 | 0 | 140 | |
| MM2 | 0.0272 | 0.0138 | 0.0092 | 0 | |
| 975 | | | | | 6877 |

Table S7: Difference Matrices.



| | MA1 | MA2 | MM1 | MM2 | |
|---|---|---|---|---|---|
| MA1 | 0 | 149 | 0 | 66 | |
| MA2 | 0.0217 | 0 | 149 | 139 | |
| MM1 | 0.0000 | 0.0217 | 0 | 66 | |
| MM2 | 0.0096 | 0.0202 | 0.0096 | 0 | |
| 980 | | | | | 7516 |
| | MA1 | MA2 | MM1 | MM2 | |
| MA1 | 0 | 183 | 211 | 216 | |
| MA2 | 0.0243 | 0 | 137 | 145 | |
| MM1 | 0.0281 | 0.0182 | 0 | 84 | |
| MM2 | 0.0287 | 0.0193 | 0.0112 | 0 | |
| 988 | | | | | 10284 |
| | MA1 | MA2 | MM1 | MM2 | |
| MA1 | 0 | 244 | 244 | 264 | |
| MA2 | 0.0237 | 0 | 0 | 135 | |
| MM1 | 0.0237 | 0.0000 | 0 | 135 | |
| MM2 | 0.0257 | 0.0131 | 0.0131 | 0 | |

| 8 | | | | | 11387 |
|---|---|---|---|---|---|
| | CR1 | CR2 | MM1 | MM2 | |
| CR1 | 0 | 356 | 132 | 134 | |
| CR2 | 0.0313 | 0 | 367 | 374 | |
| MM1 | 0.0116 | 0.0322 | 0 | 88 | |
| MM2 | 0.0118 | 0.0328 | 0.0077 | 0 | |
| 16 | | | | | 10501 |
| | CR1 | CR2 | MM1 | MM2 | |
| CR1 | 0 | 333 | 129 | 159 | |
| CR2 | 0.0317 | 0 | 319 | 322 | |
| MM1 | 0.0123 | 0.0304 | 0 | 153 | |
| MM2 | 0.0151 | 0.0307 | 0.0146 | 0 | |
| 18 | | | | | 10068 |
| | CR1 | CR2 | MM1 | MM2 | |
| CR1 | 0 | 274 | 188 | 168 | |
| CR2 | 0.0272 | 0 | 253 | 271 | |
| MM1 | 0.0187 | 0.0251 | 0 | 179 | |
| MM2 | 0.0167 | 0.0269 | 0.0178 | 0 | |
| 20 | | | | | 7947 |
| | CR1 | CR2 | MM1 | MM2 | |
| CR1 | 0 | 143 | 49 | 66 | |
| CR2 | 0.0180 | 0 | 140 | 141 | |
| MM1 | 0.0062 | 0.0176 | 0 | 71 | |
| MM2 | 0.0083 | 0.0177 | 0.0089 | 0 | |
| 22 | | | | | 4936 |
| | CR1 | CR2 | MM1 | MM2 | |
| CR1 | 0 | 85 | 80 | 82 | |
| CR2 | 0.0172 | 0 | 136 | 140 | |
| MM1 | 0.0162 | 0.0276 | 0 | 47 | |
| MM2 | 0.0166 | 0.0284 | 0.0095 | 0 | |
| 26 | | | | | 12708 |
| | CR1 | CR2 | MM1 | MM2 | |
| CR1 | 0 | 307 | 141 | 186 | |
| CR2 | 0.0242 | 0 | 311 | 308 | |
| MM1 | 0.0111 | 0.0245 | 0 | 134 | |
| MM2 | 0.0146 | 0.0242 | 0.0105 | 0 | |
| 28 | | | | | 17759 |
| | CR1 | CR2 | MM1 | MM2 | |
| CR1 | 0 | 15 | 129 | 168 | |
| CR2 | 0.0008 | 0 | 116 | 155 | |
| MM1 | 0.0073 | 0.0065 | 0 | 105 | |
| MM2 | 0.0095 | 0.0087 | 0.0059 | 0 | |

| 29 | | | | | 29571 |
|---|---|---|---|---|---|
| | CR1 | CR2 | MM1 | MM2 | |
| CR1 | 0 | 903 | 713 | 859 | |
| CR2 | 0.0305 | 0 | 719 | 689 | |
| MM1 | 0.0241 | 0.0243 | 0 | 626 | |
| MM2 | 0.0290 | 0.0233 | 0.0212 | 0 | |
| 32 | | | | | 15605 |
| | CR1 | CR2 | MM1 | MM2 | |
| CR1 | 0 | 167 | 187 | 197 | |
| CR2 | 0.0107 | 0 | 222 | 235 | |
| MM1 | 0.0120 | 0.0142 | 0 | 185 | |
| MM2 | 0.0126 | 0.0151 | 0.0119 | 0 | |
| 33 | | | | | 31052 |
| | CR1 | CR2 | MM1 | MM2 | |
| CR1 | 0 | 456 | 201 | 195 | |
| CR2 | 0.0147 | 0 | 489 | 465 | |
| MM1 | 0.0065 | 0.0157 | 0 | 187 | |
| MM2 | 0.0063 | 0.0150 | 0.0060 | 0 | |
| 34 | | | | | 15338 |
| | CR1 | CR2 | MM1 | MM2 | |
| CR1 | 0 | 357 | 228 | 238 | |
| CR2 | 0.0233 | 0 | 340 | 326 | |
| MM1 | 0.0149 | 0.0222 | 0 | 187 | |
| MM2 | 0.0155 | 0.0213 | 0.0122 | 0 | |
| 35 | | | | | 8520 |
| | CR1 | CR2 | MM1 | MM2 | |
| CR1 | 0 | 190 | 116 | 116 | |
| CR2 | 0.0223 | 0 | 194 | 194 | |
| MM1 | 0.0136 | 0.0228 | 0 | 0 | |
| MM2 | 0.0136 | 0.0228 | 0.0000 | 0 | |
| 36 | | | | | 12637 |
| | CR1 | CR2 | MM1 | MM2 | |
| CR1 | 0 | 393 | 197 | 204 | |
| CR2 | 0.0311 | 0 | 396 | 379 | |
| MM1 | 0.0156 | 0.0313 | 0 | 180 | |
| MM2 | 0.0161 | 0.0300 | 0.0142 | 0 | |
| 41 | | | | | 11599 |
| | CR1 | CR2 | MM1 | MM2 | |
| CR1 | 0 | 276 | 164 | 165 | |
| CR2 | 0.0238 | 0 | 274 | 261 | |
| MM1 | 0.0141 | 0.0236 | 0 | 156 | |
| MM2 | 0.0142 | 0.0225 | 0.0134 | 0 | |

| 46 | | | | | 6594 |
|---|---|---|---|---|---|
| | CR1 | CR2 | MM1 | MM2 | |
| CR1 | 0 | 0 | 218 | 224 | |
| CR2 | 0.0000 | 0 | 218 | 224 | |
| MM1 | 0.0331 | 0.0331 | 0 | 104 | |
| MM2 | 0.0340 | 0.0340 | 0.0158 | 0 | |

| 48 | | | | | 9269 |
|---|---|---|---|---|---|
| | CR1 | CR2 | MM1 | MM2 | |
| CR1 | 0 | 142 | 223 | 232 | |
| CR2 | 0.0153 | 0 | 297 | 308 | |
| MM1 | 0.0241 | 0.0320 | 0 | 177 | |
| MM2 | 0.0250 | 0.0332 | 0.0191 | 0 | |

| 49 | | | | | 14350 |
|---|---|---|---|---|---|
| | CR1 | CR2 | MM1 | MM2 | |
| CR1 | 0 | 275 | 121 | 135 | |
| CR2 | 0.0192 | 0 | 272 | 264 | |
| MM1 | 0.0084 | 0.0190 | 0 | 134 | |
| MM2 | 0.0094 | 0.0184 | 0.0093 | 0 | |

| 52 | | | | | 7787 |
|---|---|---|---|---|---|
| | CR1 | CR2 | MM1 | MM2 | |
| CR1 | 0 | 66 | 35 | 34 | |
| CR2 | 0.0085 | 0 | 65 | 66 | |
| MM1 | 0.0045 | 0.0083 | 0 | 35 | |
| MM2 | 0.0044 | 0.0085 | 0.0045 | 0 | |

| 58 | | | | | 12482 |
|---|---|---|---|---|---|
| | CR1 | CR2 | MM1 | MM2 | |
| CR1 | 0 | 204 | 114 | 119 | |
| CR2 | 0.0163 | 0 | 204 | 192 | |
| MM1 | 0.0091 | 0.0163 | 0 | 136 | |
| MM2 | 0.0095 | 0.0154 | 0.0109 | 0 | |

| 59 | | | | | 14513 |
|---|---|---|---|---|---|
| | CR1 | CR2 | MM1 | MM2 | |
| CR1 | 0 | 273 | 110 | 140 | |
| CR2 | 0.0188 | 0 | 258 | 286 | |
| MM1 | 0.0076 | 0.0178 | 0 | 150 | |
| MM2 | 0.0096 | 0.0197 | 0.0103 | 0 | |

| 60 | | | | | 11751 |
|---|---|---|---|---|---|
| | CR1 | CR2 | MM1 | MM2 | |
| CR1 | 0 | 254 | 0 | 136 | |
| CR2 | 0.0216 | 0 | 254 | 237 | |
| MM1 | 0.0000 | 0.0216 | 0 | 136 | |
| MM2 | 0.0116 | 0.0202 | 0.0116 | 0 | |

| 62 | | | | | 8232 |
|---|---|---|---|---|---|
| | CR1 | CR2 | MM1 | MM2 | |
| CR1 | 0 | 221 | 98 | 123 | |
| CR2 | 0.0268 | 0 | 232 | 220 | |
| MM1 | 0.0119 | 0.0282 | 0 | 91 | |
| MM2 | 0.0149 | 0.0267 | 0.0111 | 0 | |
| 63 | | | | | 12360 |
| | CR1 | CR2 | MM1 | MM2 | |
| CR1 | 0 | 363 | 405 | 413 | |
| CR2 | 0.0294 | 0 | 221 | 281 | |
| MM1 | 0.0328 | 0.0179 | 0 | 246 | |
| MM2 | 0.0334 | 0.0227 | 0.0199 | 0 | |
| 64 | | | | | 19408 |
| | CR1 | CR2 | MM1 | MM2 | |
| CR1 | 0 | 227 | 343 | 345 | |
| CR2 | 0.0117 | 0 | 411 | 418 | |
| MM1 | 0.0177 | 0.0212 | 0 | 228 | |
| MM2 | 0.0178 | 0.0215 | 0.0117 | 0 | |
| 66 | | | | | 12344 |
| | CR1 | CR2 | MM1 | MM2 | |
| CR1 | 0 | 239 | 208 | 218 | |
| CR2 | 0.0194 | 0 | 324 | 300 | |
| MM1 | 0.0169 | 0.0262 | 0 | 177 | |
| MM2 | 0.0177 | 0.0243 | 0.0143 | 0 | |
| 70 | | | | | 17461 |
| | CR1 | CR2 | MM1 | MM2 | |
| CR1 | 0 | 380 | 146 | 172 | |
| CR2 | 0.0218 | 0 | 385 | 392 | |
| MM1 | 0.0084 | 0.0220 | 0 | 62 | |
| MM2 | 0.0099 | 0.0225 | 0.0036 | 0 | |
| 71 | | | | | 9708 |
| | CR1 | CR2 | MM1 | MM2 | |
| CR1 | 0 | 230 | 94 | 123 | |
| CR2 | 0.0237 | 0 | 221 | 224 | |
| MM1 | 0.0097 | 0.0228 | 0 | 109 | |
| MM2 | 0.0127 | 0.0231 | 0.0112 | 0 | |
| 72 | | | | | 17714 |
| | CR1 | CR2 | MM1 | MM2 | |
| CR1 | 0 | 318 | 187 | 200 | |
| CR2 | 0.0180 | 0 | 340 | 329 | |
| MM1 | 0.0106 | 0.0192 | 0 | 206 | |
| MM2 | 0.0113 | 0.0186 | 0.0116 | 0 | |

| 73 | | | | | 14122 |
|---|---|---|---|---|---|
| | CR1 | CR2 | MM1 | MM2 | |
| CR1 | 0 | 225 | 221 | 224 | |
| CR2 | 0.0159 | 0 | 88 | 96 | |
| MM1 | 0.0156 | 0.0062 | 0 | 93 | |
| MM2 | 0.0159 | 0.0068 | 0.0066 | 0 | |
| 76 | | | | | 8695 |
| | CR1 | CR2 | MM1 | MM2 | |
| CR1 | 0 | 185 | 204 | 211 | |
| CR2 | 0.0213 | 0 | 95 | 123 | |
| MM1 | 0.0235 | 0.0109 | 0 | 39 | |
| MM2 | 0.0243 | 0.0141 | 0.0045 | 0 | |
| 78 | | | | | 17831 |
| | CR1 | CR2 | MM1 | MM2 | |
| CR1 | 0 | 380 | 388 | 400 | |
| CR2 | 0.0213 | 0 | 178 | 203 | |
| MM1 | 0.0218 | 0.0100 | 0 | 73 | |
| MM2 | 0.0224 | 0.0114 | 0.0041 | 0 | |
| 80 | | | | | 9507 |
| | CR1 | CR2 | MM1 | MM2 | |
| CR1 | 0 | 146 | 137 | 141 | |
| CR2 | 0.0154 | 0 | 44 | 67 | |
| MM1 | 0.0144 | 0.0046 | 0 | 55 | |
| MM2 | 0.0148 | 0.0070 | 0.0058 | 0 | |
| 84 | | | | | 10425 |
| | CR1 | CR2 | MM1 | MM2 | |
| CR1 | 0 | 412 | 174 | 193 | |
| CR2 | 0.0395 | 0 | 416 | 425 | |
| MM1 | 0.0167 | 0.0399 | 0 | 150 | |
| MM2 | 0.0185 | 0.0408 | 0.0144 | 0 | |
| 86 | | | | | 9705 |
| | CR1 | CR2 | MM1 | MM2 | |
| CR1 | 0 | 330 | 217 | 217 | |
| CR2 | 0.0340 | 0 | 307 | 307 | |
| MM1 | 0.0224 | 0.0316 | 0 | 0 | |
| MM2 | 0.0224 | 0.0316 | 0.0000 | 0 | |
| 90 | | | | | 14644 |
| | CR1 | CR2 | MM1 | MM2 | |
| CR1 | 0 | 294 | 144 | 139 | |
| CR2 | 0.0201 | 0 | 326 | 332 | |
| MM1 | 0.0098 | 0.0223 | 0 | 133 | |
| MM2 | 0.0095 | 0.0227 | 0.0091 | 0 | |

| 91 | | | | | 7542 |
|---|---|---|---|---|---|
| | CR1 | CR2 | MM1 | MM2 | |
| CR1 | 0 | 43 | 150 | 151 | |
| CR2 | 0.0057 | 0 | 176 | 179 | |
| MM1 | 0.0199 | 0.0233 | 0 | 94 | |
| MM2 | 0.0200 | 0.0237 | 0.0125 | 0 | |
| 92 | | | | | 12811 |
| | CR1 | CR2 | MM1 | MM2 | |
| CR1 | 0 | 195 | 190 | 198 | |
| CR2 | 0.0152 | 0 | 84 | 82 | |
| MM1 | 0.0148 | 0.0066 | 0 | 92 | |
| MM2 | 0.0155 | 0.0064 | 0.0072 | 0 | |
| 93 | | | | | 12881 |
| | CR1 | CR2 | MM1 | MM2 | |
| CR1 | 0 | 374 | 320 | 273 | |
| CR2 | 0.0290 | 0 | 387 | 389 | |
| MM1 | 0.0248 | 0.0300 | 0 | 210 | |
| MM2 | 0.0212 | 0.0302 | 0.0163 | 0 | |
| 94 | | | | | 8340 |
| | CR1 | CR2 | MM1 | MM2 | |
| CR1 | 0 | 378 | 203 | 174 | |
| CR2 | 0.0453 | 0 | 352 | 353 | |
| MM1 | 0.0243 | 0.0422 | 0 | 216 | |
| MM2 | 0.0209 | 0.0423 | 0.0259 | 0 | |
| 97 | | | | | 9893 |
| | CR1 | CR2 | MM1 | MM2 | |
| CR1 | 0 | 114 | 187 | 236 | |
| CR2 | 0.0115 | 0 | 261 | 282 | |
| MM1 | 0.0189 | 0.0264 | 0 | 155 | |
| MM2 | 0.0239 | 0.0285 | 0.0157 | 0 | |
| 98 | | | | | 14044 |
| | CR1 | CR2 | MM1 | MM2 | |
| CR1 | 0 | 298 | 139 | 127 | |
| CR2 | 0.0212 | 0 | 302 | 294 | |
| MM1 | 0.0099 | 0.0215 | 0 | 143 | |
| MM2 | 0.0090 | 0.0209 | 0.0102 | 0 | |
| 101 | | | | | 16924 |
| | CR1 | CR2 | MM1 | MM2 | |
| CR1 | 0 | 359 | 314 | 325 | |
| CR2 | 0.0212 | 0 | 397 | 462 | |
| MM1 | 0.0186 | 0.0235 | 0 | 243 | |
| MM2 | 0.0192 | 0.0273 | 0.0144 | 0 | |

| 107 | | | | | 16111 |
|---|---|---|---|---|---|
| | CR1 | CR2 | MM1 | MM2 | |
| CR1 | 0 | 362 | 183 | 146 | |
| CR2 | 0.0225 | 0 | 347 | 334 | |
| MM1 | 0.0114 | 0.0215 | 0 | 163 | |
| MM2 | 0.0091 | 0.0207 | 0.0101 | 0 | |
| 109 | | | | | 10411 |
| | CR1 | CR2 | MM1 | MM2 | |
| CR1 | 0 | 337 | 105 | 201 | |
| CR2 | 0.0324 | 0 | 316 | 291 | |
| MM1 | 0.0101 | 0.0304 | 0 | 96 | |
| MM2 | 0.0193 | 0.0280 | 0.0092 | 0 | |
| 110 | | | | | 9137 |
| | CR1 | CR2 | MM1 | MM2 | |
| CR1 | 0 | 203 | 114 | 97 | |
| CR2 | 0.0222 | 0 | 209 | 193 | |
| MM1 | 0.0125 | 0.0229 | 0 | 113 | |
| MM2 | 0.0106 | 0.0211 | 0.0124 | 0 | |
| 113 | | | | | 10514 |
| | CR1 | CR2 | MM1 | MM2 | |
| CR1 | 0 | 190 | 184 | 183 | |
| CR2 | 0.0181 | 0 | 77 | 102 | |
| MM1 | 0.0175 | 0.0073 | 0 | 87 | |
| MM2 | 0.0174 | 0.0097 | 0.0083 | 0 | |
| 114 | | | | | 13380 |
| | CR1 | CR2 | MM1 | MM2 | |
| CR1 | 0 | 242 | 141 | 149 | |
| CR2 | 0.0181 | 0 | 242 | 231 | |
| MM1 | 0.0105 | 0.0181 | 0 | 134 | |
| MM2 | 0.0111 | 0.0173 | 0.0100 | 0 | |
| 115 | | | | | 10245 |
| | CR1 | CR2 | MM1 | MM2 | |
| CR1 | 0 | 199 | 97 | 85 | |
| CR2 | 0.0194 | 0 | 205 | 192 | |
| MM1 | 0.0095 | 0.0200 | 0 | 87 | |
| MM2 | 0.0083 | 0.0187 | 0.0085 | 0 | |
| 117 | | | | | 10130 |
| | CR1 | CR2 | MM1 | MM2 | |
| CR1 | 0 | 10 | 236 | 211 | |
| CR2 | 0.0010 | 0 | 237 | 212 | |
| MM1 | 0.0233 | 0.0234 | 0 | 147 | |
| MM2 | 0.0208 | 0.0209 | 0.0145 | 0 | |

| 120 | | | | | 10222 |
|---|---|---|---|---|---|
| | CR1 | CR2 | MM1 | MM2 | |
| CR1 | 0 | 240 | 182 | 152 | |
| CR2 | 0.0235 | 0 | 229 | 236 | |
| MM1 | 0.0178 | 0.0224 | 0 | 155 | |
| MM2 | 0.0149 | 0.0231 | 0.0152 | 0 | |
| 121 | | | | | 15870 |
| | CR1 | CR2 | MM1 | MM2 | |
| CR1 | 0 | 332 | 172 | 175 | |
| CR2 | 0.0209 | 0 | 341 | 335 | |
| MM1 | 0.0108 | 0.0215 | 0 | 171 | |
| MM2 | 0.0110 | 0.0211 | 0.0108 | 0 | |
| 124 | | | | | 11046 |
| | CR1 | CR2 | MM1 | MM2 | |
| CR1 | 0 | 183 | 86 | 128 | |
| CR2 | 0.0166 | 0 | 166 | 174 | |
| MM1 | 0.0078 | 0.0150 | 0 | 86 | |
| MM2 | 0.0116 | 0.0158 | 0.0078 | 0 | |
| 125 | | | | | 8100 |
| | CR1 | CR2 | MM1 | MM2 | |
| CR1 | 0 | 215 | 135 | 110 | |
| CR2 | 0.0265 | 0 | 258 | 239 | |
| MM1 | 0.0167 | 0.0319 | 0 | 93 | |
| MM2 | 0.0136 | 0.0295 | 0.0115 | 0 | |
| 135 | | | | | 12346 |
| | CR1 | CR2 | MM1 | MM2 | |
| CR1 | 0 | 0 | 307 | 320 | |
| CR2 | 0.0000 | 0 | 307 | 320 | |
| MM1 | 0.0249 | 0.0249 | 0 | 165 | |
| MM2 | 0.0259 | 0.0259 | 0.0134 | 0 | |
| 146 | | | | | 6794 |
| | CR1 | CR2 | MM1 | MM2 | |
| CR1 | 0 | 215 | 198 | 202 | |
| CR2 | 0.0316 | 0 | 241 | 234 | |
| MM1 | 0.0291 | 0.0355 | 0 | 123 | |
| MM2 | 0.0297 | 0.0344 | 0.0181 | 0 | |
| 150 | | | | | 9873 |
| | CR1 | CR2 | MM1 | MM2 | |
| CR1 | 0 | 264 | 249 | 263 | |
| CR2 | 0.0267 | 0 | 143 | 164 | |
| MM1 | 0.0252 | 0.0145 | 0 | 131 | |
| MM2 | 0.0266 | 0.0166 | 0.0133 | 0 | |

| 151 | | | | | 16174 |
|---|---|---|---|---|---|
| | CR1 | CR2 | MM1 | MM2 | |
| CR1 | 0 | 347 | 136 | 127 | |
| CR2 | 0.0215 | 0 | 352 | 362 | |
| MM1 | 0.0084 | 0.0218 | 0 | 156 | |
| MM2 | 0.0079 | 0.0224 | 0.0096 | 0 | |

| 152 | | | | | 11673 |
|---|---|---|---|---|---|
| | CR1 | CR2 | MM1 | MM2 | |
| CR1 | 0 | 312 | 144 | 163 | |
| CR2 | 0.0267 | 0 | 316 | 335 | |
| MM1 | 0.0123 | 0.0271 | 0 | 143 | |
| MM2 | 0.0140 | 0.0287 | 0.0123 | 0 | |

| 156 | | | | | 16144 |
|---|---|---|---|---|---|
| | CR1 | CR2 | MM1 | MM2 | |
| CR1 | 0 | 363 | 372 | 339 | |
| CR2 | 0.0225 | 0 | 245 | 206 | |
| MM1 | 0.0230 | 0.0152 | 0 | 189 | |
| MM2 | 0.0210 | 0.0128 | 0.0117 | 0 | |

| 158 | | | | | 10788 |
|---|---|---|---|---|---|
| | CR1 | CR2 | MM1 | MM2 | |
| CR1 | 0 | 232 | 74 | 14 | |
| CR2 | 0.0215 | 0 | 244 | 236 | |
| MM1 | 0.0069 | 0.0226 | 0 | 75 | |
| MM2 | 0.0013 | 0.0219 | 0.0070 | 0 | |

| 160 | | | | | 11505 |
|---|---|---|---|---|---|
| | CR1 | CR2 | MM1 | MM2 | |
| CR1 | 0 | 273 | 185 | 164 | |
| CR2 | 0.0237 | 0 | 276 | 251 | |
| MM1 | 0.0161 | 0.0240 | 0 | 174 | |
| MM2 | 0.0143 | 0.0218 | 0.0151 | 0 | |

| 164 | | | | | 14788 |
|---|---|---|---|---|---|
| | CR1 | CR2 | MM1 | MM2 | |
| CR1 | 0 | 166 | 46 | 42 | |
| CR2 | 0.0112 | 0 | 156 | 162 | |
| MM1 | 0.0031 | 0.0105 | 0 | 44 | |
| MM2 | 0.0028 | 0.0110 | 0.0030 | 0 | |

| 165 | | | | | 13463 |
|---|---|---|---|---|---|
| | CR1 | CR2 | MM1 | MM2 | |
| CR1 | 0 | 197 | 94 | 68 | |
| CR2 | 0.0146 | 0 | 187 | 198 | |
| MM1 | 0.0070 | 0.0139 | 0 | 94 | |
| MM2 | 0.0051 | 0.0147 | 0.0070 | 0 | |

| 166 | | | | | 21826 |
|---|---|---|---|---|---|
| | CR1 | CR2 | MM1 | MM2 | |
| CR1 | 0 | 444 | 205 | 191 | |
| CR2 | 0.0203 | 0 | 467 | 470 | |
| MM1 | 0.0094 | 0.0214 | 0 | 188 | |
| MM2 | 0.0088 | 0.0215 | 0.0086 | 0 | |
| 171 | | | | | 13029 |
| | CR1 | CR2 | MM1 | MM2 | |
| CR1 | 0 | 124 | 420 | 454 | |
| CR2 | 0.0095 | 0 | 383 | 399 | |
| MM1 | 0.0322 | 0.0294 | 0 | 258 | |
| MM2 | 0.0348 | 0.0306 | 0.0198 | 0 | |
| 174 | | | | | 10585 |
| | CR1 | CR2 | MM1 | MM2 | |
| CR1 | 0 | 278 | 228 | 207 | |
| CR2 | 0.0263 | 0 | 316 | 299 | |
| MM1 | 0.0215 | 0.0299 | 0 | 172 | |
| MM2 | 0.0196 | 0.0282 | 0.0162 | 0 | |
| 175 | | | | | 12795 |
| | CR1 | CR2 | MM1 | MM2 | |
| CR1 | 0 | 325 | 161 | 159 | |
| CR2 | 0.0254 | 0 | 320 | 328 | |
| MM1 | 0.0126 | 0.0250 | 0 | 133 | |
| MM2 | 0.0124 | 0.0256 | 0.0104 | 0 | |
| 179 | | | | | 13760 |
| | CR1 | CR2 | MM1 | MM2 | |
| CR1 | 0 | 202 | 108 | 88 | |
| CR2 | 0.0147 | 0 | 216 | 200 | |
| MM1 | 0.0078 | 0.0157 | 0 | 101 | |
| MM2 | 0.0064 | 0.0145 | 0.0073 | 0 | |
| 183 | | | | | 14148 |
| | CR1 | CR2 | MM1 | MM2 | |
| CR1 | 0 | 373 | 166 | 44 | |
| CR2 | 0.0264 | 0 | 380 | 372 | |
| MM1 | 0.0117 | 0.0269 | 0 | 154 | |
| MM2 | 0.0031 | 0.0263 | 0.0109 | 0 | |
| 185 | | | | | 10093 |
| | CR1 | CR2 | MM1 | MM2 | |
| CR1 | 0 | 270 | 161 | 168 | |
| CR2 | 0.0268 | 0 | 251 | 262 | |
| MM1 | 0.0160 | 0.0249 | 0 | 129 | |
| MM2 | 0.0166 | 0.0260 | 0.0128 | 0 | |

| 195 | | | | | 9895 |
|---|---|---|---|---|---|
| | CR1 | CR2 | MM1 | MM2 | |
| CR1 | 0 | 271 | 140 | 111 | |
| CR2 | 0.0274 | 0 | 288 | 281 | |
| MM1 | 0.0141 | 0.0291 | 0 | 151 | |
| MM2 | 0.0112 | 0.0284 | 0.0153 | 0 | |
| 196 | | | | | 18576 |
| | CR1 | CR2 | MM1 | MM2 | |
| CR1 | 0 | 478 | 323 | 302 | |
| CR2 | 0.0257 | 0 | 469 | 453 | |
| MM1 | 0.0174 | 0.0252 | 0 | 222 | |
| MM2 | 0.0163 | 0.0244 | 0.0120 | 0 | |
| 198 | | | | | 13241 |
| | CR1 | CR2 | MM1 | MM2 | |
| CR1 | 0 | 419 | 376 | 392 | |
| CR2 | 0.0316 | 0 | 250 | 251 | |
| MM1 | 0.0284 | 0.0189 | 0 | 202 | |
| MM2 | 0.0296 | 0.0190 | 0.0153 | 0 | |
| 209 | | | | | 10773 |
| | CR1 | CR2 | MM1 | MM2 | |
| CR1 | 0 | 239 | 152 | 156 | |
| CR2 | 0.0222 | 0 | 233 | 241 | |
| MM1 | 0.0141 | 0.0216 | 0 | 163 | |
| MM2 | 0.0145 | 0.0224 | 0.0151 | 0 | |
| 213 | | | | | 9557 |
| | CR1 | CR2 | MM1 | MM2 | |
| CR1 | 0 | 207 | 130 | 110 | |
| CR2 | 0.0217 | 0 | 219 | 209 | |
| MM1 | 0.0136 | 0.0229 | 0 | 133 | |
| MM2 | 0.0115 | 0.0219 | 0.0139 | 0 | |
| 214 | | | | | 10027 |
| | CR1 | CR2 | MM1 | MM2 | |
| CR1 | 0 | 306 | 292 | 302 | |
| CR2 | 0.0305 | 0 | 222 | 65 | |
| MM1 | 0.0291 | 0.0221 | 0 | 246 | |
| MM2 | 0.0301 | 0.0065 | 0.0245 | 0 | |
| 215 | | | | | 11372 |
| | CR1 | CR2 | MM1 | MM2 | |
| CR1 | 0 | 250 | 242 | 248 | |
| CR2 | 0.0220 | 0 | 114 | 147 | |
| MM1 | 0.0213 | 0.0100 | 0 | 166 | |
| MM2 | 0.0218 | 0.0129 | 0.0146 | 0 | |

| 219 | | | | | 9624 |
|---|---|---|---|---|---|
| | CR1 | CR2 | MM1 | MM2 | |
| CR1 | 0 | 159 | 122 | 132 | |
| CR2 | 0.0165 | 0 | 185 | 191 | |
| MM1 | 0.0127 | 0.0192 | 0 | 24 | |
| MM2 | 0.0137 | 0.0198 | 0.0025 | 0 | |
| 222 | | | | | 10036 |
| | CR1 | CR2 | MM1 | MM2 | |
| CR1 | 0 | 186 | 92 | 108 | |
| CR2 | 0.0185 | 0 | 194 | 190 | |
| MM1 | 0.0092 | 0.0193 | 0 | 86 | |
| MM2 | 0.0108 | 0.0189 | 0.0086 | 0 | |
| 223 | | | | | 13936 |
| | CR1 | CR2 | MM1 | MM2 | |
| CR1 | 0 | 218 | 231 | 222 | |
| CR2 | 0.0156 | 0 | 118 | 109 | |
| MM1 | 0.0166 | 0.0085 | 0 | 143 | |
| MM2 | 0.0159 | 0.0078 | 0.0103 | 0 | |
| 229 | | | | | 8765 |
| | CR1 | CR2 | MM1 | MM2 | |
| CR1 | 0 | 202 | 206 | 206 | |
| CR2 | 0.0230 | 0 | 89 | 70 | |
| MM1 | 0.0235 | 0.0102 | 0 | 91 | |
| MM2 | 0.0235 | 0.0080 | 0.0104 | 0 | |
| 231 | | | | | 7242 |
| | CR1 | CR2 | MM1 | MM2 | |
| CR1 | 0 | 130 | 113 | 105 | |
| CR2 | 0.0180 | 0 | 147 | 145 | |
| MM1 | 0.0156 | 0.0203 | 0 | 94 | |
| MM2 | 0.0145 | 0.0200 | 0.0130 | 0 | |
| 234 | | | | | 10953 |
| | CR1 | CR2 | MM1 | MM2 | |
| CR1 | 0 | 254 | 244 | 234 | |
| CR2 | 0.0232 | 0 | 118 | 113 | |
| MM1 | 0.0223 | 0.0108 | 0 | 93 | |
| MM2 | 0.0214 | 0.0103 | 0.0085 | 0 | |
| 239 | | | | | 11982 |
| | CR1 | CR2 | MM1 | MM2 | |
| CR1 | 0 | 235 | 239 | 243 | |
| CR2 | 0.0196 | 0 | 129 | 95 | |
| MM1 | 0.0199 | 0.0108 | 0 | 122 | |
| MM2 | 0.0203 | 0.0079 | 0.0102 | 0 | |

| 244 | | | | | 7500 |
|---|---|---|---|---|---|
| | CR1 | CR2 | MM1 | MM2 | |
| CR1 | 0 | 11 | 166 | 167 | |
| CR2 | 0.0015 | 0 | 159 | 162 | |
| MM1 | 0.0221 | 0.0212 | 0 | 69 | |
| MM2 | 0.0223 | 0.0216 | 0.0092 | 0 | |
| 247 | | | | | 10039 |
| | CR1 | CR2 | MM1 | MM2 | |
| CR1 | 0 | 228 | 214 | 217 | |
| CR2 | 0.0227 | 0 | 84 | 89 | |
| MM1 | 0.0213 | 0.0084 | 0 | 113 | |
| MM2 | 0.0216 | 0.0089 | 0.0113 | 0 | |
| 253 | | | | | 11114 |
| | CR1 | CR2 | MM1 | MM2 | |
| CR1 | 0 | 162 | 161 | 166 | |
| CR2 | 0.0146 | 0 | 1 | 80 | |
| MM1 | 0.0145 | 0.0001 | 0 | 79 | |
| MM2 | 0.0149 | 0.0072 | 0.0071 | 0 | |
| 254 | | | | | 25641 |
| | CR1 | CR2 | MM1 | MM2 | |
| CR1 | 0 | 672 | 226 | 226 | |
| CR2 | 0.0262 | 0 | 693 | 693 | |
| MM1 | 0.0088 | 0.0270 | 0 | 0 | |
| MM2 | 0.0088 | 0.0270 | 0.0000 | 0 | |
| 256 | | | | | 11104 |
| | CR1 | CR2 | MM1 | MM2 | |
| CR1 | 0 | 300 | 167 | 175 | |
| CR2 | 0.0270 | 0 | 302 | 283 | |
| MM1 | 0.0150 | 0.0272 | 0 | 154 | |
| MM2 | 0.0158 | 0.0255 | 0.0139 | 0 | |
| 262 | | | | | 7083 |
| | CR1 | CR2 | MM1 | MM2 | |
| CR1 | 0 | 6 | 25 | 37 | |
| CR2 | 0.0008 | 0 | 29 | 41 | |
| MM1 | 0.0035 | 0.0041 | 0 | 39 | |
| MM2 | 0.0052 | 0.0058 | 0.0055 | 0 | |
| 265 | | | | | 12005 |
| | CR1 | CR2 | MM1 | MM2 | |
| CR1 | 0 | 170 | 70 | 88 | |
| CR2 | 0.0142 | 0 | 155 | 167 | |
| MM1 | 0.0058 | 0.0129 | 0 | 94 | |
| MM2 | 0.0073 | 0.0139 | 0.0078 | 0 | |

| 270 | | | | | 10875 |
|---|---|---|---|---|---|
| | CR1 | CR2 | MM1 | MM2 | |
| CR1 | 0 | 183 | 148 | 141 | |
| CR2 | 0.0168 | 0 | 201 | 194 | |
| MM1 | 0.0136 | 0.0185 | 0 | 121 | |
| MM2 | 0.0130 | 0.0178 | 0.0111 | 0 | |
| 272 | | | | | 10338 |
| | CR1 | CR2 | MM1 | MM2 | |
| CR1 | 0 | 207 | 217 | 220 | |
| CR2 | 0.0200 | 0 | 89 | 113 | |
| MM1 | 0.0210 | 0.0086 | 0 | 105 | |
| MM2 | 0.0213 | 0.0109 | 0.0102 | 0 | |
| 275 | | | | | 10378 |
| | CR1 | CR2 | MM1 | MM2 | |
| CR1 | 0 | 176 | 109 | 88 | |
| CR2 | 0.0170 | 0 | 178 | 179 | |
| MM1 | 0.0105 | 0.0172 | 0 | 98 | |
| MM2 | 0.0085 | 0.0172 | 0.0094 | 0 | |
| 277 | | | | | 20480 |
| | CR1 | CR2 | MM1 | MM2 | |
| CR1 | 0 | 461 | 70 | 100 | |
| CR2 | 0.0225 | 0 | 453 | 451 | |
| MM1 | 0.0034 | 0.0221 | 0 | 152 | |
| MM2 | 0.0049 | 0.0220 | 0.0074 | 0 | |
| 281 | | | | | 11984 |
| | CR1 | CR2 | MM1 | MM2 | |
| CR1 | 0 | 346 | 173 | 173 | |
| CR2 | 0.0289 | 0 | 330 | 330 | |
| MM1 | 0.0144 | 0.0275 | 0 | 0 | |
| MM2 | 0.0144 | 0.0275 | 0.0000 | 0 | |
| 282 | | | | | 9731 |
| | CR1 | CR2 | MM1 | MM2 | |
| CR1 | 0 | 209 | 197 | 245 | |
| CR2 | 0.0215 | 0 | 115 | 185 | |
| MM1 | 0.0202 | 0.0118 | 0 | 178 | |
| MM2 | 0.0252 | 0.0190 | 0.0183 | 0 | |
| 285 | | | | | 12050 |
| | CR1 | CR2 | MM1 | MM2 | |
| CR1 | 0 | 22 | 95 | 108 | |
| CR2 | 0.0018 | 0 | 87 | 98 | |
| MM1 | 0.0079 | 0.0072 | 0 | 87 | |
| MM2 | 0.0090 | 0.0081 | 0.0072 | 0 | |

| 287 | | | | | 15768 |
|---|---|---|---|---|---|
| | CR1 | CR2 | MM1 | MM2 | |
| CR1 | 0 | 331 | 212 | 189 | |
| CR2 | 0.0210 | 0 | 360 | 326 | |
| MM1 | 0.0134 | 0.0228 | 0 | 219 | |
| MM2 | 0.0120 | 0.0207 | 0.0139 | 0 | |

| 288 | | | | | 12309 |
|---|---|---|---|---|---|
| | CR1 | CR2 | MM1 | MM2 | |
| CR1 | 0 | 228 | 135 | 142 | |
| CR2 | 0.0185 | 0 | 222 | 225 | |
| MM1 | 0.0110 | 0.0180 | 0 | 129 | |
| MM2 | 0.0115 | 0.0183 | 0.0105 | 0 | |

| 294 | | | | | 10782 |
|---|---|---|---|---|---|
| | CR1 | CR2 | MM1 | MM2 | |
| CR1 | 0 | 39 | 227 | 200 | |
| CR2 | 0.0036 | 0 | 222 | 193 | |
| MM1 | 0.0211 | 0.0206 | 0 | 170 | |
| MM2 | 0.0185 | 0.0179 | 0.0158 | 0 | |

| 295 | | | | | 6357 |
|---|---|---|---|---|---|
| | CR1 | CR2 | MM1 | MM2 | |
| CR1 | 0 | 97 | 44 | 0 | |
| CR2 | 0.0153 | 0 | 101 | 97 | |
| MM1 | 0.0069 | 0.0159 | 0 | 44 | |
| MM2 | 0.0000 | 0.0153 | 0.0069 | 0 | |

| 296 | | | | | 11939 |
|---|---|---|---|---|---|
| | CR1 | CR2 | MM1 | MM2 | |
| CR1 | 0 | 170 | 76 | 101 | |
| CR2 | 0.0142 | 0 | 176 | 182 | |
| MM1 | 0.0064 | 0.0147 | 0 | 82 | |
| MM2 | 0.0085 | 0.0152 | 0.0069 | 0 | |

| 301 | | | | | 12440 |
|---|---|---|---|---|---|
| | CR1 | CR2 | MM1 | MM2 | |
| CR1 | 0 | 1 | 203 | 167 | |
| CR2 | 0.0001 | 0 | 202 | 166 | |
| MM1 | 0.0163 | 0.0162 | 0 | 143 | |
| MM2 | 0.0134 | 0.0133 | 0.0115 | 0 | |

| 302 | | | | | 10942 |
|---|---|---|---|---|---|
| | CR1 | CR2 | MM1 | MM2 | |
| CR1 | 0 | 65 | 334 | 329 | |
| CR2 | 0.0059 | 0 | 284 | 284 | |
| MM1 | 0.0305 | 0.0260 | 0 | 190 | |
| MM2 | 0.0301 | 0.0260 | 0.0174 | 0 | |

| 304 | | | | | 9458 |
|---|---|---|---|---|---|
| | CR1 | CR2 | MM1 | MM2 | |
| CR1 | 0 | 52 | 235 | 230 | |
| CR2 | 0.0055 | 0 | 233 | 226 | |
| MM1 | 0.0248 | 0.0246 | 0 | 126 | |
| MM2 | 0.0243 | 0.0239 | 0.0133 | 0 | |
| 309 | | | | | 12657 |
| | CR1 | CR2 | MM1 | MM2 | |
| CR1 | 0 | 248 | 136 | 107 | |
| CR2 | 0.0196 | 0 | 240 | 257 | |
| MM1 | 0.0107 | 0.0190 | 0 | 85 | |
| MM2 | 0.0085 | 0.0203 | 0.0067 | 0 | |
| 322 | | | | | 9216 |
| | CR1 | CR2 | MM1 | MM2 | |
| CR1 | 0 | 229 | 91 | 101 | |
| CR2 | 0.0248 | 0 | 211 | 218 | |
| MM1 | 0.0099 | 0.0229 | 0 | 87 | |
| MM2 | 0.0110 | 0.0237 | 0.0094 | 0 | |
| 326 | | | | | 7360 |
| | CR1 | CR2 | MM1 | MM2 | |
| CR1 | 0 | 167 | 69 | 71 | |
| CR2 | 0.0227 | 0 | 171 | 179 | |
| MM1 | 0.0094 | 0.0232 | 0 | 84 | |
| MM2 | 0.0096 | 0.0243 | 0.0114 | 0 | |
| 328 | | | | | 10943 |
| | CR1 | CR2 | MM1 | MM2 | |
| CR1 | 0 | 273 | 94 | 99 | |
| CR2 | 0.0249 | 0 | 285 | 268 | |
| MM1 | 0.0086 | 0.0260 | 0 | 113 | |
| MM2 | 0.0090 | 0.0245 | 0.0103 | 0 | |
| 329 | | | | | 8201 |
| | CR1 | CR2 | MM1 | MM2 | |
| CR1 | 0 | 229 | 70 | 65 | |
| CR2 | 0.0279 | 0 | 217 | 218 | |
| MM1 | 0.0085 | 0.0265 | 0 | 62 | |
| MM2 | 0.0079 | 0.0266 | 0.0076 | 0 | |
| 330 | | | | | 7871 |
| | CR1 | CR2 | MM1 | MM2 | |
| CR1 | 0 | 324 | 383 | 392 | |
| CR2 | 0.0412 | 0 | 190 | 189 | |
| MM1 | 0.0487 | 0.0241 | 0 | 97 | |
| MM2 | 0.0498 | 0.0240 | 0.0123 | 0 | |

| 331 | | | | | 11037 |
|---|---|---|---|---|---|
| | CR1 | CR2 | MM1 | MM2 | |
| CR1 | 0 | 322 | 138 | 137 | |
| CR2 | 0.0292 | 0 | 300 | 310 | |
| MM1 | 0.0125 | 0.0272 | 0 | 138 | |
| MM2 | 0.0124 | 0.0281 | 0.0125 | 0 | |
| 333 | | | | | 9128 |
| | CR1 | CR2 | MM1 | MM2 | |
| CR1 | 0 | 156 | 80 | 85 | |
| CR2 | 0.0171 | 0 | 159 | 156 | |
| MM1 | 0.0088 | 0.0174 | 0 | 77 | |
| MM2 | 0.0093 | 0.0171 | 0.0084 | 0 | |
| 334 | | | | | 13655 |
| | CR1 | CR2 | MM1 | MM2 | |
| CR1 | 0 | 289 | 117 | 185 | |
| CR2 | 0.0212 | 0 | 296 | 289 | |
| MM1 | 0.0086 | 0.0217 | 0 | 172 | |
| MM2 | 0.0135 | 0.0212 | 0.0126 | 0 | |
| 338 | | | | | 10479 |
| | CR1 | CR2 | MM1 | MM2 | |
| CR1 | 0 | 269 | 174 | 155 | |
| CR2 | 0.0257 | 0 | 269 | 260 | |
| MM1 | 0.0166 | 0.0257 | 0 | 149 | |
| MM2 | 0.0148 | 0.0248 | 0.0142 | 0 | |
| 343 | | | | | 16022 |
| | CR1 | CR2 | MM1 | MM2 | |
| CR1 | 0 | 256 | 274 | 268 | |
| CR2 | 0.0160 | 0 | 120 | 129 | |
| MM1 | 0.0171 | 0.0075 | 0 | 115 | |
| MM2 | 0.0167 | 0.0081 | 0.0072 | 0 | |
| 345 | | | | | 11190 |
| | CR1 | CR2 | MM1 | MM2 | |
| CR1 | 0 | 271 | 104 | 127 | |
| CR2 | 0.0242 | 0 | 280 | 274 | |
| MM1 | 0.0093 | 0.0250 | 0 | 137 | |
| MM2 | 0.0113 | 0.0245 | 0.0122 | 0 | |
| 346 | | | | | 8088 |
| | CR1 | CR2 | MM1 | MM2 | |
| CR1 | 0 | 278 | 302 | 268 | |
| CR2 | 0.0344 | 0 | 149 | 148 | |
| MM1 | 0.0373 | 0.0184 | 0 | 193 | |
| MM2 | 0.0331 | 0.0183 | 0.0239 | 0 | |

Table S7: Difference Matrices.



| 348 | | | | | 11017 |
|---|---|---|---|---|---|
| | CR1 | CR2 | MM1 | MM2 | |
| CR1 | 0 | 354 | 364 | 364 | |
| CR2 | 0.0321 | 0 | 172 | 172 | |
| MM1 | 0.0330 | 0.0156 | 0 | 0 | |
| MM2 | 0.0330 | 0.0156 | 0.0000 | 0 | |
| 352 | | | | | 11430 |
| | CR1 | CR2 | MM1 | MM2 | |
| CR1 | 0 | 16 | 131 | 152 | |
| CR2 | 0.0014 | 0 | 115 | 136 | |
| MM1 | 0.0115 | 0.0101 | 0 | 73 | |
| MM2 | 0.0133 | 0.0119 | 0.0064 | 0 | |
| 353 | | | | | 11501 |
| | CR1 | CR2 | MM1 | MM2 | |
| CR1 | 0 | 0 | 347 | 347 | |
| CR2 | 0.0000 | 0 | 347 | 347 | |
| MM1 | 0.0302 | 0.0302 | 0 | 153 | |
| MM2 | 0.0302 | 0.0302 | 0.0133 | 0 | |
| 354 | | | | | 17127 |
| | CR1 | CR2 | MM1 | MM2 | |
| CR1 | 0 | 414 | 232 | 285 | |
| CR2 | 0.0242 | 0 | 401 | 383 | |
| MM1 | 0.0135 | 0.0234 | 0 | 218 | |
| MM2 | 0.0166 | 0.0224 | 0.0127 | 0 | |
| 355 | | | | | 20986 |
| | CR1 | CR2 | MM1 | MM2 | |
| CR1 | 0 | 407 | 188 | 205 | |
| CR2 | 0.0194 | 0 | 379 | 419 | |
| MM1 | 0.0090 | 0.0181 | 0 | 240 | |
| MM2 | 0.0098 | 0.0200 | 0.0114 | 0 | |
| 356 | | | | | 7532 |
| | CR1 | CR2 | MM1 | MM2 | |
| CR1 | 0 | 117 | 25 | 37 | |
| CR2 | 0.0155 | 0 | 114 | 117 | |
| MM1 | 0.0033 | 0.0151 | 0 | 26 | |
| MM2 | 0.0049 | 0.0155 | 0.0035 | 0 | |
| 358 | | | | | 6162 |
| | CR1 | CR2 | MM1 | MM2 | |
| CR1 | 0 | 140 | 69 | 86 | |
| CR2 | 0.0227 | 0 | 135 | 142 | |
| MM1 | 0.0112 | 0.0219 | 0 | 80 | |
| MM2 | 0.0140 | 0.0230 | 0.0130 | 0 | |

| 359 | | | | | 8294 |
|---|---|---|---|---|---|
| | CR1 | CR2 | MM1 | MM2 | |
| CR1 | 0 | 163 | 97 | 0 | |
| CR2 | 0.0197 | 0 | 170 | 163 | |
| MM1 | 0.0117 | 0.0205 | 0 | 97 | |
| MM2 | 0.0000 | 0.0197 | 0.0117 | 0 | |
| 368 | | | | | 13707 |
| | CR1 | CR2 | MM1 | MM2 | |
| CR1 | 0 | 255 | 273 | 250 | |
| CR2 | 0.0186 | 0 | 115 | 130 | |
| MM1 | 0.0199 | 0.0084 | 0 | 134 | |
| MM2 | 0.0182 | 0.0095 | 0.0098 | 0 | |
| 383 | | | | | 15898 |
| | CR1 | CR2 | MM1 | MM2 | |
| CR1 | 0 | 229 | 276 | 274 | |
| CR2 | 0.0144 | 0 | 168 | 136 | |
| MM1 | 0.0174 | 0.0106 | 0 | 138 | |
| MM2 | 0.0172 | 0.0086 | 0.0087 | 0 | |
| 387 | | | | | 19393 |
| | CR1 | CR2 | MM1 | MM2 | |
| CR1 | 0 | 423 | 413 | 395 | |
| CR2 | 0.0218 | 0 | 135 | 171 | |
| MM1 | 0.0213 | 0.0070 | 0 | 155 | |
| MM2 | 0.0204 | 0.0088 | 0.0080 | 0 | |
| 389 | | | | | 10214 |
| | CR1 | CR2 | MM1 | MM2 | |
| CR1 | 0 | 280 | 271 | 277 | |
| CR2 | 0.0274 | 0 | 15 | 126 | |
| MM1 | 0.0265 | 0.0015 | 0 | 123 | |
| MM2 | 0.0271 | 0.0123 | 0.0120 | 0 | |
| 390 | | | | | 10128 |
| | CR1 | CR2 | MM1 | MM2 | |
| CR1 | 0 | 254 | 138 | 127 | |
| CR2 | 0.0251 | 0 | 228 | 256 | |
| MM1 | 0.0136 | 0.0225 | 0 | 141 | |
| MM2 | 0.0125 | 0.0253 | 0.0139 | 0 | |
| 400 | | | | | 12857 |
| | CR1 | CR2 | MM1 | MM2 | |
| CR1 | 0 | 227 | 231 | 248 | |
| CR2 | 0.0177 | 0 | 72 | 118 | |
| MM1 | 0.0180 | 0.0056 | 0 | 116 | |
| MM2 | 0.0193 | 0.0092 | 0.0090 | 0 | |

| 401 | | | | | 10379 |
|---|---|---|---|---|---|
| | CR1 | CR2 | MM1 | MM2 | |
| CR1 | 0 | 178 | 197 | 189 | |
| CR2 | 0.0172 | 0 | 90 | 69 | |
| MM1 | 0.0190 | 0.0087 | 0 | 92 | |
| MM2 | 0.0182 | 0.0066 | 0.0089 | 0 | |
| 402 | | | | | 20309 |
| | CR1 | CR2 | MM1 | MM2 | |
| CR1 | 0 | 401 | 247 | 211 | |
| CR2 | 0.0197 | 0 | 374 | 384 | |
| MM1 | 0.0122 | 0.0184 | 0 | 234 | |
| MM2 | 0.0104 | 0.0189 | 0.0115 | 0 | |
| 403 | | | | | 13564 |
| | CR1 | CR2 | MM1 | MM2 | |
| CR1 | 0 | 3 | 447 | 475 | |
| CR2 | 0.0002 | 0 | 444 | 472 | |
| MM1 | 0.0330 | 0.0327 | 0 | 192 | |
| MM2 | 0.0350 | 0.0348 | 0.0142 | 0 | |
| 408 | | | | | 19578 |
| | CR1 | CR2 | MM1 | MM2 | |
| CR1 | 0 | 321 | 179 | 190 | |
| CR2 | 0.0164 | 0 | 362 | 339 | |
| MM1 | 0.0091 | 0.0185 | 0 | 194 | |
| MM2 | 0.0097 | 0.0173 | 0.0099 | 0 | |
| 411 | | | | | 8680 |
| | CR1 | CR2 | MM1 | MM2 | |
| CR1 | 0 | 37 | 191 | 194 | |
| CR2 | 0.0043 | 0 | 218 | 221 | |
| MM1 | 0.0220 | 0.0251 | 0 | 94 | |
| MM2 | 0.0224 | 0.0255 | 0.0108 | 0 | |
| 414 | | | | | 8864 |
| | CR1 | CR2 | MM1 | MM2 | |
| CR1 | 0 | 237 | 128 | 109 | |
| CR2 | 0.0267 | 0 | 235 | 245 | |
| MM1 | 0.0144 | 0.0265 | 0 | 110 | |
| MM2 | 0.0123 | 0.0276 | 0.0124 | 0 | |
| 418 | | | | | 7433 |
| | CR1 | CR2 | MM1 | MM2 | |
| CR1 | 0 | 93 | 36 | 37 | |
| CR2 | 0.0125 | 0 | 95 | 98 | |
| MM1 | 0.0048 | 0.0128 | 0 | 28 | |
| MM2 | 0.0050 | 0.0132 | 0.0038 | 0 | |

| 419 | | | | | 10035 |
|---|---|---|---|---|---|
| | CR1 | CR2 | MM1 | MM2 | |
| CR1 | 0 | 159 | 88 | 91 | |
| CR2 | 0.0158 | 0 | 160 | 160 | |
| MM1 | 0.0088 | 0.0159 | 0 | 77 | |
| MM2 | 0.0091 | 0.0159 | 0.0077 | 0 | |
| 420 | | | | | 17666 |
| | CR1 | CR2 | MM1 | MM2 | |
| CR1 | 0 | 385 | 134 | 152 | |
| CR2 | 0.0218 | 0 | 393 | 409 | |
| MM1 | 0.0076 | 0.0222 | 0 | 73 | |
| MM2 | 0.0086 | 0.0232 | 0.0041 | 0 | |
| 423 | | | | | 9879 |
| | CR1 | CR2 | MM1 | MM2 | |
| CR1 | 0 | 292 | 288 | 289 | |
| CR2 | 0.0296 | 0 | 52 | 123 | |
| MM1 | 0.0292 | 0.0053 | 0 | 97 | |
| MM2 | 0.0293 | 0.0125 | 0.0098 | 0 | |
| 427 | | | | | 12276 |
| | CR1 | CR2 | MM1 | MM2 | |
| CR1 | 0 | 253 | 165 | 167 | |
| CR2 | 0.0206 | 0 | 284 | 279 | |
| MM1 | 0.0134 | 0.0231 | 0 | 108 | |
| MM2 | 0.0136 | 0.0227 | 0.0088 | 0 | |
| 430 | | | | | 13088 |
| | CR1 | CR2 | MM1 | MM2 | |
| CR1 | 0 | 236 | 165 | 196 | |
| CR2 | 0.0180 | 0 | 318 | 307 | |
| MM1 | 0.0126 | 0.0243 | 0 | 190 | |
| MM2 | 0.0150 | 0.0235 | 0.0145 | 0 | |
| 431 | | | | | 13966 |
| | CR1 | CR2 | MM1 | MM2 | |
| CR1 | 0 | 229 | 214 | 225 | |
| CR2 | 0.0164 | 0 | 84 | 80 | |
| MM1 | 0.0153 | 0.0060 | 0 | 83 | |
| MM2 | 0.0161 | 0.0057 | 0.0059 | 0 | |
| 442 | | | | | 13390 |
| | CR1 | CR2 | MM1 | MM2 | |
| CR1 | 0 | 290 | 136 | 155 | |
| CR2 | 0.0217 | 0 | 307 | 316 | |
| MM1 | 0.0102 | 0.0229 | 0 | 174 | |
| MM2 | 0.0116 | 0.0236 | 0.0130 | 0 | |

| 443 | | | | | 8111 |
|---|---|---|---|---|---|
| | CR1 | CR2 | MM1 | MM2 | |
| CR1 | 0 | 51 | 253 | 241 | |
| CR2 | 0.0063 | 0 | 232 | 206 | |
| MM1 | 0.0312 | 0.0286 | 0 | 133 | |
| MM2 | 0.0297 | 0.0254 | 0.0164 | 0 | |
| 448 | | | | | 11735 |
| | CR1 | CR2 | MM1 | MM2 | |
| CR1 | 0 | 195 | 84 | 91 | |
| CR2 | 0.0166 | 0 | 180 | 192 | |
| MM1 | 0.0072 | 0.0153 | 0 | 70 | |
| MM2 | 0.0078 | 0.0164 | 0.0060 | 0 | |
| 450 | | | | | 11274 |
| | CR1 | CR2 | MM1 | MM2 | |
| CR1 | 0 | 111 | 42 | 48 | |
| CR2 | 0.0098 | 0 | 113 | 119 | |
| MM1 | 0.0037 | 0.0100 | 0 | 50 | |
| MM2 | 0.0043 | 0.0106 | 0.0044 | 0 | |
| 452 | | | | | 12186 |
| | CR1 | CR2 | MM1 | MM2 | |
| CR1 | 0 | 267 | 118 | 133 | |
| CR2 | 0.0219 | 0 | 257 | 260 | |
| MM1 | 0.0097 | 0.0211 | 0 | 124 | |
| MM2 | 0.0109 | 0.0213 | 0.0102 | 0 | |
| 454 | | | | | 8469 |
| | CR1 | CR2 | MM1 | MM2 | |
| CR1 | 0 | 158 | 59 | 56 | |
| CR2 | 0.0187 | 0 | 154 | 158 | |
| MM1 | 0.0070 | 0.0182 | 0 | 47 | |
| MM2 | 0.0066 | 0.0187 | 0.0055 | 0 | |
| 455 | | | | | 4511 |
| | CR1 | CR2 | MM1 | MM2 | |
| CR1 | 0 | 25 | 57 | 59 | |
| CR2 | 0.0055 | 0 | 66 | 68 | |
| MM1 | 0.0126 | 0.0146 | 0 | 48 | |
| MM2 | 0.0131 | 0.0151 | 0.0106 | 0 | |
| 460 | | | | | 10467 |
| | CR1 | CR2 | MM1 | MM2 | |
| CR1 | 0 | 149 | 30 | 15 | |
| CR2 | 0.0142 | 0 | 153 | 149 | |
| MM1 | 0.0029 | 0.0146 | 0 | 33 | |
| MM2 | 0.0014 | 0.0142 | 0.0032 | 0 | |

| 462 | | | | | 18890 |
|---|---|---|---|---|---|
| | CR1 | CR2 | MM1 | MM2 | |
| CR1 | 0 | 294 | 89 | 145 | |
| CR2 | 0.0156 | 0 | 290 | 305 | |
| MM1 | 0.0047 | 0.0154 | 0 | 128 | |
| MM2 | 0.0077 | 0.0161 | 0.0068 | 0 | |
| 465 | | | | | 17341 |
| | CR1 | CR2 | MM1 | MM2 | |
| CR1 | 0 | 349 | 366 | 383 | |
| CR2 | 0.0201 | 0 | 195 | 192 | |
| MM1 | 0.0211 | 0.0112 | 0 | 203 | |
| MM2 | 0.0221 | 0.0111 | 0.0117 | 0 | |
| 467 | | | | | 21282 |
| | CR1 | CR2 | MM1 | MM2 | |
| CR1 | 0 | 334 | 163 | 193 | |
| CR2 | 0.0157 | 0 | 374 | 382 | |
| MM1 | 0.0077 | 0.0176 | 0 | 161 | |
| MM2 | 0.0091 | 0.0179 | 0.0076 | 0 | |
| 471 | | | | | 13367 |
| | CR1 | CR2 | MM1 | MM2 | |
| CR1 | 0 | 228 | 130 | 136 | |
| CR2 | 0.0171 | 0 | 214 | 219 | |
| MM1 | 0.0097 | 0.0160 | 0 | 111 | |
| MM2 | 0.0102 | 0.0164 | 0.0083 | 0 | |
| 473 | | | | | 14424 |
| | CR1 | CR2 | MM1 | MM2 | |
| CR1 | 0 | 58 | 218 | 186 | |
| CR2 | 0.0040 | 0 | 180 | 128 | |
| MM1 | 0.0151 | 0.0125 | 0 | 168 | |
| MM2 | 0.0129 | 0.0089 | 0.0116 | 0 | |
| 475 | | | | | 11040 |
| | CR1 | CR2 | MM1 | MM2 | |
| CR1 | 0 | 279 | 156 | 156 | |
| CR2 | 0.0253 | 0 | 255 | 255 | |
| MM1 | 0.0141 | 0.0231 | 0 | 0 | |
| MM2 | 0.0141 | 0.0231 | 0.0000 | 0 | |
| 480 | | | | | 10203 |
| | CR1 | CR2 | MM1 | MM2 | |
| CR1 | 0 | 152 | 150 | 150 | |
| CR2 | 0.0149 | 0 | 108 | 102 | |
| MM1 | 0.0147 | 0.0106 | 0 | 111 | |
| MM2 | 0.0147 | 0.0100 | 0.0109 | 0 | |

| 481 | | | | | 11111 |
|---|---|---|---|---|---|
| | CR1 | CR2 | MM1 | MM2 | |
| CR1 | 0 | 272 | 175 | 177 | |
| CR2 | 0.0245 | 0 | 286 | 286 | |
| MM1 | 0.0158 | 0.0257 | 0 | 10 | |
| MM2 | 0.0159 | 0.0257 | 0.0009 | 0 | |
| 484 | | | | | 18068 |
| | CR1 | CR2 | MM1 | MM2 | |
| CR1 | 0 | 444 | 256 | 256 | |
| CR2 | 0.0246 | 0 | 468 | 468 | |
| MM1 | 0.0142 | 0.0259 | 0 | 0 | |
| MM2 | 0.0142 | 0.0259 | 0.0000 | 0 | |
| 485 | | | | | 10649 |
| | CR1 | CR2 | MM1 | MM2 | |
| CR1 | 0 | 244 | 255 | 252 | |
| CR2 | 0.0229 | 0 | 141 | 139 | |
| MM1 | 0.0239 | 0.0132 | 0 | 159 | |
| MM2 | 0.0237 | 0.0131 | 0.0149 | 0 | |
| 488 | | | | | 15140 |
| | CR1 | CR2 | MM1 | MM2 | |
| CR1 | 0 | 403 | 174 | 174 | |
| CR2 | 0.0266 | 0 | 380 | 397 | |
| MM1 | 0.0115 | 0.0251 | 0 | 193 | |
| MM2 | 0.0115 | 0.0262 | 0.0127 | 0 | |
| 489 | | | | | 10412 |
| | CR1 | CR2 | MM1 | MM2 | |
| CR1 | 0 | 255 | 267 | 254 | |
| CR2 | 0.0245 | 0 | 96 | 127 | |
| MM1 | 0.0256 | 0.0092 | 0 | 130 | |
| MM2 | 0.0244 | 0.0122 | 0.0125 | 0 | |
| 490 | | | | | 8303 |
| | CR1 | CR2 | MM1 | MM2 | |
| CR1 | 0 | 82 | 42 | 30 | |
| CR2 | 0.0099 | 0 | 87 | 82 | |
| MM1 | 0.0051 | 0.0105 | 0 | 44 | |
| MM2 | 0.0036 | 0.0099 | 0.0053 | 0 | |
| 491 | | | | | 12604 |
| | CR1 | CR2 | MM1 | MM2 | |
| CR1 | 0 | 284 | 188 | 188 | |
| CR2 | 0.0225 | 0 | 301 | 301 | |
| MM1 | 0.0149 | 0.0239 | 0 | 0 | |
| MM2 | 0.0149 | 0.0239 | 0.0000 | 0 | |

| 494 | | | | | 15117 |
|---|---|---|---|---|---|
| | CR1 | CR2 | MM1 | MM2 | |
| CR1 | 0 | 373 | 236 | 233 | |
| CR2 | 0.0247 | 0 | 378 | 381 | |
| MM1 | 0.0156 | 0.0250 | 0 | 178 | |
| MM2 | 0.0154 | 0.0252 | 0.0118 | 0 | |

| 497 | | | | | 12598 |
|---|---|---|---|---|---|
| | CR1 | CR2 | MM1 | MM2 | |
| CR1 | 0 | 297 | 150 | 154 | |
| CR2 | 0.0236 | 0 | 292 | 316 | |
| MM1 | 0.0119 | 0.0232 | 0 | 160 | |
| MM2 | 0.0122 | 0.0251 | 0.0127 | 0 | |

| 498 | | | | | 10360 |
|---|---|---|---|---|---|
| | CR1 | CR2 | MM1 | MM2 | |
| CR1 | 0 | 300 | 129 | 166 | |
| CR2 | 0.0290 | 0 | 308 | 302 | |
| MM1 | 0.0125 | 0.0297 | 0 | 139 | |
| MM2 | 0.0160 | 0.0292 | 0.0134 | 0 | |

| 499 | | | | | 17242 |
|---|---|---|---|---|---|
| | CR1 | CR2 | MM1 | MM2 | |
| CR1 | 0 | 164 | 138 | 123 | |
| CR2 | 0.0095 | 0 | 234 | 240 | |
| MM1 | 0.0080 | 0.0136 | 0 | 134 | |
| MM2 | 0.0071 | 0.0139 | 0.0078 | 0 | |

| 501 | | | | | 9410 |
|---|---|---|---|---|---|
| | CR1 | CR2 | MM1 | MM2 | |
| CR1 | 0 | 234 | 145 | 96 | |
| CR2 | 0.0249 | 0 | 251 | 227 | |
| MM1 | 0.0154 | 0.0267 | 0 | 141 | |
| MM2 | 0.0102 | 0.0241 | 0.0150 | 0 | |

| 502 | | | | | 22996 |
|---|---|---|---|---|---|
| | CR1 | CR2 | MM1 | MM2 | |
| CR1 | 0 | 449 | 274 | 280 | |
| CR2 | 0.0195 | 0 | 448 | 464 | |
| MM1 | 0.0119 | 0.0195 | 0 | 283 | |
| MM2 | 0.0122 | 0.0202 | 0.0123 | 0 | |

| 504 | | | | | 11864 |
|---|---|---|---|---|---|
| | CR1 | CR2 | MM1 | MM2 | |
| CR1 | 0 | 300 | 175 | 209 | |
| CR2 | 0.0253 | 0 | 321 | 304 | |
| MM1 | 0.0148 | 0.0271 | 0 | 208 | |
| MM2 | 0.0176 | 0.0256 | 0.0175 | 0 | |

| 505 | | | | | 8434 |
|---|---|---|---|---|---|
| | CR1 | CR2 | MM1 | MM2 | |
| CR1 | 0 | 209 | 202 | 203 | |
| CR2 | 0.0248 | 0 | 135 | 146 | |
| MM1 | 0.0240 | 0.0160 | 0 | 145 | |
| MM2 | 0.0241 | 0.0173 | 0.0172 | 0 | |
| 509 | | | | | 9248 |
| | CR1 | CR2 | MM1 | MM2 | |
| CR1 | 0 | 156 | 84 | 58 | |
| CR2 | 0.0169 | 0 | 167 | 162 | |
| MM1 | 0.0091 | 0.0181 | 0 | 79 | |
| MM2 | 0.0063 | 0.0175 | 0.0085 | 0 | |
| 511 | | | | | 15456 |
| | CR1 | CR2 | MM1 | MM2 | |
| CR1 | 0 | 280 | 141 | 158 | |
| CR2 | 0.0181 | 0 | 283 | 290 | |
| MM1 | 0.0091 | 0.0183 | 0 | 139 | |
| MM2 | 0.0102 | 0.0188 | 0.0090 | 0 | |
| 512 | | | | | 10544 |
| | CR1 | CR2 | MM1 | MM2 | |
| CR1 | 0 | 264 | 142 | 142 | |
| CR2 | 0.0250 | 0 | 230 | 237 | |
| MM1 | 0.0135 | 0.0218 | 0 | 110 | |
| MM2 | 0.0135 | 0.0225 | 0.0104 | 0 | |
| 517 | | | | | 15997 |
| | CR1 | CR2 | MM1 | MM2 | |
| CR1 | 0 | 321 | 28 | 89 | |
| CR2 | 0.0201 | 0 | 327 | 327 | |
| MM1 | 0.0018 | 0.0204 | 0 | 101 | |
| MM2 | 0.0056 | 0.0204 | 0.0063 | 0 | |
| 518 | | | | | 7783 |
| | CR1 | CR2 | MM1 | MM2 | |
| CR1 | 0 | 103 | 50 | 67 | |
| CR2 | 0.0132 | 0 | 111 | 119 | |
| MM1 | 0.0064 | 0.0143 | 0 | 53 | |
| MM2 | 0.0086 | 0.0153 | 0.0068 | 0 | |
| 524 | | | | | 15766 |
| | CR1 | CR2 | MM1 | MM2 | |
| CR1 | 0 | 332 | 28 | 3 | |
| CR2 | 0.0211 | 0 | 336 | 331 | |
| MM1 | 0.0018 | 0.0213 | 0 | 25 | |
| MM2 | 0.0002 | 0.0210 | 0.0016 | 0 | |

| 526 | | | | | 10699 |
|---|---|---|---|---|---|
| | CR1 | CR2 | MM1 | MM2 | |
| CR1 | 0 | 8 | 158 | 164 | |
| CR2 | 0.0007 | 0 | 162 | 168 | |
| MM1 | 0.0148 | 0.0151 | 0 | 58 | |
| MM2 | 0.0153 | 0.0157 | 0.0054 | 0 | |
| 527 | | | | | 24425 |
| | CR1 | CR2 | MM1 | MM2 | |
| CR1 | 0 | 282 | 241 | 224 | |
| CR2 | 0.0115 | 0 | 420 | 412 | |
| MM1 | 0.0099 | 0.0172 | 0 | 194 | |
| MM2 | 0.0092 | 0.0169 | 0.0079 | 0 | |
| 528 | | | | | 11592 |
| | CR1 | CR2 | MM1 | MM2 | |
| CR1 | 0 | 357 | 382 | 373 | |
| CR2 | 0.0308 | 0 | 184 | 206 | |
| MM1 | 0.0330 | 0.0159 | 0 | 228 | |
| MM2 | 0.0322 | 0.0178 | 0.0197 | 0 | |
| 529 | | | | | 8857 |
| | CR1 | CR2 | MM1 | MM2 | |
| CR1 | 0 | 242 | 104 | 119 | |
| CR2 | 0.0273 | 0 | 240 | 233 | |
| MM1 | 0.0117 | 0.0271 | 0 | 115 | |
| MM2 | 0.0134 | 0.0263 | 0.0130 | 0 | |
| 531 | | | | | 7832 |
| | CR1 | CR2 | MM1 | MM2 | |
| CR1 | 0 | 0 | 238 | 257 | |
| CR2 | 0.0000 | 0 | 238 | 257 | |
| MM1 | 0.0304 | 0.0304 | 0 | 141 | |
| MM2 | 0.0328 | 0.0328 | 0.0180 | 0 | |
| 532 | | | | | 12360 |
| | CR1 | CR2 | MM1 | MM2 | |
| CR1 | 0 | 281 | 222 | 218 | |
| CR2 | 0.0227 | 0 | 183 | 181 | |
| MM1 | 0.0180 | 0.0148 | 0 | 94 | |
| MM2 | 0.0176 | 0.0146 | 0.0076 | 0 | |
| 535 | | | | | 7805 |
| | CR1 | CR2 | MM1 | MM2 | |
| CR1 | 0 | 151 | 68 | 72 | |
| CR2 | 0.0193 | 0 | 146 | 136 | |
| MM1 | 0.0087 | 0.0187 | 0 | 60 | |
| MM2 | 0.0092 | 0.0174 | 0.0077 | 0 | |

| 537 | | | | | 9154 |
|---|---|---|---|---|---|
| | CR1 | CR2 | MM1 | MM2 | |
| CR1 | 0 | 307 | 159 | 207 | |
| CR2 | 0.0335 | 0 | 296 | 300 | |
| MM1 | 0.0174 | 0.0323 | 0 | 168 | |
| MM2 | 0.0226 | 0.0328 | 0.0184 | 0 | |
| 538 | | | | | 19784 |
| | CR1 | CR2 | MM1 | MM2 | |
| CR1 | 0 | 360 | 165 | 89 | |
| CR2 | 0.0182 | 0 | 356 | 373 | |
| MM1 | 0.0083 | 0.0180 | 0 | 162 | |
| MM2 | 0.0045 | 0.0189 | 0.0082 | 0 | |
| 546 | | | | | 17100 |
| | CR1 | CR2 | MM1 | MM2 | |
| CR1 | 0 | 491 | 340 | 240 | |
| CR2 | 0.0287 | 0 | 477 | 475 | |
| MM1 | 0.0199 | 0.0279 | 0 | 185 | |
| MM2 | 0.0140 | 0.0278 | 0.0108 | 0 | |
| 551 | | | | | 12174 |
| | CR1 | CR2 | MM1 | MM2 | |
| CR1 | 0 | 253 | 268 | 296 | |
| CR2 | 0.0208 | 0 | 146 | 158 | |
| MM1 | 0.0220 | 0.0120 | 0 | 184 | |
| MM2 | 0.0243 | 0.0130 | 0.0151 | 0 | |
| 552 | | | | | 9455 |
| | CR1 | CR2 | MM1 | MM2 | |
| CR1 | 0 | 245 | 245 | 276 | |
| CR2 | 0.0259 | 0 | 0 | 143 | |
| MM1 | 0.0259 | 0.0000 | 0 | 143 | |
| MM2 | 0.0292 | 0.0151 | 0.0151 | 0 | |
| 555 | | | | | 10732 |
| | CR1 | CR2 | MM1 | MM2 | |
| CR1 | 0 | 194 | 95 | 97 | |
| CR2 | 0.0181 | 0 | 204 | 190 | |
| MM1 | 0.0089 | 0.0190 | 0 | 115 | |
| MM2 | 0.0090 | 0.0177 | 0.0107 | 0 | |
| 556 | | | | | 10452 |
| | CR1 | CR2 | MM1 | MM2 | |
| CR1 | 0 | 300 | 112 | 128 | |
| CR2 | 0.0287 | 0 | 298 | 305 | |
| MM1 | 0.0107 | 0.0285 | 0 | 128 | |
| MM2 | 0.0122 | 0.0292 | 0.0122 | 0 | |

| 561 | | | | | 9721 |
|---|---|---|---|---|---|
| | CR1 | CR2 | MM1 | MM2 | |
| CR1 | 0 | 151 | 70 | 76 | |
| CR2 | 0.0155 | 0 | 150 | 154 | |
| MM1 | 0.0072 | 0.0154 | 0 | 64 | |
| MM2 | 0.0078 | 0.0158 | 0.0066 | 0 | |
| 562 | | | | | 8076 |
| | CR1 | CR2 | MM1 | MM2 | |
| CR1 | 0 | 191 | 59 | 60 | |
| CR2 | 0.0237 | 0 | 184 | 203 | |
| MM1 | 0.0073 | 0.0228 | 0 | 51 | |
| MM2 | 0.0074 | 0.0251 | 0.0063 | 0 | |
| 567 | | | | | 11206 |
| | CR1 | CR2 | MM1 | MM2 | |
| CR1 | 0 | 0 | 216 | 241 | |
| CR2 | 0.0000 | 0 | 216 | 241 | |
| MM1 | 0.0193 | 0.0193 | 0 | 205 | |
| MM2 | 0.0215 | 0.0215 | 0.0183 | 0 | |
| 572 | | | | | 10781 |
| | CR1 | CR2 | MM1 | MM2 | |
| CR1 | 0 | 243 | 162 | 211 | |
| CR2 | 0.0225 | 0 | 219 | 236 | |
| MM1 | 0.0150 | 0.0203 | 0 | 168 | |
| MM2 | 0.0196 | 0.0219 | 0.0156 | 0 | |
| 576 | | | | | 13040 |
| | CR1 | CR2 | MM1 | MM2 | |
| CR1 | 0 | 163 | 164 | 166 | |
| CR2 | 0.0125 | 0 | 76 | 77 | |
| MM1 | 0.0126 | 0.0058 | 0 | 33 | |
| MM2 | 0.0127 | 0.0059 | 0.0025 | 0 | |
| 577 | | | | | 7585 |
| | CR1 | CR2 | MM1 | MM2 | |
| CR1 | 0 | 0 | 232 | 256 | |
| CR2 | 0.0000 | 0 | 232 | 256 | |
| MM1 | 0.0306 | 0.0306 | 0 | 105 | |
| MM2 | 0.0338 | 0.0338 | 0.0138 | 0 | |
| 590 | | | | | 9830 |
| | CR1 | CR2 | MM1 | MM2 | |
| CR1 | 0 | 228 | 69 | 71 | |
| CR2 | 0.0232 | 0 | 229 | 223 | |
| MM1 | 0.0070 | 0.0233 | 0 | 89 | |
| MM2 | 0.0072 | 0.0227 | 0.0091 | 0 | |

| 597 | | | | | 9847 |
|---|---|---|---|---|---|
| | CR1 | CR2 | MM1 | MM2 | |
| CR1 | 0 | 171 | 231 | 226 | |
| CR2 | 0.0174 | 0 | 311 | 316 | |
| MM1 | 0.0235 | 0.0316 | 0 | 119 | |
| MM2 | 0.0230 | 0.0321 | 0.0121 | 0 | |
| 598 | | | | | 16051 |
| | CR1 | CR2 | MM1 | MM2 | |
| CR1 | 0 | 297 | 389 | 390 | |
| CR2 | 0.0185 | 0 | 215 | 263 | |
| MM1 | 0.0242 | 0.0134 | 0 | 300 | |
| MM2 | 0.0243 | 0.0164 | 0.0187 | 0 | |
| 600 | | | | | 16771 |
| | CR1 | CR2 | MM1 | MM2 | |
| CR1 | 0 | 173 | 159 | 180 | |
| CR2 | 0.0103 | 0 | 260 | 264 | |
| MM1 | 0.0095 | 0.0155 | 0 | 157 | |
| MM2 | 0.0107 | 0.0157 | 0.0094 | 0 | |
| 601 | | | | | 10544 |
| | CR1 | CR2 | MM1 | MM2 | |
| CR1 | 0 | 268 | 145 | 144 | |
| CR2 | 0.0254 | 0 | 262 | 280 | |
| MM1 | 0.0138 | 0.0248 | 0 | 97 | |
| MM2 | 0.0137 | 0.0266 | 0.0092 | 0 | |
| 602 | | | | | 11210 |
| | CR1 | CR2 | MM1 | MM2 | |
| CR1 | 0 | 115 | 33 | 36 | |
| CR2 | 0.0103 | 0 | 126 | 122 | |
| MM1 | 0.0029 | 0.0112 | 0 | 35 | |
| MM2 | 0.0032 | 0.0109 | 0.0031 | 0 | |
| 606 | | | | | 7019 |
| | CR1 | CR2 | MM1 | MM2 | |
| CR1 | 0 | 139 | 148 | 146 | |
| CR2 | 0.0198 | 0 | 51 | 21 | |
| MM1 | 0.0211 | 0.0073 | 0 | 42 | |
| MM2 | 0.0208 | 0.0030 | 0.0060 | 0 | |
| 608 | | | | | 14808 |
| | CR1 | CR2 | MM1 | MM2 | |
| CR1 | 0 | 357 | 173 | 144 | |
| CR2 | 0.0241 | 0 | 367 | 369 | |
| MM1 | 0.0117 | 0.0248 | 0 | 177 | |
| MM2 | 0.0097 | 0.0249 | 0.0120 | 0 | |

| 613 | | | | | 12015 |
|---|---|---|---|---|---|
| | CR1 | CR2 | MM1 | MM2 | |
| CR1 | 0 | 296 | 131 | 150 | |
| CR2 | 0.0246 | 0 | 309 | 294 | |
| MM1 | 0.0109 | 0.0257 | 0 | 181 | |
| MM2 | 0.0125 | 0.0245 | 0.0151 | 0 | |
| 614 | | | | | 8468 |
| | CR1 | CR2 | MM1 | MM2 | |
| CR1 | 0 | 108 | 111 | 116 | |
| CR2 | 0.0128 | 0 | 41 | 58 | |
| MM1 | 0.0131 | 0.0048 | 0 | 60 | |
| MM2 | 0.0137 | 0.0068 | 0.0071 | 0 | |
| 616 | | | | | 9424 |
| | CR1 | CR2 | MM1 | MM2 | |
| CR1 | 0 | 119 | 45 | 44 | |
| CR2 | 0.0126 | 0 | 121 | 111 | |
| MM1 | 0.0048 | 0.0128 | 0 | 37 | |
| MM2 | 0.0047 | 0.0118 | 0.0039 | 0 | |
| 619 | | | | | 7822 |
| | CR1 | CR2 | MM1 | MM2 | |
| CR1 | 0 | 205 | 203 | 194 | |
| CR2 | 0.0262 | 0 | 48 | 86 | |
| MM1 | 0.0260 | 0.0061 | 0 | 110 | |
| MM2 | 0.0248 | 0.0110 | 0.0141 | 0 | |
| 620 | | | | | 11687 |
| | CR1 | CR2 | MM1 | MM2 | |
| CR1 | 0 | 158 | 74 | 78 | |
| CR2 | 0.0135 | 0 | 156 | 150 | |
| MM1 | 0.0063 | 0.0133 | 0 | 74 | |
| MM2 | 0.0067 | 0.0128 | 0.0063 | 0 | |
| 621 | | | | | 12921 |
| | CR1 | CR2 | MM1 | MM2 | |
| CR1 | 0 | 300 | 109 | 78 | |
| CR2 | 0.0232 | 0 | 303 | 300 | |
| MM1 | 0.0084 | 0.0235 | 0 | 58 | |
| MM2 | 0.0060 | 0.0232 | 0.0045 | 0 | |
| 623 | | | | | 10995 |
| | CR1 | CR2 | MM1 | MM2 | |
| CR1 | 0 | 258 | 181 | 77 | |
| CR2 | 0.0235 | 0 | 285 | 276 | |
| MM1 | 0.0165 | 0.0259 | 0 | 164 | |
| MM2 | 0.0070 | 0.0251 | 0.0149 | 0 | |

| 627 | | | | | 10662 |
|---|---|---|---|---|---|
| | CR1 | CR2 | MM1 | MM2 | |
| CR1 | 0 | 204 | 86 | 83 | |
| CR2 | 0.0191 | 0 | 200 | 193 | |
| MM1 | 0.0081 | 0.0188 | 0 | 55 | |
| MM2 | 0.0078 | 0.0181 | 0.0052 | 0 | |
| 628 | | | | | 10148 |
| | CR1 | CR2 | MM1 | MM2 | |
| CR1 | 0 | 304 | 214 | 209 | |
| CR2 | 0.0300 | 0 | 282 | 277 | |
| MM1 | 0.0211 | 0.0278 | 0 | 15 | |
| MM2 | 0.0206 | 0.0273 | 0.0015 | 0 | |
| 629 | | | | | 11164 |
| | CR1 | CR2 | MM1 | MM2 | |
| CR1 | 0 | 261 | 248 | 249 | |
| CR2 | 0.0234 | 0 | 157 | 147 | |
| MM1 | 0.0222 | 0.0141 | 0 | 62 | |
| MM2 | 0.0223 | 0.0132 | 0.0056 | 0 | |
| 630 | | | | | 10567 |
| | CR1 | CR2 | MM1 | MM2 | |
| CR1 | 0 | 284 | 129 | 158 | |
| CR2 | 0.0269 | 0 | 291 | 309 | |
| MM1 | 0.0122 | 0.0275 | 0 | 130 | |
| MM2 | 0.0150 | 0.0292 | 0.0123 | 0 | |
| 633 | | | | | 21382 |
| | CR1 | CR2 | MM1 | MM2 | |
| CR1 | 0 | 491 | 216 | 195 | |
| CR2 | 0.0230 | 0 | 512 | 498 | |
| MM1 | 0.0101 | 0.0239 | 0 | 172 | |
| MM2 | 0.0091 | 0.0233 | 0.0080 | 0 | |
| 636 | | | | | 4631 |
| | CR1 | CR2 | MM1 | MM2 | |
| CR1 | 0 | 142 | 95 | 116 | |
| CR2 | 0.0307 | 0 | 157 | 167 | |
| MM1 | 0.0205 | 0.0339 | 0 | 95 | |
| MM2 | 0.0250 | 0.0361 | 0.0205 | 0 | |
| 637 | | | | | 8889 |
| | CR1 | CR2 | MM1 | MM2 | |
| CR1 | 0 | 161 | 178 | 178 | |
| CR2 | 0.0181 | 0 | 68 | 68 | |
| MM1 | 0.0200 | 0.0076 | 0 | 0 | |
| MM2 | 0.0200 | 0.0076 | 0.0000 | 0 | |

| 642 | | | | | 7163 |
|---|---|---|---|---|---|
| | CR1 | CR2 | MM1 | MM2 | |
| CR1 | 0 | 198 | 197 | 210 | |
| CR2 | 0.0276 | 0 | 39 | 88 | |
| MM1 | 0.0275 | 0.0054 | 0 | 73 | |
| MM2 | 0.0293 | 0.0123 | 0.0102 | 0 | |

| 644 | | | | | 12909 |
|---|---|---|---|---|---|
| | CR1 | CR2 | MM1 | MM2 | |
| CR1 | 0 | 261 | 254 | 232 | |
| CR2 | 0.0202 | 0 | 153 | 99 | |
| MM1 | 0.0197 | 0.0119 | 0 | 122 | |
| MM2 | 0.0180 | 0.0077 | 0.0095 | 0 | |

| 645 | | | | | 11075 |
|---|---|---|---|---|---|
| | CR1 | CR2 | MM1 | MM2 | |
| CR1 | 0 | 202 | 44 | 77 | |
| CR2 | 0.0182 | 0 | 197 | 198 | |
| MM1 | 0.0040 | 0.0178 | 0 | 72 | |
| MM2 | 0.0070 | 0.0179 | 0.0065 | 0 | |

| 647 | | | | | 10387 |
|---|---|---|---|---|---|
| | CR1 | CR2 | MM1 | MM2 | |
| CR1 | 0 | 198 | 199 | 206 | |
| CR2 | 0.0191 | 0 | 91 | 124 | |
| MM1 | 0.0192 | 0.0088 | 0 | 72 | |
| MM2 | 0.0198 | 0.0119 | 0.0069 | 0 | |

| 651 | | | | | 9782 |
|---|---|---|---|---|---|
| | CR1 | CR2 | MM1 | MM2 | |
| CR1 | 0 | 239 | 138 | 106 | |
| CR2 | 0.0244 | 0 | 250 | 229 | |
| MM1 | 0.0141 | 0.0256 | 0 | 136 | |
| MM2 | 0.0108 | 0.0234 | 0.0139 | 0 | |

| 654 | | | | | 13332 |
|---|---|---|---|---|---|
| | CR1 | CR2 | MM1 | MM2 | |
| CR1 | 0 | 271 | 175 | 140 | |
| CR2 | 0.0203 | 0 | 270 | 274 | |
| MM1 | 0.0131 | 0.0203 | 0 | 178 | |
| MM2 | 0.0105 | 0.0206 | 0.0134 | 0 | |

| 657 | | | | | 7560 |
|---|---|---|---|---|---|
| | CR1 | CR2 | MM1 | MM2 | |
| CR1 | 0 | 78 | 120 | 126 | |
| CR2 | 0.0103 | 0 | 111 | 106 | |
| MM1 | 0.0159 | 0.0147 | 0 | 67 | |
| MM2 | 0.0167 | 0.0140 | 0.0089 | 0 | |

| 658 | | | | | 15046 |
|---|---|---|---|---|---|
| | CR1 | CR2 | MM1 | MM2 | |
| CR1 | 0 | 377 | 146 | 134 | |
| CR2 | 0.0251 | 0 | 361 | 384 | |
| MM1 | 0.0097 | 0.0240 | 0 | 177 | |
| MM2 | 0.0089 | 0.0255 | 0.0118 | 0 | |
| 661 | | | | | 9874 |
| | CR1 | CR2 | MM1 | MM2 | |
| CR1 | 0 | 201 | 143 | 135 | |
| CR2 | 0.0204 | 0 | 198 | 185 | |
| MM1 | 0.0145 | 0.0201 | 0 | 132 | |
| MM2 | 0.0137 | 0.0187 | 0.0134 | 0 | |
| 662 | | | | | 8336 |
| | CR1 | CR2 | MM1 | MM2 | |
| CR1 | 0 | 188 | 104 | 98 | |
| CR2 | 0.0226 | 0 | 169 | 181 | |
| MM1 | 0.0125 | 0.0203 | 0 | 80 | |
| MM2 | 0.0118 | 0.0217 | 0.0096 | 0 | |
| 663 | | | | | 15787 |
| | CR1 | CR2 | MM1 | MM2 | |
| CR1 | 0 | 192 | 283 | 268 | |
| CR2 | 0.0122 | 0 | 143 | 136 | |
| MM1 | 0.0179 | 0.0091 | 0 | 156 | |
| MM2 | 0.0170 | 0.0086 | 0.0099 | 0 | |
| 664 | | | | | 14040 |
| | CR1 | CR2 | MM1 | MM2 | |
| CR1 | 0 | 402 | 174 | 186 | |
| CR2 | 0.0286 | 0 | 382 | 405 | |
| MM1 | 0.0124 | 0.0272 | 0 | 211 | |
| MM2 | 0.0132 | 0.0288 | 0.0150 | 0 | |
| 665 | | | | | 12663 |
| | CR1 | CR2 | MM1 | MM2 | |
| CR1 | 0 | 189 | 106 | 78 | |
| CR2 | 0.0149 | 0 | 182 | 185 | |
| MM1 | 0.0084 | 0.0144 | 0 | 92 | |
| MM2 | 0.0062 | 0.0146 | 0.0073 | 0 | |
| 669 | | | | | 12488 |
| | CR1 | CR2 | MM1 | MM2 | |
| CR1 | 0 | 233 | 125 | 106 | |
| CR2 | 0.0187 | 0 | 237 | 219 | |
| MM1 | 0.0100 | 0.0190 | 0 | 120 | |
| MM2 | 0.0085 | 0.0175 | 0.0096 | 0 | |

| 671 | | | | | 12466 |
|---|---|---|---|---|---|
| | CR1 | CR2 | MM1 | MM2 | |
| CR1 | 0 | 207 | 197 | 194 | |
| CR2 | 0.0166 | 0 | 72 | 73 | |
| MM1 | 0.0158 | 0.0058 | 0 | 44 | |
| MM2 | 0.0156 | 0.0059 | 0.0035 | 0 | |
| 672 | | | | | 10656 |
| | CR1 | CR2 | MM1 | MM2 | |
| CR1 | 0 | 172 | 73 | 80 | |
| CR2 | 0.0161 | 0 | 172 | 172 | |
| MM1 | 0.0069 | 0.0161 | 0 | 87 | |
| MM2 | 0.0075 | 0.0161 | 0.0082 | 0 | |
| 674 | | | | | 7711 |
| | CR1 | CR2 | MM1 | MM2 | |
| CR1 | 0 | 151 | 78 | 78 | |
| CR2 | 0.0196 | 0 | 141 | 141 | |
| MM1 | 0.0101 | 0.0183 | 0 | 0 | |
| MM2 | 0.0101 | 0.0183 | 0.0000 | 0 | |
| 675 | | | | | 10763 |
| | CR1 | CR2 | MM1 | MM2 | |
| CR1 | 0 | 203 | 62 | 72 | |
| CR2 | 0.0189 | 0 | 213 | 218 | |
| MM1 | 0.0058 | 0.0198 | 0 | 70 | |
| MM2 | 0.0067 | 0.0203 | 0.0065 | 0 | |
| 676 | | | | | 21441 |
| | CR1 | CR2 | MM1 | MM2 | |
| CR1 | 0 | 552 | 555 | 542 | |
| CR2 | 0.0257 | 0 | 124 | 396 | |
| MM1 | 0.0259 | 0.0058 | 0 | 374 | |
| MM2 | 0.0253 | 0.0185 | 0.0174 | 0 | |
| 678 | | | | | 9152 |
| | CR1 | CR2 | MM1 | MM2 | |
| CR1 | 0 | 275 | 184 | 211 | |
| CR2 | 0.0300 | 0 | 305 | 305 | |
| MM1 | 0.0201 | 0.0333 | 0 | 158 | |
| MM2 | 0.0231 | 0.0333 | 0.0173 | 0 | |
| 682 | | | | | 8363 |
| | CR1 | CR2 | MM1 | MM2 | |
| CR1 | 0 | 183 | 115 | 114 | |
| CR2 | 0.0219 | 0 | 185 | 195 | |
| MM1 | 0.0138 | 0.0221 | 0 | 124 | |
| MM2 | 0.0136 | 0.0233 | 0.0148 | 0 | |

| | | | | | |
|---|---|---|---|---|---|
| 686 | | | | | 13838 |
| | CR1 | CR2 | MM1 | MM2 | |
| CR1 | 0 | 104 | 195 | 175 | |
| CR2 | 0.0075 | 0 | 179 | 149 | |
| MM1 | 0.0141 | 0.0129 | 0 | 123 | |
| MM2 | 0.0126 | 0.0108 | 0.0089 | 0 | |
| 692 | | | | | 16302 |
| | CR1 | CR2 | MM1 | MM2 | |
| CR1 | 0 | 292 | 295 | 280 | |
| CR2 | 0.0179 | 0 | 110 | 93 | |
| MM1 | 0.0181 | 0.0067 | 0 | 103 | |
| MM2 | 0.0172 | 0.0057 | 0.0063 | 0 | |
| 693 | | | | | 13523 |
| | CR1 | CR2 | MM1 | MM2 | |
| CR1 | 0 | 324 | 310 | 318 | |
| CR2 | 0.0240 | 0 | 145 | 105 | |
| MM1 | 0.0229 | 0.0107 | 0 | 136 | |
| MM2 | 0.0235 | 0.0078 | 0.0101 | 0 | |
| 694 | | | | | 10238 |
| | CR1 | CR2 | MM1 | MM2 | |
| CR1 | 0 | 154 | 145 | 149 | |
| CR2 | 0.0150 | 0 | 77 | 75 | |
| MM1 | 0.0142 | 0.0075 | 0 | 65 | |
| MM2 | 0.0146 | 0.0073 | 0.0063 | 0 | |
| 695 | | | | | 10709 |
| | CR1 | CR2 | MM1 | MM2 | |
| CR1 | 0 | 310 | 161 | 162 | |
| CR2 | 0.0289 | 0 | 316 | 318 | |
| MM1 | 0.0150 | 0.0295 | 0 | 191 | |
| MM2 | 0.0151 | 0.0297 | 0.0178 | 0 | |
| 701 | | | | | 15104 |
| | CR1 | CR2 | MM1 | MM2 | |
| CR1 | 0 | 286 | 127 | 135 | |
| CR2 | 0.0189 | 0 | 307 | 296 | |
| MM1 | 0.0084 | 0.0203 | 0 | 172 | |
| MM2 | 0.0089 | 0.0196 | 0.0114 | 0 | |
| 702 | | | | | 21143 |
| | CR1 | CR2 | MM1 | MM2 | |
| CR1 | 0 | 369 | 359 | 370 | |
| CR2 | 0.0175 | 0 | 185 | 166 | |
| MM1 | 0.0170 | 0.0087 | 0 | 215 | |
| MM2 | 0.0175 | 0.0079 | 0.0102 | 0 | |

| 704 | | | | | 15102 |
|---|---|---|---|---|---|
| | CR1 | CR2 | MM1 | MM2 | |
| CR1 | 0 | 260 | 158 | 254 | |
| CR2 | 0.0172 | 0 | 310 | 333 | |
| MM1 | 0.0105 | 0.0205 | 0 | 180 | |
| MM2 | 0.0168 | 0.0221 | 0.0119 | 0 | |
| 707 | | | | | 12070 |
| | CR1 | CR2 | MM1 | MM2 | |
| CR1 | 0 | 260 | 259 | 252 | |
| CR2 | 0.0215 | 0 | 116 | 133 | |
| MM1 | 0.0215 | 0.0096 | 0 | 125 | |
| MM2 | 0.0209 | 0.0110 | 0.0104 | 0 | |
| 708 | | | | | 11536 |
| | CR1 | CR2 | MM1 | MM2 | |
| CR1 | 0 | 129 | 68 | 81 | |
| CR2 | 0.0112 | 0 | 125 | 140 | |
| MM1 | 0.0059 | 0.0108 | 0 | 79 | |
| MM2 | 0.0070 | 0.0121 | 0.0068 | 0 | |
| 712 | | | | | 10497 |
| | CR1 | CR2 | MM1 | MM2 | |
| CR1 | 0 | 155 | 67 | 68 | |
| CR2 | 0.0148 | 0 | 149 | 150 | |
| MM1 | 0.0064 | 0.0142 | 0 | 50 | |
| MM2 | 0.0065 | 0.0143 | 0.0048 | 0 | |
| 713 | | | | | 12310 |
| | CR1 | CR2 | MM1 | MM2 | |
| CR1 | 0 | 277 | 106 | 117 | |
| CR2 | 0.0225 | 0 | 281 | 280 | |
| MM1 | 0.0086 | 0.0228 | 0 | 146 | |
| MM2 | 0.0095 | 0.0227 | 0.0119 | 0 | |
| 714 | | | | | 9182 |
| | CR1 | CR2 | MM1 | MM2 | |
| CR1 | 0 | 133 | 141 | 141 | |
| CR2 | 0.0145 | 0 | 61 | 59 | |
| MM1 | 0.0154 | 0.0066 | 0 | 63 | |
| MM2 | 0.0154 | 0.0064 | 0.0069 | 0 | |
| 716 | | | | | 8885 |
| | CR1 | CR2 | MM1 | MM2 | |
| CR1 | 0 | 182 | 202 | 197 | |
| CR2 | 0.0205 | 0 | 68 | 44 | |
| MM1 | 0.0227 | 0.0077 | 0 | 80 | |
| MM2 | 0.0222 | 0.0050 | 0.0090 | 0 | |

| 719 | | | | | 10985 |
|---|---|---|---|---|---|
| | CR1 | CR2 | MM1 | MM2 | |
| CR1 | 0 | 324 | 152 | 130 | |
| CR2 | 0.0295 | 0 | 348 | 330 | |
| MM1 | 0.0138 | 0.0317 | 0 | 146 | |
| MM2 | 0.0118 | 0.0300 | 0.0133 | 0 | |
| 720 | | | | | 7592 |
| | CR1 | CR2 | MM1 | MM2 | |
| CR1 | 0 | 147 | 139 | 138 | |
| CR2 | 0.0194 | 0 | 80 | 60 | |
| MM1 | 0.0183 | 0.0105 | 0 | 61 | |
| MM2 | 0.0182 | 0.0079 | 0.0080 | 0 | |
| 721 | | | | | 11556 |
| | CR1 | CR2 | MM1 | MM2 | |
| CR1 | 0 | 225 | 112 | 148 | |
| CR2 | 0.0195 | 0 | 222 | 200 | |
| MM1 | 0.0097 | 0.0192 | 0 | 152 | |
| MM2 | 0.0128 | 0.0173 | 0.0132 | 0 | |
| 728 | | | | | 12618 |
| | CR1 | CR2 | MM1 | MM2 | |
| CR1 | 0 | 311 | 179 | 146 | |
| CR2 | 0.0246 | 0 | 330 | 301 | |
| MM1 | 0.0142 | 0.0262 | 0 | 33 | |
| MM2 | 0.0116 | 0.0239 | 0.0026 | 0 | |
| 733 | | | | | 7857 |
| | CR1 | CR2 | MM1 | MM2 | |
| CR1 | 0 | 179 | 111 | 111 | |
| CR2 | 0.0228 | 0 | 181 | 183 | |
| MM1 | 0.0141 | 0.0230 | 0 | 12 | |
| MM2 | 0.0141 | 0.0233 | 0.0015 | 0 | |
| 737 | | | | | 14103 |
| | CR1 | CR2 | MM1 | MM2 | |
| CR1 | 0 | 302 | 167 | 153 | |
| CR2 | 0.0214 | 0 | 257 | 295 | |
| MM1 | 0.0118 | 0.0182 | 0 | 156 | |
| MM2 | 0.0108 | 0.0209 | 0.0111 | 0 | |
| 738 | | | | | 13997 |
| | CR1 | CR2 | MM1 | MM2 | |
| CR1 | 0 | 370 | 184 | 160 | |
| CR2 | 0.0264 | 0 | 371 | 377 | |
| MM1 | 0.0131 | 0.0265 | 0 | 156 | |
| MM2 | 0.0114 | 0.0269 | 0.0111 | 0 | |

| 739 | | | | | 14691 |
|---|---|---|---|---|---|
| | CR1 | CR2 | MM1 | MM2 | |
| CR1 | 0 | 376 | 139 | 169 | |
| CR2 | 0.0256 | 0 | 398 | 367 | |
| MM1 | 0.0095 | 0.0271 | 0 | 161 | |
| MM2 | 0.0115 | 0.0250 | 0.0110 | 0 | |

| 740 | | | | | 11346 |
|---|---|---|---|---|---|
| | CR1 | CR2 | MM1 | MM2 | |
| CR1 | 0 | 65 | 81 | 123 | |
| CR2 | 0.0057 | 0 | 139 | 177 | |
| MM1 | 0.0071 | 0.0123 | 0 | 128 | |
| MM2 | 0.0108 | 0.0156 | 0.0113 | 0 | |

| 742 | | | | | 9985 |
|---|---|---|---|---|---|
| | CR1 | CR2 | MM1 | MM2 | |
| CR1 | 0 | 196 | 77 | 62 | |
| CR2 | 0.0196 | 0 | 191 | 192 | |
| MM1 | 0.0077 | 0.0191 | 0 | 80 | |
| MM2 | 0.0062 | 0.0192 | 0.0080 | 0 | |

| 743 | | | | | 10055 |
|---|---|---|---|---|---|
| | CR1 | CR2 | MM1 | MM2 | |
| CR1 | 0 | 192 | 160 | 147 | |
| CR2 | 0.0191 | 0 | 223 | 222 | |
| MM1 | 0.0159 | 0.0222 | 0 | 108 | |
| MM2 | 0.0146 | 0.0221 | 0.0107 | 0 | |

| 745 | | | | | 11006 |
|---|---|---|---|---|---|
| | CR1 | CR2 | MM1 | MM2 | |
| CR1 | 0 | 190 | 99 | 99 | |
| CR2 | 0.0173 | 0 | 200 | 200 | |
| MM1 | 0.0090 | 0.0182 | 0 | 0 | |
| MM2 | 0.0090 | 0.0182 | 0.0000 | 0 | |

| 746 | | | | | 11187 |
|---|---|---|---|---|---|
| | CR1 | CR2 | MM1 | MM2 | |
| CR1 | 0 | 134 | 132 | 140 | |
| CR2 | 0.0120 | 0 | 43 | 55 | |
| MM1 | 0.0118 | 0.0038 | 0 | 56 | |
| MM2 | 0.0125 | 0.0049 | 0.0050 | 0 | |

| 747 | | | | | 24512 |
|---|---|---|---|---|---|
| | CR1 | CR2 | MM1 | MM2 | |
| CR1 | 0 | 530 | 250 | 260 | |
| CR2 | 0.0216 | 0 | 497 | 522 | |
| MM1 | 0.0102 | 0.0203 | 0 | 217 | |
| MM2 | 0.0106 | 0.0213 | 0.0089 | 0 | |

| 750 | | | | | 15145 |
|---|---|---|---|---|---|
| | CR1 | CR2 | MM1 | MM2 | |
| CR1 | 0 | 221 | 175 | 34 | |
| CR2 | 0.0146 | 0 | 237 | 218 | |
| MM1 | 0.0116 | 0.0156 | 0 | 197 | |
| MM2 | 0.0022 | 0.0144 | 0.0130 | 0 | |
| 754 | | | | | 10027 |
| | CR1 | CR2 | MM1 | MM2 | |
| CR1 | 0 | 208 | 56 | 56 | |
| CR2 | 0.0207 | 0 | 218 | 218 | |
| MM1 | 0.0056 | 0.0217 | 0 | 0 | |
| MM2 | 0.0056 | 0.0217 | 0.0000 | 0 | |
| 755 | | | | | 11612 |
| | CR1 | CR2 | MM1 | MM2 | |
| CR1 | 0 | 167 | 76 | 70 | |
| CR2 | 0.0144 | 0 | 171 | 166 | |
| MM1 | 0.0065 | 0.0147 | 0 | 74 | |
| MM2 | 0.0060 | 0.0143 | 0.0064 | 0 | |
| 756 | | | | | 14462 |
| | CR1 | CR2 | MM1 | MM2 | |
| CR1 | 0 | 355 | 127 | 127 | |
| CR2 | 0.0245 | 0 | 358 | 351 | |
| MM1 | 0.0088 | 0.0248 | 0 | 125 | |
| MM2 | 0.0088 | 0.0243 | 0.0086 | 0 | |
| 763 | | | | | 12078 |
| | CR1 | CR2 | MM1 | MM2 | |
| CR1 | 0 | 152 | 56 | 62 | |
| CR2 | 0.0126 | 0 | 147 | 147 | |
| MM1 | 0.0046 | 0.0122 | 0 | 42 | |
| MM2 | 0.0051 | 0.0122 | 0.0035 | 0 | |
| 764 | | | | | 12008 |
| | CR1 | CR2 | MM1 | MM2 | |
| CR1 | 0 | 40 | 191 | 235 | |
| CR2 | 0.0033 | 0 | 210 | 257 | |
| MM1 | 0.0159 | 0.0175 | 0 | 202 | |
| MM2 | 0.0196 | 0.0214 | 0.0168 | 0 | |
| 765 | | | | | 30288 |
| | CR1 | CR2 | MM1 | MM2 | |
| CR1 | 0 | 710 | 252 | 289 | |
| CR2 | 0.0234 | 0 | 739 | 684 | |
| MM1 | 0.0083 | 0.0244 | 0 | 310 | |
| MM2 | 0.0095 | 0.0226 | 0.0102 | 0 | |

| 766 | | | | | 12364 |
|---|---|---|---|---|---|
| | CR1 | CR2 | MM1 | MM2 | |
| CR1 | 0 | 325 | 301 | 322 | |
| CR2 | 0.0263 | 0 | 118 | 139 | |
| MM1 | 0.0243 | 0.0095 | 0 | 144 | |
| MM2 | 0.0260 | 0.0112 | 0.0116 | 0 | |
| 768 | | | | | 9791 |
| | CR1 | CR2 | MM1 | MM2 | |
| CR1 | 0 | 302 | 59 | 117 | |
| CR2 | 0.0308 | 0 | 309 | 305 | |
| MM1 | 0.0060 | 0.0316 | 0 | 158 | |
| MM2 | 0.0119 | 0.0312 | 0.0161 | 0 | |
| 774 | | | | | 5683 |
| | CR1 | CR2 | MM1 | MM2 | |
| CR1 | 0 | 100 | 111 | 106 | |
| CR2 | 0.0176 | 0 | 60 | 70 | |
| MM1 | 0.0195 | 0.0106 | 0 | 61 | |
| MM2 | 0.0187 | 0.0123 | 0.0107 | 0 | |
| 775 | | | | | 14694 |
| | CR1 | CR2 | MM1 | MM2 | |
| CR1 | 0 | 283 | 283 | 275 | |
| CR2 | 0.0193 | 0 | 0 | 140 | |
| MM1 | 0.0193 | 0.0000 | 0 | 140 | |
| MM2 | 0.0187 | 0.0095 | 0.0095 | 0 | |
| 776 | | | | | 6549 |
| | CR1 | CR2 | MM1 | MM2 | |
| CR1 | 0 | 17 | 54 | 47 | |
| CR2 | 0.0026 | 0 | 69 | 64 | |
| MM1 | 0.0082 | 0.0105 | 0 | 24 | |
| MM2 | 0.0072 | 0.0098 | 0.0037 | 0 | |
| 779 | | | | | 14157 |
| | CR1 | CR2 | MM1 | MM2 | |
| CR1 | 0 | 217 | 91 | 55 | |
| CR2 | 0.0153 | 0 | 207 | 213 | |
| MM1 | 0.0064 | 0.0146 | 0 | 84 | |
| MM2 | 0.0039 | 0.0150 | 0.0059 | 0 | |
| 780 | | | | | 16707 |
| | CR1 | CR2 | MM1 | MM2 | |
| CR1 | 0 | 298 | 189 | 195 | |
| CR2 | 0.0178 | 0 | 315 | 295 | |
| MM1 | 0.0113 | 0.0189 | 0 | 197 | |
| MM2 | 0.0117 | 0.0177 | 0.0118 | 0 | |

| 781 | | | | | 11539 |
|---|---|---|---|---|---|
| | CR1 | CR2 | MM1 | MM2 | |
| CR1 | 0 | 197 | 194 | 199 | |
| CR2 | 0.0171 | 0 | 91 | 89 | |
| MM1 | 0.0168 | 0.0079 | 0 | 83 | |
| MM2 | 0.0172 | 0.0077 | 0.0072 | 0 | |
| 783 | | | | | 13113 |
| | CR1 | CR2 | MM1 | MM2 | |
| CR1 | 0 | 199 | 108 | 73 | |
| CR2 | 0.0152 | 0 | 213 | 214 | |
| MM1 | 0.0082 | 0.0162 | 0 | 111 | |
| MM2 | 0.0056 | 0.0163 | 0.0085 | 0 | |
| 785 | | | | | 11117 |
| | CR1 | CR2 | MM1 | MM2 | |
| CR1 | 0 | 195 | 206 | 199 | |
| CR2 | 0.0175 | 0 | 105 | 97 | |
| MM1 | 0.0185 | 0.0094 | 0 | 102 | |
| MM2 | 0.0179 | 0.0087 | 0.0092 | 0 | |
| 786 | | | | | 14148 |
| | CR1 | CR2 | MM1 | MM2 | |
| CR1 | 0 | 286 | 105 | 106 | |
| CR2 | 0.0202 | 0 | 296 | 277 | |
| MM1 | 0.0074 | 0.0209 | 0 | 101 | |
| MM2 | 0.0075 | 0.0196 | 0.0071 | 0 | |
| 788 | | | | | 15224 |
| | CR1 | CR2 | MM1 | MM2 | |
| CR1 | 0 | 200 | 207 | 207 | |
| CR2 | 0.0131 | 0 | 110 | 116 | |
| MM1 | 0.0136 | 0.0072 | 0 | 104 | |
| MM2 | 0.0136 | 0.0076 | 0.0068 | 0 | |
| 792 | | | | | 10318 |
| | CR1 | CR2 | MM1 | MM2 | |
| CR1 | 0 | 165 | 110 | 150 | |
| CR2 | 0.0160 | 0 | 189 | 197 | |
| MM1 | 0.0107 | 0.0183 | 0 | 93 | |
| MM2 | 0.0145 | 0.0191 | 0.0090 | 0 | |
| 795 | | | | | 9895 |
| | CR1 | CR2 | MM1 | MM2 | |
| CR1 | 0 | 215 | 217 | 206 | |
| CR2 | 0.0217 | 0 | 73 | 97 | |
| MM1 | 0.0219 | 0.0074 | 0 | 69 | |
| MM2 | 0.0208 | 0.0098 | 0.0070 | 0 | |

| 796 | | | | | 11133 |
|---|---|---|---|---|---|
| | CR1 | CR2 | MM1 | MM2 | |
| CR1 | 0 | 168 | 101 | 137 | |
| CR2 | 0.0151 | 0 | 172 | 208 | |
| MM1 | 0.0091 | 0.0154 | 0 | 91 | |
| MM2 | 0.0123 | 0.0187 | 0.0082 | 0 | |
| 801 | | | | | 10376 |
| | CR1 | CR2 | MM1 | MM2 | |
| CR1 | 0 | 183 | 123 | 132 | |
| CR2 | 0.0176 | 0 | 179 | 180 | |
| MM1 | 0.0119 | 0.0173 | 0 | 63 | |
| MM2 | 0.0127 | 0.0173 | 0.0061 | 0 | |
| 804 | | | | | 7456 |
| | CR1 | CR2 | MM1 | MM2 | |
| CR1 | 0 | 251 | 249 | 249 | |
| CR2 | 0.0337 | 0 | 122 | 122 | |
| MM1 | 0.0334 | 0.0164 | 0 | 0 | |
| MM2 | 0.0334 | 0.0164 | 0.0000 | 0 | |
| 807 | | | | | 18404 |
| | CR1 | CR2 | MM1 | MM2 | |
| CR1 | 0 | 228 | 276 | 259 | |
| CR2 | 0.0124 | 0 | 286 | 297 | |
| MM1 | 0.0150 | 0.0155 | 0 | 151 | |
| MM2 | 0.0141 | 0.0161 | 0.0082 | 0 | |
| 808 | | | | | 16972 |
| | CR1 | CR2 | MM1 | MM2 | |
| CR1 | 0 | 289 | 119 | 113 | |
| CR2 | 0.0170 | 0 | 296 | 296 | |
| MM1 | 0.0070 | 0.0174 | 0 | 62 | |
| MM2 | 0.0067 | 0.0174 | 0.0037 | 0 | |
| 813 | | | | | 9359 |
| | CR1 | CR2 | MM1 | MM2 | |
| CR1 | 0 | 100 | 47 | 42 | |
| CR2 | 0.0107 | 0 | 119 | 116 | |
| MM1 | 0.0050 | 0.0127 | 0 | 43 | |
| MM2 | 0.0045 | 0.0124 | 0.0046 | 0 | |
| 814 | | | | | 13356 |
| | CR1 | CR2 | MM1 | MM2 | |
| CR1 | 0 | 216 | 106 | 102 | |
| CR2 | 0.0162 | 0 | 227 | 237 | |
| MM1 | 0.0079 | 0.0170 | 0 | 106 | |
| MM2 | 0.0076 | 0.0177 | 0.0079 | 0 | |

| 815 | | | | | 9028 |
|---|---|---|---|---|---|
| | CR1 | CR2 | MM1 | MM2 | |
| CR1 | 0 | 55 | 183 | 182 | |
| CR2 | 0.0061 | 0 | 151 | 150 | |
| MM1 | 0.0203 | 0.0167 | 0 | 1 | |
| MM2 | 0.0202 | 0.0166 | 0.0001 | 0 | |
| 821 | | | | | 10673 |
| | CR1 | CR2 | MM1 | MM2 | |
| CR1 | 0 | 34 | 143 | 162 | |
| CR2 | 0.0032 | 0 | 140 | 152 | |
| MM1 | 0.0134 | 0.0131 | 0 | 94 | |
| MM2 | 0.0152 | 0.0142 | 0.0088 | 0 | |
| 822 | | | | | 11686 |
| | CR1 | CR2 | MM1 | MM2 | |
| CR1 | 0 | 231 | 154 | 121 | |
| CR2 | 0.0198 | 0 | 256 | 225 | |
| MM1 | 0.0132 | 0.0219 | 0 | 131 | |
| MM2 | 0.0104 | 0.0193 | 0.0112 | 0 | |
| 824 | | | | | 10830 |
| | CR1 | CR2 | MM1 | MM2 | |
| CR1 | 0 | 272 | 135 | 125 | |
| CR2 | 0.0251 | 0 | 286 | 274 | |
| MM1 | 0.0125 | 0.0264 | 0 | 134 | |
| MM2 | 0.0115 | 0.0253 | 0.0124 | 0 | |
| 826 | | | | | 12290 |
| | CR1 | CR2 | MM1 | MM2 | |
| CR1 | 0 | 325 | 172 | 193 | |
| CR2 | 0.0264 | 0 | 312 | 324 | |
| MM1 | 0.0140 | 0.0254 | 0 | 191 | |
| MM2 | 0.0157 | 0.0264 | 0.0155 | 0 | |
| 828 | | | | | 8675 |
| | CR1 | CR2 | MM1 | MM2 | |
| CR1 | 0 | 182 | 80 | 77 | |
| CR2 | 0.0210 | 0 | 187 | 175 | |
| MM1 | 0.0092 | 0.0216 | 0 | 76 | |
| MM2 | 0.0089 | 0.0202 | 0.0088 | 0 | |
| 832 | | | | | 11467 |
| | CR1 | CR2 | MM1 | MM2 | |
| CR1 | 0 | 280 | 140 | 93 | |
| CR2 | 0.0244 | 0 | 281 | 288 | |
| MM1 | 0.0122 | 0.0245 | 0 | 136 | |
| MM2 | 0.0081 | 0.0251 | 0.0119 | 0 | |

| 833 | | | | | 8710 |
|---|---|---|---|---|---|
| | CR1 | CR2 | MM1 | MM2 | |
| CR1 | 0 | 354 | 107 | 102 | |
| CR2 | 0.0406 | 0 | 357 | 350 | |
| MM1 | 0.0123 | 0.0410 | 0 | 103 | |
| MM2 | 0.0117 | 0.0402 | 0.0118 | 0 | |
| 834 | | | | | 14196 |
| | CR1 | CR2 | MM1 | MM2 | |
| CR1 | 0 | 152 | 47 | 7 | |
| CR2 | 0.0107 | 0 | 159 | 151 | |
| MM1 | 0.0033 | 0.0112 | 0 | 48 | |
| MM2 | 0.0005 | 0.0106 | 0.0034 | 0 | |
| 835 | | | | | 11057 |
| | CR1 | CR2 | MM1 | MM2 | |
| CR1 | 0 | 206 | 121 | 125 | |
| CR2 | 0.0186 | 0 | 195 | 195 | |
| MM1 | 0.0109 | 0.0176 | 0 | 10 | |
| MM2 | 0.0113 | 0.0176 | 0.0009 | 0 | |
| 836 | | | | | 8788 |
| | CR1 | CR2 | MM1 | MM2 | |
| CR1 | 0 | 182 | 104 | 61 | |
| CR2 | 0.0207 | 0 | 202 | 170 | |
| MM1 | 0.0118 | 0.0230 | 0 | 99 | |
| MM2 | 0.0069 | 0.0193 | 0.0113 | 0 | |
| 838 | | | | | 6367 |
| | CR1 | CR2 | MM1 | MM2 | |
| CR1 | 0 | 98 | 44 | 0 | |
| CR2 | 0.0154 | 0 | 102 | 98 | |
| MM1 | 0.0069 | 0.0160 | 0 | 44 | |
| MM2 | 0.0000 | 0.0154 | 0.0069 | 0 | |
| 839 | | | | | 15866 |
| | CR1 | CR2 | MM1 | MM2 | |
| CR1 | 0 | 177 | 92 | 135 | |
| CR2 | 0.0112 | 0 | 243 | 252 | |
| MM1 | 0.0058 | 0.0153 | 0 | 79 | |
| MM2 | 0.0085 | 0.0159 | 0.0050 | 0 | |
| 842 | | | | | 11716 |
| | CR1 | CR2 | MM1 | MM2 | |
| CR1 | 0 | 235 | 3 | 107 | |
| CR2 | 0.0201 | 0 | 232 | 236 | |
| MM1 | 0.0003 | 0.0198 | 0 | 104 | |
| MM2 | 0.0091 | 0.0201 | 0.0089 | 0 | |

| 843 | | | | | 18363 |
|---|---|---|---|---|---|
| | CR1 | CR2 | MM1 | MM2 | |
| CR1 | 0 | 359 | 156 | 149 | |
| CR2 | 0.0196 | 0 | 359 | 352 | |
| MM1 | 0.0085 | 0.0196 | 0 | 7 | |
| MM2 | 0.0081 | 0.0192 | 0.0004 | 0 | |
| 844 | | | | | 8972 |
| | CR1 | CR2 | MM1 | MM2 | |
| CR1 | 0 | 201 | 203 | 205 | |
| CR2 | 0.0224 | 0 | 143 | 134 | |
| MM1 | 0.0226 | 0.0159 | 0 | 133 | |
| MM2 | 0.0228 | 0.0149 | 0.0148 | 0 | |
| 849 | | | | | 7587 |
| | CR1 | CR2 | MM1 | MM2 | |
| CR1 | 0 | 157 | 6 | 67 | |
| CR2 | 0.0207 | 0 | 155 | 173 | |
| MM1 | 0.0008 | 0.0204 | 0 | 61 | |
| MM2 | 0.0088 | 0.0228 | 0.0080 | 0 | |
| 850 | | | | | 7914 |
| | CR1 | CR2 | MM1 | MM2 | |
| CR1 | 0 | 210 | 111 | 121 | |
| CR2 | 0.0265 | 0 | 210 | 229 | |
| MM1 | 0.0140 | 0.0265 | 0 | 123 | |
| MM2 | 0.0153 | 0.0289 | 0.0155 | 0 | |
| 851 | | | | | 10602 |
| | CR1 | CR2 | MM1 | MM2 | |
| CR1 | 0 | 318 | 124 | 177 | |
| CR2 | 0.0300 | 0 | 334 | 343 | |
| MM1 | 0.0117 | 0.0315 | 0 | 175 | |
| MM2 | 0.0167 | 0.0324 | 0.0165 | 0 | |
| 854 | | | | | 11028 |
| | CR1 | CR2 | MM1 | MM2 | |
| CR1 | 0 | 131 | 256 | 252 | |
| CR2 | 0.0119 | 0 | 318 | 308 | |
| MM1 | 0.0232 | 0.0288 | 0 | 174 | |
| MM2 | 0.0229 | 0.0279 | 0.0158 | 0 | |
| 858 | | | | | 11514 |
| | CR1 | CR2 | MM1 | MM2 | |
| CR1 | 0 | 251 | 191 | 189 | |
| CR2 | 0.0218 | 0 | 286 | 284 | |
| MM1 | 0.0166 | 0.0248 | 0 | 2 | |
| MM2 | 0.0164 | 0.0247 | 0.0002 | 0 | |

| 862 | | | | | 13811 |
|---|---|---|---|---|---|
| | CR1 | CR2 | MM1 | MM2 | |
| CR1 | 0 | 139 | 31 | 69 | |
| CR2 | 0.0101 | 0 | 132 | 126 | |
| MM1 | 0.0022 | 0.0096 | 0 | 72 | |
| MM2 | 0.0050 | 0.0091 | 0.0052 | 0 | |
| 863 | | | | | 18246 |
| | CR1 | CR2 | MM1 | MM2 | |
| CR1 | 0 | 408 | 188 | 169 | |
| CR2 | 0.0224 | 0 | 410 | 398 | |
| MM1 | 0.0103 | 0.0225 | 0 | 187 | |
| MM2 | 0.0093 | 0.0218 | 0.0102 | 0 | |
| 864 | | | | | 16829 |
| | CR1 | CR2 | MM1 | MM2 | |
| CR1 | 0 | 294 | 312 | 339 | |
| CR2 | 0.0175 | 0 | 154 | 187 | |
| MM1 | 0.0185 | 0.0092 | 0 | 141 | |
| MM2 | 0.0201 | 0.0111 | 0.0084 | 0 | |
| 866 | | | | | 6860 |
| | CR1 | CR2 | MM1 | MM2 | |
| CR1 | 0 | 83 | 147 | 147 | |
| CR2 | 0.0121 | 0 | 81 | 81 | |
| MM1 | 0.0214 | 0.0118 | 0 | 0 | |
| MM2 | 0.0214 | 0.0118 | 0.0000 | 0 | |
| 867 | | | | | 11851 |
| | CR1 | CR2 | MM1 | MM2 | |
| CR1 | 0 | 80 | 105 | 143 | |
| CR2 | 0.0068 | 0 | 57 | 85 | |
| MM1 | 0.0089 | 0.0048 | 0 | 76 | |
| MM2 | 0.0121 | 0.0072 | 0.0064 | 0 | |
| 871 | | | | | 10065 |
| | CR1 | CR2 | MM1 | MM2 | |
| CR1 | 0 | 92 | 193 | 193 | |
| CR2 | 0.0091 | 0 | 153 | 141 | |
| MM1 | 0.0192 | 0.0152 | 0 | 65 | |
| MM2 | 0.0192 | 0.0140 | 0.0065 | 0 | |
| 874 | | | | | 18254 |
| | CR1 | CR2 | MM1 | MM2 | |
| CR1 | 0 | 393 | 394 | 389 | |
| CR2 | 0.0215 | 0 | 182 | 83 | |
| MM1 | 0.0216 | 0.0100 | 0 | 180 | |
| MM2 | 0.0213 | 0.0045 | 0.0099 | 0 | |

| 876 | | | | | 6922 |
|---|---|---|---|---|---|
| | CR1 | CR2 | MM1 | MM2 | |
| CR1 | 0 | 137 | 97 | 98 | |
| CR2 | 0.0198 | 0 | 162 | 161 | |
| MM1 | 0.0140 | 0.0234 | 0 | 86 | |
| MM2 | 0.0142 | 0.0233 | 0.0124 | 0 | |
| 877 | | | | | 10017 |
| | CR1 | CR2 | MM1 | MM2 | |
| CR1 | 0 | 287 | 180 | 181 | |
| CR2 | 0.0287 | 0 | 238 | 232 | |
| MM1 | 0.0180 | 0.0238 | 0 | 112 | |
| MM2 | 0.0181 | 0.0232 | 0.0112 | 0 | |
| 878 | | | | | 10878 |
| | CR1 | CR2 | MM1 | MM2 | |
| CR1 | 0 | 203 | 108 | 106 | |
| CR2 | 0.0187 | 0 | 181 | 179 | |
| MM1 | 0.0099 | 0.0166 | 0 | 2 | |
| MM2 | 0.0097 | 0.0165 | 0.0002 | 0 | |
| 880 | | | | | 15372 |
| | CR1 | CR2 | MM1 | MM2 | |
| CR1 | 0 | 224 | 212 | 215 | |
| CR2 | 0.0146 | 0 | 114 | 121 | |
| MM1 | 0.0138 | 0.0074 | 0 | 57 | |
| MM2 | 0.0140 | 0.0079 | 0.0037 | 0 | |
| 882 | | | | | 23405 |
| | CR1 | CR2 | MM1 | MM2 | |
| CR1 | 0 | 509 | 524 | 510 | |
| CR2 | 0.0217 | 0 | 223 | 232 | |
| MM1 | 0.0224 | 0.0095 | 0 | 214 | |
| MM2 | 0.0218 | 0.0099 | 0.0091 | 0 | |
| 884 | | | | | 9076 |
| | CR1 | CR2 | MM1 | MM2 | |
| CR1 | 0 | 152 | 129 | 125 | |
| CR2 | 0.0167 | 0 | 69 | 68 | |
| MM1 | 0.0142 | 0.0076 | 0 | 75 | |
| MM2 | 0.0138 | 0.0075 | 0.0083 | 0 | |
| 885 | | | | | 20856 |
| | CR1 | CR2 | MM1 | MM2 | |
| CR1 | 0 | 288 | 327 | 330 | |
| CR2 | 0.0138 | 0 | 302 | 313 | |
| MM1 | 0.0157 | 0.0145 | 0 | 201 | |
| MM2 | 0.0158 | 0.0150 | 0.0096 | 0 | |

| 891 | | | | | 10748 |
|---|---|---|---|---|---|
| | CR1 | CR2 | MM1 | MM2 | |
| CR1 | 0 | 281 | 138 | 82 | |
| CR2 | 0.0261 | 0 | 288 | 279 | |
| MM1 | 0.0128 | 0.0268 | 0 | 157 | |
| MM2 | 0.0076 | 0.0260 | 0.0146 | 0 | |
| 892 | | | | | 9698 |
| | CR1 | CR2 | MM1 | MM2 | |
| CR1 | 0 | 184 | 43 | 43 | |
| CR2 | 0.0190 | 0 | 185 | 183 | |
| MM1 | 0.0044 | 0.0191 | 0 | 48 | |
| MM2 | 0.0044 | 0.0189 | 0.0049 | 0 | |
| 893 | | | | | 10108 |
| | CR1 | CR2 | MM1 | MM2 | |
| CR1 | 0 | 313 | 132 | 142 | |
| CR2 | 0.0310 | 0 | 304 | 323 | |
| MM1 | 0.0131 | 0.0301 | 0 | 85 | |
| MM2 | 0.0140 | 0.0320 | 0.0084 | 0 | |
| 894 | | | | | 9450 |
| | CR1 | CR2 | MM1 | MM2 | |
| CR1 | 0 | 208 | 90 | 76 | |
| CR2 | 0.0220 | 0 | 198 | 210 | |
| MM1 | 0.0095 | 0.0210 | 0 | 80 | |
| MM2 | 0.0080 | 0.0222 | 0.0085 | 0 | |
| 895 | | | | | 10281 |
| | CR1 | CR2 | MM1 | MM2 | |
| CR1 | 0 | 297 | 307 | 318 | |
| CR2 | 0.0289 | 0 | 177 | 179 | |
| MM1 | 0.0299 | 0.0172 | 0 | 187 | |
| MM2 | 0.0309 | 0.0174 | 0.0182 | 0 | |
| 896 | | | | | 9636 |
| | CR1 | CR2 | MM1 | MM2 | |
| CR1 | 0 | 155 | 197 | 142 | |
| CR2 | 0.0161 | 0 | 283 | 216 | |
| MM1 | 0.0204 | 0.0294 | 0 | 216 | |
| MM2 | 0.0147 | 0.0224 | 0.0224 | 0 | |
| 899 | | | | | 16575 |
| | CR1 | CR2 | MM1 | MM2 | |
| CR1 | 0 | 348 | 306 | 333 | |
| CR2 | 0.0210 | 0 | 281 | 288 | |
| MM1 | 0.0185 | 0.0170 | 0 | 204 | |
| MM2 | 0.0201 | 0.0174 | 0.0123 | 0 | |

| 900 | | | | | 8722 |
|---|---|---|---|---|---|
| | CR1 | CR2 | MM1 | MM2 | |
| CR1 | 0 | 159 | 169 | 169 | |
| CR2 | 0.0182 | 0 | 243 | 243 | |
| MM1 | 0.0194 | 0.0279 | 0 | 0 | |
| MM2 | 0.0194 | 0.0279 | 0.0000 | 0 | |

| 901 | | | | | 10172 |
|---|---|---|---|---|---|
| | CR1 | CR2 | MM1 | MM2 | |
| CR1 | 0 | 220 | 62 | 113 | |
| CR2 | 0.0216 | 0 | 216 | 240 | |
| MM1 | 0.0061 | 0.0212 | 0 | 117 | |
| MM2 | 0.0111 | 0.0236 | 0.0115 | 0 | |

| 909 | | | | | 9928 |
|---|---|---|---|---|---|
| | CR1 | CR2 | MM1 | MM2 | |
| CR1 | 0 | 276 | 274 | 275 | |
| CR2 | 0.0278 | 0 | 172 | 161 | |
| MM1 | 0.0276 | 0.0173 | 0 | 164 | |
| MM2 | 0.0277 | 0.0162 | 0.0165 | 0 | |

| 912 | | | | | 14310 |
|---|---|---|---|---|---|
| | CR1 | CR2 | MM1 | MM2 | |
| CR1 | 0 | 316 | 151 | 168 | |
| CR2 | 0.0221 | 0 | 287 | 312 | |
| MM1 | 0.0106 | 0.0201 | 0 | 137 | |
| MM2 | 0.0117 | 0.0218 | 0.0096 | 0 | |

| 913 | | | | | 17047 |
|---|---|---|---|---|---|
| | CR1 | CR2 | MM1 | MM2 | |
| CR1 | 0 | 387 | 224 | 60 | |
| CR2 | 0.0227 | 0 | 391 | 385 | |
| MM1 | 0.0131 | 0.0229 | 0 | 225 | |
| MM2 | 0.0035 | 0.0226 | 0.0132 | 0 | |

| 921 | | | | | 16190 |
|---|---|---|---|---|---|
| | CR1 | CR2 | MM1 | MM2 | |
| CR1 | 0 | 351 | 229 | 194 | |
| CR2 | 0.0217 | 0 | 342 | 356 | |
| MM1 | 0.0141 | 0.0211 | 0 | 194 | |
| MM2 | 0.0120 | 0.0220 | 0.0120 | 0 | |

| 924 | | | | | 14411 |
|---|---|---|---|---|---|
| | CR1 | CR2 | MM1 | MM2 | |
| CR1 | 0 | 418 | 148 | 164 | |
| CR2 | 0.0290 | 0 | 401 | 424 | |
| MM1 | 0.0103 | 0.0278 | 0 | 132 | |
| MM2 | 0.0114 | 0.0294 | 0.0092 | 0 | |

| 926 | | | | | 8102 |
|---|---|---|---|---|---|
| | CR1 | CR2 | MM1 | MM2 | |
| CR1 | 0 | 238 | 118 | 117 | |
| CR2 | 0.0294 | 0 | 248 | 247 | |
| MM1 | 0.0146 | 0.0306 | 0 | 1 | |
| MM2 | 0.0144 | 0.0305 | 0.0001 | 0 | |
| 933 | | | | | 11072 |
| | CR1 | CR2 | MM1 | MM2 | |
| CR1 | 0 | 324 | 217 | 198 | |
| CR2 | 0.0293 | 0 | 335 | 342 | |
| MM1 | 0.0196 | 0.0303 | 0 | 191 | |
| MM2 | 0.0179 | 0.0309 | 0.0173 | 0 | |
| 934 | | | | | 4543 |
| | CR1 | CR2 | MM1 | MM2 | |
| CR1 | 0 | 84 | 22 | 18 | |
| CR2 | 0.0185 | 0 | 83 | 82 | |
| MM1 | 0.0048 | 0.0183 | 0 | 20 | |
| MM2 | 0.0040 | 0.0180 | 0.0044 | 0 | |
| 939 | | | | | 9906 |
| | CR1 | CR2 | MM1 | MM2 | |
| CR1 | 0 | 17 | 0 | 71 | |
| CR2 | 0.0017 | 0 | 17 | 82 | |
| MM1 | 0.0000 | 0.0017 | 0 | 71 | |
| MM2 | 0.0072 | 0.0083 | 0.0072 | 0 | |
| 941 | | | | | 21459 |
| | CR1 | CR2 | MM1 | MM2 | |
| CR1 | 0 | 347 | 181 | 173 | |
| CR2 | 0.0162 | 0 | 330 | 354 | |
| MM1 | 0.0084 | 0.0154 | 0 | 160 | |
| MM2 | 0.0081 | 0.0165 | 0.0075 | 0 | |
| 943 | | | | | 10064 |
| | CR1 | CR2 | MM1 | MM2 | |
| CR1 | 0 | 228 | 123 | 114 | |
| CR2 | 0.0227 | 0 | 213 | 209 | |
| MM1 | 0.0122 | 0.0212 | 0 | 81 | |
| MM2 | 0.0113 | 0.0208 | 0.0080 | 0 | |
| 944 | | | | | 8050 |
| | CR1 | CR2 | MM1 | MM2 | |
| CR1 | 0 | 135 | 125 | 140 | |
| CR2 | 0.0168 | 0 | 44 | 57 | |
| MM1 | 0.0155 | 0.0055 | 0 | 39 | |
| MM2 | 0.0174 | 0.0071 | 0.0048 | 0 | |

| 946 | | | | | 9314 |
|---|---|---|---|---|---|
| | CR1 | CR2 | MM1 | MM2 | |
| CR1 | 0 | 288 | 118 | 125 | |
| CR2 | 0.0309 | 0 | 285 | 294 | |
| MM1 | 0.0127 | 0.0306 | 0 | 87 | |
| MM2 | 0.0134 | 0.0316 | 0.0093 | 0 | |
| 947 | | | | | 10045 |
| | CR1 | CR2 | MM1 | MM2 | |
| CR1 | 0 | 331 | 162 | 145 | |
| CR2 | 0.0330 | 0 | 332 | 301 | |
| MM1 | 0.0161 | 0.0331 | 0 | 161 | |
| MM2 | 0.0144 | 0.0300 | 0.0160 | 0 | |
| 952 | | | | | 10987 |
| | CR1 | CR2 | MM1 | MM2 | |
| CR1 | 0 | 223 | 130 | 105 | |
| CR2 | 0.0203 | 0 | 226 | 218 | |
| MM1 | 0.0118 | 0.0206 | 0 | 112 | |
| MM2 | 0.0096 | 0.0198 | 0.0102 | 0 | |
| 953 | | | | | 9372 |
| | CR1 | CR2 | MM1 | MM2 | |
| CR1 | 0 | 139 | 71 | 72 | |
| CR2 | 0.0148 | 0 | 121 | 118 | |
| MM1 | 0.0076 | 0.0129 | 0 | 49 | |
| MM2 | 0.0077 | 0.0126 | 0.0052 | 0 | |
| 954 | | | | | 11406 |
| | CR1 | CR2 | MM1 | MM2 | |
| CR1 | 0 | 218 | 110 | 90 | |
| CR2 | 0.0191 | 0 | 218 | 224 | |
| MM1 | 0.0096 | 0.0191 | 0 | 96 | |
| MM2 | 0.0079 | 0.0196 | 0.0084 | 0 | |
| 956 | | | | | 14610 |
| | CR1 | CR2 | MM1 | MM2 | |
| CR1 | 0 | 293 | 309 | 295 | |
| CR2 | 0.0201 | 0 | 142 | 145 | |
| MM1 | 0.0211 | 0.0097 | 0 | 84 | |
| MM2 | 0.0202 | 0.0099 | 0.0057 | 0 | |
| 961 | | | | | 12126 |
| | CR1 | CR2 | MM1 | MM2 | |
| CR1 | 0 | 185 | 66 | 95 | |
| CR2 | 0.0153 | 0 | 185 | 186 | |
| MM1 | 0.0054 | 0.0153 | 0 | 83 | |
| MM2 | 0.0078 | 0.0153 | 0.0068 | 0 | |

| 964 | | | | | 12650 |
|---|---|---|---|---|---|
| | CR1 | CR2 | MM1 | MM2 | |
| CR1 | 0 | 0 | 361 | 378 | |
| CR2 | 0.0000 | 0 | 361 | 378 | |
| MM1 | 0.0285 | 0.0285 | 0 | 272 | |
| MM2 | 0.0299 | 0.0299 | 0.0215 | 0 | |
| 965 | | | | | 10207 |
| | CR1 | CR2 | MM1 | MM2 | |
| CR1 | 0 | 117 | 201 | 221 | |
| CR2 | 0.0115 | 0 | 121 | 140 | |
| MM1 | 0.0197 | 0.0119 | 0 | 127 | |
| MM2 | 0.0217 | 0.0137 | 0.0124 | 0 | |
| 966 | | | | | 15590 |
| | CR1 | CR2 | MM1 | MM2 | |
| CR1 | 0 | 157 | 184 | 184 | |
| CR2 | 0.0101 | 0 | 27 | 27 | |
| MM1 | 0.0118 | 0.0017 | 0 | 0 | |
| MM2 | 0.0118 | 0.0017 | 0.0000 | 0 | |
| 970 | | | | | 10982 |
| | CR1 | CR2 | MM1 | MM2 | |
| CR1 | 0 | 266 | 124 | 129 | |
| CR2 | 0.0242 | 0 | 251 | 264 | |
| MM1 | 0.0113 | 0.0229 | 0 | 136 | |
| MM2 | 0.0117 | 0.0240 | 0.0124 | 0 | |
| 973 | | | | | 13520 |
| | CR1 | CR2 | MM1 | MM2 | |
| CR1 | 0 | 275 | 47 | 126 | |
| CR2 | 0.0203 | 0 | 277 | 275 | |
| MM1 | 0.0035 | 0.0205 | 0 | 111 | |
| MM2 | 0.0093 | 0.0203 | 0.0082 | 0 | |
| 977 | | | | | 14044 |
| | CR1 | CR2 | MM1 | MM2 | |
| CR1 | 0 | 362 | 238 | 227 | |
| CR2 | 0.0258 | 0 | 375 | 369 | |
| MM1 | 0.0169 | 0.0267 | 0 | 169 | |
| MM2 | 0.0162 | 0.0263 | 0.0120 | 0 | |
| 978 | | | | | 11812 |
| | CR1 | CR2 | MM1 | MM2 | |
| CR1 | 0 | 223 | 100 | 92 | |
| CR2 | 0.0189 | 0 | 237 | 230 | |
| MM1 | 0.0085 | 0.0201 | 0 | 90 | |
| MM2 | 0.0078 | 0.0195 | 0.0076 | 0 | |

| 982 | | | | | 10775 |
|---|---|---|---|---|---|
| | CR1 | CR2 | MM1 | MM2 | |
| CR1 | 0 | 196 | 189 | 199 | |
| CR2 | 0.0182 | 0 | 85 | 135 | |
| MM1 | 0.0175 | 0.0079 | 0 | 112 | |
| MM2 | 0.0185 | 0.0125 | 0.0104 | 0 | |
| 983 | | | | | 15181 |
| | CR1 | CR2 | MM1 | MM2 | |
| CR1 | 0 | 444 | 181 | 204 | |
| CR2 | 0.0292 | 0 | 445 | 421 | |
| MM1 | 0.0119 | 0.0293 | 0 | 231 | |
| MM2 | 0.0134 | 0.0277 | 0.0152 | 0 | |
| 985 | | | | | 21043 |
| | CR1 | CR2 | MM1 | MM2 | |
| CR1 | 0 | 352 | 160 | 163 | |
| CR2 | 0.0167 | 0 | 339 | 339 | |
| MM1 | 0.0076 | 0.0161 | 0 | 158 | |
| MM2 | 0.0077 | 0.0161 | 0.0075 | 0 | |
| 986 | | | | | 9892 |
| | CR1 | CR2 | MM1 | MM2 | |
| CR1 | 0 | 224 | 150 | 164 | |
| CR2 | 0.0226 | 0 | 222 | 239 | |
| MM1 | 0.0152 | 0.0224 | 0 | 173 | |
| MM2 | 0.0166 | 0.0242 | 0.0175 | 0 | |
| 987 | | | | | 7699 |
| | CR1 | CR2 | MM1 | MM2 | |
| CR1 | 0 | 337 | 123 | 117 | |
| CR2 | 0.0438 | 0 | 366 | 360 | |
| MM1 | 0.0160 | 0.0475 | 0 | 124 | |
| MM2 | 0.0152 | 0.0468 | 0.0161 | 0 | |
| 990 | | | | | 10415 |
| | CR1 | CR2 | MM1 | MM2 | |
| CR1 | 0 | 231 | 134 | 124 | |
| CR2 | 0.0222 | 0 | 217 | 230 | |
| MM1 | 0.0129 | 0.0208 | 0 | 76 | |
| MM2 | 0.0119 | 0.0221 | 0.0073 | 0 | |
| 991 | | | | | 7183 |
| | CR1 | CR2 | MM1 | MM2 | |
| CR1 | 0 | 0 | 255 | 130 | |
| CR2 | 0.0000 | 0 | 255 | 130 | |
| MM1 | 0.0355 | 0.0355 | 0 | 190 | |
| MM2 | 0.0181 | 0.0181 | 0.0265 | 0 | |

| 992 | | | | | 13579 |
|---|---|---|---|---|---|
| | CR1 | CR2 | MM1 | MM2 | |
| CR1 | 0 | 241 | 104 | 104 | |
| CR2 | 0.0177 | 0 | 237 | 237 | |
| MM1 | 0.0077 | 0.0175 | 0 | 0 | |
| MM2 | 0.0077 | 0.0175 | 0.0000 | 0 | |
| 996 | | | | | 20594 |
| | CR1 | CR2 | MM1 | MM2 | |
| CR1 | 0 | 465 | 274 | 320 | |
| CR2 | 0.0226 | 0 | 501 | 495 | |
| MM1 | 0.0133 | 0.0243 | 0 | 269 | |
| MM2 | 0.0155 | 0.0240 | 0.0131 | 0 | |
| 997 | | | | | 11806 |
| | CR1 | CR2 | MM1 | MM2 | |
| CR1 | 0 | 245 | 145 | 135 | |
| CR2 | 0.0208 | 0 | 223 | 227 | |
| MM1 | 0.0123 | 0.0189 | 0 | 130 | |
| MM2 | 0.0114 | 0.0192 | 0.0110 | 0 | |
| 1003 | | | | | 14569 |
| | CR1 | CR2 | MM1 | MM2 | |
| CR1 | 0 | 28 | 197 | 214 | |
| CR2 | 0.0019 | 0 | 183 | 194 | |
| MM1 | 0.0135 | 0.0126 | 0 | 133 | |
| MM2 | 0.0147 | 0.0133 | 0.0091 | 0 | |
| 1005 | | | | | 9883 |
| | CR1 | CR2 | MM1 | MM2 | |
| CR1 | 0 | 162 | 81 | 61 | |
| CR2 | 0.0164 | 0 | 163 | 159 | |
| MM1 | 0.0082 | 0.0165 | 0 | 56 | |
| MM2 | 0.0062 | 0.0161 | 0.0057 | 0 | |
| 1010 | | | | | 22736 |
| | CR1 | CR2 | MM1 | MM2 | |
| CR1 | 0 | 618 | 260 | 211 | |
| CR2 | 0.0272 | 0 | 571 | 612 | |
| MM1 | 0.0114 | 0.0251 | 0 | 284 | |
| MM2 | 0.0093 | 0.0269 | 0.0125 | 0 | |
| 1011 | | | | | 9234 |
| | CR1 | CR2 | MM1 | MM2 | |
| CR1 | 0 | 170 | 66 | 54 | |
| CR2 | 0.0184 | 0 | 180 | 177 | |
| MM1 | 0.0071 | 0.0195 | 0 | 64 | |
| MM2 | 0.0058 | 0.0192 | 0.0069 | 0 | |

| 1015 | | | | | 23371 |
|---|---|---|---|---|---|
| | CR1 | CR2 | MM1 | MM2 | |
| CR1 | 0 | 408 | 132 | 155 | |
| CR2 | 0.0175 | 0 | 408 | 422 | |
| MM1 | 0.0056 | 0.0175 | 0 | 163 | |
| MM2 | 0.0066 | 0.0181 | 0.0070 | 0 | |
| 1022 | | | | | 9868 |
| | CR1 | CR2 | MM1 | MM2 | |
| CR1 | 0 | 150 | 160 | 148 | |
| CR2 | 0.0152 | 0 | 55 | 71 | |
| MM1 | 0.0162 | 0.0056 | 0 | 69 | |
| MM2 | 0.0150 | 0.0072 | 0.0070 | 0 | |
| 1024 | | | | | 10130 |
| | CR1 | CR2 | MM1 | MM2 | |
| CR1 | 0 | 237 | 227 | 235 | |
| CR2 | 0.0234 | 0 | 112 | 115 | |
| MM1 | 0.0224 | 0.0111 | 0 | 88 | |
| MM2 | 0.0232 | 0.0114 | 0.0087 | 0 | |
| 1028 | | | | | 8220 |
| | CR1 | CR2 | MM1 | MM2 | |
| CR1 | 0 | 50 | 97 | 92 | |
| CR2 | 0.0061 | 0 | 128 | 114 | |
| MM1 | 0.0118 | 0.0156 | 0 | 108 | |
| MM2 | 0.0112 | 0.0139 | 0.0131 | 0 | |
| 1029 | | | | | 10230 |
| | CR1 | CR2 | MM1 | MM2 | |
| CR1 | 0 | 231 | 113 | 127 | |
| CR2 | 0.0226 | 0 | 234 | 235 | |
| MM1 | 0.0110 | 0.0229 | 0 | 127 | |
| MM2 | 0.0124 | 0.0230 | 0.0124 | 0 | |
| 1030 | | | | | 9022 |
| | CR1 | CR2 | MM1 | MM2 | |
| CR1 | 0 | 224 | 54 | 101 | |
| CR2 | 0.0248 | 0 | 218 | 207 | |
| MM1 | 0.0060 | 0.0242 | 0 | 85 | |
| MM2 | 0.0112 | 0.0229 | 0.0094 | 0 | |
| 1031 | | | | | 28878 |
| | CR1 | CR2 | MM1 | MM2 | |
| CR1 | 0 | 473 | 275 | 281 | |
| CR2 | 0.0164 | 0 | 462 | 452 | |
| MM1 | 0.0095 | 0.0160 | 0 | 253 | |
| MM2 | 0.0097 | 0.0157 | 0.0088 | 0 | |

| 1036 | | | | | 18254 |
|---|---|---|---|---|---|
| | CR1 | CR2 | MM1 | MM2 | |
| CR1 | 0 | 217 | 225 | 226 | |
| CR2 | 0.0119 | 0 | 123 | 118 | |
| MM1 | 0.0123 | 0.0067 | 0 | 98 | |
| MM2 | 0.0124 | 0.0065 | 0.0054 | 0 | |
| 1037 | | | | | 16421 |
| | CR1 | CR2 | MM1 | MM2 | |
| CR1 | 0 | 540 | 559 | 526 | |
| CR2 | 0.0329 | 0 | 368 | 143 | |
| MM1 | 0.0340 | 0.0224 | 0 | 326 | |
| MM2 | 0.0320 | 0.0087 | 0.0199 | 0 | |
| 1045 | | | | | 11024 |
| | CR1 | CR2 | MM1 | MM2 | |
| CR1 | 0 | 318 | 109 | 149 | |
| CR2 | 0.0288 | 0 | 323 | 311 | |
| MM1 | 0.0099 | 0.0293 | 0 | 123 | |
| MM2 | 0.0135 | 0.0282 | 0.0112 | 0 | |
| 1047 | | | | | 20449 |
| | CR1 | CR2 | MM1 | MM2 | |
| CR1 | 0 | 356 | 165 | 236 | |
| CR2 | 0.0174 | 0 | 351 | 376 | |
| MM1 | 0.0081 | 0.0172 | 0 | 214 | |
| MM2 | 0.0115 | 0.0184 | 0.0105 | 0 | |
| 1050 | | | | | 12818 |
| | CR1 | CR2 | MM1 | MM2 | |
| CR1 | 0 | 192 | 88 | 91 | |
| CR2 | 0.0150 | 0 | 210 | 206 | |
| MM1 | 0.0069 | 0.0164 | 0 | 93 | |
| MM2 | 0.0071 | 0.0161 | 0.0073 | 0 | |
| 1051 | | | | | 15223 |
| | CR1 | CR2 | MM1 | MM2 | |
| CR1 | 0 | 294 | 135 | 151 | |
| CR2 | 0.0193 | 0 | 308 | 301 | |
| MM1 | 0.0089 | 0.0202 | 0 | 156 | |
| MM2 | 0.0099 | 0.0198 | 0.0102 | 0 | |
| 1062 | | | | | 14366 |
| | CR1 | CR2 | MM1 | MM2 | |
| CR1 | 0 | 331 | 206 | 195 | |
| CR2 | 0.0230 | 0 | 334 | 345 | |
| MM1 | 0.0143 | 0.0232 | 0 | 81 | |
| MM2 | 0.0136 | 0.0240 | 0.0056 | 0 | |

| 1067 | | | | | 12199 |
|---|---|---|---|---|---|
| | CR1 | CR2 | MM1 | MM2 | |
| CR1 | 0 | 146 | 224 | 218 | |
| CR2 | 0.0120 | 0 | 282 | 295 | |
| MM1 | 0.0184 | 0.0231 | 0 | 233 | |
| MM2 | 0.0179 | 0.0242 | 0.0191 | 0 | |
| 1068 | | | | | 18532 |
| | CR1 | CR2 | MM1 | MM2 | |
| CR1 | 0 | 79 | 490 | 517 | |
| CR2 | 0.0043 | 0 | 454 | 477 | |
| MM1 | 0.0264 | 0.0245 | 0 | 177 | |
| MM2 | 0.0279 | 0.0257 | 0.0096 | 0 | |
| 1070 | | | | | 6438 |
| | CR1 | CR2 | MM1 | MM2 | |
| CR1 | 0 | 180 | 184 | 180 | |
| CR2 | 0.0280 | 0 | 90 | 111 | |
| MM1 | 0.0286 | 0.0140 | 0 | 113 | |
| MM2 | 0.0280 | 0.0172 | 0.0176 | 0 | |
| 1075 | | | | | 12763 |
| | CR1 | CR2 | MM1 | MM2 | |
| CR1 | 0 | 337 | 387 | 347 | |
| CR2 | 0.0264 | 0 | 195 | 118 | |
| MM1 | 0.0303 | 0.0153 | 0 | 199 | |
| MM2 | 0.0272 | 0.0092 | 0.0156 | 0 | |
| 1077 | | | | | 9784 |
| | CR1 | CR2 | MM1 | MM2 | |
| CR1 | 0 | 0 | 95 | 125 | |
| CR2 | 0.0000 | 0 | 95 | 125 | |
| MM1 | 0.0097 | 0.0097 | 0 | 111 | |
| MM2 | 0.0128 | 0.0128 | 0.0113 | 0 | |
| 1078 | | | | | 10276 |
| | CR1 | CR2 | MM1 | MM2 | |
| CR1 | 0 | 205 | 85 | 108 | |
| CR2 | 0.0199 | 0 | 222 | 218 | |
| MM1 | 0.0083 | 0.0216 | 0 | 97 | |
| MM2 | 0.0105 | 0.0212 | 0.0094 | 0 | |
| 1080 | | | | | 8171 |
| | CR1 | CR2 | MM1 | MM2 | |
| CR1 | 0 | 230 | 64 | 52 | |
| CR2 | 0.0281 | 0 | 220 | 228 | |
| MM1 | 0.0078 | 0.0269 | 0 | 58 | |
| MM2 | 0.0064 | 0.0279 | 0.0071 | 0 | |

| 1083 | | | | | 11112 |
|---|---|---|---|---|---|
| | CR1 | CR2 | MM1 | MM2 | |
| CR1 | 0 | 322 | 173 | 203 | |
| CR2 | 0.0290 | 0 | 318 | 290 | |
| MM1 | 0.0156 | 0.0286 | 0 | 189 | |
| MM2 | 0.0183 | 0.0261 | 0.0170 | 0 | |
| 1085 | | | | | 9735 |
| | CR1 | CR2 | MM1 | MM2 | |
| CR1 | 0 | 0 | 91 | 136 | |
| CR2 | 0.0000 | 0 | 91 | 136 | |
| MM1 | 0.0093 | 0.0093 | 0 | 129 | |
| MM2 | 0.0140 | 0.0140 | 0.0133 | 0 | |
| 1086 | | | | | 8424 |
| | CR1 | CR2 | MM1 | MM2 | |
| CR1 | 0 | 219 | 93 | 76 | |
| CR2 | 0.0260 | 0 | 210 | 231 | |
| MM1 | 0.0110 | 0.0249 | 0 | 112 | |
| MM2 | 0.0090 | 0.0274 | 0.0133 | 0 | |
| 1087 | | | | | 7465 |
| | CR1 | CR2 | MM1 | MM2 | |
| CR1 | 0 | 135 | 43 | 59 | |
| CR2 | 0.0181 | 0 | 134 | 135 | |
| MM1 | 0.0058 | 0.0180 | 0 | 57 | |
| MM2 | 0.0079 | 0.0181 | 0.0076 | 0 | |
| 1088 | | | | | 12529 |
| | CR1 | CR2 | MM1 | MM2 | |
| CR1 | 0 | 365 | 169 | 154 | |
| CR2 | 0.0291 | 0 | 361 | 366 | |
| MM1 | 0.0135 | 0.0288 | 0 | 159 | |
| MM2 | 0.0123 | 0.0292 | 0.0127 | 0 | |
| 1089 | | | | | 12057 |
| | CR1 | CR2 | MM1 | MM2 | |
| CR1 | 0 | 280 | 143 | 141 | |
| CR2 | 0.0232 | 0 | 276 | 262 | |
| MM1 | 0.0119 | 0.0229 | 0 | 123 | |
| MM2 | 0.0117 | 0.0217 | 0.0102 | 0 | |
| 1093 | | | | | 9117 |
| | CR1 | CR2 | MM1 | MM2 | |
| CR1 | 0 | 186 | 203 | 193 | |
| CR2 | 0.0204 | 0 | 149 | 135 | |
| MM1 | 0.0223 | 0.0163 | 0 | 116 | |
| MM2 | 0.0212 | 0.0148 | 0.0127 | 0 | |

| 1097 | | | | | 11537 |
|---|---|---|---|---|---|
| | CR1 | CR2 | MM1 | MM2 | |
| CR1 | 0 | 324 | 178 | 151 | |
| CR2 | 0.0281 | 0 | 323 | 334 | |
| MM1 | 0.0154 | 0.0280 | 0 | 187 | |
| MM2 | 0.0131 | 0.0290 | 0.0162 | 0 | |
| 1103 | | | | | 9910 |
| | CR1 | CR2 | MM1 | MM2 | |
| CR1 | 0 | 239 | 123 | 73 | |
| CR2 | 0.0241 | 0 | 244 | 235 | |
| MM1 | 0.0124 | 0.0246 | 0 | 138 | |
| MM2 | 0.0074 | 0.0237 | 0.0139 | 0 | |
| 1104 | | | | | 19599 |
| | CR1 | CR2 | MM1 | MM2 | |
| CR1 | 0 | 439 | 221 | 222 | |
| CR2 | 0.0224 | 0 | 421 | 422 | |
| MM1 | 0.0113 | 0.0215 | 0 | 1 | |
| MM2 | 0.0113 | 0.0215 | 0.0001 | 0 | |
| 1105 | | | | | 19582 |
| | CR1 | CR2 | MM1 | MM2 | |
| CR1 | 0 | 450 | 441 | 424 | |
| CR2 | 0.0230 | 0 | 217 | 292 | |
| MM1 | 0.0225 | 0.0111 | 0 | 235 | |
| MM2 | 0.0217 | 0.0149 | 0.0120 | 0 | |
| 1112 | | | | | 9572 |
| | CR1 | CR2 | MM1 | MM2 | |
| CR1 | 0 | 203 | 108 | 88 | |
| CR2 | 0.0212 | 0 | 178 | 194 | |
| MM1 | 0.0113 | 0.0186 | 0 | 100 | |
| MM2 | 0.0092 | 0.0203 | 0.0104 | 0 | |
| 1116 | | | | | 9828 |
| | CR1 | CR2 | MM1 | MM2 | |
| CR1 | 0 | 237 | 117 | 102 | |
| CR2 | 0.0241 | 0 | 246 | 229 | |
| MM1 | 0.0119 | 0.0250 | 0 | 106 | |
| MM2 | 0.0104 | 0.0233 | 0.0108 | 0 | |
| 1117 | | | | | 13442 |
| | CR1 | CR2 | MM1 | MM2 | |
| CR1 | 0 | 367 | 193 | 180 | |
| CR2 | 0.0273 | 0 | 386 | 365 | |
| MM1 | 0.0144 | 0.0287 | 0 | 178 | |
| MM2 | 0.0134 | 0.0272 | 0.0132 | 0 | |

| 1121 | | | | | 14393 |
|---|---|---|---|---|---|
| | CR1 | CR2 | MM1 | MM2 | |
| CR1 | 0 | 286 | 287 | 278 | |
| CR2 | 0.0199 | 0 | 115 | 116 | |
| MM1 | 0.0199 | 0.0080 | 0 | 113 | |
| MM2 | 0.0193 | 0.0081 | 0.0079 | 0 | |
| 1123 | | | | | 12864 |
| | CR1 | CR2 | MM1 | MM2 | |
| CR1 | 0 | 32 | 202 | 184 | |
| CR2 | 0.0025 | 0 | 219 | 204 | |
| MM1 | 0.0157 | 0.0170 | 0 | 103 | |
| MM2 | 0.0143 | 0.0159 | 0.0080 | 0 | |
| 1125 | | | | | 10221 |
| | CR1 | CR2 | MM1 | MM2 | |
| CR1 | 0 | 256 | 3 | 131 | |
| CR2 | 0.0250 | 0 | 257 | 251 | |
| MM1 | 0.0003 | 0.0251 | 0 | 130 | |
| MM2 | 0.0128 | 0.0246 | 0.0127 | 0 | |
| 1130 | | | | | 7425 |
| | CR1 | CR2 | MM1 | MM2 | |
| CR1 | 0 | 0 | 200 | 188 | |
| CR2 | 0.0000 | 0 | 200 | 188 | |
| MM1 | 0.0269 | 0.0269 | 0 | 98 | |
| MM2 | 0.0253 | 0.0253 | 0.0132 | 0 | |
| 1131 | | | | | 15676 |
| | CR1 | CR2 | MM1 | MM2 | |
| CR1 | 0 | 256 | 218 | 195 | |
| CR2 | 0.0163 | 0 | 391 | 375 | |
| MM1 | 0.0139 | 0.0249 | 0 | 214 | |
| MM2 | 0.0124 | 0.0239 | 0.0137 | 0 | |
| 1132 | | | | | 6200 |
| | CR1 | CR2 | MM1 | MM2 | |
| CR1 | 0 | 103 | 71 | 65 | |
| CR2 | 0.0166 | 0 | 104 | 106 | |
| MM1 | 0.0115 | 0.0168 | 0 | 61 | |
| MM2 | 0.0105 | 0.0171 | 0.0098 | 0 | |
| 1140 | | | | | 11680 |
| | CR1 | CR2 | MM1 | MM2 | |
| CR1 | 0 | 237 | 107 | 107 | |
| CR2 | 0.0203 | 0 | 236 | 236 | |
| MM1 | 0.0092 | 0.0202 | 0 | 0 | |
| MM2 | 0.0092 | 0.0202 | 0.0000 | 0 | |

| 1141 | | | | | 10272 |
|---|---|---|---|---|---|
| | CR1 | CR2 | MM1 | MM2 | |
| CR1 | 0 | 3 | 179 | 197 | |
| CR2 | 0.0003 | 0 | 176 | 194 | |
| MM1 | 0.0174 | 0.0171 | 0 | 76 | |
| MM2 | 0.0192 | 0.0189 | 0.0074 | 0 | |

| 1143 | | | | | 8625 |
|---|---|---|---|---|---|
| | CR1 | CR2 | MM1 | MM2 | |
| CR1 | 0 | 190 | 101 | 96 | |
| CR2 | 0.0220 | 0 | 183 | 181 | |
| MM1 | 0.0117 | 0.0212 | 0 | 25 | |
| MM2 | 0.0111 | 0.0210 | 0.0029 | 0 | |

| 1147 | | | | | 8005 |
|---|---|---|---|---|---|
| | CR1 | CR2 | MM1 | MM2 | |
| CR1 | 0 | 170 | 66 | 79 | |
| CR2 | 0.0212 | 0 | 179 | 177 | |
| MM1 | 0.0082 | 0.0224 | 0 | 64 | |
| MM2 | 0.0099 | 0.0221 | 0.0080 | 0 | |

| 1148 | | | | | 16447 |
|---|---|---|---|---|---|
| | CR1 | CR2 | MM1 | MM2 | |
| CR1 | 0 | 382 | 164 | 147 | |
| CR2 | 0.0232 | 0 | 369 | 364 | |
| MM1 | 0.0100 | 0.0224 | 0 | 86 | |
| MM2 | 0.0089 | 0.0221 | 0.0052 | 0 | |

| 1149 | | | | | 10051 |
|---|---|---|---|---|---|
| | CR1 | CR2 | MM1 | MM2 | |
| CR1 | 0 | 231 | 212 | 203 | |
| CR2 | 0.0230 | 0 | 104 | 100 | |
| MM1 | 0.0211 | 0.0103 | 0 | 77 | |
| MM2 | 0.0202 | 0.0099 | 0.0077 | 0 | |

| 1150 | | | | | 8578 |
|---|---|---|---|---|---|
| | CR1 | CR2 | MM1 | MM2 | |
| CR1 | 0 | 257 | 257 | 241 | |
| CR2 | 0.0300 | 0 | 111 | 113 | |
| MM1 | 0.0300 | 0.0129 | 0 | 124 | |
| MM2 | 0.0281 | 0.0132 | 0.0145 | 0 | |

| 1151 | | | | | 11064 |
|---|---|---|---|---|---|
| | CR1 | CR2 | MM1 | MM2 | |
| CR1 | 0 | 247 | 111 | 131 | |
| CR2 | 0.0223 | 0 | 244 | 253 | |
| MM1 | 0.0100 | 0.0221 | 0 | 103 | |
| MM2 | 0.0118 | 0.0229 | 0.0093 | 0 | |

| 1156 | | | | | 11945 |
|---|---|---|---|---|---|
| | CR1 | CR2 | MM1 | MM2 | |
| CR1 | 0 | 251 | 120 | 121 | |
| CR2 | 0.0210 | 0 | 263 | 264 | |
| MM1 | 0.0100 | 0.0220 | 0 | 139 | |
| MM2 | 0.0101 | 0.0221 | 0.0116 | 0 | |
| 1168 | | | | | 11757 |
| | CR1 | CR2 | MM1 | MM2 | |
| CR1 | 0 | 0 | 212 | 197 | |
| CR2 | 0.0000 | 0 | 212 | 197 | |
| MM1 | 0.0180 | 0.0180 | 0 | 114 | |
| MM2 | 0.0168 | 0.0168 | 0.0097 | 0 | |
| 1171 | | | | | 11283 |
| | CR1 | CR2 | MM1 | MM2 | |
| CR1 | 0 | 296 | 257 | 228 | |
| CR2 | 0.0262 | 0 | 359 | 344 | |
| MM1 | 0.0228 | 0.0318 | 0 | 244 | |
| MM2 | 0.0202 | 0.0305 | 0.0216 | 0 | |
| 1173 | | | | | 8351 |
| | CR1 | CR2 | MM1 | MM2 | |
| CR1 | 0 | 143 | 80 | 79 | |
| CR2 | 0.0171 | 0 | 142 | 135 | |
| MM1 | 0.0096 | 0.0170 | 0 | 7 | |
| MM2 | 0.0095 | 0.0162 | 0.0008 | 0 | |
| 1174 | | | | | 7156 |
| | CR1 | CR2 | MM1 | MM2 | |
| CR1 | 0 | 149 | 169 | 150 | |
| CR2 | 0.0208 | 0 | 64 | 49 | |
| MM1 | 0.0236 | 0.0089 | 0 | 69 | |
| MM2 | 0.0210 | 0.0068 | 0.0096 | 0 | |
| 1179 | | | | | 18420 |
| | CR1 | CR2 | MM1 | MM2 | |
| CR1 | 0 | 275 | 48 | 51 | |
| CR2 | 0.0149 | 0 | 276 | 281 | |
| MM1 | 0.0026 | 0.0150 | 0 | 59 | |
| MM2 | 0.0028 | 0.0153 | 0.0032 | 0 | |
| 1180 | | | | | 11669 |
| | CR1 | CR2 | MM1 | MM2 | |
| CR1 | 0 | 228 | 206 | 221 | |
| CR2 | 0.0195 | 0 | 91 | 77 | |
| MM1 | 0.0177 | 0.0078 | 0 | 80 | |
| MM2 | 0.0189 | 0.0066 | 0.0069 | 0 | |

| 1185 | | | | | 8195 |
|---|---|---|---|---|---|
| | CR1 | CR2 | MM1 | MM2 | |
| CR1 | 0 | 124 | 45 | 24 | |
| CR2 | 0.0151 | 0 | 118 | 120 | |
| MM1 | 0.0055 | 0.0144 | 0 | 38 | |
| MM2 | 0.0029 | 0.0146 | 0.0046 | 0 | |
| 1186 | | | | | 21367 |
| | CR1 | CR2 | MM1 | MM2 | |
| CR1 | 0 | 520 | 230 | 259 | |
| CR2 | 0.0243 | 0 | 492 | 513 | |
| MM1 | 0.0108 | 0.0230 | 0 | 255 | |
| MM2 | 0.0121 | 0.0240 | 0.0119 | 0 | |
| 1187 | | | | | 7887 |
| | CR1 | CR2 | MM1 | MM2 | |
| CR1 | 0 | 181 | 71 | 55 | |
| CR2 | 0.0229 | 0 | 188 | 192 | |
| MM1 | 0.0090 | 0.0238 | 0 | 79 | |
| MM2 | 0.0070 | 0.0243 | 0.0100 | 0 | |
| 1191 | | | | | 11399 |
| | CR1 | CR2 | MM1 | MM2 | |
| CR1 | 0 | 318 | 205 | 246 | |
| CR2 | 0.0279 | 0 | 307 | 309 | |
| MM1 | 0.0180 | 0.0269 | 0 | 199 | |
| MM2 | 0.0216 | 0.0271 | 0.0175 | 0 | |

| 6 | | | | | | | 15773 |
|---|---|---|---|---|---|---|---|
| | CR1 | CR2 | MA1 | MA2 | MM1 | MM2 | |
| CR1 | 0 | 330 | 90 | 350 | 358 | 350 | |
| CR2 | 0.0209 | 0 | 320 | 269 | 83 | 269 | |
| MA1 | 0.0057 | 0.0203 | 0 | 343 | 348 | 343 | |
| MA2 | 0.0222 | 0.0171 | 0.0217 | 0 | 261 | 0 | |
| MM1 | 0.0227 | 0.0053 | 0.0221 | 0.0165 | 0 | 261 | |
| MM2 | 0.0222 | 0.0171 | 0.0217 | 0.0000 | 0.0165 | 0 | |
| 7 | | | | | | | 12326 |
| | CR1 | CR2 | MA1 | MA2 | MM1 | MM2 | |
| CR1 | 0 | 214 | 97 | 219 | 97 | 77 | |
| CR2 | 0.0174 | 0 | 228 | 39 | 228 | 234 | |
| MA1 | 0.0079 | 0.0185 | 0 | 232 | 0 | 88 | |
| MA2 | 0.0178 | 0.0032 | 0.0188 | 0 | 232 | 239 | |
| MM1 | 0.0079 | 0.0185 | 0.0000 | 0.0188 | 0 | 88 | |
| MM2 | 0.0062 | 0.0190 | 0.0071 | 0.0194 | 0.0071 | 0 | |
| 8 | | | | | | | 18112 |
| | CR1 | CR2 | MA1 | MA2 | MM1 | MM2 | |
| CR1 | 0 | 354 | 392 | 223 | 412 | 393 | |
| CR2 | 0.0195 | 0 | 220 | 441 | 257 | 217 | |
| MA1 | 0.0216 | 0.0121 | 0 | 435 | 282 | 3 | |
| MA2 | 0.0123 | 0.0243 | 0.0240 | 0 | 456 | 438 | |
| MM1 | 0.0227 | 0.0142 | 0.0156 | 0.0252 | 0 | 280 | |
| MM2 | 0.0217 | 0.0120 | 0.0002 | 0.0242 | 0.0155 | 0 | |
| 10 | | | | | | | 12313 |
| | CR1 | CR2 | MA1 | MA2 | MM1 | MM2 | |
| CR1 | 0 | 201 | 83 | 222 | 83 | 76 | |
| CR2 | 0.0163 | 0 | 196 | 61 | 196 | 198 | |
| MA1 | 0.0067 | 0.0159 | 0 | 210 | 0 | 83 | |
| MA2 | 0.0180 | 0.0050 | 0.0171 | 0 | 210 | 210 | |
| MM1 | 0.0067 | 0.0159 | 0.0000 | 0.0171 | 0 | 83 | |
| MM2 | 0.0062 | 0.0161 | 0.0067 | 0.0171 | 0.0067 | 0 | |
| 12 | | | | | | | 13800 |
| | CR1 | CR2 | MA1 | MA2 | MM1 | MM2 | |
| CR1 | 0 | 315 | 318 | 168 | 168 | 121 | |
| CR2 | 0.0228 | 0 | 53 | 303 | 303 | 292 | |
| MA1 | 0.0230 | 0.0038 | 0 | 304 | 304 | 293 | |
| MA2 | 0.0122 | 0.0220 | 0.0220 | 0 | 0 | 139 | |
| MM1 | 0.0122 | 0.0220 | 0.0220 | 0.0000 | 0 | 139 | |
| MM2 | 0.0088 | 0.0212 | 0.0212 | 0.0101 | 0.0101 | 0 | |

| 13 | | | | | | | 16978 |
|---|---|---|---|---|---|---|---|
| | CR1 | CR2 | MA1 | MA2 | MM1 | MM2 | |
| CR1 | 0 | 378 | 365 | 96 | 365 | 387 | |
| CR2 | 0.0223 | 0 | 171 | 337 | 171 | 141 | |
| MA1 | 0.0215 | 0.0101 | 0 | 324 | 0 | 153 | |
| MA2 | 0.0057 | 0.0198 | 0.0191 | 0 | 324 | 348 | |
| MM1 | 0.0215 | 0.0101 | 0.0000 | 0.0191 | 0 | 153 | |
| MM2 | 0.0228 | 0.0083 | 0.0090 | 0.0205 | 0.0090 | 0 | |

| 14 | | | | | | | 15888 |
|---|---|---|---|---|---|---|---|
| | CR1 | CR2 | MA1 | MA2 | MM1 | MM2 | |
| CR1 | 0 | 283 | 102 | 280 | 102 | 158 | |
| CR2 | 0.0178 | 0 | 265 | 94 | 265 | 279 | |
| MA1 | 0.0064 | 0.0167 | 0 | 259 | 0 | 142 | |
| MA2 | 0.0176 | 0.0059 | 0.0163 | 0 | 259 | 270 | |
| MM1 | 0.0064 | 0.0167 | 0.0000 | 0.0163 | 0 | 142 | |
| MM2 | 0.0099 | 0.0176 | 0.0089 | 0.0170 | 0.0089 | 0 | |

| 15 | | | | | | | 11240 |
|---|---|---|---|---|---|---|---|
| | CR1 | CR2 | MA1 | MA2 | MM1 | MM2 | |
| CR1 | 0 | 80 | 211 | 119 | 55 | 126 | |
| CR2 | 0.0071 | 0 | 154 | 169 | 110 | 175 | |
| MA1 | 0.0188 | 0.0137 | 0 | 206 | 220 | 257 | |
| MA2 | 0.0106 | 0.0150 | 0.0183 | 0 | 110 | 136 | |
| MM1 | 0.0049 | 0.0098 | 0.0196 | 0.0098 | 0 | 108 | |
| MM2 | 0.0112 | 0.0156 | 0.0229 | 0.0121 | 0.0096 | 0 | |

| 18 | | | | | | | 13029 |
|---|---|---|---|---|---|---|---|
| | CR1 | CR2 | MA1 | MA2 | MM1 | MM2 | |
| CR1 | 0 | 228 | 87 | 233 | 96 | 87 | |
| CR2 | 0.0175 | 0 | 236 | 85 | 243 | 236 | |
| MA1 | 0.0067 | 0.0181 | 0 | 242 | 97 | 0 | |
| MA2 | 0.0179 | 0.0065 | 0.0186 | 0 | 245 | 242 | |
| MM1 | 0.0074 | 0.0187 | 0.0074 | 0.0188 | 0 | 97 | |
| MM2 | 0.0067 | 0.0181 | 0.0000 | 0.0186 | 0.0074 | 0 | |

| 20 | | | | | | | 14046 |
|---|---|---|---|---|---|---|---|
| | CR1 | CR2 | MA1 | MA2 | MM1 | MM2 | |
| CR1 | 0 | 191 | 91 | 188 | 91 | 112 | |
| CR2 | 0.0136 | 0 | 187 | 58 | 187 | 185 | |
| MA1 | 0.0065 | 0.0133 | 0 | 184 | 0 | 100 | |
| MA2 | 0.0134 | 0.0041 | 0.0131 | 0 | 184 | 184 | |
| MM1 | 0.0065 | 0.0133 | 0.0000 | 0.0131 | 0 | 100 | |
| MM2 | 0.0080 | 0.0132 | 0.0071 | 0.0131 | 0.0071 | 0 | |

| 23 | | | | | | | 16217 |
|---|---|---|---|---|---|---|---|
| | CR1 | CR2 | MA1 | MA2 | MM1 | MM2 | |
| CR1 | 0 | 351 | 214 | 365 | 179 | 197 | |
| CR2 | 0.0216 | 0 | 339 | 142 | 338 | 349 | |
| MA1 | 0.0132 | 0.0209 | 0 | 346 | 207 | 223 | |
| MA2 | 0.0225 | 0.0088 | 0.0213 | 0 | 346 | 354 | |
| MM1 | 0.0110 | 0.0208 | 0.0128 | 0.0213 | 0 | 210 | |
| MM2 | 0.0121 | 0.0215 | 0.0138 | 0.0218 | 0.0129 | 0 | |
| 25 | | | | | | | 20239 |
| | CR1 | CR2 | MA1 | MA2 | MM1 | MM2 | |
| CR1 | 0 | 0 | 118 | 412 | 425 | 412 | |
| CR2 | 0.0000 | 0 | 118 | 412 | 425 | 412 | |
| MA1 | 0.0058 | 0.0058 | 0 | 446 | 448 | 446 | |
| MA2 | 0.0204 | 0.0204 | 0.0220 | 0 | 209 | 0 | |
| MM1 | 0.0210 | 0.0210 | 0.0221 | 0.0103 | 0 | 209 | |
| MM2 | 0.0204 | 0.0204 | 0.0220 | 0.0000 | 0.0103 | 0 | |
| 27 | | | | | | | 11096 |
| | CR1 | CR2 | MA1 | MA2 | MM1 | MM2 | |
| CR1 | 0 | 24 | 67 | 165 | 95 | 67 | |
| CR2 | 0.0022 | 0 | 67 | 166 | 95 | 67 | |
| MA1 | 0.0060 | 0.0060 | 0 | 171 | 90 | 0 | |
| MA2 | 0.0149 | 0.0150 | 0.0154 | 0 | 177 | 171 | |
| MM1 | 0.0086 | 0.0086 | 0.0081 | 0.0160 | 0 | 90 | |
| MM2 | 0.0060 | 0.0060 | 0.0000 | 0.0154 | 0.0081 | 0 | |
| 28 | | | | | | | 12851 |
| | CR1 | CR2 | MA1 | MA2 | MM1 | MM2 | |
| CR1 | 0 | 256 | 253 | 137 | 129 | 41 | |
| CR2 | 0.0199 | 0 | 105 | 259 | 258 | 267 | |
| MA1 | 0.0197 | 0.0082 | 0 | 252 | 253 | 264 | |
| MA2 | 0.0107 | 0.0202 | 0.0196 | 0 | 132 | 124 | |
| MM1 | 0.0100 | 0.0201 | 0.0197 | 0.0103 | 0 | 100 | |
| MM2 | 0.0032 | 0.0208 | 0.0205 | 0.0096 | 0.0078 | 0 | |
| 30 | | | | | | | 13372 |
| | CR1 | CR2 | MA1 | MA2 | MM1 | MM2 | |
| CR1 | 0 | 250 | 255 | 106 | 112 | 106 | |
| CR2 | 0.0187 | 0 | 51 | 251 | 264 | 251 | |
| MA1 | 0.0191 | 0.0038 | 0 | 258 | 271 | 258 | |
| MA2 | 0.0079 | 0.0188 | 0.0193 | 0 | 106 | 0 | |
| MM1 | 0.0084 | 0.0197 | 0.0203 | 0.0079 | 0 | 106 | |
| MM2 | 0.0079 | 0.0188 | 0.0193 | 0.0000 | 0.0079 | 0 | |

| 31 | | | | | | | 9551 |
|---|---|---|---|---|---|---|---|
| | CR1 | CR2 | MA1 | MA2 | MM1 | MM2 | |
| CR1 | 0 | 136 | 21 | 137 | 132 | 137 | |
| CR2 | 0.0142 | 0 | 135 | 69 | 48 | 69 | |
| MA1 | 0.0022 | 0.0141 | 0 | 138 | 131 | 138 | |
| MA2 | 0.0143 | 0.0072 | 0.0144 | 0 | 49 | 0 | |
| MM1 | 0.0138 | 0.0050 | 0.0137 | 0.0051 | 0 | 49 | |
| MM2 | 0.0143 | 0.0072 | 0.0144 | 0.0000 | 0.0051 | 0 | |
| 33 | | | | | | | 14977 |
| | CR1 | CR2 | MA1 | MA2 | MM1 | MM2 | |
| CR1 | 0 | 133 | 339 | 95 | 91 | 95 | |
| CR2 | 0.0089 | 0 | 236 | 220 | 216 | 220 | |
| MA1 | 0.0226 | 0.0158 | 0 | 356 | 342 | 356 | |
| MA2 | 0.0063 | 0.0147 | 0.0238 | 0 | 103 | 0 | |
| MM1 | 0.0061 | 0.0144 | 0.0228 | 0.0069 | 0 | 103 | |
| MM2 | 0.0063 | 0.0147 | 0.0238 | 0.0000 | 0.0069 | 0 | |
| 34 | | | | | | | 9687 |
| | CR1 | CR2 | MA1 | MA2 | MM1 | MM2 | |
| CR1 | 0 | 157 | 89 | 155 | 89 | 82 | |
| CR2 | 0.0162 | 0 | 163 | 80 | 163 | 159 | |
| MA1 | 0.0092 | 0.0168 | 0 | 165 | 0 | 66 | |
| MA2 | 0.0160 | 0.0083 | 0.0170 | 0 | 165 | 159 | |
| MM1 | 0.0092 | 0.0168 | 0.0000 | 0.0170 | 0 | 66 | |
| MM2 | 0.0085 | 0.0164 | 0.0068 | 0.0164 | 0.0068 | 0 | |
| 36 | | | | | | | 12497 |
| | CR1 | CR2 | MA1 | MA2 | MM1 | MM2 | |
| CR1 | 0 | 126 | 188 | 187 | 221 | 207 | |
| CR2 | 0.0101 | 0 | 263 | 95 | 265 | 261 | |
| MA1 | 0.0150 | 0.0210 | 0 | 282 | 148 | 110 | |
| MA2 | 0.0150 | 0.0076 | 0.0226 | 0 | 268 | 265 | |
| MM1 | 0.0177 | 0.0212 | 0.0118 | 0.0214 | 0 | 151 | |
| MM2 | 0.0166 | 0.0209 | 0.0088 | 0.0212 | 0.0121 | 0 | |
| 37 | | | | | | | 16712 |
| | CR1 | CR2 | MA1 | MA2 | MM1 | MM2 | |
| CR1 | 0 | 185 | 83 | 176 | 83 | 81 | |
| CR2 | 0.0111 | 0 | 202 | 42 | 202 | 195 | |
| MA1 | 0.0050 | 0.0121 | 0 | 190 | 0 | 98 | |
| MA2 | 0.0105 | 0.0025 | 0.0114 | 0 | 190 | 191 | |
| MM1 | 0.0050 | 0.0121 | 0.0000 | 0.0114 | 0 | 98 | |
| MM2 | 0.0048 | 0.0117 | 0.0059 | 0.0114 | 0.0059 | 0 | |

| 38 | | | | | | | 13192 |
|---|---|---|---|---|---|---|---|
| | CR1 | CR2 | MA1 | MA2 | MM1 | MM2 | |
| CR1 | 0 | 171 | 98 | 164 | 98 | 63 | |
| CR2 | 0.0130 | 0 | 188 | 41 | 188 | 179 | |
| MA1 | 0.0074 | 0.0143 | 0 | 181 | 0 | 93 | |
| MA2 | 0.0124 | 0.0031 | 0.0137 | 0 | 181 | 172 | |
| MM1 | 0.0074 | 0.0143 | 0.0000 | 0.0137 | 0 | 93 | |
| MM2 | 0.0048 | 0.0136 | 0.0070 | 0.0130 | 0.0070 | 0 | |
| 39 | | | | | | | 11038 |
| | CR1 | CR2 | MA1 | MA2 | MM1 | MM2 | |
| CR1 | 0 | 258 | 90 | 252 | 90 | 108 | |
| CR2 | 0.0234 | 0 | 258 | 67 | 258 | 263 | |
| MA1 | 0.0082 | 0.0234 | 0 | 257 | 0 | 126 | |
| MA2 | 0.0228 | 0.0061 | 0.0233 | 0 | 257 | 260 | |
| MM1 | 0.0082 | 0.0234 | 0.0000 | 0.0233 | 0 | 126 | |
| MM2 | 0.0098 | 0.0238 | 0.0114 | 0.0236 | 0.0114 | 0 | |
| 40 | | | | | | | 13785 |
| | CR1 | CR2 | MA1 | MA2 | MM1 | MM2 | |
| CR1 | 0 | 164 | 133 | 121 | 28 | 79 | |
| CR2 | 0.0119 | 0 | 55 | 71 | 158 | 157 | |
| MA1 | 0.0096 | 0.0040 | 0 | 20 | 117 | 128 | |
| MA2 | 0.0088 | 0.0052 | 0.0015 | 0 | 103 | 124 | |
| MM1 | 0.0020 | 0.0115 | 0.0085 | 0.0075 | 0 | 79 | |
| MM2 | 0.0057 | 0.0114 | 0.0093 | 0.0090 | 0.0057 | 0 | |
| 42 | | | | | | | 17104 |
| | CR1 | CR2 | MA1 | MA2 | MM1 | MM2 | |
| CR1 | 0 | 239 | 167 | 173 | 95 | 98 | |
| CR2 | 0.0140 | 0 | 150 | 194 | 234 | 243 | |
| MA1 | 0.0098 | 0.0088 | 0 | 136 | 126 | 169 | |
| MA2 | 0.0101 | 0.0113 | 0.0080 | 0 | 124 | 184 | |
| MM1 | 0.0056 | 0.0137 | 0.0074 | 0.0072 | 0 | 106 | |
| MM2 | 0.0057 | 0.0142 | 0.0099 | 0.0108 | 0.0062 | 0 | |
| 44 | | | | | | | 14183 |
| | CR1 | CR2 | MA1 | MA2 | MM1 | MM2 | |
| CR1 | 0 | 192 | 87 | 196 | 117 | 87 | |
| CR2 | 0.0135 | 0 | 198 | 20 | 197 | 198 | |
| MA1 | 0.0061 | 0.0140 | 0 | 202 | 98 | 0 | |
| MA2 | 0.0138 | 0.0014 | 0.0142 | 0 | 201 | 202 | |
| MM1 | 0.0082 | 0.0139 | 0.0069 | 0.0142 | 0 | 98 | |
| MM2 | 0.0061 | 0.0140 | 0.0000 | 0.0142 | 0.0069 | 0 | |

| 46 | | | | | | | 12341 |
|---|---|---|---|---|---|---|---|
| | CR1 | CR2 | MA1 | MA2 | MM1 | MM2 | |
| CR1 | 0 | 204 | 60 | 202 | 211 | 202 | |
| CR2 | 0.0165 | 0 | 214 | 104 | 101 | 104 | |
| MA1 | 0.0049 | 0.0173 | 0 | 213 | 221 | 213 | |
| MA2 | 0.0164 | 0.0084 | 0.0173 | 0 | 83 | 0 | |
| MM1 | 0.0171 | 0.0082 | 0.0179 | 0.0067 | 0 | 83 | |
| MM2 | 0.0164 | 0.0084 | 0.0173 | 0.0000 | 0.0067 | 0 | |
| 47 | | | | | | | 15440 |
| | CR1 | CR2 | MA1 | MA2 | MM1 | MM2 | |
| CR1 | 0 | 266 | 275 | 151 | 80 | 151 | |
| CR2 | 0.0172 | 0 | 82 | 290 | 272 | 290 | |
| MA1 | 0.0178 | 0.0053 | 0 | 298 | 279 | 298 | |
| MA2 | 0.0098 | 0.0188 | 0.0193 | 0 | 154 | 0 | |
| MM1 | 0.0052 | 0.0176 | 0.0181 | 0.0100 | 0 | 154 | |
| MM2 | 0.0098 | 0.0188 | 0.0193 | 0.0000 | 0.0100 | 0 | |
| 49 | | | | | | | 9579 |
| | CR1 | CR2 | MA1 | MA2 | MM1 | MM2 | |
| CR1 | 0 | 71 | 122 | 150 | 43 | 90 | |
| CR2 | 0.0074 | 0 | 145 | 98 | 114 | 135 | |
| MA1 | 0.0127 | 0.0151 | 0 | 177 | 89 | 63 | |
| MA2 | 0.0157 | 0.0102 | 0.0185 | 0 | 185 | 174 | |
| MM1 | 0.0045 | 0.0119 | 0.0093 | 0.0193 | 0 | 65 | |
| MM2 | 0.0094 | 0.0141 | 0.0066 | 0.0182 | 0.0068 | 0 | |
| 50 | | | | | | | 21075 |
| | CR1 | CR2 | MA1 | MA2 | MM1 | MM2 | |
| CR1 | 0 | 607 | 627 | 300 | 658 | 628 | |
| CR2 | 0.0288 | 0 | 366 | 589 | 393 | 382 | |
| MA1 | 0.0298 | 0.0174 | 0 | 572 | 452 | 392 | |
| MA2 | 0.0142 | 0.0279 | 0.0271 | 0 | 640 | 619 | |
| MM1 | 0.0312 | 0.0186 | 0.0214 | 0.0304 | 0 | 380 | |
| MM2 | 0.0298 | 0.0181 | 0.0186 | 0.0294 | 0.0180 | 0 | |
| 51 | | | | | | | 9049 |
| | CR1 | CR2 | MA1 | MA2 | MM1 | MM2 | |
| CR1 | 0 | 107 | 31 | 105 | 34 | 40 | |
| CR2 | 0.0118 | 0 | 98 | 49 | 105 | 93 | |
| MA1 | 0.0034 | 0.0108 | 0 | 97 | 40 | 36 | |
| MA2 | 0.0116 | 0.0054 | 0.0107 | 0 | 102 | 102 | |
| MM1 | 0.0038 | 0.0116 | 0.0044 | 0.0113 | 0 | 44 | |
| MM2 | 0.0044 | 0.0103 | 0.0040 | 0.0113 | 0.0049 | 0 | |

| 52 | | | | | | | 7695 |
|---|---|---|---|---|---|---|---|
| | CR1 | CR2 | MA1 | MA2 | MM1 | MM2 | |
| CR1 | 0 | 85 | 87 | 30 | 33 | 30 | |
| CR2 | 0.0110 | 0 | 21 | 83 | 86 | 83 | |
| MA1 | 0.0113 | 0.0027 | 0 | 85 | 88 | 85 | |
| MA2 | 0.0039 | 0.0108 | 0.0110 | 0 | 23 | 0 | |
| MM1 | 0.0043 | 0.0112 | 0.0114 | 0.0030 | 0 | 23 | |
| MM2 | 0.0039 | 0.0108 | 0.0110 | 0.0000 | 0.0030 | 0 | |
| 53 | | | | | | | 11347 |
| | CR1 | CR2 | MA1 | MA2 | MM1 | MM2 | |
| CR1 | 0 | 368 | 174 | 384 | 173 | 136 | |
| CR2 | 0.0324 | 0 | 372 | 81 | 333 | 365 | |
| MA1 | 0.0153 | 0.0328 | 0 | 385 | 196 | 83 | |
| MA2 | 0.0338 | 0.0071 | 0.0339 | 0 | 348 | 380 | |
| MM1 | 0.0152 | 0.0293 | 0.0173 | 0.0307 | 0 | 113 | |
| MM2 | 0.0120 | 0.0322 | 0.0073 | 0.0335 | 0.0100 | 0 | |
| 54 | | | | | | | 16664 |
| | CR1 | CR2 | MA1 | MA2 | MM1 | MM2 | |
| CR1 | 0 | 414 | 439 | 135 | 437 | 443 | |
| CR2 | 0.0248 | 0 | 249 | 413 | 256 | 250 | |
| MA1 | 0.0263 | 0.0149 | 0 | 433 | 273 | 187 | |
| MA2 | 0.0081 | 0.0248 | 0.0260 | 0 | 435 | 440 | |
| MM1 | 0.0262 | 0.0154 | 0.0164 | 0.0261 | 0 | 260 | |
| MM2 | 0.0266 | 0.0150 | 0.0112 | 0.0264 | 0.0156 | 0 | |
| 55 | | | | | | | 13812 |
| | CR1 | CR2 | MA1 | MA2 | MM1 | MM2 | |
| CR1 | 0 | 174 | 75 | 167 | 62 | 55 | |
| CR2 | 0.0126 | 0 | 177 | 53 | 175 | 160 | |
| MA1 | 0.0054 | 0.0128 | 0 | 170 | 72 | 52 | |
| MA2 | 0.0121 | 0.0038 | 0.0123 | 0 | 168 | 153 | |
| MM1 | 0.0045 | 0.0127 | 0.0052 | 0.0122 | 0 | 53 | |
| MM2 | 0.0040 | 0.0116 | 0.0038 | 0.0111 | 0.0038 | 0 | |
| 57 | | | | | | | 13285 |
| | CR1 | CR2 | MA1 | MA2 | MM1 | MM2 | |
| CR1 | 0 | 221 | 99 | 224 | 99 | 101 | |
| CR2 | 0.0166 | 0 | 206 | 74 | 206 | 202 | |
| MA1 | 0.0075 | 0.0155 | 0 | 206 | 0 | 88 | |
| MA2 | 0.0169 | 0.0056 | 0.0155 | 0 | 206 | 205 | |
| MM1 | 0.0075 | 0.0155 | 0.0000 | 0.0155 | 0 | 88 | |
| MM2 | 0.0076 | 0.0152 | 0.0066 | 0.0154 | 0.0066 | 0 | |

| 64 | | | | | | | 11380 |
|---|---|---|---|---|---|---|---|
| | CR1 | CR2 | MA1 | MA2 | MM1 | MM2 | |
| CR1 | 0 | 5 | 207 | 151 | 220 | 230 | |
| CR2 | 0.0004 | 0 | 212 | 146 | 225 | 235 | |
| MA1 | 0.0182 | 0.0186 | 0 | 248 | 136 | 139 | |
| MA2 | 0.0133 | 0.0128 | 0.0218 | 0 | 251 | 264 | |
| MM1 | 0.0193 | 0.0198 | 0.0120 | 0.0221 | 0 | 128 | |
| MM2 | 0.0202 | 0.0207 | 0.0122 | 0.0232 | 0.0112 | 0 | |

| 65 | | | | | | | 19220 |
|---|---|---|---|---|---|---|---|
| | CR1 | CR2 | MA1 | MA2 | MM1 | MM2 | |
| CR1 | 0 | 319 | 228 | 328 | 186 | 197 | |
| CR2 | 0.0166 | 0 | 316 | 140 | 318 | 321 | |
| MA1 | 0.0119 | 0.0164 | 0 | 323 | 176 | 135 | |
| MA2 | 0.0171 | 0.0073 | 0.0168 | 0 | 325 | 330 | |
| MM1 | 0.0097 | 0.0165 | 0.0092 | 0.0169 | 0 | 162 | |
| MM2 | 0.0102 | 0.0167 | 0.0070 | 0.0172 | 0.0084 | 0 | |

| 66 | | | | | | | 12728 |
|---|---|---|---|---|---|---|---|
| | CR1 | CR2 | MA1 | MA2 | MM1 | MM2 | |
| CR1 | 0 | 244 | 88 | 248 | 88 | 81 | |
| CR2 | 0.0192 | 0 | 237 | 61 | 237 | 244 | |
| MA1 | 0.0069 | 0.0186 | 0 | 242 | 0 | 88 | |
| MA2 | 0.0195 | 0.0048 | 0.0190 | 0 | 242 | 247 | |
| MM1 | 0.0069 | 0.0186 | 0.0000 | 0.0190 | 0 | 88 | |
| MM2 | 0.0064 | 0.0192 | 0.0069 | 0.0194 | 0.0069 | 0 | |

| 68 | | | | | | | 18791 |
|---|---|---|---|---|---|---|---|
| | CR1 | CR2 | MA1 | MA2 | MM1 | MM2 | |
| CR1 | 0 | 274 | 265 | 74 | 259 | 265 | |
| CR2 | 0.0146 | 0 | 88 | 265 | 82 | 88 | |
| MA1 | 0.0141 | 0.0047 | 0 | 258 | 39 | 0 | |
| MA2 | 0.0039 | 0.0141 | 0.0137 | 0 | 254 | 258 | |
| MM1 | 0.0138 | 0.0044 | 0.0021 | 0.0135 | 0 | 39 | |
| MM2 | 0.0141 | 0.0047 | 0.0000 | 0.0137 | 0.0021 | 0 | |

| 70 | | | | | | | 12972 |
|---|---|---|---|---|---|---|---|
| | CR1 | CR2 | MA1 | MA2 | MM1 | MM2 | |
| CR1 | 0 | 200 | 204 | 129 | 129 | 119 | |
| CR2 | 0.0154 | 0 | 39 | 181 | 181 | 171 | |
| MA1 | 0.0157 | 0.0030 | 0 | 187 | 187 | 175 | |
| MA2 | 0.0099 | 0.0140 | 0.0144 | 0 | 0 | 78 | |
| MM1 | 0.0099 | 0.0140 | 0.0144 | 0.0000 | 0 | 78 | |
| MM2 | 0.0092 | 0.0132 | 0.0135 | 0.0060 | 0.0060 | 0 | |

| 71 | | | | | | | 18558 |
|---|---|---|---|---|---|---|---|
| | CR1 | CR2 | MA1 | MA2 | MM1 | MM2 | |
| CR1 | 0 | 307 | 145 | 299 | 134 | 126 | |
| CR2 | 0.0165 | 0 | 305 | 101 | 309 | 299 | |
| MA1 | 0.0078 | 0.0164 | 0 | 297 | 146 | 123 | |
| MA2 | 0.0161 | 0.0054 | 0.0160 | 0 | 295 | 288 | |
| MM1 | 0.0072 | 0.0167 | 0.0079 | 0.0159 | 0 | 127 | |
| MM2 | 0.0068 | 0.0161 | 0.0066 | 0.0155 | 0.0068 | 0 | |
| 72 | | | | | | | 11405 |
| | CR1 | CR2 | MA1 | MA2 | MM1 | MM2 | |
| CR1 | 0 | 292 | 273 | 139 | 79 | 139 | |
| CR2 | 0.0256 | 0 | 84 | 313 | 307 | 313 | |
| MA1 | 0.0239 | 0.0074 | 0 | 296 | 288 | 296 | |
| MA2 | 0.0122 | 0.0274 | 0.0260 | 0 | 120 | 0 | |
| MM1 | 0.0069 | 0.0269 | 0.0253 | 0.0105 | 0 | 120 | |
| MM2 | 0.0122 | 0.0274 | 0.0260 | 0.0000 | 0.0105 | 0 | |
| 73 | | | | | | | 18630 |
| | CR1 | CR2 | MA1 | MA2 | MM1 | MM2 | |
| CR1 | 0 | 370 | 203 | 370 | 191 | 243 | |
| CR2 | 0.0199 | 0 | 369 | 0 | 383 | 378 | |
| MA1 | 0.0109 | 0.0198 | 0 | 369 | 197 | 233 | |
| MA2 | 0.0199 | 0.0000 | 0.0198 | 0 | 383 | 378 | |
| MM1 | 0.0103 | 0.0206 | 0.0106 | 0.0206 | 0 | 245 | |
| MM2 | 0.0130 | 0.0203 | 0.0125 | 0.0203 | 0.0132 | 0 | |
| 75 | | | | | | | 9477 |
| | CR1 | CR2 | MA1 | MA2 | MM1 | MM2 | |
| CR1 | 0 | 203 | 47 | 199 | 199 | 182 | |
| CR2 | 0.0214 | 0 | 200 | 75 | 75 | 74 | |
| MA1 | 0.0050 | 0.0211 | 0 | 194 | 194 | 177 | |
| MA2 | 0.0210 | 0.0079 | 0.0205 | 0 | 0 | 73 | |
| MM1 | 0.0210 | 0.0079 | 0.0205 | 0.0000 | 0 | 73 | |
| MM2 | 0.0192 | 0.0078 | 0.0187 | 0.0077 | 0.0077 | 0 | |
| 76 | | | | | | | 8820 |
| | CR1 | CR2 | MA1 | MA2 | MM1 | MM2 | |
| CR1 | 0 | 229 | 143 | 225 | 110 | 143 | |
| CR2 | 0.0260 | 0 | 212 | 31 | 230 | 212 | |
| MA1 | 0.0162 | 0.0240 | 0 | 209 | 140 | 0 | |
| MA2 | 0.0255 | 0.0035 | 0.0237 | 0 | 226 | 209 | |
| MM1 | 0.0125 | 0.0261 | 0.0159 | 0.0256 | 0 | 140 | |
| MM2 | 0.0162 | 0.0240 | 0.0000 | 0.0237 | 0.0159 | 0 | |

| 77 | | | | | | 15195 |
|---|---|---|---|---|---|---|
| | CR1 | CR2 | MA1 | MA2 | MM1 | MM2 |
| CR1 | 0 | 109 | 310 | 191 | 226 | 191 |
| CR2 | 0.0072 | 0 | 213 | 229 | 274 | 229 |
| MA1 | 0.0204 | 0.0140 | 0 | 327 | 342 | 327 |
| MA2 | 0.0126 | 0.0151 | 0.0215 | 0 | 196 | 0 |
| MM1 | 0.0149 | 0.0180 | 0.0225 | 0.0129 | 0 | 196 |
| MM2 | 0.0126 | 0.0151 | 0.0215 | 0.0000 | 0.0129 | 0 |

| 78 | | | | | | 13466 |
|---|---|---|---|---|---|---|
| | CR1 | CR2 | MA1 | MA2 | MM1 | MM2 |
| CR1 | 0 | 367 | 191 | 326 | 191 | 220 |
| CR2 | 0.0273 | 0 | 373 | 102 | 373 | 400 |
| MA1 | 0.0142 | 0.0277 | 0 | 339 | 0 | 233 |
| MA2 | 0.0242 | 0.0076 | 0.0252 | 0 | 339 | 383 |
| MM1 | 0.0142 | 0.0277 | 0.0000 | 0.0252 | 0 | 233 |
| MM2 | 0.0163 | 0.0297 | 0.0173 | 0.0284 | 0.0173 | 0 |

| 79 | | | | | | 13694 |
|---|---|---|---|---|---|---|
| | CR1 | CR2 | MA1 | MA2 | MM1 | MM2 |
| CR1 | 0 | 180 | 104 | 177 | 82 | 87 |
| CR2 | 0.0131 | 0 | 183 | 7 | 182 | 167 |
| MA1 | 0.0076 | 0.0134 | 0 | 180 | 80 | 94 |
| MA2 | 0.0129 | 0.0005 | 0.0131 | 0 | 179 | 164 |
| MM1 | 0.0060 | 0.0133 | 0.0058 | 0.0131 | 0 | 91 |
| MM2 | 0.0064 | 0.0122 | 0.0069 | 0.0120 | 0.0066 | 0 |

| 81 | | | | | | 17906 |
|---|---|---|---|---|---|---|
| | CR1 | CR2 | MA1 | MA2 | MM1 | MM2 |
| CR1 | 0 | 338 | 184 | 375 | 184 | 152 |
| CR2 | 0.0189 | 0 | 364 | 71 | 364 | 368 |
| MA1 | 0.0103 | 0.0203 | 0 | 401 | 0 | 154 |
| MA2 | 0.0209 | 0.0040 | 0.0224 | 0 | 401 | 405 |
| MM1 | 0.0103 | 0.0203 | 0.0000 | 0.0224 | 0 | 154 |
| MM2 | 0.0085 | 0.0206 | 0.0086 | 0.0226 | 0.0086 | 0 |

| 82 | | | | | | 11012 |
|---|---|---|---|---|---|---|
| | CR1 | CR2 | MA1 | MA2 | MM1 | MM2 |
| CR1 | 0 | 265 | 272 | 159 | 159 | 134 |
| CR2 | 0.0241 | 0 | 38 | 277 | 277 | 258 |
| MA1 | 0.0247 | 0.0035 | 0 | 282 | 282 | 266 |
| MA2 | 0.0144 | 0.0252 | 0.0256 | 0 | 0 | 151 |
| MM1 | 0.0144 | 0.0252 | 0.0256 | 0.0000 | 0 | 151 |
| MM2 | 0.0122 | 0.0234 | 0.0242 | 0.0137 | 0.0137 | 0 |

| 86 | | | | | | 12347 |
|---|---|---|---|---|---|---|
| | CR1 | CR2 | MA1 | MA2 | MM1 | MM2 |
| CR1 | 0 | 181 | 90 | 188 | 90 | 87 |
| CR2 | 0.0147 | 0 | 181 | 41 | 181 | 169 |
| MA1 | 0.0073 | 0.0147 | 0 | 186 | 0 | 99 |
| MA2 | 0.0152 | 0.0033 | 0.0151 | 0 | 186 | 178 |
| MM1 | 0.0073 | 0.0147 | 0.0000 | 0.0151 | 0 | 99 |
| MM2 | 0.0070 | 0.0137 | 0.0080 | 0.0144 | 0.0080 | 0 |

| 87 | | | | | | 13357 |
|---|---|---|---|---|---|---|
| | CR1 | CR2 | MA1 | MA2 | MM1 | MM2 |
| CR1 | 0 | 264 | 137 | 255 | 132 | 137 |
| CR2 | 0.0198 | 0 | 287 | 41 | 257 | 287 |
| MA1 | 0.0103 | 0.0215 | 0 | 280 | 130 | 0 |
| MA2 | 0.0191 | 0.0031 | 0.0210 | 0 | 246 | 280 |
| MM1 | 0.0099 | 0.0192 | 0.0097 | 0.0184 | 0 | 130 |
| MM2 | 0.0103 | 0.0215 | 0.0000 | 0.0210 | 0.0097 | 0 |

| 88 | | | | | | 13194 |
|---|---|---|---|---|---|---|
| | CR1 | CR2 | MA1 | MA2 | MM1 | MM2 |
| CR1 | 0 | 265 | 93 | 260 | 144 | 93 |
| CR2 | 0.0201 | 0 | 264 | 93 | 262 | 264 |
| MA1 | 0.0070 | 0.0200 | 0 | 255 | 123 | 0 |
| MA2 | 0.0197 | 0.0070 | 0.0193 | 0 | 258 | 255 |
| MM1 | 0.0109 | 0.0199 | 0.0093 | 0.0196 | 0 | 123 |
| MM2 | 0.0070 | 0.0200 | 0.0000 | 0.0193 | 0.0093 | 0 |

| 90 | | | | | | 15614 |
|---|---|---|---|---|---|---|
| | CR1 | CR2 | MA1 | MA2 | MM1 | MM2 |
| CR1 | 0 | 351 | 168 | 351 | 146 | 161 |
| CR2 | 0.0225 | 0 | 336 | 72 | 345 | 349 |
| MA1 | 0.0108 | 0.0215 | 0 | 332 | 163 | 15 |
| MA2 | 0.0225 | 0.0046 | 0.0213 | 0 | 345 | 347 |
| MM1 | 0.0094 | 0.0221 | 0.0104 | 0.0221 | 0 | 153 |
| MM2 | 0.0103 | 0.0224 | 0.0010 | 0.0222 | 0.0098 | 0 |

| 91 | | | | | | 9860 |
|---|---|---|---|---|---|---|
| | CR1 | CR2 | MA1 | MA2 | MM1 | MM2 |
| CR1 | 0 | 121 | 152 | 79 | 139 | 152 |
| CR2 | 0.0123 | 0 | 92 | 166 | 88 | 92 |
| MA1 | 0.0154 | 0.0093 | 0 | 189 | 70 | 0 |
| MA2 | 0.0080 | 0.0168 | 0.0192 | 0 | 176 | 189 |
| MM1 | 0.0141 | 0.0089 | 0.0071 | 0.0178 | 0 | 70 |
| MM2 | 0.0154 | 0.0093 | 0.0000 | 0.0192 | 0.0071 | 0 |

| 92 | | | | | | | 11609 |
|---|---|---|---|---|---|---|---|
| | CR1 | CR2 | MA1 | MA2 | MM1 | MM2 | |
| CR1 | 0 | 192 | 103 | 181 | 126 | 96 | |
| CR2 | 0.0165 | 0 | 193 | 71 | 201 | 192 | |
| MA1 | 0.0089 | 0.0166 | 0 | 184 | 129 | 111 | |
| MA2 | 0.0156 | 0.0061 | 0.0158 | 0 | 192 | 181 | |
| MM1 | 0.0109 | 0.0173 | 0.0111 | 0.0165 | 0 | 121 | |
| MM2 | 0.0083 | 0.0165 | 0.0096 | 0.0156 | 0.0104 | 0 | |
| 94 | | | | | | | 8009 |
| | CR1 | CR2 | MA1 | MA2 | MM1 | MM2 | |
| CR1 | 0 | 119 | 78 | 118 | 78 | 83 | |
| CR2 | 0.0149 | 0 | 112 | 11 | 112 | 116 | |
| MA1 | 0.0097 | 0.0140 | 0 | 110 | 0 | 61 | |
| MA2 | 0.0147 | 0.0014 | 0.0137 | 0 | 110 | 117 | |
| MM1 | 0.0097 | 0.0140 | 0.0000 | 0.0137 | 0 | 61 | |
| MM2 | 0.0104 | 0.0145 | 0.0076 | 0.0146 | 0.0076 | 0 | |
| 96 | | | | | | | 8566 |
| | CR1 | CR2 | MA1 | MA2 | MM1 | MM2 | |
| CR1 | 0 | 116 | 112 | 100 | 106 | 107 | |
| CR2 | 0.0135 | 0 | 82 | 56 | 86 | 83 | |
| MA1 | 0.0131 | 0.0096 | 0 | 108 | 46 | 43 | |
| MA2 | 0.0117 | 0.0065 | 0.0126 | 0 | 112 | 113 | |
| MM1 | 0.0124 | 0.0100 | 0.0054 | 0.0131 | 0 | 23 | |
| MM2 | 0.0125 | 0.0097 | 0.0050 | 0.0132 | 0.0027 | 0 | |
| 97 | | | | | | | 24368 |
| | CR1 | CR2 | MA1 | MA2 | MM1 | MM2 | |
| CR1 | 0 | 468 | 238 | 237 | 209 | 212 | |
| CR2 | 0.0192 | 0 | 420 | 419 | 456 | 478 | |
| MA1 | 0.0098 | 0.0172 | 0 | 89 | 245 | 244 | |
| MA2 | 0.0097 | 0.0172 | 0.0037 | 0 | 240 | 251 | |
| MM1 | 0.0086 | 0.0187 | 0.0101 | 0.0098 | 0 | 201 | |
| MM2 | 0.0087 | 0.0196 | 0.0100 | 0.0103 | 0.0082 | 0 | |
| 98 | | | | | | | 20798 |
| | CR1 | CR2 | MA1 | MA2 | MM1 | MM2 | |
| CR1 | 0 | 452 | 218 | 461 | 224 | 192 | |
| CR2 | 0.0217 | 0 | 454 | 138 | 476 | 448 | |
| MA1 | 0.0105 | 0.0218 | 0 | 465 | 189 | 157 | |
| MA2 | 0.0222 | 0.0066 | 0.0224 | 0 | 489 | 462 | |
| MM1 | 0.0108 | 0.0229 | 0.0091 | 0.0235 | 0 | 32 | |
| MM2 | 0.0092 | 0.0215 | 0.0075 | 0.0222 | 0.0015 | 0 | |

| 99 | | | | | | | 22033 |
|---|---|---|---|---|---|---|---|
| | CR1 | CR2 | MA1 | MA2 | MM1 | MM2 | |
| CR1 | 0 | 539 | 131 | 512 | 548 | 512 | |
| CR2 | 0.0245 | 0 | 550 | 215 | 231 | 215 | |
| MA1 | 0.0059 | 0.0250 | 0 | 522 | 561 | 522 | |
| MA2 | 0.0232 | 0.0098 | 0.0237 | 0 | 198 | 0 | |
| MM1 | 0.0249 | 0.0105 | 0.0255 | 0.0090 | 0 | 198 | |
| MM2 | 0.0232 | 0.0098 | 0.0237 | 0.0000 | 0.0090 | 0 | |
| 101 | | | | | | | 8821 |
| | CR1 | CR2 | MA1 | MA2 | MM1 | MM2 | |
| CR1 | 0 | 86 | 95 | 4 | 91 | 90 | |
| CR2 | 0.0097 | 0 | 44 | 88 | 41 | 36 | |
| MA1 | 0.0108 | 0.0050 | 0 | 97 | 57 | 49 | |
| MA2 | 0.0005 | 0.0100 | 0.0110 | 0 | 93 | 92 | |
| MM1 | 0.0103 | 0.0046 | 0.0065 | 0.0105 | 0 | 53 | |
| MM2 | 0.0102 | 0.0041 | 0.0056 | 0.0104 | 0.0060 | 0 | |
| 102 | | | | | | | 17362 |
| | CR1 | CR2 | MA1 | MA2 | MM1 | MM2 | |
| CR1 | 0 | 285 | 269 | 170 | 101 | 163 | |
| CR2 | 0.0164 | 0 | 158 | 279 | 280 | 298 | |
| MA1 | 0.0155 | 0.0091 | 0 | 224 | 268 | 285 | |
| MA2 | 0.0098 | 0.0161 | 0.0129 | 0 | 155 | 162 | |
| MM1 | 0.0058 | 0.0161 | 0.0154 | 0.0089 | 0 | 158 | |
| MM2 | 0.0094 | 0.0172 | 0.0164 | 0.0093 | 0.0091 | 0 | |
| 103 | | | | | | | 10425 |
| | CR1 | CR2 | MA1 | MA2 | MM1 | MM2 | |
| CR1 | 0 | 222 | 223 | 175 | 128 | 120 | |
| CR2 | 0.0213 | 0 | 101 | 170 | 220 | 233 | |
| MA1 | 0.0214 | 0.0097 | 0 | 139 | 224 | 234 | |
| MA2 | 0.0168 | 0.0163 | 0.0133 | 0 | 183 | 95 | |
| MM1 | 0.0123 | 0.0211 | 0.0215 | 0.0176 | 0 | 143 | |
| MM2 | 0.0115 | 0.0224 | 0.0224 | 0.0091 | 0.0137 | 0 | |
| 104 | | | | | | | 15517 |
| | CR1 | CR2 | MA1 | MA2 | MM1 | MM2 | |
| CR1 | 0 | 254 | 85 | 246 | 83 | 85 | |
| CR2 | 0.0164 | 0 | 246 | 58 | 246 | 246 | |
| MA1 | 0.0055 | 0.0159 | 0 | 236 | 79 | 0 | |
| MA2 | 0.0159 | 0.0037 | 0.0152 | 0 | 238 | 236 | |
| MM1 | 0.0053 | 0.0159 | 0.0051 | 0.0153 | 0 | 79 | |
| MM2 | 0.0055 | 0.0159 | 0.0000 | 0.0152 | 0.0051 | 0 | |

| 105 | | | | | | | 10807 |
|---|---|---|---|---|---|---|---|
| | CR1 | CR2 | MA1 | MA2 | MM1 | MM2 | |
| CR1 | 0 | 159 | 37 | 150 | 165 | 150 | |
| CR2 | 0.0147 | 0 | 158 | 52 | 59 | 52 | |
| MA1 | 0.0034 | 0.0146 | 0 | 147 | 162 | 147 | |
| MA2 | 0.0139 | 0.0048 | 0.0136 | 0 | 40 | 0 | |
| MM1 | 0.0153 | 0.0055 | 0.0150 | 0.0037 | 0 | 40 | |
| MM2 | 0.0139 | 0.0048 | 0.0136 | 0.0000 | 0.0037 | 0 | |
| 110 | | | | | | | 15592 |
| | CR1 | CR2 | MA1 | MA2 | MM1 | MM2 | |
| CR1 | 0 | 293 | 186 | 277 | 164 | 191 | |
| CR2 | 0.0188 | 0 | 261 | 111 | 271 | 279 | |
| MA1 | 0.0119 | 0.0167 | 0 | 285 | 143 | 160 | |
| MA2 | 0.0178 | 0.0071 | 0.0183 | 0 | 303 | 314 | |
| MM1 | 0.0105 | 0.0174 | 0.0092 | 0.0194 | 0 | 146 | |
| MM2 | 0.0122 | 0.0179 | 0.0103 | 0.0201 | 0.0094 | 0 | |
| 111 | | | | | | | 11601 |
| | CR1 | CR2 | MA1 | MA2 | MM1 | MM2 | |
| CR1 | 0 | 329 | 116 | 308 | 116 | 155 | |
| CR2 | 0.0284 | 0 | 323 | 74 | 323 | 330 | |
| MA1 | 0.0100 | 0.0278 | 0 | 299 | 0 | 175 | |
| MA2 | 0.0265 | 0.0064 | 0.0258 | 0 | 299 | 308 | |
| MM1 | 0.0100 | 0.0278 | 0.0000 | 0.0258 | 0 | 175 | |
| MM2 | 0.0134 | 0.0284 | 0.0151 | 0.0265 | 0.0151 | 0 | |
| 112 | | | | | | | 17372 |
| | CR1 | CR2 | MA1 | MA2 | MM1 | MM2 | |
| CR1 | 0 | 366 | 369 | 250 | 242 | 232 | |
| CR2 | 0.0211 | 0 | 65 | 386 | 387 | 383 | |
| MA1 | 0.0212 | 0.0037 | 0 | 387 | 391 | 387 | |
| MA2 | 0.0144 | 0.0222 | 0.0223 | 0 | 245 | 247 | |
| MM1 | 0.0139 | 0.0223 | 0.0225 | 0.0141 | 0 | 56 | |
| MM2 | 0.0134 | 0.0220 | 0.0223 | 0.0142 | 0.0032 | 0 | |
| 114 | | | | | | | 4071 |
| | CR1 | CR2 | MA1 | MA2 | MM1 | MM2 | |
| CR1 | 0 | 64 | 30 | 64 | 30 | 25 | |
| CR2 | 0.0157 | 0 | 58 | 16 | 56 | 57 | |
| MA1 | 0.0074 | 0.0142 | 0 | 60 | 24 | 23 | |
| MA2 | 0.0157 | 0.0039 | 0.0147 | 0 | 60 | 59 | |
| MM1 | 0.0074 | 0.0138 | 0.0059 | 0.0147 | 0 | 23 | |
| MM2 | 0.0061 | 0.0140 | 0.0056 | 0.0145 | 0.0056 | 0 | |

| 115 | | | | | | | 10101 |
|---|---|---|---|---|---|---|---|
| | CR1 | CR2 | MA1 | MA2 | MM1 | MM2 | |
| CR1 | 0 | 47 | 251 | 88 | 86 | 88 | |
| CR2 | 0.0047 | 0 | 204 | 103 | 105 | 103 | |
| MA1 | 0.0248 | 0.0202 | 0 | 256 | 249 | 256 | |
| MA2 | 0.0087 | 0.0102 | 0.0253 | 0 | 74 | 0 | |
| MM1 | 0.0085 | 0.0104 | 0.0247 | 0.0073 | 0 | 74 | |
| MM2 | 0.0087 | 0.0102 | 0.0253 | 0.0000 | 0.0073 | 0 | |
| 116 | | | | | | | 9233 |
| | CR1 | CR2 | MA1 | MA2 | MM1 | MM2 | |
| CR1 | 0 | 212 | 81 | 194 | 81 | 78 | |
| CR2 | 0.0230 | 0 | 218 | 67 | 218 | 201 | |
| MA1 | 0.0088 | 0.0236 | 0 | 200 | 0 | 85 | |
| MA2 | 0.0210 | 0.0073 | 0.0217 | 0 | 200 | 182 | |
| MM1 | 0.0088 | 0.0236 | 0.0000 | 0.0217 | 0 | 85 | |
| MM2 | 0.0084 | 0.0218 | 0.0092 | 0.0197 | 0.0092 | 0 | |
| 119 | | | | | | | 12521 |
| | CR1 | CR2 | MA1 | MA2 | MM1 | MM2 | |
| CR1 | 0 | 201 | 107 | 198 | 88 | 107 | |
| CR2 | 0.0161 | 0 | 205 | 52 | 196 | 205 | |
| MA1 | 0.0085 | 0.0164 | 0 | 201 | 109 | 0 | |
| MA2 | 0.0158 | 0.0042 | 0.0161 | 0 | 197 | 201 | |
| MM1 | 0.0070 | 0.0157 | 0.0087 | 0.0157 | 0 | 109 | |
| MM2 | 0.0085 | 0.0164 | 0.0000 | 0.0161 | 0.0087 | 0 | |
| 120 | | | | | | | 23585 |
| | CR1 | CR2 | MA1 | MA2 | MM1 | MM2 | |
| CR1 | 0 | 388 | 227 | 364 | 183 | 183 | |
| CR2 | 0.0165 | 0 | 288 | 144 | 397 | 397 | |
| MA1 | 0.0096 | 0.0122 | 0 | 218 | 165 | 165 | |
| MA2 | 0.0154 | 0.0061 | 0.0092 | 0 | 383 | 383 | |
| MM1 | 0.0078 | 0.0168 | 0.0070 | 0.0162 | 0 | 0 | |
| MM2 | 0.0078 | 0.0168 | 0.0070 | 0.0162 | 0.0000 | 0 | |
| 122 | | | | | | | 11133 |
| | CR1 | CR2 | MA1 | MA2 | MM1 | MM2 | |
| CR1 | 0 | 219 | 89 | 218 | 218 | 243 | |
| CR2 | 0.0197 | 0 | 221 | 3 | 3 | 96 | |
| MA1 | 0.0080 | 0.0199 | 0 | 220 | 220 | 244 | |
| MA2 | 0.0196 | 0.0003 | 0.0198 | 0 | 0 | 93 | |
| MM1 | 0.0196 | 0.0003 | 0.0198 | 0.0000 | 0 | 93 | |
| MM2 | 0.0218 | 0.0086 | 0.0219 | 0.0084 | 0.0084 | 0 | |

| 123 | | | | | | | 11847 |
|---|---|---|---|---|---|---|---|
| | CR1 | CR2 | MA1 | MA2 | MM1 | MM2 | |
| CR1 | 0 | 211 | 152 | 207 | 109 | 156 | |
| CR2 | 0.0178 | 0 | 231 | 52 | 199 | 223 | |
| MA1 | 0.0128 | 0.0195 | 0 | 211 | 121 | 166 | |
| MA2 | 0.0175 | 0.0044 | 0.0178 | 0 | 187 | 209 | |
| MM1 | 0.0092 | 0.0168 | 0.0102 | 0.0158 | 0 | 153 | |
| MM2 | 0.0132 | 0.0188 | 0.0140 | 0.0176 | 0.0129 | 0 | |
| 124 | | | | | | | 10246 |
| | CR1 | CR2 | MA1 | MA2 | MM1 | MM2 | |
| CR1 | 0 | 240 | 117 | 193 | 107 | 117 | |
| CR2 | 0.0234 | 0 | 241 | 178 | 230 | 241 | |
| MA1 | 0.0114 | 0.0235 | 0 | 106 | 96 | 0 | |
| MA2 | 0.0188 | 0.0174 | 0.0103 | 0 | 166 | 106 | |
| MM1 | 0.0104 | 0.0224 | 0.0094 | 0.0162 | 0 | 96 | |
| MM2 | 0.0114 | 0.0235 | 0.0000 | 0.0103 | 0.0094 | 0 | |
| 125 | | | | | | | 10022 |
| | CR1 | CR2 | MA1 | MA2 | MM1 | MM2 | |
| CR1 | 0 | 219 | 49 | 205 | 197 | 205 | |
| CR2 | 0.0219 | 0 | 200 | 130 | 104 | 130 | |
| MA1 | 0.0049 | 0.0200 | 0 | 186 | 179 | 186 | |
| MA2 | 0.0205 | 0.0130 | 0.0186 | 0 | 98 | 0 | |
| MM1 | 0.0197 | 0.0104 | 0.0179 | 0.0098 | 0 | 98 | |
| MM2 | 0.0205 | 0.0130 | 0.0186 | 0.0000 | 0.0098 | 0 | |
| 126 | | | | | | | 11187 |
| | CR1 | CR2 | MA1 | MA2 | MM1 | MM2 | |
| CR1 | 0 | 342 | 324 | 278 | 156 | 160 | |
| CR2 | 0.0306 | 0 | 137 | 191 | 340 | 321 | |
| MA1 | 0.0290 | 0.0122 | 0 | 93 | 329 | 310 | |
| MA2 | 0.0249 | 0.0171 | 0.0083 | 0 | 285 | 217 | |
| MM1 | 0.0139 | 0.0304 | 0.0294 | 0.0255 | 0 | 161 | |
| MM2 | 0.0143 | 0.0287 | 0.0277 | 0.0194 | 0.0144 | 0 | |
| 128 | | | | | | | 11016 |
| | CR1 | CR2 | MA1 | MA2 | MM1 | MM2 | |
| CR1 | 0 | 129 | 129 | 54 | 54 | 61 | |
| CR2 | 0.0117 | 0 | 26 | 127 | 127 | 130 | |
| MA1 | 0.0117 | 0.0024 | 0 | 127 | 127 | 130 | |
| MA2 | 0.0049 | 0.0115 | 0.0115 | 0 | 0 | 63 | |
| MM1 | 0.0049 | 0.0115 | 0.0115 | 0.0000 | 0 | 63 | |
| MM2 | 0.0055 | 0.0118 | 0.0118 | 0.0057 | 0.0057 | 0 | |

| 129 | | | | | | | 14754 |
|---|---|---|---|---|---|---|---|
| | CR1 | CR2 | MA1 | MA2 | MM1 | MM2 | |
| CR1 | 0 | 263 | 259 | 60 | 60 | 207 | |
| CR2 | 0.0178 | 0 | 58 | 265 | 265 | 280 | |
| MA1 | 0.0176 | 0.0039 | 0 | 262 | 262 | 280 | |
| MA2 | 0.0041 | 0.0180 | 0.0178 | 0 | 0 | 231 | |
| MM1 | 0.0041 | 0.0180 | 0.0178 | 0.0000 | 0 | 231 | |
| MM2 | 0.0140 | 0.0190 | 0.0190 | 0.0157 | 0.0157 | 0 | |
| 131 | | | | | | | 18310 |
| | CR1 | CR2 | MA1 | MA2 | MM1 | MM2 | |
| CR1 | 0 | 375 | 153 | 374 | 138 | 149 | |
| CR2 | 0.0205 | 0 | 346 | 73 | 375 | 360 | |
| MA1 | 0.0084 | 0.0189 | 0 | 349 | 146 | 48 | |
| MA2 | 0.0204 | 0.0040 | 0.0191 | 0 | 370 | 365 | |
| MM1 | 0.0075 | 0.0205 | 0.0080 | 0.0202 | 0 | 104 | |
| MM2 | 0.0081 | 0.0197 | 0.0026 | 0.0199 | 0.0057 | 0 | |
| 132 | | | | | | | 14634 |
| | CR1 | CR2 | MA1 | MA2 | MM1 | MM2 | |
| CR1 | 0 | 353 | 186 | 353 | 186 | 169 | |
| CR2 | 0.0241 | 0 | 324 | 83 | 324 | 339 | |
| MA1 | 0.0127 | 0.0221 | 0 | 333 | 0 | 184 | |
| MA2 | 0.0241 | 0.0057 | 0.0228 | 0 | 333 | 345 | |
| MM1 | 0.0127 | 0.0221 | 0.0000 | 0.0228 | 0 | 184 | |
| MM2 | 0.0115 | 0.0232 | 0.0126 | 0.0236 | 0.0126 | 0 | |
| 133 | | | | | | | 17959 |
| | CR1 | CR2 | MA1 | MA2 | MM1 | MM2 | |
| CR1 | 0 | 290 | 166 | 286 | 166 | 163 | |
| CR2 | 0.0161 | 0 | 299 | 118 | 299 | 294 | |
| MA1 | 0.0092 | 0.0166 | 0 | 298 | 0 | 158 | |
| MA2 | 0.0159 | 0.0066 | 0.0166 | 0 | 298 | 290 | |
| MM1 | 0.0092 | 0.0166 | 0.0000 | 0.0166 | 0 | 158 | |
| MM2 | 0.0091 | 0.0164 | 0.0088 | 0.0161 | 0.0088 | 0 | |
| 134 | | | | | | | 13429 |
| | CR1 | CR2 | MA1 | MA2 | MM1 | MM2 | |
| CR1 | 0 | 314 | 11 | 321 | 152 | 36 | |
| CR2 | 0.0234 | 0 | 321 | 80 | 298 | 326 | |
| MA1 | 0.0008 | 0.0239 | 0 | 328 | 149 | 27 | |
| MA2 | 0.0239 | 0.0060 | 0.0244 | 0 | 303 | 333 | |
| MM1 | 0.0113 | 0.0222 | 0.0111 | 0.0226 | 0 | 122 | |
| MM2 | 0.0027 | 0.0243 | 0.0020 | 0.0248 | 0.0091 | 0 | |

137 | | | | | | 10904

| | CR1 | CR2 | MA1 | MA2 | MM1 | MM2 |
|---|---|---|---|---|---|---|
| CR1 | 0 | 71 | 36 | 111 | 36 | 52 |
| CR2 | 0.0065 | 0 | 83 | 54 | 83 | 96 |
| MA1 | 0.0033 | 0.0076 | 0 | 115 | 0 | 58 |
| MA2 | 0.0102 | 0.0050 | 0.0105 | 0 | 115 | 110 |
| MM1 | 0.0033 | 0.0076 | 0.0000 | 0.0105 | 0 | 58 |
| MM2 | 0.0048 | 0.0088 | 0.0053 | 0.0101 | 0.0053 | 0 |

138 | | | | | | 13470

| | CR1 | CR2 | MA1 | MA2 | MM1 | MM2 |
|---|---|---|---|---|---|---|
| CR1 | 0 | 292 | 150 | 298 | 298 | 298 |
| CR2 | 0.0217 | 0 | 295 | 166 | 174 | 166 |
| MA1 | 0.0111 | 0.0219 | 0 | 297 | 291 | 297 |
| MA2 | 0.0221 | 0.0123 | 0.0220 | 0 | 96 | 0 |
| MM1 | 0.0221 | 0.0129 | 0.0216 | 0.0071 | 0 | 96 |
| MM2 | 0.0221 | 0.0123 | 0.0220 | 0.0000 | 0.0071 | 0 |

139 | | | | | | 12019

| | CR1 | CR2 | MA1 | MA2 | MM1 | MM2 |
|---|---|---|---|---|---|---|
| CR1 | 0 | 263 | 158 | 264 | 189 | 158 |
| CR2 | 0.0219 | 0 | 252 | 68 | 266 | 252 |
| MA1 | 0.0131 | 0.0210 | 0 | 254 | 216 | 0 |
| MA2 | 0.0220 | 0.0057 | 0.0211 | 0 | 263 | 254 |
| MM1 | 0.0157 | 0.0221 | 0.0180 | 0.0219 | 0 | 216 |
| MM2 | 0.0131 | 0.0210 | 0.0000 | 0.0211 | 0.0180 | 0 |

140 | | | | | | 13142

| | CR1 | CR2 | MA1 | MA2 | MM1 | MM2 |
|---|---|---|---|---|---|---|
| CR1 | 0 | 301 | 177 | 306 | 178 | 155 |
| CR2 | 0.0229 | 0 | 300 | 146 | 302 | 304 |
| MA1 | 0.0135 | 0.0228 | 0 | 302 | 117 | 179 |
| MA2 | 0.0233 | 0.0111 | 0.0230 | 0 | 309 | 311 |
| MM1 | 0.0135 | 0.0230 | 0.0089 | 0.0235 | 0 | 127 |
| MM2 | 0.0118 | 0.0231 | 0.0136 | 0.0237 | 0.0097 | 0 |

141 | | | | | | 9510

| | CR1 | CR2 | MA1 | MA2 | MM1 | MM2 |
|---|---|---|---|---|---|---|
| CR1 | 0 | 144 | 218 | 49 | 217 | 236 |
| CR2 | 0.0151 | 0 | 152 | 167 | 124 | 166 |
| MA1 | 0.0229 | 0.0160 | 0 | 215 | 118 | 128 |
| MA2 | 0.0052 | 0.0176 | 0.0226 | 0 | 217 | 233 |
| MM1 | 0.0228 | 0.0130 | 0.0124 | 0.0228 | 0 | 139 |
| MM2 | 0.0248 | 0.0175 | 0.0135 | 0.0245 | 0.0146 | 0 |

| 142 | | | | | | | 10087 |
|---|---|---|---|---|---|---|---|
| | CR1 | CR2 | MA1 | MA2 | MM1 | MM2 | |
| CR1 | 0 | 182 | 82 | 169 | 82 | 84 | |
| CR2 | 0.0180 | 0 | 190 | 41 | 190 | 188 | |
| MA1 | 0.0081 | 0.0188 | 0 | 177 | 0 | 102 | |
| MA2 | 0.0168 | 0.0041 | 0.0175 | 0 | 177 | 173 | |
| MM1 | 0.0081 | 0.0188 | 0.0000 | 0.0175 | 0 | 102 | |
| MM2 | 0.0083 | 0.0186 | 0.0101 | 0.0172 | 0.0101 | 0 | |
| 143 | | | | | | | 11798 |
| | CR1 | CR2 | MA1 | MA2 | MM1 | MM2 | |
| CR1 | 0 | 229 | 109 | 217 | 124 | 109 | |
| CR2 | 0.0194 | 0 | 212 | 76 | 213 | 212 | |
| MA1 | 0.0092 | 0.0180 | 0 | 204 | 100 | 0 | |
| MA2 | 0.0184 | 0.0064 | 0.0173 | 0 | 207 | 204 | |
| MM1 | 0.0105 | 0.0181 | 0.0085 | 0.0175 | 0 | 100 | |
| MM2 | 0.0092 | 0.0180 | 0.0000 | 0.0173 | 0.0085 | 0 | |
| 144 | | | | | | | 11897 |
| | CR1 | CR2 | MA1 | MA2 | MM1 | MM2 | |
| CR1 | 0 | 325 | 316 | 143 | 302 | 309 | |
| CR2 | 0.0273 | 0 | 218 | 347 | 222 | 245 | |
| MA1 | 0.0266 | 0.0183 | 0 | 340 | 188 | 154 | |
| MA2 | 0.0120 | 0.0292 | 0.0286 | 0 | 318 | 329 | |
| MM1 | 0.0254 | 0.0187 | 0.0158 | 0.0267 | 0 | 212 | |
| MM2 | 0.0260 | 0.0206 | 0.0129 | 0.0277 | 0.0178 | 0 | |
| 146 | | | | | | | 12362 |
| | CR1 | CR2 | MA1 | MA2 | MM1 | MM2 | |
| CR1 | 0 | 141 | 222 | 75 | 215 | 213 | |
| CR2 | 0.0114 | 0 | 161 | 132 | 143 | 148 | |
| MA1 | 0.0180 | 0.0130 | 0 | 229 | 95 | 95 | |
| MA2 | 0.0061 | 0.0107 | 0.0185 | 0 | 215 | 216 | |
| MM1 | 0.0174 | 0.0116 | 0.0077 | 0.0174 | 0 | 90 | |
| MM2 | 0.0172 | 0.0120 | 0.0077 | 0.0175 | 0.0073 | 0 | |
| 147 | | | | | | | 15270 |
| | CR1 | CR2 | MA1 | MA2 | MM1 | MM2 | |
| CR1 | 0 | 251 | 245 | 52 | 245 | 252 | |
| CR2 | 0.0164 | 0 | 63 | 234 | 63 | 79 | |
| MA1 | 0.0160 | 0.0041 | 0 | 228 | 0 | 68 | |
| MA2 | 0.0034 | 0.0153 | 0.0149 | 0 | 228 | 233 | |
| MM1 | 0.0160 | 0.0041 | 0.0000 | 0.0149 | 0 | 68 | |
| MM2 | 0.0165 | 0.0052 | 0.0045 | 0.0153 | 0.0045 | 0 | |

| 148 | | | | | | | 14701 |
|---|---|---|---|---|---|---|---|
| | CR1 | CR2 | MA1 | MA2 | MM1 | MM2 | |
| CR1 | 0 | 331 | 168 | 338 | 162 | 189 | |
| CR2 | 0.0225 | 0 | 324 | 116 | 326 | 341 | |
| MA1 | 0.0114 | 0.0220 | 0 | 318 | 16 | 181 | |
| MA2 | 0.0230 | 0.0079 | 0.0216 | 0 | 334 | 347 | |
| MM1 | 0.0110 | 0.0222 | 0.0011 | 0.0227 | 0 | 165 | |
| MM2 | 0.0129 | 0.0232 | 0.0123 | 0.0236 | 0.0112 | 0 | |
| 149 | | | | | | | 14413 |
| | CR1 | CR2 | MA1 | MA2 | MM1 | MM2 | |
| CR1 | 0 | 228 | 128 | 241 | 113 | 106 | |
| CR2 | 0.0158 | 0 | 235 | 67 | 242 | 238 | |
| MA1 | 0.0089 | 0.0163 | 0 | 212 | 134 | 120 | |
| MA2 | 0.0167 | 0.0046 | 0.0147 | 0 | 252 | 246 | |
| MM1 | 0.0078 | 0.0168 | 0.0093 | 0.0175 | 0 | 98 | |
| MM2 | 0.0074 | 0.0165 | 0.0083 | 0.0171 | 0.0068 | 0 | |
| 154 | | | | | | | 9382 |
| | CR1 | CR2 | MA1 | MA2 | MM1 | MM2 | |
| CR1 | 0 | 157 | 149 | 75 | 54 | 48 | |
| CR2 | 0.0167 | 0 | 11 | 132 | 133 | 141 | |
| MA1 | 0.0159 | 0.0012 | 0 | 126 | 127 | 135 | |
| MA2 | 0.0080 | 0.0141 | 0.0134 | 0 | 53 | 37 | |
| MM1 | 0.0058 | 0.0142 | 0.0135 | 0.0056 | 0 | 24 | |
| MM2 | 0.0051 | 0.0150 | 0.0144 | 0.0039 | 0.0026 | 0 | |
| 156 | | | | | | | 12178 |
| | CR1 | CR2 | MA1 | MA2 | MM1 | MM2 | |
| CR1 | 0 | 307 | 281 | 28 | 316 | 281 | |
| CR2 | 0.0252 | 0 | 176 | 312 | 180 | 176 | |
| MA1 | 0.0231 | 0.0145 | 0 | 286 | 170 | 0 | |
| MA2 | 0.0023 | 0.0256 | 0.0235 | 0 | 321 | 286 | |
| MM1 | 0.0259 | 0.0148 | 0.0140 | 0.0264 | 0 | 170 | |
| MM2 | 0.0231 | 0.0145 | 0.0000 | 0.0235 | 0.0140 | 0 | |
| 159 | | | | | | | 12299 |
| | CR1 | CR2 | MA1 | MA2 | MM1 | MM2 | |
| CR1 | 0 | 284 | 59 | 266 | 260 | 261 | |
| CR2 | 0.0231 | 0 | 285 | 154 | 182 | 169 | |
| MA1 | 0.0048 | 0.0232 | 0 | 273 | 257 | 264 | |
| MA2 | 0.0216 | 0.0125 | 0.0222 | 0 | 146 | 148 | |
| MM1 | 0.0211 | 0.0148 | 0.0209 | 0.0119 | 0 | 157 | |
| MM2 | 0.0212 | 0.0137 | 0.0215 | 0.0120 | 0.0128 | 0 | |

| 160 | | | | | | | 9854 |
|---|---|---|---|---|---|---|---|
| | CR1 | CR2 | MA1 | MA2 | MM1 | MM2 | |
| CR1 | 0 | 85 | 56 | 23 | 23 | 27 | |
| CR2 | 0.0086 | 0 | 37 | 90 | 90 | 89 | |
| MA1 | 0.0057 | 0.0038 | 0 | 61 | 61 | 63 | |
| MA2 | 0.0023 | 0.0091 | 0.0062 | 0 | 0 | 26 | |
| MM1 | 0.0023 | 0.0091 | 0.0062 | 0.0000 | 0 | 26 | |
| MM2 | 0.0027 | 0.0090 | 0.0064 | 0.0026 | 0.0026 | 0 | |
| 161 | | | | | | | 16370 |
| | CR1 | CR2 | MA1 | MA2 | MM1 | MM2 | |
| CR1 | 0 | 371 | 384 | 160 | 386 | 391 | |
| CR2 | 0.0227 | 0 | 191 | 393 | 123 | 187 | |
| MA1 | 0.0235 | 0.0117 | 0 | 401 | 204 | 206 | |
| MA2 | 0.0098 | 0.0240 | 0.0245 | 0 | 409 | 415 | |
| MM1 | 0.0236 | 0.0075 | 0.0125 | 0.0250 | 0 | 228 | |
| MM2 | 0.0239 | 0.0114 | 0.0126 | 0.0254 | 0.0139 | 0 | |
| 165 | | | | | | | 13402 |
| | CR1 | CR2 | MA1 | MA2 | MM1 | MM2 | |
| CR1 | 0 | 311 | 155 | 311 | 312 | 307 | |
| CR2 | 0.0232 | 0 | 311 | 0 | 150 | 197 | |
| MA1 | 0.0116 | 0.0232 | 0 | 311 | 307 | 322 | |
| MA2 | 0.0232 | 0.0000 | 0.0232 | 0 | 150 | 197 | |
| MM1 | 0.0233 | 0.0112 | 0.0229 | 0.0112 | 0 | 190 | |
| MM2 | 0.0229 | 0.0147 | 0.0240 | 0.0147 | 0.0142 | 0 | |
| 167 | | | | | | | 13120 |
| | CR1 | CR2 | MA1 | MA2 | MM1 | MM2 | |
| CR1 | 0 | 300 | 0 | 297 | 106 | 0 | |
| CR2 | 0.0229 | 0 | 300 | 55 | 303 | 300 | |
| MA1 | 0.0000 | 0.0229 | 0 | 297 | 106 | 0 | |
| MA2 | 0.0226 | 0.0042 | 0.0226 | 0 | 300 | 297 | |
| MM1 | 0.0081 | 0.0231 | 0.0081 | 0.0229 | 0 | 106 | |
| MM2 | 0.0000 | 0.0229 | 0.0000 | 0.0226 | 0.0081 | 0 | |
| 168 | | | | | | | 12458 |
| | CR1 | CR2 | MA1 | MA2 | MM1 | MM2 | |
| CR1 | 0 | 305 | 207 | 307 | 189 | 228 | |
| CR2 | 0.0245 | 0 | 294 | 99 | 296 | 319 | |
| MA1 | 0.0166 | 0.0236 | 0 | 297 | 207 | 203 | |
| MA2 | 0.0246 | 0.0079 | 0.0238 | 0 | 300 | 317 | |
| MM1 | 0.0152 | 0.0238 | 0.0166 | 0.0241 | 0 | 230 | |
| MM2 | 0.0183 | 0.0256 | 0.0163 | 0.0254 | 0.0185 | 0 | |

| 170 | | | | | | | 12525 |
|---|---|---|---|---|---|---|---|
| | CR1 | CR2 | MA1 | MA2 | MM1 | MM2 | |
| CR1 | 0 | 211 | 211 | 103 | 118 | 100 | |
| CR2 | 0.0168 | 0 | 80 | 228 | 221 | 227 | |
| MA1 | 0.0168 | 0.0064 | 0 | 224 | 218 | 221 | |
| MA2 | 0.0082 | 0.0182 | 0.0179 | 0 | 129 | 91 | |
| MM1 | 0.0094 | 0.0176 | 0.0174 | 0.0103 | 0 | 138 | |
| MM2 | 0.0080 | 0.0181 | 0.0176 | 0.0073 | 0.0110 | 0 | |
| 172 | | | | | | | 14980 |
| | CR1 | CR2 | MA1 | MA2 | MM1 | MM2 | |
| CR1 | 0 | 288 | 162 | 279 | 162 | 141 | |
| CR2 | 0.0192 | 0 | 282 | 107 | 282 | 281 | |
| MA1 | 0.0108 | 0.0188 | 0 | 281 | 0 | 178 | |
| MA2 | 0.0186 | 0.0071 | 0.0188 | 0 | 281 | 276 | |
| MM1 | 0.0108 | 0.0188 | 0.0000 | 0.0188 | 0 | 178 | |
| MM2 | 0.0094 | 0.0188 | 0.0119 | 0.0184 | 0.0119 | 0 | |
| 173 | | | | | | | 11706 |
| | CR1 | CR2 | MA1 | MA2 | MM1 | MM2 | |
| CR1 | 0 | 229 | 113 | 234 | 113 | 108 | |
| CR2 | 0.0196 | 0 | 220 | 65 | 220 | 231 | |
| MA1 | 0.0097 | 0.0188 | 0 | 223 | 0 | 103 | |
| MA2 | 0.0200 | 0.0056 | 0.0191 | 0 | 223 | 234 | |
| MM1 | 0.0097 | 0.0188 | 0.0000 | 0.0191 | 0 | 103 | |
| MM2 | 0.0092 | 0.0197 | 0.0088 | 0.0200 | 0.0088 | 0 | |
| 174 | | | | | | | 11701 |
| | CR1 | CR2 | MA1 | MA2 | MM1 | MM2 | |
| CR1 | 0 | 157 | 163 | 47 | 157 | 163 | |
| CR2 | 0.0134 | 0 | 71 | 156 | 0 | 71 | |
| MA1 | 0.0139 | 0.0061 | 0 | 160 | 71 | 0 | |
| MA2 | 0.0040 | 0.0133 | 0.0137 | 0 | 156 | 160 | |
| MM1 | 0.0134 | 0.0000 | 0.0061 | 0.0133 | 0 | 71 | |
| MM2 | 0.0139 | 0.0061 | 0.0000 | 0.0137 | 0.0061 | 0 | |
| 175 | | | | | | | 12168 |
| | CR1 | CR2 | MA1 | MA2 | MM1 | MM2 | |
| CR1 | 0 | 20 | 270 | 126 | 111 | 126 | |
| CR2 | 0.0016 | 0 | 284 | 120 | 103 | 120 | |
| MA1 | 0.0222 | 0.0233 | 0 | 271 | 287 | 271 | |
| MA2 | 0.0104 | 0.0099 | 0.0223 | 0 | 104 | 0 | |
| MM1 | 0.0091 | 0.0085 | 0.0236 | 0.0085 | 0 | 104 | |
| MM2 | 0.0104 | 0.0099 | 0.0223 | 0.0000 | 0.0085 | 0 | |

| 176 | | | | | | | 14902 |
|---|---|---|---|---|---|---|---|
| | CR1 | CR2 | MA1 | MA2 | MM1 | MM2 | |
| CR1 | 0 | 8 | 174 | 198 | 200 | 216 | |
| CR2 | 0.0005 | 0 | 182 | 206 | 192 | 208 | |
| MA1 | 0.0117 | 0.0122 | 0 | 243 | 247 | 263 | |
| MA2 | 0.0133 | 0.0138 | 0.0163 | 0 | 134 | 20 | |
| MM1 | 0.0134 | 0.0129 | 0.0166 | 0.0090 | 0 | 114 | |
| MM2 | 0.0145 | 0.0140 | 0.0176 | 0.0013 | 0.0076 | 0 | |
| 177 | | | | | | | 14045 |
| | CR1 | CR2 | MA1 | MA2 | MM1 | MM2 | |
| CR1 | 0 | 165 | 178 | 161 | 142 | 161 | |
| CR2 | 0.0117 | 0 | 36 | 243 | 229 | 243 | |
| MA1 | 0.0127 | 0.0026 | 0 | 236 | 222 | 236 | |
| MA2 | 0.0115 | 0.0173 | 0.0168 | 0 | 116 | 0 | |
| MM1 | 0.0101 | 0.0163 | 0.0158 | 0.0083 | 0 | 116 | |
| MM2 | 0.0115 | 0.0173 | 0.0168 | 0.0000 | 0.0083 | 0 | |
| 178 | | | | | | | 21442 |
| | CR1 | CR2 | MA1 | MA2 | MM1 | MM2 | |
| CR1 | 0 | 445 | 192 | 465 | 260 | 245 | |
| CR2 | 0.0208 | 0 | 437 | 149 | 447 | 446 | |
| MA1 | 0.0090 | 0.0204 | 0 | 463 | 282 | 311 | |
| MA2 | 0.0217 | 0.0069 | 0.0216 | 0 | 475 | 469 | |
| MM1 | 0.0121 | 0.0208 | 0.0132 | 0.0222 | 0 | 223 | |
| MM2 | 0.0114 | 0.0208 | 0.0145 | 0.0219 | 0.0104 | 0 | |
| 179 | | | | | | | 13479 |
| | CR1 | CR2 | MA1 | MA2 | MM1 | MM2 | |
| CR1 | 0 | 285 | 151 | 276 | 151 | 137 | |
| CR2 | 0.0211 | 0 | 292 | 65 | 292 | 275 | |
| MA1 | 0.0112 | 0.0217 | 0 | 279 | 0 | 127 | |
| MA2 | 0.0205 | 0.0048 | 0.0207 | 0 | 279 | 266 | |
| MM1 | 0.0112 | 0.0217 | 0.0000 | 0.0207 | 0 | 127 | |
| MM2 | 0.0102 | 0.0204 | 0.0094 | 0.0197 | 0.0094 | 0 | |
| 180 | | | | | | | 22045 |
| | CR1 | CR2 | MA1 | MA2 | MM1 | MM2 | |
| CR1 | 0 | 554 | 239 | 354 | 239 | 240 | |
| CR2 | 0.0251 | 0 | 581 | 454 | 581 | 582 | |
| MA1 | 0.0108 | 0.0264 | 0 | 167 | 0 | 1 | |
| MA2 | 0.0161 | 0.0206 | 0.0076 | 0 | 167 | 168 | |
| MM1 | 0.0108 | 0.0264 | 0.0000 | 0.0076 | 0 | 1 | |
| MM2 | 0.0109 | 0.0264 | 0.0000 | 0.0076 | 0.0000 | 0 | |

| 181 | | | | | | | 16863 |
|---|---|---|---|---|---|---|---|
| | CR1 | CR2 | MA1 | MA2 | MM1 | MM2 | |
| CR1 | 0 | 272 | 275 | 120 | 0 | 129 | |
| CR2 | 0.0161 | 0 | 55 | 259 | 272 | 267 | |
| MA1 | 0.0163 | 0.0033 | 0 | 261 | 275 | 267 | |
| MA2 | 0.0071 | 0.0154 | 0.0155 | 0 | 120 | 120 | |
| MM1 | 0.0000 | 0.0161 | 0.0163 | 0.0071 | 0 | 129 | |
| MM2 | 0.0076 | 0.0158 | 0.0158 | 0.0071 | 0.0076 | 0 | |
| 183 | | | | | | | 9283 |
| | CR1 | CR2 | MA1 | MA2 | MM1 | MM2 | |
| CR1 | 0 | 138 | 148 | 32 | 148 | 140 | |
| CR2 | 0.0149 | 0 | 61 | 143 | 61 | 16 | |
| MA1 | 0.0159 | 0.0066 | 0 | 153 | 0 | 57 | |
| MA2 | 0.0034 | 0.0154 | 0.0165 | 0 | 153 | 145 | |
| MM1 | 0.0159 | 0.0066 | 0.0000 | 0.0165 | 0 | 57 | |
| MM2 | 0.0151 | 0.0017 | 0.0061 | 0.0156 | 0.0061 | 0 | |
| 186 | | | | | | | 15087 |
| | CR1 | CR2 | MA1 | MA2 | MM1 | MM2 | |
| CR1 | 0 | 220 | 73 | 226 | 73 | 61 | |
| CR2 | 0.0146 | 0 | 223 | 32 | 223 | 214 | |
| MA1 | 0.0048 | 0.0148 | 0 | 229 | 0 | 66 | |
| MA2 | 0.0150 | 0.0021 | 0.0152 | 0 | 229 | 220 | |
| MM1 | 0.0048 | 0.0148 | 0.0000 | 0.0152 | 0 | 66 | |
| MM2 | 0.0040 | 0.0142 | 0.0044 | 0.0146 | 0.0044 | 0 | |
| 187 | | | | | | | 15800 |
| | CR1 | CR2 | MA1 | MA2 | MM1 | MM2 | |
| CR1 | 0 | 292 | 282 | 139 | 164 | 154 | |
| CR2 | 0.0185 | 0 | 87 | 288 | 298 | 290 | |
| MA1 | 0.0178 | 0.0055 | 0 | 278 | 288 | 276 | |
| MA2 | 0.0088 | 0.0182 | 0.0176 | 0 | 153 | 160 | |
| MM1 | 0.0104 | 0.0189 | 0.0182 | 0.0097 | 0 | 160 | |
| MM2 | 0.0097 | 0.0184 | 0.0175 | 0.0101 | 0.0101 | 0 | |
| 188 | | | | | | | 12984 |
| | CR1 | CR2 | MA1 | MA2 | MM1 | MM2 | |
| CR1 | 0 | 110 | 169 | 169 | 169 | 177 | |
| CR2 | 0.0085 | 0 | 253 | 129 | 129 | 104 | |
| MA1 | 0.0130 | 0.0195 | 0 | 241 | 241 | 270 | |
| MA2 | 0.0130 | 0.0099 | 0.0186 | 0 | 0 | 139 | |
| MM1 | 0.0130 | 0.0099 | 0.0186 | 0.0000 | 0 | 139 | |
| MM2 | 0.0136 | 0.0080 | 0.0208 | 0.0107 | 0.0107 | 0 | |

| 189 | | | | | | | 12533 |
|---|---|---|---|---|---|---|---|
| | CR1 | CR2 | MA1 | MA2 | MM1 | MM2 | |
| CR1 | 0 | 161 | 82 | 162 | 82 | 60 | |
| CR2 | 0.0128 | 0 | 159 | 41 | 159 | 164 | |
| MA1 | 0.0065 | 0.0127 | 0 | 154 | 0 | 75 | |
| MA2 | 0.0129 | 0.0033 | 0.0123 | 0 | 154 | 165 | |
| MM1 | 0.0065 | 0.0127 | 0.0000 | 0.0123 | 0 | 75 | |
| MM2 | 0.0048 | 0.0131 | 0.0060 | 0.0132 | 0.0060 | 0 | |
| 197 | | | | | | | 9844 |
| | CR1 | CR2 | MA1 | MA2 | MM1 | MM2 | |
| CR1 | 0 | 203 | 206 | 84 | 103 | 84 | |
| CR2 | 0.0206 | 0 | 100 | 199 | 217 | 199 | |
| MA1 | 0.0209 | 0.0102 | 0 | 198 | 215 | 198 | |
| MA2 | 0.0085 | 0.0202 | 0.0201 | 0 | 123 | 0 | |
| MM1 | 0.0105 | 0.0220 | 0.0218 | 0.0125 | 0 | 123 | |
| MM2 | 0.0085 | 0.0202 | 0.0201 | 0.0000 | 0.0125 | 0 | |
| 198 | | | | | | | 10951 |
| | CR1 | CR2 | MA1 | MA2 | MM1 | MM2 | |
| CR1 | 0 | 195 | 83 | 194 | 52 | 83 | |
| CR2 | 0.0178 | 0 | 207 | 25 | 193 | 207 | |
| MA1 | 0.0076 | 0.0189 | 0 | 206 | 63 | 0 | |
| MA2 | 0.0177 | 0.0023 | 0.0188 | 0 | 194 | 206 | |
| MM1 | 0.0047 | 0.0176 | 0.0058 | 0.0177 | 0 | 63 | |
| MM2 | 0.0076 | 0.0189 | 0.0000 | 0.0188 | 0.0058 | 0 | |
| 199 | | | | | | | 15149 |
| | CR1 | CR2 | MA1 | MA2 | MM1 | MM2 | |
| CR1 | 0 | 310 | 309 | 139 | 169 | 141 | |
| CR2 | 0.0205 | 0 | 81 | 322 | 323 | 325 | |
| MA1 | 0.0204 | 0.0053 | 0 | 319 | 320 | 324 | |
| MA2 | 0.0092 | 0.0213 | 0.0211 | 0 | 166 | 161 | |
| MM1 | 0.0112 | 0.0213 | 0.0211 | 0.0110 | 0 | 161 | |
| MM2 | 0.0093 | 0.0215 | 0.0214 | 0.0106 | 0.0106 | 0 | |
| 201 | | | | | | | 10462 |
| | CR1 | CR2 | MA1 | MA2 | MM1 | MM2 | |
| CR1 | 0 | 176 | 63 | 180 | 63 | 84 | |
| CR2 | 0.0168 | 0 | 185 | 62 | 185 | 180 | |
| MA1 | 0.0060 | 0.0177 | 0 | 193 | 0 | 77 | |
| MA2 | 0.0172 | 0.0059 | 0.0184 | 0 | 193 | 186 | |
| MM1 | 0.0060 | 0.0177 | 0.0000 | 0.0184 | 0 | 77 | |
| MM2 | 0.0080 | 0.0172 | 0.0074 | 0.0178 | 0.0074 | 0 | |

| 203 | | | | | | | 12277 |
|---|---|---|---|---|---|---|---|
| | CR1 | CR2 | MA1 | MA2 | MM1 | MM2 | |
| CR1 | 0 | 151 | 151 | 42 | 149 | 151 | |
| CR2 | 0.0123 | 0 | 0 | 150 | 34 | 0 | |
| MA1 | 0.0123 | 0.0000 | 0 | 150 | 34 | 0 | |
| MA2 | 0.0034 | 0.0122 | 0.0122 | 0 | 149 | 150 | |
| MM1 | 0.0121 | 0.0028 | 0.0028 | 0.0121 | 0 | 34 | |
| MM2 | 0.0123 | 0.0000 | 0.0000 | 0.0122 | 0.0028 | 0 | |
| 204 | | | | | | | 10906 |
| | CR1 | CR2 | MA1 | MA2 | MM1 | MM2 | |
| CR1 | 0 | 247 | 225 | 88 | 236 | 244 | |
| CR2 | 0.0226 | 0 | 157 | 247 | 162 | 141 | |
| MA1 | 0.0206 | 0.0144 | 0 | 232 | 140 | 150 | |
| MA2 | 0.0081 | 0.0226 | 0.0213 | 0 | 238 | 241 | |
| MM1 | 0.0216 | 0.0149 | 0.0128 | 0.0218 | 0 | 128 | |
| MM2 | 0.0224 | 0.0129 | 0.0138 | 0.0221 | 0.0117 | 0 | |
| 205 | | | | | | | 6020 |
| | CR1 | CR2 | MA1 | MA2 | MM1 | MM2 | |
| CR1 | 0 | 96 | 99 | 89 | 69 | 59 | |
| CR2 | 0.0159 | 0 | 5 | 101 | 94 | 90 | |
| MA1 | 0.0164 | 0.0008 | 0 | 104 | 97 | 93 | |
| MA2 | 0.0148 | 0.0168 | 0.0173 | 0 | 50 | 46 | |
| MM1 | 0.0115 | 0.0156 | 0.0161 | 0.0083 | 0 | 14 | |
| MM2 | 0.0098 | 0.0150 | 0.0154 | 0.0076 | 0.0023 | 0 | |
| 206 | | | | | | | 11220 |
| | CR1 | CR2 | MA1 | MA2 | MM1 | MM2 | |
| CR1 | 0 | 93 | 237 | 131 | 256 | 193 | |
| CR2 | 0.0083 | 0 | 264 | 38 | 281 | 260 | |
| MA1 | 0.0211 | 0.0235 | 0 | 270 | 136 | 121 | |
| MA2 | 0.0117 | 0.0034 | 0.0241 | 0 | 288 | 263 | |
| MM1 | 0.0228 | 0.0250 | 0.0121 | 0.0257 | 0 | 133 | |
| MM2 | 0.0172 | 0.0232 | 0.0108 | 0.0234 | 0.0119 | 0 | |
| 207 | | | | | | | 9591 |
| | CR1 | CR2 | MA1 | MA2 | MM1 | MM2 | |
| CR1 | 0 | 102 | 108 | 65 | 0 | 65 | |
| CR2 | 0.0106 | 0 | 38 | 111 | 102 | 111 | |
| MA1 | 0.0113 | 0.0040 | 0 | 111 | 108 | 111 | |
| MA2 | 0.0068 | 0.0116 | 0.0116 | 0 | 65 | 0 | |
| MM1 | 0.0000 | 0.0106 | 0.0113 | 0.0068 | 0 | 65 | |
| MM2 | 0.0068 | 0.0116 | 0.0116 | 0.0000 | 0.0068 | 0 | |

| 208 | | | | | | | 4768 |
|---|---|---|---|---|---|---|---|
| | CR1 | CR2 | MA1 | MA2 | MM1 | MM2 | |
| CR1 | 0 | 70 | 33 | 74 | 36 | 31 | |
| CR2 | 0.0147 | 0 | 70 | 18 | 66 | 55 | |
| MA1 | 0.0069 | 0.0147 | 0 | 74 | 34 | 45 | |
| MA2 | 0.0155 | 0.0038 | 0.0155 | 0 | 70 | 59 | |
| MM1 | 0.0076 | 0.0138 | 0.0071 | 0.0147 | 0 | 47 | |
| MM2 | 0.0065 | 0.0115 | 0.0094 | 0.0124 | 0.0099 | 0 | |
| 209 | | | | | | | 20956 |
| | CR1 | CR2 | MA1 | MA2 | MM1 | MM2 | |
| CR1 | 0 | 219 | 222 | 65 | 65 | 75 | |
| CR2 | 0.0105 | 0 | 50 | 210 | 210 | 206 | |
| MA1 | 0.0106 | 0.0024 | 0 | 213 | 213 | 209 | |
| MA2 | 0.0031 | 0.0100 | 0.0102 | 0 | 0 | 80 | |
| MM1 | 0.0031 | 0.0100 | 0.0102 | 0.0000 | 0 | 80 | |
| MM2 | 0.0036 | 0.0098 | 0.0100 | 0.0038 | 0.0038 | 0 | |
| 210 | | | | | | | 6858 |
| | CR1 | CR2 | MA1 | MA2 | MM1 | MM2 | |
| CR1 | 0 | 45 | 39 | 17 | 17 | 10 | |
| CR2 | 0.0066 | 0 | 16 | 46 | 46 | 45 | |
| MA1 | 0.0057 | 0.0023 | 0 | 34 | 34 | 37 | |
| MA2 | 0.0025 | 0.0067 | 0.0050 | 0 | 0 | 15 | |
| MM1 | 0.0025 | 0.0067 | 0.0050 | 0.0000 | 0 | 15 | |
| MM2 | 0.0015 | 0.0066 | 0.0054 | 0.0022 | 0.0022 | 0 | |
| 211 | | | | | | | 12335 |
| | CR1 | CR2 | MA1 | MA2 | MM1 | MM2 | |
| CR1 | 0 | 294 | 132 | 291 | 119 | 123 | |
| CR2 | 0.0238 | 0 | 256 | 89 | 289 | 287 | |
| MA1 | 0.0107 | 0.0208 | 0 | 233 | 142 | 130 | |
| MA2 | 0.0236 | 0.0072 | 0.0189 | 0 | 288 | 284 | |
| MM1 | 0.0096 | 0.0234 | 0.0115 | 0.0233 | 0 | 113 | |
| MM2 | 0.0100 | 0.0233 | 0.0105 | 0.0230 | 0.0092 | 0 | |
| 212 | | | | | | | 11850 |
| | CR1 | CR2 | MA1 | MA2 | MM1 | MM2 | |
| CR1 | 0 | 254 | 195 | 188 | 158 | 130 | |
| CR2 | 0.0214 | 0 | 127 | 121 | 237 | 228 | |
| MA1 | 0.0165 | 0.0107 | 0 | 180 | 179 | 162 | |
| MA2 | 0.0159 | 0.0102 | 0.0152 | 0 | 202 | 175 | |
| MM1 | 0.0133 | 0.0200 | 0.0151 | 0.0170 | 0 | 132 | |
| MM2 | 0.0110 | 0.0192 | 0.0137 | 0.0148 | 0.0111 | 0 | |

| 213 | | | | | | | 23936 |
|---|---|---|---|---|---|---|---|
| | CR1 | CR2 | MA1 | MA2 | MM1 | MM2 | |
| CR1 | 0 | 529 | 298 | 549 | 299 | 335 | |
| CR2 | 0.0221 | 0 | 510 | 138 | 511 | 516 | |
| MA1 | 0.0124 | 0.0213 | 0 | 520 | 1 | 320 | |
| MA2 | 0.0229 | 0.0058 | 0.0217 | 0 | 521 | 527 | |
| MM1 | 0.0125 | 0.0213 | 0.0000 | 0.0218 | 0 | 319 | |
| MM2 | 0.0140 | 0.0216 | 0.0134 | 0.0220 | 0.0133 | 0 | |
| 216 | | | | | | | 12934 |
| | CR1 | CR2 | MA1 | MA2 | MM1 | MM2 | |
| CR1 | 0 | 135 | 99 | 189 | 200 | 158 | |
| CR2 | 0.0104 | 0 | 178 | 84 | 267 | 230 | |
| MA1 | 0.0077 | 0.0138 | 0 | 133 | 204 | 147 | |
| MA2 | 0.0146 | 0.0065 | 0.0103 | 0 | 271 | 242 | |
| MM1 | 0.0155 | 0.0206 | 0.0158 | 0.0210 | 0 | 213 | |
| MM2 | 0.0122 | 0.0178 | 0.0114 | 0.0187 | 0.0165 | 0 | |
| 217 | | | | | | | 11135 |
| | CR1 | CR2 | MA1 | MA2 | MM1 | MM2 | |
| CR1 | 0 | 273 | 48 | 266 | 141 | 48 | |
| CR2 | 0.0245 | 0 | 262 | 86 | 255 | 262 | |
| MA1 | 0.0043 | 0.0235 | 0 | 259 | 138 | 0 | |
| MA2 | 0.0239 | 0.0077 | 0.0233 | 0 | 250 | 259 | |
| MM1 | 0.0127 | 0.0229 | 0.0124 | 0.0225 | 0 | 138 | |
| MM2 | 0.0043 | 0.0235 | 0.0000 | 0.0233 | 0.0124 | 0 | |
| 218 | | | | | | | 14214 |
| | CR1 | CR2 | MA1 | MA2 | MM1 | MM2 | |
| CR1 | 0 | 298 | 283 | 74 | 301 | 287 | |
| CR2 | 0.0210 | 0 | 73 | 299 | 116 | 134 | |
| MA1 | 0.0199 | 0.0051 | 0 | 276 | 127 | 131 | |
| MA2 | 0.0052 | 0.0210 | 0.0194 | 0 | 294 | 282 | |
| MM1 | 0.0212 | 0.0082 | 0.0089 | 0.0207 | 0 | 132 | |
| MM2 | 0.0202 | 0.0094 | 0.0092 | 0.0198 | 0.0093 | 0 | |
| 220 | | | | | | | 14620 |
| | CR1 | CR2 | MA1 | MA2 | MM1 | MM2 | |
| CR1 | 0 | 307 | 0 | 305 | 0 | 123 | |
| CR2 | 0.0210 | 0 | 307 | 73 | 307 | 302 | |
| MA1 | 0.0000 | 0.0210 | 0 | 305 | 0 | 123 | |
| MA2 | 0.0209 | 0.0050 | 0.0209 | 0 | 305 | 301 | |
| MM1 | 0.0000 | 0.0210 | 0.0000 | 0.0209 | 0 | 123 | |
| MM2 | 0.0084 | 0.0207 | 0.0084 | 0.0206 | 0.0084 | 0 | |

| 222 | | | | | | | 12476 |
|---|---|---|---|---|---|---|---|
| | CR1 | CR2 | MA1 | MA2 | MM1 | MM2 | |
| CR1 | 0 | 156 | 162 | 0 | 162 | 156 | |
| CR2 | 0.0125 | 0 | 95 | 156 | 95 | 97 | |
| MA1 | 0.0130 | 0.0076 | 0 | 162 | 0 | 107 | |
| MA2 | 0.0000 | 0.0125 | 0.0130 | 0 | 162 | 156 | |
| MM1 | 0.0130 | 0.0076 | 0.0000 | 0.0130 | 0 | 107 | |
| MM2 | 0.0125 | 0.0078 | 0.0086 | 0.0125 | 0.0086 | 0 | |
| 223 | | | | | | | 12325 |
| | CR1 | CR2 | MA1 | MA2 | MM1 | MM2 | |
| CR1 | 0 | 288 | 141 | 280 | 163 | 139 | |
| CR2 | 0.0234 | 0 | 272 | 111 | 290 | 264 | |
| MA1 | 0.0114 | 0.0221 | 0 | 266 | 156 | 113 | |
| MA2 | 0.0227 | 0.0090 | 0.0216 | 0 | 279 | 262 | |
| MM1 | 0.0132 | 0.0235 | 0.0127 | 0.0226 | 0 | 148 | |
| MM2 | 0.0113 | 0.0214 | 0.0092 | 0.0213 | 0.0120 | 0 | |
| 224 | | | | | | | 12473 |
| | CR1 | CR2 | MA1 | MA2 | MM1 | MM2 | |
| CR1 | 0 | 195 | 209 | 125 | 148 | 125 | |
| CR2 | 0.0156 | 0 | 72 | 198 | 192 | 198 | |
| MA1 | 0.0168 | 0.0058 | 0 | 210 | 198 | 210 | |
| MA2 | 0.0100 | 0.0159 | 0.0168 | 0 | 159 | 0 | |
| MM1 | 0.0119 | 0.0154 | 0.0159 | 0.0127 | 0 | 159 | |
| MM2 | 0.0100 | 0.0159 | 0.0168 | 0.0000 | 0.0127 | 0 | |
| 227 | | | | | | | 10819 |
| | CR1 | CR2 | MA1 | MA2 | MM1 | MM2 | |
| CR1 | 0 | 267 | 283 | 0 | 146 | 136 | |
| CR2 | 0.0247 | 0 | 59 | 267 | 274 | 273 | |
| MA1 | 0.0262 | 0.0055 | 0 | 283 | 291 | 289 | |
| MA2 | 0.0000 | 0.0247 | 0.0262 | 0 | 146 | 136 | |
| MM1 | 0.0135 | 0.0253 | 0.0269 | 0.0135 | 0 | 101 | |
| MM2 | 0.0126 | 0.0252 | 0.0267 | 0.0126 | 0.0093 | 0 | |
| 228 | | | | | | | 13188 |
| | CR1 | CR2 | MA1 | MA2 | MM1 | MM2 | |
| CR1 | 0 | 244 | 101 | 242 | 116 | 101 | |
| CR2 | 0.0185 | 0 | 245 | 83 | 238 | 245 | |
| MA1 | 0.0077 | 0.0186 | 0 | 241 | 101 | 0 | |
| MA2 | 0.0184 | 0.0063 | 0.0183 | 0 | 236 | 241 | |
| MM1 | 0.0088 | 0.0180 | 0.0077 | 0.0179 | 0 | 101 | |
| MM2 | 0.0077 | 0.0186 | 0.0000 | 0.0183 | 0.0077 | 0 | |

| 231 | | | | | | | 10881 |
|---|---|---|---|---|---|---|---|
| | CR1 | CR2 | MA1 | MA2 | MM1 | MM2 | |
| CR1 | 0 | 196 | 203 | 204 | 38 | 118 | |
| CR2 | 0.0180 | 0 | 55 | 53 | 189 | 185 | |
| MA1 | 0.0187 | 0.0051 | 0 | 4 | 194 | 191 | |
| MA2 | 0.0187 | 0.0049 | 0.0004 | 0 | 195 | 190 | |
| MM1 | 0.0035 | 0.0174 | 0.0178 | 0.0179 | 0 | 116 | |
| MM2 | 0.0108 | 0.0170 | 0.0176 | 0.0175 | 0.0107 | 0 | |

| 232 | | | | | | | 11762 |
|---|---|---|---|---|---|---|---|
| | CR1 | CR2 | MA1 | MA2 | MM1 | MM2 | |
| CR1 | 0 | 294 | 188 | 312 | 188 | 121 | |
| CR2 | 0.0250 | 0 | 293 | 161 | 293 | 304 | |
| MA1 | 0.0160 | 0.0249 | 0 | 303 | 0 | 169 | |
| MA2 | 0.0265 | 0.0137 | 0.0258 | 0 | 303 | 312 | |
| MM1 | 0.0160 | 0.0249 | 0.0000 | 0.0258 | 0 | 169 | |
| MM2 | 0.0103 | 0.0258 | 0.0144 | 0.0265 | 0.0144 | 0 | |

| 233 | | | | | | | 13283 |
|---|---|---|---|---|---|---|---|
| | CR1 | CR2 | MA1 | MA2 | MM1 | MM2 | |
| CR1 | 0 | 247 | 96 | 241 | 37 | 96 | |
| CR2 | 0.0186 | 0 | 236 | 46 | 245 | 236 | |
| MA1 | 0.0072 | 0.0178 | 0 | 230 | 97 | 0 | |
| MA2 | 0.0181 | 0.0035 | 0.0173 | 0 | 239 | 230 | |
| MM1 | 0.0028 | 0.0184 | 0.0073 | 0.0180 | 0 | 97 | |
| MM2 | 0.0072 | 0.0178 | 0.0000 | 0.0173 | 0.0073 | 0 | |

| 235 | | | | | | | 15877 |
|---|---|---|---|---|---|---|---|
| | CR1 | CR2 | MA1 | MA2 | MM1 | MM2 | |
| CR1 | 0 | 360 | 346 | 207 | 171 | 137 | |
| CR2 | 0.0227 | 0 | 98 | 341 | 349 | 372 | |
| MA1 | 0.0218 | 0.0062 | 0 | 340 | 338 | 360 | |
| MA2 | 0.0130 | 0.0215 | 0.0214 | 0 | 210 | 224 | |
| MM1 | 0.0108 | 0.0220 | 0.0213 | 0.0132 | 0 | 160 | |
| MM2 | 0.0086 | 0.0234 | 0.0227 | 0.0141 | 0.0101 | 0 | |

| 237 | | | | | | | 11297 |
|---|---|---|---|---|---|---|---|
| | CR1 | CR2 | MA1 | MA2 | MM1 | MM2 | |
| CR1 | 0 | 234 | 77 | 238 | 233 | 230 | |
| CR2 | 0.0207 | 0 | 228 | 74 | 91 | 74 | |
| MA1 | 0.0068 | 0.0202 | 0 | 244 | 229 | 225 | |
| MA2 | 0.0211 | 0.0066 | 0.0216 | 0 | 95 | 87 | |
| MM1 | 0.0206 | 0.0081 | 0.0203 | 0.0084 | 0 | 87 | |
| MM2 | 0.0204 | 0.0066 | 0.0199 | 0.0077 | 0.0077 | 0 | |

| 239 | | | | | | | 14851 |
|---|---|---|---|---|---|---|---|
| | CR1 | CR2 | MA1 | MA2 | MM1 | MM2 | |
| CR1 | 0 | 365 | 180 | 352 | 190 | 155 | |
| CR2 | 0.0246 | 0 | 379 | 133 | 387 | 366 | |
| MA1 | 0.0121 | 0.0255 | 0 | 365 | 191 | 169 | |
| MA2 | 0.0237 | 0.0090 | 0.0246 | 0 | 374 | 353 | |
| MM1 | 0.0128 | 0.0261 | 0.0129 | 0.0252 | 0 | 147 | |
| MM2 | 0.0104 | 0.0246 | 0.0114 | 0.0238 | 0.0099 | 0 | |
| 240 | | | | | | | 12500 |
| | CR1 | CR2 | MA1 | MA2 | MM1 | MM2 | |
| CR1 | 0 | 248 | 124 | 241 | 148 | 151 | |
| CR2 | 0.0198 | 0 | 227 | 68 | 240 | 262 | |
| MA1 | 0.0099 | 0.0182 | 0 | 222 | 130 | 139 | |
| MA2 | 0.0193 | 0.0054 | 0.0178 | 0 | 234 | 259 | |
| MM1 | 0.0118 | 0.0192 | 0.0104 | 0.0187 | 0 | 148 | |
| MM2 | 0.0121 | 0.0210 | 0.0111 | 0.0207 | 0.0118 | 0 | |
| 242 | | | | | | | 26124 |
| | CR1 | CR2 | MA1 | MA2 | MM1 | MM2 | |
| CR1 | 0 | 612 | 602 | 397 | 346 | 379 | |
| CR2 | 0.0234 | 0 | 201 | 607 | 593 | 608 | |
| MA1 | 0.0230 | 0.0077 | 0 | 617 | 600 | 613 | |
| MA2 | 0.0152 | 0.0232 | 0.0236 | 0 | 418 | 409 | |
| MM1 | 0.0132 | 0.0227 | 0.0230 | 0.0160 | 0 | 327 | |
| MM2 | 0.0145 | 0.0233 | 0.0235 | 0.0157 | 0.0125 | 0 | |
| 243 | | | | | | | 16373 |
| | CR1 | CR2 | MA1 | MA2 | MM1 | MM2 | |
| CR1 | 0 | 291 | 172 | 288 | 172 | 175 | |
| CR2 | 0.0178 | 0 | 286 | 86 | 286 | 278 | |
| MA1 | 0.0105 | 0.0175 | 0 | 292 | 0 | 45 | |
| MA2 | 0.0176 | 0.0053 | 0.0178 | 0 | 292 | 284 | |
| MM1 | 0.0105 | 0.0175 | 0.0000 | 0.0178 | 0 | 45 | |
| MM2 | 0.0107 | 0.0170 | 0.0027 | 0.0173 | 0.0027 | 0 | |
| 245 | | | | | | | 12089 |
| | CR1 | CR2 | MA1 | MA2 | MM1 | MM2 | |
| CR1 | 0 | 235 | 254 | 149 | 142 | 149 | |
| CR2 | 0.0194 | 0 | 87 | 252 | 248 | 252 | |
| MA1 | 0.0210 | 0.0072 | 0 | 257 | 263 | 257 | |
| MA2 | 0.0123 | 0.0208 | 0.0213 | 0 | 133 | 0 | |
| MM1 | 0.0117 | 0.0205 | 0.0218 | 0.0110 | 0 | 133 | |
| MM2 | 0.0123 | 0.0208 | 0.0213 | 0.0000 | 0.0110 | 0 | |

| 246 | | | | | | | 18758 |
|---|---|---|---|---|---|---|---|
| | CR1 | CR2 | MA1 | MA2 | MM1 | MM2 | |
| CR1 | 0 | 492 | 264 | 483 | 246 | 264 | |
| CR2 | 0.0262 | 0 | 473 | 163 | 480 | 473 | |
| MA1 | 0.0141 | 0.0252 | 0 | 422 | 306 | 0 | |
| MA2 | 0.0257 | 0.0087 | 0.0225 | 0 | 500 | 422 | |
| MM1 | 0.0131 | 0.0256 | 0.0163 | 0.0267 | 0 | 306 | |
| MM2 | 0.0141 | 0.0252 | 0.0000 | 0.0225 | 0.0163 | 0 | |
| 247 | | | | | | | 13639 |
| | CR1 | CR2 | MA1 | MA2 | MM1 | MM2 | |
| CR1 | 0 | 267 | 245 | 47 | 274 | 255 | |
| CR2 | 0.0196 | 0 | 95 | 258 | 122 | 107 | |
| MA1 | 0.0180 | 0.0070 | 0 | 239 | 95 | 87 | |
| MA2 | 0.0034 | 0.0189 | 0.0175 | 0 | 266 | 246 | |
| MM1 | 0.0201 | 0.0089 | 0.0070 | 0.0195 | 0 | 100 | |
| MM2 | 0.0187 | 0.0078 | 0.0064 | 0.0180 | 0.0073 | 0 | |
| 248 | | | | | | | 12426 |
| | CR1 | CR2 | MA1 | MA2 | MM1 | MM2 | |
| CR1 | 0 | 209 | 93 | 231 | 99 | 93 | |
| CR2 | 0.0168 | 0 | 202 | 79 | 215 | 202 | |
| MA1 | 0.0075 | 0.0163 | 0 | 229 | 84 | 0 | |
| MA2 | 0.0186 | 0.0064 | 0.0184 | 0 | 238 | 229 | |
| MM1 | 0.0080 | 0.0173 | 0.0068 | 0.0192 | 0 | 84 | |
| MM2 | 0.0075 | 0.0163 | 0.0000 | 0.0184 | 0.0068 | 0 | |
| 249 | | | | | | | 11825 |
| | CR1 | CR2 | MA1 | MA2 | MM1 | MM2 | |
| CR1 | 0 | 156 | 166 | 65 | 166 | 166 | |
| CR2 | 0.0132 | 0 | 102 | 161 | 88 | 102 | |
| MA1 | 0.0140 | 0.0086 | 0 | 171 | 102 | 0 | |
| MA2 | 0.0055 | 0.0136 | 0.0145 | 0 | 168 | 171 | |
| MM1 | 0.0140 | 0.0074 | 0.0086 | 0.0142 | 0 | 102 | |
| MM2 | 0.0140 | 0.0086 | 0.0000 | 0.0145 | 0.0086 | 0 | |
| 250 | | | | | | | 14740 |
| | CR1 | CR2 | MA1 | MA2 | MM1 | MM2 | |
| CR1 | 0 | 60 | 112 | 281 | 112 | 119 | |
| CR2 | 0.0041 | 0 | 124 | 261 | 124 | 127 | |
| MA1 | 0.0076 | 0.0084 | 0 | 283 | 18 | 103 | |
| MA2 | 0.0191 | 0.0177 | 0.0192 | 0 | 283 | 291 | |
| MM1 | 0.0076 | 0.0084 | 0.0012 | 0.0192 | 0 | 85 | |
| MM2 | 0.0081 | 0.0086 | 0.0070 | 0.0197 | 0.0058 | 0 | |

| 251 | | | | | | | 13127 |
|---|---|---|---|---|---|---|---|
| | CR1 | CR2 | MA1 | MA2 | MM1 | MM2 | |
| CR1 | 0 | 262 | 145 | 272 | 155 | 150 | |
| CR2 | 0.0200 | 0 | 284 | 82 | 291 | 290 | |
| MA1 | 0.0110 | 0.0216 | 0 | 297 | 127 | 133 | |
| MA2 | 0.0207 | 0.0062 | 0.0226 | 0 | 304 | 304 | |
| MM1 | 0.0118 | 0.0222 | 0.0097 | 0.0232 | 0 | 85 | |
| MM2 | 0.0114 | 0.0221 | 0.0101 | 0.0232 | 0.0065 | 0 | |
| 252 | | | | | | | 6201 |
| | CR1 | CR2 | MA1 | MA2 | MM1 | MM2 | |
| CR1 | 0 | 100 | 29 | 98 | 109 | 108 | |
| CR2 | 0.0161 | 0 | 95 | 66 | 85 | 66 | |
| MA1 | 0.0047 | 0.0153 | 0 | 93 | 104 | 103 | |
| MA2 | 0.0158 | 0.0106 | 0.0150 | 0 | 69 | 65 | |
| MM1 | 0.0176 | 0.0137 | 0.0168 | 0.0111 | 0 | 78 | |
| MM2 | 0.0174 | 0.0106 | 0.0166 | 0.0105 | 0.0126 | 0 | |
| 253 | | | | | | | 11706 |
| | CR1 | CR2 | MA1 | MA2 | MM1 | MM2 | |
| CR1 | 0 | 152 | 141 | 55 | 143 | 142 | |
| CR2 | 0.0130 | 0 | 88 | 159 | 82 | 81 | |
| MA1 | 0.0120 | 0.0075 | 0 | 146 | 80 | 79 | |
| MA2 | 0.0047 | 0.0136 | 0.0125 | 0 | 146 | 145 | |
| MM1 | 0.0122 | 0.0070 | 0.0068 | 0.0125 | 0 | 1 | |
| MM2 | 0.0121 | 0.0069 | 0.0067 | 0.0124 | 0.0001 | 0 | |
| 256 | | | | | | | 10706 |
| | CR1 | CR2 | MA1 | MA2 | MM1 | MM2 | |
| CR1 | 0 | 174 | 71 | 177 | 61 | 71 | |
| CR2 | 0.0163 | 0 | 179 | 21 | 183 | 179 | |
| MA1 | 0.0066 | 0.0167 | 0 | 182 | 84 | 0 | |
| MA2 | 0.0165 | 0.0020 | 0.0170 | 0 | 186 | 182 | |
| MM1 | 0.0057 | 0.0171 | 0.0078 | 0.0174 | 0 | 84 | |
| MM2 | 0.0066 | 0.0167 | 0.0000 | 0.0170 | 0.0078 | 0 | |
| 257 | | | | | | | 9720 |
| | CR1 | CR2 | MA1 | MA2 | MM1 | MM2 | |
| CR1 | 0 | 202 | 152 | 180 | 148 | 152 | |
| CR2 | 0.0208 | 0 | 150 | 118 | 166 | 150 | |
| MA1 | 0.0156 | 0.0154 | 0 | 195 | 123 | 0 | |
| MA2 | 0.0185 | 0.0121 | 0.0201 | 0 | 202 | 195 | |
| MM1 | 0.0152 | 0.0171 | 0.0127 | 0.0208 | 0 | 123 | |
| MM2 | 0.0156 | 0.0154 | 0.0000 | 0.0201 | 0.0127 | 0 | |

| 258 | | | | | | | 16555 |
|---|---|---|---|---|---|---|---|
| | CR1 | CR2 | MA1 | MA2 | MM1 | MM2 | |
| CR1 | 0 | 324 | 325 | 156 | 139 | 0 | |
| CR2 | 0.0196 | 0 | 56 | 315 | 307 | 324 | |
| MA1 | 0.0196 | 0.0034 | 0 | 317 | 306 | 325 | |
| MA2 | 0.0094 | 0.0190 | 0.0191 | 0 | 147 | 156 | |
| MM1 | 0.0084 | 0.0185 | 0.0185 | 0.0089 | 0 | 139 | |
| MM2 | 0.0000 | 0.0196 | 0.0196 | 0.0094 | 0.0084 | 0 | |
| 259 | | | | | | | 9510 |
| | CR1 | CR2 | MA1 | MA2 | MM1 | MM2 | |
| CR1 | 0 | 288 | 260 | 161 | 149 | 199 | |
| CR2 | 0.0303 | 0 | 136 | 283 | 291 | 298 | |
| MA1 | 0.0273 | 0.0143 | 0 | 237 | 272 | 271 | |
| MA2 | 0.0169 | 0.0298 | 0.0249 | 0 | 209 | 215 | |
| MM1 | 0.0157 | 0.0306 | 0.0286 | 0.0220 | 0 | 162 | |
| MM2 | 0.0209 | 0.0313 | 0.0285 | 0.0226 | 0.0170 | 0 | |
| 261 | | | | | | | 9311 |
| | CR1 | CR2 | MA1 | MA2 | MM1 | MM2 | |
| CR1 | 0 | 146 | 77 | 138 | 78 | 78 | |
| CR2 | 0.0157 | 0 | 122 | 75 | 129 | 136 | |
| MA1 | 0.0083 | 0.0131 | 0 | 116 | 7 | 89 | |
| MA2 | 0.0148 | 0.0081 | 0.0125 | 0 | 123 | 132 | |
| MM1 | 0.0084 | 0.0139 | 0.0008 | 0.0132 | 0 | 90 | |
| MM2 | 0.0084 | 0.0146 | 0.0096 | 0.0142 | 0.0097 | 0 | |
| 262 | | | | | | | 10784 |
| | CR1 | CR2 | MA1 | MA2 | MM1 | MM2 | |
| CR1 | 0 | 149 | 62 | 153 | 55 | 62 | |
| CR2 | 0.0138 | 0 | 159 | 54 | 153 | 159 | |
| MA1 | 0.0057 | 0.0147 | 0 | 162 | 63 | 0 | |
| MA2 | 0.0142 | 0.0050 | 0.0150 | 0 | 154 | 162 | |
| MM1 | 0.0051 | 0.0142 | 0.0058 | 0.0143 | 0 | 63 | |
| MM2 | 0.0057 | 0.0147 | 0.0000 | 0.0150 | 0.0058 | 0 | |
| 264 | | | | | | | 13931 |
| | CR1 | CR2 | MA1 | MA2 | MM1 | MM2 | |
| CR1 | 0 | 146 | 47 | 133 | 47 | 52 | |
| CR2 | 0.0105 | 0 | 144 | 25 | 144 | 145 | |
| MA1 | 0.0034 | 0.0103 | 0 | 131 | 0 | 54 | |
| MA2 | 0.0095 | 0.0018 | 0.0094 | 0 | 131 | 132 | |
| MM1 | 0.0034 | 0.0103 | 0.0000 | 0.0094 | 0 | 54 | |
| MM2 | 0.0037 | 0.0104 | 0.0039 | 0.0095 | 0.0039 | 0 | |

| 265 | | | | | | | 8721 |
|---|---|---|---|---|---|---|---|
| | CR1 | CR2 | MA1 | MA2 | MM1 | MM2 | |
| CR1 | 0 | 131 | 158 | 249 | 125 | 122 | |
| CR2 | 0.0150 | 0 | 186 | 124 | 191 | 162 | |
| MA1 | 0.0181 | 0.0213 | 0 | 246 | 161 | 78 | |
| MA2 | 0.0286 | 0.0142 | 0.0282 | 0 | 234 | 242 | |
| MM1 | 0.0143 | 0.0219 | 0.0185 | 0.0268 | 0 | 143 | |
| MM2 | 0.0140 | 0.0186 | 0.0089 | 0.0277 | 0.0164 | 0 | |

| 266 | | | | | | | 16146 |
|---|---|---|---|---|---|---|---|
| | CR1 | CR2 | MA1 | MA2 | MM1 | MM2 | |
| CR1 | 0 | 210 | 215 | 50 | 227 | 215 | |
| CR2 | 0.0130 | 0 | 23 | 231 | 63 | 23 | |
| MA1 | 0.0133 | 0.0014 | 0 | 236 | 68 | 0 | |
| MA2 | 0.0031 | 0.0143 | 0.0146 | 0 | 248 | 236 | |
| MM1 | 0.0141 | 0.0039 | 0.0042 | 0.0154 | 0 | 68 | |
| MM2 | 0.0133 | 0.0014 | 0.0000 | 0.0146 | 0.0042 | 0 | |

| 267 | | | | | | | 17862 |
|---|---|---|---|---|---|---|---|
| | CR1 | CR2 | MA1 | MA2 | MM1 | MM2 | |
| CR1 | 0 | 412 | 439 | 202 | 193 | 184 | |
| CR2 | 0.0231 | 0 | 54 | 427 | 427 | 426 | |
| MA1 | 0.0246 | 0.0030 | 0 | 450 | 452 | 449 | |
| MA2 | 0.0113 | 0.0239 | 0.0252 | 0 | 190 | 78 | |
| MM1 | 0.0108 | 0.0239 | 0.0253 | 0.0106 | 0 | 190 | |
| MM2 | 0.0103 | 0.0238 | 0.0251 | 0.0044 | 0.0106 | 0 | |

| 268 | | | | | | | 11333 |
|---|---|---|---|---|---|---|---|
| | CR1 | CR2 | MA1 | MA2 | MM1 | MM2 | |
| CR1 | 0 | 85 | 42 | 90 | 42 | 33 | |
| CR2 | 0.0075 | 0 | 88 | 27 | 88 | 81 | |
| MA1 | 0.0037 | 0.0078 | 0 | 93 | 0 | 33 | |
| MA2 | 0.0079 | 0.0024 | 0.0082 | 0 | 93 | 86 | |
| MM1 | 0.0037 | 0.0078 | 0.0000 | 0.0082 | 0 | 33 | |
| MM2 | 0.0029 | 0.0071 | 0.0029 | 0.0076 | 0.0029 | 0 | |

| 270 | | | | | | | 11444 |
|---|---|---|---|---|---|---|---|
| | CR1 | CR2 | MA1 | MA2 | MM1 | MM2 | |
| CR1 | 0 | 121 | 39 | 124 | 59 | 39 | |
| CR2 | 0.0106 | 0 | 127 | 32 | 126 | 127 | |
| MA1 | 0.0034 | 0.0111 | 0 | 128 | 52 | 0 | |
| MA2 | 0.0108 | 0.0028 | 0.0112 | 0 | 127 | 128 | |
| MM1 | 0.0052 | 0.0110 | 0.0045 | 0.0111 | 0 | 52 | |
| MM2 | 0.0034 | 0.0111 | 0.0000 | 0.0112 | 0.0045 | 0 | |

| 273 | | | | | | | 12242 |
|---|---|---|---|---|---|---|---|
| | CR1 | CR2 | MA1 | MA2 | MM1 | MM2 | |
| CR1 | 0 | 214 | 95 | 212 | 95 | 85 | |
| CR2 | 0.0175 | 0 | 214 | 40 | 214 | 184 | |
| MA1 | 0.0078 | 0.0175 | 0 | 201 | 0 | 80 | |
| MA2 | 0.0173 | 0.0033 | 0.0164 | 0 | 201 | 184 | |
| MM1 | 0.0078 | 0.0175 | 0.0000 | 0.0164 | 0 | 80 | |
| MM2 | 0.0069 | 0.0150 | 0.0065 | 0.0150 | 0.0065 | 0 | |
| 275 | | | | | | | 11688 |
| | CR1 | CR2 | MA1 | MA2 | MM1 | MM2 | |
| CR1 | 0 | 286 | 156 | 286 | 149 | 156 | |
| CR2 | 0.0245 | 0 | 290 | 0 | 284 | 290 | |
| MA1 | 0.0133 | 0.0248 | 0 | 290 | 165 | 0 | |
| MA2 | 0.0245 | 0.0000 | 0.0248 | 0 | 284 | 290 | |
| MM1 | 0.0127 | 0.0243 | 0.0141 | 0.0243 | 0 | 165 | |
| MM2 | 0.0133 | 0.0248 | 0.0000 | 0.0248 | 0.0141 | 0 | |
| 276 | | | | | | | 9082 |
| | CR1 | CR2 | MA1 | MA2 | MM1 | MM2 | |
| CR1 | 0 | 102 | 109 | 13 | 109 | 113 | |
| CR2 | 0.0112 | 0 | 25 | 102 | 25 | 24 | |
| MA1 | 0.0120 | 0.0028 | 0 | 109 | 0 | 29 | |
| MA2 | 0.0014 | 0.0112 | 0.0120 | 0 | 109 | 113 | |
| MM1 | 0.0120 | 0.0028 | 0.0000 | 0.0120 | 0 | 29 | |
| MM2 | 0.0124 | 0.0026 | 0.0032 | 0.0124 | 0.0032 | 0 | |
| 277 | | | | | | | 10843 |
| | CR1 | CR2 | MA1 | MA2 | MM1 | MM2 | |
| CR1 | 0 | 183 | 16 | 183 | 69 | 16 | |
| CR2 | 0.0169 | 0 | 187 | 48 | 186 | 187 | |
| MA1 | 0.0015 | 0.0172 | 0 | 185 | 73 | 0 | |
| MA2 | 0.0169 | 0.0044 | 0.0171 | 0 | 182 | 185 | |
| MM1 | 0.0064 | 0.0172 | 0.0067 | 0.0168 | 0 | 73 | |
| MM2 | 0.0015 | 0.0172 | 0.0000 | 0.0171 | 0.0067 | 0 | |
| 278 | | | | | | | 15048 |
| | CR1 | CR2 | MA1 | MA2 | MM1 | MM2 | |
| CR1 | 0 | 339 | 278 | 286 | 194 | 193 | |
| CR2 | 0.0225 | 0 | 227 | 194 | 331 | 321 | |
| MA1 | 0.0185 | 0.0151 | 0 | 129 | 287 | 259 | |
| MA2 | 0.0190 | 0.0129 | 0.0086 | 0 | 281 | 264 | |
| MM1 | 0.0129 | 0.0220 | 0.0191 | 0.0187 | 0 | 187 | |
| MM2 | 0.0128 | 0.0213 | 0.0172 | 0.0175 | 0.0124 | 0 | |

| 279 | | | | | | | 12895 |
|---|---|---|---|---|---|---|---|
| | CR1 | CR2 | MA1 | MA2 | MM1 | MM2 | |
| CR1 | 0 | 264 | 264 | 93 | 254 | 248 | |
| CR2 | 0.0205 | 0 | 155 | 277 | 125 | 138 | |
| MA1 | 0.0205 | 0.0120 | 0 | 279 | 130 | 153 | |
| MA2 | 0.0072 | 0.0215 | 0.0216 | 0 | 265 | 269 | |
| MM1 | 0.0197 | 0.0097 | 0.0101 | 0.0206 | 0 | 150 | |
| MM2 | 0.0192 | 0.0107 | 0.0119 | 0.0209 | 0.0116 | 0 | |
| 280 | | | | | | | 9542 |
| | CR1 | CR2 | MA1 | MA2 | MM1 | MM2 | |
| CR1 | 0 | 146 | 78 | 153 | 62 | 49 | |
| CR2 | 0.0153 | 0 | 116 | 33 | 157 | 144 | |
| MA1 | 0.0082 | 0.0122 | 0 | 111 | 101 | 40 | |
| MA2 | 0.0160 | 0.0035 | 0.0116 | 0 | 164 | 151 | |
| MM1 | 0.0065 | 0.0165 | 0.0106 | 0.0172 | 0 | 76 | |
| MM2 | 0.0051 | 0.0151 | 0.0042 | 0.0158 | 0.0080 | 0 | |
| 281 | | | | | | | 16255 |
| | CR1 | CR2 | MA1 | MA2 | MM1 | MM2 | |
| CR1 | 0 | 393 | 145 | 401 | 137 | 190 | |
| CR2 | 0.0242 | 0 | 369 | 150 | 374 | 401 | |
| MA1 | 0.0089 | 0.0227 | 0 | 377 | 170 | 194 | |
| MA2 | 0.0247 | 0.0092 | 0.0232 | 0 | 385 | 408 | |
| MM1 | 0.0084 | 0.0230 | 0.0105 | 0.0237 | 0 | 208 | |
| MM2 | 0.0117 | 0.0247 | 0.0119 | 0.0251 | 0.0128 | 0 | |
| 282 | | | | | | | 15093 |
| | CR1 | CR2 | MA1 | MA2 | MM1 | MM2 | |
| CR1 | 0 | 275 | 265 | 85 | 273 | 269 | |
| CR2 | 0.0182 | 0 | 158 | 284 | 99 | 123 | |
| MA1 | 0.0176 | 0.0105 | 0 | 275 | 145 | 140 | |
| MA2 | 0.0056 | 0.0188 | 0.0182 | 0 | 279 | 276 | |
| MM1 | 0.0181 | 0.0066 | 0.0096 | 0.0185 | 0 | 108 | |
| MM2 | 0.0178 | 0.0081 | 0.0093 | 0.0183 | 0.0072 | 0 | |
| 283 | | | | | | | 16951 |
| | CR1 | CR2 | MA1 | MA2 | MM1 | MM2 | |
| CR1 | 0 | 471 | 471 | 413 | 325 | 340 | |
| CR2 | 0.0278 | 0 | 180 | 232 | 444 | 430 | |
| MA1 | 0.0278 | 0.0106 | 0 | 170 | 404 | 431 | |
| MA2 | 0.0244 | 0.0137 | 0.0100 | 0 | 262 | 363 | |
| MM1 | 0.0192 | 0.0262 | 0.0238 | 0.0155 | 0 | 300 | |
| MM2 | 0.0201 | 0.0254 | 0.0254 | 0.0214 | 0.0177 | 0 | |

| 284 | | | | | | | 19766 |
|---|---|---|---|---|---|---|---|
| | CR1 | CR2 | MA1 | MA2 | MM1 | MM2 | |
| CR1 | 0 | 207 | 319 | 119 | 316 | 325 | |
| CR2 | 0.0105 | 0 | 212 | 257 | 194 | 227 | |
| MA1 | 0.0161 | 0.0107 | 0 | 332 | 172 | 137 | |
| MA2 | 0.0060 | 0.0130 | 0.0168 | 0 | 331 | 339 | |
| MM1 | 0.0160 | 0.0098 | 0.0087 | 0.0167 | 0 | 174 | |
| MM2 | 0.0164 | 0.0115 | 0.0069 | 0.0172 | 0.0088 | 0 | |
| 287 | | | | | | | 22865 |
| | CR1 | CR2 | MA1 | MA2 | MM1 | MM2 | |
| CR1 | 0 | 539 | 250 | 537 | 251 | 240 | |
| CR2 | 0.0236 | 0 | 541 | 162 | 569 | 554 | |
| MA1 | 0.0109 | 0.0237 | 0 | 535 | 261 | 242 | |
| MA2 | 0.0235 | 0.0071 | 0.0234 | 0 | 562 | 550 | |
| MM1 | 0.0110 | 0.0249 | 0.0114 | 0.0246 | 0 | 267 | |
| MM2 | 0.0105 | 0.0242 | 0.0106 | 0.0241 | 0.0117 | 0 | |
| 290 | | | | | | | 8590 |
| | CR1 | CR2 | MA1 | MA2 | MM1 | MM2 | |
| CR1 | 0 | 161 | 38 | 35 | 74 | 81 | |
| CR2 | 0.0187 | 0 | 132 | 132 | 146 | 156 | |
| MA1 | 0.0044 | 0.0154 | 0 | 9 | 82 | 94 | |
| MA2 | 0.0041 | 0.0154 | 0.0010 | 0 | 85 | 91 | |
| MM1 | 0.0086 | 0.0170 | 0.0095 | 0.0099 | 0 | 71 | |
| MM2 | 0.0094 | 0.0182 | 0.0109 | 0.0106 | 0.0083 | 0 | |
| 291 | | | | | | | 11158 |
| | CR1 | CR2 | MA1 | MA2 | MM1 | MM2 | |
| CR1 | 0 | 340 | 292 | 77 | 356 | 338 | |
| CR2 | 0.0305 | 0 | 54 | 322 | 209 | 6 | |
| MA1 | 0.0262 | 0.0048 | 0 | 268 | 245 | 52 | |
| MA2 | 0.0069 | 0.0289 | 0.0240 | 0 | 340 | 320 | |
| MM1 | 0.0319 | 0.0187 | 0.0220 | 0.0305 | 0 | 205 | |
| MM2 | 0.0303 | 0.0005 | 0.0047 | 0.0287 | 0.0184 | 0 | |
| 297 | | | | | | | 13382 |
| | CR1 | CR2 | MA1 | MA2 | MM1 | MM2 | |
| CR1 | 0 | 292 | 100 | 294 | 100 | 83 | |
| CR2 | 0.0218 | 0 | 290 | 89 | 290 | 290 | |
| MA1 | 0.0075 | 0.0217 | 0 | 293 | 0 | 142 | |
| MA2 | 0.0220 | 0.0067 | 0.0219 | 0 | 293 | 293 | |
| MM1 | 0.0075 | 0.0217 | 0.0000 | 0.0219 | 0 | 142 | |
| MM2 | 0.0062 | 0.0217 | 0.0106 | 0.0219 | 0.0106 | 0 | |

| 300 | | | | | | | 12407 |
|---|---|---|---|---|---|---|---|
| | CR1 | CR2 | MA1 | MA2 | MM1 | MM2 | |
| CR1 | 0 | 123 | 159 | 167 | 0 | 99 | |
| CR2 | 0.0099 | 0 | 134 | 144 | 123 | 154 | |
| MA1 | 0.0128 | 0.0108 | 0 | 45 | 159 | 121 | |
| MA2 | 0.0135 | 0.0116 | 0.0036 | 0 | 167 | 127 | |
| MM1 | 0.0000 | 0.0099 | 0.0128 | 0.0135 | 0 | 99 | |
| MM2 | 0.0080 | 0.0124 | 0.0098 | 0.0102 | 0.0080 | 0 | |
| 302 | | | | | | | 11498 |
| | CR1 | CR2 | MA1 | MA2 | MM1 | MM2 | |
| CR1 | 0 | 224 | 142 | 223 | 142 | 137 | |
| CR2 | 0.0195 | 0 | 225 | 63 | 225 | 208 | |
| MA1 | 0.0123 | 0.0196 | 0 | 220 | 0 | 115 | |
| MA2 | 0.0194 | 0.0055 | 0.0191 | 0 | 220 | 208 | |
| MM1 | 0.0123 | 0.0196 | 0.0000 | 0.0191 | 0 | 115 | |
| MM2 | 0.0119 | 0.0181 | 0.0100 | 0.0181 | 0.0100 | 0 | |
| 303 | | | | | | | 9258 |
| | CR1 | CR2 | MA1 | MA2 | MM1 | MM2 | |
| CR1 | 0 | 137 | 27 | 132 | 27 | 53 | |
| CR2 | 0.0148 | 0 | 128 | 48 | 128 | 128 | |
| MA1 | 0.0029 | 0.0138 | 0 | 123 | 0 | 46 | |
| MA2 | 0.0143 | 0.0052 | 0.0133 | 0 | 123 | 127 | |
| MM1 | 0.0029 | 0.0138 | 0.0000 | 0.0133 | 0 | 46 | |
| MM2 | 0.0057 | 0.0138 | 0.0050 | 0.0137 | 0.0050 | 0 | |
| 304 | | | | | | | 17409 |
| | CR1 | CR2 | MA1 | MA2 | MM1 | MM2 | |
| CR1 | 0 | 361 | 198 | 355 | 200 | 198 | |
| CR2 | 0.0207 | 0 | 325 | 44 | 346 | 329 | |
| MA1 | 0.0114 | 0.0187 | 0 | 319 | 178 | 18 | |
| MA2 | 0.0204 | 0.0025 | 0.0183 | 0 | 341 | 323 | |
| MM1 | 0.0115 | 0.0199 | 0.0102 | 0.0196 | 0 | 180 | |
| MM2 | 0.0114 | 0.0189 | 0.0010 | 0.0186 | 0.0103 | 0 | |
| 308 | | | | | | | 13935 |
| | CR1 | CR2 | MA1 | MA2 | MM1 | MM2 | |
| CR1 | 0 | 195 | 104 | 141 | 31 | 86 | |
| CR2 | 0.0140 | 0 | 193 | 107 | 201 | 205 | |
| MA1 | 0.0075 | 0.0139 | 0 | 102 | 98 | 24 | |
| MA2 | 0.0101 | 0.0077 | 0.0073 | 0 | 147 | 126 | |
| MM1 | 0.0022 | 0.0144 | 0.0070 | 0.0105 | 0 | 81 | |
| MM2 | 0.0062 | 0.0147 | 0.0017 | 0.0090 | 0.0058 | 0 | |

| 311 | | | | | | | 10664 |
|---|---|---|---|---|---|---|---|
| | CR1 | CR2 | MA1 | MA2 | MM1 | MM2 | |
| CR1 | 0 | 144 | 48 | 148 | 49 | 48 | |
| CR2 | 0.0135 | 0 | 135 | 53 | 132 | 135 | |
| MA1 | 0.0045 | 0.0127 | 0 | 138 | 21 | 0 | |
| MA2 | 0.0139 | 0.0050 | 0.0129 | 0 | 135 | 138 | |
| MM1 | 0.0046 | 0.0124 | 0.0020 | 0.0127 | 0 | 21 | |
| MM2 | 0.0045 | 0.0127 | 0.0000 | 0.0129 | 0.0020 | 0 | |
| 312 | | | | | | | 12618 |
| | CR1 | CR2 | MA1 | MA2 | MM1 | MM2 | |
| CR1 | 0 | 168 | 70 | 171 | 70 | 75 | |
| CR2 | 0.0133 | 0 | 155 | 35 | 155 | 153 | |
| MA1 | 0.0055 | 0.0123 | 0 | 160 | 0 | 49 | |
| MA2 | 0.0136 | 0.0028 | 0.0127 | 0 | 160 | 156 | |
| MM1 | 0.0055 | 0.0123 | 0.0000 | 0.0127 | 0 | 49 | |
| MM2 | 0.0059 | 0.0121 | 0.0039 | 0.0124 | 0.0039 | 0 | |
| 313 | | | | | | | 11017 |
| | CR1 | CR2 | MA1 | MA2 | MM1 | MM2 | |
| CR1 | 0 | 87 | 145 | 162 | 221 | 238 | |
| CR2 | 0.0079 | 0 | 195 | 144 | 239 | 251 | |
| MA1 | 0.0132 | 0.0177 | 0 | 74 | 141 | 199 | |
| MA2 | 0.0147 | 0.0131 | 0.0067 | 0 | 215 | 243 | |
| MM1 | 0.0201 | 0.0217 | 0.0128 | 0.0195 | 0 | 78 | |
| MM2 | 0.0216 | 0.0228 | 0.0181 | 0.0221 | 0.0071 | 0 | |
| 315 | | | | | | | 14756 |
| | CR1 | CR2 | MA1 | MA2 | MM1 | MM2 | |
| CR1 | 0 | 177 | 59 | 175 | 48 | 59 | |
| CR2 | 0.0120 | 0 | 170 | 25 | 172 | 170 | |
| MA1 | 0.0040 | 0.0115 | 0 | 168 | 57 | 0 | |
| MA2 | 0.0119 | 0.0017 | 0.0114 | 0 | 170 | 168 | |
| MM1 | 0.0033 | 0.0117 | 0.0039 | 0.0115 | 0 | 57 | |
| MM2 | 0.0040 | 0.0115 | 0.0000 | 0.0114 | 0.0039 | 0 | |
| 316 | | | | | | | 14582 |
| | CR1 | CR2 | MA1 | MA2 | MM1 | MM2 | |
| CR1 | 0 | 184 | 54 | 183 | 194 | 183 | |
| CR2 | 0.0126 | 0 | 198 | 94 | 82 | 94 | |
| MA1 | 0.0037 | 0.0136 | 0 | 195 | 208 | 195 | |
| MA2 | 0.0125 | 0.0064 | 0.0134 | 0 | 89 | 0 | |
| MM1 | 0.0133 | 0.0056 | 0.0143 | 0.0061 | 0 | 89 | |
| MM2 | 0.0125 | 0.0064 | 0.0134 | 0.0000 | 0.0061 | 0 | |

| 317 | | | | | | | 12769 |
|---|---|---|---|---|---|---|---|
| | CR1 | CR2 | MA1 | MA2 | MM1 | MM2 | |
| CR1 | 0 | 297 | 152 | 303 | 165 | 152 | |
| CR2 | 0.0233 | 0 | 295 | 88 | 315 | 295 | |
| MA1 | 0.0119 | 0.0231 | 0 | 310 | 152 | 0 | |
| MA2 | 0.0237 | 0.0069 | 0.0243 | 0 | 324 | 310 | |
| MM1 | 0.0129 | 0.0247 | 0.0119 | 0.0254 | 0 | 152 | |
| MM2 | 0.0119 | 0.0231 | 0.0000 | 0.0243 | 0.0119 | 0 | |
| 318 | | | | | | | 13424 |
| | CR1 | CR2 | MA1 | MA2 | MM1 | MM2 | |
| CR1 | 0 | 271 | 250 | 51 | 266 | 280 | |
| CR2 | 0.0202 | 0 | 122 | 263 | 120 | 114 | |
| MA1 | 0.0186 | 0.0091 | 0 | 236 | 142 | 36 | |
| MA2 | 0.0038 | 0.0196 | 0.0176 | 0 | 258 | 272 | |
| MM1 | 0.0198 | 0.0089 | 0.0106 | 0.0192 | 0 | 128 | |
| MM2 | 0.0209 | 0.0085 | 0.0027 | 0.0203 | 0.0095 | 0 | |
| 319 | | | | | | | 14205 |
| | CR1 | CR2 | MA1 | MA2 | MM1 | MM2 | |
| CR1 | 0 | 175 | 176 | 95 | 95 | 87 | |
| CR2 | 0.0123 | 0 | 31 | 156 | 176 | 165 | |
| MA1 | 0.0124 | 0.0022 | 0 | 149 | 177 | 170 | |
| MA2 | 0.0067 | 0.0110 | 0.0105 | 0 | 97 | 21 | |
| MM1 | 0.0067 | 0.0124 | 0.0125 | 0.0068 | 0 | 87 | |
| MM2 | 0.0061 | 0.0116 | 0.0120 | 0.0015 | 0.0061 | 0 | |
| 320 | | | | | | | 10719 |
| | CR1 | CR2 | MA1 | MA2 | MM1 | MM2 | |
| CR1 | 0 | 130 | 93 | 151 | 91 | 89 | |
| CR2 | 0.0121 | 0 | 127 | 61 | 140 | 153 | |
| MA1 | 0.0087 | 0.0118 | 0 | 143 | 80 | 83 | |
| MA2 | 0.0141 | 0.0057 | 0.0133 | 0 | 158 | 165 | |
| MM1 | 0.0085 | 0.0131 | 0.0075 | 0.0147 | 0 | 71 | |
| MM2 | 0.0083 | 0.0143 | 0.0077 | 0.0154 | 0.0066 | 0 | |
| 322 | | | | | | | 10461 |
| | CR1 | CR2 | MA1 | MA2 | MM1 | MM2 | |
| CR1 | 0 | 164 | 173 | 101 | 101 | 116 | |
| CR2 | 0.0157 | 0 | 39 | 172 | 172 | 170 | |
| MA1 | 0.0165 | 0.0037 | 0 | 175 | 175 | 177 | |
| MA2 | 0.0097 | 0.0164 | 0.0167 | 0 | 0 | 58 | |
| MM1 | 0.0097 | 0.0164 | 0.0167 | 0.0000 | 0 | 58 | |
| MM2 | 0.0111 | 0.0163 | 0.0169 | 0.0055 | 0.0055 | 0 | |

324 10252

| | CR1 | CR2 | MA1 | MA2 | MM1 | MM2 |
|---|---|---|---|---|---|---|
| CR1 | 0 | 217 | 197 | 134 | 162 | 134 |
| CR2 | 0.0212 | 0 | 71 | 198 | 212 | 198 |
| MA1 | 0.0192 | 0.0069 | 0 | 174 | 188 | 174 |
| MA2 | 0.0131 | 0.0193 | 0.0170 | 0 | 128 | 0 |
| MM1 | 0.0158 | 0.0207 | 0.0183 | 0.0125 | 0 | 128 |
| MM2 | 0.0131 | 0.0193 | 0.0170 | 0.0000 | 0.0125 | 0 |

326 15107

| | CR1 | CR2 | MA1 | MA2 | MM1 | MM2 |
|---|---|---|---|---|---|---|
| CR1 | 0 | 201 | 67 | 200 | 199 | 200 |
| CR2 | 0.0133 | 0 | 186 | 72 | 73 | 72 |
| MA1 | 0.0044 | 0.0123 | 0 | 189 | 188 | 189 |
| MA2 | 0.0132 | 0.0048 | 0.0125 | 0 | 81 | 0 |
| MM1 | 0.0132 | 0.0048 | 0.0124 | 0.0054 | 0 | 81 |
| MM2 | 0.0132 | 0.0048 | 0.0125 | 0.0000 | 0.0054 | 0 |

329 17384

| | CR1 | CR2 | MA1 | MA2 | MM1 | MM2 |
|---|---|---|---|---|---|---|
| CR1 | 0 | 357 | 213 | 353 | 231 | 226 |
| CR2 | 0.0205 | 0 | 337 | 100 | 346 | 344 |
| MA1 | 0.0123 | 0.0194 | 0 | 331 | 102 | 175 |
| MA2 | 0.0203 | 0.0058 | 0.0190 | 0 | 335 | 344 |
| MM1 | 0.0133 | 0.0199 | 0.0059 | 0.0193 | 0 | 176 |
| MM2 | 0.0130 | 0.0198 | 0.0101 | 0.0198 | 0.0101 | 0 |

334 6212

| | CR1 | CR2 | MA1 | MA2 | MM1 | MM2 |
|---|---|---|---|---|---|---|
| CR1 | 0 | 150 | 122 | 74 | 71 | 66 |
| CR2 | 0.0241 | 0 | 72 | 127 | 132 | 144 |
| MA1 | 0.0196 | 0.0116 | 0 | 95 | 86 | 112 |
| MA2 | 0.0119 | 0.0204 | 0.0153 | 0 | 9 | 68 |
| MM1 | 0.0114 | 0.0212 | 0.0138 | 0.0014 | 0 | 61 |
| MM2 | 0.0106 | 0.0232 | 0.0180 | 0.0109 | 0.0098 | 0 |

335 11776

| | CR1 | CR2 | MA1 | MA2 | MM1 | MM2 |
|---|---|---|---|---|---|---|
| CR1 | 0 | 227 | 218 | 47 | 205 | 222 |
| CR2 | 0.0193 | 0 | 141 | 226 | 132 | 146 |
| MA1 | 0.0185 | 0.0120 | 0 | 220 | 124 | 143 |
| MA2 | 0.0040 | 0.0192 | 0.0187 | 0 | 199 | 224 |
| MM1 | 0.0174 | 0.0112 | 0.0105 | 0.0169 | 0 | 109 |
| MM2 | 0.0189 | 0.0124 | 0.0121 | 0.0190 | 0.0093 | 0 |

| 336 | | | | | | | 8843 |
|---|---|---|---|---|---|---|---|
| | CR1 | CR2 | MA1 | MA2 | MM1 | MM2 | |
| CR1 | 0 | 118 | 17 | 117 | 2 | 1 | |
| CR2 | 0.0133 | 0 | 120 | 33 | 116 | 117 | |
| MA1 | 0.0019 | 0.0136 | 0 | 119 | 17 | 16 | |
| MA2 | 0.0132 | 0.0037 | 0.0135 | 0 | 115 | 116 | |
| MM1 | 0.0002 | 0.0131 | 0.0019 | 0.0130 | 0 | 3 | |
| MM2 | 0.0001 | 0.0132 | 0.0018 | 0.0131 | 0.0003 | 0 | |

| 337 | | | | | | | 12346 |
|---|---|---|---|---|---|---|---|
| | CR1 | CR2 | MA1 | MA2 | MM1 | MM2 | |
| CR1 | 0 | 126 | 173 | 97 | 165 | 173 | |
| CR2 | 0.0102 | 0 | 67 | 201 | 106 | 69 | |
| MA1 | 0.0140 | 0.0054 | 0 | 190 | 97 | 64 | |
| MA2 | 0.0079 | 0.0163 | 0.0154 | 0 | 192 | 199 | |
| MM1 | 0.0134 | 0.0086 | 0.0079 | 0.0156 | 0 | 113 | |
| MM2 | 0.0140 | 0.0056 | 0.0052 | 0.0161 | 0.0092 | 0 | |

| 342 | | | | | | | 12842 |
|---|---|---|---|---|---|---|---|
| | CR1 | CR2 | MA1 | MA2 | MM1 | MM2 | |
| CR1 | 0 | 277 | 172 | 271 | 164 | 185 | |
| CR2 | 0.0216 | 0 | 283 | 10 | 290 | 302 | |
| MA1 | 0.0134 | 0.0220 | 0 | 277 | 163 | 207 | |
| MA2 | 0.0211 | 0.0008 | 0.0216 | 0 | 284 | 296 | |
| MM1 | 0.0128 | 0.0226 | 0.0127 | 0.0221 | 0 | 152 | |
| MM2 | 0.0144 | 0.0235 | 0.0161 | 0.0230 | 0.0118 | 0 | |

| 343 | | | | | | | 15883 |
|---|---|---|---|---|---|---|---|
| | CR1 | CR2 | MA1 | MA2 | MM1 | MM2 | |
| CR1 | 0 | 359 | 179 | 358 | 167 | 179 | |
| CR2 | 0.0226 | 0 | 370 | 96 | 372 | 370 | |
| MA1 | 0.0113 | 0.0233 | 0 | 369 | 172 | 0 | |
| MA2 | 0.0225 | 0.0060 | 0.0232 | 0 | 373 | 369 | |
| MM1 | 0.0105 | 0.0234 | 0.0108 | 0.0235 | 0 | 172 | |
| MM2 | 0.0113 | 0.0233 | 0.0000 | 0.0232 | 0.0108 | 0 | |

| 344 | | | | | | | 20715 |
|---|---|---|---|---|---|---|---|
| | CR1 | CR2 | MA1 | MA2 | MM1 | MM2 | |
| CR1 | 0 | 538 | 511 | 144 | 493 | 510 | |
| CR2 | 0.0260 | 0 | 320 | 519 | 293 | 297 | |
| MA1 | 0.0247 | 0.0154 | 0 | 490 | 224 | 288 | |
| MA2 | 0.0070 | 0.0251 | 0.0237 | 0 | 475 | 495 | |
| MM1 | 0.0238 | 0.0141 | 0.0108 | 0.0229 | 0 | 279 | |
| MM2 | 0.0246 | 0.0143 | 0.0139 | 0.0239 | 0.0135 | 0 | |

| 346 | | | | | | | 23486 |
|---|---|---|---|---|---|---|---|
| | CR1 | CR2 | MA1 | MA2 | MM1 | MM2 | |
| CR1 | 0 | 487 | 288 | 483 | 300 | 288 | |
| CR2 | 0.0207 | 0 | 475 | 88 | 456 | 475 | |
| MA1 | 0.0123 | 0.0202 | 0 | 473 | 274 | 0 | |
| MA2 | 0.0206 | 0.0037 | 0.0201 | 0 | 453 | 473 | |
| MM1 | 0.0128 | 0.0194 | 0.0117 | 0.0193 | 0 | 274 | |
| MM2 | 0.0123 | 0.0202 | 0.0000 | 0.0201 | 0.0117 | 0 | |
| 348 | | | | | | | 9735 |
| | CR1 | CR2 | MA1 | MA2 | MM1 | MM2 | |
| CR1 | 0 | 133 | 136 | 57 | 44 | 51 | |
| CR2 | 0.0137 | 0 | 36 | 130 | 130 | 132 | |
| MA1 | 0.0140 | 0.0037 | 0 | 129 | 128 | 131 | |
| MA2 | 0.0059 | 0.0134 | 0.0133 | 0 | 46 | 58 | |
| MM1 | 0.0045 | 0.0134 | 0.0131 | 0.0047 | 0 | 48 | |
| MM2 | 0.0052 | 0.0136 | 0.0135 | 0.0060 | 0.0049 | 0 | |
| 349 | | | | | | | 10583 |
| | CR1 | CR2 | MA1 | MA2 | MM1 | MM2 | |
| CR1 | 0 | 157 | 91 | 159 | 90 | 82 | |
| CR2 | 0.0148 | 0 | 157 | 64 | 164 | 165 | |
| MA1 | 0.0086 | 0.0148 | 0 | 157 | 85 | 106 | |
| MA2 | 0.0150 | 0.0060 | 0.0148 | 0 | 165 | 167 | |
| MM1 | 0.0085 | 0.0155 | 0.0080 | 0.0156 | 0 | 101 | |
| MM2 | 0.0077 | 0.0156 | 0.0100 | 0.0158 | 0.0095 | 0 | |
| 352 | | | | | | | 16727 |
| | CR1 | CR2 | MA1 | MA2 | MM1 | MM2 | |
| CR1 | 0 | 288 | 139 | 285 | 141 | 139 | |
| CR2 | 0.0172 | 0 | 289 | 111 | 295 | 289 | |
| MA1 | 0.0083 | 0.0173 | 0 | 278 | 136 | 0 | |
| MA2 | 0.0170 | 0.0066 | 0.0166 | 0 | 278 | 278 | |
| MM1 | 0.0084 | 0.0176 | 0.0081 | 0.0166 | 0 | 136 | |
| MM2 | 0.0083 | 0.0173 | 0.0000 | 0.0166 | 0.0081 | 0 | |
| 354 | | | | | | | 13162 |
| | CR1 | CR2 | MA1 | MA2 | MM1 | MM2 | |
| CR1 | 0 | 360 | 100 | 144 | 361 | 345 | |
| CR2 | 0.0274 | 0 | 335 | 306 | 196 | 182 | |
| MA1 | 0.0076 | 0.0255 | 0 | 140 | 311 | 326 | |
| MA2 | 0.0109 | 0.0232 | 0.0106 | 0 | 259 | 285 | |
| MM1 | 0.0274 | 0.0149 | 0.0236 | 0.0197 | 0 | 177 | |
| MM2 | 0.0262 | 0.0138 | 0.0248 | 0.0217 | 0.0134 | 0 | |

| 356 | | | | | | | 3160 |
|---|---|---|---|---|---|---|---|
| | CR1 | CR2 | MA1 | MA2 | MM1 | MM2 | |
| CR1 | 0 | 33 | 16 | 35 | 18 | 19 | |
| CR2 | 0.0104 | 0 | 33 | 8 | 33 | 32 | |
| MA1 | 0.0051 | 0.0104 | 0 | 35 | 14 | 11 | |
| MA2 | 0.0111 | 0.0025 | 0.0111 | 0 | 35 | 34 | |
| MM1 | 0.0057 | 0.0104 | 0.0044 | 0.0111 | 0 | 9 | |
| MM2 | 0.0060 | 0.0101 | 0.0035 | 0.0108 | 0.0028 | 0 | |
| 358 | | | | | | | 10976 |
| | CR1 | CR2 | MA1 | MA2 | MM1 | MM2 | |
| CR1 | 0 | 127 | 79 | 87 | 56 | 70 | |
| CR2 | 0.0116 | 0 | 121 | 103 | 114 | 124 | |
| MA1 | 0.0072 | 0.0110 | 0 | 22 | 70 | 67 | |
| MA2 | 0.0079 | 0.0094 | 0.0020 | 0 | 76 | 77 | |
| MM1 | 0.0051 | 0.0104 | 0.0064 | 0.0069 | 0 | 64 | |
| MM2 | 0.0064 | 0.0113 | 0.0061 | 0.0070 | 0.0058 | 0 | |
| 361 | | | | | | | 10100 |
| | CR1 | CR2 | MA1 | MA2 | MM1 | MM2 | |
| CR1 | 0 | 208 | 197 | 158 | 197 | 228 | |
| CR2 | 0.0206 | 0 | 86 | 203 | 86 | 157 | |
| MA1 | 0.0195 | 0.0085 | 0 | 145 | 0 | 144 | |
| MA2 | 0.0156 | 0.0201 | 0.0144 | 0 | 145 | 192 | |
| MM1 | 0.0195 | 0.0085 | 0.0000 | 0.0144 | 0 | 144 | |
| MM2 | 0.0226 | 0.0155 | 0.0143 | 0.0190 | 0.0143 | 0 | |
| 362 | | | | | | | 6892 |
| | CR1 | CR2 | MA1 | MA2 | MM1 | MM2 | |
| CR1 | 0 | 100 | 79 | 99 | 47 | 48 | |
| CR2 | 0.0145 | 0 | 78 | 49 | 105 | 105 | |
| MA1 | 0.0115 | 0.0113 | 0 | 53 | 54 | 74 | |
| MA2 | 0.0144 | 0.0071 | 0.0077 | 0 | 106 | 108 | |
| MM1 | 0.0068 | 0.0152 | 0.0078 | 0.0154 | 0 | 36 | |
| MM2 | 0.0070 | 0.0152 | 0.0107 | 0.0157 | 0.0052 | 0 | |
| 364 | | | | | | | 2705 |
| | CR1 | CR2 | MA1 | MA2 | MM1 | MM2 | |
| CR1 | 0 | 0 | 58 | 23 | 24 | 14 | |
| CR2 | 0.0000 | 0 | 58 | 23 | 24 | 14 | |
| MA1 | 0.0214 | 0.0214 | 0 | 57 | 60 | 54 | |
| MA2 | 0.0085 | 0.0085 | 0.0211 | 0 | 27 | 15 | |
| MM1 | 0.0089 | 0.0089 | 0.0222 | 0.0100 | 0 | 22 | |
| MM2 | 0.0052 | 0.0052 | 0.0200 | 0.0055 | 0.0081 | 0 | |

| 366 | | | | | | | 16893 |
|---|---|---|---|---|---|---|---|
| | CR1 | CR2 | MA1 | MA2 | MM1 | MM2 | |
| CR1 | 0 | 250 | 85 | 261 | 261 | 257 | |
| CR2 | 0.0148 | 0 | 263 | 110 | 110 | 67 | |
| MA1 | 0.0050 | 0.0156 | 0 | 277 | 277 | 268 | |
| MA2 | 0.0155 | 0.0065 | 0.0164 | 0 | 0 | 122 | |
| MM1 | 0.0155 | 0.0065 | 0.0164 | 0.0000 | 0 | 122 | |
| MM2 | 0.0152 | 0.0040 | 0.0159 | 0.0072 | 0.0072 | 0 | |
| 367 | | | | | | | 19894 |
| | CR1 | CR2 | MA1 | MA2 | MM1 | MM2 | |
| CR1 | 0 | 409 | 173 | 401 | 184 | 197 | |
| CR2 | 0.0206 | 0 | 392 | 56 | 444 | 443 | |
| MA1 | 0.0087 | 0.0197 | 0 | 378 | 184 | 202 | |
| MA2 | 0.0202 | 0.0028 | 0.0190 | 0 | 430 | 431 | |
| MM1 | 0.0092 | 0.0223 | 0.0092 | 0.0216 | 0 | 215 | |
| MM2 | 0.0099 | 0.0223 | 0.0102 | 0.0217 | 0.0108 | 0 | |
| 368 | | | | | | | 17637 |
| | CR1 | CR2 | MA1 | MA2 | MM1 | MM2 | |
| CR1 | 0 | 493 | 275 | 502 | 200 | 269 | |
| CR2 | 0.0280 | 0 | 489 | 109 | 516 | 484 | |
| MA1 | 0.0156 | 0.0277 | 0 | 510 | 252 | 237 | |
| MA2 | 0.0285 | 0.0062 | 0.0289 | 0 | 528 | 496 | |
| MM1 | 0.0113 | 0.0293 | 0.0143 | 0.0299 | 0 | 277 | |
| MM2 | 0.0153 | 0.0274 | 0.0134 | 0.0281 | 0.0157 | 0 | |
| 370 | | | | | | | 20855 |
| | CR1 | CR2 | MA1 | MA2 | MM1 | MM2 | |
| CR1 | 0 | 368 | 194 | 373 | 194 | 193 | |
| CR2 | 0.0176 | 0 | 397 | 126 | 397 | 374 | |
| MA1 | 0.0093 | 0.0190 | 0 | 405 | 0 | 169 | |
| MA2 | 0.0179 | 0.0060 | 0.0194 | 0 | 405 | 387 | |
| MM1 | 0.0093 | 0.0190 | 0.0000 | 0.0194 | 0 | 169 | |
| MM2 | 0.0093 | 0.0179 | 0.0081 | 0.0186 | 0.0081 | 0 | |
| 371 | | | | | | | 8672 |
| | CR1 | CR2 | MA1 | MA2 | MM1 | MM2 | |
| CR1 | 0 | 116 | 108 | 25 | 108 | 108 | |
| CR2 | 0.0134 | 0 | 48 | 107 | 48 | 48 | |
| MA1 | 0.0125 | 0.0055 | 0 | 99 | 34 | 0 | |
| MA2 | 0.0029 | 0.0123 | 0.0114 | 0 | 99 | 99 | |
| MM1 | 0.0125 | 0.0055 | 0.0039 | 0.0114 | 0 | 34 | |
| MM2 | 0.0125 | 0.0055 | 0.0000 | 0.0114 | 0.0039 | 0 | |

| 372 | | | | | | | 9299 |
|---|---|---|---|---|---|---|---|
| | CR1 | CR2 | MA1 | MA2 | MM1 | MM2 | |
| CR1 | 0 | 100 | 100 | 32 | 32 | 32 | |
| CR2 | 0.0108 | 0 | 31 | 101 | 101 | 101 | |
| MA1 | 0.0108 | 0.0033 | 0 | 100 | 100 | 100 | |
| MA2 | 0.0034 | 0.0109 | 0.0108 | 0 | 0 | 0 | |
| MM1 | 0.0034 | 0.0109 | 0.0108 | 0.0000 | 0 | 0 | |
| MM2 | 0.0034 | 0.0109 | 0.0108 | 0.0000 | 0.0000 | 0 | |
| 373 | | | | | | | 11126 |
| | CR1 | CR2 | MA1 | MA2 | MM1 | MM2 | |
| CR1 | 0 | 163 | 73 | 149 | 73 | 76 | |
| CR2 | 0.0147 | 0 | 140 | 55 | 140 | 132 | |
| MA1 | 0.0066 | 0.0126 | 0 | 152 | 0 | 47 | |
| MA2 | 0.0134 | 0.0049 | 0.0137 | 0 | 152 | 148 | |
| MM1 | 0.0066 | 0.0126 | 0.0000 | 0.0137 | 0 | 47 | |
| MM2 | 0.0068 | 0.0119 | 0.0042 | 0.0133 | 0.0042 | 0 | |
| 374 | | | | | | | 9168 |
| | CR1 | CR2 | MA1 | MA2 | MM1 | MM2 | |
| CR1 | 0 | 185 | 9 | 180 | 9 | 110 | |
| CR2 | 0.0202 | 0 | 184 | 55 | 184 | 189 | |
| MA1 | 0.0010 | 0.0201 | 0 | 179 | 0 | 105 | |
| MA2 | 0.0196 | 0.0060 | 0.0195 | 0 | 179 | 188 | |
| MM1 | 0.0010 | 0.0201 | 0.0000 | 0.0195 | 0 | 105 | |
| MM2 | 0.0120 | 0.0206 | 0.0115 | 0.0205 | 0.0115 | 0 | |
| 376 | | | | | | | 13235 |
| | CR1 | CR2 | MA1 | MA2 | MM1 | MM2 | |
| CR1 | 0 | 42 | 263 | 154 | 263 | 251 | |
| CR2 | 0.0032 | 0 | 277 | 112 | 277 | 275 | |
| MA1 | 0.0199 | 0.0209 | 0 | 296 | 0 | 120 | |
| MA2 | 0.0116 | 0.0085 | 0.0224 | 0 | 296 | 293 | |
| MM1 | 0.0199 | 0.0209 | 0.0000 | 0.0224 | 0 | 120 | |
| MM2 | 0.0190 | 0.0208 | 0.0091 | 0.0221 | 0.0091 | 0 | |
| 378 | | | | | | | 26248 |
| | CR1 | CR2 | MA1 | MA2 | MM1 | MM2 | |
| CR1 | 0 | 502 | 111 | 481 | 490 | 481 | |
| CR2 | 0.0191 | 0 | 506 | 225 | 239 | 225 | |
| MA1 | 0.0042 | 0.0193 | 0 | 485 | 488 | 485 | |
| MA2 | 0.0183 | 0.0086 | 0.0185 | 0 | 242 | 0 | |
| MM1 | 0.0187 | 0.0091 | 0.0186 | 0.0092 | 0 | 242 | |
| MM2 | 0.0183 | 0.0086 | 0.0185 | 0.0000 | 0.0092 | 0 | |

| 379 | | | | | | 11889 |
|---|---|---|---|---|---|---|
| | CR1 | CR2 | MA1 | MA2 | MM1 | MM2 |
| CR1 | 0 | 275 | 174 | 280 | 280 | 282 |
| CR2 | 0.0231 | 0 | 229 | 205 | 205 | 186 |
| MA1 | 0.0146 | 0.0193 | 0 | 313 | 313 | 312 |
| MA2 | 0.0236 | 0.0172 | 0.0263 | 0 | 0 | 210 |
| MM1 | 0.0236 | 0.0172 | 0.0263 | 0.0000 | 0 | 210 |
| MM2 | 0.0237 | 0.0156 | 0.0262 | 0.0177 | 0.0177 | 0 |

| 380 | | | | | | 15239 |
|---|---|---|---|---|---|---|
| | CR1 | CR2 | MA1 | MA2 | MM1 | MM2 |
| CR1 | 0 | 230 | 115 | 227 | 79 | 115 |
| CR2 | 0.0151 | 0 | 226 | 67 | 219 | 226 |
| MA1 | 0.0075 | 0.0148 | 0 | 221 | 78 | 0 |
| MA2 | 0.0149 | 0.0044 | 0.0145 | 0 | 214 | 221 |
| MM1 | 0.0052 | 0.0144 | 0.0051 | 0.0140 | 0 | 78 |
| MM2 | 0.0075 | 0.0148 | 0.0000 | 0.0145 | 0.0051 | 0 |

| 382 | | | | | | 13353 |
|---|---|---|---|---|---|---|
| | CR1 | CR2 | MA1 | MA2 | MM1 | MM2 |
| CR1 | 0 | 233 | 109 | 222 | 0 | 119 |
| CR2 | 0.0174 | 0 | 248 | 65 | 233 | 251 |
| MA1 | 0.0082 | 0.0186 | 0 | 236 | 109 | 143 |
| MA2 | 0.0166 | 0.0049 | 0.0177 | 0 | 222 | 239 |
| MM1 | 0.0000 | 0.0174 | 0.0082 | 0.0166 | 0 | 119 |
| MM2 | 0.0089 | 0.0188 | 0.0107 | 0.0179 | 0.0089 | 0 |

| 384 | | | | | | 11135 |
|---|---|---|---|---|---|---|
| | CR1 | CR2 | MA1 | MA2 | MM1 | MM2 |
| CR1 | 0 | 243 | 229 | 75 | 120 | 79 |
| CR2 | 0.0218 | 0 | 83 | 236 | 227 | 247 |
| MA1 | 0.0206 | 0.0075 | 0 | 227 | 212 | 234 |
| MA2 | 0.0067 | 0.0212 | 0.0204 | 0 | 108 | 102 |
| MM1 | 0.0108 | 0.0204 | 0.0190 | 0.0097 | 0 | 141 |
| MM2 | 0.0071 | 0.0222 | 0.0210 | 0.0092 | 0.0127 | 0 |

| 385 | | | | | | 11660 |
|---|---|---|---|---|---|---|
| | CR1 | CR2 | MA1 | MA2 | MM1 | MM2 |
| CR1 | 0 | 151 | 62 | 157 | 62 | 75 |
| CR2 | 0.0130 | 0 | 154 | 61 | 154 | 147 |
| MA1 | 0.0053 | 0.0132 | 0 | 160 | 0 | 69 |
| MA2 | 0.0135 | 0.0052 | 0.0137 | 0 | 160 | 153 |
| MM1 | 0.0053 | 0.0132 | 0.0000 | 0.0137 | 0 | 69 |
| MM2 | 0.0064 | 0.0126 | 0.0059 | 0.0131 | 0.0059 | 0 |

| 386 | | | | | | | 9402 |
|---|---|---|---|---|---|---|---|
| | CR1 | CR2 | MA1 | MA2 | MM1 | MM2 | |
| CR1 | 0 | 268 | 204 | 281 | 189 | 188 | |
| CR2 | 0.0285 | 0 | 308 | 99 | 303 | 302 | |
| MA1 | 0.0217 | 0.0328 | 0 | 310 | 191 | 190 | |
| MA2 | 0.0299 | 0.0105 | 0.0330 | 0 | 310 | 309 | |
| MM1 | 0.0201 | 0.0322 | 0.0203 | 0.0330 | 0 | 1 | |
| MM2 | 0.0200 | 0.0321 | 0.0202 | 0.0329 | 0.0001 | 0 | |

| 387 | | | | | | | 15361 |
|---|---|---|---|---|---|---|---|
| | CR1 | CR2 | MA1 | MA2 | MM1 | MM2 | |
| CR1 | 0 | 269 | 270 | 297 | 387 | 387 | |
| CR2 | 0.0175 | 0 | 15 | 456 | 252 | 252 | |
| MA1 | 0.0176 | 0.0010 | 0 | 457 | 245 | 245 | |
| MA2 | 0.0193 | 0.0297 | 0.0298 | 0 | 452 | 452 | |
| MM1 | 0.0252 | 0.0164 | 0.0159 | 0.0294 | 0 | 0 | |
| MM2 | 0.0252 | 0.0164 | 0.0159 | 0.0294 | 0.0000 | 0 | |

| 388 | | | | | | | 20079 |
|---|---|---|---|---|---|---|---|
| | CR1 | CR2 | MA1 | MA2 | MM1 | MM2 | |
| CR1 | 0 | 350 | 156 | 352 | 156 | 217 | |
| CR2 | 0.0174 | 0 | 317 | 73 | 317 | 347 | |
| MA1 | 0.0078 | 0.0158 | 0 | 318 | 0 | 164 | |
| MA2 | 0.0175 | 0.0036 | 0.0158 | 0 | 318 | 345 | |
| MM1 | 0.0078 | 0.0158 | 0.0000 | 0.0158 | 0 | 164 | |
| MM2 | 0.0108 | 0.0173 | 0.0082 | 0.0172 | 0.0082 | 0 | |

| 391 | | | | | | | 11422 |
|---|---|---|---|---|---|---|---|
| | CR1 | CR2 | MA1 | MA2 | MM1 | MM2 | |
| CR1 | 0 | 325 | 232 | 242 | 166 | 169 | |
| CR2 | 0.0285 | 0 | 194 | 206 | 319 | 321 | |
| MA1 | 0.0203 | 0.0170 | 0 | 301 | 137 | 227 | |
| MA2 | 0.0212 | 0.0180 | 0.0264 | 0 | 164 | 223 | |
| MM1 | 0.0145 | 0.0279 | 0.0120 | 0.0144 | 0 | 143 | |
| MM2 | 0.0148 | 0.0281 | 0.0199 | 0.0195 | 0.0125 | 0 | |

| 392 | | | | | | | 14525 |
|---|---|---|---|---|---|---|---|
| | CR1 | CR2 | MA1 | MA2 | MM1 | MM2 | |
| CR1 | 0 | 258 | 159 | 227 | 163 | 109 | |
| CR2 | 0.0178 | 0 | 257 | 151 | 248 | 257 | |
| MA1 | 0.0109 | 0.0177 | 0 | 184 | 163 | 152 | |
| MA2 | 0.0156 | 0.0104 | 0.0127 | 0 | 211 | 231 | |
| MM1 | 0.0112 | 0.0171 | 0.0112 | 0.0145 | 0 | 157 | |
| MM2 | 0.0075 | 0.0177 | 0.0105 | 0.0159 | 0.0108 | 0 | |

| 393 | | | | | | | 18341 |
|---|---|---|---|---|---|---|---|
| | CR1 | CR2 | MA1 | MA2 | MM1 | MM2 | |
| CR1 | 0 | 349 | 329 | 122 | 343 | 350 | |
| CR2 | 0.0190 | 0 | 174 | 336 | 219 | 203 | |
| MA1 | 0.0179 | 0.0095 | 0 | 317 | 216 | 202 | |
| MA2 | 0.0067 | 0.0183 | 0.0173 | 0 | 331 | 338 | |
| MM1 | 0.0187 | 0.0119 | 0.0118 | 0.0180 | 0 | 203 | |
| MM2 | 0.0191 | 0.0111 | 0.0110 | 0.0184 | 0.0111 | 0 | |
| 394 | | | | | | | 11952 |
| | CR1 | CR2 | MA1 | MA2 | MM1 | MM2 | |
| CR1 | 0 | 228 | 211 | 102 | 101 | 118 | |
| CR2 | 0.0191 | 0 | 84 | 224 | 223 | 221 | |
| MA1 | 0.0177 | 0.0070 | 0 | 206 | 206 | 206 | |
| MA2 | 0.0085 | 0.0187 | 0.0172 | 0 | 113 | 110 | |
| MM1 | 0.0085 | 0.0187 | 0.0172 | 0.0095 | 0 | 109 | |
| MM2 | 0.0099 | 0.0185 | 0.0172 | 0.0092 | 0.0091 | 0 | |
| 395 | | | | | | | 15585 |
| | CR1 | CR2 | MA1 | MA2 | MM1 | MM2 | |
| CR1 | 0 | 264 | 263 | 48 | 263 | 298 | |
| CR2 | 0.0169 | 0 | 104 | 256 | 104 | 154 | |
| MA1 | 0.0169 | 0.0067 | 0 | 256 | 0 | 143 | |
| MA2 | 0.0031 | 0.0164 | 0.0164 | 0 | 256 | 291 | |
| MM1 | 0.0169 | 0.0067 | 0.0000 | 0.0164 | 0 | 143 | |
| MM2 | 0.0191 | 0.0099 | 0.0092 | 0.0187 | 0.0092 | 0 | |
| 396 | | | | | | | 16656 |
| | CR1 | CR2 | MA1 | MA2 | MM1 | MM2 | |
| CR1 | 0 | 276 | 288 | 47 | 288 | 287 | |
| CR2 | 0.0166 | 0 | 121 | 279 | 121 | 132 | |
| MA1 | 0.0173 | 0.0073 | 0 | 293 | 0 | 105 | |
| MA2 | 0.0028 | 0.0168 | 0.0176 | 0 | 293 | 290 | |
| MM1 | 0.0173 | 0.0073 | 0.0000 | 0.0176 | 0 | 105 | |
| MM2 | 0.0172 | 0.0079 | 0.0063 | 0.0174 | 0.0063 | 0 | |
| 397 | | | | | | | 11106 |
| | CR1 | CR2 | MA1 | MA2 | MM1 | MM2 | |
| CR1 | 0 | 160 | 106 | 187 | 269 | 254 | |
| CR2 | 0.0144 | 0 | 136 | 223 | 178 | 165 | |
| MA1 | 0.0095 | 0.0122 | 0 | 109 | 210 | 200 | |
| MA2 | 0.0168 | 0.0201 | 0.0098 | 0 | 183 | 181 | |
| MM1 | 0.0242 | 0.0160 | 0.0189 | 0.0165 | 0 | 68 | |
| MM2 | 0.0229 | 0.0149 | 0.0180 | 0.0163 | 0.0061 | 0 | |

| 398 | | | | | | | 9683 |
|---|---|---|---|---|---|---|---|
| | CR1 | CR2 | MA1 | MA2 | MM1 | MM2 | |
| CR1 | 0 | 170 | 102 | 171 | 90 | 102 | |
| CR2 | 0.0176 | 0 | 163 | 17 | 171 | 163 | |
| MA1 | 0.0105 | 0.0168 | 0 | 162 | 79 | 0 | |
| MA2 | 0.0177 | 0.0018 | 0.0167 | 0 | 172 | 162 | |
| MM1 | 0.0093 | 0.0177 | 0.0082 | 0.0178 | 0 | 79 | |
| MM2 | 0.0105 | 0.0168 | 0.0000 | 0.0167 | 0.0082 | 0 | |

| 400 | | | | | | | 11622 |
|---|---|---|---|---|---|---|---|
| | CR1 | CR2 | MA1 | MA2 | MM1 | MM2 | |
| CR1 | 0 | 240 | 80 | 228 | 82 | 80 | |
| CR2 | 0.0207 | 0 | 238 | 69 | 227 | 238 | |
| MA1 | 0.0069 | 0.0205 | 0 | 226 | 95 | 0 | |
| MA2 | 0.0196 | 0.0059 | 0.0194 | 0 | 217 | 226 | |
| MM1 | 0.0071 | 0.0195 | 0.0082 | 0.0187 | 0 | 95 | |
| MM2 | 0.0069 | 0.0205 | 0.0000 | 0.0194 | 0.0082 | 0 | |

| 402 | | | | | | | 16736 |
|---|---|---|---|---|---|---|---|
| | CR1 | CR2 | MA1 | MA2 | MM1 | MM2 | |
| CR1 | 0 | 279 | 277 | 274 | 324 | 344 | |
| CR2 | 0.0167 | 0 | 201 | 212 | 194 | 186 | |
| MA1 | 0.0166 | 0.0120 | 0 | 19 | 66 | 229 | |
| MA2 | 0.0164 | 0.0127 | 0.0011 | 0 | 85 | 248 | |
| MM1 | 0.0194 | 0.0116 | 0.0039 | 0.0051 | 0 | 206 | |
| MM2 | 0.0206 | 0.0111 | 0.0137 | 0.0148 | 0.0123 | 0 | |

| 404 | | | | | | | 9283 |
|---|---|---|---|---|---|---|---|
| | CR1 | CR2 | MA1 | MA2 | MM1 | MM2 | |
| CR1 | 0 | 142 | 52 | 183 | 36 | 85 | |
| CR2 | 0.0153 | 0 | 150 | 117 | 134 | 157 | |
| MA1 | 0.0056 | 0.0162 | 0 | 171 | 16 | 77 | |
| MA2 | 0.0197 | 0.0126 | 0.0184 | 0 | 187 | 194 | |
| MM1 | 0.0039 | 0.0144 | 0.0017 | 0.0201 | 0 | 71 | |
| MM2 | 0.0092 | 0.0169 | 0.0083 | 0.0209 | 0.0076 | 0 | |

| 407 | | | | | | | 12028 |
|---|---|---|---|---|---|---|---|
| | CR1 | CR2 | MA1 | MA2 | MM1 | MM2 | |
| CR1 | 0 | 151 | 284 | 126 | 121 | 126 | |
| CR2 | 0.0126 | 0 | 199 | 187 | 170 | 187 | |
| MA1 | 0.0236 | 0.0165 | 0 | 296 | 284 | 296 | |
| MA2 | 0.0105 | 0.0155 | 0.0246 | 0 | 92 | 0 | |
| MM1 | 0.0101 | 0.0141 | 0.0236 | 0.0076 | 0 | 92 | |
| MM2 | 0.0105 | 0.0155 | 0.0246 | 0.0000 | 0.0076 | 0 | |

| 408 | | | | | | | 13526 |
|---|---|---|---|---|---|---|---|
| | CR1 | CR2 | MA1 | MA2 | MM1 | MM2 | |
| CR1 | 0 | 277 | 56 | 251 | 251 | 238 | |
| CR2 | 0.0205 | 0 | 278 | 127 | 127 | 127 | |
| MA1 | 0.0041 | 0.0206 | 0 | 251 | 251 | 239 | |
| MA2 | 0.0186 | 0.0094 | 0.0186 | 0 | 0 | 95 | |
| MM1 | 0.0186 | 0.0094 | 0.0186 | 0.0000 | 0 | 95 | |
| MM2 | 0.0176 | 0.0094 | 0.0177 | 0.0070 | 0.0070 | 0 | |

| 409 | | | | | | | 17532 |
|---|---|---|---|---|---|---|---|
| | CR1 | CR2 | MA1 | MA2 | MM1 | MM2 | |
| CR1 | 0 | 288 | 278 | 94 | 294 | 293 | |
| CR2 | 0.0164 | 0 | 128 | 294 | 124 | 130 | |
| MA1 | 0.0159 | 0.0073 | 0 | 282 | 116 | 131 | |
| MA2 | 0.0054 | 0.0168 | 0.0161 | 0 | 299 | 298 | |
| MM1 | 0.0168 | 0.0071 | 0.0066 | 0.0171 | 0 | 116 | |
| MM2 | 0.0167 | 0.0074 | 0.0075 | 0.0170 | 0.0066 | 0 | |

| 410 | | | | | | | 13582 |
|---|---|---|---|---|---|---|---|
| | CR1 | CR2 | MA1 | MA2 | MM1 | MM2 | |
| CR1 | 0 | 228 | 377 | 77 | 350 | 368 | |
| CR2 | 0.0168 | 0 | 257 | 251 | 256 | 259 | |
| MA1 | 0.0278 | 0.0189 | 0 | 368 | 165 | 196 | |
| MA2 | 0.0057 | 0.0185 | 0.0271 | 0 | 339 | 361 | |
| MM1 | 0.0258 | 0.0188 | 0.0121 | 0.0250 | 0 | 185 | |
| MM2 | 0.0271 | 0.0191 | 0.0144 | 0.0266 | 0.0136 | 0 | |

| 413 | | | | | | | 11643 |
|---|---|---|---|---|---|---|---|
| | CR1 | CR2 | MA1 | MA2 | MM1 | MM2 | |
| CR1 | 0 | 173 | 167 | 75 | 176 | 182 | |
| CR2 | 0.0149 | 0 | 90 | 178 | 64 | 84 | |
| MA1 | 0.0143 | 0.0077 | 0 | 179 | 92 | 111 | |
| MA2 | 0.0064 | 0.0153 | 0.0154 | 0 | 181 | 187 | |
| MM1 | 0.0151 | 0.0055 | 0.0079 | 0.0155 | 0 | 93 | |
| MM2 | 0.0156 | 0.0072 | 0.0095 | 0.0161 | 0.0080 | 0 | |

| 414 | | | | | | | 5846 |
|---|---|---|---|---|---|---|---|
| | CR1 | CR2 | MA1 | MA2 | MM1 | MM2 | |
| CR1 | 0 | 125 | 54 | 128 | 141 | 128 | |
| CR2 | 0.0214 | 0 | 135 | 52 | 79 | 52 | |
| MA1 | 0.0092 | 0.0231 | 0 | 137 | 150 | 137 | |
| MA2 | 0.0219 | 0.0089 | 0.0234 | 0 | 86 | 0 | |
| MM1 | 0.0241 | 0.0135 | 0.0257 | 0.0147 | 0 | 86 | |
| MM2 | 0.0219 | 0.0089 | 0.0234 | 0.0000 | 0.0147 | 0 | |

| 417 | | | | | | 10057 |
|---|---|---|---|---|---|---|
| | CR1 | CR2 | MA1 | MA2 | MM1 | MM2 |
| CR1 | 0 | 184 | 173 | 51 | 66 | 51 |
| CR2 | 0.0183 | 0 | 31 | 190 | 197 | 190 |
| MA1 | 0.0172 | 0.0031 | 0 | 179 | 186 | 179 |
| MA2 | 0.0051 | 0.0189 | 0.0178 | 0 | 67 | 0 |
| MM1 | 0.0066 | 0.0196 | 0.0185 | 0.0067 | 0 | 67 |
| MM2 | 0.0051 | 0.0189 | 0.0178 | 0.0000 | 0.0067 | 0 |

| 418 | | | | | | 13107 |
|---|---|---|---|---|---|---|
| | CR1 | CR2 | MA1 | MA2 | MM1 | MM2 |
| CR1 | 0 | 220 | 105 | 211 | 135 | 127 |
| CR2 | 0.0168 | 0 | 231 | 70 | 233 | 237 |
| MA1 | 0.0080 | 0.0176 | 0 | 225 | 153 | 128 |
| MA2 | 0.0161 | 0.0053 | 0.0172 | 0 | 229 | 226 |
| MM1 | 0.0103 | 0.0178 | 0.0117 | 0.0175 | 0 | 145 |
| MM2 | 0.0097 | 0.0181 | 0.0098 | 0.0172 | 0.0111 | 0 |

| 422 | | | | | | 14783 |
|---|---|---|---|---|---|---|
| | CR1 | CR2 | MA1 | MA2 | MM1 | MM2 |
| CR1 | 0 | 292 | 136 | 295 | 174 | 141 |
| CR2 | 0.0198 | 0 | 311 | 68 | 309 | 305 |
| MA1 | 0.0092 | 0.0210 | 0 | 314 | 134 | 97 |
| MA2 | 0.0200 | 0.0046 | 0.0212 | 0 | 312 | 310 |
| MM1 | 0.0118 | 0.0209 | 0.0091 | 0.0211 | 0 | 130 |
| MM2 | 0.0095 | 0.0206 | 0.0066 | 0.0210 | 0.0088 | 0 |

| 424 | | | | | | 10954 |
|---|---|---|---|---|---|---|
| | CR1 | CR2 | MA1 | MA2 | MM1 | MM2 |
| CR1 | 0 | 295 | 189 | 253 | 145 | 139 |
| CR2 | 0.0269 | 0 | 316 | 159 | 313 | 298 |
| MA1 | 0.0173 | 0.0288 | 0 | 211 | 174 | 176 |
| MA2 | 0.0231 | 0.0145 | 0.0193 | 0 | 257 | 245 |
| MM1 | 0.0132 | 0.0286 | 0.0159 | 0.0235 | 0 | 131 |
| MM2 | 0.0127 | 0.0272 | 0.0161 | 0.0224 | 0.0120 | 0 |

| 427 | | | | | | 15243 |
|---|---|---|---|---|---|---|
| | CR1 | CR2 | MA1 | MA2 | MM1 | MM2 |
| CR1 | 0 | 296 | 152 | 286 | 150 | 150 |
| CR2 | 0.0194 | 0 | 281 | 114 | 293 | 293 |
| MA1 | 0.0100 | 0.0184 | 0 | 278 | 132 | 130 |
| MA2 | 0.0188 | 0.0075 | 0.0182 | 0 | 287 | 287 |
| MM1 | 0.0098 | 0.0192 | 0.0087 | 0.0188 | 0 | 2 |
| MM2 | 0.0098 | 0.0192 | 0.0085 | 0.0188 | 0.0001 | 0 |

| 428 | | | | | | | 16934 |
|---|---|---|---|---|---|---|---|
| | CR1 | CR2 | MA1 | MA2 | MM1 | MM2 | |
| CR1 | 0 | 304 | 303 | 33 | 163 | 27 | |
| CR2 | 0.0180 | 0 | 33 | 304 | 292 | 306 | |
| MA1 | 0.0179 | 0.0019 | 0 | 303 | 293 | 305 | |
| MA2 | 0.0019 | 0.0180 | 0.0179 | 0 | 158 | 10 | |
| MM1 | 0.0096 | 0.0172 | 0.0173 | 0.0093 | 0 | 164 | |
| MM2 | 0.0016 | 0.0181 | 0.0180 | 0.0006 | 0.0097 | 0 | |
| 429 | | | | | | | 13366 |
| | CR1 | CR2 | MA1 | MA2 | MM1 | MM2 | |
| CR1 | 0 | 276 | 275 | 90 | 280 | 280 | |
| CR2 | 0.0206 | 0 | 128 | 286 | 142 | 142 | |
| MA1 | 0.0206 | 0.0096 | 0 | 282 | 153 | 153 | |
| MA2 | 0.0067 | 0.0214 | 0.0211 | 0 | 286 | 286 | |
| MM1 | 0.0209 | 0.0106 | 0.0114 | 0.0214 | 0 | 0 | |
| MM2 | 0.0209 | 0.0106 | 0.0114 | 0.0214 | 0.0000 | 0 | |
| 432 | | | | | | | 11131 |
| | CR1 | CR2 | MA1 | MA2 | MM1 | MM2 | |
| CR1 | 0 | 112 | 102 | 58 | 32 | 41 | |
| CR2 | 0.0101 | 0 | 34 | 88 | 116 | 117 | |
| MA1 | 0.0092 | 0.0031 | 0 | 96 | 104 | 99 | |
| MA2 | 0.0052 | 0.0079 | 0.0086 | 0 | 54 | 33 | |
| MM1 | 0.0029 | 0.0104 | 0.0093 | 0.0049 | 0 | 33 | |
| MM2 | 0.0037 | 0.0105 | 0.0089 | 0.0030 | 0.0030 | 0 | |
| 433 | | | | | | | 11833 |
| | CR1 | CR2 | MA1 | MA2 | MM1 | MM2 | |
| CR1 | 0 | 172 | 80 | 172 | 98 | 80 | |
| CR2 | 0.0145 | 0 | 172 | 0 | 203 | 172 | |
| MA1 | 0.0068 | 0.0145 | 0 | 172 | 100 | 0 | |
| MA2 | 0.0145 | 0.0000 | 0.0145 | 0 | 203 | 172 | |
| MM1 | 0.0083 | 0.0172 | 0.0085 | 0.0172 | 0 | 100 | |
| MM2 | 0.0068 | 0.0145 | 0.0000 | 0.0145 | 0.0085 | 0 | |
| 435 | | | | | | | 29510 |
| | CR1 | CR2 | MA1 | MA2 | MM1 | MM2 | |
| CR1 | 0 | 633 | 625 | 176 | 620 | 625 | |
| CR2 | 0.0215 | 0 | 299 | 639 | 320 | 299 | |
| MA1 | 0.0212 | 0.0101 | 0 | 634 | 314 | 0 | |
| MA2 | 0.0060 | 0.0217 | 0.0215 | 0 | 630 | 634 | |
| MM1 | 0.0210 | 0.0108 | 0.0106 | 0.0213 | 0 | 314 | |
| MM2 | 0.0212 | 0.0101 | 0.0000 | 0.0215 | 0.0106 | 0 | |

436 10400

| | CR1 | CR2 | MA1 | MA2 | MM1 | MM2 |
|---|---|---|---|---|---|---|
| CR1 | 0 | 228 | 220 | 45 | 210 | 225 |
| CR2 | 0.0219 | 0 | 129 | 223 | 116 | 127 |
| MA1 | 0.0212 | 0.0124 | 0 | 217 | 111 | 128 |
| MA2 | 0.0043 | 0.0214 | 0.0209 | 0 | 205 | 220 |
| MM1 | 0.0202 | 0.0112 | 0.0107 | 0.0197 | 0 | 100 |
| MM2 | 0.0216 | 0.0122 | 0.0123 | 0.0212 | 0.0096 | 0 |

437 16654

| | CR1 | CR2 | MA1 | MA2 | MM1 | MM2 |
|---|---|---|---|---|---|---|
| CR1 | 0 | 243 | 156 | 97 | 245 | 244 |
| CR2 | 0.0146 | 0 | 165 | 208 | 99 | 117 |
| MA1 | 0.0094 | 0.0099 | 0 | 119 | 129 | 179 |
| MA2 | 0.0058 | 0.0125 | 0.0071 | 0 | 180 | 210 |
| MM1 | 0.0147 | 0.0059 | 0.0077 | 0.0108 | 0 | 111 |
| MM2 | 0.0147 | 0.0070 | 0.0107 | 0.0126 | 0.0067 | 0 |

441 8680

| | CR1 | CR2 | MA1 | MA2 | MM1 | MM2 |
|---|---|---|---|---|---|---|
| CR1 | 0 | 246 | 134 | 115 | 114 | 104 |
| CR2 | 0.0283 | 0 | 182 | 199 | 235 | 235 |
| MA1 | 0.0154 | 0.0210 | 0 | 35 | 77 | 97 |
| MA2 | 0.0132 | 0.0229 | 0.0040 | 0 | 42 | 78 |
| MM1 | 0.0131 | 0.0271 | 0.0089 | 0.0048 | 0 | 63 |
| MM2 | 0.0120 | 0.0271 | 0.0112 | 0.0090 | 0.0073 | 0 |

442 16573

| | CR1 | CR2 | MA1 | MA2 | MM1 | MM2 |
|---|---|---|---|---|---|---|
| CR1 | 0 | 293 | 289 | 239 | 186 | 175 |
| CR2 | 0.0177 | 0 | 42 | 194 | 304 | 304 |
| MA1 | 0.0174 | 0.0025 | 0 | 172 | 302 | 282 |
| MA2 | 0.0144 | 0.0117 | 0.0104 | 0 | 264 | 110 |
| MM1 | 0.0112 | 0.0183 | 0.0182 | 0.0159 | 0 | 219 |
| MM2 | 0.0106 | 0.0183 | 0.0170 | 0.0066 | 0.0132 | 0 |

445 14604

| | CR1 | CR2 | MA1 | MA2 | MM1 | MM2 |
|---|---|---|---|---|---|---|
| CR1 | 0 | 338 | 198 | 344 | 197 | 221 |
| CR2 | 0.0231 | 0 | 292 | 52 | 301 | 297 |
| MA1 | 0.0136 | 0.0200 | 0 | 298 | 184 | 176 |
| MA2 | 0.0236 | 0.0036 | 0.0204 | 0 | 309 | 299 |
| MM1 | 0.0135 | 0.0206 | 0.0126 | 0.0212 | 0 | 164 |
| MM2 | 0.0151 | 0.0203 | 0.0121 | 0.0205 | 0.0112 | 0 |

| 447 | | | | | | | 16392 |
|---|---|---|---|---|---|---|---|
| | CR1 | CR2 | MA1 | MA2 | MM1 | MM2 | |
| CR1 | 0 | 346 | 110 | 346 | 144 | 111 | |
| CR2 | 0.0211 | 0 | 330 | 0 | 341 | 341 | |
| MA1 | 0.0067 | 0.0201 | 0 | 330 | 139 | 113 | |
| MA2 | 0.0211 | 0.0000 | 0.0201 | 0 | 341 | 341 | |
| MM1 | 0.0088 | 0.0208 | 0.0085 | 0.0208 | 0 | 126 | |
| MM2 | 0.0068 | 0.0208 | 0.0069 | 0.0208 | 0.0077 | 0 | |
| 448 | | | | | | | 10630 |
| | CR1 | CR2 | MA1 | MA2 | MM1 | MM2 | |
| CR1 | 0 | 259 | 240 | 123 | 242 | 240 | |
| CR2 | 0.0244 | 0 | 120 | 238 | 106 | 120 | |
| MA1 | 0.0226 | 0.0113 | 0 | 215 | 95 | 0 | |
| MA2 | 0.0116 | 0.0224 | 0.0202 | 0 | 220 | 215 | |
| MM1 | 0.0228 | 0.0100 | 0.0089 | 0.0207 | 0 | 95 | |
| MM2 | 0.0226 | 0.0113 | 0.0000 | 0.0202 | 0.0089 | 0 | |
| 452 | | | | | | | 7486 |
| | CR1 | CR2 | MA1 | MA2 | MM1 | MM2 | |
| CR1 | 0 | 164 | 166 | 105 | 105 | 88 | |
| CR2 | 0.0219 | 0 | 21 | 169 | 169 | 143 | |
| MA1 | 0.0222 | 0.0028 | 0 | 170 | 170 | 144 | |
| MA2 | 0.0140 | 0.0226 | 0.0227 | 0 | 0 | 99 | |
| MM1 | 0.0140 | 0.0226 | 0.0227 | 0.0000 | 0 | 99 | |
| MM2 | 0.0118 | 0.0191 | 0.0192 | 0.0132 | 0.0132 | 0 | |
| 453 | | | | | | | 12819 |
| | CR1 | CR2 | MA1 | MA2 | MM1 | MM2 | |
| CR1 | 0 | 342 | 237 | 349 | 225 | 219 | |
| CR2 | 0.0267 | 0 | 309 | 54 | 333 | 314 | |
| MA1 | 0.0185 | 0.0241 | 0 | 316 | 263 | 248 | |
| MA2 | 0.0272 | 0.0042 | 0.0247 | 0 | 339 | 322 | |
| MM1 | 0.0176 | 0.0260 | 0.0205 | 0.0264 | 0 | 159 | |
| MM2 | 0.0171 | 0.0245 | 0.0193 | 0.0251 | 0.0124 | 0 | |
| 454 | | | | | | | 12570 |
| | CR1 | CR2 | MA1 | MA2 | MM1 | MM2 | |
| CR1 | 0 | 205 | 120 | 215 | 114 | 120 | |
| CR2 | 0.0163 | 0 | 250 | 53 | 245 | 250 | |
| MA1 | 0.0095 | 0.0199 | 0 | 254 | 103 | 0 | |
| MA2 | 0.0171 | 0.0042 | 0.0202 | 0 | 252 | 254 | |
| MM1 | 0.0091 | 0.0195 | 0.0082 | 0.0200 | 0 | 103 | |
| MM2 | 0.0095 | 0.0199 | 0.0000 | 0.0202 | 0.0082 | 0 | |

| 456 | | | | | | | 10232 |
|---|---|---|---|---|---|---|---|
| | CR1 | CR2 | MA1 | MA2 | MM1 | MM2 | |
| CR1 | 0 | 230 | 143 | 252 | 143 | 127 | |
| CR2 | 0.0225 | 0 | 205 | 97 | 205 | 201 | |
| MA1 | 0.0140 | 0.0200 | 0 | 226 | 0 | 134 | |
| MA2 | 0.0246 | 0.0095 | 0.0221 | 0 | 226 | 223 | |
| MM1 | 0.0140 | 0.0200 | 0.0000 | 0.0221 | 0 | 134 | |
| MM2 | 0.0124 | 0.0196 | 0.0131 | 0.0218 | 0.0131 | 0 | |
| 457 | | | | | | | 6287 |
| | CR1 | CR2 | MA1 | MA2 | MM1 | MM2 | |
| CR1 | 0 | 124 | 59 | 90 | 82 | 90 | |
| CR2 | 0.0197 | 0 | 83 | 74 | 88 | 74 | |
| MA1 | 0.0094 | 0.0132 | 0 | 112 | 121 | 112 | |
| MA2 | 0.0143 | 0.0118 | 0.0178 | 0 | 44 | 0 | |
| MM1 | 0.0130 | 0.0140 | 0.0192 | 0.0070 | 0 | 44 | |
| MM2 | 0.0143 | 0.0118 | 0.0178 | 0.0000 | 0.0070 | 0 | |
| 458 | | | | | | | 15000 |
| | CR1 | CR2 | MA1 | MA2 | MM1 | MM2 | |
| CR1 | 0 | 321 | 308 | 88 | 310 | 308 | |
| CR2 | 0.0214 | 0 | 122 | 329 | 129 | 122 | |
| MA1 | 0.0205 | 0.0081 | 0 | 318 | 65 | 0 | |
| MA2 | 0.0059 | 0.0219 | 0.0212 | 0 | 320 | 318 | |
| MM1 | 0.0207 | 0.0086 | 0.0043 | 0.0213 | 0 | 65 | |
| MM2 | 0.0205 | 0.0081 | 0.0000 | 0.0212 | 0.0043 | 0 | |
| 459 | | | | | | | 11549 |
| | CR1 | CR2 | MA1 | MA2 | MM1 | MM2 | |
| CR1 | 0 | 283 | 151 | 274 | 161 | 141 | |
| CR2 | 0.0245 | 0 | 270 | 53 | 287 | 287 | |
| MA1 | 0.0131 | 0.0234 | 0 | 263 | 189 | 138 | |
| MA2 | 0.0237 | 0.0046 | 0.0228 | 0 | 281 | 277 | |
| MM1 | 0.0139 | 0.0249 | 0.0164 | 0.0243 | 0 | 191 | |
| MM2 | 0.0122 | 0.0249 | 0.0119 | 0.0240 | 0.0165 | 0 | |
| 461 | | | | | | | 11337 |
| | CR1 | CR2 | MA1 | MA2 | MM1 | MM2 | |
| CR1 | 0 | 257 | 113 | 259 | 107 | 124 | |
| CR2 | 0.0227 | 0 | 236 | 36 | 264 | 259 | |
| MA1 | 0.0100 | 0.0208 | 0 | 236 | 28 | 131 | |
| MA2 | 0.0228 | 0.0032 | 0.0208 | 0 | 264 | 259 | |
| MM1 | 0.0094 | 0.0233 | 0.0025 | 0.0233 | 0 | 103 | |
| MM2 | 0.0109 | 0.0228 | 0.0116 | 0.0228 | 0.0091 | 0 | |

| 462 | | | | | | | 12233 |
|---|---|---|---|---|---|---|---|
| | CR1 | CR2 | MA1 | MA2 | MM1 | MM2 | |
| CR1 | 0 | 322 | 282 | 132 | 292 | 293 | |
| CR2 | 0.0263 | 0 | 150 | 318 | 133 | 141 | |
| MA1 | 0.0231 | 0.0123 | 0 | 281 | 132 | 124 | |
| MA2 | 0.0108 | 0.0260 | 0.0230 | 0 | 286 | 284 | |
| MM1 | 0.0239 | 0.0109 | 0.0108 | 0.0234 | 0 | 148 | |
| MM2 | 0.0240 | 0.0115 | 0.0101 | 0.0232 | 0.0121 | 0 | |
| 464 | | | | | | | 14277 |
| | CR1 | CR2 | MA1 | MA2 | MM1 | MM2 | |
| CR1 | 0 | 218 | 214 | 88 | 218 | 226 | |
| CR2 | 0.0153 | 0 | 87 | 237 | 0 | 68 | |
| MA1 | 0.0150 | 0.0061 | 0 | 228 | 87 | 19 | |
| MA2 | 0.0062 | 0.0166 | 0.0160 | 0 | 237 | 247 | |
| MM1 | 0.0153 | 0.0000 | 0.0061 | 0.0166 | 0 | 68 | |
| MM2 | 0.0158 | 0.0048 | 0.0013 | 0.0173 | 0.0048 | 0 | |
| 465 | | | | | | | 15674 |
| | CR1 | CR2 | MA1 | MA2 | MM1 | MM2 | |
| CR1 | 0 | 221 | 216 | 76 | 76 | 85 | |
| CR2 | 0.0141 | 0 | 49 | 217 | 217 | 209 | |
| MA1 | 0.0138 | 0.0031 | 0 | 211 | 211 | 204 | |
| MA2 | 0.0048 | 0.0138 | 0.0135 | 0 | 0 | 83 | |
| MM1 | 0.0048 | 0.0138 | 0.0135 | 0.0000 | 0 | 83 | |
| MM2 | 0.0054 | 0.0133 | 0.0130 | 0.0053 | 0.0053 | 0 | |
| 467 | | | | | | | 13327 |
| | CR1 | CR2 | MA1 | MA2 | MM1 | MM2 | |
| CR1 | 0 | 179 | 134 | 161 | 107 | 106 | |
| CR2 | 0.0134 | 0 | 135 | 80 | 209 | 215 | |
| MA1 | 0.0101 | 0.0101 | 0 | 83 | 154 | 158 | |
| MA2 | 0.0121 | 0.0060 | 0.0062 | 0 | 198 | 201 | |
| MM1 | 0.0080 | 0.0157 | 0.0116 | 0.0149 | 0 | 61 | |
| MM2 | 0.0080 | 0.0161 | 0.0119 | 0.0151 | 0.0046 | 0 | |
| 472 | | | | | | | 13996 |
| | CR1 | CR2 | MA1 | MA2 | MM1 | MM2 | |
| CR1 | 0 | 262 | 250 | 124 | 124 | 129 | |
| CR2 | 0.0187 | 0 | 68 | 260 | 260 | 262 | |
| MA1 | 0.0179 | 0.0049 | 0 | 247 | 247 | 251 | |
| MA2 | 0.0089 | 0.0186 | 0.0176 | 0 | 0 | 149 | |
| MM1 | 0.0089 | 0.0186 | 0.0176 | 0.0000 | 0 | 149 | |
| MM2 | 0.0092 | 0.0187 | 0.0179 | 0.0106 | 0.0106 | 0 | |

| 473 | | | | | | | 7669 |
|---|---|---|---|---|---|---|---|
| | CR1 | CR2 | MA1 | MA2 | MM1 | MM2 | |
| CR1 | 0 | 102 | 101 | 6 | 99 | 96 | |
| CR2 | 0.0133 | 0 | 33 | 100 | 19 | 24 | |
| MA1 | 0.0132 | 0.0043 | 0 | 99 | 28 | 23 | |
| MA2 | 0.0008 | 0.0130 | 0.0129 | 0 | 97 | 94 | |
| MM1 | 0.0129 | 0.0025 | 0.0037 | 0.0126 | 0 | 9 | |
| MM2 | 0.0125 | 0.0031 | 0.0030 | 0.0123 | 0.0012 | 0 | |
| 475 | | | | | | | 8551 |
| | CR1 | CR2 | MA1 | MA2 | MM1 | MM2 | |
| CR1 | 0 | 100 | 34 | 178 | 180 | 189 | |
| CR2 | 0.0117 | 0 | 126 | 118 | 118 | 133 | |
| MA1 | 0.0040 | 0.0147 | 0 | 185 | 187 | 196 | |
| MA2 | 0.0208 | 0.0138 | 0.0216 | 0 | 6 | 63 | |
| MM1 | 0.0211 | 0.0138 | 0.0219 | 0.0007 | 0 | 67 | |
| MM2 | 0.0221 | 0.0156 | 0.0229 | 0.0074 | 0.0078 | 0 | |
| 476 | | | | | | | 12824 |
| | CR1 | CR2 | MA1 | MA2 | MM1 | MM2 | |
| CR1 | 0 | 239 | 154 | 241 | 174 | 167 | |
| CR2 | 0.0186 | 0 | 290 | 95 | 287 | 257 | |
| MA1 | 0.0120 | 0.0226 | 0 | 285 | 174 | 181 | |
| MA2 | 0.0188 | 0.0074 | 0.0222 | 0 | 280 | 250 | |
| MM1 | 0.0136 | 0.0224 | 0.0136 | 0.0218 | 0 | 156 | |
| MM2 | 0.0130 | 0.0200 | 0.0141 | 0.0195 | 0.0122 | 0 | |
| 477 | | | | | | | 20695 |
| | CR1 | CR2 | MA1 | MA2 | MM1 | MM2 | |
| CR1 | 0 | 284 | 132 | 261 | 139 | 128 | |
| CR2 | 0.0137 | 0 | 280 | 109 | 266 | 292 | |
| MA1 | 0.0064 | 0.0135 | 0 | 224 | 138 | 119 | |
| MA2 | 0.0126 | 0.0053 | 0.0108 | 0 | 259 | 247 | |
| MM1 | 0.0067 | 0.0129 | 0.0067 | 0.0125 | 0 | 155 | |
| MM2 | 0.0062 | 0.0141 | 0.0058 | 0.0119 | 0.0075 | 0 | |
| 478 | | | | | | | 19160 |
| | CR1 | CR2 | MA1 | MA2 | MM1 | MM2 | |
| CR1 | 0 | 401 | 394 | 158 | 394 | 393 | |
| CR2 | 0.0209 | 0 | 166 | 410 | 166 | 144 | |
| MA1 | 0.0206 | 0.0087 | 0 | 399 | 0 | 125 | |
| MA2 | 0.0082 | 0.0214 | 0.0208 | 0 | 399 | 400 | |
| MM1 | 0.0206 | 0.0087 | 0.0000 | 0.0208 | 0 | 125 | |
| MM2 | 0.0205 | 0.0075 | 0.0065 | 0.0209 | 0.0065 | 0 | |

| 480 | | | | | | | 7398 |
|---|---|---|---|---|---|---|---|
| | CR1 | CR2 | MA1 | MA2 | MM1 | MM2 | |
| CR1 | 0 | 75 | 20 | 76 | 18 | 20 | |
| CR2 | 0.0101 | 0 | 72 | 19 | 74 | 72 | |
| MA1 | 0.0027 | 0.0097 | 0 | 73 | 26 | 0 | |
| MA2 | 0.0103 | 0.0026 | 0.0099 | 0 | 74 | 73 | |
| MM1 | 0.0024 | 0.0100 | 0.0035 | 0.0100 | 0 | 26 | |
| MM2 | 0.0027 | 0.0097 | 0.0000 | 0.0099 | 0.0035 | 0 | |
| 481 | | | | | | | 11860 |
| | CR1 | CR2 | MA1 | MA2 | MM1 | MM2 | |
| CR1 | 0 | 149 | 348 | 206 | 373 | 348 | |
| CR2 | 0.0126 | 0 | 298 | 330 | 320 | 298 | |
| MA1 | 0.0293 | 0.0251 | 0 | 416 | 244 | 0 | |
| MA2 | 0.0174 | 0.0278 | 0.0351 | 0 | 438 | 416 | |
| MM1 | 0.0315 | 0.0270 | 0.0206 | 0.0369 | 0 | 244 | |
| MM2 | 0.0293 | 0.0251 | 0.0000 | 0.0351 | 0.0206 | 0 | |
| 482 | | | | | | | 20094 |
| | CR1 | CR2 | MA1 | MA2 | MM1 | MM2 | |
| CR1 | 0 | 372 | 326 | 383 | 326 | 305 | |
| CR2 | 0.0185 | 0 | 444 | 206 | 449 | 444 | |
| MA1 | 0.0162 | 0.0221 | 0 | 439 | 221 | 33 | |
| MA2 | 0.0191 | 0.0103 | 0.0218 | 0 | 438 | 439 | |
| MM1 | 0.0162 | 0.0223 | 0.0110 | 0.0218 | 0 | 188 | |
| MM2 | 0.0152 | 0.0221 | 0.0016 | 0.0218 | 0.0094 | 0 | |
| 483 | | | | | | | 9877 |
| | CR1 | CR2 | MA1 | MA2 | MM1 | MM2 | |
| CR1 | 0 | 140 | 159 | 92 | 166 | 159 | |
| CR2 | 0.0142 | 0 | 81 | 188 | 92 | 81 | |
| MA1 | 0.0161 | 0.0082 | 0 | 203 | 87 | 0 | |
| MA2 | 0.0093 | 0.0190 | 0.0206 | 0 | 206 | 203 | |
| MM1 | 0.0168 | 0.0093 | 0.0088 | 0.0209 | 0 | 87 | |
| MM2 | 0.0161 | 0.0082 | 0.0000 | 0.0206 | 0.0088 | 0 | |
| 484 | | | | | | | 12197 |
| | CR1 | CR2 | MA1 | MA2 | MM1 | MM2 | |
| CR1 | 0 | 128 | 36 | 124 | 36 | 32 | |
| CR2 | 0.0105 | 0 | 129 | 32 | 129 | 127 | |
| MA1 | 0.0030 | 0.0106 | 0 | 125 | 0 | 40 | |
| MA2 | 0.0102 | 0.0026 | 0.0102 | 0 | 125 | 125 | |
| MM1 | 0.0030 | 0.0106 | 0.0000 | 0.0102 | 0 | 40 | |
| MM2 | 0.0026 | 0.0104 | 0.0033 | 0.0102 | 0.0033 | 0 | |

| 485 | | | | | | 10128 |
|---|---|---|---|---|---|---|
| | CR1 | CR2 | MA1 | MA2 | MM1 | MM2 |
| CR1 | 0 | 184 | 95 | 187 | 199 | 187 |
| CR2 | 0.0182 | 0 | 193 | 78 | 90 | 78 |
| MA1 | 0.0094 | 0.0191 | 0 | 198 | 203 | 198 |
| MA2 | 0.0185 | 0.0077 | 0.0195 | 0 | 106 | 0 |
| MM1 | 0.0196 | 0.0089 | 0.0200 | 0.0105 | 0 | 106 |
| MM2 | 0.0185 | 0.0077 | 0.0195 | 0.0000 | 0.0105 | 0 |

| 486 | | | | | | 14553 |
|---|---|---|---|---|---|---|
| | CR1 | CR2 | MA1 | MA2 | MM1 | MM2 |
| CR1 | 0 | 265 | 322 | 199 | 325 | 312 |
| CR2 | 0.0182 | 0 | 166 | 358 | 179 | 177 |
| MA1 | 0.0221 | 0.0114 | 0 | 363 | 195 | 201 |
| MA2 | 0.0137 | 0.0246 | 0.0249 | 0 | 340 | 345 |
| MM1 | 0.0223 | 0.0123 | 0.0134 | 0.0234 | 0 | 186 |
| MM2 | 0.0214 | 0.0122 | 0.0138 | 0.0237 | 0.0128 | 0 |

| 487 | | | | | | 12026 |
|---|---|---|---|---|---|---|
| | CR1 | CR2 | MA1 | MA2 | MM1 | MM2 |
| CR1 | 0 | 310 | 227 | 256 | 148 | 190 |
| CR2 | 0.0258 | 0 | 256 | 212 | 303 | 308 |
| MA1 | 0.0189 | 0.0213 | 0 | 58 | 221 | 206 |
| MA2 | 0.0213 | 0.0176 | 0.0048 | 0 | 258 | 243 |
| MM1 | 0.0123 | 0.0252 | 0.0184 | 0.0215 | 0 | 184 |
| MM2 | 0.0158 | 0.0256 | 0.0171 | 0.0202 | 0.0153 | 0 |

| 488 | | | | | | 16102 |
|---|---|---|---|---|---|---|
| | CR1 | CR2 | MA1 | MA2 | MM1 | MM2 |
| CR1 | 0 | 331 | 185 | 316 | 185 | 188 |
| CR2 | 0.0206 | 0 | 303 | 111 | 303 | 306 |
| MA1 | 0.0115 | 0.0188 | 0 | 271 | 0 | 3 |
| MA2 | 0.0196 | 0.0069 | 0.0168 | 0 | 271 | 274 |
| MM1 | 0.0115 | 0.0188 | 0.0000 | 0.0168 | 0 | 3 |
| MM2 | 0.0117 | 0.0190 | 0.0002 | 0.0170 | 0.0002 | 0 |

| 491 | | | | | | 10487 |
|---|---|---|---|---|---|---|
| | CR1 | CR2 | MA1 | MA2 | MM1 | MM2 |
| CR1 | 0 | 246 | 220 | 107 | 248 | 253 |
| CR2 | 0.0235 | 0 | 127 | 235 | 98 | 112 |
| MA1 | 0.0210 | 0.0121 | 0 | 209 | 29 | 117 |
| MA2 | 0.0102 | 0.0224 | 0.0199 | 0 | 237 | 238 |
| MM1 | 0.0236 | 0.0093 | 0.0028 | 0.0226 | 0 | 89 |
| MM2 | 0.0241 | 0.0107 | 0.0112 | 0.0227 | 0.0085 | 0 |

| 492 | | | | | | | 12510 |
|---|---|---|---|---|---|---|---|
| | CR1 | CR2 | MA1 | MA2 | MM1 | MM2 | |
| CR1 | 0 | 290 | 169 | 276 | 163 | 179 | |
| CR2 | 0.0232 | 0 | 285 | 85 | 290 | 279 | |
| MA1 | 0.0135 | 0.0228 | 0 | 275 | 128 | 169 | |
| MA2 | 0.0221 | 0.0068 | 0.0220 | 0 | 281 | 274 | |
| MM1 | 0.0130 | 0.0232 | 0.0102 | 0.0225 | 0 | 208 | |
| MM2 | 0.0143 | 0.0223 | 0.0135 | 0.0219 | 0.0166 | 0 | |
| 493 | | | | | | | 19008 |
| | CR1 | CR2 | MA1 | MA2 | MM1 | MM2 | |
| CR1 | 0 | 135 | 102 | 143 | 164 | 102 | |
| CR2 | 0.0071 | 0 | 217 | 68 | 218 | 217 | |
| MA1 | 0.0054 | 0.0114 | 0 | 233 | 135 | 0 | |
| MA2 | 0.0075 | 0.0036 | 0.0123 | 0 | 232 | 233 | |
| MM1 | 0.0086 | 0.0115 | 0.0071 | 0.0122 | 0 | 135 | |
| MM2 | 0.0054 | 0.0114 | 0.0000 | 0.0123 | 0.0071 | 0 | |
| 494 | | | | | | | 11484 |
| | CR1 | CR2 | MA1 | MA2 | MM1 | MM2 | |
| CR1 | 0 | 263 | 119 | 214 | 119 | 155 | |
| CR2 | 0.0229 | 0 | 272 | 164 | 272 | 276 | |
| MA1 | 0.0104 | 0.0237 | 0 | 169 | 0 | 141 | |
| MA2 | 0.0186 | 0.0143 | 0.0147 | 0 | 169 | 205 | |
| MM1 | 0.0104 | 0.0237 | 0.0000 | 0.0147 | 0 | 141 | |
| MM2 | 0.0135 | 0.0240 | 0.0123 | 0.0179 | 0.0123 | 0 | |
| 496 | | | | | | | 17913 |
| | CR1 | CR2 | MA1 | MA2 | MM1 | MM2 | |
| CR1 | 0 | 77 | 238 | 427 | 434 | 457 | |
| CR2 | 0.0043 | 0 | 207 | 470 | 458 | 485 | |
| MA1 | 0.0133 | 0.0116 | 0 | 478 | 455 | 488 | |
| MA2 | 0.0238 | 0.0262 | 0.0267 | 0 | 282 | 248 | |
| MM1 | 0.0242 | 0.0256 | 0.0254 | 0.0157 | 0 | 272 | |
| MM2 | 0.0255 | 0.0271 | 0.0272 | 0.0138 | 0.0152 | 0 | |
| 500 | | | | | | | 10444 |
| | CR1 | CR2 | MA1 | MA2 | MM1 | MM2 | |
| CR1 | 0 | 0 | 158 | 211 | 158 | 102 | |
| CR2 | 0.0000 | 0 | 158 | 211 | 158 | 102 | |
| MA1 | 0.0151 | 0.0151 | 0 | 222 | 0 | 147 | |
| MA2 | 0.0202 | 0.0202 | 0.0213 | 0 | 222 | 209 | |
| MM1 | 0.0151 | 0.0151 | 0.0000 | 0.0213 | 0 | 147 | |
| MM2 | 0.0098 | 0.0098 | 0.0141 | 0.0200 | 0.0141 | 0 | |

| 501 | | | | | | | 13769 |
|---|---|---|---|---|---|---|---|
| | CR1 | CR2 | MA1 | MA2 | MM1 | MM2 | |
| CR1 | 0 | 324 | 132 | 329 | 102 | 134 | |
| CR2 | 0.0235 | 0 | 332 | 73 | 314 | 327 | |
| MA1 | 0.0096 | 0.0241 | 0 | 337 | 126 | 142 | |
| MA2 | 0.0239 | 0.0053 | 0.0245 | 0 | 321 | 332 | |
| MM1 | 0.0074 | 0.0228 | 0.0092 | 0.0233 | 0 | 126 | |
| MM2 | 0.0097 | 0.0237 | 0.0103 | 0.0241 | 0.0092 | 0 | |
| 502 | | | | | | | 12465 |
| | CR1 | CR2 | MA1 | MA2 | MM1 | MM2 | |
| CR1 | 0 | 211 | 214 | 49 | 211 | 214 | |
| CR2 | 0.0169 | 0 | 95 | 205 | 66 | 95 | |
| MA1 | 0.0172 | 0.0076 | 0 | 208 | 105 | 0 | |
| MA2 | 0.0039 | 0.0164 | 0.0167 | 0 | 205 | 208 | |
| MM1 | 0.0169 | 0.0053 | 0.0084 | 0.0164 | 0 | 105 | |
| MM2 | 0.0172 | 0.0076 | 0.0000 | 0.0167 | 0.0084 | 0 | |
| 503 | | | | | | | 13598 |
| | CR1 | CR2 | MA1 | MA2 | MM1 | MM2 | |
| CR1 | 0 | 254 | 12 | 165 | 12 | 96 | |
| CR2 | 0.0187 | 0 | 250 | 125 | 250 | 266 | |
| MA1 | 0.0009 | 0.0184 | 0 | 161 | 0 | 96 | |
| MA2 | 0.0121 | 0.0092 | 0.0118 | 0 | 161 | 205 | |
| MM1 | 0.0009 | 0.0184 | 0.0000 | 0.0118 | 0 | 96 | |
| MM2 | 0.0071 | 0.0196 | 0.0071 | 0.0151 | 0.0071 | 0 | |
| 505 | | | | | | | 16410 |
| | CR1 | CR2 | MA1 | MA2 | MM1 | MM2 | |
| CR1 | 0 | 283 | 261 | 82 | 261 | 292 | |
| CR2 | 0.0172 | 0 | 144 | 288 | 144 | 129 | |
| MA1 | 0.0159 | 0.0088 | 0 | 266 | 0 | 129 | |
| MA2 | 0.0050 | 0.0176 | 0.0162 | 0 | 266 | 294 | |
| MM1 | 0.0159 | 0.0088 | 0.0000 | 0.0162 | 0 | 129 | |
| MM2 | 0.0178 | 0.0079 | 0.0079 | 0.0179 | 0.0079 | 0 | |
| 507 | | | | | | | 12734 |
| | CR1 | CR2 | MA1 | MA2 | MM1 | MM2 | |
| CR1 | 0 | 1 | 237 | 237 | 252 | 218 | |
| CR2 | 0.0001 | 0 | 238 | 238 | 251 | 219 | |
| MA1 | 0.0186 | 0.0187 | 0 | 0 | 227 | 216 | |
| MA2 | 0.0186 | 0.0187 | 0.0000 | 0 | 227 | 216 | |
| MM1 | 0.0198 | 0.0197 | 0.0178 | 0.0178 | 0 | 235 | |
| MM2 | 0.0171 | 0.0172 | 0.0170 | 0.0170 | 0.0185 | 0 | |

| 510 | | | | | | | 15156 |
|---|---|---|---|---|---|---|---|
| | CR1 | CR2 | MA1 | MA2 | MM1 | MM2 | |
| CR1 | 0 | 375 | 211 | 368 | 154 | 238 | |
| CR2 | 0.0247 | 0 | 376 | 45 | 370 | 381 | |
| MA1 | 0.0139 | 0.0248 | 0 | 369 | 139 | 177 | |
| MA2 | 0.0243 | 0.0030 | 0.0243 | 0 | 365 | 376 | |
| MM1 | 0.0102 | 0.0244 | 0.0092 | 0.0241 | 0 | 212 | |
| MM2 | 0.0157 | 0.0251 | 0.0117 | 0.0248 | 0.0140 | 0 | |

| 511 | | | | | | | 15769 |
|---|---|---|---|---|---|---|---|
| | CR1 | CR2 | MA1 | MA2 | MM1 | MM2 | |
| CR1 | 0 | 184 | 64 | 189 | 63 | 56 | |
| CR2 | 0.0117 | 0 | 174 | 19 | 187 | 178 | |
| MA1 | 0.0041 | 0.0110 | 0 | 179 | 51 | 60 | |
| MA2 | 0.0120 | 0.0012 | 0.0114 | 0 | 192 | 183 | |
| MM1 | 0.0040 | 0.0119 | 0.0032 | 0.0122 | 0 | 59 | |
| MM2 | 0.0036 | 0.0113 | 0.0038 | 0.0116 | 0.0037 | 0 | |

| 512 | | | | | | | 17190 |
|---|---|---|---|---|---|---|---|
| | CR1 | CR2 | MA1 | MA2 | MM1 | MM2 | |
| CR1 | 0 | 356 | 280 | 295 | 201 | 210 | |
| CR2 | 0.0207 | 0 | 329 | 219 | 375 | 362 | |
| MA1 | 0.0163 | 0.0191 | 0 | 203 | 272 | 109 | |
| MA2 | 0.0172 | 0.0127 | 0.0118 | 0 | 306 | 202 | |
| MM1 | 0.0117 | 0.0218 | 0.0158 | 0.0178 | 0 | 195 | |
| MM2 | 0.0122 | 0.0211 | 0.0063 | 0.0118 | 0.0113 | 0 | |

| 515 | | | | | | | 11212 |
|---|---|---|---|---|---|---|---|
| | CR1 | CR2 | MA1 | MA2 | MM1 | MM2 | |
| CR1 | 0 | 258 | 76 | 255 | 153 | 149 | |
| CR2 | 0.0230 | 0 | 261 | 74 | 290 | 269 | |
| MA1 | 0.0068 | 0.0233 | 0 | 259 | 161 | 163 | |
| MA2 | 0.0227 | 0.0066 | 0.0231 | 0 | 295 | 264 | |
| MM1 | 0.0136 | 0.0259 | 0.0144 | 0.0263 | 0 | 172 | |
| MM2 | 0.0133 | 0.0240 | 0.0145 | 0.0235 | 0.0153 | 0 | |

| 516 | | | | | | | 14612 |
|---|---|---|---|---|---|---|---|
| | CR1 | CR2 | MA1 | MA2 | MM1 | MM2 | |
| CR1 | 0 | 128 | 55 | 114 | 56 | 52 | |
| CR2 | 0.0088 | 0 | 132 | 40 | 133 | 138 | |
| MA1 | 0.0038 | 0.0090 | 0 | 118 | 1 | 59 | |
| MA2 | 0.0078 | 0.0027 | 0.0081 | 0 | 119 | 124 | |
| MM1 | 0.0038 | 0.0091 | 0.0001 | 0.0081 | 0 | 58 | |
| MM2 | 0.0036 | 0.0094 | 0.0040 | 0.0085 | 0.0040 | 0 | |

| 517 | | | | | | | 11185 |
|---|---|---|---|---|---|---|---|
| | CR1 | CR2 | MA1 | MA2 | MM1 | MM2 | |
| CR1 | 0 | 248 | 118 | 240 | 0 | 132 | |
| CR2 | 0.0222 | 0 | 249 | 87 | 248 | 255 | |
| MA1 | 0.0105 | 0.0223 | 0 | 240 | 118 | 114 | |
| MA2 | 0.0215 | 0.0078 | 0.0215 | 0 | 240 | 244 | |
| MM1 | 0.0000 | 0.0222 | 0.0105 | 0.0215 | 0 | 132 | |
| MM2 | 0.0118 | 0.0228 | 0.0102 | 0.0218 | 0.0118 | 0 | |
| 519 | | | | | | | 13392 |
| | CR1 | CR2 | MA1 | MA2 | MM1 | MM2 | |
| CR1 | 0 | 271 | 228 | 177 | 172 | 52 | |
| CR2 | 0.0202 | 0 | 185 | 282 | 283 | 264 | |
| MA1 | 0.0170 | 0.0138 | 0 | 132 | 212 | 200 | |
| MA2 | 0.0132 | 0.0211 | 0.0099 | 0 | 167 | 159 | |
| MM1 | 0.0128 | 0.0211 | 0.0158 | 0.0125 | 0 | 153 | |
| MM2 | 0.0039 | 0.0197 | 0.0149 | 0.0119 | 0.0114 | 0 | |
| 520 | | | | | | | 19499 |
| | CR1 | CR2 | MA1 | MA2 | MM1 | MM2 | |
| CR1 | 0 | 445 | 186 | 445 | 186 | 361 | |
| CR2 | 0.0228 | 0 | 421 | 0 | 421 | 499 | |
| MA1 | 0.0095 | 0.0216 | 0 | 421 | 0 | 327 | |
| MA2 | 0.0228 | 0.0000 | 0.0216 | 0 | 421 | 499 | |
| MM1 | 0.0095 | 0.0216 | 0.0000 | 0.0216 | 0 | 327 | |
| MM2 | 0.0185 | 0.0256 | 0.0168 | 0.0256 | 0.0168 | 0 | |
| 521 | | | | | | | 12330 |
| | CR1 | CR2 | MA1 | MA2 | MM1 | MM2 | |
| CR1 | 0 | 175 | 173 | 52 | 174 | 173 | |
| CR2 | 0.0142 | 0 | 79 | 177 | 1 | 79 | |
| MA1 | 0.0140 | 0.0064 | 0 | 176 | 78 | 0 | |
| MA2 | 0.0042 | 0.0144 | 0.0143 | 0 | 176 | 176 | |
| MM1 | 0.0141 | 0.0001 | 0.0063 | 0.0143 | 0 | 78 | |
| MM2 | 0.0140 | 0.0064 | 0.0000 | 0.0143 | 0.0063 | 0 | |
| 522 | | | | | | | 13020 |
| | CR1 | CR2 | MA1 | MA2 | MM1 | MM2 | |
| CR1 | 0 | 164 | 75 | 163 | 75 | 68 | |
| CR2 | 0.0126 | 0 | 155 | 19 | 155 | 167 | |
| MA1 | 0.0058 | 0.0119 | 0 | 154 | 0 | 49 | |
| MA2 | 0.0125 | 0.0015 | 0.0118 | 0 | 154 | 166 | |
| MM1 | 0.0058 | 0.0119 | 0.0000 | 0.0118 | 0 | 49 | |
| MM2 | 0.0052 | 0.0128 | 0.0038 | 0.0127 | 0.0038 | 0 | |

| 524 | | | | | | | 14371 |
|---|---|---|---|---|---|---|---|
| | CR1 | CR2 | MA1 | MA2 | MM1 | MM2 | |
| CR1 | 0 | 213 | 124 | 205 | 59 | 124 | |
| CR2 | 0.0148 | 0 | 212 | 34 | 225 | 212 | |
| MA1 | 0.0086 | 0.0148 | 0 | 204 | 118 | 0 | |
| MA2 | 0.0143 | 0.0024 | 0.0142 | 0 | 217 | 204 | |
| MM1 | 0.0041 | 0.0157 | 0.0082 | 0.0151 | 0 | 118 | |
| MM2 | 0.0086 | 0.0148 | 0.0000 | 0.0142 | 0.0082 | 0 | |
| 525 | | | | | | | 17359 |
| | CR1 | CR2 | MA1 | MA2 | MM1 | MM2 | |
| CR1 | 0 | 135 | 421 | 193 | 202 | 158 | |
| CR2 | 0.0078 | 0 | 337 | 276 | 285 | 264 | |
| MA1 | 0.0243 | 0.0194 | 0 | 403 | 410 | 413 | |
| MA2 | 0.0111 | 0.0159 | 0.0232 | 0 | 189 | 208 | |
| MM1 | 0.0116 | 0.0164 | 0.0236 | 0.0109 | 0 | 176 | |
| MM2 | 0.0091 | 0.0152 | 0.0238 | 0.0120 | 0.0101 | 0 | |
| 526 | | | | | | | 9808 |
| | CR1 | CR2 | MA1 | MA2 | MM1 | MM2 | |
| CR1 | 0 | 78 | 93 | 64 | 64 | 46 | |
| CR2 | 0.0080 | 0 | 55 | 98 | 98 | 78 | |
| MA1 | 0.0095 | 0.0056 | 0 | 107 | 107 | 89 | |
| MA2 | 0.0065 | 0.0100 | 0.0109 | 0 | 0 | 60 | |
| MM1 | 0.0065 | 0.0100 | 0.0109 | 0.0000 | 0 | 60 | |
| MM2 | 0.0047 | 0.0080 | 0.0091 | 0.0061 | 0.0061 | 0 | |
| 527 | | | | | | | 9752 |
| | CR1 | CR2 | MA1 | MA2 | MM1 | MM2 | |
| CR1 | 0 | 234 | 219 | 141 | 144 | 123 | |
| CR2 | 0.0240 | 0 | 88 | 246 | 253 | 241 | |
| MA1 | 0.0225 | 0.0090 | 0 | 236 | 242 | 227 | |
| MA2 | 0.0145 | 0.0252 | 0.0242 | 0 | 142 | 151 | |
| MM1 | 0.0148 | 0.0259 | 0.0248 | 0.0146 | 0 | 129 | |
| MM2 | 0.0126 | 0.0247 | 0.0233 | 0.0155 | 0.0132 | 0 | |
| 529 | | | | | | | 12887 |
| | CR1 | CR2 | MA1 | MA2 | MM1 | MM2 | |
| CR1 | 0 | 258 | 278 | 81 | 274 | 277 | |
| CR2 | 0.0200 | 0 | 157 | 252 | 174 | 167 | |
| MA1 | 0.0216 | 0.0122 | 0 | 271 | 151 | 183 | |
| MA2 | 0.0063 | 0.0196 | 0.0210 | 0 | 268 | 271 | |
| MM1 | 0.0213 | 0.0135 | 0.0117 | 0.0208 | 0 | 151 | |
| MM2 | 0.0215 | 0.0130 | 0.0142 | 0.0210 | 0.0117 | 0 | |

| 530 | | | | | | | 13427 |
|---|---|---|---|---|---|---|---|
| | CR1 | CR2 | MA1 | MA2 | MM1 | MM2 | |
| CR1 | 0 | 190 | 65 | 213 | 202 | 211 | |
| CR2 | 0.0142 | 0 | 193 | 75 | 77 | 86 | |
| MA1 | 0.0048 | 0.0144 | 0 | 216 | 202 | 205 | |
| MA2 | 0.0159 | 0.0056 | 0.0161 | 0 | 98 | 104 | |
| MM1 | 0.0150 | 0.0057 | 0.0150 | 0.0073 | 0 | 33 | |
| MM2 | 0.0157 | 0.0064 | 0.0153 | 0.0077 | 0.0025 | 0 | |
